# A single DPE core promoter motif contributes to *in vivo* transcriptional regulation and affects cardiac function

**DOI:** 10.1101/2023.06.11.544490

**Authors:** Anna Sloutskin, Dekel Itzhak, Georg Vogler, Diana Ideses, Hadar Alter, Hadar Shachar, Tirza Doniger, Manfred Frasch, Rolf Bodmer, Sascha H Duttke, Tamar Juven-Gershon

**Affiliations:** The Mina and Everard Goodman Faculty of Life Sciences, Bar-Ilan University, Ramat Gan, Israel; Development, Aging and Regeneration Program, Sanford Burnham Prebys Medical Discovery Institute, La Jolla, CA, USA; Division of Developmental Biology, Department of Biology, Friedrich-Alexander University of Erlangen-Nürnberg, Erlangen, Germany; School of Molecular Biosciences, College of Veterinary Medicine, Washington State University, Pullman, WA, USA

**Keywords:** Core Promoter, RNA Polymerase II transcription, DPE, Gene Expression, nascent transcription, *Drosophila*, heart development

## Abstract

Transcription is initiated at the core promoter, which confers specific functions depending on the unique combination of core promoter elements. The downstream core promoter element (DPE) is found in many genes related to heart and mesodermal development. However, the function of these core promoter elements has thus far been studied primarily in isolated, *in vitro* or reporter gene settings. *tinman* (*tin*) encodes a key transcription factor that regulates the formation of the dorsal musculature and heart. Pioneering a novel approach utilizing both CRISPR and nascent transcriptomics, we show that a substitution mutation of the functional *tin* DPE motif within the natural context of the core promoter results in a massive perturbation of Tinman’s regulatory network orchestrating dorsal musculature and heart formation. Mutation of endogenous *tin* DPE reduced the expression of *tin* and distinct target genes, resulting in significantly reduced viability and an overall decrease in adult heart function. We demonstrate the feasibility and importance of characterizing DNA sequence elements *in vivo* in their natural context, and accentuate the critical impact a single DPE motif has during *Drosophila* embryogenesis and functional heart formation.

## Introduction

Transcription initiation by RNA Polymerase II (Pol II) occurs at the core promoter region (–40 to +40 relative to the transcription start site (TSS)), which is often referred to as the “gateway to transcription” (Heintzman and Ren 2007; Juven-Gershon et al. 2008b). Although the core promoter was previously regarded as a universal component that functions by a similar mechanism for all the protein-coding genes, it is nowadays appreciated that core promoters are divergent in their composition and function, with a growing body of evidence indicating that each individual core promoter is rather unique. Core promoters may contain one or more short DNA sequence motifs, termed core promoter elements or motifs, which confer specific properties to the core promoter (Goldberg 1979; Smale and Baltimore 1989; Burke and Kadonaga 1996; Lo and Smale 1996; Burke and Kadonaga 1997; Lagrange et al. 1998; Kutach and Kadonaga 2000; Ohler et al. 2002; Lim et al. 2004; Deng and Roberts 2005; Tokusumi et al. 2007; Hendrix et al. 2008; Anish et al. 2009; Parry et al. 2010; Theisen et al. 2010; Vo Ngoc et al. 2017; Wang et al. 2017; Vo Ngoc et al. 2020). Interestingly, distinct core promoter compositions were demonstrated to result in various transcriptional outputs and to be associated with specific gene regulatory networks (Danino et al. 2015; Haberle and Stark 2018; Vo Ngoc et al. 2019; Sloutskin et al. 2021).

The downstream core promoter element (DPE) is enriched in the promoters of developmentally-regulated genes, including most homeotic (*Hox*) genes (Juven-Gershon et al. 2008a) and those regulating dorsal-ventral patterning (Zehavi et al. 2014a; Zehavi et al. 2014b). Interestingly, the DPE is also found in the promoters of many genes involved in heart and mesodermal development (Sloutskin et al. 2015), including *tinman* (*tin*). *tinman* encodes an extensively studied transcription factor that orchestrates the formation of the heart and heart-associated tissues during *Drosophila* embryonic development (Azpiazu and Frasch 1993; Bodmer 1993). Tinman plays a key role in early mesoderm patterning and is essential for the formation of all dorsal mesodermal derivatives, which in addition to working cardioblasts, valve cardioblasts and pericardial cells, include visceral and specific somatic muscles (Cripps and Olson 2002; Zaffran et al. 2006; Bryantsev and Cripps 2009; Reim and Frasch 2010; Rotstein and Paululat 2016). Genome-wide approaches have identified Tinman binding sites, which led to the discovery of additional Tinman-responsive enhancers and Tinman target genes (Liu et al. 2009; Jin et al. 2013).

Substitution mutations of the DPE element in isolated *tinman* core promoters has previously been shown to significantly reduce transcriptional output in reporter transfection assays and *in vitro* transcription analysis with embryonic extracts (Zehavi et al. 2014a). However, in the genome, mutations in regulatory elements are often buffered, in part due to a diversity of sequence-specific transcription factors and the functional redundancy of regulatory motifs (Jin et al. 2013; Spivakov 2014; Osterwalder et al. 2018). To explore a possible function of the DPE motif in regulating heart and mesodermal development, as well as to promote the study of core promoter elements within their native genomic context *in vivo*, we mutated the DPE motif of the *tin* core promoter (*tin^mDPE^*) using a CRISPR-based strategy (Levi et al. 2020). We found that mutation of the DPE motif is sufficient to reduce *tin* expression, at both the RNA and protein levels, with no accompanied changes detected in *tin* expression patterns. Although the dorsal vessel is formed in *tin^mDPE^* homozygous embryos, both alleles are required for survival, with one copy of *tin^mDPE^* being unable to fully compensate for the loss of a *tinman* allele. Importantly, major defects in adult heart physiology were observed. Nascent transcription analysis of *tin^WT^* and *tin^mDPE^* embryos detected differential expression of *tin* target genes, many of which are implicated in heart development and tube formation. Moreover, DPE-like motifs are significantly enriched among the differentially regulated peaks.

Altogether, our results demonstrate the feasibility and importance of studying core promoter elements in their native genomic context, demonstrate the function of the DPE motif of *tin* in dorsal vessel specification in *Drosophila*, and highlight the importance a single core promoter element can have in development, viability and functional heart formation.

## Results

### Reduced expression levels of endogenous *tinman* in the mDPE strains

To investigate the contribution of the DPE motif to the regulation of the *tin* gene *in vivo*, we utilized the co-CRISPR approach to substitute the endogenous DPE sequence (AGACACG) with CTCATGT ((Levi et al. 2020), Fig. 1A). Using *in vitro* transcription and reporter gene analysis, this 7bp mutation was previously shown to reduce expression of DPE-containing promoters, including *tin* (Burke and Kadonaga 1997; Kutach and Kadonaga 2000; Zehavi et al. 2014a). Two independent *tin^mDPE^ Drosophila melanogaster* strains, namely F3 and M6, were extensively characterized in this study, and compared to the injected strain, Cas9, which is referred to as *tin^WT^*. Quantification of endogenous *tin* RNA levels in *tin^WT^* and *tin^mDPE^* embryos within 1h windows during the first 8 hours of embryonic development (up to Bownes developmental stage 12 embryos, Fig. 1B) revealed a marked reduction of endogenous *tin* expression levels in *tin^mDPE^* embryos. Differences in *tin* expression levels were evident starting from the earliest tested timepoint (0-1h), and were most substantial at 3-4h, when Tinman activity becomes critical for mesoderm development (Yin et al. 1997; Zaffran et al. 2006). Later in development (6-7h and 7-8h, stage 11-12), *tin* levels were indistinguishable between *tin^mDPE^* and *tin^WT^* embryos. Despite differences in *tin* expression levels at 0-6h, no apparent differences in the *tin* expression pattern were detected in early *tin^mDPE^* and *tin^WT^*embryos by *in situ* hybridization (Supplemental Fig. S1). Both *tin^mDPE^*strains and *tin^WT^* exhibit *tin* expression pattern that highly matches the reported one (Bodmer et al. 1990; Azpiazu and Frasch 1993; Bodmer 1993).

**Figure 1.**
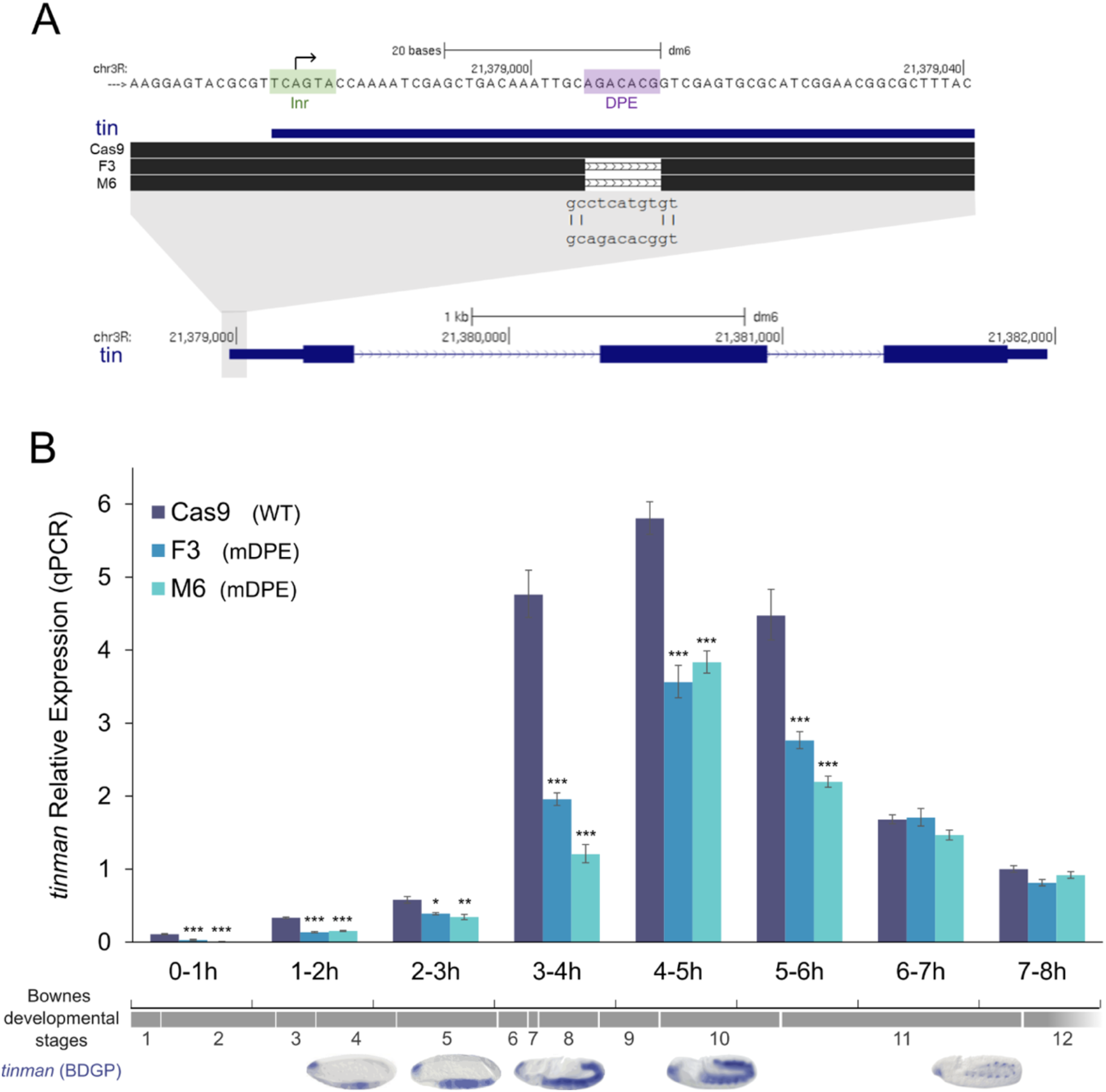
Endogenous *tinman* RNA levels are reduced in *tin^mDPE^ Drosophila melanogaster* embryos. (*A*) Graphical summary of the *tinman* core promoter and gene. The Inr and DPE motifs are annotated, along with the motif sequences in the generated flies. Cas9 - WT strain; F3, M6 - mDPE *tinman* strains. Note the different scales for the top and bottom parts. (*B*) Cas9, F3, and M6 embryos were collected at 1h intervals during the first 8h of development. RNA was purified and reverse transcribed. Endogenous *tinman* expression levels were measured by RT-qPCR and analyzed using the StepOnePlus software. Each qPCR experiment was performed in triplicates. Error bars represent 95% confidence interval, n≥3 for each timepoint. *p<0.05, **p<0.01, ***p<0.001 one-way ANOVA followed by Tukey’s post hoc test. Only comparisons to the Cas9 samples are presented. Bownes developmental stages (Interactive Fly) and the expected *tinman* expression patterns are indicated below the relevant time points (*in-situ* hybridization patterns obtained with permission from Berkeley *Drosophila* Genome Project, https://insitu.fruitfly.org/cgi-bin/ex/insitu.pl) (Tomancak et al. 2007).

To examine whether the DPE mutation impacts Tinman protein levels, we performed western blot analysis. While at 2-4h Tinman protein levels were below detectability, 4-6h and 8-10h *tin^mDPE^* embryos showed about a 2-fold decrease compared to *tin^WT^* (Fig. 2A, B, Supplemental Fig. S2). Staining the embryos with antibodies against Tinman revealed a normal staining pattern in the dorsal mesoderm for both *tin^mDPE^* and *tin^WT^* embryos (Fig. 2C), recapitulating the observations at the RNA level. Thus, mutation of the endogenous *tin* DPE results in reduced *tin* RNA and protein expression levels, whereas the spatial localization of *tin* RNA and protein are not affected.

**Figure 2.**
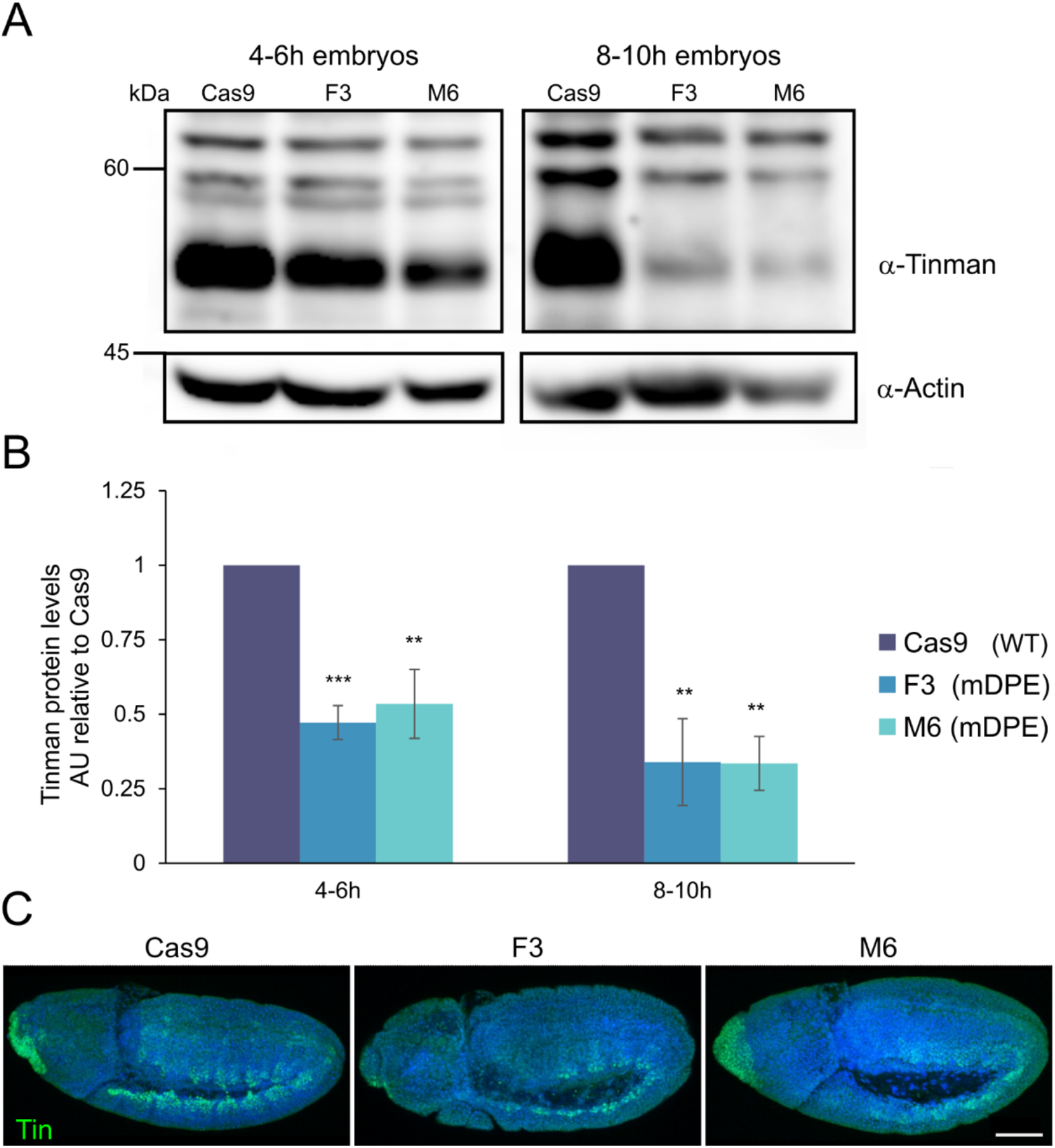
Tinman protein expression levels are reduced in *tinman* mDPE embryos, whereas the Tinman expression pattern is retained. (*A*) Representative western blot images of embryos collected at 4-6h or 8-10h time intervals. (*B*) Quantification of Tinman protein levels, n=4. AU - arbitrary units. Error bars represent the SEM. **p ≤ 0.01, ***p < 0.001, two-tailed Students t-test; comparison to Cas9. (*C*) Embryos at a developmental window of 4-6h (Bownes stage ∼10) were stained using anti-Tinman antibodies (green) and counterstained with a nuclear dye (Hoechst, blue). The observed expression pattern is similar between *tin^mDPE^* and *tin^WT^* embryos. Z-stack maximal projections are shown; anterior to the left, dorsal side up. Scale bar = 50µm.

### Functional consequences of reduced Tinman expression

*tinman* encodes a homeodomain transcription factor and is a master regulator of mesoderm and heart development (Bryantsev and Cripps 2009). We therefore tested the endogenous expression levels of the following Tinman target genes in 4-6h *tin^WT^* and *tin^mDPE^*embryos (stages 8-10)- *Dorsocross 2* (*Doc2)*, *seven up* (*svp*), *Myocyte enhancer factor 2* (*Mef2*) and *even skipped* (*eve*), along with *tin* expression (Fig. 3A). *Doc2* and *svp* are two genes required to generate the developing heart that are regulated by Tinman (Lo and Frasch 2003; Reim and Frasch 2005; Ryan et al. 2007). Indeed, both *Doc2* and *svp* are downregulated in *tin^mDPE^* strains at the 4-6h timepoint. Mef2 and Eve were shown to be perturbed in classical *tin* knockouts (Azpiazu and Frasch 1993; Bodmer 1993; Gajewski et al. 1997). Although *Mef2* and *eve* RNA levels were slightly elevated in early *tin^mDPE^* embryos (stage 8-10), the formation of an apparently normal dorsal vessel in late *tin^mDPE^* embryos (stage ≥13) was evident, based on either Mef2 or Eve protein localization (Fig. 3B,C). The use of LacZ combined with Odd-skipped (Odd), a key marker of pericardial cells (Ward and Skeath 2000), enabled the distinction between homozygous and heterozygous *tin^null^* embryos (Supplemental Fig. S3). *tin* null homozygotes display no overall Tinman staining and dorsal-vessel specific Odd staining, while in *tin* null heterozygotes a dorsal vessel is detected by the presence of Tinman and Odd staining. Misexpression of Odd, specifically in *tin^mDPE^* embryos, was detected, presenting an ectopic Odd pattern almost completely masking the Odd-positive pericardial cells (Supplemental Fig. S4). Nevertheless, based on Tin and Odd staining patterns, dorsal vessel formation is evident in homozygous *tin^mDPE^* embryos.

**Figure 3.**
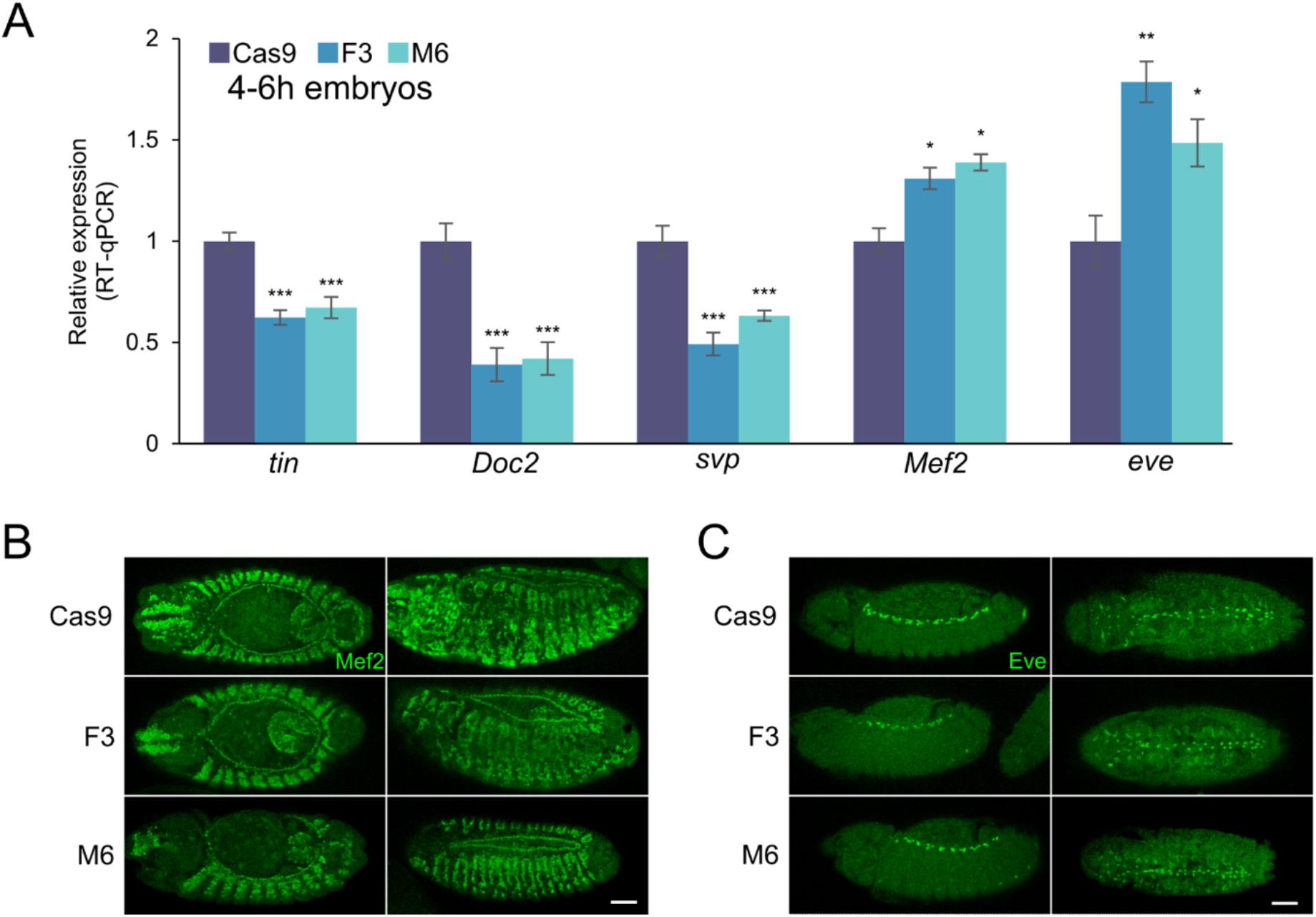
The expression of distinct Tinman target genes is affected in early *tin^mDPE^* embryos, yet dorsal vessel formation, as evident from Mef2 and Eve protein expression patterns at later stages, appears intact in *tin^mDPE^* embryos. (*A*) Endogenous *tin*, *Doc2*, *svp*, *Mef2* and *eve* RNA levels were quantified at 4-6h embryos (Bownes stage ∼10) using RT-qPCR and analyzed using the StepOnePlus software. Expression of all genes in Cas9 is defined to be 1. Each qPCR experiment was performed in triplicates. Error bars represent 95% confidence interval. *p<0.05, **p<0.01, ***p<0.001 one-way ANOVA followed by Tukey’s post hoc test. Only comparisons to Cas9 samples are presented, n≥3. In addition, embryos were stained using antibodies against (*B*) Mef2 and (*C*) Eve of late embryos (Bownes stage >13). Z-stack maximal projections are shown, scale bar = 50µm.

### Viability and adult heart function are reduced upon mutation of endogenous *tinman* DPE

In contrast to *tin* null homozygous flies (Azpiazu and Frasch 1993), *tin^mDPE^* homozygous flies were viable but appeared frail. We therefore tested whether a single *tin^mDPE^* allele can compensate for the loss of *tin* by crossing *tin^WT^* and *tin^mDPE^* flies to flies with a *tin* null allele maintained over a balancer (*tin^346^*/TM3 *eve*-*lacZ*). Eclosed flies were scored for the Stubble (Sb) phenotype (shortened and thick bristles; indicative of the TM3 balancer), which enables the distinction between the examined allele over *tin^346^* (non-Sb) or over a WT (balancer, Sb) allele. Thus, a non-Sb/Sb ratio reflects the presence of *tin^WT^* or *tin^mDPE^ in trans* to *tin* null allele. For full compensation by the WT allele, we expect the non-Sb/Sb ratio to be equal to 1, *i.e.*, same numbers of rescued *tin^346^*/*tin^WT^* flies and *tin^346^*/TM3. Strikingly, only half of the number of viable flies are produced in *tin^mDPE^/tin^346^* as compared to *tin^WT^/tin^346^* (Fig. 4). These data indicate a substantial decrease in viability when *tin^mDPE^* is present as a single copy, in contrast to a single copy of the *tin* WT allele when tested in trans to a null allele. These results strongly support the notion that the reduced function of *tin^mDPE^* is due to its decreased mRNA expression levels. We next tested whether *tin* DPE is necessary for heart function and analyzed the hearts of *tin^mDPE^*homozygous adult flies for defects using semi-automated heart analysis (SOHA, (Fink et al. 2009)). Adult WT and mDPE females were dissected, and their hearts were imaged using high-speed video recording, followed by determination of spatial and temporal parameters (*e.g.*, heart diameters during diastole (DD), systolic intervals (SI) and stroke volume). Both *tin^mDPE^*alleles have smaller diastolic diameters (Fig. 5A) and reduced contractility (measure by fractional shortening, FS, Fig. 5B), resulting in a reduced stroke volume (Fig. 5C). Furthermore, *tin^mDPE^*mutant hearts show longer systolic intervals (SI, Fig. 5D) indicating prolonged contraction intervals. This indicates that the *tin* DPE is required to establish proper heart physiology.

**Figure 4.**
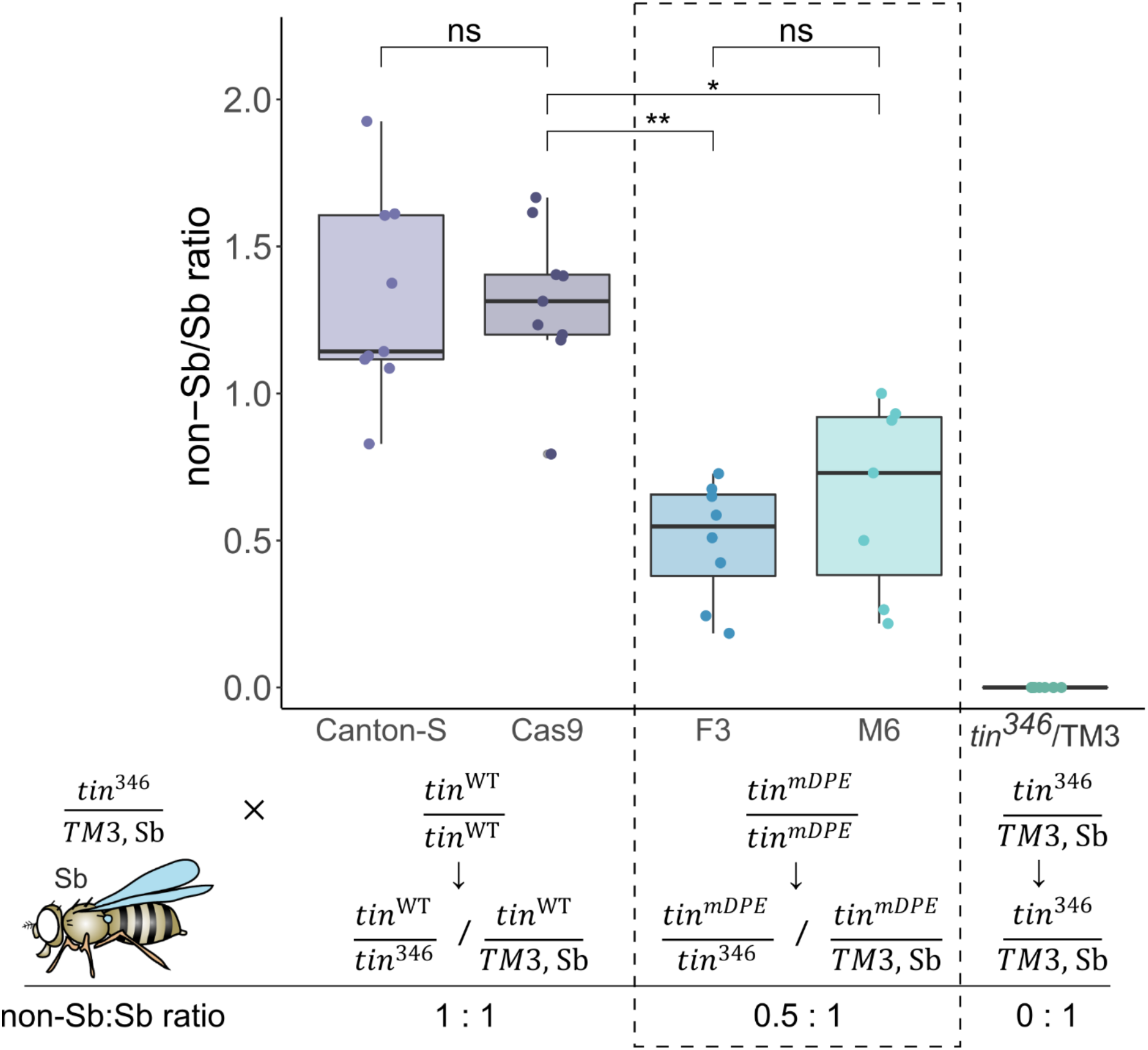
Mutation of *tinman* endogenous DPE reduces viability of mature flies when tested in trans to *tin* null mutation. *tin^WT^* (Canton-S, Cas9) and *tin^mDPE^*(F3, M6) strains were crossed to *tin^346^*/TM3, Sb flies. The Sb phenotype of hatched flies was used to distinguish between *tin^mDPE^*over *tin^346^* (null) or WT allele, *tin^346^* was also scored as a background. * p< 0.05, ** p<0.01 One-way nested ANOVA followed by TukeyHSD post hoc test. Schematic representation of the relevant cross and the expected non-Sb/Sb ratio is indicated. Schematic fly image is from (Roote and Prokop 2013). The data and genotypes of the *tin^mDPE^* flies that were crossed with *tin* null heterozygote are marked by a dashed frame.

**Figure 5.**
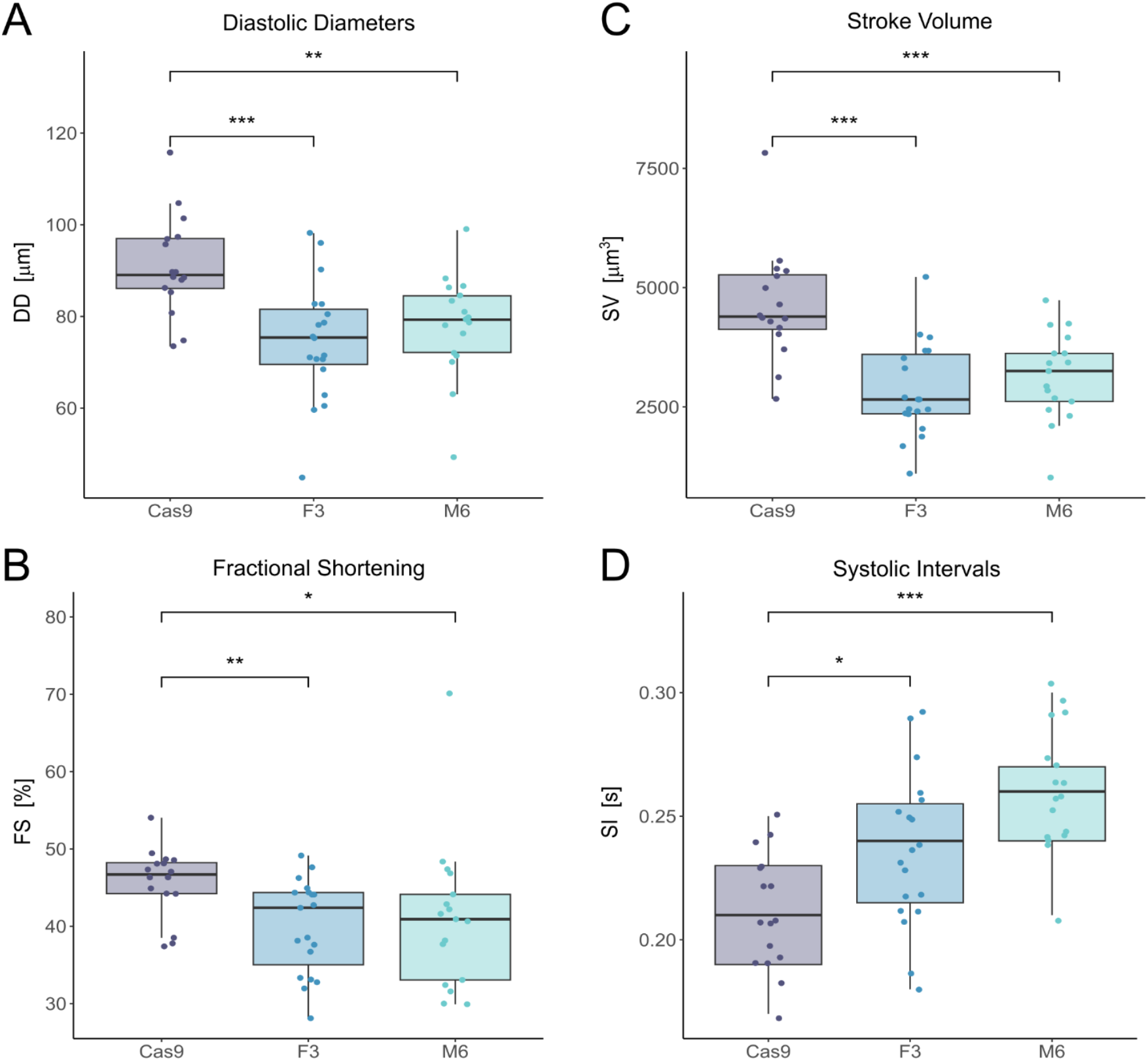
Mutation of *tinman* endogenous DPE affects adult heart size and function. *tin^mDPE^*hearts (F3, M6) have smaller diastolic diameters (*A*, DD) and reduced fractional shortening (*B*, FS), resulting in reduced stroke volume (*C*, SV). *Tin* DPE mutant hearts also exhibit prolonged systolic intervals (*D*, SI). In all panels significance was tested using Wilcoxon tests; ***<0.001, **<0.01, *<0.05 significance levels.

### Nascent transcription analysis following *tinman* DPE mutation reveals significant changes in muscle and heart transcriptomes, preferentially among genes with DPE motifs

The reduced viability and functional heart parameters of *tin^mDPE^* flies indicated that this 7bp substitution mutation within a single promoter results in major transcriptional changes. To capture and quantify these changes genome-wide, we captured active or ‘nascent’ transcription, which offers a high-resolution of impacted genomic loci and the underlying gene regulatory programs (Wissink et al. 2019). In addition, combining nascent assays with 5’Cap selection enables the precise determination of the transcription initiation position and consequently the detection of any alternative initiation sites (Policastro and Zentner 2021). Nuclei isolation as needed for most nascent methods (Wissink et al. 2019) results in significant bias when digesting tissues (Lepage et al. 2021) or embryos, while mechanical tissue homogenization results in preferential loss of fragile cells. We therefore performed capped small (cs)RNA-seq, which similar to GRO-cap accurately captures nascent transcription start sites (TSSs) from total RNA (Duttke et al. 2019; Yao et al. 2022). Analysis of *tin^mDPE^* and *tin^WT^* embryos collected at 0-2h, 2-4h, 4-6h and 6-8h time intervals confirmed markedly reduced *tin* levels in *tin^mDPE^*, especially at 2-4h (Fig.6A). No alternative TSSs were detected (Supplemental Fig. S5).

**Figure 6.**
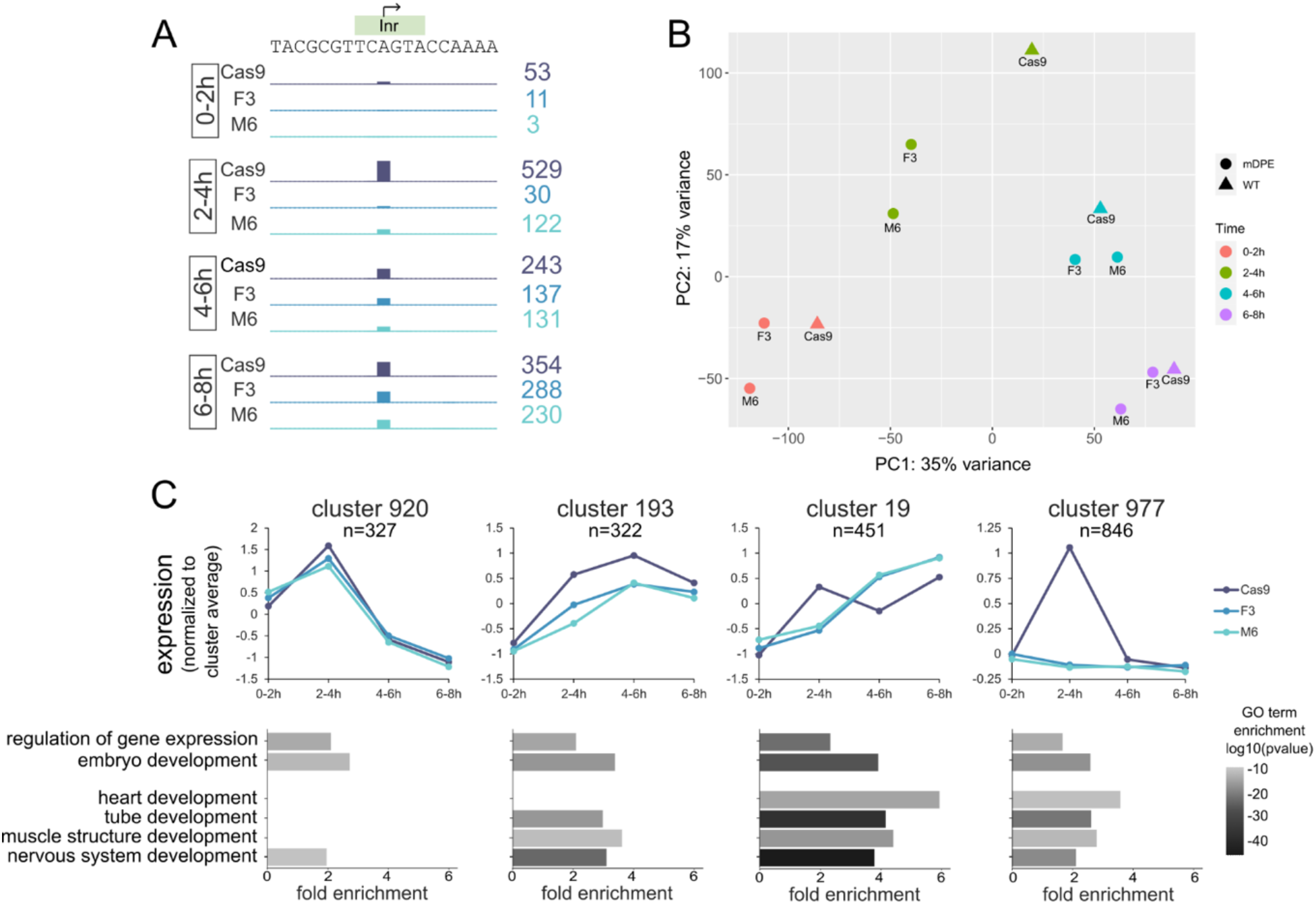
Nascent transcription of developmental genes is altered in *tin^mDPE^* embryos. csRNA-seq analysis was performed on 0-2h, 2-4h, 4-6h and 6-8h embryos for both *tin^WT^* (Cas9) and *tin^mDPE^* (F3, M6) embryos. (*A*) Nascent transcription profile of the *tinman* locus, ±10bp relative to the TSS. Scale is 0 to 530 for all tracks. (*B*) PCA analysis of the csRNAseq samples. While samples are separated by developmental time, a difference between *tin^mDPE^* and *tin^WT^* strains is clearly evident for each time point. (*C*) Expression and selected GO terms of representative clusters from HOMER’s analyzeClusters.pl script. (top) Mean expression of the cluster is shown for each strain at each timepoint. (bottom) Same selected GO terms are shown for each cluster, with the associated fold enrichment and p-value over genomic background. Note the strong association of cluster 19 with heart and tube development. See Supplemental Fig. S7 and Supplemental File S1 for complete data.

Principal component analysis (PCA) shows that samples first cluster by time, and then by mDPE versus WT alleles (Fig. 6B). In accordance with *tin* RT-qPCR results, the main overall difference between the mDPE and WT alleles is at the 2-4h timepoint. Within each timepoint, the F3 and M6 samples clustered apart from Cas9 samples, indicating some impairment in overall transcription and developmental programs.

Peaks were annotated based on the EPDnew database (Supplemental Fig. S6), which identifies TSSs based on the 5’ cap of transcripts and quantifies the expression levels of individual transcripts based on experimental data (Meylan et al. 2020). HOMER (Heinz et al. 2010) was used to cluster the csRNA-seq peaks according to their expression, and analyze the groups for enriched motifs and GO terms (Supplemental Fig. S7, Supplemental File S1). The resulting clusters present either similar or differential *tin^mDPE^* and *tin^WT^*expression (Supplemental File S2). The differential expression encompasses several modes, for example *tin^mDPE^* expression being lower than *tin^WT^* and alternating expression patterns (Fig. 6C, top, clusters #193 and #19, respectively). All the depicted clusters are similarly enriched for general GO terms, such as regulation of gene expression and embryo development, while differentially expressed clusters (#193, #19 and #977) are specifically enriched for heart and muscle structure formation GO terms (Fig. 6C, bottom). Interestingly, nervous system development is also associated with the differentially expressed clusters.

In each time interval, differentially expressed peaks were enriched for DPE-like motifs (Fig. 7A). Given that the differentially expressed peaks for each timepoint are not overlapping (Supplemental Fig. S8), the similarity of the over-represented motifs is striking. While the exact motif composition and enrichment varies across the examined timepoints, combining the peaks that are differentially expressed at least in one timepoint revealed a common DPE-like motif. This common motif is significantly enriched in all differentially expressed peaks (Fig. 7B). As expected from a canonical DPE motif, all the DPE-like motifs are strictly positioned at the csRNA-seq peaks center (Supplemental Fig. S9). Taken together, we demonstrate the *in vivo* importance of a single core promoter element, the DPE motif (Fig. 7C). This 7bp change within the *tinman* promoter was sufficient to decrease Tinman levels resulting in marked changes of nascent transcription patterns, specifically of DPE-containing genes required for muscle and heart formation. Moreover, the insufficiency of a single *tin^mDPE^* copy to support viability in a deficient background highlights its critical role in establishing the adequate Tinman levels required for functional heart formation.

**Figure 7.**
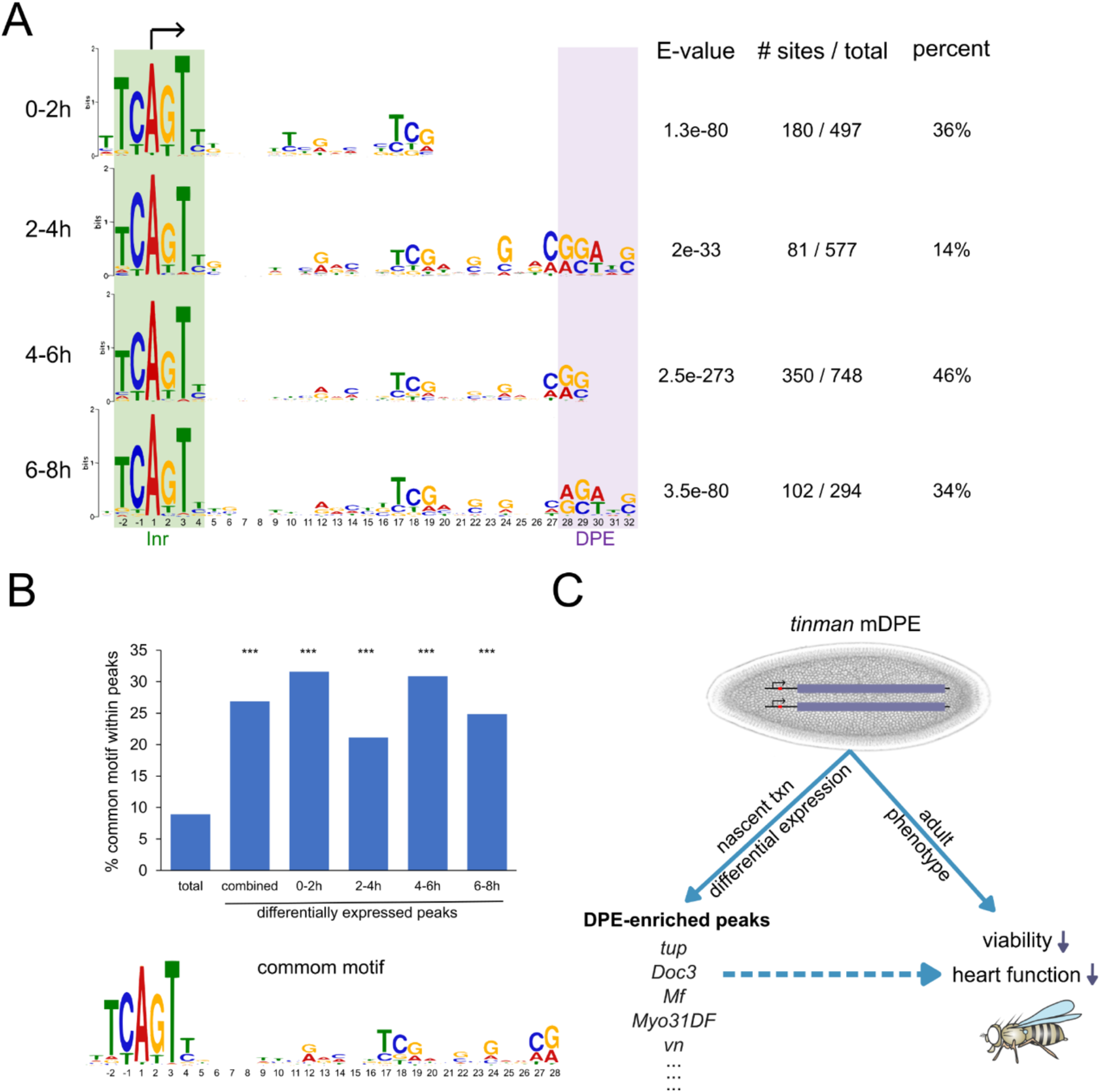
The DPE motif is enriched in differentially expressed genes following *tinman* DPE mutation. (*A*) A significant Inr+DPE resembling motif is evident for each tested time point following MEME analysis. Both raw numbers and percentages are presented. Inr and DPE motif locations are highlighted, arrow represents transcription start site. (*B*) The common motif enriched among all the differentially expressed promoters (bottom) is enriched above the expected background (total peaks) in all differentially expressed peaks (combined), as well as in each individual time interval (0-2h, 2-4h, 4-6h, 6-8h). ***p< 0.001, one proportion Z-test. (*C*) Schematic summary. Endogenous *tinman* promoter was genetically edited using CRISPR to change the 7bp encompassing the DPE motif of the *tinman* promoter. This genomic editing resulted in viable homozygotes, which present major changes in nascent transcription (txn) of muscle and developmental genes, enriched for DPE-like motifs in their core promoter. In addition, the *tinman* mDPE homozygotes present impaired viability and heart function parameters, demonstrating the *in vivo* importance of a single core promoter element, the DPE motif. Schematic fly image is from (Roote and Prokop 2013).

## Discussion

### Mutation of *tinman* DPE alters transcriptional programs regulating heart development

In this study, we present a detailed characterization of flies harboring a mutation in the endogenous sequence of a downstream core promoter element, the DPE motif. The 7bp substitution in the 5’UTR of the *tin* gene resulted in reduced expression of both RNA and protein levels in the mDPE embryos (Figs. 1-3; Supplemental Fig. S2), albeit with no apparent change in expression patterns detected using RNA *in situ* hybridization or antibody staining (Fig. 2C, Supplemental Fig. S1). Interestingly, Mef2 and Eve protein expression patterns are not affected by the *tin* DPE mutation, while the *Doc2* and *svp* RNA levels are markedly reduced in 4-6h mDPE embryos (Fig. 3). In the *Drosophila* heart, Tinman activates *svp* (Ryan et al. 2007), which encodes a repressor that acts through both DNA-binding competition and protein-protein interactions (Zelhof et al. 1995). *eve* expression in pericardial cells is *Doc2*-independent (Reim and Frasch 2005), suggesting that the reduction of Tinman levels following the DPE mutation differentially affects its target genes. Notably, the specification of Svp-expressing cells is additionally regulated by Hox genes, establishing the anterio-posterior polarity of the dorsal tube (Lo and Frasch 2003). While the majority of the *Hox* genes and mesodermal targets are DPE dependent (Juven-Gershon and Kadonaga 2010; Sloutskin et al. 2021), it remains to be discovered whether Hox-driven patterning of the dorsal tube is mediated via the DPE.

Strikingly, both *tin^mDPE^* independent fly lines have an overall decrease in adult heart function as compared to the WT. The functional reduction was demonstrated through structural and temporal aspects, wherein mDPE strains exhibited diminished diastolic diameters and decreased contractility, resulting in elongated contraction intervals and reduced stroke volume (Fig. 5). These results highlight the *in vivo* importance of the DPE. The functional effects manifested in the adult heart may result from reduced expression of Tinman target genes required for normal heart physiology in the adult heart. The observed diminished heart function may also be due to subtle morphological alterations in the embryonic heart.

Developmental programs are largely executed by transcription of the relevant genes. Capped-small RNA-seq (csRNA-seq) was developed in order to accurately quantify changes in transcription initiation during dynamic processes (Duttke et al. 2019). We applied csRNA-seq to study transcriptional dynamics at 2h resolution, comparing *tin^WT^* and *tin^mDPE^* embryos from 0 to 8h of development. Detected peaks were assigned to genes and clustered based on similar expression pattern. Reassuringly, development-related GO terms were enriched in most clusters, while heart-related and tube formation GO terms were associated with differentially regulated genes (Fig. 6). Together, our results further demonstrate the specificity of the generated *tin* DPE mutation, leading to a reduction of *tin* levels and specifically altering transcriptional programs governing the functional heart formation.

### Functional compensation in *tinman* mutant DPE embryos

Remarkably, while homozygous *tin^mDPE^* flies are viable, they cannot fully compensate for the loss of *tin* (Fig. 4). Nevertheless, despite the reduced Tinman levels in the *tin^mDPE^* embryos, a dorsal vessel with a normal pattern of four Tinman-expressing cardioblasts is formed, presenting a very similar pattern to *tin* null heterozygotes. The viable phenotype of the *tin^mDPE^* flies strongly suggests the existence of a compensatory mechanism that ensures that heart development is resistant to small-scale perturbations. In fact, several key mesodermal transcription factors, mainly Tinman, Pannier and Doc2, can bind the same genomic loci and define cardiac enhancers (Zinzen et al. 2009; Junion et al. 2012; Jin et al. 2013). Moreover, it was suggested that functional flexibility exists, where the jointly bound transcription factors (TFs) cooperate to recruit the relevant TF to lower-affinity binding sites (Frasch 2016). In such a case, it is more plausible that the target enhancer will be activated even in the presence of lower levels of the Tinman protein, resulting from the perturbation to the *tin* DPE sequence motif. Consistent with the existence of a compensatory mechanism, not all Tinman target genes are affected by the reduction of Tinman levels in *tin* mDPE flies. Our findings provide further evidence for the complexity of the mesodermal regulatory network, which contains some redundant connections that are challenging to detect with standard genetic experiments. This redundancy may support fine-tuning of expression circuits, which in turn generates a gene regulatory network that is more resistant to disruption. In this regard, the *tinman* DPE can be thought of as fine-tuning its expression.

### Potential impact of the DPE motif on transcriptional regulation and dynamics *in vivo*

Compatibility between specific promoters and enhancers was demonstrated to be functional in different organisms (Butler and Kadonaga 2001; Zabidi et al. 2015; Lim and Levine 2021; Martinez-Ara et al. 2022). During development, *tin* is expressed in the head, trunk mesoderm, dorsal mesoderm and in cardioblasts. Each expression pattern represents a specific developmental stage, and is explicitly controlled by a distinct enhancer integrating the relevant regulatory signals (Yin et al. 1997). These characterized enhancers were cloned and used to demonstrate that early Tinman expression is sufficient for dorsal vessel development and mesoderm specification (Zaffran et al. 2006). Remarkably, *tin* levels show the most pronounced decrease during the 3-4-hour developmental phase (Fig. 1B), precisely when *twist* activates *tin*. These results suggest that the interaction between the twist-dependent intronic enhancer (Yin et al. 1997) and the *tin* promoter is highly sensitive to the presence of a functional DPE motif. However, it remains to be determined whether the function of a specific enhancer is disturbed in the *tin^mDPE^* embryos.

In *Drosophila* embryos, core promoter composition affects transcriptional dynamics profiles, detected with MS2-based reporters and a shared enhancer (Fukaya et al. 2016). Furthermore, the spacing of enhancer-promoter pair modulates gene activity by changing the temporal and quantitative parameters of transcriptional bursts in the developing *Drosophila* embryo (Yokoshi et al. 2020). Direct examination of TATA-box and Initiator elements using synthetic constructs in *Drosophila* revealed differences in the modes of action of these motifs (Pimmett et al. 2021). The essential role of native TATA box and DPE motifs within the *fushi tarazu (ftz)* promoter in transcriptional dynamics regulation was recently demonstrated *in vivo* (Yokoshi et al. 2022). While the proper expression of *ftz* requires both motifs, the DPE was found to regulate transcriptional onset, and the TATA-box to affect overall intensity. Our findings demonstrate that in the *tin* promoter, which lacks a natural TATA box, mutation of the DPE motif is sufficient to reduce overall nascent RNA levels (Fig. 6A). A comprehensive analysis of the regulation of transcriptional dynamics by endogenous promoter motifs will help to fully elucidate their fascinating roles during embryonic development.

### Prospective relevance of the DPE motif for human cardiogenesis

The vertebrate *tinman* homolog, Nkx2-5, is required for heart specification and is expressed in early cardial progenitors (Harvey 1996). The molecular mechanisms controlling heart formation are highly conserved in evolution from flies to humans (reviewed in (Bodmer and Venkatesh 1998; Cripps and Olson 2002; Rotstein and Paululat 2016)). Vertebrate Nkx2-5 mutants are able to properly specify cardiac progenitor cells, however the final organization of the heart is disturbed (Lyons et al. 1995; Targoff et al. 2008; Targoff et al. 2013). Interestingly, additional Nkx2-family members cannot compensate for the specific loss of Nkx2-5, demonstrating the strong specific requirement of Nkx2-5 and possibly its cofactors (Stutt et al. 2022).

The DPE motif was originally discovered as conserved from flies to human (Burke and Kadonaga 1997), yet for many years, only a few human genes were experimentally shown to contain a functional DPE (Burke and Kadonaga 1997; Zhou and Chiang 2001; Duttke 2014). Recently, machine learning models were used to define the downstream core promoter region (DPR) in human and *Drosophila* (Vo Ngoc et al. 2020; Vo Ngoc et al. 2023). In parallel, preferred downstream positions required for proper transcriptional output were identified (PDP, (Dreos et al. 2021)). Interestingly, the core promoter of the human Nkx2-5 contains both the DPR and PDP motifs. It remains to be determined whether Nkx2-5 levels are controlled by its core promoter composition. If so, this would suggest that the regulatory function of the DPE during heart formation is not limited to *Drosophila* but is rather conserved, along with many components of the gene regulatory network. Notably, multiple cardiac pathologies (*e.g.*, septal openings and conduction defects) result from mutations in the coding region of Nkx2-5 (Schott et al. 1998; Benson et al. 1999; Elliott et al. 2003; McElhinney et al. 2003). Thus, it is conceivable that, in addition to mutations in the protein coding region of Nkx2-5, homozygous or heterozygous mutations in downstream core promoter motifs of Nkx2-5 could likewise be responsible for congenital heart defects.

In summary, we demonstrated the *in vivo* contribution of a single core promoter element, namely the DPE motif, to the regulation of the *tin* gene and its developmental gene regulatory network. This exemplifies the contribution of the endogenous core promoter to transcriptional regulation during *Drosophila melanogaster* embryogenesis, thus paving the way for further exciting discoveries related to transcriptional regulation of developmental genes via their core promoter.

## Materials and Methods

### Fly culture and stocks

Flies were cultured and crossed on standard media (cornmeal, yeast, molasses, and agar) at 25°C, 60% relative humidity and under a 12h light/12h dark cycle. All the described embryonic development was performed at 25°C. F3 and M6 *tin^mDPE^* strains were generated based on a *white* co-conversion approach (Ge et al. 2016) using ssODN, as detailed in (Levi et al. 2020). Cas9 is shorthand for the injected strain that was used as *tin^WT^* control in all the experiments. w; *tin^346^*/TM3, *eve-lacZ* is a balanced null allele described in (Azpiazu and Frasch 1993).

### RNA extraction and Real Time PCR analysis

0-8h embryos were collected and aged at 25°C as indicated. For each time point, WT (Cas9) and two independent mDPE (F3, M6) strains were collected and processed in parallel. Total RNA was extracted from dechorionated embryos using the TRI Reagent (Sigma-Merck) according to the manufacturer’s protocol, followed by ethanol precipitation for further purification. 1 μg RNA was further used for cDNA synthesis (qScript cDNA Synthesis Kit, Quantabio). Quantitative PCR using SYBR green (qPCRBIO SyGreen Blue Mix, PCR Biosystems) was performed using a StepOnePlus Real-Time PCR machine. Control reactions lacking reverse transcriptase were also performed to ensure that the levels of contaminating genomic DNA were negligible. Transcript levels were analyzed by the ΔΔCT method using Polr2F (RpII18) as an internal control. Each sample was run in triplicates. Statistical analysis was performed using ‘HH’ R package (https://CRAN.R-project.org/package=HH), with mean and standard deviation values exported from StepOnePlus software. Primer sequences are provided in Supplemental File S4.

### Western blot analysis

Protein extracts from dechorionated embryos were prepared in 2X DTT-based sample buffer at a final concentration of ∼0.5mg embryos/µl. 10µl of the sample was analyzed using 10% SDS-PAGE gel, followed by rabbit anti-Tinman polyclonal antibodies (1:1000 in 3% BSA, (Yin et al. 1997)) and then by goat-anti-rabbit IgG-HRP (1:5000 in 5% milk, Jackson ImmunoResearch). HRP signal was detected using EZ-ECL kit (Biological Industries), and imaged using ImageQuant LAS 4000 instrument (GE). The use of the Tinman antibody results in background bands, however the major band is above the 45kDa size marker, as predicted. The same membrane was stripped (ST010, Gene Bio-Application) and re-blotted with mouse anti-Actin monoclonal antibodies (1:1000 in 3% BSA, Abcam ab8227) to ensure proper gel loading. Images were quantified using the “Gels” analysis feature of ImageJ software; each sample was normalized to the detected Actin levels. Statistics was calculated with two-tailed Students t-test function.

### Immunostaining and staging *Drosophila* embryos

Dechorionated embryos were fixed in freshly prepared 1:1 mixture of heptane and 3.7% paraformaldehyde solution (diluted 1:10 in PBS) for 20 min with vigorous shaking. Devitellinization was performed in heptane:methanol 1:1 solution, and embryos were stored in methanol at -20°C. Before staining, embryos were washed 3 times in PBST (0.1% Tween-20 in PBS) and blocked in 2% BSA supplemented with 0.2% fetal calf serum. Embryos were incubated overnight at 4°C with the following primary antibodies as indicated: rabbit anti-Tinman (1:800), anti-Eve (1:800), anti-Mef2 (1:800), guinea pig anti-Odd (1:200, #805 from Asian Distribution Center for Segmentation Antibodies, distributed by Prof. Zeev Paroush) and mouse anti-LacZ (1:1000, Promega z3781). Detection was performed using mainly goat anti-rabbit IgG H&L (DyLight® 488) (1:500, Abcam ab96883), Cy3-donkey anti guinea pig IgG H+L (1:1000, JacksonImmunoResearch 706-165-148) and Cy5-goat anti-mouse IgG H+L (1:000, JacksonImmunoResearch 115-175-166). Embryos were counter-stained with Hoechst 33342 (Sigma-Aldrich), and mounted in n-propyl gallate based anti-fade mounting medium (5% w/v n-propyl gallate dissolved in 0.1M Tris pH 9 and glycerol (1:9 ratio)). Images were acquired with a Leica SP8 confocal microscope, using oil immersion objectives. Z-stack maximal projections are shown. Quantification of imaged embryos was performed using CGPfunctions R package (https://github.com/ibecav/CGPfunctions).

Bownes developmental stages were used for embryo development classification (after (José A. Campos-Ortega 1985) and https://www.sdbonline.org/sites/fly/aimain/2stages.htm). *tinman* in-situ images were obtained from Berkeley *Drosophila* Genome Project (Tomancak et al. 2002; Tomancak et al. 2007) via FlyExpress website (Kumar et al. 2011).

### Viability testing

Cas9, F3 and M6 virgins were crosses to *tin^346^*/TM3 (Sb), *eve-lacZ* in triplicates (biological replicates), and each vial was flipped 3 times (technical replicates). Parental flies were discarded, and F1 flies were anaesthetized, separated based on Sb phenotype, counted in groups of 5, and then discarded. Each vial was counted twice, ensuring most of the eclosed flies are scored. For analysis, non-Sb to Sb ratios were log2-transformed. One-way nested ANOVA was performed to test the effect of strain on non-Sb/Sb ratios. Specifically, a linear mixed effect model was performed, and the ANOVA was performed on the resulting model. Post hoc analysis was performed as pairwise comparisons using Tukey’s method.

### Adult *Drosophila* heart assay

All dissection steps were done using artificial hemolymph. In brief, 3-week-old female flies were anesthetized with FlyNap (Carolina Biological), transferred to a petroleum jelly-coated Petri dish, and dissected as described (Vogler and Ocorr 2009). The dissected hearts were equilibrated for 15 min at room temperature under constant oxygenation. High-speed movies were taken on an Olympus BX61WI microscope with a 10x immersion objective, using a Hamamatsu Orca Flash4 CMOS digital camera and HCI image capture software (Hamamatsu). Movies were then analyzed with custom-designed software (Ocorr et al. 2009).

### csRNA-seq samples and processing

F3, M6 and Cas9 embryos were aged at 25°C and collected at 0-2h, 2-4h, 4-6h and 6-8h time points. Total RNA was extracted from dechorionated embryos using the TRI Reagent (Sigma-Merck) according to the manufacturer’s protocol, followed by ethanol precipitation for further purification. Reduction of *tin* levels was verified using RT-qPCR, and samples were subjected to csRNA-seq analysis protocol version 5.2 (Duttke et al. 2022). Briefly, RNA was heat denatured and short RNAs (18-65nt) purified by 15% UREA-PAGE. A small fraction (5%) of these short RNAs was used to generate input libraries (conventional small RNA-seq) and the remainder cap-selected with 5′ monophosphate-dependent exonuclease (TER51020) followed by two phosphatase (CIP) treatments. Sequencing libraries for sRNA-seq and csRNA-seq were generated using the NEB sRNA kit, but with addition of RppH for decapping (Hetzel et al. 2016).

Sequencing data was analyzed using HOMER csRNAseq module (http://homer.ucsd.edu/homer/ngs/csRNAseq/index.html, (Duttke et al. 2019)) and R custom scripts. 3′ adapter sequences of the reads were trimmed using HOMER (Heinz et al. 2010) and aligned to *dm6* genome using STAR (version 2.7.10a) (Dobin et al. 2013). Reads were visualized as strand-specific bedGraph using HOMER `makeUCSCfile` command with `-style` tss parameter. Peak calling was performed using the `findcsRNATSS.pl` function in HOMER (Duttke et al. 2019), with input RNA-seq used as background to eliminate transcripts from degraded and high-abundance RNAs in csRNAseq. HOMER `annotatePeaks.pl` command was used with `-rlog` parameter for calculating the normalized expression values for each peak used in downstream analyses. It was also used for generating transcription profile plots, for example `annotatePeaks.pl tss dm6 -size 400 -hist 10 -pc 3`. For differential expression, `getDiffExpression.pl -edgeR -simpleNorm -dispersion 0.05 -AvsA` was used on raw counts. ComplexHeatmap R package (Gu et al. 2016) was used for hierarchal clustering. HOMER `analyzeClusters.pl` was used for motifs and GO terms enrichment analysis in the identified cluster. plotPCA function from DESeq2 package was used with parameter *ntop = 40000.* Raw sequence data were deposited in the NCBI GEO database under the following accession number GSE221852.

### Motif enrichment analysis

For each time point, the list of peaks with pAdj < 0.1 for both F3 and M6 vs. Cas9 was extracted based on differential expression analysis (above). The “combined” list comprises the unique list of differentially expressed peaks within at least one time point. Peak coordinates were used for construction of BED files, and sequences were extracted based on dm6 genome. MEME analysis (Bailey and Elkan 1994) was performed on each list separately (Supplemental File S3). For the analysis, only peaks with “promoter” annotation were used, however similar results were obtained when using all the differentially expressed peaks. Over-represented motifs were converted to HOMER format, which was then used to scan the relevant BED files with the motifs of interest.

## Supporting information

SupFile1

SupFile2

SupFile3

SupFile4

## Acknowledgments

We thank Nati Malachi for technical assistance with embryo collection at early stages of the project. We thank Prof. Zeev Paroush and his lab members, Dr. Shaked Bar-Cohen and Dr. Tanya Kushnir for teaching and guidance of the *in situ* hybridization protocol, followed by many fruitful discussions and suggestions. We also thank Prof. Adi Salzberg, Prof. Galit Shohat-Ophir, Prof. Ron Wides, Dr. Mali Levi and Dr. Adel Avetisyan for sharing their expertise and reagents during different stages of the project. We thank Orit Adato for critical reading of the manuscript.

The study was partially supported by the German-Israeli Foundation (GIF grant I-1220-363.13/2012 to Tamar Juven-Gershon and Eileen Furlong), Yad Hanadiv (to Tamar Juven-Gershon) and the NIH (NIGMS R00-GM135515 to Sascha Duttke and R01 HL054732 to Rolf Bodmer). Anna Sloutskin was also supported by the Nehemia Levzion Scholarship and Bar-Ilan University President’s Scholarship.

## Author Contributions

A.S., G.V., M.F., R.B., S.H.D. and T.J-G. conceived and designed the experiments. A.S., D.Y., G.V., D.I., H.A., H.S. and S.H.D. performed laboratory experiments. A.S. and T.D. performed bioinformatics analysis. A.S. and G.V. performed statistical analysis. A.S., G.V., D.I., M.F., R.B., S.H.D. and T.J-G. interpreted the experiments. A.S., D.Y. and G.V. prepared the figures and tables. A.S., G.V., M.F., R.B., S.H.D. and T.J-G. wrote the manuscript with input from all authors. M.F., R.B., S.H.D. and T.J-G. supervised the project.

## Supplementary Information

**Supplemental Figure S1.**
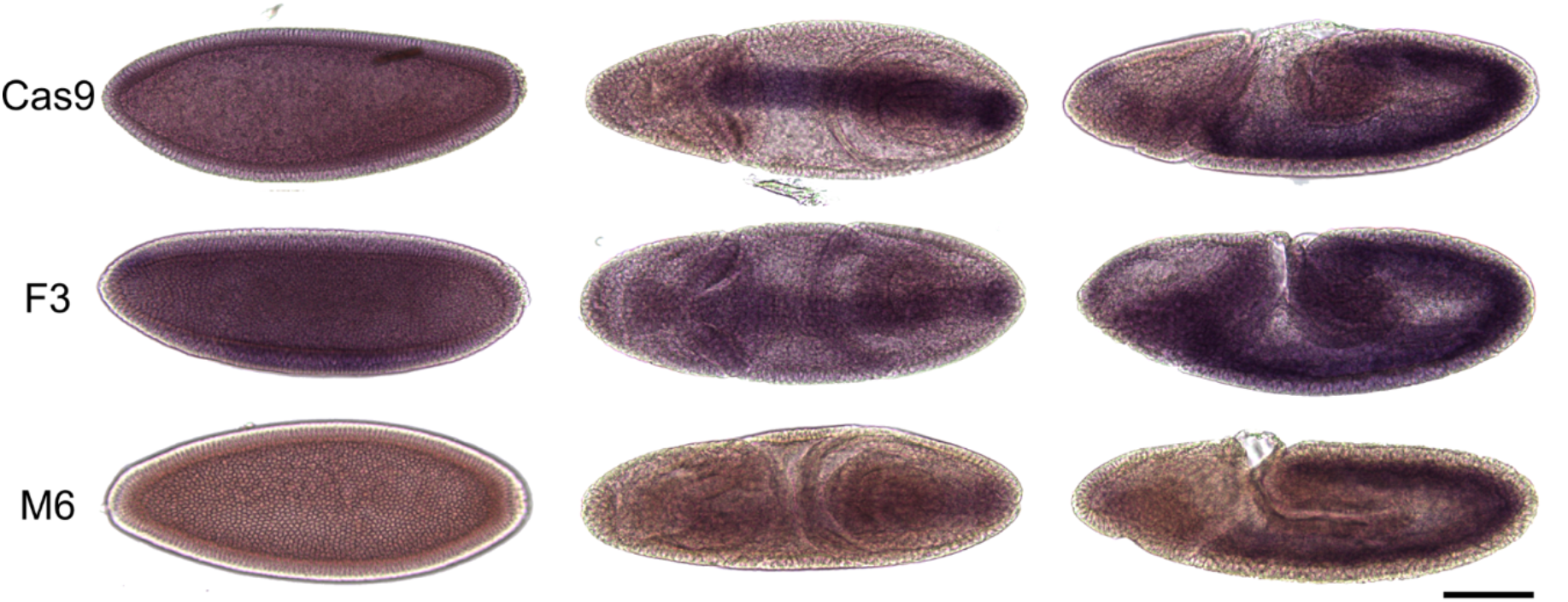
*tinman* expression pattern is similar in early (<stage 8) *tin^WT^* and *tin^mDPE^* embryos. *In situ* RNA hybridization using a DIG-labeled *tinman* probe resulted in no detectable difference of the staining pattern for 0-4h embryos (Bownes stage <8). Embryos were dechorionated in bleach and fixed in 4% formaldehyde/PBS/heptane for 20 minutes. pFlc-1-tin plasmid RE01329 (DGRC Stock 7896; https://dgrc.bio.indiana.edu//stock/7896; RRID:DGRC_7896) was linearized with NotI and transcribed in the presence of DIG labeling mix (Roche) to create a DIG-UTP labeled antisense RNA probe. Expression of *tin* was visualized by whole-mount *in situ* hybridization using the DIG-labeled probe detected by anti-DIG antibodies conjugated to alkaline phosphatase (Roche). Whole-mount *in situ* hybridization procedure was carried out essentially as described in (Wilk et al. 2010). Specimens were mounted in 70% glycerol/PBS, and imaged within a week. DIC microscope images were acquired with a Leica LMD7 microscope using a 10X objective lens. All embryos were imaged under the same conditions. Scale bar = 100µm.

**Supplemental Figure S2.**
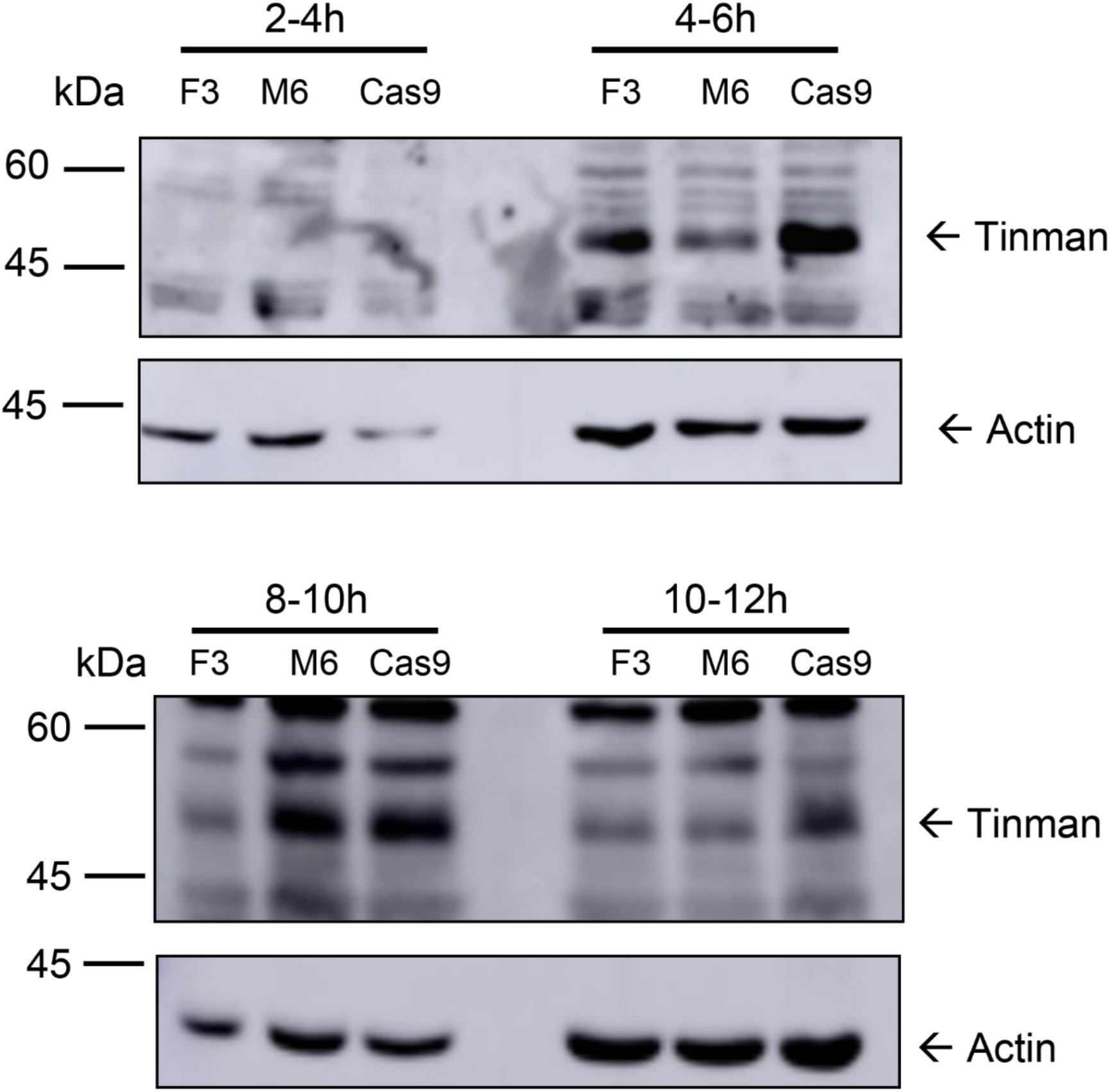
Overall Tinman levels are markedly reduced at 2-4h and 10-12h of embryonic development. Representative western blot images of embryos collected at the indicated time interval. For each membrane, embryos from the same fly population cage were collected. Each membrane was first blotted with rabbit α-Tinman antibodies and following chemiluminescent detection, stripped and re-blotted using mouse α-Actin.

**Supplemental Figure S3.**
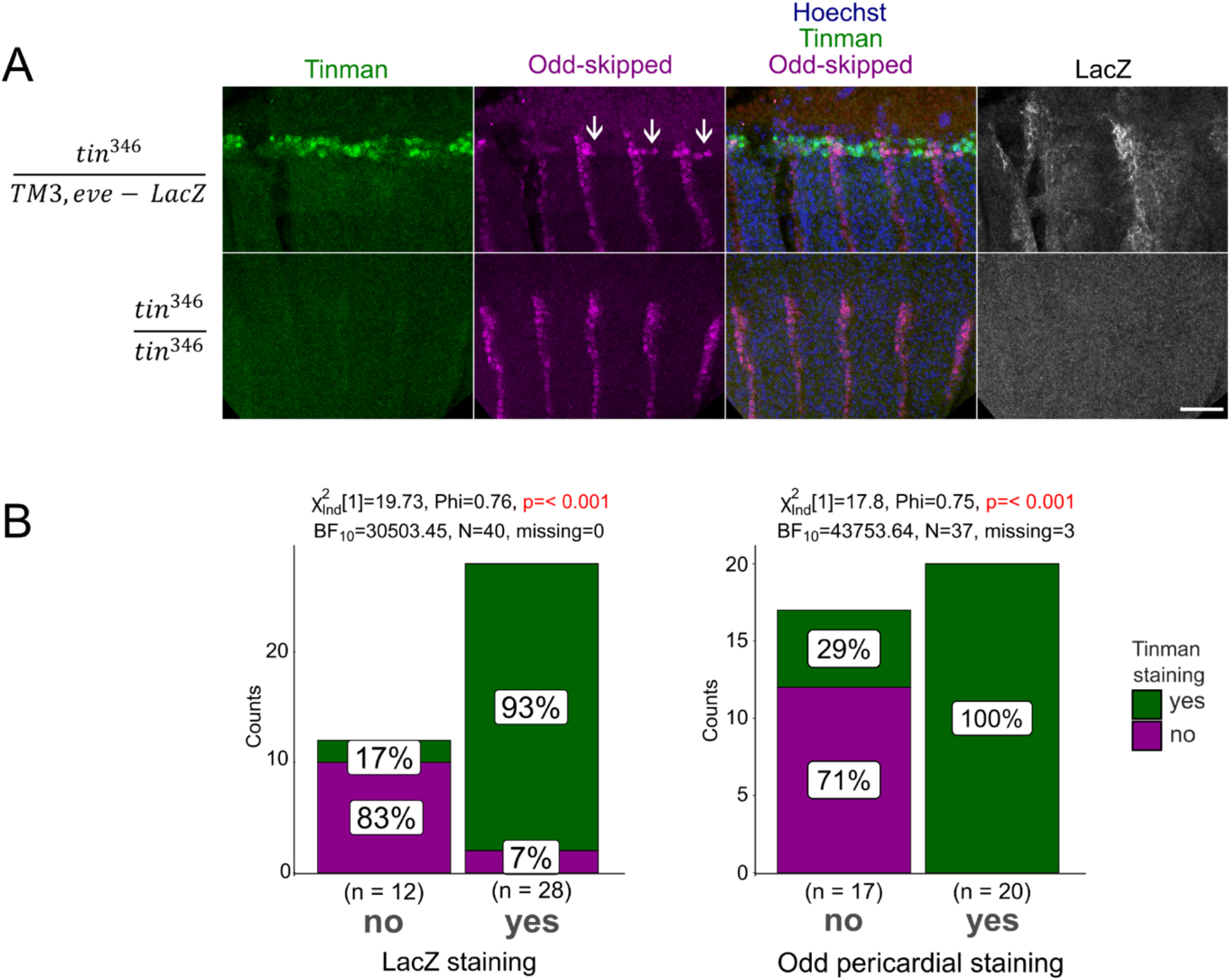
*tin* null heterozygote present both Tinman and pericardial Odd staining, indicating normal heart development, as opposed to homozygote embryos. (*A*) *tin^346^*/TM3, *eve-lacZ* were stained for Tinman (green), Odd-skipped (magenta) and LacZ (white). The Odd pericardial precursor cells (marked by white arrows) are present in embryos stained for both Tinman and LacZ. Segmental (non-pericardial) Odd staining is detected in both *tin* null heterozygotes and homozygotes. Z-stack maximal projections are shown. Scale bar = 25µm. Quantification of imaged embryos using CGPfunctions R package. (*B*) Quantification of LacZ (indicating *tin^346^*/TM3 genotype) and of Odd-expressing pericardial cell staining as a function of Tinman staining. Both LacZ and Odd pericardial cell staining are significantly correlated with Tinman staining, indicating normal heart development in *tin* null heterozygous embryos.

**Supplemental Figure S4.**
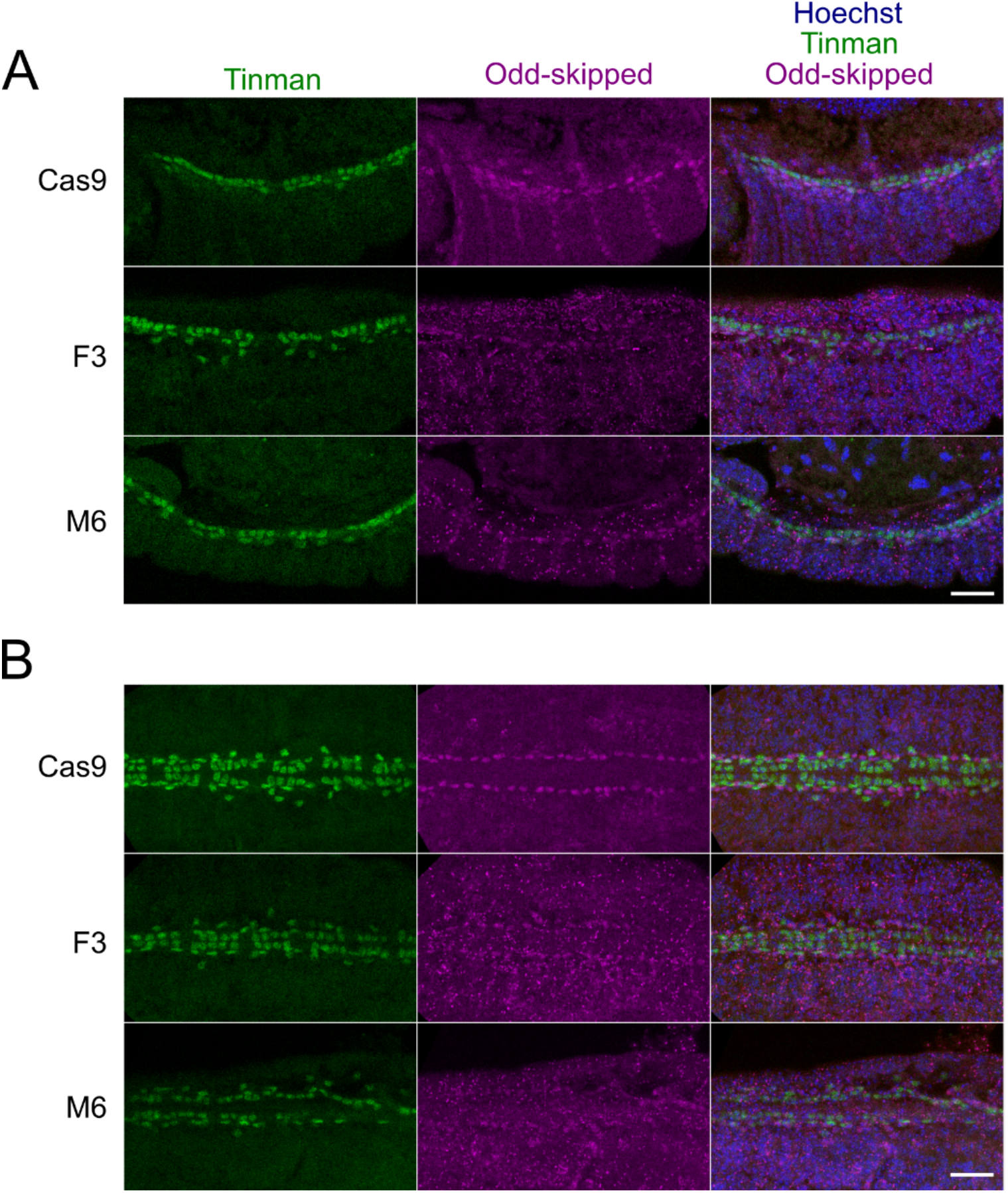
Tinman and Odd staining of the mDPE embryos. Embryos from *tin^WT^* (Cas9), *tin^mDPE^* (F3, M6) were stained for Tinman (green) and Odd-skipped (magenta). (*A*) Stage 13 and (*B*) Stage 17 embryos. Z-stack maximal projections are shown, scale bar = 25µm.

**Supplemental Figure S5.**
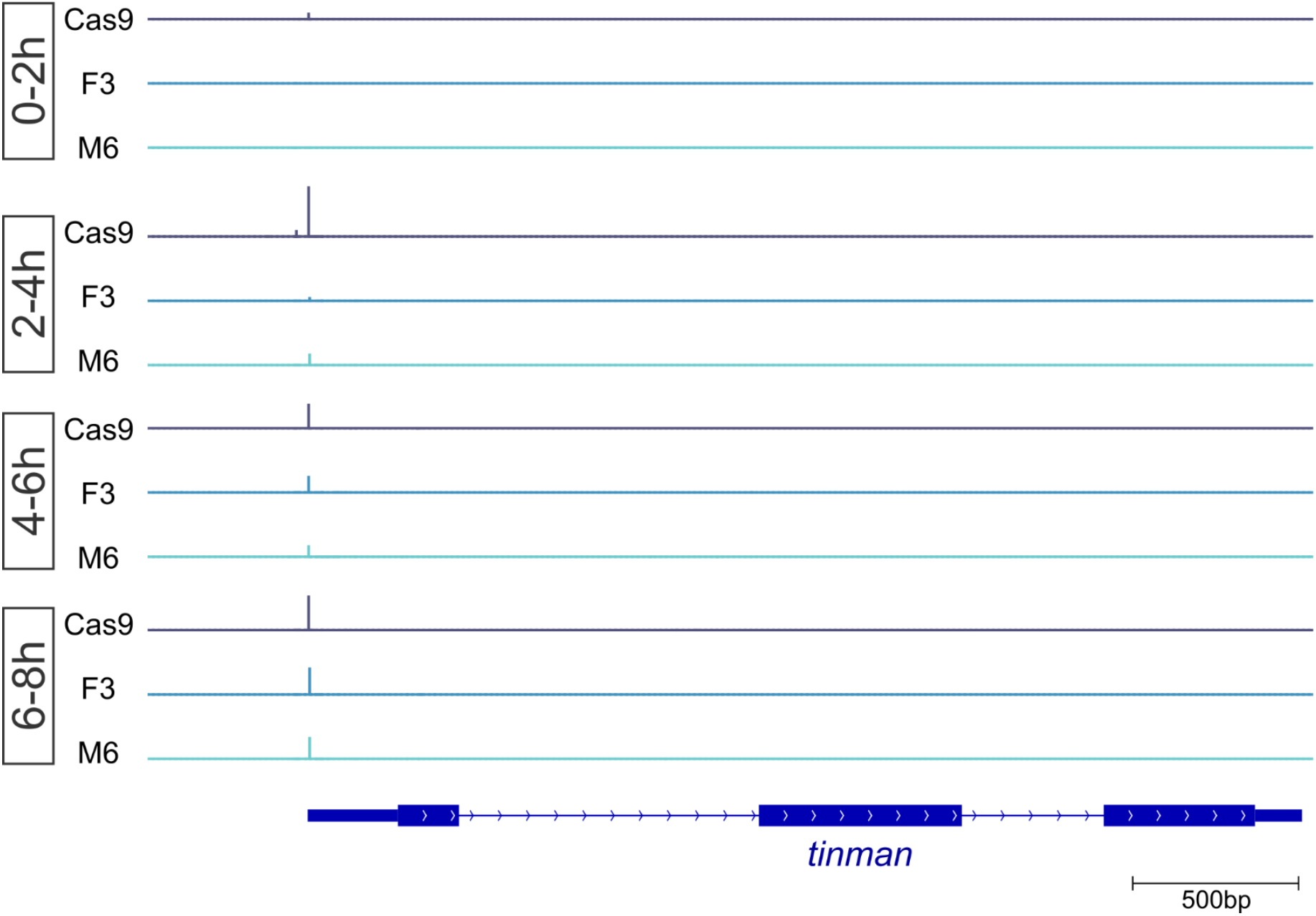
The tin mDPE allele does not result in additional TSSs. Nascent transcription profile of the whole *tinman* locus. Depicted region is Chr3R:21378500-218200 (dm6), scale is 0 to 530 for all tracks.

**Supplemental Figure S6.**
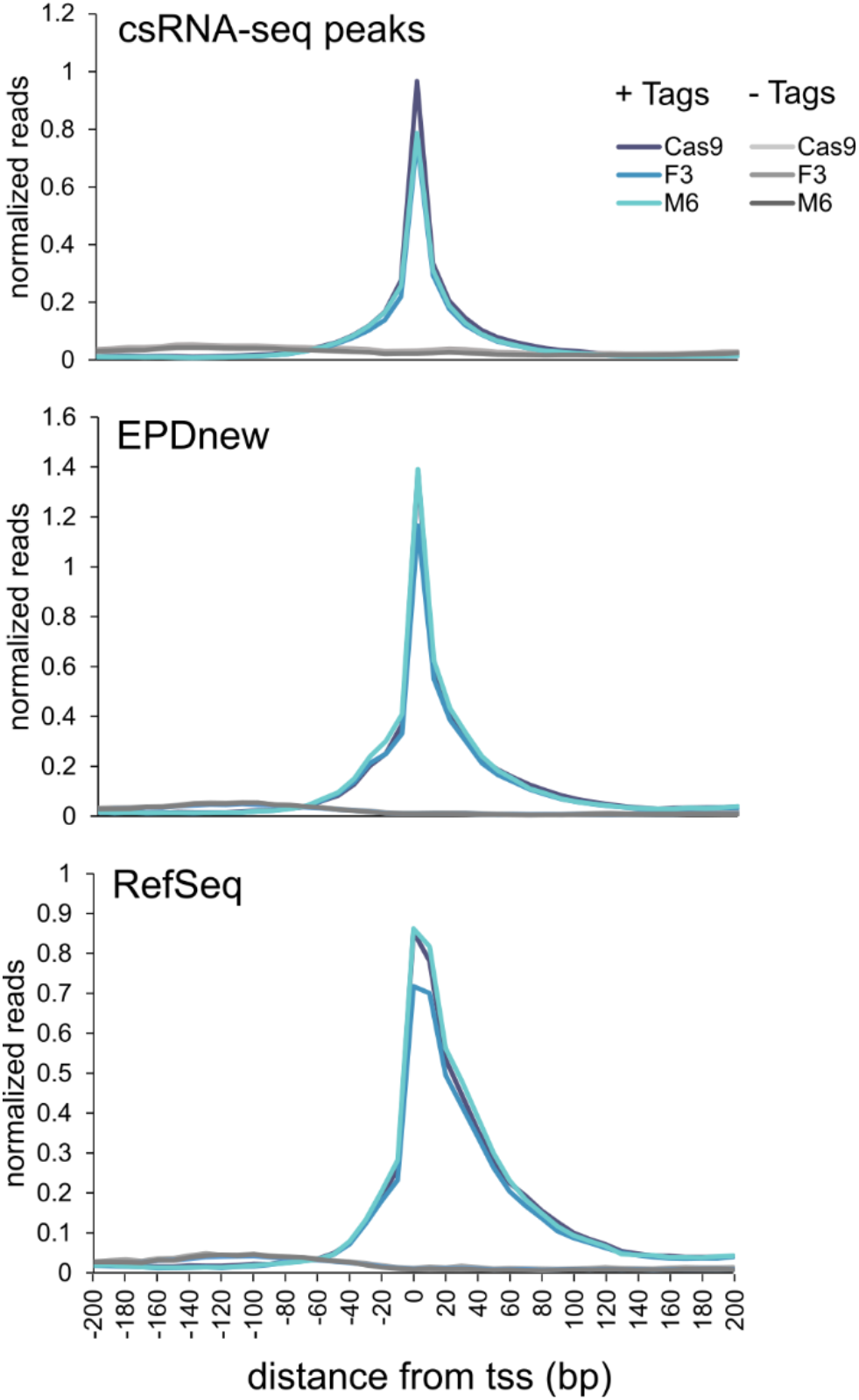
The choice of TSSs source result in slightly different nascent transcription profiles. The same csRNA-seq data (2-4h embryos of Cas9, F3 and M6 strains) was mapped using called csRNA-seq peaks, EPDnew or RefSeq. Note that csRNA-seq reads exhibited rather symmetrical distribution around EPDnew TSSs as compared to HOMER’s default (RefSeq). Transcription initiation seems to be detectable at sequences downstream of RefSeq-defined TSSs. In contrast, EPDnew-based TSSs exhibit a more symmetric distribution, similar to the csRNAseq-based TSSs. Divergent transcription (- tags) is barely detectable, consistent with previous reports that divergent transcription is predominantly absent in *Drosophila melanogaster* (Core et al. 2012; Meers et al. 2018).

**Supplemental Figure S7.**
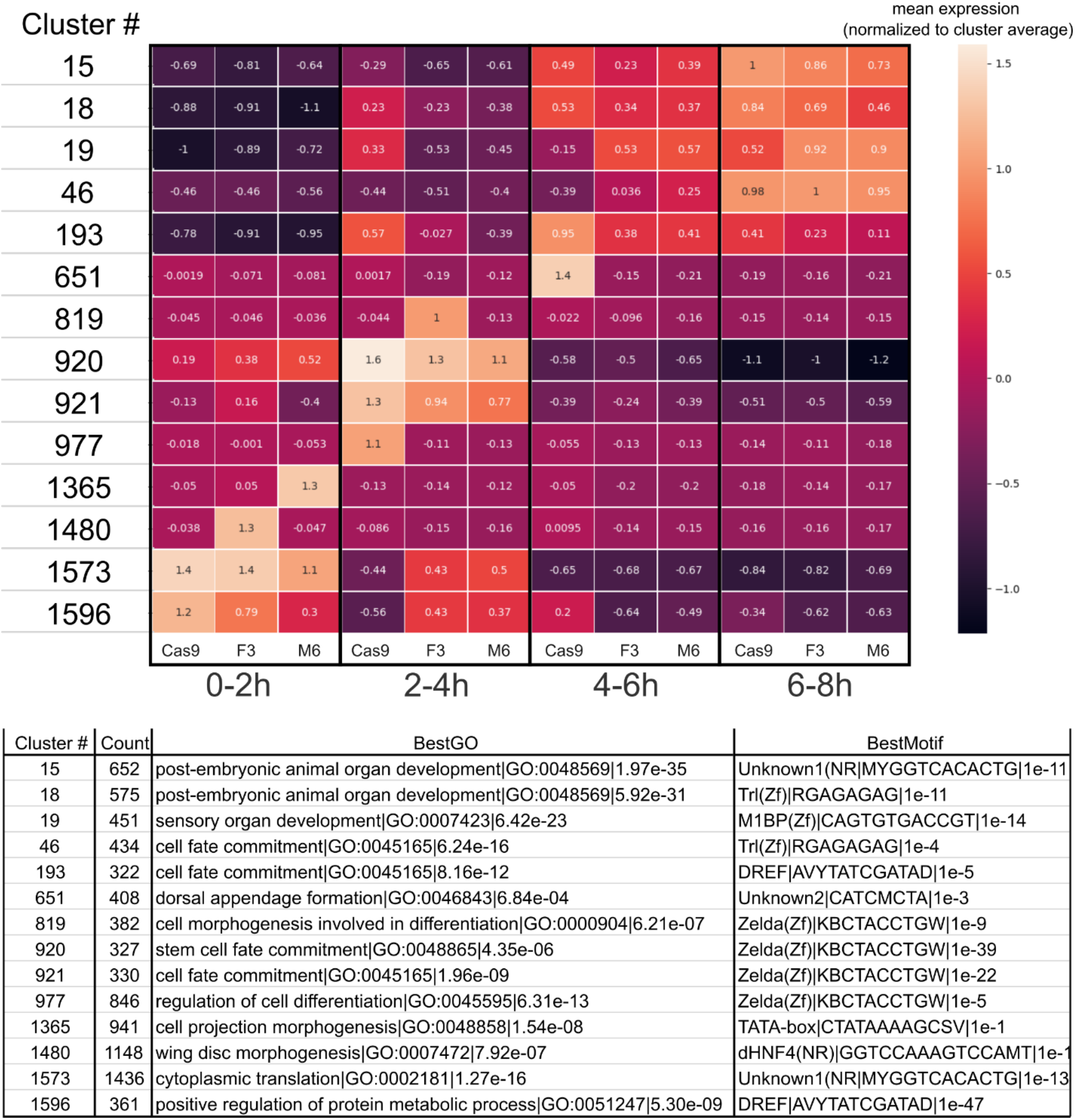
Full output of the analyzeClusters.pl script. Each cluster is annotated with cluster number, number of consisting peaks (n), top enriched HOMER motif and GO term. For each sample, the mean average expression of peaks in the cluster was normalized by the average expression per cluster (derived from all the samples). The following command was used: *analyzeClusters.pl -i all.rlog.byTime.txt -o ./cluster.allRlog.c300.t8/ -peaks -minDiff 1 -center -genome dm6 -size -100,100 -thresh -0.8 -min 300 -cpu 30*

**Supplemental Figure S8.**
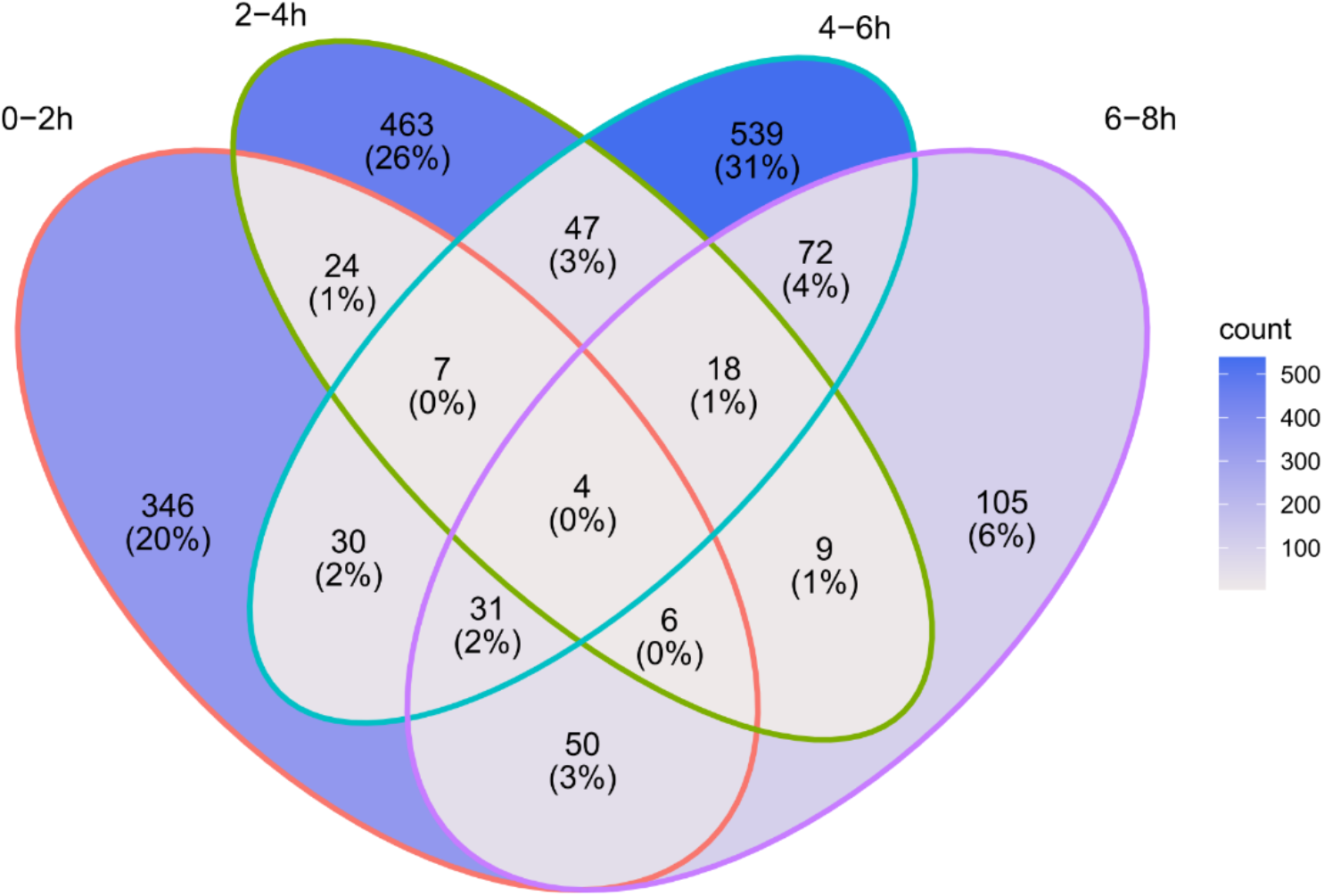
The majority of differentially-expressed csRNA-seq peaks do not overlap across the tested time points. The overlap between differentially-expressed peaks for each time point was examined using ggVennDiagram R package (Gao et al. 2021). Each section is colored according to the number of peaks.

**Supplemental Figure S9.**
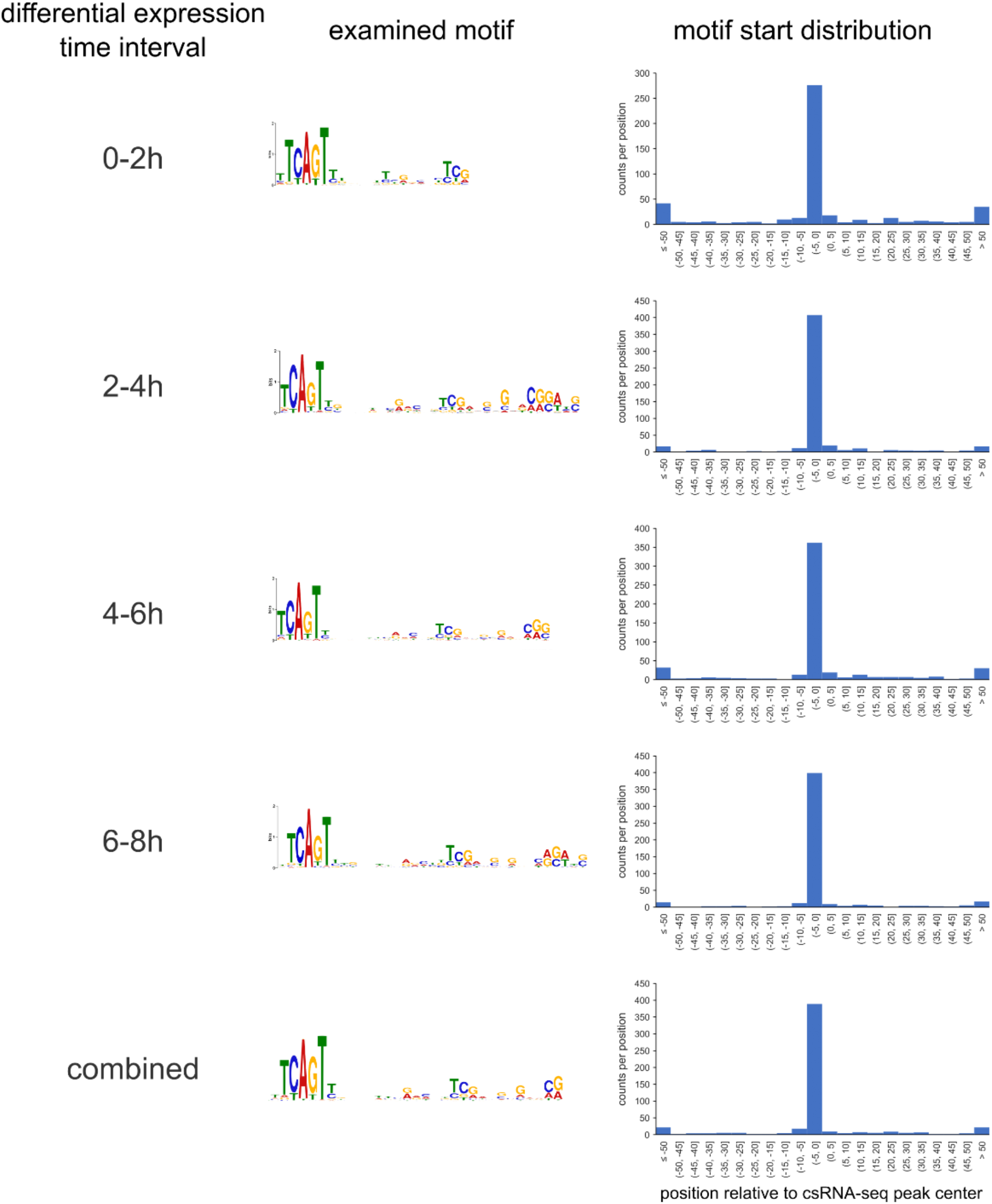
The detected Inr+DPE motifs are strictly positioned relative to the differentially expressed csRNA-seq peaks. The lists of differentially expressed peaks at each time point, and at all time points combined, were scanned using the Inr+DPE-like motifs detected for each time point (shown on the right, same as in Figure 6A). A specific enrichment at positions ±5bp is evident for all motifs. The indicated positions (x-axis) are of the motif start relative to the detected csRNA-seq peak center.

**SupFile1_ClusterSum_exp_GO.xlsx**

Summary of expression profiles and selected GO terms for each cluster, used for Figure 6.

**SupFile2_cluster.allRlog.c300.t8.zip**

Full output of the HOMER-based analyzeClusters.pl script with the following command: *analyzeClusters.pl -i all.rlog.byTime.txt -o ./cluster.allRlog.c300.t8/ -peaks -minDiff 1 -center -genome dm6 -size -100,100 -thresh -0.8 -min 300 -cpu 30*

The script performs clustering based on expression, then each cluster is analyzed for both GO terms and TF motifs enrichment. The output contains both summary files an individual cluster analysis (in separate folders).

**SupFile3_MEMEoutput.zip**

Complete MEME output of differentially expressed csRNA-seq peaks at individual and combined timepoints. BED files used to extract the sequences are provided as well.

**SupFile4_primers.xlsx**

RT-qPCR primers used in the manuscript.

## References

Anish R, Hossain MB, Jacobson RH, Takada S. 2009. Characterization of transcription from TATA-less promoters: identification of a new core promoter element XCPE2 and analysis of factor requirements. PloS one 4: e5103.

Azpiazu N, Frasch M. 1993. tinman and bagpipe: two homeo box genes that determine cell fates in the dorsal mesoderm of Drosophila. Genes Dev 7: 1325–1340.

Bailey TL, Elkan C. 1994. Fitting a mixture model by expectation maximization to discover motifs in biopolymers. Proc Int Conf Intell Syst Mol Biol 2: 28–36.

Benson DW, Silberbach GM, Kavanaugh-McHugh A, Cottrill C, Zhang Y, Riggs S, Smalls O, Johnson MC, Watson MS, Seidman JG et al. 1999. Mutations in the cardiac transcription factor NKX2.5 affect diverse cardiac developmental pathways. J Clin Invest 104: 1567–1573.

Bodmer R. 1993. The gene tinman is required for specification of the heart and visceral muscles in Drosophila. Development 118: 719–729.

Bodmer R, Jan LY, Jan YN. 1990. A new homeobox-containing gene, msh-2, is transiently expressed early during mesoderm formation of Drosophila. Development 110: 661–669.

Bodmer R, Venkatesh TV. 1998. Heart development in Drosophila and vertebrates: conservation of molecular mechanisms. Dev Genet 22: 181–186.

Bryantsev AL, Cripps RM. 2009. Cardiac gene regulatory networks in Drosophila. Biochimica et biophysica acta 1789: 343–353.

Burke TW, Kadonaga JT. 1996. Drosophila TFIID binds to a conserved downstream basal promoter element that is present in many TATA-box-deficient promoters. Genes & Development 10: 711–724.

Burke TW, Kadonaga JT. 1997. The downstream core promoter element, DPE, is conserved from Drosophila to humans and is recognized by TAF(II)60 of Drosophila. Genes & Development 11: 3020-3031.

Butler JE, Kadonaga JT. 2001. Enhancer-promoter specificity mediated by DPE or TATA core promoter motifs. Genes Dev 15: 2515–2519.

Cripps RM, Olson EN. 2002. Control of cardiac development by an evolutionarily conserved transcriptional network. Developmental biology 246: 14–28.

Danino YM, Even D, Ideses D, Juven-Gershon T. 2015. The core promoter: At the heart of gene expression. Biochimica et biophysica acta 1849: 1116–1131.

Deng W, Roberts SG. 2005. A core promoter element downstream of the TATA box that is recognized by TFIIB. Genes Dev 19: 2418–2423.

Dobin A, Davis CA, Schlesinger F, Drenkow J, Zaleski C, Jha S, Batut P, Chaisson M, Gingeras TR. 2013. STAR: ultrafast universal RNA-seq aligner. Bioinformatics 29: 15–21.

Dreos R, Sloutskin A, Malachi N, Ideses D, Bucher P, Juven-Gershon T. 2021. Computational identification and experimental characterization of preferred downstream positions in human core promoters. PLoS Comput Biol 17: e1009256.

Duttke SH. 2014. RNA polymerase III accurately initiates transcription from RNA polymerase II promoters in vitro. The Journal of biological chemistry 289: 20396–20404.

Duttke SH, Beyhan S, Singh R, Neal S, Viriyakosol S, Fierer J, Kirkland TN, Stajich JE, Benner C, Carlin AF. 2022. Decoding Transcription Regulatory Mechanisms Associated with Coccidioides immitis Phase Transition Using Total RNA. mSystems 7: e0140421.

Duttke SH, Chang MW, Heinz S, Benner C. 2019. Identification and dynamic quantification of regulatory elements using total RNA. Genome Res 29: 1836–1846.

Elliott DA, Kirk EP, Yeoh T, Chandar S, McKenzie F, Taylor P, Grossfeld P, Fatkin D, Jones O, Hayes P et al. 2003. Cardiac homeobox gene NKX2-5 mutations and congenital heart disease: associations with atrial septal defect and hypoplastic left heart syndrome. J Am Coll Cardiol 41: 2072–2076.

Fink M, Callol-Massot C, Chu A, Ruiz-Lozano P, Izpisua Belmonte JC, Giles W, Bodmer R, Ocorr K. 2009. A new method for detection and quantification of heartbeat parameters in Drosophila, zebrafish, and embryonic mouse hearts. Biotechniques 46: 101–113.

Frasch M. 2016. Genome-Wide Approaches to Drosophila Heart Development. Journal of cardiovascular development and disease 3.

Fukaya T, Lim B, Levine M. 2016. Enhancer Control of Transcriptional Bursting. Cell 166: 358–368.

Gajewski K, Kim Y, Lee YM, Olson EN, Schulz RA. 1997. D-mef2 is a target for Tinman activation during Drosophila heart development. EMBO J 16: 515–522.

Ge DT, Tipping C, Brodsky MH, Zamore PD. 2016. Rapid Screening for CRISPR-Directed Editing of the Drosophila Genome Using white Coconversion. G3 6: 3197–3206.

Goldberg ML. 1979. Sequence analysis of Drosophila histone genes. Ph.D. Thesis, Stanford University

Gu Z, Eils R, Schlesner M. 2016. Complex heatmaps reveal patterns and correlations in multidimensional genomic data. Bioinformatics 32: 2847–2849.

Haberle V, Stark A. 2018. Eukaryotic core promoters and the functional basis of transcription initiation. Nat Rev Mol Cell Biol 19: 621–637.

Harvey RP. 1996. NK-2 homeobox genes and heart development. Developmental biology 178: 203–216.

Heintzman ND, Ren B. 2007. The gateway to transcription: identifying, characterizing and understanding promoters in the eukaryotic genome. Cell Mol Life Sci 64: 386–400.

Heinz S, Benner C, Spann N, Bertolino E, Lin YC, Laslo P, Cheng JX, Murre C, Singh H, Glass CK. 2010. Simple combinations of lineage-determining transcription factors prime cis-regulatory elements required for macrophage and B cell identities. Mol Cell 38: 576–589.

Hendrix DA, Hong JW, Zeitlinger J, Rokhsar DS, Levine MS. 2008. Promoter elements associated with RNA Pol II stalling in the Drosophila embryo. Proceedings of the National Academy of Sciences of the United States of America 105: 7762–7767.

Hetzel J, Duttke SH, Benner C, Chory J. 2016. Nascent RNA sequencing reveals distinct features in plant transcription. Proceedings of the National Academy of Sciences of the United States of America 113: 12316–12321.

Jin H, Stojnic R, Adryan B, Ozdemir A, Stathopoulos A, Frasch M. 2013. Genome-wide screens for in vivo Tinman binding sites identify cardiac enhancers with diverse functional architectures. PLoS genetics 9: e1003195.

José A. Campos-Ortega VH. 1985. The Embryonic Development of Drosophila melanogaster. Springer Berlin, Heidelberg.

Junion G, Spivakov M, Girardot C, Braun M, Gustafson EH, Birney E, Furlong EE. 2012. A transcription factor collective defines cardiac cell fate and reflects lineage history. Cell 148: 473–486.

Juven-Gershon T, Hsu J-Y, Kadonaga JT. 2008a. Caudal, a key developmental regulator, is a DPE-specific transcriptional factor. Genes & Development 22: 2823–2830.

Juven-Gershon T, Hsu JY, Theisen JW, Kadonaga JT. 2008b. The RNA polymerase II core promoter - the gateway to transcription. Current opinion in cell biology 20: 253–259.

Juven-Gershon T, Kadonaga JT. 2010. Regulation of gene expression via the core promoter and the basal transcriptional machinery. Developmental biology 339: 225–229.

Kumar S, Konikoff C, Van Emden B, Busick C, Davis KT, Ji S, Wu LW, Ramos H, Brody T, Panchanathan S et al. 2011. FlyExpress: visual mining of spatiotemporal patterns for genes and publications in Drosophila embryogenesis. Bioinformatics 27: 3319–3320.

Kutach AK, Kadonaga JT. 2000. The downstream promoter element DPE appears to be as widely used as the TATA box in Drosophila core promoters. Molecular and cellular biology 20: 4754–4764.

Lagrange T, Kapanidis AN, Tang H, Reinberg D, Ebright RH. 1998. New core promoter element in RNA polymerase II-dependent transcription: sequence-specific DNA binding by transcription factor IIB. Genes Dev 12: 34–44.

Lepage SIM, Sharma R, Dukoff D, Stalker L, LaMarre J, Koch TG. 2021. Gene Expression Profile Is Different between Intact and Enzymatically Digested Equine Articular Cartilage. Cartilage 12: 222–225.

Levi T, Sloutskin A, Kalifa R, Juven-Gershon T, Gerlitz O. 2020. Efficient In Vivo Introduction of Point Mutations Using ssODN and a Co-CRISPR Approach. Biological procedures online 22: 14.

Lim B, Levine MS. 2021. Enhancer-promoter communication: hubs or loops? Curr Opin Genet Dev 67: 5–9.

Lim CY, Santoso B, Boulay T, Dong E, Ohler U, Kadonaga JT. 2004. The MTE, a new core promoter element for transcription by RNA polymerase II. Genes & Development 18: 1606–1617.

Liu YH, Jakobsen JS, Valentin G, Amarantos I, Gilmour DT, Furlong EE. 2009. A systematic analysis of Tinman function reveals Eya and JAK-STAT signaling as essential regulators of muscle development. Developmental cell 16: 280–291.

Lo K, Smale ST. 1996. Generality of a functional initiator consensus sequence. Gene 182: 13–22.

Lo PC, Frasch M. 2003. Establishing A-P polarity in the embryonic heart tube: a conserved function of Hox genes in Drosophila and vertebrates? Trends in cardiovascular medicine 13: 182–187.

Lyons I, Parsons LM, Hartley L, Li R, Andrews JE, Robb L, Harvey RP. 1995. Myogenic and morphogenetic defects in the heart tubes of murine embryos lacking the homeo box gene Nkx2-5. Genes Dev 9: 1654–1666.

Martinez-Ara M, Comoglio F, van Arensbergen J, van Steensel B. 2022. Systematic analysis of intrinsic enhancer-promoter compatibility in the mouse genome. Mol Cell 82: 2519–2531 e2516.

McElhinney DB, Geiger E, Blinder J, Benson DW, Goldmuntz E. 2003. NKX2.5 mutations in patients with congenital heart disease. J Am Coll Cardiol 42: 1650–1655.

Meylan P, Dreos R, Ambrosini G, Groux R, Bucher P. 2020. EPD in 2020: enhanced data visualization and extension to ncRNA promoters. Nucleic Acids Res 48: D65–D69.

Ocorr K, Fink M, Cammarato A, Bernstein S, Bodmer R. 2009. Semi-automated Optical Heartbeat Analysis of small hearts. J Vis Exp.

Ohler U, Liao GC, Niemann H, Rubin GM. 2002. Computational analysis of core promoters in the Drosophila genome. Genome biology 3: RESEARCH0087.

Osterwalder M, Barozzi I, Tissieres V, Fukuda-Yuzawa Y, Mannion BJ, Afzal SY, Lee EA, Zhu Y, Plajzer-Frick I, Pickle CS et al. 2018. Enhancer redundancy provides phenotypic robustness in mammalian development. Nature 554: 239–243.

Parry TJ, Theisen JWM, Hsu J-Y, Wang Y-L, Corcoran DL, Eustice M, Ohler U, Kadonaga JT. 2010. The TCT motif, a key component of an RNA polymerase II transcription system for the translational machinery. Genes & Development 24: 2013–2018.

Pimmett VL, Dejean M, Fernandez C, Trullo A, Bertrand E, Radulescu O, Lagha M. 2021. Quantitative imaging of transcription in living Drosophila embryos reveals the impact of core promoter motifs on promoter state dynamics. Nat Commun 12: 4504.

Policastro RA, Zentner GE. 2021. Global approaches for profiling transcription initiation. Cell Rep Methods 1.

Reim I, Frasch M. 2005. The Dorsocross T-box genes are key components of the regulatory network controlling early cardiogenesis in Drosophila. Development 132: 4911–4925.

Reim I, Frasch M. 2010. Genetic and genomic dissection of cardiogenesis in the Drosophila model. Pediatric cardiology 31: 325–334.

Rotstein B, Paululat A. 2016. On the Morphology of the Drosophila Heart. Journal of cardiovascular development and disease 3.

Ryan KM, Hendren JD, Helander LA, Cripps RM. 2007. The NK homeodomain transcription factor Tinman is a direct activator of seven-up in the Drosophila dorsal vessel. Developmental biology 302: 694–702.

Schott JJ, Benson DW, Basson CT, Pease W, Silberbach GM, Moak JP, Maron BJ, Seidman CE, Seidman JG. 1998. Congenital heart disease caused by mutations in the transcription factor NKX2-5. Science 281: 108–111.

Sloutskin A, Danino YM, Orenstein Y, Zehavi Y, Doniger T, Shamir R, Juven-Gershon T. 2015. ElemeNT: a computational tool for detecting core promoter elements. Transcription 6: 41–50.

Sloutskin A, Shir-Shapira H, Freiman RN, Juven-Gershon T. 2021. The Core Promoter Is a Regulatory Hub for Developmental Gene Expression. Front Cell Dev Biol 9: 666508.

Smale ST, Baltimore D. 1989. The "initiator" as a transcription control element. Cell 57: 103–113.

Spivakov M. 2014. Spurious transcription factor binding: non-functional or genetically redundant? Bioessays 36: 798-806.

Stutt N, Song M, Wilson MD, Scott IC. 2022. Cardiac specification during gastrulation - The Yellow Brick Road leading to Tinman. Semin Cell Dev Biol 127: 46–58.

Targoff KL, Colombo S, George V, Schell T, Kim S-H, Solnica-Krezel L, Yelon D. 2013. Nkx genes are essential for maintenance of ventricular identity. Development 140: 4203–4213.

Targoff KL, Schell T, Yelon D. 2008. Nkx genes regulate heart tube extension and exert differential effects on ventricular and atrial cell number. Developmental biology 322: 314–321.

Theisen JW, Lim CY, Kadonaga JT. 2010. Three key subregions contribute to the function of the downstream RNA polymerase II core promoter. Molecular and cellular biology 30: 3471–3479.

Tokusumi Y, Ma Y, Song X, Jacobson RH, Takada S. 2007. The new core promoter element XCPE1 (X Core Promoter Element 1) directs activator-, mediator-, and TATA-binding protein-dependent but TFIID-independent RNA polymerase II transcription from TATA-less promoters. Molecular and cellular biology 27: 1844–1858.

Tomancak P, Beaton A, Weiszmann R, Kwan E, Shu S, Lewis SE, Richards S, Ashburner M, Hartenstein V, Celniker SE et al. 2002. Systematic determination of patterns of gene expression during Drosophila embryogenesis. Genome biology 3: RESEARCH0088.

Tomancak P, Berman BP, Beaton A, Weiszmann R, Kwan E, Hartenstein V, Celniker SE, Rubin GM. 2007. Global analysis of patterns of gene expression during Drosophila embryogenesis. Genome biology 8: R145.

Vo Ngoc L, Cassidy CJ, Huang CY, Duttke SH, Kadonaga JT. 2017. The human initiator is a distinct and abundant element that is precisely positioned in focused core promoters. Genes Dev 31: 6–11.

Vo Ngoc L, Huang CY, Cassidy CJ, Medrano C, Kadonaga JT. 2020. Identification of the human DPR core promoter element using machine learning. Nature 585: 459–463.

Vo Ngoc L, Kassavetis GA, Kadonaga JT. 2019. The RNA Polymerase II Core Promoter in Drosophila. Genetics 212: 13–24.

Vo Ngoc L, Rhyne TE, Kadonaga JT. 2023. Analysis of the Drosophila and human DPR elements reveals a distinct human variant whose specificity can be enhanced by machine learning. Genes Dev 37: 377–382.

Vogler G, Ocorr K. 2009. Visualizing the beating heart in Drosophila. J Vis Exp.

Wang J, Zhao S, He W, Wei Y, Zhang Y, Pegg H, Shore P, Roberts SGE, Deng W. 2017. A transcription factor IIA-binding site differentially regulates RNA polymerase II-mediated transcription in a promoter context-dependent manner. The Journal of biological chemistry 292: 11873–11885.

Ward EJ, Skeath JB. 2000. Characterization of a novel subset of cardiac cells and their progenitors in the Drosophila embryo. Development 127: 4959–4969.

Wissink EM, Vihervaara A, Tippens ND, Lis JT. 2019. Nascent RNA analyses: tracking transcription and its regulation. Nat Rev Genet 20: 705–723.

Yao L, Liang J, Ozer A, Leung AK, Lis JT, Yu H. 2022. A comparison of experimental assays and analytical methods for genome-wide identification of active enhancers. Nat Biotechnol 40: 1056–1065.

Yin Z, Xu XL, Frasch M. 1997. Regulation of the twist target gene tinman by modular cis-regulatory elements during early mesoderm development. Development 124: 4971–4982.

Yokoshi M, Kawasaki K, Cambon M, Fukaya T. 2022. Dynamic modulation of enhancer responsiveness by core promoter elements in living Drosophila embryos. Nucleic Acids Res 50: 92–107.

Yokoshi M, Segawa K, Fukaya T. 2020. Visualizing the Role of Boundary Elements in Enhancer-Promoter Communication. Mol Cell 78: 224–235 e225.

Zabidi MA, Arnold CD, Schernhuber K, Pagani M, Rath M, Frank O, Stark A. 2015. Enhancer-core-promoter specificity separates developmental and housekeeping gene regulation. Nature 518: 556–559.

Zaffran S, Reim I, Qian L, Lo PC, Bodmer R, Frasch M. 2006. Cardioblast-intrinsic Tinman activity controls proper diversification and differentiation of myocardial cells in Drosophila. Development 133: 4073–4083.

Zehavi Y, Kuznetsov O, Ovadia-Shochat A, Juven-Gershon T. 2014a. Core promoter functions in the regulation of gene expression of Drosophila dorsal target genes. The Journal of biological chemistry 289: 11993–12004.

Zehavi Y, Sloutskin A, Kuznetsov O, Juven-Gershon T. 2014b. The core promoter composition establishes a new dimension in developmental gene networks. Nucleus 5.

Zelhof AC, Yao TP, Chen JD, Evans RM, McKeown M. 1995. Seven-up inhibits ultraspiracle-based signaling pathways in vitro and in vivo. Molecular and cellular biology 15: 6736–6745.

Zhou T, Chiang CM. 2001. The intronless and TATA-less human TAF(II)55 gene contains a functional initiator and a downstream promoter element. The Journal of biological chemistry 276: 25503–25511.

Zinzen RP, Girardot C, Gagneur J, Braun M, Furlong EE. 2009. Combinatorial binding predicts spatio-temporal cis-regulatory activity. Nature 462: 65–70.

## References

Core LJ, Waterfall JJ, Gilchrist DA, Fargo DC, Kwak H, Adelman K, Lis JT. 2012. Defining the status of RNA polymerase at promoters. Cell Rep 2: 1025–1035.

Gao CH, Yu G, Cai P. 2021. ggVennDiagram: An Intuitive, Easy-to-Use, and Highly Customizable R Package to Generate Venn Diagram. Front Genet 12: 706907.

Meers MP, Adelman K, Duronio RJ, Strahl BD, McKay DJ, Matera AG. 2018. Transcription start site profiling uncovers divergent transcription and enhancer-associated RNAs in Drosophila melanogaster. BMC Genomics 19: 157.

Roote J, Prokop A. 2013. How to design a genetic mating scheme: a basic training package for Drosophila genetics. G3 3: 353–358.

Wilk R, Murthy SUM, Yan. H, Krause HM. 2010. In Situ Hybridization: Fruit Fly Embryos and Tissues. Curr Protoc Essential Lab Tech 4: 9.3.1-9.3.24.

