## Supplementary material for "A single DPE core promoter motif contributes to *in vivo* transcriptional regulation and affects cardiac function": SupFile2: geneOntology.html

Gene Ontology Results

### Gene Ontology Enrichment Results

Homer *de novo* Motif Enrichment Results  
Known Motif Enrichment Results  

###### Text file version of complete results (i.e. open with Excel) - biological process: Functional groupings of proteins (Gene Ontology) - molecular function: Mechanistic actions of proteins (Gene Ontology) - cellular component: Protein localization (Gene Ontology) - chromosome location: Genes with similar chromosome localization (NCBI Gene) - KEGG pathways: Groups of proteins in the same pathways (KEGG) - protein interactions: "Proteins interacting with a common protein (BIND, EcoCyc, HPRD)" (NCBI Gene) - interpro domains: Proteins with similar domains and features (Interpro) - pfam domains: Proteins with similar domains and features (Pfam) - smart domains: Proteins with similar domains and features (SMART) - gene3d domains: Proteins with similar domains and features (Gene3D) - prosite domains: Proteins with similar domains and features (Prosite) - prints domains: Proteins with similar domains and features (PRINTS) - MSigDB lists: "Genes sets for pathways, factor/miRNA target predictions, expression patterns, etc." (MSigDB) - BIOCYC pathways: Groups of proteins in the same pathways (BIOCYC) - COSMIC cancer mutations: Genes mutated in similar cancers (COSMIC) - GWAS genes: Genes mutated in similar diseases (GWAS Catalog) - Lipid Maps pathways: Groups of proteins in the same lipid pathways (Lipid Maps/Biosystems) - Pathway Interaction DB: Groups of proteins in the same pathways (Pathway Interaction Database) - REACTOME pathways: Groups of proteins in the same pathways (REACTOME) - SMPDB pathways: Groups of proteins in the same pathways (SMPDB) - WikiPathways: Groups of proteins in the same pathways (Wikipathways) Enriched Categories | | | | | | | | | | | | --- | --- | --- | --- | --- | --- | --- | --- | --- | --- | | P-value | ln(P) | Term | GO Tree | GO ID | # of Genes in Term | # of Target Genes in Term | # of Total Genes | # of Target Genes | Common Genes | | 8.324e-52 | -117.62 | system development | biological process | GO:0048731 | 1808 | 194 | 12345 | 442 | nuf,Ptp4E,frc,Atg17,siz,Dl,cv-d,Dronc,H,emc,mop,hyx,fz,sns,tkv,flfl,Mef2,tai,ey,Sox14,ft,Pvf3,edl,Drak,vn,bowl,Fkbp14,Bsg,scrib,comm2,betaTub60D,Ppcs,zfh2,beat-IIa,RhoGAP19D,unc-119,InR,nub,sick,Egfr,LanA,cact,al,glec,aop,p130CAS,spdo,chn,rg,path,Scr,fz2,spi,ttk,otk,Eph,Kdm2,Arf79F,Actbeta,Gli,E(Pc),Nrk,Snoo,fl(2)d,mam,hth,h,disco-r,jar,boi,htl,Dad,crp,kon,Rab5,cic,Tom,14-3-3epsilon,kay,eff,Poxn,arm,chic,CG6191,Cdk4,corto,gukh,Psc,ed,kn,fuss,ena,step,wgn,sli,f,C1GalTA,bou,dpy,Src42A,ss,Sema1b,pum,PI31,abd-A,moody,Inx2,Cbl,ds,Sema1a,rictor,sqz,baz,Abd-B,Mpcp2,CG34401,robo1,Syx1A,Abl,Adf1,pnr,Cks30A,AP-1gamma,Cip4,Tollo,ap,sens-2,cindr,N,drl,dock,Mob2,eyg,ec,tara,SMC3,sina,spen,B-H1,rdx,dome,shg,tsh,Fas1,Dg,wnd,fs(1)h,sens,RhoL,SCAR,imd,Irk3,Hr39,Lmx1a,unc-5,d,eya,rib,Rab1,Raf,nmo,Pdk1,crol,Alh,IA-2,E(spl)m5-HLH,yrt,Diap1,CtBP,Pen,Socs36E,shot,Pvr,px,pdm3,dlp,dve,klu,jeb,ph-p,S,fend,grn,CG1129,msi,sim,esg,cora,sd,bab2,Sxl,lilli,klar,jumu | | 9.319e-50 | -112.90 | anatomical structure morphogenesis | biological process | GO:0009653 | 1579 | 178 | 12345 | 442 | nmo,crol,scyl,CtBP,Dp,Diap1,yrt,fs(1)h,sens,Dg,wnd,SCAR,Hr39,Raf,unc-5,d,eya,rib,msi,cora,sd,sim,esg,bab2,lilli,jumu,klar,Socs36E,Pvr,shot,Pen,pdm3,px,klu,jeb,dlp,dve,grn,ph-p,S,fend,Abd-B,rictor,baz,sqz,Atg16,Syx1A,robo1,Abl,Cip4,Tollo,pnr,Cks30A,abd-A,pum,Sema1b,Inx2,Cbl,Myo10A,Sema1a,ds,SMC3,ec,rdx,B-H1,sina,spen,CenB1A,dome,Fas1,shg,tsh,orb2,ap,drl,N,cindr,eyg,dock,Mob2,boi,htl,Dad,jar,Rab5,kon,cic,Tom,kay,14-3-3epsilon,Poxn,arm,eff,Arf79F,trbl,otk,Eph,Actbeta,Gli,Nrk,Snoo,E(Pc),h,hth,disco-r,mam,wgn,ena,step,f,C1GalTA,sli,dpy,Src42A,ss,RhoU,chic,corto,CG6191,kn,Psc,gukh,ed,fuss,Orct2,tkv,Mef2,ey,Sox14,ft,Shrm,tai,Atg17,Ptp4E,frc,siz,nuf,Dl,hyx,emc,Dronc,cv-d,H,yuri,fz,sns,LanA,cact,Egfr,sick,aop,al,rg,spdo,chn,ttk,path,Scr,fz2,spi,Bsg,neo,Pgk,bowl,Drak,vn,Fkbp14,zfh2,beat-IIa,RhoGAP19D,scrib,betaTub60D,Ppcs,comm2,14-3-3zeta,chrb,nub,InR | | 7.659e-47 | -106.19 | animal organ development | biological process | GO:0048513 | 1267 | 155 | 12345 | 442 | Poxn,arm,eff,kay,14-3-3epsilon,Rab5,kon,Tom,cic,crp,htl,boi,Dad,jar,hth,h,disco-r,mam,Nrk,fl(2)d,Gli,Kdm2,Arf79F,otk,Eph,Src42A,ss,dpy,f,sli,ena,step,fuss,kn,Psc,ed,corto,CG6191,chic,Pvf3,ey,ft,Mef2,tkv,sns,fz,emc,hyx,mop,Dronc,cv-d,H,Dl,Atg17,frc,siz,nuf,ttk,path,Scr,spi,fz2,rg,spdo,chn,aop,al,LanA,cact,Egfr,InR,nub,zfh2,RhoGAP19D,scrib,betaTub60D,Ppcs,edl,Drak,bowl,vn,Fkbp14,CtBP,yrt,Diap1,Alh,crol,nmo,Raf,unc-5,rib,eya,d,Hr39,Irk3,RhoL,SCAR,sens,Dg,lilli,klar,jumu,Sxl,bab2,cora,sd,sim,esg,CG1129,msi,grn,S,klu,jeb,dlp,dve,px,Socs36E,Pvr,shot,Pen,Cip4,pnr,AP-1gamma,Cks30A,Abl,Syx1A,robo1,Abd-B,Mpcp2,baz,Sema1a,ds,Cbl,moody,abd-A,PI31,Sema1b,shg,tsh,dome,B-H1,rdx,sina,spen,tara,SMC3,ec,eyg,Mob2,drl,cindr,N,sens-2,ap | | 2.138e-45 | -102.86 | animal organ morphogenesis | biological process | GO:0009887 | 782 | 120 | 12345 | 442 | fuss,S,grn,dlp,ed,dve,klu,kn,jeb,px,CG6191,corto,chic,Socs36E,shot,Pvr,lilli,Src42A,ss,jumu,klar,bab2,dpy,sli,esg,f,cora,sd,ena,step,msi,unc-5,mam,eya,rib,d,hth,h,Raf,disco-r,Nrk,Hr39,SCAR,Gli,Dg,otk,Arf79F,sens,eff,Diap1,yrt,Poxn,CtBP,arm,kay,crol,Rab5,cic,Tom,jar,boi,Dad,nmo,Mob2,eyg,scrib,Ppcs,cindr,N,zfh2,drl,RhoGAP19D,bowl,vn,Drak,ap,Fkbp14,shg,Scr,path,tsh,fz2,spi,ttk,spdo,chn,dome,rg,sina,al,spen,aop,rdx,B-H1,Egfr,ec,LanA,SMC3,sns,fz,ds,Dronc,H,cv-d,Cbl,emc,hyx,Dl,nuf,frc,Atg17,siz,pnr,Cks30A,Cip4,ey,ft,robo1,Abd-B,tkv | | 6.316e-45 | -101.77 | regulation of cellular process | biological process | GO:0050794 | 3560 | 267 | 12345 | 442 | nmo,Hr4,Pdk1,Alh,fz3,E(spl)m5-HLH,CtBP,Dp,hng3,lqf,sens,Tis11,CG12769,Lmx1a,unc-5,eya,Rab1,CG5522,Raf,CG34376,msi,esg,REPTOR,bin3,Sxl,jumu,Socs36E,pxb,pdm3,Lk6,dlp,H2.0,Spn42De,S,ph-p,hfw,rho-4,grn,Ddr,rictor,baz,morgue,Atg16,Syx1A,wdb,Cip4,Cals,Tollo,rgr,moody,Cbl,SERCA,ds,hang,Sema1a,ec,CG11617,tara,312,spen,Kdm4B,B-H1,rdx,shg,otp,dmrt11E,Jupiter,pita,CG11360,drl,Mob2,eyg,jar,boi,Dad,crp,Rab5,kon,cic,14-3-3epsilon,eff,Poxn,arm,trbl,apolpp,Gli,Pat1,Nrk,Snoo,mam,h,disco-r,ena,wgn,mrt,MEP-1,lncRNA:bxd,dpy,CG8312,Cdk4,corto,ed,kek5,Pitslre,tkv,Dcr-2,Mef2,tai,ft,Sox14,Ptp4E,CycG,hyx,emc,CycB3,Egfr,LanA,cact,Pez,Scr,spi,ttk,RhoGEF3,vn,bowl,su(sable),Ir21a,Elba2,scrib,zfh2,chrb,hppy,Wdr62,nub,InR,mim,scyl,crol,Diap1,Dg,wnd,fs(1)h,CG9784,imd,SCAR,ITP,Hr39,d,Dyrk2,rib,HmgZ,ZnT41F,sim,sd,bab2,lilli,klar,Pvr,shot,bol,dve,klu,jeb,CG42788,Abd-B,CG34401,robo1,SNF4Agamma,Abl,Adf1,pnr,Cks30A,pum,Sema1b,PI31,abd-A,Rgl,LRR,Ocrl,CG10348,sina,CG10321,CG13204,dome,Su(z)2,tsh,ap,sens-2,orb2,N,cindr,FBXO11,dock,lt,CG4896,htl,ogre,Tom,HLH3B,kay,otk,Eph,Kdm2,Arf79F,cue,Ziz,Actbeta,Tet,fl(2)d,hth,step,sli,C1GalTA,Nmda1,Src42A,ss,chic,lncRNA:Hsromega,Rm62,RhoU,CG6191,tna,bru2,Psc,kn,Cerk,fuss,Orct2,CG1218,CG42684,ey,Tob,Pvf3,Cadps,RapGAP1,lin-28,nuf,CG6891,Atg17,lili,ps,siz,Hel89B,RhoGEF4,Dl,H,cv-d,Fem-1,Dronc,mop,fz,Ric,CG34172,al,aop,spdo,chn,Prdm13,fz2,edl,Fkbp14,Pgk,neo,14-3-3zeta,Magi,nAChRalpha4,RhoGAP19D | | 1.099e-44 | -101.22 | regulation of biological process | biological process | GO:0050789 | 3862 | 279 | 12345 | 442 | Src42A,Nmda1,ss,C1GalTA,sli,step,CG1218,fuss,Cerk,Orct2,kn,Psc,bru2,CG6191,tna,RhoU,lncRNA:Hsromega,Rm62,chic,kay,HLH3B,ogre,Tom,htl,CG4896,hth,Tet,fl(2)d,Actbeta,Arf79F,cue,Ziz,Kdm2,otk,Eph,fz2,Prdm13,spdo,chn,aop,al,CG34172,RhoGAP19D,nAChRalpha4,14-3-3zeta,Magi,neo,Pgk,edl,Fkbp14,RapGAP1,Pvf3,Cadps,ey,Tob,CG42684,Ric,fz,mop,cv-d,Fem-1,H,Dronc,RhoGEF4,Dl,Hel89B,Him,lili,Atg17,siz,ps,lin-28,nuf,CG6891,lilli,klar,bab2,sd,ZnT41F,sim,klu,CG42788,jeb,dve,bol,shot,Pvr,tipE,vig,Diap1,IA-2,crol,scyl,mld,mim,HmgZ,Dyrk2,d,rib,Hr39,ITP,CG2765,imd,RhoL,SCAR,CG9784,fs(1)h,Dg,wnd,tsh,CG10321,CG13204,dome,Su(z)2,sina,Ocrl,CG10348,lt,dock,FBXO11,DPCoAC,N,cindr,sens-2,orb2,ap,pnr,Cks30A,Adf1,SNF4Agamma,Abl,CG34401,robo1,Abd-B,LRR,PI31,Rgl,abd-A,pum,Sema1b,CG8312,MEP-1,lncRNA:bxd,dpy,mrt,wgn,ena,kek5,Pitslre,ed,Cdk4,corto,Poxn,arm,eff,14-3-3epsilon,Rab5,kon,cic,crp,boi,Dad,jar,h,disco-r,mam,Snoo,Nrk,Pat1,Gli,apolpp,trbl,ttk,RhoGEF3,path,CG32795,Scr,spi,Pez,LanA,cact,Egfr,Wdr62,nub,InR,hppy,chrb,zfh2,scrib,Elba2,su(sable),Ir21a,vn,bowl,Sox14,ft,tai,Mef2,tkv,Dcr-2,CycB3,hyx,emc,Ptp4E,CycG,jumu,Sxl,bin3,REPTOR,esg,CG34376,msi,hfw,grn,rho-4,ph-p,S,Spn42De,dlp,H2.0,Lk6,pdm3,Socs36E,pxb,CtBP,Dp,E(spl)m5-HLH,Alh,fz3,Pdk1,Hr4,nmo,wbl,CG5522,Rab1,Raf,Lmx1a,unc-5,eya,CG12769,Tis11,sens,lqf,hng3,Jupiter,shg,dmrt11E,otp,B-H1,rdx,Kdm4B,spen,jigr1,tara,312,ec,CG11617,eyg,Mob2,CG11360,drl,pita,Cip4,Cals,Tollo,wdb,Atg16,Syx1A,morgue,Ddr,rictor,baz,hang,Sema1a,SERCA,ds,Cbl,moody,rgr | | 2.027e-43 | -98.30 | biological regulation | biological process | GO:0065007 | 4350 | 296 | 12345 | 442 | ec,CG11617,tara,jigr1,312,Kdm4B,spen,rdx,B-H1,shg,dmrt11E,otp,Jupiter,pita,CG11360,drl,CG3149,Mob2,eyg,baz,Ddr,rictor,morgue,Atg16,Syx1A,wdb,Cip4,Cals,Tollo,rgr,moody,Cbl,SERCA,ds,hang,Sema1a,CG34376,msi,REPTOR,esg,cora,bin3,Sxl,jumu,Socs36E,pxb,Mid1,Lk6,pdm3,dlp,H2.0,Spn42De,ph-p,S,hfw,rho-4,grn,wbl,nmo,Hr4,Pdk1,Alh,fz3,E(spl)m5-HLH,Dp,CtBP,lqf,hng3,sens,Tis11,CG12769,unc-5,Lmx1a,eya,CG5522,Rab1,Raf,Egfr,LanA,cact,Pez,p130CAS,path,Scr,CG32795,spi,ttk,RhoGEF3,vn,bowl,su(sable),Ir21a,scrib,Elba2,Ppcs,zfh2,hppy,chrb,InR,Wdr62,nub,tkv,Dcr-2,Mef2,tai,Sox14,ft,Ptp4E,CycG,emc,hyx,CycB3,ena,wgn,Atox1,mrt,lncRNA:bxd,MEP-1,dpy,CG8312,Cdk4,corto,ed,kek5,Pitslre,Vps13,jar,boi,Dad,crp,kon,Rab5,cic,14-3-3epsilon,eff,Poxn,arm,trbl,apolpp,Pat1,Gli,Snoo,Nrk,mam,Esp,h,disco-r,CG10348,ATPCL,Ocrl,sina,CG10321,CG13204,CenB1A,Su(z)2,dome,tsh,ap,sens-2,orb2,DPCoAC,cindr,N,FBXO11,dock,lt,Abd-B,CG34401,robo1,SNF4Agamma,Abl,Adf1,pnr,Cks30A,pum,Sema1b,Rgl,PI31,abd-A,LRR,sim,ZnT41F,sd,bab2,lilli,klar,tipE,shot,Pvr,bol,dve,klu,CG42788,jeb,mim,mld,crol,scyl,IA-2,Diap1,yrt,vig,Dg,wnd,CG9784,fs(1)h,SCAR,imd,RhoL,CG17883,CG2765,Irk3,ITP,Hr39,Dyrk2,d,rib,HmgZ,CG34172,al,aop,spdo,chn,Prdm13,fz2,edl,Fkbp14,neo,Pgk,14-3-3zeta,Magi,drongo,nAChRalpha4,RhoGAP19D,CG42684,TBC1D5,Tob,ey,Pvf3,Cadps,RapGAP1,lin-28,CG6891,nuf,lili,Him,Atg17,siz,ps,Hel89B,RhoGEF4,Dl,Fem-1,cv-d,Dronc,H,mop,fz,Ric,step,sli,C1GalTA,bou,Nmda1,Src42A,ss,chic,Rm62,RhoU,lncRNA:Hsromega,tna,CG6191,bru2,Psc,kn,Cerk,fuss,Orct2,CG1218,CG4896,htl,ogre,Tom,HLH3B,kay,otk,Eph,cue,Arf79F,Kdm2,Ziz,CG7394,Actbeta,Tet,fl(2)d,hth | | 3.643e-41 | -93.11 | multicellular organism development | biological process | GO:0007275 | 2496 | 213 | 12345 | 442 | Irk3,imd,RhoL,SCAR,sens,fs(1)h,wnd,Dg,Raf,Rab1,d,eya,rib,unc-5,Lmx1a,Hr39,crol,scyl,Pdk1,nmo,mld,wbl,Dp,CtBP,yrt,Diap1,E(spl)m5-HLH,IA-2,Alh,pdm3,px,shot,Pvr,Socs36E,Pen,grn,hfw,fend,S,ph-p,jeb,klu,dve,dlp,sd,cora,sim,esg,CG1129,msi,klar,jumu,lilli,Sxl,bab2,Inx2,moody,abd-A,PI31,Sema1b,pum,Sema1a,ds,Cbl,Myo10A,robo1,Syx1A,Atg16,CG34401,Mpcp2,Abd-B,sqz,rictor,baz,Tollo,Cip4,Cks30A,AP-1gamma,larp,pnr,Adf1,wdb,Abl,drl,N,cindr,orb2,sens-2,ap,lt,eyg,dock,Mob2,B-H1,rdx,spen,sina,tara,SMC3,ec,Fas1,tsh,shg,dome,Pat1,Gli,Actbeta,Arf79F,Kdm2,Eph,otk,trbl,disco-r,hth,h,mam,fl(2)d,E(Pc),Snoo,Nrk,cic,Tom,kon,Rab5,crp,Dad,htl,boi,jar,arm,Poxn,eff,14-3-3epsilon,kay,corto,Cdk4,CG6191,chic,Orct2,fuss,kn,ed,gukh,Psc,C1GalTA,f,sli,wgn,step,ena,ss,Src42A,dpy,bou,Dl,siz,Atg17,frc,Ptp4E,nuf,alpha-Man-Ia,yuri,CycB3,fz,sns,mop,hyx,emc,H,cv-d,Dronc,flfl,Mef2,tkv,Pvf3,ft,Sox14,ey,tai,RhoGAP19D,beat-IIa,zfh2,betaTub60D,14-3-3zeta,comm2,Ppcs,scrib,Bsg,Fkbp14,Drak,vn,bowl,edl,nub,InR,unc-119,chrb,aop,glec,al,cact,LanA,Egfr,sick,ttk,fz2,spi,Scr,path,rg,chn,spdo,p130CAS | | 1.340e-39 | -89.51 | tissue development | biological process | GO:0009888 | 1113 | 135 | 12345 | 442 | px,Socs36E,Pvr,shot,S,ph-p,Npc2g,grn,dlp,dve,jeb,sim,esg,cora,sd,meso18E,lilli,jumu,bab2,Sxl,RhoL,Irk3,Dg,eya,rib,d,Raf,Hr39,crol,mim,nmo,yrt,Diap1,Dp,CtBP,Alh,cindr,N,drl,ap,eyg,sina,spen,B-H1,rdx,ec,tara,SMC3,shg,tsh,dome,Inx2,Sema1b,Dgat2,abd-A,SERCA,ds,Sema1a,Myo10A,Cbl,Atg16,Syx1A,robo1,baz,Mpcp2,Abd-B,pnr,Cks30A,Cip4,Tollo,Abl,CG6191,corto,chic,fuss,ed,kn,sli,f,ena,step,Src42A,ss,dpy,Gli,otk,mam,hth,disco-r,Rab5,cic,jar,htl,boi,Dad,Poxn,arm,14-3-3epsilon,kay,scrib,Ppcs,zfh2,RhoGAP19D,Ptpmeg2,bowl,vn,Drak,Fkbp14,neo,InR,al,aop,Egfr,LanA,cact,path,spi,ttk,Dl,nuf,frc,Ptp4E,CycG,fz,H,cv-d,emc,hyx,mop,Mef2,tkv,tai,Shrm,ey,ft | | 1.581e-39 | -89.34 | developmental process | biological process | GO:0032502 | 2988 | 233 | 12345 | 442 | Kdm2,Arf79F,Eph,trbl,otk,Pat1,Gli,Actbeta,fl(2)d,Snoo,E(Pc),Tet,Nrk,disco-r,hth,h,mam,Dad,htl,boi,jar,Tom,cic,kon,Rab5,crp,kay,14-3-3epsilon,His2Av,arm,Poxn,eff,lncRNA:Hsromega,RhoU,chic,corto,Cdk4,CG6191,kn,ed,gukh,Psc,Orct2,fuss,wgn,step,ena,C1GalTA,f,sli,dpy,bou,ss,Src42A,siz,CycG,frc,Atg17,Ptp4E,nuf,lin-28,Dl,hyx,mop,emc,cv-d,H,Dronc,alpha-Man-Ia,yuri,CycB3,sns,fz,tkv,Mef2,flfl,Sox14,ft,ey,Shrm,tai,Pvf3,Pgk,neo,Bsg,Fkbp14,Ptpmeg2,Drak,vn,bowl,edl,RhoGAP19D,zfh2,beat-IIa,betaTub60D,comm2,Ppcs,14-3-3zeta,scrib,unc-119,chrb,nub,InR,cact,LanA,sick,Egfr,aop,glec,al,rg,chn,p130CAS,spdo,ttk,fz2,spi,Scr,path,sens,fs(1)h,wnd,Dg,Irk3,SCAR,RhoL,imd,Hr39,Rab1,Raf,rib,d,eya,unc-5,Lmx1a,nmo,mld,wbl,mim,scyl,Pdk1,crol,E(spl)m5-HLH,IA-2,Pex10,Alh,Dp,CtBP,Diap1,yrt,shot,Pvr,Socs36E,Pen,pdm3,bol,px,jeb,klu,dve,dlp,grn,hfw,Npc2g,fend,ph-p,S,CG1129,msi,sd,bin3,cora,sim,esg,bab2,Sxl,jumu,klar,lilli,meso18E,PI31,abd-A,Sema1b,Dgat2,pum,Inx2,moody,Cbl,Myo10A,Sema1a,ds,SERCA,Mpcp2,Abd-B,Ddr,rictor,sqz,baz,robo1,Syx1A,Atg16,CG34401,Adf1,wdb,Abl,Tollo,Cip4,AP-1gamma,Cks30A,larp,pnr,orb2,sens-2,ap,drl,cindr,N,eyg,lt,dock,Mob2,tara,SMC3,ec,B-H1,rdx,spen,sina,dome,CenB1A,Fas1,tsh,shg | | 1.855e-39 | -89.18 | anatomical structure development | biological process | GO:0048856 | 2924 | 230 | 12345 | 442 | Mpcp2,Abd-B,baz,sqz,rictor,CG34401,Atg16,robo1,Syx1A,Adf1,wdb,Abl,Cip4,Tollo,larp,pnr,AP-1gamma,Cks30A,PI31,abd-A,Dgat2,pum,Sema1b,moody,Inx2,Cbl,Myo10A,Sema1a,SERCA,ds,SMC3,tara,ec,B-H1,rdx,sina,spen,CenB1A,dome,Fas1,shg,tsh,sens-2,orb2,ap,drl,cindr,N,eyg,lt,dock,Mob2,nmo,mld,mim,wbl,Pdk1,scyl,crol,Pex10,IA-2,E(spl)m5-HLH,Alh,Dp,CtBP,Diap1,yrt,fs(1)h,sens,Dg,wnd,Irk3,RhoL,SCAR,imd,Hr39,Rab1,Raf,unc-5,Lmx1a,rib,d,eya,msi,CG1129,cora,sd,sim,esg,Sxl,bab2,lilli,jumu,klar,meso18E,Socs36E,Pvr,shot,Pen,pdm3,bol,px,klu,jeb,dlp,dve,hfw,grn,ph-p,S,Npc2g,fend,tkv,flfl,Mef2,ey,ft,Sox14,tai,Shrm,Pvf3,Ptp4E,Atg17,frc,siz,CycG,lin-28,nuf,Dl,mop,emc,hyx,cv-d,H,Dronc,yuri,alpha-Man-Ia,sns,fz,CycB3,LanA,cact,Egfr,sick,glec,aop,al,rg,p130CAS,spdo,chn,ttk,Scr,path,spi,fz2,Pgk,Bsg,neo,vn,Drak,Ptpmeg2,edl,bowl,Fkbp14,zfh2,beat-IIa,RhoGAP19D,scrib,betaTub60D,14-3-3zeta,Ppcs,comm2,unc-119,chrb,nub,InR,htl,boi,Dad,jar,kon,Rab5,Tom,cic,crp,14-3-3epsilon,kay,Poxn,arm,eff,Kdm2,Arf79F,trbl,otk,Eph,Actbeta,Gli,Pat1,E(Pc),Tet,Snoo,Nrk,fl(2)d,h,hth,disco-r,mam,wgn,ena,step,f,C1GalTA,sli,dpy,bou,Src42A,ss,lncRNA:Hsromega,RhoU,chic,Cdk4,corto,CG6191,kn,gukh,Psc,ed,fuss,Orct2 | | 1.763e-38 | -86.93 | generation of neurons | biological process | GO:0048699 | 880 | 118 | 12345 | 442 | robo1,flfl,CG34401,baz,sqz,rictor,Pvf3,Tollo,Cip4,Abl,tai,Adf1,Sox14,ft,ey,Dl,Sema1b,pum,abd-A,Ptp4E,ds,fz,Sema1a,Dronc,emc,mop,Cbl,spen,al,sina,B-H1,aop,sick,Egfr,SMC3,LanA,fz2,spi,shg,path,Fas1,ttk,chn,p130CAS,rg,comm2,N,betaTub60D,scrib,drl,beat-IIa,ap,edl,Bsg,dock,Mob2,nub,InR,Rab5,jar,Dad,boi,yrt,eff,arm,E(spl)m5-HLH,14-3-3epsilon,kay,SCAR,Gli,Actbeta,wnd,Eph,Dg,otk,sens,Arf79F,fs(1)h,eya,Lmx1a,unc-5,mam,Raf,h,Rab1,hth,Snoo,Nrk,E(Pc),sim,esg,sli,sd,cora,f,msi,ena,wgn,klar,ss,Src42A,lilli,pdm3,chic,shot,Socs36E,fend,ph-p,fuss,S,grn,ed,gukh,Psc,dlp,kn,jeb,klu | | 7.869e-38 | -85.44 | neuron differentiation | biological process | GO:0030182 | 710 | 105 | 12345 | 442 | Rab5,boi,Dad,eff,yrt,arm,14-3-3epsilon,kay,SCAR,Actbeta,Gli,otk,Dg,wnd,Eph,fs(1)h,sens,Lmx1a,mam,unc-5,eya,hth,Raf,Nrk,Snoo,E(Pc),sli,sim,cora,f,ena,msi,wgn,lilli,Src42A,klar,ss,pdm3,chic,shot,fuss,S,ph-p,fend,grn,gukh,dlp,Psc,klu,jeb,kn,robo1,sqz,baz,rictor,Pvf3,Cip4,Tollo,Abl,tai,ey,Sox14,ft,Dl,Sema1b,pum,Ptp4E,fz,ds,Sema1a,Dronc,Cbl,emc,mop,sina,al,spen,B-H1,aop,sick,Egfr,SMC3,LanA,path,shg,fz2,spi,ttk,Fas1,p130CAS,chn,rg,scrib,comm2,N,betaTub60D,beat-IIa,drl,edl,ap,Bsg,dock,Mob2,nub,InR | | 8.271e-38 | -85.39 | neurogenesis | biological process | GO:0022008 | 923 | 120 | 12345 | 442 | msi,ena,wgn,sim,esg,sli,sd,cora,f,klar,ss,Src42A,lilli,chic,shot,Socs36E,pdm3,ed,Psc,gukh,dlp,jeb,kn,klu,fend,S,fuss,ph-p,grn,jar,Dad,boi,htl,Rab5,E(spl)m5-HLH,kay,14-3-3epsilon,yrt,eff,arm,wnd,Eph,otk,Dg,Arf79F,sens,fs(1)h,SCAR,Gli,Actbeta,E(Pc),Snoo,Nrk,eya,mam,Lmx1a,unc-5,h,Raf,hth,Rab1,Egfr,sick,LanA,SMC3,spen,al,sina,aop,B-H1,chn,p130CAS,rg,spi,fz2,path,shg,Fas1,ttk,ap,edl,Bsg,N,betaTub60D,comm2,scrib,drl,beat-IIa,Mob2,dock,InR,nub,sqz,rictor,baz,robo1,flfl,CG34401,tai,Abl,ft,Sox14,Adf1,ey,Pvf3,Tollo,Cip4,Sema1b,pum,abd-A,Ptp4E,Dl,moody,Dronc,mop,emc,Cbl,ds,fz,Sema1a | | 9.319e-38 | -85.27 | nervous system development | biological process | GO:0007399 | 1108 | 132 | 12345 | 442 | tai,ey,ft,Sox14,Pvf3,flfl,Dronc,emc,mop,fz,Ptp4E,siz,Dl,spdo,p130CAS,chn,rg,path,spi,fz2,ttk,Egfr,sick,LanA,cact,al,glec,aop,unc-119,InR,nub,edl,vn,Bsg,scrib,comm2,betaTub60D,beat-IIa,zfh2,14-3-3epsilon,kay,eff,arm,jar,htl,boi,Dad,Rab5,Nrk,E(Pc),Snoo,mam,hth,h,otk,Eph,Arf79F,Actbeta,Gli,Src42A,ss,ena,wgn,sli,f,C1GalTA,Psc,gukh,ed,kn,fuss,chic,Abl,Adf1,Cip4,Tollo,baz,rictor,sqz,CG34401,robo1,Cbl,ds,Sema1a,Sema1b,pum,abd-A,moody,shg,Fas1,SMC3,sina,spen,B-H1,dock,Mob2,ap,N,drl,E(spl)m5-HLH,yrt,Pdk1,unc-5,Lmx1a,rib,eya,Raf,Rab1,Dg,wnd,fs(1)h,sens,SCAR,imd,lilli,klar,msi,esg,sim,cora,sd,dlp,klu,jeb,ph-p,S,fend,grn,Socs36E,Pvr,shot,pdm3 | | 8.036e-37 | -83.11 | epithelium development | biological process | GO:0060429 | 1020 | 125 | 12345 | 442 | ey,ft,tai,Shrm,Abl,Cip4,Tollo,pnr,Cks30A,Mpcp2,Abd-B,tkv,baz,Atg16,robo1,Syx1A,Mef2,Cbl,emc,hyx,mop,Myo10A,H,cv-d,fz,SERCA,ds,frc,Ptp4E,CycG,abd-A,nuf,Inx2,Dl,dome,ttk,path,shg,tsh,spi,SMC3,tara,cact,Egfr,ec,aop,rdx,B-H1,sina,spen,al,eyg,InR,neo,Ptpmeg2,Drak,vn,bowl,ap,Fkbp14,zfh2,RhoGAP19D,drl,scrib,N,cindr,Ppcs,14-3-3epsilon,kay,Alh,Poxn,arm,CtBP,Dp,Diap1,yrt,boi,Dad,nmo,mim,jar,crol,Rab5,cic,hth,Raf,disco-r,mam,d,rib,eya,otk,Dg,Gli,Irk3,RhoL,dpy,Sxl,bab2,lilli,Src42A,jumu,ss,ena,step,cora,f,sd,sli,sim,esg,kn,dlp,ed,dve,ph-p,fuss,S,Socs36E,shot,Pvr,chic,corto,px,CG6191 | | 1.532e-36 | -82.47 | tube development | biological process | GO:0035295 | 747 | 106 | 12345 | 442 | Fkbp14,ap,vn,bowl,Drak,Ppcs,N,scrib,RhoGAP19D,drl,zfh2,eyg,InR,Egfr,SMC3,tara,al,spen,B-H1,dome,tsh,spi,shg,Scr,path,ttk,nuf,abd-A,Ptp4E,frc,Dl,Inx2,cv-d,H,hyx,emc,Cbl,ds,fz,baz,tkv,Abd-B,Mpcp2,Syx1A,robo1,Mef2,ft,ey,Cks30A,pnr,Cip4,chic,shot,Pvr,Socs36E,px,CG6191,corto,ed,dve,dlp,kn,S,fuss,step,esg,sim,sli,sd,f,cora,bab2,Sxl,dpy,ss,jumu,klar,Src42A,lilli,Dg,otk,Gli,Irk3,eya,rib,d,mam,disco-r,h,Raf,hth,jar,nmo,Dad,boi,cic,Rab5,crol,Alh,14-3-3epsilon,kay,IA-2,Diap1,arm,CtBP,Poxn | | 1.590e-35 | -80.13 | imaginal disc development | biological process | GO:0007444 | 591 | 93 | 12345 | 442 | Dl,nuf,abd-A,frc,ds,fz,cv-d,H,emc,hyx,Cbl,Syx1A,tkv,Mpcp2,Abd-B,Cks30A,Cip4,ft,ey,N,Ppcs,scrib,RhoGAP19D,drl,zfh2,ap,Fkbp14,vn,Drak,bowl,InR,eyg,al,spen,B-H1,Egfr,tara,SMC3,spi,tsh,shg,path,dome,Irk3,Gli,otk,Dg,d,eya,mam,disco-r,hth,Raf,cic,Rab5,crol,jar,Dad,nmo,boi,Diap1,arm,CtBP,Poxn,Alh,14-3-3epsilon,kay,CG6191,px,corto,chic,shot,Pvr,Socs36E,fuss,S,ed,dve,dlp,kn,esg,sim,sd,f,cora,step,jumu,ss,lilli,Src42A,bab2,Sxl,dpy | | 1.777e-35 | -80.02 | tissue morphogenesis | biological process | GO:0048729 | 644 | 97 | 12345 | 442 | abd-A,frc,Ptp4E,Dl,Inx2,cv-d,H,Myo10A,hyx,emc,Cbl,ds,fz,baz,tkv,Shrm,Abl,ft,ey,Cks30A,pnr,Tollo,Cip4,ap,Fkbp14,Drak,bowl,vn,Ppcs,N,scrib,RhoGAP19D,zfh2,ec,Egfr,SMC3,al,spen,aop,rdx,B-H1,dome,spi,tsh,shg,path,ttk,otk,Dg,Gli,eya,rib,d,mam,disco-r,hth,Raf,jar,Dad,nmo,htl,cic,Rab5,crol,kay,yrt,Diap1,arm,Dp,Poxn,chic,shot,Pvr,Socs36E,CG6191,px,corto,ed,dve,dlp,jeb,kn,fuss,ph-p,S,step,ena,esg,sli,sd,cora,f,bab2,dpy,ss,jumu,Src42A | | 1.972e-35 | -79.91 | post-embryonic animal organ development | biological process | GO:0048569 | 481 | 84 | 12345 | 442 | ft,ey,Cip4,Cks30A,tkv,emc,hyx,Cbl,H,cv-d,ds,sns,fz,frc,Dl,chn,dome,tsh,spi,shg,path,SMC3,Egfr,B-H1,al,spen,ap,Fkbp14,vn,Drak,bowl,RhoGAP19D,zfh2,Ppcs,N,scrib,kay,Alh,arm,Poxn,Diap1,nmo,Dad,htl,jar,cic,Rab5,crol,crp,disco-r,hth,Raf,eya,d,mam,Kdm2,Dg,otk,Gli,dpy,bab2,jumu,klar,ss,Src42A,step,CG1129,sd,f,cora,esg,kn,dve,ed,dlp,S,fuss,Pvr,shot,Socs36E,chic,corto,CG6191,px | | 2.277e-35 | -79.77 | cellular developmental process | biological process | GO:0048869 | 1818 | 169 | 12345 | 442 | Pvf3,tai,Shrm,ey,Sox14,ft,flfl,Mef2,tkv,fz,sns,yuri,H,Dronc,emc,mop,Dl,lin-28,nuf,Ptp4E,Atg17,siz,path,spi,fz2,ttk,p130CAS,chn,rg,al,aop,sick,Egfr,LanA,cact,nub,InR,scrib,14-3-3zeta,Ppcs,betaTub60D,comm2,beat-IIa,vn,Ptpmeg2,edl,Pgk,Bsg,neo,eff,arm,kay,14-3-3epsilon,Rab5,kon,Tom,cic,jar,boi,htl,Dad,mam,h,hth,disco-r,Tet,Snoo,E(Pc),Nrk,Actbeta,Gli,Pat1,otk,Eph,Arf79F,Src42A,ss,sli,f,C1GalTA,ena,wgn,fuss,Psc,gukh,ed,kn,chic,RhoU,lncRNA:Hsromega,pnr,Cip4,Tollo,Abl,Adf1,Atg16,CG34401,robo1,rictor,baz,Ddr,sqz,Abd-B,SERCA,ds,Sema1a,Cbl,moody,Inx2,Sema1b,pum,abd-A,shg,tsh,Fas1,CenB1A,dome,sina,spen,B-H1,ec,SMC3,dock,Mob2,cindr,N,drl,ap,orb2,yrt,Diap1,CtBP,Dp,Pex10,E(spl)m5-HLH,mim,wbl,unc-5,Lmx1a,eya,Raf,Rab1,SCAR,RhoL,Dg,wnd,fs(1)h,sens,lilli,klar,Sxl,sim,esg,cora,sd,msi,ph-p,S,fend,grn,dlp,klu,jeb,pdm3,bol,Pen,Socs36E,shot,Pvr | | 5.213e-35 | -78.94 | cell differentiation | biological process | GO:0030154 | 1773 | 166 | 12345 | 442 | chic,lncRNA:Hsromega,fuss,Psc,gukh,ed,kn,sli,f,C1GalTA,ena,wgn,Src42A,ss,Actbeta,Gli,Pat1,otk,Eph,Arf79F,mam,h,hth,disco-r,Nrk,Snoo,E(Pc),Tet,kon,Rab5,Tom,cic,jar,htl,boi,Dad,eff,arm,kay,14-3-3epsilon,scrib,14-3-3zeta,comm2,betaTub60D,Ppcs,beat-IIa,edl,vn,Ptpmeg2,neo,Pgk,Bsg,nub,InR,al,aop,Egfr,sick,LanA,cact,path,spi,fz2,ttk,p130CAS,chn,rg,Dl,lin-28,nuf,Ptp4E,Atg17,siz,sns,fz,yuri,Dronc,H,emc,mop,Mef2,flfl,tkv,Pvf3,tai,ey,Sox14,ft,bol,pdm3,Pen,Socs36E,shot,Pvr,ph-p,S,fend,grn,dlp,klu,jeb,esg,sim,cora,sd,msi,lilli,klar,Sxl,RhoL,SCAR,Dg,wnd,fs(1)h,sens,Lmx1a,unc-5,eya,Rab1,Raf,wbl,mim,yrt,Diap1,Dp,CtBP,Pex10,E(spl)m5-HLH,N,cindr,drl,ap,orb2,dock,Mob2,sina,spen,B-H1,ec,SMC3,shg,tsh,Fas1,dome,moody,Inx2,Sema1b,pum,abd-A,SERCA,ds,Sema1a,Cbl,CG34401,Atg16,robo1,Ddr,baz,sqz,rictor,Abd-B,pnr,Cip4,Tollo,Abl,Adf1 | | 1.627e-33 | -75.50 | morphogenesis of an epithelium | biological process | GO:0002009 | 625 | 93 | 12345 | 442 | ena,step,esg,f,cora,sd,bab2,dpy,Src42A,ss,jumu,chic,Socs36E,shot,Pvr,CG6191,px,corto,dlp,ed,dve,kn,fuss,ph-p,S,jar,Dad,nmo,Rab5,crol,cic,kay,yrt,Diap1,Poxn,Dp,arm,otk,Dg,Gli,mam,eya,d,rib,hth,Raf,disco-r,Egfr,ec,SMC3,spen,al,B-H1,rdx,aop,dome,path,shg,spi,tsh,ttk,bowl,Drak,vn,ap,Fkbp14,scrib,Ppcs,N,zfh2,RhoGAP19D,baz,tkv,Abl,Shrm,ey,ft,pnr,Cks30A,Cip4,Tollo,frc,Ptp4E,Dl,Inx2,Myo10A,cv-d,H,Cbl,hyx,emc,fz,ds | | 4.471e-33 | -74.49 | imaginal disc morphogenesis | biological process | GO:0007560 | 420 | 76 | 12345 | 442 | hth,Raf,disco-r,mam,d,eya,otk,Dg,Gli,kay,Poxn,arm,Diap1,Dad,nmo,jar,Rab5,crol,cic,kn,dlp,ed,dve,fuss,S,Socs36E,Pvr,shot,chic,corto,CG6191,px,dpy,bab2,Src42A,jumu,ss,step,cora,f,sd,esg,Cbl,emc,hyx,H,cv-d,fz,ds,frc,Dl,ey,ft,Cip4,Cks30A,tkv,bowl,Drak,vn,Fkbp14,ap,zfh2,RhoGAP19D,scrib,Ppcs,N,dome,path,shg,spi,tsh,SMC3,Egfr,B-H1,spen,al | | 4.471e-33 | -74.49 | post-embryonic animal organ morphogenesis | biological process | GO:0048563 | 420 | 76 | 12345 | 442 | Dl,frc,fz,ds,Cbl,emc,hyx,H,cv-d,tkv,Cip4,Cks30A,ey,ft,zfh2,RhoGAP19D,scrib,N,Ppcs,Drak,vn,bowl,Fkbp14,ap,B-H1,al,spen,SMC3,Egfr,path,shg,tsh,spi,dome,Gli,otk,Dg,Raf,hth,disco-r,mam,eya,d,crol,Rab5,cic,nmo,Dad,jar,Poxn,arm,Diap1,kay,corto,px,CG6191,Socs36E,Pvr,shot,chic,fuss,S,kn,dlp,ed,dve,f,cora,sd,esg,step,Src42A,ss,jumu,dpy,bab2 | | 4.527e-32 | -72.17 | instar larval or pupal morphogenesis | biological process | GO:0048707 | 470 | 79 | 12345 | 442 | kay,Diap1,arm,Poxn,jar,nmo,Dad,cic,crol,Rab5,eya,d,mam,disco-r,hth,Raf,Dg,otk,Gli,bab2,dpy,ss,jumu,Src42A,step,esg,sd,cora,f,dve,ed,dlp,kn,S,fuss,chic,shot,Pvr,Socs36E,CG6191,px,corto,ft,ey,Cks30A,Cip4,tkv,Atg16,cv-d,Dronc,H,emc,hyx,Cbl,ds,fz,Atg17,frc,Dl,dome,tsh,spi,path,shg,Egfr,SMC3,al,spen,B-H1,ap,Fkbp14,Drak,vn,bowl,N,Ppcs,scrib,RhoGAP19D,zfh2 | | 5.567e-32 | -71.97 | positive regulation of cellular process | biological process | GO:0048522 | 1464 | 144 | 12345 | 442 | Egfr,Pez,aop,chn,spdo,spi,Scr,ttk,Fkbp14,edl,vn,bowl,Pgk,14-3-3zeta,zfh2,hppy,InR,nub,Dcr-2,tkv,Mef2,tai,ft,Sox14,ey,Pvf3,Cadps,lin-28,siz,ps,CycG,lili,Atg17,Dl,RhoGEF4,cv-d,Dronc,hyx,mop,fz,step,ena,sli,ss,CG8312,Src42A,chic,RhoU,lncRNA:Hsromega,tna,Cdk4,ed,kn,Pitslre,Orct2,jar,htl,kon,kay,14-3-3epsilon,arm,trbl,Arf79F,apolpp,Gli,Actbeta,Tet,mam,hth,ec,tara,spen,sina,B-H1,rdx,Su(z)2,dome,CG13204,dmrt11E,tsh,shg,Jupiter,ap,orb2,N,cindr,Ddr,rictor,Abd-B,morgue,robo1,Abl,wdb,SNF4Agamma,Adf1,Cks30A,pnr,Cals,Sema1b,pum,PI31,abd-A,Cbl,ds,Sema1a,esg,REPTOR,sim,sd,Sxl,lilli,Pvr,shot,Socs36E,bol,Lk6,dlp,jeb,klu,S,rho-4,nmo,crol,Pdk1,Diap1,Dp,CtBP,wnd,hng3,lqf,sens,fs(1)h,CG9784,imd,SCAR,Tis11,Hr39,CG12769,Dyrk2,eya,unc-5,Lmx1a,Raf,Rab1 | | 3.003e-31 | -70.28 | epithelial tube morphogenesis | biological process | GO:0060562 | 495 | 80 | 12345 | 442 | ap,Fkbp14,Drak,vn,bowl,RhoGAP19D,zfh2,N,Ppcs,scrib,dome,ttk,tsh,spi,path,shg,SMC3,Egfr,B-H1,spen,al,hyx,emc,Cbl,cv-d,H,ds,fz,frc,Ptp4E,Dl,ft,ey,Cip4,Cks30A,tkv,baz,kn,ed,dve,dlp,fuss,S,shot,Pvr,Socs36E,chic,corto,CG6191,px,dpy,bab2,ss,jumu,Src42A,step,sd,cora,f,esg,disco-r,hth,Raf,d,eya,rib,mam,Dg,otk,Gli,kay,arm,Poxn,Diap1,Dad,nmo,jar,cic,crol,Rab5 | | 3.712e-31 | -70.07 | post-embryonic animal morphogenesis | biological process | GO:0009886 | 484 | 79 | 12345 | 442 | Dg,otk,Gli,mam,d,eya,Raf,hth,disco-r,jar,Dad,nmo,Rab5,crol,cic,kay,Diap1,Poxn,arm,chic,Socs36E,shot,Pvr,CG6191,px,corto,dlp,dve,ed,kn,fuss,S,step,esg,f,cora,sd,bab2,dpy,Src42A,jumu,ss,frc,Atg17,Dl,cv-d,H,Dronc,Cbl,hyx,emc,fz,ds,tkv,Atg16,ey,ft,Cks30A,Cip4,Drak,vn,bowl,ap,Fkbp14,scrib,N,Ppcs,zfh2,RhoGAP19D,Egfr,SMC3,spen,al,B-H1,dome,path,shg,tsh,spi | | 4.014e-31 | -69.99 | metamorphosis | biological process | GO:0007552 | 497 | 80 | 12345 | 442 | Atg17,frc,Dl,Dronc,H,cv-d,Cbl,emc,hyx,fz,ds,tkv,Atg16,ey,Sox14,ft,Cks30A,Cip4,vn,bowl,Drak,Fkbp14,ap,scrib,N,Ppcs,zfh2,RhoGAP19D,Egfr,SMC3,spen,al,B-H1,dome,shg,path,tsh,spi,Dg,otk,Gli,mam,d,eya,hth,Raf,disco-r,jar,Dad,nmo,crol,Rab5,cic,kay,Diap1,Poxn,arm,chic,Socs36E,shot,Pvr,CG6191,px,corto,dlp,dve,ed,kn,fuss,S,step,esg,cora,f,sd,bab2,dpy,Src42A,jumu,ss | | 4.555e-31 | -69.86 | tube morphogenesis | biological process | GO:0035239 | 536 | 83 | 12345 | 442 | Diap1,Poxn,arm,kay,Rab5,crol,cic,jar,nmo,Dad,mam,rib,eya,d,hth,h,Raf,disco-r,Gli,otk,Dg,Src42A,ss,jumu,klar,bab2,dpy,esg,cora,f,sd,step,fuss,S,dlp,ed,dve,kn,px,CG6191,corto,chic,Socs36E,Pvr,shot,Cks30A,Cip4,ey,ft,baz,tkv,fz,ds,cv-d,H,Cbl,emc,hyx,Dl,Inx2,Ptp4E,frc,shg,path,spi,tsh,ttk,dome,al,spen,B-H1,Egfr,SMC3,scrib,N,Ppcs,zfh2,RhoGAP19D,bowl,Drak,vn,ap,Fkbp14 | | 5.680e-31 | -69.64 | positive regulation of biological process | biological process | GO:0048518 | 1642 | 152 | 12345 | 442 | lilli,Sxl,REPTOR,sim,esg,sd,S,rho-4,dlp,jeb,klu,Lk6,bol,shot,Pvr,Socs36E,Diap1,CtBP,Dp,Hr4,crol,Pdk1,nmo,mld,eya,Dyrk2,d,unc-5,Lmx1a,Raf,Rab1,CG12769,Hr39,SCAR,RhoL,imd,Tis11,wnd,lqf,hng3,sens,CG9784,fs(1)h,dmrt11E,tsh,shg,Jupiter,dome,Su(z)2,CG13204,spen,sina,rdx,B-H1,ec,tara,cindr,N,DPCoAC,ap,orb2,Cks30A,pnr,Cals,Abl,wdb,SNF4Agamma,Adf1,morgue,robo1,rictor,Ddr,Abd-B,ds,SERCA,Sema1a,Cbl,pum,Sema1b,PI31,abd-A,ss,Src42A,CG8312,sli,step,ena,Pitslre,Orct2,ed,kn,tna,Cdk4,chic,lncRNA:Hsromega,RhoU,arm,14-3-3epsilon,kay,kon,jar,htl,mam,hth,Tet,Gli,Actbeta,trbl,Arf79F,cue,apolpp,spi,Scr,CG32795,ttk,chn,spdo,aop,Pez,Egfr,nub,InR,hppy,14-3-3zeta,zfh2,Fkbp14,vn,edl,bowl,Pgk,Cadps,Pvf3,tai,ft,Sox14,ey,Mef2,tkv,Dcr-2,fz,Dronc,cv-d,hyx,mop,Dl,RhoGEF4,lin-28,siz,CycG,ps,Atg17,lili | | 1.168e-30 | -68.92 | wing disc development | biological process | GO:0035220 | 407 | 72 | 12345 | 442 | jumu,Src42A,lilli,dpy,esg,sd,cora,f,step,fuss,S,ed,dve,dlp,kn,px,CG6191,corto,shot,Socs36E,arm,CtBP,Poxn,kay,14-3-3epsilon,cic,crol,Rab5,Dad,nmo,boi,d,mam,disco-r,Raf,hth,Irk3,Gli,Dg,spi,tsh,path,shg,dome,spen,Egfr,SMC3,tara,eyg,Ppcs,N,scrib,zfh2,ap,Fkbp14,bowl,Drak,vn,Cip4,ft,tkv,Mpcp2,ds,fz,cv-d,H,hyx,emc,Cbl,Dl,nuf,frc | | 2.313e-30 | -68.24 | post-embryonic development | biological process | GO:0009791 | 642 | 90 | 12345 | 442 | tkv,Atg16,ey,ft,Cks30A,Cip4,Atg17,frc,Dl,H,Dronc,cv-d,Cbl,hyx,emc,fz,sns,ds,Egfr,SMC3,spen,al,B-H1,chn,dome,path,shg,tsh,fz2,spi,vn,bowl,Drak,Fkbp14,ap,scrib,N,Ppcs,zfh2,RhoGAP19D,jar,htl,Dad,mld,nmo,crp,Rab5,crol,cic,Alh,kay,Diap1,Poxn,arm,Dg,otk,Kdm2,Gli,mam,eya,d,hth,Raf,disco-r,CG1129,step,esg,f,cora,sd,bab2,dpy,Src42A,ss,jumu,klar,chic,Socs36E,Pvr,shot,px,CG6191,corto,dlp,dve,ed,kn,S,fuss,hfw | | 2.707e-30 | -68.08 | appendage development | biological process | GO:0048736 | 366 | 68 | 12345 | 442 | Alh,Poxn,arm,Diap1,nmo,Dad,crol,cic,Raf,hth,disco-r,mam,d,Dg,Gli,dpy,Src42A,jumu,ss,step,f,cora,sd,kn,dlp,dve,ed,S,fuss,Socs36E,shot,corto,px,CG6191,ft,Cip4,tkv,Syx1A,Cbl,hyx,emc,H,cv-d,fz,ds,frc,Dl,shg,path,spi,tsh,SMC3,Egfr,al,spen,eyg,nub,vn,bowl,Drak,Fkbp14,ap,zfh2,drl,RhoGAP19D,scrib,Ppcs,N | | 9.613e-30 | -66.81 | imaginal disc-derived appendage development | biological process | GO:0048737 | 362 | 67 | 12345 | 442 | shot,Socs36E,px,CG6191,corto,dve,ed,dlp,kn,S,fuss,step,sd,f,cora,dpy,ss,jumu,Src42A,Dg,Gli,d,mam,disco-r,hth,Raf,nmo,Dad,cic,crol,Alh,Diap1,arm,Poxn,Fkbp14,ap,bowl,vn,Drak,N,Ppcs,scrib,RhoGAP19D,drl,zfh2,eyg,Egfr,SMC3,al,spen,spi,tsh,path,shg,frc,Dl,cv-d,H,emc,hyx,Cbl,ds,fz,tkv,Syx1A,ft,Cip4 | | 1.266e-29 | -66.54 | sensory organ development | biological process | GO:0007423 | 496 | 78 | 12345 | 442 | frc,siz,nuf,Dl,Cbl,emc,hyx,Dronc,H,fz,sns,ds,tkv,ey,ft,Abl,pnr,AP-1gamma,edl,Fkbp14,scrib,Ppcs,cindr,N,eyg,Mob2,Egfr,ec,aop,B-H1,rdx,sina,al,spen,rg,spdo,dome,chn,ttk,shg,Scr,tsh,spi,sens,Arf79F,SCAR,Nrk,fl(2)d,hth,Raf,eya,d,boi,nmo,Tom,cic,kay,Poxn,CtBP,arm,eff,Diap1,yrt,shot,klu,jeb,dlp,ed,S,ena,msi,f,sd,lilli,Src42A,jumu,klar,ss | | 2.118e-29 | -66.02 | cell development | biological process | GO:0048468 | 1436 | 138 | 12345 | 442 | Abl,Tollo,Cip4,baz,rictor,sqz,robo1,Atg16,Cbl,ds,SERCA,Sema1a,Sema1b,pum,abd-A,Inx2,moody,dome,shg,Fas1,ec,spen,Mob2,dock,ap,orb2,cindr,N,drl,Pex10,yrt,Diap1,Dp,wbl,mim,eya,unc-5,Raf,wnd,Dg,sens,fs(1)h,RhoL,SCAR,Sxl,klar,lilli,msi,sim,sd,cora,dlp,jeb,klu,fend,S,ph-p,grn,Pen,Pvr,shot,Socs36E,bol,pdm3,tai,Sox14,ft,Pvf3,tkv,Mef2,Dronc,emc,mop,sns,fz,yuri,lin-28,Atg17,Ptp4E,Dl,chn,p130CAS,rg,fz2,spi,path,ttk,Egfr,sick,cact,LanA,al,aop,InR,nub,Ptpmeg2,edl,neo,Bsg,comm2,betaTub60D,Ppcs,14-3-3zeta,scrib,beat-IIa,kay,14-3-3epsilon,eff,arm,jar,Dad,htl,cic,kon,Rab5,Snoo,Nrk,E(Pc),Tet,mam,Eph,otk,Pat1,Gli,Actbeta,ss,Src42A,ena,wgn,sli,C1GalTA,ed,Psc,gukh,kn,fuss,chic,lncRNA:Hsromega | | 8.644e-29 | -64.62 | instar larval or pupal development | biological process | GO:0002165 | 576 | 83 | 12345 | 442 | bowl,Drak,vn,Fkbp14,ap,zfh2,RhoGAP19D,scrib,Ppcs,N,dome,shg,path,tsh,spi,fz2,SMC3,Egfr,B-H1,spen,al,Cbl,emc,hyx,Dronc,cv-d,H,fz,ds,frc,Atg17,Dl,ey,ft,Cip4,Cks30A,tkv,Atg16,kn,dlp,ed,dve,hfw,fuss,S,Socs36E,shot,Pvr,chic,corto,px,CG6191,dpy,bab2,Src42A,ss,jumu,step,cora,f,sd,esg,Raf,hth,disco-r,mam,eya,d,Dg,otk,Gli,kay,Alh,Poxn,arm,Diap1,Dad,mld,nmo,jar,crol,Rab5,cic | | 1.600e-28 | -64.00 | protein binding | molecular function | GO:0005515 | 2410 | 179 | 12176 | 414 | Abl,Diap1,hyx,bab2,vig,CG6891,Ziz,ena,eya,Pex14,HmgZ,arm,Atg17,RhoGEF3,Slbp,CG5522,spi,CG1218,Dl,chn,Unc-76,ec,step,ed,crp,14-3-3epsilon,cv-d,tna,hng3,fuss,Dronc,Dyb,CG17883,pita,ey,kon,Syx1A,Cdk4,morgue,rg,CG30156,flfl,Nrk,d,cindr,Tom,eff,Nlp,Dg,Socs36E,Raf,Pvf3,yrt,Mob2,otk,fz3,Magi,Hr39,CycG,kay,Pvr,shg,rdx,SMC3,fz2,fl(2)d,Actbeta,Pp2C1,f,Scr,TBC1D5,nuf,htl,InR,His3.3A,vn,Gli,Fas1,drl,baz,Shrm,14-3-3zeta,HLH3B,Dad,RhoGEF4,orb2,corto,RhoU,SERCA,tkv,cic,gukh,dlp,ss,REPTOR,edl,scrib,sim,His2Av,SCAR,sns,tsh,wbl,chic,aop,LanA,Cip4,spdo,h,SNF4Agamma,wgn,imd,ft,bin3,fz,boi,Rm62,dock,His3.3B,Myo10A,cue,Dp,ttk,CG4645,Sema1a,Pez,Mef2,ds,emc,Jupiter,mim,pnr,Rab1,siz,jar,trbl,CG13204,sli,kn,Lk6,subdued,klar,CG2765,cact,robo1,sens,Cbl,hth,sd,ITP,tai,CG31457,Tob,Snoo,Cks30A,N,l(2)37Cg,Ric,Sxl,ph-p,CycB3,Src42A,Sema1b,apolpp,shot,H,CtBP,Egfr,cora,Bsg,Rab5,Dcr-2,lilli,Rgl,E(spl)m5-HLH,MEP-1,dome,Mfe2,sina | | 2.369e-28 | -63.61 | appendage morphogenesis | biological process | GO:0035107 | 358 | 65 | 12345 | 442 | dpy,ss,jumu,Src42A,step,sd,f,cora,kn,ed,dve,dlp,S,fuss,shot,Socs36E,corto,px,CG6191,arm,Poxn,Diap1,Dad,nmo,cic,crol,disco-r,Raf,hth,d,mam,Dg,Gli,spi,tsh,path,shg,SMC3,Egfr,al,spen,eyg,nub,Fkbp14,ap,vn,Drak,bowl,RhoGAP19D,zfh2,Ppcs,N,scrib,ft,Cip4,tkv,emc,hyx,Cbl,H,cv-d,ds,fz,frc,Dl | | 9.807e-28 | -62.19 | imaginal disc-derived appendage morphogenesis | biological process | GO:0035114 | 355 | 64 | 12345 | 442 | Dg,Gli,d,mam,disco-r,Raf,hth,Dad,nmo,cic,crol,Diap1,arm,Poxn,shot,Socs36E,CG6191,px,corto,ed,dve,dlp,kn,fuss,S,step,sd,cora,f,dpy,jumu,ss,Src42A,frc,Dl,cv-d,H,hyx,emc,Cbl,ds,fz,tkv,ft,Cip4,Fkbp14,ap,bowl,Drak,vn,N,Ppcs,scrib,RhoGAP19D,zfh2,eyg,Egfr,SMC3,spen,al,tsh,spi,shg,path | | 1.086e-27 | -62.09 | cellular component morphogenesis | biological process | GO:0032989 | 667 | 88 | 12345 | 442 | LanA,cact,Egfr,sick,spen,CenB1A,chn,ttk,Fas1,shg,path,fz2,neo,Bsg,ap,beat-IIa,drl,scrib,Ppcs,betaTub60D,N,comm2,nub,InR,Mob2,dock,tkv,sqz,rictor,baz,robo1,ft,Sox14,Shrm,tai,Abl,Ptp4E,Sema1b,pum,Dl,emc,Dronc,Sema1a,fz,ds,wgn,ena,sli,sim,Src42A,shot,RhoU,chic,pdm3,jeb,kn,Psc,dlp,gukh,ed,grn,S,ph-p,fend,htl,Dad,kon,Rab5,cic,kay,14-3-3epsilon,Dp,arm,fs(1)h,sens,otk,Dg,wnd,Eph,Actbeta,SCAR,Nrk,E(Pc),Snoo,Raf,h,unc-5,eya | | 1.386e-27 | -61.84 | regulation of developmental process | biological process | GO:0050793 | 757 | 94 | 12345 | 442 | Egfr,ec,LanA,cact,sina,aop,chn,shg,fz2,tsh,spi,ttk,vn,neo,scrib,N,Mob2,InR,nub,baz,rictor,Abd-B,tkv,morgue,CG34401,Atg16,Mef2,Syx1A,robo1,tai,Abl,ey,Adf1,Sox14,ft,Cip4,Cals,pum,Sema1b,Him,abd-A,Dl,Dronc,Cbl,emc,mop,fz,ds,Sema1a,ena,step,sli,esg,C1GalTA,sd,bab2,Src42A,klar,ss,chic,Socs36E,Pvr,shot,pxb,dlp,ed,klu,kn,fuss,Orct2,wbl,htl,nmo,Dad,Hr4,Pdk1,kay,E(spl)m5-HLH,eff,Diap1,Dp,otk,lqf,Arf79F,SCAR,Gli,Snoo,rib,eya,d,h,Rab1,Raf,hth | | 1.544e-27 | -61.74 | multicellular organismal process | biological process | GO:0032501 | 3844 | 246 | 12345 | 442 | Hr39,Rab1,Raf,unc-5,Lmx1a,Dyrk2,rib,d,eya,fs(1)h,sens,Dg,wnd,Irk3,imd,RhoL,SCAR,IA-2,Pex10,E(spl)m5-HLH,Alh,Dp,CtBP,Diap1,yrt,nmo,mld,wbl,mim,His3.3A,crol,Pdk1,scyl,subdued,His3.3B,klu,jeb,dlp,dve,hfw,grn,ph-p,S,fend,Socs36E,Pvr,shot,pxb,tipE,Pen,bol,pdm3,px,bab2,Sxl,lilli,jumu,klar,msi,CG1129,cora,sd,esg,sim,Cbl,Myo10A,lncRNA:CR44420,hang,Sema1a,SERCA,ds,PI31,abd-A,Sema1b,pum,moody,Inx2,Adf1,wdb,Abl,Cip4,Tollo,pnr,larp,Cks30A,AP-1gamma,Abd-B,Mpcp2,baz,rictor,sqz,CG34401,Atg16,robo1,Syx1A,mura,eyg,lt,dock,Mob2,sens-2,orb2,ap,drl,N,cindr,dome,Fas1,shg,dmrt11E,tsh,SMC3,tara,ec,rdx,B-H1,sina,spen,Tet,Nrk,Snoo,E(Pc),fl(2)d,h,hth,disco-r,mam,Esp,Arf79F,Kdm2,otk,trbl,Eph,Actbeta,Gli,Pat1,14-3-3epsilon,kay,His2Av,Poxn,arm,eff,htl,boi,Dad,jar,Rab5,kon,ogre,Tom,cic,crp,kn,Psc,gukh,ed,Cerk,fuss,Orct2,lncRNA:Hsromega,CG30344,chic,Cdk4,Bacc,corto,CG6191,dpy,bou,Src42A,ss,wgn,Gr93c,ena,step,f,C1GalTA,sli,Tmhs,hyx,mop,emc,Dronc,H,cv-d,yuri,alpha-Man-Ia,sns,fz,CycB3,Ptp4E,frc,Atg17,siz,CycG,lin-28,nuf,Dl,ey,Tob,ft,Sox14,tai,Acp63F,Pvf3,tkv,flfl,Mef2,unc-119,chrb,hppy,nub,InR,Bsg,Ir21a,Ptpmeg2,edl,vn,Drak,bowl,Fkbp14,zfh2,beat-IIa,nAChRalpha4,RhoGAP19D,scrib,comm2,Ppcs,betaTub60D,14-3-3zeta,rg,SKIP,p130CAS,spdo,chn,ttk,Scr,path,fz2,spi,LanA,cact,Egfr,sick,glec,aop,al | | 1.773e-27 | -61.60 | cell morphogenesis | biological process | GO:0000902 | 533 | 78 | 12345 | 442 | arm,kay,14-3-3epsilon,Rab5,htl,Dad,h,unc-5,eya,Nrk,Snoo,E(Pc),Actbeta,SCAR,fs(1)h,sens,Dg,otk,Eph,wnd,Src42A,sli,sim,wgn,ena,grn,ph-p,fend,kn,jeb,Psc,gukh,dlp,pdm3,shot,RhoU,chic,Sox14,ft,Abl,tai,Shrm,robo1,tkv,sqz,rictor,baz,Sema1a,fz,ds,Dronc,Dl,Ptp4E,Sema1b,pum,Fas1,path,shg,fz2,CenB1A,chn,spen,LanA,sick,InR,nub,Mob2,dock,beat-IIa,drl,scrib,betaTub60D,N,Ppcs,comm2,Bsg,neo,ap | | 4.846e-27 | -60.59 | wing disc morphogenesis | biological process | GO:0007472 | 308 | 59 | 12345 | 442 | arm,Poxn,Cip4,ft,cic,crol,Dad,tkv,nmo,ds,d,mam,fz,disco-r,Raf,hth,H,cv-d,hyx,emc,Cbl,Dl,Gli,Dg,frc,tsh,spi,shg,path,jumu,Src42A,dome,dpy,spen,sd,f,cora,step,Egfr,SMC3,S,fuss,dve,ed,dlp,kn,N,CG6191,px,Ppcs,scrib,corto,zfh2,Fkbp14,ap,vn,Drak,shot,Socs36E | | 5.738e-27 | -60.42 | imaginal disc-derived wing morphogenesis | biological process | GO:0007476 | 298 | 58 | 12345 | 442 | dpy,Src42A,jumu,path,shg,tsh,spi,SMC3,Egfr,step,f,cora,sd,spen,kn,dlp,dve,ed,S,fuss,Socs36E,shot,vn,Drak,ap,Fkbp14,zfh2,corto,scrib,Ppcs,N,px,CG6191,ft,Cip4,Poxn,arm,nmo,tkv,Dad,crol,cic,Cbl,emc,hyx,cv-d,H,Raf,hth,disco-r,mam,fz,d,ds,frc,Dg,Dl,Gli | | 6.323e-27 | -60.33 | neuron development | biological process | GO:0048666 | 584 | 81 | 12345 | 442 | ena,wgn,sim,sli,klar,ss,lilli,Src42A,chic,shot,pdm3,dlp,gukh,Psc,kn,jeb,fend,ph-p,fuss,grn,Dad,Rab5,14-3-3epsilon,kay,yrt,eff,Eph,wnd,otk,Dg,sens,fs(1)h,SCAR,Gli,Actbeta,Snoo,E(Pc),Nrk,eya,unc-5,Raf,sick,LanA,spen,al,chn,p130CAS,rg,fz2,shg,path,Fas1,ttk,ap,edl,Bsg,N,comm2,betaTub60D,drl,beat-IIa,Mob2,dock,InR,nub,baz,sqz,rictor,robo1,tai,Abl,Sox14,Pvf3,Tollo,Sema1b,pum,Ptp4E,Dronc,mop,fz,Sema1a | | 8.532e-27 | -60.03 | post-embryonic appendage morphogenesis | biological process | GO:0035120 | 345 | 62 | 12345 | 442 | Fkbp14,ap,bowl,vn,Drak,shot,Socs36E,px,CG6191,Ppcs,N,scrib,corto,RhoGAP19D,zfh2,dve,ed,dlp,kn,S,fuss,step,Egfr,SMC3,al,spen,sd,cora,f,dpy,tsh,spi,shg,path,ss,jumu,Src42A,Dg,frc,Gli,Dl,H,cv-d,hyx,emc,Cbl,ds,d,fz,mam,disco-r,hth,Raf,nmo,Dad,tkv,cic,crol,ft,arm,Cip4,Poxn | | 9.849e-27 | -59.88 | regulation of nitrogen compound metabolic process | biological process | GO:0051171 | 1870 | 156 | 12345 | 442 | bol,Lk6,pdm3,Pvr,Socs36E,grn,ph-p,Spn42De,jeb,klu,dve,H2.0,sd,bin3,esg,REPTOR,sim,msi,CG34376,jumu,lilli,bab2,Sxl,imd,Tis11,sens,fs(1)h,wnd,hng3,Dg,HmgZ,Raf,rib,eya,Lmx1a,CG12769,Hr39,crol,Pdk1,Hr4,CtBP,Dp,Diap1,E(spl)m5-HLH,Alh,FBXO11,CG11360,N,pita,orb2,sens-2,ap,eyg,B-H1,rdx,Kdm4B,spen,sina,CG10348,312,tara,CG11617,tsh,otp,dmrt11E,Su(z)2,CG13204,CG10321,PI31,abd-A,pum,rgr,Cbl,Abd-B,rictor,Tollo,Cks30A,pnr,Adf1,wdb,SNF4Agamma,corto,bru2,tna,lncRNA:Hsromega,Rm62,CG1218,kek5,fuss,kn,Psc,step,ss,Src42A,CG8312,dpy,lncRNA:bxd,MEP-1,Actbeta,Kdm2,trbl,hth,h,mam,fl(2)d,Tet,Snoo,cic,crp,Dad,CG4896,arm,Poxn,14-3-3epsilon,kay,HLH3B,zfh2,14-3-3zeta,scrib,Elba2,su(sable),edl,vn,bowl,nub,InR,hppy,aop,al,cact,Egfr,ttk,spi,Scr,Prdm13,chn,Hel89B,CycG,ps,Atg17,lili,lin-28,CycB3,hyx,emc,cv-d,Dronc,Fem-1,H,Mef2,Dcr-2,tkv,Sox14,Tob,ey,tai | | 1.261e-26 | -59.64 | regulation of primary metabolic process | biological process | GO:0080090 | 1894 | 157 | 12345 | 442 | sim,esg,REPTOR,sd,bin3,msi,CG34376,jumu,lilli,bab2,Sxl,bol,pdm3,Lk6,Pvr,Socs36E,ph-p,grn,dve,H2.0,jeb,Spn42De,klu,Hr4,crol,Pdk1,Diap1,CtBP,Dp,Alh,E(spl)m5-HLH,Tis11,CG2765,wnd,hng3,Dg,sens,fs(1)h,eya,rib,Lmx1a,HmgZ,Raf,CG12769,Hr39,spen,Kdm4B,sina,B-H1,rdx,CG11617,CG10348,312,tara,dmrt11E,tsh,otp,Su(z)2,CG13204,CG10321,N,pita,FBXO11,CG11360,ap,orb2,sens-2,eyg,rictor,Abd-B,Cks30A,pnr,Tollo,wdb,SNF4Agamma,Adf1,pum,rgr,abd-A,PI31,Cbl,step,ss,CG8312,Src42A,Nmda1,dpy,lncRNA:bxd,MEP-1,tna,corto,bru2,lncRNA:Hsromega,Rm62,kek5,fuss,CG1218,Psc,kn,crp,cic,CG4896,Dad,arm,Poxn,HLH3B,kay,14-3-3epsilon,Actbeta,trbl,Kdm2,mam,hth,h,fl(2)d,Tet,Snoo,al,aop,Egfr,cact,spi,Scr,ttk,chn,Prdm13,14-3-3zeta,Elba2,scrib,zfh2,vn,edl,bowl,su(sable),InR,nub,hppy,Mef2,Dcr-2,tkv,tai,Sox14,Tob,ey,Hel89B,lin-28,ps,CycG,lili,Atg17,CycB3,Dronc,cv-d,Fem-1,H,hyx,emc | | 2.107e-26 | -59.12 | binding | molecular function | GO:0005488 | 5743 | 300 | 12176 | 414 | eya,ena,Ziz,klu,vig,bab2,hyx,unc-119,Abl,CG14441,msi,Pex5,spi,Pdk1,Lmx1a,CG7763,arm,CG32772,Dronc,l(1)G0196,Dyrk2,Gfat2,tna,14-3-3epsilon,crp,ed,ec,step,chn,Dl,CG1218,CG3165,Sox14,CG10321,Syx1A,ey,CG17883,CG7536,Socs36E,cindr,Tom,d,rgr,Nrk,Su(z)2,CG12769,abd-A,morgue,rg,SmD1,CenB1A,Elba2,Pp2C1,fl(2)d,SMC3,glec,rdx,Pitslre,Drak,Pvr,fz3,Magi,Fkbp14,RhoGEF4,CG13895,14-3-3zeta,Hr4,Shrm,drl,His3.3A,vn,Gli,TBC1D5,htl,rib,edl,sim,REPTOR,Pgk,CG3107,jigr1,ss,cic,gukh,dlp,tkv,SERCA,aop,mura,tsh,CG17683,sns,Psa,CG14506,imd,Tet,wgn,CG4896,spdo,Cals,hang,dock,Rm62,boi,sick,betaTub60D,ft,jumu,ds,Arf79F,CG11360,Poxn,Psc,kn,CG13204,siz,jar,rho-4,disco-r,pnr,mim,dmrt11E,mld,Jupiter,Cks30A,Npc2g,ITP,tai,sd,H2.0,wnd,hth,robo1,CG13481,Cyp6a16,klar,CG2765,shot,nmo,CG11414,Adf1,CalpC,CG10348,Sxl,Ric,l(2)37Cg,sina,Mfe2,MEP-1,CG17646,Rab5,cora,CtBP,Pex14,hppy,CG6891,bru2,Pex10,bowl,Diap1,B-H1,crol,CG1316,CG5522,CG2991,Slbp,Abd-B,RhoGEF3,Atg17,esg,HmgZ,fuss,hng3,cv-d,al,Eph,ps,Unc-76,alpha-Man-Ia,CG31121,Cdk4,kon,GluProRS,pita,Dyb,eyg,Raf,Nlp,Dg,CG4935,eff,CG5853,flfl,CG30156,grn,mrt,Scr,f,CG11617,Actbeta,fz2,Atox1,sens-2,kay,CycG,shg,Hr39,otk,Mob2,yrt,Pvf3,corto,orb2,CG34376,Tis11,CG11247,HLH3B,Hel89B,Dad,Ddr,baz,lin-28,Fas1,su(sable),InR,nuf,scrib,dve,CG11122,RhoU,312,spen,LanA,chic,wbl,CG4467,His2Av,SCAR,Prdm13,nub,SNF4Agamma,CG2017,Cip4,h,cue,CG7560,Myo10A,His3.3B,bol,fz,bin3,emc,Mef2,Cadps,Pez,Sema1a,CG4645,ttk,Dp,Kdm2,Lk6,CG45050,sli,CG10417,pdm3,trbl,Rab1,pum,Snoo,Tob,CG31457,CG3436,Cbl,sens,cact,subdued,H,zfh2,apolpp,Sema1b,Src42A,CycB3,ph-p,CG34401,ATPCL,N,CG11069,sqz,ap,dome,FBXO11,E(spl)m5-HLH,Rgl,lt,lilli,Dcr-2,otp,Bsg,dpy,Egfr | | 3.706e-26 | -58.56 | pattern specification process | biological process | GO:0007389 | 544 | 77 | 12345 | 442 | Pat1,Gli,sens,Kdm2,fs(1)h,Dg,hth,Raf,h,unc-5,Tom,cic,nmo,Dad,boi,wbl,arm,CtBP,Dp,E(spl)m5-HLH,14-3-3epsilon,corto,Cdk4,chic,S,fuss,kn,klu,ed,dve,dlp,cora,f,sli,step,ena,ss,lilli,Dl,abd-A,frc,pum,nuf,ds,fz,emc,Cbl,H,cv-d,robo1,tkv,Mpcp2,Abd-B,baz,pnr,ft,N,14-3-3zeta,scrib,ap,edl,vn,bowl,nub,rdx,B-H1,spen,cact,tara,ec,Egfr,spi,fz2,tsh,shg,Scr,dome | | 8.410e-26 | -57.74 | regulation of cellular metabolic process | biological process | GO:0031323 | 1987 | 160 | 12345 | 442 | h,hth,mam,fl(2)d,Snoo,Tet,Actbeta,Kdm2,trbl,arm,Poxn,14-3-3epsilon,kay,HLH3B,cic,crp,Dad,CG4896,CG1218,kek5,fuss,kn,Psc,corto,bru2,tna,lncRNA:Hsromega,Rm62,ss,CG8312,Src42A,Nmda1,dpy,MEP-1,lncRNA:bxd,step,CycB3,hyx,emc,cv-d,H,Dronc,Fem-1,Hel89B,ps,CycG,Atg17,lili,lin-28,Sox14,Tob,ey,tai,Mef2,tkv,Dcr-2,nub,InR,hppy,zfh2,14-3-3zeta,scrib,Elba2,Pgk,su(sable),bowl,edl,vn,ttk,spi,Scr,Prdm13,chn,aop,al,cact,CG34172,Egfr,HmgZ,Raf,rib,eya,Lmx1a,CG12769,Hr39,imd,Tis11,sens,fs(1)h,wnd,Dg,hng3,Dp,CtBP,Diap1,E(spl)m5-HLH,Alh,Pdk1,crol,Hr4,grn,ph-p,Spn42De,jeb,klu,dve,H2.0,Lk6,bol,pdm3,Pvr,Socs36E,jumu,lilli,Sxl,bab2,sd,bin3,esg,REPTOR,sim,msi,CG34376,Cbl,abd-A,PI31,pum,rgr,Tollo,Cks30A,pnr,Adf1,wdb,SNF4Agamma,Atg16,Abd-B,rictor,eyg,FBXO11,CG11360,N,pita,orb2,sens-2,ap,dmrt11E,tsh,otp,Su(z)2,CG13204,CG10321,rdx,B-H1,Kdm4B,spen,sina,312,CG10348,tara,CG11617 | | 5.029e-25 | -55.95 | regulation of macromolecule metabolic process | biological process | GO:0060255 | 2019 | 160 | 12345 | 442 | Dcr-2,tkv,Mef2,Tob,ey,Sox14,ft,tai,Atg17,lili,CycG,ps,lin-28,Hel89B,emc,hyx,cv-d,H,Fem-1,Dronc,CycB3,cact,Egfr,aop,al,Prdm13,chn,ttk,Scr,spi,su(sable),bowl,vn,edl,zfh2,scrib,Elba2,14-3-3zeta,hppy,nub,InR,Dad,CG4896,cic,crp,kay,14-3-3epsilon,HLH3B,Poxn,arm,Kdm2,trbl,Actbeta,Tet,Snoo,fl(2)d,h,hth,mam,step,MEP-1,lncRNA:bxd,dpy,mrt,CG8312,Src42A,ss,lncRNA:Hsromega,Rm62,bru2,corto,tna,kn,Psc,CG1218,kek5,fuss,Abd-B,rictor,Adf1,SNF4Agamma,wdb,Tollo,pnr,Cks30A,PI31,abd-A,rgr,pum,Cbl,tara,jigr1,CG10348,312,CG11617,B-H1,rdx,sina,Kdm4B,spen,CG13204,CG10321,Su(z)2,otp,dmrt11E,tsh,sens-2,orb2,ap,CG11360,FBXO11,pita,N,eyg,nmo,crol,Pdk1,Hr4,E(spl)m5-HLH,Alh,vig,Dp,CtBP,Diap1,fs(1)h,sens,hng3,Dg,wnd,Tis11,CG12769,Hr39,Raf,HmgZ,Lmx1a,eya,rib,CG34376,msi,bin3,sd,sim,esg,REPTOR,Sxl,bab2,lilli,jumu,Socs36E,Pvr,Lk6,pdm3,bol,klu,jeb,Spn42De,H2.0,dve,grn,ph-p | | 1.650e-24 | -54.76 | regulation of metabolic process | biological process | GO:0019222 | 2165 | 166 | 12345 | 442 | aop,al,cact,CG34172,Egfr,ttk,spi,Scr,Prdm13,chn,zfh2,14-3-3zeta,Elba2,scrib,su(sable),Pgk,edl,bowl,vn,nub,InR,hppy,Mef2,Dcr-2,tkv,Sox14,ft,ey,Tob,tai,Hel89B,CycG,ps,Atg17,lili,lin-28,CycB3,hyx,emc,Dronc,Fem-1,H,cv-d,step,ss,CG8312,Nmda1,Src42A,dpy,MEP-1,lncRNA:bxd,mrt,corto,bru2,tna,Rm62,lncRNA:Hsromega,CG1218,kek5,fuss,kn,Psc,cic,crp,Dad,CG4896,arm,Poxn,kay,14-3-3epsilon,HLH3B,Actbeta,Kdm2,trbl,h,hth,mam,fl(2)d,Tet,Snoo,B-H1,rdx,Kdm4B,spen,sina,312,CG10348,jigr1,tara,CG11617,otp,dmrt11E,tsh,Su(z)2,CG13204,CG10321,FBXO11,CG11360,N,pita,orb2,sens-2,ap,eyg,Atg16,Abd-B,rictor,Tollo,Cks30A,pnr,Adf1,wdb,SNF4Agamma,abd-A,PI31,pum,rgr,Cbl,sd,bin3,REPTOR,esg,sim,msi,CG34376,jumu,lilli,Sxl,bab2,pdm3,Lk6,bol,Pvr,Socs36E,grn,ph-p,Spn42De,jeb,klu,H2.0,dve,Pdk1,crol,Hr4,nmo,Dp,CtBP,vig,Diap1,E(spl)m5-HLH,Alh,CG2765,imd,Tis11,sens,fs(1)h,wnd,Dg,hng3,HmgZ,Raf,rib,eya,Lmx1a,CG12769,Hr39 | | 1.750e-24 | -54.70 | cell morphogenesis involved in differentiation | biological process | GO:0000904 | 468 | 69 | 12345 | 442 | chn,fz2,path,shg,Fas1,sick,LanA,spen,dock,nub,InR,ap,Bsg,neo,N,Ppcs,betaTub60D,comm2,drl,beat-IIa,Abl,tai,Sox14,ft,sqz,baz,rictor,robo1,Dronc,ds,fz,Sema1a,pum,Sema1b,Ptp4E,Dl,Src42A,ena,wgn,sim,sli,gukh,Psc,dlp,jeb,kn,fend,ph-p,grn,chic,shot,pdm3,14-3-3epsilon,kay,Dad,Rab5,Snoo,E(Pc),Nrk,eya,unc-5,Eph,wnd,otk,Dg,sens,fs(1)h,SCAR,Actbeta | | 1.843e-24 | -54.65 | locomotion | biological process | GO:0040011 | 664 | 83 | 12345 | 442 | Dad,nmo,htl,mim,jar,Rab5,14-3-3epsilon,kay,Diap1,sens,Eph,otk,Dg,SCAR,RhoL,E(Pc),Nrk,Raf,eya,rib,unc-5,wgn,ena,sim,esg,sli,klar,Src42A,shot,Pvr,RhoU,Socs36E,chic,Pen,pdm3,jeb,Psc,dlp,gukh,grn,fend,Cerk,tkv,Abd-B,sqz,baz,robo1,ey,Shrm,tai,Abl,Pvf3,abd-A,Rim2,Ptp4E,pum,Sema1b,Dl,Sema1a,yuri,fz,LanA,Egfr,aop,spen,dome,p130CAS,Fas1,fz2,shg,orb2,Bsg,ap,Ptpmeg2,drl,beat-IIa,betaTub60D,cindr,comm2,N,Ppcs,InR,dock | | 2.039e-24 | -54.55 | negative regulation of cellular process | biological process | GO:0048523 | 1291 | 121 | 12345 | 442 | Kdm4B,al,Pez,B-H1,aop,rdx,Egfr,312,cact,LanA,tsh,spi,ttk,Su(z)2,chn,spdo,Prdm13,N,scrib,Elba2,FBXO11,zfh2,bowl,edl,orb2,su(sable),Mob2,dock,eyg,nub,InR,chrb,hppy,CG42684,baz,Dcr-2,Abd-B,pnr,RapGAP1,Cip4,Abl,wdb,ft,Tob,Dl,Sema1b,pum,lin-28,abd-A,CycG,Ptp4E,ds,Sema1a,H,Dronc,emc,Cbl,LRR,esg,C1GalTA,sd,bin3,msi,ena,wgn,ss,klar,Src42A,CG8312,Sxl,MEP-1,lncRNA:bxd,bol,Lk6,corto,bru2,chic,shot,pxb,Pvr,Socs36E,fuss,ph-p,kek5,ed,dve,Psc,jeb,Spn42De,klu,Hr4,Tom,cic,Pdk1,Rab5,scyl,crol,mim,Dad,nmo,Diap1,eff,arm,CtBP,E(spl)m5-HLH,14-3-3epsilon,Tis11,trbl,otk,lqf,cue,Kdm2,sens,fs(1)h,d,eya,disco-r,h,Raf,fl(2)d,Snoo | | 2.340e-24 | -54.41 | cell projection organization | biological process | GO:0030030 | 637 | 81 | 12345 | 442 | Nrk,E(Pc),Snoo,unc-5,eya,otk,Dg,wnd,Eph,fs(1)h,sens,SCAR,Actbeta,Gli,kay,14-3-3epsilon,mim,htl,Dad,Rab5,gukh,Psc,dlp,kn,jeb,fend,grn,chic,Pvr,shot,pdm3,Src42A,ena,wgn,sli,sim,cora,f,Myo10A,Dronc,LRR,fz,ds,Sema1a,yuri,Sema1b,pum,Ptp4E,Abl,tai,ft,Sox14,Cip4,sqz,rictor,robo1,unc-119,dock,InR,nub,vn,ap,Bsg,orb2,scrib,betaTub60D,N,comm2,beat-IIa,drl,p130CAS,chn,shg,path,fz2,Fas1,sick,Egfr,LanA,SMC3,spen | | 2.764e-24 | -54.25 | compound eye development | biological process | GO:0048749 | 370 | 61 | 12345 | 442 | spi,tsh,shg,klar,ss,jumu,lilli,Src42A,ttk,dome,chn,rg,spen,sina,sd,B-H1,aop,rdx,ec,msi,Egfr,ena,Mob2,S,eyg,ed,dlp,jeb,klu,N,cindr,scrib,AP-1gamma,eff,arm,Abl,ft,kay,ey,nmo,tkv,boi,d,ds,eya,sns,fz,hth,Raf,Dronc,fl(2)d,Nrk,hyx,emc,Cbl,SCAR,Dl,Arf79F,siz,sens,frc | | 4.915e-24 | -53.67 | regionalization | biological process | GO:0003002 | 503 | 71 | 12345 | 442 | Pat1,Gli,sens,Kdm2,fs(1)h,Dg,h,hth,Raf,unc-5,cic,Tom,nmo,Dad,boi,wbl,CtBP,Dp,arm,E(spl)m5-HLH,14-3-3epsilon,corto,Cdk4,chic,S,fuss,kn,klu,dve,dlp,cora,f,sli,step,ena,ss,lilli,Dl,abd-A,frc,pum,nuf,ds,fz,emc,Cbl,cv-d,H,robo1,tkv,Mpcp2,Abd-B,baz,pnr,ft,N,scrib,ap,bowl,vn,rdx,B-H1,spen,cact,tara,Egfr,fz2,tsh,shg,Scr,dome | | 6.795e-24 | -53.35 | sensory system development | biological process | GO:0048880 | 401 | 63 | 12345 | 442 | Abl,ft,ey,kay,AP-1gamma,yrt,eff,arm,tkv,nmo,boi,cic,fl(2)d,Dronc,Nrk,emc,hyx,Cbl,ds,eya,d,sns,fz,hth,Raf,siz,sens,Arf79F,frc,SCAR,Dl,chn,dome,rg,tsh,spi,shg,klar,ss,jumu,ttk,lilli,Src42A,msi,ec,ena,Egfr,spen,sina,rdx,sd,aop,B-H1,ed,dlp,jeb,klu,Mob2,S,eyg,cindr,N,scrib | | 6.795e-24 | -53.35 | eye development | biological process | GO:0001654 | 401 | 63 | 12345 | 442 | scrib,N,cindr,eyg,S,Mob2,klu,jeb,dlp,ed,rdx,aop,B-H1,sd,sina,spen,Egfr,ena,msi,ec,lilli,Src42A,ttk,jumu,ss,klar,shg,tsh,spi,rg,chn,dome,Dl,SCAR,frc,siz,sens,Arf79F,hth,Raf,fz,sns,eya,d,ds,Cbl,emc,hyx,Nrk,Dronc,fl(2)d,cic,boi,nmo,tkv,arm,eff,AP-1gamma,yrt,kay,ey,ft,Abl | | 6.795e-24 | -53.35 | visual system development | biological process | GO:0150063 | 401 | 63 | 12345 | 442 | ey,kay,ft,Abl,arm,eff,AP-1gamma,yrt,boi,nmo,tkv,cic,Cbl,hyx,emc,Nrk,Dronc,fl(2)d,hth,Raf,sns,fz,eya,d,ds,frc,Arf79F,sens,siz,Dl,SCAR,rg,dome,chn,Src42A,lilli,ttk,klar,ss,jumu,shg,tsh,spi,Egfr,ena,msi,ec,aop,rdx,sd,B-H1,sina,spen,klu,jeb,dlp,ed,eyg,S,Mob2,scrib,cindr,N | | 1.486e-23 | -52.56 | embryo development | biological process | GO:0009790 | 568 | 75 | 12345 | 442 | robo1,tkv,Abd-B,baz,Pvf3,Cks30A,pnr,larp,Abl,Inx2,abd-A,frc,pum,Sema1b,CycB3,sns,fz,emc,Dronc,Myo10A,rdx,aop,Egfr,fz2,spi,tsh,shg,chn,dome,N,14-3-3zeta,scrib,edl,bowl,InR,chrb,cic,scyl,Rab5,Dad,htl,wbl,jar,arm,yrt,kay,Pat1,RhoL,fs(1)h,Dg,trbl,hth,Raf,h,rib,mam,cora,sli,step,ena,jumu,lilli,Src42A,Cdk4,Pvr,shot,chic,Orct2,S,ph-p,kn,jeb,ed,dlp,Psc | | 4.176e-23 | -51.53 | regulation of RNA metabolic process | biological process | GO:0051252 | 1281 | 118 | 12345 | 442 | hyx,emc,cv-d,H,lili,abd-A,ps,lin-28,rgr,pum,Hel89B,ey,Tob,Adf1,Sox14,tai,pnr,Abd-B,Dcr-2,tkv,rictor,Mef2,nub,eyg,sens-2,su(sable),edl,bowl,ap,zfh2,CG11360,FBXO11,Elba2,pita,scrib,14-3-3zeta,N,Prdm13,CG13204,CG10321,Su(z)2,chn,ttk,Scr,otp,dmrt11E,tsh,tara,cact,CG10348,CG11617,B-H1,aop,Kdm4B,al,spen,Hr39,CG12769,Tet,Snoo,fl(2)d,h,hth,HmgZ,Lmx1a,mam,rib,eya,fs(1)h,sens,Dg,hng3,Tis11,kay,14-3-3epsilon,E(spl)m5-HLH,HLH3B,Alh,Poxn,CtBP,Dp,arm,Dad,CG4896,crol,cic,Hr4,crp,klu,kn,Psc,H2.0,dve,grn,kek5,ph-p,fuss,Rm62,bru2,bol,corto,pdm3,tna,MEP-1,lncRNA:bxd,dpy,bab2,Sxl,CG8312,lilli,jumu,ss,CG34376,bin3,sd,sim,esg,REPTOR | | 5.256e-23 | -51.30 | regulation of multicellular organismal development | biological process | GO:2000026 | 594 | 76 | 12345 | 442 | Cip4,Cals,Adf1,ft,Sox14,tai,Abl,Syx1A,robo1,Mef2,CG34401,morgue,tkv,Abd-B,rictor,baz,Sema1a,ds,fz,mop,emc,Cbl,Dronc,Dl,abd-A,Him,pum,Sema1b,ttk,tsh,spi,shg,chn,aop,sina,LanA,ec,Egfr,nub,InR,Mob2,N,scrib,neo,vn,Diap1,eff,E(spl)m5-HLH,kay,Hr4,Dad,nmo,hth,Raf,Rab1,h,d,Gli,Arf79F,otk,lqf,ss,klar,Src42A,sd,C1GalTA,sli,ena,kn,klu,ed,dlp,shot,Pvr,Socs36E,chic | | 5.866e-23 | -51.19 | regulation of signal transduction | biological process | GO:0009966 | 745 | 86 | 12345 | 442 | step,Src42A,Sxl,Lk6,Cdk4,lncRNA:Hsromega,Pvr,Socs36E,Pitslre,fuss,kek5,S,rho-4,ed,dlp,jeb,Hr4,Tom,scyl,Pdk1,crol,Dad,nmo,Diap1,eff,CtBP,14-3-3epsilon,Actbeta,wnd,otk,lqf,trbl,Ziz,cue,apolpp,eya,Dyrk2,d,CG5522,Raf,hth,Snoo,spen,Pez,aop,rdx,Egfr,cact,spi,RhoGEF3,spdo,N,14-3-3zeta,scrib,Fkbp14,vn,edl,dock,InR,chrb,hppy,CG42684,rictor,Dcr-2,tkv,RapGAP1,wdb,ft,ey,Dl,RhoGEF4,pum,lin-28,CycG,siz,Rgl,Ptp4E,lili,ds,Dronc,H,cv-d,hyx,mop,LRR,Cbl | | 7.781e-23 | -50.91 | regulation of response to stimulus | biological process | GO:0048583 | 958 | 99 | 12345 | 442 | spdo,RhoGEF3,CG32795,spi,cact,Egfr,rdx,Pez,aop,spen,hppy,chrb,InR,nub,dock,edl,vn,Fkbp14,DPCoAC,scrib,14-3-3zeta,N,ey,ft,wdb,RapGAP1,pnr,Dcr-2,tkv,rictor,CG34401,CG42684,Cbl,LRR,hyx,mop,Dronc,cv-d,H,Sema1a,fz,ds,lili,Ptp4E,Rgl,CycG,siz,lin-28,Sema1b,pum,Dl,RhoGEF4,Sxl,Src42A,step,sli,jeb,dlp,ed,rho-4,fuss,kek5,S,Pitslre,Socs36E,Pvr,lncRNA:Hsromega,Rm62,chic,Cdk4,Lk6,14-3-3epsilon,CtBP,eff,Diap1,htl,Dad,nmo,crol,Pdk1,scyl,Tom,Hr4,Snoo,hth,CG5522,Raf,d,Dyrk2,eya,apolpp,cue,Ziz,lqf,otk,trbl,wnd,Actbeta,imd | | 1.142e-22 | -50.52 | cell morphogenesis involved in neuron differentiation | biological process | GO:0048667 | 422 | 63 | 12345 | 442 | Rab5,robo1,baz,sqz,rictor,Dad,tai,Abl,kay,14-3-3epsilon,Sox14,SCAR,Actbeta,Dg,otk,Eph,pum,Sema1b,wnd,Ptp4E,fs(1)h,sens,unc-5,fz,eya,Sema1a,Snoo,E(Pc),Nrk,Dronc,sli,sim,spen,ena,sick,LanA,wgn,shg,path,fz2,Src42A,Fas1,chn,betaTub60D,comm2,N,beat-IIa,pdm3,drl,chic,ap,Bsg,shot,dock,fend,nub,InR,grn,gukh,Psc,dlp,kn,jeb | | 1.748e-22 | -50.10 | cellular component organization | biological process | GO:0016043 | 2830 | 192 | 12345 | 442 | ttk,RhoGEF3,path,fz2,rg,Spc105R,p130CAS,chn,aop,LanA,cact,Egfr,sick,CG34172,Wdr62,InR,nub,unc-119,hppy,beat-IIa,scrib,Elba2,14-3-3zeta,comm2,betaTub60D,Magi,Ppcs,Bsg,neo,vn,Pvf3,Tob,ft,Sox14,Shrm,tai,tkv,Dcr-2,ReepA,yuri,sns,fz,Drep4,emc,hyx,mop,Dronc,Pex5,Dl,Hel89B,Atg17,Ptp4E,Rim2,siz,CycG,nuf,CG6891,Src42A,Pex14,bou,f,C1GalTA,sli,wgn,ena,kn,Psc,gukh,ed,Cdk4,bru2,corto,tna,RhoU,lncRNA:Hsromega,chic,arm,eff,Slbp,kay,14-3-3epsilon,His2Av,kon,Rab5,cic,htl,Dad,jar,Pex7,h,Nrk,Snoo,E(Pc),fl(2)d,Actbeta,Gli,Pat1,CG7394,Arf79F,Kdm2,trbl,otk,Eph,Jupiter,Fas1,shg,CenB1A,Su(z)2,B-H1,Kdm4B,spen,SMC3,tara,lt,Mob2,dock,CG14562,FBXO11,CG3149,drl,N,cindr,orb2,ap,Cip4,larp,AP-1gamma,Cks30A,Adf1,Abl,wdb,Atg16,Nlp,Syx1A,robo1,rictor,baz,sqz,Sema1a,ds,LRR,Myo10A,moody,PI31,Sema1b,Dgat2,pum,lilli,klar,jumu,Sxl,bin3,cora,sim,grn,S,ph-p,fend,jeb,dlp,dve,pdm3,Pvr,shot,Pen,CG17683,vig,CtBP,Dp,Diap1,yrt,Pex10,Ppn,Pdk1,crol,His3.3B,mim,His3.3A,SmD1,Raf,Rab1,HmgZ,unc-5,rib,eya,Hr39,SCAR,fs(1)h,sens,Dg,CG33474,wnd | | 2.127e-22 | -49.90 | regulation of cell communication | biological process | GO:0010646 | 855 | 92 | 12345 | 442 | rictor,Dcr-2,tkv,CG42684,wdb,ft,ey,RapGAP1,Cals,pum,lin-28,Rgl,CycG,siz,lili,Ptp4E,Dl,RhoGEF4,Dronc,cv-d,H,hyx,mop,LRR,Cbl,ds,Egfr,cact,spen,Pez,rdx,aop,spdo,spi,RhoGEF3,Fkbp14,vn,edl,Ir21a,14-3-3zeta,N,scrib,chrb,hppy,dock,InR,nmo,Dad,Hr4,Tom,crol,Pdk1,scyl,14-3-3epsilon,Diap1,eff,CtBP,wnd,lqf,trbl,Dg,otk,cue,Ziz,apolpp,Gli,Actbeta,Snoo,eya,Dyrk2,d,hth,Rab1,Raf,CG5522,step,wgn,Sxl,Src42A,lncRNA:Hsromega,Pvr,Socs36E,Lk6,Cdk4,ed,dlp,jeb,Pitslre,S,fuss,kek5,rho-4 | | 2.127e-22 | -49.90 | regulation of signaling | biological process | GO:0023051 | 855 | 92 | 12345 | 442 | Gli,Actbeta,wnd,lqf,otk,trbl,Dg,Ziz,cue,apolpp,Dyrk2,d,eya,hth,CG5522,Rab1,Raf,Snoo,Hr4,Tom,scyl,crol,Pdk1,Dad,nmo,Diap1,eff,CtBP,14-3-3epsilon,Lk6,Cdk4,Pvr,lncRNA:Hsromega,Socs36E,Pitslre,S,fuss,kek5,rho-4,ed,dlp,jeb,step,wgn,Src42A,Sxl,RhoGEF4,Dl,pum,lin-28,siz,CycG,Rgl,lili,Ptp4E,ds,Dronc,cv-d,H,mop,hyx,LRR,Cbl,CG42684,rictor,tkv,Dcr-2,RapGAP1,Cals,wdb,ft,ey,14-3-3zeta,N,scrib,Fkbp14,edl,vn,Ir21a,dock,InR,chrb,hppy,spen,aop,Pez,rdx,Egfr,cact,spi,RhoGEF3,spdo | | 2.469e-22 | -49.75 | cell projection morphogenesis | biological process | GO:0048858 | 428 | 63 | 12345 | 442 | chn,Fas1,Src42A,fz2,path,shg,wgn,LanA,ena,sick,spen,sim,sli,kn,jeb,gukh,Psc,dlp,grn,nub,InR,dock,fend,shot,Bsg,ap,chic,drl,pdm3,beat-IIa,N,betaTub60D,comm2,Sox14,kay,14-3-3epsilon,Abl,tai,Dad,htl,rictor,sqz,robo1,Rab5,Dronc,Snoo,E(Pc),Nrk,Sema1a,eya,unc-5,fz,sens,fs(1)h,Ptp4E,Sema1b,Eph,wnd,pum,otk,Dg,Actbeta,SCAR | | 3.134e-22 | -49.51 | plasma membrane bounded cell projection organization | biological process | GO:0120036 | 626 | 77 | 12345 | 442 | Bsg,orb2,ap,beat-IIa,drl,scrib,comm2,N,betaTub60D,unc-119,nub,InR,dock,SMC3,LanA,sick,spen,p130CAS,chn,Fas1,path,shg,fz2,Ptp4E,pum,Sema1b,LRR,Myo10A,Dronc,yuri,Sema1a,fz,ds,rictor,sqz,robo1,ft,Sox14,Abl,tai,Cip4,shot,chic,pdm3,kn,jeb,dlp,gukh,Psc,grn,fend,wgn,ena,f,cora,sli,sim,Src42A,fs(1)h,sens,Dg,otk,wnd,Eph,Actbeta,Gli,SCAR,Snoo,Nrk,E(Pc),unc-5,eya,Dad,mim,Rab5,kay,14-3-3epsilon | | 3.623e-22 | -49.37 | regulation of multicellular organismal process | biological process | GO:0051239 | 781 | 87 | 12345 | 442 | chn,shg,spi,tsh,fz2,ttk,Egfr,ec,LanA,cact,sina,aop,Mob2,InR,nub,vn,neo,scrib,N,tai,Abl,ey,ft,Sox14,Adf1,Cip4,Cals,rictor,baz,Abd-B,tkv,morgue,CG34401,robo1,Mef2,Syx1A,Dronc,Cbl,emc,mop,fz,ds,Sema1a,pum,Sema1b,Him,abd-A,Dl,Src42A,ss,klar,ena,step,sli,sd,C1GalTA,dlp,ed,klu,kn,Orct2,chic,Socs36E,pxb,shot,Pvr,kay,E(spl)m5-HLH,eff,Diap1,Dp,wbl,nmo,mld,Dad,Hr4,Pdk1,d,Raf,hth,h,Rab1,otk,lqf,cue,Arf79F,Gli | | 3.926e-22 | -49.29 | regulation of nucleobase-containing compound metabolic process | biological process | GO:0019219 | 1315 | 118 | 12345 | 442 | FBXO11,zfh2,CG11360,N,14-3-3zeta,pita,Elba2,scrib,su(sable),sens-2,ap,edl,bowl,nub,eyg,B-H1,aop,al,spen,Kdm4B,CG10348,cact,tara,CG11617,ttk,tsh,otp,dmrt11E,Scr,Prdm13,Su(z)2,chn,CG10321,CG13204,Hel89B,abd-A,ps,lili,pum,rgr,lin-28,hyx,emc,cv-d,H,Mef2,tkv,Dcr-2,Abd-B,rictor,pnr,Adf1,Sox14,Tob,ey,tai,pdm3,corto,bol,bru2,tna,Rm62,grn,ph-p,fuss,kek5,kn,klu,H2.0,dve,Psc,sd,bin3,REPTOR,sim,esg,CG34376,ss,jumu,CG8312,lilli,dpy,MEP-1,lncRNA:bxd,bab2,Sxl,Tis11,sens,fs(1)h,hng3,Dg,HmgZ,h,hth,rib,eya,Lmx1a,mam,Hr39,CG12769,fl(2)d,Tet,Snoo,cic,crol,Hr4,crp,Dad,CG4896,arm,CtBP,Dp,Poxn,E(spl)m5-HLH,kay,14-3-3epsilon,HLH3B,Alh | | 6.059e-22 | -48.86 | movement of cell or subcellular component | biological process | GO:0006928 | 724 | 83 | 12345 | 442 | Sema1a,fz,Dl,Ptp4E,abd-A,pum,Sema1b,Pvf3,ey,Abl,tai,Shrm,robo1,Abd-B,tkv,sqz,baz,InR,dock,beat-IIa,drl,Ppcs,N,cindr,betaTub60D,comm2,Bsg,neo,orb2,Ptpmeg2,ap,Fas1,shg,fz2,p130CAS,dome,aop,spen,LanA,Egfr,Raf,Rab1,unc-5,rib,eya,Nrk,E(Pc),Pat1,SCAR,RhoL,sens,Dg,otk,Eph,Diap1,14-3-3epsilon,kay,Rab5,htl,Dad,jar,mim,grn,Cerk,fend,jeb,dlp,Psc,gukh,pdm3,Socs36E,RhoU,Pvr,shot,chic,Src42A,klar,sli,sim,esg,wgn,Unc-76,ena | | 6.373e-22 | -48.80 | eye morphogenesis | biological process | GO:0048592 | 310 | 53 | 12345 | 442 | sina,sd,aop,rdx,B-H1,ec,msi,Egfr,spi,shg,klar,ss,ttk,lilli,chn,dome,rg,N,cindr,scrib,Mob2,S,eyg,ed,jeb,klu,cic,nmo,tkv,boi,yrt,eff,arm,ft,ey,kay,SCAR,Dl,Arf79F,sens,siz,frc,eya,ds,d,sns,fz,hth,Raf,Nrk,hyx,emc,Cbl | | 6.373e-22 | -48.80 | sensory organ morphogenesis | biological process | GO:0090596 | 310 | 53 | 12345 | 442 | ft,kay,ey,yrt,eff,arm,nmo,tkv,boi,cic,Nrk,emc,hyx,Cbl,ds,eya,d,fz,sns,Raf,hth,sens,Arf79F,siz,frc,SCAR,Dl,dome,chn,rg,spi,shg,ss,klar,ttk,lilli,ec,msi,Egfr,sina,aop,sd,B-H1,rdx,ed,jeb,klu,Mob2,S,eyg,cindr,N,scrib | | 7.644e-22 | -48.62 | cell part morphogenesis | biological process | GO:0032990 | 437 | 63 | 12345 | 442 | beat-IIa,pdm3,drl,comm2,N,betaTub60D,Bsg,shot,chic,ap,nub,InR,grn,fend,dock,kn,jeb,Psc,gukh,dlp,sli,sim,spen,LanA,wgn,ena,sick,Src42A,Fas1,shg,path,fz2,chn,Actbeta,SCAR,Ptp4E,fs(1)h,sens,otk,Dg,wnd,Eph,Sema1b,pum,Sema1a,fz,unc-5,eya,Snoo,E(Pc),Nrk,Dronc,Rab5,robo1,htl,Dad,rictor,sqz,14-3-3epsilon,kay,Sox14,tai,Abl | | 8.909e-22 | -48.47 | neuron projection morphogenesis | biological process | GO:0048812 | 425 | 62 | 12345 | 442 | unc-5,fz,eya,Sema1a,E(Pc),Nrk,Snoo,Dronc,SCAR,Actbeta,otk,Dg,Eph,pum,wnd,Sema1b,Ptp4E,fs(1)h,sens,tai,Abl,14-3-3epsilon,kay,Sox14,Rab5,robo1,sqz,rictor,Dad,dock,fend,InR,nub,grn,Psc,gukh,dlp,jeb,kn,comm2,betaTub60D,N,beat-IIa,drl,pdm3,ap,chic,Bsg,shot,shg,path,fz2,Src42A,Fas1,chn,sli,spen,sim,ena,sick,wgn,LanA | | 1.010e-21 | -48.34 | plasma membrane bounded cell projection morphogenesis | biological process | GO:0120039 | 426 | 62 | 12345 | 442 | beat-IIa,pdm3,drl,N,betaTub60D,comm2,Bsg,shot,ap,chic,nub,InR,grn,fend,dock,kn,jeb,dlp,Psc,gukh,sli,spen,sim,wgn,LanA,sick,ena,Src42A,Fas1,path,shg,fz2,chn,Actbeta,SCAR,Ptp4E,fs(1)h,sens,otk,Dg,wnd,Eph,Sema1b,pum,Sema1a,fz,unc-5,eya,Snoo,E(Pc),Nrk,Dronc,Rab5,robo1,Dad,sqz,rictor,14-3-3epsilon,kay,Sox14,Abl,tai | | 1.088e-21 | -48.27 | compound eye morphogenesis | biological process | GO:0001745 | 290 | 51 | 12345 | 442 | scrib,cindr,N,eyg,S,Mob2,klu,jeb,ed,B-H1,rdx,aop,sd,sina,Egfr,msi,ec,lilli,ttk,ss,klar,shg,spi,rg,dome,chn,Dl,SCAR,frc,Arf79F,sens,siz,Raf,hth,fz,sns,eya,ds,d,Cbl,emc,hyx,Nrk,boi,tkv,nmo,arm,eff,ey,kay,ft | | 1.186e-21 | -48.18 | negative regulation of biological process | biological process | GO:0048519 | 1502 | 127 | 12345 | 442 | Su(z)2,chn,spdo,Prdm13,spi,tsh,ttk,Egfr,312,cact,LanA,al,Kdm4B,rdx,aop,Pez,B-H1,hppy,chrb,Mob2,dock,InR,nub,eyg,bowl,edl,orb2,su(sable),14-3-3zeta,N,Elba2,scrib,FBXO11,zfh2,wdb,Abl,ft,Tob,pnr,RapGAP1,Cip4,baz,tkv,Dcr-2,Abd-B,CG42684,Dronc,H,emc,Cbl,LRR,ds,Sema1a,Sema1b,pum,lin-28,CycG,abd-A,Him,Ptp4E,Dl,mrt,Sxl,lncRNA:bxd,MEP-1,ss,klar,CG8312,Src42A,msi,ena,wgn,esg,sd,C1GalTA,bin3,ed,dve,Psc,Spn42De,jeb,klu,fuss,kek5,ph-p,chic,Pvr,Rm62,pxb,shot,Socs36E,corto,bol,Lk6,bru2,E(spl)m5-HLH,14-3-3epsilon,Diap1,eff,CtBP,arm,vig,mim,Dad,nmo,Hr4,cic,Tom,crol,scyl,Rab5,Pdk1,fl(2)d,Snoo,eya,d,disco-r,Raf,h,lqf,trbl,otk,cue,sens,Kdm2,fs(1)h,Tis11 | | 1.297e-21 | -48.09 | regulation of cell differentiation | biological process | GO:0045595 | 428 | 62 | 12345 | 442 | Snoo,Dronc,Cbl,mop,emc,fz,eya,hth,h,Rab1,Raf,Sema1a,lqf,Sema1b,Arf79F,Dl,Gli,tai,Abl,kay,Adf1,E(spl)m5-HLH,Sox14,eff,Diap1,Dp,rictor,Abd-B,tkv,morgue,CG34401,Atg16,robo1,Syx1A,ed,klu,kn,fuss,InR,nub,vn,chic,Socs36E,shot,Pvr,N,chn,bab2,shg,spi,ttk,Src42A,klar,ss,Egfr,ec,cact,sina,esg,sd,aop,C1GalTA | | 1.736e-21 | -47.80 | regulation of gene expression | biological process | GO:0010468 | 1547 | 129 | 12345 | 442 | h,hth,mam,fl(2)d,Tet,Snoo,cic,crp,Dad,CG4896,arm,Poxn,14-3-3epsilon,kay,HLH3B,corto,bru2,tna,Rm62,fuss,kek5,kn,Psc,ss,CG8312,dpy,MEP-1,lncRNA:bxd,mrt,Hel89B,ps,lili,lin-28,hyx,emc,cv-d,H,Mef2,tkv,Dcr-2,ft,Sox14,Tob,ey,tai,zfh2,14-3-3zeta,Elba2,scrib,su(sable),bowl,edl,nub,aop,al,cact,Egfr,ttk,spi,Scr,Prdm13,chn,Tis11,sens,fs(1)h,hng3,Dg,HmgZ,eya,rib,Lmx1a,CG12769,Hr39,crol,Hr4,nmo,CtBP,Dp,vig,Diap1,E(spl)m5-HLH,Alh,bol,pdm3,grn,ph-p,klu,H2.0,dve,sd,bin3,esg,sim,REPTOR,msi,CG34376,jumu,lilli,Sxl,bab2,abd-A,pum,rgr,Abd-B,rictor,pnr,Adf1,FBXO11,CG11360,N,pita,orb2,sens-2,ap,eyg,B-H1,Kdm4B,spen,312,CG10348,tara,jigr1,CG11617,tsh,dmrt11E,otp,Su(z)2,CG13204,CG10321 | | 1.977e-21 | -47.67 | regulation of RNA biosynthetic process | biological process | GO:2001141 | 1158 | 108 | 12345 | 442 | lili,abd-A,rgr,pum,Hel89B,emc,hyx,cv-d,H,Abd-B,tkv,Dcr-2,rictor,Mef2,ey,Tob,Sox14,Adf1,tai,pnr,su(sable),sens-2,bowl,edl,ap,zfh2,FBXO11,scrib,pita,Elba2,N,14-3-3zeta,eyg,nub,tara,CG10348,cact,CG11617,B-H1,aop,Kdm4B,al,spen,Prdm13,CG10321,CG13204,chn,Su(z)2,ttk,Scr,dmrt11E,tsh,otp,fs(1)h,sens,hng3,Hr39,CG12769,Tet,Snoo,fl(2)d,hth,h,HmgZ,Lmx1a,mam,rib,eya,Dad,crol,cic,crp,Hr4,kay,14-3-3epsilon,E(spl)m5-HLH,HLH3B,Alh,Poxn,arm,CtBP,Dp,pdm3,corto,tna,klu,kn,Psc,H2.0,dve,grn,ph-p,kek5,fuss,CG34376,bin3,sd,sim,esg,REPTOR,MEP-1,lncRNA:bxd,dpy,bab2,CG8312,lilli,jumu,ss | | 1.977e-21 | -47.67 | regulation of nucleic acid-templated transcription | biological process | GO:1903506 | 1158 | 108 | 12345 | 442 | CtBP,Dp,arm,Poxn,Alh,HLH3B,E(spl)m5-HLH,14-3-3epsilon,kay,crp,Hr4,cic,crol,Dad,eya,rib,mam,Lmx1a,HmgZ,h,hth,fl(2)d,Tet,Snoo,CG12769,Hr39,hng3,sens,fs(1)h,jumu,ss,CG8312,lilli,bab2,dpy,MEP-1,lncRNA:bxd,sim,REPTOR,esg,sd,bin3,CG34376,ph-p,kek5,fuss,grn,H2.0,dve,Psc,kn,klu,tna,pdm3,corto,pnr,tai,Sox14,Adf1,ey,Tob,Mef2,rictor,Dcr-2,tkv,Abd-B,H,cv-d,emc,hyx,Hel89B,pum,rgr,abd-A,lili,otp,dmrt11E,tsh,Scr,ttk,chn,Su(z)2,CG13204,CG10321,Prdm13,Kdm4B,spen,al,B-H1,aop,CG11617,CG10348,cact,tara,nub,eyg,14-3-3zeta,N,scrib,Elba2,pita,FBXO11,zfh2,ap,bowl,edl,sens-2,su(sable) | | 2.569e-21 | -47.41 | regulation of transcription, DNA-templated | biological process | GO:0006355 | 1144 | 107 | 12345 | 442 | hng3,fs(1)h,sens,mam,Lmx1a,rib,eya,h,hth,HmgZ,Snoo,Tet,fl(2)d,Hr39,CG12769,crp,Hr4,crol,cic,Dad,Poxn,Dp,CtBP,arm,HLH3B,Alh,14-3-3epsilon,kay,E(spl)m5-HLH,tna,corto,pdm3,fuss,ph-p,kek5,grn,Psc,H2.0,dve,klu,kn,REPTOR,esg,sim,bin3,sd,CG34376,lilli,CG8312,ss,jumu,bab2,lncRNA:bxd,MEP-1,dpy,Hel89B,rgr,pum,lili,abd-A,cv-d,H,hyx,emc,Mef2,rictor,Abd-B,Dcr-2,tkv,pnr,tai,ey,Sox14,Adf1,scrib,pita,Elba2,14-3-3zeta,N,zfh2,FBXO11,bowl,edl,ap,su(sable),sens-2,nub,eyg,al,spen,Kdm4B,B-H1,aop,CG11617,tara,CG10348,cact,Scr,tsh,otp,dmrt11E,ttk,CG10321,CG13204,Su(z)2,chn,Prdm13 | | 2.586e-21 | -47.40 | neuron projection development | biological process | GO:0031175 | 447 | 63 | 12345 | 442 | p130CAS,chn,shg,path,fz2,Src42A,Fas1,sick,ena,wgn,LanA,sli,spen,sim,dlp,Psc,gukh,jeb,kn,dock,fend,InR,nub,grn,chic,ap,Bsg,shot,comm2,betaTub60D,N,beat-IIa,pdm3,drl,Abl,tai,14-3-3epsilon,kay,Sox14,rictor,sqz,Dad,Rab5,robo1,Nrk,E(Pc),Snoo,Dronc,unc-5,fz,eya,Sema1a,Dg,otk,wnd,pum,Eph,Sema1b,fs(1)h,Ptp4E,sens,SCAR,Actbeta | | 2.889e-21 | -47.29 | cell | cellular component | GO:0005623 | 7811 | 357 | 11940 | 417 | scyl,h,ey,flfl,Pex5,UQCR-11,Adf1,InR,Cdk4,bou,tkv,Fas1,Jupiter,lt,robo1,Pdk1,msi,Pex10,jigr1,Snoo,ReepA,Sox14,betaTub60D,CG13481,sim,IntS3,CtBP,Pp2C1,CG32772,ttk,DPCoAC,orb2,Tis11,klu,Syx1A,CG17883,Ddr,sd,bol,boi,mRpL41,step,Psa,path,Pvr,klar,lncRNA:bxd,CycG,l(2)37Cg,Abd-B,morgue,CG33082,fs(1)h,Vps13,Hel89B,Sema1b,Pex7,Dronc,Dad,CG17683,chn,Pat1,HLH3B,Abl,Nrk,Lk6,frc,nAChRalpha4,Pgant5,Ir21a,N,glec,CG3436,CG5853,Mfe2,otk,14-3-3zeta,Tsp96F,l(1)G0196,RhoU,Esp,cue,kn,fz3,Cadps,jar,Bsg,aop,Ric,yuri,siz,RhoGEF4,Kr-h2,htl,fuss,312,CG14562,rg,gukh,dock,CG31211,Shrm,cact,sens,Dcr-2,kon,jumu,RhoGAP19D,su(sable),Drak,nmo,emp,chrb,fl(2)d,PI31,CalpC,lin-28,Eph,rgr,ps,p130CAS,tai,abd-A,CG7560,His3.3A,ogre,fz,Pitslre,S,Unc-76,spen,CG31457,cic,H,Dp,SmD1,Dyrk2,ft,corto,bowl,CG14441,Wdr62,grn,pnr,CG8312,ATPCL,Tpc1,dmrt11E,dlp,spdo,alpha-Man-Ia,tna,Drep4,His3.3B,al,GluProRS,Ptpmeg2,LanA,Mob2,ap,Cks30A,Egfr,ena,hth,CG10321,Rab5,CG9220,sqz,koi,Slbp,CG31121,CG4935,rho-4,AP-1gamma,sina,pdgy,Pez,CG12769,bab2,wbl,crol,Gli,Alh,CG12402,Dg,hang,tsh,pita,CG6465,kay,SNF4Agamma,CG7536,CG6656,CG17646,Dyb,CG34172,Raf,CG9784,lawc,pum,Magi,crp,Elba2,E(Pc),Sxl,Atg16,drongo,mam,larp,CG42788,Rm62,Hr4,drl,Src42A,CycB3,wnd,snRNA:U5:38ABb,ed,Tob,spi,Pex14,Cbl,shg,Tom,baz,bru2,Dl,rdx,CG13204,Gr93c,Pgk,CG3107,CG6891,SERCA,Kdm4B,mura,CG4645,REPTOR,shot,Scr,chic,cora,RapGAP1,dome,trbl,Poxn,Psc,Pen,emc,CG14464,Bacc,CG2765,CG11069,bin3,Mpcp2,arm,CG34376,Arf79F,stx,CG30156,Diap1,GlcAT-I,mld,eff,CG42684,eyg,CG11360,HmgZ,CG11617,ZnT41F,Rim2,CG8306,edl,vig,dve,lilli,beat-IIa,Su(z)2,cindr,CG4896,mrt,hppy,CG4467,sns,rictor,Hr39,kek5,sens-2,SCAR,apolpp,CG3149,Ocrl,Rab1,eya,ph-p,lncRNA:Hsromega,jeb,fz2,Mef2,Inx2,zfh2,Prdm13,ds,Tpst,CG1218,moody,CG12910,wgn,TBC1D5,Myo10A,mop,Ptp4E,MEP-1,scrib,esg,Tet,f,Atg17,ss,Mid1,CG11247,px,muc,tipE,CG7394,Fkbp14,rib,yrt,CG33474,Atox1,Cals,B-H1,CG6340,Cip4,Pgant6,Nlp,lqf,CG1673,SMC3,Spc105R,nuf,CG34401,Lmx1a,dpy,d,Kdm2,hng3,wdb,His2Av,14-3-3epsilon,CG6983,CG11030,CG10417,hyx,Cerk,FBXO11,E(spl)m5-HLH | | 2.889e-21 | -47.29 | cell part | cellular component | GO:0044464 | 7811 | 357 | 11940 | 417 | yuri,siz,Ric,aop,312,RhoGEF4,Kr-h2,htl,fuss,dock,CG14562,rg,gukh,RhoGAP19D,kon,jumu,su(sable),Drak,nmo,CG31211,cact,Shrm,Dcr-2,sens,emp,CalpC,PI31,Eph,lin-28,rgr,chrb,fl(2)d,abd-A,CG7560,His3.3A,ogre,ps,p130CAS,tai,cic,H,SmD1,Dp,Dyrk2,fz,Pitslre,S,Unc-76,spen,CG31457,pnr,CG8312,ft,grn,bowl,corto,Wdr62,CG14441,tna,alpha-Man-Ia,His3.3B,Drep4,al,GluProRS,Ptpmeg2,ATPCL,Tpc1,dmrt11E,dlp,spdo,Mob2,Cks30A,Egfr,ap,hth,ena,LanA,sqz,koi,CG10321,Rab5,CG9220,rho-4,AP-1gamma,sina,Slbp,CG31121,CG4935,pdgy,Pez,CG12769,CG12402,Alh,Dg,hang,bab2,wbl,Gli,crol,CG7536,CG6656,tsh,pita,CG6465,kay,SNF4Agamma,scyl,h,Pex5,UQCR-11,Adf1,ey,flfl,Jupiter,lt,robo1,InR,Cdk4,bou,Fas1,tkv,Pex10,jigr1,Sox14,ReepA,Snoo,Pdk1,msi,sim,IntS3,CtBP,CG13481,betaTub60D,Pp2C1,CG32772,ttk,orb2,DPCoAC,klu,Syx1A,Tis11,boi,Ddr,sd,CG17883,bol,mRpL41,klar,lncRNA:bxd,step,Psa,Pvr,path,Abd-B,morgue,fs(1)h,CG33082,CycG,l(2)37Cg,Dronc,Dad,Vps13,Hel89B,Sema1b,Pex7,Pat1,Abl,HLH3B,Lk6,Nrk,CG17683,chn,N,frc,nAChRalpha4,Pgant5,Ir21a,Tsp96F,otk,14-3-3zeta,Esp,RhoU,l(1)G0196,glec,CG3436,Mfe2,CG5853,Cadps,fz3,Bsg,jar,cue,kn,Rab1,eya,ph-p,kek5,sens-2,Hr39,SCAR,CG3149,apolpp,Ocrl,fz2,lncRNA:Hsromega,jeb,Prdm13,ds,Tpst,CG1218,Mef2,Inx2,zfh2,wgn,TBC1D5,moody,CG12910,MEP-1,scrib,Tet,esg,Myo10A,Ptp4E,mop,Mid1,ss,CG11247,px,f,Atg17,CG7394,muc,tipE,yrt,Fkbp14,rib,Cals,B-H1,CG33474,Atox1,Nlp,lqf,CG6340,Cip4,Pgant6,Spc105R,nuf,SMC3,CG1673,dpy,CG34401,Lmx1a,hng3,Kdm2,wdb,d,CG11030,CG10417,His2Av,CG6983,14-3-3epsilon,hyx,Cerk,E(spl)m5-HLH,FBXO11,Raf,CG9784,CG17646,Dyb,CG34172,crp,Magi,Elba2,lawc,pum,drongo,mam,larp,E(Pc),Sxl,Atg16,Src42A,wnd,CycB3,Rm62,Hr4,CG42788,drl,Tob,spi,Pex14,Cbl,snRNA:U5:38ABb,ed,bru2,shg,Tom,baz,Gr93c,Pgk,CG3107,Dl,rdx,CG13204,CG6891,SERCA,mura,CG4645,Kdm4B,Scr,chic,cora,RapGAP1,REPTOR,shot,Psc,dome,trbl,Poxn,CG14464,Bacc,CG11069,CG2765,Pen,emc,arm,CG34376,bin3,Mpcp2,Diap1,GlcAT-I,mld,eff,Arf79F,stx,CG30156,CG42684,eyg,CG11360,HmgZ,dve,beat-IIa,lilli,Su(z)2,cindr,CG4896,ZnT41F,CG11617,CG8306,Rim2,edl,vig,CG4467,sns,rictor,mrt,hppy | | 4.964e-21 | -46.75 | regulation of biosynthetic process | biological process | GO:0009889 | 1374 | 119 | 12345 | 442 | nub,eyg,FBXO11,zfh2,N,14-3-3zeta,pita,scrib,Elba2,orb2,su(sable),sens-2,ap,edl,bowl,ttk,tsh,otp,dmrt11E,Scr,Prdm13,chn,Su(z)2,CG10321,CG13204,aop,B-H1,al,spen,Kdm4B,CG10348,312,cact,tara,CG11617,hyx,emc,Cbl,H,cv-d,Hel89B,abd-A,lili,pum,rgr,Tollo,pnr,Sox14,Adf1,ey,Tob,tai,Mef2,tkv,Dcr-2,Abd-B,rictor,grn,fuss,ph-p,kek5,kn,klu,dve,H2.0,Psc,pdm3,corto,bol,bru2,tna,jumu,ss,CG8312,lilli,dpy,MEP-1,lncRNA:bxd,Sxl,bab2,sd,bin3,sim,REPTOR,esg,msi,CG34376,HmgZ,hth,h,eya,rib,Lmx1a,mam,CG12769,Hr39,fl(2)d,Tet,Snoo,CG2765,imd,Tis11,sens,fs(1)h,hng3,CtBP,arm,Dp,Poxn,E(spl)m5-HLH,kay,14-3-3epsilon,Alh,HLH3B,cic,crol,crp,Hr4,Dad | | 7.990e-21 | -46.28 | regulation of cellular macromolecule biosynthetic process | biological process | GO:2000112 | 1307 | 115 | 12345 | 442 | hyx,emc,H,cv-d,Hel89B,abd-A,lili,pum,rgr,Tollo,pnr,Sox14,Adf1,ey,tai,Mef2,tkv,Dcr-2,Abd-B,rictor,nub,eyg,FBXO11,zfh2,N,14-3-3zeta,Elba2,pita,scrib,orb2,su(sable),sens-2,ap,edl,bowl,ttk,tsh,dmrt11E,otp,Scr,Prdm13,chn,Su(z)2,CG13204,CG10321,B-H1,aop,al,Kdm4B,spen,CG10348,cact,312,tara,CG11617,HmgZ,h,hth,eya,rib,mam,Lmx1a,Hr39,CG12769,fl(2)d,Tet,Snoo,Tis11,sens,fs(1)h,hng3,arm,Dp,CtBP,Poxn,E(spl)m5-HLH,kay,14-3-3epsilon,Alh,HLH3B,cic,crol,crp,Hr4,Dad,grn,kek5,ph-p,fuss,kn,klu,dve,H2.0,Psc,bol,corto,pdm3,bru2,tna,jumu,ss,CG8312,lilli,dpy,MEP-1,lncRNA:bxd,bab2,Sxl,sd,bin3,esg,sim,REPTOR,msi,CG34376 | | 8.190e-21 | -46.25 | regulation of macromolecule biosynthetic process | biological process | GO:0010556 | 1326 | 116 | 12345 | 442 | Hel89B,lili,abd-A,rgr,pum,hyx,emc,cv-d,H,Mef2,Abd-B,tkv,Dcr-2,rictor,Tollo,pnr,ey,Tob,Sox14,Adf1,tai,zfh2,FBXO11,Elba2,scrib,pita,N,14-3-3zeta,su(sable),sens-2,orb2,bowl,edl,ap,nub,eyg,B-H1,aop,al,Kdm4B,spen,tara,CG10348,cact,312,CG11617,ttk,Scr,dmrt11E,tsh,otp,Prdm13,CG13204,CG10321,chn,Su(z)2,Tis11,fs(1)h,sens,hng3,h,hth,HmgZ,mam,Lmx1a,rib,eya,CG12769,Hr39,Snoo,Tet,fl(2)d,crol,cic,Hr4,crp,Dad,Poxn,arm,Dp,CtBP,kay,14-3-3epsilon,E(spl)m5-HLH,Alh,HLH3B,bru2,corto,bol,pdm3,tna,grn,kek5,ph-p,fuss,klu,kn,Psc,dve,H2.0,bin3,sd,sim,REPTOR,esg,CG34376,msi,lilli,CG8312,jumu,ss,lncRNA:bxd,MEP-1,dpy,Sxl,bab2 | | 1.297e-20 | -45.79 | regulation of cellular biosynthetic process | biological process | GO:0031326 | 1371 | 118 | 12345 | 442 | msi,CG34376,esg,sim,REPTOR,sd,bin3,bab2,Sxl,dpy,MEP-1,lncRNA:bxd,jumu,ss,CG8312,lilli,tna,corto,bol,pdm3,bru2,H2.0,dve,Psc,kn,klu,ph-p,fuss,kek5,grn,Dad,Hr4,crp,cic,crol,Alh,HLH3B,E(spl)m5-HLH,14-3-3epsilon,kay,CtBP,Dp,arm,Poxn,hng3,sens,fs(1)h,imd,Tis11,fl(2)d,Tet,Snoo,Hr39,CG12769,rib,eya,mam,Lmx1a,HmgZ,hth,h,CG11617,312,CG10348,cact,tara,al,Kdm4B,spen,B-H1,aop,Su(z)2,chn,CG10321,CG13204,Prdm13,dmrt11E,tsh,otp,Scr,ttk,ap,edl,bowl,orb2,su(sable),sens-2,14-3-3zeta,N,Elba2,pita,scrib,FBXO11,zfh2,nub,eyg,rictor,tkv,Dcr-2,Abd-B,Mef2,tai,Adf1,Sox14,ey,Tob,pnr,Tollo,pum,rgr,abd-A,lili,Hel89B,cv-d,H,hyx,emc,Cbl | | 2.988e-20 | -44.96 | cellular component organization or biogenesis | biological process | GO:0071840 | 2975 | 193 | 12345 | 442 | ds,Sema1a,Myo10A,LRR,moody,Dgat2,pum,Sema1b,PI31,Cks30A,AP-1gamma,larp,Cip4,Abl,wdb,Adf1,Syx1A,Nlp,robo1,Atg16,baz,sqz,rictor,Mob2,dock,lt,CG14562,cindr,N,CG3149,drl,FBXO11,ap,orb2,shg,Fas1,Jupiter,CG11030,Su(z)2,CenB1A,spen,Kdm4B,B-H1,SMC3,tara,rib,eya,unc-5,HmgZ,Rab1,Raf,Hr39,SCAR,wnd,CG33474,Dg,sens,fs(1)h,Diap1,yrt,CtBP,Dp,vig,CG17683,Ppn,Pex10,His3.3B,crol,Pdk1,SmD1,mim,His3.3A,fend,ph-p,S,grn,dve,dlp,jeb,pdm3,Pen,Pvr,shot,jumu,klar,lilli,Sxl,sim,bin3,cora,sns,fz,yuri,Dronc,hyx,emc,mop,Drep4,Hel89B,Dl,Pex5,CG6891,nuf,CycG,siz,Ptp4E,Atg17,Rim2,Pvf3,Shrm,tai,Sox14,ft,Tob,ReepA,Dcr-2,tkv,nub,InR,Wdr62,hppy,unc-119,comm2,Ppcs,14-3-3zeta,betaTub60D,Magi,Elba2,scrib,beat-IIa,vn,Bsg,neo,fz2,path,RhoGEF3,ttk,chn,p130CAS,rg,Spc105R,aop,CG34172,Egfr,sick,cact,LanA,h,fl(2)d,E(Pc),Nrk,Snoo,CG7394,Gli,Pat1,Actbeta,Eph,otk,trbl,Kdm2,Arf79F,Slbp,eff,arm,His2Av,14-3-3epsilon,kay,cic,Rab5,kon,jar,Pex7,Dad,htl,ed,gukh,Psc,kn,tna,corto,bru2,Cdk4,chic,lncRNA:Hsromega,RhoU,Src42A,bou,Pex14,sli,C1GalTA,f,ena,wgn | | 1.408e-19 | -43.41 | embryonic morphogenesis | biological process | GO:0048598 | 251 | 45 | 12345 | 442 | Inx2,trbl,pum,abd-A,mam,rib,Raf,Myo10A,Dronc,emc,Rab5,scyl,jar,baz,htl,Dad,tkv,pnr,yrt,arm,Abl,kay,scrib,N,bowl,chic,Pvr,shot,S,Orct2,InR,ed,chrb,jeb,sli,cora,aop,ena,Egfr,step,shg,tsh,spi,Src42A,jumu | | 1.647e-19 | -43.25 | cell migration | biological process | GO:0016477 | 287 | 48 | 12345 | 442 | htl,Abd-B,tkv,Dad,mim,jar,baz,Rab5,ey,14-3-3epsilon,kay,Abl,tai,Shrm,Pvf3,Diap1,abd-A,pum,Sema1b,Dl,SCAR,RhoL,Raf,Sema1a,unc-5,fz,eya,rib,ena,Egfr,aop,sli,esg,p130CAS,dome,Src42A,shg,fz2,Socs36E,Pvr,RhoU,Ptpmeg2,cindr,N,Ppcs,jeb,dlp,Cerk | | 3.791e-19 | -42.42 | Egfr (Epidermal growth factor receptor) | protein interactions | 37455 | 158 | 40 | 9204 | 432 | dome,Egfr,nmo,hppy,spen,arm,fz,rg,vn,eya,tkv,ed,scrib,S,Src42A,kay,htl,H,spi,pum,Dl,abd-A,shg,step,emc,Pvr,B-H1,Cbl,aop,klu,Raf,kn,sli,Abl,px,N,lilli,sens,dock,cora | | 4.977e-19 | -42.14 | H (Hairless) | protein interactions | 42445 | 143 | 38 | 9204 | 432 | ey,14-3-3zeta,Diap1,lilli,mam,trbl,baz,eyg,px,N,dve,klu,aop,esg,Gli,emc,Dl,muc,sd,cv-d,kay,ft,CycB3,S,CtBP,wdb,tkv,ap,vn,ed,CycG,fl(2)d,Dad,rg,Hr4,nmo,spen,Egfr | | 4.706e-18 | -39.90 | axon development | biological process | GO:0061564 | 336 | 50 | 12345 | 442 | 14-3-3epsilon,kay,Abl,tai,Dad,robo1,Rab5,Nrk,Sema1a,eya,fz,unc-5,sens,Ptp4E,wnd,Eph,Sema1b,Dg,otk,SCAR,p130CAS,Fas1,Src42A,fz2,shg,wgn,LanA,sick,ena,spen,sim,sli,jeb,gukh,dlp,Psc,grn,InR,dock,fend,shot,Bsg,chic,ap,pdm3,drl,beat-IIa,betaTub60D,N,comm2 | | 6.809e-18 | -39.53 | axonogenesis | biological process | GO:0007409 | 326 | 49 | 12345 | 442 | fz2,shg,Fas1,Src42A,ena,sick,wgn,LanA,spen,sim,sli,Psc,gukh,dlp,jeb,fend,dock,grn,InR,ap,chic,shot,Bsg,betaTub60D,comm2,N,drl,pdm3,beat-IIa,Abl,tai,kay,14-3-3epsilon,Dad,robo1,Rab5,Nrk,eya,fz,unc-5,Sema1a,wnd,Sema1b,Eph,otk,Dg,sens,Ptp4E,SCAR | | 9.183e-18 | -39.23 | anatomical structure formation involved in morphogenesis | biological process | GO:0048646 | 553 | 65 | 12345 | 442 | spi,shg,Scr,ttk,B-H1,Egfr,cact,dock,InR,N,14-3-3zeta,Ppcs,orb2,Bsg,Pgk,Cip4,Abl,ft,robo1,Mef2,Syx1A,baz,tkv,ds,sns,fz,yuri,H,Dronc,emc,Dl,Inx2,nuf,abd-A,siz,ss,klar,sli,ena,ph-p,S,ed,jeb,klu,Pen,shot,Dp,arm,CtBP,14-3-3epsilon,cic,Tom,Rab5,kon,jar,htl,eya,unc-5,mam,Raf,hth,SCAR,Dg,trbl,sens | | 1.191e-17 | -38.97 | neuron projection guidance | biological process | GO:0097485 | 280 | 45 | 12345 | 442 | SCAR,otk,Dg,Sema1b,Eph,Ptp4E,sens,unc-5,eya,Sema1a,Nrk,E(Pc),robo1,sqz,Dad,Abl,14-3-3epsilon,betaTub60D,N,comm2,beat-IIa,pdm3,drl,ap,chic,Bsg,shot,dock,fend,InR,grn,dlp,Psc,gukh,jeb,sli,sim,spen,ena,LanA,wgn,shg,fz2,Src42A,Fas1 | | 1.294e-17 | -38.89 | regulation of cell development | biological process | GO:0060284 | 344 | 50 | 12345 | 442 | klar,ss,ttk,Src42A,spi,shg,ec,Egfr,aop,sd,sina,kn,klu,ed,nub,InR,Pvr,shot,Socs36E,chic,vn,N,E(spl)m5-HLH,Sox14,Adf1,kay,tai,Abl,Dp,Diap1,eff,rictor,Syx1A,robo1,CG34401,morgue,mop,emc,Cbl,Dronc,Sema1a,h,Raf,Rab1,hth,fz,Arf79F,Sema1b,Gli,Dl | | 4.809e-17 | -37.57 | positive regulation of response to stimulus | biological process | GO:0048584 | 481 | 59 | 12345 | 442 | 14-3-3epsilon,ft,wdb,CtBP,pnr,Diap1,htl,Dcr-2,tkv,rictor,Pdk1,mop,hyx,cv-d,Dronc,hth,Raf,fz,eya,ds,Dyrk2,lili,apolpp,CycG,siz,lqf,lin-28,wnd,Actbeta,RhoGEF4,imd,spdo,Sxl,Src42A,CG32795,spi,Egfr,step,Pez,rdx,sli,spen,jeb,dlp,ed,hppy,InR,rho-4,S,Pitslre,lncRNA:Hsromega,Pvr,edl,vn,Fkbp14,chic,Cdk4,DPCoAC,14-3-3zeta | | 5.270e-17 | -37.48 | positive regulation of multicellular organismal process | biological process | GO:0051240 | 266 | 43 | 12345 | 442 | ey,kay,Sox14,tai,Cals,Dp,Diap1,mld,tkv,nmo,rictor,Pdk1,morgue,Hr4,Cbl,mop,Dronc,hth,Rab1,Raf,fz,d,abd-A,cue,Arf79F,Gli,ttk,Src42A,shg,spi,tsh,Egfr,step,ec,aop,klu,InR,Orct2,Pvr,shot,vn,chic,N | | 5.735e-17 | -37.40 | DNA-binding transcription factor activity | molecular function | GO:0003700 | 463 | 56 | 12176 | 414 | sd,Abd-B,Scr,nub,esg,h,sens,kay,Lmx1a,hth,Hr39,pdm3,CG13204,tsh,bowl,bab2,eyg,aop,kn,klu,B-H1,grn,CG12769,abd-A,rgr,dmrt11E,pnr,Alh,emc,CG10321,Mef2,E(spl)m5-HLH,sim,edl,dve,ap,Sox14,jumu,REPTOR,dpy,ss,ttk,Dp,cic,lilli,ey,hng3,su(sable),rib,chn,Hr4,al,HLH3B,Adf1,Dad,crp | | 6.266e-17 | -37.31 | chemotaxis | biological process | GO:0006935 | 292 | 45 | 12345 | 442 | Psc,dlp,gukh,jeb,fend,dock,grn,InR,chic,ap,shot,Bsg,comm2,betaTub60D,N,drl,pdm3,beat-IIa,fz2,shg,Fas1,Src42A,ena,wgn,LanA,sim,spen,sli,E(Pc),Nrk,eya,unc-5,Sema1a,Eph,Sema1b,Dg,otk,sens,Ptp4E,SCAR,Abl,14-3-3epsilon,sqz,Dad,robo1 | | 6.520e-17 | -37.27 | cell motility | biological process | GO:0048870 | 344 | 49 | 12345 | 442 | esg,sli,aop,Egfr,ena,fz2,shg,Src42A,dome,p130CAS,Ppcs,N,cindr,Ptpmeg2,Pvr,RhoU,orb2,Socs36E,Cerk,dlp,jeb,Rab5,mim,jar,baz,Dad,tkv,Abd-B,htl,Diap1,Pvf3,Abl,tai,Shrm,ey,kay,14-3-3epsilon,SCAR,RhoL,Dl,Sema1b,pum,abd-A,eya,rib,fz,unc-5,Sema1a,Raf | | 8.163e-17 | -37.04 | embryonic development via the syncytial blastoderm | biological process | GO:0001700 | 199 | 37 | 12345 | 442 | Abl,kay,pnr,yrt,arm,jar,tkv,scyl,Rab5,Myo10A,Dronc,emc,sns,rib,pum,Sema1b,chn,shg,tsh,spi,Src42A,jumu,Egfr,ena,step,cora,aop,chrb,ed,kn,Orct2,InR,chic,Pvr,shot,scrib,N | | 1.069e-16 | -36.77 | axon guidance | biological process | GO:0007411 | 271 | 43 | 12345 | 442 | shg,fz2,Src42A,Fas1,ena,wgn,LanA,sli,sim,spen,Psc,dlp,gukh,jeb,dock,fend,InR,grn,ap,chic,Bsg,shot,betaTub60D,N,comm2,beat-IIa,drl,pdm3,Abl,14-3-3epsilon,Dad,robo1,Nrk,unc-5,eya,Sema1a,Dg,otk,Eph,Sema1b,Ptp4E,sens,SCAR | | 1.106e-16 | -36.74 | positive regulation of developmental process | biological process | GO:0051094 | 259 | 42 | 12345 | 442 | spi,tsh,shg,ttk,Src42A,ec,step,Egfr,aop,klu,Orct2,InR,chic,vn,Pvr,shot,N,tai,Sox14,kay,ey,Diap1,Dp,Cals,rictor,nmo,tkv,Abd-B,Hr4,morgue,Pdk1,Dronc,mop,Cbl,d,fz,Raf,hth,Rab1,abd-A,Arf79F,Gli | | 1.342e-16 | -36.55 | epithelial cell differentiation | biological process | GO:0030855 | 419 | 54 | 12345 | 442 | Rab5,Atg16,Mef2,cic,baz,mim,jar,Abd-B,tkv,yrt,Diap1,Cip4,Dp,tai,Abl,kay,ft,RhoL,Gli,Dl,fz,SERCA,mam,ds,eya,Raf,Cbl,mop,emc,sina,spen,f,cora,aop,ena,Egfr,SMC3,cact,shg,tsh,ttk,dome,scrib,cindr,N,Ppcs,Ptpmeg2,Socs36E,neo,Pvr,ph-p,S,InR,ed | | 1.503e-16 | -36.43 | localization of cell | biological process | GO:0051674 | 351 | 49 | 12345 | 442 | p130CAS,dome,Src42A,shg,fz2,Egfr,ena,aop,sli,esg,jeb,dlp,Cerk,Socs36E,RhoU,Pvr,orb2,Ptpmeg2,Ppcs,cindr,N,14-3-3epsilon,kay,ey,Shrm,Abl,tai,Diap1,Pvf3,htl,Abd-B,tkv,Dad,mim,jar,baz,Rab5,Raf,Sema1a,fz,unc-5,rib,eya,abd-A,pum,Sema1b,Dl,RhoL,SCAR | | 1.800e-16 | -36.25 | embryo development ending in birth or egg hatching | biological process | GO:0009792 | 215 | 38 | 12345 | 442 | sns,rib,Myo10A,Dronc,emc,pum,Sema1b,pnr,yrt,arm,Abl,kay,Rab5,scyl,jar,wbl,tkv,Orct2,InR,ed,chrb,kn,scrib,N,chic,shot,Pvr,shg,tsh,spi,Src42A,jumu,chn,cora,aop,ena,Egfr,step | | 2.308e-16 | -36.00 | transcription regulator activity | molecular function | GO:0140110 | 605 | 64 | 12176 | 414 | crp,al,chn,tna,hng3,ttk,Dp,ey,Sox14,jumu,Mef2,Alh,emc,CG10321,B-H1,aop,klu,eya,tsh,bowl,bab2,mam,arm,Lmx1a,h,esg,nub,Abd-B,Hr4,Adf1,HLH3B,Dad,rib,su(sable),H,cic,lilli,ss,CtBP,dpy,ap,REPTOR,dve,sim,edl,E(spl)m5-HLH,dmrt11E,pnr,rgr,grn,abd-A,CG12769,kn,eyg,pdm3,CG13204,Hr39,sens,kay,hth,tai,Tob,Scr,sd | | 2.507e-16 | -35.92 | positive regulation of signal transduction | biological process | GO:0009967 | 369 | 50 | 12345 | 442 | Actbeta,RhoGEF4,apolpp,lili,siz,CycG,lqf,lin-28,wnd,hth,Raf,Dyrk2,ds,hyx,mop,Dronc,cv-d,Pdk1,Dcr-2,tkv,rictor,CtBP,Diap1,14-3-3epsilon,ft,wdb,Cdk4,14-3-3zeta,lncRNA:Hsromega,Pvr,vn,edl,Fkbp14,InR,rho-4,S,Pitslre,jeb,dlp,ed,hppy,Pez,rdx,spen,Egfr,step,Src42A,spi,spdo,Sxl | | 4.917e-16 | -35.25 | positive regulation of signaling | biological process | GO:0023056 | 403 | 52 | 12345 | 442 | jeb,dlp,hppy,ed,InR,rho-4,S,Pitslre,Pvr,lncRNA:Hsromega,edl,vn,Fkbp14,Cdk4,14-3-3zeta,spdo,Sxl,Src42A,spi,Egfr,step,rdx,Pez,spen,hyx,mop,cv-d,Dronc,Raf,hth,Rab1,ds,Dyrk2,apolpp,lili,CycG,siz,lin-28,lqf,wnd,Actbeta,RhoGEF4,14-3-3epsilon,ft,wdb,Cals,CtBP,Diap1,Dcr-2,tkv,rictor,Pdk1 | | 4.917e-16 | -35.25 | positive regulation of cell communication | biological process | GO:0010647 | 403 | 52 | 12345 | 442 | 14-3-3zeta,Cdk4,Fkbp14,edl,vn,lncRNA:Hsromega,Pvr,Pitslre,S,rho-4,InR,ed,hppy,dlp,jeb,spen,rdx,Pez,step,Egfr,spi,Src42A,Sxl,spdo,RhoGEF4,Actbeta,wnd,lin-28,lqf,siz,CycG,apolpp,lili,Dyrk2,ds,Raf,Rab1,hth,cv-d,Dronc,mop,hyx,Pdk1,rictor,tkv,Dcr-2,Diap1,CtBP,Cals,wdb,ft,14-3-3epsilon | | 5.331e-16 | -35.17 | positive regulation of cell motility | biological process | GO:2000147 | 53 | 20 | 12345 | 442 | ttk,Diap1,Pvf3,spi,shg,kay,tai,robo1,aop,sli,Egfr,ena,Sema1a,unc-5,Cbl,Gli,Pvr,CG9784,Sema1b,vn | | 7.470e-16 | -34.83 | taxis | biological process | GO:0042330 | 365 | 49 | 12345 | 442 | sli,sim,esg,spen,ena,wgn,LanA,shg,fz2,Src42A,Fas1,betaTub60D,comm2,N,beat-IIa,pdm3,drl,ap,Pen,chic,Bsg,shot,fend,dock,InR,grn,dlp,Psc,gukh,jeb,robo1,sqz,nmo,Dad,Abl,14-3-3epsilon,SCAR,otk,Dg,Eph,Sema1b,Ptp4E,sens,unc-5,eya,yuri,Sema1a,Nrk,E(Pc) | | 1.058e-15 | -34.48 | cell fate commitment | biological process | GO:0045165 | 301 | 44 | 12345 | 442 | jeb,kn,klu,S,nub,ap,vn,shot,N,14-3-3zeta,scrib,spi,ttk,msi,Egfr,spen,esg,sim,sina,aop,H,emc,Cbl,mam,fz,Raf,hth,nuf,abd-A,sens,Dl,14-3-3epsilon,kay,ey,pnr,CtBP,arm,jar,baz,tkv,htl,Abd-B,Tom,flfl | | 1.140e-15 | -34.41 | positive regulation of locomotion | biological process | GO:0040017 | 69 | 22 | 12345 | 442 | sli,robo1,aop,ena,Egfr,shg,spi,Diap1,Pvf3,ttk,tai,kay,Gli,vn,chic,Sema1b,CG9784,Pvr,fz,unc-5,Sema1a,Cbl | | 1.565e-15 | -34.09 | photoreceptor cell differentiation | biological process | GO:0046530 | 183 | 34 | 12345 | 442 | rg,spi,ttk,lilli,ss,klar,Egfr,msi,sina,B-H1,aop,klu,jeb,S,Mob2,edl,Bsg,scrib,N,kay,ey,eff,yrt,arm,baz,boi,Nrk,Cbl,emc,fz,eya,hth,Raf,sens | | 1.599e-15 | -34.07 | positive regulation of cell migration | biological process | GO:0030335 | 49 | 19 | 12345 | 442 | Gli,Sema1b,vn,Pvr,unc-5,Sema1a,Cbl,sli,robo1,aop,Egfr,ena,Diap1,Pvf3,spi,shg,ttk,tai,kay | | 2.255e-15 | -33.73 | positive regulation of cellular component movement | biological process | GO:0051272 | 71 | 22 | 12345 | 442 | Egfr,ena,sli,aop,robo1,tai,kay,shg,Pvf3,Diap1,spi,ttk,vn,chic,Sema1b,CG9784,Pvr,Gli,Cbl,fz,unc-5,Sema1a | | 1.339e-14 | -31.94 | regulation of transcription by RNA polymerase II | biological process | GO:0006357 | 741 | 71 | 12345 | 442 | ss,lilli,CG8312,MEP-1,bab2,sd,esg,sim,CG34376,grn,kek5,fuss,kn,klu,dve,tna,arm,Dp,CtBP,E(spl)m5-HLH,kay,HLH3B,cic,crol,crp,Dad,HmgZ,h,hth,eya,Lmx1a,mam,CG12769,Hr39,Snoo,Tet,sens,fs(1)h,ttk,tsh,dmrt11E,Scr,Prdm13,chn,Su(z)2,aop,B-H1,al,nub,eyg,zfh2,N,scrib,ap,edl,bowl,pnr,Sox14,Adf1,ey,tai,Mef2,tkv,Abd-B,emc,hyx,cv-d,H,Hel89B,abd-A,lili | | 1.436e-14 | -31.87 | imaginal disc pattern formation | biological process | GO:0007447 | 112 | 26 | 12345 | 442 | Egfr,boi,tara,Abd-B,Mpcp2,tkv,Dad,nmo,spen,B-H1,cic,14-3-3epsilon,ft,tsh,lilli,vn,nuf,ap,frc,abd-A,N,Dl,dlp,H,ds,hth | | 1.534e-14 | -31.81 | regulation of animal organ morphogenesis | biological process | GO:2000027 | 94 | 24 | 12345 | 442 | Socs36E,otk,N,klu,ed,Dronc,hth,ds,tkv,Abd-B,ec,baz,Egfr,aop,sd,morgue,sina,chn,ss,ttk,Src42A,Diap1,tsh,eff | | 1.543e-14 | -31.80 | regulation of growth | biological process | GO:0040008 | 310 | 43 | 12345 | 442 | nmo,Dad,tkv,wbl,rictor,Pdk1,scyl,cic,Hr4,14-3-3epsilon,ey,ft,tai,Abl,Cip4,CycG,pum,Sema1b,mop,Raf,Sema1a,fz,d,LanA,ena,Egfr,step,Src42A,lilli,shg,path,shot,chic,Cdk4,Lk6,scrib,N,dlp,chrb,hppy,InR,Mob2,Orct2 | | 1.614e-14 | -31.76 | nucleus | cellular component | GO:0005634 | 2809 | 167 | 11940 | 417 | mura,Kdm4B,sd,bol,klu,Tis11,rdx,CG13204,bru2,Pp2C1,CG32772,ttk,orb2,Tob,sim,IntS3,CtBP,snRNA:U5:38ABb,CG13481,jigr1,Snoo,Sox14,CycB3,Hr4,Rm62,Pdk1,msi,Jupiter,mam,Cdk4,E(Pc),Sxl,crp,Elba2,Adf1,ey,pum,flfl,CG9784,h,kn,mrt,14-3-3zeta,dve,lilli,Su(z)2,CG4896,CG11617,edl,glec,CG3436,vig,N,eyg,HmgZ,CG11360,Diap1,HLH3B,Lk6,mld,chn,stx,arm,CG34376,bin3,Hel89B,Abd-B,fs(1)h,Bacc,CG33082,Pen,CycG,l(2)37Cg,emc,klar,Psc,lncRNA:bxd,trbl,Poxn,Scr,REPTOR,cic,yrt,H,SmD1,Dp,Pitslre,rib,spen,CG31457,abd-A,ps,tai,ss,rgr,CG11247,px,fl(2)d,MEP-1,Tet,esg,jumu,Drak,nmo,su(sable),CG31211,Dcr-2,sens,Prdm13,dock,CG1218,Mef2,zfh2,fz2,RhoGEF4,Kr-h2,lncRNA:Hsromega,yuri,eya,ph-p,Hr39,sens-2,SCAR,aop,CG6656,E(spl)m5-HLH,tsh,pita,SNF4Agamma,kay,hyx,Alh,hang,bab2,crol,CG11030,His2Av,CG12769,14-3-3epsilon,Kdm2,hng3,wdb,sina,Slbp,CG4935,sqz,dpy,koi,CG10321,Lmx1a,ap,hth,tna,Drep4,His3.3B,al,Nlp,lqf,CG6340,dmrt11E,pnr,CG8312,B-H1,CG14441,grn,corto,bowl | | 1.649e-14 | -31.74 | Ras85D (Ras oncogene at 85D) | protein interactions | 41140 | 241 | 43 | 9204 | 432 | Cbl,B-H1,aop,Pvr,step,Rgl,pum,spi,dock,chic,cindr,lilli,mam,baz,jeb,wgn,eyg,Dg,N,14-3-3epsilon,klu,Raf,sli,vn,eya,ed,edl,rg,Hr4,spen,fz,dome,Egfr,IntS3,htl,ttk,kay,imd,Lk6,S,Src42A,cic,scrib | | 1.805e-14 | -31.65 | regulation of cellular component movement | biological process | GO:0051270 | 113 | 26 | 12345 | 442 | CG9784,Pvr,vn,chic,Sema1b,Pat1,Gli,Cbl,Rab1,Sema1a,unc-5,fz,LanA,Egfr,ena,CG34401,aop,robo1,sli,kay,tai,ttk,shg,spi,Pvf3,Diap1 | | 2.037e-14 | -31.52 | regulation of cell motility | biological process | GO:2000145 | 78 | 22 | 12345 | 442 | Sema1b,vn,Pvr,CG9784,Gli,Cbl,unc-5,Sema1a,Rab1,ena,Egfr,LanA,sli,robo1,aop,tai,kay,Diap1,Pvf3,spi,shg,ttk | | 2.094e-14 | -31.50 | cell surface receptor signaling pathway | biological process | GO:0007166 | 340 | 45 | 12345 | 442 | InR,jeb,drl,N,Pvr,Ir21a,vn,Src42A,ttk,fz2,spi,tsh,dome,aop,sli,sina,spen,wgn,cact,Egfr,Raf,Sema1a,mam,fz,unc-5,Nrk,Actbeta,Dl,imd,otk,Sema1b,Eph,arm,pnr,Pvf3,fz3,Pdk1,cic,robo1,boi,htl,tkv,Dad,Ddr,rictor | | 2.948e-14 | -31.16 | positive regulation of macromolecule metabolic process | biological process | GO:0010604 | 858 | 77 | 12345 | 442 | Raf,hth,eya,Lmx1a,mam,Hr39,CG12769,Tet,Actbeta,Tis11,sens,fs(1)h,wnd,hng3,trbl,CtBP,arm,Dp,Diap1,kay,Pdk1,jeb,kn,bol,Lk6,tna,Pvr,lncRNA:Hsromega,Socs36E,ss,CG8312,Src42A,lilli,sd,REPTOR,sim,esg,step,hyx,Dronc,PI31,ps,abd-A,Atg17,pum,lin-28,Cks30A,pnr,Adf1,ey,tai,Mef2,Abd-B,rictor,InR,nub,hppy,zfh2,N,orb2,ap,vn,bowl,ttk,tsh,spi,dmrt11E,Scr,chn,Su(z)2,CG13204,B-H1,rdx,aop,sina,tara,Egfr | | 3.927e-14 | -30.87 | positive regulation of nitrogen compound metabolic process | biological process | GO:0051173 | 827 | 75 | 12345 | 442 | tna,Lk6,bol,lncRNA:Hsromega,Pvr,Socs36E,kn,jeb,REPTOR,esg,sim,sd,step,ss,lilli,CG8312,Src42A,imd,Tis11,Actbeta,wnd,trbl,hng3,sens,fs(1)h,eya,Lmx1a,mam,Raf,hth,Tet,CG12769,Hr39,Pdk1,Diap1,arm,CtBP,kay,N,zfh2,ap,vn,bowl,orb2,InR,nub,hppy,sina,rdx,B-H1,Egfr,tara,spi,tsh,dmrt11E,Scr,ttk,Su(z)2,chn,CG13204,pum,abd-A,ps,PI31,Atg17,Dronc,hyx,Mef2,rictor,Abd-B,Cks30A,pnr,tai,Adf1,ey | | 4.000e-14 | -30.85 | intracellular part | cellular component | GO:0044424 | 6762 | 309 | 11940 | 417 | lncRNA:bxd,klar,path,Psa,step,mRpL41,Dad,Dronc,Sema1b,Pex7,Hel89B,Vps13,fs(1)h,morgue,CG33082,Abd-B,l(2)37Cg,CycG,N,Pgant5,frc,Lk6,HLH3B,Abl,Pat1,chn,CG17683,jar,Bsg,Cadps,kn,l(1)G0196,RhoU,14-3-3zeta,Mfe2,CG3436,glec,Adf1,UQCR-11,Pex5,flfl,ey,h,scyl,Sox14,Snoo,ReepA,jigr1,Pex10,msi,Pdk1,lt,Jupiter,tkv,Cdk4,DPCoAC,orb2,ttk,Pp2C1,CG32772,IntS3,CtBP,sim,betaTub60D,CG13481,bol,CG17883,sd,Syx1A,klu,Tis11,Ptpmeg2,GluProRS,His3.3B,Drep4,al,tna,alpha-Man-Ia,spdo,dmrt11E,ATPCL,Tpc1,CG8312,pnr,Wdr62,bowl,CG14441,corto,grn,koi,sqz,CG9220,Rab5,CG10321,hth,ena,ap,Cks30A,Mob2,CG12769,Pez,pdgy,sina,rho-4,AP-1gamma,CG4935,Slbp,CG6656,CG7536,SNF4Agamma,kay,CG6465,pita,tsh,hang,Dg,CG12402,Alh,crol,bab2,wbl,312,fuss,Kr-h2,RhoGEF4,siz,yuri,aop,Drak,nmo,su(sable),jumu,kon,RhoGAP19D,sens,Dcr-2,Shrm,cact,CG31211,dock,gukh,rg,CG14562,rgr,lin-28,CalpC,fl(2)d,chrb,emp,Dyrk2,SmD1,Dp,H,cic,spen,CG31457,Unc-76,S,fz,Pitslre,His3.3A,CG7560,abd-A,tai,p130CAS,ps,Psc,Poxn,trbl,RapGAP1,chic,cora,Scr,shot,REPTOR,CG34376,arm,Mpcp2,bin3,CG2765,Bacc,CG14464,emc,Pen,CG11360,HmgZ,eyg,CG42684,mld,eff,GlcAT-I,Diap1,CG30156,stx,Arf79F,rictor,CG4467,hppy,mrt,cindr,Su(z)2,CG4896,lilli,dve,vig,CG8306,edl,Rim2,CG11617,Magi,Elba2,crp,pum,lawc,CG9784,Raf,CG34172,CycB3,Src42A,CG42788,Hr4,Rm62,larp,mam,drongo,Sxl,Atg16,E(Pc),bru2,Tom,baz,shg,Cbl,Pex14,spi,Tob,ed,snRNA:U5:38ABb,mura,Kdm4B,CG4645,SERCA,CG6891,CG3107,Pgk,CG13204,rdx,Dl,lqf,Nlp,Pgant6,CG6340,B-H1,Atox1,CG33474,dpy,CG34401,Lmx1a,nuf,Spc105R,CG1673,SMC3,CG10417,CG11030,CG6983,14-3-3epsilon,His2Av,wdb,hng3,Kdm2,d,E(spl)m5-HLH,FBXO11,hyx,fz2,jeb,lncRNA:Hsromega,ph-p,eya,Rab1,Ocrl,CG3149,apolpp,SCAR,Hr39,sens-2,TBC1D5,CG12910,CG1218,Tpst,Prdm13,zfh2,Mef2,px,CG11247,ss,Atg17,f,esg,Tet,scrib,MEP-1,mop,Myo10A,yrt,rib,Fkbp14,CG7394,muc | | 4.000e-14 | -30.85 | intracellular | cellular component | GO:0005622 | 6762 | 309 | 11940 | 417 | Adf1,UQCR-11,Pex5,flfl,ey,h,scyl,Sox14,Snoo,ReepA,jigr1,Pex10,msi,Pdk1,Jupiter,lt,tkv,Cdk4,DPCoAC,orb2,ttk,Pp2C1,CG32772,IntS3,CtBP,sim,betaTub60D,CG13481,bol,sd,CG17883,Syx1A,klu,Tis11,lncRNA:bxd,klar,path,step,Psa,mRpL41,Dad,Dronc,Pex7,Sema1b,Hel89B,Vps13,morgue,CG33082,fs(1)h,Abd-B,l(2)37Cg,CycG,N,Pgant5,frc,Lk6,HLH3B,Abl,Pat1,CG17683,chn,Bsg,jar,Cadps,kn,l(1)G0196,RhoU,14-3-3zeta,Mfe2,CG3436,glec,312,fuss,Kr-h2,RhoGEF4,siz,yuri,aop,su(sable),Drak,nmo,RhoGAP19D,jumu,kon,sens,Dcr-2,cact,Shrm,CG31211,dock,gukh,rg,CG14562,rgr,lin-28,CalpC,fl(2)d,chrb,emp,Dyrk2,Dp,SmD1,H,cic,spen,CG31457,Unc-76,S,fz,Pitslre,His3.3A,CG7560,abd-A,tai,p130CAS,ps,Ptpmeg2,GluProRS,al,His3.3B,Drep4,tna,alpha-Man-Ia,spdo,dmrt11E,Tpc1,ATPCL,CG8312,pnr,Wdr62,bowl,CG14441,corto,grn,koi,sqz,CG9220,Rab5,CG10321,ena,hth,Cks30A,ap,Mob2,CG12769,Pez,pdgy,sina,AP-1gamma,rho-4,CG4935,Slbp,CG7536,CG6656,SNF4Agamma,kay,CG6465,tsh,pita,hang,Dg,CG12402,Alh,crol,bab2,wbl,Magi,crp,Elba2,pum,lawc,CG9784,Raf,CG34172,CycB3,Src42A,Hr4,Rm62,CG42788,larp,mam,drongo,Atg16,Sxl,E(Pc),bru2,baz,Tom,shg,Cbl,Pex14,spi,Tob,ed,snRNA:U5:38ABb,mura,CG4645,Kdm4B,SERCA,CG6891,CG3107,Pgk,rdx,CG13204,Dl,Psc,Poxn,trbl,RapGAP1,cora,chic,Scr,shot,REPTOR,CG34376,arm,Mpcp2,bin3,CG2765,Bacc,CG14464,emc,Pen,CG11360,HmgZ,eyg,CG42684,eff,mld,GlcAT-I,Diap1,CG30156,stx,Arf79F,rictor,CG4467,hppy,mrt,Su(z)2,CG4896,cindr,lilli,dve,vig,CG8306,Rim2,edl,CG11617,fz2,lncRNA:Hsromega,jeb,ph-p,eya,Rab1,Ocrl,CG3149,apolpp,SCAR,Hr39,sens-2,TBC1D5,CG12910,CG1218,Tpst,Prdm13,zfh2,Mef2,px,CG11247,ss,Atg17,f,Tet,esg,scrib,MEP-1,mop,Myo10A,yrt,rib,Fkbp14,CG7394,muc,lqf,Nlp,Pgant6,CG6340,B-H1,Atox1,CG33474,dpy,Lmx1a,CG34401,nuf,Spc105R,SMC3,CG1673,CG10417,CG11030,CG6983,14-3-3epsilon,His2Av,wdb,hng3,Kdm2,d,E(spl)m5-HLH,FBXO11,hyx | | 5.064e-14 | -30.61 | negative regulation of nitrogen compound metabolic process | biological process | GO:0051172 | 625 | 63 | 12345 | 442 | 312,msi,aop,sd,B-H1,bin3,al,Kdm4B,esg,Prdm13,lncRNA:bxd,MEP-1,chn,Sxl,Su(z)2,ss,ttk,CG8312,tsh,orb2,Socs36E,su(sable),bowl,FBXO11,corto,bol,zfh2,bru2,N,scrib,Elba2,Spn42De,klu,dve,Psc,eyg,nub,ph-p,Dcr-2,cic,crol,Hr4,E(spl)m5-HLH,Tob,wdb,arm,CtBP,Diap1,pnr,sens,CycG,Kdm2,abd-A,fs(1)h,pum,trbl,lin-28,Tis11,emc,Cbl,H,fl(2)d,h | | 5.500e-14 | -30.53 | morphogenesis of embryonic epithelium | biological process | GO:0016331 | 128 | 27 | 12345 | 442 | ed,Myo10A,emc,rib,chic,Pvr,shot,N,scrib,Inx2,Abl,kay,spi,yrt,pnr,jumu,arm,Src42A,step,Egfr,ena,jar,baz,tkv,aop,Rab5,cora | | 6.771e-14 | -30.32 | negative regulation of cell communication | biological process | GO:0010648 | 351 | 45 | 12345 | 442 | N,Lk6,edl,Socs36E,dock,kek5,fuss,InR,ed,chrb,hppy,jeb,rdx,aop,Egfr,cact,wgn,Src42A,spdo,Dl,pum,trbl,otk,CycG,cue,Ptp4E,ds,eya,d,Raf,H,Snoo,Cbl,LRR,Hr4,CG42684,Tom,scyl,crol,nmo,Dad,eff,CtBP,RapGAP1,wdb | | 7.038e-14 | -30.28 | regulation of cell migration | biological process | GO:0030334 | 74 | 21 | 12345 | 442 | Gli,Sema1b,vn,Pvr,unc-5,Sema1a,Rab1,Cbl,sli,aop,robo1,Egfr,ena,LanA,Diap1,Pvf3,spi,shg,ttk,tai,kay | | 7.513e-14 | -30.22 | negative regulation of signaling | biological process | GO:0023057 | 352 | 45 | 12345 | 442 | InR,dock,kek5,fuss,jeb,hppy,chrb,ed,Lk6,N,Socs36E,edl,Src42A,spdo,rdx,aop,cact,wgn,Egfr,Raf,ds,d,eya,LRR,Cbl,H,Snoo,Dl,cue,CycG,Ptp4E,pum,otk,trbl,RapGAP1,CtBP,eff,wdb,Tom,crol,scyl,Hr4,CG42684,Dad,nmo | | 8.089e-14 | -30.15 | negative regulation of gene expression | biological process | GO:0010629 | 582 | 60 | 12345 | 442 | arm,CtBP,vig,Diap1,pnr,E(spl)m5-HLH,ft,cic,crol,Hr4,tkv,Dcr-2,h,emc,fl(2)d,H,Tis11,abd-A,sens,fs(1)h,pum,lin-28,ss,ttk,spi,tsh,Prdm13,MEP-1,lncRNA:bxd,chn,mrt,Sxl,Su(z)2,aop,sd,B-H1,bin3,Kdm4B,al,esg,312,msi,Egfr,nub,eyg,ph-p,klu,dve,Psc,FBXO11,corto,zfh2,bru2,N,scrib,Elba2,Rm62,orb2,su(sable),bowl | | 9.011e-14 | -30.04 | negative regulation of cellular biosynthetic process | biological process | GO:0031327 | 457 | 52 | 12345 | 442 | arm,CtBP,pnr,E(spl)m5-HLH,Tob,cic,crol,Hr4,Dcr-2,h,emc,Cbl,H,fl(2)d,Tis11,abd-A,sens,fs(1)h,pum,ss,ttk,tsh,Prdm13,lncRNA:bxd,MEP-1,chn,Su(z)2,Sxl,sd,aop,B-H1,bin3,al,Kdm4B,esg,312,msi,eyg,nub,ph-p,klu,dve,Psc,corto,FBXO11,bru2,zfh2,Elba2,scrib,orb2,su(sable),bowl | | 9.011e-14 | -30.04 | negative regulation of biosynthetic process | biological process | GO:0009890 | 457 | 52 | 12345 | 442 | pnr,arm,CtBP,Tob,E(spl)m5-HLH,Hr4,crol,cic,Dcr-2,h,H,fl(2)d,Cbl,emc,Tis11,pum,fs(1)h,sens,abd-A,tsh,ttk,ss,chn,Sxl,Su(z)2,lncRNA:bxd,MEP-1,Prdm13,al,Kdm4B,esg,bin3,aop,sd,B-H1,msi,312,ph-p,nub,eyg,Psc,dve,klu,scrib,Elba2,zfh2,bru2,FBXO11,corto,bowl,su(sable),orb2 | | 9.833e-14 | -29.95 | regulation of locomotion | biological process | GO:0040012 | 111 | 25 | 12345 | 442 | Gli,Sema1b,chic,vn,Pvr,CG9784,unc-5,fz,Sema1a,Rab1,Cbl,sli,aop,robo1,CG34401,ena,Egfr,LanA,Diap1,spi,Pvf3,shg,ttk,tai,kay | | 9.833e-14 | -29.95 | dorsal closure | biological process | GO:0007391 | 111 | 25 | 12345 | 442 | pnr,yrt,spi,Src42A,jumu,arm,Abl,kay,Rab5,cora,aop,Egfr,ena,jar,step,tkv,rib,Myo10A,ed,emc,scrib,N,chic,Pvr,shot | | 9.840e-14 | -29.95 | negative regulation of RNA metabolic process | biological process | GO:0051253 | 398 | 48 | 12345 | 442 | E(spl)m5-HLH,Tob,pnr,CtBP,arm,Dcr-2,Hr4,cic,crol,H,fl(2)d,emc,h,pum,lin-28,sens,abd-A,fs(1)h,Su(z)2,chn,Sxl,Prdm13,lncRNA:bxd,MEP-1,tsh,ss,ttk,Kdm4B,esg,al,aop,B-H1,sd,bin3,dve,Psc,klu,ph-p,eyg,nub,bowl,su(sable),Elba2,scrib,corto,bol,FBXO11,zfh2 | | 1.049e-13 | -29.89 | negative regulation of developmental process | biological process | GO:0051093 | 222 | 35 | 12345 | 442 | chn,tsh,klar,ttk,Src42A,Egfr,ena,cact,LanA,esg,sd,aop,C1GalTA,ed,Mob2,fuss,InR,Socs36E,N,scrib,Abl,ft,Diap1,Cip4,Dad,tkv,Abd-B,Snoo,emc,eya,Sema1a,Sema1b,lqf,Him,Dl | | 1.229e-13 | -29.73 | membrane-bounded organelle | cellular component | GO:0043227 | 4716 | 238 | 11940 | 417 | CG4935,Slbp,wdb,sina,hng3,Kdm2,AP-1gamma,rho-4,CG12769,14-3-3epsilon,pdgy,His2Av,CG11030,crol,wbl,bab2,hang,Alh,hyx,kay,SNF4Agamma,pita,tsh,E(spl)m5-HLH,CG6656,CG7536,grn,CG14441,bowl,corto,CG33474,B-H1,CG8312,pnr,spdo,Cip4,Pgant6,dmrt11E,CG6340,Tpc1,lqf,Nlp,al,Drep4,His3.3B,tna,alpha-Man-Ia,CG1673,nuf,ena,hth,ap,Lmx1a,CG9220,Rab5,CG10321,koi,dpy,sqz,mop,emp,esg,Tet,MEP-1,fl(2)d,px,rgr,ss,CG11247,tai,ps,muc,CG7394,abd-A,spen,CG31457,S,rib,Fkbp14,fz,Pitslre,SmD1,Dp,H,yrt,cic,Ocrl,aop,apolpp,CG3149,SCAR,sens-2,Hr39,ph-p,siz,eya,Rab1,yuri,jeb,lncRNA:Hsromega,Kr-h2,RhoGEF4,312,fz2,zfh2,Mef2,CG1218,dock,Tpst,Prdm13,Dcr-2,sens,CG31211,Drak,nmo,su(sable),TBC1D5,jumu,CG30156,chn,stx,Arf79F,mld,Lk6,GlcAT-I,HLH3B,Diap1,HmgZ,CG11360,Pgant5,eyg,frc,N,Mfe2,vig,CG3436,glec,CG8306,Rim2,edl,CG11617,Su(z)2,CG4896,lilli,dve,14-3-3zeta,kn,mrt,jar,Cadps,REPTOR,Scr,mRpL41,trbl,path,Poxn,step,lncRNA:bxd,klar,Psc,l(2)37Cg,emc,Pen,CycG,fs(1)h,CG33082,Bacc,Abd-B,CG14464,Mpcp2,Pex7,Hel89B,bin3,Vps13,CG34376,arm,Dronc,ed,CG13481,snRNA:U5:38ABb,CtBP,IntS3,Pex14,spi,Tob,sim,orb2,DPCoAC,shg,ttk,Pp2C1,CG32772,bru2,rdx,CG13204,Tis11,Dl,CG3107,Syx1A,klu,bol,sd,CG17883,CG4645,Kdm4B,mura,SERCA,CG34172,h,CG9784,pum,flfl,ey,Adf1,UQCR-11,Elba2,crp,Pex5,Sxl,Atg16,tkv,E(Pc),Cdk4,larp,mam,robo1,lt,Jupiter,msi,Pdk1,Hr4,Rm62,CycB3,Sox14,ReepA,Snoo,jigr1,Pex10 | | 1.296e-13 | -29.67 | formation of anatomical boundary | biological process | GO:0048859 | 46 | 17 | 12345 | 442 | N,sens,nuf,hth,ds,unc-5,fz,klu,H,robo1,Tom,B-H1,sli,ena,CtBP,Scr,ft | | 1.311e-13 | -29.66 | negative regulation of macromolecule biosynthetic process | biological process | GO:0010558 | 446 | 51 | 12345 | 442 | fs(1)h,abd-A,sens,pum,Tis11,emc,H,fl(2)d,h,Dcr-2,crol,cic,Hr4,Tob,E(spl)m5-HLH,CtBP,arm,pnr,su(sable),orb2,bowl,zfh2,bru2,FBXO11,corto,Elba2,scrib,klu,Psc,dve,eyg,nub,ph-p,312,msi,bin3,sd,B-H1,aop,esg,al,Kdm4B,lncRNA:bxd,MEP-1,Prdm13,Sxl,Su(z)2,chn,ttk,ss,tsh | | 1.525e-13 | -29.51 | regulation of anatomical structure morphogenesis | biological process | GO:0022603 | 250 | 37 | 12345 | 442 | tkv,Abd-B,htl,baz,CG34401,morgue,Abl,Diap1,eff,Sema1b,otk,SCAR,Dronc,Sema1a,hth,Rab1,ds,rib,fz,ec,Egfr,aop,sd,sina,chn,ss,Src42A,ttk,tsh,shot,neo,Socs36E,chic,N,kn,klu,ed | | 2.041e-13 | -29.22 | organelle | cellular component | GO:0043226 | 5470 | 264 | 11940 | 417 | kon,jumu,su(sable),Drak,nmo,Shrm,CG31211,Dcr-2,sens,dock,gukh,312,Kr-h2,RhoGEF4,yuri,siz,aop,H,cic,Dyrk2,Dp,SmD1,S,Pitslre,fz,spen,CG31457,His3.3A,abd-A,ps,tai,rgr,fl(2)d,emp,sqz,koi,CG9220,CG10321,Rab5,ap,ena,hth,His3.3B,Drep4,al,alpha-Man-Ia,tna,Tpc1,spdo,dmrt11E,CG8312,pnr,grn,corto,bowl,CG14441,Wdr62,CG6656,CG7536,tsh,pita,kay,SNF4Agamma,Alh,hang,Dg,bab2,wbl,crol,pdgy,CG12769,Pez,rho-4,AP-1gamma,sina,Slbp,CG4935,Pex10,Sox14,ReepA,Snoo,jigr1,Pdk1,msi,Jupiter,lt,robo1,Cdk4,tkv,UQCR-11,Pex5,Adf1,ey,flfl,h,CG17883,sd,bol,klu,Syx1A,Tis11,ttk,Pp2C1,CG32772,DPCoAC,orb2,sim,CtBP,IntS3,betaTub60D,CG13481,Dronc,Hel89B,Vps13,Pex7,fs(1)h,CG33082,Abd-B,CycG,l(2)37Cg,klar,lncRNA:bxd,step,path,mRpL41,Bsg,jar,Cadps,kn,14-3-3zeta,RhoU,glec,Mfe2,CG3436,N,frc,Pgant5,HLH3B,Pat1,Lk6,chn,TBC1D5,CG12910,Prdm13,CG1218,Tpst,Mef2,zfh2,fz2,jeb,lncRNA:Hsromega,Rab1,ph-p,eya,SCAR,sens-2,Hr39,Ocrl,CG3149,apolpp,yrt,rib,Fkbp14,CG7394,muc,px,ss,CG11247,f,MEP-1,esg,Tet,scrib,mop,Myo10A,dpy,Lmx1a,nuf,Spc105R,SMC3,CG1673,lqf,Nlp,Pgant6,Cip4,CG6340,B-H1,CG33474,E(spl)m5-HLH,hyx,CG11030,His2Av,14-3-3epsilon,wdb,hng3,Kdm2,d,CycB3,CG42788,Hr4,Rm62,larp,mam,E(Pc),Sxl,Atg16,Magi,crp,Elba2,lawc,pum,CG9784,CG34172,CG6891,mura,Kdm4B,CG4645,SERCA,CG3107,Pgk,Dl,CG13204,rdx,bru2,shg,spi,Tob,Pex14,ed,snRNA:U5:38ABb,CG34376,arm,bin3,Mpcp2,Bacc,CG14464,Pen,emc,Psc,trbl,Poxn,Scr,cora,REPTOR,shot,mrt,dve,Su(z)2,CG4896,lilli,Rim2,edl,CG8306,CG11617,vig,HmgZ,CG11360,eyg,GlcAT-I,Diap1,mld,Arf79F,CG30156,stx | | 2.212e-13 | -29.14 | developmental process involved in reproduction | biological process | GO:0003006 | 928 | 79 | 12345 | 442 | wbl,mim,jar,htl,cic,Pdk1,Rab5,14-3-3epsilon,kay,Pex10,Diap1,eff,arm,Dp,Dg,otk,RhoL,SCAR,Pat1,Gli,fl(2)d,Tet,rib,eya,mam,Raf,msi,ena,sli,Sxl,bab2,chic,Pen,lncRNA:Hsromega,Pvr,Socs36E,bol,ed,dve,S,ph-p,baz,tkv,Abd-B,Mef2,robo1,tai,Abl,Cip4,pum,lin-28,PI31,abd-A,Inx2,Dl,Dronc,emc,mop,Cbl,fz,SERCA,sns,yuri,Egfr,cact,aop,dome,fz2,Scr,shg,ttk,Ptpmeg2,orb2,N,14-3-3zeta,cindr,Ppcs,scrib,InR | | 2.467e-13 | -29.03 | positive regulation of cellular metabolic process | biological process | GO:0031325 | 875 | 76 | 12345 | 442 | lncRNA:Hsromega,Pvr,Socs36E,tna,Lk6,bol,kn,jeb,step,REPTOR,esg,sim,sd,ss,Src42A,CG8312,lilli,wnd,hng3,trbl,sens,fs(1)h,imd,Tis11,Actbeta,Tet,CG12769,Hr39,eya,Lmx1a,mam,Raf,hth,Pdk1,kay,Diap1,CtBP,arm,ap,bowl,vn,orb2,Pgk,N,zfh2,hppy,nub,InR,Egfr,tara,sina,B-H1,rdx,Su(z)2,chn,CG13204,tsh,dmrt11E,spi,Scr,ttk,pum,ps,PI31,abd-A,Atg17,Dronc,hyx,rictor,Abd-B,Mef2,tai,Adf1,ey,Cks30A,pnr | | 2.481e-13 | -29.03 | negative regulation of cellular metabolic process | biological process | GO:0031324 | 681 | 65 | 12345 | 442 | Diap1,CtBP,arm,E(spl)m5-HLH,Hr4,crol,cic,Raf,h,fl(2)d,Tis11,trbl,fs(1)h,sens,Kdm2,CG8312,ss,Sxl,lncRNA:bxd,MEP-1,esg,bin3,sd,msi,ph-p,Psc,dve,klu,Spn42De,bru2,bol,corto,Socs36E,pnr,wdb,Tob,Dcr-2,H,Cbl,emc,lin-28,pum,abd-A,CycG,tsh,ttk,chn,Su(z)2,Prdm13,al,Kdm4B,aop,B-H1,312,eyg,InR,nub,Elba2,scrib,N,zfh2,FBXO11,bowl,su(sable),orb2 | | 2.487e-13 | -29.02 | negative regulation of cellular macromolecule biosynthetic process | biological process | GO:2000113 | 438 | 50 | 12345 | 442 | E(spl)m5-HLH,pnr,arm,CtBP,Dcr-2,Hr4,crol,cic,H,fl(2)d,emc,h,pum,fs(1)h,abd-A,sens,Tis11,Sxl,chn,Su(z)2,MEP-1,lncRNA:bxd,Prdm13,tsh,ttk,ss,msi,312,Kdm4B,al,esg,bin3,aop,B-H1,sd,Psc,dve,klu,ph-p,eyg,nub,bowl,su(sable),orb2,scrib,Elba2,bru2,zfh2,FBXO11,corto | | 3.305e-13 | -28.74 | positive regulation of cell development | biological process | GO:0010720 | 148 | 28 | 12345 | 442 | Dronc,Cbl,klu,mop,fz,InR,Rab1,Raf,vn,chic,Arf79F,Pvr,shot,N,Gli,tai,kay,Sox14,shg,Diap1,spi,ttk,Src42A,Dp,Egfr,ec,morgue,aop | | 3.689e-13 | -28.63 | positive regulation of metabolic process | biological process | GO:0009893 | 937 | 79 | 12345 | 442 | ss,CG8312,lilli,Src42A,REPTOR,sim,esg,sd,step,kn,jeb,tna,bol,Lk6,Pvr,lncRNA:Hsromega,Socs36E,Diap1,Dp,CtBP,arm,kay,Pdk1,eya,Lmx1a,mam,Raf,hth,Tet,Hr39,CG12769,imd,Tis11,Actbeta,wnd,hng3,trbl,sens,fs(1)h,tsh,dmrt11E,spi,Scr,ttk,Su(z)2,chn,CG13204,sina,aop,rdx,B-H1,Egfr,tara,nub,InR,hppy,N,zfh2,ap,bowl,vn,orb2,Pgk,Cks30A,pnr,tai,Adf1,ey,Mef2,rictor,Abd-B,Dronc,hyx,pum,lin-28,PI31,ps,abd-A,Atg17 | | 4.130e-13 | -28.52 | negative regulation of signal transduction | biological process | GO:0009968 | 326 | 42 | 12345 | 442 | Tom,crol,scyl,Hr4,CG42684,nmo,Dad,CtBP,RapGAP1,eff,wdb,Dl,CycG,cue,Ptp4E,pum,trbl,otk,Raf,eya,d,ds,Cbl,LRR,H,Snoo,aop,rdx,cact,Egfr,Src42A,spdo,Lk6,N,Socs36E,edl,dock,kek5,fuss,chrb,ed,hppy | | 4.163e-13 | -28.51 | response to stimulus | biological process | GO:0050896 | 2655 | 160 | 12345 | 442 | Rab1,Raf,Dyrk2,eya,unc-5,Hr39,ITP,imd,SCAR,RhoL,sens,wnd,Dg,Dp,fz3,Pdk1,scyl,Hr4,subdued,nmo,grn,hfw,koi,fend,S,ph-p,CG42788,jeb,dlp,Lk6,pdm3,pxb,shot,Pvr,Socs36E,Pen,tipE,jumu,Sxl,sd,REPTOR,esg,ZnT41F,sim,msi,Sema1a,hang,CG13481,Cbl,LRR,moody,Rgl,CG7536,Sema1b,pum,Tollo,pnr,Adf1,Abl,SNF4Agamma,robo1,Atg16,rictor,Ddr,sqz,lt,mura,dock,drl,N,DPCoAC,ap,Fas1,tsh,shg,Tpc1,dome,rdx,spen,sina,Ocrl,h,mam,E(Pc),Nrk,Actbeta,Arf79F,Eph,otk,arm,Poxn,14-3-3epsilon,kay,cic,ogre,Rab5,Dad,htl,boi,CG1218,Cerk,kn,ed,Psc,gukh,Bacc,Rm62,RhoU,chic,CG30344,ss,Src42A,Atox1,sli,wgn,Gr93c,ena,yuri,alpha-Man-Ia,Ric,fz,Dronc,Dl,Pex5,CycG,Ptp4E,Atg17,Pvf3,tai,flfl,Mef2,Dcr-2,tkv,ReepA,InR,CG30156,hppy,chrb,RhoGAP19D,nAChRalpha4,beat-IIa,comm2,14-3-3zeta,betaTub60D,Magi,scrib,Ir21a,Bsg,vn,ttk,RhoGEF3,spi,fz2,CG32795,rg,aop,cact,LanA,Egfr,sick | | 4.442e-13 | -28.44 | negative regulation of macromolecule metabolic process | biological process | GO:0010605 | 777 | 70 | 12345 | 442 | pum,lin-28,abd-A,CycG,H,emc,Cbl,Dcr-2,tkv,wdb,ft,Tob,pnr,bowl,orb2,su(sable),N,scrib,Elba2,FBXO11,zfh2,nub,eyg,Egfr,312,al,Kdm4B,aop,B-H1,chn,Su(z)2,Prdm13,tsh,spi,ttk,trbl,sens,Kdm2,fs(1)h,Tis11,fl(2)d,h,Hr4,cic,crol,E(spl)m5-HLH,Diap1,CtBP,arm,vig,Rm62,Socs36E,bol,corto,bru2,dve,Psc,Spn42De,klu,ph-p,msi,esg,sd,bin3,Sxl,mrt,lncRNA:bxd,MEP-1,ss,CG8312 | | 4.806e-13 | -28.36 | regulation of nervous system development | biological process | GO:0051960 | 356 | 44 | 12345 | 442 | CG34401,robo1,rictor,nmo,tkv,Dad,eff,Cals,Cip4,Abl,Adf1,Sox14,E(spl)m5-HLH,Dl,pum,Sema1b,Arf79F,fz,Raf,hth,Rab1,h,Sema1a,mop,emc,sina,sd,aop,ena,LanA,spi,ttk,Src42A,ss,N,chic,Socs36E,shot,Mob2,nub,InR,ed,klu,kn | | 5.181e-13 | -28.29 | intracellular organelle | cellular component | GO:0043229 | 5391 | 260 | 11940 | 417 | nuf,Spc105R,CG1673,SMC3,dpy,Lmx1a,B-H1,CG33474,lqf,Nlp,Pgant6,CG6340,hyx,E(spl)m5-HLH,wdb,Kdm2,hng3,d,CG11030,His2Av,14-3-3epsilon,Prdm13,CG1218,Tpst,Mef2,zfh2,TBC1D5,CG12910,Rab1,ph-p,eya,SCAR,sens-2,Hr39,Ocrl,CG3149,apolpp,fz2,jeb,lncRNA:Hsromega,CG7394,muc,yrt,rib,Fkbp14,MEP-1,Tet,esg,mop,Myo10A,px,CG11247,ss,f,Bacc,CG14464,Pen,emc,CG34376,arm,bin3,Mpcp2,Scr,cora,REPTOR,shot,Psc,trbl,Poxn,dve,Su(z)2,CG4896,lilli,edl,CG8306,Rim2,CG11617,vig,mrt,GlcAT-I,Diap1,mld,Arf79F,CG30156,stx,CG11360,HmgZ,eyg,larp,mam,E(Pc),Sxl,Atg16,CycB3,CG42788,Hr4,Rm62,CG9784,CG34172,crp,Elba2,lawc,pum,CG3107,Pgk,Dl,CG13204,rdx,CG6891,Kdm4B,CG4645,mura,SERCA,spi,Tob,Pex14,ed,snRNA:U5:38ABb,bru2,shg,ap,hth,ena,sqz,koi,CG9220,Rab5,CG10321,CG8312,pnr,corto,grn,bowl,CG14441,Wdr62,al,His3.3B,Drep4,tna,alpha-Man-Ia,Tpc1,spdo,dmrt11E,Alh,hang,Dg,wbl,bab2,crol,CG7536,CG6656,pita,tsh,kay,SNF4Agamma,rho-4,AP-1gamma,sina,Slbp,CG4935,pdgy,CG12769,Pez,dock,gukh,kon,jumu,su(sable),Drak,nmo,Shrm,CG31211,sens,Dcr-2,yuri,siz,aop,312,Kr-h2,RhoGEF4,His3.3A,abd-A,ps,tai,H,cic,Dyrk2,SmD1,Dp,S,Pitslre,fz,spen,CG31457,emp,rgr,fl(2)d,CG33082,fs(1)h,Abd-B,CycG,l(2)37Cg,Dronc,Hel89B,Vps13,Pex7,mRpL41,klar,lncRNA:bxd,step,path,14-3-3zeta,RhoU,glec,Mfe2,CG3436,jar,Bsg,Cadps,kn,HLH3B,Pat1,Lk6,chn,N,frc,Pgant5,Jupiter,lt,Cdk4,tkv,Pex10,ReepA,Snoo,Sox14,jigr1,Pdk1,msi,h,UQCR-11,Pex5,Adf1,ey,flfl,klu,Syx1A,Tis11,sd,CG17883,bol,sim,CtBP,IntS3,betaTub60D,CG13481,ttk,Pp2C1,CG32772,orb2,DPCoAC | | 5.712e-13 | -28.19 | negative regulation of nucleobase-containing compound metabolic process | biological process | GO:0045934 | 417 | 48 | 12345 | 442 | Tob,E(spl)m5-HLH,CtBP,arm,pnr,Dcr-2,crol,cic,Hr4,emc,fl(2)d,H,h,fs(1)h,sens,abd-A,lin-28,pum,lncRNA:bxd,MEP-1,Prdm13,chn,Su(z)2,Sxl,ttk,ss,tsh,bin3,aop,sd,B-H1,Kdm4B,al,esg,klu,Psc,dve,eyg,nub,ph-p,su(sable),bowl,zfh2,FBXO11,corto,bol,scrib,Elba2 | | 6.101e-13 | -28.13 | columnar/cuboidal epithelial cell differentiation | biological process | GO:0002065 | 344 | 43 | 12345 | 442 | Atg16,Rab5,cic,Mef2,mim,jar,baz,Abd-B,tkv,Diap1,Cip4,Dp,tai,kay,RhoL,Dl,mam,fz,SERCA,eya,Raf,Cbl,mop,emc,sina,aop,Egfr,ena,cact,shg,ttk,dome,scrib,N,Ppcs,cindr,Ptpmeg2,Socs36E,Pvr,ph-p,S,InR,ed | | 6.847e-13 | -28.01 | negative regulation of response to stimulus | biological process | GO:0048585 | 389 | 46 | 12345 | 442 | Snoo,H,LRR,Cbl,d,ds,eya,Raf,Sema1a,trbl,otk,pum,Sema1b,Ptp4E,cue,CycG,Dl,wdb,eff,CtBP,RapGAP1,Dad,nmo,CG42684,Hr4,scyl,crol,Tom,chrb,ed,hppy,kek5,fuss,dock,nub,edl,Socs36E,Pvr,N,Lk6,spdo,Src42A,Egfr,cact,rdx,aop | | 7.700e-13 | -27.89 | DNA-binding transcription factor activity, RNA polymerase II-specific | molecular function | GO:0000981 | 277 | 37 | 12176 | 414 | klu,kn,aop,bowl,tsh,pnr,dmrt11E,B-H1,CG12769,abd-A,grn,Scr,nub,sd,hth,Lmx1a,kay,sens,Hr39,esg,h,HLH3B,Adf1,Hr4,al,chn,edl,sim,Sox14,ap,Mef2,E(spl)m5-HLH,ss,cic,ey,Dp,ttk | | 8.286e-13 | -27.82 | negative regulation of metabolic process | biological process | GO:0009892 | 823 | 72 | 12345 | 442 | Tob,ft,wdb,pnr,Dcr-2,tkv,Cbl,emc,H,CycG,abd-A,lin-28,pum,Prdm13,chn,Su(z)2,ttk,spi,tsh,312,Egfr,aop,B-H1,Kdm4B,al,eyg,nub,InR,su(sable),orb2,bowl,zfh2,FBXO11,Elba2,scrib,N,E(spl)m5-HLH,vig,arm,CtBP,Diap1,crol,cic,Hr4,fl(2)d,h,Raf,fs(1)h,sens,Kdm2,trbl,Tis11,MEP-1,lncRNA:bxd,mrt,Sxl,CG8312,ss,msi,bin3,sd,esg,klu,Spn42De,Psc,dve,ph-p,Socs36E,Rm62,bru2,bol,corto | | 9.209e-13 | -27.71 | pbl (pebble) | protein interactions | 38879 | 147 | 31 | 9204 | 432 | htl,shot,ttk,Diap1,Lk6,imd,Cerk,trbl,CycB3,CenB1A,fs(1)h,mura,Ptp4E,stx,Mob2,wdb,Cdk4,Vps13,chrb,Dad,RapGAP1,flfl,Pdk1,His2Av,Sema1a,arm,Pitslre,crol,nuf,pum,spi | | 9.378e-13 | -27.70 | ameboidal-type cell migration | biological process | GO:0001667 | 177 | 30 | 12345 | 442 | Sema1a,Raf,rib,fz,jeb,dlp,Dl,N,RhoL,cindr,Ppcs,Pvr,Socs36E,Sema1b,Ptpmeg2,Diap1,shg,kay,dome,tai,aop,Rab5,esg,sli,htl,jar,ena,baz,Egfr,mim | | 1.075e-12 | -27.56 | negative regulation of nucleic acid-templated transcription | biological process | GO:1903507 | 379 | 45 | 12345 | 442 | pum,sens,abd-A,fs(1)h,H,fl(2)d,emc,h,Dcr-2,Hr4,cic,crol,E(spl)m5-HLH,Tob,pnr,CtBP,arm,bowl,su(sable),scrib,Elba2,FBXO11,corto,zfh2,dve,Psc,klu,ph-p,eyg,nub,esg,al,Kdm4B,aop,B-H1,sd,bin3,Su(z)2,chn,Prdm13,lncRNA:bxd,MEP-1,tsh,ss,ttk | | 1.075e-12 | -27.56 | negative regulation of RNA biosynthetic process | biological process | GO:1902679 | 379 | 45 | 12345 | 442 | su(sable),bowl,zfh2,FBXO11,corto,scrib,Elba2,klu,Psc,dve,eyg,nub,ph-p,bin3,B-H1,sd,aop,al,esg,Kdm4B,lncRNA:bxd,MEP-1,Prdm13,Su(z)2,chn,ttk,ss,tsh,fs(1)h,sens,abd-A,pum,emc,H,fl(2)d,h,Dcr-2,crol,cic,Hr4,Tob,E(spl)m5-HLH,arm,CtBP,pnr | | 1.280e-12 | -27.38 | sequence-specific DNA binding | molecular function | GO:0043565 | 515 | 52 | 12176 | 414 | zfh2,CG34376,CG11247,HLH3B,al,Hr4,crp,ph-p,su(sable),chn,sim,dve,edl,jumu,ap,Sox14,CG10321,E(spl)m5-HLH,Mef2,otp,ey,cic,pita,Dp,ttk,eyg,klu,Psc,aop,bowl,bab2,rgr,pnr,dmrt11E,B-H1,CG12769,abd-A,Prdm13,grn,Scr,Elba2,nub,sd,Abd-B,H2.0,Lmx1a,hth,kay,sens,Hr39,esg,h | | 1.285e-12 | -27.38 | positive regulation of cell differentiation | biological process | GO:0045597 | 156 | 28 | 12345 | 442 | Sox14,kay,tai,Dp,Src42A,ttk,Diap1,spi,shg,ec,Egfr,aop,morgue,mop,Cbl,klu,Dronc,Rab1,InR,Raf,fz,shot,Pvr,Arf79F,chic,vn,Gli,N | | 1.322e-12 | -27.35 | RNA polymerase II regulatory region sequence-specific DNA binding | molecular function | GO:0000977 | 254 | 35 | 12176 | 414 | klu,aop,eyg,bowl,pnr,dmrt11E,rgr,abd-A,Prdm13,CG12769,grn,B-H1,Scr,Abd-B,sd,Hr39,Lmx1a,kay,h,esg,zfh2,CG11247,CG34376,HLH3B,Hr4,al,su(sable),ap,sim,dve,Mef2,E(spl)m5-HLH,cic,ey,ttk | | 1.949e-12 | -26.96 | regulation of localization | biological process | GO:0032879 | 478 | 51 | 12345 | 442 | S,lt,InR,dlp,scrib,cindr,vn,tipE,chic,Pvr,shg,spi,ttk,lilli,klar,sli,ZnT41F,rdx,aop,Egfr,ena,step,LanA,SERCA,unc-5,fz,Rab1,Sema1a,Cbl,mop,Pat1,Gli,trbl,Sema1b,CG9784,Arf79F,Pvf3,Cadps,Diap1,wdb,tai,Abl,IA-2,kay,ft,Rab5,CG34401,Syx1A,robo1,mim,jar | | 2.031e-12 | -26.92 | negative regulation of transcription, DNA-templated | biological process | GO:0045892 | 371 | 44 | 12345 | 442 | pum,fs(1)h,abd-A,sens,h,fl(2)d,H,emc,Hr4,crol,cic,Dcr-2,pnr,arm,CtBP,E(spl)m5-HLH,scrib,Elba2,zfh2,FBXO11,corto,bowl,su(sable),ph-p,nub,eyg,Psc,dve,klu,Kdm4B,esg,al,bin3,sd,aop,B-H1,tsh,ttk,ss,Su(z)2,chn,MEP-1,lncRNA:bxd,Prdm13 | | 2.059e-12 | -26.91 | animal organ formation | biological process | GO:0048645 | 46 | 16 | 12345 | 442 | N,nuf,sens,fz,unc-5,hth,H,klu,sli,B-H1,robo1,Tom,Scr,CtBP,arm,ss | | 2.641e-12 | -26.66 | establishment of planar polarity | biological process | GO:0001736 | 97 | 22 | 12345 | 442 | ed,ds,d,S,fz,otk,Dl,N,scrib,ft,kay,Shrm,spi,shg,nmo,SMC3,ec,Egfr,baz,aop,rdx,spen | | 2.641e-12 | -26.66 | establishment of tissue polarity | biological process | GO:0007164 | 97 | 22 | 12345 | 442 | shg,spi,kay,ft,Shrm,rdx,aop,spen,SMC3,nmo,baz,Egfr,ec,fz,S,ds,d,ed,Dl,scrib,N,otk | | 2.769e-12 | -26.61 | N (Notch) | protein interactions | 31293 | 301 | 45 | 9204 | 432 | nmo,arm,spen,fz,msi,Egfr,Ocrl,Sxl,jigr1,rictor,ed,ap,vn,tkv,CycG,ena,S,CtBP,sd,Lk6,shg,Gli,emc,Rgl,InR,H,Dl,B-H1,aop,esg,spdo,Abl,N,px,eyg,klu,chn,ey,orb2,lqf,lilli,mam,SERCA,frc,trbl | | 2.948e-12 | -26.55 | RNA polymerase II regulatory region DNA binding | molecular function | GO:0001012 | 261 | 35 | 12176 | 414 | ey,cic,ttk,E(spl)m5-HLH,Mef2,ap,dve,sim,su(sable),HLH3B,al,Hr4,CG11247,CG34376,zfh2,h,esg,Hr39,Lmx1a,kay,Abd-B,sd,Scr,abd-A,Prdm13,CG12769,grn,B-H1,pnr,dmrt11E,rgr,bowl,klu,aop,eyg | | 3.180e-12 | -26.47 | epithelial cell development | biological process | GO:0002064 | 361 | 43 | 12345 | 442 | Gli,Dl,RhoL,Cbl,mop,emc,Raf,mam,SERCA,fz,eya,tkv,mim,jar,baz,Rab5,cic,Mef2,kay,Abl,tai,Cip4,Dp,Diap1,Socs36E,neo,Pvr,Ptpmeg2,scrib,N,Ppcs,cindr,ed,InR,S,ph-p,cact,Egfr,ena,aop,dome,ttk,shg | | 3.382e-12 | -26.41 | response to chemical | biological process | GO:0042221 | 1191 | 90 | 12345 | 442 | dome,Tpc1,rg,shg,spi,fz2,Fas1,Egfr,LanA,spen,aop,rdx,hppy,CG30156,dock,InR,mura,vn,ap,Bsg,Ir21a,scrib,N,comm2,betaTub60D,14-3-3zeta,beat-IIa,drl,Abl,tai,Adf1,pnr,sqz,rictor,tkv,robo1,LRR,hang,alpha-Man-Ia,Sema1a,pum,Sema1b,Ptp4E,moody,Pex5,Src42A,ss,ena,msi,Gr93c,wgn,sli,esg,REPTOR,ZnT41F,sim,dlp,Psc,gukh,jeb,S,ph-p,fend,hfw,grn,CG30344,chic,pxb,shot,Pvr,Bacc,pdm3,14-3-3epsilon,Poxn,htl,Dad,Hr4,Pdk1,E(Pc),Nrk,Hr39,unc-5,Dyrk2,eya,Raf,Dg,otk,Eph,sens,SCAR | | 3.408e-12 | -26.41 | morphogenesis of a polarized epithelium | biological process | GO:0001738 | 108 | 23 | 12345 | 442 | shg,spi,yrt,Shrm,kay,ft,spen,rdx,aop,baz,Egfr,ec,SMC3,nmo,fz,S,ds,d,ed,scrib,N,Dl,otk | | 4.374e-12 | -26.16 | cell fate determination | biological process | GO:0001709 | 141 | 26 | 12345 | 442 | H,klu,S,mam,fz,nub,nuf,sens,shot,abd-A,14-3-3zeta,N,14-3-3epsilon,spi,pnr,arm,CtBP,msi,baz,jar,Egfr,htl,spen,esg,Tom,aop | | 4.573e-12 | -26.11 | tissue migration | biological process | GO:0090130 | 164 | 28 | 12345 | 442 | jeb,dlp,Raf,fz,rib,Socs36E,Pvr,Ptpmeg2,Dl,cindr,Ppcs,RhoL,N,kay,tai,dome,shg,Diap1,htl,baz,Egfr,mim,jar,ena,Rab5,aop,sli,esg | | 4.575e-12 | -26.11 | arm (armadillo) | protein interactions | 31151 | 119 | 27 | 9204 | 432 | crol,Egfr,SCAR,spi,dlp,nmo,hyx,spen,shg,jar,fz,ds,Dad,tkv,aop,tsh,klu,Ptp4E,baz,ft,Abl,Src42A,Magi,N,CtBP,lilli,mam | | 5.053e-12 | -26.01 | rho (rhomboid) | protein interactions | 38168 | 64 | 20 | 9204 | 432 | N,px,Src42A,S,ft,lilli,Rab5,hth,abd-A,spi,Egfr,H,emc,shg,arm,aop,CycG,ed,vn,tkv | | 5.081e-12 | -26.01 | sex differentiation | biological process | GO:0007548 | 100 | 22 | 12345 | 442 | Abd-B,htl,Egfr,ena,robo1,aop,Rab5,Pdk1,sli,Sxl,bab2,fz2,Scr,shg,Pvr,PI31,abd-A,otk,N,InR,rib,eya | | 5.328e-12 | -25.96 | eye photoreceptor cell differentiation | biological process | GO:0001754 | 165 | 28 | 12345 | 442 | Nrk,jeb,emc,klu,Cbl,eya,Mob2,S,fz,Raf,hth,sens,N,scrib,kay,rg,yrt,eff,klar,ss,ttk,lilli,msi,Egfr,boi,sina,aop,B-H1 | | 5.472e-12 | -25.93 | response to external stimulus | biological process | GO:0009605 | 987 | 79 | 12345 | 442 | fz2,CG32795,shg,Fas1,RhoGEF3,dome,spen,sick,cact,LanA,dock,InR,lt,N,comm2,betaTub60D,DPCoAC,drl,beat-IIa,ap,Bsg,Tollo,Abl,SNF4Agamma,robo1,Mef2,Atg16,rictor,sqz,Dcr-2,yuri,Sema1a,Ric,Dronc,Sema1b,CG7536,Atg17,Ptp4E,jumu,Src42A,esg,sim,REPTOR,sli,ena,wgn,fend,Cerk,grn,dlp,Psc,gukh,jeb,kn,pdm3,chic,Pen,Rm62,Pvr,shot,14-3-3epsilon,kay,subdued,ogre,Dad,nmo,htl,eya,unc-5,Raf,Nrk,E(Pc),SCAR,imd,RhoL,Eph,otk,Dg,sens | | 6.066e-12 | -25.83 | DNA binding | molecular function | GO:0003677 | 984 | 76 | 12176 | 414 | B-H1,His2Av,Prdm13,aop,klu,tsh,bowl,bab2,Lmx1a,HmgZ,esg,h,nub,Tet,Abd-B,al,crp,chn,CG32772,His3.3B,hng3,pita,ttk,Dp,ey,Kdm2,Poxn,Sox14,jumu,CG10321,Mef2,rgr,dmrt11E,Su(z)2,pnr,grn,CG12769,abd-A,CG45050,eyg,kn,Psc,CG13204,pdm3,kay,sens,hth,Hr39,Scr,mrt,Elba2,sd,CG11617,H2.0,Hr4,Hel89B,Adf1,HLH3B,ph-p,His3.3A,su(sable),rib,l(2)37Cg,zfh2,CG34376,CG13895,CG11247,lilli,otp,cic,sim,edl,dve,ap,jigr1,E(spl)m5-HLH | | 6.404e-12 | -25.77 | regulation of intracellular signal transduction | biological process | GO:1902531 | 339 | 41 | 12345 | 442 | ft,14-3-3epsilon,wdb,RapGAP1,rictor,Pdk1,scyl,CG42684,LRR,Cbl,Raf,CG5522,d,ds,siz,CycG,Rgl,Ziz,wnd,trbl,RhoGEF4,Actbeta,Src42A,RhoGEF3,spi,step,Egfr,Pez,rdx,jeb,ed,chrb,hppy,InR,Pvr,lncRNA:Hsromega,vn,Lk6,Cdk4,14-3-3zeta,N | | 7.907e-12 | -25.56 | establishment or maintenance of cell polarity | biological process | GO:0007163 | 217 | 32 | 12345 | 442 | scrib,14-3-3zeta,N,S,ed,cora,rdx,aop,Egfr,ec,shg,spi,p130CAS,Dl,Dg,fz,d,rib,ds,Raf,Cbl,Rab5,rictor,baz,nmo,yrt,Dp,fz3,Shrm,14-3-3epsilon,kay,ft | | 8.117e-12 | -25.54 | transcription regulatory region sequence-specific DNA binding | molecular function | GO:0000976 | 329 | 39 | 12176 | 414 | su(sable),Hr4,al,HLH3B,CG11247,CG34376,zfh2,ttk,ey,cic,Mef2,E(spl)m5-HLH,CG10321,ap,jumu,sim,dve,grn,Prdm13,CG12769,abd-A,B-H1,dmrt11E,pnr,rgr,bowl,aop,klu,eyg,h,esg,Hr39,kay,hth,Lmx1a,Abd-B,sd,nub,Scr | | 8.199e-12 | -25.53 | ovarian follicle cell development | biological process | GO:0030707 | 327 | 40 | 12345 | 442 | shg,ttk,dome,aop,Egfr,ena,cact,ph-p,S,InR,ed,scrib,cindr,N,Ppcs,Ptpmeg2,Socs36E,Pvr,Diap1,Cip4,Dp,tai,kay,Rab5,cic,Mef2,mim,baz,jar,tkv,SERCA,fz,mam,eya,Raf,Cbl,mop,emc,RhoL,Dl | | 8.405e-12 | -25.50 | S (Star) | protein interactions | 33281 | 52 | 18 | 9204 | 432 | Src42A,px,N,Raf,chic,su(sable),shg,step,sina,H,rho-4,Egfr,spi,Dl,aop,rg,edl,ec | | 9.036e-12 | -25.43 | columnar/cuboidal epithelial cell development | biological process | GO:0002066 | 328 | 40 | 12345 | 442 | aop,ena,Egfr,cact,shg,ttk,dome,Ppcs,cindr,N,scrib,Ptpmeg2,Pvr,Socs36E,ph-p,S,InR,ed,cic,Mef2,Rab5,mim,jar,baz,tkv,Diap1,Dp,Cip4,tai,kay,RhoL,Dl,eya,SERCA,fz,mam,Raf,mop,emc,Cbl | | 9.126e-12 | -25.42 | cellular process involved in reproduction in multicellular organism | biological process | GO:0022412 | 997 | 79 | 12345 | 442 | dome,shg,ttk,Egfr,cact,LanA,aop,InR,Ptpmeg2,orb2,14-3-3zeta,cindr,Ppcs,N,scrib,Abl,tai,Cks30A,larp,Cip4,baz,tkv,Abd-B,Mef2,Dronc,emc,mop,Cbl,sns,fz,SERCA,yuri,pum,lin-28,PI31,abd-A,CycG,Inx2,Dl,Sxl,msi,ena,ed,S,ph-p,Cerk,Pen,chic,lncRNA:Hsromega,Pvr,Socs36E,bol,kay,14-3-3epsilon,Pex10,Diap1,eff,arm,Dp,His3.3A,wbl,jar,mim,htl,His3.3B,cic,Rab5,Tet,Hr39,eya,mam,Rab1,Raf,Dg,Arf79F,RhoL,SCAR,Pat1,Gli | | 1.012e-11 | -25.32 | reproductive system development | biological process | GO:0061458 | 84 | 20 | 12345 | 442 | sli,robo1,aop,Rab5,Egfr,ena,Abd-B,htl,fz2,shg,bab2,N,otk,Pvr,abd-A,PI31,eya,rib,InR,dve | | 1.012e-11 | -25.32 | reproductive structure development | biological process | GO:0048608 | 84 | 20 | 12345 | 442 | fz2,shg,bab2,robo1,aop,Rab5,sli,Abd-B,htl,ena,Egfr,InR,eya,rib,dve,N,PI31,abd-A,Pvr,otk | | 1.226e-11 | -25.12 | cell communication | biological process | GO:0007154 | 1239 | 91 | 12345 | 442 | Src42A,sd,REPTOR,sli,wgn,hfw,Cerk,CG42788,kn,jeb,pdm3,Lk6,RhoU,shot,Pvr,Socs36E,arm,Dp,fz3,cic,ogre,Pdk1,Rab5,Hr4,nmo,Dad,boi,htl,Raf,Rab1,unc-5,mam,Hr39,ITP,Nrk,Actbeta,imd,Arf79F,wnd,Eph,otk,lqf,ttk,tsh,spi,fz2,dome,aop,spen,sina,cact,Ocrl,Egfr,InR,lt,hppy,drl,RhoGAP19D,nAChRalpha4,N,Magi,Ir21a,Pgk,vn,Tollo,Cadps,Pvf3,pnr,Abl,SNF4Agamma,robo1,Syx1A,Atg16,tkv,rictor,Ddr,Ric,Sema1a,hang,fz,SERCA,Cbl,Dronc,Dl,Inx2,moody,Rgl,CG7536,Atg17,pum,Sema1b | | 1.302e-11 | -25.06 | regulation of neurogenesis | biological process | GO:0050767 | 261 | 35 | 12345 | 442 | Sema1b,Arf79F,Dl,emc,mop,fz,Sema1a,Raf,hth,h,Rab1,rictor,robo1,CG34401,Abl,Adf1,E(spl)m5-HLH,Sox14,eff,chic,shot,Socs36E,N,ed,kn,klu,InR,nub,sina,sd,aop,spi,ss,ttk,Src42A | | 1.334e-11 | -25.04 | formation of animal organ boundary | biological process | GO:0010160 | 37 | 14 | 12345 | 442 | CtBP,hth,unc-5,Scr,fz,klu,H,Tom,B-H1,robo1,N,sli,sens,nuf | | 1.614e-11 | -24.85 | developmental growth involved in morphogenesis | biological process | GO:0060560 | 86 | 20 | 12345 | 442 | arm,Tollo,ttk,fz2,shg,path,kay,Abl,tai,Rab5,al,tkv,Dad,LanA,baz,disco-r,fz,SCAR,wnd,otk | | 1.858e-11 | -24.71 | regulatory region nucleic acid binding | molecular function | GO:0001067 | 369 | 41 | 12176 | 414 | B-H1,abd-A,CG12769,Prdm13,grn,rgr,pnr,dmrt11E,bowl,eyg,klu,aop,esg,h,Lmx1a,hth,kay,Hr39,sd,Abd-B,mrt,Scr,nub,su(sable),HLH3B,Hr4,al,CG34376,CG11247,zfh2,ey,cic,ttk,CG10321,Mef2,E(spl)m5-HLH,jigr1,sim,dve,jumu,ap | | 1.858e-11 | -24.71 | transcription regulatory region DNA binding | molecular function | GO:0044212 | 369 | 41 | 12176 | 414 | klu,aop,eyg,bowl,pnr,dmrt11E,rgr,CG12769,Prdm13,abd-A,grn,B-H1,nub,mrt,Scr,Abd-B,sd,Hr39,Lmx1a,hth,kay,h,esg,zfh2,CG11247,CG34376,HLH3B,al,Hr4,su(sable),jumu,ap,sim,dve,E(spl)m5-HLH,Mef2,jigr1,CG10321,cic,ey,ttk | | 2.018e-11 | -24.63 | plasma membrane | cellular component | GO:0005886 | 1265 | 90 | 11940 | 417 | Gli,CG5853,ZnT41F,glec,RhoU,Esp,Dg,beat-IIa,Cerk,cindr,otk,Tsp96F,14-3-3zeta,cue,sns,CG7536,fz3,Bsg,CG31121,d,Arf79F,Nrk,rho-4,Abl,Ir21a,14-3-3epsilon,nAChRalpha4,N,LanA,CG11069,nuf,Mob2,Egfr,Rab5,arm,ft,cora,Cals,dlp,spdo,Cip4,Pvr,path,dome,step,tipE,Dl,p130CAS,ogre,Syx1A,fz,S,Ddr,CG6891,yrt,Ptp4E,ed,emp,scrib,Cbl,Tob,baz,orb2,shg,Mid1,Eph,Inx2,tkv,Fas1,gukh,InR,bou,robo1,ds,drl,moody,Shrm,Pdk1,Src42A,wgn,Dyb,Ric,CG17646,kek5,Raf,Rab1,CG9784,htl,fz2,Magi | | 2.685e-11 | -24.34 | localization | biological process | GO:0051179 | 2206 | 135 | 12345 | 442 | Diap1,Pex10,subdued,Tpst,CG13384,wbl,mim,Rab1,Raf,eya,rib,d,unc-5,CG17883,Irk3,RhoL,SCAR,lqf,Dg,klar,CG5853,cora,ZnT41F,esg,koi,Npc2g,S,jeb,dve,dlp,Mid1,Pvr,Socs36E,Pen,tipE,Cip4,AP-1gamma,larp,Abl,SNF4Agamma,Syx1A,CG31121,Abd-B,Mpcp2,baz,rictor,Sema1a,ds,SERCA,Myo10A,Inx2,abd-A,CG7536,pum,Sema1b,shg,dome,Tpc1,Ocrl,lt,CG3149,CG4645,cindr,N,DPCoAC,orb2,arm,Slbp,kay,14-3-3epsilon,His2Av,CG11069,Rab5,ogre,Dad,htl,boi,jar,hth,Esp,Pat1,Gli,CG7394,Arf79F,apolpp,Src42A,Pex14,Atox1,sli,Unc-76,ena,CG4991,Vps13,Orct2,Cerk,gukh,lncRNA:Hsromega,RhoU,chic,CG8034,CG30344,Pvf3,Cadps,ft,ey,tai,Shrm,flfl,Mef2,TBC1D5,tkv,Dcr-2,fz,mop,cv-d,Dl,Pex5,frc,Rim2,nuf,fz2,path,CG17646,p130CAS,aop,cact,Egfr,unc-119,nAChRalpha4,14-3-3zeta,Ppcs,scrib,Bsg,Ptpmeg2,edl | | 2.704e-11 | -24.33 | gamete generation | biological process | GO:0007276 | 1097 | 83 | 12345 | 442 | Sxl,ena,msi,S,ph-p,Cerk,ed,bol,Pen,chic,Socs36E,Pvr,lncRNA:Hsromega,eff,Diap1,Dp,arm,Pex10,kay,14-3-3epsilon,His3.3B,Pdk1,Rab5,cic,His3.3A,wbl,mim,jar,htl,mam,eya,Raf,Rab1,Tet,Hr39,RhoL,SCAR,Pat1,Gli,Dg,wnd,Arf79F,shg,dmrt11E,ttk,dome,aop,Egfr,LanA,cact,InR,scrib,Ppcs,cindr,14-3-3zeta,N,Ptpmeg2,orb2,larp,Cks30A,Cip4,Abl,tai,Mef2,baz,Abd-B,tkv,fz,sns,SERCA,yuri,lncRNA:CR44420,Dronc,Cbl,emc,mop,Inx2,Dl,lin-28,pum,abd-A,PI31,CycG | | 3.052e-11 | -24.21 | sequence-specific double-stranded DNA binding | molecular function | GO:1990837 | 359 | 40 | 12176 | 414 | nub,Scr,Abd-B,sd,Hr39,hth,Lmx1a,kay,h,esg,klu,aop,eyg,bab2,bowl,pnr,dmrt11E,rgr,abd-A,Prdm13,CG12769,grn,B-H1,jumu,ap,sim,dve,Mef2,E(spl)m5-HLH,CG10321,cic,ey,ttk,zfh2,CG11247,CG34376,HLH3B,al,Hr4,su(sable) | | 3.228e-11 | -24.16 | establishment of ommatidial planar polarity | biological process | GO:0042067 | 62 | 17 | 12345 | 442 | spi,shg,ft,kay,aop,rdx,nmo,ec,Egfr,ds,d,fz,S,ed,Dl,N,scrib | | 3.386e-11 | -24.11 | segmentation | biological process | GO:0035282 | 256 | 34 | 12345 | 442 | arm,pnr,cic,Abd-B,baz,Raf,hth,h,fz,Pat1,Dl,Kdm2,frc,fs(1)h,pum,Dg,ss,lilli,tsh,fz2,shg,dome,rdx,spen,Egfr,kn,dve,dlp,Cdk4,N,scrib,chic,ap,bowl | | 3.493e-11 | -24.08 | gland development | biological process | GO:0048732 | 178 | 28 | 12345 | 442 | htl,Abd-B,CG1129,Egfr,robo1,sli,Src42A,tsh,fz2,shg,pnr,Scr,Pvr,Atg17,chic,Pen,drl,N,jeb,Dronc,Raf,hth,eyg,h,eya,rib,fz,unc-5 | | 3.765e-11 | -24.00 | growth | biological process | GO:0040007 | 191 | 29 | 12345 | 442 | arm,Tollo,ttk,fz2,shg,path,ft,kay,Pex10,Sxl,tai,Abl,Rab5,Pdk1,al,Dad,tkv,htl,LanA,baz,disco-r,InR,fz,jeb,SCAR,N,wnd,otk,Dg | | 3.765e-11 | -24.00 | developmental growth | biological process | GO:0048589 | 191 | 29 | 12345 | 442 | kay,Pex10,ft,tai,Sxl,Abl,ttk,Tollo,arm,path,shg,fz2,htl,LanA,Dad,tkv,baz,Pdk1,Rab5,al,jeb,InR,disco-r,fz,Dg,otk,wnd,N,SCAR | | 4.043e-11 | -23.93 | salivary gland development | biological process | GO:0007431 | 121 | 23 | 12345 | 442 | Src42A,Scr,shg,fz2,tsh,htl,Abd-B,Egfr,robo1,sli,jeb,Dronc,eyg,h,hth,fz,unc-5,rib,eya,Atg17,Pvr,chic,drl | | 4.043e-11 | -23.93 | exocrine system development | biological process | GO:0035272 | 121 | 23 | 12345 | 442 | Abd-B,htl,Egfr,robo1,sli,Src42A,tsh,fz2,Scr,shg,Pvr,Atg17,chic,drl,jeb,Dronc,eyg,h,hth,rib,eya,unc-5,fz | | 4.079e-11 | -23.92 | epithelial cell migration | biological process | GO:0010631 | 155 | 26 | 12345 | 442 | Raf,fz,rib,dlp,Dl,RhoL,N,cindr,Ppcs,Socs36E,Pvr,Ptpmeg2,shg,Diap1,kay,dome,tai,Rab5,aop,sli,esg,baz,jar,mim,ena,Egfr | | 4.223e-11 | -23.89 | cell periphery | cellular component | GO:0071944 | 1409 | 96 | 11940 | 417 | Raf,Rab1,CG9784,Ric,Dyb,CG17646,kek5,Magi,fz2,htl,dock,robo1,drongo,ds,Inx2,Fas1,tkv,gukh,InR,bou,wgn,Src42A,drl,moody,Pdk1,Shrm,scrib,Cbl,Tob,emp,ed,Ptp4E,Mid1,Eph,orb2,baz,shg,ogre,Syx1A,tipE,p130CAS,Dl,yrt,CG6891,fz,Ddr,S,chic,cora,Cals,ft,dlp,Cip4,Pvr,path,spdo,dome,step,nuf,CG11069,Mob2,Egfr,LanA,arm,Rab5,Nrk,rho-4,Abl,CG31121,d,Arf79F,N,Pez,14-3-3epsilon,Ir21a,CG42684,nAChRalpha4,beat-IIa,Dg,Cerk,Esp,RhoU,cindr,otk,14-3-3zeta,Tsp96F,Gli,CG5853,ZnT41F,glec,fz3,CG7536,sns,Bsg,jar,cue | | 4.589e-11 | -23.80 | regulation of neuron differentiation | biological process | GO:0045664 | 180 | 28 | 12345 | 442 | N,Arf79F,shot,Socs36E,chic,Sema1b,Sema1a,InR,Raf,hth,Rab1,fz,emc,kn,mop,ed,aop,robo1,CG34401,sina,rictor,ss,ttk,Src42A,eff,Sox14,Adf1,Abl | | 4.620e-11 | -23.80 | neuron fate commitment | biological process | GO:0048663 | 72 | 18 | 12345 | 442 | sina,aop,Egfr,msi,spi,ttk,ey,kay,scrib,N,Dl,S,mam,fz,Raf,hth,Cbl,emc | | 4.736e-11 | -23.77 | epithelium migration | biological process | GO:0090132 | 156 | 26 | 12345 | 442 | baz,jar,ena,mim,Egfr,sli,esg,Rab5,aop,dome,tai,kay,shg,Diap1,Ptpmeg2,Socs36E,Pvr,Ppcs,RhoL,N,cindr,Dl,dlp,fz,rib,Raf | | 5.490e-11 | -23.63 | compound eye photoreceptor cell differentiation | biological process | GO:0001751 | 157 | 26 | 12345 | 442 | Egfr,msi,boi,sina,aop,B-H1,kay,eff,lilli,ttk,ss,klar,sens,scrib,N,Nrk,Cbl,klu,emc,jeb,fz,S,eya,Mob2,hth,Raf | | 8.946e-11 | -23.14 | muscle structure development | biological process | GO:0061061 | 224 | 31 | 12345 | 442 | chn,Alh,klar,ttk,htl,crp,sd,aop,C1GalTA,Mef2,cora,kon,ed,kn,jeb,dock,eya,sns,hth,InR,ap,Dg,shot,siz,abd-A,Kdm2,Pgk,Bsg,SCAR,N,drl | | 9.149e-11 | -23.11 | cellular process | biological process | GO:0009987 | 7485 | 331 | 12345 | 442 | IA-2,vig,CG17683,yrt,Diap1,Tpst,mld,mim,scyl,crol,Hr39,ITP,HmgZ,d,rib,Dyrk2,fs(1)h,CG9784,wnd,CG33474,Dg,SCAR,imd,RhoL,bab2,klar,lilli,sd,muc,sim,CG42788,jeb,klu,dve,koi,shot,Pvr,tipE,bol,Pp2C1,CG12402,Adf1,Abl,SNF4Agamma,Cks30A,pnr,larp,Abd-B,sqz,robo1,CG34401,CG13481,LRR,GluProRS,Rgl,abd-A,PI31,pum,Dgat2,Sema1b,IntS3,dome,Su(z)2,CenB1A,Fas1,CG11030,tsh,CG31522,ATPCL,Ocrl,CG6656,sina,CG14562,lt,dock,orb2,ap,FBXO11,CG12910,N,cindr,kay,htl,Pex7,Tom,ogre,fl(2)d,Tet,E(Pc),hth,Arf79F,Kdm2,Eph,otk,Actbeta,CG7394,bou,ss,Src42A,Unc-76,C1GalTA,sli,kn,Psc,CG1218,Cerk,fuss,lncRNA:Hsromega,RhoU,Rm62,chic,bru2,tna,Tob,ey,Pvf3,Cadps,ReepA,flfl,mop,Drep4,Fem-1,H,Dronc,yuri,Ric,fz,sns,siz,UQCR-11,Rim2,Atg17,nuf,CG6891,lin-28,Dl,Pex5,Hel89B,Spc105R,rg,chn,CG17646,spdo,fz2,CG34172,sick,aop,al,CG30156,Pgk,neo,Bsg,Fkbp14,edl,Ptpmeg2,Drak,nAChRalpha4,RhoGAP19D,Magi,14-3-3zeta,E(spl)m5-HLH,Pex10,Ppn,fz3,Dp,CtBP,nmo,SmD1,His3.3A,wbl,Pdk1,Hr4,His3.3B,CG8306,Rab1,Raf,eya,Lmx1a,unc-5,pdgy,sens,lqf,Tis11,CG4935,Gfat2,Sxl,jumu,l(2)37Cg,msi,cora,bin3,REPTOR,esg,CG1673,l(1)G0196,dlp,rho-4,CG33082,grn,CG3119,hfw,fend,S,ph-p,Socs36E,Pen,stx,Lk6,pdm3,Mid1,wdb,Tollo,Cip4,AP-1gamma,baz,Ddr,rictor,Nlp,Syx1A,CG42327,Atg16,morgue,Cbl,Myo10A,Sema1a,Psa,hang,ds,SERCA,CG7536,Inx2,moody,Ubc87F,CG10417,Jupiter,shg,CG3436,CG7560,SMC3,tara,ec,rdx,B-H1,Kdm4B,spen,CG9220,mura,Mob2,GlcAT-I,CG11414,drl,CG3149,pita,14-3-3epsilon,His2Av,arm,Slbp,eff,Dad,CG3961,boi,jar,cic,kon,Rab5,CG2017,Nrk,Snoo,disco-r,h,mam,CG34057,trbl,Pat1,Gli,Mfe2,Pex14,mrt,wgn,ena,f,Atox1,ed,gukh,Vps13,Pitslre,CG6465,corto,Cdk4,CG9485,Sox14,ft,Shrm,tai,tkv,Dcr-2,Mef2,emc,hyx,CG7460,alpha-Man-Ia,CycB3,CycG,Ptp4E,p130CAS,RhoGEF3,ttk,spi,path,cact,LanA,Egfr,Pez,CG4467,unc-119,mRpL41,chrb,hppy,snRNA:U5:38ABb,Wdr62,nub,InR,Ir21a,su(sable),vn,beat-IIa,betaTub60D,comm2,Ppcs,scrib,Elba2 | | 9.397e-11 | -23.09 | pnr (pannier) | protein interactions | 44849 | 107 | 24 | 9204 | 432 | Scr,CG43759,chrb,esg,Dad,Hr39,Orct2,Abd-B,Mef2,Hr4,emc,spi,pum,abd-A,tara,muc,lin-28,sd,ps,tai,glec,Drak,CtBP,chn | | 9.883e-11 | -23.04 | signal transduction | biological process | GO:0007165 | 968 | 75 | 12345 | 442 | imd,Actbeta,Eph,wnd,otk,Arf79F,mam,unc-5,Raf,Rab1,ITP,Nrk,Hr39,Hr4,cic,ogre,Rab5,Pdk1,nmo,Dad,boi,htl,Dp,arm,fz3,Lk6,Pvr,RhoU,Socs36E,Cerk,hfw,jeb,CG42788,REPTOR,sli,sd,wgn,Src42A,Dl,moody,Sema1b,Rgl,fz,Sema1a,Ric,hang,Dronc,Cbl,robo1,rictor,Ddr,tkv,Pvf3,pnr,Tollo,Magi,N,RhoGAP19D,drl,nAChRalpha4,vn,Ir21a,InR,hppy,spen,sina,aop,Egfr,Ocrl,cact,fz2,tsh,spi,ttk,dome | | 1.038e-10 | -22.99 | negative regulation of multicellular organismal process | biological process | GO:0051241 | 212 | 30 | 12345 | 442 | Egfr,ena,Abd-B,LanA,tkv,Dad,aop,sd,C1GalTA,chn,Abl,ft,Diap1,tsh,Src42A,Cip4,ttk,klar,lqf,Sema1b,Socs36E,Him,scrib,N,Dl,ed,emc,Mob2,InR,Sema1a | | 1.109e-10 | -22.92 | regulation of biological quality | biological process | GO:0065008 | 1352 | 94 | 12345 | 442 | hppy,InR,Mob2,neo,Pgk,Fkbp14,nAChRalpha4,CG3149,scrib,Ppcs,N,14-3-3zeta,p130CAS,ttk,shg,path,LanA,cact,ATPCL,Egfr,rdx,Cbl,H,Sema1a,SERCA,fz,ds,Atg17,CG6891,Sema1b,pum,ey,ft,SNF4Agamma,wdb,tai,Abl,Cip4,Cals,Pvf3,Cadps,Abd-B,tkv,rictor,Atg16,Syx1A,Mef2,robo1,ed,Vps13,shot,Pvr,RhoU,chic,bol,Mid1,CG6191,dpy,bou,Sxl,lilli,Src42A,klar,wgn,ena,cora,Atox1,ZnT41F,esg,Nrk,Raf,Rab1,rib,Esp,Arf79F,cue,lqf,trbl,Dg,wnd,Gli,Irk3,Tis11,SCAR,14-3-3epsilon,eff,yrt,htl,boi,nmo,Dad,mld,Pdk1,Rab5 | | 1.131e-10 | -22.90 | reproductive process | biological process | GO:0022414 | 1436 | 98 | 12345 | 442 | Rab1,Raf,mam,rib,Esp,eya,Hr39,Tet,fl(2)d,Gli,Pat1,RhoL,SCAR,Arf79F,Dg,otk,wnd,Poxn,arm,Dp,eff,Diap1,Pex10,14-3-3epsilon,kay,Rab5,Pdk1,cic,His3.3B,htl,His3.3A,jar,mim,wbl,ph-p,Cerk,S,dve,ed,bol,Socs36E,lncRNA:Hsromega,Pvr,tipE,chic,Pen,ss,Sxl,bab2,sli,Gr93c,ena,msi,yuri,sns,fz,SERCA,Cbl,emc,mop,lncRNA:CR44420,Dronc,Dl,Inx2,PI31,abd-A,CycG,lin-28,pum,Cip4,larp,Cks30A,tai,Abl,robo1,Mef2,Nlp,Abd-B,tkv,baz,InR,scrib,N,Ppcs,14-3-3zeta,cindr,orb2,Ptpmeg2,ttk,shg,Scr,dmrt11E,fz2,dome,aop,LanA,cact,Egfr | | 1.153e-10 | -22.88 | double-stranded DNA binding | molecular function | GO:0003690 | 391 | 41 | 12176 | 414 | nub,Scr,Abd-B,sd,Hr39,Lmx1a,hth,kay,h,esg,klu,aop,eyg,bowl,bab2,pnr,dmrt11E,rgr,abd-A,Prdm13,CG12769,grn,B-H1,jumu,ap,sim,dve,E(spl)m5-HLH,Mef2,CG10321,lilli,ey,cic,ttk,zfh2,CG11247,CG34376,HLH3B,Hr4,al,su(sable) | | 1.632e-10 | -22.54 | regulation of phosphorylation | biological process | GO:0042325 | 271 | 34 | 12345 | 442 | InR,jeb,hppy,Lk6,N,Pvr,lncRNA:Hsromega,Pgk,Socs36E,vn,edl,Src42A,CG8312,spi,rdx,step,CG34172,Egfr,Raf,CycB3,Cbl,Actbeta,CycG,Atg17,wnd,Dg,trbl,Cks30A,kay,wdb,SNF4Agamma,Pdk1,Dad,rictor | | 1.652e-10 | -22.52 | intracellular membrane-bounded organelle | cellular component | GO:0043231 | 4479 | 219 | 11940 | 417 | TBC1D5,Drak,su(sable),nmo,jumu,Dcr-2,sens,CG31211,dock,Tpst,CG1218,Prdm13,zfh2,Mef2,312,fz2,lncRNA:Hsromega,RhoGEF4,Kr-h2,eya,siz,ph-p,yuri,Rab1,aop,CG3149,sens-2,Hr39,SCAR,SmD1,Dp,yrt,cic,H,CG31457,spen,Fkbp14,Pitslre,rib,S,abd-A,CG7394,tai,ps,muc,CG11247,ss,rgr,px,fl(2)d,esg,Tet,MEP-1,emp,dpy,koi,sqz,Lmx1a,CG10321,CG9220,ena,hth,ap,CG1673,Nlp,lqf,tna,alpha-Man-Ia,al,Drep4,His3.3B,CG6340,dmrt11E,Pgant6,Tpc1,B-H1,pnr,CG8312,corto,bowl,grn,CG14441,CG33474,E(spl)m5-HLH,CG6656,CG7536,kay,SNF4Agamma,tsh,pita,hang,hyx,Alh,crol,wbl,bab2,CG11030,14-3-3epsilon,CG12769,His2Av,pdgy,hng3,Kdm2,wdb,sina,rho-4,AP-1gamma,CG4935,Slbp,jigr1,ReepA,Sox14,Snoo,CycB3,Pex10,msi,Hr4,Rm62,Pdk1,mam,larp,Jupiter,Sxl,Atg16,Cdk4,E(Pc),Adf1,crp,Pex5,Elba2,UQCR-11,flfl,pum,ey,h,CG9784,CG34172,SERCA,mura,Kdm4B,CG4645,bol,sd,CG17883,CG3107,klu,rdx,CG13204,Tis11,bru2,orb2,DPCoAC,Pp2C1,CG32772,ttk,IntS3,CtBP,Pex14,sim,Tob,spi,snRNA:U5:38ABb,CG13481,arm,Dronc,CG34376,Pex7,Mpcp2,bin3,Hel89B,CG14464,Abd-B,CG33082,Bacc,fs(1)h,l(2)37Cg,emc,Pen,CycG,lncRNA:bxd,Psc,klar,Poxn,trbl,path,step,mRpL41,Scr,REPTOR,kn,mrt,lilli,Su(z)2,CG4896,14-3-3zeta,dve,CG3436,vig,Mfe2,CG11617,Rim2,glec,edl,CG8306,N,eyg,Pgant5,HmgZ,CG11360,frc,Lk6,mld,Diap1,GlcAT-I,HLH3B,chn,stx,CG30156,Arf79F | | 1.899e-10 | -22.38 | regulation of cellular component organization | biological process | GO:0051128 | 791 | 65 | 12345 | 442 | Sema1b,pum,CG6891,RhoGEF4,Cbl,Sema1a,fz,tkv,Ddr,rictor,baz,CG34401,robo1,Syx1A,ey,Adf1,Abl,Cals,Cip4,sens-2,neo,scrib,N,cindr,lt,InR,Mob2,LanA,Ocrl,Su(z)2,ttk,Jupiter,Arf79F,Kdm2,trbl,wnd,SCAR,Rab1,rib,htl,nmo,Dad,mim,jar,Pdk1,Rab5,kon,crol,cic,14-3-3epsilon,shot,RhoU,chic,Cdk4,kn,ed,Orct2,ena,step,C1GalTA,mrt,Src42A,lilli,ss,klar | | 1.920e-10 | -22.37 | eya (eyes absent) | protein interactions | 33916 | 35 | 14 | 9204 | 432 | nmo,Abl,Egfr,Socs36E,Tis11,Raf,abd-A,eya,Scr,ey,Lmx1a,aop,CG42788,Abd-B | | 1.957e-10 | -22.35 | photoreceptor cell fate commitment | biological process | GO:0046552 | 60 | 16 | 12345 | 442 | msi,Egfr,aop,sina,kay,ey,ttk,spi,N,scrib,emc,Cbl,Raf,hth,S,fz | | 2.017e-10 | -22.32 | enzyme linked receptor protein signaling pathway | biological process | GO:0007167 | 154 | 25 | 12345 | 442 | cic,aop,Pdk1,spen,sina,tkv,Dad,htl,Egfr,Ddr,rictor,Src42A,ttk,spi,Pvf3,pnr,drl,Actbeta,Pvr,Eph,vn,InR,Raf,jeb,Nrk | | 2.653e-10 | -22.05 | digestive tract development | biological process | GO:0048565 | 110 | 21 | 12345 | 442 | Abd-B,Dad,Egfr,Mef2,sli,IA-2,dome,shg,Scr,tsh,spi,Pvr,abd-A,vn,bowl,Inx2,N,emc,dve,S,rib | | 2.653e-10 | -22.05 | digestive system development | biological process | GO:0055123 | 110 | 21 | 12345 | 442 | sli,Mef2,Egfr,Dad,Abd-B,spi,tsh,shg,Scr,dome,IA-2,N,Inx2,bowl,vn,Pvr,abd-A,rib,S,dve,emc | | 2.725e-10 | -22.02 | cell junction | cellular component | GO:0030054 | 160 | 25 | 11940 | 417 | Shrm,moody,Src42A,kon,sns,yrt,arm,Gli,Inx2,p130CAS,Dl,LRR,ena,ogre,otk,baz,shg,step,N,Magi,ed,shot,cora,scrib,Abl | | 2.774e-10 | -22.01 | signaling | biological process | GO:0023052 | 1169 | 84 | 12345 | 442 | Syx1A,robo1,rictor,Ddr,tkv,Cadps,Pvf3,pnr,Tollo,Abl,Dl,moody,Sema1b,pum,Rgl,SERCA,fz,Ric,Sema1a,hang,Dronc,Cbl,spen,sina,aop,Egfr,Ocrl,cact,spi,fz2,tsh,ttk,dome,N,Magi,RhoGAP19D,drl,nAChRalpha4,vn,Ir21a,Pgk,InR,hppy,Hr4,cic,ogre,Rab5,Pdk1,nmo,Dad,boi,htl,Dp,arm,fz3,imd,Actbeta,Eph,wnd,lqf,otk,Arf79F,unc-5,mam,Rab1,Raf,ITP,Nrk,Hr39,REPTOR,sli,sd,wgn,Src42A,Lk6,Pvr,shot,RhoU,Socs36E,Cerk,hfw,kn,CG42788,jeb | | 2.782e-10 | -22.00 | female gamete generation | biological process | GO:0007292 | 761 | 63 | 12345 | 442 | shg,ttk,dome,Sxl,aop,Egfr,ena,LanA,cact,ph-p,S,InR,ed,scrib,cindr,N,Ppcs,14-3-3zeta,Ptpmeg2,Pen,chic,Socs36E,lncRNA:Hsromega,Pvr,eff,Cks30A,Diap1,Cip4,Dp,arm,tai,Abl,14-3-3epsilon,kay,Rab5,Pdk1,cic,Mef2,mim,wbl,jar,baz,tkv,fz,SERCA,mam,eya,Raf,Dronc,Cbl,Hr39,emc,mop,RhoL,SCAR,Gli,Pat1,Inx2,Dl,lin-28,Dg,pum,CycG | | 2.885e-10 | -21.97 | development of primary sexual characteristics | biological process | GO:0045137 | 53 | 15 | 12345 | 442 | ena,Egfr,htl,Abd-B,PI31,abd-A,sli,robo1,aop,bab2,shg,fz2,eya,rib,InR | | 2.885e-10 | -21.97 | gonad development | biological process | GO:0008406 | 53 | 15 | 12345 | 442 | bab2,eya,rib,fz2,shg,InR,Egfr,ena,abd-A,PI31,htl,Abd-B,sli,aop,robo1 | | 3.056e-10 | -21.91 | open tracheal system development | biological process | GO:0007424 | 235 | 31 | 12345 | 442 | Raf,InR,h,fz,rib,ds,dlp,ed,Cdk4,Gli,N,Ptp4E,shot,Src42A,ttk,klar,arm,shg,spi,ft,dpy,bou,dome,cora,sli,esg,LanA,Abd-B,tkv,Egfr,baz | | 3.769e-10 | -21.70 | transmembrane receptor protein tyrosine kinase signaling pathway | biological process | GO:0007169 | 112 | 21 | 12345 | 442 | Raf,InR,jeb,Nrk,drl,Pvr,Eph,vn,ttk,Src42A,spi,Pvf3,cic,aop,Pdk1,spen,sina,htl,rictor,Ddr,Egfr | | 3.898e-10 | -21.67 | regulation of epithelial cell migration | biological process | GO:0010632 | 46 | 14 | 12345 | 442 | aop,robo1,Gli,sli,Pvr,vn,Egfr,ttk,spi,Diap1,shg,kay,Cbl,tai | | 4.033e-10 | -21.63 | negative regulation of cell differentiation | biological process | GO:0045596 | 135 | 23 | 12345 | 442 | Abl,chn,Diap1,klar,ttk,Egfr,cact,Abd-B,esg,aop,C1GalTA,sd,ed,Snoo,emc,eya,fuss,Sema1a,Sema1b,lqf,Socs36E,N,Dl | | 4.252e-10 | -21.58 | stem cell differentiation | biological process | GO:0048863 | 72 | 17 | 12345 | 442 | spen,sqz,Egfr,htl,spi,pnr,arm,N,scrib,Dl,Sema1b,abd-A,SERCA,S,Sema1a,nub,klu | | 4.358e-10 | -21.55 | ovarian follicle cell migration | biological process | GO:0007297 | 124 | 22 | 12345 | 442 | Diap1,shg,kay,tai,dome,aop,Rab5,jar,mim,Egfr,baz,ena,Raf,fz,Dl,RhoL,cindr,Ppcs,N,Pvr,Socs36E,Ptpmeg2 | | 5.184e-10 | -21.38 | leg disc development | biological process | GO:0035218 | 103 | 20 | 12345 | 442 | vn,bowl,Drak,ap,N,Dl,RhoGAP19D,dve,hyx,ds,d,hth,Egfr,al,B-H1,sd,bab2,tsh,Poxn,ss | | 5.900e-10 | -21.25 | spi (spitz) | protein interactions | 35253 | 73 | 19 | 9204 | 432 | Lk6,CG14441,ph-p,stx,fs(1)h,Src42A,S,pnr,ec,edl,vn,abd-A,Pgant6,Socs36E,rho-4,Egfr,nmo,spen,arm | | 7.877e-10 | -20.96 | respiratory system development | biological process | GO:0060541 | 244 | 31 | 12345 | 442 | LanA,Abd-B,tkv,Egfr,baz,cora,sli,esg,ft,dpy,bou,dome,ttk,Src42A,arm,klar,shg,spi,Ptp4E,shot,Cdk4,Gli,N,dlp,ed,InR,h,Raf,fz,ds,rib | | 8.054e-10 | -20.94 | Ubx (Ultrabithorax) | protein interactions | 42034 | 118 | 24 | 9204 | 432 | 14-3-3epsilon,N,kn,hth,al,Hel89B,drl,arm,nmo,Egfr,Alh,Dl,abd-A,Syx1A,vn,ap,tkv,tsh,fl(2)d,h,yuri,pnr,Abd-B,Mef2 | | 8.784e-10 | -20.85 | multi-organism reproductive process | biological process | GO:0044703 | 1300 | 89 | 12345 | 442 | Abl,tai,larp,Cks30A,Cip4,baz,Abd-B,tkv,Mef2,Nlp,Dronc,lncRNA:CR44420,Cbl,mop,emc,fz,sns,SERCA,yuri,lin-28,pum,PI31,abd-A,CycG,Dl,Inx2,dome,shg,dmrt11E,ttk,Egfr,LanA,cact,aop,InR,Ptpmeg2,orb2,scrib,cindr,N,14-3-3zeta,Ppcs,kay,Pex10,14-3-3epsilon,eff,Diap1,Poxn,arm,Dp,jar,mim,His3.3A,wbl,htl,His3.3B,Rab5,Pdk1,cic,Tet,Hr39,mam,eya,Esp,Raf,Rab1,Dg,wnd,Arf79F,SCAR,RhoL,Gli,Pat1,Sxl,ss,ena,msi,Gr93c,ed,Cerk,ph-p,S,tipE,chic,Pen,Socs36E,lncRNA:Hsromega,Pvr,bol | | 8.784e-10 | -20.85 | sexual reproduction | biological process | GO:0019953 | 1300 | 89 | 12345 | 442 | ena,msi,Gr93c,ss,Sxl,bol,tipE,Pen,chic,Socs36E,lncRNA:Hsromega,Pvr,S,Cerk,ph-p,ed,His3.3B,Rab5,Pdk1,cic,mim,His3.3A,wbl,jar,htl,eff,Diap1,Poxn,arm,Dp,Pex10,kay,14-3-3epsilon,RhoL,SCAR,Pat1,Gli,Dg,wnd,Arf79F,mam,eya,Esp,Rab1,Raf,Tet,Hr39,aop,Egfr,LanA,cact,shg,dmrt11E,ttk,dome,scrib,N,Ppcs,14-3-3zeta,cindr,Ptpmeg2,orb2,InR,Mef2,Nlp,baz,Abd-B,tkv,larp,Cks30A,Cip4,tai,Abl,Dl,Inx2,lin-28,pum,CycG,PI31,abd-A,SERCA,sns,fz,yuri,lncRNA:CR44420,Dronc,Cbl,emc,mop | | 9.211e-10 | -20.81 | organic cyclic compound binding | molecular function | GO:0097159 | 3086 | 160 | 12176 | 414 | l(1)G0196,CG32772,Dyrk2,hng3,al,crp,Eph,ps,chn,CG3165,Sox14,CG31121,CG10321,Cdk4,pita,ey,GluProRS,hppy,klu,vig,bru2,bowl,bab2,Abl,CG14441,B-H1,msi,crol,CG1316,Pdk1,Abd-B,Slbp,Lmx1a,HmgZ,esg,orb2,Tis11,CG34376,CG13895,CG11247,Hr4,HLH3B,Hel89B,drl,lin-28,Ddr,His3.3A,su(sable),htl,rib,InR,sim,edl,dve,Pgk,CG11122,jigr1,SERCA,tkv,cic,RhoU,eyg,Raf,d,rgr,Su(z)2,CG5853,Nrk,grn,abd-A,CG12769,SmD1,Scr,mrt,Elba2,CG11617,Pitslre,SMC3,kay,sens-2,Pvr,Drak,otk,fz3,Hr39,CG7560,hang,Rm62,His3.3B,Myo10A,bol,sick,betaTub60D,bin3,jumu,Mef2,Arf79F,ttk,Dp,CG11360,Kdm2,Poxn,spen,aop,tsh,Prdm13,His2Av,CG14506,nub,Tet,CG2017,SNF4Agamma,CG4896,h,zfh2,nmo,apolpp,Src42A,CG11414,Adf1,ph-p,CG10348,ATPCL,Ric,Sxl,l(2)37Cg,CG11069,sqz,ap,CG17646,MEP-1,E(spl)m5-HLH,otp,Rab5,Dcr-2,lilli,CtBP,Egfr,CG45050,Lk6,kn,Psc,trbl,CG13204,pdm3,jar,Rab1,disco-r,dmrt11E,pnr,pum,mld,Npc2g,sd,CG3436,H2.0,sens,hth,wnd,Cyp6a16 | | 9.942e-10 | -20.73 | regulation of epidermal growth factor receptor signaling pathway | biological process | GO:0042058 | 49 | 14 | 12345 | 442 | rho-4,Src42A,kek5,S,mop,Cbl,hppy,ed,scrib,Ptp4E,Socs36E,step,pum,edl | | 9.942e-10 | -20.73 | regulation of ERBB signaling pathway | biological process | GO:1901184 | 49 | 14 | 12345 | 442 | step,pum,edl,Ptp4E,Socs36E,scrib,hppy,ed,mop,Cbl,kek5,S,rho-4,Src42A | | 1.026e-09 | -20.70 | positive regulation of border follicle cell migration | biological process | GO:1903688 | 34 | 12 | 12345 | 442 | Cbl,kay,tai,ttk,Diap1,spi,shg,Pvr,Egfr,vn,Gli,aop | | 1.104e-09 | -20.62 | multicellular organismal reproductive process | biological process | GO:0048609 | 1285 | 88 | 12345 | 442 | dmrt11E,shg,ttk,dome,aop,Egfr,cact,LanA,InR,14-3-3zeta,N,Ppcs,cindr,scrib,Ptpmeg2,orb2,Cks30A,larp,Cip4,Abl,tai,Mef2,baz,tkv,Abd-B,sns,fz,SERCA,yuri,lncRNA:CR44420,Dronc,emc,mop,Cbl,Dl,Inx2,pum,lin-28,PI31,abd-A,CycG,ss,Sxl,msi,ena,Gr93c,Cerk,ph-p,S,ed,bol,chic,Pen,tipE,lncRNA:Hsromega,Pvr,Socs36E,Diap1,eff,Dp,arm,Poxn,kay,Pex10,14-3-3epsilon,His3.3B,cic,Pdk1,Rab5,wbl,jar,His3.3A,mim,htl,eya,Esp,mam,Raf,Rab1,Tet,Hr39,SCAR,RhoL,Pat1,Gli,wnd,Dg,Arf79F | | 1.152e-09 | -20.58 | heterocyclic compound binding | molecular function | GO:1901363 | 3069 | 159 | 12176 | 414 | tsh,spen,aop,Prdm13,His2Av,CG14506,Tet,nub,CG4896,h,SNF4Agamma,CG2017,sick,bol,hang,CG7560,Rm62,His3.3B,Myo10A,betaTub60D,bin3,Mef2,jumu,Kdm2,CG11360,Poxn,Arf79F,Dp,ttk,trbl,pdm3,CG13204,jar,CG45050,Lk6,kn,Psc,pum,mld,disco-r,Rab1,dmrt11E,pnr,sd,H2.0,CG3436,Cyp6a16,sens,wnd,hth,Src42A,CG11414,zfh2,nmo,apolpp,ATPCL,Ric,Sxl,l(2)37Cg,Adf1,ph-p,CG10348,CG17646,MEP-1,E(spl)m5-HLH,CG11069,ap,sqz,CtBP,Egfr,otp,Dcr-2,Rab5,lilli,bru2,vig,bab2,bowl,hppy,klu,B-H1,CG14441,msi,crol,Abl,Abd-B,Slbp,CG1316,Pdk1,HmgZ,esg,Lmx1a,Dyrk2,hng3,l(1)G0196,CG32772,ps,chn,al,crp,Eph,CG10321,Cdk4,CG3165,Sox14,CG31121,pita,ey,GluProRS,eyg,Raf,grn,CG12769,abd-A,d,rgr,Nrk,Su(z)2,CG5853,CG11617,Pitslre,SMC3,SmD1,Scr,mrt,Elba2,Pvr,kay,sens-2,Drak,fz3,otk,Hr39,Tis11,CG34376,CG13895,CG11247,orb2,His3.3A,su(sable),htl,rib,InR,Hr4,HLH3B,Hel89B,drl,lin-28,Ddr,CG11122,Pgk,jigr1,dve,edl,sim,RhoU,tkv,SERCA,cic | | 1.156e-09 | -20.58 | wg (wingless) | protein interactions | 34009 | 120 | 24 | 9204 | 432 | mam,ey,sd,Raf,ft,Dg,N,CtBP,ds,pnr,ap,Dronc,tsh,nub,dome,fz3,spi,dlp,shg,spen,arm,fz2,apolpp,fz | | 1.319e-09 | -20.45 | positive regulation of oogenesis | biological process | GO:1905881 | 42 | 13 | 12345 | 442 | Pvr,Egfr,vn,Gli,aop,Cbl,kay,tai,ttk,Dp,shg,Diap1,spi | | 1.511e-09 | -20.31 | positive regulation of epithelial cell migration | biological process | GO:0010634 | 35 | 12 | 12345 | 442 | aop,Gli,Pvr,vn,Egfr,ttk,Diap1,spi,shg,kay,Cbl,tai | | 1.511e-09 | -20.31 | cell fate commitment involved in pattern specification | biological process | GO:0060581 | 35 | 12 | 12345 | 442 | CtBP,fz,klu,H,Tom,scrib,N,sens,abd-A,Egfr,vn,nuf | | 1.511e-09 | -20.31 | regulation of border follicle cell migration | biological process | GO:1903684 | 35 | 12 | 12345 | 442 | tai,Cbl,kay,shg,spi,Diap1,ttk,Egfr,vn,Pvr,aop,Gli | | 1.541e-09 | -20.29 | oogenesis | biological process | GO:0048477 | 682 | 57 | 12345 | 442 | pum,lin-28,Dg,Dl,Pat1,Inx2,RhoL,SCAR,mop,emc,Cbl,Dronc,Raf,eya,SERCA,fz,mam,tkv,mim,wbl,baz,jar,cic,Mef2,Rab5,kay,14-3-3epsilon,tai,Abl,Dp,arm,Cip4,Diap1,eff,lncRNA:Hsromega,Pvr,Socs36E,chic,Pen,Ptpmeg2,cindr,Ppcs,14-3-3zeta,N,scrib,ed,InR,ph-p,S,cact,ena,Egfr,aop,Sxl,dome,ttk,shg | | 1.618e-09 | -20.24 | mam (mastermind) | protein interactions | 36555 | 69 | 18 | 9204 | 432 | Scr,Abd-B,emc,arm,fz2,pum,Dl,abd-A,tara,H,Lk6,Mfe2,lilli,N,Raf,wdb,Fas1,klu | | 1.621e-09 | -20.24 | cell-cell junction | cellular component | GO:0005911 | 101 | 19 | 11940 | 417 | Inx2,Gli,Dl,ena,ogre,otk,moody,Shrm,arm,sns,yrt,ed,scrib,cora,Abl,baz,step,shg,N | | 1.736e-09 | -20.17 | maintenance of cell number | biological process | GO:0098727 | 99 | 19 | 12345 | 442 | His2Av,arm,lilli,shg,eff,Dad,tkv,msi,Egfr,esg,klu,H,InR,mam,Socs36E,chic,Dl,Inx2,N | | 1.736e-09 | -20.17 | stem cell population maintenance | biological process | GO:0019827 | 99 | 19 | 12345 | 442 | esg,Egfr,msi,Dad,tkv,shg,eff,lilli,arm,His2Av,N,Dl,Inx2,chic,Socs36E,mam,InR,H,klu | | 1.751e-09 | -20.16 | border follicle cell migration | biological process | GO:0007298 | 110 | 20 | 12345 | 442 | RhoL,cindr,N,Dl,Ptpmeg2,Socs36E,Pvr,fz,Raf,Rab5,aop,Egfr,baz,ena,mim,jar,shg,Diap1,dome,tai | | 1.896e-09 | -20.08 | Mad (Mothers against dpp) | protein interactions | 33529 | 104 | 22 | 9204 | 432 | Actbeta,scrib,bowl,Snoo,fuss,CG45050,Tob,CG6891,Rab5,lqf,rib,abd-A,nmo,shg,arm,pnr,Abd-B,Dad,CycG,tkv,eya,CG14657 | | 1.951e-09 | -20.05 | cell adhesion | biological process | GO:0007155 | 171 | 25 | 12345 | 442 | beat-IIa,Gli,scrib,SCAR,Bsg,otk,fz,sns,ds,mop,ed,glec,cora,crol,LanA,baz,Ddr,Egfr,Cals,Src42A,arm,Fas1,shg,ft,p130CAS | | 2.001e-09 | -20.03 | aop (anterior open) | protein interactions | 33392 | 62 | 17 | 9204 | 432 | aop,eya,B-H1,ed,edl,hppy,arm,spen,shg,Syx1A,H,Egfr,ttk,lilli,N,S,Raf | | 2.184e-09 | -19.94 | germ cell development | biological process | GO:0007281 | 820 | 64 | 12345 | 442 | Dl,Inx2,lin-28,pum,SERCA,sns,fz,yuri,Dronc,Cbl,mop,emc,Mef2,baz,tkv,Cip4,tai,Abl,scrib,N,Ppcs,cindr,14-3-3zeta,Ptpmeg2,orb2,InR,aop,Egfr,cact,shg,ttk,dome,SCAR,RhoL,Pat1,Dg,mam,eya,Raf,Tet,Rab5,cic,wbl,mim,jar,eff,Diap1,arm,Dp,14-3-3epsilon,Pex10,kay,bol,chic,Pen,Socs36E,Pvr,lncRNA:Hsromega,S,ph-p,ed,ena,msi,Sxl | | 2.242e-09 | -19.92 | shg (shotgun) | protein interactions | 37386 | 87 | 20 | 9204 | 432 | RhoL,lilli,cindr,ey,N,Src42A,S,ft,baz,Myo10A,ec,ds,aop,Inx2,tkv,Egfr,step,arm,nmo,jar | | 2.444e-09 | -19.83 | wing disc pattern formation | biological process | GO:0035222 | 80 | 17 | 12345 | 442 | Dl,N,nuf,ap,ds,H,dlp,cic,spen,nmo,tkv,Mpcp2,tara,boi,lilli,tsh,14-3-3epsilon | | 2.471e-09 | -19.82 | heart development | biological process | GO:0007507 | 101 | 19 | 12345 | 442 | betaTub60D,Gli,abd-A,mam,fz,sli,sd,Mef2,robo1,cora,tkv,LanA,Abd-B,htl,shg,pnr,arm,jumu,spdo | | 2.676e-09 | -19.74 | positive regulation of RNA metabolic process | biological process | GO:0051254 | 513 | 47 | 12345 | 442 | Tet,Hr39,CG12769,hyx,Lmx1a,mam,eya,hth,hng3,pum,fs(1)h,abd-A,ps,sens,Tis11,tai,kay,ey,Adf1,pnr,arm,CtBP,rictor,Abd-B,Mef2,kn,nub,bowl,ap,tna,N,zfh2,CG13204,chn,Scr,dmrt11E,tsh,lilli,CG8312,ttk,ss,tara,esg,REPTOR,sim,B-H1,sd | | 2.730e-09 | -19.72 | blastoderm segmentation | biological process | GO:0007350 | 214 | 28 | 12345 | 442 | dome,shg,pnr,tsh,fz2,lilli,arm,Egfr,baz,Abd-B,cic,rdx,dlp,kn,fz,Raf,h,hth,bowl,Dg,chic,pum,frc,fs(1)h,scrib,N,Cdk4,Pat1 | | 2.744e-09 | -19.71 | regulation of developmental growth | biological process | GO:0048638 | 228 | 29 | 12345 | 442 | pum,chic,Sema1b,shot,scrib,dlp,Orct2,Mob2,d,fz,Sema1a,InR,Raf,step,rictor,wbl,Egfr,ena,nmo,tkv,Dad,LanA,Hr4,Pdk1,Abl,ft,ey,Cip4,Src42A | | 2.756e-09 | -19.71 | multi-organism process | biological process | GO:0051704 | 1653 | 104 | 12345 | 442 | aop,spen,cact,LanA,sick,Egfr,ttk,RhoGEF3,dmrt11E,CG32795,shg,dome,N,14-3-3zeta,Ppcs,cindr,DPCoAC,scrib,orb2,Ptpmeg2,InR,Nlp,Mef2,tkv,Dcr-2,Abd-B,baz,Tollo,Cip4,Cks30A,larp,Abl,tai,Inx2,Dl,CycG,abd-A,PI31,pum,lin-28,yuri,fz,SERCA,sns,mop,emc,Cbl,lncRNA:CR44420,Dronc,Gr93c,msi,ena,ss,jumu,Src42A,Sxl,bol,Mid1,Pvr,lncRNA:Hsromega,Rm62,Socs36E,chic,Pen,tipE,Cerk,ph-p,S,kn,ed,cic,Pdk1,Rab5,His3.3B,subdued,htl,His3.3A,jar,mim,wbl,arm,Dp,Poxn,Diap1,eff,14-3-3epsilon,Pex10,kay,Gli,Pat1,SCAR,RhoL,imd,Arf79F,wnd,Eph,Dg,Raf,Rab1,Esp,eya,mam,Hr39,Tet | | 2.789e-09 | -19.70 | haltere development | biological process | GO:0007482 | 13 | 8 | 12345 | 442 | Syx1A,drl,Socs36E,Egfr,vn,ap,hth,Alh | | 2.821e-09 | -19.69 | biological adhesion | biological process | GO:0022610 | 174 | 25 | 12345 | 442 | beat-IIa,Gli,scrib,SCAR,Bsg,otk,sns,fz,ds,mop,ed,glec,cora,crol,LanA,baz,Ddr,Egfr,Src42A,Cals,Fas1,arm,shg,ft,p130CAS | | 2.929e-09 | -19.65 | regulation of phosphate metabolic process | biological process | GO:0019220 | 302 | 34 | 12345 | 442 | Atg17,CycG,Dg,trbl,wnd,Actbeta,Cbl,Raf,CycB3,Dad,rictor,Pdk1,kay,SNF4Agamma,wdb,Cks30A,Socs36E,Pgk,lncRNA:Hsromega,Pvr,vn,edl,Lk6,N,jeb,hppy,InR,Egfr,CG34172,step,rdx,CG8312,Src42A,spi | | 2.929e-09 | -19.65 | regulation of phosphorus metabolic process | biological process | GO:0051174 | 302 | 34 | 12345 | 442 | Pdk1,Dad,rictor,Cks30A,kay,SNF4Agamma,wdb,Actbeta,Atg17,CycG,Dg,trbl,wnd,Raf,CycB3,Cbl,rdx,Egfr,CG34172,step,Src42A,CG8312,spi,Lk6,N,Pgk,Socs36E,lncRNA:Hsromega,Pvr,vn,edl,InR,jeb,hppy | | 2.938e-09 | -19.65 | cell-cell adhesion | biological process | GO:0098609 | 102 | 19 | 12345 | 442 | Fas1,arm,Src42A,Cals,shg,ft,p130CAS,glec,Egfr,baz,ds,fz,sns,mop,ed,Gli,beat-IIa,scrib,Bsg | | 3.040e-09 | -19.61 | regulation of cell population proliferation | biological process | GO:0042127 | 229 | 29 | 12345 | 442 | fz,Raf,InR,nub,hth,Dronc,klu,Cbl,scrib,N,Dl,lin-28,vn,Pvr,pxb,fz2,spi,tsh,dome,Abl,ey,ft,esg,Rab5,Pez,sd,Ddr,Egfr,htl | | 3.095e-09 | -19.59 | ion binding | molecular function | GO:0043167 | 2058 | 117 | 12176 | 414 | otk,robo1,Hr39,shg,Atox1,Pvr,Drak,Cbl,wnd,Cyp6a16,CG13481,Pp2C1,CenB1A,CG31457,Pitslre,SMC3,Su(z)2,CG5853,Nrk,pnr,mim,d,Rab1,grn,mld,CG10417,sli,Psc,CG45050,Raf,Lk6,jar,rho-4,trbl,Dg,tkv,SERCA,Dcr-2,Rab5,lt,Egfr,RhoU,dpy,FBXO11,CG11069,sina,CG3107,Pgk,CG17646,drl,ph-p,lin-28,baz,Ddr,Hr4,CalpC,Hel89B,N,htl,InR,ATPCL,su(sable),vn,Ric,CG34401,apolpp,Fkbp14,shot,nmo,CG11247,Src42A,Tis11,CG11414,CG2017,Cip4,Tet,Pdk1,CG2991,Abl,Diap1,CG4467,Psa,CG14441,mura,hppy,Pex10,CG7536,pita,GluProRS,Arf79F,Sema1a,Dyb,CG11360,Kdm2,alpha-Man-Ia,CG31121,Cdk4,Cadps,ds,CG10321,Eph,betaTub60D,Dl,CG1218,ft,step,Rm62,l(1)G0196,Myo10A,hang,CG7560,Cals,cue,cv-d,tna,Dyrk2,sick | | 3.142e-09 | -19.58 | positive regulation of epithelial cell differentiation | biological process | GO:0030858 | 37 | 12 | 12345 | 442 | aop,Gli,Pvr,vn,Egfr,ttk,spi,Diap1,shg,kay,Cbl,tai | | 3.235e-09 | -19.55 | negative regulation of animal organ morphogenesis | biological process | GO:0110111 | 30 | 11 | 12345 | 442 | N,aop,Egfr,Abd-B,Socs36E,tkv,Diap1,tsh,ttk,chn,ed | | 3.252e-09 | -19.54 | Dl (Delta) | protein interactions | 42313 | 126 | 24 | 9204 | 432 | sens,Fkbp14,lqf,mam,S,N,px,eyg,Dg,cic,boi,ed,ap,tsh,CycG,rg,Dad,CG34376,nmo,emc,fz,Egfr,H,Dl | | 3.365e-09 | -19.51 | embryonic pattern specification | biological process | GO:0009880 | 230 | 29 | 12345 | 442 | Cdk4,Pat1,scrib,N,fs(1)h,frc,Dg,edl,bowl,pum,chic,h,Raf,hth,fz,kn,dlp,rdx,cic,Abd-B,baz,Egfr,lilli,arm,shg,pnr,tsh,fz2,dome | | 3.706e-09 | -19.41 | positive regulation of gene expression | biological process | GO:0010628 | 571 | 50 | 12345 | 442 | nub,kn,tna,N,zfh2,bol,bowl,ap,orb2,Scr,dmrt11E,tsh,lilli,CG8312,ttk,ss,chn,REPTOR,esg,sim,B-H1,sd,aop,Egfr,tara,mam,Lmx1a,eya,hth,Tet,hyx,CG12769,Hr39,lin-28,pum,fs(1)h,sens,abd-A,ps,pnr,arm,Dp,CtBP,tai,kay,ey,Adf1,Mef2,rictor,Abd-B | | 4.125e-09 | -19.31 | circulatory system development | biological process | GO:0072359 | 104 | 19 | 12345 | 442 | sli,cora,sd,Mef2,robo1,LanA,Abd-B,htl,tkv,shg,pnr,arm,jumu,spdo,betaTub60D,Gli,abd-A,fz,mam | | 4.552e-09 | -19.21 | positive regulation of intracellular signal transduction | biological process | GO:1902533 | 178 | 25 | 12345 | 442 | ed,hppy,jeb,ds,Raf,InR,vn,wnd,Pvr,CycG,siz,lncRNA:Hsromega,14-3-3zeta,Cdk4,RhoGEF4,14-3-3epsilon,ft,spi,Src42A,rictor,Egfr,step,Pdk1,Pez,rdx | | 4.704e-09 | -19.17 | dendrite morphogenesis | biological process | GO:0048813 | 140 | 22 | 12345 | 442 | Dronc,E(Pc),Snoo,kn,Sema1a,nub,pum,shot,fs(1)h,drl,Actbeta,Abl,chn,Sox14,kay,path,sqz,rictor,ena,sli,robo1,Rab5 | | 4.818e-09 | -19.15 | Pc (Polycomb) | protein interactions | 40358 | 109 | 22 | 9204 | 432 | His2Av,lawc,Abd-B,Scr,dome,eff,tara,Adf1,Rm62,SMC3,tna,Kdm2,sens,corto,sd,ph-p,rdx,Psc,stx,Su(z)2,E(Pc),fs(1)h | | 4.982e-09 | -19.12 | positive regulation of nucleobase-containing compound metabolic process | biological process | GO:0045935 | 523 | 47 | 12345 | 442 | Tis11,hng3,pum,fs(1)h,sens,abd-A,ps,mam,Lmx1a,eya,hth,Tet,CG12769,Hr39,hyx,Mef2,rictor,Abd-B,pnr,CtBP,arm,tai,ey,kay,Adf1,tna,N,zfh2,bowl,ap,nub,kn,esg,sim,REPTOR,B-H1,sd,tara,Scr,dmrt11E,tsh,ttk,CG8312,lilli,ss,CG13204,chn | | 5.507e-09 | -19.02 | adherens junction | cellular component | GO:0005912 | 97 | 18 | 11940 | 417 | arm,yrt,Src42A,kon,Shrm,ena,LRR,p130CAS,Dl,Magi,N,shg,step,baz,Abl,scrib,shot,ed | | 5.507e-09 | -19.02 | anchoring junction | cellular component | GO:0070161 | 97 | 18 | 11940 | 417 | step,shg,baz,Magi,N,shot,ed,Abl,scrib,Shrm,yrt,arm,Src42A,kon,p130CAS,Dl,ena,LRR | | 6.304e-09 | -18.88 | haltere disc development | biological process | GO:0035216 | 14 | 8 | 12345 | 442 | Syx1A,drl,Socs36E,ap,vn,Egfr,hth,Alh | | 6.349e-09 | -18.87 | MAPK signaling pathway - fly | KEGG pathways | ko04013 | 93 | 18 | 3198 | 123 | Raf,sina,aop,RhoL,vn,Src42A,ttk,Mef2,chic,Egfr,kay,edl,eff,tkv,14-3-3zeta,cic,wgn,spi | | 6.349e-09 | -18.87 | MAPK signaling pathway - fly | KEGG pathways | dme04013 | 93 | 18 | 3198 | 123 | tkv,eff,edl,spi,wgn,cic,14-3-3zeta,Src42A,ttk,chic,Mef2,vn,RhoL,aop,Raf,sina,kay,Egfr | | 7.174e-09 | -18.75 | aos (argos) | protein interactions | 39833 | 59 | 16 | 9204 | 432 | rg,ed,Cbl,vn,Egfr,H,Dl,spi,arm,nmo,fz,Diap1,Raf,Dcr-2,S,Dg | | 8.077e-09 | -18.63 | peripheral nervous system development | biological process | GO:0007422 | 66 | 15 | 12345 | 442 | chn,spdo,spi,Tollo,ttk,hth,Eph,Egfr,vn,sens,abd-A,spen,N,Gli,Dl | | 8.162e-09 | -18.62 | cellular response to stimulus | biological process | GO:0051716 | 1467 | 94 | 12345 | 442 | vn,Ir21a,Magi,N,RhoGAP19D,drl,nAChRalpha4,hppy,CG30156,lt,InR,Egfr,Ocrl,cact,sina,spen,aop,rdx,dome,fz2,spi,tsh,ttk,Sema1b,CG7536,Atg17,Rgl,CycG,moody,Dl,Dronc,Cbl,CG13481,fz,hang,Ric,Sema1a,rictor,Ddr,ReepA,tkv,Atg16,robo1,flfl,SNF4Agamma,tai,pnr,Pvf3,Tollo,tipE,Socs36E,Pvr,RhoU,Lk6,CG42788,jeb,Cerk,koi,hfw,CG1218,msi,wgn,sli,REPTOR,sd,Src42A,ss,otk,Eph,wnd,Arf79F,imd,Actbeta,Nrk,ITP,Hr39,mam,unc-5,Rab1,Raf,boi,htl,nmo,Dad,Hr4,Pdk1,ogre,Rab5,cic,fz3,kay,14-3-3epsilon,Dp,arm | | 8.713e-09 | -18.56 | positive regulation of reproductive process | biological process | GO:2000243 | 57 | 14 | 12345 | 442 | ttk,Dp,shg,spi,Diap1,Cbl,kay,tai,Gli,aop,CG9784,Pvr,vn,Egfr | | 8.951e-09 | -18.53 | regulation of protein modification process | biological process | GO:0031399 | 347 | 36 | 12345 | 442 | InR,CG1218,jeb,hppy,Lk6,N,sens-2,Socs36E,lncRNA:Hsromega,Pvr,vn,Src42A,CG8312,spi,Su(z)2,rdx,Egfr,step,Raf,CycB3,Cbl,Fem-1,Actbeta,Atg17,Kdm2,CycG,trbl,wnd,Tollo,Diap1,Cks30A,kay,SNF4Agamma,wdb,Pdk1,rictor | | 9.188e-09 | -18.51 | dendrite development | biological process | GO:0016358 | 145 | 22 | 12345 | 442 | robo1,Rab5,sli,ena,rictor,sqz,path,Sox14,kay,chn,Abl,drl,Actbeta,shot,fs(1)h,pum,Sema1a,nub,kn,Dronc,E(Pc),Snoo | | 1.176e-08 | -18.26 | regulation of epithelial cell differentiation | biological process | GO:0030856 | 41 | 12 | 12345 | 442 | aop,Gli,Pvr,Egfr,vn,ttk,spi,Diap1,shg,kay,Cbl,tai | | 1.190e-08 | -18.25 | Su(H) (Suppressor of Hairless) | protein interactions | 34881 | 69 | 17 | 9204 | 432 | mam,lilli,Elba2,Tom,CtBP,N,eyg,Src42A,Abd-B,rg,aop,Scr,vn,Dl,abd-A,H,arm | | 1.285e-08 | -18.17 | positive regulation of transcription by RNA polymerase II | biological process | GO:0045944 | 368 | 37 | 12345 | 442 | bowl,ap,N,kn,nub,sim,esg,B-H1,sd,chn,Scr,tsh,dmrt11E,CG8312,ttk,lilli,ss,fs(1)h,abd-A,sens,Tet,CG12769,Hr39,hyx,mam,Lmx1a,eya,hth,Abd-B,Mef2,tai,ey,kay,Adf1,pnr,arm,CtBP | | 1.405e-08 | -18.08 | transmembrane receptor protein kinase activity | molecular function | GO:0019199 | 28 | 10 | 12176 | 414 | otk,tkv,drl,Nrk,Eph,Ddr,Pvr,htl,Egfr,InR | | 1.438e-08 | -18.06 | regulation of protein phosphorylation | biological process | GO:0001932 | 245 | 29 | 12345 | 442 | wnd,vn,trbl,CycG,lncRNA:Hsromega,Pvr,Atg17,Socs36E,N,Lk6,Actbeta,hppy,jeb,Cbl,CycB3,InR,Raf,step,Egfr,rictor,rdx,Pdk1,wdb,SNF4Agamma,kay,spi,Cks30A,CG8312,Src42A | | 1.485e-08 | -18.03 | organelle organization | biological process | GO:0006996 | 1885 | 112 | 12345 | 442 | 14-3-3epsilon,Pex10,His2Av,vig,CtBP,eff,yrt,Diap1,Slbp,jar,Pex7,His3.3A,crol,Rab5,kon,His3.3B,Hr39,E(Pc),Nrk,fl(2)d,Rab1,HmgZ,rib,eya,Kdm2,Arf79F,CG33474,Dg,wnd,Gli,CG7394,SCAR,Pex14,Sxl,Src42A,klar,jumu,ena,cora,bin3,f,Psc,gukh,ed,dve,ph-p,lncRNA:Hsromega,shot,RhoU,chic,Cdk4,corto,tna,ft,wdb,Shrm,larp,Cks30A,AP-1gamma,Dcr-2,tkv,rictor,baz,ReepA,Atg16,Syx1A,Nlp,LRR,mop,hyx,yuri,sns,ds,Rim2,Atg17,siz,CycG,PI31,nuf,Dgat2,Pex5,moody,Dl,Hel89B,Spc105R,p130CAS,Su(z)2,RhoGEF3,Jupiter,tara,SMC3,LanA,sick,CG34172,B-H1,Kdm4B,CG14562,unc-119,hppy,Wdr62,lt,nub,Mob2,Bsg,orb2,FBXO11,Elba2,14-3-3zeta,cindr,betaTub60D,Ppcs,N | | 1.508e-08 | -18.01 | positive regulation of cell population proliferation | biological process | GO:0008284 | 79 | 16 | 12345 | 442 | ey,dome,spi,tsh,htl,Egfr,Ddr,klu,InR,hth,Raf,Pvr,vn,lin-28,Dl,N | | 1.959e-08 | -17.75 | learning or memory | biological process | GO:0007611 | 164 | 23 | 12345 | 442 | mura,Mob2,drl,scrib,14-3-3zeta,N,Socs36E,orb2,pxb,vn,wnd,pum,lilli,arm,ss,spi,rg,Tob,Adf1,dome,Abl,mld,Egfr | | 1.959e-08 | -17.75 | cognition | biological process | GO:0050890 | 164 | 23 | 12345 | 442 | mld,Egfr,rg,Tob,Adf1,dome,Abl,lilli,ss,arm,spi,Socs36E,orb2,pxb,vn,wnd,pum,drl,scrib,14-3-3zeta,N,mura,Mob2 | | 2.073e-08 | -17.69 | regulation of anatomical structure size | biological process | GO:0090066 | 249 | 29 | 12345 | 442 | Gli,SCAR,CG6191,RhoU,shot,Pvr,trbl,CG6891,Sema1b,chic,InR,Sema1a,fz,rib,ds,H,hppy,cora,Pdk1,ena,rictor,Cip4,lilli,ttk,path,Pvf3,dpy,ft,Abl | | 2.085e-08 | -17.69 | positive regulation of nucleic acid-templated transcription | biological process | GO:1903508 | 494 | 44 | 12345 | 442 | hth,eya,Lmx1a,mam,hyx,Hr39,CG12769,Tet,abd-A,sens,fs(1)h,hng3,CtBP,arm,pnr,Adf1,kay,ey,tai,Mef2,Abd-B,rictor,nub,kn,zfh2,tna,N,ap,bowl,ss,ttk,CG8312,lilli,tsh,dmrt11E,Scr,chn,CG13204,B-H1,sd,sim,REPTOR,esg,tara | | 2.085e-08 | -17.69 | positive regulation of RNA biosynthetic process | biological process | GO:1902680 | 494 | 44 | 12345 | 442 | bowl,ap,tna,N,zfh2,kn,nub,tara,REPTOR,sim,esg,sd,B-H1,CG13204,chn,Scr,dmrt11E,tsh,ttk,lilli,CG8312,ss,hng3,fs(1)h,sens,abd-A,Tet,Hr39,CG12769,hyx,mam,Lmx1a,eya,hth,rictor,Abd-B,Mef2,tai,kay,ey,Adf1,pnr,arm,CtBP | | 2.140e-08 | -17.66 | eye-antennal disc morphogenesis | biological process | GO:0007455 | 43 | 12 | 12345 | 442 | bab2,ey,eya,Diap1,ss,Poxn,Egfr,N,al,Dl,B-H1,f | | 2.199e-08 | -17.63 | chaeta development | biological process | GO:0022416 | 81 | 16 | 12345 | 442 | hyx,emc,Dronc,sens,Fkbp14,Dl,N,Ppcs,CtBP,pnr,Diap1,msi,f,B-H1,al,spen | | 2.199e-08 | -17.63 | leg disc morphogenesis | biological process | GO:0007478 | 81 | 16 | 12345 | 442 | bab2,Poxn,ss,tsh,sd,al,hyx,dve,hth,ds,Drak,bowl,ap,RhoGAP19D,Dl,N | | 2.301e-08 | -17.59 | proximal/distal pattern formation | biological process | GO:0009954 | 28 | 10 | 12345 | 442 | tsh,hth,ap,vn,Egfr,spen,N,Gli,B-H1,cora | | 2.336e-08 | -17.57 | eye-antennal disc development | biological process | GO:0035214 | 71 | 15 | 12345 | 442 | eyg,Poxn,ss,tsh,eya,Diap1,ey,bab2,f,B-H1,Dl,al,N,Dad,Egfr | | 2.336e-08 | -17.57 | regulation of oogenesis | biological process | GO:1905879 | 71 | 15 | 12345 | 442 | Diap1,spi,shg,Dp,klar,ttk,tai,Cbl,kay,aop,Gli,Syx1A,vn,Egfr,Pvr | | 2.720e-08 | -17.42 | positive regulation of macromolecule biosynthetic process | biological process | GO:0010557 | 552 | 47 | 12345 | 442 | CG13204,chn,Scr,dmrt11E,tsh,ttk,lilli,CG8312,ss,tara,REPTOR,sim,esg,B-H1,sd,kn,nub,bowl,ap,orb2,tna,N,zfh2,bol,tai,kay,ey,Adf1,pnr,arm,CtBP,rictor,Abd-B,Mef2,Tet,Hr39,hyx,CG12769,Lmx1a,mam,eya,hth,hng3,pum,fs(1)h,sens,abd-A | | 2.981e-08 | -17.33 | transmembrane receptor protein tyrosine kinase activity | molecular function | GO:0004714 | 23 | 9 | 12176 | 414 | Pvr,Ddr,Nrk,Eph,otk,drl,Egfr,InR,htl | | 3.234e-08 | -17.25 | protein tyrosine kinase activity | molecular function | GO:0004713 | 38 | 11 | 12176 | 414 | Src42A,Nrk,Ddr,Eph,drl,Abl,otk,Pvr,InR,Egfr,htl | | 4.035e-08 | -17.03 | peptidyl-tyrosine modification | biological process | GO:0018212 | 37 | 11 | 12345 | 442 | InR,Src42A,Nrk,Abl,drl,Ddr,Ptpmeg2,Eph,htl,Tpst,Pvr | | 4.214e-08 | -16.98 | actin cytoskeleton organization | biological process | GO:0030036 | 242 | 28 | 12345 | 442 | nuf,chic,Dg,siz,shot,RhoU,Bsg,SCAR,Ppcs,cindr,N,Dl,moody,ed,gukh,sns,rictor,jar,sick,ena,cora,f,kon,Shrm,p130CAS,yrt,Src42A,RhoGEF3 | | 4.433e-08 | -16.93 | reproduction | biological process | GO:0000003 | 1651 | 100 | 12345 | 442 | Sxl,bab2,ss,ena,msi,Gr93c,sli,dve,ed,ph-p,S,Cerk,tipE,Pen,chic,Socs36E,lncRNA:Hsromega,Pvr,bol,kay,14-3-3epsilon,Pex10,eff,Diap1,Poxn,arm,Dp,wbl,His3.3A,mim,jar,htl,His3.3B,scyl,Pdk1,Rab5,cic,Tet,fl(2)d,Hr39,mam,rib,eya,Esp,Raf,Rab1,otk,Dg,wnd,Arf79F,SCAR,RhoL,Pat1,Gli,dome,shg,Scr,dmrt11E,fz2,ttk,Egfr,LanA,cact,aop,InR,Ptpmeg2,orb2,scrib,Ppcs,14-3-3zeta,cindr,N,tai,Abl,larp,Acp63F,Cks30A,Cip4,baz,Abd-B,tkv,robo1,Nlp,Mef2,Dronc,lncRNA:CR44420,Cbl,mop,emc,fz,SERCA,sns,yuri,lin-28,pum,abd-A,PI31,CycG,Inx2,Dl | | 4.854e-08 | -16.84 | protein dimerization activity | molecular function | GO:0046983 | 390 | 36 | 12176 | 414 | h,subdued,otk,shg,kay,rdx,SMC3,tai,Scr,His2Av,pnr,mim,wbl,trbl,kn,CtBP,ttk,ss,E(spl)m5-HLH,Mef2,emc,dome,Mfe2,REPTOR,sim,l(2)37Cg,His3.3A,step,crp,drl,14-3-3epsilon,HLH3B,14-3-3zeta,corto,Dronc,His3.3B | | 4.869e-08 | -16.84 | positive regulation of transcription, DNA-templated | biological process | GO:0045893 | 473 | 42 | 12345 | 442 | Hr39,hyx,CG12769,Tet,hth,Lmx1a,mam,eya,fs(1)h,abd-A,sens,ey,kay,Adf1,tai,arm,CtBP,pnr,Abd-B,rictor,Mef2,kn,nub,bowl,ap,zfh2,N,tna,chn,ttk,lilli,CG8312,ss,Scr,tsh,dmrt11E,tara,B-H1,sd,REPTOR,sim,esg | | 5.075e-08 | -16.80 | dgo (diego) | protein interactions | 36139 | 24 | 10 | 9204 | 432 | shg,nmo,baz,fz,Dl,kn,ed,aop,ds,pnr | | 5.126e-08 | -16.79 | larval development | biological process | GO:0002164 | 159 | 22 | 12345 | 442 | scrib,N,Pvr,Kdm2,ap,chic,Raf,fz,sns,eya,kn,crp,htl,Dad,jar,CG1129,arm,klar,fz2,ft,Alh,chn | | 6.430e-08 | -16.56 | actin filament-based process | biological process | GO:0030029 | 262 | 29 | 12345 | 442 | RhoGEF3,Src42A,yrt,p130CAS,Shrm,f,kon,cora,rictor,sick,jar,ena,sns,gukh,ed,moody,Dl,Ppcs,cindr,SCAR,N,Bsg,neo,RhoU,shot,siz,Dg,nuf,chic | | 6.598e-08 | -16.53 | Pkinase\_Tyr | pfam domains | PF07714 | 51 | 13 | 10480 | 419 | wnd,Abl,InR,Egfr,Raf,Pvr,Ddr,Eph,Nrk,otk,drl,Src42A,htl | | 6.697e-08 | -16.52 | transcription factor binding | molecular function | GO:0008134 | 184 | 23 | 12176 | 414 | eya,tna,hyx,14-3-3zeta,bin3,14-3-3epsilon,pnr,Scr,edl,tai,sd,emc,hth,sens,kay,cact,cic,arm,lilli,Dp,HmgZ,CtBP,h | | 6.866e-08 | -16.49 | TYRKINASE | prints domains | PR00109 | 32 | 11 | 2185 | 101 | Egfr,drl,otk,htl,wnd,Src42A,InR,Abl,Eph,Nrk,Ddr | | 6.915e-08 | -16.49 | upd1 (unpaired 1) | protein interactions | 32813 | 77 | 17 | 9204 | 432 | Dl,Socs36E,Egfr,dome,tkv,His2Av,Dad,esg,CtBP,N,eyg,px,Cdk4,Mob2,Cip4,trbl,lilli | | 7.262e-08 | -16.44 | GO\_EPITHELIAL\_CELL\_DEVELOPMENT | MSigDB lists | GO\_EPITHELIAL\_CELL\_DEVELOPMENT | 38 | 11 | 4161 | 156 | Cdk4,Dl,CG17883,C1GalTA,Fem-1,N,Pdk1,sns,arm,Mfe2,Rab1 | | 7.396e-08 | -16.42 | imaginal disc-derived leg morphogenesis | biological process | GO:0007480 | 77 | 15 | 12345 | 442 | ap,Drak,bowl,RhoGAP19D,Dl,sd,al,N,hyx,dve,ss,hth,Poxn,ds,tsh | | 7.482e-08 | -16.41 | positive regulation of multi-organism process | biological process | GO:0043902 | 149 | 21 | 12345 | 442 | Pitslre,mop,Cbl,Gli,imd,DPCoAC,Pvr,vn,Dp,ttk,spi,Diap1,pnr,CG32795,shg,kay,tai,aop,Dcr-2,htl,Egfr | | 7.518e-08 | -16.40 | Ser-Thr/Tyr\_kinase\_cat\_dom | interpro domains | IPR001245 | 52 | 13 | 11383 | 450 | Eph,htl,Pvr,InR,Raf,Src42A,otk,Egfr,Nrk,drl,Ddr,Abl,wnd | | 7.751e-08 | -16.37 | dorsal/ventral pattern formation, imaginal disc | biological process | GO:0007450 | 57 | 13 | 12345 | 442 | 14-3-3epsilon,H,dlp,lilli,tsh,nmo,tara,Mpcp2,ap,nuf,cic,Dl,N | | 8.277e-08 | -16.31 | mesoderm development | biological process | GO:0007498 | 100 | 17 | 12345 | 442 | sli,Mef2,htl,LanA,Abd-B,meso18E,shg,Cip4,RhoL,N,Dl,Dgat2,abd-A,Npc2g,eya,mam,jeb | | 8.736e-08 | -16.25 | lilli (lilliputian) | protein interactions | 33496 | 38 | 12 | 9204 | 432 | Raf,H,Egfr,N,sina,arm,shg,Pvr,mam,aop,ttk,sens | | 8.861e-08 | -16.24 | urogenital system development | biological process | GO:0001655 | 78 | 15 | 12345 | 442 | hth,tsh,spi,rib,pnr,sns,S,N,scrib,Dad,tkv,Pvr,htl,baz,Egfr | | 8.861e-08 | -16.24 | renal system development | biological process | GO:0072001 | 78 | 15 | 12345 | 442 | hth,rib,spi,tsh,sns,pnr,S,tkv,Pvr,Dad,htl,Egfr,baz,N,scrib | | 9.135e-08 | -16.21 | nmo (nemo) | protein interactions | 38890 | 53 | 14 | 9204 | 432 | Dl,spi,Egfr,H,N,px,arm,nmo,fz2,shg,ec,Dad,eya,tkv | | 9.241e-08 | -16.20 | negative regulation of transcription by RNA polymerase II | biological process | GO:0000122 | 221 | 26 | 12345 | 442 | arm,ss,CtBP,ttk,tsh,pnr,Prdm13,E(spl)m5-HLH,MEP-1,chn,Su(z)2,B-H1,cic,aop,esg,eyg,h,emc,klu,dve,H,zfh2,scrib,sens,abd-A,bowl | | 9.668e-08 | -16.15 | positive regulation of biosynthetic process | biological process | GO:0009891 | 594 | 48 | 12345 | 442 | Tet,hyx,CG12769,Hr39,Lmx1a,mam,eya,hth,hng3,pum,fs(1)h,abd-A,sens,imd,tai,ey,kay,Adf1,pnr,arm,CtBP,rictor,Abd-B,Mef2,kn,nub,bowl,ap,orb2,N,tna,zfh2,bol,CG13204,chn,Scr,tsh,dmrt11E,CG8312,lilli,ttk,ss,tara,sim,esg,REPTOR,B-H1,sd | | 9.668e-08 | -16.15 | positive regulation of cellular biosynthetic process | biological process | GO:0031328 | 594 | 48 | 12345 | 442 | zfh2,bol,tna,N,orb2,bowl,ap,nub,kn,sd,B-H1,REPTOR,esg,sim,tara,lilli,CG8312,ttk,ss,Scr,dmrt11E,tsh,CG13204,chn,imd,fs(1)h,sens,abd-A,hng3,pum,hth,mam,Lmx1a,eya,CG12769,Hr39,hyx,Tet,Mef2,Abd-B,rictor,arm,CtBP,pnr,ey,kay,Adf1,tai | | 9.899e-08 | -16.13 | regulation of cell fate commitment | biological process | GO:0010453 | 40 | 11 | 12345 | 442 | lqf,Abd-B,Socs36E,N,aop,chn,ed,eya,hth,ttk,Src42A | | 1.082e-07 | -16.04 | positive regulation of phosphorylation | biological process | GO:0042327 | 139 | 20 | 12345 | 442 | hppy,jeb,Raf,InR,wnd,vn,Pvr,lncRNA:Hsromega,Pgk,Atg17,Lk6,Actbeta,Cks30A,spi,Src42A,step,Egfr,rictor,rdx,Pdk1 | | 1.093e-07 | -16.03 | sensory organ boundary specification | biological process | GO:0008052 | 25 | 9 | 12345 | 442 | klu,H,CtBP,fz,sens,nuf,B-H1,Tom,N | | 1.334e-07 | -15.83 | dorsal/ventral pattern formation | biological process | GO:0009953 | 154 | 21 | 12345 | 442 | Raf,S,Cbl,dlp,H,Dl,N,nuf,ap,lilli,Dp,tsh,14-3-3epsilon,cic,tara,Mpcp2,tkv,cact,nmo,wbl,Egfr | | 1.348e-07 | -15.82 | genital disc development | biological process | GO:0035215 | 50 | 12 | 12345 | 442 | otk,Egfr,abd-A,Pvr,tkv,Abd-B,N,sim,esg,Rab5,spi,S | | 1.463e-07 | -15.74 | sd (scalloped) | protein interactions | 32536 | 47 | 13 | 9204 | 432 | sens,Dp,kay,pnr,Mef2,jar,N,H,ph-p,Psc,kn,Dcr-2,sli | | 1.497e-07 | -15.71 | cell growth | biological process | GO:0016049 | 81 | 15 | 12345 | 442 | fz2,shg,fz,path,arm,InR,tai,Pex10,kay,SCAR,Rab5,wnd,tkv,Dad,htl | | 1.615e-07 | -15.64 | regulation of cellular protein metabolic process | biological process | GO:0032268 | 623 | 49 | 12345 | 442 | Spn42De,jeb,hppy,CG1218,InR,Socs36E,sens-2,Pvr,orb2,lncRNA:Hsromega,vn,bru2,bol,Lk6,N,Su(z)2,Sxl,CG8312,Src42A,spi,312,Egfr,msi,step,bin3,rdx,sina,Cbl,Dronc,Fem-1,Raf,CycB3,Atg17,Kdm2,PI31,CycG,trbl,pum,wnd,Actbeta,Tis11,kay,SNF4Agamma,wdb,Tollo,Diap1,Cks30A,rictor,Pdk1 | | 1.675e-07 | -15.60 | Homeobox-like\_sf | interpro domains | IPR009057 | 142 | 21 | 11383 | 450 | rib,CG13895,bab2,otp,nub,zfh2,B-H1,eyg,Scr,al,Poxn,Lmx1a,ap,ey,hth,pdm3,abd-A,Abd-B,H2.0,CG11617,dve | | 1.700e-07 | -15.59 | plasma membrane part | cellular component | GO:0044459 | 820 | 58 | 11940 | 417 | cindr,beat-IIa,ogre,Syx1A,Esp,Dg,otk,tipE,Gli,Dl,yrt,Ddr,fz,Nrk,scrib,Abl,rho-4,Tob,Ptp4E,emp,ed,Arf79F,d,N,Eph,orb2,Ir21a,baz,shg,nAChRalpha4,nuf,robo1,ds,Egfr,gukh,tkv,Inx2,bou,LanA,InR,Src42A,wgn,arm,drl,Shrm,Rab1,Cals,Dyb,ft,kek5,Magi,fz2,Pvr,Cip4,dlp,htl,step,dome | | 1.752e-07 | -15.56 | positive regulation of protein phosphorylation | biological process | GO:0001934 | 130 | 19 | 12345 | 442 | hppy,jeb,Raf,InR,vn,wnd,Atg17,lncRNA:Hsromega,Pvr,Actbeta,Lk6,spi,Cks30A,Src42A,rictor,Egfr,step,Pdk1,rdx | | 1.912e-07 | -15.47 | muscle cell differentiation | biological process | GO:0042692 | 118 | 18 | 12345 | 442 | C1GalTA,aop,sd,Mef2,kon,htl,dock,sns,InR,kn,jeb,SCAR,N,Dg,abd-A,siz,Bsg,Pgk | | 1.912e-07 | -15.47 | positive regulation of growth | biological process | GO:0045927 | 118 | 18 | 12345 | 442 | Pdk1,Hr4,nmo,tkv,Egfr,rictor,step,ey,14-3-3epsilon,tai,Cdk4,shot,CycG,chic,InR,fz,d,Orct2 | | 1.986e-07 | -15.43 | mesenchyme development | biological process | GO:0060485 | 14 | 7 | 12345 | 442 | shg,Sema1a,disco-r,Sema1b,abd-A,Abd-B,htl | | 2.148e-07 | -15.35 | morphogenesis of follicular epithelium | biological process | GO:0016333 | 52 | 12 | 12345 | 442 | scrib,Ppcs,N,Dg,Egfr,jar,shg,ph-p,Raf,Cip4,Dp,Cbl | | 2.235e-07 | -15.31 | regulation of photoreceptor cell differentiation | biological process | GO:0046532 | 43 | 11 | 12345 | 442 | ss,ttk,Raf,Src42A,eff,mop,ed,aop,N,sina,Socs36E | | 2.241e-07 | -15.31 | pnt (pointed) | protein interactions | 42757 | 41 | 12 | 9204 | 432 | spen,N,Egfr,cic,abd-A,Raf,ed,B-H1,Scr,aop,edl,Abd-B | | 2.267e-07 | -15.30 | behavior | biological process | GO:0007610 | 630 | 49 | 12345 | 442 | arm,Poxn,Adf1,ey,kay,Tob,Abl,Mef2,ogre,mld,Dcr-2,Abd-B,hang,Esp,Dyrk2,SERCA,ITP,moody,wnd,pum,ss,lilli,spi,rg,dome,muc,sim,tara,Gr93c,Egfr,mura,InR,Mob2,S,klu,hppy,drl,Bacc,bol,betaTub60D,N,14-3-3zeta,Mid1,scrib,pxb,orb2,Socs36E,vn,tipE | | 2.467e-07 | -15.22 | cell fate specification | biological process | GO:0001708 | 84 | 15 | 12345 | 442 | arm,ttk,jeb,kn,ey,N,esg,scrib,Dl,Tom,vn,Egfr,abd-A,tkv,htl | | 2.565e-07 | -15.18 | CycE (Cyclin E) | protein interactions | 34924 | 168 | 25 | 9204 | 432 | Cdk4,Poxn,scrib,E(Pc),N,Abl,ft,glec,Dp,hth,ttk,spi,abd-A,crol,Egfr,emc,sina,spen,arm,shg,sim,lawc,Mef2,Dad,Scr | | 2.587e-07 | -15.17 | GO\_EPITHELIAL\_CELL\_DIFFERENTIATION | MSigDB lists | GO\_EPITHELIAL\_CELL\_DIFFERENTIATION | 94 | 16 | 4161 | 156 | Rab1,sns,N,otk,Dl,Mef2,Cdk4,arm,Mfe2,Pgk,Pdk1,fz,LanA,Fem-1,C1GalTA,CG17883 | | 2.687e-07 | -15.13 | wing disc dorsal/ventral pattern formation | biological process | GO:0048190 | 53 | 12 | 12345 | 442 | lilli,H,dlp,14-3-3epsilon,N,Dl,cic,ap,nuf,nmo,Mpcp2,tara | | 2.918e-07 | -15.05 | central nervous system development | biological process | GO:0007417 | 281 | 29 | 12345 | 442 | C1GalTA,esg,sim,htl,shg,ey,rg,Abl,drl,moody,N,shot,siz,Pvr,Ptp4E,chic,Eph,ap,vn,hth,nub,eya,rib,mam,ph-p,mop,kn,Dronc,Psc | | 2.929e-07 | -15.04 | Hippo signaling pathway - fly | KEGG pathways | ko04391 | 61 | 13 | 3198 | 123 | scrib,d,vn,ed,sd,ds,wgn,14-3-3epsilon,14-3-3zeta,baz,hth,ft,tsh | | 2.929e-07 | -15.04 | Hippo signaling pathway - fly | KEGG pathways | dme04391 | 61 | 13 | 3198 | 123 | d,scrib,sd,ed,ds,vn,wgn,14-3-3zeta,14-3-3epsilon,baz,ft,hth,tsh | | 3.112e-07 | -14.98 | nucleic acid binding | molecular function | GO:0003676 | 2082 | 111 | 12176 | 414 | sim,dve,edl,sqz,ap,CG11122,MEP-1,jigr1,E(spl)m5-HLH,Dcr-2,otp,cic,lilli,zfh2,orb2,Tis11,CG11414,CG34376,CG13895,CG11247,Hr4,Adf1,HLH3B,Hel89B,CG10348,ph-p,lin-28,His3.3A,su(sable),Sxl,rib,l(2)37Cg,SmD1,Scr,mrt,Elba2,sd,CG11617,CG3436,H2.0,kay,sens-2,sens,hth,fz3,Hr39,CG45050,eyg,kn,Psc,pdm3,CG13204,rgr,disco-r,dmrt11E,Su(z)2,pnr,pum,grn,CG12769,mld,abd-A,CG3165,Sox14,jumu,CG10321,Mef2,pita,ttk,Dp,ey,Kdm2,CG11360,Poxn,hang,Rm62,His3.3B,CG32772,bol,hng3,al,crp,bin3,ps,chn,CG14506,CG1316,Tet,nub,Slbp,Abd-B,Lmx1a,CG2017,HmgZ,CG4896,esg,h,spen,aop,klu,vig,bru2,tsh,bab2,bowl,CG14441,B-H1,msi,crol,Prdm13,His2Av | | 3.180e-07 | -14.96 | rl (rolled) | protein interactions | 3354888 | 85 | 17 | 9204 | 432 | rg,aop,eya,vn,ed,pum,Egfr,step,sina,Lk6,lilli,kay,corto,Raf,cic,Src42A,jeb | | 3.358e-07 | -14.91 | regulation of cell fate specification | biological process | GO:0042659 | 28 | 9 | 12345 | 442 | eya,ttk,hth,chn,ed,N,aop,lqf,Abd-B | | 3.373e-07 | -14.90 | sty (sprouty) | protein interactions | 38424 | 105 | 19 | 9204 | 432 | aop,LanA,ed,Cbl,Pvr,mop,fz,sina,spi,H,Egfr,htl,Ppn,kay,robo1,N,S,Raf,Mob2 | | 3.387e-07 | -14.90 | drk (downstream of receptor kinase) | protein interactions | 36497 | 76 | 16 | 9204 | 432 | Pez,moody,cindr,14-3-3zeta,Raf,mim,Src42A,14-3-3epsilon,Pvr,mop,ena,CG42788,Cbl,B-H1,pum,Egfr | | 3.445e-07 | -14.88 | peptidyl-tyrosine phosphorylation | biological process | GO:0018108 | 36 | 10 | 12345 | 442 | Eph,Ptpmeg2,Ddr,Pvr,htl,drl,Abl,Nrk,InR,Src42A | | 3.449e-07 | -14.88 | positive regulation of phosphate metabolic process | biological process | GO:0045937 | 149 | 20 | 12345 | 442 | step,Egfr,rictor,rdx,Pdk1,Src42A,spi,Cks30A,Pvr,lncRNA:Hsromega,Atg17,Pgk,wnd,vn,Lk6,Actbeta,jeb,hppy,Raf,InR | | 3.449e-07 | -14.88 | positive regulation of phosphorus metabolic process | biological process | GO:0010562 | 149 | 20 | 12345 | 442 | Raf,InR,jeb,hppy,Actbeta,Lk6,Atg17,Pgk,lncRNA:Hsromega,Pvr,vn,wnd,Src42A,Cks30A,spi,Pdk1,rdx,rictor,Egfr,step | | 3.609e-07 | -14.83 | compound eye cone cell differentiation | biological process | GO:0042675 | 15 | 7 | 12345 | 442 | rg,emc,Egfr,ttk,Dl,eya,N | | 3.786e-07 | -14.79 | determination of genital disc primordium | biological process | GO:0035225 | 10 | 6 | 12345 | 442 | Egfr,abd-A,Abd-B,spi,sim,S | | 3.900e-07 | -14.76 | fz (frizzled) | protein interactions | 45307 | 59 | 14 | 9204 | 432 | ed,Rab5,ds,mop,kay,drl,baz,ft,fz2,arm,N,Egfr,Dl,otk | | 4.274e-07 | -14.67 | multicellular organism reproduction | biological process | GO:0032504 | 1503 | 90 | 12345 | 442 | dome,ttk,shg,dmrt11E,LanA,cact,Egfr,aop,InR,orb2,Ptpmeg2,scrib,cindr,Ppcs,14-3-3zeta,N,tai,Abl,Cip4,larp,Acp63F,Cks30A,Abd-B,tkv,baz,Mef2,Cbl,emc,mop,lncRNA:CR44420,Dronc,yuri,SERCA,fz,sns,abd-A,PI31,CycG,lin-28,pum,Dl,Inx2,Sxl,ss,Gr93c,ena,msi,ed,S,ph-p,Cerk,Socs36E,Pvr,lncRNA:Hsromega,tipE,Pen,chic,bol,14-3-3epsilon,kay,Pex10,Poxn,Dp,arm,eff,Diap1,htl,mim,wbl,His3.3A,jar,scyl,Pdk1,Rab5,cic,His3.3B,Hr39,Tet,Rab1,Raf,mam,eya,Esp,Arf79F,Dg,wnd,Pat1,Gli,RhoL,SCAR | | 4.494e-07 | -14.62 | Homeobox\_dom | interpro domains | IPR001356 | 102 | 17 | 11383 | 450 | otp,nub,zfh2,B-H1,eyg,Scr,al,pdm3,ap,hth,ey,Lmx1a,abd-A,H2.0,Abd-B,dve,CG11617 | | 4.514e-07 | -14.61 | GO\_KIDNEY\_EPITHELIUM\_DEVELOPMENT | MSigDB lists | GO\_KIDNEY\_EPITHELIUM\_DEVELOPMENT | 36 | 10 | 4161 | 156 | robo1,Mef2,Dl,ds,sli,ft,LanA,N,sns,arm | | 4.663e-07 | -14.58 | dpp (decapentaplegic) | protein interactions | 33432 | 97 | 18 | 9204 | 432 | eya,tkv,pnr,ds,nmo,spen,arm,Tollo,spi,Egfr,H,ey,lilli,cv-d,N,Snoo,Dg,ph-p | | 4.716e-07 | -14.57 | regulation of cell-cell adhesion | biological process | GO:0022407 | 29 | 9 | 12345 | 442 | arm,disco-r,aop,robo1,RhoL,N,sli,tkv,baz | | 5.071e-07 | -14.49 | GO\_EPITHELIUM\_DEVELOPMENT | MSigDB lists | GO\_EPITHELIUM\_DEVELOPMENT | 193 | 23 | 4161 | 156 | Rab1,N,Actbeta,sns,ap,Cdk4,Mef2,robo1,Dl,otk,fz2,arm,Mfe2,fz,Pdk1,Pgk,ft,ds,CG17883,sli,C1GalTA,Fem-1,LanA | | 5.102e-07 | -14.49 | imaginal disc-derived wing vein specification | biological process | GO:0007474 | 56 | 12 | 12345 | 442 | Raf,S,fuss,kn,cv-d,corto,cic,N,nmo,Dg,Egfr,step | | 5.210e-07 | -14.47 | sensory organ precursor cell fate determination | biological process | GO:0016360 | 22 | 8 | 12345 | 442 | CtBP,fz,klu,H,Tom,N,sens,nuf | | 5.210e-07 | -14.47 | cell fate determination involved in pattern specification | biological process | GO:0060582 | 22 | 8 | 12345 | 442 | klu,H,CtBP,fz,sens,nuf,Tom,N | | 5.260e-07 | -14.46 | regulation of protein metabolic process | biological process | GO:0051246 | 648 | 49 | 12345 | 442 | Pdk1,rictor,Cks30A,Diap1,Tollo,wdb,SNF4Agamma,kay,Tis11,Actbeta,wnd,pum,trbl,CycG,Kdm2,PI31,Atg17,CycB3,Raf,Fem-1,Dronc,Cbl,sina,rdx,bin3,msi,step,Egfr,312,spi,CG8312,Src42A,Su(z)2,Sxl,N,Lk6,bol,bru2,vn,lncRNA:Hsromega,Pvr,orb2,Socs36E,sens-2,CG1218,InR,hppy,Spn42De,jeb | | 5.325e-07 | -14.45 | Src42A (Src oncogene at 42A) | protein interactions | 35524 | 52 | 13 | 9204 | 432 | dock,14-3-3zeta,Pvr,baz,l(2)37Cg,shg,arm,S,Src42A,bowl,Egfr,spi,Raf | | 5.758e-07 | -14.37 | ci (cubitus interruptus) | protein interactions | 43767 | 187 | 26 | 9204 | 432 | cv-d,ey,hth,zfh2,sd,Su(z)2,ph-p,rdx,Psc,kn,sli,boi,px,RapGAP1,bin3,tsh,dome,fz3,eff,Egfr,Dl,Sxl,hyx,arm,CG34401,IA-2 | | 5.825e-07 | -14.36 | germ-line stem cell population maintenance | biological process | GO:0030718 | 78 | 14 | 12345 | 442 | His2Av,InR,mam,eff,shg,Socs36E,tkv,Dad,Egfr,chic,Inx2,Dl,N,esg | | 5.842e-07 | -14.35 | apical junction complex | cellular component | GO:0043296 | 58 | 12 | 11940 | 417 | Dl,Gli,ed,scrib,cora,moody,shg,Shrm,baz,arm,yrt,N | | 5.909e-07 | -14.34 | protein kinase binding | molecular function | GO:0019901 | 82 | 14 | 12176 | 414 | trbl,dome,Pp2C1,dock,Cks30A,Lk6,RhoU,Mob2,SNF4Agamma,drl,Atg17,CycG,CycB3,Cbl | | 6.219e-07 | -14.29 | regulation of protein kinase B signaling | biological process | GO:0051896 | 16 | 7 | 12345 | 442 | wdb,trbl,rictor,Pvr,CycG,InR,Pdk1 | | 6.364e-07 | -14.27 | Tyr\_kinase\_AS | interpro domains | IPR008266 | 35 | 10 | 11383 | 450 | Eph,htl,Pvr,InR,Src42A,otk,Egfr,Nrk,drl,Abl | | 6.379e-07 | -14.27 | fng (fringe) | protein interactions | 40314 | 18 | 8 | 9204 | 432 | ap,ey,spdo,eyg,N,InR,Egfr,Dl | | 6.393e-07 | -14.26 | membrane | cellular component | GO:0016020 | 2722 | 138 | 11940 | 417 | CenB1A,gukh,rg,Inx2,ds,Shrm,moody,Irk3,wgn,kek5,Ocrl,CG3149,Ric,Rab1,siz,Kr-h2,htl,fz2,CG8034,p130CAS,tipE,CG7394,ogre,S,fz,yrt,neo,Ptp4E,emp,scrib,Atg17,unc-5,Eph,Mid1,LanA,Egfr,Mob2,nuf,CG9220,Rab5,ft,CG33474,Cals,Tpc1,Cip4,spdo,dlp,alpha-Man-Ia,Gli,Cerk,Dg,CG7536,d,CG31121,AP-1gamma,CG30344,rho-4,14-3-3epsilon,bou,InR,tkv,Atg16,Fas1,lt,drongo,robo1,Pdk1,drl,Orct2,CG34057,Src42A,Pex10,Sema1a,CG17646,CG34172,Dyb,CG3119,CG9784,Raf,UQCR-11,Pex5,Magi,Dl,Gr93c,Syx1A,CG17883,Ddr,CG6891,CG4645,SERCA,Pvf3,ed,spi,Tob,Cbl,pes,Pex14,shg,DPCoAC,baz,orb2,Socs36E,CG11069,Vps13,C1GalTA,Mpcp2,arm,shot,cora,step,dome,Pvr,path,klar,Rim2,glec,ZnT41F,CG5853,Tsp96F,otk,14-3-3zeta,cindr,Esp,RhoU,beat-IIa,cue,Bsg,fz3,sns,Arf79F,CG30156,GlcAT-I,Abl,CG32795,Nrk,nAChRalpha4,Ir21a,N | | 6.526e-07 | -14.24 | ommatidial rotation | biological process | GO:0016318 | 30 | 9 | 12345 | 442 | nmo,Egfr,ec,fz,S,shg,spi,ds,ed | | 7.468e-07 | -14.11 | germ cell migration | biological process | GO:0008354 | 48 | 11 | 12345 | 442 | mim,pum,Abd-B,htl,tkv,abd-A,14-3-3epsilon,Cerk,shg,eya,Diap1 | | 7.674e-07 | -14.08 | kinase binding | molecular function | GO:0019900 | 96 | 15 | 12176 | 414 | trbl,Cks30A,Lk6,dock,Pp2C1,dome,Mob2,RhoU,Cbl,CycB3,CycG,Atg17,arm,SNF4Agamma,drl | | 8.077e-07 | -14.03 | determination of imaginal disc primordium | biological process | GO:0007445 | 11 | 6 | 12345 | 442 | S,spi,sim,Abd-B,abd-A,Egfr | | 8.527e-07 | -13.97 | abd-A (abdominal A) | protein interactions | 42037 | 54 | 13 | 9204 | 432 | Egfr,abd-A,spi,kn,mam,Abd-B,Mef2,pnr,ap,eya,tsh,CycG,hth | | 8.529e-07 | -13.97 | vg (vestigial) | protein interactions | 36421 | 31 | 10 | 9204 | 432 | Diap1,Mef2,tkv,ap,sd,Egfr,sli,spi,ft,N | | 8.819e-07 | -13.94 | Rho1 (Rho1) | protein interactions | 36775 | 91 | 17 | 9204 | 432 | H,Sema1a,shg,step,fz,Dad,ena,tkv,SNF4Agamma,baz,Abl,robo1,cindr,lilli,Drak,wnd,Rab5 | | 9.307e-07 | -13.89 | cell-cell adherens junction | cellular component | GO:0005913 | 24 | 8 | 11940 | 417 | baz,shg,step,N,arm,yrt,ed,scrib | | 9.323e-07 | -13.89 | imaginal disc-derived wing hair organization | biological process | GO:0035317 | 49 | 11 | 12345 | 442 | scrib,N,spen,f,cora,Gli,SMC3,fz,ds,Cip4,ft | | 9.609e-07 | -13.86 | negative regulation of nervous system development | biological process | GO:0051961 | 118 | 17 | 12345 | 442 | ena,Dad,LanA,aop,sd,Abl,ttk,Src42A,Cip4,Sema1b,Socs36E,N,Dl,ed,emc,Mob2,Sema1a | | 1.047e-06 | -13.77 | regulation of multi-organism process | biological process | GO:0043900 | 250 | 26 | 12345 | 442 | vn,Rm62,Pvr,DPCoAC,imd,Gli,Cbl,mop,Pitslre,nub,Egfr,htl,Dcr-2,cact,Syx1A,aop,tai,kay,shg,pnr,CG32795,Diap1,spi,ttk,Dp,klar | | 1.070e-06 | -13.75 | Antp (Antennapedia) | protein interactions | 40835 | 55 | 13 | 9204 | 432 | Scr,sens,ey,hth,tsh,tna,mam,pnr,Mef2,spen,N,ss,kn | | 1.147e-06 | -13.68 | mys (myospheroid) | protein interactions | 44885 | 47 | 12 | 9204 | 432 | Dl,sli,crol,kon,Sema1a,mop,p130CAS,mam,shot,Bsg,vn,tkv | | 1.149e-06 | -13.68 | Wnt-protein binding | molecular function | GO:0017147 | 18 | 7 | 12176 | 414 | Nrk,dlp,drl,fz3,otk,fz,fz2 | | 1.157e-06 | -13.67 | regulation of cell adhesion | biological process | GO:0030155 | 50 | 11 | 12345 | 442 | baz,Ddr,tkv,LanA,N,RhoL,sli,aop,robo1,arm,disco-r | | 1.157e-06 | -13.67 | non-sensory hair organization | biological process | GO:0035316 | 50 | 11 | 12345 | 442 | f,cora,Gli,scrib,N,spen,SMC3,Cip4,fz,ds,ft | | 1.157e-06 | -13.67 | hair cell differentiation | biological process | GO:0035315 | 50 | 11 | 12345 | 442 | ft,Cip4,fz,ds,SMC3,cora,f,Gli,scrib,spen,N | | 1.223e-06 | -13.61 | memory | biological process | GO:0007613 | 120 | 17 | 12345 | 442 | rg,Tob,Adf1,dome,arm,ss,mld,mura,Mob2,Socs36E,orb2,pxb,pum,wnd,drl,scrib,N | | 1.281e-06 | -13.57 | positive regulation of protein modification process | biological process | GO:0031401 | 176 | 21 | 12345 | 442 | step,rictor,Egfr,rdx,Pdk1,Su(z)2,Diap1,spi,Cks30A,Src42A,wnd,vn,Pvr,lncRNA:Hsromega,Atg17,Lk6,Actbeta,hppy,jeb,Raf,InR | | 1.294e-06 | -13.56 | establishment or maintenance of bipolar cell polarity | biological process | GO:0061245 | 41 | 10 | 12345 | 442 | Dg,Egfr,baz,scrib,Rab5,ft,yrt,rib,ds,fz | | 1.294e-06 | -13.56 | establishment or maintenance of apical/basal cell polarity | biological process | GO:0035088 | 41 | 10 | 12345 | 442 | yrt,ds,rib,fz,ft,scrib,Rab5,baz,Egfr,Dg | | 1.322e-06 | -13.54 | MODULE\_27 | MSigDB lists | MODULE\_27 | 40 | 10 | 4161 | 156 | Nrk,htl,tkv,trbl,nAChRalpha4,fz2,otk,Tollo,Eph,Ddr | | 1.353e-06 | -13.51 | zonula adherens | cellular component | GO:0005915 | 12 | 6 | 11940 | 417 | N,yrt,arm,baz,ed,shg | | 1.354e-06 | -13.51 | regulation of plasma membrane bounded cell projection organization | biological process | GO:0120035 | 148 | 19 | 12345 | 442 | shot,chic,Sema1b,SCAR,N,kn,Rab1,Sema1a,fz,ena,baz,rictor,kon,CG34401,robo1,C1GalTA,Adf1,Abl,ss | | 1.400e-06 | -13.48 | positive regulation of cellular protein metabolic process | biological process | GO:0032270 | 303 | 29 | 12345 | 442 | Socs36E,Atg17,Pvr,lncRNA:Hsromega,PI31,orb2,vn,trbl,wnd,pum,Actbeta,Lk6,bol,jeb,Dronc,hppy,Raf,InR,rictor,Egfr,step,Pdk1,rdx,sina,Su(z)2,Src42A,Cks30A,Diap1,spi | | 1.443e-06 | -13.45 | Snr1 (Snf5-related 1) | protein interactions | 40657 | 137 | 21 | 9204 | 432 | sens,corto,CG12581,Kdm2,S,N,H2.0,px,robo1,SKIP,Npc2g,ed,tkv,CG2017,lawc,jar,Egfr,H,Tet,Dl,HmgZ | | 1.488e-06 | -13.42 | CTTTGCA\_MIR527 | MSigDB lists | CTTTGCA\_MIR527 | 60 | 12 | 4161 | 156 | Kdm2,eff,wdb,Vps13,sd,CG2812,Lk6,Ocrl,AP-1gamma,frc,rg,Mef2 | | 1.502e-06 | -13.41 | regulation of cell projection organization | biological process | GO:0031344 | 149 | 19 | 12345 | 442 | SCAR,N,Sema1b,chic,shot,fz,Rab1,Sema1a,kn,kon,CG34401,robo1,C1GalTA,ena,baz,rictor,ss,Abl,Adf1 | | 1.567e-06 | -13.37 | establishment of apical/basal cell polarity | biological process | GO:0035089 | 12 | 6 | 12345 | 442 | scrib,fz,ds,yrt,ft,baz | | 1.599e-06 | -13.35 | negative regulation of neuron differentiation | biological process | GO:0045665 | 33 | 9 | 12345 | 442 | ttk,Sema1a,emc,ed,Abl,aop,N,Socs36E,Sema1b | | 1.601e-06 | -13.35 | regulation of cellular component biogenesis | biological process | GO:0044087 | 305 | 29 | 12345 | 442 | C1GalTA,Syx1A,kon,tkv,Dad,312,nmo,Ocrl,LanA,baz,rictor,ena,Jupiter,klar,Cip4,Cals,Src42A,Abl,RhoGEF4,SCAR,N,RhoU,shot,CG6891,chic,pum,lt,Mob2,fz | | 1.603e-06 | -13.34 | pole cell migration | biological process | GO:0007280 | 25 | 8 | 12345 | 442 | abd-A,htl,Abd-B,pum,eya,Cerk,shg,14-3-3epsilon | | 1.620e-06 | -13.33 | positive regulation of MAPK cascade | biological process | GO:0043410 | 73 | 13 | 12345 | 442 | step,wnd,Egfr,vn,Pvr,lncRNA:Hsromega,rdx,hppy,jeb,spi,Src42A,Raf,InR | | 1.625e-06 | -13.33 | positive regulation of hippo signaling | biological process | GO:0035332 | 18 | 7 | 12345 | 442 | ed,hppy,14-3-3epsilon,ft,ds,14-3-3zeta,Pez | | 1.727e-06 | -13.27 | plasma membrane region | cellular component | GO:0098590 | 314 | 29 | 11940 | 417 | Egfr,ds,cindr,nuf,Syx1A,Dg,ogre,LanA,gukh,Inx2,yrt,arm,Shrm,Rab1,Cals,Tob,scrib,ft,Arf79F,d,Ptp4E,Magi,step,nAChRalpha4,dome,orb2,baz,Cip4,Ir21a | | 1.754e-06 | -13.25 | eye photoreceptor cell fate commitment | biological process | GO:0042706 | 52 | 11 | 12345 | 442 | msi,scrib,sina,N,aop,Cbl,kay,fz,hth,ttk,Raf | | 1.754e-06 | -13.25 | compound eye photoreceptor fate commitment | biological process | GO:0001752 | 52 | 11 | 12345 | 442 | msi,N,sina,scrib,aop,kay,Cbl,fz,hth,Raf,ttk | | 1.889e-06 | -13.18 | Abl (Abl tyrosine kinase) | protein interactions | 45821 | 41 | 11 | 9204 | 432 | N,robo1,arm,Abl,sli,Egfr,chic,eya,cindr,ena,Pvr | | 1.889e-06 | -13.18 | sli (slit) | protein interactions | 36746 | 41 | 11 | 9204 | 432 | Egfr,siz,sli,Abl,robo1,sim,LanA,shot,dock,Atg16,sd | | 1.889e-06 | -13.18 | Sos (Son of sevenless) | protein interactions | 34790 | 41 | 11 | 9204 | 432 | B-H1,dock,Fem-1,aop,Egfr,sli,pum,Raf,N,Src42A,robo1 | | 1.902e-06 | -13.17 | long-term memory | biological process | GO:0007616 | 74 | 13 | 12345 | 442 | N,Socs36E,orb2,pxb,mld,pum,wnd,mura,arm,Mob2,Tob,Adf1,dome | | 1.917e-06 | -13.16 | cytoskeleton organization | biological process | GO:0007010 | 580 | 44 | 12345 | 442 | ena,sick,cora,f,p130CAS,RhoGEF3,Src42A,Jupiter,klar,chic,Bsg,orb2,shot,RhoU,Ppcs,14-3-3zeta,cindr,N,betaTub60D,gukh,ed,Wdr62,jar,baz,rictor,kon,wdb,Shrm,14-3-3epsilon,ft,larp,Cks30A,yrt,Dg,nuf,siz,SCAR,moody,Dl,mop,sns,rib,ds,yuri | | 1.987e-06 | -13.13 | protein domain specific binding | molecular function | GO:0019904 | 78 | 13 | 12176 | 414 | ena,imd,Dronc,aop,sd,fz2,14-3-3zeta,Cbl,cindr,mim,14-3-3epsilon,ttk,Egfr | | 1.988e-06 | -13.13 | negative regulation of cell development | biological process | GO:0010721 | 86 | 14 | 12345 | 442 | Abl,ed,emc,Diap1,ttk,klar,Sema1a,Egfr,Sema1b,Socs36E,N,aop,sd,Dl | | 2.098e-06 | -13.07 | signaling receptor binding | molecular function | GO:0005102 | 256 | 25 | 12176 | 414 | Actbeta,tai,ITP,spi,scrib,Pvf3,spdo,otk,arm,Sema1a,Cbl,Src42A,Sema1b,sli,apolpp,LanA,dock,Socs36E,Pex14,cue,Dl,htl,vn,Gli,ed | | 2.106e-06 | -13.07 | negative regulation of epidermal growth factor receptor signaling pathway | biological process | GO:0042059 | 34 | 9 | 12345 | 442 | kek5,Src42A,hppy,ed,Cbl,edl,pum,Ptp4E,Socs36E | | 2.106e-06 | -13.07 | regulation of compound eye photoreceptor cell differentiation | biological process | GO:0110116 | 34 | 9 | 12345 | 442 | ed,ttk,Src42A,ss,eff,Socs36E,aop,sina,N | | 2.106e-06 | -13.07 | negative regulation of ERBB signaling pathway | biological process | GO:1901185 | 34 | 9 | 12345 | 442 | edl,pum,Ptp4E,Socs36E,kek5,Src42A,ed,hppy,Cbl | | 2.143e-06 | -13.05 | epidermal cell differentiation | biological process | GO:0009913 | 53 | 11 | 12345 | 442 | cora,f,Gli,scrib,spen,N,SMC3,Cip4,fz,ds,ft | | 2.151e-06 | -13.05 | GO\_PROTEIN\_AUTOPHOSPHORYLATION | MSigDB lists | GO\_PROTEIN\_AUTOPHOSPHORYLATION | 42 | 10 | 4161 | 156 | nmo,Drak,Pdk1,Nrk,Lk6,htl,InR,Src42A,Ddr,Eph | | 2.182e-06 | -13.04 | regulation of MAPK cascade | biological process | GO:0043408 | 125 | 17 | 12345 | 442 | lncRNA:Hsromega,Pvr,vn,trbl,wnd,Actbeta,N,Cbl,jeb,hppy,Raf,InR,Egfr,step,rdx,Src42A,spi | | 2.210e-06 | -13.02 | PH-like\_dom\_sf | interpro domains | IPR011993 | 139 | 19 | 11383 | 450 | ena,Myo10A,RhoGEF3,yrt,CG8312,CG12581,LRR,step,CG5522,CenB1A,CG34383,Pez,siz,Cadps,flfl,Ziz,Pdk1,cora,RhoGEF4 | | 2.217e-06 | -13.02 | Rbfox1 (RNA-binding Fox protein 1) | protein interactions | 39198 | 15 | 7 | 9204 | 432 | sens,N,H,kn,pum,spdo,mir-980 | | 2.240e-06 | -13.01 | spen (split ends) | protein interactions | 44205 | 27 | 9 | 9204 | 432 | Egfr,H,Raf,spi,arm,N,aop,tsh,lqf | | 2.314e-06 | -12.98 | epidermis development | biological process | GO:0008544 | 64 | 12 | 12345 | 442 | Gli,f,cora,N,spen,scrib,SMC3,Cip4,tsh,ds,fz,ft | | 2.424e-06 | -12.93 | GO\_TISSUE\_DEVELOPMENT | MSigDB lists | GO\_TISSUE\_DEVELOPMENT | 292 | 28 | 4161 | 156 | hyx,LanA,Fem-1,C1GalTA,ds,sli,CG17883,Ddr,ft,CG9220,tkv,Pgk,Pdk1,fz,Mfe2,arm,fz2,Dl,otk,robo1,Cdk4,Mef2,ap,htl,sns,Actbeta,N,Rab1 | | 2.452e-06 | -12.92 | peb (pebbled) | protein interactions | 31391 | 68 | 14 | 9204 | 432 | ed,cact,tkv,LanA,aop,chic,ps,px,InR,Egfr,dve,Dl,kn,chn | | 2.746e-06 | -12.81 | segment specification | biological process | GO:0007379 | 35 | 9 | 12345 | 442 | spen,Abd-B,Kdm2,shg,tsh,hth,lilli,dlp,kn | | 2.822e-06 | -12.78 | positive regulation of ERBB signaling pathway | biological process | GO:1901186 | 13 | 6 | 12345 | 442 | step,edl,mop,S,rho-4,Src42A | | 2.822e-06 | -12.78 | positive regulation of epidermal growth factor receptor signaling pathway | biological process | GO:0045742 | 13 | 6 | 12345 | 442 | mop,edl,step,Src42A,rho-4,S | | 2.836e-06 | -12.77 | anterior/posterior pattern specification | biological process | GO:0009952 | 200 | 22 | 12345 | 442 | ena,baz,tkv,Dad,Abd-B,cic,E(spl)m5-HLH,shg,Scr,lilli,chic,pum,Dg,bowl,abd-A,fs(1)h,N,scrib,Pat1,Dl,Raf,h | | 2.869e-06 | -12.76 | GO\_NEPHRON\_EPITHELIUM\_DEVELOPMENT | MSigDB lists | GO\_NEPHRON\_EPITHELIUM\_DEVELOPMENT | 26 | 8 | 4161 | 156 | arm,N,sns,ds,ft,LanA,Mef2,Dl | | 2.900e-06 | -12.75 | enzyme binding | molecular function | GO:0019899 | 481 | 37 | 12176 | 414 | sina,dome,RhoU,Rgl,CG4645,CG17883,RhoGEF4,dock,ec,step,nuf,TBC1D5,CycB3,drl,ph-p,CG5522,RhoGEF3,rdx,Cks30A,CG31457,Pp2C1,Mob2,Cbl,CycG,Atg17,arm,Magi,SNF4Agamma,trbl,siz,eff,hyx,Lk6,Ziz,morgue,Tom,Diap1 | | 2.935e-06 | -12.74 | scrib (scribble) | protein interactions | 44448 | 51 | 12 | 9204 | 432 | baz,gukh,CG11158,Egfr,scrib,dome,Raf,Socs36E,tkv,hth,Rab5,kay | | 3.063e-06 | -12.70 | polytene chromosome | cellular component | GO:0005700 | 206 | 22 | 11940 | 417 | mam,His3.3A,vig,E(Pc),bab2,CycG,su(sable),jumu,sens,Rm62,pita,CtBP,MEP-1,corto,Adf1,Nlp,His3.3B,Psc,Elba2,ttk,lawc,His2Av | | 3.361e-06 | -12.60 | cell division | biological process | GO:0051301 | 266 | 26 | 12345 | 442 | N,bol,Dl,lin-28,chic,pum,Arf79F,orb2,mam,fz,CycB3,Rab1,InR,esg,Syx1A,flfl,jar,baz,msi,Ocrl,tkv,eff,shg,Diap1,spdo,Pex10 | | 3.483e-06 | -12.57 | MODULE\_199 | MSigDB lists | MODULE\_199 | 13 | 6 | 4161 | 156 | Ddr,Eph,otk,htl,tkv,Nrk | | 3.483e-06 | -12.57 | MODULE\_85 | MSigDB lists | MODULE\_85 | 13 | 6 | 4161 | 156 | Nrk,otk,Eph,htl,tkv,Ddr | | 3.696e-06 | -12.51 | positive regulation of protein metabolic process | biological process | GO:0051247 | 318 | 29 | 12345 | 442 | Egfr,rictor,step,sina,Pdk1,rdx,Su(z)2,Cks30A,spi,Diap1,Src42A,vn,trbl,pum,wnd,Atg17,Socs36E,orb2,Pvr,PI31,lncRNA:Hsromega,Actbeta,bol,Lk6,Dronc,hppy,jeb,InR,Raf | | 3.720e-06 | -12.50 | somatic stem cell population maintenance | biological process | GO:0035019 | 20 | 7 | 12345 | 442 | His2Av,H,chic,arm,shg,mam,esg | | 3.720e-06 | -12.50 | negative regulation of cell fate commitment | biological process | GO:0010454 | 20 | 7 | 12345 | 442 | Socs36E,Abd-B,chn,ed,lqf,aop,eya | | 3.740e-06 | -12.50 | cindr (CIN85 and CD2AP related) | protein interactions | 43654 | 61 | 13 | 9204 | 432 | rictor,mim,siz,rdx,shg,Abl,imd,Pdk1,ena,ds,chic,tsh,Cbl | | 3.783e-06 | -12.49 | pb (proboscipedia) | protein interactions | 40826 | 16 | 7 | 9204 | 432 | H,abd-A,Abd-B,sens,Scr,ey,eyg | | 3.783e-06 | -12.49 | px (plexus) | protein interactions | 37573 | 16 | 7 | 9204 | 432 | Dl,H,Egfr,N,emc,S,nmo | | 3.783e-06 | -12.49 | Caf1-105 (Chromatin assembly factor 1, p105 subunit) | protein interactions | 36107 | 16 | 7 | 9204 | 432 | His3.3A,His3.3B,eyg,N,H,mam,Dl | | 3.947e-06 | -12.44 | eve (even skipped) | protein interactions | 36039 | 22 | 8 | 9204 | 432 | unc-5,Abd-B,Scr,kn,grn,Dl,abd-A,nmo | | 3.977e-06 | -12.43 | tkv (thickveins) | protein interactions | 33753 | 101 | 17 | 9204 | 432 | Dad,tkv,H,kek5,Egfr,lili,Gli,shg,Tollo,nmo,arm,Rab5,ey,Raf,scrib,Snoo,N | | 4.006e-06 | -12.43 | head involution | biological process | GO:0008258 | 46 | 10 | 12345 | 442 | pum,ena,scyl,emc,Dronc,chrb,rib,tsh,yrt,shg | | 4.197e-06 | -12.38 | positive regulation of nervous system development | biological process | GO:0051962 | 131 | 17 | 12345 | 442 | chic,shot,Arf79F,N,klu,mop,fz,Raf,InR,Rab1,rictor,tkv,nmo,Sox14,spi,Src42A,Cals | | 4.259e-06 | -12.37 | Homeodomain | pfam domains | PF00046 | 94 | 15 | 10480 | 419 | B-H1,Scr,nub,dve,zfh2,pdm3,eyg,al,ap,abd-A,H2.0,Abd-B,otp,Lmx1a,ey | | 4.358e-06 | -12.34 | CtBP (C-terminal Binding Protein) | protein interactions | 41602 | 81 | 15 | 9204 | 432 | h,tsh,ttk,ey,sens,imd,pnr,esg,Diap1,N,CtBP,arm,MEP-1,CG10348,H | | 4.400e-06 | -12.33 | cno (canoe) | protein interactions | 40620 | 29 | 9 | 9204 | 432 | kn,H,scrib,robo1,N,S,shg,ena,ed | | 4.583e-06 | -12.29 | apicolateral plasma membrane | cellular component | GO:0016327 | 21 | 7 | 11940 | 417 | Magi,ds,scrib,d,ft,Inx2,baz | | 4.656e-06 | -12.28 | photoreceptor cell development | biological process | GO:0042461 | 105 | 15 | 12345 | 442 | rg,jeb,Nrk,Raf,ttk,lilli,klar,ss,Mob2,yrt,eya,Bsg,sens,edl,baz | | 5.196e-06 | -12.17 | dock (dreadlocks) | protein interactions | 33262 | 37 | 10 | 9204 | 432 | sli,SCAR,Egfr,InR,sick,Src42A,sns,robo1,Pvr,dock | | 5.281e-06 | -12.15 | renal tubule development | biological process | GO:0061326 | 69 | 12 | 12345 | 442 | hth,S,rib,tsh,spi,Pvr,tkv,Dad,baz,Egfr,scrib,N | | 5.281e-06 | -12.15 | Malpighian tubule development | biological process | GO:0072002 | 69 | 12 | 12345 | 442 | scrib,N,Egfr,baz,Pvr,tkv,Dad,S,tsh,spi,rib,hth | | 5.408e-06 | -12.13 | regulation of R7 cell differentiation | biological process | GO:0045676 | 21 | 7 | 12345 | 442 | sina,eff,aop,ss,ttk,Src42A,Socs36E | | 5.432e-06 | -12.12 | cell junction organization | biological process | GO:0034330 | 81 | 13 | 12345 | 442 | cora,moody,Gli,scrib,cindr,Magi,jar,baz,Src42A,arm,sns,shg,bou | | 5.450e-06 | -12.12 | PROTEIN\_KINASE\_TYR | prosite domains | PS00109 | 34 | 10 | 6243 | 325 | drl,Abl,htl,Nrk,otk,InR,Src42A,Egfr,Eph,Pvr | | 5.718e-06 | -12.07 | Raf (Raf oncogene) | protein interactions | 31221 | 149 | 21 | 9204 | 432 | step,spen,morgue,Egfr,aop,B-H1,eya,tkv,betaTub60D,Psa,14-3-3epsilon,Src42A,ft,S,scrib,14-3-3zeta,Lk6,mam,SERCA,lilli,Gfat2 | | 5.805e-06 | -12.06 | fj (four-jointed) | protein interactions | 37089 | 23 | 8 | 9204 | 432 | ena,SERCA,ds,N,ft,baz,ph-p,d | | 5.811e-06 | -12.06 | regulation of cellular component size | biological process | GO:0032535 | 178 | 20 | 12345 | 442 | path,Pvf3,Cip4,lilli,Abl,Pdk1,rictor,ena,fz,InR,Sema1a,hppy,SCAR,trbl,chic,Sema1b,CG6891,Pvr,RhoU,shot | | 5.877e-06 | -12.04 | positive regulation of developmental growth | biological process | GO:0048639 | 94 | 14 | 12345 | 442 | Egfr,rictor,chic,step,nmo,shot,tkv,Hr4,Pdk1,ey,fz,d,Orct2,InR | | 5.907e-06 | -12.04 | VART\_KSHV\_INFECTION\_ANGIOGENIC\_MARKERS\_DN | MSigDB lists | VART\_KSHV\_INFECTION\_ANGIOGENIC\_MARKERS\_DN | 14 | 6 | 4161 | 156 | Dl,htl,N,fz,LanA,fz2 | | 5.907e-06 | -12.04 | GO\_TRANSMEMBRANE\_RECEPTOR\_PROTEIN\_KINASE\_ACTIVITY | MSigDB lists | GO\_TRANSMEMBRANE\_RECEPTOR\_PROTEIN\_KINASE\_ACTIVITY | 14 | 6 | 4161 | 156 | Ddr,Nrk,tkv,InR,htl,Eph | | 6.028e-06 | -12.02 | tsh (teashirt) | protein interactions | 35430 | 30 | 9 | 9204 | 432 | abd-A,Dl,CtBP,arm,spen,Abd-B,cindr,hth,Scr | | 6.073e-06 | -12.01 | cell cortex | cellular component | GO:0005938 | 183 | 20 | 11940 | 417 | baz,spdo,Pez,step,shg,CG42684,Cbl,chic,scrib,Abl,ed,jar,Bsg,CG6891,Shrm,Rab5,cindr,nuf,drongo,Dl | | 6.170e-06 | -12.00 | cell-cell adhesion via plasma-membrane adhesion molecules | biological process | GO:0098742 | 70 | 12 | 12345 | 442 | baz,Bsg,glec,beat-IIa,ed,ft,fz,sns,shg,ds,Cals,Fas1 | | 6.172e-06 | -12.00 | net (net) | protein interactions | 45339 | 17 | 7 | 9204 | 432 | Egfr,H,Dl,nmo,S,emc,CycG | | 6.285e-06 | -11.98 | regulation of reproductive process | biological process | GO:2000241 | 121 | 16 | 12345 | 442 | kay,tai,ttk,klar,Dp,shg,spi,Diap1,Egfr,aop,Syx1A,Cbl,CG9784,Pvr,vn,Gli | | 6.410e-06 | -11.96 | Tyr\_kinase\_cat\_dom | interpro domains | IPR020635 | 27 | 8 | 11383 | 450 | Eph,htl,InR,Src42A,Egfr,otk,Nrk,Abl | | 6.435e-06 | -11.95 | neuron fate determination | biological process | GO:0048664 | 9 | 5 | 12345 | 442 | Egfr,S,mam,spi,N | | 6.484e-06 | -11.95 | developmental cell growth | biological process | GO:0048588 | 59 | 11 | 12345 | 442 | wnd,htl,SCAR,Rab5,tai,kay,Pex10,fz2,fz,path,InR | | 6.584e-06 | -11.93 | ct (cut) | protein interactions | 44540 | 94 | 16 | 9204 | 432 | ss,shg,sqz,aop,h,path,rgr,kn,Psc,Su(z)2,N,fs(1)h,mam,chic,14-3-3zeta,ey | | 6.601e-06 | -11.93 | dsh (dishevelled) | protein interactions | 32078 | 127 | 19 | 9204 | 432 | H,Dl,fz2,arm,shg,fz,ds,CG2017,ed,tkv,aop,otk,Abl,baz,N,mam,kay,ey,Rab5 | | 6.855e-06 | -11.89 | hid (head involution defective) | protein interactions | 40009 | 105 | 17 | 9204 | 432 | Egfr,eff,pum,spi,arm,fz2,nmo,Hr4,morgue,Cbl,Dronc,scrib,klu,Raf,lncRNA:Hsromega,Diap1,sens | | 6.973e-06 | -11.87 | CUI\_TCF21\_TARGETS\_2\_DN | MSigDB lists | CUI\_TCF21\_TARGETS\_2\_DN | 177 | 20 | 4161 | 156 | Mef2,cact,Pgant6,Hel89B,Vps13,trbl,l(1)G0196,shot,CenB1A,nmo,hyx,Tollo,Eph,Src42A,Ids,Kr-h2,rdx,jar,CG13384,bru2 | | 7.079e-06 | -11.86 | protein kinase activity | molecular function | GO:0004672 | 274 | 25 | 12176 | 414 | trbl,Src42A,Dyrk2,Raf,hppy,nmo,Lk6,InR,htl,CycB3,Ddr,Nrk,Eph,Abl,drl,Cdk4,Pitslre,Pdk1,Egfr,wnd,Drak,CycG,Pvr,tkv,otk | | 7.335e-06 | -11.82 | periodic partitioning | biological process | GO:0007365 | 49 | 10 | 12345 | 442 | rdx,frc,Egfr,arm,lilli,h,fz2,fz,kn,dlp | | 7.498e-06 | -11.80 | Krn (Keren) | protein interactions | 326198 | 12 | 6 | 9204 | 432 | unc-119,S,vn,spi,rho-4,Egfr | | 7.689e-06 | -11.78 | proximal/distal pattern formation, imaginal disc | biological process | GO:0007449 | 22 | 7 | 12345 | 442 | tsh,N,B-H1,hth,ap,Egfr,vn | | 7.689e-06 | -11.78 | epidermal growth factor receptor signaling pathway | biological process | GO:0007173 | 22 | 7 | 12345 | 442 | vn,Egfr,spi,spen,Raf,cic,aop | | 7.689e-06 | -11.78 | ERBB signaling pathway | biological process | GO:0038127 | 22 | 7 | 12345 | 442 | spi,spen,Raf,cic,aop,Egfr,vn | | 7.689e-06 | -11.78 | positive regulation of animal organ morphogenesis | biological process | GO:0110110 | 22 | 7 | 12345 | 442 | Src42A,hth,tsh,morgue,klu,ec,Dronc | | 7.741e-06 | -11.77 | R3/R4 cell differentiation | biological process | GO:0048056 | 15 | 6 | 12345 | 442 | fz,scrib,N,aop,kay,emc | | 7.741e-06 | -11.77 | antennal morphogenesis | biological process | GO:0048800 | 15 | 6 | 12345 | 442 | al,Diap1,f,Poxn,ss,Dl | | 7.741e-06 | -11.77 | negative regulation of cell fate specification | biological process | GO:0009996 | 15 | 6 | 12345 | 442 | Abd-B,chn,ed,lqf,aop,eya | | 7.741e-06 | -11.77 | establishment of monopolar cell polarity | biological process | GO:0061162 | 15 | 6 | 12345 | 442 | ft,baz,yrt,ds,scrib,fz | | 7.741e-06 | -11.77 | establishment or maintenance of monopolar cell polarity | biological process | GO:0061339 | 15 | 6 | 12345 | 442 | ds,yrt,fz,scrib,ft,baz | | 7.755e-06 | -11.77 | MODULE\_259 | MSigDB lists | MODULE\_259 | 9 | 5 | 4161 | 156 | Ddr,htl,otk,Eph,Nrk | | 7.818e-06 | -11.76 | Diap1 (Death-associated inhibitor of apoptosis 1) | protein interactions | 39753 | 106 | 17 | 9204 | 432 | dome,eff,H,pum,hppy,morgue,sina,Pvr,Dronc,ap,mir-1000,Dcr-2,CG32772,CtBP,lncRNA:Hsromega,Diap1,chic | | 7.879e-06 | -11.75 | baz (bazooka) | protein interactions | 32703 | 65 | 13 | 9204 | 432 | 14-3-3zeta,ed,Arf79F,Magi,14-3-3epsilon,fz,Src42A,shg,arm,baz,Dcr-2,scrib,H | | 8.213e-06 | -11.71 | somatic muscle development | biological process | GO:0007525 | 84 | 13 | 12345 | 442 | eya,hth,InR,Alh,chn,kn,crp,cora,sd,ap,htl,Pgk,Kdm2 | | 8.353e-06 | -11.69 | Abi (Abelson interacting protein) | protein interactions | 41718 | 24 | 8 | 9204 | 432 | ena,Cip4,orb2,Egfr,SCAR,CG14818,Abl,CG42353 | | 8.551e-06 | -11.67 | salivary gland morphogenesis | biological process | GO:0007435 | 97 | 14 | 12345 | 442 | eyg,Src42A,h,fz2,eya,rib,fz,shg,jeb,Dronc,drl,Pvr,Atg17,chic | | 8.551e-06 | -11.67 | gland morphogenesis | biological process | GO:0022612 | 97 | 14 | 12345 | 442 | shg,fz,fz2,eya,rib,Src42A,h,eyg,Dronc,jeb,drl,chic,Atg17,Pvr | | 9.159e-06 | -11.60 | enhancer binding | molecular function | GO:0035326 | 42 | 9 | 12176 | 414 | abd-A,ttk,ey,hth,Abd-B,Mef2,sd,aop,klu | | 9.238e-06 | -11.59 | GO\_APOPTOTIC\_PROCESS\_INVOLVED\_IN\_DEVELOPMENT | MSigDB lists | GO\_APOPTOTIC\_PROCESS\_INVOLVED\_IN\_DEVELOPMENT | 5 | 4 | 4161 | 156 | robo1,N,sli,fz2 | | 9.454e-06 | -11.57 | GO\_NEPHRON\_DEVELOPMENT | MSigDB lists | GO\_NEPHRON\_DEVELOPMENT | 30 | 8 | 4161 | 156 | LanA,ft,ds,Dl,Mef2,arm,sns,N | | 9.522e-06 | -11.56 | GO\_LOCOMOTION | MSigDB lists | GO\_LOCOMOTION | 212 | 22 | 4161 | 156 | tkv,sina,Tsp96F,Pdk1,arm,Eph,LanA,emp,ds,sli,Unc-76,ap,dock,N,14-3-3epsilon,shot,Rab1,unc-5,otk,robo1,Mef2,Pex7 | | 9.655e-06 | -11.55 | regulation of actin cytoskeleton organization | biological process | GO:0032956 | 98 | 14 | 12345 | 442 | Ocrl,RhoU,ena,jar,rictor,step,chic,CG6891,RhoGEF4,SCAR,ed,Abl,Cip4,fz | | 9.675e-06 | -11.55 | regulation of ERK1 and ERK2 cascade | biological process | GO:0070372 | 31 | 8 | 12345 | 442 | vn,Egfr,step,Pvr,spi,Src42A,Raf,Cbl | | 9.675e-06 | -11.55 | establishment or maintenance of epithelial cell apical/basal polarity | biological process | GO:0045197 | 31 | 8 | 12345 | 442 | ft,rib,ds,yrt,fz,baz,Dg,scrib | | 9.888e-06 | -11.52 | Signaling by Robo receptor | REACTOME pathways | R-DME-376176 | 13 | 6 | 3923 | 176 | sli,dlp,robo1,Abl,dock,ena | | 9.994e-06 | -11.51 | dom (domino) | protein interactions | 45655 | 97 | 16 | 9204 | 432 | lilli,mam,Hel89B,Lk6,E(Pc),N,ph-p,wdb,CG31211,His2Av,Sema1a,emc,pum,jigr1,Dl,tara | | 1.052e-05 | -11.46 | DNA-binding transcription factor binding | molecular function | GO:0140297 | 65 | 11 | 12176 | 414 | Scr,edl,tai,sens,kay,cact,cic,arm,lilli,pnr,CtBP | | 1.071e-05 | -11.44 | positive regulation of ERK1 and ERK2 cascade | biological process | GO:0070374 | 23 | 7 | 12345 | 442 | Pvr,Egfr,vn,step,Raf,Src42A,spi | | 1.087e-05 | -11.43 | fra (frazzled) | protein interactions | 36377 | 32 | 9 | 9204 | 432 | ena,drl,unc-5,siz,sli,Abl,N,robo1,Src42A | | 1.103e-05 | -11.42 | protein phosphorylation | biological process | GO:0006468 | 284 | 26 | 12345 | 442 | Nrk,hppy,Raf,InR,Pitslre,CycB3,Dyrk2,CycG,Pvr,otk,Ptpmeg2,trbl,Drak,wnd,Eph,Cdk4,Lk6,drl,Abl,Src42A,Cks30A,htl,tkv,nmo,Ddr,Pdk1 | | 1.109e-05 | -11.41 | Cortactin (cortactin) | protein interactions | 42491 | 8 | 5 | 9204 | 432 | ft,Src42A,d,Pvr,mim | | 1.114e-05 | -11.40 | Psn (Presenilin) | protein interactions | 40260 | 40 | 10 | 9204 | 432 | arm,S,N,H,Dl,eya,mam,Diap1,spdo,dpy | | 1.148e-05 | -11.38 | yki (yorkie) | protein interactions | 37851 | 109 | 17 | 9204 | 432 | tkv,cact,ed,Dronc,mop,Pen,ft,CG30089,14-3-3epsilon,scrib,ZnT41F,hth,14-3-3zeta,sd,Diap1,Pez,tai | | 1.201e-05 | -11.33 | regulation of muscle organ development | biological process | GO:0048634 | 16 | 6 | 12345 | 442 | abd-A,tkv,Dad,Abd-B,Him,Mef2 | | 1.225e-05 | -11.31 | regulation of actin filament-based process | biological process | GO:0032970 | 100 | 14 | 12345 | 442 | SCAR,RhoGEF4,CG6891,chic,step,rictor,ena,jar,RhoU,Ocrl,fz,Cip4,ed,Abl | | 1.234e-05 | -11.30 | GO\_PEPTIDYL\_TYROSINE\_MODIFICATION | MSigDB lists | GO\_PEPTIDYL\_TYROSINE\_MODIFICATION | 31 | 8 | 4161 | 156 | Eph,Src42A,Ddr,htl,InR,Nrk,Dyrk2,Tpst | | 1.249e-05 | -11.29 | polarized epithelial cell differentiation | biological process | GO:0030859 | 10 | 5 | 12345 | 442 | ft,ds,yrt,fz,scrib | | 1.249e-05 | -11.29 | compound eye retinal cell programmed cell death | biological process | GO:0046667 | 10 | 5 | 12345 | 442 | ec,arm,Dl,S,N | | 1.249e-05 | -11.29 | establishment of epithelial cell apical/basal polarity | biological process | GO:0045198 | 10 | 5 | 12345 | 442 | scrib,fz,yrt,ds,ft | | 1.249e-05 | -11.29 | positive regulation of protein kinase B signaling | biological process | GO:0051897 | 10 | 5 | 12345 | 442 | Pvr,CycG,rictor,InR,Pdk1 | | 1.250e-05 | -11.29 | homeostasis of number of cells | biological process | GO:0048872 | 32 | 8 | 12345 | 442 | Pvr,esg,N,Atg16,H,SERCA,Src42A,InR | | 1.274e-05 | -11.27 | cellular component assembly | biological process | GO:0022607 | 1109 | 67 | 12345 | 442 | kay,Dp,arm,CG17683,yrt,SmD1,mim,His3.3A,cic,Pdk1,kon,His3.3B,Raf,Dg,Gli,Pat1,SCAR,bou,Src42A,ena,f,jeb,ed,S,Pvr,chic,bru2,Adf1,Tob,wdb,Cks30A,larp,tkv,Dcr-2,rictor,baz,Nlp,Atg16,emc,mop,hyx,LRR,Dronc,Myo10A,Sema1a,sns,PI31,Atg17,moody,Spc105R,ttk,shg,cact,CG34172,Egfr,aop,unc-119,InR,orb2,Bsg,vn,drl,FBXO11,N,Ppcs,cindr,scrib | | 1.300e-05 | -11.25 | GO\_ORGAN\_MORPHOGENESIS | MSigDB lists | GO\_ORGAN\_MORPHOGENESIS | 169 | 19 | 4161 | 156 | Eph,CG17883,ds,ft,LanA,Fem-1,fz,CG9220,tkv,arm,fz2,Mef2,Raf,Dl,otk,Pex7,N,Actbeta,htl | | 1.337e-05 | -11.22 | l(1)sc (lethal of scute) | protein interactions | 30983 | 13 | 6 | 9204 | 432 | jumu,h,emc,H,mam,Dl | | 1.337e-05 | -11.22 | siz (schizo) | protein interactions | 40327 | 13 | 6 | 9204 | 432 | sli,cindr,Rgl,CG31211,Arf79F,robo1 | | 1.371e-05 | -11.20 | cell adhesion molecule binding | molecular function | GO:0050839 | 44 | 9 | 12176 | 414 | dock,siz,ds,cindr,arm,otk,Fas1,ed,ft | | 1.388e-05 | -11.19 | chaeta morphogenesis | biological process | GO:0008407 | 42 | 9 | 12345 | 442 | emc,hyx,pnr,sens,Dl,B-H1,f,N,spen | | 1.425e-05 | -11.16 | GO\_BIOLOGICAL\_ADHESION | MSigDB lists | GO\_BIOLOGICAL\_ADHESION | 155 | 18 | 4161 | 156 | Pdk1,Tsp96F,fz2,arm,Eph,Ddr,ft,ds,LanA,sns,dock,shot,Rab1,Cdk4,Raf,robo1,Cals,otk | | 1.504e-05 | -11.10 | GO\_TRANSMEMBRANE\_RECEPTOR\_PROTEIN\_TYROSINE\_KINASE\_ACTIVITY | MSigDB lists | GO\_TRANSMEMBRANE\_RECEPTOR\_PROTEIN\_TYROSINE\_KINASE\_ACTIVITY | 10 | 5 | 4161 | 156 | InR,htl,Eph,Nrk,Ddr | | 1.538e-05 | -11.08 | regulation of programmed cell death | biological process | GO:0043067 | 222 | 22 | 12345 | 442 | morgue,Pdk1,rdx,Egfr,ec,Abd-B,spi,Diap1,Dp,ft,N,Actbeta,FBXO11,Pvr,lncRNA:Hsromega,sens,eya,Raf,hppy,Dronc,klu,Cbl | | 1.599e-05 | -11.04 | regulation of filopodium assembly | biological process | GO:0051489 | 33 | 8 | 12345 | 442 | shot,chic,ena,baz,C1GalTA,kon,SCAR,N | | 1.633e-05 | -11.02 | emc (extra macrochaetae) | protein interactions | 38091 | 26 | 8 | 9204 | 432 | N,px,Egfr,H,Dl,vn,mam,pnr | | 1.669e-05 | -11.00 | HOMEOBOX\_2 | prosite domains | PS50071 | 102 | 17 | 6243 | 325 | B-H1,ap,pdm3,CG11617,eyg,Scr,dve,Abd-B,H2.0,abd-A,hth,al,nub,Lmx1a,otp,ey,zfh2 | | 1.704e-05 | -10.98 | GO\_UROGENITAL\_SYSTEM\_DEVELOPMENT | MSigDB lists | GO\_UROGENITAL\_SYSTEM\_DEVELOPMENT | 75 | 12 | 4161 | 156 | arm,sns,N,LanA,Fem-1,ds,C1GalTA,sli,ft,Dl,Mef2,robo1 | | 1.752e-05 | -10.95 | regulation of protein kinase activity | biological process | GO:0045859 | 131 | 16 | 12345 | 442 | Pdk1,rictor,step,Cks30A,kay,SNF4Agamma,Atg17,Socs36E,CycG,trbl,wnd,Raf,CycB3,Cbl,jeb,hppy | | 1.774e-05 | -10.94 | gastrulation | biological process | GO:0007369 | 90 | 13 | 12345 | 442 | Abl,jeb,spi,shg,mam,arm,Raf,trbl,baz,tkv,abd-A,htl,sli | | 1.777e-05 | -10.94 | ed (echinoid) | protein interactions | 33619 | 42 | 10 | 9204 | 432 | aop,ed,Dl,Egfr,H,sina,N,fz,jar,baz | | 1.791e-05 | -10.93 | orb2 (orb2) | protein interactions | 39018 | 80 | 14 | 9204 | 432 | ps,Tob,CG6891,CG42684,bol,ReepA,path,orb2,CG12769,MEP-1,wdb,CG1316,mura,N | | 1.838e-05 | -10.90 | gro (groucho) | protein interactions | 43162 | 70 | 13 | 9204 | 432 | klu,H,Egfr,cic,CtBP,E(spl)m5-HLH,N,H2.0,arm,al,h,ey,Him | | 1.840e-05 | -10.90 | GO\_RENAL\_TUBULE\_DEVELOPMENT | MSigDB lists | GO\_RENAL\_TUBULE\_DEVELOPMENT | 24 | 7 | 4161 | 156 | Dl,N,Mef2,LanA,arm,ds,ft | | 1.849e-05 | -10.90 | molecular function regulator | molecular function | GO:0098772 | 563 | 39 | 12176 | 414 | RhoGEF3,Ocrl,CG5522,Actbeta,ITP,rictor,wdb,Pdk1,CenB1A,spi,Cks30A,Pvf3,Atg17,CycG,siz,trbl,Ziz,Socs36E,RhoGAP19D,Ppn,Diap1,Spn42De,tipE,CG7394,CG17883,Rgl,Sema1a,RhoGEF4,Sema1b,RapGAP1,Dl,htl,TBC1D5,drongo,vn,step,jeb,CycB3,14-3-3zeta | | 1.869e-05 | -10.89 | ap (apterous) | protein interactions | 35509 | 34 | 9 | 9204 | 432 | vn,Scr,Abd-B,Diap1,N,sqz,abd-A,Dl,H | | 1.869e-05 | -10.89 | vn (vein) | protein interactions | 38657 | 34 | 9 | 9204 | 432 | boi,spi,Egfr,H,N,emc,Pvr,CycG,ap | | 1.872e-05 | -10.89 | Wnt-activated receptor activity | molecular function | GO:0042813 | 6 | 4 | 12176 | 414 | fz,fz2,fz3,drl | | 1.953e-05 | -10.84 | regulation of apoptotic process | biological process | GO:0042981 | 193 | 20 | 12345 | 442 | spi,Diap1,Dp,ft,morgue,rdx,Pdk1,Egfr,Abd-B,eya,Raf,Dronc,hppy,Cbl,N,FBXO11,Actbeta,Pvr,sens,lncRNA:Hsromega | | 1.974e-05 | -10.83 | stem cell development | biological process | GO:0048864 | 25 | 7 | 12345 | 442 | N,arm,Sema1a,Sema1b,sqz,abd-A,klu | | 2.070e-05 | -10.79 | regulation of plasma membrane bounded cell projection assembly | biological process | GO:0120032 | 44 | 9 | 12345 | 442 | SCAR,N,C1GalTA,kon,chic,baz,ena,shot,Abl | | 2.070e-05 | -10.79 | homophilic cell adhesion via plasma membrane adhesion molecules | biological process | GO:0007156 | 44 | 9 | 12345 | 442 | ed,ft,ds,fz,sns,shg,Fas1,Cals,Bsg | | 2.070e-05 | -10.79 | regulation of cell projection assembly | biological process | GO:0060491 | 44 | 9 | 12345 | 442 | Abl,C1GalTA,kon,N,SCAR,shot,chic,baz,ena | | 2.109e-05 | -10.77 | PTPase\_domain | interpro domains | IPR000242 | 16 | 6 | 11383 | 450 | IA-2,CG42327,mop,Ptpmeg2,Ptp4E,Pez | | 2.134e-05 | -10.76 | Prot\_kinase\_dom | interpro domains | IPR000719 | 221 | 23 | 11383 | 450 | drl,Ddr,tkv,nmo,wnd,trbl,Lk6,Src42A,otk,Raf,Eph,hppy,Cdk4,Pdk1,Abl,Egfr,Nrk,InR,Pitslre,Dyrk2,htl,Pvr,Drak | | 2.146e-05 | -10.75 | protein homodimerization activity | molecular function | GO:0042803 | 140 | 16 | 12176 | 414 | CtBP,shg,otk,ttk,rdx,E(spl)m5-HLH,Scr,Mfe2,step,14-3-3zeta,drl,mim,trbl,wbl,corto,Dronc | | 2.194e-05 | -10.73 | synapse | cellular component | GO:0045202 | 285 | 25 | 11940 | 417 | Syx1A,ena,Dg,LanA,Gli,gukh,tkv,Cadps,Bsg,ReepA,CG31211,Rab5,cact,Pdk1,Tob,Cals,scrib,Magi,nAChRalpha4,dlp,pum,Cip4,jeb,orb2,Ir21a | | 2.219e-05 | -10.72 | ventral cord development | biological process | GO:0007419 | 79 | 12 | 12345 | 442 | Psc,Abl,mop,kn,eya,nub,ap,htl,Pvr,N,sim,moody | | 2.220e-05 | -10.72 | Scr (Sex combs reduced) | protein interactions | 40833 | 43 | 10 | 9204 | 432 | tsh,hth,eya,Scr,ap,Mef2,pnr,mam,tna,kn | | 2.223e-05 | -10.71 | negative regulation of compound eye photoreceptor cell differentiation | biological process | GO:0110118 | 11 | 5 | 12345 | 442 | ed,Socs36E,N,aop,ttk | | 2.226e-05 | -10.71 | neur (neuralized) | protein interactions | 41085 | 27 | 8 | 9204 | 432 | H,eff,Dl,Tom,CG34401,koi,N,lqf | | 2.243e-05 | -10.71 | Y\_phosphatase | pfam domains | PF00102 | 16 | 6 | 10480 | 419 | IA-2,CG42327,mop,Ptp4E,Pez,Ptpmeg2 | | 2.248e-05 | -10.70 | MESR4 (Misexpression suppressor of ras 4) | protein interactions | 36986 | 14 | 6 | 9204 | 432 | pnr,Syx1A,Pvr,H,N,Sema1a | | 2.280e-05 | -10.69 | EGF-like\_CS | interpro domains | IPR013032 | 61 | 11 | 11383 | 450 | LanA,nahoda,shg,vn,dpy,spi,cue,N,ft,sli,Dl | | 2.337e-05 | -10.66 | GCANCTGNY\_MYOD\_Q6 | MSigDB lists | GCANCTGNY\_MYOD\_Q6 | 176 | 19 | 4161 | 156 | LanA,Cadps,sli,hyx,unc-119,rg,Ric,lili,rdx,betaTub60D,CG2812,Nrk,Dl,shot,14-3-3epsilon,Psa,InR,N,14-3-3zeta | | 2.398e-05 | -10.64 | tin (tinman) | protein interactions | 42536 | 9 | 5 | 9204 | 432 | pnr,abd-A,Abd-B,Tom,Scr | | 2.410e-05 | -10.63 | asymmetric cell division | biological process | GO:0008356 | 106 | 14 | 12345 | 442 | InR,shg,fz,mam,eff,Diap1,spdo,flfl,Dl,esg,N,orb2,jar,baz | | 2.527e-05 | -10.59 | chromatin organization involved in regulation of transcription | biological process | GO:0034401 | 80 | 12 | 12345 | 442 | nub,Psc,Su(z)2,fl(2)d,crol,bin3,corto,FBXO11,Elba2,tna,tara,Dcr-2 | | 2.540e-05 | -10.58 | dorsal appendage formation | biological process | GO:0046843 | 56 | 10 | 12345 | 442 | Raf,ttk,S,emc,ed,cic,Ppcs,N,cact,Egfr | | 2.620e-05 | -10.55 | branch fusion, open tracheal system | biological process | GO:0035147 | 18 | 6 | 12345 | 442 | ed,shot,esg,shg,arm,ttk | | 2.620e-05 | -10.55 | regulation of compound eye retinal cell programmed cell death | biological process | GO:0046669 | 18 | 6 | 12345 | 442 | morgue,Diap1,Dronc,ec,Egfr,klu | | 2.620e-05 | -10.55 | adherens junction assembly | biological process | GO:0034333 | 18 | 6 | 12345 | 442 | arm,shg,sns,scrib,cindr,baz | | 2.620e-05 | -10.55 | tube fusion | biological process | GO:0035146 | 18 | 6 | 12345 | 442 | ed,shot,esg,shg,arm,ttk | | 2.620e-05 | -10.55 | leg disc proximal/distal pattern formation | biological process | GO:0007479 | 18 | 6 | 12345 | 442 | Egfr,vn,ap,tsh,hth,B-H1 | | 2.674e-05 | -10.53 | LI\_WILMS\_TUMOR\_VS\_FETAL\_KIDNEY\_2\_DN | MSigDB lists | LI\_WILMS\_TUMOR\_VS\_FETAL\_KIDNEY\_2\_DN | 11 | 5 | 4161 | 156 | dlp,Ddr,Actbeta,fz,InR | | 2.674e-05 | -10.53 | GO\_PEROXISOMAL\_TRANSPORT | MSigDB lists | GO\_PEROXISOMAL\_TRANSPORT | 11 | 5 | 4161 | 156 | Pex10,CG45050,Pex7,Pex5,Pex14 | | 2.690e-05 | -10.52 | GO\_HIPPO\_SIGNALING | MSigDB lists | GO\_HIPPO\_SIGNALING | 6 | 4 | 4161 | 156 | 14-3-3epsilon,ft,ds,sd | | 2.808e-05 | -10.48 | regulation of kinase activity | biological process | GO:0043549 | 136 | 16 | 12345 | 442 | CycG,Socs36E,Atg17,wnd,trbl,Raf,CycB3,jeb,Cbl,hppy,Pdk1,step,rictor,Cks30A,kay,SNF4Agamma | | 2.827e-05 | -10.47 | regulation of transferase activity | biological process | GO:0051338 | 151 | 17 | 12345 | 442 | Pdk1,step,rictor,Cks30A,SNF4Agamma,kay,wnd,trbl,CycG,Atg17,Socs36E,CycB3,Raf,hppy,Fem-1,jeb,Cbl | | 2.984e-05 | -10.42 | regulation of cell size | biological process | GO:0008361 | 108 | 14 | 12345 | 442 | Sema1b,chic,trbl,rictor,Pvr,shot,SCAR,Pdk1,hppy,Pvf3,fz,Sema1a,lilli,InR | | 3.019e-05 | -10.41 | R7 cell differentiation | biological process | GO:0045466 | 46 | 9 | 12345 | 442 | aop,N,sina,emc,klu,Cbl,ss,ttk,Raf | | 3.081e-05 | -10.39 | Signal Transduction | REACTOME pathways | R-DME-162582 | 717 | 54 | 3923 | 176 | htl,CG31431,dock,Dronc,Ptp4E,Actbeta,p130CAS,wdb,Cbl,Dl,Egfr,tkv,RhoU,14-3-3zeta,Lk6,arm,RhoGEF4,baz,Dad,Dp,lqf,Rgl,muc,Ocrl,boi,fz,CG4935,CG42684,vn,RhoGAP19D,Cip4,N,PI31,rdx,rictor,Pdk1,moody,CtBP,14-3-3epsilon,Sox14,Diap1,Abl,trbl,Snoo,hyx,eff,SCAR,fz2,Raf,cact,SNF4Agamma,cindr,Pvr,nmo | | 3.085e-05 | -10.39 | Ser (Serrate) | protein interactions | 43275 | 36 | 9 | 9204 | 432 | ap,tsh,mam,Src42A,eyg,N,H,fz3,Dl | | 3.132e-05 | -10.37 | jing (jing) | protein interactions | 35555 | 21 | 7 | 9204 | 432 | sli,sim,spi,kay,Egfr,hth,S | | 3.132e-05 | -10.37 | robo2 (roundabout 2) | protein interactions | 44522 | 21 | 7 | 9204 | 432 | Pvr,ena,sli,Atg16,Abl,robo1,Dg | | 3.150e-05 | -10.37 | brm (brahma) | protein interactions | 39744 | 106 | 16 | 9204 | 432 | N,ph-p,Psc,Raf,sd,tna,arm,shg,Egfr,H,tara,Dl,rictor,vn,tkv,CG2017 | | 3.171e-05 | -10.36 | adherens junction organization | biological process | GO:0034332 | 36 | 8 | 12345 | 442 | Src42A,arm,shg,sns,scrib,cindr,Magi,baz | | 3.171e-05 | -10.36 | midgut development | biological process | GO:0007494 | 36 | 8 | 12345 | 442 | Mef2,vn,Abd-B,abd-A,Scr,tsh,dve,emc | | 3.203e-05 | -10.35 | muscle organ development | biological process | GO:0007517 | 95 | 13 | 12345 | 442 | ed,kn,jeb,sns,klar,ap,Dg,shot,htl,N,drl,Mef2,kon | | 3.239e-05 | -10.34 | GO\_NEGATIVE\_REGULATION\_OF\_HEMOPOIESIS | MSigDB lists | GO\_NEGATIVE\_REGULATION\_OF\_HEMOPOIESIS | 35 | 8 | 4161 | 156 | N,Actbeta,arm,Cdk4,hyx,Tollo,Dl,cact | | 3.239e-05 | -10.34 | GO\_MORPHOGENESIS\_OF\_A\_BRANCHING\_STRUCTURE | MSigDB lists | GO\_MORPHOGENESIS\_OF\_A\_BRANCHING\_STRUCTURE | 35 | 8 | 4161 | 156 | LanA,Fem-1,sli,ds,ft,arm,fz2,N | | 3.282e-05 | -10.32 | DACOSTA\_UV\_RESPONSE\_VIA\_ERCC3\_TTD\_DN | MSigDB lists | DACOSTA\_UV\_RESPONSE\_VIA\_ERCC3\_TTD\_DN | 26 | 7 | 4161 | 156 | Pex14,IntS3,Hel89B,shot,CenB1A,Ziz,sd | | 3.329e-05 | -10.31 | enhancer sequence-specific DNA binding | molecular function | GO:0001158 | 38 | 8 | 12176 | 414 | ey,ttk,hth,abd-A,aop,Mef2,Abd-B,sd | | 3.376e-05 | -10.30 | zinc ion binding | molecular function | GO:0008270 | 516 | 36 | 12176 | 414 | hang,tna,CG11247,CG11414,ph-p,lin-28,Hr4,CG1218,CG34401,FBXO11,sina,CG10321,pita,lt,Dyb,dpy,CG11360,Kdm2,mura,Psc,CG45050,Pex10,pnr,CG4467,Su(z)2,Diap1,grn,mld,Psa,CG14441,Tet,CG31457,CG2991,Hr39,Cbl,CG13481 | | 3.427e-05 | -10.28 | antennal development | biological process | GO:0007469 | 27 | 7 | 12345 | 442 | ss,Dl,eyg,Poxn,f,al,Diap1 | | 3.486e-05 | -10.26 | Rap2l (Ras-associated protein 2-like) | protein interactions | 37850 | 64 | 12 | 9204 | 432 | spdo,CG42788,ey,Fem-1,Raf,emp,Rgl,rdx,scrib,Magi,Kr-h2,IA-2 | | 3.491e-05 | -10.26 | behavioral response to ethanol | biological process | GO:0048149 | 58 | 10 | 12345 | 442 | hppy,spi,S,hang,mura,pum,Egfr,pxb,Bacc,moody | | 3.517e-05 | -10.26 | regulation of protein serine/threonine kinase activity | biological process | GO:0071900 | 70 | 11 | 12345 | 442 | hppy,SNF4Agamma,kay,CycB3,Cks30A,Raf,wnd,step,rictor,trbl,CycG | | 3.586e-05 | -10.24 | positive regulation of neurogenesis | biological process | GO:0050769 | 96 | 13 | 12345 | 442 | Sox14,mop,klu,Raf,InR,Src42A,Rab1,spi,fz,shot,Arf79F,chic,N | | 3.618e-05 | -10.23 | imaginal disc-derived wing vein morphogenesis | biological process | GO:0008586 | 47 | 9 | 12345 | 442 | Socs36E,vn,Egfr,spen,px,hyx,emc,H,d | | 3.676e-05 | -10.21 | negative regulation of growth | biological process | GO:0045926 | 110 | 14 | 12345 | 442 | Sema1a,Cip4,Src42A,Mob2,ft,Abl,chrb,cic,scyl,scrib,Dad,LanA,Sema1b,ena | | 3.699e-05 | -10.20 | retinal cell programmed cell death | biological process | GO:0046666 | 12 | 5 | 12345 | 442 | arm,Dl,N,S,ec | | 3.699e-05 | -10.20 | histoblast morphogenesis | biological process | GO:0007488 | 12 | 5 | 12345 | 442 | chic,tkv,Dad,Cks30A,esg | | 3.699e-05 | -10.20 | germarium-derived female germ-line cyst encapsulation | biological process | GO:0030708 | 12 | 5 | 12345 | 442 | Cip4,N,Ppcs,shg,ph-p | | 3.699e-05 | -10.20 | salivary gland boundary specification | biological process | GO:0007432 | 12 | 5 | 12345 | 442 | Scr,unc-5,sli,robo1,hth | | 3.699e-05 | -10.20 | female germ-line cyst encapsulation | biological process | GO:0048139 | 12 | 5 | 12345 | 442 | shg,ph-p,Ppcs,N,Cip4 | | 3.714e-05 | -10.20 | regulation of retinal cell programmed cell death | biological process | GO:0046668 | 19 | 6 | 12345 | 442 | Egfr,ec,Dronc,klu,Diap1,morgue | | 3.725e-05 | -10.20 | - | gene3d domains | 2.30.29.30 | 139 | 19 | 4929 | 241 | Cadps,CG5522,yrt,Pdk1,CG8312,siz,LRR,cora,Myo10A,ena,step,RhoGEF4,CG12581,CenB1A,flfl,Pez,RhoGEF3,Ziz,CG34383 | | 3.909e-05 | -10.15 | Dad (Daughters against dpp) | protein interactions | 42059 | 37 | 9 | 9204 | 432 | pnr,tkv,Dl,H,Dg,lili,Gli,arm,nmo | | 3.909e-05 | -10.15 | robo1 (roundabout 1) | protein interactions | 37603 | 37 | 9 | 9204 | 432 | Abl,robo1,Dg,siz,sli,dock,shot,ena,unc-5 | | 3.920e-05 | -10.15 | regulation of hippo signaling | biological process | GO:0035330 | 37 | 8 | 12345 | 442 | Pez,14-3-3zeta,ds,d,ft,14-3-3epsilon,ed,hppy | | 3.920e-05 | -10.15 | heart morphogenesis | biological process | GO:0003007 | 37 | 8 | 12345 | 442 | sli,cora,Gli,robo1,mam,shg,arm,spdo | | 3.945e-05 | -10.14 | gcm (glial cells missing) | protein interactions | 34277 | 55 | 11 | 9204 | 432 | dome,crol,MEP-1,chn,robo1,Dg,lilli,esg,Pen,Hr39,ttk | | 4.071e-05 | -10.11 | actin filament organization | biological process | GO:0007015 | 111 | 14 | 12345 | 442 | ena,sick,jar,chic,RhoU,cindr,SCAR,N,f,Dl,p130CAS,Shrm,ed,Src42A | | 4.164e-05 | -10.09 | stau (staufen) | protein interactions | 37065 | 86 | 14 | 9204 | 432 | Dl,nub,Kdm4B,baz,sqz,Adf1,spen,CG4788,klar,CG43736,Tob,ps,flfl,vn | | 4.292e-05 | -10.06 | AGCATTA\_MIR155 | MSigDB lists | AGCATTA\_MIR155 | 27 | 7 | 4161 | 156 | CG1673,FBXO11,14-3-3zeta,hyx,14-3-3epsilon,bru2,Rab1 | | 4.430e-05 | -10.02 | regulation of stem cell differentiation | biological process | GO:2000736 | 28 | 7 | 12345 | 442 | N,C1GalTA,lqf,vn,bab2,Abd-B,Arf79F | | 4.447e-05 | -10.02 | GO\_REGULATION\_OF\_SYNAPSE\_ASSEMBLY | MSigDB lists | GO\_REGULATION\_OF\_SYNAPSE\_ASSEMBLY | 12 | 5 | 4161 | 156 | Mef2,Nrk,robo1,Cals,Eph | | 4.555e-05 | -10.00 | RNA polymerase II transcription factor binding | molecular function | GO:0001085 | 13 | 5 | 12176 | 414 | sd,emc,tna,pnr,bin3 | | 4.560e-05 | -10.00 | maintenance of imaginal disc-derived wing hair orientation | biological process | GO:0035321 | 3 | 3 | 12345 | 442 | spen,cora,Gli | | 4.560e-05 | -10.00 | negative regulation of muscle organ development | biological process | GO:0048635 | 3 | 3 | 12345 | 442 | Him,Abd-B,Dad | | 4.623e-05 | -9.98 | PRTYPHPHTASE | prints domains | PR00700 | 16 | 6 | 2185 | 101 | Pez,mop,Ptp4E,CG42327,Ptpmeg2,IA-2 | | 4.812e-05 | -9.94 | protein autophosphorylation | biological process | GO:0046777 | 38 | 8 | 12345 | 442 | Lk6,Eph,wnd,nmo,InR,Src42A,Raf,Abl | | 4.912e-05 | -9.92 | bw (brown) | protein interactions | 37724 | 38 | 9 | 9204 | 432 | Kdm4B,crol,CG13895,HmgZ,His3.3B,pita,Rm62,tai,Hr39 | | 4.995e-05 | -9.90 | immune system process | biological process | GO:0002376 | 328 | 27 | 12345 | 442 | Mef2,LanA,htl,Dcr-2,cact,CG1129,RhoGEF3,Tollo,pnr,shg,Pvf3,kay,dome,imd,N,RhoL,sens-2,Rm62,Pvr,Eph,Pen,Raf,alpha-Man-Ia,eya,LRR,kn,Dronc | | 5.149e-05 | -9.87 | GO\_RESPONSE\_TO\_EXTERNAL\_STIMULUS | MSigDB lists | GO\_RESPONSE\_TO\_EXTERNAL\_STIMULUS | 380 | 30 | 4161 | 156 | robo1,CG6891,Mef2,Cdk4,CG11069,cact,Tpc1,N,ap,dock,Unc-76,unc-5,SmD1,Rab1,hyx,unc-119,Eph,Tollo,sli,AP-1gamma,ReepA,CG4645,sina,fz,CG9220,subdued,tkv,CG9485,arm,fz2 | | 5.154e-05 | -9.87 | HOX | smart domains | SM00389 | 102 | 17 | 5086 | 290 | zfh2,Lmx1a,H2.0,eyg,ap,ey,dve,pdm3,otp,abd-A,al,B-H1,Abd-B,CG11617,nub,Scr,hth | | 5.207e-05 | -9.86 | second mitotic wave involved in compound eye morphogenesis | biological process | GO:0016330 | 7 | 4 | 12345 | 442 | Dl,N,kay,Egfr | | 5.207e-05 | -9.86 | protein localization involved in establishment of planar polarity | biological process | GO:0090251 | 7 | 4 | 12345 | 442 | ds,d,Shrm,ft | | 5.207e-05 | -9.86 | axis elongation | biological process | GO:0003401 | 7 | 4 | 12345 | 442 | Abl,otk,baz,Tollo | | 5.222e-05 | -9.86 | gene silencing | biological process | GO:0016458 | 143 | 16 | 12345 | 442 | vig,mrt,Su(z)2,bin3,crol,Dcr-2,ph-p,nub,fl(2)d,Psc,Elba2,Tis11,FBXO11,corto,pum,Rm62 | | 5.429e-05 | -9.82 | GO\_CELL\_DEVELOPMENT | MSigDB lists | GO\_CELL\_DEVELOPMENT | 289 | 25 | 4161 | 156 | Pdk1,sina,tkv,CG9220,fz2,arm,Mfe2,Eph,ft,sli,CG17883,C1GalTA,Fem-1,LanA,N,Actbeta,sns,Unc-76,ap,unc-5,Rab1,Mef2,Cdk4,robo1,Dl | | 5.545e-05 | -9.80 | GSE22886\_CD8\_VS\_CD4\_NAIVE\_TCELL\_DN | MSigDB lists | GSE22886\_CD8\_VS\_CD4\_NAIVE\_TCELL\_DN | 28 | 7 | 4161 | 156 | lili,Rab5,step,ReepA,unc-119,sns,trbl | | 5.589e-05 | -9.79 | crb (crumbs) | protein interactions | 42896 | 57 | 11 | 9204 | 432 | rib,mam,yrt,tai,baz,arm,shg,N,SCAR,scrib,Dl | | 5.661e-05 | -9.78 | intestinal stem cell homeostasis | biological process | GO:0036335 | 29 | 7 | 12345 | 442 | InR,Src42A,N,esg,SERCA,Pvr,H | | 5.686e-05 | -9.77 | regulation of cell morphogenesis | biological process | GO:0022604 | 144 | 16 | 12345 | 442 | CG34401,htl,ss,ttk,Abl,SCAR,chic,Sema1b,shot,neo,rib,fz,Sema1a,Rab1,ed,kn | | 5.834e-05 | -9.75 | negative regulation of cell-cell adhesion | biological process | GO:0022408 | 13 | 5 | 12345 | 442 | baz,N,disco-r,arm,aop | | 5.919e-05 | -9.73 | regulation of molecular function | biological process | GO:0065009 | 523 | 37 | 12345 | 442 | edl,Socs36E,14-3-3zeta,drongo,RhoGAP19D,hppy,Spn42De,jeb,step,Ocrl,cact,CenB1A,ttk,trbl,wnd,Atg17,CycG,Rgl,CG7394,CG17883,Fem-1,Dronc,Cbl,emc,SERCA,CycB3,Raf,rictor,CG42684,Pdk1,TBC1D5,SNF4Agamma,14-3-3epsilon,kay,Diap1,Cks30A,RapGAP1 | | 5.945e-05 | -9.73 | regulation of cytoskeleton organization | biological process | GO:0051493 | 160 | 17 | 12345 | 442 | Ocrl,step,ena,rictor,jar,Jupiter,lilli,Cip4,Abl,RhoGEF4,SCAR,shot,RhoU,CG6891,chic,fz,ed | | 5.998e-05 | -9.72 | GO\_CELL\_MORPHOGENESIS\_INVOLVED\_IN\_DIFFERENTIATION | MSigDB lists | GO\_CELL\_MORPHOGENESIS\_INVOLVED\_IN\_DIFFERENTIATION | 112 | 14 | 4161 | 156 | N,sina,Unc-76,tkv,ap,unc-5,arm,Rab1,Mef2,robo1,Eph,CG17883,sli,LanA | | 6.042e-05 | -9.71 | GO\_NEGATIVE\_REGULATION\_OF\_IMMUNE\_SYSTEM\_PROCESS | MSigDB lists | GO\_NEGATIVE\_REGULATION\_OF\_IMMUNE\_SYSTEM\_PROCESS | 72 | 11 | 4161 | 156 | arm,Pdk1,N,Actbeta,Spn42De,sli,Tollo,Dl,cact,hyx,Cdk4 | | 6.093e-05 | -9.71 | DACOSTA\_UV\_RESPONSE\_VIA\_ERCC3\_DN | MSigDB lists | DACOSTA\_UV\_RESPONSE\_VIA\_ERCC3\_DN | 291 | 25 | 4161 | 156 | shot,CenB1A,l(1)G0196,dock,Psa,Pgant5,InR,sd,Vps13,CtBP,Hel89B,step,eff,Tpst,Pat1,rdx,Pex14,CG9220,SMC3,wdb,Rgl,Ziz,sli,IntS3,Ddr | | 6.094e-05 | -9.71 | GO\_RETINAL\_GANGLION\_CELL\_AXON\_GUIDANCE | MSigDB lists | GO\_RETINAL\_GANGLION\_CELL\_AXON\_GUIDANCE | 7 | 4 | 4161 | 156 | sli,tkv,Eph,robo1 | | 6.110e-05 | -9.70 | endocytic vesicle | cellular component | GO:0030139 | 30 | 7 | 11940 | 417 | apolpp,Rab5,Dl,nuf,N,arm,jar | | 6.125e-05 | -9.70 | Dfd (Deformed) | protein interactions | 40832 | 39 | 9 | 9204 | 432 | fs(1)h,N,Cdk4,CycG,hth,tsh,LanA,orb2,Diap1 | | 6.134e-05 | -9.70 | cnk (connector enhancer of ksr) | protein interactions | 36952 | 48 | 10 | 9204 | 432 | Raf,H,Src42A,step,14-3-3epsilon,jeb,kay,cindr,14-3-3zeta,aop | | 6.241e-05 | -9.68 | GO\_PROTEIN\_KINASE\_ACTIVITY | MSigDB lists | GO\_PROTEIN\_KINASE\_ACTIVITY | 157 | 17 | 4161 | 156 | otk,Raf,Cdk4,hppy,InR,htl,trbl,nmo,Drak,Eph,Ddr,Src42A,tkv,Lk6,Nrk,Pdk1,Dyrk2 | | 6.252e-05 | -9.68 | Homeobox\_CS | interpro domains | IPR017970 | 92 | 13 | 11383 | 450 | nub,Scr,al,otp,eyg,B-H1,abd-A,Abd-B,H2.0,ap,Lmx1a,ey,zfh2 | | 6.320e-05 | -9.67 | WOO\_LIVER\_CANCER\_RECURRENCE\_UP | MSigDB lists | WOO\_LIVER\_CANCER\_RECURRENCE\_UP | 20 | 6 | 4161 | 156 | fz,Mef2,sli,dlp,jar,LanA | | 6.518e-05 | -9.64 | Bx (Beadex) | protein interactions | 32846 | 68 | 12 | 9204 | 432 | nuf,cic,H,nub,Dl,tara,N,Mpcp2,lilli,SERCA,pnr,ap | | 6.617e-05 | -9.62 | GO\_MORPHOGENESIS\_OF\_AN\_EPITHELIUM | MSigDB lists | GO\_MORPHOGENESIS\_OF\_AN\_EPITHELIUM | 99 | 13 | 4161 | 156 | Mef2,Dl,otk,ft,sli,ds,Fem-1,LanA,fz,N,ap,fz2,arm | | 6.659e-05 | -9.62 | ast (asteroid) | protein interactions | 33282 | 6 | 4 | 9204 | 432 | dpy,Egfr,px,S | | 6.730e-05 | -9.61 | axon | cellular component | GO:0030424 | 233 | 21 | 11940 | 417 | path,jeb,orb2,pum,Magi,Ptp4E,shot,Abl,CG9784,drl,Unc-76,wnd,jar,Bsg,arm,gukh,InR,robo1,ena,Syx1A,Gr93c | | 6.778e-05 | -9.60 | regulation of Notch signaling pathway | biological process | GO:0008593 | 75 | 11 | 12345 | 442 | Cbl,ed,H,spdo,Dl,Tom,scrib,CycG,cue,Fkbp14,lqf | | 7.008e-05 | -9.57 | MODULE\_438 | MSigDB lists | MODULE\_438 | 13 | 5 | 4161 | 156 | Syx1A,step,LanA,Dyb,betaTub60D | | 7.081e-05 | -9.56 | VECCHI\_GASTRIC\_CANCER\_ADVANCED\_VS\_EARLY\_UP | MSigDB lists | VECCHI\_GASTRIC\_CANCER\_ADVANCED\_VS\_EARLY\_UP | 29 | 7 | 4161 | 156 | Fkbp14,Actbeta,CG1673,tkv,sli,Ddr,Gfat2 | | 7.081e-05 | -9.56 | GO\_PROTEIN\_TYROSINE\_KINASE\_ACTIVITY | MSigDB lists | GO\_PROTEIN\_TYROSINE\_KINASE\_ACTIVITY | 29 | 7 | 4161 | 156 | Dyrk2,Ddr,Src42A,htl,InR,Eph,Nrk | | 7.107e-05 | -9.55 | segment polarity determination | biological process | GO:0007367 | 40 | 8 | 12345 | 442 | fz,fz2,arm,dlp,kn,rdx,Egfr,frc | | 7.159e-05 | -9.54 | male sex differentiation | biological process | GO:0046661 | 30 | 7 | 12345 | 442 | otk,Abd-B,Pvr,PI31,N,eya,Rab5 | | 7.287e-05 | -9.53 | cell-cell junction organization | biological process | GO:0045216 | 63 | 10 | 12345 | 442 | cora,moody,Gli,scrib,baz,jar,arm,sns,shg,bou | | 7.444e-05 | -9.51 | positive regulation of programmed cell death | biological process | GO:0043068 | 89 | 12 | 12345 | 442 | rdx,morgue,N,lncRNA:Hsromega,Abd-B,ec,Dp,ft,Cbl,klu,hppy,Dronc | | 7.451e-05 | -9.50 | GO\_REGULATION\_OF\_OSSIFICATION | MSigDB lists | GO\_REGULATION\_OF\_OSSIFICATION | 39 | 8 | 4161 | 156 | arm,N,fz,CG9220,tkv,Ddr,Cdk4,Mef2 | | 7.513e-05 | -9.50 | regulation of gene expression, epigenetic | biological process | GO:0040029 | 163 | 17 | 12345 | 442 | bin3,crol,tara,Dcr-2,vig,Sxl,Su(z)2,Tis11,Elba2,tna,FBXO11,corto,pum,Rm62,nub,Psc,fl(2)d | | 7.674e-05 | -9.48 | negative regulation of neurogenesis | biological process | GO:0050768 | 76 | 11 | 12345 | 442 | ttk,Sema1a,ed,Abl,emc,N,sd,Dl,aop,Sema1b,Socs36E | | 7.799e-05 | -9.46 | Fmr1 (Fmr1) | protein interactions | 37528 | 114 | 16 | 9204 | 432 | pnr,Cbl,cact,dlp,rictor,SCAR,d,Rm62,SmD1,Diap1,chic,dock,orb2,vig,Abl,baz | | 8.126e-05 | -9.42 | kinase activity | molecular function | GO:0016301 | 375 | 28 | 12176 | 414 | Pdk1,Pgk,Cdk4,Pitslre,wnd,Drak,CycG,Pvr,tkv,otk,Egfr,Raf,nmo,Lk6,hppy,l(1)G0196,trbl,Src42A,Dyrk2,CycB3,Eph,Ddr,Nrk,Abl,drl,Cerk,InR,htl | | 8.129e-05 | -9.42 | Vav (Vav guanine nucleotide exchange factor) | protein interactions | 32920 | 11 | 5 | 9204 | 432 | cindr,Egfr,Pvr,spi,shg | | 8.129e-05 | -9.42 | kek1 (kekkon 1) | protein interactions | 34688 | 11 | 5 | 9204 | 432 | Dg,ed,spi,kek5,Egfr | | 8.212e-05 | -9.41 | kuz (kuzbanian) | protein interactions | 34772 | 17 | 6 | 9204 | 432 | N,robo1,Dl,sli,SNF4Agamma,H | | 8.212e-05 | -9.41 | Hem (HEM-protein) | protein interactions | 40462 | 17 | 6 | 9204 | 432 | Abl,Cip4,dock,ft,SCAR,Arf79F | | 8.227e-05 | -9.41 | AP2\_Q6 | MSigDB lists | AP2\_Q6 | 62 | 10 | 4161 | 156 | ap,FBXO11,wdb,Kdm2,dlp,fz2,Eph,Fkbp14,hyx,Pgant6 | | 8.247e-05 | -9.40 | sl (small wing) | protein interactions | 32601 | 24 | 7 | 9204 | 432 | vn,S,Pvr,InR,Egfr,spi,Dcr-2 | | 8.247e-05 | -9.40 | stan (starry night) | protein interactions | 36125 | 24 | 7 | 9204 | 432 | Dl,ed,S,baz,fz,SMC3,Dg | | 8.383e-05 | -9.39 | striated muscle cell differentiation | biological process | GO:0051146 | 104 | 13 | 12345 | 442 | SCAR,Mef2,sd,aop,kon,Dg,siz,Pgk,htl,Bsg,dock,sns,InR | | 8.557e-05 | -9.37 | regulation of cell division | biological process | GO:0051302 | 41 | 8 | 12345 | 442 | klu,Sxl,jumu,Ocrl,Egfr,nuf,N,cindr | | 8.630e-05 | -9.36 | macromolecule localization | biological process | GO:0033036 | 868 | 53 | 12345 | 442 | CG7394,Pex5,Pat1,Dg,apolpp,Arf79F,rib,d,ds,hth,Rab1,Myo10A,cv-d,Rab5,flfl,Mef2,Syx1A,TBC1D5,wbl,jar,rictor,baz,boi,Dcr-2,Tpst,AP-1gamma,Slbp,Cip4,arm,SNF4Agamma,Shrm,His2Av,14-3-3epsilon,Pex10,ft,scrib,14-3-3zeta,CG3149,Pen,chic,Bsg,lncRNA:Hsromega,Npc2g,lt,Vps13,gukh,unc-119,cora,cact,shg,klar,p130CAS,Pex14 | | 8.687e-05 | -9.35 | TyrKc | smart domains | SM00219 | 27 | 8 | 5086 | 290 | otk,Abl,htl,InR,Nrk,Eph,Src42A,Egfr | | 8.794e-05 | -9.34 | receptor complex | cellular component | GO:0043235 | 79 | 11 | 11940 | 417 | Ddr,nAChRalpha4,Pvr,drl,htl,Eph,InR,tkv,Egfr,Nrk,scrib | | 8.809e-05 | -9.34 | negative regulation of photoreceptor cell differentiation | biological process | GO:0046533 | 14 | 5 | 12345 | 442 | ed,Socs36E,N,ttk,aop | | 8.809e-05 | -9.34 | R3/R4 cell fate commitment | biological process | GO:0007464 | 14 | 5 | 12345 | 442 | kay,aop,N,scrib,fz | | 8.949e-05 | -9.32 | GO\_NEGATIVE\_REGULATION\_OF\_MYELOID\_CELL\_DIFFERENTIATION | MSigDB lists | GO\_NEGATIVE\_REGULATION\_OF\_MYELOID\_CELL\_DIFFERENTIATION | 30 | 7 | 4161 | 156 | cact,Dl,Tollo,hyx,Actbeta,Cdk4,arm | | 8.949e-05 | -9.32 | GO\_REGULATION\_OF\_OSTEOBLAST\_DIFFERENTIATION | MSigDB lists | GO\_REGULATION\_OF\_OSTEOBLAST\_DIFFERENTIATION | 30 | 7 | 4161 | 156 | tkv,fz,Cdk4,Mef2,N,Ddr,arm | | 8.965e-05 | -9.32 | actin cytoskeleton reorganization | biological process | GO:0031532 | 31 | 7 | 12345 | 442 | SCAR,RhoGEF3,Dg,rictor,p130CAS,nuf,RhoU | | 8.965e-05 | -9.32 | larval somatic muscle development | biological process | GO:0007526 | 31 | 7 | 12345 | 442 | Alh,ap,chn,Kdm2,kn,crp,eya | | 8.966e-05 | -9.32 | Ig-like\_fold | interpro domains | IPR013783 | 195 | 20 | 11383 | 450 | ed,Ppn,kn,Eph,Dg,otk,Bsg,dome,sns,InR,boi,Pvr,robo1,htl,vn,unc-5,Ptp4E,CG31431,Ocrl,kek5 | | 9.174e-05 | -9.30 | cell cycle | biological process | GO:0007049 | 678 | 44 | 12345 | 442 | Egfr,Ocrl,LanA,CG3436,Sxl,Spc105R,N,14-3-3zeta,betaTub60D,pita,corto,bol,chic,CG11414,orb2,ph-p,Wdr62,Psc,His3.3B,Syx1A,Rab5,baz,His3.3A,jar,Slbp,Cks30A,larp,eff,Dp,wdb,14-3-3epsilon,kay,Pex10,SCAR,Dl,Gli,pum,Arf79F,CycG,PI31,CycB3,Rab1,Hr39,mop | | 9.314e-05 | -9.28 | hop (hopscotch) | protein interactions | 32080 | 41 | 9 | 9204 | 432 | kay,pxb,eya,dome,Cdk4,Dl,Socs36E,Raf,baz | | 9.320e-05 | -9.28 | leg disc pattern formation | biological process | GO:0035223 | 22 | 6 | 12345 | 442 | tsh,hth,B-H1,Egfr,vn,ap | | 9.450e-05 | -9.27 | ptc (patched) | protein interactions | 35851 | 153 | 19 | 9204 | 432 | LanA,vn,pnr,hyx,fz2,apolpp,sina,crol,Pgant6,Sxl,dlp,CG9784,Ppn,sns,N,Ziz,kn,MEP-1,boi | | 9.759e-05 | -9.23 | TYR\_PHOSPHATASE\_PTP | prosite domains | PS50055 | 16 | 6 | 6243 | 325 | Ptp4E,mop,CG42327,Pez,Ptpmeg2,IA-2 | | 9.821e-05 | -9.23 | GO\_REPRODUCTIVE\_SYSTEM\_DEVELOPMENT | MSigDB lists | GO\_REPRODUCTIVE\_SYSTEM\_DEVELOPMENT | 89 | 12 | 4161 | 156 | tkv,Actbeta,N,CG7394,arm,Mfe2,fz2,Tollo,robo1,Fem-1,sli,CG17883 | | 1.009e-04 | -9.20 | hep (hemipterous) | protein interactions | 32256 | 71 | 12 | 9204 | 432 | chic,wnd,tkv,kay,ena,Pvr,Src42A,arm,Raf,scrib,InR,H | | 1.012e-04 | -9.20 | non-canonical Wnt signaling pathway | biological process | GO:0035567 | 8 | 4 | 12345 | 442 | fz2,fz,fz3,otk | | 1.012e-04 | -9.20 | gonadal mesoderm development | biological process | GO:0007506 | 8 | 4 | 12345 | 442 | htl,Abd-B,abd-A,shg | | 1.018e-04 | -9.19 | identical protein binding | molecular function | GO:0042802 | 159 | 16 | 12176 | 414 | CtBP,ttk,otk,shg,E(spl)m5-HLH,rdx,Mfe2,Scr,step,mim,drl,14-3-3zeta,wbl,trbl,Dronc,corto | | 1.027e-04 | -9.18 | tio (tiptop) | protein interactions | 44272 | 3 | 3 | 9204 | 432 | hth,CtBP,tsh | | 1.047e-04 | -9.16 | GO\_SIGNAL\_TRANSDUCER\_ACTIVITY | MSigDB lists | GO\_SIGNAL\_TRANSDUCER\_ACTIVITY | 196 | 19 | 4161 | 156 | hppy,otk,Raf,robo1,nmo,unc-5,dlp,InR,htl,Arf79F,Ddr,Tollo,Eph,fz2,arm,tkv,nAChRalpha4,fz,Nrk | | 1.086e-04 | -9.13 | kis (kismet) | protein interactions | 33185 | 33 | 8 | 9204 | 432 | tna,pnr,Syx1A,Raf,rictor,Sema1a,N,Dg | | 1.100e-04 | -9.12 | Nrg (Neuroglian) | protein interactions | 31792 | 25 | 7 | 9204 | 432 | htl,ed,cora,unc-5,Arf79F,yrt,Egfr | | 1.100e-04 | -9.12 | step (steppke) | protein interactions | 35425 | 25 | 7 | 9204 | 432 | shg,S,step,alpha-Man-Ia,ena,Egfr,Raf | | 1.100e-04 | -9.12 | response to ethanol | biological process | GO:0045471 | 79 | 11 | 12345 | 442 | moody,Pdk1,Bacc,Egfr,pum,pxb,S,spi,hang,mura,hppy | | 1.105e-04 | -9.11 | GO\_PROTEIN\_PHOSPHORYLATION | MSigDB lists | GO\_PROTEIN\_PHOSPHORYLATION | 248 | 22 | 4161 | 156 | Drak,nmo,trbl,htl,InR,sns,CG4935,hppy,Cdk4,Raf,Mef2,otk,Dyrk2,Pdk1,Nrk,Lk6,tkv,Src42A,Ddr,PI31,Tollo,Eph | | 1.130e-04 | -9.09 | adenyl ribonucleotide binding | molecular function | GO:0032559 | 655 | 41 | 12176 | 414 | d,CG5853,Nrk,Abl,Raf,Lk6,hppy,trbl,jar,Drak,wnd,Pvr,SNF4Agamma,otk,Pdk1,SMC3,Pitslre,Hel89B,Eph,Ddr,drl,ATPCL,InR,htl,nmo,Myo10A,Rm62,l(1)G0196,Src42A,sick,Dyrk2,GluProRS,Dcr-2,SERCA,tkv,Egfr,CG11069,CG31121,Pgk,CG17646,Cdk4 | | 1.142e-04 | -9.08 | purine ribonucleotide binding | molecular function | GO:0032555 | 811 | 48 | 12176 | 414 | otk,CG2017,SNF4Agamma,wnd,Drak,Pvr,SMC3,Pitslre,Pdk1,CG5853,Nrk,Abl,Rab1,d,jar,trbl,Raf,Lk6,hppy,Egfr,RhoU,GluProRS,Rab5,Dcr-2,SERCA,tkv,Arf79F,Cdk4,CG17646,Pgk,CG31121,CG11069,InR,htl,Ric,ATPCL,Ddr,Eph,drl,Hel89B,betaTub60D,Dyrk2,sick,Src42A,Myo10A,l(1)G0196,Rm62,nmo | | 1.183e-04 | -9.04 | GO\_TUBE\_DEVELOPMENT | MSigDB lists | GO\_TUBE\_DEVELOPMENT | 134 | 15 | 4161 | 156 | Dl,otk,Mef2,robo1,LanA,ft,ds,CG17883,C1GalTA,sli,ap,fz,N,alpha-Man-Ia,arm | | 1.183e-04 | -9.04 | elav (embryonic lethal abnormal vision) | protein interactions | 31000 | 18 | 6 | 9204 | 432 | robo1,msi,Fas1,sli,Sxl,pum | | 1.183e-04 | -9.04 | GO\_PROTEIN\_IMPORT\_INTO\_PEROXISOME\_MATRIX | MSigDB lists | GO\_PROTEIN\_IMPORT\_INTO\_PEROXISOME\_MATRIX | 8 | 4 | 4161 | 156 | Pex10,Pex7,Pex5,Pex14 | | 1.183e-04 | -9.04 | BIOCARTA\_PS1\_PATHWAY | MSigDB lists | BIOCARTA\_PS1\_PATHWAY | 8 | 4 | 4161 | 156 | arm,fz,N,Dl | | 1.223e-04 | -9.01 | negative chemotaxis | biological process | GO:0050919 | 23 | 6 | 12345 | 442 | drl,Sema1a,sli,unc-5,fz2,Sema1b | | 1.233e-04 | -9.00 | purine ribonucleoside triphosphate binding | molecular function | GO:0035639 | 791 | 47 | 12176 | 414 | Cdk4,Pgk,CG17646,CG31121,CG11069,Egfr,RhoU,GluProRS,Dcr-2,Rab5,tkv,SERCA,Arf79F,Dyrk2,sick,Src42A,Myo10A,l(1)G0196,Rm62,nmo,InR,htl,Ric,ATPCL,Ddr,Eph,drl,betaTub60D,Hel89B,SMC3,Pitslre,Pdk1,otk,CG2017,wnd,Drak,Pvr,jar,trbl,Lk6,Raf,hppy,Nrk,CG5853,Abl,Rab1,d | | 1.233e-04 | -9.00 | Role of Abl in Robo-Slit signaling | REACTOME pathways | R-DME-428890 | 7 | 4 | 3923 | 176 | Abl,sli,dlp,robo1 | | 1.233e-04 | -9.00 | Znf\_C2H2\_type | interpro domains | IPR013087 | 297 | 26 | 11383 | 450 | hang,CG14441,chn,CG12769,tsh,CG10348,sens,disco-r,crol,sqz,Prdm13,CG32772,klu,mld,ttk,sens-2,CG10321,rgr,MEP-1,bowl,CG11414,CG11122,pita,esg,zfh2,CG11247 | | 1.245e-04 | -8.99 | maintenance of location | biological process | GO:0051235 | 67 | 10 | 12345 | 442 | Vps13,SNF4Agamma,p130CAS,14-3-3epsilon,14-3-3zeta,Mef2,Dg,chic,boi,cact | | 1.289e-04 | -8.96 | adenyl nucleotide binding | molecular function | GO:0030554 | 659 | 41 | 12176 | 414 | Egfr,Dcr-2,GluProRS,tkv,SERCA,CG17646,Pgk,Cdk4,CG11069,CG31121,ATPCL,InR,htl,Hel89B,Ddr,Eph,drl,Src42A,sick,Dyrk2,nmo,Myo10A,l(1)G0196,Rm62,Drak,wnd,Pvr,otk,SNF4Agamma,SMC3,Pitslre,Pdk1,d,Nrk,CG5853,Abl,trbl,jar,Lk6,Raf,hppy | | 1.310e-04 | -8.94 | embryonic organ development | biological process | GO:0048568 | 55 | 9 | 12345 | 442 | robo1,N,RhoL,sli,Pvr,jumu,Pvf3,shg,pnr | | 1.314e-04 | -8.94 | phosphotransferase activity, alcohol group as acceptor | molecular function | GO:0016773 | 346 | 26 | 12176 | 414 | Cdk4,Pitslre,Pdk1,Egfr,Pvr,CycG,wnd,Drak,tkv,otk,Src42A,trbl,Dyrk2,hppy,Lk6,Raf,nmo,Cerk,htl,InR,CycB3,drl,Abl,Ddr,Eph,Nrk | | 1.372e-04 | -8.89 | ATP binding | molecular function | GO:0005524 | 639 | 40 | 12176 | 414 | Dcr-2,GluProRS,tkv,SERCA,Egfr,CG31121,CG11069,Cdk4,CG17646,Pgk,Eph,Ddr,drl,Hel89B,InR,htl,ATPCL,Myo10A,l(1)G0196,Rm62,nmo,Dyrk2,sick,Src42A,otk,wnd,Drak,Pvr,Pdk1,SMC3,Pitslre,CG5853,Nrk,Abl,d,Raf,Lk6,hppy,jar,trbl | | 1.377e-04 | -8.89 | cation binding | molecular function | GO:0043169 | 1072 | 59 | 12176 | 414 | Cyp6a16,CG13481,Hr39,Cbl,Atox1,shg,CG2991,CG31457,Tet,Pp2C1,mld,Psa,grn,CG14441,Su(z)2,Diap1,CG4467,pnr,rho-4,Pex10,Dg,Psc,CG10417,sli,mura,CG45050,CG11360,Kdm2,dpy,Dyb,tkv,pita,lt,ds,CG3107,Cadps,CG10321,alpha-Man-Ia,FBXO11,sina,N,CG1218,Dl,CG34401,su(sable),ft,lin-28,ph-p,CalpC,Hr4,CG11247,tna,CG11414,Tis11,shot,Cals,cue,hang,Fkbp14 | | 1.389e-04 | -8.88 | GSE28726\_NAIVE\_CD4\_TCELL\_VS\_NAIVE\_NKTCELL\_DN | MSigDB lists | GSE28726\_NAIVE\_CD4\_TCELL\_VS\_NAIVE\_NKTCELL\_DN | 32 | 7 | 4161 | 156 | DPCoAC,Ocrl,step,Cadps,Kdm2,htl,trbl | | 1.409e-04 | -8.87 | regulation of cell death | biological process | GO:0010941 | 257 | 22 | 12345 | 442 | ec,Egfr,Abd-B,morgue,rdx,Pdk1,ft,spi,Diap1,Dp,sens,Pvr,lncRNA:Hsromega,N,FBXO11,Actbeta,Dronc,hppy,Cbl,klu,eya,Raf | | 1.411e-04 | -8.87 | supramolecular fiber organization | biological process | GO:0097435 | 188 | 18 | 12345 | 442 | ed,cindr,N,SCAR,Dl,chic,Dg,shot,RhoU,Bsg,Src42A,Shrm,p130CAS,f,kon,ena,sick,jar | | 1.412e-04 | -8.87 | hematopoietic or lymphoid organ development | biological process | GO:0048534 | 95 | 12 | 12345 | 442 | N,RhoL,htl,sens-2,cact,Pvr,CG1129,Pen,Raf,pnr,shg,Pvf3 | | 1.412e-04 | -8.87 | immune system development | biological process | GO:0002520 | 95 | 12 | 12345 | 442 | Raf,Pvf3,pnr,shg,Pvr,cact,sens-2,htl,CG1129,Pen,RhoL,N | | 1.423e-04 | -8.86 | GO\_REGULATION\_OF\_MULTICELLULAR\_ORGANISMAL\_DEVELOPMENT | MSigDB lists | GO\_REGULATION\_OF\_MULTICELLULAR\_ORGANISMAL\_DEVELOPMENT | 325 | 26 | 4161 | 156 | Cals,cact,otk,Dl,robo1,Mef2,Cdk4,Raf,Arf79F,Actbeta,N,dlp,Eph,Tollo,hyx,PI31,LanA,Ddr,ft,sli,tkv,nAChRalpha4,fz,Nrk,fz2,arm | | 1.435e-04 | -8.85 | EGF-like\_dom | interpro domains | IPR000742 | 74 | 11 | 11383 | 450 | cue,spi,sli,ft,Dl,N,nahoda,shg,LanA,dpy,vn | | 1.439e-04 | -8.85 | Developmental Biology | REACTOME pathways | R-DME-1266738 | 211 | 22 | 3923 | 176 | ena,Pvr,eyg,sina,Raf,sli,Abl,shot,dlp,14-3-3zeta,PI31,Egfr,vn,wdb,Eph,dock,CG42684,boi,CG4935,robo1,Syx1A,htl | | 1.445e-04 | -8.84 | Mef2 (Myocyte enhancer factor 2) | protein interactions | 36032 | 26 | 7 | 9204 | 432 | abd-A,Mef2,Abd-B,pnr,Him,Scr,sd | | 1.445e-04 | -8.84 | dally (division abnormally delayed) | protein interactions | 39013 | 26 | 7 | 9204 | 432 | apolpp,fz2,arm,tkv,Sema1a,dlp,Dad | | 1.445e-04 | -8.84 | pyd (polychaetoid) | protein interactions | 41062 | 26 | 7 | 9204 | 432 | Dl,ena,N,h,emc,tkv,shg | | 1.445e-04 | -8.84 | bun (bunched) | protein interactions | 34665 | 26 | 7 | 9204 | 432 | aop,N,vn,S,pnr,Egfr,InR | | 1.445e-04 | -8.84 | maintenance of protein location | biological process | GO:0045185 | 44 | 8 | 12345 | 442 | Vps13,14-3-3epsilon,p130CAS,14-3-3zeta,boi,cact,Dg,chic | | 1.449e-04 | -8.84 | SH3 domain binding | molecular function | GO:0017124 | 9 | 4 | 12176 | 414 | cindr,Cbl,ena,mim | | 1.461e-04 | -8.83 | programmed cell death | biological process | GO:0012501 | 172 | 17 | 12345 | 442 | Drep4,chrb,Dronc,hppy,S,abd-A,Atg17,Dl,N,Dp,arm,Diap1,Dcr-2,wgn,ec,Atg16,scyl | | 1.470e-04 | -8.83 | neuron part | cellular component | GO:0097458 | 540 | 36 | 11940 | 417 | arm,wnd,ReepA,CG31211,Rab5,drl,ena,robo1,LanA,InR,rg,tkv,gukh,Magi,pum,jeb,path,Rab1,CG9784,chic,shot,Cadps,jar,Bsg,Unc-76,bol,Gr93c,RhoU,Syx1A,Eph,nAChRalpha4,orb2,Abl,scrib,Myo10A,Ptp4E | | 1.496e-04 | -8.81 | miple1 (midkine and pleiotrophin 1) | protein interactions | 38047 | 7 | 4 | 9204 | 432 | Egfr,shg,nmo,arm | | 1.496e-04 | -8.81 | tyn (trynity) | protein interactions | 3354987 | 7 | 4 | 9204 | 432 | pnr,arm,shg,neo | | 1.499e-04 | -8.81 | anion binding | molecular function | GO:0043168 | 1099 | 60 | 12176 | 414 | Abl,Nrk,mim,CG5853,d,Rab1,sli,Lk6,hppy,Raf,jar,trbl,CG2017,otk,robo1,Pvr,wnd,Drak,Cip4,CenB1A,Pdk1,Pitslre,SMC3,drl,baz,Eph,Ddr,betaTub60D,Hel89B,htl,InR,ATPCL,vn,step,Ric,Rm62,l(1)G0196,Myo10A,apolpp,CG7560,nmo,cv-d,Dyrk2,sick,Src42A,SERCA,tkv,CG7536,Rab5,Dcr-2,GluProRS,Arf79F,Sema1a,Egfr,RhoU,CG31121,CG11069,Cdk4,CG17646,Pgk | | 1.506e-04 | -8.80 | Rab6 (Rab6) | protein interactions | 34636 | 85 | 13 | 9204 | 432 | N,Dg,CenB1A,CG30344,Unc-76,Ocrl,ttk,Rab5,Rab1,TBC1D5,SERCA,flfl,Arf79F | | 1.516e-04 | -8.79 | DNA-binding transcription activator activity | molecular function | GO:0001216 | 86 | 11 | 12176 | 414 | CG12769,B-H1,dmrt11E,Adf1,sens,kay,Mef2,nub,kn,Scr,sim | | 1.516e-04 | -8.79 | DNA-binding transcription activator activity, RNA polymerase II-specific | molecular function | GO:0001228 | 86 | 11 | 12176 | 414 | dmrt11E,Adf1,sens,kay,CG12769,B-H1,nub,kn,Scr,sim,Mef2 | | 1.530e-04 | -8.79 | purine nucleotide binding | molecular function | GO:0017076 | 821 | 48 | 12176 | 414 | Arf79F,Rab5,GluProRS,Dcr-2,SERCA,tkv,RhoU,Egfr,CG11069,CG31121,CG17646,Pgk,Cdk4,betaTub60D,Hel89B,Eph,Ddr,drl,Ric,ATPCL,InR,htl,nmo,Myo10A,l(1)G0196,Rm62,Src42A,sick,Dyrk2,wnd,Drak,Pvr,CG2017,otk,SNF4Agamma,Pdk1,SMC3,Pitslre,Rab1,d,CG5853,Nrk,Abl,Lk6,Raf,hppy,trbl,jar | | 1.575e-04 | -8.76 | ribonucleotide binding | molecular function | GO:0032553 | 822 | 48 | 12176 | 414 | wnd,Drak,Pvr,SNF4Agamma,otk,CG2017,SMC3,Pitslre,Pdk1,Rab1,d,Nrk,CG5853,Abl,trbl,jar,Raf,Lk6,hppy,RhoU,Egfr,Arf79F,GluProRS,Rab5,Dcr-2,SERCA,tkv,Pgk,CG17646,Cdk4,CG11069,CG31121,Ric,ATPCL,InR,htl,Hel89B,betaTub60D,Ddr,Eph,drl,sick,Src42A,Dyrk2,nmo,Myo10A,l(1)G0196,Rm62 | | 1.601e-04 | -8.74 | positive regulation of apoptotic process | biological process | GO:0043065 | 69 | 10 | 12345 | 442 | rdx,N,morgue,lncRNA:Hsromega,Abd-B,Dp,ft,Cbl,Dronc,hppy | | 1.607e-04 | -8.74 | PTPc | smart domains | SM00194 | 16 | 6 | 5086 | 290 | IA-2,Pez,Ptp4E,mop,Ptpmeg2,CG42327 | | 1.621e-04 | -8.73 | axis specification | biological process | GO:0009798 | 207 | 19 | 12345 | 442 | fs(1)h,bowl,Dg,chic,pum,Pat1,scrib,14-3-3zeta,N,Cbl,Raf,cact,wbl,baz,Egfr,cic,14-3-3epsilon,Dp,shg | | 1.661e-04 | -8.70 | Hipk (Homeodomain interacting protein kinase) | protein interactions | 38070 | 19 | 6 | 9204 | 432 | N,fz2,arm,spen,ey,Dl | | 1.661e-04 | -8.70 | Dscam1 (Down syndrome cell adhesion molecule 1) | protein interactions | 35652 | 19 | 6 | 9204 | 432 | sli,Src42A,robo1,dock,Abl,wnd | | 1.675e-04 | -8.69 | Ranbp9 (Ranbp9) | protein interactions | 41367 | 64 | 11 | 9204 | 432 | Raf,C1GalTA,rictor,Kr-h2,Mpcp2,GluProRS,ds,Gfat2,Pdk1,SERCA,Rab5 | | 1.681e-04 | -8.69 | GO\_SEX\_DIFFERENTIATION | MSigDB lists | GO\_SEX\_DIFFERENTIATION | 55 | 9 | 4161 | 156 | robo1,Tollo,CG17883,sli,CG7394,Actbeta,tkv,arm,Mfe2 | | 1.695e-04 | -8.68 | stg (string) | protein interactions | 43466 | 35 | 8 | 9204 | 432 | ttk,aop,trbl,pnr,shg,arm,N,crol | | 1.695e-04 | -8.68 | ds (dachsous) | protein interactions | 33245 | 35 | 8 | 9204 | 432 | arm,ft,shg,fz,Mpcp2,d,cindr,SERCA | | 1.728e-04 | -8.66 | regulation of cell growth | biological process | GO:0001558 | 83 | 11 | 12345 | 442 | cic,Pdk1,Cdk4,shot,chic,Sema1b,Sema1a,InR,Orct2,fz,ey | | 1.757e-04 | -8.65 | carbohydrate derivative binding | molecular function | GO:0097367 | 1011 | 56 | 12176 | 414 | robo1,CG2017,SNF4Agamma,otk,Drak,wnd,Pvr,SMC3,Pitslre,Pdk1,Nrk,CG5853,Abl,Rab1,d,jar,trbl,sli,Lk6,Raf,hppy,Egfr,RhoU,GluProRS,Rab5,Dcr-2,SERCA,tkv,Sema1a,Arf79F,Cdk4,CG17646,Pgk,CG31121,CG11069,InR,Dl,CG1218,htl,Ric,ATPCL,vn,Ddr,Eph,drl,betaTub60D,Hel89B,Gfat2,Dyrk2,cv-d,Src42A,sick,Myo10A,l(1)G0196,Rm62,nmo | | 1.770e-04 | -8.64 | positive regulation of compound eye retinal cell programmed cell death | biological process | GO:0046672 | 9 | 4 | 12345 | 442 | morgue,klu,Dronc,ec | | 1.770e-04 | -8.64 | notum development | biological process | GO:0007477 | 9 | 4 | 12345 | 442 | Socs36E,Egfr,vn,eyg | | 1.770e-04 | -8.64 | zonula adherens assembly | biological process | GO:0045186 | 9 | 4 | 12345 | 442 | scrib,shg,arm,baz | | 1.770e-04 | -8.64 | positive regulation of retinal cell programmed cell death | biological process | GO:0046670 | 9 | 4 | 12345 | 442 | ec,Dronc,klu,morgue | | 1.775e-04 | -8.64 | cytoplasmic sequestering of transcription factor | biological process | GO:0042994 | 4 | 3 | 12345 | 442 | 14-3-3zeta,cact,14-3-3epsilon | | 1.775e-04 | -8.64 | outflow tract morphogenesis | biological process | GO:0003151 | 4 | 3 | 12345 | 442 | robo1,sli,shg | | 1.775e-04 | -8.64 | cytoplasmic sequestering of protein | biological process | GO:0051220 | 4 | 3 | 12345 | 442 | 14-3-3epsilon,cact,14-3-3zeta | | 1.810e-04 | -8.62 | GO\_TUBE\_MORPHOGENESIS | MSigDB lists | GO\_TUBE\_MORPHOGENESIS | 81 | 11 | 4161 | 156 | Mef2,Dl,otk,sli,ds,ft,LanA,N,fz,ap,arm | | 1.811e-04 | -8.62 | negative regulation of cell adhesion | biological process | GO:0007162 | 16 | 5 | 12345 | 442 | baz,N,aop,disco-r,arm | | 1.811e-04 | -8.62 | establishment of epithelial cell polarity | biological process | GO:0090162 | 16 | 5 | 12345 | 442 | ft,ds,yrt,fz,scrib | | 1.829e-04 | -8.61 | Myc (Myc) | protein interactions | 31310 | 174 | 20 | 9204 | 432 | hang,tkv,Mef2,sim,ds,larp,shg,hyx,Sxl,crol,Rab5,Dp,beat-IIa,cindr,Diap1,N,ft,Atg17,Poxn,Su(z)2 | | 1.937e-04 | -8.55 | Adaptive Immune System | REACTOME pathways | R-DME-1280218 | 186 | 20 | 3923 | 176 | vn,Egfr,htl,mura,pes,CG4467,Dronc,wdb,Raf,cact,sina,PI31,Pdk1,rictor,14-3-3zeta,RapGAP1,FBXO11,trbl,eff,Psa | | 1.937e-04 | -8.55 | GENTILE\_UV\_RESPONSE\_CLUSTER\_D4 | MSigDB lists | GENTILE\_UV\_RESPONSE\_CLUSTER\_D4 | 24 | 6 | 4161 | 156 | sina,CG9220,Pen,Rab1,Tpst,CG3436 | | 1.937e-04 | -8.55 | GO\_SKIN\_EPIDERMIS\_DEVELOPMENT | MSigDB lists | GO\_SKIN\_EPIDERMIS\_DEVELOPMENT | 24 | 6 | 4161 | 156 | arm,LanA,Actbeta,N,ap,Dl | | 1.976e-04 | -8.53 | ft (fat) | protein interactions | 33627 | 76 | 12 | 9204 | 432 | arm,ft,shg,Sema1a,fz,nub,H,d,Raf,cora,ena,ds | | 1.994e-04 | -8.52 | Dcp-1 (Death caspase-1) | protein interactions | 37729 | 45 | 9 | 9204 | 432 | Dl,Egfr,SNF4Agamma,eff,lncRNA:Hsromega,Diap1,esg,aop,Dronc | | 2.019e-04 | -8.51 | imaginal disc-derived leg segmentation | biological process | GO:0036011 | 25 | 6 | 12345 | 442 | Dl,ss,N,dve,ap,bowl | | 2.019e-04 | -8.51 | appendage segmentation | biological process | GO:0035285 | 25 | 6 | 12345 | 442 | ss,Dl,N,bowl,ap,dve | | 2.019e-04 | -8.51 | axon midline choice point recognition | biological process | GO:0016199 | 25 | 6 | 12345 | 442 | shot,Abl,robo1,Sema1a,drl,sli | | 2.026e-04 | -8.50 | ovarian nurse cell to oocyte transport | biological process | GO:0007300 | 35 | 7 | 12345 | 442 | ena,jar,Pen,chic,Abl,SCAR,Ppcs | | 2.039e-04 | -8.50 | cytoplasmic region | cellular component | GO:0099568 | 252 | 21 | 11940 | 417 | nuf,cindr,drongo,Dl,CG6891,jar,Bsg,Rab5,Shrm,chic,scrib,Cbl,Abl,ed,Pez,pum,baz,spdo,CG42684,step,shg | | 2.040e-04 | -8.50 | imaginal disc-derived wing margin morphogenesis | biological process | GO:0008587 | 71 | 10 | 12345 | 442 | N,Dl,Fkbp14,shot,mam,arm,dlp,H,hyx,dpy | | 2.040e-04 | -8.50 | digestive tract morphogenesis | biological process | GO:0048546 | 71 | 10 | 12345 | 442 | Inx2,N,Dad,Pvr,Egfr,bowl,S,rib,spi,dome | | 2.043e-04 | -8.50 | Cdk1 (Cyclin-dependent kinase 1) | protein interactions | 34411 | 55 | 10 | 9204 | 432 | Atg16,CycG,CG34376,shg,arm,CycB3,Abl,Cks30A,Psc,tara | | 2.068e-04 | -8.48 | MODULE\_533 | MSigDB lists | MODULE\_533 | 9 | 4 | 4161 | 156 | Dyb,betaTub60D,Syx1A,step | | 2.084e-04 | -8.48 | GSE5099\_CLASSICAL\_M1\_VS\_ALTERNATIVE\_M2\_MACROPHAGE\_DN | MSigDB lists | GSE5099\_CLASSICAL\_M1\_VS\_ALTERNATIVE\_M2\_MACROPHAGE\_DN | 34 | 7 | 4161 | 156 | fz,Cdk4,nAChRalpha4,CG9220,muc,shot,CG13893 | | 2.084e-04 | -8.48 | ISRE\_01 | MSigDB lists | ISRE\_01 | 34 | 7 | 4161 | 156 | unc-5,lqf,Hel89B,Mef2,Pgk,Cdk4,FBXO11 | | 2.084e-04 | -8.48 | GSE43863\_TH1\_VS\_TFH\_MEMORY\_CD4\_TCELL\_DN | MSigDB lists | GSE43863\_TH1\_VS\_TFH\_MEMORY\_CD4\_TCELL\_DN | 34 | 7 | 4161 | 156 | Cerk,Drak,Kdm2,wbl,emp,trbl,CG2017 | | 2.094e-04 | -8.47 | sca (scabrous) | protein interactions | 36411 | 13 | 5 | 9204 | 432 | N,nmo,shg,Dl,mam | | 2.094e-04 | -8.47 | Gli (Gliotactin) | protein interactions | 34927 | 13 | 5 | 9204 | 432 | N,tkv,His2Av,H,Dad | | 2.099e-04 | -8.47 | positive regulation of cell death | biological process | GO:0010942 | 99 | 12 | 12345 | 442 | ec,Abd-B,lncRNA:Hsromega,morgue,N,rdx,hppy,Dronc,klu,Cbl,ft,Dp | | 2.136e-04 | -8.45 | Rac1 (Rac1) | protein interactions | 38146 | 124 | 16 | 9204 | 432 | robo1,fs(1)h,N,Raf,kn,scrib,SNF4Agamma,chic,muc,Diap1,RhoGEF3,lilli,nmo,shg,tkv,Dronc | | 2.141e-04 | -8.45 | negative regulation of developmental growth | biological process | GO:0048640 | 85 | 11 | 12345 | 442 | Abl,ft,Mob2,Sema1a,Cip4,Src42A,Sema1b,ena,Dad,LanA,scrib | | 2.148e-04 | -8.45 | neuron recognition | biological process | GO:0008038 | 114 | 13 | 12345 | 442 | p130CAS,Abl,Fas1,Sema1a,fz2,Pvf3,Bsg,shot,robo1,Gli,drl,sli,SCAR | | 2.148e-04 | -8.45 | regulation of neuron projection development | biological process | GO:0010975 | 114 | 13 | 12345 | 442 | Rab1,Sema1a,ss,fz,kn,Adf1,Abl,CG34401,robo1,shot,rictor,Sema1b,chic | | 2.170e-04 | -8.44 | WALLACE\_PROSTATE\_CANCER\_RACE\_UP | MSigDB lists | WALLACE\_PROSTATE\_CANCER\_RACE\_UP | 16 | 5 | 4161 | 156 | CG6891,Pex14,arm,bru2,Drak | | 2.179e-04 | -8.43 | GO\_SIGNALING\_RECEPTOR\_ACTIVITY | MSigDB lists | GO\_SIGNALING\_RECEPTOR\_ACTIVITY | 126 | 14 | 4161 | 156 | Nrk,fz,nAChRalpha4,tkv,InR,htl,dlp,unc-5,fz2,robo1,Tollo,Eph,otk,Ddr | | 2.201e-04 | -8.42 | CAGCTG\_AP4\_Q5 | MSigDB lists | CAGCTG\_AP4\_Q5 | 278 | 23 | 4161 | 156 | rdx,Pat1,Kdm2,CG2017,Nrk,wdb,betaTub60D,LanA,Cadps,Ziz,rg,unc-119,N,l(2)37Cg,14-3-3zeta,ap,InR,CG4935,Pex7,Mef2,Dl,otk,Cals | | 2.211e-04 | -8.42 | Chc (Clathrin heavy chain) | protein interactions | 32537 | 112 | 15 | 9204 | 432 | lqf,Rab5,Rab1,AP-1gamma,Gfat2,Pdk1,SERCA,Arf79F,Mpcp2,N,Psa,Kr-h2,GluProRS,Sxl,rictor | | 2.223e-04 | -8.41 | Dref (DNA replication-related element factor) | protein interactions | 34328 | 66 | 11 | 9204 | 432 | betaTub60D,l(1)G0196,Pdk1,Diap1,ds,fs(1)h,E(Pc),S,Raf,dome,crol | | 2.279e-04 | -8.39 | regulation of immune system process | biological process | GO:0002682 | 230 | 20 | 12345 | 442 | nub,Pitslre,fz,mop,kn,LRR,imd,N,DPCoAC,Rm62,pxb,Pvr,Arf79F,fz2,pnr,CG32795,C1GalTA,cact,Dcr-2,htl | | 2.280e-04 | -8.39 | dia (diaphanous) | protein interactions | 35340 | 20 | 6 | 9204 | 432 | ena,spi,Sema1a,Cip4,Abl,chic | | 2.281e-04 | -8.39 | branching involved in open tracheal system development | biological process | GO:0060446 | 59 | 9 | 12345 | 442 | shg,arm,ttk,Raf,ed,esg,baz,shot,Ptp4E | | 2.281e-04 | -8.39 | branching morphogenesis of an epithelial tube | biological process | GO:0048754 | 59 | 9 | 12345 | 442 | baz,Ptp4E,shot,esg,ed,shg,ttk,Raf,arm | | 2.281e-04 | -8.39 | morphogenesis of a branching structure | biological process | GO:0001763 | 59 | 9 | 12345 | 442 | esg,baz,Ptp4E,shot,shg,ttk,Raf,arm,ed | | 2.281e-04 | -8.39 | morphogenesis of a branching epithelium | biological process | GO:0061138 | 59 | 9 | 12345 | 442 | baz,Ptp4E,shot,esg,ed,shg,Raf,ttk,arm | | 2.310e-04 | -8.37 | germarium-derived egg chamber formation | biological process | GO:0007293 | 100 | 12 | 12345 | 442 | 14-3-3epsilon,shg,ph-p,Cip4,Dg,baz,Pen,N,Ppcs,14-3-3zeta,Inx2,Dl | | 2.323e-04 | -8.37 | zf-C2H2 | pfam domains | PF00096 | 207 | 20 | 10480 | 419 | sens-2,CG10321,sqz,CG11247,sens,pita,CG32772,klu,ttk,hang,esg,chn,bowl,tsh,zfh2,mld,CG10348,Prdm13,crol,rgr | | 2.337e-04 | -8.36 | GO\_REGULATION\_OF\_PHOSPHORUS\_METABOLIC\_PROCESS | MSigDB lists | GO\_REGULATION\_OF\_PHOSPHORUS\_METABOLIC\_PROCESS | 335 | 26 | 4161 | 156 | nAChRalpha4,Tsp96F,Nrk,Pdk1,fz,wdb,CycG,wbl,arm,fz2,Eph,Tollo,unc-119,sli,Ddr,trbl,dock,InR,htl,Arf79F,Actbeta,14-3-3epsilon,Raf,Mef2,hppy,muc | | 2.344e-04 | -8.36 | KEGG\_PATHWAYS\_IN\_CANCER | MSigDB lists | KEGG\_PATHWAYS\_IN\_CANCER | 70 | 10 | 4161 | 156 | cact,Raf,Cdk4,LanA,InR,htl,CtBP,fz,fz2,arm | | 2.366e-04 | -8.35 | regulation of synapse structure or activity | biological process | GO:0050803 | 179 | 17 | 12345 | 442 | LanA,tkv,Dad,nmo,ena,rictor,Rab5,Abl,Cals,Src42A,Cip4,Dg,wnd,pum,scrib,Rab1,Mob2 | | 2.378e-04 | -8.34 | eye photoreceptor cell development | biological process | GO:0042462 | 86 | 11 | 12345 | 442 | klar,lilli,ttk,Raf,yrt,eya,Mob2,jeb,rg,Nrk,sens | | 2.397e-04 | -8.34 | faf (fat facets) | protein interactions | 43749 | 28 | 7 | 9204 | 432 | 14-3-3zeta,lqf,aop,wnd,imd,rictor,eff | | 2.402e-04 | -8.33 | metal ion binding | molecular function | GO:0046872 | 1047 | 57 | 12176 | 414 | Diap1,pnr,CG4467,Su(z)2,mld,Psa,grn,CG14441,Psc,mura,sli,CG45050,rho-4,Dg,Pex10,Hr39,Cbl,shg,Atox1,Cyp6a16,CG13481,Tet,CG31457,CG2991,lin-28,ph-p,CalpC,Hr4,N,CG1218,Dl,CG34401,su(sable),ft,cue,shot,Cals,hang,Fkbp14,CG11247,tna,CG11414,Tis11,pita,tkv,lt,CG11360,Kdm2,Dyb,dpy,alpha-Man-Ia,FBXO11,sina,ds,CG3107,Cadps,CG10321 | | 2.402e-04 | -8.33 | RNA polymerase II distal enhancer sequence-specific DNA binding | molecular function | GO:0000980 | 27 | 6 | 12176 | 414 | ey,aop,ttk,sd,abd-A,Abd-B | | 2.431e-04 | -8.32 | regulation of transport | biological process | GO:0051049 | 267 | 22 | 12345 | 442 | klar,lilli,Cadps,IA-2,Abl,Syx1A,rdx,Rab5,step,jar,mim,InR,Rab1,lt,SERCA,S,mop,Cbl,cindr,scrib,Arf79F,tipE | | 2.438e-04 | -8.32 | regulation of insulin receptor signaling pathway | biological process | GO:0046626 | 36 | 7 | 12345 | 442 | dock,step,wdb,trbl,rictor,lin-28,CycG | | 2.438e-04 | -8.32 | regulation of cellular response to insulin stimulus | biological process | GO:1900076 | 36 | 7 | 12345 | 442 | CycG,wdb,step,trbl,rictor,lin-28,dock | | 2.491e-04 | -8.30 | syncytial blastoderm mitotic cell cycle | biological process | GO:0035186 | 17 | 5 | 12345 | 442 | Psc,larp,ph-p,Cks30A,CycB3 | | 2.524e-04 | -8.28 | GSE37533\_PPARG1\_FOXP3\_VS\_PPARG2\_FOXP3\_TRANSDUCED\_CD4\_TCELL\_PIOGLITAZONE\_TREATED\_DN | MSigDB lists | GSE37533\_PPARG1\_FOXP3\_VS\_PPARG2\_FOXP3\_TRANSDUCED\_CD4\_TCELL\_PIOGLITAZONE\_TREATED\_DN | 35 | 7 | 4161 | 156 | Tollo,trbl,fz2,CG3961,ft,step,Dyrk2 | | 2.524e-04 | -8.28 | BOQUEST\_STEM\_CELL\_UP | MSigDB lists | BOQUEST\_STEM\_CELL\_UP | 35 | 7 | 4161 | 156 | Actbeta,bru2,Gfat2,Ddr,CG9485,dlp,sli | | 2.527e-04 | -8.28 | GO\_REGULATION\_OF\_MYELOID\_CELL\_DIFFERENTIATION | MSigDB lists | GO\_REGULATION\_OF\_MYELOID\_CELL\_DIFFERENTIATION | 46 | 8 | 4161 | 156 | arm,Actbeta,cact,Tollo,Dl,Mef2,Cdk4,hyx | | 2.527e-04 | -8.28 | GO\_NEURON\_PROJECTION\_GUIDANCE | MSigDB lists | GO\_NEURON\_PROJECTION\_GUIDANCE | 46 | 8 | 4161 | 156 | sli,Eph,robo1,unc-5,tkv,Unc-76,ap,sina | | 2.529e-04 | -8.28 | cell death | biological process | GO:0008219 | 180 | 17 | 12345 | 442 | arm,Dp,Diap1,Atg16,scyl,wgn,Dcr-2,ec,S,Drep4,Dronc,hppy,chrb,Dl,N,Atg17,abd-A | | 2.546e-04 | -8.28 | establishment of imaginal disc-derived wing hair orientation | biological process | GO:0001737 | 26 | 6 | 12345 | 442 | ft,SMC3,ds,N,scrib,fz | | 2.550e-04 | -8.27 | ena (enabled) | protein interactions | 37201 | 67 | 11 | 9204 | 432 | SCAR,SKIP,Abl,ft,N,step,robo1,ena,cindr,chic,CycG | | 2.557e-04 | -8.27 | GSE17186\_NAIVE\_VS\_CD21LOW\_TRANSITIONAL\_BCELL\_CORD\_BLOOD\_UP | MSigDB lists | GSE17186\_NAIVE\_VS\_CD21LOW\_TRANSITIONAL\_BCELL\_CORD\_BLOOD\_UP | 58 | 9 | 4161 | 156 | dock,sina,Pgk,nmo,alpha-Man-Ia,CG6465,step,Pex5,Pgant6 | | 2.560e-04 | -8.27 | Imp (IGF-II mRNA-binding protein) | protein interactions | 32009 | 37 | 8 | 9204 | 432 | H,jigr1,rictor,kn,msi,His2Av,orb2,chic | | 2.575e-04 | -8.26 | motor neuron axon guidance | biological process | GO:0008045 | 73 | 10 | 12345 | 442 | grn,Sema1a,unc-5,fend,fz2,Abl,beat-IIa,robo1,N,Ptp4E | | 2.575e-04 | -8.26 | negative regulation of gene expression, epigenetic | biological process | GO:0045814 | 73 | 10 | 12345 | 442 | FBXO11,corto,crol,bin3,Elba2,Dcr-2,nub,fl(2)d,Su(z)2,Psc | | 2.575e-04 | -8.26 | chromatin organization involved in negative regulation of transcription | biological process | GO:0097549 | 73 | 10 | 12345 | 442 | nub,Psc,fl(2)d,Su(z)2,Elba2,bin3,crol,corto,FBXO11,Dcr-2 | | 2.575e-04 | -8.26 | chromatin silencing | biological process | GO:0006342 | 73 | 10 | 12345 | 442 | Psc,fl(2)d,Su(z)2,nub,Dcr-2,crol,bin3,corto,FBXO11,Elba2 | | 2.577e-04 | -8.26 | GO\_TISSUE\_MORPHOGENESIS | MSigDB lists | GO\_TISSUE\_MORPHOGENESIS | 128 | 14 | 4161 | 156 | N,Actbeta,fz,ap,arm,fz2,Mef2,Dl,otk,sli,ds,ft,LanA,Fem-1 | | 2.592e-04 | -8.26 | Frizzled\_dom\_sf | interpro domains | IPR036790 | 9 | 4 | 11383 | 450 | fz,fz3,fz2,Nrk | | 2.605e-04 | -8.25 | PRC1 complex | cellular component | GO:0035102 | 10 | 4 | 11940 | 417 | ph-p,Su(z)2,Psc,corto | | 2.606e-04 | -8.25 | PIP3 activates AKT signaling | REACTOME pathways | R-DME-1257604 | 39 | 8 | 3923 | 176 | Pdk1,htl,rictor,wdb,Dronc,trbl,Egfr,vn | | 2.641e-04 | -8.24 | negative regulation of cellular component organization | biological process | GO:0051129 | 164 | 16 | 12345 | 442 | ena,baz,mim,Dad,LanA,Abl,klar,Src42A,Cip4,chic,Sema1b,trbl,shot,Kdm2,Mob2,Sema1a | | 2.704e-04 | -8.22 | Fz | pfam domains | PF01392 | 9 | 4 | 10480 | 419 | Nrk,fz2,fz3,fz | | 2.714e-04 | -8.21 | regulation of multicellular organism growth | biological process | GO:0040014 | 48 | 8 | 12345 | 442 | wbl,step,Hr4,Pdk1,ey,Orct2,InR,Raf | | 2.714e-04 | -8.21 | positive regulation of cell growth | biological process | GO:0030307 | 48 | 8 | 12345 | 442 | ey,Orct2,fz,InR,chic,shot,Cdk4,Pdk1 | | 2.776e-04 | -8.19 | RNA polymerase II proximal promoter sequence-specific DNA binding | molecular function | GO:0000978 | 92 | 11 | 12176 | 414 | sim,zfh2,CG34376,kay,Hr4,ttk,Hr39,pnr,h,grn,CG12769 | | 2.844e-04 | -8.16 | EGF\_1 | prosite domains | PS00022 | 52 | 10 | 6243 | 325 | shg,vn,N,sli,LanA,spi,ft,cue,nahoda,Dl | | 2.872e-04 | -8.16 | Pvr (PDGF- and VEGF-receptor related) | protein interactions | 34127 | 79 | 12 | 9204 | 432 | vn,Cbl,dock,bol,CG10321,Arf79F,Diap1,lilli,Src42A,Abl,Egfr,CG8034 | | 2.877e-04 | -8.15 | GO\_KINASE\_ACTIVITY | MSigDB lists | GO\_KINASE\_ACTIVITY | 229 | 20 | 4161 | 156 | Drak,nmo,l(1)G0196,htl,InR,trbl,hppy,otk,Cdk4,Raf,Cerk,Dyrk2,tkv,Lk6,Pgk,Pdk1,Nrk,Ddr,Src42A,Eph | | 2.882e-04 | -8.15 | CG10916 (uncharacterized protein) | protein interactions | 37081 | 8 | 4 | 9204 | 432 | InR,Egfr,Pvr,N | | 2.882e-04 | -8.15 | ihog (interference hedgehog) | protein interactions | 33972 | 8 | 4 | 9204 | 432 | Dl,dlp,boi,vn | | 2.921e-04 | -8.14 | DNA-binding transcription repressor activity | molecular function | GO:0001217 | 64 | 9 | 12176 | 414 | aop,klu,tsh,E(spl)m5-HLH,ttk,cic,h,chn,esg | | 2.921e-04 | -8.14 | DNA-binding transcription repressor activity, RNA polymerase II-specific | molecular function | GO:0001227 | 64 | 9 | 12176 | 414 | E(spl)m5-HLH,tsh,klu,aop,esg,chn,h,cic,ttk | | 2.952e-04 | -8.13 | protein localization | biological process | GO:0008104 | 693 | 43 | 12345 | 442 | CG7394,Pex5,Dg,Arf79F,ds,d,rib,hth,Rab1,Myo10A,Rab5,flfl,Syx1A,TBC1D5,wbl,baz,jar,rictor,boi,Tpst,AP-1gamma,Cip4,arm,His2Av,Shrm,Pex10,14-3-3epsilon,ft,scrib,14-3-3zeta,chic,Pen,Bsg,lncRNA:Hsromega,lt,Vps13,gukh,unc-119,cora,cact,shg,p130CAS,Pex14 | | 2.955e-04 | -8.13 | hindgut development | biological process | GO:0061525 | 61 | 9 | 12345 | 442 | Dad,Pvr,bowl,Egfr,N,dome,S,rib,spi | | 2.955e-04 | -8.13 | negative regulation of cellular amide metabolic process | biological process | GO:0034249 | 61 | 9 | 12345 | 442 | nub,Sxl,bru2,bin3,Tis11,312,orb2,msi,pum | | 2.955e-04 | -8.13 | hindgut morphogenesis | biological process | GO:0007442 | 61 | 9 | 12345 | 442 | rib,spi,S,dome,N,Pvr,Dad,Egfr,bowl | | 2.955e-04 | -8.13 | cell junction assembly | biological process | GO:0034329 | 61 | 9 | 12345 | 442 | arm,shg,sns,bou,moody,Gli,scrib,cindr,baz | | 2.982e-04 | -8.12 | PID\_PS1\_PATHWAY | MSigDB lists | PID\_PS1\_PATHWAY | 17 | 5 | 4161 | 156 | N,fz,Dl,arm,nmo | | 3.011e-04 | -8.11 | molecular transducer activity | molecular function | GO:0060089 | 447 | 30 | 12176 | 414 | boi,nAChRalpha4,fz,cv-d,unc-5,Eph,Nrk,Ddr,moody,drl,Dad,Hr4,Gr93c,InR,htl,N,Gli,dome,Ptp4E,Ir21a,Pitslre,Cdk4,fz2,Hr39,wgn,tkv,otk,fz3,Pvr,Egfr | | 3.033e-04 | -8.10 | if (inflated) | protein interactions | 32661 | 29 | 7 | 9204 | 432 | px,14-3-3epsilon,Sema1a,Raf,sli,H,crol | | 3.033e-04 | -8.10 | fz2 (frizzled 2) | protein interactions | 40090 | 29 | 7 | 9204 | 432 | Pen,otk,mam,fz,nmo,fz2,LanA | | 3.036e-04 | -8.10 | GO\_BONE\_DEVELOPMENT | MSigDB lists | GO\_BONE\_DEVELOPMENT | 36 | 7 | 4161 | 156 | ds,ft,Pex7,Mef2,CG9220,tkv,htl | | 3.063e-04 | -8.09 | Growth\_fac\_rcpt\_cys\_sf | interpro domains | IPR009030 | 34 | 7 | 11383 | 450 | Egfr,Dl,sli,dpy,N,Eph,InR | | 3.067e-04 | -8.09 | tou (toutatis) | protein interactions | 36241 | 21 | 6 | 9204 | 432 | pnr,kn,ph-p,Pvr,CtBP,Sema1a | | 3.112e-04 | -8.08 | da (daughterless) | protein interactions | 34413 | 38 | 8 | 9204 | 432 | E(spl)m5-HLH,emc,arm,Sxl,Egfr,ey,HLH3B,sens | | 3.114e-04 | -8.07 | GO\_REGULATION\_OF\_EMBRYONIC\_DEVELOPMENT | MSigDB lists | GO\_REGULATION\_OF\_EMBRYONIC\_DEVELOPMENT | 26 | 6 | 4161 | 156 | Dl,otk,fz,N,LanA,arm | | 3.132e-04 | -8.07 | Zyx (Zyxin) | protein interactions | 317824 | 14 | 5 | 9204 | 432 | ena,d,ds,H2.0,ft | | 3.135e-04 | -8.07 | PI3K/AKT activation | REACTOME pathways | R-DME-198203 | 40 | 8 | 3923 | 176 | vn,Egfr,Pdk1,rictor,htl,Dronc,trbl,wdb | | 3.176e-04 | -8.05 | photoreceptor cell axon guidance | biological process | GO:0072499 | 27 | 6 | 12345 | 442 | wgn,otk,Nrk,Abl,robo1,eya | | 3.176e-04 | -8.05 | axon choice point recognition | biological process | GO:0016198 | 27 | 6 | 12345 | 442 | robo1,Sema1a,drl,sli,shot,Abl | | 3.176e-04 | -8.05 | ectoderm development | biological process | GO:0007398 | 27 | 6 | 12345 | 442 | pnr,mam,N,sim,spi,Dl | | 3.214e-04 | -8.04 | Stat92E (Signal-transducer and activator of transcription protein at 92E) | protein interactions | 42428 | 58 | 10 | 9204 | 432 | ttk,Diap1,baz,Src42A,Egfr,scrib,dome,Cdk4,Sxl,Socs36E | | 3.224e-04 | -8.04 | instar larval development | biological process | GO:0002168 | 75 | 10 | 12345 | 442 | arm,Raf,fz2,fz,ft,Alh,scrib,Dad,chic,jar | | 3.224e-04 | -8.04 | positive regulation of neuron differentiation | biological process | GO:0045666 | 75 | 10 | 12345 | 442 | shot,Arf79F,chic,Src42A,InR,Rab1,Raf,fz,Sox14,mop | | 3.225e-04 | -8.04 | regulation of tube architecture, open tracheal system | biological process | GO:0035152 | 89 | 11 | 12345 | 442 | shot,Egfr,cora,Gli,ft,bou,ttk,klar,fz,shg,ds | | 3.238e-04 | -8.04 | ttk (tramtrack) | protein interactions | 48317 | 80 | 12 | 9204 | 432 | Pex5,MEP-1,eff,otp,msi,CtBP,sina,Pex7,lilli,ttk,aop,hang | | 3.293e-04 | -8.02 | cell recognition | biological process | GO:0008037 | 119 | 13 | 12345 | 442 | Pvf3,fz2,Fas1,Sema1a,Abl,p130CAS,SCAR,sli,robo1,drl,Gli,shot,Bsg | | 3.332e-04 | -8.01 | GO\_POSITIVE\_REGULATION\_OF\_NERVOUS\_SYSTEM\_DEVELOPMENT | MSigDB lists | GO\_POSITIVE\_REGULATION\_OF\_NERVOUS\_SYSTEM\_DEVELOPMENT | 73 | 10 | 4161 | 156 | sli,Eph,otk,Cals,Mef2,Raf,robo1,arm,N,Arf79F | | 3.341e-04 | -8.00 | regulation of small GTPase mediated signal transduction | biological process | GO:0051056 | 104 | 12 | 12345 | 442 | Rgl,Ziz,siz,step,Lk6,RhoGEF4,CG42684,14-3-3zeta,14-3-3epsilon,CG5522,RhoGEF3,RapGAP1 | | 3.343e-04 | -8.00 | TGCCAAR\_NF1\_Q6 | MSigDB lists | TGCCAAR\_NF1\_Q6 | 147 | 15 | 4161 | 156 | shot,rdx,GlcAT-I,Ppn,CG2812,wdb,Nrk,hppy,PI31,Rab5,Pex7,Dyb,otk,Cdk4,CG1129 | | 3.346e-04 | -8.00 | PID\_ECADHERIN\_STABILIZATION\_PATHWAY | MSigDB lists | PID\_ECADHERIN\_STABILIZATION\_PATHWAY | 10 | 4 | 4161 | 156 | arm,jar,dock,InR | | 3.348e-04 | -8.00 | heart process | biological process | GO:0003015 | 18 | 5 | 12345 | 442 | Egfr,cora,SERCA,pnr,spi | | 3.352e-04 | -8.00 | AACTTT\_UNKNOWN | MSigDB lists | AACTTT\_UNKNOWN | 421 | 30 | 4161 | 156 | Eph,hyx,Syx1A,Cadps,Ddr,CG45050,Kdm2,Ric,Pat1,fz2,arm,Cals,cact,Dl,Cdk4,Mef2,CG1129,hppy,Gfat2,InR,sd,Tmhs,ap,CtBP,N,Actbeta,nmo,Rab1,alpha-Man-Ia,dlp | | 3.377e-04 | -7.99 | rpr (reaper) | protein interactions | 40015 | 92 | 13 | 9204 | 432 | C1GalTA,Raf,eff,klu,14-3-3epsilon,lncRNA:Hsromega,nmo,S,Hr4,morgue,lilli,Diap1,Dronc | | 3.383e-04 | -7.99 | TTTGCAC\_MIR19A\_MIR19B | MSigDB lists | TTTGCAC\_MIR19A\_MIR19B | 116 | 13 | 4161 | 156 | Kdm2,eff,subdued,CG2812,wdb,Rgl,AP-1gamma,Hel89B,CG5522,Ocrl,rg,robo1,Atg16 | | 3.442e-04 | -7.97 | DAZARD\_RESPONSE\_TO\_UV\_NHEK\_DN | MSigDB lists | DAZARD\_RESPONSE\_TO\_UV\_NHEK\_DN | 87 | 11 | 4161 | 156 | Fem-1,Ziz,step,dock,Vps13,CtBP,Rim2,Pat1,shot,l(1)G0196,rdx | | 3.465e-04 | -7.97 | formation of primary germ layer | biological process | GO:0001704 | 38 | 7 | 12345 | 442 | htl,tkv,abd-A,jeb,sli,mam,spi | | 3.480e-04 | -7.96 | GO\_POSITIVE\_REGULATION\_OF\_RESPONSE\_TO\_STIMULUS | MSigDB lists | GO\_POSITIVE\_REGULATION\_OF\_RESPONSE\_TO\_STIMULUS | 402 | 29 | 4161 | 156 | eff,wbl,fz2,arm,tkv,subdued,CG9220,Pdk1,Kr-h2,Tsp96F,sina,Syx1A,PI31,AP-1gamma,sli,Tollo,hyx,14-3-3epsilon,htl,dock,14-3-3zeta,CtBP,N,Actbeta,hppy,cact,Dl,Mef2,Raf | | 3.559e-04 | -7.94 | Ig\_sub2 | interpro domains | IPR003598 | 109 | 13 | 11383 | 450 | CG31431,unc-5,kek5,otk,Bsg,sns,ed,Ppn,boi,robo1,Pvr,vn,htl | | 3.601e-04 | -7.93 | TGTTTAC\_MIR30A5P\_MIR30C\_MIR30D\_MIR30B\_MIR30E5P | MSigDB lists | TGTTTAC\_MIR30A5P\_MIR30C\_MIR30D\_MIR30B\_MIR30E5P | 148 | 15 | 4161 | 156 | lili,unc-5,lin-28,bru2,CenB1A,alpha-Man-Ia,eff,14-3-3zeta,Pgant5,Src42A,Rgl,Cadps,CG5522,Gfat2,kn | | 3.620e-04 | -7.92 | GO\_POSITIVE\_REGULATION\_OF\_TRANSCRIPTION\_FROM\_RNA\_POLYMERASE\_II\_PROMOTER | MSigDB lists | GO\_POSITIVE\_REGULATION\_OF\_TRANSCRIPTION\_FROM\_RNA\_POLYMERASE\_II\_PROMOTER | 181 | 17 | 4161 | 156 | ap,dock,sd,Actbeta,N,CtBP,Dl,cact,Mef2,jar,arm,fz2,tkv,Tsp96F,Tollo,hyx,kn | | 3.628e-04 | -7.92 | GSE13522\_CTRL\_VS\_T\_CRUZI\_Y\_STRAIN\_INF\_SKIN\_IFNAR\_KO\_DN | MSigDB lists | GSE13522\_CTRL\_VS\_T\_CRUZI\_Y\_STRAIN\_INF\_SKIN\_IFNAR\_KO\_DN | 37 | 7 | 4161 | 156 | Cerk,frc,unc-119,Tsp96F,Cdk4,cact,CG9220 | | 3.689e-04 | -7.90 | bsk (basket) | protein interactions | 44801 | 130 | 16 | 9204 | 432 | wnd,shot,cindr,kay,Src42A,N,scrib,aop,Pvr,ec,Gli,hppy,arm,nmo,fz,H | | 3.708e-04 | -7.90 | Ral GTPase binding | molecular function | GO:0017160 | 5 | 3 | 12176 | 414 | Rgl,Magi,CG5522 | | 3.708e-04 | -7.90 | SMAD binding | molecular function | GO:0046332 | 5 | 3 | 12176 | 414 | Snoo,tkv,fuss | | 3.711e-04 | -7.90 | Hrs (Hepatocyte growth factor regulated tyrosine kinase substrate) | protein interactions | 33458 | 59 | 10 | 9204 | 432 | mop,lilli,cact,hth,stx,FBXO11,rictor,ft,fz,CtBP | | 3.711e-04 | -7.90 | pan (pangolin) | protein interactions | 43769 | 59 | 10 | 9204 | 432 | Abd-B,aop,CycG,Cdk4,shg,arm,nmo,N,Src42A,eyg | | 3.729e-04 | -7.89 | Dsor1 (Downstream of raf1) | protein interactions | 31872 | 105 | 14 | 9204 | 432 | aop,14-3-3zeta,CG1129,CG12128,SERCA,cindr,mam,14-3-3epsilon,larp,N,Src42A,arm,Raf,Egfr | | 3.748e-04 | -7.89 | Role of LAT2/NTAL/LAB on calcium mobilization | REACTOME pathways | R-DME-2730905 | 41 | 8 | 3923 | 176 | Pdk1,rictor,htl,trbl,Dronc,wdb,vn,Egfr | | 3.765e-04 | -7.88 | GO\_REGULATION\_OF\_CELLULAR\_COMPONENT\_BIOGENESIS | MSigDB lists | GO\_REGULATION\_OF\_CELLULAR\_COMPONENT\_BIOGENESIS | 216 | 19 | 4161 | 156 | Eph,Syx1A,sli,CG17883,Nrk,Pex14,fz2,arm,Cals,Mef2,robo1,Raf,Pex5,sns,Unc-76,dock,lt,Arf79F,N | | 3.781e-04 | -7.88 | GO\_REGULATION\_OF\_HEMOPOIESIS | MSigDB lists | GO\_REGULATION\_OF\_HEMOPOIESIS | 61 | 9 | 4161 | 156 | Actbeta,N,arm,Mef2,hyx,Cdk4,Dl,Tollo,cact | | 3.781e-04 | -7.88 | GO\_LEUKOCYTE\_ACTIVATION | MSigDB lists | GO\_LEUKOCYTE\_ACTIVATION | 61 | 9 | 4161 | 156 | arm,fz2,dock,Raf,Cdk4,Mef2,Dl,Eph,Tollo | | 3.783e-04 | -7.88 | chic (chickadee) | protein interactions | 33834 | 70 | 11 | 9204 | 432 | Nlp,rictor,Jupiter,Bacc,S,Abl,Diap1,ena,cindr,Pgk,Fkbp14 | | 3.808e-04 | -7.87 | ATGTACA\_MIR493 | MSigDB lists | ATGTACA\_MIR493 | 88 | 11 | 4161 | 156 | bru2,CenB1A,dlp,Pdk1,Ziz,Hel89B,CG5522,Dl,rg,hyx,Mef2 | | 3.881e-04 | -7.85 | GO\_MESONEPHROS\_DEVELOPMENT | MSigDB lists | GO\_MESONEPHROS\_DEVELOPMENT | 27 | 6 | 4161 | 156 | sli,ds,arm,ft,LanA,robo1 | | 3.881e-04 | -7.85 | GO\_REGULATION\_OF\_REGULATED\_SECRETORY\_PATHWAY | MSigDB lists | GO\_REGULATION\_OF\_REGULATED\_SECRETORY\_PATHWAY | 27 | 6 | 4161 | 156 | Syx1A,AP-1gamma,Pdk1,N,Raf,Arf79F | | 3.921e-04 | -7.84 | positive regulation of plasma membrane bounded cell projection assembly | biological process | GO:0120034 | 28 | 6 | 12345 | 442 | shot,ena,chic,Abl,kon,SCAR | | 3.921e-04 | -7.84 | positive regulation of multicellular organism growth | biological process | GO:0040018 | 28 | 6 | 12345 | 442 | ey,step,Pdk1,InR,Hr4,Orct2 | | 4.005e-04 | -7.82 | GO\_BONE\_MORPHOGENESIS | MSigDB lists | GO\_BONE\_MORPHOGENESIS | 18 | 5 | 4161 | 156 | Mef2,tkv,htl,CG9220,Pex7 | | 4.005e-04 | -7.82 | GO\_ENDOTHELIAL\_CELL\_DIFFERENTIATION | MSigDB lists | GO\_ENDOTHELIAL\_CELL\_DIFFERENTIATION | 18 | 5 | 4161 | 156 | Dl,N,LanA,Rab1,arm | | 4.033e-04 | -7.82 | proximal promoter sequence-specific DNA binding | molecular function | GO:0000987 | 96 | 11 | 12176 | 414 | grn,h,CG12769,Hr4,kay,ttk,pnr,Hr39,CG34376,sim,zfh2 | | 4.052e-04 | -7.81 | jigr1 (jing interacting gene regulatory 1) | protein interactions | 43093 | 22 | 6 | 9204 | 432 | N,Pen,jigr1,kay,emp,drongo | | 4.095e-04 | -7.80 | stem cell fate commitment | biological process | GO:0048865 | 39 | 7 | 12345 | 442 | Egfr,abd-A,N,spi,spen,S,nub | | 4.095e-04 | -7.80 | Wnt signaling pathway | biological process | GO:0016055 | 39 | 7 | 12345 | 442 | fz2,tsh,fz,drl,arm,fz3,otk | | 4.134e-04 | -7.79 | HLH\_DNA-bd\_sf | interpro domains | IPR036638 | 58 | 9 | 11383 | 450 | emc,h,ss,sim,E(spl)m5-HLH,kn,tai,crp,HLH3B | | 4.168e-04 | -7.78 | SCHAEFFER\_PROSTATE\_DEVELOPMENT\_48HR\_DN | MSigDB lists | SCHAEFFER\_PROSTATE\_DEVELOPMENT\_48HR\_DN | 75 | 10 | 4161 | 156 | Fem-1,sli,Ddr,CG1673,Eph,robo1,kn,rg,Tmhs,Actbeta | | 4.185e-04 | -7.78 | Frizzled\_dom | interpro domains | IPR020067 | 10 | 4 | 11383 | 450 | Nrk,fz,fz3,fz2 | | 4.271e-04 | -7.76 | Abd-B (Abdominal B) | protein interactions | 47763 | 60 | 10 | 9204 | 432 | abd-A,mam,pnr,Mef2,eya,ap,hth,CycG,tsh,corto | | 4.309e-04 | -7.75 | GO\_DEVELOPMENTAL\_MATURATION | MSigDB lists | GO\_DEVELOPMENTAL\_MATURATION | 38 | 7 | 4161 | 156 | htl,Fem-1,AP-1gamma,ds,arm,ft,fz2 | | 4.320e-04 | -7.75 | regulation of cardioblast cell fate specification | biological process | GO:0042686 | 5 | 3 | 12345 | 442 | Abd-B,lqf,N | | 4.320e-04 | -7.75 | regulation of cardiac cell fate specification | biological process | GO:2000043 | 5 | 3 | 12345 | 442 | Abd-B,lqf,N | | 4.320e-04 | -7.75 | dorsal/ventral lineage restriction, imaginal disc | biological process | GO:0007451 | 5 | 3 | 12345 | 442 | ap,Dl,N | | 4.322e-04 | -7.75 | synapse part | cellular component | GO:0044456 | 211 | 18 | 11940 | 417 | Cals,Tob,scrib,Magi,nAChRalpha4,jeb,Ir21a,orb2,pum,Syx1A,ena,LanA,gukh,tkv,Bsg,Cadps,ReepA,CG31211 | | 4.354e-04 | -7.74 | cellular component biogenesis | biological process | GO:0044085 | 1270 | 68 | 12345 | 442 | Atg17,PI31,moody,Myo10A,Dronc,LRR,mop,emc,hyx,sns,Sema1a,baz,rictor,tkv,Dcr-2,Atg16,Nlp,wdb,Tob,Adf1,larp,Cks30A,vn,Bsg,orb2,scrib,N,Ppcs,cindr,drl,FBXO11,unc-119,InR,Egfr,CG34172,cact,aop,Spc105R,shg,CG11030,ttk,Dg,SCAR,Pat1,Gli,Raf,His3.3A,mim,SmD1,His3.3B,Pdk1,kon,cic,kay,yrt,CG17683,arm,Dp,chic,Pvr,bru2,ed,jeb,S,ena,f,bou,Src42A | | 4.378e-04 | -7.73 | cell elongation involved in imaginal disc-derived wing morphogenesis | biological process | GO:0090254 | 11 | 4 | 12345 | 442 | tkv,Dad,arm,shg | | 4.378e-04 | -7.73 | regulation of cellular response to transforming growth factor beta stimulus | biological process | GO:1903844 | 11 | 4 | 12345 | 442 | fuss,Snoo,Dad,tkv | | 4.378e-04 | -7.73 | oenocyte differentiation | biological process | GO:0001742 | 11 | 4 | 12345 | 442 | abd-A,Egfr,hth,spi | | 4.378e-04 | -7.73 | epithelial cell proliferation involved in Malpighian tubule morphogenesis | biological process | GO:0061331 | 11 | 4 | 12345 | 442 | N,spi,S,Egfr | | 4.378e-04 | -7.73 | regulation of transforming growth factor beta receptor signaling pathway | biological process | GO:0017015 | 11 | 4 | 12345 | 442 | Snoo,Dad,tkv,fuss | | 4.378e-04 | -7.73 | negative regulation of DNA-binding transcription factor activity | biological process | GO:0043433 | 11 | 4 | 12345 | 442 | 14-3-3zeta,edl,cact,14-3-3epsilon | | 4.378e-04 | -7.73 | epithelial cell proliferation involved in renal tubule morphogenesis | biological process | GO:2001013 | 11 | 4 | 12345 | 442 | Egfr,N,spi,S | | 4.378e-04 | -7.73 | epithelial cell proliferation | biological process | GO:0050673 | 11 | 4 | 12345 | 442 | N,spi,S,Egfr | | 4.378e-04 | -7.73 | delamination | biological process | GO:0060232 | 11 | 4 | 12345 | 442 | aop,arm,disco-r,baz | | 4.455e-04 | -7.72 | GAB1 signalosome | REACTOME pathways | R-DME-180292 | 42 | 8 | 3923 | 176 | Egfr,vn,wdb,trbl,Dronc,rictor,Pdk1,htl | | 4.491e-04 | -7.71 | synaptic membrane | cellular component | GO:0097060 | 66 | 9 | 11940 | 417 | scrib,Syx1A,Cals,Tob,gukh,LanA,orb2,Ir21a,nAChRalpha4 | | 4.519e-04 | -7.70 | klu (klumpfuss) | protein interactions | 39228 | 15 | 5 | 9204 | 432 | H,mam,Egfr,arm,N | | 4.519e-04 | -7.70 | rst (roughest) | protein interactions | 31290 | 15 | 5 | 9204 | 432 | dock,morgue,sns,cindr,Dl | | 4.519e-04 | -7.70 | Acsl (Acyl-CoA synthetase long-chain) | protein interactions | 46068 | 15 | 5 | 9204 | 432 | Dad,N,Mpcp2,tkv,CG31431 | | 4.541e-04 | -7.70 | GO\_REGULATION\_OF\_CELL\_DEATH | MSigDB lists | GO\_REGULATION\_OF\_CELL\_DEATH | 330 | 25 | 4161 | 156 | CG4896,CG3961,Dl,cact,Mef2,Raf,14-3-3epsilon,Drak,unc-5,dock,InR,Actbeta,N,14-3-3zeta,Fem-1,sli,Eph,Tollo,wbl,arm,Nmda1,tkv,CG45050,sina,Pdk1 | | 4.576e-04 | -7.69 | EGF\_2 | prosite domains | PS01186 | 55 | 10 | 6243 | 325 | dpy,Dl,cue,spi,LanA,ft,nahoda,shg,sli,N | | 4.578e-04 | -7.69 | GSE1460\_INTRATHYMIC\_T\_PROGENITOR\_VS\_THYMIC\_STROMAL\_CELL\_DN | MSigDB lists | GSE1460\_INTRATHYMIC\_T\_PROGENITOR\_VS\_THYMIC\_STROMAL\_CELL\_DN | 50 | 8 | 4161 | 156 | Tpst,Rab1,jar,dlp,CG9220,Rgl,CG31522,sli | | 4.645e-04 | -7.67 | hpo (hippo) | protein interactions | 37247 | 95 | 13 | 9204 | 432 | Dronc,ed,sd,ds,Pez,Arf79F,Diap1,SERCA,ena,klar,Egfr,dome,wdb | | 4.645e-04 | -7.67 | BUYTAERT\_PHOTODYNAMIC\_THERAPY\_STRESS\_UP | MSigDB lists | BUYTAERT\_PHOTODYNAMIC\_THERAPY\_STRESS\_UP | 168 | 16 | 4161 | 156 | Fem-1,Syx1A,Rgl,CG2017,CG2812,lili,Pat1,Kdm2,cact,hppy,Actbeta,N,Rim2,InR,shot,Drak | | 4.711e-04 | -7.66 | WASp (WASp) | protein interactions | 43402 | 31 | 7 | 9204 | 432 | H,SCAR,cindr,dock,Cip4,chic,N | | 4.736e-04 | -7.66 | eggshell chorion assembly | biological process | GO:0007306 | 93 | 11 | 12345 | 442 | Egfr,cact,N,Ppcs,cic,ed,emc,S,Raf,ttk,Dp | | 4.736e-04 | -7.66 | response to antibiotic | biological process | GO:0046677 | 93 | 11 | 12345 | 442 | Pdk1,moody,Bacc,Egfr,pum,pxb,S,spi,hang,mura,hppy | | 4.736e-04 | -7.66 | regulation of Wnt signaling pathway | biological process | GO:0030111 | 93 | 11 | 12345 | 442 | CtBP,Diap1,Dyrk2,hyx,dlp,aop,crol,spen,nmo,apolpp,otk | | 4.779e-04 | -7.65 | calcium ion binding | molecular function | GO:0005509 | 200 | 17 | 12176 | 414 | dpy,Cbl,shg,ds,Cadps,alpha-Man-Ia,N,Dl,ft,CalpC,rho-4,Dg,sli,Cals,cue,shot,Fkbp14 | | 4.788e-04 | -7.64 | ACCTGTTG\_UNKNOWN | MSigDB lists | ACCTGTTG\_UNKNOWN | 28 | 6 | 4161 | 156 | Ocrl,shot,Rab5,N,Nrk,Actbeta | | 4.788e-04 | -7.64 | MODULE\_63 | MSigDB lists | MODULE\_63 | 28 | 6 | 4161 | 156 | otk,Eph,nAChRalpha4,tkv,htl,Ddr | | 4.796e-04 | -7.64 | glial cell migration | biological process | GO:0008347 | 29 | 6 | 12345 | 442 | Dl,sli,unc-5,N,ey,htl | | 4.796e-04 | -7.64 | establishment or maintenance of polarity of follicular epithelium | biological process | GO:0016334 | 29 | 6 | 12345 | 442 | Dp,Raf,scrib,Cbl,Egfr,Dg | | 4.840e-04 | -7.63 | LINDGREN\_BLADDER\_CANCER\_CLUSTER\_2B | MSigDB lists | LINDGREN\_BLADDER\_CANCER\_CLUSTER\_2B | 63 | 9 | 4161 | 156 | 14-3-3zeta,CG9220,Tpst,bru2,Cerk,CG6891,cact,Ddr,Rgl | | 4.891e-04 | -7.62 | GO\_BONE\_MATURATION | MSigDB lists | GO\_BONE\_MATURATION | 5 | 3 | 4161 | 156 | ds,ft,htl | | 4.891e-04 | -7.62 | GO\_TYPE\_B\_PANCREATIC\_CELL\_DEVELOPMENT | MSigDB lists | GO\_TYPE\_B\_PANCREATIC\_CELL\_DEVELOPMENT | 5 | 3 | 4161 | 156 | Cdk4,Pdk1,Dl | | 4.891e-04 | -7.62 | GO\_REGULATION\_OF\_CHROMATIN\_BINDING | MSigDB lists | GO\_REGULATION\_OF\_CHROMATIN\_BINDING | 5 | 3 | 4161 | 156 | Hel89B,arm,CtBP | | 4.891e-04 | -7.62 | GO\_PROXIMAL\_DISTAL\_PATTERN\_FORMATION | MSigDB lists | GO\_PROXIMAL\_DISTAL\_PATTERN\_FORMATION | 5 | 3 | 4161 | 156 | Dl,CG9220,arm | | 4.891e-04 | -7.62 | GO\_EPITHELIAL\_CELL\_DIFFERENTIATION\_INVOLVED\_IN\_KIDNEY\_DEVELOPMENT | MSigDB lists | GO\_EPITHELIAL\_CELL\_DIFFERENTIATION\_INVOLVED\_IN\_KIDNEY\_DEVELOPMENT | 5 | 3 | 4161 | 156 | sns,Mef2,N | | 4.891e-04 | -7.62 | GO\_ORGAN\_MATURATION | MSigDB lists | GO\_ORGAN\_MATURATION | 5 | 3 | 4161 | 156 | htl,ds,ft | | 4.898e-04 | -7.62 | Cdc42 (Cdc42) | protein interactions | 32981 | 61 | 10 | 9204 | 432 | N,robo1,SCAR,Cip4,muc,tkv,chic,lilli,Pvr,bol | | 4.995e-04 | -7.60 | otk (off-track) | protein interactions | 36283 | 9 | 4 | 9204 | 432 | otk,fz,fz2,Sema1a | | 4.995e-04 | -7.60 | Csp (Cysteine string protein) | protein interactions | 40459 | 9 | 4 | 9204 | 432 | Syx1A,Dl,pum,kn | | 4.995e-04 | -7.60 | Myo31DF (Myosin 31DF) | protein interactions | 34445 | 9 | 4 | 9204 | 432 | ds,Dronc,arm,shg | | 4.995e-04 | -7.60 | Wnt2 (Wnt oncogene analog 2) | protein interactions | 35975 | 9 | 4 | 9204 | 432 | fz3,otk,fz2,fz | | 4.995e-04 | -7.60 | qsm (quasimodo) | protein interactions | 47065 | 9 | 4 | 9204 | 432 | H,Ptp4E,pnr,CG11247 | | 5.088e-04 | -7.58 | GSE16450\_CTRL\_VS\_IFNA\_6H\_STIM\_IMMATURE\_NEURON\_CELL\_LINE\_UP | MSigDB lists | GSE16450\_CTRL\_VS\_IFNA\_6H\_STIM\_IMMATURE\_NEURON\_CELL\_LINE\_UP | 39 | 7 | 4161 | 156 | Rim2,robo1,N,Cdk4,Tollo,CenB1A,CycG | | 5.088e-04 | -7.58 | GSE29617\_CTRL\_VS\_TIV\_FLU\_VACCINE\_PBMC\_2008\_UP | MSigDB lists | GSE29617\_CTRL\_VS\_TIV\_FLU\_VACCINE\_PBMC\_2008\_UP | 39 | 7 | 4161 | 156 | dlp,sli,arm,Rab1,sina,Raf,kn | | 5.106e-04 | -7.58 | GO\_ENDOCHONDRAL\_BONE\_MORPHOGENESIS | MSigDB lists | GO\_ENDOCHONDRAL\_BONE\_MORPHOGENESIS | 11 | 4 | 4161 | 156 | Mef2,htl,tkv,Pex7 | | 5.106e-04 | -7.58 | GO\_NUCLEOTIDE\_TRANSMEMBRANE\_TRANSPORTER\_ACTIVITY | MSigDB lists | GO\_NUCLEOTIDE\_TRANSMEMBRANE\_TRANSPORTER\_ACTIVITY | 11 | 4 | 4161 | 156 | Rim2,Tpc1,frc,DPCoAC | | 5.106e-04 | -7.58 | GO\_EXTRINSIC\_APOPTOTIC\_SIGNALING\_PATHWAY | MSigDB lists | GO\_EXTRINSIC\_APOPTOTIC\_SIGNALING\_PATHWAY | 11 | 4 | 4161 | 156 | Actbeta,Pdk1,htl,Tollo | | 5.106e-04 | -7.58 | PID\_IGF1\_PATHWAY | MSigDB lists | PID\_IGF1\_PATHWAY | 11 | 4 | 4161 | 156 | InR,Pdk1,14-3-3zeta,14-3-3epsilon | | 5.106e-04 | -7.58 | YAMASHITA\_LIVER\_CANCER\_STEM\_CELL\_UP | MSigDB lists | YAMASHITA\_LIVER\_CANCER\_STEM\_CELL\_UP | 11 | 4 | 4161 | 156 | robo1,14-3-3zeta,htl,fz2 | | 5.110e-04 | -7.58 | GO\_REGULATION\_OF\_APOPTOTIC\_SIGNALING\_PATHWAY | MSigDB lists | GO\_REGULATION\_OF\_APOPTOTIC\_SIGNALING\_PATHWAY | 91 | 11 | 4161 | 156 | CG3961,Fem-1,14-3-3zeta,sina,Actbeta,tkv,dock,arm,Nmda1,14-3-3epsilon,wbl | | 5.247e-04 | -7.55 | cadherin binding | molecular function | GO:0045296 | 12 | 4 | 12176 | 414 | siz,ds,ft,arm | | 5.247e-04 | -7.55 | translation repressor activity, mRNA regulatory element binding | molecular function | GO:0000900 | 12 | 4 | 12176 | 414 | Sxl,pum,msi,orb2 | | 5.259e-04 | -7.55 | GSE24671\_CTRL\_VS\_BAKIMULC\_INFECTED\_MOUSE\_SPLENOCYTES\_UP | MSigDB lists | GSE24671\_CTRL\_VS\_BAKIMULC\_INFECTED\_MOUSE\_SPLENOCYTES\_UP | 51 | 8 | 4161 | 156 | unc-119,Ids,hppy,step,lili,wbl,bru2,jar | | 5.267e-04 | -7.55 | lolal (lola like) | protein interactions | 44703 | 23 | 6 | 9204 | 432 | ph-p,H,CG6891,ttk,rib,N | | 5.267e-04 | -7.55 | ac (achaete) | protein interactions | 30981 | 23 | 6 | 9204 | 432 | klu,Egfr,ttk,emc,h,sens | | 5.270e-04 | -7.55 | GO\_REGULATION\_OF\_TRANSPORT | MSigDB lists | GO\_REGULATION\_OF\_TRANSPORT | 412 | 29 | 4161 | 156 | Psa,lt,sns,Arf79F,N,Actbeta,14-3-3zeta,14-3-3epsilon,GlcAT-I,Dl,cact,Raf,Mef2,CG11069,Rab5,subdued,nAChRalpha4,Pdk1,fz,wbl,jar,arm,fz2,Tollo,unc-119,Syx1A,ReepA,AP-1gamma,CG17883 | | 5.273e-04 | -7.55 | KEGG\_PROSTATE\_CANCER | MSigDB lists | KEGG\_PROSTATE\_CANCER | 19 | 5 | 4161 | 156 | Pdk1,Raf,cact,InR,arm | | 5.276e-04 | -7.55 | Cadherin-like\_sf | interpro domains | IPR015919 | 18 | 5 | 11383 | 450 | shg,Cals,ft,ds,Dg | | 5.373e-04 | -7.53 | Rala (Ras-like protein A) | protein interactions | 31332 | 41 | 8 | 9204 | 432 | kay,ey,Dronc,lqf,Rgl,Dl,N,Magi | | 5.387e-04 | -7.53 | regulation of transmembrane receptor protein serine/threonine kinase signaling pathway | biological process | GO:0090092 | 66 | 9 | 12345 | 442 | lili,tkv,Dad,Actbeta,Snoo,cv-d,hth,fuss,kek5 | | 5.462e-04 | -7.51 | hemopoiesis | biological process | GO:0030097 | 80 | 10 | 12345 | 442 | Pvf3,shg,Raf,RhoL,N,CG1129,cact,Pvr,sens-2,htl | | 5.473e-04 | -7.51 | cell population proliferation | biological process | GO:0008283 | 175 | 16 | 12345 | 442 | ft,spi,lilli,Egfr,jar,baz,esg,flfl,sd,Cbl,S,ds,scrib,N,Cdk4,Inx2 | | 5.601e-04 | -7.49 | mbl (muscleblind) | protein interactions | 36945 | 62 | 10 | 9204 | 432 | Dcr-2,crp,CG3386,jumu,Dg,Pen,SERCA,Diap1,Elba2,Dp | | 5.627e-04 | -7.48 | olfactory learning | biological process | GO:0008355 | 41 | 7 | 12345 | 442 | rg,Adf1,Egfr,vn,drl,14-3-3zeta,spi | | 5.690e-04 | -7.47 | regulation of supramolecular fiber organization | biological process | GO:1902903 | 95 | 11 | 12345 | 442 | shot,RhoU,chic,CG6891,ena,RhoGEF4,SCAR,Abl,Jupiter,Cip4,fz | | 5.709e-04 | -7.47 | cell cycle comprising mitosis without cytokinesis | biological process | GO:0033301 | 20 | 5 | 12345 | 442 | Psc,Cks30A,CycB3,ph-p,larp | | 5.709e-04 | -7.47 | germ-line cyst encapsulation | biological process | GO:0048138 | 20 | 5 | 12345 | 442 | Cip4,ph-p,shg,N,Ppcs | | 5.709e-04 | -7.47 | neuroblast development | biological process | GO:0014019 | 20 | 5 | 12345 | 442 | arm,N,abd-A,klu,sqz | | 5.750e-04 | -7.46 | repressing transcription factor binding | molecular function | GO:0070491 | 21 | 5 | 12176 | 414 | edl,kay,cic,pnr,CtBP | | 5.782e-04 | -7.46 | - | gene3d domains | 1.10.2000.10 | 9 | 4 | 4929 | 241 | fz2,Nrk,fz3,fz | | 5.783e-04 | -7.46 | Frizzled/SFRP | interpro domains | IPR015526 | 5 | 3 | 11383 | 450 | fz,fz3,fz2 | | 5.783e-04 | -7.46 | Frizzled/Smoothened\_TM | interpro domains | IPR000539 | 5 | 3 | 11383 | 450 | fz2,fz,fz3 | | 5.790e-04 | -7.45 | Axn (Axin) | protein interactions | 43565 | 32 | 7 | 9204 | 432 | Ppn,crol,N,tsh,aop,nmo,arm | | 5.790e-04 | -7.45 | dome (domeless) | protein interactions | 32976 | 32 | 7 | 9204 | 432 | Socs36E,Egfr,scrib,dome,Diap1,Rab5,Dronc | | 5.807e-04 | -7.45 | filopodium | cellular component | GO:0030175 | 12 | 4 | 11940 | 417 | Myo10A,shot,shg,ena | | 5.816e-04 | -7.45 | mitotic cell cycle, embryonic | biological process | GO:0045448 | 30 | 6 | 12345 | 442 | 14-3-3zeta,CycB3,Cks30A,larp,ph-p,Psc | | 5.962e-04 | -7.42 | GO\_ANATOMICAL\_STRUCTURE\_FORMATION\_INVOLVED\_IN\_MORPHOGENESIS | MSigDB lists | GO\_ANATOMICAL\_STRUCTURE\_FORMATION\_INVOLVED\_IN\_MORPHOGENESIS | 224 | 19 | 4161 | 156 | sns,ap,Actbeta,N,Rab1,otk,Dl,Mef2,fz,fz2,arm,Eph,hyx,LanA,ds,C1GalTA,CG17883,sli,Ocrl | | 5.975e-04 | -7.42 | Frizzled | pfam domains | PF01534 | 5 | 3 | 10480 | 419 | fz,fz3,fz2 | | 5.976e-04 | -7.42 | IRF\_Q6 | MSigDB lists | IRF\_Q6 | 40 | 7 | 4161 | 156 | Hel89B,Kdm2,CG3961,Psa,Dl,Nrk,Cdk4 | | 6.030e-04 | -7.41 | positive regulation of protein kinase activity | biological process | GO:0045860 | 67 | 9 | 12345 | 442 | jeb,hppy,Raf,Cks30A,Atg17,wnd,step,rictor,Pdk1 | | 6.158e-04 | -7.39 | EGF\_3 | prosite domains | PS50026 | 57 | 10 | 6243 | 325 | dpy,Dl,nahoda,cue,ft,spi,sli,N,vn,shg | | 6.170e-04 | -7.39 | AATGTGA\_MIR23A\_MIR23B | MSigDB lists | AATGTGA\_MIR23A\_MIR23B | 93 | 11 | 4161 | 156 | dlp,lin-28,bru2,CycG,wdb,sd,FBXO11,Hel89B,Mef2,robo1,kn | | 6.186e-04 | -7.39 | aPKC (atypical protein kinase C) | protein interactions | 47594 | 52 | 9 | 9204 | 432 | arm,baz,fz,Magi,nuf,Pex5,orb2,yrt,Myo10A | | 6.186e-04 | -7.39 | ey (eyeless) | protein interactions | 43812 | 52 | 9 | 9204 | 432 | tkv,eya,ey,hth,shg,eyg,N,CtBP,H | | 6.186e-04 | -7.39 | dlg1 (discs large 1) | protein interactions | 32083 | 52 | 9 | 9204 | 432 | ph-p,scrib,Su(z)2,Socs36E,pum,shg,arm,baz,gukh | | 6.203e-04 | -7.39 | embryonic hindgut morphogenesis | biological process | GO:0048619 | 54 | 8 | 12345 | 442 | N,Dad,Pvr,Egfr,bowl,spi,rib,S | | 6.224e-04 | -7.38 | regulation of Ras protein signal transduction | biological process | GO:0046578 | 96 | 11 | 12345 | 442 | Lk6,RhoGEF4,CG42684,14-3-3zeta,siz,Ziz,Rgl,step,CG5522,RhoGEF3,14-3-3epsilon | | 6.298e-04 | -7.37 | GO\_MOLECULAR\_FUNCTION\_REGULATOR | MSigDB lists | GO\_MOLECULAR\_FUNCTION\_REGULATOR | 280 | 22 | 4161 | 156 | GlcAT-I,14-3-3epsilon,CenB1A,N,Arf79F,CG3107,htl,trbl,Ppn,dock,step,CG5522,wdb,Pdk1,CG17883,Ocrl,sli,PI31,Spn42De,Ziz,Syx1A,Rgl | | 6.298e-04 | -7.37 | GO\_MOVEMENT\_OF\_CELL\_OR\_SUBCELLULAR\_COMPONENT | MSigDB lists | GO\_MOVEMENT\_OF\_CELL\_OR\_SUBCELLULAR\_COMPONENT | 280 | 22 | 4161 | 156 | Rab1,shot,14-3-3epsilon,unc-5,ap,dock,Unc-76,N,Pex7,otk,robo1,Mef2,jar,arm,Pex14,tkv,sina,Pdk1,LanA,sli,ds,Eph | | 6.321e-04 | -7.37 | hbs (hibris) | protein interactions | 44129 | 16 | 5 | 9204 | 432 | cindr,dock,CG3386,N,sns | | 6.382e-04 | -7.36 | basal protein localization | biological process | GO:0045175 | 12 | 4 | 12345 | 442 | flfl,scrib,jar,baz | | 6.547e-04 | -7.33 | cell surface receptor signaling pathway involved in cell-cell signaling | biological process | GO:1905114 | 42 | 7 | 12345 | 442 | otk,fz3,arm,drl,fz,tsh,fz2 | | 6.674e-04 | -7.31 | response to alcohol | biological process | GO:0097305 | 128 | 13 | 12345 | 442 | Pdk1,moody,Bacc,pxb,Egfr,pum,hang,hfw,mura,ph-p,S,spi,hppy | | 6.734e-04 | -7.30 | regulation of cellular response to growth factor stimulus | biological process | GO:0090287 | 68 | 9 | 12345 | 442 | hth,kek5,fuss,cv-d,dlp,Snoo,Dad,tkv,lili | | 6.748e-04 | -7.30 | Cbl (Cbl proto-oncogene) | protein interactions | 38961 | 24 | 6 | 9204 | 432 | Socs36E,Pvr,mop,cindr,Egfr,Cbl | | 6.799e-04 | -7.29 | stem cell division | biological process | GO:0017145 | 97 | 11 | 12345 | 442 | InR,shg,eff,flfl,esg,tkv,baz,jar,lin-28,pum,msi | | 6.819e-04 | -7.29 | GO\_REGULATION\_OF\_SYNAPSE\_ORGANIZATION | MSigDB lists | GO\_REGULATION\_OF\_SYNAPSE\_ORGANIZATION | 20 | 5 | 4161 | 156 | Nrk,Mef2,robo1,Eph,Cals | | 6.819e-04 | -7.29 | GO\_ENDOTHELIUM\_DEVELOPMENT | MSigDB lists | GO\_ENDOTHELIUM\_DEVELOPMENT | 20 | 5 | 4161 | 156 | Dl,N,Rab1,LanA,arm | | 6.930e-04 | -7.27 | Tyr\_Pase\_cat | interpro domains | IPR003595 | 19 | 5 | 11383 | 450 | Pez,Ptp4E,Ptpmeg2,CG42327,IA-2 | | 6.930e-04 | -7.27 | Laminin\_G | interpro domains | IPR001791 | 19 | 5 | 11383 | 450 | ft,sli,kon,shg,LanA | | 6.935e-04 | -7.27 | puc (puckered) | protein interactions | 40958 | 75 | 11 | 9204 | 432 | hppy,arm,Gli,Src42A,rdx,H,tkv,sd,Rab5,kay,Pvr | | 7.056e-04 | -7.26 | bs (blistered) | protein interactions | 37890 | 33 | 7 | 9204 | 432 | vn,px,emc,N,H,Egfr,Dl | | 7.056e-04 | -7.26 | gbb (glass bottom boat) | protein interactions | 37778 | 33 | 7 | 9204 | 432 | Dad,kek5,cv-d,H2.0,robo1,tkv,HLH3B | | 7.102e-04 | -7.25 | system process | biological process | GO:0003008 | 636 | 39 | 12345 | 442 | Mob2,mura,N,14-3-3zeta,scrib,pdm3,drl,nAChRalpha4,vn,CG30344,orb2,Ir21a,pxb,Socs36E,spi,ss,lilli,dome,SKIP,rg,Tmhs,f,cora,Egfr,Gr93c,Dyrk2,sns,SERCA,Myo10A,wnd,pum,Atg17,pnr,arm,Abl,Adf1,Tob,subdued,mld | | 7.148e-04 | -7.24 | DCTN1-p150 (Dynactin 1, p150 subunit) | protein interactions | 39536 | 53 | 9 | 9204 | 432 | h,klar,betaTub60D,LRR,S,spi,Dl,nuf,Egfr | | 7.148e-04 | -7.24 | Iswi (Imitation SWI) | protein interactions | 36390 | 53 | 9 | 9204 | 432 | rictor,eff,CtBP,lncRNA:Hsromega,arm,Sema1a,Pen,pnr,ttk | | 7.238e-04 | -7.23 | intrinsic component of plasma membrane | cellular component | GO:0031226 | 480 | 31 | 11940 | 417 | robo1,Esp,beat-IIa,ds,Egfr,otk,tkv,Gli,tipE,bou,InR,wgn,drl,Ddr,fz,scrib,Nrk,Cals,rho-4,emp,ed,ft,kek5,N,Eph,fz2,Pvr,dlp,htl,shg,nAChRalpha4 | | 7.257e-04 | -7.23 | Rbf (Retinoblastoma-family protein) | protein interactions | 31027 | 64 | 10 | 9204 | 432 | N,arm,pum,tara,Cdk4,Dp,tkv,Cbl,lilli,mam | | 7.275e-04 | -7.23 | circulatory system process | biological process | GO:0003013 | 21 | 5 | 12345 | 442 | cora,spi,SERCA,pnr,Egfr | | 7.275e-04 | -7.23 | peroxisome organization | biological process | GO:0007031 | 21 | 5 | 12345 | 442 | Pex7,CG33474,Pex10,Pex14,Pex5 | | 7.275e-04 | -7.23 | genitalia morphogenesis | biological process | GO:0035112 | 21 | 5 | 12345 | 442 | Abd-B,Pvr,otk,Rab5,N | | 7.302e-04 | -7.22 | GO\_REGULATION\_OF\_IMMUNE\_SYSTEM\_PROCESS | MSigDB lists | GO\_REGULATION\_OF\_IMMUNE\_SYSTEM\_PROCESS | 302 | 23 | 4161 | 156 | hyx,Tollo,sli,ReepA,Spn42De,CG4645,PI31,AP-1gamma,Tsp96F,Pdk1,fz2,arm,eff,Mef2,Cdk4,cact,Dl,Tpc1,N,Arf79F,Actbeta,dock,SmD1 | | 7.410e-04 | -7.21 | protein tyrosine phosphatase activity | molecular function | GO:0004725 | 45 | 7 | 12176 | 414 | Ptp4E,eya,CG42327,Ptpmeg2,Pez,IA-2,mop | | 7.436e-04 | -7.20 | GO\_ORGANOPHOSPHATE\_ESTER\_TRANSMEMBRANE\_TRANSPORTER\_ACTIVITY | MSigDB lists | GO\_ORGANOPHOSPHATE\_ESTER\_TRANSMEMBRANE\_TRANSPORTER\_ACTIVITY | 12 | 4 | 4161 | 156 | Rim2,Tpc1,frc,DPCoAC | | 7.436e-04 | -7.20 | GO\_CHONDROCYTE\_DIFFERENTIATION | MSigDB lists | GO\_CHONDROCYTE\_DIFFERENTIATION | 12 | 4 | 4161 | 156 | Mef2,CG9220,htl,tkv | | 7.436e-04 | -7.20 | GO\_NUCLEOTIDE\_TRANSPORT | MSigDB lists | GO\_NUCLEOTIDE\_TRANSPORT | 12 | 4 | 4161 | 156 | frc,Tpc1,DPCoAC,Rim2 | | 7.436e-04 | -7.20 | GO\_HOMOPHILIC\_CELL\_ADHESION\_VIA\_PLASMA\_MEMBRANE\_ADHESION\_MOLECULES | MSigDB lists | GO\_HOMOPHILIC\_CELL\_ADHESION\_VIA\_PLASMA\_MEMBRANE\_ADHESION\_MOLECULES | 12 | 4 | 4161 | 156 | Cals,robo1,ds,ft | | 7.436e-04 | -7.20 | GO\_PHOSPHATE\_TRANSMEMBRANE\_TRANSPORTER\_ACTIVITY | MSigDB lists | GO\_PHOSPHATE\_TRANSMEMBRANE\_TRANSPORTER\_ACTIVITY | 12 | 4 | 4161 | 156 | Tpc1,frc,DPCoAC,Rim2 | | 7.436e-04 | -7.20 | GO\_GLANDULAR\_EPITHELIAL\_CELL\_DIFFERENTIATION | MSigDB lists | GO\_GLANDULAR\_EPITHELIAL\_CELL\_DIFFERENTIATION | 12 | 4 | 4161 | 156 | N,Pdk1,Cdk4,Dl | | 7.504e-04 | -7.19 | positive regulation of kinase activity | biological process | GO:0033674 | 69 | 9 | 12345 | 442 | Pdk1,Atg17,rictor,wnd,step,Raf,Cks30A,jeb,hppy | | 7.583e-04 | -7.18 | negative regulation of Notch signaling pathway | biological process | GO:0045746 | 43 | 7 | 12345 | 442 | H,spdo,CycG,cue,Cbl,Tom,Dl | | 7.583e-04 | -7.18 | positive regulation of Wnt signaling pathway | biological process | GO:0030177 | 43 | 7 | 12345 | 442 | dlp,hyx,apolpp,Dyrk2,spen,Diap1,CtBP | | 7.583e-04 | -7.18 | positive regulation of cell size | biological process | GO:0045793 | 43 | 7 | 12345 | 442 | Pvf3,Pdk1,InR,rictor,trbl,hppy,Pvr | | 7.740e-04 | -7.16 | Protein\_kinase\_ATP\_BS | interpro domains | IPR017441 | 164 | 16 | 11383 | 450 | Lk6,Src42A,nmo,tkv,Pdk1,Cdk4,hppy,Eph,Raf,Egfr,Nrk,Abl,htl,Dyrk2,Pvr,InR | | 7.744e-04 | -7.16 | Endocytosis | KEGG pathways | ko04144 | 124 | 13 | 3198 | 123 | nuf,siz,Arf79F,InR,htl,Cbl,Pvr,CenB1A,baz,Egfr,Rab5,step,lqf | | 7.744e-04 | -7.16 | Endocytosis | KEGG pathways | dme04144 | 124 | 13 | 3198 | 123 | Rab5,step,lqf,Egfr,baz,CenB1A,Pvr,htl,Cbl,InR,siz,Arf79F,nuf | | 7.942e-04 | -7.14 | regulation of catalytic activity | biological process | GO:0050790 | 450 | 30 | 12345 | 442 | kay,SNF4Agamma,CenB1A,RapGAP1,Cks30A,Diap1,Ocrl,rictor,step,Pdk1,TBC1D5,CG42684,Cbl,Spn42De,jeb,Dronc,hppy,Fem-1,Raf,CycB3,Socs36E,Atg17,Rgl,CycG,trbl,wnd,drongo,CG17883,RhoGAP19D,CG7394 | | 7.957e-04 | -7.14 | negative regulation of translation | biological process | GO:0017148 | 56 | 8 | 12345 | 442 | Sxl,msi,pum,orb2,312,Tis11,bru2,bin3 | | 8.018e-04 | -7.13 | sv (shaven) | protein interactions | 43825 | 10 | 4 | 9204 | 432 | ttk,rg,Poxn,abd-A | | 8.018e-04 | -7.13 | Ptp10D (Protein tyrosine phosphatase 10D) | protein interactions | 32115 | 10 | 4 | 9204 | 432 | Ptp4E,ena,Egfr,Sema1a | | 8.018e-04 | -7.13 | chm (chameau) | protein interactions | 43928 | 10 | 4 | 9204 | 432 | kay,ph-p,Abd-B,sd | | 8.018e-04 | -7.13 | Amun (amun) | protein interactions | 32123 | 10 | 4 | 9204 | 432 | Dl,H,Pvr,N | | 8.018e-04 | -7.13 | nvy (nervy) | protein interactions | 37886 | 10 | 4 | 9204 | 432 | sens,Sema1a,N,Dl | | 8.018e-04 | -7.13 | stet (stem cell tumor) | protein interactions | 38169 | 10 | 4 | 9204 | 432 | spi,Raf,S,tkv | | 8.018e-04 | -7.13 | ec (echinus) | protein interactions | 31339 | 10 | 4 | 9204 | 432 | spi,nmo,shg,S | | 8.018e-04 | -7.13 | YME1L (YME1 like ATPase) | protein interactions | 37636 | 10 | 4 | 9204 | 432 | E(Pc),CycG,Dronc,Diap1 | | 8.019e-04 | -7.13 | CTTTGA\_LEF1\_Q2 | MSigDB lists | CTTTGA\_LEF1\_Q2 | 285 | 22 | 4161 | 156 | Rab5,hppy,CG31522,Gfat2,CG3961,Cals,Mef2,CenB1A,nmo,dlp,htl,ap,Ppn,Actbeta,PI31,Rgl,Cadps,Syx1A,Ddr,ds,rg,unc-119 | | 8.120e-04 | -7.12 | GSE37605\_FOXP3\_FUSION\_GFP\_VS\_IRES\_GFP\_TREG\_C57BL6\_DN | MSigDB lists | GSE37605\_FOXP3\_FUSION\_GFP\_VS\_IRES\_GFP\_TREG\_C57BL6\_DN | 42 | 7 | 4161 | 156 | Mfe2,CG9485,Ocrl,hppy,Unc-76,sd,Dl | | 8.174e-04 | -7.11 | cell-cell signaling | biological process | GO:0007267 | 311 | 23 | 12345 | 442 | kn,fz,SERCA,mam,Pgk,Arf79F,shot,Ir21a,lqf,otk,pum,Dl,nAChRalpha4,drl,N,Abl,fz3,arm,tsh,fz2,Cadps,tkv,Syx1A | | 8.225e-04 | -7.10 | E2f1 (E2F transcription factor 1) | protein interactions | 42550 | 65 | 10 | 9204 | 432 | crol,Su(z)2,eff,tara,arm,E(Pc),Diap1,His2Av,Dp,sd | | 8.229e-04 | -7.10 | shi (shibire) | protein interactions | 45928 | 54 | 9 | 9204 | 432 | Cip4,Cbl,Dronc,lqf,ttk,mam,Sema1a,ft,sli | | 8.293e-04 | -7.09 | Znf\_C2H2\_sf | interpro domains | IPR036236 | 265 | 22 | 11383 | 450 | mld,ttk,sens-2,CG10321,klu,crol,CG32772,Prdm13,sqz,sens,CG10348,tsh,chn,CG12769,CG14441,hang,CG11247,esg,zfh2,pita,bowl,rgr | | 8.355e-04 | -7.09 | muscle cell cellular homeostasis | biological process | GO:0046716 | 32 | 6 | 12345 | 442 | robo1,N,Pgk,Fkbp14,Dg,Nrk | | 8.410e-04 | -7.08 | neuron migration | biological process | GO:0001764 | 6 | 3 | 12345 | 442 | SCAR,sli,Abl | | 8.410e-04 | -7.08 | regulation of cardioblast differentiation | biological process | GO:0051890 | 6 | 3 | 12345 | 442 | N,Abd-B,lqf | | 8.410e-04 | -7.08 | photoreceptor cell fate determination | biological process | GO:0043703 | 6 | 3 | 12345 | 442 | Egfr,S,spi | | 8.410e-04 | -7.08 | mesenchymal cell differentiation | biological process | GO:0048762 | 6 | 3 | 12345 | 442 | Sema1b,Sema1a,disco-r | | 8.525e-04 | -7.07 | GO\_BRANCHING\_MORPHOGENESIS\_OF\_AN\_EPITHELIAL\_TUBE | MSigDB lists | GO\_BRANCHING\_MORPHOGENESIS\_OF\_AN\_EPITHELIAL\_TUBE | 31 | 6 | 4161 | 156 | N,LanA,ft,sli,ds,arm | | 8.532e-04 | -7.07 | shot (short stop) | protein interactions | 36542 | 25 | 6 | 9204 | 432 | kay,sli,betaTub60D,wnd,shot,robo1 | | 8.538e-04 | -7.07 | negative regulation of cellular protein metabolic process | biological process | GO:0032269 | 218 | 18 | 12345 | 442 | Spn42De,Cbl,Kdm2,CycG,orb2,Socs36E,pum,trbl,bru2,N,Tis11,wdb,Sxl,CG8312,Diap1,312,msi,bin3 | | 8.572e-04 | -7.06 | response to toxic substance | biological process | GO:0009636 | 165 | 15 | 12345 | 442 | spi,S,mura,hang,hppy,LRR,esg,ZnT41F,Bacc,Pdk1,moody,pum,CG30344,Egfr,pxb | | 8.613e-04 | -7.06 | Sdc (Syndecan) | protein interactions | 37447 | 17 | 5 | 9204 | 432 | sli,dlp,kay,robo1,Dg | | 8.613e-04 | -7.06 | Apc2 (Adenomatous polyposis coli 2) | protein interactions | 42871 | 17 | 5 | 9204 | 432 | arm,chic,CtBP,klu,Pex5 | | 8.613e-04 | -7.06 | amos (absent MD neurons and olfactory sensilla) | protein interactions | 35110 | 17 | 5 | 9204 | 432 | Dl,H,emc,N,h | | 8.615e-04 | -7.06 | vesicle | cellular component | GO:0031982 | 422 | 28 | 11940 | 417 | N,fz2,jeb,path,Cip4,spdo,shg,Rab1,AP-1gamma,mop,Ocrl,ed,apolpp,jar,CG6656,arm,Cadps,S,Vps13,Rab5,fz,nuf,Syx1A,robo1,dock,lt,tkv,Dl | | 8.682e-04 | -7.05 | mitotic cell cycle | biological process | GO:0000278 | 432 | 29 | 12345 | 442 | 14-3-3epsilon,kay,Cks30A,Slbp,eff,larp,CG3436,Dp,baz,jar,Egfr,Ocrl,Syx1A,Rab5,Psc,CycB3,ph-p,Wdr62,Rab1,pum,chic,CG11414,Arf79F,CycG,betaTub60D,14-3-3zeta,N,Dl,corto | | 8.682e-04 | -7.05 | LXR\_DR4\_Q3 | MSigDB lists | LXR\_DR4\_Q3 | 21 | 5 | 4161 | 156 | rdx,Kdm2,14-3-3zeta,Mef2,otk | | 8.688e-04 | -7.05 | HOMEOBOX\_1 | prosite domains | PS00027 | 92 | 13 | 6243 | 325 | Lmx1a,nub,al,B-H1,ap,H2.0,Abd-B,abd-A,zfh2,otp,eyg,Scr,ey | | 8.696e-04 | -7.05 | hh (hedgehog) | protein interactions | 42737 | 77 | 11 | 9204 | 432 | Psc,Egfr,boi,Cdk4,Dl,dlp,Raf,kn,apolpp,mam,ey | | 8.696e-04 | -7.05 | Hsp83 (Heat shock protein 83) | protein interactions | 38389 | 77 | 11 | 9204 | 432 | Gli,Src42A,sina,Egfr,InR,Nlp,Dcr-2,chn,Raf,Arf79F,SERCA | | 8.745e-04 | -7.04 | negative regulation of neuromuscular junction development | biological process | GO:1904397 | 44 | 7 | 12345 | 442 | Abl,ena,Dad,LanA,Mob2,Src42A,Cip4 | | 8.745e-04 | -7.04 | peptidyl-tyrosine dephosphorylation | biological process | GO:0035335 | 44 | 7 | 12345 | 442 | Ptpmeg2,mop,IA-2,Ptp4E,eya,Pez,CG42327 | | 8.745e-04 | -7.04 | negative regulation of synaptic growth at neuromuscular junction | biological process | GO:0045886 | 44 | 7 | 12345 | 442 | Abl,ena,Dad,LanA,Mob2,Src42A,Cip4 | | 8.754e-04 | -7.04 | GO\_GOLGI\_APPARATUS\_PART | MSigDB lists | GO\_GOLGI\_APPARATUS\_PART | 268 | 21 | 4161 | 156 | GlcAT-I,dlp,Rab1,alpha-Man-Ia,N,Arf79F,Pgant5,dock,lt,Pgant6,step,Cals,fz2,Tpst,CG9220,Ocrl,C1GalTA,CG17883,AP-1gamma,rg,Tollo | | 8.810e-04 | -7.03 | tsr (twinstar) | protein interactions | 37841 | 44 | 8 | 9204 | 432 | fl(2)d,ey,imd,cindr,Sema1a,Jupiter,scrib,sick | | 8.810e-04 | -7.03 | sev (sevenless) | protein interactions | 32039 | 44 | 8 | 9204 | 432 | kay,B-H1,Cbl,Socs36E,Actbeta,Egfr,Src42A,S | | 8.854e-04 | -7.03 | GO\_ANGIOGENESIS | MSigDB lists | GO\_ANGIOGENESIS | 55 | 8 | 4161 | 156 | C1GalTA,sli,LanA,Eph,Dl,fz2,arm,N | | 8.959e-04 | -7.02 | positive regulation of insulin receptor signaling pathway | biological process | GO:0046628 | 13 | 4 | 12345 | 442 | rictor,lin-28,step,CycG | | 8.959e-04 | -7.02 | positive regulation of cellular response to insulin stimulus | biological process | GO:1900078 | 13 | 4 | 12345 | 442 | step,lin-28,rictor,CycG | | 8.963e-04 | -7.02 | FRIZZLED | prints domains | PR00489 | 5 | 3 | 2185 | 101 | fz,fz2,fz3 | | 8.972e-04 | -7.02 | cell-cell junction assembly | biological process | GO:0007043 | 57 | 8 | 12345 | 442 | baz,scrib,moody,Gli,bou,sns,shg,arm | | 9.003e-04 | -7.01 | negative regulation of protein metabolic process | biological process | GO:0051248 | 219 | 18 | 12345 | 442 | bru2,Tis11,N,Socs36E,orb2,Kdm2,CycG,trbl,pum,Cbl,Spn42De,bin3,312,msi,CG8312,Diap1,wdb,Sxl | | 9.040e-04 | -7.01 | regulation of synapse assembly | biological process | GO:0051963 | 116 | 12 | 12345 | 442 | LanA,Dad,nmo,tkv,rictor,ena,pum,Cip4,Src42A,Cals,Mob2,Abl | | 9.073e-04 | -7.01 | cytoskeletal protein binding | molecular function | GO:0008092 | 330 | 23 | 12176 | 414 | shg,yrt,cora,Dyb,CG2765,klar,f,Pez,d,Shrm,mim,crp,Unc-76,SCAR,nuf,Jupiter,shot,ena,Myo10A,apolpp,CG6891,chic,jar | | 9.141e-04 | -7.00 | germ-band shortening | biological process | GO:0007390 | 22 | 5 | 12345 | 442 | ena,Egfr,Orct2,yrt,InR | | 9.182e-04 | -6.99 | GO\_CELL\_JUNCTION | MSigDB lists | GO\_CELL\_JUNCTION | 307 | 23 | 4161 | 156 | otk,Dl,htl,sns,dock,14-3-3zeta,CtBP,N,Arf79F,14-3-3epsilon,shot,unc-5,Cadps,Syx1A,Ddr,emp,nAChRalpha4,fz,Nrk,Tsp96F,Pdk1,fz2,arm | | 9.220e-04 | -6.99 | fn3 | pfam domains | PF00041 | 40 | 7 | 10480 | 419 | Eph,robo1,boi,ed,sns,dome,Ptp4E | | 9.252e-04 | -6.99 | homeostatic process | biological process | GO:0042592 | 475 | 31 | 12345 | 442 | InR,Rab1,Raf,SERCA,hppy,H,Nrk,N,Ppcs,Mid1,Pvr,cue,Pgk,Atg17,Fkbp14,Dg,trbl,Src42A,yrt,shg,ey,tai,SNF4Agamma,robo1,Atg16,esg,Atox1,ZnT41F,ATPCL,wgn,Egfr | | 9.296e-04 | -6.98 | ebi (ebi) | protein interactions | 33212 | 66 | 10 | 9204 | 432 | CG2017,Diap1,ttk,sd,Egfr,Dl,Dcr-2,chn,N,sina | | 9.436e-04 | -6.97 | cell maturation | biological process | GO:0048469 | 184 | 16 | 12345 | 442 | mop,Dronc,Pat1,scrib,N,14-3-3zeta,Dg,chic,eff,shg,14-3-3epsilon,Sox14,cic,wbl,baz,Egfr | | 9.513e-04 | -6.96 | GO\_NEGATIVE\_REGULATION\_OF\_EPITHELIAL\_CELL\_DIFFERENTIATION | MSigDB lists | GO\_NEGATIVE\_REGULATION\_OF\_EPITHELIAL\_CELL\_DIFFERENTIATION | 6 | 3 | 4161 | 156 | arm,N,Dl | | 9.513e-04 | -6.96 | FINETTI\_BREAST\_CANCERS\_KINOME\_BLUE | MSigDB lists | FINETTI\_BREAST\_CANCERS\_KINOME\_BLUE | 6 | 3 | 4161 | 156 | htl,tkv,InR | | 9.513e-04 | -6.96 | GO\_NEGATIVE\_REGULATION\_OF\_MYELOID\_LEUKOCYTE\_DIFFERENTIATION | MSigDB lists | GO\_NEGATIVE\_REGULATION\_OF\_MYELOID\_LEUKOCYTE\_DIFFERENTIATION | 6 | 3 | 4161 | 156 | Tollo,Actbeta,arm | | 9.513e-04 | -6.96 | MODULE\_80 | MSigDB lists | MODULE\_80 | 6 | 3 | 4161 | 156 | Nrk,Eph,Ddr | | 9.513e-04 | -6.96 | GO\_GLANDULAR\_EPITHELIAL\_CELL\_DEVELOPMENT | MSigDB lists | GO\_GLANDULAR\_EPITHELIAL\_CELL\_DEVELOPMENT | 6 | 3 | 4161 | 156 | Pdk1,Cdk4,Dl | | 9.513e-04 | -6.96 | MODULE\_51 | MSigDB lists | MODULE\_51 | 6 | 3 | 4161 | 156 | Ddr,Nrk,Eph | | 9.513e-04 | -6.96 | GO\_EPITHELIAL\_CELL\_DIFFERENTIATION\_INVOLVED\_IN\_PROSTATE\_GLAND\_DEVELOPMENT | MSigDB lists | GO\_EPITHELIAL\_CELL\_DIFFERENTIATION\_INVOLVED\_IN\_PROSTATE\_GLAND\_DEVELOPMENT | 6 | 3 | 4161 | 156 | arm,Fem-1,N | | 9.513e-04 | -6.96 | KEGG\_O\_GLYCAN\_BIOSYNTHESIS | MSigDB lists | KEGG\_O\_GLYCAN\_BIOSYNTHESIS | 6 | 3 | 4161 | 156 | C1GalTA,Pgant6,Pgant5 | | 9.513e-04 | -6.96 | BIOCARTA\_GSK3\_PATHWAY | MSigDB lists | BIOCARTA\_GSK3\_PATHWAY | 6 | 3 | 4161 | 156 | fz,Pdk1,arm | | 9.567e-04 | -6.95 | Grip (Glutamate receptor binding protein) | protein interactions | 50391 | 5 | 3 | 9204 | 432 | kon,fz2,ed | | 9.567e-04 | -6.95 | edl (ETS-domain lacking) | protein interactions | 37149 | 5 | 3 | 9204 | 432 | S,aop,spi | | 9.567e-04 | -6.95 | chas (chascon) | protein interactions | 32748 | 5 | 3 | 9204 | 432 | dpy,Pex5,N | | 9.567e-04 | -6.95 | CG5986 (uncharacterized protein) | protein interactions | 192506 | 5 | 3 | 9204 | 432 | Diap1,H,Sema1a | | 9.567e-04 | -6.95 | otk2 (off-track2) | protein interactions | 36284 | 5 | 3 | 9204 | 432 | fz,fz2,otk | | 9.567e-04 | -6.95 | slp2 (sloppy paired 2) | protein interactions | 33608 | 5 | 3 | 9204 | 432 | Scr,abd-A,Abd-B | | 9.567e-04 | -6.95 | bap (bagpipe) | protein interactions | 42537 | 5 | 3 | 9204 | 432 | abd-A,Abd-B,Scr | | 9.578e-04 | -6.95 | positive regulation of cellular component biogenesis | biological process | GO:0044089 | 133 | 13 | 12345 | 442 | tkv,shot,nmo,RhoU,ena,rictor,chic,kon,RhoGEF4,SCAR,Abl,Cals,Jupiter | | 9.645e-04 | -6.94 | GO\_CELL\_ACTIVATION | MSigDB lists | GO\_CELL\_ACTIVATION | 98 | 11 | 4161 | 156 | Cdk4,Mef2,Raf,Eph,Dl,Tollo,fz2,arm,14-3-3zeta,Pdk1,dock | | 9.686e-04 | -6.94 | GO\_INTRACELLULAR\_SIGNAL\_TRANSDUCTION | MSigDB lists | GO\_INTRACELLULAR\_SIGNAL\_TRANSDUCTION | 387 | 27 | 4161 | 156 | Tollo,ds,ft,PI31,Rgl,Ziz,Pdk1,Lk6,Ric,SERCA,jar,Mef2,Raf,cact,CG4935,CG5522,Rab5,hppy,Arf79F,InR,sns,htl,sd,Rab1,Drak,14-3-3epsilon,nmo | | 9.863e-04 | -6.92 | GO\_NEUROGENESIS | MSigDB lists | GO\_NEUROGENESIS | 328 | 24 | 4161 | 156 | Pex7,Cdk4,robo1,Raf,Mef2,otk,Dl,unc-5,14-3-3epsilon,Actbeta,N,Arf79F,Unc-76,ap,ft,ds,sli,Eph,fz2,arm,fz,sina,tkv,nAChRalpha4 | | 9.908e-04 | -6.92 | regulation of chemotaxis | biological process | GO:0050920 | 33 | 6 | 12345 | 442 | chic,Sema1b,sli,fz,CG34401,Sema1a | | 9.951e-04 | -6.91 | membrane part | cellular component | GO:0044425 | 2068 | 97 | 11940 | 417 | cindr,Esp,Dg,beat-IIa,otk,Tsp96F,Gli,glec,ZnT41F,fz3,CG7536,Nrk,CG32795,Abl,AP-1gamma,rho-4,CG30344,CG30156,Arf79F,d,N,Ir21a,nAChRalpha4,nuf,Egfr,LanA,arm,Mpcp2,Vps13,C1GalTA,Cals,ft,CG33474,klar,Cip4,Pvr,dlp,Tpc1,step,dome,ogre,Syx1A,CG7394,Gr93c,tipE,CG8034,Dl,SERCA,yrt,Ddr,S,CG17883,fz,Cbl,scrib,spi,Tob,emp,Ptp4E,ed,neo,unc-5,Eph,Socs36E,Atg17,baz,DPCoAC,orb2,shg,robo1,ds,lt,gukh,tkv,rg,Inx2,bou,InR,Src42A,wgn,Orct2,Pex10,Irk3,drl,Shrm,moody,Rab1,CG34172,Dyb,CG3149,Sema1a,kek5,UQCR-11,Magi,fz2,htl,Kr-h2 | | 9.969e-04 | -6.91 | protein modification process | biological process | GO:0036211 | 1215 | 64 | 12345 | 442 | hppy,mura,InR,Drak,Ptpmeg2,FBXO11,drl,Su(z)2,CG10417,ec,sina,Kdm4B,Pez,rdx,Fem-1,Cbl,CG13481,hyx,mop,ds,CycB3,Ptp4E,CycG,Ubc87F,Abl,wdb,ft,Cks30A,Ddr,tkv,morgue,Atg16,CG42327,Pitslre,CG3119,CG1218,Pvr,CG12402,tna,Cdk4,Lk6,Pp2C1,Gfat2,Src42A,C1GalTA,E(Pc),Nrk,Dyrk2,eya,Raf,otk,trbl,wnd,Eph,Kdm2,CG34057,Pex10,IA-2,eff,Diap1,htl,Tpst,nmo,Pdk1 | | 9.969e-04 | -6.91 | cellular protein modification process | biological process | GO:0006464 | 1215 | 64 | 12345 | 442 | Fem-1,hyx,mop,CG13481,Cbl,CycB3,ds,CycG,Ptp4E,Ubc87F,Abl,wdb,ft,Cks30A,Ddr,tkv,morgue,CG42327,Atg16,hppy,InR,mura,Drak,Ptpmeg2,FBXO11,drl,Su(z)2,CG10417,ec,Kdm4B,sina,rdx,Pez,Nrk,E(Pc),eya,Dyrk2,Raf,Eph,wnd,trbl,otk,CG34057,Kdm2,Pex10,IA-2,Diap1,eff,Tpst,nmo,htl,Pdk1,Pitslre,CG1218,CG3119,Pvr,tna,CG12402,Pp2C1,Lk6,Cdk4,Gfat2,Src42A,C1GalTA | | 1.001e-03 | -6.91 | GO\_LYMPHOCYTE\_ACTIVATION | MSigDB lists | GO\_LYMPHOCYTE\_ACTIVATION | 56 | 8 | 4161 | 156 | Cdk4,Mef2,Raf,Eph,Dl,dock,fz2,arm | | 1.004e-03 | -6.90 | negative regulation of synapse assembly | biological process | GO:0051964 | 45 | 7 | 12345 | 442 | Mob2,Src42A,Cip4,ena,Abl,LanA,Dad | | 1.004e-03 | -6.90 | negative regulation of synapse organization | biological process | GO:1905809 | 45 | 7 | 12345 | 442 | Abl,ena,Dad,LanA,Mob2,Src42A,Cip4 | | 1.017e-03 | -6.89 | GO\_MESENCHYME\_DEVELOPMENT | MSigDB lists | GO\_MESENCHYME\_DEVELOPMENT | 32 | 6 | 4161 | 156 | ds,arm,LanA,Mef2,N,otk | | 1.024e-03 | -6.88 | hth (homothorax) | protein interactions | 41273 | 35 | 7 | 9204 | 432 | scrib,Sxl,Abd-B,abd-A,ey,Scr,tsh | | 1.024e-03 | -6.88 | tor (torso) | protein interactions | 35717 | 35 | 7 | 9204 | 432 | Raf,lilli,14-3-3epsilon,bowl,Pex14,14-3-3zeta,tkv | | 1.025e-03 | -6.88 | regulation of actin filament organization | biological process | GO:0110053 | 72 | 9 | 12345 | 442 | fz,Cip4,Abl,SCAR,RhoGEF4,chic,CG6891,ena,RhoU | | 1.028e-03 | -6.88 | numb (numb) | protein interactions | 34263 | 45 | 8 | 9204 | 432 | N,Rm62,fz,baz,H,Dronc,AP-1gamma,spdo | | 1.038e-03 | -6.87 | transferase activity, transferring phosphorus-containing groups | molecular function | GO:0016772 | 482 | 30 | 12176 | 414 | CycB3,CG1129,drl,Abl,Eph,Ddr,Nrk,Cerk,htl,InR,l(2)37Cg,Raf,Lk6,nmo,hppy,l(1)G0196,Src42A,trbl,Dyrk2,Pvr,CycG,Drak,wnd,tkv,otk,Egfr,Pdk1,Pgk,Pitslre,Cdk4 | | 1.043e-03 | -6.87 | GO\_PANCREAS\_DEVELOPMENT | MSigDB lists | GO\_PANCREAS\_DEVELOPMENT | 13 | 4 | 4161 | 156 | arm,Pdk1,Cdk4,Dl | | 1.043e-03 | -6.87 | GO\_CHONDROITIN\_SULFATE\_PROTEOGLYCAN\_METABOLIC\_PROCESS | MSigDB lists | GO\_CHONDROITIN\_SULFATE\_PROTEOGLYCAN\_METABOLIC\_PROCESS | 13 | 4 | 4161 | 156 | CG9220,Ids,GlcAT-I,frc | | 1.043e-03 | -6.87 | ACAWNRNSRCGG\_UNKNOWN | MSigDB lists | ACAWNRNSRCGG\_UNKNOWN | 13 | 4 | 4161 | 156 | 14-3-3epsilon,sli,trbl,Arf79F | | 1.043e-03 | -6.87 | REACTOME\_CHONDROITIN\_SULFATE\_DERMATAN\_SULFATE\_METABOLISM | MSigDB lists | REACTOME\_CHONDROITIN\_SULFATE\_DERMATAN\_SULFATE\_METABOLISM | 13 | 4 | 4161 | 156 | CG9220,dlp,GlcAT-I,Ids | | 1.048e-03 | -6.86 | csw (corkscrew) | protein interactions | 45278 | 67 | 10 | 9204 | 432 | Pvr,tkv,vn,14-3-3zeta,Egfr,Dl,spi,Raf,S,Src42A | | 1.056e-03 | -6.85 | kinase regulator activity | molecular function | GO:0019207 | 76 | 9 | 12176 | 414 | jeb,Atg17,14-3-3zeta,CycG,CycB3,Pdk1,Socs36E,Cks30A,trbl | | 1.074e-03 | -6.84 | PROTEIN\_KINASE\_DOM | prosite domains | PS50011 | 221 | 23 | 6243 | 325 | Cdk4,Drak,hppy,tkv,nmo,Nrk,Dyrk2,Pdk1,InR,drl,wnd,Abl,Lk6,trbl,htl,Ddr,Pitslre,Eph,Raf,Pvr,Src42A,Egfr,otk | | 1.079e-03 | -6.83 | ph-p (polyhomeotic proximal) | protein interactions | 31181 | 56 | 9 | 9204 | 432 | corto,sd,Su(z)2,ph-p,eff,Psc,spi,tara,CG32772 | | 1.083e-03 | -6.83 | PIT1\_Q6 | MSigDB lists | PIT1\_Q6 | 44 | 7 | 4161 | 156 | ds,nmo,Raf,Nrk,Actbeta,htl,sd | | 1.105e-03 | -6.81 | signaling receptor activity | molecular function | GO:0038023 | 419 | 27 | 12176 | 414 | Eph,Ddr,Nrk,drl,moody,Gr93c,Hr4,InR,htl,N,Gli,boi,nAChRalpha4,fz,cv-d,unc-5,Hr39,otk,fz3,tkv,wgn,Pvr,Egfr,dome,Ptp4E,Ir21a,fz2 | | 1.111e-03 | -6.80 | muscle tendon junction | cellular component | GO:0005927 | 14 | 4 | 11940 | 417 | Abl,kon,LRR,shot | | 1.121e-03 | -6.79 | Fc epsilon receptor (FCERI) signaling | REACTOME pathways | R-DME-2454202 | 167 | 17 | 3923 | 176 | 14-3-3zeta,Pdk1,rictor,PI31,trbl,eff,Raf,cact,Pvr,htl,kay,CG4935,Dronc,CG42684,wdb,vn,Egfr | | 1.134e-03 | -6.78 | R7 cell fate commitment | biological process | GO:0007465 | 23 | 5 | 12345 | 442 | Cbl,sina,Raf,ttk,aop | | 1.134e-03 | -6.78 | positive regulation of filopodium assembly | biological process | GO:0051491 | 23 | 5 | 12345 | 442 | chic,ena,shot,SCAR,kon | | 1.134e-03 | -6.78 | male genitalia development | biological process | GO:0030539 | 23 | 5 | 12345 | 442 | N,Rab5,otk,Pvr,Abd-B | | 1.136e-03 | -6.78 | FERM\_domain | interpro domains | IPR000299 | 21 | 5 | 11383 | 450 | yrt,Pez,cora,CG42788,Myo10A | | 1.147e-03 | -6.77 | Jon99Ciii (Jonah 99Ciii) | protein interactions | 43543 | 18 | 5 | 9204 | 432 | CG12581,Sxl,Pez,al,Tpc1 | | 1.147e-03 | -6.77 | salm (spalt major) | protein interactions | 34569 | 18 | 5 | 9204 | 432 | Abd-B,abd-A,spi,ph-p,Scr | | 1.147e-03 | -6.77 | trr (trithorax-related) | protein interactions | 31149 | 18 | 5 | 9204 | 432 | sd,N,Sema1a,His3.3A,spen | | 1.149e-03 | -6.77 | syncytium formation by plasma membrane fusion | biological process | GO:0000768 | 46 | 7 | 12345 | 442 | Mef2,InR,dock,SCAR,sns,siz,Pgk | | 1.149e-03 | -6.77 | cell-cell signaling by wnt | biological process | GO:0198738 | 46 | 7 | 12345 | 442 | arm,drl,fz,tsh,fz2,otk,fz3 | | 1.149e-03 | -6.77 | myoblast fusion | biological process | GO:0007520 | 46 | 7 | 12345 | 442 | sns,dock,SCAR,InR,Mef2,Pgk,siz | | 1.149e-03 | -6.77 | apical junction assembly | biological process | GO:0043297 | 46 | 7 | 12345 | 442 | baz,bou,arm,Gli,moody,shg,scrib | | 1.149e-03 | -6.77 | syncytium formation | biological process | GO:0006949 | 46 | 7 | 12345 | 442 | Mef2,InR,dock,SCAR,sns,siz,Pgk | | 1.149e-03 | -6.77 | cell-cell fusion | biological process | GO:0140253 | 46 | 7 | 12345 | 442 | InR,Mef2,sns,dock,SCAR,Pgk,siz | | 1.153e-03 | -6.77 | Ral guanyl-nucleotide exchange factor activity | molecular function | GO:0008321 | 2 | 2 | 12176 | 414 | CG5522,Rgl | | 1.153e-03 | -6.77 | structural constituent of chromatin | molecular function | GO:0030527 | 2 | 2 | 12176 | 414 | His3.3B,His3.3A | | 1.160e-03 | -6.76 | Laminin\_G\_1 | pfam domains | PF00054 | 6 | 3 | 10480 | 419 | sli,ft,LanA | | 1.167e-03 | -6.75 | genitalia development | biological process | GO:0048806 | 34 | 6 | 12345 | 442 | N,Rab5,otk,Pvr,abd-A,Abd-B | | 1.171e-03 | -6.75 | enzyme activator activity | molecular function | GO:0008047 | 143 | 13 | 12176 | 414 | Pdk1,wdb,CenB1A,rictor,Cks30A,RapGAP1,Ocrl,CG7394,jeb,CG17883,drongo,RhoGAP19D,TBC1D5 | | 1.179e-03 | -6.74 | FZ | prosite domains | PS50038 | 10 | 4 | 6243 | 325 | fz2,Nrk,fz,fz3 | | 1.184e-03 | -6.74 | GO\_SENSORY\_ORGAN\_DEVELOPMENT | MSigDB lists | GO\_SENSORY\_ORGAN\_DEVELOPMENT | 116 | 12 | 4161 | 156 | fz2,arm,tkv,ap,N,Actbeta,ft,CG17883,ds,Dl,otk,Eph | | 1.185e-03 | -6.74 | regulation of synapse organization | biological process | GO:0050807 | 153 | 14 | 12345 | 442 | Mob2,Cals,Cip4,Rab1,Src42A,Abl,rictor,ena,pum,wnd,LanA,nmo,tkv,Dad | | 1.193e-03 | -6.73 | FERM\_M | pfam domains | PF00373 | 21 | 5 | 10480 | 419 | cora,yrt,Pez,CG42788,Myo10A | | 1.195e-03 | -6.73 | msk (moleskin) | protein interactions | 44747 | 46 | 8 | 9204 | 432 | rictor,Mpcp2,Dg,GluProRS,fl(2)d,CG12769,sens,CG10417 | | 1.197e-03 | -6.73 | apical part of cell | cellular component | GO:0045177 | 157 | 14 | 11940 | 417 | arm,dome,CG42684,Shrm,Pez,baz,Abl,nuf,cindr,Dl,Arf79F,SCAR,Inx2,Ptp4E | | 1.198e-03 | -6.73 | - | gene3d domains | 2.60.40.10 | 195 | 20 | 4929 | 241 | CG31431,vn,unc-5,ed,Bsg,robo1,otk,Eph,boi,dome,Ocrl,kek5,sns,htl,InR,kn,Dg,Ptp4E,Ppn,Pvr | | 1.202e-03 | -6.72 | cleavage furrow | cellular component | GO:0032154 | 35 | 6 | 11940 | 417 | cindr,nuf,Rab1,Cip4,step,Arf79F | | 1.202e-03 | -6.72 | basal part of cell | cellular component | GO:0045178 | 35 | 6 | 11940 | 417 | apolpp,SCAR,scrib,Dg,jar,Egfr | | 1.205e-03 | -6.72 | DACOSTA\_UV\_RESPONSE\_VIA\_ERCC3\_XPCS\_DN | MSigDB lists | DACOSTA\_UV\_RESPONSE\_VIA\_ERCC3\_XPCS\_DN | 33 | 6 | 4161 | 156 | shot,CenB1A,Ziz,sli,IntS3,InR | | 1.205e-03 | -6.72 | GO\_STEM\_CELL\_DIFFERENTIATION | MSigDB lists | GO\_STEM\_CELL\_DIFFERENTIATION | 33 | 6 | 4161 | 156 | arm,LanA,fz,Cdk4,Mef2,N | | 1.205e-03 | -6.72 | GO\_COLUMNAR\_CUBOIDAL\_EPITHELIAL\_CELL\_DIFFERENTIATION | MSigDB lists | GO\_COLUMNAR\_CUBOIDAL\_EPITHELIAL\_CELL\_DIFFERENTIATION | 33 | 6 | 4161 | 156 | C1GalTA,Dl,N,Mef2,Cdk4,Pdk1 | | 1.207e-03 | -6.72 | Notch signaling pathway | KEGG pathways | ko04330 | 22 | 5 | 3198 | 123 | N,mam,H,CtBP,Dl | | 1.207e-03 | -6.72 | Notch signaling pathway | KEGG pathways | dme04330 | 22 | 5 | 3198 | 123 | H,mam,N,Dl,CtBP | | 1.213e-03 | -6.71 | mew (multiple edematous wings) | protein interactions | 32275 | 11 | 4 | 9204 | 432 | 14-3-3epsilon,tkv,sli,H | | 1.213e-03 | -6.71 | Pvf1 (PDGF- and VEGF-related factor 1) | protein interactions | 32876 | 11 | 4 | 9204 | 432 | kay,Egfr,tai,Pvr | | 1.213e-03 | -6.71 | p120ctn (Adherens junction protein p120) | protein interactions | 3355143 | 11 | 4 | 9204 | 432 | shg,arm,Sxl,ena | | 1.213e-03 | -6.71 | sno (strawberry notch) | protein interactions | 32273 | 11 | 4 | 9204 | 432 | Dl,mam,H,N | | 1.215e-03 | -6.71 | male gamete generation | biological process | GO:0048232 | 360 | 25 | 12345 | 442 | larp,eff,dmrt11E,Diap1,Pex10,Pdk1,His3.3B,jar,His3.3A,Rab1,Raf,InR,yuri,sns,eya,lncRNA:CR44420,Dronc,Inx2,bol,Gli,orb2,Arf79F,PI31,wnd,Pen | | 1.217e-03 | -6.71 | Slit-Robo signaling complex | cellular component | GO:0071666 | 2 | 2 | 11940 | 417 | sli,robo1 | | 1.219e-03 | -6.71 | cell-cell adhesion mediated by cadherin | biological process | GO:0044331 | 14 | 4 | 12345 | 442 | Cals,shg,ds,ft | | 1.219e-03 | -6.71 | muscle cell fate commitment | biological process | GO:0042693 | 14 | 4 | 12345 | 442 | jeb,kn,abd-A,N | | 1.236e-03 | -6.70 | endosome | cellular component | GO:0005768 | 212 | 17 | 11940 | 417 | S,fz,Vps13,Rab5,tkv,Dl,nuf,lt,spdo,path,shg,N,fz2,Ocrl,ed,mop,AP-1gamma | | 1.262e-03 | -6.67 | cellular component assembly involved in morphogenesis | biological process | GO:0010927 | 154 | 14 | 12345 | 442 | ed,emc,S,Dp,ttk,Raf,Dg,Egfr,cact,Bsg,N,Ppcs,cic,kon | | 1.267e-03 | -6.67 | GO\_TELENCEPHALON\_DEVELOPMENT | MSigDB lists | GO\_TELENCEPHALON\_DEVELOPMENT | 58 | 8 | 4161 | 156 | ap,Actbeta,14-3-3epsilon,arm,Cdk4,robo1,sli,ft | | 1.277e-03 | -6.66 | GO\_DEVELOPMENTAL\_PROCESS\_INVOLVED\_IN\_REPRODUCTION | MSigDB lists | GO\_DEVELOPMENTAL\_PROCESS\_INVOLVED\_IN\_REPRODUCTION | 117 | 12 | 4161 | 156 | robo1,Tollo,CG17883,sli,Fem-1,N,CG7394,Actbeta,tkv,arm,Mfe2,fz2 | | 1.279e-03 | -6.66 | negative regulation of fusion cell fate specification | biological process | GO:0035157 | 2 | 2 | 12345 | 442 | aop,ed | | 1.279e-03 | -6.66 | haltere morphogenesis | biological process | GO:0048735 | 2 | 2 | 12345 | 442 | ap,hth | | 1.279e-03 | -6.66 | regulation of glycolytic process | biological process | GO:0006110 | 2 | 2 | 12345 | 442 | Dg,N | | 1.279e-03 | -6.66 | positive regulation of cell proliferation involved in compound eye morphogenesis | biological process | GO:2000497 | 2 | 2 | 12345 | 442 | hth,tsh | | 1.279e-03 | -6.66 | regulation of hematopoietic stem cell differentiation | biological process | GO:1902036 | 2 | 2 | 12345 | 442 | C1GalTA,Arf79F | | 1.279e-03 | -6.66 | regulation of hematopoietic progenitor cell differentiation | biological process | GO:1901532 | 2 | 2 | 12345 | 442 | C1GalTA,Arf79F | | 1.279e-03 | -6.66 | stomatogastric nervous system development | biological process | GO:0007421 | 2 | 2 | 12345 | 442 | S,spi | | 1.291e-03 | -6.65 | Mcad (Medium-chain acyl-CoA dehydrogenase) | protein interactions | 38864 | 93 | 12 | 9204 | 432 | GluProRS,Mpcp2,Kr-h2,Egfr,Raf,Sxl,Rab5,Arf79F,SERCA,Gfat2,Pdk1,Rab1 | | 1.296e-03 | -6.65 | GO\_REGULATION\_OF\_NERVOUS\_SYSTEM\_DEVELOPMENT | MSigDB lists | GO\_REGULATION\_OF\_NERVOUS\_SYSTEM\_DEVELOPMENT | 150 | 14 | 4161 | 156 | fz,Nrk,Arf79F,N,nAChRalpha4,arm,robo1,Raf,Mef2,Cals,otk,Eph,Dl,sli | | 1.304e-03 | -6.64 | ACTGAAA\_MIR30A3P\_MIR30E3P | MSigDB lists | ACTGAAA\_MIR30A3P\_MIR30E3P | 72 | 9 | 4161 | 156 | wdb,sina,Pgant5,CG1316,14-3-3epsilon,bru2,Mef2,AP-1gamma,step | | 1.327e-03 | -6.62 | nej (nejire) | protein interactions | 43856 | 106 | 13 | 9204 | 432 | lilli,ena,eya,Scr,ey,H,Psc,Egfr,spi,MEP-1,shg,arm,lncRNA:Hsromega | | 1.333e-03 | -6.62 | intercalary heterochromatin | cellular component | GO:0005725 | 7 | 3 | 11940 | 417 | Su(z)2,Psc,E(Pc) | | 1.351e-03 | -6.61 | BIOCARTA\_MAPK\_PATHWAY | MSigDB lists | BIOCARTA\_MAPK\_PATHWAY | 23 | 5 | 4161 | 156 | hppy,Raf,Mef2,cact,Lk6 | | 1.351e-03 | -6.61 | GO\_PROTEIN\_LOCALIZATION\_TO\_VACUOLE | MSigDB lists | GO\_PROTEIN\_LOCALIZATION\_TO\_VACUOLE | 23 | 5 | 4161 | 156 | CenB1A,emp,lt,Tsp96F,Vps13 | | 1.351e-03 | -6.61 | GO\_MOLTING\_CYCLE | MSigDB lists | GO\_MOLTING\_CYCLE | 23 | 5 | 4161 | 156 | LanA,arm,ap,N,Actbeta | | 1.351e-03 | -6.61 | PID\_VEGFR1\_2\_PATHWAY | MSigDB lists | PID\_VEGFR1\_2\_PATHWAY | 23 | 5 | 4161 | 156 | arm,dock,Raf,Arf79F,Pdk1 | | 1.351e-03 | -6.61 | GO\_CAMERA\_TYPE\_EYE\_MORPHOGENESIS | MSigDB lists | GO\_CAMERA\_TYPE\_EYE\_MORPHOGENESIS | 23 | 5 | 4161 | 156 | Dl,Eph,CG17883,arm,fz2 | | 1.351e-03 | -6.61 | GO\_PEROXISOME\_ORGANIZATION | MSigDB lists | GO\_PEROXISOME\_ORGANIZATION | 23 | 5 | 4161 | 156 | Pex7,Pex5,Pex14,Pex10,CG45050 | | 1.353e-03 | -6.61 | GO\_POSITIVE\_REGULATION\_OF\_CELL\_DEATH | MSigDB lists | GO\_POSITIVE\_REGULATION\_OF\_CELL\_DEATH | 134 | 13 | 4161 | 156 | arm,unc-5,14-3-3epsilon,Drak,Actbeta,sina,N,14-3-3zeta,dock,tkv,sli,Mef2,Tollo | | 1.358e-03 | -6.60 | Kinase-like\_dom\_sf | interpro domains | IPR011009 | 293 | 23 | 11383 | 450 | Egfr,Nrk,Abl,Dyrk2,htl,Pvr,Drak,InR,Pitslre,trbl,Lk6,Src42A,otk,tkv,drl,Ddr,nmo,wnd,hppy,Eph,Cdk4,Pdk1,Raf | | 1.361e-03 | -6.60 | transcription factor complex | cellular component | GO:0005667 | 177 | 15 | 11940 | 417 | arm,Dp,jigr1,Dad,Sox14,N,Hr4,sd,kay,sim,hyx,abd-A,mam,hth,Hr39 | | 1.365e-03 | -6.60 | LAM\_G\_DOMAIN | prosite domains | PS50025 | 17 | 5 | 6243 | 325 | shg,sli,kon,LanA,ft | | 1.382e-03 | -6.58 | wdb (widerborst) | protein interactions | 43312 | 47 | 8 | 9204 | 432 | H,orb2,Atg16,fl(2)d,CycG,mam,p130CAS,esg | | 1.391e-03 | -6.58 | anterior/posterior pattern specification, imaginal disc | biological process | GO:0007448 | 24 | 5 | 12345 | 442 | Dl,Abd-B,abd-A,tkv,Dad | | 1.396e-03 | -6.57 | Trl (Trithorax-like) | protein interactions | 2768981 | 58 | 9 | 9204 | 432 | ph-p,MEP-1,Adf1,fs(1)h,Abd-B,ttk,sd,corto,bab2 | | 1.397e-03 | -6.57 | GO\_REGULATION\_OF\_METANEPHROS\_DEVELOPMENT | MSigDB lists | GO\_REGULATION\_OF\_METANEPHROS\_DEVELOPMENT | 2 | 2 | 4161 | 156 | arm,ft | | 1.397e-03 | -6.57 | MODULE\_457 | MSigDB lists | MODULE\_457 | 2 | 2 | 4161 | 156 | SmD1,Pen | | 1.397e-03 | -6.57 | REACTOME\_RIP\_MEDIATED\_NFKB\_ACTIVATION\_VIA\_DAI | MSigDB lists | REACTOME\_RIP\_MEDIATED\_NFKB\_ACTIVATION\_VIA\_DAI | 2 | 2 | 4161 | 156 | Tollo,cact | | 1.397e-03 | -6.57 | GO\_POSITIVE\_REGULATION\_OF\_ENDOTHELIAL\_CELL\_DIFFERENTIATION | MSigDB lists | GO\_POSITIVE\_REGULATION\_OF\_ENDOTHELIAL\_CELL\_DIFFERENTIATION | 2 | 2 | 4161 | 156 | N,arm | | 1.397e-03 | -6.57 | GO\_REGULATION\_OF\_INTERFERON\_GAMMA\_BIOSYNTHETIC\_PROCESS | MSigDB lists | GO\_REGULATION\_OF\_INTERFERON\_GAMMA\_BIOSYNTHETIC\_PROCESS | 2 | 2 | 4161 | 156 | Tollo,Actbeta | | 1.397e-03 | -6.57 | GO\_DETECTION\_OF\_TEMPERATURE\_STIMULUS\_INVOLVED\_IN\_SENSORY\_PERCEPTION | MSigDB lists | GO\_DETECTION\_OF\_TEMPERATURE\_STIMULUS\_INVOLVED\_IN\_SENSORY\_PERCEPTION | 2 | 2 | 4161 | 156 | Eph,subdued | | 1.397e-03 | -6.57 | GO\_NEGATIVE\_REGULATION\_OF\_INTERLEUKIN\_10\_PRODUCTION | MSigDB lists | GO\_NEGATIVE\_REGULATION\_OF\_INTERLEUKIN\_10\_PRODUCTION | 2 | 2 | 4161 | 156 | Dl,trbl | | 1.397e-03 | -6.57 | GO\_LENS\_MORPHOGENESIS\_IN\_CAMERA\_TYPE\_EYE | MSigDB lists | GO\_LENS\_MORPHOGENESIS\_IN\_CAMERA\_TYPE\_EYE | 2 | 2 | 4161 | 156 | CG17883,arm | | 1.403e-03 | -6.57 | nucleotide binding | molecular function | GO:0000166 | 955 | 50 | 12176 | 414 | Pdk1,SMC3,Pitslre,SNF4Agamma,otk,CG2017,wnd,Drak,Pvr,Lk6,Raf,hppy,jar,trbl,Nrk,CG5853,Abl,Rab1,d,CG31121,CG11069,Cdk4,Pgk,CG17646,Rab5,GluProRS,Dcr-2,SERCA,tkv,Arf79F,Egfr,CtBP,RhoU,Myo10A,Rm62,l(1)G0196,nmo,CG7560,Dyrk2,Src42A,sick,Eph,Ddr,drl,Hel89B,betaTub60D,InR,htl,Ric,ATPCL | | 1.403e-03 | -6.57 | nucleoside phosphate binding | molecular function | GO:1901265 | 955 | 50 | 12176 | 414 | SMC3,Pitslre,Pdk1,SNF4Agamma,otk,CG2017,wnd,Drak,Pvr,jar,trbl,hppy,Lk6,Raf,CG5853,Nrk,Abl,Rab1,d,Cdk4,Pgk,CG17646,CG31121,CG11069,Egfr,CtBP,RhoU,GluProRS,Rab5,Dcr-2,tkv,SERCA,Arf79F,Dyrk2,Src42A,sick,Myo10A,Rm62,l(1)G0196,nmo,CG7560,InR,htl,Ric,ATPCL,Eph,Ddr,drl,Hel89B,betaTub60D | | 1.416e-03 | -6.56 | GO\_RECEPTOR\_ACTIVITY | MSigDB lists | GO\_RECEPTOR\_ACTIVITY | 186 | 16 | 4161 | 156 | emp,Ddr,Tollo,Eph,fz2,Nrk,fz,nAChRalpha4,tkv,robo1,otk,dlp,unc-5,N,htl,InR | | 1.418e-03 | -6.56 | GO\_ENDODERM\_DEVELOPMENT | MSigDB lists | GO\_ENDODERM\_DEVELOPMENT | 14 | 4 | 4161 | 156 | arm,Actbeta,N,hyx | | 1.418e-03 | -6.56 | GO\_REGULATION\_OF\_NEUROTRANSMITTER\_SECRETION | MSigDB lists | GO\_REGULATION\_OF\_NEUROTRANSMITTER\_SECRETION | 14 | 4 | 4161 | 156 | Syx1A,nAChRalpha4,Mef2,Raf | | 1.418e-03 | -6.56 | HALLMARK\_NOTCH\_SIGNALING | MSigDB lists | HALLMARK\_NOTCH\_SIGNALING | 14 | 4 | 4161 | 156 | fz2,Dl,N,fz | | 1.419e-03 | -6.56 | CycB (Cyclin B) | protein interactions | 37618 | 94 | 12 | 9204 | 432 | CG31211,chic,pum,CG7763,Sxl,H,crol,Psc,msi,CG32772,Cks30A,CycB3 | | 1.419e-03 | -6.56 | GSE4590\_PRE\_BCELL\_VS\_SMALL\_PRE\_BCELL\_UP | MSigDB lists | GSE4590\_PRE\_BCELL\_VS\_SMALL\_PRE\_BCELL\_UP | 34 | 6 | 4161 | 156 | Tollo,Unc-76,CG2017,Fkbp14,Tsp96F,CG6465 | | 1.420e-03 | -6.56 | GSE11961\_FOLLICULAR\_BCELL\_VS\_MARGINAL\_ZONE\_BCELL\_UP | MSigDB lists | GSE11961\_FOLLICULAR\_BCELL\_VS\_MARGINAL\_ZONE\_BCELL\_UP | 46 | 7 | 4161 | 156 | hppy,Rab1,Ddr,trbl,CG9220,dock,Mef2 | | 1.420e-03 | -6.56 | GSE39556\_CD8A\_DC\_VS\_NK\_CELL\_MOUSE\_3H\_POST\_POLYIC\_INJ\_DN | MSigDB lists | GSE39556\_CD8A\_DC\_VS\_NK\_CELL\_MOUSE\_3H\_POST\_POLYIC\_INJ\_DN | 46 | 7 | 4161 | 156 | Unc-76,Lk6,trbl,CG4645,step,Ziz,LanA | | 1.420e-03 | -6.56 | GO\_DEVELOPMENT\_OF\_PRIMARY\_SEXUAL\_CHARACTERISTICS | MSigDB lists | GO\_DEVELOPMENT\_OF\_PRIMARY\_SEXUAL\_CHARACTERISTICS | 46 | 7 | 4161 | 156 | Tollo,tkv,Actbeta,robo1,CG17883,Mfe2,sli | | 1.423e-03 | -6.56 | FERM\_2 | interpro domains | IPR035963 | 22 | 5 | 11383 | 450 | cora,Myo10A,CG42788,Pez,yrt | | 1.423e-03 | -6.56 | FERM\_central | interpro domains | IPR019748 | 22 | 5 | 11383 | 450 | CG42788,Myo10A,cora,yrt,Pez | | 1.429e-03 | -6.55 | Ig\_sub | interpro domains | IPR003599 | 111 | 12 | 11383 | 450 | ed,Ppn,boi,htl,Pvr,robo1,CG31431,unc-5,sns,kek5,otk,Bsg | | 1.433e-03 | -6.55 | regulation of cardiocyte differentiation | biological process | GO:1905207 | 7 | 3 | 12345 | 442 | Abd-B,lqf,N | | 1.433e-03 | -6.55 | muscle cell fate specification | biological process | GO:0042694 | 7 | 3 | 12345 | 442 | jeb,abd-A,kn | | 1.433e-03 | -6.55 | negative regulation of transforming growth factor beta receptor signaling pathway | biological process | GO:0030512 | 7 | 3 | 12345 | 442 | fuss,Snoo,Dad | | 1.433e-03 | -6.55 | negative regulation of R7 cell differentiation | biological process | GO:0045677 | 7 | 3 | 12345 | 442 | Socs36E,ttk,aop | | 1.433e-03 | -6.55 | gonad morphogenesis | biological process | GO:0035262 | 7 | 3 | 12345 | 442 | shg,ena,bab2 | | 1.433e-03 | -6.55 | negative regulation of cellular response to transforming growth factor beta stimulus | biological process | GO:1903845 | 7 | 3 | 12345 | 442 | Snoo,Dad,fuss | | 1.446e-03 | -6.54 | par-1 (par-1) | protein interactions | 2768852 | 37 | 7 | 9204 | 432 | wdb,ft,baz,N,sd,14-3-3epsilon,14-3-3zeta | | 1.446e-03 | -6.54 | chb (chromosome bows) | protein interactions | 43901 | 37 | 7 | 9204 | 432 | aop,shot,Abl,wnd,trbl,dve,Su(z)2 | | 1.450e-03 | -6.54 | GO\_REGULATION\_OF\_SECRETION | MSigDB lists | GO\_REGULATION\_OF\_SECRETION | 135 | 13 | 4161 | 156 | Syx1A,AP-1gamma,Raf,Mef2,jar,wbl,sns,subdued,nAChRalpha4,Pdk1,Arf79F,N,Actbeta | | 1.459e-03 | -6.53 | protein heterodimerization activity | molecular function | GO:0046982 | 221 | 17 | 12176 | 414 | His3.3A,His2Av,14-3-3zeta,pnr,14-3-3epsilon,drl,trbl,His3.3B,kay,ss,otk,emc,SMC3,E(spl)m5-HLH,sim,REPTOR,dome | | 1.460e-03 | -6.53 | Signaling by BMP | REACTOME pathways | R-DME-201451 | 12 | 4 | 3923 | 176 | eff,tkv,Dad,Snoo | | 1.460e-03 | -6.53 | Costimulation by the CD28 family | REACTOME pathways | R-DME-388841 | 12 | 4 | 3923 | 176 | rictor,Pdk1,wdb,trbl | | 1.477e-03 | -6.52 | Kap-alpha3 (karyopherin alpha3) | protein interactions | 41158 | 82 | 11 | 9204 | 432 | Mpcp2,N,Kr-h2,GluProRS,Dl,rictor,fl(2)d,sd,Lmx1a,Myo10A,Pen | | 1.486e-03 | -6.51 | Malpighian tubule morphogenesis | biological process | GO:0007443 | 48 | 7 | 12345 | 442 | S,spi,rib,N,Pvr,Dad,Egfr | | 1.486e-03 | -6.51 | renal tubule morphogenesis | biological process | GO:0061333 | 48 | 7 | 12345 | 442 | Egfr,Pvr,Dad,S,spi,N,rib | | 1.493e-03 | -6.51 | GO\_CELL\_MORPHOGENESIS\_INVOLVED\_IN\_NEURON\_DIFFERENTIATION | MSigDB lists | GO\_CELL\_MORPHOGENESIS\_INVOLVED\_IN\_NEURON\_DIFFERENTIATION | 88 | 10 | 4161 | 156 | N,sina,ap,tkv,Unc-76,unc-5,Mef2,robo1,Eph,sli | | 1.497e-03 | -6.50 | gl (glass) | protein interactions | 42210 | 19 | 5 | 9204 | 432 | h,aop,N,sina,tkv | | 1.497e-03 | -6.50 | Jon99Cii (Jonah 99Cii) | protein interactions | 43544 | 19 | 5 | 9204 | 432 | al,Tpc1,Pez,Sxl,CG12581 | | 1.500e-03 | -6.50 | SH3\_domain | interpro domains | IPR001452 | 69 | 9 | 11383 | 450 | SKIP,dock,cindr,shot,Cip4,RhoGEF3,Src42A,Abl,p130CAS | | 1.515e-03 | -6.49 | Signaling by SCF-KIT | REACTOME pathways | R-DME-1433557 | 142 | 15 | 3923 | 176 | Egfr,vn,Cbl,Raf,Pvr,CG4935,rictor,htl,PI31,Pdk1,14-3-3zeta,wdb,trbl,CG42684,Dronc | | 1.537e-03 | -6.48 | fly chr70D5-70D5\|3-42 cM | chromosome location | fly chr70D5-70D5\|3-42 cM | 3 | 2 | 25010 | 571 | lncRNA:CR45025,CG13481 | | 1.537e-03 | -6.48 | fly chr70D5-70D5\|3-42 | chromosome location | fly chr70D5-70D5\|3-42 | 3 | 2 | 25010 | 571 | lncRNA:CR45025,CG13481 | | 1.537e-03 | -6.48 | fly chr42C3-42C3\|2-55 | chromosome location | fly chr42C3-42C3\|2-55 | 3 | 2 | 25010 | 571 | Adf1,lncRNA:CR45128 | | 1.537e-03 | -6.48 | fly chr42C3-42C3\|2-55 cM | chromosome location | fly chr42C3-42C3\|2-55 cM | 3 | 2 | 25010 | 571 | Adf1,lncRNA:CR45128 | | 1.539e-03 | -6.48 | GGGCGGR\_SP1\_Q6 | MSigDB lists | GGGCGGR\_SP1\_Q6 | 882 | 49 | 4161 | 156 | Fkbp14,Raf,Mef2,CG1129,CG6465,cact,Pex5,CG4896,CG31522,Arf79F,l(2)37Cg,14-3-3zeta,trbl,InR,dlp,alpha-Man-Ia,14-3-3epsilon,nmo,frc,LanA,CG13893,Rgl,AP-1gamma,Cadps,CG4645,CG45050,CG2017,FBXO11,lili,Pat1,Dyb,Cals,ATPCL,Gfat2,hppy,N,ap,Psa,Unc-76,GlcAT-I,Rab1,CenB1A,lqf,Syx1A,His2Av,wdb,Pex14,fz2,Kdm2 | | 1.546e-03 | -6.47 | Wnt signaling | KEGG pathways | M00677 | 14 | 4 | 3198 | 123 | fz3,fz,arm,fz2 | | 1.546e-03 | -6.47 | Wnt signaling | KEGG pathways | dme\_M00677 | 14 | 4 | 3198 | 123 | fz2,fz3,arm,fz | | 1.552e-03 | -6.47 | positive regulation of molecular function | biological process | GO:0044093 | 287 | 21 | 12345 | 442 | Raf,SERCA,jeb,Dronc,hppy,RhoGAP19D,CG17883,drongo,CG7394,Rgl,Atg17,wnd,RapGAP1,ttk,Cks30A,CenB1A,TBC1D5,Pdk1,Ocrl,step,rictor | | 1.559e-03 | -6.46 | RTP801-like\_C\_sf | interpro domains | IPR038281 | 2 | 2 | 11383 | 450 | scyl,chrb | | 1.559e-03 | -6.46 | SEM1A\_sema\_dom | interpro domains | IPR042068 | 2 | 2 | 11383 | 450 | Sema1a,Sema1b | | 1.559e-03 | -6.46 | SCAR/WAVE\_fam | interpro domains | IPR028288 | 2 | 2 | 11383 | 450 | SCAR,gukh | | 1.559e-03 | -6.46 | 14-3-3\_CS | interpro domains | IPR023409 | 2 | 2 | 11383 | 450 | 14-3-3zeta,14-3-3epsilon | | 1.559e-03 | -6.46 | 14-3-3 | interpro domains | IPR000308 | 2 | 2 | 11383 | 450 | 14-3-3epsilon,14-3-3zeta | | 1.559e-03 | -6.46 | Tscrpt\_reg\_SKI\_SnoN | interpro domains | IPR023216 | 2 | 2 | 11383 | 450 | fuss,Snoo | | 1.559e-03 | -6.46 | 14-3-3\_domain | interpro domains | IPR023410 | 2 | 2 | 11383 | 450 | 14-3-3epsilon,14-3-3zeta | | 1.559e-03 | -6.46 | c-SKI\_SMAD4-bd\_dom | interpro domains | IPR014890 | 2 | 2 | 11383 | 450 | Snoo,fuss | | 1.559e-03 | -6.46 | 14-3-3\_dom\_sf | interpro domains | IPR036815 | 2 | 2 | 11383 | 450 | 14-3-3zeta,14-3-3epsilon | | 1.559e-03 | -6.46 | RTP801-like | interpro domains | IPR012918 | 2 | 2 | 11383 | 450 | scyl,chrb | | 1.571e-03 | -6.46 | muscle cell development | biological process | GO:0055001 | 62 | 8 | 12345 | 442 | kon,sd,Mef2,aop,C1GalTA,Dg,Bsg,htl | | 1.577e-03 | -6.45 | - | gene3d domains | 4.10.280.10 | 57 | 9 | 4929 | 241 | E(spl)m5-HLH,tai,HLH3B,kn,h,ss,sim,crp,emc | | 1.587e-03 | -6.45 | PENG\_GLUCOSE\_DEPRIVATION\_DN | MSigDB lists | PENG\_GLUCOSE\_DEPRIVATION\_DN | 60 | 8 | 4161 | 156 | CG18549,betaTub60D,Pgk,His2Av,CycG,Rab1,shot,CG9485 | | 1.587e-03 | -6.45 | GATA1\_01 | MSigDB lists | GATA1\_01 | 60 | 8 | 4161 | 156 | ATPCL,Eph,otk,rdx,eff,Ric,14-3-3epsilon,Unc-76 | | 1.587e-03 | -6.45 | GSE17812\_WT\_VS\_THPOK\_KO\_MEMORY\_CD8\_TCELL\_UP | MSigDB lists | GSE17812\_WT\_VS\_THPOK\_KO\_MEMORY\_CD8\_TCELL\_UP | 60 | 8 | 4161 | 156 | Pgk,dock,alpha-Man-Ia,nmo,CG6465,Ocrl,Pgant6,step | | 1.592e-03 | -6.44 | Csk (C-terminal Src kinase) | protein interactions | 41398 | 48 | 8 | 9204 | 432 | Cdk4,Dcr-2,SCAR,Src42A,shg,Gli,ena,CycG | | 1.592e-03 | -6.44 | phyl (phyllopod) | protein interactions | 36606 | 48 | 8 | 9204 | 432 | Raf,chn,sina,N,lilli,esg,ttk,14-3-3zeta | | 1.592e-03 | -6.44 | CG34168 (uncharacterized protein) | protein interactions | 50457 | 48 | 8 | 9204 | 432 | CG34401,sina,eyg,nub,Mob2,Poxn,esg,CG34376 | | 1.592e-03 | -6.44 | Rap1 (Rap1 GTPase) | protein interactions | 38244 | 48 | 8 | 9204 | 432 | Raf,Rgl,Rab1,RhoL,SERCA,RapGAP1,Rab5,CG42788 | | 1.595e-03 | -6.44 | 14-3-3 | pfam domains | PF00244 | 2 | 2 | 10480 | 419 | 14-3-3epsilon,14-3-3zeta | | 1.595e-03 | -6.44 | c-SKI\_SMAD\_bind | pfam domains | PF08782 | 2 | 2 | 10480 | 419 | fuss,Snoo | | 1.595e-03 | -6.44 | RTP801\_C | pfam domains | PF07809 | 2 | 2 | 10480 | 419 | scyl,chrb | | 1.606e-03 | -6.43 | Activation of Rac | REACTOME pathways | R-DME-428540 | 6 | 3 | 3923 | 176 | sli,robo1,dock | | 1.611e-03 | -6.43 | ato (atonal) | protein interactions | 40975 | 28 | 6 | 9204 | 432 | Dl,Raf,Egfr,lilli,h,ed | | 1.616e-03 | -6.43 | visceral muscle development | biological process | GO:0007522 | 15 | 4 | 12345 | 442 | jeb,htl,klar,sns | | 1.616e-03 | -6.43 | cytoplasmic transport, nurse cell to oocyte | biological process | GO:0007303 | 15 | 4 | 12345 | 442 | Ppcs,jar,ena,Abl | | 1.616e-03 | -6.43 | dorsal/ventral axis specification, ovarian follicular epithelium | biological process | GO:0008069 | 15 | 4 | 12345 | 442 | Raf,Dp,Cbl,Egfr | | 1.619e-03 | -6.43 | DORN\_ADENOVIRUS\_INFECTION\_48HR\_DN | MSigDB lists | DORN\_ADENOVIRUS\_INFECTION\_48HR\_DN | 7 | 3 | 4161 | 156 | cact,Rab1,step | | 1.619e-03 | -6.43 | GO\_CELL\_DIFFERENTIATION\_INVOLVED\_IN\_KIDNEY\_DEVELOPMENT | MSigDB lists | GO\_CELL\_DIFFERENTIATION\_INVOLVED\_IN\_KIDNEY\_DEVELOPMENT | 7 | 3 | 4161 | 156 | N,Mef2,sns | | 1.619e-03 | -6.43 | GO\_ENTEROENDOCRINE\_CELL\_DIFFERENTIATION | MSigDB lists | GO\_ENTEROENDOCRINE\_CELL\_DIFFERENTIATION | 7 | 3 | 4161 | 156 | Cdk4,Pdk1,Dl | | 1.619e-03 | -6.43 | GO\_BETA\_CATENIN\_DESTRUCTION\_COMPLEX\_DISASSEMBLY | MSigDB lists | GO\_BETA\_CATENIN\_DESTRUCTION\_COMPLEX\_DISASSEMBLY | 7 | 3 | 4161 | 156 | arm,fz2,fz | | 1.619e-03 | -6.43 | GO\_NEGATIVE\_REGULATION\_OF\_CHEMOTAXIS | MSigDB lists | GO\_NEGATIVE\_REGULATION\_OF\_CHEMOTAXIS | 7 | 3 | 4161 | 156 | sli,robo1,N | | 1.619e-03 | -6.43 | BROWNE\_HCMV\_INFECTION\_8HR\_DN | MSigDB lists | BROWNE\_HCMV\_INFECTION\_8HR\_DN | 7 | 3 | 4161 | 156 | shot,Fem-1,trbl | | 1.619e-03 | -6.43 | MODULE\_200 | MSigDB lists | MODULE\_200 | 7 | 3 | 4161 | 156 | Ddr,Nrk,Eph | | 1.619e-03 | -6.43 | GO\_ENDOCRINE\_PANCREAS\_DEVELOPMENT | MSigDB lists | GO\_ENDOCRINE\_PANCREAS\_DEVELOPMENT | 7 | 3 | 4161 | 156 | Dl,Cdk4,Pdk1 | | 1.619e-03 | -6.43 | BROWNE\_INTERFERON\_RESPONSIVE\_GENES | MSigDB lists | BROWNE\_INTERFERON\_RESPONSIVE\_GENES | 7 | 3 | 4161 | 156 | Tollo,Rgl,step | | 1.623e-03 | -6.42 | tight junction | cellular component | GO:0070160 | 37 | 6 | 11940 | 417 | Gli,Shrm,moody,scrib,cora,yrt | | 1.628e-03 | -6.42 | GO\_VASCULATURE\_DEVELOPMENT | MSigDB lists | GO\_VASCULATURE\_DEVELOPMENT | 89 | 10 | 4161 | 156 | arm,fz2,N,C1GalTA,sli,LanA,Mef2,Eph,Dl,otk | | 1.634e-03 | -6.42 | mRNA 3'-UTR binding | molecular function | GO:0003730 | 38 | 6 | 12176 | 414 | orb2,Dcr-2,Tis11,bol,Sxl,pum | | 1.651e-03 | -6.41 | early endosome | cellular component | GO:0005769 | 64 | 8 | 11940 | 417 | Dl,ed,tkv,Ocrl,Rab5,spdo,fz2,N | | 1.656e-03 | -6.40 | GO\_CELL\_SUBSTRATE\_ADHESION | MSigDB lists | GO\_CELL\_SUBSTRATE\_ADHESION | 24 | 5 | 4161 | 156 | Pdk1,Eph,arm,Rab1,LanA | | 1.656e-03 | -6.40 | HALLMARK\_ALLOGRAFT\_REJECTION | MSigDB lists | HALLMARK\_ALLOGRAFT\_REJECTION | 24 | 5 | 4161 | 156 | Tollo,Pgant5,CG1673,dock,Actbeta | | 1.656e-03 | -6.40 | SETLUR\_PROSTATE\_CANCER\_TMPRSS2\_ERG\_FUSION\_UP | MSigDB lists | SETLUR\_PROSTATE\_CANCER\_TMPRSS2\_ERG\_FUSION\_UP | 24 | 5 | 4161 | 156 | Cadps,jar,tkv,Pex10,otk | | 1.656e-03 | -6.40 | WNT\_SIGNALING | MSigDB lists | WNT\_SIGNALING | 24 | 5 | 4161 | 156 | fz,CtBP,nmo,fz2,arm | | 1.660e-03 | -6.40 | GSE19825\_NAIVE\_VS\_IL2RAHIGH\_DAY3\_EFF\_CD8\_TCELL\_UP | MSigDB lists | GSE19825\_NAIVE\_VS\_IL2RAHIGH\_DAY3\_EFF\_CD8\_TCELL\_UP | 35 | 6 | 4161 | 156 | LanA,unc-5,Ids,Eph,betaTub60D,Cals | | 1.681e-03 | -6.39 | myotube differentiation | biological process | GO:0014902 | 49 | 7 | 12345 | 442 | Pgk,siz,InR,Mef2,sns,dock,SCAR | | 1.684e-03 | -6.39 | Frizzled | smart domains | SM01330 | 5 | 3 | 5086 | 290 | fz3,fz,fz2 | | 1.688e-03 | -6.38 | female sex differentiation | biological process | GO:0046660 | 25 | 5 | 12345 | 442 | Abd-B,bab2,otk,InR,fz2 | | 1.688e-03 | -6.38 | cell fate commitment involved in formation of primary germ layer | biological process | GO:0060795 | 25 | 5 | 12345 | 442 | mam,spi,htl,tkv,abd-A | | 1.690e-03 | -6.38 | GGGAGGRR\_MAZ\_Q6 | MSigDB lists | GGGAGGRR\_MAZ\_Q6 | 506 | 32 | 4161 | 156 | Psa,Tmhs,N,Rab5,hppy,Kdm2,fz2,wdb,lqf,Ddr,shot,nmo,14-3-3epsilon,dlp,dock,trbl,InR,14-3-3zeta,l(2)37Cg,otk,cact,Cdk4,CycG,betaTub60D,FBXO11,CG2017,SMC3,LanA,Rgl,Cadps,ds,unc-119 | | 1.710e-03 | -6.37 | GO\_POSITIVE\_REGULATION\_OF\_KINASE\_ACTIVITY | MSigDB lists | GO\_POSITIVE\_REGULATION\_OF\_KINASE\_ACTIVITY | 105 | 11 | 4161 | 156 | Tsp96F,Pdk1,nAChRalpha4,htl,fz2,wbl,Raf,unc-119,Tollo,Ddr,hppy | | 1.732e-03 | -6.36 | translation repressor activity | molecular function | GO:0030371 | 16 | 4 | 12176 | 414 | orb2,msi,Sxl,pum | | 1.742e-03 | -6.35 | SH3\_1 | pfam domains | PF00018 | 33 | 6 | 10480 | 419 | Cip4,Abl,RhoGEF3,Src42A,p130CAS,dock | | 1.745e-03 | -6.35 | regulation of organ growth | biological process | GO:0046620 | 63 | 8 | 12345 | 442 | ft,dlp,InR,d,tkv,Egfr,Pdk1,scrib | | 1.753e-03 | -6.35 | nau (nautilus) | protein interactions | 42799 | 12 | 4 | 9204 | 432 | kn,abd-A,Abd-B,Scr | | 1.753e-03 | -6.35 | rg (rugose) | protein interactions | 44531 | 12 | 4 | 9204 | 432 | Egfr,H,Dl,S | | 1.753e-03 | -6.35 | ro (rough) | protein interactions | 43234 | 12 | 4 | 9204 | 432 | sens,H,lilli,Dl | | 1.753e-03 | -6.35 | Sulf1 (Sulfated) | protein interactions | 53437 | 12 | 4 | 9204 | 432 | vn,Egfr,Ddr,dlp | | 1.760e-03 | -6.34 | Band\_41\_domain | interpro domains | IPR019749 | 23 | 5 | 11383 | 450 | cora,CG42788,Myo10A,yrt,Pez | | 1.770e-03 | -6.34 | GO\_REGIONALIZATION | MSigDB lists | GO\_REGIONALIZATION | 61 | 8 | 4161 | 156 | fz2,arm,N,tkv,CG9220,ap,Mef2,Dl | | 1.784e-03 | -6.33 | grk (gurken) | protein interactions | 34171 | 60 | 9 | 9204 | 432 | Gfat2,Pvr,CG34376,Cbl,vn,cact,rho-4,S,sina | | 1.789e-03 | -6.33 | regulation of synaptic growth at neuromuscular junction | biological process | GO:0008582 | 109 | 11 | 12345 | 442 | Mob2,Src42A,Cip4,Abl,rictor,ena,pum,LanA,tkv,nmo,Dad | | 1.817e-03 | -6.31 | positive regulation of catalytic activity | biological process | GO:0043085 | 252 | 19 | 12345 | 442 | TBC1D5,Pdk1,step,rictor,Ocrl,Cks30A,RapGAP1,CenB1A,CG7394,CG17883,RhoGAP19D,drongo,wnd,Rgl,Atg17,Raf,Dronc,hppy,jeb | | 1.818e-03 | -6.31 | intercellular bridge | cellular component | GO:0045171 | 26 | 5 | 11940 | 417 | dock,cindr,14-3-3zeta,jar,14-3-3epsilon | | 1.820e-03 | -6.31 | positive regulation of transferase activity | biological process | GO:0051347 | 78 | 9 | 12345 | 442 | rictor,step,wnd,Atg17,Pdk1,hppy,jeb,Cks30A,Raf | | 1.820e-03 | -6.31 | gliogenesis | biological process | GO:0042063 | 78 | 9 | 12345 | 442 | htl,sli,spen,N,moody,cora,Dl,ey,unc-5 | | 1.831e-03 | -6.30 | Mucin type O-glycan biosynthesis | KEGG pathways | dme00512 | 24 | 5 | 3198 | 123 | CG34057,Pgant6,C1GalTA,CG3119,Pgant5 | | 1.831e-03 | -6.30 | Mucin type O-glycan biosynthesis | KEGG pathways | ko00512 | 24 | 5 | 3198 | 123 | CG3119,Pgant5,CG34057,C1GalTA,Pgant6 | | 1.835e-03 | -6.30 | GO\_REGULATION\_OF\_SYNAPSE\_STRUCTURE\_OR\_ACTIVITY | MSigDB lists | GO\_REGULATION\_OF\_SYNAPSE\_STRUCTURE\_OR\_ACTIVITY | 48 | 7 | 4161 | 156 | Cals,Eph,Nrk,robo1,Raf,Arf79F,Mef2 | | 1.835e-03 | -6.30 | GAUSSMANN\_MLL\_AF4\_FUSION\_TARGETS\_C\_UP | MSigDB lists | GAUSSMANN\_MLL\_AF4\_FUSION\_TARGETS\_C\_UP | 48 | 7 | 4161 | 156 | dock,14-3-3zeta,Fkbp14,CG6891,CG4896,GlcAT-I,ft | | 1.838e-03 | -6.30 | GO\_REGULATION\_OF\_INTRACELLULAR\_SIGNAL\_TRANSDUCTION | MSigDB lists | GO\_REGULATION\_OF\_INTRACELLULAR\_SIGNAL\_TRANSDUCTION | 343 | 24 | 4161 | 156 | dock,trbl,InR,htl,N,Actbeta,nmo,cact,Raf,CG5522,step,hppy,CG45050,sina,Tsp96F,Pdk1,wbl,arm,Nmda1,fz2,Eph,Tollo,sli,Ocrl | | 1.843e-03 | -6.30 | positive regulation of canonical Wnt signaling pathway | biological process | GO:0090263 | 37 | 6 | 12345 | 442 | CtBP,Diap1,spen,apolpp,hyx,dlp | | 1.843e-03 | -6.30 | neuron projection extension | biological process | GO:1990138 | 37 | 6 | 12345 | 442 | tai,Rab5,fz,path,fz2,SCAR | | 1.847e-03 | -6.29 | gkt (glaikit) | protein interactions | 33530 | 6 | 3 | 9204 | 432 | Abd-B,drl,orb2 | | 1.847e-03 | -6.29 | Nlg4 (Neuroligin 4) | protein interactions | 42402 | 6 | 3 | 9204 | 432 | Dad,dpy,tkv | | 1.876e-03 | -6.28 | integral component of plasma membrane | cellular component | GO:0005887 | 466 | 29 | 11940 | 417 | Esp,robo1,beat-IIa,otk,Egfr,ds,Gli,tipE,tkv,InR,wgn,drl,fz,Ddr,scrib,Nrk,rho-4,Cals,emp,ed,kek5,ft,N,fz2,Eph,htl,Pvr,nAChRalpha4,shg | | 1.878e-03 | -6.28 | GGGATGC\_MIR3245P | MSigDB lists | GGGATGC\_MIR3245P | 15 | 4 | 4161 | 156 | FBXO11,AP-1gamma,hppy,unc-5 | | 1.878e-03 | -6.28 | GO\_CELL\_CELL\_ADHESION\_VIA\_PLASMA\_MEMBRANE\_ADHESION\_MOLECULES | MSigDB lists | GO\_CELL\_CELL\_ADHESION\_VIA\_PLASMA\_MEMBRANE\_ADHESION\_MOLECULES | 15 | 4 | 4161 | 156 | ft,ds,robo1,Cals | | 1.878e-03 | -6.28 | GTACAGG\_MIR486 | MSigDB lists | GTACAGG\_MIR486 | 15 | 4 | 4161 | 156 | Hel89B,bru2,SmD1,unc-5 | | 1.891e-03 | -6.27 | bHLH\_dom | interpro domains | IPR011598 | 58 | 8 | 11383 | 450 | h,emc,ss,E(spl)m5-HLH,sim,crp,HLH3B,tai | | 1.909e-03 | -6.26 | actin binding | molecular function | GO:0003779 | 133 | 12 | 12176 | 414 | d,Shrm,mim,cora,SCAR,shot,ena,Myo10A,f,CG6891,chic,jar | | 1.920e-03 | -6.26 | AP-2alpha (Adaptor Protein complex 2, alpha subunit) | protein interactions | 33211 | 20 | 5 | 9204 | 432 | Dl,step,lqf,N,ft | | 1.924e-03 | -6.25 | semaphorin receptor binding | molecular function | GO:0030215 | 8 | 3 | 12176 | 414 | Sema1a,Sema1b,otk | | 1.927e-03 | -6.25 | anatomical structure homeostasis | biological process | GO:0060249 | 161 | 14 | 12345 | 442 | Atg16,robo1,N,Atg17,Pgk,wgn,Egfr,Dg,Fkbp14,InR,shg,yrt,Nrk,tai | | 1.932e-03 | -6.25 | YAGI\_AML\_WITH\_T\_9\_11\_TRANSLOCATION | MSigDB lists | YAGI\_AML\_WITH\_T\_9\_11\_TRANSLOCATION | 36 | 6 | 4161 | 156 | frc,jar,IntS3,Mef2,Nrk,Rim2 | | 1.933e-03 | -6.25 | establishment of cell polarity | biological process | GO:0030010 | 64 | 8 | 12345 | 442 | ft,yrt,ds,fz,Dg,rictor,baz,scrib | | 1.933e-03 | -6.25 | synapse assembly | biological process | GO:0007416 | 64 | 8 | 12345 | 442 | baz,Gli,drl,Pdk1,Adf1,jeb,kay,Sema1a | | 1.952e-03 | -6.24 | Traf4 (TNF-receptor-associated factor 4) | protein interactions | 33638 | 29 | 6 | 9204 | 432 | shg,wgn,arm,baz,scrib,Diap1 | | 1.955e-03 | -6.24 | macromolecule modification | biological process | GO:0043412 | 1320 | 67 | 12345 | 442 | Ddr,tkv,morgue,CG42327,Atg16,Abl,wdb,ft,Cks30A,CycG,Ptp4E,Ubc87F,Fem-1,CG13481,mop,hyx,Cbl,ds,CycB3,ec,Kdm4B,sina,Pez,rdx,Su(z)2,CG10417,Ptpmeg2,Drak,drl,FBXO11,hppy,InR,mura,nmo,Tpst,htl,Pdk1,Pex10,IA-2,Diap1,eff,wnd,Eph,trbl,otk,CG34057,Kdm2,fl(2)d,Tet,Nrk,E(Pc),eya,Dyrk2,Raf,C1GalTA,bin3,Gfat2,Src42A,Pvr,tna,CG12402,Pp2C1,Lk6,Cdk4,Pitslre,CG1218,CG3119 | | 1.962e-03 | -6.23 | GO\_TRANSFERASE\_ACTIVITY\_TRANSFERRING\_PHOSPHORUS\_CONTAINING\_GROUPS | MSigDB lists | GO\_TRANSFERASE\_ACTIVITY\_TRANSFERRING\_PHOSPHORUS\_CONTAINING\_GROUPS | 305 | 22 | 4161 | 156 | l(1)G0196,Drak,nmo,l(2)37Cg,trbl,htl,InR,hppy,CG1129,Cdk4,Raf,otk,Dyrk2,Cerk,Pdk1,Nrk,Pgk,Lk6,tkv,Src42A,Ddr,Eph | | 1.969e-03 | -6.23 | AP2\_Q6\_01 | MSigDB lists | AP2\_Q6\_01 | 62 | 8 | 4161 | 156 | ap,Kdm2,CycG,hyx,Fkbp14,otk,Dyb,Eph | | 1.969e-03 | -6.23 | LIM\_MAMMARY\_STEM\_CELL\_UP | MSigDB lists | LIM\_MAMMARY\_STEM\_CELL\_UP | 62 | 8 | 4161 | 156 | ds,sli,kn,Mef2,Dl,Eph,shot,Tpst | | 1.973e-03 | -6.23 | GO\_REGULATION\_OF\_MAPK\_CASCADE | MSigDB lists | GO\_REGULATION\_OF\_MAPK\_CASCADE | 123 | 12 | 4161 | 156 | hppy,Raf,Tollo,Eph,arm,fz2,wbl,Actbeta,Tsp96F,trbl,htl,InR | | 1.981e-03 | -6.22 | head development | biological process | GO:0060322 | 144 | 13 | 12345 | 442 | drl,sim,shot,Eph,chic,vn,hth,h,tsh,shg,kn,rg,ey | | 1.992e-03 | -6.22 | nito (spenito) | protein interactions | 35756 | 39 | 7 | 9204 | 432 | fl(2)d,larp,arm,spen,CG31121,Sxl,H | | 1.992e-03 | -6.22 | Kr (Kruppel) | protein interactions | 38012 | 39 | 7 | 9204 | 432 | crol,abd-A,Abd-B,S,CtBP,Psa,emc | | 2.008e-03 | -6.21 | KCCGNSWTTT\_UNKNOWN | MSigDB lists | KCCGNSWTTT\_UNKNOWN | 25 | 5 | 4161 | 156 | 14-3-3epsilon,Kdm2,AP-1gamma,ATPCL,SMC3 | | 2.008e-03 | -6.21 | GO\_LYMPHOCYTE\_DIFFERENTIATION | MSigDB lists | GO\_LYMPHOCYTE\_DIFFERENTIATION | 25 | 5 | 4161 | 156 | arm,fz2,Cdk4,Raf,Dl | | 2.008e-03 | -6.21 | msn (misshapen) | protein interactions | 44030 | 61 | 9 | 9204 | 432 | Pdk1,tkv,dock,H,Egfr,rictor,ft,nmo,N | | 2.008e-03 | -6.21 | kay (kayak) | protein interactions | 3772082 | 61 | 9 | 9204 | 432 | sd,Fkbp14,wnd,shot,jigr1,H,Egfr,scrib,fz | | 2.029e-03 | -6.20 | regulation of extent of cell growth | biological process | GO:0061387 | 26 | 5 | 12345 | 442 | chic,Sema1b,shot,fz,Sema1a | | 2.029e-03 | -6.20 | regulation of axon extension | biological process | GO:0030516 | 26 | 5 | 12345 | 442 | Sema1a,fz,shot,chic,Sema1b | | 2.041e-03 | -6.19 | LamG | smart domains | SM00282 | 17 | 5 | 5086 | 290 | LanA,ft,kon,shg,sli | | 2.075e-03 | -6.18 | KEGG\_WNT\_SIGNALING\_PATHWAY | MSigDB lists | KEGG\_WNT\_SIGNALING\_PATHWAY | 49 | 7 | 4161 | 156 | CtBP,wdb,fz,sina,fz2,arm,nmo | | 2.075e-03 | -6.18 | GSE360\_L\_MAJOR\_VS\_B\_MALAYI\_LOW\_DOSE\_DC\_UP | MSigDB lists | GSE360\_L\_MAJOR\_VS\_B\_MALAYI\_LOW\_DOSE\_DC\_UP | 49 | 7 | 4161 | 156 | betaTub60D,Pgk,Actbeta,Syx1A,bru2,step,arm | | 2.075e-03 | -6.18 | GSE37301\_PRO\_BCELL\_VS\_RAG2\_KO\_NK\_CELL\_DN | MSigDB lists | GSE37301\_PRO\_BCELL\_VS\_RAG2\_KO\_NK\_CELL\_DN | 49 | 7 | 4161 | 156 | CG5522,shot,CG1316,Cerk,CG9220,Cdk4,Tsp96F | | 2.094e-03 | -6.17 | ovarian follicle cell stalk formation | biological process | GO:0030713 | 16 | 4 | 12345 | 442 | N,Ppcs,ph-p,Dl | | 2.094e-03 | -6.17 | canonical Wnt signaling pathway | biological process | GO:0060070 | 16 | 4 | 12345 | 442 | arm,fz,fz2,fz3 | | 2.116e-03 | -6.16 | 1433ZETA | prints domains | PR00305 | 2 | 2 | 2185 | 101 | 14-3-3epsilon,14-3-3zeta | | 2.138e-03 | -6.15 | immune response | biological process | GO:0006955 | 199 | 16 | 12345 | 442 | eya,kn,LRR,Dronc,RhoL,imd,Pvr,Rm62,Eph,Tollo,RhoGEF3,kay,dome,Mef2,cact,LanA | | 2.158e-03 | -6.14 | Ig-like\_dom | interpro domains | IPR007110 | 132 | 13 | 11383 | 450 | sns,otk,kek5,Bsg,unc-5,CG31431,vn,htl,Pvr,robo1,Ppn,ed,boi | | 2.184e-03 | -6.13 | GTGCAAT\_MIR25\_MIR32\_MIR92\_MIR363\_MIR367 | MSigDB lists | GTGCAAT\_MIR25\_MIR32\_MIR92\_MIR363\_MIR367 | 63 | 8 | 4161 | 156 | Ziz,Gfat2,robo1,nmo,lili,Ppcs,lin-28,sd | | 2.198e-03 | -6.12 | COX4L (Cytochrome c oxidase subunit 4-like) | protein interactions | 35491 | 2 | 2 | 9204 | 432 | ZnT41F,CG30089 | | 2.198e-03 | -6.12 | HDAC4 (Histone deacetylase 4) | protein interactions | 32278 | 2 | 2 | 9204 | 432 | Dl,14-3-3zeta | | 2.198e-03 | -6.12 | ND-B14.5B (NADH dehydrogenase (ubiquinone) B14.5 B subunit) | protein interactions | 33528 | 2 | 2 | 9204 | 432 | 14-3-3zeta,Myo10A | | 2.198e-03 | -6.12 | IRSp53 (Insulin receptor substrate 53 kDa) | protein interactions | 39258 | 2 | 2 | 9204 | 432 | ena,Dl | | 2.198e-03 | -6.12 | CG13377 (uncharacterized protein) | protein interactions | 30972 | 2 | 2 | 9204 | 432 | Jupiter,CG6891 | | 2.232e-03 | -6.11 | morphogenesis of larval imaginal disc epithelium | biological process | GO:0016335 | 8 | 3 | 12345 | 442 | scrib,ft,Dad | | 2.232e-03 | -6.11 | protein import into peroxisome matrix | biological process | GO:0016558 | 8 | 3 | 12345 | 442 | Pex5,Pex10,Pex14 | | 2.232e-03 | -6.11 | development of primary male sexual characteristics | biological process | GO:0046546 | 8 | 3 | 12345 | 442 | eya,Abd-B,PI31 | | 2.232e-03 | -6.11 | male gonad development | biological process | GO:0008584 | 8 | 3 | 12345 | 442 | PI31,Abd-B,eya | | 2.232e-03 | -6.11 | regulation of salivary gland boundary specification | biological process | GO:0045704 | 8 | 3 | 12345 | 442 | Abd-B,tkv,tsh | | 2.232e-03 | -6.11 | filopodium assembly | biological process | GO:0046847 | 8 | 3 | 12345 | 442 | mim,ena,Myo10A | | 2.232e-03 | -6.11 | negative regulation of salivary gland boundary specification | biological process | GO:0045705 | 8 | 3 | 12345 | 442 | tsh,tkv,Abd-B | | 2.235e-03 | -6.10 | GSE26928\_EFF\_MEMORY\_VS\_CXCR5\_POS\_CD4\_TCELL\_DN | MSigDB lists | GSE26928\_EFF\_MEMORY\_VS\_CXCR5\_POS\_CD4\_TCELL\_DN | 37 | 6 | 4161 | 156 | InR,Dl,Pgant5,Ppcs,Pex7,Kdm2 | | 2.264e-03 | -6.09 | Signaling by EGFR | REACTOME pathways | R-DME-177929 | 178 | 17 | 3923 | 176 | lqf,cindr,Pvr,Raf,trbl,PI31,rictor,14-3-3zeta,Pdk1,vn,Cbl,Egfr,CG42684,Dronc,wdb,htl,CG4935 | | 2.264e-03 | -6.09 | JOHNSTONE\_PARVB\_TARGETS\_2\_DN | MSigDB lists | JOHNSTONE\_PARVB\_TARGETS\_2\_DN | 125 | 12 | 4161 | 156 | Kr-h2,Vps13,SMC3,wdb,CtBP,trbl,Tpst,shot,Drak,Cdk4,C1GalTA,Ziz | | 2.273e-03 | -6.09 | transcription coactivator activity | molecular function | GO:0003713 | 69 | 8 | 12176 | 414 | CtBP,arm,CG13204,hng3,mam,tna,eya,tai | | 2.275e-03 | -6.09 | GO\_REGULATION\_OF\_IMMUNE\_EFFECTOR\_PROCESS | MSigDB lists | GO\_REGULATION\_OF\_IMMUNE\_EFFECTOR\_PROCESS | 93 | 10 | 4161 | 156 | Spn42De,Tpc1,CG4645,AP-1gamma,ReepA,Tollo,SmD1,fz2,Pdk1,Arf79F | | 2.283e-03 | -6.08 | NUYTTEN\_EZH2\_TARGETS\_UP | MSigDB lists | NUYTTEN\_EZH2\_TARGETS\_UP | 213 | 17 | 4161 | 156 | Pgant5,dock,Ppn,Actbeta,14-3-3zeta,Rim2,cact,Gfat2,step,CG9220,fz2,Tollo,PI31,Rgl,emp,sli,Ids | | 2.309e-03 | -6.07 | Ig-like\_dom\_sf | interpro domains | IPR036179 | 133 | 13 | 11383 | 450 | boi,ed,Ppn,robo1,Pvr,htl,vn,unc-5,CG31431,otk,Bsg,kek5,sns | | 2.332e-03 | -6.06 | Axon guidance | REACTOME pathways | R-DME-422475 | 194 | 18 | 3923 | 176 | sina,ena,Pvr,Raf,sli,Abl,PI31,14-3-3zeta,dlp,vn,Egfr,dock,CG42684,Eph,wdb,htl,CG4935,robo1 | | 2.338e-03 | -6.06 | AP4\_Q6\_01 | MSigDB lists | AP4\_Q6\_01 | 50 | 7 | 4161 | 156 | rdx,Kdm2,Cadps,N,14-3-3zeta,betaTub60D,InR | | 2.338e-03 | -6.06 | CAGCTTT\_MIR320 | MSigDB lists | CAGCTTT\_MIR320 | 50 | 7 | 4161 | 156 | FBXO11,InR,Cdk4,14-3-3zeta,CG1316,14-3-3epsilon,arm | | 2.338e-03 | -6.06 | CTTGTAT\_MIR381 | MSigDB lists | CTTGTAT\_MIR381 | 50 | 7 | 4161 | 156 | CG3436,nmo,Pat1,CG5522,rg,sina,cact | | 2.338e-03 | -6.06 | KRAS.600\_UP.V1\_UP | MSigDB lists | KRAS.600\_UP.V1\_UP | 50 | 7 | 4161 | 156 | Syx1A,ReepA,Actbeta,Cals,subdued,trbl,nAChRalpha4 | | 2.345e-03 | -6.06 | trio (trio) | protein interactions | 43974 | 30 | 6 | 9204 | 432 | dock,Abl,N,ena,SCAR,Dl | | 2.359e-03 | -6.05 | - | gene3d domains | 3.90.190.10 | 29 | 6 | 4929 | 241 | CG42327,Pez,Ptpmeg2,mop,Ptp4E,IA-2 | | 2.361e-03 | -6.05 | regulation of cell cycle | biological process | GO:0051726 | 317 | 22 | 12345 | 442 | scrib,CG6191,cindr,N,Dl,trbl,pum,nuf,CycG,CycB3,Wdr62,emc,esg,crol,Egfr,htl,Ocrl,Cks30A,Dp,SNF4Agamma,kay,14-3-3epsilon | | 2.381e-03 | -6.04 | - | gene3d domains | 1.20.190.20 | 2 | 2 | 4929 | 241 | 14-3-3epsilon,14-3-3zeta | | 2.381e-03 | -6.04 | - | gene3d domains | 1.10.490.100 | 2 | 2 | 4929 | 241 | scyl,chrb | | 2.386e-03 | -6.04 | regulation of neuromuscular junction development | biological process | GO:1904396 | 113 | 11 | 12345 | 442 | pum,ena,rictor,nmo,Dad,tkv,LanA,Mob2,Cip4,Src42A,Abl | | 2.388e-03 | -6.04 | associative learning | biological process | GO:0008306 | 52 | 7 | 12345 | 442 | rg,Adf1,vn,Egfr,drl,spi,14-3-3zeta | | 2.395e-03 | -6.03 | rau (rau) | protein interactions | 33814 | 87 | 11 | 9204 | 432 | CG3386,pita,Pp2C1,rgr,CG32795,cora,hang,Elba2,CG42788,Pez,Ric | | 2.397e-03 | -6.03 | CTTTGT\_LEF1\_Q2 | MSigDB lists | CTTTGT\_LEF1\_Q2 | 453 | 29 | 4161 | 156 | FBXO11,CG2812,betaTub60D,tkv,CG2017,fz,arm,rdx,lili,crol,ds,Psa,trbl,htl,sd,InR,N,14-3-3zeta,shot,nmo,14-3-3epsilon,dlp,Dl,otk,Mef2,Raf,Cdk4,CG6465,Gfat2 | | 2.404e-03 | -6.03 | GO\_CELL\_MOTILITY | MSigDB lists | GO\_CELL\_MOTILITY | 160 | 14 | 4161 | 156 | N,Pdk1,dock,arm,14-3-3epsilon,Rab1,shot,Mef2,Eph,otk,sli,ds,Pex7,LanA | | 2.412e-03 | -6.03 | GO\_PROTEOGLYCAN\_METABOLIC\_PROCESS | MSigDB lists | GO\_PROTEOGLYCAN\_METABOLIC\_PROCESS | 26 | 5 | 4161 | 156 | CG9220,tkv,Ids,GlcAT-I,frc | | 2.417e-03 | -6.03 | cell cortex part | cellular component | GO:0044448 | 116 | 11 | 11940 | 417 | CG6891,Bsg,spdo,baz,CG42684,Shrm,step,cindr,nuf,Abl,Dl | | 2.418e-03 | -6.02 | heterophilic cell-cell adhesion via plasma membrane cell adhesion molecules | biological process | GO:0007157 | 27 | 5 | 12345 | 442 | beat-IIa,glec,sns,ds,ft | | 2.418e-03 | -6.02 | embryonic hemopoiesis | biological process | GO:0035162 | 27 | 5 | 12345 | 442 | shg,Pvf3,N,RhoL,Pvr | | 2.419e-03 | -6.02 | HATADA\_METHYLATED\_IN\_LUNG\_CANCER\_UP | MSigDB lists | HATADA\_METHYLATED\_IN\_LUNG\_CANCER\_UP | 64 | 8 | 4161 | 156 | sli,Cadps,hppy,Gfat2,kn,CG1673,Dyb,Tsp96F | | 2.419e-03 | -6.02 | GSE43700\_UNTREATED\_VS\_IL10\_TREATED\_PBMC\_DN | MSigDB lists | GSE43700\_UNTREATED\_VS\_IL10\_TREATED\_PBMC\_DN | 64 | 8 | 4161 | 156 | Drep4,CG2017,sd,arm,Ric,eff,CG1129,Rgl | | 2.425e-03 | -6.02 | Su(dx) (Suppressor of deltex) | protein interactions | 33379 | 21 | 5 | 9204 | 432 | H,Pez,mop,Dl,N | | 2.425e-03 | -6.02 | RpS15Ab (Ribosomal protein S15Ab) | protein interactions | 36142 | 21 | 5 | 9204 | 432 | Bsg,His3.3B,Pex14,Arf79F,Pex5 | | 2.425e-03 | -6.02 | gt (giant) | protein interactions | 31227 | 21 | 5 | 9204 | 432 | ttk,tsh,CtBP,REPTOR,kay | | 2.425e-03 | -6.02 | Deactivation of the beta-catenin transactivating complex | REACTOME pathways | R-DME-3769402 | 22 | 5 | 3923 | 176 | Diap1,14-3-3zeta,Sox14,CtBP,arm | | 2.431e-03 | -6.02 | GO\_NUCLEOBASE\_CONTAINING\_COMPOUND\_TRANSMEMBRANE\_TRANSPORTER\_ACTIVITY | MSigDB lists | GO\_NUCLEOBASE\_CONTAINING\_COMPOUND\_TRANSMEMBRANE\_TRANSPORTER\_ACTIVITY | 16 | 4 | 4161 | 156 | Tpc1,frc,DPCoAC,Rim2 | | 2.431e-03 | -6.02 | BOYAULT\_LIVER\_CANCER\_SUBCLASS\_G123\_UP | MSigDB lists | BOYAULT\_LIVER\_CANCER\_SUBCLASS\_G123\_UP | 16 | 4 | 4161 | 156 | ATPCL,SmD1,hppy,Pen | | 2.431e-03 | -6.02 | GO\_BRANCHING\_INVOLVED\_IN\_URETERIC\_BUD\_MORPHOGENESIS | MSigDB lists | GO\_BRANCHING\_INVOLVED\_IN\_URETERIC\_BUD\_MORPHOGENESIS | 16 | 4 | 4161 | 156 | LanA,ft,ds,arm | | 2.431e-03 | -6.02 | GO\_MESONEPHRIC\_TUBULE\_MORPHOGENESIS | MSigDB lists | GO\_MESONEPHRIC\_TUBULE\_MORPHOGENESIS | 16 | 4 | 4161 | 156 | LanA,arm,ds,ft | | 2.431e-03 | -6.02 | WEIGEL\_OXIDATIVE\_STRESS\_BY\_HNE\_AND\_TBH | MSigDB lists | WEIGEL\_OXIDATIVE\_STRESS\_BY\_HNE\_AND\_TBH | 16 | 4 | 4161 | 156 | Raf,Drep4,InR,shot | | 2.431e-03 | -6.02 | GO\_POSITIVE\_REGULATION\_OF\_OSTEOBLAST\_DIFFERENTIATION | MSigDB lists | GO\_POSITIVE\_REGULATION\_OF\_OSTEOBLAST\_DIFFERENTIATION | 16 | 4 | 4161 | 156 | arm,Ddr,Mef2,tkv | | 2.431e-03 | -6.02 | GO\_NEGATIVE\_REGULATION\_OF\_OSSIFICATION | MSigDB lists | GO\_NEGATIVE\_REGULATION\_OF\_OSSIFICATION | 16 | 4 | 4161 | 156 | Mef2,Cdk4,N,CG9220 | | 2.435e-03 | -6.02 | positive regulation of protein serine/threonine kinase activity | biological process | GO:0071902 | 39 | 6 | 12345 | 442 | Cks30A,Raf,rictor,step,wnd,hppy | | 2.439e-03 | -6.02 | CLIP-190 (Cytoplasmic linker protein 190) | protein interactions | 35042 | 13 | 4 | 9204 | 432 | h,jar,Sema1a,Alh | | 2.439e-03 | -6.02 | Doc2 (Dorsocross2) | protein interactions | 39038 | 13 | 4 | 9204 | 432 | pnr,abd-A,Abd-B,Scr | | 2.439e-03 | -6.02 | ab (abrupt) | protein interactions | 34560 | 13 | 4 | 9204 | 432 | spi,kn,tai,drongo | | 2.439e-03 | -6.02 | tgo (tango) | protein interactions | 41084 | 13 | 4 | 9204 | 432 | mRpL41,ss,Myo10A,sim | | 2.439e-03 | -6.02 | dan (distal antenna) | protein interactions | 43023 | 13 | 4 | 9204 | 432 | Cdk4,ss,CtBP,ey | | 2.520e-03 | -5.98 | GO\_REPLACEMENT\_OSSIFICATION | MSigDB lists | GO\_REPLACEMENT\_OSSIFICATION | 8 | 3 | 4161 | 156 | Pex7,Mef2,htl | | 2.520e-03 | -5.98 | GO\_SMOOTH\_MUSCLE\_CELL\_DIFFERENTIATION | MSigDB lists | GO\_SMOOTH\_MUSCLE\_CELL\_DIFFERENTIATION | 8 | 3 | 4161 | 156 | Mef2,N,arm | | 2.520e-03 | -5.98 | GO\_ACETYLGALACTOSAMINYLTRANSFERASE\_ACTIVITY | MSigDB lists | GO\_ACETYLGALACTOSAMINYLTRANSFERASE\_ACTIVITY | 8 | 3 | 4161 | 156 | CG9220,Pgant5,Pgant6 | | 2.520e-03 | -5.98 | PID\_PI3KCI\_AKT\_PATHWAY | MSigDB lists | PID\_PI3KCI\_AKT\_PATHWAY | 8 | 3 | 4161 | 156 | 14-3-3epsilon,Pdk1,14-3-3zeta | | 2.520e-03 | -5.98 | DORN\_ADENOVIRUS\_INFECTION\_32HR\_DN | MSigDB lists | DORN\_ADENOVIRUS\_INFECTION\_32HR\_DN | 8 | 3 | 4161 | 156 | cact,step,Rab1 | | 2.520e-03 | -5.98 | COLIN\_PILOCYTIC\_ASTROCYTOMA\_VS\_GLIOBLASTOMA\_DN | MSigDB lists | COLIN\_PILOCYTIC\_ASTROCYTOMA\_VS\_GLIOBLASTOMA\_DN | 8 | 3 | 4161 | 156 | arm,Pgk,CG45050 | | 2.520e-03 | -5.98 | GO\_ENDODERMAL\_CELL\_DIFFERENTIATION | MSigDB lists | GO\_ENDODERMAL\_CELL\_DIFFERENTIATION | 8 | 3 | 4161 | 156 | hyx,Actbeta,arm | | 2.520e-03 | -5.98 | GO\_CHONDROITIN\_SULFATE\_PROTEOGLYCAN\_BIOSYNTHETIC\_PROCESS | MSigDB lists | GO\_CHONDROITIN\_SULFATE\_PROTEOGLYCAN\_BIOSYNTHETIC\_PROCESS | 8 | 3 | 4161 | 156 | frc,GlcAT-I,CG9220 | | 2.523e-03 | -5.98 | AGO2 (Argonaute 2) | protein interactions | 39683 | 63 | 9 | 9204 | 432 | mir-1000,Dcr-2,pum,rictor,SmD1,vig,Rm62,Pdk1,Diap1 | | 2.544e-03 | -5.97 | distal axon | cellular component | GO:0150034 | 100 | 10 | 11940 | 417 | jeb,orb2,pum,jar,Magi,gukh,shot,Syx1A,robo1,ena | | 2.553e-03 | -5.97 | GO\_REGULATION\_OF\_PROTEIN\_MODIFICATION\_PROCESS | MSigDB lists | GO\_REGULATION\_OF\_PROTEIN\_MODIFICATION\_PROCESS | 413 | 27 | 4161 | 156 | CycG,wbl,arm,fz2,nAChRalpha4,Tsp96F,Nrk,Pdk1,wdb,fz,PI31,Fem-1,sli,Ddr,Tollo,Eph,unc-119,Rab1,14-3-3epsilon,trbl,dock,InR,htl,Actbeta,hppy,Raf,Mef2 | | 2.565e-03 | -5.97 | PcG protein complex | cellular component | GO:0031519 | 28 | 5 | 11940 | 417 | Kdm2,Su(z)2,ph-p,Psc,corto | | 2.573e-03 | -5.96 | GO\_POSITIVE\_REGULATION\_OF\_APOPTOTIC\_SIGNALING\_PATHWAY | MSigDB lists | GO\_POSITIVE\_REGULATION\_OF\_APOPTOTIC\_SIGNALING\_PATHWAY | 38 | 6 | 4161 | 156 | 14-3-3zeta,sina,Actbeta,tkv,dock,14-3-3epsilon | | 2.573e-03 | -5.96 | GO\_SKIN\_DEVELOPMENT | MSigDB lists | GO\_SKIN\_DEVELOPMENT | 38 | 6 | 4161 | 156 | arm,LanA,Actbeta,N,Dl,ap | | 2.573e-03 | -5.96 | GSE40273\_GATA1\_KO\_VS\_WT\_TREG\_DN | MSigDB lists | GSE40273\_GATA1\_KO\_VS\_WT\_TREG\_DN | 38 | 6 | 4161 | 156 | lt,CG9220,Mef2,hppy,step,CenB1A | | 2.579e-03 | -5.96 | chorion-containing eggshell formation | biological process | GO:0007304 | 131 | 12 | 12345 | 442 | ed,emc,S,ttk,Raf,Dp,Egfr,cact,tkv,N,Ppcs,cic | | 2.579e-03 | -5.96 | compound eye photoreceptor development | biological process | GO:0042051 | 82 | 9 | 12345 | 442 | Mob2,eya,Raf,lilli,ttk,klar,Nrk,jeb,sens | | 2.614e-03 | -5.95 | meiotic nuclear division | biological process | GO:0140013 | 203 | 16 | 12345 | 442 | Rab5,His3.3B,LanA,His3.3A,Cks30A,eff,larp,Pex10,Sxl,Gli,PI31,orb2,Arf79F,CycG,Rab1,Hr39 | | 2.622e-03 | -5.94 | EGF-like\_Ca-bd\_dom | interpro domains | IPR001881 | 36 | 6 | 11383 | 450 | sli,dpy,ft,Dl,N,cue | | 2.641e-03 | -5.94 | cytoplasm | cellular component | GO:0005737 | 4500 | 185 | 11940 | 417 | Rab5,CG9220,koi,CG1673,hth,ena,nuf,Mob2,Pgant6,spdo,ATPCL,Tpc1,GluProRS,Nlp,lqf,Ptpmeg2,alpha-Man-Ia,Drep4,Atox1,corto,CG33474,kay,SNF4Agamma,FBXO11,tsh,CG6465,CG7536,wbl,Dg,Pez,14-3-3epsilon,CG6983,pdgy,CG10417,d,Slbp,wdb,sina,AP-1gamma,rho-4,Dcr-2,cact,Shrm,TBC1D5,nmo,RhoGAP19D,kon,rg,CG14562,Tpst,dock,jeb,fuss,RhoGEF4,Kr-h2,312,fz2,apolpp,CG3149,aop,Ocrl,SCAR,eya,siz,Rab1,yuri,Unc-76,Fkbp14,fz,S,SmD1,Dyrk2,cic,ps,muc,p130CAS,CG7394,CG7560,Atg17,chrb,f,ss,CalpC,lin-28,mop,emp,scrib,Pex7,Sema1b,Mpcp2,Vps13,Dronc,arm,emc,CycG,Pen,CG2765,trbl,path,Psa,step,klar,shot,REPTOR,chic,cora,RapGAP1,mRpL41,hppy,Cadps,CG4467,Bsg,jar,vig,Mfe2,CG8306,Rim2,l(1)G0196,RhoU,cindr,14-3-3zeta,dve,eyg,Pgant5,frc,CG42684,N,CG17683,CG30156,Arf79F,Lk6,Diap1,GlcAT-I,Abl,Hr4,CG42788,Rm62,Pdk1,Snoo,CycB3,ReepA,Pex10,Src42A,Sxl,tkv,Atg16,larp,Jupiter,lt,drongo,flfl,pum,Magi,Pex5,UQCR-11,CG34172,scyl,Raf,CG9784,bol,CG17883,SERCA,CG4645,CG6891,rdx,Dl,Tis11,Pgk,Syx1A,CG3107,orb2,baz,Tom,DPCoAC,Pp2C1,shg,bru2,betaTub60D,ed,Pex14,IntS3,Cbl,sim,Tob,spi | | 2.662e-03 | -5.93 | sevenless signaling pathway | biological process | GO:0045500 | 17 | 4 | 12345 | 442 | aop,ttk,Raf,sina | | 2.662e-03 | -5.93 | terminal region determination | biological process | GO:0007362 | 17 | 4 | 12345 | 442 | bowl,fs(1)h,Raf,cic | | 2.662e-03 | -5.93 | regulation of ATP metabolic process | biological process | GO:1903578 | 17 | 4 | 12345 | 442 | CG34172,Dg,Pgk,N | | 2.669e-03 | -5.93 | skeletal muscle organ development | biological process | GO:0060538 | 53 | 7 | 12345 | 442 | shot,kn,Dg,drl,Mef2,kon,N | | 2.669e-03 | -5.93 | regulation of tube size | biological process | GO:0035150 | 53 | 7 | 12345 | 442 | ttk,cora,Gli,fz,rib,ds,ft | | 2.672e-03 | -5.92 | GO\_TAXIS | MSigDB lists | GO\_TAXIS | 65 | 8 | 4161 | 156 | robo1,Eph,sli,sina,Unc-76,tkv,ap,unc-5 | | 2.696e-03 | -5.92 | fly chr93F14-93F14\|3-73 cM | chromosome location | fly chr93F14-93F14\|3-73 cM | 26 | 4 | 25010 | 571 | CG6656,CG31431,Gr93c,glec | | 2.696e-03 | -5.92 | fly chr93F14-93F14\|3-73 | chromosome location | fly chr93F14-93F14\|3-73 | 26 | 4 | 25010 | 571 | glec,Gr93c,CG6656,CG31431 | | 2.702e-03 | -5.91 | 1433\_2 | prosite domains | PS00797 | 2 | 2 | 6243 | 325 | 14-3-3zeta,14-3-3epsilon | | 2.702e-03 | -5.91 | 1433\_1 | prosite domains | PS00796 | 2 | 2 | 6243 | 325 | 14-3-3zeta,14-3-3epsilon | | 2.719e-03 | -5.91 | RHO GTPases activate PKNs | REACTOME pathways | R-DME-5625740 | 7 | 3 | 3923 | 176 | 14-3-3epsilon,14-3-3zeta,Pdk1 | | 2.719e-03 | -5.91 | CD28 dependent PI3K/Akt signaling | REACTOME pathways | R-DME-389357 | 7 | 3 | 3923 | 176 | trbl,Pdk1,rictor | | 2.719e-03 | -5.91 | Rap1 signalling | REACTOME pathways | R-DME-392517 | 7 | 3 | 3923 | 176 | 14-3-3zeta,Raf,RapGAP1 | | 2.727e-03 | -5.90 | protein kinase regulator activity | molecular function | GO:0019887 | 71 | 8 | 12176 | 414 | 14-3-3zeta,CycG,CycB3,jeb,Atg17,trbl,Cks30A,Pdk1 | | 2.732e-03 | -5.90 | GO\_REGULATION\_OF\_KINASE\_ACTIVITY | MSigDB lists | GO\_REGULATION\_OF\_KINASE\_ACTIVITY | 180 | 15 | 4161 | 156 | trbl,nAChRalpha4,dock,htl,InR,Tsp96F,Pdk1,wbl,CycG,fz2,Tollo,Raf,unc-119,hppy,Ddr | | 2.759e-03 | -5.89 | heparin binding | molecular function | GO:0008201 | 18 | 4 | 12176 | 414 | Sema1a,sli,robo1,vn | | 2.780e-03 | -5.89 | heterochromatin organization | biological process | GO:0070828 | 40 | 6 | 12345 | 442 | Dcr-2,fl(2)d,crol,nub,bin3,vig | | 2.794e-03 | -5.88 | Hrb98DE (Heterogeneous nuclear ribonucleoprotein at 98DE) | protein interactions | 43385 | 31 | 6 | 9204 | 432 | lncRNA:Hsromega,CG31431,shg,pum,Arf79F,InR | | 2.798e-03 | -5.88 | VEGFA-VEGFR2 Pathway | REACTOME pathways | R-DME-4420097 | 151 | 15 | 3923 | 176 | Pvr,vn,Raf,Egfr,trbl,CG42684,dock,wdb,Pdk1,htl,14-3-3zeta,rictor,PI31,CG4935,arm | | 2.798e-03 | -5.88 | Signaling by VEGF | REACTOME pathways | R-DME-194138 | 151 | 15 | 3923 | 176 | Egfr,Raf,vn,Pvr,arm,CG4935,Pdk1,14-3-3zeta,rictor,htl,PI31,wdb,dock,CG42684,trbl | | 2.811e-03 | -5.87 | GO\_POSITIVE\_REGULATION\_OF\_MULTICELLULAR\_ORGANISMAL\_PROCESS | MSigDB lists | GO\_POSITIVE\_REGULATION\_OF\_MULTICELLULAR\_ORGANISMAL\_PROCESS | 236 | 18 | 4161 | 156 | l(2)37Cg,N,Arf79F,Actbeta,Cals,otk,Dl,robo1,Raf,Mef2,tkv,SERCA,fz2,arm,Eph,Tollo,Ddr,sli | | 2.813e-03 | -5.87 | epidermal growth factor receptor binding | molecular function | GO:0005154 | 9 | 3 | 12176 | 414 | ed,vn,spi | | 2.813e-03 | -5.87 | coreceptor activity | molecular function | GO:0015026 | 9 | 3 | 12176 | 414 | otk,boi,cv-d | | 2.851e-03 | -5.86 | positive regulation of cell projection organization | biological process | GO:0031346 | 68 | 8 | 12345 | 442 | shot,ena,chic,kon,SCAR,Abl,Rab1,fz | | 2.854e-03 | -5.86 | fly chr102F8-102F8\|4-0 cM | chromosome location | fly chr102F8-102F8\|4-0 cM | 13 | 3 | 25010 | 571 | Actbeta,Cadps,Cals | | 2.854e-03 | -5.86 | fly chr102F8-102F8\|4-0 | chromosome location | fly chr102F8-102F8\|4-0 | 13 | 3 | 25010 | 571 | Cadps,Actbeta,Cals | | 2.854e-03 | -5.86 | fly chr22D1-22D1\|2-5 | chromosome location | fly chr22D1-22D1\|2-5 | 13 | 3 | 25010 | 571 | CG34172,lncRNA:CR43751,aop | | 2.854e-03 | -5.86 | fly chr22D1-22D1\|2-5 cM | chromosome location | fly chr22D1-22D1\|2-5 cM | 13 | 3 | 25010 | 571 | lncRNA:CR43751,CG34172,aop | | 2.859e-03 | -5.86 | positive regulation of transmembrane receptor protein serine/threonine kinase signaling pathway | biological process | GO:0090100 | 28 | 5 | 12345 | 442 | lili,tkv,cv-d,hth,Actbeta | | 2.859e-03 | -5.86 | imaginal disc-derived genitalia development | biological process | GO:0007484 | 28 | 5 | 12345 | 442 | Abd-B,Pvr,otk,Rab5,N | | 2.872e-03 | -5.85 | LXR\_Q3 | MSigDB lists | LXR\_Q3 | 27 | 5 | 4161 | 156 | Kdm2,rdx,cact,CG6465,hyx | | 2.872e-03 | -5.85 | NRL\_DN.V1\_UP | MSigDB lists | NRL\_DN.V1\_UP | 27 | 5 | 4161 | 156 | Lk6,Mef2,CG11030,Drak,Pgant6 | | 2.918e-03 | -5.84 | GO\_POSITIVE\_REGULATION\_OF\_CELL\_COMMUNICATION | MSigDB lists | GO\_POSITIVE\_REGULATION\_OF\_CELL\_COMMUNICATION | 355 | 24 | 4161 | 156 | dock,htl,Actbeta,N,CtBP,14-3-3zeta,14-3-3epsilon,Dl,Cals,Raf,hppy,subdued,CG9220,tkv,sina,Kr-h2,Tsp96F,wbl,arm,fz2,Tollo,hyx,Syx1A,PI31 | | 2.941e-03 | -5.83 | GSE7509\_DC\_VS\_MONOCYTE\_DN | MSigDB lists | GSE7509\_DC\_VS\_MONOCYTE\_DN | 52 | 7 | 4161 | 156 | Tpst,CG4896,CG3436,Actbeta,dock,Pgant5,cact | | 2.949e-03 | -5.83 | KRAS.600.LUNG.BREAST\_UP.V1\_UP | MSigDB lists | KRAS.600.LUNG.BREAST\_UP.V1\_UP | 39 | 6 | 4161 | 156 | Syx1A,sli,subdued,Cals,sns,Actbeta | | 2.949e-03 | -5.83 | GSE30153\_LUPUS\_VS\_HEALTHY\_DONOR\_BCELL\_DN | MSigDB lists | GSE30153\_LUPUS\_VS\_HEALTHY\_DONOR\_BCELL\_DN | 39 | 6 | 4161 | 156 | Pdk1,Mef2,Nrk,Kdm2,Gfat2,Ric | | 2.949e-03 | -5.83 | GSE24210\_CTRL\_VS\_IL35\_TREATED\_TCONV\_CD4\_TCELL\_UP | MSigDB lists | GSE24210\_CTRL\_VS\_IL35\_TREATED\_TCONV\_CD4\_TCELL\_UP | 39 | 6 | 4161 | 156 | Raf,trbl,lt,step,bru2,Tpst | | 2.949e-03 | -5.83 | GO\_LEUKOCYTE\_DIFFERENTIATION | MSigDB lists | GO\_LEUKOCYTE\_DIFFERENTIATION | 39 | 6 | 4161 | 156 | Cdk4,Mef2,Raf,Dl,arm,fz2 | | 2.977e-03 | -5.82 | GO\_PHOSPHORYLATION | MSigDB lists | GO\_PHOSPHORYLATION | 376 | 25 | 4161 | 156 | htl,InR,sns,trbl,Drak,nmo,l(1)G0196,otk,Raf,Mef2,Cdk4,hppy,CG4935,tkv,Lk6,Nrk,Pdk1,Pgk,Cerk,Dyrk2,Tollo,Eph,PI31,Ddr,Src42A | | 2.998e-03 | -5.81 | GO\_APICAL\_PART\_OF\_CELL | MSigDB lists | GO\_APICAL\_PART\_OF\_CELL | 81 | 9 | 4161 | 156 | Tsp96F,N,subdued,arm,jar,CG11069,Dl,ft,ds | | 3.002e-03 | -5.81 | GO\_POSITIVE\_REGULATION\_OF\_PHOSPHORUS\_METABOLIC\_PROCESS | MSigDB lists | GO\_POSITIVE\_REGULATION\_OF\_PHOSPHORUS\_METABOLIC\_PROCESS | 200 | 16 | 4161 | 156 | wbl,fz2,arm,nAChRalpha4,fz,Tsp96F,Pdk1,Nrk,Ddr,Tollo,unc-119,htl,Actbeta,hppy,Mef2,Raf | | 3.003e-03 | -5.81 | Cadherin-like\_dom | interpro domains | IPR002126 | 16 | 4 | 11383 | 450 | ds,ft,Cals,shg | | 3.019e-03 | -5.80 | PlexA (Plexin A) | protein interactions | 43832 | 22 | 5 | 9204 | 432 | Sema1a,14-3-3epsilon,rho-4,Sema1b,otk | | 3.019e-03 | -5.80 | alpha-Cat (alpha Catenin) | protein interactions | 40517 | 22 | 5 | 9204 | 432 | SCAR,N,shg,arm,baz | | 3.020e-03 | -5.80 | adult behavior | biological process | GO:0030534 | 169 | 14 | 12345 | 442 | Egfr,pum,pxb,sim,moody,Bacc,hppy,ey,SERCA,S,spi,hang,Poxn,mura | | 3.026e-03 | -5.80 | GO\_CIRCULATORY\_SYSTEM\_DEVELOPMENT | MSigDB lists | GO\_CIRCULATORY\_SYSTEM\_DEVELOPMENT | 164 | 14 | 4161 | 156 | Mef2,otk,Eph,Dl,ft,C1GalTA,sli,ds,LanA,fz,N,nAChRalpha4,fz2,arm | | 3.026e-03 | -5.80 | developmental maturation | biological process | GO:0021700 | 206 | 16 | 12345 | 442 | Dg,chic,Pat1,scrib,14-3-3zeta,N,mop,Dronc,Egfr,wbl,baz,cic,14-3-3epsilon,Sox14,shg,eff | | 3.028e-03 | -5.80 | fly chr82D6-82D6\|3-47.1 cM | chromosome location | fly chr82D6-82D6\|3-47.1 cM | 4 | 2 | 25010 | 571 | CG1129,CG14657 | | 3.028e-03 | -5.80 | fly chr94E1-94E1\|3-78 cM | chromosome location | fly chr94E1-94E1\|3-78 cM | 4 | 2 | 25010 | 571 | CG31457,CenB1A | | 3.028e-03 | -5.80 | fly chr56C4-56C4\|2-88 | chromosome location | fly chr56C4-56C4\|2-88 | 4 | 2 | 25010 | 571 | cora,wbl | | 3.028e-03 | -5.80 | fly chr61C9-61C9\|3-0.5 | chromosome location | fly chr61C9-61C9\|3-0.5 | 4 | 2 | 25010 | 571 | hng3,emc | | 3.028e-03 | -5.80 | fly chr61C9-61C9\|3-0.5 cM | chromosome location | fly chr61C9-61C9\|3-0.5 cM | 4 | 2 | 25010 | 571 | emc,hng3 | | 3.028e-03 | -5.80 | fly chr94E1-94E1\|3-78 | chromosome location | fly chr94E1-94E1\|3-78 | 4 | 2 | 25010 | 571 | CenB1A,CG31457 | | 3.028e-03 | -5.80 | fly chr56C4-56C4\|2-88 cM | chromosome location | fly chr56C4-56C4\|2-88 cM | 4 | 2 | 25010 | 571 | cora,wbl | | 3.028e-03 | -5.80 | fly chr82D6-82D6\|3-47.1 | chromosome location | fly chr82D6-82D6\|3-47.1 | 4 | 2 | 25010 | 571 | CG1129,CG14657 | | 3.036e-03 | -5.80 | cytoneme | cellular component | GO:0035230 | 9 | 3 | 11940 | 417 | tkv,boi,Egfr | | 3.047e-03 | -5.79 | presynapse | cellular component | GO:0098793 | 155 | 13 | 11940 | 417 | CG31211,jeb,orb2,pum,Bsg,Magi,Cadps,ReepA,LanA,gukh,tkv,ena,Syx1A | | 3.055e-03 | -5.79 | dx (deltex) | protein interactions | 31589 | 77 | 10 | 9204 | 432 | N,Src42A,koi,CG4896,H,klu,Dl,CycG,mam,esg | | 3.070e-03 | -5.79 | GO\_REGULATION\_OF\_CELLULAR\_LOCALIZATION | MSigDB lists | GO\_REGULATION\_OF\_CELLULAR\_LOCALIZATION | 316 | 22 | 4161 | 156 | fz,Pdk1,subdued,nAChRalpha4,fz2,arm,wbl,Tollo,CG17883,ReepA,Syx1A,AP-1gamma,14-3-3zeta,N,Arf79F,Actbeta,Psa,GlcAT-I,14-3-3epsilon,Mef2,Raf,cact | | 3.080e-03 | -5.78 | phosphorylation | biological process | GO:0016310 | 470 | 29 | 12345 | 442 | Abl,Src42A,Cks30A,nmo,tkv,htl,Ddr,Pdk1,hppy,Nrk,Raf,InR,Dyrk2,CycB3,Pitslre,UQCR-11,CycG,Pvr,Pgk,Socs36E,Eph,wnd,trbl,Drak,otk,Ptpmeg2,Lk6,drl,Cdk4 | | 3.087e-03 | -5.78 | GO\_COLUMNAR\_CUBOIDAL\_EPITHELIAL\_CELL\_DEVELOPMENT | MSigDB lists | GO\_COLUMNAR\_CUBOIDAL\_EPITHELIAL\_CELL\_DEVELOPMENT | 17 | 4 | 4161 | 156 | C1GalTA,Dl,Cdk4,Pdk1 | | 3.087e-03 | -5.78 | GO\_REGULATION\_OF\_CALCIUM\_ION\_DEPENDENT\_EXOCYTOSIS | MSigDB lists | GO\_REGULATION\_OF\_CALCIUM\_ION\_DEPENDENT\_EXOCYTOSIS | 17 | 4 | 4161 | 156 | Arf79F,Raf,N,Syx1A | | 3.087e-03 | -5.78 | KYNG\_DNA\_DAMAGE\_BY\_GAMMA\_RADIATION | MSigDB lists | KYNG\_DNA\_DAMAGE\_BY\_GAMMA\_RADIATION | 17 | 4 | 4161 | 156 | Arf79F,Ric,Pat1,AP-1gamma | | 3.087e-03 | -5.78 | GTATGAT\_MIR154\_MIR487 | MSigDB lists | GTATGAT\_MIR154\_MIR487 | 17 | 4 | 4161 | 156 | trbl,sd,nmo,CG3436 | | 3.087e-03 | -5.78 | CAHOY\_NEURONAL | MSigDB lists | CAHOY\_NEURONAL | 17 | 4 | 4161 | 156 | Mfe2,Syx1A,Mef2,Cals | | 3.087e-03 | -5.78 | GO\_REGULATION\_OF\_NEUROTRANSMITTER\_TRANSPORT | MSigDB lists | GO\_REGULATION\_OF\_NEUROTRANSMITTER\_TRANSPORT | 17 | 4 | 4161 | 156 | Mef2,Raf,nAChRalpha4,Syx1A | | 3.096e-03 | -5.78 | sc (scute) | protein interactions | 30982 | 42 | 7 | 9204 | 432 | chn,Sxl,esg,emc,h,sens,ed | | 3.108e-03 | -5.77 | Pten (Phosphatase and tensin homolog) | protein interactions | 43991 | 103 | 12 | 9204 | 432 | Rm62,larp,baz,hppy,rictor,wdb,MEP-1,Egfr,InR,14-3-3zeta,trbl,lilli | | 3.111e-03 | -5.77 | anterior/posterior axis specification, embryo | biological process | GO:0008595 | 134 | 12 | 12345 | 442 | cic,Pat1,N,scrib,fs(1)h,chic,pum,Dg,baz,bowl,Raf,shg | | 3.111e-03 | -5.77 | tripartite regional subdivision | biological process | GO:0007351 | 134 | 12 | 12345 | 442 | Raf,shg,cic,Pat1,scrib,N,fs(1)h,bowl,Dg,baz,pum,chic | | 3.111e-03 | -5.77 | eggshell formation | biological process | GO:0030703 | 134 | 12 | 12345 | 442 | ed,emc,S,Raf,ttk,Dp,Egfr,tkv,cact,Ppcs,N,cic | | 3.120e-03 | -5.77 | MED21 (Mediator complex subunit 21) | protein interactions | 3354977 | 7 | 3 | 9204 | 432 | N,arm,pnr | | 3.120e-03 | -5.77 | ems (empty spiracles) | protein interactions | 41697 | 7 | 3 | 9204 | 432 | Abd-B,abd-A,Scr | | 3.120e-03 | -5.77 | Brd (Bearded) | protein interactions | 39620 | 7 | 3 | 9204 | 432 | Dl,H,N | | 3.120e-03 | -5.77 | Atg4b (Autophagy-related 4b) | protein interactions | 41868 | 7 | 3 | 9204 | 432 | N,H,Dl | | 3.120e-03 | -5.77 | amn (amnesiac) | protein interactions | 32999 | 7 | 3 | 9204 | 432 | H,pnr,orb2 | | 3.120e-03 | -5.77 | robo3 (roundabout 3) | protein interactions | 33314 | 7 | 3 | 9204 | 432 | sli,Abl,robo1 | | 3.120e-03 | -5.77 | CG3651 (uncharacterized protein) | protein interactions | 35440 | 7 | 3 | 9204 | 432 | ReepA,morgue,Raf | | 3.120e-03 | -5.77 | bib (big brain) | protein interactions | 34330 | 7 | 3 | 9204 | 432 | N,pnr,Dl | | 3.125e-03 | -5.77 | Laminin\_G\_2 | pfam domains | PF02210 | 16 | 4 | 10480 | 419 | kon,shg,ft,LanA | | 3.137e-03 | -5.76 | 14-3-3zeta (14-3-3zeta) | protein interactions | 36059 | 90 | 11 | 9204 | 432 | 14-3-3zeta,rictor,Raf,Syx1A,H,REPTOR,InR,Src42A,14-3-3epsilon,baz,Sema1a | | 3.160e-03 | -5.76 | sulfur compound biosynthetic process | biological process | GO:0044272 | 41 | 6 | 12345 | 442 | ATPCL,CG7560,pdgy,CG9220,muc,GlcAT-I | | 3.219e-03 | -5.74 | FN3\_dom | interpro domains | IPR003961 | 63 | 8 | 11383 | 450 | Ptp4E,sns,dome,InR,boi,ed,Eph,robo1 | | 3.219e-03 | -5.74 | FN3\_sf | interpro domains | IPR036116 | 63 | 8 | 11383 | 450 | ed,boi,InR,robo1,Eph,Ptp4E,dome,sns | | 3.239e-03 | -5.73 | Pi3K92E (Pi3K92E) | protein interactions | 42446 | 117 | 13 | 9204 | 432 | Dronc,ap,lilli,Pdk1,Pvr,Arf79F,larp,Rm62,vig,hppy,Sema1a,rictor,InR | | 3.241e-03 | -5.73 | 14\_3\_3 | smart domains | SM00101 | 2 | 2 | 5086 | 290 | 14-3-3epsilon,14-3-3zeta | | 3.241e-03 | -5.73 | c-SKI\_SMAD\_bind | smart domains | SM01046 | 2 | 2 | 5086 | 290 | fuss,Snoo | | 3.241e-03 | -5.73 | GO\_BLOOD\_VESSEL\_MORPHOGENESIS | MSigDB lists | GO\_BLOOD\_VESSEL\_MORPHOGENESIS | 67 | 8 | 4161 | 156 | fz2,arm,N,sli,C1GalTA,LanA,Eph,Dl | | 3.259e-03 | -5.73 | protein localization to peroxisome | biological process | GO:0072662 | 9 | 3 | 12345 | 442 | Pex5,Pex10,Pex14 | | 3.259e-03 | -5.73 | determination of muscle attachment site | biological process | GO:0016204 | 9 | 3 | 12345 | 442 | drl,shot,kn | | 3.259e-03 | -5.73 | enteroendocrine cell differentiation | biological process | GO:0035883 | 9 | 3 | 12345 | 442 | sina,ttk,Atg16 | | 3.259e-03 | -5.73 | imaginal disc lineage restriction | biological process | GO:0035161 | 9 | 3 | 12345 | 442 | Dl,N,ap | | 3.259e-03 | -5.73 | regulation of animal organ formation | biological process | GO:0003156 | 9 | 3 | 12345 | 442 | Abd-B,tkv,tsh | | 3.259e-03 | -5.73 | epithelial cell-cell adhesion | biological process | GO:0090136 | 9 | 3 | 12345 | 442 | baz,mop,Src42A | | 3.259e-03 | -5.73 | larval visceral muscle development | biological process | GO:0007523 | 9 | 3 | 12345 | 442 | htl,sns,klar | | 3.259e-03 | -5.73 | glandular epithelial cell differentiation | biological process | GO:0002067 | 9 | 3 | 12345 | 442 | sina,ttk,Atg16 | | 3.259e-03 | -5.73 | protein targeting to peroxisome | biological process | GO:0006625 | 9 | 3 | 12345 | 442 | Pex5,Pex14,Pex10 | | 3.259e-03 | -5.73 | establishment of protein localization to peroxisome | biological process | GO:0072663 | 9 | 3 | 12345 | 442 | Pex14,Pex10,Pex5 | | 3.284e-03 | -5.72 | GSE26030\_UNSTIM\_VS\_RESTIM\_TH17\_DAY15\_POST\_POLARIZATION\_DN | MSigDB lists | GSE26030\_UNSTIM\_VS\_RESTIM\_TH17\_DAY15\_POST\_POLARIZATION\_DN | 53 | 7 | 4161 | 156 | CenB1A,Ric,Ids,ft,emp,arm,CG6891 | | 3.290e-03 | -5.72 | SRm160 (SRm160) | protein interactions | 39483 | 14 | 4 | 9204 | 432 | Dronc,dpy,CG31211,Diap1 | | 3.290e-03 | -5.72 | Mmp2 (Matrix metalloproteinase 2) | protein interactions | 35997 | 14 | 4 | 9204 | 432 | arm,shg,Sema1a,dlp | | 3.290e-03 | -5.72 | CG3557 (uncharacterized protein) | protein interactions | 33426 | 14 | 4 | 9204 | 432 | rdx,Mob2,crp,CG4896 | | 3.290e-03 | -5.72 | Rab8 (Rab8) | protein interactions | 40168 | 14 | 4 | 9204 | 432 | kay,dome,pdm3,tkv | | 3.290e-03 | -5.72 | mir-8 (mir-8 stem loop) | protein interactions | 12798320 | 14 | 4 | 9204 | 432 | arm,SmD1,ena,spi | | 3.290e-03 | -5.72 | Fak (Focal adhesion kinase) | protein interactions | 37233 | 14 | 4 | 9204 | 432 | kon,Sema1a,Pen,Egfr | | 3.306e-03 | -5.71 | Arl8 (ADP ribosylation factor-like 8) | protein interactions | 40961 | 32 | 6 | 9204 | 432 | Rab5,CG14818,Arf79F,SERCA,Rab1,RhoL | | 3.306e-03 | -5.71 | svp (seven up) | protein interactions | 41491 | 32 | 6 | 9204 | 432 | Raf,abd-A,Abd-B,Egfr,kay,aop | | 3.307e-03 | -5.71 | oocyte axis specification | biological process | GO:0007309 | 135 | 12 | 12345 | 442 | baz,Egfr,Dg,wbl,chic,Pat1,cic,scrib,N,14-3-3zeta,14-3-3epsilon,shg | | 3.326e-03 | -5.71 | ribonucleoside bisphosphate biosynthetic process | biological process | GO:0034030 | 18 | 4 | 12345 | 442 | Ppcs,muc,ATPCL,pdgy | | 3.326e-03 | -5.71 | purine nucleoside bisphosphate biosynthetic process | biological process | GO:0034033 | 18 | 4 | 12345 | 442 | Ppcs,muc,pdgy,ATPCL | | 3.326e-03 | -5.71 | nucleoside bisphosphate biosynthetic process | biological process | GO:0033866 | 18 | 4 | 12345 | 442 | muc,Ppcs,ATPCL,pdgy | | 3.326e-03 | -5.71 | imaginal disc-derived male genitalia morphogenesis | biological process | GO:0048803 | 18 | 4 | 12345 | 442 | otk,Pvr,N,Rab5 | | 3.355e-03 | -5.70 | hemocyte differentiation | biological process | GO:0042386 | 29 | 5 | 12345 | 442 | Dronc,htl,N,RhoL,Raf | | 3.364e-03 | -5.69 | GSE19512\_NAUTRAL\_VS\_INDUCED\_TREG\_DN | MSigDB lists | GSE19512\_NAUTRAL\_VS\_INDUCED\_TREG\_DN | 40 | 6 | 4161 | 156 | dlp,14-3-3epsilon,Rgl,Cdk4,Tollo,InR | | 3.364e-03 | -5.69 | NFKB\_C | MSigDB lists | NFKB\_C | 40 | 6 | 4161 | 156 | Rgl,nmo,wdb,Cdk4,cact,trbl | | 3.364e-03 | -5.69 | GO\_RESPIRATORY\_SYSTEM\_DEVELOPMENT | MSigDB lists | GO\_RESPIRATORY\_SYSTEM\_DEVELOPMENT | 40 | 6 | 4161 | 156 | arm,alpha-Man-Ia,LanA,N,fz,otk | | 3.382e-03 | -5.69 | mitogen-activated protein kinase binding | molecular function | GO:0051019 | 3 | 2 | 12176 | 414 | Lk6,Pp2C1 | | 3.392e-03 | -5.69 | GO\_CONNECTIVE\_TISSUE\_DEVELOPMENT | MSigDB lists | GO\_CONNECTIVE\_TISSUE\_DEVELOPMENT | 28 | 5 | 4161 | 156 | N,Mef2,CG9220,tkv,htl | | 3.392e-03 | -5.69 | MEF2\_01 | MSigDB lists | MEF2\_01 | 28 | 5 | 4161 | 156 | Mef2,Eph,Tmhs,dlp,CenB1A | | 3.413e-03 | -5.68 | GO\_REGULATION\_OF\_CELL\_DIFFERENTIATION | MSigDB lists | GO\_REGULATION\_OF\_CELL\_DIFFERENTIATION | 279 | 20 | 4161 | 156 | cact,Dl,otk,Cdk4,robo1,Mef2,Raf,trbl,Arf79F,N,Actbeta,Tollo,hyx,ft,Ddr,sli,tkv,nAChRalpha4,fz,arm | | 3.435e-03 | -5.67 | CAGGTG\_E12\_Q6 | MSigDB lists | CAGGTG\_E12\_Q6 | 464 | 29 | 4161 | 156 | CG31522,hppy,ATPCL,Dl,Dyb,Mef2,Rab1,shot,nmo,trbl,Psa,InR,Vps13,l(2)37Cg,14-3-3zeta,LanA,Syx1A,PI31,sli,hyx,rg,Ric,CG9485,fz2,CG2812,Pex10,tkv,CG45050,Nrk | | 3.447e-03 | -5.67 | phosphate-containing compound metabolic process | biological process | GO:0006796 | 900 | 48 | 12345 | 442 | CG6656,muc,Pez,ATPCL,Ocrl,Src42A,Gfat2,CG10417,Ppcs,Cdk4,Pp2C1,Lk6,drl,Drak,Ptpmeg2,Socs36E,Pgk,Pvr,Pitslre,InR,l(1)G0196,hppy,Pdk1,CG42327,Ddr,htl,tkv,nmo,Cks30A,wdb,Abl,IA-2,trbl,otk,wnd,Eph,CG9784,Ptp4E,UQCR-11,CycG,pdgy,CycB3,eya,Dyrk2,CG8306,Raf,Nrk,mop | | 3.464e-03 | -5.67 | GO\_POSITIVE\_REGULATION\_OF\_CATALYTIC\_ACTIVITY | MSigDB lists | GO\_POSITIVE\_REGULATION\_OF\_CATALYTIC\_ACTIVITY | 401 | 26 | 4161 | 156 | CG5522,hppy,step,Raf,Mef2,GlcAT-I,Rab1,CenB1A,CG3107,Arf79F,dock,htl,Ocrl,CG17883,Ddr,Rgl,Ziz,PI31,unc-119,Tollo,arm,fz2,wbl,Pdk1,Tsp96F,nAChRalpha4 | | 3.475e-03 | -5.66 | regulation of response to external stimulus | biological process | GO:0032101 | 228 | 17 | 12345 | 442 | sli,CG34401,cact,Dcr-2,htl,pnr,CG32795,imd,DPCoAC,chic,Sema1b,Rm62,Pitslre,fz,Sema1a,nub,mop | | 3.476e-03 | -5.66 | Prot-tyrosine\_phosphatase-like | interpro domains | IPR029021 | 38 | 6 | 11383 | 450 | mop,CG42327,Ptpmeg2,IA-2,Pez,Ptp4E | | 3.488e-03 | -5.66 | GO\_REGULATION\_OF\_CELL\_PROLIFERATION | MSigDB lists | GO\_REGULATION\_OF\_CELL\_PROLIFERATION | 260 | 19 | 4161 | 156 | N,Actbeta,CtBP,dock,InR,htl,Cdk4,Mef2,Raf,Dl,cact,Tsp96F,tkv,arm,fz2,hyx,Src42A,Ddr,LanA | | 3.490e-03 | -5.66 | PTPc\_motif | smart domains | SM00404 | 19 | 5 | 5086 | 290 | IA-2,Pez,Ptp4E,Ptpmeg2,CG42327 | | 3.503e-03 | -5.65 | intracellular vesicle | cellular component | GO:0097708 | 378 | 24 | 11940 | 417 | AP-1gamma,apolpp,ed,Ocrl,mop,fz2,N,shg,jeb,spdo,path,lt,Syx1A,dock,nuf,Dl,tkv,Cadps,arm,jar,Rab5,fz,Vps13,S | | 3.503e-03 | -5.65 | cytoplasmic vesicle | cellular component | GO:0031410 | 378 | 24 | 11940 | 417 | jeb,spdo,path,shg,N,fz2,mop,Ocrl,ed,apolpp,AP-1gamma,S,Vps13,fz,Rab5,jar,Cadps,arm,tkv,Dl,nuf,dock,Syx1A,lt | | 3.506e-03 | -5.65 | late endosome | cellular component | GO:0005770 | 72 | 8 | 11940 | 417 | N,fz2,spdo,S,Vps13,lt,tkv,Ocrl | | 3.510e-03 | -5.65 | response to stress | biological process | GO:0006950 | 1136 | 58 | 12345 | 442 | shot,Pvr,chic,tipE,N,DPCoAC,CG30156,kn,hppy,ed,chrb,CG1218,lt,InR,koi,LanA,msi,sick,rdx,Atox1,ZnT41F,REPTOR,spen,Sxl,dome,jumu,RhoGEF3,Src42A,CG32795,shg,CycG,CG7536,Atg17,Eph,wnd,imd,RhoL,CG13481,Dronc,Ric,Raf,h,hang,eya,Dcr-2,htl,ReepA,rictor,flfl,Atg16,scyl,Hr4,subdued,14-3-3epsilon,kay,SNF4Agamma,Tollo,Dp | | 3.514e-03 | -5.65 | embryonic axis specification | biological process | GO:0000578 | 136 | 12 | 12345 | 442 | N,scrib,cic,Pat1,pum,chic,baz,Dg,bowl,fs(1)h,shg,Raf | | 3.544e-03 | -5.64 | CADHERIN\_2 | prosite domains | PS50268 | 13 | 4 | 6243 | 325 | shg,ds,Cals,ft | | 3.552e-03 | -5.64 | Vap33 (VAMP-associated protein 33kDa) | protein interactions | 31349 | 43 | 7 | 9204 | 432 | Dl,rictor,Eph,d,ft,ReepA,tkv | | 3.552e-03 | -5.64 | Cp1 (Cysteine proteinase-1) | protein interactions | 36546 | 43 | 7 | 9204 | 432 | C1GalTA,Raf,imd,Arf79F,SERCA,Gfat2,Rab5 | | 3.552e-03 | -5.64 | MEP-1 (MEP-1) | protein interactions | 38327 | 43 | 7 | 9204 | 432 | rictor,MEP-1,Pat1,CtBP,Rm62,ttk,orb2 | | 3.559e-03 | -5.64 | GO\_OSSIFICATION | MSigDB lists | GO\_OSSIFICATION | 68 | 8 | 4161 | 156 | Mef2,Pex7,ds,Ddr,ft,htl,fz,Mfe2 | | 3.566e-03 | -5.64 | CSL-Notch-Mastermind transcription factor complex | cellular component | GO:1990433 | 3 | 2 | 11940 | 417 | mam,N | | 3.566e-03 | -5.64 | protein-carbohydrate complex | cellular component | GO:0032992 | 3 | 2 | 11940 | 417 | robo1,sli | | 3.571e-03 | -5.63 | olfactory behavior | biological process | GO:0042048 | 86 | 9 | 12345 | 442 | rg,Adf1,spi,Dyrk2,Egfr,vn,drl,scrib,14-3-3zeta | | 3.571e-03 | -5.63 | fly chr6E4-6E4\|1-18 | chromosome location | fly chr6E4-6E4\|1-18 | 14 | 3 | 25010 | 571 | ogre,bou,Inx2 | | 3.571e-03 | -5.63 | fly chr6E4-6E4\|1-18 cM | chromosome location | fly chr6E4-6E4\|1-18 cM | 14 | 3 | 25010 | 571 | Inx2,ogre,bou | | 3.576e-03 | -5.63 | regulation of vesicle-mediated transport | biological process | GO:0060627 | 119 | 11 | 12345 | 442 | step,jar,mim,Arf79F,cindr,scrib,Syx1A,Rab5,Cbl,Cadps,lt | | 3.576e-03 | -5.63 | REN\_ALVEOLAR\_RHABDOMYOSARCOMA\_DN | MSigDB lists | REN\_ALVEOLAR\_RHABDOMYOSARCOMA\_DN | 132 | 12 | 4161 | 156 | wbl,Tpst,Rab1,Pgant5,Actbeta,Tsp96F,Arf79F,fz,CG31522,Rab5,Ids,Fkbp14 | | 3.578e-03 | -5.63 | regulation of imaginal disc growth | biological process | GO:0045570 | 42 | 6 | 12345 | 442 | tkv,ft,Egfr,dlp,d,scrib | | 3.622e-03 | -5.62 | Netrin-1 signaling | REACTOME pathways | R-DME-373752 | 15 | 4 | 3923 | 176 | robo1,sli,dock,sina | | 3.636e-03 | -5.62 | axon part | cellular component | GO:0033267 | 105 | 10 | 11940 | 417 | gukh,shot,ena,Syx1A,robo1,orb2,jeb,pum,jar,Magi | | 3.657e-03 | -5.61 | GO\_ENZYME\_INHIBITOR\_ACTIVITY | MSigDB lists | GO\_ENZYME\_INHIBITOR\_ACTIVITY | 54 | 7 | 4161 | 156 | sli,Spn42De,PI31,N,Ppn,trbl,dock | | 3.657e-03 | -5.61 | ATGAAGG\_MIR205 | MSigDB lists | ATGAAGG\_MIR205 | 54 | 7 | 4161 | 156 | sina,Actbeta,Ppn,Lk6,sd,Src42A,AP-1gamma | | 3.657e-03 | -5.61 | SP3\_Q3 | MSigDB lists | SP3\_Q3 | 54 | 7 | 4161 | 156 | Eph,ap,wdb,hyx,Cdk4,14-3-3epsilon,nmo | | 3.663e-03 | -5.61 | regulation of endocytosis | biological process | GO:0030100 | 56 | 7 | 12345 | 442 | step,mim,Arf79F,Cbl,cindr,scrib,Rab5 | | 3.676e-03 | -5.61 | GO\_REGULATION\_OF\_CYTOKINE\_BIOSYNTHETIC\_PROCESS | MSigDB lists | GO\_REGULATION\_OF\_CYTOKINE\_BIOSYNTHETIC\_PROCESS | 9 | 3 | 4161 | 156 | Actbeta,Tollo,trbl | | 3.676e-03 | -5.61 | GO\_HEART\_VALVE\_DEVELOPMENT | MSigDB lists | GO\_HEART\_VALVE\_DEVELOPMENT | 9 | 3 | 4161 | 156 | N,Mef2,ds | | 3.676e-03 | -5.61 | MORF\_ZNF10 | MSigDB lists | MORF\_ZNF10 | 9 | 3 | 4161 | 156 | nAChRalpha4,lt,Drep4 | | 3.676e-03 | -5.61 | REACTOME\_TCR\_SIGNALING | MSigDB lists | REACTOME\_TCR\_SIGNALING | 9 | 3 | 4161 | 156 | cact,dock,Pdk1 | | 3.676e-03 | -5.61 | GO\_ADHERENS\_JUNCTION\_ASSEMBLY | MSigDB lists | GO\_ADHERENS\_JUNCTION\_ASSEMBLY | 9 | 3 | 4161 | 156 | Pdk1,arm,LanA | | 3.676e-03 | -5.61 | GO\_POSITIVE\_REGULATION\_OF\_SKELETAL\_MUSCLE\_TISSUE\_DEVELOPMENT | MSigDB lists | GO\_POSITIVE\_REGULATION\_OF\_SKELETAL\_MUSCLE\_TISSUE\_DEVELOPMENT | 9 | 3 | 4161 | 156 | Dl,Mef2,arm | | 3.676e-03 | -5.61 | GO\_GLOMERULUS\_DEVELOPMENT | MSigDB lists | GO\_GLOMERULUS\_DEVELOPMENT | 9 | 3 | 4161 | 156 | sns,N,Mef2 | | 3.676e-03 | -5.61 | PID\_PI3K\_PLC\_TRK\_PATHWAY | MSigDB lists | PID\_PI3K\_PLC\_TRK\_PATHWAY | 9 | 3 | 4161 | 156 | 14-3-3epsilon,14-3-3zeta,Pdk1 | | 3.676e-03 | -5.61 | GO\_ENDODERM\_FORMATION | MSigDB lists | GO\_ENDODERM\_FORMATION | 9 | 3 | 4161 | 156 | arm,hyx,Actbeta | | 3.676e-03 | -5.61 | GO\_REGULATION\_OF\_INTERFERON\_GAMMA\_PRODUCTION | MSigDB lists | GO\_REGULATION\_OF\_INTERFERON\_GAMMA\_PRODUCTION | 9 | 3 | 4161 | 156 | fz2,Actbeta,Tollo | | 3.676e-03 | -5.61 | DORN\_ADENOVIRUS\_INFECTION\_24HR\_DN | MSigDB lists | DORN\_ADENOVIRUS\_INFECTION\_24HR\_DN | 9 | 3 | 4161 | 156 | Rab1,step,cact | | 3.676e-03 | -5.61 | GO\_SPROUTING\_ANGIOGENESIS | MSigDB lists | GO\_SPROUTING\_ANGIOGENESIS | 9 | 3 | 4161 | 156 | N,Dl,sli | | 3.676e-03 | -5.61 | REACTOME\_O\_LINKED\_GLYCOSYLATION\_OF\_MUCINS | MSigDB lists | REACTOME\_O\_LINKED\_GLYCOSYLATION\_OF\_MUCINS | 9 | 3 | 4161 | 156 | Pgant5,Pgant6,C1GalTA | | 3.676e-03 | -5.61 | KEGG\_MELANOMA | MSigDB lists | KEGG\_MELANOMA | 9 | 3 | 4161 | 156 | Cdk4,Raf,InR | | 3.711e-03 | -5.60 | Socs36E (Suppressor of cytokine signaling at 36E) | protein interactions | 35085 | 23 | 5 | 9204 | 432 | eya,Cbl,spi,scrib,dome | | 3.711e-03 | -5.60 | tup (tailup) | protein interactions | 35147 | 23 | 5 | 9204 | 432 | pnr,Abd-B,abd-A,ap,Scr | | 3.711e-03 | -5.60 | lqfR (liquid facets-Related) | protein interactions | 42632 | 23 | 5 | 9204 | 432 | InR,rictor,Drep4,shg,arm | | 3.711e-03 | -5.60 | Arp3 (Actin-related protein 3) | protein interactions | 38898 | 23 | 5 | 9204 | 432 | cindr,CG14478,arm,Cip4,chic | | 3.711e-03 | -5.60 | Btk29A (Btk family kinase at 29A) | protein interactions | 34132 | 23 | 5 | 9204 | 432 | Pvr,Egfr,Src42A,N,chic | | 3.711e-03 | -5.60 | RacGAP84C (Rac GTPase activating protein at 84C) | protein interactions | 117503 | 23 | 5 | 9204 | 432 | muc,hppy,CG2017,SNF4Agamma,cindr | | 3.711e-03 | -5.60 | E(spl)mbeta-HLH (Enhancer of split mbeta, helix-loop-helix) | protein interactions | 43152 | 23 | 5 | 9204 | 432 | E(spl)m5-HLH,N,emc,nmo,tkv | | 3.728e-03 | -5.59 | Nsf2 (N-ethylmaleimide-sensitive factor 2) | protein interactions | 41694 | 92 | 11 | 9204 | 432 | N,l(2)37Cg,Gli,Dl,Bacc,Syx1A,fend,spi,CG7492,Rab5,LanA | | 3.743e-03 | -5.59 | Dorso-ventral axis formation | KEGG pathways | ko04320 | 28 | 5 | 3198 | 123 | Egfr,orb2,N,Raf,aop | | 3.743e-03 | -5.59 | Dorso-ventral axis formation | KEGG pathways | dme04320 | 28 | 5 | 3198 | 123 | aop,Raf,N,orb2,Egfr | | 3.746e-03 | -5.59 | positive regulation of ARF protein signal transduction | biological process | GO:0032014 | 3 | 2 | 12345 | 442 | step,siz | | 3.746e-03 | -5.59 | protein import into peroxisome matrix, docking | biological process | GO:0016560 | 3 | 2 | 12345 | 442 | Pex14,Pex5 | | 3.746e-03 | -5.59 | regulation of production of miRNAs involved in gene silencing by miRNA | biological process | GO:1903798 | 3 | 2 | 12345 | 442 | lin-28,Dg | | 3.746e-03 | -5.59 | peptide cross-linking | biological process | GO:0018149 | 3 | 2 | 12345 | 442 | ft,ds | | 3.746e-03 | -5.59 | endodermal cell differentiation | biological process | GO:0035987 | 3 | 2 | 12345 | 442 | htl,tkv | | 3.746e-03 | -5.59 | myoblast fate specification | biological process | GO:0048626 | 3 | 2 | 12345 | 442 | ttk,htl | | 3.746e-03 | -5.59 | pyrimidine-containing compound transmembrane transport | biological process | GO:0072531 | 3 | 2 | 12345 | 442 | Tpc1,Rim2 | | 3.746e-03 | -5.59 | microtubule cytoskeleton organization involved in establishment of planar polarity | biological process | GO:0090176 | 3 | 2 | 12345 | 442 | ft,ds | | 3.746e-03 | -5.59 | haltere disc morphogenesis | biological process | GO:0007481 | 3 | 2 | 12345 | 442 | ap,hth | | 3.746e-03 | -5.59 | endodermal cell fate commitment | biological process | GO:0001711 | 3 | 2 | 12345 | 442 | htl,tkv | | 3.746e-03 | -5.59 | Roundabout signaling pathway | biological process | GO:0035385 | 3 | 2 | 12345 | 442 | robo1,sli | | 3.746e-03 | -5.59 | glial cell fate determination | biological process | GO:0007403 | 3 | 2 | 12345 | 442 | N,spen | | 3.746e-03 | -5.59 | collateral sprouting of injured axon | biological process | GO:0048674 | 3 | 2 | 12345 | 442 | kay,wnd | | 3.746e-03 | -5.59 | glial cell fate commitment | biological process | GO:0021781 | 3 | 2 | 12345 | 442 | N,spen | | 3.746e-03 | -5.59 | collateral sprouting | biological process | GO:0048668 | 3 | 2 | 12345 | 442 | kay,wnd | | 3.779e-03 | -5.58 | apoptotic process | biological process | GO:0006915 | 103 | 10 | 12345 | 442 | scyl,wgn,abd-A,Dcr-2,Diap1,Dp,chrb,Dronc,hppy,Drep4 | | 3.780e-03 | -5.58 | FERM\_3 | prosite domains | PS50057 | 21 | 5 | 6243 | 325 | yrt,cora,Myo10A,CG42788,Pez | | 3.784e-03 | -5.58 | nervous system process | biological process | GO:0050877 | 564 | 33 | 12345 | 442 | mld,subdued,Tob,Adf1,Abl,arm,wnd,pum,Myo10A,Dyrk2,Gr93c,Egfr,f,Tmhs,rg,SKIP,dome,lilli,ss,spi,Socs36E,orb2,pxb,Ir21a,vn,nAChRalpha4,pdm3,drl,scrib,14-3-3zeta,N,mura,Mob2 | | 3.784e-03 | -5.58 | cell cycle process | biological process | GO:0022402 | 564 | 33 | 12345 | 442 | Pex10,14-3-3epsilon,wdb,larp,eff,Cks30A,Slbp,baz,jar,His3.3A,Rab5,Syx1A,His3.3B,Hr39,mop,Rab1,CycB3,CycG,Arf79F,PI31,Gli,SCAR,Spc105R,Sxl,LanA,Ocrl,Wdr62,ph-p,orb2,chic,bol,corto,pita | | 3.815e-03 | -5.57 | ZINC\_FINGER\_C2H2\_1 | prosite domains | PS00028 | 287 | 26 | 6243 | 325 | hang,Prdm13,mld,CG10321,CG32772,crol,ttk,esg,CG11414,sens-2,CG10348,CG11247,zfh2,disco-r,sens,rgr,chn,CG11122,sqz,klu,CG12769,CG14441,pita,tsh,bowl,MEP-1 | | 3.821e-03 | -5.57 | CATGTAA\_MIR496 | MSigDB lists | CATGTAA\_MIR496 | 41 | 6 | 4161 | 156 | Mef2,Kr-h2,14-3-3zeta,rg,Rim2,Rab5 | | 3.821e-03 | -5.57 | MCBRYAN\_PUBERTAL\_TGFB1\_TARGETS\_UP | MSigDB lists | MCBRYAN\_PUBERTAL\_TGFB1\_TARGETS\_UP | 41 | 6 | 4161 | 156 | step,Rab5,muc,subdued,Mef2,CG6891 | | 3.855e-03 | -5.56 | REACTOME\_NETRIN1\_SIGNALING | MSigDB lists | REACTOME\_NETRIN1\_SIGNALING | 18 | 4 | 4161 | 156 | dock,sina,unc-5,sli | | 3.855e-03 | -5.56 | LEE\_NEURAL\_CREST\_STEM\_CELL\_DN | MSigDB lists | LEE\_NEURAL\_CREST\_STEM\_CELL\_DN | 18 | 4 | 4161 | 156 | fz2,sli,ap,N | | 3.855e-03 | -5.56 | GO\_ORGAN\_GROWTH | MSigDB lists | GO\_ORGAN\_GROWTH | 18 | 4 | 4161 | 156 | Ddr,htl,Dl,N | | 3.855e-03 | -5.56 | KEGG\_CHRONIC\_MYELOID\_LEUKEMIA | MSigDB lists | KEGG\_CHRONIC\_MYELOID\_LEUKEMIA | 18 | 4 | 4161 | 156 | Cdk4,Raf,CtBP,cact | | 3.855e-03 | -5.56 | GO\_REGULATION\_OF\_EPITHELIAL\_CELL\_DIFFERENTIATION | MSigDB lists | GO\_REGULATION\_OF\_EPITHELIAL\_CELL\_DIFFERENTIATION | 18 | 4 | 4161 | 156 | N,Dl,ft,arm | | 3.855e-03 | -5.56 | MORF\_DCC | MSigDB lists | MORF\_DCC | 18 | 4 | 4161 | 156 | nAChRalpha4,lt,Drep4,Src42A | | 3.855e-03 | -5.56 | GTTATAT\_MIR410 | MSigDB lists | GTTATAT\_MIR410 | 18 | 4 | 4161 | 156 | Atg16,sd,Rab1,Ziz | | 3.855e-03 | -5.56 | GO\_PROTEOGLYCAN\_BIOSYNTHETIC\_PROCESS | MSigDB lists | GO\_PROTEOGLYCAN\_BIOSYNTHETIC\_PROCESS | 18 | 4 | 4161 | 156 | GlcAT-I,frc,CG9220,tkv | | 3.865e-03 | -5.56 | Jra (Jun-related antigen) | protein interactions | 36057 | 67 | 9 | 9204 | 432 | nmo,fz,CG14818,Egfr,Raf,tkv,sd,aop,kay | | 3.883e-03 | -5.55 | ksr (kinase suppressor of ras) | protein interactions | 40660 | 33 | 6 | 9204 | 432 | ttk,aop,Raf,Gfat2,cindr,SERCA | | 3.883e-03 | -5.55 | Sce (Sex combs extra) | protein interactions | 43327 | 33 | 6 | 9204 | 432 | Psc,dome,ph-p,Su(z)2,Kdm2,Socs36E | | 3.884e-03 | -5.55 | meiotic cell cycle process | biological process | GO:1903046 | 250 | 18 | 12345 | 442 | Cks30A,eff,larp,Sxl,Pex10,His3.3B,Rab5,His3.3A,LanA,Rab1,Hr39,bol,Gli,chic,orb2,Arf79F,PI31,CycG | | 3.893e-03 | -5.55 | CHARAFE\_BREAST\_CANCER\_LUMINAL\_VS\_MESENCHYMAL\_DN | MSigDB lists | CHARAFE\_BREAST\_CANCER\_LUMINAL\_VS\_MESENCHYMAL\_DN | 100 | 10 | 4161 | 156 | SmD1,Actbeta,CG9220,Unc-76,sd,sli,ft,Gfat2,Cdk4,CG1673 | | 3.899e-03 | -5.55 | GO\_POSITIVE\_REGULATION\_OF\_MOLECULAR\_FUNCTION | MSigDB lists | GO\_POSITIVE\_REGULATION\_OF\_MOLECULAR\_FUNCTION | 468 | 29 | 4161 | 156 | PI31,Ziz,Rgl,CG17883,Ocrl,Ddr,Tollo,unc-119,wbl,arm,fz2,nAChRalpha4,Tsp96F,Pdk1,fz,CG5522,step,hppy,cact,Raf,Mef2,Rab1,CenB1A,GlcAT-I,dock,htl,CG3107,Arf79F,CtBP | | 3.909e-03 | -5.54 | positive regulation of smoothened signaling pathway | biological process | GO:0045880 | 30 | 5 | 12345 | 442 | dlp,wdb,Sxl,apolpp,hyx | | 3.909e-03 | -5.54 | regulation of axon guidance | biological process | GO:1902667 | 30 | 5 | 12345 | 442 | fz,Sema1a,CG34401,chic,Sema1b | | 3.909e-03 | -5.54 | heterochromatin organization involved in chromatin silencing | biological process | GO:0070868 | 30 | 5 | 12345 | 442 | bin3,crol,nub,Dcr-2,fl(2)d | | 3.957e-03 | -5.53 | oocyte construction | biological process | GO:0007308 | 138 | 12 | 12345 | 442 | scrib,N,14-3-3zeta,cic,Pat1,Egfr,baz,Dg,wbl,chic,shg,14-3-3epsilon | | 3.960e-03 | -5.53 | CADHERIN | prints domains | PR00205 | 15 | 4 | 2185 | 101 | ds,ft,shg,Cals | | 3.977e-03 | -5.53 | TAXCREB\_01 | MSigDB lists | TAXCREB\_01 | 29 | 5 | 4161 | 156 | Pat1,Kdm2,nmo,muc,Raf | | 3.977e-03 | -5.53 | ZHANG\_RESPONSE\_TO\_IKK\_INHIBITOR\_AND\_TNF\_UP | MSigDB lists | ZHANG\_RESPONSE\_TO\_IKK\_INHIBITOR\_AND\_TNF\_UP | 29 | 5 | 4161 | 156 | Actbeta,cact,dock,Kdm2,step | | 3.977e-03 | -5.53 | GO\_DIGESTIVE\_SYSTEM\_DEVELOPMENT | MSigDB lists | GO\_DIGESTIVE\_SYSTEM\_DEVELOPMENT | 29 | 5 | 4161 | 156 | ds,arm,C1GalTA,ft,N | | 3.977e-03 | -5.53 | GO\_REGULATION\_OF\_LEUKOCYTE\_MEDIATED\_IMMUNITY | MSigDB lists | GO\_REGULATION\_OF\_LEUKOCYTE\_MEDIATED\_IMMUNITY | 29 | 5 | 4161 | 156 | Pdk1,Tollo,fz2,Spn42De,AP-1gamma | | 4.014e-03 | -5.52 | cell division site part | cellular component | GO:0032155 | 44 | 6 | 11940 | 417 | Arf79F,step,Cip4,Rab1,nuf,cindr | | 4.057e-03 | -5.51 | response to biotic stimulus | biological process | GO:0009607 | 415 | 26 | 12345 | 442 | kay,dome,jumu,Tollo,RhoGEF3,Src42A,CG32795,shg,cact,Dcr-2,LanA,htl,sick,Mef2,spen,subdued,kn,Dronc,eya,Pvr,Rm62,Eph,RhoL,N,imd,DPCoAC | | 4.057e-03 | -5.51 | response to external biotic stimulus | biological process | GO:0043207 | 415 | 26 | 12345 | 442 | kn,Dronc,eya,Rm62,Pvr,Eph,DPCoAC,imd,N,RhoL,kay,dome,Src42A,RhoGEF3,Tollo,jumu,shg,CG32795,LanA,htl,cact,Dcr-2,sick,Mef2,subdued,spen | | 4.057e-03 | -5.51 | response to other organism | biological process | GO:0051707 | 415 | 26 | 12345 | 442 | spen,subdued,Mef2,sick,Dcr-2,cact,LanA,htl,shg,CG32795,Tollo,jumu,RhoGEF3,Src42A,dome,kay,N,imd,RhoL,DPCoAC,Eph,Rm62,Pvr,eya,Dronc,kn | | 4.087e-03 | -5.50 | GO\_MAINTENANCE\_OF\_CELL\_POLARITY | MSigDB lists | GO\_MAINTENANCE\_OF\_CELL\_POLARITY | 3 | 2 | 4161 | 156 | shot,ap | | 4.087e-03 | -5.50 | GO\_HETEROPHILIC\_CELL\_CELL\_ADHESION\_VIA\_PLASMA\_MEMBRANE\_CELL\_ADHESION\_MOLECULES | MSigDB lists | GO\_HETEROPHILIC\_CELL\_CELL\_ADHESION\_VIA\_PLASMA\_MEMBRANE\_CELL\_ADHESION\_MOLECULES | 3 | 2 | 4161 | 156 | ft,ds | | 4.087e-03 | -5.50 | GO\_POSITIVE\_REGULATION\_OF\_B\_CELL\_PROLIFERATION | MSigDB lists | GO\_POSITIVE\_REGULATION\_OF\_B\_CELL\_PROLIFERATION | 3 | 2 | 4161 | 156 | Mef2,Tsp96F | | 4.087e-03 | -5.50 | GO\_APOPTOTIC\_PROCESS\_INVOLVED\_IN\_MORPHOGENESIS | MSigDB lists | GO\_APOPTOTIC\_PROCESS\_INVOLVED\_IN\_MORPHOGENESIS | 3 | 2 | 4161 | 156 | N,fz2 | | 4.087e-03 | -5.50 | GO\_LIPASE\_ACTIVATOR\_ACTIVITY | MSigDB lists | GO\_LIPASE\_ACTIVATOR\_ACTIVITY | 3 | 2 | 4161 | 156 | Pdk1,Arf79F | | 4.087e-03 | -5.50 | GO\_REGULATION\_OF\_B\_CELL\_PROLIFERATION | MSigDB lists | GO\_REGULATION\_OF\_B\_CELL\_PROLIFERATION | 3 | 2 | 4161 | 156 | Tsp96F,Mef2 | | 4.087e-03 | -5.50 | GO\_NEGATIVE\_REGULATION\_OF\_CYTOKINE\_BIOSYNTHETIC\_PROCESS | MSigDB lists | GO\_NEGATIVE\_REGULATION\_OF\_CYTOKINE\_BIOSYNTHETIC\_PROCESS | 3 | 2 | 4161 | 156 | trbl,Actbeta | | 4.087e-03 | -5.50 | MODULE\_76 | MSigDB lists | MODULE\_76 | 3 | 2 | 4161 | 156 | Tpst,Tollo | | 4.087e-03 | -5.50 | GO\_ROUNDABOUT\_SIGNALING\_PATHWAY | MSigDB lists | GO\_ROUNDABOUT\_SIGNALING\_PATHWAY | 3 | 2 | 4161 | 156 | sli,robo1 | | 4.087e-03 | -5.50 | GENTLES\_LEUKEMIC\_STEM\_CELL\_UP | MSigDB lists | GENTLES\_LEUKEMIC\_STEM\_CELL\_UP | 3 | 2 | 4161 | 156 | kn,Mef2 | | 4.087e-03 | -5.50 | GO\_NEGATIVE\_REGULATION\_OF\_EPIDERMAL\_CELL\_DIFFERENTIATION | MSigDB lists | GO\_NEGATIVE\_REGULATION\_OF\_EPIDERMAL\_CELL\_DIFFERENTIATION | 3 | 2 | 4161 | 156 | Dl,N | | 4.087e-03 | -5.50 | ONO\_AML1\_TARGETS\_DN | MSigDB lists | ONO\_AML1\_TARGETS\_DN | 3 | 2 | 4161 | 156 | Cdk4,Tsp96F | | 4.087e-03 | -5.50 | GO\_DETECTION\_OF\_TEMPERATURE\_STIMULUS | MSigDB lists | GO\_DETECTION\_OF\_TEMPERATURE\_STIMULUS | 3 | 2 | 4161 | 156 | Eph,subdued | | 4.087e-03 | -5.50 | GO\_REGULATION\_OF\_INTERLEUKIN\_10\_PRODUCTION | MSigDB lists | GO\_REGULATION\_OF\_INTERLEUKIN\_10\_PRODUCTION | 3 | 2 | 4161 | 156 | trbl,Dl | | 4.087e-03 | -5.50 | BIOCARTA\_AKT\_PATHWAY | MSigDB lists | BIOCARTA\_AKT\_PATHWAY | 3 | 2 | 4161 | 156 | Pdk1,cact | | 4.087e-03 | -5.50 | SHIPP\_DLBCL\_CURED\_VS\_FATAL\_UP | MSigDB lists | SHIPP\_DLBCL\_CURED\_VS\_FATAL\_UP | 3 | 2 | 4161 | 156 | nAChRalpha4,SERCA | | 4.087e-03 | -5.50 | GO\_DEVELOPMENTAL\_PROGRAMMED\_CELL\_DEATH | MSigDB lists | GO\_DEVELOPMENTAL\_PROGRAMMED\_CELL\_DEATH | 3 | 2 | 4161 | 156 | N,fz2 | | 4.087e-03 | -5.50 | GO\_REGULATION\_OF\_PROTEIN\_HOMOOLIGOMERIZATION | MSigDB lists | GO\_REGULATION\_OF\_PROTEIN\_HOMOOLIGOMERIZATION | 3 | 2 | 4161 | 156 | Pex5,Pex14 | | 4.087e-03 | -5.50 | SARTIPY\_BLUNTED\_BY\_INSULIN\_RESISTANCE\_UP | MSigDB lists | SARTIPY\_BLUNTED\_BY\_INSULIN\_RESISTANCE\_UP | 3 | 2 | 4161 | 156 | Rgl,Cerk | | 4.087e-03 | -5.50 | GO\_HEART\_VALVE\_FORMATION | MSigDB lists | GO\_HEART\_VALVE\_FORMATION | 3 | 2 | 4161 | 156 | N,ds | | 4.087e-03 | -5.50 | GO\_CARDIAC\_CHAMBER\_FORMATION | MSigDB lists | GO\_CARDIAC\_CHAMBER\_FORMATION | 3 | 2 | 4161 | 156 | Mef2,N | | 4.087e-03 | -5.50 | GO\_CELL\_MIGRATION\_INVOLVED\_IN\_HEART\_DEVELOPMENT | MSigDB lists | GO\_CELL\_MIGRATION\_INVOLVED\_IN\_HEART\_DEVELOPMENT | 3 | 2 | 4161 | 156 | N,ds | | 4.087e-03 | -5.50 | SCHOEN\_NFKB\_SIGNALING | MSigDB lists | SCHOEN\_NFKB\_SIGNALING | 3 | 2 | 4161 | 156 | Tpst,cact | | 4.087e-03 | -5.50 | KORKOLA\_EMBRYONIC\_CARCINOMA\_VS\_SEMINOMA\_UP | MSigDB lists | KORKOLA\_EMBRYONIC\_CARCINOMA\_VS\_SEMINOMA\_UP | 3 | 2 | 4161 | 156 | CG1673,dlp | | 4.087e-03 | -5.50 | DORN\_ADENOVIRUS\_INFECTION\_12HR\_UP | MSigDB lists | DORN\_ADENOVIRUS\_INFECTION\_12HR\_UP | 3 | 2 | 4161 | 156 | step,Rab1 | | 4.096e-03 | -5.50 | imaginal disc-derived genitalia morphogenesis | biological process | GO:0048805 | 19 | 4 | 12345 | 442 | Rab5,N,Pvr,otk | | 4.096e-03 | -5.50 | ventral furrow formation | biological process | GO:0007370 | 19 | 4 | 12345 | 442 | arm,shg,trbl,Abl | | 4.096e-03 | -5.50 | male anatomical structure morphogenesis | biological process | GO:0090598 | 19 | 4 | 12345 | 442 | otk,Pvr,N,Rab5 | | 4.096e-03 | -5.50 | male genitalia morphogenesis | biological process | GO:0048808 | 19 | 4 | 12345 | 442 | otk,Pvr,N,Rab5 | | 4.096e-03 | -5.50 | post-embryonic genitalia morphogenesis | biological process | GO:0035126 | 19 | 4 | 12345 | 442 | otk,Pvr,N,Rab5 | | 4.156e-03 | -5.48 | cell cortex region | cellular component | GO:0099738 | 74 | 8 | 11940 | 417 | Dl,cindr,nuf,Abl,baz,CG42684,step,Bsg | | 4.156e-03 | -5.48 | cell surface | cellular component | GO:0009986 | 74 | 8 | 11940 | 417 | Cals,Dl,Gli,ed,wgn,N,boi,dlp | | 4.165e-03 | -5.48 | GO\_CELL\_CELL\_ADHESION | MSigDB lists | GO\_CELL\_CELL\_ADHESION | 85 | 9 | 4161 | 156 | dock,arm,fz2,Cals,Cdk4,Raf,robo1,ds,ft | | 4.203e-03 | -5.47 | GO\_ANCHORING\_JUNCTION | MSigDB lists | GO\_ANCHORING\_JUNCTION | 170 | 14 | 4161 | 156 | shot,14-3-3epsilon,arm,htl,N,Arf79F,Tsp96F,Pdk1,14-3-3zeta,fz,emp,Ddr,Dl,otk | | 4.225e-03 | -5.47 | sperm individualization complex | cellular component | GO:0070864 | 10 | 3 | 11940 | 417 | PI31,yuri,jar | | 4.242e-03 | -5.46 | GO\_REGULATION\_OF\_RESPONSE\_TO\_STRESS | MSigDB lists | GO\_REGULATION\_OF\_RESPONSE\_TO\_STRESS | 407 | 26 | 4161 | 156 | dock,InR,Unc-76,Arf79F,14-3-3epsilon,SmD1,cact,Cdk4,Mef2,Raf,Tpc1,Pdk1,Kr-h2,wbl,eff,Nmda1,fz2,Tollo,Eph,Fem-1,Spn42De,CG4645,PI31,ReepA,AP-1gamma,sli | | 4.300e-03 | -5.45 | TGCTGAY\_UNKNOWN | MSigDB lists | TGCTGAY\_UNKNOWN | 135 | 12 | 4161 | 156 | FBXO11,trbl,sns,sd,Arf79F,Rab1,Kdm2,rdx,Dyb,Hel89B,AP-1gamma,Pgant6 | | 4.318e-03 | -5.45 | GO\_REGULATION\_OF\_VESICLE\_MEDIATED\_TRANSPORT | MSigDB lists | GO\_REGULATION\_OF\_VESICLE\_MEDIATED\_TRANSPORT | 118 | 11 | 4161 | 156 | lt,Arf79F,N,Pdk1,Rab5,Syx1A,AP-1gamma,CG17883,Dl,Raf,unc-119 | | 4.322e-03 | -5.44 | B-H1 (BarH1) | protein interactions | 32724 | 15 | 4 | 9204 | 432 | Raf,Egfr,aop,N | | 4.322e-03 | -5.44 | upd3 (unpaired 3) | protein interactions | 3346149 | 15 | 4 | 9204 | 432 | N,Egfr,dome,Socs36E | | 4.322e-03 | -5.44 | twi (twist) | protein interactions | 37655 | 15 | 4 | 9204 | 432 | Scr,Mef2,abd-A,Abd-B | | 4.322e-03 | -5.44 | IKKepsilon (I-kappaB kinase epsilon) | protein interactions | 35329 | 15 | 4 | 9204 | 432 | f,Dronc,nuf,Diap1 | | 4.322e-03 | -5.44 | pr (purple) | protein interactions | 35292 | 15 | 4 | 9204 | 432 | Pez,drongo,Bacc,su(sable) | | 4.322e-03 | -5.44 | comm (commissureless) | protein interactions | 39717 | 15 | 4 | 9204 | 432 | Abl,robo1,ena,sli | | 4.322e-03 | -5.44 | tara (taranis) | protein interactions | 53562 | 15 | 4 | 9204 | 432 | pnr,ph-p,mam,tna | | 4.322e-03 | -5.44 | GSE41176\_UNSTIM\_VS\_ANTI\_IGM\_STIM\_BCELL\_1H\_UP | MSigDB lists | GSE41176\_UNSTIM\_VS\_ANTI\_IGM\_STIM\_BCELL\_1H\_UP | 42 | 6 | 4161 | 156 | cact,Actbeta,jar,Tpst,CG3961,Nmda1 | | 4.322e-03 | -5.44 | GSE17186\_BLOOD\_VS\_CORD\_BLOOD\_NAIVE\_BCELL\_DN | MSigDB lists | GSE17186\_BLOOD\_VS\_CORD\_BLOOD\_NAIVE\_BCELL\_DN | 42 | 6 | 4161 | 156 | Pgant6,alpha-Man-Ia,nmo,sina,Pgk,CG1673 | | 4.322e-03 | -5.44 | GSE8921\_3H\_VS\_24H\_TLR1\_2\_STIM\_MONOCYTE\_DN | MSigDB lists | GSE8921\_3H\_VS\_24H\_TLR1\_2\_STIM\_MONOCYTE\_DN | 42 | 6 | 4161 | 156 | Cdk4,cact,Psa,CG4935,step,CG13893 | | 4.361e-03 | -5.43 | GO\_NEGATIVE\_REGULATION\_OF\_MULTICELLULAR\_ORGANISMAL\_PROCESS | MSigDB lists | GO\_NEGATIVE\_REGULATION\_OF\_MULTICELLULAR\_ORGANISMAL\_PROCESS | 189 | 15 | 4161 | 156 | hyx,Cdk4,Mef2,robo1,CG11069,Tollo,Dl,cact,sli,Actbeta,N,trbl,CG9220,arm,SERCA | | 4.386e-03 | -5.43 | Synthesis of PIPs at the plasma membrane | REACTOME pathways | R-DME-1660499 | 25 | 5 | 3923 | 176 | CG9784,Ocrl,Rab5,Arf79F,CG34383 | | 4.437e-03 | -5.42 | GO\_NEURON\_DIFFERENTIATION | MSigDB lists | GO\_NEURON\_DIFFERENTIATION | 208 | 16 | 4161 | 156 | fz,sina,tkv,fz2,arm,Eph,ft,sli,Actbeta,N,Unc-76,ap,unc-5,robo1,Mef2,Dl | | 4.490e-03 | -5.41 | signaling receptor activator activity | molecular function | GO:0030546 | 111 | 10 | 12176 | 414 | Sema1a,jeb,vn,Pvf3,Dl,htl,spi,Sema1b,ITP,Actbeta | | 4.494e-03 | -5.40 | programmed cell death involved in cell development | biological process | GO:0010623 | 89 | 9 | 12345 | 442 | Dronc,S,Dp,arm,ec,Atg17,N,Atg16,Dl | | 4.498e-03 | -5.40 | cell division site | cellular component | GO:0032153 | 45 | 6 | 11940 | 417 | Cip4,Arf79F,step,nuf,cindr,Rab1 | | 4.506e-03 | -5.40 | GO\_SKELETAL\_SYSTEM\_DEVELOPMENT | MSigDB lists | GO\_SKELETAL\_SYSTEM\_DEVELOPMENT | 86 | 9 | 4161 | 156 | ft,ds,Pex7,frc,Mef2,arm,tkv,htl,CG9220 | | 4.510e-03 | -5.40 | SCAR (SCAR) | protein interactions | 34519 | 24 | 5 | 9204 | 432 | SCAR,ena,dock,arm,Cip4 | | 4.510e-03 | -5.40 | CG42709 (uncharacterized protein) | protein interactions | 39453 | 24 | 5 | 9204 | 432 | N,Acp63F,H,scyl,Lk6 | | 4.510e-03 | -5.40 | Dsp1 (Dorsal switch protein 1) | protein interactions | 117294 | 24 | 5 | 9204 | 432 | H,pnr,Scr,fl(2)d,Rm62 | | 4.510e-03 | -5.40 | Map60 (Microtubule-associated protein 60) | protein interactions | 35991 | 24 | 5 | 9204 | 432 | emp,jigr1,rictor,Pen,Sxl | | 4.526e-03 | -5.40 | mesoderm formation | biological process | GO:0001707 | 31 | 5 | 12345 | 442 | abd-A,jeb,htl,sli,mam | | 4.533e-03 | -5.40 | para (paralytic) | protein interactions | 32619 | 34 | 6 | 9204 | 432 | Rab5,LanA,tipE,pum,chn,esg | | 4.533e-03 | -5.40 | germ-line stem-cell niche homeostasis | biological process | GO:0060250 | 10 | 3 | 12345 | 442 | InR,N,tai | | 4.533e-03 | -5.40 | regulation of stem cell division | biological process | GO:2000035 | 10 | 3 | 12345 | 442 | Sxl,klu,N | | 4.533e-03 | -5.40 | peroxisomal transport | biological process | GO:0043574 | 10 | 3 | 12345 | 442 | Pex5,Pex10,Pex14 | | 4.533e-03 | -5.40 | foregut morphogenesis | biological process | GO:0007440 | 10 | 3 | 12345 | 442 | N,Inx2,bowl | | 4.533e-03 | -5.40 | neuronal stem cell population maintenance | biological process | GO:0097150 | 10 | 3 | 12345 | 442 | N,lilli,klu | | 4.533e-03 | -5.40 | regulation of axon extension involved in axon guidance | biological process | GO:0048841 | 10 | 3 | 12345 | 442 | Sema1a,chic,Sema1b | | 4.533e-03 | -5.40 | peroxisomal membrane transport | biological process | GO:0015919 | 10 | 3 | 12345 | 442 | Pex14,Pex10,Pex5 | | 4.534e-03 | -5.40 | activation of protein kinase activity | biological process | GO:0032147 | 44 | 6 | 12345 | 442 | wnd,hppy,jeb,Atg17,Pdk1,Raf | | 4.554e-03 | -5.39 | polytene chromosome chromocenter | cellular component | GO:0005701 | 20 | 4 | 11940 | 417 | jumu,His2Av,E(Pc),bab2 | | 4.556e-03 | -5.39 | Znf\_CXXC | interpro domains | IPR002857 | 3 | 2 | 11383 | 450 | Tet,Kdm2 | | 4.556e-03 | -5.39 | Tyr-kin\_ephrin\_A/B\_rcpt-like | interpro domains | IPR011641 | 3 | 2 | 11383 | 450 | LanA,Eph | | 4.556e-03 | -5.39 | Inosine/uridine\_hydrolase\_dom | interpro domains | IPR001910 | 3 | 2 | 11383 | 450 | CG11158,CG12177 | | 4.556e-03 | -5.39 | Ski\_DNA-bd\_sf | interpro domains | IPR037000 | 3 | 2 | 11383 | 450 | fuss,Snoo | | 4.556e-03 | -5.39 | Vitellinogen\_open\_b-sht | interpro domains | IPR015255 | 3 | 2 | 11383 | 450 | apolpp,cv-d | | 4.556e-03 | -5.39 | Ribo\_hydro-like | interpro domains | IPR036452 | 3 | 2 | 11383 | 450 | CG12177,CG11158 | | 4.556e-03 | -5.39 | SAND-like\_dom\_sf | interpro domains | IPR010919 | 3 | 2 | 11383 | 450 | Snoo,fuss | | 4.556e-03 | -5.39 | SKI/SNO/DAC | interpro domains | IPR003380 | 3 | 2 | 11383 | 450 | Snoo,fuss | | 4.592e-03 | -5.38 | GO\_CELL\_DEATH | MSigDB lists | GO\_CELL\_DEATH | 247 | 18 | 4161 | 156 | unc-5,Drak,14-3-3epsilon,N,Actbeta,htl,Mef2,robo1,cact,fz2,SERCA,Pdk1,sina,Drep4,CG45050,sli,Fem-1,Tollo | | 4.600e-03 | -5.38 | transition metal ion binding | molecular function | GO:0046914 | 720 | 38 | 12176 | 414 | Hr39,Cbl,Atox1,CG13481,Cyp6a16,CG31457,Tet,CG2991,Diap1,pnr,Su(z)2,CG4467,Psa,mld,grn,CG14441,Psc,mura,CG45050,Pex10,pita,lt,CG11360,Kdm2,dpy,Dyb,FBXO11,sina,CG10321,lin-28,ph-p,Hr4,CG1218,CG34401,hang,CG11247,tna,CG11414 | | 4.616e-03 | -5.38 | trx (trithorax) | protein interactions | 41737 | 45 | 7 | 9204 | 432 | pnr,Kdm2,Abd-B,stx,ph-p,fs(1)h,sd | | 4.627e-03 | -5.38 | transmembrane signaling receptor activity | molecular function | GO:0004888 | 353 | 22 | 12176 | 414 | htl,N,InR,Gr93c,drl,moody,Ddr,Eph,Nrk,unc-5,fz,nAChRalpha4,Egfr,Pvr,tkv,wgn,otk,fz3,fz2,Ir21a,dome,Ptp4E | | 4.630e-03 | -5.38 | GO\_EYE\_MORPHOGENESIS | MSigDB lists | GO\_EYE\_MORPHOGENESIS | 30 | 5 | 4161 | 156 | Eph,Dl,arm,CG17883,fz2 | | 4.630e-03 | -5.38 | KESHELAVA\_MULTIPLE\_DRUG\_RESISTANCE | MSigDB lists | KESHELAVA\_MULTIPLE\_DRUG\_RESISTANCE | 30 | 5 | 4161 | 156 | step,Ziz,shot,FBXO11,Pex10 | | 4.630e-03 | -5.38 | GSE24671\_CTRL\_VS\_SENDAI\_VIRUS\_INFECTED\_MOUSE\_SPLENOCYTES\_UP | MSigDB lists | GSE24671\_CTRL\_VS\_SENDAI\_VIRUS\_INFECTED\_MOUSE\_SPLENOCYTES\_UP | 30 | 5 | 4161 | 156 | InR,lt,Dl,CG45050,bru2 | | 4.657e-03 | -5.37 | Ski\_Sno | pfam domains | PF02437 | 3 | 2 | 10480 | 419 | fuss,Snoo | | 4.657e-03 | -5.37 | DUF1943 | pfam domains | PF09172 | 3 | 2 | 10480 | 419 | apolpp,cv-d | | 4.657e-03 | -5.37 | IU\_nuc\_hydro | pfam domains | PF01156 | 3 | 2 | 10480 | 419 | CG12177,CG11158 | | 4.662e-03 | -5.37 | VEGFR2 mediated vascular permeability | REACTOME pathways | R-DME-5218920 | 16 | 4 | 3923 | 176 | rictor,Pdk1,arm,trbl | | 4.742e-03 | -5.35 | GO\_POSITIVE\_REGULATION\_OF\_MUSCLE\_TISSUE\_DEVELOPMENT | MSigDB lists | GO\_POSITIVE\_REGULATION\_OF\_MUSCLE\_TISSUE\_DEVELOPMENT | 19 | 4 | 4161 | 156 | Mef2,N,Dl,arm | | 4.742e-03 | -5.35 | HALLMARK\_WNT\_BETA\_CATENIN\_SIGNALING | MSigDB lists | HALLMARK\_WNT\_BETA\_CATENIN\_SIGNALING | 19 | 4 | 4161 | 156 | arm,N,fz,Dl | | 4.742e-03 | -5.35 | VART\_KSHV\_INFECTION\_ANGIOGENIC\_MARKERS\_UP | MSigDB lists | VART\_KSHV\_INFECTION\_ANGIOGENIC\_MARKERS\_UP | 19 | 4 | 4161 | 156 | Eph,Dl,htl,LanA | | 4.742e-03 | -5.35 | GO\_REGULATION\_OF\_FIBROBLAST\_PROLIFERATION | MSigDB lists | GO\_REGULATION\_OF\_FIBROBLAST\_PROLIFERATION | 19 | 4 | 4161 | 156 | Ddr,arm,Cdk4,hyx | | 4.742e-03 | -5.35 | GO\_REGULATION\_OF\_LEUKOCYTE\_PROLIFERATION | MSigDB lists | GO\_REGULATION\_OF\_LEUKOCYTE\_PROLIFERATION | 19 | 4 | 4161 | 156 | Tsp96F,Mef2,dock,arm | | 4.742e-03 | -5.35 | MORF\_FRK | MSigDB lists | MORF\_FRK | 19 | 4 | 4161 | 156 | Drep4,lt,nAChRalpha4,Src42A | | 4.818e-03 | -5.34 | Ocrl (Oculocerebrorenal syndrome of Lowe) | protein interactions | 31157 | 8 | 3 | 9204 | 432 | Rab5,N,Rab1 | | 4.818e-03 | -5.34 | Vrp1 (Verprolin 1) | protein interactions | 37521 | 8 | 3 | 9204 | 432 | SCAR,dock,sns | | 4.818e-03 | -5.34 | Ndg (Nidogen/entactin) | protein interactions | 36089 | 8 | 3 | 9204 | 432 | LanA,N,Myo10A | | 4.818e-03 | -5.34 | unc-5 (unc-5) | protein interactions | 36703 | 8 | 3 | 9204 | 432 | sns,robo1,grn | | 4.818e-03 | -5.34 | CG12084 (uncharacterized protein) | protein interactions | 192509 | 8 | 3 | 9204 | 432 | pnr,H,N | | 4.818e-03 | -5.34 | CRMP (Collapsin Response Mediator Protein) | protein interactions | 40675 | 8 | 3 | 9204 | 432 | N,pnr,Dl | | 4.818e-03 | -5.34 | nesd (nessun dorma) | protein interactions | 35298 | 8 | 3 | 9204 | 432 | 14-3-3epsilon,14-3-3zeta,Raf | | 4.818e-03 | -5.34 | exp (expansion) | protein interactions | 36276 | 8 | 3 | 9204 | 432 | Egfr,Sxl,Raf | | 4.818e-03 | -5.34 | tutl (turtle) | protein interactions | 46015 | 8 | 3 | 9204 | 432 | Actbeta,H,ed | | 4.818e-03 | -5.34 | wap (wings apart) | protein interactions | 33131 | 8 | 3 | 9204 | 432 | ft,ds,C1GalTA | | 4.818e-03 | -5.34 | Madm (MLF1-adaptor molecule) | protein interactions | 40710 | 8 | 3 | 9204 | 432 | Egfr,InR,vn | | 4.818e-03 | -5.34 | NHP2 (NHP2) | protein interactions | 44005 | 8 | 3 | 9204 | 432 | tsh,rictor,Sxl | | 4.827e-03 | -5.33 | Nfs1 (Nfs1 cysteine desulfurase) | protein interactions | 34613 | 57 | 8 | 9204 | 432 | ds,SERCA,Gfat2,Pdk1,Rab5,Raf,Mpcp2,GluProRS | | 4.833e-03 | -5.33 | response to organic substance | biological process | GO:0010033 | 529 | 31 | 12345 | 442 | hppy,CG30156,Hr39,ph-p,S,InR,hang,hfw,Raf,mura,pum,pxb,Pvr,Bacc,moody,tai,dome,spi,pnr,Egfr,rictor,Dad,tkv,wgn,Gr93c,htl,REPTOR,Hr4,rdx,aop,Pdk1 | | 4.841e-03 | -5.33 | Pp2A-29B (Protein phosphatase 2A at 29B) | protein interactions | 2768940 | 82 | 10 | 9204 | 432 | CycG,Gfat2,Pdk1,SERCA,GluProRS,baz,Mpcp2,Kr-h2,REPTOR,wdb | | 4.871e-03 | -5.32 | GSE26928\_CENTR\_MEMORY\_VS\_CXCR5\_POS\_CD4\_TCELL\_DN | MSigDB lists | GSE26928\_CENTR\_MEMORY\_VS\_CXCR5\_POS\_CD4\_TCELL\_DN | 43 | 6 | 4161 | 156 | lt,Dl,Hel89B,Kdm2,CG2006,CG9485 | | 4.871e-03 | -5.32 | IKEDA\_MIR30\_TARGETS\_UP | MSigDB lists | IKEDA\_MIR30\_TARGETS\_UP | 43 | 6 | 4161 | 156 | eff,Gfat2,alpha-Man-Ia,bru2,Cdk4,14-3-3zeta | | 4.871e-03 | -5.32 | GSE33374\_CD8\_ALPHAALPHA\_VS\_ALPHABETA\_CD161\_HIGH\_TCELL\_UP | MSigDB lists | GSE33374\_CD8\_ALPHAALPHA\_VS\_ALPHABETA\_CD161\_HIGH\_TCELL\_UP | 43 | 6 | 4161 | 156 | CG6465,N,Raf,Lk6,Ddr,Tpc1 | | 4.871e-03 | -5.32 | GSE360\_L\_DONOVANI\_VS\_L\_MAJOR\_MAC\_DN | MSigDB lists | GSE360\_L\_DONOVANI\_VS\_L\_MAJOR\_MAC\_DN | 43 | 6 | 4161 | 156 | Pex5,step,AP-1gamma,frc,Dyrk2,cact | | 4.871e-03 | -5.32 | MASSARWEH\_TAMOXIFEN\_RESISTANCE\_DN | MSigDB lists | MASSARWEH\_TAMOXIFEN\_RESISTANCE\_DN | 43 | 6 | 4161 | 156 | CycG,Src42A,sd,InR,l(2)37Cg,fz | | 4.871e-03 | -5.32 | WAMUNYOKOLI\_OVARIAN\_CANCER\_LMP\_DN | MSigDB lists | WAMUNYOKOLI\_OVARIAN\_CANCER\_LMP\_DN | 43 | 6 | 4161 | 156 | Rgl,bru2,Cerk,Ddr,sli,wdb | | 4.894e-03 | -5.32 | positive regulation of response to external stimulus | biological process | GO:0032103 | 124 | 11 | 12345 | 442 | CG32795,pnr,fz,Pitslre,mop,sli,DPCoAC,imd,chic,htl,Dcr-2 | | 4.970e-03 | -5.30 | GSE17721\_CTRL\_VS\_POLYIC\_0.5H\_BMDC\_DN | MSigDB lists | GSE17721\_CTRL\_VS\_POLYIC\_0.5H\_BMDC\_DN | 57 | 7 | 4161 | 156 | CG17883,jar,Syx1A,rg,Dgat2,sina,betaTub60D | | 4.970e-03 | -5.30 | ACTGCCT\_MIR34B | MSigDB lists | ACTGCCT\_MIR34B | 57 | 7 | 4161 | 156 | N,Cdk4,14-3-3zeta,Dl,otk,Ocrl,bru2 | | 4.971e-03 | -5.30 | fly chr68B4-68B4\|3-35 | chromosome location | fly chr68B4-68B4\|3-35 | 5 | 2 | 25010 | 571 | lncRNA:CR44715,lncRNA:CR44713 | | 4.971e-03 | -5.30 | fly chr68B4-68B4\|3-35 cM | chromosome location | fly chr68B4-68B4\|3-35 cM | 5 | 2 | 25010 | 571 | lncRNA:CR44715,lncRNA:CR44713 | | 4.977e-03 | -5.30 | hemocyte migration | biological process | GO:0035099 | 20 | 4 | 12345 | 442 | Pvr,Pvf3,RhoL,shg | | 4.977e-03 | -5.30 | regulation of establishment of planar polarity | biological process | GO:0090175 | 20 | 4 | 12345 | 442 | ds,aop,otk,baz | | 5.077e-03 | -5.28 | neuroblast differentiation | biological process | GO:0014016 | 45 | 6 | 12345 | 442 | N,spen,arm,sqz,abd-A,klu | | 5.077e-03 | -5.28 | germ-line stem cell division | biological process | GO:0042078 | 45 | 6 | 12345 | 442 | tkv,pum,msi,InR,shg,eff | | 5.077e-03 | -5.28 | GSE26290\_WT\_VS\_PDK1\_KO\_ANTI\_CD3\_AND\_IL2\_STIM\_CD8\_TCELL\_DN | MSigDB lists | GSE26290\_WT\_VS\_PDK1\_KO\_ANTI\_CD3\_AND\_IL2\_STIM\_CD8\_TCELL\_DN | 72 | 8 | 4161 | 156 | DPCoAC,Rab5,CG18549,Fkbp14,Mef2,CG9485,Pdk1,CG2017 | | 5.108e-03 | -5.28 | WU\_HBX\_TARGETS\_2\_UP | MSigDB lists | WU\_HBX\_TARGETS\_2\_UP | 10 | 3 | 4161 | 156 | 14-3-3epsilon,Rab5,CycG | | 5.108e-03 | -5.28 | GO\_PROSTATE\_GLAND\_DEVELOPMENT | MSigDB lists | GO\_PROSTATE\_GLAND\_DEVELOPMENT | 10 | 3 | 4161 | 156 | arm,Fem-1,N | | 5.108e-03 | -5.28 | GO\_POSITIVE\_REGULATION\_OF\_REGULATED\_SECRETORY\_PATHWAY | MSigDB lists | GO\_POSITIVE\_REGULATION\_OF\_REGULATED\_SECRETORY\_PATHWAY | 10 | 3 | 4161 | 156 | Arf79F,AP-1gamma,Syx1A | | 5.108e-03 | -5.28 | GO\_EXCRETION | MSigDB lists | GO\_EXCRETION | 10 | 3 | 4161 | 156 | CG11069,sns,nAChRalpha4 | | 5.108e-03 | -5.28 | GO\_NEGATIVE\_REGULATION\_OF\_LEUKOCYTE\_DIFFERENTIATION | MSigDB lists | GO\_NEGATIVE\_REGULATION\_OF\_LEUKOCYTE\_DIFFERENTIATION | 10 | 3 | 4161 | 156 | arm,Actbeta,Tollo | | 5.108e-03 | -5.28 | MODULE\_284 | MSigDB lists | MODULE\_284 | 10 | 3 | 4161 | 156 | InR,frc,l(1)G0196 | | 5.108e-03 | -5.28 | GO\_VENTRICULAR\_SEPTUM\_DEVELOPMENT | MSigDB lists | GO\_VENTRICULAR\_SEPTUM\_DEVELOPMENT | 10 | 3 | 4161 | 156 | fz,N,otk | | 5.108e-03 | -5.28 | GO\_OLFACTORY\_LOBE\_DEVELOPMENT | MSigDB lists | GO\_OLFACTORY\_LOBE\_DEVELOPMENT | 10 | 3 | 4161 | 156 | sli,robo1,ap | | 5.175e-03 | -5.26 | ScsbetaG (Succinyl-coenzyme A synthetase beta subunit, GDP-forming) | protein interactions | 44001 | 96 | 11 | 9204 | 432 | Mob2,GluProRS,Kr-h2,Mpcp2,Arf79F,Gfat2,Pdk1,SERCA,Rab1,Pen,Rab5 | | 5.177e-03 | -5.26 | SH3-like\_dom\_sf | interpro domains | IPR036028 | 68 | 8 | 11383 | 450 | Cip4,cindr,Src42A,RhoGEF3,Abl,p130CAS,SKIP,dock | | 5.192e-03 | -5.26 | FN3 | smart domains | SM00060 | 48 | 8 | 5086 | 290 | dome,sns,InR,boi,Eph,Ptp4E,ed,robo1 | | 5.230e-03 | -5.25 | lola (longitudinals lacking) | protein interactions | 44548 | 46 | 7 | 9204 | 432 | kn,sli,H,mam,robo1,rib,N | | 5.236e-03 | -5.25 | phosphorus metabolic process | biological process | GO:0006793 | 920 | 48 | 12345 | 442 | Eph,wnd,otk,trbl,UQCR-11,pdgy,CycG,Ptp4E,CG9784,eya,Dyrk2,CycB3,Raf,CG8306,Nrk,mop,CG42327,Pdk1,Ddr,nmo,tkv,htl,Cks30A,Abl,wdb,IA-2,Ppcs,Lk6,Pp2C1,drl,Cdk4,Drak,Ptpmeg2,Pvr,Socs36E,Pgk,Pitslre,InR,hppy,l(1)G0196,CG6656,Pez,muc,Ocrl,ATPCL,Src42A,CG10417,Gfat2 | | 5.237e-03 | -5.25 | TP53 Regulates Metabolic Genes | REACTOME pathways | R-DME-5628897 | 26 | 5 | 3923 | 176 | SNF4Agamma,scyl,14-3-3epsilon,14-3-3zeta,chrb | | 5.252e-03 | -5.25 | glycoprotein-N-acetylgalactosamine 3-beta-galactosyltransferase activity | molecular function | GO:0016263 | 11 | 3 | 12176 | 414 | C1GalTA,CG34057,CG3119 | | 5.258e-03 | -5.25 | sax (saxophone) | protein interactions | 35731 | 35 | 6 | 9204 | 432 | fuss,tkv,Pvr,H,Dad,kek5 | | 5.280e-03 | -5.24 | Gp93 (Glycoprotein 93) | protein interactions | 43354 | 83 | 10 | 9204 | 432 | rictor,InR,Mpcp2,Kr-h2,Sema1a,GluProRS,Arf79F,SERCA,Gfat2,Rab5 | | 5.302e-03 | -5.24 | neuromuscular junction | cellular component | GO:0031594 | 77 | 8 | 11940 | 417 | Dg,scrib,dlp,pum,Cip4,cact,Pdk1,Magi | | 5.370e-03 | -5.23 | septate junction | cellular component | GO:0005918 | 33 | 5 | 11940 | 417 | moody,Gli,yrt,cora,scrib | | 5.375e-03 | -5.23 | Ssu72 (Ssu72 CTD phosphatase) | protein interactions | 32973 | 58 | 8 | 9204 | 432 | cora,LanA,CG34376,Gli,Rm62,IA-2,siz,Pp2C1 | | 5.399e-03 | -5.22 | regulation of actin polymerization or depolymerization | biological process | GO:0008064 | 60 | 7 | 12345 | 442 | SCAR,Cip4,CG6891,chic,Abl,ena,RhoU | | 5.399e-03 | -5.22 | lymph gland development | biological process | GO:0048542 | 60 | 7 | 12345 | 442 | htl,Pvr,CG1129,Pen,Raf,pnr,N | | 5.399e-03 | -5.22 | regulation of actin filament length | biological process | GO:0030832 | 60 | 7 | 12345 | 442 | chic,CG6891,Abl,ena,RhoU,SCAR,Cip4 | | 5.422e-03 | -5.22 | receptor regulator activity | molecular function | GO:0030545 | 114 | 10 | 12176 | 414 | Dl,Pvf3,htl,vn,jeb,Sema1a,Actbeta,ITP,Sema1b,spi | | 5.444e-03 | -5.21 | GO\_CELL\_CELL\_SIGNALING | MSigDB lists | GO\_CELL\_CELL\_SIGNALING | 139 | 12 | 4161 | 156 | Syx1A,Cadps,unc-119,Mef2,Dl,arm,Ric,Rab1,fz,Actbeta,htl,nAChRalpha4 | | 5.450e-03 | -5.21 | enzyme regulator activity | molecular function | GO:0030234 | 358 | 22 | 12176 | 414 | Ocrl,CG7394,CenB1A,Pdk1,wdb,rictor,Cks30A,Atg17,CG17883,CycG,trbl,Socs36E,RapGAP1,drongo,RhoGAP19D,TBC1D5,jeb,Diap1,Ppn,14-3-3zeta,CycB3,Spn42De | | 5.469e-03 | -5.21 | GCAAAAA\_MIR129 | MSigDB lists | GCAAAAA\_MIR129 | 44 | 6 | 4161 | 156 | Eph,Psa,Pgant5,sina,bru2,Kdm2 | | 5.469e-03 | -5.21 | GO\_PALLIUM\_DEVELOPMENT | MSigDB lists | GO\_PALLIUM\_DEVELOPMENT | 44 | 6 | 4161 | 156 | ap,Cdk4,14-3-3epsilon,ft,arm,sli | | 5.469e-03 | -5.21 | GSE41867\_DAY6\_EFFECTOR\_VS\_DAY30\_MEMORY\_CD8\_TCELL\_LCMV\_ARMSTRONG\_DN | MSigDB lists | GSE41867\_DAY6\_EFFECTOR\_VS\_DAY30\_MEMORY\_CD8\_TCELL\_LCMV\_ARMSTRONG\_DN | 44 | 6 | 4161 | 156 | Lk6,CG45050,wbl,Cerk,step,hppy | | 5.469e-03 | -5.21 | GSE17721\_CTRL\_VS\_POLYIC\_8H\_BMDC\_DN | MSigDB lists | GSE17721\_CTRL\_VS\_POLYIC\_8H\_BMDC\_DN | 44 | 6 | 4161 | 156 | trbl,N,CG2017,Rgl,step,lili | | 5.473e-03 | -5.21 | microbody membrane | cellular component | GO:0031903 | 21 | 4 | 11940 | 417 | CG33474,Pex14,Pex10,Pex5 | | 5.473e-03 | -5.21 | peroxisomal membrane | cellular component | GO:0005778 | 21 | 4 | 11940 | 417 | CG33474,Pex10,Pex5,Pex14 | | 5.473e-03 | -5.21 | peroxisomal part | cellular component | GO:0044439 | 21 | 4 | 11940 | 417 | CG33474,Pex14,Pex5,Pex10 | | 5.473e-03 | -5.21 | microbody part | cellular component | GO:0044438 | 21 | 4 | 11940 | 417 | CG33474,Pex10,Pex5,Pex14 | | 5.489e-03 | -5.21 | sgg (shaggy) | protein interactions | 31248 | 183 | 17 | 9204 | 432 | Dronc,ec,pnr,CG11122,fz,apolpp,nmo,shg,arm,crol,wnd,dpy,mam,kay,N,eyg,ft | | 5.552e-03 | -5.19 | vap (vacuolar peduncle) | protein interactions | 32569 | 16 | 4 | 9204 | 432 | htl,Rab5,Egfr,Pvr | | 5.552e-03 | -5.19 | Apc (APC-like) | protein interactions | 44642 | 16 | 4 | 9204 | 432 | CG13895,Src42A,N,arm | | 5.552e-03 | -5.19 | l(2)gd1 (lethal (2) giant discs 1) | protein interactions | 34543 | 16 | 4 | 9204 | 432 | tkv,Sema1a,ft,N | | 5.552e-03 | -5.19 | fmt (fiery mountain) | protein interactions | 38714 | 16 | 4 | 9204 | 432 | scrib,Pdk1,Cerk,Raf | | 5.552e-03 | -5.19 | nkd (naked cuticle) | protein interactions | 44843 | 16 | 4 | 9204 | 432 | ds,fz,shg,arm | | 5.552e-03 | -5.19 | alphaTub84D (alpha-Tubulin at 84D) | protein interactions | 40904 | 16 | 4 | 9204 | 432 | Myo10A,klar,imd,shot | | 5.552e-03 | -5.19 | Crk (Crk oncogene) | protein interactions | 43775 | 16 | 4 | 9204 | 432 | sns,Egfr,scrib,Pvr | | 5.562e-03 | -5.19 | anterior/posterior axis specification | biological process | GO:0009948 | 144 | 12 | 12345 | 442 | fs(1)h,baz,bowl,Dg,chic,pum,Pat1,cic,scrib,N,Raf,shg | | 5.566e-03 | -5.19 | CREIGHTON\_ENDOCRINE\_THERAPY\_RESISTANCE\_3 | MSigDB lists | CREIGHTON\_ENDOCRINE\_THERAPY\_RESISTANCE\_3 | 194 | 15 | 4161 | 156 | C1GalTA,Ziz,CG4645,Rgl,CG18549,Atg16,CG1673,fz2,dlp,Drak,CG2006,Tpst,Rab1,Pdk1,Pgant5 | | 5.594e-03 | -5.19 | regulation of cell morphogenesis involved in differentiation | biological process | GO:0010769 | 92 | 9 | 12345 | 442 | CG34401,shot,Sema1b,chic,Rab1,ss,Sema1a,fz,kn | | 5.632e-03 | -5.18 | FN3 | prosite domains | PS50853 | 53 | 8 | 6243 | 325 | InR,Ptp4E,Eph,boi,dome,robo1,ed,sns | | 5.653e-03 | -5.18 | protein dephosphorylation | biological process | GO:0006470 | 109 | 10 | 12345 | 442 | wdb,mop,IA-2,CG10417,eya,Ptpmeg2,Ptp4E,CG42327,Pez,Pp2C1 | | 5.679e-03 | -5.17 | dorsal/ventral axis specification | biological process | GO:0009950 | 76 | 8 | 12345 | 442 | Raf,Dp,Cbl,cic,N,cact,wbl,Egfr | | 5.685e-03 | -5.17 | ZnF\_C2H2 | smart domains | SM00355 | 271 | 26 | 5086 | 290 | klu,Prdm13,CG12769,sens,bowl,rgr,MEP-1,CG14441,chn,CG10348,zfh2,CG11414,esg,CG32772,sqz,disco-r,CG10321,CG11122,crol,CG11247,ttk,tsh,pita,hang,mld,sens-2 | | 5.757e-03 | -5.16 | GO\_POSITIVE\_REGULATION\_OF\_OSSIFICATION | MSigDB lists | GO\_POSITIVE\_REGULATION\_OF\_OSSIFICATION | 20 | 4 | 4161 | 156 | Mef2,tkv,Ddr,arm | | 5.757e-03 | -5.16 | GO\_T\_CELL\_DIFFERENTIATION | MSigDB lists | GO\_T\_CELL\_DIFFERENTIATION | 20 | 4 | 4161 | 156 | Raf,Cdk4,arm,fz2 | | 5.757e-03 | -5.16 | GO\_KIDNEY\_MORPHOGENESIS | MSigDB lists | GO\_KIDNEY\_MORPHOGENESIS | 20 | 4 | 4161 | 156 | LanA,ds,arm,ft | | 5.757e-03 | -5.16 | GO\_OVULATION\_CYCLE\_PROCESS | MSigDB lists | GO\_OVULATION\_CYCLE\_PROCESS | 20 | 4 | 4161 | 156 | Actbeta,robo1,tkv,sli | | 5.757e-03 | -5.16 | GSE42021\_TREG\_PLN\_VS\_TREG\_PRECURSORS\_THYMUS\_DN | MSigDB lists | GSE42021\_TREG\_PLN\_VS\_TREG\_PRECURSORS\_THYMUS\_DN | 20 | 4 | 4161 | 156 | emp,step,CG2017,Tollo | | 5.757e-03 | -5.16 | GENTILE\_UV\_LOW\_DOSE\_DN | MSigDB lists | GENTILE\_UV\_LOW\_DOSE\_DN | 20 | 4 | 4161 | 156 | Pex14,sli,Tpst,InR | | 5.806e-03 | -5.15 | response to drug | biological process | GO:0042493 | 240 | 17 | 12345 | 442 | Bacc,moody,Pex5,pum,pxb,S,alpha-Man-Ia,mura,InR,hang,hppy,esg,Pdk1,Egfr,Gr93c,spi,Tpc1 | | 5.828e-03 | -5.15 | Activation of AKT2 | REACTOME pathways | R-DME-165158 | 3 | 2 | 3923 | 176 | trbl,Pdk1 | | 5.828e-03 | -5.15 | GRB7 events in ERBB2 signaling | REACTOME pathways | R-DME-1306955 | 3 | 2 | 3923 | 176 | vn,Egfr | | 5.828e-03 | -5.15 | GP1b-IX-V activation signalling | REACTOME pathways | R-DME-430116 | 3 | 2 | 3923 | 176 | 14-3-3zeta,Raf | | 5.828e-03 | -5.15 | PI3K events in ERBB4 signaling | REACTOME pathways | R-DME-1250342 | 3 | 2 | 3923 | 176 | vn,Egfr | | 5.828e-03 | -5.15 | Interactions of neurexins and neuroligins at synapses | REACTOME pathways | R-DME-6794361 | 3 | 2 | 3923 | 176 | yrt,cora | | 5.828e-03 | -5.15 | Trafficking of dietary sterols | REACTOME pathways | R-DME-265473 | 3 | 2 | 3923 | 176 | CG11069,CG31121 | | 5.868e-03 | -5.14 | BLALOCK\_ALZHEIMERS\_DISEASE\_UP | MSigDB lists | BLALOCK\_ALZHEIMERS\_DISEASE\_UP | 417 | 26 | 4161 | 156 | CG11030,PI31,Ddr,Kdm2,jar,rdx,lili,FBXO11,His2Av,CG4896,Gfat2,CG5522,step,Pgant6,cact,CG6891,shot,CenB1A,dlp,dock,Pgant5,htl,InR,Unc-76,CtBP,l(2)37Cg | | 5.903e-03 | -5.13 | Grasp65 (Grasp65) | protein interactions | 40177 | 47 | 7 | 9204 | 432 | msi,mura,ed,Unc-76,Bacc,yrt,otp | | 5.904e-03 | -5.13 | GO\_RECEPTOR\_BINDING | MSigDB lists | GO\_RECEPTOR\_BINDING | 253 | 18 | 4161 | 156 | Tsp96F,Pdk1,fz,arm,Mfe2,Pex14,LanA,ReepA,Fem-1,Src42A,sli,dock,InR,N,Actbeta,CtBP,shot,Dl | | 5.916e-03 | -5.13 | striated muscle cell development | biological process | GO:0055002 | 61 | 7 | 12345 | 442 | Bsg,htl,Dg,kon,sd,aop,Mef2 | | 5.916e-03 | -5.13 | regulation of smoothened signaling pathway | biological process | GO:0008589 | 61 | 7 | 12345 | 442 | eff,rdx,dlp,Sxl,wdb,apolpp,hyx | | 5.916e-03 | -5.13 | cellular response to growth factor stimulus | biological process | GO:0071363 | 61 | 7 | 12345 | 442 | Dad,tkv,Pvr,htl,pnr,aop,Raf | | 5.939e-03 | -5.13 | TYR\_PHOSPHATASE\_dom | interpro domains | IPR000387 | 30 | 5 | 11383 | 450 | Pez,Ptp4E,IA-2,CG42327,Ptpmeg2 | | 5.971e-03 | -5.12 | wts (warts) | protein interactions | 43651 | 59 | 8 | 9204 | 432 | baz,jar,hppy,ft,d,Rab5,sd,SERCA | | 5.971e-03 | -5.12 | sina (seven in absentia) | protein interactions | 39884 | 59 | 8 | 9204 | 432 | Diap1,lilli,ed,ttk,eff,S,msi,sina | | 5.977e-03 | -5.12 | imaginal disc growth | biological process | GO:0007446 | 21 | 4 | 12345 | 442 | ft,Sxl,InR,N | | 5.977e-03 | -5.12 | calcium-dependent cell-cell adhesion via plasma membrane cell adhesion molecules | biological process | GO:0016339 | 21 | 4 | 12345 | 442 | Cals,shg,ds,ft | | 5.983e-03 | -5.12 | DEATH-like\_dom\_sf | interpro domains | IPR011029 | 10 | 3 | 11383 | 450 | imd,unc-5,Dronc | | 5.983e-03 | -5.12 | Paired\_dom | interpro domains | IPR001523 | 10 | 3 | 11383 | 450 | ey,eyg,Poxn | | 6.002e-03 | -5.12 | BOQUEST\_STEM\_CELL\_CULTURED\_VS\_FRESH\_UP | MSigDB lists | BOQUEST\_STEM\_CELL\_CULTURED\_VS\_FRESH\_UP | 74 | 8 | 4161 | 156 | cact,Tollo,CG3961,ATPCL,subdued,Pgk,Actbeta,bru2 | | 6.021e-03 | -5.11 | GO\_REGULATION\_OF\_INTRACELLULAR\_TRANSPORT | MSigDB lists | GO\_REGULATION\_OF\_INTRACELLULAR\_TRANSPORT | 177 | 14 | 4161 | 156 | Psa,Arf79F,Pdk1,fz,14-3-3zeta,14-3-3epsilon,GlcAT-I,fz2,Tollo,cact,Raf,Syx1A,ReepA,CG17883 | | 6.025e-03 | -5.11 | MODULE\_19 | MSigDB lists | MODULE\_19 | 59 | 7 | 4161 | 156 | Mef2,trbl,sli,CG9485,Mfe2,Ddr,Rab5 | | 6.025e-03 | -5.11 | GSE3337\_4H\_VS\_16H\_IFNG\_IN\_CD8POS\_DC\_DN | MSigDB lists | GSE3337\_4H\_VS\_16H\_IFNG\_IN\_CD8POS\_DC\_DN | 59 | 7 | 4161 | 156 | Mef2,CtBP,betaTub60D,lqf,Nmda1,wbl,hppy | | 6.055e-03 | -5.11 | meiotic cell cycle | biological process | GO:0051321 | 261 | 18 | 12345 | 442 | Arf79F,PI31,CycG,orb2,chic,Gli,bol,Hr39,Rab1,LanA,His3.3A,Rab5,His3.3B,Pex10,Sxl,eff,larp,Cks30A | | 6.068e-03 | -5.10 | acyl-CoA biosynthetic process | biological process | GO:0071616 | 11 | 3 | 12345 | 442 | ATPCL,pdgy,muc | | 6.068e-03 | -5.10 | O-glycan processing, core 1 | biological process | GO:0016267 | 11 | 3 | 12345 | 442 | C1GalTA,CG3119,CG34057 | | 6.068e-03 | -5.10 | thioester biosynthetic process | biological process | GO:0035384 | 11 | 3 | 12345 | 442 | pdgy,ATPCL,muc | | 6.068e-03 | -5.10 | specification of segmental identity, head | biological process | GO:0007380 | 11 | 3 | 12345 | 442 | kn,tsh,hth | | 6.068e-03 | -5.10 | negative regulation of stem cell differentiation | biological process | GO:2000737 | 11 | 3 | 12345 | 442 | C1GalTA,Abd-B,lqf | | 6.068e-03 | -5.10 | mesoderm migration involved in gastrulation | biological process | GO:0007509 | 11 | 3 | 12345 | 442 | jeb,htl,sli | | 6.068e-03 | -5.10 | positive regulation of cell-cell adhesion | biological process | GO:0022409 | 11 | 3 | 12345 | 442 | sli,RhoL,robo1 | | 6.104e-03 | -5.10 | Translocation of GLUT4 to the plasma membrane | REACTOME pathways | R-DME-1445148 | 9 | 3 | 3923 | 176 | 14-3-3zeta,14-3-3epsilon,CG4467 | | 6.104e-03 | -5.10 | CD28 co-stimulation | REACTOME pathways | R-DME-389356 | 9 | 3 | 3923 | 176 | Pdk1,rictor,trbl | | 6.119e-03 | -5.10 | GSE7768\_OVA\_ALONE\_VS\_OVA\_WITH\_MPL\_IMMUNIZED\_MOUSE\_WHOLE\_SPLEEN\_6H\_DN | MSigDB lists | GSE7768\_OVA\_ALONE\_VS\_OVA\_WITH\_MPL\_IMMUNIZED\_MOUSE\_WHOLE\_SPLEEN\_6H\_DN | 45 | 6 | 4161 | 156 | Tollo,Psa,Unc-76,Raf,CG2017,Tpst | | 6.119e-03 | -5.10 | GSE14769\_UNSTIM\_VS\_240MIN\_LPS\_BMDM\_DN | MSigDB lists | GSE14769\_UNSTIM\_VS\_240MIN\_LPS\_BMDM\_DN | 45 | 6 | 4161 | 156 | Unc-76,otk,dock,fz,Rgl,CG2006 | | 6.119e-03 | -5.10 | GO\_SKELETAL\_SYSTEM\_MORPHOGENESIS | MSigDB lists | GO\_SKELETAL\_SYSTEM\_MORPHOGENESIS | 45 | 6 | 4161 | 156 | Pex7,arm,CG9220,tkv,htl,Mef2 | | 6.119e-03 | -5.10 | GSE24142\_DN2\_VS\_DN3\_THYMOCYTE\_DN | MSigDB lists | GSE24142\_DN2\_VS\_DN3\_THYMOCYTE\_DN | 45 | 6 | 4161 | 156 | Tpst,frc,Lk6,trbl,N,Pdk1 | | 6.150e-03 | -5.09 | Ada2b (transcriptional Adaptor 2b) | protein interactions | 40966 | 112 | 12 | 9204 | 432 | dve,chn,Alh,Bacc,mam,kay,pnr,Mef2,zfh2,Dp,bab2,rib | | 6.160e-03 | -5.09 | GSE8921\_UNSTIM\_VS\_TLR1\_2\_STIM\_MONOCYTE\_6H\_DN | MSigDB lists | GSE8921\_UNSTIM\_VS\_TLR1\_2\_STIM\_MONOCYTE\_6H\_DN | 32 | 5 | 4161 | 156 | Tpc1,CG3961,muc,Pen,CG9220 | | 6.160e-03 | -5.09 | KEGG\_AXON\_GUIDANCE | MSigDB lists | KEGG\_AXON\_GUIDANCE | 32 | 5 | 4161 | 156 | unc-5,sli,robo1,Eph,dock | | 6.160e-03 | -5.09 | GSE24142\_EARLY\_THYMIC\_PROGENITOR\_VS\_DN2\_THYMOCYTE\_ADULT\_UP | MSigDB lists | GSE24142\_EARLY\_THYMIC\_PROGENITOR\_VS\_DN2\_THYMOCYTE\_ADULT\_UP | 32 | 5 | 4161 | 156 | rdx,CtBP,Rim2,Mef2,Tsp96F | | 6.160e-03 | -5.09 | GSE34156\_UNTREATED\_VS\_24H\_NOD2\_LIGAND\_TREATED\_MONOCYTE\_UP | MSigDB lists | GSE34156\_UNTREATED\_VS\_24H\_NOD2\_LIGAND\_TREATED\_MONOCYTE\_UP | 32 | 5 | 4161 | 156 | Drak,muc,Dyb,Vps13,Nrk | | 6.160e-03 | -5.09 | KEGG\_NEUROTROPHIN\_SIGNALING\_PATHWAY | MSigDB lists | KEGG\_NEUROTROPHIN\_SIGNALING\_PATHWAY | 32 | 5 | 4161 | 156 | 14-3-3epsilon,cact,14-3-3zeta,Pdk1,Raf | | 6.160e-03 | -5.09 | GSE43863\_LY6C\_INT\_CXCR5POS\_VS\_LY6C\_LOW\_CXCR5NEG\_EFFECTOR\_CD4\_TCELL\_UP | MSigDB lists | GSE43863\_LY6C\_INT\_CXCR5POS\_VS\_LY6C\_LOW\_CXCR5NEG\_EFFECTOR\_CD4\_TCELL\_UP | 32 | 5 | 4161 | 156 | Cdk4,Kdm2,Drak,Cerk,step | | 6.160e-03 | -5.09 | GSE16266\_CTRL\_VS\_LPS\_STIM\_MEF\_UP | MSigDB lists | GSE16266\_CTRL\_VS\_LPS\_STIM\_MEF\_UP | 32 | 5 | 4161 | 156 | Ids,Cerk,Tpc1,N,cact | | 6.162e-03 | -5.09 | small molecule binding | molecular function | GO:0036094 | 1078 | 52 | 12176 | 414 | otk,CG2017,SNF4Agamma,Pvr,wnd,Drak,Pdk1,Pitslre,SMC3,Abl,Nrk,CG5853,d,Rab1,hppy,Lk6,Raf,jar,trbl,tkv,SERCA,CG7536,Dcr-2,Rab5,GluProRS,Arf79F,CtBP,Egfr,RhoU,CG31121,CG11069,Cdk4,Pgk,CG17646,drl,Eph,Ddr,betaTub60D,Hel89B,htl,InR,ATPCL,Ric,Rm62,l(1)G0196,Myo10A,apolpp,CG7560,nmo,Dyrk2,sick,Src42A | | 6.172e-03 | -5.09 | FERM\_C | pfam domains | PF09380 | 10 | 3 | 10480 | 419 | Pez,yrt,cora | | 6.172e-03 | -5.09 | PAX | pfam domains | PF00292 | 10 | 3 | 10480 | 419 | ey,Poxn,eyg | | 6.198e-03 | -5.08 | Negative regulation of the PI3K/AKT network | REACTOME pathways | R-DME-199418 | 27 | 5 | 3923 | 176 | wdb,trbl,Egfr,vn,htl | | 6.198e-03 | -5.08 | Heparan sulfate/heparin (HS-GAG) metabolism | REACTOME pathways | R-DME-1638091 | 27 | 5 | 3923 | 176 | dlp,frc,kon,Ids,GlcAT-I | | 6.209e-03 | -5.08 | RNA polymerase II-specific DNA-binding transcription factor binding | molecular function | GO:0061629 | 35 | 5 | 12176 | 414 | cact,pnr,arm,lilli,tai | | 6.271e-03 | -5.07 | cellular localization | biological process | GO:0051641 | 1000 | 51 | 12345 | 442 | Ppcs,Mid1,scrib,chic,Pen,edl,Bsg,koi,Npc2g,S,Vps13,lt,cora,ena,Unc-76,klar,p130CAS,Pex14,CG7394,Pat1,Gli,CG17883,Pex5,nuf,lqf,Dg,Arf79F,Rim2,d,ds,SERCA,Rab1,hth,Myo10A,mop,Syx1A,TBC1D5,Rab5,jar,baz,rictor,Cadps,AP-1gamma,larp,arm,Cip4,His2Av,Abl,Shrm,ft,Pex10 | | 6.290e-03 | -5.07 | TGAATGT\_MIR181A\_MIR181B\_MIR181C\_MIR181D | MSigDB lists | TGAATGT\_MIR181A\_MIR181B\_MIR181C\_MIR181D | 124 | 11 | 4161 | 156 | AP-1gamma,robo1,hyx,l(1)G0196,lin-28,CenB1A,nmo,CG45050,subdued,Psa,FBXO11 | | 6.310e-03 | -5.07 | ubiquitin ligase complex | cellular component | GO:0000151 | 150 | 12 | 11940 | 417 | Psc,CG34401,FBXO11,eff,sina,ph-p,Su(z)2,Kdm2,Cks30A,CG12402,morgue,corto | | 6.319e-03 | -5.06 | regulation of organelle organization | biological process | GO:0033043 | 408 | 25 | 12345 | 442 | crol,Ocrl,step,rictor,ena,jar,Jupiter,Cip4,lilli,14-3-3epsilon,mrt,Abl,Su(z)2,RhoGEF4,SCAR,Kdm2,shot,RhoU,sens-2,chic,CG6891,trbl,lt,fz,ed | | 6.326e-03 | -5.06 | GO\_POSITIVE\_REGULATION\_OF\_DEVELOPMENTAL\_PROCESS | MSigDB lists | GO\_POSITIVE\_REGULATION\_OF\_DEVELOPMENTAL\_PROCESS | 178 | 14 | 4161 | 156 | Raf,robo1,Mef2,Cals,Dl,Eph,otk,Ddr,sli,N,Actbeta,Arf79F,tkv,arm | | 6.389e-03 | -5.05 | bonsai (bonsai) | protein interactions | 37587 | 3 | 2 | 9204 | 432 | rictor,Dp | | 6.389e-03 | -5.05 | GPHR (Golgi pH Regulator) | protein interactions | 36682 | 3 | 2 | 9204 | 432 | mam,N | | 6.389e-03 | -5.05 | CG5599 (uncharacterized protein) | protein interactions | 32441 | 3 | 2 | 9204 | 432 | Dyb,d | | 6.389e-03 | -5.05 | ckd (cracked) | protein interactions | 38402 | 3 | 2 | 9204 | 432 | H,pnr | | 6.389e-03 | -5.05 | kud (kuduk) | protein interactions | 39882 | 3 | 2 | 9204 | 432 | koi,klar | | 6.389e-03 | -5.05 | CalpA (Calpain-A) | protein interactions | 37232 | 3 | 2 | 9204 | 432 | Atg16,cact | | 6.389e-03 | -5.05 | Eogt (EGF-domain O-GlcNAc transferase) | protein interactions | 33424 | 3 | 2 | 9204 | 432 | N,dpy | | 6.389e-03 | -5.05 | spz6 (spatzle 6) | protein interactions | 37956 | 3 | 2 | 9204 | 432 | CG32772,Sxl | | 6.389e-03 | -5.05 | CG6567 (uncharacterized protein) | protein interactions | 41304 | 3 | 2 | 9204 | 432 | Myo10A,Fem-1 | | 6.389e-03 | -5.05 | app (approximated) | protein interactions | 39399 | 3 | 2 | 9204 | 432 | d,ft | | 6.389e-03 | -5.05 | lbe (ladybird early) | protein interactions | 42542 | 3 | 2 | 9204 | 432 | abd-A,Abd-B | | 6.389e-03 | -5.05 | Non2 (Novel nucleolar protein 2) | protein interactions | 38341 | 3 | 2 | 9204 | 432 | rictor,mam | | 6.389e-03 | -5.05 | Inx3 (Innexin 3) | protein interactions | 44266 | 3 | 2 | 9204 | 432 | shg,Inx2 | | 6.389e-03 | -5.05 | Gmd (GDP-mannose 4,6-dehydratase) | protein interactions | 33716 | 3 | 2 | 9204 | 432 | N,Syx1A | | 6.389e-03 | -5.05 | Mo25 (Mo25) | protein interactions | 39858 | 3 | 2 | 9204 | 432 | N,Dl | | 6.389e-03 | -5.05 | CG14742 (uncharacterized protein) | protein interactions | 12797890 | 3 | 2 | 9204 | 432 | jumu,Poxn | | 6.418e-03 | -5.05 | negative regulation of intracellular signal transduction | biological process | GO:1902532 | 111 | 10 | 12345 | 442 | ds,d,RapGAP1,wdb,Cbl,LRR,N,CG42684,Lk6,trbl | | 6.450e-03 | -5.04 | defense response to other organism | biological process | GO:0098542 | 345 | 22 | 12345 | 442 | Dronc,kn,eya,Eph,Pvr,DPCoAC,imd,RhoL,N,dome,CG32795,shg,RhoGEF3,Src42A,Tollo,jumu,sick,LanA,htl,Dcr-2,subdued,spen | | 6.462e-03 | -5.04 | kibra (kibra) | protein interactions | 41783 | 26 | 5 | 9204 | 432 | ed,pnr,H,Pez,Diap1 | | 6.462e-03 | -5.04 | Pdhb (Pyruvate dehydrogenase E1 beta subunit) | protein interactions | 43437 | 26 | 5 | 9204 | 432 | pes,step,Kr-h2,fl(2)d,jigr1 | | 6.462e-03 | -5.04 | Lis-1 (Lissencephaly-1) | protein interactions | 36791 | 26 | 5 | 9204 | 432 | Egfr,Sema1b,h,shg,tkv | | 6.469e-03 | -5.04 | regulation of cell shape | biological process | GO:0008360 | 62 | 7 | 12345 | 442 | ttk,SCAR,rib,neo,htl,ed,Abl | | 6.469e-03 | -5.04 | response to growth factor | biological process | GO:0070848 | 62 | 7 | 12345 | 442 | Pvr,Dad,tkv,htl,aop,Raf,pnr | | 6.571e-03 | -5.03 | BAND41 | prints domains | PR00935 | 9 | 3 | 2185 | 101 | cora,yrt,Pez | | 6.612e-03 | -5.02 | chemorepellent activity | molecular function | GO:0045499 | 4 | 2 | 12176 | 414 | Sema1a,Sema1b | | 6.612e-03 | -5.02 | hedgehog family protein binding | molecular function | GO:0097108 | 4 | 2 | 12176 | 414 | dlp,boi | | 6.615e-03 | -5.02 | 14-3-3epsilon (14-3-3epsilon) | protein interactions | 42186 | 60 | 8 | 9204 | 432 | 14-3-3zeta,nuf,Nlp,REPTOR,rictor,Raf,baz,14-3-3epsilon | | 6.639e-03 | -5.01 | fu (fused) | protein interactions | 32855 | 48 | 7 | 9204 | 432 | Cip4,Gli,tkv,Sxl,kn,l(1)G0196,Dad | | 6.788e-03 | -4.99 | zip (zipper) | protein interactions | 38001 | 86 | 10 | 9204 | 432 | Shrm,Sema1a,baz,nmo,S,step,fz,Drak,rib,chic | | 6.788e-03 | -4.99 | CCT6 (Chaperonin containing TCP1 subunit 6) | protein interactions | 32518 | 86 | 10 | 9204 | 432 | GluProRS,Kr-h2,Mpcp2,Psa,Rab5,Pdk1,Gfat2,SERCA,Arf79F,imd | | 6.790e-03 | -4.99 | mesoderm morphogenesis | biological process | GO:0048332 | 34 | 5 | 12345 | 442 | mam,sli,htl,abd-A,jeb | | 6.824e-03 | -4.99 | OCT1\_03 | MSigDB lists | OCT1\_03 | 46 | 6 | 4161 | 156 | AP-1gamma,Mef2,cact,FBXO11,CG3149,CG2812 | | 6.824e-03 | -4.99 | GSE17721\_CTRL\_VS\_LPS\_2H\_BMDC\_DN | MSigDB lists | GSE17721\_CTRL\_VS\_LPS\_2H\_BMDC\_DN | 46 | 6 | 4161 | 156 | cact,Dyb,betaTub60D,Rgl,Pat1,fz2 | | 6.824e-03 | -4.99 | GSE3691\_CONVENTIONAL\_VS\_PLASMACYTOID\_DC\_SPLEEN\_UP | MSigDB lists | GSE3691\_CONVENTIONAL\_VS\_PLASMACYTOID\_DC\_SPLEEN\_UP | 46 | 6 | 4161 | 156 | trbl,Unc-76,fz,Tpc1,jar,nmo | | 6.824e-03 | -4.99 | GO\_PROTEINACEOUS\_EXTRACELLULAR\_MATRIX | MSigDB lists | GO\_PROTEINACEOUS\_EXTRACELLULAR\_MATRIX | 46 | 6 | 4161 | 156 | Ppn,SMC3,shot,LanA,sli,dlp | | 6.824e-03 | -4.99 | GSE20366\_EX\_VIVO\_VS\_HOMEOSTATIC\_CONVERSION\_NAIVE\_CD4\_TCELL\_DN | MSigDB lists | GSE20366\_EX\_VIVO\_VS\_HOMEOSTATIC\_CONVERSION\_NAIVE\_CD4\_TCELL\_DN | 46 | 6 | 4161 | 156 | Tollo,ap,CG1673,Pen,Tsp96F,l(2)37Cg | | 6.824e-03 | -4.99 | RUTELLA\_RESPONSE\_TO\_HGF\_VS\_CSF2RB\_AND\_IL4\_DN | MSigDB lists | RUTELLA\_RESPONSE\_TO\_HGF\_VS\_CSF2RB\_AND\_IL4\_DN | 46 | 6 | 4161 | 156 | Drak,bru2,step,fz2,lili,Actbeta | | 6.824e-03 | -4.99 | GSE39556\_CD8A\_DC\_VS\_NK\_CELL\_DN | MSigDB lists | GSE39556\_CD8A\_DC\_VS\_NK\_CELL\_DN | 46 | 6 | 4161 | 156 | step,CG4645,CG5522,LanA,CG9485,trbl | | 6.832e-03 | -4.99 | GO\_WNT\_SIGNALING\_PATHWAY\_CALCIUM\_MODULATING\_PATHWAY | MSigDB lists | GO\_WNT\_SIGNALING\_PATHWAY\_CALCIUM\_MODULATING\_PATHWAY | 11 | 3 | 4161 | 156 | arm,fz2,nmo | | 6.832e-03 | -4.99 | GO\_REGULATION\_OF\_ENDOPLASMIC\_RETICULUM\_STRESS\_INDUCED\_INTRINSIC\_APOPTOTIC\_SIGNALING\_PATHWAY | MSigDB lists | GO\_REGULATION\_OF\_ENDOPLASMIC\_RETICULUM\_STRESS\_INDUCED\_INTRINSIC\_APOPTOTIC\_SIGNALING\_PATHWAY | 11 | 3 | 4161 | 156 | wbl,Nmda1,dock | | 6.832e-03 | -4.99 | GARCIA\_TARGETS\_OF\_FLI1\_AND\_DAX1\_UP | MSigDB lists | GARCIA\_TARGETS\_OF\_FLI1\_AND\_DAX1\_UP | 11 | 3 | 4161 | 156 | jar,lili,Fkbp14 | | 6.832e-03 | -4.99 | GO\_CELL\_SUBSTRATE\_JUNCTION\_ASSEMBLY | MSigDB lists | GO\_CELL\_SUBSTRATE\_JUNCTION\_ASSEMBLY | 11 | 3 | 4161 | 156 | shot,LanA,Pdk1 | | 6.832e-03 | -4.99 | GO\_REGULATION\_OF\_B\_CELL\_ACTIVATION | MSigDB lists | GO\_REGULATION\_OF\_B\_CELL\_ACTIVATION | 11 | 3 | 4161 | 156 | Actbeta,Mef2,Tsp96F | | 6.832e-03 | -4.99 | GO\_O\_GLYCAN\_PROCESSING | MSigDB lists | GO\_O\_GLYCAN\_PROCESSING | 11 | 3 | 4161 | 156 | Pgant5,C1GalTA,Pgant6 | | 6.832e-03 | -4.99 | BASSO\_CD40\_SIGNALING\_UP | MSigDB lists | BASSO\_CD40\_SIGNALING\_UP | 11 | 3 | 4161 | 156 | unc-119,Mef2,cact | | 6.832e-03 | -4.99 | BIOCARTA\_WNT\_PATHWAY | MSigDB lists | BIOCARTA\_WNT\_PATHWAY | 11 | 3 | 4161 | 156 | fz,nmo,arm | | 6.832e-03 | -4.99 | ST\_PHOSPHOINOSITIDE\_3\_KINASE\_PATHWAY | MSigDB lists | ST\_PHOSPHOINOSITIDE\_3\_KINASE\_PATHWAY | 11 | 3 | 4161 | 156 | 14-3-3epsilon,14-3-3zeta,Pdk1 | | 6.832e-03 | -4.99 | REACTOME\_NEGATIVE\_REGULATION\_OF\_FGFR\_SIGNALING | MSigDB lists | REACTOME\_NEGATIVE\_REGULATION\_OF\_FGFR\_SIGNALING | 11 | 3 | 4161 | 156 | Lk6,htl,Raf | | 6.832e-03 | -4.99 | GO\_POSITIVE\_REGULATION\_OF\_PEPTIDYL\_TYROSINE\_PHOSPHORYLATION | MSigDB lists | GO\_POSITIVE\_REGULATION\_OF\_PEPTIDYL\_TYROSINE\_PHOSPHORYLATION | 11 | 3 | 4161 | 156 | unc-119,Tsp96F,htl | | 6.832e-03 | -4.99 | REACTOME\_FATTY\_ACYL\_COA\_BIOSYNTHESIS | MSigDB lists | REACTOME\_FATTY\_ACYL\_COA\_BIOSYNTHESIS | 11 | 3 | 4161 | 156 | ATPCL,CG3961,CG31522 | | 6.832e-03 | -4.99 | GO\_REGULATION\_OF\_MYOBLAST\_DIFFERENTIATION | MSigDB lists | GO\_REGULATION\_OF\_MYOBLAST\_DIFFERENTIATION | 11 | 3 | 4161 | 156 | N,Mef2,Dl | | 6.832e-03 | -4.99 | GO\_CELL\_FATE\_DETERMINATION | MSigDB lists | GO\_CELL\_FATE\_DETERMINATION | 11 | 3 | 4161 | 156 | Mef2,Dl,arm | | 6.832e-03 | -4.99 | GO\_POSITIVE\_REGULATION\_OF\_LEUKOCYTE\_PROLIFERATION | MSigDB lists | GO\_POSITIVE\_REGULATION\_OF\_LEUKOCYTE\_PROLIFERATION | 11 | 3 | 4161 | 156 | dock,Tsp96F,Mef2 | | 6.832e-03 | -4.99 | SCHAEFFER\_SOX9\_TARGETS\_IN\_PROSTATE\_DEVELOPMENT\_DN | MSigDB lists | SCHAEFFER\_SOX9\_TARGETS\_IN\_PROSTATE\_DEVELOPMENT\_DN | 11 | 3 | 4161 | 156 | sli,14-3-3zeta,Actbeta | | 6.832e-03 | -4.99 | GO\_ANATOMICAL\_STRUCTURE\_MATURATION | MSigDB lists | GO\_ANATOMICAL\_STRUCTURE\_MATURATION | 11 | 3 | 4161 | 156 | ft,ds,htl | | 6.877e-03 | -4.98 | Signalling by NGF | REACTOME pathways | R-DME-166520 | 182 | 16 | 3923 | 176 | Egfr,vn,CG4935,htl,wdb,Dronc,CG42684,cact,Raf,Rgl,Pvr,PI31,Pdk1,rictor,14-3-3zeta,trbl | | 6.890e-03 | -4.98 | male meiotic nuclear division | biological process | GO:0007140 | 95 | 9 | 12345 | 442 | His3.3A,orb2,PI31,Arf79F,His3.3B,Pex10,eff,larp,Rab1 | | 6.908e-03 | -4.98 | GO\_OVULATION\_CYCLE | MSigDB lists | GO\_OVULATION\_CYCLE | 21 | 4 | 4161 | 156 | Actbeta,robo1,tkv,sli | | 6.908e-03 | -4.98 | GO\_AXIS\_SPECIFICATION | MSigDB lists | GO\_AXIS\_SPECIFICATION | 21 | 4 | 4161 | 156 | fz2,arm,Dl,N | | 6.908e-03 | -4.98 | GO\_CANONICAL\_WNT\_SIGNALING\_PATHWAY | MSigDB lists | GO\_CANONICAL\_WNT\_SIGNALING\_PATHWAY | 21 | 4 | 4161 | 156 | fz,otk,fz2,arm | | 6.908e-03 | -4.98 | GO\_CARDIAC\_VENTRICLE\_DEVELOPMENT | MSigDB lists | GO\_CARDIAC\_VENTRICLE\_DEVELOPMENT | 21 | 4 | 4161 | 156 | otk,N,Mef2,fz | | 6.908e-03 | -4.98 | GSE20727\_ROS\_INH\_VS\_ROS\_INH\_AND\_DNFB\_ALLERGEN\_TREATED\_DC\_DN | MSigDB lists | GSE20727\_ROS\_INH\_VS\_ROS\_INH\_AND\_DNFB\_ALLERGEN\_TREATED\_DC\_DN | 21 | 4 | 4161 | 156 | nAChRalpha4,Tollo,fz,sli | | 6.913e-03 | -4.97 | - | gene3d domains | 3.10.390.10 | 3 | 2 | 4929 | 241 | Snoo,fuss | | 6.913e-03 | -4.97 | - | gene3d domains | 3.10.260.20 | 3 | 2 | 4929 | 241 | fuss,Snoo | | 6.913e-03 | -4.97 | - | gene3d domains | 3.90.245.10 | 3 | 2 | 4929 | 241 | CG11158,CG12177 | | 6.954e-03 | -4.97 | GO\_INTRACELLULAR\_PROTEIN\_TRANSPORT | MSigDB lists | GO\_INTRACELLULAR\_PROTEIN\_TRANSPORT | 359 | 23 | 4161 | 156 | cact,Pex5,Pex7,Rab5,14-3-3zeta,Vps13,Pen,lt,14-3-3epsilon,Rab1,emp,ReepA,AP-1gamma,Syx1A,CG45050,CG7394,Pex10,Pex14,arm,jar,Ric,Pat1,wbl | | 6.967e-03 | -4.97 | spot adherens junction | cellular component | GO:0005914 | 4 | 2 | 11940 | 417 | baz,shg | | 6.967e-03 | -4.97 | catenin complex | cellular component | GO:0016342 | 4 | 2 | 11940 | 417 | arm,shg | | 6.974e-03 | -4.97 | GO\_RESPONSE\_TO\_BIOTIC\_STIMULUS | MSigDB lists | GO\_RESPONSE\_TO\_BIOTIC\_STIMULUS | 180 | 14 | 4161 | 156 | Tpc1,ReepA,CG4645,hyx,Mef2,CG6891,Cdk4,cact,Tollo,fz2,SmD1,Rab1,N,dock | | 6.978e-03 | -4.97 | Hsp60D (Heat shock protein 60D) | protein interactions | 34763 | 9 | 3 | 9204 | 432 | Dronc,Diap1,trbl | | 6.978e-03 | -4.97 | cas (castor) | protein interactions | 44018 | 9 | 3 | 9204 | 432 | Dcr-2,abd-A,drl | | 6.978e-03 | -4.97 | lgs (legless) | protein interactions | 43791 | 9 | 3 | 9204 | 432 | hyx,arm,imd | | 6.978e-03 | -4.97 | E(spl)m4-BFM (Enhancer of split m4, Bearded family member) | protein interactions | 43157 | 9 | 3 | 9204 | 432 | N,H,Bacc | | 6.978e-03 | -4.97 | Thiolase (thiolase) | protein interactions | 37784 | 9 | 3 | 9204 | 432 | ft,morgue,ds | | 6.978e-03 | -4.97 | krimp (krimper) | protein interactions | 36804 | 9 | 3 | 9204 | 432 | rdx,Pat1,CG4896 | | 6.978e-03 | -4.97 | scny (scrawny) | protein interactions | 38648 | 9 | 3 | 9204 | 432 | Diap1,imd,Dronc | | 6.978e-03 | -4.97 | Uvrag (UV-resistance associated gene) | protein interactions | 34735 | 9 | 3 | 9204 | 432 | Rab5,N,mam | | 6.978e-03 | -4.97 | mirr (mirror) | protein interactions | 39441 | 9 | 3 | 9204 | 432 | cic,Dl,ttk | | 6.978e-03 | -4.97 | Ythdc1 (YTH domain containing 1) | protein interactions | 38420 | 9 | 3 | 9204 | 432 | fl(2)d,d,Sxl | | 6.978e-03 | -4.97 | Strica (Ser/Thr-rich caspase) | protein interactions | 35528 | 9 | 3 | 9204 | 432 | Dronc,Pdk1,Diap1 | | 6.978e-03 | -4.97 | O-fut1 (O-fucosyltransferase 1) | protein interactions | 36564 | 9 | 3 | 9204 | 432 | N,Dl,CG9784 | | 6.978e-03 | -4.97 | sgl (sugarless) | protein interactions | 38760 | 9 | 3 | 9204 | 432 | nmo,htl,Sema1a | | 6.993e-03 | -4.96 | regulation of lipid localization | biological process | GO:1905952 | 48 | 6 | 12345 | 442 | SERCA,lilli,InR,wdb,trbl,Arf79F | | 6.995e-03 | -4.96 | HnRNP-K (Heterogeneous nuclear ribonucleoprotein K) | protein interactions | 43862 | 17 | 4 | 9204 | 432 | N,ps,Diap1,Pdk1 | | 6.995e-03 | -4.96 | Su(Tpl) (Su(Tpl)) | protein interactions | 40171 | 17 | 4 | 9204 | 432 | sina,N,aop,lilli | | 6.995e-03 | -4.96 | Pak3 (Pak3) | protein interactions | 41995 | 17 | 4 | 9204 | 432 | scrib,SCAR,rictor,kn | | 6.995e-03 | -4.96 | sdt (stardust) | protein interactions | 44861 | 17 | 4 | 9204 | 432 | baz,shg,arm,scrib | | 7.032e-03 | -4.96 | TTGCACT\_MIR130A\_MIR301\_MIR130B | MSigDB lists | TTGCACT\_MIR130A\_MIR301\_MIR130B | 92 | 9 | 4161 | 156 | CG2812,eff,Kdm2,robo1,Atg16,rg,Dl,Hel89B,AP-1gamma | | 7.045e-03 | -4.96 | GO\_MALE\_SEX\_DIFFERENTIATION | MSigDB lists | GO\_MALE\_SEX\_DIFFERENTIATION | 33 | 5 | 4161 | 156 | Tollo,Actbeta,arm,CG17883,Mfe2 | | 7.045e-03 | -4.96 | GO\_POSITIVE\_REGULATION\_OF\_CELL\_MORPHOGENESIS\_INVOLVED\_IN\_DIFFERENTIATION | MSigDB lists | GO\_POSITIVE\_REGULATION\_OF\_CELL\_MORPHOGENESIS\_INVOLVED\_IN\_DIFFERENTIATION | 33 | 5 | 4161 | 156 | arm,sli,Raf,N,robo1 | | 7.063e-03 | -4.95 | organelle fission | biological process | GO:0048285 | 306 | 20 | 12345 | 442 | CG33474,orb2,CycG,Arf79F,PI31,corto,Gli,Hr39,ph-p,Rab1,His3.3A,LanA,His3.3B,Rab5,Sxl,Pex10,larp,eff,Cks30A,Slbp | | 7.085e-03 | -4.95 | GO\_CALCIUM\_ION\_BINDING | MSigDB lists | GO\_CALCIUM\_ION\_BINDING | 126 | 11 | 4161 | 156 | sli,ds,ft,Dl,Cals,Fkbp14,Raf,shot,alpha-Man-Ia,SERCA,N | | 7.102e-03 | -4.95 | head segmentation | biological process | GO:0035287 | 22 | 4 | 12345 | 442 | kn,tsh,h,hth | | 7.102e-03 | -4.95 | cardiocyte differentiation | biological process | GO:0035051 | 22 | 4 | 12345 | 442 | htl,abd-A,pnr,sd | | 7.102e-03 | -4.95 | genital disc morphogenesis | biological process | GO:0007483 | 22 | 4 | 12345 | 442 | Rab5,N,Pvr,otk | | 7.102e-03 | -4.95 | negative regulation of cell projection organization | biological process | GO:0031345 | 22 | 4 | 12345 | 442 | baz,Sema1b,Abl,Sema1a | | 7.102e-03 | -4.95 | regulation of apoptotic signaling pathway | biological process | GO:2001233 | 22 | 4 | 12345 | 442 | Raf,eya,Dronc,Egfr | | 7.163e-03 | -4.94 | regulation of canonical Wnt signaling pathway | biological process | GO:0060828 | 79 | 8 | 12345 | 442 | nmo,apolpp,otk,spen,hyx,dlp,CtBP,Diap1 | | 7.238e-03 | -4.93 | ER\_Q6 | MSigDB lists | ER\_Q6 | 61 | 7 | 4161 | 156 | nmo,Kdm2,PI31,Gfat2,dlp,cact,Raf | | 7.277e-03 | -4.92 | Signaling by FGFR1 | REACTOME pathways | R-DME-5654736 | 28 | 5 | 3923 | 176 | CG31431,Lk6,htl,Raf,Cbl | | 7.306e-03 | -4.92 | EGFR downregulation | REACTOME pathways | R-DME-182971 | 18 | 4 | 3923 | 176 | lqf,cindr,Egfr,Cbl | | 7.313e-03 | -4.92 | Ig\_I-set | interpro domains | IPR013098 | 72 | 8 | 11383 | 450 | Ppn,vn,Pvr,robo1,unc-5,sns,otk,kek5 | | 7.315e-03 | -4.92 | histone H2A ubiquitination | biological process | GO:0033522 | 4 | 2 | 12345 | 442 | Kdm2,Su(z)2 | | 7.315e-03 | -4.92 | neural crest cell migration | biological process | GO:0001755 | 4 | 2 | 12345 | 442 | Sema1a,Sema1b | | 7.315e-03 | -4.92 | sex determination, primary response to X:A ratio | biological process | GO:0007541 | 4 | 2 | 12345 | 442 | Sxl,emc | | 7.315e-03 | -4.92 | sprouting of injured axon | biological process | GO:0048682 | 4 | 2 | 12345 | 442 | kay,wnd | | 7.315e-03 | -4.92 | protein localization to cell junction | biological process | GO:1902414 | 4 | 2 | 12345 | 442 | Shrm,arm | | 7.315e-03 | -4.92 | neural crest cell differentiation | biological process | GO:0014033 | 4 | 2 | 12345 | 442 | Sema1a,Sema1b | | 7.315e-03 | -4.92 | myoblast fate commitment | biological process | GO:0048625 | 4 | 2 | 12345 | 442 | htl,ttk | | 7.315e-03 | -4.92 | negative regulation of cardioblast differentiation | biological process | GO:0051892 | 4 | 2 | 12345 | 442 | lqf,Abd-B | | 7.315e-03 | -4.92 | regulation of growth rate | biological process | GO:0040009 | 4 | 2 | 12345 | 442 | lilli,hppy | | 7.315e-03 | -4.92 | regulation of cell proliferation involved in compound eye morphogenesis | biological process | GO:2000495 | 4 | 2 | 12345 | 442 | tsh,hth | | 7.315e-03 | -4.92 | protein localization to adherens junction | biological process | GO:0071896 | 4 | 2 | 12345 | 442 | Shrm,arm | | 7.315e-03 | -4.92 | specification of animal organ identity | biological process | GO:0010092 | 4 | 2 | 12345 | 442 | hth,ss | | 7.315e-03 | -4.92 | primary spermatocyte growth | biological process | GO:0007285 | 4 | 2 | 12345 | 442 | InR,Pex10 | | 7.315e-03 | -4.92 | mesenchymal cell development | biological process | GO:0014031 | 4 | 2 | 12345 | 442 | Sema1a,Sema1b | | 7.315e-03 | -4.92 | neural crest cell development | biological process | GO:0014032 | 4 | 2 | 12345 | 442 | Sema1a,Sema1b | | 7.315e-03 | -4.92 | negative regulation of cardiocyte differentiation | biological process | GO:1905208 | 4 | 2 | 12345 | 442 | Abd-B,lqf | | 7.315e-03 | -4.92 | formation of a compartment boundary | biological process | GO:0060288 | 4 | 2 | 12345 | 442 | ena,N | | 7.315e-03 | -4.92 | negative regulation of cardioblast cell fate specification | biological process | GO:0009997 | 4 | 2 | 12345 | 442 | lqf,Abd-B | | 7.315e-03 | -4.92 | negative regulation of cardiac cell fate specification | biological process | GO:2000044 | 4 | 2 | 12345 | 442 | Abd-B,lqf | | 7.344e-03 | -4.91 | fly chr99C2-99C2\|3-99 | chromosome location | fly chr99C2-99C2\|3-99 | 6 | 2 | 25010 | 571 | Atg16,neo | | 7.344e-03 | -4.91 | fly chr53C9-53C9\|2-80 cM | chromosome location | fly chr53C9-53C9\|2-80 cM | 6 | 2 | 25010 | 571 | CG8306,Cdk4 | | 7.344e-03 | -4.91 | fly chr12B2-12B2\|1-45 cM | chromosome location | fly chr12B2-12B2\|1-45 cM | 6 | 2 | 25010 | 571 | CG11158,CG12177 | | 7.344e-03 | -4.91 | fly chr12B2-12B2\|1-45 | chromosome location | fly chr12B2-12B2\|1-45 | 6 | 2 | 25010 | 571 | CG12177,CG11158 | | 7.344e-03 | -4.91 | fly chr99C2-99C2\|3-99 cM | chromosome location | fly chr99C2-99C2\|3-99 cM | 6 | 2 | 25010 | 571 | Atg16,neo | | 7.344e-03 | -4.91 | fly chr98C4-98C4\|3-95 cM | chromosome location | fly chr98C4-98C4\|3-95 cM | 6 | 2 | 25010 | 571 | larp,Gfat2 | | 7.344e-03 | -4.91 | fly chr61D4-61D4\|3-0.5 cM | chromosome location | fly chr61D4-61D4\|3-0.5 cM | 6 | 2 | 25010 | 571 | lncRNA:CR46200,CG34057 | | 7.344e-03 | -4.91 | fly chr98C4-98C4\|3-95 | chromosome location | fly chr98C4-98C4\|3-95 | 6 | 2 | 25010 | 571 | Gfat2,larp | | 7.344e-03 | -4.91 | fly chr53C9-53C9\|2-80 | chromosome location | fly chr53C9-53C9\|2-80 | 6 | 2 | 25010 | 571 | Cdk4,CG8306 | | 7.344e-03 | -4.91 | fly chr61D4-61D4\|3-0.5 | chromosome location | fly chr61D4-61D4\|3-0.5 | 6 | 2 | 25010 | 571 | CG34057,lncRNA:CR46200 | | 7.344e-03 | -4.91 | fly chr75C2-75C2\|3-45 cM | chromosome location | fly chr75C2-75C2\|3-45 cM | 6 | 2 | 25010 | 571 | lncRNA:CR45935,CG7320 | | 7.344e-03 | -4.91 | fly chr75C2-75C2\|3-45 | chromosome location | fly chr75C2-75C2\|3-45 | 6 | 2 | 25010 | 571 | CG7320,lncRNA:CR45935 | | 7.353e-03 | -4.91 | GO\_ENZYME\_REGULATOR\_ACTIVITY | MSigDB lists | GO\_ENZYME\_REGULATOR\_ACTIVITY | 200 | 15 | 4161 | 156 | GlcAT-I,CenB1A,wdb,CG3107,Arf79F,N,Pdk1,dock,Ppn,trbl,CG17883,Ocrl,sli,Spn42De,PI31 | | 7.441e-03 | -4.90 | eIF4E1 (eukaryotic translation initiation factor 4E1) | protein interactions | 45525 | 49 | 7 | 9204 | 432 | fl(2)d,path,Sxl,Lk6,pum,Dcr-2,Diap1 | | 7.441e-03 | -4.90 | Tor (Target of rapamycin) | protein interactions | 47396 | 49 | 7 | 9204 | 432 | rictor,Socs36E,REPTOR,Sema1a,hppy,path,hyx | | 7.492e-03 | -4.89 | Arf79F (ADP ribosylation factor at 79F) | protein interactions | 40506 | 74 | 9 | 9204 | 432 | AP-1gamma,Rab1,Pvr,SERCA,Rab5,rictor,siz,CG4896,baz | | 7.511e-03 | -4.89 | GO\_NEGATIVE\_REGULATION\_OF\_CELL\_PROLIFERATION | MSigDB lists | GO\_NEGATIVE\_REGULATION\_OF\_CELL\_PROLIFERATION | 127 | 11 | 4161 | 156 | arm,fz2,N,Actbeta,CtBP,tkv,Src42A,Cdk4,Mef2,hyx,Dl | | 7.586e-03 | -4.88 | GO\_REGULATION\_OF\_CELLULAR\_RESPONSE\_TO\_GROWTH\_FACTOR\_STIMULUS | MSigDB lists | GO\_REGULATION\_OF\_CELLULAR\_RESPONSE\_TO\_GROWTH\_FACTOR\_STIMULUS | 47 | 6 | 4161 | 156 | arm,sli,Dl,N,Pdk1,fz | | 7.586e-03 | -4.88 | GSE20366\_CD103\_KLRG1\_DP\_VS\_DN\_TREG\_DN | MSigDB lists | GSE20366\_CD103\_KLRG1\_DP\_VS\_DN\_TREG\_DN | 47 | 6 | 4161 | 156 | Nmda1,fz2,CycG,Tpc1,rg,CG9220 | | 7.623e-03 | -4.88 | GO\_EYE\_DEVELOPMENT | MSigDB lists | GO\_EYE\_DEVELOPMENT | 77 | 8 | 4161 | 156 | ap,tkv,Actbeta,arm,fz2,Eph,Dl,CG17883 | | 7.651e-03 | -4.87 | GO\_INTRACELLULAR\_VESICLE | MSigDB lists | GO\_INTRACELLULAR\_VESICLE | 341 | 22 | 4161 | 156 | Ocrl,Cadps,Syx1A,AP-1gamma,CG4645,Kr-h2,Pdk1,fz2,wbl,Pat1,jar,Dl,Rab5,Vps13,N,Arf79F,14-3-3zeta,lt,htl,shot,Rab1,14-3-3epsilon | | 7.665e-03 | -4.87 | germline ring canal | cellular component | GO:0045172 | 23 | 4 | 11940 | 417 | jar,14-3-3zeta,dock,14-3-3epsilon | | 7.694e-03 | -4.87 | positive regulation of secretion by cell | biological process | GO:1903532 | 35 | 5 | 12345 | 442 | jar,mop,S,Cadps,Rab1 | | 7.694e-03 | -4.87 | axon extension | biological process | GO:0048675 | 35 | 5 | 12345 | 442 | SCAR,fz2,fz,Rab5,tai | | 7.734e-03 | -4.86 | regulation of tube size, open tracheal system | biological process | GO:0035151 | 49 | 6 | 12345 | 442 | ft,fz,ds,cora,ttk,Gli | | 7.752e-03 | -4.86 | ZINC\_FINGER\_C2H2\_2 | prosite domains | PS50157 | 273 | 24 | 6243 | 325 | CG32772,CG10321,esg,crol,ttk,hang,Prdm13,mld,zfh2,CG11247,disco-r,sens,sens-2,CG10348,CG11122,rgr,chn,bowl,klu,CG12769,sqz,tsh,CG14441,pita | | 7.789e-03 | -4.86 | I-set | pfam domains | PF07679 | 72 | 8 | 10480 | 419 | unc-5,robo1,kek5,Pvr,otk,vn,sns,Ppn | | 7.827e-03 | -4.85 | ZF\_CXXC | prosite domains | PS51058 | 3 | 2 | 6243 | 325 | Kdm2,Tet | | 7.849e-03 | -4.85 | phosphoprotein phosphatase activity | molecular function | GO:0004721 | 102 | 9 | 12176 | 414 | mop,Ptpmeg2,IA-2,Pez,CG10417,Pp2C1,Ptp4E,eya,CG42327 | | 7.879e-03 | -4.84 | primary sex determination, soma | biological process | GO:0007539 | 12 | 3 | 12345 | 442 | emc,fl(2)d,Sxl | | 7.879e-03 | -4.84 | lateral inhibition | biological process | GO:0046331 | 12 | 3 | 12345 | 442 | Dl,mam,N | | 7.879e-03 | -4.84 | negative regulation of apoptotic signaling pathway | biological process | GO:2001234 | 12 | 3 | 12345 | 442 | eya,Raf,Egfr | | 7.879e-03 | -4.84 | retinal ganglion cell axon guidance | biological process | GO:0031290 | 12 | 3 | 12345 | 442 | otk,Dad,fz2 | | 7.909e-03 | -4.84 | GO\_ACTIVATION\_OF\_PROTEIN\_KINASE\_ACTIVITY | MSigDB lists | GO\_ACTIVATION\_OF\_PROTEIN\_KINASE\_ACTIVITY | 62 | 7 | 4161 | 156 | Raf,Pdk1,Tsp96F,nAChRalpha4,Tollo,hppy,wbl | | 7.946e-03 | -4.84 | exd (extradenticle) | protein interactions | 32567 | 38 | 6 | 9204 | 432 | abd-A,Abd-B,tsh,hth,ey,Scr | | 7.973e-03 | -4.83 | GO\_CHONDROCYTE\_DEVELOPMENT | MSigDB lists | GO\_CHONDROCYTE\_DEVELOPMENT | 4 | 2 | 4161 | 156 | CG9220,tkv | | 7.973e-03 | -4.83 | GO\_BETA\_CATENIN\_DESTRUCTION\_COMPLEX | MSigDB lists | GO\_BETA\_CATENIN\_DESTRUCTION\_COMPLEX | 4 | 2 | 4161 | 156 | arm,sina | | 7.973e-03 | -4.83 | GO\_REGULATION\_OF\_EPITHELIAL\_CELL\_DIFFERENTIATION\_INVOLVED\_IN\_KIDNEY\_DEVELOPMENT | MSigDB lists | GO\_REGULATION\_OF\_EPITHELIAL\_CELL\_DIFFERENTIATION\_INVOLVED\_IN\_KIDNEY\_DEVELOPMENT | 4 | 2 | 4161 | 156 | arm,ft | | 7.973e-03 | -4.83 | GO\_WNT\_SIGNALOSOME | MSigDB lists | GO\_WNT\_SIGNALOSOME | 4 | 2 | 4161 | 156 | fz,arm | | 7.973e-03 | -4.83 | GO\_PROSTATE\_GLAND\_MORPHOGENESIS | MSigDB lists | GO\_PROSTATE\_GLAND\_MORPHOGENESIS | 4 | 2 | 4161 | 156 | N,Fem-1 | | 7.973e-03 | -4.83 | DAVICIONI\_PAX\_FOXO1\_SIGNATURE\_IN\_ARMS\_DN | MSigDB lists | DAVICIONI\_PAX\_FOXO1\_SIGNATURE\_IN\_ARMS\_DN | 4 | 2 | 4161 | 156 | SERCA,trbl | | 7.973e-03 | -4.83 | GO\_REGULATION\_OF\_ENDOTHELIAL\_CELL\_DIFFERENTIATION | MSigDB lists | GO\_REGULATION\_OF\_ENDOTHELIAL\_CELL\_DIFFERENTIATION | 4 | 2 | 4161 | 156 | arm,N | | 7.973e-03 | -4.83 | GO\_REGULATION\_OF\_ESTABLISHMENT\_OF\_PLANAR\_POLARITY\_INVOLVED\_IN\_NEURAL\_TUBE\_CLOSURE | MSigDB lists | GO\_REGULATION\_OF\_ESTABLISHMENT\_OF\_PLANAR\_POLARITY\_INVOLVED\_IN\_NEURAL\_TUBE\_CLOSURE | 4 | 2 | 4161 | 156 | fz,otk | | 7.973e-03 | -4.83 | GO\_NEGATIVE\_REGULATION\_OF\_EPIDERMIS\_DEVELOPMENT | MSigDB lists | GO\_NEGATIVE\_REGULATION\_OF\_EPIDERMIS\_DEVELOPMENT | 4 | 2 | 4161 | 156 | Dl,N | | 7.973e-03 | -4.83 | GO\_DORSAL\_VENTRAL\_AXIS\_SPECIFICATION | MSigDB lists | GO\_DORSAL\_VENTRAL\_AXIS\_SPECIFICATION | 4 | 2 | 4161 | 156 | fz2,arm | | 7.973e-03 | -4.83 | GO\_MORPHOGENESIS\_OF\_AN\_ENDOTHELIUM | MSigDB lists | GO\_MORPHOGENESIS\_OF\_AN\_ENDOTHELIUM | 4 | 2 | 4161 | 156 | N,arm | | 7.973e-03 | -4.83 | NIKOLSKY\_BREAST\_CANCER\_15Q26\_AMPLICON | MSigDB lists | NIKOLSKY\_BREAST\_CANCER\_15Q26\_AMPLICON | 4 | 2 | 4161 | 156 | CG9220,InR | | 7.973e-03 | -4.83 | BERENJENO\_TRANSFORMED\_BY\_RHOA\_REVERSIBLY\_DN | MSigDB lists | BERENJENO\_TRANSFORMED\_BY\_RHOA\_REVERSIBLY\_DN | 4 | 2 | 4161 | 156 | Unc-76,Actbeta | | 7.973e-03 | -4.83 | REACTOME\_NEPHRIN\_INTERACTIONS | MSigDB lists | REACTOME\_NEPHRIN\_INTERACTIONS | 4 | 2 | 4161 | 156 | sns,dock | | 7.973e-03 | -4.83 | GO\_POSITIVE\_REGULATION\_OF\_STAT\_CASCADE | MSigDB lists | GO\_POSITIVE\_REGULATION\_OF\_STAT\_CASCADE | 4 | 2 | 4161 | 156 | htl,N | | 7.973e-03 | -4.83 | GO\_ENDOCARDIAL\_CUSHION\_FORMATION | MSigDB lists | GO\_ENDOCARDIAL\_CUSHION\_FORMATION | 4 | 2 | 4161 | 156 | N,ds | | 7.973e-03 | -4.83 | GO\_REGULATION\_OF\_SPROUTING\_ANGIOGENESIS | MSigDB lists | GO\_REGULATION\_OF\_SPROUTING\_ANGIOGENESIS | 4 | 2 | 4161 | 156 | N,Dl | | 7.973e-03 | -4.83 | GO\_NEGATIVE\_REGULATION\_OF\_PROTEIN\_OLIGOMERIZATION | MSigDB lists | GO\_NEGATIVE\_REGULATION\_OF\_PROTEIN\_OLIGOMERIZATION | 4 | 2 | 4161 | 156 | Pex5,Pex14 | | 7.973e-03 | -4.83 | GO\_VENTRICULAR\_SEPTUM\_MORPHOGENESIS | MSigDB lists | GO\_VENTRICULAR\_SEPTUM\_MORPHOGENESIS | 4 | 2 | 4161 | 156 | fz,N | | 7.973e-03 | -4.83 | TIAN\_TNF\_SIGNALING\_VIA\_NFKB | MSigDB lists | TIAN\_TNF\_SIGNALING\_VIA\_NFKB | 4 | 2 | 4161 | 156 | cact,Gfat2 | | 7.973e-03 | -4.83 | GO\_NEGATIVE\_REGULATION\_OF\_OLIGODENDROCYTE\_DIFFERENTIATION | MSigDB lists | GO\_NEGATIVE\_REGULATION\_OF\_OLIGODENDROCYTE\_DIFFERENTIATION | 4 | 2 | 4161 | 156 | N,arm | | 7.973e-03 | -4.83 | SCHEIDEREIT\_IKK\_TARGETS | MSigDB lists | SCHEIDEREIT\_IKK\_TARGETS | 4 | 2 | 4161 | 156 | cact,arm | | 7.973e-03 | -4.83 | GO\_NEGATIVE\_REGULATION\_OF\_OSTEOCLAST\_DIFFERENTIATION | MSigDB lists | GO\_NEGATIVE\_REGULATION\_OF\_OSTEOCLAST\_DIFFERENTIATION | 4 | 2 | 4161 | 156 | arm,Tollo | | 7.973e-03 | -4.83 | GO\_POSITIVE\_REGULATION\_OF\_SYNAPSE\_ASSEMBLY | MSigDB lists | GO\_POSITIVE\_REGULATION\_OF\_SYNAPSE\_ASSEMBLY | 4 | 2 | 4161 | 156 | Cals,Eph | | 7.973e-03 | -4.83 | GO\_ALPHA\_LINOLENIC\_ACID\_METABOLIC\_PROCESS | MSigDB lists | GO\_ALPHA\_LINOLENIC\_ACID\_METABOLIC\_PROCESS | 4 | 2 | 4161 | 156 | Mfe2,CG31522 | | 7.973e-03 | -4.83 | GO\_BONE\_GROWTH | MSigDB lists | GO\_BONE\_GROWTH | 4 | 2 | 4161 | 156 | Ddr,htl | | 7.973e-03 | -4.83 | GO\_RESPONSE\_TO\_FOLLICLE\_STIMULATING\_HORMONE | MSigDB lists | GO\_RESPONSE\_TO\_FOLLICLE\_STIMULATING\_HORMONE | 4 | 2 | 4161 | 156 | N,Actbeta | | 7.987e-03 | -4.83 | FERM\_PH-like\_C | interpro domains | IPR018980 | 11 | 3 | 11383 | 450 | cora,Pez,yrt | | 8.015e-03 | -4.83 | MANALO\_HYPOXIA\_UP | MSigDB lists | MANALO\_HYPOXIA\_UP | 34 | 5 | 4161 | 156 | trbl,CG9220,Actbeta,bru2,ds | | 8.015e-03 | -4.83 | GO\_MICROBODY\_MEMBRANE | MSigDB lists | GO\_MICROBODY\_MEMBRANE | 34 | 5 | 4161 | 156 | Arf79F,Pex10,Mfe2,Pex5,Pex14 | | 8.015e-03 | -4.83 | GSE39110\_UNTREATED\_VS\_IL2\_TREATED\_CD8\_TCELL\_DAY6\_POST\_IMMUNIZATION\_DN | MSigDB lists | GSE39110\_UNTREATED\_VS\_IL2\_TREATED\_CD8\_TCELL\_DAY6\_POST\_IMMUNIZATION\_DN | 34 | 5 | 4161 | 156 | trbl,InR,Tsp96F,CG6891,eff | | 8.015e-03 | -4.83 | GSE23502\_WT\_VS\_HDC\_KO\_MYELOID\_DERIVED\_SUPPRESSOR\_CELL\_COLON\_TUMOR\_DN | MSigDB lists | GSE23502\_WT\_VS\_HDC\_KO\_MYELOID\_DERIVED\_SUPPRESSOR\_CELL\_COLON\_TUMOR\_DN | 34 | 5 | 4161 | 156 | Fkbp14,Eph,Tollo,CG9485,Gfat2 | | 8.015e-03 | -4.83 | GO\_CEREBRAL\_CORTEX\_DEVELOPMENT | MSigDB lists | GO\_CEREBRAL\_CORTEX\_DEVELOPMENT | 34 | 5 | 4161 | 156 | 14-3-3epsilon,arm,sli,ft,ap | | 8.015e-03 | -4.83 | GAL\_LEUKEMIC\_STEM\_CELL\_UP | MSigDB lists | GAL\_LEUKEMIC\_STEM\_CELL\_UP | 34 | 5 | 4161 | 156 | InR,DPCoAC,Pex5,sli,shot | | 8.018e-03 | -4.83 | intracellular signal transduction | biological process | GO:0035556 | 373 | 23 | 12345 | 442 | CG42788,hppy,Ric,hfw,Raf,hang,Rab1,Rgl,Arf79F,RhoU,Socs36E,wnd,vn,RhoGAP19D,Lk6,imd,Dp,nmo,rictor,sd,Pdk1,Rab5,REPTOR | | 8.059e-03 | -4.82 | Sin3A (Sin3A) | protein interactions | 36382 | 62 | 8 | 9204 | 432 | N,sina,shg,H,aop,pnr,mam,Diap1 | | 8.079e-03 | -4.82 | GO\_NEURON\_PROJECTION\_MORPHOGENESIS | MSigDB lists | GO\_NEURON\_PROJECTION\_MORPHOGENESIS | 94 | 9 | 4161 | 156 | sli,Eph,robo1,unc-5,Unc-76,tkv,ap,N,sina | | 8.128e-03 | -4.81 | dephosphorylation | biological process | GO:0016311 | 170 | 13 | 12345 | 442 | IA-2,CG10417,mop,wdb,eya,CG9784,Ptp4E,Ocrl,Ptpmeg2,Pp2C1,CG6656,CG42327,Pez | | 8.158e-03 | -4.81 | growth factor receptor binding | molecular function | GO:0070851 | 24 | 4 | 12176 | 414 | vn,ed,Pvf3,spi | | 8.193e-03 | -4.80 | SH3 | prosite domains | PS50002 | 68 | 9 | 6243 | 325 | shot,Cip4,SKIP,Src42A,cindr,p130CAS,RhoGEF3,dock,Abl | | 8.201e-03 | -4.80 | GO\_FEMALE\_SEX\_DIFFERENTIATION | MSigDB lists | GO\_FEMALE\_SEX\_DIFFERENTIATION | 22 | 4 | 4161 | 156 | robo1,Actbeta,tkv,sli | | 8.201e-03 | -4.80 | GSE2706\_R848\_VS\_LPS\_8H\_STIM\_DC\_UP | MSigDB lists | GSE2706\_R848\_VS\_LPS\_8H\_STIM\_DC\_UP | 22 | 4 | 4161 | 156 | Ric,CG7914,ap,sns | | 8.201e-03 | -4.80 | GO\_CARTILAGE\_DEVELOPMENT | MSigDB lists | GO\_CARTILAGE\_DEVELOPMENT | 22 | 4 | 4161 | 156 | Mef2,CG9220,htl,tkv | | 8.201e-03 | -4.80 | GSE19198\_1H\_VS\_6H\_IL21\_TREATED\_TCELL\_UP | MSigDB lists | GSE19198\_1H\_VS\_6H\_IL21\_TREATED\_TCELL\_UP | 22 | 4 | 4161 | 156 | CG5522,Rgl,bru2,Lk6 | | 8.201e-03 | -4.80 | GO\_SIGNAL\_SEQUENCE\_BINDING | MSigDB lists | GO\_SIGNAL\_SEQUENCE\_BINDING | 22 | 4 | 4161 | 156 | Pex5,Pex7,cact,Pen | | 8.237e-03 | -4.80 | hEGF | pfam domains | PF12661 | 11 | 3 | 10480 | 419 | sli,Dl,N | | 8.291e-03 | -4.79 | FRI | smart domains | SM00063 | 8 | 3 | 5086 | 290 | fz3,fz,fz2 | | 8.319e-03 | -4.79 | positive regulation of cellular component organization | biological process | GO:0051130 | 290 | 19 | 12345 | 442 | Cals,Jupiter,Abl,Su(z)2,kon,rictor,ena,tkv,nmo,fz,Rab1,Cbl,SCAR,cindr,RhoGEF4,chic,shot,Arf79F,RhoU | | 8.323e-03 | -4.79 | ConA-like\_dom\_sf | interpro domains | IPR013320 | 59 | 7 | 11383 | 450 | Cals,LanA,shg,rg,kon,ft,sli | | 8.359e-03 | -4.78 | unidimensional cell growth | biological process | GO:0009826 | 23 | 4 | 12345 | 442 | shg,arm,tkv,Dad | | 8.374e-03 | -4.78 | B41 | smart domains | SM00295 | 23 | 5 | 5086 | 290 | CG42788,yrt,Myo10A,cora,Pez | | 8.404e-03 | -4.78 | GOBERT\_OLIGODENDROCYTE\_DIFFERENTIATION\_DN | MSigDB lists | GOBERT\_OLIGODENDROCYTE\_DIFFERENTIATION\_DN | 242 | 17 | 4161 | 156 | N,GlcAT-I,shot,CG4935,Pex5,CG5522,step,Tsp96F,His2Av,betaTub60D,CycG,jar,Cerk,emp,DPCoAC,Ziz,PI31 | | 8.407e-03 | -4.78 | GO\_REGULATION\_OF\_EXOCYTOSIS | MSigDB lists | GO\_REGULATION\_OF\_EXOCYTOSIS | 48 | 6 | 4161 | 156 | Syx1A,AP-1gamma,N,Pdk1,Arf79F,Raf | | 8.407e-03 | -4.78 | TITF1\_Q3 | MSigDB lists | TITF1\_Q3 | 48 | 6 | 4161 | 156 | Ric,Rab1,otk,wdb,CG1129,Mef2 | | 8.407e-03 | -4.78 | GO\_CELLULAR\_PROCESS\_INVOLVED\_IN\_REPRODUCTION\_IN\_MULTICELLULAR\_ORGANISM | MSigDB lists | GO\_CELLULAR\_PROCESS\_INVOLVED\_IN\_REPRODUCTION\_IN\_MULTICELLULAR\_ORGANISM | 48 | 6 | 4161 | 156 | sli,CG17883,arm,tkv,robo1,Actbeta | | 8.407e-03 | -4.78 | KEGG\_PEROXISOME | MSigDB lists | KEGG\_PEROXISOME | 48 | 6 | 4161 | 156 | Pex7,CG3961,Pex5,Pex14,Mfe2,Pex10 | | 8.407e-03 | -4.78 | GSE21360\_NAIVE\_VS\_TERTIARY\_MEMORY\_CD8\_TCELL\_UP | MSigDB lists | GSE21360\_NAIVE\_VS\_TERTIARY\_MEMORY\_CD8\_TCELL\_UP | 48 | 6 | 4161 | 156 | CG6891,CG1129,Tpst,Cerk,ds,CG3961 | | 8.420e-03 | -4.78 | catalytic activity, acting on a protein | molecular function | GO:0140096 | 1481 | 67 | 12176 | 414 | Dyrk2,tna,Tpst,Dronc,ec,CG43336,Eph,CG6465,Cdk4,Pez,Ubc87F,Kdm2,Pex10,CG42327,eya,hppy,mura,Psa,CG4467,Ppn,Diap1,CG12402,Abl,Nepl21,IA-2,Pdk1,Ptp4E,C1GalTA,Src42A,mop,nmo,muc,InR,htl,CalpC,CycB3,Ddr,Pgant6,drl,CG3107,sina,FBXO11,Egfr,CG3119,tkv,trbl,Pgant5,rho-4,eff,Lk6,Raf,CG34057,CG10417,Kdm4B,morgue,Nrk,Pitslre,Ptpmeg2,Fem-1,Pp2C1,CG13481,wnd,Drak,Cbl,CycG,Pvr,otk | | 8.424e-03 | -4.78 | GO\_EMBRYONIC\_MORPHOGENESIS | MSigDB lists | GO\_EMBRYONIC\_MORPHOGENESIS | 129 | 11 | 4161 | 156 | LanA,Dl,otk,hyx,Mef2,fz2,arm,ap,fz,Actbeta,N | | 8.480e-03 | -4.77 | Chondroitin sulfate/dermatan sulfate metabolism | REACTOME pathways | R-DME-1793185 | 29 | 5 | 3923 | 176 | Ids,kon,dlp,CG9220,GlcAT-I | | 8.564e-03 | -4.76 | PH | pfam domains | PF00169 | 45 | 6 | 10480 | 419 | step,CenB1A,Ziz,Cadps,CG5522,CG34383 | | 8.577e-03 | -4.76 | IGc2 | smart domains | SM00408 | 109 | 13 | 5086 | 290 | CG31431,ed,kek5,Bsg,robo1,vn,Pvr,otk,sns,htl,Ppn,boi,unc-5 | | 8.625e-03 | -4.75 | YYCATTCAWW\_UNKNOWN | MSigDB lists | YYCATTCAWW\_UNKNOWN | 63 | 7 | 4161 | 156 | Ddr,Rab5,CG3436,Actbeta,trbl,InR,htl | | 8.646e-03 | -4.75 | GO\_FOREBRAIN\_DEVELOPMENT | MSigDB lists | GO\_FOREBRAIN\_DEVELOPMENT | 95 | 9 | 4161 | 156 | 14-3-3epsilon,arm,ap,N,Actbeta,sli,ft,robo1,Cdk4 | | 8.653e-03 | -4.75 | UQCR-C2 (Ubiquinol-cytochrome c reductase core protein 2) | protein interactions | 39846 | 117 | 12 | 9204 | 432 | Kr-h2,Mpcp2,Sema1a,UQCR-11,rictor,InR,Rab5,Gfat2,SERCA,Pdk1,Arf79F,ds | | 8.665e-03 | -4.75 | insc (inscuteable) | protein interactions | 37355 | 18 | 4 | 9204 | 432 | baz,N,H,Pen | | 8.678e-03 | -4.75 | cuticle pattern formation | biological process | GO:0035017 | 36 | 5 | 12345 | 442 | boi,fz2,fz,arm,f | | 8.702e-03 | -4.74 | GO\_REGULATION\_OF\_TRANSFERASE\_ACTIVITY | MSigDB lists | GO\_REGULATION\_OF\_TRANSFERASE\_ACTIVITY | 283 | 19 | 4161 | 156 | Raf,hppy,htl,InR,dock,trbl,Rab1,Tollo,unc-119,PI31,Fem-1,Ddr,nAChRalpha4,Tsp96F,Pdk1,CycG,wbl,fz2,arm | | 8.746e-03 | -4.74 | GO\_PROTEIN\_COMPLEX\_BINDING | MSigDB lists | GO\_PROTEIN\_COMPLEX\_BINDING | 243 | 17 | 4161 | 156 | jar,arm,Pex14,SMC3,Pdk1,Tsp96F,LanA,PI31,Ddr,Eph,hyx,shot,14-3-3epsilon,lt,InR,CG4896,cact | | 8.861e-03 | -4.73 | REACTOME\_SIGNALING\_BY\_ROBO\_RECEPTOR | MSigDB lists | REACTOME\_SIGNALING\_BY\_ROBO\_RECEPTOR | 12 | 3 | 4161 | 156 | sli,dock,robo1 | | 8.861e-03 | -4.73 | GO\_ADHERENS\_JUNCTION\_ORGANIZATION | MSigDB lists | GO\_ADHERENS\_JUNCTION\_ORGANIZATION | 12 | 3 | 4161 | 156 | LanA,arm,Pdk1 | | 8.861e-03 | -4.73 | MA\_MYELOID\_DIFFERENTIATION\_DN | MSigDB lists | MA\_MYELOID\_DIFFERENTIATION\_DN | 12 | 3 | 4161 | 156 | N,CG10417,CycG | | 8.861e-03 | -4.73 | NAKAYAMA\_SOFT\_TISSUE\_TUMORS\_PCA1\_DN | MSigDB lists | NAKAYAMA\_SOFT\_TISSUE\_TUMORS\_PCA1\_DN | 12 | 3 | 4161 | 156 | dlp,ap,htl | | 8.861e-03 | -4.73 | WILCOX\_RESPONSE\_TO\_PROGESTERONE\_DN | MSigDB lists | WILCOX\_RESPONSE\_TO\_PROGESTERONE\_DN | 12 | 3 | 4161 | 156 | Actbeta,InR,shot | | 8.861e-03 | -4.73 | WANG\_TARGETS\_OF\_MLL\_CBP\_FUSION\_UP | MSigDB lists | WANG\_TARGETS\_OF\_MLL\_CBP\_FUSION\_UP | 12 | 3 | 4161 | 156 | CG2017,sina,14-3-3zeta | | 8.861e-03 | -4.73 | BIOCARTA\_ALK\_PATHWAY | MSigDB lists | BIOCARTA\_ALK\_PATHWAY | 12 | 3 | 4161 | 156 | fz,Mef2,arm | | 8.861e-03 | -4.73 | ZUCCHI\_METASTASIS\_DN | MSigDB lists | ZUCCHI\_METASTASIS\_DN | 12 | 3 | 4161 | 156 | cact,CG10417,14-3-3epsilon | | 8.861e-03 | -4.73 | TONKS\_TARGETS\_OF\_RUNX1\_RUNX1T1\_FUSION\_GRANULOCYTE\_UP | MSigDB lists | TONKS\_TARGETS\_OF\_RUNX1\_RUNX1T1\_FUSION\_GRANULOCYTE\_UP | 12 | 3 | 4161 | 156 | CG1673,Mef2,Cdk4 | | 8.861e-03 | -4.73 | BRUECKNER\_TARGETS\_OF\_MIRLET7A3\_DN | MSigDB lists | BRUECKNER\_TARGETS\_OF\_MIRLET7A3\_DN | 12 | 3 | 4161 | 156 | Pgant6,Pgant5,CG6891 | | 8.861e-03 | -4.73 | PID\_PI3KCI\_PATHWAY | MSigDB lists | PID\_PI3KCI\_PATHWAY | 12 | 3 | 4161 | 156 | Pdk1,Arf79F,step | | 8.873e-03 | -4.72 | Lipovitellin\_superhlx\_dom | interpro domains | IPR011030 | 4 | 2 | 11383 | 450 | apolpp,cv-d | | 8.873e-03 | -4.72 | PAS\_fold\_3 | interpro domains | IPR013655 | 4 | 2 | 11383 | 450 | ss,sim | | 8.873e-03 | -4.72 | BEN\_domain | interpro domains | IPR018379 | 4 | 2 | 11383 | 450 | Elba2,pre-mod(mdg4)-C | | 8.873e-03 | -4.72 | Lipid\_transp\_b-sht\_shell | interpro domains | IPR015819 | 4 | 2 | 11383 | 450 | apolpp,cv-d | | 8.873e-03 | -4.72 | Rcpt\_L-dom | interpro domains | IPR000494 | 4 | 2 | 11383 | 450 | InR,Egfr | | 8.873e-03 | -4.72 | Lipid\_transpt\_N | interpro domains | IPR001747 | 4 | 2 | 11383 | 450 | apolpp,cv-d | | 8.873e-03 | -4.72 | Vitellinogen\_b-sht\_N | interpro domains | IPR015816 | 4 | 2 | 11383 | 450 | apolpp,cv-d | | 8.873e-03 | -4.72 | Furin-like\_Cys-rich\_dom | interpro domains | IPR006211 | 4 | 2 | 11383 | 450 | Egfr,InR | | 8.873e-03 | -4.72 | RAWUL | interpro domains | IPR032443 | 4 | 2 | 11383 | 450 | Psc,Su(z)2 | | 8.873e-03 | -4.72 | Rcpt\_L-dom\_sf | interpro domains | IPR036941 | 4 | 2 | 11383 | 450 | Egfr,InR | | 8.874e-03 | -4.72 | AAGCCAT\_MIR135A\_MIR135B | MSigDB lists | AAGCCAT\_MIR135A\_MIR135B | 79 | 8 | 4161 | 156 | Lk6,CG9220,sina,Actbeta,rg,Mef2,Rgl,Src42A | | 8.874e-03 | -4.72 | GO\_POSITIVE\_REGULATION\_OF\_MAPK\_CASCADE | MSigDB lists | GO\_POSITIVE\_REGULATION\_OF\_MAPK\_CASCADE | 79 | 8 | 4161 | 156 | Tollo,Raf,hppy,htl,Tsp96F,wbl,arm,fz2 | | 8.874e-03 | -4.72 | Znf\_RING | interpro domains | IPR001841 | 122 | 11 | 11383 | 450 | CG11414,mura,Pex10,CG11360,CG13481,Su(z)2,Cbl,lt,sina,Psc,Diap1 | | 8.909e-03 | -4.72 | GARY\_CD5\_TARGETS\_UP | MSigDB lists | GARY\_CD5\_TARGETS\_UP | 148 | 12 | 4161 | 156 | nmo,14-3-3epsilon,CycG,CG32795,rdx,Nmda1,Pgk,Cadps,ATPCL,cact,CG6465,Raf | | 8.909e-03 | -4.72 | SENESE\_HDAC3\_TARGETS\_UP | MSigDB lists | SENESE\_HDAC3\_TARGETS\_UP | 148 | 12 | 4161 | 156 | emp,Ids,Gfat2,Spn42De,Syx1A,Cdk4,cact,shot,alpha-Man-Ia,jar,14-3-3zeta,sd | | 8.936e-03 | -4.72 | Negative regulation of FGFR3 signaling | REACTOME pathways | R-DME-5654732 | 19 | 4 | 3923 | 176 | Lk6,htl,Raf,Cbl | | 8.936e-03 | -4.72 | ABC transporters in lipid homeostasis | REACTOME pathways | R-DME-1369062 | 19 | 4 | 3923 | 176 | CG17646,CG11069,CG5853,CG31121 | | 8.936e-03 | -4.72 | Negative regulation of FGFR2 signaling | REACTOME pathways | R-DME-5654727 | 19 | 4 | 3923 | 176 | htl,Lk6,Cbl,Raf | | 8.936e-03 | -4.72 | Negative regulation of FGFR4 signaling | REACTOME pathways | R-DME-5654733 | 19 | 4 | 3923 | 176 | Cbl,Raf,htl,Lk6 | | 8.939e-03 | -4.72 | CG10650 (uncharacterized protein) | protein interactions | 35162 | 28 | 5 | 9204 | 432 | rdx,CG31211,H2.0,Acp63F,CG34401 | | 8.939e-03 | -4.72 | Cks85A (Cyclin-dependent kinase subunit 85A) | protein interactions | 41034 | 28 | 5 | 9204 | 432 | Cks30A,CycB3,CG31211,ena,CG6191 | | 8.939e-03 | -4.72 | pnut (peanut) | protein interactions | 35801 | 28 | 5 | 9204 | 432 | Syx1A,Diap1,sina,S,ft | | 8.939e-03 | -4.72 | kni (knirps) | protein interactions | 40287 | 28 | 5 | 9204 | 432 | Abd-B,abd-A,Egfr,CtBP,Scr | | 8.978e-03 | -4.71 | SH2\_dom\_sf | interpro domains | IPR036860 | 33 | 5 | 11383 | 450 | Src42A,dock,Socs36E,Abl,Cbl | | 9.029e-03 | -4.71 | Mbs (Myosin binding subunit) | protein interactions | 49070 | 39 | 6 | 9204 | 432 | rictor,Egfr,Dad,rdx,kay,baz | | 9.029e-03 | -4.71 | msi (musashi) | protein interactions | 43087 | 39 | 6 | 9204 | 432 | ps,sina,ttk,N,msi,CG2233 | | 9.029e-03 | -4.71 | SsRbeta (Signal sequence receptor beta) | protein interactions | 39768 | 39 | 6 | 9204 | 432 | Rab5,SERCA,Pdk1,Arf79F,rictor,Rab1 | | 9.052e-03 | -4.70 | neuron projection | cellular component | GO:0043005 | 431 | 25 | 11940 | 417 | Eph,Magi,orb2,jeb,path,pum,nAChRalpha4,chic,Abl,CG9784,Ptp4E,shot,wnd,jar,Bsg,arm,Unc-76,drl,ena,Syx1A,robo1,RhoU,Gr93c,gukh,InR | | 9.069e-03 | -4.70 | Vitellogenin\_N | pfam domains | PF01347 | 4 | 2 | 10480 | 419 | cv-d,apolpp | | 9.069e-03 | -4.70 | PAS\_3 | pfam domains | PF08447 | 4 | 2 | 10480 | 419 | sim,ss | | 9.069e-03 | -4.70 | Furin-like | pfam domains | PF00757 | 4 | 2 | 10480 | 419 | Egfr,InR | | 9.069e-03 | -4.70 | Recep\_L\_domain | pfam domains | PF01030 | 4 | 2 | 10480 | 419 | Egfr,InR | | 9.069e-03 | -4.70 | RAWUL | pfam domains | PF16207 | 4 | 2 | 10480 | 419 | Psc,Su(z)2 | | 9.074e-03 | -4.70 | GSE45739\_UNSTIM\_VS\_ACD3\_ACD28\_STIM\_NRAS\_KO\_CD4\_TCELL\_DN | MSigDB lists | GSE45739\_UNSTIM\_VS\_ACD3\_ACD28\_STIM\_NRAS\_KO\_CD4\_TCELL\_DN | 35 | 5 | 4161 | 156 | ft,jar,Tollo,CG9220,Pgant5 | | 9.074e-03 | -4.70 | GSE25087\_FETAL\_VS\_ADULT\_TREG\_DN | MSigDB lists | GSE25087\_FETAL\_VS\_ADULT\_TREG\_DN | 35 | 5 | 4161 | 156 | Ppn,Atg16,bru2,CG3961,Ids | | 9.074e-03 | -4.70 | GSE41867\_DAY8\_VS\_DAY15\_LCMV\_CLONE13\_EFFECTOR\_CD8\_TCELL\_UP | MSigDB lists | GSE41867\_DAY8\_VS\_DAY15\_LCMV\_CLONE13\_EFFECTOR\_CD8\_TCELL\_UP | 35 | 5 | 4161 | 156 | Lk6,CG9220,Atg16,Rim2,bru2 | | 9.074e-03 | -4.70 | GO\_POSITIVE\_REGULATION\_OF\_CELL\_ACTIVATION | MSigDB lists | GO\_POSITIVE\_REGULATION\_OF\_CELL\_ACTIVATION | 35 | 5 | 4161 | 156 | AP-1gamma,dock,Tsp96F,Pdk1,Mef2 | | 9.233e-03 | -4.69 | nuclear division | biological process | GO:0000280 | 293 | 19 | 12345 | 442 | corto,Gli,PI31,orb2,Arf79F,CycG,Rab1,ph-p,Hr39,Rab5,His3.3B,LanA,His3.3A,Slbp,Cks30A,eff,larp,Pex10,Sxl | | 9.291e-03 | -4.68 | HALLMARK\_PEROXISOME | MSigDB lists | HALLMARK\_PEROXISOME | 49 | 6 | 4161 | 156 | sina,Pex5,CG3961,Pex14,Mfe2,Tpc1 | | 9.291e-03 | -4.68 | GSE16385\_MONOCYTE\_VS\_12H\_IL4\_TREATED\_MACROPHAGE\_DN | MSigDB lists | GSE16385\_MONOCYTE\_VS\_12H\_IL4\_TREATED\_MACROPHAGE\_DN | 49 | 6 | 4161 | 156 | eff,wbl,CG5522,step,Rab5,CG6891 | | 9.291e-03 | -4.68 | BROWNE\_HCMV\_INFECTION\_24HR\_UP | MSigDB lists | BROWNE\_HCMV\_INFECTION\_24HR\_UP | 49 | 6 | 4161 | 156 | cact,Dyb,Rab5,fz2,CG10417,CG9485 | | 9.291e-03 | -4.68 | GO\_HEART\_MORPHOGENESIS | MSigDB lists | GO\_HEART\_MORPHOGENESIS | 49 | 6 | 4161 | 156 | ds,ft,N,Mef2,fz,Dl | | 9.329e-03 | -4.67 | transferase activity | molecular function | GO:0016740 | 1384 | 63 | 12176 | 414 | Pex10,trbl,Pgant5,eff,Lk6,hppy,Raf,CG34057,mura,morgue,Diap1,Nrk,Abl,CG12402,Pitslre,Fem-1,Pdk1,CG13481,Cbl,wnd,Atg16,Drak,CycG,Pvr,C1GalTA,otk,CG12910,Src42A,Dyrk2,Gfat2,tna,Tpst,nmo,CG31522,Dgat2,GlcAT-I,muc,l(1)G0196,Cerk,ATPCL,l(2)37Cg,InR,htl,CycB3,Ddr,bin3,Eph,Pgant6,drl,CG1129,Pgk,Cdk4,sina,Ubc87F,FBXO11,CG1673,Kdm2,Egfr,CG9220,CG3119,E(Pc),CG9485,tkv | | 9.355e-03 | -4.67 | Ephrin\_rec\_like | smart domains | SM01411 | 3 | 2 | 5086 | 290 | Eph,LanA | | 9.355e-03 | -4.67 | DUF1943 | smart domains | SM01169 | 3 | 2 | 5086 | 290 | apolpp,cv-d | | 9.372e-03 | -4.67 | GO\_CELLULAR\_MACROMOLECULE\_LOCALIZATION | MSigDB lists | GO\_CELLULAR\_MACROMOLECULE\_LOCALIZATION | 498 | 29 | 4161 | 156 | cact,Raf,Rab5,Pex7,Pex5,lt,Pen,Vps13,14-3-3zeta,Rab1,14-3-3epsilon,CenB1A,LanA,AP-1gamma,Syx1A,ReepA,emp,Pex10,Nrk,CG45050,Tsp96F,CG7394,Kr-h2,Ric,wbl,Pat1,jar,arm,Pex14 | | 9.383e-03 | -4.67 | regulation of dendrite development | biological process | GO:0050773 | 51 | 6 | 12345 | 442 | rictor,Adf1,kn,robo1,ss,Rab1 | | 9.406e-03 | -4.67 | cell body | cellular component | GO:0044297 | 158 | 12 | 11940 | 417 | bol,orb2,path,Rab5,Ptp4E,rg,Myo10A,robo1,CG9784,Rab1,Tob,Gr93c | | 9.442e-03 | -4.66 | NGF signalling via TRKA from the plasma membrane | REACTOME pathways | R-DME-187037 | 172 | 15 | 3923 | 176 | wdb,trbl,CG42684,Dronc,CG4935,rictor,PI31,htl,14-3-3zeta,Pdk1,Pvr,Rgl,Egfr,vn,Raf | | 9.524e-03 | -4.65 | chromatin organization | biological process | GO:0006325 | 466 | 27 | 12345 | 442 | vig,CtBP,jumu,His2Av,Su(z)2,Rab5,bin3,crol,Nlp,His3.3B,Kdm4B,tara,Dcr-2,His3.3A,nub,HmgZ,eya,hyx,Psc,E(Pc),fl(2)d,dve,FBXO11,corto,Elba2,tna,Kdm2 | | 9.611e-03 | -4.64 | drug binding | molecular function | GO:0008144 | 828 | 41 | 12176 | 414 | CG11069,CG31121,Pgk,CG17646,Cdk4,tkv,CG7536,SERCA,GluProRS,Dcr-2,Egfr,nmo,Rm62,l(1)G0196,Myo10A,sick,Src42A,Dyrk2,Hel89B,drl,Ddr,Eph,ATPCL,htl,InR,Pdk1,Pitslre,SMC3,Pvr,wnd,Drak,otk,Raf,hppy,Lk6,trbl,jar,d,Abl,CG5853,Nrk | | 9.625e-03 | -4.64 | CG14490 (uncharacterized protein) | protein interactions | 37041 | 10 | 3 | 9204 | 432 | CG31211,CG34383,crp | | 9.625e-03 | -4.64 | uif (uninflatable) | protein interactions | 33983 | 10 | 3 | 9204 | 432 | Dl,Dg,N | | 9.625e-03 | -4.64 | PDZ-GEF (PDZ domain-containing guanine nucleotide exchange factor) | protein interactions | 33881 | 10 | 3 | 9204 | 432 | shg,Egfr,Raf | | 9.625e-03 | -4.64 | Ube3a (Ubiquitin protein ligase E3A) | protein interactions | 39266 | 10 | 3 | 9204 | 432 | drl,eff,tkv | | 9.625e-03 | -4.64 | PIP4K (Phosphatidylinositol 5-phosphate 4-kinase) | protein interactions | 3885565 | 10 | 3 | 9204 | 432 | ft,fs(1)h,Rab5 | | 9.625e-03 | -4.64 | DCAF12 (DDB1 and CUL4 associated factor 12) | protein interactions | 41460 | 10 | 3 | 9204 | 432 | scrib,Diap1,Dronc | | 9.625e-03 | -4.64 | lab (labial) | protein interactions | 40817 | 10 | 3 | 9204 | 432 | Scr,hth,abd-A | | 9.625e-03 | -4.64 | CG5791 (uncharacterized protein) | protein interactions | 50055 | 10 | 3 | 9204 | 432 | H2.0,nub,fend | | 9.625e-03 | -4.64 | Ptp99A (Protein tyrosine phosphatase 99A) | protein interactions | 43469 | 10 | 3 | 9204 | 432 | nmo,Abl,InR | | 9.625e-03 | -4.64 | CG18135 (uncharacterized protein) | protein interactions | 40073 | 10 | 3 | 9204 | 432 | Socs36E,Ppn,Myo10A | | 9.625e-03 | -4.64 | sha (shavenoid) | protein interactions | 36213 | 10 | 3 | 9204 | 432 | sina,fz,Sema1a | | 9.625e-03 | -4.64 | mRpS30 (mitochondrial ribosomal protein S30) | protein interactions | 32530 | 10 | 3 | 9204 | 432 | CycG,14-3-3zeta,Sxl | | 9.625e-03 | -4.64 | PVRAP (PVR adaptor protein) | protein interactions | 38677 | 10 | 3 | 9204 | 432 | Pvr,CG9784,Jupiter | | 9.625e-03 | -4.64 | Dlish (Dachs ligand with SH3s) | protein interactions | 37014 | 10 | 3 | 9204 | 432 | ft,d,ds | | 9.643e-03 | -4.64 | GCACCTT\_MIR18A\_MIR18B | MSigDB lists | GCACCTT\_MIR18A\_MIR18B | 23 | 4 | 4161 | 156 | trbl,alpha-Man-Ia,Kdm2,Rab5 | | 9.643e-03 | -4.64 | HALLMARK\_INFLAMMATORY\_RESPONSE | MSigDB lists | HALLMARK\_INFLAMMATORY\_RESPONSE | 23 | 4 | 4161 | 156 | fz2,Actbeta,cact,Tollo | | 9.643e-03 | -4.64 | GO\_EMBRYONIC\_HEART\_TUBE\_DEVELOPMENT | MSigDB lists | GO\_EMBRYONIC\_HEART\_TUBE\_DEVELOPMENT | 23 | 4 | 4161 | 156 | N,Mef2,Dl,arm | | 9.643e-03 | -4.64 | KEGG\_PATHOGENIC\_ESCHERICHIA\_COLI\_INFECTION | MSigDB lists | KEGG\_PATHOGENIC\_ESCHERICHIA\_COLI\_INFECTION | 23 | 4 | 4161 | 156 | dock,betaTub60D,14-3-3zeta,arm | | 9.643e-03 | -4.64 | CTCAAGA\_MIR526B | MSigDB lists | CTCAAGA\_MIR526B | 23 | 4 | 4161 | 156 | Rab1,PI31,CG1129,Cdk4 | | 9.643e-03 | -4.64 | ACAGGGT\_MIR10A\_MIR10B | MSigDB lists | ACAGGGT\_MIR10A\_MIR10B | 23 | 4 | 4161 | 156 | Pgant5,step,bru2,Pat1 | | 9.643e-03 | -4.64 | CARD\_MIR302A\_TARGETS | MSigDB lists | CARD\_MIR302A\_TARGETS | 23 | 4 | 4161 | 156 | Pen,Dl,Cdk4,Rab5 | | 9.643e-03 | -4.64 | GO\_MUCOPOLYSACCHARIDE\_METABOLIC\_PROCESS | MSigDB lists | GO\_MUCOPOLYSACCHARIDE\_METABOLIC\_PROCESS | 23 | 4 | 4161 | 156 | CG9220,GlcAT-I,Ids,frc | | 9.727e-03 | -4.63 | BHLH | prosite domains | PS50888 | 58 | 8 | 6243 | 325 | sim,HLH3B,tai,emc,crp,h,ss,E(spl)m5-HLH | | 9.728e-03 | -4.63 | Pi3K68D (Phosphotidylinositol 3 kinase 68D) | protein interactions | 39329 | 64 | 8 | 9204 | 432 | Rab5,14-3-3zeta,Gfat2,SERCA,GluProRS,N,Mpcp2,Egfr | | 9.728e-03 | -4.63 | Cyp12d1-p (Cyp12d1-p) | protein interactions | 246648 | 64 | 8 | 9204 | 432 | Mpcp2,Kr-h2,GluProRS,Rab5,Pen,Rab1,Gfat2,SERCA | | 9.746e-03 | -4.63 | positive regulation of secretion | biological process | GO:0051047 | 37 | 5 | 12345 | 442 | mop,jar,Rab1,Cadps,S | | 9.752e-03 | -4.63 | dendrite guidance | biological process | GO:0070983 | 24 | 4 | 12345 | 442 | E(Pc),sqz,robo1,drl | | 9.752e-03 | -4.63 | striated muscle tissue development | biological process | GO:0014706 | 24 | 4 | 12345 | 442 | Mef2,sd,N,abd-A | | 9.752e-03 | -4.63 | organ growth | biological process | GO:0035265 | 24 | 4 | 12345 | 442 | ft,Sxl,InR,N | | 9.752e-03 | -4.63 | embryonic heart tube development | biological process | GO:0035050 | 24 | 4 | 12345 | 442 | sli,pnr,robo1,jumu | | 9.752e-03 | -4.63 | regulation of carbohydrate metabolic process | biological process | GO:0006109 | 24 | 4 | 12345 | 442 | Cbl,Dg,Nmda1,N | | 9.813e-03 | -4.62 | oocyte development | biological process | GO:0048599 | 155 | 12 | 12345 | 442 | shg,14-3-3epsilon,N,14-3-3zeta,scrib,cic,Pat1,chic,Dg,baz,wbl,Egfr | | 9.828e-03 | -4.62 | stem cell proliferation | biological process | GO:0072089 | 67 | 7 | 12345 | 442 | scrib,esg,N,flfl,sd,baz,jar | | 9.973e-03 | -4.61 | glycogen catabolic process | biological process | GO:0005980 | 13 | 3 | 12345 | 442 | Atg17,CG9485,Atg16 | | 9.973e-03 | -4.61 | muscle fiber development | biological process | GO:0048747 | 13 | 3 | 12345 | 442 | aop,Mef2,htl | | 9.973e-03 | -4.61 | female gonad development | biological process | GO:0008585 | 13 | 3 | 12345 | 442 | InR,fz2,bab2 | | 9.973e-03 | -4.61 | development of primary female sexual characteristics | biological process | GO:0046545 | 13 | 3 | 12345 | 442 | bab2,fz2,InR | | 9.973e-03 | -4.61 | cardioblast differentiation | biological process | GO:0010002 | 13 | 3 | 12345 | 442 | pnr,htl,abd-A | | 9.973e-03 | -4.61 | glucan catabolic process | biological process | GO:0009251 | 13 | 3 | 12345 | 442 | CG9485,Atg16,Atg17 | | 9.973e-03 | -4.61 | cell-cell signaling involved in cell fate commitment | biological process | GO:0045168 | 13 | 3 | 12345 | 442 | mam,N,Dl | | 9.973e-03 | -4.61 | nephrocyte differentiation | biological process | GO:0061319 | 13 | 3 | 12345 | 442 | sns,pnr,htl | | 9.973e-03 | -4.61 | positive regulation of neuron maturation | biological process | GO:0014042 | 13 | 3 | 12345 | 442 | Arf79F,Sox14,InR | | 9.973e-03 | -4.61 | wing and notum subfield formation | biological process | GO:0035309 | 13 | 3 | 12345 | 442 | Raf,Egfr,vn | | 9.973e-03 | -4.61 | renal filtration cell differentiation | biological process | GO:0061318 | 13 | 3 | 12345 | 442 | pnr,sns,htl | | 9.973e-03 | -4.61 | positive regulation of neuron remodeling | biological process | GO:1904801 | 13 | 3 | 12345 | 442 | Arf79F,Sox14,InR | | 9.973e-03 | -4.61 | imaginal disc-derived leg joint morphogenesis | biological process | GO:0016348 | 13 | 3 | 12345 | 442 | bowl,dve,N | | 1.000e-02 | -4.60 | receptor ligand activity | molecular function | GO:0048018 | 106 | 9 | 12176 | 414 | Actbeta,ITP,Sema1b,spi,Pvf3,htl,Dl,vn,Sema1a | | 1.013e-02 | -4.59 | fly chr38A4-38A4\|2-54 | chromosome location | fly chr38A4-38A4\|2-54 | 7 | 2 | 25010 | 571 | snRNA:U5:38ABb,sick | | 1.013e-02 | -4.59 | fly chr2B14-2B14\|1-0.2 cM | chromosome location | fly chr2B14-2B14\|1-0.2 cM | 7 | 2 | 25010 | 571 | arm,CG14818 | | 1.013e-02 | -4.59 | fly chr63D2-63D2\|3-8 cM | chromosome location | fly chr63D2-63D2\|3-8 cM | 7 | 2 | 25010 | 571 | lncRNA:CR45737,Ids | | 1.013e-02 | -4.59 | fly chr38A4-38A4\|2-54 cM | chromosome location | fly chr38A4-38A4\|2-54 cM | 7 | 2 | 25010 | 571 | sick,snRNA:U5:38ABb | | 1.013e-02 | -4.59 | fly chr61C7-61C7\|3-0.5 cM | chromosome location | fly chr61C7-61C7\|3-0.5 cM | 7 | 2 | 25010 | 571 | lncRNA:CR43334,CG43337 | | 1.013e-02 | -4.59 | fly chr61C7-61C7\|3-0.5 | chromosome location | fly chr61C7-61C7\|3-0.5 | 7 | 2 | 25010 | 571 | lncRNA:CR43334,CG43337 | | 1.013e-02 | -4.59 | fly chr2B14-2B14\|1-0.2 | chromosome location | fly chr2B14-2B14\|1-0.2 | 7 | 2 | 25010 | 571 | CG14818,arm | | 1.013e-02 | -4.59 | fly chr63D2-63D2\|3-8 | chromosome location | fly chr63D2-63D2\|3-8 | 7 | 2 | 25010 | 571 | lncRNA:CR45737,Ids | | 1.020e-02 | -4.59 | EGR1\_01 | MSigDB lists | EGR1\_01 | 65 | 7 | 4161 | 156 | nmo,14-3-3epsilon,Kdm2,AP-1gamma,Mef2,fz,Eph | | 1.023e-02 | -4.58 | MCCLUNG\_CREB1\_TARGETS\_UP | MSigDB lists | MCCLUNG\_CREB1\_TARGETS\_UP | 36 | 5 | 4161 | 156 | Syx1A,Cadps,Cerk,CG1673,Mef2 | | 1.023e-02 | -4.58 | GSE46606\_IRF4\_KO\_VS\_WT\_CD40L\_IL2\_IL5\_3DAY\_STIMULATED\_BCELL\_UP | MSigDB lists | GSE46606\_IRF4\_KO\_VS\_WT\_CD40L\_IL2\_IL5\_3DAY\_STIMULATED\_BCELL\_UP | 36 | 5 | 4161 | 156 | lili,InR,Rim2,Vps13,CG3107 | | 1.023e-02 | -4.58 | GSE36476\_YOUNG\_VS\_OLD\_DONOR\_MEMORY\_CD4\_TCELL\_40H\_TSST\_ACT\_UP | MSigDB lists | GSE36476\_YOUNG\_VS\_OLD\_DONOR\_MEMORY\_CD4\_TCELL\_40H\_TSST\_ACT\_UP | 36 | 5 | 4161 | 156 | Atg16,sns,CG1673,unc-5,Gfat2 | | 1.023e-02 | -4.58 | STAT4\_01 | MSigDB lists | STAT4\_01 | 36 | 5 | 4161 | 156 | arm,Mef2,l(2)37Cg,otk,Unc-76 | | 1.024e-02 | -4.58 | GSE360\_L\_MAJOR\_VS\_M\_TUBERCULOSIS\_DC\_UP | MSigDB lists | GSE360\_L\_MAJOR\_VS\_M\_TUBERCULOSIS\_DC\_UP | 50 | 6 | 4161 | 156 | frc,bru2,Dyb,Mef2,Actbeta,His2Av | | 1.024e-02 | -4.58 | GO\_EPIDERMIS\_DEVELOPMENT | MSigDB lists | GO\_EPIDERMIS\_DEVELOPMENT | 50 | 6 | 4161 | 156 | ap,Dl,Actbeta,N,LanA,arm | | 1.024e-02 | -4.58 | PEREZ\_TP63\_TARGETS | MSigDB lists | PEREZ\_TP63\_TARGETS | 50 | 6 | 4161 | 156 | InR,fz,step,bru2,Gfat2,sli | | 1.024e-02 | -4.58 | GSE3920\_IFNA\_VS\_IFNG\_TREATED\_FIBROBLAST\_DN | MSigDB lists | GSE3920\_IFNA\_VS\_IFNG\_TREATED\_FIBROBLAST\_DN | 50 | 6 | 4161 | 156 | Cerk,nmo,CG4896,InR,trbl,ap | | 1.024e-02 | -4.58 | GO\_MUSCLE\_ORGAN\_DEVELOPMENT | MSigDB lists | GO\_MUSCLE\_ORGAN\_DEVELOPMENT | 50 | 6 | 4161 | 156 | Dl,sns,Mef2,N,fz,LanA | | 1.026e-02 | -4.58 | cellular response to starvation | biological process | GO:0009267 | 84 | 8 | 12345 | 442 | Atg16,rictor,CG7536,Atg17,Raf,InR,lt,SNF4Agamma | | 1.027e-02 | -4.58 | GSE39820\_TGFBETA1\_VS\_TGFBETA3\_IN\_IL6\_TREATED\_CD4\_TCELL\_UP | MSigDB lists | GSE39820\_TGFBETA1\_VS\_TGFBETA3\_IN\_IL6\_TREATED\_CD4\_TCELL\_UP | 81 | 8 | 4161 | 156 | Syx1A,step,Hel89B,CG4896,14-3-3zeta,Arf79F,alpha-Man-Ia,arm | | 1.028e-02 | -4.58 | UQCR-C1 (Ubiquinol-cytochrome c reductase core protein 1) | protein interactions | 41800 | 52 | 7 | 9204 | 432 | p130CAS,UQCR-11,Sxl,ft,Sema1a,bab2,msi | | 1.028e-02 | -4.58 | Khc (Kinesin heavy chain) | protein interactions | 36810 | 52 | 7 | 9204 | 432 | ena,nuf,Unc-76,klar,tkv,Pat1,sina | | 1.034e-02 | -4.57 | FERM\_N | interpro domains | IPR018979 | 12 | 3 | 11383 | 450 | cora,Pez,yrt | | 1.034e-02 | -4.57 | Fringe-like | interpro domains | IPR003378 | 12 | 3 | 11383 | 450 | CG34057,C1GalTA,CG3119 | | 1.039e-02 | -4.57 | EGF | pfam domains | PF00008 | 22 | 4 | 10480 | 419 | ft,N,sli,Dl | | 1.039e-02 | -4.57 | HMG\_box | pfam domains | PF00505 | 22 | 4 | 10480 | 419 | HmgZ,Sox14,CG12104,cic | | 1.040e-02 | -4.57 | Agpat3 (1-Acylglycerol-3-phosphate O-acyltransferase 3) | protein interactions | 39820 | 29 | 5 | 9204 | 432 | SERCA,GluProRS,Rab5,Kr-h2,Mpcp2 | | 1.040e-02 | -4.57 | ovo (ovo) | protein interactions | 31429 | 29 | 5 | 9204 | 432 | arm,Sxl,spi,Egfr,crol | | 1.043e-02 | -4.56 | GO\_POSITIVE\_REGULATION\_OF\_TRANSPORT | MSigDB lists | GO\_POSITIVE\_REGULATION\_OF\_TRANSPORT | 208 | 15 | 4161 | 156 | 14-3-3zeta,Actbeta,Pdk1,Arf79F,Psa,subdued,GlcAT-I,fz2,14-3-3epsilon,cact,Tollo,Dl,CG17883,Syx1A,AP-1gamma | | 1.051e-02 | -4.56 | GO\_POSITIVE\_REGULATION\_OF\_CELL\_DIFFERENTIATION | MSigDB lists | GO\_POSITIVE\_REGULATION\_OF\_CELL\_DIFFERENTIATION | 133 | 11 | 4161 | 156 | otk,Raf,robo1,Mef2,Ddr,sli,tkv,Actbeta,Arf79F,N,arm | | 1.053e-02 | -4.55 | GO\_POSITIVE\_REGULATION\_OF\_INTRACELLULAR\_TRANSPORT | MSigDB lists | GO\_POSITIVE\_REGULATION\_OF\_INTRACELLULAR\_TRANSPORT | 98 | 9 | 4161 | 156 | Arf79F,Pdk1,14-3-3zeta,Psa,GlcAT-I,fz2,14-3-3epsilon,Tollo,CG17883 | | 1.056e-02 | -4.55 | MADF\_DNA\_bdg | pfam domains | PF10545 | 47 | 6 | 10480 | 419 | CG3386,Adf1,rgr,jigr1,CG13204,hng3 | | 1.058e-02 | -4.55 | CG7611 (uncharacterized protein) | protein interactions | 40386 | 19 | 4 | 9204 | 432 | H2.0,jar,CG14506,rdx | | 1.058e-02 | -4.55 | Patronin (patronin) | protein interactions | 36978 | 19 | 4 | 9204 | 432 | CG14478,Fem-1,shot,His2Av | | 1.058e-02 | -4.55 | slbo (slow border cells) | protein interactions | 37889 | 19 | 4 | 9204 | 432 | trbl,aop,jar,Abl | | 1.058e-02 | -4.55 | Ras64B (Ras oncogene at 64B) | protein interactions | 38494 | 19 | 4 | 9204 | 432 | Raf,Rgl,14-3-3epsilon,tkv | | 1.058e-02 | -4.55 | Idh (Isocitrate dehydrogenase) | protein interactions | 44291 | 19 | 4 | 9204 | 432 | Diap1,Dronc,l(2)37Cg,14-3-3zeta | | 1.058e-02 | -4.55 | lap (like-AP180) | protein interactions | 40863 | 19 | 4 | 9204 | 432 | pnr,jigr1,nuf,H | | 1.058e-02 | -4.55 | run (runt) | protein interactions | 33059 | 19 | 4 | 9204 | 432 | lilli,H,CtBP,ttk | | 1.058e-02 | -4.55 | Lar (Leukocyte-antigen-related-like) | protein interactions | 35259 | 19 | 4 | 9204 | 432 | Abl,ena,InR,dlp | | 1.058e-02 | -4.55 | zfh1 (Zn finger homeodomain 1) | protein interactions | 43650 | 19 | 4 | 9204 | 432 | Scr,abd-A,Abd-B,grn | | 1.058e-02 | -4.55 | Tnks (tankyrase) | protein interactions | 43095 | 19 | 4 | 9204 | 432 | PI31,Myo10A,kn,spi | | 1.065e-02 | -4.54 | CycA (Cyclin A) | protein interactions | 39340 | 65 | 8 | 9204 | 432 | ena,Cks30A,His3.3A,CycB3,arm,dome,eff,tara | | 1.066e-02 | -4.54 | FERM\_N | pfam domains | PF09379 | 12 | 3 | 10480 | 419 | cora,yrt,Pez | | 1.066e-02 | -4.54 | Fringe | pfam domains | PF02434 | 12 | 3 | 10480 | 419 | C1GalTA,CG34057,CG3119 | | 1.068e-02 | -4.54 | defense response | biological process | GO:0006952 | 361 | 22 | 12345 | 442 | LanA,htl,Dcr-2,sick,subdued,spen,dome,Src42A,RhoGEF3,jumu,Tollo,shg,CG32795,Pvr,Eph,DPCoAC,imd,N,RhoL,kn,Dronc,eya | | 1.074e-02 | -4.53 | demethylase activity | molecular function | GO:0032451 | 14 | 3 | 12176 | 414 | Tet,Kdm2,Kdm4B | | 1.077e-02 | -4.53 | transcription coactivator binding | molecular function | GO:0001223 | 5 | 2 | 12176 | 414 | sd,hth | | 1.077e-02 | -4.53 | translation activator activity | molecular function | GO:0008494 | 5 | 2 | 12176 | 414 | bol,orb2 | | 1.077e-02 | -4.53 | histone demethylase activity (H3-K36 specific) | molecular function | GO:0051864 | 5 | 2 | 12176 | 414 | Kdm2,Kdm4B | | 1.077e-02 | -4.53 | DNA binding, bending | molecular function | GO:0008301 | 5 | 2 | 12176 | 414 | HmgZ,Sox14 | | 1.079e-02 | -4.53 | Negative regulation of FGFR1 signaling | REACTOME pathways | R-DME-5654726 | 20 | 4 | 3923 | 176 | Cbl,Raf,Lk6,htl | | 1.088e-02 | -4.52 | galactosyltransferase activity | molecular function | GO:0008378 | 26 | 4 | 12176 | 414 | CG3119,CG12910,CG34057,C1GalTA | | 1.092e-02 | -4.52 | CA | smart domains | SM00112 | 16 | 4 | 5086 | 290 | ft,ds,Cals,shg | | 1.097e-02 | -4.51 | apical cortex | cellular component | GO:0045179 | 54 | 6 | 11940 | 417 | baz,Dl,CG42684,cindr,nuf,Abl | | 1.101e-02 | -4.51 | PROTEIN\_KINASE\_ATP | prosite domains | PS00107 | 164 | 16 | 6243 | 325 | InR,Pdk1,Dyrk2,nmo,Nrk,tkv,hppy,Cdk4,Egfr,Src42A,Raf,Pvr,Eph,htl,Lk6,Abl | | 1.103e-02 | -4.51 | GO\_AMIDE\_BINDING | MSigDB lists | GO\_AMIDE\_BINDING | 82 | 8 | 4161 | 156 | fz2,Psa,InR,Pen,Actbeta,Pex7,Pex5,cact | | 1.106e-02 | -4.50 | RIGGI\_EWING\_SARCOMA\_PROGENITOR\_UP | MSigDB lists | RIGGI\_EWING\_SARCOMA\_PROGENITOR\_UP | 66 | 7 | 4161 | 156 | Atg16,Cals,trbl,Eph,ft,sli,Src42A | | 1.106e-02 | -4.50 | GSE16522\_MEMORY\_VS\_NAIVE\_ANTI\_CD3CD28\_STIM\_CD8\_TCELL\_UP | MSigDB lists | GSE16522\_MEMORY\_VS\_NAIVE\_ANTI\_CD3CD28\_STIM\_CD8\_TCELL\_UP | 66 | 7 | 4161 | 156 | CtBP,CG6891,CG7394,Lk6,Pgant5,emp,Ocrl | | 1.110e-02 | -4.50 | GO\_POSITIVE\_REGULATION\_OF\_GENE\_EXPRESSION | MSigDB lists | GO\_POSITIVE\_REGULATION\_OF\_GENE\_EXPRESSION | 395 | 24 | 4161 | 156 | wbl,jar,arm,fz2,tkv,Tsp96F,Nrk,fz,Ddr,Tollo,kn,hyx,dock,ap,sd,Actbeta,N,l(2)37Cg,CtBP,Dl,cact,Raf,Cdk4,Mef2 | | 1.110e-02 | -4.50 | MADF-dom | interpro domains | IPR006578 | 48 | 6 | 11383 | 450 | rgr,CG13204,CG3386,jigr1,hng3,Adf1 | | 1.120e-02 | -4.49 | oocyte differentiation | biological process | GO:0009994 | 177 | 13 | 12345 | 442 | wbl,baz,Dg,Egfr,chic,Pat1,cic,scrib,14-3-3zeta,N,14-3-3epsilon,Sxl,shg | | 1.121e-02 | -4.49 | ATCMNTCCGY\_UNKNOWN | MSigDB lists | ATCMNTCCGY\_UNKNOWN | 13 | 3 | 4161 | 156 | AP-1gamma,eff,N | | 1.121e-02 | -4.49 | MODULE\_122 | MSigDB lists | MODULE\_122 | 13 | 3 | 4161 | 156 | LanA,Ddr,otk | | 1.121e-02 | -4.49 | GO\_REGULATION\_OF\_SKELETAL\_MUSCLE\_TISSUE\_DEVELOPMENT | MSigDB lists | GO\_REGULATION\_OF\_SKELETAL\_MUSCLE\_TISSUE\_DEVELOPMENT | 13 | 3 | 4161 | 156 | arm,Dl,Mef2 | | 1.121e-02 | -4.49 | KEGG\_NON\_SMALL\_CELL\_LUNG\_CANCER | MSigDB lists | KEGG\_NON\_SMALL\_CELL\_LUNG\_CANCER | 13 | 3 | 4161 | 156 | Cdk4,Raf,Pdk1 | | 1.121e-02 | -4.49 | chr7q11 | MSigDB lists | chr7q11 | 13 | 3 | 4161 | 156 | Syx1A,CG32795,Tpst | | 1.121e-02 | -4.49 | PID\_MAPK\_TRK\_PATHWAY | MSigDB lists | PID\_MAPK\_TRK\_PATHWAY | 13 | 3 | 4161 | 156 | Raf,Mef2,Ric | | 1.121e-02 | -4.49 | WAMUNYOKOLI\_OVARIAN\_CANCER\_GRADES\_1\_2\_DN | MSigDB lists | WAMUNYOKOLI\_OVARIAN\_CANCER\_GRADES\_1\_2\_DN | 13 | 3 | 4161 | 156 | bru2,Rgl,sli | | 1.121e-02 | -4.49 | GO\_THYMOCYTE\_AGGREGATION | MSigDB lists | GO\_THYMOCYTE\_AGGREGATION | 13 | 3 | 4161 | 156 | Cdk4,fz2,arm | | 1.122e-02 | -4.49 | SUMOylation of RNA binding proteins | REACTOME pathways | R-DME-4570464 | 11 | 3 | 3923 | 176 | Psc,ph-p,Su(z)2 | | 1.122e-02 | -4.49 | A tetrasaccharide linker sequence is required for GAG synthesis | REACTOME pathways | R-DME-1971475 | 11 | 3 | 3923 | 176 | kon,dlp,GlcAT-I | | 1.122e-02 | -4.49 | Programmed Cell Death | REACTOME pathways | R-DME-5357801 | 11 | 3 | 3923 | 176 | Diap1,Drep4,shot | | 1.122e-02 | -4.49 | Proton-coupled neutral amino acid transporters | REACTOME pathways | R-DME-428559 | 11 | 3 | 3923 | 176 | path,CG13384,CG4991 | | 1.124e-02 | -4.49 | GO\_MESENCHYMAL\_CELL\_DIFFERENTIATION | MSigDB lists | GO\_MESENCHYMAL\_CELL\_DIFFERENTIATION | 24 | 4 | 4161 | 156 | LanA,arm,Mef2,N | | 1.124e-02 | -4.49 | ESC\_V6.5\_UP\_EARLY.V1\_DN | MSigDB lists | ESC\_V6.5\_UP\_EARLY.V1\_DN | 24 | 4 | 4161 | 156 | Dgat2,Actbeta,Cadps,fz2 | | 1.124e-02 | -4.49 | PTEN\_DN.V2\_DN | MSigDB lists | PTEN\_DN.V2\_DN | 24 | 4 | 4161 | 156 | Ric,Dyrk2,htl,unc-119 | | 1.124e-02 | -4.49 | REACTOME\_GOLGI\_ASSOCIATED\_VESICLE\_BIOGENESIS | MSigDB lists | REACTOME\_GOLGI\_ASSOCIATED\_VESICLE\_BIOGENESIS | 24 | 4 | 4161 | 156 | Ocrl,Rab5,AP-1gamma,Arf79F | | 1.124e-02 | -4.49 | CTAGGAA\_MIR384 | MSigDB lists | CTAGGAA\_MIR384 | 24 | 4 | 4161 | 156 | FBXO11,bru2,nmo,unc-5 | | 1.124e-02 | -4.49 | GO\_DORSAL\_VENTRAL\_PATTERN\_FORMATION | MSigDB lists | GO\_DORSAL\_VENTRAL\_PATTERN\_FORMATION | 24 | 4 | 4161 | 156 | arm,fz2,ap,tkv | | 1.124e-02 | -4.49 | GO\_DEVELOPMENTAL\_GROWTH\_INVOLVED\_IN\_MORPHOGENESIS | MSigDB lists | GO\_DEVELOPMENTAL\_GROWTH\_INVOLVED\_IN\_MORPHOGENESIS | 24 | 4 | 4161 | 156 | sli,otk,ap,N | | 1.125e-02 | -4.49 | GSE19401\_PLN\_VS\_PEYERS\_PATCH\_FOLLICULAR\_DC\_UP | MSigDB lists | GSE19401\_PLN\_VS\_PEYERS\_PATCH\_FOLLICULAR\_DC\_UP | 51 | 6 | 4161 | 156 | Mfe2,arm,IntS3,frc,Pat1,trbl | | 1.125e-02 | -4.49 | TFIIA\_Q6 | MSigDB lists | TFIIA\_Q6 | 51 | 6 | 4161 | 156 | lili,dlp,unc-5,14-3-3zeta,betaTub60D,FBXO11 | | 1.125e-02 | -4.49 | MARTORIATI\_MDM4\_TARGETS\_FETAL\_LIVER\_UP | MSigDB lists | MARTORIATI\_MDM4\_TARGETS\_FETAL\_LIVER\_UP | 51 | 6 | 4161 | 156 | Pgk,Tsp96F,otk,CG4935,Rab5,CycG | | 1.127e-02 | -4.49 | regulation of BMP signaling pathway | biological process | GO:0030510 | 53 | 6 | 12345 | 442 | fuss,kek5,hth,Snoo,cv-d,lili | | 1.129e-02 | -4.48 | positive regulation of organ growth | biological process | GO:0046622 | 25 | 4 | 12345 | 442 | Egfr,d,InR,Pdk1 | | 1.129e-02 | -4.48 | imaginal disc-derived male genitalia development | biological process | GO:0007485 | 25 | 4 | 12345 | 442 | Rab5,N,Pvr,otk | | 1.129e-02 | -4.48 | mitochondrion localization | biological process | GO:0051646 | 25 | 4 | 12345 | 442 | Bsg,Rim2,jar,larp | | 1.129e-02 | -4.48 | regulation of morphogenesis of an epithelium | biological process | GO:1905330 | 25 | 4 | 12345 | 442 | baz,otk,aop,ds | | 1.131e-02 | -4.48 | Inactivation of Cdc42 and Rac | REACTOME pathways | R-DME-428543 | 4 | 2 | 3923 | 176 | sli,robo1 | | 1.134e-02 | -4.48 | investment cone | cellular component | GO:0070865 | 5 | 2 | 11940 | 417 | yuri,jar | | 1.144e-02 | -4.47 | FoxO signaling pathway | KEGG pathways | dme04068 | 65 | 7 | 3198 | 123 | Raf,SNF4Agamma,CycB3,nmo,Egfr,InR,Pdk1 | | 1.146e-02 | -4.47 | GO\_PROTEIN\_LOCALIZATION | MSigDB lists | GO\_PROTEIN\_LOCALIZATION | 664 | 36 | 4161 | 156 | Ric,Pex14,arm,CG7394,Nrk,Syx1A,CenB1A,Rab1,N,Rab5,jar,wbl,Pat1,Pex10,CG45050,Tsp96F,Kr-h2,AP-1gamma,Cadps,ReepA,LanA,emp,unc-119,rg,14-3-3epsilon,sns,Pen,lt,14-3-3zeta,Vps13,Arf79F,Pex7,Pex5,cact,Raf,Atg16 | | 1.147e-02 | -4.47 | GSE40666\_WT\_VS\_STAT4\_KO\_CD8\_TCELL\_WITH\_IFNA\_STIM\_90MIN\_UP | MSigDB lists | GSE40666\_WT\_VS\_STAT4\_KO\_CD8\_TCELL\_WITH\_IFNA\_STIM\_90MIN\_UP | 37 | 5 | 4161 | 156 | dock,trbl,CG45050,CenB1A,Ids | | 1.147e-02 | -4.47 | ACACTGG\_MIR199A\_MIR199B | MSigDB lists | ACACTGG\_MIR199A\_MIR199B | 37 | 5 | 4161 | 156 | nmo,AP-1gamma,bru2,Pat1,hyx | | 1.150e-02 | -4.47 | Dg (Dystroglycan) | protein interactions | 36773 | 41 | 6 | 9204 | 432 | Dl,Dad,Pgk,robo1,cora,Sema1a | | 1.150e-02 | -4.47 | Dll (Distal-less) | protein interactions | 37973 | 41 | 6 | 9204 | 432 | Abd-B,abd-A,Su(z)2,ss,hth,Scr | | 1.154e-02 | -4.46 | GO\_POSITIVE\_REGULATION\_OF\_TRANSFERASE\_ACTIVITY | MSigDB lists | GO\_POSITIVE\_REGULATION\_OF\_TRANSFERASE\_ACTIVITY | 191 | 14 | 4161 | 156 | unc-119,Raf,Tollo,Ddr,PI31,hppy,Tsp96F,Pdk1,htl,nAChRalpha4,fz2,arm,Rab1,wbl | | 1.155e-02 | -4.46 | cyclin-dependent protein kinase holoenzyme complex | cellular component | GO:0000307 | 14 | 3 | 11940 | 417 | CycG,CycB3,Cks30A | | 1.155e-02 | -4.46 | subapical complex | cellular component | GO:0035003 | 14 | 3 | 11940 | 417 | baz,Dl,N | | 1.164e-02 | -4.45 | PHDP (Putative homeodomain protein) | protein interactions | 37788 | 66 | 8 | 9204 | 432 | CG32772,Cdk4,Bacc,CG7492,CycG,chic,orb2,Lk6 | | 1.164e-02 | -4.45 | Src64B (Src oncogene at 64B) | protein interactions | 48973 | 66 | 8 | 9204 | 432 | shg,sina,Src42A,kek5,Egfr,CG12496,Socs36E,drl | | 1.164e-02 | -4.45 | hale (hale-bopp) | protein interactions | 40183 | 66 | 8 | 9204 | 432 | CG14478,sns,mura,CG44838,CG34401,Poxn,drongo,nub | | 1.164e-02 | -4.45 | transcription coregulator activity | molecular function | GO:0003712 | 147 | 11 | 12176 | 414 | CtBP,arm,tna,hng3,mam,CG13204,emc,tai,Tob,eya,H | | 1.169e-02 | -4.45 | GO\_REGULATION\_OF\_TRANSCRIPTION\_FROM\_RNA\_POLYMERASE\_II\_PROMOTER | MSigDB lists | GO\_REGULATION\_OF\_TRANSCRIPTION\_FROM\_RNA\_POLYMERASE\_II\_PROMOTER | 312 | 20 | 4161 | 156 | Mef2,cact,Dl,CtBP,N,Actbeta,sd,ap,dock,hyx,kn,Tollo,Src42A,fz,Tsp96F,tkv,fz2,arm,jar,eff | | 1.173e-02 | -4.45 | perinuclear region of cytoplasm | cellular component | GO:0048471 | 71 | 7 | 11940 | 417 | klar,nuf,koi,chic,S,FBXO11,orb2 | | 1.174e-02 | -4.44 | HMG\_box\_dom | interpro domains | IPR009071 | 23 | 4 | 11383 | 450 | cic,Sox14,HmgZ,CG12104 | | 1.175e-02 | -4.44 | wound healing | biological process | GO:0042060 | 86 | 8 | 12345 | 442 | Sxl,ed,kay,shg,RhoGEF3,chic,shot,Pvr | | 1.190e-02 | -4.43 | mesodermal cell fate determination | biological process | GO:0007500 | 5 | 2 | 12345 | 442 | mam,htl | | 1.190e-02 | -4.43 | negative regulation of protein kinase B signaling | biological process | GO:0051898 | 5 | 2 | 12345 | 442 | wdb,trbl | | 1.190e-02 | -4.43 | compound eye cone cell fate commitment | biological process | GO:0042676 | 5 | 2 | 12345 | 442 | Dl,N | | 1.190e-02 | -4.43 | symmetric stem cell division | biological process | GO:0098724 | 5 | 2 | 12345 | 442 | lin-28,shg | | 1.190e-02 | -4.43 | histone H3-K36 demethylation | biological process | GO:0070544 | 5 | 2 | 12345 | 442 | Kdm4B,Kdm2 | | 1.190e-02 | -4.43 | establishment or maintenance of actin cytoskeleton polarity | biological process | GO:0030950 | 5 | 2 | 12345 | 442 | rictor,Shrm | | 1.190e-02 | -4.43 | myoblast differentiation | biological process | GO:0045445 | 5 | 2 | 12345 | 442 | htl,ttk | | 1.190e-02 | -4.43 | notum cell fate specification | biological process | GO:0035310 | 5 | 2 | 12345 | 442 | Egfr,vn | | 1.190e-02 | -4.43 | positive regulation of exocytosis | biological process | GO:0045921 | 5 | 2 | 12345 | 442 | jar,Cadps | | 1.190e-02 | -4.43 | wing disc proximal/distal pattern formation | biological process | GO:0007473 | 5 | 2 | 12345 | 442 | N,tsh | | 1.190e-02 | -4.43 | axon regeneration | biological process | GO:0031103 | 5 | 2 | 12345 | 442 | kay,wnd | | 1.190e-02 | -4.43 | epidermal growth factor receptor ligand maturation | biological process | GO:0038004 | 5 | 2 | 12345 | 442 | S,rho-4 | | 1.190e-02 | -4.43 | regulation of rhodopsin gene expression | biological process | GO:0007468 | 5 | 2 | 12345 | 442 | ss,sens | | 1.190e-02 | -4.43 | negative regulation of axon guidance | biological process | GO:1902668 | 5 | 2 | 12345 | 442 | Sema1b,Sema1a | | 1.190e-02 | -4.43 | negative regulation of axon extension involved in axon guidance | biological process | GO:0048843 | 5 | 2 | 12345 | 442 | Sema1b,Sema1a | | 1.190e-02 | -4.43 | regulation of compound eye cone cell fate specification | biological process | GO:0042682 | 5 | 2 | 12345 | 442 | chn,ttk | | 1.190e-02 | -4.43 | positive regulation of phosphatidylinositol 3-kinase signaling | biological process | GO:0014068 | 5 | 2 | 12345 | 442 | step,InR | | 1.192e-02 | -4.43 | GO\_GOLGI\_MEMBRANE | MSigDB lists | GO\_GOLGI\_MEMBRANE | 231 | 16 | 4161 | 156 | Tollo,C1GalTA,CG17883,AP-1gamma,CG9220,fz2,Tpst,Cals,Pgant6,step,N,Arf79F,Pgant5,GlcAT-I,Rab1,alpha-Man-Ia | | 1.197e-02 | -4.43 | GSE37416\_CTRL\_VS\_48H\_F\_TULARENSIS\_LVS\_NEUTROPHIL\_DN | MSigDB lists | GSE37416\_CTRL\_VS\_48H\_F\_TULARENSIS\_LVS\_NEUTROPHIL\_DN | 67 | 7 | 4161 | 156 | l(2)37Cg,CG2017,PI31,CG13384,Pat1,Hel89B,arm | | 1.197e-02 | -4.43 | GSE39820\_TGFBETA1\_IL6\_VS\_TGFBETA1\_IL6\_IL23A\_TREATED\_CD4\_TCELL\_UP | MSigDB lists | GSE39820\_TGFBETA1\_IL6\_VS\_TGFBETA1\_IL6\_IL23A\_TREATED\_CD4\_TCELL\_UP | 67 | 7 | 4161 | 156 | Gfat2,Syx1A,arm,Pdk1,Arf79F,sina,N | | 1.197e-02 | -4.43 | AGCACTT\_MIR93\_MIR302A\_MIR302B\_MIR302C\_MIR302D\_MIR372\_MIR373\_MIR520E\_MIR520A\_MIR526B\_MIR520B\_MIR520C\_MIR520D | MSigDB lists | AGCACTT\_MIR93\_MIR302A\_MIR302B\_MIR302C\_MIR302D\_MIR372\_MIR373\_MIR520E\_MIR520A\_MIR526B\_MIR520B\_MIR520C\_MIR520D | 67 | 7 | 4161 | 156 | rdx,Rgl,bru2,14-3-3zeta,Mef2,Pen,FBXO11 | | 1.201e-02 | -4.42 | sens (senseless) | protein interactions | 45328 | 30 | 5 | 9204 | 432 | Dl,Egfr,lilli,sd,CtBP | | 1.210e-02 | -4.41 | endomembrane system | cellular component | GO:0012505 | 1113 | 53 | 11940 | 417 | Arf79F,mop,CG30156,ed,spi,GlcAT-I,rho-4,AP-1gamma,Lk6,IntS3,shg,frc,Pgant5,N,wbl,CG8306,Dl,14-3-3zeta,Syx1A,S,CG17883,fz,Fkbp14,yrt,Cadps,CG7536,CG4645,SERCA,Ocrl,CG3149,yuri,Rab1,siz,Kr-h2,step,jeb,Pgant6,path,spdo,pum,klar,alpha-Man-Ia,fz2,tkv,lt,nuf,Tpst,CG9220,Rab5,Vps13,koi,ReepA,TBC1D5 | | 1.215e-02 | -4.41 | encapsulation of foreign target | biological process | GO:0035010 | 39 | 5 | 12345 | 442 | alpha-Man-Ia,RhoGEF3,RhoL,LanA,Eph | | 1.215e-02 | -4.41 | brain development | biological process | GO:0007420 | 122 | 10 | 12345 | 442 | shot,Eph,chic,vn,drl,sim,ey,rg,hth,shg | | 1.218e-02 | -4.41 | innate immune response | biological process | GO:0045087 | 104 | 9 | 12345 | 442 | LanA,Pvr,Eph,RhoL,kn,Dronc,RhoGEF3,Tollo,eya | | 1.220e-02 | -4.41 | cell-substrate junction | cellular component | GO:0030055 | 40 | 5 | 11940 | 417 | shot,p130CAS,LRR,kon,Abl | | 1.220e-02 | -4.41 | cell-substrate adherens junction | cellular component | GO:0005924 | 40 | 5 | 11940 | 417 | shot,p130CAS,LRR,kon,Abl | | 1.229e-02 | -4.40 | w (white) | protein interactions | 31271 | 80 | 9 | 9204 | 432 | Diap1,lt,Su(z)2,ph-p,Psc,Dcr-2,Alh,E(Pc),Rm62 | | 1.232e-02 | -4.40 | GO\_CELL\_SURFACE | MSigDB lists | GO\_CELL\_SURFACE | 118 | 10 | 4161 | 156 | sli,Tollo,Cals,robo1,wbl,dlp,fz2,htl,N,fz | | 1.234e-02 | -4.40 | GO\_SENSORY\_ORGAN\_MORPHOGENESIS | MSigDB lists | GO\_SENSORY\_ORGAN\_MORPHOGENESIS | 52 | 6 | 4161 | 156 | arm,CG17883,fz2,otk,Eph,Dl | | 1.234e-02 | -4.40 | GO\_MICROBODY\_PART | MSigDB lists | GO\_MICROBODY\_PART | 52 | 6 | 4161 | 156 | Pex10,Arf79F,Pex7,Pex14,Pex5,Mfe2 | | 1.234e-02 | -4.40 | FOXO4\_01 | MSigDB lists | FOXO4\_01 | 52 | 6 | 4161 | 156 | His2Av,Actbeta,Eph,lqf,Cadps,shot | | 1.234e-02 | -4.40 | GSE28237\_FOLLICULAR\_VS\_LATE\_GC\_BCELL\_DN | MSigDB lists | GSE28237\_FOLLICULAR\_VS\_LATE\_GC\_BCELL\_DN | 52 | 6 | 4161 | 156 | Mfe2,CG13384,Rab1,wdb,Cdk4,CG9220 | | 1.236e-02 | -4.39 | chromosomal region | cellular component | GO:0098687 | 204 | 14 | 11940 | 417 | jumu,Elba2,Psc,Adf1,Nlp,lawc,Rm62,His2Av,flfl,mam,Spc105R,E(Pc),bab2,vig | | 1.236e-02 | -4.39 | positive regulation of photoreceptor cell differentiation | biological process | GO:0046534 | 14 | 3 | 12345 | 442 | mop,Src42A,Raf | | 1.236e-02 | -4.39 | cellular polysaccharide catabolic process | biological process | GO:0044247 | 14 | 3 | 12345 | 442 | Atg17,CG9485,Atg16 | | 1.236e-02 | -4.39 | neuron fate specification | biological process | GO:0048665 | 14 | 3 | 12345 | 442 | ey,N,Dl | | 1.236e-02 | -4.39 | demethylation | biological process | GO:0070988 | 14 | 3 | 12345 | 442 | Kdm4B,Tet,Kdm2 | | 1.236e-02 | -4.39 | polysaccharide catabolic process | biological process | GO:0000272 | 14 | 3 | 12345 | 442 | Atg16,CG9485,Atg17 | | 1.236e-02 | -4.39 | O-glycan processing | biological process | GO:0016266 | 14 | 3 | 12345 | 442 | CG3119,C1GalTA,CG34057 | | 1.237e-02 | -4.39 | learning | biological process | GO:0007612 | 70 | 7 | 12345 | 442 | Adf1,rg,Egfr,vn,drl,spi,14-3-3zeta | | 1.238e-02 | -4.39 | CG10082 (uncharacterized protein) | protein interactions | 37465 | 4 | 2 | 9204 | 432 | kn,Pvr | | 1.238e-02 | -4.39 | CG8207 (uncharacterized protein) | protein interactions | 36730 | 4 | 2 | 9204 | 432 | CG1129,step | | 1.238e-02 | -4.39 | Asap (ArfGAP with SH3 domain, ankyrin repeat and PH domain) | protein interactions | 35783 | 4 | 2 | 9204 | 432 | baz,cindr | | 1.238e-02 | -4.39 | Optix (optix) | protein interactions | 44108 | 4 | 2 | 9204 | 432 | eya,ey | | 1.238e-02 | -4.39 | Nhe2 (Na[+]/H[+] hydrogen exchanger 2) | protein interactions | 35376 | 4 | 2 | 9204 | 432 | fz,Raf | | 1.238e-02 | -4.39 | Slob (Slowpoke binding protein) | protein interactions | 34038 | 4 | 2 | 9204 | 432 | 14-3-3zeta,Fem-1 | | 1.238e-02 | -4.39 | CG31637 (uncharacterized protein) | protein interactions | 33914 | 4 | 2 | 9204 | 432 | unc-119,imd | | 1.238e-02 | -4.39 | CG13344 (uncharacterized protein) | protein interactions | 36541 | 4 | 2 | 9204 | 432 | eff,Cdk4 | | 1.238e-02 | -4.39 | Burs (Bursicon) | protein interactions | 42560 | 4 | 2 | 9204 | 432 | vn,Egfr | | 1.238e-02 | -4.39 | lin-28 (lin-28) | protein interactions | 38639 | 4 | 2 | 9204 | 432 | InR,pnr | | 1.238e-02 | -4.39 | CG3760 (uncharacterized protein) | protein interactions | 37987 | 4 | 2 | 9204 | 432 | ft,Syx1A | | 1.238e-02 | -4.39 | CG11779 (uncharacterized protein) | protein interactions | 42298 | 4 | 2 | 9204 | 432 | rictor,H | | 1.238e-02 | -4.39 | Asator (asator) | protein interactions | 43794 | 4 | 2 | 9204 | 432 | Sema1a,CycG | | 1.238e-02 | -4.39 | CG17556 (uncharacterized protein) | protein interactions | 42046 | 4 | 2 | 9204 | 432 | CG14153,CG32772 | | 1.238e-02 | -4.39 | mir-2b-1 (mir-2b-1 stem loop) | protein interactions | 12798306 | 4 | 2 | 9204 | 432 | SmD1,vig | | 1.238e-02 | -4.39 | CG6791 (uncharacterized protein) | protein interactions | 41391 | 4 | 2 | 9204 | 432 | spi,Rgl | | 1.238e-02 | -4.39 | CG6227 (uncharacterized protein) | protein interactions | 32464 | 4 | 2 | 9204 | 432 | rictor,Sema1a | | 1.241e-02 | -4.39 | Signaling by Rho GTPases | REACTOME pathways | R-DME-194315 | 114 | 11 | 3923 | 176 | RhoU,Cip4,RhoGAP19D,dock,RhoGEF4,Abl,SCAR,Ocrl,Pdk1,14-3-3zeta,14-3-3epsilon | | 1.267e-02 | -4.37 | GO\_POSITIVE\_REGULATION\_OF\_CELL\_DEVELOPMENT | MSigDB lists | GO\_POSITIVE\_REGULATION\_OF\_CELL\_DEVELOPMENT | 84 | 8 | 4161 | 156 | arm,Arf79F,N,sli,otk,Mef2,Raf,robo1 | | 1.267e-02 | -4.37 | GSE20727\_CTRL\_VS\_ROS\_INH\_AND\_DNFB\_ALLERGEN\_TREATED\_DC\_DN | MSigDB lists | GSE20727\_CTRL\_VS\_ROS\_INH\_AND\_DNFB\_ALLERGEN\_TREATED\_DC\_DN | 84 | 8 | 4161 | 156 | Cdk4,rg,Kr-h2,Pgant5,Pen,alpha-Man-Ia,CycG,CG1316 | | 1.270e-02 | -4.37 | aub (aubergine) | protein interactions | 34524 | 67 | 8 | 9204 | 432 | shg,Rm62,eyg,Sxl,Cbl,tkv,Pgk,CG4991 | | 1.274e-02 | -4.36 | EbpIII (Ejaculatory bulb protein III) | protein interactions | 49821 | 20 | 4 | 9204 | 432 | CG32795,14-3-3zeta,apolpp,Pen | | 1.274e-02 | -4.36 | Tlk (Tousled-like kinase) | protein interactions | 318206 | 20 | 4 | 9204 | 432 | pnr,H,Pvr,N | | 1.274e-02 | -4.36 | srp (serpent) | protein interactions | 41944 | 20 | 4 | 9204 | 432 | Scr,Adf1,abd-A,Abd-B | | 1.274e-02 | -4.36 | mats (mob as tumor suppressor) | protein interactions | 42634 | 20 | 4 | 9204 | 432 | SERCA,ft,hppy,Mpcp2 | | 1.274e-02 | -4.36 | DAAM (Dishevelled Associated Activator of Morphogenesis) | protein interactions | 31075 | 20 | 4 | 9204 | 432 | ena,Src42A,fz,chic | | 1.274e-02 | -4.36 | wrd (well-rounded) | protein interactions | 42169 | 20 | 4 | 9204 | 432 | CycG,Scr,baz,wdb | | 1.274e-02 | -4.36 | Ptp61F (Protein tyrosine phosphatase 61F) | protein interactions | 38160 | 20 | 4 | 9204 | 432 | Abl,dock,InR,Egfr | | 1.276e-02 | -4.36 | PAIRED\_2 | prosite domains | PS51057 | 10 | 3 | 6243 | 325 | ey,Poxn,eyg | | 1.278e-02 | -4.36 | scb (scab) | protein interactions | 36692 | 11 | 3 | 9204 | 432 | sli,LanA,tkv | | 1.278e-02 | -4.36 | bgcn (benign gonial cell neoplasm) | protein interactions | 47873 | 11 | 3 | 9204 | 432 | Sxl,pum,shg | | 1.278e-02 | -4.36 | caup (caupolican) | protein interactions | 39440 | 11 | 3 | 9204 | 432 | Diap1,Dl,N | | 1.278e-02 | -4.36 | Usp2 (Ubiquitin specific protease 2) | protein interactions | 33132 | 11 | 3 | 9204 | 432 | imd,CycG,Dronc | | 1.278e-02 | -4.36 | pygo (pygopus) | protein interactions | 43718 | 11 | 3 | 9204 | 432 | arm,N,CtBP | | 1.278e-02 | -4.36 | cbt (cabut) | protein interactions | 33224 | 11 | 3 | 9204 | 432 | tkv,pnr,pum | | 1.278e-02 | -4.36 | CkIalpha (Casein kinase Ialpha) | protein interactions | 32221 | 11 | 3 | 9204 | 432 | arm,H,Pdk1 | | 1.278e-02 | -4.36 | Aldh (Aldehyde dehydrogenase) | protein interactions | 34256 | 11 | 3 | 9204 | 432 | rictor,Raf,Pdk1 | | 1.278e-02 | -4.36 | TwdlW (TweedleW) | protein interactions | 42075 | 11 | 3 | 9204 | 432 | nuf,Pdk1,CG4991 | | 1.278e-02 | -4.36 | Dab (Disabled) | protein interactions | 39866 | 11 | 3 | 9204 | 432 | Abl,N,ena | | 1.278e-02 | -4.36 | Wnt4 (Wnt oncogene analog 4) | protein interactions | 34007 | 11 | 3 | 9204 | 432 | fz2,fz,otk | | 1.282e-02 | -4.36 | YAGI\_AML\_SURVIVAL | MSigDB lists | YAGI\_AML\_SURVIVAL | 38 | 5 | 4161 | 156 | cact,CG1673,Cdk4,Pat1,Hel89B | | 1.282e-02 | -4.36 | GSE29949\_CD8\_POS\_DC\_SPLEEN\_VS\_DC\_BRAIN\_DN | MSigDB lists | GSE29949\_CD8\_POS\_DC\_SPLEEN\_VS\_DC\_BRAIN\_DN | 38 | 5 | 4161 | 156 | Eph,Dyb,Pex10,Rgl,dlp | | 1.282e-02 | -4.36 | GSE19941\_UNSTIM\_VS\_LPS\_STIM\_IL10\_KO\_MACROPHAGE\_UP | MSigDB lists | GSE19941\_UNSTIM\_VS\_LPS\_STIM\_IL10\_KO\_MACROPHAGE\_UP | 38 | 5 | 4161 | 156 | Gfat2,CycG,ATPCL,CG17883,Nrk | | 1.282e-02 | -4.36 | CTCTGGA\_MIR520A\_MIR525 | MSigDB lists | CTCTGGA\_MIR520A\_MIR525 | 38 | 5 | 4161 | 156 | bru2,eff,ds,wdb,Atg16 | | 1.290e-02 | -4.35 | O-linked glycosylation of mucins | REACTOME pathways | R-DME-913709 | 32 | 5 | 3923 | 176 | C1GalTA,Pgant6,CG34057,CG3119,Pgant5 | | 1.290e-02 | -4.35 | Psc (Posterior sex combs) | protein interactions | 36431 | 42 | 6 | 9204 | 432 | Kdm2,Su(z)2,Psc,ph-p,fs(1)h,sd | | 1.290e-02 | -4.35 | bam (bag of marbles) | protein interactions | 43038 | 42 | 6 | 9204 | 432 | tkv,lili,pum,Socs36E,trbl,Sxl | | 1.295e-02 | -4.35 | Galactose-bd-like\_sf | interpro domains | IPR008979 | 36 | 5 | 11383 | 450 | LanA,Ddr,nahoda,Eph,koi | | 1.296e-02 | -4.35 | GO\_LEFT\_RIGHT\_AXIS\_SPECIFICATION | MSigDB lists | GO\_LEFT\_RIGHT\_AXIS\_SPECIFICATION | 5 | 2 | 4161 | 156 | N,Dl | | 1.296e-02 | -4.35 | GO\_EPITHELIAL\_CELL\_FATE\_COMMITMENT | MSigDB lists | GO\_EPITHELIAL\_CELL\_FATE\_COMMITMENT | 5 | 2 | 4161 | 156 | Dl,N | | 1.296e-02 | -4.35 | GO\_A\_BAND | MSigDB lists | GO\_A\_BAND | 5 | 2 | 4161 | 156 | shot,SERCA | | 1.296e-02 | -4.35 | GO\_CENTRAL\_NERVOUS\_SYSTEM\_PROJECTION\_NEURON\_AXONOGENESIS | MSigDB lists | GO\_CENTRAL\_NERVOUS\_SYSTEM\_PROJECTION\_NEURON\_AXONOGENESIS | 5 | 2 | 4161 | 156 | Eph,sli | | 1.296e-02 | -4.35 | GO\_PROTEIN\_LOCALIZATION\_TO\_LYSOSOME | MSigDB lists | GO\_PROTEIN\_LOCALIZATION\_TO\_LYSOSOME | 5 | 2 | 4161 | 156 | Tsp96F,emp | | 1.296e-02 | -4.35 | GO\_NEGATIVE\_REGULATION\_OF\_ENDOTHELIAL\_CELL\_MIGRATION | MSigDB lists | GO\_NEGATIVE\_REGULATION\_OF\_ENDOTHELIAL\_CELL\_MIGRATION | 5 | 2 | 4161 | 156 | sli,N | | 1.296e-02 | -4.35 | GO\_NEGATIVE\_REGULATION\_OF\_GLIAL\_CELL\_DIFFERENTIATION | MSigDB lists | GO\_NEGATIVE\_REGULATION\_OF\_GLIAL\_CELL\_DIFFERENTIATION | 5 | 2 | 4161 | 156 | N,arm | | 1.296e-02 | -4.35 | MODULE\_234 | MSigDB lists | MODULE\_234 | 5 | 2 | 4161 | 156 | htl,Actbeta | | 1.296e-02 | -4.35 | ZHAN\_MULTIPLE\_MYELOMA\_MS\_UP | MSigDB lists | ZHAN\_MULTIPLE\_MYELOMA\_MS\_UP | 5 | 2 | 4161 | 156 | htl,sd | | 1.296e-02 | -4.35 | GO\_HEPARAN\_SULFATE\_PROTEOGLYCAN\_BINDING | MSigDB lists | GO\_HEPARAN\_SULFATE\_PROTEOGLYCAN\_BINDING | 5 | 2 | 4161 | 156 | sli,dlp | | 1.296e-02 | -4.35 | FUKUSHIMA\_TNFSF11\_TARGETS | MSigDB lists | FUKUSHIMA\_TNFSF11\_TARGETS | 5 | 2 | 4161 | 156 | Dl,N | | 1.296e-02 | -4.35 | GO\_ATRIOVENTRICULAR\_VALVE\_MORPHOGENESIS | MSigDB lists | GO\_ATRIOVENTRICULAR\_VALVE\_MORPHOGENESIS | 5 | 2 | 4161 | 156 | N,ds | | 1.296e-02 | -4.35 | GO\_NEGATIVE\_REGULATION\_OF\_MYOBLAST\_DIFFERENTIATION | MSigDB lists | GO\_NEGATIVE\_REGULATION\_OF\_MYOBLAST\_DIFFERENTIATION | 5 | 2 | 4161 | 156 | Dl,N | | 1.296e-02 | -4.35 | RIZ\_ERYTHROID\_DIFFERENTIATION\_12HR | MSigDB lists | RIZ\_ERYTHROID\_DIFFERENTIATION\_12HR | 5 | 2 | 4161 | 156 | sd,N | | 1.296e-02 | -4.35 | GO\_INSULIN\_RECEPTOR\_BINDING | MSigDB lists | GO\_INSULIN\_RECEPTOR\_BINDING | 5 | 2 | 4161 | 156 | InR,Pdk1 | | 1.296e-02 | -4.35 | LIU\_IL13\_MEMORY\_MODEL\_UP | MSigDB lists | LIU\_IL13\_MEMORY\_MODEL\_UP | 5 | 2 | 4161 | 156 | Lk6,unc-119 | | 1.296e-02 | -4.35 | GO\_OLFACTORY\_BULB\_INTERNEURON\_DEVELOPMENT | MSigDB lists | GO\_OLFACTORY\_BULB\_INTERNEURON\_DEVELOPMENT | 5 | 2 | 4161 | 156 | robo1,sli | | 1.296e-02 | -4.35 | GO\_WNT\_ACTIVATED\_RECEPTOR\_ACTIVITY | MSigDB lists | GO\_WNT\_ACTIVATED\_RECEPTOR\_ACTIVITY | 5 | 2 | 4161 | 156 | fz,fz2 | | 1.296e-02 | -4.35 | GO\_POSITIVE\_REGULATION\_OF\_INTERFERON\_GAMMA\_PRODUCTION | MSigDB lists | GO\_POSITIVE\_REGULATION\_OF\_INTERFERON\_GAMMA\_PRODUCTION | 5 | 2 | 4161 | 156 | fz2,Tollo | | 1.296e-02 | -4.35 | REACTOME\_DOWNSTREAM\_TCR\_SIGNALING | MSigDB lists | REACTOME\_DOWNSTREAM\_TCR\_SIGNALING | 5 | 2 | 4161 | 156 | cact,Pdk1 | | 1.296e-02 | -4.35 | GO\_OLFACTORY\_BULB\_INTERNEURON\_DIFFERENTIATION | MSigDB lists | GO\_OLFACTORY\_BULB\_INTERNEURON\_DIFFERENTIATION | 5 | 2 | 4161 | 156 | sli,robo1 | | 1.296e-02 | -4.35 | GO\_POSITIVE\_REGULATION\_OF\_NEUROBLAST\_PROLIFERATION | MSigDB lists | GO\_POSITIVE\_REGULATION\_OF\_NEUROBLAST\_PROLIFERATION | 5 | 2 | 4161 | 156 | N,arm | | 1.296e-02 | -4.35 | GO\_POSITIVE\_REGULATION\_OF\_GLYCOPROTEIN\_METABOLIC\_PROCESS | MSigDB lists | GO\_POSITIVE\_REGULATION\_OF\_GLYCOPROTEIN\_METABOLIC\_PROCESS | 5 | 2 | 4161 | 156 | Rab1,arm | | 1.296e-02 | -4.35 | chr6q13 | MSigDB lists | chr6q13 | 5 | 2 | 4161 | 156 | Dl,jar | | 1.296e-02 | -4.35 | GO\_REGULATION\_OF\_DEVELOPMENT\_HETEROCHRONIC | MSigDB lists | GO\_REGULATION\_OF\_DEVELOPMENT\_HETEROCHRONIC | 5 | 2 | 4161 | 156 | Dl,N | | 1.296e-02 | -4.35 | GO\_ENDODERMAL\_CELL\_FATE\_COMMITMENT | MSigDB lists | GO\_ENDODERMAL\_CELL\_FATE\_COMMITMENT | 5 | 2 | 4161 | 156 | arm,hyx | | 1.296e-02 | -4.35 | GO\_POLYPEPTIDE\_N\_ACETYLGALACTOSAMINYLTRANSFERASE\_ACTIVITY | MSigDB lists | GO\_POLYPEPTIDE\_N\_ACETYLGALACTOSAMINYLTRANSFERASE\_ACTIVITY | 5 | 2 | 4161 | 156 | Pgant6,Pgant5 | | 1.296e-02 | -4.35 | GO\_CELL\_SURFACE\_RECEPTOR\_SIGNALING\_PATHWAY\_INVOLVED\_IN\_HEART\_DEVELOPMENT | MSigDB lists | GO\_CELL\_SURFACE\_RECEPTOR\_SIGNALING\_PATHWAY\_INVOLVED\_IN\_HEART\_DEVELOPMENT | 5 | 2 | 4161 | 156 | arm,N | | 1.296e-02 | -4.35 | GO\_LUNG\_ALVEOLUS\_DEVELOPMENT | MSigDB lists | GO\_LUNG\_ALVEOLUS\_DEVELOPMENT | 5 | 2 | 4161 | 156 | fz,alpha-Man-Ia | | 1.296e-02 | -4.35 | GO\_EMBRYONIC\_HINDLIMB\_MORPHOGENESIS | MSigDB lists | GO\_EMBRYONIC\_HINDLIMB\_MORPHOGENESIS | 5 | 2 | 4161 | 156 | N,arm | | 1.296e-02 | -4.35 | GO\_NEGATIVE\_REGULATION\_OF\_TOLL\_LIKE\_RECEPTOR\_SIGNALING\_PATHWAY | MSigDB lists | GO\_NEGATIVE\_REGULATION\_OF\_TOLL\_LIKE\_RECEPTOR\_SIGNALING\_PATHWAY | 5 | 2 | 4161 | 156 | Tollo,Pdk1 | | 1.296e-02 | -4.35 | GO\_RESPONSE\_TO\_MURAMYL\_DIPEPTIDE | MSigDB lists | GO\_RESPONSE\_TO\_MURAMYL\_DIPEPTIDE | 5 | 2 | 4161 | 156 | N,cact | | 1.296e-02 | -4.35 | GO\_ARF\_GUANYL\_NUCLEOTIDE\_EXCHANGE\_FACTOR\_ACTIVITY | MSigDB lists | GO\_ARF\_GUANYL\_NUCLEOTIDE\_EXCHANGE\_FACTOR\_ACTIVITY | 5 | 2 | 4161 | 156 | step,dock | | 1.297e-02 | -4.35 | regulation of hemopoiesis | biological process | GO:1903706 | 26 | 4 | 12345 | 442 | Arf79F,kn,N,C1GalTA | | 1.297e-02 | -4.35 | regulation of imaginal disc-derived wing size | biological process | GO:0044719 | 26 | 4 | 12345 | 442 | dpy,H,path,CG6191 | | 1.300e-02 | -4.34 | MORF\_EPHA7 | MSigDB lists | MORF\_EPHA7 | 25 | 4 | 4161 | 156 | Drep4,nAChRalpha4,lt,Src42A | | 1.300e-02 | -4.34 | GO\_CARBOHYDRATE\_DERIVATIVE\_TRANSPORT | MSigDB lists | GO\_CARBOHYDRATE\_DERIVATIVE\_TRANSPORT | 25 | 4 | 4161 | 156 | CG3149,Tpc1,frc,DPCoAC | | 1.300e-02 | -4.34 | GO\_RECEPTOR\_SIGNALING\_PROTEIN\_SERINE\_THREONINE\_KINASE\_ACTIVITY | MSigDB lists | GO\_RECEPTOR\_SIGNALING\_PROTEIN\_SERINE\_THREONINE\_KINASE\_ACTIVITY | 25 | 4 | 4161 | 156 | tkv,Raf,hppy,nmo | | 1.300e-02 | -4.34 | GO\_ENDOCRINE\_SYSTEM\_DEVELOPMENT | MSigDB lists | GO\_ENDOCRINE\_SYSTEM\_DEVELOPMENT | 25 | 4 | 4161 | 156 | Dl,Pdk1,Cdk4,Raf | | 1.300e-02 | -4.34 | GO\_SYNAPTIC\_VESICLE\_LOCALIZATION | MSigDB lists | GO\_SYNAPTIC\_VESICLE\_LOCALIZATION | 25 | 4 | 4161 | 156 | Arf79F,arm,Cadps,Syx1A | | 1.300e-02 | -4.34 | CATRRAGC\_UNKNOWN | MSigDB lists | CATRRAGC\_UNKNOWN | 25 | 4 | 4161 | 156 | nmo,14-3-3epsilon,bru2,fz | | 1.300e-02 | -4.34 | GO\_GLYCOPROTEIN\_BINDING | MSigDB lists | GO\_GLYCOPROTEIN\_BINDING | 25 | 4 | 4161 | 156 | tkv,sli,dlp,Syx1A | | 1.312e-02 | -4.33 | GO\_REGULATION\_OF\_DEFENSE\_RESPONSE | MSigDB lists | GO\_REGULATION\_OF\_DEFENSE\_RESPONSE | 194 | 14 | 4161 | 156 | Tollo,cact,Mef2,Tpc1,ReepA,CG4645,AP-1gamma,Spn42De,PI31,sli,Pdk1,Arf79F,eff,SmD1 | | 1.326e-02 | -4.32 | GO\_NEGATIVE\_REGULATION\_OF\_DEVELOPMENTAL\_PROCESS | MSigDB lists | GO\_NEGATIVE\_REGULATION\_OF\_DEVELOPMENTAL\_PROCESS | 156 | 12 | 4161 | 156 | htl,trbl,Actbeta,N,arm,cact,Tollo,Dl,Cdk4,hyx,robo1,sli | | 1.330e-02 | -4.32 | fly chr47A11-47A11\|2-62 | chromosome location | fly chr47A11-47A11\|2-62 | 8 | 2 | 25010 | 571 | snoRNA:lola-a,asRNA:CR45134 | | 1.330e-02 | -4.32 | fly chr47A11-47A11\|2-62 cM | chromosome location | fly chr47A11-47A11\|2-62 cM | 8 | 2 | 25010 | 571 | snoRNA:lola-a,asRNA:CR45134 | | 1.330e-02 | -4.32 | fly chr62A3-62A3\|3-0.5 | chromosome location | fly chr62A3-62A3\|3-0.5 | 8 | 2 | 25010 | 571 | Psa,cue | | 1.330e-02 | -4.32 | fly chr62A3-62A3\|3-0.5 cM | chromosome location | fly chr62A3-62A3\|3-0.5 cM | 8 | 2 | 25010 | 571 | cue,Psa | | 1.331e-02 | -4.32 | regulation of humoral immune response | biological process | GO:0002920 | 71 | 7 | 12345 | 442 | mop,Pvr,cact,Pitslre,imd,pnr,nub | | 1.338e-02 | -4.31 | - | gene3d domains | 2.30.230.10 | 4 | 2 | 4929 | 241 | cv-d,apolpp | | 1.338e-02 | -4.31 | - | gene3d domains | 1.25.10.20 | 4 | 2 | 4929 | 241 | apolpp,cv-d | | 1.338e-02 | -4.31 | - | gene3d domains | 3.80.20.20 | 4 | 2 | 4929 | 241 | InR,Egfr | | 1.338e-02 | -4.31 | cellular response to nutrient levels | biological process | GO:0031669 | 88 | 8 | 12345 | 442 | rictor,Atg17,CG7536,Atg16,SNF4Agamma,lt,InR,Raf | | 1.338e-02 | -4.31 | regulation of immune effector process | biological process | GO:0002697 | 88 | 8 | 12345 | 442 | nub,pnr,Pitslre,mop,imd,Rm62,cact,Dcr-2 | | 1.348e-02 | -4.31 | regulation of cell maturation | biological process | GO:1903429 | 40 | 5 | 12345 | 442 | klar,Syx1A,InR,Sox14,Arf79F | | 1.349e-02 | -4.31 | GSE17721\_CPG\_VS\_GARDIQUIMOD\_0.5H\_BMDC\_DN | MSigDB lists | GSE17721\_CPG\_VS\_GARDIQUIMOD\_0.5H\_BMDC\_DN | 53 | 6 | 4161 | 156 | 14-3-3epsilon,lqf,arm,GlcAT-I,Lk6,N | | 1.349e-02 | -4.31 | CACCAGC\_MIR138 | MSigDB lists | CACCAGC\_MIR138 | 53 | 6 | 4161 | 156 | Arf79F,CG6891,rg,Tmhs,CG5522,PI31 | | 1.349e-02 | -4.31 | SOX9\_B1 | MSigDB lists | SOX9\_B1 | 53 | 6 | 4161 | 156 | Cadps,sli,arm,sd,trbl,Actbeta | | 1.349e-02 | -4.31 | GO\_POSITIVE\_REGULATION\_OF\_NEURON\_DIFFERENTIATION | MSigDB lists | GO\_POSITIVE\_REGULATION\_OF\_NEURON\_DIFFERENTIATION | 53 | 6 | 4161 | 156 | sli,otk,Mef2,Arf79F,robo1,Raf | | 1.349e-02 | -4.31 | GSE30962\_ACUTE\_VS\_CHRONIC\_LCMV\_PRIMARY\_INF\_CD8\_TCELL\_DN | MSigDB lists | GSE30962\_ACUTE\_VS\_CHRONIC\_LCMV\_PRIMARY\_INF\_CD8\_TCELL\_DN | 53 | 6 | 4161 | 156 | Cdk4,Mef2,Tsp96F,Pen,CG1673,bru2 | | 1.362e-02 | -4.30 | Wnt signaling pathway | KEGG pathways | ko04310 | 100 | 9 | 3198 | 123 | CtBP,dlp,sina,RhoL,arm,fz,nmo,fz3,fz2 | | 1.362e-02 | -4.30 | Wnt signaling pathway | KEGG pathways | dme04310 | 100 | 9 | 3198 | 123 | fz2,fz3,nmo,arm,fz,RhoL,dlp,sina,CtBP | | 1.378e-02 | -4.28 | ox (oxen) | protein interactions | 45401 | 31 | 5 | 9204 | 432 | Rab5,Kr-h2,Arf79F,SERCA,UQCR-11 | | 1.378e-02 | -4.28 | tws (twins) | protein interactions | 47877 | 31 | 5 | 9204 | 432 | shg,arm,baz,Cdk4,Egfr | | 1.378e-02 | -4.28 | pk (prickle) | protein interactions | 45343 | 31 | 5 | 9204 | 432 | fz,N,nmo,ds,d | | 1.382e-02 | -4.28 | Akap200 (A kinase anchor protein 200) | protein interactions | 34170 | 68 | 8 | 9204 | 432 | Pdk1,imd,pnr,Fkbp14,Nlp,rictor,N,Src42A | | 1.383e-02 | -4.28 | Cdk2 (Cyclin-dependent kinase 2) | protein interactions | 42453 | 55 | 7 | 9204 | 432 | wdb,Cdk4,Sxl,CG6191,eyg,CycG,Cks30A | | 1.388e-02 | -4.28 | GSE41867\_LCMV\_ARMSTRONG\_VS\_CLONE13\_DAY15\_EFFECTOR\_CD8\_TCELL\_DN | MSigDB lists | GSE41867\_LCMV\_ARMSTRONG\_VS\_CLONE13\_DAY15\_EFFECTOR\_CD8\_TCELL\_DN | 14 | 3 | 4161 | 156 | CG7914,CenB1A,LanA | | 1.388e-02 | -4.28 | CTIP\_DN.V1\_DN | MSigDB lists | CTIP\_DN.V1\_DN | 14 | 3 | 4161 | 156 | ReepA,frc,subdued | | 1.388e-02 | -4.28 | GO\_ASTROCYTE\_DIFFERENTIATION | MSigDB lists | GO\_ASTROCYTE\_DIFFERENTIATION | 14 | 3 | 4161 | 156 | Dl,Cdk4,N | | 1.388e-02 | -4.28 | DUTERTRE\_ESTRADIOL\_RESPONSE\_6HR\_DN | MSigDB lists | DUTERTRE\_ESTRADIOL\_RESPONSE\_6HR\_DN | 14 | 3 | 4161 | 156 | rg,Cdk4,Pgant6 | | 1.388e-02 | -4.28 | KEGG\_BASAL\_CELL\_CARCINOMA | MSigDB lists | KEGG\_BASAL\_CELL\_CARCINOMA | 14 | 3 | 4161 | 156 | arm,fz2,fz | | 1.388e-02 | -4.28 | GO\_CLATHRIN\_COATED\_ENDOCYTIC\_VESICLE | MSigDB lists | GO\_CLATHRIN\_COATED\_ENDOCYTIC\_VESICLE | 14 | 3 | 4161 | 156 | AP-1gamma,jar,fz2 | | 1.388e-02 | -4.28 | GO\_REGULATION\_OF\_BIOMINERAL\_TISSUE\_DEVELOPMENT | MSigDB lists | GO\_REGULATION\_OF\_BIOMINERAL\_TISSUE\_DEVELOPMENT | 14 | 3 | 4161 | 156 | Mef2,tkv,Ddr | | 1.388e-02 | -4.28 | KEGG\_ENDOMETRIAL\_CANCER | MSigDB lists | KEGG\_ENDOMETRIAL\_CANCER | 14 | 3 | 4161 | 156 | Pdk1,Raf,arm | | 1.388e-02 | -4.28 | GO\_CELL\_FATE\_SPECIFICATION | MSigDB lists | GO\_CELL\_FATE\_SPECIFICATION | 14 | 3 | 4161 | 156 | arm,N,Dl | | 1.388e-02 | -4.28 | GCM\_PTPRD | MSigDB lists | GCM\_PTPRD | 14 | 3 | 4161 | 156 | nmo,Ziz,emp | | 1.388e-02 | -4.28 | AGTCTTA\_MIR499 | MSigDB lists | AGTCTTA\_MIR499 | 14 | 3 | 4161 | 156 | Rab5,bru2,lin-28 | | 1.388e-02 | -4.28 | GO\_REGULATION\_OF\_MYELOID\_LEUKOCYTE\_DIFFERENTIATION | MSigDB lists | GO\_REGULATION\_OF\_MYELOID\_LEUKOCYTE\_DIFFERENTIATION | 14 | 3 | 4161 | 156 | arm,Actbeta,Tollo | | 1.388e-02 | -4.28 | YAO\_TEMPORAL\_RESPONSE\_TO\_PROGESTERONE\_CLUSTER\_8 | MSigDB lists | YAO\_TEMPORAL\_RESPONSE\_TO\_PROGESTERONE\_CLUSTER\_8 | 14 | 3 | 4161 | 156 | CG1673,fz,CycG | | 1.388e-02 | -4.28 | GO\_REGULATION\_OF\_STAT\_CASCADE | MSigDB lists | GO\_REGULATION\_OF\_STAT\_CASCADE | 14 | 3 | 4161 | 156 | htl,N,nmo | | 1.388e-02 | -4.28 | BHAT\_ESR1\_TARGETS\_NOT\_VIA\_AKT1\_DN | MSigDB lists | BHAT\_ESR1\_TARGETS\_NOT\_VIA\_AKT1\_DN | 14 | 3 | 4161 | 156 | Cdk4,Dl,Rgl | | 1.390e-02 | -4.28 | GO\_ENZYME\_LINKED\_RECEPTOR\_PROTEIN\_SIGNALING\_PATHWAY | MSigDB lists | GO\_ENZYME\_LINKED\_RECEPTOR\_PROTEIN\_SIGNALING\_PATHWAY | 157 | 12 | 4161 | 156 | Src42A,Ddr,Eph,nmo,nAChRalpha4,dock,htl,tkv,InR,Nrk,Actbeta,Pdk1 | | 1.395e-02 | -4.27 | GO\_PROTEIN\_IMPORT | MSigDB lists | GO\_PROTEIN\_IMPORT | 69 | 7 | 4161 | 156 | Pex10,cact,Pen,arm,Pex14,Pex5,Pex7 | | 1.395e-02 | -4.27 | GO\_REGULATION\_OF\_MAP\_KINASE\_ACTIVITY | MSigDB lists | GO\_REGULATION\_OF\_MAP\_KINASE\_ACTIVITY | 69 | 7 | 4161 | 156 | wbl,hppy,fz2,trbl,InR,Raf,Tsp96F | | 1.395e-02 | -4.27 | GSE43955\_1H\_VS\_10H\_ACT\_CD4\_TCELL\_UP | MSigDB lists | GSE43955\_1H\_VS\_10H\_ACT\_CD4\_TCELL\_UP | 69 | 7 | 4161 | 156 | Tsp96F,CG1129,CG2017,sd,Rab1,step,Fem-1 | | 1.397e-02 | -4.27 | TBPH (TAR DNA-binding protein-43 homolog) | protein interactions | 37781 | 110 | 11 | 9204 | 432 | nub,Syx1A,Raf,CalpC,N,lncRNA:Hsromega,Diap1,Myo10A,Pen,sens,Fem-1 | | 1.401e-02 | -4.27 | - | gene3d domains | 2.60.120.260 | 30 | 5 | 4929 | 241 | nahoda,Ddr,LanA,Eph,koi | | 1.405e-02 | -4.26 | GO\_POSITIVE\_REGULATION\_OF\_PROTEIN\_MODIFICATION\_PROCESS | MSigDB lists | GO\_POSITIVE\_REGULATION\_OF\_PROTEIN\_MODIFICATION\_PROCESS | 276 | 18 | 4161 | 156 | Rab1,htl,Actbeta,hppy,Raf,Mef2,wbl,fz2,arm,nAChRalpha4,fz,Tsp96F,Nrk,Pdk1,PI31,Ddr,Tollo,unc-119 | | 1.426e-02 | -4.25 | regulation of secretion | biological process | GO:0051046 | 89 | 8 | 12345 | 442 | jar,Syx1A,mop,IA-2,InR,Rab1,Cadps,S | | 1.428e-02 | -4.25 | XU\_GH1\_AUTOCRINE\_TARGETS\_DN | MSigDB lists | XU\_GH1\_AUTOCRINE\_TARGETS\_DN | 39 | 5 | 4161 | 156 | rg,l(1)G0196,unc-5,Mfe2,LanA | | 1.428e-02 | -4.25 | GSE26488\_CTRL\_VS\_PEPTIDE\_INJECTION\_OT2\_THYMOCYTE\_DN | MSigDB lists | GSE26488\_CTRL\_VS\_PEPTIDE\_INJECTION\_OT2\_THYMOCYTE\_DN | 39 | 5 | 4161 | 156 | Cerk,C1GalTA,trbl,Mef2,SMC3 | | 1.428e-02 | -4.25 | LI\_WILMS\_TUMOR\_VS\_FETAL\_KIDNEY\_1\_UP | MSigDB lists | LI\_WILMS\_TUMOR\_VS\_FETAL\_KIDNEY\_1\_UP | 39 | 5 | 4161 | 156 | htl,Tpst,shot,Rgl,Pex7 | | 1.428e-02 | -4.25 | GO\_RECEPTOR\_SIGNALING\_PROTEIN\_ACTIVITY | MSigDB lists | GO\_RECEPTOR\_SIGNALING\_PROTEIN\_ACTIVITY | 39 | 5 | 4161 | 156 | Raf,Arf79F,tkv,nmo,hppy | | 1.432e-02 | -4.25 | capt (capulet) | protein interactions | 45233 | 82 | 9 | 9204 | 432 | orb2,chic,Fkbp14,Nlp,Bacc,sli,ft,Abl,S | | 1.432e-02 | -4.25 | DAP12 interactions | REACTOME pathways | R-DME-2172127 | 164 | 14 | 3923 | 176 | Egfr,Raf,vn,Pvr,CG4935,htl,rictor,PI31,Pdk1,14-3-3zeta,wdb,Dronc,CG42684,trbl | | 1.432e-02 | -4.25 | Downstream signal transduction | REACTOME pathways | R-DME-186763 | 164 | 14 | 3923 | 176 | Egfr,vn,Raf,Pvr,CG4935,14-3-3zeta,Pdk1,htl,rictor,PI31,wdb,trbl,CG42684,Dronc | | 1.432e-02 | -4.25 | DAP12 signaling | REACTOME pathways | R-DME-2424491 | 164 | 14 | 3923 | 176 | Pvr,Raf,vn,Egfr,Dronc,CG42684,trbl,wdb,PI31,htl,rictor,Pdk1,14-3-3zeta,CG4935 | | 1.432e-02 | -4.25 | Signaling by PDGF | REACTOME pathways | R-DME-186797 | 164 | 14 | 3923 | 176 | wdb,CG42684,trbl,Dronc,CG4935,PI31,rictor,Pdk1,14-3-3zeta,htl,Pvr,Egfr,vn,Raf | | 1.440e-02 | -4.24 | Sec7\_C\_sf | interpro domains | IPR023394 | 5 | 2 | 11383 | 450 | siz,step | | 1.440e-02 | -4.24 | Transcription\_factor\_GATA | interpro domains | IPR039355 | 5 | 2 | 11383 | 450 | pnr,grn | | 1.440e-02 | -4.24 | Sec7\_dom\_sf | interpro domains | IPR035999 | 5 | 2 | 11383 | 450 | step,siz | | 1.440e-02 | -4.24 | POU\_dom | interpro domains | IPR000327 | 5 | 2 | 11383 | 450 | nub,pdm3 | | 1.440e-02 | -4.24 | Semaphorin | interpro domains | IPR027231 | 5 | 2 | 11383 | 450 | Sema1b,Sema1a | | 1.440e-02 | -4.24 | POU | interpro domains | IPR013847 | 5 | 2 | 11383 | 450 | nub,pdm3 | | 1.442e-02 | -4.24 | Poxm (Pox meso) | protein interactions | 40990 | 43 | 6 | 9204 | 432 | CG6656,abd-A,Abd-B,His2Av,CG14478,Scr | | 1.442e-02 | -4.24 | Ret (Ret oncogene) | protein interactions | 43875 | 43 | 6 | 9204 | 432 | Dl,spi,scrib,Src42A,px,S | | 1.448e-02 | -4.23 | RUTELLA\_RESPONSE\_TO\_CSF2RB\_AND\_IL4\_UP | MSigDB lists | RUTELLA\_RESPONSE\_TO\_CSF2RB\_AND\_IL4\_UP | 86 | 8 | 4161 | 156 | step,Rgl,Actbeta,Tsp96F,fz2,lili,bru2,Tpst | | 1.448e-02 | -4.23 | FOSTER\_TOLERANT\_MACROPHAGE\_DN | MSigDB lists | FOSTER\_TOLERANT\_MACROPHAGE\_DN | 86 | 8 | 4161 | 156 | Tpst,fz2,ap,Unc-76,N,fz,Dl,Tollo | | 1.460e-02 | -4.23 | HLH | smart domains | SM00353 | 57 | 8 | 5086 | 290 | ss,tai,sim,emc,E(spl)m5-HLH,crp,h,HLH3B | | 1.472e-02 | -4.22 | Pou | pfam domains | PF00157 | 5 | 2 | 10480 | 419 | pdm3,nub | | 1.472e-02 | -4.22 | Sec7 | pfam domains | PF01369 | 5 | 2 | 10480 | 419 | step,siz | | 1.472e-02 | -4.22 | GO\_MAINTENANCE\_OF\_CELL\_NUMBER | MSigDB lists | GO\_MAINTENANCE\_OF\_CELL\_NUMBER | 54 | 6 | 4161 | 156 | hyx,SMC3,N,Raf,Dl,arm | | 1.472e-02 | -4.22 | E12\_Q6 | MSigDB lists | E12\_Q6 | 54 | 6 | 4161 | 156 | CG31522,LanA,Pex10,Psa,Dl,l(2)37Cg | | 1.472e-02 | -4.22 | GSE45365\_NK\_CELL\_VS\_BCELL\_MCMV\_INFECTION\_DN | MSigDB lists | GSE45365\_NK\_CELL\_VS\_BCELL\_MCMV\_INFECTION\_DN | 54 | 6 | 4161 | 156 | lili,Drak,CG4896,Tpst,CG6465,cact | | 1.472e-02 | -4.22 | GSE17721\_12H\_VS\_24H\_POLYIC\_BMDC\_UP | MSigDB lists | GSE17721\_12H\_VS\_24H\_POLYIC\_BMDC\_UP | 54 | 6 | 4161 | 156 | dock,Tsp96F,SERCA,Tpst,Pat1,Ids | | 1.481e-02 | -4.21 | gravitaxis | biological process | GO:0042332 | 27 | 4 | 12345 | 442 | esg,yuri,Pen,nmo | | 1.481e-02 | -4.21 | muscle tissue development | biological process | GO:0060537 | 27 | 4 | 12345 | 442 | sd,Mef2,N,abd-A | | 1.481e-02 | -4.21 | response to gravity | biological process | GO:0009629 | 27 | 4 | 12345 | 442 | nmo,Pen,yuri,esg | | 1.492e-02 | -4.21 | regulation of lipid storage | biological process | GO:0010883 | 41 | 5 | 12345 | 442 | Arf79F,wdb,trbl,InR,SERCA | | 1.492e-02 | -4.21 | muscle attachment | biological process | GO:0016203 | 41 | 5 | 12345 | 442 | kn,shot,Dg,kon,drl | | 1.492e-02 | -4.20 | GO\_AMINOGLYCAN\_BIOSYNTHETIC\_PROCESS | MSigDB lists | GO\_AMINOGLYCAN\_BIOSYNTHETIC\_PROCESS | 26 | 4 | 4161 | 156 | GlcAT-I,dlp,frc,CG9220 | | 1.492e-02 | -4.20 | HALLMARK\_KRAS\_SIGNALING\_DN | MSigDB lists | HALLMARK\_KRAS\_SIGNALING\_DN | 26 | 4 | 4161 | 156 | sns,htl,tkv,Dyb | | 1.492e-02 | -4.20 | GSE13484\_UNSTIM\_VS\_YF17D\_VACCINE\_STIM\_PBMC\_DN | MSigDB lists | GSE13484\_UNSTIM\_VS\_YF17D\_VACCINE\_STIM\_PBMC\_DN | 26 | 4 | 4161 | 156 | Actbeta,CG2017,Tollo,C1GalTA | | 1.492e-02 | -4.20 | GSE13547\_2H\_VS\_12\_H\_ANTI\_IGM\_STIM\_ZFX\_KO\_BCELL\_UP | MSigDB lists | GSE13547\_2H\_VS\_12\_H\_ANTI\_IGM\_STIM\_ZFX\_KO\_BCELL\_UP | 26 | 4 | 4161 | 156 | dock,CycG,shot,CG9485 | | 1.492e-02 | -4.20 | GO\_ION\_CHANNEL\_BINDING | MSigDB lists | GO\_ION\_CHANNEL\_BINDING | 26 | 4 | 4161 | 156 | Nmda1,arm,Syx1A,14-3-3epsilon | | 1.492e-02 | -4.20 | BROWNE\_HCMV\_INFECTION\_12HR\_UP | MSigDB lists | BROWNE\_HCMV\_INFECTION\_12HR\_UP | 26 | 4 | 4161 | 156 | step,Rab5,Raf,Drep4 | | 1.492e-02 | -4.20 | EGF\_CA | smart domains | SM00179 | 36 | 6 | 5086 | 290 | Dl,N,ft,dpy,cue,sli | | 1.501e-02 | -4.20 | ScsbetaA (Succinyl-coenzyme A synthetase beta subunit, ADP-forming) | protein interactions | 41067 | 69 | 8 | 9204 | 432 | Rab1,SERCA,Pdk1,Gfat2,Rab5,Raf,Cdk4,Kr-h2 | | 1.503e-02 | -4.20 | GO\_NON\_CANONICAL\_WNT\_SIGNALING\_PATHWAY | MSigDB lists | GO\_NON\_CANONICAL\_WNT\_SIGNALING\_PATHWAY | 70 | 7 | 4161 | 156 | PI31,nmo,arm,dlp,fz2,otk,fz | | 1.503e-02 | -4.20 | GSE27670\_CTRL\_VS\_LMP1\_TRANSDUCED\_GC\_BCELL\_DN | MSigDB lists | GSE27670\_CTRL\_VS\_LMP1\_TRANSDUCED\_GC\_BCELL\_DN | 70 | 7 | 4161 | 156 | Cerk,Rgl,Ric,Pex5,lqf,Dl,Rim2 | | 1.503e-02 | -4.20 | LINDGREN\_BLADDER\_CANCER\_CLUSTER\_3\_DN | MSigDB lists | LINDGREN\_BLADDER\_CANCER\_CLUSTER\_3\_DN | 70 | 7 | 4161 | 156 | htl,InR,N,Tsp96F,Dyrk2,CG32795,shot | | 1.503e-02 | -4.20 | GO\_REGULATION\_OF\_CELL\_ACTIVATION | MSigDB lists | GO\_REGULATION\_OF\_CELL\_ACTIVATION | 70 | 7 | 4161 | 156 | arm,AP-1gamma,Tsp96F,Pdk1,Actbeta,Mef2,dock | | 1.505e-02 | -4.20 | negative regulation of humoral immune response | biological process | GO:0002921 | 15 | 3 | 12345 | 442 | cact,Pvr,nub | | 1.505e-02 | -4.20 | male germ-line stem cell population maintenance | biological process | GO:0036098 | 15 | 3 | 12345 | 442 | Inx2,His2Av,chic | | 1.505e-02 | -4.20 | positive regulation of cell maturation | biological process | GO:1903431 | 15 | 3 | 12345 | 442 | Sox14,Arf79F,InR | | 1.505e-02 | -4.20 | plasmatocyte differentiation | biological process | GO:0042387 | 15 | 3 | 12345 | 442 | Raf,N,htl | | 1.505e-02 | -4.20 | oocyte localization involved in germarium-derived egg chamber formation | biological process | GO:0030720 | 15 | 3 | 12345 | 442 | Dl,shg,N | | 1.505e-02 | -4.20 | positive regulation of axon extension | biological process | GO:0045773 | 15 | 3 | 12345 | 442 | shot,chic,fz | | 1.505e-02 | -4.20 | GO\_SYSTEM\_PROCESS | MSigDB lists | GO\_SYSTEM\_PROCESS | 278 | 18 | 4161 | 156 | unc-119,Eph,sli,ReepA,CG7394,Nrk,subdued,FBXO11,nAChRalpha4,SERCA,jar,Mef2,Dgat2,Raf,CG11069,Dl,Actbeta,sns | | 1.508e-02 | -4.19 | negative regulation of molecular function | biological process | GO:0044092 | 145 | 11 | 12345 | 442 | Diap1,14-3-3epsilon,Cbl,Spn42De,emc,14-3-3zeta,Socs36E,CycG,cact,trbl,edl | | 1.512e-02 | -4.19 | VITELLOGENIN | prosite domains | PS51211 | 4 | 2 | 6243 | 325 | apolpp,cv-d | | 1.512e-02 | -4.19 | BEN | prosite domains | PS51457 | 4 | 2 | 6243 | 325 | pre-mod(mdg4)-C,Elba2 | | 1.517e-02 | -4.19 | hng1 (hinge1) | protein interactions | 37412 | 21 | 4 | 9204 | 432 | CG13204,hng3,tsh,hth | | 1.517e-02 | -4.19 | Tip60 (Tat interactive protein 60kDa) | protein interactions | 31362 | 21 | 4 | 9204 | 432 | His2Av,Bacc,Dp,E(Pc) | | 1.517e-02 | -4.19 | mira (miranda) | protein interactions | 42379 | 21 | 4 | 9204 | 432 | shot,jar,baz,flfl | | 1.517e-02 | -4.19 | shn (schnurri) | protein interactions | 36171 | 21 | 4 | 9204 | 432 | tkv,CtBP,pnr,pum | | 1.517e-02 | -4.19 | cg (combgap) | protein interactions | 36571 | 21 | 4 | 9204 | 432 | CG44838,eya,Psc,ph-p | | 1.518e-02 | -4.19 | Mer (Merlin) | protein interactions | 32979 | 172 | 15 | 9204 | 432 | yrt,CG42699,lncRNA:CR42862,CycG,ed,cora,cact,MEP-1,Pex5,chn,Dyb,apolpp,E(Pc),jar,ft | | 1.518e-02 | -4.19 | cellular response to extracellular stimulus | biological process | GO:0031668 | 90 | 8 | 12345 | 442 | Atg16,CG7536,Atg17,rictor,InR,lt,Raf,SNF4Agamma | | 1.519e-02 | -4.19 | kn (knot) | protein interactions | 45318 | 56 | 7 | 9204 | 432 | Scr,sd,unc-119,Egfr,disco-r,abd-A,kn | | 1.532e-02 | -4.18 | response to wounding | biological process | GO:0009611 | 108 | 9 | 12345 | 442 | wnd,chic,Pvr,shot,Sxl,ed,kay,shg,RhoGEF3 | | 1.545e-02 | -4.17 | CATTGTYY\_SOX9\_B1 | MSigDB lists | CATTGTYY\_SOX9\_B1 | 87 | 8 | 4161 | 156 | ft,sd,htl,FBXO11,trbl,14-3-3zeta,Actbeta,Rab1 | | 1.545e-02 | -4.17 | GO\_PATTERN\_SPECIFICATION\_PROCESS | MSigDB lists | GO\_PATTERN\_SPECIFICATION\_PROCESS | 87 | 8 | 4161 | 156 | Dl,Mef2,tkv,ap,CG9220,N,fz2,arm | | 1.546e-02 | -4.17 | chromocenter | cellular component | GO:0010369 | 28 | 4 | 11940 | 417 | jumu,His2Av,bab2,E(Pc) | | 1.546e-02 | -4.17 | nuclear ubiquitin ligase complex | cellular component | GO:0000152 | 28 | 4 | 11940 | 417 | corto,Psc,ph-p,Su(z)2 | | 1.546e-02 | -4.17 | actin-based cell projection | cellular component | GO:0098858 | 28 | 4 | 11940 | 417 | ena,shg,shot,Myo10A | | 1.555e-02 | -4.16 | chromatin binding | molecular function | GO:0003682 | 237 | 15 | 12176 | 414 | SMC3,tsh,MEP-1,Nlp,Psc,Elba2,corto,His3.3B,Prdm13,N,Su(z)2,Hr39,ttk,pita,ph-p | | 1.555e-02 | -4.16 | axon terminus | cellular component | GO:0043679 | 75 | 7 | 11940 | 417 | ena,Syx1A,Magi,pum,gukh,jeb,orb2 | | 1.572e-02 | -4.15 | beta-1,3-galactosyltransferase activity | molecular function | GO:0048531 | 16 | 3 | 12176 | 414 | C1GalTA,CG34057,CG3119 | | 1.572e-02 | -4.15 | Graf (GTPase regulator associated with FAK) | protein interactions | 32522 | 32 | 5 | 9204 | 432 | Sema1a,koi,Egfr,imd,spi | | 1.577e-02 | -4.15 | positive regulation of transport | biological process | GO:0051050 | 127 | 10 | 12345 | 442 | Rab1,Cadps,SERCA,S,mop,Cbl,Abl,cindr,Arf79F,jar | | 1.580e-02 | -4.15 | gap junction channel activity | molecular function | GO:0005243 | 6 | 2 | 12176 | 414 | ogre,Inx2 | | 1.580e-02 | -4.15 | microfilament motor activity | molecular function | GO:0000146 | 6 | 2 | 12176 | 414 | jar,d | | 1.580e-02 | -4.15 | phosphatidylinositol-4,5-bisphosphate 5-phosphatase activity | molecular function | GO:0004439 | 6 | 2 | 12176 | 414 | Ocrl,CG9784 | | 1.580e-02 | -4.15 | promoter-specific chromatin binding | molecular function | GO:1990841 | 6 | 2 | 12176 | 414 | ttk,Su(z)2 | | 1.580e-02 | -4.15 | phosphatidylinositol phosphate 5-phosphatase activity | molecular function | GO:0034595 | 6 | 2 | 12176 | 414 | Ocrl,CG9784 | | 1.581e-02 | -4.15 | Signalling to RAS | REACTOME pathways | R-DME-167044 | 118 | 11 | 3923 | 176 | CG4935,PI31,14-3-3zeta,htl,wdb,CG42684,Egfr,Raf,vn,Pvr,Rgl | | 1.581e-02 | -4.15 | FCERI mediated MAPK activation | REACTOME pathways | R-DME-2871796 | 118 | 11 | 3923 | 176 | Egfr,Raf,vn,Pvr,kay,CG4935,PI31,htl,14-3-3zeta,wdb,CG42684 | | 1.581e-02 | -4.15 | Signalling to ERKs | REACTOME pathways | R-DME-187687 | 118 | 11 | 3923 | 176 | Raf,vn,Egfr,Rgl,Pvr,PI31,htl,14-3-3zeta,CG4935,CG42684,wdb | | 1.583e-02 | -4.15 | GSE30971\_CTRL\_VS\_LPS\_STIM\_MACROPHAGE\_WBP7\_KO\_4H\_UP | MSigDB lists | GSE30971\_CTRL\_VS\_LPS\_STIM\_MACROPHAGE\_WBP7\_KO\_4H\_UP | 40 | 5 | 4161 | 156 | CG3961,Gfat2,cact,Dl,CG1673 | | 1.583e-02 | -4.15 | GSE26559\_TCF1\_KO\_VS\_WT\_LIN\_NEG\_CELL\_DN | MSigDB lists | GSE26559\_TCF1\_KO\_VS\_WT\_LIN\_NEG\_CELL\_DN | 40 | 5 | 4161 | 156 | Actbeta,Mef2,Pdk1,Gfat2,jar | | 1.583e-02 | -4.15 | GSE15930\_NAIVE\_VS\_48H\_IN\_VITRO\_STIM\_CD8\_TCELL\_UP | MSigDB lists | GSE15930\_NAIVE\_VS\_48H\_IN\_VITRO\_STIM\_CD8\_TCELL\_UP | 40 | 5 | 4161 | 156 | sina,betaTub60D,Nmda1,Pgant6,hppy | | 1.583e-02 | -4.15 | GSE17721\_LPS\_VS\_POLYIC\_16H\_BMDC\_DN | MSigDB lists | GSE17721\_LPS\_VS\_POLYIC\_16H\_BMDC\_DN | 40 | 5 | 4161 | 156 | emp,Nmda1,CG1129,CG1673,Psa | | 1.583e-02 | -4.15 | GSE24634\_NAIVE\_CD4\_TCELL\_VS\_DAY5\_IL4\_CONV\_TREG\_UP | MSigDB lists | GSE24634\_NAIVE\_CD4\_TCELL\_VS\_DAY5\_IL4\_CONV\_TREG\_UP | 40 | 5 | 4161 | 156 | step,Tpst,Rim2,Cdk4,cact | | 1.583e-02 | -4.15 | GO\_CELL\_FATE\_COMMITMENT | MSigDB lists | GO\_CELL\_FATE\_COMMITMENT | 40 | 5 | 4161 | 156 | arm,Dl,N,hyx,Mef2 | | 1.583e-02 | -4.15 | GSE41176\_UNSTIM\_VS\_ANTI\_IGM\_STIM\_BCELL\_6H\_DN | MSigDB lists | GSE41176\_UNSTIM\_VS\_ANTI\_IGM\_STIM\_BCELL\_6H\_DN | 40 | 5 | 4161 | 156 | C1GalTA,jar,Raf,Atg16,CG9220 | | 1.583e-02 | -4.15 | AAAGGAT\_MIR501 | MSigDB lists | AAAGGAT\_MIR501 | 40 | 5 | 4161 | 156 | bru2,Rgl,wdb,robo1,Mef2 | | 1.597e-02 | -4.14 | GO\_CELL\_SUBSTRATE\_JUNCTION | MSigDB lists | GO\_CELL\_SUBSTRATE\_JUNCTION | 160 | 12 | 4161 | 156 | otk,emp,Ddr,Arf79F,Pdk1,Tsp96F,14-3-3zeta,fz,htl,arm,shot,14-3-3epsilon | | 1.597e-02 | -4.14 | GO\_EMBRYO\_DEVELOPMENT\_ENDING\_IN\_BIRTH\_OR\_EGG\_HATCHING | MSigDB lists | GO\_EMBRYO\_DEVELOPMENT\_ENDING\_IN\_BIRTH\_OR\_EGG\_HATCHING | 160 | 12 | 4161 | 156 | arm,fz2,ap,N,fz,frc,ds,Ocrl,lqf,otk,Dl,Mef2 | | 1.603e-02 | -4.13 | GSE39820\_TGFBETA3\_IL6\_VS\_TGFBETA3\_IL6\_IL23A\_TREATED\_CD4\_TCELL\_UP | MSigDB lists | GSE39820\_TGFBETA3\_IL6\_VS\_TGFBETA3\_IL6\_IL23A\_TREATED\_CD4\_TCELL\_UP | 55 | 6 | 4161 | 156 | arm,Syx1A,CG18549,FBXO11,Pgant5,InR | | 1.603e-02 | -4.13 | GSE27786\_CD4\_VS\_CD8\_TCELL\_UP | MSigDB lists | GSE27786\_CD4\_VS\_CD8\_TCELL\_UP | 55 | 6 | 4161 | 156 | CG2017,InR,Ids,rdx,CenB1A,hppy | | 1.603e-02 | -4.13 | GTTTGTT\_MIR495 | MSigDB lists | GTTTGTT\_MIR495 | 55 | 6 | 4161 | 156 | ap,Cdk4,Pat1,bru2,AP-1gamma,SmD1 | | 1.614e-02 | -4.13 | Cadherin\_CS | interpro domains | IPR020894 | 14 | 3 | 11383 | 450 | shg,ds,ft | | 1.614e-02 | -4.13 | Myosin\_head\_motor\_dom | interpro domains | IPR001609 | 14 | 3 | 11383 | 450 | d,jar,Myo10A | | 1.614e-02 | -4.13 | circadian rhythm | biological process | GO:0007623 | 91 | 8 | 12345 | 442 | ITP,dome,kay,emc,InR,tara,Mid1,Mef2 | | 1.617e-02 | -4.12 | LIU\_PROSTATE\_CANCER\_DN | MSigDB lists | LIU\_PROSTATE\_CANCER\_DN | 71 | 7 | 4161 | 156 | Tpst,shot,Cerk,Pgant6,Ddr,Pex14,subdued | | 1.617e-02 | -4.12 | GSE23321\_CD8\_STEM\_CELL\_MEMORY\_VS\_EFFECTOR\_MEMORY\_CD8\_TCELL\_UP | MSigDB lists | GSE23321\_CD8\_STEM\_CELL\_MEMORY\_VS\_EFFECTOR\_MEMORY\_CD8\_TCELL\_UP | 71 | 7 | 4161 | 156 | frc,PI31,IntS3,Pex14,Psa,Pgant5,Vps13 | | 1.617e-02 | -4.12 | GO\_REGULATION\_OF\_BINDING | MSigDB lists | GO\_REGULATION\_OF\_BINDING | 71 | 7 | 4161 | 156 | ap,cact,Mef2,CtBP,Hel89B,arm,Pex14 | | 1.617e-02 | -4.12 | Homeobox\_metazoa | interpro domains | IPR020479 | 38 | 5 | 11383 | 450 | Scr,B-H1,abd-A,Abd-B,H2.0 | | 1.619e-02 | -4.12 | positive regulation of immune system process | biological process | GO:0002684 | 109 | 9 | 12345 | 442 | DPCoAC,N,imd,htl,Dcr-2,CG32795,pnr,Pitslre,mop | | 1.632e-02 | -4.12 | PAX | smart domains | SM00351 | 10 | 3 | 5086 | 290 | eyg,Poxn,ey | | 1.645e-02 | -4.11 | DCX-EMAP (Doublecortin-domain-containing echinoderm-microtubule-associated protein) | protein interactions | 39617 | 12 | 3 | 9204 | 432 | ed,RapGAP1,chn | | 1.645e-02 | -4.11 | l(1)G0320 (lethal (1) G0320) | protein interactions | 326234 | 12 | 3 | 9204 | 432 | N,Rab1,Pdk1 | | 1.645e-02 | -4.11 | Fbxl7 (F-box and leucine-rich repeat protein 7) | protein interactions | 41935 | 12 | 3 | 9204 | 432 | ft,cindr,d | | 1.645e-02 | -4.11 | Jafrac2 (Thioredoxin peroxidase 2) | protein interactions | 53577 | 12 | 3 | 9204 | 432 | Mpcp2,Pez,Diap1 | | 1.645e-02 | -4.11 | ush (u-shaped) | protein interactions | 33225 | 12 | 3 | 9204 | 432 | pnr,htl,CtBP | | 1.645e-02 | -4.11 | mid (midline) | protein interactions | 33770 | 12 | 3 | 9204 | 432 | Abd-B,abd-A,pnr | | 1.645e-02 | -4.11 | B4 (B4) | protein interactions | 34767 | 12 | 3 | 9204 | 432 | InR,14-3-3zeta,14-3-3epsilon | | 1.645e-02 | -4.11 | Pmp70 (Peroxisomal Membrane Protein 70 kDa) | protein interactions | 32992 | 12 | 3 | 9204 | 432 | CG2017,C1GalTA,eyg | | 1.645e-02 | -4.11 | Mical (Molecule interacting with CasL) | protein interactions | 41225 | 12 | 3 | 9204 | 432 | InR,Sox14,Abl | | 1.645e-02 | -4.11 | shv (shriveled) | protein interactions | 33220 | 12 | 3 | 9204 | 432 | Pdk1,Raf,shg | | 1.645e-02 | -4.11 | dac (dachshund) | protein interactions | 34982 | 12 | 3 | 9204 | 432 | Egfr,eya,ey | | 1.647e-02 | -4.11 | CATTTCA\_MIR203 | MSigDB lists | CATTTCA\_MIR203 | 88 | 8 | 4161 | 156 | rg,Mef2,kn,AP-1gamma,hppy,CG45050,nmo,CycG | | 1.647e-02 | -4.11 | SENGUPTA\_NASOPHARYNGEAL\_CARCINOMA\_WITH\_LMP1\_UP | MSigDB lists | SENGUPTA\_NASOPHARYNGEAL\_CARCINOMA\_WITH\_LMP1\_UP | 88 | 8 | 4161 | 156 | Cadps,jar,lin-28,Pgant5,subdued,Rim2,CG45050,SMC3 | | 1.662e-02 | -4.10 | Myosin\_head | pfam domains | PF00063 | 14 | 3 | 10480 | 419 | jar,d,Myo10A | | 1.662e-02 | -4.10 | Cadherin | pfam domains | PF00028 | 14 | 3 | 10480 | 419 | ds,shg,ft | | 1.663e-02 | -4.10 | type Is terminal bouton | cellular component | GO:0061177 | 6 | 2 | 11940 | 417 | jeb,pum | | 1.664e-02 | -4.10 | neuron projection terminus | cellular component | GO:0044306 | 76 | 7 | 11940 | 417 | gukh,orb2,jeb,pum,Magi,Syx1A,ena | | 1.664e-02 | -4.10 | extrinsic component of membrane | cellular component | GO:0019898 | 76 | 7 | 11940 | 417 | arm,Socs36E,Src42A,Abl,Vps13,shg,Atg17 | | 1.666e-02 | -4.09 | Immune System | REACTOME pathways | R-DME-168256 | 744 | 45 | 3923 | 176 | moody,rictor,Pdk1,PI31,SCAR,eff,trbl,Abl,l(2)37Cg,CG9485,cact,Atox1,IA-2,Raf,Pvr,CG6656,CG4935,ATPCL,CG42684,Src42A,betaTub60D,vn,FBXO11,shot,arm,14-3-3zeta,RapGAP1,Psa,apolpp,sina,kay,pes,htl,Syx1A,mura,wdb,CG4467,Ptp4E,dock,Dronc,Egfr,Rab5,Cbl,CG6891,CG42788 | | 1.673e-02 | -4.09 | GGGTGGRR\_PAX4\_03 | MSigDB lists | GGGTGGRR\_PAX4\_03 | 240 | 16 | 4161 | 156 | trbl,14-3-3zeta,14-3-3epsilon,Rab1,dlp,otk,Cdk4,CG1129,hppy,Rab5,CG2812,betaTub60D,rdx,Eph,AP-1gamma,ds | | 1.680e-02 | -4.09 | positive regulation of TORC1 signaling | biological process | GO:1904263 | 28 | 4 | 12345 | 442 | InR,Cdk4,Pvr,hppy | | 1.680e-02 | -4.09 | regulation of cyclin-dependent protein serine/threonine kinase activity | biological process | GO:0000079 | 28 | 4 | 12345 | 442 | kay,CycG,CycB3,Cks30A | | 1.680e-02 | -4.09 | zygotic determination of anterior/posterior axis, embryo | biological process | GO:0007354 | 28 | 4 | 12345 | 442 | bowl,fs(1)h,cic,Raf | | 1.680e-02 | -4.09 | regulation of generation of precursor metabolites and energy | biological process | GO:0043467 | 28 | 4 | 12345 | 442 | N,Pgk,Dg,CG34172 | | 1.680e-02 | -4.09 | regulation of cyclin-dependent protein kinase activity | biological process | GO:1904029 | 28 | 4 | 12345 | 442 | Cks30A,CycB3,kay,CycG | | 1.684e-02 | -4.08 | fly chr11F1-11F1\|1-43 | chromosome location | fly chr11F1-11F1\|1-43 | 9 | 2 | 25010 | 571 | lncRNA:CR44621,dmrt11E | | 1.684e-02 | -4.08 | fly chr11F1-11F1\|1-43 cM | chromosome location | fly chr11F1-11F1\|1-43 cM | 9 | 2 | 25010 | 571 | lncRNA:CR44621,dmrt11E | | 1.688e-02 | -4.08 | KEGG\_NOTCH\_SIGNALING\_PATHWAY | MSigDB lists | KEGG\_NOTCH\_SIGNALING\_PATHWAY | 15 | 3 | 4161 | 156 | N,CtBP,Dl | | 1.688e-02 | -4.08 | MIKKELSEN\_MEF\_LCP\_WITH\_H3K4ME3 | MSigDB lists | MIKKELSEN\_MEF\_LCP\_WITH\_H3K4ME3 | 15 | 3 | 4161 | 156 | CG31522,CG45050,Actbeta | | 1.688e-02 | -4.08 | PID\_BETA\_CATENIN\_NUC\_PATHWAY | MSigDB lists | PID\_BETA\_CATENIN\_NUC\_PATHWAY | 15 | 3 | 4161 | 156 | arm,14-3-3epsilon,14-3-3zeta | | 1.688e-02 | -4.08 | KRAS.PROSTATE\_UP.V1\_UP | MSigDB lists | KRAS.PROSTATE\_UP.V1\_UP | 15 | 3 | 4161 | 156 | ReepA,nAChRalpha4,rg | | 1.688e-02 | -4.08 | SIG\_INSULIN\_RECEPTOR\_PATHWAY\_IN\_CARDIAC\_MYOCYTES | MSigDB lists | SIG\_INSULIN\_RECEPTOR\_PATHWAY\_IN\_CARDIAC\_MYOCYTES | 15 | 3 | 4161 | 156 | 14-3-3epsilon,14-3-3zeta,Pdk1 | | 1.688e-02 | -4.08 | BHAT\_ESR1\_TARGETS\_VIA\_AKT1\_DN | MSigDB lists | BHAT\_ESR1\_TARGETS\_VIA\_AKT1\_DN | 15 | 3 | 4161 | 156 | Pgant6,Dl,Cdk4 | | 1.688e-02 | -4.08 | PID\_A6B1\_A6B4\_INTEGRIN\_PATHWAY | MSigDB lists | PID\_A6B1\_A6B4\_INTEGRIN\_PATHWAY | 15 | 3 | 4161 | 156 | 14-3-3zeta,LanA,14-3-3epsilon | | 1.688e-02 | -4.08 | GO\_NUCLEOTIDE\_SUGAR\_BIOSYNTHETIC\_PROCESS | MSigDB lists | GO\_NUCLEOTIDE\_SUGAR\_BIOSYNTHETIC\_PROCESS | 15 | 3 | 4161 | 156 | CG1129,frc,Gfat2 | | 1.688e-02 | -4.08 | CCACACA\_MIR147 | MSigDB lists | CCACACA\_MIR147 | 15 | 3 | 4161 | 156 | ATPCL,Pdk1,robo1 | | 1.688e-02 | -4.08 | nuclear transcriptional repressor complex | cellular component | GO:0090568 | 16 | 3 | 11940 | 417 | Elba2,H,MEP-1 | | 1.692e-02 | -4.08 | SERCA (Sarco/endoplasmic reticulum Ca(2+)-ATPase) | protein interactions | 49297 | 128 | 12 | 9204 | 432 | Mpcp2,Kr-h2,N,Raf,CG30344,Rab5,Rab1,imd,Arf79F,ds,Pdk1,Gfat2 | | 1.702e-02 | -4.07 | GO\_FATTY\_ACYL\_COA\_METABOLIC\_PROCESS | MSigDB lists | GO\_FATTY\_ACYL\_COA\_METABOLIC\_PROCESS | 27 | 4 | 4161 | 156 | CG3961,ATPCL,Mfe2,CG31522 | | 1.702e-02 | -4.07 | REACTOME\_TRANS\_GOLGI\_NETWORK\_VESICLE\_BUDDING | MSigDB lists | REACTOME\_TRANS\_GOLGI\_NETWORK\_VESICLE\_BUDDING | 27 | 4 | 4161 | 156 | Arf79F,Rab5,AP-1gamma,Ocrl | | 1.702e-02 | -4.07 | GO\_NEURON\_MIGRATION | MSigDB lists | GO\_NEURON\_MIGRATION | 27 | 4 | 4161 | 156 | Pex7,14-3-3epsilon,arm,Mef2 | | 1.702e-02 | -4.07 | GSE6259\_33D1\_POS\_VS\_DEC205\_POS\_SPLENIC\_DC\_UP | MSigDB lists | GSE6259\_33D1\_POS\_VS\_DEC205\_POS\_SPLENIC\_DC\_UP | 27 | 4 | 4161 | 156 | Mef2,Rim2,nmo,rdx | | 1.702e-02 | -4.07 | GO\_REGULATION\_OF\_JNK\_CASCADE | MSigDB lists | GO\_REGULATION\_OF\_JNK\_CASCADE | 27 | 4 | 4161 | 156 | fz2,Eph,Tollo,InR | | 1.702e-02 | -4.07 | GSE4590\_PRE\_BCELL\_VS\_LARGE\_PRE\_BCELL\_UP | MSigDB lists | GSE4590\_PRE\_BCELL\_VS\_LARGE\_PRE\_BCELL\_UP | 27 | 4 | 4161 | 156 | Dyb,Tollo,Fkbp14,shot | | 1.705e-02 | -4.07 | BRUINS\_UVC\_RESPONSE\_VIA\_TP53\_GROUP\_B | MSigDB lists | BRUINS\_UVC\_RESPONSE\_VIA\_TP53\_GROUP\_B | 106 | 9 | 4161 | 156 | CG31522,otk,kn,jar,dlp,unc-5,InR,subdued,Actbeta | | 1.705e-02 | -4.07 | GO\_REGULATION\_OF\_PROTEIN\_SERINE\_THREONINE\_KINASE\_ACTIVITY | MSigDB lists | GO\_REGULATION\_OF\_PROTEIN\_SERINE\_THREONINE\_KINASE\_ACTIVITY | 106 | 9 | 4161 | 156 | Raf,Tollo,hppy,Tsp96F,InR,trbl,fz2,wbl,CycG | | 1.709e-02 | -4.07 | cytoskeleton | cellular component | GO:0005856 | 525 | 28 | 11940 | 417 | Dyrk2,CycB3,CG6891,yrt,jar,Bsg,CG42788,Shrm,RhoU,nuf,Jupiter,gukh,klar,Pez,14-3-3epsilon,spdo,f,Lk6,cora,Pat1,yuri,corto,Myo10A,betaTub60D,Wdr62,shot,d,SCAR | | 1.736e-02 | -4.05 | YKACATTT\_UNKNOWN | MSigDB lists | YKACATTT\_UNKNOWN | 72 | 7 | 4161 | 156 | Nrk,Mef2,dock,lt,unc-5,Mfe2,crol | | 1.741e-02 | -4.05 | GO\_REGULATION\_OF\_DEFENSE\_RESPONSE\_TO\_VIRUS | MSigDB lists | GO\_REGULATION\_OF\_DEFENSE\_RESPONSE\_TO\_VIRUS | 56 | 6 | 4161 | 156 | SmD1,CG4645,AP-1gamma,Tpc1,ReepA,Arf79F | | 1.741e-02 | -4.05 | GSE8921\_UNSTIM\_VS\_TLR1\_2\_STIM\_MONOCYTE\_6H\_UP | MSigDB lists | GSE8921\_UNSTIM\_VS\_TLR1\_2\_STIM\_MONOCYTE\_6H\_UP | 56 | 6 | 4161 | 156 | Raf,InR,trbl,Syx1A,CG13384,frc | | 1.741e-02 | -4.05 | AAGCACT\_MIR520F | MSigDB lists | AAGCACT\_MIR520F | 56 | 6 | 4161 | 156 | bru2,Rgl,lili,Ocrl,Mef2,Arf79F | | 1.741e-02 | -4.05 | GSE3039\_NKT\_CELL\_VS\_ALPHAALPHA\_CD8\_TCELL\_UP | MSigDB lists | GSE3039\_NKT\_CELL\_VS\_ALPHAALPHA\_CD8\_TCELL\_UP | 56 | 6 | 4161 | 156 | step,CG1316,arm,InR,Pdk1,CG6465 | | 1.741e-02 | -4.05 | GSE27786\_NKCELL\_VS\_NKTCELL\_UP | MSigDB lists | GSE27786\_NKCELL\_VS\_NKTCELL\_UP | 56 | 6 | 4161 | 156 | CG9485,ATPCL,hppy,Arf79F,14-3-3zeta,Lk6 | | 1.743e-02 | -4.05 | establishment of body hair or bristle planar orientation | biological process | GO:0048104 | 6 | 2 | 12345 | 442 | ft,spen | | 1.743e-02 | -4.05 | response to nicotine | biological process | GO:0035094 | 6 | 2 | 12345 | 442 | moody,esg | | 1.743e-02 | -4.05 | response to symbiont | biological process | GO:0009608 | 6 | 2 | 12345 | 442 | N,kn | | 1.743e-02 | -4.05 | R1/R6 cell differentiation | biological process | GO:0048052 | 6 | 2 | 12345 | 442 | N,ttk | | 1.743e-02 | -4.05 | protein heterooligomerization | biological process | GO:0051291 | 6 | 2 | 12345 | 442 | Dronc,InR | | 1.743e-02 | -4.05 | equator specification | biological process | GO:0045317 | 6 | 2 | 12345 | 442 | ft,ds | | 1.743e-02 | -4.05 | signal maturation | biological process | GO:0035638 | 6 | 2 | 12345 | 442 | rho-4,S | | 1.743e-02 | -4.05 | negative regulation of chemotaxis | biological process | GO:0050922 | 6 | 2 | 12345 | 442 | Sema1a,Sema1b | | 1.743e-02 | -4.05 | maternal specification of dorsal/ventral axis, oocyte, soma encoded | biological process | GO:0007313 | 6 | 2 | 12345 | 442 | wbl,cic | | 1.743e-02 | -4.05 | ovarian follicle cell-cell adhesion | biological process | GO:0007299 | 6 | 2 | 12345 | 442 | mop,baz | | 1.743e-02 | -4.05 | acetyl-CoA biosynthetic process | biological process | GO:0006085 | 6 | 2 | 12345 | 442 | muc,ATPCL | | 1.743e-02 | -4.05 | small RNA loading onto RISC | biological process | GO:0070922 | 6 | 2 | 12345 | 442 | FBXO11,Dcr-2 | | 1.743e-02 | -4.05 | regulation of nucleotide metabolic process | biological process | GO:0006140 | 6 | 2 | 12345 | 442 | N,Dg | | 1.743e-02 | -4.05 | regulation of purine nucleotide metabolic process | biological process | GO:1900542 | 6 | 2 | 12345 | 442 | N,Dg | | 1.743e-02 | -4.05 | semaphorin-plexin signaling pathway | biological process | GO:0071526 | 6 | 2 | 12345 | 442 | Sema1b,Sema1a | | 1.743e-02 | -4.05 | actin filament reorganization | biological process | GO:0090527 | 6 | 2 | 12345 | 442 | p130CAS,SCAR | | 1.743e-02 | -4.05 | regulation of carbohydrate catabolic process | biological process | GO:0043470 | 6 | 2 | 12345 | 442 | Dg,N | | 1.743e-02 | -4.05 | establishment or maintenance of polarity of larval imaginal disc epithelium | biological process | GO:0016336 | 6 | 2 | 12345 | 442 | ft,scrib | | 1.743e-02 | -4.05 | regulation of production of small RNA involved in gene silencing by RNA | biological process | GO:0070920 | 6 | 2 | 12345 | 442 | lin-28,Dg | | 1.750e-02 | -4.05 | GSE40666\_UNTREATED\_VS\_IFNA\_STIM\_STAT4\_KO\_EFFECTOR\_CD8\_TCELL\_90MIN\_DN | MSigDB lists | GSE40666\_UNTREATED\_VS\_IFNA\_STIM\_STAT4\_KO\_EFFECTOR\_CD8\_TCELL\_90MIN\_DN | 41 | 5 | 4161 | 156 | frc,Tpst,rg,Raf,Mef2 | | 1.750e-02 | -4.05 | RODWELL\_AGING\_KIDNEY\_NO\_BLOOD\_UP | MSigDB lists | RODWELL\_AGING\_KIDNEY\_NO\_BLOOD\_UP | 41 | 5 | 4161 | 156 | Unc-76,nAChRalpha4,l(2)37Cg,Actbeta,GlcAT-I | | 1.750e-02 | -4.05 | PID\_ERBB1\_DOWNSTREAM\_PATHWAY | MSigDB lists | PID\_ERBB1\_DOWNSTREAM\_PATHWAY | 41 | 5 | 4161 | 156 | 14-3-3epsilon,Mef2,Raf,Pdk1,14-3-3zeta | | 1.750e-02 | -4.05 | GO\_REGULATION\_OF\_I\_KAPPAB\_KINASE\_NF\_KAPPAB\_SIGNALING | MSigDB lists | GO\_REGULATION\_OF\_I\_KAPPAB\_KINASE\_NF\_KAPPAB\_SIGNALING | 41 | 5 | 4161 | 156 | Tollo,cact,CG45050,Pdk1,arm | | 1.750e-02 | -4.05 | MCBRYAN\_PUBERTAL\_BREAST\_5\_6WK\_DN | MSigDB lists | MCBRYAN\_PUBERTAL\_BREAST\_5\_6WK\_DN | 41 | 5 | 4161 | 156 | Rab5,step,ATPCL,kn,Mef2 | | 1.750e-02 | -4.05 | FOX\_Q2 | MSigDB lists | FOX\_Q2 | 41 | 5 | 4161 | 156 | Cadps,FBXO11,His2Av,Fkbp14,Mef2 | | 1.750e-02 | -4.05 | GSE23505\_UNTREATED\_VS\_4DAY\_IL6\_IL1\_IL23\_TREATED\_CD4\_TCELL\_UP | MSigDB lists | GSE23505\_UNTREATED\_VS\_4DAY\_IL6\_IL1\_IL23\_TREATED\_CD4\_TCELL\_UP | 41 | 5 | 4161 | 156 | Drak,hppy,step,Dyb,Raf | | 1.750e-02 | -4.05 | GSE3994\_WT\_VS\_PAC1\_KO\_ACTIVATED\_MAST\_CELL\_UP | MSigDB lists | GSE3994\_WT\_VS\_PAC1\_KO\_ACTIVATED\_MAST\_CELL\_UP | 41 | 5 | 4161 | 156 | Fkbp14,sd,DPCoAC,Syx1A,Rgl | | 1.769e-02 | -4.03 | regulation of immune response | biological process | GO:0050776 | 168 | 12 | 12345 | 442 | htl,Dcr-2,cact,Pvr,DPCoAC,imd,LRR,mop,nub,CG32795,pnr,Pitslre | | 1.774e-02 | -4.03 | sulfur compound binding | molecular function | GO:1901681 | 45 | 5 | 12176 | 414 | cv-d,vn,sli,robo1,Sema1a | | 1.775e-02 | -4.03 | VEGFR2 mediated cell proliferation | REACTOME pathways | R-DME-5218921 | 120 | 11 | 3923 | 176 | CG4935,htl,PI31,Pdk1,14-3-3zeta,wdb,CG42684,Egfr,Raf,vn,Pvr | | 1.783e-02 | -4.03 | dco (discs overgrown) | protein interactions | 43673 | 33 | 5 | 9204 | 432 | fz,ft,fz2,d,Pvr | | 1.787e-02 | -4.02 | TwdlL (TweedleL) | protein interactions | 43203 | 22 | 4 | 9204 | 432 | CG6891,imd,msi,sns | | 1.787e-02 | -4.02 | ph-d (polyhomeotic distal) | protein interactions | 44889 | 22 | 4 | 9204 | 432 | ph-p,Psc,Su(z)2,corto | | 1.787e-02 | -4.02 | Marcal1 (Marcal1) | protein interactions | 33709 | 22 | 4 | 9204 | 432 | E(Pc),ph-p,Psc,Su(z)2 | | 1.787e-02 | -4.02 | Hrb87F (Heterogeneous nuclear ribonucleoprotein at 87F) | protein interactions | 48535 | 22 | 4 | 9204 | 432 | Src42A,lncRNA:Hsromega,orb2,Raf | | 1.787e-02 | -4.02 | eIF3i (eukaryotic translation initiation factor 3 subunit i) | protein interactions | 33710 | 22 | 4 | 9204 | 432 | Pdk1,rictor,imd,orb2 | | 1.801e-02 | -4.02 | LPD\_N | smart domains | SM00638 | 4 | 2 | 5086 | 290 | apolpp,cv-d | | 1.803e-02 | -4.02 | GO\_RESPONSE\_TO\_GROWTH\_FACTOR | MSigDB lists | GO\_RESPONSE\_TO\_GROWTH\_FACTOR | 107 | 9 | 4161 | 156 | Mef2,tkv,htl,fz,Pdk1,N,nmo,CenB1A,arm | | 1.804e-02 | -4.02 | positive regulation of stem cell proliferation | biological process | GO:2000648 | 16 | 3 | 12345 | 442 | N,lin-28,klu | | 1.804e-02 | -4.02 | positive regulation of chemotaxis | biological process | GO:0050921 | 16 | 3 | 12345 | 442 | chic,fz,sli | | 1.804e-02 | -4.02 | regulation of neuron maturation | biological process | GO:0014041 | 16 | 3 | 12345 | 442 | Arf79F,Sox14,InR | | 1.804e-02 | -4.02 | regulation of neuron remodeling | biological process | GO:1904799 | 16 | 3 | 12345 | 442 | InR,Sox14,Arf79F | | 1.804e-02 | -4.02 | larval lymph gland hemocyte differentiation | biological process | GO:0035168 | 16 | 3 | 12345 | 442 | htl,Raf,N | | 1.805e-02 | -4.01 | TYR\_PHOSPHATASE\_2 | prosite domains | PS50056 | 30 | 5 | 6243 | 325 | IA-2,Ptpmeg2,Pez,CG42327,Ptp4E | | 1.807e-02 | -4.01 | Znf\_NHR/GATA | interpro domains | IPR013088 | 26 | 4 | 11383 | 450 | Hr39,Hr4,pnr,grn | | 1.808e-02 | -4.01 | melanization defense response | biological process | GO:0035006 | 43 | 5 | 12345 | 442 | RhoL,RhoGEF3,Dronc,Eph,LanA | | 1.808e-02 | -4.01 | gastrulation with mouth forming first | biological process | GO:0001703 | 43 | 5 | 12345 | 442 | trbl,baz,Abl,arm,shg | | 1.808e-02 | -4.01 | multi-organism metabolic process | biological process | GO:0044033 | 43 | 5 | 12345 | 442 | RhoGEF3,RhoL,LanA,Eph,Dronc | | 1.808e-02 | -4.01 | gastrulation involving germ band extension | biological process | GO:0010004 | 43 | 5 | 12345 | 442 | trbl,baz,Abl,shg,arm | | 1.814e-02 | -4.01 | cell projection assembly | biological process | GO:0030031 | 149 | 11 | 12345 | 442 | shg,unc-119,LRR,Myo10A,SCAR,Pvr,orb2,vn,mim,ena,Egfr | | 1.816e-02 | -4.01 | transport | biological process | GO:0006810 | 1659 | 75 | 12345 | 442 | Pex10,Slbp,Tpst,CG13384,jar,wbl,ogre,Rab5,CG11069,subdued,Rab1,Esp,apolpp,Arf79F,lqf,Irk3,Gli,CG17883,CG7394,SCAR,Pex14,Src42A,klar,Unc-76,ena,CG4991,cora,CG5853,ZnT41F,Atox1,dve,Vps13,S,Npc2g,Orct2,koi,RhoU,tipE,CG30344,chic,Pen,CG8034,Mid1,Abl,AP-1gamma,Cadps,Mpcp2,Dcr-2,CG31121,Syx1A,TBC1D5,mop,Myo10A,cv-d,SERCA,CG7536,frc,Rim2,nuf,Pex5,Inx2,CG17646,Tpc1,path,Ocrl,aop,unc-119,lt,edl,CG3149,nAChRalpha4,DPCoAC,scrib,Ppcs,CG4645 | | 1.819e-02 | -4.01 | eIF4A (eukaryotic translation initiation factor 4A) | protein interactions | 33835 | 58 | 7 | 9204 | 432 | Pdk1,rictor,Raf,Shrm,trbl,Dp,N | | 1.819e-02 | -4.01 | small GTPase mediated signal transduction | biological process | GO:0007264 | 93 | 8 | 12345 | 442 | Ric,Rab1,Raf,RhoGAP19D,Rab5,Arf79F,Rgl,RhoU | | 1.829e-02 | -4.00 | - | gene3d domains | 3.30.505.10 | 32 | 5 | 4929 | 241 | dock,Cbl,Socs36E,Src42A,Abl | | 1.830e-02 | -4.00 | CTLA4 inhibitory signaling | REACTOME pathways | R-DME-389513 | 5 | 2 | 3923 | 176 | Pdk1,wdb | | 1.830e-02 | -4.00 | GRB2 events in ERBB2 signaling | REACTOME pathways | R-DME-1963640 | 5 | 2 | 3923 | 176 | Egfr,vn | | 1.831e-02 | -4.00 | GO\_SINGLE\_ORGANISM\_CELLULAR\_LOCALIZATION | MSigDB lists | GO\_SINGLE\_ORGANISM\_CELLULAR\_LOCALIZATION | 369 | 22 | 4161 | 156 | Syx1A,ReepA,Cadps,LanA,CG17883,emp,SERCA,Pex14,arm,Pex10,CG7394,CG45050,Nrk,Pex7,Pex5,cact,Raf,Rab1,Pen,lt,Arf79F,Vps13 | | 1.850e-02 | -3.99 | HLH | pfam domains | PF00010 | 53 | 6 | 10480 | 419 | h,HLH3B,emc,sim,E(spl)m5-HLH,crp | | 1.851e-02 | -3.99 | tissue homeostasis | biological process | GO:0001894 | 59 | 6 | 12345 | 442 | shg,yrt,N,Atg16,Egfr,Atg17 | | 1.862e-02 | -3.98 | STAT1\_03 | MSigDB lists | STAT1\_03 | 73 | 7 | 4161 | 156 | lt,Dl,FBXO11,Cdk4,crol,nmo,Mfe2 | | 1.866e-02 | -3.98 | GO\_DEVELOPMENTAL\_GROWTH | MSigDB lists | GO\_DEVELOPMENTAL\_GROWTH | 90 | 8 | 4161 | 156 | htl,tkv,ap,N,otk,Dl,Ddr,sli | | 1.878e-02 | -3.97 | Interleukin-3, 5 and GM-CSF signaling | REACTOME pathways | R-DME-512988 | 121 | 11 | 3923 | 176 | CG42684,wdb,14-3-3zeta,htl,PI31,CG4935,Pvr,Raf,Cbl,vn,Egfr | | 1.885e-02 | -3.97 | GO\_CELL\_PROJECTION\_ORGANIZATION | MSigDB lists | GO\_CELL\_PROJECTION\_ORGANIZATION | 223 | 15 | 4161 | 156 | LanA,ft,Ocrl,sli,Eph,robo1,Rab1,unc-5,Unc-76,tkv,ap,dock,Arf79F,sina,N | | 1.888e-02 | -3.97 | GSE3337\_CTRL\_VS\_16H\_IFNG\_IN\_CD8POS\_DC\_UP | MSigDB lists | GSE3337\_CTRL\_VS\_16H\_IFNG\_IN\_CD8POS\_DC\_UP | 57 | 6 | 4161 | 156 | Mfe2,Rgl,Spn42De,fz,Nrk,Pgant5 | | 1.888e-02 | -3.97 | E2A\_Q2 | MSigDB lists | E2A\_Q2 | 57 | 6 | 4161 | 156 | otk,Vps13,Nrk,N,sli,rdx | | 1.888e-02 | -3.97 | GO\_PIGMENT\_GRANULE | MSigDB lists | GO\_PIGMENT\_GRANULE | 57 | 6 | 4161 | 156 | wbl,Rab1,14-3-3epsilon,Rab5,Kr-h2,14-3-3zeta | | 1.888e-02 | -3.97 | GO\_RECEPTOR\_COMPLEX | MSigDB lists | GO\_RECEPTOR\_COMPLEX | 57 | 6 | 4161 | 156 | nAChRalpha4,tkv,InR,N,Nrk,CG11069 | | 1.895e-02 | -3.97 | regulation of MAP kinase activity | biological process | GO:0043405 | 29 | 4 | 12345 | 442 | Raf,trbl,wnd,hppy | | 1.895e-02 | -3.97 | vascular endothelial growth factor receptor signaling pathway | biological process | GO:0048010 | 29 | 4 | 12345 | 442 | Pvr,Raf,aop,Pvf3 | | 1.897e-02 | -3.96 | REACTOME\_SIGNALLING\_TO\_P38\_VIA\_RIT\_AND\_RIN | MSigDB lists | REACTOME\_SIGNALLING\_TO\_P38\_VIA\_RIT\_AND\_RIN | 6 | 2 | 4161 | 156 | Ric,Raf | | 1.897e-02 | -3.96 | REACTOME\_CTNNB1\_PHOSPHORYLATION\_CASCADE | MSigDB lists | REACTOME\_CTNNB1\_PHOSPHORYLATION\_CASCADE | 6 | 2 | 4161 | 156 | wdb,arm | | 1.897e-02 | -3.96 | PID\_WNT\_CANONICAL\_PATHWAY | MSigDB lists | PID\_WNT\_CANONICAL\_PATHWAY | 6 | 2 | 4161 | 156 | fz2,arm | | 1.897e-02 | -3.96 | GO\_ATRIOVENTRICULAR\_VALVE\_DEVELOPMENT | MSigDB lists | GO\_ATRIOVENTRICULAR\_VALVE\_DEVELOPMENT | 6 | 2 | 4161 | 156 | N,ds | | 1.897e-02 | -3.96 | GO\_MITOGEN\_ACTIVATED\_PROTEIN\_KINASE\_KINASE\_BINDING | MSigDB lists | GO\_MITOGEN\_ACTIVATED\_PROTEIN\_KINASE\_KINASE\_BINDING | 6 | 2 | 4161 | 156 | Raf,trbl | | 1.897e-02 | -3.96 | REACTOME\_MYOGENESIS | MSigDB lists | REACTOME\_MYOGENESIS | 6 | 2 | 4161 | 156 | Mef2,arm | | 1.897e-02 | -3.96 | GRAHAM\_NORMAL\_QUIESCENT\_VS\_NORMAL\_DIVIDING\_UP | MSigDB lists | GRAHAM\_NORMAL\_QUIESCENT\_VS\_NORMAL\_DIVIDING\_UP | 6 | 2 | 4161 | 156 | shot,Actbeta | | 1.897e-02 | -3.96 | GO\_REGULATION\_OF\_NEUROBLAST\_PROLIFERATION | MSigDB lists | GO\_REGULATION\_OF\_NEUROBLAST\_PROLIFERATION | 6 | 2 | 4161 | 156 | arm,N | | 1.897e-02 | -3.96 | GO\_ENDOCARDIAL\_CUSHION\_MORPHOGENESIS | MSigDB lists | GO\_ENDOCARDIAL\_CUSHION\_MORPHOGENESIS | 6 | 2 | 4161 | 156 | ds,N | | 1.897e-02 | -3.96 | GO\_CELLULAR\_RESPONSE\_TO\_GONADOTROPIN\_STIMULUS | MSigDB lists | GO\_CELLULAR\_RESPONSE\_TO\_GONADOTROPIN\_STIMULUS | 6 | 2 | 4161 | 156 | Actbeta,N | | 1.897e-02 | -3.96 | GO\_PROTEOGLYCAN\_BINDING | MSigDB lists | GO\_PROTEOGLYCAN\_BINDING | 6 | 2 | 4161 | 156 | dlp,sli | | 1.897e-02 | -3.96 | GO\_REGULATION\_OF\_GLYCOPROTEIN\_METABOLIC\_PROCESS | MSigDB lists | GO\_REGULATION\_OF\_GLYCOPROTEIN\_METABOLIC\_PROCESS | 6 | 2 | 4161 | 156 | arm,Rab1 | | 1.897e-02 | -3.96 | BIOCARTA\_PITX2\_PATHWAY | MSigDB lists | BIOCARTA\_PITX2\_PATHWAY | 6 | 2 | 4161 | 156 | fz,arm | | 1.897e-02 | -3.96 | GO\_RESPONSE\_TO\_EXOGENOUS\_DSRNA | MSigDB lists | GO\_RESPONSE\_TO\_EXOGENOUS\_DSRNA | 6 | 2 | 4161 | 156 | cact,Tollo | | 1.897e-02 | -3.96 | GO\_REGULATION\_OF\_CELL\_PROLIFERATION\_INVOLVED\_IN\_HEART\_MORPHOGENESIS | MSigDB lists | GO\_REGULATION\_OF\_CELL\_PROLIFERATION\_INVOLVED\_IN\_HEART\_MORPHOGENESIS | 6 | 2 | 4161 | 156 | N,arm | | 1.897e-02 | -3.96 | GO\_DEVELOPMENTAL\_INDUCTION | MSigDB lists | GO\_DEVELOPMENTAL\_INDUCTION | 6 | 2 | 4161 | 156 | fz2,arm | | 1.897e-02 | -3.96 | GO\_CELL\_FATE\_COMMITMENT\_INVOLVED\_IN\_FORMATION\_OF\_PRIMARY\_GERM\_LAYER | MSigDB lists | GO\_CELL\_FATE\_COMMITMENT\_INVOLVED\_IN\_FORMATION\_OF\_PRIMARY\_GERM\_LAYER | 6 | 2 | 4161 | 156 | hyx,arm | | 1.897e-02 | -3.96 | REACTOME\_A\_TETRASACCHARIDE\_LINKER\_SEQUENCE\_IS\_REQUIRED\_FOR\_GAG\_SYNTHESIS | MSigDB lists | REACTOME\_A\_TETRASACCHARIDE\_LINKER\_SEQUENCE\_IS\_REQUIRED\_FOR\_GAG\_SYNTHESIS | 6 | 2 | 4161 | 156 | dlp,GlcAT-I | | 1.897e-02 | -3.96 | NIELSEN\_LIPOSARCOMA\_DN | MSigDB lists | NIELSEN\_LIPOSARCOMA\_DN | 6 | 2 | 4161 | 156 | fz,Actbeta | | 1.897e-02 | -3.96 | GO\_ACETYL\_COA\_BIOSYNTHETIC\_PROCESS | MSigDB lists | GO\_ACETYL\_COA\_BIOSYNTHETIC\_PROCESS | 6 | 2 | 4161 | 156 | ATPCL,muc | | 1.897e-02 | -3.96 | FARDIN\_HYPOXIA\_11 | MSigDB lists | FARDIN\_HYPOXIA\_11 | 6 | 2 | 4161 | 156 | Pgk,InR | | 1.897e-02 | -3.96 | GO\_NEGATIVE\_REGULATION\_OF\_BMP\_SIGNALING\_PATHWAY | MSigDB lists | GO\_NEGATIVE\_REGULATION\_OF\_BMP\_SIGNALING\_PATHWAY | 6 | 2 | 4161 | 156 | fz,N | | 1.897e-02 | -3.96 | ZWANG\_CLASS\_2\_TRANSIENTLY\_INDUCED\_BY\_EGF | MSigDB lists | ZWANG\_CLASS\_2\_TRANSIENTLY\_INDUCED\_BY\_EGF | 6 | 2 | 4161 | 156 | Actbeta,Gfat2 | | 1.897e-02 | -3.96 | GO\_SULFATION | MSigDB lists | GO\_SULFATION | 6 | 2 | 4161 | 156 | Tpst,CG9220 | | 1.897e-02 | -3.96 | GO\_MHC\_PROTEIN\_COMPLEX\_BINDING | MSigDB lists | GO\_MHC\_PROTEIN\_COMPLEX\_BINDING | 6 | 2 | 4161 | 156 | Tsp96F,14-3-3epsilon | | 1.897e-02 | -3.96 | GO\_REGULATION\_OF\_CYTOKINE\_PRODUCTION\_INVOLVED\_IN\_IMMUNE\_RESPONSE | MSigDB lists | GO\_REGULATION\_OF\_CYTOKINE\_PRODUCTION\_INVOLVED\_IN\_IMMUNE\_RESPONSE | 6 | 2 | 4161 | 156 | Tollo,fz2 | | 1.897e-02 | -3.96 | MATTIOLI\_MULTIPLE\_MYELOMA\_WITH\_14Q32\_TRANSLOCATIONS | MSigDB lists | MATTIOLI\_MULTIPLE\_MYELOMA\_WITH\_14Q32\_TRANSLOCATIONS | 6 | 2 | 4161 | 156 | htl,Pgant6 | | 1.897e-02 | -3.96 | GO\_MHC\_CLASS\_II\_PROTEIN\_COMPLEX\_BINDING | MSigDB lists | GO\_MHC\_CLASS\_II\_PROTEIN\_COMPLEX\_BINDING | 6 | 2 | 4161 | 156 | Tsp96F,14-3-3epsilon | | 1.897e-02 | -3.96 | GO\_VIRAL\_GENOME\_REPLICATION | MSigDB lists | GO\_VIRAL\_GENOME\_REPLICATION | 6 | 2 | 4161 | 156 | CtBP,Atg16 | | 1.897e-02 | -3.96 | GO\_CYTOKINE\_SECRETION | MSigDB lists | GO\_CYTOKINE\_SECRETION | 6 | 2 | 4161 | 156 | Rab1,N | | 1.897e-02 | -3.96 | GO\_POSITIVE\_REGULATION\_OF\_STEM\_CELL\_DIFFERENTIATION | MSigDB lists | GO\_POSITIVE\_REGULATION\_OF\_STEM\_CELL\_DIFFERENTIATION | 6 | 2 | 4161 | 156 | arm,N | | 1.897e-02 | -3.96 | GO\_POSITIVE\_REGULATION\_OF\_EPITHELIAL\_TO\_MESENCHYMAL\_TRANSITION | MSigDB lists | GO\_POSITIVE\_REGULATION\_OF\_EPITHELIAL\_TO\_MESENCHYMAL\_TRANSITION | 6 | 2 | 4161 | 156 | N,arm | | 1.897e-02 | -3.96 | GO\_NOTCH\_BINDING | MSigDB lists | GO\_NOTCH\_BINDING | 6 | 2 | 4161 | 156 | Dl,N | | 1.897e-02 | -3.96 | GO\_MORPHOGENESIS\_OF\_A\_POLARIZED\_EPITHELIUM | MSigDB lists | GO\_MORPHOGENESIS\_OF\_A\_POLARIZED\_EPITHELIUM | 6 | 2 | 4161 | 156 | LanA,otk | | 1.897e-02 | -3.96 | KEGG\_BLADDER\_CANCER | MSigDB lists | KEGG\_BLADDER\_CANCER | 6 | 2 | 4161 | 156 | htl,Raf | | 1.897e-02 | -3.96 | GO\_SKELETAL\_MUSCLE\_CELL\_DIFFERENTIATION | MSigDB lists | GO\_SKELETAL\_MUSCLE\_CELL\_DIFFERENTIATION | 6 | 2 | 4161 | 156 | N,Mef2 | | 1.897e-02 | -3.96 | GO\_HYPEROSMOTIC\_RESPONSE | MSigDB lists | GO\_HYPEROSMOTIC\_RESPONSE | 6 | 2 | 4161 | 156 | Tollo,Pdk1 | | 1.897e-02 | -3.96 | GRAHAM\_CML\_QUIESCENT\_VS\_NORMAL\_DIVIDING\_UP | MSigDB lists | GRAHAM\_CML\_QUIESCENT\_VS\_NORMAL\_DIVIDING\_UP | 6 | 2 | 4161 | 156 | Actbeta,SMC3 | | 1.897e-02 | -3.96 | GO\_SENSORY\_PERCEPTION\_OF\_TEMPERATURE\_STIMULUS | MSigDB lists | GO\_SENSORY\_PERCEPTION\_OF\_TEMPERATURE\_STIMULUS | 6 | 2 | 4161 | 156 | Eph,subdued | | 1.897e-02 | -3.96 | PLASARI\_NFIC\_TARGETS\_BASAL\_DN | MSigDB lists | PLASARI\_NFIC\_TARGETS\_BASAL\_DN | 6 | 2 | 4161 | 156 | sli,kn | | 1.897e-02 | -3.96 | PHONG\_TNF\_TARGETS\_UP | MSigDB lists | PHONG\_TNF\_TARGETS\_UP | 6 | 2 | 4161 | 156 | cact,Actbeta | | 1.897e-02 | -3.96 | BIOCARTA\_ARAP\_PATHWAY | MSigDB lists | BIOCARTA\_ARAP\_PATHWAY | 6 | 2 | 4161 | 156 | Arf79F,step | | 1.897e-02 | -3.96 | GO\_NEGATIVE\_REGULATION\_OF\_CALCIUM\_ION\_DEPENDENT\_EXOCYTOSIS | MSigDB lists | GO\_NEGATIVE\_REGULATION\_OF\_CALCIUM\_ION\_DEPENDENT\_EXOCYTOSIS | 6 | 2 | 4161 | 156 | Raf,N | | 1.897e-02 | -3.96 | GO\_HINDLIMB\_MORPHOGENESIS | MSigDB lists | GO\_HINDLIMB\_MORPHOGENESIS | 6 | 2 | 4161 | 156 | arm,N | | 1.918e-02 | -3.95 | O-linked glycosylation | REACTOME pathways | R-DME-5173105 | 48 | 6 | 3923 | 176 | CG34057,Pgant5,CG3119,C1GalTA,Pgant6,Dg | | 1.928e-02 | -3.95 | CAIRO\_HEPATOBLASTOMA\_CLASSES\_DN | MSigDB lists | CAIRO\_HEPATOBLASTOMA\_CLASSES\_DN | 42 | 5 | 4161 | 156 | Kdm2,ft,CG3961,cact,Tsp96F | | 1.928e-02 | -3.95 | GSE3920\_IFNB\_VS\_IFNG\_TREATED\_ENDOTHELIAL\_CELL\_UP | MSigDB lists | GSE3920\_IFNB\_VS\_IFNG\_TREATED\_ENDOTHELIAL\_CELL\_UP | 42 | 5 | 4161 | 156 | Pgk,trbl,InR,shot,Cerk | | 1.928e-02 | -3.95 | GSE15930\_NAIVE\_VS\_24H\_IN\_VITRO\_STIM\_INFAB\_CD8\_TCELL\_UP | MSigDB lists | GSE15930\_NAIVE\_VS\_24H\_IN\_VITRO\_STIM\_INFAB\_CD8\_TCELL\_UP | 42 | 5 | 4161 | 156 | sina,unc-119,Nmda1,CG17883,hppy | | 1.928e-02 | -3.95 | ATATGCA\_MIR448 | MSigDB lists | ATATGCA\_MIR448 | 42 | 5 | 4161 | 156 | InR,Mef2,rg,Pat1,Ziz | | 1.928e-02 | -3.95 | GO\_MORPHOGENESIS\_OF\_EMBRYONIC\_EPITHELIUM | MSigDB lists | GO\_MORPHOGENESIS\_OF\_EMBRYONIC\_EPITHELIUM | 42 | 5 | 4161 | 156 | ap,otk,fz,LanA,arm | | 1.928e-02 | -3.95 | GO\_TUBE\_FORMATION | MSigDB lists | GO\_TUBE\_FORMATION | 42 | 5 | 4161 | 156 | arm,fz,N,otk,ap | | 1.928e-02 | -3.95 | GSE22443\_IL2\_VS\_IL12\_TREATED\_ACT\_CD8\_TCELL\_UP | MSigDB lists | GSE22443\_IL2\_VS\_IL12\_TREATED\_ACT\_CD8\_TCELL\_UP | 42 | 5 | 4161 | 156 | Pgant6,Pex5,Gfat2,PI31,Rgl | | 1.928e-02 | -3.95 | GSE15930\_NAIVE\_VS\_24H\_IN\_VITRO\_STIM\_IL12\_CD8\_TCELL\_UP | MSigDB lists | GSE15930\_NAIVE\_VS\_24H\_IN\_VITRO\_STIM\_IL12\_CD8\_TCELL\_UP | 42 | 5 | 4161 | 156 | hppy,CG17883,Nmda1,unc-119,sina | | 1.928e-02 | -3.95 | GO\_CENTRAL\_NERVOUS\_SYSTEM\_NEURON\_DIFFERENTIATION | MSigDB lists | GO\_CENTRAL\_NERVOUS\_SYSTEM\_NEURON\_DIFFERENTIATION | 42 | 5 | 4161 | 156 | robo1,Actbeta,fz,Eph,sli | | 1.928e-02 | -3.95 | GSE22886\_DAY1\_VS\_DAY7\_MONOCYTE\_IN\_CULTURE\_UP | MSigDB lists | GSE22886\_DAY1\_VS\_DAY7\_MONOCYTE\_IN\_CULTURE\_UP | 42 | 5 | 4161 | 156 | cact,unc-119,Actbeta,step,CG3961 | | 1.928e-02 | -3.95 | NFKAPPAB65\_01 | MSigDB lists | NFKAPPAB65\_01 | 42 | 5 | 4161 | 156 | 14-3-3zeta,Cdk4,cact,trbl,nmo | | 1.928e-02 | -3.95 | MULLIGHAN\_NPM1\_MUTATED\_SIGNATURE\_2\_UP | MSigDB lists | MULLIGHAN\_NPM1\_MUTATED\_SIGNATURE\_2\_UP | 42 | 5 | 4161 | 156 | Mfe2,shot,CtBP,Pdk1,CG45050 | | 1.929e-02 | -3.95 | rhythmic process | biological process | GO:0048511 | 94 | 8 | 12345 | 442 | Mid1,Mef2,tara,InR,dome,ITP,kay,emc | | 1.929e-02 | -3.95 | GO\_REGULATION\_OF\_MUSCLE\_TISSUE\_DEVELOPMENT | MSigDB lists | GO\_REGULATION\_OF\_MUSCLE\_TISSUE\_DEVELOPMENT | 28 | 4 | 4161 | 156 | Dl,Mef2,N,arm | | 1.929e-02 | -3.95 | GSE37534\_UNTREATED\_VS\_PIOGLITAZONE\_TREATED\_CD4\_TCELL\_PPARG1\_AND\_FOXP3\_TRASDUCED\_DN | MSigDB lists | GSE37534\_UNTREATED\_VS\_PIOGLITAZONE\_TREATED\_CD4\_TCELL\_PPARG1\_AND\_FOXP3\_TRASDUCED\_DN | 28 | 4 | 4161 | 156 | Tollo,trbl,CG1673,CG2017 | | 1.929e-02 | -3.95 | DAZARD\_RESPONSE\_TO\_UV\_SCC\_DN | MSigDB lists | DAZARD\_RESPONSE\_TO\_UV\_SCC\_DN | 28 | 4 | 4161 | 156 | sina,rdx,CG11030,Kdm2 | | 1.929e-02 | -3.95 | GO\_THIOESTER\_BIOSYNTHETIC\_PROCESS | MSigDB lists | GO\_THIOESTER\_BIOSYNTHETIC\_PROCESS | 28 | 4 | 4161 | 156 | muc,ATPCL,CG3961,CG31522 | | 1.929e-02 | -3.95 | SARRIO\_EPITHELIAL\_MESENCHYMAL\_TRANSITION\_DN | MSigDB lists | SARRIO\_EPITHELIAL\_MESENCHYMAL\_TRANSITION\_DN | 28 | 4 | 4161 | 156 | CycG,shot,dock,cact | | 1.929e-02 | -3.95 | RAY\_TUMORIGENESIS\_BY\_ERBB2\_CDC25A\_DN | MSigDB lists | RAY\_TUMORIGENESIS\_BY\_ERBB2\_CDC25A\_DN | 28 | 4 | 4161 | 156 | CG31522,dlp,Ppcs,N | | 1.929e-02 | -3.95 | FOXD3\_01 | MSigDB lists | FOXD3\_01 | 28 | 4 | 4161 | 156 | Eph,FBXO11,Mef2,sina | | 1.931e-02 | -3.95 | GATAZNFINGER | prints domains | PR00619 | 5 | 2 | 2185 | 101 | grn,pnr | | 1.931e-02 | -3.95 | POUDOMAIN | prints domains | PR00028 | 5 | 2 | 2185 | 101 | pdm3,nub | | 1.940e-02 | -3.94 | IG\_LIKE | prosite domains | PS50835 | 132 | 13 | 6243 | 325 | boi,Bsg,robo1,kek5,Ppn,sns,vn,ed,CG31431,unc-5,htl,Pvr,otk | | 1.959e-02 | -3.93 | ABC\_2\_trans | interpro domains | IPR013525 | 15 | 3 | 11383 | 450 | CG31121,CG5853,CG17646 | | 1.970e-02 | -3.93 | Tctp (Translationally controlled tumor protein) | protein interactions | 41341 | 46 | 6 | 9204 | 432 | Nlp,rdx,C1GalTA,14-3-3zeta,chic,14-3-3epsilon | | 1.970e-02 | -3.93 | Past1 (Putative Achaete Scute Target 1) | protein interactions | 41569 | 46 | 6 | 9204 | 432 | H,Syx1A,Raf,rictor,Kr-h2,N | | 1.982e-02 | -3.92 | neuron remodeling | biological process | GO:0016322 | 44 | 5 | 12345 | 442 | eff,Sox14,mop,chic,Dronc | | 1.982e-02 | -3.92 | multi-organism cellular process | biological process | GO:0044764 | 44 | 5 | 12345 | 442 | Dronc,Eph,LanA,RhoL,RhoGEF3 | | 1.982e-02 | -3.92 | larval lymph gland hemopoiesis | biological process | GO:0035167 | 44 | 5 | 12345 | 442 | CG1129,htl,Pvr,N,Raf | | 1.982e-02 | -3.92 | protein complex oligomerization | biological process | GO:0051259 | 44 | 5 | 12345 | 442 | yrt,aop,InR,Dronc,Tob | | 1.982e-02 | -3.92 | post-embryonic hemopoiesis | biological process | GO:0035166 | 44 | 5 | 12345 | 442 | N,Raf,CG1129,Pvr,htl | | 1.982e-02 | -3.92 | glial cell differentiation | biological process | GO:0010001 | 44 | 5 | 12345 | 442 | htl,cora,moody,N,spen | | 1.983e-02 | -3.92 | GENTILE\_UV\_HIGH\_DOSE\_DN | MSigDB lists | GENTILE\_UV\_HIGH\_DOSE\_DN | 91 | 8 | 4161 | 156 | CG9220,dock,trbl,Pen,sina,Rab1,Tpst,CG3436 | | 1.983e-02 | -3.92 | WTGAAAT\_UNKNOWN | MSigDB lists | WTGAAAT\_UNKNOWN | 127 | 10 | 4161 | 156 | Ocrl,AP-1gamma,frc,Hel89B,rg,Mef2,Cdk4,wdb,sd,lt | | 2.000e-02 | -3.91 | CG34163 (uncharacterized protein) | protein interactions | 5740407 | 5 | 2 | 9204 | 432 | al,lin-28 | | 2.000e-02 | -3.91 | Eip78C (Ecdysone-induced protein 78C) | protein interactions | 40345 | 5 | 2 | 9204 | 432 | H,Diap1 | | 2.000e-02 | -3.91 | CG5110 (uncharacterized protein) | protein interactions | 35055 | 5 | 2 | 9204 | 432 | Rm62,corto | | 2.000e-02 | -3.91 | CG17612 (uncharacterized protein) | protein interactions | 33639 | 5 | 2 | 9204 | 432 | mam,ey | | 2.000e-02 | -3.91 | CG4266 (uncharacterized protein) | protein interactions | 37411 | 5 | 2 | 9204 | 432 | Pvr,rictor | | 2.000e-02 | -3.91 | asrij (asrij) | protein interactions | 37637 | 5 | 2 | 9204 | 432 | Arf79F,kn | | 2.000e-02 | -3.91 | 2mit (2mit) | protein interactions | 41616 | 5 | 2 | 9204 | 432 | arm,apolpp | | 2.000e-02 | -3.91 | mir-7 (mir-7 stem loop) | protein interactions | 12798133 | 5 | 2 | 9204 | 432 | aop,Dl | | 2.000e-02 | -3.91 | CG3402 (uncharacterized protein) | protein interactions | 38077 | 5 | 2 | 9204 | 432 | arm,CG34383 | | 2.000e-02 | -3.91 | Impbeta11 (Importin beta11) | protein interactions | 36732 | 5 | 2 | 9204 | 432 | tkv,fz2 | | 2.000e-02 | -3.91 | JMJD4 (Jumonji domain containing 4) | protein interactions | 35089 | 5 | 2 | 9204 | 432 | ttk,Sxl | | 2.000e-02 | -3.91 | chrb (charybde) | protein interactions | 39284 | 5 | 2 | 9204 | 432 | pnr,Sema1a | | 2.000e-02 | -3.91 | meru (meru) | protein interactions | 39727 | 5 | 2 | 9204 | 432 | baz,fz | | 2.000e-02 | -3.91 | CG9281 (uncharacterized protein) | protein interactions | 32508 | 5 | 2 | 9204 | 432 | Pdk1,N | | 2.000e-02 | -3.91 | mir-2b-2 (mir-2b-2 stem loop) | protein interactions | 12798018 | 5 | 2 | 9204 | 432 | SmD1,vig | | 2.000e-02 | -3.91 | Usp15-31 (Ubiquitin specific protease 15/31) | protein interactions | 37954 | 5 | 2 | 9204 | 432 | Dronc,Diap1 | | 2.000e-02 | -3.91 | Socs16D (Suppressor of Cytokine Signaling at 16D) | protein interactions | 32760 | 5 | 2 | 9204 | 432 | spi,Myo10A | | 2.000e-02 | -3.91 | Rat1 (Rat1 5'-3' exoribonuclease) | protein interactions | 33964 | 5 | 2 | 9204 | 432 | Myo10A,Sxl | | 2.000e-02 | -3.91 | Gel (Gelsolin) | protein interactions | 46008 | 5 | 2 | 9204 | 432 | HLH3B,Cdk4 | | 2.000e-02 | -3.91 | CG43373 (uncharacterized protein) | protein interactions | 39835 | 5 | 2 | 9204 | 432 | Fem-1,klar | | 2.000e-02 | -3.91 | DopEcR (Dopamine/Ecdysteroid receptor) | protein interactions | 38539 | 5 | 2 | 9204 | 432 | mld,Egfr | | 2.000e-02 | -3.91 | Roc2 (Regulator of cullins 2) | protein interactions | 36246 | 5 | 2 | 9204 | 432 | baz,CG7763 | | 2.000e-02 | -3.91 | RhoGEF64C (Rho guanine nucleotide exchange factor at 64C) | protein interactions | 38578 | 5 | 2 | 9204 | 432 | pnr,robo1 | | 2.000e-02 | -3.91 | CG3776 (uncharacterized protein) | protein interactions | 37996 | 5 | 2 | 9204 | 432 | pnr,SmD1 | | 2.003e-02 | -3.91 | regulation of GTPase activity | biological process | GO:0043087 | 113 | 9 | 12345 | 442 | RapGAP1,CenB1A,CG42684,TBC1D5,CG17883,RhoGAP19D,drongo,Rgl,Ocrl | | 2.009e-02 | -3.91 | awd (abnormal wing discs) | protein interactions | 43739 | 116 | 11 | 9204 | 432 | Myo10A,Fkbp14,chic,Rab5,Pgk,14-3-3zeta,Bacc,Jupiter,Nlp,14-3-3epsilon,Sema1a | | 2.012e-02 | -3.91 | Cenp-C (Centromeric protein-C) | protein interactions | 41014 | 34 | 5 | 9204 | 432 | 14-3-3zeta,14-3-3epsilon,Spc105R,flfl,jigr1 | | 2.012e-02 | -3.91 | E(spl)m7-HLH (Enhancer of split m7, helix-loop-helix) | protein interactions | 43160 | 34 | 5 | 9204 | 432 | CG12496,N,E(spl)m5-HLH,CG3386,CG34401 | | 2.012e-02 | -3.91 | Gug (Grunge) | protein interactions | 46156 | 34 | 5 | 9204 | 432 | Scr,ft,aop,d,al | | 2.012e-02 | -3.91 | Tbp (TATA binding protein) | protein interactions | 37476 | 34 | 5 | 9204 | 432 | N,Hel89B,ph-p,Psc,Mef2 | | 2.012e-02 | -3.91 | E(spl)m8-HLH (Enhancer of split m8, helix-loop-helix) | protein interactions | 43161 | 34 | 5 | 9204 | 432 | H,sens,h,N,E(spl)m5-HLH | | 2.013e-02 | -3.91 | GO\_GROWTH | MSigDB lists | GO\_GROWTH | 109 | 9 | 4161 | 156 | htl,tkv,ap,N,Actbeta,Ddr,sli,otk,Dl | | 2.013e-02 | -3.91 | GO\_SIGNAL\_TRANSDUCTION\_BY\_PROTEIN\_PHOSPHORYLATION | MSigDB lists | GO\_SIGNAL\_TRANSDUCTION\_BY\_PROTEIN\_PHOSPHORYLATION | 109 | 9 | 4161 | 156 | nmo,tkv,htl,sns,CG4935,PI31,hppy,Mef2,Raf | | 2.014e-02 | -3.90 | regulation of mRNA metabolic process | biological process | GO:1903311 | 132 | 10 | 12345 | 442 | fl(2)d,Sxl,bru2,CG11360,bol,Tis11,Rm62,ps,CG4896,pum | | 2.017e-02 | -3.90 | ABC2\_membrane | pfam domains | PF01061 | 15 | 3 | 10480 | 419 | CG31121,CG17646,CG5853 | | 2.021e-02 | -3.90 | GO\_MEMBRANE\_REGION | MSigDB lists | GO\_MEMBRANE\_REGION | 266 | 17 | 4161 | 156 | Cals,Dl,CG11069,robo1,shot,Arf79F,N,Ocrl,lqf,Eph,jar,arm,subdued,nAChRalpha4,Tsp96F,Pgk,Nrk | | 2.021e-02 | -3.90 | GTTAAAG\_MIR302B | MSigDB lists | GTTAAAG\_MIR302B | 16 | 3 | 4161 | 156 | nmo,AP-1gamma,robo1 | | 2.021e-02 | -3.90 | GCTGAGT\_MIR5125P | MSigDB lists | GCTGAGT\_MIR5125P | 16 | 3 | 4161 | 156 | CG5522,wbl,arm | | 2.021e-02 | -3.90 | GO\_ESTABLISHMENT\_OF\_PROTEIN\_LOCALIZATION\_TO\_VACUOLE | MSigDB lists | GO\_ESTABLISHMENT\_OF\_PROTEIN\_LOCALIZATION\_TO\_VACUOLE | 16 | 3 | 4161 | 156 | emp,lt,Vps13 | | 2.021e-02 | -3.90 | FONTAINE\_FOLLICULAR\_THYROID\_ADENOMA\_DN | MSigDB lists | FONTAINE\_FOLLICULAR\_THYROID\_ADENOMA\_DN | 16 | 3 | 4161 | 156 | Pen,N,Gfat2 | | 2.021e-02 | -3.90 | MORF\_RAB3A | MSigDB lists | MORF\_RAB3A | 16 | 3 | 4161 | 156 | lt,nAChRalpha4,SERCA | | 2.021e-02 | -3.90 | GSE7509\_UNSTIM\_VS\_IFNA\_STIM\_IMMATURE\_DC\_DN | MSigDB lists | GSE7509\_UNSTIM\_VS\_IFNA\_STIM\_IMMATURE\_DC\_DN | 16 | 3 | 4161 | 156 | N,CG9220,cact | | 2.021e-02 | -3.90 | KEGG\_GLIOMA | MSigDB lists | KEGG\_GLIOMA | 16 | 3 | 4161 | 156 | InR,Cdk4,Raf | | 2.021e-02 | -3.90 | GO\_POSITIVE\_REGULATION\_OF\_AXONOGENESIS | MSigDB lists | GO\_POSITIVE\_REGULATION\_OF\_AXONOGENESIS | 16 | 3 | 4161 | 156 | sli,Raf,robo1 | | 2.021e-02 | -3.90 | BIOCARTA\_CHREBP2\_PATHWAY | MSigDB lists | BIOCARTA\_CHREBP2\_PATHWAY | 16 | 3 | 4161 | 156 | 14-3-3epsilon,wdb,14-3-3zeta | | 2.021e-02 | -3.90 | GO\_RESPONSE\_TO\_DSRNA | MSigDB lists | GO\_RESPONSE\_TO\_DSRNA | 16 | 3 | 4161 | 156 | lin-28,Tollo,cact | | 2.021e-02 | -3.90 | GO\_REGULATION\_OF\_DNA\_TEMPLATED\_TRANSCRIPTION\_IN\_RESPONSE\_TO\_STRESS | MSigDB lists | GO\_REGULATION\_OF\_DNA\_TEMPLATED\_TRANSCRIPTION\_IN\_RESPONSE\_TO\_STRESS | 16 | 3 | 4161 | 156 | dock,N,eff | | 2.021e-02 | -3.90 | GO\_FOREBRAIN\_GENERATION\_OF\_NEURONS | MSigDB lists | GO\_FOREBRAIN\_GENERATION\_OF\_NEURONS | 16 | 3 | 4161 | 156 | sli,Actbeta,robo1 | | 2.021e-02 | -3.90 | PID\_TGFBR\_PATHWAY | MSigDB lists | PID\_TGFBR\_PATHWAY | 16 | 3 | 4161 | 156 | 14-3-3epsilon,arm,Pdk1 | | 2.028e-02 | -3.90 | GO\_CELLULAR\_COMPONENT\_MORPHOGENESIS | MSigDB lists | GO\_CELLULAR\_COMPONENT\_MORPHOGENESIS | 225 | 15 | 4161 | 156 | tkv,Unc-76,ap,N,sina,Rab1,arm,unc-5,Eph,Mef2,robo1,LanA,Ocrl,CG17883,sli | | 2.038e-02 | -3.89 | response to endogenous stimulus | biological process | GO:0009719 | 212 | 14 | 12345 | 442 | Hr39,tai,hfw,Raf,InR,pnr,ph-p,tkv,Dad,htl,rictor,aop,Pdk1,Hr4 | | 2.042e-02 | -3.89 | cellular response to organic substance | biological process | GO:0071310 | 275 | 17 | 12345 | 442 | wgn,htl,tkv,Dad,rictor,Pdk1,rdx,aop,Hr4,tai,dome,pnr,Pvr,Hr39,CG30156,Raf,InR | | 2.042e-02 | -3.89 | GO\_GUANYL\_NUCLEOTIDE\_EXCHANGE\_FACTOR\_ACTIVITY | MSigDB lists | GO\_GUANYL\_NUCLEOTIDE\_EXCHANGE\_FACTOR\_ACTIVITY | 58 | 6 | 4161 | 156 | CG5522,Ziz,step,Rgl,dock,htl | | 2.042e-02 | -3.89 | GSE26030\_TH1\_VS\_TH17\_RESTIMULATED\_DAY5\_POST\_POLARIZATION\_DN | MSigDB lists | GSE26030\_TH1\_VS\_TH17\_RESTIMULATED\_DAY5\_POST\_POLARIZATION\_DN | 58 | 6 | 4161 | 156 | Pgant5,CG2017,Pdk1,crol,emp,Nmda1 | | 2.042e-02 | -3.89 | GO\_EXTRACELLULAR\_MATRIX | MSigDB lists | GO\_EXTRACELLULAR\_MATRIX | 58 | 6 | 4161 | 156 | SMC3,Ppn,dlp,sli,shot,LanA | | 2.042e-02 | -3.89 | GSE22611\_NOD2\_VS\_CTRL\_TRANSDUCED\_HEK293T\_CELL\_UP | MSigDB lists | GSE22611\_NOD2\_VS\_CTRL\_TRANSDUCED\_HEK293T\_CELL\_UP | 58 | 6 | 4161 | 156 | sd,InR,Pen,Lk6,rdx,CG3436 | | 2.042e-02 | -3.89 | AP2\_Q3 | MSigDB lists | AP2\_Q3 | 58 | 6 | 4161 | 156 | InR,CG9220,Eph,ap,rdx,frc | | 2.042e-02 | -3.89 | GSE25123\_CTRL\_VS\_ROSIGLITAZONE\_STIM\_MACROPHAGE\_UP | MSigDB lists | GSE25123\_CTRL\_VS\_ROSIGLITAZONE\_STIM\_MACROPHAGE\_UP | 58 | 6 | 4161 | 156 | fz2,wbl,eff,Kdm2,Rab5,step | | 2.044e-02 | -3.89 | Mec2 (Mec2) | protein interactions | 32905 | 87 | 9 | 9204 | 432 | Mpcp2,Kr-h2,sns,Rab1,SERCA,Gfat2,Pdk1,Arf79F,Rab5 | | 2.044e-02 | -3.89 | CG33303 (uncharacterized protein) | protein interactions | 2768916 | 87 | 9 | 9204 | 432 | Rab5,Arf79F,SERCA,Gfat2,imd,Rab1,GluProRS,Mpcp2,Kr-h2 | | 2.058e-02 | -3.88 | PI5P, PP2A and IER3 Regulate PI3K/AKT Signaling | REACTOME pathways | R-DME-6811558 | 24 | 4 | 3923 | 176 | Egfr,htl,vn,wdb | | 2.064e-02 | -3.88 | Downstream signaling events of B Cell Receptor (BCR) | REACTOME pathways | R-DME-1168372 | 107 | 10 | 3923 | 176 | trbl,Dronc,wdb,Pdk1,rictor,htl,PI31,vn,Egfr,cact | | 2.066e-02 | -3.88 | kirre (kin of irre) | protein interactions | 31292 | 13 | 3 | 9204 | 432 | dock,sns,siz | | 2.066e-02 | -3.88 | path (pathetic) | protein interactions | 39106 | 13 | 3 | 9204 | 432 | hppy,orb2,pum | | 2.066e-02 | -3.88 | Uro (Urate oxidase) | protein interactions | 34060 | 13 | 3 | 9204 | 432 | Pex5,step,Sox14 | | 2.066e-02 | -3.88 | Daxx (Daxx-like protein) | protein interactions | 33906 | 13 | 3 | 9204 | 432 | His3.3A,His3.3B,Diap1 | | 2.066e-02 | -3.88 | Shc (SHC-adaptor protein) | protein interactions | 44052 | 13 | 3 | 9204 | 432 | Pvr,Egfr,Nrk | | 2.066e-02 | -3.88 | mop (myopic) | protein interactions | 39610 | 13 | 3 | 9204 | 432 | fz,Rab5,Cbl | | 2.066e-02 | -3.88 | bowl (brother of odd with entrails limited) | protein interactions | 33602 | 13 | 3 | 9204 | 432 | Src42A,Dcr-2,Pex5 | | 2.066e-02 | -3.88 | boi (brother of ihog) | protein interactions | 31229 | 13 | 3 | 9204 | 432 | boi,Dl,vn | | 2.066e-02 | -3.88 | nerfin-1 (nervous fingers 1) | protein interactions | 44786 | 13 | 3 | 9204 | 432 | Dl,sd,N | | 2.066e-02 | -3.88 | raptor (raptor) | protein interactions | 31543 | 13 | 3 | 9204 | 432 | hppy,REPTOR,Socs36E | | 2.066e-02 | -3.88 | ps (pasilla) | protein interactions | 44258 | 13 | 3 | 9204 | 432 | orb2,msi,pnr | | 2.066e-02 | -3.88 | inc (insomniac) | protein interactions | 31110 | 13 | 3 | 9204 | 432 | Sxl,Myo10A,Fem-1 | | 2.066e-02 | -3.88 | l(2)efl (lethal (2) essential for life) | protein interactions | 37744 | 13 | 3 | 9204 | 432 | koi,CycG,Myo10A | | 2.084e-02 | -3.87 | GO\_NEGATIVE\_REGULATION\_OF\_CELL\_DIFFERENTIATION | MSigDB lists | GO\_NEGATIVE\_REGULATION\_OF\_CELL\_DIFFERENTIATION | 128 | 10 | 4161 | 156 | sli,cact,Tollo,Dl,hyx,Cdk4,arm,trbl,N,Actbeta | | 2.085e-02 | -3.87 | HmgD (High mobility group protein D) | protein interactions | 37481 | 23 | 4 | 9204 | 432 | chic,Nlp,Pvr,Bacc | | 2.085e-02 | -3.87 | Ku80 (Ku80) | protein interactions | 34930 | 23 | 4 | 9204 | 432 | ttk,N,al,otp | | 2.085e-02 | -3.87 | hppy (happyhour) | protein interactions | 37203 | 23 | 4 | 9204 | 432 | Diap1,Egfr,aop,path | | 2.085e-02 | -3.87 | corto (corto) | protein interactions | 40616 | 23 | 4 | 9204 | 432 | corto,CycG,ph-p,Abd-B | | 2.085e-02 | -3.87 | cora (coracle) | protein interactions | 37205 | 23 | 4 | 9204 | 432 | Egfr,rictor,ft,Dg | | 2.085e-02 | -3.87 | RYBP (Ring and YY1 Binding Protein) | protein interactions | 37601 | 23 | 4 | 9204 | 432 | Diap1,Su(z)2,Psc,Kdm2 | | 2.085e-02 | -3.87 | LanA (Laminin A) | protein interactions | 38723 | 23 | 4 | 9204 | 432 | Sema1a,fz2,Dronc,sli | | 2.104e-02 | -3.86 | Znf\_GATA | interpro domains | IPR000679 | 6 | 2 | 11383 | 450 | pnr,grn | | 2.104e-02 | -3.86 | Ez/rad/moesin-like | interpro domains | IPR000798 | 6 | 2 | 11383 | 450 | yrt,cora | | 2.104e-02 | -3.86 | FERM\_CS | interpro domains | IPR019747 | 6 | 2 | 11383 | 450 | cora,yrt | | 2.104e-02 | -3.86 | FERM-adjacent | interpro domains | IPR014847 | 6 | 2 | 11383 | 450 | yrt,cora | | 2.104e-02 | -3.86 | IPPc | interpro domains | IPR000300 | 6 | 2 | 11383 | 450 | Ocrl,CG9784 | | 2.104e-02 | -3.86 | DNA-bd\_dom\_put\_sf | interpro domains | IPR009061 | 6 | 2 | 11383 | 450 | fuss,Snoo | | 2.104e-02 | -3.86 | Sec7\_dom | interpro domains | IPR000904 | 6 | 2 | 11383 | 450 | step,siz | | 2.104e-02 | -3.86 | VWF\_type-D | interpro domains | IPR001846 | 6 | 2 | 11383 | 450 | apolpp,cv-d | | 2.106e-02 | -3.86 | GO\_ENZYME\_BINDING | MSigDB lists | GO\_ENZYME\_BINDING | 530 | 29 | 4161 | 156 | rg,hyx,Cadps,Syx1A,Spn42De,AP-1gamma,Ocrl,emp,CG17883,Pdk1,Drep4,rdx,fz2,arm,cact,Raf,CG6891,Mef2,Pex7,Pex5,Pen,dock,lt,trbl,14-3-3zeta,N,CenB1A,14-3-3epsilon,nmo | | 2.107e-02 | -3.86 | Diap2 (Death-associated inhibitor of apoptosis 2) | protein interactions | 36748 | 179 | 15 | 9204 | 432 | larp,CtBP,Mpcp2,pum,Jupiter,MEP-1,cact,Dronc,Pvr,Diap1,SERCA,cindr,kay,imd,betaTub60D | | 2.118e-02 | -3.85 | ERB2\_UP.V1\_DN | MSigDB lists | ERB2\_UP.V1\_DN | 43 | 5 | 4161 | 156 | CG9485,hppy,SMC3,InR,Psa | | 2.118e-02 | -3.85 | GO\_THIOESTER\_METABOLIC\_PROCESS | MSigDB lists | GO\_THIOESTER\_METABOLIC\_PROCESS | 43 | 5 | 4161 | 156 | muc,ATPCL,Mfe2,CG3961,CG31522 | | 2.118e-02 | -3.85 | GO\_LEUKOCYTE\_CELL\_CELL\_ADHESION | MSigDB lists | GO\_LEUKOCYTE\_CELL\_CELL\_ADHESION | 43 | 5 | 4161 | 156 | Cdk4,Raf,dock,arm,fz2 | | 2.118e-02 | -3.85 | IK2\_01 | MSigDB lists | IK2\_01 | 43 | 5 | 4161 | 156 | CG11069,Cdk4,InR,dlp,nmo | | 2.118e-02 | -3.85 | GSE28726\_NAIVE\_VS\_ACTIVATED\_VA24NEG\_NKTCELL\_DN | MSigDB lists | GSE28726\_NAIVE\_VS\_ACTIVATED\_VA24NEG\_NKTCELL\_DN | 43 | 5 | 4161 | 156 | Cadps,jar,DPCoAC,Ocrl,trbl | | 2.118e-02 | -3.85 | MYOD\_01 | MSigDB lists | MYOD\_01 | 43 | 5 | 4161 | 156 | shot,rdx,otk,l(2)37Cg,Nrk | | 2.118e-02 | -3.85 | GSE40666\_NAIVE\_VS\_EFFECTOR\_CD8\_TCELL\_WITH\_IFNA\_STIM\_90MIN\_UP | MSigDB lists | GSE40666\_NAIVE\_VS\_EFFECTOR\_CD8\_TCELL\_WITH\_IFNA\_STIM\_90MIN\_UP | 43 | 5 | 4161 | 156 | trbl,CG6891,l(2)37Cg,jar,fz2 | | 2.118e-02 | -3.85 | GSE360\_T\_GONDII\_VS\_B\_MALAYI\_LOW\_DOSE\_DC\_UP | MSigDB lists | GSE360\_T\_GONDII\_VS\_B\_MALAYI\_LOW\_DOSE\_DC\_UP | 43 | 5 | 4161 | 156 | Kdm2,ReepA,bru2,sina,cact | | 2.124e-02 | -3.85 | GO\_REGULATION\_OF\_CELL\_ADHESION | MSigDB lists | GO\_REGULATION\_OF\_CELL\_ADHESION | 110 | 9 | 4161 | 156 | LanA,step,Raf,Cdk4,Dl,arm,Pdk1,N,dock | | 2.124e-02 | -3.85 | GO\_HEART\_DEVELOPMENT | MSigDB lists | GO\_HEART\_DEVELOPMENT | 110 | 9 | 4161 | 156 | fz,N,nAChRalpha4,arm,Mef2,Dl,otk,ft,ds | | 2.126e-02 | -3.85 | positive regulation of JNK cascade | biological process | GO:0046330 | 30 | 4 | 12345 | 442 | hppy,lncRNA:Hsromega,Pvr,rdx | | 2.126e-02 | -3.85 | melanotic encapsulation of foreign target | biological process | GO:0035011 | 30 | 4 | 12345 | 442 | LanA,Eph,RhoGEF3,RhoL | | 2.126e-02 | -3.85 | triglyceride homeostasis | biological process | GO:0070328 | 30 | 4 | 12345 | 442 | hppy,cue,Ppcs,InR | | 2.133e-02 | -3.85 | guanyl-nucleotide exchange factor activity | molecular function | GO:0005085 | 82 | 7 | 12176 | 414 | Rgl,Ziz,RhoGEF4,step,CG5522,RhoGEF3,siz | | 2.133e-02 | -3.85 | negative regulation of small GTPase mediated signal transduction | biological process | GO:0051058 | 17 | 3 | 12345 | 442 | Lk6,RapGAP1,CG42684 | | 2.133e-02 | -3.85 | regulation of lamellipodium assembly | biological process | GO:0010591 | 17 | 3 | 12345 | 442 | SCAR,ena,Abl | | 2.133e-02 | -3.85 | primary sex determination | biological process | GO:0007538 | 17 | 3 | 12345 | 442 | emc,Sxl,fl(2)d | | 2.133e-02 | -3.85 | regulation of lamellipodium organization | biological process | GO:1902743 | 17 | 3 | 12345 | 442 | SCAR,Abl,ena | | 2.133e-02 | -3.85 | tracheal outgrowth, open tracheal system | biological process | GO:0007426 | 17 | 3 | 12345 | 442 | Egfr,rib,ttk | | 2.133e-02 | -3.85 | regulation of actomyosin structure organization | biological process | GO:0110020 | 17 | 3 | 12345 | 442 | RhoGEF4,step,Ocrl | | 2.133e-02 | -3.85 | positive regulation of BMP signaling pathway | biological process | GO:0030513 | 17 | 3 | 12345 | 442 | hth,lili,cv-d | | 2.133e-02 | -3.85 | photoreceptor cell morphogenesis | biological process | GO:0008594 | 17 | 3 | 12345 | 442 | baz,Bsg,sens | | 2.133e-02 | -3.85 | trunk segmentation | biological process | GO:0035290 | 17 | 3 | 12345 | 442 | Abd-B,tsh,h | | 2.133e-02 | -3.85 | cell migration involved in gastrulation | biological process | GO:0042074 | 17 | 3 | 12345 | 442 | jeb,htl,sli | | 2.140e-02 | -3.84 | SH3 | smart domains | SM00326 | 61 | 8 | 5086 | 290 | RhoGEF3,Src42A,dock,Cip4,cindr,SKIP,Abl,p130CAS | | 2.141e-02 | -3.84 | Ras protein signal transduction | biological process | GO:0007265 | 78 | 7 | 12345 | 442 | Raf,Rab5,Rab1,Ric,RhoGAP19D,RhoU,Arf79F | | 2.149e-02 | -3.84 | GATA | pfam domains | PF00320 | 6 | 2 | 10480 | 419 | pnr,grn | | 2.149e-02 | -3.84 | FA | pfam domains | PF08736 | 6 | 2 | 10480 | 419 | yrt,cora | | 2.150e-02 | -3.84 | regulation of axonogenesis | biological process | GO:0050770 | 61 | 6 | 12345 | 442 | Sema1b,chic,shot,fz,Sema1a,CG34401 | | 2.150e-02 | -3.84 | FERM\_C | smart domains | SM01196 | 11 | 3 | 5086 | 290 | Pez,yrt,cora | | 2.155e-02 | -3.84 | Hippo signaling pathway -multiple species | KEGG pathways | dme04392 | 16 | 3 | 3198 | 123 | sd,ds,ft | | 2.155e-02 | -3.84 | Hippo signaling pathway -multiple species | KEGG pathways | ko04392 | 16 | 3 | 3198 | 123 | sd,ds,ft | | 2.158e-02 | -3.84 | smo (smoothened) | protein interactions | 33196 | 60 | 7 | 9204 | 432 | Kdm4B,pnr,boi,wdb,Dl,Sxl,tkv | | 2.158e-02 | -3.84 | Rab1 (Rab1) | protein interactions | 42524 | 60 | 7 | 9204 | 432 | SERCA,Arf79F,Ocrl,cact,Kr-h2,apolpp,Rab5 | | 2.158e-02 | -3.84 | - | gene3d domains | 1.10.1000.11 | 5 | 2 | 4929 | 241 | siz,step | | 2.159e-02 | -3.84 | IRS-mediated signalling | REACTOME pathways | R-DME-112399 | 156 | 13 | 3923 | 176 | Pvr,Egfr,SNF4Agamma,Raf,vn,wdb,trbl,CG42684,CG4935,htl,Pdk1,14-3-3zeta,PI31 | | 2.159e-02 | -3.84 | IRS-related events triggered by IGF1R | REACTOME pathways | R-DME-2428928 | 156 | 13 | 3923 | 176 | PI31,14-3-3zeta,htl,Pdk1,CG4935,CG42684,trbl,wdb,Raf,vn,Egfr,SNF4Agamma,Pvr | | 2.159e-02 | -3.84 | Signaling by Type 1 Insulin-like Growth Factor 1 Receptor (IGF1R) | REACTOME pathways | R-DME-2404192 | 156 | 13 | 3923 | 176 | Pvr,Egfr,SNF4Agamma,vn,Raf,wdb,trbl,CG42684,CG4935,htl,PI31,Pdk1,14-3-3zeta | | 2.159e-02 | -3.84 | Signaling by Interleukins | REACTOME pathways | R-DME-449147 | 156 | 13 | 3923 | 176 | Pvr,Egfr,vn,Raf,Cbl,wdb,CG42684,Dronc,CG4935,14-3-3zeta,PI31,htl,Syx1A | | 2.159e-02 | -3.84 | Insulin receptor signalling cascade | REACTOME pathways | R-DME-74751 | 156 | 13 | 3923 | 176 | trbl,CG42684,wdb,htl,Pdk1,14-3-3zeta,PI31,CG4935,Pvr,vn,Raf,Egfr,SNF4Agamma | | 2.159e-02 | -3.84 | IGF1R signaling cascade | REACTOME pathways | R-DME-2428924 | 156 | 13 | 3923 | 176 | CG4935,htl,PI31,Pdk1,14-3-3zeta,wdb,CG42684,trbl,SNF4Agamma,Egfr,Raf,vn,Pvr | | 2.162e-02 | -3.83 | cellular response to external stimulus | biological process | GO:0071496 | 96 | 8 | 12345 | 442 | Raf,InR,lt,SNF4Agamma,Atg16,rictor,Atg17,CG7536 | | 2.162e-02 | -3.83 | cytokine binding | molecular function | GO:0019955 | 7 | 2 | 12176 | 414 | cv-d,tkv | | 2.162e-02 | -3.83 | transcription cofactor binding | molecular function | GO:0001221 | 7 | 2 | 12176 | 414 | sd,hth | | 2.162e-02 | -3.83 | phosphatidylinositol-4,5-bisphosphate phosphatase activity | molecular function | GO:0106019 | 7 | 2 | 12176 | 414 | CG9784,Ocrl | | 2.162e-02 | -3.83 | wide pore channel activity | molecular function | GO:0022829 | 7 | 2 | 12176 | 414 | ogre,Inx2 | | 2.162e-02 | -3.83 | ARF guanyl-nucleotide exchange factor activity | molecular function | GO:0005086 | 7 | 2 | 12176 | 414 | siz,step | | 2.162e-02 | -3.83 | GTPase activating protein binding | molecular function | GO:0032794 | 7 | 2 | 12176 | 414 | Cip4,cindr | | 2.170e-02 | -3.83 | SMID\_BREAST\_CANCER\_BASAL\_DN | MSigDB lists | SMID\_BREAST\_CANCER\_BASAL\_DN | 167 | 12 | 4161 | 156 | Pgant6,CG5522,unc-119,rg,Cals,CG9485,Pat1,jar,subdued,tkv,htl,InR | | 2.172e-02 | -3.83 | PH\_domain | interpro domains | IPR001849 | 71 | 7 | 11383 | 450 | CG34383,CenB1A,CG5522,step,RhoGEF4,Cadps,Ziz | | 2.172e-02 | -3.83 | Vha68-2 (Vacuolar H[+] ATPase 68 kDa subunit 2) | protein interactions | 45012 | 47 | 6 | 9204 | 432 | Gfat2,SERCA,Dl,Sema1a,Rab5,koi | | 2.172e-02 | -3.83 | Dcr-2 (Dicer-2) | protein interactions | 36993 | 47 | 6 | 9204 | 432 | Diap1,Dcr-2,SmD1,baz,sd,bowl | | 2.172e-02 | -3.83 | Tektin-A (Tektin A) | protein interactions | 34934 | 47 | 6 | 9204 | 432 | 14-3-3zeta,koi,CG14153,Sema1a,CG12496,esg | | 2.172e-02 | -3.83 | gus (gustavus) | protein interactions | 35478 | 47 | 6 | 9204 | 432 | CycG,Adf1,CG10348,Cdk4,lilli,Egfr | | 2.174e-02 | -3.83 | JAZAG\_TGFB1\_SIGNALING\_VIA\_SMAD4\_UP | MSigDB lists | JAZAG\_TGFB1\_SIGNALING\_VIA\_SMAD4\_UP | 29 | 4 | 4161 | 156 | ap,bru2,14-3-3epsilon,SERCA | | 2.174e-02 | -3.83 | TGCAAAC\_MIR452 | MSigDB lists | TGCAAAC\_MIR452 | 29 | 4 | 4161 | 156 | Ziz,Eph,CG45050,wdb | | 2.174e-02 | -3.83 | GSE9650\_GP33\_VS\_GP276\_LCMV\_SPECIFIC\_EXHAUSTED\_CD8\_TCELL\_UP | MSigDB lists | GSE9650\_GP33\_VS\_GP276\_LCMV\_SPECIFIC\_EXHAUSTED\_CD8\_TCELL\_UP | 29 | 4 | 4161 | 156 | CG1129,CG6891,Src42A,Cadps | | 2.174e-02 | -3.83 | GO\_NEGATIVE\_REGULATION\_OF\_CELLULAR\_RESPONSE\_TO\_GROWTH\_FACTOR\_STIMULUS | MSigDB lists | GO\_NEGATIVE\_REGULATION\_OF\_CELLULAR\_RESPONSE\_TO\_GROWTH\_FACTOR\_STIMULUS | 29 | 4 | 4161 | 156 | fz,N,Pdk1,sli | | 2.174e-02 | -3.83 | BMI1\_DN.V1\_UP | MSigDB lists | BMI1\_DN.V1\_UP | 29 | 4 | 4161 | 156 | Actbeta,fz2,Gfat2,Syx1A | | 2.174e-02 | -3.83 | GSE20366\_EX\_VIVO\_VS\_DEC205\_CONVERSION\_NAIVE\_CD4\_TCELL\_DN | MSigDB lists | GSE20366\_EX\_VIVO\_VS\_DEC205\_CONVERSION\_NAIVE\_CD4\_TCELL\_DN | 29 | 4 | 4161 | 156 | Tsp96F,kn,Rgl,CG1316 | | 2.174e-02 | -3.83 | GSE37301\_CD4\_TCELL\_VS\_GRANULOCYTE\_MONOCYTE\_PROGENITOR\_UP | MSigDB lists | GSE37301\_CD4\_TCELL\_VS\_GRANULOCYTE\_MONOCYTE\_PROGENITOR\_UP | 29 | 4 | 4161 | 156 | rdx,CG7914,Pgant6,nmo | | 2.174e-02 | -3.83 | DAVICIONI\_MOLECULAR\_ARMS\_VS\_ERMS\_DN | MSigDB lists | DAVICIONI\_MOLECULAR\_ARMS\_VS\_ERMS\_DN | 29 | 4 | 4161 | 156 | trbl,ft,dlp,Rgl | | 2.174e-02 | -3.83 | HALLMARK\_KRAS\_SIGNALING\_UP | MSigDB lists | HALLMARK\_KRAS\_SIGNALING\_UP | 29 | 4 | 4161 | 156 | Gfat2,subdued,trbl,Actbeta | | 2.174e-02 | -3.83 | GO\_ORGANOPHOSPHATE\_ESTER\_TRANSPORT | MSigDB lists | GO\_ORGANOPHOSPHATE\_ESTER\_TRANSPORT | 29 | 4 | 4161 | 156 | DPCoAC,Tpc1,frc,Rim2 | | 2.174e-02 | -3.83 | ATAGGAA\_MIR202 | MSigDB lists | ATAGGAA\_MIR202 | 29 | 4 | 4161 | 156 | Tmhs,CtBP,robo1,Pat1 | | 2.178e-02 | -3.83 | pre-mRNA binding | molecular function | GO:0036002 | 18 | 3 | 12176 | 414 | ps,Sxl,Slbp | | 2.178e-02 | -3.83 | Notch binding | molecular function | GO:0005112 | 18 | 3 | 12176 | 414 | cue,spdo,Dl | | 2.180e-02 | -3.83 | GO\_EMBRYO\_DEVELOPMENT | MSigDB lists | GO\_EMBRYO\_DEVELOPMENT | 227 | 15 | 4161 | 156 | lqf,ds,Ocrl,LanA,frc,hyx,Mef2,otk,Dl,arm,fz2,Actbeta,N,fz,ap | | 2.201e-02 | -3.82 | GO\_HEAD\_DEVELOPMENT | MSigDB lists | GO\_HEAD\_DEVELOPMENT | 187 | 13 | 4161 | 156 | N,Actbeta,fz,ap,unc-5,arm,14-3-3epsilon,robo1,Raf,Cdk4,Dl,sli,ft | | 2.202e-02 | -3.82 | GO\_PROTEIN\_TARGETING | MSigDB lists | GO\_PROTEIN\_TARGETING | 207 | 14 | 4161 | 156 | Pen,Pex10,lt,14-3-3zeta,CG45050,CG7394,Vps13,14-3-3epsilon,Pex14,arm,cact,Pex7,Pex5,emp | | 2.205e-02 | -3.81 | GSE39022\_LN\_VS\_SPLEEN\_DC\_UP | MSigDB lists | GSE39022\_LN\_VS\_SPLEEN\_DC\_UP | 59 | 6 | 4161 | 156 | CG3961,ap,Unc-76,Pen,Fkbp14,Cdk4 | | 2.205e-02 | -3.81 | GSE17721\_POLYIC\_VS\_GARDIQUIMOD\_1H\_BMDC\_DN | MSigDB lists | GSE17721\_POLYIC\_VS\_GARDIQUIMOD\_1H\_BMDC\_DN | 59 | 6 | 4161 | 156 | muc,ATPCL,Drak,Kr-h2,dock,Lk6 | | 2.212e-02 | -3.81 | Dys (Dystrophin) | protein interactions | 42327 | 74 | 8 | 9204 | 432 | Pen,Dad,sli,Dl,Nrk,Dg,robo1,Sema1a | | 2.213e-02 | -3.81 | GO\_POSITIVE\_REGULATION\_OF\_CELLULAR\_COMPONENT\_ORGANIZATION | MSigDB lists | GO\_POSITIVE\_REGULATION\_OF\_CELLULAR\_COMPONENT\_ORGANIZATION | 311 | 19 | 4161 | 156 | arm,fz2,Pdk1,sli,Ddr,Eph,Rab1,14-3-3epsilon,dock,Psa,sns,Arf79F,N,14-3-3zeta,otk,Dl,Cals,robo1,Raf | | 2.234e-02 | -3.80 | Peroxisome | KEGG pathways | ko04146 | 91 | 8 | 3198 | 123 | Mfe2,CG8306,Pex14,CG33474,Pex10,Pex7,Pex5,CG3961 | | 2.234e-02 | -3.80 | Peroxisome | KEGG pathways | dme04146 | 91 | 8 | 3198 | 123 | Pex14,CG33474,Mfe2,CG8306,Pex5,CG3961,Pex10,Pex7 | | 2.239e-02 | -3.80 | GO\_GLYCOPROTEIN\_METABOLIC\_PROCESS | MSigDB lists | GO\_GLYCOPROTEIN\_METABOLIC\_PROCESS | 111 | 9 | 4161 | 156 | frc,Ids,Pgant6,C1GalTA,tkv,Pgant5,CG9220,alpha-Man-Ia,GlcAT-I | | 2.242e-02 | -3.80 | RAF activation | REACTOME pathways | R-DME-5673000 | 14 | 3 | 3923 | 176 | 14-3-3zeta,Raf,wdb | | 2.244e-02 | -3.80 | CycG (Cyclin G) | protein interactions | 43724 | 103 | 10 | 9204 | 432 | corto,vn,Abd-B,ena,N,abd-A,Cdk4,Dl,wdb,H | | 2.260e-02 | -3.79 | cpa (capping protein alpha) | protein interactions | 37387 | 35 | 5 | 9204 | 432 | Diap1,cindr,LRR,Sema1a,CG14506 | | 2.260e-02 | -3.79 | eff (effete) | protein interactions | 41785 | 35 | 5 | 9204 | 432 | Diap1,ph-p,Dronc,sina,ttk | | 2.274e-02 | -3.78 | sarcoplasmic reticulum | cellular component | GO:0016529 | 7 | 2 | 11940 | 417 | SERCA,yuri | | 2.283e-02 | -3.78 | fly chr98A6-98A8\|3-94 | chromosome location | fly chr98A6-98A8\|3-94 | 1 | 1 | 25010 | 571 | wdb | | 2.283e-02 | -3.78 | fly chr30A2-30A6\|2-35 | chromosome location | fly chr30A2-30A6\|2-35 | 1 | 1 | 25010 | 571 | tai | | 2.283e-02 | -3.78 | fly chr28F4-28F5\|2-30 | chromosome location | fly chr28F4-28F5\|2-30 | 1 | 1 | 25010 | 571 | Pvr | | 2.283e-02 | -3.78 | fly chr46E6-46E8\|2-62 cM | chromosome location | fly chr46E6-46E8\|2-62 cM | 1 | 1 | 25010 | 571 | 14-3-3zeta | | 2.283e-02 | -3.78 | fly chr40A5-40A5\|2-54 | chromosome location | fly chr40A5-40A5\|2-54 | 1 | 1 | 25010 | 571 | tsh | | 2.283e-02 | -3.78 | fly chr83B8-83B9\|3-47.5 cM | chromosome location | fly chr83B8-83B9\|3-47.5 cM | 1 | 1 | 25010 | 571 | Atg17 | | 2.283e-02 | -3.78 | fly chr78A5-78B1\|3-47 | chromosome location | fly chr78A5-78B1\|3-47 | 1 | 1 | 25010 | 571 | siz | | 2.283e-02 | -3.78 | fly chr27B4-27B4\|2-22 | chromosome location | fly chr27B4-27B4\|2-22 | 1 | 1 | 25010 | 571 | sens-2 | | 2.283e-02 | -3.78 | fly chr6F3-6F5\|1-19 | chromosome location | fly chr6F3-6F5\|1-19 | 1 | 1 | 25010 | 571 | Sxl | | 2.283e-02 | -3.78 | fly chr85D16-85D17\|3-48 | chromosome location | fly chr85D16-85D17\|3-48 | 1 | 1 | 25010 | 571 | ps | | 2.283e-02 | -3.78 | fly chr41F-41F8\|2-55 cM | chromosome location | fly chr41F-41F8\|2-55 cM | 1 | 1 | 25010 | 571 | ap | | 2.283e-02 | -3.78 | fly chr88A1-88A4\|3-54 cM | chromosome location | fly chr88A1-88A4\|3-54 cM | 1 | 1 | 25010 | 571 | rdx | | 2.283e-02 | -3.78 | fly chr8C11-8C13\|1-26 | chromosome location | fly chr8C11-8C13\|1-26 | 1 | 1 | 25010 | 571 | fend | | 2.283e-02 | -3.78 | fly chr33A1-33A2\|2-45 | chromosome location | fly chr33A1-33A2\|2-45 | 1 | 1 | 25010 | 571 | crol | | 2.283e-02 | -3.78 | fly chr89A5-89A6\|3-58 cM | chromosome location | fly chr89A5-89A6\|3-58 cM | 1 | 1 | 25010 | 571 | cv-d | | 2.283e-02 | -3.78 | fly chr67E5-67E6\|3-33 | chromosome location | fly chr67E5-67E6\|3-33 | 1 | 1 | 25010 | 571 | CG32066 | | 2.283e-02 | -3.78 | fly chr61B3-61C1\|3-1 cM | chromosome location | fly chr61B3-61C1\|3-1 cM | 1 | 1 | 25010 | 571 | RhoGEF3 | | 2.283e-02 | -3.78 | fly chr56C1-56C4\|2-88 cM | chromosome location | fly chr56C1-56C4\|2-88 cM | 1 | 1 | 25010 | 571 | hppy | | 2.283e-02 | -3.78 | fly chr85D27-85E1\|3-49 | chromosome location | fly chr85D27-85E1\|3-49 | 1 | 1 | 25010 | 571 | CG8312 | | 2.283e-02 | -3.78 | fly chr53D8-53D9\|2-80 cM | chromosome location | fly chr53D8-53D9\|2-80 cM | 1 | 1 | 25010 | 571 | CG5522 | | 2.283e-02 | -3.78 | fly chr22B1-22B2\|2-3 cM | chromosome location | fly chr22B1-22B2\|2-3 cM | 1 | 1 | 25010 | 571 | CG17646 | | 2.283e-02 | -3.78 | fly chr11B9-11B10\|1-39 | chromosome location | fly chr11B9-11B10\|1-39 | 1 | 1 | 25010 | 571 | Tis11 | | 2.283e-02 | -3.78 | fly chr88E1-88E2\|3-56 cM | chromosome location | fly chr88E1-88E2\|3-56 cM | 1 | 1 | 25010 | 571 | CG42788 | | 2.283e-02 | -3.78 | fly chr24F4-25A1\|2-12 cM | chromosome location | fly chr24F4-25A1\|2-12 cM | 1 | 1 | 25010 | 571 | dpy | | 2.283e-02 | -3.78 | fly chr18B9-18C1\|1-62 | chromosome location | fly chr18B9-18C1\|1-62 | 1 | 1 | 25010 | 571 | CG8034 | | 2.283e-02 | -3.78 | fly chr80D5-80E3\|3-47 cM | chromosome location | fly chr80D5-80E3\|3-47 cM | 1 | 1 | 25010 | 571 | nAChRalpha4 | | 2.283e-02 | -3.78 | fly chr57C2-57C3\|2-93 | chromosome location | fly chr57C2-57C3\|2-93 | 1 | 1 | 25010 | 571 | Magi | | 2.283e-02 | -3.78 | fly chr56F5-56F5\|2-90 cM | chromosome location | fly chr56F5-56F5\|2-90 cM | 1 | 1 | 25010 | 571 | lncRNA:CR44630 | | 2.283e-02 | -3.78 | fly chr85C3-85C4\|3-48 | chromosome location | fly chr85C3-85C4\|3-48 | 1 | 1 | 25010 | 571 | Kdm2 | | 2.283e-02 | -3.78 | fly chr95F6-95F8\|3-84 | chromosome location | fly chr95F6-95F8\|3-84 | 1 | 1 | 25010 | 571 | jar | | 2.283e-02 | -3.78 | fly chr89A5-89A6\|3-58 | chromosome location | fly chr89A5-89A6\|3-58 | 1 | 1 | 25010 | 571 | cv-d | | 2.283e-02 | -3.78 | fly chr23C1-23C3\|2-8 | chromosome location | fly chr23C1-23C3\|2-8 | 1 | 1 | 25010 | 571 | lilli | | 2.283e-02 | -3.78 | fly chr7D3-7D5\|1-22 | chromosome location | fly chr7D3-7D5\|1-22 | 1 | 1 | 25010 | 571 | fs(1)h | | 2.283e-02 | -3.78 | fly chr21-60\|2-54 | chromosome location | fly chr21-60\|2-54 | 1 | 1 | 25010 | 571 | Hr39 | | 2.283e-02 | -3.78 | fly chr7D18-7F1\|1-23 cM | chromosome location | fly chr7D18-7F1\|1-23 cM | 1 | 1 | 25010 | 571 | lawc | | 2.283e-02 | -3.78 | fly chr94E7-94E8\|3-78 | chromosome location | fly chr94E7-94E8\|3-78 | 1 | 1 | 25010 | 571 | CG4467 | | 2.283e-02 | -3.78 | fly chr90E2-90E2\|3-62 cM | chromosome location | fly chr90E2-90E2\|3-62 cM | 1 | 1 | 25010 | 571 | htl | | 2.283e-02 | -3.78 | fly chr83C5-83C6\|3-47.5 cM | chromosome location | fly chr83C5-83C6\|3-47.5 cM | 1 | 1 | 25010 | 571 | CG2017 | | 2.283e-02 | -3.78 | fly chr14F1-14F2\|1-54 cM | chromosome location | fly chr14F1-14F2\|1-54 cM | 1 | 1 | 25010 | 571 | CG9784 | | 2.283e-02 | -3.78 | fly chr68B4-68C1\|3-35 cM | chromosome location | fly chr68B4-68C1\|3-35 cM | 1 | 1 | 25010 | 571 | scyl | | 2.283e-02 | -3.78 | fly chr29E1-29E3\|2-33 | chromosome location | fly chr29E1-29E3\|2-33 | 1 | 1 | 25010 | 571 | Sema1a | | 2.283e-02 | -3.78 | fly chr53D11-53D12\|2-80 cM | chromosome location | fly chr53D11-53D12\|2-80 cM | 1 | 1 | 25010 | 571 | lncRNA:CR43650 | | 2.283e-02 | -3.78 | fly chr76A1-76A1\|3-45 cM | chromosome location | fly chr76A1-76A1\|3-45 cM | 1 | 1 | 25010 | 571 | lncRNA:CR45941 | | 2.283e-02 | -3.78 | fly chr89A13-89B1\|3-58 cM | chromosome location | fly chr89A13-89B1\|3-58 cM | 1 | 1 | 25010 | 571 | pnr | | 2.283e-02 | -3.78 | fly chr96A7-96A9\|3-85 cM | chromosome location | fly chr96A7-96A9\|3-85 cM | 1 | 1 | 25010 | 571 | REPTOR | | 2.283e-02 | -3.78 | fly chr8C9-8C10\|1-26 cM | chromosome location | fly chr8C9-8C10\|1-26 cM | 1 | 1 | 25010 | 571 | AP-1gamma | | 2.283e-02 | -3.78 | fly chr90A2-90A3\|3-60 cM | chromosome location | fly chr90A2-90A3\|3-60 cM | 1 | 1 | 25010 | 571 | beat-IIa | | 2.283e-02 | -3.78 | fly chr89A1-89A2\|3-58 | chromosome location | fly chr89A1-89A2\|3-58 | 1 | 1 | 25010 | 571 | pxb | | 2.283e-02 | -3.78 | fly chr75B2-75B2\|3-45 | chromosome location | fly chr75B2-75B2\|3-45 | 1 | 1 | 25010 | 571 | snoRNA:Me28S-A30 | | 2.283e-02 | -3.78 | fly chr52C9-52D2\|2-75 | chromosome location | fly chr52C9-52D2\|2-75 | 1 | 1 | 25010 | 571 | sli | | 2.283e-02 | -3.78 | fly chr70F4-70F5\|3-42 | chromosome location | fly chr70F4-70F5\|3-42 | 1 | 1 | 25010 | 571 | mop | | 2.283e-02 | -3.78 | fly chr92A1-92A2\|3-66 cM | chromosome location | fly chr92A1-92A2\|3-66 cM | 1 | 1 | 25010 | 571 | Dl | | 2.283e-02 | -3.78 | fly chr42A6-42A7\|2-55 cM | chromosome location | fly chr42A6-42A7\|2-55 cM | 1 | 1 | 25010 | 571 | Src42A | | 2.283e-02 | -3.78 | fly chr44D4-31F4\|2-59 cM | chromosome location | fly chr44D4-31F4\|2-59 cM | 1 | 1 | 25010 | 571 | rgr | | 2.283e-02 | -3.78 | fly chr46E6-46E8\|2-62 | chromosome location | fly chr46E6-46E8\|2-62 | 1 | 1 | 25010 | 571 | 14-3-3zeta | | 2.283e-02 | -3.78 | fly chr66A4-66A5\|3-21 | chromosome location | fly chr66A4-66A5\|3-21 | 1 | 1 | 25010 | 571 | lqf | | 2.283e-02 | -3.78 | fly chr61B1-61B2\|3-1 | chromosome location | fly chr61B1-61B2\|3-1 | 1 | 1 | 25010 | 571 | p130CAS | | 2.283e-02 | -3.78 | fly chr17F3-18A2\|1-61 cM | chromosome location | fly chr17F3-18A2\|1-61 cM | 1 | 1 | 25010 | 571 | CG43759 | | 2.283e-02 | -3.78 | fly chr29E1-29E3\|2-33 cM | chromosome location | fly chr29E1-29E3\|2-33 cM | 1 | 1 | 25010 | 571 | Sema1a | | 2.283e-02 | -3.78 | fly chr60A11-60A11\|2-105 | chromosome location | fly chr60A11-60A11\|2-105 | 1 | 1 | 25010 | 571 | CG2812 | | 2.283e-02 | -3.78 | fly chr17F3-18A2\|1-61 | chromosome location | fly chr17F3-18A2\|1-61 | 1 | 1 | 25010 | 571 | CG43759 | | 2.283e-02 | -3.78 | fly chr96B2-96B4\|3-86 | chromosome location | fly chr96B2-96B4\|3-86 | 1 | 1 | 25010 | 571 | CG31121 | | 2.283e-02 | -3.78 | fly chr70F4-70F5\|3-42 cM | chromosome location | fly chr70F4-70F5\|3-42 cM | 1 | 1 | 25010 | 571 | mop | | 2.283e-02 | -3.78 | fly chr17D4-17D5\|1-60 cM | chromosome location | fly chr17D4-17D5\|1-60 cM | 1 | 1 | 25010 | 571 | CG6891 | | 2.283e-02 | -3.78 | fly chr66E4-66E4\|3-27 | chromosome location | fly chr66E4-66E4\|3-27 | 1 | 1 | 25010 | 571 | orb2 | | 2.283e-02 | -3.78 | fly chr43E9-43E11\|2-57 | chromosome location | fly chr43E9-43E11\|2-57 | 1 | 1 | 25010 | 571 | LRR | | 2.283e-02 | -3.78 | fly chr10C1-10C5\|1-35 cM | chromosome location | fly chr10C1-10C5\|1-35 cM | 1 | 1 | 25010 | 571 | Drak | | 2.283e-02 | -3.78 | fly chr60D14-60D16\|2-107 cM | chromosome location | fly chr60D14-60D16\|2-107 cM | 1 | 1 | 25010 | 571 | Mid1 | | 2.283e-02 | -3.78 | fly chr66A22-66B3\|3-21 cM | chromosome location | fly chr66A22-66B3\|3-21 cM | 1 | 1 | 25010 | 571 | nmo | | 2.283e-02 | -3.78 | fly chr89B8-89B9\|3-58 | chromosome location | fly chr89B8-89B9\|3-58 | 1 | 1 | 25010 | 571 | tara | | 2.283e-02 | -3.78 | fly chr16C5-16C6\|1-58 | chromosome location | fly chr16C5-16C6\|1-58 | 1 | 1 | 25010 | 571 | CG42684 | | 2.283e-02 | -3.78 | fly chr56B5-56C1\|2-88 cM | chromosome location | fly chr56B5-56C1\|2-88 cM | 1 | 1 | 25010 | 571 | ena | | 2.283e-02 | -3.78 | fly chr18D13-18E1\|1-64 | chromosome location | fly chr18D13-18E1\|1-64 | 1 | 1 | 25010 | 571 | dome | | 2.283e-02 | -3.78 | fly chr21D3-21D3\|2-1 | chromosome location | fly chr21D3-21D3\|2-1 | 1 | 1 | 25010 | 571 | lncRNA:CR46259 | | 2.283e-02 | -3.78 | fly chr59A2-59A3\|2-101 cM | chromosome location | fly chr59A2-59A3\|2-101 cM | 1 | 1 | 25010 | 571 | robo1 | | 2.283e-02 | -3.78 | fly chr56F5-56F5\|2-90 | chromosome location | fly chr56F5-56F5\|2-90 | 1 | 1 | 25010 | 571 | lncRNA:CR44630 | | 2.283e-02 | -3.78 | fly chr6C7-6C9\|1-18 cM | chromosome location | fly chr6C7-6C9\|1-18 cM | 1 | 1 | 25010 | 571 | CG14441 | | 2.283e-02 | -3.78 | fly chr54D4-54D5\|2-83 cM | chromosome location | fly chr54D4-54D5\|2-83 cM | 1 | 1 | 25010 | 571 | Sema1b | | 2.283e-02 | -3.78 | fly chr65A8-65A9\|3-16 cM | chromosome location | fly chr65A8-65A9\|3-16 cM | 1 | 1 | 25010 | 571 | LanA | | 2.283e-02 | -3.78 | fly chr39A1-39A1 | chromosome location | fly chr39A1-39A1 | 1 | 1 | 25010 | 571 | snoRNA:Me28S-C2789 | | 2.283e-02 | -3.78 | fly chr2D2-2D3\|1-0.6 | chromosome location | fly chr2D2-2D3\|1-0.6 | 1 | 1 | 25010 | 571 | ph-p | | 2.283e-02 | -3.78 | fly chr43E9-43E11\|2-57 cM | chromosome location | fly chr43E9-43E11\|2-57 cM | 1 | 1 | 25010 | 571 | LRR | | 2.283e-02 | -3.78 | fly chr85E14-85E14\|3-49 cM | chromosome location | fly chr85E14-85E14\|3-49 cM | 1 | 1 | 25010 | 571 | lncRNA:CR45029 | | 2.283e-02 | -3.78 | fly chr66C12-66C12\|3-24 cM | chromosome location | fly chr66C12-66C12\|3-24 cM | 1 | 1 | 25010 | 571 | Cbl | | 2.283e-02 | -3.78 | fly chr91E2-91E3\|3-65 | chromosome location | fly chr91E2-91E3\|3-65 | 1 | 1 | 25010 | 571 | gukh | | 2.283e-02 | -3.78 | fly chr61F2-61F3\|3-0 | chromosome location | fly chr61F2-61F3\|3-0 | 1 | 1 | 25010 | 571 | bab2 | | 2.283e-02 | -3.78 | fly chr96E2-96E4\|3-88 cM | chromosome location | fly chr96E2-96E4\|3-88 cM | 1 | 1 | 25010 | 571 | msi | | 2.283e-02 | -3.78 | fly chr50F1-50F6\|2-70 cM | chromosome location | fly chr50F1-50F6\|2-70 cM | 1 | 1 | 25010 | 571 | Shrm | | 2.283e-02 | -3.78 | fly chr28A1-28A4\|2-27 cM | chromosome location | fly chr28A1-28A4\|2-27 cM | 1 | 1 | 25010 | 571 | Ziz | | 2.283e-02 | -3.78 | fly chr85D27-85E1\|3-49 cM | chromosome location | fly chr85D27-85E1\|3-49 cM | 1 | 1 | 25010 | 571 | CG8312 | | 2.283e-02 | -3.78 | fly chr10A4-10A6\|1-33 cM | chromosome location | fly chr10A4-10A6\|1-33 cM | 1 | 1 | 25010 | 571 | CG11122 | | 2.283e-02 | -3.78 | fly chr7D18-7F1\|1-23 | chromosome location | fly chr7D18-7F1\|1-23 | 1 | 1 | 25010 | 571 | lawc | | 2.283e-02 | -3.78 | fly chr52C2-52C4\|2-75 cM | chromosome location | fly chr52C2-52C4\|2-75 cM | 1 | 1 | 25010 | 571 | CG30089 | | 2.283e-02 | -3.78 | fly chr102C3-102C4\|4-0 | chromosome location | fly chr102C3-102C4\|4-0 | 1 | 1 | 25010 | 571 | zfh2 | | 2.283e-02 | -3.78 | fly chr22B1-22B2\|2-3 | chromosome location | fly chr22B1-22B2\|2-3 | 1 | 1 | 25010 | 571 | CG17646 | | 2.283e-02 | -3.78 | fly chr88E12-88E12\|3-56 cM | chromosome location | fly chr88E12-88E12\|3-56 cM | 1 | 1 | 25010 | 571 | CG45218 | | 2.283e-02 | -3.78 | fly chr48D6-48D7\|2-65 cM | chromosome location | fly chr48D6-48D7\|2-65 cM | 1 | 1 | 25010 | 571 | otk | | 2.283e-02 | -3.78 | fly chr51E1-51E2\|2-74 | chromosome location | fly chr51E1-51E2\|2-74 | 1 | 1 | 25010 | 571 | chn | | 2.283e-02 | -3.78 | fly chr41B3-41B3\|2-55 cM | chromosome location | fly chr41B3-41B3\|2-55 cM | 1 | 1 | 25010 | 571 | CG17883 | | 2.283e-02 | -3.78 | fly chr26A5-26B2\|2-18 cM | chromosome location | fly chr26A5-26B2\|2-18 cM | 1 | 1 | 25010 | 571 | chic | | 2.283e-02 | -3.78 | fly chr19D1-19D2\|1-64 | chromosome location | fly chr19D1-19D2\|1-64 | 1 | 1 | 25010 | 571 | RhoGAP19D | | 2.283e-02 | -3.78 | fly chr41C3-41C3\|2-55 | chromosome location | fly chr41C3-41C3\|2-55 | 1 | 1 | 25010 | 571 | CG14464 | | 2.283e-02 | -3.78 | fly chr64E12-64F2\|3-15 cM | chromosome location | fly chr64E12-64F2\|3-15 cM | 1 | 1 | 25010 | 571 | vn | | 2.283e-02 | -3.78 | fly chr60A11-60A12\|2-105 | chromosome location | fly chr60A11-60A12\|2-105 | 1 | 1 | 25010 | 571 | SERCA | | 2.283e-02 | -3.78 | fly chr89B14-89B15\|3-58 | chromosome location | fly chr89B14-89B15\|3-58 | 1 | 1 | 25010 | 571 | ss | | 2.283e-02 | -3.78 | fly chr26E1-26E2\|2-21 cM | chromosome location | fly chr26E1-26E2\|2-21 cM | 1 | 1 | 25010 | 571 | eya | | 2.283e-02 | -3.78 | fly chr97B9-97C1\|3-91 | chromosome location | fly chr97B9-97C1\|3-91 | 1 | 1 | 25010 | 571 | scrib | | 2.283e-02 | -3.78 | fly chr96A9-96A12\|3-85 | chromosome location | fly chr96A9-96A12\|3-85 | 1 | 1 | 25010 | 571 | mld | | 2.283e-02 | -3.78 | fly chr87F12-87F13\|3-54 | chromosome location | fly chr87F12-87F13\|3-54 | 1 | 1 | 25010 | 571 | CG34383 | | 2.283e-02 | -3.78 | fly chr69B1-69B2\|3-38 | chromosome location | fly chr69B1-69B2\|3-38 | 1 | 1 | 25010 | 571 | Lmx1a | | 2.283e-02 | -3.78 | fly chr26C1-26C2\|2-19 cM | chromosome location | fly chr26C1-26C2\|2-19 cM | 1 | 1 | 25010 | 571 | Ddr | | 2.283e-02 | -3.78 | fly chr44D4-31F4\|2-59 | chromosome location | fly chr44D4-31F4\|2-59 | 1 | 1 | 25010 | 571 | rgr | | 2.283e-02 | -3.78 | fly chr7D3-7D3\|1-22 | chromosome location | fly chr7D3-7D3\|1-22 | 1 | 1 | 25010 | 571 | CG2233 | | 2.283e-02 | -3.78 | fly chr6E1-6E2\|1-18 | chromosome location | fly chr6E1-6E2\|1-18 | 1 | 1 | 25010 | 571 | CG43736 | | 2.283e-02 | -3.78 | fly chr78A5-78B1\|3-47 cM | chromosome location | fly chr78A5-78B1\|3-47 cM | 1 | 1 | 25010 | 571 | siz | | 2.283e-02 | -3.78 | fly chr24D4-24D6\|2-11 | chromosome location | fly chr24D4-24D6\|2-11 | 1 | 1 | 25010 | 571 | ed | | 2.283e-02 | -3.78 | fly chr17D4-17D5\|1-60 | chromosome location | fly chr17D4-17D5\|1-60 | 1 | 1 | 25010 | 571 | CG6891 | | 2.283e-02 | -3.78 | fly chr91E2-91E3\|3-65 cM | chromosome location | fly chr91E2-91E3\|3-65 cM | 1 | 1 | 25010 | 571 | gukh | | 2.283e-02 | -3.78 | fly chr66F5-67A1\|3-28 cM | chromosome location | fly chr66F5-67A1\|3-28 cM | 1 | 1 | 25010 | 571 | bol | | 2.283e-02 | -3.78 | fly chr93D4-93D5\|3-71 | chromosome location | fly chr93D4-93D5\|3-71 | 1 | 1 | 25010 | 571 | lncRNA:Hsromega | | 2.283e-02 | -3.78 | fly chr57E9-57F1\|2-95 | chromosome location | fly chr57E9-57F1\|2-95 | 1 | 1 | 25010 | 571 | Egfr | | 2.283e-02 | -3.78 | fly chr21D3-21D3\|2-1 cM | chromosome location | fly chr21D3-21D3\|2-1 cM | 1 | 1 | 25010 | 571 | lncRNA:CR46259 | | 2.283e-02 | -3.78 | fly chr49F10-49F11\|2-68 | chromosome location | fly chr49F10-49F11\|2-68 | 1 | 1 | 25010 | 571 | Dp | | 2.283e-02 | -3.78 | fly chr3A1-3A1\|1-0.9 | chromosome location | fly chr3A1-3A1\|1-0.9 | 1 | 1 | 25010 | 571 | Raf | | 2.283e-02 | -3.78 | fly chr68C7-68C7\|3-36 cM | chromosome location | fly chr68C7-68C7\|3-36 cM | 1 | 1 | 25010 | 571 | CG7560 | | 2.283e-02 | -3.78 | fly chr83B8-83B9\|3-47.5 | chromosome location | fly chr83B8-83B9\|3-47.5 | 1 | 1 | 25010 | 571 | Atg17 | | 2.283e-02 | -3.78 | fly chr96B2-96B4\|3-86 cM | chromosome location | fly chr96B2-96B4\|3-86 cM | 1 | 1 | 25010 | 571 | CG31121 | | 2.283e-02 | -3.78 | fly chr66A22-66B3\|3-21 | chromosome location | fly chr66A22-66B3\|3-21 | 1 | 1 | 25010 | 571 | nmo | | 2.283e-02 | -3.78 | fly chr53D11-53D12\|2-80 | chromosome location | fly chr53D11-53D12\|2-80 | 1 | 1 | 25010 | 571 | lncRNA:CR43650 | | 2.283e-02 | -3.78 | fly chr89B8-89B9\|3-58 cM | chromosome location | fly chr89B8-89B9\|3-58 cM | 1 | 1 | 25010 | 571 | tara | | 2.283e-02 | -3.78 | fly chr75E3-75E4\|3-45 cM | chromosome location | fly chr75E3-75E4\|3-45 cM | 1 | 1 | 25010 | 571 | CG3961 | | 2.283e-02 | -3.78 | fly chr87F12-87F13\|3-54 cM | chromosome location | fly chr87F12-87F13\|3-54 cM | 1 | 1 | 25010 | 571 | CG34383 | | 2.283e-02 | -3.78 | fly chr18B6-18B6\|1-62 cM | chromosome location | fly chr18B6-18B6\|1-62 cM | 1 | 1 | 25010 | 571 | rictor | | 2.283e-02 | -3.78 | fly chr23B6-23B7\|2-7 cM | chromosome location | fly chr23B6-23B7\|2-7 cM | 1 | 1 | 25010 | 571 | CG2991 | | 2.283e-02 | -3.78 | fly chr52C2-52C4\|2-75 | chromosome location | fly chr52C2-52C4\|2-75 | 1 | 1 | 25010 | 571 | CG30089 | | 2.283e-02 | -3.78 | fly chr50C23-50D3\|2-68 | chromosome location | fly chr50C23-50D3\|2-68 | 1 | 1 | 25010 | 571 | mam | | 2.283e-02 | -3.78 | fly chr65A14-65B1\|3-16 cM | chromosome location | fly chr65A14-65B1\|3-16 cM | 1 | 1 | 25010 | 571 | CG13300 | | 2.283e-02 | -3.78 | fly chr24C3-24C4\|2-11 | chromosome location | fly chr24C3-24C4\|2-11 | 1 | 1 | 25010 | 571 | bowl | | 2.283e-02 | -3.78 | fly chr86C1-86C3\|3-50 cM | chromosome location | fly chr86C1-86C3\|3-50 cM | 1 | 1 | 25010 | 571 | hth | | 2.283e-02 | -3.78 | fly chr75F9-76A3\|3-45 cM | chromosome location | fly chr75F9-76A3\|3-45 cM | 1 | 1 | 25010 | 571 | fz2 | | 2.283e-02 | -3.78 | fly chr60D14-60D16\|2-107 | chromosome location | fly chr60D14-60D16\|2-107 | 1 | 1 | 25010 | 571 | Mid1 | | 2.283e-02 | -3.78 | fly chr95F6-95F8\|3-84 cM | chromosome location | fly chr95F6-95F8\|3-84 cM | 1 | 1 | 25010 | 571 | jar | | 2.283e-02 | -3.78 | fly chr26C1-26C2\|2-19 | chromosome location | fly chr26C1-26C2\|2-19 | 1 | 1 | 25010 | 571 | Ddr | | 2.283e-02 | -3.78 | fly chr24D4-24D6\|2-11 cM | chromosome location | fly chr24D4-24D6\|2-11 cM | 1 | 1 | 25010 | 571 | ed | | 2.283e-02 | -3.78 | fly chr83D1-83D2\|3-47.5 | chromosome location | fly chr83D1-83D2\|3-47.5 | 1 | 1 | 25010 | 571 | Rm62 | | 2.283e-02 | -3.78 | fly chr66E4-66E4\|3-27 cM | chromosome location | fly chr66E4-66E4\|3-27 cM | 1 | 1 | 25010 | 571 | orb2 | | 2.283e-02 | -3.78 | fly chr2B16-2C1\|1-0.2 | chromosome location | fly chr2B16-2C1\|1-0.2 | 1 | 1 | 25010 | 571 | Hr4 | | 2.283e-02 | -3.78 | fly chr81F5-81F6\|3-47.1 cM | chromosome location | fly chr81F5-81F6\|3-47.1 cM | 1 | 1 | 25010 | 571 | CG12581 | | 2.283e-02 | -3.78 | fly chr3A1-3A1\|1-0.9 cM | chromosome location | fly chr3A1-3A1\|1-0.9 cM | 1 | 1 | 25010 | 571 | Raf | | 2.283e-02 | -3.78 | fly chr49E6-49E6\|2-67 | chromosome location | fly chr49E6-49E6\|2-67 | 1 | 1 | 25010 | 571 | Psc | | 2.283e-02 | -3.78 | fly chr6C12-6C12\|1-18 cM | chromosome location | fly chr6C12-6C12\|1-18 cM | 1 | 1 | 25010 | 571 | lncRNA:CR43496 | | 2.283e-02 | -3.78 | fly chr66F5-67A1\|3-28 | chromosome location | fly chr66F5-67A1\|3-28 | 1 | 1 | 25010 | 571 | bol | | 2.283e-02 | -3.78 | fly chr13F1-13F5\|1-52 | chromosome location | fly chr13F1-13F5\|1-52 | 1 | 1 | 25010 | 571 | sd | | 2.283e-02 | -3.78 | fly chr48D6-48D7\|2-65 | chromosome location | fly chr48D6-48D7\|2-65 | 1 | 1 | 25010 | 571 | otk | | 2.283e-02 | -3.78 | fly chr2C10-2D1\|1-0.5 | chromosome location | fly chr2C10-2D1\|1-0.5 | 1 | 1 | 25010 | 571 | Unc-76 | | 2.283e-02 | -3.78 | fly chr62A11-62A12\|3-0.5 | chromosome location | fly chr62A11-62A12\|3-0.5 | 1 | 1 | 25010 | 571 | CG12011 | | 2.283e-02 | -3.78 | fly chr71C1-71C1\|3-43 cM | chromosome location | fly chr71C1-71C1\|3-43 cM | 1 | 1 | 25010 | 571 | Tollo | | 2.283e-02 | -3.78 | fly chr93E4-93E9\|3-72 cM | chromosome location | fly chr93E4-93E9\|3-72 cM | 1 | 1 | 25010 | 571 | InR | | 2.283e-02 | -3.78 | fly chr70D3-70D4\|3-42 | chromosome location | fly chr70D3-70D4\|3-42 | 1 | 1 | 25010 | 571 | nuf | | 2.283e-02 | -3.78 | fly chr29B4-29C1\|2-31 cM | chromosome location | fly chr29B4-29C1\|2-31 cM | 1 | 1 | 25010 | 571 | CG13384 | | 2.283e-02 | -3.78 | fly chr40A5-40A5\|2-54 cM | chromosome location | fly chr40A5-40A5\|2-54 cM | 1 | 1 | 25010 | 571 | tsh | | 2.283e-02 | -3.78 | fly chr23A4-23A3\|2-7 cM | chromosome location | fly chr23A4-23A3\|2-7 cM | 1 | 1 | 25010 | 571 | Pgk | | 2.283e-02 | -3.78 | fly chr51E1-51E2\|2-74 cM | chromosome location | fly chr51E1-51E2\|2-74 cM | 1 | 1 | 25010 | 571 | chn | | 2.283e-02 | -3.78 | fly chr62A11-62A12\|3-0.5 cM | chromosome location | fly chr62A11-62A12\|3-0.5 cM | 1 | 1 | 25010 | 571 | CG12011 | | 2.283e-02 | -3.78 | fly chr4F2-4F3\|1-10 | chromosome location | fly chr4F2-4F3\|1-10 | 1 | 1 | 25010 | 571 | rg | | 2.283e-02 | -3.78 | fly chr9B6-9B7\|1-31 cM | chromosome location | fly chr9B6-9B7\|1-31 cM | 1 | 1 | 25010 | 571 | alpha-Man-Ia | | 2.283e-02 | -3.78 | fly chr36F5-36F6\|2-53 | chromosome location | fly chr36F5-36F6\|2-53 | 1 | 1 | 25010 | 571 | kon | | 2.283e-02 | -3.78 | fly chr13D4-13E1\|1-51 cM | chromosome location | fly chr13D4-13E1\|1-51 cM | 1 | 1 | 25010 | 571 | CG9220 | | 2.283e-02 | -3.78 | fly chr82B1-82B2\|3-47.1 cM | chromosome location | fly chr82B1-82B2\|3-47.1 cM | 1 | 1 | 25010 | 571 | CG31522 | | 2.283e-02 | -3.78 | fly chr28B1-28B2\|2-27 | chromosome location | fly chr28B1-28B2\|2-27 | 1 | 1 | 25010 | 571 | RapGAP1 | | 2.283e-02 | -3.78 | fly chr68C7-68C7\|3-36 | chromosome location | fly chr68C7-68C7\|3-36 | 1 | 1 | 25010 | 571 | CG7560 | | 2.283e-02 | -3.78 | fly chr85D22-85D24\|3-48 cM | chromosome location | fly chr85D22-85D24\|3-48 cM | 1 | 1 | 25010 | 571 | mura | | 2.283e-02 | -3.78 | fly chr85C4-85D1\|3-48 cM | chromosome location | fly chr85C4-85D1\|3-48 cM | 1 | 1 | 25010 | 571 | pum | | 2.283e-02 | -3.78 | fly chr28F4-28F5\|2-30 cM | chromosome location | fly chr28F4-28F5\|2-30 cM | 1 | 1 | 25010 | 571 | Pvr | | 2.283e-02 | -3.78 | fly chr51C2-51C3\|2-72 cM | chromosome location | fly chr51C2-51C3\|2-72 cM | 1 | 1 | 25010 | 571 | kn | | 2.283e-02 | -3.78 | fly chr74F4-75A1\|3-45 | chromosome location | fly chr74F4-75A1\|3-45 | 1 | 1 | 25010 | 571 | CG7460 | | 2.283e-02 | -3.78 | fly chr27B4-27B4\|2-22 cM | chromosome location | fly chr27B4-27B4\|2-22 cM | 1 | 1 | 25010 | 571 | sens-2 | | 2.283e-02 | -3.78 | fly chr14B1-14B1\|1-53 | chromosome location | fly chr14B1-14B1\|1-53 | 1 | 1 | 25010 | 571 | disco-r | | 2.283e-02 | -3.78 | fly chr16B6-16B7\|1-57 cM | chromosome location | fly chr16B6-16B7\|1-57 cM | 1 | 1 | 25010 | 571 | CG12991 | | 2.283e-02 | -3.78 | fly chr45A12-45A13\|2-60 cM | chromosome location | fly chr45A12-45A13\|2-60 cM | 1 | 1 | 25010 | 571 | CG30344 | | 2.283e-02 | -3.78 | fly chr50C23-50D3\|2-68 cM | chromosome location | fly chr50C23-50D3\|2-68 cM | 1 | 1 | 25010 | 571 | mam | | 2.283e-02 | -3.78 | fly chr12A6-12A7\|1-44 cM | chromosome location | fly chr12A6-12A7\|1-44 cM | 1 | 1 | 25010 | 571 | Tpst | | 2.283e-02 | -3.78 | fly chr98D3-98D4\|3-96 | chromosome location | fly chr98D3-98D4\|3-96 | 1 | 1 | 25010 | 571 | Ppn | | 2.283e-02 | -3.78 | fly chr89D5-89D6\|3-59 cM | chromosome location | fly chr89D5-89D6\|3-59 cM | 1 | 1 | 25010 | 571 | Fas1 | | 2.283e-02 | -3.78 | fly chr18C3-18C7\|1-62 cM | chromosome location | fly chr18C3-18C7\|1-62 cM | 1 | 1 | 25010 | 571 | kek5 | | 2.283e-02 | -3.78 | fly chr85D16-85D17\|3-48 cM | chromosome location | fly chr85D16-85D17\|3-48 cM | 1 | 1 | 25010 | 571 | ps | | 2.283e-02 | -3.78 | fly chr45A12-45A13\|2-60 | chromosome location | fly chr45A12-45A13\|2-60 | 1 | 1 | 25010 | 571 | CG30344 | | 2.283e-02 | -3.78 | fly chr70D3-70D4\|3-42 cM | chromosome location | fly chr70D3-70D4\|3-42 cM | 1 | 1 | 25010 | 571 | nuf | | 2.283e-02 | -3.78 | fly chr10A4-10A6\|1-33 | chromosome location | fly chr10A4-10A6\|1-33 | 1 | 1 | 25010 | 571 | CG11122 | | 2.283e-02 | -3.78 | fly chr96A9-96A12\|3-85 cM | chromosome location | fly chr96A9-96A12\|3-85 cM | 1 | 1 | 25010 | 571 | mld | | 2.283e-02 | -3.78 | fly chr66C13-66C13\|3-24 cM | chromosome location | fly chr66C13-66C13\|3-24 cM | 1 | 1 | 25010 | 571 | CG6983 | | 2.283e-02 | -3.78 | fly chr38F6-39A1\|2-54 cM | chromosome location | fly chr38F6-39A1\|2-54 cM | 1 | 1 | 25010 | 571 | CG42238 | | 2.283e-02 | -3.78 | fly chr6F3-6F5\|1-19 cM | chromosome location | fly chr6F3-6F5\|1-19 cM | 1 | 1 | 25010 | 571 | Sxl | | 2.283e-02 | -3.78 | fly chr26D8-26D8\|2-20 cM | chromosome location | fly chr26D8-26D8\|2-20 cM | 1 | 1 | 25010 | 571 | lncRNA:CR45463 | | 2.283e-02 | -3.78 | fly chr47F6-47F7\|2-64 cM | chromosome location | fly chr47F6-47F7\|2-64 cM | 1 | 1 | 25010 | 571 | CG13204 | | 2.283e-02 | -3.78 | fly chr94A2-94A4\|3-74 | chromosome location | fly chr94A2-94A4\|3-74 | 1 | 1 | 25010 | 571 | SKIP | | 2.283e-02 | -3.78 | fly chr24C3-24C4\|2-11 cM | chromosome location | fly chr24C3-24C4\|2-11 cM | 1 | 1 | 25010 | 571 | bowl | | 2.283e-02 | -3.78 | fly chr4F2-4F3\|1-10 cM | chromosome location | fly chr4F2-4F3\|1-10 cM | 1 | 1 | 25010 | 571 | rg | | 2.283e-02 | -3.78 | fly chr18E2-18E3\|1-64 | chromosome location | fly chr18E2-18E3\|1-64 | 1 | 1 | 25010 | 571 | meso18E | | 2.283e-02 | -3.78 | fly chr93D5-93D6\|3-71 | chromosome location | fly chr93D5-93D6\|3-71 | 1 | 1 | 25010 | 571 | CG16791 | | 2.283e-02 | -3.78 | fly chr47A12-47A13\|2-62 | chromosome location | fly chr47A12-47A13\|2-62 | 1 | 1 | 25010 | 571 | asRNA:CR45135 | | 2.283e-02 | -3.78 | fly chr86E17-86E18\|3-51 cM | chromosome location | fly chr86E17-86E18\|3-51 cM | 1 | 1 | 25010 | 571 | Lk6 | | 2.283e-02 | -3.78 | fly chr8C11-8C11\|1-26 cM | chromosome location | fly chr8C11-8C11\|1-26 cM | 1 | 1 | 25010 | 571 | lncRNA:CR45539 | | 2.283e-02 | -3.78 | fly chr47F6-47F7\|2-64 | chromosome location | fly chr47F6-47F7\|2-64 | 1 | 1 | 25010 | 571 | CG13204 | | 2.283e-02 | -3.78 | fly chr98D3-98D4\|3-96 cM | chromosome location | fly chr98D3-98D4\|3-96 cM | 1 | 1 | 25010 | 571 | Ppn | | 2.283e-02 | -3.78 | fly chr16C5-16C6\|1-58 cM | chromosome location | fly chr16C5-16C6\|1-58 cM | 1 | 1 | 25010 | 571 | CG42684 | | 2.283e-02 | -3.78 | fly chr57B15-57B16\|2-92 cM | chromosome location | fly chr57B15-57B16\|2-92 cM | 1 | 1 | 25010 | 571 | shg | | 2.283e-02 | -3.78 | fly chr2B16-2C1\|1-0.2 cM | chromosome location | fly chr2B16-2C1\|1-0.2 cM | 1 | 1 | 25010 | 571 | Hr4 | | 2.283e-02 | -3.78 | fly chr99B10-99C1\|3-99 | chromosome location | fly chr99B10-99C1\|3-99 | 1 | 1 | 25010 | 571 | kay | | 2.283e-02 | -3.78 | fly chr85E1-85E1 | chromosome location | fly chr85E1-85E1 | 1 | 1 | 25010 | 571 | sisRNA:CR46357 | | 2.283e-02 | -3.78 | fly chr88A1-88A4\|3-54 | chromosome location | fly chr88A1-88A4\|3-54 | 1 | 1 | 25010 | 571 | rdx | | 2.283e-02 | -3.78 | fly chr65A8-65A9\|3-16 | chromosome location | fly chr65A8-65A9\|3-16 | 1 | 1 | 25010 | 571 | LanA | | 2.283e-02 | -3.78 | fly chr21-60\|2-54 cM | chromosome location | fly chr21-60\|2-54 cM | 1 | 1 | 25010 | 571 | Hr39 | | 2.283e-02 | -3.78 | fly chr28E4-28E6\|2-30 | chromosome location | fly chr28E4-28E6\|2-30 | 1 | 1 | 25010 | 571 | Bsg | | 2.283e-02 | -3.78 | fly chr16A4-16A5\|1-57 | chromosome location | fly chr16A4-16A5\|1-57 | 1 | 1 | 25010 | 571 | B-H1 | | 2.283e-02 | -3.78 | fly chr52D9-52D11\|2-76 | chromosome location | fly chr52D9-52D11\|2-76 | 1 | 1 | 25010 | 571 | ATPCL | | 2.283e-02 | -3.78 | fly chr11F1-11F3\|1-43 cM | chromosome location | fly chr11F1-11F3\|1-43 cM | 1 | 1 | 25010 | 571 | CG1673 | | 2.283e-02 | -3.78 | fly chr97B9-97C1\|3-91 cM | chromosome location | fly chr97B9-97C1\|3-91 cM | 1 | 1 | 25010 | 571 | scrib | | 2.283e-02 | -3.78 | fly chr102D4-102D5\|4-0 cM | chromosome location | fly chr102D4-102D5\|4-0 cM | 1 | 1 | 25010 | 571 | ey | | 2.283e-02 | -3.78 | fly chr4C12-4C13\|1-8 | chromosome location | fly chr4C12-4C13\|1-8 | 1 | 1 | 25010 | 571 | Pp2C1 | | 2.283e-02 | -3.78 | fly chr28E4-28E6\|2-30 cM | chromosome location | fly chr28E4-28E6\|2-30 cM | 1 | 1 | 25010 | 571 | Bsg | | 2.283e-02 | -3.78 | fly chr51F7-51F11\|2-74 cM | chromosome location | fly chr51F7-51F11\|2-74 cM | 1 | 1 | 25010 | 571 | unc-5 | | 2.283e-02 | -3.78 | fly chr65A14-65B1\|3-16 | chromosome location | fly chr65A14-65B1\|3-16 | 1 | 1 | 25010 | 571 | CG13300 | | 2.283e-02 | -3.78 | fly chr2C10-2D1\|1-0.5 cM | chromosome location | fly chr2C10-2D1\|1-0.5 cM | 1 | 1 | 25010 | 571 | Unc-76 | | 2.283e-02 | -3.78 | fly chr61C1-61C3\|3-0.5 | chromosome location | fly chr61C1-61C3\|3-0.5 | 1 | 1 | 25010 | 571 | klar | | 2.283e-02 | -3.78 | fly chr53D8-53D9\|2-80 | chromosome location | fly chr53D8-53D9\|2-80 | 1 | 1 | 25010 | 571 | CG5522 | | 2.283e-02 | -3.78 | fly chr41B3-41B3\|2-55 | chromosome location | fly chr41B3-41B3\|2-55 | 1 | 1 | 25010 | 571 | CG17883 | | 2.283e-02 | -3.78 | fly chr11D1-11D2\|1-41 cM | chromosome location | fly chr11D1-11D2\|1-41 cM | 1 | 1 | 25010 | 571 | CG4645 | | 2.283e-02 | -3.78 | fly chr26B2-26B3\|2-18 | chromosome location | fly chr26B2-26B3\|2-18 | 1 | 1 | 25010 | 571 | H2.0 | | 2.283e-02 | -3.78 | fly chr48E1-48E2\|2-66 | chromosome location | fly chr48E1-48E2\|2-66 | 1 | 1 | 25010 | 571 | jeb | | 2.283e-02 | -3.78 | fly chr24F4-25A1\|2-12 | chromosome location | fly chr24F4-25A1\|2-12 | 1 | 1 | 25010 | 571 | dpy | | 2.283e-02 | -3.78 | fly chr49E6-49E6\|2-67 cM | chromosome location | fly chr49E6-49E6\|2-67 cM | 1 | 1 | 25010 | 571 | Psc | | 2.283e-02 | -3.78 | fly chr70E5-70E7\|3-42 cM | chromosome location | fly chr70E5-70E7\|3-42 cM | 1 | 1 | 25010 | 571 | dlp | | 2.283e-02 | -3.78 | fly chr75E3-75E4\|3-45 | chromosome location | fly chr75E3-75E4\|3-45 | 1 | 1 | 25010 | 571 | CG3961 | | 2.283e-02 | -3.78 | fly chr46D4-46D6\|2-61 cM | chromosome location | fly chr46D4-46D6\|2-61 cM | 1 | 1 | 25010 | 571 | CG12128 | | 2.283e-02 | -3.78 | fly chr9B6-9B7\|1-31 | chromosome location | fly chr9B6-9B7\|1-31 | 1 | 1 | 25010 | 571 | alpha-Man-Ia | | 2.283e-02 | -3.78 | fly chr86E17-86E18\|3-51 | chromosome location | fly chr86E17-86E18\|3-51 | 1 | 1 | 25010 | 571 | Lk6 | | 2.283e-02 | -3.78 | fly chr21B3-21B4\|2-0.5 | chromosome location | fly chr21B3-21B4\|2-0.5 | 1 | 1 | 25010 | 571 | spen | | 2.283e-02 | -3.78 | fly chr87F10-87F11\|3-54 | chromosome location | fly chr87F10-87F11\|3-54 | 1 | 1 | 25010 | 571 | flfl | | 2.283e-02 | -3.78 | fly chr60E10-60E11\|2-107 | chromosome location | fly chr60E10-60E11\|2-107 | 1 | 1 | 25010 | 571 | CG2765 | | 2.283e-02 | -3.78 | fly chr86E13-86E15\|3-51 | chromosome location | fly chr86E13-86E15\|3-51 | 1 | 1 | 25010 | 571 | CG42327 | | 2.283e-02 | -3.78 | fly chr60A11-60A11\|2-105 cM | chromosome location | fly chr60A11-60A11\|2-105 cM | 1 | 1 | 25010 | 571 | CG2812 | | 2.283e-02 | -3.78 | fly chr18C3-18C7\|1-62 | chromosome location | fly chr18C3-18C7\|1-62 | 1 | 1 | 25010 | 571 | kek5 | | 2.283e-02 | -3.78 | fly chr42A6-42A7\|2-55 | chromosome location | fly chr42A6-42A7\|2-55 | 1 | 1 | 25010 | 571 | Src42A | | 2.283e-02 | -3.78 | fly chr25C4-25C6\|2-15 cM | chromosome location | fly chr25C4-25C6\|2-15 cM | 1 | 1 | 25010 | 571 | Pgant5 | | 2.283e-02 | -3.78 | fly chr52E2-52E4\|2-77 | chromosome location | fly chr52E2-52E4\|2-77 | 1 | 1 | 25010 | 571 | Dg | | 2.283e-02 | -3.78 | fly chr88E12-88E12\|3-56 | chromosome location | fly chr88E12-88E12\|3-56 | 1 | 1 | 25010 | 571 | CG45218 | | 2.283e-02 | -3.78 | fly chr92A1-92A2\|3-66 | chromosome location | fly chr92A1-92A2\|3-66 | 1 | 1 | 25010 | 571 | Dl | | 2.283e-02 | -3.78 | fly chr96F5-96F5\|3-89 cM | chromosome location | fly chr96F5-96F5\|3-89 cM | 1 | 1 | 25010 | 571 | Tsp96F | | 2.283e-02 | -3.78 | fly chr89A13-89B1\|3-58 | chromosome location | fly chr89A13-89B1\|3-58 | 1 | 1 | 25010 | 571 | pnr | | 2.283e-02 | -3.78 | fly chr82F11-83A1\|3-47.5 | chromosome location | fly chr82F11-83A1\|3-47.5 | 1 | 1 | 25010 | 571 | Cerk | | 2.283e-02 | -3.78 | fly chr6E1-6E2\|1-18 cM | chromosome location | fly chr6E1-6E2\|1-18 cM | 1 | 1 | 25010 | 571 | CG43736 | | 2.283e-02 | -3.78 | fly chr60E10-60E11\|2-107 cM | chromosome location | fly chr60E10-60E11\|2-107 cM | 1 | 1 | 25010 | 571 | CG2765 | | 2.283e-02 | -3.78 | fly chr16A4-16A5\|1-57 cM | chromosome location | fly chr16A4-16A5\|1-57 cM | 1 | 1 | 25010 | 571 | B-H1 | | 2.283e-02 | -3.78 | fly chr81F5-81F6\|3-47.1 | chromosome location | fly chr81F5-81F6\|3-47.1 | 1 | 1 | 25010 | 571 | CG12581 | | 2.283e-02 | -3.78 | fly chr3F2-3F3\|1-5 cM | chromosome location | fly chr3F2-3F3\|1-5 cM | 1 | 1 | 25010 | 571 | ec | | 2.283e-02 | -3.78 | fly chr77E6-77E8\|3-47 | chromosome location | fly chr77E6-77E8\|3-47 | 1 | 1 | 25010 | 571 | Spc105R | | 2.283e-02 | -3.78 | fly chr47A12-47A13\|2-62 cM | chromosome location | fly chr47A12-47A13\|2-62 cM | 1 | 1 | 25010 | 571 | asRNA:CR45135 | | 2.283e-02 | -3.78 | fly chr53D9-53D10\|2-80 cM | chromosome location | fly chr53D9-53D10\|2-80 cM | 1 | 1 | 25010 | 571 | lncRNA:CR44465 | | 2.283e-02 | -3.78 | fly chr89E2-89E4\|3-59 | chromosome location | fly chr89E2-89E4\|3-59 | 1 | 1 | 25010 | 571 | lncRNA:iab8 | | 2.283e-02 | -3.78 | fly chr72D1-72D5\|3-44 | chromosome location | fly chr72D1-72D5\|3-44 | 1 | 1 | 25010 | 571 | Diap1 | | 2.283e-02 | -3.78 | fly chr84B6-84C3\|3-48 | chromosome location | fly chr84B6-84C3\|3-48 | 1 | 1 | 25010 | 571 | Alh | | 2.283e-02 | -3.78 | fly chr93C1-93C\|3-71 | chromosome location | fly chr93C1-93C\|3-71 | 1 | 1 | 25010 | 571 | SNF4Agamma | | 2.283e-02 | -3.78 | fly chr96E2-96E4\|3-88 | chromosome location | fly chr96E2-96E4\|3-88 | 1 | 1 | 25010 | 571 | msi | | 2.283e-02 | -3.78 | fly chr67E5-67E6\|3-33 cM | chromosome location | fly chr67E5-67E6\|3-33 cM | 1 | 1 | 25010 | 571 | CG32066 | | 2.283e-02 | -3.78 | fly chr94E7-94E8\|3-78 cM | chromosome location | fly chr94E7-94E8\|3-78 cM | 1 | 1 | 25010 | 571 | CG4467 | | 2.283e-02 | -3.78 | fly chr40A1-40A2\|2-54 cM | chromosome location | fly chr40A1-40A2\|2-54 cM | 1 | 1 | 25010 | 571 | step | | 2.283e-02 | -3.78 | fly chr6C7-6C9\|1-18 | chromosome location | fly chr6C7-6C9\|1-18 | 1 | 1 | 25010 | 571 | CG14441 | | 2.283e-02 | -3.78 | fly chr17F3-17F3\|1-61 cM | chromosome location | fly chr17F3-17F3\|1-61 cM | 1 | 1 | 25010 | 571 | lncRNA:CR44708 | | 2.283e-02 | -3.78 | fly chr64B1-64B2\|3-12 | chromosome location | fly chr64B1-64B2\|3-12 | 1 | 1 | 25010 | 571 | Cip4 | | 2.283e-02 | -3.78 | fly chr7D3-7D3\|1-22 cM | chromosome location | fly chr7D3-7D3\|1-22 cM | 1 | 1 | 25010 | 571 | CG2233 | | 2.283e-02 | -3.78 | fly chr89E2-89E4\|3-59 cM | chromosome location | fly chr89E2-89E4\|3-59 cM | 1 | 1 | 25010 | 571 | lncRNA:iab8 | | 2.283e-02 | -3.78 | fly chr25C1-25C3\|2-15 cM | chromosome location | fly chr25C1-25C3\|2-15 cM | 1 | 1 | 25010 | 571 | CG9121 | | 2.283e-02 | -3.78 | fly chr23B6-23B7\|2-7 | chromosome location | fly chr23B6-23B7\|2-7 | 1 | 1 | 25010 | 571 | CG2991 | | 2.283e-02 | -3.78 | fly chr86E13-86E15\|3-51 cM | chromosome location | fly chr86E13-86E15\|3-51 cM | 1 | 1 | 25010 | 571 | CG42327 | | 2.283e-02 | -3.78 | fly chr11D1-11D2\|1-41 | chromosome location | fly chr11D1-11D2\|1-41 | 1 | 1 | 25010 | 571 | CG4645 | | 2.283e-02 | -3.78 | fly chr93E5-93E5\|3-72 | chromosome location | fly chr93E5-93E5\|3-72 | 1 | 1 | 25010 | 571 | lncRNA:CR43653 | | 2.283e-02 | -3.78 | fly chr61C1-61C3\|3-0.5 cM | chromosome location | fly chr61C1-61C3\|3-0.5 cM | 1 | 1 | 25010 | 571 | klar | | 2.283e-02 | -3.78 | fly chr96A7-96A9\|3-85 | chromosome location | fly chr96A7-96A9\|3-85 | 1 | 1 | 25010 | 571 | REPTOR | | 2.283e-02 | -3.78 | fly chr75B2-75B2\|3-45 cM | chromosome location | fly chr75B2-75B2\|3-45 cM | 1 | 1 | 25010 | 571 | snoRNA:Me28S-A30 | | 2.283e-02 | -3.78 | fly chr48E1-48E2\|2-66 cM | chromosome location | fly chr48E1-48E2\|2-66 cM | 1 | 1 | 25010 | 571 | jeb | | 2.283e-02 | -3.78 | fly chr36F5-36F6\|2-53 cM | chromosome location | fly chr36F5-36F6\|2-53 cM | 1 | 1 | 25010 | 571 | kon | | 2.283e-02 | -3.78 | fly chr3C7-3C9\|1-2 | chromosome location | fly chr3C7-3C9\|1-2 | 1 | 1 | 25010 | 571 | N | | 2.283e-02 | -3.78 | fly chr69D3-69D3\|3-38 cM | chromosome location | fly chr69D3-69D3\|3-38 cM | 1 | 1 | 25010 | 571 | SmD1 | | 2.283e-02 | -3.78 | fly chr46C4-46C7\|2-61 | chromosome location | fly chr46C4-46C7\|2-61 | 1 | 1 | 25010 | 571 | Mef2 | | 2.283e-02 | -3.78 | fly chr87D8-87D9\|3-53 cM | chromosome location | fly chr87D8-87D9\|3-53 cM | 1 | 1 | 25010 | 571 | CtBP | | 2.283e-02 | -3.78 | fly chr50A1-50A3\|2-68 | chromosome location | fly chr50A1-50A3\|2-68 | 1 | 1 | 25010 | 571 | lncRNA:CR44206 | | 2.283e-02 | -3.78 | fly chr71C1-71C1\|3-43 | chromosome location | fly chr71C1-71C1\|3-43 | 1 | 1 | 25010 | 571 | Tollo | | 2.283e-02 | -3.78 | fly chr85D22-85D24\|3-48 | chromosome location | fly chr85D22-85D24\|3-48 | 1 | 1 | 25010 | 571 | mura | | 2.283e-02 | -3.78 | fly chr14B13-14B13\|1-53 cM | chromosome location | fly chr14B13-14B13\|1-53 cM | 1 | 1 | 25010 | 571 | CalpC | | 2.283e-02 | -3.78 | fly chr9B1-9B2\|1-31 cM | chromosome location | fly chr9B1-9B2\|1-31 cM | 1 | 1 | 25010 | 571 | RhoU | | 2.283e-02 | -3.78 | fly chr53D9-53D10\|2-80 | chromosome location | fly chr53D9-53D10\|2-80 | 1 | 1 | 25010 | 571 | lncRNA:CR44465 | | 2.283e-02 | -3.78 | fly chr68C10-68C10\|3-36 cM | chromosome location | fly chr68C10-68C10\|3-36 cM | 1 | 1 | 25010 | 571 | chrb | | 2.283e-02 | -3.78 | fly chr58E4-58E8\|2-100 cM | chromosome location | fly chr58E4-58E8\|2-100 cM | 1 | 1 | 25010 | 571 | px | | 2.283e-02 | -3.78 | fly chr91F3-91F4\|3-66 cM | chromosome location | fly chr91F3-91F4\|3-66 cM | 1 | 1 | 25010 | 571 | Ppcs | | 2.283e-02 | -3.78 | fly chr77E6-77E8\|3-47 cM | chromosome location | fly chr77E6-77E8\|3-47 cM | 1 | 1 | 25010 | 571 | Spc105R | | 2.283e-02 | -3.78 | fly chr72D1-72D5\|3-44 cM | chromosome location | fly chr72D1-72D5\|3-44 cM | 1 | 1 | 25010 | 571 | Diap1 | | 2.283e-02 | -3.78 | fly chr23A4-23A3\|2-7 | chromosome location | fly chr23A4-23A3\|2-7 | 1 | 1 | 25010 | 571 | Pgk | | 2.283e-02 | -3.78 | fly chr8C11-8C11\|1-26 | chromosome location | fly chr8C11-8C11\|1-26 | 1 | 1 | 25010 | 571 | lncRNA:CR45539 | | 2.283e-02 | -3.78 | fly chr52C9-52D2\|2-75 cM | chromosome location | fly chr52C9-52D2\|2-75 cM | 1 | 1 | 25010 | 571 | sli | | 2.283e-02 | -3.78 | fly chr41F-41F8\|2-55 | chromosome location | fly chr41F-41F8\|2-55 | 1 | 1 | 25010 | 571 | ap | | 2.283e-02 | -3.78 | fly chr89E4-89E5\|3-59 | chromosome location | fly chr89E4-89E5\|3-59 | 1 | 1 | 25010 | 571 | Abd-B | | 2.283e-02 | -3.78 | fly chr4C12-4C13\|1-8 cM | chromosome location | fly chr4C12-4C13\|1-8 cM | 1 | 1 | 25010 | 571 | Pp2C1 | | 2.283e-02 | -3.78 | fly chr43D7-43E1\|2-57 | chromosome location | fly chr43D7-43E1\|2-57 | 1 | 1 | 25010 | 571 | Vps13 | | 2.283e-02 | -3.78 | fly chr18D13-18E1\|1-64 cM | chromosome location | fly chr18D13-18E1\|1-64 cM | 1 | 1 | 25010 | 571 | dome | | 2.283e-02 | -3.78 | fly chr50B3-50B3\|2-68 cM | chromosome location | fly chr50B3-50B3\|2-68 cM | 1 | 1 | 25010 | 571 | lncRNA:CR44366 | | 2.283e-02 | -3.78 | fly chr8C11-8C13\|1-26 cM | chromosome location | fly chr8C11-8C13\|1-26 cM | 1 | 1 | 25010 | 571 | fend | | 2.283e-02 | -3.78 | fly chr25D1-25D2\|2-16 cM | chromosome location | fly chr25D1-25D2\|2-16 cM | 1 | 1 | 25010 | 571 | tkv | | 2.283e-02 | -3.78 | fly chr28B1-28B2\|2-27 cM | chromosome location | fly chr28B1-28B2\|2-27 cM | 1 | 1 | 25010 | 571 | RapGAP1 | | 2.283e-02 | -3.78 | fly chr9B1-9B2\|1-31 | chromosome location | fly chr9B1-9B2\|1-31 | 1 | 1 | 25010 | 571 | RhoU | | 2.283e-02 | -3.78 | fly chr53F4-53F6\|2-81 cM | chromosome location | fly chr53F4-53F6\|2-81 cM | 1 | 1 | 25010 | 571 | CG30460 | | 2.283e-02 | -3.78 | fly chr73B1-73B4\|3-44 | chromosome location | fly chr73B1-73B4\|3-44 | 1 | 1 | 25010 | 571 | Abl | | 2.283e-02 | -3.78 | fly chr80D5-80E3\|3-47 | chromosome location | fly chr80D5-80E3\|3-47 | 1 | 1 | 25010 | 571 | nAChRalpha4 | | 2.283e-02 | -3.78 | fly chr2D2-2D3\|1-0.6 cM | chromosome location | fly chr2D2-2D3\|1-0.6 cM | 1 | 1 | 25010 | 571 | ph-p | | 2.283e-02 | -3.78 | fly chr62F6-63A1\|3-4 | chromosome location | fly chr62F6-63A1\|3-4 | 1 | 1 | 25010 | 571 | Tet | | 2.283e-02 | -3.78 | fly chr12A6-12A7\|1-44 | chromosome location | fly chr12A6-12A7\|1-44 | 1 | 1 | 25010 | 571 | Tpst | | 2.283e-02 | -3.78 | fly chr3C7-3C9\|1-2 cM | chromosome location | fly chr3C7-3C9\|1-2 cM | 1 | 1 | 25010 | 571 | N | | 2.283e-02 | -3.78 | fly chr41F10-41F11\|2-55 cM | chromosome location | fly chr41F10-41F11\|2-55 cM | 1 | 1 | 25010 | 571 | ZnT41F | | 2.283e-02 | -3.78 | fly chr8F2-8F6\|1-29 | chromosome location | fly chr8F2-8F6\|1-29 | 1 | 1 | 25010 | 571 | Ptpmeg2 | | 2.283e-02 | -3.78 | fly chr83C5-83C6\|3-47.5 | chromosome location | fly chr83C5-83C6\|3-47.5 | 1 | 1 | 25010 | 571 | CG2017 | | 2.283e-02 | -3.78 | fly chr85E9-85E10\|3-49 cM | chromosome location | fly chr85E9-85E10\|3-49 cM | 1 | 1 | 25010 | 571 | FBXO11 | | 2.283e-02 | -3.78 | fly chr68C10-68C10\|3-36 | chromosome location | fly chr68C10-68C10\|3-36 | 1 | 1 | 25010 | 571 | chrb | | 2.283e-02 | -3.78 | fly chr61F2-61F3\|3-0 cM | chromosome location | fly chr61F2-61F3\|3-0 cM | 1 | 1 | 25010 | 571 | bab2 | | 2.283e-02 | -3.78 | fly chr56B5-56C1\|2-88 | chromosome location | fly chr56B5-56C1\|2-88 | 1 | 1 | 25010 | 571 | ena | | 2.283e-02 | -3.78 | fly chr11B9-11B10\|1-39 cM | chromosome location | fly chr11B9-11B10\|1-39 cM | 1 | 1 | 25010 | 571 | Tis11 | | 2.283e-02 | -3.78 | fly chr85C3-85C4\|3-48 cM | chromosome location | fly chr85C3-85C4\|3-48 cM | 1 | 1 | 25010 | 571 | Kdm2 | | 2.283e-02 | -3.78 | fly chr5C2-5C2\|1-13 | chromosome location | fly chr5C2-5C2\|1-13 | 1 | 1 | 25010 | 571 | CG42699 | | 2.283e-02 | -3.78 | fly chr18E2-18E3\|1-64 cM | chromosome location | fly chr18E2-18E3\|1-64 cM | 1 | 1 | 25010 | 571 | meso18E | | 2.283e-02 | -3.78 | fly chr90E2-90E2\|3-62 | chromosome location | fly chr90E2-90E2\|3-62 | 1 | 1 | 25010 | 571 | htl | | 2.283e-02 | -3.78 | fly chr47F13-47F14\|2-64 | chromosome location | fly chr47F13-47F14\|2-64 | 1 | 1 | 25010 | 571 | E(Pc) | | 2.283e-02 | -3.78 | fly chr96B4-96B4\|3-86 cM | chromosome location | fly chr96B4-96B4\|3-86 cM | 1 | 1 | 25010 | 571 | CG11069 | | 2.283e-02 | -3.78 | fly chr25C4-25C6\|2-15 | chromosome location | fly chr25C4-25C6\|2-15 | 1 | 1 | 25010 | 571 | Pgant5 | | 2.283e-02 | -3.78 | fly chr57E9-57F1\|2-95 cM | chromosome location | fly chr57E9-57F1\|2-95 cM | 1 | 1 | 25010 | 571 | Egfr | | 2.283e-02 | -3.78 | fly chr28A1-28A4\|2-27 | chromosome location | fly chr28A1-28A4\|2-27 | 1 | 1 | 25010 | 571 | Ziz | | 2.283e-02 | -3.78 | fly chr70E5-70E7\|3-42 | chromosome location | fly chr70E5-70E7\|3-42 | 1 | 1 | 25010 | 571 | dlp | | 2.283e-02 | -3.78 | fly chr11F1-11F3\|1-43 | chromosome location | fly chr11F1-11F3\|1-43 | 1 | 1 | 25010 | 571 | CG1673 | | 2.283e-02 | -3.78 | fly chr87F10-87F11\|3-54 cM | chromosome location | fly chr87F10-87F11\|3-54 cM | 1 | 1 | 25010 | 571 | flfl | | 2.283e-02 | -3.78 | fly chr53F4-53F6\|2-81 | chromosome location | fly chr53F4-53F6\|2-81 | 1 | 1 | 25010 | 571 | CG30460 | | 2.283e-02 | -3.78 | fly chr60A11-60A12\|2-105 cM | chromosome location | fly chr60A11-60A12\|2-105 cM | 1 | 1 | 25010 | 571 | SERCA | | 2.283e-02 | -3.78 | fly chr58E4-58E8\|2-100 | chromosome location | fly chr58E4-58E8\|2-100 | 1 | 1 | 25010 | 571 | px | | 2.283e-02 | -3.78 | fly chr46C4-46C7\|2-61 cM | chromosome location | fly chr46C4-46C7\|2-61 cM | 1 | 1 | 25010 | 571 | Mef2 | | 2.283e-02 | -3.78 | fly chr52E2-52E4\|2-77 cM | chromosome location | fly chr52E2-52E4\|2-77 cM | 1 | 1 | 25010 | 571 | Dg | | 2.283e-02 | -3.78 | fly chr96B4-96B4\|3-86 | chromosome location | fly chr96B4-96B4\|3-86 | 1 | 1 | 25010 | 571 | CG11069 | | 2.283e-02 | -3.78 | fly chr83D1-83D2\|3-47.5 cM | chromosome location | fly chr83D1-83D2\|3-47.5 cM | 1 | 1 | 25010 | 571 | Rm62 | | 2.283e-02 | -3.78 | fly chr70D4-70D5\|3-42 cM | chromosome location | fly chr70D4-70D5\|3-42 cM | 1 | 1 | 25010 | 571 | fz | | 2.283e-02 | -3.78 | fly chr14F1-14F2\|1-54 | chromosome location | fly chr14F1-14F2\|1-54 | 1 | 1 | 25010 | 571 | CG9784 | | 2.283e-02 | -3.78 | fly chr31A1-31A2\|2-39 | chromosome location | fly chr31A1-31A2\|2-39 | 1 | 1 | 25010 | 571 | Pen | | 2.283e-02 | -3.78 | fly chr26E1-26E2\|2-21 | chromosome location | fly chr26E1-26E2\|2-21 | 1 | 1 | 25010 | 571 | eya | | 2.283e-02 | -3.78 | fly chr66C13-66C13\|3-24 | chromosome location | fly chr66C13-66C13\|3-24 | 1 | 1 | 25010 | 571 | CG6983 | | 2.283e-02 | -3.78 | fly chr82F11-83A1\|3-47.5 cM | chromosome location | fly chr82F11-83A1\|3-47.5 cM | 1 | 1 | 25010 | 571 | Cerk | | 2.283e-02 | -3.78 | fly chr66A4-66A5\|3-21 cM | chromosome location | fly chr66A4-66A5\|3-21 cM | 1 | 1 | 25010 | 571 | lqf | | 2.283e-02 | -3.78 | fly chr86E11-86E13\|3-51 | chromosome location | fly chr86E11-86E13\|3-51 | 1 | 1 | 25010 | 571 | Jupiter | | 2.283e-02 | -3.78 | fly chr50B3-50B3\|2-68 | chromosome location | fly chr50B3-50B3\|2-68 | 1 | 1 | 25010 | 571 | lncRNA:CR44366 | | 2.283e-02 | -3.78 | fly chr94A2-94A4\|3-74 cM | chromosome location | fly chr94A2-94A4\|3-74 cM | 1 | 1 | 25010 | 571 | SKIP | | 2.283e-02 | -3.78 | fly chr23C1-23C3\|2-8 cM | chromosome location | fly chr23C1-23C3\|2-8 cM | 1 | 1 | 25010 | 571 | lilli | | 2.283e-02 | -3.78 | fly chr22B4-22B6\|2-3 cM | chromosome location | fly chr22B4-22B6\|2-3 cM | 1 | 1 | 25010 | 571 | Wdr62 | | 2.283e-02 | -3.78 | fly chr14B1-14B1\|1-53 cM | chromosome location | fly chr14B1-14B1\|1-53 cM | 1 | 1 | 25010 | 571 | disco-r | | 2.283e-02 | -3.78 | fly chr31A1-31A2\|2-39 cM | chromosome location | fly chr31A1-31A2\|2-39 cM | 1 | 1 | 25010 | 571 | Pen | | 2.283e-02 | -3.78 | fly chr61B3-61C1\|3-1 | chromosome location | fly chr61B3-61C1\|3-1 | 1 | 1 | 25010 | 571 | RhoGEF3 | | 2.283e-02 | -3.78 | fly chr57C2-57C3\|2-93 cM | chromosome location | fly chr57C2-57C3\|2-93 cM | 1 | 1 | 25010 | 571 | Magi | | 2.283e-02 | -3.78 | fly chr25C1-25C3\|2-15 | chromosome location | fly chr25C1-25C3\|2-15 | 1 | 1 | 25010 | 571 | CG9121 | | 2.283e-02 | -3.78 | fly chr95F12-95F12\|3-84 | chromosome location | fly chr95F12-95F12\|3-84 | 1 | 1 | 25010 | 571 | asRNA:CR46093 | | 2.283e-02 | -3.78 | fly chr85E9-85E10\|3-49 | chromosome location | fly chr85E9-85E10\|3-49 | 1 | 1 | 25010 | 571 | FBXO11 | | 2.283e-02 | -3.78 | fly chr50C6-50C9\|2-68 | chromosome location | fly chr50C6-50C9\|2-68 | 1 | 1 | 25010 | 571 | shot | | 2.283e-02 | -3.78 | fly chr33A1-33A2\|2-45 cM | chromosome location | fly chr33A1-33A2\|2-45 cM | 1 | 1 | 25010 | 571 | crol | | 2.283e-02 | -3.78 | fly chr93C1-93C\|3-71 cM | chromosome location | fly chr93C1-93C\|3-71 cM | 1 | 1 | 25010 | 571 | SNF4Agamma | | 2.283e-02 | -3.78 | fly chr64B1-64B2\|3-12 cM | chromosome location | fly chr64B1-64B2\|3-12 cM | 1 | 1 | 25010 | 571 | Cip4 | | 2.283e-02 | -3.78 | fly chr27E1-27E2\|2-25 cM | chromosome location | fly chr27E1-27E2\|2-25 cM | 1 | 1 | 25010 | 571 | Pvf3 | | 2.283e-02 | -3.78 | fly chr76A1-76A1\|3-45 | chromosome location | fly chr76A1-76A1\|3-45 | 1 | 1 | 25010 | 571 | lncRNA:CR45941 | | 2.283e-02 | -3.78 | fly chr68C12-68C13\|3-36 cM | chromosome location | fly chr68C12-68C13\|3-36 cM | 1 | 1 | 25010 | 571 | Mob2 | | 2.283e-02 | -3.78 | fly chr2B5-2B6\|1-0.2 | chromosome location | fly chr2B5-2B6\|1-0.2 | 1 | 1 | 25010 | 571 | hfw | | 2.283e-02 | -3.78 | fly chr84B6-84C3\|3-48 cM | chromosome location | fly chr84B6-84C3\|3-48 cM | 1 | 1 | 25010 | 571 | Alh | | 2.283e-02 | -3.78 | fly chr29B4-29C1\|2-31 | chromosome location | fly chr29B4-29C1\|2-31 | 1 | 1 | 25010 | 571 | CG13384 | | 2.283e-02 | -3.78 | fly chr40A1-40A2\|2-54 | chromosome location | fly chr40A1-40A2\|2-54 | 1 | 1 | 25010 | 571 | step | | 2.283e-02 | -3.78 | fly chr10C1-10C5\|1-35 | chromosome location | fly chr10C1-10C5\|1-35 | 1 | 1 | 25010 | 571 | Drak | | 2.283e-02 | -3.78 | fly chr66C12-66C12\|3-24 | chromosome location | fly chr66C12-66C12\|3-24 | 1 | 1 | 25010 | 571 | Cbl | | 2.283e-02 | -3.78 | fly chr91F3-91F4\|3-66 | chromosome location | fly chr91F3-91F4\|3-66 | 1 | 1 | 25010 | 571 | Ppcs | | 2.283e-02 | -3.78 | fly chr13D4-13E1\|1-51 | chromosome location | fly chr13D4-13E1\|1-51 | 1 | 1 | 25010 | 571 | CG9220 | | 2.283e-02 | -3.78 | fly chr5C2-5C2\|1-13 cM | chromosome location | fly chr5C2-5C2\|1-13 cM | 1 | 1 | 25010 | 571 | CG42699 | | 2.283e-02 | -3.78 | fly chr61B1-61B2\|3-1 cM | chromosome location | fly chr61B1-61B2\|3-1 cM | 1 | 1 | 25010 | 571 | p130CAS | | 2.283e-02 | -3.78 | fly chr75F9-76A3\|3-45 | chromosome location | fly chr75F9-76A3\|3-45 | 1 | 1 | 25010 | 571 | fz2 | | 2.283e-02 | -3.78 | fly chr25F3-25F4\|2-17 cM | chromosome location | fly chr25F3-25F4\|2-17 cM | 1 | 1 | 25010 | 571 | CG11030 | | 2.283e-02 | -3.78 | fly chr86C1-86C3\|3-50 | chromosome location | fly chr86C1-86C3\|3-50 | 1 | 1 | 25010 | 571 | hth | | 2.283e-02 | -3.78 | fly chr52D9-52D11\|2-76 cM | chromosome location | fly chr52D9-52D11\|2-76 cM | 1 | 1 | 25010 | 571 | ATPCL | | 2.283e-02 | -3.78 | fly chr64E12-64F2\|3-15 | chromosome location | fly chr64E12-64F2\|3-15 | 1 | 1 | 25010 | 571 | vn | | 2.283e-02 | -3.78 | fly chr87D8-87D9\|3-53 | chromosome location | fly chr87D8-87D9\|3-53 | 1 | 1 | 25010 | 571 | CtBP | | 2.283e-02 | -3.78 | fly chr102D4-102D5\|4-0 | chromosome location | fly chr102D4-102D5\|4-0 | 1 | 1 | 25010 | 571 | ey | | 2.283e-02 | -3.78 | fly chr66D9-66D10\|3-26 cM | chromosome location | fly chr66D9-66D10\|3-26 cM | 1 | 1 | 25010 | 571 | Pex7 | | 2.283e-02 | -3.78 | fly chr22B4-22B6\|2-3 | chromosome location | fly chr22B4-22B6\|2-3 | 1 | 1 | 25010 | 571 | Wdr62 | | 2.283e-02 | -3.78 | fly chr16B6-16B7\|1-57 | chromosome location | fly chr16B6-16B7\|1-57 | 1 | 1 | 25010 | 571 | CG12991 | | 2.283e-02 | -3.78 | fly chr7A7-7A7\|1-20 | chromosome location | fly chr7A7-7A7\|1-20 | 1 | 1 | 25010 | 571 | CG1958 | | 2.283e-02 | -3.78 | fly chr3F2-3F3\|1-5 | chromosome location | fly chr3F2-3F3\|1-5 | 1 | 1 | 25010 | 571 | ec | | 2.283e-02 | -3.78 | fly chr19D1-19D2\|1-64 cM | chromosome location | fly chr19D1-19D2\|1-64 cM | 1 | 1 | 25010 | 571 | RhoGAP19D | | 2.283e-02 | -3.78 | fly chr62F6-63A1\|3-4 cM | chromosome location | fly chr62F6-63A1\|3-4 cM | 1 | 1 | 25010 | 571 | Tet | | 2.283e-02 | -3.78 | fly chr38F6-39A1\|2-54 | chromosome location | fly chr38F6-39A1\|2-54 | 1 | 1 | 25010 | 571 | CG42238 | | 2.283e-02 | -3.78 | fly chr89A1-89A2\|3-58 cM | chromosome location | fly chr89A1-89A2\|3-58 cM | 1 | 1 | 25010 | 571 | pxb | | 2.283e-02 | -3.78 | fly chr89E4-89E5\|3-59 cM | chromosome location | fly chr89E4-89E5\|3-59 cM | 1 | 1 | 25010 | 571 | Abd-B | | 2.283e-02 | -3.78 | fly chr88E1-88E2\|3-56 | chromosome location | fly chr88E1-88E2\|3-56 | 1 | 1 | 25010 | 571 | CG42788 | | 2.283e-02 | -3.78 | fly chr26A5-26B2\|2-18 | chromosome location | fly chr26A5-26B2\|2-18 | 1 | 1 | 25010 | 571 | chic | | 2.283e-02 | -3.78 | fly chr6D7-6D8\|1-18 cM | chromosome location | fly chr6D7-6D8\|1-18 cM | 1 | 1 | 25010 | 571 | lncRNA:CR45531 | | 2.283e-02 | -3.78 | fly chr18B6-18B6\|1-62 | chromosome location | fly chr18B6-18B6\|1-62 | 1 | 1 | 25010 | 571 | rictor | | 2.283e-02 | -3.78 | fly chr93D5-93D6\|3-71 cM | chromosome location | fly chr93D5-93D6\|3-71 cM | 1 | 1 | 25010 | 571 | CG16791 | | 2.283e-02 | -3.78 | fly chr43D7-43E1\|2-57 cM | chromosome location | fly chr43D7-43E1\|2-57 cM | 1 | 1 | 25010 | 571 | Vps13 | | 2.283e-02 | -3.78 | fly chr96F5-96F5\|3-89 | chromosome location | fly chr96F5-96F5\|3-89 | 1 | 1 | 25010 | 571 | Tsp96F | | 2.283e-02 | -3.78 | fly chr82B1-82B2\|3-47.1 | chromosome location | fly chr82B1-82B2\|3-47.1 | 1 | 1 | 25010 | 571 | CG31522 | | 2.283e-02 | -3.78 | fly chr8C9-8C10\|1-26 | chromosome location | fly chr8C9-8C10\|1-26 | 1 | 1 | 25010 | 571 | AP-1gamma | | 2.283e-02 | -3.78 | fly chr54D4-54D5\|2-83 | chromosome location | fly chr54D4-54D5\|2-83 | 1 | 1 | 25010 | 571 | Sema1b | | 2.283e-02 | -3.78 | fly chr6D7-6D8\|1-18 | chromosome location | fly chr6D7-6D8\|1-18 | 1 | 1 | 25010 | 571 | lncRNA:CR45531 | | 2.283e-02 | -3.78 | fly chr51C2-51C3\|2-72 | chromosome location | fly chr51C2-51C3\|2-72 | 1 | 1 | 25010 | 571 | kn | | 2.283e-02 | -3.78 | fly chr5A13-5A14\|1-12 | chromosome location | fly chr5A13-5A14\|1-12 | 1 | 1 | 25010 | 571 | CG3149 | | 2.283e-02 | -3.78 | fly chr25D1-25D2\|2-16 | chromosome location | fly chr25D1-25D2\|2-16 | 1 | 1 | 25010 | 571 | tkv | | 2.283e-02 | -3.78 | fly chr99B10-99C1\|3-99 cM | chromosome location | fly chr99B10-99C1\|3-99 cM | 1 | 1 | 25010 | 571 | kay | | 2.283e-02 | -3.78 | fly chr85E14-85E14\|3-49 | chromosome location | fly chr85E14-85E14\|3-49 | 1 | 1 | 25010 | 571 | lncRNA:CR45029 | | 2.283e-02 | -3.78 | fly chr89D5-89D6\|3-59 | chromosome location | fly chr89D5-89D6\|3-59 | 1 | 1 | 25010 | 571 | Fas1 | | 2.283e-02 | -3.78 | fly chr95F12-95F12\|3-84 cM | chromosome location | fly chr95F12-95F12\|3-84 cM | 1 | 1 | 25010 | 571 | asRNA:CR46093 | | 2.283e-02 | -3.78 | fly chr89B14-89B15\|3-58 cM | chromosome location | fly chr89B14-89B15\|3-58 cM | 1 | 1 | 25010 | 571 | ss | | 2.283e-02 | -3.78 | fly chr47F13-47F14\|2-64 cM | chromosome location | fly chr47F13-47F14\|2-64 cM | 1 | 1 | 25010 | 571 | E(Pc) | | 2.283e-02 | -3.78 | fly chr19C1-19C4\|1-64 | chromosome location | fly chr19C1-19C4\|1-64 | 1 | 1 | 25010 | 571 | CG1504 | | 2.283e-02 | -3.78 | fly chr46D4-46D6\|2-61 | chromosome location | fly chr46D4-46D6\|2-61 | 1 | 1 | 25010 | 571 | CG12128 | | 2.283e-02 | -3.78 | fly chr30A2-30A6\|2-35 cM | chromosome location | fly chr30A2-30A6\|2-35 cM | 1 | 1 | 25010 | 571 | tai | | 2.283e-02 | -3.78 | fly chr93E5-93E5\|3-72 cM | chromosome location | fly chr93E5-93E5\|3-72 cM | 1 | 1 | 25010 | 571 | lncRNA:CR43653 | | 2.283e-02 | -3.78 | fly chr87D11-87D11\|3-53 | chromosome location | fly chr87D11-87D11\|3-53 | 1 | 1 | 25010 | 571 | sim | | 2.283e-02 | -3.78 | fly chr82E7-82E7\|3-47.1 cM | chromosome location | fly chr82E7-82E7\|3-47.1 cM | 1 | 1 | 25010 | 571 | corto | | 2.283e-02 | -3.78 | fly chr102C3-102C4\|4-0 cM | chromosome location | fly chr102C3-102C4\|4-0 cM | 1 | 1 | 25010 | 571 | zfh2 | | 2.283e-02 | -3.78 | fly chr74F4-75A1\|3-45 cM | chromosome location | fly chr74F4-75A1\|3-45 cM | 1 | 1 | 25010 | 571 | CG7460 | | 2.283e-02 | -3.78 | fly chr17F3-17F3\|1-61 | chromosome location | fly chr17F3-17F3\|1-61 | 1 | 1 | 25010 | 571 | lncRNA:CR44708 | | 2.283e-02 | -3.78 | fly chr13F1-13F5\|1-52 cM | chromosome location | fly chr13F1-13F5\|1-52 cM | 1 | 1 | 25010 | 571 | sd | | 2.283e-02 | -3.78 | fly chr94A14-94A15\|3-74 cM | chromosome location | fly chr94A14-94A15\|3-74 cM | 1 | 1 | 25010 | 571 | CG34376 | | 2.283e-02 | -3.78 | fly chr73B1-73B4\|3-44 cM | chromosome location | fly chr73B1-73B4\|3-44 cM | 1 | 1 | 25010 | 571 | Abl | | 2.283e-02 | -3.78 | fly chr68C12-68C13\|3-36 | chromosome location | fly chr68C12-68C13\|3-36 | 1 | 1 | 25010 | 571 | Mob2 | | 2.283e-02 | -3.78 | fly chr50F1-50F6\|2-70 | chromosome location | fly chr50F1-50F6\|2-70 | 1 | 1 | 25010 | 571 | Shrm | | 2.283e-02 | -3.78 | fly chr98A6-98A8\|3-94 cM | chromosome location | fly chr98A6-98A8\|3-94 cM | 1 | 1 | 25010 | 571 | wdb | | 2.283e-02 | -3.78 | fly chr5A13-5A14\|1-12 cM | chromosome location | fly chr5A13-5A14\|1-12 cM | 1 | 1 | 25010 | 571 | CG3149 | | 2.283e-02 | -3.78 | fly chr50C6-50C9\|2-68 cM | chromosome location | fly chr50C6-50C9\|2-68 cM | 1 | 1 | 25010 | 571 | shot | | 2.283e-02 | -3.78 | fly chr41F10-41F11\|2-55 | chromosome location | fly chr41F10-41F11\|2-55 | 1 | 1 | 25010 | 571 | ZnT41F | | 2.283e-02 | -3.78 | fly chr59A2-59A3\|2-101 | chromosome location | fly chr59A2-59A3\|2-101 | 1 | 1 | 25010 | 571 | robo1 | | 2.283e-02 | -3.78 | fly chr27E1-27E2\|2-25 | chromosome location | fly chr27E1-27E2\|2-25 | 1 | 1 | 25010 | 571 | Pvf3 | | 2.283e-02 | -3.78 | fly chr85C4-85D1\|3-48 | chromosome location | fly chr85C4-85D1\|3-48 | 1 | 1 | 25010 | 571 | pum | | 2.283e-02 | -3.78 | fly chr17C5-17C6\|1-60 cM | chromosome location | fly chr17C5-17C6\|1-60 cM | 1 | 1 | 25010 | 571 | wgn | | 2.283e-02 | -3.78 | fly chr14B13-14B13\|1-53 | chromosome location | fly chr14B13-14B13\|1-53 | 1 | 1 | 25010 | 571 | CalpC | | 2.283e-02 | -3.78 | fly chr70D4-70D5\|3-42 | chromosome location | fly chr70D4-70D5\|3-42 | 1 | 1 | 25010 | 571 | fz | | 2.283e-02 | -3.78 | fly chr7D3-7D5\|1-22 cM | chromosome location | fly chr7D3-7D5\|1-22 cM | 1 | 1 | 25010 | 571 | fs(1)h | | 2.283e-02 | -3.78 | fly chr7A7-7A7\|1-20 cM | chromosome location | fly chr7A7-7A7\|1-20 cM | 1 | 1 | 25010 | 571 | CG1958 | | 2.283e-02 | -3.78 | fly chr93E4-93E9\|3-72 | chromosome location | fly chr93E4-93E9\|3-72 | 1 | 1 | 25010 | 571 | InR | | 2.283e-02 | -3.78 | fly chr94A14-94A15\|3-74 | chromosome location | fly chr94A14-94A15\|3-74 | 1 | 1 | 25010 | 571 | CG34376 | | 2.283e-02 | -3.78 | fly chr8F2-8F6\|1-29 cM | chromosome location | fly chr8F2-8F6\|1-29 cM | 1 | 1 | 25010 | 571 | Ptpmeg2 | | 2.283e-02 | -3.78 | fly chr69B1-69B2\|3-38 cM | chromosome location | fly chr69B1-69B2\|3-38 cM | 1 | 1 | 25010 | 571 | Lmx1a | | 2.283e-02 | -3.78 | fly chr68B4-68C1\|3-35 | chromosome location | fly chr68B4-68C1\|3-35 | 1 | 1 | 25010 | 571 | scyl | | 2.283e-02 | -3.78 | fly chr6C12-6C12\|1-18 | chromosome location | fly chr6C12-6C12\|1-18 | 1 | 1 | 25010 | 571 | lncRNA:CR43496 | | 2.283e-02 | -3.78 | fly chr69D3-69D3\|3-38 | chromosome location | fly chr69D3-69D3\|3-38 | 1 | 1 | 25010 | 571 | SmD1 | | 2.283e-02 | -3.78 | fly chr66D9-66D10\|3-26 | chromosome location | fly chr66D9-66D10\|3-26 | 1 | 1 | 25010 | 571 | Pex7 | | 2.283e-02 | -3.78 | fly chr25F3-25F4\|2-17 | chromosome location | fly chr25F3-25F4\|2-17 | 1 | 1 | 25010 | 571 | CG11030 | | 2.283e-02 | -3.78 | fly chr18B9-18C1\|1-62 cM | chromosome location | fly chr18B9-18C1\|1-62 cM | 1 | 1 | 25010 | 571 | CG8034 | | 2.283e-02 | -3.78 | fly chr57B15-57B16\|2-92 | chromosome location | fly chr57B15-57B16\|2-92 | 1 | 1 | 25010 | 571 | shg | | 2.283e-02 | -3.78 | fly chr56C1-56C4\|2-88 | chromosome location | fly chr56C1-56C4\|2-88 | 1 | 1 | 25010 | 571 | hppy | | 2.283e-02 | -3.78 | fly chr93D4-93D5\|3-71 cM | chromosome location | fly chr93D4-93D5\|3-71 cM | 1 | 1 | 25010 | 571 | lncRNA:Hsromega | | 2.283e-02 | -3.78 | fly chr19C1-19C4\|1-64 cM | chromosome location | fly chr19C1-19C4\|1-64 cM | 1 | 1 | 25010 | 571 | CG1504 | | 2.283e-02 | -3.78 | fly chr87D11-87D11\|3-53 cM | chromosome location | fly chr87D11-87D11\|3-53 cM | 1 | 1 | 25010 | 571 | sim | | 2.283e-02 | -3.78 | fly chr26B2-26B3\|2-18 cM | chromosome location | fly chr26B2-26B3\|2-18 cM | 1 | 1 | 25010 | 571 | H2.0 | | 2.283e-02 | -3.78 | fly chr49F10-49F11\|2-68 cM | chromosome location | fly chr49F10-49F11\|2-68 cM | 1 | 1 | 25010 | 571 | Dp | | 2.283e-02 | -3.78 | fly chr21B3-21B4\|2-0.5 cM | chromosome location | fly chr21B3-21B4\|2-0.5 cM | 1 | 1 | 25010 | 571 | spen | | 2.283e-02 | -3.78 | fly chr41C3-41C3\|2-55 cM | chromosome location | fly chr41C3-41C3\|2-55 cM | 1 | 1 | 25010 | 571 | CG14464 | | 2.283e-02 | -3.78 | fly chr17C5-17C6\|1-60 | chromosome location | fly chr17C5-17C6\|1-60 | 1 | 1 | 25010 | 571 | wgn | | 2.283e-02 | -3.78 | fly chr2B5-2B6\|1-0.2 cM | chromosome location | fly chr2B5-2B6\|1-0.2 cM | 1 | 1 | 25010 | 571 | hfw | | 2.283e-02 | -3.78 | fly chr86E11-86E13\|3-51 cM | chromosome location | fly chr86E11-86E13\|3-51 cM | 1 | 1 | 25010 | 571 | Jupiter | | 2.283e-02 | -3.78 | fly chr82E7-82E7\|3-47.1 | chromosome location | fly chr82E7-82E7\|3-47.1 | 1 | 1 | 25010 | 571 | corto | | 2.283e-02 | -3.78 | fly chr26D8-26D8\|2-20 | chromosome location | fly chr26D8-26D8\|2-20 | 1 | 1 | 25010 | 571 | lncRNA:CR45463 | | 2.283e-02 | -3.78 | fly chr51F7-51F11\|2-74 | chromosome location | fly chr51F7-51F11\|2-74 | 1 | 1 | 25010 | 571 | unc-5 | | 2.283e-02 | -3.78 | fly chr90A2-90A3\|3-60 | chromosome location | fly chr90A2-90A3\|3-60 | 1 | 1 | 25010 | 571 | beat-IIa | | 2.283e-02 | -3.78 | fly chr50A1-50A3\|2-68 cM | chromosome location | fly chr50A1-50A3\|2-68 cM | 1 | 1 | 25010 | 571 | lncRNA:CR44206 | | 2.311e-02 | -3.77 | regulation of antimicrobial peptide production | biological process | GO:0002784 | 62 | 6 | 12345 | 442 | cact,mop,nub,Pitslre,imd,pnr | | 2.319e-02 | -3.76 | GO\_AMINOGLYCAN\_METABOLIC\_PROCESS | MSigDB lists | GO\_AMINOGLYCAN\_METABOLIC\_PROCESS | 44 | 5 | 4161 | 156 | CG9220,frc,Ids,GlcAT-I,dlp | | 2.319e-02 | -3.76 | HORIUCHI\_WTAP\_TARGETS\_UP | MSigDB lists | HORIUCHI\_WTAP\_TARGETS\_UP | 44 | 5 | 4161 | 156 | CG3961,Kdm2,lt,Tollo,trbl | | 2.319e-02 | -3.76 | GSE41176\_UNSTIM\_VS\_ANTI\_IGM\_STIM\_BCELL\_1H\_DN | MSigDB lists | GSE41176\_UNSTIM\_VS\_ANTI\_IGM\_STIM\_BCELL\_1H\_DN | 44 | 5 | 4161 | 156 | hppy,bru2,Ids,C1GalTA,emp | | 2.319e-02 | -3.76 | MEF2\_03 | MSigDB lists | MEF2\_03 | 44 | 5 | 4161 | 156 | Mef2,Nrk,rg,FBXO11,dlp | | 2.319e-02 | -3.76 | SMID\_BREAST\_CANCER\_NORMAL\_LIKE\_UP | MSigDB lists | SMID\_BREAST\_CANCER\_NORMAL\_LIKE\_UP | 44 | 5 | 4161 | 156 | bru2,shot,Dyb,CG6891,Mef2 | | 2.319e-02 | -3.76 | GSE17721\_CTRL\_VS\_LPS\_1H\_BMDC\_DN | MSigDB lists | GSE17721\_CTRL\_VS\_LPS\_1H\_BMDC\_DN | 44 | 5 | 4161 | 156 | Tpst,CycG,Kr-h2,Actbeta,Pdk1 | | 2.319e-02 | -3.76 | BOUDOUKHA\_BOUND\_BY\_IGF2BP2 | MSigDB lists | BOUDOUKHA\_BOUND\_BY\_IGF2BP2 | 44 | 5 | 4161 | 156 | rdx,SERCA,CycG,eff,Tsp96F | | 2.319e-02 | -3.76 | GSE19198\_CTRL\_VS\_IL21\_TREATED\_TCELL\_24H\_DN | MSigDB lists | GSE19198\_CTRL\_VS\_IL21\_TREATED\_TCELL\_24H\_DN | 44 | 5 | 4161 | 156 | Pgant6,Pdk1,unc-119,Pgant5,lt | | 2.319e-02 | -3.76 | ACCATTT\_MIR522 | MSigDB lists | ACCATTT\_MIR522 | 44 | 5 | 4161 | 156 | bru2,CenB1A,alpha-Man-Ia,subdued,14-3-3zeta | | 2.319e-02 | -3.76 | GSE360\_T\_GONDII\_VS\_B\_MALAYI\_HIGH\_DOSE\_DC\_UP | MSigDB lists | GSE360\_T\_GONDII\_VS\_B\_MALAYI\_HIGH\_DOSE\_DC\_UP | 44 | 5 | 4161 | 156 | cact,sns,betaTub60D,ReepA,Kdm2 | | 2.333e-02 | -3.76 | IG | smart domains | SM00409 | 111 | 12 | 5086 | 290 | CG31431,ed,robo1,kek5,Bsg,sns,Pvr,otk,unc-5,boi,htl,Ppn | | 2.334e-02 | -3.76 | phosphatase activity | molecular function | GO:0016791 | 184 | 12 | 12176 | 414 | Pp2C1,CG10417,Ptp4E,CG42327,eya,CG6656,Ocrl,mop,Pez,IA-2,Ptpmeg2,CG9784 | | 2.342e-02 | -3.75 | AA\_transpt\_TM | interpro domains | IPR013057 | 16 | 3 | 11383 | 450 | path,CG13384,CG4991 | | 2.342e-02 | -3.75 | Cyclin\_N | interpro domains | IPR006671 | 16 | 3 | 11383 | 450 | CG6191,CycB3,CycG | | 2.342e-02 | -3.75 | PAS | interpro domains | IPR000014 | 16 | 3 | 11383 | 450 | sim,ss,tai | | 2.342e-02 | -3.75 | PAS-like\_dom\_sf | interpro domains | IPR035965 | 16 | 3 | 11383 | 450 | sim,tai,ss | | 2.342e-02 | -3.75 | CG14715 (uncharacterized protein) | protein interactions | 41403 | 61 | 7 | 9204 | 432 | Jupiter,Bacc,Nlp,meso18E,Pgk,chic,Fkbp14 | | 2.359e-02 | -3.75 | GO\_REGULATION\_OF\_NEURON\_DIFFERENTIATION | MSigDB lists | GO\_REGULATION\_OF\_NEURON\_DIFFERENTIATION | 112 | 9 | 4161 | 156 | nAChRalpha4,Arf79F,N,otk,Dl,robo1,Raf,Mef2,sli | | 2.362e-02 | -3.75 | negative regulation of Wnt signaling pathway | biological process | GO:0030178 | 46 | 5 | 12345 | 442 | otk,nmo,aop,CtBP,crol | | 2.362e-02 | -3.75 | positive regulation of small GTPase mediated signal transduction | biological process | GO:0051057 | 46 | 5 | 12345 | 442 | RhoGEF4,14-3-3zeta,siz,14-3-3epsilon,step | | 2.362e-02 | -3.75 | positive regulation of Ras protein signal transduction | biological process | GO:0046579 | 46 | 5 | 12345 | 442 | 14-3-3zeta,RhoGEF4,step,14-3-3epsilon,siz | | 2.362e-02 | -3.75 | neuron maturation | biological process | GO:0042551 | 46 | 5 | 12345 | 442 | Sox14,mop,chic,Dronc,eff | | 2.366e-02 | -3.74 | Signaling by FGFR3 | REACTOME pathways | R-DME-5654741 | 25 | 4 | 3923 | 176 | Raf,Cbl,htl,Lk6 | | 2.366e-02 | -3.74 | Signaling by FGFR4 | REACTOME pathways | R-DME-5654743 | 25 | 4 | 3923 | 176 | htl,Lk6,Cbl,Raf | | 2.374e-02 | -3.74 | larval chitin-based cuticle development | biological process | GO:0008363 | 31 | 4 | 12345 | 442 | jar,fz2,fz,arm | | 2.374e-02 | -3.74 | acylglycerol homeostasis | biological process | GO:0055090 | 31 | 4 | 12345 | 442 | InR,Ppcs,cue,hppy | | 2.374e-02 | -3.74 | stem cell fate determination | biological process | GO:0048867 | 31 | 4 | 12345 | 442 | abd-A,nub,spen,N | | 2.377e-02 | -3.74 | MYOD\_Q6\_01 | MSigDB lists | MYOD\_Q6\_01 | 60 | 6 | 4161 | 156 | fz2,rdx,shot,l(2)37Cg,Vps13,Cals | | 2.377e-02 | -3.74 | SOX5\_01 | MSigDB lists | SOX5\_01 | 60 | 6 | 4161 | 156 | arm,dlp,Actbeta,Eph,FBXO11,betaTub60D | | 2.377e-02 | -3.74 | WAKABAYASHI\_ADIPOGENESIS\_PPARG\_RXRA\_BOUND\_WITH\_H4K20ME1\_MARK | MSigDB lists | WAKABAYASHI\_ADIPOGENESIS\_PPARG\_RXRA\_BOUND\_WITH\_H4K20ME1\_MARK | 60 | 6 | 4161 | 156 | wbl,step,Cerk,emp,Pex5,Kr-h2 | | 2.377e-02 | -3.74 | GO\_ANTIGEN\_RECEPTOR\_MEDIATED\_SIGNALING\_PATHWAY | MSigDB lists | GO\_ANTIGEN\_RECEPTOR\_MEDIATED\_SIGNALING\_PATHWAY | 60 | 6 | 4161 | 156 | PI31,eff,Pdk1,Mef2,cact,dock | | 2.377e-02 | -3.74 | GSE17721\_0.5H\_VS\_24H\_LPS\_BMDC\_DN | MSigDB lists | GSE17721\_0.5H\_VS\_24H\_LPS\_BMDC\_DN | 60 | 6 | 4161 | 156 | Tsp96F,CG2017,Kr-h2,Nmda1,Pat1,AP-1gamma | | 2.377e-02 | -3.74 | THEILGAARD\_NEUTROPHIL\_AT\_SKIN\_WOUND\_DN | MSigDB lists | THEILGAARD\_NEUTROPHIL\_AT\_SKIN\_WOUND\_DN | 60 | 6 | 4161 | 156 | Psa,Unc-76,bru2,step,ATPCL,Ids | | 2.384e-02 | -3.74 | neuron projection regeneration | biological process | GO:0031102 | 7 | 2 | 12345 | 442 | kay,wnd | | 2.384e-02 | -3.74 | heart contraction | biological process | GO:0060047 | 7 | 2 | 12345 | 442 | SERCA,pnr | | 2.384e-02 | -3.74 | negative regulation of protein tyrosine kinase activity | biological process | GO:0061099 | 7 | 2 | 12345 | 442 | Socs36E,Cbl | | 2.384e-02 | -3.74 | symmetric cell division | biological process | GO:0098725 | 7 | 2 | 12345 | 442 | lin-28,shg | | 2.384e-02 | -3.74 | lamellocyte differentiation | biological process | GO:0035171 | 7 | 2 | 12345 | 442 | N,Raf | | 2.384e-02 | -3.74 | negative regulation of axon extension | biological process | GO:0030517 | 7 | 2 | 12345 | 442 | Sema1b,Sema1a | | 2.384e-02 | -3.74 | specification of segmental identity, trunk | biological process | GO:0035292 | 7 | 2 | 12345 | 442 | Abd-B,tsh | | 2.384e-02 | -3.74 | endoderm formation | biological process | GO:0001706 | 7 | 2 | 12345 | 442 | htl,tkv | | 2.384e-02 | -3.74 | hemocyte development | biological process | GO:0007516 | 7 | 2 | 12345 | 442 | RhoL,Dronc | | 2.384e-02 | -3.74 | hippo signaling | biological process | GO:0035329 | 7 | 2 | 12345 | 442 | CG42788,sd | | 2.384e-02 | -3.74 | regulation of ARF protein signal transduction | biological process | GO:0032012 | 7 | 2 | 12345 | 442 | step,siz | | 2.384e-02 | -3.74 | negative regulation of compound eye retinal cell programmed cell death | biological process | GO:0046673 | 7 | 2 | 12345 | 442 | Diap1,Egfr | | 2.384e-02 | -3.74 | histone ubiquitination | biological process | GO:0016574 | 7 | 2 | 12345 | 442 | Kdm2,Su(z)2 | | 2.384e-02 | -3.74 | negative regulation of retinal cell programmed cell death | biological process | GO:0046671 | 7 | 2 | 12345 | 442 | Egfr,Diap1 | | 2.384e-02 | -3.74 | positive regulation of muscle organ development | biological process | GO:0048636 | 7 | 2 | 12345 | 442 | abd-A,tkv | | 2.388e-02 | -3.73 | KRAS.AMP.LUNG\_UP.V1\_UP | MSigDB lists | KRAS.AMP.LUNG\_UP.V1\_UP | 17 | 3 | 4161 | 156 | CG11069,sns,Cals | | 2.388e-02 | -3.73 | CCAATNNSNNNGCG\_UNKNOWN | MSigDB lists | CCAATNNSNNNGCG\_UNKNOWN | 17 | 3 | 4161 | 156 | Mef2,ap,dock | | 2.388e-02 | -3.73 | MORF\_IL9 | MSigDB lists | MORF\_IL9 | 17 | 3 | 4161 | 156 | lt,nAChRalpha4,Src42A | | 2.388e-02 | -3.73 | GO\_PALATE\_DEVELOPMENT | MSigDB lists | GO\_PALATE\_DEVELOPMENT | 17 | 3 | 4161 | 156 | fz,Actbeta,Mef2 | | 2.388e-02 | -3.73 | WYAAANNRNNNGCG\_UNKNOWN | MSigDB lists | WYAAANNRNNNGCG\_UNKNOWN | 17 | 3 | 4161 | 156 | SMC3,wdb,Kdm2 | | 2.388e-02 | -3.73 | SIG\_PIP3\_SIGNALING\_IN\_CARDIAC\_MYOCTES | MSigDB lists | SIG\_PIP3\_SIGNALING\_IN\_CARDIAC\_MYOCTES | 17 | 3 | 4161 | 156 | 14-3-3epsilon,Pdk1,14-3-3zeta | | 2.388e-02 | -3.73 | chr6q21 | MSigDB lists | chr6q21 | 17 | 3 | 4161 | 156 | lin-28,Src42A,Pex7 | | 2.388e-02 | -3.73 | MODULE\_46 | MSigDB lists | MODULE\_46 | 17 | 3 | 4161 | 156 | Tsp96F,cact,CG1673 | | 2.388e-02 | -3.73 | GO\_TOLL\_LIKE\_RECEPTOR\_SIGNALING\_PATHWAY | MSigDB lists | GO\_TOLL\_LIKE\_RECEPTOR\_SIGNALING\_PATHWAY | 17 | 3 | 4161 | 156 | eff,Tollo,cact | | 2.388e-02 | -3.73 | MODULE\_456 | MSigDB lists | MODULE\_456 | 17 | 3 | 4161 | 156 | cact,otk,trbl | | 2.388e-02 | -3.73 | GO\_REGULATION\_OF\_PHOSPHOLIPID\_METABOLIC\_PROCESS | MSigDB lists | GO\_REGULATION\_OF\_PHOSPHOLIPID\_METABOLIC\_PROCESS | 17 | 3 | 4161 | 156 | htl,Arf79F,Tsp96F | | 2.388e-02 | -3.73 | POMEROY\_MEDULLOBLASTOMA\_DESMOPLASIC\_VS\_CLASSIC\_UP | MSigDB lists | POMEROY\_MEDULLOBLASTOMA\_DESMOPLASIC\_VS\_CLASSIC\_UP | 17 | 3 | 4161 | 156 | ap,Dyrk2,ATPCL | | 2.391e-02 | -3.73 | Sema1a (Semaphorin 1a) | protein interactions | 34192 | 150 | 13 | 9204 | 432 | apolpp,Dg,CG3107,ft,CG7320,otk,InR,mld,14-3-3zeta,chrb,LanA,CG11122,Hel89B | | 2.400e-02 | -3.73 | GTPase activator activity | molecular function | GO:0005096 | 84 | 7 | 12176 | 414 | TBC1D5,drongo,Ocrl,RhoGAP19D,CG17883,CenB1A,RapGAP1 | | 2.406e-02 | -3.73 | MYOSINHEAVY | prints domains | PR00193 | 14 | 3 | 2185 | 101 | Myo10A,jar,d | | 2.407e-02 | -3.73 | foxo (forkhead box, sub-group O) | protein interactions | 41709 | 182 | 15 | 9204 | 432 | Rgl,InR,wdb,rictor,arm,larp,14-3-3epsilon,vig,Rm62,cindr,trbl,path,ey,cora,14-3-3zeta | | 2.411e-02 | -3.73 | Cyclin\_N | pfam domains | PF00134 | 16 | 3 | 10480 | 419 | CycG,CycB3,CG6191 | | 2.411e-02 | -3.73 | Aa\_trans | pfam domains | PF01490 | 16 | 3 | 10480 | 419 | CG4991,path,CG13384 | | 2.412e-02 | -3.72 | AP-2mu (Adaptor Protein complex 2, mu subunit) | protein interactions | 42642 | 24 | 4 | 9204 | 432 | ed,ttk,Pdk1,lt | | 2.412e-02 | -3.72 | TwdlB (TweedleB) | protein interactions | 43202 | 24 | 4 | 9204 | 432 | msi,sns,CG6891,imd | | 2.412e-02 | -3.72 | Cda5 (Chitin deacetylase-like 5) | protein interactions | 33158 | 24 | 4 | 9204 | 432 | sns,eyg,Sxl,Abd-B | | 2.412e-02 | -3.72 | elfless (elfless) | protein interactions | 35076 | 24 | 4 | 9204 | 432 | eff,Dronc,His3.3B,CG34401 | | 2.412e-02 | -3.72 | betaTub60D (beta-Tubulin at 60D) | protein interactions | 37888 | 24 | 4 | 9204 | 432 | Raf,rictor,Pdk1,shot | | 2.414e-02 | -3.72 | negative regulation of protein modification process | biological process | GO:0031400 | 98 | 8 | 12345 | 442 | trbl,Kdm2,CycG,Socs36E,N,wdb,Cbl,CG8312 | | 2.424e-02 | -3.72 | negative regulation of immune system process | biological process | GO:0002683 | 80 | 7 | 12345 | 442 | cact,pxb,Pvr,LRR,N,C1GalTA,nub | | 2.428e-02 | -3.72 | GO\_REGULATION\_OF\_ORGAN\_MORPHOGENESIS | MSigDB lists | GO\_REGULATION\_OF\_ORGAN\_MORPHOGENESIS | 77 | 7 | 4161 | 156 | dlp,arm,fz2,PI31,N,fz,otk | | 2.433e-02 | -3.72 | GATA\_ZN\_FINGER\_2 | prosite domains | PS50114 | 5 | 2 | 6243 | 325 | pnr,grn | | 2.433e-02 | -3.72 | POU\_1 | prosite domains | PS00035 | 5 | 2 | 6243 | 325 | pdm3,nub | | 2.433e-02 | -3.72 | SEC7 | prosite domains | PS50190 | 5 | 2 | 6243 | 325 | siz,step | | 2.433e-02 | -3.72 | GATA\_ZN\_FINGER\_1 | prosite domains | PS00344 | 5 | 2 | 6243 | 325 | pnr,grn | | 2.433e-02 | -3.72 | FERM\_2 | prosite domains | PS00661 | 5 | 2 | 6243 | 325 | yrt,cora | | 2.433e-02 | -3.72 | POU\_2 | prosite domains | PS00465 | 5 | 2 | 6243 | 325 | pdm3,nub | | 2.433e-02 | -3.72 | POU\_3 | prosite domains | PS51179 | 5 | 2 | 6243 | 325 | pdm3,nub | | 2.435e-02 | -3.72 | transcriptional repressor complex | cellular component | GO:0017053 | 32 | 4 | 11940 | 417 | Elba2,MEP-1,H,ttk | | 2.437e-02 | -3.71 | PID\_MTOR\_4PATHWAY | MSigDB lists | PID\_MTOR\_4PATHWAY | 30 | 4 | 4161 | 156 | 14-3-3zeta,Pdk1,Raf,14-3-3epsilon | | 2.437e-02 | -3.71 | GO\_SKELETAL\_MUSCLE\_ORGAN\_DEVELOPMENT | MSigDB lists | GO\_SKELETAL\_MUSCLE\_ORGAN\_DEVELOPMENT | 30 | 4 | 4161 | 156 | Mef2,N,Dl,sns | | 2.437e-02 | -3.71 | GSE7768\_OVA\_ALONE\_VS\_OVA\_WITH\_LPS\_IMMUNIZED\_MOUSE\_WHOLE\_SPLEEN\_6H\_DN | MSigDB lists | GSE7768\_OVA\_ALONE\_VS\_OVA\_WITH\_LPS\_IMMUNIZED\_MOUSE\_WHOLE\_SPLEEN\_6H\_DN | 30 | 4 | 4161 | 156 | Rgl,CG2006,cact,dock | | 2.437e-02 | -3.71 | TOOKER\_GEMCITABINE\_RESISTANCE\_UP | MSigDB lists | TOOKER\_GEMCITABINE\_RESISTANCE\_UP | 30 | 4 | 4161 | 156 | CG3961,CG31522,Mef2,Tsp96F | | 2.437e-02 | -3.71 | GO\_NOTCH\_SIGNALING\_PATHWAY | MSigDB lists | GO\_NOTCH\_SIGNALING\_PATHWAY | 30 | 4 | 4161 | 156 | N,Cdk4,Dl,lqf | | 2.437e-02 | -3.71 | GSE34156\_UNTREATED\_VS\_6H\_NOD2\_LIGAND\_TREATED\_MONOCYTE\_DN | MSigDB lists | GSE34156\_UNTREATED\_VS\_6H\_NOD2\_LIGAND\_TREATED\_MONOCYTE\_DN | 30 | 4 | 4161 | 156 | Mef2,CG2017,Tollo,Tpst | | 2.437e-02 | -3.71 | GSE2770\_UNTREATED\_VS\_TGFB\_AND\_IL4\_TREATED\_ACT\_CD4\_TCELL\_4H\_DN | MSigDB lists | GSE2770\_UNTREATED\_VS\_TGFB\_AND\_IL4\_TREATED\_ACT\_CD4\_TCELL\_4H\_DN | 30 | 4 | 4161 | 156 | CtBP,Raf,shot,ATPCL | | 2.437e-02 | -3.71 | GO\_CARDIAC\_CHAMBER\_DEVELOPMENT | MSigDB lists | GO\_CARDIAC\_CHAMBER\_DEVELOPMENT | 30 | 4 | 4161 | 156 | otk,fz,Mef2,N | | 2.460e-02 | -3.71 | GO\_REGULATION\_OF\_ANATOMICAL\_STRUCTURE\_MORPHOGENESIS | MSigDB lists | GO\_REGULATION\_OF\_ANATOMICAL\_STRUCTURE\_MORPHOGENESIS | 210 | 14 | 4161 | 156 | CG17883,sli,PI31,Mef2,Raf,robo1,otk,Dl,arm,dlp,fz2,N,fz,nAChRalpha4 | | 2.480e-02 | -3.70 | regulation of production of molecular mediator of immune response | biological process | GO:0002700 | 63 | 6 | 12345 | 442 | nub,pnr,Pitslre,imd,mop,cact | | 2.480e-02 | -3.70 | locomotor rhythm | biological process | GO:0045475 | 63 | 6 | 12345 | 442 | Mef2,Mid1,kay,tara,dome,ITP | | 2.485e-02 | -3.69 | nuclear part | cellular component | GO:0044428 | 1416 | 63 | 11940 | 417 | CG4896,Su(z)2,lilli,hyx,14-3-3zeta,abd-A,CG3436,glec,Kdm4B,SmD1,Dp,H,yrt,Kdm2,IntS3,Lk6,CtBP,MEP-1,snRNA:U5:38ABb,CG4935,stx,Slbp,N,px,rgr,CG11030,14-3-3epsilon,HmgZ,His2Av,dock,hth,mam,Bacc,CG33082,Prdm13,emc,l(2)37Cg,E(Pc),Sox14,su(sable),koi,jumu,arm,msi,CG31211,Rm62,ph-p,yuri,CG8312,corto,aop,Hr39,lqf,Nlp,Drep4,tna,klar,Elba2,Psc,lncRNA:Hsromega,pum,CG6340,Kr-h2 | | 2.493e-02 | -3.69 | negative regulation of neuron projection development | biological process | GO:0010977 | 18 | 3 | 12345 | 442 | Sema1a,Abl,Sema1b | | 2.497e-02 | -3.69 | fly chr37A1-37A1\|2-54 | chromosome location | fly chr37A1-37A1\|2-54 | 11 | 2 | 25010 | 571 | Irk3,CG10348 | | 2.497e-02 | -3.69 | fly chr37A1-37A1\|2-54 cM | chromosome location | fly chr37A1-37A1\|2-54 cM | 11 | 2 | 25010 | 571 | Irk3,CG10348 | | 2.523e-02 | -3.68 | catalytic activity, acting on a glycoprotein | molecular function | GO:0140103 | 19 | 3 | 12176 | 414 | CG3119,C1GalTA,CG34057 | | 2.523e-02 | -3.68 | cytokine receptor binding | molecular function | GO:0005126 | 19 | 3 | 12176 | 414 | Socs36E,Pvf3,Actbeta | | 2.524e-02 | -3.68 | GO\_REGULATION\_OF\_CELLULAR\_COMPONENT\_MOVEMENT | MSigDB lists | GO\_REGULATION\_OF\_CELLULAR\_COMPONENT\_MOVEMENT | 132 | 10 | 4161 | 156 | sli,Ddr,LanA,Cdk4,Raf,unc-5,N,Pdk1,Tsp96F,InR | | 2.527e-02 | -3.68 | Irbp (Inverted repeat-binding protein) | protein interactions | 117419 | 36 | 5 | 9204 | 432 | His3.3A,ttk,otp,CG16791,al | | 2.527e-02 | -3.68 | lid (little imaginal discs) | protein interactions | 33837 | 36 | 5 | 9204 | 432 | Kdm2,H,N,CtBP,shg | | 2.533e-02 | -3.68 | HOXA4\_Q2 | MSigDB lists | HOXA4\_Q2 | 45 | 5 | 4161 | 156 | tkv,betaTub60D,CG45050,Mef2,arm | | 2.533e-02 | -3.68 | PAX4\_04 | MSigDB lists | PAX4\_04 | 45 | 5 | 4161 | 156 | nmo,CycG,InR,FBXO11,Mef2 | | 2.533e-02 | -3.68 | CHOP\_01 | MSigDB lists | CHOP\_01 | 45 | 5 | 4161 | 156 | FBXO11,cact,Mef2,Cdk4,frc | | 2.533e-02 | -3.68 | PBX1\_01 | MSigDB lists | PBX1\_01 | 45 | 5 | 4161 | 156 | tkv,FBXO11,14-3-3zeta,nmo,dlp | | 2.533e-02 | -3.68 | GO\_POSITIVE\_REGULATION\_OF\_NEURON\_PROJECTION\_DEVELOPMENT | MSigDB lists | GO\_POSITIVE\_REGULATION\_OF\_NEURON\_PROJECTION\_DEVELOPMENT | 45 | 5 | 4161 | 156 | sli,otk,Arf79F,Raf,robo1 | | 2.533e-02 | -3.68 | MARTORIATI\_MDM4\_TARGETS\_NEUROEPITHELIUM\_UP | MSigDB lists | MARTORIATI\_MDM4\_TARGETS\_NEUROEPITHELIUM\_UP | 45 | 5 | 4161 | 156 | Pgk,Cdk4,Tsp96F,otk,CycG | | 2.535e-02 | -3.68 | protein kinase complex | cellular component | GO:1902911 | 48 | 5 | 11940 | 417 | Atg17,InR,CycG,CycB3,Cks30A | | 2.538e-02 | -3.67 | egr (eiger) | protein interactions | 36054 | 62 | 7 | 9204 | 432 | Dronc,arm,wgn,lncRNA:Hsromega,Rab5,scrib,Diap1 | | 2.540e-02 | -3.67 | Tim8 (Tim8) | protein interactions | 32081 | 14 | 3 | 9204 | 432 | al,CG31211,B-H1 | | 2.540e-02 | -3.67 | Kat60 (Katanin 60) | protein interactions | 53566 | 14 | 3 | 9204 | 432 | ap,shot,Myo10A | | 2.540e-02 | -3.67 | Cisd2 (CDGSH iron sulfur domain 2) | protein interactions | 43459 | 14 | 3 | 9204 | 432 | Rab1,CG12769,Kr-h2 | | 2.540e-02 | -3.67 | CG8478 (uncharacterized protein) | protein interactions | 41194 | 14 | 3 | 9204 | 432 | Spc105R,sens,Pdk1 | | 2.540e-02 | -3.67 | Spt-I (Serine palmitoyltransferase subunit I) | protein interactions | 36448 | 14 | 3 | 9204 | 432 | Rab1,Rab5,Kr-h2 | | 2.540e-02 | -3.67 | SdhB (Succinate dehydrogenase, subunit B (iron-sulfur)) | protein interactions | 35590 | 14 | 3 | 9204 | 432 | CG42788,CG9784,Mob2 | | 2.540e-02 | -3.67 | sug (sugarbabe) | protein interactions | 36424 | 14 | 3 | 9204 | 432 | CG2233,CtBP,Cdk4 | | 2.540e-02 | -3.67 | Khc-73 (Kinesin heavy chain 73) | protein interactions | 36718 | 14 | 3 | 9204 | 432 | Rab5,14-3-3zeta,14-3-3epsilon | | 2.540e-02 | -3.67 | ck (crinkled) | protein interactions | 34882 | 14 | 3 | 9204 | 432 | fz,Dronc,spen | | 2.540e-02 | -3.67 | stum (stumble) | protein interactions | 37473 | 14 | 3 | 9204 | 432 | ATPCL,Sema1a,CG14506 | | 2.542e-02 | -3.67 | PILON\_KLF1\_TARGETS\_DN | MSigDB lists | PILON\_KLF1\_TARGETS\_DN | 701 | 36 | 4161 | 156 | IntS3,CG11030,Fem-1,eff,CG2006,bru2,wdb,ATPCL,Rab1,Drak,CG3436,CenB1A,N,Psa,emp,AP-1gamma,CG18549,CG9485,Mfe2,rdx,Pat1,Pdk1,FBXO11,CG3961,CG4896,Pex7,CG31522,Mef2,CG6891,Cdk4,14-3-3epsilon,CtBP,Pgant5,dock,trbl,InR | | 2.543e-02 | -3.67 | synapse organization | biological process | GO:0050808 | 177 | 12 | 12345 | 442 | dlp,jeb,Adf1,kay,rg,fz2,Sema1a,baz,Gli,drl,C1GalTA,Pdk1 | | 2.547e-02 | -3.67 | negative regulation of apoptotic process | biological process | GO:0043066 | 99 | 8 | 12345 | 442 | eya,Diap1,spi,Raf,Pdk1,Egfr,sens,Pvr | | 2.547e-02 | -3.67 | positive regulation of immune response | biological process | GO:0050778 | 99 | 8 | 12345 | 442 | mop,pnr,CG32795,Pitslre,htl,Dcr-2,DPCoAC,imd | | 2.551e-02 | -3.67 | spermatogenesis | biological process | GO:0007283 | 304 | 18 | 12345 | 442 | Pex10,dmrt11E,Diap1,jar,Pdk1,lncRNA:CR44420,Dronc,yuri,Raf,InR,eya,sns,orb2,wnd,Pen,bol,Gli,Inx2 | | 2.554e-02 | -3.67 | SkpA (SKP1-related A) | protein interactions | 31016 | 76 | 8 | 9204 | 432 | kay,ds,Diap1,wnd,MEP-1,FBXO11,InR,morgue | | 2.557e-02 | -3.67 | GSE26727\_WT\_VS\_KLF2\_KO\_LPS\_STIM\_MACROPHAGE\_DN | MSigDB lists | GSE26727\_WT\_VS\_KLF2\_KO\_LPS\_STIM\_MACROPHAGE\_DN | 61 | 6 | 4161 | 156 | Fem-1,14-3-3epsilon,Ids,CG2812,CG9220,Unc-76 | | 2.557e-02 | -3.67 | GO\_REGULATION\_OF\_RESPONSE\_TO\_BIOTIC\_STIMULUS | MSigDB lists | GO\_REGULATION\_OF\_RESPONSE\_TO\_BIOTIC\_STIMULUS | 61 | 6 | 4161 | 156 | Arf79F,ReepA,CG4645,AP-1gamma,Tpc1,SmD1 | | 2.557e-02 | -3.67 | CEBPGAMMA\_Q6 | MSigDB lists | CEBPGAMMA\_Q6 | 61 | 6 | 4161 | 156 | otk,CtBP,Raf,Mef2,Cdk4,bru2 | | 2.557e-02 | -3.67 | GSE42021\_TREG\_PLN\_VS\_CD24LO\_TREG\_THYMUS\_UP | MSigDB lists | GSE42021\_TREG\_PLN\_VS\_CD24LO\_TREG\_THYMUS\_UP | 61 | 6 | 4161 | 156 | tkv,sd,unc-119,alpha-Man-Ia,Cerk,Src42A | | 2.557e-02 | -3.67 | GSE37301\_HEMATOPOIETIC\_STEM\_CELL\_VS\_MULTIPOTENT\_PROGENITOR\_UP | MSigDB lists | GSE37301\_HEMATOPOIETIC\_STEM\_CELL\_VS\_MULTIPOTENT\_PROGENITOR\_UP | 61 | 6 | 4161 | 156 | bru2,Pex14,htl,Mef2,Drep4,fz | | 2.568e-02 | -3.66 | melanin metabolic process | biological process | GO:0006582 | 47 | 5 | 12345 | 442 | RhoL,RhoGEF3,Dronc,Eph,LanA | | 2.591e-02 | -3.65 | GO\_REGULATION\_OF\_ORGAN\_FORMATION | MSigDB lists | GO\_REGULATION\_OF\_ORGAN\_FORMATION | 7 | 2 | 4161 | 156 | arm,N | | 2.591e-02 | -3.65 | ACCAATC\_MIR509 | MSigDB lists | ACCAATC\_MIR509 | 7 | 2 | 4161 | 156 | nmo,Rab5 | | 2.591e-02 | -3.65 | REACTOME\_CTLA4\_INHIBITORY\_SIGNALING | MSigDB lists | REACTOME\_CTLA4\_INHIBITORY\_SIGNALING | 7 | 2 | 4161 | 156 | Pdk1,wdb | | 2.591e-02 | -3.65 | GO\_REGULATION\_OF\_EPIDERMAL\_CELL\_DIFFERENTIATION | MSigDB lists | GO\_REGULATION\_OF\_EPIDERMAL\_CELL\_DIFFERENTIATION | 7 | 2 | 4161 | 156 | N,Dl | | 2.591e-02 | -3.65 | HOX13\_01 | MSigDB lists | HOX13\_01 | 7 | 2 | 4161 | 156 | FBXO11,unc-5 | | 2.591e-02 | -3.65 | GO\_NEGATIVE\_REGULATION\_OF\_DNA\_BINDING | MSigDB lists | GO\_NEGATIVE\_REGULATION\_OF\_DNA\_BINDING | 7 | 2 | 4161 | 156 | ap,cact | | 2.591e-02 | -3.65 | GO\_POSITIVE\_REGULATION\_OF\_EPITHELIAL\_CELL\_DIFFERENTIATION | MSigDB lists | GO\_POSITIVE\_REGULATION\_OF\_EPITHELIAL\_CELL\_DIFFERENTIATION | 7 | 2 | 4161 | 156 | arm,N | | 2.591e-02 | -3.65 | GO\_DIGESTIVE\_TRACT\_MORPHOGENESIS | MSigDB lists | GO\_DIGESTIVE\_TRACT\_MORPHOGENESIS | 7 | 2 | 4161 | 156 | arm,N | | 2.591e-02 | -3.65 | AMIT\_SERUM\_RESPONSE\_60\_MCF10A | MSigDB lists | AMIT\_SERUM\_RESPONSE\_60\_MCF10A | 7 | 2 | 4161 | 156 | Actbeta,frc | | 2.591e-02 | -3.65 | GO\_POSITIVE\_REGULATION\_OF\_B\_CELL\_ACTIVATION | MSigDB lists | GO\_POSITIVE\_REGULATION\_OF\_B\_CELL\_ACTIVATION | 7 | 2 | 4161 | 156 | Mef2,Tsp96F | | 2.591e-02 | -3.65 | GO\_ASTROCYTE\_DEVELOPMENT | MSigDB lists | GO\_ASTROCYTE\_DEVELOPMENT | 7 | 2 | 4161 | 156 | Cdk4,Dl | | 2.591e-02 | -3.65 | GO\_DENDRITIC\_SPINE\_ORGANIZATION | MSigDB lists | GO\_DENDRITIC\_SPINE\_ORGANIZATION | 7 | 2 | 4161 | 156 | Eph,Arf79F | | 2.591e-02 | -3.65 | GO\_PIGMENT\_CELL\_DIFFERENTIATION | MSigDB lists | GO\_PIGMENT\_CELL\_DIFFERENTIATION | 7 | 2 | 4161 | 156 | Mef2,AP-1gamma | | 2.591e-02 | -3.65 | GO\_REGULATION\_OF\_SULFUR\_METABOLIC\_PROCESS | MSigDB lists | GO\_REGULATION\_OF\_SULFUR\_METABOLIC\_PROCESS | 7 | 2 | 4161 | 156 | muc,arm | | 2.591e-02 | -3.65 | TAGAACC\_MIR182 | MSigDB lists | TAGAACC\_MIR182 | 7 | 2 | 4161 | 156 | Ziz,Pgant5 | | 2.591e-02 | -3.65 | GO\_REGULATION\_OF\_SYNAPTIC\_VESICLE\_EXOCYTOSIS | MSigDB lists | GO\_REGULATION\_OF\_SYNAPTIC\_VESICLE\_EXOCYTOSIS | 7 | 2 | 4161 | 156 | Syx1A,Raf | | 2.591e-02 | -3.65 | TACGGGT\_MIR99A\_MIR100\_MIR99B | MSigDB lists | TACGGGT\_MIR99A\_MIR100\_MIR99B | 7 | 2 | 4161 | 156 | trbl,htl | | 2.591e-02 | -3.65 | GO\_CELL\_SUBSTRATE\_ADHERENS\_JUNCTION\_ASSEMBLY | MSigDB lists | GO\_CELL\_SUBSTRATE\_ADHERENS\_JUNCTION\_ASSEMBLY | 7 | 2 | 4161 | 156 | Pdk1,LanA | | 2.591e-02 | -3.65 | ZHENG\_IL22\_SIGNALING\_UP | MSigDB lists | ZHENG\_IL22\_SIGNALING\_UP | 7 | 2 | 4161 | 156 | fz,Actbeta | | 2.591e-02 | -3.65 | BIOCARTA\_DEATH\_PATHWAY | MSigDB lists | BIOCARTA\_DEATH\_PATHWAY | 7 | 2 | 4161 | 156 | Drep4,cact | | 2.591e-02 | -3.65 | GO\_REGULATION\_OF\_ER\_TO\_GOLGI\_VESICLE\_MEDIATED\_TRANSPORT | MSigDB lists | GO\_REGULATION\_OF\_ER\_TO\_GOLGI\_VESICLE\_MEDIATED\_TRANSPORT | 7 | 2 | 4161 | 156 | Arf79F,CG17883 | | 2.591e-02 | -3.65 | GO\_POSITIVE\_REGULATION\_OF\_CALCIUM\_ION\_DEPENDENT\_EXOCYTOSIS | MSigDB lists | GO\_POSITIVE\_REGULATION\_OF\_CALCIUM\_ION\_DEPENDENT\_EXOCYTOSIS | 7 | 2 | 4161 | 156 | Syx1A,Arf79F | | 2.591e-02 | -3.65 | KEGG\_THYROID\_CANCER | MSigDB lists | KEGG\_THYROID\_CANCER | 7 | 2 | 4161 | 156 | Raf,arm | | 2.591e-02 | -3.65 | PID\_VEGFR1\_PATHWAY | MSigDB lists | PID\_VEGFR1\_PATHWAY | 7 | 2 | 4161 | 156 | dock,Pdk1 | | 2.591e-02 | -3.65 | GO\_NUCLEOTIDE\_TRANSMEMBRANE\_TRANSPORT | MSigDB lists | GO\_NUCLEOTIDE\_TRANSMEMBRANE\_TRANSPORT | 7 | 2 | 4161 | 156 | DPCoAC,frc | | 2.591e-02 | -3.65 | WANG\_NFKB\_TARGETS | MSigDB lists | WANG\_NFKB\_TARGETS | 7 | 2 | 4161 | 156 | SERCA,N | | 2.591e-02 | -3.65 | GO\_PEPTIDYL\_TYROSINE\_AUTOPHOSPHORYLATION | MSigDB lists | GO\_PEPTIDYL\_TYROSINE\_AUTOPHOSPHORYLATION | 7 | 2 | 4161 | 156 | Src42A,InR | | 2.591e-02 | -3.65 | SIMBULAN\_PARP1\_TARGETS\_UP | MSigDB lists | SIMBULAN\_PARP1\_TARGETS\_UP | 7 | 2 | 4161 | 156 | CycG,Tsp96F | | 2.591e-02 | -3.65 | PID\_HIV\_NEF\_PATHWAY | MSigDB lists | PID\_HIV\_NEF\_PATHWAY | 7 | 2 | 4161 | 156 | cact,Drep4 | | 2.591e-02 | -3.65 | GO\_NEURONAL\_STEM\_CELL\_POPULATION\_MAINTENANCE | MSigDB lists | GO\_NEURONAL\_STEM\_CELL\_POPULATION\_MAINTENANCE | 7 | 2 | 4161 | 156 | Dl,N | | 2.591e-02 | -3.65 | REACTOME\_CREB\_PHOSPHORYLATION\_THROUGH\_THE\_ACTIVATION\_OF\_RAS | MSigDB lists | REACTOME\_CREB\_PHOSPHORYLATION\_THROUGH\_THE\_ACTIVATION\_OF\_RAS | 7 | 2 | 4161 | 156 | Raf,Pdk1 | | 2.591e-02 | -3.65 | GO\_RESPONSE\_TO\_GONADOTROPIN | MSigDB lists | GO\_RESPONSE\_TO\_GONADOTROPIN | 7 | 2 | 4161 | 156 | Actbeta,N | | 2.591e-02 | -3.65 | GO\_REGULATION\_OF\_PROTEIN\_OLIGOMERIZATION | MSigDB lists | GO\_REGULATION\_OF\_PROTEIN\_OLIGOMERIZATION | 7 | 2 | 4161 | 156 | Pex14,Pex5 | | 2.591e-02 | -3.65 | GO\_CORECEPTOR\_ACTIVITY | MSigDB lists | GO\_CORECEPTOR\_ACTIVITY | 7 | 2 | 4161 | 156 | dlp,otk | | 2.591e-02 | -3.65 | TAKEDA\_TARGETS\_OF\_NUP98\_HOXA9\_FUSION\_6HR\_DN | MSigDB lists | TAKEDA\_TARGETS\_OF\_NUP98\_HOXA9\_FUSION\_6HR\_DN | 7 | 2 | 4161 | 156 | trbl,Actbeta | | 2.591e-02 | -3.65 | SHIPP\_DLBCL\_VS\_FOLLICULAR\_LYMPHOMA\_DN | MSigDB lists | SHIPP\_DLBCL\_VS\_FOLLICULAR\_LYMPHOMA\_DN | 7 | 2 | 4161 | 156 | trbl,Mef2 | | 2.591e-02 | -3.65 | REACTOME\_HS\_GAG\_DEGRADATION | MSigDB lists | REACTOME\_HS\_GAG\_DEGRADATION | 7 | 2 | 4161 | 156 | Ids,dlp | | 2.591e-02 | -3.65 | GO\_REGULATION\_OF\_NF\_KAPPAB\_IMPORT\_INTO\_NUCLEUS | MSigDB lists | GO\_REGULATION\_OF\_NF\_KAPPAB\_IMPORT\_INTO\_NUCLEUS | 7 | 2 | 4161 | 156 | cact,Tollo | | 2.591e-02 | -3.65 | KEGG\_CYTOKINE\_CYTOKINE\_RECEPTOR\_INTERACTION | MSigDB lists | KEGG\_CYTOKINE\_CYTOKINE\_RECEPTOR\_INTERACTION | 7 | 2 | 4161 | 156 | tkv,Actbeta | | 2.591e-02 | -3.65 | GO\_SUBSTRATE\_DEPENDENT\_CELL\_MIGRATION | MSigDB lists | GO\_SUBSTRATE\_DEPENDENT\_CELL\_MIGRATION | 7 | 2 | 4161 | 156 | dock,sli | | 2.591e-02 | -3.65 | GO\_CHONDROITIN\_SULFATE\_BIOSYNTHETIC\_PROCESS | MSigDB lists | GO\_CHONDROITIN\_SULFATE\_BIOSYNTHETIC\_PROCESS | 7 | 2 | 4161 | 156 | frc,CG9220 | | 2.591e-02 | -3.65 | MIKKELSEN\_MEF\_ICP\_WITH\_H3K4ME3\_AND\_H3K27ME3 | MSigDB lists | MIKKELSEN\_MEF\_ICP\_WITH\_H3K4ME3\_AND\_H3K27ME3 | 7 | 2 | 4161 | 156 | lin-28,robo1 | | 2.591e-02 | -3.65 | PID\_P38\_MK2\_PATHWAY | MSigDB lists | PID\_P38\_MK2\_PATHWAY | 7 | 2 | 4161 | 156 | 14-3-3zeta,14-3-3epsilon | | 2.591e-02 | -3.65 | NABA\_BASEMENT\_MEMBRANES | MSigDB lists | NABA\_BASEMENT\_MEMBRANES | 7 | 2 | 4161 | 156 | Ppn,LanA | | 2.591e-02 | -3.65 | BIOCARTA\_HDAC\_PATHWAY | MSigDB lists | BIOCARTA\_HDAC\_PATHWAY | 7 | 2 | 4161 | 156 | Mef2,InR | | 2.591e-02 | -3.65 | GO\_ESTABLISHMENT\_OF\_ENDOTHELIAL\_BARRIER | MSigDB lists | GO\_ESTABLISHMENT\_OF\_ENDOTHELIAL\_BARRIER | 7 | 2 | 4161 | 156 | arm,Rab1 | | 2.607e-02 | -3.65 | regulation of response to stress | biological process | GO:0080134 | 283 | 17 | 12345 | 442 | hppy,eya,nub,chic,lncRNA:Hsromega,Rm62,Pvr,DPCoAC,imd,N,pnr,CG32795,Egfr,htl,Dcr-2,cact,rdx | | 2.617e-02 | -3.64 | Tyr\_Pase\_AS | interpro domains | IPR016130 | 29 | 4 | 11383 | 450 | Ptpmeg2,CG42327,Ptp4E,Pez | | 2.617e-02 | -3.64 | FERM/acyl-CoA-bd\_prot\_sf | interpro domains | IPR014352 | 29 | 4 | 11383 | 450 | Pez,yrt,cora,CG42788 | | 2.617e-02 | -3.64 | Mi-2 (Mi-2) | protein interactions | 40170 | 49 | 6 | 9204 | 432 | ttk,Rm62,CtBP,Egfr,MEP-1,rictor | | 2.621e-02 | -3.64 | CAGGTA\_AREB6\_01 | MSigDB lists | CAGGTA\_AREB6\_01 | 152 | 11 | 4161 | 156 | CG3961,Ziz,Cdk4,rg,otk,Dyb,dlp,Rab1,N,Kr-h2,CG9220 | | 2.639e-02 | -3.63 | cuticle development involved in chitin-based cuticle molting cycle | biological process | GO:0042337 | 32 | 4 | 12345 | 442 | jar,arm,fz,fz2 | | 2.639e-02 | -3.63 | maintenance of protein location in cell | biological process | GO:0032507 | 32 | 4 | 12345 | 442 | Vps13,chic,p130CAS,Dg | | 2.639e-02 | -3.63 | positive regulation of stress-activated MAPK cascade | biological process | GO:0032874 | 32 | 4 | 12345 | 442 | hppy,lncRNA:Hsromega,Pvr,rdx | | 2.639e-02 | -3.63 | positive regulation of stress-activated protein kinase signaling cascade | biological process | GO:0070304 | 32 | 4 | 12345 | 442 | rdx,hppy,Pvr,lncRNA:Hsromega | | 2.645e-02 | -3.63 | GO\_EARLY\_ENDOSOME | MSigDB lists | GO\_EARLY\_ENDOSOME | 96 | 8 | 4161 | 156 | lt,sina,Rab1,fz2,Eph,Tollo,Rab5,Ocrl | | 2.657e-02 | -3.63 | regulation of protein polymerization | biological process | GO:0032271 | 64 | 6 | 12345 | 442 | chic,CG6891,RhoU,SCAR,Cip4,Jupiter | | 2.657e-02 | -3.63 | antimicrobial humoral response | biological process | GO:0019730 | 64 | 6 | 12345 | 442 | kay,Rm62,cact,Pvr,imd,Mef2 | | 2.665e-02 | -3.63 | SHC1 events in ERBB2 signaling | REACTOME pathways | R-DME-1250196 | 6 | 2 | 3923 | 176 | vn,Egfr | | 2.665e-02 | -3.63 | SHC1 events in ERBB4 signaling | REACTOME pathways | R-DME-1250347 | 6 | 2 | 3923 | 176 | vn,Egfr | | 2.665e-02 | -3.63 | Regulation of KIT signaling | REACTOME pathways | R-DME-1433559 | 6 | 2 | 3923 | 176 | Cbl,Pvr | | 2.665e-02 | -3.63 | ERBB2 Regulates Cell Motility | REACTOME pathways | R-DME-6785631 | 6 | 2 | 3923 | 176 | Egfr,vn | | 2.665e-02 | -3.63 | PI3K events in ERBB2 signaling | REACTOME pathways | R-DME-1963642 | 6 | 2 | 3923 | 176 | Egfr,vn | | 2.685e-02 | -3.62 | positive regulation of GTPase activity | biological process | GO:0043547 | 100 | 8 | 12345 | 442 | Ocrl,Rgl,drongo,CG17883,TBC1D5,RhoGAP19D,CenB1A,RapGAP1 | | 2.687e-02 | -3.62 | GO\_MACROMOLECULAR\_COMPLEX\_BINDING | MSigDB lists | GO\_MACROMOLECULAR\_COMPLEX\_BINDING | 361 | 21 | 4161 | 156 | CG4896,cact,Mef2,14-3-3epsilon,shot,InR,lt,ap,CtBP,N,PI31,LanA,Ddr,Eph,hyx,jar,Pex14,arm,SMC3,Tsp96F,Pdk1 | | 2.704e-02 | -3.61 | nuclear heterochromatin | cellular component | GO:0005720 | 19 | 3 | 11940 | 417 | Su(z)2,Psc,E(Pc) | | 2.712e-02 | -3.61 | Integrin cell surface interactions | REACTOME pathways | R-DME-216083 | 15 | 3 | 3923 | 176 | Bsg,Pvr,Dg | | 2.712e-02 | -3.61 | Spry regulation of FGF signaling | REACTOME pathways | R-DME-1295596 | 15 | 3 | 3923 | 176 | Lk6,Cbl,Raf | | 2.719e-02 | -3.60 | GO\_PROTEIN\_TRANSMEMBRANE\_TRANSPORT | MSigDB lists | GO\_PROTEIN\_TRANSMEMBRANE\_TRANSPORT | 31 | 4 | 4161 | 156 | Pex14,Pex5,Pex7,Pex10 | | 2.719e-02 | -3.60 | GSE13762\_CTRL\_VS\_125\_VITAMIND\_DAY12\_DC\_DN | MSigDB lists | GSE13762\_CTRL\_VS\_125\_VITAMIND\_DAY12\_DC\_DN | 31 | 4 | 4161 | 156 | rg,Src42A,dlp,IntS3 | | 2.719e-02 | -3.60 | REACTOME\_GLYCOSAMINOGLYCAN\_METABOLISM | MSigDB lists | REACTOME\_GLYCOSAMINOGLYCAN\_METABOLISM | 31 | 4 | 4161 | 156 | CG9220,dlp,GlcAT-I,Ids | | 2.719e-02 | -3.60 | PEREZ\_TP53\_AND\_TP63\_TARGETS | MSigDB lists | PEREZ\_TP53\_AND\_TP63\_TARGETS | 31 | 4 | 4161 | 156 | bru2,sli,InR,fz | | 2.719e-02 | -3.60 | GSE35825\_UNTREATED\_VS\_IFNA\_STIM\_MACROPHAGE\_DN | MSigDB lists | GSE35825\_UNTREATED\_VS\_IFNA\_STIM\_MACROPHAGE\_DN | 31 | 4 | 4161 | 156 | Tpst,CG2006,fz2,robo1 | | 2.719e-02 | -3.60 | GSE20484\_MCSG\_VS\_CXCL4\_MONOCYTE\_DERIVED\_MACROPHAGE\_DN | MSigDB lists | GSE20484\_MCSG\_VS\_CXCL4\_MONOCYTE\_DERIVED\_MACROPHAGE\_DN | 31 | 4 | 4161 | 156 | CG2812,InR,CG3961,Hel89B | | 2.719e-02 | -3.60 | SMIRNOV\_RESPONSE\_TO\_IR\_6HR\_UP | MSigDB lists | SMIRNOV\_RESPONSE\_TO\_IR\_6HR\_UP | 31 | 4 | 4161 | 156 | N,Rab1,CycG,frc | | 2.719e-02 | -3.60 | GSE42021\_CD24INT\_VS\_CD24LOW\_TCONV\_THYMUS\_DN | MSigDB lists | GSE42021\_CD24INT\_VS\_CD24LOW\_TCONV\_THYMUS\_DN | 31 | 4 | 4161 | 156 | emp,hppy,step,CG2017 | | 2.722e-02 | -3.60 | establishment of localization | biological process | GO:0051234 | 1717 | 76 | 12345 | 442 | nuf,Rim2,CG7536,frc,Pex5,Inx2,Myo10A,cv-d,mop,SERCA,baz,Mpcp2,Dcr-2,CG31121,TBC1D5,Syx1A,Abl,AP-1gamma,Cadps,edl,DPCoAC,scrib,Ppcs,CG4645,CG3149,nAChRalpha4,unc-119,lt,Ocrl,aop,CG17646,Tpc1,path,lqf,apolpp,Arf79F,SCAR,CG7394,Gli,Irk3,CG17883,Esp,Rab1,CG13384,jar,wbl,Tpst,subdued,ogre,Rab5,CG11069,Pex10,Slbp,CG30344,tipE,chic,Pen,CG8034,RhoU,Mid1,dve,S,Npc2g,Orct2,koi,Vps13,ena,CG4991,Unc-76,Atox1,ZnT41F,cora,CG5853,Pex14,Src42A,klar | | 2.745e-02 | -3.60 | Ldh (Lactate dehydrogenase) | protein interactions | 45880 | 63 | 7 | 9204 | 432 | Rab5,N,Kr-h2,Arf79F,SERCA,Tis11,Rab1 | | 2.746e-02 | -3.59 | GSE4748\_CTRL\_VS\_CYANOBACTERIUM\_LPSLIKE\_STIM\_DC\_1H\_DN | MSigDB lists | GSE4748\_CTRL\_VS\_CYANOBACTERIUM\_LPSLIKE\_STIM\_DC\_1H\_DN | 62 | 6 | 4161 | 156 | Raf,sd,Pex14,AP-1gamma,14-3-3epsilon,CycG | | 2.746e-02 | -3.59 | GSE17721\_0.5H\_VS\_12H\_GARDIQUIMOD\_BMDC\_DN | MSigDB lists | GSE17721\_0.5H\_VS\_12H\_GARDIQUIMOD\_BMDC\_DN | 62 | 6 | 4161 | 156 | cact,ap,Tollo,Dyb,alpha-Man-Ia,CG3961 | | 2.746e-02 | -3.59 | GSE3982\_MAC\_VS\_BCELL\_UP | MSigDB lists | GSE3982\_MAC\_VS\_BCELL\_UP | 62 | 6 | 4161 | 156 | shot,jar,Ocrl,fz2,Pen,Actbeta | | 2.746e-02 | -3.59 | GTACTGT\_MIR101 | MSigDB lists | GTACTGT\_MIR101 | 62 | 6 | 4161 | 156 | eff,Rab1,nmo,rdx,robo1,Cdk4 | | 2.751e-02 | -3.59 | GSE28726\_NAIVE\_VS\_ACTIVATED\_NKTCELL\_DN | MSigDB lists | GSE28726\_NAIVE\_VS\_ACTIVATED\_NKTCELL\_DN | 79 | 7 | 4161 | 156 | Syx1A,14-3-3epsilon,CG2812,CG9220,Unc-76,Arf79F,Tsp96F | | 2.759e-02 | -3.59 | LEF1\_Q2 | MSigDB lists | LEF1\_Q2 | 46 | 5 | 4161 | 156 | CG31522,Rgl,ds,CG3961,Ddr | | 2.759e-02 | -3.59 | GSE5589\_UNSTIM\_VS\_45MIN\_LPS\_AND\_IL10\_STIM\_MACROPHAGE\_UP | MSigDB lists | GSE5589\_UNSTIM\_VS\_45MIN\_LPS\_AND\_IL10\_STIM\_MACROPHAGE\_UP | 46 | 5 | 4161 | 156 | GlcAT-I,CG11030,unc-119,Pdk1,N | | 2.759e-02 | -3.59 | GSE3039\_B2\_VS\_B1\_BCELL\_DN | MSigDB lists | GSE3039\_B2\_VS\_B1\_BCELL\_DN | 46 | 5 | 4161 | 156 | CG1673,otk,Pen,CG6891,GlcAT-I | | 2.759e-02 | -3.59 | MCBRYAN\_PUBERTAL\_BREAST\_3\_4WK\_UP | MSigDB lists | MCBRYAN\_PUBERTAL\_BREAST\_3\_4WK\_UP | 46 | 5 | 4161 | 156 | unc-119,fz,subdued,muc,ATPCL | | 2.759e-02 | -3.59 | GSE9988\_ANTI\_TREM1\_VS\_LPS\_MONOCYTE\_UP | MSigDB lists | GSE9988\_ANTI\_TREM1\_VS\_LPS\_MONOCYTE\_UP | 46 | 5 | 4161 | 156 | CG9220,Syx1A,hppy,emp,Pex5 | | 2.759e-02 | -3.59 | TGTGTGA\_MIR377 | MSigDB lists | TGTGTGA\_MIR377 | 46 | 5 | 4161 | 156 | sd,Psa,rg,bru2,eff | | 2.759e-02 | -3.59 | GSE24574\_BCL6\_HIGH\_VS\_LOW\_TFH\_CD4\_TCELL\_UP | MSigDB lists | GSE24574\_BCL6\_HIGH\_VS\_LOW\_TFH\_CD4\_TCELL\_UP | 46 | 5 | 4161 | 156 | Cadps,step,Kdm2,14-3-3zeta,trbl | | 2.762e-02 | -3.59 | HTH\_Psq | interpro domains | IPR007889 | 17 | 3 | 11383 | 450 | CG13895,rib,bab2 | | 2.768e-02 | -3.59 | PCB (Pyruvate carboxylase) | protein interactions | 36020 | 25 | 4 | 9204 | 432 | Gfat2,SERCA,Mpcp2,GluProRS | | 2.768e-02 | -3.59 | grh (grainy head) | protein interactions | 37038 | 25 | 4 | 9204 | 432 | abd-A,Kdm2,fz,Dg | | 2.768e-02 | -3.59 | alph (alphabet) | protein interactions | 43481 | 25 | 4 | 9204 | 432 | rdx,Raf,wnd,aop | | 2.768e-02 | -3.59 | trol (terribly reduced optic lobes) | protein interactions | 45320 | 25 | 4 | 9204 | 432 | Dg,Sema1a,Egfr,Rgl | | 2.768e-02 | -3.59 | hook (hook) | protein interactions | 35169 | 25 | 4 | 9204 | 432 | CG5522,hyx,baz,Rab5 | | 2.768e-02 | -3.59 | dgrn (degringolade) | protein interactions | 40725 | 25 | 4 | 9204 | 432 | eff,h,N,E(spl)m5-HLH | | 2.773e-02 | -3.59 | TGCCTTA\_MIR124A | MSigDB lists | TGCCTTA\_MIR124A | 173 | 12 | 4161 | 156 | CG6891,kn,Cdk4,C1GalTA,CG5522,Cadps,wdb,CG9220,ap,sd,lin-28,dlp | | 2.785e-02 | -3.58 | carbohydrate homeostasis | biological process | GO:0033500 | 48 | 5 | 12345 | 442 | Rab1,InR,ey,ATPCL,trbl | | 2.788e-02 | -3.58 | GO\_POSITIVE\_REGULATION\_OF\_IMMUNE\_EFFECTOR\_PROCESS | MSigDB lists | GO\_POSITIVE\_REGULATION\_OF\_IMMUNE\_EFFECTOR\_PROCESS | 18 | 3 | 4161 | 156 | Pdk1,AP-1gamma,fz2 | | 2.788e-02 | -3.58 | GO\_EMBRYONIC\_PATTERN\_SPECIFICATION | MSigDB lists | GO\_EMBRYONIC\_PATTERN\_SPECIFICATION | 18 | 3 | 4161 | 156 | Dl,fz2,arm | | 2.788e-02 | -3.58 | SCHUETZ\_BREAST\_CANCER\_DUCTAL\_INVASIVE\_DN | MSigDB lists | SCHUETZ\_BREAST\_CANCER\_DUCTAL\_INVASIVE\_DN | 18 | 3 | 4161 | 156 | shot,Pex7,jar | | 2.788e-02 | -3.58 | KEGG\_P53\_SIGNALING\_PATHWAY | MSigDB lists | KEGG\_P53\_SIGNALING\_PATHWAY | 18 | 3 | 4161 | 156 | CycG,Cdk4,sina | | 2.788e-02 | -3.58 | MODULE\_75 | MSigDB lists | MODULE\_75 | 18 | 3 | 4161 | 156 | Tsp96F,CG1673,cact | | 2.788e-02 | -3.58 | KRAS.LUNG\_UP.V1\_UP | MSigDB lists | KRAS.LUNG\_UP.V1\_UP | 18 | 3 | 4161 | 156 | Actbeta,subdued,sns | | 2.788e-02 | -3.58 | CAAGGAT\_MIR362 | MSigDB lists | CAAGGAT\_MIR362 | 18 | 3 | 4161 | 156 | Psa,CG9220,Dl | | 2.788e-02 | -3.58 | CERVERA\_SDHB\_TARGETS\_1\_UP | MSigDB lists | CERVERA\_SDHB\_TARGETS\_1\_UP | 18 | 3 | 4161 | 156 | Actbeta,ap,Dyrk2 | | 2.788e-02 | -3.58 | GO\_CELL\_MATRIX\_ADHESION | MSigDB lists | GO\_CELL\_MATRIX\_ADHESION | 18 | 3 | 4161 | 156 | Pdk1,arm,LanA | | 2.810e-02 | -3.57 | ERMFAMILY | prints domains | PR00661 | 6 | 2 | 2185 | 101 | yrt,cora | | 2.813e-02 | -3.57 | Lk6 (Lk6 kinase) | protein interactions | 44672 | 37 | 5 | 9204 | 432 | N,mam,spi,Raf,Lk6 | | 2.813e-02 | -3.57 | Pex5 (Peroxin 5) | protein interactions | 31141 | 37 | 5 | 9204 | 432 | His3.3B,ttk,Pex14,bowl,CG34376 | | 2.819e-02 | -3.57 | MAP kinase kinase kinase activity | molecular function | GO:0004709 | 8 | 2 | 12176 | 414 | wnd,Raf | | 2.825e-02 | -3.57 | cellular protein localization | biological process | GO:0034613 | 558 | 29 | 12345 | 442 | scrib,CG7394,Pex5,Dg,chic,Pen,Bsg,Arf79F,ds,d,lt,Rab1,hth,Vps13,Myo10A,Rab5,cora,Syx1A,TBC1D5,rictor,AP-1gamma,Cip4,arm,p130CAS,Shrm,His2Av,Pex14,Pex10,ft | | 2.829e-02 | -3.57 | positive regulation of response to biotic stimulus | biological process | GO:0002833 | 101 | 8 | 12345 | 442 | Pitslre,CG32795,pnr,mop,imd,DPCoAC,Dcr-2,htl | | 2.842e-02 | -3.56 | regulation of TOR signaling | biological process | GO:0032006 | 65 | 6 | 12345 | 442 | scyl,Cdk4,InR,chrb,hppy,Pvr | | 2.848e-02 | -3.56 | HOMEOBOX | prints domains | PR00024 | 38 | 5 | 2185 | 101 | B-H1,H2.0,Scr,abd-A,Abd-B | | 2.861e-02 | -3.55 | Gfat2 (Glutamine:fructose-6-phosphate aminotransferase 2) | protein interactions | 43373 | 50 | 6 | 9204 | 432 | Mpcp2,Rab5,GluProRS,Raf,Pdk1,SERCA | | 2.870e-02 | -3.55 | OAR\_dom | interpro domains | IPR003654 | 7 | 2 | 11383 | 450 | al,otp | | 2.870e-02 | -3.55 | PAC | interpro domains | IPR001610 | 7 | 2 | 11383 | 450 | ss,sim | | 2.870e-02 | -3.55 | Cyclin | interpro domains | IPR039361 | 7 | 2 | 11383 | 450 | CycG,CycB3 | | 2.870e-02 | -3.55 | Furin\_repeat | interpro domains | IPR006212 | 7 | 2 | 11383 | 450 | InR,Egfr | | 2.870e-02 | -3.55 | Pointed\_dom | interpro domains | IPR003118 | 7 | 2 | 11383 | 450 | aop,edl | | 2.870e-02 | -3.55 | PAS\_fold | interpro domains | IPR013767 | 7 | 2 | 11383 | 450 | ss,sim | | 2.870e-02 | -3.55 | Homeobox\_KN\_domain | interpro domains | IPR008422 | 7 | 2 | 11383 | 450 | hth,CG11617 | | 2.870e-02 | -3.55 | Semap\_dom\_sf | interpro domains | IPR036352 | 7 | 2 | 11383 | 450 | Sema1a,Sema1b | | 2.870e-02 | -3.55 | Semap\_dom | interpro domains | IPR001627 | 7 | 2 | 11383 | 450 | Sema1b,Sema1a | | 2.884e-02 | -3.55 | positive regulation of transcription elongation from RNA polymerase II promoter | biological process | GO:0032968 | 19 | 3 | 12345 | 442 | Mef2,fs(1)h,hyx | | 2.884e-02 | -3.55 | Notch signaling pathway | biological process | GO:0007219 | 19 | 3 | 12345 | 442 | Dl,N,mam | | 2.888e-02 | -3.54 | GO\_ESTABLISHMENT\_OF\_PROTEIN\_LOCALIZATION | MSigDB lists | GO\_ESTABLISHMENT\_OF\_PROTEIN\_LOCALIZATION | 567 | 30 | 4161 | 156 | LanA,AP-1gamma,ReepA,Syx1A,Cadps,emp,unc-119,Ric,wbl,Pat1,jar,arm,Pex14,Pex10,CG45050,CG7394,Pex7,Rab5,Pex5,cact,Raf,Atg16,Rab1,14-3-3epsilon,lt,Pen,Arf79F,Vps13,N,14-3-3zeta | | 2.888e-02 | -3.54 | SAM/pointed\_sf | interpro domains | IPR013761 | 44 | 5 | 11383 | 450 | ph-p,aop,SKIP,edl,Eph | | 2.889e-02 | -3.54 | POU | smart domains | SM00352 | 5 | 2 | 5086 | 290 | pdm3,nub | | 2.895e-02 | -3.54 | cyclin-dependent protein serine/threonine kinase regulator activity | molecular function | GO:0016538 | 20 | 3 | 12176 | 414 | CycG,CycB3,Cks30A | | 2.896e-02 | -3.54 | lipid localization | biological process | GO:0010876 | 83 | 7 | 12345 | 442 | apolpp,cv-d,SNF4Agamma,Mef2,CG3149,klar,Npc2g | | 2.896e-02 | -3.54 | GO\_POSITIVE\_REGULATION\_OF\_CELL\_PROLIFERATION | MSigDB lists | GO\_POSITIVE\_REGULATION\_OF\_CELL\_PROLIFERATION | 135 | 10 | 4161 | 156 | arm,htl,InR,dock,Tsp96F,N,Ddr,Dl,Cdk4,Mef2 | | 2.898e-02 | -3.54 | RAF/MAP kinase cascade | REACTOME pathways | R-DME-5673001 | 113 | 10 | 3923 | 176 | Raf,vn,Egfr,Pvr,14-3-3zeta,PI31,htl,CG4935,CG42684,wdb | | 2.898e-02 | -3.54 | SHC1 events in EGFR signaling | REACTOME pathways | R-DME-180336 | 113 | 10 | 3923 | 176 | CG4935,14-3-3zeta,PI31,htl,wdb,CG42684,Egfr,Raf,vn,Pvr | | 2.898e-02 | -3.54 | Frs2-mediated activation | REACTOME pathways | R-DME-170968 | 113 | 10 | 3923 | 176 | CG42684,wdb,htl,14-3-3zeta,PI31,CG4935,Pvr,vn,Raf,Egfr | | 2.898e-02 | -3.54 | ARMS-mediated activation | REACTOME pathways | R-DME-170984 | 113 | 10 | 3923 | 176 | Pvr,Raf,vn,Egfr,CG42684,wdb,14-3-3zeta,htl,PI31,CG4935 | | 2.898e-02 | -3.54 | RET signaling | REACTOME pathways | R-DME-8853659 | 113 | 10 | 3923 | 176 | Egfr,Raf,vn,Pvr,CG4935,PI31,htl,14-3-3zeta,wdb,CG42684 | | 2.898e-02 | -3.54 | Interleukin receptor SHC signaling | REACTOME pathways | R-DME-912526 | 113 | 10 | 3923 | 176 | wdb,CG42684,CG4935,PI31,14-3-3zeta,htl,Pvr,Egfr,Raf,vn | | 2.898e-02 | -3.54 | Prolonged ERK activation events | REACTOME pathways | R-DME-169893 | 113 | 10 | 3923 | 176 | CG4935,htl,PI31,14-3-3zeta,wdb,CG42684,Egfr,Raf,vn,Pvr | | 2.898e-02 | -3.54 | Signaling by the B Cell Receptor (BCR) | REACTOME pathways | R-DME-983705 | 113 | 10 | 3923 | 176 | vn,cact,Egfr,Dronc,trbl,wdb,PI31,Pdk1,rictor,htl | | 2.898e-02 | -3.54 | GRB2 events in EGFR signaling | REACTOME pathways | R-DME-179812 | 113 | 10 | 3923 | 176 | CG4935,14-3-3zeta,htl,PI31,wdb,CG42684,Egfr,vn,Raf,Pvr | | 2.898e-02 | -3.54 | SOS-mediated signalling | REACTOME pathways | R-DME-112412 | 113 | 10 | 3923 | 176 | CG4935,14-3-3zeta,htl,PI31,wdb,CG42684,Egfr,Raf,vn,Pvr | | 2.898e-02 | -3.54 | Signalling to p38 via RIT and RIN | REACTOME pathways | R-DME-187706 | 113 | 10 | 3923 | 176 | CG42684,wdb,PI31,htl,14-3-3zeta,CG4935,Pvr,Raf,vn,Egfr | | 2.907e-02 | -3.54 | toe (twin of eyg) | protein interactions | 39418 | 6 | 2 | 9204 | 432 | N,eyg | | 2.907e-02 | -3.54 | CG5003 (uncharacterized protein) | protein interactions | 43344 | 6 | 2 | 9204 | 432 | rib,CycG | | 2.907e-02 | -3.54 | TMEM216 (Transmembrane protein 216) | protein interactions | 41035 | 6 | 2 | 9204 | 432 | CG14506,pnr | | 2.907e-02 | -3.54 | Clect27 (C-type lectin 27kD) | protein interactions | 33678 | 6 | 2 | 9204 | 432 | msi,Sema1a | | 2.907e-02 | -3.54 | Lnk (Lnk) | protein interactions | 43130 | 6 | 2 | 9204 | 432 | Egfr,InR | | 2.907e-02 | -3.54 | alpha4GT1 (alpha1,4-galactosyltransferase 1) | protein interactions | 33512 | 6 | 2 | 9204 | 432 | Tom,N | | 2.907e-02 | -3.54 | rho-5 (rhomboid-5) | protein interactions | 2768944 | 6 | 2 | 9204 | 432 | Egfr,spi | | 2.907e-02 | -3.54 | heix (heixuedian) | protein interactions | 34961 | 6 | 2 | 9204 | 432 | Kr-h2,H | | 2.907e-02 | -3.54 | lili (lilipod) | protein interactions | 42956 | 6 | 2 | 9204 | 432 | Dad,tkv | | 2.907e-02 | -3.54 | rumi (rumi) | protein interactions | 326122 | 6 | 2 | 9204 | 432 | Dl,N | | 2.907e-02 | -3.54 | CG5746 (uncharacterized protein) | protein interactions | 42903 | 6 | 2 | 9204 | 432 | Pat1,Tet | | 2.907e-02 | -3.54 | boss (bride of sevenless) | protein interactions | 43146 | 6 | 2 | 9204 | 432 | H,Src42A | | 2.907e-02 | -3.54 | NetB (Netrin-B) | protein interactions | 32400 | 6 | 2 | 9204 | 432 | spi,siz | | 2.907e-02 | -3.54 | gwl (greatwall) | protein interactions | 45969 | 6 | 2 | 9204 | 432 | 14-3-3zeta,14-3-3epsilon | | 2.907e-02 | -3.54 | Vps8 (Vacuolar protein sorting 8) | protein interactions | 38735 | 6 | 2 | 9204 | 432 | Rab5,lt | | 2.907e-02 | -3.54 | mir-305 (mir-305 stem loop) | protein interactions | 12798300 | 6 | 2 | 9204 | 432 | H,InR | | 2.907e-02 | -3.54 | Xpc (Xeroderma pigmentosum, complementation group C) | protein interactions | 36697 | 6 | 2 | 9204 | 432 | otp,Sema1a | | 2.907e-02 | -3.54 | CG5144 (uncharacterized protein) | protein interactions | 39040 | 6 | 2 | 9204 | 432 | CG14478,bab2 | | 2.907e-02 | -3.54 | BCAS2 (BCAS2) | protein interactions | 43348 | 6 | 2 | 9204 | 432 | CycG,Raf | | 2.907e-02 | -3.54 | Eip63E (Ecdysone-induced protein 63E) | protein interactions | 38433 | 6 | 2 | 9204 | 432 | 14-3-3epsilon,Raf | | 2.907e-02 | -3.54 | shf (shifted) | protein interactions | 31617 | 6 | 2 | 9204 | 432 | fz2,boi | | 2.907e-02 | -3.54 | upd2 (unpaired 2) | protein interactions | 32805 | 6 | 2 | 9204 | 432 | N,dome | | 2.907e-02 | -3.54 | side (sidestep) | protein interactions | 43300 | 6 | 2 | 9204 | 432 | beat-IIa,Pex5 | | 2.907e-02 | -3.54 | sfl (sulfateless) | protein interactions | 38736 | 6 | 2 | 9204 | 432 | Sema1a,htl | | 2.913e-02 | -3.54 | HMG\_BOX\_2 | prosite domains | PS50118 | 23 | 4 | 6243 | 325 | Sox14,cic,CG12104,HmgZ | | 2.921e-02 | -3.53 | apposition of dorsal and ventral imaginal disc-derived wing surfaces | biological process | GO:0007475 | 33 | 4 | 12345 | 442 | Dl,mam,dpy,shot | | 2.921e-02 | -3.53 | transmembrane receptor protein serine/threonine kinase signaling pathway | biological process | GO:0007178 | 33 | 4 | 12345 | 442 | Actbeta,pnr,Dad,tkv | | 2.921e-02 | -3.53 | regulation of DNA-binding transcription factor activity | biological process | GO:0051090 | 33 | 4 | 12345 | 442 | edl,14-3-3epsilon,cact,14-3-3zeta | | 2.923e-02 | -3.53 | GO\_SINGLE\_ORGANISM\_CELL\_ADHESION | MSigDB lists | GO\_SINGLE\_ORGANISM\_CELL\_ADHESION | 80 | 7 | 4161 | 156 | arm,fz2,Rab1,LanA,Cdk4,Raf,dock | | 2.928e-02 | -3.53 | EGF\_Ca-bd\_CS | interpro domains | IPR018097 | 30 | 4 | 11383 | 450 | dpy,sli,Dl,N | | 2.928e-02 | -3.53 | SH2 | interpro domains | IPR000980 | 30 | 4 | 11383 | 450 | Socs36E,Abl,Src42A,dock | | 2.928e-02 | -3.53 | F-box\_dom | interpro domains | IPR001810 | 30 | 4 | 11383 | 450 | morgue,FBXO11,Kdm2,CG12402 | | 2.928e-02 | -3.53 | EGF-type\_Asp/Asn\_hydroxyl\_site | interpro domains | IPR000152 | 30 | 4 | 11383 | 450 | dpy,sli,Dl,N | | 2.930e-02 | -3.53 | Homeobox\_KN | pfam domains | PF05920 | 7 | 2 | 10480 | 419 | CG11617,hth | | 2.930e-02 | -3.53 | Sema | pfam domains | PF01403 | 7 | 2 | 10480 | 419 | Sema1b,Sema1a | | 2.930e-02 | -3.53 | PAS | pfam domains | PF00989 | 7 | 2 | 10480 | 419 | sim,ss | | 2.930e-02 | -3.53 | Death | pfam domains | PF00531 | 7 | 2 | 10480 | 419 | imd,unc-5 | | 2.930e-02 | -3.53 | SAM\_PNT | pfam domains | PF02198 | 7 | 2 | 10480 | 419 | aop,edl | | 2.945e-02 | -3.53 | GSE5589\_LPS\_VS\_LPS\_AND\_IL6\_STIM\_IL6\_KO\_MACROPHAGE\_45MIN\_DN | MSigDB lists | GSE5589\_LPS\_VS\_LPS\_AND\_IL6\_STIM\_IL6\_KO\_MACROPHAGE\_45MIN\_DN | 63 | 6 | 4161 | 156 | unc-119,jar,CG17883,ATPCL,SmD1,emp | | 2.951e-02 | -3.52 | fly chr96B1-96B1\|3-86 cM | chromosome location | fly chr96B1-96B1\|3-86 cM | 12 | 2 | 25010 | 571 | CycB3,Esp | | 2.951e-02 | -3.52 | fly chr96B1-96B1\|3-86 | chromosome location | fly chr96B1-96B1\|3-86 | 12 | 2 | 25010 | 571 | Esp,CycB3 | | 2.958e-02 | -3.52 | cellular response to endogenous stimulus | biological process | GO:0071495 | 181 | 12 | 12345 | 442 | Hr39,tai,Raf,InR,pnr,Dad,tkv,htl,rictor,aop,Pdk1,Hr4 | | 2.961e-02 | -3.52 | chromosome | cellular component | GO:0005694 | 696 | 34 | 11940 | 417 | Dp,pita,His3.3A,Su(z)2,lilli,bab2,vig,crol,ttk,His2Av,HmgZ,14-3-3epsilon,MEP-1,CtBP,jumu,su(sable),Rm62,sens,mam,Spc105R,E(Pc),SMC3,CycG,Drep4,His3.3B,Psc,tna,Elba2,Adf1,Nlp,lawc,lncRNA:Hsromega,flfl,corto | | 2.962e-02 | -3.52 | ewg (erect wing) | protein interactions | 30975 | 64 | 7 | 9204 | 432 | Cbl,CycB3,CtBP,rdx,esg,CG4788,CG6340 | | 2.963e-02 | -3.52 | type Ib terminal bouton | cellular component | GO:0061176 | 8 | 2 | 11940 | 417 | pum,jeb | | 2.963e-02 | -3.52 | gap junction | cellular component | GO:0005921 | 8 | 2 | 11940 | 417 | ogre,Inx2 | | 2.965e-02 | -3.52 | GO\_IDENTICAL\_PROTEIN\_BINDING | MSigDB lists | GO\_IDENTICAL\_PROTEIN\_BINDING | 321 | 19 | 4161 | 156 | CG4896,Pex7,cact,Atg16,robo1,Raf,shot,lt,InR,Actbeta,CtBP,14-3-3zeta,sli,Tollo,CG1673,wbl,SERCA,Mfe2,sina | | 2.976e-02 | -3.51 | GO\_INTRINSIC\_COMPONENT\_OF\_PLASMA\_MEMBRANE | MSigDB lists | GO\_INTRINSIC\_COMPONENT\_OF\_PLASMA\_MEMBRANE | 257 | 16 | 4161 | 156 | Dl,otk,sns,htl,InR,dlp,Eph,Tollo,Ddr,Syx1A,ReepA,Tsp96F,Nrk,nAChRalpha4,tkv,SERCA | | 2.997e-02 | -3.51 | GSE42724\_NAIVE\_VS\_MEMORY\_BCELL\_DN | MSigDB lists | GSE42724\_NAIVE\_VS\_MEMORY\_BCELL\_DN | 47 | 5 | 4161 | 156 | Pex5,l(1)G0196,CG3961,14-3-3zeta,ap | | 2.997e-02 | -3.51 | GO\_EAR\_DEVELOPMENT | MSigDB lists | GO\_EAR\_DEVELOPMENT | 47 | 5 | 4161 | 156 | otk,Dl,N,ft,ds | | 2.997e-02 | -3.51 | GSE17721\_LPS\_VS\_POLYIC\_8H\_BMDC\_DN | MSigDB lists | GSE17721\_LPS\_VS\_POLYIC\_8H\_BMDC\_DN | 47 | 5 | 4161 | 156 | ap,Lk6,alpha-Man-Ia,step,emp | | 2.997e-02 | -3.51 | GSE22935\_UNSTIM\_VS\_24H\_MBOVIS\_BCG\_STIM\_MACROPHAGE\_DN | MSigDB lists | GSE22935\_UNSTIM\_VS\_24H\_MBOVIS\_BCG\_STIM\_MACROPHAGE\_DN | 47 | 5 | 4161 | 156 | muc,CG13893,Pat1,Raf,Tollo | | 2.997e-02 | -3.51 | GSE42724\_NAIVE\_VS\_MEMORY\_BCELL\_UP | MSigDB lists | GSE42724\_NAIVE\_VS\_MEMORY\_BCELL\_UP | 47 | 5 | 4161 | 156 | Rgl,PI31,hyx,Atg16,dock | | 2.997e-02 | -3.51 | GSE40666\_WT\_VS\_STAT1\_KO\_CD8\_TCELL\_WITH\_IFNA\_STIM\_90MIN\_UP | MSigDB lists | GSE40666\_WT\_VS\_STAT1\_KO\_CD8\_TCELL\_WITH\_IFNA\_STIM\_90MIN\_UP | 47 | 5 | 4161 | 156 | jar,fz2,dock,unc-119,l(2)37Cg | | 2.997e-02 | -3.51 | GSE17721\_LPS\_VS\_POLYIC\_0.5H\_BMDC\_DN | MSigDB lists | GSE17721\_LPS\_VS\_POLYIC\_0.5H\_BMDC\_DN | 47 | 5 | 4161 | 156 | CG17883,14-3-3epsilon,jar,Syx1A,sina | | 2.997e-02 | -3.51 | GSE15930\_NAIVE\_VS\_48H\_IN\_VITRO\_STIM\_IFNAB\_CD8\_TCELL\_UP | MSigDB lists | GSE15930\_NAIVE\_VS\_48H\_IN\_VITRO\_STIM\_IFNAB\_CD8\_TCELL\_UP | 47 | 5 | 4161 | 156 | sina,Lk6,Pgant6,Spn42De,hppy | | 2.997e-02 | -3.51 | GATA6\_01 | MSigDB lists | GATA6\_01 | 47 | 5 | 4161 | 156 | Dgat2,Mef2,Ddr,rdx,Spn42De | | 2.997e-02 | -3.51 | NFKB\_Q6 | MSigDB lists | NFKB\_Q6 | 47 | 5 | 4161 | 156 | cact,trbl,Cdk4,Rgl,nmo | | 2.997e-02 | -3.51 | GSE37301\_CD4\_TCELL\_VS\_RAG2\_KO\_NK\_CELL\_DN | MSigDB lists | GSE37301\_CD4\_TCELL\_VS\_RAG2\_KO\_NK\_CELL\_DN | 47 | 5 | 4161 | 156 | CG4645,ATPCL,Pgant5,Dyb,Raf | | 2.999e-02 | -3.51 | GO\_CELL\_PART\_MORPHOGENESIS | MSigDB lists | GO\_CELL\_PART\_MORPHOGENESIS | 175 | 12 | 4161 | 156 | Unc-76,tkv,ap,N,sina,Rab1,unc-5,Eph,robo1,LanA,Ocrl,sli | | 3.013e-02 | -3.50 | synaptic target recognition | biological process | GO:0008039 | 49 | 5 | 12345 | 442 | sli,fz2,Pvf3,Gli,Sema1a | | 3.020e-02 | -3.50 | GSE15733\_BM\_VS\_SPLEEN\_MEMORY\_CD4\_TCELL\_UP | MSigDB lists | GSE15733\_BM\_VS\_SPLEEN\_MEMORY\_CD4\_TCELL\_UP | 32 | 4 | 4161 | 156 | Src42A,Cerk,CG9220,nAChRalpha4 | | 3.020e-02 | -3.50 | GSE22432\_MULTIPOTENT\_PROGENITOR\_VS\_CDC\_UP | MSigDB lists | GSE22432\_MULTIPOTENT\_PROGENITOR\_VS\_CDC\_UP | 32 | 4 | 4161 | 156 | Tollo,Psa,rg,Tpst | | 3.020e-02 | -3.50 | RAPA\_EARLY\_UP.V1\_UP | MSigDB lists | RAPA\_EARLY\_UP.V1\_UP | 32 | 4 | 4161 | 156 | ap,CtBP,alpha-Man-Ia,DPCoAC | | 3.020e-02 | -3.50 | GSE6674\_UNSTIM\_VS\_CPG\_STIM\_BCELL\_UP | MSigDB lists | GSE6674\_UNSTIM\_VS\_CPG\_STIM\_BCELL\_UP | 32 | 4 | 4161 | 156 | Tpc1,lin-28,ATPCL,CG6891 | | 3.020e-02 | -3.50 | GO\_NEGATIVE\_REGULATION\_OF\_BINDING | MSigDB lists | GO\_NEGATIVE\_REGULATION\_OF\_BINDING | 32 | 4 | 4161 | 156 | Pex14,Hel89B,cact,ap | | 3.020e-02 | -3.50 | GSE2405\_0H\_VS\_12H\_A\_PHAGOCYTOPHILUM\_STIM\_NEUTROPHIL\_UP | MSigDB lists | GSE2405\_0H\_VS\_12H\_A\_PHAGOCYTOPHILUM\_STIM\_NEUTROPHIL\_UP | 32 | 4 | 4161 | 156 | trbl,CG6891,jar,Pgant6 | | 3.020e-02 | -3.50 | GSE23925\_LIGHT\_ZONE\_VS\_NAIVE\_BCELL\_UP | MSigDB lists | GSE23925\_LIGHT\_ZONE\_VS\_NAIVE\_BCELL\_UP | 32 | 4 | 4161 | 156 | l(2)37Cg,CG1129,Drak,Rab1 | | 3.020e-02 | -3.50 | KRAS.KIDNEY\_UP.V1\_UP | MSigDB lists | KRAS.KIDNEY\_UP.V1\_UP | 32 | 4 | 4161 | 156 | trbl,betaTub60D,Mef2,Ric | | 3.020e-02 | -3.50 | GSE36392\_EOSINOPHIL\_VS\_MAC\_IL25\_TREATED\_LUNG\_UP | MSigDB lists | GSE36392\_EOSINOPHIL\_VS\_MAC\_IL25\_TREATED\_LUNG\_UP | 32 | 4 | 4161 | 156 | Gfat2,Rab1,FBXO11,Atg16 | | 3.020e-02 | -3.50 | GO\_CELLULAR\_RESPONSE\_TO\_BIOTIC\_STIMULUS | MSigDB lists | GO\_CELLULAR\_RESPONSE\_TO\_BIOTIC\_STIMULUS | 32 | 4 | 4161 | 156 | cact,Mef2,hyx,fz2 | | 3.033e-02 | -3.50 | SH2 | pfam domains | PF00017 | 30 | 4 | 10480 | 419 | Abl,Socs36E,dock,Src42A | | 3.038e-02 | -3.49 | chromosome organization | biological process | GO:0051276 | 680 | 34 | 12345 | 442 | Hel89B,Kdm2,eya,HmgZ,fl(2)d,E(Pc),hyx,His3.3B,Nlp,Rab5,crol,His3.3A,Dcr-2,Slbp,eff,CtBP,vig,His2Av,tna,Elba2,corto,FBXO11,chic,ph-p,nub,dve,Psc,Kdm4B,bin3,SMC3,tara,jumu,Su(z)2,Spc105R | | 3.044e-02 | -3.49 | fly chr35F1-35F1\|2-52 cM | chromosome location | fly chr35F1-35F1\|2-52 cM | 30 | 3 | 25010 | 571 | CG4935,cact,crp | | 3.044e-02 | -3.49 | fly chr35F1-35F1\|2-52 | chromosome location | fly chr35F1-35F1\|2-52 | 30 | 3 | 25010 | 571 | CG4935,cact,crp | | 3.047e-02 | -3.49 | protein ubiquitination | biological process | GO:0016567 | 245 | 15 | 12345 | 442 | Kdm2,FBXO11,rdx,Ubc87F,morgue,sina,CG12402,CG13481,Cbl,Pex10,Su(z)2,Fem-1,mura,Diap1,eff | | 3.057e-02 | -3.49 | MAPK1/MAPK3 signaling | REACTOME pathways | R-DME-5684996 | 114 | 10 | 3923 | 176 | PI31,htl,14-3-3zeta,CG4935,CG42684,wdb,vn,Raf,Egfr,Pvr | | 3.057e-02 | -3.49 | Signaling by Leptin | REACTOME pathways | R-DME-2586552 | 114 | 10 | 3923 | 176 | CG42684,wdb,PI31,htl,14-3-3zeta,CG4935,Pvr,vn,Raf,Egfr | | 3.057e-02 | -3.49 | Interleukin-2 signaling | REACTOME pathways | R-DME-451927 | 114 | 10 | 3923 | 176 | wdb,CG42684,CG4935,PI31,14-3-3zeta,htl,Pvr,Egfr,Raf,vn | | 3.066e-02 | -3.48 | Bap170 (Brahma associated protein 170kD) | protein interactions | 35560 | 15 | 3 | 9204 | 432 | Egfr,Raf,vn | | 3.066e-02 | -3.48 | Nup75 (Nucleoporin 75kD) | protein interactions | 37078 | 15 | 3 | 9204 | 432 | ph-p,cact,jar | | 3.066e-02 | -3.48 | bur (burgundy) | protein interactions | 45830 | 15 | 3 | 9204 | 432 | CG7824,sina,CtBP | | 3.066e-02 | -3.48 | Nedd4 (Nedd4) | protein interactions | 39958 | 15 | 3 | 9204 | 432 | N,cindr,Myo10A | | 3.066e-02 | -3.48 | kra (krasavietz) | protein interactions | 40680 | 15 | 3 | 9204 | 432 | robo1,shot,sli | | 3.066e-02 | -3.48 | slpr (slipper) | protein interactions | 44111 | 15 | 3 | 9204 | 432 | kay,Pvr,aop | | 3.066e-02 | -3.48 | Br140 (Bromodomain-containing protein, 140kD) | protein interactions | 35648 | 15 | 3 | 9204 | 432 | Psc,ph-p,Su(z)2 | | 3.066e-02 | -3.48 | slou (slouch) | protein interactions | 42547 | 15 | 3 | 9204 | 432 | Scr,abd-A,Abd-B | | 3.066e-02 | -3.48 | Sox21a (Sox21a) | protein interactions | 39567 | 15 | 3 | 9204 | 432 | spi,Egfr,kay | | 3.066e-02 | -3.48 | Pabp2 (Pabp2) | protein interactions | 35788 | 15 | 3 | 9204 | 432 | orb2,d,mam | | 3.066e-02 | -3.48 | Pop2 (Pop2) | protein interactions | 39366 | 15 | 3 | 9204 | 432 | Tob,Tis11,pum | | 3.066e-02 | -3.48 | PQBP1 (Poly-glutamine tract binding protein 1) | protein interactions | 43004 | 15 | 3 | 9204 | 432 | CG34376,hth,CG11247 | | 3.066e-02 | -3.48 | CadN (Cadherin-N) | protein interactions | 35070 | 15 | 3 | 9204 | 432 | siz,arm,shg | | 3.070e-02 | -3.48 | GO\_POSITIVE\_REGULATION\_OF\_PROTEIN\_METABOLIC\_PROCESS | MSigDB lists | GO\_POSITIVE\_REGULATION\_OF\_PROTEIN\_METABOLIC\_PROCESS | 366 | 21 | 4161 | 156 | cact,Raf,Mef2,hppy,trbl,dock,htl,Actbeta,Rab1,Tollo,unc-119,PI31,Ddr,nAChRalpha4,Tsp96F,Nrk,Pdk1,fz,wbl,arm,fz2 | | 3.085e-02 | -3.48 | GO\_GOLGI\_APPARATUS | MSigDB lists | GO\_GOLGI\_APPARATUS | 433 | 24 | 4161 | 156 | Cals,step,Pgant6,lt,dock,Pgant5,htl,Arf79F,N,alpha-Man-Ia,Rab1,dlp,GlcAT-I,Tollo,rg,AP-1gamma,CG17883,C1GalTA,Ocrl,CG9220,Tpst,jar,Nmda1,fz2 | | 3.102e-02 | -3.47 | GO\_REGULATION\_OF\_NEURON\_PROJECTION\_DEVELOPMENT | MSigDB lists | GO\_REGULATION\_OF\_NEURON\_PROJECTION\_DEVELOPMENT | 81 | 7 | 4161 | 156 | sli,Raf,robo1,Mef2,Arf79F,otk,nAChRalpha4 | | 3.102e-02 | -3.47 | SMTTTTGT\_UNKNOWN | MSigDB lists | SMTTTTGT\_UNKNOWN | 81 | 7 | 4161 | 156 | fz2,lqf,unc-5,ATPCL,Kdm2,14-3-3epsilon,Gfat2 | | 3.104e-02 | -3.47 | positive regulation of DNA metabolic process | biological process | GO:0051054 | 8 | 2 | 12345 | 442 | Tet,eya | | 3.104e-02 | -3.47 | dorsal trunk growth, open tracheal system | biological process | GO:0035001 | 8 | 2 | 12345 | 442 | ttk,LanA | | 3.104e-02 | -3.47 | protein localization to Golgi apparatus | biological process | GO:0034067 | 8 | 2 | 12345 | 442 | Vps13,Rab1 | | 3.104e-02 | -3.47 | regulation of striated muscle tissue development | biological process | GO:0016202 | 8 | 2 | 12345 | 442 | Abd-B,Mef2 | | 3.104e-02 | -3.47 | negative regulation of peptidyl-tyrosine phosphorylation | biological process | GO:0050732 | 8 | 2 | 12345 | 442 | Cbl,Socs36E | | 3.104e-02 | -3.47 | oenocyte development | biological process | GO:0007438 | 8 | 2 | 12345 | 442 | spi,abd-A | | 3.104e-02 | -3.47 | imaginal disc-derived female genitalia development | biological process | GO:0007486 | 8 | 2 | 12345 | 442 | otk,Abd-B | | 3.104e-02 | -3.47 | negative regulation of antimicrobial peptide production | biological process | GO:0002785 | 8 | 2 | 12345 | 442 | cact,nub | | 3.104e-02 | -3.47 | cell fate specification involved in pattern specification | biological process | GO:0060573 | 8 | 2 | 12345 | 442 | vn,Egfr | | 3.104e-02 | -3.47 | chondroitin sulfate metabolic process | biological process | GO:0030204 | 8 | 2 | 12345 | 442 | CG9220,GlcAT-I | | 3.104e-02 | -3.47 | progression of morphogenetic furrow involved in compound eye morphogenesis | biological process | GO:0007458 | 8 | 2 | 12345 | 442 | emc,chn | | 3.104e-02 | -3.47 | chondroitin sulfate biosynthetic process | biological process | GO:0030206 | 8 | 2 | 12345 | 442 | GlcAT-I,CG9220 | | 3.104e-02 | -3.47 | genital disc anterior/posterior pattern formation | biological process | GO:0035224 | 8 | 2 | 12345 | 442 | Abd-B,abd-A | | 3.104e-02 | -3.47 | imaginal disc-derived wing expansion | biological process | GO:0048526 | 8 | 2 | 12345 | 442 | disco-r,arm | | 3.104e-02 | -3.47 | establishment or maintenance of polarity of embryonic epithelium | biological process | GO:0016332 | 8 | 2 | 12345 | 442 | scrib,baz | | 3.104e-02 | -3.47 | anterior head development | biological process | GO:0097065 | 8 | 2 | 12345 | 442 | kn,hth | | 3.104e-02 | -3.47 | peptidyl-tyrosine autophosphorylation | biological process | GO:0038083 | 8 | 2 | 12345 | 442 | Src42A,Abl | | 3.104e-02 | -3.47 | genital disc pattern formation | biological process | GO:0035221 | 8 | 2 | 12345 | 442 | Abd-B,abd-A | | 3.104e-02 | -3.47 | DNA replication-independent nucleosome assembly | biological process | GO:0006336 | 8 | 2 | 12345 | 442 | His3.3B,His3.3A | | 3.104e-02 | -3.47 | regulation of muscle tissue development | biological process | GO:1901861 | 8 | 2 | 12345 | 442 | Mef2,Abd-B | | 3.104e-02 | -3.47 | positive regulation of apoptotic process involved in development | biological process | GO:1904747 | 8 | 2 | 12345 | 442 | Abd-B,Dp | | 3.104e-02 | -3.47 | periodic partitioning by pair rule gene | biological process | GO:0007366 | 8 | 2 | 12345 | 442 | h,lilli | | 3.104e-02 | -3.47 | cytokine-mediated signaling pathway | biological process | GO:0019221 | 8 | 2 | 12345 | 442 | dome,wgn | | 3.104e-02 | -3.47 | anterior head segmentation | biological process | GO:0035288 | 8 | 2 | 12345 | 442 | hth,kn | | 3.104e-02 | -3.47 | cellular response to cytokine stimulus | biological process | GO:0071345 | 8 | 2 | 12345 | 442 | wgn,dome | | 3.105e-02 | -3.47 | terminal bouton | cellular component | GO:0043195 | 68 | 6 | 11940 | 417 | Magi,ena,Syx1A,gukh,jeb,pum | | 3.116e-02 | -3.47 | ubiquitin protein ligase binding | molecular function | GO:0031625 | 52 | 5 | 12176 | 414 | Tom,Diap1,rdx,eff,morgue | | 3.118e-02 | -3.47 | Ubi-p63E (Ubiquitin-63E) | protein interactions | 38456 | 38 | 5 | 9204 | 432 | d,rdx,lqf,morgue,Dronc | | 3.125e-02 | -3.47 | regulation of response to biotic stimulus | biological process | GO:0002831 | 162 | 11 | 12345 | 442 | nub,Pitslre,pnr,CG32795,mop,imd,DPCoAC,cact,Rm62,Dcr-2,htl | | 3.130e-02 | -3.46 | intrinsic component of membrane | cellular component | GO:0031224 | 1460 | 64 | 11940 | 417 | fz,S,Ddr,SERCA,Dl,CG8034,tipE,Gr93c,Syx1A,ogre,shg,DPCoAC,Eph,unc-5,neo,ed,emp,spi,scrib,moody,drl,Irk3,Pex10,wgn,Orct2,InR,bou,rg,Inx2,tkv,ds,robo1,Kr-h2,htl,fz2,Sema1a,kek5,CG3149,fz3,CG7536,ZnT41F,glec,Gli,otk,Tsp96F,Esp,beat-IIa,nAChRalpha4,Ir21a,N,CG30344,rho-4,Nrk,CG32795,C1GalTA,Mpcp2,Egfr,Tpc1,dlp,Pvr,klar,ft,CG33474,Cals | | 3.137e-02 | -3.46 | plasma membrane bounded cell projection | cellular component | GO:0120025 | 626 | 31 | 11940 | 417 | Magi,Eph,pum,orb2,path,jeb,nAChRalpha4,shg,chic,Abl,CG9784,Myo10A,shot,Ptp4E,wnd,boi,arm,Bsg,jar,drl,Unc-76,RhoU,robo1,Syx1A,ena,Mob2,Gr93c,Egfr,tkv,gukh,InR | | 3.152e-02 | -3.46 | GSE17721\_LPS\_VS\_PAM3CSK4\_24H\_BMDC\_UP | MSigDB lists | GSE17721\_LPS\_VS\_PAM3CSK4\_24H\_BMDC\_UP | 64 | 6 | 4161 | 156 | Kr-h2,Tsp96F,Pgant6,rdx,Tpst,CG4896 | | 3.152e-02 | -3.46 | GSE360\_DC\_VS\_MAC\_UP | MSigDB lists | GSE360\_DC\_VS\_MAC\_UP | 64 | 6 | 4161 | 156 | SMC3,sina,Lk6,ATPCL,bru2,AP-1gamma | | 3.153e-02 | -3.46 | Ubi-p5E (Ubiquitin-5E) | protein interactions | 326237 | 26 | 4 | 9204 | 432 | rdx,d,Dronc,morgue | | 3.153e-02 | -3.46 | ftz (fushi tarazu) | protein interactions | 40834 | 26 | 4 | 9204 | 432 | hth,h,CG14506,tna | | 3.153e-02 | -3.46 | Rbp6 (RNA-binding protein 6) | protein interactions | 39919 | 26 | 4 | 9204 | 432 | CycG,orb2,Cdk4,yuri | | 3.153e-02 | -3.46 | Vang (Van Gogh) | protein interactions | 35922 | 26 | 4 | 9204 | 432 | shg,nmo,fz,scrib | | 3.164e-02 | -3.45 | PI Metabolism | REACTOME pathways | R-DME-1483255 | 40 | 5 | 3923 | 176 | CG34383,Arf79F,Rab5,Ocrl,CG9784 | | 3.167e-02 | -3.45 | GTPase regulator activity | molecular function | GO:0030695 | 89 | 7 | 12176 | 414 | RapGAP1,CenB1A,CG17883,RhoGAP19D,Ocrl,drongo,TBC1D5 | | 3.177e-02 | -3.45 | MODULE\_11 | MSigDB lists | MODULE\_11 | 118 | 9 | 4161 | 156 | sli,Syx1A,CG5522,Gfat2,Mef2,Dyb,bru2,htl,ap | | 3.177e-02 | -3.45 | RAO\_BOUND\_BY\_SALL4\_ISOFORM\_B | MSigDB lists | RAO\_BOUND\_BY\_SALL4\_ISOFORM\_B | 118 | 9 | 4161 | 156 | Fem-1,hppy,arm,l(1)G0196,shot,CG45050,Rim2,dock,sd | | 3.191e-02 | -3.44 | lwr (lesswright) | protein interactions | 33226 | 65 | 7 | 9204 | 432 | tna,jumu,sina,ttk,HLH3B,cact,orb2 | | 3.197e-02 | -3.44 | basolateral plasma membrane | cellular component | GO:0016323 | 51 | 5 | 11940 | 417 | yrt,Egfr,ogre,Dg,scrib | | 3.208e-02 | -3.44 | PEREZ\_TP53\_TARGETS | MSigDB lists | PEREZ\_TP53\_TARGETS | 197 | 13 | 4161 | 156 | jar,bru2,Kdm2,lili,dlp,InR,htl,CG2812,fz,Ziz,Src42A,Pgant6,unc-119 | | 3.219e-02 | -3.44 | Cyclin-like | interpro domains | IPR013763 | 18 | 3 | 11383 | 450 | CycG,CG6191,CycB3 | | 3.220e-02 | -3.44 | positive regulation of Notch signaling pathway | biological process | GO:0045747 | 34 | 4 | 12345 | 442 | spdo,lqf,ed,Fkbp14 | | 3.222e-02 | -3.44 | NCAM signaling for neurite out-growth | REACTOME pathways | R-DME-375165 | 115 | 10 | 3923 | 176 | CG4935,htl,PI31,14-3-3zeta,wdb,CG42684,Egfr,Raf,vn,Pvr | | 3.222e-02 | -3.44 | PID\_FOXO\_PATHWAY | MSigDB lists | PID\_FOXO\_PATHWAY | 19 | 3 | 4161 | 156 | 14-3-3zeta,14-3-3epsilon,arm | | 3.222e-02 | -3.44 | GO\_REGULATION\_OF\_NOTCH\_SIGNALING\_PATHWAY | MSigDB lists | GO\_REGULATION\_OF\_NOTCH\_SIGNALING\_PATHWAY | 19 | 3 | 4161 | 156 | N,cact,Dl | | 3.222e-02 | -3.44 | BEGUM\_TARGETS\_OF\_PAX3\_FOXO1\_FUSION\_UP | MSigDB lists | BEGUM\_TARGETS\_OF\_PAX3\_FOXO1\_FUSION\_UP | 19 | 3 | 4161 | 156 | Drak,sli,Cals | | 3.222e-02 | -3.44 | GO\_GLAND\_MORPHOGENESIS | MSigDB lists | GO\_GLAND\_MORPHOGENESIS | 19 | 3 | 4161 | 156 | N,LanA,Fem-1 | | 3.222e-02 | -3.44 | NABA\_ECM\_GLYCOPROTEINS | MSigDB lists | NABA\_ECM\_GLYCOPROTEINS | 19 | 3 | 4161 | 156 | LanA,sli,Ppn | | 3.222e-02 | -3.44 | GO\_REGULATION\_OF\_CHEMOTAXIS | MSigDB lists | GO\_REGULATION\_OF\_CHEMOTAXIS | 19 | 3 | 4161 | 156 | robo1,N,sli | | 3.222e-02 | -3.44 | MIKKELSEN\_MCV6\_LCP\_WITH\_H3K4ME3 | MSigDB lists | MIKKELSEN\_MCV6\_LCP\_WITH\_H3K4ME3 | 19 | 3 | 4161 | 156 | CG31522,CG4896,CG45050 | | 3.222e-02 | -3.44 | REACTOME\_TRIGLYCERIDE\_BIOSYNTHESIS | MSigDB lists | REACTOME\_TRIGLYCERIDE\_BIOSYNTHESIS | 19 | 3 | 4161 | 156 | CG3961,ATPCL,CG31522 | | 3.222e-02 | -3.44 | GO\_HEMATOPOIETIC\_PROGENITOR\_CELL\_DIFFERENTIATION | MSigDB lists | GO\_HEMATOPOIETIC\_PROGENITOR\_CELL\_DIFFERENTIATION | 19 | 3 | 4161 | 156 | Actbeta,Raf,Cdk4 | | 3.222e-02 | -3.44 | MORI\_SMALL\_PRE\_BII\_LYMPHOCYTE\_UP | MSigDB lists | MORI\_SMALL\_PRE\_BII\_LYMPHOCYTE\_UP | 19 | 3 | 4161 | 156 | C1GalTA,Cerk,wbl | | 3.222e-02 | -3.44 | WIELAND\_UP\_BY\_HBV\_INFECTION | MSigDB lists | WIELAND\_UP\_BY\_HBV\_INFECTION | 19 | 3 | 4161 | 156 | Pen,CG6891,Rgl | | 3.222e-02 | -3.44 | REACTOME\_HEPARAN\_SULFATE\_HEPARIN\_HS\_GAG\_METABOLISM | MSigDB lists | REACTOME\_HEPARAN\_SULFATE\_HEPARIN\_HS\_GAG\_METABOLISM | 19 | 3 | 4161 | 156 | dlp,GlcAT-I,Ids | | 3.222e-02 | -3.44 | AAGGGAT\_MIR188 | MSigDB lists | AAGGGAT\_MIR188 | 19 | 3 | 4161 | 156 | rg,Mef2,FBXO11 | | 3.222e-02 | -3.44 | GO\_BASEMENT\_MEMBRANE | MSigDB lists | GO\_BASEMENT\_MEMBRANE | 19 | 3 | 4161 | 156 | SMC3,LanA,shot | | 3.230e-02 | -3.43 | Negative regulation of MET activity | REACTOME pathways | R-DME-6807004 | 16 | 3 | 3923 | 176 | Cbl,Ptp4E,cindr | | 3.237e-02 | -3.43 | asymmetric stem cell division | biological process | GO:0098722 | 67 | 6 | 12345 | 442 | flfl,InR,esg,eff,jar,baz | | 3.237e-02 | -3.43 | multicellular organismal homeostasis | biological process | GO:0048871 | 67 | 6 | 12345 | 442 | Egfr,Atg17,shg,yrt,N,Atg16 | | 3.237e-02 | -3.43 | regulation of antimicrobial humoral response | biological process | GO:0002759 | 67 | 6 | 12345 | 442 | nub,pnr,Pitslre,imd,cact,mop | | 3.240e-02 | -3.43 | Glycosaminoglycan metabolism | REACTOME pathways | R-DME-1630316 | 54 | 6 | 3923 | 176 | GlcAT-I,kon,frc,dlp,Ids,CG9220 | | 3.242e-02 | -3.43 | regulation of secretion by cell | biological process | GO:1903530 | 85 | 7 | 12345 | 442 | Syx1A,Rab1,InR,Cadps,S,mop,jar | | 3.247e-02 | -3.43 | GSE26023\_PHD3\_KO\_VS\_WT\_NEUTROPHIL\_HYPOXIA\_UP | MSigDB lists | GSE26023\_PHD3\_KO\_VS\_WT\_NEUTROPHIL\_HYPOXIA\_UP | 48 | 5 | 4161 | 156 | fz2,fz,Nrk,Eph,CG1673 | | 3.247e-02 | -3.43 | GSE18893\_CTRL\_VS\_TNF\_TREATED\_TCONV\_24H\_DN | MSigDB lists | GSE18893\_CTRL\_VS\_TNF\_TREATED\_TCONV\_24H\_DN | 48 | 5 | 4161 | 156 | Atg16,Dgat2,FBXO11,fz2,CG13384 | | 3.247e-02 | -3.43 | CREL\_01 | MSigDB lists | CREL\_01 | 48 | 5 | 4161 | 156 | nmo,14-3-3zeta,Cdk4,cact,trbl | | 3.247e-02 | -3.43 | GSE10325\_MYELOID\_VS\_LUPUS\_MYELOID\_DN | MSigDB lists | GSE10325\_MYELOID\_VS\_LUPUS\_MYELOID\_DN | 48 | 5 | 4161 | 156 | hyx,CG2017,Nmda1,Rgl,PI31 | | 3.247e-02 | -3.43 | GO\_CLATHRIN\_COATED\_VESICLE | MSigDB lists | GO\_CLATHRIN\_COATED\_VESICLE | 48 | 5 | 4161 | 156 | lt,fz2,Ocrl,jar,AP-1gamma | | 3.247e-02 | -3.43 | GSE22611\_MUTANT\_NOD2\_VS\_CTRL\_TRANSDUCED\_HEK293T\_CELL\_DN | MSigDB lists | GSE22611\_MUTANT\_NOD2\_VS\_CTRL\_TRANSDUCED\_HEK293T\_CELL\_DN | 48 | 5 | 4161 | 156 | Drep4,14-3-3zeta,FBXO11,unc-5,CG2006 | | 3.247e-02 | -3.43 | GO\_RESPONSE\_TO\_MOLECULE\_OF\_BACTERIAL\_ORIGIN | MSigDB lists | GO\_RESPONSE\_TO\_MOLECULE\_OF\_BACTERIAL\_ORIGIN | 48 | 5 | 4161 | 156 | Mef2,hyx,N,cact,fz2 | | 3.247e-02 | -3.43 | SMAD\_Q6 | MSigDB lists | SMAD\_Q6 | 48 | 5 | 4161 | 156 | ds,rdx,Cadps,Rab5,sd | | 3.247e-02 | -3.43 | GSE22886\_NAIVE\_CD4\_TCELL\_VS\_DC\_UP | MSigDB lists | GSE22886\_NAIVE\_CD4\_TCELL\_VS\_DC\_UP | 48 | 5 | 4161 | 156 | Drak,Ziz,step,CycG,Atg16 | | 3.254e-02 | -3.43 | establishment or maintenance of cytoskeleton polarity | biological process | GO:0030952 | 50 | 5 | 12345 | 442 | 14-3-3epsilon,Shrm,rictor,baz,14-3-3zeta | | 3.271e-02 | -3.42 | EGF | smart domains | SM00181 | 66 | 8 | 5086 | 290 | dpy,ft,N,Dl,spi,LanA,sli,cue | | 3.276e-02 | -3.42 | GO\_NEGATIVE\_REGULATION\_OF\_TRANSPORT | MSigDB lists | GO\_NEGATIVE\_REGULATION\_OF\_TRANSPORT | 100 | 8 | 4161 | 156 | cact,unc-119,CG11069,Raf,fz,Actbeta,N,wbl | | 3.305e-02 | -3.41 | somatic sex determination | biological process | GO:0018993 | 20 | 3 | 12345 | 442 | fl(2)d,Sxl,emc | | 3.305e-02 | -3.41 | regulation of cytokinesis | biological process | GO:0032465 | 20 | 3 | 12345 | 442 | cindr,Ocrl,nuf | | 3.305e-02 | -3.41 | negative regulation of cell growth | biological process | GO:0030308 | 20 | 3 | 12345 | 442 | Sema1b,cic,Sema1a | | 3.305e-02 | -3.41 | regulation of hemocyte differentiation | biological process | GO:0045610 | 20 | 3 | 12345 | 442 | cact,kn,N | | 3.305e-02 | -3.41 | chitin-based larval cuticle pattern formation | biological process | GO:0035293 | 20 | 3 | 12345 | 442 | arm,fz2,fz | | 3.305e-02 | -3.41 | regulation of defense response to virus | biological process | GO:0050688 | 20 | 3 | 12345 | 442 | imd,Rm62,Dcr-2 | | 3.305e-02 | -3.41 | mesodermal cell fate commitment | biological process | GO:0001710 | 20 | 3 | 12345 | 442 | htl,abd-A,mam | | 3.305e-02 | -3.41 | mesodermal cell differentiation | biological process | GO:0048333 | 20 | 3 | 12345 | 442 | abd-A,htl,mam | | 3.305e-02 | -3.41 | mesodermal cell migration | biological process | GO:0008078 | 20 | 3 | 12345 | 442 | sli,jeb,htl | | 3.305e-02 | -3.41 | cellular carbohydrate catabolic process | biological process | GO:0044275 | 20 | 3 | 12345 | 442 | Atg17,CG9485,Atg16 | | 3.313e-02 | -3.41 | F-box-like | pfam domains | PF12937 | 18 | 3 | 10480 | 419 | morgue,Kdm2,FBXO11 | | 3.317e-02 | -3.41 | CADHERIN\_1 | prosite domains | PS00232 | 14 | 3 | 6243 | 325 | ft,shg,ds | | 3.317e-02 | -3.41 | MYOSIN\_MOTOR | prosite domains | PS51456 | 14 | 3 | 6243 | 325 | d,Myo10A,jar | | 3.317e-02 | -3.41 | cell projection | cellular component | GO:0042995 | 629 | 31 | 11940 | 417 | Egfr,Gr93c,Mob2,RhoU,ena,Syx1A,robo1,InR,gukh,tkv,Bsg,jar,arm,wnd,boi,drl,Unc-76,CG9784,Abl,chic,Ptp4E,shot,Myo10A,Eph,Magi,shg,nAChRalpha4,orb2,jeb,path,pum | | 3.327e-02 | -3.40 | SH3DOMAIN | prints domains | PR00452 | 27 | 4 | 2185 | 101 | cindr,dock,Src42A,Abl | | 3.330e-02 | -3.40 | neuronal cell body | cellular component | GO:0043025 | 147 | 10 | 11940 | 417 | bol,path,Rab5,Ptp4E,Myo10A,rg,robo1,CG9784,Rab1,Gr93c | | 3.339e-02 | -3.40 | GO\_REGULATION\_OF\_MUSCLE\_CELL\_DIFFERENTIATION | MSigDB lists | GO\_REGULATION\_OF\_MUSCLE\_CELL\_DIFFERENTIATION | 33 | 4 | 4161 | 156 | Dl,Mef2,N,arm | | 3.339e-02 | -3.40 | GO\_NEGATIVE\_REGULATION\_OF\_SECRETION | MSigDB lists | GO\_NEGATIVE\_REGULATION\_OF\_SECRETION | 33 | 4 | 4161 | 156 | wbl,Actbeta,Raf,N | | 3.339e-02 | -3.40 | GSE360\_CTRL\_VS\_L\_DONOVANI\_DC\_DN | MSigDB lists | GSE360\_CTRL\_VS\_L\_DONOVANI\_DC\_DN | 33 | 4 | 4161 | 156 | emp,Rgl,sina,cact | | 3.339e-02 | -3.40 | ICSBP\_Q6 | MSigDB lists | ICSBP\_Q6 | 33 | 4 | 4161 | 156 | lqf,Hel89B,Kdm2,Cdk4 | | 3.339e-02 | -3.40 | GSE20715\_0H\_VS\_48H\_OZONE\_TLR4\_KO\_LUNG\_UP | MSigDB lists | GSE20715\_0H\_VS\_48H\_OZONE\_TLR4\_KO\_LUNG\_UP | 33 | 4 | 4161 | 156 | Mef2,lili,Drak,Tpst | | 3.339e-02 | -3.40 | GSE30971\_CTRL\_VS\_LPS\_STIM\_MACROPHAGE\_WBP7\_KO\_2H\_UP | MSigDB lists | GSE30971\_CTRL\_VS\_LPS\_STIM\_MACROPHAGE\_WBP7\_KO\_2H\_UP | 33 | 4 | 4161 | 156 | cact,Dl,CG1673,CG3961 | | 3.339e-02 | -3.40 | GSE25085\_FETAL\_LIVER\_VS\_FETAL\_BM\_SP4\_THYMIC\_IMPLANT\_DN | MSigDB lists | GSE25085\_FETAL\_LIVER\_VS\_FETAL\_BM\_SP4\_THYMIC\_IMPLANT\_DN | 33 | 4 | 4161 | 156 | Tpst,Actbeta,nAChRalpha4,Dyb | | 3.339e-02 | -3.40 | GO\_EARLY\_ENDOSOME\_MEMBRANE | MSigDB lists | GO\_EARLY\_ENDOSOME\_MEMBRANE | 33 | 4 | 4161 | 156 | Eph,Rab5,Ocrl,fz2 | | 3.339e-02 | -3.40 | GSE23114\_PERITONEAL\_CAVITY\_B1A\_BCELL\_VS\_SPLEEN\_BCELL\_IN\_SLE2C1\_MOUSE\_UP | MSigDB lists | GSE23114\_PERITONEAL\_CAVITY\_B1A\_BCELL\_VS\_SPLEEN\_BCELL\_IN\_SLE2C1\_MOUSE\_UP | 33 | 4 | 4161 | 156 | Ocrl,jar,Cadps,Kdm2 | | 3.339e-02 | -3.40 | HALLMARK\_UV\_RESPONSE\_DN | MSigDB lists | HALLMARK\_UV\_RESPONSE\_DN | 33 | 4 | 4161 | 156 | InR,rdx,Pex14,bru2 | | 3.355e-02 | -3.39 | HMG | smart domains | SM00398 | 22 | 4 | 5086 | 290 | Sox14,cic,CG12104,HmgZ | | 3.369e-02 | -3.39 | GO\_REGULATION\_OF\_CELL\_MORPHOGENESIS\_INVOLVED\_IN\_DIFFERENTIATION | MSigDB lists | GO\_REGULATION\_OF\_CELL\_MORPHOGENESIS\_INVOLVED\_IN\_DIFFERENTIATION | 65 | 6 | 4161 | 156 | nAChRalpha4,robo1,N,Raf,sli,arm | | 3.369e-02 | -3.39 | MODULE\_2 | MSigDB lists | MODULE\_2 | 65 | 6 | 4161 | 156 | sli,LanA,shot,Cadps,Rgl,trbl | | 3.369e-02 | -3.39 | ZF5\_B | MSigDB lists | ZF5\_B | 65 | 6 | 4161 | 156 | htl,trbl,ap,nmo,CG4896,shot | | 3.369e-02 | -3.39 | TAGCTTT\_MIR9 | MSigDB lists | TAGCTTT\_MIR9 | 65 | 6 | 4161 | 156 | bru2,nmo,Mef2,Vps13,14-3-3zeta,sd | | 3.369e-02 | -3.39 | GSE20198\_UNTREATED\_VS\_IL12\_IL18\_TREATED\_ACT\_CD4\_TCELL\_DN | MSigDB lists | GSE20198\_UNTREATED\_VS\_IL12\_IL18\_TREATED\_ACT\_CD4\_TCELL\_DN | 65 | 6 | 4161 | 156 | rg,nmo,Rab5,CG11030,alpha-Man-Ia,CycG | | 3.369e-02 | -3.39 | GSE17721\_CTRL\_VS\_PAM3CSK4\_8H\_BMDC\_DN | MSigDB lists | GSE17721\_CTRL\_VS\_PAM3CSK4\_8H\_BMDC\_DN | 65 | 6 | 4161 | 156 | Gfat2,Rgl,Dyb,nAChRalpha4,N,Raf | | 3.369e-02 | -3.39 | GSE20152\_SPHK1\_KO\_VS\_WT\_HTNFA\_OVERXPRESS\_ANKLE\_DN | MSigDB lists | GSE20152\_SPHK1\_KO\_VS\_WT\_HTNFA\_OVERXPRESS\_ANKLE\_DN | 65 | 6 | 4161 | 156 | SMC3,Pgant6,Ppcs,muc,Pex14,CG4645 | | 3.369e-02 | -3.39 | GSE20500\_CTRL\_VS\_RETINOIC\_ACID\_TREATED\_CD4\_TCELL\_UP | MSigDB lists | GSE20500\_CTRL\_VS\_RETINOIC\_ACID\_TREATED\_CD4\_TCELL\_UP | 65 | 6 | 4161 | 156 | wbl,Pex7,CG4645,Ppcs,Pex14,Unc-76 | | 3.370e-02 | -3.39 | GO\_ORGAN\_FORMATION | MSigDB lists | GO\_ORGAN\_FORMATION | 8 | 2 | 4161 | 156 | Mef2,arm | | 3.370e-02 | -3.39 | MA\_PITUITARY\_FETAL\_VS\_ADULT\_UP | MSigDB lists | MA\_PITUITARY\_FETAL\_VS\_ADULT\_UP | 8 | 2 | 4161 | 156 | CycG,arm | | 3.370e-02 | -3.39 | BIOCARTA\_RACCYCD\_PATHWAY | MSigDB lists | BIOCARTA\_RACCYCD\_PATHWAY | 8 | 2 | 4161 | 156 | Cdk4,cact | | 3.370e-02 | -3.39 | GO\_CRANIAL\_NERVE\_DEVELOPMENT | MSigDB lists | GO\_CRANIAL\_NERVE\_DEVELOPMENT | 8 | 2 | 4161 | 156 | arm,Eph | | 3.370e-02 | -3.39 | GO\_REGULATION\_OF\_PROTEIN\_INSERTION\_INTO\_MITOCHONDRIAL\_MEMBRANE\_INVOLVED\_IN\_APOPTOTIC\_SIGNALING\_PATHWAY | MSigDB lists | GO\_REGULATION\_OF\_PROTEIN\_INSERTION\_INTO\_MITOCHONDRIAL\_MEMBRANE\_INVOLVED\_IN\_APOPTOTIC\_SIGNALING\_PATHWAY | 8 | 2 | 4161 | 156 | 14-3-3epsilon,14-3-3zeta | | 3.370e-02 | -3.39 | MODULE\_563 | MSigDB lists | MODULE\_563 | 8 | 2 | 4161 | 156 | Syx1A,Cadps | | 3.370e-02 | -3.39 | REACTOME\_ACTIVATION\_OF\_RAC | MSigDB lists | REACTOME\_ACTIVATION\_OF\_RAC | 8 | 2 | 4161 | 156 | sli,dock | | 3.370e-02 | -3.39 | GO\_NEGATIVE\_REGULATION\_OF\_EXOCYTOSIS | MSigDB lists | GO\_NEGATIVE\_REGULATION\_OF\_EXOCYTOSIS | 8 | 2 | 4161 | 156 | Raf,N | | 3.370e-02 | -3.39 | PID\_WNT\_SIGNALING\_PATHWAY | MSigDB lists | PID\_WNT\_SIGNALING\_PATHWAY | 8 | 2 | 4161 | 156 | fz2,fz | | 3.370e-02 | -3.39 | REACTOME\_SYNTHESIS\_OF\_VERY\_LONG\_CHAIN\_FATTY\_ACYL\_COAS | MSigDB lists | REACTOME\_SYNTHESIS\_OF\_VERY\_LONG\_CHAIN\_FATTY\_ACYL\_COAS | 8 | 2 | 4161 | 156 | CG31522,CG3961 | | 3.370e-02 | -3.39 | GO\_REGULATION\_OF\_TOLL\_LIKE\_RECEPTOR\_SIGNALING\_PATHWAY | MSigDB lists | GO\_REGULATION\_OF\_TOLL\_LIKE\_RECEPTOR\_SIGNALING\_PATHWAY | 8 | 2 | 4161 | 156 | Tollo,Pdk1 | | 3.370e-02 | -3.39 | PID\_ARF6\_PATHWAY | MSigDB lists | PID\_ARF6\_PATHWAY | 8 | 2 | 4161 | 156 | CenB1A,dock | | 3.370e-02 | -3.39 | GO\_ENDOTHELIAL\_CELL\_DEVELOPMENT | MSigDB lists | GO\_ENDOTHELIAL\_CELL\_DEVELOPMENT | 8 | 2 | 4161 | 156 | arm,Rab1 | | 3.370e-02 | -3.39 | GO\_REGULATION\_OF\_ERYTHROCYTE\_DIFFERENTIATION | MSigDB lists | GO\_REGULATION\_OF\_ERYTHROCYTE\_DIFFERENTIATION | 8 | 2 | 4161 | 156 | Actbeta,Cdk4 | | 3.370e-02 | -3.39 | GO\_DEVELOPMENTAL\_PIGMENTATION | MSigDB lists | GO\_DEVELOPMENTAL\_PIGMENTATION | 8 | 2 | 4161 | 156 | AP-1gamma,Mef2 | | 3.370e-02 | -3.39 | PID\_INTEGRIN1\_PATHWAY | MSigDB lists | PID\_INTEGRIN1\_PATHWAY | 8 | 2 | 4161 | 156 | Tsp96F,LanA | | 3.370e-02 | -3.39 | ZHENG\_GLIOBLASTOMA\_PLASTICITY\_DN | MSigDB lists | ZHENG\_GLIOBLASTOMA\_PLASTICITY\_DN | 8 | 2 | 4161 | 156 | Eph,Cadps | | 3.370e-02 | -3.39 | GO\_CENTRAL\_NERVOUS\_SYSTEM\_NEURON\_AXONOGENESIS | MSigDB lists | GO\_CENTRAL\_NERVOUS\_SYSTEM\_NEURON\_AXONOGENESIS | 8 | 2 | 4161 | 156 | Eph,sli | | 3.370e-02 | -3.39 | GO\_ESTABLISHMENT\_OR\_MAINTENANCE\_OF\_EPITHELIAL\_CELL\_APICAL\_BASAL\_POLARITY | MSigDB lists | GO\_ESTABLISHMENT\_OR\_MAINTENANCE\_OF\_EPITHELIAL\_CELL\_APICAL\_BASAL\_POLARITY | 8 | 2 | 4161 | 156 | otk,ap | | 3.370e-02 | -3.39 | GO\_POSITIVE\_REGULATION\_OF\_MITOCHONDRIAL\_OUTER\_MEMBRANE\_PERMEABILIZATION\_INVOLVED\_IN\_APOPTOTIC\_SIGNALING\_PATHWAY | MSigDB lists | GO\_POSITIVE\_REGULATION\_OF\_MITOCHONDRIAL\_OUTER\_MEMBRANE\_PERMEABILIZATION\_INVOLVED\_IN\_APOPTOTIC\_SIGNALING\_PATHWAY | 8 | 2 | 4161 | 156 | 14-3-3zeta,14-3-3epsilon | | 3.370e-02 | -3.39 | GO\_NEURONAL\_POSTSYNAPTIC\_DENSITY | MSigDB lists | GO\_NEURONAL\_POSTSYNAPTIC\_DENSITY | 8 | 2 | 4161 | 156 | Cals,Pdk1 | | 3.370e-02 | -3.39 | KEGG\_GLYCOSAMINOGLYCAN\_BIOSYNTHESIS\_CHONDROITIN\_SULFATE | MSigDB lists | KEGG\_GLYCOSAMINOGLYCAN\_BIOSYNTHESIS\_CHONDROITIN\_SULFATE | 8 | 2 | 4161 | 156 | GlcAT-I,CG9220 | | 3.370e-02 | -3.39 | GO\_POSITIVE\_REGULATION\_OF\_LYMPHOCYTE\_MEDIATED\_IMMUNITY | MSigDB lists | GO\_POSITIVE\_REGULATION\_OF\_LYMPHOCYTE\_MEDIATED\_IMMUNITY | 8 | 2 | 4161 | 156 | AP-1gamma,fz2 | | 3.370e-02 | -3.39 | MUELLER\_COMMON\_TARGETS\_OF\_AML\_FUSIONS\_DN | MSigDB lists | MUELLER\_COMMON\_TARGETS\_OF\_AML\_FUSIONS\_DN | 8 | 2 | 4161 | 156 | ds,Lk6 | | 3.370e-02 | -3.39 | GO\_REGULATION\_OF\_CARDIAC\_MUSCLE\_CELL\_PROLIFERATION | MSigDB lists | GO\_REGULATION\_OF\_CARDIAC\_MUSCLE\_CELL\_PROLIFERATION | 8 | 2 | 4161 | 156 | N,Mef2 | | 3.370e-02 | -3.39 | GO\_REGULATION\_OF\_CHONDROCYTE\_DIFFERENTIATION | MSigDB lists | GO\_REGULATION\_OF\_CHONDROCYTE\_DIFFERENTIATION | 8 | 2 | 4161 | 156 | arm,tkv | | 3.370e-02 | -3.39 | GO\_NEGATIVE\_REGULATION\_OF\_GLIOGENESIS | MSigDB lists | GO\_NEGATIVE\_REGULATION\_OF\_GLIOGENESIS | 8 | 2 | 4161 | 156 | N,arm | | 3.370e-02 | -3.39 | SYED\_ESTRADIOL\_RESPONSE | MSigDB lists | SYED\_ESTRADIOL\_RESPONSE | 8 | 2 | 4161 | 156 | wdb,Pen | | 3.370e-02 | -3.39 | GO\_ENDOCARDIAL\_CUSHION\_DEVELOPMENT | MSigDB lists | GO\_ENDOCARDIAL\_CUSHION\_DEVELOPMENT | 8 | 2 | 4161 | 156 | N,ds | | 3.370e-02 | -3.39 | GO\_EMBRYONIC\_AXIS\_SPECIFICATION | MSigDB lists | GO\_EMBRYONIC\_AXIS\_SPECIFICATION | 8 | 2 | 4161 | 156 | arm,fz2 | | 3.370e-02 | -3.39 | GO\_WNT\_PROTEIN\_BINDING | MSigDB lists | GO\_WNT\_PROTEIN\_BINDING | 8 | 2 | 4161 | 156 | fz,fz2 | | 3.370e-02 | -3.39 | GO\_POSITIVE\_REGULATION\_OF\_NEURAL\_PRECURSOR\_CELL\_PROLIFERATION | MSigDB lists | GO\_POSITIVE\_REGULATION\_OF\_NEURAL\_PRECURSOR\_CELL\_PROLIFERATION | 8 | 2 | 4161 | 156 | N,arm | | 3.370e-02 | -3.39 | GO\_POSITIVE\_REGULATION\_OF\_CARDIAC\_MUSCLE\_CELL\_PROLIFERATION | MSigDB lists | GO\_POSITIVE\_REGULATION\_OF\_CARDIAC\_MUSCLE\_CELL\_PROLIFERATION | 8 | 2 | 4161 | 156 | N,Mef2 | | 3.370e-02 | -3.39 | MODULE\_121 | MSigDB lists | MODULE\_121 | 8 | 2 | 4161 | 156 | CG1673,Tsp96F | | 3.370e-02 | -3.39 | GO\_REGULATION\_OF\_OLIGODENDROCYTE\_DIFFERENTIATION | MSigDB lists | GO\_REGULATION\_OF\_OLIGODENDROCYTE\_DIFFERENTIATION | 8 | 2 | 4161 | 156 | N,arm | | 3.370e-02 | -3.39 | GO\_REGULATION\_OF\_HEART\_MORPHOGENESIS | MSigDB lists | GO\_REGULATION\_OF\_HEART\_MORPHOGENESIS | 8 | 2 | 4161 | 156 | arm,N | | 3.370e-02 | -3.39 | PID\_BETA\_CATENIN\_DEG\_PATHWAY | MSigDB lists | PID\_BETA\_CATENIN\_DEG\_PATHWAY | 8 | 2 | 4161 | 156 | fz2,arm | | 3.370e-02 | -3.39 | chr1p33 | MSigDB lists | chr1p33 | 8 | 2 | 4161 | 156 | Lk6,CG4645 | | 3.370e-02 | -3.39 | GO\_NEGATIVE\_REGULATION\_OF\_REGULATED\_SECRETORY\_PATHWAY | MSigDB lists | GO\_NEGATIVE\_REGULATION\_OF\_REGULATED\_SECRETORY\_PATHWAY | 8 | 2 | 4161 | 156 | Raf,N | | 3.371e-02 | -3.39 | MADF | prosite domains | PS51029 | 47 | 6 | 6243 | 325 | rgr,jigr1,CG13204,CG3386,hng3,Adf1 | | 3.377e-02 | -3.39 | nucleoside-triphosphatase regulator activity | molecular function | GO:0060589 | 110 | 8 | 12176 | 414 | CenB1A,RapGAP1,Ocrl,CG7394,CG17883,TBC1D5,drongo,RhoGAP19D | | 3.392e-02 | -3.38 | p115 (p115) | protein interactions | 31698 | 52 | 6 | 9204 | 432 | eyg,shot,Dcr-2,Mob2,CG31211,CG34383 | | 3.400e-02 | -3.38 | amylo-alpha-1,6-glucosidase activity | molecular function | GO:0004135 | 1 | 1 | 12176 | 414 | CG9485 | | 3.400e-02 | -3.38 | peroxisome targeting sequence binding | molecular function | GO:0000268 | 1 | 1 | 12176 | 414 | Pex5 | | 3.400e-02 | -3.38 | S-acetyltransferase activity | molecular function | GO:0016418 | 1 | 1 | 12176 | 414 | muc | | 3.400e-02 | -3.38 | transforming growth factor beta binding | molecular function | GO:0050431 | 1 | 1 | 12176 | 414 | tkv | | 3.400e-02 | -3.38 | NEDD8 ligase activity | molecular function | GO:0061663 | 1 | 1 | 12176 | 414 | Diap1 | | 3.400e-02 | -3.38 | Golgi transport complex binding | molecular function | GO:0140164 | 1 | 1 | 12176 | 414 | Rab1 | | 3.400e-02 | -3.38 | myosin VI light chain binding | molecular function | GO:0070856 | 1 | 1 | 12176 | 414 | jar | | 3.400e-02 | -3.38 | peroxisome matrix targeting signal-1 binding | molecular function | GO:0005052 | 1 | 1 | 12176 | 414 | Pex5 | | 3.400e-02 | -3.38 | epidermal growth factor-activated receptor activity | molecular function | GO:0005006 | 1 | 1 | 12176 | 414 | Egfr | | 3.400e-02 | -3.38 | protein kinase C inhibitor activity | molecular function | GO:0008426 | 1 | 1 | 12176 | 414 | 14-3-3zeta | | 3.400e-02 | -3.38 | vascular endothelial growth factor-activated receptor activity | molecular function | GO:0005021 | 1 | 1 | 12176 | 414 | Pvr | | 3.400e-02 | -3.38 | mechanosensitive cation channel activity | molecular function | GO:0140135 | 1 | 1 | 12176 | 414 | Mid1 | | 3.400e-02 | -3.38 | alpha-catenin binding | molecular function | GO:0045294 | 1 | 1 | 12176 | 414 | arm | | 3.400e-02 | -3.38 | glycogen debranching enzyme activity | molecular function | GO:0004133 | 1 | 1 | 12176 | 414 | CG9485 | | 3.400e-02 | -3.38 | pyrimidine nucleotide transmembrane transporter activity | molecular function | GO:0015218 | 1 | 1 | 12176 | 414 | Rim2 | | 3.400e-02 | -3.38 | oxidative DNA demethylase activity | molecular function | GO:0035516 | 1 | 1 | 12176 | 414 | Tet | | 3.400e-02 | -3.38 | vascular endothelial growth factor binding | molecular function | GO:0038085 | 1 | 1 | 12176 | 414 | Pvr | | 3.400e-02 | -3.38 | iduronate-2-sulfatase activity | molecular function | GO:0004423 | 1 | 1 | 12176 | 414 | Ids | | 3.400e-02 | -3.38 | inositol hexakisphosphate kinase activity | molecular function | GO:0000828 | 1 | 1 | 12176 | 414 | l(1)G0196 | | 3.400e-02 | -3.38 | protein tyrosine phosphatase activity, metal-dependent | molecular function | GO:0030946 | 1 | 1 | 12176 | 414 | eya | | 3.400e-02 | -3.38 | transforming growth factor beta receptor, inhibitory cytoplasmic mediator activity | molecular function | GO:0030617 | 1 | 1 | 12176 | 414 | Dad | | 3.400e-02 | -3.38 | branched-chain-amino-acid transaminase activity | molecular function | GO:0004084 | 1 | 1 | 12176 | 414 | CG1673 | | 3.400e-02 | -3.38 | diphosphoinositol-pentakisphosphate kinase activity | molecular function | GO:0033857 | 1 | 1 | 12176 | 414 | l(1)G0196 | | 3.400e-02 | -3.38 | protein-tyrosine sulfotransferase activity | molecular function | GO:0008476 | 1 | 1 | 12176 | 414 | Tpst | | 3.400e-02 | -3.38 | phosphopantothenate--cysteine ligase activity | molecular function | GO:0004632 | 1 | 1 | 12176 | 414 | Ppcs | | 3.400e-02 | -3.38 | inositol heptakisphosphate kinase activity | molecular function | GO:0000829 | 1 | 1 | 12176 | 414 | l(1)G0196 | | 3.400e-02 | -3.38 | dihydrolipoyllysine-residue acetyltransferase activity | molecular function | GO:0004742 | 1 | 1 | 12176 | 414 | muc | | 3.400e-02 | -3.38 | 3-phosphoinositide-dependent protein kinase activity | molecular function | GO:0004676 | 1 | 1 | 12176 | 414 | Pdk1 | | 3.400e-02 | -3.38 | co-SMAD binding | molecular function | GO:0070410 | 1 | 1 | 12176 | 414 | fuss | | 3.400e-02 | -3.38 | retinol binding | molecular function | GO:0019841 | 1 | 1 | 12176 | 414 | apolpp | | 3.400e-02 | -3.38 | ATP citrate synthase activity | molecular function | GO:0003878 | 1 | 1 | 12176 | 414 | ATPCL | | 3.400e-02 | -3.38 | histone pre-mRNA stem-loop binding | molecular function | GO:0071207 | 1 | 1 | 12176 | 414 | Slbp | | 3.400e-02 | -3.38 | stretch-activated, cation-selective, calcium channel activity | molecular function | GO:0015275 | 1 | 1 | 12176 | 414 | Mid1 | | 3.400e-02 | -3.38 | netrin receptor activity involved in chemorepulsion | molecular function | GO:0005043 | 1 | 1 | 12176 | 414 | unc-5 | | 3.400e-02 | -3.38 | methylenetetrahydrofolate reductase (NAD(P)H) activity | molecular function | GO:0004489 | 1 | 1 | 12176 | 414 | CG7560 | | 3.400e-02 | -3.38 | DNA demethylase activity | molecular function | GO:0035514 | 1 | 1 | 12176 | 414 | Tet | | 3.400e-02 | -3.38 | SOCS family protein binding | molecular function | GO:0097678 | 1 | 1 | 12176 | 414 | dome | | 3.400e-02 | -3.38 | inositol hexakisphosphate 5-kinase activity | molecular function | GO:0000832 | 1 | 1 | 12176 | 414 | l(1)G0196 | | 3.400e-02 | -3.38 | insulin-activated receptor activity | molecular function | GO:0005009 | 1 | 1 | 12176 | 414 | InR | | 3.400e-02 | -3.38 | 17-beta-hydroxysteroid dehydrogenase (NAD+) activity | molecular function | GO:0044594 | 1 | 1 | 12176 | 414 | Mfe2 | | 3.400e-02 | -3.38 | 3R-hydroxyacyl-CoA dehydratase activity | molecular function | GO:0080023 | 1 | 1 | 12176 | 414 | Mfe2 | | 3.400e-02 | -3.38 | inositol-1,3,4,5,6-pentakisphosphate kinase activity | molecular function | GO:0000827 | 1 | 1 | 12176 | 414 | l(1)G0196 | | 3.400e-02 | -3.38 | 4-alpha-glucanotransferase activity | molecular function | GO:0004134 | 1 | 1 | 12176 | 414 | CG9485 | | 3.426e-02 | -3.37 | PAS | smart domains | SM00091 | 13 | 3 | 5086 | 290 | ss,tai,sim | | 3.431e-02 | -3.37 | cib (ciboulot) | protein interactions | 31359 | 66 | 7 | 9204 | 432 | Bacc,Npc2g,Jupiter,Nlp,CG9121,Fkbp14,chic | | 3.436e-02 | -3.37 | fly chr77E6-77E6\|3-47 | chromosome location | fly chr77E6-77E6\|3-47 | 13 | 2 | 25010 | 571 | Pex14,Pitslre | | 3.436e-02 | -3.37 | fly chr77E6-77E6\|3-47 cM | chromosome location | fly chr77E6-77E6\|3-47 cM | 13 | 2 | 25010 | 571 | Pex14,Pitslre | | 3.437e-02 | -3.37 | GO\_REGULATION\_OF\_HYDROLASE\_ACTIVITY | MSigDB lists | GO\_REGULATION\_OF\_HYDROLASE\_ACTIVITY | 283 | 17 | 4161 | 156 | wdb,Pdk1,Ocrl,CG17883,sli,PI31,Ziz,Rgl,Spn42De,htl,Ppn,dock,14-3-3epsilon,CenB1A,Mef2,step,CG5522 | | 3.444e-02 | -3.37 | RasGAP1 (Ras GTPase activating protein 1) | protein interactions | 39158 | 39 | 5 | 9204 | 432 | larp,aop,htl,Egfr,cindr | | 3.444e-02 | -3.37 | CG15098 (uncharacterized protein) | protein interactions | 37174 | 39 | 5 | 9204 | 432 | Rab5,tsh,Kr-h2,al,SERCA | | 3.444e-02 | -3.37 | kermit (kermit) | protein interactions | 46027 | 39 | 5 | 9204 | 432 | CG7492,Shrm,Sema1a,jar,fz | | 3.444e-02 | -3.37 | Sirt1 (Sirtuin 1) | protein interactions | 34708 | 39 | 5 | 9204 | 432 | h,N,Diap1,H,Dl | | 3.448e-02 | -3.37 | negative regulation of protein phosphorylation | biological process | GO:0001933 | 68 | 6 | 12345 | 442 | trbl,wdb,Socs36E,Cbl,N,CG8312 | | 3.448e-02 | -3.37 | glycogen degradation II | BIOCYC pathways | FLY\_PWY-5941 | 1 | 1 | 29 | 1 | CG9485 | | 3.468e-02 | -3.36 | protein modification by small protein conjugation | biological process | GO:0032446 | 271 | 16 | 12345 | 442 | Diap1,eff,Su(z)2,Pex10,morgue,sina,rdx,mura,Fem-1,CG13481,Cbl,tna,CG12402,FBXO11,Ubc87F,Kdm2 | | 3.483e-02 | -3.36 | STAT1\_02 | MSigDB lists | STAT1\_02 | 83 | 7 | 4161 | 156 | CG9485,Mfe2,crol,Cdk4,lt,Dl,FBXO11 | | 3.492e-02 | -3.35 | mitochondrial pyruvate dehydrogenase complex | cellular component | GO:0005967 | 1 | 1 | 11940 | 417 | muc | | 3.492e-02 | -3.35 | dystrobrevin complex | cellular component | GO:0016014 | 1 | 1 | 11940 | 417 | Dyb | | 3.492e-02 | -3.35 | cell leading edge cell cortex | cellular component | GO:1904269 | 1 | 1 | 11940 | 417 | step | | 3.492e-02 | -3.35 | citrate lyase complex | cellular component | GO:0009346 | 1 | 1 | 11940 | 417 | ATPCL | | 3.492e-02 | -3.35 | hemidesmosome | cellular component | GO:0030056 | 1 | 1 | 11940 | 417 | shot | | 3.492e-02 | -3.35 | insulin receptor complex | cellular component | GO:0005899 | 1 | 1 | 11940 | 417 | InR | | 3.511e-02 | -3.35 | MODULE\_64 | MSigDB lists | MODULE\_64 | 49 | 5 | 4161 | 156 | Nrk,Actbeta,Eph,Tollo,Ddr | | 3.511e-02 | -3.35 | GSE21360\_SECONDARY\_VS\_QUATERNARY\_MEMORY\_CD8\_TCELL\_DN | MSigDB lists | GSE21360\_SECONDARY\_VS\_QUATERNARY\_MEMORY\_CD8\_TCELL\_DN | 49 | 5 | 4161 | 156 | Nmda1,Drak,Tsp96F,Pgk,CG9220 | | 3.511e-02 | -3.35 | GSE17721\_LPS\_VS\_PAM3CSK4\_12H\_BMDC\_UP | MSigDB lists | GSE17721\_LPS\_VS\_PAM3CSK4\_12H\_BMDC\_UP | 49 | 5 | 4161 | 156 | emp,rdx,step,CG2017,cact | | 3.511e-02 | -3.35 | GSE21927\_SPLENIC\_VS\_TUMOR\_MONOCYTES\_FROM\_C26GM\_TUMOROUS\_MICE\_BALBC\_DN | MSigDB lists | GSE21927\_SPLENIC\_VS\_TUMOR\_MONOCYTES\_FROM\_C26GM\_TUMOROUS\_MICE\_BALBC\_DN | 49 | 5 | 4161 | 156 | alpha-Man-Ia,CycG,trbl,Unc-76,CG45050 | | 3.511e-02 | -3.35 | GSE17721\_CTRL\_VS\_CPG\_8H\_BMDC\_DN | MSigDB lists | GSE17721\_CTRL\_VS\_CPG\_8H\_BMDC\_DN | 49 | 5 | 4161 | 156 | cact,ap,CG2017,Raf,Pat1 | | 3.511e-02 | -3.35 | MTF1\_Q4 | MSigDB lists | MTF1\_Q4 | 49 | 5 | 4161 | 156 | Cdk4,His2Av,Dl,dlp,Cadps | | 3.511e-02 | -3.35 | GSE37416\_0H\_VS\_6H\_F\_TULARENSIS\_LVS\_NEUTROPHIL\_DN | MSigDB lists | GSE37416\_0H\_VS\_6H\_F\_TULARENSIS\_LVS\_NEUTROPHIL\_DN | 49 | 5 | 4161 | 156 | Cerk,CG13384,Rab5,Raf,cact | | 3.511e-02 | -3.35 | GSE15930\_NAIVE\_VS\_48H\_IN\_VITRO\_STIM\_IL12\_CD8\_TCELL\_UP | MSigDB lists | GSE15930\_NAIVE\_VS\_48H\_IN\_VITRO\_STIM\_IL12\_CD8\_TCELL\_UP | 49 | 5 | 4161 | 156 | Nmda1,CG17883,Pgant6,hppy,sina | | 3.512e-02 | -3.35 | somatodendritic compartment | cellular component | GO:0036477 | 212 | 13 | 11940 | 417 | bol,orb2,path,drl,Rab5,Ptp4E,Myo10A,rg,robo1,ena,CG9784,Rab1,Gr93c | | 3.525e-02 | -3.35 | PNT | prosite domains | PS51433 | 6 | 2 | 6243 | 325 | edl,aop | | 3.525e-02 | -3.35 | VWFD | prosite domains | PS51233 | 6 | 2 | 6243 | 325 | cv-d,apolpp | | 3.529e-02 | -3.34 | Glycosaminoglycan biosynthesis - chondroitin sulfate / dermatan sulfate | KEGG pathways | dme00532 | 8 | 2 | 3198 | 123 | GlcAT-I,CG9220 | | 3.529e-02 | -3.34 | Glycosaminoglycan biosynthesis - chondroitin sulfate / dermatan sulfate | KEGG pathways | ko00532 | 8 | 2 | 3198 | 123 | GlcAT-I,CG9220 | | 3.529e-02 | -3.34 | Notch signaling | KEGG pathways | dme\_M00682 | 8 | 2 | 3198 | 123 | N,Dl | | 3.529e-02 | -3.34 | Notch signaling | KEGG pathways | M00682 | 8 | 2 | 3198 | 123 | Dl,N | | 3.536e-02 | -3.34 | oocyte fate determination | biological process | GO:0030716 | 35 | 4 | 12345 | 442 | 14-3-3zeta,14-3-3epsilon,shot,baz | | 3.536e-02 | -3.34 | positive regulation of protein transport | biological process | GO:0051222 | 35 | 4 | 12345 | 442 | Abl,mop,S,Rab1 | | 3.541e-02 | -3.34 | regulation of cellular amide metabolic process | biological process | GO:0034248 | 186 | 12 | 12345 | 442 | Sxl,pnr,nub,msi,pum,orb2,312,Tis11,imd,bru2,bin3,bol | | 3.544e-02 | -3.34 | nuclear receptor transcription coactivator activity | molecular function | GO:0030374 | 9 | 2 | 12176 | 414 | tai,tna | | 3.544e-02 | -3.34 | long-chain fatty acid-CoA ligase activity | molecular function | GO:0004467 | 9 | 2 | 12176 | 414 | pdgy,CG3961 | | 3.544e-02 | -3.34 | non-membrane spanning protein tyrosine kinase activity | molecular function | GO:0004715 | 9 | 2 | 12176 | 414 | Src42A,Abl | | 3.544e-02 | -3.34 | kinesin binding | molecular function | GO:0019894 | 9 | 2 | 12176 | 414 | Unc-76,klar | | 3.544e-02 | -3.34 | nuclear hormone receptor binding | molecular function | GO:0035257 | 9 | 2 | 12176 | 414 | arm,tai | | 3.544e-02 | -3.34 | protein phosphorylated amino acid binding | molecular function | GO:0045309 | 9 | 2 | 12176 | 414 | 14-3-3epsilon,Cbl | | 3.544e-02 | -3.34 | protein tyrosine kinase binding | molecular function | GO:1990782 | 9 | 2 | 12176 | 414 | Cbl,dome | | 3.567e-02 | -3.33 | GO\_REGULATION\_OF\_RESPONSE\_TO\_EXTERNAL\_STIMULUS | MSigDB lists | GO\_REGULATION\_OF\_RESPONSE\_TO\_EXTERNAL\_STIMULUS | 200 | 13 | 4161 | 156 | ReepA,Tpc1,AP-1gamma,CG4645,sli,Tollo,Raf,robo1,fz2,SmD1,Unc-76,N,Arf79F | | 3.569e-02 | -3.33 | Atg5 (Autophagy-related 5) | protein interactions | 31666 | 27 | 4 | 9204 | 432 | Poxn,wdb,Fem-1,Atg16 | | 3.569e-02 | -3.33 | D (Dichaete) | protein interactions | 39570 | 27 | 4 | 9204 | 432 | nub,sim,ttk,N | | 3.569e-02 | -3.33 | Pcl (Polycomblike) | protein interactions | 37069 | 27 | 4 | 9204 | 432 | Abd-B,Scr,sd,corto | | 3.569e-02 | -3.33 | Tl (Toll) | protein interactions | 43222 | 27 | 4 | 9204 | 432 | cact,Rab1,pnr,Dcr-2 | | 3.570e-02 | -3.33 | - | gene3d domains | 3.30.50.10 | 26 | 4 | 4929 | 241 | grn,Hr4,pnr,Hr39 | | 3.570e-02 | -3.33 | - | gene3d domains | 1.10.30.10 | 26 | 4 | 4929 | 241 | HmgZ,cic,Sox14,CG12104 | | 3.572e-02 | -3.33 | MAPK family signaling cascades | REACTOME pathways | R-DME-5683057 | 117 | 10 | 3923 | 176 | Pvr,Egfr,Raf,vn,wdb,CG42684,CG4935,14-3-3zeta,htl,PI31 | | 3.580e-02 | -3.33 | positive regulation of calcium ion-dependent exocytosis | biological process | GO:0045956 | 1 | 1 | 12345 | 442 | Cadps | | 3.580e-02 | -3.33 | RNA destabilization | biological process | GO:0050779 | 1 | 1 | 12345 | 442 | Tis11 | | 3.580e-02 | -3.33 | positive regulation of cAMP-dependent protein kinase activity | biological process | GO:2000481 | 1 | 1 | 12345 | 442 | step | | 3.580e-02 | -3.33 | carbohydrate localization | biological process | GO:0052575 | 1 | 1 | 12345 | 442 | Mef2 | | 3.580e-02 | -3.33 | neurotransmitter receptor transport to plasma membrane | biological process | GO:0098877 | 1 | 1 | 12345 | 442 | scrib | | 3.580e-02 | -3.33 | regulation of negative chemotaxis | biological process | GO:0050923 | 1 | 1 | 12345 | 442 | sli | | 3.580e-02 | -3.33 | regulation of lysosomal protein catabolic process | biological process | GO:1905165 | 1 | 1 | 12345 | 442 | Socs36E | | 3.580e-02 | -3.33 | female gonad morphogenesis | biological process | GO:0061040 | 1 | 1 | 12345 | 442 | bab2 | | 3.580e-02 | -3.33 | negative regulation of antifungal peptide production | biological process | GO:0002789 | 1 | 1 | 12345 | 442 | cact | | 3.580e-02 | -3.33 | deoxynucleotide transport | biological process | GO:0030302 | 1 | 1 | 12345 | 442 | Rim2 | | 3.580e-02 | -3.33 | regulation of DNA demethylation | biological process | GO:1901535 | 1 | 1 | 12345 | 442 | Tet | | 3.580e-02 | -3.33 | glycolipid translocation | biological process | GO:0034203 | 1 | 1 | 12345 | 442 | CG3149 | | 3.580e-02 | -3.33 | neurotransmitter receptor transport, endosome to postsynaptic membrane | biological process | GO:0098887 | 1 | 1 | 12345 | 442 | scrib | | 3.580e-02 | -3.33 | positive regulation of protein export from nucleus | biological process | GO:0046827 | 1 | 1 | 12345 | 442 | Abl | | 3.580e-02 | -3.33 | negative regulation of intracellular steroid hormone receptor signaling pathway | biological process | GO:0033144 | 1 | 1 | 12345 | 442 | Hr4 | | 3.580e-02 | -3.33 | negative regulation of cell-cell adhesion mediated by cadherin | biological process | GO:2000048 | 1 | 1 | 12345 | 442 | N | | 3.580e-02 | -3.33 | positive regulation of ATPase-coupled calcium transmembrane transporter activity | biological process | GO:1901896 | 1 | 1 | 12345 | 442 | SERCA | | 3.580e-02 | -3.33 | pyrimidine nucleotide import into mitochondrion | biological process | GO:1990519 | 1 | 1 | 12345 | 442 | Rim2 | | 3.580e-02 | -3.33 | negative regulation of ecdysone receptor-mediated signaling pathway | biological process | GO:0120143 | 1 | 1 | 12345 | 442 | Hr4 | | 3.580e-02 | -3.33 | protein localization to endosome | biological process | GO:0036010 | 1 | 1 | 12345 | 442 | scrib | | 3.580e-02 | -3.33 | maternal determination of dorsal/ventral axis, ovarian follicular epithelium, soma encoded | biological process | GO:0008071 | 1 | 1 | 12345 | 442 | Egfr | | 3.580e-02 | -3.33 | neurotransmitter receptor transport, endosome to plasma membrane | biological process | GO:0099639 | 1 | 1 | 12345 | 442 | scrib | | 3.580e-02 | -3.33 | branched-chain amino acid biosynthetic process | biological process | GO:0009082 | 1 | 1 | 12345 | 442 | CG1673 | | 3.580e-02 | -3.33 | negative regulation of intracellular mRNA localization | biological process | GO:1904581 | 1 | 1 | 12345 | 442 | klar | | 3.580e-02 | -3.33 | positive regulation of fat cell proliferation | biological process | GO:0070346 | 1 | 1 | 12345 | 442 | InR | | 3.580e-02 | -3.33 | negative regulation of hematopoietic stem cell differentiation | biological process | GO:1902037 | 1 | 1 | 12345 | 442 | C1GalTA | | 3.580e-02 | -3.33 | maintenance of imaginal histoblast diploidy | biological process | GO:0007489 | 1 | 1 | 12345 | 442 | esg | | 3.580e-02 | -3.33 | pigment cell development | biological process | GO:0070285 | 1 | 1 | 12345 | 442 | eya | | 3.580e-02 | -3.33 | 5-methylcytosine catabolic process | biological process | GO:0006211 | 1 | 1 | 12345 | 442 | Tet | | 3.580e-02 | -3.33 | mRNA destabilization | biological process | GO:0061157 | 1 | 1 | 12345 | 442 | Tis11 | | 3.580e-02 | -3.33 | regulation of extrinsic apoptotic signaling pathway in absence of ligand | biological process | GO:2001239 | 1 | 1 | 12345 | 442 | eya | | 3.580e-02 | -3.33 | positive regulation of protein catabolic process in the vacuole | biological process | GO:1904352 | 1 | 1 | 12345 | 442 | Socs36E | | 3.580e-02 | -3.33 | neurotransmitter receptor transport to postsynaptic membrane | biological process | GO:0098969 | 1 | 1 | 12345 | 442 | scrib | | 3.580e-02 | -3.33 | positive regulation of lysosomal protein catabolic process | biological process | GO:1905167 | 1 | 1 | 12345 | 442 | Socs36E | | 3.580e-02 | -3.33 | induction of negative chemotaxis | biological process | GO:0050929 | 1 | 1 | 12345 | 442 | sli | | 3.580e-02 | -3.33 | clathrin-coated vesicle cargo loading | biological process | GO:0035652 | 1 | 1 | 12345 | 442 | AP-1gamma | | 3.580e-02 | -3.33 | melanosome organization | biological process | GO:0032438 | 1 | 1 | 12345 | 442 | Shrm | | 3.580e-02 | -3.33 | limb joint morphogenesis | biological process | GO:0036022 | 1 | 1 | 12345 | 442 | nub | | 3.580e-02 | -3.33 | protein sulfation | biological process | GO:0006477 | 1 | 1 | 12345 | 442 | Tpst | | 3.580e-02 | -3.33 | endodermal cell fate determination | biological process | GO:0007493 | 1 | 1 | 12345 | 442 | htl | | 3.580e-02 | -3.33 | histone dephosphorylation | biological process | GO:0016576 | 1 | 1 | 12345 | 442 | eya | | 3.580e-02 | -3.33 | negative regulation of response to alcohol | biological process | GO:1901420 | 1 | 1 | 12345 | 442 | Hr4 | | 3.580e-02 | -3.33 | establishment of thoracic bristle planar orientation | biological process | GO:0048106 | 1 | 1 | 12345 | 442 | spen | | 3.580e-02 | -3.33 | vascular endothelial growth factor signaling pathway | biological process | GO:0038084 | 1 | 1 | 12345 | 442 | Pvr | | 3.580e-02 | -3.33 | detection of virus | biological process | GO:0009597 | 1 | 1 | 12345 | 442 | Dcr-2 | | 3.580e-02 | -3.33 | DNA demethylation | biological process | GO:0080111 | 1 | 1 | 12345 | 442 | Tet | | 3.580e-02 | -3.33 | negative regulation of extrinsic apoptotic signaling pathway in absence of ligand | biological process | GO:2001240 | 1 | 1 | 12345 | 442 | eya | | 3.580e-02 | -3.33 | oxidative DNA demethylation | biological process | GO:0035511 | 1 | 1 | 12345 | 442 | Tet | | 3.580e-02 | -3.33 | negative regulation of cysteine-type endopeptidase activity involved in execution phase of apoptosis | biological process | GO:2001271 | 1 | 1 | 12345 | 442 | Diap1 | | 3.580e-02 | -3.33 | limb morphogenesis | biological process | GO:0035108 | 1 | 1 | 12345 | 442 | nub | | 3.580e-02 | -3.33 | negative regulation of ribosome biogenesis | biological process | GO:0090071 | 1 | 1 | 12345 | 442 | 312 | | 3.580e-02 | -3.33 | negative regulation of oocyte development | biological process | GO:0060283 | 1 | 1 | 12345 | 442 | klar | | 3.580e-02 | -3.33 | phospholipid homeostasis | biological process | GO:0055091 | 1 | 1 | 12345 | 442 | Ppcs | | 3.580e-02 | -3.33 | male pigmentation | biological process | GO:0048094 | 1 | 1 | 12345 | 442 | Abd-B | | 3.580e-02 | -3.33 | establishment of blood-nerve barrier | biological process | GO:0008065 | 1 | 1 | 12345 | 442 | Gli | | 3.580e-02 | -3.33 | neurotransmitter receptor transport postsynaptic membrane to endosome | biological process | GO:0098968 | 1 | 1 | 12345 | 442 | scrib | | 3.580e-02 | -3.33 | peptidyl-serine ADP-ribosylation | biological process | GO:0018312 | 1 | 1 | 12345 | 442 | CG1218 | | 3.580e-02 | -3.33 | reticulophagy | biological process | GO:0061709 | 1 | 1 | 12345 | 442 | Atg17 | | 3.580e-02 | -3.33 | negative regulation of embryonic pattern specification | biological process | GO:1902876 | 1 | 1 | 12345 | 442 | klar | | 3.580e-02 | -3.33 | branchiomotor neuron axon guidance | biological process | GO:0021785 | 1 | 1 | 12345 | 442 | Abl | | 3.580e-02 | -3.33 | cell-cell adhesion involved in establishment of planar polarity | biological process | GO:0090250 | 1 | 1 | 12345 | 442 | baz | | 3.580e-02 | -3.33 | regulation of actin cytoskeleton organization by cell-cell adhesion | biological process | GO:0090138 | 1 | 1 | 12345 | 442 | ed | | 3.580e-02 | -3.33 | cytoplasmic sequestering of NF-kappaB | biological process | GO:0007253 | 1 | 1 | 12345 | 442 | cact | | 3.580e-02 | -3.33 | calcium ion transport from cytosol to endoplasmic reticulum | biological process | GO:1903515 | 1 | 1 | 12345 | 442 | SERCA | | 3.580e-02 | -3.33 | valine biosynthetic process | biological process | GO:0009099 | 1 | 1 | 12345 | 442 | CG1673 | | 3.580e-02 | -3.33 | negative regulation of muscle tissue development | biological process | GO:1901862 | 1 | 1 | 12345 | 442 | Abd-B | | 3.580e-02 | -3.33 | synaptic vesicle to endosome fusion | biological process | GO:0016189 | 1 | 1 | 12345 | 442 | Rab5 | | 3.580e-02 | -3.33 | compound eye corneal lens morphogenesis | biological process | GO:0048750 | 1 | 1 | 12345 | 442 | ttk | | 3.580e-02 | -3.33 | specification of segmental identity, intercalary segment | biological process | GO:0035291 | 1 | 1 | 12345 | 442 | kn | | 3.580e-02 | -3.33 | positive regulation of DNA demethylation | biological process | GO:1901537 | 1 | 1 | 12345 | 442 | Tet | | 3.580e-02 | -3.33 | negative regulation of non-canonical Wnt signaling pathway | biological process | GO:2000051 | 1 | 1 | 12345 | 442 | aop | | 3.580e-02 | -3.33 | calcium-independent cell-cell adhesion via plasma membrane cell-adhesion molecules | biological process | GO:0016338 | 1 | 1 | 12345 | 442 | Fas1 | | 3.580e-02 | -3.33 | negative regulation of signal transduction in absence of ligand | biological process | GO:1901099 | 1 | 1 | 12345 | 442 | eya | | 3.580e-02 | -3.33 | protein localization to early endosome | biological process | GO:1902946 | 1 | 1 | 12345 | 442 | scrib | | 3.580e-02 | -3.33 | 5-methylcytosine metabolic process | biological process | GO:0019857 | 1 | 1 | 12345 | 442 | Tet | | 3.580e-02 | -3.33 | 3'-UTR-mediated mRNA destabilization | biological process | GO:0061158 | 1 | 1 | 12345 | 442 | Tis11 | | 3.580e-02 | -3.33 | R1/R6 development | biological process | GO:0048053 | 1 | 1 | 12345 | 442 | ttk | | 3.580e-02 | -3.33 | positive regulation of compound eye retinal cell apoptotic process | biological process | GO:1901694 | 1 | 1 | 12345 | 442 | morgue | | 3.580e-02 | -3.33 | negative regulation of cellular response to alcohol | biological process | GO:1905958 | 1 | 1 | 12345 | 442 | Hr4 | | 3.580e-02 | -3.33 | behavioral response to nicotine | biological process | GO:0035095 | 1 | 1 | 12345 | 442 | moody | | 3.580e-02 | -3.33 | establishment of protein localization to postsynaptic membrane | biological process | GO:1903540 | 1 | 1 | 12345 | 442 | scrib | | 3.580e-02 | -3.33 | regulation of protein catabolic process in the vacuole | biological process | GO:1904350 | 1 | 1 | 12345 | 442 | Socs36E | | 3.580e-02 | -3.33 | regulation of R8 cell differentiation | biological process | GO:0045679 | 1 | 1 | 12345 | 442 | ed | | 3.580e-02 | -3.33 | negative regulation of pole plasm oskar mRNA localization | biological process | GO:0045855 | 1 | 1 | 12345 | 442 | klar | | 3.580e-02 | -3.33 | lipoprotein localization | biological process | GO:0044872 | 1 | 1 | 12345 | 442 | unc-119 | | 3.580e-02 | -3.33 | negative regulation of embryonic development | biological process | GO:0045992 | 1 | 1 | 12345 | 442 | klar | | 3.580e-02 | -3.33 | negative regulation of striated muscle tissue development | biological process | GO:0045843 | 1 | 1 | 12345 | 442 | Abd-B | | 3.580e-02 | -3.33 | leucine biosynthetic process | biological process | GO:0009098 | 1 | 1 | 12345 | 442 | CG1673 | | 3.580e-02 | -3.33 | ARF protein signal transduction | biological process | GO:0032011 | 1 | 1 | 12345 | 442 | Arf79F | | 3.580e-02 | -3.33 | regulation of Ral protein signal transduction | biological process | GO:0032485 | 1 | 1 | 12345 | 442 | CG5522 | | 3.580e-02 | -3.33 | establishment of centrosome localization | biological process | GO:0051660 | 1 | 1 | 12345 | 442 | baz | | 3.580e-02 | -3.33 | vacuole fusion | biological process | GO:0097576 | 1 | 1 | 12345 | 442 | lt | | 3.580e-02 | -3.33 | regulation of cyclin-dependent protein serine/threonine kinase activity involved in G2/M transition of mitotic cell cycle | biological process | GO:0031660 | 1 | 1 | 12345 | 442 | kay | | 3.580e-02 | -3.33 | establishment of body hair planar orientation | biological process | GO:0048105 | 1 | 1 | 12345 | 442 | ft | | 3.580e-02 | -3.33 | negative regulation of R8 cell differentiation | biological process | GO:0045680 | 1 | 1 | 12345 | 442 | ed | | 3.580e-02 | -3.33 | cellular response to vascular endothelial growth factor stimulus | biological process | GO:0035924 | 1 | 1 | 12345 | 442 | Pvr | | 3.580e-02 | -3.33 | regulation of protein ADP-ribosylation | biological process | GO:0010835 | 1 | 1 | 12345 | 442 | CG1218 | | 3.580e-02 | -3.33 | peptidyl-tyrosine sulfation | biological process | GO:0006478 | 1 | 1 | 12345 | 442 | Tpst | | 3.580e-02 | -3.33 | negative regulation of hematopoietic progenitor cell differentiation | biological process | GO:1901533 | 1 | 1 | 12345 | 442 | C1GalTA | | 3.580e-02 | -3.33 | vacuole fusion, non-autophagic | biological process | GO:0042144 | 1 | 1 | 12345 | 442 | lt | | 3.580e-02 | -3.33 | external genitalia morphogenesis | biological process | GO:0035261 | 1 | 1 | 12345 | 442 | Abd-B | | 3.580e-02 | -3.33 | pyrimidine nucleotide transport | biological process | GO:0006864 | 1 | 1 | 12345 | 442 | Rim2 | | 3.580e-02 | -3.33 | positive regulation of negative chemotaxis | biological process | GO:0050924 | 1 | 1 | 12345 | 442 | sli | | 3.580e-02 | -3.33 | carbohydrate storage | biological process | GO:0052576 | 1 | 1 | 12345 | 442 | Mef2 | | 3.580e-02 | -3.33 | glycolipid transport | biological process | GO:0046836 | 1 | 1 | 12345 | 442 | CG3149 | | 3.580e-02 | -3.33 | lipoprotein transport | biological process | GO:0042953 | 1 | 1 | 12345 | 442 | unc-119 | | 3.580e-02 | -3.33 | suture of dorsal opening | biological process | GO:0007396 | 1 | 1 | 12345 | 442 | ena | | 3.580e-02 | -3.33 | transforming growth factor beta receptor complex assembly | biological process | GO:0007181 | 1 | 1 | 12345 | 442 | tkv | | 3.587e-02 | -3.33 | osk (oskar) | protein interactions | 41066 | 96 | 9 | 9204 | 432 | tkv,cue,orb2,14-3-3zeta,esg,klar,Myo10A,jar,larp | | 3.591e-02 | -3.33 | ubiquitin-like protein ligase binding | molecular function | GO:0044389 | 54 | 5 | 12176 | 414 | morgue,rdx,eff,Diap1,Tom | | 3.595e-02 | -3.33 | PAX4\_01 | MSigDB lists | PAX4\_01 | 66 | 6 | 4161 | 156 | 14-3-3zeta,CtBP,hyx,Atg16,cact,Pgant6 | | 3.595e-02 | -3.33 | GSE1791\_CTRL\_VS\_NEUROMEDINU\_IN\_T\_CELL\_LINE\_6H\_DN | MSigDB lists | GSE1791\_CTRL\_VS\_NEUROMEDINU\_IN\_T\_CELL\_LINE\_6H\_DN | 66 | 6 | 4161 | 156 | Mfe2,frc,alpha-Man-Ia,Tpst,His2Av,Pgant5 | | 3.595e-02 | -3.33 | GO\_EMBRYONIC\_ORGAN\_MORPHOGENESIS | MSigDB lists | GO\_EMBRYONIC\_ORGAN\_MORPHOGENESIS | 66 | 6 | 4161 | 156 | Dl,otk,N,Mef2,arm,fz2 | | 3.595e-02 | -3.33 | GSE1740\_MCSF\_VS\_MCSF\_AND\_IFNG\_DAY2\_DERIVED\_MACROPHAGE\_DN | MSigDB lists | GSE1740\_MCSF\_VS\_MCSF\_AND\_IFNG\_DAY2\_DERIVED\_MACROPHAGE\_DN | 66 | 6 | 4161 | 156 | Pgant5,His2Av,Tsp96F,Tpst,frc,Mfe2 | | 3.615e-02 | -3.32 | regulation of alternative mRNA splicing, via spliceosome | biological process | GO:0000381 | 87 | 7 | 12345 | 442 | fl(2)d,Sxl,CG4896,Rm62,ps,CG11360,bru2 | | 3.615e-02 | -3.32 | actomyosin structure organization | biological process | GO:0031032 | 87 | 7 | 12345 | 442 | yrt,SCAR,kon,cora,Dg,chic,Bsg | | 3.622e-02 | -3.32 | CREB phosphorylation through the activation of Ras | REACTOME pathways | R-DME-442742 | 7 | 2 | 3923 | 176 | Raf,Pdk1 | | 3.622e-02 | -3.32 | Chk1/Chk2(Cds1) mediated inactivation of Cyclin B:Cdk1 complex | REACTOME pathways | R-DME-75035 | 7 | 2 | 3923 | 176 | 14-3-3zeta,14-3-3epsilon | | 3.646e-02 | -3.31 | Wnt5 (Wnt oncogene analog 5) | protein interactions | 32838 | 16 | 3 | 9204 | 432 | fz,fz2,drl | | 3.646e-02 | -3.31 | CycB3 (Cyclin B3) | protein interactions | 42971 | 16 | 3 | 9204 | 432 | Cks30A,H,Hr39 | | 3.646e-02 | -3.31 | Smr (Smrter) | protein interactions | 32225 | 16 | 3 | 9204 | 432 | kn,rictor,H | | 3.646e-02 | -3.31 | TfIIB (Transcription factor IIB) | protein interactions | 34430 | 16 | 3 | 9204 | 432 | Scr,Abd-B,abd-A | | 3.646e-02 | -3.31 | hb (hunchback) | protein interactions | 41032 | 16 | 3 | 9204 | 432 | Adf1,cindr,pum | | 3.646e-02 | -3.31 | spz (spatzle) | protein interactions | 43256 | 16 | 3 | 9204 | 432 | cic,Egfr,cact | | 3.652e-02 | -3.31 | GO\_TRANSMEMBRANE\_RECEPTOR\_PROTEIN\_TYROSINE\_KINASE\_SIGNALING\_PATHWAY | MSigDB lists | GO\_TRANSMEMBRANE\_RECEPTOR\_PROTEIN\_TYROSINE\_KINASE\_SIGNALING\_PATHWAY | 121 | 9 | 4161 | 156 | dock,nAChRalpha4,htl,InR,Pdk1,Nrk,Src42A,Ddr,Eph | | 3.677e-02 | -3.30 | GO\_ANTERIOR\_POSTERIOR\_PATTERN\_SPECIFICATION | MSigDB lists | GO\_ANTERIOR\_POSTERIOR\_PATTERN\_SPECIFICATION | 34 | 4 | 4161 | 156 | Dl,N,fz2,arm | | 3.677e-02 | -3.30 | MEK\_UP.V1\_DN | MSigDB lists | MEK\_UP.V1\_DN | 34 | 4 | 4161 | 156 | InR,Psa,alpha-Man-Ia,CG9485 | | 3.677e-02 | -3.30 | GSE12845\_IGD\_NEG\_BLOOD\_VS\_NAIVE\_TONSIL\_BCELL\_DN | MSigDB lists | GSE12845\_IGD\_NEG\_BLOOD\_VS\_NAIVE\_TONSIL\_BCELL\_DN | 34 | 4 | 4161 | 156 | trbl,N,Kdm2,lili | | 3.677e-02 | -3.30 | GSE19923\_WT\_VS\_HEB\_KO\_DP\_THYMOCYTE\_UP | MSigDB lists | GSE19923\_WT\_VS\_HEB\_KO\_DP\_THYMOCYTE\_UP | 34 | 4 | 4161 | 156 | Ddr,frc,Tsp96F,cact | | 3.677e-02 | -3.30 | GSE1112\_HY\_CD8AB\_VS\_HY\_CD8AA\_THYMOCYTE\_RTOC\_CULTURE\_DN | MSigDB lists | GSE1112\_HY\_CD8AB\_VS\_HY\_CD8AA\_THYMOCYTE\_RTOC\_CULTURE\_DN | 34 | 4 | 4161 | 156 | Drak,Tpst,CG1673,Eph | | 3.677e-02 | -3.30 | GSE37301\_MULTIPOTENT\_PROGENITOR\_VS\_LYMPHOID\_PRIMED\_MPP\_DN | MSigDB lists | GSE37301\_MULTIPOTENT\_PROGENITOR\_VS\_LYMPHOID\_PRIMED\_MPP\_DN | 34 | 4 | 4161 | 156 | shot,jar,CG9220,cact | | 3.677e-02 | -3.30 | GSE11057\_NAIVE\_VS\_CENT\_MEMORY\_CD4\_TCELL\_UP | MSigDB lists | GSE11057\_NAIVE\_VS\_CENT\_MEMORY\_CD4\_TCELL\_UP | 34 | 4 | 4161 | 156 | InR,rg,sina,Ocrl | | 3.677e-02 | -3.30 | GAGACTG\_MIR452 | MSigDB lists | GAGACTG\_MIR452 | 34 | 4 | 4161 | 156 | FBXO11,Psa,Ocrl,CG10417 | | 3.679e-02 | -3.30 | pum (pumilio) | protein interactions | 41094 | 53 | 6 | 9204 | 432 | path,Diap1,Egfr,mam,pnr,AP-1gamma | | 3.679e-02 | -3.30 | Pglym78 (Phosphoglyceromutase 78) | protein interactions | 43447 | 53 | 6 | 9204 | 432 | Bacc,Nlp,Fkbp14,Pgk,CycG,apolpp | | 3.679e-02 | -3.30 | CG9572 (uncharacterized protein) | protein interactions | 33013 | 53 | 6 | 9204 | 432 | CG34376,Rgl,esg,nub,Mob2,H2.0 | | 3.684e-02 | -3.30 | spidey (spidey) | protein interactions | 31703 | 67 | 7 | 9204 | 432 | Rab5,Mpcp2,Kr-h2,Rab1,Arf79F,SERCA,Gfat2 | | 3.684e-02 | -3.30 | Sec61alpha (Sec61 alpha subunit) | protein interactions | 33905 | 67 | 7 | 9204 | 432 | GluProRS,Kr-h2,Mpcp2,Rab5,SERCA,lt,H | | 3.684e-02 | -3.30 | brk (brinker) | protein interactions | 31665 | 67 | 7 | 9204 | 432 | CtBP,tsh,ey,tkv,Abd-B,Hr39,Dad | | 3.689e-02 | -3.30 | GO\_CELL\_MATURATION | MSigDB lists | GO\_CELL\_MATURATION | 20 | 3 | 4161 | 156 | Fem-1,arm,fz2 | | 3.689e-02 | -3.30 | GO\_REGULATION\_OF\_ENDOTHELIAL\_CELL\_MIGRATION | MSigDB lists | GO\_REGULATION\_OF\_ENDOTHELIAL\_CELL\_MIGRATION | 20 | 3 | 4161 | 156 | sli,Pdk1,N | | 3.689e-02 | -3.30 | GO\_REGULATION\_OF\_LYMPHOCYTE\_MEDIATED\_IMMUNITY | MSigDB lists | GO\_REGULATION\_OF\_LYMPHOCYTE\_MEDIATED\_IMMUNITY | 20 | 3 | 4161 | 156 | fz2,AP-1gamma,Spn42De | | 3.689e-02 | -3.30 | NAKAYAMA\_SOFT\_TISSUE\_TUMORS\_PCA2\_UP | MSigDB lists | NAKAYAMA\_SOFT\_TISSUE\_TUMORS\_PCA2\_UP | 20 | 3 | 4161 | 156 | htl,Actbeta,fz | | 3.689e-02 | -3.30 | GO\_CARDIAC\_CHAMBER\_MORPHOGENESIS | MSigDB lists | GO\_CARDIAC\_CHAMBER\_MORPHOGENESIS | 20 | 3 | 4161 | 156 | N,Mef2,fz | | 3.689e-02 | -3.30 | GSE16385\_UNTREATED\_VS\_12H\_ROSIGLITAZONE\_IFNG\_TNF\_TREATED\_MACROPHAGE\_UP | MSigDB lists | GSE16385\_UNTREATED\_VS\_12H\_ROSIGLITAZONE\_IFNG\_TNF\_TREATED\_MACROPHAGE\_UP | 20 | 3 | 4161 | 156 | CG1129,GlcAT-I,Drak | | 3.689e-02 | -3.30 | GO\_EMBRYONIC\_HEART\_TUBE\_MORPHOGENESIS | MSigDB lists | GO\_EMBRYONIC\_HEART\_TUBE\_MORPHOGENESIS | 20 | 3 | 4161 | 156 | Mef2,N,Dl | | 3.689e-02 | -3.30 | INGRAM\_SHH\_TARGETS\_UP | MSigDB lists | INGRAM\_SHH\_TARGETS\_UP | 20 | 3 | 4161 | 156 | otk,InR,dlp | | 3.689e-02 | -3.30 | GCM\_MAP1B | MSigDB lists | GCM\_MAP1B | 20 | 3 | 4161 | 156 | emp,Pat1,nmo | | 3.689e-02 | -3.30 | LEIN\_MIDBRAIN\_MARKERS | MSigDB lists | LEIN\_MIDBRAIN\_MARKERS | 20 | 3 | 4161 | 156 | Eph,CG1673,kn | | 3.689e-02 | -3.30 | GO\_CARDIAC\_SEPTUM\_DEVELOPMENT | MSigDB lists | GO\_CARDIAC\_SEPTUM\_DEVELOPMENT | 20 | 3 | 4161 | 156 | N,fz,otk | | 3.689e-02 | -3.30 | GSE1112\_OT1\_CD8AB\_VS\_HY\_CD8AA\_THYMOCYTE\_RTOC\_CULTURE\_UP | MSigDB lists | GSE1112\_OT1\_CD8AB\_VS\_HY\_CD8AA\_THYMOCYTE\_RTOC\_CULTURE\_UP | 20 | 3 | 4161 | 156 | Eph,CG7394,ft | | 3.698e-02 | -3.30 | response to starvation | biological process | GO:0042594 | 146 | 10 | 12345 | 442 | Ric,InR,lt,Raf,SNF4Agamma,Atg16,REPTOR,Atg17,CG7536,rictor | | 3.721e-02 | -3.29 | protein kinase activator activity | molecular function | GO:0030295 | 22 | 3 | 12176 | 414 | Cks30A,Pdk1,jeb | | 3.721e-02 | -3.29 | kinase activator activity | molecular function | GO:0019209 | 22 | 3 | 12176 | 414 | jeb,Pdk1,Cks30A | | 3.723e-02 | -3.29 | dystrophin-associated glycoprotein complex | cellular component | GO:0016010 | 9 | 2 | 11940 | 417 | Dg,Dyb | | 3.723e-02 | -3.29 | glycoprotein complex | cellular component | GO:0090665 | 9 | 2 | 11940 | 417 | Dyb,Dg | | 3.723e-02 | -3.29 | RISC complex | cellular component | GO:0016442 | 9 | 2 | 11940 | 417 | vig,Dcr-2 | | 3.723e-02 | -3.29 | RNAi effector complex | cellular component | GO:0031332 | 9 | 2 | 11940 | 417 | vig,Dcr-2 | | 3.726e-02 | -3.29 | Pi3K21B (Pi3K21B) | protein interactions | 33203 | 112 | 10 | 9204 | 432 | Egfr,dome,InR,Unc-76,14-3-3epsilon,larp,Psa,Pvr,tkv,14-3-3zeta | | 3.728e-02 | -3.29 | Innexin | interpro domains | IPR000990 | 8 | 2 | 11383 | 450 | Inx2,ogre | | 3.728e-02 | -3.29 | Small\_GTPase\_Rho | interpro domains | IPR003578 | 8 | 2 | 11383 | 450 | RhoL,RhoU | | 3.728e-02 | -3.29 | PP2C\_BS | interpro domains | IPR000222 | 8 | 2 | 11383 | 450 | CG10417,Pp2C1 | | 3.728e-02 | -3.29 | Lambda\_DNA-bd\_dom\_sf | interpro domains | IPR010982 | 8 | 2 | 11383 | 450 | pdm3,nub | | 3.728e-02 | -3.29 | BOULE/DAZ | interpro domains | IPR037366 | 8 | 2 | 11383 | 450 | bol,msi | | 3.728e-02 | -3.29 | Death\_domain | interpro domains | IPR000488 | 8 | 2 | 11383 | 450 | imd,unc-5 | | 3.728e-02 | -3.29 | Histone\_H2A/H2B/H3 | interpro domains | IPR007125 | 8 | 2 | 11383 | 450 | His2Av,His3.3A | | 3.728e-02 | -3.29 | ARFGAP/RecO | interpro domains | IPR037278 | 8 | 2 | 11383 | 450 | drongo,CenB1A | | 3.728e-02 | -3.29 | ArfGAP\_dom\_sf | interpro domains | IPR038508 | 8 | 2 | 11383 | 450 | drongo,CenB1A | | 3.728e-02 | -3.29 | ArfGAP\_dom | interpro domains | IPR001164 | 8 | 2 | 11383 | 450 | CenB1A,drongo | | 3.729e-02 | -3.29 | apical plasma membrane | cellular component | GO:0016324 | 71 | 6 | 11940 | 417 | arm,Ptp4E,baz,Inx2,Shrm,dome | | 3.749e-02 | -3.28 | ZAIDI\_OSTEOBLAST\_TRANSCRIPTION\_FACTORS | MSigDB lists | ZAIDI\_OSTEOBLAST\_TRANSCRIPTION\_FACTORS | 1 | 1 | 4161 | 156 | arm | | 3.749e-02 | -3.28 | REACTOME\_ADHERENS\_JUNCTIONS\_INTERACTIONS | MSigDB lists | REACTOME\_ADHERENS\_JUNCTIONS\_INTERACTIONS | 1 | 1 | 4161 | 156 | arm | | 3.749e-02 | -3.28 | GO\_NEGATIVE\_REGULATION\_OF\_LEUKOCYTE\_CHEMOTAXIS | MSigDB lists | GO\_NEGATIVE\_REGULATION\_OF\_LEUKOCYTE\_CHEMOTAXIS | 1 | 1 | 4161 | 156 | sli | | 3.749e-02 | -3.28 | TERAO\_AOX4\_TARGETS\_HG\_DN | MSigDB lists | TERAO\_AOX4\_TARGETS\_HG\_DN | 1 | 1 | 4161 | 156 | CG31522 | | 3.749e-02 | -3.28 | REACTOME\_GLYCOPROTEIN\_HORMONES | MSigDB lists | REACTOME\_GLYCOPROTEIN\_HORMONES | 1 | 1 | 4161 | 156 | Actbeta | | 3.749e-02 | -3.28 | JONES\_TCOF1\_TARGETS | MSigDB lists | JONES\_TCOF1\_TARGETS | 1 | 1 | 4161 | 156 | CycG | | 3.749e-02 | -3.28 | GO\_POSITIVE\_REGULATION\_OF\_INTERFERON\_GAMMA\_BIOSYNTHETIC\_PROCESS | MSigDB lists | GO\_POSITIVE\_REGULATION\_OF\_INTERFERON\_GAMMA\_BIOSYNTHETIC\_PROCESS | 1 | 1 | 4161 | 156 | Tollo | | 3.749e-02 | -3.28 | GILMORE\_CORE\_NFKB\_PATHWAY | MSigDB lists | GILMORE\_CORE\_NFKB\_PATHWAY | 1 | 1 | 4161 | 156 | cact | | 3.749e-02 | -3.28 | RICKMAN\_HEAD\_AND\_NECK\_CANCER\_D | MSigDB lists | RICKMAN\_HEAD\_AND\_NECK\_CANCER\_D | 1 | 1 | 4161 | 156 | Dyb | | 3.749e-02 | -3.28 | CHOI\_ATL\_ACUTE\_STAGE | MSigDB lists | CHOI\_ATL\_ACUTE\_STAGE | 1 | 1 | 4161 | 156 | ap | | 3.749e-02 | -3.28 | MODULE\_382 | MSigDB lists | MODULE\_382 | 1 | 1 | 4161 | 156 | Actbeta | | 3.749e-02 | -3.28 | GO\_NEGATIVE\_REGULATION\_OF\_CELLULAR\_SENESCENCE | MSigDB lists | GO\_NEGATIVE\_REGULATION\_OF\_CELLULAR\_SENESCENCE | 1 | 1 | 4161 | 156 | Cdk4 | | 3.749e-02 | -3.28 | GO\_NEGATIVE\_REGULATION\_OF\_DIGESTIVE\_SYSTEM\_PROCESS | MSigDB lists | GO\_NEGATIVE\_REGULATION\_OF\_DIGESTIVE\_SYSTEM\_PROCESS | 1 | 1 | 4161 | 156 | CG11069 | | 3.749e-02 | -3.28 | GO\_ACROSOME\_ASSEMBLY | MSigDB lists | GO\_ACROSOME\_ASSEMBLY | 1 | 1 | 4161 | 156 | CG17883 | | 3.749e-02 | -3.28 | MATZUK\_LUTEAL\_GENES | MSigDB lists | MATZUK\_LUTEAL\_GENES | 1 | 1 | 4161 | 156 | Psa | | 3.749e-02 | -3.28 | REACTOME\_SIGNALING\_BY\_FGFR3\_MUTANTS | MSigDB lists | REACTOME\_SIGNALING\_BY\_FGFR3\_MUTANTS | 1 | 1 | 4161 | 156 | htl | | 3.749e-02 | -3.28 | GO\_INSULIN\_LIKE\_GROWTH\_FACTOR\_BINDING | MSigDB lists | GO\_INSULIN\_LIKE\_GROWTH\_FACTOR\_BINDING | 1 | 1 | 4161 | 156 | InR | | 3.749e-02 | -3.28 | REACTOME\_IKK\_COMPLEX\_RECRUITMENT\_MEDIATED\_BY\_RIP1 | MSigDB lists | REACTOME\_IKK\_COMPLEX\_RECRUITMENT\_MEDIATED\_BY\_RIP1 | 1 | 1 | 4161 | 156 | Tollo | | 3.749e-02 | -3.28 | TURJANSKI\_MAPK11\_TARGETS | MSigDB lists | TURJANSKI\_MAPK11\_TARGETS | 1 | 1 | 4161 | 156 | Mef2 | | 3.749e-02 | -3.28 | GO\_POSITIVE\_REGULATION\_OF\_TYROSINE\_PHOSPHORYLATION\_OF\_STAT1\_PROTEIN | MSigDB lists | GO\_POSITIVE\_REGULATION\_OF\_TYROSINE\_PHOSPHORYLATION\_OF\_STAT1\_PROTEIN | 1 | 1 | 4161 | 156 | htl | | 3.749e-02 | -3.28 | AMUNDSON\_GAMMA\_RADIATION\_RESISTANCE | MSigDB lists | AMUNDSON\_GAMMA\_RADIATION\_RESISTANCE | 1 | 1 | 4161 | 156 | InR | | 3.749e-02 | -3.28 | MATZUK\_CUMULUS\_EXPANSION | MSigDB lists | MATZUK\_CUMULUS\_EXPANSION | 1 | 1 | 4161 | 156 | tkv | | 3.749e-02 | -3.28 | NIKOLSKY\_OVERCONNECTED\_IN\_BREAST\_CANCER | MSigDB lists | NIKOLSKY\_OVERCONNECTED\_IN\_BREAST\_CANCER | 1 | 1 | 4161 | 156 | Mef2 | | 3.749e-02 | -3.28 | HINATA\_NFKB\_IMMU\_INF | MSigDB lists | HINATA\_NFKB\_IMMU\_INF | 1 | 1 | 4161 | 156 | cact | | 3.749e-02 | -3.28 | GO\_REGULATION\_OF\_CHEMOKINE\_BIOSYNTHETIC\_PROCESS | MSigDB lists | GO\_REGULATION\_OF\_CHEMOKINE\_BIOSYNTHETIC\_PROCESS | 1 | 1 | 4161 | 156 | Tollo | | 3.749e-02 | -3.28 | NIKOLSKY\_BREAST\_CANCER\_19Q13.1\_AMPLICON | MSigDB lists | NIKOLSKY\_BREAST\_CANCER\_19Q13.1\_AMPLICON | 1 | 1 | 4161 | 156 | crol | | 3.749e-02 | -3.28 | REACTOME\_PEPTIDE\_HORMONE\_BIOSYNTHESIS | MSigDB lists | REACTOME\_PEPTIDE\_HORMONE\_BIOSYNTHESIS | 1 | 1 | 4161 | 156 | Actbeta | | 3.749e-02 | -3.28 | GO\_MICROGLIAL\_CELL\_ACTIVATION | MSigDB lists | GO\_MICROGLIAL\_CELL\_ACTIVATION | 1 | 1 | 4161 | 156 | Tollo | | 3.749e-02 | -3.28 | GO\_RENAL\_SYSTEM\_VASCULATURE\_DEVELOPMENT | MSigDB lists | GO\_RENAL\_SYSTEM\_VASCULATURE\_DEVELOPMENT | 1 | 1 | 4161 | 156 | N | | 3.749e-02 | -3.28 | GO\_SEGMENT\_SPECIFICATION | MSigDB lists | GO\_SEGMENT\_SPECIFICATION | 1 | 1 | 4161 | 156 | Dl | | 3.749e-02 | -3.28 | GO\_POSITIVE\_REGULATION\_OF\_KERATINOCYTE\_DIFFERENTIATION | MSigDB lists | GO\_POSITIVE\_REGULATION\_OF\_KERATINOCYTE\_DIFFERENTIATION | 1 | 1 | 4161 | 156 | N | | 3.749e-02 | -3.28 | VANLOO\_SP3\_TARGETS\_UP | MSigDB lists | VANLOO\_SP3\_TARGETS\_UP | 1 | 1 | 4161 | 156 | CG1316 | | 3.749e-02 | -3.28 | GO\_TRIPARTITE\_REGIONAL\_SUBDIVISION | MSigDB lists | GO\_TRIPARTITE\_REGIONAL\_SUBDIVISION | 1 | 1 | 4161 | 156 | fz2 | | 3.749e-02 | -3.28 | GO\_MATURE\_B\_CELL\_DIFFERENTIATION | MSigDB lists | GO\_MATURE\_B\_CELL\_DIFFERENTIATION | 1 | 1 | 4161 | 156 | Dl | | 3.749e-02 | -3.28 | MIKKELSEN\_NPC\_ICP\_WITH\_H3K27ME3 | MSigDB lists | MIKKELSEN\_NPC\_ICP\_WITH\_H3K27ME3 | 1 | 1 | 4161 | 156 | lin-28 | | 3.749e-02 | -3.28 | MODULE\_424 | MSigDB lists | MODULE\_424 | 1 | 1 | 4161 | 156 | Tollo | | 3.749e-02 | -3.28 | HUMMERICH\_MALIGNANT\_SKIN\_TUMOR\_DN | MSigDB lists | HUMMERICH\_MALIGNANT\_SKIN\_TUMOR\_DN | 1 | 1 | 4161 | 156 | SERCA | | 3.749e-02 | -3.28 | PID\_IL23\_PATHWAY | MSigDB lists | PID\_IL23\_PATHWAY | 1 | 1 | 4161 | 156 | cact | | 3.749e-02 | -3.28 | GO\_POSITIVE\_REGULATION\_OF\_SPROUTING\_ANGIOGENESIS | MSigDB lists | GO\_POSITIVE\_REGULATION\_OF\_SPROUTING\_ANGIOGENESIS | 1 | 1 | 4161 | 156 | Dl | | 3.749e-02 | -3.28 | GO\_MATURE\_B\_CELL\_DIFFERENTIATION\_INVOLVED\_IN\_IMMUNE\_RESPONSE | MSigDB lists | GO\_MATURE\_B\_CELL\_DIFFERENTIATION\_INVOLVED\_IN\_IMMUNE\_RESPONSE | 1 | 1 | 4161 | 156 | Dl | | 3.749e-02 | -3.28 | RODRIGUES\_NTN1\_TARGETS\_UP | MSigDB lists | RODRIGUES\_NTN1\_TARGETS\_UP | 1 | 1 | 4161 | 156 | FBXO11 | | 3.749e-02 | -3.28 | GO\_EPHRIN\_RECEPTOR\_ACTIVITY | MSigDB lists | GO\_EPHRIN\_RECEPTOR\_ACTIVITY | 1 | 1 | 4161 | 156 | Eph | | 3.749e-02 | -3.28 | SHIN\_B\_CELL\_LYMPHOMA\_CLUSTER\_3 | MSigDB lists | SHIN\_B\_CELL\_LYMPHOMA\_CLUSTER\_3 | 1 | 1 | 4161 | 156 | fz | | 3.749e-02 | -3.28 | GO\_REGULATION\_OF\_MONONUCLEAR\_CELL\_MIGRATION | MSigDB lists | GO\_REGULATION\_OF\_MONONUCLEAR\_CELL\_MIGRATION | 1 | 1 | 4161 | 156 | sli | | 3.749e-02 | -3.28 | GO\_ACTIVATION\_OF\_MAPKKK\_ACTIVITY | MSigDB lists | GO\_ACTIVATION\_OF\_MAPKKK\_ACTIVITY | 1 | 1 | 4161 | 156 | hppy | | 3.749e-02 | -3.28 | ALONSO\_METASTASIS\_DN | MSigDB lists | ALONSO\_METASTASIS\_DN | 1 | 1 | 4161 | 156 | Ids | | 3.749e-02 | -3.28 | GO\_TRANSCRIPTIONAL\_ACTIVATOR\_ACTIVITY\_RNA\_POLYMERASE\_II\_DISTAL\_ENHANCER\_SEQUENCE\_SPECIFIC\_BINDING | MSigDB lists | GO\_TRANSCRIPTIONAL\_ACTIVATOR\_ACTIVITY\_RNA\_POLYMERASE\_II\_DISTAL\_ENHANCER\_SEQUENCE\_SPECIFIC\_BINDING | 1 | 1 | 4161 | 156 | Mef2 | | 3.749e-02 | -3.28 | GO\_REGULATION\_OF\_ENDOTHELIAL\_CELL\_CHEMOTAXIS | MSigDB lists | GO\_REGULATION\_OF\_ENDOTHELIAL\_CELL\_CHEMOTAXIS | 1 | 1 | 4161 | 156 | N | | 3.749e-02 | -3.28 | TURJANSKI\_MAPK14\_TARGETS | MSigDB lists | TURJANSKI\_MAPK14\_TARGETS | 1 | 1 | 4161 | 156 | Mef2 | | 3.749e-02 | -3.28 | HUMMEL\_BURKITTS\_LYMPHOMA\_DN | MSigDB lists | HUMMEL\_BURKITTS\_LYMPHOMA\_DN | 1 | 1 | 4161 | 156 | cact | | 3.749e-02 | -3.28 | TURJANSKI\_MAPK7\_TARGETS | MSigDB lists | TURJANSKI\_MAPK7\_TARGETS | 1 | 1 | 4161 | 156 | Mef2 | | 3.749e-02 | -3.28 | HUMMERICH\_BENIGN\_SKIN\_TUMOR\_DN | MSigDB lists | HUMMERICH\_BENIGN\_SKIN\_TUMOR\_DN | 1 | 1 | 4161 | 156 | SERCA | | 3.749e-02 | -3.28 | GO\_WHITE\_FAT\_CELL\_DIFFERENTIATION | MSigDB lists | GO\_WHITE\_FAT\_CELL\_DIFFERENTIATION | 1 | 1 | 4161 | 156 | CtBP | | 3.749e-02 | -3.28 | REACTOME\_APOPTOTIC\_CLEAVAGE\_OF\_CELL\_ADHESION\_PROTEINS | MSigDB lists | REACTOME\_APOPTOTIC\_CLEAVAGE\_OF\_CELL\_ADHESION\_PROTEINS | 1 | 1 | 4161 | 156 | arm | | 3.749e-02 | -3.28 | RIZ\_ERYTHROID\_DIFFERENTIATION\_APOBEC2 | MSigDB lists | RIZ\_ERYTHROID\_DIFFERENTIATION\_APOBEC2 | 1 | 1 | 4161 | 156 | Mef2 | | 3.749e-02 | -3.28 | GO\_NEGATIVE\_REGULATION\_OF\_ENDOTHELIAL\_CELL\_APOPTOTIC\_PROCESS | MSigDB lists | GO\_NEGATIVE\_REGULATION\_OF\_ENDOTHELIAL\_CELL\_APOPTOTIC\_PROCESS | 1 | 1 | 4161 | 156 | Raf | | 3.749e-02 | -3.28 | TSUTSUMI\_FBXW8\_TARGETS | MSigDB lists | TSUTSUMI\_FBXW8\_TARGETS | 1 | 1 | 4161 | 156 | SERCA | | 3.749e-02 | -3.28 | GO\_POSITIVE\_REGULATION\_OF\_NATURAL\_KILLER\_CELL\_ACTIVATION | MSigDB lists | GO\_POSITIVE\_REGULATION\_OF\_NATURAL\_KILLER\_CELL\_ACTIVATION | 1 | 1 | 4161 | 156 | AP-1gamma | | 3.749e-02 | -3.28 | TSAI\_DNAJB4\_TARGETS\_DN | MSigDB lists | TSAI\_DNAJB4\_TARGETS\_DN | 1 | 1 | 4161 | 156 | CG1673 | | 3.749e-02 | -3.28 | GO\_REGULATION\_OF\_PRI\_MIRNA\_TRANSCRIPTION\_FROM\_RNA\_POLYMERASE\_II\_PROMOTER | MSigDB lists | GO\_REGULATION\_OF\_PRI\_MIRNA\_TRANSCRIPTION\_FROM\_RNA\_POLYMERASE\_II\_PROMOTER | 1 | 1 | 4161 | 156 | sd | | 3.749e-02 | -3.28 | CGGTGTG\_MIR220 | MSigDB lists | CGGTGTG\_MIR220 | 1 | 1 | 4161 | 156 | ap | | 3.749e-02 | -3.28 | RASHI\_NFKB1\_TARGETS | MSigDB lists | RASHI\_NFKB1\_TARGETS | 1 | 1 | 4161 | 156 | cact | | 3.749e-02 | -3.28 | GO\_POSITIVE\_REGULATION\_OF\_PRI\_MIRNA\_TRANSCRIPTION\_FROM\_RNA\_POLYMERASE\_II\_PROMOTER | MSigDB lists | GO\_POSITIVE\_REGULATION\_OF\_PRI\_MIRNA\_TRANSCRIPTION\_FROM\_RNA\_POLYMERASE\_II\_PROMOTER | 1 | 1 | 4161 | 156 | sd | | 3.749e-02 | -3.28 | BIOCARTA\_TNFR2\_PATHWAY | MSigDB lists | BIOCARTA\_TNFR2\_PATHWAY | 1 | 1 | 4161 | 156 | cact | | 3.749e-02 | -3.28 | GO\_REGULATION\_OF\_MONOCYTE\_CHEMOTAXIS | MSigDB lists | GO\_REGULATION\_OF\_MONOCYTE\_CHEMOTAXIS | 1 | 1 | 4161 | 156 | sli | | 3.756e-02 | -3.28 | imaginal disc fusion | biological process | GO:0046528 | 21 | 3 | 12345 | 442 | Src42A,kay,Pvr | | 3.756e-02 | -3.28 | positive regulation of MAP kinase activity | biological process | GO:0043406 | 21 | 3 | 12345 | 442 | Raf,hppy,wnd | | 3.756e-02 | -3.28 | positive regulation of protein secretion | biological process | GO:0050714 | 21 | 3 | 12345 | 442 | Rab1,S,mop | | 3.756e-02 | -3.28 | imaginal disc fusion, thorax closure | biological process | GO:0046529 | 21 | 3 | 12345 | 442 | Src42A,Pvr,kay | | 3.756e-02 | -3.28 | positive regulation of cell adhesion | biological process | GO:0045785 | 21 | 3 | 12345 | 442 | RhoL,sli,robo1 | | 3.769e-02 | -3.28 | purine nucleoside bisphosphate metabolic process | biological process | GO:0034032 | 52 | 5 | 12345 | 442 | CG8306,muc,Ppcs,ATPCL,pdgy | | 3.769e-02 | -3.28 | nucleoside bisphosphate metabolic process | biological process | GO:0033865 | 52 | 5 | 12345 | 442 | muc,CG8306,Ppcs,pdgy,ATPCL | | 3.769e-02 | -3.28 | ribonucleoside bisphosphate metabolic process | biological process | GO:0033875 | 52 | 5 | 12345 | 442 | pdgy,ATPCL,Ppcs,CG8306,muc | | 3.787e-02 | -3.27 | GSE5542\_UNTREATED\_VS\_IFNA\_TREATED\_EPITHELIAL\_CELLS\_6H\_DN | MSigDB lists | GSE5542\_UNTREATED\_VS\_IFNA\_TREATED\_EPITHELIAL\_CELLS\_6H\_DN | 50 | 5 | 4161 | 156 | InR,rg,Drak,crol,C1GalTA | | 3.787e-02 | -3.27 | GO\_REGULATION\_OF\_EPITHELIAL\_CELL\_PROLIFERATION | MSigDB lists | GO\_REGULATION\_OF\_EPITHELIAL\_CELL\_PROLIFERATION | 50 | 5 | 4161 | 156 | Cdk4,N,hyx,Mef2,arm | | 3.787e-02 | -3.27 | TAATGTG\_MIR323 | MSigDB lists | TAATGTG\_MIR323 | 50 | 5 | 4161 | 156 | Fem-1,Rab1,Hel89B,sd,kn | | 3.787e-02 | -3.27 | LI\_INDUCED\_T\_TO\_NATURAL\_KILLER\_UP | MSigDB lists | LI\_INDUCED\_T\_TO\_NATURAL\_KILLER\_UP | 50 | 5 | 4161 | 156 | CG9220,CG6891,CycG,CG32795,CG31522 | | 3.787e-02 | -3.27 | GSE24726\_WT\_VS\_E2\_2\_KO\_PDC\_DAY4\_POST\_DELETION\_UP | MSigDB lists | GSE24726\_WT\_VS\_E2\_2\_KO\_PDC\_DAY4\_POST\_DELETION\_UP | 50 | 5 | 4161 | 156 | Arf79F,Psa,Nmda1,muc,Rgl | | 3.787e-02 | -3.27 | MEK\_UP.V1\_UP | MSigDB lists | MEK\_UP.V1\_UP | 50 | 5 | 4161 | 156 | CG6891,Mef2,otk,htl,Rab5 | | 3.787e-02 | -3.27 | GSE4984\_GALECTIN1\_VS\_LPS\_STIM\_DC\_UP | MSigDB lists | GSE4984\_GALECTIN1\_VS\_LPS\_STIM\_DC\_UP | 50 | 5 | 4161 | 156 | Vps13,Dyb,GlcAT-I,Pex5,step | | 3.787e-02 | -3.27 | GSE20715\_WT\_VS\_TLR4\_KO\_24H\_OZONE\_LUNG\_UP | MSigDB lists | GSE20715\_WT\_VS\_TLR4\_KO\_24H\_OZONE\_LUNG\_UP | 50 | 5 | 4161 | 156 | Tpst,lt,subdued,ap,CG3107 | | 3.787e-02 | -3.27 | GSE12507\_PDC\_CELL\_LINE\_VS\_IMMATUE\_T\_CELL\_LINE\_UP | MSigDB lists | GSE12507\_PDC\_CELL\_LINE\_VS\_IMMATUE\_T\_CELL\_LINE\_UP | 50 | 5 | 4161 | 156 | CycG,CG1673,Pen,Cdk4,Pgk | | 3.787e-02 | -3.27 | GSE17186\_MEMORY\_VS\_NAIVE\_BCELL\_UP | MSigDB lists | GSE17186\_MEMORY\_VS\_NAIVE\_BCELL\_UP | 50 | 5 | 4161 | 156 | sina,Pgk,Pgant6,nmo,alpha-Man-Ia | | 3.790e-02 | -3.27 | Vps4 (Vacuolar protein sorting 4) | protein interactions | 32777 | 40 | 5 | 9204 | 432 | Pdk1,Gfat2,Mpcp2,GluProRS,Dronc | | 3.796e-02 | -3.27 | locomotory behavior | biological process | GO:0007626 | 188 | 12 | 12345 | 442 | Dcr-2,tara,Mef2,sim,Mid1,Adf1,kay,ey,ITP,dome,InR,SERCA | | 3.803e-02 | -3.27 | regulation of protein localization | biological process | GO:0032880 | 107 | 8 | 12345 | 442 | rdx,jar,Rab1,S,mop,ft,dlp,Abl | | 3.803e-02 | -3.27 | negative regulation of programmed cell death | biological process | GO:0043069 | 107 | 8 | 12345 | 442 | Raf,spi,eya,Diap1,Pvr,sens,Egfr,Pdk1 | | 3.805e-02 | -3.27 | ArfGap | pfam domains | PF01412 | 8 | 2 | 10480 | 419 | CenB1A,drongo | | 3.805e-02 | -3.27 | Innexin | pfam domains | PF00876 | 8 | 2 | 10480 | 419 | Inx2,ogre | | 3.805e-02 | -3.27 | HTH\_psq | pfam domains | PF05225 | 8 | 2 | 10480 | 419 | bab2,rib | | 3.805e-02 | -3.27 | Histone | pfam domains | PF00125 | 8 | 2 | 10480 | 419 | His2Av,His3.3A | | 3.807e-02 | -3.27 | GTPase binding | molecular function | GO:0051020 | 198 | 12 | 12176 | 414 | nuf,TBC1D5,step,Magi,CG17883,Rgl,CG4645,siz,RhoGEF3,CG5522,RhoGEF4,Ziz | | 3.812e-02 | -3.27 | negative regulation of cell population proliferation | biological process | GO:0008285 | 88 | 7 | 12345 | 442 | scrib,Pez,sd,Rab5,Dronc,pxb,ft | | 3.831e-02 | -3.26 | MODULE\_48 | MSigDB lists | MODULE\_48 | 67 | 6 | 4161 | 156 | sina,N,Unc-76,tkv,dlp,14-3-3epsilon | | 3.831e-02 | -3.26 | YAMAZAKI\_TCEB3\_TARGETS\_DN | MSigDB lists | YAMAZAKI\_TCEB3\_TARGETS\_DN | 67 | 6 | 4161 | 156 | fz2,ATPCL,Pen,InR,CtBP,CG45050 | | 3.831e-02 | -3.26 | KEGG\_MAPK\_SIGNALING\_PATHWAY | MSigDB lists | KEGG\_MAPK\_SIGNALING\_PATHWAY | 67 | 6 | 4161 | 156 | nmo,hppy,Mef2,Raf,htl,Lk6 | | 3.831e-02 | -3.26 | GO\_CYTOSOLIC\_TRANSPORT | MSigDB lists | GO\_CYTOSOLIC\_TRANSPORT | 67 | 6 | 4161 | 156 | Cadps,AP-1gamma,Syx1A,arm,Vps13,Arf79F | | 3.837e-02 | -3.26 | Cytokine Signaling in Immune system | REACTOME pathways | R-DME-1280215 | 169 | 13 | 3923 | 176 | Pvr,Egfr,vn,Cbl,Raf,wdb,CG42684,Dronc,CG4935,14-3-3zeta,PI31,htl,Syx1A | | 3.869e-02 | -3.25 | epithelial cell migration, open tracheal system | biological process | GO:0007427 | 36 | 4 | 12345 | 442 | dlp,rib,esg,sli | | 3.869e-02 | -3.25 | septate junction assembly | biological process | GO:0019991 | 36 | 4 | 12345 | 442 | scrib,Gli,moody,bou | | 3.869e-02 | -3.25 | positive regulation of establishment of protein localization | biological process | GO:1904951 | 36 | 4 | 12345 | 442 | S,Rab1,Abl,mop | | 3.869e-02 | -3.25 | maintenance of location in cell | biological process | GO:0051651 | 36 | 4 | 12345 | 442 | Vps13,p130CAS,Dg,chic | | 3.894e-02 | -3.25 | negative regulation of phosphorylation | biological process | GO:0042326 | 70 | 6 | 12345 | 442 | wdb,trbl,Cbl,Socs36E,N,CG8312 | | 3.897e-02 | -3.24 | negative regulation of production of molecular mediator of immune response | biological process | GO:0002701 | 9 | 2 | 12345 | 442 | nub,cact | | 3.897e-02 | -3.24 | ventral midline development | biological process | GO:0007418 | 9 | 2 | 12345 | 442 | sim,mam | | 3.897e-02 | -3.24 | regulation of actin nucleation | biological process | GO:0051125 | 9 | 2 | 12345 | 442 | SCAR,Cip4 | | 3.897e-02 | -3.24 | activation of MAPKK activity | biological process | GO:0000186 | 9 | 2 | 12345 | 442 | Raf,wnd | | 3.897e-02 | -3.24 | endoderm development | biological process | GO:0007492 | 9 | 2 | 12345 | 442 | tkv,htl | | 3.897e-02 | -3.24 | positive regulation of lamellipodium assembly | biological process | GO:0010592 | 9 | 2 | 12345 | 442 | Abl,SCAR | | 3.897e-02 | -3.24 | female genitalia development | biological process | GO:0030540 | 9 | 2 | 12345 | 442 | Abd-B,otk | | 3.897e-02 | -3.24 | regulation of protein tyrosine kinase activity | biological process | GO:0061097 | 9 | 2 | 12345 | 442 | Cbl,Socs36E | | 3.897e-02 | -3.24 | negative regulation of axonogenesis | biological process | GO:0050771 | 9 | 2 | 12345 | 442 | Sema1b,Sema1a | | 3.897e-02 | -3.24 | regulation of lamellocyte differentiation | biological process | GO:0035203 | 9 | 2 | 12345 | 442 | N,kn | | 3.897e-02 | -3.24 | regulation of plasmatocyte differentiation | biological process | GO:0045613 | 9 | 2 | 12345 | 442 | kn,N | | 3.897e-02 | -3.24 | DNA replication-independent nucleosome organization | biological process | GO:0034724 | 9 | 2 | 12345 | 442 | His3.3B,His3.3A | | 3.897e-02 | -3.24 | dorsal closure, spreading of leading edge cells | biological process | GO:0007395 | 9 | 2 | 12345 | 442 | Src42A,Egfr | | 3.897e-02 | -3.24 | negative regulation of neuromuscular synaptic transmission | biological process | GO:1900074 | 9 | 2 | 12345 | 442 | wgn,jeb | | 3.897e-02 | -3.24 | positive regulation of lamellipodium organization | biological process | GO:1902745 | 9 | 2 | 12345 | 442 | Abl,SCAR | | 3.897e-02 | -3.24 | compartment pattern specification | biological process | GO:0007386 | 9 | 2 | 12345 | 442 | N,ena | | 3.897e-02 | -3.24 | intercellular transport | biological process | GO:0010496 | 9 | 2 | 12345 | 442 | Inx2,ogre | | 3.912e-02 | -3.24 | BONOME\_OVARIAN\_CANCER\_SURVIVAL\_SUBOPTIMAL\_DEBULKING | MSigDB lists | BONOME\_OVARIAN\_CANCER\_SURVIVAL\_SUBOPTIMAL\_DEBULKING | 142 | 10 | 4161 | 156 | ft,muc,ds,lqf,Dyb,Mef2,Fkbp14,Cerk,shot,rdx | | 3.929e-02 | -3.24 | GO\_NEURON\_DEVELOPMENT | MSigDB lists | GO\_NEURON\_DEVELOPMENT | 162 | 11 | 4161 | 156 | Eph,robo1,Mef2,ft,sli,tkv,Unc-76,ap,N,sina,unc-5 | | 3.946e-02 | -3.23 | klg (klingon) | protein interactions | 42707 | 7 | 2 | 9204 | 432 | apolpp,N | | 3.946e-02 | -3.23 | Scox (Synthesis of cytochrome c oxidase) | protein interactions | 33711 | 7 | 2 | 9204 | 432 | Diap1,Pdk1 | | 3.946e-02 | -3.23 | Nab2 (Nuclear polyadenosine RNA-binding 2) | protein interactions | 42897 | 7 | 2 | 9204 | 432 | H,pnr | | 3.946e-02 | -3.23 | Pngl (PNGase-like) | protein interactions | 35527 | 7 | 2 | 9204 | 432 | N,tkv | | 3.946e-02 | -3.23 | nAChRalpha6 (nicotinic Acetylcholine Receptor alpha6) | protein interactions | 34304 | 7 | 2 | 9204 | 432 | Dg,Fem-1 | | 3.946e-02 | -3.23 | lin (lines) | protein interactions | 45325 | 7 | 2 | 9204 | 432 | bowl,Abd-B | | 3.946e-02 | -3.23 | puf (puffyeye) | protein interactions | 42935 | 7 | 2 | 9204 | 432 | Sema1a,Diap1 | | 3.946e-02 | -3.23 | CG16904 (uncharacterized protein) | protein interactions | 41212 | 7 | 2 | 9204 | 432 | nub,rdx | | 3.946e-02 | -3.23 | pad (poils au dos) | protein interactions | 41981 | 7 | 2 | 9204 | 432 | h,tkv | | 3.946e-02 | -3.23 | CG8360 (uncharacterized protein) | protein interactions | 34122 | 7 | 2 | 9204 | 432 | rdx,nub | | 3.946e-02 | -3.23 | CG7239 (uncharacterized protein) | protein interactions | 33809 | 7 | 2 | 9204 | 432 | CG30344,Adf1 | | 3.946e-02 | -3.23 | Su(var)2-HP2 (Su(var)2-HP2) | protein interactions | 36621 | 7 | 2 | 9204 | 432 | Sema1a,vig | | 3.946e-02 | -3.23 | mib1 (mind bomb 1) | protein interactions | 39750 | 7 | 2 | 9204 | 432 | Dl,N | | 3.946e-02 | -3.23 | CG30183 (uncharacterized protein) | protein interactions | 37749 | 7 | 2 | 9204 | 432 | CtBP,nmo | | 3.946e-02 | -3.23 | CG15803 (uncharacterized protein) | protein interactions | 42206 | 7 | 2 | 9204 | 432 | CG4896,ed | | 3.946e-02 | -3.23 | atms (antimeros) | protein interactions | 40593 | 7 | 2 | 9204 | 432 | hyx,CG12769 | | 3.946e-02 | -3.23 | Mmp1 (Matrix metalloproteinase 1) | protein interactions | 37949 | 7 | 2 | 9204 | 432 | shg,Sema1a | | 3.946e-02 | -3.23 | CG14589 (uncharacterized protein) | protein interactions | 35541 | 7 | 2 | 9204 | 432 | Kr-h2,lncRNA:CR42862 | | 3.948e-02 | -3.23 | Fkbp14 (FK506-binding protein 14) | protein interactions | 37449 | 68 | 7 | 9204 | 432 | Pgk,chic,Jupiter,Dl,Bacc,Nlp,kay | | 3.949e-02 | -3.23 | fly chr100D1-100D1\|3-102 | chromosome location | fly chr100D1-100D1\|3-102 | 14 | 2 | 25010 | 571 | lncRNA:CR44091,ttk | | 3.949e-02 | -3.23 | fly chr100D1-100D1\|3-102 cM | chromosome location | fly chr100D1-100D1\|3-102 cM | 14 | 2 | 25010 | 571 | lncRNA:CR44091,ttk | | 3.953e-02 | -3.23 | postsynapse | cellular component | GO:0098794 | 72 | 6 | 11940 | 417 | nAChRalpha4,Ir21a,pum,Cals,Magi,scrib | | 3.953e-02 | -3.23 | AGL/Gdb1 | interpro domains | IPR010401 | 1 | 1 | 11383 | 450 | CG9485 | | 3.953e-02 | -3.23 | VPS41 | interpro domains | IPR016902 | 1 | 1 | 11383 | 450 | lt | | 3.953e-02 | -3.23 | Adaptor\_Cbl | interpro domains | IPR024162 | 1 | 1 | 11383 | 450 | Cbl | | 3.953e-02 | -3.23 | Crust\_neurohorm | interpro domains | IPR031098 | 1 | 1 | 11383 | 450 | ITP | | 3.953e-02 | -3.23 | PFN\_sf | interpro domains | IPR036140 | 1 | 1 | 11383 | 450 | chic | | 3.953e-02 | -3.23 | AGL\_glucanoTrfase | interpro domains | IPR032792 | 1 | 1 | 11383 | 450 | CG9485 | | 3.953e-02 | -3.23 | TF\_DP\_C\_sf | interpro domains | IPR038168 | 1 | 1 | 11383 | 450 | Dp | | 3.953e-02 | -3.23 | Crust\_neurhormone\_CS | interpro domains | IPR018251 | 1 | 1 | 11383 | 450 | ITP | | 3.953e-02 | -3.23 | RBM45\_RRM3 | interpro domains | IPR034207 | 1 | 1 | 11383 | 450 | CG1316 | | 3.953e-02 | -3.23 | MYSc\_Myo15 | interpro domains | IPR036057 | 1 | 1 | 11383 | 450 | Myo10A | | 3.953e-02 | -3.23 | Myosin-VI\_CBD | interpro domains | IPR032412 | 1 | 1 | 11383 | 450 | jar | | 3.953e-02 | -3.23 | ABL1 | interpro domains | IPR033221 | 1 | 1 | 11383 | 450 | Abl | | 3.953e-02 | -3.23 | Tyr\_kinase\_EGF/ERB/XmrK\_rcpt | interpro domains | IPR016245 | 1 | 1 | 11383 | 450 | Egfr | | 3.953e-02 | -3.23 | BTG-like\_sf | interpro domains | IPR036054 | 1 | 1 | 11383 | 450 | Tob | | 3.953e-02 | -3.23 | Neuregulin | interpro domains | IPR040180 | 1 | 1 | 11383 | 450 | vn | | 3.953e-02 | -3.23 | PNISR | interpro domains | IPR031937 | 1 | 1 | 11383 | 450 | CG31211 | | 3.953e-02 | -3.23 | BRD4\_CDT | interpro domains | IPR031354 | 1 | 1 | 11383 | 450 | fs(1)h | | 3.953e-02 | -3.23 | DUF625 | interpro domains | IPR006887 | 1 | 1 | 11383 | 450 | flfl | | 3.953e-02 | -3.23 | Epsin-2\_metazoa | interpro domains | IPR027319 | 1 | 1 | 11383 | 450 | lqf | | 3.953e-02 | -3.23 | BCAT\_family | interpro domains | IPR033939 | 1 | 1 | 11383 | 450 | CG1673 | | 3.953e-02 | -3.23 | Sxl | interpro domains | IPR006546 | 1 | 1 | 11383 | 450 | Sxl | | 3.953e-02 | -3.23 | Shroom\_fam | interpro domains | IPR027685 | 1 | 1 | 11383 | 450 | Shrm | | 3.953e-02 | -3.23 | RFT1 | interpro domains | IPR007594 | 1 | 1 | 11383 | 450 | CG3149 | | 3.953e-02 | -3.23 | NCK | interpro domains | IPR017304 | 1 | 1 | 11383 | 450 | dock | | 3.953e-02 | -3.23 | 2OGFeDO\_noxygenase\_dom | interpro domains | IPR024779 | 1 | 1 | 11383 | 450 | Tet | | 3.953e-02 | -3.23 | OCRL1/INPP5B\_INPP5c | interpro domains | IPR037793 | 1 | 1 | 11383 | 450 | Ocrl | | 3.953e-02 | -3.23 | EYA\_dom\_sf | interpro domains | IPR038102 | 1 | 1 | 11383 | 450 | eya | | 3.953e-02 | -3.23 | ATP-citrate\_synthase | interpro domains | IPR014608 | 1 | 1 | 11383 | 450 | ATPCL | | 3.953e-02 | -3.23 | Cyclin\_B3 | interpro domains | IPR015452 | 1 | 1 | 11383 | 450 | CycB3 | | 3.953e-02 | -3.23 | CREBRF | interpro domains | IPR039165 | 1 | 1 | 11383 | 450 | REPTOR | | 3.953e-02 | -3.23 | RING-HC\_ZNF598/Hel2 | interpro domains | IPR041888 | 1 | 1 | 11383 | 450 | CG11414 | | 3.953e-02 | -3.23 | HPS3 | interpro domains | IPR017216 | 1 | 1 | 11383 | 450 | CG14562 | | 3.953e-02 | -3.23 | Mehydrof\_redctse | interpro domains | IPR003171 | 1 | 1 | 11383 | 450 | CG7560 | | 3.953e-02 | -3.23 | Pianissimo\_fam | interpro domains | IPR028268 | 1 | 1 | 11383 | 450 | rictor | | 3.953e-02 | -3.23 | TEF\_metazoa | interpro domains | IPR016361 | 1 | 1 | 11383 | 450 | sd | | 3.953e-02 | -3.23 | Pro-tRNA-ligase\_IIa\_arc-type | interpro domains | IPR004499 | 1 | 1 | 11383 | 450 | GluProRS | | 3.953e-02 | -3.23 | COE\_HLH | interpro domains | IPR032201 | 1 | 1 | 11383 | 450 | kn | | 3.953e-02 | -3.23 | VIP1\_N | interpro domains | IPR040557 | 1 | 1 | 11383 | 450 | l(1)G0196 | | 3.953e-02 | -3.23 | SERCA1/2 | interpro domains | IPR030332 | 1 | 1 | 11383 | 450 | SERCA | | 3.953e-02 | -3.23 | Tc1-like\_DDE\_dom | interpro domains | IPR038717 | 1 | 1 | 11383 | 450 | CG14478 | | 3.953e-02 | -3.23 | CLP | interpro domains | IPR030502 | 1 | 1 | 11383 | 450 | CG6891 | | 3.953e-02 | -3.23 | Nav/unc-53 | interpro domains | IPR039041 | 1 | 1 | 11383 | 450 | sick | | 3.953e-02 | -3.23 | Profilin\_CS | interpro domains | IPR027310 | 1 | 1 | 11383 | 450 | chic | | 3.953e-02 | -3.23 | Autophagy-rel\_prot\_16 | interpro domains | IPR013923 | 1 | 1 | 11383 | 450 | Atg16 | | 3.953e-02 | -3.23 | VASP\_tetra | interpro domains | IPR014885 | 1 | 1 | 11383 | 450 | ena | | 3.953e-02 | -3.23 | Notch\_NODP\_dom | interpro domains | IPR011656 | 1 | 1 | 11383 | 450 | N | | 3.953e-02 | -3.23 | UNC119\_homologue\_A | interpro domains | IPR032977 | 1 | 1 | 11383 | 450 | unc-119 | | 3.953e-02 | -3.23 | TPST | interpro domains | IPR026634 | 1 | 1 | 11383 | 450 | Tpst | | 3.953e-02 | -3.23 | Fe\_hydrogenase\_lsu\_C | interpro domains | IPR004108 | 1 | 1 | 11383 | 450 | CG17683 | | 3.953e-02 | -3.23 | HJURP\_C | interpro domains | IPR022102 | 1 | 1 | 11383 | 450 | Mef2 | | 3.953e-02 | -3.23 | Band\_4 | interpro domains | IPR008379 | 1 | 1 | 11383 | 450 | cora | | 3.953e-02 | -3.23 | Hyperglycemic | interpro domains | IPR001166 | 1 | 1 | 11383 | 450 | ITP | | 3.953e-02 | -3.23 | AGL\_central | interpro domains | IPR032788 | 1 | 1 | 11383 | 450 | CG9485 | | 3.953e-02 | -3.23 | STX1 | interpro domains | IPR028669 | 1 | 1 | 11383 | 450 | Syx1A | | 3.953e-02 | -3.23 | MaoC-like\_dehydrat\_N | interpro domains | IPR039569 | 1 | 1 | 11383 | 450 | Mfe2 | | 3.953e-02 | -3.23 | Fe\_hydrogenase | interpro domains | IPR009016 | 1 | 1 | 11383 | 450 | CG17683 | | 3.953e-02 | -3.23 | MYSc\_Myo20 | interpro domains | IPR036029 | 1 | 1 | 11383 | 450 | d | | 3.953e-02 | -3.23 | Transcrpt\_fac\_DP | interpro domains | IPR015648 | 1 | 1 | 11383 | 450 | Dp | | 3.953e-02 | -3.23 | Adaptor\_Cbl\_EF\_hand-like | interpro domains | IPR014741 | 1 | 1 | 11383 | 450 | Cbl | | 3.953e-02 | -3.23 | ERp29 | interpro domains | IPR016855 | 1 | 1 | 11383 | 450 | wbl | | 3.953e-02 | -3.23 | Citrate-bd | interpro domains | IPR032263 | 1 | 1 | 11383 | 450 | ATPCL | | 3.953e-02 | -3.23 | Rictor\_IV | interpro domains | IPR029453 | 1 | 1 | 11383 | 450 | rictor | | 3.953e-02 | -3.23 | SHARP\_RRM2 | interpro domains | IPR034173 | 1 | 1 | 11383 | 450 | spen | | 3.953e-02 | -3.23 | CARMIL\_C | interpro domains | IPR031943 | 1 | 1 | 11383 | 450 | LRR | | 3.953e-02 | -3.23 | AHR/AHRR | interpro domains | IPR039091 | 1 | 1 | 11383 | 450 | ss | | 3.953e-02 | -3.23 | C2\_Dock-D | interpro domains | IPR037809 | 1 | 1 | 11383 | 450 | Ziz | | 3.953e-02 | -3.23 | Neuroggenic\_mastermind-like\_N | interpro domains | IPR019082 | 1 | 1 | 11383 | 450 | mam | | 3.953e-02 | -3.23 | Distrobrevin | interpro domains | IPR017432 | 1 | 1 | 11383 | 450 | Dyb | | 3.953e-02 | -3.23 | Notch | interpro domains | IPR008297 | 1 | 1 | 11383 | 450 | N | | 3.953e-02 | -3.23 | RICTOR\_V | interpro domains | IPR029452 | 1 | 1 | 11383 | 450 | rictor | | 3.953e-02 | -3.23 | MADS\_MEF2-like | interpro domains | IPR033896 | 1 | 1 | 11383 | 450 | Mef2 | | 3.953e-02 | -3.23 | Tyr\_kinase\_ephrin\_rcpt | interpro domains | IPR016257 | 1 | 1 | 11383 | 450 | Eph | | 3.953e-02 | -3.23 | ARF7EP\_C | interpro domains | IPR029264 | 1 | 1 | 11383 | 450 | CG14464 | | 3.953e-02 | -3.23 | SMC3\_ABC\_euk | interpro domains | IPR041741 | 1 | 1 | 11383 | 450 | SMC3 | | 3.953e-02 | -3.23 | INPP5B\_PH\_dom | interpro domains | IPR031896 | 1 | 1 | 11383 | 450 | Ocrl | | 3.953e-02 | -3.23 | GDE\_C | interpro domains | IPR032790 | 1 | 1 | 11383 | 450 | CG9485 | | 3.953e-02 | -3.23 | NosD\_dom | interpro domains | IPR007742 | 1 | 1 | 11383 | 450 | FBXO11 | | 3.953e-02 | -3.23 | CoaB-like\_sf | interpro domains | IPR035929 | 1 | 1 | 11383 | 450 | Ppcs | | 3.953e-02 | -3.23 | RBM45\_RRM2 | interpro domains | IPR034206 | 1 | 1 | 11383 | 450 | CG1316 | | 3.953e-02 | -3.23 | Notch\_NOD\_dom | interpro domains | IPR010660 | 1 | 1 | 11383 | 450 | N | | 3.953e-02 | -3.23 | Yip5/YIPF1/YIPF2 | interpro domains | IPR039765 | 1 | 1 | 11383 | 450 | CG4645 | | 3.953e-02 | -3.23 | SLBP\_RNA\_bind\_sf | interpro domains | IPR038294 | 1 | 1 | 11383 | 450 | Slbp | | 3.953e-02 | -3.23 | CAPS | interpro domains | IPR033227 | 1 | 1 | 11383 | 450 | Cadps | | 3.953e-02 | -3.23 | Pumilio\_RNA-bd | interpro domains | IPR033712 | 1 | 1 | 11383 | 450 | pum | | 3.953e-02 | -3.23 | Int\_cplx\_su3 | interpro domains | IPR019333 | 1 | 1 | 11383 | 450 | IntS3 | | 3.953e-02 | -3.23 | Transc\_factor\_DP\_C | interpro domains | IPR014889 | 1 | 1 | 11383 | 450 | Dp | | 3.953e-02 | -3.23 | DAZ\_BOULE\_RRM | interpro domains | IPR034988 | 1 | 1 | 11383 | 450 | bol | | 3.953e-02 | -3.23 | CEP162-like | interpro domains | IPR038774 | 1 | 1 | 11383 | 450 | CG42699 | | 3.953e-02 | -3.23 | EYA\_dom | interpro domains | IPR006545 | 1 | 1 | 11383 | 450 | eya | | 3.953e-02 | -3.23 | DFF40 | interpro domains | IPR039729 | 1 | 1 | 11383 | 450 | Drep4 | | 3.953e-02 | -3.23 | VASP\_sf | interpro domains | IPR038023 | 1 | 1 | 11383 | 450 | ena | | 3.953e-02 | -3.23 | FAM49 | interpro domains | IPR039789 | 1 | 1 | 11383 | 450 | CG32066 | | 3.953e-02 | -3.23 | CSPG\_rpt | interpro domains | IPR039005 | 1 | 1 | 11383 | 450 | kon | | 3.953e-02 | -3.23 | RICTOR\_M | interpro domains | IPR029451 | 1 | 1 | 11383 | 450 | rictor | | 3.953e-02 | -3.23 | SEA\_DG\_dom | interpro domains | IPR030398 | 1 | 1 | 11383 | 450 | Dg | | 3.953e-02 | -3.23 | TEA/ATTS\_dom | interpro domains | IPR000818 | 1 | 1 | 11383 | 450 | sd | | 3.953e-02 | -3.23 | Abdominal-A | interpro domains | IPR022132 | 1 | 1 | 11383 | 450 | abd-A | | 3.953e-02 | -3.23 | Cbl\_PTB | interpro domains | IPR024159 | 1 | 1 | 11383 | 450 | Cbl | | 3.953e-02 | -3.23 | PKNOX/Meis\_N | interpro domains | IPR032453 | 1 | 1 | 11383 | 450 | hth | | 3.953e-02 | -3.23 | LAT1 | interpro domains | IPR006257 | 1 | 1 | 11383 | 450 | muc | | 3.953e-02 | -3.23 | Ribosomal\_L27/L41\_mit | interpro domains | IPR019189 | 1 | 1 | 11383 | 450 | mRpL41 | | 3.953e-02 | -3.23 | Tyr\_kinase\_rcpt\_ROR | interpro domains | IPR016247 | 1 | 1 | 11383 | 450 | Nrk | | 3.953e-02 | -3.23 | PTS1R\_family | interpro domains | IPR024111 | 1 | 1 | 11383 | 450 | Pex5 | | 3.953e-02 | -3.23 | COE\_DBD\_sf | interpro domains | IPR038173 | 1 | 1 | 11383 | 450 | kn | | 3.953e-02 | -3.23 | CARMIL\_PH | interpro domains | IPR041245 | 1 | 1 | 11383 | 450 | LRR | | 3.953e-02 | -3.23 | Transcription\_factor\_COE | interpro domains | IPR003523 | 1 | 1 | 11383 | 450 | kn | | 3.953e-02 | -3.23 | DM15 | interpro domains | IPR006607 | 1 | 1 | 11383 | 450 | larp | | 3.953e-02 | -3.23 | IDS | interpro domains | IPR035874 | 1 | 1 | 11383 | 450 | Ids | | 3.953e-02 | -3.23 | LIMR | interpro domains | IPR008075 | 1 | 1 | 11383 | 450 | lili | | 3.953e-02 | -3.23 | Serine\_rich\_dom | interpro domains | IPR014928 | 1 | 1 | 11383 | 450 | p130CAS | | 3.953e-02 | -3.23 | SATB\_ULD\_sf | interpro domains | IPR038224 | 1 | 1 | 11383 | 450 | dve | | 3.953e-02 | -3.23 | DUF1088 | interpro domains | IPR010508 | 1 | 1 | 11383 | 450 | rg | | 3.953e-02 | -3.23 | PTPN14/21\_FERM\_C | interpro domains | IPR041782 | 1 | 1 | 11383 | 450 | Pez | | 3.953e-02 | -3.23 | Para\_beta\_helix\_rpt-2 | interpro domains | IPR022441 | 1 | 1 | 11383 | 450 | FBXO11 | | 3.953e-02 | -3.23 | FGF\_rcpt\_fam | interpro domains | IPR016248 | 1 | 1 | 11383 | 450 | htl | | 3.953e-02 | -3.23 | Transcription\_factor\_COE\_CS | interpro domains | IPR018350 | 1 | 1 | 11383 | 450 | kn | | 3.953e-02 | -3.23 | F-actin\_binding | interpro domains | IPR015015 | 1 | 1 | 11383 | 450 | Abl | | 3.953e-02 | -3.23 | MAP3K12\_MAP3K13 | interpro domains | IPR017419 | 1 | 1 | 11383 | 450 | wnd | | 3.953e-02 | -3.23 | PDK1-typ\_PH | interpro domains | IPR033931 | 1 | 1 | 11383 | 450 | Pdk1 | | 3.953e-02 | -3.23 | DNA-bd\_prot-inh | interpro domains | IPR026052 | 1 | 1 | 11383 | 450 | emc | | 3.953e-02 | -3.23 | SHARP\_RRM3 | interpro domains | IPR034174 | 1 | 1 | 11383 | 450 | spen | | 3.953e-02 | -3.23 | ATG11 | interpro domains | IPR040040 | 1 | 1 | 11383 | 450 | Atg17 | | 3.953e-02 | -3.23 | SLBP\_RNA\_bind | interpro domains | IPR029344 | 1 | 1 | 11383 | 450 | Slbp | | 3.953e-02 | -3.23 | RNAP\_AC19 | interpro domains | IPR033898 | 1 | 1 | 11383 | 450 | l(2)37Cg | | 3.953e-02 | -3.23 | Pectin\_lyas\_fold | interpro domains | IPR012334 | 1 | 1 | 11383 | 450 | FBXO11 | | 3.953e-02 | -3.23 | COE\_DBD | interpro domains | IPR032200 | 1 | 1 | 11383 | 450 | kn | | 3.953e-02 | -3.23 | FEZ | interpro domains | IPR011680 | 1 | 1 | 11383 | 450 | Unc-76 | | 3.953e-02 | -3.23 | UPF0687 | interpro domains | IPR026794 | 1 | 1 | 11383 | 450 | CG6983 | | 3.953e-02 | -3.23 | Alpha-dystroglycan\_domain\_2 | interpro domains | IPR027468 | 1 | 1 | 11383 | 450 | Dg | | 3.953e-02 | -3.23 | Sm\_D1 | interpro domains | IPR034102 | 1 | 1 | 11383 | 450 | SmD1 | | 3.953e-02 | -3.23 | Carb-bd\_sugar\_hydrolysis-dom | interpro domains | IPR006633 | 1 | 1 | 11383 | 450 | FBXO11 | | 3.953e-02 | -3.23 | Adaptor\_Cbl\_SH2-like | interpro domains | IPR014742 | 1 | 1 | 11383 | 450 | Cbl | | 3.953e-02 | -3.23 | TMPIT | interpro domains | IPR012926 | 1 | 1 | 11383 | 450 | CG32795 | | 3.953e-02 | -3.23 | RBM45\_RRM1 | interpro domains | IPR034203 | 1 | 1 | 11383 | 450 | CG1316 | | 3.953e-02 | -3.23 | CABLES1/2 | interpro domains | IPR012388 | 1 | 1 | 11383 | 450 | CG6191 | | 3.953e-02 | -3.23 | Pianissimo\_N | interpro domains | IPR028267 | 1 | 1 | 11383 | 450 | rictor | | 3.953e-02 | -3.23 | PFN | interpro domains | IPR005455 | 1 | 1 | 11383 | 450 | chic | | 3.953e-02 | -3.23 | CBL-B\_RING-HC | interpro domains | IPR039520 | 1 | 1 | 11383 | 450 | Cbl | | 3.953e-02 | -3.23 | UPA | interpro domains | IPR033772 | 1 | 1 | 11383 | 450 | unc-5 | | 3.953e-02 | -3.23 | TF\_AF4/FMR2 | interpro domains | IPR007797 | 1 | 1 | 11383 | 450 | lilli | | 3.953e-02 | -3.23 | APPBP2 | interpro domains | IPR042476 | 1 | 1 | 11383 | 450 | Pat1 | | 3.953e-02 | -3.23 | Tribbles/Ser\_Thr\_kinase\_40 | interpro domains | IPR024104 | 1 | 1 | 11383 | 450 | trbl | | 3.953e-02 | -3.23 | Nuclear\_rcpt\_coactivator | interpro domains | IPR017426 | 1 | 1 | 11383 | 450 | tai | | 3.953e-02 | -3.23 | DFF40\_C | interpro domains | IPR015311 | 1 | 1 | 11383 | 450 | Drep4 | | 3.953e-02 | -3.23 | Eph\_TM | interpro domains | IPR027936 | 1 | 1 | 11383 | 450 | Eph | | 3.953e-02 | -3.23 | Aminotrans\_IV\_CS | interpro domains | IPR018300 | 1 | 1 | 11383 | 450 | CG1673 | | 3.953e-02 | -3.23 | WTAP | interpro domains | IPR033757 | 1 | 1 | 11383 | 450 | fl(2)d | | 3.953e-02 | -3.23 | Adaptor\_Cbl\_N\_dom\_sf | interpro domains | IPR036537 | 1 | 1 | 11383 | 450 | Cbl | | 3.953e-02 | -3.23 | Beta-catenin | interpro domains | IPR013284 | 1 | 1 | 11383 | 450 | arm | | 3.953e-02 | -3.23 | Yuri\_gagarin | interpro domains | IPR031843 | 1 | 1 | 11383 | 450 | yuri | | 3.953e-02 | -3.23 | TMEM33/Pom33 | interpro domains | IPR005344 | 1 | 1 | 11383 | 450 | Kr-h2 | | 3.953e-02 | -3.23 | PDPK1 | interpro domains | IPR039046 | 1 | 1 | 11383 | 450 | Pdk1 | | 3.953e-02 | -3.23 | Plakin\_repeat\_sf | interpro domains | IPR035915 | 1 | 1 | 11383 | 450 | shot | | 3.953e-02 | -3.23 | Serine\_rich\_sf | interpro domains | IPR038319 | 1 | 1 | 11383 | 450 | p130CAS | | 3.953e-02 | -3.23 | Tob1/2 | interpro domains | IPR015676 | 1 | 1 | 11383 | 450 | Tob | | 3.953e-02 | -3.23 | AP1\_complex\_gsu | interpro domains | IPR017107 | 1 | 1 | 11383 | 450 | AP-1gamma | | 3.953e-02 | -3.23 | UNC5 | interpro domains | IPR037936 | 1 | 1 | 11383 | 450 | unc-5 | | 3.953e-02 | -3.23 | YBD | interpro domains | IPR041086 | 1 | 1 | 11383 | 450 | sd | | 3.953e-02 | -3.23 | Par3/HAL\_N | interpro domains | IPR021922 | 1 | 1 | 11383 | 450 | baz | | 3.953e-02 | -3.23 | Rpr2 | interpro domains | IPR007175 | 1 | 1 | 11383 | 450 | CG33082 | | 3.953e-02 | -3.23 | Atg11\_C | interpro domains | IPR019460 | 1 | 1 | 11383 | 450 | Atg17 | | 3.953e-02 | -3.23 | Tyr\_kinase\_rcpt\_V\_CS | interpro domains | IPR001426 | 1 | 1 | 11383 | 450 | Eph | | 3.953e-02 | -3.23 | Laminin\_aI | interpro domains | IPR009254 | 1 | 1 | 11383 | 450 | LanA | | 3.953e-02 | -3.23 | Peptidase\_M16C\_assoc | interpro domains | IPR013578 | 1 | 1 | 11383 | 450 | CG3107 | | 3.953e-02 | -3.23 | PKHA4/5/6/7\_PH | interpro domains | IPR040392 | 1 | 1 | 11383 | 450 | CG34383 | | 3.953e-02 | -3.23 | Desmoplakin\_SH3 | interpro domains | IPR041615 | 1 | 1 | 11383 | 450 | shot | | 3.953e-02 | -3.23 | ARL14EP | interpro domains | IPR026515 | 1 | 1 | 11383 | 450 | CG14464 | | 3.953e-02 | -3.23 | ERp29\_N | interpro domains | IPR012883 | 1 | 1 | 11383 | 450 | wbl | | 3.953e-02 | -3.23 | Beta\_helix | interpro domains | IPR039448 | 1 | 1 | 11383 | 450 | FBXO11 | | 3.953e-02 | -3.23 | HPS3\_central\_dom | interpro domains | IPR028167 | 1 | 1 | 11383 | 450 | CG14562 | | 3.953e-02 | -3.23 | Pokkuri | interpro domains | IPR033077 | 1 | 1 | 11383 | 450 | aop | | 3.953e-02 | -3.23 | Receptor\_IA-2\_dom | interpro domains | IPR021613 | 1 | 1 | 11383 | 450 | IA-2 | | 3.953e-02 | -3.23 | Spot\_14 | interpro domains | IPR009786 | 1 | 1 | 11383 | 450 | CG2765 | | 3.953e-02 | -3.23 | DUF3535 | interpro domains | IPR022707 | 1 | 1 | 11383 | 450 | Hel89B | | 3.953e-02 | -3.23 | SLBP1/SLBP2 | interpro domains | IPR026502 | 1 | 1 | 11383 | 450 | Slbp | | 3.953e-02 | -3.23 | SMAP\_dom | interpro domains | IPR028124 | 1 | 1 | 11383 | 450 | CG6340 | | 3.953e-02 | -3.23 | Receptor\_IA-2\_ectodomain\_sf | interpro domains | IPR038112 | 1 | 1 | 11383 | 450 | IA-2 | | 3.953e-02 | -3.23 | WTAP/Mum2 | interpro domains | IPR029732 | 1 | 1 | 11383 | 450 | fl(2)d | | 3.953e-02 | -3.23 | Glu-tRNA-synth\_arc/euk | interpro domains | IPR004526 | 1 | 1 | 11383 | 450 | GluProRS | | 3.953e-02 | -3.23 | MYSc\_Myo6 | interpro domains | IPR036114 | 1 | 1 | 11383 | 450 | jar | | 3.953e-02 | -3.23 | Aminotrans\_IV | interpro domains | IPR001544 | 1 | 1 | 11383 | 450 | CG1673 | | 3.953e-02 | -3.23 | DUF4819 | interpro domains | IPR032147 | 1 | 1 | 11383 | 450 | cic | | 3.953e-02 | -3.23 | Znf\_SWIM | interpro domains | IPR007527 | 1 | 1 | 11383 | 450 | CG34401 | | 3.953e-02 | -3.23 | WDR47 | interpro domains | IPR040067 | 1 | 1 | 11383 | 450 | CG2812 | | 3.953e-02 | -3.23 | P-type\_ATPase\_IIA | interpro domains | IPR005782 | 1 | 1 | 11383 | 450 | SERCA | | 3.953e-02 | -3.23 | Crust\_neurohorm\_sf | interpro domains | IPR035957 | 1 | 1 | 11383 | 450 | ITP | | 3.953e-02 | -3.23 | Anc-1-like | interpro domains | IPR039906 | 1 | 1 | 11383 | 450 | klar | | 3.953e-02 | -3.23 | Ephrin\_rcpt\_lig-bd\_dom | interpro domains | IPR001090 | 1 | 1 | 11383 | 450 | Eph | | 3.953e-02 | -3.23 | GUCD1 | interpro domains | IPR018616 | 1 | 1 | 11383 | 450 | CG13760 | | 3.953e-02 | -3.23 | ABL\_SH2 | interpro domains | IPR035837 | 1 | 1 | 11383 | 450 | Abl | | 3.953e-02 | -3.23 | Adaptor\_Cbl\_N\_hlx | interpro domains | IPR003153 | 1 | 1 | 11383 | 450 | Cbl | | 3.953e-02 | -3.23 | Plectin\_repeat | interpro domains | IPR001101 | 1 | 1 | 11383 | 450 | shot | | 3.953e-02 | -3.23 | Fe\_hydrogenase\_ssu | interpro domains | IPR003149 | 1 | 1 | 11383 | 450 | CG17683 | | 3.953e-02 | -3.23 | ProRS\_core\_arch\_euk | interpro domains | IPR033721 | 1 | 1 | 11383 | 450 | GluProRS | | 3.953e-02 | -3.23 | AGL\_euk\_N | interpro domains | IPR029436 | 1 | 1 | 11383 | 450 | CG9485 | | 3.953e-02 | -3.23 | Pro-tRNA\_ligase\_II\_C | interpro domains | IPR016061 | 1 | 1 | 11383 | 450 | GluProRS | | 3.953e-02 | -3.23 | CCNG2 | interpro domains | IPR028895 | 1 | 1 | 11383 | 450 | CycG | | 3.953e-02 | -3.23 | SKICH | interpro domains | IPR041611 | 1 | 1 | 11383 | 450 | CG9784 | | 3.953e-02 | -3.23 | ERp29\_C | interpro domains | IPR011679 | 1 | 1 | 11383 | 450 | wbl | | 3.953e-02 | -3.23 | IQSEC\_PH | interpro domains | IPR033742 | 1 | 1 | 11383 | 450 | siz | | 3.953e-02 | -3.23 | SHARP\_RRM4 | interpro domains | IPR034175 | 1 | 1 | 11383 | 450 | spen | | 3.953e-02 | -3.23 | Enhancer\_polycomb | interpro domains | IPR024943 | 1 | 1 | 11383 | 450 | E(Pc) | | 3.953e-02 | -3.23 | ULD | interpro domains | IPR032392 | 1 | 1 | 11383 | 450 | dve | | 3.953e-02 | -3.23 | Iojap/RsfS/C7orf30 | interpro domains | IPR004394 | 1 | 1 | 11383 | 450 | 312 | | 3.953e-02 | -3.23 | HPF1 | interpro domains | IPR019361 | 1 | 1 | 11383 | 450 | CG1218 | | 3.953e-02 | -3.23 | DUF3498 | interpro domains | IPR021887 | 1 | 1 | 11383 | 450 | CG42684 | | 3.953e-02 | -3.23 | ASD2\_dom | interpro domains | IPR014799 | 1 | 1 | 11383 | 450 | Shrm | | 3.953e-02 | -3.23 | Anti\_prolifrtn | interpro domains | IPR002087 | 1 | 1 | 11383 | 450 | Tob | | 3.953e-02 | -3.23 | DUF1084 | interpro domains | IPR009457 | 1 | 1 | 11383 | 450 | CG2991 | | 3.953e-02 | -3.23 | Pro-tRNA\_synth\_II | interpro domains | IPR017449 | 1 | 1 | 11383 | 450 | GluProRS | | 3.953e-02 | -3.23 | TEA/ATTS\_sf | interpro domains | IPR038096 | 1 | 1 | 11383 | 450 | sd | | 3.953e-02 | -3.23 | PEX14 | interpro domains | IPR025655 | 1 | 1 | 11383 | 450 | Pex14 | | 3.953e-02 | -3.23 | EYA\_dom\_metazoan | interpro domains | IPR042577 | 1 | 1 | 11383 | 450 | eya | | 3.953e-02 | -3.23 | Shroom | interpro domains | IPR030493 | 1 | 1 | 11383 | 450 | Shrm | | 3.953e-02 | -3.23 | ERp29\_C\_sf | interpro domains | IPR036356 | 1 | 1 | 11383 | 450 | wbl | | 3.953e-02 | -3.23 | Pex14\_N | interpro domains | IPR006785 | 1 | 1 | 11383 | 450 | Pex14 | | 3.953e-02 | -3.23 | HPS3\_N | interpro domains | IPR029437 | 1 | 1 | 11383 | 450 | CG14562 | | 3.953e-02 | -3.23 | Laminin\_dom\_II | interpro domains | IPR010307 | 1 | 1 | 11383 | 450 | LanA | | 3.953e-02 | -3.23 | Glycogen\_debranch\_met | interpro domains | IPR006421 | 1 | 1 | 11383 | 450 | CG9485 | | 3.953e-02 | -3.23 | Aminotransferase-like | interpro domains | IPR036038 | 1 | 1 | 11383 | 450 | CG1673 | | 3.953e-02 | -3.23 | CAS\_fam | interpro domains | IPR037362 | 1 | 1 | 11383 | 450 | p130CAS | | 3.953e-02 | -3.23 | FAM43A/B\_PTB | interpro domains | IPR033930 | 1 | 1 | 11383 | 450 | CG8312 | | 3.953e-02 | -3.23 | CAS\_C | interpro domains | IPR021901 | 1 | 1 | 11383 | 450 | p130CAS | | 3.953e-02 | -3.23 | B\_amino\_transII | interpro domains | IPR005786 | 1 | 1 | 11383 | 450 | CG1673 | | 3.953e-02 | -3.23 | His\_Pase\_VIP1 | interpro domains | IPR037446 | 1 | 1 | 11383 | 450 | l(1)G0196 | | 3.953e-02 | -3.23 | Hexapep\_transf\_CS | interpro domains | IPR018357 | 1 | 1 | 11383 | 450 | CG1129 | | 3.953e-02 | -3.23 | COE\_IPT | interpro domains | IPR038006 | 1 | 1 | 11383 | 450 | kn | | 3.953e-02 | -3.23 | DAG1\_C | interpro domains | IPR008465 | 1 | 1 | 11383 | 450 | Dg | | 3.953e-02 | -3.23 | TMEM72 | interpro domains | IPR032055 | 1 | 1 | 11383 | 450 | CG14086 | | 3.953e-02 | -3.23 | EYA\_fam | interpro domains | IPR028472 | 1 | 1 | 11383 | 450 | eya | | 3.953e-02 | -3.23 | Calsyntenin | interpro domains | IPR026914 | 1 | 1 | 11383 | 450 | Cals | | 3.953e-02 | -3.23 | Put\_MeTrfase | interpro domains | IPR003750 | 1 | 1 | 11383 | 450 | CG12128 | | 3.953e-02 | -3.23 | FBX11/DRE-1 | interpro domains | IPR029799 | 1 | 1 | 11383 | 450 | FBXO11 | | 3.953e-02 | -3.23 | TET1/2/3 | interpro domains | IPR040175 | 1 | 1 | 11383 | 450 | Tet | | 3.953e-02 | -3.23 | Midnolin | interpro domains | IPR039336 | 1 | 1 | 11383 | 450 | stx | | 3.953e-02 | -3.23 | RBM45\_RRM4 | interpro domains | IPR034208 | 1 | 1 | 11383 | 450 | CG1316 | | 3.953e-02 | -3.23 | MyoXV\_FERM\_C | interpro domains | IPR041795 | 1 | 1 | 11383 | 450 | Myo10A | | 3.953e-02 | -3.23 | AMPKG-2 | interpro domains | IPR039170 | 1 | 1 | 11383 | 450 | SNF4Agamma | | 3.953e-02 | -3.23 | SATB1/SATB2 | interpro domains | IPR039673 | 1 | 1 | 11383 | 450 | dve | | 3.953e-02 | -3.23 | DNA/pantothenate-metab\_flavo\_C | interpro domains | IPR007085 | 1 | 1 | 11383 | 450 | Ppcs | | 3.953e-02 | -3.23 | DUF3454\_notch | interpro domains | IPR024600 | 1 | 1 | 11383 | 450 | N | | 3.981e-02 | -3.22 | presynaptic membrane | cellular component | GO:0042734 | 22 | 3 | 11940 | 417 | Syx1A,LanA,gukh | | 3.982e-02 | -3.22 | brat (brain tumor) | protein interactions | 35197 | 54 | 6 | 9204 | 432 | arm,orb2,Sxl,pum,klu,Mfe2 | | 3.982e-02 | -3.22 | CG1416 (uncharacterized protein) | protein interactions | 35426 | 54 | 6 | 9204 | 432 | SERCA,Pdk1,Arf79F,Rab1,Psa,Rab5 | | 3.988e-02 | -3.22 | HMG\_box\_dom\_sf | interpro domains | IPR036910 | 33 | 4 | 11383 | 450 | cic,Sox14,HmgZ,CG12104 | | 3.995e-02 | -3.22 | GO\_NEURON\_PROJECTION\_DEVELOPMENT | MSigDB lists | GO\_NEURON\_PROJECTION\_DEVELOPMENT | 123 | 9 | 4161 | 156 | robo1,Eph,sli,N,sina,Unc-76,tkv,ap,unc-5 | | 3.998e-02 | -3.22 | UPF0121 | pfam domains | PF03661 | 1 | 1 | 10480 | 419 | Kr-h2 | | 3.998e-02 | -3.22 | Ints3 | pfam domains | PF10189 | 1 | 1 | 10480 | 419 | IntS3 | | 3.998e-02 | -3.22 | Laminin\_I | pfam domains | PF06008 | 1 | 1 | 10480 | 419 | LanA | | 3.998e-02 | -3.22 | MamL-1 | pfam domains | PF09596 | 1 | 1 | 10480 | 419 | mam | | 3.998e-02 | -3.22 | SH3\_10 | pfam domains | PF17902 | 1 | 1 | 10480 | 419 | shot | | 3.998e-02 | -3.22 | ATG16 | pfam domains | PF08614 | 1 | 1 | 10480 | 419 | Atg16 | | 3.998e-02 | -3.22 | BRD4\_CDT | pfam domains | PF17105 | 1 | 1 | 10480 | 419 | fs(1)h | | 3.998e-02 | -3.22 | DUF1088 | pfam domains | PF06469 | 1 | 1 | 10480 | 419 | rg | | 3.998e-02 | -3.22 | LRRCT | pfam domains | PF01463 | 1 | 1 | 10480 | 419 | sli | | 3.998e-02 | -3.22 | Fe\_hyd\_lg\_C | pfam domains | PF02906 | 1 | 1 | 10480 | 419 | CG17683 | | 3.998e-02 | -3.22 | COE1\_DBD | pfam domains | PF16422 | 1 | 1 | 10480 | 419 | kn | | 3.998e-02 | -3.22 | Cbl\_N3 | pfam domains | PF02762 | 1 | 1 | 10480 | 419 | Cbl | | 3.998e-02 | -3.22 | MaoC\_dehydrat\_N | pfam domains | PF13452 | 1 | 1 | 10480 | 419 | Mfe2 | | 3.998e-02 | -3.22 | DUF3498 | pfam domains | PF12004 | 1 | 1 | 10480 | 419 | CG42684 | | 3.998e-02 | -3.22 | hGDE\_N | pfam domains | PF14699 | 1 | 1 | 10480 | 419 | CG9485 | | 3.998e-02 | -3.22 | Rft-1 | pfam domains | PF04506 | 1 | 1 | 10480 | 419 | CG3149 | | 3.998e-02 | -3.22 | NosD | pfam domains | PF05048 | 1 | 1 | 10480 | 419 | FBXO11 | | 3.998e-02 | -3.22 | TEA | pfam domains | PF01285 | 1 | 1 | 10480 | 419 | sd | | 3.998e-02 | -3.22 | SKICH | pfam domains | PF17751 | 1 | 1 | 10480 | 419 | CG9784 | | 3.998e-02 | -3.22 | HJURP\_C | pfam domains | PF12347 | 1 | 1 | 10480 | 419 | Mef2 | | 3.998e-02 | -3.22 | DAG1 | pfam domains | PF05454 | 1 | 1 | 10480 | 419 | Dg | | 3.998e-02 | -3.22 | ARF7EP\_C | pfam domains | PF14949 | 1 | 1 | 10480 | 419 | CG14464 | | 3.998e-02 | -3.22 | Abdominal-A | pfam domains | PF12407 | 1 | 1 | 10480 | 419 | abd-A | | 3.998e-02 | -3.22 | Pex14\_N | pfam domains | PF04695 | 1 | 1 | 10480 | 419 | Pex14 | | 3.998e-02 | -3.22 | ERp29 | pfam domains | PF07749 | 1 | 1 | 10480 | 419 | wbl | | 3.998e-02 | -3.22 | DUF4819 | pfam domains | PF16090 | 1 | 1 | 10480 | 419 | cic | | 3.998e-02 | -3.22 | ULD | pfam domains | PF16534 | 1 | 1 | 10480 | 419 | dve | | 3.998e-02 | -3.22 | PPIP5K2\_N | pfam domains | PF18086 | 1 | 1 | 10480 | 419 | l(1)G0196 | | 3.998e-02 | -3.22 | SWIM | pfam domains | PF04434 | 1 | 1 | 10480 | 419 | CG34401 | | 3.998e-02 | -3.22 | ProRS-C\_1 | pfam domains | PF09180 | 1 | 1 | 10480 | 419 | GluProRS | | 3.998e-02 | -3.22 | RICTOR\_V | pfam domains | PF14668 | 1 | 1 | 10480 | 419 | rictor | | 3.998e-02 | -3.22 | DFP | pfam domains | PF04127 | 1 | 1 | 10480 | 419 | Ppcs | | 3.998e-02 | -3.22 | HPS3\_Mid | pfam domains | PF14762 | 1 | 1 | 10480 | 419 | CG14562 | | 3.998e-02 | -3.22 | TNFR\_c6 | pfam domains | PF00020 | 1 | 1 | 10480 | 419 | wgn | | 3.998e-02 | -3.22 | CARMIL\_C | pfam domains | PF16000 | 1 | 1 | 10480 | 419 | LRR | | 3.998e-02 | -3.22 | TMEM72 | pfam domains | PF16054 | 1 | 1 | 10480 | 419 | CG14086 | | 3.998e-02 | -3.22 | AF-4 | pfam domains | PF05110 | 1 | 1 | 10480 | 419 | lilli | | 3.998e-02 | -3.22 | IQ\_SEC7\_PH | pfam domains | PF16453 | 1 | 1 | 10480 | 419 | siz | | 3.998e-02 | -3.22 | PH\_3 | pfam domains | PF14593 | 1 | 1 | 10480 | 419 | Pdk1 | | 3.998e-02 | -3.22 | TMPIT | pfam domains | PF07851 | 1 | 1 | 10480 | 419 | CG32795 | | 3.998e-02 | -3.22 | zf-CXXC | pfam domains | PF02008 | 1 | 1 | 10480 | 419 | Kdm2 | | 3.998e-02 | -3.22 | PID\_2 | pfam domains | PF14719 | 1 | 1 | 10480 | 419 | CG8312 | | 3.998e-02 | -3.22 | hDGE\_amylase | pfam domains | PF14701 | 1 | 1 | 10480 | 419 | CG9485 | | 3.998e-02 | -3.22 | Yuri\_gagarin | pfam domains | PF15934 | 1 | 1 | 10480 | 419 | yuri | | 3.998e-02 | -3.22 | M16C\_assoc | pfam domains | PF08367 | 1 | 1 | 10480 | 419 | CG3107 | | 3.998e-02 | -3.22 | Myosin-VI\_CBD | pfam domains | PF16521 | 1 | 1 | 10480 | 419 | jar | | 3.998e-02 | -3.22 | DUF3454 | pfam domains | PF11936 | 1 | 1 | 10480 | 419 | N | | 3.998e-02 | -3.22 | DUF2228 | pfam domains | PF10228 | 1 | 1 | 10480 | 419 | CG1218 | | 3.998e-02 | -3.22 | DDE\_3 | pfam domains | PF13358 | 1 | 1 | 10480 | 419 | CG14478 | | 3.998e-02 | -3.22 | CAS\_C | pfam domains | PF12026 | 1 | 1 | 10480 | 419 | p130CAS | | 3.998e-02 | -3.22 | Tet\_JBP | pfam domains | PF12851 | 1 | 1 | 10480 | 419 | Tet | | 3.998e-02 | -3.22 | hGDE\_central | pfam domains | PF14702 | 1 | 1 | 10480 | 419 | CG9485 | | 3.998e-02 | -3.22 | Receptor\_IA-2 | pfam domains | PF11548 | 1 | 1 | 10480 | 419 | IA-2 | | 3.998e-02 | -3.22 | Meis\_PKNOX\_N | pfam domains | PF16493 | 1 | 1 | 10480 | 419 | hth | | 3.998e-02 | -3.22 | RasGEF\_N\_2 | pfam domains | PF14663 | 1 | 1 | 10480 | 419 | rictor | | 3.998e-02 | -3.22 | VASP\_tetra | pfam domains | PF08776 | 1 | 1 | 10480 | 419 | ena | | 3.998e-02 | -3.22 | DP | pfam domains | PF08781 | 1 | 1 | 10480 | 419 | Dp | | 3.998e-02 | -3.22 | MTHFR | pfam domains | PF02219 | 1 | 1 | 10480 | 419 | CG7560 | | 3.998e-02 | -3.22 | Laminin\_II | pfam domains | PF06009 | 1 | 1 | 10480 | 419 | LanA | | 3.998e-02 | -3.22 | Carm\_PH | pfam domains | PF17888 | 1 | 1 | 10480 | 419 | LRR | | 3.998e-02 | -3.22 | FEZ | pfam domains | PF07763 | 1 | 1 | 10480 | 419 | Unc-76 | | 3.998e-02 | -3.22 | SLBP\_RNA\_bind | pfam domains | PF15247 | 1 | 1 | 10480 | 419 | Slbp | | 3.998e-02 | -3.22 | Rpr2 | pfam domains | PF04032 | 1 | 1 | 10480 | 419 | CG33082 | | 3.998e-02 | -3.22 | Spot\_14 | pfam domains | PF07084 | 1 | 1 | 10480 | 419 | CG2765 | | 3.998e-02 | -3.22 | DUF3535 | pfam domains | PF12054 | 1 | 1 | 10480 | 419 | Hel89B | | 3.998e-02 | -3.22 | Methyltrn\_RNA\_3 | pfam domains | PF02598 | 1 | 1 | 10480 | 419 | CG12128 | | 3.998e-02 | -3.22 | EphA2\_TM | pfam domains | PF14575 | 1 | 1 | 10480 | 419 | Eph | | 3.998e-02 | -3.22 | Crust\_neurohorm | pfam domains | PF01147 | 1 | 1 | 10480 | 419 | ITP | | 3.998e-02 | -3.22 | SMAP | pfam domains | PF15477 | 1 | 1 | 10480 | 419 | CG6340 | | 3.998e-02 | -3.22 | MRP-L27 | pfam domains | PF09809 | 1 | 1 | 10480 | 419 | mRpL41 | | 3.998e-02 | -3.22 | DUF1084 | pfam domains | PF06454 | 1 | 1 | 10480 | 419 | CG2991 | | 3.998e-02 | -3.22 | Ephrin\_lbd | pfam domains | PF01404 | 1 | 1 | 10480 | 419 | Eph | | 3.998e-02 | -3.22 | PNISR | pfam domains | PF15996 | 1 | 1 | 10480 | 419 | CG31211 | | 3.998e-02 | -3.22 | RICTOR\_M | pfam domains | PF14666 | 1 | 1 | 10480 | 419 | rictor | | 3.998e-02 | -3.22 | DFF40 | pfam domains | PF09230 | 1 | 1 | 10480 | 419 | Drep4 | | 3.998e-02 | -3.22 | DUF4517 | pfam domains | PF15006 | 1 | 1 | 10480 | 419 | CG6983 | | 3.998e-02 | -3.22 | INPP5B\_PH | pfam domains | PF16776 | 1 | 1 | 10480 | 419 | Ocrl | | 3.998e-02 | -3.22 | JUPITER | pfam domains | PF17054 | 1 | 1 | 10480 | 419 | Jupiter | | 3.998e-02 | -3.22 | Wtap | pfam domains | PF17098 | 1 | 1 | 10480 | 419 | fl(2)d | | 3.998e-02 | -3.22 | HPS3\_N | pfam domains | PF14761 | 1 | 1 | 10480 | 419 | CG14562 | | 3.998e-02 | -3.22 | NODP | pfam domains | PF07684 | 1 | 1 | 10480 | 419 | N | | 3.998e-02 | -3.22 | PHD\_4 | pfam domains | PF16866 | 1 | 1 | 10480 | 419 | Kdm2 | | 3.998e-02 | -3.22 | F\_actin\_bind | pfam domains | PF08919 | 1 | 1 | 10480 | 419 | Abl | | 3.998e-02 | -3.22 | SMK-1 | pfam domains | PF04802 | 1 | 1 | 10480 | 419 | flfl | | 3.998e-02 | -3.22 | 4\_1\_CTD | pfam domains | PF05902 | 1 | 1 | 10480 | 419 | cora | | 3.998e-02 | -3.22 | YBD | pfam domains | PF17725 | 1 | 1 | 10480 | 419 | sd | | 3.998e-02 | -3.22 | GDE\_C | pfam domains | PF06202 | 1 | 1 | 10480 | 419 | CG9485 | | 3.998e-02 | -3.22 | Cbl\_N | pfam domains | PF02262 | 1 | 1 | 10480 | 419 | Cbl | | 3.998e-02 | -3.22 | Cbl\_N2 | pfam domains | PF02761 | 1 | 1 | 10480 | 419 | Cbl | | 3.998e-02 | -3.22 | UPA | pfam domains | PF17217 | 1 | 1 | 10480 | 419 | unc-5 | | 3.998e-02 | -3.22 | Beta\_helix | pfam domains | PF13229 | 1 | 1 | 10480 | 419 | FBXO11 | | 3.998e-02 | -3.22 | RICTOR\_N | pfam domains | PF14664 | 1 | 1 | 10480 | 419 | rictor | | 3.998e-02 | -3.22 | NOD | pfam domains | PF06816 | 1 | 1 | 10480 | 419 | N | | 3.998e-02 | -3.22 | COE1\_HLH | pfam domains | PF16423 | 1 | 1 | 10480 | 419 | kn | | 3.998e-02 | -3.22 | Fe\_hyd\_SSU | pfam domains | PF02256 | 1 | 1 | 10480 | 419 | CG17683 | | 3.998e-02 | -3.22 | Par3\_HAL\_N\_term | pfam domains | PF12053 | 1 | 1 | 10480 | 419 | baz | | 3.998e-02 | -3.22 | ASD2 | pfam domains | PF08687 | 1 | 1 | 10480 | 419 | Shrm | | 3.998e-02 | -3.22 | Aminotran\_4 | pfam domains | PF01063 | 1 | 1 | 10480 | 419 | CG1673 | | 3.998e-02 | -3.22 | ERp29\_N | pfam domains | PF07912 | 1 | 1 | 10480 | 419 | wbl | | 3.998e-02 | -3.22 | Citrate\_bind | pfam domains | PF16114 | 1 | 1 | 10480 | 419 | ATPCL | | 3.998e-02 | -3.22 | Serine\_rich | pfam domains | PF08824 | 1 | 1 | 10480 | 419 | p130CAS | | 3.998e-02 | -3.22 | BTG | pfam domains | PF07742 | 1 | 1 | 10480 | 419 | Tob | | 3.998e-02 | -3.22 | Profilin | pfam domains | PF00235 | 1 | 1 | 10480 | 419 | chic | | 3.998e-02 | -3.22 | Guanylate\_cyc\_2 | pfam domains | PF09778 | 1 | 1 | 10480 | 419 | CG13760 | | 3.998e-02 | -3.22 | Plectin | pfam domains | PF00681 | 1 | 1 | 10480 | 419 | shot | | 3.998e-02 | -3.22 | ATG11 | pfam domains | PF10377 | 1 | 1 | 10480 | 419 | Atg17 | | 4.006e-02 | -3.22 | TCF dependent signaling in response to WNT | REACTOME pathways | R-DME-201681 | 103 | 9 | 3923 | 176 | PI31,14-3-3zeta,fz,arm,CtBP,Sox14,Diap1,hyx,fz2 | | 4.015e-02 | -3.22 | CG6770 (uncharacterized protein) | protein interactions | 34621 | 28 | 4 | 9204 | 432 | scrib,CG7763,ey,CG32772 | | 4.015e-02 | -3.22 | Arf102F (ADP ribosylation factor at 102F) | protein interactions | 43823 | 28 | 4 | 9204 | 432 | CG4896,Kr-h2,AP-1gamma,Arf79F | | 4.016e-02 | -3.21 | negative regulation of phosphorus metabolic process | biological process | GO:0010563 | 89 | 7 | 12345 | 442 | wdb,trbl,CycG,Cbl,Socs36E,N,CG8312 | | 4.016e-02 | -3.21 | negative regulation of phosphate metabolic process | biological process | GO:0045936 | 89 | 7 | 12345 | 442 | N,CG8312,wdb,trbl,CycG,Cbl,Socs36E | | 4.035e-02 | -3.21 | GSE13306\_TREG\_RA\_VS\_TCONV\_RA\_DN | MSigDB lists | GSE13306\_TREG\_RA\_VS\_TCONV\_RA\_DN | 35 | 4 | 4161 | 156 | Pex5,Gfat2,Ppn,subdued | | 4.035e-02 | -3.21 | GSE360\_CTRL\_VS\_L\_MAJOR\_DC\_DN | MSigDB lists | GSE360\_CTRL\_VS\_L\_MAJOR\_DC\_DN | 35 | 4 | 4161 | 156 | Vps13,bru2,Rgl,Ocrl | | 4.035e-02 | -3.21 | FREAC7\_01 | MSigDB lists | FREAC7\_01 | 35 | 4 | 4161 | 156 | lqf,FBXO11,Pgant5,Raf | | 4.035e-02 | -3.21 | GSE22935\_UNSTIM\_VS\_12H\_MBOVIS\_BCG\_STIM\_MYD88\_KO\_MACROPHAGE\_UP | MSigDB lists | GSE22935\_UNSTIM\_VS\_12H\_MBOVIS\_BCG\_STIM\_MYD88\_KO\_MACROPHAGE\_UP | 35 | 4 | 4161 | 156 | Tollo,InR,Atg16,CG2017 | | 4.035e-02 | -3.21 | GO\_CELL\_JUNCTION\_ASSEMBLY | MSigDB lists | GO\_CELL\_JUNCTION\_ASSEMBLY | 35 | 4 | 4161 | 156 | Pdk1,arm,shot,LanA | | 4.035e-02 | -3.21 | NIELSEN\_GIST | MSigDB lists | NIELSEN\_GIST | 35 | 4 | 4161 | 156 | jar,CG17883,subdued,Tsp96F | | 4.044e-02 | -3.21 | anatomical structure maturation | biological process | GO:0071695 | 53 | 5 | 12345 | 442 | eff,Sox14,mop,chic,Dronc | | 4.044e-02 | -3.21 | protein localization to cell periphery | biological process | GO:1990778 | 53 | 5 | 12345 | 442 | Bsg,Arf79F,scrib,Rab1,arm | | 4.044e-02 | -3.21 | positive regulation of innate immune response | biological process | GO:0045089 | 53 | 5 | 12345 | 442 | Dcr-2,htl,imd,CG32795,DPCoAC | | 4.075e-02 | -3.20 | GO\_REGULATION\_OF\_CYTOPLASMIC\_TRANSPORT | MSigDB lists | GO\_REGULATION\_OF\_CYTOPLASMIC\_TRANSPORT | 143 | 10 | 4161 | 156 | fz2,Psa,fz,Pdk1,Arf79F,Syx1A,CG17883,cact,Tollo,Raf | | 4.077e-02 | -3.20 | KEGG\_FOCAL\_ADHESION | MSigDB lists | KEGG\_FOCAL\_ADHESION | 51 | 5 | 4161 | 156 | InR,Pdk1,Raf,LanA,arm | | 4.077e-02 | -3.20 | GSE16385\_ROSIGLITAZONE\_IL4\_VS\_IFNG\_TNF\_STIM\_MACROPHAGE\_UP | MSigDB lists | GSE16385\_ROSIGLITAZONE\_IL4\_VS\_IFNG\_TNF\_STIM\_MACROPHAGE\_UP | 51 | 5 | 4161 | 156 | Kdm2,Drak,Ids,Pgant6,wdb | | 4.077e-02 | -3.20 | GO\_NEURAL\_TUBE\_DEVELOPMENT | MSigDB lists | GO\_NEURAL\_TUBE\_DEVELOPMENT | 51 | 5 | 4161 | 156 | ds,ap,otk,N,fz | | 4.077e-02 | -3.20 | GSE37534\_UNTREATED\_VS\_PIOGLITAZONE\_TREATED\_CD4\_TCELL\_PPARG1\_AND\_FOXP3\_TRASDUCED\_UP | MSigDB lists | GSE37534\_UNTREATED\_VS\_PIOGLITAZONE\_TREATED\_CD4\_TCELL\_PPARG1\_AND\_FOXP3\_TRASDUCED\_UP | 51 | 5 | 4161 | 156 | Vps13,rg,nmo,Drak,dlp | | 4.077e-02 | -3.20 | GSE9988\_ANTI\_TREM1\_VS\_LOW\_LPS\_MONOCYTE\_UP | MSigDB lists | GSE9988\_ANTI\_TREM1\_VS\_LOW\_LPS\_MONOCYTE\_UP | 51 | 5 | 4161 | 156 | CG9220,Pex5,emp,hppy,Syx1A | | 4.077e-02 | -3.20 | GSE12198\_NK\_VS\_NK\_ACT\_EXPANSION\_SYSTEM\_DERIVED\_NK\_CELL\_UP | MSigDB lists | GSE12198\_NK\_VS\_NK\_ACT\_EXPANSION\_SYSTEM\_DERIVED\_NK\_CELL\_UP | 51 | 5 | 4161 | 156 | nmo,ap,trbl,Rim2,CG18549 | | 4.077e-02 | -3.20 | chr3p21 | MSigDB lists | chr3p21 | 51 | 5 | 4161 | 156 | Cadps,arm,CG3149,CG1129,CG6465 | | 4.077e-02 | -3.20 | GSE37605\_C57BL6\_VS\_NOD\_FOXP3\_IRES\_GFP\_TREG\_DN | MSigDB lists | GSE37605\_C57BL6\_VS\_NOD\_FOXP3\_IRES\_GFP\_TREG\_DN | 51 | 5 | 4161 | 156 | otk,14-3-3zeta,CycG,CG4896,step | | 4.077e-02 | -3.20 | GSE25088\_ROSIGLITAZONE\_VS\_IL4\_AND\_ROSIGLITAZONE\_STIM\_MACROPHAGE\_DAY10\_DN | MSigDB lists | GSE25088\_ROSIGLITAZONE\_VS\_IL4\_AND\_ROSIGLITAZONE\_STIM\_MACROPHAGE\_DAY10\_DN | 51 | 5 | 4161 | 156 | Raf,frc,CG32795,Rgl,rdx | | 4.077e-02 | -3.20 | DAZARD\_UV\_RESPONSE\_CLUSTER\_G6 | MSigDB lists | DAZARD\_UV\_RESPONSE\_CLUSTER\_G6 | 51 | 5 | 4161 | 156 | Rim2,CtBP,Ziz,Pat1,rdx | | 4.077e-02 | -3.20 | GSE5679\_CTRL\_VS\_PPARG\_LIGAND\_ROSIGLITAZONE\_AND\_RARA\_AGONIST\_AM580\_TREATED\_DC\_UP | MSigDB lists | GSE5679\_CTRL\_VS\_PPARG\_LIGAND\_ROSIGLITAZONE\_AND\_RARA\_AGONIST\_AM580\_TREATED\_DC\_UP | 51 | 5 | 4161 | 156 | Rab5,Pgant6,Actbeta,N,Nrk | | 4.077e-02 | -3.20 | GSE42021\_CD24LO\_TREG\_VS\_CD24LO\_TCONV\_THYMUS\_UP | MSigDB lists | GSE42021\_CD24LO\_TREG\_VS\_CD24LO\_TCONV\_THYMUS\_UP | 68 | 6 | 4161 | 156 | Unc-76,fz,CG6465,Cerk,LanA,Ids | | 4.077e-02 | -3.20 | GSE9960\_GRAM\_POS\_VS\_GRAM\_NEG\_AND\_POS\_SEPSIS\_PBMC\_UP | MSigDB lists | GSE9960\_GRAM\_POS\_VS\_GRAM\_NEG\_AND\_POS\_SEPSIS\_PBMC\_UP | 68 | 6 | 4161 | 156 | IntS3,Rgl,CG3436,hppy,CtBP,SMC3 | | 4.077e-02 | -3.20 | GSE17580\_UNINFECTED\_VS\_S\_MANSONI\_INF\_TEFF\_UP | MSigDB lists | GSE17580\_UNINFECTED\_VS\_S\_MANSONI\_INF\_TEFF\_UP | 68 | 6 | 4161 | 156 | Mef2,nAChRalpha4,Pgant6,Syx1A,Cerk,14-3-3epsilon | | 4.077e-02 | -3.20 | ATGTTAA\_MIR302C | MSigDB lists | ATGTTAA\_MIR302C | 68 | 6 | 4161 | 156 | rdx,dlp,Hel89B,wdb,Kr-h2,sd | | 4.108e-02 | -3.19 | actin filament binding | molecular function | GO:0051015 | 56 | 5 | 12176 | 414 | jar,CG6891,f,d,Shrm | | 4.110e-02 | -3.19 | GO\_POSITIVE\_REGULATION\_OF\_DEFENSE\_RESPONSE | MSigDB lists | GO\_POSITIVE\_REGULATION\_OF\_DEFENSE\_RESPONSE | 86 | 7 | 4161 | 156 | Mef2,Pdk1,cact,Tollo,PI31,AP-1gamma,eff | | 4.110e-02 | -3.19 | GSE15330\_LYMPHOID\_MULTIPOTENT\_VS\_GRANULOCYTE\_MONOCYTE\_PROGENITOR\_IKAROS\_KO\_UP | MSigDB lists | GSE15330\_LYMPHOID\_MULTIPOTENT\_VS\_GRANULOCYTE\_MONOCYTE\_PROGENITOR\_IKAROS\_KO\_UP | 86 | 7 | 4161 | 156 | InR,cact,ap,FBXO11,SMC3,Drep4,CG3961 | | 4.110e-02 | -3.19 | RGAGGAARY\_PU1\_Q6 | MSigDB lists | RGAGGAARY\_PU1\_Q6 | 86 | 7 | 4161 | 156 | CtBP,14-3-3zeta,Raf,otk,FBXO11,muc,Rab5 | | 4.131e-02 | -3.19 | circadian behavior | biological process | GO:0048512 | 71 | 6 | 12345 | 442 | tara,kay,dome,ITP,Mef2,Mid1 | | 4.137e-02 | -3.19 | RRM\_dom | interpro domains | IPR000504 | 135 | 10 | 11383 | 450 | CG4896,orb2,spen,Sxl,bru2,msi,bol,fz3,CG14506,CG1316 | | 4.151e-02 | -3.18 | TTGTTT\_FOXO4\_01 | MSigDB lists | TTGTTT\_FOXO4\_01 | 423 | 23 | 4161 | 156 | Ziz,ft,FBXO11,Nrk,Drep4,CG2017,His2Av,Pat1,Ric,Kdm2,CG9485,arm,lili,Dl,Cals,Cdk4,Mef2,Ppn,sd,InR,Arf79F,Actbeta,shot | | 4.157e-02 | -3.18 | STUB1 (STIP1 homology and U-box containing protein 1) | protein interactions | 34433 | 41 | 5 | 9204 | 432 | Pgant6,CG5522,InR,sqz,cact | | 4.171e-02 | -3.18 | Rab5 (Rab5) | protein interactions | 33418 | 130 | 11 | 9204 | 432 | fz,Ocrl,scrib,dome,tkv,Dronc,Arf79F,mop,Gfat2,SERCA,Rab1 | | 4.171e-02 | -3.18 | flw (flapwing) | protein interactions | 44289 | 130 | 11 | 9204 | 432 | CG14441,Drak,dpy,ec,ena,N,Pitslre,Abl,GluProRS,Dcr-2,CG31457 | | 4.172e-02 | -3.18 | IPPc | smart domains | SM00128 | 6 | 2 | 5086 | 290 | Ocrl,CG9784 | | 4.172e-02 | -3.18 | FA | smart domains | SM01195 | 6 | 2 | 5086 | 290 | yrt,cora | | 4.172e-02 | -3.18 | FU | smart domains | SM00261 | 6 | 2 | 5086 | 290 | InR,Egfr | | 4.172e-02 | -3.18 | Sec7 | smart domains | SM00222 | 6 | 2 | 5086 | 290 | siz,step | | 4.172e-02 | -3.18 | ZnF\_GATA | smart domains | SM00401 | 6 | 2 | 5086 | 290 | grn,pnr | | 4.175e-02 | -3.18 | GO\_REGULATION\_OF\_PROTEIN\_COMPLEX\_ASSEMBLY | MSigDB lists | GO\_REGULATION\_OF\_PROTEIN\_COMPLEX\_ASSEMBLY | 124 | 9 | 4161 | 156 | arm,Pex14,dock,lt,sns,Arf79F,Syx1A,sli,Pex5 | | 4.181e-02 | -3.17 | MYSc | smart domains | SM00242 | 14 | 3 | 5086 | 290 | Myo10A,d,jar | | 4.189e-02 | -3.17 | GCCNNNWTAAR\_UNKNOWN | MSigDB lists | GCCNNNWTAAR\_UNKNOWN | 21 | 3 | 4161 | 156 | trbl,lili,Pex5 | | 4.189e-02 | -3.17 | BARIS\_THYROID\_CANCER\_DN | MSigDB lists | BARIS\_THYROID\_CANCER\_DN | 21 | 3 | 4161 | 156 | CG6465,InR,step | | 4.189e-02 | -3.17 | GO\_CARBOHYDRATE\_TRANSPORT | MSigDB lists | GO\_CARBOHYDRATE\_TRANSPORT | 21 | 3 | 4161 | 156 | CG3149,Raf,frc | | 4.189e-02 | -3.17 | GSE12003\_4D\_VS\_8D\_CULTURE\_MIR223\_KO\_BM\_PROGENITOR\_UP | MSigDB lists | GSE12003\_4D\_VS\_8D\_CULTURE\_MIR223\_KO\_BM\_PROGENITOR\_UP | 21 | 3 | 4161 | 156 | Nmda1,Dyrk2,Cals | | 4.189e-02 | -3.17 | GO\_PROTEIN\_SECRETION | MSigDB lists | GO\_PROTEIN\_SECRETION | 21 | 3 | 4161 | 156 | Rab1,wbl,N | | 4.189e-02 | -3.17 | AR\_Q2 | MSigDB lists | AR\_Q2 | 21 | 3 | 4161 | 156 | Pat1,CG2812,Actbeta | | 4.189e-02 | -3.17 | PID\_NFAT\_3PATHWAY | MSigDB lists | PID\_NFAT\_3PATHWAY | 21 | 3 | 4161 | 156 | 14-3-3epsilon,Pen,14-3-3zeta | | 4.189e-02 | -3.17 | HALLMARK\_TNFA\_SIGNALING\_VIA\_NFKB | MSigDB lists | HALLMARK\_TNFA\_SIGNALING\_VIA\_NFKB | 21 | 3 | 4161 | 156 | Actbeta,cact,Gfat2 | | 4.189e-02 | -3.17 | GO\_CENTRAL\_NERVOUS\_SYSTEM\_NEURON\_DEVELOPMENT | MSigDB lists | GO\_CENTRAL\_NERVOUS\_SYSTEM\_NEURON\_DEVELOPMENT | 21 | 3 | 4161 | 156 | sli,Eph,robo1 | | 4.189e-02 | -3.17 | GO\_PEPTIDYL\_THREONINE\_MODIFICATION | MSigDB lists | GO\_PEPTIDYL\_THREONINE\_MODIFICATION | 21 | 3 | 4161 | 156 | Pdk1,Pgant5,nmo | | 4.189e-02 | -3.17 | GO\_EXTRACELLULAR\_MATRIX\_COMPONENT | MSigDB lists | GO\_EXTRACELLULAR\_MATRIX\_COMPONENT | 21 | 3 | 4161 | 156 | shot,LanA,SMC3 | | 4.189e-02 | -3.17 | NABA\_CORE\_MATRISOME | MSigDB lists | NABA\_CORE\_MATRISOME | 21 | 3 | 4161 | 156 | Ppn,LanA,sli | | 4.189e-02 | -3.17 | GO\_REGULATION\_OF\_DNA\_BINDING | MSigDB lists | GO\_REGULATION\_OF\_DNA\_BINDING | 21 | 3 | 4161 | 156 | arm,ap,cact | | 4.189e-02 | -3.17 | KEGG\_ADHERENS\_JUNCTION | MSigDB lists | KEGG\_ADHERENS\_JUNCTION | 21 | 3 | 4161 | 156 | InR,arm,nmo | | 4.195e-02 | -3.17 | transferase complex | cellular component | GO:1990234 | 496 | 25 | 11940 | 417 | CycG,InR,E(Pc),l(2)37Cg,Atg16,hyx,Prdm13,CG12402,Cks30A,morgue,Su(z)2,FBXO11,CG34401,Kdm4B,CycB3,corto,Kdm2,sina,ph-p,eff,fl(2)d,Atg17,Psc,Socs36E,rgr | | 4.220e-02 | -3.17 | female meiosis chromosome segregation | biological process | GO:0016321 | 37 | 4 | 12345 | 442 | LanA,Hr39,Rab5,Gli | | 4.224e-02 | -3.16 | Atg1 (Autophagy-related 1) | protein interactions | 39454 | 69 | 7 | 9204 | 432 | Dronc,wnd,Atg17,klar,Unc-76,mam,Diap1 | | 4.224e-02 | -3.16 | Cp190 (Centrosomal protein 190kD) | protein interactions | 41848 | 69 | 7 | 9204 | 432 | imd,Pen,Dl,fs(1)h,Dp,Rm62,pita | | 4.228e-02 | -3.16 | GO\_DOPAMINERGIC\_NEURON\_DIFFERENTIATION | MSigDB lists | GO\_DOPAMINERGIC\_NEURON\_DIFFERENTIATION | 9 | 2 | 4161 | 156 | arm,fz | | 4.228e-02 | -3.16 | ALONSO\_METASTASIS\_EMT\_UP | MSigDB lists | ALONSO\_METASTASIS\_EMT\_UP | 9 | 2 | 4161 | 156 | Rab1,fz | | 4.228e-02 | -3.16 | PID\_INSULIN\_GLUCOSE\_PATHWAY | MSigDB lists | PID\_INSULIN\_GLUCOSE\_PATHWAY | 9 | 2 | 4161 | 156 | 14-3-3zeta,14-3-3epsilon | | 4.228e-02 | -3.16 | GO\_COCHLEA\_DEVELOPMENT | MSigDB lists | GO\_COCHLEA\_DEVELOPMENT | 9 | 2 | 4161 | 156 | ds,otk | | 4.228e-02 | -3.16 | GO\_INTESTINAL\_ABSORPTION | MSigDB lists | GO\_INTESTINAL\_ABSORPTION | 9 | 2 | 4161 | 156 | CG11069,Dgat2 | | 4.228e-02 | -3.16 | BURTON\_ADIPOGENESIS\_10 | MSigDB lists | BURTON\_ADIPOGENESIS\_10 | 9 | 2 | 4161 | 156 | CG45050,Ziz | | 4.228e-02 | -3.16 | GO\_ESTABLISHMENT\_OR\_MAINTENANCE\_OF\_BIPOLAR\_CELL\_POLARITY | MSigDB lists | GO\_ESTABLISHMENT\_OR\_MAINTENANCE\_OF\_BIPOLAR\_CELL\_POLARITY | 9 | 2 | 4161 | 156 | ap,otk | | 4.228e-02 | -3.16 | GO\_LEFT\_RIGHT\_PATTERN\_FORMATION | MSigDB lists | GO\_LEFT\_RIGHT\_PATTERN\_FORMATION | 9 | 2 | 4161 | 156 | Dl,N | | 4.228e-02 | -3.16 | REACTOME\_SPRY\_REGULATION\_OF\_FGF\_SIGNALING | MSigDB lists | REACTOME\_SPRY\_REGULATION\_OF\_FGF\_SIGNALING | 9 | 2 | 4161 | 156 | Raf,Lk6 | | 4.228e-02 | -3.16 | BIOCARTA\_TOLL\_PATHWAY | MSigDB lists | BIOCARTA\_TOLL\_PATHWAY | 9 | 2 | 4161 | 156 | cact,Tollo | | 4.228e-02 | -3.16 | GO\_REGULATION\_OF\_OSTEOCLAST\_DIFFERENTIATION | MSigDB lists | GO\_REGULATION\_OF\_OSTEOCLAST\_DIFFERENTIATION | 9 | 2 | 4161 | 156 | arm,Tollo | | 4.228e-02 | -3.16 | GO\_CORONARY\_VASCULATURE\_DEVELOPMENT | MSigDB lists | GO\_CORONARY\_VASCULATURE\_DEVELOPMENT | 9 | 2 | 4161 | 156 | N,otk | | 4.228e-02 | -3.16 | GO\_REGULATION\_OF\_TRANSCRIPTION\_REGULATORY\_REGION\_DNA\_BINDING | MSigDB lists | GO\_REGULATION\_OF\_TRANSCRIPTION\_REGULATORY\_REGION\_DNA\_BINDING | 9 | 2 | 4161 | 156 | arm,ap | | 4.228e-02 | -3.16 | chr18q12 | MSigDB lists | chr18q12 | 9 | 2 | 4161 | 156 | Ric,Pgant5 | | 4.228e-02 | -3.16 | GO\_EPIDERMIS\_MORPHOGENESIS | MSigDB lists | GO\_EPIDERMIS\_MORPHOGENESIS | 9 | 2 | 4161 | 156 | N,arm | | 4.228e-02 | -3.16 | WIEDERSCHAIN\_TARGETS\_OF\_BMI1\_AND\_PCGF2 | MSigDB lists | WIEDERSCHAIN\_TARGETS\_OF\_BMI1\_AND\_PCGF2 | 9 | 2 | 4161 | 156 | Cdk4,Actbeta | | 4.228e-02 | -3.16 | BONOME\_OVARIAN\_CANCER\_POOR\_SURVIVAL\_UP | MSigDB lists | BONOME\_OVARIAN\_CANCER\_POOR\_SURVIVAL\_UP | 9 | 2 | 4161 | 156 | ft,Cerk | | 4.228e-02 | -3.16 | GO\_MYD88\_INDEPENDENT\_TOLL\_LIKE\_RECEPTOR\_SIGNALING\_PATHWAY | MSigDB lists | GO\_MYD88\_INDEPENDENT\_TOLL\_LIKE\_RECEPTOR\_SIGNALING\_PATHWAY | 9 | 2 | 4161 | 156 | eff,Tollo | | 4.228e-02 | -3.16 | GO\_REGULATION\_OF\_TRANSCRIPTION\_FROM\_RNA\_POLYMERASE\_II\_PROMOTER\_IN\_RESPONSE\_TO\_HYPOXIA | MSigDB lists | GO\_REGULATION\_OF\_TRANSCRIPTION\_FROM\_RNA\_POLYMERASE\_II\_PROMOTER\_IN\_RESPONSE\_TO\_HYPOXIA | 9 | 2 | 4161 | 156 | N,eff | | 4.228e-02 | -3.16 | GO\_REGULATION\_OF\_DENDRITIC\_SPINE\_DEVELOPMENT | MSigDB lists | GO\_REGULATION\_OF\_DENDRITIC\_SPINE\_DEVELOPMENT | 9 | 2 | 4161 | 156 | Arf79F,Mef2 | | 4.228e-02 | -3.16 | GO\_POSITIVE\_REGULATION\_OF\_RESPONSE\_TO\_WOUNDING | MSigDB lists | GO\_POSITIVE\_REGULATION\_OF\_RESPONSE\_TO\_WOUNDING | 9 | 2 | 4161 | 156 | Tollo,Raf | | 4.228e-02 | -3.16 | ST\_GAQ\_PATHWAY | MSigDB lists | ST\_GAQ\_PATHWAY | 9 | 2 | 4161 | 156 | cact,Pdk1 | | 4.228e-02 | -3.16 | GO\_REGULATION\_OF\_LEUKOCYTE\_DEGRANULATION | MSigDB lists | GO\_REGULATION\_OF\_LEUKOCYTE\_DEGRANULATION | 9 | 2 | 4161 | 156 | Pdk1,AP-1gamma | | 4.228e-02 | -3.16 | GO\_REGULATION\_OF\_NATURAL\_KILLER\_CELL\_MEDIATED\_IMMUNITY | MSigDB lists | GO\_REGULATION\_OF\_NATURAL\_KILLER\_CELL\_MEDIATED\_IMMUNITY | 9 | 2 | 4161 | 156 | Spn42De,AP-1gamma | | 4.228e-02 | -3.16 | PLASARI\_TGFB1\_SIGNALING\_VIA\_NFIC\_10HR\_UP | MSigDB lists | PLASARI\_TGFB1\_SIGNALING\_VIA\_NFIC\_10HR\_UP | 9 | 2 | 4161 | 156 | subdued,unc-5 | | 4.228e-02 | -3.16 | GO\_NEGATIVE\_REGULATION\_OF\_EPITHELIAL\_CELL\_MIGRATION | MSigDB lists | GO\_NEGATIVE\_REGULATION\_OF\_EPITHELIAL\_CELL\_MIGRATION | 9 | 2 | 4161 | 156 | sli,N | | 4.228e-02 | -3.16 | MORF\_PDCD1 | MSigDB lists | MORF\_PDCD1 | 9 | 2 | 4161 | 156 | fz2,hyx | | 4.228e-02 | -3.16 | GO\_GENITALIA\_DEVELOPMENT | MSigDB lists | GO\_GENITALIA\_DEVELOPMENT | 9 | 2 | 4161 | 156 | CG7394,arm | | 4.228e-02 | -3.16 | REACTOME\_NEF\_MEDIATES\_DOWN\_MODULATION\_OF\_CELL\_SURFACE\_RECEPTORS\_BY\_RECRUITING\_THEM\_TO\_CLATHRIN\_ADAPTERS | MSigDB lists | REACTOME\_NEF\_MEDIATES\_DOWN\_MODULATION\_OF\_CELL\_SURFACE\_RECEPTORS\_BY\_RECRUITING\_THEM\_TO\_CLATHRIN\_ADAPTERS | 9 | 2 | 4161 | 156 | Arf79F,AP-1gamma | | 4.228e-02 | -3.16 | GO\_POSITIVE\_REGULATION\_OF\_LEUKOCYTE\_MEDIATED\_IMMUNITY | MSigDB lists | GO\_POSITIVE\_REGULATION\_OF\_LEUKOCYTE\_MEDIATED\_IMMUNITY | 9 | 2 | 4161 | 156 | fz2,AP-1gamma | | 4.228e-02 | -3.16 | WAGNER\_APO2\_SENSITIVITY | MSigDB lists | WAGNER\_APO2\_SENSITIVITY | 9 | 2 | 4161 | 156 | Pgant6,Pgant5 | | 4.228e-02 | -3.16 | WU\_HBX\_TARGETS\_1\_DN | MSigDB lists | WU\_HBX\_TARGETS\_1\_DN | 9 | 2 | 4161 | 156 | Rab5,14-3-3epsilon | | 4.228e-02 | -3.16 | YRCCAKNNGNCGC\_UNKNOWN | MSigDB lists | YRCCAKNNGNCGC\_UNKNOWN | 9 | 2 | 4161 | 156 | fz2,ap | | 4.237e-02 | -3.16 | response to fibroblast growth factor | biological process | GO:0071774 | 22 | 3 | 12345 | 442 | aop,Raf,htl | | 4.237e-02 | -3.16 | fibroblast growth factor receptor signaling pathway | biological process | GO:0008543 | 22 | 3 | 12345 | 442 | aop,Raf,htl | | 4.237e-02 | -3.16 | regulation of post-embryonic development | biological process | GO:0048580 | 22 | 3 | 12345 | 442 | Hr4,N,kn | | 4.237e-02 | -3.16 | cellular response to fibroblast growth factor stimulus | biological process | GO:0044344 | 22 | 3 | 12345 | 442 | htl,aop,Raf | | 4.237e-02 | -3.16 | actin filament bundle assembly | biological process | GO:0051017 | 22 | 3 | 12345 | 442 | Src42A,f,ed | | 4.248e-02 | -3.16 | - | gene3d domains | 2.60.210.10 | 7 | 2 | 4929 | 241 | rdx,sina | | 4.251e-02 | -3.16 | cortical cytoskeleton | cellular component | GO:0030863 | 38 | 4 | 11940 | 417 | CG6891,Bsg,Shrm,spdo | | 4.277e-02 | -3.15 | CG5445 (uncharacterized protein) | protein interactions | 32722 | 17 | 3 | 9204 | 432 | CG9784,Pat1,mura | | 4.277e-02 | -3.15 | Nrt (Neurotactin) | protein interactions | 39873 | 17 | 3 | 9204 | 432 | sli,Abl,N | | 4.277e-02 | -3.15 | Pkc98E (Protein C kinase 98E) | protein interactions | 43428 | 17 | 3 | 9204 | 432 | B-H1,hth,N | | 4.277e-02 | -3.15 | east (enhanced adult sensory threshold) | protein interactions | 46006 | 17 | 3 | 9204 | 432 | imd,fs(1)h,ttk | | 4.277e-02 | -3.15 | CG12081 (uncharacterized protein) | protein interactions | 31799 | 17 | 3 | 9204 | 432 | bin3,lili,LRR | | 4.277e-02 | -3.15 | Rgl (Ral guanine nucleotide dissociation stimulator-like) | protein interactions | 44115 | 17 | 3 | 9204 | 432 | N,siz,Ppn | | 4.277e-02 | -3.15 | Grip71 (Grip71) | protein interactions | 35130 | 17 | 3 | 9204 | 432 | CG32772,CG14478,p130CAS | | 4.277e-02 | -3.15 | Pp4-19C (Protein phosphatase 19C) | protein interactions | 45031 | 17 | 3 | 9204 | 432 | Cdk4,flfl,ttk | | 4.277e-02 | -3.15 | Acn (Acinus) | protein interactions | 35173 | 17 | 3 | 9204 | 432 | rictor,Egfr,N | | 4.288e-02 | -3.15 | MRP (Multidrug-Resistance like Protein 1) | protein interactions | 34686 | 84 | 8 | 9204 | 432 | Rab5,Pen,SERCA,Gfat2,Mpcp2,Kr-h2,GluProRS,rictor | | 4.319e-02 | -3.14 | chemosensory behavior | biological process | GO:0007635 | 150 | 10 | 12345 | 442 | Dyrk2,spi,Adf1,rg,drl,14-3-3zeta,scrib,Gr93c,vn,Egfr | | 4.322e-02 | -3.14 | GRAESSMANN\_APOPTOSIS\_BY\_DOXORUBICIN\_UP | MSigDB lists | GRAESSMANN\_APOPTOSIS\_BY\_DOXORUBICIN\_UP | 335 | 19 | 4161 | 156 | DPCoAC,frc,Rgl,PI31,Tollo,Nmda1,Mfe2,lili,fz2,CycG,Pdk1,sina,Drep4,CG5522,CG6891,Ppcs,unc-5,Drak,ap | | 4.331e-02 | -3.14 | regulation of protein stability | biological process | GO:0031647 | 54 | 5 | 12345 | 442 | rdx,eff,14-3-3zeta,Sxl,Abl | | 4.331e-02 | -3.14 | sperm individualization | biological process | GO:0007291 | 54 | 5 | 12345 | 442 | orb2,Pen,Dronc,jar,yuri | | 4.332e-02 | -3.14 | phosphate ion transmembrane transporter activity | molecular function | GO:0015114 | 10 | 2 | 12176 | 414 | Mpcp2,CG7536 | | 4.332e-02 | -3.14 | histone demethylase activity | molecular function | GO:0032452 | 10 | 2 | 12176 | 414 | Kdm2,Kdm4B | | 4.332e-02 | -3.14 | protein kinase inhibitor activity | molecular function | GO:0004860 | 10 | 2 | 12176 | 414 | trbl,14-3-3zeta | | 4.332e-02 | -3.14 | magnesium-dependent protein serine/threonine phosphatase activity | molecular function | GO:0004724 | 10 | 2 | 12176 | 414 | Pp2C1,CG10417 | | 4.332e-02 | -3.14 | hormone receptor binding | molecular function | GO:0051427 | 10 | 2 | 12176 | 414 | tai,arm | | 4.332e-02 | -3.14 | phosphatidylinositol bisphosphate phosphatase activity | molecular function | GO:0034593 | 10 | 2 | 12176 | 414 | Ocrl,CG9784 | | 4.332e-02 | -3.14 | kinase inhibitor activity | molecular function | GO:0019210 | 10 | 2 | 12176 | 414 | trbl,14-3-3zeta | | 4.332e-02 | -3.14 | GO\_MICROBODY | MSigDB lists | GO\_MICROBODY | 69 | 6 | 4161 | 156 | Pex5,Pex14,Mfe2,Pex7,Arf79F,Pex10 | | 4.332e-02 | -3.14 | GSE13484\_UNSTIM\_VS\_12H\_YF17D\_VACCINE\_STIM\_PBMC\_UP | MSigDB lists | GSE13484\_UNSTIM\_VS\_12H\_YF17D\_VACCINE\_STIM\_PBMC\_UP | 69 | 6 | 4161 | 156 | Mef2,Rim2,trbl,Mfe2,Hel89B,CG11030 | | 4.332e-02 | -3.14 | TORCHIA\_TARGETS\_OF\_EWSR1\_FLI1\_FUSION\_DN | MSigDB lists | TORCHIA\_TARGETS\_OF\_EWSR1\_FLI1\_FUSION\_DN | 69 | 6 | 4161 | 156 | bru2,Ziz,Mef2,Tsp96F,CtBP,trbl | | 4.332e-02 | -3.14 | GO\_REGULATION\_OF\_ACTIN\_FILAMENT\_BASED\_PROCESS | MSigDB lists | GO\_REGULATION\_OF\_ACTIN\_FILAMENT\_BASED\_PROCESS | 69 | 6 | 4161 | 156 | Arf79F,Raf,Mef2,dock,sns,sli | | 4.335e-02 | -3.14 | GO\_SULFUR\_COMPOUND\_BIOSYNTHETIC\_PROCESS | MSigDB lists | GO\_SULFUR\_COMPOUND\_BIOSYNTHETIC\_PROCESS | 87 | 7 | 4161 | 156 | CG9220,muc,ATPCL,CG3961,GlcAT-I,frc,CG31522 | | 4.360e-02 | -3.13 | GO\_PROTEIN\_SERINE\_THREONINE\_KINASE\_ACTIVITY | MSigDB lists | GO\_PROTEIN\_SERINE\_THREONINE\_KINASE\_ACTIVITY | 125 | 9 | 4161 | 156 | Dyrk2,Drak,nmo,Pdk1,Lk6,tkv,hppy,Raf,Cdk4 | | 4.365e-02 | -3.13 | cellularization | biological process | GO:0007349 | 110 | 8 | 12345 | 442 | Cip4,yuri,Dronc,Rab5,Syx1A,jar,Pen,orb2 | | 4.379e-02 | -3.13 | GSE29617\_CTRL\_VS\_DAY3\_TIV\_FLU\_VACCINE\_PBMC\_2008\_UP | MSigDB lists | GSE29617\_CTRL\_VS\_DAY3\_TIV\_FLU\_VACCINE\_PBMC\_2008\_UP | 52 | 5 | 4161 | 156 | kn,sina,Raf,sli,dlp | | 4.379e-02 | -3.13 | GSE36009\_UNSTIM\_VS\_LPS\_STIM\_DC\_DN | MSigDB lists | GSE36009\_UNSTIM\_VS\_LPS\_STIM\_DC\_DN | 52 | 5 | 4161 | 156 | CG17883,C1GalTA,Cerk,Drak,Pex10 | | 4.379e-02 | -3.13 | GSE2770\_UNTREATED\_VS\_IL4\_TREATED\_ACT\_CD4\_TCELL\_6H\_DN | MSigDB lists | GSE2770\_UNTREATED\_VS\_IL4\_TREATED\_ACT\_CD4\_TCELL\_6H\_DN | 52 | 5 | 4161 | 156 | Pex7,SmD1,Dyb,l(2)37Cg,unc-119 | | 4.379e-02 | -3.13 | GSE6092\_B\_BURGDOFERI\_VS\_B\_BURGDORFERI\_AND\_IFNG\_STIM\_ENDOTHELIAL\_CELL\_DN | MSigDB lists | GSE6092\_B\_BURGDOFERI\_VS\_B\_BURGDORFERI\_AND\_IFNG\_STIM\_ENDOTHELIAL\_CELL\_DN | 52 | 5 | 4161 | 156 | dock,Raf,CG2017,Rgl,CG3961 | | 4.379e-02 | -3.13 | GSE36078\_WT\_VS\_IL1R\_KO\_LUNG\_DC\_AFTER\_AD5\_INF\_DN | MSigDB lists | GSE36078\_WT\_VS\_IL1R\_KO\_LUNG\_DC\_AFTER\_AD5\_INF\_DN | 52 | 5 | 4161 | 156 | Tsp96F,InR,trbl,CG9485,jar | | 4.379e-02 | -3.13 | GSE5589\_WT\_VS\_IL10\_KO\_LPS\_AND\_IL6\_STIM\_MACROPHAGE\_45MIN\_UP | MSigDB lists | GSE5589\_WT\_VS\_IL10\_KO\_LPS\_AND\_IL6\_STIM\_MACROPHAGE\_45MIN\_UP | 52 | 5 | 4161 | 156 | wdb,sina,jar,Fem-1,Gfat2 | | 4.379e-02 | -3.13 | GSE9960\_GRAM\_NEG\_VS\_GRAM\_POS\_SEPSIS\_PBMC\_DN | MSigDB lists | GSE9960\_GRAM\_NEG\_VS\_GRAM\_POS\_SEPSIS\_PBMC\_DN | 52 | 5 | 4161 | 156 | lt,wbl,Dyrk2,CG2006,hppy | | 4.379e-02 | -3.13 | GSE5542\_IFNA\_VS\_IFNA\_AND\_IFNG\_TREATED\_EPITHELIAL\_CELLS\_6H\_UP | MSigDB lists | GSE5542\_IFNA\_VS\_IFNA\_AND\_IFNG\_TREATED\_EPITHELIAL\_CELLS\_6H\_UP | 52 | 5 | 4161 | 156 | CenB1A,GlcAT-I,CG3961,CG6891,Tsp96F | | 4.379e-02 | -3.13 | GSE1460\_DP\_VS\_CD4\_THYMOCYTE\_DN | MSigDB lists | GSE1460\_DP\_VS\_CD4\_THYMOCYTE\_DN | 52 | 5 | 4161 | 156 | Eph,CG6891,eff,emp,Pgant6 | | 4.379e-02 | -3.13 | GSE3039\_ALPHAALPHA\_CD8\_TCELL\_VS\_B1\_BCELL\_DN | MSigDB lists | GSE3039\_ALPHAALPHA\_CD8\_TCELL\_VS\_B1\_BCELL\_DN | 52 | 5 | 4161 | 156 | Pat1,Syx1A,kn,Mef2,Dyb | | 4.379e-02 | -3.13 | GO\_REGULATION\_OF\_NEUROTRANSMITTER\_LEVELS | MSigDB lists | GO\_REGULATION\_OF\_NEUROTRANSMITTER\_LEVELS | 52 | 5 | 4161 | 156 | Syx1A,Cadps,nAChRalpha4,Mef2,Raf | | 4.379e-02 | -3.13 | GSE29949\_CD8\_NEG\_DC\_SPLEEN\_VS\_MONOCYTE\_BONE\_MARROW\_UP | MSigDB lists | GSE29949\_CD8\_NEG\_DC\_SPLEEN\_VS\_MONOCYTE\_BONE\_MARROW\_UP | 52 | 5 | 4161 | 156 | Unc-76,betaTub60D,14-3-3zeta,Kdm2,frc | | 4.409e-02 | -3.12 | MAP2K and MAPK activation | REACTOME pathways | R-DME-5674135 | 18 | 3 | 3923 | 176 | CG4935,Raf,14-3-3zeta | | 4.411e-02 | -3.12 | GSE15624\_3H\_VS\_6H\_HALOFUGINONE\_TREATED\_CD4\_TCELL\_DN | MSigDB lists | GSE15624\_3H\_VS\_6H\_HALOFUGINONE\_TREATED\_CD4\_TCELL\_DN | 36 | 4 | 4161 | 156 | Pgant6,CG10417,Pat1,Kr-h2 | | 4.411e-02 | -3.12 | GSE25087\_TREG\_VS\_TCONV\_ADULT\_UP | MSigDB lists | GSE25087\_TREG\_VS\_TCONV\_ADULT\_UP | 36 | 4 | 4161 | 156 | CG2812,CG45050,Mef2,Ids | | 4.411e-02 | -3.12 | GSE41867\_DAY6\_VS\_DAY8\_LCMV\_CLONE13\_EFFECTOR\_CD8\_TCELL\_UP | MSigDB lists | GSE41867\_DAY6\_VS\_DAY8\_LCMV\_CLONE13\_EFFECTOR\_CD8\_TCELL\_UP | 36 | 4 | 4161 | 156 | Actbeta,CG2017,Eph,sd | | 4.411e-02 | -3.12 | GO\_REGULATION\_OF\_EXTRINSIC\_APOPTOTIC\_SIGNALING\_PATHWAY | MSigDB lists | GO\_REGULATION\_OF\_EXTRINSIC\_APOPTOTIC\_SIGNALING\_PATHWAY | 36 | 4 | 4161 | 156 | CG3961,Fem-1,Actbeta,tkv | | 4.411e-02 | -3.12 | REACTOME\_SIGNALING\_BY\_FGFR | MSigDB lists | REACTOME\_SIGNALING\_BY\_FGFR | 36 | 4 | 4161 | 156 | Pdk1,Raf,Lk6,htl | | 4.411e-02 | -3.12 | PASQUALUCCI\_LYMPHOMA\_BY\_GC\_STAGE\_DN | MSigDB lists | PASQUALUCCI\_LYMPHOMA\_BY\_GC\_STAGE\_DN | 36 | 4 | 4161 | 156 | bru2,CG6891,CG1673,trbl | | 4.411e-02 | -3.12 | GSE29615\_DAY3\_VS\_DAY7\_LAIV\_FLU\_VACCINE\_PBMC\_DN | MSigDB lists | GSE29615\_DAY3\_VS\_DAY7\_LAIV\_FLU\_VACCINE\_PBMC\_DN | 36 | 4 | 4161 | 156 | sd,Dyb,arm,crol | | 4.411e-02 | -3.12 | CEBP\_C | MSigDB lists | CEBP\_C | 36 | 4 | 4161 | 156 | cact,Ddr,CG3961,Rab1 | | 4.411e-02 | -3.12 | CAGCCTC\_MIR4855P | MSigDB lists | CAGCCTC\_MIR4855P | 36 | 4 | 4161 | 156 | rg,fz,lin-28,ds | | 4.415e-02 | -3.12 | YTATTTTNR\_MEF2\_02 | MSigDB lists | YTATTTTNR\_MEF2\_02 | 145 | 10 | 4161 | 156 | Rgl,lqf,Pex5,Dyb,Raf,rg,nmo,subdued,FBXO11,Nrk | | 4.447e-02 | -3.11 | Clathrin-mediated endocytosis | REACTOME pathways | R-DME-8856828 | 105 | 9 | 3923 | 176 | pes,Ocrl,Egfr,Rab5,Cbl,Cip4,drongo,lqf,cindr | | 4.474e-02 | -3.11 | Znf\_RING/FYVE/PHD | interpro domains | IPR013083 | 214 | 14 | 11383 | 450 | Kdm2,tna,Psc,Diap1,sina,CG13481,Cbl,Su(z)2,CG11360,CG2991,CG11414,Alh,Pex10,mura | | 4.483e-02 | -3.10 | Ank\_2 | pfam domains | PF12796 | 65 | 6 | 10480 | 419 | CG9121,N,CenB1A,Fem-1,f,cact | | 4.486e-02 | -3.10 | LRR FLII-interacting protein 1 (LRRFIP1) activates type I IFN production | REACTOME pathways | R-DME-3134973 | 1 | 1 | 3923 | 176 | arm | | 4.486e-02 | -3.10 | Interleukin-1 processing | REACTOME pathways | R-DME-448706 | 1 | 1 | 3923 | 176 | Dronc | | 4.486e-02 | -3.10 | Keratan sulfate biosynthesis | REACTOME pathways | R-DME-2022854 | 1 | 1 | 3923 | 176 | frc | | 4.487e-02 | -3.10 | ZF\_RING\_2 | prosite domains | PS50089 | 119 | 11 | 6243 | 325 | Diap1,Pex10,CG13481,mura,sina,Psc,Su(z)2,lt,CG11360,Cbl,CG11414 | | 4.491e-02 | -3.10 | sog (short gastrulation) | protein interactions | 32498 | 29 | 4 | 9204 | 432 | LanA,tkv,rdx,kek5 | | 4.491e-02 | -3.10 | Rok (Rho kinase) | protein interactions | 43916 | 29 | 4 | 9204 | 432 | H,Shrm,Rab5,fz | | 4.491e-02 | -3.10 | Arc1 (Activity-regulated cytoskeleton associated protein 1) | protein interactions | 36595 | 29 | 4 | 9204 | 432 | Raf,Src42A,CG1129,CG1218 | | 4.491e-02 | -3.10 | bru1 (bruno 1) | protein interactions | 34648 | 29 | 4 | 9204 | 432 | S,msi,Lk6,Ppn | | 4.491e-02 | -3.10 | Galphao (G protein alpha o subunit) | protein interactions | 36104 | 29 | 4 | 9204 | 432 | fz2,Rab5,fz,RapGAP1 | | 4.491e-02 | -3.10 | pros (prospero) | protein interactions | 41363 | 29 | 4 | 9204 | 432 | Abl,sina,aop,esg | | 4.491e-02 | -3.10 | mask (multiple ankyrin repeats single KH domain) | protein interactions | 50070 | 29 | 4 | 9204 | 432 | rictor,Pdk1,CG9121,Sema1a | | 4.491e-02 | -3.10 | betaTub85D (beta-Tubulin at 85D) | protein interactions | 41124 | 29 | 4 | 9204 | 432 | Arf79F,Pex5,His3.3B,Pex14 | | 4.491e-02 | -3.10 | Dap160 (Dynamin associated protein 160) | protein interactions | 35378 | 29 | 4 | 9204 | 432 | Cbl,tkv,Dad,drongo | | 4.499e-02 | -3.10 | integral component of membrane | cellular component | GO:0016021 | 1438 | 62 | 11940 | 417 | Kr-h2,Tpc1,htl,Pvr,klar,fz2,kek5,Sema1a,ft,CG33474,CG3149,Cals,moody,C1GalTA,drl,Mpcp2,Irk3,Pex10,Orct2,wgn,InR,Inx2,rg,tkv,Egfr,ds,robo1,shg,nAChRalpha4,DPCoAC,Ir21a,Eph,unc-5,N,neo,ed,emp,CG30344,rho-4,spi,CG32795,scrib,Nrk,fz,Ddr,S,fz3,CG7536,SERCA,ZnT41F,Dl,glec,CG8034,Gli,tipE,Tsp96F,otk,Gr93c,Syx1A,beat-IIa,Esp,ogre | | 4.514e-02 | -3.10 | fly chr60D4-60D5\|2-107 | chromosome location | fly chr60D4-60D5\|2-107 | 2 | 1 | 25010 | 571 | ITP | | 4.514e-02 | -3.10 | fly chr29F2-29F2\|2-34 cM | chromosome location | fly chr29F2-29F2\|2-34 cM | 2 | 1 | 25010 | 571 | lncRNA:CR45365 | | 4.514e-02 | -3.10 | fly chr33F1-33F1\|2-47 cM | chromosome location | fly chr33F1-33F1\|2-47 cM | 2 | 1 | 25010 | 571 | nub | | 4.514e-02 | -3.10 | fly chr4F2-4F2\|1-11 cM | chromosome location | fly chr4F2-4F2\|1-11 cM | 2 | 1 | 25010 | 571 | lncRNA:CR44834 | | 4.514e-02 | -3.10 | fly chr62F4-62F4\|3-4 cM | chromosome location | fly chr62F4-62F4\|3-4 cM | 2 | 1 | 25010 | 571 | Pgant6 | | 4.514e-02 | -3.10 | fly chr57F9-57F10\|2-96 | chromosome location | fly chr57F9-57F10\|2-96 | 2 | 1 | 25010 | 571 | HmgZ | | 4.514e-02 | -3.10 | fly chr50B3-50B4\|2-68 | chromosome location | fly chr50B3-50B4\|2-68 | 2 | 1 | 25010 | 571 | CG6191 | | 4.514e-02 | -3.10 | fly chr35A1-35A2\|2-50 cM | chromosome location | fly chr35A1-35A2\|2-50 cM | 2 | 1 | 25010 | 571 | Dyrk2 | | 4.514e-02 | -3.10 | fly chr14A9-14A9\|1-52 cM | chromosome location | fly chr14A9-14A9\|1-52 cM | 2 | 1 | 25010 | 571 | CG42353 | | 4.514e-02 | -3.10 | fly chr27F2-27F3\|2-26 | chromosome location | fly chr27F2-27F3\|2-26 | 2 | 1 | 25010 | 571 | asRNA:CR44063 | | 4.514e-02 | -3.10 | fly chr4B1-4B1\|1-7 | chromosome location | fly chr4B1-4B1\|1-7 | 2 | 1 | 25010 | 571 | GlcAT-I | | 4.514e-02 | -3.10 | fly chr35A1-35A2\|2-50 | chromosome location | fly chr35A1-35A2\|2-50 | 2 | 1 | 25010 | 571 | Dyrk2 | | 4.514e-02 | -3.10 | fly chr14F1-14F1\|1-54 | chromosome location | fly chr14F1-14F1\|1-54 | 2 | 1 | 25010 | 571 | SMC3 | | 4.514e-02 | -3.10 | fly chr21A5-21B1\|2-0 | chromosome location | fly chr21A5-21B1\|2-0 | 2 | 1 | 25010 | 571 | Ir21a | | 4.514e-02 | -3.10 | fly chr18A7-18A7\|1-62 cM | chromosome location | fly chr18A7-18A7\|1-62 cM | 2 | 1 | 25010 | 571 | CG7914 | | 4.514e-02 | -3.10 | fly chr89A11-89A11\|3-58 cM | chromosome location | fly chr89A11-89A11\|3-58 cM | 2 | 1 | 25010 | 571 | Hel89B | | 4.514e-02 | -3.10 | fly chr67B9-67B9\|3-29 | chromosome location | fly chr67B9-67B9\|3-29 | 2 | 1 | 25010 | 571 | CG44838 | | 4.514e-02 | -3.10 | fly chr95F8-95F8\|3-84 cM | chromosome location | fly chr95F8-95F8\|3-84 cM | 2 | 1 | 25010 | 571 | Orct2 | | 4.514e-02 | -3.10 | fly chr102F7-102F8\|4-0 cM | chromosome location | fly chr102F7-102F8\|4-0 cM | 2 | 1 | 25010 | 571 | apolpp | | 4.514e-02 | -3.10 | fly chr57F9-57F10\|2-96 cM | chromosome location | fly chr57F9-57F10\|2-96 cM | 2 | 1 | 25010 | 571 | HmgZ | | 4.514e-02 | -3.10 | fly chr57E6-57E6\|2-95 cM | chromosome location | fly chr57E6-57E6\|2-95 cM | 2 | 1 | 25010 | 571 | Fkbp14 | | 4.514e-02 | -3.10 | fly chr4F2-4F2\|1-11 | chromosome location | fly chr4F2-4F2\|1-11 | 2 | 1 | 25010 | 571 | lncRNA:CR44834 | | 4.514e-02 | -3.10 | fly chr55E5-55E5\|2-86 | chromosome location | fly chr55E5-55E5\|2-86 | 2 | 1 | 25010 | 571 | CG42306 | | 4.514e-02 | -3.10 | fly chr43B3-43C1\|2-56 cM | chromosome location | fly chr43B3-43C1\|2-56 cM | 2 | 1 | 25010 | 571 | CG11145 | | 4.514e-02 | -3.10 | fly chr57E6-57E6\|2-95 | chromosome location | fly chr57E6-57E6\|2-95 | 2 | 1 | 25010 | 571 | Fkbp14 | | 4.514e-02 | -3.10 | fly chr5A9-5A9\|1-12 cM | chromosome location | fly chr5A9-5A9\|1-12 cM | 2 | 1 | 25010 | 571 | OtopLa | | 4.514e-02 | -3.10 | fly chr60D4-60D5\|2-107 cM | chromosome location | fly chr60D4-60D5\|2-107 cM | 2 | 1 | 25010 | 571 | ITP | | 4.514e-02 | -3.10 | fly chr89A11-89A11\|3-58 | chromosome location | fly chr89A11-89A11\|3-58 | 2 | 1 | 25010 | 571 | Hel89B | | 4.514e-02 | -3.10 | fly chr74B5-74C1\|3-45 | chromosome location | fly chr74B5-74C1\|3-45 | 2 | 1 | 25010 | 571 | frc | | 4.514e-02 | -3.10 | fly chr62B9-62B9\|3-1.0 | chromosome location | fly chr62B9-62B9\|3-1.0 | 2 | 1 | 25010 | 571 | Tmhs | | 4.514e-02 | -3.10 | fly chrh44-h46\|2-55 cM | chromosome location | fly chrh44-h46\|2-55 cM | 2 | 1 | 25010 | 571 | IntS3 | | 4.514e-02 | -3.10 | fly chr27F2-27F3\|2-26 cM | chromosome location | fly chr27F2-27F3\|2-26 cM | 2 | 1 | 25010 | 571 | asRNA:CR44063 | | 4.514e-02 | -3.10 | fly chr102F7-102F8\|4-0 | chromosome location | fly chr102F7-102F8\|4-0 | 2 | 1 | 25010 | 571 | apolpp | | 4.514e-02 | -3.10 | fly chr62A7-62A7\|3-0.5 cM | chromosome location | fly chr62A7-62A7\|3-0.5 cM | 2 | 1 | 25010 | 571 | CG12104 | | 4.514e-02 | -3.10 | fly chr14F1-14F1\|1-54 cM | chromosome location | fly chr14F1-14F1\|1-54 cM | 2 | 1 | 25010 | 571 | SMC3 | | 4.514e-02 | -3.10 | fly chr5A9-5A9\|1-12 | chromosome location | fly chr5A9-5A9\|1-12 | 2 | 1 | 25010 | 571 | OtopLa | | 4.514e-02 | -3.10 | fly chr66D10-66D10\|3-26 cM | chromosome location | fly chr66D10-66D10\|3-26 cM | 2 | 1 | 25010 | 571 | h | | 4.514e-02 | -3.10 | fly chr61F7-61F7\|3-0 | chromosome location | fly chr61F7-61F7\|3-0 | 2 | 1 | 25010 | 571 | 312 | | 4.514e-02 | -3.10 | fly chr43B3-43C1\|2-56 | chromosome location | fly chr43B3-43C1\|2-56 | 2 | 1 | 25010 | 571 | CG11145 | | 4.514e-02 | -3.10 | fly chr68C3-68C3\|3-36 cM | chromosome location | fly chr68C3-68C3\|3-36 cM | 2 | 1 | 25010 | 571 | lncRNA:CR45250 | | 4.514e-02 | -3.10 | fly chr61F7-61F7\|3-0 cM | chromosome location | fly chr61F7-61F7\|3-0 cM | 2 | 1 | 25010 | 571 | 312 | | 4.514e-02 | -3.10 | fly chr71A3-71A3\|3-42 cM | chromosome location | fly chr71A3-71A3\|3-42 cM | 2 | 1 | 25010 | 571 | Tom | | 4.514e-02 | -3.10 | fly chr2C7-2C8\|1-0.5 cM | chromosome location | fly chr2C7-2C8\|1-0.5 cM | 2 | 1 | 25010 | 571 | moody | | 4.514e-02 | -3.10 | fly chr68C3-68C3\|3-36 | chromosome location | fly chr68C3-68C3\|3-36 | 2 | 1 | 25010 | 571 | lncRNA:CR45250 | | 4.514e-02 | -3.10 | fly chr3B6-3B6\|1-1.5 cM | chromosome location | fly chr3B6-3B6\|1-1.5 cM | 2 | 1 | 25010 | 571 | CG32795 | | 4.514e-02 | -3.10 | fly chr10C10-10C10\|1-35 | chromosome location | fly chr10C10-10C10\|1-35 | 2 | 1 | 25010 | 571 | rho-4 | | 4.514e-02 | -3.10 | fly chrh44-h46\|2-55 | chromosome location | fly chrh44-h46\|2-55 | 2 | 1 | 25010 | 571 | IntS3 | | 4.514e-02 | -3.10 | fly chr66D10-66D10\|3-26 | chromosome location | fly chr66D10-66D10\|3-26 | 2 | 1 | 25010 | 571 | h | | 4.514e-02 | -3.10 | fly chr15F1-15F1\|1-57 cM | chromosome location | fly chr15F1-15F1\|1-57 cM | 2 | 1 | 25010 | 571 | baz | | 4.514e-02 | -3.10 | fly chr23B3-23B3\|2-7 | chromosome location | fly chr23B3-23B3\|2-7 | 2 | 1 | 25010 | 571 | CG3119 | | 4.514e-02 | -3.10 | fly chr60A16-60A16\|2-105 cM | chromosome location | fly chr60A16-60A16\|2-105 cM | 2 | 1 | 25010 | 571 | Sox14 | | 4.514e-02 | -3.10 | fly chr74B5-74C1\|3-45 cM | chromosome location | fly chr74B5-74C1\|3-45 cM | 2 | 1 | 25010 | 571 | frc | | 4.514e-02 | -3.10 | fly chr2C7-2C8\|1-0.5 | chromosome location | fly chr2C7-2C8\|1-0.5 | 2 | 1 | 25010 | 571 | moody | | 4.514e-02 | -3.10 | fly chr21A5-21B1\|2-0 cM | chromosome location | fly chr21A5-21B1\|2-0 cM | 2 | 1 | 25010 | 571 | Ir21a | | 4.514e-02 | -3.10 | fly chr10C10-10C10\|1-35 cM | chromosome location | fly chr10C10-10C10\|1-35 cM | 2 | 1 | 25010 | 571 | rho-4 | | 4.514e-02 | -3.10 | fly chr3B6-3B6\|1-1.5 | chromosome location | fly chr3B6-3B6\|1-1.5 | 2 | 1 | 25010 | 571 | CG32795 | | 4.514e-02 | -3.10 | fly chr15F1-15F1\|1-57 | chromosome location | fly chr15F1-15F1\|1-57 | 2 | 1 | 25010 | 571 | baz | | 4.514e-02 | -3.10 | fly chr62A7-62A7\|3-0.5 | chromosome location | fly chr62A7-62A7\|3-0.5 | 2 | 1 | 25010 | 571 | CG12104 | | 4.514e-02 | -3.10 | fly chr52C7-52C7\|2-75 | chromosome location | fly chr52C7-52C7\|2-75 | 2 | 1 | 25010 | 571 | Poxn | | 4.514e-02 | -3.10 | fly chr79F4-79F4\|3-47 | chromosome location | fly chr79F4-79F4\|3-47 | 2 | 1 | 25010 | 571 | lncRNA:CR43270 | | 4.514e-02 | -3.10 | fly chr29F2-29F2\|2-34 | chromosome location | fly chr29F2-29F2\|2-34 | 2 | 1 | 25010 | 571 | lncRNA:CR45365 | | 4.514e-02 | -3.10 | fly chr15E2-15E3\|1-57 cM | chromosome location | fly chr15E2-15E3\|1-57 cM | 2 | 1 | 25010 | 571 | CG4991 | | 4.514e-02 | -3.10 | fly chr60A16-60A16\|2-105 | chromosome location | fly chr60A16-60A16\|2-105 | 2 | 1 | 25010 | 571 | Sox14 | | 4.514e-02 | -3.10 | fly chr95F8-95F8\|3-84 | chromosome location | fly chr95F8-95F8\|3-84 | 2 | 1 | 25010 | 571 | Orct2 | | 4.514e-02 | -3.10 | fly chr73D2-73D3\|3-45 | chromosome location | fly chr73D2-73D3\|3-45 | 2 | 1 | 25010 | 571 | sina | | 4.514e-02 | -3.10 | fly chr71A3-71A3\|3-42 | chromosome location | fly chr71A3-71A3\|3-42 | 2 | 1 | 25010 | 571 | Tom | | 4.514e-02 | -3.10 | fly chr14A9-14A9\|1-52 | chromosome location | fly chr14A9-14A9\|1-52 | 2 | 1 | 25010 | 571 | CG42353 | | 4.514e-02 | -3.10 | fly chr15E2-15E3\|1-57 | chromosome location | fly chr15E2-15E3\|1-57 | 2 | 1 | 25010 | 571 | CG4991 | | 4.514e-02 | -3.10 | fly chr33F1-33F1\|2-47 | chromosome location | fly chr33F1-33F1\|2-47 | 2 | 1 | 25010 | 571 | nub | | 4.514e-02 | -3.10 | fly chr49D6-49D7\|2-67 | chromosome location | fly chr49D6-49D7\|2-67 | 2 | 1 | 25010 | 571 | Nmda1 | | 4.514e-02 | -3.10 | fly chr4D7-4D7\|1-9 | chromosome location | fly chr4D7-4D7\|1-9 | 2 | 1 | 25010 | 571 | Ptp4E | | 4.514e-02 | -3.10 | fly chr23B3-23B3\|2-7 cM | chromosome location | fly chr23B3-23B3\|2-7 cM | 2 | 1 | 25010 | 571 | CG3119 | | 4.514e-02 | -3.10 | fly chr50B3-50B4\|2-68 cM | chromosome location | fly chr50B3-50B4\|2-68 cM | 2 | 1 | 25010 | 571 | CG6191 | | 4.514e-02 | -3.10 | fly chr79F4-79F4\|3-47 cM | chromosome location | fly chr79F4-79F4\|3-47 cM | 2 | 1 | 25010 | 571 | lncRNA:CR43270 | | 4.514e-02 | -3.10 | fly chr11B9-11B9\|1-39 | chromosome location | fly chr11B9-11B9\|1-39 | 2 | 1 | 25010 | 571 | mir-9377 | | 4.514e-02 | -3.10 | fly chr52C7-52C7\|2-75 cM | chromosome location | fly chr52C7-52C7\|2-75 cM | 2 | 1 | 25010 | 571 | Poxn | | 4.514e-02 | -3.10 | fly chr49E7-49E7\|2-67 cM | chromosome location | fly chr49E7-49E7\|2-67 cM | 2 | 1 | 25010 | 571 | Su(z)2 | | 4.514e-02 | -3.10 | fly chr49E7-49E7\|2-67 | chromosome location | fly chr49E7-49E7\|2-67 | 2 | 1 | 25010 | 571 | Su(z)2 | | 4.514e-02 | -3.10 | fly chr49D6-49D7\|2-67 cM | chromosome location | fly chr49D6-49D7\|2-67 cM | 2 | 1 | 25010 | 571 | Nmda1 | | 4.514e-02 | -3.10 | fly chr4D7-4D7\|1-9 cM | chromosome location | fly chr4D7-4D7\|1-9 cM | 2 | 1 | 25010 | 571 | Ptp4E | | 4.514e-02 | -3.10 | fly chr73D2-73D3\|3-45 cM | chromosome location | fly chr73D2-73D3\|3-45 cM | 2 | 1 | 25010 | 571 | sina | | 4.514e-02 | -3.10 | fly chr35B10-35C1\|2-50 | chromosome location | fly chr35B10-35C1\|2-50 | 2 | 1 | 25010 | 571 | vig | | 4.514e-02 | -3.10 | fly chr55E5-55E5\|2-86 cM | chromosome location | fly chr55E5-55E5\|2-86 cM | 2 | 1 | 25010 | 571 | CG42306 | | 4.514e-02 | -3.10 | fly chr11B9-11B9\|1-39 cM | chromosome location | fly chr11B9-11B9\|1-39 cM | 2 | 1 | 25010 | 571 | mir-9377 | | 4.514e-02 | -3.10 | fly chr18A7-18A7\|1-62 | chromosome location | fly chr18A7-18A7\|1-62 | 2 | 1 | 25010 | 571 | CG7914 | | 4.514e-02 | -3.10 | fly chr35B10-35C1\|2-50 cM | chromosome location | fly chr35B10-35C1\|2-50 cM | 2 | 1 | 25010 | 571 | vig | | 4.514e-02 | -3.10 | fly chr62F4-62F4\|3-4 | chromosome location | fly chr62F4-62F4\|3-4 | 2 | 1 | 25010 | 571 | Pgant6 | | 4.514e-02 | -3.10 | fly chr4B1-4B1\|1-7 cM | chromosome location | fly chr4B1-4B1\|1-7 cM | 2 | 1 | 25010 | 571 | GlcAT-I | | 4.514e-02 | -3.10 | fly chr67B9-67B9\|3-29 cM | chromosome location | fly chr67B9-67B9\|3-29 cM | 2 | 1 | 25010 | 571 | CG44838 | | 4.514e-02 | -3.10 | fly chr62B9-62B9\|3-1.0 cM | chromosome location | fly chr62B9-62B9\|3-1.0 cM | 2 | 1 | 25010 | 571 | Tmhs | | 4.544e-02 | -3.09 | su(Hw) (suppressor of Hairy wing) | protein interactions | 41740 | 42 | 5 | 9204 | 432 | Mfe2,abd-A,f,ap,fs(1)h | | 4.544e-02 | -3.09 | ex (expanded) | protein interactions | 33218 | 42 | 5 | 9204 | 432 | mop,tai,ed,ft,sd | | 4.544e-02 | -3.09 | ftz-f1 (ftz transcription factor 1) | protein interactions | 40045 | 42 | 5 | 9204 | 432 | Pgk,14-3-3zeta,cic,ps,Hr39 | | 4.544e-02 | -3.09 | bip2 (bip2) | protein interactions | 43793 | 42 | 5 | 9204 | 432 | Ddr,chn,abd-A,koi,ttk | | 4.548e-02 | -3.09 | intrinsic component of peroxisomal membrane | cellular component | GO:0031231 | 10 | 2 | 11940 | 417 | Pex10,CG33474 | | 4.548e-02 | -3.09 | integral component of peroxisomal membrane | cellular component | GO:0005779 | 10 | 2 | 11940 | 417 | Pex10,CG33474 | | 4.548e-02 | -3.09 | sarcoplasm | cellular component | GO:0016528 | 10 | 2 | 11940 | 417 | yuri,SERCA | | 4.556e-02 | -3.09 | GO\_NEGATIVE\_REGULATION\_OF\_CELL\_COMMUNICATION | MSigDB lists | GO\_NEGATIVE\_REGULATION\_OF\_CELL\_COMMUNICATION | 293 | 17 | 4161 | 156 | Raf,cact,Arf79F,Actbeta,N,Unc-76,InR,dock,nmo,Tollo,lqf,sli,PI31,fz,Pdk1,Nmda1,arm | | 4.567e-02 | -3.09 | GO\_RESPONSE\_TO\_BACTERIUM | MSigDB lists | GO\_RESPONSE\_TO\_BACTERIUM | 88 | 7 | 4161 | 156 | cact,Tollo,N,hyx,Mef2,Rab1,fz2 | | 4.567e-02 | -3.09 | REACTOME\_POST\_TRANSLATIONAL\_PROTEIN\_MODIFICATION | MSigDB lists | REACTOME\_POST\_TRANSLATIONAL\_PROTEIN\_MODIFICATION | 88 | 7 | 4161 | 156 | Pgant5,CG3149,CG1129,Gfat2,alpha-Man-Ia,Pgant6,C1GalTA | | 4.588e-02 | -3.08 | tight junction organization | biological process | GO:0120193 | 38 | 4 | 12345 | 442 | moody,Gli,scrib,bou | | 4.588e-02 | -3.08 | tight junction assembly | biological process | GO:0120192 | 38 | 4 | 12345 | 442 | bou,scrib,moody,Gli | | 4.592e-02 | -3.08 | GO\_REGULATION\_OF\_GTPASE\_ACTIVITY | MSigDB lists | GO\_REGULATION\_OF\_GTPASE\_ACTIVITY | 146 | 10 | 4161 | 156 | CenB1A,htl,dock,Ocrl,CG17883,sli,Ziz,step,Rgl,CG5522 | | 4.597e-02 | -3.08 | GSE20198\_IL12\_VS\_IL12\_IL18\_TREATED\_ACT\_CD4\_TCELL\_DN | MSigDB lists | GSE20198\_IL12\_VS\_IL12\_IL18\_TREATED\_ACT\_CD4\_TCELL\_DN | 70 | 6 | 4161 | 156 | Tsp96F,CG2006,Ids,Nmda1,CG10417,Mfe2 | | 4.597e-02 | -3.08 | GSE32423\_MEMORY\_VS\_NAIVE\_CD8\_TCELL\_IL7\_IL4\_DN | MSigDB lists | GSE32423\_MEMORY\_VS\_NAIVE\_CD8\_TCELL\_IL7\_IL4\_DN | 70 | 6 | 4161 | 156 | alpha-Man-Ia,Hel89B,hppy,l(1)G0196,CG2812,CG3107 | | 4.597e-02 | -3.08 | GSE13411\_NAIVE\_VS\_MEMORY\_BCELL\_DN | MSigDB lists | GSE13411\_NAIVE\_VS\_MEMORY\_BCELL\_DN | 70 | 6 | 4161 | 156 | Pgk,Fkbp14,Pgant6,CG10417,PI31,CycG | | 4.598e-02 | -3.08 | transcription corepressor activity | molecular function | GO:0003714 | 40 | 4 | 12176 | 414 | CtBP,emc,Tob,H | | 4.598e-02 | -3.08 | glycosaminoglycan binding | molecular function | GO:0005539 | 40 | 4 | 12176 | 414 | Sema1a,sli,robo1,vn | | 4.611e-02 | -3.08 | serine/threonine protein kinase complex | cellular component | GO:1902554 | 39 | 4 | 11940 | 417 | Atg17,CycG,CycB3,Cks30A | | 4.622e-02 | -3.07 | HYPRGLYCEMIC | prints domains | PR00550 | 1 | 1 | 2185 | 101 | ITP | | 4.622e-02 | -3.07 | PROFILINPLNT | prints domains | PR01640 | 1 | 1 | 2185 | 101 | chic | | 4.622e-02 | -3.07 | ANTIPRLFBTG1 | prints domains | PR00310 | 1 | 1 | 2185 | 101 | Tob | | 4.622e-02 | -3.07 | PROFILIN | prints domains | PR00392 | 1 | 1 | 2185 | 101 | chic | | 4.622e-02 | -3.07 | LNOTCHREPEAT | prints domains | PR01452 | 1 | 1 | 2185 | 101 | N | | 4.622e-02 | -3.07 | TEADOMAIN | prints domains | PR00065 | 1 | 1 | 2185 | 101 | sd | | 4.622e-02 | -3.07 | BCATNINFAMLY | prints domains | PR01869 | 1 | 1 | 2185 | 101 | arm | | 4.622e-02 | -3.07 | LIPOCALINIMR | prints domains | PR01692 | 1 | 1 | 2185 | 101 | lili | | 4.630e-02 | -3.07 | cytosolic transport | biological process | GO:0016482 | 55 | 5 | 12345 | 442 | Ppcs,jar,ena,Abl,mop | | 4.630e-02 | -3.07 | somatic stem cell division | biological process | GO:0048103 | 55 | 5 | 12345 | 442 | baz,jar,shg,esg,flfl | | 4.631e-02 | -3.07 | regulation of peptide transport | biological process | GO:0090087 | 73 | 6 | 12345 | 442 | S,Rab1,InR,rdx,Abl,mop | | 4.634e-02 | -3.07 | Rcd5 (Reduction in Cnn dots 5) | protein interactions | 38478 | 56 | 6 | 9204 | 432 | Pitslre,eyg,CG42684,esg,CG31211,CG12496 | | 4.659e-02 | -3.07 | Nucleotide-diphossugar\_trans | interpro domains | IPR029044 | 50 | 5 | 11383 | 450 | CG1129,Pgant5,GlcAT-I,Pgant6,CG9220 | | 4.669e-02 | -3.06 | His\_Pase\_clade-2 | interpro domains | IPR000560 | 9 | 2 | 11383 | 450 | l(1)G0196,CG6656 | | 4.669e-02 | -3.06 | Ras-like\_GEF | interpro domains | IPR008937 | 9 | 2 | 11383 | 450 | Rgl,CG5522 | | 4.669e-02 | -3.06 | RASGEF\_cat\_dom\_sf | interpro domains | IPR036964 | 9 | 2 | 11383 | 450 | Rgl,CG5522 | | 4.689e-02 | -3.06 | Chondroitin sulfate biosynthesis | REACTOME pathways | R-DME-2022870 | 8 | 2 | 3923 | 176 | CG9220,kon | | 4.689e-02 | -3.06 | Signaling by NOTCH4 | REACTOME pathways | R-DME-1980150 | 8 | 2 | 3923 | 176 | N,Dl | | 4.689e-02 | -3.06 | DNA Double Strand Break Response | REACTOME pathways | R-DME-5693606 | 8 | 2 | 3923 | 176 | Abl,Pen | | 4.689e-02 | -3.06 | Signaling by NOTCH3 | REACTOME pathways | R-DME-1980148 | 8 | 2 | 3923 | 176 | Dl,N | | 4.689e-02 | -3.06 | Signaling by NOTCH2 | REACTOME pathways | R-DME-1980145 | 8 | 2 | 3923 | 176 | Dl,N | | 4.689e-02 | -3.06 | Binding and Uptake of Ligands by Scavenger Receptors | REACTOME pathways | R-DME-2173782 | 8 | 2 | 3923 | 176 | apolpp,pes | | 4.689e-02 | -3.06 | Nephrin interactions | REACTOME pathways | R-DME-373753 | 8 | 2 | 3923 | 176 | dock,sns | | 4.689e-02 | -3.06 | NOTCH2 Activation and Transmission of Signal to the Nucleus | REACTOME pathways | R-DME-2979096 | 8 | 2 | 3923 | 176 | N,Dl | | 4.689e-02 | -3.06 | Scavenging by Class B Receptors | REACTOME pathways | R-DME-3000471 | 8 | 2 | 3923 | 176 | apolpp,pes | | 4.689e-02 | -3.06 | G2/M DNA damage checkpoint | REACTOME pathways | R-DME-69473 | 8 | 2 | 3923 | 176 | 14-3-3epsilon,14-3-3zeta | | 4.689e-02 | -3.06 | Ephrin signaling | REACTOME pathways | R-DME-3928664 | 8 | 2 | 3923 | 176 | Eph,dock | | 4.694e-02 | -3.06 | Tbh (Tyramine beta hydroxylase) | protein interactions | 31718 | 1 | 1 | 9204 | 432 | Bacc | | 4.694e-02 | -3.06 | CG31908 (uncharacterized protein) | protein interactions | 261613 | 1 | 1 | 9204 | 432 | Sxl | | 4.694e-02 | -3.06 | CG8740 (uncharacterized protein) | protein interactions | 35866 | 1 | 1 | 9204 | 432 | Sema1a | | 4.694e-02 | -3.06 | 5-HT2A (5-hydroxytryptamine (serotonin) receptor 2A) | protein interactions | 40575 | 1 | 1 | 9204 | 432 | Mob2 | | 4.694e-02 | -3.06 | Spn28B (Serpin 28B) | protein interactions | 34035 | 1 | 1 | 9204 | 432 | CG32772 | | 4.694e-02 | -3.06 | lncRNA:CR32660 (long non-coding RNA:CR32660) | protein interactions | 3771836 | 1 | 1 | 9204 | 432 | CG32772 | | 4.694e-02 | -3.06 | CG3698 (uncharacterized protein) | protein interactions | 40298 | 1 | 1 | 9204 | 432 | AP-1gamma | | 4.694e-02 | -3.06 | CG1909 (uncharacterized protein) | protein interactions | 43800 | 1 | 1 | 9204 | 432 | Tpc1 | | 4.694e-02 | -3.06 | gprs (gprs) | protein interactions | 36862 | 1 | 1 | 9204 | 432 | scrib | | 4.694e-02 | -3.06 | snRNA:U6:96Ab (small nuclear RNA U6 at 96A b) | protein interactions | 3772330 | 1 | 1 | 9204 | 432 | SmD1 | | 4.694e-02 | -3.06 | Hml (Hemolectin) | protein interactions | 39529 | 1 | 1 | 9204 | 432 | C1GalTA | | 4.694e-02 | -3.06 | MIB1 (mindbomb E3 ubiquitin protein ligase 1) | protein interactions | 57534 | 1 | 1 | 9204 | 432 | Dl | | 4.694e-02 | -3.06 | CG4041 (uncharacterized protein) | protein interactions | 31422 | 1 | 1 | 9204 | 432 | Rab5 | | 4.694e-02 | -3.06 | CG34384 (uncharacterized protein) | protein interactions | 5740362 | 1 | 1 | 9204 | 432 | Dp | | 4.694e-02 | -3.06 | SclB (Sarcolamban B) | protein interactions | 19834948 | 1 | 1 | 9204 | 432 | SERCA | | 4.694e-02 | -3.06 | ocn (ocnus) | protein interactions | 43567 | 1 | 1 | 9204 | 432 | CG34401 | | 4.694e-02 | -3.06 | CG15879 (uncharacterized protein) | protein interactions | 38274 | 1 | 1 | 9204 | 432 | Pex5 | | 4.694e-02 | -3.06 | CadN2 (Cadherin-N2) | protein interactions | 35071 | 1 | 1 | 9204 | 432 | Sema1a | | 4.694e-02 | -3.06 | CG4210 (uncharacterized protein) | protein interactions | 41832 | 1 | 1 | 9204 | 432 | tsh | | 4.694e-02 | -3.06 | mir-318 (mir-318 stem loop) | protein interactions | 12797881 | 1 | 1 | 9204 | 432 | ttk | | 4.694e-02 | -3.06 | CG15249 (uncharacterized protein) | protein interactions | 31924 | 1 | 1 | 9204 | 432 | Atg16 | | 4.694e-02 | -3.06 | Exd2 (Exonuclease 3'-5' domain-containing 2) | protein interactions | 41374 | 1 | 1 | 9204 | 432 | Pex14 | | 4.694e-02 | -3.06 | GluRIA (Glutamate receptor IA) | protein interactions | 38742 | 1 | 1 | 9204 | 432 | mir-1000 | | 4.694e-02 | -3.06 | CG17470 (uncharacterized protein) | protein interactions | 35321 | 1 | 1 | 9204 | 432 | Ddr | | 4.694e-02 | -3.06 | CG6834 (uncharacterized protein) | protein interactions | 41409 | 1 | 1 | 9204 | 432 | Myo10A | | 4.694e-02 | -3.06 | CG43759 (uncharacterized protein) | protein interactions | 32889 | 1 | 1 | 9204 | 432 | pnr | | 4.694e-02 | -3.06 | CG8051 (uncharacterized protein) | protein interactions | 32925 | 1 | 1 | 9204 | 432 | pnr | | 4.694e-02 | -3.06 | CG31644 (uncharacterized protein) | protein interactions | 318868 | 1 | 1 | 9204 | 432 | Jupiter | | 4.694e-02 | -3.06 | CG7110 (uncharacterized protein) | protein interactions | 34757 | 1 | 1 | 9204 | 432 | Syx1A | | 4.694e-02 | -3.06 | CG6409 (uncharacterized protein) | protein interactions | 39220 | 1 | 1 | 9204 | 432 | CG32772 | | 4.694e-02 | -3.06 | EGFR (epidermal growth factor receptor) | protein interactions | 1956 | 1 | 1 | 9204 | 432 | Cbl | | 4.694e-02 | -3.06 | CG13869 (uncharacterized protein) | protein interactions | 37283 | 1 | 1 | 9204 | 432 | Sxl | | 4.694e-02 | -3.06 | Jon25Bii (Jonah 25Bii) | protein interactions | 33707 | 1 | 1 | 9204 | 432 | CG12581 | | 4.694e-02 | -3.06 | HSPA8 (heat shock protein family A (Hsp70) member 8) | protein interactions | 3312 | 1 | 1 | 9204 | 432 | lqf | | 4.694e-02 | -3.06 | CG13325 (uncharacterized protein) | protein interactions | 36437 | 1 | 1 | 9204 | 432 | Mpcp2 | | 4.694e-02 | -3.06 | PARP1 (poly(ADP-ribose) polymerase 1) | protein interactions | 142 | 1 | 1 | 9204 | 432 | CG1218 | | 4.694e-02 | -3.06 | CG6277 (uncharacterized protein) | protein interactions | 43251 | 1 | 1 | 9204 | 432 | Ppn | | 4.694e-02 | -3.06 | Mctp (Multiple C2 domain and transmembrane region protein) | protein interactions | 37165 | 1 | 1 | 9204 | 432 | ey | | 4.694e-02 | -3.06 | CG32373 (uncharacterized protein) | protein interactions | 318000 | 1 | 1 | 9204 | 432 | nmo | | 4.694e-02 | -3.06 | Cbp53E (Calbindin 53E) | protein interactions | 36905 | 1 | 1 | 9204 | 432 | orb2 | | 4.694e-02 | -3.06 | CG16854 (uncharacterized protein) | protein interactions | 34539 | 1 | 1 | 9204 | 432 | 14-3-3zeta | | 4.694e-02 | -3.06 | rk (rickets) | protein interactions | 34819 | 1 | 1 | 9204 | 432 | vn | | 4.694e-02 | -3.06 | CG2100 (uncharacterized protein) | protein interactions | 40707 | 1 | 1 | 9204 | 432 | Sema1a | | 4.694e-02 | -3.06 | Dci (Dodecenoyl-CoA delta-isomerase) | protein interactions | 38100 | 1 | 1 | 9204 | 432 | Pex5 | | 4.694e-02 | -3.06 | CG16852 (uncharacterized protein) | protein interactions | 34769 | 1 | 1 | 9204 | 432 | sina | | 4.694e-02 | -3.06 | Mp (Multiplexin) | protein interactions | 38769 | 1 | 1 | 9204 | 432 | sli | | 4.694e-02 | -3.06 | CG4269 (uncharacterized protein) | protein interactions | 37578 | 1 | 1 | 9204 | 432 | Cdk4 | | 4.694e-02 | -3.06 | CG1090 (uncharacterized protein) | protein interactions | 40551 | 1 | 1 | 9204 | 432 | Pvr | | 4.694e-02 | -3.06 | Usp1 (Ubiquitin specific protease 1) | protein interactions | 43462 | 1 | 1 | 9204 | 432 | Dronc | | 4.694e-02 | -3.06 | Pask (PAS kinase) | protein interactions | 37824 | 1 | 1 | 9204 | 432 | CtBP | | 4.694e-02 | -3.06 | mir-311 (mir-311 stem loop) | protein interactions | 12798127 | 1 | 1 | 9204 | 432 | esg | | 4.694e-02 | -3.06 | CG14357 (uncharacterized protein) | protein interactions | 41716 | 1 | 1 | 9204 | 432 | CtBP | | 4.694e-02 | -3.06 | Asph (Aspartyl beta-hydroxylase) | protein interactions | 36778 | 1 | 1 | 9204 | 432 | Pex5 | | 4.694e-02 | -3.06 | CG7091 (uncharacterized protein) | protein interactions | 41590 | 1 | 1 | 9204 | 432 | pita | | 4.694e-02 | -3.06 | CG4752 (uncharacterized protein) | protein interactions | 37569 | 1 | 1 | 9204 | 432 | kn | | 4.694e-02 | -3.06 | f-cup (flyers-cup) | protein interactions | 41677 | 1 | 1 | 9204 | 432 | orb2 | | 4.694e-02 | -3.06 | fit (female-specific independent of transformer) | protein interactions | 42578 | 1 | 1 | 9204 | 432 | InR | | 4.694e-02 | -3.06 | CG11210 (uncharacterized protein) | protein interactions | 35779 | 1 | 1 | 9204 | 432 | baz | | 4.694e-02 | -3.06 | dcc.L (DCC netrin 1 receptor L homeolog) | protein interactions | 378529 | 1 | 1 | 9204 | 432 | sina | | 4.694e-02 | -3.06 | CG11362 (uncharacterized protein) | protein interactions | 37574 | 1 | 1 | 9204 | 432 | ena | | 4.694e-02 | -3.06 | CG4942 (uncharacterized protein) | protein interactions | 39044 | 1 | 1 | 9204 | 432 | Inx2 | | 4.694e-02 | -3.06 | snRNA:U6:96Ac (small nuclear RNA U6 at 96A c) | protein interactions | 3772628 | 1 | 1 | 9204 | 432 | SmD1 | | 4.694e-02 | -3.06 | CG14120 (uncharacterized protein) | protein interactions | 39463 | 1 | 1 | 9204 | 432 | His3.3A | | 4.694e-02 | -3.06 | CG12056 (uncharacterized protein) | protein interactions | 31855 | 1 | 1 | 9204 | 432 | Ppn | | 4.694e-02 | -3.06 | CG7714 (uncharacterized protein) | protein interactions | 42250 | 1 | 1 | 9204 | 432 | Dl | | 4.694e-02 | -3.06 | CG8630 (uncharacterized protein) | protein interactions | 41629 | 1 | 1 | 9204 | 432 | Arf79F | | 4.694e-02 | -3.06 | CG16903 (uncharacterized protein) | protein interactions | 31178 | 1 | 1 | 9204 | 432 | koi | | 4.694e-02 | -3.06 | CG18278 (uncharacterized protein) | protein interactions | 36487 | 1 | 1 | 9204 | 432 | CycG | | 4.694e-02 | -3.06 | CG12862 (uncharacterized protein) | protein interactions | 36625 | 1 | 1 | 9204 | 432 | Tpc1 | | 4.694e-02 | -3.06 | Nepl20 (Neprilysin-like 20) | protein interactions | 43412 | 1 | 1 | 9204 | 432 | N | | 4.694e-02 | -3.06 | CG1951 (uncharacterized protein) | protein interactions | 43423 | 1 | 1 | 9204 | 432 | baz | | 4.694e-02 | -3.06 | CG32055 (uncharacterized protein) | protein interactions | 39188 | 1 | 1 | 9204 | 432 | sina | | 4.694e-02 | -3.06 | CG32521 (uncharacterized protein) | protein interactions | 33098 | 1 | 1 | 9204 | 432 | H | | 4.694e-02 | -3.06 | CG10907 (uncharacterized protein) | protein interactions | 39331 | 1 | 1 | 9204 | 432 | rictor | | 4.694e-02 | -3.06 | FDY (flagrante delicto Y) | protein interactions | 8674077 | 1 | 1 | 9204 | 432 | Pdk1 | | 4.694e-02 | -3.06 | CG33098 (uncharacterized protein) | protein interactions | 326253 | 1 | 1 | 9204 | 432 | orb2 | | 4.694e-02 | -3.06 | S-Lap7 (Sperm-Leucylaminopeptidase 7) | protein interactions | 36524 | 1 | 1 | 9204 | 432 | Ptp4E | | 4.694e-02 | -3.06 | lbm (late bloomer) | protein interactions | 35623 | 1 | 1 | 9204 | 432 | kay | | 4.694e-02 | -3.06 | CG7320 (uncharacterized protein) | protein interactions | 40010 | 1 | 1 | 9204 | 432 | Sema1a | | 4.694e-02 | -3.06 | Root (Rootletin) | protein interactions | 42860 | 1 | 1 | 9204 | 432 | orb2 | | 4.694e-02 | -3.06 | Prosalpha4T1 (Proteasome alpha4 subunit, Testis-specific 1) | protein interactions | 42457 | 1 | 1 | 9204 | 432 | arm | | 4.694e-02 | -3.06 | CSN1b (COP9 signalosome subunit 1b) | protein interactions | 40063 | 1 | 1 | 9204 | 432 | cic | | 4.694e-02 | -3.06 | CG5819 (uncharacterized protein) | protein interactions | 37547 | 1 | 1 | 9204 | 432 | CG1504 | | 4.694e-02 | -3.06 | CG13896 (uncharacterized protein) | protein interactions | 38089 | 1 | 1 | 9204 | 432 | ey | | 4.694e-02 | -3.06 | Hex-C (Hexokinase C) | protein interactions | 44008 | 1 | 1 | 9204 | 432 | IA-2 | | 4.694e-02 | -3.06 | CG4594 (uncharacterized protein) | protein interactions | 34316 | 1 | 1 | 9204 | 432 | dpy | | 4.694e-02 | -3.06 | mir-2c (mir-2c stem loop) | protein interactions | 12798412 | 1 | 1 | 9204 | 432 | N | | 4.694e-02 | -3.06 | Dbp73D (Dead box protein 73D) | protein interactions | 39871 | 1 | 1 | 9204 | 432 | kn | | 4.694e-02 | -3.06 | CG32109 (uncharacterized protein) | protein interactions | 39444 | 1 | 1 | 9204 | 432 | ds | | 4.694e-02 | -3.06 | CG5397 (uncharacterized protein) | protein interactions | 33313 | 1 | 1 | 9204 | 432 | cact | | 4.694e-02 | -3.06 | CG10357 (uncharacterized protein) | protein interactions | 38435 | 1 | 1 | 9204 | 432 | CG11247 | | 4.694e-02 | -3.06 | CG42671 (uncharacterized protein) | protein interactions | 39268 | 1 | 1 | 9204 | 432 | Sema1a | | 4.694e-02 | -3.06 | CG32040 (uncharacterized protein) | protein interactions | 317835 | 1 | 1 | 9204 | 432 | dpy | | 4.694e-02 | -3.06 | CG3328 (uncharacterized protein) | protein interactions | 37871 | 1 | 1 | 9204 | 432 | mam | | 4.694e-02 | -3.06 | Or2a (Odorant receptor 2a) | protein interactions | 31207 | 1 | 1 | 9204 | 432 | CG32772 | | 4.694e-02 | -3.06 | CG17734 (uncharacterized protein) | protein interactions | 41359 | 1 | 1 | 9204 | 432 | CG13204 | | 4.694e-02 | -3.06 | Uxt (Uxt prefoldin-like subunit) | protein interactions | 34893 | 1 | 1 | 9204 | 432 | CG13204 | | 4.694e-02 | -3.06 | rempA (reduced mechanoreceptor potential A) | protein interactions | 33230 | 1 | 1 | 9204 | 432 | Sema1a | | 4.694e-02 | -3.06 | CG1339 (uncharacterized protein) | protein interactions | 35705 | 1 | 1 | 9204 | 432 | cindr | | 4.694e-02 | -3.06 | CG6511 (uncharacterized protein) | protein interactions | 38997 | 1 | 1 | 9204 | 432 | Pat1 | | 4.694e-02 | -3.06 | Cpr76Bb (Cuticular protein 76Bb) | protein interactions | 40121 | 1 | 1 | 9204 | 432 | crp | | 4.694e-02 | -3.06 | Lrt (Leucine-rich tendon-specific protein) | protein interactions | 37342 | 1 | 1 | 9204 | 432 | robo1 | | 4.694e-02 | -3.06 | CG10859 (uncharacterized protein) | protein interactions | 34756 | 1 | 1 | 9204 | 432 | Mfe2 | | 4.694e-02 | -3.06 | Dyrk2 (Dual-specificity tyrosine phosphorylation-regulated kinase 2) | protein interactions | 34831 | 1 | 1 | 9204 | 432 | Dyrk2 | | 4.694e-02 | -3.06 | DCC (DCC netrin 1 receptor) | protein interactions | 1630 | 1 | 1 | 9204 | 432 | sina | | 4.694e-02 | -3.06 | Ts (Thymidylate synthase) | protein interactions | 33499 | 1 | 1 | 9204 | 432 | H2.0 | | 4.694e-02 | -3.06 | Hacd2 (3-hydroxyacyl-CoA dehydratase 2) | protein interactions | 34762 | 1 | 1 | 9204 | 432 | drongo | | 4.694e-02 | -3.06 | CG9346 (uncharacterized protein) | protein interactions | 37381 | 1 | 1 | 9204 | 432 | CG31211 | | 4.694e-02 | -3.06 | CG13579 (uncharacterized protein) | protein interactions | 37905 | 1 | 1 | 9204 | 432 | ZnT41F | | 4.694e-02 | -3.06 | snRNA:U6:96Aa (small nuclear RNA U6 at 96A a) | protein interactions | 3772327 | 1 | 1 | 9204 | 432 | SmD1 | | 4.694e-02 | -3.06 | CG7840 (uncharacterized protein) | protein interactions | 34136 | 1 | 1 | 9204 | 432 | unc-119 | | 4.694e-02 | -3.06 | Nadsyn (NAD synthetase) | protein interactions | 32328 | 1 | 1 | 9204 | 432 | Rab5 | | 4.694e-02 | -3.06 | EXTL3 (exostosin like glycosyltransferase 3) | protein interactions | 2137 | 1 | 1 | 9204 | 432 | Diap1 | | 4.694e-02 | -3.06 | Tsp42Ee (Tetraspanin 42Ee) | protein interactions | 35614 | 1 | 1 | 9204 | 432 | sns | | 4.694e-02 | -3.06 | hfw (halfway) | protein interactions | 31120 | 1 | 1 | 9204 | 432 | Fem-1 | | 4.694e-02 | -3.06 | CG8270 (uncharacterized protein) | protein interactions | 38734 | 1 | 1 | 9204 | 432 | Rab5 | | 4.694e-02 | -3.06 | shams (shams) | protein interactions | 43008 | 1 | 1 | 9204 | 432 | N | | 4.694e-02 | -3.06 | CG11158 (uncharacterized protein) | protein interactions | 32327 | 1 | 1 | 9204 | 432 | scrib | | 4.694e-02 | -3.06 | CG34126 (uncharacterized protein) | protein interactions | 318872 | 1 | 1 | 9204 | 432 | Rab5 | | 4.694e-02 | -3.06 | EOGT (EGF domain specific O-linked N-acetylglucosamine transferase) | protein interactions | 285203 | 1 | 1 | 9204 | 432 | N | | 4.694e-02 | -3.06 | CG42342 (uncharacterized protein) | protein interactions | 7354466 | 1 | 1 | 9204 | 432 | Rgl | | 4.694e-02 | -3.06 | CG5693 (uncharacterized protein) | protein interactions | 50463 | 1 | 1 | 9204 | 432 | Mob2 | | 4.694e-02 | -3.06 | CG15431 (uncharacterized protein) | protein interactions | 33646 | 1 | 1 | 9204 | 432 | scrib | | 4.694e-02 | -3.06 | CG9920 (uncharacterized protein) | protein interactions | 41712 | 1 | 1 | 9204 | 432 | Sxl | | 4.694e-02 | -3.06 | CG5989 (uncharacterized protein) | protein interactions | 38984 | 1 | 1 | 9204 | 432 | CG31211 | | 4.694e-02 | -3.06 | Kat2b (K(lysine) acetyltransferase 2B) | protein interactions | 18519 | 1 | 1 | 9204 | 432 | N | | 4.694e-02 | -3.06 | CG9497 (uncharacterized protein) | protein interactions | 33884 | 1 | 1 | 9204 | 432 | esg | | 4.694e-02 | -3.06 | CG14253 (uncharacterized protein) | protein interactions | 43243 | 1 | 1 | 9204 | 432 | ATPCL | | 4.694e-02 | -3.06 | lmd (lame duck) | protein interactions | 42717 | 1 | 1 | 9204 | 432 | abd-A | | 4.694e-02 | -3.06 | mex1 (midgut expression 1) | protein interactions | 39677 | 1 | 1 | 9204 | 432 | ZnT41F | | 4.694e-02 | -3.06 | pinta (prolonged depolarization afterpotential (PDA) is not apparent) | protein interactions | 42635 | 1 | 1 | 9204 | 432 | nub | | 4.694e-02 | -3.06 | CG12730 (uncharacterized protein) | protein interactions | 31466 | 1 | 1 | 9204 | 432 | trbl | | 4.694e-02 | -3.06 | CG10702 (uncharacterized protein) | protein interactions | 35181 | 1 | 1 | 9204 | 432 | baz | | 4.694e-02 | -3.06 | CG12560 (uncharacterized protein) | protein interactions | 34092 | 1 | 1 | 9204 | 432 | Tpc1 | | 4.694e-02 | -3.06 | Lrr47 (Leucine-rich repeat 47) | protein interactions | 34449 | 1 | 1 | 9204 | 432 | Sxl | | 4.694e-02 | -3.06 | Mst57Da (Male-specific RNA 57Da) | protein interactions | 47954 | 1 | 1 | 9204 | 432 | Pdk1 | | 4.694e-02 | -3.06 | CG1632 (uncharacterized protein) | protein interactions | 31763 | 1 | 1 | 9204 | 432 | Pat1 | | 4.694e-02 | -3.06 | CG3061 (uncharacterized protein) | protein interactions | 41707 | 1 | 1 | 9204 | 432 | Ddr | | 4.695e-02 | -3.06 | GSE37532\_WT\_VS\_PPARG\_KO\_VISCERAL\_ADIPOSE\_TISSUE\_TREG\_UP | MSigDB lists | GSE37532\_WT\_VS\_PPARG\_KO\_VISCERAL\_ADIPOSE\_TISSUE\_TREG\_UP | 53 | 5 | 4161 | 156 | Pex5,Pen,CG2812,14-3-3zeta,His2Av | | 4.695e-02 | -3.06 | GSE11924\_TH1\_VS\_TH2\_CD4\_TCELL\_DN | MSigDB lists | GSE11924\_TH1\_VS\_TH2\_CD4\_TCELL\_DN | 53 | 5 | 4161 | 156 | Gfat2,Rab1,Hel89B,Tpc1,hppy | | 4.695e-02 | -3.06 | GSE17721\_CTRL\_VS\_GARDIQUIMOD\_1H\_BMDC\_DN | MSigDB lists | GSE17721\_CTRL\_VS\_GARDIQUIMOD\_1H\_BMDC\_DN | 53 | 5 | 4161 | 156 | Kr-h2,Actbeta,Lk6,cact,CG9485 | | 4.695e-02 | -3.06 | GSE4748\_CTRL\_VS\_LPS\_AND\_CYANOBACTERIUM\_LPSLIKE\_STIM\_DC\_3H\_UP | MSigDB lists | GSE4748\_CTRL\_VS\_LPS\_AND\_CYANOBACTERIUM\_LPSLIKE\_STIM\_DC\_3H\_UP | 53 | 5 | 4161 | 156 | CG4896,Gfat2,Fem-1,sli,trbl | | 4.695e-02 | -3.06 | DR4\_Q2 | MSigDB lists | DR4\_Q2 | 53 | 5 | 4161 | 156 | CtBP,CG6465,otk,Psa,cact | | 4.695e-02 | -3.06 | WANG\_SMARCE1\_TARGETS\_UP | MSigDB lists | WANG\_SMARCE1\_TARGETS\_UP | 53 | 5 | 4161 | 156 | Spn42De,fz,trbl,Dl,CG1673 | | 4.695e-02 | -3.06 | HADDAD\_B\_LYMPHOCYTE\_PROGENITOR | MSigDB lists | HADDAD\_B\_LYMPHOCYTE\_PROGENITOR | 53 | 5 | 4161 | 156 | Vps13,Mef2,CtBP,trbl,frc | | 4.695e-02 | -3.06 | GSE17721\_CPG\_VS\_GARDIQUIMOD\_24H\_BMDC\_UP | MSigDB lists | GSE17721\_CPG\_VS\_GARDIQUIMOD\_24H\_BMDC\_UP | 53 | 5 | 4161 | 156 | Ppn,FBXO11,Dyb,eff,CG4896 | | 4.695e-02 | -3.06 | CTGAGCC\_MIR24 | MSigDB lists | CTGAGCC\_MIR24 | 53 | 5 | 4161 | 156 | Rab5,CG10417,dlp,lili,htl | | 4.695e-02 | -3.06 | GSE20366\_TREG\_VS\_NAIVE\_CD4\_TCELL\_HOMEOSTATIC\_CONVERSION\_UP | MSigDB lists | GSE20366\_TREG\_VS\_NAIVE\_CD4\_TCELL\_HOMEOSTATIC\_CONVERSION\_UP | 53 | 5 | 4161 | 156 | eff,frc,Syx1A,14-3-3zeta,Pen | | 4.695e-02 | -3.06 | AP2GAMMA\_01 | MSigDB lists | AP2GAMMA\_01 | 53 | 5 | 4161 | 156 | otk,ap,hyx,Fkbp14,Kdm2 | | 4.721e-02 | -3.05 | GO\_POSITIVE\_REGULATION\_OF\_EXOCYTOSIS | MSigDB lists | GO\_POSITIVE\_REGULATION\_OF\_EXOCYTOSIS | 22 | 3 | 4161 | 156 | Syx1A,AP-1gamma,Arf79F | | 4.721e-02 | -3.05 | REACTOME\_NFKB\_AND\_MAP\_KINASES\_ACTIVATION\_MEDIATED\_BY\_TLR4\_SIGNALING\_REPERTOIRE | MSigDB lists | REACTOME\_NFKB\_AND\_MAP\_KINASES\_ACTIVATION\_MEDIATED\_BY\_TLR4\_SIGNALING\_REPERTOIRE | 22 | 3 | 4161 | 156 | cact,Tollo,Mef2 | | 4.721e-02 | -3.05 | ZHANG\_RESPONSE\_TO\_IKK\_INHIBITOR\_AND\_TNF\_DN | MSigDB lists | ZHANG\_RESPONSE\_TO\_IKK\_INHIBITOR\_AND\_TNF\_DN | 22 | 3 | 4161 | 156 | C1GalTA,kn,SMC3 | | 4.721e-02 | -3.05 | REACTOME\_BIOSYNTHESIS\_OF\_THE\_N\_GLYCAN\_PRECURSOR\_DOLICHOL\_LIPID\_LINKED\_OLIGOSACCHARIDE\_LLO\_AND\_TRANSFER\_TO\_A\_NASCENT\_PROTEIN | MSigDB lists | REACTOME\_BIOSYNTHESIS\_OF\_THE\_N\_GLYCAN\_PRECURSOR\_DOLICHOL\_LIPID\_LINKED\_OLIGOSACCHARIDE\_LLO\_AND\_TRANSFER\_TO\_A\_NASCENT\_PROTEIN | 22 | 3 | 4161 | 156 | CG3149,CG1129,Gfat2 | | 4.721e-02 | -3.05 | STAMBOLSKY\_RESPONSE\_TO\_VITAMIN\_D3\_UP | MSigDB lists | STAMBOLSKY\_RESPONSE\_TO\_VITAMIN\_D3\_UP | 22 | 3 | 4161 | 156 | CtBP,trbl,tkv | | 4.721e-02 | -3.05 | TCCAGAT\_MIR5165P | MSigDB lists | TCCAGAT\_MIR5165P | 22 | 3 | 4161 | 156 | Eph,trbl,muc | | 4.721e-02 | -3.05 | MODULE\_183 | MSigDB lists | MODULE\_183 | 22 | 3 | 4161 | 156 | bru2,rdx,SmD1 | | 4.721e-02 | -3.05 | KEGG\_SMALL\_CELL\_LUNG\_CANCER | MSigDB lists | KEGG\_SMALL\_CELL\_LUNG\_CANCER | 22 | 3 | 4161 | 156 | cact,Cdk4,LanA | | 4.727e-02 | -3.05 | PAS | prosite domains | PS50112 | 16 | 3 | 6243 | 325 | ss,sim,tai | | 4.745e-02 | -3.05 | GO\_POSITIVE\_REGULATION\_OF\_IMMUNE\_RESPONSE | MSigDB lists | GO\_POSITIVE\_REGULATION\_OF\_IMMUNE\_RESPONSE | 127 | 9 | 4161 | 156 | dock,Pdk1,eff,fz2,cact,Tollo,Mef2,AP-1gamma,PI31 | | 4.747e-02 | -3.05 | glucan metabolic process | biological process | GO:0044042 | 23 | 3 | 12345 | 442 | Atg17,Atg16,CG9485 | | 4.747e-02 | -3.05 | spiracle morphogenesis, open tracheal system | biological process | GO:0035277 | 23 | 3 | 12345 | 442 | Egfr,Abd-B,spi | | 4.747e-02 | -3.05 | carbohydrate derivative transport | biological process | GO:1901264 | 23 | 3 | 12345 | 442 | CG3149,Rim2,frc | | 4.747e-02 | -3.05 | glycogen metabolic process | biological process | GO:0005977 | 23 | 3 | 12345 | 442 | Atg17,Atg16,CG9485 | | 4.747e-02 | -3.05 | negative regulation of insulin receptor signaling pathway | biological process | GO:0046627 | 23 | 3 | 12345 | 442 | trbl,wdb,dock | | 4.747e-02 | -3.05 | regeneration | biological process | GO:0031099 | 23 | 3 | 12345 | 442 | wnd,Sxl,kay | | 4.747e-02 | -3.05 | negative regulation of cellular response to insulin stimulus | biological process | GO:1900077 | 23 | 3 | 12345 | 442 | dock,trbl,wdb | | 4.747e-02 | -3.05 | regulation of transcription elongation from RNA polymerase II promoter | biological process | GO:0034243 | 23 | 3 | 12345 | 442 | fs(1)h,hyx,Mef2 | | 4.747e-02 | -3.05 | regulation of receptor-mediated endocytosis | biological process | GO:0048259 | 23 | 3 | 12345 | 442 | cindr,Cbl,mim | | 4.747e-02 | -3.05 | energy reserve metabolic process | biological process | GO:0006112 | 23 | 3 | 12345 | 442 | Atg17,Atg16,CG9485 | | 4.747e-02 | -3.05 | cellular glucan metabolic process | biological process | GO:0006073 | 23 | 3 | 12345 | 442 | Atg16,CG9485,Atg17 | | 4.758e-02 | -3.05 | histone demethylation | biological process | GO:0016577 | 10 | 2 | 12345 | 442 | Kdm4B,Kdm2 | | 4.758e-02 | -3.05 | dedifferentiation | biological process | GO:0043696 | 10 | 2 | 12345 | 442 | Socs36E,N | | 4.758e-02 | -3.05 | histone lysine demethylation | biological process | GO:0070076 | 10 | 2 | 12345 | 442 | Kdm4B,Kdm2 | | 4.758e-02 | -3.05 | response to axon injury | biological process | GO:0048678 | 10 | 2 | 12345 | 442 | kay,wnd | | 4.758e-02 | -3.05 | cardiac muscle tissue development | biological process | GO:0048738 | 10 | 2 | 12345 | 442 | sd,abd-A | | 4.758e-02 | -3.05 | chondroitin sulfate proteoglycan metabolic process | biological process | GO:0050654 | 10 | 2 | 12345 | 442 | CG9220,GlcAT-I | | 4.758e-02 | -3.05 | chondroitin sulfate proteoglycan biosynthetic process | biological process | GO:0050650 | 10 | 2 | 12345 | 442 | CG9220,GlcAT-I | | 4.758e-02 | -3.05 | pericardial nephrocyte differentiation | biological process | GO:0061320 | 10 | 2 | 12345 | 442 | htl,pnr | | 4.758e-02 | -3.05 | chromatin-mediated maintenance of transcription | biological process | GO:0048096 | 10 | 2 | 12345 | 442 | tara,tna | | 4.758e-02 | -3.05 | blood circulation | biological process | GO:0008015 | 10 | 2 | 12345 | 442 | SERCA,pnr | | 4.758e-02 | -3.05 | cell dedifferentiation | biological process | GO:0043697 | 10 | 2 | 12345 | 442 | Socs36E,N | | 4.758e-02 | -3.05 | organic cation transport | biological process | GO:0015695 | 10 | 2 | 12345 | 442 | Tpc1,Orct2 | | 4.758e-02 | -3.05 | regulation of peptidyl-tyrosine phosphorylation | biological process | GO:0050730 | 10 | 2 | 12345 | 442 | Cbl,Socs36E | | 4.758e-02 | -3.05 | regulation of sevenless signaling pathway | biological process | GO:0045501 | 10 | 2 | 12345 | 442 | Src42A,Socs36E | | 4.758e-02 | -3.05 | epithelial cell type specification, open tracheal system | biological process | GO:0035153 | 10 | 2 | 12345 | 442 | N,arm | | 4.758e-02 | -3.05 | sensory organ precursor cell division | biological process | GO:0045035 | 10 | 2 | 12345 | 442 | Diap1,spdo | | 4.765e-02 | -3.04 | SOCS\_box | pfam domains | PF07525 | 9 | 2 | 10480 | 419 | CG9121,Socs36E | | 4.765e-02 | -3.04 | His\_Phos\_2 | pfam domains | PF00328 | 9 | 2 | 10480 | 419 | l(1)G0196,CG6656 | | 4.765e-02 | -3.04 | Ig\_2 | pfam domains | PF13895 | 9 | 2 | 10480 | 419 | ed,htl | | 4.765e-02 | -3.04 | Arm | pfam domains | PF00514 | 9 | 2 | 10480 | 419 | Pen,arm | | 4.768e-02 | -3.04 | OAR | prosite domains | PS50803 | 7 | 2 | 6243 | 325 | otp,al | | 4.768e-02 | -3.04 | SEMA | prosite domains | PS51004 | 7 | 2 | 6243 | 325 | Sema1b,Sema1a | | 4.768e-02 | -3.04 | DEATH\_DOMAIN | prosite domains | PS50017 | 7 | 2 | 6243 | 325 | unc-5,imd | | 4.790e-02 | -3.04 | GO\_VACUOLAR\_PART | MSigDB lists | GO\_VACUOLAR\_PART | 230 | 14 | 4161 | 156 | jar,CenB1A,fz2,dlp,lt,AP-1gamma,Rab5,Ids,Ocrl,emp,CG4935,Eph,Tollo,Atg16 | | 4.800e-02 | -3.04 | GO\_DEFENSE\_RESPONSE | MSigDB lists | GO\_DEFENSE\_RESPONSE | 188 | 12 | 4161 | 156 | Tollo,CG6891,CG4645,ReepA,Tpc1,Src42A,tkv,N,Actbeta,Rab1,Tpst,SmD1 | | 4.807e-02 | -3.04 | GSE24142\_EARLY\_THYMIC\_PROGENITOR\_VS\_DN3\_THYMOCYTE\_ADULT\_UP | MSigDB lists | GSE24142\_EARLY\_THYMIC\_PROGENITOR\_VS\_DN3\_THYMOCYTE\_ADULT\_UP | 37 | 4 | 4161 | 156 | cact,Tsp96F,Drak,rdx | | 4.807e-02 | -3.04 | GTTRYCATRR\_UNKNOWN | MSigDB lists | GTTRYCATRR\_UNKNOWN | 37 | 4 | 4161 | 156 | betaTub60D,CG2812,14-3-3zeta,14-3-3epsilon | | 4.807e-02 | -3.04 | GO\_SYNAPSE\_ORGANIZATION | MSigDB lists | GO\_SYNAPSE\_ORGANIZATION | 37 | 4 | 4161 | 156 | fz,Nrk,fz2,arm | | 4.807e-02 | -3.04 | GSE21360\_NAIVE\_VS\_PRIMARY\_MEMORY\_CD8\_TCELL\_UP | MSigDB lists | GSE21360\_NAIVE\_VS\_PRIMARY\_MEMORY\_CD8\_TCELL\_UP | 37 | 4 | 4161 | 156 | Nmda1,Mfe2,Pgk,CG9220 | | 4.807e-02 | -3.04 | GSE28726\_NAIVE\_VS\_ACTIVATED\_CD4\_TCELL\_UP | MSigDB lists | GSE28726\_NAIVE\_VS\_ACTIVATED\_CD4\_TCELL\_UP | 37 | 4 | 4161 | 156 | trbl,Cadps,Kdm2,Ocrl | | 4.807e-02 | -3.04 | GSE6259\_FLT3L\_INDUCED\_33D1\_POS\_DC\_VS\_CD4\_TCELL\_UP | MSigDB lists | GSE6259\_FLT3L\_INDUCED\_33D1\_POS\_DC\_VS\_CD4\_TCELL\_UP | 37 | 4 | 4161 | 156 | Ocrl,shot,CG18549,InR | | 4.807e-02 | -3.04 | RACTNNRTTTNC\_UNKNOWN | MSigDB lists | RACTNNRTTTNC\_UNKNOWN | 37 | 4 | 4161 | 156 | Lk6,lt,crol,Mfe2 | | 4.807e-02 | -3.04 | GO\_GLIAL\_CELL\_DIFFERENTIATION | MSigDB lists | GO\_GLIAL\_CELL\_DIFFERENTIATION | 37 | 4 | 4161 | 156 | N,Cdk4,Dl,arm | | 4.807e-02 | -3.04 | GSE13229\_IMM\_VS\_MATURE\_NKCELL\_DN | MSigDB lists | GSE13229\_IMM\_VS\_MATURE\_NKCELL\_DN | 37 | 4 | 4161 | 156 | Pex10,Arf79F,Cerk,ft | | 4.807e-02 | -3.04 | GSE7348\_LPS\_VS\_TOLERIZED\_AND\_LPS\_STIM\_MACROPHAGE\_DN | MSigDB lists | GSE7348\_LPS\_VS\_TOLERIZED\_AND\_LPS\_STIM\_MACROPHAGE\_DN | 37 | 4 | 4161 | 156 | Atg16,CG2017,Tollo,Rgl | | 4.807e-02 | -3.04 | GSE19923\_E2A\_KO\_VS\_HEB\_AND\_E2A\_KO\_DP\_THYMOCYTE\_DN | MSigDB lists | GSE19923\_E2A\_KO\_VS\_HEB\_AND\_E2A\_KO\_DP\_THYMOCYTE\_DN | 37 | 4 | 4161 | 156 | Atg16,Cdk4,Ddr,Cadps | | 4.807e-02 | -3.04 | GSE19772\_HCMV\_INFL\_VS\_HCMV\_INF\_MONOCYTES\_AND\_PI3K\_INHIBITION\_UP | MSigDB lists | GSE19772\_HCMV\_INFL\_VS\_HCMV\_INF\_MONOCYTES\_AND\_PI3K\_INHIBITION\_UP | 37 | 4 | 4161 | 156 | Ids,Ddr,CG6465,N | | 4.807e-02 | -3.04 | XBP1\_01 | MSigDB lists | XBP1\_01 | 37 | 4 | 4161 | 156 | Cadps,Rab1,Fkbp14,trbl | | 4.807e-02 | -3.04 | GO\_XENOPHAGY | MSigDB lists | GO\_XENOPHAGY | 37 | 4 | 4161 | 156 | SmD1,ReepA,CG4645,Tpc1 | | 4.807e-02 | -3.04 | GSE30971\_CTRL\_VS\_LPS\_STIM\_MACROPHAGE\_WBP7\_HET\_2H\_UP | MSigDB lists | GSE30971\_CTRL\_VS\_LPS\_STIM\_MACROPHAGE\_WBP7\_HET\_2H\_UP | 37 | 4 | 4161 | 156 | CG3961,CG1673,Dl,cact | | 4.807e-02 | -3.04 | GSE10211\_UV\_INACT\_SENDAI\_VS\_LIVE\_SENDAI\_VIRUS\_TRACHEAL\_EPITHELIAL\_CELLS\_DN | MSigDB lists | GSE10211\_UV\_INACT\_SENDAI\_VS\_LIVE\_SENDAI\_VIRUS\_TRACHEAL\_EPITHELIAL\_CELLS\_DN | 37 | 4 | 4161 | 156 | Cerk,CG1673,Dyb,CG6891 | | 4.807e-02 | -3.04 | GSE36476\_CTRL\_VS\_TSST\_ACT\_72H\_MEMORY\_CD4\_TCELL\_OLD\_UP | MSigDB lists | GSE36476\_CTRL\_VS\_TSST\_ACT\_72H\_MEMORY\_CD4\_TCELL\_OLD\_UP | 37 | 4 | 4161 | 156 | Tollo,emp,Ids,Gfat2 | | 4.807e-02 | -3.04 | GO\_MUSCLE\_STRUCTURE\_DEVELOPMENT | MSigDB lists | GO\_MUSCLE\_STRUCTURE\_DEVELOPMENT | 89 | 7 | 4161 | 156 | N,Mef2,fz,Dl,sns,arm,LanA | | 4.811e-02 | -3.03 | Cyclin-like\_sf | interpro domains | IPR036915 | 21 | 3 | 11383 | 450 | CycG,CycB3,CG6191 | | 4.819e-02 | -3.03 | CG10638 (uncharacterized protein) | protein interactions | 39424 | 86 | 8 | 9204 | 432 | chic,Fkbp14,Pgk,14-3-3epsilon,Bacc,Slbp,Jupiter,Nlp | | 4.832e-02 | -3.03 | MODULE\_94 | MSigDB lists | MODULE\_94 | 108 | 8 | 4161 | 156 | ap,Pex14,bru2,Tpst,CG6465,Eph,Pex5,Pgant6 | | 4.866e-02 | -3.02 | WH-like\_DNA-bd\_sf | interpro domains | IPR036388 | 102 | 8 | 11383 | 450 | jumu,Dp,Poxn,eyg,larp,aop,ey,Pex14 | | 4.873e-02 | -3.02 | GSE17721\_4H\_VS\_24H\_POLYIC\_BMDC\_UP | MSigDB lists | GSE17721\_4H\_VS\_24H\_POLYIC\_BMDC\_UP | 71 | 6 | 4161 | 156 | ATPCL,unc-119,Mef2,CG6891,Pgk,Pgant5 | | 4.873e-02 | -3.02 | GSE17721\_POLYIC\_VS\_GARDIQUIMOD\_4H\_BMDC\_DN | MSigDB lists | GSE17721\_POLYIC\_VS\_GARDIQUIMOD\_4H\_BMDC\_DN | 71 | 6 | 4161 | 156 | Pat1,frc,ATPCL,CG3961,dock,Raf | | 4.873e-02 | -3.02 | GSE5589\_LPS\_VS\_LPS\_AND\_IL10\_STIM\_MACROPHAGE\_180MIN\_DN | MSigDB lists | GSE5589\_LPS\_VS\_LPS\_AND\_IL10\_STIM\_MACROPHAGE\_180MIN\_DN | 71 | 6 | 4161 | 156 | cact,Arf79F,CG7394,Rgl,arm,Src42A | | 4.889e-02 | -3.02 | - | gene3d domains | 1.20.140.80 | 1 | 1 | 4929 | 241 | Dp | | 4.889e-02 | -3.02 | - | gene3d domains | 3.30.70.1040 | 1 | 1 | 4929 | 241 | Dg | | 4.889e-02 | -3.02 | - | gene3d domains | 1.20.120.830 | 1 | 1 | 4929 | 241 | p130CAS | | 4.889e-02 | -3.02 | - | gene3d domains | 1.10.2010.10 | 1 | 1 | 4929 | 241 | ITP | | 4.889e-02 | -3.02 | - | gene3d domains | 3.40.50.12350 | 1 | 1 | 4929 | 241 | eya | | 4.889e-02 | -3.02 | - | gene3d domains | 3.40.50.10300 | 1 | 1 | 4929 | 241 | Ppcs | | 4.889e-02 | -3.02 | - | gene3d domains | 1.20.1150.12 | 1 | 1 | 4929 | 241 | wbl | | 4.889e-02 | -3.02 | - | gene3d domains | 1.20.120.1120 | 1 | 1 | 4929 | 241 | Tob | | 4.889e-02 | -3.02 | - | gene3d domains | 3.30.70.2470 | 1 | 1 | 4929 | 241 | IA-2 | | 4.889e-02 | -3.02 | - | gene3d domains | 2.60.40.3180 | 1 | 1 | 4929 | 241 | kn | | 4.889e-02 | -3.02 | - | gene3d domains | 1.20.930.20 | 1 | 1 | 4929 | 241 | Cbl | | 4.889e-02 | -3.02 | - | gene3d domains | 2.160.20.10 | 1 | 1 | 4929 | 241 | FBXO11 | | 4.889e-02 | -3.02 | - | gene3d domains | 1.10.10.1610 | 1 | 1 | 4929 | 241 | sd | | 4.889e-02 | -3.02 | - | gene3d domains | 1.10.8.1120 | 1 | 1 | 4929 | 241 | Slbp | | 4.889e-02 | -3.02 | - | gene3d domains | 3.30.110.30 | 1 | 1 | 4929 | 241 | GluProRS | | 4.889e-02 | -3.02 | - | gene3d domains | 3.90.1290.10 | 1 | 1 | 4929 | 241 | shot | | 4.889e-02 | -3.02 | - | gene3d domains | 3.10.20.710 | 1 | 1 | 4929 | 241 | dve | | 4.894e-02 | -3.02 | rhythmic behavior | biological process | GO:0007622 | 74 | 6 | 12345 | 442 | dome,ITP,kay,tara,Mid1,Mef2 | | 4.938e-02 | -3.01 | REVINTRACTNG | prints domains | PR00405 | 8 | 2 | 2185 | 101 | drongo,CenB1A | | 4.938e-02 | -3.01 | INNEXIN | prints domains | PR01262 | 8 | 2 | 2185 | 101 | ogre,Inx2 | | 4.938e-02 | -3.01 | PAIREDBOX | prints domains | PR00027 | 8 | 2 | 2185 | 101 | Poxn,ey | | 4.941e-02 | -3.01 | regulation of actin filament polymerization | biological process | GO:0030833 | 56 | 5 | 12345 | 442 | Cip4,SCAR,RhoU,CG6891,chic | | 4.947e-02 | -3.01 | GO\_REGULATION\_OF\_MULTI\_ORGANISM\_PROCESS | MSigDB lists | GO\_REGULATION\_OF\_MULTI\_ORGANISM\_PROCESS | 128 | 9 | 4161 | 156 | CG17883,ReepA,CG4645,AP-1gamma,Tpc1,Arf79F,Actbeta,N,SmD1 | | 4.952e-02 | -3.01 | pont (pontin) | protein interactions | 53439 | 43 | 5 | 9204 | 432 | E(Pc),shg,arm,imd,His2Av | | 4.952e-02 | -3.01 | Rm62 (Rm62) | protein interactions | 40739 | 43 | 5 | 9204 | 432 | Rm62,SmD1,MEP-1,d,Pdk1 | | 4.952e-02 | -3.01 | EndoGI (Endonuclease G inhibitor) | protein interactions | 34953 | 43 | 5 | 9204 | 432 | Abl,Fkbp14,14-3-3epsilon,mam,Nlp | | 4.960e-02 | -3.00 | Ccp84Aa (Ccp84Aa) | protein interactions | 40825 | 18 | 3 | 9204 | 432 | CycG,sina,bol | | 4.960e-02 | -3.00 | lqf (liquid facets) | protein interactions | 38846 | 18 | 3 | 9204 | 432 | Dl,spen,N | | 4.960e-02 | -3.00 | Fs(2)Ket (Female sterile (2) Ketel) | protein interactions | 35336 | 18 | 3 | 9204 | 432 | Jupiter,Pen,Dl | | 4.960e-02 | -3.00 | CG2540 (uncharacterized protein) | protein interactions | 32216 | 18 | 3 | 9204 | 432 | LanA,CG14478,CG1218 | | 4.960e-02 | -3.00 | CG13731 (uncharacterized protein) | protein interactions | 39938 | 18 | 3 | 9204 | 432 | CG14153,14-3-3zeta,msi | | 4.960e-02 | -3.00 | Unr (Upstream of N-ras) | protein interactions | 38963 | 18 | 3 | 9204 | 432 | Rgl,fend,Sxl | | 4.960e-02 | -3.00 | GluRIIA (Glutamate receptor IIA) | protein interactions | 33788 | 18 | 3 | 9204 | 432 | Eph,pum,cora | | 4.960e-02 | -3.00 | RFeSP (Rieske iron-sulfur protein) | protein interactions | 44390 | 18 | 3 | 9204 | 432 | UQCR-11,Egfr,CG9121 | | 4.960e-02 | -3.00 | spri (sprint) | protein interactions | 31987 | 18 | 3 | 9204 | 432 | Pvr,Cbl,Rab5 | | 4.960e-02 | -3.00 | E(spl)m3-HLH (Enhancer of split m3, helix-loop-helix) | protein interactions | 43156 | 18 | 3 | 9204 | 432 | Rim2,E(spl)m5-HLH,h | | 4.960e-02 | -3.00 | fry (furry) | protein interactions | 39122 | 18 | 3 | 9204 | 432 | CycG,sina,Cdk4 | | 4.960e-02 | -3.00 | htk (hat-trick) | protein interactions | 32877 | 18 | 3 | 9204 | 432 | N,Dl,imd | | 4.960e-02 | -3.00 | Ccp84Ab (Ccp84Ab) | protein interactions | 40824 | 18 | 3 | 9204 | 432 | bol,sina,CycG | | 4.960e-02 | -3.00 | Ack (Activated Cdc42 kinase) | protein interactions | 38489 | 18 | 3 | 9204 | 432 | tkv,dock,Magi | | 4.960e-02 | -3.00 | CG14767 (uncharacterized protein) | protein interactions | 50250 | 18 | 3 | 9204 | 432 | Pvr,mam,CG8034 | | 4.971e-02 | -3.00 | GO\_NEGATIVE\_REGULATION\_OF\_CELL\_DEATH | MSigDB lists | GO\_NEGATIVE\_REGULATION\_OF\_CELL\_DEATH | 210 | 13 | 4161 | 156 | Dl,cact,Mef2,Raf,14-3-3epsilon,Nmda1,arm,dock,InR,Pdk1,N,CG45050,14-3-3zeta | | 4.974e-02 | -3.00 | positive regulation of synapse assembly | biological process | GO:0051965 | 39 | 4 | 12345 | 442 | Cals,rictor,nmo,tkv | | 4.974e-02 | -3.00 | positive regulation of cell morphogenesis involved in differentiation | biological process | GO:0010770 | 39 | 4 | 12345 | 442 | fz,Rab1,chic,shot | | 4.984e-02 | -3.00 | grim (grim) | protein interactions | 40014 | 57 | 6 | 9204 | 432 | Diap1,klu,Dronc,Hr4,morgue,lncRNA:Hsromega | | 4.987e-02 | -3.00 | extrinsic component of plasma membrane | cellular component | GO:0019897 | 40 | 4 | 11940 | 417 | Abl,Src42A,arm,shg | | 4.997e-02 | -3.00 | Nup50 (Nucleoporin 50kD) | protein interactions | 35784 | 30 | 4 | 9204 | 432 | Raf,Pen,Pdk1,ft | | 4.997e-02 | -3.00 | Dp1 (Dodeca-satellite-binding protein 1) | protein interactions | 37116 | 30 | 4 | 9204 | 432 | Dronc,ft,rictor,Pdk1 | | 4.997e-02 | -3.00 | esc (extra sexcombs) | protein interactions | 34595 | 30 | 4 | 9204 | 432 | corto,sens,ph-p,Psc |
