## Supplementary material for "A single DPE core promoter motif contributes to *in vivo* transcriptional regulation and affects cardiac function": SupFile2: knownResults.html

./cluster.allRlog.c300.t8//15.cluster.txt.motifs/ - Homer Known Motif Enrichment Results


### Homer Known Motif Enrichment Results (./cluster.allRlog.c300.t8//15.cluster.txt.motifs/)

Homer *de novo* Motif Results  
Gene Ontology Enrichment Results  
Known Motif Enrichment Results (txt file)  
Total Target Sequences = 652, Total Background Sequences = 26454

|  |  |  |  |  |  |  |  |  |  |  |  |
| --- | --- | --- | --- | --- | --- | --- | --- | --- | --- | --- | --- |
| Rank | Motif | Name | P-value | log P-pvalue | q-value (Benjamini) | # Target Sequences with Motif | % of Targets Sequences with Motif | # Background Sequences with Motif | % of Background Sequences with Motif | Motif File | SVG |
| 1 | T G C A A G T C C T A G A C T G A G C T G T A C T C G A A G T C C G T A A G T C G A C T C T A G | Unknown1(NR/Ini-like)/Drosophila-Promoters/Homer | 1e-11 | -2.623e+01 | 0.0000 | 49.0 | 7.52% | 629.6 | 2.38% | motif file (matrix) | svg |
| 2 | G A T C C T G A C T A G G C A T C T A G C A G T C A T G T C G A G T A C A G T C T C A G A G C T | M1BP(Zf)/S2R+-M1BP-ChIP-Seq(GSE49842)/Homer | 1e-10 | -2.468e+01 | 0.0000 | 49.0 | 7.52% | 658.2 | 2.49% | motif file (matrix) | svg |
| 3 | C G T A T C G A G A T C G C A T C G T A A C G T G T A C T C A G G T C A G A C T C G T A C T A G | DREF/Drosophila-Promoters/Homer | 1e-6 | -1.536e+01 | 0.0000 | 31.0 | 4.75% | 431.2 | 1.63% | motif file (matrix) | svg |
| 4 | G C A T A C G T A T G C C T G A C T A G G A C T G A T C A C T G | Initiator/Drosophila-Promoters/Homer | 1e-6 | -1.454e+01 | 0.0000 | 209.0 | 32.06% | 6225.3 | 23.56% | motif file (matrix) | svg |
| 5 | C T A G T C A G C T G A T C A G T G C A A C T G T C G A T C A G | Trl(Zf)/S2-GAGAfactor-ChIP-Seq(GSE40646)/Homer | 1e-4 | -1.105e+01 | 0.0001 | 182.0 | 27.91% | 5544.4 | 20.98% | motif file (matrix) | svg |
| 6 | C T G A C G T A C G T A G C T A C T G A A C G T G T C A A G T C A G T C C T G A G T C A G C T A | Unknown4/Drosophila-Promoters/Homer | 1e-4 | -1.016e+01 | 0.0001 | 23.0 | 3.53% | 355.7 | 1.35% | motif file (matrix) | svg |
| 7 | A G T C C G T A C G A T A G T C G T C A A G T C A C G T C T G A | Unknown2/Drosophila-Promoters/Homer | 1e-3 | -7.158e+00 | 0.0019 | 61.0 | 9.36% | 1617.2 | 6.12% | motif file (matrix) | svg |
| 8 | T C G A G T A C T C G A T C G A C A T G A T G C A C G T A C T G A C T G A G T C C G T A C T A G A G T C A T C G A G T C | Unknown3/Drosophila-Promoters/Homer | 1e-2 | -5.861e+00 | 0.0061 | 18.0 | 2.76% | 345.8 | 1.31% | motif file (matrix) | svg |
| 9 | C T A G A G T C A G C T A C T G C G T A C A G T C G T A C T G A T A G C T G A C | Unknown5/Drosophila-Promoters/Homer | 1e-2 | -5.786e+00 | 0.0061 | 76.0 | 11.66% | 2236.5 | 8.46% | motif file (matrix) | svg |
