## Supplementary material for "A single DPE core promoter motif contributes to *in vivo* transcriptional regulation and affects cardiac function": SupFile2: geneOntology.html

Gene Ontology Results

### Gene Ontology Enrichment Results

Homer *de novo* Motif Enrichment Results  
Known Motif Enrichment Results  

###### Text file version of complete results (i.e. open with Excel) - biological process: Functional groupings of proteins (Gene Ontology) - molecular function: Mechanistic actions of proteins (Gene Ontology) - cellular component: Protein localization (Gene Ontology) - chromosome location: Genes with similar chromosome localization (NCBI Gene) - KEGG pathways: Groups of proteins in the same pathways (KEGG) - protein interactions: "Proteins interacting with a common protein (BIND, EcoCyc, HPRD)" (NCBI Gene) - interpro domains: Proteins with similar domains and features (Interpro) - pfam domains: Proteins with similar domains and features (Pfam) - smart domains: Proteins with similar domains and features (SMART) - gene3d domains: Proteins with similar domains and features (Gene3D) - prosite domains: Proteins with similar domains and features (Prosite) - prints domains: Proteins with similar domains and features (PRINTS) - MSigDB lists: "Genes sets for pathways, factor/miRNA target predictions, expression patterns, etc." (MSigDB) - BIOCYC pathways: Groups of proteins in the same pathways (BIOCYC) - COSMIC cancer mutations: Genes mutated in similar cancers (COSMIC) - GWAS genes: Genes mutated in similar diseases (GWAS Catalog) - Lipid Maps pathways: Groups of proteins in the same lipid pathways (Lipid Maps/Biosystems) - Pathway Interaction DB: Groups of proteins in the same pathways (Pathway Interaction Database) - REACTOME pathways: Groups of proteins in the same pathways (REACTOME) - SMPDB pathways: Groups of proteins in the same pathways (SMPDB) - WikiPathways: Groups of proteins in the same pathways (Wikipathways) Enriched Categories | | | | | | | | | | | | --- | --- | --- | --- | --- | --- | --- | --- | --- | --- | | P-value | ln(P) | Term | GO Tree | GO ID | # of Genes in Term | # of Target Genes in Term | # of Total Genes | # of Target Genes | Common Genes | | 4.331e-50 | -113.66 | system development | biological process | GO:0048731 | 1808 | 184 | 12345 | 414 | drk,eya,ft,bab1,wb,pnt,trn,slp1,Atg17,E(spl)m4-BFM,Doa,egr,Syx1A,Mmp2,ttk,HGTX,Prosap,Utx,CG43658,lbe,nrv2,S6kII,ds,wnd,disco,cas,S,aop,Fas2,Rbfox1,Ptp99A,nw,scb,bowl,Rho1,Acf,Fas3,Fur1,Ten-m,CadN,bdg,ci,psq,Snx3,pAbp,betaTub56D,shn,Lpin,Acsl,NetA,sm,osa,RhoGAP15B,Ptpmeg,pio,cnc,exex,Doc2,sno,Notum,Dr,edl,dve,E(spl)mdelta-HLH,ytr,Pex1,Stat92E,l(3)mbt,mmy,Nhe2,hh,par-1,dpp,stwl,dally,wake,beat-IIIc,fax,Cdk5alpha,ras,RhoGAP16F,Mrp4,sens-2,aos,Sin3A,Pcl,rau,Kr,Lim1,spi,Lac,N,crb,ckn,flfl,salm,PsGEF,NiPp1,uzip,robo2,nuf,Pvf3,jing,MFS14,ec,Toll-7,NAT1,Dmtn,bou,sbb,Hipk,phyl,lz,tou,dlp,mle,Egfr,hts,otk2,sty,ush,Tl,lama,Ser,slow,CtBP,Hph,heph,CG13366,nudE,pon,shd,Dad,Bx,nrm,if,B-H2,twi,Sox21a,elav,Ubx,Inx2,slou,sdt,tkv,Dscam1,Imp,stg,enok,ham,Zif,TfAP-2,CycE,oc,RhoGAP93B,kni,ap,comm,sas,RhoGEF2,ocm,flz,Smox,pbl,Doc1,NetB,EcR,mbt,gus,wech,tin,spen,esn,bi,ey,Fbxl7,Con,fj,ImpL2,mam,Rac2,so,tara,Psc | | 6.556e-43 | -97.13 | cellular developmental process | biological process | GO:0048869 | 1818 | 174 | 12345 | 414 | flfl,salm,N,crb,ckn,Kr,Lim1,spi,wun2,nuf,robo2,uzip,NiPp1,Cdk5alpha,CG46339,fax,Arp2,Sox102F,rau,aos,ras,l(3)mbt,mmy,par-1,hh,Stat92E,E(spl)mdelta-HLH,FoxL1,ytr,Pex1,edl,Syp,Mos,beat-IIIc,dally,wake,stwl,dpp,Doc2,exex,cnc,Pak3,Ptpmeg,Dr,qsm,sno,bora,tsl,pAbp,ci,bdg,psq,CadN,osa,sm,Acsl,NetA,shn,Pat1,scb,Ptp99A,Rbfox1,CG41099,Fas2,goe,Ten-m,Fur1,Rho1,Fas3,Utx,lbe,Prosap,Mmp2,HGTX,ttk,egr,S,cas,aop,ds,spoon,wnd,S6kII,nrv2,Myb,pnt,trn,slp1,wb,ft,drk,eya,Doa,baf,E(spl)m4-BFM,Atg17,neo,mam,ImpL2,Rac2,fj,Hydr2,Con,ey,Psc,so,pbl,tyn,ocm,Smox,bi,esn,tin,spen,Doc1,EcR,NetB,gus,mbt,CycE,oc,RhoGAP93B,Zif,RhoGEF2,AGO2,yps,sas,ap,comm,slou,Inx2,elav,cta,Ubx,ham,tkv,sdt,Imp,Dscam1,enok,Sox21a,twi,Npc1a,B-H2,if,nrm,pon,Dad,shd,CtBP,slow,Ser,lama,ush,Tl,nudE,Tet,CG13366,heph,bbg,Mitofilin,phyl,JIL-1,sty,dlp,mle,Egfr,hts,lz,jing,ec,Pvf3,sbb,Toll-7 | | 8.369e-42 | -94.58 | cell differentiation | biological process | GO:0030154 | 1773 | 170 | 12345 | 414 | sno,Dr,Pak3,Ptpmeg,Doc2,exex,cnc,stwl,dpp,Syp,Mos,wake,dally,beat-IIIc,E(spl)mdelta-HLH,ytr,FoxL1,Pex1,edl,l(3)mbt,mmy,hh,par-1,Stat92E,Sox102F,rau,aos,ras,fax,Cdk5alpha,NiPp1,nuf,robo2,uzip,N,crb,ckn,Kr,Lim1,wun2,spi,flfl,salm,E(spl)m4-BFM,Atg17,neo,Doa,baf,ft,eya,drk,trn,pnt,slp1,wb,nrv2,S6kII,Myb,S,cas,aop,ds,spoon,wnd,Mmp2,HGTX,ttk,egr,Utx,lbe,Prosap,Rho1,Fas3,Ten-m,Fur1,Ptp99A,Rbfox1,CG41099,Fas2,goe,scb,Acsl,NetA,Pat1,shn,osa,sm,bdg,ci,psq,CadN,tsl,bora,pAbp,tkv,sdt,enok,Imp,Dscam1,ham,Inx2,cta,elav,Ubx,slou,sas,comm,ap,RhoGEF2,AGO2,yps,Zif,CycE,RhoGAP93B,oc,Doc1,NetB,EcR,gus,mbt,esn,bi,tin,spen,ocm,tyn,Smox,pbl,so,Psc,Con,Hydr2,ey,ImpL2,mam,Rac2,fj,Toll-7,sbb,Pvf3,jing,ec,lz,sty,mle,dlp,Egfr,hts,JIL-1,bbg,phyl,heph,nudE,Tet,CG13366,lama,ush,Tl,CtBP,slow,Ser,nrm,pon,shd,Dad,Npc1a,Sox21a,twi,B-H2,if | | 1.287e-40 | -91.85 | anatomical structure morphogenesis | biological process | GO:0009653 | 1579 | 158 | 12345 | 414 | Atg17,neo,Doa,eya,drk,ft,bab1,wb,pnt,trn,slp1,Myb,S6kII,nrv2,spoon,ds,wnd,disco,S,aop,Syx1A,Mmp2,ttk,Prosap,CG43658,Utx,lbe,Rho1,Fas3,Ten-m,Fas2,Ptp99A,Rbfox1,CG41099,nw,scb,bowl,shn,Lpin,NetA,Acsl,sm,osa,RhoGAP15B,CadN,Myo10A,ci,bdg,psq,sno,qsm,Dr,Ptpmeg,Pak3,pio,cnc,exex,dpp,beat-IIIc,dally,Syp,Inx3,dve,E(spl)mdelta-HLH,FoxL1,Stat92E,Nhe2,mmy,par-1,hh,ras,RhoGAP16F,Arp2,aos,rau,fax,Cdk5alpha,NiPp1,robo2,uzip,nuf,Kr,Lim1,spi,Lac,N,crb,ckn,salm,Toll-7,sbb,PhKgamma,jing,ec,Dtg,lz,mle,Egfr,dlp,otk2,hts,sty,Hipk,Mitofilin,phyl,Hph,heph,CG13366,nudE,ush,lama,Ser,CtBP,slow,Dad,shd,Bx,if,B-H2,Npc1a,twi,tkv,sdt,stg,Dscam1,enok,ham,cta,Ubx,Inx2,slou,ap,comm,sas,AGO2,RhoGEF2,TfAP-2,CycE,RhoGAP93B,oc,EcR,NetB,gus,mbt,tin,spen,bi,tyn,flz,Smox,pbl,so,Psc,ey,fj,ImpL2,mam,Rac2 | | 2.205e-40 | -91.31 | developmental process | biological process | GO:0032502 | 2988 | 224 | 12345 | 414 | Tlk,Prosap,Utx,CG43658,lbe,egr,bin3,Syx1A,Mmp2,HGTX,ttk,spoon,ds,wnd,S,disco,cas,aop,Myb,S6kII,nrv2,wb,pnt,trn,slp1,eya,Chro,drk,ft,bab1,Doa,baf,Atg17,neo,E(spl)m4-BFM,pAbp,Snx3,betaTub56D,bora,tsl,CadN,Myo10A,ci,bdg,psq,sm,osa,RhoGAP15B,Tif-IA,shn,Pat1,Lpin,NetA,Acsl,scb,bowl,Fas2,goe,Ptp99A,Rbfox1,CG41099,nw,Fur1,Ten-m,Acf,Rho1,Fas3,Stat92E,Nhe2,l(3)mbt,mmy,hh,par-1,edl,Inx3,dve,ytr,E(spl)mdelta-HLH,FoxL1,Pex1,dally,beat-IIIc,wake,Mos,Syp,dpp,stwl,pio,cnc,exex,Doc2,Ptpmeg,Pak3,Dr,sno,Notum,qsm,flfl,salm,Kr,Lac,Lim1,spi,wun2,N,crb,ckn,uzip,robo2,nuf,PsGEF,meso18E,NiPp1,Cdk5alpha,cpo,fax,CG46339,mgl,ras,RhoGAP16F,Mrp4,Sox102F,Arp2,sens-2,Pcl,aos,Sin3A,rau,Mitofilin,phyl,bbg,Hipk,JIL-1,dlp,mle,Egfr,hts,otk2,sty,Dtg,lz,tou,jing,MFS14,ec,Pvf3,PhKgamma,sbb,NAT1,Toll-7,Dmtn,bou,if,B-H2,Npc1a,twi,Sox21a,pon,shd,Dad,Bx,nrm,VepD,Ser,CtBP,slow,ush,Tl,lama,Tet,CG13366,nudE,Hph,heph,CycE,oc,RhoGAP93B,kni,Zif,TfAP-2,AGO2,yps,RhoGEF2,comm,ap,sas,slou,cta,elav,Ubx,Inx2,ham,sdt,tkv,Imp,Dscam1,stg,enok,fj,ImpL2,mam,Rac2,Fbxl7,ey,Hydr2,Con,Psc,so,tara,pbl,ocm,tyn,flz,Smox,tin,spen,bi,esn,NetB,EcR,Doc1,gus,mbt,wech | | 1.712e-39 | -89.26 | animal organ development | biological process | GO:0048513 | 1267 | 139 | 12345 | 414 | tkv,stg,Dscam1,enok,slou,Ubx,RhoGEF2,ap,CycE,oc,TfAP-2,Zif,bi,tin,spen,wech,Doc1,NetB,EcR,gus,mbt,pbl,ocm,Smox,flz,Psc,tara,so,mam,Rac2,fj,Fbxl7,ey,sbb,NAT1,Dmtn,jing,MFS14,ec,Pvf3,sty,dlp,Egfr,otk2,lz,phyl,Hipk,CG13366,heph,CtBP,slow,Ser,lama,ush,Tl,twi,B-H2,if,Bx,pon,Dad,Dr,Notum,sno,Doc2,pio,exex,Ptpmeg,dally,stwl,dpp,Nhe2,hh,par-1,Stat92E,ytr,E(spl)mdelta-HLH,edl,dve,sens-2,Sin3A,rau,aos,Pcl,Mrp4,RhoGAP16F,nuf,robo2,PsGEF,salm,crb,N,Kr,Lac,Lim1,spi,Doa,E(spl)m4-BFM,Atg17,pnt,trn,slp1,wb,ft,bab1,eya,drk,disco,S,cas,aop,ds,S6kII,nrv2,Utx,CG43658,lbe,Prosap,Mmp2,Syx1A,ttk,Ten-m,Rho1,bowl,scb,Rbfox1,nw,Fas2,osa,RhoGAP15B,NetA,shn,Lpin,betaTub56D,Snx3,bdg,ci,psq,CadN | | 4.323e-39 | -88.34 | regulation of cellular process | biological process | GO:0050794 | 3560 | 245 | 12345 | 414 | CtBP,Ser,tna,Tsp26A,ush,CG4325,Tl,ps,nudE,Tet,heph,Hph,grnd,Sox21a,Pits,twi,if,B-H2,Socs16D,CG31875,Bx,Dad,CaMKII,Hip1,KP78a,CG34172,jing,ec,PhKgamma,CG5522,Pvf3,CG11486,sbb,Toll-7,phyl,Hipk,JIL-1,l(3)neo38,CG5815,sty,hiw,Gbs-70E,dlp,Egfr,mle,hts,otk2,lz,RapGAP1,tou,pbl,tyn,ocm,Smox,mof,bi,tin,spen,phtf,EcR,Doc1,NetB,gus,Rhp,mbt,Lrt,mam,ImpL2,Taf1,Rac2,CG18766,Hydr2,Fbxl7,ey,lili,Psc,tara,so,slou,Tudor-SN,mub,RhoGAP18B,elav,Ubx,cta,Cirl,CG42684,ham,sdt,tkv,stg,wcy,Imp,Dscam1,enok,kni,CycE,RhoGAP93B,oc,CG31612,ctrip,CG12099,TfAP-2,fne,Kah,Zif,RhoGEF2,AGO2,Ank,CG44774,sas,ap,comm,bowl,smg,scb,CG8485,Rbfox1,Fas2,goe,Ten-m,CG5830,RhoGAP102A,lncRNA:roX1,Rho1,Acf,CG8441,tsl,bora,pAbp,Snx3,ci,CG5728,psq,CadN,osa,PEK,RhoGAP15B,NetA,Acsl,Usp2,shn,Pat1,Lpin,corn,pnt,slp1,smal,CCKLR-17D3,ft,bab1,drk,eya,Chro,Doa,Gyf,nAChRalpha4,E(spl)m4-BFM,Atg17,neo,CG43658,CG9650,Utx,lbe,HmgZ,Tlk,Prosap,Mmp2,bin3,CG32758,Syx1A,NFAT,HGTX,ttk,egr,S,CG31224,disco,cas,aop,ds,spoon,wnd,S6kII,Myb,Cdk5alpha,cpo,Haspin,mgl,Arp2,sens-2,Sox102F,aos,Pcl,Sin3A,rau,RhoGAP16F,salm,TfIIS,crb,CG34384,N,ckn,hang,Kr,Hsc70-3,spi,Lim1,nuf,robo2,NiPp1,PsGEF,ITP,Doc2,CG4360,exex,udd,cnc,Pak3,Mondo,Fili,Ptpmeg,Dr,Notum,sno,mmy,l(3)mbt,hh,par-1,MESR3,Pli,Stat92E,FoxL1,E(spl)mdelta-HLH,edl,Inx3,dve,Syp,Mos,Cdk12,dally,wake,stwl,Cnx99A,dpp | | 7.445e-39 | -87.79 | multicellular organism development | biological process | GO:0007275 | 2496 | 200 | 12345 | 414 | slou,Ubx,elav,cta,Inx2,ham,enok,stg,Imp,Dscam1,tkv,sdt,RhoGAP93B,oc,CycE,kni,Zif,TfAP-2,AGO2,RhoGEF2,ap,comm,sas,pbl,Smox,flz,ocm,spen,tin,bi,esn,mbt,gus,Doc1,EcR,NetB,wech,fj,Rac2,mam,ImpL2,ey,Fbxl7,Con,Psc,so,tara,ec,MFS14,jing,Pvf3,PhKgamma,sbb,Dmtn,bou,Toll-7,NAT1,phyl,Hipk,hts,otk2,dlp,mle,Egfr,sty,tou,lz,Dtg,Ser,slow,CtBP,Tl,ush,lama,CG13366,nudE,Hph,heph,if,B-H2,twi,Npc1a,Sox21a,shd,Dad,pon,nrm,Bx,VepD,exex,cnc,pio,Doc2,Ptpmeg,Dr,sno,Notum,Stat92E,par-1,hh,Nhe2,l(3)mbt,mmy,Inx3,dve,edl,Pex1,E(spl)mdelta-HLH,ytr,wake,dally,beat-IIIc,Syp,dpp,stwl,Cdk5alpha,fax,mgl,RhoGAP16F,Mrp4,ras,aos,Sin3A,rau,Pcl,sens-2,salm,flfl,Lac,spi,Lim1,Kr,ckn,crb,N,robo2,uzip,nuf,PsGEF,NiPp1,wb,slp1,pnt,trn,eya,drk,Chro,bab1,ft,Doa,Atg17,E(spl)m4-BFM,Prosap,Tlk,lbe,Utx,CG43658,egr,HGTX,ttk,Mmp2,Syx1A,wnd,spoon,ds,aop,S,disco,cas,S6kII,nrv2,scb,bowl,Fas2,CG41099,nw,Rbfox1,Ptp99A,Fur1,Ten-m,Fas3,Rho1,Acf,pAbp,Snx3,tsl,betaTub56D,Myo10A,CadN,psq,ci,bdg,sm,RhoGAP15B,osa,Lpin,Pat1,shn,Acsl,NetA | | 1.895e-38 | -86.86 | anatomical structure development | biological process | GO:0048856 | 2924 | 218 | 12345 | 414 | Ser,slow,CtBP,ush,Tl,lama,Tet,CG13366,nudE,Hph,heph,if,B-H2,Sox21a,twi,Npc1a,pon,Dad,shd,Bx,nrm,VepD,jing,MFS14,ec,Pvf3,PhKgamma,sbb,Toll-7,NAT1,bou,Dmtn,Mitofilin,phyl,bbg,JIL-1,Hipk,dlp,Egfr,mle,otk2,hts,sty,lz,Dtg,tou,pbl,tyn,ocm,Smox,flz,tin,spen,bi,esn,Doc1,EcR,NetB,mbt,gus,wech,fj,ImpL2,mam,Rac2,Fbxl7,ey,Hydr2,Con,Psc,so,tara,slou,elav,Ubx,cta,Inx2,ham,sdt,tkv,stg,Dscam1,enok,Imp,CycE,oc,RhoGAP93B,kni,Zif,TfAP-2,yps,AGO2,RhoGEF2,comm,ap,sas,scb,bowl,Fas2,goe,Rbfox1,Ptp99A,nw,CG41099,Fur1,Ten-m,Acf,Rho1,Fas3,pAbp,Snx3,betaTub56D,tsl,CadN,Myo10A,ci,bdg,psq,sm,osa,RhoGAP15B,Pat1,shn,Lpin,Acsl,NetA,wb,pnt,trn,slp1,eya,drk,Chro,ft,bab1,Doa,baf,Atg17,neo,E(spl)m4-BFM,Tlk,Prosap,CG43658,Utx,lbe,egr,Syx1A,Mmp2,ttk,HGTX,spoon,ds,wnd,cas,disco,S,aop,Myb,S6kII,nrv2,Cdk5alpha,fax,mgl,ras,RhoGAP16F,Mrp4,sens-2,Arp2,rau,aos,Sin3A,Pcl,flfl,salm,Kr,wun2,Lac,Lim1,spi,N,crb,ckn,uzip,robo2,nuf,PsGEF,meso18E,NiPp1,pio,cnc,exex,Doc2,Ptpmeg,Pak3,Dr,sno,qsm,Notum,Stat92E,mmy,Nhe2,l(3)mbt,par-1,hh,edl,dve,Inx3,FoxL1,E(spl)mdelta-HLH,ytr,Pex1,dally,beat-IIIc,wake,Syp,Mos,dpp,stwl | | 1.157e-37 | -85.05 | nervous system development | biological process | GO:0007399 | 1108 | 127 | 12345 | 414 | scb,Ser,slow,Fas2,Tl,Ptp99A,Rbfox1,Fur1,Ten-m,nudE,Acf,Rho1,Fas3,B-H2,if,pAbp,CadN,pon,shd,Dad,bdg,nrm,ci,sm,osa,shn,Acsl,NetA,wb,jing,pnt,trn,Pvf3,drk,eya,ft,sbb,Doa,Toll-7,Dmtn,phyl,Prosap,lbe,egr,Mmp2,ttk,HGTX,mle,ds,dlp,Egfr,wnd,hts,S,cas,disco,sty,aop,lz,tou,nrv2,S6kII,Cdk5alpha,pbl,fax,Smox,tin,spen,esn,NetB,EcR,ras,mbt,Pcl,rau,aos,fj,mam,ImpL2,flfl,Rac2,salm,Kr,ey,Lim1,spi,crb,N,Con,ckn,uzip,robo2,Psc,so,PsGEF,NiPp1,cnc,exex,elav,Ptpmeg,ham,Dr,sno,sdt,Notum,enok,Dscam1,Imp,CycE,RhoGAP93B,oc,Stat92E,l(3)mbt,mmy,par-1,hh,edl,Zif,E(spl)mdelta-HLH,Pex1,beat-IIIc,wake,dally,RhoGEF2,comm,ap,sas | | 1.335e-37 | -84.91 | regulation of biological process | biological process | GO:0050789 | 3862 | 254 | 12345 | 414 | pbl,jigr1,mof,Smox,tyn,ocm,ScpX,spen,tin,bi,gus,mbt,Rhp,NetB,Doc1,EcR,wech,phtf,CG18766,Rac2,Taf1,ImpL2,mam,Lrt,ey,Fbxl7,Hydr2,Psc,lili,so,tara,RhoGAP18B,mub,Tudor-SN,Etl1,slou,Ubx,CG42684,cta,elav,Cirl,ham,stg,Imp,Dscam1,enok,wcy,sdt,tkv,oc,ctrip,CG31612,RhoGAP93B,CycE,kni,Zif,Kah,fne,TfAP-2,CG12099,Ank,AGO2,RhoGEF2,ap,comm,sas,CG44774,Ser,CtBP,Tl,ps,CG4325,ush,tna,Tsp26A,Tet,nudE,Hph,heph,Socs16D,B-H2,if,twi,Sox21a,Pits,Npc1a,grnd,Dad,CG31875,Bx,KP78a,Hip1,CaMKII,CG34172,NKAIN,ec,jing,CG11486,CG5522,Pvf3,PhKgamma,sbb,Toll-7,phyl,l(3)neo38,JIL-1,Hipk,hts,otk2,mle,Egfr,dlp,Gbs-70E,sty,hiw,CG5815,RapGAP1,tou,lz,cpo,Cdk5alpha,CG17265,mgl,Haspin,RhoGAP16F,Pcl,rau,aos,Sin3A,sens-2,Sox102F,Arp2,TfIIS,salm,Lim1,spi,Hsc70-3,Kr,hang,ckn,CG34384,N,crb,CG17168,robo2,nuf,ITP,PsGEF,NiPp1,cnc,udd,exex,CG4360,Doc2,Ptpmeg,Fili,Mondo,Pak3,Dr,sno,Notum,Stat92E,Pli,MESR3,hh,par-1,l(3)mbt,mmy,Inx3,dve,edl,E(spl)mdelta-HLH,FoxL1,wake,dally,Cdk12,Syp,Mos,dpp,Cnx99A,stwl,CG8485,scb,smg,bowl,Fas2,goe,Rbfox1,RhoGAP102A,Ten-m,CG5830,CG8441,Acf,Rho1,lncRNA:roX1,pAbp,Snx3,bora,tsl,CadN,CG5728,psq,ci,RhoGAP15B,PEK,osa,Lpin,Pat1,shn,Usp2,Acsl,NetA,slp1,CCKLR-17D3,smal,pnt,corn,Chro,eya,drk,bab1,ft,nAChRalpha4,Doa,Gyf,neo,Atg17,E(spl)m4-BFM,Prosap,Tlk,HmgZ,lbe,CG9650,Utx,CG43658,egr,ttk,HGTX,NFAT,Mmp2,Syx1A,CG32758,bin3,wnd,ds,spoon,aop,cas,disco,S,cv,CG31224,Myb,S6kII | | 1.868e-37 | -84.57 | tissue development | biological process | GO:0009888 | 1113 | 127 | 12345 | 414 | neo,sbb,bab1,ft,drk,eya,ec,slp1,MFS14,pnt,trn,jing,wb,S6kII,Myb,lz,aop,sty,disco,S,otk2,dlp,Egfr,spoon,ds,ttk,Syx1A,Hipk,Mmp2,lbe,bbg,CG43658,phyl,heph,Fas3,Rho1,Hph,nudE,CG13366,nw,CG41099,Rbfox1,lama,Fas2,ush,slow,CtBP,bowl,Ser,scb,NetA,Lpin,shn,RhoGAP15B,osa,psq,ci,bdg,Bx,Dad,shd,Myo10A,CadN,pon,Sox21a,twi,Snx3,B-H2,if,stg,tkv,sdt,Notum,sno,Dr,Inx2,cta,Ubx,slou,Doc2,exex,pio,ap,dpp,RhoGEF2,Syp,dally,TfAP-2,dve,Inx3,par-1,hh,mmy,Stat92E,oc,CycE,Sin3A,aos,gus,Mrp4,RhoGAP16F,EcR,Doc1,NetB,bi,spen,tin,Smox,tyn,pbl,tara,meso18E,so,nuf,robo2,crb,N,spi,Lim1,Lac,Kr,Fbxl7,ey,salm,Rac2,mam,fj | | 2.891e-37 | -84.13 | neurogenesis | biological process | GO:0022008 | 923 | 115 | 12345 | 414 | Acsl,NetA,shn,osa,sm,nrm,bdg,ci,pon,CadN,Dad,if,B-H2,pAbp,Rho1,Fas3,Ten-m,nudE,Fur1,Ptp99A,Tl,Fas2,slow,scb,Ser,S6kII,nrv2,lz,S,cas,sty,aop,mle,Egfr,ds,dlp,hts,wnd,Mmp2,ttk,HGTX,egr,lbe,Prosap,phyl,Toll-7,Doa,sbb,ft,Pvf3,drk,eya,pnt,jing,trn,wb,NiPp1,so,uzip,robo2,Psc,crb,N,Con,ckn,Kr,ey,Lim1,spi,mam,flfl,ImpL2,Rac2,salm,fj,aos,rau,NetB,ras,EcR,mbt,esn,spen,fax,Smox,Cdk5alpha,pbl,sas,ap,comm,RhoGEF2,dally,beat-IIIc,wake,E(spl)mdelta-HLH,Pex1,edl,Zif,mmy,l(3)mbt,hh,par-1,CycE,RhoGAP93B,oc,sdt,Dscam1,enok,Imp,sno,ham,Dr,Ptpmeg,cnc,exex | | 3.313e-37 | -84.00 | epithelium development | biological process | GO:0060429 | 1020 | 121 | 12345 | 414 | osa,RhoGAP15B,shn,Lpin,if,B-H2,Snx3,twi,Sox21a,pon,CadN,Dad,shd,Myo10A,Bx,ci,bdg,psq,CG13366,nudE,Hph,Rho1,Fas3,heph,scb,Ser,bowl,slow,CtBP,ush,Fas2,lama,Rbfox1,nw,CG41099,dlp,spoon,Egfr,ds,otk2,disco,S,sty,aop,lz,Myb,S6kII,phyl,bbg,CG43658,lbe,Mmp2,Syx1A,Hipk,ttk,sbb,neo,pnt,trn,jing,MFS14,ec,drk,eya,ft,bab1,robo2,nuf,so,tara,fj,mam,Rac2,salm,Fbxl7,Kr,ey,Lac,Lim1,spi,crb,N,tin,spen,bi,Doc1,EcR,Mrp4,RhoGAP16F,gus,Sin3A,aos,pbl,tyn,Smox,dally,Syp,RhoGEF2,ap,dpp,CycE,Stat92E,oc,mmy,hh,par-1,Inx3,dve,TfAP-2,Dr,sno,sdt,tkv,Notum,stg,pio,exex,Doc2,Ubx,cta,Inx2 | | 5.232e-37 | -83.54 | tube development | biological process | GO:0035295 | 747 | 103 | 12345 | 414 | pbl,flz,Smox,bi,spen,tin,aos,Sin3A,RhoGAP16F,Mrp4,gus,Doc1,EcR,salm,Rac2,mam,fj,N,crb,Lim1,spi,Kr,ey,Fbxl7,nuf,robo2,tara,so,Doc2,cnc,pio,Inx2,Ubx,Dr,stg,Notum,tkv,sno,hh,par-1,mmy,oc,Stat92E,CycE,TfAP-2,dve,RhoGEF2,dally,ap,dpp,CtBP,slow,bowl,Ser,scb,nw,lama,Rbfox1,Fas2,CG13366,heph,Rho1,Hph,twi,Sox21a,Snx3,if,B-H2,psq,ci,Bx,Dad,shd,pon,RhoGAP15B,osa,Lpin,shn,MFS14,pnt,trn,jing,wb,bab1,ft,eya,drk,sbb,lbe,CG43658,ttk,Hipk,Mmp2,Syx1A,sty,disco,S,otk2,dlp,Egfr,ds,S6kII,lz | | 5.921e-36 | -81.11 | biological regulation | biological process | GO:0065007 | 4350 | 269 | 12345 | 414 | EcR,Rhp,phtf,tin,mof,flz,pbl,so,tara,mam,ImpL2,Lrt,tkv,enok,elav,scaf6,cta,CG42684,mub,slou,comm,ap,AGO2,Ank,RhoGEF2,Zif,CG12099,TfAP-2,ctrip,RhoGAP93B,CG31612,oc,heph,Tet,ush,ps,Ser,slow,CtBP,CG34172,NKAIN,Hip1,CaMKII,shd,CG31875,if,Socs16D,grnd,Toll-7,sbb,PhKgamma,lz,RapGAP1,hts,hiw,Gbs-70E,l(3)neo38,phyl,Arp2,Sox102F,sens-2,Sin3A,aos,CG46339,Trpm,cpo,PsGEF,CG13784,NiPp1,robo2,Hsc70-3,Kr,Lim1,spi,crb,ckn,TfIIS,salm,sno,Mondo,Fili,cnc,exex,CG4360,wake,Mos,edl,dve,FoxL1,Pli,MESR3,Rho1,CG8441,Fur1,RhoGAP102A,Fas2,CG8485,shn,Usp2,RhoGAP15B,CadN,ci,CG5728,psq,pAbp,bora,Gyf,Doa,pnt,corn,CCKLR-17D3,nrv2,ds,S,cas,CG31224,aop,Mmp2,bin3,Syx1A,CG32758,NFAT,ttk,CG9650,CG43658,lbe,NetB,Doc1,mbt,gus,wech,ScpX,spen,bi,tyn,ocm,Smox,jigr1,Psc,lili,ey,Fbxl7,Hydr2,CG18766,fj,Taf1,Rac2,sdt,Dscam1,wcy,stg,Imp,ham,Ubx,Cirl,Tudor-SN,RhoGAP18B,Etl1,CG44774,sas,fne,Kah,CycE,kni,Hph,nudE,CG4325,Tl,Tsp26A,tna,KP78a,Dad,Bx,B-H2,Pits,twi,Sox21a,Npc1a,bou,CG5522,Pvf3,CG11486,jing,ec,tou,dlp,Egfr,mle,otk2,CG5815,sty,Hipk,JIL-1,RhoGAP16F,rau,Pcl,CG17265,Haspin,mgl,Cdk5alpha,ITP,CG17168,nuf,Lac,N,CG34384,hang,Notum,Dr,Ptpmeg,Pak3,udd,Doc2,dpp,CG8531,stwl,Cnx99A,dally,Syp,Cdk12,Inx3,E(spl)mdelta-HLH,Stat92E,Nhe2,l(3)mbt,mmy,hh,par-1,Acf,lncRNA:roX1,Ten-m,CG5830,goe,Rbfox1,nw,smg,scb,bowl,Pat1,Lpin,Acsl,NetA,osa,PEK,Snx3,tsl,Atg17,neo,E(spl)m4-BFM,nAChRalpha4,Chro,drk,eya,ft,bab1,smal,slp1,Myb,S6kII,spoon,wnd,disco,cv,egr,HGTX,Tlk,HmgZ,Prosap,Utx | | 1.203e-35 | -80.41 | tissue morphogenesis | biological process | GO:0048729 | 644 | 94 | 12345 | 414 | CG43658,lbe,Hipk,Mmp2,ttk,ds,dlp,spoon,Egfr,otk2,S,sty,aop,S6kII,trn,jing,pnt,ec,slp1,eya,drk,ft,bab1,sbb,if,B-H2,twi,CadN,shd,Dad,Myo10A,Bx,bdg,psq,osa,RhoGAP15B,shn,Lpin,scb,Ser,bowl,slow,ush,Fas2,Rbfox1,nw,CG13366,Hph,Rho1,heph,CycE,Stat92E,oc,mmy,par-1,hh,dve,Inx3,TfAP-2,dally,RhoGEF2,ap,dpp,pio,slou,Ubx,cta,Inx2,Dr,sno,tkv,sdt,stg,fj,mam,Rac2,salm,Kr,ey,Lac,Lim1,spi,crb,N,so,pbl,Smox,tin,spen,bi,EcR,gus,RhoGAP16F,aos | | 1.220e-35 | -80.39 | generation of neurons | biological process | GO:0048699 | 880 | 110 | 12345 | 414 | Rho1,Fas3,nudE,Ten-m,Fur1,Ptp99A,Fas2,Tl,slow,scb,NetA,Acsl,shn,osa,sm,nrm,ci,bdg,pon,CadN,Dad,B-H2,if,pAbp,Toll-7,Doa,sbb,ft,Pvf3,drk,eya,trn,pnt,jing,wb,S6kII,lz,S,cas,aop,sty,dlp,Egfr,mle,ds,hts,wnd,Mmp2,HGTX,ttk,lbe,phyl,Prosap,rau,aos,NetB,ras,EcR,mbt,esn,spen,fax,Smox,Cdk5alpha,pbl,NiPp1,robo2,uzip,Psc,N,crb,Con,ckn,ey,Kr,spi,Lim1,flfl,ImpL2,mam,salm,Rac2,fj,sdt,enok,Dscam1,Imp,sno,ham,Dr,Ptpmeg,cnc,exex,sas,comm,ap,RhoGEF2,wake,beat-IIIc,dally,E(spl)mdelta-HLH,edl,Zif,mmy,l(3)mbt,par-1,hh,CycE,RhoGAP93B,oc | | 3.757e-35 | -79.27 | morphogenesis of an epithelium | biological process | GO:0002009 | 625 | 92 | 12345 | 414 | lbe,CG43658,ttk,Mmp2,Hipk,otk2,Egfr,dlp,ds,spoon,sty,aop,S,S6kII,ec,jing,trn,pnt,eya,drk,bab1,ft,sbb,if,B-H2,twi,shd,Myo10A,Dad,CadN,psq,bdg,Bx,RhoGAP15B,osa,Lpin,shn,Ser,scb,slow,bowl,Fas2,ush,nw,Rbfox1,CG13366,Hph,heph,Rho1,oc,Stat92E,CycE,par-1,hh,mmy,dve,Inx3,TfAP-2,dally,RhoGEF2,ap,dpp,pio,cta,Ubx,Inx2,Dr,sno,stg,sdt,tkv,fj,Rac2,salm,mam,Lac,Lim1,spi,Kr,ey,crb,N,so,pbl,Smox,spen,tin,bi,RhoGAP16F,gus,EcR,aos | | 2.213e-34 | -77.49 | neuron differentiation | biological process | GO:0030182 | 710 | 97 | 12345 | 414 | NiPp1,robo2,Psc,uzip,ckn,Con,N,crb,Lim1,spi,Kr,ey,salm,Rac2,mam,ImpL2,fj,rau,mbt,EcR,ras,NetB,esn,spen,Smox,fax,pbl,Cdk5alpha,sas,comm,ap,dally,beat-IIIc,E(spl)mdelta-HLH,edl,par-1,hh,mmy,RhoGAP93B,oc,CycE,Imp,enok,Dscam1,sdt,sno,ham,Ptpmeg,cnc,exex,Acsl,NetA,shn,osa,sm,nrm,ci,bdg,Dad,CadN,B-H2,if,Fas3,Rho1,nudE,Ten-m,Fur1,Ptp99A,Fas2,Tl,slow,scb,lz,aop,S,wnd,hts,Egfr,ds,mle,dlp,HGTX,ttk,Mmp2,Prosap,phyl,Toll-7,Doa,sbb,ft,eya,drk,Pvf3,jing,pnt,trn,wb | | 5.083e-33 | -74.36 | animal organ morphogenesis | biological process | GO:0009887 | 782 | 100 | 12345 | 414 | Dad,CadN,psq,ci,Bx,bdg,if,B-H2,twi,Lpin,shn,NetA,RhoGAP15B,osa,Fas2,nw,Rbfox1,Ser,slow,CtBP,bowl,heph,Rho1,CG13366,Ten-m,ttk,Mmp2,Hipk,Prosap,phyl,lbe,Utx,CG43658,lz,S6kII,nrv2,otk2,Egfr,ds,dlp,sty,aop,S,eya,drk,bab1,ft,wb,slp1,ec,jing,trn,pnt,Atg17,sbb,Doa,Lac,spi,Lim1,ey,N,crb,fj,Rac2,salm,mam,so,robo2,nuf,flz,Smox,RhoGAP16F,gus,mbt,EcR,NetB,rau,aos,spen,tin,bi,dve,TfAP-2,E(spl)mdelta-HLH,oc,Stat92E,CycE,hh,par-1,Nhe2,ap,dpp,dally,RhoGEF2,Ubx,pio,sno,stg,tkv,Dr | | 5.917e-31 | -69.60 | post-embryonic animal organ development | biological process | GO:0048569 | 481 | 76 | 12345 | 414 | eya,drk,ft,bab1,pnt,jing,sbb,Hipk,Mmp2,CG43658,lbe,lz,S6kII,ds,Egfr,dlp,otk2,S,sty,ush,Fas2,Tl,Rbfox1,nw,scb,Ser,bowl,slow,Rho1,heph,CG13366,Dad,Bx,psq,B-H2,if,shn,Lpin,osa,RhoGAP15B,Ubx,pio,sno,tkv,stg,Dr,dve,Zif,TfAP-2,CycE,Stat92E,oc,hh,par-1,ap,dpp,stwl,dally,RhoGEF2,ocm,Smox,EcR,RhoGAP16F,gus,aos,Sin3A,spen,bi,ey,spi,Lim1,N,fj,mam,salm,so | | 7.715e-31 | -69.34 | tube morphogenesis | biological process | GO:0035239 | 536 | 80 | 12345 | 414 | Dr,sno,stg,tkv,pio,Ubx,Inx2,dally,RhoGEF2,dpp,ap,oc,Stat92E,CycE,par-1,hh,mmy,dve,TfAP-2,spen,tin,bi,RhoGAP16F,gus,EcR,aos,pbl,flz,Smox,so,fj,salm,Rac2,mam,Lim1,spi,ey,Kr,crb,N,sbb,trn,pnt,jing,drk,eya,bab1,ft,otk2,Egfr,dlp,ds,sty,S,S6kII,lbe,CG43658,ttk,Mmp2,Hipk,CG13366,Hph,heph,Rho1,Ser,slow,bowl,Fas2,nw,Rbfox1,RhoGAP15B,osa,Lpin,shn,B-H2,if,twi,Dad,psq,Bx | | 4.240e-30 | -67.63 | epithelial tube morphogenesis | biological process | GO:0060562 | 495 | 76 | 12345 | 414 | slow,bowl,Ser,nw,Rbfox1,Fas2,CG13366,heph,Rho1,Hph,twi,if,B-H2,psq,Bx,Dad,RhoGAP15B,osa,Lpin,shn,pnt,trn,jing,bab1,ft,drk,eya,sbb,lbe,CG43658,ttk,Hipk,Mmp2,sty,S,otk2,dlp,Egfr,ds,S6kII,pbl,Smox,bi,spen,aos,RhoGAP16F,gus,EcR,salm,mam,fj,N,crb,spi,Lim1,Kr,ey,so,pio,Ubx,Dr,stg,tkv,sno,hh,par-1,mmy,Stat92E,oc,CycE,TfAP-2,dve,RhoGEF2,dally,dpp,ap | | 4.355e-30 | -67.61 | imaginal disc development | biological process | GO:0007444 | 591 | 83 | 12345 | 414 | Fas2,lama,Rbfox1,nw,Ser,bowl,CtBP,slow,Rho1,heph,CG13366,Dad,ci,Bx,psq,B-H2,if,Snx3,shn,Lpin,osa,RhoGAP15B,drk,eya,ft,bab1,pnt,jing,MFS14,sbb,Syx1A,Mmp2,Hipk,CG43658,lbe,lz,S6kII,ds,Egfr,dlp,otk2,disco,S,Smox,Doc1,EcR,gus,RhoGAP16F,Mrp4,Sin3A,aos,spen,bi,ey,Fbxl7,Lim1,spi,N,crb,fj,mam,salm,so,tara,nuf,Ubx,pio,sno,tkv,Notum,stg,Dr,dve,TfAP-2,CycE,oc,Stat92E,par-1,hh,dpp,ap,dally,RhoGEF2 | | 3.807e-29 | -65.44 | cell development | biological process | GO:0048468 | 1436 | 132 | 12345 | 414 | twi,Npc1a,if,nrm,shd,Dad,nudE,Tet,CG13366,heph,slow,lama,Tl,sty,Egfr,dlp,mle,hts,lz,bbg,JIL-1,sbb,Toll-7,jing,ec,Pvf3,Psc,mam,ImpL2,Rac2,Hydr2,Con,esn,spen,EcR,NetB,gus,pbl,tyn,Smox,RhoGEF2,AGO2,yps,sas,comm,ap,CycE,RhoGAP93B,oc,ham,sdt,tkv,Imp,enok,Dscam1,Inx2,elav,cta,sm,Acsl,NetA,shn,Pat1,tsl,pAbp,ci,psq,CadN,Ten-m,Fur1,Rho1,Fas3,scb,Rbfox1,Ptp99A,CG41099,Fas2,goe,S,cas,aop,ds,spoon,wnd,nrv2,Myb,Utx,Prosap,Mmp2,ttk,Doa,baf,Atg17,neo,trn,pnt,wb,ft,eya,drk,uzip,robo2,NiPp1,salm,crb,N,ckn,Kr,wun2,spi,rau,ras,Cdk5alpha,fax,Syp,Mos,dally,beat-IIIc,stwl,dpp,l(3)mbt,mmy,par-1,hh,Stat92E,Pex1,edl,Dr,sno,cnc,exex,Ptpmeg | | 6.029e-29 | -64.98 | post-embryonic development | biological process | GO:0009791 | 642 | 85 | 12345 | 414 | heph,Rho1,CG13366,Fas2,Tl,ush,nw,Rbfox1,Ser,scb,slow,bowl,Lpin,shn,RhoGAP15B,osa,Dad,shd,psq,Bx,if,B-H2,Npc1a,tsl,Atg17,sbb,drk,eya,bab1,ft,pnt,trn,jing,lz,S6kII,otk2,ds,Egfr,dlp,sty,S,cas,Hipk,Mmp2,lbe,CG43658,Utx,gus,RhoGAP16F,EcR,aos,Sin3A,wech,spen,bi,Smox,ocm,so,spi,Lim1,ey,N,fj,salm,mam,sno,stg,tkv,Dr,Ubx,pio,ap,dpp,sas,stwl,dally,RhoGEF2,dve,Zif,TfAP-2,Stat92E,oc,CycE,hh,par-1 | | 7.606e-29 | -64.75 | pattern specification process | biological process | GO:0007389 | 544 | 78 | 12345 | 414 | Rho1,heph,Ser,bowl,CtBP,Tl,Rbfox1,osa,Pat1,shn,Lpin,Acsl,B-H2,pAbp,tsl,twi,CadN,Dad,ci,Bx,psq,sbb,Doa,pnt,jing,MFS14,slp1,ec,ft,spoon,Egfr,ds,dlp,S,disco,sty,lz,S6kII,phyl,tin,spen,bi,NetB,EcR,gus,Mrp4,aos,robo2,nuf,tara,fj,salm,Fbxl7,Kr,Lim1,spi,crb,N,Dr,sno,Notum,tkv,cnc,Ubx,AGO2,dally,Syp,ap,dpp,Stat92E,oc,par-1,kni,hh,edl,dve,E(spl)mdelta-HLH,TfAP-2 | | 8.487e-28 | -62.33 | negative regulation of cellular process | biological process | GO:0048523 | 1291 | 122 | 12345 | 414 | CG31875,psq,CadN,Dad,Sox21a,B-H2,pAbp,Acsl,Usp2,shn,Lpin,osa,CaMKII,Rbfox1,ush,Tl,goe,Fas2,bowl,CtBP,smg,scb,Ser,lncRNA:roX1,Acf,Rho1,heph,CG8441,Hipk,Mmp2,bin3,JIL-1,NFAT,ttk,HGTX,egr,Utx,CG9650,lbe,Tlk,phyl,S6kII,lz,RapGAP1,CG31224,cas,aop,hiw,sty,spoon,ds,Egfr,hts,ft,PhKgamma,Chro,eya,pnt,jing,slp1,E(spl)m4-BFM,Doa,sbb,crb,N,Kr,Lim1,spi,Lrt,ImpL2,salm,NiPp1,so,Psc,ocm,Smox,mof,pbl,Sox102F,Pcl,Sin3A,aos,EcR,Doc1,bi,tin,E(spl)mdelta-HLH,TfAP-2,edl,Zif,dve,l(3)mbt,hh,MESR3,par-1,kni,CycE,Pli,oc,Stat92E,ctrip,stwl,CG44774,sas,dpp,Mos,Syp,AGO2,dally,CG42684,Ptpmeg,Ubx,elav,Doc2,slou,CG4360,Tudor-SN,Notum,stg,wcy,enok,ham | | 1.673e-27 | -61.66 | post-embryonic animal organ morphogenesis | biological process | GO:0048563 | 420 | 67 | 12345 | 414 | RhoGAP15B,osa,Lpin,shn,if,B-H2,Dad,psq,Bx,CG13366,heph,Rho1,Ser,slow,bowl,Fas2,nw,Rbfox1,otk2,dlp,Egfr,ds,S,S6kII,lbe,CG43658,Hipk,Mmp2,sbb,jing,pnt,drk,eya,bab1,ft,so,fj,salm,mam,spi,Lim1,ey,N,spen,bi,gus,RhoGAP16F,EcR,aos,Smox,dally,RhoGEF2,ap,dpp,Stat92E,oc,CycE,hh,par-1,dve,TfAP-2,Dr,sno,stg,tkv,pio,Ubx | | 1.673e-27 | -61.66 | imaginal disc morphogenesis | biological process | GO:0007560 | 420 | 67 | 12345 | 414 | Dr,sno,tkv,stg,pio,Ubx,dally,RhoGEF2,dpp,ap,CycE,oc,Stat92E,hh,par-1,dve,TfAP-2,spen,bi,EcR,gus,RhoGAP16F,aos,Smox,so,fj,mam,salm,ey,Lim1,spi,N,sbb,jing,pnt,eya,drk,ft,bab1,Egfr,ds,dlp,otk2,S,S6kII,CG43658,lbe,Hipk,Mmp2,CG13366,Rho1,heph,Ser,bowl,slow,Fas2,Rbfox1,nw,osa,RhoGAP15B,shn,Lpin,if,B-H2,Dad,Bx,psq | | 3.689e-27 | -60.86 | regulation of developmental process | biological process | GO:0050793 | 757 | 90 | 12345 | 414 | stg,Imp,enok,Dscam1,tkv,Dr,ham,Ubx,cta,Ptpmeg,Pak3,cnc,comm,dpp,dally,Syp,RhoGEF2,E(spl)mdelta-HLH,Stat92E,oc,CycE,hh,par-1,mmy,l(3)mbt,mbt,EcR,NetB,Sin3A,aos,Arp2,mgl,Smox,tyn,cpo,pbl,robo2,spi,Fbxl7,ey,Kr,crb,N,salm,Rac2,ImpL2,neo,sbb,eya,bab1,ft,ec,pnt,lz,S6kII,otk2,ds,dlp,Egfr,hiw,sty,aop,cas,cv,ttk,Hipk,Syx1A,Prosap,lbe,Rho1,nudE,Fas2,goe,Tl,ush,Ser,scb,shn,Usp2,Acsl,NetA,osa,CaMKII,Dad,CadN,ci,pAbp,tsl,Sox21a,twi | | 1.236e-26 | -59.66 | negative regulation of biological process | biological process | GO:0048519 | 1502 | 131 | 12345 | 414 | l(3)mbt,par-1,MESR3,hh,Pli,Stat92E,E(spl)mdelta-HLH,edl,dve,Mos,Syp,dally,stwl,dpp,Doc2,CG4360,Ptpmeg,Dr,Notum,salm,N,crb,Kr,Hsc70-3,spi,Lim1,CG17168,NiPp1,CG17265,Sox102F,Sin3A,Pcl,aos,Utx,CG9650,lbe,Tlk,Mmp2,bin3,HGTX,ttk,NFAT,egr,CG31224,cas,aop,ds,spoon,S6kII,pnt,slp1,ft,bab1,eya,Chro,Doa,E(spl)m4-BFM,pAbp,psq,CadN,osa,Acsl,Usp2,shn,Lpin,bowl,scb,smg,Rbfox1,goe,Fas2,Rho1,lncRNA:roX1,Acf,CG8441,kni,CycE,oc,ctrip,TfAP-2,Zif,AGO2,CG44774,sas,slou,Tudor-SN,elav,Ubx,CG42684,ham,tkv,enok,stg,wcy,Lrt,ImpL2,Fbxl7,Psc,so,pbl,ocm,Smox,mof,bi,tin,Doc1,EcR,phyl,Hipk,JIL-1,hiw,sty,Egfr,hts,lz,RapGAP1,jing,PhKgamma,CG11486,sbb,Sox21a,twi,B-H2,CG31875,Dad,CaMKII,CtBP,Ser,ush,Tl,heph | | 1.292e-26 | -59.61 | positive regulation of biological process | biological process | GO:0048518 | 1642 | 138 | 12345 | 414 | Pak3,Doc2,udd,cnc,Notum,sno,FoxL1,edl,hh,mmy,Pli,Stat92E,dpp,Cdk12,Mos,wake,dally,Cdk5alpha,aos,Pcl,rau,Arp2,Sox102F,ckn,crb,N,spi,Lim1,PsGEF,robo2,ft,drk,eya,corn,pnt,Atg17,Doa,NFAT,HGTX,ttk,Mmp2,egr,CG43658,Utx,Prosap,Myb,aop,S,cv,disco,wnd,ds,Rbfox1,bowl,smg,lncRNA:roX1,Rho1,Ten-m,ci,bora,tsl,Snx3,pAbp,Usp2,Acsl,Lpin,shn,PEK,osa,Ubx,slou,Etl1,stg,Imp,Dscam1,enok,wcy,tkv,sdt,CG12099,TfAP-2,ctrip,oc,CG31612,CycE,ap,comm,RhoGEF2,ocm,mof,mbt,gus,EcR,Doc1,bi,spen,ScpX,tin,ey,Fbxl7,Rac2,Taf1,ImpL2,mam,CG18766,tara,so,lili,CG11486,Pvf3,ec,jing,Toll-7,JIL-1,Hipk,phyl,tou,lz,dlp,Egfr,mle,Tsp26A,tna,ps,Tl,ush,CG4325,CtBP,Hph,nudE,Tet,Bx,Pits,twi,grnd,Socs16D,B-H2,Hip1 | | 1.771e-26 | -59.30 | cell fate commitment | biological process | GO:0045165 | 301 | 56 | 12345 | 414 | ttk,Mmp2,Zif,egr,lbe,hh,par-1,oc,phyl,CycE,lz,dpp,ap,aop,cas,S,hts,wake,Egfr,drk,Ubx,pnt,slou,Doc2,exex,E(spl)m4-BFM,sdt,tkv,Dr,ham,N,bdg,spi,pon,Kr,ey,bora,salm,tsl,twi,mam,flfl,nuf,osa,robo2,ocm,ush,CtBP,Ser,Sox102F,EcR,Doc1,nudE,bi,spen,tin | | 4.030e-26 | -58.47 | metamorphosis | biological process | GO:0007552 | 497 | 71 | 12345 | 414 | dally,RhoGEF2,ap,dpp,CycE,oc,Stat92E,par-1,hh,dve,TfAP-2,Dr,sno,tkv,stg,pio,Ubx,so,fj,mam,salm,ey,spi,Lim1,N,tin,spen,bi,EcR,gus,RhoGAP16F,aos,Smox,ds,Egfr,dlp,otk2,S,S6kII,CG43658,Utx,lbe,Mmp2,Hipk,sbb,Atg17,jing,pnt,Chro,eya,drk,ft,bab1,osa,RhoGAP15B,shn,Lpin,if,B-H2,Dad,Bx,psq,CG13366,Rho1,heph,Ser,bowl,slow,Fas2,Rbfox1,nw | | 4.232e-26 | -58.42 | instar larval or pupal morphogenesis | biological process | GO:0048707 | 470 | 69 | 12345 | 414 | N,ey,Lim1,spi,mam,salm,fj,so,Smox,aos,EcR,RhoGAP16F,gus,bi,spen,TfAP-2,dve,par-1,hh,CycE,Stat92E,oc,ap,dpp,RhoGEF2,dally,Ubx,pio,tkv,stg,sno,Dr,Bx,psq,Dad,if,B-H2,shn,Lpin,osa,RhoGAP15B,Rbfox1,nw,Fas2,bowl,slow,Ser,Rho1,heph,CG13366,Mmp2,Hipk,Utx,CG43658,lbe,S6kII,S,dlp,ds,Egfr,otk2,ft,bab1,eya,drk,jing,pnt,Atg17,sbb | | 4.437e-26 | -58.38 | post-embryonic animal morphogenesis | biological process | GO:0009886 | 484 | 70 | 12345 | 414 | CG13366,heph,Rho1,Ser,slow,bowl,Fas2,nw,Rbfox1,RhoGAP15B,osa,Lpin,shn,if,B-H2,Dad,psq,Bx,sbb,Atg17,pnt,jing,trn,drk,eya,bab1,ft,otk2,dlp,Egfr,ds,S,S6kII,lbe,CG43658,Utx,Hipk,Mmp2,spen,bi,RhoGAP16F,gus,EcR,aos,Smox,so,fj,salm,mam,spi,Lim1,ey,N,Dr,sno,stg,tkv,pio,Ubx,dally,RhoGEF2,ap,dpp,Stat92E,oc,CycE,par-1,hh,dve,TfAP-2 | | 8.486e-26 | -57.73 | regionalization | biological process | GO:0003002 | 503 | 71 | 12345 | 414 | ap,dpp,dally,AGO2,Syp,dve,TfAP-2,E(spl)mdelta-HLH,Stat92E,oc,hh,par-1,kni,sno,tkv,Notum,Dr,Ubx,cnc,tara,robo2,nuf,Lim1,Fbxl7,Kr,crb,N,fj,salm,gus,Mrp4,NetB,EcR,aos,spen,bi,lz,S6kII,dlp,Egfr,ds,spoon,sty,S,disco,phyl,sbb,Doa,ft,slp1,MFS14,jing,pnt,Lpin,shn,Pat1,Acsl,osa,Dad,psq,Bx,ci,pAbp,B-H2,twi,tsl,Tl,Rbfox1,Ser,CtBP,bowl | | 1.528e-25 | -57.14 | regulation of macromolecule metabolic process | biological process | GO:0060255 | 2019 | 154 | 12345 | 414 | Cdk5alpha,CG17265,sens-2,Sox102F,aos,Sin3A,Pcl,TfIIS,salm,Hsc70-3,Kr,Lim1,spi,crb,N,CG17168,NiPp1,exex,udd,cnc,Doc2,CG4360,Mondo,Dr,sno,Stat92E,l(3)mbt,mmy,hh,MESR3,par-1,edl,dve,FoxL1,E(spl)mdelta-HLH,dally,Syp,Mos,Cdk12,dpp,stwl,smg,bowl,Rbfox1,CG5830,Rho1,lncRNA:roX1,Acf,CG8441,pAbp,bora,ci,CG5728,psq,PEK,osa,shn,Lpin,Acsl,Usp2,pnt,slp1,drk,eya,ft,bab1,Doa,Atg17,HmgZ,Tlk,Utx,CG9650,lbe,bin3,ttk,HGTX,NFAT,wnd,cas,disco,CG31224,aop,Myb,jigr1,mof,ocm,Smox,tin,spen,bi,EcR,Doc1,gus,mbt,phtf,CG18766,mam,Taf1,ey,Psc,lili,so,tara,mub,Tudor-SN,slou,Ubx,elav,ham,tkv,enok,CycE,ctrip,oc,CG31612,kni,fne,Kah,Zif,TfAP-2,CG12099,AGO2,ap,comm,CtBP,ush,Tl,ps,tna,Tet,heph,B-H2,grnd,twi,Pits,Sox21a,Dad,Bx,CG31875,jing,CG11486,sbb,phyl,Hipk,JIL-1,l(3)neo38,Egfr,mle,CG5815,Gbs-70E,hiw,sty,lz,tou | | 2.167e-25 | -56.79 | regulation of gene expression | biological process | GO:0010468 | 1547 | 131 | 12345 | 414 | Mondo,cnc,udd,exex,CG4360,Doc2,sno,Dr,dve,edl,E(spl)mdelta-HLH,FoxL1,Stat92E,hh,MESR3,l(3)mbt,dpp,stwl,dally,Cdk12,Syp,Sin3A,Pcl,sens-2,Sox102F,CG17265,Lim1,spi,Kr,Hsc70-3,N,TfIIS,salm,NiPp1,CG17168,eya,bab1,ft,slp1,pnt,Doa,NFAT,ttk,HGTX,bin3,HmgZ,Tlk,lbe,Utx,CG9650,Myb,aop,disco,cas,CG31224,Rbfox1,smg,bowl,lncRNA:roX1,Acf,CG5830,psq,CG5728,ci,pAbp,Lpin,shn,Acsl,osa,PEK,Ubx,elav,Tudor-SN,mub,slou,enok,tkv,ham,Zif,fne,Kah,TfAP-2,CG12099,oc,kni,ap,AGO2,mof,Smox,ocm,jigr1,EcR,Doc1,phtf,spen,tin,bi,ey,CG18766,Taf1,mam,so,tara,Psc,lili,CG11486,jing,sbb,l(3)neo38,JIL-1,Hipk,tou,lz,mle,Egfr,hiw,CG5815,ps,Tl,ush,tna,CtBP,heph,Tet,Dad,Bx,CG31875,B-H2,Pits,Sox21a,twi | | 2.521e-25 | -56.64 | neuron development | biological process | GO:0048666 | 584 | 76 | 12345 | 414 | Smox,fax,pbl,Cdk5alpha,NetB,ras,EcR,rau,spen,esn,Kr,Con,ckn,N,crb,Rac2,ImpL2,NiPp1,uzip,Psc,robo2,Ptpmeg,exex,cnc,sno,enok,Imp,Dscam1,sdt,ham,edl,RhoGAP93B,oc,hh,mmy,comm,ap,sas,beat-IIIc,dally,Tl,Fas2,Ptp99A,scb,slow,Fas3,Rho1,Fur1,nudE,Ten-m,Dad,CadN,ci,nrm,if,shn,NetA,Acsl,sm,eya,Pvf3,wb,jing,pnt,trn,Toll-7,sbb,Doa,ttk,Mmp2,Prosap,lz,wnd,hts,mle,dlp | | 3.374e-25 | -56.35 | regulation of metabolic process | biological process | GO:0019222 | 2165 | 160 | 12345 | 414 | pAbp,bora,ci,psq,CG5728,osa,PEK,shn,Lpin,Acsl,Usp2,smg,bowl,Rbfox1,CG5830,lncRNA:roX1,Rho1,Acf,CG8441,Tlk,HmgZ,CG9650,Utx,lbe,bin3,HGTX,NFAT,ttk,wnd,cas,CG31224,disco,aop,Myb,pnt,slp1,drk,eya,ft,bab1,Doa,Gyf,Atg17,TfIIS,salm,Hsc70-3,Kr,spi,Lim1,N,crb,CG17168,NiPp1,Cdk5alpha,CG17265,Sox102F,sens-2,Sin3A,Pcl,aos,Stat92E,Pli,l(3)mbt,mmy,par-1,hh,MESR3,edl,dve,FoxL1,E(spl)mdelta-HLH,dally,Mos,Syp,Cdk12,dpp,stwl,udd,cnc,exex,Doc2,CG4360,Mondo,Dr,sno,B-H2,grnd,Sox21a,twi,Pits,Dad,CG31875,Bx,CG34172,CtBP,ush,CG4325,ps,Tl,tna,Tet,heph,phyl,Hipk,JIL-1,l(3)neo38,mle,Egfr,CG5815,hiw,sty,Gbs-70E,lz,tou,jing,CG11486,sbb,Toll-7,CG18766,mam,Taf1,ey,Psc,lili,so,tara,jigr1,mof,ocm,Smox,tin,spen,bi,Doc1,EcR,mbt,gus,phtf,CycE,CG31612,oc,ctrip,kni,fne,Kah,Zif,TfAP-2,CG12099,AGO2,comm,ap,mub,Tudor-SN,slou,elav,Ubx,ham,tkv,wcy,enok | | 5.302e-25 | -55.90 | instar larval or pupal development | biological process | GO:0002165 | 576 | 75 | 12345 | 414 | psq,Bx,shd,Dad,Npc1a,tsl,if,B-H2,Lpin,shn,RhoGAP15B,osa,nw,Rbfox1,Fas2,slow,bowl,Ser,heph,Rho1,CG13366,Hipk,Mmp2,lbe,CG43658,Utx,S6kII,S,otk2,ds,Egfr,dlp,bab1,ft,drk,eya,jing,trn,pnt,Atg17,sbb,N,spi,Lim1,ey,salm,mam,fj,so,Smox,aos,wech,RhoGAP16F,gus,EcR,bi,spen,TfAP-2,dve,par-1,hh,Stat92E,oc,CycE,sas,dpp,ap,RhoGEF2,dally,Ubx,pio,stg,tkv,sno,Dr | | 5.609e-25 | -55.84 | cellular component morphogenesis | biological process | GO:0032989 | 667 | 81 | 12345 | 414 | CadN,Dad,ci,if,shn,Acsl,NetA,sm,Fas2,Ptp99A,lama,CG41099,scb,Rho1,Fas3,CG13366,nudE,Ten-m,Mmp2,ttk,Mitofilin,Prosap,Myb,spoon,ds,dlp,mle,Egfr,wnd,hts,S,eya,ft,wb,jing,trn,Toll-7,neo,sbb,Kr,N,crb,ckn,ImpL2,Rac2,NiPp1,Psc,uzip,robo2,fax,tyn,Smox,Cdk5alpha,EcR,ras,NetB,gus,Arp2,spen,CycE,RhoGAP93B,Stat92E,oc,hh,comm,dpp,ap,sas,dally,beat-IIIc,Syp,RhoGEF2,Ptpmeg,cta,cnc,exex,qsm,tkv,Dscam1,enok,ham | | 7.259e-25 | -55.58 | regulation of transcription, DNA-templated | biological process | GO:0006355 | 1144 | 109 | 12345 | 414 | enok,tkv,sno,Dr,ham,Ubx,Mondo,CG4360,slou,Doc2,exex,cnc,udd,stwl,ap,dpp,Cdk12,dally,AGO2,TfAP-2,CG12099,FoxL1,E(spl)mdelta-HLH,dve,Zif,edl,Kah,MESR3,kni,l(3)mbt,Stat92E,oc,Pcl,Sin3A,sens-2,phtf,Sox102F,Doc1,EcR,bi,spen,tin,Smox,ocm,mof,tara,so,lili,Psc,N,Lim1,Kr,ey,Taf1,salm,mam,TfIIS,CG18766,sbb,bab1,eya,slp1,pnt,jing,tou,Myb,lz,hiw,aop,cas,CG31224,disco,CG5815,mle,HGTX,l(3)neo38,ttk,NFAT,JIL-1,bin3,lbe,CG9650,Utx,HmgZ,Tlk,Acf,lncRNA:roX1,CG5830,Tet,tna,Rbfox1,Tl,ush,CtBP,bowl,smg,Acsl,Lpin,shn,osa,psq,Bx,ci,CG31875,Dad,twi,Pits,Sox21a,B-H2 | | 7.526e-25 | -55.55 | regulation of multicellular organismal development | biological process | GO:2000026 | 594 | 76 | 12345 | 414 | cv,sty,aop,hiw,ds,dlp,Egfr,otk2,S6kII,lz,lbe,Prosap,Syx1A,ttk,sbb,neo,pnt,ec,ft,bab1,osa,CaMKII,Acsl,NetA,Usp2,shn,tsl,twi,pAbp,ci,CadN,Dad,nudE,Rho1,scb,Ser,Fas2,Tl,RhoGEF2,Syp,dally,comm,dpp,l(3)mbt,mmy,par-1,hh,oc,E(spl)mdelta-HLH,ham,tkv,Dscam1,Imp,stg,cnc,Pak3,Ubx,Ptpmeg,cta,robo2,salm,Rac2,N,crb,Kr,Fbxl7,spi,mgl,Arp2,aos,EcR,NetB,mbt,pbl,tyn,Smox | | 8.906e-25 | -55.38 | regulation of cellular metabolic process | biological process | GO:0031323 | 1987 | 151 | 12345 | 414 | bowl,smg,Rbfox1,CG5830,Rho1,Acf,lncRNA:roX1,CG8441,bora,pAbp,ci,psq,CG5728,osa,PEK,Acsl,Usp2,shn,Lpin,pnt,slp1,bab1,eya,drk,Doa,Gyf,Atg17,CG9650,Utx,lbe,HmgZ,Tlk,bin3,HGTX,ttk,NFAT,CG31224,cas,disco,aop,wnd,Myb,Cdk5alpha,Sox102F,sens-2,Pcl,aos,Sin3A,salm,TfIIS,N,crb,Kr,Lim1,spi,NiPp1,Doc2,CG4360,exex,cnc,udd,Mondo,Dr,sno,l(3)mbt,mmy,hh,MESR3,par-1,Stat92E,Pli,E(spl)mdelta-HLH,FoxL1,edl,dve,Syp,Mos,Cdk12,dally,stwl,dpp,CtBP,tna,CG4325,ush,Tl,ps,Tet,heph,grnd,Sox21a,Pits,twi,B-H2,Bx,CG31875,Dad,CG34172,jing,sbb,Toll-7,phyl,JIL-1,l(3)neo38,CG5815,Gbs-70E,hiw,sty,mle,Egfr,lz,tou,ocm,Smox,mof,bi,tin,spen,phtf,Doc1,EcR,mbt,mam,Taf1,CG18766,ey,lili,Psc,tara,so,slou,mub,elav,Ubx,ham,tkv,enok,wcy,kni,CycE,oc,CG31612,ctrip,CG12099,TfAP-2,fne,Kah,Zif,AGO2,comm,ap | | 9.791e-25 | -55.28 | multicellular organismal process | biological process | GO:0032501 | 3844 | 228 | 12345 | 414 | CG41099,nw,Ptp99A,Rbfox1,goe,Fas2,bowl,scb,Fas3,Rho1,Acf,Ten-m,Fur1,psq,ci,bdg,Myo10A,CadN,tsl,betaTub56D,pAbp,Snx3,Acsl,NetA,Lpin,Tif-IA,Pat1,shn,RhoGAP15B,osa,sm,bab1,ft,Chro,drk,eya,slp1,CCKLR-17D3,trn,pnt,wb,E(spl)m4-BFM,Atg17,baf,nAChRalpha4,Doa,HGTX,ttk,Mmp2,Syx1A,egr,lbe,CG43658,Utx,Prosap,Tlk,S6kII,nrv2,Myb,aop,disco,cv,S,cas,wnd,ds,spoon,CG46339,fax,Cdk5alpha,aos,Pcl,rau,Sin3A,sens-2,RhoGAP16F,Mrp4,ras,mgl,hang,ckn,N,crb,Lac,spi,Lim1,wun2,Kr,Hsc70-3,salm,flfl,NiPp1,PsGEF,nuf,robo2,uzip,Ptpmeg,Doc2,exex,cnc,pio,Notum,sno,Dr,Pex1,E(spl)mdelta-HLH,ytr,Inx3,dve,edl,par-1,hh,mmy,Nhe2,l(3)mbt,Stat92E,Cnx99A,stwl,dpp,Mos,Syp,beat-IIIc,dally,wake,lama,Tl,ush,CtBP,slow,Ser,heph,Hph,nudE,CG13366,Tet,nrm,Bx,Dad,shd,pon,Sox21a,twi,Npc1a,if,B-H2,VepD,CaMKII,PhKgamma,Pvf3,ec,MFS14,jing,bou,Dmtn,Toll-7,NAT1,sbb,Hipk,JIL-1,bbg,phyl,CG31680,tou,lz,Dtg,hiw,sty,Gbs-70E,otk2,hts,Egfr,dlp,mle,flz,Smox,ocm,dbr,pbl,wech,gus,mbt,EcR,NetB,Doc1,bi,esn,spen,dpr7,tin,Con,Hydr2,Fbxl7,ey,Rac2,mam,ImpL2,fj,tara,so,Psc,Inx2,Ubx,elav,Cirl,cta,slou,Tudor-SN,stg,Dscam1,enok,wcy,Imp,tkv,sdt,ham,TfAP-2,Zif,fne,kni,RhoGAP93B,oc,CycE,sas,ap,comm,RhoGEF2,yps,AGO2 | | 1.855e-24 | -54.64 | regulation of cellular macromolecule biosynthetic process | biological process | GO:2000112 | 1307 | 117 | 12345 | 414 | mle,aop,hiw,disco,cas,CG5815,CG31224,Myb,tou,lz,Tlk,HmgZ,lbe,CG9650,Utx,HGTX,l(3)neo38,ttk,NFAT,bin3,JIL-1,sbb,slp1,jing,pnt,eya,bab1,PEK,osa,Lpin,shn,Acsl,pAbp,B-H2,twi,Pits,Sox21a,Dad,psq,ci,CG31875,Bx,Tet,CG5830,heph,Acf,lncRNA:roX1,smg,CtBP,bowl,Tl,ush,tna,Rbfox1,dally,AGO2,Cdk12,Syp,dpp,ap,stwl,Stat92E,CG31612,oc,CycE,kni,MESR3,mmy,l(3)mbt,dve,Zif,Kah,edl,TfAP-2,CG12099,FoxL1,E(spl)mdelta-HLH,Dr,ham,sno,enok,tkv,udd,cnc,exex,CG4360,slou,Doc2,Ubx,elav,Mondo,Psc,lili,so,tara,TfIIS,CG18766,Taf1,salm,mam,Lim1,ey,Kr,N,spen,tin,bi,EcR,Doc1,Sin3A,Pcl,sens-2,phtf,Sox102F,mof,Smox,ocm | | 1.994e-24 | -54.57 | regulation of nucleic acid-templated transcription | biological process | GO:1903506 | 1158 | 109 | 12345 | 414 | eya,bab1,pnt,jing,slp1,sbb,bin3,JIL-1,l(3)neo38,ttk,HGTX,NFAT,Tlk,HmgZ,CG9650,Utx,lbe,lz,Myb,tou,mle,disco,CG31224,cas,CG5815,aop,hiw,ush,Tl,Rbfox1,tna,smg,bowl,CtBP,Acf,lncRNA:roX1,Tet,CG5830,Dad,Bx,ci,CG31875,psq,B-H2,Pits,twi,Sox21a,shn,Lpin,Acsl,osa,Mondo,Ubx,cnc,udd,exex,slou,Doc2,CG4360,sno,tkv,enok,ham,Dr,Kah,edl,Zif,dve,E(spl)mdelta-HLH,FoxL1,TfAP-2,CG12099,Stat92E,oc,l(3)mbt,kni,MESR3,ap,dpp,stwl,AGO2,dally,Cdk12,mof,ocm,Smox,EcR,Doc1,phtf,sens-2,Sox102F,Sin3A,Pcl,tin,spen,bi,ey,Kr,Lim1,N,CG18766,TfIIS,mam,salm,Taf1,so,tara,Psc,lili | | 1.994e-24 | -54.57 | regulation of RNA biosynthetic process | biological process | GO:2001141 | 1158 | 109 | 12345 | 414 | ocm,Smox,mof,bi,tin,spen,sens-2,phtf,Sox102F,Pcl,Sin3A,Doc1,EcR,mam,Taf1,salm,CG18766,TfIIS,N,Kr,ey,Lim1,lili,Psc,tara,so,slou,Doc2,CG4360,cnc,exex,udd,Mondo,Ubx,ham,Dr,tkv,enok,sno,l(3)mbt,MESR3,kni,oc,Stat92E,E(spl)mdelta-HLH,FoxL1,CG12099,TfAP-2,Kah,edl,Zif,dve,Cdk12,AGO2,dally,stwl,ap,dpp,bowl,CtBP,smg,Rbfox1,tna,ush,Tl,CG5830,Tet,Acf,lncRNA:roX1,Sox21a,Pits,twi,B-H2,Bx,CG31875,ci,psq,Dad,osa,Acsl,shn,Lpin,pnt,jing,slp1,bab1,eya,sbb,Utx,CG9650,lbe,HmgZ,Tlk,bin3,JIL-1,ttk,HGTX,NFAT,l(3)neo38,cas,CG31224,disco,CG5815,hiw,aop,mle,lz,Myb,tou | | 4.172e-24 | -53.83 | regulation of RNA metabolic process | biological process | GO:0051252 | 1281 | 115 | 12345 | 414 | Sox102F,sens-2,phtf,Sin3A,Pcl,EcR,Doc1,bi,tin,spen,ocm,Smox,mof,tara,so,lili,Psc,N,ey,Kr,Lim1,mam,Taf1,salm,CG18766,TfIIS,tkv,enok,sno,ham,Dr,Mondo,elav,Ubx,Doc2,slou,CG4360,mub,exex,udd,cnc,stwl,ap,dpp,Cdk12,AGO2,dally,E(spl)mdelta-HLH,FoxL1,CG12099,TfAP-2,Kah,fne,edl,Zif,dve,l(3)mbt,kni,MESR3,oc,Stat92E,lncRNA:roX1,Acf,CG5830,Tet,Rbfox1,tna,ush,Tl,ps,bowl,CtBP,smg,Acsl,shn,Lpin,osa,ci,Bx,CG31875,CG5728,psq,Dad,Sox21a,twi,Pits,B-H2,Doa,sbb,bab1,eya,pnt,jing,slp1,lz,Myb,tou,CG31224,CG5815,disco,cas,hiw,aop,mle,bin3,JIL-1,l(3)neo38,ttk,HGTX,NFAT,CG9650,Utx,lbe,Tlk,HmgZ | | 5.259e-24 | -53.60 | regulation of nitrogen compound metabolic process | biological process | GO:0051171 | 1870 | 144 | 12345 | 414 | AGO2,ap,kni,oc,ctrip,CycE,CG12099,TfAP-2,Zif,fne,Kah,ham,enok,tkv,slou,mub,Ubx,elav,lili,Psc,tara,so,Taf1,mam,CG18766,ey,bi,spen,tin,phtf,gus,mbt,Doc1,EcR,Smox,ocm,mof,hiw,sty,CG5815,mle,Egfr,tou,lz,phyl,l(3)neo38,JIL-1,sbb,jing,twi,Sox21a,Pits,grnd,B-H2,Bx,CG31875,Dad,Tet,heph,CtBP,tna,ps,Tl,CG4325,ush,Cdk12,Syp,Mos,dally,stwl,dpp,hh,MESR3,par-1,l(3)mbt,mmy,Stat92E,E(spl)mdelta-HLH,FoxL1,dve,edl,Dr,sno,CG4360,Doc2,exex,cnc,udd,Mondo,NiPp1,salm,TfIIS,crb,N,spi,Lim1,Kr,Pcl,aos,Sin3A,sens-2,Sox102F,Cdk5alpha,aop,CG31224,cas,disco,wnd,Myb,lbe,CG9650,Utx,Tlk,HmgZ,ttk,NFAT,HGTX,bin3,Doa,Atg17,slp1,pnt,bab1,drk,eya,PEK,osa,Usp2,Acsl,Lpin,shn,bora,pAbp,psq,CG5728,ci,CG5830,CG8441,lncRNA:roX1,Acf,Rho1,bowl,smg,Rbfox1 | | 5.912e-24 | -53.49 | cell morphogenesis involved in differentiation | biological process | GO:0000904 | 468 | 66 | 12345 | 414 | sas,comm,ap,RhoGEF2,dally,beat-IIIc,hh,oc,RhoGAP93B,Stat92E,enok,Dscam1,ham,cta,Ptpmeg,exex,cnc,NiPp1,robo2,uzip,Psc,N,crb,ckn,Kr,ImpL2,Rac2,ras,NetB,EcR,spen,tyn,fax,Smox,Cdk5alpha,mle,ds,dlp,hts,wnd,Mmp2,Prosap,Toll-7,neo,sbb,ft,eya,trn,jing,wb,NetA,Acsl,shn,sm,ci,CadN,Dad,if,Rho1,Fas3,Ten-m,nudE,Ptp99A,CG41099,Fas2,scb | | 6.266e-24 | -53.43 | imaginal disc-derived appendage development | biological process | GO:0048737 | 362 | 58 | 12345 | 414 | sno,tkv,Dr,sbb,drk,Ubx,ft,pio,pnt,jing,ap,lz,dpp,S6kII,dally,dlp,Egfr,ds,disco,S,RhoGEF2,dve,TfAP-2,Hipk,Syx1A,Stat92E,CycE,par-1,hh,lbe,CG43658,RhoGAP16F,EcR,heph,aos,Rho1,spen,bi,Smox,nw,Rbfox1,Ser,slow,bowl,Lpin,shn,RhoGAP15B,osa,Lim1,Dad,spi,psq,N,Bx,fj,if,salm,mam | | 6.513e-24 | -53.39 | regulation of macromolecule biosynthetic process | biological process | GO:0010556 | 1326 | 117 | 12345 | 414 | sbb,eya,bab1,slp1,jing,pnt,tou,Myb,lz,mle,aop,hiw,CG31224,disco,CG5815,cas,NFAT,l(3)neo38,HGTX,ttk,JIL-1,bin3,Tlk,HmgZ,lbe,CG9650,Utx,heph,Acf,lncRNA:roX1,Tet,CG5830,Tl,ush,tna,Rbfox1,smg,CtBP,bowl,Lpin,shn,Acsl,osa,PEK,Dad,psq,Bx,CG31875,ci,pAbp,B-H2,Sox21a,Pits,twi,sno,enok,tkv,Dr,ham,Ubx,elav,Mondo,cnc,exex,udd,CG4360,slou,Doc2,dpp,ap,stwl,dally,AGO2,Cdk12,Syp,Zif,dve,edl,Kah,CG12099,TfAP-2,E(spl)mdelta-HLH,FoxL1,CG31612,Stat92E,oc,CycE,kni,MESR3,mmy,l(3)mbt,EcR,Doc1,Pcl,Sin3A,sens-2,phtf,Sox102F,spen,tin,bi,mof,Smox,ocm,so,tara,Psc,lili,Lim1,Kr,ey,N,TfIIS,CG18766,Taf1,salm,mam | | 7.059e-24 | -53.31 | positive regulation of cellular process | biological process | GO:0048522 | 1464 | 124 | 12345 | 414 | sno,Notum,tkv,sdt,enok,wcy,stg,Imp,Dscam1,Pak3,cnc,udd,slou,Doc2,ap,dpp,comm,dally,RhoGEF2,Mos,Cdk12,edl,TfAP-2,CG12099,CycE,CG31612,Pli,Stat92E,oc,mmy,Doc1,EcR,mbt,Arp2,Sox102F,rau,aos,Pcl,tin,spen,bi,mof,ocm,Cdk5alpha,PsGEF,so,tara,robo2,lili,Fbxl7,ey,spi,Lim1,crb,N,ckn,CG18766,mam,ImpL2,Taf1,Atg17,Toll-7,Doa,Pvf3,eya,CG11486,drk,ft,pnt,corn,jing,ec,lz,tou,Myb,Egfr,dlp,mle,ds,wnd,S,disco,aop,Hipk,JIL-1,NFAT,HGTX,ttk,phyl,Prosap,Utx,CG43658,Hph,lncRNA:roX1,Rho1,Tet,nudE,ush,CG4325,ps,Tl,Rbfox1,Tsp26A,tna,smg,bowl,CtBP,shn,Lpin,Usp2,Hip1,osa,PEK,ci,Bx,B-H2,Socs16D,Snx3,pAbp,grnd,twi,Pits,tsl,bora | | 9.119e-24 | -53.05 | locomotion | biological process | GO:0040011 | 664 | 79 | 12345 | 414 | cta,Ptpmeg,tkv,Dscam1,enok,kni,hh,par-1,RhoGAP93B,Stat92E,sas,ap,comm,dpp,beat-IIIc,dally,Smox,Cdk5alpha,Trpm,pbl,Sin3A,ras,NetB,EcR,gus,mbt,tin,spen,N,Con,ckn,Kr,ey,wun2,Lrt,ImpL2,Rac2,salm,NiPp1,so,robo2,Psc,uzip,Pvf3,drk,eya,CG6330,trn,pnt,jing,wb,Toll-7,sbb,Hipk,Mmp2,bbg,lbe,hiw,aop,spoon,l(3)L1231,dlp,Egfr,hts,Ptp99A,Tl,Fas2,scb,Rho1,Fas3,Ten-m,nudE,CadN,Dad,shd,if,Acsl,NetA,sm | | 1.114e-23 | -52.85 | appendage development | biological process | GO:0048736 | 366 | 58 | 12345 | 414 | dve,Syx1A,Hipk,TfAP-2,CycE,Stat92E,CG43658,par-1,lbe,hh,dpp,ap,lz,S6kII,ds,Egfr,dlp,dally,S,RhoGEF2,disco,drk,Ubx,ft,pio,jing,pnt,sno,tkv,sbb,Dr,spi,Lim1,Dad,N,Bx,psq,if,fj,mam,salm,shn,Lpin,osa,RhoGAP15B,Rbfox1,Smox,nw,Ser,bowl,slow,EcR,RhoGAP16F,Rho1,heph,aos,spen,bi | | 1.415e-23 | -52.61 | regulation of multicellular organismal process | biological process | GO:0051239 | 781 | 86 | 12345 | 414 | cnc,Pak3,Ubx,Ptpmeg,cta,ham,Dscam1,Imp,stg,tkv,par-1,hh,l(3)mbt,mmy,oc,TfAP-2,E(spl)mdelta-HLH,Syp,RhoGEF2,wake,dally,dpp,comm,pbl,cpo,Smox,tyn,mgl,Sin3A,aos,Arp2,Sox102F,mbt,EcR,NetB,Rac2,salm,crb,N,spi,Kr,ey,Fbxl7,robo2,ec,pnt,bab1,ft,Doa,sbb,neo,lbe,Prosap,ttk,Syx1A,Hipk,egr,hiw,sty,aop,cv,otk2,ds,dlp,Egfr,S6kII,lz,Ser,scb,Tl,Fas2,ush,nudE,Rho1,twi,tsl,pAbp,ci,Dad,CadN,osa,CaMKII,Usp2,Acsl,NetA,shn | | 1.429e-23 | -52.60 | imaginal disc-derived appendage morphogenesis | biological process | GO:0035114 | 355 | 57 | 12345 | 414 | TfAP-2,Hipk,dve,par-1,lbe,hh,CG43658,Stat92E,CycE,S6kII,dpp,lz,ap,RhoGEF2,disco,S,dally,dlp,ds,Egfr,ft,drk,Ubx,pnt,jing,pio,tkv,sno,Dr,sbb,psq,Bx,N,Dad,spi,Lim1,salm,mam,fj,if,Lpin,shn,RhoGAP15B,osa,Smox,nw,Rbfox1,slow,bowl,Ser,aos,heph,Rho1,RhoGAP16F,EcR,bi,spen | | 1.438e-23 | -52.60 | wing disc development | biological process | GO:0035220 | 407 | 61 | 12345 | 414 | if,fj,Snx3,mam,salm,spi,Dad,N,crb,Bx,ci,psq,osa,nuf,shn,Lpin,tara,Ser,bowl,CtBP,slow,Rbfox1,Smox,nw,spen,CG13366,bi,EcR,Doc1,Mrp4,gus,Rho1,heph,Sin3A,aos,CycE,Stat92E,hh,par-1,dve,Hipk,dlp,ds,Egfr,dally,S,RhoGEF2,dpp,ap,S6kII,pio,jing,pnt,MFS14,drk,ft,sbb,Dr,sno,tkv,Notum | | 1.910e-23 | -52.31 | regulation of primary metabolic process | biological process | GO:0080090 | 1894 | 144 | 12345 | 414 | bi,spen,tin,phtf,gus,mbt,Doc1,EcR,Smox,ocm,mof,lili,Psc,tara,so,Taf1,mam,CG18766,ey,ham,enok,tkv,slou,mub,elav,Ubx,AGO2,ap,kni,ctrip,oc,CycE,TfAP-2,CG12099,Zif,Kah,fne,Tet,heph,CtBP,tna,Tl,ps,ush,Sox21a,Pits,twi,grnd,B-H2,CG31875,Bx,Dad,sbb,jing,hiw,sty,Gbs-70E,CG5815,Egfr,mle,tou,lz,phyl,l(3)neo38,JIL-1,Sin3A,aos,Pcl,Sox102F,sens-2,Cdk5alpha,NiPp1,salm,TfIIS,N,crb,Lim1,spi,Kr,Dr,sno,CG4360,Doc2,exex,udd,cnc,Mondo,Cdk12,Syp,Mos,dally,stwl,dpp,MESR3,hh,par-1,l(3)mbt,mmy,Stat92E,FoxL1,E(spl)mdelta-HLH,dve,edl,CG5830,CG8441,lncRNA:roX1,Rho1,Acf,bowl,smg,Rbfox1,osa,PEK,Usp2,Acsl,Lpin,shn,bora,pAbp,psq,CG5728,ci,Doa,Atg17,slp1,pnt,bab1,drk,eya,aop,disco,cas,CG31224,wnd,Myb,lbe,Utx,CG9650,HmgZ,Tlk,ttk,HGTX,NFAT,bin3 | | 2.204e-23 | -52.17 | appendage morphogenesis | biological process | GO:0035107 | 358 | 57 | 12345 | 414 | CG43658,hh,lbe,par-1,CycE,Stat92E,Hipk,TfAP-2,dve,RhoGEF2,disco,S,ds,Egfr,dlp,dally,S6kII,dpp,ap,lz,jing,pnt,pio,ft,drk,Ubx,sbb,Dr,tkv,sno,mam,salm,if,fj,N,Bx,psq,spi,Lim1,Dad,osa,RhoGAP15B,shn,Lpin,bowl,slow,Ser,Rbfox1,Smox,nw,bi,spen,Rho1,heph,aos,EcR,RhoGAP16F | | 3.369e-23 | -51.74 | regulation of cellular biosynthetic process | biological process | GO:0031326 | 1371 | 118 | 12345 | 414 | elav,Ubx,Mondo,udd,exex,cnc,CG4360,Doc2,slou,sno,enok,tkv,Dr,ham,dve,Zif,Kah,edl,CG12099,TfAP-2,E(spl)mdelta-HLH,FoxL1,CG31612,oc,Stat92E,CycE,kni,MESR3,l(3)mbt,mmy,ap,dpp,stwl,dally,AGO2,Cdk12,Syp,mof,Smox,ocm,Doc1,EcR,Sin3A,Pcl,sens-2,phtf,Sox102F,spen,tin,bi,Lim1,ey,Kr,N,TfIIS,CG18766,salm,Taf1,mam,so,tara,Psc,lili,eya,bab1,slp1,pnt,jing,sbb,HGTX,l(3)neo38,ttk,NFAT,bin3,JIL-1,Tlk,HmgZ,lbe,Utx,CG9650,tou,Myb,lz,mle,aop,hiw,CG31224,CG5815,cas,disco,Tl,CG4325,ush,tna,Rbfox1,smg,CtBP,bowl,heph,Acf,lncRNA:roX1,Tet,CG5830,Dad,psq,Bx,CG31875,ci,pAbp,B-H2,Pits,Sox21a,twi,Lpin,shn,Acsl,osa,PEK | | 3.890e-23 | -51.60 | regulation of nucleobase-containing compound metabolic process | biological process | GO:0019219 | 1315 | 115 | 12345 | 414 | CtBP,bowl,smg,tna,Rbfox1,Tl,ps,ush,CG5830,Tet,lncRNA:roX1,Acf,Pits,twi,Sox21a,B-H2,CG5728,psq,Bx,ci,CG31875,Dad,osa,Acsl,Lpin,shn,slp1,pnt,jing,bab1,eya,Doa,sbb,lbe,Utx,CG9650,Tlk,HmgZ,ttk,HGTX,NFAT,l(3)neo38,bin3,JIL-1,hiw,aop,CG31224,disco,cas,CG5815,mle,tou,Myb,lz,Smox,ocm,mof,bi,spen,tin,Sin3A,Pcl,phtf,Sox102F,sens-2,EcR,Doc1,salm,Taf1,mam,TfIIS,CG18766,N,Lim1,Kr,ey,lili,Psc,tara,so,CG4360,slou,Doc2,exex,udd,cnc,mub,elav,Ubx,Mondo,Dr,ham,enok,tkv,sno,kni,MESR3,l(3)mbt,Stat92E,oc,TfAP-2,CG12099,FoxL1,E(spl)mdelta-HLH,dve,Zif,fne,edl,Kah,Cdk12,dally,AGO2,stwl,dpp,ap | | 4.068e-23 | -51.56 | regulation of biosynthetic process | biological process | GO:0009889 | 1374 | 118 | 12345 | 414 | lili,Psc,tara,so,Taf1,salm,mam,TfIIS,CG18766,N,Lim1,ey,Kr,bi,spen,tin,Pcl,Sin3A,phtf,sens-2,Sox102F,EcR,Doc1,Smox,ocm,mof,Cdk12,Syp,dally,AGO2,stwl,ap,dpp,kni,MESR3,l(3)mbt,mmy,oc,CG31612,Stat92E,CycE,TfAP-2,CG12099,FoxL1,E(spl)mdelta-HLH,dve,Zif,Kah,edl,Dr,ham,enok,tkv,sno,CG4360,Doc2,slou,cnc,udd,exex,elav,Ubx,Mondo,PEK,osa,Acsl,Lpin,shn,Sox21a,Pits,twi,pAbp,B-H2,psq,CG31875,ci,Bx,Dad,CG5830,Tet,heph,lncRNA:roX1,Acf,CtBP,bowl,smg,tna,Rbfox1,Tl,ush,CG4325,aop,hiw,CG31224,disco,CG5815,cas,mle,tou,Myb,lz,lbe,Utx,CG9650,HmgZ,Tlk,NFAT,l(3)neo38,HGTX,ttk,bin3,JIL-1,sbb,slp1,pnt,jing,bab1,eya | | 7.661e-23 | -50.92 | cell morphogenesis | biological process | GO:0000902 | 533 | 69 | 12345 | 414 | neo,Toll-7,sbb,ft,eya,jing,trn,wb,wnd,hts,mle,dlp,ds,Mmp2,Prosap,Fas3,Rho1,nudE,Ten-m,CG41099,Ptp99A,Fas2,scb,Acsl,NetA,shn,sm,ci,Dad,CadN,if,enok,Dscam1,qsm,tkv,ham,cta,Ptpmeg,cnc,exex,sas,comm,ap,RhoGEF2,beat-IIIc,dally,hh,Stat92E,oc,RhoGAP93B,Arp2,ras,NetB,EcR,spen,Smox,tyn,fax,Cdk5alpha,NiPp1,robo2,uzip,Psc,ckn,crb,N,Kr,Rac2,ImpL2 | | 1.320e-22 | -50.38 | post-embryonic appendage morphogenesis | biological process | GO:0035120 | 345 | 55 | 12345 | 414 | dve,TfAP-2,Hipk,Stat92E,CycE,hh,par-1,lbe,CG43658,dpp,ap,S6kII,dally,dlp,Egfr,ds,RhoGEF2,S,drk,Ubx,ft,pio,pnt,jing,sno,tkv,Dr,sbb,Lim1,spi,Dad,psq,N,Bx,fj,if,salm,mam,Lpin,shn,RhoGAP15B,osa,nw,Smox,Rbfox1,Ser,slow,bowl,RhoGAP16F,EcR,heph,aos,Rho1,spen,bi | | 5.119e-22 | -49.02 | chemotaxis | biological process | GO:0006935 | 292 | 50 | 12345 | 414 | Mmp2,hh,RhoGAP93B,sas,dpp,comm,ap,dlp,hts,beat-IIIc,dally,Ptpmeg,eya,trn,jing,wb,enok,Dscam1,Toll-7,sbb,N,Con,ckn,CadN,Kr,wun2,Dad,Lrt,ImpL2,Rac2,if,Acsl,NetA,NiPp1,uzip,Psc,robo2,sm,Ptp99A,Smox,Fas2,Tl,scb,Cdk5alpha,Rho1,Fas3,ras,NetB,Ten-m,spen | | 1.388e-21 | -48.03 | axon development | biological process | GO:0061564 | 336 | 53 | 12345 | 414 | Acsl,NetA,NiPp1,uzip,Psc,robo2,sm,CadN,Kr,Dad,N,Con,ckn,if,ImpL2,Rac2,ras,NetB,Rho1,Fas3,spen,nudE,Ten-m,Fas2,Ptp99A,fax,Smox,scb,Cdk5alpha,pbl,ap,comm,sas,mle,dlp,dally,wnd,beat-IIIc,hts,Mmp2,RhoGAP93B,mmy,hh,Toll-7,Dscam1,enok,sbb,eya,Ptpmeg,exex,wb,jing,trn | | 1.807e-21 | -47.76 | imaginal disc pattern formation | biological process | GO:0007447 | 112 | 32 | 12345 | 414 | disco,dally,Egfr,dlp,ds,ap,dpp,hh,Dr,sbb,tkv,Notum,sno,MFS14,jing,pnt,ft,nuf,osa,tara,shn,B-H2,N,ci,crb,Lim1,Dad,Fbxl7,bi,spen,Mrp4,Ser | | 2.103e-21 | -47.61 | neuron projection development | biological process | GO:0031175 | 447 | 61 | 12345 | 414 | Prosap,oc,RhoGAP93B,mmy,hh,Mmp2,mle,dlp,hts,wnd,beat-IIIc,dally,comm,ap,sas,wb,cnc,exex,jing,trn,eya,Ptpmeg,sbb,ham,Toll-7,enok,Dscam1,if,ImpL2,Rac2,CadN,Kr,Dad,ci,N,crb,Con,ckn,uzip,Psc,sm,robo2,shn,Acsl,NetA,NiPp1,scb,Cdk5alpha,pbl,Fas2,Ptp99A,fax,Smox,spen,Ten-m,nudE,EcR,NetB,ras,Rho1,Fas3 | | 9.497e-21 | -46.10 | movement of cell or subcellular component | biological process | GO:0006928 | 724 | 78 | 12345 | 414 | spen,tin,mbt,gus,NetB,EcR,ras,Sin3A,pbl,Cdk5alpha,Smox,tyn,uzip,Psc,robo2,so,NiPp1,Rac2,salm,ImpL2,Lrt,wun2,Kr,ey,ckn,crb,N,CG14535,enok,Dscam1,tkv,cta,Ptpmeg,beat-IIIc,dally,RhoGEF2,ap,dpp,comm,sas,RhoGAP93B,Stat92E,kni,hh,par-1,nudE,Ten-m,Fas3,Rho1,scb,Fas2,Ptp99A,sm,Pat1,NetA,Acsl,if,Dad,shd,CadN,sbb,neo,Toll-7,wb,jing,pnt,trn,drk,eya,Pvf3,hts,dlp,Egfr,spoon,aop,lbe,bbg,Mmp2 | | 9.802e-21 | -46.07 | regulation of transcription by RNA polymerase II | biological process | GO:0006357 | 741 | 79 | 12345 | 414 | enok,tkv,sno,Ubx,Mondo,CG4360,slou,Doc2,cnc,ap,dpp,Cdk12,dally,TfAP-2,CG12099,FoxL1,E(spl)mdelta-HLH,Zif,dve,edl,kni,MESR3,l(3)mbt,oc,Stat92E,Sox102F,phtf,EcR,Doc1,bi,tin,ocm,mof,so,lili,N,Lim1,Kr,ey,Taf1,mam,sbb,bab1,eya,slp1,pnt,jing,tou,Myb,lz,aop,hiw,CG31224,disco,mle,HGTX,NFAT,ttk,lbe,Utx,HmgZ,CG5830,Tet,tna,Tl,ush,CtBP,bowl,Acsl,Lpin,shn,osa,psq,ci,Dad,twi,Pits,Sox21a,B-H2 | | 1.206e-20 | -45.86 | taxis | biological process | GO:0042330 | 365 | 54 | 12345 | 414 | Rho1,Fas3,NetB,ras,Ten-m,spen,Ptp99A,Smox,Tl,Fas2,Cdk5alpha,scb,Trpm,Acsl,NetA,NiPp1,uzip,sm,Psc,robo2,N,ckn,Con,CadN,Kr,Dad,wun2,Lrt,ImpL2,Rac2,if,Dscam1,enok,Toll-7,sbb,CG6330,eya,Ptpmeg,trn,jing,wb,sas,ap,dpp,comm,dlp,l(3)L1231,dally,hts,beat-IIIc,Hipk,Mmp2,hh,RhoGAP93B | | 1.372e-20 | -45.74 | embryo development | biological process | GO:0009790 | 568 | 68 | 12345 | 414 | AGO2,dally,RhoGEF2,dpp,Stat92E,oc,l(3)mbt,mmy,kni,hh,par-1,edl,Inx3,Dr,tkv,sdt,stg,cnc,slou,elav,cta,Ubx,Inx2,robo2,Psc,mam,Rac2,Kr,spi,Lac,N,crb,tin,Doc1,EcR,gus,pbl,dlp,Egfr,S,aop,lz,Dtg,Tlk,lbe,Mmp2,Doa,jing,pnt,slp1,Pvf3,PhKgamma,shn,Pat1,if,twi,tsl,pon,Dad,Myo10A,shd,ci,psq,Rho1,scb,bowl,ush,CG41099 | | 1.592e-20 | -45.59 | cell morphogenesis involved in neuron differentiation | biological process | GO:0048667 | 422 | 58 | 12345 | 414 | sm,Psc,uzip,robo2,shn,NiPp1,Acsl,NetA,if,Rac2,ImpL2,Dad,Kr,CadN,ckn,crb,N,ci,spen,nudE,Ten-m,EcR,NetB,ras,Fas3,Rho1,Cdk5alpha,scb,Fas2,Smox,fax,Ptp99A,wnd,hts,beat-IIIc,dally,mle,dlp,ap,comm,sas,Prosap,oc,RhoGAP93B,hh,Mmp2,sbb,ham,Toll-7,enok,Dscam1,exex,wb,cnc,jing,trn,eya,Ptpmeg | | 1.766e-20 | -45.48 | cell part morphogenesis | biological process | GO:0032990 | 437 | 59 | 12345 | 414 | Dad,Kr,CadN,ckn,crb,ci,N,if,Rac2,ImpL2,shn,NiPp1,Acsl,NetA,sm,Psc,robo2,uzip,Fas2,Smox,fax,Ptp99A,scb,Cdk5alpha,ras,NetB,EcR,Fas3,Rho1,spen,Ten-m,nudE,Mmp2,Prosap,oc,RhoGAP93B,Mitofilin,hh,ap,comm,sas,dally,beat-IIIc,hts,wnd,dlp,mle,eya,Ptpmeg,cnc,wb,exex,trn,jing,Toll-7,Dscam1,enok,ham,sbb | | 2.265e-20 | -45.23 | neuron projection morphogenesis | biological process | GO:0048812 | 425 | 58 | 12345 | 414 | Ten-m,nudE,spen,Fas3,Rho1,ras,EcR,NetB,Cdk5alpha,scb,Smox,Ptp99A,fax,Fas2,robo2,sm,Psc,uzip,NiPp1,Acsl,NetA,shn,Rac2,ImpL2,if,ckn,crb,ci,N,Dad,Kr,CadN,ham,sbb,enok,Dscam1,Toll-7,trn,jing,wb,cnc,exex,Ptpmeg,eya,hts,dally,beat-IIIc,wnd,dlp,mle,sas,ap,comm,hh,Prosap,oc,RhoGAP93B,Mmp2 | | 2.545e-20 | -45.12 | plasma membrane bounded cell projection morphogenesis | biological process | GO:0120039 | 426 | 58 | 12345 | 414 | Psc,sm,uzip,robo2,Acsl,NetA,NiPp1,shn,ImpL2,Rac2,if,crb,ci,N,ckn,Kr,CadN,Dad,nudE,Ten-m,spen,Rho1,Fas3,ras,EcR,NetB,scb,Cdk5alpha,Ptp99A,fax,Smox,Fas2,mle,dlp,dally,hts,beat-IIIc,wnd,sas,comm,ap,hh,RhoGAP93B,oc,Prosap,Mmp2,ham,sbb,enok,Dscam1,Toll-7,jing,trn,cnc,exex,wb,Ptpmeg,eya | | 3.211e-20 | -44.89 | cell projection morphogenesis | biological process | GO:0048858 | 428 | 58 | 12345 | 414 | scb,Cdk5alpha,Fas2,Ptp99A,fax,Smox,spen,nudE,Ten-m,EcR,ras,NetB,Rho1,Fas3,if,ImpL2,Rac2,CadN,Kr,Dad,N,ci,crb,ckn,Psc,uzip,robo2,sm,shn,NetA,Acsl,NiPp1,cnc,wb,exex,jing,trn,eya,Ptpmeg,ham,sbb,Toll-7,Dscam1,enok,RhoGAP93B,Prosap,oc,hh,Mmp2,dlp,mle,beat-IIIc,hts,dally,wnd,ap,comm,sas | | 3.620e-20 | -44.77 | wing disc morphogenesis | biological process | GO:0007472 | 308 | 49 | 12345 | 414 | ap,dpp,S6kII,Egfr,dlp,ds,dally,S,RhoGEF2,dve,Hipk,CycE,Stat92E,par-1,hh,sno,tkv,sbb,Dr,drk,ft,pio,jing,pnt,shn,Lpin,osa,spi,Dad,Bx,N,psq,if,fj,mam,salm,EcR,gus,Rho1,heph,aos,spen,CG13366,bi,Rbfox1,Smox,nw,Ser,slow | | 4.892e-20 | -44.46 | protein binding | molecular function | GO:0005515 | 2410 | 156 | 12176 | 396 | enok,Gbs-70E,crb,Doc1,ush,CtBP,hh,kni,Pvf3,corn,Prosap,scb,Mhcl,RhoGEF2,cta,EcR,Atg17,Taf1,tou,rau,Stat92E,Lrt,ImpL2,Sin3A,hts,CG17778,ttk,pnt,sas,shn,CG4538,nudE,PhKgamma,CG43658,Chro,dpp,egr,Tl,twi,PsGEF,l(3)mbt,Doc2,CG42748,CG5815,CG8441,Hipk,AGO2,Fas2,NiPp1,Pak3,HmgZ,Lac,oc,Tlk,ft,smg,mbt,Dad,tsl,Mitofilin,mof,bi,bab1,bin3,eya,Mondo,Pli,so,ey,Ubx,otk2,Ten-m,Pcl,phyl,N,spi,S6kII,CG11486,ec,robo2,pbl,ITP,if,CG31875,Dscam1,Socs16D,Df31,CG10283,CG14535,CadN,cnc,dlp,E(spl)mdelta-HLH,Syx1A,Myo10A,aos,Gmap,spoon,Tif-IA,Rbfox1,NKAIN,ds,Ptpmeg,ocm,dally,bora,RhoGAP18B,NFAT,HGTX,hiw,Ser,flfl,Bx,yps,Arp2,Tudor-SN,edl,psq,tna,gus,Rho1,Ank,tkv,phtf,comm,Imp,wake,Tpr2,E(spl)m4-BFM,nuf,aop,Hip1,Rac2,sdt,Cdk12,Hsc70-3,Cnx99A,CaMKII,ci,drk,Acf,Smox,Svil,Cyp1,Ykt6,Egfr,DnaJ-H,CG5522,JIL-1,CycE,Utx,fj,ckn,slow,pAbp,wech | | 7.440e-20 | -44.04 | axonogenesis | biological process | GO:0007409 | 326 | 50 | 12345 | 414 | Psc,sm,uzip,robo2,Acsl,NetA,NiPp1,ImpL2,Rac2,if,N,ckn,CadN,Kr,Dad,nudE,Ten-m,spen,Rho1,Fas3,ras,NetB,scb,Cdk5alpha,Ptp99A,fax,Smox,Fas2,dlp,mle,beat-IIIc,wnd,hts,dally,sas,comm,ap,hh,RhoGAP93B,Mmp2,sbb,Dscam1,enok,Toll-7,jing,trn,exex,wb,Ptpmeg,eya | | 1.858e-19 | -43.13 | binding | molecular function | GO:0005488 | 5743 | 274 | 12176 | 396 | sba,lz,Sin3A,salm,Hph,exex,osa,sas,Tet,ap,shn,heph,nudE,Chro,jing,Arl4,CG41099,enok,crb,Gbs-70E,ush,Doc1,CtBP,sens-2,mub,Pvf3,corn,Haspin,Prosap,scb,CG8097,RhoGEF2,cta,Doa,CG3919,Atg17,Taf1,Thd1,tou,slp1,Stat92E,ctrip,smg,tsl,CG12010,bi,CG17168,bab1,CG18446,ytr,hang,sbb,Mondo,ey,ham,betaTub56D,CG6051,otk2,Ten-m,CG9855,Tl,PsGEF,ScsbetaG,l(3)mbt,CG42748,CG8441,Lim1,Hipk,AGO2,Dr,Fas2,Lac,RhoGAP15B,oc,dbr,E(spl)mdelta-HLH,NAT1,Syx1A,aos,slou,NKAIN,scaf6,Toll-7,Ptpmeg,ocm,bora,par-1,hiw,Psc,Myb,flfl,Bx,yps,Pcl,B-H2,phyl,N,spi,S6kII,ec,robo2,pbl,CG4325,Syp,ITP,Df31,Acf,CG32758,Svil,Cyp1,Ykt6,TfIIS,CG9281,DnaJ-H,CG6752,ckn,mgl,Arp2,lbe,Tudor-SN,CG32772,CG12099,cpo,gus,Rho1,CG32369,bowl,Ank,tkv,phtf,comm,wake,KP78a,E(spl)m4-BFM,Zif,Rac2,Hip1,CG4360,CaMKII,ci,TfAP-2,ImpL2,mle,hts,CG17778,Mmp2,ttk,Cirl,pnt,LRP1,CG4538,PhKgamma,CG43658,Cyt-c1,Snx3,hh,kni,spen,PEK,Mhcl,Cyp4p2,l(3)neo38,elav,CG5728,rush,EcR,rau,Lrt,stwl,mbt,Dad,mof,dve,Mitofilin,CG2162,baf,CG9541,CG13894,bin3,eya,Pli,so,CG3097,Ubx,CG13675,CG31224,dpp,Etl1,CG11247,Mrp4,egr,twi,MESR3,Doc2,CG5815,cas,Kr,Usp2,NiPp1,Pak3,HmgZ,Tlk,ft,CadN,CG14535,cnc,dlp,Myo10A,FoxL1,Gmap,spoon,Tif-IA,Rbfox1,CG4199,CG2991,ds,Mos,dally,tin,CG8485,NFAT,RhoGAP18B,HGTX,Ser,CG31612,fne,smal,CG45050,jigr1,CG1142,CG11486,RpS4,wnd,ps,if,CG31875,Dscam1,Socs16D,CG10283,Smox,CG9650,sm,Egfr,Cad87A,CG5522,JIL-1,Utx,CycE,fj,BtbVII,Pex1,pAbp,slow,wech,shd,Asator,edl,tna,psq,disco,Imp,bor,esn,Tpr2,LManII,nuf,aop,Pits,Kah,sdt,Cdk12,Hsc70-3,Cnx99A,drk | | 3.232e-19 | -42.58 | imaginal disc-derived wing morphogenesis | biological process | GO:0007476 | 298 | 47 | 12345 | 414 | RhoGEF2,S,dally,Egfr,dlp,ds,S6kII,dpp,ap,par-1,hh,Stat92E,CycE,Hipk,dve,Dr,sbb,tkv,sno,jing,pnt,pio,ft,drk,osa,Lpin,shn,salm,mam,fj,if,psq,N,Bx,spi,Dad,bi,spen,heph,aos,Rho1,EcR,slow,Ser,nw,Smox,Rbfox1 | | 4.225e-19 | -42.31 | positive regulation of macromolecule metabolic process | biological process | GO:0010604 | 858 | 83 | 12345 | 414 | Myb,tou,lz,aop,disco,wnd,Egfr,mle,NFAT,ttk,HGTX,Hipk,JIL-1,Utx,phyl,Atg17,drk,eya,jing,pnt,Usp2,Lpin,shn,osa,PEK,Bx,ci,bora,Pits,twi,grnd,pAbp,B-H2,Rho1,lncRNA:roX1,Tet,tna,Rbfox1,Tl,ps,CtBP,bowl,smg,dpp,ap,comm,Cdk12,Mos,CG12099,TfAP-2,FoxL1,mmy,ctrip,CG31612,Stat92E,oc,enok,sno,Ubx,Doc2,slou,udd,cnc,tara,so,N,Lim1,spi,ey,Taf1,mam,CG18766,Pcl,Sox102F,gus,mbt,Doc1,EcR,bi,tin,ocm,mof,Cdk5alpha | | 6.983e-19 | -41.81 | positive regulation of metabolic process | biological process | GO:0009893 | 937 | 87 | 12345 | 414 | ci,Bx,pAbp,B-H2,Pits,bora,twi,grnd,Lpin,shn,Usp2,osa,PEK,ps,Tl,ush,CG4325,tna,Rbfox1,smg,CtBP,bowl,lncRNA:roX1,Rho1,Tet,HGTX,ttk,NFAT,JIL-1,Hipk,phyl,Utx,tou,Myb,lz,wnd,mle,Egfr,aop,disco,drk,eya,pnt,jing,Atg17,Toll-7,Doa,spi,Lim1,ey,N,CG18766,Taf1,mam,so,tara,mof,ocm,Cdk5alpha,gus,mbt,EcR,Doc1,Pcl,Sox102F,tin,bi,CG12099,TfAP-2,FoxL1,CG31612,Stat92E,ctrip,oc,mmy,dpp,ap,comm,Cdk12,Mos,Ubx,udd,cnc,slou,Doc2,sno,enok | | 1.528e-18 | -41.02 | axon guidance | biological process | GO:0007411 | 271 | 44 | 12345 | 414 | Cdk5alpha,scb,Smox,Ptp99A,Fas2,Ten-m,spen,Fas3,Rho1,NetB,ras,Rac2,ImpL2,if,ckn,N,Dad,Kr,CadN,uzip,Psc,robo2,sm,NiPp1,Acsl,NetA,jing,trn,wb,Ptpmeg,eya,sbb,enok,Dscam1,Toll-7,RhoGAP93B,Mmp2,beat-IIIc,dally,hts,dlp,sas,ap,comm | | 4.165e-18 | -40.02 | negative regulation of gene expression | biological process | GO:0010629 | 582 | 65 | 12345 | 414 | smg,CtBP,bowl,ush,Rbfox1,heph,Acf,B-H2,twi,psq,shn,slp1,jing,CG11486,ft,sbb,Tlk,lbe,ttk,HGTX,bin3,Egfr,hiw,aop,cas,CG31224,lz,mof,ocm,tin,CG17265,bi,Doc1,EcR,Sin3A,Pcl,Sox102F,Lim1,spi,Kr,Hsc70-3,N,Psc,CG17168,so,NiPp1,Tudor-SN,CG4360,Doc2,slou,Ubx,elav,Dr,enok,tkv,oc,kni,l(3)mbt,dve,Zif,TfAP-2,E(spl)mdelta-HLH,AGO2,dpp,stwl | | 5.553e-18 | -39.73 | neuron projection guidance | biological process | GO:0097485 | 280 | 44 | 12345 | 414 | sbb,enok,Dscam1,Toll-7,jing,trn,wb,eya,Ptpmeg,dlp,dally,beat-IIIc,hts,sas,ap,comm,RhoGAP93B,Mmp2,Ten-m,spen,Rho1,Fas3,NetB,ras,Cdk5alpha,scb,Ptp99A,Smox,Fas2,Psc,sm,uzip,robo2,Acsl,NetA,NiPp1,ImpL2,Rac2,if,N,ckn,Kr,CadN,Dad | | 6.513e-18 | -39.57 | positive regulation of gene expression | biological process | GO:0010628 | 571 | 64 | 12345 | 414 | Taf1,mam,CG18766,N,Lim1,ey,tara,so,ocm,mof,bi,tin,Pcl,Sox102F,EcR,Doc1,Stat92E,oc,CG12099,TfAP-2,FoxL1,Cdk12,ap,dpp,slou,Doc2,cnc,udd,Ubx,enok,sno,twi,Pits,pAbp,B-H2,ci,Bx,osa,Lpin,shn,CtBP,bowl,tna,Rbfox1,ps,Tl,Tet,lncRNA:roX1,Utx,NFAT,ttk,HGTX,JIL-1,Hipk,aop,disco,Egfr,mle,Myb,tou,lz,pnt,jing,eya | | 7.613e-18 | -39.42 | positive regulation of nitrogen compound metabolic process | biological process | GO:0051173 | 827 | 79 | 12345 | 414 | lncRNA:roX1,Rho1,Tet,Rbfox1,tna,CG4325,ush,ps,Tl,bowl,CtBP,smg,Usp2,shn,Lpin,osa,PEK,Bx,ci,grnd,twi,bora,Pits,B-H2,pAbp,Atg17,drk,eya,pnt,jing,lz,Myb,tou,disco,mle,Egfr,wnd,JIL-1,ttk,HGTX,NFAT,Utx,phyl,Sox102F,Pcl,EcR,Doc1,mbt,gus,bi,tin,ocm,mof,Cdk5alpha,tara,so,N,ey,spi,Lim1,mam,Taf1,CG18766,enok,sno,slou,Doc2,cnc,udd,ap,dpp,Mos,Cdk12,CG12099,TfAP-2,mmy,Stat92E,ctrip,oc | | 1.762e-17 | -38.58 | positive regulation of cellular metabolic process | biological process | GO:0031325 | 875 | 81 | 12345 | 414 | cnc,udd,slou,Doc2,sno,enok,TfAP-2,CG12099,oc,Stat92E,CG31612,mmy,dpp,ap,comm,Cdk12,Mos,mof,ocm,Cdk5alpha,mbt,EcR,Doc1,Pcl,Sox102F,tin,bi,spi,Lim1,ey,N,CG18766,Taf1,mam,so,tara,eya,drk,pnt,jing,Atg17,Toll-7,Doa,ttk,NFAT,HGTX,JIL-1,phyl,Utx,Myb,tou,lz,wnd,Egfr,mle,disco,ps,Tl,CG4325,ush,tna,Rbfox1,smg,CtBP,bowl,Rho1,lncRNA:roX1,Tet,ci,Bx,pAbp,B-H2,Pits,twi,bora,grnd,Lpin,shn,Usp2,osa,PEK | | 1.879e-17 | -38.51 | regulation of cell differentiation | biological process | GO:0045595 | 428 | 54 | 12345 | 414 | ush,Fas2,goe,pbl,Ser,EcR,NetB,mbt,Rho1,aos,nudE,CadN,Kr,spi,N,ci,pAbp,salm,Sox21a,Rac2,tsl,NetA,osa,Ubx,Ptpmeg,eya,cnc,pnt,ec,tkv,enok,Dscam1,Imp,ham,Dr,E(spl)mdelta-HLH,Syx1A,ttk,CycE,Stat92E,Prosap,oc,mmy,l(3)mbt,lbe,par-1,dpp,comm,lz,S6kII,Egfr,RhoGEF2,sty,aop | | 1.941e-17 | -38.48 | embryonic morphogenesis | biological process | GO:0048598 | 251 | 41 | 12345 | 414 | Dr,tkv,sdt,stg,slou,pnt,slp1,Inx2,PhKgamma,Ubx,cta,S,RhoGEF2,aop,Egfr,dpp,Dtg,mmy,lbe,Mmp2,Inx3,tin,Rho1,EcR,bowl,scb,pbl,CG41099,ush,shn,mam,twi,Rac2,N,crb,Kr,Dad,Lac,shd,spi,Myo10A | | 2.775e-17 | -38.12 | cell projection organization | biological process | GO:0030030 | 637 | 67 | 12345 | 414 | Rho1,Fas3,Ten-m,nudE,Fas2,Ptp99A,scb,shn,NetA,Acsl,sm,CadN,Dad,Myo10A,ci,if,Toll-7,sbb,eya,ft,wb,jing,trn,dlp,Egfr,mle,ds,wnd,hts,Mmp2,Prosap,EcR,NetB,ras,spen,fax,Smox,Cdk5alpha,pbl,NiPp1,robo2,Psc,uzip,Kr,crb,N,ckn,Con,fj,ImpL2,Rac2,enok,Dscam1,ham,Ptpmeg,cnc,exex,ap,comm,sas,dally,beat-IIIc,RhoGAP93B,oc,mmy,hh,par-1 | | 3.924e-17 | -37.78 | sensory organ development | biological process | GO:0007423 | 496 | 58 | 12345 | 414 | shn,so,NetA,nuf,Lim1,spi,CadN,ey,Kr,crb,bdg,ci,Bx,N,fj,B-H2,salm,mbt,NetB,aos,rau,Rho1,spen,Ten-m,bi,Fas2,ush,Ser,CtBP,lz,dpp,dally,ds,Egfr,dlp,aop,sty,S,edl,ttk,E(spl)mdelta-HLH,Hipk,Stat92E,phyl,oc,CycE,hh,Utx,Nhe2,NAT1,tkv,E(spl)m4-BFM,Doa,eya,drk,ft,ec,pnt | | 4.401e-17 | -37.66 | plasma membrane bounded cell projection organization | biological process | GO:0120036 | 626 | 66 | 12345 | 414 | exex,cnc,Ptpmeg,ham,Dscam1,enok,par-1,hh,mmy,RhoGAP93B,oc,dally,beat-IIIc,sas,ap,comm,pbl,Cdk5alpha,Smox,fax,spen,ras,NetB,EcR,Rac2,ImpL2,fj,ckn,Con,N,crb,Kr,robo2,Psc,uzip,NiPp1,trn,jing,wb,ft,eya,sbb,Toll-7,Prosap,Mmp2,wnd,hts,ds,mle,dlp,scb,Ptp99A,Fas2,Ten-m,nudE,Fas3,Rho1,if,ci,Myo10A,Dad,CadN,sm,Acsl,NetA,shn | | 6.108e-17 | -37.33 | wing disc pattern formation | biological process | GO:0035222 | 80 | 24 | 12345 | 414 | hh,N,ci,crb,ds,dlp,dally,osa,nuf,shn,dpp,ap,tara,Ser,pnt,MFS14,sbb,Dr,spen,bi,sno,Mrp4,tkv,Notum | | 8.325e-17 | -37.02 | positive regulation of transcription, DNA-templated | biological process | GO:0045893 | 473 | 56 | 12345 | 414 | Utx,oc,Stat92E,JIL-1,TfAP-2,ttk,NFAT,CG12099,HGTX,disco,Cdk12,mle,lz,ap,tou,Myb,slou,pnt,Doc2,jing,udd,cnc,eya,enok,sno,mam,twi,Taf1,Pits,CG18766,B-H2,ci,N,Bx,ey,Lim1,osa,tara,shn,so,Lpin,bowl,CtBP,ocm,Rbfox1,tna,Tl,mof,bi,Tet,tin,lncRNA:roX1,Sox102F,Pcl,EcR,Doc1 | | 8.342e-17 | -37.02 | cell migration | biological process | GO:0016477 | 287 | 43 | 12345 | 414 | Mmp2,par-1,kni,lbe,hh,bbg,Stat92E,dpp,aop,dally,spoon,Egfr,dlp,cta,drk,eya,Pvf3,pnt,jing,trn,tkv,N,Dad,wun2,shd,ey,salm,Rac2,Lrt,if,NetA,so,robo2,pbl,scb,Sin3A,Rho1,gus,mbt,EcR,NetB,nudE,tin | | 9.102e-17 | -36.94 | compound eye development | biological process | GO:0048749 | 370 | 49 | 12345 | 414 | Ser,ush,spen,Ten-m,bi,NetB,mbt,Rho1,rau,aos,B-H2,fj,salm,CadN,ey,Kr,Lim1,spi,bdg,N,crb,shn,so,NetA,pnt,ec,eya,drk,ft,Doa,tkv,CycE,oc,Stat92E,phyl,Nhe2,hh,Hipk,E(spl)mdelta-HLH,ttk,ds,Egfr,dlp,dally,S,aop,sty,lz,dpp | | 1.140e-16 | -36.71 | sensory system development | biological process | GO:0048880 | 401 | 51 | 12345 | 414 | S,aop,sty,Egfr,ds,dlp,dally,lz,dpp,Nhe2,hh,CycE,Stat92E,oc,phyl,Hipk,E(spl)mdelta-HLH,ttk,Doa,tkv,pnt,ec,ft,drk,eya,NetA,so,shn,salm,B-H2,fj,N,ci,crb,bdg,CadN,Kr,ey,Lim1,spi,Ten-m,bi,spen,Rho1,rau,aos,NetB,mbt,Ser,ush,Fas2 | | 1.140e-16 | -36.71 | eye development | biological process | GO:0001654 | 401 | 51 | 12345 | 414 | crb,N,ci,bdg,ey,CadN,Kr,Lim1,spi,salm,B-H2,fj,NetA,so,shn,ush,Fas2,Ser,Rho1,rau,aos,NetB,mbt,Ten-m,bi,spen,E(spl)mdelta-HLH,Hipk,ttk,Nhe2,hh,CycE,oc,phyl,Stat92E,lz,dpp,S,sty,aop,ds,dlp,Egfr,dally,ft,eya,drk,pnt,ec,tkv,Doa | | 1.140e-16 | -36.71 | visual system development | biological process | GO:0150063 | 401 | 51 | 12345 | 414 | salm,B-H2,fj,N,ci,crb,bdg,ey,Kr,CadN,spi,Lim1,NetA,so,shn,Ser,ush,Fas2,bi,Ten-m,spen,Rho1,rau,aos,NetB,mbt,Nhe2,hh,CycE,oc,Stat92E,phyl,E(spl)mdelta-HLH,Hipk,ttk,S,sty,aop,Egfr,ds,dlp,dally,dpp,lz,pnt,ec,ft,drk,eya,Doa,tkv | | 1.211e-16 | -36.65 | negative regulation of macromolecule metabolic process | biological process | GO:0010605 | 777 | 74 | 12345 | 414 | AGO2,dpp,stwl,oc,ctrip,CycE,kni,hh,l(3)mbt,Zif,dve,TfAP-2,E(spl)mdelta-HLH,Dr,enok,tkv,Tudor-SN,CG4360,slou,Doc2,Ubx,elav,Psc,CG17168,so,NiPp1,spi,Lim1,Kr,Hsc70-3,N,crb,CG17265,tin,bi,Doc1,EcR,aos,Sin3A,Pcl,Sox102F,mof,ocm,Egfr,sty,hiw,aop,CG31224,cas,lz,Tlk,lbe,Utx,ttk,HGTX,bin3,sbb,slp1,jing,CG11486,ft,shn,B-H2,twi,psq,CG31875,heph,CG8441,Acf,smg,CtBP,bowl,ush,Rbfox1 | | 1.881e-16 | -36.21 | positive regulation of RNA metabolic process | biological process | GO:0051254 | 513 | 58 | 12345 | 414 | slou,jing,Doc2,pnt,cnc,udd,eya,enok,sno,Utx,Stat92E,oc,JIL-1,CG12099,TfAP-2,ttk,HGTX,NFAT,disco,Cdk12,mle,ap,lz,tou,Myb,bowl,CtBP,smg,ocm,Rbfox1,tna,Tl,ps,mof,bi,tin,Tet,Sox102F,lncRNA:roX1,Pcl,EcR,Doc1,mam,twi,Pits,Taf1,B-H2,CG18766,Bx,ci,N,ey,Lim1,osa,tara,shn,so,Lpin | | 1.914e-16 | -36.19 | digestive tract development | biological process | GO:0048565 | 110 | 27 | 12345 | 414 | Stat92E,if,twi,hh,Sox21a,dve,spi,shd,Dad,Kr,N,crb,Egfr,S,shn,dpp,pbl,wb,cnc,scb,pnt,bowl,Ubx,Inx2,tin,EcR,Rho1 | | 1.914e-16 | -36.19 | digestive system development | biological process | GO:0055123 | 110 | 27 | 12345 | 414 | Inx2,Ubx,bowl,pnt,scb,wb,cnc,pbl,Rho1,EcR,tin,crb,N,Kr,spi,shd,Dad,dve,Sox21a,hh,twi,if,Stat92E,dpp,shn,S,Egfr | | 2.438e-16 | -35.95 | negative regulation of metabolic process | biological process | GO:0009892 | 823 | 76 | 12345 | 414 | tkv,enok,wcy,Dr,elav,Ubx,Tudor-SN,Doc2,slou,CG4360,dpp,stwl,AGO2,dve,Zif,E(spl)mdelta-HLH,TfAP-2,CycE,ctrip,oc,l(3)mbt,kni,hh,EcR,Doc1,Sox102F,aos,Pcl,Sin3A,tin,CG17265,bi,mof,ocm,Smox,so,NiPp1,CG17168,Psc,Kr,Hsc70-3,Lim1,spi,N,crb,sbb,CG11486,ft,jing,slp1,lz,Egfr,cas,CG31224,hiw,sty,aop,bin3,ttk,HGTX,Tlk,Utx,lbe,Acf,heph,CG8441,ush,Rbfox1,smg,bowl,CtBP,shn,CG31875,psq,B-H2,twi | | 3.338e-16 | -35.64 | leg disc development | biological process | GO:0035218 | 103 | 26 | 12345 | 414 | osa,disco,RhoGAP15B,Egfr,ds,dally,ap,dpp,CG43658,lbe,hh,B-H2,fj,N,Bx,TfAP-2,Lim1,dve,Rho1,RhoGAP16F,jing,bowl,pnt,Ser,bab1,Ubx | | 3.784e-16 | -35.51 | transcription regulator activity | molecular function | GO:0140110 | 605 | 62 | 12176 | 396 | kni,CtBP,twi,B-H2,Doc1,tna,edl,Pcl,lbe,Lim1,Doc2,bowl,MESR3,Taf1,cas,Kr,EcR,phtf,ci,CG4360,oc,Kah,aop,Pits,Stat92E,CG31875,Zif,slp1,FoxL1,dve,mam,Smox,E(spl)mdelta-HLH,Dad,Sin3A,cnc,lz,TfAP-2,pnt,CG9650,ttk,slou,Tif-IA,bab1,bi,Lpin,Mondo,Sox21a,tin,ocm,sbb,eya,shn,ap,Myb,CG31224,HGTX,Ubx,ham,NFAT,ey,so | | 4.570e-16 | -35.32 | positive regulation of nucleobase-containing compound metabolic process | biological process | GO:0045935 | 523 | 58 | 12345 | 414 | Myb,tou,ap,lz,mle,Cdk12,disco,HGTX,CG12099,NFAT,ttk,TfAP-2,JIL-1,Stat92E,oc,Utx,sno,enok,eya,cnc,udd,slou,jing,pnt,Doc2,Lpin,shn,so,tara,osa,Lim1,ey,Bx,N,ci,B-H2,CG18766,twi,Taf1,Pits,mam,Doc1,EcR,Pcl,lncRNA:roX1,Sox102F,tin,Tet,bi,mof,ps,Tl,tna,ocm,Rbfox1,smg,CtBP,bowl | | 4.862e-16 | -35.26 | arm (armadillo) | protein interactions | 31151 | 119 | 31 | 9204 | 412 | Egfr,Mmp2,dally,ci,dpp,Hipk,sdt,spi,spen,crb,Dad,CG3402,ft,mam,N,ds,Ubx,pbl,stg,tkv,dlp,mbt,aos,Fas3,egr,CtBP,CG9541,aop,tyn,CycE,CadN | | 5.877e-16 | -35.07 | positive regulation of RNA biosynthetic process | biological process | GO:1902680 | 494 | 56 | 12345 | 414 | HGTX,ttk,TfAP-2,NFAT,CG12099,JIL-1,Utx,Stat92E,oc,tou,Myb,ap,lz,Cdk12,disco,mle,eya,jing,Doc2,slou,pnt,cnc,udd,enok,sno,ci,N,Bx,Lim1,ey,twi,Taf1,Pits,mam,B-H2,CG18766,tara,Lpin,shn,so,osa,tna,ocm,Rbfox1,Tl,mof,CtBP,bowl,Pcl,lncRNA:roX1,Sox102F,EcR,Doc1,bi,Tet,tin | | 5.877e-16 | -35.07 | positive regulation of nucleic acid-templated transcription | biological process | GO:1903508 | 494 | 56 | 12345 | 414 | Doc1,EcR,lncRNA:roX1,Sox102F,Pcl,tin,Tet,bi,mof,Tl,ocm,Rbfox1,tna,bowl,CtBP,so,shn,Lpin,tara,osa,ey,Lim1,ci,Bx,N,CG18766,B-H2,mam,twi,Taf1,Pits,sno,enok,eya,cnc,udd,slou,pnt,Doc2,jing,lz,ap,Myb,tou,mle,disco,Cdk12,JIL-1,TfAP-2,ttk,HGTX,CG12099,NFAT,oc,Stat92E,Utx | | 6.683e-16 | -34.94 | DNA-binding transcription factor activity | molecular function | GO:0003700 | 463 | 53 | 12176 | 396 | Doc2,Lim1,MESR3,bowl,edl,Doc1,twi,B-H2,kni,lbe,Pcl,oc,CG4360,ci,Zif,CG31875,slp1,Stat92E,Pits,aop,Kah,phtf,cas,EcR,Kr,CG9650,ttk,pnt,bi,bab1,slou,E(spl)mdelta-HLH,Smox,dve,FoxL1,TfAP-2,cnc,lz,Dad,CG31224,Myb,so,NFAT,ey,Ubx,HGTX,ham,ocm,tin,Sox21a,Mondo,ap,shn | | 1.359e-15 | -34.23 | positive regulation of macromolecule biosynthetic process | biological process | GO:0010557 | 552 | 59 | 12345 | 414 | oc,CG31612,Stat92E,mmy,Utx,JIL-1,CG12099,HGTX,NFAT,ttk,TfAP-2,mle,disco,Cdk12,ap,lz,tou,Myb,cnc,udd,jing,Doc2,pnt,slou,eya,sno,enok,B-H2,CG18766,pAbp,mam,Pits,Taf1,twi,ey,Lim1,Bx,N,ci,osa,shn,so,Lpin,tara,bowl,CtBP,mof,Tl,Rbfox1,ocm,tna,Tet,tin,bi,Doc1,EcR,lncRNA:roX1,Sox102F,Pcl | | 2.629e-15 | -33.57 | open tracheal system development | biological process | GO:0007424 | 235 | 37 | 12345 | 414 | scb,Fas2,flz,Hph,Rho1,fj,if,salm,Rac2,spi,Lac,crb,N,ci,pio,jing,trn,pnt,drk,Ubx,ft,bou,tkv,Stat92E,kni,hh,mmy,ttk,Mmp2,dally,Egfr,ds,dlp,sty,RhoGEF2,dpp,nrv2 | | 2.758e-15 | -33.52 | positive regulation of cellular biosynthetic process | biological process | GO:0031328 | 594 | 61 | 12345 | 414 | mmy,Utx,oc,CG31612,Stat92E,HGTX,NFAT,CG12099,ttk,TfAP-2,JIL-1,Cdk12,disco,mle,tou,Myb,lz,ap,slou,jing,Doc2,pnt,cnc,udd,eya,enok,sno,Pits,twi,Taf1,mam,pAbp,CG18766,B-H2,Bx,N,ci,Lim1,ey,osa,tara,Lpin,shn,so,CtBP,bowl,tna,ocm,Rbfox1,mof,Tl,CG4325,ush,bi,Tet,tin,Pcl,Sox102F,lncRNA:roX1,Doc1,EcR | | 2.758e-15 | -33.52 | positive regulation of biosynthetic process | biological process | GO:0009891 | 594 | 61 | 12345 | 414 | CtBP,bowl,tna,Rbfox1,ocm,mof,Tl,ush,CG4325,bi,Tet,tin,Pcl,lncRNA:roX1,Sox102F,Doc1,EcR,twi,Taf1,Pits,mam,pAbp,CG18766,B-H2,N,ci,Bx,Lim1,ey,osa,tara,Lpin,shn,so,slou,pnt,jing,Doc2,cnc,udd,eya,enok,sno,mmy,Utx,Stat92E,oc,CG31612,ttk,TfAP-2,NFAT,HGTX,CG12099,JIL-1,Cdk12,disco,mle,tou,Myb,ap,lz | | 3.437e-15 | -33.30 | regulation of response to stimulus | biological process | GO:0048583 | 958 | 81 | 12345 | 414 | Tsp26A,ush,CG4325,Tl,Fas2,CtBP,Ser,Rho1,ci,CadN,Dad,grnd,Socs16D,Acsl,NetA,Usp2,Lpin,PEK,ft,CG5522,eya,drk,pnt,E(spl)m4-BFM,Toll-7,Doa,Hipk,Mmp2,NFAT,CG43658,Prosap,phyl,S6kII,RapGAP1,S,hiw,aop,sty,ds,spoon,dlp,Egfr,wnd,mof,pbl,aos,rau,Sin3A,NetB,mbt,spen,crb,N,ckn,Fbxl7,ey,spi,Lrt,ImpL2,Rac2,PsGEF,lili,CG42684,Ptpmeg,Etl1,Notum,tkv,Imp,wcy,sno,edl,mmy,par-1,MESR3,CycE,Pli,Stat92E,dpp,RhoGEF2,Mos,dally | | 3.685e-15 | -33.23 | regulation of cellular component organization | biological process | GO:0051128 | 791 | 72 | 12345 | 414 | Mos,RhoGEF2,Syp,comm,dpp,kni,par-1,mmy,Stat92E,ctrip,CycE,ham,Imp,Dscam1,sdt,tkv,cnc,Pak3,cta,PsGEF,Rac2,N,ey,mgl,Pcl,sens-2,Arp2,NetB,pbl,tyn,mof,hiw,CG5815,wnd,hts,mle,spoon,S6kII,Myb,lbe,Utx,CG43658,Prosap,Tlk,ttk,l(3)neo38,JIL-1,Syx1A,Mmp2,neo,CG11486,drk,Chro,CaMKII,Hip1,NetA,Acsl,tsl,bora,if,psq,Dad,CadN,Ten-m,nudE,Rho1,lncRNA:roX1,Acf,Hph,scb,Fas2,Tl | | 5.320e-15 | -32.87 | negative regulation of RNA biosynthetic process | biological process | GO:1902679 | 379 | 47 | 12345 | 414 | ush,mof,ocm,bowl,CtBP,EcR,Doc1,Sox102F,Acf,Pcl,Sin3A,tin,bi,Kr,Lim1,CG31875,psq,B-H2,shn,so,Psc,Ubx,slou,jing,Doc2,CG4360,slp1,enok,sbb,Zif,dve,E(spl)mdelta-HLH,bin3,HGTX,ttk,TfAP-2,Tlk,oc,l(3)mbt,lbe,kni,lz,stwl,AGO2,cas,CG31224,aop | | 5.320e-15 | -32.87 | negative regulation of nucleic acid-templated transcription | biological process | GO:1903507 | 379 | 47 | 12345 | 414 | Psc,shn,so,B-H2,CG31875,psq,Kr,Lim1,bi,tin,Acf,Sox102F,Sin3A,Pcl,Doc1,EcR,bowl,CtBP,ocm,ush,mof,cas,CG31224,aop,AGO2,stwl,lz,l(3)mbt,kni,lbe,Tlk,oc,E(spl)mdelta-HLH,bin3,HGTX,ttk,TfAP-2,dve,Zif,sbb,enok,slou,jing,Doc2,slp1,CG4360,Ubx | | 6.415e-15 | -32.68 | regulation of signal transduction | biological process | GO:0009966 | 745 | 69 | 12345 | 414 | PEK,Acsl,Usp2,Lpin,grnd,Socs16D,ci,Dad,Rho1,CtBP,Ser,Tsp26A,Fas2,Tl,S,aop,hiw,sty,spoon,ds,dlp,Egfr,wnd,S6kII,RapGAP1,CG43658,Prosap,phyl,Mmp2,Hipk,NFAT,Doa,E(spl)m4-BFM,pnt,ft,CG5522,eya,drk,lili,PsGEF,ImpL2,N,crb,ckn,Fbxl7,ey,spi,spen,rau,Sin3A,aos,mbt,pbl,RhoGEF2,Mos,dally,dpp,MESR3,par-1,CycE,Pli,edl,tkv,Notum,Imp,wcy,sno,CG42684,Ptpmeg | | 7.948e-15 | -32.47 | negative regulation of RNA metabolic process | biological process | GO:0051253 | 398 | 48 | 12345 | 414 | elav,Ubx,jing,slou,Doc2,slp1,CG4360,enok,sbb,bin3,E(spl)mdelta-HLH,HGTX,ttk,TfAP-2,Zif,dve,l(3)mbt,lbe,kni,Tlk,oc,stwl,lz,CG31224,cas,aop,AGO2,ocm,ush,mof,bowl,CtBP,Acf,Sox102F,Sin3A,Pcl,EcR,Doc1,bi,tin,CG31875,psq,Kr,Lim1,B-H2,shn,so,Psc | | 7.981e-15 | -32.46 | cellular component organization | biological process | GO:0016043 | 2830 | 164 | 12345 | 414 | twi,B-H2,if,Dad,CaMKII,Hip1,KP78a,CG34172,CtBP,lama,tna,Tl,nudE,CG13366,heph,Mitofilin,JIL-1,Mocs2B,CG5815,hiw,mle,dlp,Egfr,hts,tou,jing,CG42748,Pvf3,sbb,Toll-7,bou,ImpL2,Taf1,Rac2,fj,Con,bor,Psc,tara,pbl,tyn,Smox,mof,dpr7,spen,NetB,EcR,gus,mbt,CycE,oc,RhoGAP93B,RhoGEF2,AGO2,Ank,DIP-epsilon,sas,comm,ap,Tim17b,Etl1,RhoGAP18B,cta,ham,sdt,tkv,Dscam1,Imp,stg,enok,wcy,betaTub56D,pAbp,ci,psq,CadN,Myo10A,osa,sm,Acsl,NetA,Pat1,shn,rush,scb,Ptp99A,CG41099,Fas2,Ten-m,Fur1,Rho1,lncRNA:roX1,Acf,Fas3,CG43658,Utx,HmgZ,Tlk,Prosap,Mmp2,bin3,Syx1A,Svil,ttk,egr,S,Gmap,aop,spoon,ds,wnd,S6kII,nrv2,Myb,trn,wb,ft,eya,drk,Chro,Doa,baf,Atg17,neo,N,crb,CG34384,ckn,Kr,Lac,nuf,uzip,robo2,Df31,NiPp1,CG13784,PsGEF,Cdk5alpha,Trpm,fax,Haspin,Ykt6,Arp2,Pcl,ras,mmy,hh,par-1,CG11498,Stat92E,Pex1,dve,Syp,beat-IIIc,dally,wake,stwl,CG8531,dpp,exex,cnc,Pak3,Ptpmeg,qsm | | 9.010e-15 | -32.34 | respiratory system development | biological process | GO:0060541 | 244 | 37 | 12345 | 414 | ttk,Mmp2,Stat92E,hh,kni,mmy,dpp,nrv2,dally,dlp,ds,Egfr,sty,RhoGEF2,Ubx,drk,ft,pio,jing,trn,pnt,bou,tkv,Lac,spi,crb,ci,N,fj,if,salm,Rac2,Fas2,flz,scb,Hph,Rho1 | | 9.446e-15 | -32.29 | negative regulation of macromolecule biosynthetic process | biological process | GO:0010558 | 446 | 51 | 12345 | 414 | mof,ush,ocm,Rbfox1,smg,CtBP,bowl,Doc1,EcR,heph,Pcl,Sin3A,Sox102F,Acf,tin,bi,Lim1,Kr,psq,CG31875,B-H2,shn,so,Psc,elav,Ubx,slp1,CG4360,jing,slou,Doc2,enok,sbb,dve,Zif,HGTX,ttk,TfAP-2,E(spl)mdelta-HLH,bin3,oc,Tlk,kni,lbe,l(3)mbt,lz,stwl,AGO2,aop,cas,CG31224 | | 1.068e-14 | -32.17 | negative regulation of transcription, DNA-templated | biological process | GO:0045892 | 371 | 46 | 12345 | 414 | stwl,lz,aop,CG31224,cas,AGO2,ttk,TfAP-2,HGTX,E(spl)mdelta-HLH,bin3,dve,Zif,kni,lbe,l(3)mbt,oc,Tlk,enok,sbb,Ubx,CG4360,slp1,Doc2,slou,jing,so,shn,Psc,psq,Lim1,Kr,B-H2,Sin3A,Pcl,Acf,Sox102F,Doc1,EcR,bi,tin,ocm,mof,ush,CtBP,bowl | | 1.896e-14 | -31.60 | negative regulation of cellular macromolecule biosynthetic process | biological process | GO:2000113 | 438 | 50 | 12345 | 414 | Sin3A,Pcl,heph,Sox102F,Acf,Doc1,EcR,bi,tin,ocm,Rbfox1,mof,ush,CtBP,bowl,smg,shn,so,Psc,psq,Lim1,Kr,B-H2,enok,sbb,elav,Ubx,CG4360,slp1,jing,Doc2,slou,stwl,lz,aop,cas,CG31224,AGO2,HGTX,ttk,TfAP-2,E(spl)mdelta-HLH,bin3,Zif,dve,lbe,kni,l(3)mbt,oc,Tlk | | 2.151e-14 | -31.47 | anatomical structure formation involved in morphogenesis | biological process | GO:0048646 | 553 | 57 | 12345 | 414 | so,nuf,robo2,N,ci,Kr,spi,mam,Npc1a,twi,salm,Rac2,B-H2,if,Rho1,heph,EcR,NetB,gus,nudE,tin,CG13366,Rbfox1,lama,flz,CtBP,pbl,dpp,lz,Myb,Syp,RhoGEF2,S,ds,spoon,Egfr,AGO2,hts,dally,Mmp2,Syx1A,ttk,mmy,par-1,lbe,hh,CycE,Stat92E,phyl,tkv,ft,Pak3,Inx2,Ubx,eya,slou,slp1 | | 2.302e-14 | -31.40 | DNA binding | molecular function | GO:0003677 | 984 | 79 | 12176 | 396 | CG31224,Utx,so,ey,Ubx,BtbVII,Mondo,CG13894,CG9650,bi,baf,bab1,Smox,dve,Acf,stwl,oc,CG4360,ci,Zif,Pits,aop,Kah,Cdk12,HmgZ,Kr,Dr,cas,Doc2,disco,CG5815,Lim1,MESR3,l(3)mbt,bowl,edl,CG32772,psq,CG12099,twi,lbe,Etl1,CG11247,Psc,Myb,NFAT,HGTX,tin,ocm,Tet,ap,ttk,osa,pnt,slou,Rbfox1,exex,E(spl)mdelta-HLH,FoxL1,sba,TfAP-2,lz,mle,cnc,CG31875,slp1,Stat92E,Thd1,tou,Taf1,CG3919,EcR,Doc1,B-H2,kni,CG45050,Pcl,jigr1,enok | | 2.530e-14 | -31.31 | negative regulation of biosynthetic process | biological process | GO:0009890 | 457 | 51 | 12345 | 414 | Doc2,slou,jing,slp1,CG4360,elav,Ubx,sbb,enok,l(3)mbt,kni,lbe,Tlk,oc,E(spl)mdelta-HLH,bin3,ttk,TfAP-2,HGTX,Zif,dve,CG31224,cas,aop,AGO2,stwl,lz,bowl,CtBP,smg,ocm,Rbfox1,ush,mof,bi,tin,Acf,Sox102F,heph,Pcl,Sin3A,Doc1,EcR,B-H2,CG31875,psq,Kr,Lim1,Psc,so,shn | | 2.530e-14 | -31.31 | negative regulation of cellular biosynthetic process | biological process | GO:0031327 | 457 | 51 | 12345 | 414 | stwl,lz,aop,cas,CG31224,AGO2,ttk,TfAP-2,HGTX,bin3,E(spl)mdelta-HLH,Zif,dve,kni,lbe,l(3)mbt,oc,Tlk,enok,sbb,elav,Ubx,slp1,CG4360,Doc2,slou,jing,shn,so,Psc,psq,CG31875,Lim1,Kr,B-H2,Sin3A,Pcl,heph,Sox102F,Acf,Doc1,EcR,bi,tin,Rbfox1,ocm,mof,ush,CtBP,bowl,smg | | 2.869e-14 | -31.18 | negative regulation of nitrogen compound metabolic process | biological process | GO:0051172 | 625 | 61 | 12345 | 414 | bowl,CtBP,smg,ocm,Rbfox1,ush,mof,bi,tin,Sox102F,Acf,Sin3A,Pcl,aos,CG8441,heph,EcR,Doc1,B-H2,CG31875,crb,N,psq,Kr,Lim1,Psc,NiPp1,so,shn,slou,jing,Doc2,slp1,CG4360,Ubx,elav,sbb,enok,Utx,l(3)mbt,hh,kni,lbe,CycE,Tlk,oc,ctrip,bin3,E(spl)mdelta-HLH,TfAP-2,HGTX,ttk,dve,Zif,CG31224,cas,aop,sty,AGO2,stwl,lz | | 2.897e-14 | -31.17 | cell fate specification | biological process | GO:0001708 | 84 | 22 | 12345 | 414 | lbe,twi,hh,salm,CycE,N,ttk,ey,robo2,Egfr,dpp,slou,pnt,Doc2,exex,ocm,Ubx,bi,tin,E(spl)m4-BFM,tkv,Doc1 | | 3.548e-14 | -30.97 | positive regulation of transcription by RNA polymerase II | biological process | GO:0045944 | 368 | 45 | 12345 | 414 | mle,disco,Cdk12,ap,lz,tou,Myb,Stat92E,oc,ttk,TfAP-2,CG12099,NFAT,HGTX,sno,enok,cnc,Doc2,pnt,slou,jing,eya,shn,so,Lpin,B-H2,mam,twi,Taf1,Pits,ey,Lim1,ci,N,Tet,tin,bi,EcR,Doc1,Sox102F,bowl,CtBP,Tl,mof,ocm | | 3.682e-14 | -30.93 | sensory organ morphogenesis | biological process | GO:0090596 | 310 | 41 | 12345 | 414 | shn,so,B-H2,fj,salm,ey,CadN,spi,crb,N,ci,bdg,Ten-m,bi,mbt,Rho1,rau,aos,Ser,Fas2,ds,Egfr,S,sty,aop,lz,dpp,CycE,oc,phyl,Nhe2,hh,E(spl)mdelta-HLH,ttk,Doa,tkv,pnt,ec,drk,eya,ft | | 3.682e-14 | -30.93 | eye morphogenesis | biological process | GO:0048592 | 310 | 41 | 12345 | 414 | E(spl)mdelta-HLH,ttk,Nhe2,hh,CycE,oc,phyl,dpp,lz,S,aop,sty,ds,Egfr,ft,eya,drk,pnt,ec,tkv,Doa,ci,crb,bdg,N,CadN,ey,spi,salm,B-H2,fj,so,shn,Fas2,Ser,Rho1,rau,aos,mbt,Ten-m,bi | | 4.277e-14 | -30.78 | neuron recognition | biological process | GO:0008038 | 114 | 25 | 12345 | 414 | Pvf3,Tl,Fas2,Ptp99A,pbl,slow,NetB,Rho1,Fas3,Dscam1,Fur1,esn,Ten-m,CadN,Mmp2,nrm,Con,if,RhoGAP93B,ImpL2,mmy,comm,uzip,robo2,beat-IIIc | | 4.823e-14 | -30.66 | negative regulation of nucleobase-containing compound metabolic process | biological process | GO:0045934 | 417 | 48 | 12345 | 414 | Doc1,EcR,Acf,Sox102F,Pcl,Sin3A,tin,bi,ush,mof,ocm,bowl,CtBP,so,shn,Psc,Kr,Lim1,CG31875,psq,B-H2,enok,sbb,Ubx,elav,Doc2,slou,jing,CG4360,slp1,lz,stwl,AGO2,cas,CG31224,aop,dve,Zif,E(spl)mdelta-HLH,bin3,HGTX,TfAP-2,ttk,Tlk,oc,l(3)mbt,lbe,kni | | 6.319e-14 | -30.39 | cell motility | biological process | GO:0048870 | 344 | 43 | 12345 | 414 | pbl,scb,gus,mbt,EcR,NetB,Sin3A,Rho1,tin,nudE,Dad,wun2,shd,ey,N,if,salm,Rac2,Lrt,so,NetA,robo2,drk,cta,eya,Pvf3,trn,jing,pnt,tkv,Mmp2,Stat92E,par-1,kni,hh,lbe,bbg,dpp,dally,Egfr,spoon,dlp,aop | | 6.378e-14 | -30.38 | regulation of signaling | biological process | GO:0023051 | 855 | 73 | 12345 | 414 | PEK,Acsl,Usp2,Lpin,grnd,Snx3,Socs16D,ci,Dad,Rho1,CtBP,Ser,Tsp26A,Fas2,Tl,S,sty,aop,hiw,ds,Egfr,dlp,spoon,wnd,S6kII,RapGAP1,CG43658,CG9650,phyl,Prosap,Mmp2,Hipk,NFAT,egr,Doa,E(spl)m4-BFM,pnt,ft,CG5522,drk,eya,lili,PsGEF,ImpL2,crb,N,ckn,Fbxl7,ey,spi,spen,Sin3A,aos,rau,mbt,pbl,Syp,RhoGEF2,Mos,dally,dpp,MESR3,par-1,CycE,Pli,edl,tkv,Notum,wcy,Imp,sno,CG42684,Ptpmeg | | 6.378e-14 | -30.38 | regulation of cell communication | biological process | GO:0010646 | 855 | 73 | 12345 | 414 | grnd,Snx3,Socs16D,ci,Dad,PEK,Acsl,Usp2,Lpin,CtBP,Ser,Tsp26A,Tl,Fas2,Rho1,CG43658,CG9650,Prosap,phyl,Mmp2,Hipk,NFAT,egr,S,aop,hiw,sty,dlp,ds,Egfr,spoon,wnd,S6kII,RapGAP1,pnt,ft,CG5522,drk,eya,Doa,E(spl)m4-BFM,ImpL2,N,crb,ckn,ey,Fbxl7,spi,lili,PsGEF,pbl,spen,Sin3A,aos,rau,mbt,par-1,MESR3,CycE,Pli,edl,RhoGEF2,Syp,Mos,dally,dpp,Ptpmeg,CG42684,tkv,Notum,wcy,Imp,sno | | 8.570e-14 | -30.09 | negative regulation of developmental process | biological process | GO:0051093 | 222 | 34 | 12345 | 414 | Stat92E,ttk,Egfr,Syp,sty,aop,hiw,dpp,S6kII,pnt,Ptpmeg,eya,ft,bab1,tkv,enok,Sox21a,salm,CadN,Fbxl7,Dad,N,crb,osa,shn,Acsl,scb,Ser,ush,Fas2,Tl,goe,EcR,aos | | 9.291e-14 | -30.01 | compound eye morphogenesis | biological process | GO:0001745 | 290 | 39 | 12345 | 414 | ft,drk,eya,pnt,ec,tkv,Doa,E(spl)mdelta-HLH,ttk,Nhe2,hh,CycE,phyl,oc,dpp,lz,S,aop,sty,Egfr,ds,Ser,Rho1,rau,aos,mbt,bi,Ten-m,bdg,crb,N,ey,CadN,spi,salm,B-H2,fj,so,shn | | 1.219e-13 | -29.74 | cell recognition | biological process | GO:0008037 | 119 | 25 | 12345 | 414 | Ten-m,esn,Fur1,Rho1,Dscam1,Fas3,NetB,slow,pbl,Ptp99A,Pvf3,Tl,Fas2,robo2,uzip,beat-IIIc,comm,mmy,ImpL2,if,RhoGAP93B,Mmp2,nrm,Con,CadN | | 1.221e-13 | -29.73 | negative regulation of cellular metabolic process | biological process | GO:0031324 | 681 | 63 | 12345 | 414 | sbb,wcy,enok,CG4360,slp1,Doc2,jing,slou,Ubx,elav,AGO2,sty,aop,CG31224,cas,lz,stwl,ctrip,oc,CycE,Tlk,lbe,kni,hh,l(3)mbt,Utx,Zif,dve,ttk,HGTX,TfAP-2,bin3,E(spl)mdelta-HLH,tin,bi,EcR,Doc1,CG8441,Sin3A,aos,Pcl,heph,Acf,Sox102F,smg,CtBP,bowl,mof,ush,Smox,Rbfox1,ocm,Psc,shn,so,NiPp1,B-H2,Lim1,Kr,psq,N,CG31875,crb | | 1.285e-13 | -29.68 | localization of cell | biological process | GO:0051674 | 351 | 43 | 12345 | 414 | dpp,aop,dally,spoon,dlp,Egfr,Mmp2,lbe,hh,kni,par-1,bbg,Stat92E,tkv,drk,eya,cta,Pvf3,pnt,trn,jing,NetA,so,robo2,N,wun2,shd,Dad,ey,salm,Rac2,Lrt,if,Sin3A,Rho1,mbt,gus,NetB,EcR,nudE,tin,pbl,scb | | 1.363e-13 | -29.62 | cell fate determination | biological process | GO:0001709 | 141 | 27 | 12345 | 414 | CtBP,slou,Ubx,ush,nudE,spen,Dr,tin,tsl,twi,par-1,bora,mam,phyl,CycE,N,spi,egr,Kr,aop,nuf,cas,S,hts,Egfr,dpp,lz | | 1.680e-13 | -29.41 | cellular component organization or biogenesis | biological process | GO:0071840 | 2975 | 166 | 12345 | 414 | S6kII,nrv2,Myb,aop,Gmap,S,wnd,spoon,ds,Svil,ttk,Syx1A,bin3,Mmp2,egr,CG43658,Utx,Prosap,Tlk,HmgZ,neo,Atg17,baf,Doa,ft,Chro,eya,drk,trn,wb,NetA,Acsl,Pat1,shn,osa,sm,psq,ci,Myo10A,CadN,betaTub56D,pAbp,Fas3,CG11030,Acf,lncRNA:roX1,Rho1,Ten-m,Fur1,CG41099,Ptp99A,Fas2,rush,scb,stwl,CG8531,dpp,Syp,beat-IIIc,wake,dally,Pex1,dve,hh,par-1,mmy,Stat92E,CG11498,qsm,Pak3,Ptpmeg,cnc,exex,NiPp1,Df31,CG13784,PsGEF,nuf,uzip,robo2,ckn,N,CG34384,crb,Lac,Kr,Pcl,Arp2,ras,Ykt6,Haspin,fax,Trpm,Cdk5alpha,tou,hiw,CG5815,hts,Egfr,mle,dlp,Mocs2B,JIL-1,Mitofilin,bou,Toll-7,sbb,Pvf3,CG42748,jing,CG34172,CaMKII,KP78a,Hip1,Dad,twi,if,B-H2,heph,nudE,CG13366,tna,lama,Tl,CtBP,sas,Tim17b,comm,ap,RhoGEF2,Ank,DIP-epsilon,AGO2,RhoGAP93B,oc,CycE,Imp,wcy,Dscam1,enok,stg,tkv,sdt,ham,cta,Etl1,RhoGAP18B,tara,Nopp140,bor,Psc,Con,Taf1,Rac2,ImpL2,fj,gus,mbt,NetB,EcR,spen,dpr7,Smox,tyn,mof,pbl | | 1.690e-13 | -29.41 | nucleus | cellular component | GO:0005634 | 2809 | 161 | 11940 | 407 | ap,bin3,ham,lncRNA:roX1,Asator,slp1,l(3)L1231,ps,udd,CG6700,CG3919,bbg,Stat92E,oc,shn,tou,CG11030,Rbfox1,mle,bab1,Taf1,MESR3,slou,EcR,cas,dbr,twi,hh,sbb,NFAT,Kah,CG31224,mub,CG32772,pAbp,cnc,Cdk12,CG9650,Utx,cpo,disco,CG31612,pnt,Kr,Tudor-SN,Thd1,sm,crb,Psc,stwl,hiw,eya,bowl,elav,bora,Tlk,Mocs2B,Myb,CG17168,Nopp140,so,Lim1,psq,CG18766,HmgZ,baf,Cyp1,Syp,ttk,hang,Zif,Tif-IA,kni,Sox21a,ci,bi,Bx,drk,N,PhKgamma,Doa,spen,IntS6,Pcl,dve,ytr,l(3)mbt,Chro,Acf,Doc1,yps,TfIIS,gus,mof,jigr1,Sin3A,CG1142,pbl,phtf,flfl,Hsc70-3,E(spl)mdelta-HLH,ocm,Lpin,tna,enok,Tet,ctrip,Df31,B-H2,Smox,wcy,ey,Hsc70Cb,edl,CG5728,CG2162,Ubx,CG8531,JIL-1,CG11247,CG31712,Dr,osa,CtBP,sens-2,CG8441,FoxL1,Hipk,lz,salm,Doc2,ush,Etl1,sno,Imp,mam,CG8485,nudE,exex,Pits,Haspin,CG5815,NiPp1,lbe,tin,CG44774,Fas3,heph,AGO2,TfAP-2,CycE,HGTX,jing,CG4360,Hph,CG31875,stg,phyl,aop,Mondo | | 1.850e-13 | -29.32 | dorsal/ventral pattern formation | biological process | GO:0009953 | 154 | 28 | 12345 | 414 | Tl,pnt,MFS14,cnc,tkv,Notum,aos,sno,Mrp4,sbb,Dr,N,crb,twi,pAbp,tara,shn,dpp,ap,osa,Syp,S,sty,nuf,spoon,Egfr,dlp,dally | | 4.435e-13 | -28.44 | cell part | cellular component | GO:0044464 | 7811 | 331 | 11940 | 407 | Psc,NTPase,crb,sm,pio,CG31612,pnt,RhoGAP18B,kdn,bowl,Ugt49C1,eya,Fas2,ft,hiw,stwl,Karl,so,Arl4,CG2201,sas,l(3)neo38,CG5830,PsGEF,ci,kni,Con,Tif-IA,baf,Syp,ttk,psq,Arp2,Lim1,slow,CG32758,wun2,Asator,Mos,lncRNA:roX1,Fbxl7,ham,egr,bin3,S6kII,shn,Ptr,tou,Stat92E,oc,Cnx99A,CG3919,Lrt,fax,Tsp39D,Dtg,if,CG13784,scb,cas,comm,EcR,CG13029,Rac2,mbt,slou,MFS14,bab1,CG11030,disco,CG9650,LRP1,mub,sdt,goe,CadN,sbb,hh,RhoGAP93B,dally,Ser,Ubx,CG8531,Pli,CG6051,edl,DIP-epsilon,Ptp99A,Hip1,ey,Hsc70Cb,wcy,retm,betaTub56D,Hipk,smg,sens-2,CG8441,Ppa,CtBP,CG11247,CG1271,RpS4,Pat1,CG5815,spi,exex,CG14646,CG17778,Pits,nAChRalpha4,Usp2,dlp,mam,sno,Ptpmeg,Imp,CG13366,Etl1,ush,lz,Mondo,phyl,CG31875,rush,Hph,HGTX,Fas3,heph,CG1492,dve,CG41128,CG14535,Pcl,RhoGAP16F,spen,Mitofilin,Acsl,IntS6,Doa,N,PhKgamma,drk,Bx,wnd,CG9220,hts,Pex1,ckn,CG3253,CG6330,yps,wake,nuf,ras,CG34384,l(3)mbt,Lpin,tna,flfl,robo2,Cdk5alpha,lawc,CG11498,Atg17,Smox,Pak3,Myo10A,CG6752,corn,CG41099,Tet,enok,Ykt6,Tl,Thd1,Kr,Tudor-SN,tyn,bou,RhoGEF2,sty,CG17224,bor,LManII,otk2,Rho1,nrm,CG17168,Nopp140,Ten-m,Lac,Tlk,Mocs2B,Myb,bora,elav,dpp,smal,bi,Sox21a,hang,Zif,CD98hc,HmgZ,Cyp1,ScpX,CG18766,NetB,shd,l(3)L1231,ps,slp1,Snx3,ap,bbg,CG6700,udd,emp,twi,ds,Mhcl,dbr,Cyt-c1,Taf1,MESR3,KP78a,mle,par-1,Rbfox1,Tsp26A,CG4538,cpo,Utx,mgl,SamDC,Cdk12,CG32772,fne,pAbp,cnc,CG31224,Kah,NFAT,Inx2,pon,CG9855,cta,CG5728,NetA,CG2162,tkv,Cirl,FoxL1,osa,Dr,RhoGAP15B,CG31712,JIL-1,S,Syx1A,CG43313,Haspin,dpr7,CG11486,Egfr,fj,CG10283,Dad,nudE,grnd,ND-B14.5AL,CG8485,Toll-7,CaMKII,salm,Doc2,DnaJ-H,stg,aop,spoon,CG4360,jing,Dscam1,Ank,AGO2,TfAP-2,CycE,RapGAP1,CG1344,CG44774,NiPp1,lbe,tin,nrv2,Fur1,CG42684,Dmtn,mof,gus,ImpL2,Nhe2,CG42748,PEK,CG34172,TfIIS,Chro,Acf,Doc1,Gmap,ytr,CG11752,ocm,Hsc70-3,E(spl)mdelta-HLH,phtf,Inx3,Mmp2,Cad87A,Hydr2,Prosap,pbl,CG1142,jigr1,Sin3A,ScsbetaG,Tim17b,B-H2,Df31,ctrip,uzip | | 4.435e-13 | -28.44 | cell | cellular component | GO:0005623 | 7811 | 331 | 11940 | 407 | AGO2,CycE,RapGAP1,TfAP-2,CG1344,CG44774,NiPp1,tin,lbe,stg,DnaJ-H,aop,spoon,jing,CG4360,Dscam1,Ank,ND-B14.5AL,CG8485,Toll-7,CaMKII,salm,Doc2,Haspin,dpr7,CG11486,Egfr,CG10283,fj,Dad,grnd,nudE,Dr,RhoGAP15B,CG31712,S,JIL-1,Syx1A,CG43313,FoxL1,osa,tkv,Cirl,cta,CG5728,NetA,CG2162,ctrip,uzip,ScsbetaG,Tim17b,B-H2,Df31,Cad87A,CG1142,Prosap,Hydr2,pbl,jigr1,Sin3A,ocm,Hsc70-3,E(spl)mdelta-HLH,phtf,Inx3,Mmp2,TfIIS,Acf,Chro,Doc1,Gmap,ytr,CG11752,Dmtn,mof,gus,Nhe2,CG42748,ImpL2,PEK,CG34172,Fur1,CG42684,nrv2,HmgZ,CD98hc,Cyp1,ScpX,CG18766,NetB,shd,smal,bi,Sox21a,hang,Zif,bora,elav,dpp,CG17168,Ten-m,Nopp140,Lac,Mocs2B,Tlk,Myb,bor,LManII,nrm,otk2,Rho1,sty,CG17224,Kr,Tudor-SN,tyn,RhoGEF2,bou,Tl,Thd1,Kah,NFAT,Inx2,pon,CG9855,Utx,Tsp26A,CG4538,cpo,mgl,Cdk12,SamDC,pAbp,fne,CG32772,cnc,CG31224,Taf1,MESR3,KP78a,mle,par-1,Rbfox1,twi,ds,Mhcl,dbr,Cyt-c1,CG6700,udd,emp,bbg,Snx3,ap,ps,l(3)L1231,slp1,Fas3,heph,CG1492,phyl,Mondo,Hph,rush,CG31875,HGTX,dlp,mam,Ptpmeg,sno,Imp,CG13366,Etl1,ush,lz,CG5815,spi,CG14646,exex,CG17778,Pits,nAChRalpha4,Usp2,CG1271,CG11247,RpS4,Pat1,betaTub56D,Hipk,smg,sens-2,CG8441,Ppa,CtBP,CG6051,DIP-epsilon,edl,Ptp99A,ey,Hip1,Hsc70Cb,wcy,retm,Ser,Ubx,CG8531,Pli,CG41099,Tet,enok,Ykt6,Smox,Pak3,Myo10A,CG6752,corn,lawc,Cdk5alpha,CG11498,Atg17,Lpin,tna,flfl,robo2,ckn,CG3253,CG6330,yps,wake,nuf,CG34384,ras,l(3)mbt,CG9220,hts,Pex1,Mitofilin,spen,IntS6,Acsl,Doa,N,PhKgamma,drk,Bx,wnd,CG41128,dve,CG14535,Pcl,RhoGAP16F,baf,Syp,ttk,Arp2,psq,Lim1,ci,kni,Con,Tif-IA,CG5830,PsGEF,Karl,so,Arl4,CG2201,sas,l(3)neo38,Fas2,ft,hiw,stwl,Ugt49C1,bowl,eya,CG31612,pnt,RhoGAP18B,kdn,Psc,NTPase,sm,crb,pio,goe,sdt,CadN,sbb,hh,RhoGAP93B,dally,disco,CG9650,LRP1,mub,bab1,MFS14,CG11030,scb,CG13784,cas,comm,CG13029,EcR,Rac2,mbt,slou,Lrt,fax,Dtg,Tsp39D,if,S6kII,shn,tou,Ptr,Stat92E,oc,Cnx99A,CG3919,Asator,lncRNA:roX1,Mos,ham,Fbxl7,egr,bin3,wun2,slow,CG32758 | | 6.568e-13 | -28.05 | cell communication | biological process | GO:0007154 | 1239 | 90 | 12345 | 414 | Rho1,RhoGAP102A,Ten-m,Tl,CG4325,Ser,CG8485,Acsl,Lpin,shn,RhoGAP15B,osa,PEK,CaMKII,KP78a,ci,Dad,CadN,twi,tsl,grnd,pAbp,Socs16D,Toll-7,Atg17,nAChRalpha4,Doa,sbb,PhKgamma,drk,Pvf3,smal,CCKLR-17D3,pnt,aop,hiw,otk2,wnd,Egfr,ttk,Syx1A,CG32758,Mmp2,JIL-1,egr,phyl,Tlk,RhoGAP16F,mbt,Rhp,gus,EcR,Haspin,spen,Smox,mof,cpo,Cdk5alpha,ITP,Hydr2,hang,ckn,CG34384,crb,N,spi,Hsc70-3,Rac2,Lrt,mam,fj,tkv,sno,Inx2,cta,Cirl,RhoGAP18B,exex,Cnx99A,stwl,dpp,Ank,dally,wake,Inx3,par-1,hh,RhoGAP93B,Stat92E,Pli | | 7.301e-13 | -27.95 | positive regulation of response to stimulus | biological process | GO:0048584 | 481 | 50 | 12345 | 414 | PsGEF,Lpin,PEK,lili,Fbxl7,spi,ci,crb,ckn,Socs16D,grnd,ImpL2,Rac2,mbt,Rho1,rau,spen,ush,CG4325,Tl,Tsp26A,CtBP,dpp,dlp,Egfr,ds,dally,wnd,RhoGEF2,S,Mos,edl,Hipk,Mmp2,CycE,Pli,Prosap,mmy,CG43658,Toll-7,sno,tkv,Notum,Imp,wcy,Doa,eya,drk,ft,Etl1 | | 7.361e-13 | -27.94 | H (Hairless) | protein interactions | 42445 | 143 | 30 | 9204 | 412 | Sin3A,ey,dve,S,Egfr,so,dpp,sno,Rac2,Bx,E(spl)m4-BFM,Rho1,ap,spen,if,Dad,tkv,aos,mam,ft,qsm,N,Ser,aop,lz,sty,Tlk,CtBP,Rbfox1,Imp | | 8.004e-13 | -27.85 | central nervous system development | biological process | GO:0007417 | 281 | 37 | 12345 | 414 | Dscam1,enok,Notum,Dmtn,Dr,Ptpmeg,elav,eya,jing,exex,ap,disco,cas,Pex1,HGTX,Mmp2,lbe,hh,mmy,oc,Prosap,CycE,Pcl,mbt,EcR,tin,Smox,Fas2,PsGEF,Psc,robo2,N,shd,Kr,ey,mam,if | | 1.118e-12 | -27.52 | digestive tract morphogenesis | biological process | GO:0048546 | 71 | 19 | 12345 | 414 | Kr,spi,Dad,crb,N,Stat92E,twi,hh,shn,dpp,Egfr,S,Inx2,pbl,bowl,pnt,EcR,Rho1,tin | | 1.468e-12 | -27.25 | leg disc morphogenesis | biological process | GO:0007478 | 81 | 20 | 12345 | 414 | Ubx,bab1,Ser,bowl,RhoGAP16F,Rho1,dve,Lim1,N,Bx,TfAP-2,fj,CG43658,hh,lbe,ap,ds,dally,osa,RhoGAP15B | | 2.130e-12 | -26.87 | regulation of localization | biological process | GO:0032879 | 478 | 49 | 12345 | 414 | scb,mgl,ScpX,Sin3A,Rho1,NetB,Rac2,Npc1a,tsl,Lrt,ImpL2,Snx3,crb,spi,CadN,Fbxl7,CaMKII,Hip1,robo2,Usp2,NetA,NKAIN,Acsl,Pat1,corn,cnc,ft,Ptpmeg,drk,Pvf3,Dscam1,sdt,par-1,hh,mmy,CG9650,Prosap,ttk,NFAT,Syx1A,aop,S,dally,Egfr,dlp,Cnx99A,CG44774,S6kII,comm | | 2.453e-12 | -26.73 | heterocyclic compound binding | molecular function | GO:1901363 | 3069 | 162 | 12176 | 396 | Arl4,Cyt-c1,jing,PhKgamma,heph,shn,ap,Tet,pnt,osa,ttk,exex,salm,lz,mle,sba,TfAP-2,Stat92E,slp1,tou,Taf1,Thd1,CG3919,EcR,cta,Doa,CG5728,elav,CG8097,Cyp4p2,l(3)neo38,PEK,Mhcl,Haspin,mub,sens-2,spen,kni,CtBP,ush,Doc1,enok,CG31224,betaTub56D,Ubx,ham,ey,so,Mondo,sbb,hang,bin3,CG13894,CG9541,ytr,CG18446,bab1,baf,bi,CG17168,CG2162,mof,dve,CG12010,mbt,stwl,smg,Tlk,oc,HmgZ,Pak3,NiPp1,Kr,Dr,cas,AGO2,Lim1,Hipk,CG8441,CG5815,Doc2,l(3)mbt,ScsbetaG,MESR3,twi,Mrp4,CG11247,Etl1,yps,fne,Myb,Psc,CG31612,HGTX,par-1,CG8485,NFAT,ocm,tin,Mos,scaf6,CG4199,slou,Rbfox1,spoon,FoxL1,NAT1,Myo10A,E(spl)mdelta-HLH,cnc,CG14535,CG31875,ps,Syp,wnd,RpS4,CG11486,CG1142,S6kII,B-H2,jigr1,Pcl,CG45050,shd,pAbp,Pex1,BtbVII,Utx,JIL-1,Egfr,CG9281,TfIIS,sm,CG9650,Smox,Acf,ci,CaMKII,CG4360,Hsc70-3,Cdk12,Kah,Pits,Rac2,aop,Zif,KP78a,bor,Imp,tkv,disco,bowl,Rho1,cpo,CG32772,CG12099,psq,edl,Asator,Tudor-SN,Arp2,lbe | | 2.548e-12 | -26.70 | photoreceptor cell differentiation | biological process | GO:0046530 | 183 | 29 | 12345 | 414 | aop,S,osa,Egfr,lz,salm,hh,phyl,oc,B-H2,ttk,crb,bdg,N,E(spl)mdelta-HLH,spi,edl,ey,CadN,Ten-m,Doa,rau,sdt,mbt,EcR,sno,pnt,drk,eya | | 3.326e-12 | -26.43 | genital disc development | biological process | GO:0035215 | 50 | 16 | 12345 | 414 | dpp,lz,S,Egfr,otk2,ci,N,TfAP-2,spi,lbe,hh,tkv,aos,Dr,Fas2,pnt | | 3.683e-12 | -26.33 | epithelial cell differentiation | biological process | GO:0030855 | 419 | 45 | 12345 | 414 | Fas3,Rho1,gus,EcR,nudE,spen,tin,CG41099,tyn,lama,ush,crb,N,ci,shd,Rac2,Sox21a,mam,fj,if,tkv,neo,ft,cta,eya,drk,jing,exex,Myb,dpp,sty,aop,RhoGEF2,S,Syp,spoon,ds,Egfr,ttk,hh,par-1,bbg,Stat92E,phyl,CycE | | 3.906e-12 | -26.27 | negative regulation of multicellular organismal process | biological process | GO:0051241 | 212 | 31 | 12345 | 414 | tkv,Sin3A,aos,EcR,Doa,ft,bab1,Tl,Ptpmeg,Fas2,pnt,scb,Ser,S6kII,Acsl,shn,dpp,Syp,osa,CaMKII,hiw,aop,sty,Egfr,crb,N,ttk,CadN,Fbxl7,Dad,salm | | 3.976e-12 | -26.25 | organic cyclic compound binding | molecular function | GO:0097159 | 3086 | 162 | 12176 | 396 | scaf6,Mos,ocm,tin,NFAT,CG8485,HGTX,par-1,Psc,CG31612,fne,Myb,yps,CG14535,cnc,E(spl)mdelta-HLH,NAT1,Myo10A,FoxL1,spoon,Rbfox1,slou,CG4199,wnd,Syp,ps,CG31875,CG45050,Pcl,jigr1,B-H2,S6kII,CG1142,CG11486,RpS4,CG9281,Egfr,JIL-1,Utx,BtbVII,Pex1,pAbp,shd,Acf,Smox,CG9650,sm,TfIIS,tkv,Imp,bor,KP78a,Zif,Pits,Rac2,aop,Cdk12,Kah,Hsc70-3,CG4360,CaMKII,ci,lbe,Arp2,Asator,Tudor-SN,edl,CG32772,psq,CG12099,cpo,Rho1,bowl,disco,Tet,ap,shn,heph,PhKgamma,jing,Cyt-c1,Arl4,sba,TfAP-2,lz,mle,salm,exex,ttk,osa,pnt,CG5728,Doa,cta,CG3919,EcR,Taf1,tou,Thd1,slp1,Stat92E,enok,Doc1,ush,CtBP,kni,sens-2,spen,mub,Haspin,PEK,Mhcl,l(3)neo38,CG8097,Cyp4p2,elav,CG18446,ytr,hang,CG13894,CG9541,bin3,sbb,Mondo,so,ey,Ubx,ham,betaTub56D,CG31224,smg,stwl,mbt,CG12010,mof,dve,CG2162,baf,CG17168,bi,bab1,AGO2,Dr,Kr,cas,NiPp1,Pak3,HmgZ,oc,Tlk,Etl1,CG11247,Mrp4,twi,ScsbetaG,MESR3,l(3)mbt,Doc2,CG5815,CG8441,Hipk,Lim1 | | 4.684e-12 | -26.09 | DNA-binding transcription factor activity, RNA polymerase II-specific | molecular function | GO:0000981 | 277 | 35 | 12176 | 396 | E(spl)mdelta-HLH,FoxL1,TfAP-2,ttk,pnt,bi,Sox21a,ocm,tin,ap,shn,CG31224,Myb,ey,so,HGTX,Doc1,edl,kni,B-H2,twi,lbe,Doc2,Lim1,bowl,EcR,Kr,CG4360,oc,ci,Stat92E,Zif,slp1,Pits,aop | | 5.229e-12 | -25.98 | regulation of cell development | biological process | GO:0060284 | 344 | 40 | 12345 | 414 | NetA,osa,spi,CadN,Kr,N,pAbp,tsl,Rac2,mbt,EcR,NetB,aos,Rho1,nudE,Fas2,pbl,lz,comm,S6kII,Egfr,aop,sty,RhoGEF2,ttk,Syx1A,E(spl)mdelta-HLH,Prosap,oc,CycE,lbe,par-1,mmy,Imp,Dscam1,ham,Ptpmeg,cnc,ec,pnt | | 6.973e-12 | -25.69 | renal system development | biological process | GO:0072001 | 78 | 19 | 12345 | 414 | S,Egfr,shn,dpp,twi,crb,N,Kr,spi,Dad,tin,Rho1,tkv,Doc1,EcR,pnt,Doc2,pbl,Ubx | | 6.973e-12 | -25.69 | urogenital system development | biological process | GO:0001655 | 78 | 19 | 12345 | 414 | crb,N,Kr,spi,Dad,twi,dpp,shn,S,Egfr,Ubx,Doc2,pnt,pbl,Rho1,tkv,EcR,Doc1,tin | | 7.141e-12 | -25.67 | Malpighian tubule development | biological process | GO:0072002 | 69 | 18 | 12345 | 414 | Rho1,tkv,Doc1,EcR,Ubx,Doc2,pnt,pbl,dpp,shn,S,Egfr,crb,N,spi,Dad,Kr,twi | | 7.141e-12 | -25.67 | renal tubule development | biological process | GO:0061326 | 69 | 18 | 12345 | 414 | Ubx,pnt,Doc2,pbl,tkv,Rho1,Doc1,EcR,crb,N,Kr,spi,Dad,twi,dpp,shn,S,Egfr | | 8.430e-12 | -25.50 | N (Notch) | protein interactions | 31293 | 301 | 43 | 9204 | 412 | jigr1,Bx,E(spl)m4-BFM,ap,crb,ey,ham,PEK,aop,CycE,Acsl,par-1,CtBP,tkv,sm,N,Ubx,E(spl)mdelta-HLH,CG41099,Utx,fj,spen,enok,Hipk,Sin3A,CG9281,Egfr,S,sno,dpp,TfAP-2,Ser,Tlk,sty,bi,Hip1,Toll-7,Rbfox1,ban,phyl,pnt,mam,stg | | 8.586e-12 | -25.48 | hindgut development | biological process | GO:0061525 | 61 | 17 | 12345 | 414 | S,Egfr,shn,dpp,hh,twi,Stat92E,N,crb,spi,Dad,Kr,Rho1,EcR,pnt,bowl,pbl | | 8.586e-12 | -25.48 | hindgut morphogenesis | biological process | GO:0007442 | 61 | 17 | 12345 | 414 | bowl,pnt,pbl,Rho1,EcR,N,crb,spi,Dad,Kr,twi,hh,Stat92E,shn,dpp,S,Egfr | | 1.036e-11 | -25.29 | sequence-specific DNA binding | molecular function | GO:0043565 | 515 | 49 | 12176 | 396 | l(3)mbt,bowl,MESR3,Lim1,Doc2,disco,Pcl,CG11247,lbe,B-H2,kni,edl,Doc1,psq,Pits,aop,Zif,slp1,Stat92E,ci,oc,CG4360,Kr,Dr,EcR,Taf1,bab1,slou,exex,bi,pnt,ttk,CG9650,lz,TfAP-2,FoxL1,dve,E(spl)mdelta-HLH,Ubx,HGTX,Utx,so,ey,NFAT,CG31224,Psc,ap,ocm,tin | | 1.047e-11 | -25.28 | signal transduction | biological process | GO:0007165 | 968 | 74 | 12345 | 414 | wake,dally,Ank,dpp,Cnx99A,Stat92E,Pli,RhoGAP93B,par-1,hh,Inx3,sno,tkv,RhoGAP18B,Cirl,cta,ITP,Rac2,mam,Lrt,spi,Hsc70-3,hang,Hydr2,ckn,N,crb,CG34384,spen,Haspin,RhoGAP16F,mbt,gus,Rhp,EcR,Cdk5alpha,mof,Smox,otk2,wnd,Egfr,aop,hiw,phyl,Tlk,egr,ttk,JIL-1,CG32758,nAChRalpha4,Doa,Toll-7,CCKLR-17D3,smal,pnt,drk,Pvf3,PhKgamma,KP78a,RhoGAP15B,PEK,osa,shn,Socs16D,tsl,grnd,Dad,ci,RhoGAP102A,Rho1,CG8485,Ser,Tl,CG4325 | | 1.075e-11 | -25.26 | CycE (Cyclin E) | protein interactions | 34924 | 168 | 31 | 9204 | 412 | Imp,ttk,ban,sty,CadN,shn,N,pbl,stg,ft,hh,phyl,Smox,pnt,crb,Dad,spen,enok,osa,Rho1,scb,Rac2,PEK,ci,dally,dpp,NFAT,Egfr,drk,spi,lawc | | 1.207e-11 | -25.14 | wg (wingless) | protein interactions | 34009 | 120 | 26 | 9204 | 412 | ds,N,hh,ft,mam,egr,Stat92E,dlp,jing,CtBP,Snx3,bi,CycE,Ser,slp1,sno,ci,dally,dpp,Mmp2,twi,spi,ey,Hipk,spen,ap | | 1.367e-11 | -25.02 | regulation of anatomical structure morphogenesis | biological process | GO:0022603 | 250 | 33 | 12345 | 414 | oc,mmy,lbe,par-1,ttk,Egfr,ds,otk2,RhoGEF2,aop,sty,dpp,lz,S6kII,ec,cta,neo,tkv,Dscam1,pAbp,tsl,Rac2,CadN,N,shn,NetA,pbl,tyn,Smox,nudE,NetB,mbt,Rho1 | | 1.432e-11 | -24.97 | heart development | biological process | GO:0007507 | 101 | 21 | 12345 | 414 | Mmp2,ci,Lac,pon,twi,lbe,hh,mam,nrv2,dpp,robo2,Tl,Ubx,ush,slp1,Doc2,scb,tkv,Doc1,EcR,tin | | 1.550e-11 | -24.89 | regulation of nervous system development | biological process | GO:0051960 | 356 | 40 | 12345 | 414 | mbt,EcR,NetB,aos,Rho1,Arp2,nudE,Fas2,pbl,scb,Acsl,NetA,CaMKII,osa,Dad,spi,CadN,Kr,N,pAbp,Rac2,Imp,Dscam1,tkv,ham,Pak3,pnt,comm,S6kII,sty,hiw,aop,RhoGEF2,Syp,ttk,E(spl)mdelta-HLH,oc,Prosap,lbe,mmy | | 1.806e-11 | -24.74 | transcription regulatory region DNA binding | molecular function | GO:0044212 | 369 | 40 | 12176 | 396 | pnt,ttk,slou,Rbfox1,bi,dve,E(spl)mdelta-HLH,lz,TfAP-2,CG31224,HGTX,Ubx,so,Utx,NFAT,ey,ocm,tin,ap,Lim1,Doc2,disco,bowl,MESR3,B-H2,Doc1,CG12099,CG11247,jigr1,ci,CG4360,Pits,aop,Cdk12,CG31875,slp1,Stat92E,Taf1,EcR,Dr | | 1.806e-11 | -24.74 | regulatory region nucleic acid binding | molecular function | GO:0001067 | 369 | 40 | 12176 | 396 | tin,ocm,ap,CG31224,Utx,so,ey,NFAT,Ubx,HGTX,E(spl)mdelta-HLH,dve,TfAP-2,lz,ttk,pnt,bi,slou,Rbfox1,Taf1,Dr,EcR,CG4360,ci,slp1,CG31875,Stat92E,aop,Pits,Cdk12,Doc1,CG12099,B-H2,jigr1,CG11247,Doc2,disco,Lim1,MESR3,bowl | | 1.856e-11 | -24.71 | shg (shotgun) | protein interactions | 37386 | 87 | 22 | 9204 | 412 | crb,Rho1,Egfr,Mmp2,S,ey,sdt,Sin3A,Inx2,tyn,CycE,Inx3,CadN,aop,ec,N,ds,stg,ft,Myo10A,Fas3,tkv | | 2.155e-11 | -24.56 | signaling | biological process | GO:0023052 | 1169 | 83 | 12345 | 414 | Dad,ci,Socs16D,pAbp,grnd,tsl,twi,shn,Acsl,KP78a,CaMKII,osa,PEK,RhoGAP15B,CG4325,Tl,CG8485,Ser,Rho1,Ten-m,RhoGAP102A,egr,Syx1A,JIL-1,Mmp2,CG32758,ttk,Tlk,phyl,Egfr,otk2,wnd,hiw,aop,Pvf3,drk,PhKgamma,pnt,smal,CCKLR-17D3,Toll-7,Doa,nAChRalpha4,Hsc70-3,spi,CG34384,crb,N,hang,ckn,Hydr2,fj,Lrt,mam,Rac2,ITP,mof,Smox,Cdk5alpha,cpo,EcR,gus,RhoGAP16F,Rhp,mbt,spen,Haspin,Inx3,RhoGAP93B,Stat92E,Pli,par-1,hh,dpp,Cnx99A,wake,Ank,dally,Cirl,cta,RhoGAP18B,sno,tkv | | 2.587e-11 | -24.38 | circulatory system development | biological process | GO:0072359 | 104 | 21 | 12345 | 414 | ush,Ubx,Tl,scb,Doc2,slp1,Doc1,EcR,tkv,tin,pon,Lac,ci,Mmp2,mam,hh,twi,lbe,dpp,nrv2,robo2 | | 2.677e-11 | -24.34 | proximal/distal pattern formation, imaginal disc | biological process | GO:0007449 | 22 | 11 | 12345 | 414 | N,Fbxl7,Lim1,jing,pnt,hh,B-H2,ap,dpp,disco,Egfr | | 3.152e-11 | -24.18 | dorsal/ventral pattern formation, imaginal disc | biological process | GO:0007450 | 57 | 16 | 12345 | 414 | MFS14,pnt,Notum,Mrp4,sno,Dr,sbb,N,crb,tara,ap,shn,nuf,osa,dally,dlp | | 3.233e-11 | -24.15 | regulation of cell fate specification | biological process | GO:0042659 | 28 | 12 | 12345 | 414 | S6kII,Dr,aop,ush,eya,Ubx,N,ttk,l(3)mbt,pnt,salm,lbe | | 3.233e-11 | -24.15 | proximal/distal pattern formation | biological process | GO:0009954 | 28 | 12 | 12345 | 414 | Fbxl7,Lim1,N,B-H2,jing,pnt,hh,dpp,ap,Egfr,spen,disco | | 3.371e-11 | -24.11 | nucleic acid binding | molecular function | GO:0003676 | 2082 | 120 | 12176 | 396 | pAbp,CG31224,BtbVII,Ubx,ham,ey,Utx,so,Mondo,sbb,bin3,hang,CG13894,ytr,CG18446,TfIIS,sm,CG9650,bab1,bi,CG17168,baf,CG2162,mof,dve,Smox,stwl,Acf,smg,ci,CG4360,oc,Kah,Cdk12,aop,Pits,Zif,HmgZ,NiPp1,Dr,Imp,Kr,cas,AGO2,Lim1,CG8441,CG5815,disco,Doc2,bowl,l(3)mbt,MESR3,twi,cpo,CG12099,CG32772,psq,edl,Tudor-SN,CG11247,lbe,Etl1,yps,fne,Myb,CG31612,Psc,jing,HGTX,NFAT,heph,ocm,tin,shn,scaf6,ap,Tet,pnt,osa,ttk,Rbfox1,slou,exex,spoon,FoxL1,NAT1,salm,E(spl)mdelta-HLH,mle,cnc,lz,TfAP-2,sba,Stat92E,CG31875,slp1,ps,tou,Thd1,Taf1,Syp,EcR,CG3919,CG5728,RpS4,elav,l(3)neo38,CG11486,CG1142,mub,spen,sens-2,kni,B-H2,Doc1,ush,enok,jigr1,Pcl,CG45050 | | 3.495e-11 | -24.08 | pnr (pannier) | protein interactions | 44849 | 107 | 24 | 9204 | 412 | Ubx,qsm,ush,stg,Pits,tin,CtBP,Doc2,shn,tyn,Tlk,wech,Tl,dpp,NFAT,tou,tara,Sin3A,spi,osa,Dad,Bx,ras,ps | | 3.980e-11 | -23.95 | negative regulation of cell communication | biological process | GO:0010648 | 351 | 39 | 12345 | 414 | MESR3,par-1,CG9650,Pli,phyl,NFAT,Hipk,Mmp2,edl,egr,aop,hiw,sty,dally,Egfr,ds,S6kII,RapGAP1,pnt,Ptpmeg,eya,CG42684,Notum,E(spl)m4-BFM,ImpL2,crb,N,Dad,Usp2,Acsl,Lpin,CtBP,pbl,Ser,Tl,Fas2,aos,Sin3A,Rho1 | | 4.209e-11 | -23.89 | gland development | biological process | GO:0048732 | 178 | 27 | 12345 | 414 | Ser,scb,trn,pnt,eya,ush,ocm,tin,EcR,Atg17,NetB,Rho1,Stat92E,if,Rac2,twi,Utx,Zif,crb,N,Egfr,robo2,sty,lz,so,dpp,stwl | | 4.340e-11 | -23.86 | negative regulation of signaling | biological process | GO:0023057 | 352 | 39 | 12345 | 414 | Pli,phyl,par-1,MESR3,CG9650,egr,edl,NFAT,Hipk,Mmp2,dally,ds,Egfr,hiw,aop,sty,RapGAP1,S6kII,pnt,Ptpmeg,eya,CG42684,E(spl)m4-BFM,Notum,ImpL2,Dad,N,crb,Lpin,Usp2,Acsl,pbl,Ser,CtBP,Fas2,Tl,Sin3A,aos,Rho1 | | 4.748e-11 | -23.77 | positive regulation of signal transduction | biological process | GO:0009967 | 369 | 40 | 12345 | 414 | dpp,S,RhoGEF2,Mos,dlp,ds,Egfr,wnd,dally,Hipk,edl,CG43658,CycE,Prosap,Pli,Notum,tkv,Imp,wcy,sno,Doa,ft,drk,PsGEF,Lpin,PEK,lili,ci,crb,ckn,Fbxl7,spi,grnd,ImpL2,Socs16D,rau,mbt,spen,Tsp26A,CtBP | | 5.293e-11 | -23.66 | imaginal disc-derived leg morphogenesis | biological process | GO:0007480 | 77 | 18 | 12345 | 414 | RhoGAP16F,Rho1,Ubx,Ser,bowl,ap,ds,dally,osa,RhoGAP15B,dve,Lim1,N,Bx,TfAP-2,fj,CG43658,lbe | | 5.939e-11 | -23.55 | negative regulation of transcription by RNA polymerase II | biological process | GO:0000122 | 221 | 30 | 12345 | 414 | l(3)mbt,lbe,kni,B-H2,oc,E(spl)mdelta-HLH,ttk,HGTX,TfAP-2,Kr,Zif,dve,CG31224,aop,lz,shn,jing,bowl,slou,Doc2,slp1,CG4360,CtBP,ocm,ush,Ubx,bi,sbb,Doc1,EcR | | 6.037e-11 | -23.53 | cell surface receptor signaling pathway | biological process | GO:0007166 | 340 | 38 | 12345 | 414 | phyl,Pli,Stat92E,hh,egr,ttk,Egfr,dally,otk2,aop,hiw,dpp,pnt,smal,Pvf3,Cirl,drk,Toll-7,sno,tkv,Socs16D,Lrt,grnd,mam,tsl,Rac2,spi,Dad,crb,N,ci,ckn,osa,shn,Ser,Tl,Smox,spen | | 6.672e-11 | -23.43 | gliogenesis | biological process | GO:0042063 | 78 | 18 | 12345 | 414 | cas,sty,so,NetA,nrv2,salm,hh,egr,ey,N,Pex1,Dr,spen,NetB,Rho1,rau,Ser,jing | | 6.995e-11 | -23.38 | negative regulation of signal transduction | biological process | GO:0009968 | 326 | 37 | 12345 | 414 | Lpin,Usp2,Acsl,Dad,N,crb,ImpL2,Sin3A,aos,Rho1,Tl,Fas2,Ser,pbl,CtBP,RapGAP1,S6kII,dally,ds,Egfr,aop,sty,hiw,edl,NFAT,Hipk,Mmp2,Pli,phyl,par-1,MESR3,Notum,E(spl)m4-BFM,eya,Ptpmeg,CG42684,pnt | | 8.602e-11 | -23.18 | abd-A (abdominal A) | protein interactions | 42037 | 54 | 17 | 9204 | 412 | pnt,lbe,mam,Ubx,Doc2,CycE,slou,kni,tin,Kr,twi,spi,Egfr,cas,salm,eya,ap | | 8.780e-11 | -23.16 | response to stimulus | biological process | GO:0050896 | 2655 | 145 | 12345 | 414 | Mrp4,RhoGAP16F,ras,Pcl,Haspin,Trpm,Cdk5alpha,ITP,NiPp1,robo2,uzip,spi,wun2,Hsc70-3,Kr,ckn,hang,crb,N,CG34384,flfl,SamDC,sno,Ptpmeg,Mondo,cnc,Drat,dpp,Cnx99A,stwl,beat-IIIc,wake,dally,Inx3,Stat92E,Pli,hh,par-1,Fas3,Rho1,RhoGAP102A,Ten-m,Fas2,Ptp99A,CG8485,scb,Lpin,shn,Acsl,NetA,sm,RhoGAP15B,PEK,osa,CadN,ci,tsl,Atg17,Thd1,nAChRalpha4,Doa,eya,drk,wb,CCKLR-17D3,smal,pnt,trn,wnd,spoon,aop,S,egr,NFAT,ttk,CG32758,Mmp2,Tlk,Utx,Rhp,mbt,gus,EcR,NetB,spen,mof,Smox,pbl,so,Psc,Hydr2,Con,Rac2,ImpL2,Lrt,mam,Dscam1,enok,Imp,tkv,Cirl,cta,RhoGAP18B,mub,Etl1,ap,comm,sas,Ank,AGO2,RhoGEF2,CG11873,ctrip,RhoGAP93B,Karl,Hph,Tl,CG4325,Ser,KP78a,CaMKII,Dad,Bx,Socs16D,if,grnd,Toll-7,sbb,CG6330,Pvf3,PhKgamma,MFS14,jing,lz,otk2,hts,l(3)L1231,dlp,Egfr,Gbs-70E,hiw,JIL-1,Hipk,phyl,bbg | | 9.150e-11 | -23.11 | cellular process | biological process | GO:0009987 | 7485 | 312 | 12345 | 414 | smg,scb,Ptp99A,CG3961,Rbfox1,goe,Ten-m,CG5830,lncRNA:roX1,Acf,Fas3,betaTub56D,tsl,bdg,Myo10A,PEK,osa,sm,NetA,Acsl,Pat1,Lpin,trn,slp1,smal,wb,ft,bab1,CG1271,Chro,eya,drk,baf,nAChRalpha4,E(spl)m4-BFM,Atg17,neo,Utx,HmgZ,Tlk,Prosap,HGTX,Svil,egr,CG3253,Tpr2,spoon,wnd,S6kII,Myb,Cdk5alpha,fax,Haspin,Ykt6,CG17265,rau,Pcl,ras,RhoGAP16F,SamDC,N,CG34384,hang,wun2,Lac,nuf,CG17168,uzip,CG13029,Df31,ITP,Doc2,Pak3,CG6752,Ptpmeg,Cyp1,Dr,Notum,mmy,Nhe2,l(3)mbt,hh,par-1,Stat92E,E(spl)mdelta-HLH,Pex1,Inx3,Syp,Cdk12,dally,stwl,Cnx99A,dpp,CG8531,IntS6,lama,tna,CG4325,Tl,nudE,CG13366,Hph,Npc1a,Sox21a,twi,B-H2,Ppa,pon,Dad,KP78a,jing,ec,Pvf3,CG11486,bou,Mitofilin,JIL-1,Hipk,Mocs2B,RpS4,Asator,CG5815,sty,CG9855,dlp,Egfr,mle,otk2,tou,CG1492,tyn,ocm,Smox,bi,spen,wech,CG2201,NetB,Doc1,gus,mbt,Taf1,Rac2,fj,Con,CG4538,Hydr2,Fbxl7,ey,Psc,Etl1,Tudor-SN,RhoGAP18B,Cirl,Ubx,CG14535,ham,sdt,wcy,stg,Dscam1,Imp,CG1344,kni,CycE,Cyt-c1,yps,sas,Tim17b,rush,CG8485,CG41099,Fas2,RhoGAP102A,Fur1,Rho1,CG10082,CG11030,bora,pAbp,ci,CG5728,psq,CadN,RhoGAP15B,DnaJ-H,Usp2,Tif-IA,shn,pnt,CCKLR-17D3,CG31637,Doa,Gyf,CG14291,Thd1,CG14646,CG43658,lbe,Syx1A,CG32758,bin3,Mmp2,ttk,Gmap,cas,S,aop,ds,nrv2,Trpm,cpo,CG46339,Arp2,Sox102F,aos,Sin3A,flfl,salm,TfIIS,CG9541,crb,ckn,Hsc70-3,Kr,spi,Lim1,robo2,NiPp1,CG13784,PsGEF,Drat,cnc,exex,qsm,sno,CG11498,Pli,FoxL1,ytr,edl,dve,Mos,kdn,wake,beat-IIIc,slow,CtBP,Ser,ush,CG9220,Tet,CG13675,heph,grnd,if,Socs16D,nrm,shd,CaMKII,Hip1,CG34172,PhKgamma,CG42748,ScsbetaG,CG6330,sbb,CG17224,Toll-7,NAT1,bbg,phyl,hiw,hts,lz,LManII,pbl,mof,esn,dpr7,tin,EcR,Rhp,ImpL2,mam,CG8097,ND-B14.5AL,Lrt,bor,tara,Nopp140,so,slou,Inx2,elav,cta,scaf6,CG17778,tkv,enok,oc,ctrip,RhoGAP93B,CG12099,Zif,CG11873,RhoGEF2,AGO2,DIP-epsilon,Ank,comm,ap | | 1.156e-10 | -22.88 | wing disc dorsal/ventral pattern formation | biological process | GO:0048190 | 53 | 15 | 12345 | 414 | dally,dlp,sbb,nuf,osa,Mrp4,ap,shn,sno,Notum,tara,MFS14,pnt,N,crb | | 1.197e-10 | -22.85 | segmentation | biological process | GO:0035282 | 256 | 32 | 12345 | 414 | AGO2,dlp,Egfr,dally,dpp,ap,kni,par-1,hh,oc,Stat92E,TfAP-2,dve,Doa,jing,slp1,cnc,Ubx,osa,Acsl,Pat1,tsl,fj,ci,Bx,N,psq,Kr,spen,gus,bowl,Ser | | 1.217e-10 | -22.83 | eye-antennal disc development | biological process | GO:0035214 | 71 | 17 | 12345 | 414 | ey,Dad,N,B-H2,oc,Stat92E,salm,lz,dpp,so,Egfr,disco,eya,bab1,Ser,pnt,aos | | 1.298e-10 | -22.76 | Ras85D (Ras oncogene at 85D) | protein interactions | 41140 | 241 | 36 | 9204 | 412 | Rho1,spen,osa,spi,twi,Sin3A,dpp,S,Egfr,sty,Ptpmeg,egr,Stat92E,aos,pnt,phyl,stg,S6kII,mam,hh,eya,Sox21a,MESR3,NFAT,cnc,drk,rau,Taf1,CycE,aop,lz,RhoGEF2,ttk,edl,jing,N | | 1.552e-10 | -22.59 | embryonic hindgut morphogenesis | biological process | GO:0048619 | 54 | 15 | 12345 | 414 | pbl,pnt,bowl,twi,Kr,spi,Dad,crb,N,Egfr,S,EcR,shn,dpp,Rho1 | | 1.562e-10 | -22.58 | cell population proliferation | biological process | GO:0008283 | 175 | 26 | 12345 | 414 | ds,Egfr,wake,hts,S,shn,Myb,CycE,Stat92E,flfl,l(3)mbt,lbe,Sox21a,egr,pon,Zif,spi,N,Rho1,sdt,enok,stg,Ser,pnt,ft,Inx2 | | 1.823e-10 | -22.43 | response to external stimulus | biological process | GO:0009605 | 987 | 72 | 12345 | 414 | Pcl,ras,NetB,EcR,spen,Smox,Trpm,Cdk5alpha,NiPp1,so,uzip,Psc,robo2,ckn,Con,N,wun2,Kr,Rac2,ImpL2,SamDC,Lrt,Imp,Dscam1,enok,Ptpmeg,Etl1,sas,stwl,comm,dpp,ap,beat-IIIc,dally,AGO2,Inx3,par-1,hh,Stat92E,RhoGAP93B,Pli,Fas3,Rho1,Ten-m,Ptp99A,Tl,Fas2,scb,Acsl,NetA,Lpin,CaMKII,sm,Dad,CadN,if,Toll-7,Atg17,sbb,CG6330,eya,trn,jing,wb,hts,dlp,l(3)L1231,Mmp2,Hipk,egr,bbg,Tlk | | 1.948e-10 | -22.36 | positive regulation of signaling | biological process | GO:0023056 | 403 | 41 | 12345 | 414 | CtBP,Tsp26A,spen,rau,mbt,grnd,ImpL2,Snx3,Socs16D,crb,ci,ckn,Fbxl7,spi,lili,PEK,PsGEF,Lpin,ft,drk,Doa,Notum,tkv,wcy,Imp,sno,CG43658,CycE,Pli,Prosap,Hipk,edl,RhoGEF2,S,Mos,dlp,ds,Egfr,dally,wnd,dpp | | 1.948e-10 | -22.36 | positive regulation of cell communication | biological process | GO:0010647 | 403 | 41 | 12345 | 414 | Lpin,PsGEF,lili,PEK,ckn,crb,ci,spi,Fbxl7,ImpL2,grnd,Socs16D,Snx3,rau,mbt,spen,Tsp26A,CtBP,dpp,Mos,S,RhoGEF2,wnd,dally,Egfr,ds,dlp,Hipk,edl,CG43658,Pli,Prosap,CycE,Imp,wcy,tkv,Notum,sno,Doa,ft,drk | | 2.374e-10 | -22.16 | morphogenesis of embryonic epithelium | biological process | GO:0016331 | 128 | 22 | 12345 | 414 | Rac2,mmy,shd,Lac,spi,Myo10A,Inx3,Mmp2,crb,N,Egfr,aop,RhoGEF2,shn,dpp,scb,ush,Inx2,tin,tkv,sdt,Rho1 | | 2.947e-10 | -21.95 | regulation of neuron differentiation | biological process | GO:0045664 | 180 | 26 | 12345 | 414 | nudE,ham,Rho1,Dscam1,Imp,NetB,EcR,mbt,pnt,pbl,Fas2,RhoGEF2,sty,aop,S6kII,NetA,comm,mmy,lbe,Rac2,oc,pAbp,Prosap,N,ttk,CadN | | 3.117e-10 | -21.89 | Malpighian tubule morphogenesis | biological process | GO:0007443 | 48 | 14 | 12345 | 414 | EcR,shn,dpp,Rho1,Egfr,S,spi,Dad,Kr,N,crb,pbl,twi,pnt | | 3.117e-10 | -21.89 | renal tubule morphogenesis | biological process | GO:0061333 | 48 | 14 | 12345 | 414 | Egfr,S,dpp,shn,EcR,Rho1,pbl,twi,pnt,spi,Dad,Kr,crb,N | | 7.338e-10 | -21.03 | ventral cord development | biological process | GO:0007419 | 79 | 17 | 12345 | 414 | Mmp2,N,Kr,lbe,oc,CycE,if,ap,cas,robo2,Psc,eya,Dscam1,Pcl,Notum,Dr,tin | | 8.520e-10 | -20.88 | negative regulation of response to stimulus | biological process | GO:0048585 | 389 | 39 | 12345 | 414 | hiw,sty,aop,dally,ds,Egfr,S6kII,RapGAP1,MESR3,par-1,Pli,phyl,Stat92E,NFAT,Mmp2,Hipk,edl,Notum,E(spl)m4-BFM,pnt,eya,Ptpmeg,CG42684,Usp2,Acsl,Lpin,ImpL2,Lrt,crb,N,Dad,Sin3A,aos,Rho1,CtBP,Ser,pbl,Fas2,Tl | | 8.525e-10 | -20.88 | regulation of neurogenesis | biological process | GO:0050767 | 261 | 31 | 12345 | 414 | Fas2,pbl,pnt,mbt,NetB,EcR,Dscam1,aos,Imp,Rho1,ham,nudE,spi,Kr,CadN,ttk,E(spl)mdelta-HLH,N,pAbp,oc,Prosap,lbe,Rac2,mmy,comm,S6kII,NetA,aop,sty,RhoGEF2,osa | | 9.035e-10 | -20.82 | hemopoiesis | biological process | GO:0030097 | 80 | 17 | 12345 | 414 | Zif,ytr,N,if,Stat92E,Rac2,dpp,lz,stwl,sty,ush,Pvf3,ocm,Ser,pnt,sens-2,Rho1 | | 9.200e-10 | -20.81 | fng (fringe) | protein interactions | 40314 | 18 | 10 | 9204 | 412 | ap,CycE,heph,Ser,Bx,pbl,N,Egfr,ey,Hipk | | 9.221e-10 | -20.80 | regulation of cellular component movement | biological process | GO:0051270 | 113 | 20 | 12345 | 414 | Pvf3,Ptpmeg,scb,NetB,Rho1,Dscam1,Sin3A,CadN,spi,ttk,mmy,Lrt,hh,tsl,Pat1,NetA,CG44774,robo2,Egfr,aop | | 1.002e-09 | -20.72 | plasma membrane | cellular component | GO:0005886 | 1265 | 84 | 11940 | 407 | sty,S,Syx1A,Fas2,otk2,nrm,ft,Rho1,hiw,Pli,crb,Tl,cta,Ser,tkv,Cirl,bou,RhoGEF2,DIP-epsilon,Ptp99A,Hip1,tyn,Dscam1,Ank,smal,Con,CG1492,CD98hc,Fas3,fj,grnd,sas,dpr7,Ten-m,Lac,Egfr,nAChRalpha4,Arl4,CaMKII,dlp,CG5830,Toll-7,Stat92E,ImpL2,Nhe2,CG42748,Cnx99A,Dmtn,hts,Ptr,Tsp39D,emp,nuf,if,Lrt,fax,wun2,nrv2,N,Fur1,drk,egr,Tsp26A,mgl,CadN,Inx2,uzip,hh,dally,sdt,goe,Rac2,Inx3,Mmp2,mbt,ds,scb,comm,Cad87A,Prosap,Hydr2,pbl,par-1,MFS14 | | 1.057e-09 | -20.67 | ameboidal-type cell migration | biological process | GO:0001667 | 177 | 25 | 12345 | 414 | if,Stat92E,bbg,hh,Rac2,par-1,salm,kni,shd,N,spoon,Egfr,dlp,dally,aop,dpp,pbl,pnt,jing,drk,cta,nudE,EcR,Rho1,Sin3A | | 1.162e-09 | -20.57 | Antp (Antennapedia) | protein interactions | 40835 | 55 | 16 | 9204 | 412 | spen,osa,salm,tna,ci,Kr,ey,twi,slou,Doc2,CycE,mam,Ubx,N,pnt,Pcl | | 1.212e-09 | -20.53 | dpp (decapentaplegic) | protein interactions | 33432 | 97 | 21 | 9204 | 412 | ci,dally,Tl,Egfr,so,spi,ey,Kr,spen,osa,eya,N,ds,Ubx,hh,Stat92E,tkv,cv,bi,CycE,shn | | 1.274e-09 | -20.48 | transcription factor binding | molecular function | GO:0008134 | 184 | 25 | 12176 | 396 | CG31875,so,Utx,NFAT,HGTX,Cdk12,eya,Taf1,tou,ocm,HmgZ,Mondo,EcR,shn,bin3,Doc2,Hipk,bi,Rbfox1,edl,tna,ush,Doc1,CtBP,kni | | 1.355e-09 | -20.42 | developmental process involved in reproduction | biological process | GO:0003006 | 928 | 67 | 12345 | 414 | gus,EcR,tin,Hydr2,N,wun2,Rac2,salm,mam,robo2,Inx2,elav,cnc,tkv,Dr,TfAP-2,dve,hh,par-1,l(3)mbt,Stat92E,CycE,stwl,dpp,Syp,RhoGEF2,Mos,dally,yps,AGO2,lama,Rbfox1,Fas2,goe,scb,heph,Fas3,Rho1,nudE,Tet,psq,ci,shd,tsl,Npc1a,pAbp,if,Pat1,bab1,eya,drk,jing,wb,baf,Doa,ttk,JIL-1,bbg,Myb,lz,aop,sty,S,hts,otk2,Egfr,spoon | | 1.572e-09 | -20.27 | cell periphery | cellular component | GO:0071944 | 1409 | 90 | 11940 | 407 | goe,sdt,hh,dally,CadN,Inx2,pon,uzip,Tsp26A,mgl,corn,MFS14,par-1,Prosap,Hydr2,pbl,Cad87A,comm,scb,ds,Inx3,mbt,Mmp2,Rac2,fax,Lrt,if,nuf,Tsp39D,emp,Dmtn,Ptr,hts,Cnx99A,gus,Stat92E,Nhe2,ImpL2,CG42748,drk,CG42684,egr,N,Fur1,nrv2,wun2,Fas3,CD98hc,Syp,CG1492,Arp2,Con,smal,Ank,Dscam1,Toll-7,dlp,CG5830,CaMKII,Lac,Egfr,Arl4,nAChRalpha4,dpr7,Ten-m,sas,grnd,fj,Rho1,otk2,nrm,ft,hiw,Fas2,Syx1A,S,sty,tyn,Hip1,DIP-epsilon,Ptp99A,Cirl,RhoGEF2,bou,tkv,Ser,cta,Tl,Pli,crb | | 1.644e-09 | -20.23 | MAPK signaling pathway - fly | KEGG pathways | dme04013 | 93 | 17 | 3198 | 100 | aos,spi,egr,sty,drk,Egfr,pnt,edl,phyl,aop,ttk,lz,tkv,tsl,Rac2,bi,dpp | | 1.644e-09 | -20.23 | MAPK signaling pathway - fly | KEGG pathways | ko04013 | 93 | 17 | 3198 | 100 | phyl,aop,edl,Egfr,pnt,drk,spi,egr,sty,aos,dpp,bi,tsl,Rac2,tkv,ttk,lz | | 1.726e-09 | -20.18 | regulation of animal organ morphogenesis | biological process | GO:2000027 | 94 | 18 | 12345 | 414 | ec,Smox,mbt,tkv,oc,pAbp,par-1,N,ttk,ds,Egfr,otk2,aop,sty,lz,shn,dpp,S6kII | | 1.800e-09 | -20.14 | muscle structure development | biological process | GO:0061061 | 224 | 28 | 12345 | 414 | if,Stat92E,betaTub56D,lbe,Rac2,salm,twi,Kr,N,E(spl)mdelta-HLH,ttk,hts,sty,aop,ap,pnt,slou,slow,eya,Ubx,Tl,Pak3,tin,CG13366,Dr,Rho1,wech,Sin3A | | 2.043e-09 | -20.01 | spi (spitz) | protein interactions | 35253 | 73 | 18 | 9204 | 412 | Sox21a,spen,Prosap,salm,dpp,Egfr,S,MFS14,NetB,CG13784,sty,CycE,ec,pbl,aos,Socs16D,edl,jing | | 2.064e-09 | -20.00 | hematopoietic or lymphoid organ development | biological process | GO:0048534 | 95 | 18 | 12345 | 414 | tin,Rho1,sens-2,pnt,Ser,ocm,ush,Pvf3,sty,stwl,lz,dpp,Rac2,if,Stat92E,N,ytr,Zif | | 2.064e-09 | -20.00 | immune system development | biological process | GO:0002520 | 95 | 18 | 12345 | 414 | if,Stat92E,Rac2,Zif,ytr,N,sty,lz,dpp,stwl,Ser,pnt,Pvf3,ush,ocm,tin,Rho1,sens-2 | | 2.092e-09 | -19.99 | animal organ formation | biological process | GO:0048645 | 46 | 13 | 12345 | 414 | lz,NetB,dally,robo2,nuf,Ubx,N,ci,phyl,B-H2,hh,CtBP,slp1 | | 2.712e-09 | -19.73 | morphogenesis of a polarized epithelium | biological process | GO:0001738 | 108 | 19 | 12345 | 414 | Rho1,sdt,aos,spen,ft,pnt,ec,Egfr,ds,otk2,S,aop,sty,CadN,spi,N,bdg,fj,par-1 | | 2.933e-09 | -19.65 | establishment of planar polarity | biological process | GO:0001736 | 97 | 18 | 12345 | 414 | S,aop,sty,ds,Egfr,otk2,par-1,fj,N,bdg,CadN,spi,spen,Rho1,aos,pnt,ec,ft | | 2.933e-09 | -19.65 | establishment of tissue polarity | biological process | GO:0007164 | 97 | 18 | 12345 | 414 | ft,ec,pnt,aos,Rho1,spen,spi,CadN,N,bdg,fj,par-1,otk2,Egfr,ds,sty,aop,S | | 2.979e-09 | -19.63 | regulation of cell population proliferation | biological process | GO:0042127 | 229 | 28 | 12345 | 414 | EcR,stg,Sin3A,ham,Doa,Tl,ft,Smox,cpo,pnt,shn,dpp,Myb,Egfr,osa,cas,ey,Kr,spi,Mmp2,N,crb,E(spl)mdelta-HLH,CycE,Stat92E,l(3)mbt,Utx,hh | | 3.110e-09 | -19.59 | larval development | biological process | GO:0002164 | 159 | 23 | 12345 | 414 | hh,Stat92E,N,Zif,Dad,sty,stwl,sas,ap,dpp,lz,pnt,trn,scb,Ser,ft,ocm,ush,eya,Tl,wech,Sin3A,EcR | | 3.595e-09 | -19.44 | vg (vestigial) | protein interactions | 36421 | 31 | 12 | 9204 | 412 | ap,salm,Ser,Bx,Egfr,ft,dpp,ci,N,sno,tkv,spi | | 3.749e-09 | -19.40 | RNA polymerase II regulatory region sequence-specific DNA binding | molecular function | GO:0000977 | 254 | 29 | 12176 | 396 | CG4360,CG31224,ci,ey,NFAT,Stat92E,Utx,HGTX,Ubx,aop,Pits,ocm,tin,ap,Dr,EcR,Doc2,ttk,Lim1,MESR3,bi,bowl,slou,Doc1,E(spl)mdelta-HLH,B-H2,dve,TfAP-2,CG11247 | | 4.119e-09 | -19.31 | regulation of cell fate commitment | biological process | GO:0010453 | 40 | 12 | 12345 | 414 | aop,Dr,S6kII,pnt,l(3)mbt,salm,lbe,N,ttk,ush,Ubx,eya | | 4.367e-09 | -19.25 | regulation of locomotion | biological process | GO:0040012 | 111 | 19 | 12345 | 414 | Ptpmeg,Pvf3,scb,NetB,Dscam1,Sin3A,Rho1,Ten-m,spi,CadN,ttk,hh,mmy,Lrt,NetA,CG44774,robo2,Egfr,aop | | 4.544e-09 | -19.21 | epithelial cell development | biological process | GO:0002064 | 361 | 36 | 12345 | 414 | gus,EcR,Fas3,Rho1,nudE,CG41099,lama,tyn,shd,crb,N,ci,if,Rac2,mam,neo,tkv,drk,cta,eya,jing,Myb,dpp,spoon,Egfr,aop,sty,RhoGEF2,Syp,S,ttk,Stat92E,CycE,hh,par-1,bbg | | 5.269e-09 | -19.06 | developmental growth | biological process | GO:0048589 | 191 | 25 | 12345 | 414 | RhoGEF2,otk2,wnd,mle,dpp,Rac2,par-1,hh,fj,Prosap,ttk,N,shd,Dad,CadN,Ten-m,bi,Imp,Dscam1,Rho1,tkv,jing,scb,ft,Ptpmeg | | 5.269e-09 | -19.06 | growth | biological process | GO:0040007 | 191 | 25 | 12345 | 414 | bi,Ten-m,Dscam1,Imp,tkv,Rho1,jing,scb,ft,Ptpmeg,RhoGEF2,wnd,otk2,mle,dpp,par-1,hh,Rac2,Prosap,fj,ttk,N,shd,Dad,CadN | | 5.729e-09 | -18.98 | tissue migration | biological process | GO:0090130 | 164 | 23 | 12345 | 414 | EcR,Rho1,Sin3A,nudE,drk,pbl,jing,pnt,dpp,dlp,spoon,Egfr,dally,aop,shd,N,if,Stat92E,bbg,kni,hh,salm,par-1 | | 6.717e-09 | -18.82 | Egfr (Epidermal growth factor receptor) | protein interactions | 37455 | 158 | 26 | 9204 | 412 | Sox21a,spen,psq,eya,dpp,ci,S,drk,Egfr,spi,gus,kni,Fas2,CG33993,Ptpmeg,sty,CycE,aop,Ubx,N,hh,Stat92E,aos,jing,pnt,tkv | | 7.007e-09 | -18.78 | RNA polymerase II regulatory region DNA binding | molecular function | GO:0001012 | 261 | 29 | 12176 | 396 | ocm,tin,Dr,EcR,ap,ci,CG4360,CG31224,HGTX,Pits,Ubx,aop,NFAT,Stat92E,ey,Utx,B-H2,dve,Doc1,E(spl)mdelta-HLH,CG11247,TfAP-2,Lim1,ttk,Doc2,slou,bowl,MESR3,bi | | 9.857e-09 | -18.44 | epithelial cell migration | biological process | GO:0010631 | 155 | 22 | 12345 | 414 | N,shd,par-1,hh,kni,salm,bbg,Stat92E,if,dpp,aop,dally,spoon,dlp,Egfr,drk,pnt,jing,Sin3A,Rho1,EcR,nudE | | 1.032e-08 | -18.39 | eye-antennal disc morphogenesis | biological process | GO:0007455 | 43 | 12 | 12345 | 414 | Egfr,aos,so,dpp,pnt,B-H2,oc,Stat92E,N,bab1,ey,eya | | 1.077e-08 | -18.35 | regulation of intracellular signal transduction | biological process | GO:1902531 | 339 | 34 | 12345 | 414 | PsGEF,Lpin,PEK,Fbxl7,spi,ci,crb,N,Socs16D,ImpL2,grnd,mbt,aos,pbl,dpp,RapGAP1,S6kII,spoon,ds,Egfr,wnd,RhoGEF2,Mos,sty,Hipk,NFAT,CG43658,par-1,MESR3,wcy,CG5522,CG42684,drk,ft | | 1.083e-08 | -18.34 | GO\_MORPHOGENESIS\_OF\_AN\_EPITHELIUM | MSigDB lists | GO\_MORPHOGENESIS\_OF\_AN\_EPITHELIUM | 99 | 18 | 4161 | 153 | Rho1,sdt,ds,Utx,ap,N,Smox,NetB,Ser,Lim1,dpp,drk,hh,so,TfAP-2,cta,ft,wech | | 1.113e-08 | -18.31 | epithelium migration | biological process | GO:0090132 | 156 | 22 | 12345 | 414 | pnt,jing,drk,nudE,Rho1,Sin3A,EcR,bbg,kni,hh,salm,par-1,if,Stat92E,N,shd,aop,Egfr,spoon,dlp,dally,dpp | | 1.192e-08 | -18.25 | stem cell differentiation | biological process | GO:0048863 | 72 | 15 | 12345 | 414 | N,Kr,spi,CycE,EcR,dpp,cas,S,osa,Egfr,tin,robo2,ham,spen,Dr | | 1.192e-08 | -18.25 | neuron fate commitment | biological process | GO:0048663 | 72 | 15 | 12345 | 414 | drk,spi,ey,ttk,bdg,N,exex,phyl,oc,CycE,pnt,mam,Egfr,aop,S | | 1.206e-08 | -18.23 | membrane-bounded organelle | cellular component | GO:0043227 | 4716 | 216 | 11940 | 407 | B-H2,Df31,ScsbetaG,Smox,Tim17b,enok,Ykt6,ctrip,CG41099,Tet,flfl,Hsc70-3,E(spl)mdelta-HLH,phtf,Lpin,tna,ocm,pbl,CG11498,CG1142,jigr1,Sin3A,gus,Pex1,CG34172,CG9220,mof,Gmap,CG34384,nuf,CG11752,ytr,l(3)mbt,TfIIS,CG3253,Acf,Chro,yps,Doc1,CG41128,dve,Pcl,N,Fur1,PhKgamma,Bx,drk,spen,Mitofilin,Acsl,IntS6,Doa,spoon,CG4360,jing,HGTX,stg,phyl,aop,Mondo,CG31875,Hph,rush,CG44774,NiPp1,lbe,tin,AGO2,TfAP-2,CycE,Fas3,heph,nudE,Haspin,CG5815,spi,exex,CG14646,Pits,Etl1,ush,lz,salm,Doc2,ND-B14.5AL,CG8485,mam,Ptpmeg,sno,Imp,sens-2,CG8441,osa,CtBP,Hipk,FoxL1,JIL-1,S,Syx1A,CG43313,Dr,CG1271,RhoGAP15B,CG11247,CG31712,CG5728,CG2162,Ubx,CG8531,tkv,wcy,retm,Cirl,edl,CG6051,ey,Hip1,Hsc70Cb,CG32772,pAbp,cnc,CG31224,mub,Utx,CG4538,cpo,disco,Cdk12,CG9650,sbb,CG9855,hh,Kah,NFAT,CG13029,EcR,Rac2,slou,Cyt-c1,CG13784,twi,cas,comm,dbr,mle,Rbfox1,CG11030,Taf1,MESR3,bab1,Stat92E,oc,bbg,CG3919,Cnx99A,tou,shn,Ptr,Dtg,emp,CG6700,udd,l(3)L1231,ps,slp1,CG32758,ham,ap,bin3,Asator,lncRNA:roX1,Snx3,hang,Tif-IA,Zif,ci,bi,kni,Sox21a,psq,ScpX,CG18766,shd,Lim1,HmgZ,baf,ttk,Syp,Cyp1,Tlk,Mocs2B,Myb,CG17168,so,Nopp140,dpp,bora,elav,bowl,Ugt49C1,eya,stwl,bor,LManII,Rho1,hiw,Thd1,crb,sm,Psc,Tl,kdn,Kr,Tudor-SN,CG31612,pnt | | 1.224e-08 | -18.22 | establishment of ommatidial planar polarity | biological process | GO:0042067 | 62 | 14 | 12345 | 414 | pnt,ec,fj,ft,bdg,N,CadN,spi,S,aop,sty,ds,Egfr,aos | | 1.255e-08 | -18.19 | compound eye photoreceptor cell differentiation | biological process | GO:0001751 | 157 | 22 | 12345 | 414 | B-H2,oc,phyl,salm,hh,CadN,crb,E(spl)mdelta-HLH,bdg,N,ttk,Egfr,S,aop,lz,pnt,drk,eya,Doa,Ten-m,mbt,rau | | 1.315e-08 | -18.15 | head development | biological process | GO:0060322 | 144 | 21 | 12345 | 414 | Smox,Fas2,Ptpmeg,slp1,jing,cnc,enok,Dscam1,Dmtn,mbt,EcR,Dr,ey,hh,oc,Prosap,PsGEF,dpp,cas,disco,robo2 | | 1.559e-08 | -17.98 | columnar/cuboidal epithelial cell differentiation | biological process | GO:0002065 | 344 | 34 | 12345 | 414 | lama,tin,nudE,gus,EcR,Fas3,Rho1,if,Sox21a,Rac2,mam,shd,N,ci,exex,jing,eya,drk,tkv,phyl,Stat92E,CycE,hh,par-1,bbg,ttk,spoon,Egfr,aop,sty,S,Syp,Myb,dpp | | 1.697e-08 | -17.89 | establishment or maintenance of cell polarity | biological process | GO:0007163 | 217 | 26 | 12345 | 414 | cnc,ec,pnt,cta,ft,nudE,heph,aos,sdt,Rho1,fj,par-1,Zif,spi,CadN,crb,bdg,N,wake,KP78a,spoon,ds,Egfr,sty,aop,S | | 2.152e-08 | -17.65 | positive regulation of multicellular organismal process | biological process | GO:0051240 | 266 | 29 | 12345 | 414 | nudE,stg,Imp,Arp2,tkv,Sox102F,mbt,EcR,ec,pnt,Pak3,Ubx,aop,cv,wake,Egfr,Usp2,dpp,lz,tsl,mmy,Prosap,oc,ttk,Hipk,N,spi,egr,ey | | 2.228e-08 | -17.62 | negative regulation of cell differentiation | biological process | GO:0045596 | 135 | 20 | 12345 | 414 | pnt,Ser,ush,Ptpmeg,eya,Fas2,goe,aos,enok,EcR,salm,Sox21a,Stat92E,N,ttk,CadN,osa,aop,Egfr,S6kII | | 2.370e-08 | -17.56 | GO\_TISSUE\_MORPHOGENESIS | MSigDB lists | GO\_TISSUE\_MORPHOGENESIS | 128 | 20 | 4161 | 153 | so,hh,drk,wech,ft,cta,TfAP-2,sdt,fj,Rho1,Lim1,Ser,dpp,Smox,N,NetB,ds,ap,Utx,if | | 2.390e-08 | -17.55 | R7 cell differentiation | biological process | GO:0045466 | 46 | 12 | 12345 | 414 | rau,lz,aop,Ten-m,ttk,N,E(spl)mdelta-HLH,drk,CadN,salm,pnt,phyl | | 2.575e-08 | -17.48 | Scr (Sex combs reduced) | protein interactions | 40833 | 43 | 13 | 9204 | 412 | salm,Doc2,eya,CycE,ap,slou,tin,kni,pnt,Pcl,twi,mam,tna | | 2.811e-08 | -17.39 | formation of primary germ layer | biological process | GO:0001704 | 38 | 11 | 12345 | 414 | dpp,tkv,tin,Ubx,spi,pbl,slp1,lbe,twi,slou,mam | | 2.959e-08 | -17.34 | transcription regulatory region sequence-specific DNA binding | molecular function | GO:0000976 | 329 | 32 | 12176 | 396 | MESR3,bowl,Doc2,disco,Lim1,CG11247,Doc1,B-H2,slp1,Stat92E,aop,Pits,CG4360,ci,EcR,Dr,bi,slou,ttk,TfAP-2,E(spl)mdelta-HLH,dve,so,Utx,NFAT,ey,Ubx,HGTX,CG31224,ap,ocm,tin | | 3.162e-08 | -17.27 | eye photoreceptor cell differentiation | biological process | GO:0001754 | 165 | 22 | 12345 | 414 | rau,mbt,Doa,Ten-m,drk,eya,pnt,lz,S,aop,Egfr,crb,N,bdg,E(spl)mdelta-HLH,ttk,CadN,salm,hh,B-H2,oc,phyl | | 3.197e-08 | -17.26 | sex differentiation | biological process | GO:0007548 | 100 | 17 | 12345 | 414 | tin,Dr,Doa,eya,Fas2,bab1,wb,lz,dpp,robo2,Egfr,otk2,aop,N,TfAP-2,hh,salm | | 3.456e-08 | -17.18 | upd1 (unpaired 1) | protein interactions | 32813 | 77 | 17 | 9204 | 412 | crb,Dad,Rho1,Ten-m,dally,NFAT,dpp,pAbp,Egfr,cpo,Imp,CtBP,Ser,N,hh,Stat92E,tkv | | 3.459e-08 | -17.18 | Snr1 (Snf5-related 1) | protein interactions | 40657 | 137 | 23 | 9204 | 412 | fax,CycE,CG1737,l(3)L1231,Tet,aos,Stat92E,tkv,CG43658,cv,HmgZ,N,bab1,osa,lawc,Zif,Ank,esn,ci,dpp,sbb,Egfr,S | | 3.747e-08 | -17.10 | germ-line stem cell population maintenance | biological process | GO:0030718 | 78 | 15 | 12345 | 414 | fax,N,CG46339,Inx2,Dad,mam,Ser,Stat92E,stwl,tkv,enok,Tif-IA,dpp,Egfr,dally | | 3.799e-08 | -17.09 | stem cell fate commitment | biological process | GO:0048865 | 39 | 11 | 12345 | 414 | cas,S,osa,Dr,spen,ham,Egfr,CycE,N,spi,Kr | | 4.233e-08 | -16.98 | gastrulation | biological process | GO:0007369 | 90 | 16 | 12345 | 414 | tkv,Rho1,stg,tin,cta,Ubx,slou,slp1,pbl,dpp,Dtg,RhoGEF2,spi,mam,lbe,twi | | 4.550e-08 | -16.91 | positive regulation of developmental process | biological process | GO:0051094 | 259 | 28 | 12345 | 414 | Prosap,oc,mmy,ImpL2,tsl,ey,spi,Hipk,ci,N,ttk,Egfr,cv,aop,lz,dpp,Usp2,pnt,ec,Ubx,Pak3,nudE,EcR,mbt,tkv,Arp2,Imp,stg | | 4.873e-08 | -16.84 | leg disc proximal/distal pattern formation | biological process | GO:0007479 | 18 | 8 | 12345 | 414 | jing,pnt,B-H2,Lim1,disco,Egfr,ap,dpp | | 4.892e-08 | -16.83 | response to chemical | biological process | GO:0042221 | 1191 | 75 | 12345 | 414 | beat-IIIc,dally,Cnx99A,sas,comm,dpp,ap,hh,Stat92E,RhoGAP93B,Karl,enok,Dscam1,tkv,Drat,Ptpmeg,Mondo,robo2,Psc,uzip,NiPp1,Rac2,Lrt,ImpL2,Hydr2,Con,ckn,hang,N,wun2,spi,Hsc70-3,Kr,spen,ras,EcR,NetB,Cdk5alpha,Smox,aop,hiw,Gbs-70E,S,hts,dlp,Egfr,Utx,Mmp2,sbb,Toll-7,pnt,jing,trn,wb,eya,drk,PEK,sm,NetA,Acsl,Lpin,shn,grnd,if,Bx,Dad,CadN,Ten-m,Fas3,Rho1,Hph,scb,Ptp99A,Tl,Fas2 | | 5.299e-08 | -16.75 | cellular response to stimulus | biological process | GO:0051716 | 1467 | 87 | 12345 | 414 | egr,JIL-1,CG32758,ttk,Tlk,phyl,Utx,Egfr,otk2,wnd,hiw,aop,Pvf3,drk,PhKgamma,pnt,smal,CCKLR-17D3,Atg17,Toll-7,Thd1,Doa,nAChRalpha4,Dad,ci,Socs16D,grnd,tsl,shn,Lpin,KP78a,osa,PEK,CaMKII,RhoGAP15B,CG4325,Tl,CG8485,Ser,Hph,Rho1,RhoGAP102A,Inx3,Pli,RhoGAP93B,ctrip,Stat92E,par-1,hh,dpp,stwl,Cnx99A,dally,wake,Ank,CG11873,Cirl,cta,Drat,cnc,RhoGAP18B,sno,tkv,Hsc70-3,wun2,spi,CG34384,crb,N,ckn,Hydr2,hang,mam,Lrt,flfl,Rac2,ITP,mof,Smox,Cdk5alpha,EcR,Rhp,mbt,gus,RhoGAP16F,spen,Haspin | | 5.354e-08 | -16.74 | double-stranded DNA binding | molecular function | GO:0003690 | 391 | 35 | 12176 | 396 | E(spl)mdelta-HLH,dve,TfAP-2,mle,ttk,bi,slou,bab1,tin,ocm,ap,CG31224,NFAT,ey,Utx,so,HGTX,Ubx,Doc1,B-H2,CG11247,disco,Doc2,Lim1,MESR3,bowl,Thd1,EcR,Dr,CG4360,ci,Stat92E,slp1,Pits,aop | | 5.912e-08 | -16.64 | anterior/posterior pattern specification | biological process | GO:0009952 | 200 | 24 | 12345 | 414 | hh,kni,tsl,par-1,oc,E(spl)mdelta-HLH,N,ci,psq,Kr,Dad,Pat1,shn,dpp,bowl,slp1,Ser,cnc,Ubx,bi,tkv,aos,EcR,gus | | 6.238e-08 | -16.59 | cell fate commitment involved in formation of primary germ layer | biological process | GO:0060795 | 25 | 9 | 12345 | 414 | spi,Ubx,slou,mam,twi,lbe,dpp,tkv,tin | | 6.824e-08 | -16.50 | GO\_EAR\_DEVELOPMENT | MSigDB lists | GO\_EAR\_DEVELOPMENT | 47 | 12 | 4161 | 153 | Rac2,so,lbk,hh,ft,NetB,N,if,ds,TfAP-2,dpp,Ser | | 6.987e-08 | -16.48 | sequence-specific double-stranded DNA binding | molecular function | GO:1990837 | 359 | 33 | 12176 | 396 | Dr,EcR,ci,CG4360,aop,Pits,slp1,Stat92E,B-H2,Doc1,CG11247,Lim1,Doc2,disco,bowl,MESR3,tin,ocm,ap,CG31224,Ubx,HGTX,Utx,so,NFAT,ey,dve,E(spl)mdelta-HLH,TfAP-2,ttk,bab1,slou,bi | | 7.152e-08 | -16.45 | AACTTT\_UNKNOWN | MSigDB lists | AACTTT\_UNKNOWN | 421 | 38 | 4161 | 153 | ras,DIP-epsilon,Tlk,Imp,S6kII,Trpm,CaMKII,Arl4,smal,CG32758,CG45050,LRP1,Syx1A,N,Smox,Ser,CG34384,dlp,CtBP,TfAP-2,cta,NKAIN,CG5830,CG9281,SamDC,JIL-1,Ank,sdt,fne,Pat1,Utx,ap,NetB,pbl,Dscam1,Mhcl,mmy,Lim1 | | 1.003e-07 | -16.12 | plasma membrane region | cellular component | GO:0098590 | 314 | 31 | 11940 | 407 | Stat92E,mgl,hts,if,nuf,Syx1A,dally,Inx2,Fas3,sdt,Rho1,ft,Fas2,sas,crb,Arl4,nAChRalpha4,Egfr,Ser,scb,Tl,cta,ds,dpr7,Ten-m,par-1,CaMKII,RhoGEF2,pbl,Prosap,tyn | | 1.057e-07 | -16.06 | reproductive structure development | biological process | GO:0048608 | 84 | 15 | 12345 | 414 | Egfr,tin,robo2,Dr,otk2,aop,lz,wb,hh,salm,eya,dve,Fas2,N,TfAP-2 | | 1.057e-07 | -16.06 | reproductive system development | biological process | GO:0061458 | 84 | 15 | 12345 | 414 | dve,eya,Fas2,TfAP-2,N,wb,salm,hh,lz,otk2,Dr,tin,robo2,Egfr,aop | | 1.122e-07 | -16.00 | cell junction | cellular component | GO:0030054 | 160 | 21 | 11940 | 407 | if,CadN,Inx2,Rho1,Fas3,sdt,otk2,Lrt,ckn,wech,bbg,smal,Fbxl7,N,Inx3,mbt,crb,Lac,nrv2,slow,scb | | 1.233e-07 | -15.91 | spen (split ends) | protein interactions | 44205 | 27 | 10 | 9204 | 412 | spi,pnt,smash,Hipk,CG2865,dpp,N,Egfr,CycE,aop | | 1.244e-07 | -15.90 | organelle | cellular component | GO:0043226 | 5470 | 238 | 11940 | 407 | slou,Cyt-c1,Rac2,CG13029,EcR,dbr,comm,cas,Mhcl,CG13784,twi,Rbfox1,CG11030,mle,MESR3,Taf1,bab1,mub,CG31224,cnc,pAbp,CG32772,CG9650,Cdk12,disco,CG4538,Utx,cpo,RhoGAP93B,hh,CG9855,sbb,NFAT,Kah,slp1,ps,l(3)L1231,CG32758,bin3,ap,ham,Snx3,lncRNA:roX1,Asator,bbg,CG3919,Cnx99A,oc,Stat92E,Ptr,tou,shn,emp,Dtg,udd,CG6700,l(3)neo38,Myb,Mocs2B,Tlk,so,Nopp140,CG17168,dpp,elav,bora,Zif,Tif-IA,hang,Sox21a,kni,bi,ci,shd,Lim1,CG18766,Arp2,ScpX,psq,baf,Cyp1,ttk,Syp,HmgZ,crb,sm,Thd1,Tl,Psc,kdn,RhoGEF2,pnt,CG31612,Tudor-SN,Kr,eya,Ugt49C1,bowl,stwl,hiw,Rho1,LManII,Fas2,bor,phtf,E(spl)mdelta-HLH,Hsc70-3,flfl,ocm,tna,Lpin,Sin3A,jigr1,Hydr2,CG1142,CG11498,Prosap,pbl,lawc,Df31,B-H2,Myo10A,Tim17b,ScsbetaG,Smox,Ykt6,enok,Tet,CG41099,ctrip,Pcl,CG41128,dve,CG14535,Bx,drk,PhKgamma,Fur1,N,Doa,Acsl,IntS6,spen,Mitofilin,PEK,CG34172,gus,Pex1,mof,hts,CG9220,l(3)mbt,CG11752,CG34384,nuf,ytr,Gmap,wake,yps,Doc1,Chro,Acf,CG3253,TfIIS,nudE,Pits,CG17778,CG11486,exex,CG14646,CG5815,spi,Haspin,salm,Doc2,lz,Etl1,ush,CG13366,Imp,sno,Ptpmeg,mam,CG8485,ND-B14.5AL,Ank,jing,CG4360,HGTX,spoon,Hph,CG31875,rush,Mondo,phyl,aop,stg,lbe,tin,NiPp1,CG44774,heph,Fas3,TfAP-2,CycE,AGO2,CG2162,CG5728,CG8531,Ubx,retm,Cirl,tkv,wcy,Hsc70Cb,Hip1,ey,edl,CG6051,CtBP,osa,CG8441,sens-2,smg,FoxL1,Hipk,betaTub56D,Pat1,CG43313,Syx1A,RpS4,S,JIL-1,CG31712,CG1271,RhoGAP15B,CG11247,Dr | | 1.322e-07 | -15.84 | axonal fasciculation | biological process | GO:0007413 | 20 | 8 | 12345 | 414 | mmy,CadN,Fas2,Mmp2,Con,uzip,Fas3,Dscam1 | | 1.329e-07 | -15.83 | regulation of cell migration | biological process | GO:0030334 | 74 | 14 | 12345 | 414 | scb,Lrt,hh,Pvf3,Ptpmeg,spi,ttk,Egfr,robo2,aop,Rho1,CG44774,Sin3A,Dscam1 | | 1.348e-07 | -15.82 | ectoderm development | biological process | GO:0007398 | 27 | 9 | 12345 | 414 | N,ush,cta,spi,mam,hh,shn,dpp,Dr | | 1.412e-07 | -15.77 | GO\_SENSORY\_ORGAN\_DEVELOPMENT | MSigDB lists | GO\_SENSORY\_ORGAN\_DEVELOPMENT | 116 | 18 | 4161 | 153 | Lim1,Ser,dpp,ap,if,ds,Smox,N,NetB,fj,TfAP-2,ft,hh,lbk,so,ras,Rac2,tkv | | 1.494e-07 | -15.72 | post-embryonic hemopoiesis | biological process | GO:0035166 | 44 | 11 | 12345 | 414 | stwl,lz,dpp,sty,ocm,N,ush,Zif,pnt,Ser,Stat92E | | 1.494e-07 | -15.72 | larval lymph gland hemopoiesis | biological process | GO:0035167 | 44 | 11 | 12345 | 414 | lz,dpp,stwl,sty,Zif,ush,ocm,N,Ser,Stat92E,pnt | | 1.690e-07 | -15.59 | maintenance of cell number | biological process | GO:0098727 | 99 | 16 | 12345 | 414 | Ser,CG46339,fax,Inx2,tkv,enok,CycE,Stat92E,mam,Dad,N,Egfr,dally,dpp,Tif-IA,stwl | | 1.690e-07 | -15.59 | stem cell population maintenance | biological process | GO:0019827 | 99 | 16 | 12345 | 414 | mam,CycE,Stat92E,N,Dad,Egfr,dally,stwl,dpp,Tif-IA,Ser,CG46339,fax,Inx2,tkv,enok | | 1.895e-07 | -15.48 | immune system process | biological process | GO:0002376 | 328 | 31 | 12345 | 414 | Toll-7,sens-2,Rho1,tin,eya,Tl,ush,Pvf3,ocm,Ser,Cdk5alpha,pnt,lz,dpp,stwl,AGO2,sty,Zif,egr,ytr,N,ci,Mmp2,Stat92E,Pli,if,Tlk,Rac2,par-1,hh,bbg | | 2.058e-07 | -15.40 | intracellular part | cellular component | GO:0044424 | 6762 | 280 | 11940 | 407 | osa,FoxL1,CG43313,Syx1A,S,JIL-1,CG31712,RhoGAP15B,Dr,CG2162,NetA,CG5728,cta,Cirl,tkv,Ank,CG4360,jing,spoon,aop,stg,DnaJ-H,lbe,tin,NiPp1,CG44774,CG1344,RapGAP1,CycE,TfAP-2,AGO2,nudE,Dad,CG10283,fj,CG11486,Haspin,salm,Doc2,CaMKII,CG8485,ND-B14.5AL,PEK,CG34172,gus,mof,CG11752,ytr,Gmap,Doc1,Acf,Chro,TfIIS,CG42684,Fur1,Df31,B-H2,Tim17b,ScsbetaG,ctrip,phtf,E(spl)mdelta-HLH,Hsc70-3,ocm,Sin3A,jigr1,CG1142,pbl,sty,CG17224,Rho1,LManII,bor,Thd1,Tl,RhoGEF2,Tudor-SN,Kr,Zif,hang,Sox21a,bi,shd,NetB,CG18766,ScpX,Cyp1,HmgZ,CD98hc,Myb,Tlk,Mocs2B,Nopp140,CG17168,dpp,elav,bora,bbg,emp,udd,CG6700,slp1,l(3)L1231,ps,ap,Snx3,CG31224,cnc,pAbp,fne,CG32772,SamDC,Cdk12,mgl,CG4538,Utx,cpo,CG9855,pon,NFAT,Kah,Cyt-c1,dbr,Mhcl,twi,Rbfox1,par-1,mle,KP78a,Taf1,MESR3,CtBP,Ppa,CG8441,sens-2,smg,Hipk,betaTub56D,Pat1,RpS4,CG1271,CG11247,Pli,CG8531,Ubx,Ser,retm,wcy,Hsc70Cb,Hip1,ey,CG6051,edl,HGTX,rush,Hph,CG31875,Mondo,phyl,heph,Fas3,Usp2,Pits,CG14646,exex,CG17778,spi,CG5815,lz,Etl1,ush,CG13366,Imp,sno,Ptpmeg,mam,Pex1,hts,CG9220,l(3)mbt,CG34384,ras,nuf,wake,yps,CG6330,CG3253,ckn,RhoGAP16F,Pcl,CG41128,dve,CG14535,Bx,drk,PhKgamma,N,Doa,Acsl,IntS6,Mitofilin,spen,corn,CG6752,Myo10A,Pak3,Smox,Ykt6,enok,Tet,CG41099,flfl,tna,Lpin,Atg17,CG11498,lawc,eya,Ugt49C1,bowl,stwl,hiw,Fas2,crb,sm,NTPase,Psc,kdn,pnt,RhoGAP18B,CG31612,Tif-IA,kni,ci,Lim1,psq,Arp2,baf,ttk,Syp,l(3)neo38,CG2201,Arl4,so,Karl,PsGEF,CG3919,Cnx99A,oc,Stat92E,shn,Ptr,tou,fax,CG32758,bin3,ham,Fbxl7,Mos,lncRNA:roX1,Asator,LRP1,mub,CG9650,disco,RhoGAP93B,hh,sbb,sdt,slou,mbt,Rac2,CG13029,EcR,comm,cas,scb,CG13784,CG11030,bab1 | | 2.058e-07 | -15.40 | intracellular | cellular component | GO:0005622 | 6762 | 280 | 11940 | 407 | Taf1,MESR3,KP78a,mle,par-1,Rbfox1,twi,Mhcl,dbr,Cyt-c1,Kah,NFAT,pon,CG9855,CG4538,Utx,cpo,mgl,Cdk12,SamDC,pAbp,fne,CG32772,cnc,CG31224,Snx3,ap,ps,l(3)L1231,slp1,CG6700,udd,emp,bbg,bora,elav,dpp,CG17168,Nopp140,Tlk,Mocs2B,Myb,HmgZ,CD98hc,Cyp1,ScpX,CG18766,NetB,shd,bi,Sox21a,hang,Zif,Kr,Tudor-SN,RhoGEF2,Tl,Thd1,bor,LManII,Rho1,sty,CG17224,pbl,CG1142,jigr1,Sin3A,ocm,Hsc70-3,E(spl)mdelta-HLH,phtf,ctrip,ScsbetaG,Tim17b,B-H2,Df31,Fur1,CG42684,TfIIS,Chro,Acf,Doc1,Gmap,CG11752,ytr,mof,gus,CG34172,PEK,ND-B14.5AL,CG8485,CaMKII,salm,Doc2,Haspin,CG11486,fj,CG10283,Dad,nudE,AGO2,RapGAP1,TfAP-2,CycE,CG1344,CG44774,NiPp1,lbe,tin,DnaJ-H,stg,aop,spoon,CG4360,jing,Ank,tkv,Cirl,cta,CG5728,CG2162,NetA,Dr,RhoGAP15B,CG31712,JIL-1,S,Syx1A,CG43313,FoxL1,osa,bab1,CG11030,CG13784,scb,cas,comm,CG13029,EcR,Rac2,mbt,slou,sdt,sbb,hh,RhoGAP93B,disco,CG9650,LRP1,mub,Asator,Mos,lncRNA:roX1,Fbxl7,ham,bin3,CG32758,fax,tou,shn,Ptr,Stat92E,oc,Cnx99A,CG3919,PsGEF,Karl,so,Arl4,CG2201,l(3)neo38,ttk,baf,Syp,psq,Arp2,Lim1,ci,kni,Tif-IA,CG31612,RhoGAP18B,pnt,kdn,Psc,NTPase,crb,sm,Fas2,hiw,stwl,Ugt49C1,bowl,eya,lawc,CG11498,Atg17,Lpin,tna,flfl,CG41099,Tet,enok,Ykt6,Smox,Pak3,Myo10A,CG6752,corn,spen,Mitofilin,IntS6,Acsl,Doa,N,PhKgamma,Bx,drk,dve,CG14535,CG41128,Pcl,RhoGAP16F,ckn,CG3253,CG6330,yps,wake,CG34384,ras,nuf,l(3)mbt,CG9220,hts,Pex1,mam,Ptpmeg,sno,Imp,CG13366,Etl1,ush,lz,spi,CG5815,CG14646,exex,CG17778,Pits,Usp2,Fas3,heph,phyl,Mondo,Hph,CG31875,rush,HGTX,CG6051,edl,ey,Hip1,Hsc70Cb,wcy,retm,Ser,Ubx,CG8531,Pli,CG1271,CG11247,RpS4,Pat1,betaTub56D,Hipk,smg,sens-2,CG8441,Ppa,CtBP | | 2.074e-07 | -15.39 | neuron projection fasciculation | biological process | GO:0106030 | 21 | 8 | 12345 | 414 | Con,Mmp2,Fas2,CadN,mmy,Dscam1,Fas3,uzip | | 2.089e-07 | -15.38 | enzyme linked receptor protein signaling pathway | biological process | GO:0007167 | 154 | 20 | 12345 | 414 | smal,pnt,drk,Pvf3,Smox,spen,tkv,Socs16D,phyl,tsl,Dad,spi,ttk,ckn,dally,Egfr,hiw,aop,dpp,shn | | 2.310e-07 | -15.28 | regulation of biological quality | biological process | GO:0065008 | 1352 | 80 | 12345 | 414 | fj,ImpL2,Rac2,Hsc70-3,ey,Lac,crb,N,PsGEF,CG13784,pbl,Trpm,CG46339,tyn,flz,EcR,mbt,Arp2,Sin3A,CycE,mmy,Nhe2,Ank,dally,Syp,RhoGEF2,dpp,Cnx99A,cnc,Mondo,Cirl,cta,scaf6,Pak3,tkv,stg,if,Snx3,Npc1a,CadN,shd,Dad,CaMKII,Lpin,Acsl,smg,scb,slow,ush,Fas2,nw,Fur1,Ten-m,Rho1,Prosap,CG9650,egr,Syx1A,JIL-1,Hipk,NFAT,ttk,spoon,Egfr,ds,wnd,hts,disco,hiw,Myb,nrv2,S6kII,Pvf3,drk,ft,Gyf,nAChRalpha4,Atg17,neo,bou | | 2.446e-07 | -15.22 | formation of anatomical boundary | biological process | GO:0048859 | 46 | 11 | 12345 | 414 | ft,N,CtBP,phyl,B-H2,NetB,lz,nuf,dally,ds,robo2 | | 2.491e-07 | -15.21 | intracellular organelle | cellular component | GO:0043229 | 5391 | 234 | 11940 | 407 | Ubx,CG8531,CG5728,CG2162,Hip1,ey,Hsc70Cb,edl,CG6051,Cirl,retm,wcy,tkv,FoxL1,smg,betaTub56D,Hipk,osa,CtBP,sens-2,CG8441,CG1271,CG11247,RhoGAP15B,CG31712,Dr,Syx1A,CG43313,Pat1,S,JIL-1,RpS4,CG14646,exex,CG11486,CG17778,Pits,Haspin,CG5815,spi,nudE,sno,Ptpmeg,Imp,ND-B14.5AL,mam,CG8485,lz,Doc2,salm,CG13366,ush,Etl1,rush,Hph,CG31875,stg,phyl,Mondo,aop,Ank,spoon,jing,CG4360,HGTX,Fas3,heph,AGO2,CycE,TfAP-2,NiPp1,tin,lbe,CG44774,Pcl,CG41128,CG14535,dve,Doa,spen,Mitofilin,Acsl,IntS6,Bx,drk,N,PhKgamma,Fur1,CG9220,mof,hts,PEK,CG34172,Pex1,gus,Chro,Acf,yps,Doc1,TfIIS,CG3253,CG11752,ytr,nuf,CG34384,l(3)mbt,wake,Gmap,ocm,Lpin,tna,phtf,Hsc70-3,flfl,E(spl)mdelta-HLH,lawc,jigr1,Sin3A,CG11498,CG1142,pbl,Myo10A,Smox,ScsbetaG,Tim17b,Df31,B-H2,CG41099,Tet,ctrip,enok,Ykt6,Psc,Tl,Thd1,sm,crb,CG31612,pnt,Kr,Tudor-SN,kdn,RhoGEF2,eya,Ugt49C1,bowl,Rho1,hiw,bor,LManII,stwl,CG17168,so,Nopp140,l(3)neo38,Tlk,Mocs2B,Myb,elav,bora,dpp,kni,Sox21a,ci,bi,hang,Tif-IA,Zif,HmgZ,Syp,ttk,baf,Cyp1,Lim1,shd,ScpX,Arp2,psq,CG18766,CG32758,slp1,ps,l(3)L1231,Snx3,lncRNA:roX1,Asator,ap,bin3,ham,tou,shn,Ptr,bbg,CG3919,Cnx99A,Stat92E,oc,udd,CG6700,emp,comm,cas,dbr,CG13784,twi,Mhcl,slou,Cyt-c1,EcR,CG13029,Rac2,Taf1,MESR3,bab1,Rbfox1,CG11030,mle,Cdk12,CG9650,cpo,CG4538,Utx,disco,CG31224,mub,CG32772,pAbp,cnc,Kah,NFAT,CG9855,RhoGAP93B,hh,sbb | | 2.640e-07 | -15.15 | regulation of cell motility | biological process | GO:2000145 | 78 | 14 | 12345 | 414 | Pvf3,Ptpmeg,spi,ttk,scb,Lrt,hh,CG44774,Rho1,Dscam1,Sin3A,Egfr,robo2,aop | | 2.688e-07 | -15.13 | intracellular membrane-bounded organelle | cellular component | GO:0043231 | 4479 | 202 | 11940 | 407 | elav,bora,Mocs2B,Tlk,Myb,CG17168,so,Nopp140,Lim1,shd,ScpX,psq,CG18766,HmgZ,baf,Syp,Cyp1,ttk,hang,Zif,Tif-IA,kni,Sox21a,ci,bi,kdn,CG31612,pnt,Kr,Tudor-SN,Thd1,crb,sm,Psc,stwl,Rho1,hiw,bor,LManII,eya,bowl,Ugt49C1,Rbfox1,CG11030,mle,bab1,Taf1,MESR3,slou,Cyt-c1,EcR,CG13029,cas,comm,dbr,twi,CG13784,hh,CG9855,sbb,NFAT,Kah,CG31224,mub,pAbp,CG32772,cnc,CG9650,Cdk12,cpo,Utx,CG4538,disco,ap,bin3,ham,lncRNA:roX1,Asator,slp1,l(3)L1231,ps,emp,udd,CG6700,Cnx99A,CG3919,bbg,Stat92E,oc,shn,tou,lz,salm,Doc2,Etl1,ush,Ptpmeg,sno,Imp,ND-B14.5AL,CG8485,mam,nudE,CG14646,exex,Pits,Haspin,spi,CG5815,NiPp1,tin,lbe,CG44774,Fas3,heph,AGO2,TfAP-2,CycE,spoon,HGTX,CG4360,jing,CG31875,Hph,stg,phyl,Mondo,aop,retm,wcy,ey,Hip1,Hsc70Cb,edl,CG5728,CG2162,Ubx,CG8531,CG43313,S,JIL-1,RhoGAP15B,CG1271,CG11247,CG31712,Dr,CtBP,osa,sens-2,CG8441,FoxL1,Hipk,jigr1,Sin3A,CG11498,CG1142,pbl,phtf,flfl,Hsc70-3,E(spl)mdelta-HLH,ocm,Lpin,tna,enok,Ykt6,CG41099,Tet,ctrip,Df31,B-H2,ScsbetaG,Smox,Tim17b,Bx,drk,N,PhKgamma,Fur1,Doa,spen,Mitofilin,Acsl,IntS6,Pcl,dve,CG41128,ytr,CG11752,l(3)mbt,Gmap,Acf,Chro,yps,Doc1,TfIIS,CG3253,CG34172,Pex1,gus,CG9220,mof | | 2.714e-07 | -15.12 | hemocyte differentiation | biological process | GO:0042386 | 29 | 9 | 12345 | 414 | lz,dpp,sty,N,ytr,ush,pnt,Ser,Stat92E | | 2.714e-07 | -15.12 | regulation of cell-cell adhesion | biological process | GO:0022407 | 29 | 9 | 12345 | 414 | par-1,N,robo2,Ten-m,aop,mbt,wech,tkv,aos | | 2.735e-07 | -15.11 | Abd-B (Abdominal B) | protein interactions | 47763 | 60 | 14 | 9204 | 412 | Kr,pnt,twi,Pcl,lbe,mam,Ubx,Doc2,salm,eya,ap,slou,tin,kni | | 3.164e-07 | -14.97 | leg disc pattern formation | biological process | GO:0035223 | 22 | 8 | 12345 | 414 | Lim1,jing,pnt,B-H2,ap,dpp,disco,Egfr | | 3.693e-07 | -14.81 | apical part of cell | cellular component | GO:0045177 | 157 | 20 | 11940 | 407 | wun2,scb,Ser,sas,pio,crb,tyn,CG42684,RhoGEF2,Fbxl7,mgl,bbg,corn,Stat92E,sty,Rho1,sdt,nuf,dally,Inx2 | | 3.751e-07 | -14.80 | ttk (tramtrack) | protein interactions | 48317 | 80 | 16 | 9204 | 412 | hang,pbl,stg,phyl,Pits,Stat92E,CtBP,ttk,lz,aop,ocm,CycE,Doc1,lama,Rac2,CG17168 | | 3.763e-07 | -14.79 | ommatidial rotation | biological process | GO:0016318 | 30 | 9 | 12345 | 414 | aos,Egfr,ds,S,sty,CadN,spi,pnt,ec | | 3.763e-07 | -14.79 | negative regulation of animal organ morphogenesis | biological process | GO:0110111 | 30 | 9 | 12345 | 414 | N,ttk,dpp,shn,S6kII,tkv,Egfr,sty,aop | | 3.993e-07 | -14.73 | wing disc anterior/posterior pattern formation | biological process | GO:0048100 | 16 | 7 | 12345 | 414 | hh,bi,Ser,ci,tkv,dpp,shn | | 3.993e-07 | -14.73 | larval lymph gland hemocyte differentiation | biological process | GO:0035168 | 16 | 7 | 12345 | 414 | N,ush,dpp,lz,pnt,sty,Stat92E | | 4.224e-07 | -14.68 | aop (anterior open) | protein interactions | 33392 | 62 | 14 | 9204 | 412 | Egfr,S,Syx1A,N,stg,Sin3A,pnt,edl,spen,ttk,aop,lz,sty,eya | | 4.880e-07 | -14.53 | synaptic target recognition | biological process | GO:0008039 | 49 | 11 | 12345 | 414 | nrm,Con,Pvf3,Tl,slow,Fas3,NetB,Ten-m,Fur1,robo2,beat-IIIc | | 5.617e-07 | -14.39 | lymph gland development | biological process | GO:0048542 | 60 | 12 | 12345 | 414 | lz,dpp,stwl,tin,sty,ush,Zif,N,ocm,Stat92E,Ser,pnt | | 5.617e-07 | -14.39 | photoreceptor cell fate commitment | biological process | GO:0046552 | 60 | 12 | 12345 | 414 | bdg,N,ttk,ey,spi,drk,pnt,phyl,oc,S,aop,Egfr | | 5.689e-07 | -14.38 | ovarian follicle cell development | biological process | GO:0030707 | 327 | 30 | 12345 | 414 | lama,drk,eya,jing,Rho1,tkv,Fas3,EcR,gus,nudE,ci,N,ttk,shd,mam,bbg,Rac2,par-1,hh,if,CycE,Stat92E,dpp,Myb,S,Syp,aop,sty,Egfr,spoon | | 5.999e-07 | -14.33 | kinase activity | molecular function | GO:0016301 | 375 | 32 | 12176 | 396 | JIL-1,Egfr,Mos,CG9541,CG34384,PhKgamma,par-1,CycE,fj,CG8485,mbt,KP78a,Pak3,Taf1,CG1271,tkv,wnd,CG10082,Doa,CaMKII,Tlk,Cdk12,sdt,CG2201,Asator,smal,CG11486,Hipk,Haspin,CG1344,PEK,S6kII | | 6.071e-07 | -14.31 | columnar/cuboidal epithelial cell development | biological process | GO:0002066 | 328 | 30 | 12345 | 414 | shd,ci,N,ttk,CycE,if,Stat92E,mam,bbg,par-1,hh,Rac2,dpp,Myb,spoon,Egfr,S,Syp,aop,sty,drk,eya,lama,jing,EcR,gus,Rho1,tkv,Fas3,nudE | | 6.271e-07 | -14.28 | brain development | biological process | GO:0007420 | 122 | 17 | 12345 | 414 | Dr,mbt,Dmtn,EcR,enok,Dscam1,Fas2,Ptpmeg,Smox,robo2,disco,cas,PsGEF,Prosap,oc,hh,ey | | 6.859e-07 | -14.19 | anterior/posterior pattern specification, imaginal disc | biological process | GO:0007448 | 24 | 8 | 12345 | 414 | shn,dpp,tkv,bi,Dad,ci,Ser,hh | | 7.780e-07 | -14.07 | motor neuron axon guidance | biological process | GO:0008045 | 73 | 13 | 12345 | 414 | beat-IIIc,dally,Ten-m,NetB,NetA,Rho1,Cdk5alpha,Rac2,trn,Fas2,N,Mmp2,Ptp99A | | 7.813e-07 | -14.06 | kn (knot) | protein interactions | 45318 | 56 | 13 | 9204 | 412 | heph,Imp,Rbfox1,Pak3,Tpr2,CG10082,hh,Egfr,cnc,ci,tou,Ubx,ctrip | | 7.920e-07 | -14.05 | sty (sprouty) | protein interactions | 38424 | 105 | 18 | 9204 | 412 | Kr,spi,S,drk,Egfr,CG46339,Ppa,Pits,Socs16D,aos,N,aop,Ser,disco,rau,CycE,LRP1,Fas2 | | 8.126e-07 | -14.02 | embryonic pattern specification | biological process | GO:0009880 | 230 | 24 | 12345 | 414 | Ubx,cnc,slp1,bowl,jing,gus,Doa,edl,Kr,psq,N,ci,oc,Stat92E,kni,hh,par-1,tsl,dpp,Pat1,dally,Egfr,AGO2,dlp | | 8.224e-07 | -14.01 | blastoderm segmentation | biological process | GO:0007350 | 214 | 23 | 12345 | 414 | gus,Doa,Ubx,cnc,jing,bowl,slp1,Pat1,dpp,dlp,Egfr,AGO2,dally,Kr,N,ci,psq,Stat92E,oc,hh,par-1,tsl,kni | | 8.437e-07 | -13.99 | dorsal closure | biological process | GO:0007391 | 111 | 16 | 12345 | 414 | scb,ush,Rho1,tkv,mmy,Rac2,spi,shd,Myo10A,Inx3,N,crb,Egfr,aop,shn,dpp | | 8.842e-07 | -13.94 | embryonic development via the syncytial blastoderm | biological process | GO:0001700 | 199 | 22 | 12345 | 414 | crb,N,Myo10A,Inx3,spi,shd,mmy,l(3)mbt,Rac2,oc,shn,dpp,RhoGEF2,aop,AGO2,Egfr,ush,scb,Rho1,tkv,EcR,Doc1 | | 8.928e-07 | -13.93 | embryo development ending in birth or egg hatching | biological process | GO:0009792 | 215 | 23 | 12345 | 414 | elav,ush,scb,Doc1,EcR,Rho1,tkv,Myo10A,spi,Inx3,shd,crb,N,oc,Rac2,l(3)mbt,mmy,dpp,shn,AGO2,Egfr,aop,RhoGEF2 | | 9.079e-07 | -13.91 | regulation of imaginal disc growth | biological process | GO:0045570 | 42 | 10 | 12345 | 414 | Fbxl7,Ubx,ft,crb,dpp,tkv,dlp,Egfr,sbb,dally | | 9.111e-07 | -13.91 | plasma membrane part | cellular component | GO:0044459 | 820 | 55 | 11940 | 407 | Dscam1,smal,mgl,Tsp26A,Inx2,CadN,dally,Fas3,sdt,Mmp2,Inx3,sas,Ten-m,dpr7,ds,scb,Arl4,Egfr,nAChRalpha4,Cad87A,pbl,Prosap,par-1,CaMKII,dlp,Toll-7,CG42748,sty,Stat92E,hts,emp,Tsp39D,Syx1A,if,nuf,Fas2,otk2,Rho1,ft,crb,Pli,wun2,Tl,cta,Ser,nrv2,N,tkv,egr,RhoGEF2,bou,Cirl,DIP-epsilon,Hip1,tyn | | 9.154e-07 | -13.90 | nej (nejire) | protein interactions | 43856 | 106 | 18 | 9204 | 412 | CG2865,Psc,RhoGAP93B,Ubx,heph,sty,kni,EcR,Kr,ey,spi,Myb,so,Egfr,dpp,ci,eya,Utx | | 9.793e-07 | -13.84 | appendage segmentation | biological process | GO:0035285 | 25 | 8 | 12345 | 414 | dve,Bx,N,TfAP-2,Ser,fj,bowl,ap | | 9.793e-07 | -13.84 | imaginal disc-derived leg segmentation | biological process | GO:0036011 | 25 | 8 | 12345 | 414 | N,Bx,TfAP-2,dve,bowl,fj,Ser,ap | | 9.853e-07 | -13.83 | regulation of anatomical structure size | biological process | GO:0090066 | 249 | 25 | 12345 | 414 | Pvf3,Fas2,drk,ft,nw,pbl,slow,EcR,mbt,Rho1,Arp2,stg,CadN,Lac,ttk,CycE,fj,mmy,PsGEF,dpp,Myb,nrv2,ds,spoon,hts | | 1.005e-06 | -13.81 | DNA-binding transcription factor binding | molecular function | GO:0140297 | 65 | 12 | 12176 | 396 | EcR,shn,edl,ush,Doc1,CtBP,ocm,kni,bi,Utx,HGTX,Doc2 | | 1.063e-06 | -13.75 | protein phosphorylation | biological process | GO:0006468 | 284 | 27 | 12345 | 414 | JIL-1,Hipk,egr,Asator,Taf1,Rac2,par-1,CycE,Tlk,fj,S6kII,PEK,Mos,CaMKII,Cdk12,KP78a,wnd,PhKgamma,Pak3,CG11486,Ptpmeg,smal,CG8485,tkv,mbt,Haspin,Doa | | 1.107e-06 | -13.71 | mesoderm development | biological process | GO:0007498 | 100 | 15 | 12345 | 414 | eya,Ubx,cta,N,pbl,wb,slou,mam,hh,twi,lbe,slp1,meso18E,dpp,tin | | 1.131e-06 | -13.69 | regulation of tube size | biological process | GO:0035150 | 53 | 11 | 12345 | 414 | ds,nrv2,Rho1,dpp,EcR,mmy,fj,ttk,ft,Lac,Fas2 | | 1.185e-06 | -13.65 | pnt (pointed) | protein interactions | 42757 | 41 | 11 | 9204 | 412 | Egfr,N,stg,Ubx,aos,edl,spen,Bx,aop,CycE,rau | | 1.187e-06 | -13.64 | EGF\_1 | prosite domains | PS00022 | 52 | 13 | 6243 | 316 | spi,crb,mgl,Ser,NetA,Ten-m,slow,NetB,CadN,wb,N,ft,LRP1 | | 1.222e-06 | -13.62 | regulation of embryonic development | biological process | GO:0045995 | 88 | 14 | 12345 | 414 | Rho1,stg,neo,RhoGEF2,Syx1A,tyn,ttk,Tl,cta,l(3)mbt,par-1,Rac2,tsl,cnc | | 1.241e-06 | -13.60 | GO\_RENAL\_TUBULE\_DEVELOPMENT | MSigDB lists | GO\_RENAL\_TUBULE\_DEVELOPMENT | 24 | 8 | 4161 | 153 | hh,so,Ser,Lim1,dpp,ds,N,ft | | 1.267e-06 | -13.58 | germ cell development | biological process | GO:0007281 | 820 | 54 | 12345 | 414 | Pat1,Rac2,Npc1a,tsl,mam,pAbp,if,psq,Hydr2,ci,N,wun2,shd,nudE,Tet,heph,Fas3,Rho1,gus,EcR,scb,lama,Rbfox1,goe,aop,sty,Syp,Mos,S,dally,hts,spoon,Egfr,yps,stwl,Myb,dpp,par-1,hh,bbg,l(3)mbt,Stat92E,CycE,ttk,JIL-1,baf,Doa,tkv,jing,cnc,Inx2,eya,elav,drk | | 1.312e-06 | -13.54 | GO\_CELL\_DEVELOPMENT | MSigDB lists | GO\_CELL\_DEVELOPMENT | 289 | 28 | 4161 | 153 | Fas2,nudE,TfAP-2,ft,CaMKII,DIP-epsilon,hh,CG9220,drk,tkv,Rac2,dpp,Ser,Lim1,l(3)mbt,if,ap,Dscam1,NetB,Ptr,N,Smox,trn,JIL-1,Ank,mbt,sdt,Hydr2 | | 1.392e-06 | -13.48 | GGGTGGRR\_PAX4\_03 | MSigDB lists | GGGTGGRR\_PAX4\_03 | 240 | 25 | 4161 | 153 | Snx3,LRP1,Tpr2,fne,trn,LManII,Nhe2,CG12547,Mhcl,mgl,NetB,ds,Utx,kdn,Imp,hh,dlp,Rac2,PhKgamma,Tsp26A,Lac,hts,CG2812,sm,S6kII | | 1.406e-06 | -13.47 | regulation of tube architecture, open tracheal system | biological process | GO:0035152 | 89 | 14 | 12345 | 414 | Egfr,ds,dpp,bou,Rho1,nrv2,if,fj,mmy,Lac,Fas2,ft,ttk,flz | | 1.434e-06 | -13.46 | GO\_POSITIVE\_REGULATION\_OF\_DEVELOPMENTAL\_PROCESS | MSigDB lists | GO\_POSITIVE\_REGULATION\_OF\_DEVELOPMENTAL\_PROCESS | 178 | 21 | 4161 | 153 | CG12099,N,Smox,pbl,NetB,Ser,Lim1,dpp,tkv,Rac2,CycE,CG9855,so,hh,CG11367,S6kII,TfAP-2,nudE,Tif-IA,smal,Lac | | 1.458e-06 | -13.44 | Homeobox-like\_sf | interpro domains | IPR009057 | 142 | 19 | 11383 | 428 | slou,exex,lbe,Dr,tin,ey,dve,CG13894,oc,HGTX,so,Myb,ap,B-H2,psq,BtbVII,Ubx,Lim1,bab1 | | 1.588e-06 | -13.35 | GO\_GROWTH | MSigDB lists | GO\_GROWTH | 109 | 16 | 4161 | 153 | Smox,N,NTPase,Utx,ap,Lim1,dpp,bor,mbt,CaMKII,hts,nudE,smal,tkv,CycE,hh | | 1.600e-06 | -13.35 | cell fate commitment involved in pattern specification | biological process | GO:0060581 | 35 | 9 | 12345 | 414 | EcR,lz,Egfr,nuf,Ubx,N,phyl,salm,CtBP | | 1.612e-06 | -13.34 | Acsl (Acyl-CoA synthetase long-chain) | protein interactions | 46068 | 15 | 7 | 9204 | 412 | tkv,Kr,fax,Cnx99A,N,Dad,kni | | 1.725e-06 | -13.27 | axis specification | biological process | GO:0009798 | 207 | 22 | 12345 | 414 | Tl,slp1,bowl,cnc,aos,heph,gus,psq,N,Kr,hh,kni,par-1,twi,tsl,oc,Pat1,dpp,sty,Syp,spoon,Egfr | | 1.765e-06 | -13.25 | GO\_NEUROGENESIS | MSigDB lists | GO\_NEUROGENESIS | 328 | 30 | 4161 | 153 | Lac,nudE,Tif-IA,Fas2,TfAP-2,CG5830,ft,hh,DIP-epsilon,so,drk,nAChRalpha4,tkv,Rac2,Lim1,Ser,dpp,ap,ds,N,pbl,NetB,Dscam1,LRP1,CG12099,Ank,sdt,mbt,JIL-1,trn | | 1.833e-06 | -13.21 | shn (schnurri) | protein interactions | 36171 | 21 | 8 | 9204 | 412 | Sin3A,tkv,dpp,pbl,CycE,Bx,CtBP,Fas2 | | 1.875e-06 | -13.19 | TAATTA\_CHX10\_01 | MSigDB lists | TAATTA\_CHX10\_01 | 166 | 20 | 4161 | 153 | Nhe2,Pli,kdn,Utx,if,mgl,Cnx99A,Dscam1,Doa,fne,ScsbetaG,sdt,SamDC,CG45050,phtf,TfAP-2,cta,so,CG16890,tkv | | 2.018e-06 | -13.11 | imaginal disc-derived wing vein specification | biological process | GO:0007474 | 56 | 11 | 12345 | 414 | S,Egfr,S6kII,aos,dpp,Lpin,jing,hh,fj,N,Rbfox1 | | 2.134e-06 | -13.06 | cell chemotaxis | biological process | GO:0060326 | 8 | 5 | 12345 | 414 | Lrt,hh,dpp,wun2,Dad | | 2.239e-06 | -13.01 | regulation of plasma membrane bounded cell projection organization | biological process | GO:0120035 | 148 | 18 | 12345 | 414 | Prosap,Rac2,mmy,CadN,N,RhoGEF2,CaMKII,comm,PsGEF,NetA,pbl,ham,nudE,NetB,Imp,Dscam1,Arp2,Rho1 | | 2.302e-06 | -12.98 | GO\_CELL\_MORPHOGENESIS\_INVOLVED\_IN\_DIFFERENTIATION | MSigDB lists | GO\_CELL\_MORPHOGENESIS\_INVOLVED\_IN\_DIFFERENTIATION | 112 | 16 | 4161 | 153 | N,NetB,Dscam1,if,ap,Lim1,dpp,Ank,trn,JIL-1,nudE,Fas2,Rac2,tkv,drk,hh | | 2.470e-06 | -12.91 | regulation of cell projection organization | biological process | GO:0031344 | 149 | 18 | 12345 | 414 | mmy,Rac2,Prosap,N,CadN,RhoGEF2,CaMKII,NetA,comm,PsGEF,pbl,nudE,ham,Arp2,Rho1,Dscam1,Imp,NetB | | 2.476e-06 | -12.91 | GO\_NEPHRON\_EPITHELIUM\_DEVELOPMENT | MSigDB lists | GO\_NEPHRON\_EPITHELIUM\_DEVELOPMENT | 26 | 8 | 4161 | 153 | so,hh,dpp,Lim1,Ser,ft,N,ds | | 2.477e-06 | -12.91 | regulation of developmental growth | biological process | GO:0048638 | 228 | 23 | 12345 | 414 | scb,ft,Fas2,Tl,Ubx,sbb,tkv,Arp2,tsl,crb,Hipk,Dad,Fbxl7,CadN,ey,hiw,CaMKII,Syp,dally,Egfr,dlp,Acsl,dpp | | 2.656e-06 | -12.84 | formation of animal organ boundary | biological process | GO:0010160 | 37 | 9 | 12345 | 414 | N,CtBP,phyl,B-H2,NetB,lz,nuf,dally,robo2 | | 2.716e-06 | -12.82 | Raf (Raf oncogene) | protein interactions | 31221 | 149 | 21 | 9204 | 412 | eya,CG9775,spen,if,S,drk,Egfr,Gyf,sty,aop,RhoGEF2,aos,pnt,NAT1,Nhe2,phyl,tkv,S6kII,mam,ft,hh | | 2.723e-06 | -12.81 | exocrine system development | biological process | GO:0035272 | 121 | 16 | 12345 | 414 | trn,scb,Ser,eya,tin,Rho1,EcR,NetB,Atg17,Utx,twi,Rac2,if,crb,Egfr,robo2 | | 2.723e-06 | -12.81 | salivary gland development | biological process | GO:0007431 | 121 | 16 | 12345 | 414 | robo2,Egfr,if,Utx,twi,Rac2,crb,tin,NetB,EcR,Atg17,Rho1,scb,Ser,trn,eya | | 2.783e-06 | -12.79 | GO\_CELL\_MORPHOGENESIS\_INVOLVED\_IN\_NEURON\_DIFFERENTIATION | MSigDB lists | GO\_CELL\_MORPHOGENESIS\_INVOLVED\_IN\_NEURON\_DIFFERENTIATION | 88 | 14 | 4161 | 153 | trn,JIL-1,drk,Ank,Rac2,tkv,hh,ap,NetB,Dscam1,N,Lim1,Fas2,nudE | | 3.023e-06 | -12.71 | GO\_MORPHOGENESIS\_OF\_A\_BRANCHING\_STRUCTURE | MSigDB lists | GO\_MORPHOGENESIS\_OF\_A\_BRANCHING\_STRUCTURE | 35 | 9 | 4161 | 153 | drk,hh,so,cta,ds,ft,N,dpp,Lim1 | | 3.153e-06 | -12.67 | cell-cell adhesion via plasma-membrane adhesion molecules | biological process | GO:0098742 | 70 | 12 | 12345 | 414 | Fas3,beat-IIIc,ds,Lac,Fas2,CadN,Con,ft,scb,if,Cad87A,ImpL2 | | 3.238e-06 | -12.64 | protein kinase activity | molecular function | GO:0004672 | 274 | 25 | 12176 | 396 | CG11486,Hipk,Haspin,PEK,S6kII,Asator,mbt,smal,CaMKII,Tlk,PhKgamma,par-1,Cdk12,CycE,CG8485,fj,KP78a,JIL-1,Egfr,Mos,Taf1,Pak3,tkv,Doa,wnd | | 3.454e-06 | -12.58 | glial cell migration | biological process | GO:0008347 | 29 | 8 | 12345 | 414 | so,NetB,NetA,Rho1,jing,hh,ey,N | | 3.508e-06 | -12.56 | GO\_EMBRYONIC\_ORGAN\_MORPHOGENESIS | MSigDB lists | GO\_EMBRYONIC\_ORGAN\_MORPHOGENESIS | 66 | 12 | 4161 | 153 | so,hh,lbk,Rac2,Lim1,NetB,Dscam1,N,Smox,if,Utx,TfAP-2 | | 3.670e-06 | -12.52 | GO\_EMBRYONIC\_ORGAN\_DEVELOPMENT | MSigDB lists | GO\_EMBRYONIC\_ORGAN\_DEVELOPMENT | 90 | 14 | 4161 | 153 | Lim1,dpp,N,Smox,NetB,Dscam1,TfAP-2,if,Utx,so,lbk,hh,Rac2,drk | | 3.905e-06 | -12.45 | GO\_KIDNEY\_EPITHELIUM\_DEVELOPMENT | MSigDB lists | GO\_KIDNEY\_EPITHELIUM\_DEVELOPMENT | 36 | 9 | 4161 | 153 | hh,so,ds,ft,Smox,N,dpp,Ser,Lim1 | | 3.916e-06 | -12.45 | rho (rhomboid) | protein interactions | 38168 | 64 | 13 | 9204 | 412 | tkv,aos,spi,ft,Egfr,S,N,dpp,Ser,aop,sty,Fas2,kni | | 4.110e-06 | -12.40 | regulation of tube size, open tracheal system | biological process | GO:0035151 | 49 | 10 | 12345 | 414 | dpp,Rho1,nrv2,ds,Fas2,Lac,ft,ttk,fj,mmy | | 4.169e-06 | -12.39 | regulation of cellular component biogenesis | biological process | GO:0044087 | 305 | 27 | 12345 | 414 | PsGEF,Acsl,hts,mle,spoon,hiw,CG5815,CaMKII,Syp,Dad,N,Mmp2,Syx1A,Stat92E,if,par-1,CG43658,Acf,Arp2,tkv,Rho1,Fas2,CG11486,Pak3,cnc,pbl,scb | | 4.337e-06 | -12.35 | apical plasma membrane | cellular component | GO:0016324 | 71 | 12 | 11940 | 407 | crb,Stat92E,sas,mgl,scb,Ser,Inx2,RhoGEF2,dally,tyn,Rho1,sdt | | 4.391e-06 | -12.34 | GO\_NEURON\_PROJECTION\_GUIDANCE | MSigDB lists | GO\_NEURON\_PROJECTION\_GUIDANCE | 46 | 10 | 4161 | 153 | Lim1,Fas2,ap,NetB,hh,drk,Ank,JIL-1,Rac2,tkv | | 4.433e-06 | -12.33 | lilli (lilliputian) | protein interactions | 33496 | 38 | 10 | 9204 | 412 | aop,Bx,Rho1,ttk,phyl,gus,mam,Egfr,dpp,N | | 4.456e-06 | -12.32 | oogenesis | biological process | GO:0048477 | 682 | 46 | 12345 | 414 | dally,hts,Egfr,yps,spoon,sty,aop,Mos,S,Syp,Myb,dpp,Stat92E,CycE,hh,par-1,bbg,ttk,JIL-1,baf,Doa,tkv,cnc,jing,drk,eya,Inx2,Pat1,pAbp,if,Rac2,mam,shd,psq,N,ci,nudE,gus,EcR,Fas3,heph,Rho1,scb,goe,Rbfox1,lama | | 4.668e-06 | -12.27 | neuron fate determination | biological process | GO:0048664 | 9 | 5 | 12345 | 414 | Egfr,mam,S,spi,N | | 4.874e-06 | -12.23 | cellular response to growth factor stimulus | biological process | GO:0071363 | 61 | 11 | 12345 | 414 | dpp,Toll-7,shn,tkv,dally,aop,hiw,Dad,drk,Smox,pnt | | 4.997e-06 | -12.21 | cardiocyte differentiation | biological process | GO:0035051 | 22 | 7 | 12345 | 414 | robo2,tin,Doc2,EcR,dpp,Doc1,Ubx | | 5.258e-06 | -12.16 | GO\_POSITIVE\_REGULATION\_OF\_CELL\_DIFFERENTIATION | MSigDB lists | GO\_POSITIVE\_REGULATION\_OF\_CELL\_DIFFERENTIATION | 133 | 17 | 4161 | 153 | S6kII,Lac,Tif-IA,nudE,smal,CycE,tkv,Rac2,hh,CG11367,Smox,N,pbl,NetB,Ser,Lim1,dpp | | 5.274e-06 | -12.15 | GO\_ARTERY\_DEVELOPMENT | MSigDB lists | GO\_ARTERY\_DEVELOPMENT | 14 | 6 | 4161 | 153 | hh,LRP1,Ser,so,mgl,N | | 5.284e-06 | -12.15 | developmental growth involved in morphogenesis | biological process | GO:0060560 | 86 | 13 | 12345 | 414 | scb,Rac2,Dad,Ptpmeg,CadN,ttk,wnd,otk2,mle,RhoGEF2,Dscam1,Rho1,tkv | | 5.309e-06 | -12.15 | negative regulation of cell fate specification | biological process | GO:0009996 | 15 | 6 | 12345 | 414 | S6kII,ush,eya,pnt,salm,aop | | 5.698e-06 | -12.08 | Sulf1 (Sulfated) | protein interactions | 53437 | 12 | 6 | 9204 | 412 | rau,Stat92E,Notum,dlp,dally,Egfr | | 5.698e-06 | -12.08 | dac (dachshund) | protein interactions | 34982 | 12 | 6 | 9204 | 412 | ey,CycE,eya,Egfr,hh,dpp | | 5.749e-06 | -12.07 | response to growth factor | biological process | GO:0070848 | 62 | 11 | 12345 | 414 | Smox,Dad,drk,pnt,tkv,Toll-7,dpp,shn,hiw,aop,dally | | 5.817e-06 | -12.05 | GO\_TUBE\_MORPHOGENESIS | MSigDB lists | GO\_TUBE\_MORPHOGENESIS | 81 | 13 | 4161 | 153 | hh,so,Lim1,dpp,wech,ap,Utx,cta,ds,N,Smox,NetB,ft | | 5.984e-06 | -12.03 | mesoderm formation | biological process | GO:0001707 | 31 | 8 | 12345 | 414 | Ubx,mam,slou,lbe,slp1,twi,pbl,tin | | 6.033e-06 | -12.02 | GGGCGGR\_SP1\_Q6 | MSigDB lists | GGGCGGR\_SP1\_Q6 | 882 | 56 | 4161 | 153 | CG43313,ras,CycE,Hsc70Cb,Mocs2B,Imp,Tlk,hts,sm,CaMKII,Arl4,S6kII,Arp2,Fas2,CG6752,CG45050,LManII,CG32758,RpS4,LRP1,N,Cnx99A,Smox,Notum,Syx1A,Tim17b,Nhe2,Ser,Sin3A,drk,dlp,TfIIS,Hsc70-3,Taf1,CG5830,mle,NKAIN,nudE,Tsp26A,Hydr2,Pat1,EcR,mbt,fne,sdt,Tpr2,Ppa,NetB,pAbp,mgl,kdn,Utx,ap,lili,dpp,Mhcl | | 6.135e-06 | -12.00 | yki (yorkie) | protein interactions | 37851 | 109 | 17 | 9204 | 412 | Utx,osa,Hsc70-3,ci,l(3)mbt,Hipk,RhoGAP15B,ImpL2,EcR,ban,par-1,pbl,ft,CG30089,tkv,CG43658,Smox | | 6.227e-06 | -11.99 | GO\_NEURON\_PROJECTION\_MORPHOGENESIS | MSigDB lists | GO\_NEURON\_PROJECTION\_MORPHOGENESIS | 94 | 14 | 4161 | 153 | Rac2,tkv,JIL-1,trn,Ank,drk,hh,NetB,Dscam1,N,ap,Fas2,nudE,Lim1 | | 6.759e-06 | -11.90 | regulation of organ growth | biological process | GO:0046620 | 63 | 11 | 12345 | 414 | dlp,Egfr,sbb,dally,dpp,tkv,Fbxl7,Ubx,ft,Hipk,crb | | 6.910e-06 | -11.88 | GO\_REGULATION\_OF\_CELL\_DIFFERENTIATION | MSigDB lists | GO\_REGULATION\_OF\_CELL\_DIFFERENTIATION | 279 | 26 | 4161 | 153 | hh,CG11367,so,CycE,tkv,heph,Rac2,nAChRalpha4,Lac,smal,Tif-IA,nudE,TfAP-2,S6kII,ft,CG5830,CG12099,LRP1,EcR,dpp,Ser,Lim1,pbl,NetB,Smox,N | | 6.978e-06 | -11.87 | spiracle morphogenesis, open tracheal system | biological process | GO:0035277 | 23 | 7 | 12345 | 414 | Egfr,salm,hh,RhoGEF2,spi,ci,Rho1 | | 7.092e-06 | -11.86 | trr (trithorax-related) | protein interactions | 31149 | 18 | 7 | 9204 | 412 | N,dpp,hh,Utx,spen,EcR,Ser | | 7.341e-06 | -11.82 | polytene chromosome | cellular component | GO:0005700 | 206 | 21 | 11940 | 407 | JIL-1,l(3)mbt,stwl,ttk,Chro,cnc,Df31,CtBP,mof,tou,Utx,Cdk12,mle,Sin3A,mam,lawc,Myb,Mocs2B,EcR,Psc,Pits | | 7.400e-06 | -11.81 | svp (seven up) | protein interactions | 41491 | 32 | 9 | 9204 | 412 | sty,psq,aop,Ubx,Egfr,drk,phyl,EcR,Kr | | 7.406e-06 | -11.81 | Signal Transduction | REACTOME pathways | R-DME-162582 | 717 | 49 | 3923 | 148 | RhoGAP18B,Fur1,if,CG42684,Rtnl2,S6kII,Tl,Dad,dpp,stg,drk,N,NFAT,Ser,RhoGAP15B,nrv2,Egfr,dally,Stat92E,Cirl,JIL-1,RhoGAP102A,Arp2,esn,Notum,cta,Fas2,par-1,Rhp,CG34384,pbl,Rho1,lbk,Snx3,sty,Ptpmeg,Rbfox1,heph,hh,CaMKII,Sox102F,RhoGEF2,tkv,RhoGAP93B,ci,Rac2,CtBP,scb,Smox | | 7.857e-06 | -11.75 | cell-cell junction | cellular component | GO:0005911 | 101 | 14 | 11940 | 407 | Inx2,CadN,Fbxl7,N,sdt,otk2,Fas3,Rho1,crb,bbg,Inx3,mbt,Lac,nrv2 | | 7.995e-06 | -11.74 | GO\_REGULATION\_OF\_OSSIFICATION | MSigDB lists | GO\_REGULATION\_OF\_OSSIFICATION | 39 | 9 | 4161 | 153 | tkv,CG11367,CG9220,TfAP-2,Smox,N,Ser,dpp,smal | | 8.182e-06 | -11.71 | GO\_NEPHRON\_DEVELOPMENT | MSigDB lists | GO\_NEPHRON\_DEVELOPMENT | 30 | 8 | 4161 | 153 | so,hh,dpp,Lim1,Ser,ft,N,ds | | 8.239e-06 | -11.71 | phosphorylation | biological process | GO:0016310 | 470 | 35 | 12345 | 414 | KP78a,CaMKII,PEK,CG34384,fj,Socs16D,ND-B14.5AL,Rac2,Taf1,mbt,CG2201,CG10082,Haspin,CG8485,S6kII,wnd,Mos,Cdk12,Cyt-c1,egr,Asator,Hipk,JIL-1,Tlk,CycE,par-1,CG1344,tkv,Doa,CG11486,Ptpmeg,CG1271,PhKgamma,Pak3,smal | | 8.256e-06 | -11.70 | regulation of muscle organ development | biological process | GO:0048634 | 16 | 6 | 12345 | 414 | tkv,dpp,Dad,Ubx,cv,twi | | 8.258e-06 | -11.70 | positive regulation of intracellular signal transduction | biological process | GO:1902533 | 178 | 19 | 12345 | 414 | RhoGEF2,PEK,Mos,wnd,Egfr,ds,Lpin,PsGEF,CG43658,grnd,Socs16D,ci,crb,spi,Fbxl7,wcy,mbt,ft,drk | | 8.531e-06 | -11.67 | phosphotransferase activity, alcohol group as acceptor | molecular function | GO:0016773 | 346 | 28 | 12176 | 396 | smal,Asator,mbt,S6kII,Haspin,PEK,CG11486,Hipk,CG1271,tkv,wnd,CG34384,Doa,Egfr,Taf1,Mos,Pak3,KP78a,JIL-1,CycE,CG2201,fj,CG8485,PhKgamma,par-1,Cdk12,Tlk,CaMKII | | 9.079e-06 | -11.61 | negative regulation of cell motility | biological process | GO:2000146 | 10 | 5 | 12345 | 414 | Ptpmeg,Sin3A,CG44774,scb,Lrt | | 9.079e-06 | -11.61 | foregut morphogenesis | biological process | GO:0007440 | 10 | 5 | 12345 | 414 | Inx2,N,dpp,hh,bowl | | 9.079e-06 | -11.61 | regulation of fibroblast growth factor receptor signaling pathway | biological process | GO:0040036 | 10 | 5 | 12345 | 414 | rau,Mmp2,dally,dlp,sty | | 9.079e-06 | -11.61 | negative regulation of cell migration | biological process | GO:0030336 | 10 | 5 | 12345 | 414 | Sin3A,CG44774,Ptpmeg,Lrt,scb | | 9.114e-06 | -11.61 | muscle cell differentiation | biological process | GO:0042692 | 118 | 15 | 12345 | 414 | if,twi,Rac2,lbe,slou,pnt,Ubx,Pak3,N,hts,CG13366,tin,aop,sty,Rho1 | | 9.285e-06 | -11.59 | reproductive process | biological process | GO:0022414 | 1436 | 78 | 12345 | 414 | scb,Fas2,goe,lama,Rbfox1,Tet,nudE,heph,Fas3,Rho1,pAbp,if,tsl,Npc1a,shd,psq,Bx,ci,CaMKII,Pat1,wb,jing,drk,eya,bab1,baf,Doa,bbg,ttk,JIL-1,Mmp2,wnd,otk2,hts,Egfr,mle,spoon,aop,sty,S,Myb,lz,pbl,tin,gus,EcR,salm,Rac2,mam,wun2,Hydr2,N,robo2,so,cnc,Tudor-SN,elav,Inx2,Dr,Imp,tkv,Stat92E,CycE,hh,par-1,l(3)mbt,dve,fne,TfAP-2,Pex1,dally,AGO2,yps,Syp,RhoGEF2,Mos,dpp,stwl | | 9.318e-06 | -11.58 | Abl (Abl tyrosine kinase) | protein interactions | 45821 | 41 | 10 | 9204 | 412 | comm,Dscam1,Ptp99A,Egfr,N,robo2,eya,Rho1,CycE,fax | | 9.318e-06 | -11.58 | sli (slit) | protein interactions | 36746 | 41 | 10 | 9204 | 412 | ci,Egfr,jing,Dscam1,comm,if,elav,robo2,wb,scb | | 9.756e-06 | -11.54 | Homeodomain | pfam domains | PF00046 | 94 | 14 | 10480 | 393 | exex,lbe,B-H2,dve,ap,Dr,slou,HGTX,Ubx,so,tin,Lim1,oc,ey | | 9.769e-06 | -11.54 | GO\_ORGAN\_MORPHOGENESIS | MSigDB lists | GO\_ORGAN\_MORPHOGENESIS | 169 | 19 | 4161 | 153 | ft,TfAP-2,tkv,Rac2,CG9220,so,lbk,hh,N,Smox,Dscam1,NetB,Utx,if,ds,Ser,Lim1,dpp,fj | | 1.037e-05 | -11.48 | photoreceptor cell development | biological process | GO:0042461 | 105 | 14 | 12345 | 414 | Ten-m,Doa,EcR,lz,sno,rau,sdt,oc,pnt,eya,edl,CadN,ttk,crb | | 1.040e-05 | -11.47 | RNA polymerase II activating transcription factor binding | molecular function | GO:0001102 | 17 | 6 | 12176 | 396 | Doc1,ush,Doc2,ocm,bi,shn | | 1.069e-05 | -11.45 | GO\_BRANCHING\_MORPHOGENESIS\_OF\_AN\_EPITHELIAL\_TUBE | MSigDB lists | GO\_BRANCHING\_MORPHOGENESIS\_OF\_AN\_EPITHELIAL\_TUBE | 31 | 8 | 4161 | 153 | dpp,Lim1,ds,cta,ft,N,hh,so | | 1.108e-05 | -11.41 | gro (groucho) | protein interactions | 43162 | 70 | 13 | 9204 | 412 | kni,CtBP,shn,E(spl)m4-BFM,so,Egfr,sbb,HGTX,E(spl)mdelta-HLH,slp1,N,ey,Hipk | | 1.155e-05 | -11.37 | bun (bunched) | protein interactions | 34665 | 26 | 8 | 9204 | 412 | Smox,Kr,hh,Egfr,S,CG11486,N,aop | | 1.192e-05 | -11.34 | scrib (scribble) | protein interactions | 44448 | 51 | 11 | 9204 | 412 | crb,grnd,CycE,Egfr,oc,CG42748,tkv,Pak3,sdt,egr,Stat92E | | 1.222e-05 | -11.31 | embryonic organ development | biological process | GO:0048568 | 55 | 10 | 12345 | 414 | tin,robo2,lz,Rho1,if,Rac2,pon,Pvf3,N,Mmp2 | | 1.229e-05 | -11.31 | homophilic cell adhesion via plasma membrane adhesion molecules | biological process | GO:0007156 | 44 | 9 | 12345 | 414 | Fas3,ds,Lac,Fas2,CadN,Con,ft,Cad87A,ImpL2 | | 1.229e-05 | -11.31 | glial cell differentiation | biological process | GO:0010001 | 44 | 9 | 12345 | 414 | spen,Dr,sty,cas,rau,nrv2,salm,Pex1,N | | 1.240e-05 | -11.30 | negative regulation of small GTPase mediated signal transduction | biological process | GO:0051058 | 17 | 6 | 12345 | 414 | sty,MESR3,CG42684,RapGAP1,S6kII,NFAT | | 1.253e-05 | -11.29 | Protein\_kinase\_ATP\_BS | interpro domains | IPR017441 | 164 | 19 | 11383 | 428 | Tlk,Cdk12,CaMKII,JIL-1,Egfr,PEK,S6kII,Hipk,Haspin,Asator,PhKgamma,Pak3,Doa,Mos,par-1,tkv,CG8485,mbt,KP78a | | 1.261e-05 | -11.28 | mesoderm morphogenesis | biological process | GO:0048332 | 34 | 8 | 12345 | 414 | Ubx,pbl,twi,lbe,slp1,mam,slou,tin | | 1.274e-05 | -11.27 | regulation of Wnt signaling pathway | biological process | GO:0030111 | 93 | 13 | 12345 | 414 | Prosap,phyl,pbl,CtBP,Hipk,Mmp2,dlp,spen,dally,aop,Notum,Rho1,Sin3A | | 1.291e-05 | -11.26 | sensory organ boundary specification | biological process | GO:0008052 | 25 | 7 | 12345 | 414 | dally,phyl,B-H2,nuf,CtBP,lz,N | | 1.323e-05 | -11.23 | GO\_MESONEPHRIC\_TUBULE\_MORPHOGENESIS | MSigDB lists | GO\_MESONEPHRIC\_TUBULE\_MORPHOGENESIS | 16 | 6 | 4161 | 153 | ft,ds,so,dpp,hh,Lim1 | | 1.323e-05 | -11.23 | GO\_BRANCHING\_INVOLVED\_IN\_URETERIC\_BUD\_MORPHOGENESIS | MSigDB lists | GO\_BRANCHING\_INVOLVED\_IN\_URETERIC\_BUD\_MORPHOGENESIS | 16 | 6 | 4161 | 153 | ft,ds,so,dpp,Lim1,hh | | 1.405e-05 | -11.17 | activating transcription factor binding | molecular function | GO:0033613 | 26 | 7 | 12176 | 396 | ocm,Doc2,edl,Doc1,ush,shn,bi | | 1.429e-05 | -11.16 | EGF-like\_CS | interpro domains | IPR013032 | 61 | 11 | 11383 | 428 | LRP1,crb,Ser,N,ft,Ten-m,slow,wb,spi,CadN,mgl | | 1.444e-05 | -11.15 | dorsal appendage formation | biological process | GO:0046843 | 56 | 10 | 12345 | 414 | dpp,gus,spoon,Egfr,Syp,S,lama,N,ttk,Rac2 | | 1.447e-05 | -11.14 | regulation of cellular response to growth factor stimulus | biological process | GO:0090287 | 68 | 11 | 12345 | 414 | Mmp2,Dad,rau,Acsl,tkv,Lpin,hiw,sty,lili,dally,dlp | | 1.457e-05 | -11.14 | gamete generation | biological process | GO:0007276 | 1097 | 63 | 12345 | 414 | dpp,Myb,stwl,Egfr,yps,spoon,AGO2,hts,wnd,dally,Mos,S,RhoGEF2,Syp,aop,sty,Mmp2,JIL-1,ttk,Pex1,CycE,Stat92E,l(3)mbt,bbg,par-1,hh,tkv,Imp,Doa,baf,eya,drk,elav,Inx2,Tudor-SN,cnc,jing,so,Pat1,shd,wun2,N,ci,Hydr2,psq,if,pAbp,mam,Rac2,Npc1a,tsl,EcR,gus,Rho1,Fas3,heph,Tet,tin,nudE,goe,Rbfox1,lama,scb | | 1.584e-05 | -11.05 | cellular component assembly involved in morphogenesis | biological process | GO:0010927 | 154 | 17 | 12345 | 414 | lama,CG13366,gus,Rho1,CycE,if,hh,Rac2,N,ttk,Egfr,spoon,hts,S,Syp,dpp,Myb | | 1.612e-05 | -11.04 | muscle organ development | biological process | GO:0007517 | 95 | 13 | 12345 | 414 | N,Kr,salm,lbe,slow,Rac2,betaTub56D,slou,if,Rho1,wech,ap,Dr | | 1.619e-05 | -11.03 | determination of imaginal disc primordium | biological process | GO:0007445 | 11 | 5 | 12345 | 414 | Doc1,spi,S,pnt,Egfr | | 1.619e-05 | -11.03 | positive chemotaxis | biological process | GO:0050918 | 11 | 5 | 12345 | 414 | Ten-m,hh,NetB,Fas3,Con | | 1.619e-05 | -11.03 | epithelial cell proliferation involved in renal tubule morphogenesis | biological process | GO:2001013 | 11 | 5 | 12345 | 414 | Egfr,pnt,S,spi,N | | 1.619e-05 | -11.03 | epithelial cell proliferation | biological process | GO:0050673 | 11 | 5 | 12345 | 414 | spi,N,Egfr,S,pnt | | 1.619e-05 | -11.03 | epithelial cell proliferation involved in Malpighian tubule morphogenesis | biological process | GO:0061331 | 11 | 5 | 12345 | 414 | N,spi,S,pnt,Egfr | | 1.619e-05 | -11.03 | negative regulation of stem cell differentiation | biological process | GO:2000737 | 11 | 5 | 12345 | 414 | Stat92E,pnt,goe,enok,aos | | 1.646e-05 | -11.01 | regulation of growth | biological process | GO:0040008 | 310 | 26 | 12345 | 414 | Tl,Fas2,Ubx,ft,scb,Hph,EcR,Arp2,tkv,sbb,Dad,CadN,Fbxl7,ey,crb,Hipk,N,tsl,dpp,Acsl,dally,Egfr,dlp,hiw,Syp,CaMKII | | 1.653e-05 | -11.01 | stg (string) | protein interactions | 43466 | 35 | 9 | 9204 | 412 | Myb,Smox,pnt,Sin3A,N,pbl,CycE,aop,ttk | | 1.692e-05 | -10.99 | GO\_EMBRYO\_DEVELOPMENT\_ENDING\_IN\_BIRTH\_OR\_EGG\_HATCHING | MSigDB lists | GO\_EMBRYO\_DEVELOPMENT\_ENDING\_IN\_BIRTH\_OR\_EGG\_HATCHING | 160 | 18 | 4161 | 153 | hh,so,drk,Sin3A,wech,nudE,Tif-IA,TfAP-2,cta,hts,Lim1,dpp,ds,Utx,ap,N,Smox,Dscam1 | | 1.727e-05 | -10.97 | cell adhesion | biological process | GO:0007155 | 171 | 18 | 12345 | 414 | nrv2,sas,ds,Egfr,beat-IIIc,CadN,Lac,Con,if,ImpL2,Cad87A,EcR,Fas3,Fas2,tyn,ft,scb,trn | | 1.759e-05 | -10.95 | nmo (nemo) | protein interactions | 38890 | 53 | 11 | 9204 | 412 | Dad,ec,eya,Egfr,dpp,Ubx,N,Ptp99A,tkv,spi,aos | | 1.773e-05 | -10.94 | R1/R6 cell differentiation | biological process | GO:0048052 | 6 | 4 | 12345 | 414 | E(spl)mdelta-HLH,N,ttk,phyl | | 1.773e-05 | -10.94 | synaptic target attraction | biological process | GO:0016200 | 6 | 4 | 12345 | 414 | NetB,Con,Fas3,Ten-m | | 1.813e-05 | -10.92 | Ubx (Ultrabithorax) | protein interactions | 42034 | 118 | 17 | 9204 | 412 | dpp,Syx1A,Egfr,twi,ap,N,Pcl,aos,pnt,tkv,Smox,kni,tin,Mitofilin,slou,elav,Doc2 | | 1.856e-05 | -10.89 | regulation of organelle organization | biological process | GO:0033043 | 408 | 31 | 12345 | 414 | lncRNA:roX1,sens-2,Arp2,Rho1,Acf,Pcl,pbl,CG11486,mof,cta,Chro,mle,spoon,hts,Mos,CG5815,RhoGEF2,PsGEF,Myb,if,CycE,Tlk,Stat92E,ctrip,Utx,CG43658,bora,kni,JIL-1,l(3)neo38,psq | | 1.877e-05 | -10.88 | positive regulation of cell differentiation | biological process | GO:0045597 | 156 | 17 | 12345 | 414 | EcR,mbt,Imp,nudE,pnt,ec,lz,Egfr,aop,spi,N,ci,ttk,Prosap,oc,mmy,tsl | | 1.909e-05 | -10.87 | edl (ETS-domain lacking) | protein interactions | 37149 | 5 | 4 | 9204 | 412 | aop,pnt,spi,S | | 1.981e-05 | -10.83 | epithelial cell migration, open tracheal system | biological process | GO:0007427 | 36 | 8 | 12345 | 414 | drk,pnt,hh,kni,salm,dpp,dlp,dally | | 2.011e-05 | -10.81 | GO\_CELL\_PROLIFERATION | MSigDB lists | GO\_CELL\_PROLIFERATION | 162 | 18 | 4161 | 153 | NiPp1,wech,Tif-IA,nudE,smal,hh,dlp,Imp,so,Fbxl7,ras,Rac2,dpp,N,mgl,NTPase,LRP1,mbt | | 2.060e-05 | -10.79 | DNA-binding transcription activator activity, RNA polymerase II-specific | molecular function | GO:0001228 | 86 | 12 | 12176 | 396 | EcR,Doc1,B-H2,ocm,Stat92E,bi,Pits,oc,Doc2,ci,pnt,Myb | | 2.060e-05 | -10.79 | DNA-binding transcription activator activity | molecular function | GO:0001216 | 86 | 12 | 12176 | 396 | Doc1,B-H2,ocm,EcR,oc,Doc2,Myb,pnt,ci,bi,Stat92E,Pits | | 2.084e-05 | -10.78 | GO\_EMBRYO\_DEVELOPMENT | MSigDB lists | GO\_EMBRYO\_DEVELOPMENT | 227 | 22 | 4161 | 153 | lbk,hh,so,drk,Sin3A,Rac2,wech,nudE,Tif-IA,TfAP-2,cta,hts,Lim1,dpp,if,Utx,ap,ds,Smox,N,Dscam1,NetB | | 2.117e-05 | -10.76 | Ser (Serrate) | protein interactions | 43275 | 36 | 9 | 9204 | 412 | mam,N,sno,fj,Utx,ap,crb,Bx,sty | | 2.161e-05 | -10.74 | imaginal disc-derived wing vein morphogenesis | biological process | GO:0008586 | 47 | 9 | 12345 | 414 | osa,Egfr,spen,dally,heph,dpp,shn,salm,Hipk | | 2.171e-05 | -10.74 | PAX4\_01 | MSigDB lists | PAX4\_01 | 66 | 11 | 4161 | 153 | CG8531,Hydr2,Acsl,CG12099,LRP1,CG2201,mgl,pAbp,CG5830,CtBP,Tudor-SN | | 2.180e-05 | -10.73 | GO\_GLYCOPROTEIN\_BINDING | MSigDB lists | GO\_GLYCOPROTEIN\_BINDING | 25 | 7 | 4161 | 153 | dlp,dally,Hsc70-3,hh,Syx1A,tkv,Cnx99A | | 2.190e-05 | -10.73 | biological adhesion | biological process | GO:0022610 | 174 | 18 | 12345 | 414 | Cad87A,ImpL2,if,Con,Lac,CadN,beat-IIIc,Egfr,ds,sas,nrv2,trn,scb,tyn,ft,Fas2,Fas3,EcR | | 2.300e-05 | -10.68 | cellular process involved in reproduction in multicellular organism | biological process | GO:0022412 | 997 | 58 | 12345 | 414 | goe,Rbfox1,lama,scb,gus,EcR,Fas3,heph,Rho1,Tet,tin,nudE,wun2,shd,psq,Hydr2,N,ci,pAbp,if,Npc1a,tsl,Rac2,mam,Pat1,eya,drk,elav,Inx2,cnc,jing,tkv,baf,Doa,ttk,Pex1,JIL-1,Stat92E,CycE,hh,par-1,bbg,l(3)mbt,Myb,dpp,stwl,hts,dally,AGO2,yps,Egfr,spoon,aop,sty,S,Syp,RhoGEF2,Mos | | 2.421e-05 | -10.63 | enhancer sequence-specific DNA binding | molecular function | GO:0001158 | 38 | 8 | 12176 | 396 | ey,slp1,Ubx,aop,disco,ttk,Doc1,tin | | 2.455e-05 | -10.61 | heart morphogenesis | biological process | GO:0003007 | 37 | 8 | 12345 | 414 | Lac,Mmp2,ci,mam,hh,slp1,nrv2,robo2 | | 2.517e-05 | -10.59 | GCTNWTTGK\_UNKNOWN | MSigDB lists | GCTNWTTGK\_UNKNOWN | 67 | 11 | 4161 | 153 | Smox,par-1,Tudor-SN,Lim1,CG8531,sdt,CG14646,SamDC,so,Tlk,CG12099 | | 2.587e-05 | -10.56 | regulation of cell morphogenesis | biological process | GO:0022604 | 144 | 16 | 12345 | 414 | neo,NetB,Dscam1,Rho1,nudE,cta,tyn,pbl,dpp,NetA,RhoGEF2,CadN,ttk,tsl,Rac2,mmy | | 2.637e-05 | -10.54 | Homeobox\_dom | interpro domains | IPR001356 | 102 | 14 | 11383 | 428 | tin,lbe,Dr,so,exex,slou,ey,dve,ap,HGTX,oc,Lim1,B-H2,Ubx | | 2.687e-05 | -10.52 | regulation of neuron projection development | biological process | GO:0010975 | 114 | 14 | 12345 | 414 | CadN,mmy,Rac2,Prosap,pbl,NetA,Rho1,Imp,Dscam1,NetB,comm,nudE,RhoGEF2,ham | | 2.699e-05 | -10.52 | anterior Malpighian tubule development | biological process | GO:0061327 | 12 | 5 | 12345 | 414 | Doc2,Ubx,Doc1,dpp,tkv | | 2.699e-05 | -10.52 | regulation of compound eye photoreceptor development | biological process | GO:0045314 | 12 | 5 | 12345 | 414 | oc,pAbp,sty,mbt,N | | 2.699e-05 | -10.52 | mesodermal cell fate specification | biological process | GO:0007501 | 12 | 5 | 12345 | 414 | Ubx,tin,lbe,twi,slou | | 2.751e-05 | -10.50 | B-H1 (BarH1) | protein interactions | 32724 | 15 | 6 | 9204 | 412 | B-H2,N,drk,Egfr,pnt,aop | | 2.823e-05 | -10.48 | protein serine/threonine kinase activity | molecular function | GO:0004674 | 200 | 19 | 12176 | 396 | PEK,Haspin,Hipk,S6kII,Asator,mbt,CaMKII,Tlk,CG8485,Cdk12,par-1,PhKgamma,Pak3,Taf1,JIL-1,KP78a,wnd,Doa,tkv | | 2.870e-05 | -10.46 | Growth\_fac\_rcpt\_cys\_sf | interpro domains | IPR009030 | 34 | 8 | 11383 | 428 | mgl,CadN,Egfr,N,crb,LRP1,Ser,Fur1 | | 3.045e-05 | -10.40 | ph-p (polyhomeotic proximal) | protein interactions | 31181 | 56 | 11 | 9204 | 412 | tou,ci,tara,dpp,CG32772,spi,Psc,Pli,fj,Taf1,salm | | 3.223e-05 | -10.34 | hh (hedgehog) | protein interactions | 42737 | 77 | 13 | 9204 | 412 | CycE,bi,dlp,ey,Psc,Ten-m,dally,ci,dpp,hh,Egfr,sbb,mam | | 3.273e-05 | -10.33 | regulation of chromatin organization | biological process | GO:1902275 | 116 | 14 | 12345 | 414 | CG5815,mle,sens-2,Acf,lncRNA:roX1,Pcl,Utx,Tlk,ctrip,Stat92E,JIL-1,l(3)neo38,psq,mof | | 3.326e-05 | -10.31 | regulation of gene expression, epigenetic | biological process | GO:0040029 | 163 | 17 | 12345 | 414 | CG17265,lncRNA:roX1,mof,tna,ocm,Psc,AGO2,mle,CG17168,NiPp1,tara,Tlk,Hsc70-3,Kr,psq,bin3,JIL-1 | | 3.333e-05 | -10.31 | GSE9988\_ANTI\_TREM1\_VS\_LPS\_MONOCYTE\_UP | MSigDB lists | GSE9988\_ANTI\_TREM1\_VS\_LPS\_MONOCYTE\_UP | 46 | 9 | 4161 | 153 | emp,Snx3,CG9220,Acsl,SamDC,MFS10,Npc1a,CG14291,Syx1A | | 3.382e-05 | -10.29 | mys (myospheroid) | protein interactions | 44885 | 47 | 10 | 9204 | 412 | scb,wb,if,Fas2,pio,tkv,EcR,Ptp99A,mam,CaMKII | | 3.450e-05 | -10.27 | GO\_NEURON\_PROJECTION\_DEVELOPMENT | MSigDB lists | GO\_NEURON\_PROJECTION\_DEVELOPMENT | 123 | 15 | 4161 | 153 | Rac2,tkv,trn,JIL-1,drk,mbt,Ank,hh,Dscam1,NetB,N,ap,Fas2,nudE,Lim1 | | 3.461e-05 | -10.27 | female gamete generation | biological process | GO:0007292 | 761 | 47 | 12345 | 414 | Pat1,shd,N,ci,psq,if,pAbp,mam,Rac2,EcR,gus,Rho1,heph,Fas3,nudE,goe,lama,Rbfox1,scb,dpp,Myb,spoon,yps,Egfr,dally,hts,S,Mos,Syp,aop,sty,Mmp2,JIL-1,ttk,CycE,Stat92E,bbg,par-1,hh,tkv,Doa,baf,drk,eya,Inx2,cnc,jing | | 3.492e-05 | -10.26 | cell-cell adhesion | biological process | GO:0098609 | 102 | 13 | 12345 | 414 | if,scb,Cad87A,ImpL2,Lac,Fas2,CadN,Con,ft,beat-IIIc,Egfr,ds,Fas3 | | 3.499e-05 | -10.26 | Rac1 (Rac1) | protein interactions | 38146 | 124 | 17 | 9204 | 412 | Rac2,Rho1,RhoGAP18B,crb,Pak3,CG10082,cpo,tou,PsGEF,Tlk,RhoGAP93B,tkv,aos,egr,Stat92E,N,pbl | | 3.564e-05 | -10.24 | mesodermal cell differentiation | biological process | GO:0048333 | 20 | 6 | 12345 | 414 | Ubx,mam,slou,lbe,twi,tin | | 3.564e-05 | -10.24 | negative regulation of cell fate commitment | biological process | GO:0010454 | 20 | 6 | 12345 | 414 | pnt,salm,aop,S6kII,ush,eya | | 3.564e-05 | -10.24 | mesodermal cell fate commitment | biological process | GO:0001710 | 20 | 6 | 12345 | 414 | Ubx,lbe,twi,mam,slou,tin | | 3.634e-05 | -10.22 | regulation of cell adhesion | biological process | GO:0030155 | 50 | 9 | 12345 | 414 | N,par-1,wech,tkv,aos,mbt,Ten-m,aop,robo2 | | 3.636e-05 | -10.22 | positive regulation of cell development | biological process | GO:0010720 | 148 | 16 | 12345 | 414 | spi,ttk,N,oc,Prosap,tsl,mmy,lz,Egfr,aop,ec,pnt,mbt,EcR,Imp,nudE | | 3.668e-05 | -10.21 | stet (stem cell tumor) | protein interactions | 38169 | 10 | 5 | 9204 | 412 | tkv,spi,S,dally,dpp | | 3.733e-05 | -10.20 | Homeobox\_CS | interpro domains | IPR017970 | 92 | 13 | 11383 | 428 | ey,Dr,tin,lbe,exex,so,slou,oc,HGTX,Lim1,B-H2,Ubx,ap | | 3.745e-05 | -10.19 | microtubule cytoskeleton organization involved in establishment of planar polarity | biological process | GO:0090176 | 3 | 3 | 12345 | 414 | ds,par-1,ft | | 3.785e-05 | -10.18 | GO\_MESONEPHROS\_DEVELOPMENT | MSigDB lists | GO\_MESONEPHROS\_DEVELOPMENT | 27 | 7 | 4161 | 153 | ds,Smox,ft,Lim1,hh,dpp,so | | 3.842e-05 | -10.17 | GO\_TELENCEPHALON\_DEVELOPMENT | MSigDB lists | GO\_TELENCEPHALON\_DEVELOPMENT | 58 | 10 | 4161 | 153 | hh,LRP1,Imp,CG17265,Rac2,Lim1,dpp,nudE,ap,ft | | 3.852e-05 | -10.16 | GO\_CELL\_JUNCTION | MSigDB lists | GO\_CELL\_JUNCTION | 307 | 26 | 4161 | 153 | Hsc70-3,Tlk,emp,ras,drk,Rac2,MFS10,nAChRalpha4,smal,Arp2,CtBP,cta,CaMKII,hts,LRP1,CG17265,RpS4,mbt,Ank,sdt,Ser,Syx1A,if,Dscam1,pbl,N | | 3.970e-05 | -10.13 | negative regulation of nervous system development | biological process | GO:0051961 | 118 | 14 | 12345 | 414 | scb,N,ttk,CadN,Dad,Fas2,Syp,osa,hiw,aop,S6kII,Acsl,aos,EcR | | 4.027e-05 | -10.12 | positive regulation of muscle organ development | biological process | GO:0048636 | 7 | 4 | 12345 | 414 | dpp,Ubx,tkv,cv | | 4.088e-05 | -10.10 | GO\_INNER\_EAR\_MORPHOGENESIS | MSigDB lists | GO\_INNER\_EAR\_MORPHOGENESIS | 19 | 6 | 4161 | 153 | lbk,so,if,TfAP-2,Rac2,NetB | | 4.112e-05 | -10.10 | Bx (Beadex) | protein interactions | 32846 | 68 | 12 | 9204 | 412 | sbb,Hydr2,N,trn,tara,pnt,Kr,osa,ap,nuf,Ser,shn | | 4.112e-05 | -10.10 | Syx16 (Syntaxin 16) | protein interactions | 33034 | 68 | 12 | 9204 | 412 | enok,CG3919,bi,fax,disco,Syx1A,CG11486,Acf,Sox102F,bin3,CG11873,CG43658 | | 4.209e-05 | -10.08 | regulation of phosphorylation | biological process | GO:0042325 | 271 | 23 | 12345 | 414 | Mos,PEK,sty,Egfr,wnd,CG34172,dpp,grnd,par-1,bora,CycE,Pli,N,edl,Dad,spi,Rho1,CG8441,aos,Atg17,mbt,Cdk5alpha,drk | | 4.265e-05 | -10.06 | regulation of eye photoreceptor cell development | biological process | GO:0042478 | 13 | 5 | 12345 | 414 | oc,pAbp,sty,mbt,N | | 4.265e-05 | -10.06 | cell-cell signaling involved in cell fate commitment | biological process | GO:0045168 | 13 | 5 | 12345 | 414 | hh,mam,Ser,Mmp2,N | | 4.265e-05 | -10.06 | imaginal disc-derived leg joint morphogenesis | biological process | GO:0016348 | 13 | 5 | 12345 | 414 | fj,bowl,dve,TfAP-2,N | | 4.265e-05 | -10.06 | haltere development | biological process | GO:0007482 | 13 | 5 | 12345 | 414 | Ubx,ap,aos,Syx1A,Egfr | | 4.289e-05 | -10.06 | regulation of protein modification process | biological process | GO:0031399 | 347 | 27 | 12345 | 414 | sens-2,Rho1,CG8441,aos,Pcl,Atg17,mbt,mof,drk,Cdk5alpha,NiPp1,dpp,Mos,PEK,sty,Egfr,wnd,N,spi,grnd,mmy,Utx,par-1,hh,bora,CycE,ctrip | | 4.372e-05 | -10.04 | transferase activity, transferring phosphorus-containing groups | molecular function | GO:0016772 | 482 | 33 | 12176 | 396 | Doa,CG10082,wnd,CG1271,tkv,KP78a,Taf1,Pak3,sdt,Cdk12,CG2201,CaMKII,Tlk,Asator,smal,S6kII,Hipk,CG11486,CG1344,PEK,Haspin,CG9541,CG34384,JIL-1,Mos,Egfr,PhKgamma,par-1,CG8485,fj,CycE,mbt,mmy | | 4.478e-05 | -10.01 | EGF-like\_Ca-bd\_dom | interpro domains | IPR001881 | 36 | 8 | 11383 | 428 | N,LRP1,crb,Ser,mgl,CadN,ft,slow | | 4.623e-05 | -9.98 | GO\_TISSUE\_DEVELOPMENT | MSigDB lists | GO\_TISSUE\_DEVELOPMENT | 292 | 25 | 4161 | 153 | sdt,Rho1,fj,Utx,ds,if,ap,Smox,N,NetB,Lim1,Ser,dpp,drk,Rac2,tkv,hh,CG9220,so,cta,TfAP-2,ft,CaMKII,wech,smal | | 4.728e-05 | -9.96 | neuroblast fate commitment | biological process | GO:0014017 | 30 | 7 | 12345 | 414 | Kr,N,Dr,spen,CycE,ham,osa | | 4.746e-05 | -9.96 | axon | cellular component | GO:0030424 | 233 | 21 | 11940 | 407 | Dscam1,ImpL2,Con,NetB,Syx1A,CadN,Fas2,ckn,robo2,nudE,NetA,nrv2,Ser,wnd,CaMKII,par-1,Prosap,Imp,Cdk5alpha,Acsl,Ptp99A | | 4.832e-05 | -9.94 | GO\_REGULATION\_OF\_NEURON\_DIFFERENTIATION | MSigDB lists | GO\_REGULATION\_OF\_NEURON\_DIFFERENTIATION | 112 | 14 | 4161 | 153 | Rac2,nAChRalpha4,so,hh,LRP1,NetB,pbl,N,CG5830,nudE,Tif-IA,dpp,Lac,Ser | | 4.835e-05 | -9.94 | epidermis development | biological process | GO:0008544 | 64 | 10 | 12345 | 414 | Rho1,EcR,spen,ds,N,ft,par-1,hh,lbe,fj | | 4.962e-05 | -9.91 | protein homodimerization activity | molecular function | GO:0042803 | 140 | 15 | 12176 | 396 | Lac,Dscam1,ttk,otk2,oc,Ten-m,ci,CadN,dpp,robo2,Pak3,psq,CtBP,twi,rau | | 5.043e-05 | -9.89 | MEISSNER\_BRAIN\_HCP\_WITH\_H3K4ME3\_AND\_H3K27ME3 | MSigDB lists | MEISSNER\_BRAIN\_HCP\_WITH\_H3K4ME3\_AND\_H3K27ME3 | 142 | 16 | 4161 | 153 | dpp,Ser,ap,gus,if,Notum,pbl,NetB,N,mgl,Rho1,Trpm,ft,CG5830,dally,Fbxl7 | | 5.215e-05 | -9.86 | enhancer binding | molecular function | GO:0035326 | 42 | 8 | 12176 | 396 | Doc1,tin,ttk,disco,slp1,ey,Ubx,aop | | 5.401e-05 | -9.83 | regulation of epithelial cell differentiation | biological process | GO:0030856 | 41 | 8 | 12345 | 414 | Egfr,aop,spi,Ptpmeg,ci,ttk,tsl,par-1 | | 5.436e-05 | -9.82 | mushroom body development | biological process | GO:0016319 | 78 | 11 | 12345 | 414 | ey,Fas2,Ptpmeg,Smox,robo2,cas,PsGEF,EcR,mbt,Dscam1,enok | | 5.525e-05 | -9.80 | pcx (pecanex) | protein interactions | 31204 | 6 | 4 | 9204 | 412 | Hsc70-3,CG2865,Fas2,N | | 5.525e-05 | -9.80 | rho-5 (rhomboid-5) | protein interactions | 2768944 | 6 | 4 | 9204 | 412 | Egfr,aos,sty,spi | | 5.645e-05 | -9.78 | GO\_REGULATION\_OF\_ORGAN\_FORMATION | MSigDB lists | GO\_REGULATION\_OF\_ORGAN\_FORMATION | 7 | 4 | 4161 | 153 | dpp,hh,so,N | | 5.654e-05 | -9.78 | GO\_HEART\_MORPHOGENESIS | MSigDB lists | GO\_HEART\_MORPHOGENESIS | 49 | 9 | 4161 | 153 | Ser,dpp,N,Smox,ft,Utx,ds,so,hh | | 5.664e-05 | -9.78 | GO\_POSITIVE\_REGULATION\_OF\_OSSIFICATION | MSigDB lists | GO\_POSITIVE\_REGULATION\_OF\_OSSIFICATION | 20 | 6 | 4161 | 153 | Ser,dpp,smal,TfAP-2,Smox,tkv | | 5.664e-05 | -9.78 | GO\_KIDNEY\_MORPHOGENESIS | MSigDB lists | GO\_KIDNEY\_MORPHOGENESIS | 20 | 6 | 4161 | 153 | dpp,hh,Lim1,so,ds,ft | | 5.761e-05 | -9.76 | GO\_SKELETAL\_SYSTEM\_DEVELOPMENT | MSigDB lists | GO\_SKELETAL\_SYSTEM\_DEVELOPMENT | 86 | 12 | 4161 | 153 | hh,so,CG9220,tkv,Lim1,dpp,TfAP-2,S6kII,ds,Smox,ft,Dscam1 | | 5.765e-05 | -9.76 | GO\_CELLULAR\_COMPONENT\_MORPHOGENESIS | MSigDB lists | GO\_CELLULAR\_COMPONENT\_MORPHOGENESIS | 225 | 21 | 4161 | 153 | drk,tkv,Rac2,Mrp4,hh,hts,Arp2,Fas2,nudE,trn,JIL-1,sdt,Ank,ap,if,NetB,pbl,Dscam1,N,dpp,Lim1 | | 6.038e-05 | -9.71 | GO\_REGIONALIZATION | MSigDB lists | GO\_REGIONALIZATION | 61 | 10 | 4161 | 153 | Dscam1,Smox,N,ap,Utx,dpp,Lim1,tkv,CG9220,hh | | 6.076e-05 | -9.71 | GO\_EMBRYONIC\_MORPHOGENESIS | MSigDB lists | GO\_EMBRYONIC\_MORPHOGENESIS | 129 | 15 | 4161 | 153 | N,Smox,NetB,Dscam1,TfAP-2,if,Utx,ap,Lim1,wech,Rac2,drk,so,hh,lbk | | 6.128e-05 | -9.70 | regulation of canonical Wnt signaling pathway | biological process | GO:0060828 | 79 | 11 | 12345 | 414 | Mmp2,Hipk,pbl,phyl,CtBP,Sin3A,Rho1,Notum,spen,dally,dlp | | 6.156e-05 | -9.70 | eggshell chorion assembly | biological process | GO:0007306 | 93 | 12 | 12345 | 414 | dpp,gus,Myb,Egfr,spoon,Syp,S,lama,N,ttk,CycE,Rac2 | | 6.207e-05 | -9.69 | hid (head involution defective) | protein interactions | 40009 | 105 | 15 | 9204 | 412 | NetA,Imp,ban,lz,wech,sty,CycE,Egfr,stwl,comm,pnt,aos,NetB,egr,spi | | 6.231e-05 | -9.68 | SAM\_PNT | pfam domains | PF02198 | 7 | 4 | 10480 | 393 | edl,l(3)mbt,pnt,aop | | 6.301e-05 | -9.67 | CG42788 (uncharacterized protein) | protein interactions | 41817 | 17 | 6 | 9204 | 412 | drk,CG32809,Fas2,eya,rau,fax | | 6.303e-05 | -9.67 | Pointed\_dom | interpro domains | IPR003118 | 7 | 4 | 11383 | 428 | l(3)mbt,aop,edl,pnt | | 6.345e-05 | -9.67 | peripheral nervous system development | biological process | GO:0007422 | 66 | 10 | 12345 | 414 | Rho1,Dscam1,EcR,Egfr,spen,N,ttk,Pex1,spi,CycE | | 6.452e-05 | -9.65 | haltere disc development | biological process | GO:0035216 | 14 | 5 | 12345 | 414 | ap,Ubx,Syx1A,aos,Egfr | | 6.473e-05 | -9.65 | GO\_PATTERN\_SPECIFICATION\_PROCESS | MSigDB lists | GO\_PATTERN\_SPECIFICATION\_PROCESS | 87 | 12 | 4161 | 153 | hh,so,CG9220,tkv,Lim1,dpp,Utx,cta,ap,Smox,N,Dscam1 | | 6.479e-05 | -9.64 | Vav (Vav guanine nucleotide exchange factor) | protein interactions | 32920 | 11 | 5 | 9204 | 412 | Egfr,egr,spi,aos,Rac2 | | 6.479e-05 | -9.64 | snama (something that sticks like glue) | protein interactions | 37857 | 11 | 5 | 9204 | 412 | Fas2,Tet,N,dpp,hh | | 6.481e-05 | -9.64 | epidermal growth factor receptor signaling pathway | biological process | GO:0007173 | 22 | 6 | 12345 | 414 | drk,spi,Egfr,spen,pnt,aop | | 6.481e-05 | -9.64 | ERBB signaling pathway | biological process | GO:0038127 | 22 | 6 | 12345 | 414 | drk,spi,aop,pnt,spen,Egfr | | 6.490e-05 | -9.64 | GO\_EPITHELIUM\_DEVELOPMENT | MSigDB lists | GO\_EPITHELIUM\_DEVELOPMENT | 193 | 19 | 4161 | 153 | wech,cta,TfAP-2,ft,hh,so,drk,Rac2,Ser,Lim1,dpp,Utx,ap,ds,N,Smox,NetB,Rho1,sdt | | 6.748e-05 | -9.60 | p53-like\_TF\_DNA-bd | interpro domains | IPR008967 | 20 | 6 | 11383 | 428 | Doc1,bi,NFAT,lz,Stat92E,Doc2 | | 6.880e-05 | -9.58 | CTTTGT\_LEF1\_Q2 | MSigDB lists | CTTTGT\_LEF1\_Q2 | 453 | 33 | 4161 | 153 | Hydr2,Ank,sdt,fne,Tpr2,NetB,Ppa,kdn,Utx,lili,tkv,dlp,CG2812,Taf1,CG5830,mle,cta,TfAP-2,NKAIN,trn,fj,LRP1,Smox,N,ds,Etl1,CG43313,so,Imp,Tlk,hh,Arl4,Ptpmeg | | 6.987e-05 | -9.57 | Cdc42 (Cdc42) | protein interactions | 32981 | 61 | 11 | 9204 | 412 | Pak3,tkv,CG10082,mbt,pbl,NFAT,N,RhoGEF2,Rho1,Tlk,RhoGAP18B | | 7.238e-05 | -9.53 | stem cell proliferation | biological process | GO:0072089 | 67 | 10 | 12345 | 414 | Sox21a,flfl,l(3)mbt,N,Zif,pon,wake,enok,Rho1,sdt | | 7.266e-05 | -9.53 | GO\_NEURON\_DEVELOPMENT | MSigDB lists | GO\_NEURON\_DEVELOPMENT | 162 | 17 | 4161 | 153 | Fas2,nudE,ft,DIP-epsilon,hh,tkv,Rac2,drk,Lim1,Dscam1,NetB,N,ap,trn,JIL-1,mbt,Ank | | 7.377e-05 | -9.51 | regulation of embryonic cell shape | biological process | GO:0016476 | 32 | 7 | 12345 | 414 | neo,cta,ttk,tyn,Rho1,Rac2,RhoGEF2 | | 7.439e-05 | -9.51 | GO\_UROGENITAL\_SYSTEM\_DEVELOPMENT | MSigDB lists | GO\_UROGENITAL\_SYSTEM\_DEVELOPMENT | 75 | 11 | 4161 | 153 | if,ds,TfAP-2,N,Smox,ft,Lim1,Ser,dpp,hh,so | | 7.730e-05 | -9.47 | regulation of photoreceptor cell differentiation | biological process | GO:0046532 | 43 | 8 | 12345 | 414 | N,ttk,pAbp,oc,S6kII,mbt,sty,aop | | 7.743e-05 | -9.47 | cell growth | biological process | GO:0016049 | 81 | 11 | 12345 | 414 | Rac2,scb,Ptpmeg,Dad,CadN,RhoGEF2,wnd,mle,Dscam1,tkv,Rho1 | | 7.768e-05 | -9.46 | border follicle cell migration | biological process | GO:0007298 | 110 | 13 | 12345 | 414 | Stat92E,if,par-1,jing,bbg,shd,N,spoon,Egfr,aop,nudE,EcR,Rho1 | | 7.907e-05 | -9.45 | GO\_REGULATION\_OF\_OSTEOBLAST\_DIFFERENTIATION | MSigDB lists | GO\_REGULATION\_OF\_OSTEOBLAST\_DIFFERENTIATION | 30 | 7 | 4161 | 153 | Smox,N,tkv,CG11367,Ser,dpp,smal | | 8.208e-05 | -9.41 | regulation of phosphate metabolic process | biological process | GO:0019220 | 302 | 24 | 12345 | 414 | Cdk5alpha,drk,aos,CG8441,Rho1,mbt,Atg17,bora,par-1,grnd,Pli,CycE,N,Dad,spi,edl,sty,PEK,Mos,wnd,Egfr,NiPp1,CG34172,dpp | | 8.208e-05 | -9.41 | regulation of phosphorus metabolic process | biological process | GO:0051174 | 302 | 24 | 12345 | 414 | Atg17,mbt,Rho1,CG8441,aos,drk,Cdk5alpha,CG34172,dpp,NiPp1,Egfr,wnd,PEK,Mos,sty,edl,Dad,spi,N,CycE,Pli,grnd,par-1,bora | | 8.352e-05 | -9.39 | X chromosome located dosage compensation complex, transcription activating | cellular component | GO:0016456 | 8 | 4 | 11940 | 407 | mle,JIL-1,lncRNA:roX1,mof | | 8.352e-05 | -9.39 | X chromosome | cellular component | GO:0000805 | 8 | 4 | 11940 | 407 | mof,lncRNA:roX1,JIL-1,mle | | 8.352e-05 | -9.39 | dosage compensation complex | cellular component | GO:0046536 | 8 | 4 | 11940 | 407 | mle,JIL-1,lncRNA:roX1,mof | | 8.456e-05 | -9.38 | S (Star) | protein interactions | 33281 | 52 | 10 | 9204 | 412 | ec,aop,sty,Egfr,N,Kr,jing,spi,edl,aos | | 8.456e-05 | -9.38 | ey (eyeless) | protein interactions | 43812 | 52 | 10 | 9204 | 412 | CtBP,NKAIN,eya,dpp,N,ci,hh,tkv,Hipk,ey | | 8.739e-05 | -9.35 | CTTTGA\_LEF1\_Q2 | MSigDB lists | CTTTGA\_LEF1\_Q2 | 285 | 24 | 4161 | 153 | ap,ds,Utx,kdn,Syx1A,mgl,Ser,Lim1,mmy,Tpr2,fne,wake,Ank,mbt,sdt,Acsl,Hydr2,CG9281,smal,CG34384,CG16890,CG3961,dlp,dally | | 8.907e-05 | -9.33 | NKAIN (Na,K-ATPase Interacting) | protein interactions | 37979 | 3 | 3 | 9204 | 412 | ey,esn,nrv2 | | 8.946e-05 | -9.32 | Hippo signaling pathway - fly | KEGG pathways | ko04391 | 61 | 9 | 3198 | 100 | dally,crb,ban,fj,ft,sdt,CycE,egr,ds | | 8.946e-05 | -9.32 | Hippo signaling pathway - fly | KEGG pathways | dme04391 | 61 | 9 | 3198 | 100 | ds,egr,CycE,sdt,ft,fj,ban,crb,dally | | 8.982e-05 | -9.32 | GO\_POSITIVE\_REGULATION\_OF\_TRANSCRIPTION\_FROM\_RNA\_POLYMERASE\_II\_PROMOTER | MSigDB lists | GO\_POSITIVE\_REGULATION\_OF\_TRANSCRIPTION\_FROM\_RNA\_POLYMERASE\_II\_PROMOTER | 181 | 18 | 4161 | 153 | Ser,dpp,if,ap,N,Smox,CG12099,EcR,JIL-1,CtBP,S6kII,TfAP-2,Taf1,hh,Hsc70-3,so,Sin3A,tkv | | 9.045e-05 | -9.31 | cell division | biological process | GO:0051301 | 266 | 22 | 12345 | 414 | CG17778,sdt,Rho1,tkv,pbl,wake,hts,dally,stwl,Myb,dpp,hh,mam,flfl,pAbp,Stat92E,CycE,Pex1,Syx1A,N,Zif,pon | | 9.085e-05 | -9.31 | GO\_DEVELOPMENTAL\_GROWTH | MSigDB lists | GO\_DEVELOPMENTAL\_GROWTH | 90 | 12 | 4161 | 153 | ap,Utx,N,Smox,CaMKII,hts,Lim1,nudE,smal,CycE,tkv,hh | | 9.096e-05 | -9.31 | transmembrane receptor protein serine/threonine kinase signaling pathway | biological process | GO:0007178 | 33 | 7 | 12345 | 414 | dally,hiw,shn,dpp,Dad,tkv,Smox | | 9.149e-05 | -9.30 | Cys-rich\_flank\_reg\_C | interpro domains | IPR000483 | 21 | 6 | 11383 | 428 | lbk,Toll-7,Lrt,Con,trn,Tl | | 9.207e-05 | -9.29 | EGF-like\_dom | interpro domains | IPR000742 | 74 | 11 | 11383 | 428 | spi,wb,Ten-m,slow,ft,mgl,CadN,Ser,crb,LRP1,N | | 9.211e-05 | -9.29 | GO\_SENSORY\_ORGAN\_MORPHOGENESIS | MSigDB lists | GO\_SENSORY\_ORGAN\_MORPHOGENESIS | 52 | 9 | 4161 | 153 | NetB,if,TfAP-2,Ser,Lim1,Rac2,fj,so,lbk | | 9.219e-05 | -9.29 | fly chr102C1-102C1\|4-0 | chromosome location | fly chr102C1-102C1\|4-0 | 5 | 3 | 25010 | 531 | asRNA:CR45126,Asator,CG33941 | | 9.219e-05 | -9.29 | fly chr102C1-102C1\|4-0 cM | chromosome location | fly chr102C1-102C1\|4-0 cM | 5 | 3 | 25010 | 531 | Asator,asRNA:CR45126,CG33941 | | 9.221e-05 | -9.29 | RYTTCCTG\_ETS2\_B | MSigDB lists | RYTTCCTG\_ETS2\_B | 215 | 20 | 4161 | 153 | Cyp1,Lim1,dpp,Mhcl,Smox,Cnx99A,mgl,Ppa,Utx,ap,SamDC,slow,Tudor-SN,Arp2,S6kII,Imp,dlp,Rac2,CG34384,CG3961 | | 9.222e-05 | -9.29 | ZIC3\_01 | MSigDB lists | ZIC3\_01 | 64 | 10 | 4161 | 153 | Utx,CG5830,NetB,NiPp1,Lac,ras,Rac2,LRP1,Tlk,RpS4 | | 9.337e-05 | -9.28 | salivary gland morphogenesis | biological process | GO:0007435 | 97 | 12 | 12345 | 414 | Atg17,EcR,Rho1,tin,eya,crb,Ser,if,Rac2,twi,trn,Utx | | 9.337e-05 | -9.28 | gland morphogenesis | biological process | GO:0022612 | 97 | 12 | 12345 | 414 | crb,eya,Utx,trn,Rac2,twi,if,Ser,Rho1,EcR,Atg17,tin | | 9.412e-05 | -9.27 | negative regulation of cellular component movement | biological process | GO:0051271 | 15 | 5 | 12345 | 414 | scb,Lrt,Ptpmeg,CG44774,Sin3A | | 9.412e-05 | -9.27 | R7 cell development | biological process | GO:0045467 | 15 | 5 | 12345 | 414 | Ten-m,ttk,rau,lz,CadN | | 9.412e-05 | -9.27 | crystal cell differentiation | biological process | GO:0042688 | 15 | 5 | 12345 | 414 | lz,ush,N,Ser,pnt | | 9.412e-05 | -9.27 | plasmatocyte differentiation | biological process | GO:0042387 | 15 | 5 | 12345 | 414 | N,ush,pnt,sty,Stat92E | | 9.412e-05 | -9.27 | compound eye cone cell differentiation | biological process | GO:0042675 | 15 | 5 | 12345 | 414 | Egfr,lz,eya,N,ttk | | 9.636e-05 | -9.25 | GO\_REGULATION\_OF\_BIOMINERAL\_TISSUE\_DEVELOPMENT | MSigDB lists | GO\_REGULATION\_OF\_BIOMINERAL\_TISSUE\_DEVELOPMENT | 14 | 5 | 4161 | 153 | Smox,tkv,TfAP-2,smal,dpp | | 9.713e-05 | -9.24 | EGF-type\_Asp/Asn\_hydroxyl\_site | interpro domains | IPR000152 | 30 | 7 | 11383 | 428 | slow,crb,LRP1,Ser,N,CadN,mgl | | 9.720e-05 | -9.24 | anterior/posterior axis specification | biological process | GO:0009948 | 144 | 15 | 12345 | 414 | aos,gus,dpp,Pat1,kni,hh,slp1,par-1,tsl,bowl,cnc,oc,psq,N,Kr | | 9.789e-05 | -9.23 | Gug (Grunge) | protein interactions | 46156 | 34 | 8 | 9204 | 412 | aop,Fas2,pnt,aos,Pcl,ft,ci,Ubx | | 9.855e-05 | -9.22 | GO\_REGULATION\_OF\_NERVOUS\_SYSTEM\_DEVELOPMENT | MSigDB lists | GO\_REGULATION\_OF\_NERVOUS\_SYSTEM\_DEVELOPMENT | 150 | 16 | 4161 | 153 | NetB,pbl,N,dpp,Ser,Lim1,CG12099,LRP1,CG5830,Lac,nudE,Tif-IA,Rac2,nAChRalpha4,hh,so | | 1.001e-04 | -9.21 | pum (pumilio) | protein interactions | 41094 | 53 | 10 | 9204 | 412 | stwl,pbl,mam,pAbp,Egfr,drk,shn,Rbfox1,AGO2,elav | | 1.046e-04 | -9.17 | regulation of multi-organism process | biological process | GO:0043900 | 250 | 21 | 12345 | 414 | Tl,Ptpmeg,CG4325,ush,cnc,Etl1,EcR,Toll-7,Doa,spi,ttk,Syx1A,Mmp2,Stat92E,Pli,par-1,Usp2,AGO2,Egfr,aop,CaMKII | | 1.056e-04 | -9.16 | GO\_TAXIS | MSigDB lists | GO\_TAXIS | 65 | 10 | 4161 | 153 | Fas2,Lim1,NetB,ap,hh,tkv,Rac2,drk,Ank,JIL-1 | | 1.084e-04 | -9.13 | neuroblast differentiation | biological process | GO:0014016 | 45 | 8 | 12345 | 414 | N,Kr,CycE,osa,cas,Dr,spen,ham | | 1.097e-04 | -9.12 | GO\_ENDOCARDIAL\_CUSHION\_DEVELOPMENT | MSigDB lists | GO\_ENDOCARDIAL\_CUSHION\_DEVELOPMENT | 8 | 4 | 4161 | 153 | N,ds,dpp,Ser | | 1.097e-04 | -9.12 | GO\_ARTERY\_MORPHOGENESIS | MSigDB lists | GO\_ARTERY\_MORPHOGENESIS | 8 | 4 | 4161 | 153 | Ser,LRP1,so,N | | 1.097e-04 | -9.12 | GO\_FOREBRAIN\_REGIONALIZATION | MSigDB lists | GO\_FOREBRAIN\_REGIONALIZATION | 8 | 4 | 4161 | 153 | Lim1,hh,dpp,ap | | 1.104e-04 | -9.11 | organ growth | biological process | GO:0035265 | 24 | 6 | 12345 | 414 | hh,bi,fj,ft,N,dpp | | 1.113e-04 | -9.10 | genitalia development | biological process | GO:0048806 | 34 | 7 | 12345 | 414 | salm,otk2,Dr,N,TfAP-2,lz,Fas2 | | 1.113e-04 | -9.10 | regulation of compound eye photoreceptor cell differentiation | biological process | GO:0110116 | 34 | 7 | 12345 | 414 | aop,sty,oc,pAbp,ttk,N,mbt | | 1.116e-04 | -9.10 | RNA polymerase II-specific DNA-binding transcription factor binding | molecular function | GO:0061629 | 35 | 7 | 12176 | 396 | Utx,bi,shn,Doc2,Doc1,ush,ocm | | 1.144e-04 | -9.08 | transcription factor complex | cellular component | GO:0005667 | 177 | 17 | 11940 | 407 | Smox,mof,Mondo,CG31875,Sox21a,CG3919,NFAT,udd,stwl,so,Ubx,EcR,Dad,mam,Taf1,N,jigr1 | | 1.196e-04 | -9.03 | imaginal disc-derived wing margin morphogenesis | biological process | GO:0008587 | 71 | 10 | 12345 | 414 | if,pio,Ser,mam,N,Hipk,dlp,osa,sno,heph | | 1.198e-04 | -9.03 | EGF | pfam domains | PF00008 | 22 | 6 | 10480 | 393 | Ser,ft,LRP1,N,crb,CadN | | 1.219e-04 | -9.01 | neural precursor cell proliferation | biological process | GO:0061351 | 58 | 9 | 12345 | 414 | wake,Rho1,sdt,enok,flfl,l(3)mbt,pon,Zif,N | | 1.219e-04 | -9.01 | neuroblast proliferation | biological process | GO:0007405 | 58 | 9 | 12345 | 414 | wake,sdt,Rho1,enok,flfl,l(3)mbt,pon,Zif,N | | 1.220e-04 | -9.01 | tor (torso) | protein interactions | 35717 | 35 | 8 | 9204 | 412 | tsl,sty,psq,drk,bowl,dpp,tkv,Stat92E | | 1.220e-04 | -9.01 | l(2)gl (lethal (2) giant larvae) | protein interactions | 33156 | 35 | 8 | 9204 | 412 | dpp,Stat92E,Zif,pnt,sdt,crb,nuf,CycE | | 1.229e-04 | -9.00 | GO\_MESENCHYME\_DEVELOPMENT | MSigDB lists | GO\_MESENCHYME\_DEVELOPMENT | 32 | 7 | 4161 | 153 | ds,N,Smox,dpp,hh,Ser,so | | 1.247e-04 | -8.99 | PDZ | interpro domains | IPR001478 | 64 | 10 | 11383 | 428 | CG3402,CG32758,Ptpmeg,PsGEF,Mhcl,RhoGEF2,Rhp,sdt,Prosap,bbg | | 1.254e-04 | -8.98 | learning or memory | biological process | GO:0007611 | 164 | 16 | 12345 | 414 | wnd,spoon,Egfr,hiw,CaMKII,S6kII,Stat92E,spi,N,EcR,wcy,scb,dbr,Fas2,drk,Rbfox1 | | 1.254e-04 | -8.98 | cognition | biological process | GO:0050890 | 164 | 16 | 12345 | 414 | S6kII,hiw,CaMKII,wnd,Egfr,spoon,N,spi,Stat92E,wcy,EcR,Rbfox1,drk,Fas2,dbr,scb | | 1.274e-04 | -8.97 | negative regulation of Wnt signaling pathway | biological process | GO:0030178 | 46 | 8 | 12345 | 414 | aop,Sin3A,Notum,Rho1,phyl,pbl,CtBP,Mmp2 | | 1.274e-04 | -8.97 | regulation of epithelial cell migration | biological process | GO:0010632 | 46 | 8 | 12345 | 414 | Sin3A,aop,robo2,Egfr,ttk,Ptpmeg,spi,hh | | 1.279e-04 | -8.96 | Ser/Thr\_kinase\_AS | interpro domains | IPR008271 | 147 | 16 | 11383 | 428 | Hipk,S6kII,PEK,Cdk12,Tlk,CaMKII,JIL-1,tkv,CG8485,par-1,KP78a,wnd,Pak3,PhKgamma,Mos,Doa | | 1.299e-04 | -8.95 | Rho GTPase cycle | REACTOME pathways | R-DME-194840 | 53 | 9 | 3923 | 148 | Rho1,Rac2,RhoGAP102A,RhoGAP15B,cta,RhoGAP18B,RhoGEF2,pbl,RhoGAP93B | | 1.332e-04 | -8.92 | negative regulation of locomotion | biological process | GO:0040013 | 16 | 5 | 12345 | 414 | scb,Lrt,Ptpmeg,Sin3A,CG44774 | | 1.332e-04 | -8.92 | negative regulation of Ras protein signal transduction | biological process | GO:0046580 | 16 | 5 | 12345 | 414 | CG42684,S6kII,NFAT,MESR3,sty | | 1.345e-04 | -8.91 | negative regulation of cell development | biological process | GO:0010721 | 86 | 11 | 12345 | 414 | aop,osa,Egfr,aos,S6kII,EcR,ttk,N,Ptpmeg,Fas2,CadN | | 1.351e-04 | -8.91 | regulation of tube length, open tracheal system | biological process | GO:0035159 | 35 | 7 | 12345 | 414 | ds,fj,nrv2,ft,Rho1,dpp,Fas2 | | 1.351e-04 | -8.91 | segment specification | biological process | GO:0007379 | 35 | 7 | 12345 | 414 | Ubx,spen,Stat92E,dlp,slp1,osa,jing | | 1.396e-04 | -8.88 | tissue homeostasis | biological process | GO:0001894 | 59 | 9 | 12345 | 414 | Lac,ush,crb,N,if,Atg17,Cnx99A,Egfr,disco | | 1.412e-04 | -8.87 | actin-mediated cell contraction | biological process | GO:0070252 | 25 | 6 | 12345 | 414 | Rho1,crb,tyn,cta,neo,RhoGEF2 | | 1.412e-04 | -8.87 | regulation of morphogenesis of an epithelium | biological process | GO:1905330 | 25 | 6 | 12345 | 414 | Smox,ds,otk2,aop,par-1,sty | | 1.437e-04 | -8.85 | lateral plasma membrane | cellular component | GO:0016328 | 16 | 5 | 11940 | 407 | RhoGEF2,Inx3,CG42748,Fas3,hts | | 1.437e-04 | -8.85 | external side of plasma membrane | cellular component | GO:0009897 | 16 | 5 | 11940 | 407 | dally,Mmp2,dlp,Tl,Toll-7 | | 1.461e-04 | -8.83 | labial disc development | biological process | GO:0035217 | 4 | 3 | 12345 | 414 | ci,dpp,hh | | 1.463e-04 | -8.83 | sex chromosome | cellular component | GO:0000803 | 9 | 4 | 11940 | 407 | lncRNA:roX1,mof,JIL-1,mle | | 1.469e-04 | -8.83 | cellular response to organic substance | biological process | GO:0071310 | 275 | 22 | 12345 | 414 | dpp,shn,Lpin,Cnx99A,dally,PEK,hiw,aop,Hsc70-3,Dad,Hydr2,Stat92E,grnd,Utx,Toll-7,EcR,tkv,drk,Tl,Smox,Drat,pnt | | 1.499e-04 | -8.81 | Dl (Delta) | protein interactions | 42313 | 126 | 16 | 9204 | 412 | hh,mam,N,Ubx,aos,CG9650,Ser,Egfr,S,ci,sno,Hipk,ap,crb,Dad,Bx | | 1.511e-04 | -8.80 | tkv (thickveins) | protein interactions | 33753 | 101 | 14 | 9204 | 412 | dpp,Ubx,dally,N,Egfr,tkv,ey,Dad,hiw,shn,Acsl,Rho1,scb,lili | | 1.540e-04 | -8.78 | multicellular organismal reproductive process | biological process | GO:0048609 | 1285 | 67 | 12345 | 414 | scb,goe,Rbfox1,lama,Tet,nudE,Rho1,Fas3,heph,if,pAbp,Npc1a,tsl,shd,ci,psq,CaMKII,Pat1,jing,drk,eya,Doa,baf,bbg,Mmp2,JIL-1,ttk,spoon,mle,Egfr,hts,wnd,S,sty,aop,Myb,tin,EcR,gus,mam,Rac2,wun2,N,Hydr2,so,Tudor-SN,cnc,elav,Inx2,tkv,Imp,CycE,Stat92E,l(3)mbt,hh,par-1,fne,Pex1,TfAP-2,yps,AGO2,dally,Syp,Mos,RhoGEF2,dpp,stwl | | 1.546e-04 | -8.77 | GO\_FOREBRAIN\_DEVELOPMENT | MSigDB lists | GO\_FOREBRAIN\_DEVELOPMENT | 95 | 12 | 4161 | 153 | LRP1,hh,Imp,CG17265,Rac2,Lim1,dpp,nudE,ap,mgl,N,ft | | 1.551e-04 | -8.77 | regulation of chromosome organization | biological process | GO:0033044 | 167 | 16 | 12345 | 414 | Chro,mof,Pcl,sens-2,lncRNA:roX1,Acf,Utx,ctrip,Stat92E,Tlk,l(3)neo38,psq,JIL-1,CG5815,Mos,mle | | 1.570e-04 | -8.76 | EGF\_CA | pfam domains | PF07645 | 23 | 6 | 10480 | 393 | LRP1,Ser,crb,mgl,N,slow | | 1.589e-04 | -8.75 | ptc (patched) | protein interactions | 35851 | 153 | 18 | 9204 | 412 | ci,dally,dpp,EcR,Lac,scb,N,hh,Svil,mgl,dlp,CG2865,Nhe2,Pits,LRP1,CycE,sas,hiw | | 1.598e-04 | -8.74 | AP2ALPHA\_01 | MSigDB lists | AP2ALPHA\_01 | 44 | 8 | 4161 | 153 | Dscam1,sm,pAbp,Tsp26A,Tlk,LRP1,so,Imp | | 1.599e-04 | -8.74 | tripartite regional subdivision | biological process | GO:0007351 | 134 | 14 | 12345 | 414 | N,psq,Kr,bowl,slp1,par-1,hh,tsl,kni,cnc,oc,Pat1,dpp,gus | | 1.599e-04 | -8.74 | anterior/posterior axis specification, embryo | biological process | GO:0008595 | 134 | 14 | 12345 | 414 | Kr,N,psq,cnc,oc,bowl,kni,tsl,hh,slp1,par-1,dpp,Pat1,gus | | 1.687e-04 | -8.69 | csw (corkscrew) | protein interactions | 45278 | 67 | 11 | 9204 | 412 | sty,if,aos,Stat92E,spi,tkv,pnt,dpp,drk,Egfr,S | | 1.730e-04 | -8.66 | regulation of cell cycle | biological process | GO:0051726 | 317 | 24 | 12345 | 414 | dally,Egfr,nuf,Mos,PEK,Myb,CG31612,CycE,Tlk,kni,hh,bora,l(3)mbt,ci,N,enok,Sin3A,stg,pbl,Cdk5alpha,mof,Chro,PhKgamma,Smox | | 1.733e-04 | -8.66 | Pkinase | pfam domains | PF00069 | 167 | 17 | 10480 | 393 | tkv,Pak3,mbt,CaMKII,Hipk,KP78a,Cdk12,Tlk,PhKgamma,PEK,Asator,par-1,S6kII,Mos,JIL-1,CG8485,Doa | | 1.733e-04 | -8.66 | Dorso-ventral axis formation | KEGG pathways | dme04320 | 28 | 6 | 3198 | 100 | drk,aop,Egfr,N,pnt,aos | | 1.733e-04 | -8.66 | Dorso-ventral axis formation | KEGG pathways | ko04320 | 28 | 6 | 3198 | 100 | aop,drk,aos,pnt,N,Egfr | | 1.760e-04 | -8.65 | srp (serpent) | protein interactions | 41944 | 20 | 6 | 9204 | 412 | ush,Ubx,wun2,Fas2,Kr,lz | | 1.781e-04 | -8.63 | adult behavior | biological process | GO:0030534 | 169 | 16 | 12345 | 414 | cv,S,hiw,Gbs-70E,Egfr,sm,oc,TfAP-2,hang,ey,spi,CCKLR-17D3,scb,Cdk5alpha,Fas2,Cirl | | 1.785e-04 | -8.63 | GO\_NEURON\_DIFFERENTIATION | MSigDB lists | GO\_NEURON\_DIFFERENTIATION | 208 | 19 | 4161 | 153 | ap,NetB,Dscam1,N,Lim1,Ser,trn,JIL-1,Ank,mbt,ft,Fas2,nudE,drk,tkv,Rac2,DIP-epsilon,hh,so | | 1.816e-04 | -8.61 | regulation of axonogenesis | biological process | GO:0050770 | 61 | 9 | 12345 | 414 | RhoGEF2,NetB,Dscam1,NetA,Rho1,pbl,Rac2,mmy,CadN | | 1.833e-04 | -8.60 | striated muscle cell differentiation | biological process | GO:0051146 | 104 | 12 | 12345 | 414 | Pak3,if,lbe,Rac2,twi,pnt,Rho1,CG13366,hts,tin,sty,aop | | 1.835e-04 | -8.60 | sevenless signaling pathway | biological process | GO:0045500 | 17 | 5 | 12345 | 414 | aop,pnt,phyl,ttk,drk | | 1.835e-04 | -8.60 | tracheal outgrowth, open tracheal system | biological process | GO:0007426 | 17 | 5 | 12345 | 414 | Rac2,Egfr,ttk,Mmp2,drk | | 1.850e-04 | -8.60 | Dad (Daughters against dpp) | protein interactions | 42059 | 37 | 8 | 9204 | 412 | lili,Acsl,Lpin,CycE,Rho1,pbl,dally,tkv | | 1.850e-04 | -8.60 | robo1 (roundabout 1) | protein interactions | 37603 | 37 | 8 | 9204 | 412 | sty,Rho1,robo2,elav,RhoGAP93B,comm,Dscam1,Lrt | | 1.871e-04 | -8.58 | embryonic axis specification | biological process | GO:0000578 | 136 | 14 | 12345 | 414 | tsl,kni,hh,par-1,slp1,bowl,cnc,oc,psq,N,Kr,gus,dpp,Pat1 | | 1.884e-04 | -8.58 | HOXA4\_Q2 | MSigDB lists | HOXA4\_Q2 | 45 | 8 | 4161 | 153 | tkv,CG45050,SamDC,CG16890,sdt,Nhe2,CaMKII,kdn | | 1.891e-04 | -8.57 | RNA polymerase II proximal promoter sequence-specific DNA binding | molecular function | GO:0000978 | 92 | 11 | 12176 | 396 | tin,CG31224,ttk,CG4360,bi,Utx,Stat92E,MESR3,NFAT,HGTX,slou | | 1.898e-04 | -8.57 | GO\_TELENCEPHALON\_REGIONALIZATION | MSigDB lists | GO\_TELENCEPHALON\_REGIONALIZATION | 4 | 3 | 4161 | 153 | ap,dpp,hh | | 1.898e-04 | -8.57 | GO\_REGULATION\_OF\_ODONTOGENESIS | MSigDB lists | GO\_REGULATION\_OF\_ODONTOGENESIS | 4 | 3 | 4161 | 153 | TfAP-2,hh,dpp | | 1.918e-04 | -8.56 | GO\_HEART\_VALVE\_DEVELOPMENT | MSigDB lists | GO\_HEART\_VALVE\_DEVELOPMENT | 9 | 4 | 4161 | 153 | dpp,Ser,ds,N | | 1.933e-04 | -8.55 | peb (pebbled) | protein interactions | 31391 | 68 | 11 | 9204 | 412 | tkv,dve,Egfr,dpp,ush,aop,ps,Acsl,Rho1,CycE,osa | | 1.938e-04 | -8.55 | cell surface | cellular component | GO:0009986 | 74 | 10 | 11940 | 407 | dlp,Toll-7,N,Dtg,dally,Ser,Tl,slow,Con,Mmp2 | | 2.061e-04 | -8.49 | regulation of reproductive process | biological process | GO:2000241 | 121 | 13 | 12345 | 414 | stg,Egfr,aop,Doa,Mos,CaMKII,Ptpmeg,spi,ttk,bab1,Syx1A,cnc,par-1 | | 2.063e-04 | -8.49 | regulation of cell shape | biological process | GO:0008360 | 62 | 9 | 12345 | 414 | pbl,Rac2,cta,tyn,ttk,RhoGEF2,dpp,neo,Rho1 | | 2.094e-04 | -8.47 | PDZ\_sf | interpro domains | IPR036034 | 68 | 10 | 11383 | 428 | Prosap,sdt,Rhp,bbg,CG3402,CG32758,Mhcl,RhoGEF2,Ptpmeg,PsGEF | | 2.103e-04 | -8.47 | identical protein binding | molecular function | GO:0042802 | 159 | 15 | 12176 | 396 | ci,Ten-m,ttk,oc,otk2,Dscam1,Lac,rau,twi,CtBP,Pak3,psq,robo2,CadN,dpp | | 2.130e-04 | -8.45 | dorsal/ventral axis specification | biological process | GO:0009950 | 76 | 10 | 12345 | 414 | twi,cnc,N,Tl,Syp,sty,Egfr,spoon,aos,dpp | | 2.136e-04 | -8.45 | TF\_T-box | interpro domains | IPR001699 | 9 | 4 | 11383 | 428 | Doc2,ocm,Doc1,bi | | 2.157e-04 | -8.44 | sexual reproduction | biological process | GO:0019953 | 1300 | 67 | 12345 | 414 | so,mam,Rac2,wun2,N,Hydr2,tin,EcR,gus,yps,AGO2,dally,RhoGEF2,Mos,Syp,dpp,stwl,CycE,Stat92E,l(3)mbt,par-1,hh,fne,TfAP-2,Pex1,tkv,Imp,Tudor-SN,cnc,elav,Inx2,CaMKII,Pat1,if,pAbp,Npc1a,tsl,shd,ci,psq,Tet,nudE,Rho1,Fas3,heph,scb,goe,lama,Rbfox1,Egfr,mle,spoon,wnd,hts,S,sty,aop,Myb,bbg,JIL-1,Mmp2,ttk,Doa,baf,jing,eya,drk | | 2.157e-04 | -8.44 | multi-organism reproductive process | biological process | GO:0044703 | 1300 | 67 | 12345 | 414 | CaMKII,Pat1,Npc1a,tsl,if,pAbp,ci,psq,shd,nudE,Tet,Rho1,Fas3,heph,scb,lama,Rbfox1,goe,S,sty,aop,Egfr,mle,spoon,wnd,hts,Myb,bbg,JIL-1,Mmp2,ttk,Doa,baf,jing,eya,drk,so,mam,Rac2,N,Hydr2,wun2,tin,EcR,gus,RhoGEF2,Mos,Syp,yps,AGO2,dally,stwl,dpp,l(3)mbt,hh,par-1,CycE,Stat92E,Pex1,TfAP-2,fne,tkv,Imp,Tudor-SN,cnc,Inx2,elav | | 2.207e-04 | -8.42 | ZIC1\_01 | MSigDB lists | ZIC1\_01 | 58 | 9 | 4161 | 153 | NetB,N,Utx,Lac,CG34384,mbt,CG9855,Tlk,LRP1 | | 2.221e-04 | -8.41 | GO\_EAR\_MORPHOGENESIS | MSigDB lists | GO\_EAR\_MORPHOGENESIS | 25 | 6 | 4161 | 153 | Rac2,NetB,TfAP-2,if,so,lbk | | 2.221e-04 | -8.41 | GSE16450\_CTRL\_VS\_IFNA\_12H\_STIM\_MATURE\_NEURON\_CELL\_LINE\_UP | MSigDB lists | GSE16450\_CTRL\_VS\_IFNA\_12H\_STIM\_MATURE\_NEURON\_CELL\_LINE\_UP | 25 | 6 | 4161 | 153 | Tsp26A,CG7264,EcR,Ank,nAChRalpha4,CG6330 | | 2.229e-04 | -8.41 | epithelial cell type specification, open tracheal system | biological process | GO:0035153 | 10 | 4 | 12345 | 414 | N,dpp,hh,pnt | | 2.229e-04 | -8.41 | morphogenesis of an epithelial fold | biological process | GO:0060571 | 10 | 4 | 12345 | 414 | RhoGEF2,Rho1,N,Kr | | 2.229e-04 | -8.41 | dedifferentiation | biological process | GO:0043696 | 10 | 4 | 12345 | 414 | Stat92E,pnt,CG46339,N | | 2.229e-04 | -8.41 | cell dedifferentiation | biological process | GO:0043697 | 10 | 4 | 12345 | 414 | pnt,Stat92E,N,CG46339 | | 2.229e-04 | -8.41 | determination of genital disc primordium | biological process | GO:0035225 | 10 | 4 | 12345 | 414 | S,pnt,Egfr,spi | | 2.230e-04 | -8.41 | embryonic hemopoiesis | biological process | GO:0035162 | 27 | 6 | 12345 | 414 | if,Rac2,lz,Pvf3,Rho1,N | | 2.237e-04 | -8.41 | HALLMARK\_ANDROGEN\_RESPONSE | MSigDB lists | HALLMARK\_ANDROGEN\_RESPONSE | 35 | 7 | 4161 | 153 | S6kII,Acsl,Pat1,Hydr2,tkv,Mrp4,mmy | | 2.293e-04 | -8.38 | GO\_LOCOMOTION | MSigDB lists | GO\_LOCOMOTION | 212 | 19 | 4161 | 153 | Mhcl,Lim1,NetB,N,ds,ap,JIL-1,Ank,Rho1,mbt,Fas2,nudE,Npc1a,emp,so,hh,Rac2,tkv,drk | | 2.325e-04 | -8.37 | glial cell development | biological process | GO:0021782 | 38 | 7 | 12345 | 414 | nrv2,rau,Pex1,cas,sty,salm,Dr | | 2.359e-04 | -8.35 | CtBP (C-terminal Binding Protein) | protein interactions | 41602 | 81 | 12 | 9204 | 412 | CtBP,kni,ttk,shn,disco,tou,N,E(spl)mdelta-HLH,ush,pAbp,ey,Kr | | 2.371e-04 | -8.35 | Su(fu) (Suppressor of fused) | protein interactions | 41565 | 21 | 6 | 9204 | 412 | Gyf,lili,ci,stg,Rbfox1,Gmap | | 2.371e-04 | -8.35 | cg (combgap) | protein interactions | 36571 | 21 | 6 | 9204 | 412 | Psc,EcR,CycE,eya,so,ci | | 2.468e-04 | -8.31 | regulation of cell morphogenesis involved in differentiation | biological process | GO:0010769 | 92 | 11 | 12345 | 414 | CadN,pbl,mmy,tsl,Rac2,NetB,Rho1,NetA,Dscam1,RhoGEF2,nudE | | 2.471e-04 | -8.31 | regulation of chromatin assembly or disassembly | biological process | GO:0001672 | 18 | 5 | 12345 | 414 | Stat92E,Tlk,mle,CG5815,Acf | | 2.515e-04 | -8.29 | Prot\_kinase\_dom | interpro domains | IPR000719 | 221 | 20 | 11383 | 428 | Cdk12,Tlk,CaMKII,JIL-1,Egfr,Hipk,S6kII,PEK,PhKgamma,Pak3,Asator,Haspin,Mos,Doa,tkv,CG8485,par-1,KP78a,mbt,wnd | | 2.538e-04 | -8.28 | osa (osa) | protein interactions | 42130 | 59 | 10 | 9204 | 412 | CycE,Bx,lz,osa,Kr,tara,tou,tna,dpp,bdg | | 2.538e-04 | -8.28 | fz (frizzled) | protein interactions | 45307 | 59 | 10 | 9204 | 412 | Rho1,sty,Nhe2,otk2,aos,bdg,Egfr,ft,N,ds | | 2.538e-04 | -8.28 | aos (argos) | protein interactions | 39833 | 59 | 10 | 9204 | 412 | spi,pnt,Ubx,S,Egfr,sty,salm,lz,Fas2,goe | | 2.591e-04 | -8.26 | regulation of cellular protein metabolic process | biological process | GO:0032268 | 623 | 38 | 12345 | 414 | sens-2,Rho1,heph,CG8441,aos,Pcl,mbt,Rbfox1,mof,smg,Cdk5alpha,Usp2,NiPp1,PEK,crb,N,spi,grnd,bora,pAbp,Atg17,elav,drk,dpp,Syp,Mos,sty,mle,Egfr,wnd,bin3,Utx,mmy,hh,par-1,CycE,ctrip,phyl | | 2.608e-04 | -8.25 | SAM\_PNT | smart domains | SM00251 | 7 | 4 | 5086 | 276 | pnt,edl,aop,l(3)mbt | | 2.611e-04 | -8.25 | regulation of transport | biological process | GO:0051049 | 267 | 21 | 12345 | 414 | Prosap,Snx3,CG9650,hh,Npc1a,Syx1A,crb,NFAT,Hip1,CaMKII,S,comm,NKAIN,S6kII,Cnx99A,Usp2,corn,drk,ScpX,mgl,sdt | | 2.631e-04 | -8.24 | ovarian follicle cell migration | biological process | GO:0007297 | 124 | 13 | 12345 | 414 | jing,bbg,par-1,if,Stat92E,N,shd,nudE,aop,Egfr,spoon,Rho1,EcR | | 2.633e-04 | -8.24 | GO\_REGULATION\_OF\_CELL\_DEVELOPMENT | MSigDB lists | GO\_REGULATION\_OF\_CELL\_DEVELOPMENT | 163 | 16 | 4161 | 153 | Ser,dpp,N,Smox,NetB,pbl,LRP1,CG12099,Lac,nudE,Tif-IA,CG5830,hh,so,nAChRalpha4,Rac2 | | 2.664e-04 | -8.23 | signaling receptor binding | molecular function | GO:0005102 | 256 | 20 | 12176 | 396 | crb,tsl,egr,aos,hh,dpp,Prosap,sas,spi,Pvf3,wake,ITP,cta,comm,pbl,Ser,slow,drk,Utx,Stat92E | | 2.756e-04 | -8.20 | Mi-2 (Mi-2) | protein interactions | 40170 | 49 | 9 | 9204 | 412 | CtBP,ttk,Egfr,Ubx,S6kII,Doc1,Kr,mle,EcR | | 2.759e-04 | -8.20 | imaginal disc-derived genitalia development | biological process | GO:0007484 | 28 | 6 | 12345 | 414 | otk2,Dr,N,TfAP-2,lz,Fas2 | | 2.759e-04 | -8.20 | regulation of stem cell differentiation | biological process | GO:2000736 | 28 | 6 | 12345 | 414 | goe,aos,enok,N,Stat92E,pnt | | 2.759e-04 | -8.20 | zygotic determination of anterior/posterior axis, embryo | biological process | GO:0007354 | 28 | 6 | 12345 | 414 | bowl,tsl,kni,oc,Kr,dpp | | 2.762e-04 | -8.19 | proximal promoter sequence-specific DNA binding | molecular function | GO:0000987 | 96 | 11 | 12176 | 396 | tin,CG4360,CG31224,ttk,Stat92E,MESR3,NFAT,Utx,bi,slou,HGTX | | 2.801e-04 | -8.18 | GO\_ANCHORED\_COMPONENT\_OF\_MEMBRANE | MSigDB lists | GO\_ANCHORED\_COMPONENT\_OF\_MEMBRANE | 26 | 6 | 4161 | 153 | Mmp2,Lac,DIP-epsilon,dally,dlp,Fas2 | | 2.815e-04 | -8.18 | regulation of protein metabolic process | biological process | GO:0051246 | 648 | 39 | 12345 | 414 | mof,Rbfox1,smg,Cdk5alpha,gus,mbt,sens-2,Rho1,CG8441,heph,aos,Pcl,spi,N,crb,pAbp,grnd,bora,Usp2,NiPp1,PEK,elav,drk,Atg17,bin3,CycE,phyl,ctrip,mmy,Utx,hh,par-1,dpp,Egfr,mle,wnd,Mos,Syp,sty | | 2.844e-04 | -8.17 | CG15436 (uncharacterized protein) | protein interactions | 33657 | 30 | 7 | 9204 | 412 | CG32369,Lim1,Zif,betaTub56D,CG4360,ttk,NetA | | 2.845e-04 | -8.16 | GO\_POSITIVE\_REGULATION\_OF\_NERVOUS\_SYSTEM\_DEVELOPMENT | MSigDB lists | GO\_POSITIVE\_REGULATION\_OF\_NERVOUS\_SYSTEM\_DEVELOPMENT | 73 | 10 | 4161 | 153 | hh,CG12099,Tif-IA,nudE,Lac,Lim1,dpp,N,pbl,NetB | | 2.941e-04 | -8.13 | positive regulation of cell population proliferation | biological process | GO:0008284 | 79 | 10 | 12345 | 414 | Stat92E,pnt,ey,spi,Tl,N,crb,Egfr,Myb,stg | | 2.945e-04 | -8.13 | hpo (hippo) | protein interactions | 37247 | 95 | 13 | 9204 | 412 | Smox,hh,Sox102F,Egfr,CG13366,ci,ds,disco,CG3253,CycE,CG3919,par-1,ban | | 2.957e-04 | -8.13 | pbl (pebble) | protein interactions | 38879 | 147 | 17 | 9204 | 412 | spi,CG44774,nuf,Dad,Rac2,ras,Rho1,Fur1,stg,jing,flfl,ttk,RapGAP1,shn,Tlk,Ptpmeg,CycE | | 2.979e-04 | -8.12 | Signaling by Rho GTPases | REACTOME pathways | R-DME-194315 | 114 | 13 | 3923 | 148 | RhoGAP18B,RhoGEF2,pbl,RhoGAP93B,Rho1,drk,Rac2,stg,RhoGAP102A,Arp2,RhoGAP15B,cta,Rhp | | 3.026e-04 | -8.10 | oocyte differentiation | biological process | GO:0009994 | 177 | 16 | 12345 | 414 | par-1,pAbp,N,psq,Mos,Syp,Egfr,hts,Pat1,dpp,scb,cnc,Rbfox1,nudE,heph,gus | | 3.105e-04 | -8.08 | GO\_REGULATION\_OF\_STEROL\_TRANSPORT | MSigDB lists | GO\_REGULATION\_OF\_STEROL\_TRANSPORT | 10 | 4 | 4161 | 153 | hh,ScpX,LRP1,EcR | | 3.105e-04 | -8.08 | GO\_REGULATION\_OF\_KIDNEY\_DEVELOPMENT | MSigDB lists | GO\_REGULATION\_OF\_KIDNEY\_DEVELOPMENT | 10 | 4 | 4161 | 153 | ft,Lim1,hh,so | | 3.105e-04 | -8.08 | GO\_AORTA\_DEVELOPMENT | MSigDB lists | GO\_AORTA\_DEVELOPMENT | 10 | 4 | 4161 | 153 | LRP1,Ser,so,mgl | | 3.105e-04 | -8.08 | GO\_MESENCHYME\_MORPHOGENESIS | MSigDB lists | GO\_MESENCHYME\_MORPHOGENESIS | 10 | 4 | 4161 | 153 | ds,Smox,N,dpp | | 3.126e-04 | -8.07 | fly chr61C7-61C7\|3-0.5 | chromosome location | fly chr61C7-61C7\|3-0.5 | 7 | 3 | 25010 | 531 | lncRNA:CR42719,lncRNA:CR43334,ban | | 3.126e-04 | -8.07 | fly chr61C7-61C7\|3-0.5 cM | chromosome location | fly chr61C7-61C7\|3-0.5 cM | 7 | 3 | 25010 | 531 | lncRNA:CR42719,lncRNA:CR43334,ban | | 3.231e-04 | -8.04 | behavior | biological process | GO:0007610 | 630 | 38 | 12345 | 414 | scb,Cdk5alpha,dbr,Fas2,Rbfox1,EcR,Rac2,spi,ey,hang,Bx,N,sm,CaMKII,ITP,tara,mub,CCKLR-17D3,Cirl,drk,Ptpmeg,sbb,wcy,qsm,oc,Stat92E,fne,TfAP-2,wnd,spoon,Egfr,mle,Gbs-70E,hiw,disco,S,cv,S6kII | | 3.262e-04 | -8.03 | Notch signaling pathway | biological process | GO:0007219 | 19 | 5 | 12345 | 414 | Ser,mam,sno,N,crb | | 3.262e-04 | -8.03 | dendrite self-avoidance | biological process | GO:0070593 | 19 | 5 | 12345 | 414 | Rho1,Dscam1,Fas2,ImpL2,esn | | 3.308e-04 | -8.01 | GO\_POSITIVE\_REGULATION\_OF\_MULTICELLULAR\_ORGANISMAL\_PROCESS | MSigDB lists | GO\_POSITIVE\_REGULATION\_OF\_MULTICELLULAR\_ORGANISMAL\_PROCESS | 236 | 20 | 4161 | 153 | tkv,so,hh,CG11367,CaMKII,TfAP-2,mle,nudE,Tif-IA,smal,Lac,EcR,CG12099,Smox,N,NetB,pbl,Lim1,Ser,dpp | | 3.312e-04 | -8.01 | GCACTTT\_MIR175P\_MIR20A\_MIR106A\_MIR106B\_MIR20B\_MIR519D | MSigDB lists | GCACTTT\_MIR175P\_MIR20A\_MIR106A\_MIR106B\_MIR20B\_MIR519D | 118 | 13 | 4161 | 153 | S6kII,CG41099,l(3)mbt,Ptpmeg,nudE,phtf,Hydr2,JIL-1,CG5742,fj,Rho1,Imp,CG11367 | | 3.348e-04 | -8.00 | side of membrane | cellular component | GO:0098552 | 65 | 9 | 11940 | 407 | Mmp2,Pli,Stat92E,sty,cta,Tl,dally,Toll-7,dlp | | 3.381e-04 | -7.99 | female germ-line stem cell population maintenance | biological process | GO:0036099 | 29 | 6 | 12345 | 414 | enok,N,stwl,fax,Tif-IA,dpp | | 3.388e-04 | -7.99 | negative regulation of intracellular signal transduction | biological process | GO:1902532 | 111 | 12 | 12345 | 414 | N,Hipk,NFAT,CG42684,ImpL2,MESR3,par-1,S6kII,aos,RapGAP1,sty,ds | | 3.405e-04 | -7.99 | GO\_TUBE\_DEVELOPMENT | MSigDB lists | GO\_TUBE\_DEVELOPMENT | 134 | 14 | 4161 | 153 | hh,so,dpp,wech,Ser,Lim1,cta,ap,Utx,ds,ft,NetB,Smox,N | | 3.410e-04 | -7.98 | Pc (Polycomb) | protein interactions | 40358 | 109 | 14 | 9204 | 412 | lawc,Pcl,jing,Psc,Ubx,tna,tou,tara,psq,CycE,salm,enok,osa,Utx | | 3.411e-04 | -7.98 | oenocyte differentiation | biological process | GO:0001742 | 11 | 4 | 12345 | 414 | spi,salm,pnt,Egfr | | 3.411e-04 | -7.98 | cardiac cell fate commitment | biological process | GO:0060911 | 11 | 4 | 12345 | 414 | robo2,tin,EcR,Ubx | | 3.411e-04 | -7.98 | cell elongation involved in imaginal disc-derived wing morphogenesis | biological process | GO:0090254 | 11 | 4 | 12345 | 414 | RhoGEF2,Dad,Rho1,tkv | | 3.444e-04 | -7.97 | Optix (optix) | protein interactions | 44108 | 4 | 3 | 9204 | 412 | eya,ey,Smox | | 3.444e-04 | -7.97 | Ero1L (Endoplasmic reticulum oxidoreductin-1-like) | protein interactions | 38500 | 4 | 3 | 9204 | 412 | PEK,N,cnc | | 3.504e-04 | -7.96 | Znf\_C2H2\_type | interpro domains | IPR013087 | 297 | 24 | 11383 | 428 | CG32772,shn,l(3)neo38,CG31224,CG18446,Kr,disco,ttk,sbb,Zif,ush,ci,Kah,CG31612,ham,jing,cas,CG4360,salm,sens-2,CG11247,hang,CG9650,bowl | | 3.525e-04 | -7.95 | gonad development | biological process | GO:0008406 | 53 | 8 | 12345 | 414 | aop,Egfr,robo2,tin,eya,salm,hh,wb | | 3.525e-04 | -7.95 | skeletal muscle organ development | biological process | GO:0060538 | 53 | 8 | 12345 | 414 | Rho1,wech,betaTub56D,lbe,slow,Rac2,if,N | | 3.525e-04 | -7.95 | development of primary sexual characteristics | biological process | GO:0045137 | 53 | 8 | 12345 | 414 | wb,hh,salm,eya,tin,Egfr,robo2,aop | | 3.561e-04 | -7.94 | dorsal closure, leading edge cell differentiation | biological process | GO:0046663 | 5 | 3 | 12345 | 414 | ush,dpp,Rho1 | | 3.561e-04 | -7.94 | wing disc proximal/distal pattern formation | biological process | GO:0007473 | 5 | 3 | 12345 | 414 | N,dpp,hh | | 3.561e-04 | -7.94 | leading edge cell differentiation | biological process | GO:0035026 | 5 | 3 | 12345 | 414 | Rho1,dpp,ush | | 3.575e-04 | -7.94 | cell-substrate junction | cellular component | GO:0030055 | 40 | 7 | 11940 | 407 | smal,wech,ckn,Lrt,slow,scb,if | | 3.575e-04 | -7.94 | cell-substrate adherens junction | cellular component | GO:0005924 | 40 | 7 | 11940 | 407 | smal,wech,Lrt,slow,scb,ckn,if | | 3.628e-04 | -7.92 | Lpt (Lost PHDs of trr) | protein interactions | 37795 | 15 | 5 | 9204 | 412 | EcR,tkv,rau,Utx,dpp | | 3.657e-04 | -7.91 | EGF\_2 | prosite domains | PS01186 | 55 | 10 | 6243 | 316 | spi,crb,Ser,mgl,Ten-m,slow,N,CadN,LRP1,ft | | 3.682e-04 | -7.91 | transmembrane receptor protein tyrosine kinase signaling pathway | biological process | GO:0007169 | 112 | 12 | 12345 | 414 | Socs16D,phyl,pnt,tsl,smal,Pvf3,spi,drk,ttk,Egfr,spen,aop | | 3.708e-04 | -7.90 | T3R\_Q6 | MSigDB lists | T3R\_Q6 | 62 | 9 | 4161 | 153 | Tudor-SN,baf,Utx,Cyt-c1,sm,hh,fne,JIL-1,CG3961 | | 3.726e-04 | -7.90 | BUYTAERT\_PHOTODYNAMIC\_THERAPY\_STRESS\_UP | MSigDB lists | BUYTAERT\_PHOTODYNAMIC\_THERAPY\_STRESS\_UP | 168 | 16 | 4161 | 153 | Hsc70Cb,MFS10,Npc1a,nudE,cta,par-1,CG2812,JIL-1,Pat1,dpp,sno,Cyp1,Utx,Syx1A,lili,N | | 3.751e-04 | -7.89 | multicellular organismal homeostasis | biological process | GO:0048871 | 67 | 9 | 12345 | 414 | Egfr,disco,Atg17,Cnx99A,if,Lac,ush,N,crb | | 3.799e-04 | -7.88 | establishment or maintenance of apical/basal cell polarity | biological process | GO:0035088 | 41 | 7 | 12345 | 414 | Egfr,ds,fj,Rho1,crb,sdt,ft | | 3.799e-04 | -7.88 | negative regulation of canonical Wnt signaling pathway | biological process | GO:0090090 | 41 | 7 | 12345 | 414 | Sin3A,Mmp2,Notum,Rho1,CtBP,phyl,pbl | | 3.799e-04 | -7.88 | establishment or maintenance of bipolar cell polarity | biological process | GO:0061245 | 41 | 7 | 12345 | 414 | sdt,ft,crb,Rho1,fj,ds,Egfr | | 3.826e-04 | -7.87 | GO\_CELLULAR\_RESPONSE\_TO\_INORGANIC\_SUBSTANCE | MSigDB lists | GO\_CELLULAR\_RESPONSE\_TO\_INORGANIC\_SUBSTANCE | 38 | 7 | 4161 | 153 | CaMKII,pbl,TfAP-2,Ank,Cyp1,Hsc70-3,hh | | 3.924e-04 | -7.84 | GO\_REGULATORY\_REGION\_NUCLEIC\_ACID\_BINDING | MSigDB lists | GO\_REGULATORY\_REGION\_NUCLEIC\_ACID\_BINDING | 90 | 11 | 4161 | 153 | Tif-IA,Utx,TfAP-2,N,Smox,sm,Taf1,CG12099,so,Sin3A,EcR | | 3.932e-04 | -7.84 | stem cell division | biological process | GO:0017145 | 97 | 11 | 12345 | 414 | Stat92E,hh,flfl,Zif,pon,dally,wake,dpp,tkv,sdt,stwl | | 3.945e-04 | -7.84 | membrane | cellular component | GO:0016020 | 2722 | 122 | 11940 | 407 | Snx3,Fili,Mitofilin,egr,drk,Socs16D,Fur1,N,nrv2,wun2,CG41128,fax,CG12730,Lrt,if,CG11752,nuf,Gmap,emp,Tsp39D,Pvf3,CG6231,Ptr,hts,Dmtn,CG9220,CG34172,Cnx99A,ImpL2,Nhe2,CG42748,Pex1,Stat92E,MFS14,par-1,Atg17,Hydr2,Cad87A,Prosap,pbl,CG11498,comm,neo,scb,ds,Cyt-c1,mbt,Mmp2,Inx3,Rac2,Hsc70-3,goe,sdt,dally,CG9855,hh,Ykt6,uzip,Inx2,CadN,Tim17b,mgl,Tsp26A,bdg,Hip1,tyn,Ptp99A,DIP-epsilon,CG6051,bou,RhoGEF2,Cirl,tkv,CG8531,cta,Tl,Ser,crb,Pli,hiw,nrm,otk2,Rho1,ft,LManII,Fas2,Npc1a,CG43313,Syx1A,CG45782,S,Trpm,sty,Mrp4,Toll-7,CG5830,dlp,ND-B14.5AL,CaMKII,nAChRalpha4,Arl4,Egfr,Lac,CG14646,dpr7,spi,Ten-m,grnd,sas,CCKLR-17D3,fj,CG11367,Fas3,CD98hc,CG1492,Con,smal,ci,Ank,Dscam1,MFS10,spoon | | 4.004e-04 | -7.82 | ion binding | molecular function | GO:0043167 | 2058 | 93 | 12176 | 396 | CG45050,N,kni,hh,crb,S6kII,CG8097,Cyp4p2,Mhcl,PEK,Haspin,pbl,EcR,Doa,wnd,rush,CG4325,ctrip,mle,lz,dbr,CadN,CG14535,Hph,Myo10A,Mmp2,sas,CG4199,ds,shn,LRP1,Tet,CG2991,Mos,hiw,PhKgamma,par-1,CG8485,CG41099,Snx3,Arl4,Psc,Ser,CG11247,Mrp4,Asator,CG9855,Etl1,Arp2,dpp,mgl,PsGEF,CG12099,tna,l(3)mbt,CG32369,ScsbetaG,Rho1,Hipk,esn,bor,tkv,KP78a,Pak3,Cdk12,RhoGAP15B,Rac2,Hip1,Zif,ft,CaMKII,Tlk,Cnx99A,Hsc70-3,mbt,CG12010,CG32758,TfIIS,CG9281,hang,CG9541,JIL-1,Cad87A,Egfr,betaTub56D,Utx,CG6752,CG3097,wech,shd,slow,Pex1,CG6051 | | 4.013e-04 | -7.82 | GSE23114\_WT\_VS\_SLE2C1\_MOUSE\_SPLEEN\_B1A\_BCELL\_DN | MSigDB lists | GSE23114\_WT\_VS\_SLE2C1\_MOUSE\_SPLEEN\_B1A\_BCELL\_DN | 50 | 8 | 4161 | 153 | PhKgamma,Cyp1,mub,ap,so,CycE,ras,LManII | | 4.087e-04 | -7.80 | cora (coracle) | protein interactions | 37205 | 23 | 6 | 9204 | 412 | nrv2,rau,ft,Egfr,Fas2,S6kII | | 4.090e-04 | -7.80 | negative regulation of cellular component organization | biological process | GO:0051129 | 164 | 15 | 12345 | 414 | Acsl,lncRNA:roX1,hiw,Mos,Syp,hts,spoon,JIL-1,Mmp2,Chro,Dad,CadN,Utx,ctrip,scb | | 4.109e-04 | -7.80 | negative regulation of cellular response to growth factor stimulus | biological process | GO:0090288 | 30 | 6 | 12345 | 414 | sty,hiw,Acsl,Mmp2,Dad,Lpin | | 4.167e-04 | -7.78 | MAN1 (MAN1) | protein interactions | 37838 | 9 | 4 | 9204 | 412 | Dad,dpp,baf,Smox | | 4.185e-04 | -7.78 | positive regulation of protein phosphorylation | biological process | GO:0001934 | 130 | 13 | 12345 | 414 | spi,drk,grnd,bora,Cdk5alpha,Rho1,dpp,Atg17,mbt,Mos,PEK,Egfr,wnd | | 4.212e-04 | -7.77 | localization | biological process | GO:0051179 | 2206 | 101 | 12345 | 414 | Tsp26A,CG41099,rush,scb,smg,Rho1,Hph,nudE,Fur1,bdg,Dad,Myo10A,shd,Npc1a,bora,MFS10,Snx3,if,NetA,Acsl,Pat1,ft,Chro,drk,eya,Pvf3,MFS14,trn,pnt,jing,nAChRalpha4,Doa,Syx1A,Mmp2,CG32758,egr,lbe,bbg,nrv2,CG43693,aop,S,Gmap,hts,ds,spoon,Egfr,dlp,pbl,Trpm,Sin3A,Mrp4,gus,mbt,NetB,EcR,Arl4,Ykt6,ScpX,tin,N,crb,wun2,ey,salm,Rac2,flfl,Lrt,so,CG13784,nuf,robo2,CG13029,Inx2,Cirl,cta,cnc,Dscam1,CD98hc,sdt,tkv,Pex1,CG6231,dve,Inx3,edl,Cyt-c1,kni,par-1,hh,Nhe2,l(3)mbt,Stat92E,Tim17b,dpp,comm,CG8531,Ank,dally,wake,AGO2 | | 4.258e-04 | -7.76 | cellular response to chemical stimulus | biological process | GO:0070887 | 378 | 26 | 12345 | 414 | Toll-7,Hph,EcR,tkv,drk,Tl,Smox,Drat,pnt,dpp,shn,Lpin,Cnx99A,dally,PEK,hiw,aop,Hsc70-3,Dad,wun2,Hydr2,Stat92E,grnd,Utx,Lrt,hh | | 4.313e-04 | -7.75 | GO\_RETINA\_DEVELOPMENT\_IN\_CAMERA\_TYPE\_EYE | MSigDB lists | GO\_RETINA\_DEVELOPMENT\_IN\_CAMERA\_TYPE\_EYE | 28 | 6 | 4161 | 153 | tkv,fj,ap,ras,TfAP-2,Lim1 | | 4.352e-04 | -7.74 | fra (frazzled) | protein interactions | 36377 | 32 | 7 | 9204 | 412 | comm,Dscam1,robo2,NetB,Rho1,NetA,N | | 4.352e-04 | -7.74 | Axn (Axin) | protein interactions | 43565 | 32 | 7 | 9204 | 412 | Utx,N,aop,salm,esn,Hipk,CycE | | 4.382e-04 | -7.73 | aPKC (atypical protein kinase C) | protein interactions | 47594 | 52 | 9 | 9204 | 412 | sdt,Zif,Myo10A,pbl,pon,sty,par-1,nuf,crb | | 4.382e-04 | -7.73 | Src42A (Src oncogene at 42A) | protein interactions | 35524 | 52 | 9 | 9204 | 412 | drk,Egfr,S,bowl,Stat92E,spi,smash,Dscam1,Ser | | 4.401e-04 | -7.73 | GO\_CELL\_PROJECTION\_ORGANIZATION | MSigDB lists | GO\_CELL\_PROJECTION\_ORGANIZATION | 223 | 19 | 4161 | 153 | Lim1,NetB,Dscam1,N,if,ap,JIL-1,trn,mbt,Ank,Arp2,Fas2,nudE,ft,Mrp4,hh,Rac2,tkv,drk | | 4.492e-04 | -7.71 | BENPORATH\_ES\_WITH\_H3K27ME3 | MSigDB lists | BENPORATH\_ES\_WITH\_H3K27ME3 | 154 | 15 | 4161 | 153 | ras,fne,TfIIS,dally,hh,so,Notum,Syx1A,ap,NetB,Ppa,Dscam1,mgl,Lac,Fas2 | | 4.499e-04 | -7.71 | fly chr11B9-11B9\|1-39 | chromosome location | fly chr11B9-11B9\|1-39 | 2 | 2 | 25010 | 531 | lncRNA:CR33963,mir-9377 | | 4.499e-04 | -7.71 | fly chr86E18-86E18\|3-51 cM | chromosome location | fly chr86E18-86E18\|3-51 cM | 2 | 2 | 25010 | 531 | l(3)neo38,lncRNA:CR45595 | | 4.499e-04 | -7.71 | fly chr86E18-86E18\|3-51 | chromosome location | fly chr86E18-86E18\|3-51 | 2 | 2 | 25010 | 531 | l(3)neo38,lncRNA:CR45595 | | 4.499e-04 | -7.71 | fly chr11B9-11B9\|1-39 cM | chromosome location | fly chr11B9-11B9\|1-39 cM | 2 | 2 | 25010 | 531 | mir-9377,lncRNA:CR33963 | | 4.508e-04 | -7.70 | positive regulation of nervous system development | biological process | GO:0051962 | 131 | 13 | 12345 | 414 | oc,Prosap,pnt,mmy,spi,N,Pak3,nudE,EcR,mbt,Arp2,tkv,Imp | | 4.508e-04 | -7.70 | chorion-containing eggshell formation | biological process | GO:0007304 | 131 | 13 | 12345 | 414 | CycE,Rac2,lama,N,ttk,spoon,Egfr,Syp,S,dpp,gus,Myb,tkv | | 4.614e-04 | -7.68 | GSE9988\_ANTI\_TREM1\_VS\_LOW\_LPS\_MONOCYTE\_UP | MSigDB lists | GSE9988\_ANTI\_TREM1\_VS\_LOW\_LPS\_MONOCYTE\_UP | 51 | 8 | 4161 | 153 | Npc1a,Syx1A,cta,emp,CG9220,Snx3,SamDC,Acsl | | 4.618e-04 | -7.68 | GO\_BONE\_MATURATION | MSigDB lists | GO\_BONE\_MATURATION | 5 | 3 | 4161 | 153 | dpp,ds,ft | | 4.618e-04 | -7.68 | GO\_AORTA\_MORPHOGENESIS | MSigDB lists | GO\_AORTA\_MORPHOGENESIS | 5 | 3 | 4161 | 153 | so,Ser,LRP1 | | 4.618e-04 | -7.68 | GO\_PERICARDIUM\_DEVELOPMENT | MSigDB lists | GO\_PERICARDIUM\_DEVELOPMENT | 5 | 3 | 4161 | 153 | Smox,N,dpp | | 4.618e-04 | -7.68 | CDPCR3\_01 | MSigDB lists | CDPCR3\_01 | 5 | 3 | 4161 | 153 | so,NetB,CG6330 | | 4.618e-04 | -7.68 | GO\_ORGAN\_MATURATION | MSigDB lists | GO\_ORGAN\_MATURATION | 5 | 3 | 4161 | 153 | ft,ds,dpp | | 4.618e-04 | -7.68 | GO\_REGULATION\_OF\_CHOLESTEROL\_EFFLUX | MSigDB lists | GO\_REGULATION\_OF\_CHOLESTEROL\_EFFLUX | 5 | 3 | 4161 | 153 | EcR,hh,LRP1 | | 4.618e-04 | -7.68 | GO\_ATRIOVENTRICULAR\_VALVE\_MORPHOGENESIS | MSigDB lists | GO\_ATRIOVENTRICULAR\_VALVE\_MORPHOGENESIS | 5 | 3 | 4161 | 153 | dpp,ds,N | | 4.618e-04 | -7.68 | GO\_POSITIVE\_REGULATION\_OF\_KIDNEY\_DEVELOPMENT | MSigDB lists | GO\_POSITIVE\_REGULATION\_OF\_KIDNEY\_DEVELOPMENT | 5 | 3 | 4161 | 153 | so,Lim1,hh | | 4.648e-04 | -7.67 | ex (expanded) | protein interactions | 33218 | 42 | 8 | 9204 | 412 | par-1,ban,crb,CycE,drk,l(3)mbt,ft,Kr | | 4.658e-04 | -7.67 | GO\_ANATOMICAL\_STRUCTURE\_FORMATION\_INVOLVED\_IN\_MORPHOGENESIS | MSigDB lists | GO\_ANATOMICAL\_STRUCTURE\_FORMATION\_INVOLVED\_IN\_MORPHOGENESIS | 224 | 19 | 4161 | 153 | Lim1,Ser,Smox,N,ds,if,Utx,ap,slow,fj,sdt,Arp2,wech,TfAP-2,cta,so,Mrp4,hh,drk | | 4.687e-04 | -7.67 | positive regulation of locomotion | biological process | GO:0040017 | 69 | 9 | 12345 | 414 | aop,Ten-m,Egfr,robo2,Dscam1,mmy,ttk,spi,Pvf3 | | 4.692e-04 | -7.66 | repressing transcription factor binding | molecular function | GO:0070491 | 21 | 5 | 12176 | 396 | HGTX,EcR,edl,kni,CtBP | | 4.708e-04 | -7.66 | Ig-like\_dom\_sf | interpro domains | IPR036179 | 133 | 14 | 11383 | 428 | beat-IIIc,Dscam1,Fas2,Lac,lbk,nrm,otk2,dpr7,Ptp99A,ImpL2,CG6044,Fas3,robo2,DIP-epsilon | | 4.857e-04 | -7.63 | somatic muscle development | biological process | GO:0007525 | 84 | 10 | 12345 | 414 | ap,Sin3A,hts,CG13366,Tl,eya,Ubx,E(spl)mdelta-HLH,Stat92E,salm | | 4.935e-04 | -7.61 | EGF\_3 | prosite domains | PS50026 | 57 | 10 | 6243 | 316 | slow,CadN,N,ft,LRP1,spi,crb,Ser,mgl,Ten-m | | 4.952e-04 | -7.61 | stem cell fate determination | biological process | GO:0048867 | 31 | 6 | 12345 | 414 | spen,Dr,CycE,cas,Kr,N | | 4.952e-04 | -7.61 | epithelial cell morphogenesis | biological process | GO:0003382 | 31 | 6 | 12345 | 414 | RhoGEF2,CG41099,crb,tyn,neo,cta | | 4.981e-04 | -7.60 | histoblast morphogenesis | biological process | GO:0007488 | 12 | 4 | 12345 | 414 | stg,tkv,Dad,EcR | | 4.981e-04 | -7.60 | lateral inhibition | biological process | GO:0046331 | 12 | 4 | 12345 | 414 | mam,Ser,N,Mmp2 | | 5.085e-04 | -7.58 | bi (bifid) | protein interactions | 31379 | 16 | 5 | 9204 | 412 | hh,ap,ci,N,dpp | | 5.085e-04 | -7.58 | px (plexus) | protein interactions | 37573 | 16 | 5 | 9204 | 412 | S,if,Egfr,N,ci | | 5.119e-04 | -7.58 | regulation of actin filament-based process | biological process | GO:0032970 | 100 | 11 | 12345 | 414 | cta,pbl,if,tsl,CG43658,PsGEF,Rho1,Arp2,hts,spoon,RhoGEF2 | | 5.211e-04 | -7.56 | hEGF | pfam domains | PF12661 | 11 | 4 | 10480 | 393 | Ser,N,CadN,crb | | 5.314e-04 | -7.54 | kis (kismet) | protein interactions | 33185 | 33 | 7 | 9204 | 412 | Syx1A,S6kII,pbl,N,ci,tna,rau | | 5.314e-04 | -7.54 | GSE17721\_LPS\_VS\_POLYIC\_16H\_BMDC\_DN | MSigDB lists | GSE17721\_LPS\_VS\_POLYIC\_16H\_BMDC\_DN | 40 | 7 | 4161 | 153 | LRP1,Doa,Tlk,emp,CycE,CG5830,CaMKII | | 5.323e-04 | -7.54 | sgg (shaggy) | protein interactions | 31248 | 183 | 19 | 9204 | 412 | wnd,Doa,PEK,ci,osa,CG4325,Rho1,Acf,mam,ft,S6kII,N,mgl,egr,Etl1,ec,Taf1,CadN,robo2 | | 5.400e-04 | -7.52 | calcium-dependent cell-cell adhesion via plasma membrane cell adhesion molecules | biological process | GO:0016339 | 21 | 5 | 12345 | 414 | CadN,ft,ds,scb,Cad87A | | 5.400e-04 | -7.52 | cellular response to BMP stimulus | biological process | GO:0071773 | 21 | 5 | 12345 | 414 | tkv,Dad,dpp,hiw,dally | | 5.400e-04 | -7.52 | BMP signaling pathway | biological process | GO:0030509 | 21 | 5 | 12345 | 414 | dpp,Dad,tkv,dally,hiw | | 5.400e-04 | -7.52 | imaginal disc growth | biological process | GO:0007446 | 21 | 5 | 12345 | 414 | N,ft,dpp,hh,fj | | 5.429e-04 | -7.52 | EGF\_CA | smart domains | SM00179 | 36 | 8 | 5086 | 276 | ft,mgl,CadN,Ser,crb,slow,N,LRP1 | | 5.607e-04 | -7.49 | eggshell formation | biological process | GO:0030703 | 134 | 13 | 12345 | 414 | spoon,Egfr,S,Syp,dpp,Myb,gus,tkv,CycE,Rac2,lama,N,ttk | | 5.633e-04 | -7.48 | PDZ | pfam domains | PF00595 | 51 | 8 | 10480 | 393 | Rhp,Ptpmeg,CG3402,sdt,RhoGEF2,bbg,CG32758,PsGEF | | 5.652e-04 | -7.48 | HOMEOBOX\_2 | prosite domains | PS50071 | 102 | 14 | 6243 | 316 | B-H2,exex,Lim1,ap,Dr,dve,so,lbe,HGTX,tin,ey,Ubx,oc,slou | | 5.732e-04 | -7.46 | PROTEIN\_KINASE\_ATP | prosite domains | PS00107 | 164 | 19 | 6243 | 316 | Asator,Haspin,CG8485,Tlk,Pak3,S6kII,Doa,PEK,KP78a,mbt,Cdk12,JIL-1,CaMKII,par-1,Hipk,Egfr,PhKgamma,tkv,Mos | | 5.786e-04 | -7.45 | reproduction | biological process | GO:0000003 | 1651 | 79 | 12345 | 414 | psq,Bx,ci,shd,Npc1a,tsl,pAbp,if,Pat1,CaMKII,Rbfox1,lama,Fas2,goe,scb,heph,Fas3,Rho1,nudE,Tet,ttk,Mmp2,JIL-1,bbg,CG31680,Myb,lz,sty,aop,S,otk2,wnd,hts,spoon,Egfr,mle,bab1,eya,drk,jing,wb,baf,Doa,Hydr2,N,wun2,salm,Rac2,mam,so,robo2,pbl,gus,EcR,tin,TfAP-2,Pex1,dve,fne,par-1,hh,l(3)mbt,Stat92E,CycE,stwl,dpp,Mos,RhoGEF2,Syp,dally,yps,AGO2,Inx2,elav,cnc,Tudor-SN,Imp,tkv,Dr | | 5.806e-04 | -7.45 | positive regulation of cellular component movement | biological process | GO:0051272 | 71 | 9 | 12345 | 414 | Egfr,robo2,aop,Dscam1,tsl,mmy,spi,Pvf3,ttk | | 5.814e-04 | -7.45 | apicolateral plasma membrane | cellular component | GO:0016327 | 21 | 5 | 11940 | 407 | ds,sdt,ft,crb,Inx2 | | 5.900e-04 | -7.44 | GO\_CELL\_PART\_MORPHOGENESIS | MSigDB lists | GO\_CELL\_PART\_MORPHOGENESIS | 175 | 16 | 4161 | 153 | Lim1,ap,NetB,Dscam1,N,trn,JIL-1,Ank,Fas2,Arp2,nudE,Mrp4,hh,drk,tkv,Rac2 | | 5.909e-04 | -7.43 | ASX\_HYDROXYL | prosite domains | PS00010 | 30 | 7 | 6243 | 316 | LRP1,CadN,N,Ser,mgl,slow,crb | | 5.923e-04 | -7.43 | locomotory behavior | biological process | GO:0007626 | 188 | 16 | 12345 | 414 | sbb,EcR,qsm,Cdk5alpha,CCKLR-17D3,Cirl,cv,disco,hiw,ITP,S6kII,tara,oc,ey,Bx,TfAP-2 | | 6.032e-04 | -7.41 | regulation of protein phosphorylation | biological process | GO:0001932 | 245 | 19 | 12345 | 414 | aos,CG8441,Rho1,mbt,Atg17,Cdk5alpha,drk,sty,Mos,PEK,wnd,Egfr,dpp,bora,par-1,grnd,CycE,N,spi | | 6.034e-04 | -7.41 | CACCAGC\_MIR138 | MSigDB lists | CACCAGC\_MIR138 | 53 | 8 | 4161 | 153 | Rho1,CycE,Sin3A,CG2201,DIP-epsilon,Doa,CG5522,l(3)mbt | | 6.034e-04 | -7.41 | AP2GAMMA\_01 | MSigDB lists | AP2GAMMA\_01 | 53 | 8 | 4161 | 153 | pAbp,sm,Dscam1,ap,Tsp26A,Imp,so,Tlk | | 6.173e-04 | -7.39 | GO\_HEART\_DEVELOPMENT | MSigDB lists | GO\_HEART\_DEVELOPMENT | 110 | 12 | 4161 | 153 | N,Smox,mgl,CaMKII,ft,Utx,ds,Ser,dpp,nAChRalpha4,so,hh | | 6.204e-04 | -7.39 | Rho\_GTPase\_activation\_prot | interpro domains | IPR008936 | 29 | 6 | 11383 | 428 | RhoGAP93B,RhoGAP102A,RhoGAP15B,RhoGAP16F,RhoGAP18B,CG42684 | | 6.213e-04 | -7.38 | GATA1\_02 | MSigDB lists | GATA1\_02 | 41 | 7 | 4161 | 153 | so,Pli,kdn,CycE,fne,sm,pbl | | 6.213e-04 | -7.38 | GO\_ESTABLISHMENT\_OR\_MAINTENANCE\_OF\_CELL\_POLARITY | MSigDB lists | GO\_ESTABLISHMENT\_OR\_MAINTENANCE\_OF\_CELL\_POLARITY | 41 | 7 | 4161 | 153 | Arp2,nudE,Mhcl,hh,ap,CG32758,sdt | | 6.238e-04 | -7.38 | GO\_SOMITE\_DEVELOPMENT | MSigDB lists | GO\_SOMITE\_DEVELOPMENT | 20 | 5 | 4161 | 153 | Utx,Smox,hh,Lim1,so | | 6.446e-04 | -7.35 | ebi (ebi) | protein interactions | 33212 | 66 | 10 | 9204 | 412 | Ser,crb,ttk,osa,aos,egr,phyl,N,sno,Egfr | | 6.567e-04 | -7.33 | LRRCT | smart domains | SM00082 | 21 | 6 | 5086 | 276 | Toll-7,trn,Tl,Lrt,lbk,Con | | 6.646e-04 | -7.32 | trol (terribly reduced optic lobes) | protein interactions | 45320 | 25 | 6 | 9204 | 412 | Fas2,ci,hh,Egfr,CycE,sty | | 6.678e-04 | -7.31 | HOMEOBOX\_1 | prosite domains | PS00027 | 92 | 13 | 6243 | 316 | ey,Ubx,oc,slou,B-H2,Lim1,ap,exex,Dr,lbe,HGTX,so,tin | | 6.700e-04 | -7.31 | cv-2 (crossveinless 2) | protein interactions | 45280 | 10 | 4 | 9204 | 412 | tkv,dally,dpp,pbl | | 6.798e-04 | -7.29 | response to BMP | biological process | GO:0071772 | 22 | 5 | 12345 | 414 | hiw,dally,tkv,Dad,dpp | | 6.798e-04 | -7.29 | sensory organ precursor cell fate determination | biological process | GO:0016360 | 22 | 5 | 12345 | 414 | lz,N,phyl,nuf,CtBP | | 6.798e-04 | -7.29 | cell fate determination involved in pattern specification | biological process | GO:0060582 | 22 | 5 | 12345 | 414 | N,lz,nuf,CtBP,phyl | | 6.848e-04 | -7.29 | GO\_CENTRAL\_NERVOUS\_SYSTEM\_DEVELOPMENT | MSigDB lists | GO\_CENTRAL\_NERVOUS\_SYSTEM\_DEVELOPMENT | 231 | 19 | 4161 | 153 | TfAP-2,S6kII,ft,nudE,Rac2,hh,Hsc70-3,Imp,so,if,ap,N,mgl,Dscam1,Lim1,dpp,Acsl,LRP1,CG17265 | | 6.908e-04 | -7.28 | GO\_MORPHOGENESIS\_OF\_AN\_EPITHELIAL\_SHEET | MSigDB lists | GO\_MORPHOGENESIS\_OF\_AN\_EPITHELIAL\_SHEET | 12 | 4 | 4161 | 153 | Ser,N,sdt,Rho1 | | 6.908e-04 | -7.28 | NAKAYAMA\_SOFT\_TISSUE\_TUMORS\_PCA1\_DN | MSigDB lists | NAKAYAMA\_SOFT\_TISSUE\_TUMORS\_PCA1\_DN | 12 | 4 | 4161 | 153 | dlp,so,ap,trn | | 6.908e-04 | -7.28 | GO\_NEGATIVE\_REGULATION\_OF\_MUSCLE\_CELL\_DIFFERENTIATION | MSigDB lists | GO\_NEGATIVE\_REGULATION\_OF\_MUSCLE\_CELL\_DIFFERENTIATION | 12 | 4 | 4161 | 153 | hh,dpp,N,heph | | 6.908e-04 | -7.28 | GO\_POSITIVE\_REGULATION\_OF\_BIOMINERAL\_TISSUE\_DEVELOPMENT | MSigDB lists | GO\_POSITIVE\_REGULATION\_OF\_BIOMINERAL\_TISSUE\_DEVELOPMENT | 12 | 4 | 4161 | 153 | dpp,tkv,Smox,TfAP-2 | | 6.911e-04 | -7.28 | memory | biological process | GO:0007613 | 120 | 12 | 12345 | 414 | Stat92E,scb,dbr,drk,Fas2,N,Rbfox1,wnd,spoon,hiw,CaMKII,EcR | | 6.935e-04 | -7.27 | Kinase-like\_dom\_sf | interpro domains | IPR011009 | 293 | 23 | 11383 | 428 | Egfr,JIL-1,CaMKII,Cdk12,Tlk,Hipk,S6kII,PEK,Mos,Doa,CG11486,PhKgamma,Asator,Pak3,Haspin,KP78a,mbt,wnd,CG8485,tkv,CG1344,par-1,CG2201 | | 6.944e-04 | -7.27 | positive regulation of fibroblast growth factor receptor signaling pathway | biological process | GO:0045743 | 6 | 3 | 12345 | 414 | dally,dlp,rau | | 6.944e-04 | -7.27 | heart formation | biological process | GO:0060914 | 6 | 3 | 12345 | 414 | ci,hh,slp1 | | 6.944e-04 | -7.27 | photoreceptor cell fate determination | biological process | GO:0043703 | 6 | 3 | 12345 | 414 | spi,Egfr,S | | 6.957e-04 | -7.27 | ci (cubitus interruptus) | protein interactions | 43767 | 187 | 19 | 9204 | 412 | CycE,kni,Rbfox1,LRP1,RapGAP1,bi,Stat92E,dbr,jing,CG10011,mgl,Psc,hh,bin3,ey,Doc1,dpp,ci,Egfr | | 7.005e-04 | -7.26 | negative regulation of reproductive process | biological process | GO:2000242 | 13 | 4 | 12345 | 414 | Ptpmeg,bab1,Mos,Doa | | 7.005e-04 | -7.26 | wing and notum subfield formation | biological process | GO:0035309 | 13 | 4 | 12345 | 414 | Egfr,Dr,salm,dpp | | 7.005e-04 | -7.26 | cardioblast differentiation | biological process | GO:0010002 | 13 | 4 | 12345 | 414 | EcR,dpp,robo2,tin | | 7.005e-04 | -7.26 | negative regulation of cell-cell adhesion | biological process | GO:0022408 | 13 | 4 | 12345 | 414 | aop,par-1,N,aos | | 7.005e-04 | -7.26 | muscle fiber development | biological process | GO:0048747 | 13 | 4 | 12345 | 414 | aop,twi,sty,pnt | | 7.039e-04 | -7.26 | negative regulation of neuron differentiation | biological process | GO:0045665 | 33 | 6 | 12345 | 414 | ttk,N,S6kII,Fas2,CadN,aop | | 7.039e-04 | -7.26 | actin filament-based movement | biological process | GO:0030048 | 33 | 6 | 12345 | 414 | RhoGEF2,crb,Rho1,tyn,cta,neo | | 7.052e-04 | -7.26 | negative regulation of cell population proliferation | biological process | GO:0008285 | 88 | 10 | 12345 | 414 | ham,Doa,osa,dpp,EcR,shn,l(3)mbt,Utx,Mmp2,ft | | 7.153e-04 | -7.24 | regulation of small GTPase mediated signal transduction | biological process | GO:0051056 | 104 | 11 | 12345 | 414 | CG42684,CG5522,NFAT,pbl,MESR3,CG43658,RapGAP1,PsGEF,S6kII,sty,RhoGEF2 | | 7.304e-04 | -7.22 | EGF | smart domains | SM00181 | 66 | 11 | 5086 | 276 | crb,slow,N,LRP1,spi,CadN,Ser,wb,ft,Ten-m,mgl | | 7.459e-04 | -7.20 | REACTOME\_AXON\_GUIDANCE | MSigDB lists | REACTOME\_AXON\_GUIDANCE | 68 | 9 | 4161 | 153 | NetB,S6kII,Fas2,Rac2,JIL-1,Ank,drk,Rho1,mbt | | 7.459e-04 | -7.20 | GO\_OSSIFICATION | MSigDB lists | GO\_OSSIFICATION | 68 | 9 | 4161 | 153 | ft,Smox,ds,mle,Tudor-SN,smal,dpp,hh,CG11367 | | 7.498e-04 | -7.20 | regulation of immune system process | biological process | GO:0002682 | 230 | 18 | 12345 | 414 | Usp2,lz,Rac2,Stat92E,Pli,Mmp2,N,Kr,Doa,Sin3A,EcR,Toll-7,Etl1,cpo,Ser,Tl,ush,CG4325 | | 7.513e-04 | -7.19 | EGF\_Ca-bd\_CS | interpro domains | IPR018097 | 30 | 6 | 11383 | 428 | N,mgl,slow,crb,LRP1,Ser | | 7.539e-04 | -7.19 | regulation of molecular function | biological process | GO:0065009 | 523 | 32 | 12345 | 414 | Atg17,CG42684,pnt,RhoGAP18B,nrv2,RapGAP1,CG8531,sty,wnd,mle,ttk,edl,Pli,RhoGAP93B,Prosap,CycE,CG8441,aos,Acf,Rho1,RhoGAP16F,mbt,RhoGAP102A,nudE,mof,ush,pbl,Cdk5alpha,NiPp1,RhoGAP15B,bora,fj | | 7.693e-04 | -7.17 | REACTOME\_GLYCOSAMINOGLYCAN\_METABOLISM | MSigDB lists | REACTOME\_GLYCOSAMINOGLYCAN\_METABOLISM | 31 | 6 | 4161 | 153 | CG9220,dlp,dally,CG14291,CG31637,CG43313 | | 7.742e-04 | -7.16 | eya (eyes absent) | protein interactions | 33916 | 35 | 7 | 9204 | 412 | dpp,Egfr,scaf6,so,eya,aop,ey | | 7.893e-04 | -7.14 | BENPORATH\_SUZ12\_TARGETS | MSigDB lists | BENPORATH\_SUZ12\_TARGETS | 129 | 13 | 4161 | 153 | Fas2,mgl,NetB,Dscam1,ap,so,hh,ScpX,TfIIS,dlp,dally,nAChRalpha4,ras | | 7.893e-04 | -7.14 | CAGTATT\_MIR200B\_MIR200C\_MIR429 | MSigDB lists | CAGTATT\_MIR200B\_MIR200C\_MIR429 | 129 | 13 | 4161 | 153 | CG45050,heph,Rac2,sdt,fne,Ank,LRP1,Etl1,S6kII,Syx1A,phtf,Lac,Npc1a | | 7.939e-04 | -7.14 | positive regulation of phosphorylation | biological process | GO:0042327 | 139 | 13 | 12345 | 414 | Cdk5alpha,grnd,bora,spi,drk,Egfr,wnd,PEK,Mos,Atg17,dpp,mbt,Rho1 | | 8.124e-04 | -7.12 | phosphorus metabolic process | biological process | GO:0006793 | 920 | 49 | 12345 | 414 | Acsl,KP78a,CaMKII,PEK,wun2,CG34384,Socs16D,fj,Rac2,Taf1,ND-B14.5AL,mbt,ras,CG2201,CG10082,CG5830,Haspin,Ptp99A,CG8485,S6kII,wnd,Cdk12,Mos,Asator,egr,Cyt-c1,Mocs2B,JIL-1,Hipk,CycE,Tlk,CG1344,par-1,mmy,CG17224,stg,Notum,sdt,tkv,Doa,CG11486,CG6330,eya,CG1271,Ptpmeg,ScsbetaG,PhKgamma,Pak3,smal | | 8.239e-04 | -7.10 | Developmental Biology | REACTOME pathways | R-DME-1266738 | 211 | 18 | 3923 | 148 | if,CG42684,dlp,robo2,drk,Rho1,Ank,Egfr,Syx1A,RhoGAP93B,Arp2,JIL-1,par-1,Fas2,Smox,NetA,NetB,scb | | 8.317e-04 | -7.09 | msl-2 (male-specific lethal 2) | protein interactions | 33565 | 26 | 6 | 9204 | 412 | ocm,mle,mof,AGO2,lncRNA:roX1,JIL-1 | | 8.317e-04 | -7.09 | dally (division abnormally delayed) | protein interactions | 39013 | 26 | 6 | 9204 | 412 | dlp,tkv,CycE,hh,dpp,Dad | | 8.321e-04 | -7.09 | cell projection | cellular component | GO:0042995 | 629 | 37 | 11940 | 407 | AGO2,NetB,CadN,Myo10A,Con,Pak3,Dscam1,Cdk5alpha,Imp,par-1,CaMKII,Hydr2,Prosap,dpp,Egfr,nAChRalpha4,dpr7,nudE,Mmp2,robo2,EcR,Rac2,Rho1,yps,ckn,Fas2,Syx1A,ImpL2,Ptp99A,Acsl,wnd,tkv,nrv2,Ser,cta,NetA,crb | | 8.324e-04 | -7.09 | Socs16D (Suppressor of Cytokine Signaling at 16D) | protein interactions | 32760 | 5 | 3 | 9204 | 412 | Myo10A,sty,spi | | 8.324e-04 | -7.09 | caps (capricious) | protein interactions | 39493 | 5 | 3 | 9204 | 412 | trn,Kr,Bx | | 8.367e-04 | -7.09 | neuron part | cellular component | GO:0097458 | 540 | 33 | 11940 | 407 | nAChRalpha4,dpr7,Ten-m,Mmp2,robo2,nudE,EcR,Imp,Cdk5alpha,par-1,CaMKII,Prosap,Con,Myo10A,Dscam1,AGO2,NetB,CadN,nrv2,Ser,NetA,Ptp99A,Acsl,wnd,Cirl,tkv,hts,S6kII,ImpL2,yps,Fas2,ckn,Syx1A | | 8.450e-04 | -7.08 | apical constriction | biological process | GO:0003383 | 23 | 5 | 12345 | 414 | neo,cta,crb,tyn,RhoGEF2 | | 8.450e-04 | -7.08 | R7 cell fate commitment | biological process | GO:0007465 | 23 | 5 | 12345 | 414 | ttk,drk,pnt,aop,phyl | | 8.450e-04 | -7.08 | unidimensional cell growth | biological process | GO:0009826 | 23 | 5 | 12345 | 414 | tkv,Rho1,Dad,RhoGEF2,Rac2 | | 8.491e-04 | -7.07 | dendrite morphogenesis | biological process | GO:0048813 | 140 | 13 | 12345 | 414 | ham,dally,nudE,shn,EcR,comm,NetB,Rho1,NetA,cnc,Prosap,ci,Smox | | 8.581e-04 | -7.06 | positive regulation of protein modification process | biological process | GO:0031401 | 176 | 15 | 12345 | 414 | Mos,PEK,wnd,Egfr,Pcl,Rho1,mbt,Atg17,dpp,bora,grnd,mmy,Cdk5alpha,drk,spi | | 8.601e-04 | -7.06 | brm (brahma) | protein interactions | 39744 | 106 | 13 | 9204 | 412 | Pcl,Psc,tkv,tara,N,tna,S6kII,Egfr,hh,CycE,Bx,Utx,osa | | 8.765e-04 | -7.04 | GO\_CIRCULATORY\_SYSTEM\_DEVELOPMENT | MSigDB lists | GO\_CIRCULATORY\_SYSTEM\_DEVELOPMENT | 164 | 15 | 4161 | 153 | ds,cta,Utx,Smox,N,mgl,ft,CaMKII,Ser,dpp,slow,nAChRalpha4,hh,LRP1,so | | 8.933e-04 | -7.02 | GO\_POSITIVE\_REGULATION\_OF\_CELL\_DEVELOPMENT | MSigDB lists | GO\_POSITIVE\_REGULATION\_OF\_CELL\_DEVELOPMENT | 84 | 10 | 4161 | 153 | pbl,NetB,N,Smox,nudE,Tif-IA,dpp,Lac,Rac2,hh | | 8.936e-04 | -7.02 | phosphate-containing compound metabolic process | biological process | GO:0006796 | 900 | 48 | 12345 | 414 | S6kII,wnd,Cdk12,Mos,Asator,Cyt-c1,egr,Mocs2B,JIL-1,Hipk,CycE,Tlk,par-1,CG1344,CG17224,stg,Notum,sdt,tkv,Doa,Ptpmeg,eya,CG1271,CG11486,CG6330,ScsbetaG,PhKgamma,Pak3,smal,Acsl,KP78a,PEK,CaMKII,wun2,CG34384,fj,Socs16D,Taf1,Rac2,ND-B14.5AL,mbt,ras,CG10082,CG2201,CG5830,Haspin,Ptp99A,CG8485 | | 8.945e-04 | -7.02 | RhoGAP | pfam domains | PF00620 | 21 | 5 | 10480 | 393 | RhoGAP18B,RhoGAP16F,RhoGAP93B,RhoGAP15B,RhoGAP102A | | 8.986e-04 | -7.01 | PHONG\_TNF\_TARGETS\_UP | MSigDB lists | PHONG\_TNF\_TARGETS\_UP | 6 | 3 | 4161 | 153 | fj,Utx,dpp | | 8.986e-04 | -7.01 | GO\_POSITIVE\_REGULATION\_OF\_CARTILAGE\_DEVELOPMENT | MSigDB lists | GO\_POSITIVE\_REGULATION\_OF\_CARTILAGE\_DEVELOPMENT | 6 | 3 | 4161 | 153 | Smox,tkv,dpp | | 8.986e-04 | -7.01 | GO\_ANATOMICAL\_STRUCTURE\_ARRANGEMENT | MSigDB lists | GO\_ANATOMICAL\_STRUCTURE\_ARRANGEMENT | 6 | 3 | 4161 | 153 | Hsc70-3,TfAP-2,Rac2 | | 8.986e-04 | -7.01 | GO\_POSITIVE\_REGULATION\_OF\_STEM\_CELL\_DIFFERENTIATION | MSigDB lists | GO\_POSITIVE\_REGULATION\_OF\_STEM\_CELL\_DIFFERENTIATION | 6 | 3 | 4161 | 153 | Smox,N,dpp | | 8.986e-04 | -7.01 | GO\_ATRIOVENTRICULAR\_VALVE\_DEVELOPMENT | MSigDB lists | GO\_ATRIOVENTRICULAR\_VALVE\_DEVELOPMENT | 6 | 3 | 4161 | 153 | dpp,N,ds | | 8.986e-04 | -7.01 | GO\_POSITIVE\_REGULATION\_OF\_EPITHELIAL\_TO\_MESENCHYMAL\_TRANSITION | MSigDB lists | GO\_POSITIVE\_REGULATION\_OF\_EPITHELIAL\_TO\_MESENCHYMAL\_TRANSITION | 6 | 3 | 4161 | 153 | dpp,Smox,N | | 8.986e-04 | -7.01 | LU\_TUMOR\_ANGIOGENESIS\_UP | MSigDB lists | LU\_TUMOR\_ANGIOGENESIS\_UP | 6 | 3 | 4161 | 153 | Ser,CycE,N | | 8.986e-04 | -7.01 | GO\_ENDOCARDIAL\_CUSHION\_MORPHOGENESIS | MSigDB lists | GO\_ENDOCARDIAL\_CUSHION\_MORPHOGENESIS | 6 | 3 | 4161 | 153 | N,ds,dpp | | 9.086e-04 | -7.00 | regulation of protein localization | biological process | GO:0032880 | 107 | 11 | 12345 | 414 | dally,dlp,CaMKII,S,sdt,S6kII,Snx3,hh,Fbxl7,ft,crb | | 9.096e-04 | -7.00 | synapse organization | biological process | GO:0050808 | 177 | 15 | 12345 | 414 | comm,Imp,S6kII,Arp2,DIP-epsilon,dlp,Fur1,dpr7,hiw,CaMKII,Ten-m,Fas2,Tl,Prosap,par-1 | | 9.121e-04 | -7.00 | response to organic substance | biological process | GO:0010033 | 529 | 32 | 12345 | 414 | Drat,pnt,Mondo,drk,Toll-7,tkv,Dscam1,Stat92E,Utx,Egfr,dally,S,hiw,aop,Gbs-70E,dpp,Cnx99A,scb,Fas2,Tl,Smox,EcR,grnd,Hsc70-3,Dad,spi,Bx,hang,Hydr2,PEK,shn,Lpin | | 9.138e-04 | -7.00 | ATP binding | molecular function | GO:0005524 | 639 | 36 | 12176 | 396 | CG9281,CG9541,Mos,Egfr,JIL-1,CG8485,PhKgamma,par-1,Pex1,CG14535,mbt,lz,mle,Myo10A,CG12010,Doa,wnd,tkv,bor,Pak3,KP78a,Cdk12,Hsc70-3,CaMKII,Tlk,Arp2,Etl1,Asator,Mrp4,ScsbetaG,S6kII,PEK,Mhcl,Haspin,Hipk,CG8097 | | 9.184e-04 | -6.99 | GO\_EMBRYONIC\_SKELETAL\_SYSTEM\_DEVELOPMENT | MSigDB lists | GO\_EMBRYONIC\_SKELETAL\_SYSTEM\_DEVELOPMENT | 32 | 6 | 4161 | 153 | Dscam1,Smox,TfAP-2,so,Lim1,hh | | 9.219e-04 | -6.99 | Su(H) (Suppressor of Hairless) | protein interactions | 34881 | 69 | 10 | 9204 | 412 | mam,l(3)mbt,Ubx,ci,N,Sin3A,CtBP,Rbfox1,aop,Prosap | | 9.260e-04 | -6.98 | elav (embryonic lethal abnormal vision) | protein interactions | 31000 | 18 | 5 | 9204 | 412 | Ubx,fne,elav,heph,comm | | 9.260e-04 | -6.98 | Unr (Upstream of N-ras) | protein interactions | 38963 | 18 | 5 | 9204 | 412 | lncRNA:roX1,pAbp,smg,JIL-1,mle | | 9.275e-04 | -6.98 | striated muscle cell development | biological process | GO:0055002 | 61 | 8 | 12345 | 414 | pnt,twi,if,sty,aop,hts,CG13366,Rho1 | | 9.483e-04 | -6.96 | endosome | cellular component | GO:0005768 | 212 | 17 | 11940 | 407 | S,Ykt6,nuf,CG34384,hh,Fas3,rush,tkv,N,dpp,CG6051,Snx3,CG32758,CG13784,Tl,comm,dbr | | 9.517e-04 | -6.96 | multi-organism process | biological process | GO:0051704 | 1653 | 78 | 12345 | 414 | Pat1,CaMKII,psq,ci,shd,tsl,Npc1a,pAbp,if,heph,Fas3,Rho1,nudE,Tet,lama,Rbfox1,Tl,goe,scb,Myb,sty,aop,S,wnd,hts,mle,spoon,Egfr,ttk,JIL-1,Mmp2,egr,bbg,Tlk,Toll-7,baf,Doa,drk,eya,jing,so,Hydr2,N,wun2,Rac2,SamDC,mam,Pcl,gus,EcR,spen,tin,Cdk5alpha,stwl,dpp,Mos,Syp,RhoGEF2,dally,AGO2,yps,TfAP-2,Pex1,fne,par-1,hh,l(3)mbt,Stat92E,Pli,CycE,Imp,Dscam1,tkv,Inx2,elav,Etl1,cnc,Tudor-SN | | 9.549e-04 | -6.95 | negative regulation of photoreceptor cell differentiation | biological process | GO:0046533 | 14 | 4 | 12345 | 414 | aop,ttk,N,S6kII | | 9.549e-04 | -6.95 | Bolwig's organ morphogenesis | biological process | GO:0001746 | 14 | 4 | 12345 | 414 | hh,so,Fas2,eya | | 9.549e-04 | -6.95 | regulation of chromatin assembly | biological process | GO:0010847 | 14 | 4 | 12345 | 414 | CG5815,Stat92E,mle,Acf | | 9.549e-04 | -6.95 | R3/R4 cell fate commitment | biological process | GO:0007464 | 14 | 4 | 12345 | 414 | bdg,N,aop,pnt | | 9.549e-04 | -6.95 | Bolwig's organ development | biological process | GO:0055034 | 14 | 4 | 12345 | 414 | Fas2,eya,so,hh | | 9.549e-04 | -6.95 | cell-cell adhesion mediated by cadherin | biological process | GO:0044331 | 14 | 4 | 12345 | 414 | Cad87A,ds,ft,CadN | | 9.549e-04 | -6.95 | neuron fate specification | biological process | GO:0048665 | 14 | 4 | 12345 | 414 | ey,N,CycE,exex | | 9.587e-04 | -6.95 | negative regulation of neurogenesis | biological process | GO:0050768 | 76 | 9 | 12345 | 414 | Fas2,CadN,ttk,N,EcR,aos,S6kII,aop,osa | | 9.618e-04 | -6.95 | Laminin\_N | pfam domains | PF00055 | 6 | 3 | 10480 | 393 | NetA,wb,NetB | | 9.637e-04 | -6.94 | Rho GTPase binding | molecular function | GO:0017048 | 49 | 7 | 12176 | 396 | Pak3,PsGEF,RhoGAP18B,RhoGEF2,mbt,CG43658,pbl | | 9.654e-04 | -6.94 | GO\_PALLIUM\_DEVELOPMENT | MSigDB lists | GO\_PALLIUM\_DEVELOPMENT | 44 | 7 | 4161 | 153 | ap,Rac2,ft,LRP1,nudE,Imp,CG17265 | | 9.679e-04 | -6.94 | Release of Hh-Np from the secreting cell | REACTOME pathways | R-DME-5362798 | 6 | 3 | 3923 | 148 | dally,Notum,hh | | 9.679e-04 | -6.94 | p75NTR regulates axonogenesis | REACTOME pathways | R-DME-193697 | 6 | 3 | 3923 | 148 | Rtnl2,Rho1,RhoGEF2 | | 9.679e-04 | -6.94 | Axonal growth inhibition (RHOA activation) | REACTOME pathways | R-DME-193634 | 6 | 3 | 3923 | 148 | Rho1,RhoGEF2,Rtnl2 | | 9.694e-04 | -6.94 | GO\_REGULATION\_OF\_NEURAL\_PRECURSOR\_CELL\_PROLIFERATION | MSigDB lists | GO\_REGULATION\_OF\_NEURAL\_PRECURSOR\_CELL\_PROLIFERATION | 13 | 4 | 4161 | 153 | wech,Lim1,hh,N | | 9.694e-04 | -6.94 | REACTOME\_CHONDROITIN\_SULFATE\_DERMATAN\_SULFATE\_METABOLISM | MSigDB lists | REACTOME\_CHONDROITIN\_SULFATE\_DERMATAN\_SULFATE\_METABOLISM | 13 | 4 | 4161 | 153 | CG9220,dally,dlp,CG43313 | | 9.699e-04 | -6.94 | Laminin\_N\_sf | interpro domains | IPR038684 | 6 | 3 | 11383 | 428 | NetB,wb,NetA | | 9.699e-04 | -6.94 | Laminin\_N | interpro domains | IPR008211 | 6 | 3 | 11383 | 428 | NetB,wb,NetA | | 9.762e-04 | -6.93 | GO\_POSITIVE\_REGULATION\_OF\_GENE\_EXPRESSION | MSigDB lists | GO\_POSITIVE\_REGULATION\_OF\_GENE\_EXPRESSION | 395 | 27 | 4161 | 153 | Ser,Lim1,dpp,Smox,N,Etl1,if,Utx,ap,RpS4,mof,CG12099,Ank,EcR,JIL-1,Tif-IA,smal,Taf1,CtBP,S6kII,TfAP-2,so,hh,Hsc70-3,tkv,Sin3A,CycE | | 9.816e-04 | -6.93 | GO\_REGULATION\_OF\_CELLULAR\_COMPONENT\_MOVEMENT | MSigDB lists | GO\_REGULATION\_OF\_CELLULAR\_COMPONENT\_MOVEMENT | 132 | 13 | 4161 | 153 | Rac2,Hydr2,Hsc70-3,LRP1,hh,cta,NetB,CaMKII,Smox,N,dpp,Ser,smal | | 9.819e-04 | -6.93 | response to wounding | biological process | GO:0009611 | 108 | 11 | 12345 | 414 | lz,Rho1,wnd,hiw,RhoGEF2,Tl,Mmp2,pbl,scb,Utx,jing | | 9.930e-04 | -6.91 | GO\_RECEPTOR\_COMPLEX | MSigDB lists | GO\_RECEPTOR\_COMPLEX | 57 | 8 | 4161 | 153 | LRP1,tkv,nAChRalpha4,dpp,Smox,N,mgl,if | | 9.939e-04 | -6.91 | regulation of MAPK cascade | biological process | GO:0043408 | 125 | 12 | 12345 | 414 | grnd,N,spi,drk,sty,PEK,Mos,wnd,Egfr,aos,mbt,dpp | | 1.016e-03 | -6.89 | Dab (Disabled) | protein interactions | 39866 | 11 | 4 | 9204 | 412 | drk,N,Fas2,CG33993 | | 1.016e-03 | -6.89 | Gprk2 (G protein-coupled receptor kinase 2) | protein interactions | 49045 | 11 | 4 | 9204 | 412 | Tl,RapGAP1,cta,hh | | 1.016e-03 | -6.89 | veli (veli) | protein interactions | 43003 | 11 | 4 | 9204 | 412 | sdt,grnd,crb,Egfr | | 1.017e-03 | -6.89 | germ cell migration | biological process | GO:0008354 | 48 | 7 | 12345 | 414 | tin,hh,gus,wun2,eya,dpp,tkv | | 1.018e-03 | -6.89 | Dcr-2 (Dicer-2) | protein interactions | 36993 | 47 | 8 | 9204 | 412 | CG44774,aos,Tl,lncRNA:roX1,bowl,cas,osp,AGO2 | | 1.034e-03 | -6.87 | muscle cell development | biological process | GO:0055001 | 62 | 8 | 12345 | 414 | Rho1,aop,sty,hts,CG13366,twi,pnt,if | | 1.038e-03 | -6.87 | positive regulation of mitotic cell cycle phase transition | biological process | GO:1901992 | 24 | 5 | 12345 | 414 | CycE,ci,N,stg,Myb | | 1.038e-03 | -6.87 | Znf\_C2H2\_sf | interpro domains | IPR036236 | 265 | 21 | 11383 | 428 | Zif,ush,ci,CG31612,Kah,ttk,Kr,l(3)neo38,CG31224,CG32772,CG34283,shn,CG9650,bowl,sens-2,salm,CG11247,hang,jing,CG4360,ham | | 1.060e-03 | -6.85 | basal part of cell | cellular component | GO:0045178 | 35 | 6 | 11940 | 407 | Egfr,Syp,scb,RhoGEF2,if,pon | | 1.066e-03 | -6.84 | Signal transduction by L1 | REACTOME pathways | R-DME-445144 | 13 | 4 | 3923 | 148 | scb,Fas2,Egfr,if | | 1.072e-03 | -6.84 | regulation of cell death | biological process | GO:0010941 | 257 | 19 | 12345 | 414 | Egfr,lz,dpp,sas,pAbp,CycE,Utx,grnd,spi,Hipk,N,EcR,aos,ec,pnt,Fili,mof,eya,ft | | 1.084e-03 | -6.83 | cytokine binding | molecular function | GO:0019955 | 7 | 3 | 12176 | 396 | Imp,tkv,Tl | | 1.089e-03 | -6.82 | GO\_POSITIVE\_REGULATION\_OF\_CELL\_MORPHOGENESIS\_INVOLVED\_IN\_DIFFERENTIATION | MSigDB lists | GO\_POSITIVE\_REGULATION\_OF\_CELL\_MORPHOGENESIS\_INVOLVED\_IN\_DIFFERENTIATION | 33 | 6 | 4161 | 153 | nudE,dpp,N,Smox,NetB,Rac2 | | 1.093e-03 | -6.82 | cytoskeleton organization | biological process | GO:0007010 | 580 | 34 | 12345 | 414 | RhoGAP18B,cnc,ft,Pak3,cta,Chro,stg,CG43658,par-1,Svil,RhoGEF2,ds,hts,wake,Ank,Myb,pbl,tyn,nudE,Ten-m,CG13366,Rho1,Arp2,heph,mbt,betaTub56D,Rac2,twi,if,crb,N,nuf,Hip1,KP78a | | 1.093e-03 | -6.82 | Ig-like\_fold | interpro domains | IPR013783 | 195 | 17 | 11383 | 428 | Fas3,nrm,NFAT,Lac,beat-IIIc,wake,ImpL2,DIP-epsilon,robo2,CG6044,otk2,dpr7,lbk,Ptp99A,Fas2,sas,Dscam1 | | 1.094e-03 | -6.82 | HOX | smart domains | SM00389 | 102 | 14 | 5086 | 276 | HGTX,ey,so,Dr,lbe,Lim1,tin,ap,B-H2,oc,slou,Ubx,dve,exex | | 1.109e-03 | -6.80 | GO\_SKELETAL\_SYSTEM\_MORPHOGENESIS | MSigDB lists | GO\_SKELETAL\_SYSTEM\_MORPHOGENESIS | 45 | 7 | 4161 | 153 | TfAP-2,Smox,Dscam1,tkv,Lim1,so,CG9220 | | 1.116e-03 | -6.80 | PDZ | prosite domains | PS50106 | 63 | 10 | 6243 | 316 | PsGEF,Mhcl,bbg,CG3402,Prosap,sdt,RhoGEF2,Rhp,Ptpmeg,CG32758 | | 1.117e-03 | -6.80 | E4BP4\_01 | MSigDB lists | E4BP4\_01 | 58 | 8 | 4161 | 153 | Lac,Pli,S6kII,Utx,CG2812,Smox,ras,sdt | | 1.117e-03 | -6.80 | AP2\_Q3 | MSigDB lists | AP2\_Q3 | 58 | 8 | 4161 | 153 | Acsl,Tpr2,CG9220,Ppa,CG5830,ap,S6kII,CG9281 | | 1.122e-03 | -6.79 | positive regulation of eye photoreceptor cell development | biological process | GO:0042479 | 2 | 2 | 12345 | 414 | mbt,oc | | 1.122e-03 | -6.79 | stomatogastric nervous system development | biological process | GO:0007421 | 2 | 2 | 12345 | 414 | S,spi | | 1.122e-03 | -6.79 | positive regulation of compound eye photoreceptor development | biological process | GO:0045315 | 2 | 2 | 12345 | 414 | oc,mbt | | 1.122e-03 | -6.79 | negative regulation of epithelial cell migration | biological process | GO:0010633 | 2 | 2 | 12345 | 414 | Ptpmeg,Sin3A | | 1.122e-03 | -6.79 | regulation of cell-cell adhesion mediated by cadherin | biological process | GO:2000047 | 2 | 2 | 12345 | 414 | N,mbt | | 1.122e-03 | -6.79 | negative regulation of terminal cell fate specification, open tracheal system | biological process | GO:0035155 | 2 | 2 | 12345 | 414 | aop,salm | | 1.122e-03 | -6.79 | oenocyte delamination | biological process | GO:0060233 | 2 | 2 | 12345 | 414 | aop,aos | | 1.137e-03 | -6.78 | midgut development | biological process | GO:0007494 | 36 | 6 | 12345 | 414 | dve,Ubx,shd,shn,if,scb | | 1.137e-03 | -6.78 | RhoGAP\_dom | interpro domains | IPR000198 | 22 | 5 | 11383 | 428 | RhoGAP16F,RhoGAP18B,RhoGAP93B,RhoGAP15B,RhoGAP102A | | 1.153e-03 | -6.77 | regulation of ERBB signaling pathway | biological process | GO:1901184 | 49 | 7 | 12345 | 414 | S,sty,rau,aos,edl,Fas2,Ptpmeg | | 1.153e-03 | -6.77 | positive regulation of cell migration | biological process | GO:0030335 | 49 | 7 | 12345 | 414 | Dscam1,ttk,spi,Pvf3,aop,Egfr,robo2 | | 1.153e-03 | -6.77 | imaginal disc-derived wing hair organization | biological process | GO:0035317 | 49 | 7 | 12345 | 414 | par-1,spen,fj,ds,ft,N,Rho1 | | 1.153e-03 | -6.77 | regulation of epidermal growth factor receptor signaling pathway | biological process | GO:0042058 | 49 | 7 | 12345 | 414 | S,sty,edl,Fas2,Ptpmeg,rau,aos | | 1.161e-03 | -6.76 | SAM/pointed\_sf | interpro domains | IPR013761 | 44 | 7 | 11383 | 428 | smg,edl,aop,l(3)mbt,ckn,CG34384,pnt | | 1.176e-03 | -6.75 | dendrite development | biological process | GO:0016358 | 145 | 13 | 12345 | 414 | ci,Smox,cnc,Prosap,NetB,comm,EcR,shn,NetA,Rho1,ham,dally,nudE | | 1.180e-03 | -6.74 | neuromuscular junction | cellular component | GO:0031594 | 77 | 9 | 11940 | 407 | Con,S6kII,Ten-m,par-1,Prosap,Fas3,dlp,Fas2,Syp | | 1.185e-03 | -6.74 | second mitotic wave involved in compound eye morphogenesis | biological process | GO:0016330 | 7 | 3 | 12345 | 414 | N,Ser,Egfr | | 1.185e-03 | -6.74 | axon extension involved in axon guidance | biological process | GO:0048846 | 7 | 3 | 12345 | 414 | CadN,Ptpmeg,Dscam1 | | 1.185e-03 | -6.74 | lamellocyte differentiation | biological process | GO:0035171 | 7 | 3 | 12345 | 414 | sty,Stat92E,N | | 1.185e-03 | -6.74 | protein localization involved in establishment of planar polarity | biological process | GO:0090251 | 7 | 3 | 12345 | 414 | ds,ft,Rho1 | | 1.185e-03 | -6.74 | neuron projection extension involved in neuron projection guidance | biological process | GO:1902284 | 7 | 3 | 12345 | 414 | Ptpmeg,CadN,Dscam1 | | 1.185e-03 | -6.74 | negative regulation of protein ubiquitination | biological process | GO:0031397 | 7 | 3 | 12345 | 414 | CycE,ctrip,hh | | 1.196e-03 | -6.73 | GO\_MOVEMENT\_OF\_CELL\_OR\_SUBCELLULAR\_COMPONENT | MSigDB lists | GO\_MOVEMENT\_OF\_CELL\_OR\_SUBCELLULAR\_COMPONENT | 280 | 21 | 4161 | 153 | Fas2,Arp2,nudE,CaMKII,cta,so,hh,Rac2,tkv,drk,Mhcl,Lim1,NetB,N,ap,ds,betaTub56D,JIL-1,Rho1,Ank,mbt | | 1.205e-03 | -6.72 | GGGAGGRR\_MAZ\_Q6 | MSigDB lists | GGGAGGRR\_MAZ\_Q6 | 506 | 32 | 4161 | 153 | heph,Sin3A,dlp,TfIIS,Taf1,NKAIN,JIL-1,Rho1,fne,Snx3,kdn,Utx,dpp,Lim1,CG43313,ras,CycE,so,Imp,Tlk,hh,CG31637,sm,Fas2,smal,NiPp1,PhKgamma,trn,LRP1,N,ds,Ser | | 1.211e-03 | -6.72 | Ras64B (Ras oncogene at 64B) | protein interactions | 38494 | 19 | 5 | 9204 | 412 | dpp,rau,aos,sty,tkv | | 1.211e-03 | -6.72 | gl (glass) | protein interactions | 42210 | 19 | 5 | 9204 | 412 | N,hh,so,aop,tkv | | 1.217e-03 | -6.71 | GO\_HEAD\_DEVELOPMENT | MSigDB lists | GO\_HEAD\_DEVELOPMENT | 187 | 16 | 4161 | 153 | Rac2,Imp,hh,Hsc70-3,ft,nudE,Acsl,CG17265,LRP1,N,mgl,Dscam1,if,ap,Lim1,dpp | | 1.229e-03 | -6.70 | GO\_POSITIVE\_REGULATION\_OF\_BIOSYNTHETIC\_PROCESS | MSigDB lists | GO\_POSITIVE\_REGULATION\_OF\_BIOSYNTHETIC\_PROCESS | 401 | 27 | 4161 | 153 | RpS4,CG12099,mof,EcR,Acsl,JIL-1,dpp,Lim1,Ser,Smox,N,ap,if,Etl1,so,Hsc70-3,hh,tkv,Rac2,Sin3A,CycE,smal,Tif-IA,Taf1,TfAP-2,S6kII,CtBP | | 1.235e-03 | -6.70 | CAGGTG\_E12\_Q6 | MSigDB lists | CAGGTG\_E12\_Q6 | 464 | 30 | 4161 | 153 | CG45050,CG8531,Ank,trn,SamDC,Ser,Mhcl,pAbp,Smox,Cnx99A,Dscam1,Syx1A,Imp,CG9855,hh,Tlk,tkv,Sin3A,drk,CG16890,Ptpmeg,Fas2,Lac,Arl4,CG2812,CG31637,S6kII,CG10663,TfAP-2,cta | | 1.262e-03 | -6.68 | faf (fat facets) | protein interactions | 43749 | 28 | 6 | 9204 | 412 | par-1,wnd,pbl,aop,pnt,hiw | | 1.262e-03 | -6.68 | kni (knirps) | protein interactions | 40287 | 28 | 6 | 9204 | 412 | Acsl,salm,ci,CtBP,Ubx,Egfr | | 1.263e-03 | -6.67 | pole cell migration | biological process | GO:0007280 | 25 | 5 | 12345 | 414 | eya,wun2,gus,hh,tin | | 1.263e-03 | -6.67 | imaginal disc-derived male genitalia development | biological process | GO:0007485 | 25 | 5 | 12345 | 414 | Fas2,TfAP-2,N,Dr,otk2 | | 1.263e-03 | -6.67 | dosage compensation | biological process | GO:0007549 | 25 | 5 | 12345 | 414 | ocm,lncRNA:roX1,mof,AGO2,mle | | 1.268e-03 | -6.67 | GSE4142\_NAIVE\_BCELL\_VS\_PLASMA\_CELL\_UP | MSigDB lists | GSE4142\_NAIVE\_BCELL\_VS\_PLASMA\_CELL\_UP | 46 | 7 | 4161 | 153 | CG34384,Syx1A,CG45050,PhKgamma,dally,emp,Pli | | 1.268e-03 | -6.67 | GSE20715\_WT\_VS\_TLR4\_KO\_48H\_OZONE\_LUNG\_UP | MSigDB lists | GSE20715\_WT\_VS\_TLR4\_KO\_48H\_OZONE\_LUNG\_UP | 46 | 7 | 4161 | 153 | if,sdt,rush,Arl4,Lim1,CG9281,nudE | | 1.268e-03 | -6.67 | regulation of tube diameter | biological process | GO:0035296 | 15 | 4 | 12345 | 414 | nrv2,EcR,dpp,Fas2 | | 1.268e-03 | -6.67 | dosage compensation by hyperactivation of X chromosome | biological process | GO:0009047 | 15 | 4 | 12345 | 414 | AGO2,mle,lncRNA:roX1,mof | | 1.268e-03 | -6.67 | R3/R4 cell differentiation | biological process | GO:0048056 | 15 | 4 | 12345 | 414 | aop,pnt,bdg,N | | 1.278e-03 | -6.66 | establishment of cell polarity | biological process | GO:0030010 | 64 | 8 | 12345 | 414 | KP78a,wake,ds,nudE,fj,par-1,cta,ft | | 1.283e-03 | -6.66 | GO\_CEREBRAL\_CORTEX\_DEVELOPMENT | MSigDB lists | GO\_CEREBRAL\_CORTEX\_DEVELOPMENT | 34 | 6 | 4161 | 153 | Rac2,ft,ap,nudE,CG17265,LRP1 | | 1.283e-03 | -6.66 | GO\_ANTERIOR\_POSTERIOR\_PATTERN\_SPECIFICATION | MSigDB lists | GO\_ANTERIOR\_POSTERIOR\_PATTERN\_SPECIFICATION | 34 | 6 | 4161 | 153 | dpp,hh,Lim1,N,Smox,Utx | | 1.284e-03 | -6.66 | 14-3-3epsilon (14-3-3epsilon) | protein interactions | 42186 | 60 | 9 | 9204 | 412 | EcR,drk,Ubx,S6kII,Df31,par-1,nudE,nuf,if | | 1.303e-03 | -6.64 | non-sensory hair organization | biological process | GO:0035316 | 50 | 7 | 12345 | 414 | N,ft,Rho1,spen,fj,ds,par-1 | | 1.303e-03 | -6.64 | hair cell differentiation | biological process | GO:0035315 | 50 | 7 | 12345 | 414 | Rho1,N,ft,par-1,ds,fj,spen | | 1.320e-03 | -6.63 | positive regulation of epithelial cell differentiation | biological process | GO:0030858 | 37 | 6 | 12345 | 414 | ttk,ci,spi,tsl,aop,Egfr | | 1.320e-03 | -6.63 | regulation of hippo signaling | biological process | GO:0035330 | 37 | 6 | 12345 | 414 | ds,par-1,Fbxl7,ft,crb,Hipk | | 1.323e-03 | -6.63 | regulation of programmed cell death | biological process | GO:0043067 | 222 | 17 | 12345 | 414 | grnd,Utx,CycE,Hipk,N,spi,Egfr,lz,dpp,ec,pnt,ft,Fili,mof,eya,aos,EcR | | 1.331e-03 | -6.62 | fly chr43C5-43C5\|2-56 cM | chromosome location | fly chr43C5-43C5\|2-56 cM | 3 | 2 | 25010 | 531 | asRNA:CR43723,wech | | 1.331e-03 | -6.62 | fly chr43C5-43C5\|2-56 | chromosome location | fly chr43C5-43C5\|2-56 | 3 | 2 | 25010 | 531 | wech,asRNA:CR43723 | | 1.331e-03 | -6.62 | fly chr32E1-32E1\|2-44 | chromosome location | fly chr32E1-32E1\|2-44 | 3 | 2 | 25010 | 531 | lncRNA:CR43859,CG4788 | | 1.331e-03 | -6.62 | fly chr32E1-32E1\|2-44 cM | chromosome location | fly chr32E1-32E1\|2-44 cM | 3 | 2 | 25010 | 531 | CG4788,lncRNA:CR43859 | | 1.345e-03 | -6.61 | actin filament-based process | biological process | GO:0030029 | 262 | 19 | 12345 | 414 | if,twi,Rac2,CG43658,crb,N,hts,Hip1,nuf,RhoGEF2,pbl,RhoGAP18B,cta,Pak3,tyn,CG13366,neo,Rho1,Arp2 | | 1.348e-03 | -6.61 | - | gene3d domains | 2.60.120.1490 | 6 | 3 | 4929 | 208 | NetB,NetA,wb | | 1.351e-03 | -6.61 | pzg (putzig) | protein interactions | 40351 | 49 | 8 | 9204 | 412 | Chro,CycE,mof,JIL-1,ci,N,S6kII,EcR | | 1.360e-03 | -6.60 | rl (rolled) | protein interactions | 3354888 | 85 | 11 | 9204 | 412 | osa,Fas2,aop,eya,Lpin,Svil,Egfr,S6kII,pnt,comm,aos | | 1.383e-03 | -6.58 | Ig-like\_dom | interpro domains | IPR007110 | 132 | 13 | 11383 | 428 | beat-IIIc,Fas2,Dscam1,Lac,otk2,nrm,lbk,dpr7,ImpL2,DIP-epsilon,Fas3,robo2,CG6044 | | 1.390e-03 | -6.58 | positive regulation of neurogenesis | biological process | GO:0050769 | 96 | 10 | 12345 | 414 | mbt,EcR,Imp,nudE,spi,N,Prosap,oc,pnt,mmy | | 1.390e-03 | -6.58 | regulation of Ras protein signal transduction | biological process | GO:0046578 | 96 | 10 | 12345 | 414 | RhoGEF2,sty,S6kII,PsGEF,CG43658,MESR3,pbl,NFAT,CG5522,CG42684 | | 1.401e-03 | -6.57 | GSE17721\_LPS\_VS\_GARDIQUIMOD\_8H\_BMDC\_UP | MSigDB lists | GSE17721\_LPS\_VS\_GARDIQUIMOD\_8H\_BMDC\_UP | 60 | 8 | 4161 | 153 | Hsc70-3,ras,Tpr2,nAChRalpha4,CG43313,smal,cta,rush | | 1.403e-03 | -6.57 | RRM\_8 | pfam domains | PF11835 | 2 | 2 | 10480 | 393 | heph,sm | | 1.403e-03 | -6.57 | SERTA | pfam domains | PF06031 | 2 | 2 | 10480 | 393 | CG2865,tara | | 1.403e-03 | -6.57 | WSD | pfam domains | PF15613 | 2 | 2 | 10480 | 393 | tou,Acf | | 1.403e-03 | -6.57 | Glypican | pfam domains | PF01153 | 2 | 2 | 10480 | 393 | dlp,dally | | 1.403e-03 | -6.57 | NTR | pfam domains | PF01759 | 2 | 2 | 10480 | 393 | NetA,NetB | | 1.411e-03 | -6.56 | HnRNP-L/PTB | interpro domains | IPR006536 | 2 | 2 | 11383 | 428 | heph,sm | | 1.411e-03 | -6.56 | Netrin\_module\_non-TIMP | interpro domains | IPR018933 | 2 | 2 | 11383 | 428 | NetB,NetA | | 1.411e-03 | -6.56 | WHIM2\_dom | interpro domains | IPR028941 | 2 | 2 | 11383 | 428 | Acf,tou | | 1.411e-03 | -6.56 | Glypican | interpro domains | IPR001863 | 2 | 2 | 11383 | 428 | dally,dlp | | 1.411e-03 | -6.56 | SERTA\_dom | interpro domains | IPR009263 | 2 | 2 | 11383 | 428 | tara,CG2865 | | 1.411e-03 | -6.56 | PTBP1\_RRM | interpro domains | IPR021790 | 2 | 2 | 11383 | 428 | sm,heph | | 1.411e-03 | -6.56 | Thioredoxin-like\_fold | interpro domains | IPR012336 | 2 | 2 | 11383 | 428 | fax,CG12547 | | 1.413e-03 | -6.56 | adenyl ribonucleotide binding | molecular function | GO:0032559 | 655 | 36 | 12176 | 396 | Pex1,PhKgamma,par-1,CG8485,JIL-1,Egfr,Mos,CG9541,CG9281,CG12010,Myo10A,mle,lz,mbt,CG14535,CaMKII,Tlk,Hsc70-3,Cdk12,KP78a,Pak3,bor,tkv,wnd,Doa,CG8097,Hipk,Haspin,Mhcl,PEK,S6kII,ScsbetaG,Asator,Mrp4,Etl1,Arp2 | | 1.414e-03 | -6.56 | RHO GTPases Activate Rhotekin and Rhophilins | REACTOME pathways | R-DME-5666185 | 2 | 2 | 3923 | 148 | Rhp,Rho1 | | 1.445e-03 | -6.54 | GSE17721\_LPS\_VS\_POLYIC\_0.5H\_BMDC\_DN | MSigDB lists | GSE17721\_LPS\_VS\_POLYIC\_0.5H\_BMDC\_DN | 47 | 7 | 4161 | 153 | Fas2,Ptpmeg,PhKgamma,hh,Ykt6,Syx1A,Mmp2 | | 1.445e-03 | -6.54 | CREBP1\_01 | MSigDB lists | CREBP1\_01 | 47 | 7 | 4161 | 153 | Lac,Pli,ras,S6kII,sdt,CG2812,Smox | | 1.445e-03 | -6.54 | Cadherin | pfam domains | PF00028 | 14 | 4 | 10480 | 393 | CadN,Cad87A,ft,ds | | 1.448e-03 | -6.54 | msn (misshapen) | protein interactions | 44030 | 61 | 9 | 9204 | 412 | egr,tkv,N,S6kII,ft,Egfr,drk,Mocs2B,rau | | 1.454e-03 | -6.53 | plasma membrane bounded cell projection | cellular component | GO:0120025 | 626 | 36 | 11940 | 407 | crb,NetA,Ser,cta,nrv2,tkv,wnd,Ptp99A,Acsl,ImpL2,Syx1A,Fas2,ckn,yps,Rho1,EcR,Mmp2,robo2,nudE,dpr7,nAChRalpha4,Egfr,Prosap,dpp,Hydr2,par-1,CaMKII,Imp,Cdk5alpha,Dscam1,Pak3,Con,Myo10A,CadN,NetB,AGO2 | | 1.461e-03 | -6.53 | Cadherin\_CS | interpro domains | IPR020894 | 14 | 4 | 11383 | 428 | Cad87A,CadN,ft,ds | | 1.470e-03 | -6.52 | Ptp69D (Protein tyrosine phosphatase 69D) | protein interactions | 39443 | 12 | 4 | 9204 | 412 | robo2,sas,Dscam1,Ptp99A | | 1.470e-03 | -6.52 | mid (midline) | protein interactions | 33770 | 12 | 4 | 9204 | 412 | ci,tin,Ubx,HGTX | | 1.470e-03 | -6.52 | fog (folded gastrulation) | protein interactions | 33148 | 12 | 4 | 9204 | 412 | tsl,cta,twi,RhoGEF2 | | 1.475e-03 | -6.52 | GO\_BIOLOGICAL\_ADHESION | MSigDB lists | GO\_BIOLOGICAL\_ADHESION | 155 | 14 | 4161 | 153 | DIP-epsilon,hh,Rac2,trn,slow,Fas2,smal,Lac,Dscam1,NetB,ft,Smox,if,ds | | 1.480e-03 | -6.52 | determination of adult lifespan | biological process | GO:0008340 | 167 | 14 | 12345 | 414 | Sin3A,S6kII,Mrp4,EcR,aop,Egfr,sm,mle,Smox,N,ImpL2,pnt,cnc,Cdk5alpha | | 1.495e-03 | -6.51 | cell cortex | cellular component | GO:0005938 | 183 | 15 | 11940 | 407 | corn,gus,Ser,CG42684,par-1,nuf,RhoGEF2,Arp2,pbl,pon,Rho1,Hip1,sdt,CD98hc,Syp | | 1.502e-03 | -6.50 | regulation of mitotic cell cycle | biological process | GO:0007346 | 186 | 15 | 12345 | 414 | Smox,PhKgamma,ci,N,Chro,bora,hh,kni,l(3)mbt,CycE,stg,Sin3A,Myb,PEK,Egfr | | 1.509e-03 | -6.50 | positive regulation of phosphorus metabolic process | biological process | GO:0010562 | 149 | 13 | 12345 | 414 | Egfr,wnd,PEK,Mos,dpp,Atg17,mbt,Rho1,Cdk5alpha,grnd,bora,drk,spi | | 1.509e-03 | -6.50 | positive regulation of phosphate metabolic process | biological process | GO:0045937 | 149 | 13 | 12345 | 414 | Egfr,wnd,PEK,Mos,Atg17,dpp,mbt,Rho1,Cdk5alpha,grnd,bora,drk,spi | | 1.516e-03 | -6.49 | chaeta development | biological process | GO:0022416 | 81 | 9 | 12345 | 414 | ush,N,Bx,B-H2,CtBP,NAT1,dally,spen,Ten-m | | 1.516e-03 | -6.49 | cell junction organization | biological process | GO:0034330 | 81 | 9 | 12345 | 414 | Rho1,nrv2,sdt,bou,Mmp2,crb,CG42748,Lac,Rac2 | | 1.518e-03 | -6.49 | Kr (Kruppel) | protein interactions | 38012 | 39 | 7 | 9204 | 412 | dpp,CtBP,S,osa,Acsl,sty,Bx | | 1.518e-03 | -6.49 | GSE8835\_CD4\_VS\_CD8\_TCELL\_DN | MSigDB lists | GSE8835\_CD4\_VS\_CD8\_TCELL\_DN | 24 | 5 | 4161 | 153 | TfAP-2,Fbxl7,CG12099,CG5522,DIP-epsilon | | 1.518e-03 | -6.49 | GO\_DEVELOPMENTAL\_GROWTH\_INVOLVED\_IN\_MORPHOGENESIS | MSigDB lists | GO\_DEVELOPMENTAL\_GROWTH\_INVOLVED\_IN\_MORPHOGENESIS | 24 | 5 | 4161 | 153 | N,ap,nudE,hh,Lim1 | | 1.518e-03 | -6.49 | GO\_EMBRYONIC\_SKELETAL\_SYSTEM\_MORPHOGENESIS | MSigDB lists | GO\_EMBRYONIC\_SKELETAL\_SYSTEM\_MORPHOGENESIS | 24 | 5 | 4161 | 153 | so,Lim1,Dscam1,Smox,TfAP-2 | | 1.518e-03 | -6.49 | GO\_DORSAL\_VENTRAL\_PATTERN\_FORMATION | MSigDB lists | GO\_DORSAL\_VENTRAL\_PATTERN\_FORMATION | 24 | 5 | 4161 | 153 | tkv,Dscam1,ap,hh,Lim1 | | 1.521e-03 | -6.49 | neuroblast fate determination | biological process | GO:0007400 | 26 | 5 | 12345 | 414 | Kr,N,Dr,spen,CycE | | 1.521e-03 | -6.49 | establishment of imaginal disc-derived wing hair orientation | biological process | GO:0001737 | 26 | 5 | 12345 | 414 | fj,ds,par-1,ft,N | | 1.523e-03 | -6.49 | protein autophosphorylation | biological process | GO:0046777 | 38 | 6 | 12345 | 414 | Pak3,JIL-1,wnd,Tlk,Doa,CaMKII | | 1.525e-03 | -6.49 | GO\_REGULATION\_OF\_MULTICELLULAR\_ORGANISMAL\_DEVELOPMENT | MSigDB lists | GO\_REGULATION\_OF\_MULTICELLULAR\_ORGANISMAL\_DEVELOPMENT | 325 | 23 | 4161 | 153 | TfAP-2,CG5830,ft,Lac,nudE,Tif-IA,smal,nAChRalpha4,tkv,Rac2,hh,CG11367,dlp,so,N,Smox,pbl,NetB,Ser,Lim1,dpp,LRP1,CG12099 | | 1.530e-03 | -6.48 | GO\_POSITIVE\_REGULATION\_OF\_GLIAL\_CELL\_DIFFERENTIATION | MSigDB lists | GO\_POSITIVE\_REGULATION\_OF\_GLIAL\_CELL\_DIFFERENTIATION | 7 | 3 | 4161 | 153 | hh,dpp,N | | 1.530e-03 | -6.48 | REACTOME\_HS\_GAG\_DEGRADATION | MSigDB lists | REACTOME\_HS\_GAG\_DEGRADATION | 7 | 3 | 4161 | 153 | dlp,dally,CG14291 | | 1.530e-03 | -6.48 | GO\_POSITIVE\_REGULATION\_OF\_STEROL\_TRANSPORT | MSigDB lists | GO\_POSITIVE\_REGULATION\_OF\_STEROL\_TRANSPORT | 7 | 3 | 4161 | 153 | ScpX,LRP1,EcR | | 1.530e-03 | -6.48 | GO\_NEURONAL\_STEM\_CELL\_POPULATION\_MAINTENANCE | MSigDB lists | GO\_NEURONAL\_STEM\_CELL\_POPULATION\_MAINTENANCE | 7 | 3 | 4161 | 153 | N,Ser,Imp | | 1.530e-03 | -6.48 | GO\_DIGESTIVE\_TRACT\_MORPHOGENESIS | MSigDB lists | GO\_DIGESTIVE\_TRACT\_MORPHOGENESIS | 7 | 3 | 4161 | 153 | Smox,N,hh | | 1.530e-03 | -6.48 | GO\_THYROID\_GLAND\_DEVELOPMENT | MSigDB lists | GO\_THYROID\_GLAND\_DEVELOPMENT | 7 | 3 | 4161 | 153 | Smox,hh,so | | 1.532e-03 | -6.48 | bru1 (bruno 1) | protein interactions | 34648 | 29 | 6 | 9204 | 412 | smg,Rbfox1,ci,enok,S,mub | | 1.532e-03 | -6.48 | cno (canoe) | protein interactions | 40620 | 29 | 6 | 9204 | 412 | S,drk,N,ci,Ser,mbt | | 1.532e-03 | -6.48 | Rok (Rho kinase) | protein interactions | 43916 | 29 | 6 | 9204 | 412 | RhoGEF2,egr,Rho1,hts,pbl,crb | | 1.532e-03 | -6.48 | CG42797 (uncharacterized protein) | protein interactions | 31905 | 29 | 6 | 9204 | 412 | CG3919,Fas2,phyl,betaTub56D,Hsc70-3,CG1142 | | 1.556e-03 | -6.47 | msl-1 (male-specific lethal 1) | protein interactions | 35121 | 20 | 5 | 9204 | 412 | JIL-1,lncRNA:roX1,mle,ocm,mof | | 1.563e-03 | -6.46 | ZF5\_01 | MSigDB lists | ZF5\_01 | 61 | 8 | 4161 | 153 | Dscam1,NetB,pAbp,CycE,heph,Tlk,CG12099,Imp | | 1.564e-03 | -6.46 | regulation of transmembrane receptor protein serine/threonine kinase signaling pathway | biological process | GO:0090092 | 66 | 8 | 12345 | 414 | dally,lili,hiw,dpp,Lpin,tkv,Acsl,Dad | | 1.570e-03 | -6.46 | adenyl nucleotide binding | molecular function | GO:0030554 | 659 | 36 | 12176 | 396 | Myo10A,CG12010,mbt,mle,lz,CG14535,JIL-1,Mos,Egfr,CG9281,CG9541,Pex1,par-1,PhKgamma,CG8485,Mrp4,Asator,Arp2,Etl1,Hipk,CG8097,Mhcl,PEK,Haspin,ScsbetaG,S6kII,KP78a,Pak3,bor,Doa,wnd,tkv,Tlk,CaMKII,Hsc70-3,Cdk12 | | 1.570e-03 | -6.46 | GO\_NEGATIVE\_REGULATION\_OF\_DEVELOPMENTAL\_PROCESS | MSigDB lists | GO\_NEGATIVE\_REGULATION\_OF\_DEVELOPMENTAL\_PROCESS | 156 | 14 | 4161 | 153 | Ser,dpp,Smox,CG5830,N,NetB,so,CG9855,LRP1,CG11367,hh,CG12099,heph,EcR | | 1.585e-03 | -6.45 | RNA polymerase II distal enhancer sequence-specific DNA binding | molecular function | GO:0000980 | 27 | 5 | 12176 | 396 | Ubx,aop,ey,tin,ttk | | 1.586e-03 | -6.45 | dsh (dishevelled) | protein interactions | 32078 | 127 | 14 | 9204 | 412 | osa,RhoGEF2,aop,CycE,Rho1,rau,bdg,mam,ds,N,Ubx,ey,tkv,egr | | 1.609e-03 | -6.43 | Snoo (Sno oncogene) | protein interactions | 5740414 | 6 | 3 | 9204 | 412 | tkv,Smox,dpp | | 1.627e-03 | -6.42 | regulation of actin cytoskeleton organization | biological process | GO:0032956 | 98 | 10 | 12345 | 414 | pbl,if,CG43658,cta,hts,spoon,RhoGEF2,PsGEF,Rho1,Arp2 | | 1.646e-03 | -6.41 | negative regulation of cell adhesion | biological process | GO:0007162 | 16 | 4 | 12345 | 414 | aop,par-1,N,aos | | 1.646e-03 | -6.41 | initiation of dorsal closure | biological process | GO:0007392 | 16 | 4 | 12345 | 414 | ush,dpp,Rho1,Rac2 | | 1.647e-03 | -6.41 | eye photoreceptor cell fate commitment | biological process | GO:0042706 | 52 | 7 | 12345 | 414 | aop,pnt,phyl,ttk,bdg,N,drk | | 1.647e-03 | -6.41 | compound eye photoreceptor fate commitment | biological process | GO:0001752 | 52 | 7 | 12345 | 414 | drk,ttk,bdg,N,phyl,aop,pnt | | 1.648e-03 | -6.41 | - | gene3d domains | 1.10.150.50 | 42 | 7 | 4929 | 208 | l(3)mbt,edl,pnt,smg,ckn,aop,CG34384 | | 1.653e-03 | -6.40 | compound eye photoreceptor development | biological process | GO:0042051 | 82 | 9 | 12345 | 414 | Doa,Ten-m,rau,lz,oc,ttk,crb,eya,CadN | | 1.695e-03 | -6.38 | anchoring junction | cellular component | GO:0070161 | 97 | 10 | 11940 | 407 | mbt,smal,scb,slow,if,N,sdt,wech,ckn,Lrt | | 1.695e-03 | -6.38 | adherens junction | cellular component | GO:0005912 | 97 | 10 | 11940 | 407 | smal,scb,slow,mbt,wech,ckn,Lrt,sdt,N,if | | 1.705e-03 | -6.37 | GO\_SEQUENCE\_SPECIFIC\_DNA\_BINDING | MSigDB lists | GO\_SEQUENCE\_SPECIFIC\_DNA\_BINDING | 107 | 11 | 4161 | 153 | Sin3A,EcR,so,Smox,N,Taf1,ap,TfAP-2,Utx,Tif-IA,Lim1 | | 1.721e-03 | -6.36 | neuron projection | cellular component | GO:0043005 | 431 | 27 | 11940 | 407 | NetA,EcR,Mmp2,robo2,nudE,dpr7,Ser,nAChRalpha4,nrv2,Prosap,wnd,par-1,CaMKII,Acsl,Ptp99A,Imp,Cdk5alpha,ImpL2,Dscam1,Con,CadN,NetB,Syx1A,Fas2,AGO2,ckn,yps | | 1.739e-03 | -6.35 | GO\_POSITIVE\_REGULATION\_OF\_LOCOMOTION | MSigDB lists | GO\_POSITIVE\_REGULATION\_OF\_LOCOMOTION | 62 | 8 | 4161 | 153 | CaMKII,NetB,Smox,N,dpp,smal,Rac2,Hsc70-3 | | 1.747e-03 | -6.35 | BIOCARTA\_CREB\_PATHWAY | MSigDB lists | BIOCARTA\_CREB\_PATHWAY | 15 | 4 | 4161 | 153 | JIL-1,drk,CaMKII,Rac2 | | 1.748e-03 | -6.35 | GO\_CELL\_GROWTH | MSigDB lists | GO\_CELL\_GROWTH | 36 | 6 | 4161 | 153 | ap,mbt,NTPase,CaMKII,bor,nudE | | 1.748e-03 | -6.35 | CAGCCTC\_MIR4855P | MSigDB lists | CAGCCTC\_MIR4855P | 36 | 6 | 4161 | 153 | Hydr2,Rac2,CG43313,ds,wake,mbt | | 1.748e-03 | -6.35 | GO\_BONE\_DEVELOPMENT | MSigDB lists | GO\_BONE\_DEVELOPMENT | 36 | 6 | 4161 | 153 | dpp,CG9220,TfAP-2,ds,tkv,ft | | 1.748e-03 | -6.35 | COULOUARN\_TEMPORAL\_TGFB1\_SIGNATURE\_DN | MSigDB lists | COULOUARN\_TEMPORAL\_TGFB1\_SIGNATURE\_DN | 36 | 6 | 4161 | 153 | NTPase,Acsl,nudE,Ser,LRP1,ScpX | | 1.753e-03 | -6.35 | multicellular organism aging | biological process | GO:0010259 | 170 | 14 | 12345 | 414 | aop,sm,Egfr,mle,Sin3A,S6kII,Mrp4,EcR,ImpL2,pnt,cnc,Cdk5alpha,Smox,N | | 1.771e-03 | -6.34 | slmb (supernumerary limbs) | protein interactions | 42504 | 75 | 10 | 9204 | 412 | par-1,crb,dpp,tara,ci,ds,tou,EcR,Hipk,dbr | | 1.816e-03 | -6.31 | positive regulation of cell cycle phase transition | biological process | GO:1901989 | 27 | 5 | 12345 | 414 | CycE,Myb,ci,N,stg | | 1.816e-03 | -6.31 | heterophilic cell-cell adhesion via plasma membrane cell adhesion molecules | biological process | GO:0007157 | 27 | 5 | 12345 | 414 | scb,ds,if,beat-IIIc,ft | | 1.841e-03 | -6.30 | GO\_EYE\_DEVELOPMENT | MSigDB lists | GO\_EYE\_DEVELOPMENT | 77 | 9 | 4161 | 153 | hh,tkv,ras,fj,Ser,Lim1,Smox,TfAP-2,ap | | 1.842e-03 | -6.30 | GSE45365\_BCELL\_VS\_CD8\_TCELL\_UP | MSigDB lists | GSE45365\_BCELL\_VS\_CD8\_TCELL\_UP | 25 | 5 | 4161 | 153 | CG4538,Pli,so,DIP-epsilon,Dscam1 | | 1.843e-03 | -6.30 | mre11 (meiotic recombination 11) | protein interactions | 34565 | 30 | 6 | 9204 | 412 | NiPp1,flfl,NAT1,tou,oc,CG5728 | | 1.843e-03 | -6.30 | tsh (teashirt) | protein interactions | 35430 | 30 | 6 | 9204 | 412 | Ser,betaTub56D,ci,CtBP,Ubx,spen | | 1.844e-03 | -6.30 | positive regulation of cell motility | biological process | GO:2000147 | 53 | 7 | 12345 | 414 | Dscam1,ttk,spi,Pvf3,aop,Egfr,robo2 | | 1.844e-03 | -6.30 | epidermal cell differentiation | biological process | GO:0009913 | 53 | 7 | 12345 | 414 | spen,fj,ds,par-1,Rho1,N,ft | | 1.849e-03 | -6.29 | tracheal pit formation in open tracheal system | biological process | GO:0035202 | 8 | 3 | 12345 | 414 | salm,Mmp2,Rho1 | | 1.849e-03 | -6.29 | regulation of salivary gland boundary specification | biological process | GO:0045704 | 8 | 3 | 12345 | 414 | tkv,dpp,shn | | 1.849e-03 | -6.29 | genital disc pattern formation | biological process | GO:0035221 | 8 | 3 | 12345 | 414 | hh,ci,dpp | | 1.849e-03 | -6.29 | negative regulation of salivary gland boundary specification | biological process | GO:0045705 | 8 | 3 | 12345 | 414 | shn,dpp,tkv | | 1.849e-03 | -6.29 | morphogenesis of larval imaginal disc epithelium | biological process | GO:0016335 | 8 | 3 | 12345 | 414 | hh,ft,Dad | | 1.849e-03 | -6.29 | genital disc anterior/posterior pattern formation | biological process | GO:0035224 | 8 | 3 | 12345 | 414 | ci,dpp,hh | | 1.865e-03 | -6.28 | - | gene3d domains | 1.10.555.10 | 22 | 5 | 4929 | 208 | RhoGAP16F,RhoGAP102A,RhoGAP18B,RhoGAP15B,RhoGAP93B | | 1.915e-03 | -6.26 | regulation of synapse organization | biological process | GO:0050807 | 153 | 13 | 12345 | 414 | Pak3,Dad,Fas2,scb,Arp2,S6kII,tkv,Acsl,Syp,CaMKII,Ten-m,hiw,wnd | | 1.949e-03 | -6.24 | collagen-containing extracellular matrix | cellular component | GO:0062023 | 27 | 5 | 11940 | 407 | dlp,NetA,wb,NetB,dally | | 1.968e-03 | -6.23 | coro (coro) | protein interactions | 35596 | 21 | 5 | 9204 | 412 | Syx1A,pbl,dpp,tkv,PhKgamma | | 1.968e-03 | -6.23 | jing (jing) | protein interactions | 35555 | 21 | 5 | 9204 | 412 | pbl,ci,S,Egfr,spi | | 1.968e-03 | -6.23 | Zn72D (Zinc-finger protein at 72D) | protein interactions | 39764 | 21 | 5 | 9204 | 412 | CycE,S6kII,Ubx,CG9775,Fas2 | | 1.968e-03 | -6.23 | Mtor (Megator) | protein interactions | 36264 | 21 | 5 | 9204 | 412 | Chro,EcR,mof,Psc,Asator | | 1.968e-03 | -6.23 | hpRNA:CR18854 (hairpin RNA:CR18854) | protein interactions | 10178800 | 21 | 5 | 9204 | 412 | l(3)L1231,Sin3A,CG32772,AGO2,CG7264 | | 1.999e-03 | -6.22 | CG10300 (uncharacterized protein) | protein interactions | 42798 | 2 | 2 | 9204 | 412 | sty,CG11247 | | 1.999e-03 | -6.22 | CG30460 (uncharacterized protein) | protein interactions | 36924 | 2 | 2 | 9204 | 412 | Fas2,CG2865 | | 1.999e-03 | -6.22 | CG17344 (uncharacterized protein) | protein interactions | 35185 | 2 | 2 | 9204 | 412 | Dmtn,CG9650 | | 1.999e-03 | -6.22 | CG14339 (uncharacterized protein) | protein interactions | 33286 | 2 | 2 | 9204 | 412 | Fas2,Gbs-70E | | 1.999e-03 | -6.22 | CG7115 (uncharacterized protein) | protein interactions | 34069 | 2 | 2 | 9204 | 412 | CG18766,Bx | | 1.999e-03 | -6.22 | CG9302 (uncharacterized protein) | protein interactions | 34750 | 2 | 2 | 9204 | 412 | AGO2,CG32772 | | 1.999e-03 | -6.22 | Ssadh (Succinic semialdehyde dehydrogenase) | protein interactions | 43092 | 2 | 2 | 9204 | 412 | Drat,HmgZ | | 1.999e-03 | -6.22 | Oamb (Octopamine receptor in mushroom bodies) | protein interactions | 43982 | 2 | 2 | 9204 | 412 | CaMKII,Mmp2 | | 2.002e-03 | -6.21 | segment polarity determination | biological process | GO:0007367 | 40 | 6 | 12345 | 414 | AGO2,dlp,Egfr,dally,hh,ci | | 2.005e-03 | -6.21 | dlg1 (discs large 1) | protein interactions | 32083 | 52 | 8 | 9204 | 412 | Fas2,crb,par-1,wake,CycE,CaMKII,hts,Syp | | 2.012e-03 | -6.21 | zf-C2H2 | pfam domains | PF00096 | 207 | 17 | 10480 | 393 | CG31224,ham,CG4360,Kah,hang,Kr,CG11247,ttk,bowl,Zif,CG32772,shn,ci,salm,sens-2,l(3)neo38,CG9650 | | 2.029e-03 | -6.20 | intrinsic component of plasma membrane | cellular component | GO:0031226 | 480 | 29 | 11940 | 407 | smal,Tsp26A,Dscam1,ft,otk2,emp,Tsp39D,CadN,dally,if,dpr7,Ten-m,wun2,ds,scb,Tl,nAChRalpha4,nrv2,Egfr,sas,dlp,DIP-epsilon,N,Prosap,Cad87A,tkv,bou,egr,Cirl | | 2.043e-03 | -6.19 | Rbf (Retinoblastoma-family protein) | protein interactions | 31027 | 64 | 9 | 9204 | 412 | Myb,tkv,EcR,tara,N,mam,CycE,ban,Utx | | 2.043e-03 | -6.19 | Pi3K68D (Phosphotidylinositol 3 kinase 68D) | protein interactions | 39329 | 64 | 9 | 9204 | 412 | Ank,aos,bor,ScsbetaG,Egfr,drk,N,fax,Cnx99A | | 2.049e-03 | -6.19 | hyd (hyperplastic discs) | protein interactions | 41181 | 13 | 4 | 9204 | 412 | pAbp,hh,ci,CycE | | 2.049e-03 | -6.19 | mir-14 (mir-14 stem loop) | protein interactions | 12798384 | 13 | 4 | 9204 | 412 | EcR,Bx,pbl,hh | | 2.049e-03 | -6.19 | sca (scabrous) | protein interactions | 36411 | 13 | 4 | 9204 | 412 | hh,mam,N,pbl | | 2.049e-03 | -6.19 | Shc (SHC-adaptor protein) | protein interactions | 44052 | 13 | 4 | 9204 | 412 | Egfr,drk,sty,CycE | | 2.049e-03 | -6.19 | ab (abrupt) | protein interactions | 34560 | 13 | 4 | 9204 | 412 | spi,EcR,bi,so | | 2.054e-03 | -6.19 | hop (hopscotch) | protein interactions | 32080 | 41 | 7 | 9204 | 412 | Ser,Rho1,CycE,eya,Myb,Stat92E,ush | | 2.077e-03 | -6.18 | chromatin organization | biological process | GO:0006325 | 466 | 28 | 12345 | 414 | AGO2,Psc,CG5815,osa,tou,Df31,stwl,tara,HmgZ,Tlk,Taf1,Utx,dve,Kr,psq,Mocs2B,bin3,JIL-1,Haspin,enok,wcy,Pcl,Acf,CtBP,Etl1,mof,eya,tna | | 2.092e-03 | -6.17 | AFP1\_Q6 | MSigDB lists | AFP1\_Q6 | 50 | 7 | 4161 | 153 | Pli,so,dlp,CG8531,pbl,Dscam1,fne | | 2.096e-03 | -6.17 | trunk segmentation | biological process | GO:0035290 | 17 | 4 | 12345 | 414 | Ubx,Kr,kni,hh | | 2.096e-03 | -6.17 | transforming growth factor beta receptor signaling pathway | biological process | GO:0007179 | 17 | 4 | 12345 | 414 | shn,dpp,tkv,Smox | | 2.101e-03 | -6.17 | PDZ | smart domains | SM00228 | 64 | 10 | 5086 | 276 | CG3402,RhoGEF2,PsGEF,Rhp,Ptpmeg,CG32758,bbg,sdt,Prosap,Mhcl | | 2.127e-03 | -6.15 | negative regulation of developmental growth | biological process | GO:0048640 | 85 | 9 | 12345 | 414 | scb,Fbxl7,Dad,Tl,ft,crb,Syp,hiw,Acsl | | 2.140e-03 | -6.15 | ACTTTAT\_MIR1425P | MSigDB lists | ACTTTAT\_MIR1425P | 64 | 8 | 4161 | 153 | ds,mgl,dpp,Ptpmeg,JIL-1,Rho1,Tpr2,fne | | 2.150e-03 | -6.14 | oocyte development | biological process | GO:0048599 | 155 | 13 | 12345 | 414 | heph,gus,dpp,Pat1,Mos,Syp,Egfr,psq,N,par-1,cnc,pAbp,scb | | 2.164e-03 | -6.14 | sinah (sina homologue) | protein interactions | 39885 | 77 | 10 | 9204 | 412 | esn,phyl,slow,CG8929,CG17265,Gyf,slou,Cnx99A,CG11030,ttk | | 2.207e-03 | -6.12 | PH-like\_dom\_sf | interpro domains | IPR011993 | 139 | 13 | 11383 | 428 | RhoGEF2,PsGEF,Ptpmeg,CG5522,CG34383,osp,Myo10A,pbl,MESR3,GramD1B,CG34384,flfl,rush | | 2.214e-03 | -6.11 | GSE27291\_0H\_VS\_7D\_STIM\_GAMMADELTA\_TCELL\_DN | MSigDB lists | GSE27291\_0H\_VS\_7D\_STIM\_GAMMADELTA\_TCELL\_DN | 26 | 5 | 4161 | 153 | rush,JIL-1,Smox,CG45050,phtf | | 2.214e-03 | -6.11 | GO\_ESTABLISHMENT\_OF\_CELL\_POLARITY | MSigDB lists | GO\_ESTABLISHMENT\_OF\_CELL\_POLARITY | 26 | 5 | 4161 | 153 | hh,Mhcl,nudE,sdt,CG32758 | | 2.214e-03 | -6.11 | GO\_PROTEOGLYCAN\_METABOLIC\_PROCESS | MSigDB lists | GO\_PROTEOGLYCAN\_METABOLIC\_PROCESS | 26 | 5 | 4161 | 153 | tkv,CG43313,CG9220,dpp,CG14291 | | 2.214e-03 | -6.11 | GO\_AMINOGLYCAN\_BIOSYNTHETIC\_PROCESS | MSigDB lists | GO\_AMINOGLYCAN\_BIOSYNTHETIC\_PROCESS | 26 | 5 | 4161 | 153 | CG9220,dally,dlp,CG43313,CG31637 | | 2.246e-03 | -6.10 | cellular protein modification process | biological process | GO:0006464 | 1215 | 59 | 12345 | 414 | wnd,CG9855,ds,hiw,Cdk12,Mos,S6kII,Pli,ctrip,CycE,Tlk,par-1,Utx,Asator,CG3253,egr,Mocs2B,CG12099,JIL-1,Hipk,Doa,CG14646,enok,wcy,stg,Notum,tkv,ec,smal,eya,CG11486,Ptpmeg,Cyp1,CG6752,Pak3,PhKgamma,ft,KP78a,CG13029,CaMKII,PEK,Usp2,Ppa,fj,Taf1,Rac2,Fbxl7,CG5830,Haspin,mbt,Hph,wech,Acf,CG8485,LManII,mof,CG4325,tna,Ptp99A | | 2.246e-03 | -6.10 | protein modification process | biological process | GO:0036211 | 1215 | 59 | 12345 | 414 | Mos,Cdk12,hiw,ds,CG9855,wnd,S6kII,Utx,par-1,CycE,Tlk,Pli,ctrip,JIL-1,Hipk,CG12099,Mocs2B,egr,CG3253,Asator,Doa,tkv,Notum,enok,wcy,stg,CG14646,smal,ec,ft,PhKgamma,Pak3,CG6752,eya,Ptpmeg,Cyp1,CG11486,PEK,CaMKII,CG13029,KP78a,Usp2,Taf1,Rac2,Ppa,fj,Fbxl7,CG5830,Haspin,Acf,wech,Hph,mbt,LManII,CG8485,Ptp99A,tna,CG4325,mof | | 2.264e-03 | -6.09 | REACTOME\_G\_ALPHA1213\_SIGNALLING\_EVENTS | MSigDB lists | REACTOME\_G\_ALPHA1213\_SIGNALLING\_EVENTS | 16 | 4 | 4161 | 153 | Rho1,cta,pbl,Rac2 | | 2.264e-03 | -6.09 | REACTOME\_SIGNAL\_TRANSDUCTION\_BY\_L1 | MSigDB lists | REACTOME\_SIGNAL\_TRANSDUCTION\_BY\_L1 | 16 | 4 | 4161 | 153 | Rac2,S6kII,JIL-1,Fas2 | | 2.264e-03 | -6.09 | GO\_REGULATION\_OF\_DNA\_TEMPLATED\_TRANSCRIPTION\_IN\_RESPONSE\_TO\_STRESS | MSigDB lists | GO\_REGULATION\_OF\_DNA\_TEMPLATED\_TRANSCRIPTION\_IN\_RESPONSE\_TO\_STRESS | 16 | 4 | 4161 | 153 | N,Sin3A,S6kII,Hsc70-3 | | 2.264e-03 | -6.09 | GO\_POSITIVE\_REGULATION\_OF\_OSTEOBLAST\_DIFFERENTIATION | MSigDB lists | GO\_POSITIVE\_REGULATION\_OF\_OSTEOBLAST\_DIFFERENTIATION | 16 | 4 | 4161 | 153 | tkv,smal,Ser,dpp | | 2.264e-03 | -6.09 | GO\_REGULATION\_OF\_LIPASE\_ACTIVITY | MSigDB lists | GO\_REGULATION\_OF\_LIPASE\_ACTIVITY | 16 | 4 | 4161 | 153 | Rho1,cta,EcR,LRP1 | | 2.264e-03 | -6.09 | GO\_NEGATIVE\_REGULATION\_OF\_OSSIFICATION | MSigDB lists | GO\_NEGATIVE\_REGULATION\_OF\_OSSIFICATION | 16 | 4 | 4161 | 153 | Smox,N,CG11367,CG9220 | | 2.271e-03 | -6.09 | Iswi (Imitation SWI) | protein interactions | 36390 | 53 | 8 | 9204 | 412 | Acf,tou,mle,EcR,Sin3A,CtBP,ttk,mof | | 2.280e-03 | -6.08 | muscle attachment | biological process | GO:0016203 | 41 | 6 | 12345 | 414 | Rho1,wech,betaTub56D,slow,Rac2,if | | 2.292e-03 | -6.08 | PNT | prosite domains | PS51433 | 6 | 3 | 6243 | 316 | pnt,edl,aop | | 2.292e-03 | -6.08 | LAMININ\_NTER | prosite domains | PS51117 | 6 | 3 | 6243 | 316 | wb,NetA,NetB | | 2.293e-03 | -6.08 | somatic stem cell division | biological process | GO:0048103 | 55 | 7 | 12345 | 414 | wake,Stat92E,flfl,hh,pon,Zif,sdt | | 2.298e-03 | -6.08 | aging | biological process | GO:0007568 | 175 | 14 | 12345 | 414 | EcR,Mrp4,S6kII,Sin3A,mle,sm,Egfr,aop,N,Smox,Cdk5alpha,cnc,pnt,ImpL2 | | 2.307e-03 | -6.07 | eye photoreceptor cell development | biological process | GO:0042462 | 86 | 9 | 12345 | 414 | oc,ttk,crb,eya,CadN,Ten-m,Doa,rau,lz | | 2.308e-03 | -6.07 | Axon guidance | REACTOME pathways | R-DME-422475 | 194 | 16 | 3923 | 148 | RhoGAP93B,NetA,par-1,Fas2,scb,NetB,Arp2,JIL-1,robo2,if,CG42684,dlp,Egfr,Ank,drk,Rho1 | | 2.311e-03 | -6.07 | YGCGYRCGC\_UNKNOWN | MSigDB lists | YGCGYRCGC\_UNKNOWN | 95 | 10 | 4161 | 153 | JIL-1,sdt,CycE,hh,RpS4,Mitofilin,so,sm,CG6752,Nhe2 | | 2.311e-03 | -6.07 | GO\_IN\_UTERO\_EMBRYONIC\_DEVELOPMENT | MSigDB lists | GO\_IN\_UTERO\_EMBRYONIC\_DEVELOPMENT | 95 | 10 | 4161 | 153 | drk,Sin3A,Smox,N,hts,cta,Utx,nudE,Tif-IA,dpp | | 2.322e-03 | -6.07 | fly chr22D1-22D1\|2-5 cM | chromosome location | fly chr22D1-22D1\|2-5 cM | 13 | 3 | 25010 | 531 | CG34172,lncRNA:CR43751,aop | | 2.322e-03 | -6.07 | fly chr22D1-22D1\|2-5 | chromosome location | fly chr22D1-22D1\|2-5 | 13 | 3 | 25010 | 531 | CG34172,aop,lncRNA:CR43751 | | 2.332e-03 | -6.06 | GSE2770\_IL12\_VS\_IL4\_TREATED\_ACT\_CD4\_TCELL\_48H\_DN | MSigDB lists | GSE2770\_IL12\_VS\_IL4\_TREATED\_ACT\_CD4\_TCELL\_48H\_DN | 38 | 6 | 4161 | 153 | S6kII,NTPase,Arl4,sno,Hsc70Cb,Nhe2 | | 2.351e-03 | -6.05 | GO\_NEGATIVE\_REGULATION\_OF\_CELL\_DIFFERENTIATION | MSigDB lists | GO\_NEGATIVE\_REGULATION\_OF\_CELL\_DIFFERENTIATION | 128 | 12 | 4161 | 153 | dpp,Ser,NetB,N,CG5830,Smox,CG12099,LRP1,hh,CG11367,EcR,heph | | 2.351e-03 | -6.05 | FOXO1\_02 | MSigDB lists | FOXO1\_02 | 51 | 7 | 4161 | 153 | Utx,Sin3A,NetB,dpp,Ser,Pli,CG4538 | | 2.365e-03 | -6.05 | E2F1\_Q3\_01 | MSigDB lists | E2F1\_Q3\_01 | 65 | 8 | 4161 | 153 | CG5830,NetB,ap,Pli,Fas2,CG8531,JIL-1,Tlk | | 2.383e-03 | -6.04 | SHARMA\_PILOCYTIC\_ASTROCYTOMA\_LOCATION\_UP | MSigDB lists | SHARMA\_PILOCYTIC\_ASTROCYTOMA\_LOCATION\_UP | 8 | 3 | 4161 | 153 | ap,so,sno | | 2.383e-03 | -6.04 | GO\_POSITIVE\_REGULATION\_OF\_NEURAL\_PRECURSOR\_CELL\_PROLIFERATION | MSigDB lists | GO\_POSITIVE\_REGULATION\_OF\_NEURAL\_PRECURSOR\_CELL\_PROLIFERATION | 8 | 3 | 4161 | 153 | N,Lim1,hh | | 2.383e-03 | -6.04 | GO\_REGULATION\_OF\_HEART\_MORPHOGENESIS | MSigDB lists | GO\_REGULATION\_OF\_HEART\_MORPHOGENESIS | 8 | 3 | 4161 | 153 | N,dpp,so | | 2.383e-03 | -6.04 | GO\_RETINA\_LAYER\_FORMATION | MSigDB lists | GO\_RETINA\_LAYER\_FORMATION | 8 | 3 | 4161 | 153 | fj,TfAP-2,Lim1 | | 2.383e-03 | -6.04 | GO\_SMOOTH\_MUSCLE\_CELL\_DIFFERENTIATION | MSigDB lists | GO\_SMOOTH\_MUSCLE\_CELL\_DIFFERENTIATION | 8 | 3 | 4161 | 153 | so,if,N | | 2.383e-03 | -6.04 | SA\_B\_CELL\_RECEPTOR\_COMPLEXES | MSigDB lists | SA\_B\_CELL\_RECEPTOR\_COMPLEXES | 8 | 3 | 4161 | 153 | Rac2,drk,S6kII | | 2.386e-03 | -6.04 | CADHERIN | prints domains | PR00205 | 15 | 4 | 2185 | 88 | Cad87A,ds,ft,CadN | | 2.455e-03 | -6.01 | crol (crooked legs) | protein interactions | 34592 | 22 | 5 | 9204 | 412 | Kr,CycE,if,pbl,stg | | 2.455e-03 | -6.01 | jigr1 (jing interacting gene regulatory 1) | protein interactions | 43093 | 22 | 5 | 9204 | 412 | jigr1,N,emp,Imp,Chro | | 2.455e-03 | -6.01 | cdc14 (cell division cycle 14) | protein interactions | 34067 | 22 | 5 | 9204 | 412 | pbl,CG11486,heph,EcR,gus | | 2.472e-03 | -6.00 | regulation of cellular localization | biological process | GO:0060341 | 139 | 12 | 12345 | 414 | sdt,S6kII,Cnx99A,dlp,dally,ScpX,crb,Syx1A,NFAT,cnc,par-1,hh | | 2.476e-03 | -6.00 | Laminin\_G\_2 | pfam domains | PF02210 | 16 | 4 | 10480 | 393 | ft,CadN,crb,wb | | 2.477e-03 | -6.00 | Dsor1 (Downstream of raf1) | protein interactions | 31872 | 105 | 12 | 9204 | 412 | if,RhoGAP18B,Imp,aop,mam,hh,Egfr,S6kII,N,pnt,ScsbetaG,aos | | 2.478e-03 | -6.00 | epidermal growth factor receptor binding | molecular function | GO:0005154 | 9 | 3 | 12176 | 396 | drk,aos,spi | | 2.502e-03 | -5.99 | Cadherin-like\_dom | interpro domains | IPR002126 | 16 | 4 | 11383 | 428 | ft,ds,Cad87A,CadN | | 2.506e-03 | -5.99 | regulation of oogenesis | biological process | GO:1905879 | 71 | 8 | 12345 | 414 | aop,Egfr,par-1,cnc,Syx1A,ttk,spi,Ptpmeg | | 2.540e-03 | -5.98 | chromatin remodeling | biological process | GO:0006338 | 104 | 10 | 12345 | 414 | CtBP,Etl1,HmgZ,Mocs2B,dve,CG5815,osa,Psc,Acf,tou | | 2.546e-03 | -5.97 | T-box | pfam domains | PF00907 | 8 | 3 | 10480 | 393 | Doc1,Doc2,bi | | 2.546e-03 | -5.97 | HTH\_psq | pfam domains | PF05225 | 8 | 3 | 10480 | 393 | bab1,BtbVII,psq | | 2.546e-03 | -5.97 | LRR\_1 | pfam domains | PF00560 | 8 | 3 | 10480 | 393 | Fili,Toll-7,Tl | | 2.553e-03 | -5.97 | regulation of cell cycle process | biological process | GO:0010564 | 177 | 14 | 12345 | 414 | Myb,stg,Egfr,PEK,Mos,nuf,Chro,N,ci,CycE,CG31612,pbl,kni,bora | | 2.554e-03 | -5.97 | SERTA | prosite domains | PS51053 | 2 | 2 | 6243 | 316 | CG2865,tara | | 2.563e-03 | -5.97 | Chondroitin sulfate biosynthesis | REACTOME pathways | R-DME-2022870 | 8 | 3 | 3923 | 148 | CG9220,CG31637,CG43313 | | 2.567e-03 | -5.97 | T-box\_sf | interpro domains | IPR036960 | 8 | 3 | 11383 | 428 | Doc2,bi,Doc1 | | 2.567e-03 | -5.97 | TF\_T-box\_CS | interpro domains | IPR018186 | 8 | 3 | 11383 | 428 | Doc2,Doc1,bi | | 2.583e-03 | -5.96 | protein phosphatase binding | molecular function | GO:0019903 | 30 | 5 | 12176 | 396 | ec,NiPp1,CG42748,Gbs-70E,ckn | | 2.625e-03 | -5.94 | positive regulation of hippo signaling | biological process | GO:0035332 | 18 | 4 | 12345 | 414 | ft,crb,Fbxl7,ds | | 2.627e-03 | -5.94 | BENPORATH\_PRC2\_TARGETS | MSigDB lists | BENPORATH\_PRC2\_TARGETS | 81 | 9 | 4161 | 153 | so,hh,TfIIS,dally,mgl,Dscam1,NetB,ap,Fas2 | | 2.635e-03 | -5.94 | GSE33425\_CD8\_ALPHAALPHA\_VS\_ALPHABETA\_CD161\_HIGH\_TCELL\_UP | MSigDB lists | GSE33425\_CD8\_ALPHAALPHA\_VS\_ALPHABETA\_CD161\_HIGH\_TCELL\_UP | 52 | 7 | 4161 | 153 | EcR,nudE,Fas2,Cyp1,Mitofilin,phtf,Doa | | 2.635e-03 | -5.94 | GTGCCAT\_MIR183 | MSigDB lists | GTGCCAT\_MIR183 | 52 | 7 | 4161 | 153 | Ptpmeg,l(3)mbt,CG5742,Sin3A,S6kII,SamDC,CG5830 | | 2.635e-03 | -5.94 | GSE2770\_UNTREATED\_VS\_IL4\_TREATED\_ACT\_CD4\_TCELL\_6H\_DN | MSigDB lists | GSE2770\_UNTREATED\_VS\_IL4\_TREATED\_ACT\_CD4\_TCELL\_6H\_DN | 52 | 7 | 4161 | 153 | JIL-1,S6kII,cta,drk,pbl,l(3)mbt,Nhe2 | | 2.638e-03 | -5.94 | SHEPARD\_CRUSH\_AND\_BURN\_MUTANT\_DN | MSigDB lists | SHEPARD\_CRUSH\_AND\_BURN\_MUTANT\_DN | 27 | 5 | 4161 | 153 | Lim1,Ser,Smox,pbl,Utx | | 2.638e-03 | -5.94 | GSE3982\_BCELL\_VS\_NKCELL\_DN | MSigDB lists | GSE3982\_BCELL\_VS\_NKCELL\_DN | 27 | 5 | 4161 | 153 | CG14291,retm,emp,Fas2,Fbxl7 | | 2.638e-03 | -5.94 | GO\_LYSOSOMAL\_LUMEN | MSigDB lists | GO\_LYSOSOMAL\_LUMEN | 27 | 5 | 4161 | 153 | emp,dlp,dally,CG14291,LManII | | 2.705e-03 | -5.91 | regulation of crystal cell differentiation | biological process | GO:0042689 | 9 | 3 | 12345 | 414 | ush,lz,N | | 2.705e-03 | -5.91 | regulation of animal organ formation | biological process | GO:0003156 | 9 | 3 | 12345 | 414 | tkv,dpp,shn | | 2.705e-03 | -5.91 | imaginal disc lineage restriction | biological process | GO:0035161 | 9 | 3 | 12345 | 414 | hh,ap,N | | 2.705e-03 | -5.91 | positive regulation of G1/S transition of mitotic cell cycle | biological process | GO:1900087 | 9 | 3 | 12345 | 414 | CycE,ci,N | | 2.705e-03 | -5.91 | epithelial tube formation | biological process | GO:0072175 | 9 | 3 | 12345 | 414 | mmy,Kr,Mmp2 | | 2.705e-03 | -5.91 | ventral midline development | biological process | GO:0007418 | 9 | 3 | 12345 | 414 | hh,mam,jing | | 2.710e-03 | -5.91 | 6-blade\_b-propeller\_TolB-like | interpro domains | IPR011042 | 38 | 6 | 11383 | 428 | CG12547,mgl,yellow-e3,wech,Ten-m,LRP1 | | 2.710e-03 | -5.91 | Homeobox\_metazoa | interpro domains | IPR020479 | 38 | 6 | 11383 | 428 | Dr,exex,slou,HGTX,Ubx,B-H2 | | 2.723e-03 | -5.91 | Deaf1 (Deformed epidermal autoregulatory factor-1) | protein interactions | 40164 | 7 | 3 | 9204 | 412 | Hph,Pat1,MFS10 | | 2.723e-03 | -5.91 | lncRNA:roX2 (long non-coding RNA on the X 2) | protein interactions | 3772415 | 7 | 3 | 9204 | 412 | mof,mle,lncRNA:roX1 | | 2.723e-03 | -5.91 | miple1 (midkine and pleiotrophin 1) | protein interactions | 38047 | 7 | 3 | 9204 | 412 | Ptp99A,aos,Egfr | | 2.723e-03 | -5.91 | CG9932 (uncharacterized protein) | protein interactions | 34701 | 7 | 3 | 9204 | 412 | Fas2,CycE,Bx | | 2.723e-03 | -5.91 | Fsn (F-box synaptic protein) | protein interactions | 36460 | 7 | 3 | 9204 | 412 | hiw,gus,wnd | | 2.726e-03 | -5.90 | Ret (Ret oncogene) | protein interactions | 43875 | 43 | 7 | 9204 | 412 | crb,stg,drk,hh,S,spi,Sin3A | | 2.768e-03 | -5.89 | Dd (Dullard) | protein interactions | 33107 | 14 | 4 | 9204 | 412 | Dad,dpp,Lpin,tkv | | 2.768e-03 | -5.89 | CG5783 (uncharacterized protein) | protein interactions | 35088 | 14 | 4 | 9204 | 412 | Doa,Pvf3,Pits,CG43658 | | 2.768e-03 | -5.89 | mir-8 (mir-8 stem loop) | protein interactions | 12798320 | 14 | 4 | 9204 | 412 | spi,Ser,ush,AGO2 | | 2.768e-03 | -5.89 | RhoGAP18B (Rho GTPase activating protein at 18B) | protein interactions | 32898 | 14 | 4 | 9204 | 412 | Pli,S6kII,drk,Rho1 | | 2.797e-03 | -5.88 | LamNT | smart domains | SM00136 | 6 | 3 | 5086 | 276 | wb,NetB,NetA | | 2.819e-03 | -5.87 | Pits (Protein interacting with Ttk69 and Sin3A) | protein interactions | 32205 | 67 | 9 | 9204 | 412 | heph,CG32369,sty,Rac2,Gbs-70E,ttk,Chro,CG2865,Sin3A | | 2.819e-03 | -5.87 | brk (brinker) | protein interactions | 31665 | 67 | 9 | 9204 | 412 | tkv,ey,cv,CtBP,Dad,ban,sty,CG32369,shn | | 2.833e-03 | -5.87 | cytoneme | cellular component | GO:0035230 | 9 | 3 | 11940 | 407 | Egfr,tkv,dpp | | 2.846e-03 | -5.86 | Glycosaminoglycan biosynthesis, chondroitin sulfate backbone | KEGG pathways | dme\_M00058 | 3 | 2 | 3198 | 100 | CG43313,CG9220 | | 2.846e-03 | -5.86 | Glycosaminoglycan biosynthesis, chondroitin sulfate backbone | KEGG pathways | M00058 | 3 | 2 | 3198 | 100 | CG43313,CG9220 | | 2.876e-03 | -5.85 | GO\_METANEPHROS\_DEVELOPMENT | MSigDB lists | GO\_METANEPHROS\_DEVELOPMENT | 17 | 4 | 4161 | 153 | if,Lim1,hh,so | | 2.876e-03 | -5.85 | GO\_MECHANORECEPTOR\_DIFFERENTIATION | MSigDB lists | GO\_MECHANORECEPTOR\_DIFFERENTIATION | 17 | 4 | 4161 | 153 | Ser,N,Rac2,ft | | 2.876e-03 | -5.85 | GSE13485\_DAY1\_VS\_DAY7\_YF17D\_VACCINE\_PBMC\_UP | MSigDB lists | GSE13485\_DAY1\_VS\_DAY7\_YF17D\_VACCINE\_PBMC\_UP | 17 | 4 | 4161 | 153 | NKAIN,Trpm,CG7264,CG14291 | | 2.891e-03 | -5.85 | Sema1a (Semaphorin 1a) | protein interactions | 34192 | 150 | 15 | 9204 | 412 | tou,dally,Ten-m,pbl,ft,Mmp2,Asator,RhoGAP15B,Fas2,if,CG42671,Rho1,Taf1,RhoGEF2,Cyp1 | | 2.897e-03 | -5.84 | cytokine receptor binding | molecular function | GO:0005126 | 19 | 4 | 12176 | 396 | egr,Pvf3,dpp,Stat92E | | 2.909e-03 | -5.84 | fly chr6E4-6E4\|1-18 | chromosome location | fly chr6E4-6E4\|1-18 | 14 | 3 | 25010 | 531 | lncRNA:CR43837,bou,Inx2 | | 2.909e-03 | -5.84 | fly chr6E4-6E4\|1-18 cM | chromosome location | fly chr6E4-6E4\|1-18 cM | 14 | 3 | 25010 | 531 | Inx2,bou,lncRNA:CR43837 | | 2.921e-03 | -5.84 | positive regulation of programmed cell death | biological process | GO:0043068 | 89 | 9 | 12345 | 414 | aos,lz,EcR,N,ft,grnd,Utx,ec,CycE | | 2.922e-03 | -5.84 | asymmetric cell division | biological process | GO:0008356 | 106 | 10 | 12345 | 414 | flfl,mam,CycE,N,Zif,pon,hts,wake,dally,sdt | | 2.935e-03 | -5.83 | C345C | smart domains | SM00643 | 2 | 2 | 5086 | 276 | NetA,NetB | | 2.944e-03 | -5.83 | G alpha (12/13) signalling events | REACTOME pathways | R-DME-416482 | 27 | 5 | 3923 | 148 | RhoGEF2,Rho1,Rac2,cta,pbl | | 2.944e-03 | -5.83 | PROTEIN\_KINASE\_ST | prosite domains | PS00108 | 147 | 16 | 6243 | 316 | Tlk,Pak3,Doa,S6kII,wnd,CG8485,CaMKII,Hipk,par-1,tkv,PhKgamma,Mos,PEK,Cdk12,KP78a,JIL-1 | | 2.944e-03 | -5.83 | DR4\_Q2 | MSigDB lists | DR4\_Q2 | 53 | 7 | 4161 | 153 | fne,SamDC,JIL-1,Acsl,Utx,CtBP,NKAIN | | 2.944e-03 | -5.83 | GSE17721\_12H\_VS\_24H\_PAM3CSK4\_BMDC\_UP | MSigDB lists | GSE17721\_12H\_VS\_24H\_PAM3CSK4\_BMDC\_UP | 53 | 7 | 4161 | 153 | Imp,Hydr2,heph,cta,CG11030,Rho1,CG14646 | | 2.951e-03 | -5.83 | male sex differentiation | biological process | GO:0046661 | 30 | 5 | 12345 | 414 | otk2,salm,Fas2,eya,N | | 2.988e-03 | -5.81 | positive regulation of MAPK cascade | biological process | GO:0043410 | 73 | 8 | 12345 | 414 | drk,spi,grnd,mbt,PEK,Mos,Egfr,wnd | | 3.001e-03 | -5.81 | anatomical structure homeostasis | biological process | GO:0060249 | 161 | 13 | 12345 | 414 | Egfr,disco,Gyf,Atg17,Cnx99A,if,egr,ush,Lac,crb,JIL-1,N,CG46339 | | 3.023e-03 | -5.80 | fj (four-jointed) | protein interactions | 37089 | 23 | 5 | 9204 | 412 | fj,ft,N,ds,Ser | | 3.023e-03 | -5.80 | Btk29A (Btk family kinase at 29A) | protein interactions | 34132 | 23 | 5 | 9204 | 412 | rau,CycE,sty,N,Egfr | | 3.023e-03 | -5.80 | Pax (Paxillin) | protein interactions | 35215 | 23 | 5 | 9204 | 412 | ttk,CG10663,EcR,Rho1,sty | | 3.023e-03 | -5.80 | tup (tailup) | protein interactions | 35147 | 23 | 5 | 9204 | 412 | CG32369,Bx,ush,Ubx,ap | | 3.052e-03 | -5.79 | GO\_CELL\_FATE\_COMMITMENT | MSigDB lists | GO\_CELL\_FATE\_COMMITMENT | 40 | 6 | 4161 | 153 | Ser,hh,dpp,so,N,Dscam1 | | 3.053e-03 | -5.79 | TBOX | prints domains | PR00937 | 8 | 3 | 2185 | 88 | Doc2,bi,Doc1 | | 3.070e-03 | -5.79 | DENR (Density regulated protein) | protein interactions | 32679 | 33 | 6 | 9204 | 412 | CG2865,EcR,Sin3A,salm,hang,trn | | 3.073e-03 | -5.79 | ct (cut) | protein interactions | 44540 | 94 | 11 | 9204 | 412 | mam,N,sno,Psc,ey,osa,aop,Lim1,shn,heph,CycE | | 3.097e-03 | -5.78 | uridine phosphorylase activity | molecular function | GO:0004850 | 3 | 2 | 12176 | 396 | CG17224,CG6330 | | 3.117e-03 | -5.77 | MODULE\_63 | MSigDB lists | MODULE\_63 | 28 | 5 | 4161 | 153 | tkv,mgl,nAChRalpha4,LRP1,smal | | 3.117e-03 | -5.77 | REACTOME\_CELL\_CELL\_COMMUNICATION | MSigDB lists | REACTOME\_CELL\_CELL\_COMMUNICATION | 28 | 5 | 4161 | 153 | Dscam1,NetB,Rac2,drk,sdt | | 3.117e-03 | -5.77 | GO\_REGULATION\_OF\_MUSCLE\_TISSUE\_DEVELOPMENT | MSigDB lists | GO\_REGULATION\_OF\_MUSCLE\_TISSUE\_DEVELOPMENT | 28 | 5 | 4161 | 153 | N,Smox,dpp,hh,so | | 3.118e-03 | -5.77 | behavioral response to ethanol | biological process | GO:0048149 | 58 | 7 | 12345 | 414 | Egfr,scb,S,Gbs-70E,spi,Fas2,hang | | 3.123e-03 | -5.77 | MARSON\_BOUND\_BY\_FOXP3\_UNSTIMULATED | MSigDB lists | MARSON\_BOUND\_BY\_FOXP3\_UNSTIMULATED | 406 | 26 | 4161 | 153 | MFS10,CG5742,drk,Hsc70Cb,CaMKII,hts,sm,Arl4,CG5830,S6kII,cta,Arp2,Npc1a,CG14291,phtf,betaTub56D,Hydr2,CG45050,JIL-1,Snx3,NTPase,pbl,Smox,Utx,Syx1A,CG12547 | | 3.124e-03 | -5.77 | Chi (Chip) | protein interactions | 37837 | 68 | 9 | 9204 | 412 | ush,Hydr2,tou,N,Lim1,lz,Bx,osa,ap | | 3.132e-03 | -5.77 | cellular response to endogenous stimulus | biological process | GO:0071495 | 181 | 14 | 12345 | 414 | pnt,Utx,Smox,Hydr2,drk,Dad,hiw,aop,dally,tkv,Lpin,shn,dpp,EcR | | 3.165e-03 | -5.76 | endocytic vesicle | cellular component | GO:0030139 | 30 | 5 | 11940 | 407 | comm,Hip1,hh,nuf,N | | 3.176e-03 | -5.75 | ZP\_dom | interpro domains | IPR001507 | 17 | 4 | 11383 | 428 | pio,neo,tyn,qsm | | 3.178e-03 | -5.75 | organelle organization | biological process | GO:0006996 | 1885 | 84 | 12345 | 414 | CG34172,KP78a,Hip1,osa,psq,pAbp,if,B-H2,twi,betaTub56D,heph,Rho1,Acf,CG13366,Ten-m,nudE,tna,rush,CtBP,tou,Myb,wnd,hts,mle,ds,spoon,CG5815,Gmap,Mocs2B,Svil,JIL-1,Syx1A,bin3,Prosap,Mitofilin,Tlk,HmgZ,Utx,CG43658,Atg17,baf,Doa,eya,Chro,ft,Df31,tara,Psc,nuf,bor,Kr,N,crb,Rac2,Taf1,mbt,Pcl,Arp2,Haspin,mof,tyn,pbl,dpp,CG8531,Tim17b,stwl,wake,Ank,AGO2,RhoGEF2,dve,Pex1,oc,CG11498,par-1,enok,stg,wcy,tkv,cta,Pak3,cnc,RhoGAP18B,Etl1 | | 3.198e-03 | -5.75 | Fas2 (Fasciclin 2) | protein interactions | 31364 | 294 | 24 | 9204 | 412 | tou,CG6700,baf,Egfr,RpS4,sdt,betaTub56D,bbg,Hsc70-3,CG5815,CG6695,CG42748,Myo10A,aos,Tudor-SN,CG43658,Psc,LRP1,Fas2,pio,par-1,sty,shn,LManII | | 3.200e-03 | -5.74 | CADHERIN\_2 | prosite domains | PS50268 | 13 | 4 | 6243 | 316 | ft,Cad87A,ds,CadN | | 3.237e-03 | -5.73 | response to transforming growth factor beta | biological process | GO:0071559 | 19 | 4 | 12345 | 414 | shn,dpp,tkv,Smox | | 3.237e-03 | -5.73 | cellular response to transforming growth factor beta stimulus | biological process | GO:0071560 | 19 | 4 | 12345 | 414 | Smox,tkv,shn,dpp | | 3.254e-03 | -5.73 | long-term memory | biological process | GO:0007616 | 74 | 8 | 12345 | 414 | N,Stat92E,dbr,EcR,spoon,wnd,CaMKII,hiw | | 3.283e-03 | -5.72 | - | gene3d domains | 2.60.40.10 | 195 | 17 | 4929 | 208 | ImpL2,Fas2,wake,lbk,NFAT,Dscam1,sas,otk2,dpr7,Ptp99A,beat-IIIc,Lac,DIP-epsilon,CG6044,Fas3,nrm,robo2 | | 3.291e-03 | -5.72 | actin cytoskeleton organization | biological process | GO:0030036 | 242 | 17 | 12345 | 414 | hts,Hip1,nuf,RhoGEF2,if,twi,Rac2,CG43658,N,crb,CG13366,Rho1,Arp2,RhoGAP18B,pbl,Pak3,tyn | | 3.291e-03 | -5.72 | peptide cross-linking | biological process | GO:0018149 | 3 | 2 | 12345 | 414 | ft,ds | | 3.291e-03 | -5.72 | glial cell fate commitment | biological process | GO:0021781 | 3 | 2 | 12345 | 414 | N,spen | | 3.291e-03 | -5.72 | ectoderm formation | biological process | GO:0001705 | 3 | 2 | 12345 | 414 | spi,dpp | | 3.291e-03 | -5.72 | negative regulation of developmental pigmentation | biological process | GO:0048086 | 3 | 2 | 12345 | 414 | S6kII,bab1 | | 3.291e-03 | -5.72 | glial cell proliferation | biological process | GO:0014009 | 3 | 2 | 12345 | 414 | egr,Ser | | 3.291e-03 | -5.72 | glial cell fate determination | biological process | GO:0007403 | 3 | 2 | 12345 | 414 | spen,N | | 3.291e-03 | -5.72 | leading edge cell fate commitment | biological process | GO:0035027 | 3 | 2 | 12345 | 414 | ush,dpp | | 3.291e-03 | -5.72 | cell chemotaxis involved in Malpighian tubule morphogenesis | biological process | GO:0061352 | 3 | 2 | 12345 | 414 | dpp,Dad | | 3.291e-03 | -5.72 | epithelial cell fate commitment | biological process | GO:0072148 | 3 | 2 | 12345 | 414 | ush,dpp | | 3.291e-03 | -5.72 | regulation of sterol transport | biological process | GO:0032371 | 3 | 2 | 12345 | 414 | Npc1a,ScpX | | 3.291e-03 | -5.72 | spectrosome organization | biological process | GO:0030721 | 3 | 2 | 12345 | 414 | dpp,tkv | | 3.291e-03 | -5.72 | assembly of actomyosin apparatus involved in mitotic cytokinesis | biological process | GO:1902407 | 3 | 2 | 12345 | 414 | pbl,Rho1 | | 3.291e-03 | -5.72 | ectodermal cell differentiation | biological process | GO:0010668 | 3 | 2 | 12345 | 414 | dpp,spi | | 3.291e-03 | -5.72 | ectodermal cell fate commitment | biological process | GO:0001712 | 3 | 2 | 12345 | 414 | dpp,spi | | 3.291e-03 | -5.72 | leading edge cell fate determination | biological process | GO:0035028 | 3 | 2 | 12345 | 414 | dpp,ush | | 3.291e-03 | -5.72 | embryonic heart tube anterior/posterior pattern specification | biological process | GO:0035054 | 3 | 2 | 12345 | 414 | Ubx,EcR | | 3.291e-03 | -5.72 | dorsal closure, leading edge cell fate commitment | biological process | GO:0035029 | 3 | 2 | 12345 | 414 | ush,dpp | | 3.291e-03 | -5.72 | regulation of cholesterol transport | biological process | GO:0032374 | 3 | 2 | 12345 | 414 | Npc1a,ScpX | | 3.291e-03 | -5.72 | dorsal closure, leading edge cell fate determination | biological process | GO:0007393 | 3 | 2 | 12345 | 414 | dpp,ush | | 3.291e-03 | -5.72 | compound eye corneal lens development | biological process | GO:0048058 | 3 | 2 | 12345 | 414 | ttk,Ten-m | | 3.291e-03 | -5.72 | regulation of non-canonical Wnt signaling pathway | biological process | GO:2000050 | 3 | 2 | 12345 | 414 | aop,Prosap | | 3.291e-03 | -5.72 | mitotic actomyosin contractile ring assembly | biological process | GO:1903475 | 3 | 2 | 12345 | 414 | Rho1,pbl | | 3.291e-03 | -5.72 | BMP signaling pathway involved in Malpighian tubule cell chemotaxis | biological process | GO:0061353 | 3 | 2 | 12345 | 414 | Dad,dpp | | 3.291e-03 | -5.72 | cell rearrangement involved in Malpighian tubule morphogenesis | biological process | GO:0061334 | 3 | 2 | 12345 | 414 | dpp,Dad | | 3.347e-03 | -5.70 | RHOGAP | prosite domains | PS50238 | 21 | 5 | 6243 | 316 | RhoGAP18B,RhoGAP15B,RhoGAP102A,RhoGAP16F,RhoGAP93B | | 3.392e-03 | -5.69 | EGF\_CA | prosite domains | PS01187 | 30 | 6 | 6243 | 316 | N,LRP1,crb,Ser,mgl,slow | | 3.399e-03 | -5.68 | CSL-Notch-Mastermind transcription factor complex | cellular component | GO:1990433 | 3 | 2 | 11940 | 407 | N,mam | | 3.409e-03 | -5.68 | apical junction complex | cellular component | GO:0043296 | 58 | 7 | 11940 | 407 | nrv2,Lac,Fas3,sdt,crb,N,bbg | | 3.423e-03 | -5.68 | regulation of ERK1 and ERK2 cascade | biological process | GO:0070372 | 31 | 5 | 12345 | 414 | Egfr,sty,spi,drk,aos | | 3.423e-03 | -5.68 | larval somatic muscle development | biological process | GO:0007526 | 31 | 5 | 12345 | 414 | Sin3A,eya,Tl,ap,Stat92E | | 3.423e-03 | -5.68 | establishment or maintenance of epithelial cell apical/basal polarity | biological process | GO:0045197 | 31 | 5 | 12345 | 414 | Rho1,sdt,ft,fj,ds | | 3.423e-03 | -5.68 | actin cytoskeleton reorganization | biological process | GO:0031532 | 31 | 5 | 12345 | 414 | pbl,nuf,Rac2,RhoGEF2,Rho1 | | 3.454e-03 | -5.67 | mam (mastermind) | protein interactions | 36555 | 69 | 9 | 9204 | 412 | Ser,Mrp4,crb,Sin3A,EcR,hh,N,sno,tara | | 3.455e-03 | -5.67 | calmodulin-dependent protein kinase activity | molecular function | GO:0004683 | 10 | 3 | 12176 | 396 | PEK,CaMKII,PhKgamma | | 3.462e-03 | -5.67 | Kap-alpha3 (karyopherin alpha3) | protein interactions | 41158 | 82 | 10 | 9204 | 412 | Myo10A,cnc,S6kII,N,ci,Smox,ScsbetaG,Cnx99A,Ser,rau | | 3.469e-03 | -5.66 | SRY\_02 | MSigDB lists | SRY\_02 | 41 | 6 | 4161 | 153 | Pli,Mhcl,LRP1,Smox,ds,fne | | 3.469e-03 | -5.66 | GO\_OSTEOBLAST\_DIFFERENTIATION | MSigDB lists | GO\_OSTEOBLAST\_DIFFERENTIATION | 41 | 6 | 4161 | 153 | Tudor-SN,hh,CG11367,dpp,Smox,mle | | 3.476e-03 | -5.66 | L1CAM interactions | REACTOME pathways | R-DME-373760 | 28 | 5 | 3923 | 148 | Fas2,Egfr,scb,if,Ank | | 3.478e-03 | -5.66 | GO\_COCHLEA\_DEVELOPMENT | MSigDB lists | GO\_COCHLEA\_DEVELOPMENT | 9 | 3 | 4161 | 153 | Rac2,ds,so | | 3.478e-03 | -5.66 | PID\_BMP\_PATHWAY | MSigDB lists | PID\_BMP\_PATHWAY | 9 | 3 | 4161 | 153 | dpp,CG5830,tkv | | 3.478e-03 | -5.66 | REACTOME\_DSCAM\_INTERACTIONS | MSigDB lists | REACTOME\_DSCAM\_INTERACTIONS | 9 | 3 | 4161 | 153 | Rac2,Dscam1,NetB | | 3.478e-03 | -5.66 | GO\_POSITIVE\_REGULATION\_OF\_EMBRYONIC\_DEVELOPMENT | MSigDB lists | GO\_POSITIVE\_REGULATION\_OF\_EMBRYONIC\_DEVELOPMENT | 9 | 3 | 4161 | 153 | hh,Lim1,so | | 3.478e-03 | -5.66 | GO\_REGULATION\_OF\_MORPHOGENESIS\_OF\_A\_BRANCHING\_STRUCTURE | MSigDB lists | GO\_REGULATION\_OF\_MORPHOGENESIS\_OF\_A\_BRANCHING\_STRUCTURE | 9 | 3 | 4161 | 153 | so,hh,Lim1 | | 3.478e-03 | -5.66 | HEDENFALK\_BREAST\_CANCER\_BRACX\_UP | MSigDB lists | HEDENFALK\_BREAST\_CANCER\_BRACX\_UP | 9 | 3 | 4161 | 153 | RpS4,CG43313,Utx | | 3.478e-03 | -5.66 | GO\_GLOMERULUS\_DEVELOPMENT | MSigDB lists | GO\_GLOMERULUS\_DEVELOPMENT | 9 | 3 | 4161 | 153 | N,Lim1,Ser | | 3.511e-03 | -5.65 | HOMEOBOX | prints domains | PR00024 | 38 | 6 | 2185 | 88 | Ubx,slou,HGTX,exex,Dr,B-H2 | | 3.539e-03 | -5.64 | positive regulation of neuron differentiation | biological process | GO:0045666 | 75 | 8 | 12345 | 414 | mbt,EcR,Imp,nudE,oc,Prosap,pnt,mmy | | 3.544e-03 | -5.64 | - | gene3d domains | 2.60.40.820 | 8 | 3 | 4929 | 208 | Doc1,bi,Doc2 | | 3.544e-03 | -5.64 | purine ribonucleoside triphosphate binding | molecular function | GO:0035639 | 791 | 40 | 12176 | 396 | CG9541,CG9281,JIL-1,Egfr,Mos,PhKgamma,par-1,betaTub56D,CG8485,Arl4,Pex1,lz,mle,mbt,CG14535,CG12010,Myo10A,bor,tkv,Doa,wnd,KP78a,Pak3,Rac2,Cdk12,Tlk,CaMKII,Hsc70-3,Mrp4,Asator,Etl1,Arp2,Rho1,ScsbetaG,S6kII,CG8097,Hipk,Haspin,Mhcl,PEK | | 3.579e-03 | -5.63 | positive regulation of immune system process | biological process | GO:0002684 | 109 | 10 | 12345 | 414 | lz,Toll-7,Doa,N,Mmp2,Tl,CG4325,ush,Etl1,Pli | | 3.590e-03 | -5.63 | cell cortex region | cellular component | GO:0099738 | 74 | 8 | 11940 | 407 | Ser,Rho1,sdt,Syp,RhoGEF2,nuf,CG42684,pon | | 3.590e-03 | -5.63 | vn (vein) | protein interactions | 38657 | 34 | 6 | 9204 | 412 | ap,Egfr,Ubx,N,spi,aos | | 3.590e-03 | -5.63 | ap (apterous) | protein interactions | 35509 | 34 | 6 | 9204 | 412 | Ubx,Dr,N,bi,Ser,Bx | | 3.593e-03 | -5.63 | GO\_APICAL\_JUNCTION\_COMPLEX | MSigDB lists | GO\_APICAL\_JUNCTION\_COMPLEX | 18 | 4 | 4161 | 153 | CG17265,pbl,Ank,sdt | | 3.593e-03 | -5.63 | KANG\_IMMORTALIZED\_BY\_TERT\_UP | MSigDB lists | KANG\_IMMORTALIZED\_BY\_TERT\_UP | 18 | 4 | 4161 | 153 | nAChRalpha4,Pat1,TfAP-2,mle | | 3.593e-03 | -5.63 | GO\_ORGAN\_GROWTH | MSigDB lists | GO\_ORGAN\_GROWTH | 18 | 4 | 4161 | 153 | N,CaMKII,hh,smal | | 3.593e-03 | -5.63 | GO\_EMBRYONIC\_PATTERN\_SPECIFICATION | MSigDB lists | GO\_EMBRYONIC\_PATTERN\_SPECIFICATION | 18 | 4 | 4161 | 153 | hh,Lim1,Utx,Smox | | 3.593e-03 | -5.63 | GO\_REGULATION\_OF\_LIPID\_TRANSPORT | MSigDB lists | GO\_REGULATION\_OF\_LIPID\_TRANSPORT | 18 | 4 | 4161 | 153 | ScpX,hh,LRP1,EcR | | 3.643e-03 | -5.61 | tara (taranis) | protein interactions | 53562 | 15 | 4 | 9204 | 412 | Bx,mam,osa,tna | | 3.643e-03 | -5.61 | Sox21a (Sox21a) | protein interactions | 39567 | 15 | 4 | 9204 | 412 | Mmp2,Egfr,spi,Stat92E | | 3.643e-03 | -5.61 | Nedd4 (Nedd4) | protein interactions | 39958 | 15 | 4 | 9204 | 412 | Myo10A,N,comm,sty | | 3.643e-03 | -5.61 | upd3 (unpaired 3) | protein interactions | 3346149 | 15 | 4 | 9204 | 412 | Stat92E,Sox21a,N,Egfr | | 3.645e-03 | -5.61 | GO\_RIBONUCLEOPROTEIN\_GRANULE | MSigDB lists | GO\_RIBONUCLEOPROTEIN\_GRANULE | 55 | 7 | 4161 | 153 | Imp,RpS4,wech,pAbp,Rac2,sm,mle | | 3.645e-03 | -5.61 | GSE32034\_UNTREATED\_VS\_ROSIGLIZATONE\_TREATED\_LY6C\_HIGH\_MONOCYTE\_UP | MSigDB lists | GSE32034\_UNTREATED\_VS\_ROSIGLIZATONE\_TREATED\_LY6C\_HIGH\_MONOCYTE\_UP | 55 | 7 | 4161 | 153 | CG9281,Tim17b,phtf,Npc1a,dpp,CycE,Ank | | 3.657e-03 | -5.61 | NELSON\_RESPONSE\_TO\_ANDROGEN\_UP | MSigDB lists | NELSON\_RESPONSE\_TO\_ANDROGEN\_UP | 29 | 5 | 4161 | 153 | mmy,Mrp4,Acsl,Hydr2,Pat1 | | 3.657e-03 | -5.61 | GO\_DIGESTIVE\_SYSTEM\_DEVELOPMENT | MSigDB lists | GO\_DIGESTIVE\_SYSTEM\_DEVELOPMENT | 29 | 5 | 4161 | 153 | ds,ft,Smox,N,hh | | 3.657e-03 | -5.61 | GO\_VACUOLAR\_LUMEN | MSigDB lists | GO\_VACUOLAR\_LUMEN | 29 | 5 | 4161 | 153 | emp,CG14291,dally,dlp,LManII | | 3.657e-03 | -5.61 | GO\_MESODERM\_DEVELOPMENT | MSigDB lists | GO\_MESODERM\_DEVELOPMENT | 29 | 5 | 4161 | 153 | if,Utx,ap,Smox,hh | | 3.681e-03 | -5.60 | Rac2 (Rac2) | protein interactions | 38831 | 24 | 5 | 9204 | 412 | ttk,pbl,CycE,Pak3,Pits | | 3.681e-03 | -5.60 | Nipped-B (Nipped-B) | protein interactions | 3355136 | 24 | 5 | 9204 | 412 | hh,scaf6,mam,N,S6kII | | 3.681e-03 | -5.60 | sl (small wing) | protein interactions | 32601 | 24 | 5 | 9204 | 412 | S,Egfr,drk,spi,aos | | 3.691e-03 | -5.60 | germ-line stem cell division | biological process | GO:0042078 | 45 | 6 | 12345 | 414 | tkv,stwl,dpp,hh,dally,Stat92E | | 3.719e-03 | -5.59 | GO\_CELL\_SURFACE | MSigDB lists | GO\_CELL\_SURFACE | 118 | 11 | 4161 | 153 | Ank,Rho1,hh,dlp,Hsc70-3,if,N,pbl,Dscam1,dpp,Fas2 | | 3.719e-03 | -5.59 | CTTTGTA\_MIR524 | MSigDB lists | CTTTGTA\_MIR524 | 118 | 11 | 4161 | 153 | Arl4,Smox,pbl,hts,Etl1,nudE,l(3)mbt,NiPp1,Npc1a,Rac2,fne | | 3.746e-03 | -5.59 | macromolecule modification | biological process | GO:0043412 | 1320 | 62 | 12345 | 414 | ctrip,Pli,CycE,Tlk,par-1,Utx,Asator,egr,CG3253,Mocs2B,CG12099,Hipk,JIL-1,bin3,wnd,CG9855,ds,Cdk12,hiw,Mos,S6kII,smal,ec,Cyp1,Ptpmeg,eya,CG11486,CG6752,Pak3,PhKgamma,ft,Doa,CG14646,wcy,stg,enok,tkv,Notum,fj,Ppa,Rac2,Taf1,Fbxl7,KP78a,CG13029,CaMKII,PEK,Nopp140,Usp2,CG8485,LManII,mof,CG4325,tna,Ptp99A,Tet,CG5830,Haspin,mbt,Hph,wech,Acf | | 3.768e-03 | -5.58 | negative regulation of protein modification by small protein conjugation or removal | biological process | GO:1903321 | 10 | 3 | 12345 | 414 | hh,ctrip,CycE | | 3.768e-03 | -5.58 | establishment of epithelial cell apical/basal polarity | biological process | GO:0045198 | 10 | 3 | 12345 | 414 | fj,ds,ft | | 3.768e-03 | -5.58 | compound eye retinal cell programmed cell death | biological process | GO:0046667 | 10 | 3 | 12345 | 414 | S,ec,N | | 3.768e-03 | -5.58 | pericardial nephrocyte differentiation | biological process | GO:0061320 | 10 | 3 | 12345 | 414 | dpp,tin,pnt | | 3.768e-03 | -5.58 | polarized epithelial cell differentiation | biological process | GO:0030859 | 10 | 3 | 12345 | 414 | ft,fj,ds | | 3.768e-03 | -5.58 | central complex development | biological process | GO:0048036 | 10 | 3 | 12345 | 414 | robo2,Ptpmeg,ey | | 3.770e-03 | -5.58 | Glycosaminoglycan metabolism | REACTOME pathways | R-DME-1630316 | 54 | 7 | 3923 | 148 | CG43313,CG31637,Mrp4,dlp,dally,CG14291,CG9220 | | 3.782e-03 | -5.58 | regulation of neural precursor cell proliferation | biological process | GO:2000177 | 60 | 7 | 12345 | 414 | osa,cas,pnt,N,E(spl)mdelta-HLH,Kr,EcR | | 3.823e-03 | -5.57 | negative regulation of growth | biological process | GO:0045926 | 110 | 10 | 12345 | 414 | ft,crb,Fbxl7,Tl,Dad,scb,Acsl,EcR,Syp,hiw | | 3.909e-03 | -5.54 | MODULE\_23 | MSigDB lists | MODULE\_23 | 102 | 10 | 4161 | 153 | Taf1,ds,Syx1A,Tim17b,CG43313,Fbxl7,LManII,CG7264,ScpX,LRP1 | | 3.927e-03 | -5.54 | TCCATTKW\_UNKNOWN | MSigDB lists | TCCATTKW\_UNKNOWN | 42 | 6 | 4161 | 153 | CG12547,Imp,dpp,Mhcl,pAbp,fne | | 3.927e-03 | -5.54 | GO\_REGULATION\_OF\_TRANSMEMBRANE\_RECEPTOR\_PROTEIN\_SERINE\_THREONINE\_KINASE\_SIGNALING\_PATHWAY | MSigDB lists | GO\_REGULATION\_OF\_TRANSMEMBRANE\_RECEPTOR\_PROTEIN\_SERINE\_THREONINE\_KINASE\_SIGNALING\_PATHWAY | 42 | 6 | 4161 | 153 | Smox,N,if,hh,Hsc70-3,dpp | | 3.927e-03 | -5.54 | WANG\_MLL\_TARGETS | MSigDB lists | WANG\_MLL\_TARGETS | 42 | 6 | 4161 | 153 | lbk,ScpX,Lac,Pli,Notum,Fbxl7 | | 3.927e-03 | -5.54 | GO\_TUBE\_FORMATION | MSigDB lists | GO\_TUBE\_FORMATION | 42 | 6 | 4161 | 153 | wech,hh,so,ap,Utx,N | | 3.933e-03 | -5.54 | GO\_ORGAN\_INDUCTION | MSigDB lists | GO\_ORGAN\_INDUCTION | 3 | 2 | 4161 | 153 | so,dpp | | 3.933e-03 | -5.54 | GO\_PARASYMPATHETIC\_NERVOUS\_SYSTEM\_DEVELOPMENT | MSigDB lists | GO\_PARASYMPATHETIC\_NERVOUS\_SYSTEM\_DEVELOPMENT | 3 | 2 | 4161 | 153 | so,TfAP-2 | | 3.933e-03 | -5.54 | WANG\_LSD1\_TARGETS\_UP | MSigDB lists | WANG\_LSD1\_TARGETS\_UP | 3 | 2 | 4161 | 153 | so,CycE | | 3.933e-03 | -5.54 | GO\_REGULATION\_OF\_SKELETAL\_MUSCLE\_CELL\_PROLIFERATION | MSigDB lists | GO\_REGULATION\_OF\_SKELETAL\_MUSCLE\_CELL\_PROLIFERATION | 3 | 2 | 4161 | 153 | so,hh | | 3.933e-03 | -5.54 | GO\_CELL\_MIGRATION\_INVOLVED\_IN\_HEART\_DEVELOPMENT | MSigDB lists | GO\_CELL\_MIGRATION\_INVOLVED\_IN\_HEART\_DEVELOPMENT | 3 | 2 | 4161 | 153 | ds,N | | 3.933e-03 | -5.54 | GO\_HEART\_VALVE\_FORMATION | MSigDB lists | GO\_HEART\_VALVE\_FORMATION | 3 | 2 | 4161 | 153 | N,ds | | 3.933e-03 | -5.54 | YAGUE\_PRETUMOR\_DRUG\_RESISTANCE\_DN | MSigDB lists | YAGUE\_PRETUMOR\_DRUG\_RESISTANCE\_DN | 3 | 2 | 4161 | 153 | ds,DIP-epsilon | | 3.933e-03 | -5.54 | GO\_REGULATION\_OF\_MESONEPHROS\_DEVELOPMENT | MSigDB lists | GO\_REGULATION\_OF\_MESONEPHROS\_DEVELOPMENT | 3 | 2 | 4161 | 153 | Lim1,so | | 3.933e-03 | -5.54 | GO\_OSTEOBLAST\_DEVELOPMENT | MSigDB lists | GO\_OSTEOBLAST\_DEVELOPMENT | 3 | 2 | 4161 | 153 | hh,Smox | | 3.933e-03 | -5.54 | GO\_REGULATION\_OF\_CELL\_COMMUNICATION\_BY\_ELECTRICAL\_COUPLING | MSigDB lists | GO\_REGULATION\_OF\_CELL\_COMMUNICATION\_BY\_ELECTRICAL\_COUPLING | 3 | 2 | 4161 | 153 | Ank,CaMKII | | 3.933e-03 | -5.54 | GO\_HETEROPHILIC\_CELL\_CELL\_ADHESION\_VIA\_PLASMA\_MEMBRANE\_CELL\_ADHESION\_MOLECULES | MSigDB lists | GO\_HETEROPHILIC\_CELL\_CELL\_ADHESION\_VIA\_PLASMA\_MEMBRANE\_CELL\_ADHESION\_MOLECULES | 3 | 2 | 4161 | 153 | ft,ds | | 3.933e-03 | -5.54 | GO\_NEGATIVE\_REGULATION\_OF\_TRANSCRIPTION\_REGULATORY\_REGION\_DNA\_BINDING | MSigDB lists | GO\_NEGATIVE\_REGULATION\_OF\_TRANSCRIPTION\_REGULATORY\_REGION\_DNA\_BINDING | 3 | 2 | 4161 | 153 | Sin3A,ap | | 3.933e-03 | -5.54 | GO\_REGULATION\_OF\_EXTRACELLULAR\_MATRIX\_ASSEMBLY | MSigDB lists | GO\_REGULATION\_OF\_EXTRACELLULAR\_MATRIX\_ASSEMBLY | 3 | 2 | 4161 | 153 | N,Smox | | 3.933e-03 | -5.54 | GO\_AXON\_INITIAL\_SEGMENT | MSigDB lists | GO\_AXON\_INITIAL\_SEGMENT | 3 | 2 | 4161 | 153 | Ank,CaMKII | | 3.933e-03 | -5.54 | GO\_REGULATION\_OF\_CARDIAC\_MUSCLE\_CELL\_ACTION\_POTENTIAL | MSigDB lists | GO\_REGULATION\_OF\_CARDIAC\_MUSCLE\_CELL\_ACTION\_POTENTIAL | 3 | 2 | 4161 | 153 | CaMKII,Ank | | 3.933e-03 | -5.54 | GO\_POSITIVE\_REGULATION\_OF\_MESONEPHROS\_DEVELOPMENT | MSigDB lists | GO\_POSITIVE\_REGULATION\_OF\_MESONEPHROS\_DEVELOPMENT | 3 | 2 | 4161 | 153 | Lim1,so | | 3.933e-03 | -5.54 | GO\_DISTAL\_TUBULE\_DEVELOPMENT | MSigDB lists | GO\_DISTAL\_TUBULE\_DEVELOPMENT | 3 | 2 | 4161 | 153 | Ser,N | | 3.933e-03 | -5.54 | GO\_POSITIVE\_REGULATION\_OF\_ASTROCYTE\_DIFFERENTIATION | MSigDB lists | GO\_POSITIVE\_REGULATION\_OF\_ASTROCYTE\_DIFFERENTIATION | 3 | 2 | 4161 | 153 | dpp,N | | 3.933e-03 | -5.54 | GO\_REGULATION\_OF\_CHOLESTEROL\_HOMEOSTASIS | MSigDB lists | GO\_REGULATION\_OF\_CHOLESTEROL\_HOMEOSTASIS | 3 | 2 | 4161 | 153 | EcR,Npc1a | | 3.933e-03 | -5.54 | BRUNEAU\_HEART\_GREAT\_VESSELS\_AND\_VALVULOGENESIS | MSigDB lists | BRUNEAU\_HEART\_GREAT\_VESSELS\_AND\_VALVULOGENESIS | 3 | 2 | 4161 | 153 | Ser,N | | 3.936e-03 | -5.54 | EGFR downregulation | REACTOME pathways | R-DME-182971 | 18 | 4 | 3923 | 148 | sty,Ptpmeg,drk,Egfr | | 3.941e-03 | -5.54 | hemocyte migration | biological process | GO:0035099 | 20 | 4 | 12345 | 414 | if,Rac2,Pvf3,Rho1 | | 3.941e-03 | -5.54 | somatic stem cell population maintenance | biological process | GO:0035019 | 20 | 4 | 12345 | 414 | mam,CycE,Stat92E,CG46339 | | 3.941e-03 | -5.54 | regulation of hemocyte differentiation | biological process | GO:0045610 | 20 | 4 | 12345 | 414 | Ser,lz,ush,N | | 3.941e-03 | -5.54 | positive regulation of response to wounding | biological process | GO:1903036 | 20 | 4 | 12345 | 414 | Egfr,Rac2,Imp,Rho1 | | 3.942e-03 | -5.54 | purine nucleotide binding | molecular function | GO:0017076 | 821 | 41 | 12176 | 396 | CG12010,Myo10A,CG14535,mle,lz,mbt,Pex1,Arl4,CG8485,par-1,PhKgamma,betaTub56D,Egfr,Mos,JIL-1,CG9541,CG9281,Haspin,Mhcl,PEK,CG8097,Hipk,Rho1,ScsbetaG,S6kII,Arp2,Etl1,Asator,Mrp4,Hsc70-3,CaMKII,Tlk,Rac2,Cdk12,Pak3,KP78a,tkv,cta,Doa,wnd,bor | | 3.964e-03 | -5.53 | Cadherin-like\_sf | interpro domains | IPR015919 | 18 | 4 | 11383 | 428 | ds,ft,CadN,Cad87A | | 3.995e-03 | -5.52 | cell division site | cellular component | GO:0032153 | 45 | 6 | 11940 | 407 | Arl4,Rho1,Tl,RhoGEF2,nuf,pbl | | 4.040e-03 | -5.51 | SENESE\_HDAC1\_AND\_HDAC2\_TARGETS\_UP | MSigDB lists | SENESE\_HDAC1\_AND\_HDAC2\_TARGETS\_UP | 56 | 7 | 4161 | 153 | Cyp1,phtf,dpp,heph,CycE,cta,mle | | 4.040e-03 | -5.51 | GSE17721\_0.5H\_VS\_12H\_POLYIC\_BMDC\_DN | MSigDB lists | GSE17721\_0.5H\_VS\_12H\_POLYIC\_BMDC\_DN | 56 | 7 | 4161 | 153 | Rhp,CaMKII,Pat1,ap,rush,CycE,Etl1 | | 4.040e-03 | -5.51 | MODULE\_1 | MSigDB lists | MODULE\_1 | 56 | 7 | 4161 | 153 | LRP1,ScpX,smal,Fas2,Fbxl7,LManII,CG43313 | | 4.044e-03 | -5.51 | Stat92E (Signal-transducer and activator of transcription protein at 92E) | protein interactions | 42428 | 58 | 8 | 9204 | 412 | dpp,Tl,ci,Egfr,Taf1,ttk,tsl,Sox21a | | 4.075e-03 | -5.50 | Chondroitin sulfate/dermatan sulfate metabolism | REACTOME pathways | R-DME-1793185 | 29 | 5 | 3923 | 148 | CG9220,CG31637,CG43313,dlp,dally | | 4.103e-03 | -5.50 | GO\_REGULATION\_OF\_CELL\_DEATH | MSigDB lists | GO\_REGULATION\_OF\_CELL\_DEATH | 330 | 22 | 4161 | 153 | Mhcl,dpp,bor,Cyp1,N,Smox,pbl,LRP1,CG45050,Npc1a,Tif-IA,S6kII,TfAP-2,CaMKII,hh,Hsc70-3,so,Sin3A,CG5742,CG3961,tkv,Rac2 | | 4.104e-03 | -5.50 | CHGN | pfam domains | PF05679 | 3 | 2 | 10480 | 393 | CG43313,CG9220 | | 4.124e-03 | -5.49 | enzyme binding | molecular function | GO:0019899 | 481 | 27 | 12176 | 396 | crb,Gbs-70E,Mitofilin,PsGEF,Gmap,mbt,CG42748,CG8441,Rho1,Tif-IA,NiPp1,Pak3,CG5522,bora,rau,E(spl)m4-BFM,RhoGEF2,ec,Atg17,pbl,ckn,CG10283,CycE,nuf,RhoGAP18B,Rac2,CG43658 | | 4.126e-03 | -5.49 | Chond\_GalNAc | interpro domains | IPR008428 | 3 | 2 | 11383 | 428 | CG9220,CG43313 | | 4.126e-03 | -5.49 | PAK\_BD | interpro domains | IPR033923 | 3 | 2 | 11383 | 428 | mbt,Pak3 | | 4.126e-03 | -5.49 | Uridine\_phosphorylase\_euk | interpro domains | IPR010059 | 3 | 2 | 11383 | 428 | CG6330,CG17224 | | 4.126e-03 | -5.49 | Netrin\_domain | interpro domains | IPR001134 | 3 | 2 | 11383 | 428 | NetB,NetA | | 4.126e-03 | -5.49 | ELAV/Hu\_RRM1 | interpro domains | IPR034775 | 3 | 2 | 11383 | 428 | elav,fne | | 4.126e-03 | -5.49 | TIMP-like\_OB-fold | interpro domains | IPR008993 | 3 | 2 | 11383 | 428 | NetB,NetA | | 4.126e-03 | -5.49 | ELAD\_HU\_SF | interpro domains | IPR006548 | 3 | 2 | 11383 | 428 | elav,fne | | 4.129e-03 | -5.49 | TGCCTTA\_MIR124A | MSigDB lists | TGCCTTA\_MIR124A | 173 | 14 | 4161 | 153 | dlp,Mrp4,CG9220,ScsbetaG,CG14646,heph,CG5522,phtf,Lac,Ser,ap,CG41099,cta,Etl1 | | 4.172e-03 | -5.48 | hth (homothorax) | protein interactions | 41273 | 35 | 6 | 9204 | 412 | Ubx,ci,cnc,CycE,jing,ey | | 4.172e-03 | -5.48 | sax (saxophone) | protein interactions | 35731 | 35 | 6 | 9204 | 412 | Mhcl,Dad,dpp,tkv,CycE,sty | | 4.196e-03 | -5.47 | hep (hemipterous) | protein interactions | 32256 | 71 | 9 | 9204 | 412 | tkv,egr,wnd,bdg,PEK,dpp,grnd,bi,Sox21a | | 4.213e-03 | -5.47 | CG10916 (uncharacterized protein) | protein interactions | 37081 | 8 | 3 | 9204 | 412 | Egfr,N,CG13323 | | 4.213e-03 | -5.47 | CG12084 (uncharacterized protein) | protein interactions | 192509 | 8 | 3 | 9204 | 412 | CycE,Fas2,N | | 4.213e-03 | -5.47 | Tollo (tollo) | protein interactions | 44497 | 8 | 3 | 9204 | 412 | dpp,Toll-7,tkv | | 4.213e-03 | -5.47 | eg (eagle) | protein interactions | 40428 | 8 | 3 | 9204 | 412 | robo2,CtBP,kni | | 4.213e-03 | -5.47 | Cortactin (cortactin) | protein interactions | 42491 | 8 | 3 | 9204 | 412 | Ptpmeg,drk,ft | | 4.213e-03 | -5.47 | trk (trunk) | protein interactions | 34376 | 8 | 3 | 9204 | 412 | sty,Fur1,tsl | | 4.213e-03 | -5.47 | milt (milton) | protein interactions | 45683 | 8 | 3 | 9204 | 412 | ey,Myo10A,par-1 | | 4.260e-03 | -5.46 | REACTOME\_L1CAM\_INTERACTIONS | MSigDB lists | REACTOME\_L1CAM\_INTERACTIONS | 30 | 5 | 4161 | 153 | Fas2,S6kII,JIL-1,Ank,Rac2 | | 4.260e-03 | -5.46 | BMI1\_DN\_MEL18\_DN.V1\_UP | MSigDB lists | BMI1\_DN\_MEL18\_DN.V1\_UP | 30 | 5 | 4161 | 153 | Syx1A,fj,Arl4,DIP-epsilon,Ser | | 4.268e-03 | -5.46 | Notch signaling pathway | KEGG pathways | dme04330 | 22 | 4 | 3198 | 100 | CtBP,mam,Ser,N | | 4.268e-03 | -5.46 | Notch signaling pathway | KEGG pathways | ko04330 | 22 | 4 | 3198 | 100 | N,Ser,mam,CtBP | | 4.303e-03 | -5.45 | CADHERIN\_1 | prosite domains | PS00232 | 14 | 4 | 6243 | 316 | CadN,ft,ds,Cad87A | | 4.312e-03 | -5.45 | fly chr93E4-93E4\|3-72 | chromosome location | fly chr93E4-93E4\|3-72 | 5 | 2 | 25010 | 531 | CG15498,slou | | 4.312e-03 | -5.45 | fly chr102B1-102B1\|4-0 cM | chromosome location | fly chr102B1-102B1\|4-0 cM | 5 | 2 | 25010 | 531 | dpr7,RhoGAP102A | | 4.312e-03 | -5.45 | fly chr102B1-102B1\|4-0 | chromosome location | fly chr102B1-102B1\|4-0 | 5 | 2 | 25010 | 531 | RhoGAP102A,dpr7 | | 4.312e-03 | -5.45 | fly chr93E4-93E4\|3-72 cM | chromosome location | fly chr93E4-93E4\|3-72 cM | 5 | 2 | 25010 | 531 | slou,CG15498 | | 4.332e-03 | -5.44 | regulation of response to external stimulus | biological process | GO:0032101 | 228 | 16 | 12345 | 414 | Usp2,NetA,Mmp2,CadN,mmy,Lrt,Pli,Stat92E,Imp,NetB,Toll-7,Doa,Tl,CG4325,ush,Etl1 | | 4.337e-03 | -5.44 | fly chr61C8-61C8\|3-0.5 cM | chromosome location | fly chr61C8-61C8\|3-0.5 cM | 16 | 3 | 25010 | 531 | CG3386,CG3402,CG13894 | | 4.337e-03 | -5.44 | fly chr61C8-61C8\|3-0.5 | chromosome location | fly chr61C8-61C8\|3-0.5 | 16 | 3 | 25010 | 531 | CG13894,CG3386,CG3402 | | 4.350e-03 | -5.44 | GO\_POSITIVE\_REGULATION\_OF\_CELLULAR\_COMPONENT\_ORGANIZATION | MSigDB lists | GO\_POSITIVE\_REGULATION\_OF\_CELLULAR\_COMPONENT\_ORGANIZATION | 311 | 21 | 4161 | 153 | dpp,Cyp1,N,Smox,NetB,Ank,EcR,JIL-1,Acsl,Pcl,Lac,nudE,Tif-IA,Arp2,smal,CG41099,Taf1,CG9855,Sin3A,drk,Rac2 | | 4.366e-03 | -5.43 | protein dimerization activity | molecular function | GO:0046983 | 390 | 23 | 12176 | 396 | ttk,scb,Sin3A,cnc,dpp,CadN,twi,CtBP,E(spl)mdelta-HLH,psq,Dscam1,Lac,ci,otk2,oc,Ten-m,EcR,robo2,phtf,rau,if,Mondo,Pak3 | | 4.367e-03 | -5.43 | positive regulation of multi-organism process | biological process | GO:0043902 | 149 | 12 | 12345 | 414 | Etl1,Pli,ttk,Mmp2,Tl,spi,ush,CG4325,aop,Doa,Egfr,Toll-7 | | 4.423e-03 | -5.42 | GO\_REGULATION\_OF\_NOTCH\_SIGNALING\_PATHWAY | MSigDB lists | GO\_REGULATION\_OF\_NOTCH\_SIGNALING\_PATHWAY | 19 | 4 | 4161 | 153 | N,slow,Tsp26A,Ser | | 4.423e-03 | -5.42 | GO\_EMBRYONIC\_CRANIAL\_SKELETON\_MORPHOGENESIS | MSigDB lists | GO\_EMBRYONIC\_CRANIAL\_SKELETON\_MORPHOGENESIS | 19 | 4 | 4161 | 153 | Smox,TfAP-2,so,Lim1 | | 4.423e-03 | -5.42 | GO\_REGULATION\_OF\_GLIOGENESIS | MSigDB lists | GO\_REGULATION\_OF\_GLIOGENESIS | 19 | 4 | 4161 | 153 | hh,dpp,CG12099,N | | 4.423e-03 | -5.42 | CCATCCA\_MIR432 | MSigDB lists | CCATCCA\_MIR432 | 19 | 4 | 4161 | 153 | Doa,Tlk,S6kII,Sin3A | | 4.428e-03 | -5.42 | GSE4142\_NAIVE\_VS\_GC\_BCELL\_UP | MSigDB lists | GSE4142\_NAIVE\_VS\_GC\_BCELL\_UP | 43 | 6 | 4161 | 153 | CG2812,NetB,Fbxl7,Snx3,PhKgamma,CG2201 | | 4.428e-03 | -5.42 | GSE13306\_TREG\_VS\_TCONV\_SPLEEN\_UP | MSigDB lists | GSE13306\_TREG\_VS\_TCONV\_SPLEEN\_UP | 43 | 6 | 4161 | 153 | NetB,CaMKII,Dscam1,mub,CG34384,TfIIS | | 4.435e-03 | -5.42 | Nrg (Neuroglian) | protein interactions | 31792 | 25 | 5 | 9204 | 412 | Fas2,elav,Egfr,Fas3,Ank | | 4.435e-03 | -5.42 | pcs (parcas) | protein interactions | 36674 | 25 | 5 | 9204 | 412 | baf,Pat1,betaTub56D,esn,eya | | 4.435e-03 | -5.42 | geminin (geminin) | protein interactions | 35563 | 25 | 5 | 9204 | 412 | CG10663,Tlk,Egfr,osa,mub | | 4.517e-03 | -5.40 | GO\_REGULATION\_OF\_TRANSCRIPTION\_FROM\_RNA\_POLYMERASE\_II\_PROMOTER | MSigDB lists | GO\_REGULATION\_OF\_TRANSCRIPTION\_FROM\_RNA\_POLYMERASE\_II\_PROMOTER | 312 | 21 | 4161 | 153 | CG12099,JIL-1,EcR,Ser,dpp,Smox,N,if,ap,so,hh,Hsc70-3,TfIIS,tkv,Sin3A,CG5830,Taf1,CaMKII,S6kII,TfAP-2,CtBP | | 4.527e-03 | -5.40 | pole cell development | biological process | GO:0007277 | 33 | 5 | 12345 | 414 | RhoGEF2,tsl,AGO2,CycE,wun2 | | 4.527e-03 | -5.40 | regulation of chemotaxis | biological process | GO:0050920 | 33 | 5 | 12345 | 414 | Lrt,mmy,NetB,CadN,NetA | | 4.536e-03 | -5.40 | phosphatase binding | molecular function | GO:0019902 | 34 | 5 | 12176 | 396 | ckn,CG42748,Gbs-70E,NiPp1,ec | | 4.549e-03 | -5.39 | negative regulation of multi-organism process | biological process | GO:0043901 | 62 | 7 | 12345 | 414 | EcR,Ptpmeg,Usp2,AGO2,Stat92E,Pli,Doa | | 4.597e-03 | -5.38 | DNA-binding transcription repressor activity, RNA polymerase II-specific | molecular function | GO:0001227 | 64 | 7 | 12176 | 396 | CG4360,ttk,E(spl)mdelta-HLH,CG31224,kni,Zif,aop | | 4.597e-03 | -5.38 | DNA-binding transcription repressor activity | molecular function | GO:0001217 | 64 | 7 | 12176 | 396 | kni,CG31224,E(spl)mdelta-HLH,ttk,CG4360,aop,Zif | | 4.600e-03 | -5.38 | positive regulation of cell cycle process | biological process | GO:0090068 | 47 | 6 | 12345 | 414 | Myb,ci,N,stg,CycE,CG31612 | | 4.633e-03 | -5.37 | regulation of neuromuscular junction development | biological process | GO:1904396 | 113 | 10 | 12345 | 414 | Fas2,Dad,scb,Acsl,S6kII,Arp2,tkv,hiw,Syp,CaMKII | | 4.636e-03 | -5.37 | poly(U) RNA binding | molecular function | GO:0008266 | 11 | 3 | 12176 | 396 | fne,pAbp,elav | | 4.678e-03 | -5.36 | - | gene3d domains | 2.120.10.30 | 38 | 6 | 4929 | 208 | yellow-e3,mgl,CG12547,wech,Ten-m,LRP1 | | 4.688e-03 | -5.36 | pb (proboscipedia) | protein interactions | 40826 | 16 | 4 | 9204 | 412 | ey,Ubx,ci,hh | | 4.688e-03 | -5.36 | dlp (dally-like) | protein interactions | 39596 | 16 | 4 | 9204 | 412 | dally,Mmp2,hh,Notum | | 4.688e-03 | -5.36 | vap (vacuolar peduncle) | protein interactions | 32569 | 16 | 4 | 9204 | 412 | sty,Dscam1,Egfr,drk | | 4.688e-03 | -5.36 | htl (heartless) | protein interactions | 42160 | 16 | 4 | 9204 | 412 | Egfr,pbl,ush,sty | | 4.688e-03 | -5.36 | l(2)gd1 (lethal (2) giant discs 1) | protein interactions | 34543 | 16 | 4 | 9204 | 412 | tkv,Ser,N,ft | | 4.748e-03 | -5.35 | SH2 | pfam domains | PF00017 | 30 | 5 | 10480 | 393 | CG33993,Socs16D,Stat92E,CG10479,drk | | 4.808e-03 | -5.34 | SH2 | interpro domains | IPR000980 | 30 | 5 | 11383 | 428 | Socs16D,drk,Stat92E,CG33993,CG10479 | | 4.818e-03 | -5.34 | REACTOME\_DEVELOPMENTAL\_BIOLOGY | MSigDB lists | REACTOME\_DEVELOPMENTAL\_BIOLOGY | 105 | 10 | 4161 | 153 | Ank,mbt,drk,Rho1,JIL-1,Rac2,Fas2,S6kII,Smox,NetB | | 4.822e-03 | -5.33 | lid (little imaginal discs) | protein interactions | 33837 | 36 | 6 | 9204 | 412 | CycE,Stat92E,Sin3A,N,ci,CtBP | | 4.836e-03 | -5.33 | GO\_REGULATION\_OF\_EXTRACELLULAR\_MATRIX\_ORGANIZATION | MSigDB lists | GO\_REGULATION\_OF\_EXTRACELLULAR\_MATRIX\_ORGANIZATION | 10 | 3 | 4161 | 153 | smal,Smox,N | | 4.836e-03 | -5.33 | GCM\_SMARCC1 | MSigDB lists | GCM\_SMARCC1 | 10 | 3 | 4161 | 153 | Tlk,mle,sm | | 4.836e-03 | -5.33 | GO\_NEGATIVE\_REGULATION\_OF\_OSTEOBLAST\_DIFFERENTIATION | MSigDB lists | GO\_NEGATIVE\_REGULATION\_OF\_OSTEOBLAST\_DIFFERENTIATION | 10 | 3 | 4161 | 153 | Smox,N,CG11367 | | 4.836e-03 | -5.33 | KEGG\_PRION\_DISEASES | MSigDB lists | KEGG\_PRION\_DISEASES | 10 | 3 | 4161 | 153 | N,Hsc70-3,Fas2 | | 4.836e-03 | -5.33 | GO\_POSITIVE\_REGULATION\_OF\_GLIOGENESIS | MSigDB lists | GO\_POSITIVE\_REGULATION\_OF\_GLIOGENESIS | 10 | 3 | 4161 | 153 | hh,dpp,N | | 4.848e-03 | -5.33 | septate junction | cellular component | GO:0005918 | 33 | 5 | 11940 | 407 | Fas3,Lac,sdt,nrv2,bbg | | 4.874e-03 | -5.32 | Laminin\_G | interpro domains | IPR001791 | 19 | 4 | 11383 | 428 | CadN,ft,crb,wb | | 4.875e-03 | -5.32 | response to ethanol | biological process | GO:0045471 | 79 | 8 | 12345 | 414 | Drat,scb,spi,Fas2,hang,Egfr,S,Gbs-70E | | 4.893e-03 | -5.32 | stau (staufen) | protein interactions | 37065 | 86 | 10 | 9204 | 412 | spen,CG4788,Ppa,l(3)neo38,ps,Syp,CG5830,oc,flfl,CG44774 | | 4.926e-03 | -5.31 | AHR\_Q5 | MSigDB lists | AHR\_Q5 | 58 | 7 | 4161 | 153 | Mitofilin,Snx3,Nhe2,CG6752,sm,CaMKII,sdt | | 4.926e-03 | -5.31 | GO\_EXTRACELLULAR\_MATRIX | MSigDB lists | GO\_EXTRACELLULAR\_MATRIX | 58 | 7 | 4161 | 153 | CG3097,hh,dlp,dally,NetB,Mmp2,slow | | 4.932e-03 | -5.31 | RP58\_01 | MSigDB lists | RP58\_01 | 31 | 5 | 4161 | 153 | SamDC,if,CG8531,NetB,Tudor-SN | | 4.932e-03 | -5.31 | AREB6\_01 | MSigDB lists | AREB6\_01 | 31 | 5 | 4161 | 153 | TfAP-2,Syx1A,Imp,hh,Tlk | | 4.932e-03 | -5.31 | PPARA\_02 | MSigDB lists | PPARA\_02 | 31 | 5 | 4161 | 153 | Cyt-c1,CG5830,LManII,Tpr2,PhKgamma | | 4.956e-03 | -5.31 | WAKABAYASHI\_ADIPOGENESIS\_PPARG\_RXRA\_BOUND\_8D | MSigDB lists | WAKABAYASHI\_ADIPOGENESIS\_PPARG\_RXRA\_BOUND\_8D | 294 | 20 | 4161 | 153 | Acsl,betaTub56D,Mitofilin,rush,kdn,Cnx99A,NetB,ras,CG6330,heph,Mrp4,Hsc70-3,Mocs2B,emp,CG9855,hiw,sm,Lac,Npc1a,nudE | | 4.961e-03 | -5.31 | nucleoplasm | cellular component | GO:0005654 | 557 | 31 | 11940 | 407 | CG18766,CG44774,enok,stwl,NiPp1,baf,ctrip,CG31712,jing,gus,osa,mof,aop,Utx,Cdk12,Hph,mle,pbl,CG11030,Rbfox1,Sin3A,drk,mam,elav,lncRNA:roX1,Taf1,sm,Mocs2B,Tlk,Pits,Pcl | | 4.974e-03 | -5.30 | cell-cell junction organization | biological process | GO:0045216 | 63 | 7 | 12345 | 414 | CG42748,Lac,bou,nrv2,crb,sdt,Mmp2 | | 4.974e-03 | -5.30 | STAT5A\_03 | MSigDB lists | STAT5A\_03 | 44 | 6 | 4161 | 153 | Pli,dlp,LRP1,Utx,SamDC,mbt | | 4.974e-03 | -5.30 | GO\_AMINOGLYCAN\_METABOLIC\_PROCESS | MSigDB lists | GO\_AMINOGLYCAN\_METABOLIC\_PROCESS | 44 | 6 | 4161 | 153 | CG9220,dlp,dally,CG14291,CG31637,CG43313 | | 4.974e-03 | -5.30 | GSE5455\_HEALTHY\_VS\_TUMOR\_BEARING\_MOUSE\_SPLEEN\_MONOCYTE\_DN | MSigDB lists | GSE5455\_HEALTHY\_VS\_TUMOR\_BEARING\_MOUSE\_SPLEEN\_MONOCYTE\_DN | 44 | 6 | 4161 | 153 | Pat1,ras,Syx1A,CG1271,l(3)mbt,CG11367 | | 4.995e-03 | -5.30 | Pak (p21-activated kinase) | protein interactions | 44039 | 60 | 8 | 9204 | 412 | E(spl)mdelta-HLH,N,EcR,Dscam1,Pak3,crb,Rac2,Rho1 | | 5.004e-03 | -5.30 | PH\_domain | interpro domains | IPR001849 | 71 | 8 | 11383 | 428 | CG43658,rush,CG34384,RhoGAP15B,osp,CG5522,CG34383,RhoGEF2 | | 5.054e-03 | -5.29 | regulation of tube diameter, open tracheal system | biological process | GO:0035158 | 11 | 3 | 12345 | 414 | nrv2,Fas2,dpp | | 5.054e-03 | -5.29 | negative regulation of compound eye photoreceptor cell differentiation | biological process | GO:0110118 | 11 | 3 | 12345 | 414 | ttk,N,aop | | 5.054e-03 | -5.29 | histone phosphorylation | biological process | GO:0016572 | 11 | 3 | 12345 | 414 | Haspin,Tlk,JIL-1 | | 5.054e-03 | -5.29 | negative regulation of DNA-binding transcription factor activity | biological process | GO:0043433 | 11 | 3 | 12345 | 414 | pnt,edl,ush | | 5.054e-03 | -5.29 | adherens junction maintenance | biological process | GO:0034334 | 11 | 3 | 12345 | 414 | Rho1,crb,Rac2 | | 5.054e-03 | -5.29 | negative regulation of post-embryonic development | biological process | GO:0048581 | 11 | 3 | 12345 | 414 | N,Ser,sty | | 5.054e-03 | -5.29 | delamination | biological process | GO:0060232 | 11 | 3 | 12345 | 414 | par-1,aop,aos | | 5.095e-03 | -5.28 | protein kinase binding | molecular function | GO:0019901 | 82 | 8 | 12176 | 396 | CG8441,Rac2,Rho1,CycE,Mitofilin,bora,crb,Atg17 | | 5.112e-03 | -5.28 | regulation of lipid localization | biological process | GO:1905952 | 48 | 6 | 12345 | 414 | Acsl,Rac2,Npc1a,ImpL2,CaMKII,ScpX | | 5.143e-03 | -5.27 | toy (twin of eyeless) | protein interactions | 43833 | 48 | 7 | 9204 | 412 | Doa,dpp,ytr,Stat92E,gus,ey,Hipk | | 5.143e-03 | -5.27 | phyl (phyllopod) | protein interactions | 36606 | 48 | 7 | 9204 | 412 | N,ttk,CG11486,CycE,dbr,phyl,betaTub56D | | 5.164e-03 | -5.27 | negative regulation of ERBB signaling pathway | biological process | GO:1901185 | 34 | 5 | 12345 | 414 | edl,Fas2,Ptpmeg,aos,sty | | 5.164e-03 | -5.27 | negative regulation of epidermal growth factor receptor signaling pathway | biological process | GO:0042059 | 34 | 5 | 12345 | 414 | edl,Fas2,Ptpmeg,aos,sty | | 5.184e-03 | -5.26 | fly chr3-47.1 | chromosome location | fly chr3-47.1 | 17 | 3 | 25010 | 531 | CG41128,CG41099,Tim17b | | 5.184e-03 | -5.26 | fly chr3-47.1 cM | chromosome location | fly chr3-47.1 cM | 17 | 3 | 25010 | 531 | CG41128,CG41099,Tim17b | | 5.256e-03 | -5.25 | cytoplasmic vesicle | cellular component | GO:0031410 | 378 | 23 | 11940 | 407 | Rac2,dbr,comm,Tl,CG13784,CG32758,Cirl,tkv,dpp,N,Snx3,Hip1,CG6051,rush,Ptr,hh,CG34384,Syx1A,nuf,Ykt6,S,hiw,Fas3 | | 5.256e-03 | -5.25 | intracellular vesicle | cellular component | GO:0097708 | 378 | 23 | 11940 | 407 | rush,Ptr,hh,CG34384,nuf,Syx1A,Ykt6,S,hiw,Fas3,Rac2,dbr,comm,CG13784,CG32758,Tl,Cirl,tkv,N,dpp,Hip1,Snx3,CG6051 | | 5.263e-03 | -5.25 | chromatin organization involved in regulation of transcription | biological process | GO:0034401 | 80 | 8 | 12345 | 414 | Tlk,bin3,tna,psq,Kr,Psc,AGO2,tara | | 5.289e-03 | -5.24 | anchored component of plasma membrane | cellular component | GO:0046658 | 11 | 3 | 11940 | 407 | bou,dally,dlp | | 5.289e-03 | -5.24 | MSL complex | cellular component | GO:0072487 | 11 | 3 | 11940 | 407 | lncRNA:roX1,mof,mle | | 5.293e-03 | -5.24 | emc (extra macrochaetae) | protein interactions | 38091 | 26 | 5 | 9204 | 412 | N,mam,Egfr,CycE,Kr | | 5.293e-03 | -5.24 | RhoGEF2 (Rho guanine nucleotide exchange factor 2) | protein interactions | 36915 | 26 | 5 | 9204 | 412 | so,cta,drk,tsl,Rho1 | | 5.293e-03 | -5.24 | Mef2 (Myocyte enhancer factor 2) | protein interactions | 36032 | 26 | 5 | 9204 | 412 | Ubx,Svil,twi,CycE,TfIIS | | 5.293e-03 | -5.24 | pyd (polychaetoid) | protein interactions | 41062 | 26 | 5 | 9204 | 412 | drk,N,tkv,rau,CycE | | 5.294e-03 | -5.24 | positive regulation of protein metabolic process | biological process | GO:0051247 | 318 | 20 | 12345 | 414 | mbt,gus,Atg17,Pcl,Rho1,drk,Cdk5alpha,dpp,Usp2,wnd,Egfr,PEK,Mos,spi,phyl,ctrip,pAbp,bora,mmy,grnd | | 5.328e-03 | -5.23 | transferase activity | molecular function | GO:0016740 | 1384 | 62 | 12176 | 396 | Ugt49C1,hiw,par-1,PhKgamma,CG8485,fj,CycE,CG6752,wech,Fbxl7,bin3,Mocs2B,CG9541,ScpX,CG1492,CG34384,Pli,JIL-1,Mos,Egfr,CG17224,Ykt6,mbt,Acf,mmy,mof,CG43313,Cdk12,sdt,ctrip,CG2201,CG31637,CG3253,Tlk,CaMKII,CG10082,Doa,wnd,CG13029,tkv,CG1271,KP78a,Pak3,Taf1,CG4325,CG14646,S6kII,kdn,Hipk,Ppa,CG11486,CG1344,PEK,Haspin,enok,Asator,CG9220,CG9855,smal,CG12099,CG6330,tna | | 5.351e-03 | -5.23 | I-set | pfam domains | PF07679 | 72 | 8 | 10480 | 393 | Fas2,lbk,ImpL2,DIP-epsilon,Dscam1,Lac,otk2,robo2 | | 5.369e-03 | -5.23 | DUTERTRE\_ESTRADIOL\_RESPONSE\_24HR\_DN | MSigDB lists | DUTERTRE\_ESTRADIOL\_RESPONSE\_24HR\_DN | 90 | 9 | 4161 | 153 | Acsl,Ank,Rho1,MFS10,TfAP-2,Utx,rush,ft,Smox | | 5.373e-03 | -5.23 | GCM\_MAP1B | MSigDB lists | GCM\_MAP1B | 20 | 4 | 4161 | 153 | Pat1,Acsl,Fas2,emp | | 5.373e-03 | -5.23 | GO\_NEUROEPITHELIAL\_CELL\_DIFFERENTIATION | MSigDB lists | GO\_NEUROEPITHELIAL\_CELL\_DIFFERENTIATION | 20 | 4 | 4161 | 153 | Ser,dpp,N,Rac2 | | 5.373e-03 | -5.23 | GO\_POSITIVE\_REGULATION\_OF\_EPITHELIAL\_CELL\_PROLIFERATION | MSigDB lists | GO\_POSITIVE\_REGULATION\_OF\_EPITHELIAL\_CELL\_PROLIFERATION | 20 | 4 | 4161 | 153 | hh,dpp,N,slow | | 5.416e-03 | -5.22 | purine ribonucleotide binding | molecular function | GO:0032555 | 811 | 40 | 12176 | 396 | Mos,Egfr,JIL-1,CG9281,CG9541,Pex1,Arl4,CG8485,betaTub56D,par-1,PhKgamma,Myo10A,CG12010,CG14535,mbt,mle,lz,Pak3,KP78a,Doa,wnd,tkv,bor,Hsc70-3,CaMKII,Tlk,Cdk12,Rac2,Arp2,Etl1,Asator,Mrp4,Mhcl,PEK,Haspin,Hipk,CG8097,S6kII,ScsbetaG,Rho1 | | 5.421e-03 | -5.22 | MAF\_Q6 | MSigDB lists | MAF\_Q6 | 59 | 7 | 4161 | 153 | Arp2,so,Tlk,dlp,Lac,S6kII,Utx | | 5.421e-03 | -5.22 | GO\_MUSCLE\_CELL\_DIFFERENTIATION | MSigDB lists | GO\_MUSCLE\_CELL\_DIFFERENTIATION | 59 | 7 | 4161 | 153 | CaMKII,N,if,so,dpp,hh,Ser | | 5.450e-03 | -5.21 | Ig\_I-set | interpro domains | IPR013098 | 72 | 8 | 11383 | 428 | Dscam1,Fas2,Lac,lbk,otk2,ImpL2,DIP-epsilon,robo2 | | 5.460e-03 | -5.21 | - | gene3d domains | 2.30.29.30 | 139 | 13 | 4929 | 208 | GramD1B,Ptpmeg,CG34384,flfl,CG34383,rush,CG5522,MESR3,RhoGEF2,PsGEF,Myo10A,osp,pbl | | 5.480e-03 | -5.21 | GGGYGTGNY\_UNKNOWN | MSigDB lists | GGGYGTGNY\_UNKNOWN | 160 | 13 | 4161 | 153 | sm,Cnx99A,Arl4,S6kII,Syx1A,Fas2,Ser,Hydr2,trn,JIL-1,mbt,Hsc70-3,dlp | | 5.503e-03 | -5.20 | ZINC\_FINGER\_C2H2\_2 | prosite domains | PS50157 | 273 | 24 | 6243 | 316 | salm,Kah,disco,ttk,shn,bowl,Zif,sens-2,l(3)neo38,ham,CG9650,ci,CG4360,hang,CG31224,ush,CG32772,Kr,sbb,CG11247,cas,CG18446,CG31612,jing | | 5.560e-03 | -5.19 | negative regulation of protein modification process | biological process | GO:0031400 | 98 | 9 | 12345 | 414 | NiPp1,CG8441,aos,sty,N,CycE,ctrip,Utx,hh | | 5.569e-03 | -5.19 | LEF1\_Q6 | MSigDB lists | LEF1\_Q6 | 45 | 6 | 4161 | 153 | dlp,so,fne,mle,Utx,Hydr2 | | 5.569e-03 | -5.19 | GATA\_C | MSigDB lists | GATA\_C | 45 | 6 | 4161 | 153 | so,lbk,CG31637,sdt,CG10663,Fbxl7 | | 5.569e-03 | -5.19 | REACTOME\_PLATELET\_ACTIVATION\_SIGNALING\_AND\_AGGREGATION | MSigDB lists | REACTOME\_PLATELET\_ACTIVATION\_SIGNALING\_AND\_AGGREGATION | 45 | 6 | 4161 | 153 | Hsc70-3,Mrp4,cta,CG34384,drk,Rac2 | | 5.569e-03 | -5.19 | GSE36527\_CD62L\_HIGH\_CD69\_NEG\_VS\_CD62L\_LOW\_CD69\_POS\_TREG\_KLRG1\_NEG\_DN | MSigDB lists | GSE36527\_CD62L\_HIGH\_CD69\_NEG\_VS\_CD62L\_LOW\_CD69\_POS\_TREG\_KLRG1\_NEG\_DN | 45 | 6 | 4161 | 153 | pAbp,TfAP-2,CycE,RpS4,Ptpmeg,NiPp1 | | 5.571e-03 | -5.19 | RhoGAP | smart domains | SM00324 | 22 | 5 | 5086 | 276 | RhoGAP102A,RhoGAP15B,RhoGAP16F,RhoGAP93B,RhoGAP18B | | 5.573e-03 | -5.19 | regulation of synapse assembly | biological process | GO:0051963 | 116 | 10 | 12345 | 414 | CaMKII,Syp,hiw,Arp2,Acsl,tkv,scb,Fas2,Dad,Pak3 | | 5.575e-03 | -5.19 | beta-Oxidation, acyl-CoA synthesis | KEGG pathways | M00086 | 4 | 2 | 3198 | 100 | Acsl,CG3961 | | 5.575e-03 | -5.19 | beta-Oxidation, acyl-CoA synthesis | KEGG pathways | dme\_M00086 | 4 | 2 | 3198 | 100 | Acsl,CG3961 | | 5.600e-03 | -5.19 | ZWANG\_TRANSIENTLY\_UP\_BY\_2ND\_EGF\_PULSE\_ONLY | MSigDB lists | ZWANG\_TRANSIENTLY\_UP\_BY\_2ND\_EGF\_PULSE\_ONLY | 179 | 14 | 4161 | 153 | Npc1a,Lac,Cyp1,Notum,mub,CG31637,Dscam1,retm,Mrp4,DIP-epsilon,hh,Fbxl7,fj,wake | | 5.644e-03 | -5.18 | negative regulation of cell size | biological process | GO:0045792 | 22 | 4 | 12345 | 414 | mbt,Myb,stg,pbl | | 5.644e-03 | -5.18 | germ-band shortening | biological process | GO:0007390 | 22 | 4 | 12345 | 414 | EcR,ush,Egfr,Rac2 | | 5.644e-03 | -5.18 | regulation of apoptotic signaling pathway | biological process | GO:2001233 | 22 | 4 | 12345 | 414 | Egfr,grnd,pnt,eya | | 5.644e-03 | -5.18 | genital disc morphogenesis | biological process | GO:0007483 | 22 | 4 | 12345 | 414 | Fas2,N,otk2,lbe | | 5.644e-03 | -5.18 | head segmentation | biological process | GO:0035287 | 22 | 4 | 12345 | 414 | oc,jing,slp1,hh | | 5.644e-03 | -5.18 | positive regulation of animal organ morphogenesis | biological process | GO:0110110 | 22 | 4 | 12345 | 414 | oc,ec,mbt,lz | | 5.644e-03 | -5.18 | regulation of post-embryonic development | biological process | GO:0048580 | 22 | 4 | 12345 | 414 | Ser,sty,N,Smox | | 5.665e-03 | -5.17 | periodic partitioning | biological process | GO:0007365 | 49 | 6 | 12345 | 414 | ci,Egfr,AGO2,dlp,dally,hh | | 5.675e-03 | -5.17 | KEGG\_NEUROTROPHIN\_SIGNALING\_PATHWAY | MSigDB lists | KEGG\_NEUROTROPHIN\_SIGNALING\_PATHWAY | 32 | 5 | 4161 | 153 | CaMKII,Rac2,drk,JIL-1,S6kII | | 5.675e-03 | -5.17 | LTE2\_UP.V1\_DN | MSigDB lists | LTE2\_UP.V1\_DN | 32 | 5 | 4161 | 153 | Ser,Tif-IA,TfAP-2,S6kII,N | | 5.789e-03 | -5.15 | cytoplasmic region | cellular component | GO:0099568 | 252 | 17 | 11940 | 407 | pbl,RhoGEF2,par-1,CG42684,Hip1,Ser,pon,Arp2,nuf,Syp,CD98hc,AGO2,yps,sdt,Rho1,gus,corn | | 5.815e-03 | -5.15 | multicellular organism reproduction | biological process | GO:0032504 | 1503 | 68 | 12345 | 414 | Doa,baf,jing,drk,eya,S,aop,sty,spoon,Egfr,mle,wnd,hts,CG31680,Myb,bbg,JIL-1,Mmp2,ttk,nudE,Tet,Rho1,Fas3,heph,scb,lama,Rbfox1,goe,CaMKII,Pat1,tsl,Npc1a,if,pAbp,ci,psq,shd,tkv,Imp,Tudor-SN,cnc,Inx2,elav,Syp,RhoGEF2,Mos,AGO2,yps,dally,stwl,dpp,l(3)mbt,hh,par-1,CycE,Stat92E,Pex1,TfAP-2,fne,tin,EcR,gus,so,mam,Rac2,N,Hydr2,wun2 | | 5.819e-03 | -5.15 | CG11134 (uncharacterized protein) | protein interactions | 32334 | 3 | 2 | 9204 | 412 | CG30089,l(3)L1231 | | 5.819e-03 | -5.15 | CG11399 (uncharacterized protein) | protein interactions | 40295 | 3 | 2 | 9204 | 412 | N,Bx | | 5.819e-03 | -5.15 | Vps13 (Vacuolar protein sorting 13) | protein interactions | 35693 | 3 | 2 | 9204 | 412 | Fas2,pbl | | 5.819e-03 | -5.15 | Gmd (GDP-mannose 4,6-dehydratase) | protein interactions | 33716 | 3 | 2 | 9204 | 412 | Syx1A,N | | 5.819e-03 | -5.15 | ogre (optic ganglion reduced) | protein interactions | 45382 | 3 | 2 | 9204 | 412 | Pli,Fas2 | | 5.819e-03 | -5.15 | CG5762 (uncharacterized protein) | protein interactions | 42907 | 3 | 2 | 9204 | 412 | bdg,tara | | 5.819e-03 | -5.15 | GPHR (Golgi pH Regulator) | protein interactions | 36682 | 3 | 2 | 9204 | 412 | mam,N | | 5.819e-03 | -5.15 | Edem2 (ER degradation enhancer, mannosidase alpha-like 2) | protein interactions | 34714 | 3 | 2 | 9204 | 412 | aos,sty | | 5.819e-03 | -5.15 | ClC-b (Chloride channel-b) | protein interactions | 36381 | 3 | 2 | 9204 | 412 | par-1,CG32772 | | 5.819e-03 | -5.15 | sll (slalom) | protein interactions | 42115 | 3 | 2 | 9204 | 412 | dally,hh | | 5.819e-03 | -5.15 | E(spl)malpha-BFM (Enhancer of split malpha, Bearded family member) | protein interactions | 43153 | 3 | 2 | 9204 | 412 | N,E(spl)m4-BFM | | 5.819e-03 | -5.15 | CG14441 (uncharacterized protein) | protein interactions | 31611 | 3 | 2 | 9204 | 412 | spi,Fas2 | | 5.819e-03 | -5.15 | Brms1 (BRMS1) | protein interactions | 32243 | 3 | 2 | 9204 | 412 | Sin3A,EcR | | 5.854e-03 | -5.14 | Ank | pfam domains | PF00023 | 20 | 4 | 10480 | 393 | CG41099,Ank,CG10011,N | | 5.862e-03 | -5.14 | regulation of border follicle cell migration | biological process | GO:1903684 | 35 | 5 | 12345 | 414 | ttk,Ptpmeg,spi,aop,Egfr | | 5.908e-03 | -5.13 | anion binding | molecular function | GO:0043168 | 1099 | 51 | 12176 | 396 | Hsc70-3,CaMKII,Tlk,Hip1,Rac2,Cdk12,RhoGAP15B,Pak3,KP78a,rush,tkv,Doa,wnd,pbl,bor,Haspin,Mhcl,PEK,CG8097,Hipk,Rho1,ScsbetaG,S6kII,PsGEF,dpp,Arp2,Etl1,Asator,Mrp4,Pex1,Arl4,Snx3,CG8485,PhKgamma,par-1,betaTub56D,Egfr,Mos,JIL-1,CG9281,CG9541,CG4199,sas,CG32758,CG12010,Myo10A,Hph,CG14535,mle,lz,mbt | | 5.917e-03 | -5.13 | bon (bonus) | protein interactions | 44235 | 17 | 4 | 9204 | 412 | EcR,Myb,Egfr,ttk | | 5.917e-03 | -5.13 | Sdc (Syndecan) | protein interactions | 37447 | 17 | 4 | 9204 | 412 | comm,dlp,robo2,dally | | 5.917e-03 | -5.13 | kuz (kuzbanian) | protein interactions | 34772 | 17 | 4 | 9204 | 412 | sty,comm,Ser,N | | 5.917e-03 | -5.13 | amos (absent MD neurons and olfactory sensilla) | protein interactions | 35110 | 17 | 4 | 9204 | 412 | lz,hh,Dr,N | | 5.917e-03 | -5.13 | east (enhanced adult sensory threshold) | protein interactions | 46006 | 17 | 4 | 9204 | 412 | pbl,ttk,Fas2,Chro | | 5.917e-03 | -5.13 | Su(Tpl) (Su(Tpl)) | protein interactions | 40171 | 17 | 4 | 9204 | 412 | aop,Bx,N,lncRNA:roX1 | | 5.919e-03 | -5.13 | early endosome | cellular component | GO:0005769 | 64 | 7 | 11940 | 407 | CG6051,CG32758,Tl,rush,Snx3,N,tkv | | 5.941e-03 | -5.13 | positive regulation of cell death | biological process | GO:0010942 | 99 | 9 | 12345 | 414 | lz,EcR,aos,CycE,grnd,Utx,ec,ft,N | | 5.947e-03 | -5.12 | TBOX\_1 | prosite domains | PS01283 | 8 | 3 | 6243 | 316 | bi,Doc2,Doc1 | | 5.947e-03 | -5.12 | TBOX\_3 | prosite domains | PS50252 | 8 | 3 | 6243 | 316 | bi,Doc2,Doc1 | | 5.952e-03 | -5.12 | GATA1\_01 | MSigDB lists | GATA1\_01 | 60 | 7 | 4161 | 153 | sm,fne,Ank,S6kII,Mitofilin,Lim1,LRP1 | | 5.952e-03 | -5.12 | MYOD\_Q6\_01 | MSigDB lists | MYOD\_Q6\_01 | 60 | 7 | 4161 | 153 | drk,ras,Arl4,Smox,CG8531,Imp,Cyp1 | | 5.952e-03 | -5.12 | WAKABAYASHI\_ADIPOGENESIS\_PPARG\_RXRA\_BOUND\_WITH\_H4K20ME1\_MARK | MSigDB lists | WAKABAYASHI\_ADIPOGENESIS\_PPARG\_RXRA\_BOUND\_WITH\_H4K20ME1\_MARK | 60 | 7 | 4161 | 153 | Mitofilin,emp,Mocs2B,Mrp4,kdn,Acsl,rush | | 5.952e-03 | -5.12 | NKX25\_02 | MSigDB lists | NKX25\_02 | 60 | 7 | 4161 | 153 | Ank,fne,cta,mgl,N,LRP1,dlp | | 5.952e-03 | -5.12 | PXR\_Q2 | MSigDB lists | PXR\_Q2 | 60 | 7 | 4161 | 153 | Arl4,Smox,JIL-1,CG16890,so,DIP-epsilon,Mrp4 | | 5.952e-03 | -5.12 | GSE22886\_NAIVE\_TCELL\_VS\_NKCELL\_DN | MSigDB lists | GSE22886\_NAIVE\_TCELL\_VS\_NKCELL\_DN | 60 | 7 | 4161 | 153 | CtBP,N,Rac2,Hydr2,Npc1a,Fas2,CG12547 | | 6.031e-03 | -5.11 | regulation of kinase activity | biological process | GO:0043549 | 136 | 11 | 12345 | 414 | Rho1,CG8441,aos,Atg17,mbt,sty,wnd,bora,Cdk5alpha,CycE,Pli | | 6.034e-03 | -5.11 | translation repressor activity, mRNA regulatory element binding | molecular function | GO:0000900 | 12 | 3 | 12176 | 396 | smg,heph,elav | | 6.034e-03 | -5.11 | cadherin binding | molecular function | GO:0045296 | 12 | 3 | 12176 | 396 | CadN,ds,ft | | 6.051e-03 | -5.11 | puc (puckered) | protein interactions | 40958 | 75 | 9 | 9204 | 412 | egr,Sin3A,tkv,dpp,Rho1,Sox21a,crb,bi,Mrp4 | | 6.051e-03 | -5.11 | mod(mdg4) (modifier of mdg4) | protein interactions | 49228 | 75 | 9 | 9204 | 412 | Chro,ap,CG10265,Bx,lz,CycE,Smox,Pits,lawc | | 6.061e-03 | -5.11 | morphogen activity | molecular function | GO:0016015 | 4 | 2 | 12176 | 396 | dpp,hh | | 6.112e-03 | -5.10 | TpnC25D (Troponin C at 25D) | protein interactions | 33752 | 9 | 3 | 9204 | 412 | CG10265,Df31,fax | | 6.112e-03 | -5.10 | Cad99C (Cadherin 99C) | protein interactions | 43528 | 9 | 3 | 9204 | 412 | sas,hh,ci | | 6.112e-03 | -5.10 | Doc1 (Dorsocross1) | protein interactions | 39039 | 9 | 3 | 9204 | 412 | ci,ttk,Doc2 | | 6.112e-03 | -5.10 | simj (simjang) | protein interactions | 39212 | 9 | 3 | 9204 | 412 | mle,ttk,pbl | | 6.112e-03 | -5.10 | Csp (Cysteine string protein) | protein interactions | 40459 | 9 | 3 | 9204 | 412 | Syx1A,pbl,Ptp99A | | 6.112e-03 | -5.10 | Uvrag (UV-resistance associated gene) | protein interactions | 34735 | 9 | 3 | 9204 | 412 | mam,N,Stat92E | | 6.112e-03 | -5.10 | CG3386 (uncharacterized protein) | protein interactions | 38081 | 9 | 3 | 9204 | 412 | CG32758,rau,CG3386 | | 6.112e-03 | -5.10 | Ndfip (Nedd4 family interacting protein) | protein interactions | 326197 | 9 | 3 | 9204 | 412 | N,fax,Ser | | 6.112e-03 | -5.10 | CG6441 (uncharacterized protein) | protein interactions | 34029 | 9 | 3 | 9204 | 412 | slow,disco,pAbp | | 6.112e-03 | -5.10 | Socs44A (Suppressor of Cytokine Signaling at 44A) | protein interactions | 35786 | 9 | 3 | 9204 | 412 | aos,eya,Stat92E | | 6.112e-03 | -5.10 | CG7849 (uncharacterized protein) | protein interactions | 35537 | 9 | 3 | 9204 | 412 | betaTub56D,CG11486,Hip1 | | 6.135e-03 | -5.09 | GO\_CALCIUM\_ION\_BINDING | MSigDB lists | GO\_CALCIUM\_ION\_BINDING | 126 | 11 | 4161 | 153 | mgl,Cnx99A,N,ft,ds,Ser,Mmp2,slow,hh,LRP1,Hsc70-3 | | 6.135e-03 | -5.09 | GO\_TRANSCRIPTION\_FACTOR\_ACTIVITY\_PROTEIN\_BINDING | MSigDB lists | GO\_TRANSCRIPTION\_FACTOR\_ACTIVITY\_PROTEIN\_BINDING | 126 | 11 | 4161 | 153 | ap,CtBP,TfAP-2,Taf1,N,Smox,Lim1,Tudor-SN,EcR,Sin3A,CycE | | 6.214e-03 | -5.08 | GO\_PROTEINACEOUS\_EXTRACELLULAR\_MATRIX | MSigDB lists | GO\_PROTEINACEOUS\_EXTRACELLULAR\_MATRIX | 46 | 6 | 4161 | 153 | NetB,Mmp2,CG3097,hh,dlp,dally | | 6.214e-03 | -5.08 | GATA3\_01 | MSigDB lists | GATA3\_01 | 46 | 6 | 4161 | 153 | Trpm,sm,Arl4,Mhcl,LRP1,so | | 6.214e-03 | -5.08 | GSE9988\_ANTI\_TREM1\_VS\_VEHICLE\_TREATED\_MONOCYTES\_UP | MSigDB lists | GSE9988\_ANTI\_TREM1\_VS\_VEHICLE\_TREATED\_MONOCYTES\_UP | 46 | 6 | 4161 | 153 | CG9220,Npc1a,mmy,MFS10,Acsl,cta | | 6.214e-03 | -5.08 | OCT1\_03 | MSigDB lists | OCT1\_03 | 46 | 6 | 4161 | 153 | Ser,Snx3,sdt,CG16890,SamDC,CG2812 | | 6.214e-03 | -5.08 | LEF1\_Q2 | MSigDB lists | LEF1\_Q2 | 46 | 6 | 4161 | 153 | mmy,smal,Ank,ds,CG16890,CG3961 | | 6.223e-03 | -5.08 | cell cortex part | cellular component | GO:0044448 | 116 | 10 | 11940 | 407 | RhoGEF2,CG42684,nuf,pon,Arp2,Hip1,sdt,Rho1,Syp,Ser | | 6.260e-03 | -5.07 | establishment or maintenance of cytoskeleton polarity | biological process | GO:0030952 | 50 | 6 | 12345 | 414 | cnc,par-1,nudE,cta,heph,Rho1 | | 6.261e-03 | -5.07 | Tl (Toll) | protein interactions | 43222 | 27 | 5 | 9204 | 412 | Tl,dpp,Fas3,Stat92E,Myb | | 6.261e-03 | -5.07 | LanB1 (LanB1) | protein interactions | 34068 | 27 | 5 | 9204 | 412 | Cnx99A,LRP1,ci,CG43658,sty | | 6.261e-03 | -5.07 | D (Dichaete) | protein interactions | 39570 | 27 | 5 | 9204 | 412 | CG2865,N,ttk,dpp,Sox21a | | 6.261e-03 | -5.07 | hiw (highwire) | protein interactions | 32429 | 27 | 5 | 9204 | 412 | tkv,betaTub56D,Dscam1,Syp,wnd | | 6.283e-03 | -5.07 | GSE22886\_NAIVE\_CD4\_TCELL\_VS\_NKCELL\_DN | MSigDB lists | GSE22886\_NAIVE\_CD4\_TCELL\_VS\_NKCELL\_DN | 76 | 8 | 4161 | 153 | Fas2,Npc1a,NiPp1,CtBP,CG45050,Hydr2,Rac2,Acsl | | 6.340e-03 | -5.06 | bw (brown) | protein interactions | 37724 | 38 | 6 | 9204 | 412 | l(3)neo38,HmgZ,psq,Etl1,Acf,ban | | 6.377e-03 | -5.05 | fly chr68B3-68B3\|3-35 | chromosome location | fly chr68B3-68B3\|3-35 | 6 | 2 | 25010 | 531 | lncRNA:CR44719,CG42671 | | 6.377e-03 | -5.05 | fly chr39E3-39E3\|2-54 cM | chromosome location | fly chr39E3-39E3\|2-54 cM | 6 | 2 | 25010 | 531 | CG2201,Df31 | | 6.377e-03 | -5.05 | fly chr39E3-39E3\|2-54 | chromosome location | fly chr39E3-39E3\|2-54 | 6 | 2 | 25010 | 531 | CG2201,Df31 | | 6.377e-03 | -5.05 | fly chr68B3-68B3\|3-35 cM | chromosome location | fly chr68B3-68B3\|3-35 cM | 6 | 2 | 25010 | 531 | lncRNA:CR44719,CG42671 | | 6.384e-03 | -5.05 | ConA-like\_dom\_sf | interpro domains | IPR013320 | 59 | 7 | 11383 | 428 | CadN,gus,Cnx99A,wb,ft,CG6752,crb | | 6.436e-03 | -5.05 | embryonic crystal cell differentiation | biological process | GO:0035165 | 4 | 2 | 12345 | 414 | lz,N | | 6.436e-03 | -5.05 | immune response-regulating cell surface receptor signaling pathway | biological process | GO:0002768 | 4 | 2 | 12345 | 414 | Rac2,Toll-7 | | 6.436e-03 | -5.05 | ectodermal placode development | biological process | GO:0071696 | 4 | 2 | 12345 | 414 | eya,so | | 6.436e-03 | -5.05 | positive regulation of G2/M transition of mitotic cell cycle | biological process | GO:0010971 | 4 | 2 | 12345 | 414 | Myb,stg | | 6.436e-03 | -5.05 | ectodermal placode formation | biological process | GO:0060788 | 4 | 2 | 12345 | 414 | so,eya | | 6.436e-03 | -5.05 | regulation of meiotic cell cycle phase transition | biological process | GO:1901993 | 4 | 2 | 12345 | 414 | Mos,stg | | 6.436e-03 | -5.05 | regulation of Malpighian tubule size | biological process | GO:0035298 | 4 | 2 | 12345 | 414 | EcR,Rho1 | | 6.436e-03 | -5.05 | positive regulation of cell cycle G2/M phase transition | biological process | GO:1902751 | 4 | 2 | 12345 | 414 | stg,Myb | | 6.436e-03 | -5.05 | negative regulation of fibroblast growth factor receptor signaling pathway | biological process | GO:0040037 | 4 | 2 | 12345 | 414 | sty,Mmp2 | | 6.436e-03 | -5.05 | ectodermal placode morphogenesis | biological process | GO:0071697 | 4 | 2 | 12345 | 414 | so,eya | | 6.436e-03 | -5.05 | cellular response to lipopolysaccharide | biological process | GO:0071222 | 4 | 2 | 12345 | 414 | EcR,Stat92E | | 6.436e-03 | -5.05 | optic lobe placode formation | biological process | GO:0001744 | 4 | 2 | 12345 | 414 | eya,so | | 6.436e-03 | -5.05 | posterior midgut invagination | biological process | GO:0007374 | 4 | 2 | 12345 | 414 | RhoGEF2,Rho1 | | 6.436e-03 | -5.05 | embryonic hemocyte differentiation | biological process | GO:0035163 | 4 | 2 | 12345 | 414 | lz,N | | 6.450e-03 | -5.04 | HALLMARK\_TNFA\_SIGNALING\_VIA\_NFKB | MSigDB lists | HALLMARK\_TNFA\_SIGNALING\_VIA\_NFKB | 21 | 4 | 4161 | 153 | Ser,dpp,Smox,fj | | 6.450e-03 | -5.04 | GO\_CRANIAL\_SKELETAL\_SYSTEM\_DEVELOPMENT | MSigDB lists | GO\_CRANIAL\_SKELETAL\_SYSTEM\_DEVELOPMENT | 21 | 4 | 4161 | 153 | TfAP-2,Smox,Lim1,so | | 6.450e-03 | -5.04 | GO\_AXIS\_SPECIFICATION | MSigDB lists | GO\_AXIS\_SPECIFICATION | 21 | 4 | 4161 | 153 | N,Utx,Lim1,hh | | 6.450e-03 | -5.04 | AZARE\_NEOPLASTIC\_TRANSFORMATION\_BY\_STAT3\_UP | MSigDB lists | AZARE\_NEOPLASTIC\_TRANSFORMATION\_BY\_STAT3\_UP | 21 | 4 | 4161 | 153 | Ser,fne,mle,Ykt6 | | 6.450e-03 | -5.04 | GCCNNNWTAAR\_UNKNOWN | MSigDB lists | GCCNNNWTAAR\_UNKNOWN | 21 | 4 | 4161 | 153 | CG5830,CaMKII,lili,hh | | 6.450e-03 | -5.04 | GO\_REGULATION\_OF\_STEM\_CELL\_DIFFERENTIATION | MSigDB lists | GO\_REGULATION\_OF\_STEM\_CELL\_DIFFERENTIATION | 21 | 4 | 4161 | 153 | Smox,N,dpp,Ser | | 6.450e-03 | -5.04 | GO\_REGULATION\_OF\_FAT\_CELL\_DIFFERENTIATION | MSigDB lists | GO\_REGULATION\_OF\_FAT\_CELL\_DIFFERENTIATION | 21 | 4 | 4161 | 153 | dpp,Ser,CG11367,Smox | | 6.471e-03 | -5.04 | SHAFFER\_IRF4\_MULTIPLE\_MYELOMA\_PROGRAM | MSigDB lists | SHAFFER\_IRF4\_MULTIPLE\_MYELOMA\_PROGRAM | 11 | 3 | 4161 | 153 | Cnx99A,smal,mmy | | 6.471e-03 | -5.04 | KEGG\_CELL\_ADHESION\_MOLECULES\_CAMS | MSigDB lists | KEGG\_CELL\_ADHESION\_MOLECULES\_CAMS | 11 | 3 | 4161 | 153 | Lac,Fas2,if | | 6.471e-03 | -5.04 | GO\_ANATOMICAL\_STRUCTURE\_MATURATION | MSigDB lists | GO\_ANATOMICAL\_STRUCTURE\_MATURATION | 11 | 3 | 4161 | 153 | ds,ft,dpp | | 6.471e-03 | -5.04 | GO\_RETINA\_MORPHOGENESIS\_IN\_CAMERA\_TYPE\_EYE | MSigDB lists | GO\_RETINA\_MORPHOGENESIS\_IN\_CAMERA\_TYPE\_EYE | 11 | 3 | 4161 | 153 | Lim1,TfAP-2,fj | | 6.471e-03 | -5.04 | GO\_APICAL\_JUNCTION\_ASSEMBLY | MSigDB lists | GO\_APICAL\_JUNCTION\_ASSEMBLY | 11 | 3 | 4161 | 153 | Rho1,sdt,pbl | | 6.471e-03 | -5.04 | GO\_REGULATION\_OF\_CARTILAGE\_DEVELOPMENT | MSigDB lists | GO\_REGULATION\_OF\_CARTILAGE\_DEVELOPMENT | 11 | 3 | 4161 | 153 | dpp,tkv,Smox | | 6.471e-03 | -5.04 | GO\_NEURAL\_RETINA\_DEVELOPMENT | MSigDB lists | GO\_NEURAL\_RETINA\_DEVELOPMENT | 11 | 3 | 4161 | 153 | fj,TfAP-2,Lim1 | | 6.471e-03 | -5.04 | LI\_WILMS\_TUMOR\_VS\_FETAL\_KIDNEY\_2\_DN | MSigDB lists | LI\_WILMS\_TUMOR\_VS\_FETAL\_KIDNEY\_2\_DN | 11 | 3 | 4161 | 153 | Lim1,dlp,smal | | 6.471e-03 | -5.04 | GO\_POSITIVE\_REGULATION\_OF\_LIPID\_TRANSPORT | MSigDB lists | GO\_POSITIVE\_REGULATION\_OF\_LIPID\_TRANSPORT | 11 | 3 | 4161 | 153 | ScpX,LRP1,EcR | | 6.471e-03 | -5.04 | GO\_CARDIAC\_CELL\_DEVELOPMENT | MSigDB lists | GO\_CARDIAC\_CELL\_DEVELOPMENT | 11 | 3 | 4161 | 153 | Ser,N,CaMKII | | 6.471e-03 | -5.04 | BIOCARTA\_P38MAPK\_PATHWAY | MSigDB lists | BIOCARTA\_P38MAPK\_PATHWAY | 11 | 3 | 4161 | 153 | Rac2,drk,JIL-1 | | 6.471e-03 | -5.04 | chrxq13 | MSigDB lists | chrxq13 | 11 | 3 | 4161 | 153 | Taf1,RpS4,Snx3 | | 6.471e-03 | -5.04 | HALMOS\_CEBPA\_TARGETS\_UP | MSigDB lists | HALMOS\_CEBPA\_TARGETS\_UP | 11 | 3 | 4161 | 153 | Arl4,dpp,mmy | | 6.521e-03 | -5.03 | NABA\_MATRISOME\_ASSOCIATED | MSigDB lists | NABA\_MATRISOME\_ASSOCIATED | 61 | 7 | 4161 | 153 | CG10663,slow,Mmp2,dpp,dlp,dally,hh | | 6.567e-03 | -5.03 | neuromuscular junction development | biological process | GO:0007528 | 83 | 8 | 12345 | 414 | Prosap,par-1,Fas2,dlp,CaMKII,Ten-m,S6kII,Imp | | 6.572e-03 | -5.02 | positive regulation of wound healing | biological process | GO:0090303 | 12 | 3 | 12345 | 414 | Egfr,Rac2,Rho1 | | 6.572e-03 | -5.02 | retinal cell programmed cell death | biological process | GO:0046666 | 12 | 3 | 12345 | 414 | N,ec,S | | 6.572e-03 | -5.02 | positive regulation of cell cycle G1/S phase transition | biological process | GO:1902808 | 12 | 3 | 12345 | 414 | N,ci,CycE | | 6.572e-03 | -5.02 | axonal defasciculation | biological process | GO:0007414 | 12 | 3 | 12345 | 414 | Ptp99A,if,pbl | | 6.572e-03 | -5.02 | maintenance of epithelial integrity, open tracheal system | biological process | GO:0035160 | 12 | 3 | 12345 | 414 | Lac,if,Egfr | | 6.572e-03 | -5.02 | establishment of apical/basal cell polarity | biological process | GO:0035089 | 12 | 3 | 12345 | 414 | ds,fj,ft | | 6.572e-03 | -5.02 | negative regulation of apoptotic signaling pathway | biological process | GO:2001234 | 12 | 3 | 12345 | 414 | pnt,Egfr,eya | | 6.572e-03 | -5.02 | regulation of heterochromatin assembly | biological process | GO:0031445 | 12 | 3 | 12345 | 414 | Acf,mle,Stat92E | | 6.600e-03 | -5.02 | SkpA (SKP1-related A) | protein interactions | 31016 | 76 | 9 | 9204 | 412 | Ppa,hiw,ci,ds,Doa,Fbxl7,wnd,CG11486,CG43658 | | 6.644e-03 | -5.01 | Sin3 complex | cellular component | GO:0016580 | 4 | 2 | 11940 | 407 | Sin3A,CG44774 | | 6.654e-03 | -5.01 | male genitalia development | biological process | GO:0030539 | 23 | 4 | 12345 | 414 | otk2,salm,Fas2,N | | 6.654e-03 | -5.01 | negative chemotaxis | biological process | GO:0050919 | 23 | 4 | 12345 | 414 | wun2,Tl,robo2,beat-IIIc | | 6.659e-03 | -5.01 | AATGTGA\_MIR23A\_MIR23B | MSigDB lists | AATGTGA\_MIR23A\_MIR23B | 93 | 9 | 4161 | 153 | Utx,gus,NTPase,Smox,Cyp1,CG32758,sdt,Pcl,dlp | | 6.669e-03 | -5.01 | Mad (Mothers against dpp) | protein interactions | 33529 | 104 | 11 | 9204 | 412 | smash,Smox,tkv,bowl,Ubx,dpp,dally,eya,shn,CG45050,Dad | | 6.741e-03 | -5.00 | negative regulation of cellular protein metabolic process | biological process | GO:0032269 | 218 | 15 | 12345 | 414 | N,Rbfox1,bin3,crb,elav,hh,Utx,ctrip,CycE,smg,aos,CG8441,NiPp1,heph,sty | | 6.765e-03 | -5.00 | ribonucleotide binding | molecular function | GO:0032553 | 822 | 40 | 12176 | 396 | Mos,Egfr,JIL-1,CG9281,CG9541,Pex1,Arl4,CG8485,betaTub56D,PhKgamma,par-1,Myo10A,CG12010,CG14535,mbt,lz,mle,Pak3,KP78a,Doa,wnd,tkv,bor,Hsc70-3,Tlk,CaMKII,Cdk12,Rac2,Arp2,Etl1,Asator,Mrp4,Mhcl,PEK,Haspin,Hipk,CG8097,S6kII,ScsbetaG,Rho1 | | 6.798e-03 | -4.99 | GO\_REGULATION\_OF\_ORGAN\_MORPHOGENESIS | MSigDB lists | GO\_REGULATION\_OF\_ORGAN\_MORPHOGENESIS | 77 | 8 | 4161 | 153 | Rac2,hh,dlp,so,TfAP-2,N,Lim1,dpp | | 6.875e-03 | -4.98 | anchored component of membrane | cellular component | GO:0031225 | 12 | 3 | 11940 | 407 | dally,bou,dlp | | 6.897e-03 | -4.98 | Laminin\_EGF | pfam domains | PF00053 | 11 | 3 | 10480 | 393 | NetA,NetB,wb | | 6.899e-03 | -4.98 | regulation of dendrite development | biological process | GO:0050773 | 51 | 6 | 12345 | 414 | nudE,ham,Prosap,Dscam1,comm,CadN | | 6.912e-03 | -4.97 | GO\_NEGATIVE\_REGULATION\_OF\_NEURON\_DIFFERENTIATION | MSigDB lists | GO\_NEGATIVE\_REGULATION\_OF\_NEURON\_DIFFERENTIATION | 47 | 6 | 4161 | 153 | NetB,CG5830,N,hh,LRP1,Ser | | 6.912e-03 | -4.97 | GSE3982\_NEUTROPHIL\_VS\_TH2\_UP | MSigDB lists | GSE3982\_NEUTROPHIL\_VS\_TH2\_UP | 47 | 6 | 4161 | 153 | par-1,ras,Arl4,CG5522,Fas2,CG9220 | | 6.912e-03 | -4.97 | OISHI\_CHOLANGIOMA\_STEM\_CELL\_LIKE\_DN | MSigDB lists | OISHI\_CHOLANGIOMA\_STEM\_CELL\_LIKE\_DN | 47 | 6 | 4161 | 153 | LRP1,Pli,smal,CG3961,CG43313,Arl4 | | 6.912e-03 | -4.97 | GSE19923\_WT\_VS\_E2A\_KO\_DP\_THYMOCYTE\_DN | MSigDB lists | GSE19923\_WT\_VS\_E2A\_KO\_DP\_THYMOCYTE\_DN | 47 | 6 | 4161 | 153 | Tudor-SN,fne,Tpr2,EcR,Pat1,heph | | 6.918e-03 | -4.97 | Znf\_RING/FYVE/PHD | interpro domains | IPR013083 | 214 | 16 | 11383 | 428 | CG12099,CG6051,hiw,rush,tna,tou,CG41099,Psc,Pcl,CG2991,Acf,CG4325,enok,CG9855,CG6752,CG32369 | | 6.939e-03 | -4.97 | GOBERT\_OLIGODENDROCYTE\_DIFFERENTIATION\_DN | MSigDB lists | GOBERT\_OLIGODENDROCYTE\_DIFFERENTIATION\_DN | 242 | 17 | 4161 | 153 | Mhcl,N,NTPase,Dscam1,Utx,Ank,LManII,SamDC,Lac,CG5522,Npc1a,Arl4,CaMKII,Hsc70Cb,emp,Tlk,MFS10 | | 6.956e-03 | -4.97 | transcription coactivator activity | molecular function | GO:0003713 | 69 | 7 | 12176 | 396 | tna,mam,eya,CtBP,Lpin,shn,EcR | | 6.977e-03 | -4.97 | regulation of antimicrobial humoral response | biological process | GO:0002759 | 67 | 7 | 12345 | 414 | Usp2,Tl,CG4325,ush,Doa,Stat92E,Pli | | 6.988e-03 | -4.96 | growth factor receptor binding | molecular function | GO:0070851 | 24 | 4 | 12176 | 396 | spi,Pvf3,aos,drk | | 7.022e-03 | -4.96 | negative regulation of protein metabolic process | biological process | GO:0051248 | 219 | 15 | 12345 | 414 | hh,Utx,ctrip,smg,CycE,Rbfox1,bin3,crb,N,elav,sty,aos,heph,NiPp1,CG8441 | | 7.099e-03 | -4.95 | regulation of response to stress | biological process | GO:0080134 | 283 | 18 | 12345 | 414 | Toll-7,Imp,Rho1,eya,mof,Tl,CG4325,ush,Etl1,Usp2,Egfr,PEK,N,Mmp2,Stat92E,Pli,Rac2,grnd | | 7.130e-03 | -4.94 | CDX2\_Q5 | MSigDB lists | CDX2\_Q5 | 62 | 7 | 4161 | 153 | Ser,mgl,Arl4,CG2812,fne,Utx,ds | | 7.203e-03 | -4.93 | regulation of cell cycle phase transition | biological process | GO:1901987 | 102 | 9 | 12345 | 414 | Egfr,Mos,PEK,Myb,stg,CycE,Chro,ci,N | | 7.214e-03 | -4.93 | LIM | pfam domains | PF00412 | 33 | 5 | 10480 | 393 | Bx,ap,Lim1,esn,smash | | 7.216e-03 | -4.93 | Mbs (Myosin binding subunit) | protein interactions | 49070 | 39 | 6 | 9204 | 412 | Rho1,Egfr,par-1,crb,pbl,Dad | | 7.216e-03 | -4.93 | RasGAP1 (Ras GTPase activating protein 1) | protein interactions | 39158 | 39 | 6 | 9204 | 412 | sty,CycE,Ptpmeg,aop,Egfr,RhoGAP18B | | 7.220e-03 | -4.93 | TBOX | smart domains | SM00425 | 8 | 3 | 5086 | 276 | Doc2,bi,Doc1 | | 7.291e-03 | -4.92 | Rho1 (Rho1) | protein interactions | 36775 | 91 | 10 | 9204 | 412 | wnd,pbl,tkv,egr,cta,RhoGAP18B,Dad,crb,RhoGEF2,CycE | | 7.303e-03 | -4.92 | SH2\_dom\_sf | interpro domains | IPR036860 | 33 | 5 | 11383 | 428 | CG10479,CG33993,drk,Socs16D,Stat92E | | 7.309e-03 | -4.92 | regulation of synapse structure or activity | biological process | GO:0050803 | 179 | 13 | 12345 | 414 | wnd,CaMKII,Syp,Ten-m,hiw,S6kII,tkv,Arp2,Acsl,scb,Fas2,Dad,Pak3 | | 7.343e-03 | -4.91 | htk (hat-trick) | protein interactions | 32877 | 18 | 4 | 9204 | 412 | N,bdg,rau,Sin3A | | 7.343e-03 | -4.91 | CSN5 (COP9 signalosome subunit 5) | protein interactions | 42000 | 18 | 4 | 9204 | 412 | Egfr,trn,EcR,CycE | | 7.343e-03 | -4.91 | vig (vasa intronic gene) | protein interactions | 34885 | 18 | 4 | 9204 | 412 | S6kII,Fas2,AGO2,Tudor-SN | | 7.343e-03 | -4.91 | Uba1 (Ubiquitin activating enzyme 1) | protein interactions | 35998 | 18 | 4 | 9204 | 412 | N,ci,egr,Stat92E | | 7.345e-03 | -4.91 | E2F1\_Q4 | MSigDB lists | E2F1\_Q4 | 78 | 8 | 4161 | 153 | CG8531,JIL-1,Sin3A,CG12547,NiPp1,CG5830,S6kII,lili | | 7.360e-03 | -4.91 | mRNA 3'-UTR binding | molecular function | GO:0003730 | 38 | 5 | 12176 | 396 | Rbfox1,Imp,smg,heph,pAbp | | 7.401e-03 | -4.91 | Rap2l (Ras-associated protein 2-like) | protein interactions | 37850 | 64 | 8 | 9204 | 412 | ey,oc,CadN,rau,RhoGEF2,Prosap,emp,Mhcl | | 7.406e-03 | -4.91 | NTR | prosite domains | PS50189 | 3 | 2 | 6243 | 316 | NetA,NetB | | 7.448e-03 | -4.90 | basolateral plasma membrane | cellular component | GO:0016323 | 51 | 6 | 11940 | 407 | dally,if,par-1,Egfr,Fas3,scb | | 7.449e-03 | -4.90 | intracellular signal transduction | biological process | GO:0035556 | 373 | 22 | 12345 | 414 | egr,JIL-1,CG34384,hang,Tlk,par-1,Rac2,KP78a,wnd,CG4325,Cirl,mof,drk,cta,PhKgamma,Cdk5alpha,CG8485,mbt,gus,Rho1,Haspin,Doa | | 7.453e-03 | -4.90 | positive regulation of canonical Wnt signaling pathway | biological process | GO:0090263 | 37 | 5 | 12345 | 414 | Hipk,dlp,dally,spen,CtBP | | 7.462e-03 | -4.90 | TGF-beta signaling pathway | KEGG pathways | ko04350 | 40 | 5 | 3198 | 100 | tkv,dpp,Rho1,Smox,Dad | | 7.462e-03 | -4.90 | TGF-beta signaling pathway | KEGG pathways | dme04350 | 40 | 5 | 3198 | 100 | tkv,Rho1,dpp,Smox,Dad | | 7.584e-03 | -4.88 | associative learning | biological process | GO:0008306 | 52 | 6 | 12345 | 414 | scb,Egfr,S6kII,drk,spi,Fas2 | | 7.639e-03 | -4.87 | chromosome organization | biological process | GO:0051276 | 680 | 35 | 12345 | 414 | nudE,Haspin,Acf,Pcl,CtBP,tna,mof,osa,Psc,Df31,tara,Taf1,psq,Kr,Doa,baf,wcy,enok,Etl1,Chro,eya,CG5815,mle,spoon,AGO2,stwl,tou,Myb,Utx,HmgZ,Tlk,JIL-1,bin3,Mocs2B,dve | | 7.657e-03 | -4.87 | H4 histone acetyltransferase activity | molecular function | GO:0010485 | 13 | 3 | 12176 | 396 | Taf1,mof,enok | | 7.657e-03 | -4.87 | RNA polymerase II transcription factor binding | molecular function | GO:0001085 | 13 | 3 | 12176 | 396 | ush,tna,bin3 | | 7.662e-03 | -4.87 | MEISSNER\_BRAIN\_HCP\_WITH\_H3K27ME3 | MSigDB lists | MEISSNER\_BRAIN\_HCP\_WITH\_H3K27ME3 | 22 | 4 | 4161 | 153 | nAChRalpha4,Imp,so,Lim1 | | 7.665e-03 | -4.87 | CACBINDINGPROTEIN\_Q6 | MSigDB lists | CACBINDINGPROTEIN\_Q6 | 48 | 6 | 4161 | 153 | mgl,ds,Fas2,Imp,Lac,LRP1 | | 7.676e-03 | -4.87 | VERRECCHIA\_RESPONSE\_TO\_TGFB1\_C4 | MSigDB lists | VERRECCHIA\_RESPONSE\_TO\_TGFB1\_C4 | 4 | 2 | 4161 | 153 | Rac2,Mmp2 | | 7.676e-03 | -4.87 | GO\_REGULATION\_OF\_EPITHELIAL\_CELL\_DIFFERENTIATION\_INVOLVED\_IN\_KIDNEY\_DEVELOPMENT | MSigDB lists | GO\_REGULATION\_OF\_EPITHELIAL\_CELL\_DIFFERENTIATION\_INVOLVED\_IN\_KIDNEY\_DEVELOPMENT | 4 | 2 | 4161 | 153 | ft,Lim1 | | 7.676e-03 | -4.87 | GO\_PROSTATE\_GLAND\_MORPHOGENESIS | MSigDB lists | GO\_PROSTATE\_GLAND\_MORPHOGENESIS | 4 | 2 | 4161 | 153 | N,hh | | 7.676e-03 | -4.87 | MODULE\_433 | MSigDB lists | MODULE\_433 | 4 | 2 | 4161 | 153 | Ser,dpp | | 7.676e-03 | -4.87 | GO\_INTERFERON\_GAMMA\_MEDIATED\_SIGNALING\_PATHWAY | MSigDB lists | GO\_INTERFERON\_GAMMA\_MEDIATED\_SIGNALING\_PATHWAY | 4 | 2 | 4161 | 153 | Fas2,CaMKII | | 7.676e-03 | -4.87 | GO\_LONG\_CHAIN\_FATTY\_ACID\_COA\_LIGASE\_ACTIVITY | MSigDB lists | GO\_LONG\_CHAIN\_FATTY\_ACID\_COA\_LIGASE\_ACTIVITY | 4 | 2 | 4161 | 153 | CG3961,Acsl | | 7.676e-03 | -4.87 | GO\_CHONDROCYTE\_DEVELOPMENT | MSigDB lists | GO\_CHONDROCYTE\_DEVELOPMENT | 4 | 2 | 4161 | 153 | CG9220,tkv | | 7.676e-03 | -4.87 | GO\_CARDIAC\_RIGHT\_VENTRICLE\_MORPHOGENESIS | MSigDB lists | GO\_CARDIAC\_RIGHT\_VENTRICLE\_MORPHOGENESIS | 4 | 2 | 4161 | 153 | N,Ser | | 7.676e-03 | -4.87 | GO\_NEGATIVE\_REGULATION\_OF\_MUSCLE\_TISSUE\_DEVELOPMENT | MSigDB lists | GO\_NEGATIVE\_REGULATION\_OF\_MUSCLE\_TISSUE\_DEVELOPMENT | 4 | 2 | 4161 | 153 | hh,dpp | | 7.676e-03 | -4.87 | REACTOME\_CHONDROITIN\_SULFATE\_BIOSYNTHESIS | MSigDB lists | REACTOME\_CHONDROITIN\_SULFATE\_BIOSYNTHESIS | 4 | 2 | 4161 | 153 | CG9220,CG43313 | | 7.676e-03 | -4.87 | REACTOME\_SIGNALING\_BY\_FGFR1\_MUTANTS | MSigDB lists | REACTOME\_SIGNALING\_BY\_FGFR1\_MUTANTS | 4 | 2 | 4161 | 153 | Mhcl,drk | | 7.676e-03 | -4.87 | YAMASHITA\_LIVER\_CANCER\_WITH\_EPCAM\_DN | MSigDB lists | YAMASHITA\_LIVER\_CANCER\_WITH\_EPCAM\_DN | 4 | 2 | 4161 | 153 | Ugt49C1,so | | 7.676e-03 | -4.87 | GO\_SMOOTH\_MUSCLE\_TISSUE\_DEVELOPMENT | MSigDB lists | GO\_SMOOTH\_MUSCLE\_TISSUE\_DEVELOPMENT | 4 | 2 | 4161 | 153 | if,so | | 7.676e-03 | -4.87 | GO\_POSITIVE\_REGULATION\_OF\_TRANSCRIPTION\_FROM\_RNA\_POLYMERASE\_II\_PROMOTER\_IN\_RESPONSE\_TO\_STRESS | MSigDB lists | GO\_POSITIVE\_REGULATION\_OF\_TRANSCRIPTION\_FROM\_RNA\_POLYMERASE\_II\_PROMOTER\_IN\_RESPONSE\_TO\_STRESS | 4 | 2 | 4161 | 153 | Hsc70-3,N | | 7.676e-03 | -4.87 | HOLLERN\_SOLID\_NODULAR\_BREAST\_TUMOR\_DN | MSigDB lists | HOLLERN\_SOLID\_NODULAR\_BREAST\_TUMOR\_DN | 4 | 2 | 4161 | 153 | CG34384,dpp | | 7.676e-03 | -4.87 | GO\_REGULATION\_OF\_OSTEOBLAST\_PROLIFERATION | MSigDB lists | GO\_REGULATION\_OF\_OSTEOBLAST\_PROLIFERATION | 4 | 2 | 4161 | 153 | dpp,Smox | | 7.676e-03 | -4.87 | REACTOME\_SIGNALING\_BY\_FGFR1\_FUSION\_MUTANTS | MSigDB lists | REACTOME\_SIGNALING\_BY\_FGFR1\_FUSION\_MUTANTS | 4 | 2 | 4161 | 153 | drk,Mhcl | | 7.676e-03 | -4.87 | GO\_POSITIVE\_REGULATION\_OF\_CHOLESTEROL\_EFFLUX | MSigDB lists | GO\_POSITIVE\_REGULATION\_OF\_CHOLESTEROL\_EFFLUX | 4 | 2 | 4161 | 153 | LRP1,EcR | | 7.676e-03 | -4.87 | GO\_ENDOCARDIAL\_CUSHION\_FORMATION | MSigDB lists | GO\_ENDOCARDIAL\_CUSHION\_FORMATION | 4 | 2 | 4161 | 153 | N,ds | | 7.676e-03 | -4.87 | GO\_POSITIVE\_REGULATION\_OF\_EXTRACELLULAR\_MATRIX\_ORGANIZATION | MSigDB lists | GO\_POSITIVE\_REGULATION\_OF\_EXTRACELLULAR\_MATRIX\_ORGANIZATION | 4 | 2 | 4161 | 153 | Smox,smal | | 7.676e-03 | -4.87 | GO\_FATTY\_ACID\_LIGASE\_ACTIVITY | MSigDB lists | GO\_FATTY\_ACID\_LIGASE\_ACTIVITY | 4 | 2 | 4161 | 153 | Acsl,CG3961 | | 7.676e-03 | -4.87 | GO\_STEROL\_TRANSPORTER\_ACTIVITY | MSigDB lists | GO\_STEROL\_TRANSPORTER\_ACTIVITY | 4 | 2 | 4161 | 153 | ScpX,Npc1a | | 7.676e-03 | -4.87 | REACTOME\_DEGRADATION\_OF\_THE\_EXTRACELLULAR\_MATRIX | MSigDB lists | REACTOME\_DEGRADATION\_OF\_THE\_EXTRACELLULAR\_MATRIX | 4 | 2 | 4161 | 153 | Mmp2,CG10663 | | 7.676e-03 | -4.87 | GO\_POSITIVE\_REGULATION\_OF\_CHONDROCYTE\_DIFFERENTIATION | MSigDB lists | GO\_POSITIVE\_REGULATION\_OF\_CHONDROCYTE\_DIFFERENTIATION | 4 | 2 | 4161 | 153 | tkv,Smox | | 7.676e-03 | -4.87 | WATANABE\_COLON\_CANCER\_MSI\_VS\_MSS\_UP | MSigDB lists | WATANABE\_COLON\_CANCER\_MSI\_VS\_MSS\_UP | 4 | 2 | 4161 | 153 | sm,TfAP-2 | | 7.676e-03 | -4.87 | GO\_NEGATIVE\_REGULATION\_OF\_GLIAL\_CELL\_PROLIFERATION | MSigDB lists | GO\_NEGATIVE\_REGULATION\_OF\_GLIAL\_CELL\_PROLIFERATION | 4 | 2 | 4161 | 153 | CG12099,N | | 7.777e-03 | -4.86 | striated muscle tissue development | biological process | GO:0014706 | 24 | 4 | 12345 | 414 | N,EcR,lbe,tin | | 7.777e-03 | -4.86 | embryonic heart tube development | biological process | GO:0035050 | 24 | 4 | 12345 | 414 | Mmp2,pon,robo2,tin | | 7.777e-03 | -4.86 | negative regulation of transmembrane receptor protein serine/threonine kinase signaling pathway | biological process | GO:0090101 | 24 | 4 | 12345 | 414 | hiw,Acsl,Lpin,Dad | | 7.779e-03 | -4.86 | GSE9988\_ANTI\_TREM1\_VS\_ANTI\_TREM1\_AND\_LPS\_MONOCYTE\_UP | MSigDB lists | GSE9988\_ANTI\_TREM1\_VS\_ANTI\_TREM1\_AND\_LPS\_MONOCYTE\_UP | 63 | 7 | 4161 | 153 | CG5830,SamDC,Snx3,emp,CG9220,Cyp1,Tsp26A | | 7.779e-03 | -4.86 | TARTE\_PLASMA\_CELL\_VS\_PLASMABLAST\_UP | MSigDB lists | TARTE\_PLASMA\_CELL\_VS\_PLASMABLAST\_UP | 63 | 7 | 4161 | 153 | CG14291,LRP1,Fas2,smal,if,Syx1A,Taf1 | | 7.779e-03 | -4.86 | GO\_NEGATIVE\_REGULATION\_OF\_NERVOUS\_SYSTEM\_DEVELOPMENT | MSigDB lists | GO\_NEGATIVE\_REGULATION\_OF\_NERVOUS\_SYSTEM\_DEVELOPMENT | 63 | 7 | 4161 | 153 | CG12099,LRP1,Ser,hh,NetB,N,CG5830 | | 7.823e-03 | -4.85 | GO\_RECEPTOR\_ACTIVITY | MSigDB lists | GO\_RECEPTOR\_ACTIVITY | 186 | 14 | 4161 | 153 | mgl,N,Ptr,Fas2,smal,Npc1a,nAChRalpha4,tkv,Hydr2,EcR,trn,emp,LRP1,dlp | | 7.823e-03 | -4.85 | RNGTGGGC\_UNKNOWN | MSigDB lists | RNGTGGGC\_UNKNOWN | 186 | 14 | 4161 | 153 | Sin3A,trn,hh,LRP1,dlp,Tlk,Imp,cta,S6kII,mgl,Arl4,sm,CG31637,Fas2 | | 7.860e-03 | -4.85 | Ig\_sub2 | interpro domains | IPR003598 | 109 | 10 | 11383 | 428 | ImpL2,robo2,DIP-epsilon,dpr7,otk2,lbk,nrm,Lac,Dscam1,Fas2 | | 7.864e-03 | -4.85 | Nsf2 (N-ethylmaleimide-sensitive factor 2) | protein interactions | 41694 | 92 | 10 | 9204 | 412 | Syx1A,cpo,N,spi,NetB,Ykt6,Ser,wech,CycE,phtf | | 7.924e-03 | -4.84 | GSE28726\_NAIVE\_CD4\_TCELL\_VS\_NAIVE\_NKTCELL\_UP | MSigDB lists | GSE28726\_NAIVE\_CD4\_TCELL\_VS\_NAIVE\_NKTCELL\_UP | 79 | 8 | 4161 | 153 | Syx1A,CG2812,mmy,nudE,Tpr2,SamDC,Acsl,Mitofilin | | 7.968e-03 | -4.83 | tight junction | cellular component | GO:0070160 | 37 | 5 | 11940 | 407 | bbg,nrv2,sdt,Lac,Fas3 | | 7.986e-03 | -4.83 | PROTEIN\_KINASE\_DOM | prosite domains | PS50011 | 221 | 20 | 6243 | 316 | CaMKII,par-1,Egfr,Hipk,PhKgamma,tkv,Mos,PEK,mbt,KP78a,Cdk12,JIL-1,Tlk,Pak3,S6kII,Doa,wnd,Asator,CG8485,Haspin | | 8.047e-03 | -4.82 | AT\_hook\_DNA-bd\_motif | interpro domains | IPR017956 | 4 | 2 | 11383 | 428 | Thd1,Taf1 | | 8.047e-03 | -4.82 | MAD\_homology\_MH1 | interpro domains | IPR013019 | 4 | 2 | 11383 | 428 | Dad,Smox | | 8.047e-03 | -4.82 | Dwarfin | interpro domains | IPR013790 | 4 | 2 | 11383 | 428 | Smox,Dad | | 8.047e-03 | -4.82 | SMAD\_MH1\_sf | interpro domains | IPR036578 | 4 | 2 | 11383 | 428 | Dad,Smox | | 8.047e-03 | -4.82 | Integrin\_alpha\_C\_CS | interpro domains | IPR018184 | 4 | 2 | 11383 | 428 | if,scb | | 8.047e-03 | -4.82 | DDT\_dom | interpro domains | IPR018501 | 4 | 2 | 11383 | 428 | tou,Acf | | 8.069e-03 | -4.82 | CREB phosphorylation | REACTOME pathways | R-DME-199920 | 4 | 2 | 3923 | 148 | S6kII,JIL-1 | | 8.158e-03 | -4.81 | positive regulation of molecular function | biological process | GO:0044093 | 287 | 18 | 12345 | 414 | RhoGAP102A,RhoGAP16F,mbt,Atg17,Rho1,pbl,RhoGAP18B,Cdk5alpha,mof,wnd,mle,RhoGAP15B,RapGAP1,CG8531,nrv2,RhoGAP93B,bora,ttk | | 8.177e-03 | -4.81 | Psn (Presenilin) | protein interactions | 40260 | 40 | 6 | 9204 | 412 | so,hh,S,mam,N,eya | | 8.177e-03 | -4.81 | CG42369 (uncharacterized protein) | protein interactions | 33894 | 40 | 6 | 9204 | 412 | Tpr2,Df31,Pak3,Sin3A,CaMKII,stg | | 8.214e-03 | -4.80 | BENPORATH\_EED\_TARGETS | MSigDB lists | BENPORATH\_EED\_TARGETS | 168 | 13 | 4161 | 153 | mgl,Dscam1,Ppa,NetB,ap,Fas2,tkv,fne,CG5742,so,hh,dally,TfIIS | | 8.304e-03 | -4.79 | Znf\_LIM | interpro domains | IPR001781 | 34 | 5 | 11383 | 428 | esn,Lim1,smash,ap,Bx | | 8.334e-03 | -4.79 | nephrocyte differentiation | biological process | GO:0061319 | 13 | 3 | 12345 | 414 | tin,pnt,dpp | | 8.334e-03 | -4.79 | axon target recognition | biological process | GO:0007412 | 13 | 3 | 12345 | 414 | sbb,exex,CadN | | 8.334e-03 | -4.79 | positive regulation of epidermal growth factor receptor signaling pathway | biological process | GO:0045742 | 13 | 3 | 12345 | 414 | rau,edl,S | | 8.334e-03 | -4.79 | renal filtration cell differentiation | biological process | GO:0061318 | 13 | 3 | 12345 | 414 | tin,pnt,dpp | | 8.334e-03 | -4.79 | positive regulation of ERBB signaling pathway | biological process | GO:1901186 | 13 | 3 | 12345 | 414 | S,rau,edl | | 8.334e-03 | -4.79 | cell junction maintenance | biological process | GO:0034331 | 13 | 3 | 12345 | 414 | Rac2,Rho1,crb | | 8.334e-03 | -4.79 | regulation of development, heterochronic | biological process | GO:0040034 | 13 | 3 | 12345 | 414 | cas,Kr,EcR | | 8.374e-03 | -4.78 | GO\_CORE\_PROMOTER\_PROXIMAL\_REGION\_DNA\_BINDING | MSigDB lists | GO\_CORE\_PROMOTER\_PROXIMAL\_REGION\_DNA\_BINDING | 35 | 5 | 4161 | 153 | so,Smox,EcR,Utx,TfAP-2 | | 8.398e-03 | -4.78 | PID\_FGF\_PATHWAY | MSigDB lists | PID\_FGF\_PATHWAY | 12 | 3 | 4161 | 153 | Fas2,mbt,drk | | 8.398e-03 | -4.78 | GO\_HOMOPHILIC\_CELL\_ADHESION\_VIA\_PLASMA\_MEMBRANE\_ADHESION\_MOLECULES | MSigDB lists | GO\_HOMOPHILIC\_CELL\_ADHESION\_VIA\_PLASMA\_MEMBRANE\_ADHESION\_MOLECULES | 12 | 3 | 4161 | 153 | ds,ft,Dscam1 | | 8.398e-03 | -4.78 | PID\_PRL\_SIGNALING\_EVENTS\_PATHWAY | MSigDB lists | PID\_PRL\_SIGNALING\_EVENTS\_PATHWAY | 12 | 3 | 4161 | 153 | Rac2,CycE,Rho1 | | 8.398e-03 | -4.78 | GO\_VASCULOGENESIS | MSigDB lists | GO\_VASCULOGENESIS | 12 | 3 | 4161 | 153 | slow,N,hh | | 8.398e-03 | -4.78 | GO\_CHONDROCYTE\_DIFFERENTIATION | MSigDB lists | GO\_CHONDROCYTE\_DIFFERENTIATION | 12 | 3 | 4161 | 153 | tkv,dpp,CG9220 | | 8.398e-03 | -4.78 | GO\_ANTERIOR\_POSTERIOR\_AXIS\_SPECIFICATION | MSigDB lists | GO\_ANTERIOR\_POSTERIOR\_AXIS\_SPECIFICATION | 12 | 3 | 4161 | 153 | Utx,Lim1,hh | | 8.398e-03 | -4.78 | GO\_CELLULAR\_RESPONSE\_TO\_CALCIUM\_ION | MSigDB lists | GO\_CELLULAR\_RESPONSE\_TO\_CALCIUM\_ION | 12 | 3 | 4161 | 153 | CaMKII,pbl,Cyp1 | | 8.398e-03 | -4.78 | MARIADASON\_RESPONSE\_TO\_BUTYRATE\_SULINDAC\_6 | MSigDB lists | MARIADASON\_RESPONSE\_TO\_BUTYRATE\_SULINDAC\_6 | 12 | 3 | 4161 | 153 | mle,CG5830,betaTub56D | | 8.398e-03 | -4.78 | GO\_POSITIVE\_REGULATION\_OF\_NOTCH\_SIGNALING\_PATHWAY | MSigDB lists | GO\_POSITIVE\_REGULATION\_OF\_NOTCH\_SIGNALING\_PATHWAY | 12 | 3 | 4161 | 153 | Ser,Tsp26A,N | | 8.398e-03 | -4.78 | GO\_POSITIVE\_REGULATION\_OF\_STEM\_CELL\_PROLIFERATION | MSigDB lists | GO\_POSITIVE\_REGULATION\_OF\_STEM\_CELL\_PROLIFERATION | 12 | 3 | 4161 | 153 | N,so,hh | | 8.398e-03 | -4.78 | GO\_CELL\_DIFFERENTIATION\_IN\_SPINAL\_CORD | MSigDB lists | GO\_CELL\_DIFFERENTIATION\_IN\_SPINAL\_CORD | 12 | 3 | 4161 | 153 | N,hh,Lim1 | | 8.444e-03 | -4.77 | cv-c (crossveinless c) | protein interactions | 41749 | 10 | 3 | 9204 | 412 | Egfr,spi,Rho1 | | 8.444e-03 | -4.77 | Dlish (Dachs ligand with SH3s) | protein interactions | 37014 | 10 | 3 | 9204 | 412 | ds,Fbxl7,ft | | 8.444e-03 | -4.77 | Mtpalpha (Mitochondrial trifunctional protein alpha subunit) | protein interactions | 34276 | 10 | 3 | 9204 | 412 | fj,ft,ds | | 8.444e-03 | -4.77 | Smyd5 (SET and MYND domain containing, class 5) | protein interactions | 42517 | 10 | 3 | 9204 | 412 | NFAT,CG3919,esn | | 8.444e-03 | -4.77 | LRP1 (LDL receptor protein 1) | protein interactions | 35799 | 10 | 3 | 9204 | 412 | ci,Fas2,sty | | 8.444e-03 | -4.77 | sv (shaven) | protein interactions | 43825 | 10 | 3 | 9204 | 412 | CycE,lz,ttk | | 8.444e-03 | -4.77 | Rtf1 (Rtf1) | protein interactions | 37554 | 10 | 3 | 9204 | 412 | N,Fas2,S6kII | | 8.444e-03 | -4.77 | CG12163 (uncharacterized protein) | protein interactions | 40628 | 10 | 3 | 9204 | 412 | Fas2,ap,Bx | | 8.444e-03 | -4.77 | Timp (Tissue inhibitor of metalloproteases) | protein interactions | 41248 | 10 | 3 | 9204 | 412 | Fas2,Sox21a,Mmp2 | | 8.472e-03 | -4.77 | E2F1\_Q4\_01 | MSigDB lists | E2F1\_Q4\_01 | 64 | 7 | 4161 | 153 | JIL-1,fne,Sin3A,CG8531,NiPp1,Lac,CG12547 | | 8.476e-03 | -4.77 | GSE41978\_WT\_VS\_ID2\_KO\_KLRG1\_LOW\_EFFECTOR\_CD8\_TCELL\_DN | MSigDB lists | GSE41978\_WT\_VS\_ID2\_KO\_KLRG1\_LOW\_EFFECTOR\_CD8\_TCELL\_DN | 49 | 6 | 4161 | 153 | nudE,Lim1,Smox,Arl4,Ppa,Mmp2 | | 8.476e-03 | -4.77 | GSE37301\_PRO\_BCELL\_VS\_RAG2\_KO\_NK\_CELL\_DN | MSigDB lists | GSE37301\_PRO\_BCELL\_VS\_RAG2\_KO\_NK\_CELL\_DN | 49 | 6 | 4161 | 153 | CaMKII,CG5522,CG11367,phtf,CG9220,CG12547 | | 8.476e-03 | -4.77 | GGCAGCT\_MIR22 | MSigDB lists | GGCAGCT\_MIR22 | 49 | 6 | 4161 | 153 | MFS10,CG5830,Tpr2,Nhe2,nudE,Tlk | | 8.481e-03 | -4.77 | Pvr (PDGF- and VEGF-receptor related) | protein interactions | 34127 | 79 | 9 | 9204 | 412 | robo2,sty,Ptpmeg,Tlk,drk,Egfr,NFAT,tou,CG10082 | | 8.481e-03 | -4.77 | Cdk4 (Cyclin-dependent kinase 4) | protein interactions | 36854 | 79 | 9 | 9204 | 412 | Stat92E,Pits,EcR,CG2865,gus,pbl,CG11486,hh,CycE | | 8.488e-03 | -4.77 | DDT | smart domains | SM00571 | 3 | 2 | 5086 | 276 | tou,Acf | | 8.560e-03 | -4.76 | vari (varicose) | protein interactions | 35343 | 29 | 5 | 9204 | 412 | bab1,egr,rau,Fas3,pio | | 8.560e-03 | -4.76 | ovo (ovo) | protein interactions | 31429 | 29 | 5 | 9204 | 412 | spi,mle,Kr,pbl,Egfr | | 8.560e-03 | -4.76 | pros (prospero) | protein interactions | 41363 | 29 | 5 | 9204 | 412 | aop,pnt,CycE,spoon,Chro | | 8.595e-03 | -4.76 | S\_TKc | smart domains | SM00220 | 167 | 17 | 5086 | 276 | wnd,Asator,Hipk,CaMKII,PEK,KP78a,Cdk12,Tlk,par-1,JIL-1,PhKgamma,Doa,tkv,CG8485,Pak3,S6kII,Mos | | 8.698e-03 | -4.74 | gene silencing | biological process | GO:0016458 | 143 | 11 | 12345 | 414 | NiPp1,CG17168,AGO2,CG17265,Psc,Hsc70-3,Kr,psq,bin3,Tlk,Tudor-SN | | 8.714e-03 | -4.74 | basal plasma membrane | cellular component | GO:0009925 | 13 | 3 | 11940 | 407 | Egfr,scb,if | | 8.803e-03 | -4.73 | fly chr26B2-26B2\|2-18 | chromosome location | fly chr26B2-26B2\|2-18 | 7 | 2 | 25010 | 531 | asRNA:CR43866,Tsp26A | | 8.803e-03 | -4.73 | fly chr26B2-26B2\|2-18 cM | chromosome location | fly chr26B2-26B2\|2-18 cM | 7 | 2 | 25010 | 531 | asRNA:CR43866,Tsp26A | | 8.815e-03 | -4.73 | positive regulation of response to external stimulus | biological process | GO:0032103 | 124 | 10 | 12345 | 414 | Doa,Imp,Toll-7,Etl1,mmy,Pli,Mmp2,Tl,ush,CG4325 | | 8.832e-03 | -4.73 | learning | biological process | GO:0007612 | 70 | 7 | 12345 | 414 | Fas2,drk,spi,wcy,S6kII,scb,Egfr | | 8.890e-03 | -4.72 | Src64B (Src oncogene at 64B) | protein interactions | 48973 | 66 | 8 | 9204 | 412 | CG33993,Rho1,pbl,Egfr,drk,udd,Sin3A,betaTub56D | | 8.896e-03 | -4.72 | Ig\_sub | interpro domains | IPR003599 | 111 | 10 | 11383 | 428 | Dscam1,Fas2,Lac,dpr7,lbk,otk2,nrm,DIP-epsilon,robo2,ImpL2 | | 8.943e-03 | -4.72 | RA | pfam domains | PF00788 | 12 | 3 | 10480 | 393 | wake,rau,CG32758 | | 8.969e-03 | -4.71 | GO\_NEGATIVE\_REGULATION\_OF\_MULTICELLULAR\_ORGANISMAL\_PROCESS | MSigDB lists | GO\_NEGATIVE\_REGULATION\_OF\_MULTICELLULAR\_ORGANISMAL\_PROCESS | 189 | 14 | 4161 | 153 | so,CG9220,CG12099,CG11367,hh,LRP1,Rac2,JIL-1,dpp,Ser,NetB,N,CG5830,Smox | | 8.978e-03 | -4.71 | Hipk (Homeodomain interacting protein kinase) | protein interactions | 38070 | 19 | 4 | 9204 | 412 | ey,Hipk,spen,N | | 8.978e-03 | -4.71 | Tnks (tankyrase) | protein interactions | 43095 | 19 | 4 | 9204 | 412 | Myo10A,spi,CG32369,Bx | | 8.978e-03 | -4.71 | lap (like-AP180) | protein interactions | 40863 | 19 | 4 | 9204 | 412 | jigr1,jing,nuf,cnc | | 8.978e-03 | -4.71 | KP78b (KP78b) | protein interactions | 41361 | 19 | 4 | 9204 | 412 | Doa,cas,CG8929,CG2865 | | 8.978e-03 | -4.71 | gish (gilgamesh) | protein interactions | 49701 | 19 | 4 | 9204 | 412 | so,hh,pbl,ey | | 8.978e-03 | -4.71 | Dscam1 (Down syndrome cell adhesion molecule 1) | protein interactions | 35652 | 19 | 4 | 9204 | 412 | Dscam1,hiw,NetB,wnd | | 9.008e-03 | -4.71 | Signaling by BMP | REACTOME pathways | R-DME-201451 | 12 | 3 | 3923 | 148 | dpp,tkv,Dad | | 9.008e-03 | -4.71 | HS-GAG degradation | REACTOME pathways | R-DME-2024096 | 12 | 3 | 3923 | 148 | dally,CG14291,dlp | | 9.014e-03 | -4.71 | GO\_EMBRYONIC\_HEART\_TUBE\_DEVELOPMENT | MSigDB lists | GO\_EMBRYONIC\_HEART\_TUBE\_DEVELOPMENT | 23 | 4 | 4161 | 153 | N,Smox,dpp,hh | | 9.014e-03 | -4.71 | BIOCARTA\_MAPK\_PATHWAY | MSigDB lists | BIOCARTA\_MAPK\_PATHWAY | 23 | 4 | 4161 | 153 | Rac2,drk,S6kII,JIL-1 | | 9.014e-03 | -4.71 | GSE15624\_3H\_VS\_6H\_HALOFUGINONE\_TREATED\_CD4\_TCELL\_UP | MSigDB lists | GSE15624\_3H\_VS\_6H\_HALOFUGINONE\_TREATED\_CD4\_TCELL\_UP | 23 | 4 | 4161 | 153 | NTPase,CG5830,JIL-1,Hsc70-3 | | 9.014e-03 | -4.71 | GO\_CAMERA\_TYPE\_EYE\_MORPHOGENESIS | MSigDB lists | GO\_CAMERA\_TYPE\_EYE\_MORPHOGENESIS | 23 | 4 | 4161 | 153 | TfAP-2,fj,Lim1,Ser | | 9.014e-03 | -4.71 | SERVITJA\_ISLET\_HNF1A\_TARGETS\_UP | MSigDB lists | SERVITJA\_ISLET\_HNF1A\_TARGETS\_UP | 23 | 4 | 4161 | 153 | Rho1,fne,Smox,DIP-epsilon | | 9.018e-03 | -4.71 | epithelial structure maintenance | biological process | GO:0010669 | 25 | 4 | 12345 | 414 | Lac,Atg17,Egfr,if | | 9.018e-03 | -4.71 | regulation of G1/S transition of mitotic cell cycle | biological process | GO:2000045 | 25 | 4 | 12345 | 414 | ci,N,Egfr,CycE | | 9.033e-03 | -4.71 | positive regulation of cellular component organization | biological process | GO:0051130 | 290 | 18 | 12345 | 414 | drk,CG11486,Pak3,nudE,Imp,Pcl,tkv,Arp2,sdt,Prosap,Stat92E,tsl,CG43658,mmy,Hip1,mle,PsGEF,comm | | 9.043e-03 | -4.71 | GO\_ANCHORING\_JUNCTION | MSigDB lists | GO\_ANCHORING\_JUNCTION | 170 | 13 | 4161 | 153 | LRP1,Hsc70-3,RpS4,emp,mbt,Rac2,Ser,Arp2,smal,if,cta,N,hts | | 9.073e-03 | -4.70 | LAM\_G\_DOMAIN | prosite domains | PS50025 | 17 | 4 | 6243 | 316 | crb,wb,CadN,ft | | 9.073e-03 | -4.70 | ZP\_2 | prosite domains | PS51034 | 17 | 4 | 6243 | 316 | pio,neo,qsm,tyn | | 9.103e-03 | -4.70 | JAK-STAT signaling | KEGG pathways | dme\_M00684 | 5 | 2 | 3198 | 100 | Socs16D,Stat92E | | 9.103e-03 | -4.70 | JAK-STAT signaling | KEGG pathways | M00684 | 5 | 2 | 3198 | 100 | Stat92E,Socs16D | | 9.119e-03 | -4.70 | cell maturation | biological process | GO:0048469 | 184 | 13 | 12345 | 414 | par-1,cnc,N,Hydr2,psq,Syp,Egfr,heph,Imp,Pat1,dpp,EcR,gus | | 9.192e-03 | -4.69 | CA | smart domains | SM00112 | 16 | 4 | 5086 | 276 | CadN,Cad87A,ft,ds | | 9.192e-03 | -4.69 | ZP | smart domains | SM00241 | 16 | 4 | 5086 | 276 | qsm,tyn,pio,neo | | 9.209e-03 | -4.69 | GO\_NEGATIVE\_REGULATION\_OF\_CELL\_DEVELOPMENT | MSigDB lists | GO\_NEGATIVE\_REGULATION\_OF\_CELL\_DEVELOPMENT | 65 | 7 | 4161 | 153 | LRP1,Ser,hh,CG12099,N,CG5830,NetB | | 9.209e-03 | -4.69 | GO\_REGULATION\_OF\_CELL\_MORPHOGENESIS\_INVOLVED\_IN\_DIFFERENTIATION | MSigDB lists | GO\_REGULATION\_OF\_CELL\_MORPHOGENESIS\_INVOLVED\_IN\_DIFFERENTIATION | 65 | 7 | 4161 | 153 | dpp,nudE,Rac2,NetB,nAChRalpha4,Smox,N | | 9.209e-03 | -4.69 | E2F\_Q3\_01 | MSigDB lists | E2F\_Q3\_01 | 65 | 7 | 4161 | 153 | Lac,NiPp1,CG12547,JIL-1,fne,Sin3A,CG8531 | | 9.210e-03 | -4.69 | GO\_KINASE\_ACTIVITY | MSigDB lists | GO\_KINASE\_ACTIVITY | 229 | 16 | 4161 | 153 | CG34384,CycE,drk,tkv,Tlk,CG1271,S6kII,par-1,CaMKII,Taf1,PhKgamma,smal,JIL-1,mbt,Doa,CG2201 | | 9.227e-03 | -4.69 | z (zeste) | protein interactions | 31230 | 41 | 6 | 9204 | 412 | ap,osa,Ubx,Pli,lz,Psc | | 9.227e-03 | -4.69 | Sos (Son of sevenless) | protein interactions | 34790 | 41 | 6 | 9204 | 412 | egr,Rho1,aop,N,Egfr,drk | | 9.267e-03 | -4.68 | regulation of immune effector process | biological process | GO:0002697 | 88 | 8 | 12345 | 414 | Doa,Toll-7,Usp2,Stat92E,Pli,Tl,ush,CG4325 | | 9.323e-03 | -4.68 | positive regulation of mitotic cell cycle | biological process | GO:0045931 | 39 | 5 | 12345 | 414 | CycE,stg,ci,N,Myb | | 9.323e-03 | -4.68 | TGCACTT\_MIR519C\_MIR519B\_MIR519A | MSigDB lists | TGCACTT\_MIR519C\_MIR519B\_MIR519A | 98 | 9 | 4161 | 153 | phtf,l(3)mbt,Ptpmeg,nudE,CG41099,CG11367,Imp,fj,Hydr2 | | 9.343e-03 | -4.67 | protein phosphatase 1 binding | molecular function | GO:0008157 | 26 | 4 | 12176 | 396 | Gbs-70E,NiPp1,CG42748,ec | | 9.347e-03 | -4.67 | KRAS.600\_UP.V1\_UP | MSigDB lists | KRAS.600\_UP.V1\_UP | 50 | 6 | 4161 | 153 | DIP-epsilon,Pli,Mmp2,Ank,Syx1A,nAChRalpha4 | | 9.347e-03 | -4.67 | GO\_REGULATION\_OF\_EPITHELIAL\_CELL\_PROLIFERATION | MSigDB lists | GO\_REGULATION\_OF\_EPITHELIAL\_CELL\_PROLIFERATION | 50 | 6 | 4161 | 153 | so,hh,dpp,N,Smox,slow | | 9.347e-03 | -4.67 | TAATGTG\_MIR323 | MSigDB lists | TAATGTG\_MIR323 | 50 | 6 | 4161 | 153 | nudE,NTPase,gus,Utx,S6kII,Etl1 | | 9.371e-03 | -4.67 | regulation of catalytic activity | biological process | GO:0050790 | 450 | 25 | 12345 | 414 | RhoGAP15B,sty,mle,wnd,nrv2,NiPp1,CG8531,RapGAP1,bora,CycE,Pli,RhoGAP93B,nudE,RhoGAP102A,Acf,Rho1,aos,CG8441,Atg17,RhoGAP16F,mbt,Cdk5alpha,pbl,RhoGAP18B,CG42684 | | 9.396e-03 | -4.67 | integral component of plasma membrane | cellular component | GO:0005887 | 466 | 26 | 11940 | 407 | otk2,ft,if,CadN,Tsp39D,emp,Tsp26A,smal,Dscam1,DIP-epsilon,Cirl,egr,Prosap,Cad87A,tkv,N,nrv2,nAChRalpha4,Egfr,ds,wun2,Tl,scb,dpr7,Ten-m,sas | | 9.418e-03 | -4.67 | Hedgehog Signaling Pathway | WikiPathways | WP492 | 10 | 3 | 499 | 24 | hh,ci,Sin3A | | 9.443e-03 | -4.66 | GSE12392\_IFNAR\_KO\_VS\_IFNB\_KO\_CD8\_NEG\_SPLEEN\_DC\_UP | MSigDB lists | GSE12392\_IFNAR\_KO\_VS\_IFNB\_KO\_CD8\_NEG\_SPLEEN\_DC\_UP | 36 | 5 | 4161 | 153 | cta,mmy,Cyp1,nudE,Tif-IA | | 9.443e-03 | -4.66 | AHRARNT\_01 | MSigDB lists | AHRARNT\_01 | 36 | 5 | 4161 | 153 | Lim1,Ser,CaMKII,fne,ds | | 9.482e-03 | -4.66 | GTGCCTT\_MIR506 | MSigDB lists | GTGCCTT\_MIR506 | 171 | 13 | 4161 | 153 | CG5830,Etl1,ap,Cyp1,Lac,Ser,MFS10,Pcl,heph,drk,Imp,CG9220,Mrp4 | | 9.514e-03 | -4.66 | Rac GTPase binding | molecular function | GO:0048365 | 14 | 3 | 12176 | 396 | Pak3,mbt,pbl | | 9.524e-03 | -4.65 | regulation of humoral immune response | biological process | GO:0002920 | 71 | 7 | 12345 | 414 | Doa,Pli,Stat92E,Usp2,CG4325,ush,Tl | | 9.550e-03 | -4.65 | GO\_NEGATIVE\_REGULATION\_OF\_CELL\_DEATH | MSigDB lists | GO\_NEGATIVE\_REGULATION\_OF\_CELL\_DEATH | 210 | 15 | 4161 | 153 | CG45050,Sin3A,so,Hsc70-3,hh,LRP1,Smox,N,S6kII,TfAP-2,Cyp1,Tif-IA,Npc1a,bor,Mhcl | | 9.568e-03 | -4.65 | axon part | cellular component | GO:0033267 | 105 | 9 | 11940 | 407 | NetA,Con,Ser,Syx1A,NetB,par-1,Prosap,Cdk5alpha,Fas2 | | 9.592e-03 | -4.65 | nucleoside-triphosphatase regulator activity | molecular function | GO:0060589 | 110 | 9 | 12176 | 396 | RhoGAP93B,CG8531,nrv2,RhoGAP16F,pbl,RhoGAP102A,RapGAP1,RhoGAP18B,RhoGAP15B | | 9.623e-03 | -4.64 | regulation of catabolic process | biological process | GO:0009894 | 227 | 15 | 12345 | 414 | Utx,ctrip,phyl,smg,Smox,crb,N,PEK,Gyf,Doa,Usp2,wcy,gus,EcR,Toll-7 | | 9.713e-03 | -4.63 | Jra (Jun-related antigen) | protein interactions | 36057 | 67 | 8 | 9204 | 412 | tkv,phyl,Sin3A,pnt,Stat92E,Egfr,Mocs2B,aop | | 9.809e-03 | -4.62 | apical cortex | cellular component | GO:0045179 | 54 | 6 | 11940 | 407 | Rho1,sdt,Ser,CG42684,nuf,RhoGEF2 | | 9.867e-03 | -4.62 | NABA\_MATRISOME | MSigDB lists | NABA\_MATRISOME | 82 | 8 | 4161 | 153 | Mmp2,slow,hh,dlp,dally,CG10663,NetB,dpp | | 9.885e-03 | -4.62 | tau-protein kinase activity | molecular function | GO:0050321 | 5 | 2 | 12176 | 396 | KP78a,par-1 | | 9.885e-03 | -4.62 | collagen binding | molecular function | GO:0005518 | 5 | 2 | 12176 | 396 | dpp,smal | | 9.894e-03 | -4.62 | programmed cell death involved in cell development | biological process | GO:0010623 | 89 | 8 | 12345 | 414 | S,Atg17,dpp,EcR,Utx,ec,wun2,N | | 9.896e-03 | -4.62 | vesicle | cellular component | GO:0031982 | 422 | 24 | 11940 | 407 | rush,Ptr,Fas3,hiw,nuf,CG34384,Syx1A,hh,S,Dtg,Ykt6,comm,dbr,CG13784,CG32758,Tl,Rac2,Snx3,Hip1,CG6051,Cirl,dpp,N,tkv | | 9.904e-03 | -4.61 | ash2 (absent, small, or homeotic discs 2) | protein interactions | 42936 | 30 | 5 | 9204 | 412 | Utx,Egfr,Ubx,kni,CycE | | 9.904e-03 | -4.61 | mxc (multi sex combs) | protein interactions | 31869 | 30 | 5 | 9204 | 412 | Pcl,Psc,Tl,dpp,hh | | 9.904e-03 | -4.61 | Chro (Chromator) | protein interactions | 40508 | 30 | 5 | 9204 | 412 | l(3)mbt,JIL-1,jigr1,mof,Pits | | 9.957e-03 | -4.61 | serine/threonine protein kinase complex | cellular component | GO:1902554 | 39 | 5 | 11940 | 407 | PhKgamma,Atg17,CycE,Cdk12,Cdk5alpha | | 1.005e-02 | -4.60 | - | gene3d domains | 3.90.520.10 | 4 | 2 | 4929 | 208 | Smox,Dad | | 1.013e-02 | -4.59 | - | gene3d domains | 3.30.505.10 | 32 | 5 | 4929 | 208 | Socs16D,CG10479,CG33993,drk,Stat92E | | 1.019e-02 | -4.59 | zinc ion binding | molecular function | GO:0008270 | 516 | 27 | 12176 | 396 | TfIIS,CG32369,l(3)mbt,Mmp2,CG12099,tna,Hph,kni,CG45050,dbr,CG9855,CG11247,Psc,wech,CG3097,Zif,CG6752,ctrip,hiw,CG4325,Tet,CG2991,EcR,bor,esn,shn,hang | | 1.028e-02 | -4.58 | GSE39152\_SPLEEN\_CD103\_NEG\_VS\_BRAIN\_CD103\_POS\_MEMORY\_CD8\_TCELL\_UP | MSigDB lists | GSE39152\_SPLEEN\_CD103\_NEG\_VS\_BRAIN\_CD103\_POS\_MEMORY\_CD8\_TCELL\_UP | 51 | 6 | 4161 | 153 | CG9281,CG4538,PhKgamma,Lim1,betaTub56D,Tpr2 | | 1.028e-02 | -4.58 | GR\_Q6 | MSigDB lists | GR\_Q6 | 51 | 6 | 4161 | 153 | Lac,so,Fas2,emp,fne,pAbp | | 1.028e-02 | -4.58 | GO\_NEURAL\_TUBE\_DEVELOPMENT | MSigDB lists | GO\_NEURAL\_TUBE\_DEVELOPMENT | 51 | 6 | 4161 | 153 | N,ds,ap,Utx,wech,hh | | 1.028e-02 | -4.58 | TFIIA\_Q6 | MSigDB lists | TFIIA\_Q6 | 51 | 6 | 4161 | 153 | so,Pli,dlp,kdn,Tpr2,lili | | 1.035e-02 | -4.57 | organ or tissue specific immune response | biological process | GO:0002251 | 14 | 3 | 12345 | 414 | bbg,hh,ci | | 1.035e-02 | -4.57 | mucosal immune response | biological process | GO:0002385 | 14 | 3 | 12345 | 414 | bbg,hh,ci | | 1.035e-02 | -4.57 | regulation of wound healing | biological process | GO:0061041 | 14 | 3 | 12345 | 414 | Rho1,Rac2,Egfr | | 1.035e-02 | -4.57 | negative regulation of hemocyte differentiation | biological process | GO:0045611 | 14 | 3 | 12345 | 414 | ush,N,Ser | | 1.035e-02 | -4.57 | dorsal closure, amnioserosa morphology change | biological process | GO:0046664 | 14 | 3 | 12345 | 414 | tkv,crb,Rho1 | | 1.035e-02 | -4.57 | muscle cell fate commitment | biological process | GO:0042693 | 14 | 3 | 12345 | 414 | slou,Ubx,N | | 1.035e-02 | -4.57 | response to molecule of bacterial origin | biological process | GO:0002237 | 14 | 3 | 12345 | 414 | Stat92E,EcR,Dscam1 | | 1.037e-02 | -4.57 | positive regulation of TOR signaling | biological process | GO:0032008 | 40 | 5 | 12345 | 414 | Socs16D,Mos,drk,wcy,ci | | 1.037e-02 | -4.57 | regulation of cell maturation | biological process | GO:1903429 | 40 | 5 | 12345 | 414 | Syx1A,EcR,Fas2,par-1,cnc | | 1.037e-02 | -4.57 | ed (echinoid) | protein interactions | 33619 | 42 | 6 | 9204 | 412 | Egfr,N,pnt,aop,aos,sty | | 1.037e-02 | -4.57 | bip2 (bip2) | protein interactions | 43793 | 42 | 6 | 9204 | 412 | BtbVII,CG18766,CG13894,CG11498,S6kII,ttk | | 1.038e-02 | -4.57 | regulation of developmental pigmentation | biological process | GO:0048070 | 26 | 4 | 12345 | 414 | mgl,bab1,S6kII,Notum | | 1.038e-02 | -4.57 | regulation of imaginal disc-derived wing size | biological process | GO:0044719 | 26 | 4 | 12345 | 414 | slow,fj,CycE,nw | | 1.038e-02 | -4.57 | positive regulation of gene expression, epigenetic | biological process | GO:0045815 | 26 | 4 | 12345 | 414 | tara,JIL-1,lncRNA:roX1,tna | | 1.038e-02 | -4.57 | regulation of embryonic pattern specification | biological process | GO:1902875 | 26 | 4 | 12345 | 414 | Syx1A,Tl,par-1,cnc | | 1.044e-02 | -4.56 | posttranscriptional regulation of gene expression | biological process | GO:0010608 | 208 | 14 | 12345 | 414 | smg,pAbp,Rbfox1,bin3,Hsc70-3,elav,PEK,Syp,CG17168,mle,CG17265,AGO2,heph,NiPp1 | | 1.047e-02 | -4.56 | Toll Like Receptor 4 (TLR4) Cascade | REACTOME pathways | R-DME-166016 | 36 | 5 | 3923 | 148 | S6kII,Pli,mle,JIL-1,Ptpmeg | | 1.049e-02 | -4.56 | notum cell fate specification | biological process | GO:0035310 | 5 | 2 | 12345 | 414 | salm,Egfr | | 1.049e-02 | -4.56 | embryonic epithelial tube formation | biological process | GO:0001838 | 5 | 2 | 12345 | 414 | Mmp2,mmy | | 1.049e-02 | -4.56 | signal transduction in response to DNA damage | biological process | GO:0042770 | 5 | 2 | 12345 | 414 | PhKgamma,mof | | 1.049e-02 | -4.56 | dosage compensation complex assembly involved in dosage compensation by hyperactivation of X chromosome | biological process | GO:0016457 | 5 | 2 | 12345 | 414 | lncRNA:roX1,mle | | 1.049e-02 | -4.56 | regulation of cardiac cell fate specification | biological process | GO:2000043 | 5 | 2 | 12345 | 414 | N,pnt | | 1.049e-02 | -4.56 | mesodermal cell fate determination | biological process | GO:0007500 | 5 | 2 | 12345 | 414 | mam,twi | | 1.049e-02 | -4.56 | regulation of photoreceptor cell axon guidance | biological process | GO:2000289 | 5 | 2 | 12345 | 414 | NetB,NetA | | 1.049e-02 | -4.56 | terminal cell fate specification, open tracheal system | biological process | GO:0035154 | 5 | 2 | 12345 | 414 | hh,pnt | | 1.049e-02 | -4.56 | muscle cell fate determination | biological process | GO:0007521 | 5 | 2 | 12345 | 414 | slou,N | | 1.049e-02 | -4.56 | positive regulation of crystal cell differentiation | biological process | GO:0042691 | 5 | 2 | 12345 | 414 | N,lz | | 1.049e-02 | -4.56 | rhabdomere morphogenesis | biological process | GO:0061541 | 5 | 2 | 12345 | 414 | oc,crb | | 1.049e-02 | -4.56 | regulation of rhodopsin gene expression | biological process | GO:0007468 | 5 | 2 | 12345 | 414 | EcR,oc | | 1.049e-02 | -4.56 | dorsal/ventral lineage restriction, imaginal disc | biological process | GO:0007451 | 5 | 2 | 12345 | 414 | ap,N | | 1.049e-02 | -4.56 | regulation of cardioblast cell fate specification | biological process | GO:0042686 | 5 | 2 | 12345 | 414 | pnt,N | | 1.049e-02 | -4.56 | regulation of compound eye cone cell fate specification | biological process | GO:0042682 | 5 | 2 | 12345 | 414 | S6kII,ttk | | 1.051e-02 | -4.56 | osk (oskar) | protein interactions | 41066 | 96 | 10 | 9204 | 412 | tkv,Myo10A,Syp,pAbp,Cad87A,heph,par-1,smg,Imp,yps | | 1.051e-02 | -4.56 | LEE\_TARGETS\_OF\_PTCH1\_AND\_SUFU\_DN | MSigDB lists | LEE\_TARGETS\_OF\_PTCH1\_AND\_SUFU\_DN | 24 | 4 | 4161 | 153 | Fas2,DIP-epsilon,Tsp26A,CaMKII | | 1.051e-02 | -4.56 | GO\_CARDIOCYTE\_DIFFERENTIATION | MSigDB lists | GO\_CARDIOCYTE\_DIFFERENTIATION | 24 | 4 | 4161 | 153 | Ser,dpp,N,CaMKII | | 1.051e-02 | -4.56 | GO\_MESENCHYMAL\_CELL\_DIFFERENTIATION | MSigDB lists | GO\_MESENCHYMAL\_CELL\_DIFFERENTIATION | 24 | 4 | 4161 | 153 | N,dpp,Ser,hh | | 1.060e-02 | -4.55 | GO\_RAS\_GUANYL\_NUCLEOTIDE\_EXCHANGE\_FACTOR\_ACTIVITY | MSigDB lists | GO\_RAS\_GUANYL\_NUCLEOTIDE\_EXCHANGE\_FACTOR\_ACTIVITY | 37 | 5 | 4161 | 153 | CG5522,Fas2,drk,CaMKII,pbl | | 1.060e-02 | -4.55 | GSE22527\_ANTI\_CD3\_INVIVO\_VS\_UNTREATED\_MOUSE\_TREG\_UP | MSigDB lists | GSE22527\_ANTI\_CD3\_INVIVO\_VS\_UNTREATED\_MOUSE\_TREG\_UP | 37 | 5 | 4161 | 153 | Cyp1,emp,CG12099,Syx1A,SamDC | | 1.063e-02 | -4.54 | ACAWNRNSRCGG\_UNKNOWN | MSigDB lists | ACAWNRNSRCGG\_UNKNOWN | 13 | 3 | 4161 | 153 | Pli,heph,mbt | | 1.063e-02 | -4.54 | GO\_SOMITOGENESIS | MSigDB lists | GO\_SOMITOGENESIS | 13 | 3 | 4161 | 153 | Lim1,Smox,Utx | | 1.063e-02 | -4.54 | GSE36095\_WT\_VS\_HDAC9\_KO\_TREG\_DN | MSigDB lists | GSE36095\_WT\_VS\_HDAC9\_KO\_TREG\_DN | 13 | 3 | 4161 | 153 | tkv,Imp,Tif-IA | | 1.063e-02 | -4.54 | GO\_REGULATION\_OF\_GLIAL\_CELL\_DIFFERENTIATION | MSigDB lists | GO\_REGULATION\_OF\_GLIAL\_CELL\_DIFFERENTIATION | 13 | 3 | 4161 | 153 | N,dpp,hh | | 1.063e-02 | -4.54 | GO\_AUDITORY\_RECEPTOR\_CELL\_DIFFERENTIATION | MSigDB lists | GO\_AUDITORY\_RECEPTOR\_CELL\_DIFFERENTIATION | 13 | 3 | 4161 | 153 | Rac2,N,Ser | | 1.063e-02 | -4.54 | GO\_POSITIVE\_REGULATION\_OF\_LIPASE\_ACTIVITY | MSigDB lists | GO\_POSITIVE\_REGULATION\_OF\_LIPASE\_ACTIVITY | 13 | 3 | 4161 | 153 | Rho1,cta,EcR | | 1.081e-02 | -4.53 | supramolecular fiber organization | biological process | GO:0097435 | 188 | 13 | 12345 | 414 | Rho1,Arp2,Ten-m,RhoGEF2,nudE,hts,CG13366,Hip1,Pak3,N,tyn,Rac2,if | | 1.081e-02 | -4.53 | subapical complex | cellular component | GO:0035003 | 14 | 3 | 11940 | 407 | sdt,crb,N | | 1.081e-02 | -4.53 | muscle tendon junction | cellular component | GO:0005927 | 14 | 3 | 11940 | 407 | wech,Lrt,slow | | 1.081e-02 | -4.53 | axonal growth cone | cellular component | GO:0044295 | 14 | 3 | 11940 | 407 | NetA,NetB,Con | | 1.081e-02 | -4.53 | ISWI-type complex | cellular component | GO:0031010 | 14 | 3 | 11940 | 407 | CtBP,tou,Acf | | 1.082e-02 | -4.53 | regulation of cellular protein localization | biological process | GO:1903827 | 56 | 6 | 12345 | 414 | dlp,dally,hh,sdt,crb,S6kII | | 1.082e-02 | -4.53 | GO\_BLOOD\_VESSEL\_MORPHOGENESIS | MSigDB lists | GO\_BLOOD\_VESSEL\_MORPHOGENESIS | 67 | 7 | 4161 | 153 | so,LRP1,Ser,hh,N,slow,cta | | 1.082e-02 | -4.53 | MODULE\_48 | MSigDB lists | MODULE\_48 | 67 | 7 | 4161 | 153 | dally,dlp,CG14291,CG3097,drk,tkv,N | | 1.082e-02 | -4.53 | GSE2770\_IL12\_VS\_TGFB\_AND\_IL12\_TREATED\_ACT\_CD4\_TCELL\_6H\_DN | MSigDB lists | GSE2770\_IL12\_VS\_TGFB\_AND\_IL12\_TREATED\_ACT\_CD4\_TCELL\_6H\_DN | 67 | 7 | 4161 | 153 | CG45050,CG2812,CG11030,drk,Mitofilin,CG12099,Ser | | 1.082e-02 | -4.53 | YAMAZAKI\_TCEB3\_TARGETS\_DN | MSigDB lists | YAMAZAKI\_TCEB3\_TARGETS\_DN | 67 | 7 | 4161 | 153 | fne,CycE,CtBP,Arl4,CG45050,pbl,betaTub56D | | 1.083e-02 | -4.53 | female germline ring canal inner rim | cellular component | GO:0035183 | 5 | 2 | 11940 | 407 | Lpin,hts | | 1.083e-02 | -4.53 | Ref1 (RNA and export factor binding protein 1) | protein interactions | 44029 | 20 | 4 | 9204 | 412 | fj,Ubx,CG9775,S6kII | | 1.083e-02 | -4.53 | Med (Medea) | protein interactions | 43725 | 20 | 4 | 9204 | 412 | dpp,Smox,tkv,hiw | | 1.083e-02 | -4.53 | CG8079 (uncharacterized protein) | protein interactions | 36689 | 20 | 4 | 9204 | 412 | ci,sty,CG32369,CG1142 | | 1.083e-02 | -4.53 | babo (baboon) | protein interactions | 35900 | 20 | 4 | 9204 | 412 | Fas2,tkv,Smox,EcR | | 1.083e-02 | -4.53 | lz (lozenge) | protein interactions | 31883 | 20 | 4 | 9204 | 412 | osa,ttk,aop,aos | | 1.090e-02 | -4.52 | response to alcohol | biological process | GO:0097305 | 128 | 10 | 12345 | 414 | Fas2,spi,hang,scb,Drat,Utx,EcR,Egfr,Gbs-70E,S | | 1.094e-02 | -4.52 | regulation of synaptic growth at neuromuscular junction | biological process | GO:0008582 | 109 | 9 | 12345 | 414 | scb,Dad,Fas2,Syp,CaMKII,hiw,tkv,Acsl,Arp2 | | 1.094e-02 | -4.52 | regulation of mRNA processing | biological process | GO:0050684 | 109 | 9 | 12345 | 414 | fne,elav,ps,Rbfox1,CG5728,mub,salm,mle,Doa | | 1.096e-02 | -4.51 | positive regulation of catalytic activity | biological process | GO:0043085 | 252 | 16 | 12345 | 414 | RhoGAP102A,Atg17,RhoGAP16F,mbt,Rho1,Cdk5alpha,pbl,RhoGAP18B,mle,wnd,RhoGAP15B,CG8531,RapGAP1,nrv2,RhoGAP93B,bora | | 1.097e-02 | -4.51 | growth cone | cellular component | GO:0030426 | 26 | 4 | 11940 | 407 | NetB,NetA,Cdk5alpha,Con | | 1.103e-02 | -4.51 | regulation of peptide transport | biological process | GO:0090087 | 73 | 7 | 12345 | 414 | Snx3,CaMKII,S,CG9650,crb,S6kII,sdt | | 1.103e-02 | -4.51 | positive regulation of cell cycle | biological process | GO:0045787 | 73 | 7 | 12345 | 414 | CG31612,CycE,Cdk5alpha,Myb,stg,N,ci | | 1.114e-02 | -4.50 | postsynapse | cellular component | GO:0098794 | 72 | 7 | 11940 | 407 | CaMKII,Prosap,nAChRalpha4,hts,Ten-m,Fas2,Syp | | 1.122e-02 | -4.49 | phosphatidylinositol phosphate binding | molecular function | GO:1901981 | 42 | 5 | 12176 | 396 | Snx3,CG32758,Hip1,RhoGAP15B,pbl | | 1.123e-02 | -4.49 | CG6664 (uncharacterized protein) | protein interactions | 39903 | 11 | 3 | 9204 | 412 | psq,flfl,CG4325 | | 1.123e-02 | -4.49 | pygo (pygopus) | protein interactions | 43718 | 11 | 3 | 9204 | 412 | N,CtBP,CycE | | 1.123e-02 | -4.49 | scb (scab) | protein interactions | 36692 | 11 | 3 | 9204 | 412 | CycE,tkv,wb | | 1.123e-02 | -4.49 | sno (strawberry notch) | protein interactions | 32273 | 11 | 3 | 9204 | 412 | N,mam,Ser | | 1.123e-02 | -4.49 | Sec15 (Secretory 15) | protein interactions | 42499 | 11 | 3 | 9204 | 412 | N,dpp,egr | | 1.123e-02 | -4.49 | seq (sequoia) | protein interactions | 3772396 | 11 | 3 | 9204 | 412 | pnt,ham,CadN | | 1.129e-02 | -4.48 | ltl (larval translucida) | protein interactions | 38845 | 4 | 2 | 9204 | 412 | Fas2,dlp | | 1.129e-02 | -4.48 | verm (vermiform) | protein interactions | 40149 | 4 | 2 | 9204 | 412 | Doa,ci | | 1.129e-02 | -4.48 | CG6695 (uncharacterized protein) | protein interactions | 42937 | 4 | 2 | 9204 | 412 | Fas2,CG2865 | | 1.129e-02 | -4.48 | CG2182 (uncharacterized protein) | protein interactions | 40697 | 4 | 2 | 9204 | 412 | spi,CG7264 | | 1.129e-02 | -4.48 | magu (magu) | protein interactions | 36048 | 4 | 2 | 9204 | 412 | dally,tkv | | 1.129e-02 | -4.48 | fz3 (frizzled 3) | protein interactions | 31023 | 4 | 2 | 9204 | 412 | ci,Ser | | 1.129e-02 | -4.48 | dlt (discs lost) | protein interactions | 46719 | 4 | 2 | 9204 | 412 | CycE,crb | | 1.129e-02 | -4.48 | CG3760 (uncharacterized protein) | protein interactions | 37987 | 4 | 2 | 9204 | 412 | ft,Syx1A | | 1.129e-02 | -4.48 | plum (plum) | protein interactions | 43197 | 4 | 2 | 9204 | 412 | Smox,EcR | | 1.129e-02 | -4.48 | CG7065 (uncharacterized protein) | protein interactions | 31844 | 4 | 2 | 9204 | 412 | S6kII,Fas2 | | 1.129e-02 | -4.48 | disp (dispatched) | protein interactions | 44274 | 4 | 2 | 9204 | 412 | dlp,hh | | 1.129e-02 | -4.48 | Irk2 (Inwardly rectifying potassium channel 2) | protein interactions | 42770 | 4 | 2 | 9204 | 412 | tkv,dpp | | 1.129e-02 | -4.48 | CG10348 (uncharacterized protein) | protein interactions | 35132 | 4 | 2 | 9204 | 412 | CtBP,gus | | 1.129e-02 | -4.48 | Ten-a (Tenascin accessory) | protein interactions | 32183 | 4 | 2 | 9204 | 412 | mam,Ten-m | | 1.129e-02 | -4.48 | CG31974 (uncharacterized protein) | protein interactions | 33174 | 4 | 2 | 9204 | 412 | CG7264,Mocs2B | | 1.138e-02 | -4.48 | CTCF (CTCF) | protein interactions | 38817 | 31 | 5 | 9204 | 412 | AGO2,enok,Chro,Myb,EcR | | 1.138e-02 | -4.48 | CycD (Cyclin D) | protein interactions | 32551 | 31 | 5 | 9204 | 412 | Usp2,CG11486,Stat92E,CG2865,EcR | | 1.139e-02 | -4.47 | Signaling by Robo receptor | REACTOME pathways | R-DME-376176 | 13 | 3 | 3923 | 148 | RhoGAP93B,robo2,dlp | | 1.139e-02 | -4.47 | RA\_dom | interpro domains | IPR000159 | 13 | 3 | 11383 | 428 | CG32758,wake,rau | | 1.139e-02 | -4.47 | Laminin\_EGF | interpro domains | IPR002049 | 13 | 3 | 11383 | 428 | NetB,wb,NetA | | 1.141e-02 | -4.47 | RhoGEF | pfam domains | PF00621 | 24 | 4 | 10480 | 393 | pbl,CG43658,RhoGEF2,PsGEF | | 1.149e-02 | -4.47 | tube formation | biological process | GO:0035148 | 41 | 5 | 12345 | 414 | Kr,flz,Rho1,Mmp2,mmy | | 1.149e-02 | -4.47 | regulation of hemocyte proliferation | biological process | GO:0035206 | 41 | 5 | 12345 | 414 | Sin3A,EcR,Kr,Tl,cpo | | 1.149e-02 | -4.47 | olfactory learning | biological process | GO:0008355 | 41 | 5 | 12345 | 414 | scb,Egfr,drk,Fas2,spi | | 1.152e-02 | -4.46 | LamG | smart domains | SM00282 | 17 | 4 | 5086 | 276 | ft,CadN,crb,wb | | 1.152e-02 | -4.46 | DH-domain | interpro domains | IPR000219 | 24 | 4 | 11383 | 428 | CG43658,RhoGEF2,PsGEF,pbl | | 1.152e-02 | -4.46 | DBL\_dom\_sf | interpro domains | IPR035899 | 24 | 4 | 11383 | 428 | pbl,PsGEF,RhoGEF2,CG43658 | | 1.157e-02 | -4.46 | fly chr10B3-10B3\|1-34 | chromosome location | fly chr10B3-10B3\|1-34 | 8 | 2 | 25010 | 531 | CG11752,CG11756 | | 1.157e-02 | -4.46 | fly chr10B3-10B3\|1-34 cM | chromosome location | fly chr10B3-10B3\|1-34 cM | 8 | 2 | 25010 | 531 | CG11756,CG11752 | | 1.159e-02 | -4.46 | CycK (Cyclin K) | protein interactions | 49816 | 83 | 9 | 9204 | 412 | heph,cta,Cdk12,CG2865,CG43658,EcR,S,CG11486,Ubx | | 1.161e-02 | -4.46 | MEP-1 (MEP-1) | protein interactions | 38327 | 43 | 6 | 9204 | 412 | Pat1,CtBP,Doc1,ttk,mle,Pits | | 1.170e-02 | -4.45 | MODULE\_38 | MSigDB lists | MODULE\_38 | 68 | 7 | 4161 | 153 | CG43313,TfAP-2,CycE,Ank,Tim17b,Ser,LRP1 | | 1.170e-02 | -4.45 | GSE17580\_UNINFECTED\_VS\_S\_MANSONI\_INF\_TEFF\_UP | MSigDB lists | GSE17580\_UNINFECTED\_VS\_S\_MANSONI\_INF\_TEFF\_UP | 68 | 7 | 4161 | 153 | nAChRalpha4,CG5830,CG43313,hts,hiw,Syx1A,Mhcl | | 1.182e-02 | -4.44 | EGF\_LAM\_1 | prosite domains | PS01248 | 10 | 3 | 6243 | 316 | NetA,NetB,wb | | 1.182e-02 | -4.44 | EGF\_LAM\_2 | prosite domains | PS50027 | 10 | 3 | 6243 | 316 | wb,NetA,NetB | | 1.186e-02 | -4.43 | HEN1\_01 | MSigDB lists | HEN1\_01 | 38 | 5 | 4161 | 153 | Sin3A,ras,Arl4,CG43313,CG31637 | | 1.186e-02 | -4.43 | GSE20727\_CTRL\_VS\_H2O2\_TREATED\_DC\_UP | MSigDB lists | GSE20727\_CTRL\_VS\_H2O2\_TREATED\_DC\_UP | 38 | 5 | 4161 | 153 | drk,rush,lili,cta,Npc1a | | 1.186e-02 | -4.43 | GO\_DEVELOPMENTAL\_MATURATION | MSigDB lists | GO\_DEVELOPMENTAL\_MATURATION | 38 | 5 | 4161 | 153 | dpp,l(3)mbt,ds,Hydr2,ft | | 1.186e-02 | -4.43 | GSE7219\_UNSTIM\_VS\_LPS\_AND\_ANTI\_CD40\_STIM\_DC\_UP | MSigDB lists | GSE7219\_UNSTIM\_VS\_LPS\_AND\_ANTI\_CD40\_STIM\_DC\_UP | 38 | 5 | 4161 | 153 | CG34384,SamDC,cta,Smox,betaTub56D | | 1.186e-02 | -4.43 | GSE17974\_IL4\_AND\_ANTI\_IL12\_VS\_UNTREATED\_4H\_ACT\_CD4\_TCELL\_UP | MSigDB lists | GSE17974\_IL4\_AND\_ANTI\_IL12\_VS\_UNTREATED\_4H\_ACT\_CD4\_TCELL\_UP | 38 | 5 | 4161 | 153 | CaMKII,Mitofilin,Ptpmeg,Rhp,CG14291 | | 1.186e-02 | -4.43 | GSE45365\_WT\_VS\_IFNAR\_KO\_CD8A\_DC\_DN | MSigDB lists | GSE45365\_WT\_VS\_IFNAR\_KO\_CD8A\_DC\_DN | 38 | 5 | 4161 | 153 | dpp,bor,Tlk,par-1,drk | | 1.187e-02 | -4.43 | muscle tissue development | biological process | GO:0060537 | 27 | 4 | 12345 | 414 | N,EcR,lbe,tin | | 1.187e-02 | -4.43 | regulation of response to wounding | biological process | GO:1903034 | 27 | 4 | 12345 | 414 | Rac2,Egfr,Rho1,Imp | | 1.195e-02 | -4.43 | Wnt signaling pathway | KEGG pathways | dme04310 | 100 | 8 | 3198 | 100 | Rho1,esn,Notum,dlp,dally,CtBP,CaMKII,Rac2 | | 1.195e-02 | -4.43 | Wnt signaling pathway | KEGG pathways | ko04310 | 100 | 8 | 3198 | 100 | Rho1,esn,Notum,dlp,CaMKII,dally,CtBP,Rac2 | | 1.215e-02 | -4.41 | GO\_CELL\_CELL\_ADHESION | MSigDB lists | GO\_CELL\_CELL\_ADHESION | 85 | 8 | 4161 | 153 | Lac,ft,NetB,Dscam1,Smox,if,ds,hh | | 1.217e-02 | -4.41 | CCCNNNNNNAAGWT\_UNKNOWN | MSigDB lists | CCCNNNNNNAAGWT\_UNKNOWN | 25 | 4 | 4161 | 153 | S6kII,Tpr2,heph,Imp | | 1.217e-02 | -4.41 | YRTCANNRCGC\_UNKNOWN | MSigDB lists | YRTCANNRCGC\_UNKNOWN | 25 | 4 | 4161 | 153 | Etl1,Pat1,Fas2,nudE | | 1.217e-02 | -4.41 | MEL18\_DN.V1\_UP | MSigDB lists | MEL18\_DN.V1\_UP | 25 | 4 | 4161 | 153 | Ser,DIP-epsilon,fj,Syx1A | | 1.217e-02 | -4.41 | GO\_ENDOCRINE\_SYSTEM\_DEVELOPMENT | MSigDB lists | GO\_ENDOCRINE\_SYSTEM\_DEVELOPMENT | 25 | 4 | 4161 | 153 | Smox,so,dpp,hh | | 1.217e-02 | -4.41 | GSE4748\_CYANOBACTERIUM\_LPSLIKE\_VS\_LPS\_AND\_CYANOBACTERIUM\_LPSLIKE\_STIM\_DC\_3H\_DN | MSigDB lists | GSE4748\_CYANOBACTERIUM\_LPSLIKE\_VS\_LPS\_AND\_CYANOBACTERIUM\_LPSLIKE\_STIM\_DC\_3H\_DN | 25 | 4 | 4161 | 153 | par-1,CG34384,betaTub56D,Arl4 | | 1.217e-02 | -4.41 | GSE17974\_0H\_VS\_6H\_IN\_VITRO\_ACT\_CD4\_TCELL\_UP | MSigDB lists | GSE17974\_0H\_VS\_6H\_IN\_VITRO\_ACT\_CD4\_TCELL\_UP | 25 | 4 | 4161 | 153 | CG12099,TfIIS,Pli,ScsbetaG | | 1.217e-02 | -4.41 | PHONG\_TNF\_RESPONSE\_VIA\_P38\_PARTIAL | MSigDB lists | PHONG\_TNF\_RESPONSE\_VIA\_P38\_PARTIAL | 25 | 4 | 4161 | 153 | dpp,Cyp1,fj,Smox | | 1.220e-02 | -4.41 | response to endogenous stimulus | biological process | GO:0009719 | 212 | 14 | 12345 | 414 | Utx,pnt,drk,Dad,Hydr2,Smox,dally,hiw,aop,shn,EcR,dpp,Lpin,tkv | | 1.223e-02 | -4.40 | actin filament organization | biological process | GO:0007015 | 111 | 9 | 12345 | 414 | Pak3,N,tyn,Rac2,Arp2,Rho1,hts,Hip1,RhoGEF2 | | 1.234e-02 | -4.39 | GATAAGR\_GATA\_C | MSigDB lists | GATAAGR\_GATA\_C | 53 | 6 | 4161 | 153 | Trpm,CG10663,kdn,CG6752,Mhcl,so | | 1.234e-02 | -4.39 | GSE360\_L\_DONOVANI\_VS\_T\_GONDII\_MAC\_UP | MSigDB lists | GSE360\_L\_DONOVANI\_VS\_T\_GONDII\_MAC\_UP | 53 | 6 | 4161 | 153 | Hsc70-3,Tlk,Cyp1,Tif-IA,ds,baf | | 1.234e-02 | -4.39 | GO\_POSITIVE\_REGULATION\_OF\_NEURON\_DIFFERENTIATION | MSigDB lists | GO\_POSITIVE\_REGULATION\_OF\_NEURON\_DIFFERENTIATION | 53 | 6 | 4161 | 153 | Tif-IA,nudE,dpp,Lac,pbl,NetB | | 1.234e-02 | -4.39 | NGFIC\_01 | MSigDB lists | NGFIC\_01 | 53 | 6 | 4161 | 153 | Imp,Smox,NetB,Sin3A,drk,S6kII | | 1.234e-02 | -4.39 | SOX9\_B1 | MSigDB lists | SOX9\_B1 | 53 | 6 | 4161 | 153 | fne,Mhcl,Fas2,so,Pli,CG9281 | | 1.235e-02 | -4.39 | Hippo signaling pathway -multiple species | KEGG pathways | dme04392 | 16 | 3 | 3198 | 100 | ft,ds,Pak3 | | 1.235e-02 | -4.39 | Hippo signaling pathway -multiple species | KEGG pathways | ko04392 | 16 | 3 | 3198 | 100 | ft,ds,Pak3 | | 1.249e-02 | -4.38 | GO\_EMBRYONIC\_HINDLIMB\_MORPHOGENESIS | MSigDB lists | GO\_EMBRYONIC\_HINDLIMB\_MORPHOGENESIS | 5 | 2 | 4161 | 153 | hh,N | | 1.249e-02 | -4.38 | GO\_PATTERNING\_OF\_BLOOD\_VESSELS | MSigDB lists | GO\_PATTERNING\_OF\_BLOOD\_VESSELS | 5 | 2 | 4161 | 153 | hh,cta | | 1.249e-02 | -4.38 | GO\_CARDIAC\_EPITHELIAL\_TO\_MESENCHYMAL\_TRANSITION | MSigDB lists | GO\_CARDIAC\_EPITHELIAL\_TO\_MESENCHYMAL\_TRANSITION | 5 | 2 | 4161 | 153 | N,dpp | | 1.249e-02 | -4.38 | GO\_NEGATIVE\_REGULATION\_OF\_KIDNEY\_DEVELOPMENT | MSigDB lists | GO\_NEGATIVE\_REGULATION\_OF\_KIDNEY\_DEVELOPMENT | 5 | 2 | 4161 | 153 | hh,so | | 1.249e-02 | -4.38 | GO\_POSITIVE\_REGULATION\_OF\_NEUROBLAST\_PROLIFERATION | MSigDB lists | GO\_POSITIVE\_REGULATION\_OF\_NEUROBLAST\_PROLIFERATION | 5 | 2 | 4161 | 153 | N,hh | | 1.249e-02 | -4.38 | GO\_COCHLEA\_MORPHOGENESIS | MSigDB lists | GO\_COCHLEA\_MORPHOGENESIS | 5 | 2 | 4161 | 153 | Rac2,so | | 1.249e-02 | -4.38 | FUKUSHIMA\_TNFSF11\_TARGETS | MSigDB lists | FUKUSHIMA\_TNFSF11\_TARGETS | 5 | 2 | 4161 | 153 | N,Ser | | 1.249e-02 | -4.38 | GO\_NEGATIVE\_REGULATION\_OF\_STRIATED\_MUSCLE\_CELL\_DIFFERENTIATION | MSigDB lists | GO\_NEGATIVE\_REGULATION\_OF\_STRIATED\_MUSCLE\_CELL\_DIFFERENTIATION | 5 | 2 | 4161 | 153 | dpp,N | | 1.249e-02 | -4.38 | REACTOME\_INTERFERON\_GAMMA\_SIGNALING | MSigDB lists | REACTOME\_INTERFERON\_GAMMA\_SIGNALING | 5 | 2 | 4161 | 153 | CaMKII,Fas2 | | 1.249e-02 | -4.38 | GO\_AUTONOMIC\_NERVOUS\_SYSTEM\_DEVELOPMENT | MSigDB lists | GO\_AUTONOMIC\_NERVOUS\_SYSTEM\_DEVELOPMENT | 5 | 2 | 4161 | 153 | so,TfAP-2 | | 1.249e-02 | -4.38 | GO\_HEPARAN\_SULFATE\_PROTEOGLYCAN\_BINDING | MSigDB lists | GO\_HEPARAN\_SULFATE\_PROTEOGLYCAN\_BINDING | 5 | 2 | 4161 | 153 | dally,dlp | | 1.249e-02 | -4.38 | GO\_EPITHELIAL\_CELL\_DIFFERENTIATION\_INVOLVED\_IN\_KIDNEY\_DEVELOPMENT | MSigDB lists | GO\_EPITHELIAL\_CELL\_DIFFERENTIATION\_INVOLVED\_IN\_KIDNEY\_DEVELOPMENT | 5 | 2 | 4161 | 153 | N,Ser | | 1.249e-02 | -4.38 | GO\_RETINOIC\_ACID\_RECEPTOR\_BINDING | MSigDB lists | GO\_RETINOIC\_ACID\_RECEPTOR\_BINDING | 5 | 2 | 4161 | 153 | CtBP,EcR | | 1.249e-02 | -4.38 | GO\_MESENCHYMAL\_CELL\_PROLIFERATION | MSigDB lists | GO\_MESENCHYMAL\_CELL\_PROLIFERATION | 5 | 2 | 4161 | 153 | dpp,hh | | 1.249e-02 | -4.38 | GO\_NEGATIVE\_REGULATION\_OF\_LIPID\_TRANSPORT | MSigDB lists | GO\_NEGATIVE\_REGULATION\_OF\_LIPID\_TRANSPORT | 5 | 2 | 4161 | 153 | EcR,hh | | 1.249e-02 | -4.38 | GO\_CRANIAL\_NERVE\_MORPHOGENESIS | MSigDB lists | GO\_CRANIAL\_NERVE\_MORPHOGENESIS | 5 | 2 | 4161 | 153 | so,TfAP-2 | | 1.249e-02 | -4.38 | GO\_LIPOPROTEIN\_LOCALIZATION | MSigDB lists | GO\_LIPOPROTEIN\_LOCALIZATION | 5 | 2 | 4161 | 153 | LRP1,mgl | | 1.249e-02 | -4.38 | GO\_LOW\_DENSITY\_LIPOPROTEIN\_RECEPTOR\_ACTIVITY | MSigDB lists | GO\_LOW\_DENSITY\_LIPOPROTEIN\_RECEPTOR\_ACTIVITY | 5 | 2 | 4161 | 153 | LRP1,mgl | | 1.249e-02 | -4.38 | GO\_METANEPHRIC\_MESENCHYME\_DEVELOPMENT | MSigDB lists | GO\_METANEPHRIC\_MESENCHYME\_DEVELOPMENT | 5 | 2 | 4161 | 153 | so,hh | | 1.249e-02 | -4.38 | GO\_POSITIVE\_REGULATION\_OF\_PHOSPHATASE\_ACTIVITY | MSigDB lists | GO\_POSITIVE\_REGULATION\_OF\_PHOSPHATASE\_ACTIVITY | 5 | 2 | 4161 | 153 | Smox,dpp | | 1.249e-02 | -4.38 | GO\_CELL\_SURFACE\_RECEPTOR\_SIGNALING\_PATHWAY\_INVOLVED\_IN\_HEART\_DEVELOPMENT | MSigDB lists | GO\_CELL\_SURFACE\_RECEPTOR\_SIGNALING\_PATHWAY\_INVOLVED\_IN\_HEART\_DEVELOPMENT | 5 | 2 | 4161 | 153 | Ser,N | | 1.249e-02 | -4.38 | GO\_REGULATION\_OF\_ASTROCYTE\_DIFFERENTIATION | MSigDB lists | GO\_REGULATION\_OF\_ASTROCYTE\_DIFFERENTIATION | 5 | 2 | 4161 | 153 | N,dpp | | 1.249e-02 | -4.38 | GO\_KIDNEY\_MESENCHYME\_DEVELOPMENT | MSigDB lists | GO\_KIDNEY\_MESENCHYME\_DEVELOPMENT | 5 | 2 | 4161 | 153 | so,hh | | 1.249e-02 | -4.38 | GO\_RESPONSE\_TO\_MURAMYL\_DIPEPTIDE | MSigDB lists | GO\_RESPONSE\_TO\_MURAMYL\_DIPEPTIDE | 5 | 2 | 4161 | 153 | N,Ser | | 1.254e-02 | -4.38 | Mitochondrial LC-Fatty Acid Beta-Oxidation | WikiPathways | WP406 | 11 | 3 | 499 | 24 | CG3961,ScpX,Acsl | | 1.254e-02 | -4.38 | site of polarized growth | cellular component | GO:0030427 | 27 | 4 | 11940 | 407 | Con,Cdk5alpha,NetA,NetB | | 1.262e-02 | -4.37 | establishment of monopolar cell polarity | biological process | GO:0061162 | 15 | 3 | 12345 | 414 | fj,ds,ft | | 1.262e-02 | -4.37 | activation of innate immune response | biological process | GO:0002218 | 15 | 3 | 12345 | 414 | Mmp2,Toll-7,Tl | | 1.262e-02 | -4.37 | dorsal/ventral axis specification, ovarian follicular epithelium | biological process | GO:0008069 | 15 | 3 | 12345 | 414 | spoon,Egfr,sty | | 1.262e-02 | -4.37 | immune response-regulating signaling pathway | biological process | GO:0002764 | 15 | 3 | 12345 | 414 | Rac2,Tl,Toll-7 | | 1.262e-02 | -4.37 | male germ-line stem cell population maintenance | biological process | GO:0036098 | 15 | 3 | 12345 | 414 | Inx2,CG46339,dpp | | 1.262e-02 | -4.37 | regulation of oskar mRNA translation | biological process | GO:0046011 | 15 | 3 | 12345 | 414 | elav,heph,Syp | | 1.262e-02 | -4.37 | synaptic target inhibition | biological process | GO:0016201 | 15 | 3 | 12345 | 414 | beat-IIIc,robo2,Tl | | 1.262e-02 | -4.37 | establishment or maintenance of monopolar cell polarity | biological process | GO:0061339 | 15 | 3 | 12345 | 414 | ft,ds,fj | | 1.263e-02 | -4.37 | GO\_REGULATION\_OF\_ACTIN\_FILAMENT\_BASED\_PROCESS | MSigDB lists | GO\_REGULATION\_OF\_ACTIN\_FILAMENT\_BASED\_PROCESS | 69 | 7 | 4161 | 153 | LRP1,Arp2,drk,Rac2,CaMKII,hts,Smox | | 1.263e-02 | -4.37 | E2F1\_Q6\_01 | MSigDB lists | E2F1\_Q6\_01 | 69 | 7 | 4161 | 153 | SamDC,Sin3A,CG8531,dpp,NiPp1,LRP1,Nhe2 | | 1.269e-02 | -4.37 | regulation of Notch signaling pathway | biological process | GO:0008593 | 75 | 7 | 12345 | 414 | phyl,Ser,Tsp26A,crb,E(spl)m4-BFM,Hipk,sno | | 1.269e-02 | -4.37 | instar larval development | biological process | GO:0002168 | 75 | 7 | 12345 | 414 | Ser,hh,trn,Dad,sas,ft,wech | | 1.269e-02 | -4.37 | regulation of protein kinase activity | biological process | GO:0045859 | 131 | 10 | 12345 | 414 | sty,wnd,aos,CG8441,Rho1,mbt,Atg17,bora,Cdk5alpha,CycE | | 1.272e-02 | -4.36 | response to antibiotic | biological process | GO:0046677 | 93 | 8 | 12345 | 414 | Drat,scb,spi,Fas2,hang,Egfr,Gbs-70E,S | | 1.273e-02 | -4.36 | MODULE\_13 | MSigDB lists | MODULE\_13 | 103 | 9 | 4161 | 153 | so,CG7264,mbt,fne,Tpr2,Ser,Cyp1,mub,Taf1 | | 1.276e-02 | -4.36 | regulation of stem cell proliferation | biological process | GO:0072091 | 58 | 6 | 12345 | 414 | pnt,osa,EcR,Mmp2,N,E(spl)mdelta-HLH | | 1.277e-02 | -4.36 | regulation of transferase activity | biological process | GO:0051338 | 151 | 11 | 12345 | 414 | aos,CG8441,Rho1,mbt,Atg17,sty,wnd,bora,Pli,CycE,Cdk5alpha | | 1.283e-02 | -4.36 | kinase binding | molecular function | GO:0019900 | 96 | 8 | 12176 | 396 | CG8441,CycE,Rho1,Rac2,crb,bora,Mitofilin,Atg17 | | 1.292e-02 | -4.35 | Ssb-c31a (Single stranded-binding protein c31A) | protein interactions | 34120 | 21 | 4 | 9204 | 412 | CG31712,Df31,S6kII,CG10265 | | 1.292e-02 | -4.35 | CG6701 (uncharacterized protein) | protein interactions | 36550 | 21 | 4 | 9204 | 412 | CycE,pbl,CG9775,Syp | | 1.295e-02 | -4.35 | Hrb27C (Heterogeneous nuclear ribonucleoprotein at 27C) | protein interactions | 33968 | 44 | 6 | 9204 | 412 | CtBP,Ubx,Imp,tna,pAbp,Syp | | 1.299e-02 | -4.34 | FOSTER\_TOLERANT\_MACROPHAGE\_DN | MSigDB lists | FOSTER\_TOLERANT\_MACROPHAGE\_DN | 86 | 8 | 4161 | 153 | CG34384,Tlk,CG9855,rush,ap,Arl4,N,CaMKII | | 1.301e-02 | -4.34 | MH1 | pfam domains | PF03165 | 5 | 2 | 10480 | 393 | Smox,Dad | | 1.301e-02 | -4.34 | zf-MYST | pfam domains | PF17772 | 5 | 2 | 10480 | 393 | enok,mof | | 1.301e-02 | -4.34 | Integrin\_alpha2 | pfam domains | PF08441 | 5 | 2 | 10480 | 393 | scb,if | | 1.301e-02 | -4.34 | Patched | pfam domains | PF02460 | 5 | 2 | 10480 | 393 | Npc1a,Ptr | | 1.301e-02 | -4.34 | MOZ\_SAS | pfam domains | PF01853 | 5 | 2 | 10480 | 393 | enok,mof | | 1.301e-02 | -4.34 | CG8301 (uncharacterized protein) | protein interactions | 41160 | 32 | 5 | 9204 | 412 | phyl,rau,nudE,CG13366,CG31224 | | 1.301e-02 | -4.34 | GstS1 (Glutathione S transferase S1) | protein interactions | 36927 | 32 | 5 | 9204 | 412 | spi,Kr,kdn,CG34172,CG6330 | | 1.301e-02 | -4.34 | CG13373 (uncharacterized protein) | protein interactions | 30988 | 32 | 5 | 9204 | 412 | Usp2,bowl,S,EcR,Pits | | 1.308e-02 | -4.34 | CRIB\_dom\_sf | interpro domains | IPR036936 | 5 | 2 | 11383 | 428 | Pak3,mbt | | 1.308e-02 | -4.34 | Integrin\_alpha | interpro domains | IPR000413 | 5 | 2 | 11383 | 428 | scb,if | | 1.308e-02 | -4.34 | Int\_alpha\_beta-p | interpro domains | IPR013519 | 5 | 2 | 11383 | 428 | if,scb | | 1.308e-02 | -4.34 | Zf-MYST | interpro domains | IPR040706 | 5 | 2 | 11383 | 428 | enok,mof | | 1.308e-02 | -4.34 | Integrin\_alpha-2 | interpro domains | IPR013649 | 5 | 2 | 11383 | 428 | scb,if | | 1.308e-02 | -4.34 | HAT\_MYST-type | interpro domains | IPR002717 | 5 | 2 | 11383 | 428 | mof,enok | | 1.308e-02 | -4.34 | MAD\_homology1\_Dwarfin-type | interpro domains | IPR003619 | 5 | 2 | 11383 | 428 | Smox,Dad | | 1.308e-02 | -4.34 | TF\_Brachyury | interpro domains | IPR002070 | 5 | 2 | 11383 | 428 | Doc2,Doc1 | | 1.308e-02 | -4.34 | Ptc/Disp | interpro domains | IPR003392 | 5 | 2 | 11383 | 428 | Ptr,Npc1a | | 1.308e-02 | -4.34 | Integrin\_alpha\_N | interpro domains | IPR028994 | 5 | 2 | 11383 | 428 | scb,if | | 1.308e-02 | -4.34 | Methyl\_CpG\_DNA-bd | interpro domains | IPR001739 | 5 | 2 | 11383 | 428 | tou,sba | | 1.310e-02 | -4.33 | crb (crumbs) | protein interactions | 42896 | 57 | 7 | 9204 | 412 | N,mam,CycE,Rho1,grnd,sdt,Ser | | 1.312e-02 | -4.33 | GRB2 events in ERBB2 signaling | REACTOME pathways | R-DME-1963640 | 5 | 2 | 3923 | 148 | drk,Egfr | | 1.312e-02 | -4.33 | MyD88 dependent cascade initiated on endosome | REACTOME pathways | R-DME-975155 | 38 | 5 | 3923 | 148 | mle,Pli,Tl,S6kII,JIL-1 | | 1.312e-02 | -4.33 | Toll Like Receptor 7/8 (TLR7/8) Cascade | REACTOME pathways | R-DME-168181 | 38 | 5 | 3923 | 148 | JIL-1,Pli,mle,S6kII,Tl | | 1.312e-02 | -4.33 | TRAF6 mediated induction of NFkB and MAP kinases upon TLR7/8 or 9 activation | REACTOME pathways | R-DME-975138 | 38 | 5 | 3923 | 148 | Tl,S6kII,mle,Pli,JIL-1 | | 1.317e-02 | -4.33 | ROSS\_AML\_WITH\_PML\_RARA\_FUSION | MSigDB lists | ROSS\_AML\_WITH\_PML\_RARA\_FUSION | 14 | 3 | 4161 | 153 | Ser,Hsc70-3,CG41099 | | 1.317e-02 | -4.33 | GO\_REGULATION\_OF\_EPITHELIAL\_TO\_MESENCHYMAL\_TRANSITION | MSigDB lists | GO\_REGULATION\_OF\_EPITHELIAL\_TO\_MESENCHYMAL\_TRANSITION | 14 | 3 | 4161 | 153 | dpp,Smox,N | | 1.317e-02 | -4.33 | GO\_REGULATION\_OF\_SODIUM\_ION\_TRANSPORT | MSigDB lists | GO\_REGULATION\_OF\_SODIUM\_ION\_TRANSPORT | 14 | 3 | 4161 | 153 | Ank,NKAIN,CaMKII | | 1.317e-02 | -4.33 | GO\_PROTEIN\_AUTOUBIQUITINATION | MSigDB lists | GO\_PROTEIN\_AUTOUBIQUITINATION | 14 | 3 | 4161 | 153 | CG9855,wech,CG12099 | | 1.317e-02 | -4.33 | GO\_ENDODERM\_DEVELOPMENT | MSigDB lists | GO\_ENDODERM\_DEVELOPMENT | 14 | 3 | 4161 | 153 | Lim1,N,Smox | | 1.317e-02 | -4.33 | WINZEN\_DEGRADED\_VIA\_KHSRP | MSigDB lists | WINZEN\_DEGRADED\_VIA\_KHSRP | 14 | 3 | 4161 | 153 | dpp,Ser,CG45050 | | 1.317e-02 | -4.33 | GO\_NEURON\_FATE\_COMMITMENT | MSigDB lists | GO\_NEURON\_FATE\_COMMITMENT | 14 | 3 | 4161 | 153 | N,hh,so | | 1.317e-02 | -4.33 | SIG\_REGULATION\_OF\_THE\_ACTIN\_CYTOSKELETON\_BY\_RHO\_GTPASES | MSigDB lists | SIG\_REGULATION\_OF\_THE\_ACTIN\_CYTOSKELETON\_BY\_RHO\_GTPASES | 14 | 3 | 4161 | 153 | Arp2,RpS4,mbt | | 1.317e-02 | -4.33 | SHAFFER\_IRF4\_TARGETS\_IN\_PLASMA\_CELL\_VS\_MATURE\_B\_LYMPHOCYTE | MSigDB lists | SHAFFER\_IRF4\_TARGETS\_IN\_PLASMA\_CELL\_VS\_MATURE\_B\_LYMPHOCYTE | 14 | 3 | 4161 | 153 | mmy,Cnx99A,pAbp | | 1.317e-02 | -4.33 | WANG\_RESPONSE\_TO\_ANDROGEN\_UP | MSigDB lists | WANG\_RESPONSE\_TO\_ANDROGEN\_UP | 14 | 3 | 4161 | 153 | Acsl,Pat1,tkv | | 1.317e-02 | -4.33 | GO\_STEM\_CELL\_PROLIFERATION | MSigDB lists | GO\_STEM\_CELL\_PROLIFERATION | 14 | 3 | 4161 | 153 | wech,dpp,hh | | 1.317e-02 | -4.33 | GO\_CELL\_FATE\_SPECIFICATION | MSigDB lists | GO\_CELL\_FATE\_SPECIFICATION | 14 | 3 | 4161 | 153 | so,hh,N | | 1.319e-02 | -4.33 | KH\_1 | pfam domains | PF00013 | 25 | 4 | 10480 | 393 | Imp,mub,ps,spoon | | 1.321e-02 | -4.33 | GSE14769\_UNSTIM\_VS\_60MIN\_LPS\_BMDM\_DN | MSigDB lists | GSE14769\_UNSTIM\_VS\_60MIN\_LPS\_BMDM\_DN | 39 | 5 | 4161 | 153 | nudE,mmy,dpp,CG2812,Acsl | | 1.321e-02 | -4.33 | YTAATTAA\_LHX3\_01 | MSigDB lists | YTAATTAA\_LHX3\_01 | 39 | 5 | 4161 | 153 | Doa,cta,TfAP-2,if,ScsbetaG | | 1.321e-02 | -4.33 | FOXM1\_01 | MSigDB lists | FOXM1\_01 | 39 | 5 | 4161 | 153 | mgl,SamDC,Fas2,Lim1,LRP1 | | 1.321e-02 | -4.33 | GSE10325\_MYELOID\_VS\_LUPUS\_MYELOID\_UP | MSigDB lists | GSE10325\_MYELOID\_VS\_LUPUS\_MYELOID\_UP | 39 | 5 | 4161 | 153 | pAbp,NetB,Fas2,RpS4,dpp | | 1.321e-02 | -4.33 | GSE21063\_WT\_VS\_NFATC1\_KO\_16H\_ANTI\_IGM\_STIM\_BCELL\_UP | MSigDB lists | GSE21063\_WT\_VS\_NFATC1\_KO\_16H\_ANTI\_IGM\_STIM\_BCELL\_UP | 39 | 5 | 4161 | 153 | pAbp,Syx1A,lili,Pli,mmy | | 1.322e-02 | -4.33 | GO\_POSITIVE\_REGULATION\_OF\_HYDROLASE\_ACTIVITY | MSigDB lists | GO\_POSITIVE\_REGULATION\_OF\_HYDROLASE\_ACTIVITY | 198 | 14 | 4161 | 153 | drk,Tpr2,Rho1,EcR,Hsc70-3,CG4538,cta,Smox,CaMKII,pbl,CG5522,dpp,nudE,Fas2 | | 1.325e-02 | -4.32 | regulation of apoptotic process | biological process | GO:0042981 | 193 | 13 | 12345 | 414 | ft,N,Fili,spi,mof,eya,grnd,pnt,CycE,aos,dpp,EcR,Egfr | | 1.330e-02 | -4.32 | protein domain specific binding | molecular function | GO:0019904 | 78 | 7 | 12176 | 396 | psq,Egfr,ttk,Tl,comm,aop,Ubx | | 1.334e-02 | -4.32 | regulation of mRNA metabolic process | biological process | GO:1903311 | 132 | 10 | 12345 | 414 | mub,smg,salm,fne,ps,elav,Rbfox1,CG5728,mle,Doa | | 1.346e-02 | -4.31 | chromosome | cellular component | GO:0005694 | 696 | 35 | 11940 | 407 | Psc,lncRNA:roX1,osa,CtBP,tou,mof,stwl,l(3)mbt,JIL-1,Acf,Chro,l(3)neo38,nudE,Myb,flfl,EcR,Tlk,Mocs2B,Pits,CG17778,tna,Sin3A,mle,lawc,mam,Df31,cnc,Cdk12,Utx,stg,CG44774,enok,ttk,HmgZ,Kah | | 1.347e-02 | -4.31 | GSE7548\_DAY7\_VS\_DAY28\_PCC\_IMMUNIZATION\_CD4\_TCELL\_UP | MSigDB lists | GSE7548\_DAY7\_VS\_DAY28\_PCC\_IMMUNIZATION\_CD4\_TCELL\_UP | 54 | 6 | 4161 | 153 | Ugt49C1,Npc1a,dpp,CG2201,CG14291,smal | | 1.347e-02 | -4.31 | SP3\_Q3 | MSigDB lists | SP3\_Q3 | 54 | 6 | 4161 | 153 | CG5830,sm,Tpr2,ap,CG9855,dpp | | 1.348e-02 | -4.31 | flw (flapwing) | protein interactions | 44289 | 130 | 12 | 9204 | 412 | ec,RhoGEF2,Lpin,NiPp1,Rho1,Mhcl,Gbs-70E,crb,CG2865,l(3)mbt,CG42748,N | | 1.349e-02 | -4.31 | positive regulation of TORC1 signaling | biological process | GO:1904263 | 28 | 4 | 12345 | 414 | Socs16D,drk,ci,wcy | | 1.349e-02 | -4.31 | regulation of generation of precursor metabolites and energy | biological process | GO:0043467 | 28 | 4 | 12345 | 414 | CG34172,EcR,N,Gbs-70E | | 1.349e-02 | -4.31 | positive regulation of transmembrane receptor protein serine/threonine kinase signaling pathway | biological process | GO:0090100 | 28 | 4 | 12345 | 414 | lili,dally,tkv,dpp | | 1.349e-02 | -4.31 | regulation of cell cycle G1/S phase transition | biological process | GO:1902806 | 28 | 4 | 12345 | 414 | ci,N,Egfr,CycE | | 1.362e-02 | -4.30 | KEGG\_PATHWAYS\_IN\_CANCER | MSigDB lists | KEGG\_PATHWAYS\_IN\_CANCER | 70 | 7 | 4161 | 153 | dpp,hh,Rac2,Smox,CtBP,drk,CycE | | 1.362e-02 | -4.30 | Znf\_FYVE\_PHD | interpro domains | IPR011011 | 68 | 7 | 11383 | 428 | Acf,tou,CG6051,enok,CG41099,Pcl,rush | | 1.373e-02 | -4.29 | positive regulation of cellular protein metabolic process | biological process | GO:0032270 | 303 | 18 | 12345 | 414 | Pcl,Rho1,mbt,Atg17,drk,Cdk5alpha,Usp2,dpp,Mos,PEK,wnd,Egfr,spi,bora,grnd,mmy,pAbp,phyl | | 1.375e-02 | -4.29 | Rho guanyl-nucleotide exchange factor activity | molecular function | GO:0005089 | 29 | 4 | 12176 | 396 | PsGEF,CG43658,pbl,RhoGEF2 | | 1.376e-02 | -4.29 | GO\_REGULATION\_OF\_CELL\_PROLIFERATION | MSigDB lists | GO\_REGULATION\_OF\_CELL\_PROLIFERATION | 260 | 17 | 4161 | 153 | wech,smal,CtBP,TfAP-2,CaMKII,hh,so,tkv,dpp,Lim1,Ser,NetB,N,Smox,CG12099,RpS4,slow | | 1.381e-02 | -4.28 | developmental cell growth | biological process | GO:0048588 | 59 | 6 | 12345 | 414 | Dscam1,Ptpmeg,CadN,wnd,mle,scb | | 1.385e-02 | -4.28 | GO\_CELL\_MOTILITY | MSigDB lists | GO\_CELL\_MOTILITY | 160 | 12 | 4161 | 153 | hh,so,mbt,drk,Rho1,Rac2,Lim1,Mhcl,nudE,ds,N,NetB | | 1.387e-02 | -4.28 | CATTGTYY\_SOX9\_B1 | MSigDB lists | CATTGTYY\_SOX9\_B1 | 87 | 8 | 4161 | 153 | Ptpmeg,CG9281,Mhcl,ft,Trpm,Imp,so,Ank | | 1.395e-02 | -4.27 | poly-pyrimidine tract binding | molecular function | GO:0008187 | 16 | 3 | 12176 | 396 | elav,pAbp,fne | | 1.395e-02 | -4.27 | translation repressor activity | molecular function | GO:0030371 | 16 | 3 | 12176 | 396 | smg,elav,heph | | 1.397e-02 | -4.27 | GO\_REGULATION\_OF\_EMBRYONIC\_DEVELOPMENT | MSigDB lists | GO\_REGULATION\_OF\_EMBRYONIC\_DEVELOPMENT | 26 | 4 | 4161 | 153 | so,Lim1,hh,N | | 1.397e-02 | -4.27 | MODULE\_123 | MSigDB lists | MODULE\_123 | 26 | 4 | 4161 | 153 | fne,ap,TfAP-2,cta | | 1.398e-02 | -4.27 | gastrulation involving germ band extension | biological process | GO:0010004 | 43 | 5 | 12345 | 414 | RhoGEF2,twi,Rho1,dpp,cta | | 1.398e-02 | -4.27 | positive regulation of Wnt signaling pathway | biological process | GO:0030177 | 43 | 5 | 12345 | 414 | Hipk,CtBP,dlp,dally,spen | | 1.398e-02 | -4.27 | gastrulation with mouth forming first | biological process | GO:0001703 | 43 | 5 | 12345 | 414 | Rho1,dpp,cta,RhoGEF2,twi | | 1.400e-02 | -4.27 | FYVE | pfam domains | PF01363 | 14 | 3 | 10480 | 393 | rush,CG6051,CG41099 | | 1.410e-02 | -4.26 | Znf\_FYVE | interpro domains | IPR000306 | 14 | 3 | 11383 | 428 | rush,CG41099,CG6051 | | 1.425e-02 | -4.25 | PcG protein complex | cellular component | GO:0031519 | 28 | 4 | 11940 | 407 | jing,psq,Pcl,Psc | | 1.425e-02 | -4.25 | focal adhesion | cellular component | GO:0005925 | 28 | 4 | 11940 | 407 | if,smal,ckn,scb | | 1.432e-02 | -4.25 | MH1 | prosite domains | PS51075 | 4 | 2 | 6243 | 316 | Smox,Dad | | 1.432e-02 | -4.25 | DDT | prosite domains | PS50827 | 4 | 2 | 6243 | 316 | Acf,tou | | 1.432e-02 | -4.25 | INTEGRIN\_ALPHA | prosite domains | PS00242 | 4 | 2 | 6243 | 316 | if,scb | | 1.434e-02 | -4.25 | adult locomotory behavior | biological process | GO:0008344 | 95 | 8 | 12345 | 414 | ey,Cirl,TfAP-2,Cdk5alpha,oc,CCKLR-17D3,cv,hiw | | 1.443e-02 | -4.24 | metal ion binding | molecular function | GO:0046872 | 1047 | 47 | 12176 | 396 | l(3)mbt,S6kII,CG32369,Cyp4p2,CG9855,CG11247,mgl,CG45050,hh,kni,N,crb,CG12099,tna,ctrip,Zif,ft,Cnx99A,EcR,esn,bor,tkv,rush,CG4325,Mmp2,TfIIS,CadN,dbr,Hph,hiw,CG3097,CG6752,Utx,slow,wech,shd,CG41099,CG6051,Ser,Psc,LRP1,ds,shn,hang,CG2991,Tet,Cad87A | | 1.448e-02 | -4.23 | Krn (Keren) | protein interactions | 326198 | 12 | 3 | 9204 | 412 | S,Egfr,spi | | 1.448e-02 | -4.23 | gpp (grappa) | protein interactions | 40793 | 12 | 3 | 9204 | 412 | hh,CG4325,Pcl | | 1.448e-02 | -4.23 | Pkn (Protein kinase N) | protein interactions | 35950 | 12 | 3 | 9204 | 412 | Rho1,drk,Fas2 | | 1.448e-02 | -4.23 | rg (rugose) | protein interactions | 44531 | 12 | 3 | 9204 | 412 | Egfr,S,aos | | 1.451e-02 | -4.23 | - | gene3d domains | 1.20.900.10 | 23 | 4 | 4929 | 208 | CG43658,PsGEF,RhoGEF2,pbl | | 1.451e-02 | -4.23 | spectrin binding | molecular function | GO:0030507 | 6 | 2 | 12176 | 396 | hts,crb | | 1.451e-02 | -4.23 | promoter-specific chromatin binding | molecular function | GO:1990841 | 6 | 2 | 12176 | 396 | ttk,Utx | | 1.451e-02 | -4.23 | gap junction channel activity | molecular function | GO:0005243 | 6 | 2 | 12176 | 396 | Inx3,Inx2 | | 1.465e-02 | -4.22 | GSE1460\_DP\_THYMOCYTE\_VS\_THYMIC\_STROMAL\_CELL\_DN | MSigDB lists | GSE1460\_DP\_THYMOCYTE\_VS\_THYMIC\_STROMAL\_CELL\_DN | 71 | 7 | 4161 | 153 | DIP-epsilon,smal,CG12547,nudE,ras,Rac2,Smox | | 1.465e-02 | -4.22 | GO\_REGULATION\_OF\_BINDING | MSigDB lists | GO\_REGULATION\_OF\_BINDING | 71 | 7 | 4161 | 153 | Sin3A,ap,CtBP,Smox,hts,dpp,Hsc70-3 | | 1.466e-02 | -4.22 | GSE1112\_OT1\_CD8AB\_VS\_HY\_CD8AA\_THYMOCYTE\_RTOC\_CULTURE\_DN | MSigDB lists | GSE1112\_OT1\_CD8AB\_VS\_HY\_CD8AA\_THYMOCYTE\_RTOC\_CULTURE\_DN | 40 | 5 | 4161 | 153 | CG12099,NiPp1,Pli,CG5742,Cnx99A | | 1.466e-02 | -4.22 | GSE3982\_BCELL\_VS\_TH2\_UP | MSigDB lists | GSE3982\_BCELL\_VS\_TH2\_UP | 40 | 5 | 4161 | 153 | CG14291,Arl4,rush,fne,CtBP | | 1.466e-02 | -4.22 | NFKB\_C | MSigDB lists | NFKB\_C | 40 | 5 | 4161 | 153 | pbl,Notum,cta,Sin3A,Etl1 | | 1.466e-02 | -4.22 | GSE41978\_ID2\_KO\_VS\_ID2\_KO\_AND\_BIM\_KO\_KLRG1\_LOW\_EFFECTOR\_CD8\_TCELL\_UP | MSigDB lists | GSE41978\_ID2\_KO\_VS\_ID2\_KO\_AND\_BIM\_KO\_KLRG1\_LOW\_EFFECTOR\_CD8\_TCELL\_UP | 40 | 5 | 4161 | 153 | N,Cnx99A,Snx3,Imp,hh | | 1.466e-02 | -4.22 | GSE360\_L\_MAJOR\_VS\_T\_GONDII\_MAC\_UP | MSigDB lists | GSE360\_L\_MAJOR\_VS\_T\_GONDII\_MAC\_UP | 40 | 5 | 4161 | 153 | Tlk,Tif-IA,Ptpmeg,Syx1A,CG43313 | | 1.466e-02 | -4.22 | GO\_INFLAMMATORY\_RESPONSE | MSigDB lists | GO\_INFLAMMATORY\_RESPONSE | 40 | 5 | 4161 | 153 | tkv,Rac2,N,JIL-1,dpp | | 1.466e-02 | -4.22 | MODULE\_27 | MSigDB lists | MODULE\_27 | 40 | 5 | 4161 | 153 | LRP1,smal,tkv,nAChRalpha4,mgl | | 1.467e-02 | -4.22 | fly chr56E4-56E4\|2-90 cM | chromosome location | fly chr56E4-56E4\|2-90 cM | 9 | 2 | 25010 | 531 | lncRNA:CR44572,Toll-7 | | 1.467e-02 | -4.22 | fly chr36F5-36F5\|2-53 cM | chromosome location | fly chr36F5-36F5\|2-53 cM | 9 | 2 | 25010 | 531 | grnd,CG10283 | | 1.467e-02 | -4.22 | fly chr36F5-36F5\|2-53 | chromosome location | fly chr36F5-36F5\|2-53 | 9 | 2 | 25010 | 531 | CG10283,grnd | | 1.467e-02 | -4.22 | fly chr66F1-66F1\|3-28 | chromosome location | fly chr66F1-66F1\|3-28 | 9 | 2 | 25010 | 531 | smg,Doc2 | | 1.467e-02 | -4.22 | fly chr66F1-66F1\|3-28 cM | chromosome location | fly chr66F1-66F1\|3-28 cM | 9 | 2 | 25010 | 531 | Doc2,smg | | 1.467e-02 | -4.22 | fly chr56E4-56E4\|2-90 | chromosome location | fly chr56E4-56E4\|2-90 | 9 | 2 | 25010 | 531 | lncRNA:CR44572,Toll-7 | | 1.468e-02 | -4.22 | ZIC2\_01 | MSigDB lists | ZIC2\_01 | 55 | 6 | 4161 | 153 | LRP1,Tlk,Rac2,Utx,ras,JIL-1 | | 1.468e-02 | -4.22 | GTTTGTT\_MIR495 | MSigDB lists | GTTTGTT\_MIR495 | 55 | 6 | 4161 | 153 | Pat1,CG5742,par-1,ap,Hsc70-3,DIP-epsilon | | 1.468e-02 | -4.22 | GSE13411\_SWITCHED\_MEMORY\_BCELL\_VS\_PLASMA\_CELL\_UP | MSigDB lists | GSE13411\_SWITCHED\_MEMORY\_BCELL\_VS\_PLASMA\_CELL\_UP | 55 | 6 | 4161 | 153 | S6kII,rush,Ppa,CG2812,Tlk,Doa | | 1.480e-02 | -4.21 | GO\_REGULATION\_OF\_HYDROLASE\_ACTIVITY | MSigDB lists | GO\_REGULATION\_OF\_HYDROLASE\_ACTIVITY | 283 | 18 | 4161 | 153 | Rho1,Tpr2,EcR,LRP1,CG4538,Smox,pbl,dpp,Cyp1,drk,CG11367,Hsc70-3,cta,S6kII,CaMKII,CG5522,nudE,Fas2 | | 1.482e-02 | -4.21 | BRACHYURY | prints domains | PR00938 | 5 | 2 | 2185 | 88 | Doc1,Doc2 | | 1.482e-02 | -4.21 | INTEGRINA | prints domains | PR01185 | 5 | 2 | 2185 | 88 | if,scb | | 1.515e-02 | -4.19 | ovarian follicle cell stalk formation | biological process | GO:0030713 | 16 | 3 | 12345 | 414 | N,hh,Stat92E | | 1.515e-02 | -4.19 | activation of immune response | biological process | GO:0002253 | 16 | 3 | 12345 | 414 | Toll-7,Tl,Mmp2 | | 1.515e-02 | -4.19 | cell-matrix adhesion | biological process | GO:0007160 | 16 | 3 | 12345 | 414 | tyn,if,scb | | 1.515e-02 | -4.19 | establishment of epithelial cell polarity | biological process | GO:0090162 | 16 | 3 | 12345 | 414 | ft,fj,ds | | 1.515e-02 | -4.19 | cellular response to ionizing radiation | biological process | GO:0071479 | 16 | 3 | 12345 | 414 | Stat92E,Utx,mof | | 1.520e-02 | -4.19 | negative regulation of organelle organization | biological process | GO:0010639 | 96 | 8 | 12345 | 414 | lncRNA:roX1,Mos,spoon,hts,JIL-1,Chro,Utx,ctrip | | 1.525e-02 | -4.18 | Marcal1 (Marcal1) | protein interactions | 33709 | 22 | 4 | 9204 | 412 | osa,Psc,Pcl,psq | | 1.525e-02 | -4.18 | CG7763 (uncharacterized protein) | protein interactions | 36235 | 22 | 4 | 9204 | 412 | CG2865,CG12182,Mhcl,CG17163 | | 1.525e-02 | -4.18 | PlexA (Plexin A) | protein interactions | 43832 | 22 | 4 | 9204 | 412 | pbl,Fas2,if,Rho1 | | 1.525e-02 | -4.18 | CG2116 (uncharacterized protein) | protein interactions | 31735 | 22 | 4 | 9204 | 412 | CG11076,phyl,esn,betaTub56D | | 1.525e-02 | -4.18 | establishment or maintenance of polarity of follicular epithelium | biological process | GO:0016334 | 29 | 4 | 12345 | 414 | crb,spoon,Egfr,sty | | 1.525e-02 | -4.18 | negative regulation of chromatin organization | biological process | GO:1905268 | 29 | 4 | 12345 | 414 | Utx,ctrip,JIL-1,lncRNA:roX1 | | 1.525e-02 | -4.18 | regulation of ATPase activity | biological process | GO:0043462 | 29 | 4 | 12345 | 414 | CG8531,Acf,nrv2,mle | | 1.525e-02 | -4.18 | peptidyl-threonine phosphorylation | biological process | GO:0018107 | 29 | 4 | 12345 | 414 | Hipk,Asator,Haspin,CaMKII | | 1.525e-02 | -4.18 | negative regulation of transferase activity | biological process | GO:0051348 | 29 | 4 | 12345 | 414 | aos,CG8441,sty,CycE | | 1.525e-02 | -4.18 | vascular endothelial growth factor receptor signaling pathway | biological process | GO:0048010 | 29 | 4 | 12345 | 414 | Socs16D,aop,drk,Pvf3 | | 1.535e-02 | -4.18 | regulation of cell projection assembly | biological process | GO:0060491 | 44 | 5 | 12345 | 414 | Arp2,Rho1,N,PsGEF,CaMKII | | 1.535e-02 | -4.18 | regulation of plasma membrane bounded cell projection assembly | biological process | GO:0120032 | 44 | 5 | 12345 | 414 | PsGEF,Arp2,N,Rho1,CaMKII | | 1.535e-02 | -4.18 | negative regulation of synaptic growth at neuromuscular junction | biological process | GO:0045886 | 44 | 5 | 12345 | 414 | Acsl,Dad,hiw,Syp,scb | | 1.535e-02 | -4.18 | negative regulation of neuromuscular junction development | biological process | GO:1904397 | 44 | 5 | 12345 | 414 | hiw,Syp,scb,Acsl,Dad | | 1.535e-02 | -4.18 | asymmetric neuroblast division | biological process | GO:0055059 | 44 | 5 | 12345 | 414 | sdt,Zif,pon,flfl,wake | | 1.539e-02 | -4.17 | positive regulation of hemocyte differentiation | biological process | GO:0045612 | 6 | 2 | 12345 | 414 | N,lz | | 1.539e-02 | -4.17 | clypeo-labral disc development | biological process | GO:0035213 | 6 | 2 | 12345 | 414 | CG13366,TfAP-2 | | 1.539e-02 | -4.17 | establishment of body hair or bristle planar orientation | biological process | GO:0048104 | 6 | 2 | 12345 | 414 | ft,spen | | 1.539e-02 | -4.17 | neuroepithelial cell differentiation | biological process | GO:0060563 | 6 | 2 | 12345 | 414 | exex,tin | | 1.539e-02 | -4.17 | neuroendocrine cell differentiation | biological process | GO:0061101 | 6 | 2 | 12345 | 414 | tin,exex | | 1.539e-02 | -4.17 | positive regulation of nervous system process | biological process | GO:0031646 | 6 | 2 | 12345 | 414 | egr,Prosap | | 1.539e-02 | -4.17 | dosage compensation complex assembly | biological process | GO:0042714 | 6 | 2 | 12345 | 414 | mle,lncRNA:roX1 | | 1.539e-02 | -4.17 | amnioserosa maintenance | biological process | GO:0046665 | 6 | 2 | 12345 | 414 | crb,ush | | 1.539e-02 | -4.17 | proboscis development | biological process | GO:0048728 | 6 | 2 | 12345 | 414 | CG13366,TfAP-2 | | 1.539e-02 | -4.17 | brain morphogenesis | biological process | GO:0048854 | 6 | 2 | 12345 | 414 | ey,Prosap | | 1.539e-02 | -4.17 | regulation of cardioblast differentiation | biological process | GO:0051890 | 6 | 2 | 12345 | 414 | N,pnt | | 1.539e-02 | -4.17 | equator specification | biological process | GO:0045317 | 6 | 2 | 12345 | 414 | ft,ds | | 1.542e-02 | -4.17 | oocyte axis specification | biological process | GO:0007309 | 135 | 10 | 12345 | 414 | Egfr,Syp,gus,dpp,Pat1,heph,cnc,par-1,psq,N | | 1.567e-02 | -4.16 | Hrs (Hepatocyte growth factor regulated tyrosine kinase substrate) | protein interactions | 33458 | 59 | 7 | 9204 | 412 | RhoGEF2,rush,hh,ft,fj,Tl,CtBP | | 1.572e-02 | -4.15 | IG\_LIKE | prosite domains | PS50835 | 132 | 13 | 6243 | 316 | CG6044,nrm,Fas2,lbk,otk2,beat-IIIc,Dscam1,robo2,ImpL2,DIP-epsilon,Lac,dpr7,Fas3 | | 1.576e-02 | -4.15 | GO\_VASCULATURE\_DEVELOPMENT | MSigDB lists | GO\_VASCULATURE\_DEVELOPMENT | 89 | 8 | 4161 | 153 | Ser,cta,mgl,N,hh,LRP1,so,slow | | 1.583e-02 | -4.15 | GO\_PROTEIN\_SERINE\_THREONINE\_KINASE\_ACTIVITY | MSigDB lists | GO\_PROTEIN\_SERINE\_THREONINE\_KINASE\_ACTIVITY | 125 | 10 | 4161 | 153 | tkv,JIL-1,mbt,Doa,Tlk,CaMKII,Taf1,par-1,S6kII,PhKgamma | | 1.587e-02 | -4.14 | Golgi cisterna membrane | cellular component | GO:0032580 | 6 | 2 | 11940 | 407 | CG43313,CG9220 | | 1.587e-02 | -4.14 | perinuclear endoplasmic reticulum | cellular component | GO:0097038 | 6 | 2 | 11940 | 407 | S,Cnx99A | | 1.589e-02 | -4.14 | polo (polo) | protein interactions | 40232 | 73 | 8 | 9204 | 412 | Acf,stg,Doc1,N,CG13380,spen,nuf,bi | | 1.595e-02 | -4.14 | LXR\_Q3 | MSigDB lists | LXR\_Q3 | 27 | 4 | 4161 | 153 | Acsl,NKAIN,SamDC,Tudor-SN | | 1.595e-02 | -4.14 | MODULE\_204 | MSigDB lists | MODULE\_204 | 27 | 4 | 4161 | 153 | TfAP-2,CG43313,CG2201,nudE | | 1.595e-02 | -4.14 | GO\_FORMATION\_OF\_PRIMARY\_GERM\_LAYER | MSigDB lists | GO\_FORMATION\_OF\_PRIMARY\_GERM\_LAYER | 27 | 4 | 4161 | 153 | Lim1,if,Utx,Smox | | 1.595e-02 | -4.14 | TCTGATA\_MIR361 | MSigDB lists | TCTGATA\_MIR361 | 27 | 4 | 4161 | 153 | DIP-epsilon,Tudor-SN,CaMKII,Rac2 | | 1.595e-02 | -4.14 | alpha-Spec (alpha Spectrin) | protein interactions | 38231 | 46 | 6 | 9204 | 412 | Syp,Ten-m,crb,S6kII,bor,rau | | 1.595e-02 | -4.14 | ZHENG\_FOXP3\_TARGETS\_IN\_THYMUS\_UP | MSigDB lists | ZHENG\_FOXP3\_TARGETS\_IN\_THYMUS\_UP | 56 | 6 | 4161 | 153 | betaTub56D,CG45050,Arl4,phtf,CG11367,l(3)mbt | | 1.595e-02 | -4.14 | GSE17721\_LPS\_VS\_CPG\_4H\_BMDC\_UP | MSigDB lists | GSE17721\_LPS\_VS\_CPG\_4H\_BMDC\_UP | 56 | 6 | 4161 | 153 | Tlk,if,ras,cta,Pat1,CG43313 | | 1.595e-02 | -4.14 | PAX4\_03 | MSigDB lists | PAX4\_03 | 56 | 6 | 4161 | 153 | Snx3,CG12547,LRP1,mgl,LManII,kdn | | 1.602e-02 | -4.13 | KEGG\_NOTCH\_SIGNALING\_PATHWAY | MSigDB lists | KEGG\_NOTCH\_SIGNALING\_PATHWAY | 15 | 3 | 4161 | 153 | N,CtBP,Ser | | 1.602e-02 | -4.13 | GO\_MYOBLAST\_DIFFERENTIATION | MSigDB lists | GO\_MYOBLAST\_DIFFERENTIATION | 15 | 3 | 4161 | 153 | N,Ser,hh | | 1.602e-02 | -4.13 | REACTOME\_NCAM\_SIGNALING\_FOR\_NEURITE\_OUT\_GROWTH | MSigDB lists | REACTOME\_NCAM\_SIGNALING\_FOR\_NEURITE\_OUT\_GROWTH | 15 | 3 | 4161 | 153 | JIL-1,drk,Fas2 | | 1.602e-02 | -4.13 | MIKKELSEN\_MEF\_LCP\_WITH\_H3K4ME3 | MSigDB lists | MIKKELSEN\_MEF\_LCP\_WITH\_H3K4ME3 | 15 | 3 | 4161 | 153 | CG3097,CG6752,CG45050 | | 1.602e-02 | -4.13 | GO\_CELL\_CELL\_ADHESION\_VIA\_PLASMA\_MEMBRANE\_ADHESION\_MOLECULES | MSigDB lists | GO\_CELL\_CELL\_ADHESION\_VIA\_PLASMA\_MEMBRANE\_ADHESION\_MOLECULES | 15 | 3 | 4161 | 153 | ds,ft,Dscam1 | | 1.602e-02 | -4.13 | GO\_SEGMENTATION | MSigDB lists | GO\_SEGMENTATION | 15 | 3 | 4161 | 153 | Smox,Utx,Lim1 | | 1.610e-02 | -4.13 | regulation of mitotic cell cycle phase transition | biological process | GO:1901990 | 97 | 8 | 12345 | 414 | CycE,Chro,N,ci,Egfr,PEK,Myb,stg | | 1.615e-02 | -4.13 | SH2 | prosite domains | PS50001 | 30 | 5 | 6243 | 316 | CG10479,CG33993,Stat92E,drk,Socs16D | | 1.616e-02 | -4.13 | fn3 | pfam domains | PF00041 | 40 | 5 | 10480 | 393 | Fas2,Dscam1,sas,Ptp99A,robo2 | | 1.621e-02 | -4.12 | GSE3994\_WT\_VS\_PAC1\_KO\_ACTIVATED\_MAST\_CELL\_UP | MSigDB lists | GSE3994\_WT\_VS\_PAC1\_KO\_ACTIVATED\_MAST\_CELL\_UP | 41 | 5 | 4161 | 153 | Arl4,Syx1A,Mmp2,retm,CG6752 | | 1.621e-02 | -4.12 | GSE7852\_THYMUS\_VS\_FAT\_TREG\_DN | MSigDB lists | GSE7852\_THYMUS\_VS\_FAT\_TREG\_DN | 41 | 5 | 4161 | 153 | Ptpmeg,nudE,Rho1,Smox,Pcl | | 1.621e-02 | -4.12 | BILD\_HRAS\_ONCOGENIC\_SIGNATURE | MSigDB lists | BILD\_HRAS\_ONCOGENIC\_SIGNATURE | 41 | 5 | 4161 | 153 | Cyp1,mmy,dpp,lbk,Syx1A | | 1.621e-02 | -4.12 | PID\_ERBB1\_DOWNSTREAM\_PATHWAY | MSigDB lists | PID\_ERBB1\_DOWNSTREAM\_PATHWAY | 41 | 5 | 4161 | 153 | Arp2,Rac2,S6kII,JIL-1,drk | | 1.621e-02 | -4.12 | GSE22886\_NAIVE\_CD8\_TCELL\_VS\_NKCELL\_DN | MSigDB lists | GSE22886\_NAIVE\_CD8\_TCELL\_VS\_NKCELL\_DN | 41 | 5 | 4161 | 153 | CtBP,Hydr2,N,Npc1a,Fas2 | | 1.629e-02 | -4.12 | - | gene3d domains | 3.90.810.10 | 5 | 2 | 4929 | 208 | Pak3,mbt | | 1.629e-02 | -4.12 | - | gene3d domains | 2.130.10.130 | 5 | 2 | 4929 | 208 | scb,if | | 1.636e-02 | -4.11 | polytene chromosome interband | cellular component | GO:0005705 | 44 | 5 | 11940 | 407 | Utx,mof,Chro,lawc,JIL-1 | | 1.636e-02 | -4.11 | cell division site part | cellular component | GO:0032155 | 44 | 5 | 11940 | 407 | Arl4,Tl,RhoGEF2,nuf,pbl | | 1.637e-02 | -4.11 | MBD | smart domains | SM00391 | 4 | 2 | 5086 | 276 | sba,tou | | 1.637e-02 | -4.11 | AT\_hook | smart domains | SM00384 | 4 | 2 | 5086 | 276 | Thd1,Taf1 | | 1.655e-02 | -4.10 | pigmentation | biological process | GO:0043473 | 79 | 7 | 12345 | 414 | bi,hh,B-H2,bab1,Cnx99A,dpp,eya | | 1.671e-02 | -4.09 | immune response | biological process | GO:0006955 | 199 | 13 | 12345 | 414 | hh,par-1,Rac2,bbg,Pli,Stat92E,Cdk5alpha,Tlk,ci,eya,Tl,egr,Rho1 | | 1.672e-02 | -4.09 | CG17265 (uncharacterized protein) | protein interactions | 33509 | 34 | 5 | 9204 | 412 | so,Mondo,CG12010,esn,CG17265 | | 1.672e-02 | -4.09 | Cul2 (Cullin 2) | protein interactions | 35420 | 34 | 5 | 9204 | 412 | EcR,tkv,CG43658,dpp,Doa | | 1.672e-02 | -4.09 | so (sine oculis) | protein interactions | 35662 | 34 | 5 | 9204 | 412 | dpp,RhoGAP93B,CG17265,RhoGEF2,eya | | 1.681e-02 | -4.09 | negative regulation of synapse assembly | biological process | GO:0051964 | 45 | 5 | 12345 | 414 | scb,Syp,hiw,Dad,Acsl | | 1.681e-02 | -4.09 | negative regulation of synapse organization | biological process | GO:1905809 | 45 | 5 | 12345 | 414 | Acsl,Dad,hiw,Syp,scb | | 1.682e-02 | -4.09 | molecular function regulator | molecular function | GO:0098772 | 563 | 28 | 12176 | 396 | RhoGEF2,RhoGAP16F,Atg17,pbl,NiPp1,Pli,nrv2,CG5522,ITP,RhoGAP18B,CycE,CG43658,RhoGAP15B,Socs16D,Ser,dpp,RhoGAP93B,tsl,PsGEF,hh,aos,CG8531,spi,Pvf3,Cdk5alpha,RhoGAP102A,RapGAP1,CG8441 | | 1.689e-02 | -4.08 | ZWANG\_CLASS\_1\_TRANSIENTLY\_INDUCED\_BY\_EGF | MSigDB lists | ZWANG\_CLASS\_1\_TRANSIENTLY\_INDUCED\_BY\_EGF | 73 | 7 | 4161 | 153 | Rhp,S6kII,JIL-1,CtBP,CaMKII,Hydr2,Smox | | 1.714e-02 | -4.07 | peptidyl-threonine modification | biological process | GO:0018210 | 30 | 4 | 12345 | 414 | CaMKII,Haspin,Hipk,Asator | | 1.714e-02 | -4.07 | neutral lipid metabolic process | biological process | GO:0006638 | 30 | 4 | 12345 | 414 | CG34384,Hydr2,Lpin,CG1271 | | 1.714e-02 | -4.07 | regulation of axon guidance | biological process | GO:1902667 | 30 | 4 | 12345 | 414 | NetB,CadN,NetA,mmy | | 1.714e-02 | -4.07 | cellular component maintenance | biological process | GO:0043954 | 30 | 4 | 12345 | 414 | crb,Rho1,Ten-m,Rac2 | | 1.714e-02 | -4.07 | positive regulation of smoothened signaling pathway | biological process | GO:0045880 | 30 | 4 | 12345 | 414 | dlp,dally,Notum,Hipk | | 1.714e-02 | -4.07 | acylglycerol metabolic process | biological process | GO:0006639 | 30 | 4 | 12345 | 414 | CG1271,Lpin,CG34384,Hydr2 | | 1.730e-02 | -4.06 | AP1\_Q2\_01 | MSigDB lists | AP1\_Q2\_01 | 57 | 6 | 4161 | 153 | CG3097,smal,nudE,ras,fne,Ank | | 1.730e-02 | -4.06 | HALLMARK\_UV\_RESPONSE\_UP | MSigDB lists | HALLMARK\_UV\_RESPONSE\_UP | 57 | 6 | 4161 | 153 | dpp,Cyp1,CycE,ap,SamDC,Ykt6 | | 1.730e-02 | -4.06 | HNF4\_01\_B | MSigDB lists | HNF4\_01\_B | 57 | 6 | 4161 | 153 | Lim1,LRP1,Tlk,sdt,mbt,CG3961 | | 1.734e-02 | -4.05 | regulation of antimicrobial peptide production | biological process | GO:0002784 | 62 | 6 | 12345 | 414 | Doa,Pli,Usp2,Tl,ush,CG4325 | | 1.742e-02 | -4.05 | KH\_dom | interpro domains | IPR004087 | 27 | 4 | 11383 | 428 | mub,spoon,ps,Imp | | 1.742e-02 | -4.05 | SMAD\_FHA\_dom\_sf | interpro domains | IPR008984 | 27 | 4 | 11383 | 428 | NiPp1,Dad,CG17168,Smox | | 1.742e-02 | -4.05 | KH\_dom\_type\_1 | interpro domains | IPR004088 | 27 | 4 | 11383 | 428 | Imp,ps,spoon,mub | | 1.747e-02 | -4.05 | cell-cell signaling | biological process | GO:0007267 | 311 | 18 | 12345 | 414 | Ten-m,nAChRalpha4,tkv,cpo,Ser,Cirl,otk2,CaMKII,osa,Acsl,pAbp,fj,mam,twi,hh,Mmp2,N,Syx1A | | 1.761e-02 | -4.04 | sd (scalloped) | protein interactions | 32536 | 47 | 6 | 9204 | 412 | Psc,Pcl,osa,par-1,N,ci | | 1.774e-02 | -4.03 | oocyte construction | biological process | GO:0007308 | 138 | 10 | 12345 | 414 | Pat1,dpp,gus,heph,Egfr,Syp,N,psq,cnc,par-1 | | 1.779e-02 | -4.03 | - | gene3d domains | 3.30.40.10 | 214 | 16 | 4929 | 208 | hiw,tna,enok,tou,Acf,rush,CG41099,Pcl,CG6752,CG9855,CG32369,CG12099,CG2991,CG6051,Psc,CG4325 | | 1.782e-02 | -4.03 | ac (achaete) | protein interactions | 30981 | 23 | 4 | 9204 | 412 | Hip1,Egfr,ttk,Bx | | 1.782e-02 | -4.03 | Dcr-1 (Dicer-1) | protein interactions | 42693 | 23 | 4 | 9204 | 412 | N,ban,AGO2,CycE | | 1.782e-02 | -4.03 | lolal (lola like) | protein interactions | 44703 | 23 | 4 | 9204 | 412 | psq,ttk,dpp,N | | 1.782e-02 | -4.03 | vkg (viking) | protein interactions | 33726 | 23 | 4 | 9204 | 412 | Myo10A,dpp,Tet,Fas2 | | 1.782e-02 | -4.03 | Vti1b (Vesicle transport through interaction with t-SNAREs 1b) | protein interactions | 42215 | 23 | 4 | 9204 | 412 | CG18766,Usp2,gus,EcR | | 1.787e-02 | -4.02 | CAIRO\_HEPATOBLASTOMA\_CLASSES\_DN | MSigDB lists | CAIRO\_HEPATOBLASTOMA\_CLASSES\_DN | 42 | 5 | 4161 | 153 | Smox,ft,if,TfAP-2,CG3961 | | 1.787e-02 | -4.02 | GO\_MORPHOGENESIS\_OF\_EMBRYONIC\_EPITHELIUM | MSigDB lists | GO\_MORPHOGENESIS\_OF\_EMBRYONIC\_EPITHELIUM | 42 | 5 | 4161 | 153 | wech,so,ap,Utx,TfAP-2 | | 1.787e-02 | -4.02 | GSE29618\_BCELL\_VS\_PDC\_DAY7\_FLU\_VACCINE\_DN | MSigDB lists | GSE29618\_BCELL\_VS\_PDC\_DAY7\_FLU\_VACCINE\_DN | 42 | 5 | 4161 | 153 | LManII,S6kII,Ser,Npc1a,emp | | 1.787e-02 | -4.02 | GSE20727\_CTRL\_VS\_H2O2\_TREATED\_DC\_DN | MSigDB lists | GSE20727\_CTRL\_VS\_H2O2\_TREATED\_DC\_DN | 42 | 5 | 4161 | 153 | Smox,hiw,Ank,Sin3A,l(3)mbt | | 1.787e-02 | -4.02 | CDPCR3HD\_01 | MSigDB lists | CDPCR3HD\_01 | 42 | 5 | 4161 | 153 | CG8531,CaMKII,drk,so,Lac | | 1.787e-02 | -4.02 | GSE21033\_CTRL\_VS\_POLYIC\_STIM\_DC\_1H\_DN | MSigDB lists | GSE21033\_CTRL\_VS\_POLYIC\_STIM\_DC\_1H\_DN | 42 | 5 | 4161 | 153 | CG12099,smal,CG9855,CG3961,nAChRalpha4 | | 1.787e-02 | -4.02 | MEF2\_Q6\_01 | MSigDB lists | MEF2\_Q6\_01 | 42 | 5 | 4161 | 153 | TfIIS,Nhe2,so,CtBP,pAbp | | 1.793e-02 | -4.02 | receptor complex | cellular component | GO:0043235 | 79 | 7 | 11940 | 407 | scb,nAChRalpha4,Egfr,EcR,Prosap,tkv,if | | 1.794e-02 | -4.02 | negative regulation of secretion by cell | biological process | GO:1903531 | 17 | 3 | 12345 | 414 | NFAT,CaMKII,CG9650 | | 1.794e-02 | -4.02 | regulation of actomyosin structure organization | biological process | GO:0110020 | 17 | 3 | 12345 | 414 | CG43658,pbl,if | | 1.794e-02 | -4.02 | apical protein localization | biological process | GO:0045176 | 17 | 3 | 12345 | 414 | egr,crb,sdt | | 1.801e-02 | -4.02 | positive regulation of immune response | biological process | GO:0050778 | 99 | 8 | 12345 | 414 | Toll-7,Doa,Mmp2,Tl,ush,CG4325,Etl1,Pli | | 1.808e-02 | -4.01 | GSE37301\_CD4\_TCELL\_VS\_RAG2\_KO\_NK\_CELL\_UP | MSigDB lists | GSE37301\_CD4\_TCELL\_VS\_RAG2\_KO\_NK\_CELL\_UP | 28 | 4 | 4161 | 153 | ScpX,Ser,Cnx99A,Dscam1 | | 1.808e-02 | -4.01 | DAZARD\_RESPONSE\_TO\_UV\_SCC\_DN | MSigDB lists | DAZARD\_RESPONSE\_TO\_UV\_SCC\_DN | 28 | 4 | 4161 | 153 | SamDC,par-1,CG11030,Tlk | | 1.808e-02 | -4.01 | GSE5589\_WT\_VS\_IL6\_KO\_LPS\_AND\_IL6\_STIM\_MACROPHAGE\_45MIN\_UP | MSigDB lists | GSE5589\_WT\_VS\_IL6\_KO\_LPS\_AND\_IL6\_STIM\_MACROPHAGE\_45MIN\_UP | 28 | 4 | 4161 | 153 | S6kII,nAChRalpha4,NetB,CG3097 | | 1.808e-02 | -4.01 | GO\_CONNECTIVE\_TISSUE\_DEVELOPMENT | MSigDB lists | GO\_CONNECTIVE\_TISSUE\_DEVELOPMENT | 28 | 4 | 4161 | 153 | N,tkv,dpp,CG9220 | | 1.808e-02 | -4.01 | GO\_PLASMA\_MEMBRANE\_RECEPTOR\_COMPLEX | MSigDB lists | GO\_PLASMA\_MEMBRANE\_RECEPTOR\_COMPLEX | 28 | 4 | 4161 | 153 | nAChRalpha4,tkv,if,dpp | | 1.808e-02 | -4.01 | WWTAAGGC\_UNKNOWN | MSigDB lists | WWTAAGGC\_UNKNOWN | 28 | 4 | 4161 | 153 | kdn,so,Fas2,Lim1 | | 1.809e-02 | -4.01 | nucleoside phosphate binding | molecular function | GO:1901265 | 955 | 43 | 12176 | 396 | CG4199,CG12010,Myo10A,mle,lz,mbt,CG14535,Arl4,Pex1,par-1,PhKgamma,betaTub56D,CG8485,JIL-1,Egfr,Mos,CG9281,CG9541,CG8097,Hipk,Haspin,Mhcl,PEK,Rho1,ScsbetaG,S6kII,CtBP,Asator,Mrp4,Etl1,Arp2,CaMKII,Tlk,Hsc70-3,Rac2,Cdk12,KP78a,Pak3,bor,tkv,Doa,cta,wnd | | 1.809e-02 | -4.01 | nucleotide binding | molecular function | GO:0000166 | 955 | 43 | 12176 | 396 | CaMKII,Tlk,Hsc70-3,Rac2,Cdk12,KP78a,Pak3,bor,tkv,cta,Doa,wnd,CG8097,Hipk,Haspin,PEK,Mhcl,Rho1,S6kII,ScsbetaG,CtBP,Asator,Mrp4,Arp2,Etl1,Arl4,Pex1,PhKgamma,par-1,betaTub56D,CG8485,JIL-1,Egfr,Mos,CG9541,CG9281,CG4199,CG12010,Myo10A,mle,lz,mbt,CG14535 | | 1.809e-02 | -4.01 | fly chr87C3-87C3\|3-52 | chromosome location | fly chr87C3-87C3\|3-52 | 10 | 2 | 25010 | 531 | Dtg,Snx3 | | 1.809e-02 | -4.01 | fly chr87C3-87C3\|3-52 cM | chromosome location | fly chr87C3-87C3\|3-52 cM | 10 | 2 | 25010 | 531 | Snx3,Dtg | | 1.810e-02 | -4.01 | EGR\_Q6 | MSigDB lists | EGR\_Q6 | 74 | 7 | 4161 | 153 | CG12099,Imp,Nhe2,Hsc70Cb,drk,Utx,sm | | 1.821e-02 | -4.01 | Myd88 (Myd88) | protein interactions | 35956 | 13 | 3 | 9204 | 412 | Pli,Tl,Toll-7 | | 1.821e-02 | -4.01 | Doc2 (Dorsocross2) | protein interactions | 39038 | 13 | 3 | 9204 | 412 | Ubx,tin,Doc1 | | 1.821e-02 | -4.01 | Pxt (Peroxinectin-like) | protein interactions | 42131 | 13 | 3 | 9204 | 412 | ci,Mhcl,l(3)mbt | | 1.821e-02 | -4.01 | Gli (Gliotactin) | protein interactions | 34927 | 13 | 3 | 9204 | 412 | tkv,Dad,N | | 1.827e-02 | -4.00 | cv-d (crossveinless d) | protein interactions | 41921 | 5 | 2 | 9204 | 412 | dpp,ci | | 1.827e-02 | -4.00 | 128up (upstream of RpIII128) | protein interactions | 36288 | 5 | 2 | 9204 | 412 | Utx,smg | | 1.827e-02 | -4.00 | CG12863 (uncharacterized protein) | protein interactions | 36623 | 5 | 2 | 9204 | 412 | CG41099,CG32369 | | 1.827e-02 | -4.00 | CG17612 (uncharacterized protein) | protein interactions | 33639 | 5 | 2 | 9204 | 412 | ey,mam | | 1.827e-02 | -4.00 | rn (rotund) | protein interactions | 40879 | 5 | 2 | 9204 | 412 | Ser,N | | 1.827e-02 | -4.00 | bab1 (bric a brac 1) | protein interactions | 38116 | 5 | 2 | 9204 | 412 | hts,FoxL1 | | 1.827e-02 | -4.00 | St3 (Sulfotransferase 3) | protein interactions | 37743 | 5 | 2 | 9204 | 412 | CG43658,Bx | | 1.827e-02 | -4.00 | CG3645 (uncharacterized protein) | protein interactions | 33190 | 5 | 2 | 9204 | 412 | Gbs-70E,ci | | 1.827e-02 | -4.00 | INPP5E (Inositol polyphosphate 5-phosphatase E) | protein interactions | 39404 | 5 | 2 | 9204 | 412 | smal,bdg | | 1.827e-02 | -4.00 | Pdk (Pyruvate dehydrogenase kinase) | protein interactions | 35970 | 5 | 2 | 9204 | 412 | Gbs-70E,Syx1A | | 1.827e-02 | -4.00 | Abp1 (Actin binding protein 1) | protein interactions | 39520 | 5 | 2 | 9204 | 412 | Arp2,CaMKII | | 1.827e-02 | -4.00 | Hip14 (Huntingtin-interacting protein 14) | protein interactions | 39747 | 5 | 2 | 9204 | 412 | sty,Dad | | 1.827e-02 | -4.00 | nemy (no extended memory) | protein interactions | 36395 | 5 | 2 | 9204 | 412 | par-1,pbl | | 1.827e-02 | -4.00 | TAF1B (TATA box-binding protein-associated factor RNA polymerase I subunit B) | protein interactions | 41242 | 5 | 2 | 9204 | 412 | Utx,udd | | 1.827e-02 | -4.00 | Snm1 (Snm1) | protein interactions | 40669 | 5 | 2 | 9204 | 412 | CadN,Acf | | 1.827e-02 | -4.00 | Zir (Zizimin-related) | protein interactions | 33165 | 5 | 2 | 9204 | 412 | Rac2,Fas2 | | 1.827e-02 | -4.00 | HERC2 (HECT and RLD domain containing protein 2) | protein interactions | 33035 | 5 | 2 | 9204 | 412 | sty,Fas2 | | 1.828e-02 | -4.00 | REACTOME\_CD28\_DEPENDENT\_VAV1\_PATHWAY | MSigDB lists | REACTOME\_CD28\_DEPENDENT\_VAV1\_PATHWAY | 6 | 2 | 4161 | 153 | Rac2,drk | | 1.828e-02 | -4.00 | GO\_REGULATION\_OF\_CELL\_PROLIFERATION\_INVOLVED\_IN\_HEART\_MORPHOGENESIS | MSigDB lists | GO\_REGULATION\_OF\_CELL\_PROLIFERATION\_INVOLVED\_IN\_HEART\_MORPHOGENESIS | 6 | 2 | 4161 | 153 | so,N | | 1.828e-02 | -4.00 | GO\_HIPPO\_SIGNALING | MSigDB lists | GO\_HIPPO\_SIGNALING | 6 | 2 | 4161 | 153 | ft,ds | | 1.828e-02 | -4.00 | GO\_REGULATION\_OF\_ACTION\_POTENTIAL | MSigDB lists | GO\_REGULATION\_OF\_ACTION\_POTENTIAL | 6 | 2 | 4161 | 153 | CaMKII,Ank | | 1.828e-02 | -4.00 | RORIE\_TARGETS\_OF\_EWSR1\_FLI1\_FUSION\_UP | MSigDB lists | RORIE\_TARGETS\_OF\_EWSR1\_FLI1\_FUSION\_UP | 6 | 2 | 4161 | 153 | dlp,TfAP-2 | | 1.828e-02 | -4.00 | PID\_HES\_HEY\_PATHWAY | MSigDB lists | PID\_HES\_HEY\_PATHWAY | 6 | 2 | 4161 | 153 | CaMKII,N | | 1.828e-02 | -4.00 | GO\_HEART\_GROWTH | MSigDB lists | GO\_HEART\_GROWTH | 6 | 2 | 4161 | 153 | CaMKII,N | | 1.828e-02 | -4.00 | GO\_RNA\_POLYMERASE\_II\_ACTIVATING\_TRANSCRIPTION\_FACTOR\_BINDING | MSigDB lists | GO\_RNA\_POLYMERASE\_II\_ACTIVATING\_TRANSCRIPTION\_FACTOR\_BINDING | 6 | 2 | 4161 | 153 | Smox,Sin3A | | 1.828e-02 | -4.00 | GO\_BILE\_ACID\_METABOLIC\_PROCESS | MSigDB lists | GO\_BILE\_ACID\_METABOLIC\_PROCESS | 6 | 2 | 4161 | 153 | ScpX,Npc1a | | 1.828e-02 | -4.00 | GO\_MESODERMAL\_CELL\_DIFFERENTIATION | MSigDB lists | GO\_MESODERMAL\_CELL\_DIFFERENTIATION | 6 | 2 | 4161 | 153 | if,Utx | | 1.828e-02 | -4.00 | CEBALLOS\_TARGETS\_OF\_TP53\_AND\_MYC\_UP | MSigDB lists | CEBALLOS\_TARGETS\_OF\_TP53\_AND\_MYC\_UP | 6 | 2 | 4161 | 153 | mmy,Acsl | | 1.828e-02 | -4.00 | GO\_NOTCH\_BINDING | MSigDB lists | GO\_NOTCH\_BINDING | 6 | 2 | 4161 | 153 | N,Ser | | 1.828e-02 | -4.00 | GO\_LIPOPROTEIN\_PARTICLE\_RECEPTOR\_ACTIVITY | MSigDB lists | GO\_LIPOPROTEIN\_PARTICLE\_RECEPTOR\_ACTIVITY | 6 | 2 | 4161 | 153 | mgl,LRP1 | | 1.828e-02 | -4.00 | GO\_HINDLIMB\_MORPHOGENESIS | MSigDB lists | GO\_HINDLIMB\_MORPHOGENESIS | 6 | 2 | 4161 | 153 | N,hh | | 1.828e-02 | -4.00 | WIKMAN\_ASBESTOS\_LUNG\_CANCER\_UP | MSigDB lists | WIKMAN\_ASBESTOS\_LUNG\_CANCER\_UP | 6 | 2 | 4161 | 153 | CtBP,hts | | 1.828e-02 | -4.00 | GO\_FORELIMB\_MORPHOGENESIS | MSigDB lists | GO\_FORELIMB\_MORPHOGENESIS | 6 | 2 | 4161 | 153 | hh,TfAP-2 | | 1.828e-02 | -4.00 | GO\_POSITIVE\_REGULATION\_OF\_MESENCHYMAL\_CELL\_PROLIFERATION | MSigDB lists | GO\_POSITIVE\_REGULATION\_OF\_MESENCHYMAL\_CELL\_PROLIFERATION | 6 | 2 | 4161 | 153 | hh,so | | 1.828e-02 | -4.00 | REACTOME\_SIGNALING\_BY\_BMP | MSigDB lists | REACTOME\_SIGNALING\_BY\_BMP | 6 | 2 | 4161 | 153 | tkv,dpp | | 1.828e-02 | -4.00 | GO\_ECTODERM\_DEVELOPMENT | MSigDB lists | GO\_ECTODERM\_DEVELOPMENT | 6 | 2 | 4161 | 153 | hh,Lim1 | | 1.828e-02 | -4.00 | GO\_POSITIVE\_REGULATION\_OF\_ADHERENS\_JUNCTION\_ORGANIZATION | MSigDB lists | GO\_POSITIVE\_REGULATION\_OF\_ADHERENS\_JUNCTION\_ORGANIZATION | 6 | 2 | 4161 | 153 | Rac2,Smox | | 1.828e-02 | -4.00 | GO\_DEVELOPMENTAL\_INDUCTION | MSigDB lists | GO\_DEVELOPMENTAL\_INDUCTION | 6 | 2 | 4161 | 153 | so,dpp | | 1.828e-02 | -4.00 | GO\_REGULATION\_OF\_MESENCHYMAL\_CELL\_PROLIFERATION | MSigDB lists | GO\_REGULATION\_OF\_MESENCHYMAL\_CELL\_PROLIFERATION | 6 | 2 | 4161 | 153 | so,hh | | 1.828e-02 | -4.00 | REACTOME\_A\_TETRASACCHARIDE\_LINKER\_SEQUENCE\_IS\_REQUIRED\_FOR\_GAG\_SYNTHESIS | MSigDB lists | REACTOME\_A\_TETRASACCHARIDE\_LINKER\_SEQUENCE\_IS\_REQUIRED\_FOR\_GAG\_SYNTHESIS | 6 | 2 | 4161 | 153 | dlp,dally | | 1.828e-02 | -4.00 | CONRAD\_STEM\_CELL | MSigDB lists | CONRAD\_STEM\_CELL | 6 | 2 | 4161 | 153 | Hsc70Cb,phtf | | 1.828e-02 | -4.00 | GO\_REGULATION\_OF\_NEUROBLAST\_PROLIFERATION | MSigDB lists | GO\_REGULATION\_OF\_NEUROBLAST\_PROLIFERATION | 6 | 2 | 4161 | 153 | N,hh | | 1.828e-02 | -4.00 | PID\_HEDGEHOG\_2PATHWAY | MSigDB lists | PID\_HEDGEHOG\_2PATHWAY | 6 | 2 | 4161 | 153 | mgl,hh | | 1.828e-02 | -4.00 | GO\_POSITIVE\_REGULATION\_OF\_TRANSCRIPTION\_FROM\_RNA\_POLYMERASE\_II\_PROMOTER\_INVOLVED\_IN\_CELLULAR\_RESPONSE\_TO\_CHEMICAL\_STIMULUS | MSigDB lists | GO\_POSITIVE\_REGULATION\_OF\_TRANSCRIPTION\_FROM\_RNA\_POLYMERASE\_II\_PROMOTER\_INVOLVED\_IN\_CELLULAR\_RESPONSE\_TO\_CHEMICAL\_STIMULUS | 6 | 2 | 4161 | 153 | N,dpp | | 1.828e-02 | -4.00 | GO\_EMBRYONIC\_FORELIMB\_MORPHOGENESIS | MSigDB lists | GO\_EMBRYONIC\_FORELIMB\_MORPHOGENESIS | 6 | 2 | 4161 | 153 | hh,TfAP-2 | | 1.828e-02 | -4.00 | VERRECCHIA\_RESPONSE\_TO\_TGFB1\_C2 | MSigDB lists | VERRECCHIA\_RESPONSE\_TO\_TGFB1\_C2 | 6 | 2 | 4161 | 153 | Rho1,LRP1 | | 1.828e-02 | -4.00 | GO\_PROTEOGLYCAN\_BINDING | MSigDB lists | GO\_PROTEOGLYCAN\_BINDING | 6 | 2 | 4161 | 153 | dally,dlp | | 1.828e-02 | -4.00 | GO\_TRANSLATION\_REGULATOR\_ACTIVITY | MSigDB lists | GO\_TRANSLATION\_REGULATOR\_ACTIVITY | 6 | 2 | 4161 | 153 | wech,Imp | | 1.829e-02 | -4.00 | membrane part | cellular component | GO:0044425 | 2068 | 87 | 11940 | 407 | Snx3,Fili,Socs16D,egr,N,Fur1,nrv2,wun2,CG41128,Lrt,CG12730,CG11752,nuf,if,Tsp39D,emp,CG6231,hts,Ptr,Cnx99A,CG34172,Stat92E,CG42748,Nhe2,MFS14,Atg17,par-1,Prosap,pbl,Cad87A,ds,scb,neo,Inx3,Cyt-c1,Mmp2,sdt,CG9855,dally,CadN,Inx2,Ykt6,Tsp26A,mgl,Tim17b,tyn,Hip1,bdg,DIP-epsilon,Cirl,bou,RhoGEF2,tkv,CG8531,Ser,Tl,cta,Pli,crb,ft,otk2,Rho1,Npc1a,Fas2,Syx1A,S,Trpm,Mrp4,sty,Toll-7,dlp,ND-B14.5AL,CaMKII,CG14646,nAChRalpha4,Egfr,Arl4,spi,Ten-m,dpr7,sas,CG11367,CCKLR-17D3,Fas3,smal,Dscam1,MFS10 | | 1.833e-02 | -4.00 | enzyme activator activity | molecular function | GO:0008047 | 143 | 10 | 12176 | 396 | RhoGAP15B,RhoGAP18B,RapGAP1,RhoGAP102A,Cdk5alpha,pbl,RhoGAP16F,nrv2,CG8531,RhoGAP93B | | 1.835e-02 | -4.00 | Hedgehog signaling | KEGG pathways | M00678 | 7 | 2 | 3198 | 100 | ci,hh | | 1.835e-02 | -4.00 | Hedgehog signaling | KEGG pathways | dme\_M00678 | 7 | 2 | 3198 | 100 | hh,ci | | 1.836e-02 | -4.00 | neuroblast division | biological process | GO:0055057 | 46 | 5 | 12345 | 414 | wake,flfl,Zif,pon,sdt | | 1.836e-02 | -4.00 | apical junction assembly | biological process | GO:0043297 | 46 | 5 | 12345 | 414 | crb,sdt,nrv2,bou,Lac | | 1.836e-02 | -4.00 | neuronal stem cell division | biological process | GO:0036445 | 46 | 5 | 12345 | 414 | wake,flfl,pon,Zif,sdt | | 1.858e-02 | -3.99 | ZINC\_FINGER\_C2H2\_1 | prosite domains | PS00028 | 287 | 23 | 6243 | 316 | CG9650,ci,CG4360,hang,CG31224,l(3)neo38,ham,CG31612,jing,Kr,CG32772,sbb,CG11247,cas,CG18446,salm,Kah,disco,bowl,sens-2,Zif,ttk,shn | | 1.858e-02 | -3.99 | kay (kayak) | protein interactions | 3772082 | 61 | 7 | 9204 | 412 | Cnx99A,Egfr,wnd,Sox21a,jigr1,jing,sty | | 1.864e-02 | -3.98 | locomotor rhythm | biological process | GO:0045475 | 63 | 6 | 12345 | 414 | disco,qsm,tara,S6kII,Bx,ITP | | 1.864e-02 | -3.98 | regulation of production of molecular mediator of immune response | biological process | GO:0002700 | 63 | 6 | 12345 | 414 | Tl,CG4325,ush,Usp2,Pli,Doa | | 1.866e-02 | -3.98 | cellular component assembly | biological process | GO:0022607 | 1109 | 50 | 12345 | 414 | pbl,Trpm,Tl,lama,CG13366,Ten-m,nudE,gus,Acf,Rho1,lncRNA:roX1,if,Rac2,Taf1,Myo10A,Lac,N,CG34384,crb,psq,Hip1,CG34172,Pat1,Df31,cnc,Atg17,bou,sdt,tkv,enok,Imp,stg,CycE,Prosap,hh,par-1,ttk,spoon,AGO2,Egfr,mle,hts,CG5815,Syp,S,aop,dpp,comm,Myb,nrv2 | | 1.873e-02 | -3.98 | GSE18804\_SPLEEN\_MACROPHAGE\_VS\_COLON\_TUMORAL\_MACROPHAGE\_DN | MSigDB lists | GSE18804\_SPLEEN\_MACROPHAGE\_VS\_COLON\_TUMORAL\_MACROPHAGE\_DN | 58 | 6 | 4161 | 153 | sdt,pAbp,Smox,phtf,CG6752,Mitofilin | | 1.873e-02 | -3.98 | GSE17721\_CTRL\_VS\_LPS\_8H\_BMDC\_DN | MSigDB lists | GSE17721\_CTRL\_VS\_LPS\_8H\_BMDC\_DN | 58 | 6 | 4161 | 153 | cta,rush,Ppa,smal,Mocs2B,nudE | | 1.873e-02 | -3.98 | OSF2\_Q6 | MSigDB lists | OSF2\_Q6 | 58 | 6 | 4161 | 153 | Tsp26A,Fas2,CG9220,Sin3A,fne,SamDC | | 1.874e-02 | -3.98 | phosphatidylinositol binding | molecular function | GO:0035091 | 65 | 6 | 12176 | 396 | Snx3,Hip1,pbl,RhoGAP15B,CG32758,rush | | 1.881e-02 | -3.97 | fru (fruitless) | protein interactions | 42226 | 35 | 5 | 9204 | 412 | ap,N,Kr,EcR,robo2 | | 1.881e-02 | -3.97 | His2A:CG31618 (histone H2A) | protein interactions | 318855 | 35 | 5 | 9204 | 412 | Acf,Fas2,Ubx,phyl,mof | | 1.881e-02 | -3.97 | B52 (B52) | protein interactions | 41670 | 35 | 5 | 9204 | 412 | EcR,Mondo,RhoGAP16F,Fur1,CG32772 | | 1.884e-02 | -3.97 | EGF\_Lam | smart domains | SM00180 | 11 | 3 | 5086 | 276 | NetA,wb,NetB | | 1.894e-02 | -3.97 | regulation of cytoskeleton organization | biological process | GO:0051493 | 160 | 11 | 12345 | 414 | cta,pbl,if,bora,CG43658,PsGEF,Rho1,Arp2,hts,spoon,RhoGEF2 | | 1.902e-02 | -3.96 | p53 (p53) | protein interactions | 2768677 | 202 | 16 | 9204 | 412 | CG42684,spoon,CycE,Cyt-c1,Imp,CG2865,Svil,Utx,if,AGO2,ap,gus,Mmp2,CG11486,tou,stwl | | 1.903e-02 | -3.96 | Laminin\_G\_1 | pfam domains | PF00054 | 6 | 2 | 10480 | 393 | crb,ft | | 1.903e-02 | -3.96 | PBD | pfam domains | PF00786 | 6 | 2 | 10480 | 393 | mbt,Pak3 | | 1.903e-02 | -3.96 | MyTH4 | pfam domains | PF00784 | 6 | 2 | 10480 | 393 | RhoGAP93B,Myo10A | | 1.903e-02 | -3.96 | MH2 | pfam domains | PF03166 | 6 | 2 | 10480 | 393 | Smox,Dad | | 1.913e-02 | -3.96 | MyTH4\_dom | interpro domains | IPR000857 | 6 | 2 | 11383 | 428 | Myo10A,RhoGAP93B | | 1.913e-02 | -3.96 | SMAD\_dom\_Dwarfin-type | interpro domains | IPR001132 | 6 | 2 | 11383 | 428 | Smox,Dad | | 1.913e-02 | -3.96 | SMAD-like\_dom\_sf | interpro domains | IPR017855 | 6 | 2 | 11383 | 428 | Smox,Dad | | 1.913e-02 | -3.96 | MyTH4\_dom\_sf | interpro domains | IPR038185 | 6 | 2 | 11383 | 428 | Myo10A,RhoGAP93B | | 1.913e-02 | -3.96 | DNA-bd\_dom\_sf | interpro domains | IPR016177 | 6 | 2 | 11383 | 428 | sba,tou | | 1.919e-02 | -3.95 | ERBB2 Regulates Cell Motility | REACTOME pathways | R-DME-6785631 | 6 | 2 | 3923 | 148 | Egfr,Rho1 | | 1.919e-02 | -3.95 | Pre-NOTCH Expression and Processing | REACTOME pathways | R-DME-1912422 | 6 | 2 | 3923 | 148 | N,Fur1 | | 1.919e-02 | -3.95 | PI3K events in ERBB2 signaling | REACTOME pathways | R-DME-1963642 | 6 | 2 | 3923 | 148 | drk,Egfr | | 1.919e-02 | -3.95 | Pre-NOTCH Processing in Golgi | REACTOME pathways | R-DME-1912420 | 6 | 2 | 3923 | 148 | Fur1,N | | 1.919e-02 | -3.95 | EGFR Transactivation by Gastrin | REACTOME pathways | R-DME-2179392 | 6 | 2 | 3923 | 148 | drk,Egfr | | 1.919e-02 | -3.95 | HuR (ELAVL1) binds and stabilizes mRNA | REACTOME pathways | R-DME-450520 | 6 | 2 | 3923 | 148 | fne,elav | | 1.919e-02 | -3.95 | SHC1 events in ERBB4 signaling | REACTOME pathways | R-DME-1250347 | 6 | 2 | 3923 | 148 | drk,Egfr | | 1.919e-02 | -3.95 | GSE5589\_WT\_VS\_IL10\_KO\_LPS\_AND\_IL10\_STIM\_MACROPHAGE\_180MIN\_DN | MSigDB lists | GSE5589\_WT\_VS\_IL10\_KO\_LPS\_AND\_IL10\_STIM\_MACROPHAGE\_180MIN\_DN | 16 | 3 | 4161 | 153 | LRP1,dally,CG7264 | | 1.919e-02 | -3.95 | NABA\_SECRETED\_FACTORS | MSigDB lists | NABA\_SECRETED\_FACTORS | 16 | 3 | 4161 | 153 | hh,dpp,slow | | 1.919e-02 | -3.95 | GO\_REGULATION\_OF\_STEM\_CELL\_PROLIFERATION | MSigDB lists | GO\_REGULATION\_OF\_STEM\_CELL\_PROLIFERATION | 16 | 3 | 4161 | 153 | so,hh,N | | 1.919e-02 | -3.95 | MODULE\_372 | MSigDB lists | MODULE\_372 | 16 | 3 | 4161 | 153 | pbl,Mrp4,CG6752 | | 1.919e-02 | -3.95 | JAZAG\_TGFB1\_SIGNALING\_VIA\_SMAD4\_DN | MSigDB lists | JAZAG\_TGFB1\_SIGNALING\_VIA\_SMAD4\_DN | 16 | 3 | 4161 | 153 | dpp,Hsc70Cb,CycE | | 1.919e-02 | -3.95 | GO\_CELL\_CELL\_JUNCTION\_ASSEMBLY | MSigDB lists | GO\_CELL\_CELL\_JUNCTION\_ASSEMBLY | 16 | 3 | 4161 | 153 | pbl,sdt,Rho1 | | 1.919e-02 | -3.95 | MODULE\_92 | MSigDB lists | MODULE\_92 | 16 | 3 | 4161 | 153 | Syx1A,dpp,Fas2 | | 1.919e-02 | -3.95 | GO\_SMAD\_BINDING | MSigDB lists | GO\_SMAD\_BINDING | 16 | 3 | 4161 | 153 | dpp,tkv,Smox | | 1.919e-02 | -3.95 | GO\_REGULATION\_OF\_STRIATED\_MUSCLE\_CELL\_DIFFERENTIATION | MSigDB lists | GO\_REGULATION\_OF\_STRIATED\_MUSCLE\_CELL\_DIFFERENTIATION | 16 | 3 | 4161 | 153 | N,hh,dpp | | 1.919e-02 | -3.95 | CTGYNNCTYTAA\_UNKNOWN | MSigDB lists | CTGYNNCTYTAA\_UNKNOWN | 16 | 3 | 4161 | 153 | Pli,fne,lili | | 1.931e-02 | -3.95 | SENESE\_HDAC3\_TARGETS\_UP | MSigDB lists | SENESE\_HDAC3\_TARGETS\_UP | 148 | 11 | 4161 | 153 | Rho1,Hsc70Cb,emp,Mrp4,CaMKII,mle,cta,Syx1A,Arp2,dpp,phtf | | 1.935e-02 | -3.94 | GTPase activator activity | molecular function | GO:0005096 | 84 | 7 | 12176 | 396 | RhoGAP93B,RapGAP1,RhoGAP102A,RhoGAP16F,RhoGAP18B,RhoGAP15B,pbl | | 1.937e-02 | -3.94 | signaling adaptor activity | molecular function | GO:0035591 | 18 | 3 | 12176 | 396 | ckn,Prosap,drk | | 1.937e-02 | -3.94 | Wnt-protein binding | molecular function | GO:0017147 | 18 | 3 | 12176 | 396 | dlp,fj,otk2 | | 1.940e-02 | -3.94 | Drep2 (DNA fragmentation factor-related protein 2) | protein interactions | 35955 | 48 | 6 | 9204 | 412 | crb,CG3386,AGO2,Con,tyn,NAT1 | | 1.940e-02 | -3.94 | fu (fused) | protein interactions | 32855 | 48 | 6 | 9204 | 412 | hh,Svil,Ten-m,ci,Dad,tkv | | 1.958e-02 | -3.93 | drug binding | molecular function | GO:0008144 | 828 | 38 | 12176 | 396 | Pak3,KP78a,Doa,wnd,tkv,bor,Hsc70-3,Tlk,CaMKII,Cdk12,Arp2,Etl1,Mrp4,Asator,PEK,Mhcl,Haspin,Hipk,CG8097,ScsbetaG,S6kII,Mos,Egfr,JIL-1,CG9541,CG9281,CG13675,Pex1,CG8485,PhKgamma,par-1,Myo10A,CG12010,CG14535,mbt,mle,lz,Cyp1 | | 1.964e-02 | -3.93 | DELYS\_THYROID\_CANCER\_DN | MSigDB lists | DELYS\_THYROID\_CANCER\_DN | 43 | 5 | 4161 | 153 | dpp,Fas2,JIL-1,mgl,Arl4 | | 1.964e-02 | -3.93 | IK2\_01 | MSigDB lists | IK2\_01 | 43 | 5 | 4161 | 153 | Ank,TfAP-2,mgl,LRP1,dlp | | 1.964e-02 | -3.93 | IRF7\_01 | MSigDB lists | IRF7\_01 | 43 | 5 | 4161 | 153 | pAbp,Trpm,fne,cta,Tlk | | 1.964e-02 | -3.93 | E47\_02 | MSigDB lists | E47\_02 | 43 | 5 | 4161 | 153 | TfAP-2,fne,Dscam1,Arl4,Mhcl | | 1.964e-02 | -3.93 | CEBP\_Q3 | MSigDB lists | CEBP\_Q3 | 43 | 5 | 4161 | 153 | Fas2,Snx3,dlp,CG34384,CG3961 | | 1.964e-02 | -3.93 | GSE33374\_CD8\_ALPHAALPHA\_VS\_ALPHABETA\_CD161\_HIGH\_TCELL\_UP | MSigDB lists | GSE33374\_CD8\_ALPHAALPHA\_VS\_ALPHABETA\_CD161\_HIGH\_TCELL\_UP | 43 | 5 | 4161 | 153 | N,EcR,Fas2,Mitofilin,smal | | 1.964e-02 | -3.93 | GSE2770\_IL12\_ACT\_VS\_ACT\_CD4\_TCELL\_2H\_UP | MSigDB lists | GSE2770\_IL12\_ACT\_VS\_ACT\_CD4\_TCELL\_2H\_UP | 43 | 5 | 4161 | 153 | CG45050,Syx1A,CG34384,Cyp1,Npc1a | | 1.964e-02 | -3.93 | GSE35435\_RESTING\_VS\_IL4\_TREATED\_MACROPHAGE\_UP | MSigDB lists | GSE35435\_RESTING\_VS\_IL4\_TREATED\_MACROPHAGE\_UP | 43 | 5 | 4161 | 153 | CG32758,sm,mof,Hsc70Cb,Fas2 | | 1.964e-02 | -3.93 | GO\_CELL\_JUNCTION\_ORGANIZATION | MSigDB lists | GO\_CELL\_JUNCTION\_ORGANIZATION | 43 | 5 | 4161 | 153 | sdt,Rho1,Smox,Rac2,pbl | | 1.964e-02 | -3.93 | GSE25123\_IL4\_VS\_IL4\_AND\_ROSIGLITAZONE\_STIM\_PPARG\_KO\_MACROPHAGE\_DAY10\_UP | MSigDB lists | GSE25123\_IL4\_VS\_IL4\_AND\_ROSIGLITAZONE\_STIM\_PPARG\_KO\_MACROPHAGE\_DAY10\_UP | 43 | 5 | 4161 | 153 | CG32758,NKAIN,ras,CG6752,Tudor-SN | | 1.964e-02 | -3.93 | GSE9988\_LOW\_LPS\_VS\_ANTI\_TREM1\_AND\_LPS\_MONOCYTE\_DN | MSigDB lists | GSE9988\_LOW\_LPS\_VS\_ANTI\_TREM1\_AND\_LPS\_MONOCYTE\_DN | 43 | 5 | 4161 | 153 | cta,Acsl,Npc1a,emp,CG9220 | | 1.964e-02 | -3.93 | GSE45365\_CD8A\_DC\_VS\_CD11B\_DC\_IFNAR\_KO\_MCMV\_INFECTION\_UP | MSigDB lists | GSE45365\_CD8A\_DC\_VS\_CD11B\_DC\_IFNAR\_KO\_MCMV\_INFECTION\_UP | 43 | 5 | 4161 | 153 | CG41099,par-1,mgl,N,Fas2 | | 1.972e-02 | -3.93 | Toll Like Receptor 9 (TLR9) Cascade | REACTOME pathways | R-DME-168138 | 42 | 5 | 3923 | 148 | S6kII,Tl,Pli,mle,JIL-1 | | 1.988e-02 | -3.92 | histone kinase activity | molecular function | GO:0035173 | 7 | 2 | 12176 | 396 | JIL-1,Haspin | | 1.988e-02 | -3.92 | wide pore channel activity | molecular function | GO:0022829 | 7 | 2 | 12176 | 396 | Inx3,Inx2 | | 1.999e-02 | -3.91 | GGATTA\_PITX2\_Q2 | MSigDB lists | GGATTA\_PITX2\_Q2 | 111 | 9 | 4161 | 153 | smal,Ptpmeg,CtBP,S6kII,Trpm,mgl,Tlk,so,ras | | 1.999e-02 | -3.91 | signal transduction by protein phosphorylation | biological process | GO:0023014 | 47 | 5 | 12345 | 414 | mbt,egr,wnd,Rac2,Doa | | 2.001e-02 | -3.91 | regulation of protein transport | biological process | GO:0051223 | 64 | 6 | 12345 | 414 | CaMKII,S,Snx3,S6kII,sdt,crb | | 2.001e-02 | -3.91 | synapse assembly | biological process | GO:0007416 | 64 | 6 | 12345 | 414 | Tl,comm,Imp,Prosap,par-1,Ten-m | | 2.008e-02 | -3.91 | positive regulation of response to biotic stimulus | biological process | GO:0002833 | 101 | 8 | 12345 | 414 | Pli,Etl1,Tl,ush,CG4325,Mmp2,Doa,Toll-7 | | 2.010e-02 | -3.91 | GO\_PLASMA\_MEMBRANE\_PROTEIN\_COMPLEX | MSigDB lists | GO\_PLASMA\_MEMBRANE\_PROTEIN\_COMPLEX | 93 | 8 | 4161 | 153 | nAChRalpha4,tkv,drk,CG9281,dpp,Syx1A,if,cta | | 2.011e-02 | -3.91 | GO\_POSITIVE\_REGULATION\_OF\_RESPONSE\_TO\_STIMULUS | MSigDB lists | GO\_POSITIVE\_REGULATION\_OF\_RESPONSE\_TO\_STIMULUS | 402 | 23 | 4161 | 153 | Rho1,JIL-1,Syx1A,if,N,Smox,pbl,Ser,dpp,Cyp1,Pli,Sin3A,drk,tkv,Rac2,hh,CG9220,CtBP,S6kII,CaMKII,Tsp26A,nudE,Arp2 | | 2.017e-02 | -3.90 | cup (cup) | protein interactions | 33934 | 62 | 7 | 9204 | 412 | yps,smg,Dad,pAbp,spoon,gus,CG2865 | | 2.017e-02 | -3.90 | Sin3A (Sin3A) | protein interactions | 36382 | 62 | 7 | 9204 | 412 | mam,stg,N,Pits,aop,CG2865,shn | | 2.024e-02 | -3.90 | GSE3982\_MAC\_VS\_TH1\_DN | MSigDB lists | GSE3982\_MAC\_VS\_TH1\_DN | 59 | 6 | 4161 | 153 | lili,Tpr2,Smox,NiPp1,Tlk,CG9220 | | 2.034e-02 | -3.90 | ENK\_UV\_RESPONSE\_EPIDERMIS\_DN | MSigDB lists | ENK\_UV\_RESPONSE\_EPIDERMIS\_DN | 130 | 10 | 4161 | 153 | mmy,Pli,nudE,cta,Utx,S6kII,Mrp4,CG4538,JIL-1,fj | | 2.034e-02 | -3.90 | GO\_ENDOCYTOSIS | MSigDB lists | GO\_ENDOCYTOSIS | 130 | 10 | 4161 | 153 | hh,LRP1,Rac2,drk,Arp2,Npc1a,mgl,Cnx99A,CG41099,lili | | 2.039e-02 | -3.89 | TGCAAAC\_MIR452 | MSigDB lists | TGCAAAC\_MIR452 | 29 | 4 | 4161 | 153 | CG45050,CG34384,Ank,Doa | | 2.039e-02 | -3.89 | GSE14415\_FOXP3\_KO\_NATURAL\_TREG\_VS\_TCONV\_UP | MSigDB lists | GSE14415\_FOXP3\_KO\_NATURAL\_TREG\_VS\_TCONV\_UP | 29 | 4 | 4161 | 153 | Lac,Ppa,sdt,Tpr2 | | 2.039e-02 | -3.89 | GGCNRNWCTTYS\_UNKNOWN | MSigDB lists | GGCNRNWCTTYS\_UNKNOWN | 29 | 4 | 4161 | 153 | S6kII,Utx,Cnx99A,Tudor-SN | | 2.039e-02 | -3.89 | GSE5589\_WT\_VS\_IL10\_KO\_LPS\_STIM\_MACROPHAGE\_180MIN\_DN | MSigDB lists | GSE5589\_WT\_VS\_IL10\_KO\_LPS\_STIM\_MACROPHAGE\_180MIN\_DN | 29 | 4 | 4161 | 153 | emp,hh,Pcl,cta | | 2.039e-02 | -3.89 | GSE19888\_CTRL\_VS\_TCELL\_MEMBRANES\_ACT\_MAST\_CELL\_PRETREAT\_A3R\_INH\_DN | MSigDB lists | GSE19888\_CTRL\_VS\_TCELL\_MEMBRANES\_ACT\_MAST\_CELL\_PRETREAT\_A3R\_INH\_DN | 29 | 4 | 4161 | 153 | CG9855,CG41099,Ppa,Rac2 | | 2.039e-02 | -3.89 | KIM\_WT1\_TARGETS\_12HR\_UP | MSigDB lists | KIM\_WT1\_TARGETS\_12HR\_UP | 29 | 4 | 4161 | 153 | sm,Smox,nAChRalpha4,kdn | | 2.039e-02 | -3.89 | ATAGGAA\_MIR202 | MSigDB lists | ATAGGAA\_MIR202 | 29 | 4 | 4161 | 153 | S6kII,CtBP,Acsl,Pat1 | | 2.039e-02 | -3.89 | GSE6259\_FLT3L\_INDUCED\_DEC205\_POS\_DC\_VS\_CD8\_TCELL\_DN | MSigDB lists | GSE6259\_FLT3L\_INDUCED\_DEC205\_POS\_DC\_VS\_CD8\_TCELL\_DN | 29 | 4 | 4161 | 153 | N,CG41099,Tudor-SN,bor | | 2.053e-02 | -3.89 | p75 NTR receptor-mediated signalling | REACTOME pathways | R-DME-193704 | 16 | 3 | 3923 | 148 | Rtnl2,Rho1,RhoGEF2 | | 2.055e-02 | -3.88 | ZnF\_C2H2 | smart domains | SM00355 | 271 | 23 | 5086 | 276 | salm,Zif,CG18446,CG32772,shn,CG4360,hang,l(3)neo38,CG31612,jing,ci,ham,cas,sens-2,disco,CG31224,Kr,ush,Kah,CG9650,CG11247,ttk,bowl | | 2.065e-02 | -3.88 | Cbl (Cbl proto-oncogene) | protein interactions | 38961 | 24 | 4 | 9204 | 412 | drk,Egfr,aos,sty | | 2.065e-02 | -3.88 | E(bx) (Enhancer of bithorax) | protein interactions | 44811 | 24 | 4 | 9204 | 412 | EcR,Ubx,Fas2,lncRNA:roX1 | | 2.074e-02 | -3.88 | distal axon | cellular component | GO:0150034 | 100 | 8 | 11940 | 407 | Con,NetA,Cdk5alpha,Fas2,NetB,Syx1A,par-1,Prosap | | 2.101e-02 | -3.86 | regulation of compound eye retinal cell programmed cell death | biological process | GO:0046669 | 18 | 3 | 12345 | 414 | lz,ec,Egfr | | 2.101e-02 | -3.86 | specification of symmetry | biological process | GO:0009799 | 18 | 3 | 12345 | 414 | Rho1,tin,S | | 2.101e-02 | -3.86 | determination of bilateral symmetry | biological process | GO:0009855 | 18 | 3 | 12345 | 414 | tin,S,Rho1 | | 2.101e-02 | -3.86 | imaginal disc-derived male genitalia morphogenesis | biological process | GO:0048803 | 18 | 3 | 12345 | 414 | Fas2,N,otk2 | | 2.101e-02 | -3.86 | negative regulation of secretion | biological process | GO:0051048 | 18 | 3 | 12345 | 414 | NFAT,CaMKII,CG9650 | | 2.101e-02 | -3.86 | adult walking behavior | biological process | GO:0007628 | 18 | 3 | 12345 | 414 | oc,hiw,ey | | 2.106e-02 | -3.86 | RpII215 (RNA polymerase II 215kD subunit) | protein interactions | 32100 | 36 | 5 | 9204 | 412 | mof,Cdk12,ci,Utx,AGO2 | | 2.106e-02 | -3.86 | Ufl1 (UFM1 specific ligase 1) | protein interactions | 40843 | 36 | 5 | 9204 | 412 | Gbs-70E,Imp,CG32758,CycE,phyl | | 2.107e-02 | -3.86 | embryonic organ morphogenesis | biological process | GO:0048562 | 7 | 2 | 12345 | 414 | tin,Mmp2 | | 2.107e-02 | -3.86 | cellular response to biotic stimulus | biological process | GO:0071216 | 7 | 2 | 12345 | 414 | EcR,Stat92E | | 2.107e-02 | -3.86 | cardioblast cell fate commitment | biological process | GO:0042684 | 7 | 2 | 12345 | 414 | tin,robo2 | | 2.107e-02 | -3.86 | lymph gland plasmatocyte differentiation | biological process | GO:0035169 | 7 | 2 | 12345 | 414 | pnt,ush | | 2.107e-02 | -3.86 | syncytial nuclear migration | biological process | GO:0035190 | 7 | 2 | 12345 | 414 | AGO2,l(3)mbt | | 2.107e-02 | -3.86 | negative regulation of R7 cell differentiation | biological process | GO:0045677 | 7 | 2 | 12345 | 414 | ttk,aop | | 2.107e-02 | -3.86 | regulation of gastrulation | biological process | GO:0010470 | 7 | 2 | 12345 | 414 | cta,tsl | | 2.107e-02 | -3.86 | regulation of histone H3-K27 methylation | biological process | GO:0061085 | 7 | 2 | 12345 | 414 | Pcl,Utx | | 2.107e-02 | -3.86 | regulation of signaling receptor activity | biological process | GO:0010469 | 7 | 2 | 12345 | 414 | Prosap,aos | | 2.107e-02 | -3.86 | cellular response to molecule of bacterial origin | biological process | GO:0071219 | 7 | 2 | 12345 | 414 | EcR,Stat92E | | 2.107e-02 | -3.86 | endoderm formation | biological process | GO:0001706 | 7 | 2 | 12345 | 414 | tkv,Ubx | | 2.107e-02 | -3.86 | response to lipopolysaccharide | biological process | GO:0032496 | 7 | 2 | 12345 | 414 | Stat92E,EcR | | 2.107e-02 | -3.86 | regulation of cardiocyte differentiation | biological process | GO:1905207 | 7 | 2 | 12345 | 414 | pnt,N | | 2.116e-02 | -3.86 | GO\_REGULATION\_OF\_ANATOMICAL\_STRUCTURE\_MORPHOGENESIS | MSigDB lists | GO\_REGULATION\_OF\_ANATOMICAL\_STRUCTURE\_MORPHOGENESIS | 210 | 14 | 4161 | 153 | Smox,N,NetB,cta,TfAP-2,nudE,Lim1,dpp,nAChRalpha4,Rac2,CG9855,so,hh,dlp | | 2.117e-02 | -3.86 | cation binding | molecular function | GO:0043169 | 1072 | 47 | 12176 | 396 | EcR,esn,bor,tkv,rush,CG4325,ctrip,Zif,ft,Cnx99A,CG9855,CG11247,mgl,CG45050,hh,kni,N,crb,CG12099,tna,l(3)mbt,CG32369,S6kII,Cyp4p2,LRP1,shn,hang,ds,Tet,CG2991,Cad87A,hiw,Utx,CG3097,CG6752,slow,CG41099,wech,shd,CG6051,Ser,Psc,CadN,dbr,Hph,Mmp2,TfIIS | | 2.123e-02 | -3.85 | fly chr1B10-1B12\|1-0 | chromosome location | fly chr1B10-1B12\|1-0 | 1 | 1 | 25010 | 531 | CG13366 | | 2.123e-02 | -3.85 | fly chr51E11-51F4\|2-74 cM | chromosome location | fly chr51E11-51F4\|2-74 cM | 1 | 1 | 25010 | 531 | Trpm | | 2.123e-02 | -3.85 | fly chr4C3-4C4\|1-8 | chromosome location | fly chr4C3-4C4\|1-8 | 1 | 1 | 25010 | 531 | bi | | 2.123e-02 | -3.85 | fly chr60E12-60F1\|2-107 | chromosome location | fly chr60E12-60F1\|2-107 | 1 | 1 | 25010 | 531 | uzip | | 2.123e-02 | -3.85 | fly chr48A2-48A3\|2-64 | chromosome location | fly chr48A2-48A3\|2-64 | 1 | 1 | 25010 | 531 | tou | | 2.123e-02 | -3.85 | fly chr13A5-13A8\|1-49 cM | chromosome location | fly chr13A5-13A8\|1-49 cM | 1 | 1 | 25010 | 531 | hiw | | 2.123e-02 | -3.85 | fly chr59A1-59A2\|2-101 cM | chromosome location | fly chr59A1-59A2\|2-101 cM | 1 | 1 | 25010 | 531 | Ppa | | 2.123e-02 | -3.85 | fly chr13C5-13C7\|1-50 cM | chromosome location | fly chr13C5-13C7\|1-50 cM | 1 | 1 | 25010 | 531 | Gmap | | 2.123e-02 | -3.85 | fly chr48A2-48A3\|2-64 cM | chromosome location | fly chr48A2-48A3\|2-64 cM | 1 | 1 | 25010 | 531 | tou | | 2.123e-02 | -3.85 | fly chr8D5-8D6\|1-27 cM | chromosome location | fly chr8D5-8D6\|1-27 cM | 1 | 1 | 25010 | 531 | lz | | 2.123e-02 | -3.85 | fly chr3F3-3F3\|1-5 cM | chromosome location | fly chr3F3-3F3\|1-5 cM | 1 | 1 | 25010 | 531 | lncRNA:roX1 | | 2.123e-02 | -3.85 | fly chr80B1-80B2\|3-47 cM | chromosome location | fly chr80B1-80B2\|3-47 cM | 1 | 1 | 25010 | 531 | Chro | | 2.123e-02 | -3.85 | fly chr43D2-43D3\|2-56 | chromosome location | fly chr43D2-43D3\|2-56 | 1 | 1 | 25010 | 531 | Drat | | 2.123e-02 | -3.85 | fly chr85D16-85D17\|3-48 cM | chromosome location | fly chr85D16-85D17\|3-48 cM | 1 | 1 | 25010 | 531 | ps | | 2.123e-02 | -3.85 | fly chr63A6-63B1\|3-5 cM | chromosome location | fly chr63A6-63B1\|3-5 cM | 1 | 1 | 25010 | 531 | CG11486 | | 2.123e-02 | -3.85 | fly chr70D6-70D7\|3-42 | chromosome location | fly chr70D6-70D7\|3-42 | 1 | 1 | 25010 | 531 | stwl | | 2.123e-02 | -3.85 | fly chr99A9-99B1\|3-98 | chromosome location | fly chr99A9-99B1\|3-98 | 1 | 1 | 25010 | 531 | Ptp99A | | 2.123e-02 | -3.85 | fly chr12F2-12F3\|1-49 | chromosome location | fly chr12F2-12F3\|1-49 | 1 | 1 | 25010 | 531 | NetB | | 2.123e-02 | -3.85 | fly chr61C3-61C5\|3-0.5 cM | chromosome location | fly chr61C3-61C5\|3-0.5 cM | 1 | 1 | 25010 | 531 | Hipk | | 2.123e-02 | -3.85 | fly chr26D4-26D5\|2-20 cM | chromosome location | fly chr26D4-26D5\|2-20 cM | 1 | 1 | 25010 | 531 | retm | | 2.123e-02 | -3.85 | fly chr42B1-42B2\|2-55 | chromosome location | fly chr42B1-42B2\|2-55 | 1 | 1 | 25010 | 531 | jing | | 2.123e-02 | -3.85 | fly chr66A18-66A19\|3-21 | chromosome location | fly chr66A18-66A19\|3-21 | 1 | 1 | 25010 | 531 | pbl | | 2.123e-02 | -3.85 | fly chr9F2-9F4\|1-33 | chromosome location | fly chr9F2-9F4\|1-33 | 1 | 1 | 25010 | 531 | Imp | | 2.123e-02 | -3.85 | fly chr58D4-58D7\|2-99 cM | chromosome location | fly chr58D4-58D7\|2-99 cM | 1 | 1 | 25010 | 531 | CG6044 | | 2.123e-02 | -3.85 | fly chr84C8-84D1\|3-48 cM | chromosome location | fly chr84C8-84D1\|3-48 cM | 1 | 1 | 25010 | 531 | sas | | 2.123e-02 | -3.85 | fly chr102E1-102E3\|4-0 | chromosome location | fly chr102E1-102E3\|4-0 | 1 | 1 | 25010 | 531 | Sox102F | | 2.123e-02 | -3.85 | fly chr57E9-57F1\|2-95 | chromosome location | fly chr57E9-57F1\|2-95 | 1 | 1 | 25010 | 531 | Egfr | | 2.123e-02 | -3.85 | fly chr15A5-15A7\|1-55 | chromosome location | fly chr15A5-15A7\|1-55 | 1 | 1 | 25010 | 531 | if | | 2.123e-02 | -3.85 | fly chr46E1-46E3\|2-62 | chromosome location | fly chr46E1-46E3\|2-62 | 1 | 1 | 25010 | 531 | egr | | 2.123e-02 | -3.85 | fly chr36C2-36C4\|2-53 cM | chromosome location | fly chr36C2-36C4\|2-53 cM | 1 | 1 | 25010 | 531 | beat-IIIc | | 2.123e-02 | -3.85 | fly chr16A5-16B1\|1-57 | chromosome location | fly chr16A5-16B1\|1-57 | 1 | 1 | 25010 | 531 | CG43658 | | 2.123e-02 | -3.85 | fly chr59A1-59A2\|2-101 | chromosome location | fly chr59A1-59A2\|2-101 | 1 | 1 | 25010 | 531 | Ppa | | 2.123e-02 | -3.85 | fly chr80C3-80C3\|3-47 cM | chromosome location | fly chr80C3-80C3\|3-47 cM | 1 | 1 | 25010 | 531 | nrm | | 2.123e-02 | -3.85 | fly chr76D2-76D2\|3-46 cM | chromosome location | fly chr76D2-76D2\|3-46 cM | 1 | 1 | 25010 | 531 | lncRNA:CR45433 | | 2.123e-02 | -3.85 | fly chr42A6-42A6\|2-55 | chromosome location | fly chr42A6-42A6\|2-55 | 1 | 1 | 25010 | 531 | mle | | 2.123e-02 | -3.85 | fly chr25F3-25F4\|2-17 | chromosome location | fly chr25F3-25F4\|2-17 | 1 | 1 | 25010 | 531 | CG11030 | | 2.123e-02 | -3.85 | fly chr62F6-63A1\|3-4 cM | chromosome location | fly chr62F6-63A1\|3-4 cM | 1 | 1 | 25010 | 531 | Tet | | 2.123e-02 | -3.85 | fly chr11B1-11B2\|1-39 cM | chromosome location | fly chr11B1-11B2\|1-39 cM | 1 | 1 | 25010 | 531 | Pits | | 2.123e-02 | -3.85 | fly chr40A2-40A3\|2-54 | chromosome location | fly chr40A2-40A3\|2-54 | 1 | 1 | 25010 | 531 | CG31612 | | 2.123e-02 | -3.85 | fly chr58A3-58A4\|2-97 cM | chromosome location | fly chr58A3-58A4\|2-97 cM | 1 | 1 | 25010 | 531 | Fili | | 2.123e-02 | -3.85 | fly chr14B9-14B12\|1-53 | chromosome location | fly chr14B9-14B12\|1-53 | 1 | 1 | 25010 | 531 | Cyp1 | | 2.123e-02 | -3.85 | fly chr35B10-35B10\|2-50 cM | chromosome location | fly chr35B10-35B10\|2-50 cM | 1 | 1 | 25010 | 531 | TfIIS | | 2.123e-02 | -3.85 | fly chr50C19-50C20\|2-68 cM | chromosome location | fly chr50C19-50C20\|2-68 cM | 1 | 1 | 25010 | 531 | CG42806 | | 2.123e-02 | -3.85 | fly chr50C23-50D3\|2-68 | chromosome location | fly chr50C23-50D3\|2-68 | 1 | 1 | 25010 | 531 | mam | | 2.123e-02 | -3.85 | fly chr70D1-70D2\|3-42 cM | chromosome location | fly chr70D1-70D2\|3-42 cM | 1 | 1 | 25010 | 531 | Pex1 | | 2.123e-02 | -3.85 | fly chr97D10-97D11\|3-92 | chromosome location | fly chr97D10-97D11\|3-92 | 1 | 1 | 25010 | 531 | CG6330 | | 2.123e-02 | -3.85 | fly chr12F1-12F2\|1-49 cM | chromosome location | fly chr12F1-12F2\|1-49 cM | 1 | 1 | 25010 | 531 | NetA | | 2.123e-02 | -3.85 | fly chr27E4-27E5\|2-25 | chromosome location | fly chr27E4-27E5\|2-25 | 1 | 1 | 25010 | 531 | CG13784 | | 2.123e-02 | -3.85 | fly chr33E9-33E9\|2-46 | chromosome location | fly chr33E9-33E9\|2-46 | 1 | 1 | 25010 | 531 | lncRNA:CR42746 | | 2.123e-02 | -3.85 | fly chr50C19-50C20\|2-68 | chromosome location | fly chr50C19-50C20\|2-68 | 1 | 1 | 25010 | 531 | CG42806 | | 2.123e-02 | -3.85 | fly chr21B3-21B4\|2-0.5 | chromosome location | fly chr21B3-21B4\|2-0.5 | 1 | 1 | 25010 | 531 | spen | | 2.123e-02 | -3.85 | fly chr5B8-5C2\|1-13 | chromosome location | fly chr5B8-5C2\|1-13 | 1 | 1 | 25010 | 531 | CG3097 | | 2.123e-02 | -3.85 | fly chr80D5-80E3\|3-47 | chromosome location | fly chr80D5-80E3\|3-47 | 1 | 1 | 25010 | 531 | nAChRalpha4 | | 2.123e-02 | -3.85 | fly chr44F12-45A1\|2-59 cM | chromosome location | fly chr44F12-45A1\|2-59 cM | 1 | 1 | 25010 | 531 | flz | | 2.123e-02 | -3.85 | fly chr67E4-67E5\|3-33 | chromosome location | fly chr67E4-67E5\|3-33 | 1 | 1 | 25010 | 531 | Rbfox1 | | 2.123e-02 | -3.85 | fly chr72C3-72D1\|3-43 | chromosome location | fly chr72C3-72D1\|3-43 | 1 | 1 | 25010 | 531 | Notum | | 2.123e-02 | -3.85 | fly chr67E1-67E2\|3-33 cM | chromosome location | fly chr67E1-67E2\|3-33 cM | 1 | 1 | 25010 | 531 | lncRNA:CR46006 | | 2.123e-02 | -3.85 | fly chr10B3-10B4\|1-34 | chromosome location | fly chr10B3-10B4\|1-34 | 1 | 1 | 25010 | 531 | CG1737 | | 2.123e-02 | -3.85 | fly chr54E10-54E10\|2-84 cM | chromosome location | fly chr54E10-54E10\|2-84 cM | 1 | 1 | 25010 | 531 | lncRNA:CR45270 | | 2.123e-02 | -3.85 | fly chr57D2-57D2\|2-94 | chromosome location | fly chr57D2-57D2\|2-94 | 1 | 1 | 25010 | 531 | Ugt49C1 | | 2.123e-02 | -3.85 | fly chr94E10-94E13\|3-78 | chromosome location | fly chr94E10-94E13\|3-78 | 1 | 1 | 25010 | 531 | pnt | | 2.123e-02 | -3.85 | fly chr24C3-24C4\|2-11 | chromosome location | fly chr24C3-24C4\|2-11 | 1 | 1 | 25010 | 531 | bowl | | 2.123e-02 | -3.85 | fly chr102E1-102E3\|4-0 cM | chromosome location | fly chr102E1-102E3\|4-0 cM | 1 | 1 | 25010 | 531 | Sox102F | | 2.123e-02 | -3.85 | fly chr99B3-99B3\|3-99 | chromosome location | fly chr99B3-99B3\|3-99 | 1 | 1 | 25010 | 531 | Dr | | 2.123e-02 | -3.85 | fly chr43D2-43D3\|2-56 cM | chromosome location | fly chr43D2-43D3\|2-56 cM | 1 | 1 | 25010 | 531 | Drat | | 2.123e-02 | -3.85 | fly chr92A10-92A11\|3-66 | chromosome location | fly chr92A10-92A11\|3-66 | 1 | 1 | 25010 | 531 | CG6231 | | 2.123e-02 | -3.85 | fly chr64C2-64C5\|3-13 cM | chromosome location | fly chr64C2-64C5\|3-13 cM | 1 | 1 | 25010 | 531 | Con | | 2.123e-02 | -3.85 | fly chr75E3-75E4\|3-45 cM | chromosome location | fly chr75E3-75E4\|3-45 cM | 1 | 1 | 25010 | 531 | CG3961 | | 2.123e-02 | -3.85 | fly chr61C5-61C6\|3-0.5 cM | chromosome location | fly chr61C5-61C6\|3-0.5 cM | 1 | 1 | 25010 | 531 | Kah | | 2.123e-02 | -3.85 | fly chr23E5-23E6\|2-9 cM | chromosome location | fly chr23E5-23E6\|2-9 cM | 1 | 1 | 25010 | 531 | GramD1B | | 2.123e-02 | -3.85 | fly chr8D10-8D10\|1-28 | chromosome location | fly chr8D10-8D10\|1-28 | 1 | 1 | 25010 | 531 | mgl | | 2.123e-02 | -3.85 | fly chr26D9-26E1\|2-21 | chromosome location | fly chr26D9-26E1\|2-21 | 1 | 1 | 25010 | 531 | CG31637 | | 2.123e-02 | -3.85 | fly chr84E1-84E3\|3-48 | chromosome location | fly chr84E1-84E3\|3-48 | 1 | 1 | 25010 | 531 | CG34384 | | 2.123e-02 | -3.85 | fly chr83E6-83E6\|3-47.5 cM | chromosome location | fly chr83E6-83E6\|3-47.5 cM | 1 | 1 | 25010 | 531 | lncRNA:CR44017 | | 2.123e-02 | -3.85 | fly chr82F3-82F8\|3-47.5 cM | chromosome location | fly chr82F3-82F8\|3-47.5 cM | 1 | 1 | 25010 | 531 | Hph | | 2.123e-02 | -3.85 | fly chr65A1-65A2\|3-16 cM | chromosome location | fly chr65A1-65A2\|3-16 cM | 1 | 1 | 25010 | 531 | CG10479 | | 2.123e-02 | -3.85 | fly chr11D11-11E1\|1-41 | chromosome location | fly chr11D11-11E1\|1-41 | 1 | 1 | 25010 | 531 | CG43313 | | 2.123e-02 | -3.85 | fly chr42B1-42B2\|2-55 cM | chromosome location | fly chr42B1-42B2\|2-55 cM | 1 | 1 | 25010 | 531 | jing | | 2.123e-02 | -3.85 | fly chr44D4-44D5\|2-59 cM | chromosome location | fly chr44D4-44D5\|2-59 cM | 1 | 1 | 25010 | 531 | Cirl | | 2.123e-02 | -3.85 | fly chr40E5-40F2\|2-54 | chromosome location | fly chr40E5-40F2\|2-54 | 1 | 1 | 25010 | 531 | CG42748 | | 2.123e-02 | -3.85 | fly chr31D1-31D4\|2-41 | chromosome location | fly chr31D1-31D4\|2-41 | 1 | 1 | 25010 | 531 | Cdk5alpha | | 2.123e-02 | -3.85 | fly chr51D6-51D6\|2-73 | chromosome location | fly chr51D6-51D6\|2-73 | 1 | 1 | 25010 | 531 | ckn | | 2.123e-02 | -3.85 | fly chr11D3-11D4\|1-41 | chromosome location | fly chr11D3-11D4\|1-41 | 1 | 1 | 25010 | 531 | fne | | 2.123e-02 | -3.85 | fly chr33E9-33E9\|2-46 cM | chromosome location | fly chr33E9-33E9\|2-46 cM | 1 | 1 | 25010 | 531 | lncRNA:CR42746 | | 2.123e-02 | -3.85 | fly chr5F2-5F4\|1-16 cM | chromosome location | fly chr5F2-5F4\|1-16 cM | 1 | 1 | 25010 | 531 | kdn | | 2.123e-02 | -3.85 | fly chr70E3-70E3\|3-42 cM | chromosome location | fly chr70E3-70E3\|3-42 cM | 1 | 1 | 25010 | 531 | HGTX | | 2.123e-02 | -3.85 | fly chr4C3-4C4\|1-8 cM | chromosome location | fly chr4C3-4C4\|1-8 cM | 1 | 1 | 25010 | 531 | bi | | 2.123e-02 | -3.85 | fly chr60A11-60A11\|2-105 cM | chromosome location | fly chr60A11-60A11\|2-105 cM | 1 | 1 | 25010 | 531 | CG2812 | | 2.123e-02 | -3.85 | fly chr17D3-17D4\|1-60 | chromosome location | fly chr17D3-17D4\|1-60 | 1 | 1 | 25010 | 531 | CCKLR-17D3 | | 2.123e-02 | -3.85 | fly chr23E5-23E6\|2-9 | chromosome location | fly chr23E5-23E6\|2-9 | 1 | 1 | 25010 | 531 | GramD1B | | 2.123e-02 | -3.85 | fly chr35B3-35B4\|2-50 cM | chromosome location | fly chr35B3-35B4\|2-50 cM | 1 | 1 | 25010 | 531 | osp | | 2.123e-02 | -3.85 | fly chr61E2-61F1\|3-0.5 | chromosome location | fly chr61E2-61F1\|3-0.5 | 1 | 1 | 25010 | 531 | bab1 | | 2.123e-02 | -3.85 | fly chr93E8-93E8\|3-72 cM | chromosome location | fly chr93E8-93E8\|3-72 cM | 1 | 1 | 25010 | 531 | lncRNA:CR44034 | | 2.123e-02 | -3.85 | fly chr7D17-7D22\|1-22 | chromosome location | fly chr7D17-7D22\|1-22 | 1 | 1 | 25010 | 531 | sdt | | 2.123e-02 | -3.85 | fly chr89D6-89D9\|3-59 cM | chromosome location | fly chr89D6-89D9\|3-59 cM | 1 | 1 | 25010 | 531 | Ubx | | 2.123e-02 | -3.85 | fly chr97E6-97E8\|3-92 cM | chromosome location | fly chr97E6-97E8\|3-92 cM | 1 | 1 | 25010 | 531 | Ser | | 2.123e-02 | -3.85 | fly chr80C3-80C3\|3-47 | chromosome location | fly chr80C3-80C3\|3-47 | 1 | 1 | 25010 | 531 | nrm | | 2.123e-02 | -3.85 | fly chr37A2-37A4\|2-54 | chromosome location | fly chr37A2-37A4\|2-54 | 1 | 1 | 25010 | 531 | ham | | 2.123e-02 | -3.85 | fly chr25D1-25D2\|2-16 | chromosome location | fly chr25D1-25D2\|2-16 | 1 | 1 | 25010 | 531 | tkv | | 2.123e-02 | -3.85 | fly chr27B1-27B2\|2-22 cM | chromosome location | fly chr27B1-27B2\|2-22 cM | 1 | 1 | 25010 | 531 | nrv2 | | 2.123e-02 | -3.85 | fly chr36F2-36F4\|2-53 cM | chromosome location | fly chr36F2-36F4\|2-53 cM | 1 | 1 | 25010 | 531 | Fas3 | | 2.123e-02 | -3.85 | fly chr50D3-50E1\|2-69 cM | chromosome location | fly chr50D3-50E1\|2-69 cM | 1 | 1 | 25010 | 531 | Prosap | | 2.123e-02 | -3.85 | fly chr70D6-70D7\|3-42 cM | chromosome location | fly chr70D6-70D7\|3-42 cM | 1 | 1 | 25010 | 531 | stwl | | 2.123e-02 | -3.85 | fly chr78D8-78E1\|3-47 cM | chromosome location | fly chr78D8-78E1\|3-47 cM | 1 | 1 | 25010 | 531 | TfAP-2 | | 2.123e-02 | -3.85 | fly chr3F1-3F1\|1-4 | chromosome location | fly chr3F1-3F1\|1-4 | 1 | 1 | 25010 | 531 | Tlk | | 2.123e-02 | -3.85 | fly chr56D9-56D11\|2-89 cM | chromosome location | fly chr56D9-56D11\|2-89 cM | 1 | 1 | 25010 | 531 | par-1 | | 2.123e-02 | -3.85 | fly chr66F2-66F3\|3-28 cM | chromosome location | fly chr66F2-66F3\|3-28 cM | 1 | 1 | 25010 | 531 | Doc1 | | 2.123e-02 | -3.85 | fly chr23B6-23B7\|2-7 cM | chromosome location | fly chr23B6-23B7\|2-7 cM | 1 | 1 | 25010 | 531 | CG2991 | | 2.123e-02 | -3.85 | fly chr58D7-58D7\|2-99 | chromosome location | fly chr58D7-58D7\|2-99 | 1 | 1 | 25010 | 531 | VepD | | 2.123e-02 | -3.85 | fly chr41F-41F8\|2-55 | chromosome location | fly chr41F-41F8\|2-55 | 1 | 1 | 25010 | 531 | ap | | 2.123e-02 | -3.85 | fly chr2F4-2F5\|1-0.9 cM | chromosome location | fly chr2F4-2F5\|1-0.9 cM | 1 | 1 | 25010 | 531 | CG2865 | | 2.123e-02 | -3.85 | fly chr7D14-7D15\|1-22 cM | chromosome location | fly chr7D14-7D15\|1-22 cM | 1 | 1 | 25010 | 531 | Smox | | 2.123e-02 | -3.85 | fly chr10E3-10E4\|1-36 | chromosome location | fly chr10E3-10E4\|1-36 | 1 | 1 | 25010 | 531 | Hsc70-3 | | 2.123e-02 | -3.85 | fly chr8D10-8D10\|1-28 cM | chromosome location | fly chr8D10-8D10\|1-28 cM | 1 | 1 | 25010 | 531 | mgl | | 2.123e-02 | -3.85 | fly chr58D4-58D7\|2-99 | chromosome location | fly chr58D4-58D7\|2-99 | 1 | 1 | 25010 | 531 | CG6044 | | 2.123e-02 | -3.85 | fly chr102A1-102A3\|4-0 | chromosome location | fly chr102A1-102A3\|4-0 | 1 | 1 | 25010 | 531 | ci | | 2.123e-02 | -3.85 | fly chr51D6-51D6\|2-73 cM | chromosome location | fly chr51D6-51D6\|2-73 cM | 1 | 1 | 25010 | 531 | ckn | | 2.123e-02 | -3.85 | fly chr18E2-18E3\|1-64 cM | chromosome location | fly chr18E2-18E3\|1-64 cM | 1 | 1 | 25010 | 531 | meso18E | | 2.123e-02 | -3.85 | fly chr50A13-50A13\|2-68 | chromosome location | fly chr50A13-50A13\|2-68 | 1 | 1 | 25010 | 531 | drk | | 2.123e-02 | -3.85 | fly chr52F5-52F5\|2-78 cM | chromosome location | fly chr52F5-52F5\|2-78 cM | 1 | 1 | 25010 | 531 | CG8441 | | 2.123e-02 | -3.85 | fly chr56D3-56D5\|2-89 | chromosome location | fly chr56D3-56D5\|2-89 | 1 | 1 | 25010 | 531 | hts | | 2.123e-02 | -3.85 | fly chr27E1-27E2\|2-25 | chromosome location | fly chr27E1-27E2\|2-25 | 1 | 1 | 25010 | 531 | Pvf3 | | 2.123e-02 | -3.85 | fly chr12A9-12B2\|1-44 cM | chromosome location | fly chr12A9-12B2\|1-44 cM | 1 | 1 | 25010 | 531 | NFAT | | 2.123e-02 | -3.85 | fly chr49B5-49B7\|2-67 | chromosome location | fly chr49B5-49B7\|2-67 | 1 | 1 | 25010 | 531 | Sin3A | | 2.123e-02 | -3.85 | fly chr49A6-49A7\|2-67 cM | chromosome location | fly chr49A6-49A7\|2-67 cM | 1 | 1 | 25010 | 531 | Lac | | 2.123e-02 | -3.85 | fly chr21D1-21E2\|2-1.5 | chromosome location | fly chr21D1-21E2\|2-1.5 | 1 | 1 | 25010 | 531 | ush | | 2.123e-02 | -3.85 | fly chr5B8-5C2\|1-13 cM | chromosome location | fly chr5B8-5C2\|1-13 cM | 1 | 1 | 25010 | 531 | CG3097 | | 2.123e-02 | -3.85 | fly chr2C9-2C10\|1-0.5 cM | chromosome location | fly chr2C9-2C10\|1-0.5 cM | 1 | 1 | 25010 | 531 | CG4199 | | 2.123e-02 | -3.85 | fly chr90C1-90C2\|3-61 cM | chromosome location | fly chr90C1-90C2\|3-61 cM | 1 | 1 | 25010 | 531 | osa | | 2.123e-02 | -3.85 | fly chr92F12-93A1\|3-69 cM | chromosome location | fly chr92F12-93A1\|3-69 cM | 1 | 1 | 25010 | 531 | Syp | | 2.123e-02 | -3.85 | fly chr41E5-41E6\|2-55 | chromosome location | fly chr41E5-41E6\|2-55 | 1 | 1 | 25010 | 531 | lncRNA:CR42646 | | 2.123e-02 | -3.85 | fly chr66E1-66E3\|3-27 cM | chromosome location | fly chr66E1-66E3\|3-27 cM | 1 | 1 | 25010 | 531 | dally | | 2.123e-02 | -3.85 | fly chr57A5-57A6\|2-91 cM | chromosome location | fly chr57A5-57A6\|2-91 cM | 1 | 1 | 25010 | 531 | qsm | | 2.123e-02 | -3.85 | fly chr7D18-7F1\|1-23 | chromosome location | fly chr7D18-7F1\|1-23 | 1 | 1 | 25010 | 531 | lawc | | 2.123e-02 | -3.85 | fly chr85D16-85D17\|3-48 | chromosome location | fly chr85D16-85D17\|3-48 | 1 | 1 | 25010 | 531 | ps | | 2.123e-02 | -3.85 | fly chr31C7-31C7\|2-40 cM | chromosome location | fly chr31C7-31C7\|2-40 cM | 1 | 1 | 25010 | 531 | Utx | | 2.123e-02 | -3.85 | fly chr65A1-65A2\|3-16 | chromosome location | fly chr65A1-65A2\|3-16 | 1 | 1 | 25010 | 531 | CG10479 | | 2.123e-02 | -3.85 | fly chr55B5-55B9\|2-84 cM | chromosome location | fly chr55B5-55B9\|2-84 cM | 1 | 1 | 25010 | 531 | pAbp | | 2.123e-02 | -3.85 | fly chr31E5-31E5\|2-42 | chromosome location | fly chr31E5-31E5\|2-42 | 1 | 1 | 25010 | 531 | LManII | | 2.123e-02 | -3.85 | fly chr5B2-5B3\|1-13 cM | chromosome location | fly chr5B2-5B3\|1-13 cM | 1 | 1 | 25010 | 531 | CG32758 | | 2.123e-02 | -3.85 | fly chr41F-41F8\|2-55 cM | chromosome location | fly chr41F-41F8\|2-55 cM | 1 | 1 | 25010 | 531 | ap | | 2.123e-02 | -3.85 | fly chr3F2-3F3\|1-5 | chromosome location | fly chr3F2-3F3\|1-5 | 1 | 1 | 25010 | 531 | ec | | 2.123e-02 | -3.85 | fly chr87F12-87F13\|3-54 | chromosome location | fly chr87F12-87F13\|3-54 | 1 | 1 | 25010 | 531 | CG34383 | | 2.123e-02 | -3.85 | fly chr69F7-69F7\|3-39 cM | chromosome location | fly chr69F7-69F7\|3-39 cM | 1 | 1 | 25010 | 531 | lncRNA:CR43652 | | 2.123e-02 | -3.85 | fly chr44B8-44C2\|2-58 cM | chromosome location | fly chr44B8-44C2\|2-58 cM | 1 | 1 | 25010 | 531 | LRP1 | | 2.123e-02 | -3.85 | fly chr18A3-18A4\|1-62 cM | chromosome location | fly chr18A3-18A4\|1-62 cM | 1 | 1 | 25010 | 531 | RhoGAP18B | | 2.123e-02 | -3.85 | fly chr98F1-98F3\|3-97 | chromosome location | fly chr98F1-98F3\|3-97 | 1 | 1 | 25010 | 531 | Doa | | 2.123e-02 | -3.85 | fly chr45F6-46A1\|2-61 cM | chromosome location | fly chr45F6-46A1\|2-61 cM | 1 | 1 | 25010 | 531 | Mmp2 | | 2.123e-02 | -3.85 | fly chr94E10-94E13\|3-78 cM | chromosome location | fly chr94E10-94E13\|3-78 cM | 1 | 1 | 25010 | 531 | pnt | | 2.123e-02 | -3.85 | fly chr55B8-55B8\|2-84 | chromosome location | fly chr55B8-55B8\|2-84 | 1 | 1 | 25010 | 531 | Pcl | | 2.123e-02 | -3.85 | fly chr55F3-55F4\|2-86 cM | chromosome location | fly chr55F3-55F4\|2-86 cM | 1 | 1 | 25010 | 531 | MFS14 | | 2.123e-02 | -3.85 | fly chr12F2-12F3\|1-49 cM | chromosome location | fly chr12F2-12F3\|1-49 cM | 1 | 1 | 25010 | 531 | NetB | | 2.123e-02 | -3.85 | fly chr87A1-87A2\|3-51 cM | chromosome location | fly chr87A1-87A2\|3-51 cM | 1 | 1 | 25010 | 531 | Cad87A | | 2.123e-02 | -3.85 | fly chr52C2-52C4\|2-75 cM | chromosome location | fly chr52C2-52C4\|2-75 cM | 1 | 1 | 25010 | 531 | CG30089 | | 2.123e-02 | -3.85 | fly chr102C4-102C5\|4-0 | chromosome location | fly chr102C4-102C5\|4-0 | 1 | 1 | 25010 | 531 | Thd1 | | 2.123e-02 | -3.85 | fly chr44F12-45A1\|2-59 | chromosome location | fly chr44F12-45A1\|2-59 | 1 | 1 | 25010 | 531 | flz | | 2.123e-02 | -3.85 | fly chr102E2-102F2\|4-0 | chromosome location | fly chr102E2-102F2\|4-0 | 1 | 1 | 25010 | 531 | CaMKII | | 2.123e-02 | -3.85 | fly chr94E7-94E7\|3-78 cM | chromosome location | fly chr94E7-94E7\|3-78 cM | 1 | 1 | 25010 | 531 | cnc | | 2.123e-02 | -3.85 | fly chr54B18-54C1\|2-82 | chromosome location | fly chr54B18-54C1\|2-82 | 1 | 1 | 25010 | 531 | nw | | 2.123e-02 | -3.85 | fly chr70D7-70E1\|3-42 cM | chromosome location | fly chr70D7-70E1\|3-42 cM | 1 | 1 | 25010 | 531 | bbg | | 2.123e-02 | -3.85 | fly chr93B10-93B11\|3-70 cM | chromosome location | fly chr93B10-93B11\|3-70 cM | 1 | 1 | 25010 | 531 | RhoGAP93B | | 2.123e-02 | -3.85 | fly chr95F10-95F11\|3-84 cM | chromosome location | fly chr95F10-95F11\|3-84 cM | 1 | 1 | 25010 | 531 | crb | | 2.123e-02 | -3.85 | fly chr17D3-17D4\|1-60 cM | chromosome location | fly chr17D3-17D4\|1-60 cM | 1 | 1 | 25010 | 531 | CCKLR-17D3 | | 2.123e-02 | -3.85 | fly chr7F10-8A1\|1-23 | chromosome location | fly chr7F10-8A1\|1-23 | 1 | 1 | 25010 | 531 | oc | | 2.123e-02 | -3.85 | fly chr89B8-89B9\|3-58 | chromosome location | fly chr89B8-89B9\|3-58 | 1 | 1 | 25010 | 531 | tara | | 2.123e-02 | -3.85 | fly chr35B3-35B4\|2-50 | chromosome location | fly chr35B3-35B4\|2-50 | 1 | 1 | 25010 | 531 | osp | | 2.123e-02 | -3.85 | fly chr50D3-50E1\|2-69 | chromosome location | fly chr50D3-50E1\|2-69 | 1 | 1 | 25010 | 531 | Prosap | | 2.123e-02 | -3.85 | fly chr63A6-63B1\|3-5 | chromosome location | fly chr63A6-63B1\|3-5 | 1 | 1 | 25010 | 531 | CG11486 | | 2.123e-02 | -3.85 | fly chr2F4-2F5\|1-0.9 | chromosome location | fly chr2F4-2F5\|1-0.9 | 1 | 1 | 25010 | 531 | CG2865 | | 2.123e-02 | -3.85 | fly chr22F1-22F3\|2-6 | chromosome location | fly chr22F1-22F3\|2-6 | 1 | 1 | 25010 | 531 | dpp | | 2.123e-02 | -3.85 | fly chr12E4-12E4\|1-48 | chromosome location | fly chr12E4-12E4\|1-48 | 1 | 1 | 25010 | 531 | l(1)G0469 | | 2.123e-02 | -3.85 | fly chr16A1-16A1\|1-57 | chromosome location | fly chr16A1-16A1\|1-57 | 1 | 1 | 25010 | 531 | B-H2 | | 2.123e-02 | -3.85 | fly chr55C2-55C4\|2-85 cM | chromosome location | fly chr55C2-55C4\|2-85 cM | 1 | 1 | 25010 | 531 | sbb | | 2.123e-02 | -3.85 | fly chr95C9-95C12\|3-81 | chromosome location | fly chr95C9-95C12\|3-81 | 1 | 1 | 25010 | 531 | sba | | 2.123e-02 | -3.85 | fly chr59C1-59C2\|2-102 | chromosome location | fly chr59C1-59C2\|2-102 | 1 | 1 | 25010 | 531 | twi | | 2.123e-02 | -3.85 | fly chr70E4-70E4\|3-42 cM | chromosome location | fly chr70E4-70E4\|3-42 cM | 1 | 1 | 25010 | 531 | shd | | 2.123e-02 | -3.85 | fly chr47D6-47D7\|2-63 | chromosome location | fly chr47D6-47D7\|2-63 | 1 | 1 | 25010 | 531 | shn | | 2.123e-02 | -3.85 | fly chr89B16-89B17\|3-58 cM | chromosome location | fly chr89B16-89B17\|3-58 cM | 1 | 1 | 25010 | 531 | Pak3 | | 2.123e-02 | -3.85 | fly chr95C5-95C8\|3-81 cM | chromosome location | fly chr95C5-95C8\|3-81 cM | 1 | 1 | 25010 | 531 | Pli | | 2.123e-02 | -3.85 | fly chr78C4-78C5\|3-47 cM | chromosome location | fly chr78C4-78C5\|3-47 cM | 1 | 1 | 25010 | 531 | CG12974 | | 2.123e-02 | -3.85 | fly chr65E5-65E5\|3-19 | chromosome location | fly chr65E5-65E5\|3-19 | 1 | 1 | 25010 | 531 | corn | | 2.123e-02 | -3.85 | fly chr83E5-83E6\|3-47.5 | chromosome location | fly chr83E5-83E6\|3-47.5 | 1 | 1 | 25010 | 531 | Dmtn | | 2.123e-02 | -3.85 | fly chr54E10-54E10\|2-84 | chromosome location | fly chr54E10-54E10\|2-84 | 1 | 1 | 25010 | 531 | lncRNA:CR45270 | | 2.123e-02 | -3.85 | fly chr90C1-90C2\|3-61 | chromosome location | fly chr90C1-90C2\|3-61 | 1 | 1 | 25010 | 531 | osa | | 2.123e-02 | -3.85 | fly chr49B5-49B7\|2-67 cM | chromosome location | fly chr49B5-49B7\|2-67 cM | 1 | 1 | 25010 | 531 | Sin3A | | 2.123e-02 | -3.85 | fly chr3C7-3C9\|1-2 | chromosome location | fly chr3C7-3C9\|1-2 | 1 | 1 | 25010 | 531 | N | | 2.123e-02 | -3.85 | fly chr67E1-67E2\|3-33 | chromosome location | fly chr67E1-67E2\|3-33 | 1 | 1 | 25010 | 531 | lncRNA:CR46006 | | 2.123e-02 | -3.85 | fly chr27E1-27E2\|2-25 cM | chromosome location | fly chr27E1-27E2\|2-25 cM | 1 | 1 | 25010 | 531 | Pvf3 | | 2.123e-02 | -3.85 | fly chr56D15-56E1\|2-90 cM | chromosome location | fly chr56D15-56E1\|2-90 cM | 1 | 1 | 25010 | 531 | sm | | 2.123e-02 | -3.85 | fly chr98F1-98F3\|3-97 cM | chromosome location | fly chr98F1-98F3\|3-97 cM | 1 | 1 | 25010 | 531 | Doa | | 2.123e-02 | -3.85 | fly chr73A2-73A2\|3-44 | chromosome location | fly chr73A2-73A2\|3-44 | 1 | 1 | 25010 | 531 | aos | | 2.123e-02 | -3.85 | fly chr44D3-44D6\|2-59 cM | chromosome location | fly chr44D3-44D6\|2-59 cM | 1 | 1 | 25010 | 531 | Acsl | | 2.123e-02 | -3.85 | fly chr47A13-47B1\|2-62 cM | chromosome location | fly chr47A13-47B1\|2-62 cM | 1 | 1 | 25010 | 531 | psq | | 2.123e-02 | -3.85 | fly chr89B16-89B17\|3-58 | chromosome location | fly chr89B16-89B17\|3-58 | 1 | 1 | 25010 | 531 | Pak3 | | 2.123e-02 | -3.85 | fly chr72E5-72F1\|3-44 cM | chromosome location | fly chr72E5-72F1\|3-44 cM | 1 | 1 | 25010 | 531 | fax | | 2.123e-02 | -3.85 | fly chr40F6-40F7\|2-54 | chromosome location | fly chr40F6-40F7\|2-54 | 1 | 1 | 25010 | 531 | Tif-IA | | 2.123e-02 | -3.85 | fly chr75E3-75E4\|3-45 | chromosome location | fly chr75E3-75E4\|3-45 | 1 | 1 | 25010 | 531 | CG3961 | | 2.123e-02 | -3.85 | fly chr44D4-44D5\|2-59 | chromosome location | fly chr44D4-44D5\|2-59 | 1 | 1 | 25010 | 531 | Cirl | | 2.123e-02 | -3.85 | fly chr56D15-56E1\|2-90 | chromosome location | fly chr56D15-56E1\|2-90 | 1 | 1 | 25010 | 531 | sm | | 2.123e-02 | -3.85 | fly chr16A5-16B1\|1-57 cM | chromosome location | fly chr16A5-16B1\|1-57 cM | 1 | 1 | 25010 | 531 | CG43658 | | 2.123e-02 | -3.85 | fly chr98D6-98D7\|3-96 | chromosome location | fly chr98D6-98D7\|3-96 | 1 | 1 | 25010 | 531 | CG10011 | | 2.123e-02 | -3.85 | fly chr61E2-61F1\|3-0.5 cM | chromosome location | fly chr61E2-61F1\|3-0.5 cM | 1 | 1 | 25010 | 531 | bab1 | | 2.123e-02 | -3.85 | fly chr66E1-66E3\|3-27 | chromosome location | fly chr66E1-66E3\|3-27 | 1 | 1 | 25010 | 531 | dally | | 2.123e-02 | -3.85 | fly chr50C23-50D3\|2-68 cM | chromosome location | fly chr50C23-50D3\|2-68 cM | 1 | 1 | 25010 | 531 | mam | | 2.123e-02 | -3.85 | fly chr79D4-79E3\|3-47 cM | chromosome location | fly chr79D4-79E3\|3-47 cM | 1 | 1 | 25010 | 531 | Ten-m | | 2.123e-02 | -3.85 | fly chr64B3-64B4\|3-12 cM | chromosome location | fly chr64B3-64B4\|3-12 cM | 1 | 1 | 25010 | 531 | slow | | 2.123e-02 | -3.85 | fly chr89B8-89B9\|3-58 cM | chromosome location | fly chr89B8-89B9\|3-58 cM | 1 | 1 | 25010 | 531 | tara | | 2.123e-02 | -3.85 | fly chr7F10-8A1\|1-23 cM | chromosome location | fly chr7F10-8A1\|1-23 cM | 1 | 1 | 25010 | 531 | oc | | 2.123e-02 | -3.85 | fly chr95C5-95C8\|3-81 | chromosome location | fly chr95C5-95C8\|3-81 | 1 | 1 | 25010 | 531 | Pli | | 2.123e-02 | -3.85 | fly chr93F9-93F10\|3-73 | chromosome location | fly chr93F9-93F10\|3-73 | 1 | 1 | 25010 | 531 | tsl | | 2.123e-02 | -3.85 | fly chr45F6-46A1\|2-61 | chromosome location | fly chr45F6-46A1\|2-61 | 1 | 1 | 25010 | 531 | Mmp2 | | 2.123e-02 | -3.85 | fly chr89E2-89E4\|3-59 | chromosome location | fly chr89E2-89E4\|3-59 | 1 | 1 | 25010 | 531 | lncRNA:iab8 | | 2.123e-02 | -3.85 | fly chr27B4-27B4\|2-22 cM | chromosome location | fly chr27B4-27B4\|2-22 cM | 1 | 1 | 25010 | 531 | sens-2 | | 2.123e-02 | -3.85 | fly chr50A1-50A3\|2-68 | chromosome location | fly chr50A1-50A3\|2-68 | 1 | 1 | 25010 | 531 | lncRNA:CR44206 | | 2.123e-02 | -3.85 | fly chr6F10-7A5\|1-19 cM | chromosome location | fly chr6F10-7A5\|1-19 cM | 1 | 1 | 25010 | 531 | CG9650 | | 2.123e-02 | -3.85 | fly chr70D3-70D4\|3-42 | chromosome location | fly chr70D3-70D4\|3-42 | 1 | 1 | 25010 | 531 | nuf | | 2.123e-02 | -3.85 | fly chr76D2-76D2\|3-46 | chromosome location | fly chr76D2-76D2\|3-46 | 1 | 1 | 25010 | 531 | lncRNA:CR45433 | | 2.123e-02 | -3.85 | fly chr78D4-78D5\|3-47 cM | chromosome location | fly chr78D4-78D5\|3-47 cM | 1 | 1 | 25010 | 531 | Cdk12 | | 2.123e-02 | -3.85 | fly chr70E3-70E3\|3-42 | chromosome location | fly chr70E3-70E3\|3-42 | 1 | 1 | 25010 | 531 | HGTX | | 2.123e-02 | -3.85 | fly chr87D8-87D9\|3-53 | chromosome location | fly chr87D8-87D9\|3-53 | 1 | 1 | 25010 | 531 | CtBP | | 2.123e-02 | -3.85 | fly chr18A3-18A4\|1-62 | chromosome location | fly chr18A3-18A4\|1-62 | 1 | 1 | 25010 | 531 | RhoGAP18B | | 2.123e-02 | -3.85 | fly chr40F6-40F7\|2-54 cM | chromosome location | fly chr40F6-40F7\|2-54 cM | 1 | 1 | 25010 | 531 | Tif-IA | | 2.123e-02 | -3.85 | fly chr36F7-36F9\|2-53 cM | chromosome location | fly chr36F7-36F9\|2-53 cM | 1 | 1 | 25010 | 531 | MESR3 | | 2.123e-02 | -3.85 | fly chr85A9-85A10\|3-48 cM | chromosome location | fly chr85A9-85A10\|3-48 cM | 1 | 1 | 25010 | 531 | CG9801 | | 2.123e-02 | -3.85 | fly chr78C4-78C5\|3-47 | chromosome location | fly chr78C4-78C5\|3-47 | 1 | 1 | 25010 | 531 | CG12974 | | 2.123e-02 | -3.85 | fly chr95F10-95F11\|3-84 | chromosome location | fly chr95F10-95F11\|3-84 | 1 | 1 | 25010 | 531 | crb | | 2.123e-02 | -3.85 | fly chr36F7-36F9\|2-53 | chromosome location | fly chr36F7-36F9\|2-53 | 1 | 1 | 25010 | 531 | MESR3 | | 2.123e-02 | -3.85 | fly chr7D17-7D22\|1-22 cM | chromosome location | fly chr7D17-7D22\|1-22 cM | 1 | 1 | 25010 | 531 | sdt | | 2.123e-02 | -3.85 | fly chr102A1-102A3\|4-0 cM | chromosome location | fly chr102A1-102A3\|4-0 cM | 1 | 1 | 25010 | 531 | ci | | 2.123e-02 | -3.85 | fly chr28B1-28B2\|2-27 cM | chromosome location | fly chr28B1-28B2\|2-27 cM | 1 | 1 | 25010 | 531 | RapGAP1 | | 2.123e-02 | -3.85 | fly chr26E1-26E2\|2-21 cM | chromosome location | fly chr26E1-26E2\|2-21 cM | 1 | 1 | 25010 | 531 | eya | | 2.123e-02 | -3.85 | fly chr59C1-59C2\|2-102 cM | chromosome location | fly chr59C1-59C2\|2-102 cM | 1 | 1 | 25010 | 531 | twi | | 2.123e-02 | -3.85 | fly chr60D7-60D8\|2-107 cM | chromosome location | fly chr60D7-60D8\|2-107 cM | 1 | 1 | 25010 | 531 | pio | | 2.123e-02 | -3.85 | fly chr12A9-12B2\|1-44 | chromosome location | fly chr12A9-12B2\|1-44 | 1 | 1 | 25010 | 531 | NFAT | | 2.123e-02 | -3.85 | fly chr97E6-97E8\|3-92 | chromosome location | fly chr97E6-97E8\|3-92 | 1 | 1 | 25010 | 531 | Ser | | 2.123e-02 | -3.85 | fly chr61C5-61C6\|3-0.5 | chromosome location | fly chr61C5-61C6\|3-0.5 | 1 | 1 | 25010 | 531 | Kah | | 2.123e-02 | -3.85 | fly chr70D7-70E1\|3-42 | chromosome location | fly chr70D7-70E1\|3-42 | 1 | 1 | 25010 | 531 | bbg | | 2.123e-02 | -3.85 | fly chr70E1-70E2\|3-42 | chromosome location | fly chr70E1-70E2\|3-42 | 1 | 1 | 25010 | 531 | Gbs-70E | | 2.123e-02 | -3.85 | fly chr7D18-7F1\|1-23 cM | chromosome location | fly chr7D18-7F1\|1-23 cM | 1 | 1 | 25010 | 531 | lawc | | 2.123e-02 | -3.85 | fly chr49E6-49E6\|2-67 | chromosome location | fly chr49E6-49E6\|2-67 | 1 | 1 | 25010 | 531 | Psc | | 2.123e-02 | -3.85 | fly chr44B4-44B5\|2-58 cM | chromosome location | fly chr44B4-44B5\|2-58 cM | 1 | 1 | 25010 | 531 | Lpin | | 2.123e-02 | -3.85 | fly chr24C6-24C6\|2-11 | chromosome location | fly chr24C6-24C6\|2-11 | 1 | 1 | 25010 | 531 | slp1 | | 2.123e-02 | -3.85 | fly chr56D7-56D8\|2-89 cM | chromosome location | fly chr56D7-56D8\|2-89 cM | 1 | 1 | 25010 | 531 | betaTub56D | | 2.123e-02 | -3.85 | fly chr3C7-3C9\|1-2 cM | chromosome location | fly chr3C7-3C9\|1-2 cM | 1 | 1 | 25010 | 531 | N | | 2.123e-02 | -3.85 | fly chr42C7-42C7\|2-55 cM | chromosome location | fly chr42C7-42C7\|2-55 cM | 1 | 1 | 25010 | 531 | phtf | | 2.123e-02 | -3.85 | fly chr9F2-9F4\|1-33 cM | chromosome location | fly chr9F2-9F4\|1-33 cM | 1 | 1 | 25010 | 531 | Imp | | 2.123e-02 | -3.85 | fly chr60B10-60B11\|2-106 cM | chromosome location | fly chr60B10-60B11\|2-106 cM | 1 | 1 | 25010 | 531 | CG3253 | | 2.123e-02 | -3.85 | fly chr62D7-62E1\|3-3 | chromosome location | fly chr62D7-62E1\|3-3 | 1 | 1 | 25010 | 531 | Svil | | 2.123e-02 | -3.85 | fly chr16C5-16C6\|1-58 cM | chromosome location | fly chr16C5-16C6\|1-58 cM | 1 | 1 | 25010 | 531 | CG42684 | | 2.123e-02 | -3.85 | fly chr70E5-70E7\|3-42 | chromosome location | fly chr70E5-70E7\|3-42 | 1 | 1 | 25010 | 531 | dlp | | 2.123e-02 | -3.85 | fly chr56D9-56D11\|2-89 | chromosome location | fly chr56D9-56D11\|2-89 | 1 | 1 | 25010 | 531 | par-1 | | 2.123e-02 | -3.85 | fly chr98F6-98F10\|3-97 cM | chromosome location | fly chr98F6-98F10\|3-97 cM | 1 | 1 | 25010 | 531 | CG11873 | | 2.123e-02 | -3.85 | fly chr1B10-1B12\|1-0 cM | chromosome location | fly chr1B10-1B12\|1-0 cM | 1 | 1 | 25010 | 531 | CG13366 | | 2.123e-02 | -3.85 | fly chr50A13-50A13\|2-68 cM | chromosome location | fly chr50A13-50A13\|2-68 cM | 1 | 1 | 25010 | 531 | drk | | 2.123e-02 | -3.85 | fly chr66A13-66A14\|3-21 cM | chromosome location | fly chr66A13-66A14\|3-21 cM | 1 | 1 | 25010 | 531 | CG32369 | | 2.123e-02 | -3.85 | fly chr32F1-32F2\|2-44 cM | chromosome location | fly chr32F1-32F2\|2-44 cM | 1 | 1 | 25010 | 531 | salm | | 2.123e-02 | -3.85 | fly chr102E2-102F2\|4-0 cM | chromosome location | fly chr102E2-102F2\|4-0 cM | 1 | 1 | 25010 | 531 | CaMKII | | 2.123e-02 | -3.85 | fly chr47A13-47B1\|2-62 | chromosome location | fly chr47A13-47B1\|2-62 | 1 | 1 | 25010 | 531 | psq | | 2.123e-02 | -3.85 | fly chr5B4-5B6\|1-13 | chromosome location | fly chr5B4-5B6\|1-13 | 1 | 1 | 25010 | 531 | IntS6 | | 2.123e-02 | -3.85 | fly chr90D1-90E1\|3-62 cM | chromosome location | fly chr90D1-90E1\|3-62 cM | 1 | 1 | 25010 | 531 | cpo | | 2.123e-02 | -3.85 | fly chr54B18-54C1\|2-82 cM | chromosome location | fly chr54B18-54C1\|2-82 cM | 1 | 1 | 25010 | 531 | nw | | 2.123e-02 | -3.85 | fly chr66A13-66A14\|3-21 | chromosome location | fly chr66A13-66A14\|3-21 | 1 | 1 | 25010 | 531 | CG32369 | | 2.123e-02 | -3.85 | fly chr24C6-24C6\|2-11 cM | chromosome location | fly chr24C6-24C6\|2-11 cM | 1 | 1 | 25010 | 531 | slp1 | | 2.123e-02 | -3.85 | fly chr87D8-87D9\|3-53 cM | chromosome location | fly chr87D8-87D9\|3-53 cM | 1 | 1 | 25010 | 531 | CtBP | | 2.123e-02 | -3.85 | fly chr97D7-97D9\|3-92 cM | chromosome location | fly chr97D7-97D9\|3-92 cM | 1 | 1 | 25010 | 531 | CG46339 | | 2.123e-02 | -3.85 | fly chr85A9-85A10\|3-48 | chromosome location | fly chr85A9-85A10\|3-48 | 1 | 1 | 25010 | 531 | CG9801 | | 2.123e-02 | -3.85 | fly chr87F12-87F13\|3-54 cM | chromosome location | fly chr87F12-87F13\|3-54 cM | 1 | 1 | 25010 | 531 | CG34383 | | 2.123e-02 | -3.85 | fly chr4B1-4B3\|1-7 cM | chromosome location | fly chr4B1-4B3\|1-7 cM | 1 | 1 | 25010 | 531 | Fas2 | | 2.123e-02 | -3.85 | fly chr89E2-89E4\|3-59 cM | chromosome location | fly chr89E2-89E4\|3-59 cM | 1 | 1 | 25010 | 531 | lncRNA:iab8 | | 2.123e-02 | -3.85 | fly chr3F3-3F3\|1-5 | chromosome location | fly chr3F3-3F3\|1-5 | 1 | 1 | 25010 | 531 | lncRNA:roX1 | | 2.123e-02 | -3.85 | fly chr44B8-44C2\|2-58 | chromosome location | fly chr44B8-44C2\|2-58 | 1 | 1 | 25010 | 531 | LRP1 | | 2.123e-02 | -3.85 | fly chr31C7-31C7\|2-40 | chromosome location | fly chr31C7-31C7\|2-40 | 1 | 1 | 25010 | 531 | Utx | | 2.123e-02 | -3.85 | fly chr66F2-66F3\|3-28 | chromosome location | fly chr66F2-66F3\|3-28 | 1 | 1 | 25010 | 531 | Doc1 | | 2.123e-02 | -3.85 | fly chr13C5-13C7\|1-50 | chromosome location | fly chr13C5-13C7\|1-50 | 1 | 1 | 25010 | 531 | Gmap | | 2.123e-02 | -3.85 | fly chr15A3-15A5\|1-55 cM | chromosome location | fly chr15A3-15A5\|1-55 cM | 1 | 1 | 25010 | 531 | goe | | 2.123e-02 | -3.85 | fly chr63D2-63D3\|3-8 cM | chromosome location | fly chr63D2-63D3\|3-8 cM | 1 | 1 | 25010 | 531 | sty | | 2.123e-02 | -3.85 | fly chr89D6-89D9\|3-59 | chromosome location | fly chr89D6-89D9\|3-59 | 1 | 1 | 25010 | 531 | Ubx | | 2.123e-02 | -3.85 | fly chr93E8-93E8\|3-72 | chromosome location | fly chr93E8-93E8\|3-72 | 1 | 1 | 25010 | 531 | lncRNA:CR44034 | | 2.123e-02 | -3.85 | fly chr8A5-8B2\|1-24 | chromosome location | fly chr8A5-8B2\|1-24 | 1 | 1 | 25010 | 531 | Lim1 | | 2.123e-02 | -3.85 | fly chr55F3-55F4\|2-86 | chromosome location | fly chr55F3-55F4\|2-86 | 1 | 1 | 25010 | 531 | MFS14 | | 2.123e-02 | -3.85 | fly chr40A2-40A3\|2-54 cM | chromosome location | fly chr40A2-40A3\|2-54 cM | 1 | 1 | 25010 | 531 | CG31612 | | 2.123e-02 | -3.85 | fly chr61C3-61C5\|3-0.5 | chromosome location | fly chr61C3-61C5\|3-0.5 | 1 | 1 | 25010 | 531 | Hipk | | 2.123e-02 | -3.85 | fly chr55B5-55B9\|2-84 | chromosome location | fly chr55B5-55B9\|2-84 | 1 | 1 | 25010 | 531 | pAbp | | 2.123e-02 | -3.85 | fly chr65E5-65E5\|3-19 cM | chromosome location | fly chr65E5-65E5\|3-19 cM | 1 | 1 | 25010 | 531 | corn | | 2.123e-02 | -3.85 | fly chr10B3-10B4\|1-34 cM | chromosome location | fly chr10B3-10B4\|1-34 cM | 1 | 1 | 25010 | 531 | CG1737 | | 2.123e-02 | -3.85 | fly chr28B1-28B2\|2-27 | chromosome location | fly chr28B1-28B2\|2-27 | 1 | 1 | 25010 | 531 | RapGAP1 | | 2.123e-02 | -3.85 | fly chr69F7-69F7\|3-39 | chromosome location | fly chr69F7-69F7\|3-39 | 1 | 1 | 25010 | 531 | lncRNA:CR43652 | | 2.123e-02 | -3.85 | fly chr5B4-5B6\|1-13 cM | chromosome location | fly chr5B4-5B6\|1-13 cM | 1 | 1 | 25010 | 531 | IntS6 | | 2.123e-02 | -3.85 | fly chr45D4-45D5\|2-60 | chromosome location | fly chr45D4-45D5\|2-60 | 1 | 1 | 25010 | 531 | wun2 | | 2.123e-02 | -3.85 | fly chr68A5-68A6\|3-34 | chromosome location | fly chr68A5-68A6\|3-34 | 1 | 1 | 25010 | 531 | JIL-1 | | 2.123e-02 | -3.85 | fly chr82F3-82F8\|3-47.5 | chromosome location | fly chr82F3-82F8\|3-47.5 | 1 | 1 | 25010 | 531 | Hph | | 2.123e-02 | -3.85 | fly chr95B7-95C1\|3-80 | chromosome location | fly chr95B7-95C1\|3-80 | 1 | 1 | 25010 | 531 | CG33111 | | 2.123e-02 | -3.85 | fly chr42F1-42F1\|2-55 cM | chromosome location | fly chr42F1-42F1\|2-55 cM | 1 | 1 | 25010 | 531 | esn | | 2.123e-02 | -3.85 | fly chr100D3-100E2\|3-103 | chromosome location | fly chr100D3-100E2\|3-103 | 1 | 1 | 25010 | 531 | heph | | 2.123e-02 | -3.85 | fly chr83E5-83E6\|3-47.5 cM | chromosome location | fly chr83E5-83E6\|3-47.5 cM | 1 | 1 | 25010 | 531 | Dmtn | | 2.123e-02 | -3.85 | fly chr15A5-15A7\|1-55 cM | chromosome location | fly chr15A5-15A7\|1-55 cM | 1 | 1 | 25010 | 531 | if | | 2.123e-02 | -3.85 | fly chr7D14-7D15\|1-22 | chromosome location | fly chr7D14-7D15\|1-22 | 1 | 1 | 25010 | 531 | Smox | | 2.123e-02 | -3.85 | fly chr64B3-64B4\|3-12 | chromosome location | fly chr64B3-64B4\|3-12 | 1 | 1 | 25010 | 531 | slow | | 2.123e-02 | -3.85 | fly chr26D9-26E1\|2-21 cM | chromosome location | fly chr26D9-26E1\|2-21 cM | 1 | 1 | 25010 | 531 | CG31637 | | 2.123e-02 | -3.85 | fly chr4B1-4B3\|1-7 | chromosome location | fly chr4B1-4B3\|1-7 | 1 | 1 | 25010 | 531 | Fas2 | | 2.123e-02 | -3.85 | fly chr97E3-97E5\|3-92 | chromosome location | fly chr97E3-97E5\|3-92 | 1 | 1 | 25010 | 531 | CG6051 | | 2.123e-02 | -3.85 | fly chr47D6-47D7\|2-63 cM | chromosome location | fly chr47D6-47D7\|2-63 cM | 1 | 1 | 25010 | 531 | shn | | 2.123e-02 | -3.85 | fly chr30E1-30E4\|2-37 cM | chromosome location | fly chr30E1-30E4\|2-37 cM | 1 | 1 | 25010 | 531 | Etl1 | | 2.123e-02 | -3.85 | fly chr70D3-70D4\|3-42 cM | chromosome location | fly chr70D3-70D4\|3-42 cM | 1 | 1 | 25010 | 531 | nuf | | 2.123e-02 | -3.85 | fly chr41E5-41E6\|2-55 cM | chromosome location | fly chr41E5-41E6\|2-55 cM | 1 | 1 | 25010 | 531 | lncRNA:CR42646 | | 2.123e-02 | -3.85 | fly chr42F1-42F1\|2-55 | chromosome location | fly chr42F1-42F1\|2-55 | 1 | 1 | 25010 | 531 | esn | | 2.123e-02 | -3.85 | fly chr87A1-87A2\|3-51 | chromosome location | fly chr87A1-87A2\|3-51 | 1 | 1 | 25010 | 531 | Cad87A | | 2.123e-02 | -3.85 | fly chr53E4-53F1\|2-81 | chromosome location | fly chr53E4-53F1\|2-81 | 1 | 1 | 25010 | 531 | RhoGEF2 | | 2.123e-02 | -3.85 | fly chr22F1-22F3\|2-6 cM | chromosome location | fly chr22F1-22F3\|2-6 cM | 1 | 1 | 25010 | 531 | dpp | | 2.123e-02 | -3.85 | fly chr80D5-80E3\|3-47 cM | chromosome location | fly chr80D5-80E3\|3-47 cM | 1 | 1 | 25010 | 531 | nAChRalpha4 | | 2.123e-02 | -3.85 | fly chr42A6-42A6\|2-55 cM | chromosome location | fly chr42A6-42A6\|2-55 cM | 1 | 1 | 25010 | 531 | mle | | 2.123e-02 | -3.85 | fly chr57D2-57D2\|2-94 cM | chromosome location | fly chr57D2-57D2\|2-94 cM | 1 | 1 | 25010 | 531 | Ugt49C1 | | 2.123e-02 | -3.85 | fly chr85E1-85E1 | chromosome location | fly chr85E1-85E1 | 1 | 1 | 25010 | 531 | sisRNA:CR46357 | | 2.123e-02 | -3.85 | fly chr64C12-64C13\|3-13 cM | chromosome location | fly chr64C12-64C13\|3-13 cM | 1 | 1 | 25010 | 531 | lama | | 2.123e-02 | -3.85 | fly chr57A5-57A6\|2-91 | chromosome location | fly chr57A5-57A6\|2-91 | 1 | 1 | 25010 | 531 | qsm | | 2.123e-02 | -3.85 | fly chr70E4-70E4\|3-42 | chromosome location | fly chr70E4-70E4\|3-42 | 1 | 1 | 25010 | 531 | shd | | 2.123e-02 | -3.85 | fly chr31D1-31D4\|2-41 cM | chromosome location | fly chr31D1-31D4\|2-41 cM | 1 | 1 | 25010 | 531 | Cdk5alpha | | 2.123e-02 | -3.85 | fly chr27E4-27E5\|2-25 cM | chromosome location | fly chr27E4-27E5\|2-25 cM | 1 | 1 | 25010 | 531 | CG13784 | | 2.123e-02 | -3.85 | fly chr94E7-94E7\|3-78 | chromosome location | fly chr94E7-94E7\|3-78 | 1 | 1 | 25010 | 531 | cnc | | 2.123e-02 | -3.85 | fly chr72C1-72C2\|3-43 | chromosome location | fly chr72C1-72C2\|3-43 | 1 | 1 | 25010 | 531 | CG5830 | | 2.123e-02 | -3.85 | fly chr4F9-4F10\|1-11 cM | chromosome location | fly chr4F9-4F10\|1-11 cM | 1 | 1 | 25010 | 531 | spoon | | 2.123e-02 | -3.85 | fly chr97D10-97D11\|3-92 cM | chromosome location | fly chr97D10-97D11\|3-92 cM | 1 | 1 | 25010 | 531 | CG6330 | | 2.123e-02 | -3.85 | fly chr60E12-60F1\|2-107 cM | chromosome location | fly chr60E12-60F1\|2-107 cM | 1 | 1 | 25010 | 531 | uzip | | 2.123e-02 | -3.85 | fly chr58D7-58D7\|2-99 cM | chromosome location | fly chr58D7-58D7\|2-99 cM | 1 | 1 | 25010 | 531 | VepD | | 2.123e-02 | -3.85 | fly chr13D4-13E1\|1-51 cM | chromosome location | fly chr13D4-13E1\|1-51 cM | 1 | 1 | 25010 | 531 | CG9220 | | 2.123e-02 | -3.85 | fly chr16C5-16C6\|1-58 | chromosome location | fly chr16C5-16C6\|1-58 | 1 | 1 | 25010 | 531 | CG42684 | | 2.123e-02 | -3.85 | fly chr62F6-63A1\|3-4 | chromosome location | fly chr62F6-63A1\|3-4 | 1 | 1 | 25010 | 531 | Tet | | 2.123e-02 | -3.85 | fly chr27B4-27B4\|2-22 | chromosome location | fly chr27B4-27B4\|2-22 | 1 | 1 | 25010 | 531 | sens-2 | | 2.123e-02 | -3.85 | fly chr70D1-70D2\|3-42 | chromosome location | fly chr70D1-70D2\|3-42 | 1 | 1 | 25010 | 531 | Pex1 | | 2.123e-02 | -3.85 | fly chr99A9-99B1\|3-98 cM | chromosome location | fly chr99A9-99B1\|3-98 cM | 1 | 1 | 25010 | 531 | Ptp99A | | 2.123e-02 | -3.85 | fly chr35B10-35B10\|2-50 | chromosome location | fly chr35B10-35B10\|2-50 | 1 | 1 | 25010 | 531 | TfIIS | | 2.123e-02 | -3.85 | fly chr102C4-102C5\|4-0 cM | chromosome location | fly chr102C4-102C5\|4-0 cM | 1 | 1 | 25010 | 531 | Thd1 | | 2.123e-02 | -3.85 | fly chr72C3-72D1\|3-43 cM | chromosome location | fly chr72C3-72D1\|3-43 cM | 1 | 1 | 25010 | 531 | Notum | | 2.123e-02 | -3.85 | fly chr46E1-46E3\|2-62 cM | chromosome location | fly chr46E1-46E3\|2-62 cM | 1 | 1 | 25010 | 531 | egr | | 2.123e-02 | -3.85 | fly chr51E10-51E11\|2-74 cM | chromosome location | fly chr51E10-51E11\|2-74 cM | 1 | 1 | 25010 | 531 | scb | | 2.123e-02 | -3.85 | fly chr78D8-78E1\|3-47 | chromosome location | fly chr78D8-78E1\|3-47 | 1 | 1 | 25010 | 531 | TfAP-2 | | 2.123e-02 | -3.85 | fly chr53E4-53F1\|2-81 cM | chromosome location | fly chr53E4-53F1\|2-81 cM | 1 | 1 | 25010 | 531 | RhoGEF2 | | 2.123e-02 | -3.85 | fly chr16A1-16A1\|1-57 cM | chromosome location | fly chr16A1-16A1\|1-57 cM | 1 | 1 | 25010 | 531 | B-H2 | | 2.123e-02 | -3.85 | fly chr70E5-70E7\|3-42 cM | chromosome location | fly chr70E5-70E7\|3-42 cM | 1 | 1 | 25010 | 531 | dlp | | 2.123e-02 | -3.85 | fly chr5B2-5B3\|1-13 | chromosome location | fly chr5B2-5B3\|1-13 | 1 | 1 | 25010 | 531 | CG32758 | | 2.123e-02 | -3.85 | fly chr12E4-12E4\|1-48 cM | chromosome location | fly chr12E4-12E4\|1-48 cM | 1 | 1 | 25010 | 531 | l(1)G0469 | | 2.123e-02 | -3.85 | fly chr83E6-83E6\|3-47.5 | chromosome location | fly chr83E6-83E6\|3-47.5 | 1 | 1 | 25010 | 531 | lncRNA:CR44017 | | 2.123e-02 | -3.85 | fly chr44B4-44B5\|2-58 | chromosome location | fly chr44B4-44B5\|2-58 | 1 | 1 | 25010 | 531 | Lpin | | 2.123e-02 | -3.85 | fly chr36F2-36F4\|2-53 | chromosome location | fly chr36F2-36F4\|2-53 | 1 | 1 | 25010 | 531 | Fas3 | | 2.123e-02 | -3.85 | fly chr99B3-99B3\|3-99 cM | chromosome location | fly chr99B3-99B3\|3-99 cM | 1 | 1 | 25010 | 531 | Dr | | 2.123e-02 | -3.85 | fly chr52C2-52C4\|2-75 | chromosome location | fly chr52C2-52C4\|2-75 | 1 | 1 | 25010 | 531 | CG30089 | | 2.123e-02 | -3.85 | fly chr60A11-60A11\|2-105 | chromosome location | fly chr60A11-60A11\|2-105 | 1 | 1 | 25010 | 531 | CG2812 | | 2.123e-02 | -3.85 | fly chr83B8-83B9\|3-47.5 cM | chromosome location | fly chr83B8-83B9\|3-47.5 cM | 1 | 1 | 25010 | 531 | Atg17 | | 2.123e-02 | -3.85 | fly chr95C9-95C12\|3-81 cM | chromosome location | fly chr95C9-95C12\|3-81 cM | 1 | 1 | 25010 | 531 | sba | | 2.123e-02 | -3.85 | fly chr5F2-5F4\|1-16 | chromosome location | fly chr5F2-5F4\|1-16 | 1 | 1 | 25010 | 531 | kdn | | 2.123e-02 | -3.85 | fly chr51E10-51E11\|2-74 | chromosome location | fly chr51E10-51E11\|2-74 | 1 | 1 | 25010 | 531 | scb | | 2.123e-02 | -3.85 | fly chr84E1-84E3\|3-48 cM | chromosome location | fly chr84E1-84E3\|3-48 cM | 1 | 1 | 25010 | 531 | CG34384 | | 2.123e-02 | -3.85 | fly chr42C7-42C7\|2-55 | chromosome location | fly chr42C7-42C7\|2-55 | 1 | 1 | 25010 | 531 | phtf | | 2.123e-02 | -3.85 | fly chr12F1-12F2\|1-49 | chromosome location | fly chr12F1-12F2\|1-49 | 1 | 1 | 25010 | 531 | NetA | | 2.123e-02 | -3.85 | fly chr6F10-7A5\|1-19 | chromosome location | fly chr6F10-7A5\|1-19 | 1 | 1 | 25010 | 531 | CG9650 | | 2.123e-02 | -3.85 | fly chr13A5-13A8\|1-49 | chromosome location | fly chr13A5-13A8\|1-49 | 1 | 1 | 25010 | 531 | hiw | | 2.123e-02 | -3.85 | fly chr60B10-60B11\|2-106 | chromosome location | fly chr60B10-60B11\|2-106 | 1 | 1 | 25010 | 531 | CG3253 | | 2.123e-02 | -3.85 | fly chr45D4-45D5\|2-60 cM | chromosome location | fly chr45D4-45D5\|2-60 cM | 1 | 1 | 25010 | 531 | wun2 | | 2.123e-02 | -3.85 | fly chr52F2-52F3\|2-78 | chromosome location | fly chr52F2-52F3\|2-78 | 1 | 1 | 25010 | 531 | lbk | | 2.123e-02 | -3.85 | fly chr11D3-11D4\|1-41 cM | chromosome location | fly chr11D3-11D4\|1-41 cM | 1 | 1 | 25010 | 531 | fne | | 2.123e-02 | -3.85 | fly chr31E5-31E5\|2-42 cM | chromosome location | fly chr31E5-31E5\|2-42 cM | 1 | 1 | 25010 | 531 | LManII | | 2.123e-02 | -3.85 | fly chr66A18-66A19\|3-21 cM | chromosome location | fly chr66A18-66A19\|3-21 cM | 1 | 1 | 25010 | 531 | pbl | | 2.123e-02 | -3.85 | fly chr25F3-25F4\|2-17 cM | chromosome location | fly chr25F3-25F4\|2-17 cM | 1 | 1 | 25010 | 531 | CG11030 | | 2.123e-02 | -3.85 | fly chr24C3-24C4\|2-11 cM | chromosome location | fly chr24C3-24C4\|2-11 cM | 1 | 1 | 25010 | 531 | bowl | | 2.123e-02 | -3.85 | fly chr10E3-10E4\|1-36 cM | chromosome location | fly chr10E3-10E4\|1-36 cM | 1 | 1 | 25010 | 531 | Hsc70-3 | | 2.123e-02 | -3.85 | fly chr85E14-85E14\|3-49 | chromosome location | fly chr85E14-85E14\|3-49 | 1 | 1 | 25010 | 531 | lncRNA:CR45029 | | 2.123e-02 | -3.85 | fly chr93B10-93B11\|3-70 | chromosome location | fly chr93B10-93B11\|3-70 | 1 | 1 | 25010 | 531 | RhoGAP93B | | 2.123e-02 | -3.85 | fly chr64C2-64C5\|3-13 | chromosome location | fly chr64C2-64C5\|3-13 | 1 | 1 | 25010 | 531 | Con | | 2.123e-02 | -3.85 | fly chr94B8-94B11\|3-75 cM | chromosome location | fly chr94B8-94B11\|3-75 cM | 1 | 1 | 25010 | 531 | wake | | 2.123e-02 | -3.85 | fly chr55C2-55C4\|2-85 | chromosome location | fly chr55C2-55C4\|2-85 | 1 | 1 | 25010 | 531 | sbb | | 2.123e-02 | -3.85 | fly chr64C12-64C13\|3-13 | chromosome location | fly chr64C12-64C13\|3-13 | 1 | 1 | 25010 | 531 | lama | | 2.123e-02 | -3.85 | fly chr3F2-3F3\|1-5 cM | chromosome location | fly chr3F2-3F3\|1-5 cM | 1 | 1 | 25010 | 531 | ec | | 2.123e-02 | -3.85 | fly chr49A6-49A7\|2-67 | chromosome location | fly chr49A6-49A7\|2-67 | 1 | 1 | 25010 | 531 | Lac | | 2.123e-02 | -3.85 | fly chr102D4-102D5\|4-0 cM | chromosome location | fly chr102D4-102D5\|4-0 cM | 1 | 1 | 25010 | 531 | ey | | 2.123e-02 | -3.85 | fly chr3F1-3F1\|1-4 cM | chromosome location | fly chr3F1-3F1\|1-4 cM | 1 | 1 | 25010 | 531 | Tlk | | 2.123e-02 | -3.85 | fly chr13D4-13E1\|1-51 | chromosome location | fly chr13D4-13E1\|1-51 | 1 | 1 | 25010 | 531 | CG9220 | | 2.123e-02 | -3.85 | fly chr4F9-4F10\|1-11 | chromosome location | fly chr4F9-4F10\|1-11 | 1 | 1 | 25010 | 531 | spoon | | 2.123e-02 | -3.85 | fly chr96D2-96E1\|3-88 | chromosome location | fly chr96D2-96E1\|3-88 | 1 | 1 | 25010 | 531 | Fur1 | | 2.123e-02 | -3.85 | fly chr55B8-55B8\|2-84 cM | chromosome location | fly chr55B8-55B8\|2-84 cM | 1 | 1 | 25010 | 531 | Pcl | | 2.123e-02 | -3.85 | fly chr98F6-98F10\|3-97 | chromosome location | fly chr98F6-98F10\|3-97 | 1 | 1 | 25010 | 531 | CG11873 | | 2.123e-02 | -3.85 | fly chr102D4-102D5\|4-0 | chromosome location | fly chr102D4-102D5\|4-0 | 1 | 1 | 25010 | 531 | ey | | 2.123e-02 | -3.85 | fly chr85E14-85E14\|3-49 cM | chromosome location | fly chr85E14-85E14\|3-49 cM | 1 | 1 | 25010 | 531 | lncRNA:CR45029 | | 2.123e-02 | -3.85 | fly chr78F4-79A4\|3-47 cM | chromosome location | fly chr78F4-79A4\|3-47 cM | 1 | 1 | 25010 | 531 | mub | | 2.123e-02 | -3.85 | fly chr53D8-53D9\|2-80 | chromosome location | fly chr53D8-53D9\|2-80 | 1 | 1 | 25010 | 531 | CG5522 | | 2.123e-02 | -3.85 | fly chr44D3-44D6\|2-59 | chromosome location | fly chr44D3-44D6\|2-59 | 1 | 1 | 25010 | 531 | Acsl | | 2.123e-02 | -3.85 | fly chr50A1-50A3\|2-68 cM | chromosome location | fly chr50A1-50A3\|2-68 cM | 1 | 1 | 25010 | 531 | lncRNA:CR44206 | | 2.123e-02 | -3.85 | fly chr26D4-26D5\|2-20 | chromosome location | fly chr26D4-26D5\|2-20 | 1 | 1 | 25010 | 531 | retm | | 2.123e-02 | -3.85 | fly chr2C9-2C10\|1-0.5 | chromosome location | fly chr2C9-2C10\|1-0.5 | 1 | 1 | 25010 | 531 | CG4199 | | 2.123e-02 | -3.85 | fly chr36C2-36C4\|2-53 | chromosome location | fly chr36C2-36C4\|2-53 | 1 | 1 | 25010 | 531 | beat-IIIc | | 2.123e-02 | -3.85 | fly chr11D11-11E1\|1-41 cM | chromosome location | fly chr11D11-11E1\|1-41 cM | 1 | 1 | 25010 | 531 | CG43313 | | 2.123e-02 | -3.85 | fly chr52F2-52F3\|2-78 cM | chromosome location | fly chr52F2-52F3\|2-78 cM | 1 | 1 | 25010 | 531 | lbk | | 2.123e-02 | -3.85 | fly chr79D4-79E3\|3-47 | chromosome location | fly chr79D4-79E3\|3-47 | 1 | 1 | 25010 | 531 | Ten-m | | 2.123e-02 | -3.85 | fly chr72E5-72F1\|3-44 | chromosome location | fly chr72E5-72F1\|3-44 | 1 | 1 | 25010 | 531 | fax | | 2.123e-02 | -3.85 | fly chr63D2-63D3\|3-8 | chromosome location | fly chr63D2-63D3\|3-8 | 1 | 1 | 25010 | 531 | sty | | 2.123e-02 | -3.85 | fly chr62D7-62E1\|3-3 cM | chromosome location | fly chr62D7-62E1\|3-3 cM | 1 | 1 | 25010 | 531 | Svil | | 2.123e-02 | -3.85 | fly chr87F10-87F11\|3-54 cM | chromosome location | fly chr87F10-87F11\|3-54 cM | 1 | 1 | 25010 | 531 | flfl | | 2.123e-02 | -3.85 | fly chr83B8-83B9\|3-47.5 | chromosome location | fly chr83B8-83B9\|3-47.5 | 1 | 1 | 25010 | 531 | Atg17 | | 2.123e-02 | -3.85 | fly chr27B1-27B2\|2-22 | chromosome location | fly chr27B1-27B2\|2-22 | 1 | 1 | 25010 | 531 | nrv2 | | 2.123e-02 | -3.85 | fly chr97E3-97E5\|3-92 cM | chromosome location | fly chr97E3-97E5\|3-92 cM | 1 | 1 | 25010 | 531 | CG6051 | | 2.123e-02 | -3.85 | fly chr80B1-80B2\|3-47 | chromosome location | fly chr80B1-80B2\|3-47 | 1 | 1 | 25010 | 531 | Chro | | 2.123e-02 | -3.85 | fly chr95B7-95C1\|3-80 cM | chromosome location | fly chr95B7-95C1\|3-80 cM | 1 | 1 | 25010 | 531 | CG33111 | | 2.123e-02 | -3.85 | fly chr97D7-97D9\|3-92 | chromosome location | fly chr97D7-97D9\|3-92 | 1 | 1 | 25010 | 531 | CG46339 | | 2.123e-02 | -3.85 | fly chr68A5-68A6\|3-34 cM | chromosome location | fly chr68A5-68A6\|3-34 cM | 1 | 1 | 25010 | 531 | JIL-1 | | 2.123e-02 | -3.85 | fly chr67E4-67E5\|3-33 cM | chromosome location | fly chr67E4-67E5\|3-33 cM | 1 | 1 | 25010 | 531 | Rbfox1 | | 2.123e-02 | -3.85 | fly chr72C1-72C2\|3-43 cM | chromosome location | fly chr72C1-72C2\|3-43 cM | 1 | 1 | 25010 | 531 | CG5830 | | 2.123e-02 | -3.85 | fly chr8A5-8B2\|1-24 cM | chromosome location | fly chr8A5-8B2\|1-24 cM | 1 | 1 | 25010 | 531 | Lim1 | | 2.123e-02 | -3.85 | fly chr73A2-73A2\|3-44 cM | chromosome location | fly chr73A2-73A2\|3-44 cM | 1 | 1 | 25010 | 531 | aos | | 2.123e-02 | -3.85 | fly chr87F10-87F11\|3-54 | chromosome location | fly chr87F10-87F11\|3-54 | 1 | 1 | 25010 | 531 | flfl | | 2.123e-02 | -3.85 | fly chr94B8-94B11\|3-75 | chromosome location | fly chr94B8-94B11\|3-75 | 1 | 1 | 25010 | 531 | wake | | 2.123e-02 | -3.85 | fly chr40E5-40F2\|2-54 cM | chromosome location | fly chr40E5-40F2\|2-54 cM | 1 | 1 | 25010 | 531 | CG42748 | | 2.123e-02 | -3.85 | fly chr18E2-18E3\|1-64 | chromosome location | fly chr18E2-18E3\|1-64 | 1 | 1 | 25010 | 531 | meso18E | | 2.123e-02 | -3.85 | fly chr84C8-84D1\|3-48 | chromosome location | fly chr84C8-84D1\|3-48 | 1 | 1 | 25010 | 531 | sas | | 2.123e-02 | -3.85 | fly chr92F12-93A1\|3-69 | chromosome location | fly chr92F12-93A1\|3-69 | 1 | 1 | 25010 | 531 | Syp | | 2.123e-02 | -3.85 | fly chr98D6-98D7\|3-96 cM | chromosome location | fly chr98D6-98D7\|3-96 cM | 1 | 1 | 25010 | 531 | CG10011 | | 2.123e-02 | -3.85 | fly chr11B1-11B2\|1-39 | chromosome location | fly chr11B1-11B2\|1-39 | 1 | 1 | 25010 | 531 | Pits | | 2.123e-02 | -3.85 | fly chr78D4-78D5\|3-47 | chromosome location | fly chr78D4-78D5\|3-47 | 1 | 1 | 25010 | 531 | Cdk12 | | 2.123e-02 | -3.85 | fly chr100D3-100E2\|3-103 cM | chromosome location | fly chr100D3-100E2\|3-103 cM | 1 | 1 | 25010 | 531 | heph | | 2.123e-02 | -3.85 | fly chr30E1-30E4\|2-37 | chromosome location | fly chr30E1-30E4\|2-37 | 1 | 1 | 25010 | 531 | Etl1 | | 2.123e-02 | -3.85 | fly chr26E1-26E2\|2-21 | chromosome location | fly chr26E1-26E2\|2-21 | 1 | 1 | 25010 | 531 | eya | | 2.123e-02 | -3.85 | fly chr56D7-56D8\|2-89 | chromosome location | fly chr56D7-56D8\|2-89 | 1 | 1 | 25010 | 531 | betaTub56D | | 2.123e-02 | -3.85 | fly chr21D1-21E2\|2-1.5 cM | chromosome location | fly chr21D1-21E2\|2-1.5 cM | 1 | 1 | 25010 | 531 | ush | | 2.123e-02 | -3.85 | fly chr8D5-8D6\|1-27 | chromosome location | fly chr8D5-8D6\|1-27 | 1 | 1 | 25010 | 531 | lz | | 2.123e-02 | -3.85 | fly chr53D8-53D9\|2-80 cM | chromosome location | fly chr53D8-53D9\|2-80 cM | 1 | 1 | 25010 | 531 | CG5522 | | 2.123e-02 | -3.85 | fly chr15A3-15A5\|1-55 | chromosome location | fly chr15A3-15A5\|1-55 | 1 | 1 | 25010 | 531 | goe | | 2.123e-02 | -3.85 | fly chr32F1-32F2\|2-44 | chromosome location | fly chr32F1-32F2\|2-44 | 1 | 1 | 25010 | 531 | salm | | 2.123e-02 | -3.85 | fly chr14B9-14B12\|1-53 cM | chromosome location | fly chr14B9-14B12\|1-53 cM | 1 | 1 | 25010 | 531 | Cyp1 | | 2.123e-02 | -3.85 | fly chr52F5-52F5\|2-78 | chromosome location | fly chr52F5-52F5\|2-78 | 1 | 1 | 25010 | 531 | CG8441 | | 2.123e-02 | -3.85 | fly chr93F9-93F10\|3-73 cM | chromosome location | fly chr93F9-93F10\|3-73 cM | 1 | 1 | 25010 | 531 | tsl | | 2.123e-02 | -3.85 | fly chr92A10-92A11\|3-66 cM | chromosome location | fly chr92A10-92A11\|3-66 cM | 1 | 1 | 25010 | 531 | CG6231 | | 2.123e-02 | -3.85 | fly chr60D7-60D8\|2-107 | chromosome location | fly chr60D7-60D8\|2-107 | 1 | 1 | 25010 | 531 | pio | | 2.123e-02 | -3.85 | fly chr23B6-23B7\|2-7 | chromosome location | fly chr23B6-23B7\|2-7 | 1 | 1 | 25010 | 531 | CG2991 | | 2.123e-02 | -3.85 | fly chr96D2-96E1\|3-88 cM | chromosome location | fly chr96D2-96E1\|3-88 cM | 1 | 1 | 25010 | 531 | Fur1 | | 2.123e-02 | -3.85 | fly chr51E11-51F4\|2-74 | chromosome location | fly chr51E11-51F4\|2-74 | 1 | 1 | 25010 | 531 | Trpm | | 2.123e-02 | -3.85 | fly chr37A2-37A4\|2-54 cM | chromosome location | fly chr37A2-37A4\|2-54 cM | 1 | 1 | 25010 | 531 | ham | | 2.123e-02 | -3.85 | fly chr78F4-79A4\|3-47 | chromosome location | fly chr78F4-79A4\|3-47 | 1 | 1 | 25010 | 531 | mub | | 2.123e-02 | -3.85 | fly chr56D3-56D5\|2-89 cM | chromosome location | fly chr56D3-56D5\|2-89 cM | 1 | 1 | 25010 | 531 | hts | | 2.123e-02 | -3.85 | fly chr25D1-25D2\|2-16 cM | chromosome location | fly chr25D1-25D2\|2-16 cM | 1 | 1 | 25010 | 531 | tkv | | 2.123e-02 | -3.85 | fly chr21B3-21B4\|2-0.5 cM | chromosome location | fly chr21B3-21B4\|2-0.5 cM | 1 | 1 | 25010 | 531 | spen | | 2.123e-02 | -3.85 | fly chr58A3-58A4\|2-97 | chromosome location | fly chr58A3-58A4\|2-97 | 1 | 1 | 25010 | 531 | Fili | | 2.123e-02 | -3.85 | fly chr70E1-70E2\|3-42 cM | chromosome location | fly chr70E1-70E2\|3-42 cM | 1 | 1 | 25010 | 531 | Gbs-70E | | 2.123e-02 | -3.85 | fly chr57E9-57F1\|2-95 cM | chromosome location | fly chr57E9-57F1\|2-95 cM | 1 | 1 | 25010 | 531 | Egfr | | 2.123e-02 | -3.85 | fly chr49E6-49E6\|2-67 cM | chromosome location | fly chr49E6-49E6\|2-67 cM | 1 | 1 | 25010 | 531 | Psc | | 2.123e-02 | -3.85 | fly chr90D1-90E1\|3-62 | chromosome location | fly chr90D1-90E1\|3-62 | 1 | 1 | 25010 | 531 | cpo | | 2.128e-02 | -3.85 | postsynaptic membrane | cellular component | GO:0045211 | 47 | 5 | 11940 | 407 | Fas2,Ten-m,nAChRalpha4,Prosap,CaMKII | | 2.134e-02 | -3.85 | regulation of Rho protein signal transduction | biological process | GO:0035023 | 32 | 4 | 12345 | 414 | PsGEF,RhoGEF2,CG43658,pbl | | 2.152e-02 | -3.84 | STAT6\_02 | MSigDB lists | STAT6\_02 | 44 | 5 | 4161 | 153 | mgl,Tpr2,if,slow,dpp | | 2.152e-02 | -3.84 | FOXO4\_02 | MSigDB lists | FOXO4\_02 | 44 | 5 | 4161 | 153 | NetB,fne,Utx,CG4538,Ser | | 2.152e-02 | -3.84 | MORF\_BCL2 | MSigDB lists | MORF\_BCL2 | 44 | 5 | 4161 | 153 | fne,mub,CG4538,mof,Ser | | 2.152e-02 | -3.84 | GSE34515\_CD16\_POS\_MONOCYTE\_VS\_DC\_DN | MSigDB lists | GSE34515\_CD16\_POS\_MONOCYTE\_VS\_DC\_DN | 44 | 5 | 4161 | 153 | Pli,dpp,N,CG32758,JIL-1 | | 2.152e-02 | -3.84 | GSE1925\_3H\_VS\_24H\_IFNG\_STIM\_MACROPHAGE\_DN | MSigDB lists | GSE1925\_3H\_VS\_24H\_IFNG\_STIM\_MACROPHAGE\_DN | 44 | 5 | 4161 | 153 | par-1,betaTub56D,hts,CG6752,Arp2 | | 2.152e-02 | -3.84 | GSE360\_T\_GONDII\_VS\_M\_TUBERCULOSIS\_DC\_DN | MSigDB lists | GSE360\_T\_GONDII\_VS\_M\_TUBERCULOSIS\_DC\_DN | 44 | 5 | 4161 | 153 | Hsc70-3,dlp,CG7264,emp,Ykt6 | | 2.152e-02 | -3.84 | GSE33292\_DN3\_THYMOCYTE\_VS\_TCELL\_LYMPHOMA\_FROM\_TCF1\_KO\_DN | MSigDB lists | GSE33292\_DN3\_THYMOCYTE\_VS\_TCELL\_LYMPHOMA\_FROM\_TCF1\_KO\_DN | 44 | 5 | 4161 | 153 | mbt,Ugt49C1,CG14291,phtf,emp | | 2.152e-02 | -3.84 | GSE24574\_BCL6\_HIGH\_TFH\_VS\_TFH\_CD4\_TCELL\_UP | MSigDB lists | GSE24574\_BCL6\_HIGH\_TFH\_VS\_TFH\_CD4\_TCELL\_UP | 44 | 5 | 4161 | 153 | dpp,Mitofilin,Hsc70Cb,if,Cyt-c1 | | 2.152e-02 | -3.84 | EGR2\_01 | MSigDB lists | EGR2\_01 | 44 | 5 | 4161 | 153 | pAbp,CG16890,Sin3A,Imp,dlp | | 2.152e-02 | -3.84 | TFIII\_Q6 | MSigDB lists | TFIII\_Q6 | 44 | 5 | 4161 | 153 | LRP1,Ser,JIL-1,Sin3A,Pat1 | | 2.152e-02 | -3.84 | GSE7548\_NAIVE\_VS\_DAY7\_PCC\_IMMUNIZATION\_CD4\_TCELL\_DN | MSigDB lists | GSE7548\_NAIVE\_VS\_DAY7\_PCC\_IMMUNIZATION\_CD4\_TCELL\_DN | 44 | 5 | 4161 | 153 | Npc1a,Tlk,CG9220,Smox,Arl4 | | 2.158e-02 | -3.84 | developmental maturation | biological process | GO:0021700 | 206 | 13 | 12345 | 414 | Pat1,EcR,dpp,gus,heph,Imp,Egfr,Syp,N,psq,Hydr2,cnc,par-1 | | 2.172e-02 | -3.83 | regulation of histone modification | biological process | GO:0031056 | 48 | 5 | 12345 | 414 | ctrip,Utx,mof,sens-2,Pcl | | 2.172e-02 | -3.83 | phagocytosis | biological process | GO:0006909 | 48 | 5 | 12345 | 414 | Rac2,scb,Dscam1,egr,EcR | | 2.173e-02 | -3.83 | actin cortical patch | cellular component | GO:0030479 | 7 | 2 | 11940 | 407 | Arp2,Hip1 | | 2.173e-02 | -3.83 | integrin complex | cellular component | GO:0008305 | 7 | 2 | 11940 | 407 | scb,if | | 2.173e-02 | -3.83 | protein complex involved in cell adhesion | cellular component | GO:0098636 | 7 | 2 | 11940 | 407 | if,scb | | 2.173e-02 | -3.83 | endocytic patch | cellular component | GO:0061645 | 7 | 2 | 11940 | 407 | Arp2,Hip1 | | 2.174e-02 | -3.83 | histone binding | molecular function | GO:0042393 | 86 | 7 | 12176 | 396 | Pcl,enok,Stat92E,mof,Chro,Df31,CG5815 | | 2.177e-02 | -3.83 | transcription coregulator activity | molecular function | GO:0003712 | 147 | 10 | 12176 | 396 | tna,sbb,mam,eya,CtBP,shn,EcR,Sin3A,CG31875,Lpin | | 2.180e-02 | -3.83 | fly chr60E5-60E5\|2-107 cM | chromosome location | fly chr60E5-60E5\|2-107 cM | 11 | 2 | 25010 | 531 | asRNA:CR43466,NKAIN | | 2.180e-02 | -3.83 | fly chr82A5-82A5\|3-47.1 | chromosome location | fly chr82A5-82A5\|3-47.1 | 11 | 2 | 25010 | 531 | CG14646,CG9855 | | 2.180e-02 | -3.83 | fly chr82A5-82A5\|3-47.1 cM | chromosome location | fly chr82A5-82A5\|3-47.1 cM | 11 | 2 | 25010 | 531 | CG14646,CG9855 | | 2.180e-02 | -3.83 | fly chr60E5-60E5\|2-107 | chromosome location | fly chr60E5-60E5\|2-107 | 11 | 2 | 25010 | 531 | NKAIN,asRNA:CR43466 | | 2.183e-02 | -3.82 | GSE20500\_RETINOIC\_ACID\_VS\_RARA\_ANTAGONIST\_TREATED\_CD4\_TCELL\_DN | MSigDB lists | GSE20500\_RETINOIC\_ACID\_VS\_RARA\_ANTAGONIST\_TREATED\_CD4\_TCELL\_DN | 60 | 6 | 4161 | 153 | CG9220,Cyp1,emp,CG12099,Smox,Acsl | | 2.183e-02 | -3.82 | GSE22033\_WT\_VS\_PPARG\_KO\_MEF\_DN | MSigDB lists | GSE22033\_WT\_VS\_PPARG\_KO\_MEF\_DN | 60 | 6 | 4161 | 153 | Trpm,Ppa,pbl,nAChRalpha4,ScpX,CG4538 | | 2.183e-02 | -3.82 | GO\_PLASMA\_MEMBRANE\_ORGANIZATION | MSigDB lists | GO\_PLASMA\_MEMBRANE\_ORGANIZATION | 60 | 6 | 4161 | 153 | ScpX,Tsp26A,sdt,Ank,ft,Rac2 | | 2.183e-02 | -3.82 | GSE2770\_UNTREATED\_VS\_TGFB\_AND\_IL4\_TREATED\_ACT\_CD4\_TCELL\_2H\_DN | MSigDB lists | GSE2770\_UNTREATED\_VS\_TGFB\_AND\_IL4\_TREATED\_ACT\_CD4\_TCELL\_2H\_DN | 60 | 6 | 4161 | 153 | Nhe2,phtf,PhKgamma,CG11367,JIL-1,drk | | 2.185e-02 | -3.82 | Smox (Smad on X) | protein interactions | 31738 | 63 | 7 | 9204 | 412 | Acf,CG12182,NetA,stg,Ubx,wb,CycE | | 2.192e-02 | -3.82 | perivitelline space | cellular component | GO:0098595 | 18 | 3 | 11940 | 407 | dlp,dally,flz | | 2.208e-02 | -3.81 | GSE41176\_WT\_VS\_TAK1\_KO\_ANTI\_IGM\_STIM\_BCELL\_24H\_UP | MSigDB lists | GSE41176\_WT\_VS\_TAK1\_KO\_ANTI\_IGM\_STIM\_BCELL\_24H\_UP | 77 | 7 | 4161 | 153 | Acsl,Rho1,betaTub56D,Hsc70-3,Ser,Cyp1,nudE | | 2.208e-02 | -3.81 | GSE5589\_LPS\_AND\_IL10\_VS\_LPS\_AND\_IL6\_STIM\_IL6\_KO\_MACROPHAGE\_45MIN\_DN | MSigDB lists | GSE5589\_LPS\_AND\_IL10\_VS\_LPS\_AND\_IL6\_STIM\_IL6\_KO\_MACROPHAGE\_45MIN\_DN | 77 | 7 | 4161 | 153 | pAbp,sm,kdn,emp,LRP1,Npc1a,Mhcl | | 2.209e-02 | -3.81 | Mer (Merlin) | protein interactions | 32979 | 172 | 14 | 9204 | 412 | fax,l(3)L1231,crb,Mhcl,MFS14,CG11873,CG18446,betaTub56D,trn,CG31224,dpp,sbb,ft,Gmap | | 2.242e-02 | -3.80 | tra (transformer) | protein interactions | 39849 | 14 | 3 | 9204 | 412 | oc,N,Doa | | 2.242e-02 | -3.80 | Kat60 (Katanin 60) | protein interactions | 53566 | 14 | 3 | 9204 | 412 | ap,Myo10A,Bx | | 2.242e-02 | -3.80 | hts (hu li tai shao) | protein interactions | 37230 | 14 | 3 | 9204 | 412 | hts,S6kII,bab1 | | 2.242e-02 | -3.80 | ssh (slingshot) | protein interactions | 42986 | 14 | 3 | 9204 | 412 | Rho1,mbt,RhoGEF2 | | 2.242e-02 | -3.80 | kto (kohtalo) | protein interactions | 44830 | 14 | 3 | 9204 | 412 | ci,EcR,CycE | | 2.242e-02 | -3.80 | jub (Ajuba LIM protein) | protein interactions | 32351 | 14 | 3 | 9204 | 412 | ft,drk,ci | | 2.242e-02 | -3.80 | TwdlJ (TweedleJ) | protein interactions | 43206 | 14 | 3 | 9204 | 412 | eya,pAbp,trn | | 2.266e-02 | -3.79 | histone acetyltransferase activity | molecular function | GO:0004402 | 50 | 5 | 12176 | 396 | enok,Mocs2B,Acf,mof,Taf1 | | 2.269e-02 | -3.79 | ALK\_DN.V1\_UP | MSigDB lists | ALK\_DN.V1\_UP | 17 | 3 | 4161 | 153 | CG7264,tkv,fj | | 2.269e-02 | -3.79 | SHETH\_LIVER\_CANCER\_VS\_TXNIP\_LOSS\_PAM5 | MSigDB lists | SHETH\_LIVER\_CANCER\_VS\_TXNIP\_LOSS\_PAM5 | 17 | 3 | 4161 | 153 | Ser,Fas2,Etl1 | | 2.269e-02 | -3.79 | SIG\_PIP3\_SIGNALING\_IN\_CARDIAC\_MYOCTES | MSigDB lists | SIG\_PIP3\_SIGNALING\_IN\_CARDIAC\_MYOCTES | 17 | 3 | 4161 | 153 | drk,mbt,S6kII | | 2.269e-02 | -3.79 | BROWNE\_HCMV\_INFECTION\_8HR\_UP | MSigDB lists | BROWNE\_HCMV\_INFECTION\_8HR\_UP | 17 | 3 | 4161 | 153 | cta,SamDC,CycE | | 2.269e-02 | -3.79 | CERVERA\_SDHB\_TARGETS\_2 | MSigDB lists | CERVERA\_SDHB\_TARGETS\_2 | 17 | 3 | 4161 | 153 | dlp,CtBP,TfAP-2 | | 2.269e-02 | -3.79 | GO\_HAIR\_CELL\_DIFFERENTIATION | MSigDB lists | GO\_HAIR\_CELL\_DIFFERENTIATION | 17 | 3 | 4161 | 153 | Rac2,N,Ser | | 2.287e-02 | -3.78 | GSE1740\_UNSTIM\_VS\_IFNA\_STIMULATED\_MCSF\_IFNG\_DERIVED\_MACROPHAGE\_UP | MSigDB lists | GSE1740\_UNSTIM\_VS\_IFNA\_STIMULATED\_MCSF\_IFNG\_DERIVED\_MACROPHAGE\_UP | 30 | 4 | 4161 | 153 | Ank,NTPase,Cyp1,Tudor-SN | | 2.287e-02 | -3.78 | PID\_CXCR4\_PATHWAY | MSigDB lists | PID\_CXCR4\_PATHWAY | 30 | 4 | 4161 | 153 | cta,if,Rho1,Rac2 | | 2.287e-02 | -3.78 | GO\_EYE\_MORPHOGENESIS | MSigDB lists | GO\_EYE\_MORPHOGENESIS | 30 | 4 | 4161 | 153 | TfAP-2,fj,Ser,Lim1 | | 2.287e-02 | -3.78 | GO\_HISTONE\_METHYLTRANSFERASE\_COMPLEX | MSigDB lists | GO\_HISTONE\_METHYLTRANSFERASE\_COMPLEX | 30 | 4 | 4161 | 153 | Utx,Pcl,Taf1,mof | | 2.287e-02 | -3.78 | GO\_LYSOSOMAL\_TRANSPORT | MSigDB lists | GO\_LYSOSOMAL\_TRANSPORT | 30 | 4 | 4161 | 153 | CG32758,CG41099,Npc1a,emp | | 2.294e-02 | -3.77 | Pdk1 (Phosphoinositide-dependent kinase 1) | protein interactions | 38017 | 124 | 11 | 9204 | 412 | pbl,S6kII,CG43658,CG9281,ScsbetaG,RhoGAP18B,AGO2,yps,l(3)neo38,Hsc70-3,CG5815 | | 2.307e-02 | -3.77 | FG\_GAP | prosite domains | PS51470 | 5 | 2 | 6243 | 316 | scb,if | | 2.307e-02 | -3.77 | MYST\_HAT | prosite domains | PS51726 | 5 | 2 | 6243 | 316 | enok,mof | | 2.307e-02 | -3.77 | MBD | prosite domains | PS50982 | 5 | 2 | 6243 | 316 | sba,tou | | 2.307e-02 | -3.77 | CAGGTA\_AREB6\_01 | MSigDB lists | CAGGTA\_AREB6\_01 | 152 | 11 | 4161 | 153 | Ser,Fas2,S6kII,TfAP-2,Utx,N,dlp,Tlk,CG9220,CG3961,CG43313 | | 2.307e-02 | -3.77 | SCHAEFFER\_PROSTATE\_DEVELOPMENT\_6HR\_DN | MSigDB lists | SCHAEFFER\_PROSTATE\_DEVELOPMENT\_6HR\_DN | 152 | 11 | 4161 | 153 | LRP1,Pat1,l(3)mbt,Tudor-SN,Tim17b,PhKgamma,ft,Taf1,sm,N,Cnx99A | | 2.311e-02 | -3.77 | protein kinase complex | cellular component | GO:1902911 | 48 | 5 | 11940 | 407 | Atg17,PhKgamma,Cdk12,Cdk5alpha,CycE | | 2.349e-02 | -3.75 | pie (pineapple eye) | protein interactions | 44315 | 37 | 5 | 9204 | 412 | gus,Pits,heph,CG14478,Gbs-70E | | 2.349e-02 | -3.75 | Lk6 (Lk6 kinase) | protein interactions | 44672 | 37 | 5 | 9204 | 412 | pbl,N,mam,spi,betaTub56D | | 2.349e-02 | -3.75 | skd (skuld) | protein interactions | 43906 | 37 | 5 | 9204 | 412 | ci,tna,EcR,Bx,CycE | | 2.349e-02 | -3.75 | Imp (IGF-II mRNA-binding protein) | protein interactions | 32009 | 37 | 5 | 9204 | 412 | jigr1,CycE,pAbp,mub,S6kII | | 2.349e-02 | -3.75 | innate immune response | biological process | GO:0045087 | 104 | 8 | 12345 | 414 | Rho1,egr,eya,Tl,bbg,Rac2,Cdk5alpha,Pli | | 2.350e-02 | -3.75 | GO\_CELL\_CELL\_JUNCTION | MSigDB lists | GO\_CELL\_CELL\_JUNCTION | 61 | 6 | 4161 | 153 | drk,Ank,sdt,CaMKII,pbl,CG17265 | | 2.350e-02 | -3.75 | GO\_MUSCLE\_TISSUE\_DEVELOPMENT | MSigDB lists | GO\_MUSCLE\_TISSUE\_DEVELOPMENT | 61 | 6 | 4161 | 153 | N,CaMKII,if,so,hh,dpp | | 2.350e-02 | -3.75 | RRAGTTGT\_UNKNOWN | MSigDB lists | RRAGTTGT\_UNKNOWN | 61 | 6 | 4161 | 153 | pbl,TfAP-2,CG32758,Utx,S6kII,Lim1 | | 2.351e-02 | -3.75 | IK3\_01 | MSigDB lists | IK3\_01 | 45 | 5 | 4161 | 153 | Ser,Trpm,Ank,TfAP-2,CG6330 | | 2.351e-02 | -3.75 | HALLMARK\_PI3K\_AKT\_MTOR\_SIGNALING | MSigDB lists | HALLMARK\_PI3K\_AKT\_MTOR\_SIGNALING | 45 | 5 | 4161 | 153 | drk,mbt,S6kII,Rac2,Arp2 | | 2.351e-02 | -3.75 | GSE369\_IFNG\_KO\_VS\_WT\_LIVER\_UP | MSigDB lists | GSE369\_IFNG\_KO\_VS\_WT\_LIVER\_UP | 45 | 5 | 4161 | 153 | CG14291,Arl4,Smox,CG45050,JIL-1 | | 2.354e-02 | -3.75 | BROWNE\_HCMV\_INFECTION\_16HR\_UP | MSigDB lists | BROWNE\_HCMV\_INFECTION\_16HR\_UP | 78 | 7 | 4161 | 153 | dpp,CG4538,cta,Ykt6,Arl4,Ppa,CG2812 | | 2.355e-02 | -3.75 | regulation of neuroblast proliferation | biological process | GO:1902692 | 49 | 5 | 12345 | 414 | EcR,N,E(spl)mdelta-HLH,osa,pnt | | 2.355e-02 | -3.75 | negative regulation of transport | biological process | GO:0051051 | 49 | 5 | 12345 | 414 | S6kII,Usp2,NFAT,CaMKII,CG9650 | | 2.366e-02 | -3.74 | germarium-derived oocyte differentiation | biological process | GO:0030706 | 33 | 4 | 12345 | 414 | Rbfox1,par-1,nudE,hts | | 2.366e-02 | -3.74 | regulation of DNA-binding transcription factor activity | biological process | GO:0051090 | 33 | 4 | 12345 | 414 | pnt,ush,edl,mof | | 2.366e-02 | -3.74 | regulation of filopodium assembly | biological process | GO:0051489 | 33 | 4 | 12345 | 414 | Arp2,N,Rho1,CaMKII | | 2.366e-02 | -3.74 | protein stabilization | biological process | GO:0050821 | 33 | 4 | 12345 | 414 | crb,Hipk,Cnx99A,Snx3 | | 2.373e-02 | -3.74 | shot (short stop) | protein interactions | 36542 | 25 | 4 | 9204 | 412 | betaTub56D,pbl,Fas2,wnd | | 2.373e-02 | -3.74 | CG15356 (uncharacterized protein) | protein interactions | 33370 | 25 | 4 | 9204 | 412 | Chro,osa,Sin3A,rau | | 2.373e-02 | -3.74 | Rop (Ras opposite) | protein interactions | 38493 | 25 | 4 | 9204 | 412 | Dad,pbl,disco,bor | | 2.373e-02 | -3.74 | trbl (tribbles) | protein interactions | 43999 | 25 | 4 | 9204 | 412 | pbl,stg,CG12730,N | | 2.373e-02 | -3.74 | e(r) (enhancer of rudimentary) | protein interactions | 43951 | 25 | 4 | 9204 | 412 | N,CG11486,CG8929,smal | | 2.376e-02 | -3.74 | - | gene3d domains | 2.60.200.10 | 6 | 2 | 4929 | 208 | Smox,Dad | | 2.376e-02 | -3.74 | - | gene3d domains | 1.25.40.530 | 6 | 2 | 4929 | 208 | RhoGAP93B,Myo10A | | 2.378e-02 | -3.74 | LIM\_DOMAIN\_1 | prosite domains | PS00478 | 33 | 5 | 6243 | 316 | smash,esn,Bx,ap,Lim1 | | 2.380e-02 | -3.74 | regulation of secretion by cell | biological process | GO:1903530 | 85 | 7 | 12345 | 414 | S,CaMKII,CG9650,corn,Snx3,Syx1A,NFAT | | 2.397e-02 | -3.73 | BMP signaling | KEGG pathways | dme\_M00679 | 8 | 2 | 3198 | 100 | dpp,tkv | | 2.397e-02 | -3.73 | BMP signaling | KEGG pathways | M00679 | 8 | 2 | 3198 | 100 | tkv,dpp | | 2.397e-02 | -3.73 | Glycosaminoglycan biosynthesis - chondroitin sulfate / dermatan sulfate | KEGG pathways | ko00532 | 8 | 2 | 3198 | 100 | CG9220,CG43313 | | 2.397e-02 | -3.73 | Glycosaminoglycan biosynthesis - chondroitin sulfate / dermatan sulfate | KEGG pathways | dme00532 | 8 | 2 | 3198 | 100 | CG9220,CG43313 | | 2.413e-02 | -3.72 | Myc (Myc) | protein interactions | 31310 | 174 | 14 | 9204 | 412 | yps,Atg17,ban,Hsc70-3,ft,cnc,hang,CaMKII,N,ds,PEK,dpp,tkv,Sin3A | | 2.424e-02 | -3.72 | Znf\_FYVE-rel | interpro domains | IPR017455 | 17 | 3 | 11383 | 428 | CG41099,rush,CG6051 | | 2.424e-02 | -3.72 | HTH\_Psq | interpro domains | IPR007889 | 17 | 3 | 11383 | 428 | psq,BtbVII,bab1 | | 2.434e-02 | -3.72 | male genitalia morphogenesis | biological process | GO:0048808 | 19 | 3 | 12345 | 414 | Fas2,N,otk2 | | 2.434e-02 | -3.72 | retina homeostasis | biological process | GO:0001895 | 19 | 3 | 12345 | 414 | crb,Cnx99A,disco | | 2.434e-02 | -3.72 | production of small RNA involved in gene silencing by RNA | biological process | GO:0070918 | 19 | 3 | 12345 | 414 | CG17168,AGO2,NiPp1 | | 2.434e-02 | -3.72 | male anatomical structure morphogenesis | biological process | GO:0090598 | 19 | 3 | 12345 | 414 | otk2,Fas2,N | | 2.434e-02 | -3.72 | post-embryonic genitalia morphogenesis | biological process | GO:0035126 | 19 | 3 | 12345 | 414 | Fas2,N,otk2 | | 2.434e-02 | -3.72 | regulation of retinal cell programmed cell death | biological process | GO:0046668 | 19 | 3 | 12345 | 414 | Egfr,ec,lz | | 2.434e-02 | -3.72 | thermosensory behavior | biological process | GO:0040040 | 19 | 3 | 12345 | 414 | spoon,mub,Ptpmeg | | 2.434e-02 | -3.72 | ventral furrow formation | biological process | GO:0007370 | 19 | 3 | 12345 | 414 | Rho1,RhoGEF2,twi | | 2.434e-02 | -3.72 | walking behavior | biological process | GO:0090659 | 19 | 3 | 12345 | 414 | ey,hiw,oc | | 2.434e-02 | -3.72 | dsRNA processing | biological process | GO:0031050 | 19 | 3 | 12345 | 414 | NiPp1,CG17168,AGO2 | | 2.434e-02 | -3.72 | imaginal disc-derived genitalia morphogenesis | biological process | GO:0048805 | 19 | 3 | 12345 | 414 | N,Fas2,otk2 | | 2.437e-02 | -3.71 | Toll-Like Receptors Cascades | REACTOME pathways | R-DME-168898 | 60 | 6 | 3923 | 148 | Ptpmeg,JIL-1,mle,Pli,Tl,S6kII | | 2.451e-02 | -3.71 | asymmetric stem cell division | biological process | GO:0098722 | 67 | 6 | 12345 | 414 | dally,wake,flfl,Zif,pon,sdt | | 2.451e-02 | -3.71 | positive regulation of protein kinase activity | biological process | GO:0045860 | 67 | 6 | 12345 | 414 | Atg17,mbt,Rho1,Cdk5alpha,wnd,bora | | 2.451e-02 | -3.71 | gene silencing by RNA | biological process | GO:0031047 | 67 | 6 | 12345 | 414 | Tudor-SN,AGO2,CG17168,CG17265,NiPp1,Hsc70-3 | | 2.451e-02 | -3.71 | regulation of establishment of protein localization | biological process | GO:0070201 | 67 | 6 | 12345 | 414 | S,CaMKII,Snx3,crb,S6kII,sdt | | 2.460e-02 | -3.70 | synaptic membrane | cellular component | GO:0097060 | 66 | 6 | 11940 | 407 | Prosap,CaMKII,Syx1A,Ten-m,Fas2,nAChRalpha4 | | 2.470e-02 | -3.70 | positive regulation of catabolic process | biological process | GO:0009896 | 125 | 9 | 12345 | 414 | Utx,phyl,ctrip,smg,Usp2,gus,Toll-7,PEK,Doa | | 2.483e-02 | -3.70 | Immune System | REACTOME pathways | R-DME-168256 | 744 | 38 | 3923 | 148 | Egfr,CG46339,NFAT,drk,smash,Pli,Tl,ctrip,S6kII,CG42684,if,Mmp2,LManII,RhoGAP18B,gus,par-1,CG32369,Fas2,Arp2,JIL-1,Cirl,Stat92E,Tsp26A,mle,Syx1A,GILT3,Ptpmeg,RapGAP1,Rho1,Tsp39D,Ppa,Cdk12,Fbxl7,scb,Trpm,bor,betaTub56D,ras | | 2.492e-02 | -3.69 | KH\_dom\_type\_1\_sf | interpro domains | IPR036612 | 30 | 4 | 11383 | 428 | spoon,mub,Imp,ps | | 2.498e-02 | -3.69 | KANNAN\_TP53\_TARGETS\_DN | MSigDB lists | KANNAN\_TP53\_TARGETS\_DN | 7 | 2 | 4161 | 153 | Arl4,CycE | | 2.498e-02 | -3.69 | KANG\_GIST\_WITH\_PDGFRA\_UP | MSigDB lists | KANG\_GIST\_WITH\_PDGFRA\_UP | 7 | 2 | 4161 | 153 | nAChRalpha4,fj | | 2.498e-02 | -3.69 | GO\_POSITIVE\_REGULATION\_OF\_CELL\_JUNCTION\_ASSEMBLY | MSigDB lists | GO\_POSITIVE\_REGULATION\_OF\_CELL\_JUNCTION\_ASSEMBLY | 7 | 2 | 4161 | 153 | Rac2,Smox | | 2.498e-02 | -3.69 | GCM\_MAX | MSigDB lists | GCM\_MAX | 7 | 2 | 4161 | 153 | baf,hiw | | 2.498e-02 | -3.69 | GO\_REGULATION\_OF\_GLIAL\_CELL\_PROLIFERATION | MSigDB lists | GO\_REGULATION\_OF\_GLIAL\_CELL\_PROLIFERATION | 7 | 2 | 4161 | 153 | CG12099,N | | 2.498e-02 | -3.69 | GO\_REGULATION\_OF\_SODIUM\_ION\_TRANSMEMBRANE\_TRANSPORTER\_ACTIVITY | MSigDB lists | GO\_REGULATION\_OF\_SODIUM\_ION\_TRANSMEMBRANE\_TRANSPORTER\_ACTIVITY | 7 | 2 | 4161 | 153 | Ank,CaMKII | | 2.498e-02 | -3.69 | GO\_NEGATIVE\_REGULATION\_OF\_DNA\_BINDING | MSigDB lists | GO\_NEGATIVE\_REGULATION\_OF\_DNA\_BINDING | 7 | 2 | 4161 | 153 | Sin3A,ap | | 2.498e-02 | -3.69 | chr2q24 | MSigDB lists | chr2q24 | 7 | 2 | 4161 | 153 | mgl,CG6330 | | 2.498e-02 | -3.69 | GGCAGAC\_MIR346 | MSigDB lists | GGCAGAC\_MIR346 | 7 | 2 | 4161 | 153 | Imp,Smox | | 2.498e-02 | -3.69 | GO\_INTERMEDIATE\_FILAMENT | MSigDB lists | GO\_INTERMEDIATE\_FILAMENT | 7 | 2 | 4161 | 153 | Tlk,CG6330 | | 2.498e-02 | -3.69 | KEGG\_CYTOKINE\_CYTOKINE\_RECEPTOR\_INTERACTION | MSigDB lists | KEGG\_CYTOKINE\_CYTOKINE\_RECEPTOR\_INTERACTION | 7 | 2 | 4161 | 153 | dpp,tkv | | 2.498e-02 | -3.69 | GO\_CELL\_DIFFERENTIATION\_INVOLVED\_IN\_KIDNEY\_DEVELOPMENT | MSigDB lists | GO\_CELL\_DIFFERENTIATION\_INVOLVED\_IN\_KIDNEY\_DEVELOPMENT | 7 | 2 | 4161 | 153 | N,Ser | | 2.498e-02 | -3.69 | GO\_ESTABLISHMENT\_OF\_EPITHELIAL\_CELL\_POLARITY | MSigDB lists | GO\_ESTABLISHMENT\_OF\_EPITHELIAL\_CELL\_POLARITY | 7 | 2 | 4161 | 153 | Mhcl,sdt | | 2.498e-02 | -3.69 | GO\_INTERMEDIATE\_FILAMENT\_BASED\_PROCESS | MSigDB lists | GO\_INTERMEDIATE\_FILAMENT\_BASED\_PROCESS | 7 | 2 | 4161 | 153 | nudE,hh | | 2.498e-02 | -3.69 | GO\_NEGATIVE\_REGULATION\_OF\_FAT\_CELL\_DIFFERENTIATION | MSigDB lists | GO\_NEGATIVE\_REGULATION\_OF\_FAT\_CELL\_DIFFERENTIATION | 7 | 2 | 4161 | 153 | Ser,Smox | | 2.498e-02 | -3.69 | PID\_S1P\_S1P3\_PATHWAY | MSigDB lists | PID\_S1P\_S1P3\_PATHWAY | 7 | 2 | 4161 | 153 | Rac2,cta | | 2.498e-02 | -3.69 | GO\_RNA\_STABILIZATION | MSigDB lists | GO\_RNA\_STABILIZATION | 7 | 2 | 4161 | 153 | mle,Imp | | 2.498e-02 | -3.69 | GARGALOVIC\_RESPONSE\_TO\_OXIDIZED\_PHOSPHOLIPIDS\_GREY\_DN | MSigDB lists | GARGALOVIC\_RESPONSE\_TO\_OXIDIZED\_PHOSPHOLIPIDS\_GREY\_DN | 7 | 2 | 4161 | 153 | TfAP-2,Arl4 | | 2.498e-02 | -3.69 | ACAACCT\_MIR453 | MSigDB lists | ACAACCT\_MIR453 | 7 | 2 | 4161 | 153 | CG32758,Utx | | 2.498e-02 | -3.69 | HOFMANN\_CELL\_LYMPHOMA\_DN | MSigDB lists | HOFMANN\_CELL\_LYMPHOMA\_DN | 7 | 2 | 4161 | 153 | Taf1,Cyt-c1 | | 2.498e-02 | -3.69 | GO\_PROTEIN\_FOLDING\_IN\_ENDOPLASMIC\_RETICULUM | MSigDB lists | GO\_PROTEIN\_FOLDING\_IN\_ENDOPLASMIC\_RETICULUM | 7 | 2 | 4161 | 153 | Cnx99A,Hsc70-3 | | 2.498e-02 | -3.69 | REACTOME\_CREB\_PHOSPHORYLATION\_THROUGH\_THE\_ACTIVATION\_OF\_RAS | MSigDB lists | REACTOME\_CREB\_PHOSPHORYLATION\_THROUGH\_THE\_ACTIVATION\_OF\_RAS | 7 | 2 | 4161 | 153 | CaMKII,S6kII | | 2.498e-02 | -3.69 | GO\_CHONDROITIN\_SULFATE\_BIOSYNTHETIC\_PROCESS | MSigDB lists | GO\_CHONDROITIN\_SULFATE\_BIOSYNTHETIC\_PROCESS | 7 | 2 | 4161 | 153 | CG43313,CG9220 | | 2.498e-02 | -3.69 | GO\_REGULATION\_OF\_ANOIKIS | MSigDB lists | GO\_REGULATION\_OF\_ANOIKIS | 7 | 2 | 4161 | 153 | N,CG5742 | | 2.498e-02 | -3.69 | GO\_POLY\_PYRIMIDINE\_TRACT\_BINDING | MSigDB lists | GO\_POLY\_PYRIMIDINE\_TRACT\_BINDING | 7 | 2 | 4161 | 153 | heph,pAbp | | 2.498e-02 | -3.69 | GO\_POSITIVE\_REGULATION\_OF\_EPITHELIAL\_CELL\_DIFFERENTIATION | MSigDB lists | GO\_POSITIVE\_REGULATION\_OF\_EPITHELIAL\_CELL\_DIFFERENTIATION | 7 | 2 | 4161 | 153 | N,Lim1 | | 2.498e-02 | -3.69 | GO\_REGULATION\_OF\_MEMBRANE\_DEPOLARIZATION | MSigDB lists | GO\_REGULATION\_OF\_MEMBRANE\_DEPOLARIZATION | 7 | 2 | 4161 | 153 | Ank,CaMKII | | 2.498e-02 | -3.69 | LUND\_SILENCED\_BY\_METHYLATION | MSigDB lists | LUND\_SILENCED\_BY\_METHYLATION | 7 | 2 | 4161 | 153 | N,SamDC | | 2.498e-02 | -3.69 | PID\_SMAD2\_3PATHWAY | MSigDB lists | PID\_SMAD2\_3PATHWAY | 7 | 2 | 4161 | 153 | Smox,CG5830 | | 2.504e-02 | -3.69 | polytene chromosome puff | cellular component | GO:0005703 | 49 | 5 | 11940 | 407 | JIL-1,cnc,mam,lawc,Cdk12 | | 2.505e-02 | -3.69 | GSE28726\_NAIVE\_VS\_ACTIVATED\_NKTCELL\_DN | MSigDB lists | GSE28726\_NAIVE\_VS\_ACTIVATED\_NKTCELL\_DN | 79 | 7 | 4161 | 153 | mmy,Npc1a,nudE,CG9220,Syx1A,CG2812,betaTub56D | | 2.505e-02 | -3.69 | AAGCCAT\_MIR135A\_MIR135B | MSigDB lists | AAGCCAT\_MIR135A\_MIR135B | 79 | 7 | 4161 | 153 | phtf,Lac,Imp,CG9220,Pli,CG34384,Ank | | 2.516e-02 | -3.68 | RA | prosite domains | PS50200 | 13 | 3 | 6243 | 316 | wake,rau,CG32758 | | 2.520e-02 | -3.68 | wound healing | biological process | GO:0042060 | 86 | 7 | 12345 | 414 | jing,Utx,scb,pbl,Rho1,Mmp2,lz | | 2.525e-02 | -3.68 | MCBRYAN\_PUBERTAL\_BREAST\_4\_5WK\_DN | MSigDB lists | MCBRYAN\_PUBERTAL\_BREAST\_4\_5WK\_DN | 62 | 6 | 4161 | 153 | NiPp1,Ser,Tsp26A,Mitofilin,S6kII,Ykt6 | | 2.525e-02 | -3.68 | BERTUCCI\_MEDULLARY\_VS\_DUCTAL\_BREAST\_CANCER\_UP | MSigDB lists | BERTUCCI\_MEDULLARY\_VS\_DUCTAL\_BREAST\_CANCER\_UP | 62 | 6 | 4161 | 153 | Hsc70Cb,Cyp1,Mrp4,Pcl,Ppa,CG3961 | | 2.525e-02 | -3.68 | AP2\_Q6\_01 | MSigDB lists | AP2\_Q6\_01 | 62 | 6 | 4161 | 153 | gus,ap,pAbp,Tlk,CG9855,Imp | | 2.539e-02 | -3.67 | basement membrane | cellular component | GO:0005604 | 19 | 3 | 11940 | 407 | NetB,wb,NetA | | 2.547e-02 | -3.67 | p38b (p38b MAP kinase) | protein interactions | 34780 | 51 | 6 | 9204 | 412 | ps,hiw,tkv,ush,dpp,bi | | 2.547e-02 | -3.67 | negative regulation of chromosome organization | biological process | GO:2001251 | 50 | 5 | 12345 | 414 | Utx,ctrip,JIL-1,lncRNA:roX1,Chro | | 2.549e-02 | -3.67 | E2f1 (E2F transcription factor 1) | protein interactions | 42550 | 65 | 7 | 9204 | 412 | CycE,l(3)mbt,pbl,tou,Doa,Imp,tara | | 2.549e-02 | -3.67 | lwr (lesswright) | protein interactions | 33226 | 65 | 7 | 9204 | 412 | tna,CaMKII,Doa,dpp,ttk,mbt,ScsbetaG | | 2.549e-02 | -3.67 | baz (bazooka) | protein interactions | 32703 | 65 | 7 | 9204 | 412 | sdt,Rho1,Stat92E,fj,par-1,pon,crb | | 2.553e-02 | -3.67 | GO\_RESPONSE\_TO\_FIBROBLAST\_GROWTH\_FACTOR | MSigDB lists | GO\_RESPONSE\_TO\_FIBROBLAST\_GROWTH\_FACTOR | 31 | 4 | 4161 | 153 | Lim1,wech,drk,heph | | 2.553e-02 | -3.67 | GSE12366\_GC\_VS\_NAIVE\_BCELL\_DN | MSigDB lists | GSE12366\_GC\_VS\_NAIVE\_BCELL\_DN | 31 | 4 | 4161 | 153 | LManII,JIL-1,CG2201,l(3)mbt | | 2.553e-02 | -3.67 | GO\_POSITIVE\_REGULATION\_OF\_CELL\_CELL\_ADHESION | MSigDB lists | GO\_POSITIVE\_REGULATION\_OF\_CELL\_CELL\_ADHESION | 31 | 4 | 4161 | 153 | hh,Ank,drk,Rac2 | | 2.553e-02 | -3.67 | GSE29617\_CTRL\_VS\_DAY7\_TIV\_FLU\_VACCINE\_PBMC\_2008\_UP | MSigDB lists | GSE29617\_CTRL\_VS\_DAY7\_TIV\_FLU\_VACCINE\_PBMC\_2008\_UP | 31 | 4 | 4161 | 153 | l(3)mbt,cta,pbl,Arl4 | | 2.553e-02 | -3.67 | MODULE\_567 | MSigDB lists | MODULE\_567 | 31 | 4 | 4161 | 153 | Fas2,CG8531,Ank,CG10663 | | 2.553e-02 | -3.67 | GSE4590\_PRE\_BCELL\_VS\_VPREB\_POS\_LARGE\_PRE\_BCELL\_UP | MSigDB lists | GSE4590\_PRE\_BCELL\_VS\_VPREB\_POS\_LARGE\_PRE\_BCELL\_UP | 31 | 4 | 4161 | 153 | mub,ap,CG6330,Arl4 | | 2.553e-02 | -3.67 | ST\_INTEGRIN\_SIGNALING\_PATHWAY | MSigDB lists | ST\_INTEGRIN\_SIGNALING\_PATHWAY | 31 | 4 | 4161 | 153 | Arp2,if,mbt,drk | | 2.562e-02 | -3.66 | GSE31082\_DN\_VS\_DP\_THYMOCYTE\_DN | MSigDB lists | GSE31082\_DN\_VS\_DP\_THYMOCYTE\_DN | 46 | 5 | 4161 | 153 | par-1,cta,EcR,rush,PhKgamma | | 2.562e-02 | -3.66 | MYB\_Q6 | MSigDB lists | MYB\_Q6 | 46 | 5 | 4161 | 153 | Ppa,Utx,Rho1,smal,Mhcl | | 2.562e-02 | -3.66 | MIYAGAWA\_TARGETS\_OF\_EWSR1\_ETS\_FUSIONS\_UP | MSigDB lists | MIYAGAWA\_TARGETS\_OF\_EWSR1\_ETS\_FUSIONS\_UP | 46 | 5 | 4161 | 153 | JIL-1,Ser,lbk,Imp,so | | 2.562e-02 | -3.66 | GSE24574\_BCL6\_HIGH\_TFH\_VS\_NAIVE\_CD4\_TCELL\_UP | MSigDB lists | GSE24574\_BCL6\_HIGH\_TFH\_VS\_NAIVE\_CD4\_TCELL\_UP | 46 | 5 | 4161 | 153 | PhKgamma,dlp,tkv,mub,ras | | 2.562e-02 | -3.66 | GO\_REGULATION\_OF\_CHROMATIN\_ORGANIZATION | MSigDB lists | GO\_REGULATION\_OF\_CHROMATIN\_ORGANIZATION | 46 | 5 | 4161 | 153 | Pcl,CaMKII,Sin3A,JIL-1,Tlk | | 2.562e-02 | -3.66 | PID\_PDGFRB\_PATHWAY | MSigDB lists | PID\_PDGFRB\_PATHWAY | 46 | 5 | 4161 | 153 | S6kII,drk,Rac2,LRP1,Arp2 | | 2.569e-02 | -3.66 | GTPase regulator activity | molecular function | GO:0030695 | 89 | 7 | 12176 | 396 | RhoGAP18B,RhoGAP16F,pbl,RhoGAP15B,RhoGAP93B,RapGAP1,RhoGAP102A | | 2.573e-02 | -3.66 | GO\_POSITIVE\_REGULATION\_OF\_CELL\_PROLIFERATION | MSigDB lists | GO\_POSITIVE\_REGULATION\_OF\_CELL\_PROLIFERATION | 135 | 10 | 4161 | 153 | N,NetB,CaMKII,Lim1,dpp,smal,slow,hh,so,RpS4 | | 2.579e-02 | -3.66 | fly chr61B3-61B3\|3-1 | chromosome location | fly chr61B3-61B3\|3-1 | 12 | 2 | 25010 | 531 | Tudor-SN,snoRNA:Psi18S-996 | | 2.579e-02 | -3.66 | fly chr23C4-23C4\|2-8 cM | chromosome location | fly chr23C4-23C4\|2-8 cM | 12 | 2 | 25010 | 531 | CG17224,CG17265 | | 2.579e-02 | -3.66 | fly chr42A13-42A13\|2-55 | chromosome location | fly chr42A13-42A13\|2-55 | 12 | 2 | 25010 | 531 | Ptr,bin3 | | 2.579e-02 | -3.66 | fly chr23C4-23C4\|2-8 | chromosome location | fly chr23C4-23C4\|2-8 | 12 | 2 | 25010 | 531 | CG17224,CG17265 | | 2.579e-02 | -3.66 | fly chr42A13-42A13\|2-55 cM | chromosome location | fly chr42A13-42A13\|2-55 cM | 12 | 2 | 25010 | 531 | bin3,Ptr | | 2.579e-02 | -3.66 | fly chr61B3-61B3\|3-1 cM | chromosome location | fly chr61B3-61B3\|3-1 cM | 12 | 2 | 25010 | 531 | snoRNA:Psi18S-996,Tudor-SN | | 2.580e-02 | -3.66 | PH | pfam domains | PF00169 | 45 | 5 | 10480 | 393 | CG34383,CG5522,rush,osp,CG34384 | | 2.594e-02 | -3.65 | POZ domain binding | molecular function | GO:0031208 | 8 | 2 | 12176 | 396 | ttk,psq | | 2.598e-02 | -3.65 | regulation of autophagy | biological process | GO:0010506 | 106 | 8 | 12345 | 414 | Smox,Utx,EcR,Toll-7,wcy,Doa,PEK,Gyf | | 2.599e-02 | -3.65 | PAN\_1 | pfam domains | PF00024 | 7 | 2 | 10480 | 393 | tyn,neo | | 2.599e-02 | -3.65 | NHL | pfam domains | PF01436 | 7 | 2 | 10480 | 393 | wech,CG12547 | | 2.606e-02 | -3.65 | regulation of immune response | biological process | GO:0050776 | 168 | 11 | 12345 | 414 | Doa,Usp2,Toll-7,Etl1,Rac2,Pli,Stat92E,Mmp2,CG4325,ush,Tl | | 2.613e-02 | -3.64 | NHL\_repeat | interpro domains | IPR001258 | 7 | 2 | 11383 | 428 | CG12547,wech | | 2.613e-02 | -3.64 | Pan\_app | interpro domains | IPR003609 | 7 | 2 | 11383 | 428 | tyn,neo | | 2.613e-02 | -3.64 | Hud\_Sxl\_RNA | interpro domains | IPR002343 | 7 | 2 | 11383 | 428 | elav,fne | | 2.613e-02 | -3.64 | Furin\_repeat | interpro domains | IPR006212 | 7 | 2 | 11383 | 428 | Egfr,Fur1 | | 2.613e-02 | -3.64 | NHL\_repeat\_subgr | interpro domains | IPR013017 | 7 | 2 | 11383 | 428 | CG12547,wech | | 2.613e-02 | -3.64 | CRIB\_dom | interpro domains | IPR000095 | 7 | 2 | 11383 | 428 | Pak3,mbt | | 2.613e-02 | -3.64 | Integrin\_dom\_sf | interpro domains | IPR032695 | 7 | 2 | 11383 | 428 | scb,if | | 2.613e-02 | -3.64 | SSD | interpro domains | IPR000731 | 7 | 2 | 11383 | 428 | Ptr,Npc1a | | 2.613e-02 | -3.64 | positive regulation of border follicle cell migration | biological process | GO:1903688 | 34 | 4 | 12345 | 414 | spi,ttk,Egfr,aop | | 2.617e-02 | -3.64 | GO\_ZINC\_ION\_BINDING | MSigDB lists | GO\_ZINC\_ION\_BINDING | 258 | 16 | 4161 | 153 | Mmp2,CG3097,CG9855,TfIIS,hh,hiw,wech,CG6752,CG45050,Pcl,EcR,LManII,CG12099,Smox,ap,Lim1 | | 2.621e-02 | -3.64 | RHO GTPases activate PKNs | REACTOME pathways | R-DME-5625740 | 7 | 2 | 3923 | 148 | Rho1,stg | | 2.621e-02 | -3.64 | CREB phosphorylation through the activation of Ras | REACTOME pathways | R-DME-442742 | 7 | 2 | 3923 | 148 | CaMKII,S6kII | | 2.621e-02 | -3.64 | Role of Abl in Robo-Slit signaling | REACTOME pathways | R-DME-428890 | 7 | 2 | 3923 | 148 | robo2,dlp | | 2.631e-02 | -3.64 | Int\_alpha | smart domains | SM00191 | 5 | 2 | 5086 | 276 | if,scb | | 2.631e-02 | -3.64 | DWA | smart domains | SM00523 | 5 | 2 | 5086 | 276 | Dad,Smox | | 2.639e-02 | -3.63 | peptide-lysine-N-acetyltransferase activity | molecular function | GO:0061733 | 52 | 5 | 12176 | 396 | Acf,Mocs2B,enok,Taf1,mof | | 2.651e-02 | -3.63 | GO\_AMINOGLYCAN\_CATABOLIC\_PROCESS | MSigDB lists | GO\_AMINOGLYCAN\_CATABOLIC\_PROCESS | 18 | 3 | 4161 | 153 | CG14291,dlp,dally | | 2.651e-02 | -3.63 | KEGG\_CHRONIC\_MYELOID\_LEUKEMIA | MSigDB lists | KEGG\_CHRONIC\_MYELOID\_LEUKEMIA | 18 | 3 | 4161 | 153 | Smox,drk,CtBP | | 2.651e-02 | -3.63 | GTTNYYNNGGTNA\_UNKNOWN | MSigDB lists | GTTNYYNNGGTNA\_UNKNOWN | 18 | 3 | 4161 | 153 | CG16890,Etl1,CG2812 | | 2.651e-02 | -3.63 | GO\_BONE\_MORPHOGENESIS | MSigDB lists | GO\_BONE\_MORPHOGENESIS | 18 | 3 | 4161 | 153 | tkv,TfAP-2,CG9220 | | 2.651e-02 | -3.63 | MODULE\_568 | MSigDB lists | MODULE\_568 | 18 | 3 | 4161 | 153 | betaTub56D,Acsl,CG9220 | | 2.651e-02 | -3.63 | SIG\_CHEMOTAXIS | MSigDB lists | SIG\_CHEMOTAXIS | 18 | 3 | 4161 | 153 | Arp2,RpS4,mbt | | 2.651e-02 | -3.63 | LEE\_NEURAL\_CREST\_STEM\_CELL\_DN | MSigDB lists | LEE\_NEURAL\_CREST\_STEM\_CELL\_DN | 18 | 3 | 4161 | 153 | ap,fj,N | | 2.651e-02 | -3.63 | GSE9946\_MATURE\_STIMULATORY\_VS\_LISTERIA\_INF\_MATURE\_DC\_DN | MSigDB lists | GSE9946\_MATURE\_STIMULATORY\_VS\_LISTERIA\_INF\_MATURE\_DC\_DN | 18 | 3 | 4161 | 153 | Ser,Hsc70-3,Dscam1 | | 2.651e-02 | -3.63 | JIANG\_HYPOXIA\_VIA\_VHL | MSigDB lists | JIANG\_HYPOXIA\_VIA\_VHL | 18 | 3 | 4161 | 153 | heph,Cnx99A,emp | | 2.651e-02 | -3.63 | GO\_PROTEOGLYCAN\_BIOSYNTHETIC\_PROCESS | MSigDB lists | GO\_PROTEOGLYCAN\_BIOSYNTHETIC\_PROCESS | 18 | 3 | 4161 | 153 | CG9220,CG43313,tkv | | 2.651e-02 | -3.63 | GO\_NEGATIVE\_REGULATION\_OF\_TRANSPORTER\_ACTIVITY | MSigDB lists | GO\_NEGATIVE\_REGULATION\_OF\_TRANSPORTER\_ACTIVITY | 18 | 3 | 4161 | 153 | Cyp1,CaMKII,Ank | | 2.651e-02 | -3.63 | GO\_VIRUS\_RECEPTOR\_ACTIVITY | MSigDB lists | GO\_VIRUS\_RECEPTOR\_ACTIVITY | 18 | 3 | 4161 | 153 | Npc1a,Fas2,emp | | 2.651e-02 | -3.63 | GO\_REGULATION\_OF\_EPITHELIAL\_CELL\_DIFFERENTIATION | MSigDB lists | GO\_REGULATION\_OF\_EPITHELIAL\_CELL\_DIFFERENTIATION | 18 | 3 | 4161 | 153 | N,ft,Lim1 | | 2.660e-02 | -3.63 | lili (lilipod) | protein interactions | 42956 | 6 | 2 | 9204 | 412 | Dad,tkv | | 2.660e-02 | -3.63 | mRpL12 (mitochondrial ribosomal protein L12) | protein interactions | 44326 | 6 | 2 | 9204 | 412 | Hph,S6kII | | 2.660e-02 | -3.63 | wb (wing blister) | protein interactions | 43946 | 6 | 2 | 9204 | 412 | scb,Smox | | 2.660e-02 | -3.63 | mtd (mustard) | protein interactions | 45467 | 6 | 2 | 9204 | 412 | mam,Fas2 | | 2.660e-02 | -3.63 | Nlg4 (Neuroligin 4) | protein interactions | 42402 | 6 | 2 | 9204 | 412 | Dad,tkv | | 2.660e-02 | -3.63 | Mnt (Mnt) | protein interactions | 31331 | 6 | 2 | 9204 | 412 | N,Sin3A | | 2.660e-02 | -3.63 | TBCD (tubulin folding cofactor D) | protein interactions | 33389 | 6 | 2 | 9204 | 412 | Fas2,Dscam1 | | 2.660e-02 | -3.63 | dbr (debra) | protein interactions | 33161 | 6 | 2 | 9204 | 412 | ci,phyl | | 2.660e-02 | -3.63 | RpL34b (Ribosomal protein L34b) | protein interactions | 41120 | 6 | 2 | 9204 | 412 | CG3386,Pli | | 2.660e-02 | -3.63 | CG42557 (uncharacterized protein) | protein interactions | 43548 | 6 | 2 | 9204 | 412 | Cyp1,baf | | 2.660e-02 | -3.63 | Kr-h1 (Kruppel homolog 1) | protein interactions | 33861 | 6 | 2 | 9204 | 412 | Kr,oc | | 2.660e-02 | -3.63 | shf (shifted) | protein interactions | 31617 | 6 | 2 | 9204 | 412 | dally,hh | | 2.660e-02 | -3.63 | stx (stuxnet) | protein interactions | 32005 | 6 | 2 | 9204 | 412 | pbl,spi | | 2.660e-02 | -3.63 | ast (asteroid) | protein interactions | 33282 | 6 | 2 | 9204 | 412 | S,Egfr | | 2.660e-02 | -3.63 | NetB (Netrin-B) | protein interactions | 32400 | 6 | 2 | 9204 | 412 | spi,Dscam1 | | 2.660e-02 | -3.63 | mtgo (miles to go) | protein interactions | 34987 | 6 | 2 | 9204 | 412 | flfl,sty | | 2.660e-02 | -3.63 | rumi (rumi) | protein interactions | 326122 | 6 | 2 | 9204 | 412 | Hsc70-3,N | | 2.660e-02 | -3.63 | upd2 (unpaired 2) | protein interactions | 32805 | 6 | 2 | 9204 | 412 | Sox21a,N | | 2.660e-02 | -3.63 | CG43867 (uncharacterized protein) | protein interactions | 31031 | 6 | 2 | 9204 | 412 | CG17265,drk | | 2.660e-02 | -3.63 | CG5746 (uncharacterized protein) | protein interactions | 42903 | 6 | 2 | 9204 | 412 | Tet,Pat1 | | 2.660e-02 | -3.63 | ACC (Acetyl-CoA carboxylase) | protein interactions | 35761 | 6 | 2 | 9204 | 412 | S6kII,Fas2 | | 2.660e-02 | -3.63 | rasp (rasp) | protein interactions | 44098 | 6 | 2 | 9204 | 412 | hh,spi | | 2.664e-02 | -3.63 | GSE3982\_MAC\_VS\_BASOPHIL\_UP | MSigDB lists | GSE3982\_MAC\_VS\_BASOPHIL\_UP | 80 | 7 | 4161 | 153 | baf,LManII,ras,Acsl,CG14291,Cyp1,CG9220 | | 2.664e-02 | -3.63 | GO\_RESPONSE\_TO\_METAL\_ION | MSigDB lists | GO\_RESPONSE\_TO\_METAL\_ION | 98 | 8 | 4161 | 153 | Npc1a,Cyp1,TfAP-2,pbl,CaMKII,Hsc70-3,hh,Ank | | 2.664e-02 | -3.63 | actomyosin structure organization | biological process | GO:0031032 | 87 | 7 | 12345 | 414 | Rho1,crb,if,pbl,CG13366,hts,twi | | 2.677e-02 | -3.62 | LIM\_DOMAIN\_2 | prosite domains | PS50023 | 34 | 5 | 6243 | 316 | smash,Bx,esn,Lim1,ap | | 2.684e-02 | -3.62 | Ras GTPase binding | molecular function | GO:0017016 | 174 | 11 | 12176 | 396 | CG43658,RhoGAP18B,nuf,pbl,mbt,RhoGEF2,PsGEF,Gmap,rau,CG5522,Pak3 | | 2.691e-02 | -3.62 | GO\_SIGNAL\_TRANSDUCER\_ACTIVITY | MSigDB lists | GO\_SIGNAL\_TRANSDUCER\_ACTIVITY | 196 | 13 | 4161 | 153 | Rho1,mbt,EcR,nAChRalpha4,Hydr2,tkv,dlp,cta,Smox,Ptr,pbl,Npc1a,smal | | 2.703e-02 | -3.61 | positive regulation of transport | biological process | GO:0051050 | 127 | 9 | 12345 | 414 | sdt,comm,S,Hip1,ScpX,drk,corn,hh,Snx3 | | 2.704e-02 | -3.61 | Leu-rich\_rpt\_typical-subtyp | interpro domains | IPR003591 | 78 | 7 | 11383 | 428 | Lrt,Toll-7,lbk,trn,Tl,Con,Fili | | 2.709e-02 | -3.61 | Eaf (ELL-associated factor) | protein interactions | 35660 | 26 | 4 | 9204 | 412 | CG3919,l(3)neo38,Socs16D,fax | | 2.709e-02 | -3.61 | fs(1)h (female sterile (1) homeotic) | protein interactions | 31722 | 26 | 4 | 9204 | 412 | pbl,cnc,spi,Psc | | 2.709e-02 | -3.61 | Lis-1 (Lissencephaly-1) | protein interactions | 36791 | 26 | 4 | 9204 | 412 | tkv,Egfr,nudE,smg | | 2.709e-02 | -3.61 | GTGCAAT\_MIR25\_MIR32\_MIR92\_MIR363\_MIR367 | MSigDB lists | GTGCAAT\_MIR25\_MIR32\_MIR92\_MIR363\_MIR367 | 63 | 6 | 4161 | 153 | lili,hiw,Dscam1,phtf,Mhcl,Rhp | | 2.712e-02 | -3.61 | slpr (slipper) | protein interactions | 44111 | 15 | 3 | 9204 | 412 | ci,aop,Rho1 | | 2.712e-02 | -3.61 | Ctf4 (Chromosome transmission fidelity 4) | protein interactions | 36551 | 15 | 3 | 9204 | 412 | ft,pbl,S6kII | | 2.712e-02 | -3.61 | Bap170 (Brahma associated protein 170kD) | protein interactions | 35560 | 15 | 3 | 9204 | 412 | pnt,Egfr,kni | | 2.712e-02 | -3.61 | kra (krasavietz) | protein interactions | 40680 | 15 | 3 | 9204 | 412 | CG8191,RpS4,Kr | | 2.712e-02 | -3.61 | CG33129 (uncharacterized protein) | protein interactions | 34514 | 15 | 3 | 9204 | 412 | CG8929,CG11076,esn | | 2.712e-02 | -3.61 | Cpr76Ba (Cuticular protein 76Ba) | protein interactions | 40120 | 15 | 3 | 9204 | 412 | CG32772,CG17163,scaf6 | | 2.712e-02 | -3.61 | klu (klumpfuss) | protein interactions | 39228 | 15 | 3 | 9204 | 412 | Egfr,mam,N | | 2.712e-02 | -3.61 | comm (commissureless) | protein interactions | 39717 | 15 | 3 | 9204 | 412 | elav,robo2,comm | | 2.712e-02 | -3.61 | twi (twist) | protein interactions | 37655 | 15 | 3 | 9204 | 412 | Ubx,twi,Df31 | | 2.712e-02 | -3.61 | PQBP1 (Poly-glutamine tract binding protein 1) | protein interactions | 43004 | 15 | 3 | 9204 | 412 | CG11247,yps,Hsc70-3 | | 2.712e-02 | -3.61 | Br140 (Bromodomain-containing protein, 140kD) | protein interactions | 35648 | 15 | 3 | 9204 | 412 | Psc,ci,enok | | 2.712e-02 | -3.61 | Rbfox1 (RNA-binding Fox protein 1) | protein interactions | 39198 | 15 | 3 | 9204 | 412 | Rbfox1,N,ci | | 2.740e-02 | -3.60 | PH | smart domains | SM00233 | 67 | 8 | 5086 | 276 | rush,CG43658,CG34384,CG34383,osp,RhoGAP15B,RhoGEF2,CG5522 | | 2.747e-02 | -3.59 | dorsal trunk growth, open tracheal system | biological process | GO:0035001 | 8 | 2 | 12345 | 414 | scb,ttk | | 2.747e-02 | -3.59 | cellular response to cytokine stimulus | biological process | GO:0071345 | 8 | 2 | 12345 | 414 | Tl,grnd | | 2.747e-02 | -3.59 | gonadal mesoderm development | biological process | GO:0007506 | 8 | 2 | 12345 | 414 | tin,hh | | 2.747e-02 | -3.59 | regulation of stress fiber assembly | biological process | GO:0051492 | 8 | 2 | 12345 | 414 | if,CG43658 | | 2.747e-02 | -3.59 | regulation of lipid transport | biological process | GO:0032368 | 8 | 2 | 12345 | 414 | ScpX,Npc1a | | 2.747e-02 | -3.59 | defasciculation of motor neuron axon | biological process | GO:0007415 | 8 | 2 | 12345 | 414 | pbl,Ptp99A | | 2.747e-02 | -3.59 | positive regulation of embryonic development | biological process | GO:0040019 | 8 | 2 | 12345 | 414 | tsl,stg | | 2.747e-02 | -3.59 | male gonad development | biological process | GO:0008584 | 8 | 2 | 12345 | 414 | salm,eya | | 2.747e-02 | -3.59 | negative regulation of ERK1 and ERK2 cascade | biological process | GO:0070373 | 8 | 2 | 12345 | 414 | aos,sty | | 2.747e-02 | -3.59 | positive regulation of DNA metabolic process | biological process | GO:0051054 | 8 | 2 | 12345 | 414 | Tet,eya | | 2.747e-02 | -3.59 | imaginal disc-derived female genitalia development | biological process | GO:0007486 | 8 | 2 | 12345 | 414 | lz,otk2 | | 2.747e-02 | -3.59 | anterior head segmentation | biological process | GO:0035288 | 8 | 2 | 12345 | 414 | oc,hh | | 2.747e-02 | -3.59 | oenocyte development | biological process | GO:0007438 | 8 | 2 | 12345 | 414 | spi,salm | | 2.747e-02 | -3.59 | progression of morphogenetic furrow involved in compound eye morphogenesis | biological process | GO:0007458 | 8 | 2 | 12345 | 414 | dpp,hh | | 2.747e-02 | -3.59 | development of primary male sexual characteristics | biological process | GO:0046546 | 8 | 2 | 12345 | 414 | eya,salm | | 2.747e-02 | -3.59 | negative regulation of lamellocyte differentiation | biological process | GO:0035204 | 8 | 2 | 12345 | 414 | N,Ser | | 2.747e-02 | -3.59 | negative regulation of oskar mRNA translation | biological process | GO:0007319 | 8 | 2 | 12345 | 414 | elav,heph | | 2.747e-02 | -3.59 | anterior head development | biological process | GO:0097065 | 8 | 2 | 12345 | 414 | hh,oc | | 2.747e-02 | -3.59 | lymph gland crystal cell differentiation | biological process | GO:0035170 | 8 | 2 | 12345 | 414 | ush,pnt | | 2.747e-02 | -3.59 | cytokine-mediated signaling pathway | biological process | GO:0019221 | 8 | 2 | 12345 | 414 | grnd,Tl | | 2.747e-02 | -3.59 | cell fate specification involved in pattern specification | biological process | GO:0060573 | 8 | 2 | 12345 | 414 | Egfr,salm | | 2.747e-02 | -3.59 | establishment or maintenance of polarity of embryonic epithelium | biological process | GO:0016332 | 8 | 2 | 12345 | 414 | crb,sdt | | 2.747e-02 | -3.59 | negative regulation of plasmatocyte differentiation | biological process | GO:0045614 | 8 | 2 | 12345 | 414 | Ser,N | | 2.747e-02 | -3.59 | hemocyte proliferation | biological process | GO:0035172 | 8 | 2 | 12345 | 414 | Stat92E,N | | 2.747e-02 | -3.59 | regulation of myosin II filament organization | biological process | GO:0043519 | 8 | 2 | 12345 | 414 | RhoGEF2,cta | | 2.776e-02 | -3.58 | Leu-rich\_rpt | interpro domains | IPR001611 | 114 | 9 | 11383 | 428 | trn,Lrt,Toll-7,Tl,Fbxl7,Con,Fili,Ppa,lbk | | 2.785e-02 | -3.58 | LMO2COM\_02 | MSigDB lists | LMO2COM\_02 | 47 | 5 | 4161 | 153 | so,Mhcl,sm,CycE,Trpm | | 2.785e-02 | -3.58 | GATA6\_01 | MSigDB lists | GATA6\_01 | 47 | 5 | 4161 | 153 | kdn,Utx,pbl,smal,so | | 2.785e-02 | -3.58 | NFAT\_Q6 | MSigDB lists | NFAT\_Q6 | 47 | 5 | 4161 | 153 | smal,CG2812,fj,Trpm,Notum | | 2.785e-02 | -3.58 | GSE6269\_E\_COLI\_VS\_STAPH\_AUREUS\_INF\_PBMC\_DN | MSigDB lists | GSE6269\_E\_COLI\_VS\_STAPH\_AUREUS\_INF\_PBMC\_DN | 47 | 5 | 4161 | 153 | ras,EcR,Rac2,Ser,Hsc70-3 | | 2.785e-02 | -3.58 | FOXO3\_01 | MSigDB lists | FOXO3\_01 | 47 | 5 | 4161 | 153 | Utx,NetB,phtf,Pli,CG4538 | | 2.785e-02 | -3.58 | SRY\_01 | MSigDB lists | SRY\_01 | 47 | 5 | 4161 | 153 | CG9855,Fas2,lili,Ank,ds | | 2.786e-02 | -3.58 | positive regulation of apoptotic process | biological process | GO:0043065 | 69 | 6 | 12345 | 414 | CycE,grnd,EcR,aos,ft,N | | 2.786e-02 | -3.58 | positive regulation of kinase activity | biological process | GO:0033674 | 69 | 6 | 12345 | 414 | wnd,Cdk5alpha,bora,mbt,Atg17,Rho1 | | 2.794e-02 | -3.58 | endoplasmic reticulum unfolded protein response | biological process | GO:0030968 | 20 | 3 | 12345 | 414 | Cnx99A,Hsc70-3,PEK | | 2.794e-02 | -3.58 | regulation of establishment of planar polarity | biological process | GO:0090175 | 20 | 3 | 12345 | 414 | aop,ds,otk2 | | 2.794e-02 | -3.58 | negative regulation of hippo signaling | biological process | GO:0035331 | 20 | 3 | 12345 | 414 | ds,par-1,Hipk | | 2.794e-02 | -3.58 | positive regulation of receptor-mediated endocytosis | biological process | GO:0048260 | 20 | 3 | 12345 | 414 | Hip1,comm,drk | | 2.794e-02 | -3.58 | chitin-based embryonic cuticle biosynthetic process | biological process | GO:0008362 | 20 | 3 | 12345 | 414 | EcR,pio,mmy | | 2.794e-02 | -3.58 | negative regulation of BMP signaling pathway | biological process | GO:0030514 | 20 | 3 | 12345 | 414 | Lpin,Acsl,hiw | | 2.809e-02 | -3.57 | Bromodomain | pfam domains | PF00439 | 18 | 3 | 10480 | 393 | Taf1,tou,Acf | | 2.820e-02 | -3.57 | Pi3K21B (Pi3K21B) | protein interactions | 33203 | 112 | 10 | 9204 | 412 | Nopp140,Syp,Egfr,mle,ImpL2,tkv,bor,yps,ras,Hsc70-3 | | 2.827e-02 | -3.57 | nuclear lumen | cellular component | GO:0031981 | 1016 | 46 | 11940 | 407 | aop,Utx,Cdk12,Hph,jing,Tif-IA,ctrip,baf,Kah,HmgZ,CG18766,CG44774,enok,NiPp1,Nopp140,CG5815,tna,Pits,Myb,Mocs2B,Tlk,mam,elav,Taf1,mle,pbl,Rbfox1,Sin3A,CG11030,tou,mof,CG8441,gus,CtBP,osa,udd,CG31712,Acf,JIL-1,stwl,Psc,Pcl,sm,CG5728,lncRNA:roX1,drk | | 2.827e-02 | -3.57 | carbohydrate derivative binding | molecular function | GO:0097367 | 1011 | 44 | 12176 | 396 | Mrp4,Asator,Etl1,Arp2,dpp,Hipk,CG8097,PEK,Mhcl,Haspin,S6kII,ScsbetaG,Rho1,KP78a,Pak3,bor,Doa,wnd,tkv,CaMKII,Tlk,Hsc70-3,Cdk12,Rac2,Myo10A,CG12010,mbt,lz,mle,CG14535,sas,JIL-1,Mos,Egfr,CG9281,CG9541,Arl4,CG13675,Pex1,Ser,betaTub56D,PhKgamma,par-1,CG8485 | | 2.829e-02 | -3.57 | CTACCTC\_LET7A\_LET7B\_LET7C\_LET7D\_LET7E\_LET7F\_MIR98\_LET7G\_LET7I | MSigDB lists | CTACCTC\_LET7A\_LET7B\_LET7C\_LET7D\_LET7E\_LET7F\_MIR98\_LET7G\_LET7I | 81 | 7 | 4161 | 153 | lbk,wech,Imp,Tim17b,Etl1,S6kII,CG41099 | | 2.830e-02 | -3.56 | Nucleotide-bd\_a/b\_plait\_sf | interpro domains | IPR012677 | 152 | 11 | 11383 | 428 | Rbfox1,heph,Syp,CG2162,cpo,elav,Imp,fne,pAbp,sm,spen | | 2.832e-02 | -3.56 | TGF-beta receptor signaling activates SMADs | REACTOME pathways | R-DME-2173789 | 18 | 3 | 3923 | 148 | Fur1,Dad,Smox | | 2.832e-02 | -3.56 | spectrosome | cellular component | GO:0045170 | 8 | 2 | 11940 | 407 | hts,Ank | | 2.832e-02 | -3.56 | MICOS complex | cellular component | GO:0061617 | 8 | 2 | 11940 | 407 | CG8531,CG41128 | | 2.832e-02 | -3.56 | gap junction | cellular component | GO:0005921 | 8 | 2 | 11940 | 407 | Inx3,Inx2 | | 2.837e-02 | -3.56 | GO\_CYTOKINESIS | MSigDB lists | GO\_CYTOKINESIS | 32 | 4 | 4161 | 153 | pbl,Rho1,Ank,Arp2 | | 2.837e-02 | -3.56 | ATCATGA\_MIR433 | MSigDB lists | ATCATGA\_MIR433 | 32 | 4 | 4161 | 153 | MFS10,mbt,drk,par-1 | | 2.837e-02 | -3.56 | GSE19941\_IL10\_KO\_VS\_IL10\_KO\_AND\_NFKBP50\_KO\_UNSTIM\_MACROPHAGE\_UP | MSigDB lists | GSE19941\_IL10\_KO\_VS\_IL10\_KO\_AND\_NFKBP50\_KO\_UNSTIM\_MACROPHAGE\_UP | 32 | 4 | 4161 | 153 | nAChRalpha4,rush,CG34384,slow | | 2.837e-02 | -3.56 | GSE16266\_CTRL\_VS\_LPS\_STIM\_MEF\_UP | MSigDB lists | GSE16266\_CTRL\_VS\_LPS\_STIM\_MEF\_UP | 32 | 4 | 4161 | 153 | mmy,Ppa,N,cta | | 2.837e-02 | -3.56 | GSE14769\_UNSTIM\_VS\_120MIN\_LPS\_BMDM\_DN | MSigDB lists | GSE14769\_UNSTIM\_VS\_120MIN\_LPS\_BMDM\_DN | 32 | 4 | 4161 | 153 | cta,nudE,Tlk,dpp | | 2.849e-02 | -3.56 | - | gene3d domains | 3.30.1370.10 | 28 | 4 | 4929 | 208 | ps,spoon,Imp,mub | | 2.850e-02 | -3.56 | GO\_PROTEIN\_KINASE\_ACTIVITY | MSigDB lists | GO\_PROTEIN\_KINASE\_ACTIVITY | 157 | 11 | 4161 | 153 | tkv,JIL-1,mbt,Doa,Tlk,CaMKII,Taf1,par-1,S6kII,smal,PhKgamma | | 2.850e-02 | -3.56 | GO\_ENZYME\_LINKED\_RECEPTOR\_PROTEIN\_SIGNALING\_PATHWAY | MSigDB lists | GO\_ENZYME\_LINKED\_RECEPTOR\_PROTEIN\_SIGNALING\_PATHWAY | 157 | 11 | 4161 | 153 | drk,JIL-1,nAChRalpha4,tkv,heph,Rac2,Smox,dpp,wech,smal,Arp2 | | 2.851e-02 | -3.56 | BROMODOMAIN | prints domains | PR00503 | 17 | 3 | 2185 | 88 | Taf1,tou,Acf | | 2.874e-02 | -3.55 | axon extension | biological process | GO:0048675 | 35 | 4 | 12345 | 414 | CadN,Ptpmeg,Dscam1,mle | | 2.874e-02 | -3.55 | positive regulation of epithelial cell migration | biological process | GO:0010634 | 35 | 4 | 12345 | 414 | ttk,spi,aop,Egfr | | 2.874e-02 | -3.55 | synaptic growth at neuromuscular junction | biological process | GO:0051124 | 35 | 4 | 12345 | 414 | Prosap,par-1,Ten-m,Imp | | 2.886e-02 | -3.55 | msi (musashi) | protein interactions | 43087 | 39 | 5 | 9204 | 412 | N,Imp,ttk,elav,ps | | 2.886e-02 | -3.55 | mle (maleless) | protein interactions | 35523 | 39 | 5 | 9204 | 412 | lncRNA:roX1,mof,Prosap,mle,ocm | | 2.901e-02 | -3.54 | GSE13306\_TREG\_VS\_TCONV\_DN | MSigDB lists | GSE13306\_TREG\_VS\_TCONV\_DN | 64 | 6 | 4161 | 153 | NTPase,heph,CG8531,kdn,Mitofilin,Tlk | | 2.901e-02 | -3.54 | GSE43955\_TH0\_VS\_TGFB\_IL6\_TH17\_ACT\_CD4\_TCELL\_60H\_DN | MSigDB lists | GSE43955\_TH0\_VS\_TGFB\_IL6\_TH17\_ACT\_CD4\_TCELL\_60H\_DN | 64 | 6 | 4161 | 153 | CG6752,N,Cnx99A,tkv,Ank,if | | 2.928e-02 | -3.53 | dom (domino) | protein interactions | 45655 | 97 | 9 | 9204 | 412 | Tudor-SN,EcR,Zif,mam,N,sno,tara,jigr1,Etl1 | | 2.952e-02 | -3.52 | HUDSXLRNA | prints domains | PR00961 | 7 | 2 | 2185 | 88 | elav,fne | | 2.954e-02 | -3.52 | ena (enabled) | protein interactions | 37201 | 67 | 7 | 9204 | 412 | Rho1,robo2,comm,N,ft,fj,drk | | 2.961e-02 | -3.52 | morphogenesis of follicular epithelium | biological process | GO:0016333 | 52 | 5 | 12345 | 414 | spoon,Egfr,sty,N,crb | | 2.963e-02 | -3.52 | CHARAFE\_BREAST\_CANCER\_LUMINAL\_VS\_MESENCHYMAL\_DN | MSigDB lists | CHARAFE\_BREAST\_CANCER\_LUMINAL\_VS\_MESENCHYMAL\_DN | 100 | 8 | 4161 | 153 | CG9220,Mrp4,Arp2,nudE,Ser,ft,Smox,pAbp | | 2.963e-02 | -3.52 | GO\_NEGATIVE\_REGULATION\_OF\_TRANSPORT | MSigDB lists | GO\_NEGATIVE\_REGULATION\_OF\_TRANSPORT | 100 | 8 | 4161 | 153 | CaMKII,N,Cyp1,Rac2,EcR,Ank,Snx3,hh | | 2.965e-02 | -3.52 | regulation of protein serine/threonine kinase activity | biological process | GO:0071900 | 70 | 6 | 12345 | 414 | wnd,CycE,Cdk5alpha,sty,mbt,Rho1 | | 2.965e-02 | -3.52 | developmental pigmentation | biological process | GO:0048066 | 70 | 6 | 12345 | 414 | Cnx99A,bab1,eya,dpp,hh,bi | | 2.970e-02 | -3.52 | regulation of secretion | biological process | GO:0051046 | 89 | 7 | 12345 | 414 | S,CG9650,corn,CaMKII,Snx3,Syx1A,NFAT | | 2.970e-02 | -3.52 | somatodendritic compartment | cellular component | GO:0036477 | 212 | 13 | 11940 | 407 | Myo10A,Ten-m,S6kII,Mmp2,nudE,Dscam1,EcR,Imp,Cdk5alpha,NetB,CaMKII,CadN,Prosap | | 2.992e-02 | -3.51 | small GTPase binding | molecular function | GO:0031267 | 177 | 11 | 12176 | 396 | RhoGEF2,pbl,mbt,Pak3,CG5522,Gmap,PsGEF,rau,nuf,RhoGAP18B,CG43658 | | 3.001e-02 | -3.51 | NMYC\_01 | MSigDB lists | NMYC\_01 | 82 | 7 | 4161 | 153 | pAbp,ds,cta,CtBP,Imp,Nhe2,bor | | 3.015e-02 | -3.50 | DCTN1-p150 (Dynactin 1, p150 subunit) | protein interactions | 39536 | 53 | 6 | 9204 | 412 | spi,betaTub56D,bbg,S,nuf,Egfr | | 3.020e-02 | -3.50 | GSE24142\_ADULT\_VS\_FETAL\_DN2\_THYMOCYTE\_DN | MSigDB lists | GSE24142\_ADULT\_VS\_FETAL\_DN2\_THYMOCYTE\_DN | 48 | 5 | 4161 | 153 | CtBP,kdn,CG9855,Fas2,Imp | | 3.020e-02 | -3.50 | GSE27786\_NKTCELL\_VS\_MONO\_MAC\_DN | MSigDB lists | GSE27786\_NKTCELL\_VS\_MONO\_MAC\_DN | 48 | 5 | 4161 | 153 | slow,LManII,Pli,Mrp4,ScpX | | 3.020e-02 | -3.50 | GSE21927\_SPLEEN\_C57BL6\_VS\_EL4\_TUMOR\_BALBC\_MONOCYTES\_UP | MSigDB lists | GSE21927\_SPLEEN\_C57BL6\_VS\_EL4\_TUMOR\_BALBC\_MONOCYTES\_UP | 48 | 5 | 4161 | 153 | CG12547,hiw,CG5830,CG45050,Ppa | | 3.020e-02 | -3.50 | GSE27786\_LIN\_NEG\_VS\_NKCELL\_DN | MSigDB lists | GSE27786\_LIN\_NEG\_VS\_NKCELL\_DN | 48 | 5 | 4161 | 153 | PhKgamma,sno,CG2201,CG5830,par-1 | | 3.020e-02 | -3.50 | DURAND\_STROMA\_S\_UP | MSigDB lists | DURAND\_STROMA\_S\_UP | 48 | 5 | 4161 | 153 | TfAP-2,ft,lbk,so,Fas2 | | 3.020e-02 | -3.50 | AAAYWAACM\_HFH4\_01 | MSigDB lists | AAAYWAACM\_HFH4\_01 | 48 | 5 | 4161 | 153 | CG9855,Fas2,Sin3A,ras,NetB | | 3.020e-02 | -3.50 | GSE20727\_ROS\_INH\_VS\_ROS\_INH\_AND\_DNFB\_ALLERGEN\_TREATED\_DC\_UP | MSigDB lists | GSE20727\_ROS\_INH\_VS\_ROS\_INH\_AND\_DNFB\_ALLERGEN\_TREATED\_DC\_UP | 48 | 5 | 4161 | 153 | pbl,betaTub56D,Rho1,ds,cta | | 3.020e-02 | -3.50 | GSE19888\_CTRL\_VS\_TCELL\_MEMBRANES\_ACT\_MAST\_CELL\_PRETREAT\_A3R\_INH\_UP | MSigDB lists | GSE19888\_CTRL\_VS\_TCELL\_MEMBRANES\_ACT\_MAST\_CELL\_PRETREAT\_A3R\_INH\_UP | 48 | 5 | 4161 | 153 | CG45050,NTPase,pAbp,CG14291,Tim17b | | 3.027e-02 | -3.50 | cleavage furrow | cellular component | GO:0032154 | 35 | 4 | 11940 | 407 | Arl4,Tl,nuf,pbl | | 3.043e-02 | -3.49 | calcium ion binding | molecular function | GO:0005509 | 200 | 12 | 12176 | 396 | ft,slow,Cnx99A,Ser,N,hh,Cad87A,crb,ds,LRP1,mgl,CadN | | 3.052e-02 | -3.49 | FN3\_sf | interpro domains | IPR036116 | 63 | 6 | 11383 | 428 | wake,Dscam1,sas,Fas2,Ptp99A,robo2 | | 3.052e-02 | -3.49 | FN3\_dom | interpro domains | IPR003961 | 63 | 6 | 11383 | 428 | robo2,wake,Ptp99A,sas,Dscam1,Fas2 | | 3.065e-02 | -3.49 | GO\_DEVELOPMENTAL\_CELL\_GROWTH | MSigDB lists | GO\_DEVELOPMENTAL\_CELL\_GROWTH | 19 | 3 | 4161 | 153 | nudE,CaMKII,ap | | 3.065e-02 | -3.49 | GO\_LIGASE\_ACTIVITY\_FORMING\_CARBON\_SULFUR\_BONDS | MSigDB lists | GO\_LIGASE\_ACTIVITY\_FORMING\_CARBON\_SULFUR\_BONDS | 19 | 3 | 4161 | 153 | Acsl,CG3961,ScsbetaG | | 3.065e-02 | -3.49 | REACTOME\_HEPARAN\_SULFATE\_HEPARIN\_HS\_GAG\_METABOLISM | MSigDB lists | REACTOME\_HEPARAN\_SULFATE\_HEPARIN\_HS\_GAG\_METABOLISM | 19 | 3 | 4161 | 153 | CG14291,dlp,dally | | 3.065e-02 | -3.49 | GO\_CELLULAR\_RESPONSE\_TO\_IONIZING\_RADIATION | MSigDB lists | GO\_CELLULAR\_RESPONSE\_TO\_IONIZING\_RADIATION | 19 | 3 | 4161 | 153 | Tlk,drk,pbl | | 3.065e-02 | -3.49 | KEGG\_HEDGEHOG\_SIGNALING\_PATHWAY | MSigDB lists | KEGG\_HEDGEHOG\_SIGNALING\_PATHWAY | 19 | 3 | 4161 | 153 | mgl,hh,dpp | | 3.065e-02 | -3.49 | GUTIERREZ\_MULTIPLE\_MYELOMA\_UP | MSigDB lists | GUTIERREZ\_MULTIPLE\_MYELOMA\_UP | 19 | 3 | 4161 | 153 | Ykt6,Cnx99A,ScsbetaG | | 3.065e-02 | -3.49 | GO\_MESODERM\_MORPHOGENESIS | MSigDB lists | GO\_MESODERM\_MORPHOGENESIS | 19 | 3 | 4161 | 153 | if,Utx,Smox | | 3.065e-02 | -3.49 | GO\_GLAND\_MORPHOGENESIS | MSigDB lists | GO\_GLAND\_MORPHOGENESIS | 19 | 3 | 4161 | 153 | N,NetB,hh | | 3.065e-02 | -3.49 | VART\_KSHV\_INFECTION\_ANGIOGENIC\_MARKERS\_UP | MSigDB lists | VART\_KSHV\_INFECTION\_ANGIOGENIC\_MARKERS\_UP | 19 | 3 | 4161 | 153 | Ser,slow,drk | | 3.065e-02 | -3.49 | GSE17974\_IL4\_AND\_ANTI\_IL12\_VS\_UNTREATED\_24H\_ACT\_CD4\_TCELL\_UP | MSigDB lists | GSE17974\_IL4\_AND\_ANTI\_IL12\_VS\_UNTREATED\_24H\_ACT\_CD4\_TCELL\_UP | 19 | 3 | 4161 | 153 | dlp,CG14646,ds | | 3.065e-02 | -3.49 | GSE40443\_INDUCED\_VS\_TOTAL\_TREG\_DN | MSigDB lists | GSE40443\_INDUCED\_VS\_TOTAL\_TREG\_DN | 19 | 3 | 4161 | 153 | CG11367,Pli,NTPase | | 3.065e-02 | -3.49 | GO\_POSITIVE\_REGULATION\_OF\_ACTIN\_FILAMENT\_POLYMERIZATION | MSigDB lists | GO\_POSITIVE\_REGULATION\_OF\_ACTIN\_FILAMENT\_POLYMERIZATION | 19 | 3 | 4161 | 153 | Arp2,Rac2,drk | | 3.065e-02 | -3.49 | GO\_REGULATION\_OF\_CHEMOTAXIS | MSigDB lists | GO\_REGULATION\_OF\_CHEMOTAXIS | 19 | 3 | 4161 | 153 | N,Smox,Rac2 | | 3.065e-02 | -3.49 | GSE37605\_TREG\_VS\_TCONV\_C57BL6\_FOXP3\_IRES\_GFP\_UP | MSigDB lists | GSE37605\_TREG\_VS\_TCONV\_C57BL6\_FOXP3\_IRES\_GFP\_UP | 19 | 3 | 4161 | 153 | so,Hsc70Cb,mbt | | 3.065e-02 | -3.49 | MORI\_MATURE\_B\_LYMPHOCYTE\_UP | MSigDB lists | MORI\_MATURE\_B\_LYMPHOCYTE\_UP | 19 | 3 | 4161 | 153 | phtf,hts,hiw | | 3.068e-02 | -3.48 | DH\_2 | prosite domains | PS50010 | 24 | 4 | 6243 | 316 | PsGEF,RhoGEF2,CG43658,pbl | | 3.071e-02 | -3.48 | Uba4 (Ubiquitin activating enzyme 4) | protein interactions | 34187 | 27 | 4 | 9204 | 412 | CG2865,Myo10A,Mocs2B,CG11486 | | 3.071e-02 | -3.48 | neur (neuralized) | protein interactions | 41085 | 27 | 4 | 9204 | 412 | E(spl)m4-BFM,sdt,Ser,N | | 3.071e-02 | -3.48 | btl (breathless) | protein interactions | 39564 | 27 | 4 | 9204 | 412 | CycE,jing,tkv,kni | | 3.096e-02 | -3.48 | RRM\_dom | interpro domains | IPR000504 | 135 | 10 | 11383 | 428 | fne,pAbp,Imp,sm,spen,Syp,Rbfox1,heph,elav,cpo | | 3.103e-02 | -3.47 | TAGCTTT\_MIR9 | MSigDB lists | TAGCTTT\_MIR9 | 65 | 6 | 4161 | 153 | CG1271,nudE,Lac,NiPp1,sdt,fne | | 3.103e-02 | -3.47 | GSE339\_EX\_VIVO\_VS\_IN\_CULTURE\_CD4CD8DN\_DC\_UP | MSigDB lists | GSE339\_EX\_VIVO\_VS\_IN\_CULTURE\_CD4CD8DN\_DC\_UP | 65 | 6 | 4161 | 153 | CG3961,Ank,ScsbetaG,Cyp1,Tudor-SN,ScpX | | 3.103e-02 | -3.47 | GSE3982\_DC\_VS\_BASOPHIL\_UP | MSigDB lists | GSE3982\_DC\_VS\_BASOPHIL\_UP | 65 | 6 | 4161 | 153 | ras,LManII,Hydr2,CG14291,Npc1a,emp | | 3.103e-02 | -3.47 | GSE1925\_CTRL\_VS\_24H\_IFNG\_STIM\_IFNG\_PRIMED\_MACROPHAGE\_DN | MSigDB lists | GSE1925\_CTRL\_VS\_24H\_IFNG\_STIM\_IFNG\_PRIMED\_MACROPHAGE\_DN | 65 | 6 | 4161 | 153 | par-1,cta,CG3961,ScsbetaG,CycE,baf | | 3.115e-02 | -3.47 | bsk (basket) | protein interactions | 44801 | 130 | 11 | 9204 | 412 | Rho1,CG32369,hiw,ec,aop,Fas2,Sox21a,egr,Sin3A,N,wnd | | 3.121e-02 | -3.47 | GO\_CHROMATIN\_BINDING | MSigDB lists | GO\_CHROMATIN\_BINDING | 101 | 8 | 4161 | 153 | Sin3A,so,Utx,ap,TfAP-2,CtBP,N,Smox | | 3.139e-02 | -3.46 | GSE26495\_NAIVE\_VS\_PD1LOW\_CD8\_TCELL\_DN | MSigDB lists | GSE26495\_NAIVE\_VS\_PD1LOW\_CD8\_TCELL\_DN | 33 | 4 | 4161 | 153 | Npc1a,Ptpmeg,Smox,CG5830 | | 3.139e-02 | -3.46 | GSE8515\_CTRL\_VS\_IL6\_4H\_STIM\_MAC\_DN | MSigDB lists | GSE8515\_CTRL\_VS\_IL6\_4H\_STIM\_MAC\_DN | 33 | 4 | 4161 | 153 | Npc1a,Lim1,CG2201,Ykt6 | | 3.139e-02 | -3.46 | GO\_REGULATION\_OF\_MUSCLE\_CELL\_DIFFERENTIATION | MSigDB lists | GO\_REGULATION\_OF\_MUSCLE\_CELL\_DIFFERENTIATION | 33 | 4 | 4161 | 153 | N,heph,hh,dpp | | 3.139e-02 | -3.46 | GSE6269\_HEALTHY\_VS\_STAPH\_PNEUMO\_INF\_PBMC\_DN | MSigDB lists | GSE6269\_HEALTHY\_VS\_STAPH\_PNEUMO\_INF\_PBMC\_DN | 33 | 4 | 4161 | 153 | drk,Pli,Cyp1,CG9220 | | 3.139e-02 | -3.46 | GSE22935\_24H\_VS\_48H\_MBOVIS\_BCG\_STIM\_MYD88\_KO\_MACROPHAGE\_DN | MSigDB lists | GSE22935\_24H\_VS\_48H\_MBOVIS\_BCG\_STIM\_MYD88\_KO\_MACROPHAGE\_DN | 33 | 4 | 4161 | 153 | S6kII,retm,CG2201,CG12547 | | 3.139e-02 | -3.46 | GSE29618\_PRE\_VS\_DAY7\_POST\_TIV\_FLU\_VACCINE\_MONOCYTE\_UP | MSigDB lists | GSE29618\_PRE\_VS\_DAY7\_POST\_TIV\_FLU\_VACCINE\_MONOCYTE\_UP | 33 | 4 | 4161 | 153 | dlp,TfAP-2,cta,kdn | | 3.139e-02 | -3.46 | GSE20715\_0H\_VS\_48H\_OZONE\_TLR4\_KO\_LUNG\_UP | MSigDB lists | GSE20715\_0H\_VS\_48H\_OZONE\_TLR4\_KO\_LUNG\_UP | 33 | 4 | 4161 | 153 | lili,mub,hts,Tsp26A | | 3.139e-02 | -3.46 | IZADPANAH\_STEM\_CELL\_ADIPOSE\_VS\_BONE\_UP | MSigDB lists | IZADPANAH\_STEM\_CELL\_ADIPOSE\_VS\_BONE\_UP | 33 | 4 | 4161 | 153 | Ser,DIP-epsilon,TfAP-2,CG3961 | | 3.139e-02 | -3.46 | GCNP\_SHH\_UP\_LATE.V1\_DN | MSigDB lists | GCNP\_SHH\_UP\_LATE.V1\_DN | 33 | 4 | 4161 | 153 | bor,mbt,LManII,Syx1A | | 3.139e-02 | -3.46 | ICSBP\_Q6 | MSigDB lists | ICSBP\_Q6 | 33 | 4 | 4161 | 153 | mgl,fne,Trpm,so | | 3.139e-02 | -3.46 | GO\_STEM\_CELL\_DIFFERENTIATION | MSigDB lists | GO\_STEM\_CELL\_DIFFERENTIATION | 33 | 4 | 4161 | 153 | dpp,Ser,hh,N | | 3.139e-02 | -3.46 | GO\_COLUMNAR\_CUBOIDAL\_EPITHELIAL\_CELL\_DIFFERENTIATION | MSigDB lists | GO\_COLUMNAR\_CUBOIDAL\_EPITHELIAL\_CELL\_DIFFERENTIATION | 33 | 4 | 4161 | 153 | N,Rac2,Ser,dpp | | 3.151e-02 | -3.46 | circadian behavior | biological process | GO:0048512 | 71 | 6 | 12345 | 414 | S6kII,tara,qsm,Bx,ITP,disco | | 3.151e-02 | -3.46 | adherens junction organization | biological process | GO:0034332 | 36 | 4 | 12345 | 414 | Rac2,crb,Rho1,sdt | | 3.180e-02 | -3.45 | GABRIELY\_MIR21\_TARGETS | MSigDB lists | GABRIELY\_MIR21\_TARGETS | 83 | 7 | 4161 | 153 | Arp2,smal,S6kII,hiw,sdt,Taf1,MFS10 | | 3.181e-02 | -3.45 | genitalia morphogenesis | biological process | GO:0035112 | 21 | 3 | 12345 | 414 | N,Fas2,otk2 | | 3.181e-02 | -3.45 | vtd (verthandi) | protein interactions | 3354896 | 40 | 5 | 9204 | 412 | egr,Psc,N,Tl,hh | | 3.182e-02 | -3.45 | regulation of BMP signaling pathway | biological process | GO:0030510 | 53 | 5 | 12345 | 414 | dally,lili,hiw,Lpin,Acsl | | 3.182e-02 | -3.45 | positive regulation of antimicrobial peptide production | biological process | GO:0002225 | 53 | 5 | 12345 | 414 | Pli,Doa,Tl,ush,CG4325 | | 3.182e-02 | -3.45 | positive regulation of antimicrobial humoral response | biological process | GO:0002760 | 53 | 5 | 12345 | 414 | Tl,CG4325,ush,Doa,Pli | | 3.182e-02 | -3.45 | positive regulation of production of molecular mediator of immune response | biological process | GO:0002702 | 53 | 5 | 12345 | 414 | Tl,ush,CG4325,Pli,Doa | | 3.182e-02 | -3.45 | positive regulation of immune effector process | biological process | GO:0002699 | 53 | 5 | 12345 | 414 | Tl,CG4325,ush,Doa,Pli | | 3.193e-02 | -3.44 | LRR\_8 | pfam domains | PF13855 | 81 | 7 | 10480 | 393 | Con,Lrt,Fili,trn,Toll-7,lbk,Tl | | 3.217e-02 | -3.44 | GO\_CELL\_SUBSTRATE\_JUNCTION | MSigDB lists | GO\_CELL\_SUBSTRATE\_JUNCTION | 160 | 11 | 4161 | 153 | cta,if,hts,Arp2,smal,mbt,Rac2,Hsc70-3,LRP1,emp,RpS4 | | 3.230e-02 | -3.43 | hb (hunchback) | protein interactions | 41032 | 16 | 3 | 9204 | 412 | Kr,cas,kni | | 3.230e-02 | -3.43 | CG3542 (uncharacterized protein) | protein interactions | 33513 | 16 | 3 | 9204 | 412 | CG31712,ytr,kdn | | 3.230e-02 | -3.43 | Smr (Smrter) | protein interactions | 32225 | 16 | 3 | 9204 | 412 | shn,EcR,Sin3A | | 3.230e-02 | -3.43 | ND-18 (NADH dehydrogenase (ubiquinone) 18 kDa subunit) | protein interactions | 32936 | 16 | 3 | 9204 | 412 | bowl,kdn,CG34172 | | 3.230e-02 | -3.43 | hbs (hibris) | protein interactions | 44129 | 16 | 3 | 9204 | 412 | CG3386,N,Hsc70-3 | | 3.230e-02 | -3.43 | PEK (pancreatic eIF-2alpha kinase) | protein interactions | 40653 | 16 | 3 | 9204 | 412 | PEK,N,CycE | | 3.230e-02 | -3.43 | mip120 (Myb-interacting protein 120) | protein interactions | 36499 | 16 | 3 | 9204 | 412 | Myb,Fas2,l(3)mbt | | 3.230e-02 | -3.43 | spz (spatzle) | protein interactions | 43256 | 16 | 3 | 9204 | 412 | Toll-7,Tl,Egfr | | 3.230e-02 | -3.43 | Art1 (Arginine methyltransferase 1) | protein interactions | 41295 | 16 | 3 | 9204 | 412 | dpp,Dad,pAbp | | 3.230e-02 | -3.43 | Wnt5 (Wnt oncogene analog 5) | protein interactions | 32838 | 16 | 3 | 9204 | 412 | CadN,uzip,pbl | | 3.235e-02 | -3.43 | - | gene3d domains | 3.40.50.1580 | 7 | 2 | 4929 | 208 | CG17224,CG6330 | | 3.238e-02 | -3.43 | receptor regulator activity | molecular function | GO:0030545 | 114 | 8 | 12176 | 396 | hh,aos,ITP,tsl,dpp,Ser,Pvf3,spi | | 3.251e-02 | -3.43 | DISTECHE\_ESCAPED\_FROM\_X\_INACTIVATION | MSigDB lists | DISTECHE\_ESCAPED\_FROM\_X\_INACTIVATION | 8 | 2 | 4161 | 153 | Utx,RpS4 | | 3.251e-02 | -3.43 | GCM\_MSN | MSigDB lists | GCM\_MSN | 8 | 2 | 4161 | 153 | mle,sm | | 3.251e-02 | -3.43 | chr2q35 | MSigDB lists | chr2q35 | 8 | 2 | 4161 | 153 | CG5830,CG43313 | | 3.251e-02 | -3.43 | GO\_TRANSFORMING\_GROWTH\_FACTOR\_BETA\_RECEPTOR\_BINDING | MSigDB lists | GO\_TRANSFORMING\_GROWTH\_FACTOR\_BETA\_RECEPTOR\_BINDING | 8 | 2 | 4161 | 153 | dpp,Smox | | 3.251e-02 | -3.43 | chr16p11 | MSigDB lists | chr16p11 | 8 | 2 | 4161 | 153 | mof,PhKgamma | | 3.251e-02 | -3.43 | PID\_EPHA2\_FWD\_PATHWAY | MSigDB lists | PID\_EPHA2\_FWD\_PATHWAY | 8 | 2 | 4161 | 153 | drk,Rac2 | | 3.251e-02 | -3.43 | HOWLIN\_CITED1\_TARGETS\_1\_DN | MSigDB lists | HOWLIN\_CITED1\_TARGETS\_1\_DN | 8 | 2 | 4161 | 153 | sdt,mmy | | 3.251e-02 | -3.43 | GO\_REGULATION\_OF\_SMOOTH\_MUSCLE\_CELL\_MIGRATION | MSigDB lists | GO\_REGULATION\_OF\_SMOOTH\_MUSCLE\_CELL\_MIGRATION | 8 | 2 | 4161 | 153 | CaMKII,LRP1 | | 3.251e-02 | -3.43 | GO\_CRANIAL\_NERVE\_DEVELOPMENT | MSigDB lists | GO\_CRANIAL\_NERVE\_DEVELOPMENT | 8 | 2 | 4161 | 153 | TfAP-2,so | | 3.251e-02 | -3.43 | GO\_RESPONSE\_TO\_PROGESTERONE | MSigDB lists | GO\_RESPONSE\_TO\_PROGESTERONE | 8 | 2 | 4161 | 153 | CycE,Hydr2 | | 3.251e-02 | -3.43 | VERRECCHIA\_EARLY\_RESPONSE\_TO\_TGFB1 | MSigDB lists | VERRECCHIA\_EARLY\_RESPONSE\_TO\_TGFB1 | 8 | 2 | 4161 | 153 | LRP1,Rho1 | | 3.251e-02 | -3.43 | GENTILE\_UV\_LOW\_DOSE\_UP | MSigDB lists | GENTILE\_UV\_LOW\_DOSE\_UP | 8 | 2 | 4161 | 153 | NiPp1,CycE | | 3.251e-02 | -3.43 | GO\_NEGATIVE\_REGULATION\_OF\_GLIOGENESIS | MSigDB lists | GO\_NEGATIVE\_REGULATION\_OF\_GLIOGENESIS | 8 | 2 | 4161 | 153 | N,CG12099 | | 3.251e-02 | -3.43 | GO\_POSITIVE\_REGULATION\_OF\_AXON\_EXTENSION | MSigDB lists | GO\_POSITIVE\_REGULATION\_OF\_AXON\_EXTENSION | 8 | 2 | 4161 | 153 | NetB,nudE | | 3.251e-02 | -3.43 | GO\_REGULATION\_OF\_CHONDROCYTE\_DIFFERENTIATION | MSigDB lists | GO\_REGULATION\_OF\_CHONDROCYTE\_DIFFERENTIATION | 8 | 2 | 4161 | 153 | tkv,Smox | | 3.251e-02 | -3.43 | GO\_REGULATION\_OF\_OLIGODENDROCYTE\_DIFFERENTIATION | MSigDB lists | GO\_REGULATION\_OF\_OLIGODENDROCYTE\_DIFFERENTIATION | 8 | 2 | 4161 | 153 | N,hh | | 3.251e-02 | -3.43 | KEGG\_GLYCOSAMINOGLYCAN\_BIOSYNTHESIS\_CHONDROITIN\_SULFATE | MSigDB lists | KEGG\_GLYCOSAMINOGLYCAN\_BIOSYNTHESIS\_CHONDROITIN\_SULFATE | 8 | 2 | 4161 | 153 | CG43313,CG9220 | | 3.251e-02 | -3.43 | GO\_CALMODULIN\_DEPENDENT\_PROTEIN\_KINASE\_ACTIVITY | MSigDB lists | GO\_CALMODULIN\_DEPENDENT\_PROTEIN\_KINASE\_ACTIVITY | 8 | 2 | 4161 | 153 | CaMKII,PhKgamma | | 3.251e-02 | -3.43 | GO\_POSITIVE\_REGULATION\_OF\_CELL\_MATRIX\_ADHESION | MSigDB lists | GO\_POSITIVE\_REGULATION\_OF\_CELL\_MATRIX\_ADHESION | 8 | 2 | 4161 | 153 | Rac2,Smox | | 3.251e-02 | -3.43 | GO\_RHO\_PROTEIN\_SIGNAL\_TRANSDUCTION | MSigDB lists | GO\_RHO\_PROTEIN\_SIGNAL\_TRANSDUCTION | 8 | 2 | 4161 | 153 | NetB,cta | | 3.251e-02 | -3.43 | REACTOME\_ACTIVATION\_OF\_RAC | MSigDB lists | REACTOME\_ACTIVATION\_OF\_RAC | 8 | 2 | 4161 | 153 | Rac2,mbt | | 3.251e-02 | -3.43 | GO\_EMBRYONIC\_AXIS\_SPECIFICATION | MSigDB lists | GO\_EMBRYONIC\_AXIS\_SPECIFICATION | 8 | 2 | 4161 | 153 | Utx,Lim1 | | 3.251e-02 | -3.43 | HOEGERKORP\_CD44\_TARGETS\_DIRECT\_UP | MSigDB lists | HOEGERKORP\_CD44\_TARGETS\_DIRECT\_UP | 8 | 2 | 4161 | 153 | CaMKII,Syx1A | | 3.251e-02 | -3.43 | GO\_CHONDROITIN\_SULFATE\_PROTEOGLYCAN\_BIOSYNTHETIC\_PROCESS | MSigDB lists | GO\_CHONDROITIN\_SULFATE\_PROTEOGLYCAN\_BIOSYNTHETIC\_PROCESS | 8 | 2 | 4161 | 153 | CG43313,CG9220 | | 3.251e-02 | -3.43 | GCM\_LTK | MSigDB lists | GCM\_LTK | 8 | 2 | 4161 | 153 | hts,EcR | | 3.251e-02 | -3.43 | GO\_SMAD\_PROTEIN\_SIGNAL\_TRANSDUCTION | MSigDB lists | GO\_SMAD\_PROTEIN\_SIGNAL\_TRANSDUCTION | 8 | 2 | 4161 | 153 | dpp,Smox | | 3.251e-02 | -3.43 | REACTOME\_SYNTHESIS\_OF\_VERY\_LONG\_CHAIN\_FATTY\_ACYL\_COAS | MSigDB lists | REACTOME\_SYNTHESIS\_OF\_VERY\_LONG\_CHAIN\_FATTY\_ACYL\_COAS | 8 | 2 | 4161 | 153 | CG3961,Acsl | | 3.251e-02 | -3.43 | GNF2\_MYD88 | MSigDB lists | GNF2\_MYD88 | 8 | 2 | 4161 | 153 | Arp2,LManII | | 3.251e-02 | -3.43 | REACTOME\_SIGNALING\_BY\_FGFR\_MUTANTS | MSigDB lists | REACTOME\_SIGNALING\_BY\_FGFR\_MUTANTS | 8 | 2 | 4161 | 153 | Mhcl,drk | | 3.251e-02 | -3.43 | GO\_MRNA\_3\_UTR\_BINDING | MSigDB lists | GO\_MRNA\_3\_UTR\_BINDING | 8 | 2 | 4161 | 153 | fne,Imp | | 3.251e-02 | -3.43 | CHANG\_IMMORTALIZED\_BY\_HPV31\_DN | MSigDB lists | CHANG\_IMMORTALIZED\_BY\_HPV31\_DN | 8 | 2 | 4161 | 153 | LRP1,CG10663 | | 3.251e-02 | -3.43 | GO\_ACETYLGALACTOSAMINYLTRANSFERASE\_ACTIVITY | MSigDB lists | GO\_ACETYLGALACTOSAMINYLTRANSFERASE\_ACTIVITY | 8 | 2 | 4161 | 153 | CG9220,CG43313 | | 3.251e-02 | -3.43 | BIOCARTA\_RACCYCD\_PATHWAY | MSigDB lists | BIOCARTA\_RACCYCD\_PATHWAY | 8 | 2 | 4161 | 153 | Rac2,CycE | | 3.252e-02 | -3.43 | virion binding | molecular function | GO:0046790 | 1 | 1 | 12176 | 396 | Toll-7 | | 3.252e-02 | -3.43 | epidermal growth factor-activated receptor activity | molecular function | GO:0005006 | 1 | 1 | 12176 | 396 | Egfr | | 3.252e-02 | -3.43 | palmitoyl-CoA ligase activity | molecular function | GO:0090433 | 1 | 1 | 12176 | 396 | Acsl | | 3.252e-02 | -3.43 | palmitoleyl hydrolase activity | molecular function | GO:1990699 | 1 | 1 | 12176 | 396 | Notum | | 3.252e-02 | -3.43 | DNA demethylase activity | molecular function | GO:0035514 | 1 | 1 | 12176 | 396 | Tet | | 3.252e-02 | -3.43 | ecdysone 20-monooxygenase activity | molecular function | GO:0004501 | 1 | 1 | 12176 | 396 | shd | | 3.252e-02 | -3.43 | filamin binding | molecular function | GO:0031005 | 1 | 1 | 12176 | 396 | Ten-m | | 3.252e-02 | -3.43 | protein tyrosine phosphatase activity, metal-dependent | molecular function | GO:0030946 | 1 | 1 | 12176 | 396 | eya | | 3.252e-02 | -3.43 | histone kinase activity (H3-T3 specific) | molecular function | GO:0072354 | 1 | 1 | 12176 | 396 | Haspin | | 3.252e-02 | -3.43 | pyrimidine-specific mismatch base pair DNA N-glycosylase activity | molecular function | GO:0008263 | 1 | 1 | 12176 | 396 | Thd1 | | 3.252e-02 | -3.43 | intracellular phosphatidylinositol-3,5-bisphosphate-sensitive cation channel activity | molecular function | GO:0097682 | 1 | 1 | 12176 | 396 | Trpm | | 3.252e-02 | -3.43 | molybdopterin synthase activity | molecular function | GO:0030366 | 1 | 1 | 12176 | 396 | Mocs2B | | 3.252e-02 | -3.43 | histone acetyltransferase activity (H3-K14 specific) | molecular function | GO:0036408 | 1 | 1 | 12176 | 396 | Taf1 | | 3.252e-02 | -3.43 | latrotoxin receptor activity | molecular function | GO:0016524 | 1 | 1 | 12176 | 396 | Cirl | | 3.252e-02 | -3.43 | mismatch base pair DNA N-glycosylase activity | molecular function | GO:0000700 | 1 | 1 | 12176 | 396 | Thd1 | | 3.252e-02 | -3.43 | oxidative DNA demethylase activity | molecular function | GO:0035516 | 1 | 1 | 12176 | 396 | Tet | | 3.252e-02 | -3.43 | RNA polymerase I general transcription initiation factor activity | molecular function | GO:0001181 | 1 | 1 | 12176 | 396 | Tif-IA | | 3.252e-02 | -3.43 | dopamine receptor binding | molecular function | GO:0050780 | 1 | 1 | 12176 | 396 | cta | | 3.252e-02 | -3.43 | lysine N-acetyltransferase activity, acting on acetyl phosphate as donor | molecular function | GO:0004468 | 1 | 1 | 12176 | 396 | mof | | 3.252e-02 | -3.43 | hydrolase activity, acting on acid sulfur-nitrogen bonds | molecular function | GO:0016826 | 1 | 1 | 12176 | 396 | CG14291 | | 3.252e-02 | -3.43 | RNA polymerase II transcription coactivator binding | molecular function | GO:0001225 | 1 | 1 | 12176 | 396 | EcR | | 3.252e-02 | -3.43 | RNA polymerase I core binding | molecular function | GO:0001042 | 1 | 1 | 12176 | 396 | Tif-IA | | 3.252e-02 | -3.43 | IMP dehydrogenase activity | molecular function | GO:0003938 | 1 | 1 | 12176 | 396 | ras | | 3.252e-02 | -3.43 | D5 dopamine receptor binding | molecular function | GO:0031752 | 1 | 1 | 12176 | 396 | cta | | 3.252e-02 | -3.43 | transforming growth factor beta receptor, inhibitory cytoplasmic mediator activity | molecular function | GO:0030617 | 1 | 1 | 12176 | 396 | Dad | | 3.252e-02 | -3.43 | histone kinase activity (H3-S10 specific) | molecular function | GO:0035175 | 1 | 1 | 12176 | 396 | JIL-1 | | 3.252e-02 | -3.43 | GKAP/Homer scaffold activity | molecular function | GO:0030160 | 1 | 1 | 12176 | 396 | Prosap | | 3.252e-02 | -3.43 | adenosylmethionine decarboxylase activity | molecular function | GO:0004014 | 1 | 1 | 12176 | 396 | SamDC | | 3.252e-02 | -3.43 | N-sulfoglucosamine sulfohydrolase activity | molecular function | GO:0016250 | 1 | 1 | 12176 | 396 | CG14291 | | 3.252e-02 | -3.43 | histone demethylase activity (H3-K27 specific) | molecular function | GO:0071558 | 1 | 1 | 12176 | 396 | Utx | | 3.252e-02 | -3.43 | elongation factor-2 kinase activity | molecular function | GO:0004686 | 1 | 1 | 12176 | 396 | PEK | | 3.252e-02 | -3.43 | UDP-N-acetylglucosamine diphosphorylase activity | molecular function | GO:0003977 | 1 | 1 | 12176 | 396 | mmy | | 3.252e-02 | -3.43 | glycogen binding | molecular function | GO:2001069 | 1 | 1 | 12176 | 396 | Gbs-70E | | 3.252e-02 | -3.43 | transforming growth factor beta binding | molecular function | GO:0050431 | 1 | 1 | 12176 | 396 | tkv | | 3.265e-02 | -3.42 | phosphatidate phosphatase activity | molecular function | GO:0008195 | 9 | 2 | 12176 | 396 | Lpin,wun2 | | 3.265e-02 | -3.42 | nuclear receptor transcription coactivator activity | molecular function | GO:0030374 | 9 | 2 | 12176 | 396 | EcR,tna | | 3.265e-02 | -3.42 | long-chain fatty acid-CoA ligase activity | molecular function | GO:0004467 | 9 | 2 | 12176 | 396 | Acsl,CG3961 | | 3.265e-02 | -3.42 | H3 histone acetyltransferase activity | molecular function | GO:0010484 | 9 | 2 | 12176 | 396 | Taf1,enok | | 3.266e-02 | -3.42 | GSE17721\_POLYIC\_VS\_GARDIQUIMOD\_12H\_BMDC\_UP | MSigDB lists | GSE17721\_POLYIC\_VS\_GARDIQUIMOD\_12H\_BMDC\_UP | 49 | 5 | 4161 | 153 | Arl4,CaMKII,rush,gus,Tsp26A | | 3.266e-02 | -3.42 | FOXJ2\_01 | MSigDB lists | FOXJ2\_01 | 49 | 5 | 4161 | 153 | NetB,ras,Fas2,phtf,Tsp26A | | 3.266e-02 | -3.42 | GSE29615\_CTRL\_VS\_DAY3\_LAIV\_IFLU\_VACCINE\_PBMC\_UP | MSigDB lists | GSE29615\_CTRL\_VS\_DAY3\_LAIV\_IFLU\_VACCINE\_PBMC\_UP | 49 | 5 | 4161 | 153 | cta,mle,Tlk,nudE,CG17265 | | 3.266e-02 | -3.42 | WHITFIELD\_CELL\_CYCLE\_G2 | MSigDB lists | WHITFIELD\_CELL\_CYCLE\_G2 | 49 | 5 | 4161 | 153 | Arl4,NTPase,betaTub56D,ScsbetaG,TfAP-2 | | 3.269e-02 | -3.42 | Bromodomain | interpro domains | IPR001487 | 19 | 3 | 11383 | 428 | tou,Taf1,Acf | | 3.299e-02 | -3.41 | circadian rhythm | biological process | GO:0007623 | 91 | 7 | 12345 | 414 | tara,Bx,qsm,S6kII,so,ITP,disco | | 3.313e-02 | -3.41 | GSE16522\_MEMORY\_VS\_NAIVE\_ANTI\_CD3CD28\_STIM\_CD8\_TCELL\_UP | MSigDB lists | GSE16522\_MEMORY\_VS\_NAIVE\_ANTI\_CD3CD28\_STIM\_CD8\_TCELL\_UP | 66 | 6 | 4161 | 153 | emp,Npc1a,mmy,hts,CtBP,drk | | 3.313e-02 | -3.41 | GSE360\_L\_DONOVANI\_VS\_B\_MALAYI\_LOW\_DOSE\_MAC\_UP | MSigDB lists | GSE360\_L\_DONOVANI\_VS\_B\_MALAYI\_LOW\_DOSE\_MAC\_UP | 66 | 6 | 4161 | 153 | Npc1a,Arp2,baf,Rho1,CG11030,Arl4 | | 3.313e-02 | -3.41 | MYB\_Q5\_01 | MSigDB lists | MYB\_Q5\_01 | 66 | 6 | 4161 | 153 | nudE,smal,Mhcl,Cnx99A,Hydr2,Rho1 | | 3.313e-02 | -3.41 | CTAWWWATA\_RSRFC4\_Q2 | MSigDB lists | CTAWWWATA\_RSRFC4\_Q2 | 66 | 6 | 4161 | 153 | TfIIS,Tlk,dlp,Nhe2,SamDC,Smox | | 3.313e-02 | -3.41 | GSE22886\_DC\_VS\_MONOCYTE\_UP | MSigDB lists | GSE22886\_DC\_VS\_MONOCYTE\_UP | 66 | 6 | 4161 | 153 | sdt,CycE,MFS10,phtf,Ser,Hsc70Cb | | 3.344e-02 | -3.40 | regulation of actin filament organization | biological process | GO:0110053 | 72 | 6 | 12345 | 414 | Arp2,PsGEF,CG43658,hts,if,spoon | | 3.346e-02 | -3.40 | MH2 | prosite domains | PS51076 | 6 | 2 | 6243 | 316 | Smox,Dad | | 3.346e-02 | -3.40 | MYTH4 | prosite domains | PS51016 | 6 | 2 | 6243 | 316 | Myo10A,RhoGAP93B | | 3.354e-02 | -3.40 | oxidative DNA demethylation | biological process | GO:0035511 | 1 | 1 | 12345 | 414 | Tet | | 3.354e-02 | -3.40 | regulation of muscle cell chemotaxis toward tendon cell | biological process | GO:2001281 | 1 | 1 | 12345 | 414 | Lrt | | 3.354e-02 | -3.40 | cell morphogenesis involved in Malpighian tubule morphogenesis | biological process | GO:0061336 | 1 | 1 | 12345 | 414 | crb | | 3.354e-02 | -3.40 | negative regulation of extracellular matrix assembly | biological process | GO:1901202 | 1 | 1 | 12345 | 414 | Mmp2 | | 3.354e-02 | -3.40 | negative regulation of histone H3-K27 trimethylation | biological process | GO:1902465 | 1 | 1 | 12345 | 414 | Utx | | 3.354e-02 | -3.40 | intein-mediated protein splicing | biological process | GO:0016539 | 1 | 1 | 12345 | 414 | hh | | 3.354e-02 | -3.40 | negative regulation of signal transduction in absence of ligand | biological process | GO:1901099 | 1 | 1 | 12345 | 414 | eya | | 3.354e-02 | -3.40 | heart rudiment formation | biological process | GO:0003315 | 1 | 1 | 12345 | 414 | Mmp2 | | 3.354e-02 | -3.40 | regulation of male germ cell proliferation | biological process | GO:2000254 | 1 | 1 | 12345 | 414 | Doa | | 3.354e-02 | -3.40 | regulation of male pigmentation | biological process | GO:0048088 | 1 | 1 | 12345 | 414 | bab1 | | 3.354e-02 | -3.40 | early endosome to recycling endosome transport | biological process | GO:0061502 | 1 | 1 | 12345 | 414 | Acsl | | 3.354e-02 | -3.40 | regulation of imaginal disc-derived wing vein specification | biological process | GO:0110107 | 1 | 1 | 12345 | 414 | sty | | 3.354e-02 | -3.40 | proboscis morphogenesis | biological process | GO:0048734 | 1 | 1 | 12345 | 414 | CG13366 | | 3.354e-02 | -3.40 | GPI anchor release | biological process | GO:0006507 | 1 | 1 | 12345 | 414 | Notum | | 3.354e-02 | -3.40 | determination of wing disc primordium | biological process | GO:0035294 | 1 | 1 | 12345 | 414 | Doc1 | | 3.354e-02 | -3.40 | regulation of ubiquitin-dependent endocytosis | biological process | GO:2000395 | 1 | 1 | 12345 | 414 | sdt | | 3.354e-02 | -3.40 | regulation of transcription involved in G2/M transition of mitotic cell cycle | biological process | GO:0000117 | 1 | 1 | 12345 | 414 | Myb | | 3.354e-02 | -3.40 | regulation of ruffle assembly | biological process | GO:1900027 | 1 | 1 | 12345 | 414 | PsGEF | | 3.354e-02 | -3.40 | defense response to oomycetes | biological process | GO:0002229 | 1 | 1 | 12345 | 414 | Tl | | 3.354e-02 | -3.40 | heart rudiment morphogenesis | biological process | GO:0003314 | 1 | 1 | 12345 | 414 | Mmp2 | | 3.354e-02 | -3.40 | cell differentiation in spinal cord | biological process | GO:0021515 | 1 | 1 | 12345 | 414 | exex | | 3.354e-02 | -3.40 | cell migration involved in heart development | biological process | GO:0060973 | 1 | 1 | 12345 | 414 | Mmp2 | | 3.354e-02 | -3.40 | positive regulation of ruffle assembly | biological process | GO:1900029 | 1 | 1 | 12345 | 414 | PsGEF | | 3.354e-02 | -3.40 | 5-methylcytosine metabolic process | biological process | GO:0019857 | 1 | 1 | 12345 | 414 | Tet | | 3.354e-02 | -3.40 | regulation of cell-cell adhesion mediated by integrin | biological process | GO:0033632 | 1 | 1 | 12345 | 414 | wech | | 3.354e-02 | -3.40 | negative regulation of imaginal disc-derived wing vein specification | biological process | GO:0110109 | 1 | 1 | 12345 | 414 | sty | | 3.354e-02 | -3.40 | heart rudiment development | biological process | GO:0003313 | 1 | 1 | 12345 | 414 | Mmp2 | | 3.354e-02 | -3.40 | heparan sulfate proteoglycan catabolic process | biological process | GO:0030200 | 1 | 1 | 12345 | 414 | CG14291 | | 3.354e-02 | -3.40 | anterior midgut invagination | biological process | GO:0007375 | 1 | 1 | 12345 | 414 | RhoGEF2 | | 3.354e-02 | -3.40 | spinal cord motor neuron differentiation | biological process | GO:0021522 | 1 | 1 | 12345 | 414 | exex | | 3.354e-02 | -3.40 | negative regulation of cell-cell adhesion mediated by cadherin | biological process | GO:2000048 | 1 | 1 | 12345 | 414 | N | | 3.354e-02 | -3.40 | regulation of preblastoderm mitotic cell cycle | biological process | GO:0007347 | 1 | 1 | 12345 | 414 | l(3)mbt | | 3.354e-02 | -3.40 | regulation of Ral protein signal transduction | biological process | GO:0032485 | 1 | 1 | 12345 | 414 | CG5522 | | 3.354e-02 | -3.40 | pH elevation | biological process | GO:0045852 | 1 | 1 | 12345 | 414 | Nhe2 | | 3.354e-02 | -3.40 | negative regulation of morphogenesis of an epithelium | biological process | GO:1905331 | 1 | 1 | 12345 | 414 | sty | | 3.354e-02 | -3.40 | negative regulation of pigment cell differentiation | biological process | GO:0050941 | 1 | 1 | 12345 | 414 | S6kII | | 3.354e-02 | -3.40 | gastric emptying | biological process | GO:0035483 | 1 | 1 | 12345 | 414 | Bx | | 3.354e-02 | -3.40 | protein splicing | biological process | GO:0030908 | 1 | 1 | 12345 | 414 | hh | | 3.354e-02 | -3.40 | pigment cell development | biological process | GO:0070285 | 1 | 1 | 12345 | 414 | eya | | 3.354e-02 | -3.40 | spinal cord motor neuron cell fate specification | biological process | GO:0021520 | 1 | 1 | 12345 | 414 | exex | | 3.354e-02 | -3.40 | negative regulation of neuron remodeling | biological process | GO:1904800 | 1 | 1 | 12345 | 414 | Fas2 | | 3.354e-02 | -3.40 | R1/R6 development | biological process | GO:0048053 | 1 | 1 | 12345 | 414 | ttk | | 3.354e-02 | -3.40 | immune response-activating cell surface receptor signaling pathway | biological process | GO:0002429 | 1 | 1 | 12345 | 414 | Toll-7 | | 3.354e-02 | -3.40 | Roundabout signaling pathway involved in muscle cell chemotaxis toward tendon cell | biological process | GO:2001283 | 1 | 1 | 12345 | 414 | Lrt | | 3.354e-02 | -3.40 | negative regulation of extrinsic apoptotic signaling pathway in absence of ligand | biological process | GO:2001240 | 1 | 1 | 12345 | 414 | eya | | 3.354e-02 | -3.40 | mitophagy by induced vacuole formation | biological process | GO:0035695 | 1 | 1 | 12345 | 414 | CG11498 | | 3.354e-02 | -3.40 | negative regulation of neuron maturation | biological process | GO:0014043 | 1 | 1 | 12345 | 414 | Fas2 | | 3.354e-02 | -3.40 | positive regulation of transcription involved in G2/M transition of mitotic cell cycle | biological process | GO:0090282 | 1 | 1 | 12345 | 414 | Myb | | 3.354e-02 | -3.40 | gastric motility | biological process | GO:0035482 | 1 | 1 | 12345 | 414 | Bx | | 3.354e-02 | -3.40 | compound eye corneal lens morphogenesis | biological process | GO:0048750 | 1 | 1 | 12345 | 414 | ttk | | 3.354e-02 | -3.40 | histone dephosphorylation | biological process | GO:0016576 | 1 | 1 | 12345 | 414 | eya | | 3.354e-02 | -3.40 | regulation of pigment cell differentiation | biological process | GO:0050932 | 1 | 1 | 12345 | 414 | S6kII | | 3.354e-02 | -3.40 | regulation of extrinsic apoptotic signaling pathway in absence of ligand | biological process | GO:2001239 | 1 | 1 | 12345 | 414 | eya | | 3.354e-02 | -3.40 | postsynaptic density assembly | biological process | GO:0097107 | 1 | 1 | 12345 | 414 | Prosap | | 3.354e-02 | -3.40 | DNA protection | biological process | GO:0042262 | 1 | 1 | 12345 | 414 | stwl | | 3.354e-02 | -3.40 | negative regulation of germ cell proliferation | biological process | GO:1905937 | 1 | 1 | 12345 | 414 | Doa | | 3.354e-02 | -3.40 | transforming growth factor beta receptor complex assembly | biological process | GO:0007181 | 1 | 1 | 12345 | 414 | tkv | | 3.354e-02 | -3.40 | negative regulation of cytokine secretion | biological process | GO:0050710 | 1 | 1 | 12345 | 414 | CaMKII | | 3.354e-02 | -3.40 | establishment of body hair planar orientation | biological process | GO:0048105 | 1 | 1 | 12345 | 414 | ft | | 3.354e-02 | -3.40 | DNA demethylation | biological process | GO:0080111 | 1 | 1 | 12345 | 414 | Tet | | 3.354e-02 | -3.40 | regulation of DNA demethylation | biological process | GO:1901535 | 1 | 1 | 12345 | 414 | Tet | | 3.354e-02 | -3.40 | histone H3-K27 demethylation | biological process | GO:0071557 | 1 | 1 | 12345 | 414 | Utx | | 3.354e-02 | -3.40 | maternal determination of dorsal/ventral axis, ovarian follicular epithelium, soma encoded | biological process | GO:0008071 | 1 | 1 | 12345 | 414 | Egfr | | 3.354e-02 | -3.40 | histone H3-T3 phosphorylation | biological process | GO:0072355 | 1 | 1 | 12345 | 414 | Haspin | | 3.354e-02 | -3.40 | excitatory synapse assembly | biological process | GO:1904861 | 1 | 1 | 12345 | 414 | Prosap | | 3.354e-02 | -3.40 | long-chain fatty-acyl-CoA biosynthetic process | biological process | GO:0035338 | 1 | 1 | 12345 | 414 | Acsl | | 3.354e-02 | -3.40 | regulation of establishment of bipolar cell polarity | biological process | GO:0061172 | 1 | 1 | 12345 | 414 | wake | | 3.354e-02 | -3.40 | cell migration to the midline involved in heart development | biological process | GO:0003318 | 1 | 1 | 12345 | 414 | Mmp2 | | 3.354e-02 | -3.40 | positive regulation of histone H3-K27 methylation | biological process | GO:0061087 | 1 | 1 | 12345 | 414 | Pcl | | 3.354e-02 | -3.40 | protein depalmitoleylation | biological process | GO:1990697 | 1 | 1 | 12345 | 414 | Notum | | 3.354e-02 | -3.40 | cell surface pattern recognition receptor signaling pathway | biological process | GO:0002752 | 1 | 1 | 12345 | 414 | Toll-7 | | 3.354e-02 | -3.40 | reticulophagy | biological process | GO:0061709 | 1 | 1 | 12345 | 414 | Atg17 | | 3.354e-02 | -3.40 | positive regulation of protein localization to cell surface | biological process | GO:2000010 | 1 | 1 | 12345 | 414 | hh | | 3.354e-02 | -3.40 | mitochondrial protein catabolic process | biological process | GO:0035694 | 1 | 1 | 12345 | 414 | CG11498 | | 3.354e-02 | -3.40 | regulation of histone H4 acetylation involved in response to DNA damage stimulus | biological process | GO:2000873 | 1 | 1 | 12345 | 414 | mof | | 3.354e-02 | -3.40 | ectodermal cell fate specification | biological process | GO:0001715 | 1 | 1 | 12345 | 414 | dpp | | 3.354e-02 | -3.40 | regulation of histone H3-K27 trimethylation | biological process | GO:1902464 | 1 | 1 | 12345 | 414 | Utx | | 3.354e-02 | -3.40 | positive regulation of DNA demethylation | biological process | GO:1901537 | 1 | 1 | 12345 | 414 | Tet | | 3.354e-02 | -3.40 | negative regulation of male germ cell proliferation | biological process | GO:2000255 | 1 | 1 | 12345 | 414 | Doa | | 3.354e-02 | -3.40 | postsynaptic specialization assembly | biological process | GO:0098698 | 1 | 1 | 12345 | 414 | Prosap | | 3.354e-02 | -3.40 | negative regulation of semaphorin-plexin signaling pathway | biological process | GO:2001261 | 1 | 1 | 12345 | 414 | dally | | 3.354e-02 | -3.40 | negative regulation of male pigmentation | biological process | GO:0048092 | 1 | 1 | 12345 | 414 | bab1 | | 3.354e-02 | -3.40 | cardioblast migration to the midline involved in heart rudiment formation | biological process | GO:0003319 | 1 | 1 | 12345 | 414 | Mmp2 | | 3.354e-02 | -3.40 | negative regulation of non-canonical Wnt signaling pathway | biological process | GO:2000051 | 1 | 1 | 12345 | 414 | aop | | 3.354e-02 | -3.40 | negative regulation of epithelial cell differentiation | biological process | GO:0030857 | 1 | 1 | 12345 | 414 | Ptpmeg | | 3.354e-02 | -3.40 | response to oomycetes | biological process | GO:0002239 | 1 | 1 | 12345 | 414 | Tl | | 3.354e-02 | -3.40 | pharynx development | biological process | GO:0060465 | 1 | 1 | 12345 | 414 | cnc | | 3.354e-02 | -3.40 | negative regulation of border follicle cell migration | biological process | GO:1903687 | 1 | 1 | 12345 | 414 | Ptpmeg | | 3.354e-02 | -3.40 | negative regulation of muscle cell chemotaxis toward tendon cell | biological process | GO:2001282 | 1 | 1 | 12345 | 414 | Lrt | | 3.354e-02 | -3.40 | dorsal vessel aortic cell fate commitment | biological process | GO:0035052 | 1 | 1 | 12345 | 414 | Ubx | | 3.354e-02 | -3.40 | cardioblast cell fate specification | biological process | GO:0042685 | 1 | 1 | 12345 | 414 | robo2 | | 3.354e-02 | -3.40 | muscle cell chemotaxis toward tendon cell | biological process | GO:0036061 | 1 | 1 | 12345 | 414 | Lrt | | 3.354e-02 | -3.40 | ventral spinal cord development | biological process | GO:0021517 | 1 | 1 | 12345 | 414 | exex | | 3.354e-02 | -3.40 | zonula adherens maintenance | biological process | GO:0045218 | 1 | 1 | 12345 | 414 | crb | | 3.354e-02 | -3.40 | intracellular pH elevation | biological process | GO:0051454 | 1 | 1 | 12345 | 414 | Nhe2 | | 3.354e-02 | -3.40 | 5-methylcytosine catabolic process | biological process | GO:0006211 | 1 | 1 | 12345 | 414 | Tet | | 3.354e-02 | -3.40 | negative regulation of rhodopsin gene expression | biological process | GO:0045871 | 1 | 1 | 12345 | 414 | oc | | 3.354e-02 | -3.40 | ocellus morphogenesis | biological process | GO:0048816 | 1 | 1 | 12345 | 414 | oc | | 3.354e-02 | -3.40 | proteoglycan catabolic process | biological process | GO:0030167 | 1 | 1 | 12345 | 414 | CG14291 | | 3.354e-02 | -3.40 | testicular fusome organization | biological process | GO:0030724 | 1 | 1 | 12345 | 414 | hts | | 3.354e-02 | -3.40 | positive regulation of excitatory postsynaptic potential | biological process | GO:2000463 | 1 | 1 | 12345 | 414 | Prosap | | 3.354e-02 | -3.40 | positive regulation of ubiquitin-dependent endocytosis | biological process | GO:2000397 | 1 | 1 | 12345 | 414 | sdt | | 3.354e-02 | -3.40 | innate immune response activating cell surface receptor signaling pathway | biological process | GO:0002220 | 1 | 1 | 12345 | 414 | Toll-7 | | 3.354e-02 | -3.40 | spinal cord development | biological process | GO:0021510 | 1 | 1 | 12345 | 414 | exex | | 3.354e-02 | -3.40 | peptidyl-threonine autophosphorylation | biological process | GO:1990443 | 1 | 1 | 12345 | 414 | CaMKII | | 3.354e-02 | -3.40 | establishment of thoracic bristle planar orientation | biological process | GO:0048106 | 1 | 1 | 12345 | 414 | spen | | 3.365e-02 | -3.39 | GSE3982\_EOSINOPHIL\_VS\_BCELL\_DN | MSigDB lists | GSE3982\_EOSINOPHIL\_VS\_BCELL\_DN | 84 | 7 | 4161 | 153 | CG9281,RpS4,Tif-IA,mub,CycE,CG11030,ScsbetaG | | 3.381e-02 | -3.39 | Ets | pfam domains | PF00178 | 8 | 2 | 10480 | 393 | aop,pnt | | 3.381e-02 | -3.39 | PNP\_UDP\_1 | pfam domains | PF01048 | 8 | 2 | 10480 | 393 | CG6330,CG17224 | | 3.381e-02 | -3.39 | Innexin | pfam domains | PF00876 | 8 | 2 | 10480 | 393 | Inx2,Inx3 | | 3.381e-02 | -3.39 | Zona\_pellucida | pfam domains | PF00100 | 8 | 2 | 10480 | 393 | pio,qsm | | 3.398e-02 | -3.38 | Innexin | interpro domains | IPR000990 | 8 | 2 | 11383 | 428 | Inx2,Inx3 | | 3.398e-02 | -3.38 | Nucleoside\_phosphorylase\_sf | interpro domains | IPR035994 | 8 | 2 | 11383 | 428 | CG17224,CG6330 | | 3.398e-02 | -3.38 | SNase\_OB-fold\_sf | interpro domains | IPR035437 | 8 | 2 | 11383 | 428 | Tudor-SN,spoon | | 3.398e-02 | -3.38 | Ets\_dom | interpro domains | IPR000418 | 8 | 2 | 11383 | 428 | pnt,aop | | 3.398e-02 | -3.38 | Small\_GTPase\_Rho | interpro domains | IPR003578 | 8 | 2 | 11383 | 428 | Rac2,Rho1 | | 3.398e-02 | -3.38 | Nucleoside\_phosphorylase\_d | interpro domains | IPR000845 | 8 | 2 | 11383 | 428 | CG6330,CG17224 | | 3.399e-02 | -3.38 | Atg1 (Autophagy-related 1) | protein interactions | 39454 | 69 | 7 | 9204 | 412 | wcy,EcR,gus,hiw,mam,wnd,Atg17 | | 3.409e-02 | -3.38 | secretory granule lumen | cellular component | GO:0034774 | 1 | 1 | 11940 | 407 | dpp | | 3.409e-02 | -3.38 | cytoplasmic vesicle lumen | cellular component | GO:0060205 | 1 | 1 | 11940 | 407 | dpp | | 3.409e-02 | -3.38 | cyclin E1-CDK2 complex | cellular component | GO:0097134 | 1 | 1 | 11940 | 407 | CycE | | 3.409e-02 | -3.38 | axolemma | cellular component | GO:0030673 | 1 | 1 | 11940 | 407 | Ser | | 3.409e-02 | -3.38 | dense core granule lumen | cellular component | GO:0098898 | 1 | 1 | 11940 | 407 | dpp | | 3.409e-02 | -3.38 | calcium- and calmodulin-dependent protein kinase complex | cellular component | GO:0005954 | 1 | 1 | 11940 | 407 | CaMKII | | 3.409e-02 | -3.38 | vesicle lumen | cellular component | GO:0031983 | 1 | 1 | 11940 | 407 | dpp | | 3.409e-02 | -3.38 | NOTCH2 Activation and Transmission of Signal to the Nucleus | REACTOME pathways | R-DME-2979096 | 8 | 2 | 3923 | 148 | Ser,N | | 3.409e-02 | -3.38 | Signaling by NOTCH2 | REACTOME pathways | R-DME-1980145 | 8 | 2 | 3923 | 148 | Ser,N | | 3.409e-02 | -3.38 | Nuclear signaling by ERBB4 | REACTOME pathways | R-DME-1251985 | 8 | 2 | 3923 | 148 | Stat92E,Egfr | | 3.414e-02 | -3.38 | regulation of protein stability | biological process | GO:0031647 | 54 | 5 | 12345 | 414 | Cnx99A,Sin3A,crb,Hipk,Snx3 | | 3.414e-02 | -3.38 | positive regulation of humoral immune response | biological process | GO:0002922 | 54 | 5 | 12345 | 414 | Pli,Doa,ush,CG4325,Tl | | 3.443e-02 | -3.37 | neuron projection extension | biological process | GO:1990138 | 37 | 4 | 12345 | 414 | Ptpmeg,CadN,Dscam1,mle | | 3.443e-02 | -3.37 | cellular response to lipid | biological process | GO:0071396 | 37 | 4 | 12345 | 414 | Utx,Stat92E,Hydr2,EcR | | 3.445e-02 | -3.37 | Ank\_2 | pfam domains | PF12796 | 65 | 6 | 10480 | 393 | CG10011,Ank,CG41099,N,Prosap,CG5742 | | 3.454e-02 | -3.37 | GO\_WOUND\_HEALING | MSigDB lists | GO\_WOUND\_HEALING | 103 | 8 | 4161 | 153 | cta,pAbp,Smox,N,hh,CG34384,Rho1,Rac2 | | 3.454e-02 | -3.37 | GO\_IMMUNE\_SYSTEM\_DEVELOPMENT | MSigDB lists | GO\_IMMUNE\_SYSTEM\_DEVELOPMENT | 103 | 8 | 4161 | 153 | Sin3A,so,hh,mof,hts,Smox,l(3)mbt,Ser | | 3.455e-02 | -3.37 | enteroendocrine cell differentiation | biological process | GO:0035883 | 9 | 2 | 12345 | 414 | ttk,phyl | | 3.455e-02 | -3.37 | positive regulation of receptor signaling pathway via STAT | biological process | GO:1904894 | 9 | 2 | 12345 | 414 | CycE,Imp | | 3.455e-02 | -3.37 | intercellular transport | biological process | GO:0010496 | 9 | 2 | 12345 | 414 | Inx2,Inx3 | | 3.455e-02 | -3.37 | regulation of lamellocyte differentiation | biological process | GO:0035203 | 9 | 2 | 12345 | 414 | N,Ser | | 3.455e-02 | -3.37 | compartment pattern specification | biological process | GO:0007386 | 9 | 2 | 12345 | 414 | hh,N | | 3.455e-02 | -3.37 | zonula adherens assembly | biological process | GO:0045186 | 9 | 2 | 12345 | 414 | crb,sdt | | 3.455e-02 | -3.37 | positive regulation of receptor signaling pathway via JAK-STAT | biological process | GO:0046427 | 9 | 2 | 12345 | 414 | CycE,Imp | | 3.455e-02 | -3.37 | endoderm development | biological process | GO:0007492 | 9 | 2 | 12345 | 414 | Ubx,tkv | | 3.455e-02 | -3.37 | dorsal closure, spreading of leading edge cells | biological process | GO:0007395 | 9 | 2 | 12345 | 414 | Egfr,Rho1 | | 3.455e-02 | -3.37 | notum development | biological process | GO:0007477 | 9 | 2 | 12345 | 414 | Egfr,CG13366 | | 3.455e-02 | -3.37 | ganglion mother cell fate determination | biological process | GO:0007402 | 9 | 2 | 12345 | 414 | Kr,cas | | 3.455e-02 | -3.37 | nuclear-transcribed mRNA poly(A) tail shortening | biological process | GO:0000289 | 9 | 2 | 12345 | 414 | CG11486,smg | | 3.455e-02 | -3.37 | mitotic cytokinetic process | biological process | GO:1902410 | 9 | 2 | 12345 | 414 | pbl,Rho1 | | 3.455e-02 | -3.37 | female genitalia development | biological process | GO:0030540 | 9 | 2 | 12345 | 414 | lz,otk2 | | 3.455e-02 | -3.37 | positive regulation of heterochromatin assembly | biological process | GO:0031453 | 9 | 2 | 12345 | 414 | mle,Stat92E | | 3.455e-02 | -3.37 | regulation of plasmatocyte differentiation | biological process | GO:0045613 | 9 | 2 | 12345 | 414 | Ser,N | | 3.455e-02 | -3.37 | positive regulation of retinal cell programmed cell death | biological process | GO:0046670 | 9 | 2 | 12345 | 414 | lz,ec | | 3.455e-02 | -3.37 | glandular epithelial cell differentiation | biological process | GO:0002067 | 9 | 2 | 12345 | 414 | ttk,phyl | | 3.455e-02 | -3.37 | positive regulation of compound eye retinal cell programmed cell death | biological process | GO:0046672 | 9 | 2 | 12345 | 414 | ec,lz | | 3.455e-02 | -3.37 | maintenance of apical/basal cell polarity | biological process | GO:0035090 | 9 | 2 | 12345 | 414 | Rho1,crb | | 3.459e-02 | -3.36 | GO\_TERPENOID\_METABOLIC\_PROCESS | MSigDB lists | GO\_TERPENOID\_METABOLIC\_PROCESS | 34 | 4 | 4161 | 153 | mgl,dlp,dally,LRP1 | | 3.459e-02 | -3.36 | GSE40277\_EOS\_AND\_LEF1\_TRANSDUCED\_VS\_CTRL\_CD4\_TCELL\_UP | MSigDB lists | GSE40277\_EOS\_AND\_LEF1\_TRANSDUCED\_VS\_CTRL\_CD4\_TCELL\_UP | 34 | 4 | 4161 | 153 | phtf,emp,hts,CG45050 | | 3.459e-02 | -3.36 | GSE2706\_R848\_VS\_R848\_AND\_LPS\_8H\_STIM\_DC\_DN | MSigDB lists | GSE2706\_R848\_VS\_R848\_AND\_LPS\_8H\_STIM\_DC\_DN | 34 | 4 | 4161 | 153 | mmy,Tif-IA,gus,rush | | 3.459e-02 | -3.36 | GSE19941\_LPS\_VS\_LPS\_AND\_IL10\_STIM\_IL10\_KO\_NFKBP50\_KO\_MACROPHAGE\_DN | MSigDB lists | GSE19941\_LPS\_VS\_LPS\_AND\_IL10\_STIM\_IL10\_KO\_NFKBP50\_KO\_MACROPHAGE\_DN | 34 | 4 | 4161 | 153 | CG11367,Syx1A,JIL-1,Rho1 | | 3.459e-02 | -3.36 | GO\_NEGATIVE\_REGULATION\_OF\_PROTEIN\_CATABOLIC\_PROCESS | MSigDB lists | GO\_NEGATIVE\_REGULATION\_OF\_PROTEIN\_CATABOLIC\_PROCESS | 34 | 4 | 4161 | 153 | Smox,hh,Tlk,Snx3 | | 3.459e-02 | -3.36 | GSE37301\_MULTIPOTENT\_PROGENITOR\_VS\_LYMPHOID\_PRIMED\_MPP\_DN | MSigDB lists | GSE37301\_MULTIPOTENT\_PROGENITOR\_VS\_LYMPHOID\_PRIMED\_MPP\_DN | 34 | 4 | 4161 | 153 | JIL-1,Arl4,ScpX,CG9220 | | 3.459e-02 | -3.36 | MORF\_DEAF1 | MSigDB lists | MORF\_DEAF1 | 34 | 4 | 4161 | 153 | CG8531,pAbp,kdn,Mitofilin | | 3.459e-02 | -3.36 | GSE5099\_CLASSICAL\_M1\_VS\_ALTERNATIVE\_M2\_MACROPHAGE\_DN | MSigDB lists | GSE5099\_CLASSICAL\_M1\_VS\_ALTERNATIVE\_M2\_MACROPHAGE\_DN | 34 | 4 | 4161 | 153 | CG9220,Pli,nAChRalpha4,pbl | | 3.459e-02 | -3.36 | GSE29949\_MICROGLIA\_BRAIN\_VS\_MONOCYTE\_BONE\_MARROW\_UP | MSigDB lists | GSE29949\_MICROGLIA\_BRAIN\_VS\_MONOCYTE\_BONE\_MARROW\_UP | 34 | 4 | 4161 | 153 | nudE,ras,mgl,betaTub56D | | 3.460e-02 | -3.36 | mof (males absent on the first) | protein interactions | 31518 | 28 | 4 | 9204 | 412 | lncRNA:roX1,Chro,mle,mof | | 3.460e-02 | -3.36 | elB (elbow B) | protein interactions | 34844 | 28 | 4 | 9204 | 412 | ci,Pits,rau,CycE | | 3.460e-02 | -3.36 | ato (atonal) | protein interactions | 40975 | 28 | 4 | 9204 | 412 | pnt,hh,Egfr,ci | | 3.460e-02 | -3.36 | Msp300 (Muscle-specific protein 300 kDa) | protein interactions | 3771968 | 28 | 4 | 9204 | 412 | Syp,Fas2,ci,sty | | 3.460e-02 | -3.36 | nuf (nuclear fallout) | protein interactions | 39572 | 28 | 4 | 9204 | 412 | Pat1,pbl,Bx,esn | | 3.468e-02 | -3.36 | LIM | smart domains | SM00132 | 34 | 5 | 5086 | 276 | ap,Bx,smash,Lim1,esn | | 3.470e-02 | -3.36 | RBD\_domain\_sf | interpro domains | IPR035979 | 157 | 11 | 11383 | 428 | spen,sm,pAbp,fne,Imp,elav,cpo,CG2162,Syp,Rbfox1,heph | | 3.495e-02 | -3.35 | actin filament binding | molecular function | GO:0051015 | 56 | 5 | 12176 | 396 | Mhcl,hts,Hip1,Arp2,Svil | | 3.495e-02 | -3.35 | Dll (Distal-less) | protein interactions | 37973 | 41 | 5 | 9204 | 412 | Ubx,CG11486,mbt,Zif,disco | | 3.495e-02 | -3.35 | Dg (Dystroglycan) | protein interactions | 36773 | 41 | 5 | 9204 | 412 | dpp,Dad,CG6700,robo2,aos | | 3.511e-02 | -3.35 | GO\_CARDIAC\_SEPTUM\_DEVELOPMENT | MSigDB lists | GO\_CARDIAC\_SEPTUM\_DEVELOPMENT | 20 | 3 | 4161 | 153 | Ser,N,mgl | | 3.511e-02 | -3.35 | PAX3\_B | MSigDB lists | PAX3\_B | 20 | 3 | 4161 | 153 | Cnx99A,fne,nudE | | 3.511e-02 | -3.35 | MODULE\_491 | MSigDB lists | MODULE\_491 | 20 | 3 | 4161 | 153 | Acsl,betaTub56D,CG9220 | | 3.511e-02 | -3.35 | GSE42021\_TREG\_PLN\_VS\_TREG\_PRECURSORS\_THYMUS\_DN | MSigDB lists | GSE42021\_TREG\_PLN\_VS\_TREG\_PRECURSORS\_THYMUS\_DN | 20 | 3 | 4161 | 153 | CG41099,Tlk,emp | | 3.511e-02 | -3.35 | WOO\_LIVER\_CANCER\_RECURRENCE\_UP | MSigDB lists | WOO\_LIVER\_CANCER\_RECURRENCE\_UP | 20 | 3 | 4161 | 153 | Ser,dlp,Tlk | | 3.511e-02 | -3.35 | GO\_EMBRYONIC\_HEART\_TUBE\_MORPHOGENESIS | MSigDB lists | GO\_EMBRYONIC\_HEART\_TUBE\_MORPHOGENESIS | 20 | 3 | 4161 | 153 | hh,Smox,N | | 3.511e-02 | -3.35 | PTEN\_DN.V1\_UP | MSigDB lists | PTEN\_DN.V1\_UP | 20 | 3 | 4161 | 153 | Ser,tkv,ap | | 3.511e-02 | -3.35 | FREAC4\_01 | MSigDB lists | FREAC4\_01 | 20 | 3 | 4161 | 153 | Sin3A,phtf,Pli | | 3.511e-02 | -3.35 | KEGG\_RENAL\_CELL\_CARCINOMA | MSigDB lists | KEGG\_RENAL\_CELL\_CARCINOMA | 20 | 3 | 4161 | 153 | mbt,drk,Rac2 | | 3.511e-02 | -3.35 | GO\_TRANSCRIPTIONAL\_ACTIVATOR\_ACTIVITY\_RNA\_POLYMERASE\_II\_CORE\_PROMOTER\_PROXIMAL\_REGION\_SEQUENCE\_SPECIFIC\_BINDING | MSigDB lists | GO\_TRANSCRIPTIONAL\_ACTIVATOR\_ACTIVITY\_RNA\_POLYMERASE\_II\_CORE\_PROMOTER\_PROXIMAL\_REGION\_SEQUENCE\_SPECIFIC\_BINDING | 20 | 3 | 4161 | 153 | so,EcR,TfAP-2 | | 3.511e-02 | -3.35 | GO\_CHROMATIN\_DNA\_BINDING | MSigDB lists | GO\_CHROMATIN\_DNA\_BINDING | 20 | 3 | 4161 | 153 | N,Smox,Utx | | 3.525e-02 | -3.35 | GO\_EPIDERMIS\_DEVELOPMENT | MSigDB lists | GO\_EPIDERMIS\_DEVELOPMENT | 50 | 5 | 4161 | 153 | ap,N,Rac2,hh,Ser | | 3.525e-02 | -3.35 | GSE42021\_TCONV\_PLN\_VS\_CD24HI\_TCONV\_THYMUS\_UP | MSigDB lists | GSE42021\_TCONV\_PLN\_VS\_CD24HI\_TCONV\_THYMUS\_UP | 50 | 5 | 4161 | 153 | Tpr2,rush,Acsl,CG3961,so | | 3.525e-02 | -3.35 | FREAC3\_01 | MSigDB lists | FREAC3\_01 | 50 | 5 | 4161 | 153 | Pli,mgl,NetB,Utx,ras | | 3.525e-02 | -3.35 | GO\_PROTEIN\_LOCALIZATION\_TO\_CELL\_PERIPHERY | MSigDB lists | GO\_PROTEIN\_LOCALIZATION\_TO\_CELL\_PERIPHERY | 50 | 5 | 4161 | 153 | Ank,sdt,Rac2,ScpX,Tsp26A | | 3.525e-02 | -3.35 | GSE37301\_PRO\_BCELL\_VS\_GRANULOCYTE\_MONOCYTE\_PROGENITOR\_DN | MSigDB lists | GSE37301\_PRO\_BCELL\_VS\_GRANULOCYTE\_MONOCYTE\_PROGENITOR\_DN | 50 | 5 | 4161 | 153 | ScpX,CG11367,drk,CG45050,Arl4 | | 3.533e-02 | -3.34 | NFY\_01 | MSigDB lists | NFY\_01 | 67 | 6 | 4161 | 153 | Lim1,Fas2,smal,CG9281,so,CG14646 | | 3.533e-02 | -3.34 | GSE17721\_LPS\_VS\_PAM3CSK4\_4H\_BMDC\_UP | MSigDB lists | GSE17721\_LPS\_VS\_PAM3CSK4\_4H\_BMDC\_UP | 67 | 6 | 4161 | 153 | ras,Pat1,Smox,DIP-epsilon,Tlk,Mocs2B | | 3.533e-02 | -3.34 | AGCACTT\_MIR93\_MIR302A\_MIR302B\_MIR302C\_MIR302D\_MIR372\_MIR373\_MIR520E\_MIR520A\_MIR526B\_MIR520B\_MIR520C\_MIR520D | MSigDB lists | AGCACTT\_MIR93\_MIR302A\_MIR302B\_MIR302C\_MIR302D\_MIR372\_MIR373\_MIR520E\_MIR520A\_MIR526B\_MIR520B\_MIR520C\_MIR520D | 67 | 6 | 4161 | 153 | Imp,JIL-1,S6kII,Rho1,CG5742,pbl | | 3.544e-02 | -3.34 | negative regulation of gene expression, epigenetic | biological process | GO:0045814 | 73 | 6 | 12345 | 414 | Kr,psq,bin3,Tlk,Psc,AGO2 | | 3.544e-02 | -3.34 | chromatin silencing | biological process | GO:0006342 | 73 | 6 | 12345 | 414 | Psc,AGO2,Tlk,Kr,psq,bin3 | | 3.544e-02 | -3.34 | chromatin organization involved in negative regulation of transcription | biological process | GO:0097549 | 73 | 6 | 12345 | 414 | Kr,psq,bin3,Tlk,Psc,AGO2 | | 3.559e-02 | -3.34 | NRF1\_Q6 | MSigDB lists | NRF1\_Q6 | 85 | 7 | 4161 | 153 | Tudor-SN,Snx3,LRP1,Ser,CG6752,sm,sdt | | 3.560e-02 | -3.34 | RNAi effector complex | cellular component | GO:0031332 | 9 | 2 | 11940 | 407 | AGO2,Tudor-SN | | 3.560e-02 | -3.34 | RISC complex | cellular component | GO:0016442 | 9 | 2 | 11940 | 407 | AGO2,Tudor-SN | | 3.577e-02 | -3.33 | late endosome | cellular component | GO:0005770 | 72 | 6 | 11940 | 407 | comm,rush,dbr,N,tkv,S | | 3.593e-02 | -3.33 | GO\_NEGATIVE\_REGULATION\_OF\_TRANSCRIPTION\_FROM\_RNA\_POLYMERASE\_II\_PROMOTER | MSigDB lists | GO\_NEGATIVE\_REGULATION\_OF\_TRANSCRIPTION\_FROM\_RNA\_POLYMERASE\_II\_PROMOTER | 123 | 9 | 4161 | 153 | dpp,CtBP,TfAP-2,Smox,N,hh,so,Sin3A,EcR | | 3.593e-02 | -3.33 | GO\_RESPONSE\_TO\_WOUNDING | MSigDB lists | GO\_RESPONSE\_TO\_WOUNDING | 123 | 9 | 4161 | 153 | cta,pAbp,Smox,N,Rho1,CG34384,Rac2,Hydr2,hh | | 3.594e-02 | -3.33 | cellular response to fibroblast growth factor stimulus | biological process | GO:0044344 | 22 | 3 | 12345 | 414 | drk,pnt,aop | | 3.594e-02 | -3.33 | Rho protein signal transduction | biological process | GO:0007266 | 22 | 3 | 12345 | 414 | Rac2,Rho1,cta | | 3.594e-02 | -3.33 | positive regulation of ATPase activity | biological process | GO:0032781 | 22 | 3 | 12345 | 414 | CG8531,nrv2,mle | | 3.594e-02 | -3.33 | response to fibroblast growth factor | biological process | GO:0071774 | 22 | 3 | 12345 | 414 | pnt,aop,drk | | 3.594e-02 | -3.33 | fibroblast growth factor receptor signaling pathway | biological process | GO:0008543 | 22 | 3 | 12345 | 414 | aop,pnt,drk | | 3.594e-02 | -3.33 | actin filament bundle assembly | biological process | GO:0051017 | 22 | 3 | 12345 | 414 | Rac2,hts,Rho1 | | 3.613e-02 | -3.32 | regulation of GTPase activity | biological process | GO:0043087 | 113 | 8 | 12345 | 414 | RhoGAP18B,RhoGAP93B,pbl,CG42684,RhoGAP15B,RhoGAP102A,RhoGAP16F,RapGAP1 | | 3.615e-02 | -3.32 | CG14829 (uncharacterized protein) | protein interactions | 38792 | 7 | 2 | 9204 | 412 | Mondo,betaTub56D | | 3.615e-02 | -3.32 | phol (pleiohomeotic like) | protein interactions | 39095 | 7 | 2 | 9204 | 412 | rau,Ubx | | 3.615e-02 | -3.32 | wun (wunen) | protein interactions | 35966 | 7 | 2 | 9204 | 412 | pbl,wun2 | | 3.615e-02 | -3.32 | CG6767 (uncharacterized protein) | protein interactions | 39132 | 7 | 2 | 9204 | 412 | phyl,Gbs-70E | | 3.615e-02 | -3.32 | janA (janus A) | protein interactions | 43569 | 7 | 2 | 9204 | 412 | CG30089,CG32809 | | 3.615e-02 | -3.32 | SoxN (SoxNeuro) | protein interactions | 44275 | 7 | 2 | 9204 | 412 | Sox21a,Egfr | | 3.615e-02 | -3.32 | mib1 (mind bomb 1) | protein interactions | 39750 | 7 | 2 | 9204 | 412 | Ser,N | | 3.615e-02 | -3.32 | ATPCL (ATP citrate lyase) | protein interactions | 36760 | 7 | 2 | 9204 | 412 | CG11486,CG13454 | | 3.615e-02 | -3.32 | CG30339 (uncharacterized protein) | protein interactions | 246549 | 7 | 2 | 9204 | 412 | Fas2,Df31 | | 3.615e-02 | -3.32 | cta (concertina) | protein interactions | 3355131 | 7 | 2 | 9204 | 412 | RhoGEF2,Rho1 | | 3.615e-02 | -3.32 | grnd (grindelwald) | protein interactions | 35103 | 7 | 2 | 9204 | 412 | crb,egr | | 3.615e-02 | -3.32 | G9a (G9a) | protein interactions | 30971 | 7 | 2 | 9204 | 412 | EcR,sba | | 3.615e-02 | -3.32 | Pitslre (pitslre) | protein interactions | 40292 | 7 | 2 | 9204 | 412 | pbl,Fas2 | | 3.615e-02 | -3.32 | fwd (four wheel drive) | protein interactions | 45374 | 7 | 2 | 9204 | 412 | pbl,ci | | 3.615e-02 | -3.32 | klg (klingon) | protein interactions | 42707 | 7 | 2 | 9204 | 412 | N,pbl | | 3.615e-02 | -3.32 | pad (poils au dos) | protein interactions | 41981 | 7 | 2 | 9204 | 412 | tkv,rau | | 3.615e-02 | -3.32 | BHD (Birt-Hogg-Dube) | protein interactions | 38793 | 7 | 2 | 9204 | 412 | Stat92E,dpp | | 3.615e-02 | -3.32 | Atg4b (Autophagy-related 4b) | protein interactions | 41868 | 7 | 2 | 9204 | 412 | N,sno | | 3.615e-02 | -3.32 | Mmp1 (Matrix metalloproteinase 1) | protein interactions | 37949 | 7 | 2 | 9204 | 412 | Mmp2,Sox21a | | 3.615e-02 | -3.32 | Rdl (Resistant to dieldrin) | protein interactions | 39054 | 7 | 2 | 9204 | 412 | Fas2,wake | | 3.615e-02 | -3.32 | tweek (tweek) | protein interactions | 35026 | 7 | 2 | 9204 | 412 | ci,CG10283 | | 3.615e-02 | -3.32 | scw (screw) | protein interactions | 46000 | 7 | 2 | 9204 | 412 | dpp,tkv | | 3.615e-02 | -3.32 | Teh4 (tipE homolog 4) | protein interactions | 38501 | 7 | 2 | 9204 | 412 | par-1,CG2865 | | 3.615e-02 | -3.32 | Brd (Bearded) | protein interactions | 39620 | 7 | 2 | 9204 | 412 | N,Ubx | | 3.615e-02 | -3.32 | CG14696 (uncharacterized protein) | protein interactions | 41317 | 7 | 2 | 9204 | 412 | Pli,phyl | | 3.615e-02 | -3.32 | CG30183 (uncharacterized protein) | protein interactions | 37749 | 7 | 2 | 9204 | 412 | CtBP,Fas2 | | 3.615e-02 | -3.32 | CG5078 (uncharacterized protein) | protein interactions | 40276 | 7 | 2 | 9204 | 412 | CG7264,CycE | | 3.615e-02 | -3.32 | Pngl (PNGase-like) | protein interactions | 35527 | 7 | 2 | 9204 | 412 | tkv,N | | 3.615e-02 | -3.32 | RapGAP1 (Rap GTPase activating protein 1) | protein interactions | 34032 | 7 | 2 | 9204 | 412 | ci,pbl | | 3.615e-02 | -3.32 | CG6758 (uncharacterized protein) | protein interactions | 37532 | 7 | 2 | 9204 | 412 | CG32772,CG17265 | | 3.615e-02 | -3.32 | Klp61F (Kinesin-like protein at 61F) | protein interactions | 38135 | 7 | 2 | 9204 | 412 | Fas2,fj | | 3.617e-02 | -3.32 | GO\_SOMATODENDRITIC\_COMPARTMENT | MSigDB lists | GO\_SOMATODENDRITIC\_COMPARTMENT | 163 | 11 | 4161 | 153 | LRP1,Imp,Ank,nAChRalpha4,tkv,nudE,CG11030,if,Cnx99A,Ykt6,CaMKII | | 3.630e-02 | -3.32 | Ankyrin\_rpt | interpro domains | IPR002110 | 83 | 7 | 11383 | 428 | CG41099,CG10011,Prosap,wake,CG5742,Ank,N | | 3.659e-02 | -3.31 | Pi3K92E (Pi3K92E) | protein interactions | 42446 | 117 | 10 | 9204 | 412 | CycE,Hsc70-3,ap,Chro,ban,yps,Syp,Trpm,CaMKII,ci | | 3.677e-02 | -3.30 | WANG\_METASTASIS\_OF\_BREAST\_CANCER | MSigDB lists | WANG\_METASTASIS\_OF\_BREAST\_CANCER | 1 | 1 | 4161 | 153 | RpS4 | | 3.677e-02 | -3.30 | chr13q22 | MSigDB lists | chr13q22 | 1 | 1 | 4161 | 153 | hiw | | 3.677e-02 | -3.30 | JI\_CARCINOGENESIS\_BY\_KRAS\_AND\_STK11\_UP | MSigDB lists | JI\_CARCINOGENESIS\_BY\_KRAS\_AND\_STK11\_UP | 1 | 1 | 4161 | 153 | TfAP-2 | | 3.677e-02 | -3.30 | GO\_POSITIVE\_REGULATION\_OF\_THYMOCYTE\_AGGREGATION | MSigDB lists | GO\_POSITIVE\_REGULATION\_OF\_THYMOCYTE\_AGGREGATION | 1 | 1 | 4161 | 153 | hh | | 3.677e-02 | -3.30 | GO\_REGULATION\_OF\_ODONTOGENESIS\_OF\_DENTIN\_CONTAINING\_TOOTH | MSigDB lists | GO\_REGULATION\_OF\_ODONTOGENESIS\_OF\_DENTIN\_CONTAINING\_TOOTH | 1 | 1 | 4161 | 153 | dpp | | 3.677e-02 | -3.30 | BEGUM\_TARGETS\_OF\_PAX3\_FOXO1\_FUSION\_AND\_PAX3 | MSigDB lists | BEGUM\_TARGETS\_OF\_PAX3\_FOXO1\_FUSION\_AND\_PAX3 | 1 | 1 | 4161 | 153 | Ser | | 3.677e-02 | -3.30 | GO\_WHITE\_FAT\_CELL\_DIFFERENTIATION | MSigDB lists | GO\_WHITE\_FAT\_CELL\_DIFFERENTIATION | 1 | 1 | 4161 | 153 | CtBP | | 3.677e-02 | -3.30 | HERNANDEZ\_MITOTIC\_ARREST\_BY\_DOCETAXEL\_1\_UP | MSigDB lists | HERNANDEZ\_MITOTIC\_ARREST\_BY\_DOCETAXEL\_1\_UP | 1 | 1 | 4161 | 153 | fne | | 3.677e-02 | -3.30 | LI\_WILMS\_TUMOR | MSigDB lists | LI\_WILMS\_TUMOR | 1 | 1 | 4161 | 153 | so | | 3.677e-02 | -3.30 | SCHAEFFER\_PROSTATE\_DEVELOPMENT\_AND\_CANCER\_BOX5\_UP | MSigDB lists | SCHAEFFER\_PROSTATE\_DEVELOPMENT\_AND\_CANCER\_BOX5\_UP | 1 | 1 | 4161 | 153 | Pli | | 3.677e-02 | -3.30 | GO\_REGULATION\_OF\_ENDOTHELIAL\_CELL\_CHEMOTAXIS | MSigDB lists | GO\_REGULATION\_OF\_ENDOTHELIAL\_CELL\_CHEMOTAXIS | 1 | 1 | 4161 | 153 | N | | 3.677e-02 | -3.30 | GO\_CD4\_POSITIVE\_OR\_CD8\_POSITIVE\_ALPHA\_BETA\_T\_CELL\_LINEAGE\_COMMITMENT | MSigDB lists | GO\_CD4\_POSITIVE\_OR\_CD8\_POSITIVE\_ALPHA\_BETA\_T\_CELL\_LINEAGE\_COMMITMENT | 1 | 1 | 4161 | 153 | hh | | 3.677e-02 | -3.30 | GO\_RENAL\_SYSTEM\_VASCULATURE\_DEVELOPMENT | MSigDB lists | GO\_RENAL\_SYSTEM\_VASCULATURE\_DEVELOPMENT | 1 | 1 | 4161 | 153 | N | | 3.677e-02 | -3.30 | GO\_NOSE\_DEVELOPMENT | MSigDB lists | GO\_NOSE\_DEVELOPMENT | 1 | 1 | 4161 | 153 | so | | 3.677e-02 | -3.30 | REACTOME\_APOBEC3G\_MEDIATED\_RESISTANCE\_TO\_HIV1\_INFECTION | MSigDB lists | REACTOME\_APOBEC3G\_MEDIATED\_RESISTANCE\_TO\_HIV1\_INFECTION | 1 | 1 | 4161 | 153 | baf | | 3.677e-02 | -3.30 | GNF2\_MATK | MSigDB lists | GNF2\_MATK | 1 | 1 | 4161 | 153 | Ptpmeg | | 3.677e-02 | -3.30 | MATZUK\_CUMULUS\_EXPANSION | MSigDB lists | MATZUK\_CUMULUS\_EXPANSION | 1 | 1 | 4161 | 153 | tkv | | 3.677e-02 | -3.30 | GO\_NEGATIVE\_REGULATION\_OF\_CARDIAC\_MUSCLE\_TISSUE\_DEVELOPMENT | MSigDB lists | GO\_NEGATIVE\_REGULATION\_OF\_CARDIAC\_MUSCLE\_TISSUE\_DEVELOPMENT | 1 | 1 | 4161 | 153 | dpp | | 3.677e-02 | -3.30 | GO\_LONG\_CHAIN\_FATTY\_ACID\_BINDING | MSigDB lists | GO\_LONG\_CHAIN\_FATTY\_ACID\_BINDING | 1 | 1 | 4161 | 153 | ScpX | | 3.677e-02 | -3.30 | GO\_T\_CELL\_LINEAGE\_COMMITMENT | MSigDB lists | GO\_T\_CELL\_LINEAGE\_COMMITMENT | 1 | 1 | 4161 | 153 | hh | | 3.677e-02 | -3.30 | GO\_RETINOID\_X\_RECEPTOR\_BINDING | MSigDB lists | GO\_RETINOID\_X\_RECEPTOR\_BINDING | 1 | 1 | 4161 | 153 | EcR | | 3.677e-02 | -3.30 | TING\_SILENCED\_BY\_DICER | MSigDB lists | TING\_SILENCED\_BY\_DICER | 1 | 1 | 4161 | 153 | Pli | | 3.677e-02 | -3.30 | GO\_NOTOCHORD\_DEVELOPMENT | MSigDB lists | GO\_NOTOCHORD\_DEVELOPMENT | 1 | 1 | 4161 | 153 | Utx | | 3.677e-02 | -3.30 | GO\_POSITIVE\_REGULATION\_OF\_BONE\_REMODELING | MSigDB lists | GO\_POSITIVE\_REGULATION\_OF\_BONE\_REMODELING | 1 | 1 | 4161 | 153 | CG11367 | | 3.677e-02 | -3.30 | GO\_NEPHRIC\_DUCT\_DEVELOPMENT | MSigDB lists | GO\_NEPHRIC\_DUCT\_DEVELOPMENT | 1 | 1 | 4161 | 153 | Lim1 | | 3.677e-02 | -3.30 | GO\_REGULATION\_OF\_MESODERM\_DEVELOPMENT | MSigDB lists | GO\_REGULATION\_OF\_MESODERM\_DEVELOPMENT | 1 | 1 | 4161 | 153 | hh | | 3.677e-02 | -3.30 | TSAI\_RESPONSE\_TO\_RADIATION\_THERAPY | MSigDB lists | TSAI\_RESPONSE\_TO\_RADIATION\_THERAPY | 1 | 1 | 4161 | 153 | Smox | | 3.677e-02 | -3.30 | GO\_EMBRYONIC\_CAMERA\_TYPE\_EYE\_FORMATION | MSigDB lists | GO\_EMBRYONIC\_CAMERA\_TYPE\_EYE\_FORMATION | 1 | 1 | 4161 | 153 | TfAP-2 | | 3.677e-02 | -3.30 | GO\_NEGATIVE\_REGULATION\_OF\_ALPHA\_BETA\_T\_CELL\_DIFFERENTIATION | MSigDB lists | GO\_NEGATIVE\_REGULATION\_OF\_ALPHA\_BETA\_T\_CELL\_DIFFERENTIATION | 1 | 1 | 4161 | 153 | hh | | 3.677e-02 | -3.30 | GO\_INTERMEDIATE\_FILAMENT\_ORGANIZATION | MSigDB lists | GO\_INTERMEDIATE\_FILAMENT\_ORGANIZATION | 1 | 1 | 4161 | 153 | hh | | 3.677e-02 | -3.30 | CHENG\_TAF7L\_TARGETS | MSigDB lists | CHENG\_TAF7L\_TARGETS | 1 | 1 | 4161 | 153 | CG3097 | | 3.677e-02 | -3.30 | MIKKELSEN\_PARTIALLY\_REPROGRAMMED\_TO\_PLURIPOTENCY | MSigDB lists | MIKKELSEN\_PARTIALLY\_REPROGRAMMED\_TO\_PLURIPOTENCY | 1 | 1 | 4161 | 153 | TfAP-2 | | 3.677e-02 | -3.30 | CHOI\_ATL\_ACUTE\_STAGE | MSigDB lists | CHOI\_ATL\_ACUTE\_STAGE | 1 | 1 | 4161 | 153 | ap | | 3.677e-02 | -3.30 | GO\_FEMALE\_GENITALIA\_DEVELOPMENT | MSigDB lists | GO\_FEMALE\_GENITALIA\_DEVELOPMENT | 1 | 1 | 4161 | 153 | Lim1 | | 3.677e-02 | -3.30 | FUJIWARA\_PARK2\_IN\_LIVER\_CANCER\_UP | MSigDB lists | FUJIWARA\_PARK2\_IN\_LIVER\_CANCER\_UP | 1 | 1 | 4161 | 153 | CycE | | 3.677e-02 | -3.30 | GO\_NEPHRIC\_DUCT\_MORPHOGENESIS | MSigDB lists | GO\_NEPHRIC\_DUCT\_MORPHOGENESIS | 1 | 1 | 4161 | 153 | Lim1 | | 3.677e-02 | -3.30 | GO\_NEGATIVE\_REGULATION\_OF\_ALPHA\_BETA\_T\_CELL\_ACTIVATION | MSigDB lists | GO\_NEGATIVE\_REGULATION\_OF\_ALPHA\_BETA\_T\_CELL\_ACTIVATION | 1 | 1 | 4161 | 153 | hh | | 3.677e-02 | -3.30 | GO\_NEGATIVE\_REGULATION\_OF\_CIRCADIAN\_RHYTHM | MSigDB lists | GO\_NEGATIVE\_REGULATION\_OF\_CIRCADIAN\_RHYTHM | 1 | 1 | 4161 | 153 | Sin3A | | 3.677e-02 | -3.30 | MIZUKAMI\_HYPOXIA\_DN | MSigDB lists | MIZUKAMI\_HYPOXIA\_DN | 1 | 1 | 4161 | 153 | LRP1 | | 3.677e-02 | -3.30 | GO\_POSITIVE\_REGULATION\_OF\_OSTEOBLAST\_PROLIFERATION | MSigDB lists | GO\_POSITIVE\_REGULATION\_OF\_OSTEOBLAST\_PROLIFERATION | 1 | 1 | 4161 | 153 | dpp | | 3.677e-02 | -3.30 | GO\_POSITIVE\_REGULATION\_OF\_KERATINOCYTE\_DIFFERENTIATION | MSigDB lists | GO\_POSITIVE\_REGULATION\_OF\_KERATINOCYTE\_DIFFERENTIATION | 1 | 1 | 4161 | 153 | N | | 3.677e-02 | -3.30 | GO\_POSITIVE\_REGULATION\_OF\_TISSUE\_REMODELING | MSigDB lists | GO\_POSITIVE\_REGULATION\_OF\_TISSUE\_REMODELING | 1 | 1 | 4161 | 153 | CG11367 | | 3.677e-02 | -3.30 | GO\_TITIN\_BINDING | MSigDB lists | GO\_TITIN\_BINDING | 1 | 1 | 4161 | 153 | CaMKII | | 3.677e-02 | -3.30 | CGGTGTG\_MIR220 | MSigDB lists | CGGTGTG\_MIR220 | 1 | 1 | 4161 | 153 | ap | | 3.677e-02 | -3.30 | JI\_CARCINOGENESIS\_BY\_KRAS\_AND\_STK11\_DN | MSigDB lists | JI\_CARCINOGENESIS\_BY\_KRAS\_AND\_STK11\_DN | 1 | 1 | 4161 | 153 | Fas2 | | 3.677e-02 | -3.30 | GO\_ACTIVATION\_OF\_CYSTEINE\_TYPE\_ENDOPEPTIDASE\_ACTIVITY\_INVOLVED\_IN\_APOPTOTIC\_SIGNALING\_PATHWAY | MSigDB lists | GO\_ACTIVATION\_OF\_CYSTEINE\_TYPE\_ENDOPEPTIDASE\_ACTIVITY\_INVOLVED\_IN\_APOPTOTIC\_SIGNALING\_PATHWAY | 1 | 1 | 4161 | 153 | Smox | | 3.677e-02 | -3.30 | GO\_URETER\_DEVELOPMENT | MSigDB lists | GO\_URETER\_DEVELOPMENT | 1 | 1 | 4161 | 153 | so | | 3.677e-02 | -3.30 | GO\_PROTEIN\_SERINE\_THREONINE\_PHOSPHATASE\_INHIBITOR\_ACTIVITY | MSigDB lists | GO\_PROTEIN\_SERINE\_THREONINE\_PHOSPHATASE\_INHIBITOR\_ACTIVITY | 1 | 1 | 4161 | 153 | NiPp1 | | 3.677e-02 | -3.30 | LOPEZ\_EPITHELIOID\_MESOTHELIOMA | MSigDB lists | LOPEZ\_EPITHELIOID\_MESOTHELIOMA | 1 | 1 | 4161 | 153 | mgl | | 3.677e-02 | -3.30 | GO\_REGULATION\_OF\_LIPOPROTEIN\_LIPASE\_ACTIVITY | MSigDB lists | GO\_REGULATION\_OF\_LIPOPROTEIN\_LIPASE\_ACTIVITY | 1 | 1 | 4161 | 153 | EcR | | 3.686e-02 | -3.30 | FYVE | smart domains | SM00064 | 14 | 3 | 5086 | 276 | CG41099,rush,CG6051 | | 3.697e-02 | -3.30 | kinase regulator activity | molecular function | GO:0019207 | 76 | 6 | 12176 | 396 | Atg17,CycE,CG8441,Pli,Socs16D,Cdk5alpha | | 3.699e-02 | -3.30 | Fatty acid biosynthesis | KEGG pathways | ko00061 | 10 | 2 | 3198 | 100 | CG3961,Acsl | | 3.699e-02 | -3.30 | Fatty acid biosynthesis | KEGG pathways | dme00061 | 10 | 2 | 3198 | 100 | CG3961,Acsl | | 3.743e-02 | -3.29 | GO\_PROTEIN\_PHOSPHORYLATION | MSigDB lists | GO\_PROTEIN\_PHOSPHORYLATION | 248 | 15 | 4161 | 153 | PhKgamma,dpp,Fas2,smal,par-1,S6kII,CaMKII,Taf1,Doa,Tlk,CycE,mbt,drk,JIL-1,tkv | | 3.746e-02 | -3.28 | MyD88 cascade initiated on plasma membrane | REACTOME pathways | R-DME-975871 | 34 | 4 | 3923 | 148 | S6kII,mle,Pli,JIL-1 | | 3.746e-02 | -3.28 | MyD88:Mal cascade initiated on plasma membrane | REACTOME pathways | R-DME-166058 | 34 | 4 | 3923 | 148 | S6kII,mle,Pli,JIL-1 | | 3.746e-02 | -3.28 | Activated TLR4 signalling | REACTOME pathways | R-DME-166054 | 34 | 4 | 3923 | 148 | mle,Pli,S6kII,JIL-1 | | 3.746e-02 | -3.28 | Toll Like Receptor TLR6:TLR2 Cascade | REACTOME pathways | R-DME-168188 | 34 | 4 | 3923 | 148 | JIL-1,mle,Pli,S6kII | | 3.746e-02 | -3.28 | Toll Like Receptor 5 (TLR5) Cascade | REACTOME pathways | R-DME-168176 | 34 | 4 | 3923 | 148 | S6kII,mle,Pli,JIL-1 | | 3.746e-02 | -3.28 | Toll Like Receptor 2 (TLR2) Cascade | REACTOME pathways | R-DME-181438 | 34 | 4 | 3923 | 148 | JIL-1,Pli,mle,S6kII | | 3.746e-02 | -3.28 | Toll Like Receptor TLR1:TLR2 Cascade | REACTOME pathways | R-DME-168179 | 34 | 4 | 3923 | 148 | JIL-1,mle,Pli,S6kII | | 3.748e-02 | -3.28 | regulation of cellular component size | biological process | GO:0032535 | 178 | 11 | 12345 | 414 | spoon,hts,Arp2,stg,PsGEF,Myb,mbt,pbl,Pvf3,CadN,drk | | 3.749e-02 | -3.28 | positive regulation of endocytosis | biological process | GO:0045807 | 38 | 4 | 12345 | 414 | Hip1,sdt,comm,drk | | 3.750e-02 | -3.28 | DUF3669 | pfam domains | PF12417 | 1 | 1 | 10480 | 393 | CG8929 | | 3.750e-02 | -3.28 | Thioredoxin\_8 | pfam domains | PF13905 | 1 | 1 | 10480 | 393 | CG12547 | | 3.750e-02 | -3.28 | Phospholip\_B | pfam domains | PF04916 | 1 | 1 | 10480 | 393 | lama | | 3.750e-02 | -3.28 | CaMKII\_AD | pfam domains | PF08332 | 1 | 1 | 10480 | 393 | CaMKII | | 3.750e-02 | -3.28 | DUF1168 | pfam domains | PF06658 | 1 | 1 | 10480 | 393 | CG8441 | | 3.750e-02 | -3.28 | Tet\_JBP | pfam domains | PF12851 | 1 | 1 | 10480 | 393 | Tet | | 3.750e-02 | -3.28 | ATG11 | pfam domains | PF10377 | 1 | 1 | 10480 | 393 | Atg17 | | 3.750e-02 | -3.28 | CCDC85 | pfam domains | PF10226 | 1 | 1 | 10480 | 393 | CG17265 | | 3.750e-02 | -3.28 | Crust\_neurohorm | pfam domains | PF01147 | 1 | 1 | 10480 | 393 | ITP | | 3.750e-02 | -3.28 | DUF3523 | pfam domains | PF12037 | 1 | 1 | 10480 | 393 | bor | | 3.750e-02 | -3.28 | NOD | pfam domains | PF06816 | 1 | 1 | 10480 | 393 | N | | 3.750e-02 | -3.28 | DUF3454 | pfam domains | PF11936 | 1 | 1 | 10480 | 393 | N | | 3.750e-02 | -3.28 | Mitofilin | pfam domains | PF09731 | 1 | 1 | 10480 | 393 | Mitofilin | | 3.750e-02 | -3.28 | SRP40\_C | pfam domains | PF05022 | 1 | 1 | 10480 | 393 | Nopp140 | | 3.750e-02 | -3.28 | DUF4801 | pfam domains | PF16059 | 1 | 1 | 10480 | 393 | ocm | | 3.750e-02 | -3.28 | IRF-2BP1\_2 | pfam domains | PF11261 | 1 | 1 | 10480 | 393 | Pits | | 3.750e-02 | -3.28 | Integrin\_alpha | pfam domains | PF00357 | 1 | 1 | 10480 | 393 | if | | 3.750e-02 | -3.28 | DUF4757 | pfam domains | PF15949 | 1 | 1 | 10480 | 393 | smash | | 3.750e-02 | -3.28 | WAC\_Acf1\_DNA\_bd | pfam domains | PF10537 | 1 | 1 | 10480 | 393 | Acf | | 3.750e-02 | -3.28 | BAF | pfam domains | PF02961 | 1 | 1 | 10480 | 393 | baf | | 3.750e-02 | -3.28 | RRN3 | pfam domains | PF05327 | 1 | 1 | 10480 | 393 | Tif-IA | | 3.750e-02 | -3.28 | AIP3 | pfam domains | PF03915 | 1 | 1 | 10480 | 393 | CG32809 | | 3.750e-02 | -3.28 | 40S\_S4\_C | pfam domains | PF16121 | 1 | 1 | 10480 | 393 | RpS4 | | 3.750e-02 | -3.28 | HH\_signal | pfam domains | PF01085 | 1 | 1 | 10480 | 393 | hh | | 3.750e-02 | -3.28 | DUF1084 | pfam domains | PF06454 | 1 | 1 | 10480 | 393 | CG2991 | | 3.750e-02 | -3.28 | Optomotor-blind | pfam domains | PF11078 | 1 | 1 | 10480 | 393 | bi | | 3.750e-02 | -3.28 | Sin3\_corepress | pfam domains | PF08295 | 1 | 1 | 10480 | 393 | Sin3A | | 3.750e-02 | -3.28 | AAA\_2 | pfam domains | PF07724 | 1 | 1 | 10480 | 393 | CG4538 | | 3.750e-02 | -3.28 | DDE\_3 | pfam domains | PF13358 | 1 | 1 | 10480 | 393 | CG14478 | | 3.750e-02 | -3.28 | CDK5\_activator | pfam domains | PF03261 | 1 | 1 | 10480 | 393 | Cdk5alpha | | 3.750e-02 | -3.28 | TBP-binding | pfam domains | PF09247 | 1 | 1 | 10480 | 393 | Taf1 | | 3.750e-02 | -3.28 | RGS-like | pfam domains | PF09128 | 1 | 1 | 10480 | 393 | RhoGEF2 | | 3.750e-02 | -3.28 | GLTSCR1 | pfam domains | PF15249 | 1 | 1 | 10480 | 393 | CG11873 | | 3.750e-02 | -3.28 | Cmyb\_C | pfam domains | PF09316 | 1 | 1 | 10480 | 393 | Myb | | 3.750e-02 | -3.28 | DUF3395 | pfam domains | PF11875 | 1 | 1 | 10480 | 393 | CG8531 | | 3.750e-02 | -3.28 | SMK-1 | pfam domains | PF04802 | 1 | 1 | 10480 | 393 | flfl | | 3.750e-02 | -3.28 | INT\_SG\_DDX\_CT\_C | pfam domains | PF15300 | 1 | 1 | 10480 | 393 | IntS6 | | 3.750e-02 | -3.28 | Fcf2 | pfam domains | PF08698 | 1 | 1 | 10480 | 393 | CG1142 | | 3.750e-02 | -3.28 | PAE | pfam domains | PF03283 | 1 | 1 | 10480 | 393 | Notum | | 3.750e-02 | -3.28 | NUDE\_C | pfam domains | PF04880 | 1 | 1 | 10480 | 393 | nudE | | 3.750e-02 | -3.28 | GST\_N\_4 | pfam domains | PF17172 | 1 | 1 | 10480 | 393 | fax | | 3.750e-02 | -3.28 | Ribosomal\_S4e | pfam domains | PF00900 | 1 | 1 | 10480 | 393 | RpS4 | | 3.750e-02 | -3.28 | Pan3\_PK | pfam domains | PF18101 | 1 | 1 | 10480 | 393 | CG11486 | | 3.750e-02 | -3.28 | Phtf-FEM1B\_bdg | pfam domains | PF12129 | 1 | 1 | 10480 | 393 | phtf | | 3.750e-02 | -3.28 | SNase | pfam domains | PF00565 | 1 | 1 | 10480 | 393 | Tudor-SN | | 3.750e-02 | -3.28 | Lipin\_mid | pfam domains | PF16876 | 1 | 1 | 10480 | 393 | Lpin | | 3.750e-02 | -3.28 | DUF4976 | pfam domains | PF16347 | 1 | 1 | 10480 | 393 | CG14291 | | 3.750e-02 | -3.28 | PAN\_4 | pfam domains | PF14295 | 1 | 1 | 10480 | 393 | tyn | | 3.750e-02 | -3.28 | MoaE | pfam domains | PF02391 | 1 | 1 | 10480 | 393 | Mocs2B | | 3.750e-02 | -3.28 | MamL-1 | pfam domains | PF09596 | 1 | 1 | 10480 | 393 | mam | | 3.750e-02 | -3.28 | TF\_AP-2 | pfam domains | PF03299 | 1 | 1 | 10480 | 393 | TfAP-2 | | 3.750e-02 | -3.28 | DUF3591 | pfam domains | PF12157 | 1 | 1 | 10480 | 393 | Taf1 | | 3.750e-02 | -3.28 | zf-CCHC\_6 | pfam domains | PF15288 | 1 | 1 | 10480 | 393 | Taf1 | | 3.750e-02 | -3.28 | MIEAP | pfam domains | PF16026 | 1 | 1 | 10480 | 393 | CG11498 | | 3.750e-02 | -3.28 | CCDC50\_N | pfam domains | PF15295 | 1 | 1 | 10480 | 393 | CG10283 | | 3.750e-02 | -3.28 | ULD | pfam domains | PF16534 | 1 | 1 | 10480 | 393 | dve | | 3.750e-02 | -3.28 | RS4NT | pfam domains | PF08071 | 1 | 1 | 10480 | 393 | RpS4 | | 3.750e-02 | -3.28 | Mtf2\_C | pfam domains | PF14061 | 1 | 1 | 10480 | 393 | Pcl | | 3.750e-02 | -3.28 | hnRNP\_Q\_AcD | pfam domains | PF18360 | 1 | 1 | 10480 | 393 | Syp | | 3.750e-02 | -3.28 | GDA1\_CD39 | pfam domains | PF01150 | 1 | 1 | 10480 | 393 | NTPase | | 3.750e-02 | -3.28 | BAF250\_C | pfam domains | PF12031 | 1 | 1 | 10480 | 393 | osa | | 3.750e-02 | -3.28 | Dscam\_C | pfam domains | PF12355 | 1 | 1 | 10480 | 393 | Dscam1 | | 3.750e-02 | -3.28 | PEX-1N | pfam domains | PF09262 | 1 | 1 | 10480 | 393 | Pex1 | | 3.750e-02 | -3.28 | STAT\_bind | pfam domains | PF02864 | 1 | 1 | 10480 | 393 | Stat92E | | 3.750e-02 | -3.28 | NKAIN | pfam domains | PF05640 | 1 | 1 | 10480 | 393 | NKAIN | | 3.750e-02 | -3.28 | NODP | pfam domains | PF07684 | 1 | 1 | 10480 | 393 | N | | 3.750e-02 | -3.28 | Lipin\_N | pfam domains | PF04571 | 1 | 1 | 10480 | 393 | Lpin | | 3.750e-02 | -3.28 | ClpB\_D2-small | pfam domains | PF10431 | 1 | 1 | 10480 | 393 | CG4538 | | 3.750e-02 | -3.28 | STAT\_alpha | pfam domains | PF01017 | 1 | 1 | 10480 | 393 | Stat92E | | 3.750e-02 | -3.28 | TRPM\_tetra | pfam domains | PF16519 | 1 | 1 | 10480 | 393 | Trpm | | 3.750e-02 | -3.28 | LSDAT\_euk | pfam domains | PF18139 | 1 | 1 | 10480 | 393 | Trpm | | 3.750e-02 | -3.28 | ARGLU | pfam domains | PF15346 | 1 | 1 | 10480 | 393 | CG31712 | | 3.750e-02 | -3.28 | SPT2 | pfam domains | PF08243 | 1 | 1 | 10480 | 393 | CG5815 | | 3.750e-02 | -3.28 | Fra10Ac1 | pfam domains | PF09725 | 1 | 1 | 10480 | 393 | CG16890 | | 3.750e-02 | -3.28 | PHAT | pfam domains | PF09246 | 1 | 1 | 10480 | 393 | smg | | 3.750e-02 | -3.28 | FAM60A | pfam domains | PF15396 | 1 | 1 | 10480 | 393 | CG44774 | | 3.750e-02 | -3.28 | C1-set | pfam domains | PF07654 | 1 | 1 | 10480 | 393 | CG6044 | | 3.750e-02 | -3.28 | DUF4588 | pfam domains | PF15251 | 1 | 1 | 10480 | 393 | CG4768 | | 3.750e-02 | -3.28 | SKA2 | pfam domains | PF16740 | 1 | 1 | 10480 | 393 | CG17778 | | 3.750e-02 | -3.28 | STAT\_int | pfam domains | PF02865 | 1 | 1 | 10480 | 393 | Stat92E | | 3.750e-02 | -3.28 | BORA\_N | pfam domains | PF15280 | 1 | 1 | 10480 | 393 | bora | | 3.750e-02 | -3.28 | RIB43A | pfam domains | PF05914 | 1 | 1 | 10480 | 393 | CG7264 | | 3.750e-02 | -3.28 | IMPDH | pfam domains | PF00478 | 1 | 1 | 10480 | 393 | ras | | 3.750e-02 | -3.28 | Pellino | pfam domains | PF04710 | 1 | 1 | 10480 | 393 | Pli | | 3.750e-02 | -3.28 | Sin3a\_C | pfam domains | PF16879 | 1 | 1 | 10480 | 393 | Sin3A | | 3.750e-02 | -3.28 | Hint | pfam domains | PF01079 | 1 | 1 | 10480 | 393 | hh | | 3.750e-02 | -3.28 | LNS2 | pfam domains | PF08235 | 1 | 1 | 10480 | 393 | Lpin | | 3.750e-02 | -3.28 | RHS\_repeat | pfam domains | PF05593 | 1 | 1 | 10480 | 393 | Ten-m | | 3.750e-02 | -3.28 | SAM\_decarbox | pfam domains | PF01536 | 1 | 1 | 10480 | 393 | SamDC | | 3.750e-02 | -3.28 | DUF3498 | pfam domains | PF12004 | 1 | 1 | 10480 | 393 | CG42684 | | 3.750e-02 | -3.28 | TNF | pfam domains | PF00229 | 1 | 1 | 10480 | 393 | egr | | 3.750e-02 | -3.28 | PAH | pfam domains | PF02671 | 1 | 1 | 10480 | 393 | Sin3A | | 3.750e-02 | -3.28 | Tmpp129 | pfam domains | PF10272 | 1 | 1 | 10480 | 393 | CG14646 | | 3.750e-02 | -3.28 | Tmemb\_cc2 | pfam domains | PF10267 | 1 | 1 | 10480 | 393 | Dmtn | | 3.750e-02 | -3.28 | SNRNP27 | pfam domains | PF08648 | 1 | 1 | 10480 | 393 | ytr | | 3.750e-02 | -3.28 | Argos | pfam domains | PF11581 | 1 | 1 | 10480 | 393 | aos | | 3.753e-02 | -3.28 | rhythmic behavior | biological process | GO:0007622 | 74 | 6 | 12345 | 414 | disco,ITP,qsm,Bx,tara,S6kII | | 3.759e-02 | -3.28 | GSE15330\_LYMPHOID\_MULTIPOTENT\_VS\_GRANULOCYTE\_MONOCYTE\_PROGENITOR\_IKAROS\_KO\_UP | MSigDB lists | GSE15330\_LYMPHOID\_MULTIPOTENT\_VS\_GRANULOCYTE\_MONOCYTE\_PROGENITOR\_IKAROS\_KO\_UP | 86 | 7 | 4161 | 153 | CG6752,Cyp1,Mitofilin,CG3961,ap,CycE,Arl4 | | 3.760e-02 | -3.28 | Ska2\_N | interpro domains | IPR042091 | 1 | 1 | 11383 | 428 | CG17778 | | 3.760e-02 | -3.28 | STAT\_N\_sf | interpro domains | IPR036535 | 1 | 1 | 11383 | 428 | Stat92E | | 3.760e-02 | -3.28 | PEX-N\_psi\_beta-barrel | interpro domains | IPR015342 | 1 | 1 | 11383 | 428 | Pex1 | | 3.760e-02 | -3.28 | S-AdoMet\_deCO2ase\_core | interpro domains | IPR016067 | 1 | 1 | 11383 | 428 | SamDC | | 3.760e-02 | -3.28 | Ribosomal\_S6\_kinase\_II | interpro domains | IPR016239 | 1 | 1 | 11383 | 428 | S6kII | | 3.760e-02 | -3.28 | MACPF | interpro domains | IPR020864 | 1 | 1 | 11383 | 428 | tsl | | 3.760e-02 | -3.28 | Staphylococal\_nuclease\_OB-fold | interpro domains | IPR016071 | 1 | 1 | 11383 | 428 | Tudor-SN | | 3.760e-02 | -3.28 | Ribosomal\_S4e\_central\_region | interpro domains | IPR013845 | 1 | 1 | 11383 | 428 | RpS4 | | 3.760e-02 | -3.28 | TF\_AP2\_C | interpro domains | IPR013854 | 1 | 1 | 11383 | 428 | TfAP-2 | | 3.760e-02 | -3.28 | SOCS7\_SOCS | interpro domains | IPR037346 | 1 | 1 | 11383 | 428 | Socs16D | | 3.760e-02 | -3.28 | Ribosomal\_S4e\_central\_sf | interpro domains | IPR038237 | 1 | 1 | 11383 | 428 | RpS4 | | 3.760e-02 | -3.28 | DUF625 | interpro domains | IPR006887 | 1 | 1 | 11383 | 428 | flfl | | 3.760e-02 | -3.28 | Ect2 | interpro domains | IPR026817 | 1 | 1 | 11383 | 428 | pbl | | 3.760e-02 | -3.28 | Notch | interpro domains | IPR008297 | 1 | 1 | 11383 | 428 | N | | 3.760e-02 | -3.28 | Hint\_dom\_sf | interpro domains | IPR036844 | 1 | 1 | 11383 | 428 | hh | | 3.760e-02 | -3.28 | C-myb\_C | interpro domains | IPR015395 | 1 | 1 | 11383 | 428 | Myb | | 3.760e-02 | -3.28 | Aurora\_borealis\_protein | interpro domains | IPR023252 | 1 | 1 | 11383 | 428 | bora | | 3.760e-02 | -3.28 | Dally | interpro domains | IPR031187 | 1 | 1 | 11383 | 428 | dally | | 3.760e-02 | -3.28 | Tc1-like\_DDE\_dom | interpro domains | IPR038717 | 1 | 1 | 11383 | 428 | CG14478 | | 3.760e-02 | -3.28 | IMP\_DH/GMP\_Rdtase\_CS | interpro domains | IPR015875 | 1 | 1 | 11383 | 428 | ras | | 3.760e-02 | -3.28 | HnRNP\_R/Q\_splicing\_fac | interpro domains | IPR006535 | 1 | 1 | 11383 | 428 | Syp | | 3.760e-02 | -3.28 | PLipase\_B-like | interpro domains | IPR007000 | 1 | 1 | 11383 | 428 | lama | | 3.760e-02 | -3.28 | PAE/NOTUM | interpro domains | IPR004963 | 1 | 1 | 11383 | 428 | Notum | | 3.760e-02 | -3.28 | RNA\_pol\_I\_trans\_ini\_fac\_RRN3 | interpro domains | IPR007991 | 1 | 1 | 11383 | 428 | Tif-IA | | 3.760e-02 | -3.28 | ATG11 | interpro domains | IPR040040 | 1 | 1 | 11383 | 428 | Atg17 | | 3.760e-02 | -3.28 | Prkrip1 | interpro domains | IPR009548 | 1 | 1 | 11383 | 428 | CG8441 | | 3.760e-02 | -3.28 | Dscam\_C | interpro domains | IPR021012 | 1 | 1 | 11383 | 428 | Dscam1 | | 3.760e-02 | -3.28 | RSK\_N | interpro domains | IPR041906 | 1 | 1 | 11383 | 428 | S6kII | | 3.760e-02 | -3.28 | CCDC50\_N | interpro domains | IPR029311 | 1 | 1 | 11383 | 428 | CG10283 | | 3.760e-02 | -3.28 | DUF4976 | interpro domains | IPR032506 | 1 | 1 | 11383 | 428 | CG14291 | | 3.760e-02 | -3.28 | Crust\_neurohorm | interpro domains | IPR031098 | 1 | 1 | 11383 | 428 | ITP | | 3.760e-02 | -3.28 | SHARP\_RRM4 | interpro domains | IPR034175 | 1 | 1 | 11383 | 428 | spen | | 3.760e-02 | -3.28 | CIP2A | interpro domains | IPR042510 | 1 | 1 | 11383 | 428 | CG12702 | | 3.760e-02 | -3.28 | DUF4588 | interpro domains | IPR029196 | 1 | 1 | 11383 | 428 | CG4768 | | 3.760e-02 | -3.28 | Phosph\_kin\_gamma | interpro domains | IPR002291 | 1 | 1 | 11383 | 428 | PhKgamma | | 3.760e-02 | -3.28 | FAM60A | interpro domains | IPR026065 | 1 | 1 | 11383 | 428 | CG44774 | | 3.760e-02 | -3.28 | Fcf2\_C | interpro domains | IPR014810 | 1 | 1 | 11383 | 428 | CG1142 | | 3.760e-02 | -3.28 | TF\_IIS-typ | interpro domains | IPR035100 | 1 | 1 | 11383 | 428 | TfIIS | | 3.760e-02 | -3.28 | PHTF1/2 | interpro domains | IPR039775 | 1 | 1 | 11383 | 428 | phtf | | 3.760e-02 | -3.28 | Ecdystd\_rcpt | interpro domains | IPR003069 | 1 | 1 | 11383 | 428 | EcR | | 3.760e-02 | -3.28 | NR\_LBD\_EcR | interpro domains | IPR041889 | 1 | 1 | 11383 | 428 | EcR | | 3.760e-02 | -3.28 | BAF250\_C | interpro domains | IPR033388 | 1 | 1 | 11383 | 428 | osa | | 3.760e-02 | -3.28 | Atg11\_C | interpro domains | IPR019460 | 1 | 1 | 11383 | 428 | Atg17 | | 3.760e-02 | -3.28 | TNF\_dom | interpro domains | IPR006052 | 1 | 1 | 11383 | 428 | egr | | 3.760e-02 | -3.28 | Tumour\_necrosis\_fac-like\_dom | interpro domains | IPR008983 | 1 | 1 | 11383 | 428 | egr | | 3.760e-02 | -3.28 | Mt-IM\_prot\_Mitofilin | interpro domains | IPR019133 | 1 | 1 | 11383 | 428 | Mitofilin | | 3.760e-02 | -3.28 | CDK5\_activator | interpro domains | IPR004944 | 1 | 1 | 11383 | 428 | Cdk5alpha | | 3.760e-02 | -3.28 | Smaug\_SAM | interpro domains | IPR037634 | 1 | 1 | 11383 | 428 | smg | | 3.760e-02 | -3.28 | STX1 | interpro domains | IPR028669 | 1 | 1 | 11383 | 428 | Syx1A | | 3.760e-02 | -3.28 | Jagged/Serrate | interpro domains | IPR026219 | 1 | 1 | 11383 | 428 | Ser | | 3.760e-02 | -3.28 | NPC1-like | interpro domains | IPR004765 | 1 | 1 | 11383 | 428 | Npc1a | | 3.760e-02 | -3.28 | DALRD3 | interpro domains | IPR037380 | 1 | 1 | 11383 | 428 | CG8097 | | 3.760e-02 | -3.28 | DnaJ-like\_C11\_C | interpro domains | IPR024586 | 1 | 1 | 11383 | 428 | CG8531 | | 3.760e-02 | -3.28 | GDA1\_CD39\_NTPase | interpro domains | IPR000407 | 1 | 1 | 11383 | 428 | NTPase | | 3.760e-02 | -3.28 | PKHA4/5/6/7\_PH | interpro domains | IPR040392 | 1 | 1 | 11383 | 428 | CG34383 | | 3.760e-02 | -3.28 | Clp\_protease\_ATP-bd\_su\_ClpX | interpro domains | IPR004487 | 1 | 1 | 11383 | 428 | CG4538 | | 3.760e-02 | -3.28 | Neural\_cell\_adh | interpro domains | IPR009138 | 1 | 1 | 11383 | 428 | Fas2 | | 3.760e-02 | -3.28 | PH\_Phafin | interpro domains | IPR037871 | 1 | 1 | 11383 | 428 | rush | | 3.760e-02 | -3.28 | FOG | interpro domains | IPR039746 | 1 | 1 | 11383 | 428 | ush | | 3.760e-02 | -3.28 | TFIID\_sub1\_DUF3591 | interpro domains | IPR022591 | 1 | 1 | 11383 | 428 | Taf1 | | 3.760e-02 | -3.28 | S-AdoMet\_deCO2ase\_CS | interpro domains | IPR018166 | 1 | 1 | 11383 | 428 | SamDC | | 3.760e-02 | -3.28 | INT\_SG\_DDX\_CT\_C | interpro domains | IPR029307 | 1 | 1 | 11383 | 428 | IntS6 | | 3.760e-02 | -3.28 | DUF3523 | interpro domains | IPR021911 | 1 | 1 | 11383 | 428 | bor | | 3.760e-02 | -3.28 | Silence\_cplx\_Nase-comp\_TudorSN | interpro domains | IPR016685 | 1 | 1 | 11383 | 428 | Tudor-SN | | 3.760e-02 | -3.28 | LIPIN | interpro domains | IPR026058 | 1 | 1 | 11383 | 428 | Lpin | | 3.760e-02 | -3.28 | Hedgehog | interpro domains | IPR001657 | 1 | 1 | 11383 | 428 | hh | | 3.760e-02 | -3.28 | BAF\_prot | interpro domains | IPR004122 | 1 | 1 | 11383 | 428 | baf | | 3.760e-02 | -3.28 | STAT\_TF\_alpha | interpro domains | IPR013800 | 1 | 1 | 11383 | 428 | Stat92E | | 3.760e-02 | -3.28 | C16orf72-like | interpro domains | IPR040308 | 1 | 1 | 11383 | 428 | CG4768 | | 3.760e-02 | -3.28 | MIEAP | interpro domains | IPR026169 | 1 | 1 | 11383 | 428 | CG11498 | | 3.760e-02 | -3.28 | WAC | interpro domains | IPR038867 | 1 | 1 | 11383 | 428 | wcy | | 3.760e-02 | -3.28 | MYO18A | interpro domains | IPR031244 | 1 | 1 | 11383 | 428 | Mhcl | | 3.760e-02 | -3.28 | MUG/TDG | interpro domains | IPR015637 | 1 | 1 | 11383 | 428 | Thd1 | | 3.760e-02 | -3.28 | Sin3-like | interpro domains | IPR039774 | 1 | 1 | 11383 | 428 | Sin3A | | 3.760e-02 | -3.28 | NUDE | interpro domains | IPR033494 | 1 | 1 | 11383 | 428 | nudE | | 3.760e-02 | -3.28 | Clp\_ATPase\_C | interpro domains | IPR019489 | 1 | 1 | 11383 | 428 | CG4538 | | 3.760e-02 | -3.28 | Chromatin\_SPT2 | interpro domains | IPR013256 | 1 | 1 | 11383 | 428 | CG5815 | | 3.760e-02 | -3.28 | RHOGAP6/36 | interpro domains | IPR037863 | 1 | 1 | 11383 | 428 | RhoGAP102A | | 3.760e-02 | -3.28 | R3HC1/R3HCL | interpro domains | IPR039884 | 1 | 1 | 11383 | 428 | CG2162 | | 3.760e-02 | -3.28 | FAX | interpro domains | IPR026928 | 1 | 1 | 11383 | 428 | fax | | 3.760e-02 | -3.28 | ULD | interpro domains | IPR032392 | 1 | 1 | 11383 | 428 | dve | | 3.760e-02 | -3.28 | Tax1-binding\_p3 | interpro domains | IPR017268 | 1 | 1 | 11383 | 428 | CG3402 | | 3.760e-02 | -3.28 | Caskin1/2\_SAM\_2 | interpro domains | IPR035498 | 1 | 1 | 11383 | 428 | ckn | | 3.760e-02 | -3.28 | Na/K-Atpase\_Interacting | interpro domains | IPR008516 | 1 | 1 | 11383 | 428 | NKAIN | | 3.760e-02 | -3.28 | MBD5/MBD6 | interpro domains | IPR037385 | 1 | 1 | 11383 | 428 | sba | | 3.760e-02 | -3.28 | Apical\_junction | interpro domains | IPR038825 | 1 | 1 | 11383 | 428 | CG42748 | | 3.760e-02 | -3.28 | Srp40\_C | interpro domains | IPR007718 | 1 | 1 | 11383 | 428 | Nopp140 | | 3.760e-02 | -3.28 | KOW\_RPS4 | interpro domains | IPR041982 | 1 | 1 | 11383 | 428 | RpS4 | | 3.760e-02 | -3.28 | BAF\_sf | interpro domains | IPR036617 | 1 | 1 | 11383 | 428 | baf | | 3.760e-02 | -3.28 | TRPM\_SLOG | interpro domains | IPR041491 | 1 | 1 | 11383 | 428 | Trpm | | 3.760e-02 | -3.28 | Pase-1\_reg-su\_3B/C/D\_met | interpro domains | IPR017434 | 1 | 1 | 11383 | 428 | Gbs-70E | | 3.760e-02 | -3.28 | WDR47 | interpro domains | IPR040067 | 1 | 1 | 11383 | 428 | CG2812 | | 3.760e-02 | -3.28 | GSCR1\_dom | interpro domains | IPR015671 | 1 | 1 | 11383 | 428 | CG11873 | | 3.760e-02 | -3.28 | MyoXV\_FERM\_C | interpro domains | IPR041795 | 1 | 1 | 11383 | 428 | Myo10A | | 3.760e-02 | -3.28 | MAP3K12\_MAP3K13 | interpro domains | IPR017419 | 1 | 1 | 11383 | 428 | wnd | | 3.760e-02 | -3.28 | Lipin\_mid | interpro domains | IPR031703 | 1 | 1 | 11383 | 428 | Lpin | | 3.760e-02 | -3.28 | Optomoror-blind\_N | interpro domains | IPR021101 | 1 | 1 | 11383 | 428 | bi | | 3.760e-02 | -3.28 | Lipin\_N | interpro domains | IPR007651 | 1 | 1 | 11383 | 428 | Lpin | | 3.760e-02 | -3.28 | TAF1\_animal | interpro domains | IPR011177 | 1 | 1 | 11383 | 428 | Taf1 | | 3.760e-02 | -3.28 | PHAT\_dom\_sf | interpro domains | IPR037093 | 1 | 1 | 11383 | 428 | smg | | 3.760e-02 | -3.28 | TAFII-230\_TBP-bd\_sf | interpro domains | IPR036741 | 1 | 1 | 11383 | 428 | Taf1 | | 3.760e-02 | -3.28 | Akap1 | interpro domains | IPR033104 | 1 | 1 | 11383 | 428 | spoon | | 3.760e-02 | -3.28 | LNS2 | interpro domains | IPR013209 | 1 | 1 | 11383 | 428 | Lpin | | 3.760e-02 | -3.28 | Notch\_NODP\_dom | interpro domains | IPR011656 | 1 | 1 | 11383 | 428 | N | | 3.760e-02 | -3.28 | PABP\_1234 | interpro domains | IPR006515 | 1 | 1 | 11383 | 428 | pAbp | | 3.760e-02 | -3.28 | SHARP\_RRM3 | interpro domains | IPR034174 | 1 | 1 | 11383 | 428 | spen | | 3.760e-02 | -3.28 | S-AdoMet\_decarboxylase | interpro domains | IPR001985 | 1 | 1 | 11383 | 428 | SamDC | | 3.760e-02 | -3.28 | Ig\_C1-set | interpro domains | IPR003597 | 1 | 1 | 11383 | 428 | CG6044 | | 3.760e-02 | -3.28 | Znf-CCHC\_6 | interpro domains | IPR041670 | 1 | 1 | 11383 | 428 | Taf1 | | 3.760e-02 | -3.28 | DUF3498 | interpro domains | IPR021887 | 1 | 1 | 11383 | 428 | CG42684 | | 3.760e-02 | -3.28 | FOX1\_RRM | interpro domains | IPR034237 | 1 | 1 | 11383 | 428 | Rbfox1 | | 3.760e-02 | -3.28 | PTPN3/4\_FERM\_C | interpro domains | IPR041783 | 1 | 1 | 11383 | 428 | Ptpmeg | | 3.760e-02 | -3.28 | Intein\_N | interpro domains | IPR006141 | 1 | 1 | 11383 | 428 | hh | | 3.760e-02 | -3.28 | CCDC50 | interpro domains | IPR039303 | 1 | 1 | 11383 | 428 | CG10283 | | 3.760e-02 | -3.28 | Crust\_neurohorm\_sf | interpro domains | IPR035957 | 1 | 1 | 11383 | 428 | ITP | | 3.760e-02 | -3.28 | TRPM\_tetra | interpro domains | IPR032415 | 1 | 1 | 11383 | 428 | Trpm | | 3.760e-02 | -3.28 | DUF3669\_Znf | interpro domains | IPR022137 | 1 | 1 | 11383 | 428 | CG8929 | | 3.760e-02 | -3.28 | AIP3-like\_C | interpro domains | IPR022782 | 1 | 1 | 11383 | 428 | CG32809 | | 3.760e-02 | -3.28 | GK5\_FGGY | interpro domains | IPR037444 | 1 | 1 | 11383 | 428 | CG1271 | | 3.760e-02 | -3.28 | STAT\_TF\_prot\_interaction | interpro domains | IPR013799 | 1 | 1 | 11383 | 428 | Stat92E | | 3.760e-02 | -3.28 | Clarin | interpro domains | IPR026748 | 1 | 1 | 11383 | 428 | CG1103 | | 3.760e-02 | -3.28 | 40S\_S4\_C | interpro domains | IPR032277 | 1 | 1 | 11383 | 428 | RpS4 | | 3.760e-02 | -3.28 | TF\_homeodomain\_male | interpro domains | IPR021980 | 1 | 1 | 11383 | 428 | phtf | | 3.760e-02 | -3.28 | Mag2/Rnf10 | interpro domains | IPR039739 | 1 | 1 | 11383 | 428 | CG12099 | | 3.760e-02 | -3.28 | DUF3454\_notch | interpro domains | IPR024600 | 1 | 1 | 11383 | 428 | N | | 3.760e-02 | -3.28 | Pan3\_PK | interpro domains | IPR041332 | 1 | 1 | 11383 | 428 | CG11486 | | 3.760e-02 | -3.28 | BICRA | interpro domains | IPR038842 | 1 | 1 | 11383 | 428 | CG11873 | | 3.760e-02 | -3.28 | MYSc\_Myo15 | interpro domains | IPR036057 | 1 | 1 | 11383 | 428 | Myo10A | | 3.760e-02 | -3.28 | SNX27\_PX | interpro domains | IPR037833 | 1 | 1 | 11383 | 428 | CG32758 | | 3.760e-02 | -3.28 | Ca/CaM-dep\_prot\_kinase-assoc | interpro domains | IPR013543 | 1 | 1 | 11383 | 428 | CaMKII | | 3.760e-02 | -3.28 | Pellino\_fam | interpro domains | IPR006800 | 1 | 1 | 11383 | 428 | Pli | | 3.760e-02 | -3.28 | TFSII | interpro domains | IPR006289 | 1 | 1 | 11383 | 428 | TfIIS | | 3.760e-02 | -3.28 | LIMR | interpro domains | IPR008075 | 1 | 1 | 11383 | 428 | lili | | 3.760e-02 | -3.28 | BAF250/Osa | interpro domains | IPR021906 | 1 | 1 | 11383 | 428 | osa | | 3.760e-02 | -3.28 | Neuroggenic\_mastermind-like\_N | interpro domains | IPR019082 | 1 | 1 | 11383 | 428 | mam | | 3.760e-02 | -3.28 | IMP\_DH | interpro domains | IPR005990 | 1 | 1 | 11383 | 428 | ras | | 3.760e-02 | -3.28 | NUDE\_dom | interpro domains | IPR006964 | 1 | 1 | 11383 | 428 | nudE | | 3.760e-02 | -3.28 | BAZ2A/B\_Bromo | interpro domains | IPR037374 | 1 | 1 | 11383 | 428 | tou | | 3.760e-02 | -3.28 | Tyr\_kinase\_EGF/ERB/XmrK\_rcpt | interpro domains | IPR016245 | 1 | 1 | 11383 | 428 | Egfr | | 3.760e-02 | -3.28 | NFAT | interpro domains | IPR008366 | 1 | 1 | 11383 | 428 | NFAT | | 3.760e-02 | -3.28 | Sin3\_C | interpro domains | IPR031693 | 1 | 1 | 11383 | 428 | Sin3A | | 3.760e-02 | -3.28 | MYSc\_Myo18 | interpro domains | IPR036064 | 1 | 1 | 11383 | 428 | Mhcl | | 3.760e-02 | -3.28 | SATB\_ULD\_sf | interpro domains | IPR038224 | 1 | 1 | 11383 | 428 | dve | | 3.760e-02 | -3.28 | SATB1/SATB2 | interpro domains | IPR039673 | 1 | 1 | 11383 | 428 | dve | | 3.760e-02 | -3.28 | MIEAP\_C | interpro domains | IPR031981 | 1 | 1 | 11383 | 428 | CG11498 | | 3.760e-02 | -3.28 | STAT | interpro domains | IPR001217 | 1 | 1 | 11383 | 428 | Stat92E | | 3.760e-02 | -3.28 | RIB43A | interpro domains | IPR008805 | 1 | 1 | 11383 | 428 | CG7264 | | 3.760e-02 | -3.28 | SOCS7\_SH2 | interpro domains | IPR035866 | 1 | 1 | 11383 | 428 | Socs16D | | 3.760e-02 | -3.28 | Pex1 | interpro domains | IPR025653 | 1 | 1 | 11383 | 428 | Pex1 | | 3.760e-02 | -3.28 | Hyperglycemic | interpro domains | IPR001166 | 1 | 1 | 11383 | 428 | ITP | | 3.760e-02 | -3.28 | STAT\_TF\_DNA-bd | interpro domains | IPR013801 | 1 | 1 | 11383 | 428 | Stat92E | | 3.760e-02 | -3.28 | RNA\_Pol\_CTD\_Phosphatase | interpro domains | IPR040078 | 1 | 1 | 11383 | 428 | CG5830 | | 3.760e-02 | -3.28 | Notch\_NOD\_dom | interpro domains | IPR010660 | 1 | 1 | 11383 | 428 | N | | 3.760e-02 | -3.28 | EYA\_dom\_metazoan | interpro domains | IPR042577 | 1 | 1 | 11383 | 428 | eya | | 3.760e-02 | -3.28 | Gastrin\_rcpt | interpro domains | IPR000314 | 1 | 1 | 11383 | 428 | CCKLR-17D3 | | 3.760e-02 | -3.28 | Fcf2/DNTTIP2 | interpro domains | IPR039883 | 1 | 1 | 11383 | 428 | CG1142 | | 3.760e-02 | -3.28 | Interferon\_reg\_fac2-bd1\_2\_Znf | interpro domains | IPR022750 | 1 | 1 | 11383 | 428 | Pits | | 3.760e-02 | -3.28 | TAF\_II\_230-bd | interpro domains | IPR009067 | 1 | 1 | 11383 | 428 | Taf1 | | 3.760e-02 | -3.28 | M-RIP\_PH | interpro domains | IPR039597 | 1 | 1 | 11383 | 428 | osp | | 3.760e-02 | -3.28 | Srp40/Nopp140 | interpro domains | IPR039191 | 1 | 1 | 11383 | 428 | Nopp140 | | 3.760e-02 | -3.28 | Mtf2\_C\_dom | interpro domains | IPR025894 | 1 | 1 | 11383 | 428 | Pcl | | 3.760e-02 | -3.28 | SIN3A | interpro domains | IPR037969 | 1 | 1 | 11383 | 428 | Sin3A | | 3.760e-02 | -3.28 | ANKFN1 | interpro domains | IPR039269 | 1 | 1 | 11383 | 428 | wake | | 3.760e-02 | -3.28 | Grb2 | interpro domains | IPR030219 | 1 | 1 | 11383 | 428 | drk | | 3.760e-02 | -3.28 | ARGLU1 | interpro domains | IPR033371 | 1 | 1 | 11383 | 428 | CG31712 | | 3.760e-02 | -3.28 | TAF1 | interpro domains | IPR040240 | 1 | 1 | 11383 | 428 | Taf1 | | 3.760e-02 | -3.28 | HDAC\_interact\_dom | interpro domains | IPR013194 | 1 | 1 | 11383 | 428 | Sin3A | | 3.760e-02 | -3.28 | DUF4801 | interpro domains | IPR032060 | 1 | 1 | 11383 | 428 | ocm | | 3.760e-02 | -3.28 | Cpo\_RRM | interpro domains | IPR034788 | 1 | 1 | 11383 | 428 | cpo | | 3.760e-02 | -3.28 | PAN3 | interpro domains | IPR030844 | 1 | 1 | 11383 | 428 | CG11486 | | 3.760e-02 | -3.28 | EYA\_dom\_sf | interpro domains | IPR038102 | 1 | 1 | 11383 | 428 | eya | | 3.760e-02 | -3.28 | Gprotein\_alpha\_12/13 | interpro domains | IPR000469 | 1 | 1 | 11383 | 428 | cta | | 3.760e-02 | -3.28 | Ribosomal\_S4e | interpro domains | IPR000876 | 1 | 1 | 11383 | 428 | RpS4 | | 3.760e-02 | -3.28 | Ribosomal\_S4e\_N | interpro domains | IPR013843 | 1 | 1 | 11383 | 428 | RpS4 | | 3.760e-02 | -3.28 | TEX28/TMCC | interpro domains | IPR019394 | 1 | 1 | 11383 | 428 | Dmtn | | 3.760e-02 | -3.28 | Shootin-1 | interpro domains | IPR024849 | 1 | 1 | 11383 | 428 | CG33111 | | 3.760e-02 | -3.28 | PHAT\_dom | interpro domains | IPR015327 | 1 | 1 | 11383 | 428 | smg | | 3.760e-02 | -3.28 | MoaE\_sf | interpro domains | IPR036563 | 1 | 1 | 11383 | 428 | Mocs2B | | 3.760e-02 | -3.28 | hnRNP\_Q\_AcD | interpro domains | IPR041337 | 1 | 1 | 11383 | 428 | Syp | | 3.760e-02 | -3.28 | ZF\_CCHC\_FOG | interpro domains | IPR034731 | 1 | 1 | 11383 | 428 | ush | | 3.760e-02 | -3.28 | EYA\_fam | interpro domains | IPR028472 | 1 | 1 | 11383 | 428 | eya | | 3.760e-02 | -3.28 | Crust\_neurhormone\_CS | interpro domains | IPR018251 | 1 | 1 | 11383 | 428 | ITP | | 3.760e-02 | -3.28 | Myb-like\_TF | interpro domains | IPR015495 | 1 | 1 | 11383 | 428 | Myb | | 3.760e-02 | -3.28 | Hint\_dom\_N | interpro domains | IPR003587 | 1 | 1 | 11383 | 428 | hh | | 3.760e-02 | -3.28 | RGS-like\_dom | interpro domains | IPR015212 | 1 | 1 | 11383 | 428 | RhoGEF2 | | 3.760e-02 | -3.28 | MPP5\_SH3 | interpro domains | IPR035601 | 1 | 1 | 11383 | 428 | sdt | | 3.760e-02 | -3.28 | ZN608/ZN609 | interpro domains | IPR040010 | 1 | 1 | 11383 | 428 | sbb | | 3.760e-02 | -3.28 | EYA\_dom | interpro domains | IPR006545 | 1 | 1 | 11383 | 428 | eya | | 3.760e-02 | -3.28 | APPBP2 | interpro domains | IPR042476 | 1 | 1 | 11383 | 428 | Pat1 | | 3.760e-02 | -3.28 | CHERP | interpro domains | IPR039037 | 1 | 1 | 11383 | 428 | scaf6 | | 3.760e-02 | -3.28 | TM129 | interpro domains | IPR018801 | 1 | 1 | 11383 | 428 | CG14646 | | 3.760e-02 | -3.28 | FAM122 | interpro domains | IPR026716 | 1 | 1 | 11383 | 428 | CG4497 | | 3.760e-02 | -3.28 | Hint\_dom\_C | interpro domains | IPR003586 | 1 | 1 | 11383 | 428 | hh | | 3.760e-02 | -3.28 | RHS\_repeat | interpro domains | IPR031325 | 1 | 1 | 11383 | 428 | Ten-m | | 3.760e-02 | -3.28 | Twist-related | interpro domains | IPR015789 | 1 | 1 | 11383 | 428 | twi | | 3.760e-02 | -3.28 | Ribosomal\_S4e\_N\_CS | interpro domains | IPR018199 | 1 | 1 | 11383 | 428 | RpS4 | | 3.760e-02 | -3.28 | 2OGFeDO\_noxygenase\_dom | interpro domains | IPR024779 | 1 | 1 | 11383 | 428 | Tet | | 3.760e-02 | -3.28 | PTBP1/2/3\_RRM2 | interpro domains | IPR034798 | 1 | 1 | 11383 | 428 | heph | | 3.760e-02 | -3.28 | WSTF\_Acf1\_Cbp146 | interpro domains | IPR013136 | 1 | 1 | 11383 | 428 | Acf | | 3.760e-02 | -3.28 | STAT\_TF\_coiled-coil | interpro domains | IPR015988 | 1 | 1 | 11383 | 428 | Stat92E | | 3.760e-02 | -3.28 | TF\_AP2 | interpro domains | IPR004979 | 1 | 1 | 11383 | 428 | TfAP-2 | | 3.760e-02 | -3.28 | ATAD3 | interpro domains | IPR039188 | 1 | 1 | 11383 | 428 | bor | | 3.760e-02 | -3.28 | Pokkuri | interpro domains | IPR033077 | 1 | 1 | 11383 | 428 | aop | | 3.760e-02 | -3.28 | Caskin1/2\_SAM\_1 | interpro domains | IPR035497 | 1 | 1 | 11383 | 428 | ckn | | 3.760e-02 | -3.28 | Hedgehog\_Hint | interpro domains | IPR001767 | 1 | 1 | 11383 | 428 | hh | | 3.760e-02 | -3.28 | SPRY\_RNF123 | interpro domains | IPR035773 | 1 | 1 | 11383 | 428 | CG6752 | | 3.760e-02 | -3.28 | UDP-sugar\_pyrophosphorylase | interpro domains | IPR039741 | 1 | 1 | 11383 | 428 | mmy | | 3.760e-02 | -3.28 | STAT\_TF\_DNA-bd\_N | interpro domains | IPR012345 | 1 | 1 | 11383 | 428 | Stat92E | | 3.760e-02 | -3.28 | Succ\_CoA\_betaG\_euk | interpro domains | IPR034722 | 1 | 1 | 11383 | 428 | ScsbetaG | | 3.760e-02 | -3.28 | RhoGAP\_ARAP | interpro domains | IPR037858 | 1 | 1 | 11383 | 428 | RhoGAP15B | | 3.760e-02 | -3.28 | Citrate\_synthase\_euk | interpro domains | IPR010109 | 1 | 1 | 11383 | 428 | kdn | | 3.760e-02 | -3.28 | CCDC85 | interpro domains | IPR019359 | 1 | 1 | 11383 | 428 | CG17265 | | 3.760e-02 | -3.28 | BIP\_NBD | interpro domains | IPR042050 | 1 | 1 | 11383 | 428 | Hsc70-3 | | 3.760e-02 | -3.28 | Arp2 | interpro domains | IPR027306 | 1 | 1 | 11383 | 428 | Arp2 | | 3.760e-02 | -3.28 | Hedgehog\_sig/DD-Pept\_Zn-bd\_sf | interpro domains | IPR009045 | 1 | 1 | 11383 | 428 | hh | | 3.760e-02 | -3.28 | DUF4757 | interpro domains | IPR031865 | 1 | 1 | 11383 | 428 | smash | | 3.760e-02 | -3.28 | IMP\_DH\_GMPRt | interpro domains | IPR001093 | 1 | 1 | 11383 | 428 | ras | | 3.760e-02 | -3.28 | Mopterin\_biosynth\_MoaE | interpro domains | IPR003448 | 1 | 1 | 11383 | 428 | Mocs2B | | 3.760e-02 | -3.28 | Hedgehog\_signalling\_dom | interpro domains | IPR000320 | 1 | 1 | 11383 | 428 | hh | | 3.760e-02 | -3.28 | Tyr\_Pase\_non-rcpt\_typ-3/4 | interpro domains | IPR012151 | 1 | 1 | 11383 | 428 | Ptpmeg | | 3.760e-02 | -3.28 | SNRNP27 | interpro domains | IPR013957 | 1 | 1 | 11383 | 428 | ytr | | 3.760e-02 | -3.28 | MPP5 | interpro domains | IPR031185 | 1 | 1 | 11383 | 428 | sdt | | 3.760e-02 | -3.28 | Argos | interpro domains | IPR021633 | 1 | 1 | 11383 | 428 | aos | | 3.760e-02 | -3.28 | SNX27\_FERM-like\_dom | interpro domains | IPR037827 | 1 | 1 | 11383 | 428 | CG32758 | | 3.760e-02 | -3.28 | SHARP\_RRM2 | interpro domains | IPR034173 | 1 | 1 | 11383 | 428 | spen | | 3.760e-02 | -3.28 | Folate-sensitive\_fs\_Fra10Ac1 | interpro domains | IPR019129 | 1 | 1 | 11383 | 428 | CG16890 | | 3.760e-02 | -3.28 | DUF1084 | interpro domains | IPR009457 | 1 | 1 | 11383 | 428 | CG2991 | | 3.760e-02 | -3.28 | ZF\_CCHC\_HIVEP | interpro domains | IPR034729 | 1 | 1 | 11383 | 428 | shn | | 3.760e-02 | -3.28 | TET1/2/3 | interpro domains | IPR040175 | 1 | 1 | 11383 | 428 | Tet | | 3.760e-02 | -3.28 | Acf1\_Bromo | interpro domains | IPR037325 | 1 | 1 | 11383 | 428 | Acf | | 3.760e-02 | -3.28 | FJX1/FJ | interpro domains | IPR024868 | 1 | 1 | 11383 | 428 | fj | | 3.760e-02 | -3.28 | MOCS2B\_euk | interpro domains | IPR028888 | 1 | 1 | 11383 | 428 | Mocs2B | | 3.761e-02 | -3.28 | GSE9006\_TYPE\_1\_DIABETES\_AT\_DX\_VS\_4MONTH\_POST\_DX\_PBMC\_UP | MSigDB lists | GSE9006\_TYPE\_1\_DIABETES\_AT\_DX\_VS\_4MONTH\_POST\_DX\_PBMC\_UP | 68 | 6 | 4161 | 153 | pAbp,Arl4,Hsc70-3,mmy,Cyp1,Arp2 | | 3.761e-02 | -3.28 | KAYO\_AGING\_MUSCLE\_UP | MSigDB lists | KAYO\_AGING\_MUSCLE\_UP | 68 | 6 | 4161 | 153 | TfAP-2,Syx1A,SamDC,fne,Fas2,smal | | 3.761e-02 | -3.28 | WGGAATGY\_TEF1\_Q6 | MSigDB lists | WGGAATGY\_TEF1\_Q6 | 68 | 6 | 4161 | 153 | Pat1,Smox,Acsl,Sin3A,Trpm,Tsp26A | | 3.773e-02 | -3.28 | Growth hormone receptor signaling | REACTOME pathways | R-DME-982772 | 1 | 1 | 3923 | 148 | Stat92E | | 3.773e-02 | -3.28 | STING mediated induction of host immune responses | REACTOME pathways | R-DME-1834941 | 1 | 1 | 3923 | 148 | Stat92E | | 3.773e-02 | -3.28 | Interleukin-4 and 13 signaling | REACTOME pathways | R-DME-6785807 | 1 | 1 | 3923 | 148 | Stat92E | | 3.773e-02 | -3.28 | Epigenetic regulation of gene expression | REACTOME pathways | R-DME-212165 | 1 | 1 | 3923 | 148 | Thd1 | | 3.773e-02 | -3.28 | Class B/2 (Secretin family receptors) | REACTOME pathways | R-DME-373080 | 1 | 1 | 3923 | 148 | Cirl | | 3.773e-02 | -3.28 | Hydrolysis of LPC | REACTOME pathways | R-DME-1483115 | 1 | 1 | 3923 | 148 | lama | | 3.773e-02 | -3.28 | PKA-mediated phosphorylation of key metabolic factors | REACTOME pathways | R-DME-163358 | 1 | 1 | 3923 | 148 | Mondo | | 3.773e-02 | -3.28 | NCAM1 interactions | REACTOME pathways | R-DME-419037 | 1 | 1 | 3923 | 148 | Fas2 | | 3.773e-02 | -3.28 | Interleukin-6 signaling | REACTOME pathways | R-DME-1059683 | 1 | 1 | 3923 | 148 | Stat92E | | 3.773e-02 | -3.28 | STAT6-mediated induction of chemokines | REACTOME pathways | R-DME-3249367 | 1 | 1 | 3923 | 148 | Stat92E | | 3.773e-02 | -3.28 | Interferon alpha/beta signaling | REACTOME pathways | R-DME-909733 | 1 | 1 | 3923 | 148 | Stat92E | | 3.773e-02 | -3.28 | Phosphate bond hydrolysis by NTPDase proteins | REACTOME pathways | R-DME-8850843 | 1 | 1 | 3923 | 148 | NTPase | | 3.773e-02 | -3.28 | mRNA Editing: C to U Conversion | REACTOME pathways | R-DME-72200 | 1 | 1 | 3923 | 148 | Syp | | 3.773e-02 | -3.28 | Formation of the Editosome | REACTOME pathways | R-DME-75094 | 1 | 1 | 3923 | 148 | Syp | | 3.773e-02 | -3.28 | Interleukin-10 signaling | REACTOME pathways | R-DME-6783783 | 1 | 1 | 3923 | 148 | Stat92E | | 3.773e-02 | -3.28 | TET1,2,3 and TDG demethylate DNA | REACTOME pathways | R-DME-5221030 | 1 | 1 | 3923 | 148 | Thd1 | | 3.773e-02 | -3.28 | Interleukin-6 family signaling | REACTOME pathways | R-DME-6783589 | 1 | 1 | 3923 | 148 | Stat92E | | 3.795e-02 | -3.27 | Pp4-19C (Protein phosphatase 19C) | protein interactions | 45031 | 17 | 3 | 9204 | 412 | ttk,Doa,flfl | | 3.795e-02 | -3.27 | Nrt (Neurotactin) | protein interactions | 39873 | 17 | 3 | 9204 | 412 | Ubx,N,ci | | 3.795e-02 | -3.27 | net (net) | protein interactions | 45339 | 17 | 3 | 9204 | 412 | osa,S,Egfr | | 3.795e-02 | -3.27 | CG8173 (uncharacterized protein) | protein interactions | 32753 | 17 | 3 | 9204 | 412 | CG17265,betaTub56D,CG11076 | | 3.795e-02 | -3.27 | Mad1 (Mitotic arrest-deficient 1) | protein interactions | 35954 | 17 | 3 | 9204 | 412 | CG7264,E(spl)mdelta-HLH,CG32369 | | 3.795e-02 | -3.27 | CG3815 (uncharacterized protein) | protein interactions | 31571 | 17 | 3 | 9204 | 412 | Sin3A,l(3)mbt,CtBP | | 3.795e-02 | -3.27 | CRIF (CR6-interacting factor) | protein interactions | 40395 | 17 | 3 | 9204 | 412 | CG9541,stg,CG17265 | | 3.795e-02 | -3.27 | Acn (Acinus) | protein interactions | 35173 | 17 | 3 | 9204 | 412 | Egfr,S6kII,N | | 3.797e-02 | -3.27 | GATA4\_Q3 | MSigDB lists | GATA4\_Q3 | 35 | 4 | 4161 | 153 | fne,SamDC,slow,pbl | | 3.797e-02 | -3.27 | GSE7831\_1H\_VS\_4H\_INFLUENZA\_STIM\_PDC\_DN | MSigDB lists | GSE7831\_1H\_VS\_4H\_INFLUENZA\_STIM\_PDC\_DN | 35 | 4 | 4161 | 153 | N,Hydr2,CG6752,CG5522 | | 3.797e-02 | -3.27 | BOQUEST\_STEM\_CELL\_UP | MSigDB lists | BOQUEST\_STEM\_CELL\_UP | 35 | 4 | 4161 | 153 | smal,LRP1,dlp,if | | 3.797e-02 | -3.27 | GSE36476\_CTRL\_VS\_TSST\_ACT\_40H\_MEMORY\_CD4\_TCELL\_YOUNG\_UP | MSigDB lists | GSE36476\_CTRL\_VS\_TSST\_ACT\_40H\_MEMORY\_CD4\_TCELL\_YOUNG\_UP | 35 | 4 | 4161 | 153 | Arl4,CG5522,Ptpmeg,nudE | | 3.797e-02 | -3.27 | GSE22886\_NAIVE\_BCELL\_VS\_BLOOD\_PLASMA\_CELL\_UP | MSigDB lists | GSE22886\_NAIVE\_BCELL\_VS\_BLOOD\_PLASMA\_CELL\_UP | 35 | 4 | 4161 | 153 | CG12547,CG5522,Smox,CG41099 | | 3.797e-02 | -3.27 | GSE37533\_PPARG1\_FOXP3\_VS\_PPARG2\_FOXP3\_TRANSDUCED\_CD4\_TCELL\_PIOGLITAZONE\_TREATED\_DN | MSigDB lists | GSE37533\_PPARG1\_FOXP3\_VS\_PPARG2\_FOXP3\_TRANSDUCED\_CD4\_TCELL\_PIOGLITAZONE\_TREATED\_DN | 35 | 4 | 4161 | 153 | ft,Ppa,CG3961,Fbxl7 | | 3.797e-02 | -3.27 | GSE21927\_BALBC\_VS\_C57BL6\_MONOCYTE\_TUMOR\_DN | MSigDB lists | GSE21927\_BALBC\_VS\_C57BL6\_MONOCYTE\_TUMOR\_DN | 51 | 5 | 4161 | 153 | rush,ScsbetaG,SamDC,CG1271,Arp2 | | 3.797e-02 | -3.27 | TAL1ALPHAE47\_01 | MSigDB lists | TAL1ALPHAE47\_01 | 51 | 5 | 4161 | 153 | trn,if,Smox,Arl4,Tudor-SN | | 3.797e-02 | -3.27 | GSE2770\_IL12\_VS\_TGFB\_AND\_IL12\_TREATED\_ACT\_CD4\_TCELL\_48H\_UP | MSigDB lists | GSE2770\_IL12\_VS\_TGFB\_AND\_IL12\_TREATED\_ACT\_CD4\_TCELL\_48H\_UP | 51 | 5 | 4161 | 153 | CG1271,l(3)mbt,Smox,Syx1A,LManII | | 3.797e-02 | -3.27 | MODULE\_6 | MSigDB lists | MODULE\_6 | 51 | 5 | 4161 | 153 | CG43313,Ank,so,LRP1,ScpX | | 3.797e-02 | -3.27 | GSE32423\_IL7\_VS\_IL4\_MEMORY\_CD8\_TCELL\_UP | MSigDB lists | GSE32423\_IL7\_VS\_IL4\_MEMORY\_CD8\_TCELL\_UP | 51 | 5 | 4161 | 153 | Snx3,CG11367,hts,S6kII,JIL-1 | | 3.797e-02 | -3.27 | GSE34006\_WT\_VS\_A2AR\_KO\_TREG\_UP | MSigDB lists | GSE34006\_WT\_VS\_A2AR\_KO\_TREG\_UP | 51 | 5 | 4161 | 153 | betaTub56D,heph,CycE,Cyp1,CG2201 | | 3.807e-02 | -3.27 | MyTH4 | smart domains | SM00139 | 6 | 2 | 5086 | 276 | Myo10A,RhoGAP93B | | 3.807e-02 | -3.27 | FU | smart domains | SM00261 | 6 | 2 | 5086 | 276 | Fur1,Egfr | | 3.807e-02 | -3.27 | DWB | smart domains | SM00524 | 6 | 2 | 5086 | 276 | Smox,Dad | | 3.811e-02 | -3.27 | REACTOME\_SIGNALING\_BY\_GPCR | MSigDB lists | REACTOME\_SIGNALING\_BY\_GPCR | 105 | 8 | 4161 | 153 | CG34384,Rho1,drk,Rac2,hh,cta,S6kII,pbl | | 3.817e-02 | -3.27 | GO\_PROTEIN\_LOCALIZATION | MSigDB lists | GO\_PROTEIN\_LOCALIZATION | 664 | 33 | 4161 | 153 | CG32758,CG14646,CG45050,ScpX,LRP1,RpS4,if,Syx1A,rush,N,Cnx99A,Tim17b,Rac2,hh,CG9855,so,Hsc70Cb,Arl4,Npc1a,Ank,Rho1,sdt,Pat1,Snx3,pbl,Ykt6,mgl,CG5742,Sin3A,Hsc70-3,emp,mle,Tsp26A | | 3.827e-02 | -3.26 | bam (bag of marbles) | protein interactions | 43038 | 42 | 5 | 9204 | 412 | udd,tkv,lili,stg,Doa | | 3.827e-02 | -3.26 | ftz-f1 (ftz transcription factor 1) | protein interactions | 40045 | 42 | 5 | 9204 | 412 | ci,ps,Arp2,EcR,gus | | 3.827e-02 | -3.26 | Psc (Posterior sex combs) | protein interactions | 36431 | 42 | 5 | 9204 | 412 | psq,Psc,ci,Fas2,hh | | 3.827e-02 | -3.26 | su(Hw) (suppressor of Hairy wing) | protein interactions | 41740 | 42 | 5 | 9204 | 412 | Bx,dbr,ap,osa,Ubx | | 3.834e-02 | -3.26 | INNEXIN | prints domains | PR01262 | 8 | 2 | 2185 | 88 | Inx2,Inx3 | | 3.834e-02 | -3.26 | ETSDOMAIN | prints domains | PR00454 | 8 | 2 | 2185 | 88 | aop,pnt | | 3.836e-02 | -3.26 | positive regulation of developmental growth | biological process | GO:0048639 | 94 | 7 | 12345 | 414 | Hipk,tkv,Arp2,ey,dpp,tsl,Egfr | | 3.836e-02 | -3.26 | rhythmic process | biological process | GO:0048511 | 94 | 7 | 12345 | 414 | disco,ITP,so,qsm,Bx,tara,S6kII | | 3.838e-02 | -3.26 | Ankyrin\_rpt-contain\_sf | interpro domains | IPR036770 | 84 | 7 | 11383 | 428 | CG41099,CG10011,Prosap,N,CG5742,wake,Ank | | 3.838e-02 | -3.26 | Ankyrin\_rpt-contain\_dom | interpro domains | IPR020683 | 84 | 7 | 11383 | 428 | Prosap,CG41099,CG10011,Ank,wake,CG5742,N | | 3.877e-02 | -3.25 | tum (tumbleweed) | protein interactions | 36538 | 29 | 4 | 9204 | 412 | pbl,CycE,aos,Rho1 | | 3.877e-02 | -3.25 | cos (costa) | protein interactions | 35653 | 29 | 4 | 9204 | 412 | ci,hh,CG13366,Gmap | | 3.877e-02 | -3.25 | Sirt4 (Sirtuin 4) | protein interactions | 31480 | 29 | 4 | 9204 | 412 | CG11486,mub,Pits,uzip | | 3.877e-02 | -3.25 | rb (ruby) | protein interactions | 31381 | 29 | 4 | 9204 | 412 | l(3)neo38,aos,CG3919,mmy | | 3.877e-02 | -3.25 | Traf4 (TNF-receptor-associated factor 4) | protein interactions | 33638 | 29 | 4 | 9204 | 412 | CG9541,RhoGEF2,CycE,E(spl)m4-BFM | | 3.877e-02 | -3.25 | fz2 (frizzled 2) | protein interactions | 40090 | 29 | 4 | 9204 | 412 | Hipk,otk2,dally,mam | | 3.877e-02 | -3.25 | sog (short gastrulation) | protein interactions | 32498 | 29 | 4 | 9204 | 412 | scb,tkv,if,dpp | | 3.877e-02 | -3.25 | how (held out wings) | protein interactions | 42596 | 29 | 4 | 9204 | 412 | stg,Hydr2,dpp,if | | 3.909e-02 | -3.24 | negative regulation of translation | biological process | GO:0017148 | 56 | 5 | 12345 | 414 | smg,elav,bin3,Rbfox1,heph | | 3.909e-02 | -3.24 | peptidyl-serine phosphorylation | biological process | GO:0018105 | 56 | 5 | 12345 | 414 | mbt,Asator,Hipk,JIL-1,Tlk | | 3.909e-02 | -3.24 | regulation of endocytosis | biological process | GO:0030100 | 56 | 5 | 12345 | 414 | sdt,comm,drk,mgl,Hip1 | | 3.946e-02 | -3.23 | TGCTGCT\_MIR15A\_MIR16\_MIR15B\_MIR195\_MIR424\_MIR497 | MSigDB lists | TGCTGCT\_MIR15A\_MIR16\_MIR15B\_MIR195\_MIR424\_MIR497 | 145 | 10 | 4161 | 153 | Pcl,CycE,CG1271,CG2812,Smox,gus,TfAP-2,Syx1A,S6kII,Arp2 | | 3.951e-02 | -3.23 | response to stress | biological process | GO:0006950 | 1136 | 49 | 12345 | 414 | Thd1,Toll-7,Atg17,Imp,Drat,cnc,jing,Etl1,MFS14,eya,PhKgamma,AGO2,wnd,RhoGEF2,CG11873,hiw,lz,stwl,Cnx99A,Karl,Stat92E,Pli,ctrip,Utx,bbg,egr,Mmp2,NFAT,spen,Hph,mbt,Mrp4,Rho1,Pcl,Cdk5alpha,scb,pbl,Tl,mof,PEK,CaMKII,Lpin,flfl,ImpL2,SamDC,Rac2,Hsc70-3,N,hang | | 3.989e-02 | -3.22 | GO\_POSITIVE\_REGULATION\_OF\_HEMOPOIESIS | MSigDB lists | GO\_POSITIVE\_REGULATION\_OF\_HEMOPOIESIS | 21 | 3 | 4161 | 153 | CG11367,hh,Ser | | 3.989e-02 | -3.22 | GSE18791\_CTRL\_VS\_NEWCASTLE\_VIRUS\_DC\_14H\_DN | MSigDB lists | GSE18791\_CTRL\_VS\_NEWCASTLE\_VIRUS\_DC\_14H\_DN | 21 | 3 | 4161 | 153 | Imp,Tlk,mgl | | 3.989e-02 | -3.22 | AR\_Q2 | MSigDB lists | AR\_Q2 | 21 | 3 | 4161 | 153 | LRP1,Pat1,CG2812 | | 3.989e-02 | -3.22 | chr1p32 | MSigDB lists | chr1p32 | 21 | 3 | 4161 | 153 | ScpX,CG5742,pAbp | | 3.989e-02 | -3.22 | GSE15750\_WT\_VS\_TRAF6KO\_DAY10\_EFF\_CD8\_TCELL\_DN | MSigDB lists | GSE15750\_WT\_VS\_TRAF6KO\_DAY10\_EFF\_CD8\_TCELL\_DN | 21 | 3 | 4161 | 153 | CG34384,Acsl,Dscam1 | | 3.989e-02 | -3.22 | TOMLINS\_PROSTATE\_CANCER\_UP | MSigDB lists | TOMLINS\_PROSTATE\_CANCER\_UP | 21 | 3 | 4161 | 153 | Cnx99A,Tudor-SN,mmy | | 3.989e-02 | -3.22 | KEGG\_PPAR\_SIGNALING\_PATHWAY | MSigDB lists | KEGG\_PPAR\_SIGNALING\_PATHWAY | 21 | 3 | 4161 | 153 | Acsl,CG3961,ScpX | | 3.989e-02 | -3.22 | GSE29614\_CTRL\_VS\_DAY3\_TIV\_FLU\_VACCINE\_PBMC\_DN | MSigDB lists | GSE29614\_CTRL\_VS\_DAY3\_TIV\_FLU\_VACCINE\_PBMC\_DN | 21 | 3 | 4161 | 153 | Hsc70-3,CG6330,CG34384 | | 3.989e-02 | -3.22 | GO\_CANONICAL\_WNT\_SIGNALING\_PATHWAY | MSigDB lists | GO\_CANONICAL\_WNT\_SIGNALING\_PATHWAY | 21 | 3 | 4161 | 153 | hh,CG11367,Utx | | 3.989e-02 | -3.22 | GSE18791\_CTRL\_VS\_NEWCASTLE\_VIRUS\_DC\_18H\_DN | MSigDB lists | GSE18791\_CTRL\_VS\_NEWCASTLE\_VIRUS\_DC\_18H\_DN | 21 | 3 | 4161 | 153 | mgl,CG34384,Utx | | 3.989e-02 | -3.22 | GO\_CARDIAC\_VENTRICLE\_DEVELOPMENT | MSigDB lists | GO\_CARDIAC\_VENTRICLE\_DEVELOPMENT | 21 | 3 | 4161 | 153 | N,mgl,Ser | | 3.989e-02 | -3.22 | FOSTER\_KDM1A\_TARGETS\_UP | MSigDB lists | FOSTER\_KDM1A\_TARGETS\_UP | 21 | 3 | 4161 | 153 | l(3)mbt,smal,ras | | 3.989e-02 | -3.22 | GSE20727\_ROS\_INH\_VS\_ROS\_INH\_AND\_DNFB\_ALLERGEN\_TREATED\_DC\_DN | MSigDB lists | GSE20727\_ROS\_INH\_VS\_ROS\_INH\_AND\_DNFB\_ALLERGEN\_TREATED\_DC\_DN | 21 | 3 | 4161 | 153 | Trpm,nAChRalpha4,Tsp26A | | 3.995e-02 | -3.22 | cell-cell adhesion mediator activity | molecular function | GO:0098632 | 10 | 2 | 12176 | 396 | Fas2,ImpL2 | | 3.995e-02 | -3.22 | cell adhesion mediator activity | molecular function | GO:0098631 | 10 | 2 | 12176 | 396 | ImpL2,Fas2 | | 3.995e-02 | -3.22 | beta-catenin binding | molecular function | GO:0008013 | 10 | 2 | 12176 | 396 | CadN,Hipk | | 3.999e-02 | -3.22 | GSE5589\_WT\_VS\_IL6\_KO\_LPS\_STIM\_MACROPHAGE\_45MIN\_UP | MSigDB lists | GSE5589\_WT\_VS\_IL6\_KO\_LPS\_STIM\_MACROPHAGE\_45MIN\_UP | 69 | 6 | 4161 | 153 | CG11030,ScsbetaG,emp,Snx3,smal,LRP1 | | 3.999e-02 | -3.22 | GCM\_RAN | MSigDB lists | GCM\_RAN | 69 | 6 | 4161 | 153 | Pat1,Acsl,CG32758,kdn,Fas2,CG5522 | | 3.999e-02 | -3.22 | GSE2770\_IL12\_ACT\_VS\_ACT\_CD4\_TCELL\_48H\_UP | MSigDB lists | GSE2770\_IL12\_ACT\_VS\_ACT\_CD4\_TCELL\_48H\_UP | 69 | 6 | 4161 | 153 | Cnx99A,ras,par-1,Tudor-SN,Npc1a,CG6752 | | 4.019e-02 | -3.21 | Equilibrium Signalling Pathway of Differentiated Cells | WikiPathways | WP3653 | 7 | 2 | 499 | 24 | hh,Stat92E | | 4.027e-02 | -3.21 | PHOSPHBKNASE | prints domains | PR01049 | 1 | 1 | 2185 | 88 | PhKgamma | | 4.027e-02 | -3.21 | AURORABORA | prints domains | PR02038 | 1 | 1 | 2185 | 88 | bora | | 4.027e-02 | -3.21 | JAGGEDFAMILY | prints domains | PR02059 | 1 | 1 | 2185 | 88 | Ser | | 4.027e-02 | -3.21 | GPROTEINA12 | prints domains | PR00440 | 1 | 1 | 2185 | 88 | cta | | 4.027e-02 | -3.21 | AP2TNSCPFCT | prints domains | PR01748 | 1 | 1 | 2185 | 88 | TfAP-2 | | 4.027e-02 | -3.21 | HYPRGLYCEMIC | prints domains | PR00550 | 1 | 1 | 2185 | 88 | ITP | | 4.027e-02 | -3.21 | NUCFACTORATC | prints domains | PR01789 | 1 | 1 | 2185 | 88 | NFAT | | 4.027e-02 | -3.21 | GASTRINR | prints domains | PR00527 | 1 | 1 | 2185 | 88 | CCKLR-17D3 | | 4.027e-02 | -3.21 | SONICHHOG | prints domains | PR00632 | 1 | 1 | 2185 | 88 | hh | | 4.027e-02 | -3.21 | LIPOCALINIMR | prints domains | PR01692 | 1 | 1 | 2185 | 88 | lili | | 4.027e-02 | -3.21 | ECDYSTEROIDR | prints domains | PR01283 | 1 | 1 | 2185 | 88 | EcR | | 4.027e-02 | -3.21 | LNOTCHREPEAT | prints domains | PR01452 | 1 | 1 | 2185 | 88 | N | | 4.027e-02 | -3.21 | 4JOINTEDBOX1 | prints domains | PR02072 | 1 | 1 | 2185 | 88 | fj | | 4.027e-02 | -3.21 | NCAMFAMILY | prints domains | PR01838 | 1 | 1 | 2185 | 88 | Fas2 | | 4.033e-02 | -3.21 | positive regulation of circadian rhythm | biological process | GO:0042753 | 23 | 3 | 12345 | 414 | EcR,ctrip,wake | | 4.033e-02 | -3.21 | positive regulation of ERK1 and ERK2 cascade | biological process | GO:0070374 | 23 | 3 | 12345 | 414 | spi,drk,Egfr | | 4.033e-02 | -3.21 | regulation of receptor-mediated endocytosis | biological process | GO:0048259 | 23 | 3 | 12345 | 414 | Hip1,comm,drk | | 4.033e-02 | -3.21 | trachea development | biological process | GO:0060438 | 23 | 3 | 12345 | 414 | crb,flz,pnt | | 4.046e-02 | -3.21 | rau (rau) | protein interactions | 33814 | 87 | 8 | 9204 | 412 | rau,sty,CG41099,CG3386,pnt,hang,Acf,Sox102F | | 4.071e-02 | -3.20 | positive regulation of protein serine/threonine kinase activity | biological process | GO:0071902 | 39 | 4 | 12345 | 414 | Rho1,mbt,wnd,Cdk5alpha | | 4.081e-02 | -3.20 | BIOCARTA\_G1\_PATHWAY | MSigDB lists | BIOCARTA\_G1\_PATHWAY | 9 | 2 | 4161 | 153 | CycE,Smox | | 4.081e-02 | -3.20 | BORLAK\_LIVER\_CANCER\_EGF\_UP | MSigDB lists | BORLAK\_LIVER\_CANCER\_EGF\_UP | 9 | 2 | 4161 | 153 | Rho1,pbl | | 4.081e-02 | -3.20 | GO\_POSITIVE\_REGULATION\_OF\_SMALL\_GTPASE\_MEDIATED\_SIGNAL\_TRANSDUCTION | MSigDB lists | GO\_POSITIVE\_REGULATION\_OF\_SMALL\_GTPASE\_MEDIATED\_SIGNAL\_TRANSDUCTION | 9 | 2 | 4161 | 153 | CaMKII,Rac2 | | 4.081e-02 | -3.20 | GO\_CEREBRAL\_CORTEX\_RADIALLY\_ORIENTED\_CELL\_MIGRATION | MSigDB lists | GO\_CEREBRAL\_CORTEX\_RADIALLY\_ORIENTED\_CELL\_MIGRATION | 9 | 2 | 4161 | 153 | nudE,Rac2 | | 4.081e-02 | -3.20 | GO\_CHANNEL\_INHIBITOR\_ACTIVITY | MSigDB lists | GO\_CHANNEL\_INHIBITOR\_ACTIVITY | 9 | 2 | 4161 | 153 | CaMKII,Syx1A | | 4.081e-02 | -3.20 | GO\_GENITALIA\_DEVELOPMENT | MSigDB lists | GO\_GENITALIA\_DEVELOPMENT | 9 | 2 | 4161 | 153 | hh,Lim1 | | 4.081e-02 | -3.20 | GO\_BICELLULAR\_TIGHT\_JUNCTION\_ASSEMBLY | MSigDB lists | GO\_BICELLULAR\_TIGHT\_JUNCTION\_ASSEMBLY | 9 | 2 | 4161 | 153 | pbl,sdt | | 4.081e-02 | -3.20 | GO\_DOPAMINERGIC\_NEURON\_DIFFERENTIATION | MSigDB lists | GO\_DOPAMINERGIC\_NEURON\_DIFFERENTIATION | 9 | 2 | 4161 | 153 | Rac2,hh | | 4.081e-02 | -3.20 | GO\_EPIDERMIS\_MORPHOGENESIS | MSigDB lists | GO\_EPIDERMIS\_MORPHOGENESIS | 9 | 2 | 4161 | 153 | N,hh | | 4.081e-02 | -3.20 | REACTOME\_NRAGE\_SIGNALS\_DEATH\_THROUGH\_JNK | MSigDB lists | REACTOME\_NRAGE\_SIGNALS\_DEATH\_THROUGH\_JNK | 9 | 2 | 4161 | 153 | pbl,Rac2 | | 4.081e-02 | -3.20 | GO\_REGULATION\_OF\_ALTERNATIVE\_MRNA\_SPLICING\_VIA\_SPLICEOSOME | MSigDB lists | GO\_REGULATION\_OF\_ALTERNATIVE\_MRNA\_SPLICING\_VIA\_SPLICEOSOME | 9 | 2 | 4161 | 153 | heph,sm | | 4.081e-02 | -3.20 | GO\_REGULATION\_OF\_TRANSCRIPTION\_REGULATORY\_REGION\_DNA\_BINDING | MSigDB lists | GO\_REGULATION\_OF\_TRANSCRIPTION\_REGULATORY\_REGION\_DNA\_BINDING | 9 | 2 | 4161 | 153 | Sin3A,ap | | 4.081e-02 | -3.20 | GO\_APOPTOTIC\_CELL\_CLEARANCE | MSigDB lists | GO\_APOPTOTIC\_CELL\_CLEARANCE | 9 | 2 | 4161 | 153 | Rac2,LRP1 | | 4.081e-02 | -3.20 | HOLLERN\_EMT\_BREAST\_TUMOR\_DN | MSigDB lists | HOLLERN\_EMT\_BREAST\_TUMOR\_DN | 9 | 2 | 4161 | 153 | Cyp1,Rhp | | 4.081e-02 | -3.20 | YRCCAKNNGNCGC\_UNKNOWN | MSigDB lists | YRCCAKNNGNCGC\_UNKNOWN | 9 | 2 | 4161 | 153 | ap,CG8531 | | 4.081e-02 | -3.20 | GO\_CORONARY\_VASCULATURE\_DEVELOPMENT | MSigDB lists | GO\_CORONARY\_VASCULATURE\_DEVELOPMENT | 9 | 2 | 4161 | 153 | N,mgl | | 4.081e-02 | -3.20 | TOMLINS\_METASTASIS\_DN | MSigDB lists | TOMLINS\_METASTASIS\_DN | 9 | 2 | 4161 | 153 | CtBP,Mrp4 | | 4.081e-02 | -3.20 | GO\_PRE\_MRNA\_BINDING | MSigDB lists | GO\_PRE\_MRNA\_BINDING | 9 | 2 | 4161 | 153 | heph,sm | | 4.081e-02 | -3.20 | GO\_SPECTRIN\_BINDING | MSigDB lists | GO\_SPECTRIN\_BINDING | 9 | 2 | 4161 | 153 | Ank,hts | | 4.081e-02 | -3.20 | CORRE\_MULTIPLE\_MYELOMA\_DN | MSigDB lists | CORRE\_MULTIPLE\_MYELOMA\_DN | 9 | 2 | 4161 | 153 | lbk,TfIIS | | 4.081e-02 | -3.20 | GO\_APOLIPOPROTEIN\_BINDING | MSigDB lists | GO\_APOLIPOPROTEIN\_BINDING | 9 | 2 | 4161 | 153 | LRP1,Cnx99A | | 4.081e-02 | -3.20 | GO\_RETROMER\_COMPLEX | MSigDB lists | GO\_RETROMER\_COMPLEX | 9 | 2 | 4161 | 153 | CG41099,CG32758 | | 4.081e-02 | -3.20 | HOQUE\_METHYLATED\_IN\_CANCER | MSigDB lists | HOQUE\_METHYLATED\_IN\_CANCER | 9 | 2 | 4161 | 153 | dpp,TfAP-2 | | 4.081e-02 | -3.20 | PID\_UPA\_UPAR\_PATHWAY | MSigDB lists | PID\_UPA\_UPAR\_PATHWAY | 9 | 2 | 4161 | 153 | Rac2,LRP1 | | 4.081e-02 | -3.20 | MODULE\_533 | MSigDB lists | MODULE\_533 | 9 | 2 | 4161 | 153 | Fas2,Syx1A | | 4.081e-02 | -3.20 | GO\_STEROID\_HORMONE\_RECEPTOR\_ACTIVITY | MSigDB lists | GO\_STEROID\_HORMONE\_RECEPTOR\_ACTIVITY | 9 | 2 | 4161 | 153 | EcR,Hydr2 | | 4.081e-02 | -3.20 | chr17q24 | MSigDB lists | chr17q24 | 9 | 2 | 4161 | 153 | cta,drk | | 4.081e-02 | -3.20 | HANN\_RESISTANCE\_TO\_BCL2\_INHIBITOR\_DN | MSigDB lists | HANN\_RESISTANCE\_TO\_BCL2\_INHIBITOR\_DN | 9 | 2 | 4161 | 153 | Rhp,CG10663 | | 4.081e-02 | -3.20 | GO\_CEREBELLAR\_PURKINJE\_CELL\_LAYER\_DEVELOPMENT | MSigDB lists | GO\_CEREBELLAR\_PURKINJE\_CELL\_LAYER\_DEVELOPMENT | 9 | 2 | 4161 | 153 | Hsc70-3,Lim1 | | 4.081e-02 | -3.20 | KEGG\_DORSO\_VENTRAL\_AXIS\_FORMATION | MSigDB lists | KEGG\_DORSO\_VENTRAL\_AXIS\_FORMATION | 9 | 2 | 4161 | 153 | drk,N | | 4.081e-02 | -3.20 | GO\_POSITIVE\_REGULATION\_OF\_SMOOTHENED\_SIGNALING\_PATHWAY | MSigDB lists | GO\_POSITIVE\_REGULATION\_OF\_SMOOTHENED\_SIGNALING\_PATHWAY | 9 | 2 | 4161 | 153 | CG9220,hh | | 4.081e-02 | -3.20 | GO\_REGULATION\_OF\_ALCOHOL\_BIOSYNTHETIC\_PROCESS | MSigDB lists | GO\_REGULATION\_OF\_ALCOHOL\_BIOSYNTHETIC\_PROCESS | 9 | 2 | 4161 | 153 | Acsl,dpp | | 4.081e-02 | -3.20 | GSE8921\_UNSTIM\_VS\_TLR1\_2\_STIM\_MONOCYTE\_12H\_UP | MSigDB lists | GSE8921\_UNSTIM\_VS\_TLR1\_2\_STIM\_MONOCYTE\_12H\_UP | 52 | 5 | 4161 | 153 | Acsl,LManII,sdt,CG14291,LRP1 | | 4.081e-02 | -3.20 | GFI1\_01 | MSigDB lists | GFI1\_01 | 52 | 5 | 4161 | 153 | CaMKII,smal,Fas2,CG9855,CG9281 | | 4.081e-02 | -3.20 | KEGG\_LYSOSOME | MSigDB lists | KEGG\_LYSOSOME | 52 | 5 | 4161 | 153 | LManII,MFS10,Npc1a,CG14291,emp | | 4.081e-02 | -3.20 | GSE6092\_B\_BURGDOFERI\_VS\_B\_BURGDORFERI\_AND\_IFNG\_STIM\_ENDOTHELIAL\_CELL\_DN | MSigDB lists | GSE6092\_B\_BURGDOFERI\_VS\_B\_BURGDORFERI\_AND\_IFNG\_STIM\_ENDOTHELIAL\_CELL\_DN | 52 | 5 | 4161 | 153 | CG3961,kdn,SamDC,CG34384,phtf | | 4.081e-02 | -3.20 | GSE46606\_DAY1\_VS\_DAY3\_CD40L\_IL2\_IL5\_STIMULATED\_IRF4HIGH\_BCELL\_DN | MSigDB lists | GSE46606\_DAY1\_VS\_DAY3\_CD40L\_IL2\_IL5\_STIMULATED\_IRF4HIGH\_BCELL\_DN | 52 | 5 | 4161 | 153 | Tpr2,hiw,Hsc70-3,nudE,Cyp1 | | 4.082e-02 | -3.20 | Pten (Phosphatase and tensin homolog) | protein interactions | 43991 | 103 | 9 | 9204 | 412 | Hsc70-3,AGO2,egr,EcR,Smox,ImpL2,CaMKII,Egfr,Syp | | 4.099e-02 | -3.19 | MODULE\_88 | MSigDB lists | MODULE\_88 | 126 | 9 | 4161 | 153 | fne,Ank,Fbxl7,ScpX,CG7264,Syx1A,TfAP-2,Arl4,Tim17b | | 4.099e-02 | -3.19 | MODULE\_55 | MSigDB lists | MODULE\_55 | 126 | 9 | 4161 | 153 | Fbxl7,Ank,fne,CG7264,ScpX,LRP1,Syx1A,TfAP-2,Tim17b | | 4.107e-02 | -3.19 | DOUGLAS\_BMI1\_TARGETS\_UP | MSigDB lists | DOUGLAS\_BMI1\_TARGETS\_UP | 146 | 10 | 4161 | 153 | Cyp1,sno,N,kdn,mle,ds,CG45050,Tpr2,ras,CG32758 | | 4.110e-02 | -3.19 | Toll Like Receptor 10 (TLR10) Cascade | REACTOME pathways | R-DME-168142 | 35 | 4 | 3923 | 148 | mle,Pli,S6kII,JIL-1 | | 4.147e-02 | -3.18 | Mpcp2 (Mitochondrial phosphate carrier protein 2) | protein interactions | 39587 | 72 | 7 | 9204 | 412 | Cnx99A,ds,Bx,bor,Acsl,ScsbetaG,Cyt-c1 | | 4.153e-02 | -3.18 | RORA2\_01 | MSigDB lists | RORA2\_01 | 36 | 4 | 4161 | 153 | hh,DIP-epsilon,Acsl,trn | | 4.153e-02 | -3.18 | AAANWWTGC\_UNKNOWN | MSigDB lists | AAANWWTGC\_UNKNOWN | 36 | 4 | 4161 | 153 | Ank,Trpm,NetB,Imp | | 4.153e-02 | -3.18 | GSE16450\_IMMATURE\_VS\_MATURE\_NEURON\_CELL\_LINE\_6H\_IFNA\_STIM\_DN | MSigDB lists | GSE16450\_IMMATURE\_VS\_MATURE\_NEURON\_CELL\_LINE\_6H\_IFNA\_STIM\_DN | 36 | 4 | 4161 | 153 | dpp,if,ras,NetB | | 4.153e-02 | -3.18 | GSE29949\_CD8\_NEG\_DC\_SPLEEN\_VS\_CD8\_POS\_DC\_SPLEEN\_UP | MSigDB lists | GSE29949\_CD8\_NEG\_DC\_SPLEEN\_VS\_CD8\_POS\_DC\_SPLEEN\_UP | 36 | 4 | 4161 | 153 | ras,betaTub56D,LRP1,Tim17b | | 4.153e-02 | -3.18 | GCM\_CALM1 | MSigDB lists | GCM\_CALM1 | 36 | 4 | 4161 | 153 | Pat1,Acsl,CG32758,Fas2 | | 4.153e-02 | -3.18 | GSE26488\_HDAC7\_KO\_VS\_VP16\_TRANSGENIC\_HDAC7\_KO\_DOUBLE\_POSITIVE\_THYMOCYTE\_UP | MSigDB lists | GSE26488\_HDAC7\_KO\_VS\_VP16\_TRANSGENIC\_HDAC7\_KO\_DOUBLE\_POSITIVE\_THYMOCYTE\_UP | 36 | 4 | 4161 | 153 | CG3097,Pli,dally,Rhp | | 4.153e-02 | -3.18 | GSE26727\_WT\_VS\_KLF2\_KO\_LPS\_STIM\_MACROPHAGE\_UP | MSigDB lists | GSE26727\_WT\_VS\_KLF2\_KO\_LPS\_STIM\_MACROPHAGE\_UP | 36 | 4 | 4161 | 153 | ras,PhKgamma,Tim17b,Cyp1 | | 4.153e-02 | -3.18 | GO\_REGULATION\_OF\_CELL\_SIZE | MSigDB lists | GO\_REGULATION\_OF\_CELL\_SIZE | 36 | 4 | 4161 | 153 | Rac2,hts,NetB,nudE | | 4.153e-02 | -3.18 | GSE21033\_1H\_VS\_12H\_POLYIC\_STIM\_DC\_UP | MSigDB lists | GSE21033\_1H\_VS\_12H\_POLYIC\_STIM\_DC\_UP | 36 | 4 | 4161 | 153 | Ppa,NTPase,Mmp2,rush | | 4.153e-02 | -3.18 | GSE6259\_FLT3L\_INDUCED\_33D1\_POS\_DC\_VS\_CD8\_TCELL\_UP | MSigDB lists | GSE6259\_FLT3L\_INDUCED\_33D1\_POS\_DC\_VS\_CD8\_TCELL\_UP | 36 | 4 | 4161 | 153 | Hydr2,pAbp,Tsp26A,CG4538 | | 4.153e-02 | -3.18 | HNF3B\_01 | MSigDB lists | HNF3B\_01 | 36 | 4 | 4161 | 153 | Pli,Sin3A,Utx,if | | 4.164e-02 | -3.18 | Signaling by TGF-beta Receptor Complex | REACTOME pathways | R-DME-170834 | 51 | 5 | 3923 | 148 | Smox,nrv2,Dad,Fur1,Rho1 | | 4.167e-02 | -3.18 | cytoskeletal protein binding | molecular function | GO:0008092 | 330 | 17 | 12176 | 396 | Ptpmeg,Hip1,nudE,nuf,Ten-m,Socs16D,smg,CG14535,Arp2,hts,crb,CG17778,Myo10A,corn,Svil,Ank,Mhcl | | 4.172e-02 | -3.18 | cell-cell junction assembly | biological process | GO:0007043 | 57 | 5 | 12345 | 414 | crb,nrv2,sdt,Lac,bou | | 4.172e-02 | -3.18 | positive regulation of reproductive process | biological process | GO:2000243 | 57 | 5 | 12345 | 414 | ttk,stg,spi,aop,Egfr | | 4.178e-02 | -3.18 | LIMK1 (LIM domain kinase 1) | protein interactions | 32207 | 43 | 5 | 9204 | 412 | RhoGAP18B,pbl,RhoGEF2,mbt,Rho1 | | 4.178e-02 | -3.18 | RhoGAP68F (Rho GTPase activating protein at 68F) | protein interactions | 39385 | 43 | 5 | 9204 | 412 | par-1,gus,Pits,heph,Fas3 | | 4.178e-02 | -3.18 | E(z) (Enhancer of zeste) | protein interactions | 39203 | 43 | 5 | 9204 | 412 | Utx,Ubx,Psc,jing,Pcl | | 4.181e-02 | -3.17 | ATGTACA\_MIR493 | MSigDB lists | ATGTACA\_MIR493 | 88 | 7 | 4161 | 153 | JIL-1,Utx,Sin3A,CG5522,dlp,CG2201,CG4538 | | 4.192e-02 | -3.17 | GO\_RESPONSE\_TO\_GROWTH\_FACTOR | MSigDB lists | GO\_RESPONSE\_TO\_GROWTH\_FACTOR | 107 | 8 | 4161 | 153 | N,Smox,wech,dpp,Lim1,heph,tkv,drk | | 4.195e-02 | -3.17 | - | gene3d domains | 2.40.50.90 | 8 | 2 | 4929 | 208 | Tudor-SN,spoon | | 4.200e-02 | -3.17 | germline ring canal | cellular component | GO:0045172 | 23 | 3 | 11940 | 407 | hts,Lpin,par-1 | | 4.201e-02 | -3.17 | fly chr91C3-91C3\|3-64 cM | chromosome location | fly chr91C3-91C3\|3-64 cM | 2 | 1 | 25010 | 531 | CG34283 | | 4.201e-02 | -3.17 | fly chr1B7-1B8\|1-0 cM | chromosome location | fly chr1B7-1B8\|1-0 cM | 2 | 1 | 25010 | 531 | elav | | 4.201e-02 | -3.17 | fly chr2C7-2C7\|1-0.5 | chromosome location | fly chr2C7-2C7\|1-0.5 | 2 | 1 | 25010 | 531 | CG4325 | | 4.201e-02 | -3.17 | fly chr95F12-95F13\|3-84 cM | chromosome location | fly chr95F12-95F13\|3-84 cM | 2 | 1 | 25010 | 531 | CG5728 | | 4.201e-02 | -3.17 | fly chr91C3-91C3\|3-64 | chromosome location | fly chr91C3-91C3\|3-64 | 2 | 1 | 25010 | 531 | CG34283 | | 4.201e-02 | -3.17 | fly chr42A9-42A12\|2-55 cM | chromosome location | fly chr42A9-42A12\|2-55 cM | 2 | 1 | 25010 | 531 | EcR | | 4.201e-02 | -3.17 | fly chr71C3-71E5\|3-43 cM | chromosome location | fly chr71C3-71E5\|3-43 cM | 2 | 1 | 25010 | 531 | AGO2 | | 4.201e-02 | -3.17 | fly chr1B7-1B8\|1-0 | chromosome location | fly chr1B7-1B8\|1-0 | 2 | 1 | 25010 | 531 | elav | | 4.201e-02 | -3.17 | fly chr16D4-16D4\|1-58 | chromosome location | fly chr16D4-16D4\|1-58 | 2 | 1 | 25010 | 531 | Socs16D | | 4.201e-02 | -3.17 | fly chr13F14-13F14\|1-52 cM | chromosome location | fly chr13F14-13F14\|1-52 cM | 2 | 1 | 25010 | 531 | Myb | | 4.201e-02 | -3.17 | fly chr36D2-36D2\|2-53 cM | chromosome location | fly chr36D2-36D2\|2-53 cM | 2 | 1 | 25010 | 531 | CadN | | 4.201e-02 | -3.17 | fly chr5A13-5A13\|1-12 cM | chromosome location | fly chr5A13-5A13\|1-12 cM | 2 | 1 | 25010 | 531 | cv | | 4.201e-02 | -3.17 | fly chr98E6-98F1\|3-97 | chromosome location | fly chr98E6-98F1\|3-97 | 2 | 1 | 25010 | 531 | Inx3 | | 4.201e-02 | -3.17 | fly chr10D5-10D5\|1-35 | chromosome location | fly chr10D5-10D5\|1-35 | 2 | 1 | 25010 | 531 | PhKgamma | | 4.201e-02 | -3.17 | fly chr27F2-27F3\|2-26 | chromosome location | fly chr27F2-27F3\|2-26 | 2 | 1 | 25010 | 531 | asRNA:CR44063 | | 4.201e-02 | -3.17 | fly chr44C2-44C2 | chromosome location | fly chr44C2-44C2 | 2 | 1 | 25010 | 531 | sisRNA:CR46360 | | 4.201e-02 | -3.17 | fly chr63B7-63B7\|3-6 | chromosome location | fly chr63B7-63B7\|3-6 | 2 | 1 | 25010 | 531 | BtbVII | | 4.201e-02 | -3.17 | fly chr42C5-42C6\|2-55 | chromosome location | fly chr42C5-42C6\|2-55 | 2 | 1 | 25010 | 531 | CG45092 | | 4.201e-02 | -3.17 | fly chr42A9-42A12\|2-55 | chromosome location | fly chr42A9-42A12\|2-55 | 2 | 1 | 25010 | 531 | EcR | | 4.201e-02 | -3.17 | fly chr43B3-43C1\|2-56 cM | chromosome location | fly chr43B3-43C1\|2-56 cM | 2 | 1 | 25010 | 531 | so | | 4.201e-02 | -3.17 | fly chr79F2-79F2\|3-47 | chromosome location | fly chr79F2-79F2\|3-47 | 2 | 1 | 25010 | 531 | ND-B14.5AL | | 4.201e-02 | -3.17 | fly chr46F1-46F2\|2-62 | chromosome location | fly chr46F1-46F2\|2-62 | 2 | 1 | 25010 | 531 | CG46319 | | 4.201e-02 | -3.17 | fly chr79F2-79F2\|3-47 cM | chromosome location | fly chr79F2-79F2\|3-47 cM | 2 | 1 | 25010 | 531 | ND-B14.5AL | | 4.201e-02 | -3.17 | fly chr7C3-7C3\|1-21 cM | chromosome location | fly chr7C3-7C3\|1-21 cM | 2 | 1 | 25010 | 531 | Ykt6 | | 4.201e-02 | -3.17 | fly chr26C4-26D1\|2-19 cM | chromosome location | fly chr26C4-26D1\|2-19 cM | 2 | 1 | 25010 | 531 | DIP-epsilon | | 4.201e-02 | -3.17 | fly chr65A1-65A1\|3-16 | chromosome location | fly chr65A1-65A1\|3-16 | 2 | 1 | 25010 | 531 | CG33993 | | 4.201e-02 | -3.17 | fly chr5C6-5C6\|1-14 | chromosome location | fly chr5C6-5C6\|1-14 | 2 | 1 | 25010 | 531 | lncRNA:CR45668 | | 4.201e-02 | -3.17 | fly chr36D2-36D2\|2-53 | chromosome location | fly chr36D2-36D2\|2-53 | 2 | 1 | 25010 | 531 | CadN | | 4.201e-02 | -3.17 | fly chr60D4-60D5\|2-107 | chromosome location | fly chr60D4-60D5\|2-107 | 2 | 1 | 25010 | 531 | ITP | | 4.201e-02 | -3.17 | fly chr92E2-92E3\|3-68 | chromosome location | fly chr92E2-92E3\|3-68 | 2 | 1 | 25010 | 531 | CG4360 | | 4.201e-02 | -3.17 | fly chr46C1-46C1\|2-61 | chromosome location | fly chr46C1-46C1\|2-61 | 2 | 1 | 25010 | 531 | CG18446 | | 4.201e-02 | -3.17 | fly chr10D5-10D5\|1-35 cM | chromosome location | fly chr10D5-10D5\|1-35 cM | 2 | 1 | 25010 | 531 | PhKgamma | | 4.201e-02 | -3.17 | fly chr7C3-7C3\|1-21 | chromosome location | fly chr7C3-7C3\|1-21 | 2 | 1 | 25010 | 531 | Ykt6 | | 4.201e-02 | -3.17 | fly chr4D6-4D6\|1-9 cM | chromosome location | fly chr4D6-4D6\|1-9 cM | 2 | 1 | 25010 | 531 | CG44774 | | 4.201e-02 | -3.17 | fly chr68A2-68A2\|3-34 cM | chromosome location | fly chr68A2-68A2\|3-34 cM | 2 | 1 | 25010 | 531 | CG43693 | | 4.201e-02 | -3.17 | fly chr4D6-4D6\|1-9 | chromosome location | fly chr4D6-4D6\|1-9 | 2 | 1 | 25010 | 531 | CG44774 | | 4.201e-02 | -3.17 | fly chr5A13-5A13\|1-12 | chromosome location | fly chr5A13-5A13\|1-12 | 2 | 1 | 25010 | 531 | cv | | 4.201e-02 | -3.17 | fly chr27F2-27F3\|2-26 cM | chromosome location | fly chr27F2-27F3\|2-26 cM | 2 | 1 | 25010 | 531 | asRNA:CR44063 | | 4.201e-02 | -3.17 | fly chr89B2-89B3\|3-58 cM | chromosome location | fly chr89B2-89B3\|3-58 cM | 2 | 1 | 25010 | 531 | Mhcl | | 4.201e-02 | -3.17 | fly chr102A3-102A3\|4-0 | chromosome location | fly chr102A3-102A3\|4-0 | 2 | 1 | 25010 | 531 | Ank | | 4.201e-02 | -3.17 | fly chr60B10-60B10\|2-106 cM | chromosome location | fly chr60B10-60B10\|2-106 cM | 2 | 1 | 25010 | 531 | enok | | 4.201e-02 | -3.17 | fly chr102A3-102A3\|4-0 cM | chromosome location | fly chr102A3-102A3\|4-0 cM | 2 | 1 | 25010 | 531 | Ank | | 4.201e-02 | -3.17 | fly chr83E8-83E8\|3-47.5 cM | chromosome location | fly chr83E8-83E8\|3-47.5 cM | 2 | 1 | 25010 | 531 | Zif | | 4.201e-02 | -3.17 | fly chr15F1-15F1\|1-57 cM | chromosome location | fly chr15F1-15F1\|1-57 cM | 2 | 1 | 25010 | 531 | lncRNA:CR44026 | | 4.201e-02 | -3.17 | fly chr43A4-43B1\|2-55 cM | chromosome location | fly chr43A4-43B1\|2-55 cM | 2 | 1 | 25010 | 531 | Dscam1 | | 4.201e-02 | -3.17 | fly chr39D1-39D1\|2-54 | chromosome location | fly chr39D1-39D1\|2-54 | 2 | 1 | 25010 | 531 | Tsp39D | | 4.201e-02 | -3.17 | fly chr65A1-65A1\|3-16 cM | chromosome location | fly chr65A1-65A1\|3-16 cM | 2 | 1 | 25010 | 531 | CG33993 | | 4.201e-02 | -3.17 | fly chr95F12-95F13\|3-84 | chromosome location | fly chr95F12-95F13\|3-84 | 2 | 1 | 25010 | 531 | CG5728 | | 4.201e-02 | -3.17 | fly chr2B6-2B7\|1-0.2 | chromosome location | fly chr2B6-2B7\|1-0.2 | 2 | 1 | 25010 | 531 | CG32809 | | 4.201e-02 | -3.17 | fly chr13F14-13F14\|1-52 | chromosome location | fly chr13F14-13F14\|1-52 | 2 | 1 | 25010 | 531 | Myb | | 4.201e-02 | -3.17 | fly chr42C5-42C6\|2-55 cM | chromosome location | fly chr42C5-42C6\|2-55 cM | 2 | 1 | 25010 | 531 | CG45092 | | 4.201e-02 | -3.17 | fly chr89B2-89B3\|3-58 | chromosome location | fly chr89B2-89B3\|3-58 | 2 | 1 | 25010 | 531 | Mhcl | | 4.201e-02 | -3.17 | fly chr98E6-98F1\|3-97 cM | chromosome location | fly chr98E6-98F1\|3-97 cM | 2 | 1 | 25010 | 531 | Inx3 | | 4.201e-02 | -3.17 | fly chr5C6-5C6\|1-14 cM | chromosome location | fly chr5C6-5C6\|1-14 cM | 2 | 1 | 25010 | 531 | lncRNA:CR45668 | | 4.201e-02 | -3.17 | fly chr39C1-39C2\|2-54 cM | chromosome location | fly chr39C1-39C2\|2-54 cM | 2 | 1 | 25010 | 531 | Mondo | | 4.201e-02 | -3.17 | fly chr26C4-26D1\|2-19 | chromosome location | fly chr26C4-26D1\|2-19 | 2 | 1 | 25010 | 531 | DIP-epsilon | | 4.201e-02 | -3.17 | fly chr2B6-2B7\|1-0.2 cM | chromosome location | fly chr2B6-2B7\|1-0.2 cM | 2 | 1 | 25010 | 531 | CG32809 | | 4.201e-02 | -3.17 | fly chr48D7-48D7\|2-65 cM | chromosome location | fly chr48D7-48D7\|2-65 cM | 2 | 1 | 25010 | 531 | otk2 | | 4.201e-02 | -3.17 | fly chr71C3-71E5\|3-43 | chromosome location | fly chr71C3-71E5\|3-43 | 2 | 1 | 25010 | 531 | AGO2 | | 4.201e-02 | -3.17 | fly chr60B10-60B10\|2-106 | chromosome location | fly chr60B10-60B10\|2-106 | 2 | 1 | 25010 | 531 | enok | | 4.201e-02 | -3.17 | fly chr57F9-57F10\|2-96 | chromosome location | fly chr57F9-57F10\|2-96 | 2 | 1 | 25010 | 531 | HmgZ | | 4.201e-02 | -3.17 | fly chr46F1-46F2\|2-62 cM | chromosome location | fly chr46F1-46F2\|2-62 cM | 2 | 1 | 25010 | 531 | CG46319 | | 4.201e-02 | -3.17 | fly chr39A2-39A3\|2-54 cM | chromosome location | fly chr39A2-39A3\|2-54 cM | 2 | 1 | 25010 | 531 | Nhe2 | | 4.201e-02 | -3.17 | fly chr43A4-43B1\|2-55 | chromosome location | fly chr43A4-43B1\|2-55 | 2 | 1 | 25010 | 531 | Dscam1 | | 4.201e-02 | -3.17 | fly chr39D1-39D1\|2-54 cM | chromosome location | fly chr39D1-39D1\|2-54 cM | 2 | 1 | 25010 | 531 | Tsp39D | | 4.201e-02 | -3.17 | fly chr43B3-43C1\|2-56 | chromosome location | fly chr43B3-43C1\|2-56 | 2 | 1 | 25010 | 531 | so | | 4.201e-02 | -3.17 | fly chr39A2-39A3\|2-54 | chromosome location | fly chr39A2-39A3\|2-54 | 2 | 1 | 25010 | 531 | Nhe2 | | 4.201e-02 | -3.17 | fly chr68A2-68A2\|3-34 | chromosome location | fly chr68A2-68A2\|3-34 | 2 | 1 | 25010 | 531 | CG43693 | | 4.201e-02 | -3.17 | fly chr83E8-83E8\|3-47.5 | chromosome location | fly chr83E8-83E8\|3-47.5 | 2 | 1 | 25010 | 531 | Zif | | 4.201e-02 | -3.17 | fly chr92E2-92E3\|3-68 cM | chromosome location | fly chr92E2-92E3\|3-68 cM | 2 | 1 | 25010 | 531 | CG4360 | | 4.201e-02 | -3.17 | fly chr16D4-16D4\|1-58 cM | chromosome location | fly chr16D4-16D4\|1-58 cM | 2 | 1 | 25010 | 531 | Socs16D | | 4.201e-02 | -3.17 | fly chr60D4-60D5\|2-107 cM | chromosome location | fly chr60D4-60D5\|2-107 cM | 2 | 1 | 25010 | 531 | ITP | | 4.201e-02 | -3.17 | fly chr57F9-57F10\|2-96 cM | chromosome location | fly chr57F9-57F10\|2-96 cM | 2 | 1 | 25010 | 531 | HmgZ | | 4.201e-02 | -3.17 | fly chr92B4-92B6\|3-67 cM | chromosome location | fly chr92B4-92B6\|3-67 cM | 2 | 1 | 25010 | 531 | CG4538 | | 4.201e-02 | -3.17 | fly chr39C1-39C2\|2-54 | chromosome location | fly chr39C1-39C2\|2-54 | 2 | 1 | 25010 | 531 | Mondo | | 4.201e-02 | -3.17 | fly chr15F1-15F1\|1-57 | chromosome location | fly chr15F1-15F1\|1-57 | 2 | 1 | 25010 | 531 | lncRNA:CR44026 | | 4.201e-02 | -3.17 | fly chr46C1-46C1\|2-61 cM | chromosome location | fly chr46C1-46C1\|2-61 cM | 2 | 1 | 25010 | 531 | CG18446 | | 4.201e-02 | -3.17 | fly chr92B4-92B6\|3-67 | chromosome location | fly chr92B4-92B6\|3-67 | 2 | 1 | 25010 | 531 | CG4538 | | 4.201e-02 | -3.17 | fly chr63B7-63B7\|3-6 cM | chromosome location | fly chr63B7-63B7\|3-6 cM | 2 | 1 | 25010 | 531 | BtbVII | | 4.201e-02 | -3.17 | fly chr2C7-2C7\|1-0.5 cM | chromosome location | fly chr2C7-2C7\|1-0.5 cM | 2 | 1 | 25010 | 531 | CG4325 | | 4.201e-02 | -3.17 | fly chr48D7-48D7\|2-65 | chromosome location | fly chr48D7-48D7\|2-65 | 2 | 1 | 25010 | 531 | otk2 | | 4.220e-02 | -3.17 | - | gene3d domains | 3.30.1380.10 | 1 | 1 | 4929 | 208 | hh | | 4.220e-02 | -3.17 | - | gene3d domains | 2.60.40.630 | 1 | 1 | 4929 | 208 | Stat92E | | 4.220e-02 | -3.17 | - | gene3d domains | 1.10.150.40 | 1 | 1 | 4929 | 208 | baf | | 4.220e-02 | -3.17 | - | gene3d domains | 1.10.532.10 | 1 | 1 | 4929 | 208 | Stat92E | | 4.220e-02 | -3.17 | - | gene3d domains | 1.10.2010.10 | 1 | 1 | 4929 | 208 | ITP | | 4.220e-02 | -3.17 | - | gene3d domains | 3.10.20.710 | 1 | 1 | 4929 | 208 | dve | | 4.220e-02 | -3.17 | - | gene3d domains | 3.90.1170.40 | 1 | 1 | 4929 | 208 | Mocs2B | | 4.220e-02 | -3.17 | - | gene3d domains | 1.25.40.170 | 1 | 1 | 4929 | 208 | smg | | 4.220e-02 | -3.17 | - | gene3d domains | 1.10.1100.10 | 1 | 1 | 4929 | 208 | Taf1 | | 4.220e-02 | -3.17 | - | gene3d domains | 2.40.50.740 | 1 | 1 | 4929 | 208 | RpS4 | | 4.220e-02 | -3.17 | - | gene3d domains | 3.40.50.12350 | 1 | 1 | 4929 | 208 | eya | | 4.220e-02 | -3.17 | - | gene3d domains | 2.60.120.40 | 1 | 1 | 4929 | 208 | egr | | 4.224e-02 | -3.16 | regulation of cellular catabolic process | biological process | GO:0031329 | 204 | 12 | 12345 | 414 | crb,Smox,Utx,smg,phyl,wcy,Usp2,EcR,Toll-7,Doa,Gyf,PEK | | 4.225e-02 | -3.16 | germ-band extension | biological process | GO:0007377 | 10 | 2 | 12345 | 414 | RhoGEF2,Rho1 | | 4.225e-02 | -3.16 | chromatin-mediated maintenance of transcription | biological process | GO:0048096 | 10 | 2 | 12345 | 414 | tara,tna | | 4.225e-02 | -3.16 | central nervous system neuron differentiation | biological process | GO:0021953 | 10 | 2 | 12345 | 414 | hh,exex | | 4.225e-02 | -3.16 | regulation of myotube differentiation | biological process | GO:0010830 | 10 | 2 | 12345 | 414 | lbe,Rho1 | | 4.225e-02 | -3.16 | positive regulation of compound eye photoreceptor cell differentiation | biological process | GO:0110117 | 10 | 2 | 12345 | 414 | oc,mbt | | 4.225e-02 | -3.16 | apical constriction involved in gastrulation | biological process | GO:0003384 | 10 | 2 | 12345 | 414 | RhoGEF2,cta | | 4.225e-02 | -3.16 | regulation of myoblast fusion | biological process | GO:1901739 | 10 | 2 | 12345 | 414 | lbe,Rho1 | | 4.225e-02 | -3.16 | guanosine-containing compound metabolic process | biological process | GO:1901068 | 10 | 2 | 12345 | 414 | sdt,ras | | 4.225e-02 | -3.16 | cardiac muscle tissue development | biological process | GO:0048738 | 10 | 2 | 12345 | 414 | EcR,tin | | 4.225e-02 | -3.16 | pseudocleavage involved in syncytial blastoderm formation | biological process | GO:0030589 | 10 | 2 | 12345 | 414 | Rho1,RhoGEF2 | | 4.225e-02 | -3.16 | response to axon injury | biological process | GO:0048678 | 10 | 2 | 12345 | 414 | wnd,hiw | | 4.225e-02 | -3.16 | regulation of syncytium formation by plasma membrane fusion | biological process | GO:0060142 | 10 | 2 | 12345 | 414 | Rho1,lbe | | 4.225e-02 | -3.16 | epithelial cell morphogenesis involved in gastrulation | biological process | GO:0003381 | 10 | 2 | 12345 | 414 | RhoGEF2,cta | | 4.225e-02 | -3.16 | positive regulation of chromatin assembly or disassembly | biological process | GO:0045799 | 10 | 2 | 12345 | 414 | Stat92E,mle | | 4.225e-02 | -3.16 | pseudocleavage | biological process | GO:0030588 | 10 | 2 | 12345 | 414 | Rho1,RhoGEF2 | | 4.225e-02 | -3.16 | germ-line stem-cell niche homeostasis | biological process | GO:0060250 | 10 | 2 | 12345 | 414 | N,CG46339 | | 4.241e-02 | -3.16 | SOCS\_box | pfam domains | PF07525 | 9 | 2 | 10480 | 393 | Socs16D,gus | | 4.241e-02 | -3.16 | Ldl\_recept\_b | pfam domains | PF00058 | 9 | 2 | 10480 | 393 | mgl,LRP1 | | 4.241e-02 | -3.16 | RINGv | pfam domains | PF12906 | 9 | 2 | 10480 | 393 | CG2991,CG9855 | | 4.246e-02 | -3.16 | NFY\_Q6 | MSigDB lists | NFY\_Q6 | 70 | 6 | 4161 | 153 | CtBP,smal,CG9281,so,Lim1,hh | | 4.246e-02 | -3.16 | GO\_NEGATIVE\_REGULATION\_OF\_WNT\_SIGNALING\_PATHWAY | MSigDB lists | GO\_NEGATIVE\_REGULATION\_OF\_WNT\_SIGNALING\_PATHWAY | 70 | 6 | 4161 | 153 | N,Notum,dpp,CG11367,hh,LRP1 | | 4.246e-02 | -3.16 | E2F\_Q4\_01 | MSigDB lists | E2F\_Q4\_01 | 70 | 6 | 4161 | 153 | CG8531,JIL-1,CG9281,CG12547,Lac,NiPp1 | | 4.247e-02 | -3.16 | Tudor | interpro domains | IPR002999 | 21 | 3 | 11383 | 428 | spoon,Tudor-SN,Pcl | | 4.262e-02 | -3.16 | SOCS\_box-like\_dom\_sf | interpro domains | IPR036036 | 9 | 2 | 11383 | 428 | gus,Socs16D | | 4.276e-02 | -3.15 | ERK/MAPK targets | REACTOME pathways | R-DME-198753 | 9 | 2 | 3923 | 148 | JIL-1,S6kII | | 4.278e-02 | -3.15 | MORF\_XRCC5 | MSigDB lists | MORF\_XRCC5 | 127 | 9 | 4161 | 153 | pAbp,hts,kdn,mle,Arp2,heph,CG8531,ras,Mitofilin | | 4.287e-02 | -3.15 | nuclear part | cellular component | GO:0044428 | 1416 | 60 | 11940 | 407 | Pits,Nopp140,tna,CG5815,nudE,Myb,Tlk,EcR,Mocs2B,Imp,elav,Taf1,mam,Sin3A,CG11030,Rbfox1,mle,pbl,Sox21a,Cdk12,Hph,aop,Utx,jing,Tif-IA,pAbp,Fas3,Syp,baf,ctrip,Kah,HmgZ,NFAT,NiPp1,CG44774,CG18766,psq,enok,Pcl,Psc,sm,CG2162,CG5728,lncRNA:roX1,IntS6,drk,N,tou,mof,CtBP,osa,CG8441,gus,CG31712,udd,yps,Acf,stwl,ytr,JIL-1 | | 4.321e-02 | -3.14 | trio (trio) | protein interactions | 43974 | 30 | 4 | 9204 | 412 | Rac2,fax,CycE,N | | 4.321e-02 | -3.14 | ND-75 (NADH dehydrogenase (ubiquinone) 75 kDa subunit) | protein interactions | 31762 | 30 | 4 | 9204 | 412 | spi,CG11752,N,smg | | 4.321e-02 | -3.14 | CG17377 (uncharacterized protein) | protein interactions | 33954 | 30 | 4 | 9204 | 412 | disco,sty,PhKgamma,mub | | 4.346e-02 | -3.14 | GO\_RECEPTOR\_BINDING | MSigDB lists | GO\_RECEPTOR\_BINDING | 253 | 15 | 4161 | 153 | Smox,N,Cnx99A,cta,CtBP,dpp,Ser,betaTub56D,EcR,CycE,CG5742,drk,hh,ScpX,LRP1 | | 4.352e-02 | -3.13 | ESC/E(Z) complex | cellular component | GO:0035098 | 10 | 2 | 11940 | 407 | Pcl,jing | | 4.360e-02 | -3.13 | MOHANKUMAR\_HOXA1\_TARGETS\_UP | MSigDB lists | MOHANKUMAR\_HOXA1\_TARGETS\_UP | 168 | 11 | 4161 | 153 | Lim1,PhKgamma,TfAP-2,kdn,RpS4,CG12099,betaTub56D,CG45050,SamDC,CG32758,drk | | 4.377e-02 | -3.13 | GSE26030\_UNSTIM\_VS\_RESTIM\_TH17\_DAY15\_POST\_POLARIZATION\_DN | MSigDB lists | GSE26030\_UNSTIM\_VS\_RESTIM\_TH17\_DAY15\_POST\_POLARIZATION\_DN | 53 | 5 | 4161 | 153 | ft,emp,Arp2,CG9855,RpS4 | | 4.377e-02 | -3.13 | ETS\_Q4 | MSigDB lists | ETS\_Q4 | 53 | 5 | 4161 | 153 | Cnx99A,SamDC,Utx,CG3961,Tudor-SN | | 4.377e-02 | -3.13 | AACTGGA\_MIR145 | MSigDB lists | AACTGGA\_MIR145 | 53 | 5 | 4161 | 153 | sdt,CG32758,gus,Smox,CG2201 | | 4.377e-02 | -3.13 | GSE37532\_WT\_VS\_PPARG\_KO\_VISCERAL\_ADIPOSE\_TISSUE\_TREG\_UP | MSigDB lists | GSE37532\_WT\_VS\_PPARG\_KO\_VISCERAL\_ADIPOSE\_TISSUE\_TREG\_UP | 53 | 5 | 4161 | 153 | CG14646,CycE,betaTub56D,pbl,CG2812 | | 4.377e-02 | -3.13 | GSE17721\_CPG\_VS\_GARDIQUIMOD\_0.5H\_BMDC\_DN | MSigDB lists | GSE17721\_CPG\_VS\_GARDIQUIMOD\_0.5H\_BMDC\_DN | 53 | 5 | 4161 | 153 | CG9855,Snx3,CG5742,N,hts | | 4.377e-02 | -3.13 | GO\_NEGATIVE\_REGULATION\_OF\_CATABOLIC\_PROCESS | MSigDB lists | GO\_NEGATIVE\_REGULATION\_OF\_CATABOLIC\_PROCESS | 53 | 5 | 4161 | 153 | Snx3,hh,Tlk,Npc1a,Smox | | 4.377e-02 | -3.13 | MORF\_TNFRSF25 | MSigDB lists | MORF\_TNFRSF25 | 53 | 5 | 4161 | 153 | CG4538,mof,Ser,mub,fne | | 4.377e-02 | -3.13 | GSE37416\_0H\_VS\_3H\_F\_TULARENSIS\_LVS\_NEUTROPHIL\_UP | MSigDB lists | GSE37416\_0H\_VS\_3H\_F\_TULARENSIS\_LVS\_NEUTROPHIL\_UP | 53 | 5 | 4161 | 153 | hts,SamDC,CtBP,Arp2,CG6752 | | 4.404e-02 | -3.12 | GO\_GLAND\_DEVELOPMENT | MSigDB lists | GO\_GLAND\_DEVELOPMENT | 89 | 7 | 4161 | 153 | Smox,N,NetB,CycE,so,hh,dpp | | 4.404e-02 | -3.12 | GO\_MUSCLE\_STRUCTURE\_DEVELOPMENT | MSigDB lists | GO\_MUSCLE\_STRUCTURE\_DEVELOPMENT | 89 | 7 | 4161 | 153 | if,N,CaMKII,hh,Ser,dpp,so | | 4.407e-02 | -3.12 | GluRIIA (Glutamate receptor IIA) | protein interactions | 33788 | 18 | 3 | 9204 | 412 | hiw,pAbp,CaMKII | | 4.407e-02 | -3.12 | CG14767 (uncharacterized protein) | protein interactions | 50250 | 18 | 3 | 9204 | 412 | mam,sty,fax | | 4.407e-02 | -3.12 | Ack (Activated Cdc42 kinase) | protein interactions | 38489 | 18 | 3 | 9204 | 412 | drk,pbl,tkv | | 4.407e-02 | -3.12 | psq (pipsqueak) | protein interactions | 36118 | 18 | 3 | 9204 | 412 | Egfr,Psc,psq | | 4.407e-02 | -3.12 | RFeSP (Rieske iron-sulfur protein) | protein interactions | 44390 | 18 | 3 | 9204 | 412 | Egfr,smg,Cyt-c1 | | 4.407e-02 | -3.12 | wnd (wallenda) | protein interactions | 40143 | 18 | 3 | 9204 | 412 | Dscam1,hiw,Rho1 | | 4.407e-02 | -3.12 | salm (spalt major) | protein interactions | 34569 | 18 | 3 | 9204 | 412 | spi,aos,kni | | 4.407e-02 | -3.12 | futsch (futsch) | protein interactions | 5740544 | 18 | 3 | 9204 | 412 | par-1,Syp,NFAT | | 4.408e-02 | -3.12 | heterochromatin organization | biological process | GO:0070828 | 40 | 4 | 12345 | 414 | stwl,bin3,Tlk,AGO2 | | 4.418e-02 | -3.12 | CG7386 (uncharacterized protein) | protein interactions | 38719 | 73 | 7 | 9204 | 412 | ttk,CG4325,CG11486,CycE,Zif,rau,CG43658 | | 4.424e-02 | -3.12 | regulation of nuclear division | biological process | GO:0051783 | 77 | 6 | 12345 | 414 | Mos,kni,bora,CycE,Chro,Myb | | 4.436e-02 | -3.12 | fly chr20C1-20C1\|1-66 cM | chromosome location | fly chr20C1-20C1\|1-66 cM | 16 | 2 | 25010 | 531 | S6kII,Usp2 | | 4.436e-02 | -3.12 | fly chr20C1-20C1\|1-66 | chromosome location | fly chr20C1-20C1\|1-66 | 16 | 2 | 25010 | 531 | Usp2,S6kII | | 4.445e-02 | -3.11 | posttranscriptional gene silencing by RNA | biological process | GO:0035194 | 58 | 5 | 12345 | 414 | CG17265,CG17168,AGO2,Hsc70-3,NiPp1 | | 4.476e-02 | -3.11 | CG15625 (uncharacterized protein) | protein interactions | 33694 | 1 | 1 | 9204 | 412 | pbl | | 4.476e-02 | -3.11 | CG14395 (uncharacterized protein) | protein interactions | 41560 | 1 | 1 | 9204 | 412 | Fas2 | | 4.476e-02 | -3.11 | CG33543 (uncharacterized protein) | protein interactions | 3346207 | 1 | 1 | 9204 | 412 | Fas2 | | 4.476e-02 | -3.11 | CG6216 (uncharacterized protein) | protein interactions | 39257 | 1 | 1 | 9204 | 412 | rau | | 4.476e-02 | -3.11 | Pask (PAS kinase) | protein interactions | 37824 | 1 | 1 | 9204 | 412 | CtBP | | 4.476e-02 | -3.11 | CG1632 (uncharacterized protein) | protein interactions | 31763 | 1 | 1 | 9204 | 412 | Pat1 | | 4.476e-02 | -3.11 | CG9733 (uncharacterized protein) | protein interactions | 43601 | 1 | 1 | 9204 | 412 | Smox | | 4.476e-02 | -3.11 | CarT (Carcinine transporter) | protein interactions | 35334 | 1 | 1 | 9204 | 412 | Fas2 | | 4.476e-02 | -3.11 | CG16800 (uncharacterized protein) | protein interactions | 34693 | 1 | 1 | 9204 | 412 | pAbp | | 4.476e-02 | -3.11 | pigs (pickled eggs) | protein interactions | 31596 | 1 | 1 | 9204 | 412 | pbl | | 4.476e-02 | -3.11 | Cda4 (Chitin deacetylase-like 4) | protein interactions | 33144 | 1 | 1 | 9204 | 412 | ci | | 4.476e-02 | -3.11 | CG8774 (uncharacterized protein) | protein interactions | 41635 | 1 | 1 | 9204 | 412 | CG9541 | | 4.476e-02 | -3.11 | CG8908 (uncharacterized protein) | protein interactions | 37293 | 1 | 1 | 9204 | 412 | Fas2 | | 4.476e-02 | -3.11 | CG30016 (uncharacterized protein) | protein interactions | 246393 | 1 | 1 | 9204 | 412 | pbl | | 4.476e-02 | -3.11 | Ilp4 (Insulin-like peptide 4) | protein interactions | 39152 | 1 | 1 | 9204 | 412 | ImpL2 | | 4.476e-02 | -3.11 | l(1)air10 (lethal aberrant immune response 10) | protein interactions | 48539 | 1 | 1 | 9204 | 412 | CycE | | 4.476e-02 | -3.11 | CG9068 (uncharacterized protein) | protein interactions | 36817 | 1 | 1 | 9204 | 412 | Fas2 | | 4.476e-02 | -3.11 | EOGT (EGF domain specific O-linked N-acetylglucosamine transferase) | protein interactions | 285203 | 1 | 1 | 9204 | 412 | N | | 4.476e-02 | -3.11 | bef (beta lobes fused) | protein interactions | 247841 | 1 | 1 | 9204 | 412 | CycE | | 4.476e-02 | -3.11 | Siah1a (siah E3 ubiquitin protein ligase 1A) | protein interactions | 20437 | 1 | 1 | 9204 | 412 | phyl | | 4.476e-02 | -3.11 | lectin-28C (lectin-28C) | protein interactions | 53542 | 1 | 1 | 9204 | 412 | Smox | | 4.476e-02 | -3.11 | CG33310 (uncharacterized protein) | protein interactions | 2768910 | 1 | 1 | 9204 | 412 | Fas2 | | 4.476e-02 | -3.11 | CG3108 (uncharacterized protein) | protein interactions | 31506 | 1 | 1 | 9204 | 412 | ci | | 4.476e-02 | -3.11 | CG32850 (uncharacterized protein) | protein interactions | 318246 | 1 | 1 | 9204 | 412 | Mhcl | | 4.476e-02 | -3.11 | CG4957 (uncharacterized protein) | protein interactions | 34374 | 1 | 1 | 9204 | 412 | Zif | | 4.476e-02 | -3.11 | Nepl20 (Neprilysin-like 20) | protein interactions | 43412 | 1 | 1 | 9204 | 412 | N | | 4.476e-02 | -3.11 | Ir76a (Ionotropic receptor 76a) | protein interactions | 40157 | 1 | 1 | 9204 | 412 | AGO2 | | 4.476e-02 | -3.11 | CG32812 (uncharacterized protein) | protein interactions | 31074 | 1 | 1 | 9204 | 412 | pbl | | 4.476e-02 | -3.11 | hpRNA:CR46343 (hairpin RNA:CR46343) | protein interactions | 12798474 | 1 | 1 | 9204 | 412 | AGO2 | | 4.476e-02 | -3.11 | CG13747 (uncharacterized protein) | protein interactions | 35907 | 1 | 1 | 9204 | 412 | CG8191 | | 4.476e-02 | -3.11 | CG7778 (uncharacterized protein) | protein interactions | 34147 | 1 | 1 | 9204 | 412 | CG8191 | | 4.476e-02 | -3.11 | CG4942 (uncharacterized protein) | protein interactions | 39044 | 1 | 1 | 9204 | 412 | Inx2 | | 4.476e-02 | -3.11 | Or2a (Odorant receptor 2a) | protein interactions | 31207 | 1 | 1 | 9204 | 412 | CG32772 | | 4.476e-02 | -3.11 | CG6959 (uncharacterized protein) | protein interactions | 41434 | 1 | 1 | 9204 | 412 | Lac | | 4.476e-02 | -3.11 | mir-100 (mir-100 stem loop) | protein interactions | 12798011 | 1 | 1 | 9204 | 412 | AGO2 | | 4.476e-02 | -3.11 | FucTC (alpha1,3-fucosyltransferase C) | protein interactions | 3355171 | 1 | 1 | 9204 | 412 | ci | | 4.476e-02 | -3.11 | CG1236 (uncharacterized protein) | protein interactions | 40708 | 1 | 1 | 9204 | 412 | Df31 | | 4.476e-02 | -3.11 | Oseg5 (Outer segment 5) | protein interactions | 35349 | 1 | 1 | 9204 | 412 | Smox | | 4.476e-02 | -3.11 | lncRNA:CR32660 (long non-coding RNA:CR32660) | protein interactions | 3771836 | 1 | 1 | 9204 | 412 | CG32772 | | 4.476e-02 | -3.11 | CG6834 (uncharacterized protein) | protein interactions | 41409 | 1 | 1 | 9204 | 412 | Myo10A | | 4.476e-02 | -3.11 | cd (cardinal) | protein interactions | 42681 | 1 | 1 | 9204 | 412 | aos | | 4.476e-02 | -3.11 | GstZ2 (Glutathione S transferase Z2) | protein interactions | 41133 | 1 | 1 | 9204 | 412 | l(3)neo38 | | 4.476e-02 | -3.11 | neo (neyo) | protein interactions | 43523 | 1 | 1 | 9204 | 412 | tyn | | 4.476e-02 | -3.11 | CG7137 (uncharacterized protein) | protein interactions | 37204 | 1 | 1 | 9204 | 412 | Fas2 | | 4.476e-02 | -3.11 | CG13896 (uncharacterized protein) | protein interactions | 38089 | 1 | 1 | 9204 | 412 | ey | | 4.476e-02 | -3.11 | CG18473 (uncharacterized protein) | protein interactions | 41116 | 1 | 1 | 9204 | 412 | Smox | | 4.476e-02 | -3.11 | CG12017 (uncharacterized protein) | protein interactions | 38404 | 1 | 1 | 9204 | 412 | cpo | | 4.476e-02 | -3.11 | Elob (elongin B) | protein interactions | 67673 | 1 | 1 | 9204 | 412 | gus | | 4.476e-02 | -3.11 | SA-2 (Stromalin-2) | protein interactions | 38167 | 1 | 1 | 9204 | 412 | Fas2 | | 4.476e-02 | -3.11 | l(1)air4 (lethal aberrant immune response 4) | protein interactions | 48281 | 1 | 1 | 9204 | 412 | CycE | | 4.476e-02 | -3.11 | mir-313 (mir-313 stem loop) | protein interactions | 12798129 | 1 | 1 | 9204 | 412 | pbl | | 4.476e-02 | -3.11 | fipi (factor of interpulse interval) | protein interactions | 33676 | 1 | 1 | 9204 | 412 | Fas2 | | 4.476e-02 | -3.11 | Gs2 (Glutamine synthetase 2) | protein interactions | 32087 | 1 | 1 | 9204 | 412 | ci | | 4.476e-02 | -3.11 | Ir67b (Ionotropic receptor 67b) | protein interactions | 39194 | 1 | 1 | 9204 | 412 | Fas2 | | 4.476e-02 | -3.11 | CG8419 (uncharacterized protein) | protein interactions | 34118 | 1 | 1 | 9204 | 412 | pbl | | 4.476e-02 | -3.11 | CG30431 (uncharacterized protein) | protein interactions | 35549 | 1 | 1 | 9204 | 412 | pAbp | | 4.476e-02 | -3.11 | CG33090 (uncharacterized protein) | protein interactions | 34835 | 1 | 1 | 9204 | 412 | so | | 4.476e-02 | -3.11 | Usp20-33 (Ubiquitin specific protease 20/33) | protein interactions | 36580 | 1 | 1 | 9204 | 412 | FoxL1 | | 4.476e-02 | -3.11 | CG4884 (uncharacterized protein) | protein interactions | 43357 | 1 | 1 | 9204 | 412 | Fas2 | | 4.476e-02 | -3.11 | mir-2c (mir-2c stem loop) | protein interactions | 12798412 | 1 | 1 | 9204 | 412 | N | | 4.476e-02 | -3.11 | COPS2 (COP9 signalosome subunit 2) | protein interactions | 9318 | 1 | 1 | 9204 | 412 | EcR | | 4.476e-02 | -3.11 | CG42534 (uncharacterized protein) | protein interactions | 43298 | 1 | 1 | 9204 | 412 | Fas2 | | 4.476e-02 | -3.11 | TMS1 (TMS1) | protein interactions | 39827 | 1 | 1 | 9204 | 412 | sty | | 4.476e-02 | -3.11 | MTPAP (mitochondrial poly(A) polymerase) | protein interactions | 31081 | 1 | 1 | 9204 | 412 | ci | | 4.476e-02 | -3.11 | mir-980 (mir-980 stem loop) | protein interactions | 12798291 | 1 | 1 | 9204 | 412 | Rbfox1 | | 4.476e-02 | -3.11 | CG9044 (uncharacterized protein) | protein interactions | 33825 | 1 | 1 | 9204 | 412 | Fas2 | | 4.476e-02 | -3.11 | Muc91C (Mucin 91C) | protein interactions | 42246 | 1 | 1 | 9204 | 412 | sty | | 4.476e-02 | -3.11 | CG10841 (uncharacterized protein) | protein interactions | 41665 | 1 | 1 | 9204 | 412 | Fas2 | | 4.476e-02 | -3.11 | CG42382 (uncharacterized protein) | protein interactions | 35938 | 1 | 1 | 9204 | 412 | gus | | 4.476e-02 | -3.11 | CG12499 (uncharacterized protein) | protein interactions | 42637 | 1 | 1 | 9204 | 412 | Fas2 | | 4.476e-02 | -3.11 | nyo (nyobe) | protein interactions | 43716 | 1 | 1 | 9204 | 412 | tyn | | 4.476e-02 | -3.11 | CG31436 (uncharacterized protein) | protein interactions | 318734 | 1 | 1 | 9204 | 412 | Smox | | 4.476e-02 | -3.11 | CG9747 (uncharacterized protein) | protein interactions | 43591 | 1 | 1 | 9204 | 412 | CG3402 | | 4.476e-02 | -3.11 | rpk (ripped pocket) | protein interactions | 40580 | 1 | 1 | 9204 | 412 | Smox | | 4.476e-02 | -3.11 | CG34124 (uncharacterized protein) | protein interactions | 4379887 | 1 | 1 | 9204 | 412 | Fas2 | | 4.476e-02 | -3.11 | Spn28B (Serpin 28B) | protein interactions | 34035 | 1 | 1 | 9204 | 412 | CG32772 | | 4.476e-02 | -3.11 | CG10357 (uncharacterized protein) | protein interactions | 38435 | 1 | 1 | 9204 | 412 | CG11247 | | 4.476e-02 | -3.11 | ich (ichor) | protein interactions | 41069 | 1 | 1 | 9204 | 412 | flz | | 4.476e-02 | -3.11 | Six4 (Six4) | protein interactions | 40297 | 1 | 1 | 9204 | 412 | so | | 4.476e-02 | -3.11 | CG33127 (uncharacterized protein) | protein interactions | 318893 | 1 | 1 | 9204 | 412 | ci | | 4.476e-02 | -3.11 | CG10375 (uncharacterized protein) | protein interactions | 42812 | 1 | 1 | 9204 | 412 | CG7264 | | 4.476e-02 | -3.11 | ND-B8 (NADH dehydrogenase (ubiquinone) B8 subunit) | protein interactions | 50178 | 1 | 1 | 9204 | 412 | smg | | 4.476e-02 | -3.11 | NKCC (sodium potassium chloride cotransporter) | protein interactions | 40663 | 1 | 1 | 9204 | 412 | qsm | | 4.476e-02 | -3.11 | CG18095 (uncharacterized protein) | protein interactions | 34823 | 1 | 1 | 9204 | 412 | Smox | | 4.476e-02 | -3.11 | CG10089 (uncharacterized protein) | protein interactions | 39517 | 1 | 1 | 9204 | 412 | Smox | | 4.476e-02 | -3.11 | Jon66Cii (Jonah 66Cii) | protein interactions | 38953 | 1 | 1 | 9204 | 412 | lama | | 4.476e-02 | -3.11 | Agpat2 (1-Acylglycerol-3-phosphate O-acyltransferase 2) | protein interactions | 34176 | 1 | 1 | 9204 | 412 | mle | | 4.476e-02 | -3.11 | CCDC151 (Coiled-coil domain containing protein 151) | protein interactions | 39345 | 1 | 1 | 9204 | 412 | Fas2 | | 4.476e-02 | -3.11 | moody (moody) | protein interactions | 31168 | 1 | 1 | 9204 | 412 | drk | | 4.476e-02 | -3.11 | CG1428 (uncharacterized protein) | protein interactions | 35433 | 1 | 1 | 9204 | 412 | Fas2 | | 4.476e-02 | -3.11 | SUMO1 (small ubiquitin like modifier 1) | protein interactions | 7341 | 1 | 1 | 9204 | 412 | Stat92E | | 4.476e-02 | -3.11 | CG5707 (uncharacterized protein) | protein interactions | 48422 | 1 | 1 | 9204 | 412 | rau | | 4.476e-02 | -3.11 | mir-318 (mir-318 stem loop) | protein interactions | 12797881 | 1 | 1 | 9204 | 412 | ttk | | 4.476e-02 | -3.11 | CG10909 (uncharacterized protein) | protein interactions | 41581 | 1 | 1 | 9204 | 412 | Gyf | | 4.476e-02 | -3.11 | CG17068 (uncharacterized protein) | protein interactions | 33024 | 1 | 1 | 9204 | 412 | Utx | | 4.476e-02 | -3.11 | CG15651 (uncharacterized protein) | protein interactions | 37375 | 1 | 1 | 9204 | 412 | CG18766 | | 4.476e-02 | -3.11 | CG9864 (uncharacterized protein) | protein interactions | 37287 | 1 | 1 | 9204 | 412 | CG2865 | | 4.476e-02 | -3.11 | hist1h4a.L (histone cluster 1, H4a L homeolog) | protein interactions | 447787 | 1 | 1 | 9204 | 412 | mof | | 4.476e-02 | -3.11 | CG7110 (uncharacterized protein) | protein interactions | 34757 | 1 | 1 | 9204 | 412 | Syx1A | | 4.476e-02 | -3.11 | AsnS (Asparagine synthetase) | protein interactions | 2768965 | 1 | 1 | 9204 | 412 | Fas2 | | 4.476e-02 | -3.11 | Ack-like (Activated Cdc42 kinase-like) | protein interactions | 36442 | 1 | 1 | 9204 | 412 | drk | | 4.476e-02 | -3.11 | CG30120 (uncharacterized protein) | protein interactions | 246467 | 1 | 1 | 9204 | 412 | Fas2 | | 4.476e-02 | -3.11 | CG6409 (uncharacterized protein) | protein interactions | 39220 | 1 | 1 | 9204 | 412 | CG32772 | | 4.476e-02 | -3.11 | CG3764 (uncharacterized protein) | protein interactions | 39902 | 1 | 1 | 9204 | 412 | ci | | 4.476e-02 | -3.11 | CG3036 (uncharacterized protein) | protein interactions | 33695 | 1 | 1 | 9204 | 412 | pbl | | 4.476e-02 | -3.11 | Mctp (Multiple C2 domain and transmembrane region protein) | protein interactions | 37165 | 1 | 1 | 9204 | 412 | ey | | 4.476e-02 | -3.11 | CG15212 (uncharacterized protein) | protein interactions | 50315 | 1 | 1 | 9204 | 412 | Smox | | 4.476e-02 | -3.11 | CG6511 (uncharacterized protein) | protein interactions | 38997 | 1 | 1 | 9204 | 412 | Pat1 | | 4.476e-02 | -3.11 | hpRNA:CR33940 (hairpin RNA:CR33940) | protein interactions | 3771790 | 1 | 1 | 9204 | 412 | AGO2 | | 4.476e-02 | -3.11 | CG10588 (uncharacterized protein) | protein interactions | 40316 | 1 | 1 | 9204 | 412 | phyl | | 4.476e-02 | -3.11 | DIP2 (DISCO Interacting Protein 2) | protein interactions | 252479 | 1 | 1 | 9204 | 412 | drk | | 4.476e-02 | -3.11 | Irp-1B (Iron regulatory protein 1B) | protein interactions | 41269 | 1 | 1 | 9204 | 412 | Fas2 | | 4.476e-02 | -3.11 | CG14357 (uncharacterized protein) | protein interactions | 41716 | 1 | 1 | 9204 | 412 | CtBP | | 4.476e-02 | -3.11 | CG33158 (uncharacterized protein) | protein interactions | 39834 | 1 | 1 | 9204 | 412 | Fas2 | | 4.476e-02 | -3.11 | Kat2b (K(lysine) acetyltransferase 2B) | protein interactions | 18519 | 1 | 1 | 9204 | 412 | N | | 4.476e-02 | -3.11 | Tsp68C (Tetraspanin 68C) | protein interactions | 117407 | 1 | 1 | 9204 | 412 | ytr | | 4.476e-02 | -3.11 | CG10734 (uncharacterized protein) | protein interactions | 36787 | 1 | 1 | 9204 | 412 | mle | | 4.476e-02 | -3.11 | CG10011 (uncharacterized protein) | protein interactions | 43387 | 1 | 1 | 9204 | 412 | ci | | 4.476e-02 | -3.11 | Mocs2A (Molybdenum cofactor synthesis 2A) | protein interactions | 8674021 | 1 | 1 | 9204 | 412 | Mocs2B | | 4.476e-02 | -3.11 | yellow-c (yellow-c) | protein interactions | 34879 | 1 | 1 | 9204 | 412 | esn | | 4.476e-02 | -3.11 | Eloc (elongin C) | protein interactions | 67923 | 1 | 1 | 9204 | 412 | gus | | 4.476e-02 | -3.11 | ng4 (new glue 4) | protein interactions | 31301 | 1 | 1 | 9204 | 412 | smal | | 4.476e-02 | -3.11 | Gr58b (Gustatory receptor 58b) | protein interactions | 37502 | 1 | 1 | 9204 | 412 | Hip1 | | 4.476e-02 | -3.11 | USP7 (ubiquitin specific peptidase 7) | protein interactions | 7874 | 1 | 1 | 9204 | 412 | ci | | 4.476e-02 | -3.11 | Sobp (Sine oculis-binding protein) | protein interactions | 36260 | 1 | 1 | 9204 | 412 | so | | 4.476e-02 | -3.11 | CG30503 (uncharacterized protein) | protein interactions | 246656 | 1 | 1 | 9204 | 412 | CG11486 | | 4.476e-02 | -3.11 | bnk (bottleneck) | protein interactions | 43687 | 1 | 1 | 9204 | 412 | Fas2 | | 4.476e-02 | -3.11 | Upf2 (Upf2) | protein interactions | 31724 | 1 | 1 | 9204 | 412 | Fas2 | | 4.476e-02 | -3.11 | CG3626 (uncharacterized protein) | protein interactions | 31377 | 1 | 1 | 9204 | 412 | Fas2 | | 4.476e-02 | -3.11 | Cyp6w1 (Cyp6w1) | protein interactions | 35543 | 1 | 1 | 9204 | 412 | rau | | 4.476e-02 | -3.11 | CG17803 (uncharacterized protein) | protein interactions | 42141 | 1 | 1 | 9204 | 412 | CG17778 | | 4.476e-02 | -3.11 | Sur (Sulfonylurea receptor) | protein interactions | 34350 | 1 | 1 | 9204 | 412 | pnt | | 4.476e-02 | -3.11 | CG32109 (uncharacterized protein) | protein interactions | 39444 | 1 | 1 | 9204 | 412 | ds | | 4.476e-02 | -3.11 | nes (nessy) | protein interactions | 40093 | 1 | 1 | 9204 | 412 | wun2 | | 4.476e-02 | -3.11 | meng (meng-po) | protein interactions | 33948 | 1 | 1 | 9204 | 412 | Smox | | 4.476e-02 | -3.11 | Osi24 (Osiris 24) | protein interactions | 40755 | 1 | 1 | 9204 | 412 | heph | | 4.476e-02 | -3.11 | goe (gone early) | protein interactions | 32660 | 1 | 1 | 9204 | 412 | aos | | 4.476e-02 | -3.11 | l(1)air16 (lethal aberrant immune response 16) | protein interactions | 49140 | 1 | 1 | 9204 | 412 | CycE | | 4.476e-02 | -3.11 | CG3328 (uncharacterized protein) | protein interactions | 37871 | 1 | 1 | 9204 | 412 | mam | | 4.476e-02 | -3.11 | CG2187 (uncharacterized protein) | protein interactions | 43743 | 1 | 1 | 9204 | 412 | Fas2 | | 4.476e-02 | -3.11 | CG8329 (uncharacterized protein) | protein interactions | 39123 | 1 | 1 | 9204 | 412 | esn | | 4.476e-02 | -3.11 | CG5597 (uncharacterized protein) | protein interactions | 37789 | 1 | 1 | 9204 | 412 | nrm | | 4.476e-02 | -3.11 | Cyp9c1 (Cytochrome P450-9c1) | protein interactions | 37941 | 1 | 1 | 9204 | 412 | Smox | | 4.476e-02 | -3.11 | CG42354 (uncharacterized protein) | protein interactions | 7354437 | 1 | 1 | 9204 | 412 | S6kII | | 4.476e-02 | -3.11 | CG17364 (uncharacterized protein) | protein interactions | 39543 | 1 | 1 | 9204 | 412 | CG8191 | | 4.476e-02 | -3.11 | blp (black pearl) | protein interactions | 41937 | 1 | 1 | 9204 | 412 | Fas2 | | 4.476e-02 | -3.11 | Eglp4 (Entomoglyceroporin 4) | protein interactions | 37739 | 1 | 1 | 9204 | 412 | Fas2 | | 4.476e-02 | -3.11 | CG31465 (uncharacterized protein) | protein interactions | 42577 | 1 | 1 | 9204 | 412 | Fas2 | | 4.476e-02 | -3.11 | yellow-e3 (yellow-e3) | protein interactions | 41651 | 1 | 1 | 9204 | 412 | Mhcl | | 4.476e-02 | -3.11 | Hyccin (hyccin) | protein interactions | 37019 | 1 | 1 | 9204 | 412 | rau | | 4.476e-02 | -3.11 | shams (shams) | protein interactions | 43008 | 1 | 1 | 9204 | 412 | N | | 4.484e-02 | -3.10 | positive regulation of growth | biological process | GO:0045927 | 118 | 8 | 12345 | 414 | Egfr,Arp2,tkv,dpp,Hph,tsl,Hipk,ey | | 4.498e-02 | -3.10 | dendrite guidance | biological process | GO:0070983 | 24 | 3 | 12345 | 414 | NetA,comm,NetB | | 4.498e-02 | -3.10 | negative regulation of organ growth | biological process | GO:0046621 | 24 | 3 | 12345 | 414 | Fbxl7,ft,crb | | 4.498e-02 | -3.10 | cell-substrate adhesion | biological process | GO:0031589 | 24 | 3 | 12345 | 414 | tyn,if,scb | | 4.498e-02 | -3.10 | postsynapse organization | biological process | GO:0099173 | 24 | 3 | 12345 | 414 | Prosap,Fur1,Ten-m | | 4.498e-02 | -3.10 | RAAGNYNNCTTY\_UNKNOWN | MSigDB lists | RAAGNYNNCTTY\_UNKNOWN | 22 | 3 | 4161 | 153 | dlp,cta,fne | | 4.498e-02 | -3.10 | GO\_POSITIVE\_REGULATION\_OF\_TRANSMEMBRANE\_RECEPTOR\_PROTEIN\_SERINE\_THREONINE\_KINASE\_SIGNALING\_PATHWAY | MSigDB lists | GO\_POSITIVE\_REGULATION\_OF\_TRANSMEMBRANE\_RECEPTOR\_PROTEIN\_SERINE\_THREONINE\_KINASE\_SIGNALING\_PATHWAY | 22 | 3 | 4161 | 153 | if,N,dpp | | 4.498e-02 | -3.10 | GO\_INTERMEDIATE\_FILAMENT\_CYTOSKELETON | MSigDB lists | GO\_INTERMEDIATE\_FILAMENT\_CYTOSKELETON | 22 | 3 | 4161 | 153 | Tlk,nudE,CG6330 | | 4.498e-02 | -3.10 | GO\_REGULATION\_OF\_ATPASE\_ACTIVITY | MSigDB lists | GO\_REGULATION\_OF\_ATPASE\_ACTIVITY | 22 | 3 | 4161 | 153 | Cyp1,CG11367,Tpr2 | | 4.498e-02 | -3.10 | REACTOME\_SIGNALING\_BY\_RHO\_GTPASES | MSigDB lists | REACTOME\_SIGNALING\_BY\_RHO\_GTPASES | 22 | 3 | 4161 | 153 | Rac2,pbl,Rho1 | | 4.498e-02 | -3.10 | KEGG\_ANTIGEN\_PROCESSING\_AND\_PRESENTATION | MSigDB lists | KEGG\_ANTIGEN\_PROCESSING\_AND\_PRESENTATION | 22 | 3 | 4161 | 153 | Cnx99A,Hsc70Cb,Hsc70-3 | | 4.498e-02 | -3.10 | TCCAGAT\_MIR5165P | MSigDB lists | TCCAGAT\_MIR5165P | 22 | 3 | 4161 | 153 | CG32758,Utx,phtf | | 4.498e-02 | -3.10 | GO\_CARTILAGE\_DEVELOPMENT | MSigDB lists | GO\_CARTILAGE\_DEVELOPMENT | 22 | 3 | 4161 | 153 | dpp,CG9220,tkv | | 4.498e-02 | -3.10 | STAMBOLSKY\_RESPONSE\_TO\_VITAMIN\_D3\_UP | MSigDB lists | STAMBOLSKY\_RESPONSE\_TO\_VITAMIN\_D3\_UP | 22 | 3 | 4161 | 153 | tkv,mle,CtBP | | 4.498e-02 | -3.10 | YTTCCNNNGGAMR\_UNKNOWN | MSigDB lists | YTTCCNNNGGAMR\_UNKNOWN | 22 | 3 | 4161 | 153 | Mhcl,Ser,Smox | | 4.498e-02 | -3.10 | GO\_FEMALE\_SEX\_DIFFERENTIATION | MSigDB lists | GO\_FEMALE\_SEX\_DIFFERENTIATION | 22 | 3 | 4161 | 153 | Lim1,tkv,CycE | | 4.498e-02 | -3.10 | GO\_NEGATIVE\_REGULATION\_OF\_TRANSMEMBRANE\_TRANSPORT | MSigDB lists | GO\_NEGATIVE\_REGULATION\_OF\_TRANSMEMBRANE\_TRANSPORT | 22 | 3 | 4161 | 153 | Ank,CaMKII,Cyp1 | | 4.498e-02 | -3.10 | GSE2706\_R848\_VS\_LPS\_2H\_STIM\_DC\_DN | MSigDB lists | GSE2706\_R848\_VS\_LPS\_2H\_STIM\_DC\_DN | 22 | 3 | 4161 | 153 | CG43313,rush,Ser | | 4.498e-02 | -3.10 | Galactose-bd-like\_sf | interpro domains | IPR008979 | 36 | 4 | 11383 | 428 | hiw,Fur1,smal,Ten-m | | 4.504e-02 | -3.10 | TGANTCA\_AP1\_C | MSigDB lists | TGANTCA\_AP1\_C | 211 | 13 | 4161 | 153 | dpp,NiPp1,S6kII,Syx1A,Tlk,LRP1,CG3097,Acsl,sdt,Rho1,fne,Ank,Hydr2 | | 4.520e-02 | -3.10 | GO\_LYTIC\_VACUOLE | MSigDB lists | GO\_LYTIC\_VACUOLE | 169 | 11 | 4161 | 153 | mgl,CG41099,Npc1a,CG14291,MFS10,LManII,Ank,emp,dally,dlp,LRP1 | | 4.528e-02 | -3.09 | WIERENGA\_STAT5A\_TARGETS\_DN | MSigDB lists | WIERENGA\_STAT5A\_TARGETS\_DN | 37 | 4 | 4161 | 153 | Syx1A,N,Cyp1,CG9855 | | 4.528e-02 | -3.09 | GSE15750\_WT\_VS\_TRAF6KO\_DAY6\_EFF\_CD8\_TCELL\_DN | MSigDB lists | GSE15750\_WT\_VS\_TRAF6KO\_DAY6\_EFF\_CD8\_TCELL\_DN | 37 | 4 | 4161 | 153 | CG8531,Sin3A,CG4538,Pli | | 4.528e-02 | -3.09 | GSE36476\_CTRL\_VS\_TSST\_ACT\_72H\_MEMORY\_CD4\_TCELL\_OLD\_UP | MSigDB lists | GSE36476\_CTRL\_VS\_TSST\_ACT\_72H\_MEMORY\_CD4\_TCELL\_OLD\_UP | 37 | 4 | 4161 | 153 | emp,mub,Trpm,Arl4 | | 4.528e-02 | -3.09 | ACACTGG\_MIR199A\_MIR199B | MSigDB lists | ACACTGG\_MIR199A\_MIR199B | 37 | 4 | 4161 | 153 | Pat1,sdt,Hsc70-3,Doa | | 4.528e-02 | -3.09 | GSE12392\_WT\_VS\_IFNB\_KO\_CD8A\_NEG\_SPLEEN\_DC\_DN | MSigDB lists | GSE12392\_WT\_VS\_IFNB\_KO\_CD8A\_NEG\_SPLEEN\_DC\_DN | 37 | 4 | 4161 | 153 | dpp,phtf,sno,hiw | | 4.528e-02 | -3.09 | GSE13485\_CTRL\_VS\_DAY3\_YF17D\_VACCINE\_PBMC\_DN | MSigDB lists | GSE13485\_CTRL\_VS\_DAY3\_YF17D\_VACCINE\_PBMC\_DN | 37 | 4 | 4161 | 153 | Pcl,CtBP,Mitofilin,emp | | 4.528e-02 | -3.09 | GO\_RESPONSE\_TO\_IONIZING\_RADIATION | MSigDB lists | GO\_RESPONSE\_TO\_IONIZING\_RADIATION | 37 | 4 | 4161 | 153 | Tlk,Doa,drk,pbl | | 4.528e-02 | -3.09 | NCX\_01 | MSigDB lists | NCX\_01 | 37 | 4 | 4161 | 153 | Ptpmeg,fne,Tpr2,Pat1 | | 4.528e-02 | -3.09 | GO\_POSITIVE\_REGULATION\_OF\_LIPID\_METABOLIC\_PROCESS | MSigDB lists | GO\_POSITIVE\_REGULATION\_OF\_LIPID\_METABOLIC\_PROCESS | 37 | 4 | 4161 | 153 | Acsl,EcR,Rac2,ScpX | | 4.528e-02 | -3.09 | GSE19772\_HCMV\_INFL\_VS\_HCMV\_INF\_MONOCYTES\_AND\_PI3K\_INHIBITION\_UP | MSigDB lists | GSE19772\_HCMV\_INFL\_VS\_HCMV\_INF\_MONOCYTES\_AND\_PI3K\_INHIBITION\_UP | 37 | 4 | 4161 | 153 | smal,Mitofilin,N,EcR | | 4.528e-02 | -3.09 | ROZANOV\_MMP14\_TARGETS\_UP | MSigDB lists | ROZANOV\_MMP14\_TARGETS\_UP | 37 | 4 | 4161 | 153 | CG9220,Fas2,Ser,lili | | 4.528e-02 | -3.09 | GSE7509\_DC\_VS\_MONOCYTE\_WITH\_FCGRIIB\_STIM\_DN | MSigDB lists | GSE7509\_DC\_VS\_MONOCYTE\_WITH\_FCGRIIB\_STIM\_DN | 37 | 4 | 4161 | 153 | CG9220,Tlk,Smox,SamDC | | 4.528e-02 | -3.09 | GO\_APPENDAGE\_DEVELOPMENT | MSigDB lists | GO\_APPENDAGE\_DEVELOPMENT | 37 | 4 | 4161 | 153 | hh,TfAP-2,tkv,N | | 4.528e-02 | -3.09 | GSE41978\_ID2\_KO\_AND\_BIM\_KO\_VS\_BIM\_KO\_KLRG1\_LOW\_EFFECTOR\_CD8\_TCELL\_DN | MSigDB lists | GSE41978\_ID2\_KO\_AND\_BIM\_KO\_VS\_BIM\_KO\_KLRG1\_LOW\_EFFECTOR\_CD8\_TCELL\_DN | 37 | 4 | 4161 | 153 | Arl4,Rac2,Tif-IA,Pli | | 4.528e-02 | -3.09 | HFH4\_01 | MSigDB lists | HFH4\_01 | 37 | 4 | 4161 | 153 | fne,NetB,phtf,Tsp26A | | 4.528e-02 | -3.09 | GSE6269\_E\_COLI\_VS\_STREP\_PNEUMO\_INF\_PBMC\_DN | MSigDB lists | GSE6269\_E\_COLI\_VS\_STREP\_PNEUMO\_INF\_PBMC\_DN | 37 | 4 | 4161 | 153 | Npc1a,Ser,ras,Hydr2 | | 4.528e-02 | -3.09 | GSE30971\_CTRL\_VS\_LPS\_STIM\_MACROPHAGE\_WBP7\_HET\_2H\_UP | MSigDB lists | GSE30971\_CTRL\_VS\_LPS\_STIM\_MACROPHAGE\_WBP7\_HET\_2H\_UP | 37 | 4 | 4161 | 153 | mmy,Npc1a,CG1271,CG3961 | | 4.528e-02 | -3.09 | GSE22045\_TREG\_VS\_TCONV\_UP | MSigDB lists | GSE22045\_TREG\_VS\_TCONV\_UP | 37 | 4 | 4161 | 153 | CG5522,Cyp1,Syx1A,Hydr2 | | 4.528e-02 | -3.09 | GO\_GLIAL\_CELL\_DIFFERENTIATION | MSigDB lists | GO\_GLIAL\_CELL\_DIFFERENTIATION | 37 | 4 | 4161 | 153 | hh,N,sdt,TfAP-2 | | 4.528e-02 | -3.09 | GSE26495\_PD1HIGH\_VS\_PD1LOW\_CD8\_TCELL\_DN | MSigDB lists | GSE26495\_PD1HIGH\_VS\_PD1LOW\_CD8\_TCELL\_DN | 37 | 4 | 4161 | 153 | CtBP,N,Hydr2,Fas2 | | 4.530e-02 | -3.09 | NHL | prosite domains | PS51125 | 7 | 2 | 6243 | 316 | CG12547,wech | | 4.530e-02 | -3.09 | CRIB | prosite domains | PS50108 | 7 | 2 | 6243 | 316 | mbt,Pak3 | | 4.530e-02 | -3.09 | PAN | prosite domains | PS50948 | 7 | 2 | 6243 | 316 | tyn,neo | | 4.530e-02 | -3.09 | SSD | prosite domains | PS50156 | 7 | 2 | 6243 | 316 | Npc1a,Ptr | | 4.544e-02 | -3.09 | Apoptosis - fly | KEGG pathways | dme04214 | 63 | 5 | 3198 | 100 | PEK,EcR,Egfr,egr,ban | | 4.544e-02 | -3.09 | Apoptosis - fly | KEGG pathways | ko04214 | 63 | 5 | 3198 | 100 | ban,egr,Egfr,EcR,PEK | | 4.548e-02 | -3.09 | sqd (squid) | protein interactions | 41666 | 44 | 5 | 9204 | 412 | Imp,S6kII,Syp,Egfr,pAbp | | 4.548e-02 | -3.09 | Scm (Sex comb on midleg) | protein interactions | 41168 | 44 | 5 | 9204 | 412 | l(3)mbt,psq,Pcl,ocm,Psc | | 4.548e-02 | -3.09 | Tsp (Thrombospondin) | protein interactions | 33941 | 44 | 5 | 9204 | 412 | if,pAbp,CG2865,PhKgamma,slow | | 4.548e-02 | -3.09 | Act57B (Actin 57B) | protein interactions | 37368 | 44 | 5 | 9204 | 412 | ci,Fas2,CG10265,Df31,Arp2 | | 4.548e-02 | -3.09 | sev (sevenless) | protein interactions | 32039 | 44 | 5 | 9204 | 412 | S6kII,S,drk,Egfr,sty | | 4.548e-02 | -3.09 | Cdk5 (Cyclin-dependent kinase 5) | protein interactions | 36727 | 44 | 5 | 9204 | 412 | CycE,CG2865,EcR,Cdk5alpha,Usp2 | | 4.597e-02 | -3.08 | HALLMARK\_MTORC1\_SIGNALING | MSigDB lists | HALLMARK\_MTORC1\_SIGNALING | 109 | 8 | 4161 | 153 | Hsc70-3,Mitofilin,Hsc70Cb,Acsl,Arp2,CG9281,Ykt6,Cnx99A | | 4.597e-02 | -3.08 | MODULE\_163 | MSigDB lists | MODULE\_163 | 109 | 8 | 4161 | 153 | tkv,drk,CG3097,dlp,dally,N,Pli,CG14291 | | 4.597e-02 | -3.08 | MODULE\_95 | MSigDB lists | MODULE\_95 | 109 | 8 | 4161 | 153 | CG3097,dally,dlp,tkv,drk,Pli,CG14291,N | | 4.623e-02 | -3.07 | regulation of response to biotic stimulus | biological process | GO:0002831 | 162 | 10 | 12345 | 414 | Mmp2,CG4325,ush,Tl,Etl1,Stat92E,Pli,Usp2,Toll-7,Doa | | 4.638e-02 | -3.07 | maternal determination of anterior/posterior axis, embryo | biological process | GO:0008358 | 98 | 7 | 12345 | 414 | cnc,par-1,tsl,gus,Pat1,psq,N | | 4.652e-02 | -3.07 | LEE\_BMP2\_TARGETS\_UP | MSigDB lists | LEE\_BMP2\_TARGETS\_UP | 129 | 9 | 4161 | 153 | gus,ds,Pli,Ser,MFS10,Ank,emp,CG2201,LRP1 | | 4.663e-02 | -3.07 | positive regulation of transferase activity | biological process | GO:0051347 | 78 | 6 | 12345 | 414 | bora,wnd,Cdk5alpha,Rho1,mbt,Atg17 | | 4.673e-02 | -3.06 | regulation of vesicle-mediated transport | biological process | GO:0060627 | 119 | 8 | 12345 | 414 | NFAT,Syx1A,drk,corn,sdt,comm,mgl,Hip1 | | 4.674e-02 | -3.06 | chromosomal region | cellular component | GO:0098687 | 204 | 12 | 11940 | 407 | JIL-1,mam,lawc,Chro,flfl,cnc,nudE,Psc,Utx,mof,CG17778,Cdk12 | | 4.679e-02 | -3.06 | lft (lowfat) | protein interactions | 34405 | 8 | 2 | 9204 | 412 | ft,ds | | 4.679e-02 | -3.06 | mahe (maheshvara) | protein interactions | 31707 | 8 | 2 | 9204 | 412 | mam,N | | 4.679e-02 | -3.06 | ncm (nucampholin) | protein interactions | 35099 | 8 | 2 | 9204 | 412 | S6kII,aos | | 4.679e-02 | -3.06 | mil (milkah) | protein interactions | 43343 | 8 | 2 | 9204 | 412 | CG11076,BtbVII | | 4.679e-02 | -3.06 | Ak6 (Adenylate kinase 6) | protein interactions | 36379 | 8 | 2 | 9204 | 412 | CG2865,CG3919 | | 4.679e-02 | -3.06 | Ndg (Nidogen/entactin) | protein interactions | 36089 | 8 | 2 | 9204 | 412 | Myo10A,N | | 4.679e-02 | -3.06 | UQCR-11 (Ubiquinol-cytochrome c reductase 11 kDa subunit) | protein interactions | 5740185 | 8 | 2 | 9204 | 412 | Cyt-c1,smg | | 4.679e-02 | -3.06 | wap (wings apart) | protein interactions | 33131 | 8 | 2 | 9204 | 412 | ft,ds | | 4.679e-02 | -3.06 | CG9609 (uncharacterized protein) | protein interactions | 32663 | 8 | 2 | 9204 | 412 | rau,nuf | | 4.679e-02 | -3.06 | ocm (over compensating males) | protein interactions | 37874 | 8 | 2 | 9204 | 412 | mle,ttk | | 4.679e-02 | -3.06 | sim (single-minded) | protein interactions | 41612 | 8 | 2 | 9204 | 412 | CycE,jing | | 4.679e-02 | -3.06 | Smg6 (Smg6) | protein interactions | 42994 | 8 | 2 | 9204 | 412 | rau,Doa | | 4.679e-02 | -3.06 | Dlic (Dynein light intermediate chain) | protein interactions | 32057 | 8 | 2 | 9204 | 412 | nuf,nudE | | 4.679e-02 | -3.06 | CG34383 (uncharacterized protein) | protein interactions | 5740131 | 8 | 2 | 9204 | 412 | Trpm,CG3402 | | 4.679e-02 | -3.06 | Ptp52F (Protein tyrosine phosphatase 52F) | protein interactions | 36790 | 8 | 2 | 9204 | 412 | CG10283,trn | | 4.679e-02 | -3.06 | D19B (D19B) | protein interactions | 38717 | 8 | 2 | 9204 | 412 | rau,spi | | 4.679e-02 | -3.06 | hang (hangover) | protein interactions | 32613 | 8 | 2 | 9204 | 412 | rau,ttk | | 4.679e-02 | -3.06 | ihog (interference hedgehog) | protein interactions | 33972 | 8 | 2 | 9204 | 412 | dlp,hh | | 4.679e-02 | -3.06 | Prosap (prosap) | protein interactions | 50225 | 8 | 2 | 9204 | 412 | mle,spi | | 4.679e-02 | -3.06 | dpr9 (defective proboscis extension response 9) | protein interactions | 2768670 | 8 | 2 | 9204 | 412 | slow,salm | | 4.679e-02 | -3.06 | CRMP (Collapsin Response Mediator Protein) | protein interactions | 40675 | 8 | 2 | 9204 | 412 | N,Rac2 | | 4.679e-02 | -3.06 | exp (expansion) | protein interactions | 36276 | 8 | 2 | 9204 | 412 | Rho1,Egfr | | 4.679e-02 | -3.06 | Mdh1 (Malate dehydrogenase 1) | protein interactions | 34414 | 8 | 2 | 9204 | 412 | Lim1,Zif | | 4.679e-02 | -3.06 | DJ-1alpha (DJ-1alpha) | protein interactions | 36543 | 8 | 2 | 9204 | 412 | Smox,ras | | 4.679e-02 | -3.06 | spdo (sanpodo) | protein interactions | 318559 | 8 | 2 | 9204 | 412 | Rbfox1,N | | 4.679e-02 | -3.06 | Rabex-5 (Rabaptin-5-associated exchange factor for Rab5) | protein interactions | 38144 | 8 | 2 | 9204 | 412 | Stat92E,Egfr | | 4.679e-02 | -3.06 | CG31802 (uncharacterized protein) | protein interactions | 318949 | 8 | 2 | 9204 | 412 | sdt,smg | | 4.679e-02 | -3.06 | Faa (Fumarylacetoacetase) | protein interactions | 45577 | 8 | 2 | 9204 | 412 | PhKgamma,CG32772 | | 4.679e-02 | -3.06 | DOR (Diabetes and obesity regulated) | protein interactions | 38543 | 8 | 2 | 9204 | 412 | CG31224,EcR | | 4.679e-02 | -3.06 | RpI135 (RNA polymerase I 135kD subunit) | protein interactions | 33210 | 8 | 2 | 9204 | 412 | udd,S6kII | | 4.679e-02 | -3.06 | Dad1 (Defender against apoptotic cell death 1) | protein interactions | 34159 | 8 | 2 | 9204 | 412 | wnd,PEK | | 4.679e-02 | -3.06 | MtnB (Metallothionein B) | protein interactions | 42424 | 8 | 2 | 9204 | 412 | esn,CG32772 | | 4.687e-02 | -3.06 | ETS2\_B | MSigDB lists | ETS2\_B | 54 | 5 | 4161 | 153 | Arp2,Cnx99A,Rac2,Utx,ras | | 4.687e-02 | -3.06 | TCF11\_01 | MSigDB lists | TCF11\_01 | 54 | 5 | 4161 | 153 | Utx,S6kII,dlp,Ser,Tsp26A | | 4.687e-02 | -3.06 | FREAC2\_01 | MSigDB lists | FREAC2\_01 | 54 | 5 | 4161 | 153 | Sin3A,Utx,dpp,CG4538,Pli | | 4.687e-02 | -3.06 | PLASARI\_TGFB1\_TARGETS\_10HR\_DN | MSigDB lists | PLASARI\_TGFB1\_TARGETS\_10HR\_DN | 54 | 5 | 4161 | 153 | ft,Fbxl7,so,CG3097,TfIIS | | 4.687e-02 | -3.06 | JIANG\_VHL\_TARGETS | MSigDB lists | JIANG\_VHL\_TARGETS | 54 | 5 | 4161 | 153 | heph,Cnx99A,cta,emp,LRP1 | | 4.687e-02 | -3.06 | E12\_Q6 | MSigDB lists | E12\_Q6 | 54 | 5 | 4161 | 153 | drk,Smox,Cnx99A,Lac,Imp | | 4.700e-02 | -3.06 | TwdlE (TweedleE) | protein interactions | 34075 | 74 | 7 | 9204 | 412 | ttk,Pat1,Doa,Imp,pAbp,Chro,Mrp4 | | 4.728e-02 | -3.05 | morphogenesis of a branching epithelium | biological process | GO:0061138 | 59 | 5 | 12345 | 414 | pnt,trn,sty,Hph,ttk | | 4.728e-02 | -3.05 | branching morphogenesis of an epithelial tube | biological process | GO:0048754 | 59 | 5 | 12345 | 414 | sty,pnt,trn,Hph,ttk | | 4.728e-02 | -3.05 | morphogenesis of a branching structure | biological process | GO:0001763 | 59 | 5 | 12345 | 414 | pnt,trn,sty,ttk,Hph | | 4.728e-02 | -3.05 | peptidyl-serine modification | biological process | GO:0018209 | 59 | 5 | 12345 | 414 | Tlk,Asator,mbt,Hipk,JIL-1 | | 4.728e-02 | -3.05 | branching involved in open tracheal system development | biological process | GO:0060446 | 59 | 5 | 12345 | 414 | pnt,trn,sty,Hph,ttk | | 4.735e-02 | -3.05 | CAGCTG\_AP4\_Q5 | MSigDB lists | CAGCTG\_AP4\_Q5 | 278 | 16 | 4161 | 153 | ras,CG43313,Hsc70-3,CG3097,CG9855,fne,LManII,trn,Pat1,LRP1,gus,ap,N,Ser,Mhcl,CG12547 | | 4.744e-02 | -3.05 | sina (seven in absentia) | protein interactions | 39884 | 59 | 6 | 9204 | 412 | Sin3A,phyl,CycE,sty,S,ttk | | 4.744e-02 | -3.05 | Sec5 (Secretory 5) | protein interactions | 33563 | 59 | 6 | 9204 | 412 | smash,hiw,egr,fj,wake,CG2991 | | 4.744e-02 | -3.05 | pan (pangolin) | protein interactions | 43769 | 59 | 6 | 9204 | 412 | Hipk,Sin3A,aop,egr,N,Ubx | | 4.752e-02 | -3.05 | HMG\_box | pfam domains | PF00505 | 22 | 3 | 10480 | 393 | Sox21a,HmgZ,Sox102F | | 4.760e-02 | -3.04 | TER94 (TER94) | protein interactions | 36040 | 106 | 9 | 9204 | 412 | Ptpmeg,fax,Df31,Fas2,Gbs-70E,Cnx99A,bor,Hsc70Cb,ft | | 4.760e-02 | -3.04 | Diap1 (Death-associated inhibitor of apoptosis 1) | protein interactions | 39753 | 106 | 9 | 9204 | 412 | CtBP,AGO2,ap,pbl,CG32772,Stat92E,egr,aos,Sin3A | | 4.761e-02 | -3.04 | regulation of TORC1 signaling | biological process | GO:1903432 | 41 | 4 | 12345 | 414 | Socs16D,wcy,ci,drk | | 4.761e-02 | -3.04 | regulation of cell division | biological process | GO:0051302 | 41 | 4 | 12345 | 414 | N,nuf,pbl,Egfr | | 4.761e-02 | -3.04 | regulation of lipid storage | biological process | GO:0010883 | 41 | 4 | 12345 | 414 | ImpL2,CaMKII,Rac2,Acsl | | 4.761e-02 | -3.04 | establishment or maintenance of microtubule cytoskeleton polarity | biological process | GO:0030951 | 41 | 4 | 12345 | 414 | heph,cnc,par-1,nudE | | 4.761e-02 | -3.04 | regulation of peptide secretion | biological process | GO:0002791 | 41 | 4 | 12345 | 414 | Snx3,S,CG9650,CaMKII | | 4.761e-02 | -3.04 | MAPK cascade | biological process | GO:0000165 | 41 | 4 | 12345 | 414 | egr,Rac2,Doa,wnd | | 4.769e-02 | -3.04 | HNF4\_DR1\_Q3 | MSigDB lists | HNF4\_DR1\_Q3 | 72 | 6 | 4161 | 153 | mbt,ds,trn,Imp,LRP1,Tlk | | 4.779e-02 | -3.04 | growth factor binding | molecular function | GO:0019838 | 11 | 2 | 12176 | 396 | tkv,ImpL2 | | 4.782e-02 | -3.04 | Ras guanyl-nucleotide exchange factor activity | molecular function | GO:0005088 | 61 | 5 | 12176 | 396 | pbl,CG43658,RhoGEF2,PsGEF,CG5522 | | 4.785e-02 | -3.04 | Bromodomain-like\_sf | interpro domains | IPR036427 | 22 | 3 | 11383 | 428 | tou,Taf1,Acf | | 4.789e-02 | -3.04 | mor (moira) | protein interactions | 41942 | 90 | 8 | 9204 | 412 | Pcl,Stat92E,tna,S6kII,CycE,Taf1,osa,Tet | | 4.792e-02 | -3.04 | Signaling by ERBB4 | REACTOME pathways | R-DME-1236394 | 22 | 3 | 3923 | 148 | Stat92E,drk,Egfr | | 4.793e-02 | -3.04 | pk (prickle) | protein interactions | 45343 | 31 | 4 | 9204 | 412 | N,ds,bdg,Rho1 | | 4.793e-02 | -3.04 | Pp1-13C (Protein phosphatase 1 at 13C) | protein interactions | 48531 | 31 | 4 | 9204 | 412 | NiPp1,ec,Gbs-70E,CG42748 | | 4.793e-02 | -3.04 | mwh (multiple wing hairs) | protein interactions | 38131 | 31 | 4 | 9204 | 412 | RhoGEF2,Rho1,wech,smg | | 4.793e-02 | -3.04 | apt (apontic) | protein interactions | 37734 | 31 | 4 | 9204 | 412 | Stat92E,hh,Ubx,pbl | | 4.793e-02 | -3.04 | Marf (Mitochondrial assembly regulatory factor) | protein interactions | 31581 | 31 | 4 | 9204 | 412 | mbt,egr,N,kdn | | 4.798e-02 | -3.04 | NUYTTEN\_EZH2\_TARGETS\_UP | MSigDB lists | NUYTTEN\_EZH2\_TARGETS\_UP | 213 | 13 | 4161 | 153 | CG9220,emp,Hsc70-3,ScpX,CG11367,sdt,CG14646,Rho1,Cyp1,mmy,S6kII,cta,CG41099 | | 4.809e-02 | -3.03 | GO\_REGULATION\_OF\_CELL\_MORPHOGENESIS | MSigDB lists | GO\_REGULATION\_OF\_CELL\_MORPHOGENESIS | 110 | 8 | 4161 | 153 | Rac2,nAChRalpha4,nudE,dpp,NetB,N,Smox,cta | | 4.846e-02 | -3.03 | small molecule binding | molecular function | GO:0036094 | 1078 | 45 | 12176 | 396 | CG9541,CG9281,Mos,Egfr,JIL-1,CG8485,betaTub56D,par-1,PhKgamma,Pex1,Arl4,CG14535,mbt,lz,mle,Myo10A,CG12010,Hph,CG4199,Doa,cta,wnd,tkv,bor,EcR,Pak3,KP78a,Cdk12,Rac2,Hsc70-3,CaMKII,Tlk,Etl1,Arp2,Asator,Mrp4,CtBP,ScsbetaG,S6kII,Rho1,PEK,Mhcl,Haspin,Hipk,CG8097 | | 4.877e-02 | -3.02 | GO\_INTRINSIC\_COMPONENT\_OF\_PLASMA\_MEMBRANE | MSigDB lists | GO\_INTRINSIC\_COMPONENT\_OF\_PLASMA\_MEMBRANE | 257 | 15 | 4161 | 153 | Mmp2,MFS10,nAChRalpha4,tkv,LRP1,dlp,dally,Syx1A,if,Ykt6,Ser,Tsp26A,dpp,Npc1a,smal | | 4.884e-02 | -3.02 | regulation of cellular amide metabolic process | biological process | GO:0034248 | 186 | 11 | 12345 | 414 | Rbfox1,bin3,ush,CG4325,elav,smg,pAbp,heph,Syp,PEK,mle | | 4.910e-02 | -3.01 | response to endoplasmic reticulum stress | biological process | GO:0034976 | 79 | 6 | 12345 | 414 | Hsc70-3,Cnx99A,cnc,Cdk5alpha,PEK,CG11873 | | 4.922e-02 | -3.01 | GSE9988\_ANTI\_TREM1\_VS\_CTRL\_TREATED\_MONOCYTES\_UP | MSigDB lists | GSE9988\_ANTI\_TREM1\_VS\_CTRL\_TREATED\_MONOCYTES\_UP | 38 | 4 | 4161 | 153 | Acsl,cta,CG9220,Npc1a | | 4.922e-02 | -3.01 | MZF1\_01 | MSigDB lists | MZF1\_01 | 38 | 4 | 4161 | 153 | LRP1,mgl,ds,SamDC | | 4.922e-02 | -3.01 | GSE23502\_BM\_VS\_COLON\_TUMOR\_HDC\_KO\_MYELOID\_DERIVED\_SUPPRESSOR\_CELL\_UP | MSigDB lists | GSE23502\_BM\_VS\_COLON\_TUMOR\_HDC\_KO\_MYELOID\_DERIVED\_SUPPRESSOR\_CELL\_UP | 38 | 4 | 4161 | 153 | nAChRalpha4,Fbxl7,DIP-epsilon,retm | | 4.922e-02 | -3.01 | GSE24634\_NAIVE\_CD4\_TCELL\_VS\_DAY7\_IL4\_CONV\_TREG\_UP | MSigDB lists | GSE24634\_NAIVE\_CD4\_TCELL\_VS\_DAY7\_IL4\_CONV\_TREG\_UP | 38 | 4 | 4161 | 153 | CG14291,Ank,CG11030,par-1 | | 4.922e-02 | -3.01 | GSE1791\_CTRL\_VS\_NEUROMEDINU\_IN\_T\_CELL\_LINE\_0.8H\_DN | MSigDB lists | GSE1791\_CTRL\_VS\_NEUROMEDINU\_IN\_T\_CELL\_LINE\_0.8H\_DN | 38 | 4 | 4161 | 153 | Cyt-c1,betaTub56D,Syx1A,TfAP-2 | | 4.922e-02 | -3.01 | AMIT\_EGF\_RESPONSE\_480\_HELA | MSigDB lists | AMIT\_EGF\_RESPONSE\_480\_HELA | 38 | 4 | 4161 | 153 | Cyp1,Rho1,Hydr2,betaTub56D | | 4.922e-02 | -3.01 | NKX3A\_01 | MSigDB lists | NKX3A\_01 | 38 | 4 | 4161 | 153 | fne,N,mgl,so | | 4.922e-02 | -3.01 | GSE15271\_CXCR4\_POS\_VS\_NEG\_GC\_BCELL\_DN | MSigDB lists | GSE15271\_CXCR4\_POS\_VS\_NEG\_GC\_BCELL\_DN | 38 | 4 | 4161 | 153 | CG45050,Hydr2,LRP1,Mhcl | | 4.922e-02 | -3.01 | GSE36888\_UNTREATED\_VS\_IL2\_TREATED\_TCELL\_17H\_DN | MSigDB lists | GSE36888\_UNTREATED\_VS\_IL2\_TREATED\_TCELL\_17H\_DN | 38 | 4 | 4161 | 153 | fj,Syx1A,betaTub56D,CG1271 | | 4.922e-02 | -3.01 | GO\_HOMEOSTASIS\_OF\_NUMBER\_OF\_CELLS | MSigDB lists | GO\_HOMEOSTASIS\_OF\_NUMBER\_OF\_CELLS | 38 | 4 | 4161 | 153 | Tif-IA,l(3)mbt,hts,Rac2 | | 4.922e-02 | -3.01 | GO\_INTERACTION\_WITH\_HOST | MSigDB lists | GO\_INTERACTION\_WITH\_HOST | 38 | 4 | 4161 | 153 | Smox,emp,Fas2,Npc1a | | 4.922e-02 | -3.01 | GO\_SKIN\_DEVELOPMENT | MSigDB lists | GO\_SKIN\_DEVELOPMENT | 38 | 4 | 4161 | 153 | ap,N,Ser,hh | | 4.922e-02 | -3.01 | RAF\_UP.V1\_DN | MSigDB lists | RAF\_UP.V1\_DN | 38 | 4 | 4161 | 153 | TfAP-2,Utx,Tif-IA,dlp | | 4.922e-02 | -3.01 | chr12q13 | MSigDB lists | chr12q13 | 38 | 4 | 4161 | 153 | lbk,LRP1,kdn,lili | | 4.951e-02 | -3.01 | import into cell | biological process | GO:0098657 | 164 | 10 | 12345 | 414 | aop,comm,EcR,CD98hc,nrv2,Rho1,rush,Nhe2,egr,CG41099 | | 4.952e-02 | -3.01 | homeostatic process | biological process | GO:0042592 | 475 | 23 | 12345 | 414 | CG46339,ush,Mondo,scaf6,cnc,Trpm,Atg17,EcR,Gyf,crb,JIL-1,N,ey,Hsc70-3,egr,Lac,Nhe2,if,nrv2,Cnx99A,Lpin,disco,Egfr | | 4.957e-02 | -3.00 | Wnt Signaling Pathway | WikiPathways | WP97 | 18 | 3 | 499 | 24 | hh,dally,ci | | 4.958e-02 | -3.00 | fly chr36D1-36D1\|2-53 | chromosome location | fly chr36D1-36D1\|2-53 | 17 | 2 | 25010 | 531 | lncRNA:CR45919,lncRNA:CR44411 | | 4.958e-02 | -3.00 | fly chr36D1-36D1\|2-53 cM | chromosome location | fly chr36D1-36D1\|2-53 cM | 17 | 2 | 25010 | 531 | lncRNA:CR44411,lncRNA:CR45919 | | 4.980e-02 | -3.00 | LIU\_LIVER\_CANCER | MSigDB lists | LIU\_LIVER\_CANCER | 10 | 2 | 4161 | 153 | smal,Dscam1 | | 4.980e-02 | -3.00 | chr3q23 | MSigDB lists | chr3q23 | 10 | 2 | 4161 | 153 | gus,CG1271 | | 4.980e-02 | -3.00 | CTIP\_DN.V1\_UP | MSigDB lists | CTIP\_DN.V1\_UP | 10 | 2 | 4161 | 153 | Nhe2,slow | | 4.980e-02 | -3.00 | GO\_T\_TUBULE | MSigDB lists | GO\_T\_TUBULE | 10 | 2 | 4161 | 153 | Ank,CaMKII | | 4.980e-02 | -3.00 | GO\_POSITIVE\_REGULATION\_OF\_ENDOTHELIAL\_CELL\_PROLIFERATION | MSigDB lists | GO\_POSITIVE\_REGULATION\_OF\_ENDOTHELIAL\_CELL\_PROLIFERATION | 10 | 2 | 4161 | 153 | dpp,slow | | 4.980e-02 | -3.00 | GO\_REGULATION\_OF\_FIBROBLAST\_MIGRATION | MSigDB lists | GO\_REGULATION\_OF\_FIBROBLAST\_MIGRATION | 10 | 2 | 4161 | 153 | smal,Rac2 | | 4.980e-02 | -3.00 | REACTOME\_HS\_GAG\_BIOSYNTHESIS | MSigDB lists | REACTOME\_HS\_GAG\_BIOSYNTHESIS | 10 | 2 | 4161 | 153 | dally,dlp | | 4.980e-02 | -3.00 | GO\_PROSTATE\_GLAND\_DEVELOPMENT | MSigDB lists | GO\_PROSTATE\_GLAND\_DEVELOPMENT | 10 | 2 | 4161 | 153 | N,hh | | 4.980e-02 | -3.00 | chr14q21 | MSigDB lists | chr14q21 | 10 | 2 | 4161 | 153 | so,Pli | | 4.980e-02 | -3.00 | GO\_HOMEOSTASIS\_OF\_NUMBER\_OF\_CELLS\_WITHIN\_A\_TISSUE | MSigDB lists | GO\_HOMEOSTASIS\_OF\_NUMBER\_OF\_CELLS\_WITHIN\_A\_TISSUE | 10 | 2 | 4161 | 153 | Rac2,hts | | 4.980e-02 | -3.00 | REACTOME\_NUCLEAR\_EVENTS\_KINASE\_AND\_TRANSCRIPTION\_FACTOR\_ACTIVATION | MSigDB lists | REACTOME\_NUCLEAR\_EVENTS\_KINASE\_AND\_TRANSCRIPTION\_FACTOR\_ACTIVATION | 10 | 2 | 4161 | 153 | JIL-1,S6kII | | 4.980e-02 | -3.00 | REACTOME\_ERK\_MAPK\_TARGETS | MSigDB lists | REACTOME\_ERK\_MAPK\_TARGETS | 10 | 2 | 4161 | 153 | S6kII,JIL-1 | | 4.980e-02 | -3.00 | GO\_CELLULAR\_RESPONSE\_TO\_INTERFERON\_GAMMA | MSigDB lists | GO\_CELLULAR\_RESPONSE\_TO\_INTERFERON\_GAMMA | 10 | 2 | 4161 | 153 | CaMKII,Fas2 | | 4.980e-02 | -3.00 | GO\_STEROL\_HOMEOSTASIS | MSigDB lists | GO\_STEROL\_HOMEOSTASIS | 10 | 2 | 4161 | 153 | EcR,Npc1a | | 4.980e-02 | -3.00 | PID\_S1P\_S1P2\_PATHWAY | MSigDB lists | PID\_S1P\_S1P2\_PATHWAY | 10 | 2 | 4161 | 153 | Rac2,cta | | 4.980e-02 | -3.00 | GO\_OLFACTORY\_LOBE\_DEVELOPMENT | MSigDB lists | GO\_OLFACTORY\_LOBE\_DEVELOPMENT | 10 | 2 | 4161 | 153 | ap,Rac2 | | 4.980e-02 | -3.00 | VERRECCHIA\_DELAYED\_RESPONSE\_TO\_TGFB1 | MSigDB lists | VERRECCHIA\_DELAYED\_RESPONSE\_TO\_TGFB1 | 10 | 2 | 4161 | 153 | Rac2,Mmp2 | | 4.980e-02 | -3.00 | PID\_NEPHRIN\_NEPH1\_PATHWAY | MSigDB lists | PID\_NEPHRIN\_NEPH1\_PATHWAY | 10 | 2 | 4161 | 153 | drk,Rac2 | | 4.980e-02 | -3.00 | MARTIN\_INTERACT\_WITH\_HDAC | MSigDB lists | MARTIN\_INTERACT\_WITH\_HDAC | 10 | 2 | 4161 | 153 | Sin3A,CaMKII | | 4.980e-02 | -3.00 | GO\_GROWTH\_FACTOR\_ACTIVITY | MSigDB lists | GO\_GROWTH\_FACTOR\_ACTIVITY | 10 | 2 | 4161 | 153 | Ser,dpp | | 4.980e-02 | -3.00 | GO\_INTERCALATED\_DISC | MSigDB lists | GO\_INTERCALATED\_DISC | 10 | 2 | 4161 | 153 | CaMKII,Ank | | 4.980e-02 | -3.00 | BIOCARTA\_SALMONELLA\_PATHWAY | MSigDB lists | BIOCARTA\_SALMONELLA\_PATHWAY | 10 | 2 | 4161 | 153 | Arp2,Rac2 | | 4.980e-02 | -3.00 | GO\_VENTRICULAR\_SEPTUM\_DEVELOPMENT | MSigDB lists | GO\_VENTRICULAR\_SEPTUM\_DEVELOPMENT | 10 | 2 | 4161 | 153 | mgl,N | | 4.980e-02 | -3.00 | REACTOME\_SEMA4D\_INDUCED\_CELL\_MIGRATION\_AND\_GROWTH\_CONE\_COLLAPSE | MSigDB lists | REACTOME\_SEMA4D\_INDUCED\_CELL\_MIGRATION\_AND\_GROWTH\_CONE\_COLLAPSE | 10 | 2 | 4161 | 153 | Rac2,Rho1 | | 4.980e-02 | -3.00 | LUI\_THYROID\_CANCER\_CLUSTER\_1 | MSigDB lists | LUI\_THYROID\_CANCER\_CLUSTER\_1 | 10 | 2 | 4161 | 153 | CtBP,Ank | | 4.980e-02 | -3.00 | REACTOME\_EXTRACELLULAR\_MATRIX\_ORGANIZATION | MSigDB lists | REACTOME\_EXTRACELLULAR\_MATRIX\_ORGANIZATION | 10 | 2 | 4161 | 153 | CG10663,Mmp2 | | 4.980e-02 | -3.00 | GO\_STEROL\_TRANSPORT | MSigDB lists | GO\_STEROL\_TRANSPORT | 10 | 2 | 4161 | 153 | Npc1a,ScpX | | 4.980e-02 | -3.00 | AGTCTAG\_MIR151 | MSigDB lists | AGTCTAG\_MIR151 | 10 | 2 | 4161 | 153 | Rhp,DIP-epsilon | | 4.980e-02 | -3.00 | GO\_PROTEIN\_KINASE\_C\_ACTIVATING\_G\_PROTEIN\_COUPLED\_RECEPTOR\_SIGNALING\_PATHWAY | MSigDB lists | GO\_PROTEIN\_KINASE\_C\_ACTIVATING\_G\_PROTEIN\_COUPLED\_RECEPTOR\_SIGNALING\_PATHWAY | 10 | 2 | 4161 | 153 | LRP1,CG34384 | | 4.980e-02 | -3.00 | BIOCARTA\_PTDINS\_PATHWAY | MSigDB lists | BIOCARTA\_PTDINS\_PATHWAY | 10 | 2 | 4161 | 153 | Ser,Rac2 | | 4.980e-02 | -3.00 | INGRAM\_SHH\_TARGETS\_DN | MSigDB lists | INGRAM\_SHH\_TARGETS\_DN | 10 | 2 | 4161 | 153 | Tudor-SN,CycE | | 4.980e-02 | -3.00 | GO\_CARDIAC\_SEPTUM\_MORPHOGENESIS | MSigDB lists | GO\_CARDIAC\_SEPTUM\_MORPHOGENESIS | 10 | 2 | 4161 | 153 | N,Ser | | 4.980e-02 | -3.00 | GO\_INTRINSIC\_COMPONENT\_OF\_MITOCHONDRIAL\_INNER\_MEMBRANE | MSigDB lists | GO\_INTRINSIC\_COMPONENT\_OF\_MITOCHONDRIAL\_INNER\_MEMBRANE | 10 | 2 | 4161 | 153 | Mitofilin,Tim17b | | 4.980e-02 | -3.00 | chr8q12 | MSigDB lists | chr8q12 | 10 | 2 | 4161 | 153 | NKAIN,CG3097 | | 4.984e-02 | -3.00 | FN3 | prosite domains | PS50853 | 53 | 6 | 6243 | 316 | robo2,Dscam1,sas,Fas2,Ptp99A,wake | | 4.988e-02 | -3.00 | positive regulation of organ growth | biological process | GO:0046622 | 25 | 3 | 12345 | 414 | dpp,Hipk,Egfr | | 4.988e-02 | -3.00 | female sex differentiation | biological process | GO:0046660 | 25 | 3 | 12345 | 414 | lz,bab1,otk2 | | 4.988e-02 | -3.00 | regulation of exocytosis | biological process | GO:0017157 | 25 | 3 | 12345 | 414 | corn,NFAT,Syx1A | | 4.988e-02 | -3.00 | actin filament bundle organization | biological process | GO:0061572 | 25 | 3 | 12345 | 414 | hts,Rac2,Rho1 | | 4.988e-02 | -3.00 | regulation of pole plasm oskar mRNA localization | biological process | GO:0007317 | 25 | 3 | 12345 | 414 | Syx1A,par-1,cnc | | 4.988e-02 | -3.00 | regulation of terminal button organization | biological process | GO:2000331 | 25 | 3 | 12345 | 414 | wnd,Ten-m,hiw | | 4.988e-02 | -3.00 | regulation of oocyte development | biological process | GO:0060281 | 25 | 3 | 12345 | 414 | par-1,cnc,Syx1A | | 4.999e-02 | -3.00 | BYSTRYKH\_HEMATOPOIESIS\_STEM\_CELL\_QTL\_TRANS | MSigDB lists | BYSTRYKH\_HEMATOPOIESIS\_STEM\_CELL\_QTL\_TRANS | 280 | 16 | 4161 | 153 | CG4538,Snx3,CG14646,Tpr2,bor,Etl1,Cnx99A,Tlk,dlp,Mmp2,ras,Tif-IA,Fas2,mle,cta,TfAP-2 |
