## Supplementary material for "A single DPE core promoter motif contributes to *in vivo* transcriptional regulation and affects cardiac function": SupFile2: knownResults.html

./cluster.allRlog.c300.t8//18.cluster.txt.motifs/ - Homer Known Motif Enrichment Results


### Homer Known Motif Enrichment Results (./cluster.allRlog.c300.t8//18.cluster.txt.motifs/)

Homer *de novo* Motif Results  
Gene Ontology Enrichment Results  
Known Motif Enrichment Results (txt file)  
Total Target Sequences = 575, Total Background Sequences = 26054

|  |  |  |  |  |  |  |  |  |  |  |  |
| --- | --- | --- | --- | --- | --- | --- | --- | --- | --- | --- | --- |
| Rank | Motif | Name | P-value | log P-pvalue | q-value (Benjamini) | # Target Sequences with Motif | % of Targets Sequences with Motif | # Background Sequences with Motif | % of Background Sequences with Motif | Motif File | SVG |
| 1 | C T A G T C A G C T G A T C A G T G C A A C T G T C G A T C A G | Trl(Zf)/S2-GAGAfactor-ChIP-Seq(GSE40646)/Homer | 1e-11 | -2.595e+01 | 0.0000 | 194.0 | 33.74% | 5570.4 | 21.38% | motif file (matrix) | svg |
| 2 | C G T A T C G A G A T C G C A T C G T A A C G T G T A C T C A G G T C A G A C T C G T A C T A G | DREF/Drosophila-Promoters/Homer | 1e-8 | -2.046e+01 | 0.0000 | 35.0 | 6.09% | 477.7 | 1.83% | motif file (matrix) | svg |
| 3 | G C A T A C G T A T G C C T G A C T A G G A C T G A T C A C T G | Initiator/Drosophila-Promoters/Homer | 1e-8 | -1.982e+01 | 0.0000 | 196.0 | 34.09% | 6056.3 | 23.24% | motif file (matrix) | svg |
| 4 | G A T C C T G A C T A G G C A T C T A G C A G T C A T G T C G A G T A C A G T C T C A G A G C T | M1BP(Zf)/S2R+-M1BP-ChIP-Seq(GSE49842)/Homer | 1e-8 | -1.848e+01 | 0.0000 | 41.0 | 7.13% | 672.6 | 2.58% | motif file (matrix) | svg |
| 5 | T G C A A G T C C T A G A C T G A G C T G T A C T C G A A G T C C G T A A G T C G A C T C T A G | Unknown1(NR/Ini-like)/Drosophila-Promoters/Homer | 1e-6 | -1.545e+01 | 0.0000 | 37.0 | 6.43% | 641.1 | 2.46% | motif file (matrix) | svg |
| 6 | A G T C C G T A C G A T A G T C G T C A A G T C A C G T C T G A | Unknown2/Drosophila-Promoters/Homer | 1e-5 | -1.162e+01 | 0.0000 | 63.0 | 10.96% | 1605.2 | 6.16% | motif file (matrix) | svg |
| 7 | C T G A C G T A C G T A G C T A C T G A A C G T G T C A A G T C A G T C C T G A G T C A G C T A | Unknown4/Drosophila-Promoters/Homer | 1e-3 | -7.693e+00 | 0.0011 | 21.0 | 3.65% | 413.4 | 1.59% | motif file (matrix) | svg |
| 8 | A C T G A T C G A G T C A C G T C G T A A G T C A G T C A C G T A C T G C G T A | Zelda(Zf)/Embryo-zld-ChIP-Seq(GSE65441)/Homer | 1e-2 | -5.872e+00 | 0.0060 | 27.0 | 4.70% | 678.6 | 2.60% | motif file (matrix) | svg |
| 9 | C G T A G C T A C G A T A C G T A G C T A G C T C T G A G T C A C G T A G C T A | Unknown6/Drosophila-Promoters/Homer | 1e-2 | -4.731e+00 | 0.0166 | 74.0 | 12.87% | 2539.4 | 9.75% | motif file (matrix) | svg |
