## Supplementary material for "A single DPE core promoter motif contributes to *in vivo* transcriptional regulation and affects cardiac function": SupFile2: geneOntology.html

Gene Ontology Results

### Gene Ontology Enrichment Results

Homer *de novo* Motif Enrichment Results  
Known Motif Enrichment Results  

###### Text file version of complete results (i.e. open with Excel) - biological process: Functional groupings of proteins (Gene Ontology) - molecular function: Mechanistic actions of proteins (Gene Ontology) - cellular component: Protein localization (Gene Ontology) - chromosome location: Genes with similar chromosome localization (NCBI Gene) - KEGG pathways: Groups of proteins in the same pathways (KEGG) - protein interactions: "Proteins interacting with a common protein (BIND, EcoCyc, HPRD)" (NCBI Gene) - interpro domains: Proteins with similar domains and features (Interpro) - pfam domains: Proteins with similar domains and features (Pfam) - smart domains: Proteins with similar domains and features (SMART) - gene3d domains: Proteins with similar domains and features (Gene3D) - prosite domains: Proteins with similar domains and features (Prosite) - prints domains: Proteins with similar domains and features (PRINTS) - MSigDB lists: "Genes sets for pathways, factor/miRNA target predictions, expression patterns, etc." (MSigDB) - BIOCYC pathways: Groups of proteins in the same pathways (BIOCYC) - COSMIC cancer mutations: Genes mutated in similar cancers (COSMIC) - GWAS genes: Genes mutated in similar diseases (GWAS Catalog) - Lipid Maps pathways: Groups of proteins in the same lipid pathways (Lipid Maps/Biosystems) - Pathway Interaction DB: Groups of proteins in the same pathways (Pathway Interaction Database) - REACTOME pathways: Groups of proteins in the same pathways (REACTOME) - SMPDB pathways: Groups of proteins in the same pathways (SMPDB) - WikiPathways: Groups of proteins in the same pathways (Wikipathways) Enriched Categories | | | | | | | | | | | | --- | --- | --- | --- | --- | --- | --- | --- | --- | --- | | P-value | ln(P) | Term | GO Tree | GO ID | # of Genes in Term | # of Target Genes in Term | # of Total Genes | # of Target Genes | Common Genes | | 2.281e-13 | -29.11 | biological regulation | biological process | GO:0065007 | 4350 | 295 | 12345 | 597 | Src42A,LpR2,Fhos,mGluR,fz2,kay,pirk,Su(H),PhKgamma,KaiR1D,Syt12,RhoGAP100F,jim,Exn,CG3530,fz4,CG42830,FoxL1,Spn88Eb,Dlic,ninaD,Flo2,Art2,l(3)neo38,trol,CG11318,lncRNA:Hsromega,luna,Stacl,Rab9Fa,trv,Myo81F,GlyP,g,shd,Antp,AANAT1,Hr96,CG14252,Atf6,Eip74EF,Vha100-5,crol,Syx1A,Ank2,Pomp,scro,numb,CG7483,Actn,Acp62F,by,CG4849,RYa,olf186-F,Cpsf160,Nlg2,bun,rdgA,GATAe,rib,CG18304,MESK2,Pld,ddbt,Qsox2,Ank,nemy,melt,EcR,Marf1,Achl,CG32006,csw,Nmdar2,CG7368,siz,CG31750,Der-1,Tk,Ste:CG33247,tld,upd3,CG3842,sky,Stlk,jus,Mondo,CG30054,CG3078,cpx,Snoo,CG32971,CG13229,Pde6,Pkc98E,RhoGEF3,CG13526,Cadps,nrv2,tyf,wbl,Pde8,Pde1c,CG41520,chinmo,toe,tin,Msp300,Raf,dnc,AstA-R1,Irk3,HDAC4,CG31224,dpp,CG17698,CG9981,can,unc-13,Elp3,hdc,Poxm,barc,E5,Fer2,Neb-cGP,Ubx,rictor,sdt,gce,CG34172,jigr1,CG12200,RhoGAP18B,ham,CG42390,SteXh:CG42398,TpnC25D,scrib,Awh,Acbp2,sno,scramb1,mid,rst,Mlp84B,kek1,raw,CG17124,et,tnc,Syn2,Sod1,Spf45,Mob2,CG4301,Set2,bnl,polybromo,msps,RhoL,sgg,fus,CG9380,Fife,CG40178,kek5,ine,Mos,Mbs,shakB,l(2)01289,sxc,hth,spri,CG9650,HPS4,Cam,Mrtf,ps,CG12071,Yeti,nuf,TfAP-2,RhoGAP1A,AGO2,Ser,CycT,Mctp,shg,step,CG17883,Obp56h,mamo,CG14182,msi,bi,CG43689,Strn-Mlck,path,ths,B-H2,TfIIFbeta,bru1,Cdep,Fad2,retn,spok,fru,woc,sws,CG18659,CG4374,Glut1,CG42797,fs(1)N,cindr,Frq2,CG34408,CG1572,MsR2,Ptp61F,REG,Cat,ems,grn,egh,CG4839,Spt20,boi,Frq1,CG6966,ACXD,Tusp,Tor,ex,Dys,CG1894,Gpb5,Khc,CG33310,Pak,Cln3,Usp1,TpnC41C,foxo,chb,dsh,qvr,CG10462,CG15073,CG5059,opa,dany,Iswi,Pdfr,mnb,Ca-beta,p130CAS,cic,ZnT63C,CG10188,Octbeta2R,bol,esg,flw,fw,CG12605,CCAP-R,Doa,Pde9,CG5522,GluRIIC,Lpin,Gprk1,JhI-26,lncRNA:bxd,Sox102F,Cep97,AGO3,CrebB,CG9514,CG13898,ITP,PGRP-LA,Sh,CG9519,CG13711,mag,Pkc53E,Maf1,MCU,wake,htk | | 3.042e-12 | -26.52 | nervous system development | biological process | GO:0007399 | 1108 | 106 | 12345 | 597 | shd,dnc,Antp,CycT,Dmtn,Ser,shg,CG5921,opa,trol,foxo,chinmo,chb,shep,Cam,dsh,tin,Raf,CG13251,Khc,jim,RhoGAP100F,ine,hth,Pak,Mbs,spri,Dlic,nrv2,Src42A,raw,boi,Syn2,Stlk,sky,Mob2,cpx,kay,fz2,app,msps,bnl,Tor,Snoo,ex,Fife,sgg,scrib,E(Pc),melt,EcR,Awh,Ptp61F,sno,Sh,ems,csw,mid,siz,klg,egh,Ptp99A,wake,grn,olf186-F,Nlg2,sqz,bun,CG42797,rib,chp,beat-IIa,sick,ham,nrm,Fer2,E5,esg,dpr12,retn,fw,CadN2,pod1,rictor,Doa,sdt,fru,sws,dsd,gukh,msi,Elp3,Ca-beta,mnb,Ank2,path,hdc,barc,esn,scro,p130CAS,ths,Poxm,numb,B-H2,haf | | 3.861e-12 | -26.28 | regulation of biological process | biological process | GO:0050789 | 3862 | 265 | 12345 | 597 | g,Antp,AANAT1,Atf6,Hr96,CG14252,l(3)neo38,trol,CG11318,lncRNA:Hsromega,luna,Rab9Fa,Stacl,trv,Myo81F,GlyP,PhKgamma,Syt12,KaiR1D,Exn,jim,RhoGAP100F,fz4,CG3530,ninaD,Dlic,CG42830,Spn88Eb,FoxL1,Art2,Flo2,Src42A,LpR2,fz2,kay,mGluR,Fhos,pirk,Su(H),Qsox2,melt,EcR,nemy,Ank,Marf1,Nmdar2,csw,CG32006,Achl,Der-1,CG31750,CG7368,siz,olf186-F,Cpsf160,Nlg2,bun,rib,CG18304,GATAe,rdgA,MESK2,Pld,CG7483,Acp62F,CG4849,RYa,Eip74EF,crol,Syx1A,Ank2,scro,Pomp,numb,dnc,AstA-R1,HDAC4,CG31224,dpp,Pde8,wbl,CG41520,Pde1c,chinmo,toe,tin,Raf,Pkc98E,RhoGEF3,Cadps,CG13526,tyf,Ste:CG33247,Tk,upd3,tld,Stlk,sky,CG30054,Mondo,cpx,CG3078,Snoo,CG32971,Pde6,CG13229,scrib,Awh,sno,mid,Mlp84B,rst,CG34172,CG12200,RhoGAP18B,jigr1,ham,TpnC25D,SteXh:CG42398,Fer2,E5,Neb-cGP,Ubx,rictor,sdt,gce,CG17698,can,Elp3,unc-13,hdc,barc,Poxm,RhoGAP1A,AGO2,CycT,Ser,shg,Mctp,step,HPS4,Cam,Mrtf,Yeti,nuf,CG12071,ps,TfAP-2,CG40178,kek5,Mos,ine,hth,Mbs,l(2)01289,shakB,sxc,spri,CG9650,et,raw,CG17124,kek1,Sod1,Syn2,Spf45,Set2,Mob2,polybromo,RhoL,msps,bnl,Fife,sgg,CG9380,fus,CG1572,MsR2,Ptp61F,Cat,REG,ems,CG4839,grn,egh,CG4374,Glut1,CG42797,fs(1)N,cindr,Frq2,CG34408,Cdep,bru1,retn,woc,fru,sws,CG18659,mamo,CG14182,CG43689,msi,bi,path,ths,TfIIFbeta,B-H2,CG15073,CG5059,opa,dany,foxo,TpnC41C,chb,dsh,qvr,CG10462,Dys,Khc,CG1894,Gpb5,Cln3,Pak,Usp1,Spt20,boi,Frq1,CG6966,ACXD,Tusp,Tor,ex,CrebB,CG13898,ITP,PGRP-LA,Sh,CG13711,Pkc53E,Maf1,wake,htk,GluRIIC,Gprk1,Lpin,Sox102F,lncRNA:bxd,Cep97,AGO3,CG10188,bol,Octbeta2R,esg,flw,CCAP-R,CG12605,Doa,Pde9,CG5522,Iswi,Pdfr,Ca-beta,mnb,cic | | 4.553e-12 | -26.12 | neuron differentiation | biological process | GO:0030182 | 710 | 78 | 12345 | 597 | Src42A,Syn2,boi,Stlk,sky,Mob2,fz2,kay,Snoo,Tor,msps,app,ex,CG13251,Khc,RhoGAP100F,jim,Mbs,Pak,hth,spri,Dlic,trol,foxo,chinmo,shep,chb,Cam,dsh,Raf,dnc,CG5921,shg,msi,dsd,gukh,Ca-beta,path,hdc,Ank2,p130CAS,Poxm,esn,scro,haf,B-H2,nrm,E5,dpr12,fw,retn,pod1,CadN2,rictor,fru,sdt,Doa,sqz,bun,chp,ham,sick,beat-IIa,scrib,E(Pc),melt,EcR,Awh,Ptp61F,Sh,sno,ems,mid,csw,grn,klg,egh,Ptp99A | | 4.703e-12 | -26.08 | neurogenesis | biological process | GO:0022008 | 923 | 93 | 12345 | 597 | sky,Stlk,boi,Syn2,Src42A,ex,sgg,msps,app,Snoo,Tor,bnl,kay,fz2,Mob2,jim,ine,RhoGAP100F,Khc,CG13251,nrv2,Dlic,spri,hth,Pak,Mbs,chb,shep,chinmo,trol,foxo,Raf,dsh,Cam,Ser,Antp,dnc,shg,CG5921,dsd,gukh,msi,numb,haf,B-H2,esn,scro,barc,ths,Poxm,p130CAS,Ank2,path,hdc,Elp3,Ca-beta,mnb,retn,fw,esg,dpr12,Fer2,E5,nrm,fru,Doa,sdt,rictor,CadN2,pod1,sqz,bun,beat-IIa,ham,sick,CG42797,chp,Ptp61F,Awh,melt,EcR,E(Pc),scrib,egh,Ptp99A,wake,grn,klg,csw,mid,ems,sno,Sh | | 7.043e-12 | -25.68 | regulation of cellular process | biological process | GO:0050794 | 3560 | 248 | 12345 | 597 | fs(1)N,rdgA,GATAe,rib,CG42797,CG18304,Pld,CG34408,Frq2,cindr,MESK2,Cpsf160,olf186-F,CG4374,bun,Glut1,Nlg2,ems,Marf1,siz,grn,CG7368,CG4839,CG31750,Der-1,Nmdar2,CG32006,csw,Qsox2,MsR2,CG1572,REG,Cat,Ptp61F,nemy,Ank,EcR,melt,Syx1A,Ank2,crol,TfIIFbeta,B-H2,numb,Pomp,ths,scro,Eip74EF,mamo,msi,bi,CG43689,CG14182,Acp62F,sws,CG18659,RYa,fru,woc,Cdep,bru1,retn,Stacl,Rab9Fa,lncRNA:Hsromega,luna,Cam,Mrtf,TfAP-2,GlyP,trv,CG12071,Myo81F,ps,nuf,Yeti,trol,HPS4,l(3)neo38,CG11318,step,CG14252,Hr96,Mctp,shg,Atf6,RhoGAP1A,Antp,g,Ser,CycT,AGO2,mGluR,Fhos,kay,fz2,Mob2,Set2,sgg,fus,CG9380,Fife,pirk,bnl,Su(H),msps,polybromo,kek1,CG17124,Src42A,raw,et,Spf45,Syn2,Sod1,spri,CG3530,fz4,l(2)01289,Mbs,sxc,shakB,hth,Flo2,CG9650,Art2,FoxL1,Spn88Eb,CG42830,ninaD,Dlic,Syt12,CG40178,KaiR1D,PhKgamma,RhoGAP100F,jim,Mos,Exn,kek5,lncRNA:bxd,Sox102F,SteXh:CG42398,TpnC25D,AGO3,ham,Cep97,CG34172,GluRIIC,Gprk1,RhoGAP18B,CG12200,Lpin,Pkc53E,Sh,sno,wake,htk,Maf1,mid,ITP,CG13898,scrib,CrebB,PGRP-LA,Awh,hdc,unc-13,mnb,Elp3,Ca-beta,cic,Poxm,barc,CG17698,Iswi,Pdfr,rictor,CG12605,CCAP-R,gce,CG5522,Doa,sdt,Pde9,E5,bol,Octbeta2R,Fer2,CG10188,Ubx,esg,flw,dsh,qvr,tin,CG10462,Raf,Pde1c,TpnC41C,CG41520,foxo,Pde8,toe,chb,chinmo,CG31224,dany,dpp,opa,CG5059,AstA-R1,CG15073,dnc,HDAC4,Tusp,ACXD,cpx,Mondo,CG30054,ex,CG13229,Pde6,Tor,Snoo,CG32971,tld,upd3,Spt20,Tk,Ste:CG33247,sky,Stlk,CG6966,Frq1,boi,CG13526,Cadps,Pak,RhoGEF3,Cln3,Usp1,tyf,Dys,Pkc98E,CG1894,Gpb5,Khc | | 1.454e-11 | -24.95 | fly chr3-47 | chromosome location | fly chr3-47 | 46 | 15 | 25010 | 845 | lncRNA:CR45177,CG40045,lncRNA:CR45159,CG40160,CG40472,scro,CG40178,CG40470,lncRNA:CR43941,CG45782,CG42598,UQCR-11,mRpS5,snoRNA:Me18S-C419,ND-AGGG | | 1.454e-11 | -24.95 | fly chr3-47 cM | chromosome location | fly chr3-47 cM | 46 | 15 | 25010 | 845 | UQCR-11,lncRNA:CR43941,CG42598,CG45782,snoRNA:Me18S-C419,ND-AGGG,mRpS5,CG40045,lncRNA:CR45177,scro,CG40178,CG40470,lncRNA:CR45159,CG40160,CG40472 | | 2.885e-11 | -24.27 | fly chr3-47.1 cM | chromosome location | fly chr3-47.1 cM | 17 | 10 | 25010 | 845 | CG41128,Myo81F,Pzl,CG40198,CG42402,CG41099,lncRNA:CR46123,spok,lncRNA:CR45220,lncRNA:CR46122 | | 2.885e-11 | -24.27 | fly chr3-47.1 | chromosome location | fly chr3-47.1 | 17 | 10 | 25010 | 845 | spok,lncRNA:CR45220,lncRNA:CR46122,lncRNA:CR46123,CG41099,CG42402,Pzl,CG40198,CG41128,Myo81F | | 2.921e-11 | -24.26 | generation of neurons | biological process | GO:0048699 | 880 | 88 | 12345 | 597 | gukh,dsd,msi,Ank2,path,hdc,Ca-beta,Elp3,mnb,haf,B-H2,numb,barc,scro,esn,Poxm,p130CAS,Fer2,E5,nrm,retn,fw,esg,dpr12,rictor,CadN2,pod1,sdt,fru,Doa,sqz,bun,CG42797,chp,beat-IIa,ham,sick,E(Pc),scrib,Ptp61F,EcR,melt,Awh,ems,sno,Sh,klg,egh,grn,Ptp99A,wake,csw,mid,Src42A,sky,Stlk,Syn2,boi,kay,fz2,Mob2,sgg,ex,app,msps,Snoo,Tor,CG13251,jim,RhoGAP100F,Khc,spri,hth,Mbs,Pak,Dlic,foxo,trol,chb,shep,chinmo,dsh,Cam,Raf,Antp,dnc,shg,CG5921 | | 7.066e-11 | -23.37 | neuron development | biological process | GO:0048666 | 584 | 66 | 12345 | 597 | haf,p130CAS,Poxm,esn,scro,path,hdc,Ank2,Ca-beta,dsd,gukh,sdt,Doa,fru,rictor,pod1,CadN2,retn,dpr12,nrm,sick,ham,beat-IIa,chp,sqz,grn,egh,klg,Ptp99A,mid,csw,ems,Sh,sno,Ptp61F,EcR,E(Pc),Tor,Snoo,msps,fz2,kay,Mob2,Stlk,sky,Syn2,Src42A,Dlic,spri,Pak,Mbs,RhoGAP100F,jim,Khc,CG13251,Raf,dsh,Cam,shep,chb,chinmo,foxo,trol,CG5921,shg,dnc | | 1.144e-10 | -22.89 | cell differentiation | biological process | GO:0030154 | 1773 | 143 | 12345 | 597 | Msp300,Raf,CG41099,dsh,tin,chinmo,chb,wbl,foxo,dpp,CG5921,dnc,Tor,Snoo,bbg,ex,boi,sky,Stlk,upd3,tld,nrv2,Pak,Rbp6,Khc,ham,sick,beat-IIa,Act88F,CG42355,Sox102F,mid,rst,Ptp99A,wake,Mlp84B,Sh,sno,Awh,scrib,Poxm,p130CAS,barc,cic,haf,DNaseII,mnb,Elp3,Ca-beta,hdc,gukh,can,Iswi,sdt,Doa,pod1,CadN2,drd,rictor,esg,flw,fw,Ubx,lncRNA:flam,nrm,E5,bol,Fer2,nuf,lncRNA:Hsromega,Mrtf,Cam,shep,trol,Hr96,shg,AGO2,Ser,shd,Antp,Su(H),bnl,msps,app,polybromo,RhoL,sgg,Mob2,Fhos,fz2,kay,Syn2,kek1,Src42A,FoxL1,Dlic,Mbs,hth,spri,RhoGAP100F,ine,Mos,jim,CG13251,cindr,GATAe,chp,Inx2,CG42797,fs(1)N,sqz,bun,CG32006,rols,csw,klg,siz,grn,egh,ems,EcR,melt,Ptp61F,E(Pc),ths,esn,scro,B-H2,numb,path,Ank2,msi,bi,dsd,Eip74EF,IP3K2,fru,Prm,dpr12,Actn,retn,CG7483,bru1 | | 2.388e-10 | -22.16 | system development | biological process | GO:0048731 | 1808 | 144 | 12345 | 597 | Lpin,beat-IIa,sick,ham,Awh,scrib,Ptp99A,wake,rst,mid,sno,Sh,gukh,cic,haf,barc,Poxm,p130CAS,hdc,Ca-beta,Elp3,mnb,Ubx,fw,flw,esg,Fer2,E5,nrm,cato,Doa,sdt,rictor,CadN2,pod1,chb,toe,chinmo,foxo,Msp300,Raf,tin,dsh,Irk3,dnc,dpp,opa,CG5921,sky,Stlk,boi,upd3,tld,ex,Snoo,Tor,cpx,Idgf6,Khc,Dys,nrv2,Pak,sqz,bun,Nlg2,olf186-F,cindr,rib,CG42797,Inx2,GATAe,chp,Cat,Ptp61F,melt,EcR,E(Pc),siz,klg,grn,egh,rols,csw,ems,dsd,bi,msi,IP3K2,Eip74EF,numb,B-H2,scro,esn,ths,Ank2,Syx1A,path,crol,retn,dpr12,sws,fru,by,shep,trol,TfAP-2,nuf,Mrtf,Cam,Dmtn,CycT,Ser,Antp,shd,step,shg,Spf45,Syn2,tnc,raw,Src42A,Fife,sgg,msps,polybromo,RhoL,app,bnl,Su(H),kay,fz2,Fhos,Set2,Mob2,jim,Exn,ine,RhoGAP100F,CG13251,Dlic,spri,hth,sxc,Mbs | | 3.545e-10 | -21.76 | cellular developmental process | biological process | GO:0048869 | 1818 | 144 | 12345 | 597 | CG42355,Sox102F,ham,sick,beat-IIa,Act88F,scrib,Awh,Sh,sno,mid,wake,Ptp99A,rst,Mlp84B,Iswi,can,gukh,mnb,Elp3,Ca-beta,hdc,p130CAS,Poxm,barc,cic,haf,DNaseII,lncRNA:flam,nrm,bol,E5,Fer2,flw,esg,fw,Ubx,pod1,CadN2,rictor,drd,sdt,Doa,wbl,foxo,chinmo,chb,dsh,tin,Raf,Msp300,CG41099,dnc,CG5921,dpp,upd3,tld,boi,Stlk,sky,Tor,Snoo,bbg,ex,Khc,Pak,Rbp6,nrv2,sqz,bun,GATAe,Inx2,chp,CG42797,fs(1)N,cindr,E(Pc),melt,EcR,Ptp61F,ems,csw,CG32006,rols,grn,egh,siz,klg,IP3K2,Eip74EF,bi,msi,dsd,path,Ank2,ths,esn,scro,B-H2,numb,CG7483,bru1,dpr12,Actn,retn,Prm,fru,trol,shep,lncRNA:Hsromega,Mrtf,Cam,nuf,shd,Antp,AGO2,Ser,Hr96,shg,kek1,raw,Src42A,Syn2,Mob2,Fhos,fz2,kay,Su(H),bnl,app,polybromo,msps,RhoL,sgg,CG13251,RhoGAP100F,ine,jim,Mos,Mbs,hth,spri,FoxL1,Dlic | | 5.157e-10 | -21.39 | animal organ morphogenesis | biological process | GO:0009887 | 782 | 78 | 12345 | 597 | Ubx,retn,fw,esg,flw,Doa,fru,CadN2,by,dsd,bi,msi,IP3K2,Eip74EF,B-H2,cic,numb,Poxm,path,hdc,crol,Cat,melt,EcR,scrib,grn,siz,klg,rst,csw,sno,bun,Lpin,cindr,rib,chp,Exn,RhoGAP100F,Khc,Dys,CG13251,nrv2,hth,Mbs,Pak,Spf45,Syn2,boi,tnc,raw,Src42A,upd3,tld,sgg,ex,app,polybromo,bnl,Su(H),kay,fz2,Fhos,Mob2,Ser,dpp,opa,step,shg,CG5921,foxo,trol,TfAP-2,nuf,Raf,tin,dsh,Cam,Mrtf | | 1.060e-09 | -20.66 | plasma membrane | cellular component | GO:0005886 | 1265 | 103 | 11940 | 538 | fz2,Pde6,dpr15,Ptp99A,sdt,Cln3,unc-13,CG30054,kek5,CCAP-R,rst,Mob2,klg,nrv2,CG10960,gukh,Pkc98E,CG11318,CG9981,LpR2,Ank2,fs(1)N,Octbeta2R,olf186-F,Cam,Raf,ine,Vha100-5,raw,chp,CG14606,D2hgdh,Dmtn,qvr,Nlg2,CG4301,step,Sh,Sk1,AstA-R1,fz4,Gprk1,Src42A,path,Syn2,Pak,sws,Dys,Fife,Syx1A,beat-IIa,dpr5,nrm,mGluR,DIP-delta,dsh,Ca-beta,Or85b,shakB,Pkc53E,Syt12,CNT1,Ank,CG33310,CG4839,Pdfr,ex,Tsp96F,shg,CG5921,Inx2,ZnT63C,cindr,DIP-lambda,Pzl,kcc,p130CAS,MsR2,mino,kek1,Or42b,Or65c,GluRIIC,Flo2,scrib,dpr12,CG13794,melt,ACXD,KaiR1D,CG13654,CadN2,Tsf1,Ser,CG3078,dpr18,CG3285,CG14182,nuf,Nmdar2,PGRP-LA,Hil,Exn | | 1.709e-09 | -20.19 | signaling | biological process | GO:0023052 | 1169 | 102 | 12345 | 597 | boi,Frq1,Stlk,sky,Tk,Src42A,et,upd3,Su(H),bnl,Tor,CG13229,Pde6,ex,sgg,Fife,Set2,CG30054,cpx,fz2,Tusp,ACXD,mGluR,Gpb5,ine,RhoGAP100F,PhKgamma,Dys,Pkc98E,KaiR1D,ninaD,shakB,Pak,Cadps,fz4,CG13526,spri,CG11318,foxo,CG41520,TpnC41C,Pde1c,Myo81F,Raf,Cam,Stacl,Rab9Fa,dsh,Ser,dnc,AstA-R1,CG5059,opa,dpp,Obp56h,Atf6,Hr96,Pdfr,Iswi,CG17698,ths,cic,Ca-beta,unc-13,Syx1A,Ank2,flw,CG5541,Octbeta2R,Cdep,Doa,Pde9,gce,RYa,CCAP-R,rictor,Gprk1,RhoGAP18B,Nlg2,bun,GluRIIC,MESK2,TpnC25D,Frq2,Pld,rdgA,fs(1)N,EcR,Ank,Cat,MsR2,CG13898,ITP,Nmdar2,csw,CG31750,Der-1,CG4839,wake,sno,Pkc53E,scramb1 | | 2.312e-09 | -19.89 | cell communication | biological process | GO:0007154 | 1239 | 106 | 12345 | 597 | CG30054,Set2,mGluR,Tusp,ACXD,cpx,fz2,bnl,Su(H),Tor,ex,sgg,Fife,Pde6,CG13229,Tk,upd3,Src42A,et,boi,Stlk,sky,Frq1,shakB,Pak,CG13526,spri,Cadps,fz4,ninaD,PhKgamma,Dys,KaiR1D,Pkc98E,Gpb5,ine,RhoGAP100F,Cam,Rab9Fa,Stacl,dsh,Raf,Myo81F,TpnC41C,Pde1c,CG41520,foxo,CG11318,Hr96,Atf6,opa,CG5059,Obp56h,dpp,dnc,AstA-R1,HDAC4,Ser,unc-13,Ca-beta,Ank2,Syx1A,ths,cic,CG17698,Iswi,Pdfr,CCAP-R,rictor,Doa,Pde9,gce,RYa,Octbeta2R,Cdep,flw,CG5541,rdgA,Inx2,fs(1)N,MESK2,Frq2,Pld,TpnC25D,GluRIIC,Lpin,Gprk1,Nlg2,RhoGAP18B,bun,sno,scramb1,Pkc53E,Nmdar2,csw,wake,CG31750,Der-1,CG4839,MsR2,ITP,CG13898,Ank,EcR,melt,Cat | | 5.780e-09 | -18.97 | cell periphery | cellular component | GO:0071944 | 1409 | 109 | 11940 | 538 | ine,Cam,Raf,app,olf186-F,Octbeta2R,fs(1)N,CG9981,Ank2,LpR2,gukh,CG11318,Pkc98E,nrv2,CG10960,klg,spri,Mob2,rst,kek5,CCAP-R,CG30054,Cln3,unc-13,dpr15,Ptp99A,sdt,Pde6,numb,fz2,dpr5,Syx1A,beat-IIa,Fife,sws,Dys,Pak,Src42A,path,Syn2,Gprk1,AstA-R1,fz4,Sk1,CG4301,Sh,step,Nlg2,Khc,qvr,Dmtn,raw,Vha100-5,D2hgdh,CG14606,chp,DIP-lambda,Pzl,kcc,cindr,CG5921,ZnT63C,Inx2,ex,Tsp96F,shg,Pdfr,CG33310,CG4839,CNT1,Mbs,Ank,Syt12,Pkc53E,Mur2B,shakB,dsh,Ca-beta,Or85b,DIP-delta,mGluR,nrm,Hil,PGRP-LA,Exn,nuf,Nmdar2,CG14182,dpr18,CG3078,CG3285,Ser,Tsf1,CG13654,CadN2,KaiR1D,melt,ACXD,CG13794,scrib,dpr12,Flo2,GluRIIC,Or65c,mino,kek1,Or42b,MsR2,p130CAS | | 8.311e-09 | -18.61 | signal transduction | biological process | GO:0007165 | 968 | 87 | 12345 | 597 | Rab9Fa,Stacl,dsh,Cam,Myo81F,Raf,foxo,CG41520,Pde1c,TpnC41C,CG11318,Atf6,Hr96,dpp,CG5059,opa,AstA-R1,dnc,Ser,fz2,mGluR,Tusp,ACXD,Set2,CG30054,CG13229,Pde6,ex,sgg,Su(H),bnl,Tor,et,Src42A,upd3,Tk,Stlk,boi,fz4,spri,CG13526,shakB,Pak,ninaD,Pkc98E,PhKgamma,RhoGAP100F,Gpb5,fs(1)N,rdgA,TpnC25D,Pld,MESK2,GluRIIC,bun,RhoGAP18B,Gprk1,Pkc53E,sno,Der-1,CG4839,CG31750,wake,csw,Nmdar2,ITP,CG13898,MsR2,EcR,Ank,Ank2,Ca-beta,unc-13,cic,ths,Iswi,CG17698,Pdfr,rictor,CCAP-R,gce,RYa,Pde9,Doa,Cdep,Octbeta2R,flw | | 1.487e-08 | -18.02 | anatomical structure morphogenesis | biological process | GO:0009653 | 1579 | 124 | 12345 | 597 | retn,Ubx,Actn,fw,esg,flw,bru1,sdt,Doa,fru,rictor,Prm,CadN2,by,pod1,dsd,gukh,bi,msi,IP3K2,Eip74EF,haf,cic,numb,B-H2,scro,Poxm,ths,Ank2,Syx1A,path,hdc,crol,Ptp61F,Cat,melt,EcR,E(Pc),scrib,Mlp84B,Ptp99A,egh,grn,siz,rst,klg,CG32006,csw,rols,mid,TwdlG,ems,sno,Sh,sqz,bun,Lpin,Act88F,Pld,beat-IIa,cindr,sick,ham,fs(1)N,rib,chp,Inx2,Exn,jim,RhoGAP100F,Khc,Dys,CG13251,PhKgamma,nrv2,Dlic,FoxL1,spri,hth,Pak,Mbs,Spf45,Stlk,tnc,boi,Syn2,Src42A,raw,tld,upd3,sgg,ex,msps,polybromo,app,Su(H),Snoo,Tor,bnl,kay,fz2,Fhos,Mob2,TwdlY,Ser,AGO2,dnc,shd,dpp,opa,step,shg,CG5921,chb,trol,foxo,TfAP-2,CG41099,nuf,Raf,dsh,tin,Mrtf,Cam | | 1.536e-08 | -17.99 | cell projection morphogenesis | biological process | GO:0048858 | 428 | 49 | 12345 | 597 | Khc,RhoGAP100F,retn,jim,fru,Dlic,pod1,Pak,Mbs,CadN2,spri,rictor,Stlk,gukh,Src42A,ths,Snoo,Poxm,bnl,Tor,scro,msps,haf,path,hdc,kay,fz2,Ank2,EcR,Ptp61F,dnc,E(Pc),mid,Ptp99A,klg,egh,grn,Sh,shg,ems,sqz,chb,foxo,trol,sick,ham,beat-IIa,Cam,dsh | | 1.751e-08 | -17.86 | cellular component morphogenesis | biological process | GO:0032989 | 667 | 66 | 12345 | 597 | Raf,CG41099,dsh,Cam,chb,foxo,trol,dpp,shg,dnc,sgg,Snoo,Tor,bnl,msps,polybromo,kay,fz2,Mob2,Stlk,Src42A,raw,Dlic,spri,Mbs,Pak,RhoGAP100F,jim,Khc,Act88F,ham,sick,beat-IIa,fs(1)N,sqz,bun,Ptp99A,klg,egh,grn,Mlp84B,mid,rols,ems,Sh,Ptp61F,EcR,E(Pc),scrib,haf,cic,ths,Poxm,scro,hdc,path,Ank2,gukh,fru,Prm,rictor,pod1,CadN2,Actn,retn,bru1 | | 1.943e-08 | -17.76 | cell development | biological process | GO:0048468 | 1436 | 115 | 12345 | 597 | Khc,Mos,jim,ine,RhoGAP100F,CG13251,Dlic,nrv2,Mbs,Pak,Rbp6,spri,Syn2,sky,Stlk,Src42A,upd3,tld,kek1,msps,polybromo,bbg,RhoL,Tor,Snoo,ex,sgg,Mob2,fz2,kay,Fhos,AGO2,shd,dnc,Antp,dpp,shg,CG5921,chinmo,chb,shep,wbl,foxo,trol,CG41099,Msp300,Raf,Mrtf,Cam,lncRNA:Hsromega,dsh,flw,dpr12,retn,Actn,nrm,CG7483,lncRNA:flam,bol,bru1,fru,Doa,sdt,CadN2,pod1,rictor,drd,Prm,gukh,dsd,can,msi,Iswi,IP3K2,Eip74EF,scro,esn,ths,p130CAS,Poxm,DNaseII,numb,haf,cic,Ca-beta,Ank2,path,hdc,EcR,Ptp61F,scrib,E(Pc),csw,rols,mid,Mlp84B,Ptp99A,egh,grn,rst,klg,sno,Sh,ems,bun,sqz,beat-IIa,sick,cindr,ham,Act88F,CG42355,chp,Inx2,fs(1)N | | 3.005e-08 | -17.32 | cell part morphogenesis | biological process | GO:0032990 | 437 | 49 | 12345 | 597 | foxo,trol,chb,sqz,Cam,dsh,beat-IIa,ham,sick,dnc,E(Pc),EcR,Ptp61F,shg,Sh,ems,mid,Ptp99A,egh,klg,grn,Src42A,gukh,Stlk,Ank2,fz2,kay,hdc,path,msps,scro,ths,Tor,Snoo,bnl,Poxm,haf,Khc,retn,jim,RhoGAP100F,CadN2,pod1,Pak,Mbs,rictor,spri,Dlic,fru | | 3.591e-08 | -17.14 | plasma membrane bounded cell projection morphogenesis | biological process | GO:0120039 | 426 | 48 | 12345 | 597 | rictor,spri,CadN2,Mbs,pod1,Pak,Dlic,fru,jim,retn,RhoGAP100F,Khc,Ank2,kay,fz2,path,hdc,haf,scro,msps,Tor,Poxm,bnl,Snoo,Src42A,gukh,Stlk,ems,shg,Sh,egh,grn,Ptp99A,klg,mid,E(Pc),dnc,Ptp61F,EcR,dsh,Cam,beat-IIa,sick,ham,foxo,trol,chb,sqz | | 6.159e-08 | -16.60 | neuron projection development | biological process | GO:0031175 | 447 | 49 | 12345 | 597 | Dlic,fru,CadN2,Mbs,Pak,pod1,rictor,spri,Khc,retn,jim,RhoGAP100F,msps,scro,Poxm,Snoo,Tor,p130CAS,haf,Ank2,kay,fz2,path,hdc,gukh,sky,Stlk,Src42A,mid,egh,klg,Ptp99A,grn,shg,Sh,ems,EcR,Ptp61F,dnc,E(Pc),beat-IIa,ham,sick,Cam,dsh,chb,sqz,foxo,trol | | 8.882e-08 | -16.24 | neuron projection morphogenesis | biological process | GO:0048812 | 425 | 47 | 12345 | 597 | chb,sqz,foxo,trol,beat-IIa,sick,ham,dsh,Cam,Ptp61F,EcR,E(Pc),dnc,Ptp99A,egh,klg,grn,mid,ems,shg,Sh,gukh,Stlk,Src42A,haf,msps,scro,Snoo,Tor,Poxm,Ank2,fz2,kay,path,hdc,jim,retn,RhoGAP100F,Khc,Dlic,fru,rictor,spri,CadN2,Mbs,pod1,Pak | | 1.281e-07 | -15.87 | plasma membrane part | cellular component | GO:0044459 | 820 | 70 | 11940 | 538 | dpr15,sdt,Pde6,fz2,CG30054,Cln3,unc-13,kek5,CCAP-R,rst,klg,nrv2,CG10960,Ank2,gukh,CG11318,olf186-F,Octbeta2R,fs(1)N,ine,CG14606,D2hgdh,chp,Nlg2,qvr,step,Sh,AstA-R1,Src42A,Syn2,Dys,Pak,Syx1A,beat-IIa,Fife,dpr5,mGluR,Ca-beta,Or85b,DIP-delta,CNT1,Pdfr,CG33310,Inx2,ZnT63C,ex,shg,DIP-lambda,kcc,cindr,kek1,Or42b,MsR2,dpr12,scrib,Flo2,Or65c,GluRIIC,ACXD,KaiR1D,Ser,CadN2,CG14182,dpr18,CG3285,PGRP-LA,Hil,Exn,nuf,Nmdar2 | | 1.310e-07 | -15.85 | regulation of cell communication | biological process | GO:0010646 | 855 | 76 | 12345 | 597 | spri,Pak,sxc,Cln3,RhoGEF3,hth,Flo2,CG9650,Dys,KaiR1D,Mos,Exn,kek5,mGluR,cpx,Fife,sgg,ex,Su(H),pirk,bnl,Snoo,Tor,kek1,tld,Src42A,raw,et,sky,Frq1,Sod1,step,Mctp,dpp,RhoGAP1A,dnc,Ser,HDAC4,qvr,lncRNA:Hsromega,Cam,Raf,foxo,trol,Pde8,chb,rictor,CG18659,CG5522,Doa,Cdep,CG10188,flw,Ank2,mnb,crol,numb,CG14182,Sh,sno,siz,scrib,Cat,Ptp61F,nemy,melt,rdgA,Frq2,CG34408,MESK2,GluRIIC,olf186-F,Glut1,Lpin,Nlg2,Gprk1 | | 1.310e-07 | -15.85 | regulation of signaling | biological process | GO:0023051 | 855 | 76 | 12345 | 597 | Exn,Mos,kek5,Dys,KaiR1D,Flo2,CG9650,spri,Pak,sxc,Cln3,RhoGEF3,hth,sky,Frq1,Sod1,tld,kek1,raw,et,Src42A,Fife,ex,sgg,bnl,Su(H),pirk,Tor,Snoo,mGluR,cpx,Ser,HDAC4,RhoGAP1A,dnc,dpp,step,Mctp,chb,foxo,trol,Pde8,Raf,qvr,lncRNA:Hsromega,Cam,flw,Cdep,CG10188,CG18659,CG5522,Doa,rictor,CG14182,numb,Ank2,mnb,crol,Cat,Ptp61F,nemy,melt,scrib,siz,Sh,sno,Glut1,Gprk1,Nlg2,Lpin,GluRIIC,olf186-F,Frq2,CG34408,MESK2,rdgA | | 1.786e-07 | -15.54 | cell morphogenesis | biological process | GO:0000902 | 533 | 54 | 12345 | 597 | CG41099,beat-IIa,sick,ham,dsh,Cam,chb,sqz,foxo,trol,egh,klg,Ptp99A,grn,mid,ems,shg,Sh,Ptp61F,EcR,E(Pc),dnc,scrib,haf,sgg,msps,scro,Snoo,ths,Poxm,bnl,Tor,fz2,Ank2,kay,path,hdc,Mob2,gukh,Stlk,raw,Src42A,Dlic,fru,rictor,spri,CadN2,Mbs,pod1,Pak,jim,retn,RhoGAP100F,Khc | | 1.875e-07 | -15.49 | cell morphogenesis involved in neuron differentiation | biological process | GO:0048667 | 422 | 46 | 12345 | 597 | Tor,Poxm,Snoo,scro,msps,haf,path,hdc,kay,Ank2,fz2,gukh,Src42A,fru,Dlic,Mbs,pod1,Pak,CadN2,spri,rictor,Khc,RhoGAP100F,jim,retn,ham,sick,beat-IIa,Cam,dsh,sqz,chb,foxo,trol,mid,egh,klg,Ptp99A,grn,Sh,shg,ems,EcR,Ptp61F,dnc,E(Pc) | | 4.059e-07 | -14.72 | animal organ development | biological process | GO:0048513 | 1267 | 100 | 12345 | 597 | CG13251,Dys,Khc,RhoGAP100F,Exn,Idgf6,Pak,Mbs,sxc,hth,nrv2,tld,upd3,raw,Src42A,boi,Syn2,tnc,Spf45,Set2,Mob2,Fhos,kay,fz2,bnl,Tor,Su(H),app,polybromo,RhoL,sgg,ex,Antp,Irk3,Ser,CycT,Dmtn,CG5921,shg,step,opa,dpp,trol,foxo,chinmo,toe,chb,Cam,Mrtf,dsh,tin,Raf,Msp300,nuf,TfAP-2,E5,flw,esg,fw,retn,Ubx,CadN2,by,fru,Doa,cato,IP3K2,Eip74EF,bi,msi,dsd,Elp3,crol,hdc,path,Syx1A,Poxm,ths,numb,B-H2,cic,scrib,Awh,EcR,melt,Cat,sno,ems,mid,rols,csw,grn,rst,klg,siz,Lpin,bun,chp,rib,cindr | | 4.237e-07 | -14.67 | regulation of transcription by RNA polymerase II | biological process | GO:0006357 | 741 | 67 | 12345 | 597 | Spt20,tld,kay,Mondo,ex,polybromo,Su(H),Snoo,CG32971,CG1894,kek5,hth,FoxL1,CG42830,foxo,toe,chinmo,tin,Mrtf,luna,TfAP-2,CG10462,CG12071,Antp,CycT,CG31224,Atf6,Hr96,dpp,dany,opa,mamo,Iswi,Eip74EF,CG43689,bi,crol,Elp3,cic,B-H2,TfIIFbeta,scro,E5,retn,Ubx,esg,CG12605,gce,woc,fru,GluRIIC,CG4374,bun,Lpin,Sox102F,GATAe,CrebB,scrib,EcR,Awh,ems,sno,CG7368,grn,htk,CG32006,mid | | 4.580e-07 | -14.60 | cell projection organization | biological process | GO:0030030 | 637 | 60 | 12345 | 597 | fru,Dlic,Pak,nompB,Mbs,pod1,CadN2,spri,rictor,Khc,fw,RhoGAP100F,retn,jim,Snoo,bnl,ths,Tor,Poxm,p130CAS,app,msps,scro,haf,hdc,path,kay,Ank2,fz2,sky,Stlk,gukh,Src42A,mid,klg,grn,Ptp99A,egh,Sh,shg,ems,EcR,Ptp61F,dnc,scrib,E(Pc),ham,IFT46,sick,beat-IIa,chp,Root,Cam,dsh,sqz,chb,CG14105,foxo,CG17387,trol | | 4.756e-07 | -14.56 | multicellular organism development | biological process | GO:0007275 | 2496 | 170 | 12345 | 597 | nrv2,Cpr67Fa1,Cln3,Pak,Idgf6,Khc,Dys,Pkc98E,ex,Tor,Snoo,cpx,sky,Stlk,stw,boi,upd3,tld,dpp,opa,CG5921,TwdlY,Irk3,dnc,CG41099,Raf,Msp300,dsh,tin,chb,toe,chinmo,foxo,wbl,cato,Doa,sdt,drd,rictor,CadN2,pod1,Ubx,fw,esg,flw,Fer2,E5,nrm,haf,cic,barc,Poxm,p130CAS,hdc,Elp3,Ca-beta,mnb,gukh,rst,Ptp99A,wake,mid,CG9519,sno,Sh,Awh,CG9514,scrib,beat-IIa,sick,ham,Lpin,Dlic,spri,hth,sxc,Mbs,jim,Exn,RhoGAP100F,ine,tamo,Cpr64Ac,CG13251,PhKgamma,Fife,sgg,msps,polybromo,RhoL,app,bnl,Su(H),fz2,kay,Fhos,Mob2,Set2,Spf45,Syn2,tnc,Sod1,raw,Src42A,step,shg,Hr96,Dmtn,CycT,Ser,AGO2,Antp,g,shd,GlyP,TfAP-2,nuf,Cam,Mrtf,luna,shep,CG17387,trol,sws,woc,fru,by,spok,retn,dpr12,CG7483,numb,B-H2,esn,scro,ths,Ank2,Syx1A,path,crol,dsd,bi,msi,IP3K2,Eip74EF,klg,siz,grn,egh,csw,rols,TwdlG,ems,Cat,Ptp61F,melt,EcR,E(Pc),Pld,cindr,fs(1)N,rib,CG42797,Inx2,GATAe,chp,bun,sqz,Nlg2,olf186-F | | 5.633e-07 | -14.39 | plasma membrane bounded cell projection organization | biological process | GO:0120036 | 626 | 59 | 12345 | 597 | ems,Sh,shg,Ptp99A,egh,klg,grn,mid,E(Pc),scrib,dnc,Ptp61F,EcR,dsh,chp,Root,Cam,sick,IFT46,ham,beat-IIa,foxo,CG17387,trol,CG14105,sqz,chb,spri,rictor,pod1,Mbs,Pak,nompB,CadN2,fru,Dlic,RhoGAP100F,fw,retn,jim,Khc,hdc,path,fz2,Ank2,kay,haf,Tor,p130CAS,Poxm,Snoo,bnl,msps,scro,app,Src42A,sky,Stlk,gukh | | 6.226e-07 | -14.29 | cell morphogenesis involved in differentiation | biological process | GO:0000904 | 468 | 48 | 12345 | 597 | klg,Ptp99A,grn,egh,mid,ems,shg,Sh,Ptp61F,EcR,E(Pc),dnc,beat-IIa,CG41099,sick,ham,dsh,Cam,sqz,chb,trol,foxo,Dlic,fru,rictor,spri,CadN2,Mbs,Pak,pod1,jim,retn,RhoGAP100F,Khc,sgg,haf,msps,scro,Tor,Poxm,Snoo,fz2,kay,Ank2,path,hdc,gukh,Src42A | | 6.419e-07 | -14.26 | developmental process | biological process | GO:0032502 | 2988 | 196 | 12345 | 597 | Pkc98E,Dys,Idgf6,Khc,Rbp6,Pak,Cpr67Fa1,Cln3,nrv2,upd3,tld,sky,Stlk,boi,stw,cpx,ex,Snoo,Tor,bbg,Irk3,dnc,TwdlY,CG17470,CG5921,dpp,opa,foxo,wbl,Pde8,toe,chb,chinmo,tin,dsh,Raf,Msp300,CG41099,bol,E5,Fer2,lncRNA:flam,nrm,fw,Ubx,flw,esg,rictor,drd,pod1,CadN2,Doa,sdt,cato,Iswi,can,gukh,hdc,unc-13,mnb,Elp3,Ca-beta,cic,haf,DNaseII,Poxm,p130CAS,barc,CG9514,scrib,Awh,CG9519,Sh,sno,rst,wake,Ptp99A,Mlp84B,Treh,mid,Lpin,CG42355,Sox102F,Act88F,ham,sick,beat-IIa,PhKgamma,CG13251,RhoGAP100F,ine,Exn,jim,Mos,Cpr64Ac,tamo,spri,Tif-IA,Mbs,sxc,hth,Spn88Eb,FoxL1,Dlic,kek1,raw,Src42A,Spf45,tnc,Syn2,Sod1,Fhos,kay,fz2,Mob2,Set2,Fife,sgg,bnl,Su(H),RhoL,app,msps,polybromo,Antp,shd,g,Ser,CycT,Dmtn,AGO2,step,Hr96,shg,CG34417,CG17387,trol,shep,luna,lncRNA:Hsromega,Mrtf,Cam,TfAP-2,GlyP,nuf,bru1,CG7483,Actn,spok,retn,dpr12,Prm,by,sws,fru,woc,Eip74EF,IP3K2,msi,bi,dsd,path,Syx1A,Ank2,crol,numb,B-H2,ths,scro,esn,E(Pc),Cat,Ptp61F,EcR,melt,ems,TwdlG,klg,siz,egh,grn,CG32006,rols,csw,olf186-F,bun,sqz,Nlg2,fs(1)N,chp,Inx2,GATAe,rib,CG42797,Pld,cindr | | 6.582e-07 | -14.23 | regulation of transport | biological process | GO:0051049 | 267 | 33 | 12345 | 597 | sdt,Flo2,CG9650,spri,Cadps,Exn,Dys,Syt12,Fife,numb,sgg,Pde6,mnb,Ca-beta,CG3078,cpx,Syx1A,sky,Frq1,LpR2,Mctp,step,Pkc53E,nemy,scrib,cindr,Pld,Frq2,Cam,CG42797,qvr,Glut1,olf186-F | | 7.097e-07 | -14.16 | DNA-binding transcription factor activity | molecular function | GO:0003700 | 463 | 47 | 12176 | 580 | FoxL1,tin,CG43689,E5,CG12071,Octbeta2R,EcR,ems,Su(H),CG15073,Ubx,B-H2,gce,ham,Eip74EF,bi,grn,CG31224,retn,TfAP-2,CG9650,kay,cic,Atf6,Mrtf,CG32006,Mondo,fru,CG12605,GATAe,opa,woc,foxo,hth,rib,bun,esg,Poxm,Awh,Antp,scro,luna,mid,Hr96,CrebB,CG10462,CG4374 | | 7.294e-07 | -14.13 | photoreceptor cell differentiation | biological process | GO:0046530 | 183 | 26 | 12345 | 597 | CG5921,hth,sno,CadN2,klg,sdt,Doa,csw,scrib,CG13251,RhoGAP100F,EcR,Khc,melt,dsh,kay,chp,Cam,Mob2,B-H2,ex,Raf,msi,bun,Syn2,boi | | 8.353e-07 | -14.00 | neuron projection extension | biological process | GO:1990138 | 37 | 11 | 12345 | 597 | Ank2,fz2,hdc,dsh,path,spri,CadN2,Sh,Mbs,dnc,RhoGAP100F | | 8.379e-07 | -13.99 | anatomical structure development | biological process | GO:0048856 | 2924 | 192 | 12345 | 597 | TwdlG,ems,csw,rols,CG32006,egh,klg,grn,siz,E(Pc),EcR,melt,Ptp61F,Cat,rib,CG42797,GATAe,Inx2,chp,fs(1)N,cindr,Pld,olf186-F,Nlg2,sqz,bun,by,Prm,woc,fru,sws,CG7483,bru1,dpr12,retn,spok,Actn,crol,Syx1A,Ank2,path,esn,scro,ths,numb,B-H2,IP3K2,Eip74EF,dsd,msi,bi,CG34417,shg,Hr96,step,g,shd,Antp,AGO2,Dmtn,CycT,Ser,Mrtf,Cam,luna,lncRNA:Hsromega,nuf,GlyP,TfAP-2,CG17387,trol,shep,hth,Mbs,sxc,spri,Dlic,Spn88Eb,FoxL1,CG13251,PhKgamma,Cpr64Ac,tamo,Mos,Exn,jim,ine,RhoGAP100F,Set2,Mob2,kay,fz2,Fhos,RhoL,app,msps,polybromo,Su(H),bnl,Fife,sgg,raw,Src42A,kek1,tnc,Syn2,Sod1,Spf45,sno,Sh,CG9519,mid,Mlp84B,wake,rst,Ptp99A,CG9514,scrib,Awh,CG42355,beat-IIa,sick,ham,Act88F,Lpin,CadN2,pod1,rictor,drd,cato,Doa,sdt,nrm,lncRNA:flam,Fer2,E5,bol,esg,flw,Ubx,fw,Ca-beta,Elp3,mnb,hdc,barc,p130CAS,Poxm,DNaseII,cic,haf,Iswi,can,gukh,CG5921,opa,dpp,dnc,Irk3,CG17470,TwdlY,dsh,tin,CG41099,Msp300,Raf,wbl,Pde8,foxo,chinmo,chb,toe,Cln3,Cpr67Fa1,Pak,Rbp6,nrv2,Dys,Pkc98E,Khc,Idgf6,cpx,bbg,Tor,Snoo,ex,tld,upd3,boi,stw,sky,Stlk | | 9.267e-07 | -13.89 | regulation of neurotransmitter secretion | biological process | GO:0046928 | 24 | 9 | 12345 | 597 | Exn,Frq1,sky,nemy,Dys,Frq2,Fife,cpx,Mctp | | 9.267e-07 | -13.89 | regulation of neurotransmitter transport | biological process | GO:0051588 | 24 | 9 | 12345 | 597 | Frq2,Fife,Mctp,cpx,nemy,Exn,Frq1,sky,Dys | | 1.081e-06 | -13.74 | DNA-binding transcription factor activity, RNA polymerase II-specific | molecular function | GO:0000981 | 277 | 33 | 12176 | 580 | Antp,Awh,esg,CG4374,CG10462,Hr96,mid,luna,CG12605,CG32006,Mrtf,hth,foxo,woc,GATAe,opa,CG31224,grn,bi,Eip74EF,gce,B-H2,cic,kay,TfAP-2,retn,CG12071,E5,CG43689,tin,FoxL1,Su(H),EcR | | 1.239e-06 | -13.60 | regulation of RNA metabolic process | biological process | GO:0051252 | 1281 | 99 | 12345 | 597 | ham,AGO3,lncRNA:bxd,GATAe,Sox102F,rib,Lpin,bun,CG4374,GluRIIC,Maf1,mid,CG32006,grn,CG7368,htk,Marf1,sno,ems,EcR,melt,Awh,CrebB,scrib,Poxm,barc,scro,TfIIFbeta,cic,B-H2,mnb,Elp3,crol,bi,CG43689,Eip74EF,Iswi,mamo,fru,Doa,woc,gce,CG12605,rictor,esg,retn,Ubx,bol,E5,bru1,Octbeta2R,Fer2,ps,CG10462,trv,CG12071,TfAP-2,luna,Mrtf,tin,chinmo,toe,HPS4,l(3)neo38,foxo,opa,dany,dpp,CG14252,Hr96,Atf6,CG31224,HDAC4,AGO2,CycT,Antp,CG15073,Snoo,Tor,Su(H),CG32971,polybromo,ex,fus,Mondo,Set2,kay,Spf45,tld,Spt20,CG42830,FoxL1,CG9650,Art2,hth,kek5,CG1894,Khc,jim | | 1.259e-06 | -13.59 | regulation of nucleobase-containing compound metabolic process | biological process | GO:0019219 | 1315 | 101 | 12345 | 597 | opa,dany,dpp,CG14252,Hr96,Atf6,CG31224,HDAC4,AGO2,CycT,Antp,CG15073,CG10462,CG12071,ps,trv,TfAP-2,luna,Mrtf,tin,chinmo,toe,HPS4,l(3)neo38,foxo,FoxL1,CG42830,Usp1,CG9650,Art2,hth,kek5,CG1894,Khc,jim,Tor,CG32971,Su(H),Snoo,polybromo,fus,ex,Mondo,Set2,kay,Spf45,tld,Spt20,Maf1,mid,CG32006,grn,CG7368,htk,Marf1,sno,ems,EcR,melt,Awh,scrib,CrebB,ham,AGO3,GATAe,lncRNA:bxd,Sox102F,rib,Lpin,CG12200,bun,CG4374,GluRIIC,Doa,fru,woc,gce,CG12605,rictor,esg,retn,Ubx,bol,bru1,E5,Octbeta2R,Fer2,Poxm,barc,scro,cic,B-H2,TfIIFbeta,mnb,Elp3,crol,bi,CG43689,Eip74EF,mamo,Iswi | | 1.363e-06 | -13.51 | positive regulation of biological process | biological process | GO:0048518 | 1642 | 120 | 12345 | 597 | dpp,CG5059,opa,step,Atf6,shg,CG14252,Hr96,CycT,HDAC4,RhoGAP1A,Antp,g,GlyP,TfAP-2,CG10462,Raf,ps,CG12071,dsh,tin,qvr,Cam,Mrtf,lncRNA:Hsromega,chb,chinmo,trol,foxo,CG41520,HPS4,Pde8,tyf,Flo2,CG42830,FoxL1,Cadps,spri,hth,sxc,Pak,Mos,Exn,CG1894,KaiR1D,sgg,ex,Fife,msps,RhoL,polybromo,bnl,Tor,Su(H),kay,Fhos,Stlk,LpR2,Src42A,tld,upd3,Tk,Der-1,wake,siz,mid,Pkc53E,ems,sno,Marf1,Sh,REG,EcR,melt,Awh,CG1572,CrebB,AGO3,Pld,cindr,MESK2,Sox102F,CG42797,GATAe,bun,Lpin,Gprk1,CG12200,Glut1,Cpsf160,CG4374,olf186-F,gce,Doa,sdt,rictor,CG4849,retn,Ubx,esg,flw,Fer2,bol,Octbeta2R,bru1,CG10188,CG7483,TfIIFbeta,B-H2,numb,barc,scro,Pomp,Ank2,crol,mnb,CG43689,bi,Iswi,Eip74EF | | 1.366e-06 | -13.50 | RNA polymerase II regulatory region sequence-specific DNA binding | molecular function | GO:0000977 | 254 | 31 | 12176 | 580 | foxo,CG15073,Su(H),ems,CG7368,EcR,opa,GATAe,tin,E5,CG43689,CG12071,CG12605,Atf6,cic,CG4374,mid,retn,TfAP-2,Hr96,CG10462,kay,Awh,bi,grn,scro,Antp,CG31224,Ubx,B-H2,esg | | 1.558e-06 | -13.37 | GO\_REGULATION\_OF\_CELL\_ADHESION | MSigDB lists | GO\_REGULATION\_OF\_CELL\_ADHESION | 110 | 17 | 4161 | 170 | Ank,step,csw,IP3K2,Tor,Ptp61F,chb,Octbeta2R,mino,Flo2,Sod1,Pkc53E,Raf,Pak,flw,Pkc98E,dpp | | 1.856e-06 | -13.20 | GO\_REGULATION\_OF\_HOMOTYPIC\_CELL\_CELL\_ADHESION | MSigDB lists | GO\_REGULATION\_OF\_HOMOTYPIC\_CELL\_CELL\_ADHESION | 47 | 11 | 4161 | 170 | Flo2,Ank,Sod1,csw,Pkc53E,IP3K2,Tor,Pak,Ptp61F,Octbeta2R,mino | | 2.026e-06 | -13.11 | PDEase | interpro domains | IPR023088 | 6 | 5 | 11383 | 585 | Pde9,Pde1c,Pde6,Pde8,dnc | | 2.026e-06 | -13.11 | PDEase\_catalytic\_dom | interpro domains | IPR002073 | 6 | 5 | 11383 | 585 | Pde6,Pde1c,Pde9,dnc,Pde8 | | 2.246e-06 | -13.01 | PDEase\_I | pfam domains | PF00233 | 6 | 5 | 10480 | 550 | Pde8,Pde1c,dnc,Pde9,Pde6 | | 2.265e-06 | -13.00 | multicellular organismal process | biological process | GO:0032501 | 3844 | 238 | 12345 | 597 | Ser,Dmtn,AANAT1,CycT,AGO2,Antp,CG34435,shd,g,Obp56h,step,Hr96,shg,geko,shep,CG17387,trol,TfAP-2,GlyP,nuf,aust,luna,lncRNA:Hsromega,Mrtf,Cam,dpr18,ine,RhoGAP100F,jim,Mos,Exn,Cpr64Ac,tamo,CG13251,PhKgamma,Dlic,spri,Tif-IA,sxc,shakB,Mbs,hth,Spf45,Sod1,Syn2,tnc,kek1,raw,Src42A,Fife,sgg,Su(H),bnl,RhoL,msps,app,polybromo,mGluR,Fhos,fz2,CG42823,kay,Set2,Mob2,Cat,Ptp61F,nemy,melt,EcR,Qsox2,E(Pc),MsR2,grn,siz,klg,egh,CG31750,Or42b,rols,Nmdar2,Gld2,csw,ems,TwdlG,bun,sqz,Nlg2,olf186-F,Pld,cindr,CG1428,fs(1)N,Inx2,GATAe,rdgA,chp,CG42797,rib,spok,retn,dpr12,Fad2,bru1,CG7483,Ebp,sws,Cht6,fru,woc,lncRNA:CR43484,Acp62F,by,bi,msi,dsd,CG16854,IP3K2,dpr15,Eip74EF,mamo,B-H2,numb,CG10000,ths,esn,scro,path,Syx1A,Ank2,crol,TwdlY,HDAC4,Irk3,dnc,ND-51L2,dany,dpp,opa,CG5921,toe,chb,chinmo,Pde1c,foxo,wbl,Msp300,Raf,CG41099,qvr,dsh,tin,Gr22c,Root,Idgf6,Khc,Pkc98E,Dys,nrv2,Or65c,lncRNA:CR43753,Pak,nompB,Cln3,Cpr67Fa1,Stlk,sky,boi,stw,upd3,tld,Tk,ex,Snoo,Tor,bbg,cpx,Gr39a,dpr5,Awh,lncRNA:CR43802,CrebB,Nha1,CG9514,scrib,Ptp99A,wake,rst,mid,Treh,MCU,CG9519,Sh,sno,JhI-26,Lpin,lncRNA:CR44585,ham,sick,Or85b,beat-IIa,CG42355,lncRNA:CR44420,fw,CG5541,Ubx,flw,esg,E5,bol,Octbeta2R,Mst87F,Fer2,lncRNA:flam,nrm,Doa,sdt,cato,rictor,drd,pod1,CadN2,CCAP-R,gukh,can,Pdfr,Iswi,Obp56e,cic,haf,DNaseII,Poxm,CG18135,p130CAS,barc,hdc,mnb,Ca-beta,Elp3 | | 2.445e-06 | -12.92 | RNA polymerase II regulatory region DNA binding | molecular function | GO:0001012 | 261 | 31 | 12176 | 580 | cic,CG4374,retn,mid,kay,CG10462,Hr96,TfAP-2,Awh,bi,CG31224,grn,Antp,scro,Ubx,B-H2,esg,foxo,CG15073,Su(H),ems,CG7368,EcR,GATAe,opa,tin,CG12071,CG12605,CG43689,E5,Atf6 | | 2.540e-06 | -12.88 | membrane | cellular component | GO:0016020 | 2722 | 168 | 11940 | 538 | ACXD,melt,scrib,dpr12,Flo2,mino,CG14024,Ir41a,nuf,ABCD,Hil,CG3285,CG3078,CG14182,Irk3,Tsf1,Ser,CG1572,PIG-O,CG3168,Cyt-b5-r,Ca-beta,dsh,mGluR,bru1,CG17600,kcc,DIP-lambda,ex,Tsp96F,Inx2,CG5921,CG4839,UQCR-11,CG7882,Pdfr,CG41128,Syt12,CG34172,fz4,Sh,CG4301,Mctp,Sk1,Stt3B,HPS4,CG14606,D2hgdh,dpr5,PIG-G,siz,CG13229,CG7720,beat-IIa,CG40472,Pak,MCU,Ir10a,ND-51L2,path,CG8468,klg,CG12121,cpx,CCAP-R,Cln3,CG30054,Pde6,sdt,Fad2,dpr15,Cam,CG45782,ND-MLRQ,ine,Octbeta2R,Ucp4A,Qsox2,LpR2,Ank2,CG9981,CG10960,CG6790,CG13794,CG32320,GluRIIC,Or65c,MsR2,kek1,Or42b,p130CAS,Nmdar2,PGRP-LA,Exn,Msp300,dpr18,CadN2,Gr22c,CG13654,g,egh,KaiR1D,Vha36-3,Pkc53E,shakB,DIP-delta,Or85b,nrm,CG6839,CG17883,cindr,Pzl,shg,CG34056,ZnT63C,CG14439,CG33310,Atf6,CG42826,ND-AGGG,Ank,CNT1,CG30036,ninaD,AstA-R1,Rab9Fa,CG33934,Gprk1,step,Der-1,qvr,Dmtn,Nlg2,chp,raw,Vha100-5,CG33090,Nha1,et,Fife,CG9109,Syx1A,CG9132,CG42235,sws,Dys,chk,Syn2,Src42A,rst,kek5,Neb-cGP,Mob2,unc-13,Gr39a,fz2,Glut1,CG5059,Ptp99A,Raf,Meltrin,fs(1)N,olf186-F,Pkc98E,CG11318,gukh,Ir56a,nrv2 | | 2.541e-06 | -12.88 | GO\_REGULATION\_OF\_CELL\_CELL\_ADHESION | MSigDB lists | GO\_REGULATION\_OF\_CELL\_CELL\_ADHESION | 58 | 12 | 4161 | 170 | Ptp61F,Pak,Octbeta2R,dpp,mino,Ank,Flo2,IP3K2,Tor,csw,Sod1,Pkc53E | | 2.553e-06 | -12.88 | eye morphogenesis | biological process | GO:0048592 | 310 | 35 | 12345 | 597 | bnl,ex,cic,B-H2,Mob2,kay,Syn2,boi,msi,Spf45,bi,Doa,hth,CadN2,Khc,RhoGAP100F,fw,CG13251,cindr,Raf,chp,Cam,dsh,bun,foxo,csw,rst,klg,siz,dpp,CG5921,shg,melt,Ser,scrib | | 2.553e-06 | -12.88 | sensory organ morphogenesis | biological process | GO:0090596 | 310 | 35 | 12345 | 597 | melt,Ser,scrib,csw,dpp,rst,siz,klg,shg,CG5921,bun,foxo,Raf,cindr,Cam,chp,dsh,Khc,RhoGAP100F,fw,CG13251,Doa,CadN2,hth,boi,Syn2,bi,Spf45,msi,bnl,B-H2,ex,cic,Mob2,kay | | 2.693e-06 | -12.82 | GO\_3\_5\_CYCLIC\_AMP\_PHOSPHODIESTERASE\_ACTIVITY | MSigDB lists | GO\_3\_5\_CYCLIC\_AMP\_PHOSPHODIESTERASE\_ACTIVITY | 4 | 4 | 4161 | 170 | Pde8,Pde1c,dnc,Pde6 | | 2.793e-06 | -12.79 | regulation of macromolecule metabolic process | biological process | GO:0060255 | 2019 | 140 | 12345 | 597 | retn,bru1,CG7483,sws,fru,woc,CG4849,Acp62F,msi,bi,CG43689,Eip74EF,mamo,B-H2,TfIIFbeta,numb,Pomp,scro,crol,Ptp61F,REG,EcR,melt,grn,CG7368,Achl,CG32006,ems,Marf1,bun,CG4374,GATAe,CG42797,rib,Mos,jim,kek5,Art2,CG9650,CG42830,Spn88Eb,FoxL1,sxc,hth,Spf45,kek1,Src42A,raw,et,fus,sgg,Su(H),bnl,polybromo,kay,Set2,CycT,AGO2,Antp,g,step,Hr96,CG14252,Atf6,HPS4,l(3)neo38,TfAP-2,GlyP,ps,trv,CG12071,Yeti,lncRNA:Hsromega,luna,Mrtf,Ubx,flw,esg,bol,Octbeta2R,E5,Fer2,gce,Doa,rictor,CG12605,can,Iswi,cic,Poxm,barc,hdc,mnb,Elp3,Awh,CrebB,scrib,htk,mid,Maf1,sno,jigr1,Lpin,CG12200,GluRIIC,SteXh:CG42398,AGO3,ham,lncRNA:bxd,Sox102F,CG1894,Khc,Usp1,tyf,Pak,Cln3,Stlk,CG6966,tld,Spt20,Ste:CG33247,ex,Tor,CG32971,Snoo,Mondo,HDAC4,CG15073,dany,dpp,opa,CG31224,toe,chinmo,foxo,CG41520,Pde8,Raf,CG10462,tin | | 3.452e-06 | -12.58 | positive regulation of gene expression | biological process | GO:0010628 | 571 | 53 | 12345 | 597 | kay,mnb,TfIIFbeta,B-H2,Su(H),Tor,scro,polybromo,barc,Eip74EF,Iswi,bi,CG43689,CG4849,rictor,hth,gce,tyf,CG42830,FoxL1,bol,Fer2,CG7483,Ubx,retn,CG1894,esg,tin,GATAe,Mrtf,Sox102F,TfAP-2,CG10462,ps,CG12071,foxo,HPS4,CG4374,Lpin,ems,Hr96,CG14252,sno,Atf6,dpp,mid,opa,Antp,CrebB,g,CycT,EcR,Awh | | 3.470e-06 | -12.57 | GO\_REGULATION\_OF\_CELL\_ACTIVATION | MSigDB lists | GO\_REGULATION\_OF\_CELL\_ACTIVATION | 70 | 13 | 4161 | 170 | Tor,IP3K2,Pkc53E,RhoGAP1A,csw,Sod1,Flo2,mino,Octbeta2R,Ptp61F,Pak,Tsp96F,scramb1 | | 3.592e-06 | -12.54 | regulation of transcription, DNA-templated | biological process | GO:0006355 | 1144 | 89 | 12345 | 597 | scrib,CrebB,EcR,melt,Awh,ems,sno,CG7368,grn,htk,CG32006,mid,Maf1,GluRIIC,CG4374,bun,Lpin,rib,Sox102F,lncRNA:bxd,GATAe,ham,Fer2,E5,Octbeta2R,Ubx,retn,esg,rictor,CG12605,gce,woc,fru,Iswi,mamo,Eip74EF,CG43689,bi,Elp3,crol,mnb,cic,TfIIFbeta,B-H2,scro,Poxm,CG15073,Antp,CycT,AGO2,HDAC4,Atf6,CG31224,Hr96,CG14252,dpp,dany,opa,foxo,l(3)neo38,HPS4,toe,chinmo,tin,Mrtf,luna,TfAP-2,CG10462,CG12071,jim,Khc,CG1894,kek5,hth,CG9650,Art2,FoxL1,CG42830,Spt20,tld,kay,Set2,Mondo,ex,polybromo,CG32971,Tor,Snoo,Su(H) | | 3.601e-06 | -12.53 | developmental cell growth | biological process | GO:0048588 | 59 | 13 | 12345 | 597 | CadN2,Sh,Mbs,fz2,Ank2,kay,path,hdc,spri,dsh,ine,RhoGAP100F,dnc | | 3.972e-06 | -12.44 | axon extension | biological process | GO:0048675 | 35 | 10 | 12345 | 597 | Mbs,Sh,CadN2,dsh,hdc,spri,Ank2,fz2,RhoGAP100F,dnc | | 4.192e-06 | -12.38 | response to stimulus | biological process | GO:0050896 | 2655 | 174 | 12345 | 597 | Iswi,CG17698,gukh,Pdfr,Ca-beta,mnb,unc-13,ZnT63C,haf,cic,Fer2,Octbeta2R,CG3556,flw,esg,rictor,CadN2,CCAP-R,pod1,gce,Pde9,Doa,GluRIIC,Lpin,Gprk1,RhoGAP18B,Ucp4A,TpnC25D,beat-IIa,Or85b,sick,ITP,CG13898,CrebB,scrib,PGRP-LA,Pkc53E,sno,Sh,wake,Ptp99A,MCU,mid,upd3,Tk,jus,Stlk,boi,Tusp,ACXD,CG7362,Gr39a,Cyp12a4,CG30054,Mondo,CG13229,Pde6,ex,bbg,Tor,Pkc98E,Idgf6,Khc,kcc,Gpb5,CG13526,Cln3,RhoGEF3,Pak,Cyp6a13,tyf,Or65c,CG41520,foxo,Ir56a,TpnC41C,Pde1c,chb,Gr22c,dsh,Root,Raf,AstA-R1,dnc,LysP,HDAC4,dpp,opa,CG5059,Ir10a,msi,Ank2,Pino,Pzl,Ir41a,Tsf1,ths,Cdep,retn,CG8492,RYa,fru,bun,sqz,fs(1)N,GATAe,rdgA,Pld,Drsl2,MESK2,E(Pc),MsR2,Ptp61F,Cat,EcR,melt,Ank,CG33934,CG31750,Der-1,CG4839,klg,egh,grn,Nmdar2,csw,Or42b,Src42A,et,St4,Spf45,Sod1,kay,fz2,mGluR,Fhos,Set2,CG9380,sgg,msps,RhoL,Su(H),pirk,bnl,Syt12,PhKgamma,Exn,ine,RhoGAP100F,eIF2Bgamma,fz4,spri,shakB,sxc,Flo2,ninaD,Spn88Eb,trol,CG11318,shep,geko,Stacl,Rab9Fa,Mrtf,Cam,GlyP,Myo81F,CycT,Ser,AGO2,Atf6,shg,Hr96,Obp56h | | 4.412e-06 | -12.33 | eye photoreceptor cell differentiation | biological process | GO:0001754 | 165 | 23 | 12345 | 597 | Syn2,boi,msi,bun,Raf,B-H2,ex,chp,Cam,Mob2,dsh,kay,Khc,melt,RhoGAP100F,scrib,CG13251,Doa,csw,klg,CG5921,hth,CadN2 | | 4.460e-06 | -12.32 | instar larval or pupal development | biological process | GO:0002165 | 576 | 53 | 12345 | 597 | Mrtf,Cam,dsh,Raf,TfAP-2,foxo,Lpin,shg,sno,step,CG9519,opa,dpp,shd,CG9514,scrib,EcR,Ser,Elp3,crol,Set2,kay,fz2,path,hdc,Fhos,polybromo,app,Tor,Su(H),cic,sgg,B-H2,Eip74EF,IP3K2,Src42A,upd3,tld,tnc,bi,by,hth,Pak,Mbs,woc,fru,Dys,flw,esg,Ubx,spok,Exn,fw | | 4.880e-06 | -12.23 | positive regulation of macromolecule metabolic process | biological process | GO:0010604 | 858 | 71 | 12345 | 597 | mnb,scro,barc,Pomp,B-H2,TfIIFbeta,Iswi,Eip74EF,CG43689,bi,rictor,CG4849,gce,CG7483,Fer2,bol,esg,retn,Ubx,CG42797,Sox102F,GATAe,AGO3,CG4374,Lpin,CG12200,sno,Marf1,ems,mid,CrebB,Awh,EcR,REG,kay,polybromo,Tor,bnl,Su(H),sgg,Src42A,Stlk,hth,Pak,CG42830,FoxL1,tyf,CG1894,Mos,Mrtf,lncRNA:Hsromega,tin,CG10462,CG12071,Raf,ps,GlyP,TfAP-2,HPS4,Pde8,foxo,CG41520,Atf6,Hr96,CG14252,step,opa,dpp,g,Antp,CycT | | 5.022e-06 | -12.20 | PDEASE\_I\_2 | prosite domains | PS51845 | 6 | 5 | 6243 | 386 | Pde1c,Pde9,Pde6,dnc,Pde8 | | 5.050e-06 | -12.20 | regulation of cellular metabolic process | biological process | GO:0031323 | 1987 | 137 | 12345 | 597 | step,Atf6,CG14252,Hr96,CycT,AGO2,Antp,GlyP,TfAP-2,Yeti,trv,CG12071,ps,Mrtf,luna,lncRNA:Hsromega,l(3)neo38,HPS4,Art2,CG9650,CG42830,FoxL1,Spn88Eb,CG3530,hth,Mos,jim,kek5,fus,sgg,polybromo,Su(H),bnl,kay,Set2,Spf45,Sod1,CG17124,et,Src42A,raw,kek1,CG7368,grn,CG32006,ems,Marf1,REG,Ptp61F,EcR,melt,rib,CG18304,GATAe,bun,Cpsf160,CG4374,sws,woc,fru,Acp62F,retn,bru1,B-H2,numb,TfIIFbeta,scro,Pomp,crol,CG43689,msi,bi,mamo,Eip74EF,dpp,dany,opa,CG31224,HDAC4,CG15073,CG10462,Raf,tin,toe,chinmo,CG41520,foxo,Pde8,tyf,Usp1,Cln3,Pak,Khc,CG1894,ex,Snoo,Tor,CG32971,Mondo,CG6966,Stlk,Spt20,tld,Ste:CG33247,htk,mid,Maf1,sno,Awh,scrib,CrebB,AGO3,SteXh:CG42398,ham,Sox102F,lncRNA:bxd,Lpin,CG12200,GluRIIC,CG34172,gce,Doa,rictor,CG12605,Ubx,esg,flw,Fer2,Octbeta2R,bol,E5,cic,barc,Poxm,Elp3,mnb,Iswi | | 5.105e-06 | -12.19 | plasma membrane region | cellular component | GO:0098590 | 314 | 33 | 11940 | 538 | KaiR1D,Ser,dpr18,Fife,CG14182,Syx1A,dpr5,nuf,Exn,Hil,Nlg2,kek1,Or42b,Or65c,GluRIIC,step,scrib,dpr12,Flo2,gukh,Ank2,ex,Inx2,ZnT63C,cindr,ine,Pde6,dpr15,sdt,Cln3,unc-13,Or85b,rst | | 5.406e-06 | -12.13 | developmental growth | biological process | GO:0048589 | 191 | 25 | 12345 | 597 | RhoGAP100F,ine,shd,dnc,Spn88Eb,dpp,shg,CadN2,Mbs,Sh,spri,Nlg2,bun,bi,raw,Tor,sgg,Elp3,cpx,kay,Ank2,fz2,path,dsh,hdc | | 5.406e-06 | -12.13 | growth | biological process | GO:0040007 | 191 | 25 | 12345 | 597 | Tor,sgg,Elp3,kay,fz2,Ank2,cpx,dsh,path,hdc,Nlg2,bun,bi,raw,Spn88Eb,dpp,CadN2,shg,Mbs,Sh,spri,ine,RhoGAP100F,shd,dnc | | 5.882e-06 | -12.04 | regulation of nucleic acid-templated transcription | biological process | GO:1903506 | 1158 | 89 | 12345 | 597 | jim,CG1894,kek5,Khc,Art2,CG9650,FoxL1,CG42830,hth,tld,Spt20,ex,Snoo,Su(H),CG32971,Tor,polybromo,kay,Mondo,Set2,CycT,HDAC4,AGO2,Antp,CG15073,dany,dpp,opa,CG14252,Hr96,Atf6,CG31224,toe,chinmo,foxo,HPS4,l(3)neo38,TfAP-2,CG10462,CG12071,tin,luna,Mrtf,retn,Ubx,esg,Octbeta2R,E5,Fer2,gce,fru,woc,rictor,CG12605,bi,CG43689,Eip74EF,mamo,Iswi,TfIIFbeta,cic,B-H2,Poxm,scro,mnb,crol,Elp3,Awh,EcR,melt,scrib,CrebB,htk,CG7368,grn,Maf1,mid,CG32006,ems,sno,bun,Lpin,GluRIIC,CG4374,ham,GATAe,lncRNA:bxd,rib,Sox102F | | 5.882e-06 | -12.04 | regulation of RNA biosynthetic process | biological process | GO:2001141 | 1158 | 89 | 12345 | 597 | Elp3,crol,mnb,scro,Poxm,cic,B-H2,TfIIFbeta,mamo,Iswi,Eip74EF,CG43689,bi,CG12605,rictor,woc,fru,gce,Fer2,Octbeta2R,E5,esg,retn,Ubx,rib,Sox102F,lncRNA:bxd,GATAe,ham,CG4374,GluRIIC,Lpin,bun,sno,ems,CG32006,Maf1,mid,htk,grn,CG7368,CrebB,scrib,EcR,melt,Awh,Set2,Mondo,kay,polybromo,Tor,Su(H),CG32971,Snoo,ex,Spt20,tld,hth,CG42830,FoxL1,Art2,CG9650,Khc,CG1894,kek5,jim,Mrtf,luna,tin,CG10462,CG12071,TfAP-2,l(3)neo38,HPS4,foxo,chinmo,toe,CG31224,Atf6,CG14252,Hr96,opa,dpp,dany,CG15073,Antp,AGO2,HDAC4,CycT | | 6.167e-06 | -12.00 | PDIESTERASE1 | prints domains | PR00387 | 6 | 5 | 2185 | 142 | dnc,Pde6,Pde9,Pde8,Pde1c | | 6.290e-06 | -11.98 | regulation of nitrogen compound metabolic process | biological process | GO:0051171 | 1870 | 130 | 12345 | 597 | retn,bru1,sws,fru,woc,Acp62F,bi,msi,CG43689,Eip74EF,mamo,TfIIFbeta,B-H2,Pomp,scro,crol,Ptp61F,REG,melt,EcR,grn,CG7368,CG32006,ems,Marf1,bun,CG4374,GATAe,CG42797,rib,Mos,jim,kek5,CG9650,Art2,CG42830,Spn88Eb,FoxL1,hth,Spf45,kek1,raw,et,Src42A,fus,sgg,Su(H),bnl,polybromo,kay,Set2,CycT,AGO2,Antp,step,Hr96,CG14252,Atf6,HPS4,l(3)neo38,TfAP-2,trv,ps,CG12071,Yeti,lncRNA:Hsromega,luna,Mrtf,Ubx,flw,esg,bol,E5,Octbeta2R,Fer2,gce,Doa,rictor,CG12605,Iswi,cic,Poxm,barc,mnb,Elp3,Awh,CrebB,scrib,htk,Maf1,mid,sno,CG12200,Lpin,GluRIIC,SteXh:CG42398,AGO3,ham,lncRNA:bxd,Sox102F,CG1894,Khc,Usp1,tyf,Pak,Cln3,Stlk,CG6966,tld,Spt20,Ste:CG33247,ex,Tor,CG32971,Snoo,Mondo,HDAC4,CG15073,dany,dpp,opa,CG31224,toe,chinmo,foxo,CG41520,Pde8,CG10462,Raf,tin | | 6.501e-06 | -11.94 | cell growth | biological process | GO:0016049 | 81 | 15 | 12345 | 597 | fz2,Ank2,kay,dsh,spri,hdc,path,shg,CadN2,Sh,Mbs,ine,RhoGAP100F,raw,dnc | | 7.441e-06 | -11.81 | regulation of primary metabolic process | biological process | GO:0080090 | 1894 | 131 | 12345 | 597 | jim,Mos,kek5,CG9650,Art2,CG42830,FoxL1,Spn88Eb,hth,Spf45,et,raw,Src42A,kek1,fus,sgg,polybromo,Su(H),bnl,kay,Set2,CycT,AGO2,Antp,step,Atf6,CG14252,Hr96,l(3)neo38,HPS4,GlyP,TfAP-2,Yeti,trv,ps,CG12071,Mrtf,lncRNA:Hsromega,luna,retn,bru1,sws,woc,fru,Acp62F,CG43689,bi,msi,mamo,Eip74EF,TfIIFbeta,B-H2,scro,Pomp,crol,REG,Ptp61F,melt,EcR,grn,CG7368,CG32006,ems,Marf1,bun,CG4374,CG42797,rib,GATAe,Khc,CG1894,tyf,Usp1,Cln3,Pak,CG6966,Stlk,Spt20,tld,Ste:CG33247,ex,CG32971,Tor,Snoo,Mondo,HDAC4,CG15073,dpp,dany,opa,CG31224,toe,chinmo,CG41520,foxo,Pde8,CG10462,Raf,tin,Ubx,esg,flw,Fer2,bol,Octbeta2R,E5,gce,Doa,rictor,CG12605,Iswi,cic,barc,Poxm,Elp3,mnb,Awh,CrebB,scrib,htk,mid,Maf1,sno,CG12200,Lpin,GluRIIC,AGO3,SteXh:CG42398,ham,Sox102F,lncRNA:bxd | | 7.462e-06 | -11.81 | modulation of chemical synaptic transmission | biological process | GO:0050804 | 124 | 19 | 12345 | 597 | KaiR1D,Dys,Cat,Exn,nemy,Sh,Mctp,Src42A,GluRIIC,Frq1,sky,Sod1,Nlg2,Ank2,cpx,qvr,mGluR,Fife,Frq2 | | 7.462e-06 | -11.81 | regulation of trans-synaptic signaling | biological process | GO:0099177 | 124 | 19 | 12345 | 597 | Sh,Mctp,nemy,Cat,Exn,Dys,KaiR1D,Fife,Frq2,cpx,Ank2,mGluR,qvr,Nlg2,Sod1,Frq1,sky,Src42A,GluRIIC | | 7.606e-06 | -11.79 | regulation of metabolic process | biological process | GO:0019222 | 2165 | 146 | 12345 | 597 | Spf45,Sod1,kek1,CG17124,raw,Src42A,et,sgg,fus,Su(H),bnl,polybromo,kay,Set2,jim,Mos,kek5,CG9650,Art2,FoxL1,Spn88Eb,CG42830,CG3530,sxc,hth,HPS4,l(3)neo38,TfAP-2,GlyP,ps,CG12071,trv,Yeti,lncRNA:Hsromega,luna,Mrtf,CycT,AGO2,Antp,g,step,CG14252,Hr96,Atf6,bi,msi,CG43689,Eip74EF,mamo,B-H2,TfIIFbeta,numb,Pomp,scro,crol,retn,bru1,CG7483,sws,fru,woc,CG4849,Acp62F,bun,Cpsf160,CG4374,GATAe,CG18304,rib,CG42797,Ptp61F,REG,melt,EcR,grn,CG7368,Achl,CG32006,ems,Marf1,Stlk,CG6966,tld,Spt20,Ste:CG33247,ex,CG32971,Tor,Snoo,Mondo,CG1894,Khc,Usp1,tyf,Pak,Cln3,toe,chinmo,foxo,CG41520,Pde8,CG10462,Raf,tin,HDAC4,CG15073,dany,dpp,opa,CG31224,can,Iswi,cic,Poxm,barc,hdc,mnb,Elp3,Ubx,flw,esg,Octbeta2R,bol,E5,Fer2,gce,Doa,rictor,CG12605,jigr1,Lpin,CG12200,GluRIIC,CG34172,SteXh:CG42398,AGO3,ham,lncRNA:bxd,Sox102F,Awh,CrebB,scrib,htk,mid,Maf1,sno | | 7.606e-06 | -11.79 | regulation of biological quality | biological process | GO:0065008 | 1352 | 100 | 12345 | 597 | Exn,ine,KaiR1D,CG40178,Dys,Dlic,CG9650,nrv2,Cln3,l(2)01289,shakB,CG33310,Pak,Cadps,Syn2,boi,Sod1,tnc,CG3842,jus,Frq1,sky,raw,Src42A,upd3,Tor,sgg,Fife,CG4301,Mob2,Mondo,cpx,Fhos,AANAT1,shd,RhoGAP1A,Irk3,CG5059,dpp,Obp56h,shg,Hr96,Mctp,chinmo,foxo,Msp300,Raf,Cam,dsh,Stacl,qvr,Fad2,esg,retn,spok,Actn,fw,bru1,bol,Octbeta2R,woc,fru,sws,by,rictor,Pdfr,CG9981,Vha100-5,p130CAS,ZnT63C,unc-13,mnb,Syx1A,Ank2,path,EcR,Acbp2,Ank,nemy,Cat,Ptp61F,ddbt,scrib,CG9514,Qsox2,MCU,egh,Sh,mag,scramb1,CG9519,Nlg2,Lpin,Glut1,bun,JhI-26,Frq2,rib,rdgA,GATAe | | 9.622e-06 | -11.55 | tube development | biological process | GO:0035295 | 747 | 63 | 12345 | 597 | Inx2,GATAe,rib,Mrtf,dsh,tin,Raf,nuf,TfAP-2,chinmo,Lpin,sno,shg,step,mid,rols,csw,opa,dpp,shd,scrib,Irk3,Antp,Awh,EcR,Ser,Set2,crol,Fhos,path,hdc,Syx1A,kay,Su(H),Tor,bnl,polybromo,app,numb,sgg,cic,B-H2,tld,upd3,Src42A,raw,tnc,boi,bi,sxc,Mbs,Pak,by,hth,fru,Dlic,Dys,flw,esg,fw,Ubx,Idgf6,Exn | | 1.176e-05 | -11.35 | transcription regulator activity | molecular function | GO:0140110 | 605 | 53 | 12176 | 580 | FoxL1,tin,CG12071,CG43689,E5,EcR,Octbeta2R,Su(H),CG15073,ems,Ubx,gce,B-H2,ham,bi,Eip74EF,Spt20,CG31224,grn,retn,kay,TfAP-2,CG9650,cic,Mrtf,Atf6,fru,mnb,Mondo,CG32006,CG12605,CG42830,CG32971,opa,GATAe,foxo,woc,rib,hth,Poxm,bun,esg,Awh,Tif-IA,scro,Antp,mid,luna,CG10462,Hr96,CrebB,Lpin,CG4374 | | 1.177e-05 | -11.35 | post-embryonic development | biological process | GO:0009791 | 642 | 56 | 12345 | 597 | tnc,bi,Eip74EF,IP3K2,upd3,tld,Src42A,ths,Poxm,Tor,Su(H),app,polybromo,sgg,B-H2,cic,Set2,Elp3,crol,Fhos,path,hdc,kay,fz2,flw,esg,fw,spok,Exn,Ubx,Dys,fru,woc,Pak,Mbs,hth,by,Lpin,foxo,Msp300,Raf,TfAP-2,Cam,Mrtf,dsh,EcR,Ser,CG9514,scrib,shd,opa,dpp,sno,shg,CG9519,step | | 1.200e-05 | -11.33 | regulation of secretion by cell | biological process | GO:1903530 | 85 | 15 | 12345 | 597 | Cadps,cpx,Syx1A,Mctp,Flo2,Frq2,Fife,CG9650,Dys,Syt12,sky,Exn,Frq1,nemy,Glut1 | | 1.305e-05 | -11.25 | Ig\_sub | interpro domains | IPR003599 | 111 | 18 | 11383 | 585 | CG17716,rst,dpr18,kek5,nrm,klg,dpr5,DIP-lambda,dpr15,dpr21,dpr12,zormin,DIP-delta,kek1,boi,trol,Strn-Mlck,side-VIII | | 1.398e-05 | -11.18 | GO\_POSITIVE\_REGULATION\_OF\_MAPK\_CASCADE | MSigDB lists | GO\_POSITIVE\_REGULATION\_OF\_MAPK\_CASCADE | 79 | 13 | 4161 | 170 | CG42672,wbl,Pkc53E,csw,Sod1,Octbeta2R,dsh,Pak,Raf,Tsp96F,dpp,fz2,Pkc98E | | 1.441e-05 | -11.15 | synapse organization | biological process | GO:0050808 | 177 | 23 | 12345 | 597 | dpr15,Nlg2,dpr21,Frq1,unc-13,Ca-beta,dpr18,Elp3,mGluR,Ank2,fz2,kay,cpx,DIP-delta,Frq2,Fife,sgg,CrebB,dpr12,RhoGAP100F,dpr5,pod1,klg | | 1.470e-05 | -11.13 | regulation of gene expression | biological process | GO:0010468 | 1547 | 110 | 12345 | 597 | scrib,CrebB,melt,EcR,Awh,Marf1,sno,ems,CG32006,mid,Achl,Maf1,CG7368,grn,htk,CG4374,GluRIIC,Lpin,jigr1,bun,rib,Sox102F,GATAe,lncRNA:bxd,ham,AGO3,CG7483,Fer2,Octbeta2R,bru1,E5,bol,esg,Ubx,retn,CG12605,rictor,CG4849,woc,Doa,fru,gce,Iswi,mamo,Eip74EF,can,CG43689,msi,bi,crol,Elp3,mnb,hdc,barc,scro,Poxm,B-H2,TfIIFbeta,cic,numb,g,CG15073,Antp,AGO2,HDAC4,CycT,CG31224,Atf6,Hr96,CG14252,opa,dpp,dany,l(3)neo38,HPS4,foxo,chinmo,toe,Mrtf,luna,tin,ps,CG10462,CG12071,trv,TfAP-2,Khc,kek5,CG1894,jim,hth,sxc,CG42830,FoxL1,tyf,CG9650,Art2,Spt20,tld,Spf45,Set2,Mondo,kay,polybromo,Tor,CG32971,Snoo,Su(H),fus,ex | | 1.482e-05 | -11.12 | membrane part | cellular component | GO:0044425 | 2068 | 131 | 11940 | 538 | Vha100-5,chp,Nlg2,qvr,Der-1,step,CG33934,AstA-R1,Src42A,chk,Syn2,sws,Dys,Syx1A,CG9132,Fife,CG33090,Nha1,Glut1,CG5059,fz2,Gr39a,unc-13,Neb-cGP,rst,kek5,nrv2,Ir56a,gukh,CG11318,olf186-F,fs(1)N,Meltrin,kek1,Or42b,MsR2,GluRIIC,Or65c,CG32320,CG13794,KaiR1D,egh,g,CG13654,CadN2,Gr22c,dpr18,Msp300,PGRP-LA,Exn,Nmdar2,Or85b,DIP-delta,Vha36-3,CNT1,ND-AGGG,CG42826,Atf6,CG33310,CG14439,ZnT63C,shg,Pzl,cindr,CG17883,D2hgdh,CG14606,Stt3B,Mctp,CG4301,Sh,CG34172,fz4,ND-51L2,Ir10a,MCU,Pak,CG40472,beat-IIa,CG13229,PIG-G,dpr5,dpr15,Fad2,sdt,Pde6,CG30054,Cln3,CCAP-R,klg,CG12121,CG8468,cpx,CG6790,CG10960,CG9981,Ank2,Qsox2,Octbeta2R,ND-MLRQ,ine,CG14024,dpr12,Flo2,scrib,ACXD,CG3168,PIG-O,CG1572,Ser,Irk3,CG14182,CG3285,ABCD,Hil,nuf,Ir41a,mGluR,Ca-beta,Cyt-b5-r,Syt12,CG41128,Pdfr,UQCR-11,CG7882,Inx2,ex,Tsp96F,DIP-lambda,kcc,CG17600 | | 1.508e-05 | -11.10 | salivary gland morphogenesis | biological process | GO:0007435 | 97 | 16 | 12345 | 597 | shg,Pak,retn,Ser,EcR,fz2,hdc,dsh,Fhos,tin,Cam,rib,Src42A,raw,IP3K2,Eip74EF | | 1.508e-05 | -11.10 | gland morphogenesis | biological process | GO:0022612 | 97 | 16 | 12345 | 597 | raw,Src42A,IP3K2,Eip74EF,fz2,hdc,Fhos,dsh,tin,Cam,rib,retn,Ser,EcR,shg,Pak | | 1.530e-05 | -11.09 | axon development | biological process | GO:0061564 | 336 | 35 | 12345 | 597 | gukh,Src42A,haf,p130CAS,msps,hdc,kay,fz2,Ank2,RhoGAP100F,retn,Khc,fru,Dlic,spri,Pak,Mbs,pod1,CadN2,chb,trol,sick,beat-IIa,dsh,Cam,Ptp61F,dnc,egh,klg,Ptp99A,grn,mid,ems,Sh,shg | | 1.650e-05 | -11.01 | GO\_NEUROGENESIS | MSigDB lists | GO\_NEUROGENESIS | 328 | 30 | 4161 | 170 | Su(H),Iswi,Sod1,Elp3,sdt,sky,eIF4E7,Frq1,Cdep,eIF2Bgamma,LRP1,Ser,dpp,Raf,LpR2,unc-13,Khc,Pak,csw,Frq2,CG42672,Awh,Tor,TfAP-2,Ank,CG33090,fz2,Tif-IA,AstA-R1,Dys | | 1.664e-05 | -11.00 | positive regulation of cellular process | biological process | GO:0048522 | 1464 | 105 | 12345 | 597 | GATAe,Sox102F,Pld,AGO3,MESK2,cindr,Cpsf160,olf186-F,CG4374,bun,Glut1,CG12200,Lpin,Gprk1,ems,Pkc53E,Marf1,sno,siz,Der-1,mid,CrebB,CG1572,REG,EcR,melt,Awh,Ank2,mnb,crol,TfIIFbeta,B-H2,numb,Pomp,barc,scro,Eip74EF,Iswi,bi,CG43689,rictor,gce,sdt,Doa,Octbeta2R,bru1,bol,Fer2,CG10188,retn,esg,flw,tin,dsh,lncRNA:Hsromega,Mrtf,TfAP-2,GlyP,CG10462,Raf,ps,CG12071,trol,CG41520,foxo,Pde8,chb,chinmo,step,CG14252,Hr96,Atf6,shg,dpp,opa,CG5059,Antp,RhoGAP1A,CycT,Fhos,kay,ex,Fife,sgg,Tor,bnl,Su(H),polybromo,msps,tld,upd3,Src42A,Stlk,spri,Cadps,sxc,Pak,hth,Flo2,tyf,CG42830,KaiR1D,Exn,Mos,CG1894 | | 1.679e-05 | -10.99 | metamorphosis | biological process | GO:0007552 | 497 | 46 | 12345 | 597 | TfAP-2,Raf,dsh,tin,Mrtf,Cam,shep,Lpin,dpp,opa,step,shg,sno,Ser,EcR,scrib,B-H2,sgg,cic,app,polybromo,Tor,Su(H),kay,hdc,path,Fhos,crol,bi,tnc,Src42A,upd3,tld,Eip74EF,IP3K2,fru,by,hth,Pak,Mbs,Ubx,Exn,fw,esg,flw,Dys | | 1.734e-05 | -10.96 | HD/PDEase\_dom | interpro domains | IPR003607 | 8 | 5 | 11383 | 585 | Pde8,dnc,Pde1c,Pde9,Pde6 | | 1.784e-05 | -10.93 | IL2\_UP.V1\_UP | MSigDB lists | IL2\_UP.V1\_UP | 30 | 8 | 4161 | 170 | Glut1,Frq1,fus,Frq2,Meltrin,fz2,dpp,Sh | | 1.942e-05 | -10.85 | instar larval or pupal morphogenesis | biological process | GO:0048707 | 470 | 44 | 12345 | 597 | upd3,tld,Src42A,IP3K2,Eip74EF,bi,tnc,Fhos,path,hdc,kay,crol,B-H2,sgg,cic,Su(H),Tor,app,polybromo,Dys,fw,Ubx,Exn,flw,esg,Mbs,Pak,by,hth,fru,Lpin,dsh,Mrtf,Cam,TfAP-2,Raf,scrib,Ser,EcR,step,sno,shg,dpp,opa | | 1.972e-05 | -10.83 | axonogenesis | biological process | GO:0007409 | 326 | 34 | 12345 | 597 | Khc,RhoGAP100F,retn,pod1,Mbs,Pak,CadN2,spri,fru,Dlic,Src42A,gukh,hdc,Ank2,kay,fz2,msps,haf,dnc,Ptp61F,Sh,shg,ems,mid,klg,Ptp99A,egh,grn,trol,chb,Cam,dsh,sick,beat-IIa | | 2.129e-05 | -10.76 | regulation of secretion | biological process | GO:0051046 | 89 | 15 | 12345 | 597 | Dys,Syt12,sky,Frq1,Exn,nemy,Glut1,cpx,Cadps,Syx1A,Mctp,Flo2,Fife,Frq2,CG9650 | | 2.145e-05 | -10.75 | fly chr2-55 cM | chromosome location | fly chr2-55 cM | 36 | 8 | 25010 | 845 | CG17684,DIP-lambda,dpr21,lncRNA:CR44043,CG41242,Maf1,RYa,CG41520 | | 2.145e-05 | -10.75 | fly chr2-55 | chromosome location | fly chr2-55 | 36 | 8 | 25010 | 845 | CG17684,DIP-lambda,dpr21,lncRNA:CR44043,CG41242,Maf1,CG41520,RYa | | 2.203e-05 | -10.72 | tissue development | biological process | GO:0009888 | 1113 | 84 | 12345 | 597 | Awh,EcR,scrib,egh,grn,Mlp84B,mid,rols,csw,sno,bun,Lpin,cindr,fs(1)N,Inx2,GATAe,rib,fw,Ubx,flw,esg,lncRNA:flam,sdt,fru,Prm,CadN2,by,bi,dsd,numb,B-H2,cic,ths,hdc,path,Syx1A,crol,Ser,CG17470,Irk3,shd,dpp,opa,step,shg,CG34417,chinmo,trol,Pde8,TfAP-2,Msp300,Raf,CG41099,nuf,tin,dsh,Mrtf,Exn,Idgf6,Khc,Dys,Spn88Eb,Pak,Mbs,sxc,hth,tnc,boi,upd3,tld,raw,Src42A,sgg,ex,bnl,Tor,Su(H),app,RhoL,bbg,polybromo,Fhos,kay,Set2 | | 2.351e-05 | -10.66 | HALLMARK\_IL2\_STAT5\_SIGNALING | MSigDB lists | HALLMARK\_IL2\_STAT5\_SIGNALING | 40 | 9 | 4161 | 170 | scramb1,Tsp96F,dpp,CG8184,Odc1,Glut1,Frq1,Frq2,Meltrin | | 2.408e-05 | -10.63 | GO\_ENZYME\_BINDING | MSigDB lists | GO\_ENZYME\_BINDING | 530 | 41 | 4161 | 170 | Syx1A,Ptp61F,Cat,Octbeta2R,Dys,fz2,Ugt301D1,Tif-IA,Cadps,dnc,CG41099,Frq2,csw,Maf1,PhKgamma,Ca-beta,CG40045,Raf,Act87E,LRP1,kcc,Flo2,KaiR1D,Sod1,CG17883,Elp3,scramb1,dsh,Stlk,Tor,Der-1,Gas8,Pak,Pkc98E,flw,EcR,Sk1,Cpsf160,eIF4E7,Frq1,Pkc53E | | 2.589e-05 | -10.56 | 3',5'-cyclic-nucleotide phosphodiesterase activity | molecular function | GO:0004114 | 9 | 5 | 12176 | 580 | Pde1c,dnc,Pde6,Pde9,Pde8 | | 2.687e-05 | -10.52 | post-embryonic appendage morphogenesis | biological process | GO:0035120 | 345 | 35 | 12345 | 597 | sno,shg,step,dpp,scrib,EcR,Ser,Mrtf,dsh,Raf,TfAP-2,Lpin,Pak,Mbs,by,hth,fru,Dys,flw,fw,Exn,Ubx,crol,path,Fhos,Su(H),app,polybromo,sgg,cic,tld,upd3,Src42A,tnc,bi | | 2.717e-05 | -10.51 | PH-like\_dom\_sf | interpro domains | IPR011993 | 139 | 20 | 11383 | 585 | Exn,Gprk1,cert,Pld,CG5522,melt,siz,Cadps,CG9132,CG3530,ex,RhoGEF3,Cdep,CG10188,CG5004,numb,Myo81F,CG42673,by,step | | 2.767e-05 | -10.50 | behavior | biological process | GO:0007610 | 630 | 54 | 12345 | 597 | nemy,HDAC4,EcR,dnc,CrebB,scrib,MsR2,ITP,MCU,Gld2,Nmdar2,Obp56h,egh,klg,Sh,Nlg2,geko,JhI-26,shep,Pde1c,foxo,TfAP-2,Root,Gr22c,Fad2,kcc,Khc,retn,CG5541,lncRNA:flam,Ebp,bol,Pkc98E,Fer2,fru,tyf,sxc,shakB,Pdfr,jus,Tk,Su(H),Tsf1,sgg,numb,mnb,Pzl,Elp3,Gr39a,Mob2,mGluR,Pino,Ank2,kay | | 2.837e-05 | -10.47 | compound eye morphogenesis | biological process | GO:0001745 | 290 | 31 | 12345 | 597 | foxo,bun,bi,Spf45,msi,boi,Syn2,kay,dsh,Cam,Mob2,chp,B-H2,ex,Raf,cindr,bnl,scrib,Ser,RhoGAP100F,fw,melt,hth,CadN2,shg,dpp,siz,rst,klg,csw,Doa | | 2.867e-05 | -10.46 | positive regulation of metabolic process | biological process | GO:0009893 | 937 | 73 | 12345 | 597 | CycT,g,Antp,opa,dpp,Hr96,CG14252,Atf6,step,Pde8,HPS4,CG41520,foxo,CG12071,ps,Raf,CG10462,TfAP-2,GlyP,lncRNA:Hsromega,Mrtf,tin,CG1894,Mos,CG42830,FoxL1,tyf,Pak,hth,Stlk,Src42A,bnl,Su(H),Tor,polybromo,sgg,kay,Awh,EcR,REG,CrebB,mid,sno,Marf1,ems,Lpin,CG12200,CG4374,Cpsf160,AGO3,GATAe,Sox102F,CG42797,esg,Ubx,retn,CG7483,bol,Fer2,Doa,gce,CG4849,rictor,bi,CG43689,Eip74EF,Iswi,Pomp,barc,scro,B-H2,TfIIFbeta,mnb | | 2.934e-05 | -10.44 | regulation of localization | biological process | GO:0032879 | 478 | 44 | 12345 | 597 | sdt,Flo2,CG9650,spri,Cadps,Khc,Exn,Syt12,Dys,Tor,bnl,Fife,sgg,numb,Pde6,mnb,Elp3,Ca-beta,CG3078,Fhos,cpx,kay,Syx1A,sky,Frq1,LpR2,Mctp,shg,step,Pkc53E,CG13711,nemy,Ptp61F,scrib,cindr,Pld,Frq2,CG42797,Cam,qvr,dsh,Glut1,olf186-F,foxo | | 2.944e-05 | -10.43 | Signal Transduction | REACTOME pathways | R-DME-162582 | 717 | 59 | 3923 | 197 | rdgA,RhoGAP18B,Pde8,Octbeta2R,CG10188,mGluR,Raf,Strn-Mlck,Pak,Stlk,trv,CrebB,Ser,CG3091,Pkc98E,REG,csw,numb,p130CAS,CG30054,CycT,Prosbeta5R1,CG42797,Ptp61F,TfIIFbeta,Vha36-3,Pde1c,eIF4E7,Cam,Exn,Sk1,boi,rictor,esn,CG6424,flw,bnl,Mbs,fz2,dnc,sgg,Snoo,by,dpp,Gprk1,RhoGAP1A,cindr,ACXD,foxo,nrv2,RhoGAP100F,Mrtf,fus,Tor,Pkc53E,Vha100-5,Sox102F,CG41520,dsh | | 3.001e-05 | -10.41 | cellular response to stimulus | biological process | GO:0051716 | 1467 | 104 | 12345 | 597 | CG17698,Iswi,msi,Pdfr,Ank2,unc-13,Ca-beta,cic,ths,Octbeta2R,Cdep,flw,rictor,CCAP-R,gce,RYa,Doa,Pde9,GluRIIC,bun,Gprk1,RhoGAP18B,Lpin,fs(1)N,rdgA,Pld,TpnC25D,MESK2,ITP,CG13898,MsR2,Ptp61F,Cat,Ank,melt,EcR,Pkc53E,Sh,sno,wake,CG31750,CG4839,Der-1,Nmdar2,csw,upd3,Src42A,et,Tk,St4,Stlk,Spf45,Sod1,boi,Tusp,ACXD,mGluR,CG7362,fz2,kay,Set2,CG30054,sgg,ex,Pde6,CG13229,Tor,bnl,Su(H),Pkc98E,Syt12,PhKgamma,RhoGAP100F,Gpb5,kcc,eIF2Bgamma,spri,CG13526,fz4,Pak,shakB,ninaD,TpnC41C,Pde1c,CG41520,foxo,shep,CG11318,Rab9Fa,Stacl,dsh,Mrtf,Cam,Raf,Myo81F,AstA-R1,dnc,Ser,HDAC4,Hr96,Atf6,dpp,opa,CG5059 | | 3.092e-05 | -10.38 | GO\_REGULATION\_OF\_MAPK\_CASCADE | MSigDB lists | GO\_REGULATION\_OF\_MAPK\_CASCADE | 123 | 16 | 4161 | 170 | csw,CG42672,dsh,Ptp61F,Octbeta2R,CG3376,fz2,KaiR1D,Sod1,Pkc53E,wbl,Raf,Tsp96F,Pak,Pkc98E,dpp | | 3.094e-05 | -10.38 | locomotion | biological process | GO:0040011 | 664 | 56 | 12345 | 597 | CG17387,trol,shep,chb,bun,sqz,dsh,tin,rdgA,rib,Cam,Mrtf,Msp300,Raf,cindr,beat-IIa,E(Pc),shd,Ptp61F,EcR,shg,klg,Ptp99A,grn,egh,dpp,mid,opa,csw,rols,Src42A,gukh,Syn2,Pdfr,Fhos,kay,fz2,haf,ex,numb,TpnC4,p130CAS,bnl,ths,msps,bbg,RhoL,RhoGAP100F,retn,esg,Khc,Gas8,Pak,pod1,CadN2,fru | | 3.169e-05 | -10.36 | imaginal disc-derived appendage development | biological process | GO:0048737 | 362 | 36 | 12345 | 597 | Lpin,Raf,TfAP-2,Mrtf,dsh,EcR,Ser,scrib,dpp,sno,shg,step,tnc,bi,Src42A,tld,upd3,polybromo,app,Su(H),cic,sgg,crol,Syx1A,Fhos,path,flw,Exn,Ubx,fw,Dys,fru,hth,by,Mbs,Pak | | 3.313e-05 | -10.32 | PDEase\_catalytic\_dom\_sf | interpro domains | IPR036971 | 5 | 4 | 11383 | 585 | Pde8,dnc,Pde6,Pde9 | | 3.313e-05 | -10.32 | PDEase\_CS | interpro domains | IPR023174 | 5 | 4 | 11383 | 585 | dnc,Pde9,Pde1c,Pde6 | | 3.785e-05 | -10.18 | GO\_CYCLIC\_NUCLEOTIDE\_PHOSPHODIESTERASE\_ACTIVITY | MSigDB lists | GO\_CYCLIC\_NUCLEOTIDE\_PHOSPHODIESTERASE\_ACTIVITY | 6 | 4 | 4161 | 170 | Pde6,dnc,Pde8,Pde1c | | 3.796e-05 | -10.18 | Ig\_sub2 | interpro domains | IPR003598 | 109 | 17 | 11383 | 585 | dpr15,dpr5,DIP-lambda,klg,nrm,kek5,rst,CG17716,Strn-Mlck,side-VIII,trol,kek1,boi,DIP-delta,dpr12,zormin,dpr21 | | 3.800e-05 | -10.18 | regulation of multicellular organismal process | biological process | GO:0051239 | 781 | 63 | 12345 | 597 | Khc,Ubx,Octbeta2R,bru1,Doa,Dlic,Pak,Mbs,hth,spri,rictor,Syn2,Pdfr,Tk,raw,Src42A,Su(H),Tor,bnl,barc,ex,numb,CG9380,sgg,mnb,Elp3,Mob2,hdc,CG3078,kay,Syx1A,fz2,Ank2,EcR,Ser,AANAT1,Cat,PGRP-LA,scrib,CrebB,CG1572,Antp,mid,csw,rst,wake,egh,dpp,Mlp84B,Sh,shg,Pkc53E,step,bun,wbl,foxo,ham,Raf,TfAP-2,CG42797,Sox102F,qvr,dsh | | 3.901e-05 | -10.15 | imaginal disc development | biological process | GO:0007444 | 591 | 51 | 12345 | 597 | bi,boi,tnc,tld,upd3,Src42A,B-H2,sgg,cic,bnl,Su(H),app,polybromo,Fhos,path,kay,Syx1A,crol,Set2,fw,Idgf6,Ubx,Exn,esg,flw,Dys,fru,Mbs,Pak,sxc,by,hth,Lpin,chinmo,TfAP-2,Raf,nuf,dsh,Mrtf,Ser,Awh,EcR,Irk3,scrib,dpp,mid,csw,opa,step,sno,shg | | 3.960e-05 | -10.14 | regulation of cellular response to insulin stimulus | biological process | GO:1900076 | 36 | 9 | 12345 | 597 | step,rictor,sgg,Tor,foxo,Ptp61F,HDAC4,Lpin,melt | | 3.960e-05 | -10.14 | regulation of insulin receptor signaling pathway | biological process | GO:0046626 | 36 | 9 | 12345 | 597 | foxo,HDAC4,melt,Lpin,Ptp61F,step,rictor,Tor,sgg | | 3.963e-05 | -10.14 | post-embryonic animal morphogenesis | biological process | GO:0009886 | 484 | 44 | 12345 | 597 | Pak,Mbs,hth,by,fru,Dys,fw,Ubx,Exn,esg,flw,Fhos,path,hdc,kay,crol,sgg,B-H2,cic,Tor,Su(H),app,polybromo,upd3,tld,Src42A,IP3K2,Eip74EF,bi,tnc,step,shg,sno,dpp,opa,scrib,Ser,EcR,dsh,Cam,Mrtf,TfAP-2,Raf,Lpin | | 4.002e-05 | -10.13 | appendage development | biological process | GO:0048736 | 366 | 36 | 12345 | 597 | dpp,sno,shg,step,EcR,Ser,scrib,Raf,TfAP-2,Mrtf,dsh,Lpin,fru,Mbs,Pak,hth,by,flw,fw,Ubx,Exn,Dys,Su(H),polybromo,app,cic,sgg,crol,path,Fhos,Syx1A,tnc,bi,tld,upd3,Src42A | | 4.291e-05 | -10.06 | regulation of tube diameter | biological process | GO:0035296 | 15 | 6 | 12345 | 597 | tnc,EcR,dpp,nrv2,rib,raw | | 4.362e-05 | -10.04 | GSE27786\_NKCELL\_VS\_NEUTROPHIL\_DN | MSigDB lists | GSE27786\_NKCELL\_VS\_NEUTROPHIL\_DN | 43 | 9 | 4161 | 170 | sws,AstA-R1,CNT1,AGO3,EcR,KaiR1D,TfAP-2,D2hgdh,Iswi | | 4.416e-05 | -10.03 | regulation of intracellular signal transduction | biological process | GO:1902531 | 339 | 34 | 12345 | 597 | chb,Lpin,Pde8,olf186-F,CG34408,Raf,MESK2,lncRNA:Hsromega,Ptp61F,melt,RhoGAP1A,dnc,siz,dpp,step,raw,Src42A,sgg,ex,bnl,Tor,mnb,Exn,Mos,flw,Cdep,CG10188,CG18659,CG5522,spri,rictor,Pak,RhoGEF3,Cln3 | | 4.749e-05 | -9.95 | Ig-like\_dom\_sf | interpro domains | IPR036179 | 133 | 19 | 11383 | 585 | dpr21,zormin,dpr12,DIP-delta,kek1,boi,trol,Strn-Mlck,side-VIII,CG17716,rst,dpr18,kek5,nrm,klg,Ptp99A,dpr5,DIP-lambda,dpr15 | | 4.809e-05 | -9.94 | gland development | biological process | GO:0048732 | 178 | 22 | 12345 | 597 | fz2,Fhos,tin,dsh,hdc,Cam,rib,Tor,ths,Raf,Src42A,raw,Eip74EF,IP3K2,hth,shg,Pak,dpp,Antp,retn,Ser,EcR | | 4.873e-05 | -9.93 | imaginal disc-derived appendage morphogenesis | biological process | GO:0035114 | 355 | 35 | 12345 | 597 | Mrtf,dsh,Raf,TfAP-2,Lpin,shg,sno,step,dpp,scrib,EcR,Ser,crol,Fhos,path,Su(H),polybromo,app,cic,sgg,upd3,tld,Src42A,tnc,bi,Mbs,Pak,hth,by,fru,Dys,flw,fw,Exn,Ubx | | 4.976e-05 | -9.91 | cyclic-nucleotide phosphodiesterase activity | molecular function | GO:0004112 | 10 | 5 | 12176 | 580 | Pde1c,Pde9,Pde8,Pde6,dnc | | 5.015e-05 | -9.90 | regulation of phosphorylation | biological process | GO:0042325 | 271 | 29 | 12345 | 597 | Stlk,chinmo,kek1,CG34172,et,CG17124,CG41520,Src42A,raw,Pde8,Ste:CG33247,SteXh:CG42398,sgg,bnl,Tor,Raf,kay,mnb,lncRNA:Hsromega,Mos,Ptp61F,CycT,flw,sws,dpp,step,rictor,Pak,Cln3 | | 5.030e-05 | -9.90 | DNA-binding transcription activator activity | molecular function | GO:0001216 | 86 | 14 | 12176 | 580 | retn,EcR,mid,kay,CG10462,opa,foxo,Su(H),CG4374,Mrtf,B-H2,Eip74EF,bi,CG43689 | | 5.030e-05 | -9.90 | DNA-binding transcription activator activity, RNA polymerase II-specific | molecular function | GO:0001228 | 86 | 14 | 12176 | 580 | opa,kay,CG10462,mid,EcR,retn,CG4374,Su(H),foxo,B-H2,Mrtf,CG43689,bi,Eip74EF | | 5.336e-05 | -9.84 | tube morphogenesis | biological process | GO:0035239 | 536 | 47 | 12345 | 597 | Pak,Mbs,by,hth,fru,Dlic,Dys,flw,esg,fw,Ubx,Exn,crol,Fhos,hdc,path,kay,bnl,Su(H),Tor,app,polybromo,B-H2,sgg,cic,upd3,tld,Src42A,raw,tnc,bi,shg,sno,step,opa,dpp,scrib,EcR,Ser,Inx2,rib,Mrtf,dsh,tin,Raf,TfAP-2,Lpin | | 5.412e-05 | -9.82 | plasma membrane bounded cell projection | cellular component | GO:0120025 | 626 | 50 | 11940 | 538 | Ptp99A,dpr15,kl-5,Or85b,Gr39a,unc-13,Cln3,mnb,Mst87F,Mob2,pod1,IFT46,boi,cpx,klg,EcR,nrv2,Gld2,CG17122,Pdfr,Ank2,gukh,CG5921,shg,Gas8,kcc,Cam,chp,chb,Khc,Or42b,dpr12,dpp,Or65c,CG13794,path,nompB,g,Ser,Gr22c,hdc,CG14182,Syx1A,AGO2,dpr18,sky,Root,sgg,CG17387,dpr5 | | 5.458e-05 | -9.82 | regulation of protein phosphorylation | biological process | GO:0001932 | 245 | 27 | 12345 | 597 | Raf,Tor,bnl,SteXh:CG42398,sgg,lncRNA:Hsromega,mnb,kay,chinmo,Stlk,Pde8,Ste:CG33247,kek1,raw,et,CG41520,Src42A,sws,dpp,Pak,Cln3,rictor,step,flw,Ptp61F,Mos,CycT | | 5.540e-05 | -9.80 | visual system development | biological process | GO:0150063 | 401 | 38 | 12345 | 597 | foxo,bun,toe,dsh,Cam,chp,Raf,cindr,scrib,Ser,Awh,melt,shg,CG5921,dpp,klg,rst,siz,csw,Src42A,msi,bi,Spf45,Syn2,boi,kay,Mob2,cic,ex,B-H2,bnl,CG13251,fw,RhoGAP100F,Khc,hth,CadN2,Doa | | 5.540e-05 | -9.80 | sensory system development | biological process | GO:0048880 | 401 | 38 | 12345 | 597 | Doa,hth,CadN2,Khc,RhoGAP100F,fw,CG13251,bnl,B-H2,ex,cic,Mob2,kay,boi,Syn2,msi,Spf45,bi,Src42A,csw,klg,siz,rst,dpp,CG5921,shg,melt,Awh,Ser,scrib,cindr,Raf,chp,Cam,dsh,toe,bun,foxo | | 5.540e-05 | -9.80 | eye development | biological process | GO:0001654 | 401 | 38 | 12345 | 597 | Doa,CadN2,hth,fw,RhoGAP100F,Khc,CG13251,cic,B-H2,ex,bnl,kay,Mob2,msi,Spf45,bi,boi,Syn2,Src42A,dpp,siz,klg,rst,csw,shg,CG5921,Ser,melt,Awh,scrib,Raf,cindr,dsh,Cam,chp,bun,toe,foxo | | 5.790e-05 | -9.76 | cell migration | biological process | GO:0016477 | 287 | 30 | 12345 | 597 | Src42A,bun,chb,Mrtf,rib,fz2,kay,tin,dsh,Fhos,RhoL,bbg,bnl,Raf,cindr,Msp300,p130CAS,ths,ex,numb,shd,EcR,esg,shg,Pak,opa,csw,rols,dpp,egh | | 5.791e-05 | -9.76 | appendage morphogenesis | biological process | GO:0035107 | 358 | 35 | 12345 | 597 | TfAP-2,Raf,dsh,Mrtf,Lpin,dpp,step,shg,sno,Ser,EcR,scrib,cic,sgg,polybromo,app,Su(H),Fhos,path,crol,bi,tnc,Src42A,upd3,tld,fru,by,hth,Mbs,Pak,Exn,Ubx,fw,flw,Dys | | 5.899e-05 | -9.74 | GO\_POSITIVE\_REGULATION\_OF\_CELL\_ADHESION | MSigDB lists | GO\_POSITIVE\_REGULATION\_OF\_CELL\_ADHESION | 55 | 10 | 4161 | 170 | Pak,Raf,Pkc98E,mino,Ank,Flo2,IP3K2,Tor,csw,Pkc53E | | 5.931e-05 | -9.73 | developmental growth involved in morphogenesis | biological process | GO:0060560 | 86 | 14 | 12345 | 597 | path,dsh,hdc,spri,Ank2,kay,fz2,Mbs,Sh,shg,CadN2,raw,dnc,RhoGAP100F | | 5.958e-05 | -9.73 | positive regulation of nucleobase-containing compound metabolic process | biological process | GO:0045935 | 523 | 46 | 12345 | 597 | Lpin,CG12200,CG4374,foxo,CG10462,ps,CG12071,TfAP-2,AGO3,GATAe,Sox102F,Mrtf,tin,EcR,Awh,CycT,CrebB,Antp,mid,opa,CG14252,Hr96,Marf1,sno,Atf6,ems,bi,CG43689,Eip74EF,Iswi,Tor,Su(H),scro,barc,polybromo,TfIIFbeta,B-H2,mnb,kay,CG1894,esg,retn,CG42830,gce,hth,rictor | | 6.114e-05 | -9.70 | cell projection | cellular component | GO:0042995 | 629 | 50 | 11940 | 538 | Or65c,dpr12,dpp,CG13794,chp,chb,Or42b,Khc,sky,dpr18,sgg,Root,CG17387,CG14182,Syx1A,AGO2,dpr5,g,nompB,path,Gr22c,hdc,Ser,boi,Mob2,pod1,IFT46,EcR,cpx,klg,Ptp99A,dpr15,unc-13,Cln3,Mst87F,mnb,kl-5,Or85b,Gr39a,Gas8,shg,CG5921,Cam,kcc,nrv2,Gld2,gukh,Pdfr,CG17122,Ank2 | | 6.581e-05 | -9.63 | - | gene3d domains | 1.10.1300.10 | 5 | 4 | 4929 | 302 | Pde9,Pde6,Pde8,dnc | | 6.746e-05 | -9.60 | epithelial tube morphogenesis | biological process | GO:0060562 | 495 | 44 | 12345 | 597 | scrib,Ser,EcR,step,shg,sno,dpp,opa,Lpin,dsh,Mrtf,rib,TfAP-2,Raf,Dys,fw,Exn,Ubx,flw,esg,Mbs,Pak,hth,by,fru,upd3,tld,raw,Src42A,bi,tnc,path,hdc,Fhos,kay,crol,sgg,B-H2,cic,Su(H),Tor,bnl,app,polybromo | | 6.818e-05 | -9.59 | GO\_POSITIVE\_REGULATION\_OF\_MULTICELLULAR\_ORGANISMAL\_PROCESS | MSigDB lists | GO\_POSITIVE\_REGULATION\_OF\_MULTICELLULAR\_ORGANISMAL\_PROCESS | 236 | 23 | 4161 | 170 | CG42672,Tor,TfAP-2,IP3K2,mino,fz2,dnc,Tif-IA,Nha1,Dys,Octbeta2R,chb,Pkc53E,Su(H),sxc,Sod1,Pkc98E,EcR,dpp,Ser,Raf,LpR2,unc-13 | | 6.848e-05 | -9.59 | PDEASE\_I\_1 | prosite domains | PS00126 | 5 | 4 | 6243 | 386 | Pde6,dnc,Pde9,Pde1c | | 6.855e-05 | -9.59 | compound eye photoreceptor cell differentiation | biological process | GO:0001751 | 157 | 20 | 12345 | 597 | scrib,melt,RhoGAP100F,CadN2,hth,Doa,csw,klg,Syn2,boi,msi,bun,chp,Cam,Mob2,dsh,kay,Raf,B-H2,ex | | 6.906e-05 | -9.58 | imaginal disc morphogenesis | biological process | GO:0007560 | 420 | 39 | 12345 | 597 | by,hth,Mbs,Pak,fru,Dys,Exn,Ubx,fw,flw,esg,kay,Fhos,path,crol,sgg,cic,B-H2,app,polybromo,Su(H),Src42A,upd3,tld,bi,tnc,step,shg,sno,dpp,opa,scrib,Ser,EcR,dsh,Mrtf,TfAP-2,Raf,Lpin | | 6.906e-05 | -9.58 | post-embryonic animal organ morphogenesis | biological process | GO:0048563 | 420 | 39 | 12345 | 597 | Pak,Mbs,hth,by,fru,Dys,flw,esg,fw,Exn,Ubx,crol,path,Fhos,kay,Su(H),app,polybromo,B-H2,sgg,cic,tld,upd3,Src42A,tnc,bi,sno,shg,step,opa,dpp,scrib,EcR,Ser,Mrtf,dsh,Raf,TfAP-2,Lpin | | 6.963e-05 | -9.57 | salivary gland development | biological process | GO:0007431 | 121 | 17 | 12345 | 597 | Ser,retn,EcR,Pak,shg,hth,Src42A,raw,Eip74EF,IP3K2,tin,dsh,Fhos,hdc,fz2,rib,Cam | | 6.963e-05 | -9.57 | exocrine system development | biological process | GO:0035272 | 121 | 17 | 12345 | 597 | Cam,rib,dsh,Fhos,tin,hdc,fz2,Eip74EF,IP3K2,Src42A,raw,Pak,hth,shg,EcR,Ser,retn | | 7.161e-05 | -9.54 | post-embryonic animal organ development | biological process | GO:0048569 | 481 | 43 | 12345 | 597 | Dys,esg,flw,Exn,Ubx,fw,by,hth,Mbs,Pak,fru,Src42A,upd3,tld,tnc,bi,crol,kay,Fhos,path,app,polybromo,Poxm,ths,Su(H),Tor,cic,B-H2,sgg,scrib,EcR,Ser,sno,shg,step,opa,dpp,Lpin,Mrtf,dsh,Raf,Msp300,TfAP-2 | | 7.437e-05 | -9.51 | regulation of cellular macromolecule biosynthetic process | biological process | GO:2000112 | 1307 | 93 | 12345 | 597 | FoxL1,CG42830,CG9650,Art2,tyf,hth,CG1894,kek5,Khc,jim,Tor,CG32971,Su(H),Snoo,polybromo,ex,Mondo,Set2,kay,tld,Spt20,opa,dany,dpp,CG14252,Hr96,Atf6,CG31224,AGO2,HDAC4,CycT,Antp,CG15073,CG12071,CG10462,TfAP-2,luna,Mrtf,tin,chinmo,toe,HPS4,l(3)neo38,foxo,fru,woc,gce,CG12605,rictor,esg,retn,Ubx,bru1,bol,Octbeta2R,E5,Fer2,Poxm,scro,B-H2,TfIIFbeta,cic,mnb,crol,Elp3,msi,bi,CG43689,Eip74EF,Iswi,mamo,Maf1,mid,CG32006,grn,htk,CG7368,sno,ems,EcR,melt,Awh,CrebB,scrib,ham,lncRNA:bxd,GATAe,Sox102F,rib,Lpin,bun,CG4374,GluRIIC | | 7.513e-05 | -9.50 | GO\_POSITIVE\_REGULATION\_OF\_KINASE\_ACTIVITY | MSigDB lists | GO\_POSITIVE\_REGULATION\_OF\_KINASE\_ACTIVITY | 105 | 14 | 4161 | 170 | Stlk,Sod1,csw,wbl,CG17698,CG42672,LpR2,Tsp96F,Raf,Pak,dsh,Octbeta2R,fz2,dpp | | 7.513e-05 | -9.50 | maternal specification of dorsal/ventral axis, oocyte, soma encoded | biological process | GO:0007313 | 6 | 4 | 12345 | 597 | cic,dpp,wbl,tld | | 7.637e-05 | -9.48 | positive regulation of RNA metabolic process | biological process | GO:0051254 | 513 | 45 | 12345 | 597 | Mrtf,Sox102F,GATAe,tin,CG12071,CG10462,ps,AGO3,TfAP-2,CG4374,foxo,Lpin,Marf1,Atf6,sno,CG14252,Hr96,ems,opa,mid,CrebB,Antp,Awh,EcR,CycT,mnb,kay,scro,polybromo,barc,Su(H),Tor,B-H2,TfIIFbeta,Iswi,Eip74EF,CG43689,bi,hth,rictor,CG42830,gce,esg,CG1894,retn | | 7.867e-05 | -9.45 | regulatory region nucleic acid binding | molecular function | GO:0001067 | 369 | 35 | 12176 | 580 | tin,CG12071,E5,CG43689,EcR,CG15073,ems,Su(H),Ubx,B-H2,bi,CG31224,grn,retn,kay,TfAP-2,cic,Atf6,CG12605,CG7368,GATAe,opa,foxo,hth,esg,Awh,Antp,Maf1,scro,jigr1,mid,CG10462,Hr96,htk,CG4374 | | 7.867e-05 | -9.45 | transcription regulatory region DNA binding | molecular function | GO:0044212 | 369 | 35 | 12176 | 580 | cic,kay,TfAP-2,retn,CG31224,grn,bi,B-H2,Ubx,Su(H),ems,CG15073,EcR,CG12071,E5,CG43689,tin,htk,CG4374,CG10462,Hr96,jigr1,mid,Antp,Maf1,scro,Awh,esg,hth,foxo,GATAe,opa,CG7368,CG12605,Atf6 | | 8.011e-05 | -9.43 | calcium ion binding | molecular function | GO:0005509 | 200 | 23 | 12176 | 580 | Frq1,TpnC4,Cadps,LpR2,TpnC25D,Ser,unc-13,CG17698,CG13526,Frq2,CG14362,TpnC41C,Cam,Mctp,CadN2,Syt12,CG10126,LRP1,tld,shg,Actn,Scp1,CG13898 | | 8.352e-05 | -9.39 | GO\_SINGLE\_ORGANISM\_BEHAVIOR | MSigDB lists | GO\_SINGLE\_ORGANISM\_BEHAVIOR | 106 | 14 | 4161 | 170 | Pkc98E,AstA-R1,Raf,Nmdar2,LpR2,Octbeta2R,Cln3,Ca-beta,Awh,Tor,KaiR1D,eIF4E7,CG7483,Nlg2 | | 8.690e-05 | -9.35 | GO\_CELL\_DEVELOPMENT | MSigDB lists | GO\_CELL\_DEVELOPMENT | 289 | 26 | 4161 | 170 | Ank,csw,CG42672,TfAP-2,Tor,Ca-beta,Awh,AstA-R1,Dys,fz2,CG33090,dnc,Rbp6,sky,Cdep,Su(H),Sod1,CG17883,sdt,LpR2,Khc,Pak,eIF2Bgamma,AGO3,Ser,dpp | | 8.700e-05 | -9.35 | GO\_POSITIVE\_REGULATION\_OF\_TRANSPORT | MSigDB lists | GO\_POSITIVE\_REGULATION\_OF\_TRANSPORT | 208 | 21 | 4161 | 170 | Frq2,csw,Ank,CG41099,fz2,Dys,Octbeta2R,chb,AstA-R1,Syx1A,CCAP-R,CG17883,RhoGAP1A,Frq1,LRP1,CG8184,EcR,Pkc98E,Khc,MCU,unc-13 | | 8.873e-05 | -9.33 | GO\_REGULATION\_OF\_MAP\_KINASE\_ACTIVITY | MSigDB lists | GO\_REGULATION\_OF\_MAP\_KINASE\_ACTIVITY | 69 | 11 | 4161 | 170 | fz2,dpp,Raf,Tsp96F,Pak,dsh,CG3376,csw,Sod1,wbl,CG42672 | | 9.153e-05 | -9.30 | GO\_EAR\_DEVELOPMENT | MSigDB lists | GO\_EAR\_DEVELOPMENT | 47 | 9 | 4161 | 170 | Ser,dpp,dsh,Su(H),RhoGAP1A,Sod1,csw,RpL38,TfAP-2 | | 9.606e-05 | -9.25 | GO\_REGULATION\_OF\_LEUKOCYTE\_DIFFERENTIATION | MSigDB lists | GO\_REGULATION\_OF\_LEUKOCYTE\_DIFFERENTIATION | 28 | 7 | 4161 | 170 | Nha1,Ptp61F,Sod1,sxc,Pkc53E,Tor,IP3K2 | | 9.799e-05 | -9.23 | GO\_REGULATION\_OF\_SECRETION | MSigDB lists | GO\_REGULATION\_OF\_SECRETION | 135 | 16 | 4161 | 170 | Pde8,Frq1,Pkc53E,RhoGAP1A,wbl,unc-13,Raf,MCU,Pkc98E,EcR,ZnT63C,csw,Frq2,Syx1A,chb,mino | | 9.909e-05 | -9.22 | transcription regulatory region sequence-specific DNA binding | molecular function | GO:0000976 | 329 | 32 | 12176 | 580 | Ubx,B-H2,bi,CG31224,grn,retn,kay,TfAP-2,cic,tin,CG12071,E5,CG43689,EcR,Su(H),CG15073,ems,esg,Awh,scro,Antp,mid,CG10462,Hr96,CG4374,Atf6,CG12605,CG7368,GATAe,opa,foxo,hth | | 1.009e-04 | -9.20 | positive regulation of transcription by RNA polymerase II | biological process | GO:0045944 | 368 | 35 | 12345 | 597 | Lpin,CG4374,foxo,CG12071,CG10462,TfAP-2,Mrtf,Sox102F,GATAe,tin,EcR,Awh,CrebB,Antp,opa,mid,Atf6,sno,Hr96,ems,CG43689,bi,Iswi,Eip74EF,scro,Su(H),B-H2,TfIIFbeta,kay,esg,CG1894,retn,CG42830,gce,hth | | 1.051e-04 | -9.16 | cell junction | cellular component | GO:0030054 | 160 | 19 | 11940 | 538 | p130CAS,sdt,rols,bbg,rst,shakB,step,scrib,Pak,CadN2,nrv2,fw,Mbs,Src42A,by,ex,shg,Actn,Inx2 | | 1.074e-04 | -9.14 | RNA polymerase II proximal promoter sequence-specific DNA binding | molecular function | GO:0000978 | 92 | 14 | 12176 | 580 | GATAe,CG10462,kay,ems,Su(H),CG4374,Atf6,E5,grn,scro,CG12071,CG31224,tin,bi | | 1.126e-04 | -9.09 | regulation of Malpighian tubule diameter | biological process | GO:0035297 | 3 | 3 | 12345 | 597 | raw,rib,EcR | | 1.130e-04 | -9.09 | regulation of signal transduction | biological process | GO:0009966 | 745 | 59 | 12345 | 597 | tld,kek1,Src42A,et,raw,CG14182,mnb,crol,numb,ex,sgg,bnl,Tor,Snoo,Su(H),pirk,Cdep,CG10188,Mos,Exn,kek5,flw,spri,rictor,sxc,Pak,RhoGEF3,hth,Cln3,Flo2,CG18659,CG5522,Doa,GluRIIC,trol,foxo,olf186-F,Pde8,chb,Lpin,Gprk1,qvr,rdgA,lncRNA:Hsromega,Cam,CG34408,MESK2,Raf,RhoGAP1A,dnc,scrib,Ser,Ptp61F,HDAC4,melt,step,sno,siz,dpp | | 1.151e-04 | -9.07 | imaginal disc-derived wing morphogenesis | biological process | GO:0007476 | 298 | 30 | 12345 | 597 | Dys,scrib,fw,Ser,flw,EcR,step,Mbs,by,sno,hth,shg,dpp,fru,tld,upd3,Src42A,bi,Lpin,tnc,dsh,Fhos,path,Mrtf,crol,sgg,cic,Raf,Su(H),app | | 1.262e-04 | -8.98 | regulation of macromolecule biosynthetic process | biological process | GO:0010556 | 1326 | 93 | 12345 | 597 | sno,ems,CG32006,mid,Maf1,CG7368,grn,htk,scrib,CrebB,Awh,melt,EcR,rib,Sox102F,GATAe,lncRNA:bxd,ham,CG4374,GluRIIC,Lpin,bun,CG12605,rictor,woc,fru,gce,Fer2,bol,E5,Octbeta2R,bru1,esg,Ubx,retn,Elp3,crol,mnb,scro,Poxm,cic,B-H2,TfIIFbeta,Iswi,mamo,Eip74EF,CG43689,bi,msi,CG31224,Atf6,CG14252,Hr96,opa,dpp,dany,CG15073,Antp,HDAC4,AGO2,CycT,Mrtf,luna,tin,CG10462,CG12071,TfAP-2,l(3)neo38,HPS4,foxo,chinmo,toe,hth,CG42830,FoxL1,tyf,CG9650,Art2,Khc,kek5,CG1894,jim,Set2,Mondo,kay,polybromo,Su(H),Tor,CG32971,Snoo,ex,Spt20,tld | | 1.330e-04 | -8.92 | IG | smart domains | SM00409 | 111 | 18 | 5086 | 315 | dpr5,kek5,boi,kek1,Strn-Mlck,dpr15,dpr18,dpr12,CG17716,klg,nrm,rst,side-VIII,zormin,trol,dpr21,DIP-delta,DIP-lambda | | 1.368e-04 | -8.90 | GO\_POSITIVE\_REGULATION\_OF\_MAP\_KINASE\_ACTIVITY | MSigDB lists | GO\_POSITIVE\_REGULATION\_OF\_MAP\_KINASE\_ACTIVITY | 39 | 8 | 4161 | 170 | dsh,Pak,Tsp96F,dpp,fz2,wbl,csw,Sod1 | | 1.376e-04 | -8.89 | Ig-like\_dom | interpro domains | IPR007110 | 132 | 18 | 11383 | 585 | CG17716,rst,dpr18,kek5,klg,nrm,dpr5,DIP-lambda,dpr15,dpr21,dpr12,zormin,DIP-delta,kek1,boi,trol,side-VIII,Strn-Mlck | | 1.388e-04 | -8.88 | dendrite morphogenesis | biological process | GO:0048813 | 140 | 18 | 12345 | 597 | rictor,ems,Dlic,fru,E(Pc),Khc,EcR,jim,retn,kay,path,scro,Poxm,Snoo,Tor,ham,foxo,sqz | | 1.419e-04 | -8.86 | eIF5B (eukaryotic translation initiation factor 5B) | protein interactions | 44261 | 3 | 3 | 9204 | 481 | esn,rictor,CG3081 | | 1.419e-04 | -8.86 | CG1139 (uncharacterized protein) | protein interactions | 38264 | 3 | 3 | 9204 | 481 | Tor,path,foxo | | 1.419e-04 | -8.86 | CG6656 (uncharacterized protein) | protein interactions | 42576 | 3 | 3 | 9204 | 481 | Poxm,Cyp28d1,Cyp28d2 | | 1.434e-04 | -8.85 | sensory organ development | biological process | GO:0007423 | 496 | 43 | 12345 | 597 | Khc,fw,RhoGAP100F,CG13251,Doa,cato,hth,CadN2,Syn2,boi,Spf45,bi,msi,Src42A,bnl,Su(H),ex,B-H2,cic,sgg,numb,Mob2,kay,melt,Awh,Ser,scrib,csw,siz,klg,rst,dpp,CG5921,shg,toe,bun,foxo,cindr,Raf,nuf,chp,Cam,dsh | | 1.516e-04 | -8.79 | response to external stimulus | biological process | GO:0009605 | 987 | 73 | 12345 | 597 | kay,fz2,CG9380,sgg,bbg,msps,RhoL,pirk,Tor,bnl,Src42A,et,upd3,jus,RhoGEF3,shakB,Cyp6a13,Pak,tyf,Flo2,ninaD,Spn88Eb,Exn,RhoGAP100F,kcc,Khc,dsh,Root,Cam,GlyP,Raf,foxo,trol,chb,shep,shg,Hr96,dpp,LysP,HDAC4,AGO2,Pzl,haf,Tsf1,gukh,Pdfr,rictor,CadN2,pod1,CG8492,fru,retn,esg,GATAe,rdgA,Pld,beat-IIa,sick,Drsl2,sqz,Gprk1,Lpin,Sh,Ptp99A,egh,grn,klg,mid,E(Pc),PGRP-LA,Ptp61F,EcR,melt | | 1.519e-04 | -8.79 | GO\_REGULATION\_OF\_CELLULAR\_LOCALIZATION | MSigDB lists | GO\_REGULATION\_OF\_CELLULAR\_LOCALIZATION | 316 | 27 | 4161 | 170 | CG17883,RhoGAP1A,Pkc53E,wbl,sxc,Frq1,Pde8,CG8184,Pkc98E,EcR,Khc,MCU,Raf,unc-13,Tor,csw,Frq2,Ank,ZnT63C,dnc,mino,fz2,Dys,chb,dsh,Syx1A,CCAP-R | | 1.592e-04 | -8.75 | GO\_REGULATION\_OF\_MYELOID\_LEUKOCYTE\_DIFFERENTIATION | MSigDB lists | GO\_REGULATION\_OF\_MYELOID\_LEUKOCYTE\_DIFFERENTIATION | 14 | 5 | 4161 | 170 | Ptp61F,Nha1,Tor,Pkc53E,sxc | | 1.617e-04 | -8.73 | morphogenesis of an epithelium | biological process | GO:0002009 | 625 | 51 | 12345 | 597 | tin,dsh,Mrtf,rib,Inx2,TfAP-2,Raf,Lpin,step,shg,sno,dpp,egh,opa,csw,shd,scrib,Ser,EcR,kay,path,Fhos,hdc,crol,B-H2,cic,sgg,app,polybromo,Su(H),bnl,Tor,raw,Src42A,upd3,tld,bi,tnc,by,hth,CadN2,Mbs,Pak,sdt,fru,Dys,Exn,Ubx,fw,esg,flw | | 1.628e-04 | -8.72 | wing disc development | biological process | GO:0035220 | 407 | 37 | 12345 | 597 | crol,Set2,kay,path,Fhos,app,Su(H),cic,sgg,Src42A,tld,upd3,tnc,boi,bi,by,hth,sxc,Mbs,fru,Dys,flw,esg,fw,Mrtf,dsh,nuf,Raf,Lpin,sno,shg,step,dpp,scrib,Irk3,EcR,Ser | | 1.656e-04 | -8.71 | GO\_TRANSMISSION\_OF\_NERVE\_IMPULSE | MSigDB lists | GO\_TRANSMISSION\_OF\_NERVE\_IMPULSE | 8 | 4 | 4161 | 170 | Ank,KaiR1D,Ca-beta,Sod1 | | 1.656e-04 | -8.71 | GO\_SYNAPTIC\_TRANSMISSION\_GLUTAMATERGIC | MSigDB lists | GO\_SYNAPTIC\_TRANSMISSION\_GLUTAMATERGIC | 8 | 4 | 4161 | 170 | Ca-beta,unc-13,KaiR1D,Octbeta2R | | 1.669e-04 | -8.70 | Ig-like\_fold | interpro domains | IPR013783 | 195 | 23 | 11383 | 585 | rst,dpr18,CG17716,kek5,CG5445,Ptp99A,klg,nrm,dpr15,dpr5,DIP-lambda,dpr21,wake,DIP-delta,zormin,dpr12,trol,kek1,Su(H),boi,side-VIII,Strn-Mlck,CG18135 | | 1.692e-04 | -8.68 | GO\_ACTIVATION\_OF\_PROTEIN\_KINASE\_ACTIVITY | MSigDB lists | GO\_ACTIVATION\_OF\_PROTEIN\_KINASE\_ACTIVITY | 62 | 10 | 4161 | 170 | Octbeta2R,Tsp96F,Raf,dpp,Stlk,CG42672,wbl,CG17698,Sod1,csw | | 1.712e-04 | -8.67 | proximal promoter sequence-specific DNA binding | molecular function | GO:0000987 | 96 | 14 | 12176 | 580 | Atf6,E5,grn,scro,CG31224,CG12071,tin,bi,GATAe,CG10462,kay,CG4374,Su(H),ems | | 1.731e-04 | -8.66 | epithelial cell migration | biological process | GO:0010631 | 155 | 19 | 12345 | 597 | ex,bbg,RhoL,Raf,cindr,bnl,kay,dsh,rib,Mrtf,bun,dpp,egh,csw,rols,shg,esg,EcR,shd | | 1.811e-04 | -8.62 | tissue morphogenesis | biological process | GO:0048729 | 644 | 52 | 12345 | 597 | rib,Mrtf,Inx2,tin,dsh,Raf,TfAP-2,Lpin,shg,sno,step,opa,csw,dpp,egh,shd,scrib,EcR,Ser,crol,kay,path,Fhos,hdc,app,polybromo,Su(H),Tor,ths,bnl,sgg,cic,B-H2,raw,Src42A,tld,upd3,tnc,bi,CadN2,by,hth,Mbs,Pak,fru,sdt,Dys,flw,esg,Ubx,Exn,fw | | 1.843e-04 | -8.60 | positive regulation of cellular metabolic process | biological process | GO:0031325 | 875 | 66 | 12345 | 597 | Mrtf,lncRNA:Hsromega,tin,CG12071,ps,Raf,CG10462,GlyP,TfAP-2,Pde8,foxo,CG41520,Atf6,CG14252,Hr96,step,opa,dpp,Antp,CycT,kay,polybromo,Tor,bnl,Su(H),sgg,Src42A,Stlk,hth,Pak,CG42830,tyf,CG1894,Mos,Sox102F,GATAe,AGO3,CG4374,Cpsf160,Lpin,CG12200,Marf1,sno,ems,mid,CrebB,EcR,Awh,REG,mnb,barc,scro,Pomp,B-H2,TfIIFbeta,Iswi,Eip74EF,CG43689,bi,rictor,Doa,gce,Fer2,bol,esg,retn | | 1.884e-04 | -8.58 | epithelium migration | biological process | GO:0090132 | 156 | 19 | 12345 | 597 | bun,kay,dsh,rib,Mrtf,ex,RhoL,bbg,Raf,bnl,cindr,shd,esg,EcR,shg,dpp,egh,csw,rols | | 1.923e-04 | -8.56 | GO\_POSITIVE\_REGULATION\_OF\_CELL\_CELL\_ADHESION | MSigDB lists | GO\_POSITIVE\_REGULATION\_OF\_CELL\_CELL\_ADHESION | 31 | 7 | 4161 | 170 | IP3K2,Tor,mino,csw,Ank,Pak,Flo2 | | 1.941e-04 | -8.55 | GO\_REGULATION\_OF\_METAL\_ION\_TRANSPORT | MSigDB lists | GO\_REGULATION\_OF\_METAL\_ION\_TRANSPORT | 63 | 10 | 4161 | 170 | ZnT63C,Ank,Ca-beta,Dys,Khc,Octbeta2R,CCAP-R,AstA-R1,dnc,Pkc98E | | 1.961e-04 | -8.54 | GO\_RESPONSE\_TO\_ENDOGENOUS\_STIMULUS | MSigDB lists | GO\_RESPONSE\_TO\_ENDOGENOUS\_STIMULUS | 356 | 29 | 4161 | 170 | csw,Tor,CG4849,CG42672,Hr96,Uros1,mino,Ugt301D1,CG5493,dnc,TfIIFbeta,Ptp61F,CG3376,Octbeta2R,Cat,sxc,Sod1,fus,CG7483,Pkc98E,eIF2Bgamma,EcR,Ser,dpp,LpR2,Tsp96F,Raf,GlyP,Pak | | 1.980e-04 | -8.53 | positive regulation of nitrogen compound metabolic process | biological process | GO:0051173 | 827 | 63 | 12345 | 597 | REG,CycT,Awh,EcR,Antp,CrebB,dpp,opa,mid,step,ems,sno,Atf6,Marf1,CG14252,Hr96,Lpin,CG12200,CG41520,foxo,CG4374,Pde8,AGO3,TfAP-2,ps,CG10462,Raf,CG12071,tin,Sox102F,Mrtf,CG42797,lncRNA:Hsromega,GATAe,retn,Mos,esg,CG1894,bol,tyf,gce,CG42830,rictor,hth,Pak,CG43689,Stlk,bi,Src42A,Iswi,Eip74EF,B-H2,sgg,TfIIFbeta,polybromo,scro,barc,Pomp,bnl,Tor,Su(H),kay,mnb | | 2.050e-04 | -8.49 | positive regulation of transcription, DNA-templated | biological process | GO:0045893 | 473 | 41 | 12345 | 597 | ems,sno,Atf6,Hr96,CG14252,opa,mid,Antp,CrebB,CycT,EcR,Awh,tin,Sox102F,Mrtf,GATAe,TfAP-2,CG12071,CG10462,foxo,CG4374,Lpin,rictor,hth,gce,CG42830,retn,esg,CG1894,kay,mnb,B-H2,TfIIFbeta,scro,polybromo,Su(H),Tor,Iswi,Eip74EF,CG43689,bi | | 2.066e-04 | -8.48 | wing disc morphogenesis | biological process | GO:0007472 | 308 | 30 | 12345 | 597 | Lpin,tnc,bi,Src42A,upd3,tld,app,Raf,Su(H),sgg,cic,crol,Mrtf,Fhos,path,dsh,flw,EcR,fw,Ser,scrib,Dys,fru,dpp,hth,sno,by,shg,Mbs,step | | 2.106e-04 | -8.47 | memory | biological process | GO:0007613 | 120 | 16 | 12345 | 597 | mGluR,Ank2,Mob2,Su(H),Pkc98E,scrib,CrebB,dnc,CG5541,nemy,HDAC4,EcR,klg,Gld2,MCU,Nmdar2 | | 2.145e-04 | -8.45 | GO\_REGULATION\_OF\_TRANSPORT | MSigDB lists | GO\_REGULATION\_OF\_TRANSPORT | 412 | 32 | 4161 | 170 | Frq2,csw,Ca-beta,CG41099,ZnT63C,fz2,dnc,Syx1A,AstA-R1,Octbeta2R,Dys,chb,CG17883,LRP1,CG8184,Raf,unc-13,Khc,MCU,Tor,Ank,mino,Sh,CCAP-R,wbl,Pkc53E,RhoGAP1A,sxc,Frq1,Pde8,Pkc98E,EcR | | 2.158e-04 | -8.44 | regulation of cell differentiation | biological process | GO:0045595 | 428 | 38 | 12345 | 597 | PGRP-LA,Ser,EcR,Antp,dpp,csw,mid,shg,bun,Raf,ham,dsh,CG42797,Ubx,esg,Khc,bru1,Dlic,rictor,spri,hth,Pak,Mbs,Src42A,CG9380,sgg,numb,ex,barc,Tor,Su(H),Snoo,kay,Syx1A,Fhos,hdc,Elp3,mnb | | 2.172e-04 | -8.43 | dendrite development | biological process | GO:0016358 | 145 | 18 | 12345 | 597 | sqz,foxo,Poxm,ham,Tor,Snoo,scro,path,kay,jim,retn,Khc,EcR,E(Pc),fru,Dlic,ems,rictor | | 2.203e-04 | -8.42 | N (Notch) | protein interactions | 31293 | 301 | 31 | 9204 | 481 | upd3,Su(H),jigr1,htk,chinmo,bun,bi,sgg,klg,toe,shg,Ser,msi,numb,rictor,Ubx,Antp,sno,CG41099,fru,TfAP-2,Pkc98E,esg,mir-278,RpL38,flw,Pak,dsh,ham,dpp,Ank2 | | 2.226e-04 | -8.41 | jus (julius seizure) | protein interactions | 43453 | 12 | 5 | 9204 | 481 | esg,kcc,Sh,shg,shakB | | 2.256e-04 | -8.40 | sequence-specific double-stranded DNA binding | molecular function | GO:1990837 | 359 | 33 | 12176 | 580 | opa,GATAe,CG7368,hth,foxo,Atf6,CG12605,CG10462,Hr96,mid,CG4374,esg,scro,Maf1,Antp,Awh,EcR,Su(H),CG15073,ems,CG12071,E5,CG43689,tin,kay,TfAP-2,retn,cic,B-H2,Ubx,CG31224,grn,bi | | 2.272e-04 | -8.39 | eag (ether a go-go) | protein interactions | 32428 | 7 | 4 | 9204 | 481 | Cam,Khc,qvr,Sh | | 2.309e-04 | -8.37 | GO\_REGULATION\_OF\_CARBOHYDRATE\_BIOSYNTHETIC\_PROCESS | MSigDB lists | GO\_REGULATION\_OF\_CARBOHYDRATE\_BIOSYNTHETIC\_PROCESS | 15 | 5 | 4161 | 170 | Doa,Ptp61F,Tor,sxc,flw | | 2.309e-04 | -8.37 | GO\_REGULATION\_OF\_ACTIN\_FILAMENT\_BUNDLE\_ASSEMBLY | MSigDB lists | GO\_REGULATION\_OF\_ACTIN\_FILAMENT\_BUNDLE\_ASSEMBLY | 15 | 5 | 4161 | 170 | Tor,Fhos,Raf,Pak,chb | | 2.378e-04 | -8.34 | GSE28726\_NAIVE\_CD4\_TCELL\_VS\_NAIVE\_NKTCELL\_DN | MSigDB lists | GSE28726\_NAIVE\_CD4\_TCELL\_VS\_NAIVE\_NKTCELL\_DN | 32 | 7 | 4161 | 170 | Flo2,Glut1,Octbeta2R,step,Argl,Cadps,PhKgamma | | 2.420e-04 | -8.33 | positive regulation of insulin receptor signaling pathway | biological process | GO:0046628 | 13 | 5 | 12345 | 597 | rictor,step,Lpin,melt,Tor | | 2.420e-04 | -8.33 | positive regulation of cellular response to insulin stimulus | biological process | GO:1900078 | 13 | 5 | 12345 | 597 | step,rictor,Tor,Lpin,melt | | 2.436e-04 | -8.32 | compound eye development | biological process | GO:0048749 | 370 | 34 | 12345 | 597 | RhoGAP100F,fw,CadN2,hth,Doa,Src42A,Syn2,boi,Spf45,bi,msi,Mob2,kay,bnl,ex,B-H2,scrib,Awh,melt,Ser,shg,csw,dpp,rst,siz,klg,foxo,bun,toe,Cam,chp,dsh,Raf,cindr | | 2.460e-04 | -8.31 | epithelium development | biological process | GO:0060429 | 1020 | 74 | 12345 | 597 | Mbs,sxc,Pak,hth,Spn88Eb,Dys,Khc,Idgf6,Exn,Set2,Fhos,kay,Tor,bnl,Su(H),bbg,RhoL,app,polybromo,sgg,ex,upd3,tld,raw,Src42A,tnc,boi,shg,step,opa,dpp,shd,Irk3,Ser,Mrtf,tin,dsh,Raf,nuf,CG41099,TfAP-2,chinmo,by,CadN2,fru,sdt,lncRNA:flam,esg,flw,fw,Ubx,crol,hdc,path,Syx1A,numb,cic,B-H2,bi,sno,mid,csw,rols,egh,scrib,Awh,EcR,GATAe,Inx2,rib,fs(1)N,cindr,Lpin,bun | | 2.478e-04 | -8.30 | GO\_REGULATION\_OF\_INTRACELLULAR\_SIGNAL\_TRANSDUCTION | MSigDB lists | GO\_REGULATION\_OF\_INTRACELLULAR\_SIGNAL\_TRANSDUCTION | 343 | 28 | 4161 | 170 | CG3376,Cat,Octbeta2R,Dys,dsh,Ptp61F,dnc,Tif-IA,fz2,CG34408,CG42672,IP3K2,Tor,csw,step,Pak,Tsp96F,Raf,dpp,CG5522,Pkc98E,Cdep,KaiR1D,wbl,Pkc53E,RhoGAP1A,Sod1,sxc | | 2.601e-04 | -8.25 | GO\_REGULATION\_OF\_SENSORY\_PERCEPTION | MSigDB lists | GO\_REGULATION\_OF\_SENSORY\_PERCEPTION | 4 | 3 | 4161 | 170 | Tor,Nmdar2,Octbeta2R | | 2.601e-04 | -8.25 | GO\_CYCLIC\_NUCLEOTIDE\_CATABOLIC\_PROCESS | MSigDB lists | GO\_CYCLIC\_NUCLEOTIDE\_CATABOLIC\_PROCESS | 4 | 3 | 4161 | 170 | Pde8,dnc,Pde6 | | 2.624e-04 | -8.25 | GO\_POSITIVE\_REGULATION\_OF\_IMMUNE\_SYSTEM\_PROCESS | MSigDB lists | GO\_POSITIVE\_REGULATION\_OF\_IMMUNE\_SYSTEM\_PROCESS | 177 | 18 | 4161 | 170 | Ca-beta,Tor,IP3K2,csw,REG,dnc,fz2,mino,scramb1,Nha1,sxc,Pkc53E,Flo2,Ser,Act87E,Pkc98E,Pak,Tsp96F | | 2.633e-04 | -8.24 | - | gene3d domains | 2.30.29.30 | 139 | 20 | 4929 | 302 | Cadps,ex,CG5522,CG10188,numb,CG3530,cert,by,siz,CG5004,Myo81F,CG42673,Exn,Gprk1,Cdep,melt,CG9132,RhoGEF3,step,Pld | | 2.816e-04 | -8.18 | GO\_POSITIVE\_REGULATION\_OF\_DEVELOPMENTAL\_PROCESS | MSigDB lists | GO\_POSITIVE\_REGULATION\_OF\_DEVELOPMENTAL\_PROCESS | 178 | 18 | 4161 | 170 | dpp,Ser,unc-13,LpR2,Raf,sxc,Pkc53E,Su(H),Tif-IA,mino,chb,Dys,Octbeta2R,Nha1,Tor,TfAP-2,IP3K2,CG42672 | | 2.847e-04 | -8.16 | GO\_BEHAVIOR | MSigDB lists | GO\_BEHAVIOR | 133 | 15 | 4161 | 170 | Octbeta2R,LpR2,Nmdar2,AstA-R1,Raf,Pkc98E,CG7483,Nlg2,eIF4E7,KaiR1D,Tor,Awh,Ca-beta,Sod1,Cln3 | | 2.886e-04 | -8.15 | MODULE\_208 | MSigDB lists | MODULE\_208 | 9 | 4 | 4161 | 170 | Meltrin,Sh,scramb1,MCU | | 2.965e-04 | -8.12 | cell motility | biological process | GO:0048870 | 344 | 32 | 12345 | 597 | Gas8,Pak,esg,ex,numb,ths,p130CAS,bnl,bbg,RhoL,Fhos,kay,fz2,Src42A,egh,dpp,opa,rols,csw,shg,EcR,shd,Raf,Msp300,cindr,tin,dsh,rib,Mrtf,chb,bun,CG17387 | | 2.987e-04 | -8.12 | locomotory behavior | biological process | GO:0007626 | 188 | 21 | 12345 | 597 | TfAP-2,sgg,Elp3,kay,Nlg2,Pdfr,shep,Tk,foxo,tyf,Sh,sxc,shakB,nemy,EcR,Khc,CrebB,dnc,MsR2,Fer2,ITP | | 3.098e-04 | -8.08 | movement of cell or subcellular component | biological process | GO:0006928 | 724 | 56 | 12345 | 597 | CadN2,pod1,Pak,nompB,Gas8,Dlic,fru,esg,Khc,retn,RhoGAP100F,fz2,kay,Fhos,RhoL,msps,bbg,ths,bnl,p130CAS,ex,haf,numb,Src42A,gukh,shg,rols,csw,opa,mid,dpp,grn,Ptp99A,egh,klg,g,shd,E(Pc),EcR,Ptp61F,Mrtf,Cam,rib,dsh,kl-5,tin,beat-IIa,Raf,Msp300,IFT46,cindr,CG17387,trol,chb,sqz,bun | | 3.102e-04 | -8.08 | GO\_REGULATION\_OF\_GLUCOSE\_METABOLIC\_PROCESS | MSigDB lists | GO\_REGULATION\_OF\_GLUCOSE\_METABOLIC\_PROCESS | 24 | 6 | 4161 | 170 | sxc,flw,Tor,PhKgamma,Doa,Ptp61F | | 3.187e-04 | -8.05 | Post NMDA receptor activation events | REACTOME pathways | R-DME-438064 | 20 | 6 | 3923 | 197 | CrebB,Cam,Raf,CG17698,Actn,Nmdar2 | | 3.228e-04 | -8.04 | negative regulation of cellular process | biological process | GO:0048523 | 1291 | 89 | 12345 | 597 | CG12605,Acp62F,Doa,gce,sws,bru1,bol,E5,Fer2,esg,flw,retn,Ubx,mnb,crol,hdc,Ank2,cic,numb,B-H2,Iswi,CG14182,msi,bi,CG43689,Maf1,mid,csw,CG7368,htk,scrib,EcR,REG,Ptp61F,PGRP-LA,lncRNA:bxd,Sox102F,ham,Cep97,AGO3,Gprk1,Lpin,Nlg2,bun,Spn88Eb,CG9650,PhKgamma,Pkc98E,CG1894,kek5,Khc,Mos,Mob2,cpx,pirk,Tor,Su(H),Snoo,polybromo,CG9380,ex,sgg,kek1,Src42A,raw,et,sky,CG31224,dpp,dnc,g,Antp,AGO2,HDAC4,Ser,Cam,qvr,tin,CG12071,Raf,TfAP-2,Pde8,HPS4,CG41520,trol,foxo,chinmo,toe,chb | | 3.238e-04 | -8.04 | GO\_REGULATION\_OF\_KINASE\_ACTIVITY | MSigDB lists | GO\_REGULATION\_OF\_KINASE\_ACTIVITY | 180 | 18 | 4161 | 170 | csw,CG17698,Tor,CG42672,Stlk,fz2,dsh,Ptp61F,CG3376,Octbeta2R,Sod1,wbl,Elp3,dpp,LpR2,Tsp96F,Raf,Pak | | 3.272e-04 | -8.03 | presynaptic membrane | cellular component | GO:0042734 | 22 | 6 | 11940 | 538 | gukh,unc-13,Ank2,Exn,Fife,Syx1A | | 3.303e-04 | -8.02 | IGc2 | smart domains | SM00408 | 109 | 17 | 5086 | 315 | dpr15,dpr12,CG17716,klg,rst,nrm,side-VIII,zormin,trol,dpr21,DIP-delta,DIP-lambda,dpr5,kek5,boi,kek1,Strn-Mlck | | 3.310e-04 | -8.01 | GO\_REGULATION\_OF\_CARBOHYDRATE\_METABOLIC\_PROCESS | MSigDB lists | GO\_REGULATION\_OF\_CARBOHYDRATE\_METABOLIC\_PROCESS | 44 | 8 | 4161 | 170 | Ptp61F,Pkc98E,flw,Ugt301D1,Doa,sxc,PhKgamma,Tor | | 3.329e-04 | -8.01 | GO\_REGULATION\_OF\_CELL\_DIFFERENTIATION | MSigDB lists | GO\_REGULATION\_OF\_CELL\_DIFFERENTIATION | 279 | 24 | 4161 | 170 | Tif-IA,Dys,chb,Ptp61F,Nha1,CG42672,IP3K2,Tor,TfAP-2,Frq2,LRP1,dpp,Ser,EcR,Raf,unc-13,LpR2,Pkc53E,Su(H),Sod1,sxc,Frq1,Flo2,eIF4E7 | | 3.333e-04 | -8.01 | regulation of phosphorus metabolic process | biological process | GO:0051174 | 302 | 29 | 12345 | 597 | flw,CycT,Mos,Ptp61F,dpp,sws,Cln3,Pak,rictor,step,chinmo,Stlk,Ste:CG33247,Pde8,CG41520,CG17124,et,raw,Src42A,kek1,CG34172,Tor,Raf,bnl,SteXh:CG42398,sgg,lncRNA:Hsromega,mnb,kay | | 3.333e-04 | -8.01 | regulation of phosphate metabolic process | biological process | GO:0019220 | 302 | 29 | 12345 | 597 | flw,Mos,CycT,Ptp61F,Pak,Cln3,step,rictor,sws,dpp,Pde8,Ste:CG33247,kek1,CG34172,raw,et,Src42A,CG41520,CG17124,chinmo,Stlk,mnb,lncRNA:Hsromega,kay,bnl,Tor,Raf,SteXh:CG42398,sgg | | 3.533e-04 | -7.95 | B52 (B52) | protein interactions | 41670 | 35 | 8 | 9204 | 481 | bru1,hth,sgg,Mur2B,CG17716,Mondo,EcR,CG34417 | | 3.533e-04 | -7.95 | kst (karst) | protein interactions | 38418 | 35 | 8 | 9204 | 481 | sgg,foxo,bbg,flw,rst,Pak,ex,rictor | | 3.533e-04 | -7.95 | l(2)gl (lethal (2) giant larvae) | protein interactions | 33156 | 35 | 8 | 9204 | 481 | scrib,nuf,dpp,dsh,sdt,numb,Src42A,Pak | | 3.549e-04 | -7.94 | ameboidal-type cell migration | biological process | GO:0001667 | 177 | 20 | 12345 | 597 | rols,csw,egh,dpp,shg,esg,EcR,shd,Raf,ths,cindr,bnl,RhoL,bbg,ex,rib,Mrtf,dsh,kay,bun | | 3.603e-04 | -7.93 | cognition | biological process | GO:0050890 | 164 | 19 | 12345 | 597 | Sh,klg,Gld2,MCU,Nmdar2,Pkc98E,scrib,dnc,CrebB,CG5541,EcR,nemy,HDAC4,Ank2,mGluR,Mob2,mnb,sgg,Su(H) | | 3.603e-04 | -7.93 | learning or memory | biological process | GO:0007611 | 164 | 19 | 12345 | 597 | klg,MCU,Gld2,Nmdar2,Sh,CG5541,HDAC4,nemy,EcR,Pkc98E,CrebB,dnc,scrib,sgg,Su(H),mGluR,Ank2,mnb,Mob2 | | 3.603e-04 | -7.93 | tissue migration | biological process | GO:0090130 | 164 | 19 | 12345 | 597 | bun,kay,dsh,Mrtf,rib,ex,bbg,RhoL,bnl,Raf,cindr,shd,EcR,esg,shg,dpp,egh,csw,rols | | 3.648e-04 | -7.92 | GO\_HOMEOSTATIC\_PROCESS | MSigDB lists | GO\_HOMEOSTATIC\_PROCESS | 351 | 28 | 4161 | 170 | IP3K2,Ca-beta,CG4341,Der-1,csw,Cln3,ZnT63C,Ank,Tif-IA,Root,mino,CG40178,Ptp61F,Octbeta2R,Dys,AstA-R1,CG17883,Qsox2,sxc,Sod1,Act88F,kcc,KaiR1D,Act87E,Pkc98E,EcR,MCU,GlyP | | 3.670e-04 | -7.91 | hemocyte differentiation | biological process | GO:0042386 | 29 | 7 | 12345 | 597 | dpp,Ser,RhoL,Su(H),Raf,Tor,ths | | 3.700e-04 | -7.90 | GO\_DIVALENT\_INORGANIC\_CATION\_HOMEOSTASIS | MSigDB lists | GO\_DIVALENT\_INORGANIC\_CATION\_HOMEOSTASIS | 68 | 10 | 4161 | 170 | Cln3,CG4341,Ca-beta,KaiR1D,Ank,ZnT63C,Pkc98E,AstA-R1,GlyP,MCU | | 3.707e-04 | -7.90 | GO\_POSITIVE\_REGULATION\_OF\_PROTEIN\_SERINE\_THREONINE\_KINASE\_ACTIVITY | MSigDB lists | GO\_POSITIVE\_REGULATION\_OF\_PROTEIN\_SERINE\_THREONINE\_KINASE\_ACTIVITY | 56 | 9 | 4161 | 170 | wbl,csw,Sod1,Stlk,dpp,fz2,Pak,dsh,Tsp96F | | 3.795e-04 | -7.88 | neuron part | cellular component | GO:0097458 | 540 | 42 | 11940 | 538 | CG5921,kcc,Gld2,nrv2,Frq2,gukh,Ank2,Pdfr,pod1,EcR,klg,cpx,RhoGAP100F,Ptp99A,dpr15,mnb,unc-13,Gr39a,Or85b,sgg,sky,dpr18,Fife,Syx1A,AGO2,dpr5,Exn,Cadps,g,KaiR1D,path,hdc,Gr22c,Ser,Or65c,dpr12,scrib,CG13794,bol,Or42b,Khc,chb | | 3.944e-04 | -7.84 | anchoring junction | cellular component | GO:0070161 | 97 | 13 | 11940 | 538 | by,step,shg,Actn,ex,rst,scrib,Pak,CadN2,p130CAS,sdt,rols,Src42A | | 3.944e-04 | -7.84 | adherens junction | cellular component | GO:0005912 | 97 | 13 | 11940 | 538 | rst,ex,shg,Actn,step,scrib,by,p130CAS,Src42A,sdt,rols,CadN2,Pak | | 3.944e-04 | -7.84 | REACTOME\_G\_ALPHA\_S\_SIGNALLING\_EVENTS | MSigDB lists | REACTOME\_G\_ALPHA\_S\_SIGNALLING\_EVENTS | 25 | 6 | 4161 | 170 | dnc,Pde6,Octbeta2R,Pde8,CCAP-R,Pde1c | | 3.990e-04 | -7.83 | GO\_REGULATION\_OF\_PHOSPHORUS\_METABOLIC\_PROCESS | MSigDB lists | GO\_REGULATION\_OF\_PHOSPHORUS\_METABOLIC\_PROCESS | 335 | 27 | 4161 | 170 | fz2,dnc,AstA-R1,mGluR,Ptp61F,dsh,Dys,CG3376,Octbeta2R,csw,CG17698,IP3K2,Tor,CG42672,Stlk,Pkc98E,dpp,LpR2,Tsp96F,Raf,Pak,sxc,Sod1,wbl,Pkc53E,Elp3,KaiR1D | | 4.001e-04 | -7.82 | embryo development | biological process | GO:0009790 | 568 | 46 | 12345 | 597 | Src42A,raw,tld,upd3,tnc,fz2,kay,sgg,cic,numb,RhoL,msps,ths,CG7483,PhKgamma,Ubx,retn,Khc,hth,Pak,Mbs,sdt,Doa,wbl,chb,fs(1)N,dsh,tin,rib,luna,Inx2,Pld,CG41099,Raf,Antp,scrib,shd,Ptp61F,EcR,AGO2,step,ems,shg,dpp,csw,mid | | 4.058e-04 | -7.81 | regulation of cellular biosynthetic process | biological process | GO:0031326 | 1371 | 93 | 12345 | 597 | CG9650,Art2,tyf,FoxL1,CG42830,hth,jim,Khc,kek5,CG1894,ex,polybromo,Tor,CG32971,Su(H),Snoo,kay,Set2,Mondo,Spt20,tld,dpp,dany,opa,Atf6,CG31224,CG14252,Hr96,CycT,AGO2,HDAC4,CG15073,Antp,TfAP-2,CG12071,CG10462,tin,Mrtf,luna,toe,chinmo,foxo,l(3)neo38,HPS4,gce,woc,fru,rictor,CG12605,Ubx,retn,esg,Fer2,bol,bru1,Octbeta2R,E5,cic,TfIIFbeta,B-H2,scro,Poxm,Elp3,crol,mnb,CG43689,bi,msi,mamo,Iswi,Eip74EF,htk,CG7368,grn,CG32006,Maf1,mid,ems,sno,EcR,Awh,melt,scrib,CrebB,ham,rib,Sox102F,GATAe,lncRNA:bxd,bun,Lpin,GluRIIC,CG4374 | | 4.149e-04 | -7.79 | 3',5'-cyclic-AMP phosphodiesterase activity | molecular function | GO:0004115 | 4 | 3 | 12176 | 580 | Pde1c,Pde8,dnc | | 4.233e-04 | -7.77 | localization of cell | biological process | GO:0051674 | 351 | 32 | 12345 | 597 | dpp,egh,rols,opa,csw,shg,EcR,shd,Raf,Msp300,cindr,dsh,tin,rib,Mrtf,bun,chb,CG17387,Gas8,Pak,esg,numb,ex,bbg,RhoL,ths,p130CAS,bnl,fz2,kay,Fhos,Src42A | | 4.258e-04 | -7.76 | GO\_POSITIVE\_REGULATION\_OF\_PHOSPHORUS\_METABOLIC\_PROCESS | MSigDB lists | GO\_POSITIVE\_REGULATION\_OF\_PHOSPHORUS\_METABOLIC\_PROCESS | 200 | 19 | 4161 | 170 | LpR2,Tsp96F,Raf,Pak,Pkc98E,dpp,Sod1,sxc,wbl,Pkc53E,dsh,CG3376,Octbeta2R,fz2,Stlk,csw,CG17698,Tor,CG42672 | | 4.282e-04 | -7.76 | GO\_POSITIVE\_REGULATION\_OF\_CELL\_ACTIVATION | MSigDB lists | GO\_POSITIVE\_REGULATION\_OF\_CELL\_ACTIVATION | 35 | 7 | 4161 | 170 | csw,mino,IP3K2,Tor,Flo2,Tsp96F,Pak | | 4.296e-04 | -7.75 | chemotaxis | biological process | GO:0006935 | 292 | 28 | 12345 | 597 | E(Pc),Khc,retn,Ptp61F,RhoGAP100F,shg,CadN2,pod1,Pak,fru,mid,dpp,Ptp99A,klg,egh,grn,Src42A,trol,sqz,gukh,chb,Cam,fz2,dsh,beat-IIa,msps,bnl,haf | | 4.314e-04 | -7.75 | EF-hand\_7 | pfam domains | PF13499 | 54 | 10 | 10480 | 550 | Frq2,Frq1,TpnC41C,CG13526,CG10126,TpnC25D,CG13898,CG14362,TpnC4,Cam | | 4.340e-04 | -7.74 | regulation of Malpighian tubule size | biological process | GO:0035298 | 4 | 3 | 12345 | 597 | rib,raw,EcR | | 4.353e-04 | -7.74 | HDc | smart domains | SM00471 | 7 | 4 | 5086 | 315 | Pde9,Pde6,Pde8,Pde1c | | 4.369e-04 | -7.74 | regulation of biosynthetic process | biological process | GO:0009889 | 1374 | 93 | 12345 | 597 | CycT,HDAC4,AGO2,Antp,CG15073,dany,dpp,opa,Hr96,CG14252,Atf6,CG31224,toe,chinmo,foxo,HPS4,l(3)neo38,TfAP-2,CG10462,CG12071,tin,luna,Mrtf,jim,CG1894,kek5,Khc,CG9650,tyf,Art2,CG42830,FoxL1,hth,tld,Spt20,ex,Su(H),CG32971,Tor,Snoo,polybromo,kay,Mondo,Set2,EcR,Awh,melt,CrebB,scrib,CG7368,grn,htk,Maf1,mid,CG32006,ems,sno,bun,Lpin,GluRIIC,CG4374,ham,GATAe,lncRNA:bxd,Sox102F,rib,retn,Ubx,esg,bru1,Octbeta2R,bol,E5,Fer2,gce,fru,woc,rictor,CG12605,bi,msi,CG43689,Eip74EF,mamo,Iswi,TfIIFbeta,cic,B-H2,Poxm,scro,mnb,crol,Elp3 | | 4.409e-04 | -7.73 | molecular function regulator | molecular function | GO:0098772 | 563 | 45 | 12176 | 580 | CG34408,kek1,CG15369,RhoGAP1A,Mbs,CG3078,spri,Elp3,RhoGEF3,siz,Exn,Acp62F,CG41520,step,CycT,ths,Cdep,Gpb5,CG42390,Tk,Ste:CG33247,CG10188,sky,RhoGAP100F,RhoGAP18B,nrv2,rictor,bnl,Ser,upd3,CG5522,CG33310,Frq1,REG,Spn88Eb,dpp,SteXh:CG42398,ITP,CG18659,CG14298,Neb-cGP,mnb,RYa,CG17124,CG17883 | | 4.452e-04 | -7.72 | GO\_REGULATION\_OF\_MULTICELLULAR\_ORGANISM\_GROWTH | MSigDB lists | GO\_REGULATION\_OF\_MULTICELLULAR\_ORGANISM\_GROWTH | 17 | 5 | 4161 | 170 | dsd,Octbeta2R,Sod1,csw,mino | | 4.589e-04 | -7.69 | positive regulation of behavior | biological process | GO:0048520 | 30 | 7 | 12345 | 597 | wake,EcR,foxo,qvr,CrebB,mnb,Sh | | 4.704e-04 | -7.66 | GO\_REGULATION\_OF\_MEMBRANE\_POTENTIAL | MSigDB lists | GO\_REGULATION\_OF\_MEMBRANE\_POTENTIAL | 70 | 10 | 4161 | 170 | KaiR1D,Ank,CG7483,Cln3,Sod1,Ca-beta,Syx1A,Octbeta2R,Khc,Dys | | 4.752e-04 | -7.65 | GO\_POSITIVE\_REGULATION\_OF\_MOLECULAR\_FUNCTION | MSigDB lists | GO\_POSITIVE\_REGULATION\_OF\_MOLECULAR\_FUNCTION | 468 | 34 | 4161 | 170 | AstA-R1,Nmdar2,Octbeta2R,Cat,Dys,fz2,csw,CG42672,Raf,LpR2,eIF2Bgamma,Cdep,Sod1,CG17883,scramb1,dsh,REG,Ank,CG34408,Stlk,CG17698,step,Der-1,Tsp96F,Pak,EcR,Pkc98E,dpp,CG5522,sky,RhoGAP1A,wbl,Pkc53E,sxc | | 4.807e-04 | -7.64 | cGMP effects | REACTOME pathways | R-DME-418457 | 4 | 3 | 3923 | 197 | Pde1c,Pde9,Pde6 | | 4.913e-04 | -7.62 | embryonic morphogenesis | biological process | GO:0048598 | 251 | 25 | 12345 | 597 | tnc,tld,Src42A,raw,ths,Raf,CG41099,Pld,Inx2,rib,tin,dsh,kay,EcR,Ubx,Ptp61F,PhKgamma,scrib,shd,sdt,dpp,Pak,Mbs,shg,step | | 5.037e-04 | -7.59 | positive regulation of nucleic acid-templated transcription | biological process | GO:1903508 | 494 | 41 | 12345 | 597 | ems,CG14252,Hr96,sno,Atf6,mid,opa,Antp,CrebB,CycT,EcR,Awh,tin,GATAe,Mrtf,Sox102F,TfAP-2,CG10462,CG12071,foxo,CG4374,Lpin,rictor,hth,gce,CG42830,retn,CG1894,esg,kay,mnb,B-H2,TfIIFbeta,Tor,Su(H),scro,polybromo,Eip74EF,Iswi,bi,CG43689 | | 5.037e-04 | -7.59 | positive regulation of RNA biosynthetic process | biological process | GO:1902680 | 494 | 41 | 12345 | 597 | CG12071,CG10462,TfAP-2,GATAe,Sox102F,Mrtf,tin,Lpin,CG4374,foxo,mid,opa,Hr96,CG14252,Atf6,sno,ems,EcR,Awh,CycT,CrebB,Antp,Su(H),Tor,polybromo,scro,TfIIFbeta,B-H2,mnb,kay,bi,CG43689,Eip74EF,Iswi,CG42830,gce,hth,rictor,CG1894,esg,retn | | 5.116e-04 | -7.58 | GO\_POSITIVE\_REGULATION\_OF\_INTRACELLULAR\_SIGNAL\_TRANSDUCTION | MSigDB lists | GO\_POSITIVE\_REGULATION\_OF\_INTRACELLULAR\_SIGNAL\_TRANSDUCTION | 171 | 17 | 4161 | 170 | fz2,Cat,Octbeta2R,Ptp61F,dsh,CG42672,IP3K2,Tor,csw,dpp,Pkc98E,Pak,Raf,Tsp96F,Pkc53E,wbl,Sod1 | | 5.122e-04 | -7.58 | cellular component organization | biological process | GO:0016043 | 2830 | 171 | 12345 | 597 | CG33158,raw,Src42A,dpr21,Syn2,Nnf1b,fz2,kay,mGluR,Set2,Mob2,CG4301,Fife,sgg,msps,app,polybromo,bnl,jim,RhoGAP100F,spri,shakB,Mbs,cert,Art2,Dlic,CG17387,trol,HPS4,aust,Stacl,Cam,dpr18,Mrtf,lncRNA:Hsromega,luna,CG12782,DIP-delta,nuf,Yeti,RhoGAP1A,mino,g,AGO2,CG34417,shg,mamo,dpr15,Ank2,Syx1A,Strn-Mlck,path,crol,B-H2,scro,Pomp,ths,bru1,CG7483,CG6839,retn,Actn,dpr12,CG18109,CG32264,Prm,by,CG41562,sws,woc,fru,bun,sqz,Nlg2,fs(1)N,rib,CG30036,chp,Pld,Frq2,cindr,His1:CG33837,IFT46,E(Pc),ddbt,Ptp61F,EcR,Ank,ems,siz,klg,grn,egh,rols,csw,Spt20,Frq1,Stlk,sky,cpx,Tusp,spd-2,Tor,Snoo,Pkc98E,Khc,Rcd6,CG1894,RhoGEF3,Cln3,Pak,nompB,nrv2,foxo,CG41520,CG14105,chb,Act87E,chinmo,dsh,Root,Meltrin,CG40813,CG41099,Raf,Msp300,dnc,HDAC4,CG5921,dpp,CG5059,Iswi,gukh,can,CG9981,hdc,Elp3,Ca-beta,unc-13,DNaseII,cic,haf,Poxm,p130CAS,fw,flw,rictor,CadN2,pod1,sdt,Doa,Epg5,CG34172,CG12200,RhoGAP18B,CG42355,Act88F,Cep97,beat-IIa,sick,ham,CrebB,scrib,dpr5,eRF3,scramb1,Sh,Mlp84B,wake,Ptp99A,mid | | 5.154e-04 | -7.57 | photoreceptor cell development | biological process | GO:0042461 | 105 | 14 | 12345 | 597 | Doa,sdt,Raf,csw,CG5921,chp,sno,Cam,CadN2,Mob2,Syn2,EcR,RhoGAP100F,CG13251 | | 5.174e-04 | -7.57 | cell surface receptor signaling pathway | biological process | GO:0007166 | 340 | 31 | 12345 | 597 | dpp,RYa,Nmdar2,opa,csw,rictor,fz4,sno,Ser,flw,cic,ex,bnl,Su(H),Tor,ths,Raf,fz2,fs(1)N,mGluR,dsh,bun,CG11318,boi,Pdfr,Src42A,CG41520,et,foxo,upd3,GluRIIC | | 5.193e-04 | -7.56 | GPCR downstream signaling | REACTOME pathways | R-DME-388396 | 91 | 13 | 3923 | 197 | rdgA,Exn,CG30054,Octbeta2R,Pde8,mGluR,CG10188,dnc,RhoGAP1A,Gprk1,Pkc98E,Pde1c,ACXD | | 5.465e-04 | -7.51 | cell-cell adherens junction | cellular component | GO:0005913 | 24 | 6 | 11940 | 538 | CadN2,step,shg,ex,rst,scrib | | 5.531e-04 | -7.50 | GO\_REGULATION\_OF\_NEURON\_DIFFERENTIATION | MSigDB lists | GO\_REGULATION\_OF\_NEURON\_DIFFERENTIATION | 112 | 13 | 4161 | 170 | eIF4E7,Frq1,Frq2,Tor,CG42672,LpR2,unc-13,Raf,Dys,Tif-IA,dpp,Ser,LRP1 | | 5.531e-04 | -7.50 | muscle cell differentiation | biological process | GO:0042692 | 118 | 15 | 12345 | 597 | Ubx,Actn,Khc,foxo,bru1,Antp,Mlp84B,Act88F,rst,numb,siz,rols,Tor,Prm,tin | | 5.568e-04 | -7.49 | CTTTGT\_LEF1\_Q2 | MSigDB lists | CTTTGT\_LEF1\_Q2 | 453 | 33 | 4161 | 170 | TfAP-2,Odc1,Ank,app,REG,CG5493,Rbp6,Syn2,sdt,sxc,Su(H),crol,polybromo,CG6966,rdgA,flw,Pak,CG5339,CG6418,dnc,chb,mGluR,Dys,Octbeta2R,Nmdar2,Meltrin,Iswi,Rat1,KaiR1D,LRP1,Act87E,CG14480,Raf | | 5.639e-04 | -7.48 | Activation of NMDA receptor upon glutamate binding and postsynaptic events | REACTOME pathways | R-DME-442755 | 22 | 6 | 3923 | 197 | CrebB,Cam,Actn,Nmdar2,Raf,CG17698 | | 5.639e-04 | -7.48 | Platelet homeostasis | REACTOME pathways | R-DME-418346 | 22 | 6 | 3923 | 197 | Pde9,olf186-F,Pde6,Cam,CG30054,Pde1c | | 5.684e-04 | -7.47 | actin cytoskeleton reorganization | biological process | GO:0031532 | 31 | 7 | 12345 | 597 | Actn,p130CAS,nuf,rictor,CG32264,pod1,RhoGEF3 | | 5.806e-04 | -7.45 | GO\_REGULATION\_OF\_MULTICELLULAR\_ORGANISMAL\_DEVELOPMENT | MSigDB lists | GO\_REGULATION\_OF\_MULTICELLULAR\_ORGANISMAL\_DEVELOPMENT | 325 | 26 | 4161 | 170 | Tor,TfAP-2,IP3K2,CG42672,Frq2,REG,Tif-IA,fz2,chb,Ptp61F,dsh,Dys,Octbeta2R,Nha1,sxc,Sod1,Su(H),Pkc53E,Frq1,eIF4E7,dpp,Ser,LRP1,LpR2,unc-13,Raf | | 5.860e-04 | -7.44 | GO\_NEUROLOGICAL\_SYSTEM\_PROCESS | MSigDB lists | GO\_NEUROLOGICAL\_SYSTEM\_PROCESS | 173 | 17 | 4161 | 170 | Nmdar2,AstA-R1,Octbeta2R,Cln3,Tor,Awh,Ca-beta,RpL38,TfAP-2,Ank,LpR2,Raf,Sod1,RhoGAP1A,KaiR1D,Nlg2,CG7483 | | 5.861e-04 | -7.44 | cell-cell junction | cellular component | GO:0005911 | 101 | 13 | 11940 | 538 | fw,Mbs,sdt,nrv2,CadN2,Inx2,scrib,ex,rst,step,shakB,shg,bbg | | 6.098e-04 | -7.40 | neuromuscular junction development | biological process | GO:0007528 | 83 | 12 | 12345 | 597 | Nlg2,Frq1,CrebB,sgg,Frq2,unc-13,Ca-beta,Elp3,mGluR,kay,Ank2,cpx | | 6.119e-04 | -7.40 | GO\_POSITIVE\_REGULATION\_OF\_LIPID\_METABOLIC\_PROCESS | MSigDB lists | GO\_POSITIVE\_REGULATION\_OF\_LIPID\_METABOLIC\_PROCESS | 37 | 7 | 4161 | 170 | Tor,Pkc98E,EcR,ABCD,mino,Octbeta2R,Tsp96F | | 6.129e-04 | -7.40 | GO\_RESPONSE\_TO\_MOLECULE\_OF\_BACTERIAL\_ORIGIN | MSigDB lists | GO\_RESPONSE\_TO\_MOLECULE\_OF\_BACTERIAL\_ORIGIN | 48 | 8 | 4161 | 170 | RhoGAP1A,sxc,TfAP-2,Pkc98E,fz2,dnc,Ugt301D1,LpR2 | | 6.289e-04 | -7.37 | regulation of behavior | biological process | GO:0050795 | 95 | 13 | 12345 | 597 | foxo,CrebB,AANAT1,EcR,Pdfr,qvr,Sh,mnb,Mbs,wake,RYa,TfAP-2,egh | | 6.295e-04 | -7.37 | neuromuscular junction | cellular component | GO:0031594 | 77 | 11 | 11940 | 538 | unc-13,Pak,Nlg2,Ank2,cpx,Exn,sky,GluRIIC,sgg,pod1,scrib | | 6.307e-04 | -7.37 | GO\_AORTA\_MORPHOGENESIS | MSigDB lists | GO\_AORTA\_MORPHOGENESIS | 5 | 3 | 4161 | 170 | LRP1,Ser,Su(H) | | 6.307e-04 | -7.37 | GO\_NEGATIVE\_REGULATION\_OF\_STEROID\_METABOLIC\_PROCESS | MSigDB lists | GO\_NEGATIVE\_REGULATION\_OF\_STEROID\_METABOLIC\_PROCESS | 5 | 3 | 4161 | 170 | Sod1,Ugt301D1,dpp | | 6.333e-04 | -7.36 | Platelet activation, signaling and aggregation | REACTOME pathways | R-DME-76002 | 155 | 18 | 3923 | 197 | Raf,Cam,rdgA,Pkc98E,kek5,CG12547,Actn,Qsox2,CG32264,CG30054,p130CAS,fw,kek1,Tsf1,CG41520,Ptp61F,Pkc53E,ninaD | | 6.340e-04 | -7.36 | stem cell differentiation | biological process | GO:0048863 | 72 | 11 | 12345 | 597 | sqz,EcR,Antp,scrib,dpp,numb,mid,ham,Rbp6,ems,tin | | 6.626e-04 | -7.32 | REACTOME\_GPCR\_DOWNSTREAM\_SIGNALING | MSigDB lists | REACTOME\_GPCR\_DOWNSTREAM\_SIGNALING | 73 | 10 | 4161 | 170 | Pkc98E,dnc,Pde6,AstA-R1,Pde1c,CCAP-R,Octbeta2R,Pkc53E,RhoGAP1A,Pde8 | | 6.707e-04 | -7.31 | Phototransduction - fly | KEGG pathways | dme04745 | 38 | 8 | 3198 | 170 | Pkc53E,Gprk1,CG13526,Act87E,CG30054,Act88F,Cam,CG13898 | | 6.707e-04 | -7.31 | Phototransduction - fly | KEGG pathways | ko04745 | 38 | 8 | 3198 | 170 | CG13898,Act88F,Cam,CG30054,Act87E,CG13526,Pkc53E,Gprk1 | | 6.874e-04 | -7.28 | synaptic membrane | cellular component | GO:0097060 | 66 | 10 | 11940 | 538 | unc-13,gukh,Ank2,Nlg2,KaiR1D,Exn,GluRIIC,Fife,scrib,Syx1A | | 7.071e-04 | -7.25 | Spectrin/alpha-actinin | interpro domains | IPR018159 | 9 | 4 | 11383 | 585 | Dys,zormin,Msp300,Actn | | 7.071e-04 | -7.25 | Spectrin\_repeat | interpro domains | IPR002017 | 9 | 4 | 11383 | 585 | Dys,zormin,Msp300,Actn | | 7.074e-04 | -7.25 | GO\_REGULATION\_OF\_CATION\_TRANSMEMBRANE\_TRANSPORT | MSigDB lists | GO\_REGULATION\_OF\_CATION\_TRANSMEMBRANE\_TRANSPORT | 49 | 8 | 4161 | 170 | ZnT63C,Ank,Ca-beta,Dys,CCAP-R,AstA-R1,dnc,Pkc98E | | 7.085e-04 | -7.25 | GO\_REGULATION\_OF\_LIPID\_CATABOLIC\_PROCESS | MSigDB lists | GO\_REGULATION\_OF\_LIPID\_CATABOLIC\_PROCESS | 11 | 4 | 4161 | 170 | Tor,ABCD,Pkc98E,Octbeta2R | | 7.085e-04 | -7.25 | GO\_CAMP\_METABOLIC\_PROCESS | MSigDB lists | GO\_CAMP\_METABOLIC\_PROCESS | 11 | 4 | 4161 | 170 | Ca-beta,Pde6,dnc,Pde8 | | 7.085e-04 | -7.25 | MODULE\_171 | MSigDB lists | MODULE\_171 | 11 | 4 | 4161 | 170 | Sh,Meltrin,MCU,scramb1 | | 7.096e-04 | -7.25 | fly chr2-54 cM | chromosome location | fly chr2-54 cM | 22 | 5 | 25010 | 845 | Gpb5,CG17715,Spf45,mir-2490,lncRNA:CR46253 | | 7.096e-04 | -7.25 | fly chr2-54 | chromosome location | fly chr2-54 | 22 | 5 | 25010 | 845 | mir-2490,lncRNA:CR46253,Gpb5,Spf45,CG17715 | | 7.117e-04 | -7.25 | TAATAAT\_MIR126 | MSigDB lists | TAATAAT\_MIR126 | 61 | 9 | 4161 | 170 | dsh,chb,LRP1,Ser,Pkc98E,bru1,dsd,sxc,csw | | 7.188e-04 | -7.24 | GOBERT\_OLIGODENDROCYTE\_DIFFERENTIATION\_DN | MSigDB lists | GOBERT\_OLIGODENDROCYTE\_DIFFERENTIATION\_DN | 242 | 21 | 4161 | 170 | sxc,Pkc53E,Glut1,CG5522,Act87E,PIG-G,Pak,Aldh-III,Tsp96F,CG42672,step,PIG-O,ZnT63C,Stlk,pod1,dsd,Ank,app,Rbp6,dnc,KrT95D | | 7.495e-04 | -7.20 | Mhc (Myosin heavy chain) | protein interactions | 35007 | 58 | 10 | 9204 | 481 | CG5522,Act88F,Actn,bru1,TpnC41C,flw,Prm,Scp1,CG34172,Ptp61F | | 7.651e-04 | -7.18 | Spectrin | pfam domains | PF00435 | 9 | 4 | 10480 | 550 | Msp300,Dys,Actn,zormin | | 7.761e-04 | -7.16 | phosphoric diester hydrolase activity | molecular function | GO:0008081 | 33 | 7 | 12176 | 580 | Pde8,Pde9,CG18135,dnc,Pde6,Pde1c,Pld | | 8.036e-04 | -7.13 | long-term memory | biological process | GO:0007616 | 74 | 11 | 12345 | 597 | mGluR,Mob2,klg,Su(H),Gld2,Nmdar2,Pkc98E,CrebB,CG5541,HDAC4,EcR | | 8.036e-04 | -7.13 | rhythmic behavior | biological process | GO:0007622 | 74 | 11 | 12345 | 597 | Pdfr,dnc,CrebB,Fer2,ITP,foxo,numb,sgg,tyf,sxc,kay | | 8.071e-04 | -7.12 | regulation of response to stimulus | biological process | GO:0048583 | 958 | 68 | 12345 | 597 | Raf,GlyP,Cam,lncRNA:Hsromega,qvr,dsh,chb,Pde8,foxo,trol,dpp,step,HDAC4,Ser,dnc,RhoGAP1A,Tor,bnl,pirk,Su(H),Snoo,sgg,ex,CG9380,Src42A,raw,et,kek1,tld,Usp1,Flo2,RhoGEF3,hth,Cln3,Pak,sxc,spri,kek5,Mos,Exn,ine,Syt12,MESK2,CG34408,rdgA,Gprk1,Lpin,olf186-F,GluRIIC,siz,sno,melt,PGRP-LA,Ptp61F,scrib,numb,crol,Elp3,mnb,CG14182,Doa,CG5522,CG18659,rictor,flw,CG10188,Fer2,Cdep | | 8.133e-04 | -7.11 | GCM\_PTK2 | MSigDB lists | GCM\_PTK2 | 50 | 8 | 4161 | 170 | CG12121,CG33090,Frq1,Dlic,MESK2,dsd,CG42672,Frq2 | | 8.152e-04 | -7.11 | Ig\_V-set | interpro domains | IPR013106 | 31 | 7 | 11383 | 585 | dpr21,nrm,Strn-Mlck,side-VIII,dpr15,dpr5,DIP-lambda | | 8.157e-04 | -7.11 | locomotor rhythm | biological process | GO:0045475 | 63 | 10 | 12345 | 597 | Fer2,ITP,foxo,CrebB,dnc,Pdfr,kay,sxc,tyf,sgg | | 8.284e-04 | -7.10 | taxis | biological process | GO:0042330 | 365 | 32 | 12345 | 597 | dsh,Cam,rdgA,beat-IIa,trol,chb,sqz,shep,shg,dpp,egh,Ptp99A,grn,klg,mid,E(Pc),Ptp61F,fz2,haf,msps,bnl,Src42A,gukh,Pdfr,CadN2,pod1,Pak,fru,retn,RhoGAP100F,Khc,esg | | 8.503e-04 | -7.07 | sequence-specific DNA binding | molecular function | GO:0043565 | 515 | 41 | 12176 | 580 | EcR,ems,Su(H),CG15073,FoxL1,E5,CG43689,CG12071,tin,CG9650,TfAP-2,kay,retn,cic,B-H2,Ubx,grn,CG31224,bi,Eip74EF,GATAe,opa,toe,CG7368,hth,foxo,Atf6,CG12605,mamo,CG32006,Hr96,CrebB,CG10462,luna,mid,CG4374,esg,scro,Antp,Maf1,Awh | | 8.532e-04 | -7.07 | GSE19888\_ADENOSINE\_A3R\_ACT\_VS\_TCELL\_MEMBRANES\_ACT\_AND\_A3R\_INH\_PRETREAT\_IN\_MAST\_CELL\_DN | MSigDB lists | GSE19888\_ADENOSINE\_A3R\_ACT\_VS\_TCELL\_MEMBRANES\_ACT\_AND\_A3R\_INH\_PRETREAT\_IN\_MAST\_CELL\_DN | 39 | 7 | 4161 | 170 | CG17698,Frq2,rdgA,TfAP-2,CG8184,Frq1,Ucp4A | | 8.532e-04 | -7.07 | GSE39916\_B\_CELL\_SPLEEN\_VS\_PLASMA\_CELL\_BONE\_MARROW\_UP | MSigDB lists | GSE39916\_B\_CELL\_SPLEEN\_VS\_PLASMA\_CELL\_BONE\_MARROW\_UP | 39 | 7 | 4161 | 170 | Stt3B,Pgant6,sdt,app,Tsp96F,Pak,Ptp61F | | 8.807e-04 | -7.03 | negative regulation of cell communication | biological process | GO:0010648 | 351 | 31 | 12345 | 597 | CG9650,kek5,HDAC4,flw,Ser,Ptp61F,dnc,Raf,Snoo,Su(H),pirk,sgg,numb,mnb,Cam,crol,qvr,Ank2,cpx,CG14182,Lpin,Nlg2,Gprk1,sky,chb,kek1,Src42A,et,trol,foxo,raw | | 8.862e-04 | -7.03 | Hemostasis | REACTOME pathways | R-DME-109582 | 257 | 25 | 3923 | 197 | Raf,Pde6,rdgA,Cam,Pkc98E,csw,kek5,CG12547,nrv2,Qsox2,CG32264,Actn,Khc,olf186-F,Pde9,p130CAS,CG30054,Tsf1,Ptp61F,CG41520,kek1,Pde1c,fw,ninaD,Pkc53E | | 9.154e-04 | -7.00 | regulation of developmental process | biological process | GO:0050793 | 757 | 56 | 12345 | 597 | dsh,rib,CG42797,ham,Raf,foxo,wbl,bun,step,Pkc53E,shg,rst,Mlp84B,dpp,mid,csw,Antp,scrib,CG1572,Ser,Cat,PGRP-LA,EcR,Fhos,hdc,Syx1A,Ank2,fz2,kay,mnb,Elp3,Mob2,sgg,CG9380,ex,numb,bnl,Snoo,Su(H),Tor,barc,raw,Src42A,Eip74EF,Syn2,spri,rictor,Pak,Mbs,hth,Dlic,bru1,Octbeta2R,Ubx,esg,Khc | | 9.174e-04 | -6.99 | CG15422 (uncharacterized protein) | protein interactions | 33615 | 40 | 8 | 9204 | 481 | CG17600,msi,CG17777,gce,Raf,Marf1,CG41099,CG13227 | | 9.200e-04 | -6.99 | GO\_POSITIVE\_REGULATION\_OF\_CELL\_DIFFERENTIATION | MSigDB lists | GO\_POSITIVE\_REGULATION\_OF\_CELL\_DIFFERENTIATION | 133 | 14 | 4161 | 170 | CG42672,IP3K2,Tor,Pkc53E,Su(H),sxc,Dys,Raf,Nha1,LpR2,unc-13,Tif-IA,dpp,Ser | | 9.219e-04 | -6.99 | regulation of circadian sleep/wake cycle | biological process | GO:0042749 | 43 | 8 | 12345 | 597 | Sh,qvr,wake,TfAP-2,CrebB,EcR,Pdfr,AANAT1 | | 9.228e-04 | -6.99 | negative regulation of signaling | biological process | GO:0023057 | 352 | 31 | 12345 | 597 | sky,chb,CG14182,Nlg2,Lpin,Gprk1,kek1,foxo,trol,raw,et,Src42A,sgg,numb,Snoo,pirk,Su(H),Raf,qvr,Ank2,cpx,mnb,Cam,crol,Ser,Ptp61F,HDAC4,kek5,flw,dnc,CG9650 | | 9.232e-04 | -6.99 | response to chemical | biological process | GO:0042221 | 1191 | 81 | 12345 | 597 | chb,geko,Ir56a,trol,foxo,Raf,Mrtf,Cam,Gr22c,dsh,dnc,opa,dpp,Obp56h,shg,Atf6,Sod1,Tk,St4,et,Src42A,msps,Tor,bnl,sgg,ex,Set2,Cyp12a4,Gr39a,Mondo,fz2,CG7362,eIF2Bgamma,Khc,Gpb5,ine,RhoGAP100F,Pkc98E,Syt12,Or65c,Pak,Lpin,sqz,bun,beat-IIa,Or85b,EcR,Ptp61F,Cat,scrib,E(Pc),csw,mid,Or42b,Der-1,CG33934,CG31750,egh,Ptp99A,klg,grn,Sh,gukh,msi,Iswi,Ir10a,Tsf1,ths,ZnT63C,Ir41a,haf,mnb,Pino,esg,retn,CG3556,fru,gce,CadN2,pod1,rictor | | 9.315e-04 | -6.98 | GSE20366\_TREG\_VS\_NAIVE\_CD4\_TCELL\_DEC205\_CONVERSION\_UP | MSigDB lists | GSE20366\_TREG\_VS\_NAIVE\_CD4\_TCELL\_DEC205\_CONVERSION\_UP | 51 | 8 | 4161 | 170 | CG17698,Meltrin,step,Glut1,CG15093,Rbp6,CG40178,Tsp96F | | 9.346e-04 | -6.98 | stem cell population maintenance | biological process | GO:0019827 | 99 | 13 | 12345 | 597 | chb,Ser,msi,esg,foxo,bru1,dpp,sgg,Treh,Tif-IA,dsh,shg,Inx2 | | 9.346e-04 | -6.98 | maintenance of cell number | biological process | GO:0098727 | 99 | 13 | 12345 | 597 | dpp,sgg,Treh,Tif-IA,dsh,shg,Inx2,chb,msi,Ser,esg,foxo,bru1 | | 9.886e-04 | -6.92 | GO\_RESPONSE\_TO\_NITROGEN\_COMPOUND | MSigDB lists | GO\_RESPONSE\_TO\_NITROGEN\_COMPOUND | 214 | 19 | 4161 | 170 | mino,CG5493,dnc,Ptp61F,CG3376,Octbeta2R,Cat,CG4849,Tor,Uros1,Pkc98E,eIF2Bgamma,Ser,LpR2,Raf,Pak,GlyP,sxc,Sod1 | | 9.902e-04 | -6.92 | GO\_REGULATION\_OF\_CELL\_PROJECTION\_ORGANIZATION | MSigDB lists | GO\_REGULATION\_OF\_CELL\_PROJECTION\_ORGANIZATION | 119 | 13 | 4161 | 170 | sky,Frq1,Frq2,CG42672,CG17883,Tor,Raf,LpR2,unc-13,Dys,LRP1,nompB,Tif-IA | | 1.000e-03 | -6.91 | EGF-like\_CS | interpro domains | IPR013032 | 61 | 10 | 11383 | 585 | CG13654,LRP1,dsd,CadN2,Meltrin,trol,LpR2,shg,Ser,tld | | 1.004e-03 | -6.90 | stem cell development | biological process | GO:0048864 | 25 | 6 | 12345 | 597 | Rbp6,Antp,ems,sqz,numb,mid | | 1.009e-03 | -6.90 | GO\_REGULATION\_OF\_CATION\_CHANNEL\_ACTIVITY | MSigDB lists | GO\_REGULATION\_OF\_CATION\_CHANNEL\_ACTIVITY | 20 | 5 | 4161 | 170 | Dys,Ank,AstA-R1,Ca-beta,dnc | | 1.009e-03 | -6.90 | GO\_REGULATION\_OF\_CALCIUM\_ION\_IMPORT | MSigDB lists | GO\_REGULATION\_OF\_CALCIUM\_ION\_IMPORT | 20 | 5 | 4161 | 170 | CCAP-R,ZnT63C,Dys,Pkc98E,dnc | | 1.009e-03 | -6.90 | GO\_ACTION\_POTENTIAL | MSigDB lists | GO\_ACTION\_POTENTIAL | 20 | 5 | 4161 | 170 | Cln3,Ca-beta,KaiR1D,Ank,Dys | | 1.009e-03 | -6.90 | GO\_REGULATION\_OF\_PEPTIDYL\_SERINE\_PHOSPHORYLATION | MSigDB lists | GO\_REGULATION\_OF\_PEPTIDYL\_SERINE\_PHOSPHORYLATION | 20 | 5 | 4161 | 170 | dnc,sxc,Dys,Pak,Raf | | 1.016e-03 | -6.89 | GO\_MODULATION\_OF\_SYNAPTIC\_TRANSMISSION | MSigDB lists | GO\_MODULATION\_OF\_SYNAPTIC\_TRANSMISSION | 77 | 10 | 4161 | 170 | KaiR1D,ZnT63C,Nlg2,CG7483,Pkc98E,unc-13,Raf,Syx1A,mGluR,Khc | | 1.020e-03 | -6.89 | CREB phosphorylation through the activation of CaMKII | REACTOME pathways | R-DME-442729 | 10 | 4 | 3923 | 197 | Actn,Nmdar2,CrebB,Cam | | 1.022e-03 | -6.89 | double-stranded DNA binding | molecular function | GO:0003690 | 391 | 33 | 12176 | 580 | EcR,CG15073,Su(H),ems,tin,E5,CG43689,CG12071,retn,TfAP-2,kay,cic,Ubx,B-H2,bi,grn,CG31224,CG7368,opa,GATAe,foxo,hth,Atf6,CG12605,mid,Hr96,CG10462,CG4374,esg,Awh,Antp,Maf1,scro | | 1.025e-03 | -6.88 | salivary gland histolysis | biological process | GO:0035070 | 34 | 7 | 12345 | 597 | EcR,Pak,IP3K2,Eip74EF,Cam,Fhos,hdc | | 1.025e-03 | -6.88 | GO\_MEMBRANE\_REGION | MSigDB lists | GO\_MEMBRANE\_REGION | 266 | 22 | 4161 | 170 | Ank,Cln3,Frq2,CG9132,scramb1,Nmdar2,Dys,Octbeta2R,mGluR,Syn2,dnc,Sh,KaiR1D,Flo2,Nlg2,Frq1,Tsp96F,LpR2,unc-13,Pak,LRP1,Ser | | 1.028e-03 | -6.88 | mesoderm development | biological process | GO:0007498 | 100 | 13 | 12345 | 597 | Pde8,CG17470,flw,Ubx,CG34417,shg,Prm,tin,ths,Msp300,RhoL,csw,dpp | | 1.028e-03 | -6.88 | sex differentiation | biological process | GO:0007548 | 100 | 13 | 12345 | 597 | tnc,raw,dpp,TfAP-2,Doa,mid,Tor,fru,fz2,dsh,tin,shg,rib | | 1.029e-03 | -6.88 | GO\_POSITIVE\_REGULATION\_OF\_LEUKOCYTE\_DIFFERENTIATION | MSigDB lists | GO\_POSITIVE\_REGULATION\_OF\_LEUKOCYTE\_DIFFERENTIATION | 12 | 4 | 4161 | 170 | sxc,Pkc53E,IP3K2,Nha1 | | 1.029e-03 | -6.88 | GO\_POSITIVE\_REGULATION\_OF\_ACTIN\_FILAMENT\_BUNDLE\_ASSEMBLY | MSigDB lists | GO\_POSITIVE\_REGULATION\_OF\_ACTIN\_FILAMENT\_BUNDLE\_ASSEMBLY | 12 | 4 | 4161 | 170 | Fhos,Tor,Raf,Pak | | 1.031e-03 | -6.88 | Smr (Smrter) | protein interactions | 32225 | 16 | 5 | 9204 | 481 | EcR,rictor,flw,fru,Su(H) | | 1.035e-03 | -6.87 | EF\_Hand\_1\_Ca\_BS | interpro domains | IPR018247 | 83 | 12 | 11383 | 585 | TpnC25D,TpnC41C,CG10126,Scp1,CG14362,TpnC4,Cam,Frq1,Actn,CG13526,Frq2,CG13898 | | 1.039e-03 | -6.87 | REACTOME\_SIGNALING\_BY\_GPCR | MSigDB lists | REACTOME\_SIGNALING\_BY\_GPCR | 105 | 12 | 4161 | 170 | Pde8,Pkc53E,RhoGAP1A,AstA-R1,Pde1c,CCAP-R,Octbeta2R,mGluR,fz2,Pkc98E,dnc,Pde6 | | 1.043e-03 | -6.87 | Ig\_I-set | interpro domains | IPR013098 | 72 | 11 | 11383 | 585 | trol,klg,kek1,Strn-Mlck,DIP-lambda,dpr21,rst,DIP-delta,kek5,dpr12,zormin | | 1.050e-03 | -6.86 | hep (hemipterous) | protein interactions | 32256 | 71 | 11 | 9204 | 481 | scrib,sgg,bi,by,Raf,Src42A,dpp,dsh,kay,foxo,flw | | 1.053e-03 | -6.86 | negative regulation of biological process | biological process | GO:0048519 | 1502 | 98 | 12345 | 597 | E5,bru1,bol,Fer2,Ubx,retn,esg,flw,CG12605,Acp62F,sws,gce,RYa,Doa,Iswi,bi,msi,CG43689,CG14182,hdc,Ank2,mnb,crol,cic,numb,B-H2,scrib,PGRP-LA,Ptp61F,REG,EcR,melt,Marf1,htk,CG7368,Achl,Maf1,mid,csw,bun,Gprk1,Nlg2,Lpin,GATAe,lncRNA:bxd,CG42797,Sox102F,AGO3,ham,Cep97,Pkc98E,PhKgamma,Mos,kek5,CG1894,Khc,CG9650,Spn88Eb,kek1,tld,raw,Src42A,et,sky,cpx,Mob2,CG9380,ex,sgg,Pde6,pirk,Snoo,Tor,Su(H),polybromo,Antp,dnc,g,Ser,HDAC4,AGO2,CG31224,dpp,CG41520,trol,foxo,Pde8,HPS4,toe,chb,chinmo,tin,qvr,Cam,TfAP-2,GlyP,Raf,CG12071 | | 1.059e-03 | -6.85 | EFh | smart domains | SM00054 | 70 | 12 | 5086 | 315 | CG10126,Frq1,TpnC25D,Scp1,Cam,TpnC41C,CG13898,Frq2,CG14362,CG13526,TpnC4,Actn | | 1.059e-03 | -6.85 | neuron projection | cellular component | GO:0043005 | 431 | 34 | 11940 | 538 | unc-13,mnb,Gr39a,Or85b,Ptp99A,dpr15,EcR,cpx,klg,pod1,gukh,Pdfr,Ank2,nrv2,Gld2,kcc,CG5921,chb,Or42b,Khc,CG13794,Or65c,dpr12,Gr22c,hdc,Ser,g,path,dpr5,dpr18,sky,sgg,AGO2,Syx1A | | 1.063e-03 | -6.85 | GO\_POSTSYNAPTIC\_MEMBRANE | MSigDB lists | GO\_POSTSYNAPTIC\_MEMBRANE | 52 | 8 | 4161 | 170 | mGluR,Dys,Nmdar2,Nlg2,Frq1,Ank,KaiR1D,Frq2 | | 1.063e-03 | -6.85 | GO\_SENSORY\_ORGAN\_MORPHOGENESIS | MSigDB lists | GO\_SENSORY\_ORGAN\_MORPHOGENESIS | 52 | 8 | 4161 | 170 | dsh,fz2,Ser,RhoGAP1A,Sod1,RpL38,CG17883,TfAP-2 | | 1.092e-03 | -6.82 | cytoskeletal protein binding | molecular function | GO:0008092 | 330 | 29 | 12176 | 580 | Cdep,Cpsf160,Yeti,pod1,Cam,CG5921,CadN2,nuf,Fhos,by,CG18135,Actn,shg,CG13251,zormin,Gas8,CG18109,Khc,Mlp84B,Ank,Ank2,chb,CG32264,Syn2,nompB,msps,CG34417,Dys,Msp300 | | 1.094e-03 | -6.82 | neuron projection guidance | biological process | GO:0097485 | 280 | 26 | 12345 | 597 | Khc,RhoGAP100F,retn,Ptp61F,E(Pc),fru,mid,grn,Ptp99A,egh,klg,pod1,Pak,CadN2,shg,chb,gukh,sqz,trol,Src42A,msps,beat-IIa,haf,Cam,dsh,fz2 | | 1.109e-03 | -6.80 | cell part | cellular component | GO:0044464 | 7811 | 385 | 11940 | 538 | Antp,Syn2,eIF4E7,sqz,CG3594,zormin,sws,CG18659,aust,CG9132,sky,Lpin,CG10000,Fife,CG32264,dnc,Cadps,CG9451,raw,Nuak1,CG40470,Nlg2,CG11023,Act87E,Dmtn,qvr,Ptp61F,Prm,Gprk1,CrebB,loopin-1,Ste:CG33247,Rab9Fa,nrv2,Gld2,CG11318,Raf,CG6418,RpL38,CG8272,Ptp99A,CG5059,fz2,numb,CG42830,Gr39a,unc-13,flw,Mob2,Neb-cGP,Pgant6,Prosbeta5R1,rst,msi,csw,KaiR1D,egh,CG41099,GATAe,CadN2,CG9650,RhoGAP18B,Msp300,sgg,Pfdn4,PGRP-LA,E5,IP3K2,mRpL24,p130CAS,GstZ2,kek1,Or42b,CG33158,scaRNA:PsiU2-35.45,polybromo,scaRNA:MeU2-C41,Or65c,GluRIIC,CG9436,scro,CNT1,ND-AGGG,Atf6,Ugt304A1,CG33310,His1:CG33837,cert,Snoo,Pzl,rols,Odc1,wbl,nrm,CG32006,DIP-delta,Alp12,Mst87F,ham,shakB,Rat1,bbg,Gpb5,Set2,CG15093,nompB,Pomp,lncRNA:Hsromega,CG13585,beat-IIa,siz,dpr5,CG14606,dany,HPS4,chb,Stt3B,CG9449,Sk1,TfAP-2,CG31224,AANATL2,fz4,CG6790,Ank2,Usp1,Qsox2,Ucp4A,CG34417,Octbeta2R,mRpS5,Stlk,Poxm,Cam,Cpsf160,ps,Fad2,PCB,CG4374,shd,Pde6,Kyat,CG30054,Doa,kl-5,CG42390,PhKgamma,CG12782,Iswi,CG14480,IntS3,Tsf1,Ser,CG42458,CG3078,toe,CG1894,Mrtf,CG5445,ABCD,AANAT1,Pld,Cdep,bol,Rcd6,trv,dpr12,melt,CG7483,REG,hth,Fhos,Syt12,LRP1,CG15073,CG4839,CG40813,CG5921,CG12071,bi,kcc,GlyP,rib,Marf1,CG18109,RhoGAP100F,eIF2Bgamma,sno,Art2,Cyt-b5-r,Act88F,Awh,Epg5,pdgy,spd-2,Tif-IA,Src42A,CG12605,glob3,Dys,Ugt302K1,CycT,Syx1A,Actn,Nha1,CG7362,HDAC4,CG41562,Vha100-5,chp,Yeti,Su(H),foxo,Ugt301D1,dpp,Der-1,AGO3,step,Spf45,AstA-R1,B-H2,CG17122,Ipk1,gukh,Pkc98E,app,olf186-F,fs(1)N,CG7368,Mondo,CG43689,mid,CG13526,lncRNA:bxd,MESK2,Cep97,bun,tin,IFT46,Hr96,rictor,kek5,Maf1,AOX1,CG46385,CG17124,Tor,g,CG8184,ddbt,CG13654,Gr22c,hdc,htk,AGO2,dpr18,fus,Ubx,ems,Exn,Nmdar2,CG15629,MsR2,tamo,mamo,CG13794,Mbs,CG30036,Ank,Frq2,crol,Nnf1b,CG14252,ZnT63C,msps,shg,Mlp84B,cic,CG44249,CG17883,cindr,CG6839,Or85b,mnb,chinmo,jigr1,boi,TpnC25D,Pkc53E,Mur2B,EcR,path,ND-51L2,MCU,Pak,CG40472,CG13251,CG12200,PIG-G,D2hgdh,fru,Sod1,CG17698,Khc,Elp3,CG10462,Mctp,CG4301,Sh,Argl,TfIIFbeta,CG34172,l(3)neo38,CG11970,CG42797,luna,CG10960,CG9981,LpR2,Dlic,CG4849,E(Pc),ine,Mos,ND-MLRQ,esg,CG34435,dpr15,Damm,sdt,Cln3,retn,woc,CCAP-R,cpx,barc,klg,St4,spri,PIG-O,Cht6,CG32971,Eip74EF,can,Myo81F,CG14182,shep,FoxL1,mag,CG17387,Root,CG3285,Hil,nuf,mino,SteXh:CG42398,kay,scrib,Flo2,grn,ACXD,wake,Gpo3,gce,CG41128,Pdfr,UQCR-11,Inx2,ex,Tsp96F,opa,Gas8,DIP-lambda,eRF3,bru1,mGluR,Cat,dsh,Ca-beta,pod1,GstE11,Spt20,Asator,Rbp6,TTLL3A | | 1.109e-03 | -6.80 | cell | cellular component | GO:0005623 | 7811 | 385 | 11940 | 538 | Mctp,CG10462,Sh,Argl,CG4301,l(3)neo38,CG34172,TfIIFbeta,D2hgdh,Sod1,fru,Khc,CG17698,Elp3,CG13251,CG40472,CG12200,PIG-G,path,MCU,ND-51L2,Pak,CCAP-R,woc,spri,St4,barc,cpx,klg,sdt,dpr15,Damm,retn,Cln3,Dlic,E(Pc),CG4849,esg,Mos,ine,ND-MLRQ,CG34435,CG42797,CG11970,CG10960,luna,LpR2,CG9981,Flo2,scrib,kay,grn,ACXD,Gpo3,wake,mino,SteXh:CG42398,FoxL1,shep,CG14182,Myo81F,CG3285,Root,CG17387,mag,Hil,nuf,CG32971,PIG-O,Cht6,Eip74EF,can,GstE11,pod1,Asator,Spt20,Rbp6,TTLL3A,mGluR,Cat,Ca-beta,dsh,Inx2,Gas8,opa,ex,Tsp96F,DIP-lambda,bru1,eRF3,gce,CG41128,Pdfr,UQCR-11,Der-1,dpp,Ugt301D1,step,AGO3,Spf45,AstA-R1,chp,HDAC4,Vha100-5,CG41562,foxo,Su(H),Yeti,Syx1A,Actn,CG7362,Nha1,CG12605,Src42A,CycT,Dys,glob3,Ugt302K1,Hr96,IFT46,rictor,kek5,AOX1,Maf1,MESK2,lncRNA:bxd,tin,bun,Cep97,olf186-F,app,Mondo,CG7368,fs(1)N,CG43689,CG13526,mid,B-H2,CG17122,Pkc98E,Ipk1,gukh,tamo,mamo,CG13794,MsR2,CG15629,AGO2,htk,Ubx,dpr18,fus,Exn,ems,Nmdar2,CG46385,ddbt,g,CG8184,Tor,CG17124,Gr22c,hdc,CG13654,jigr1,boi,Pkc53E,TpnC25D,EcR,Mur2B,CG6839,Or85b,mnb,chinmo,ZnT63C,CG14252,shg,msps,cic,Mlp84B,cindr,CG17883,CG44249,Ank,Frq2,CG30036,Mbs,Nnf1b,crol,TfAP-2,Sk1,CG9449,CG31224,AANATL2,fz4,CG14606,dany,Stt3B,HPS4,chb,beat-IIa,siz,dpr5,Pomp,nompB,CG15093,Set2,lncRNA:Hsromega,CG13585,PhKgamma,CG42390,Iswi,CG12782,ps,Fad2,Cpsf160,Pde6,Kyat,PCB,shd,CG4374,kl-5,Doa,CG30054,CG34417,mRpS5,Octbeta2R,Stlk,Cam,Poxm,CG6790,Ank2,Ucp4A,Qsox2,Usp1,dpr12,trv,Rcd6,CG7483,melt,REG,AANAT1,Pld,Cdep,bol,toe,CG3078,CG5445,ABCD,CG1894,Mrtf,CG14480,IntS3,Ser,Tsf1,CG42458,Epg5,Act88F,Awh,pdgy,Tif-IA,spd-2,rib,Marf1,eIF2Bgamma,CG18109,RhoGAP100F,sno,Art2,Cyt-b5-r,CG5921,CG40813,kcc,GlyP,CG12071,bi,hth,Fhos,Syt12,LRP1,CG4839,CG15073,Prm,loopin-1,CrebB,Gprk1,Rab9Fa,Ste:CG33247,Nuak1,raw,CG40470,CG11023,Act87E,Nlg2,Ptp61F,qvr,Dmtn,CG9132,Lpin,Fife,CG10000,sky,CG9451,Cadps,dnc,CG32264,eIF4E7,Syn2,Antp,zormin,CG3594,sqz,sws,aust,CG18659,Neb-cGP,Mob2,rst,Pgant6,Prosbeta5R1,msi,csw,CG8272,Ptp99A,numb,fz2,CG5059,Gr39a,CG42830,unc-13,flw,Raf,RpL38,CG6418,Gld2,nrv2,CG11318,polybromo,scaRNA:MeU2-C41,GluRIIC,Or65c,CG9436,mRpL24,p130CAS,Or42b,CG33158,kek1,GstZ2,scaRNA:PsiU2-35.45,CG9650,Msp300,sgg,RhoGAP18B,PGRP-LA,Pfdn4,IP3K2,E5,egh,KaiR1D,GATAe,CG41099,CadN2,shakB,ham,Gpb5,bbg,Rat1,rols,wbl,nrm,Odc1,CG32006,Mst87F,Alp12,DIP-delta,cert,Pzl,Snoo,CNT1,scro,ND-AGGG,Atf6,His1:CG33837,CG33310,Ugt304A1 | | 1.125e-03 | -6.79 | GO\_REGULATION\_OF\_LIPID\_METABOLIC\_PROCESS | MSigDB lists | GO\_REGULATION\_OF\_LIPID\_METABOLIC\_PROCESS | 78 | 10 | 4161 | 170 | Pkc98E,EcR,mino,ABCD,Ugt301D1,dpp,Tsp96F,Octbeta2R,Sod1,Tor | | 1.131e-03 | -6.78 | cytoskeleton organization | biological process | GO:0007010 | 580 | 45 | 12345 | 597 | RhoGEF3,by,pod1,CG32264,rictor,Prm,Dlic,bru1,Khc,Rcd6,CG18109,RhoGAP100F,Actn,Elp3,Ank2,Strn-Mlck,msps,p130CAS,spd-2,sgg,Src42A,gukh,CG34417,rols,Mlp84B,wake,siz,egh,Ank,Root,rib,Cam,Mrtf,dsh,nuf,Cep97,sick,cindr,Msp300,Act88F,CG17387,foxo,RhoGAP18B,chb,Act87E | | 1.131e-03 | -6.78 | GO\_REGULATION\_OF\_PROTEIN\_SERINE\_THREONINE\_KINASE\_ACTIVITY | MSigDB lists | GO\_REGULATION\_OF\_PROTEIN\_SERINE\_THREONINE\_KINASE\_ACTIVITY | 106 | 12 | 4161 | 170 | Stlk,CG42672,csw,Sod1,wbl,dsh,Pak,CG3376,Tsp96F,Raf,dpp,fz2 | | 1.140e-03 | -6.78 | fly chr41D1-41D1\|2-55 | chromosome location | fly chr41D1-41D1\|2-55 | 2 | 2 | 25010 | 845 | lncRNA:CR44105,TpnC41C | | 1.140e-03 | -6.78 | fly chr60D4-60D5\|2-107 | chromosome location | fly chr60D4-60D5\|2-107 | 2 | 2 | 25010 | 845 | ITP,CG4622 | | 1.140e-03 | -6.78 | fly chr65D2-65D2\|3-18 cM | chromosome location | fly chr65D2-65D2\|3-18 cM | 2 | 2 | 25010 | 845 | lncRNA:CR45416,lncRNA:CR45115 | | 1.140e-03 | -6.78 | fly chr65D2-65D2\|3-18 | chromosome location | fly chr65D2-65D2\|3-18 | 2 | 2 | 25010 | 845 | lncRNA:CR45115,lncRNA:CR45416 | | 1.140e-03 | -6.78 | fly chr41D1-41D1\|2-55 cM | chromosome location | fly chr41D1-41D1\|2-55 cM | 2 | 2 | 25010 | 845 | TpnC41C,lncRNA:CR44105 | | 1.140e-03 | -6.78 | fly chr39C1-39C2\|2-54 cM | chromosome location | fly chr39C1-39C2\|2-54 cM | 2 | 2 | 25010 | 845 | Mondo,Gr39a | | 1.140e-03 | -6.78 | fly chr39C1-39C2\|2-54 | chromosome location | fly chr39C1-39C2\|2-54 | 2 | 2 | 25010 | 845 | Mondo,Gr39a | | 1.140e-03 | -6.78 | fly chr60D4-60D5\|2-107 cM | chromosome location | fly chr60D4-60D5\|2-107 cM | 2 | 2 | 25010 | 845 | ITP,CG4622 | | 1.145e-03 | -6.77 | RHO GTPases activate PAKs | REACTOME pathways | R-DME-5627123 | 17 | 5 | 3923 | 197 | Cam,Mbs,flw,Pak,Strn-Mlck | | 1.157e-03 | -6.76 | Nitric oxide stimulates guanylate cyclase | REACTOME pathways | R-DME-392154 | 5 | 3 | 3923 | 197 | Pde1c,Pde6,Pde9 | | 1.164e-03 | -6.76 | GSE7852\_LN\_VS\_THYMUS\_TREG\_UP | MSigDB lists | GSE7852\_LN\_VS\_THYMUS\_TREG\_UP | 41 | 7 | 4161 | 170 | CNT1,Glut1,Dys,Pkc98E,Pkc53E,Pgant6,D2hgdh | | 1.164e-03 | -6.76 | GO\_REGULATION\_OF\_CALCIUM\_ION\_TRANSPORT | MSigDB lists | GO\_REGULATION\_OF\_CALCIUM\_ION\_TRANSPORT | 41 | 7 | 4161 | 170 | Octbeta2R,Dys,ZnT63C,CCAP-R,dnc,Ca-beta,Pkc98E | | 1.176e-03 | -6.75 | IG\_LIKE | prosite domains | PS50835 | 132 | 18 | 6243 | 386 | nrm,side-VIII,CG17716,kek1,zormin,klg,dpr15,dpr5,DIP-lambda,trol,boi,dpr12,rst,dpr21,kek5,dpr18,DIP-delta,Strn-Mlck | | 1.182e-03 | -6.74 | Gpdh1 (Glycerol-3-phosphate dehydrogenase 1) | protein interactions | 33824 | 72 | 11 | 9204 | 481 | Tsf1,Act88F,Poxm,TpnC41C,Actn,TpnC4,GlyP,Prm,Strn-Mlck,CG34172,Scp1 | | 1.196e-03 | -6.73 | positive regulation of transport | biological process | GO:0051050 | 127 | 15 | 12345 | 597 | qvr,spri,Cadps,Pkc53E,mnb,Cam,numb,Fife,Pld,Flo2,sdt,cindr,LpR2,olf186-F,Glut1 | | 1.209e-03 | -6.72 | GO\_POSITIVE\_REGULATION\_OF\_NEURON\_DIFFERENTIATION | MSigDB lists | GO\_POSITIVE\_REGULATION\_OF\_NEURON\_DIFFERENTIATION | 53 | 8 | 4161 | 170 | Tor,CG42672,Dys,unc-13,LpR2,Raf,dpp,Tif-IA | | 1.209e-03 | -6.72 | GSE35825\_IFNA\_VS\_IFNG\_STIM\_MACROPHAGE\_DN | MSigDB lists | GSE35825\_IFNA\_VS\_IFNG\_STIM\_MACROPHAGE\_DN | 53 | 8 | 4161 | 170 | sxc,Frq2,wbl,Frq1,app,rdgA,Pak,KrT95D | | 1.215e-03 | -6.71 | fly chr80F9-80F9\|3-47 | chromosome location | fly chr80F9-80F9\|3-47 | 7 | 3 | 25010 | 845 | CG17698,AGO3,lncRNA:CR45174 | | 1.215e-03 | -6.71 | fly chr80F9-80F9\|3-47 cM | chromosome location | fly chr80F9-80F9\|3-47 cM | 7 | 3 | 25010 | 845 | lncRNA:CR45174,AGO3,CG17698 | | 1.215e-03 | -6.71 | C1\_1 | pfam domains | PF00130 | 24 | 6 | 10480 | 550 | unc-13,Pkc98E,Raf,rdgA,Pkc53E,Stacl | | 1.216e-03 | -6.71 | cellular response to endogenous stimulus | biological process | GO:0071495 | 181 | 19 | 12345 | 597 | dpp,gce,csw,opa,rictor,Sh,EcR,Gpb5,ex,Tor,bnl,ths,Raf,CG7362,Set2,bun,Lpin,foxo,Iswi | | 1.224e-03 | -6.71 | GO\_VOLTAGE\_GATED\_CALCIUM\_CHANNEL\_ACTIVITY | MSigDB lists | GO\_VOLTAGE\_GATED\_CALCIUM\_CHANNEL\_ACTIVITY | 6 | 3 | 4161 | 170 | Frq1,Frq2,Ca-beta | | 1.224e-03 | -6.71 | GO\_REGULATION\_OF\_GLYCOGEN\_METABOLIC\_PROCESS | MSigDB lists | GO\_REGULATION\_OF\_GLYCOGEN\_METABOLIC\_PROCESS | 6 | 3 | 4161 | 170 | flw,PhKgamma,Tor | | 1.224e-03 | -6.71 | MODULE\_530 | MSigDB lists | MODULE\_530 | 6 | 3 | 4161 | 170 | CG5493,Sod1,Cat | | 1.224e-03 | -6.71 | GO\_POSITIVE\_REGULATION\_OF\_MYELOID\_LEUKOCYTE\_DIFFERENTIATION | MSigDB lists | GO\_POSITIVE\_REGULATION\_OF\_MYELOID\_LEUKOCYTE\_DIFFERENTIATION | 6 | 3 | 4161 | 170 | Pkc53E,sxc,Nha1 | | 1.224e-03 | -6.71 | GO\_REGULATION\_OF\_CARDIAC\_MUSCLE\_CELL\_DIFFERENTIATION | MSigDB lists | GO\_REGULATION\_OF\_CARDIAC\_MUSCLE\_CELL\_DIFFERENTIATION | 6 | 3 | 4161 | 170 | Tor,dpp,Su(H) | | 1.229e-03 | -6.70 | regulation of synaptic plasticity | biological process | GO:0048167 | 35 | 7 | 12345 | 597 | nemy,Sod1,Fife,Cat,Mctp,Dys,Src42A | | 1.231e-03 | -6.70 | I-set | pfam domains | PF07679 | 72 | 11 | 10480 | 550 | kek1,rst,kek5,DIP-lambda,zormin,Strn-Mlck,DIP-delta,dpr21,dpr12,klg,trol | | 1.259e-03 | -6.68 | muscle structure development | biological process | GO:0061061 | 224 | 22 | 12345 | 597 | Mlp84B,rst,siz,rols,fru,Prm,hth,retn,Ubx,Actn,Khc,Dys,Antp,bru1,Act88F,numb,Tor,ths,Msp300,Poxm,tin,foxo | | 1.275e-03 | -6.67 | GO\_SYNAPTIC\_MEMBRANE | MSigDB lists | GO\_SYNAPTIC\_MEMBRANE | 66 | 9 | 4161 | 170 | Frq2,Frq1,Nlg2,Ank,KaiR1D,Dys,mGluR,unc-13,Nmdar2 | | 1.276e-03 | -6.66 | GO\_ORGAN\_MORPHOGENESIS | MSigDB lists | GO\_ORGAN\_MORPHOGENESIS | 169 | 16 | 4161 | 170 | CG7483,CG17387,Su(H),RhoGAP1A,Sod1,CG17883,Raf,dpp,Ser,RpL38,TfAP-2,Tor,Awh,Treh,dsh,fz2 | | 1.282e-03 | -6.66 | GO\_POSITIVE\_REGULATION\_OF\_HEMOPOIESIS | MSigDB lists | GO\_POSITIVE\_REGULATION\_OF\_HEMOPOIESIS | 21 | 5 | 4161 | 170 | sxc,Pkc53E,Ser,IP3K2,Nha1 | | 1.312e-03 | -6.64 | presynaptic active zone | cellular component | GO:0048786 | 28 | 6 | 11940 | 538 | unc-13,cpx,RhoGAP100F,Fife,Syx1A,KaiR1D | | 1.340e-03 | -6.62 | positive regulation of macromolecule biosynthetic process | biological process | GO:0010557 | 552 | 43 | 12345 | 597 | TfAP-2,CG10462,CG12071,tin,Sox102F,Mrtf,GATAe,Lpin,foxo,CG4374,opa,mid,ems,sno,Atf6,Hr96,CG14252,CycT,EcR,Awh,Antp,CrebB,TfIIFbeta,B-H2,scro,polybromo,Tor,Su(H),kay,mnb,CG43689,bi,Iswi,Eip74EF,tyf,gce,CG42830,rictor,hth,retn,esg,CG1894,bol | | 1.348e-03 | -6.61 | GO\_NEURON\_PROJECTION | MSigDB lists | GO\_NEURON\_PROJECTION | 254 | 21 | 4161 | 170 | Sod1,sxc,KaiR1D,sky,Frq1,CG7483,LRP1,unc-13,Raf,Pak,Khc,Frq2,Cln3,Maf1,Tor,Ank,Sh,Syx1A,mGluR,Dys,Octbeta2R | | 1.361e-03 | -6.60 | cardiac cell fate commitment | biological process | GO:0060911 | 11 | 4 | 12345 | 597 | tin,Ubx,EcR,mid | | 1.418e-03 | -6.56 | SPEC | smart domains | SM00150 | 9 | 4 | 5086 | 315 | Dys,Msp300,Actn,zormin | | 1.422e-03 | -6.56 | GO\_POSITIVE\_REGULATION\_OF\_SECRETION | MSigDB lists | GO\_POSITIVE\_REGULATION\_OF\_SECRETION | 67 | 9 | 4161 | 170 | EcR,Pkc98E,chb,MCU,unc-13,Syx1A,csw,Frq2,Frq1 | | 1.423e-03 | -6.55 | GO\_POSITIVE\_REGULATION\_OF\_CATALYTIC\_ACTIVITY | MSigDB lists | GO\_POSITIVE\_REGULATION\_OF\_CATALYTIC\_ACTIVITY | 401 | 29 | 4161 | 170 | fz2,REG,scramb1,Nmdar2,dsh,Octbeta2R,csw,step,CG17698,CG42672,CG34408,Stlk,EcR,eIF2Bgamma,Pkc98E,dpp,CG5522,LpR2,Tsp96F,Raf,Pak,Sod1,sxc,RhoGAP1A,Pkc53E,wbl,CG17883,sky,Cdep | | 1.429e-03 | -6.55 | RNA polymerase II distal enhancer sequence-specific DNA binding | molecular function | GO:0000980 | 27 | 6 | 12176 | 580 | tin,Su(H),Antp,ems,Ubx,opa | | 1.440e-03 | -6.54 | MIKKELSEN\_ES\_ICP\_WITH\_H3K4ME3\_AND\_H3K27ME3 | MSigDB lists | MIKKELSEN\_ES\_ICP\_WITH\_H3K4ME3\_AND\_H3K27ME3 | 13 | 4 | 4161 | 170 | Pde8,mGluR,Pde6,AGO3 | | 1.440e-03 | -6.54 | GCAAGAC\_MIR431 | MSigDB lists | GCAAGAC\_MIR431 | 13 | 4 | 4161 | 170 | Rbp6,Awh,bru1,Ptp61F | | 1.449e-03 | -6.54 | cytoskeleton | cellular component | GO:0005856 | 525 | 39 | 11940 | 538 | kl-5,Cep97,CG18109,TpnC25D,spd-2,TTLL3A,Act88F,pod1,IFT46,Asator,gukh,Mlp84B,CG34435,Cam,CG13526,Dlic,CG34417,ex,msps,Gas8,chb,Act87E,Khc,Cdep,wake,Prm,Rcd6,Dys,aust,Pak,nompB,nuf,Myo81F,CG13251,Fife,CG17387,sgg,Msp300,Root | | 1.455e-03 | -6.53 | Neurotransmitter Receptor Binding And Downstream Transmission In The Postsynaptic Cell | REACTOME pathways | R-DME-112314 | 66 | 10 | 3923 | 197 | Nmdar2,Actn,Raf,KaiR1D,Cam,GluRIIC,ACXD,CG17698,Pkc53E,CrebB | | 1.463e-03 | -6.53 | regulation of circadian sleep/wake cycle, sleep | biological process | GO:0045187 | 36 | 7 | 12345 | 597 | AANAT1,TfAP-2,wake,EcR,qvr,CrebB,Sh | | 1.465e-03 | -6.53 | binding | molecular function | GO:0005488 | 5743 | 309 | 12176 | 580 | Fhos,Exn,Mur89F,kl-5,CG42390,CG13898,CG12071,CG18109,shd,crol,glob3,CG5955,Mos,CG18135,CG1434,cic,CG13676,Der-1,qvr,Hil,eIF2Bgamma,Mlp84B,CG30054,gce,cato,B-H2,D2hgdh,Obp56h,Cpsf160,ITP,scaRNA:PsiU2-35.45,opa,Cyp309a1,rst,CG3530,Ptp61F,TTLL4B,sws,Hr96,fs(1)N,Cyp6a13,jim,Src42A,CG7878,Doa,Fad2,Gprk1,CG15073,CG34426,CadN2,ND-51L2,tamo,E5,Ipk1,FoxL1,Cyp12a4,CG9380,CG40813,CG9650,ham,CG31224,Ubx,CG7362,CG1239,bru1,foxo,woc,Iswi,Stlk,Pfdn4,fz4,CG12605,Syx1A,Strn-Mlck,aust,Nlg2,bnl,bol,LpR2,Tif-IA,Poxm,Nuak1,Mur2B,Cyp304a1,ems,Yeti,pod1,tin,zormin,eIF4E7,Pde6,CG10126,LRP1,TpnC4,Pld,CG40470,sick,CG44249,CG7368,mnb,Actn,CG13251,Gas8,sdt,l(3)neo38,et,unc-13,mid,Marf1,Or42b,scro,PhKgamma,CG11970,CG4988,mGluR,PCB,Gpb5,Pkc53E,dany,Ank,CG4301,csw,stw,Raf,kek1,Sod1,Rab9Fa,grn,chb,wake,Or65c,dpp,Meltrin,rib,pirk,ex,CG9981,RYa,Achl,CG8117,nompB,Frq2,CG42673,Rat1,Asator,CG1894,Syt12,Obp56e,chinmo,Cyt-b5-r,Set2,Pak,gukh,Khc,spri,Mbs,Msp300,Dys,CG14362,CG41099,Elp3,CG9519,hth,Snoo,GATAe,CG6418,CG15093,Mondo,msi,CG32006,mamo,CG18659,Cht6,Mrtf,CG41562,barc,CG42458,RhoGAP18B,polybromo,Mob2,CG13794,CG5493,CG34417,Maf1,numb,CG32264,step,CG4622,boi,EcR,Mctp,ths,CG5921,Acbp2,Cat,retn,RhoGAP1A,Kyat,Cadps,kay,loopin-1,CG42402,mRpS5,RhoGEF3,PGRP-LA,by,CG6839,toe,Cyp4p2,fru,Pde1c,Pdfr,cpx,htk,ps,jigr1,Fer2,Frq1,luna,CG10462,CG5522,Rbp6,CG9514,CG13526,esg,Syn2,trv,dsh,CycT,CG43689,Fife,ABCD,melt,TpnC25D,AGO2,rols,esn,Eip74EF,Ank2,siz,His1:CG33837,wbl,fz2,Gld2,Or85b,fus,Pkc98E,shg,AGO3,Scp1,CG4849,CAH13,AOX1,Atf6,CG4374,IP3K2,CG5445,fw,Dlic,bun,Su(H),Cdep,Idgf6,CG12782,Cam,Vha100-5,cert,CG10188,Tk,CG7483,CG7017,Tor,tld,CG17698,Spf45,TfIIFbeta,Cyp6u1,TfAP-2,bi,scrib,msps,CG3594,scaRNA:MeU2-C41,Cyp28d1,nuf,TpnC41C,Neb-cGP,tyf,CG17883,shep,sqz,sgg,spok,AstA-R1,upd3,Ser,CG4839,CrebB,Awh,cindr,Myo81F,Cyp28d2,Antp,CG12200,CG33158,CG3556,eRF3,CG14499,GlyP | | 1.470e-03 | -6.52 | DAG and IP3 signaling | REACTOME pathways | R-DME-1489509 | 26 | 6 | 3923 | 197 | Cam,Pkc53E,Gprk1,Pkc98E,ACXD,Pde1c | | 1.470e-03 | -6.52 | PLC-gamma1 signalling | REACTOME pathways | R-DME-167021 | 26 | 6 | 3923 | 197 | Pkc98E,Gprk1,Pde1c,ACXD,Cam,Pkc53E | | 1.476e-03 | -6.52 | centrosome | cellular component | GO:0005813 | 125 | 14 | 11940 | 538 | Dlic,Rcd6,CG13251,IFT46,msps,sgg,wake,spd-2,nuf,Cam,CG13526,CG18109,chb,Cep97 | | 1.479e-03 | -6.52 | cell fate commitment | biological process | GO:0045165 | 301 | 27 | 12345 | 597 | msi,bi,Sox102F,tin,kay,Raf,ths,ham,Su(H),nuf,msps,numb,scrib,Antp,EcR,melt,esg,Ser,Ubx,hth,mid,sdt,csw,klg,egh,wake,dpp | | 1.483e-03 | -6.51 | regulation of small GTPase mediated signal transduction | biological process | GO:0051056 | 104 | 13 | 12345 | 597 | RhoGEF3,step,spri,MESK2,CG5522,CG34408,CG18659,siz,CG10188,Cdep,RhoGAP1A,Exn,chb | | 1.510e-03 | -6.50 | GO\_REGULATION\_OF\_TRANSMEMBRANE\_TRANSPORT | MSigDB lists | GO\_REGULATION\_OF\_TRANSMEMBRANE\_TRANSPORT | 81 | 10 | 4161 | 170 | Tor,Ca-beta,ZnT63C,Ank,Pkc98E,Sh,dnc,CCAP-R,AstA-R1,Dys | | 1.510e-03 | -6.50 | GO\_REGULATION\_OF\_NEURON\_PROJECTION\_DEVELOPMENT | MSigDB lists | GO\_REGULATION\_OF\_NEURON\_PROJECTION\_DEVELOPMENT | 81 | 10 | 4161 | 170 | LRP1,Tif-IA,Raf,LpR2,unc-13,Dys,Frq2,CG42672,Tor,Frq1 | | 1.545e-03 | -6.47 | muscle tissue development | biological process | GO:0060537 | 27 | 6 | 12345 | 597 | tin,Msp300,EcR,Khc,flw,Mlp84B | | 1.558e-03 | -6.46 | STAT5A\_01 | MSigDB lists | STAT5A\_01 | 43 | 7 | 4161 | 170 | Ank,CG3376,scramb1,eIF4E7,CG8272,mRpL24,Sod1 | | 1.558e-03 | -6.46 | GSE28726\_ACT\_CD4\_TCELL\_VS\_ACT\_NKTCELL\_DN | MSigDB lists | GSE28726\_ACT\_CD4\_TCELL\_VS\_ACT\_NKTCELL\_DN | 43 | 7 | 4161 | 170 | Glut1,step,Cln3,RpL38,rdgA,IP3K2,dnc | | 1.558e-03 | -6.46 | GSE3982\_MAST\_CELL\_VS\_NKCELL\_DN | MSigDB lists | GSE3982\_MAST\_CELL\_VS\_NKCELL\_DN | 43 | 7 | 4161 | 170 | KaiR1D,MESK2,step,IP3K2,Pde6,AGO3,Sh | | 1.591e-03 | -6.44 | GTGCAAA\_MIR507 | MSigDB lists | GTGCAAA\_MIR507 | 32 | 6 | 4161 | 170 | flw,Pkc98E,Awh,polybromo,KaiR1D,Ptp61F | | 1.591e-03 | -6.44 | GO\_PLATELET\_ACTIVATION | MSigDB lists | GO\_PLATELET\_ACTIVATION | 32 | 6 | 4161 | 170 | Pkc53E,Pkc98E,csw,Act87E,rdgA,scramb1 | | 1.599e-03 | -6.44 | para (paralytic) | protein interactions | 32619 | 34 | 7 | 9204 | 481 | shakB,jus,Khc,ine,Sh,kcc,esg | | 1.604e-03 | -6.44 | GO\_ASSOCIATIVE\_LEARNING | MSigDB lists | GO\_ASSOCIATIVE\_LEARNING | 22 | 5 | 4161 | 170 | Tor,Cln3,Octbeta2R,CG7483,Raf | | 1.604e-03 | -6.44 | GSE19198\_1H\_VS\_6H\_IL21\_TREATED\_TCELL\_UP | MSigDB lists | GSE19198\_1H\_VS\_6H\_IL21\_TREATED\_TCELL\_UP | 22 | 5 | 4161 | 170 | CG17698,Awh,CG5522,CG6966,bru1 | | 1.609e-03 | -6.43 | EF-hand-dom\_pair | interpro domains | IPR011992 | 111 | 14 | 11383 | 585 | TpnC4,Scp1,CG14362,CG10126,TpnC41C,Dys,TpnC25D,Actn,Frq2,CG13526,CG13898,Frq1,CG34435,Cam | | 1.661e-03 | -6.40 | - | gene3d domains | 2.60.40.10 | 195 | 23 | 4929 | 302 | trol,Ptp99A,zormin,dpr21,dpr12,side-VIII,nrm,dpr15,rst,dpr5,CG17716,wake,kek5,Strn-Mlck,boi,kek1,DIP-delta,klg,DIP-lambda,CG5445,Su(H),CG18135,dpr18 | | 1.667e-03 | -6.40 | GO\_POSITIVE\_REGULATION\_OF\_PROTEIN\_MODIFICATION\_PROCESS | MSigDB lists | GO\_POSITIVE\_REGULATION\_OF\_PROTEIN\_MODIFICATION\_PROCESS | 276 | 22 | 4161 | 170 | fz2,REG,dsh,CG3376,Octbeta2R,csw,CG17698,Tor,Der-1,CG42672,Stlk,Pkc98E,dpp,AGO3,LpR2,Tsp96F,Raf,Pak,Sod1,sxc,Pkc53E,wbl | | 1.684e-03 | -6.39 | negative regulation of nitrogen compound metabolic process | biological process | GO:0051172 | 625 | 47 | 12345 | 597 | CG12605,Acp62F,sws,Spn88Eb,bol,E5,bru1,Ubx,retn,flw,Khc,esg,CG1894,crol,mnb,sgg,cic,B-H2,polybromo,Tor,Su(H),et,raw,kek1,Iswi,CG43689,bi,msi,CG31224,CG7368,mid,Maf1,Antp,scrib,Ptp61F,REG,EcR,AGO2,tin,Sox102F,lncRNA:bxd,TfAP-2,CG12071,foxo,HPS4,toe,chinmo | | 1.697e-03 | -6.38 | cell cortex region | cellular component | GO:0099738 | 74 | 10 | 11940 | 538 | cindr,nuf,app,step,Fife,Ser,dsh,unc-13,sdt,Mbs | | 1.730e-03 | -6.36 | EGF\_2 | prosite domains | PS01186 | 55 | 10 | 6243 | 386 | trol,LRP1,CadN2,LpR2,tld,CG13654,shg,Meltrin,Ser,dsd | | 1.746e-03 | -6.35 | GSE9316\_IL6\_KO\_VS\_IFNG\_KO\_INVIVO\_EXPANDED\_CD4\_TCELL\_UP | MSigDB lists | GSE9316\_IL6\_KO\_VS\_IFNG\_KO\_INVIVO\_EXPANDED\_CD4\_TCELL\_UP | 56 | 8 | 4161 | 170 | rdgA,dpp,CG33090,chb,Pak,Cat,Dys,Pgant6 | | 1.746e-03 | -6.35 | GSE29949\_MICROGLIA\_VS\_DC\_BRAIN\_UP | MSigDB lists | GSE29949\_MICROGLIA\_VS\_DC\_BRAIN\_UP | 56 | 8 | 4161 | 170 | Sk1,CG8184,Meltrin,Iswi,eRF3,eIF4E7,Flo2,CG34408 | | 1.751e-03 | -6.35 | wupA (wings up A) | protein interactions | 32794 | 54 | 9 | 9204 | 481 | TpnC4,TpnC41C,Actn,sgg,Act88F,Scp1,dpp,flw,Prm | | 1.788e-03 | -6.33 | regulation of vesicle-mediated transport | biological process | GO:0060627 | 119 | 14 | 12345 | 597 | sky,Syt12,scrib,Pld,Fife,numb,sdt,cindr,step,Cadps,Pkc53E,Syx1A,spri,mnb | | 1.799e-03 | -6.32 | pattern specification process | biological process | GO:0007389 | 544 | 42 | 12345 | 597 | fz2,B-H2,cic,numb,sgg,msps,app,Su(H),tld,bi,boi,CadN2,hth,Doa,Pkc98E,Dys,CG7483,retn,Ubx,flw,Khc,fs(1)N,dsh,tin,TfAP-2,nuf,Raf,CG17387,wbl,Lpin,step,ems,sno,shg,dpp,csw,mid,Antp,scrib,Ser,EcR,AGO2 | | 1.811e-03 | -6.31 | CG4270 (uncharacterized protein) | protein interactions | 33408 | 11 | 4 | 9204 | 481 | CG13157,Set2,CG10164,CG15198 | | 1.854e-03 | -6.29 | CG9776 (uncharacterized protein) | protein interactions | 40546 | 18 | 5 | 9204 | 481 | CG10164,Doa,CG9436,barc,TwdlG | | 1.857e-03 | -6.29 | rhythmic process | biological process | GO:0048511 | 94 | 12 | 12345 | 597 | foxo,Fer2,ITP,CrebB,dnc,Pdfr,kay,mnb,sxc,tyf,numb,sgg | | 1.866e-03 | -6.28 | enhancer sequence-specific DNA binding | molecular function | GO:0001158 | 38 | 7 | 12176 | 580 | tin,hth,Su(H),Antp,ems,Ubx,opa | | 1.880e-03 | -6.28 | AACWWCAANK\_UNKNOWN | MSigDB lists | AACWWCAANK\_UNKNOWN | 33 | 6 | 4161 | 170 | RhoGAP1A,mRpL24,app,Dys,CG3842,chb | | 1.880e-03 | -6.28 | GO\_POSITIVE\_REGULATION\_OF\_SYNAPTIC\_TRANSMISSION | MSigDB lists | GO\_POSITIVE\_REGULATION\_OF\_SYNAPTIC\_TRANSMISSION | 33 | 6 | 4161 | 170 | Khc,Raf,KaiR1D,Syx1A,unc-13,Pkc98E | | 1.880e-03 | -6.28 | GO\_NEGATIVE\_REGULATION\_OF\_SECRETION | MSigDB lists | GO\_NEGATIVE\_REGULATION\_OF\_SECRETION | 33 | 6 | 4161 | 170 | csw,wbl,RhoGAP1A,ZnT63C,Pde8,Raf | | 1.880e-03 | -6.28 | GSE43863\_TFH\_VS\_LY6C\_INT\_CXCR5POS\_EFFECTOR\_CD4\_TCELL\_DN | MSigDB lists | GSE43863\_TFH\_VS\_LY6C\_INT\_CXCR5POS\_EFFECTOR\_CD4\_TCELL\_DN | 33 | 6 | 4161 | 170 | scramb1,Spf45,Tsp96F,Awh,Pfdn4,CG4341 | | 1.880e-03 | -6.28 | GO\_REGULATION\_OF\_ION\_HOMEOSTASIS | MSigDB lists | GO\_REGULATION\_OF\_ION\_HOMEOSTASIS | 33 | 6 | 4161 | 170 | dnc,Pkc98E,Dys,Ank,ZnT63C,CCAP-R | | 1.887e-03 | -6.27 | salivary gland cell autophagic cell death | biological process | GO:0035071 | 28 | 6 | 12345 | 597 | EcR,hdc,Cam,Pak,IP3K2,Eip74EF | | 1.907e-03 | -6.26 | GO\_CHEMICAL\_HOMEOSTASIS | MSigDB lists | GO\_CHEMICAL\_HOMEOSTASIS | 209 | 18 | 4161 | 170 | ZnT63C,Ank,Ca-beta,CG4341,Der-1,csw,Cln3,Ptp61F,AstA-R1,mino,KaiR1D,CG17883,sxc,Sod1,GlyP,MCU,EcR,Pkc98E | | 1.910e-03 | -6.26 | apical part of cell | cellular component | GO:0045177 | 157 | 16 | 11940 | 538 | rst,bbg,Pde6,sdt,Cln3,kek1,dsh,ex,Inx2,ZnT63C,app,CG5921,cindr,nuf,Mbs,Ser | | 1.930e-03 | -6.25 | 3',5'-cyclic-GMP phosphodiesterase activity | molecular function | GO:0047555 | 6 | 3 | 12176 | 580 | Pde6,Pde9,Pde1c | | 1.946e-03 | -6.24 | GO\_REGULATION\_OF\_PROTEIN\_SECRETION | MSigDB lists | GO\_REGULATION\_OF\_PROTEIN\_SECRETION | 70 | 9 | 4161 | 170 | csw,Pkc53E,wbl,Pde8,EcR,Pkc98E,mino,Syx1A,MCU | | 1.946e-03 | -6.24 | GO\_COGNITION | MSigDB lists | GO\_COGNITION | 70 | 9 | 4161 | 170 | Nlg2,CG7483,Tor,Awh,Cln3,Octbeta2R,Raf,AstA-R1,LpR2 | | 1.947e-03 | -6.24 | GO\_CELL\_ACTIVATION | MSigDB lists | GO\_CELL\_ACTIVATION | 98 | 11 | 4161 | 170 | Act87E,rdgA,Pkc98E,fz2,Ptp61F,Raf,scramb1,IP3K2,Su(H),Pkc53E,csw | | 1.952e-03 | -6.24 | GO\_REGULATION\_OF\_NEUROTRANSMITTER\_SECRETION | MSigDB lists | GO\_REGULATION\_OF\_NEUROTRANSMITTER\_SECRETION | 14 | 4 | 4161 | 170 | ZnT63C,Syx1A,Raf,unc-13 | | 1.952e-03 | -6.24 | GO\_GLUTAMATE\_RECEPTOR\_SIGNALING\_PATHWAY | MSigDB lists | GO\_GLUTAMATE\_RECEPTOR\_SIGNALING\_PATHWAY | 14 | 4 | 4161 | 170 | mGluR,Nmdar2,KaiR1D,Cln3 | | 1.952e-03 | -6.24 | GO\_REGULATION\_OF\_SODIUM\_ION\_TRANSPORT | MSigDB lists | GO\_REGULATION\_OF\_SODIUM\_ION\_TRANSPORT | 14 | 4 | 4161 | 170 | Pkc98E,Dys,Ank,Octbeta2R | | 1.952e-03 | -6.24 | GO\_RETINA\_HOMEOSTASIS | MSigDB lists | GO\_RETINA\_HOMEOSTASIS | 14 | 4 | 4161 | 170 | Act87E,Act88F,Sod1,Root | | 1.952e-03 | -6.24 | KEGG\_ADIPOCYTOKINE\_SIGNALING\_PATHWAY | MSigDB lists | KEGG\_ADIPOCYTOKINE\_SIGNALING\_PATHWAY | 14 | 4 | 4161 | 170 | CG3961,CG17698,csw,Tor | | 1.952e-03 | -6.24 | GO\_POSITIVE\_REGULATION\_OF\_MYELOID\_CELL\_DIFFERENTIATION | MSigDB lists | GO\_POSITIVE\_REGULATION\_OF\_MYELOID\_CELL\_DIFFERENTIATION | 14 | 4 | 4161 | 170 | Nha1,sxc,Pkc53E,Ser | | 1.958e-03 | -6.24 | axon | cellular component | GO:0030424 | 233 | 21 | 11940 | 538 | klg,cpx,pod1,unc-13,mnb,chb,Gr39a,Khc,Ptp99A,kcc,sky,sgg,Syx1A,gukh,Gr22c,hdc,Ser,Ank2,nrv2,g,path | | 1.963e-03 | -6.23 | cytoskeletal anchoring at plasma membrane | biological process | GO:0007016 | 12 | 4 | 12345 | 597 | p130CAS,Ank,Actn,by | | 1.963e-03 | -6.23 | calcium-ion regulated exocytosis | biological process | GO:0017156 | 12 | 4 | 12345 | 597 | Syt12,Cadps,unc-13,Fife | | 1.983e-03 | -6.22 | GSE2770\_UNTREATED\_VS\_ACT\_CD4\_TCELL\_48H\_DN | MSigDB lists | GSE2770\_UNTREATED\_VS\_ACT\_CD4\_TCELL\_48H\_DN | 23 | 5 | 4161 | 170 | Hr96,Syx1A,Ank,IP3K2,dnc | | 1.983e-03 | -6.22 | ACAGGGT\_MIR10A\_MIR10B | MSigDB lists | ACAGGGT\_MIR10A\_MIR10B | 23 | 5 | 4161 | 170 | step,CG41099,dsh,bru1,mGluR | | 2.003e-03 | -6.21 | YOSHIMURA\_MAPK8\_TARGETS\_UP | MSigDB lists | YOSHIMURA\_MAPK8\_TARGETS\_UP | 298 | 23 | 4161 | 170 | CG14105,rdgA,GlyP,unc-13,Pkc53E,Sod1,Sh,dnc,CG5493,Argl,mino,Octbeta2R,Treh,mGluR,Syx1A,Nmdar2,IP3K2,CG4849,Ca-beta,CG6418,csw,Dlic,Ank | | 2.017e-03 | -6.21 | neurotransmitter receptor transport | biological process | GO:0099637 | 6 | 3 | 12345 | 597 | g,scrib,Khc | | 2.034e-03 | -6.20 | hematopoietic or lymphoid organ development | biological process | GO:0048534 | 95 | 12 | 12345 | 597 | Raf,Tor,ths,Su(H),RhoL,dpp,shg,Elp3,tin,Ser,chb,Antp | | 2.034e-03 | -6.20 | immune system development | biological process | GO:0002520 | 95 | 12 | 12345 | 597 | Ser,chb,Antp,dpp,ths,Tor,Raf,Su(H),RhoL,tin,shg,Elp3 | | 2.048e-03 | -6.19 | GO\_POSITIVE\_REGULATION\_OF\_NEURON\_PROJECTION\_DEVELOPMENT | MSigDB lists | GO\_POSITIVE\_REGULATION\_OF\_NEURON\_PROJECTION\_DEVELOPMENT | 45 | 7 | 4161 | 170 | LpR2,unc-13,Raf,Dys,Tor,Tif-IA,CG42672 | | 2.067e-03 | -6.18 | upd1 (unpaired 1) | protein interactions | 32813 | 77 | 11 | 9204 | 481 | flw,esg,jim,CG17574,dpp,Ser,spri,hdc,et,Mob2,bun | | 2.077e-03 | -6.18 | GO\_NEGATIVE\_REGULATION\_OF\_PEPTIDYL\_SERINE\_PHOSPHORYLATION | MSigDB lists | GO\_NEGATIVE\_REGULATION\_OF\_PEPTIDYL\_SERINE\_PHOSPHORYLATION | 7 | 3 | 4161 | 170 | dnc,sxc,Dys | | 2.077e-03 | -6.18 | DUNNE\_TARGETS\_OF\_AML1\_MTG8\_FUSION\_UP | MSigDB lists | DUNNE\_TARGETS\_OF\_AML1\_MTG8\_FUSION\_UP | 7 | 3 | 4161 | 170 | CG14480,Aldh-III,Cat | | 2.077e-03 | -6.18 | GO\_NEGATIVE\_REGULATION\_OF\_LYASE\_ACTIVITY | MSigDB lists | GO\_NEGATIVE\_REGULATION\_OF\_LYASE\_ACTIVITY | 7 | 3 | 4161 | 170 | Pkc53E,AstA-R1,mGluR | | 2.077e-03 | -6.18 | GO\_DENSE\_CORE\_GRANULE | MSigDB lists | GO\_DENSE\_CORE\_GRANULE | 7 | 3 | 4161 | 170 | Frq1,Frq2,Sod1 | | 2.077e-03 | -6.18 | GO\_POSITIVE\_REGULATION\_OF\_LIPID\_CATABOLIC\_PROCESS | MSigDB lists | GO\_POSITIVE\_REGULATION\_OF\_LIPID\_CATABOLIC\_PROCESS | 7 | 3 | 4161 | 170 | Octbeta2R,ABCD,Pkc98E | | 2.077e-03 | -6.18 | GO\_REGULATION\_OF\_SODIUM\_ION\_TRANSMEMBRANE\_TRANSPORTER\_ACTIVITY | MSigDB lists | GO\_REGULATION\_OF\_SODIUM\_ION\_TRANSMEMBRANE\_TRANSPORTER\_ACTIVITY | 7 | 3 | 4161 | 170 | Ank,Dys,Pkc98E | | 2.077e-03 | -6.18 | GO\_REGULATION\_OF\_POLYSACCHARIDE\_METABOLIC\_PROCESS | MSigDB lists | GO\_REGULATION\_OF\_POLYSACCHARIDE\_METABOLIC\_PROCESS | 7 | 3 | 4161 | 170 | Tor,PhKgamma,flw | | 2.085e-03 | -6.17 | circadian behavior | biological process | GO:0048512 | 71 | 10 | 12345 | 597 | kay,sxc,tyf,sgg,ITP,foxo,Fer2,CrebB,dnc,Pdfr | | 2.132e-03 | -6.15 | negative regulation of transcription, DNA-templated | biological process | GO:0045892 | 371 | 31 | 12345 | 597 | cic,TfAP-2,B-H2,polybromo,Su(H),CG12071,tin,crol,Sox102F,lncRNA:bxd,CG43689,bi,toe,foxo,Iswi,HPS4,CG7368,Maf1,mid,CG31224,CG12605,Ubx,retn,esg,Khc,EcR,CG1894,AGO2,E5,Antp,scrib | | 2.152e-03 | -6.14 | GO\_POSITIVE\_REGULATION\_OF\_CELL\_COMMUNICATION | MSigDB lists | GO\_POSITIVE\_REGULATION\_OF\_CELL\_COMMUNICATION | 355 | 26 | 4161 | 170 | Ank,csw,CG42672,Tor,IP3K2,Syx1A,Octbeta2R,Cat,Ptp61F,dsh,fz2,REG,KaiR1D,Pkc53E,Su(H),wbl,Sod1,Tsp96F,Raf,unc-13,Khc,Pak,MCU,Pkc98E,Ser,dpp | | 2.200e-03 | -6.12 | neuroblast development | biological process | GO:0014019 | 20 | 5 | 12345 | 597 | ems,Antp,numb,sqz,mid | | 2.207e-03 | -6.12 | GSE4590\_PRE\_BCELL\_VS\_SMALL\_PRE\_BCELL\_UP | MSigDB lists | GSE4590\_PRE\_BCELL\_VS\_SMALL\_PRE\_BCELL\_UP | 34 | 6 | 4161 | 170 | Frq2,Glut1,Frq1,Nmdar2,CG41099,Tsp96F | | 2.225e-03 | -6.11 | regulation of Ras protein signal transduction | biological process | GO:0046578 | 96 | 12 | 12345 | 597 | MESK2,siz,CG18659,CG5522,RhoGEF3,spri,step,chb,Exn,CG10188,Cdep,RhoGAP1A | | 2.245e-03 | -6.10 | Tyr\_Pase\_cat | interpro domains | IPR003595 | 19 | 5 | 11383 | 585 | csw,Ptp36E,CG3530,Ptp61F,Ptp99A | | 2.265e-03 | -6.09 | RNA polymerase II transcription cofactor binding | molecular function | GO:0001224 | 2 | 2 | 12176 | 580 | EcR,mid | | 2.265e-03 | -6.09 | aralkylamine N-acetyltransferase activity | molecular function | GO:0004059 | 2 | 2 | 12176 | 580 | AANATL2,AANAT1 | | 2.283e-03 | -6.08 | autophagic cell death | biological process | GO:0048102 | 29 | 6 | 12345 | 597 | EcR,hdc,Pak,Eip74EF,IP3K2,Cam | | 2.283e-03 | -6.08 | intestinal stem cell homeostasis | biological process | GO:0036335 | 29 | 6 | 12345 | 597 | Src42A,upd3,GATAe,Cat,bun,esg | | 2.287e-03 | -6.08 | EGF\_3 | prosite domains | PS50026 | 57 | 10 | 6243 | 386 | Meltrin,dsd,Ser,CadN2,LpR2,tld,LRP1,trol,shg,CG13654 | | 2.292e-03 | -6.08 | GO\_NEGATIVE\_REGULATION\_OF\_TRANSPORT | MSigDB lists | GO\_NEGATIVE\_REGULATION\_OF\_TRANSPORT | 100 | 11 | 4161 | 170 | Ank,ZnT63C,Pde8,wbl,RhoGAP1A,csw,sxc,Tor,Raf,EcR,Pkc98E | | 2.310e-03 | -6.07 | cell fate specification | biological process | GO:0001708 | 84 | 11 | 12345 | 597 | Ubx,bi,melt,esg,Antp,scrib,dpp,numb,ths,mid,tin | | 2.316e-03 | -6.07 | intracellular signal transduction | biological process | GO:0035556 | 373 | 31 | 12345 | 597 | Pde9,Doa,Pak,CG13526,Pkc53E,rictor,RhoGAP100F,PhKgamma,dnc,Cdep,Octbeta2R,Pkc98E,CG13898,Raf,Tor,Pld,TpnC25D,rdgA,unc-13,Cam,Ca-beta,Stacl,Rab9Fa,Tusp,dsh,ACXD,Pdfr,Stlk,CG11318,CG17698,TpnC41C | | 2.334e-03 | -6.06 | GO\_REGULATION\_OF\_MYELOID\_CELL\_DIFFERENTIATION | MSigDB lists | GO\_REGULATION\_OF\_MYELOID\_CELL\_DIFFERENTIATION | 46 | 7 | 4161 | 170 | Nha1,Ptp61F,sxc,Pkc53E,Tor,IP3K2,Ser | | 2.334e-03 | -6.06 | GSE21360\_TERTIARY\_VS\_QUATERNARY\_MEMORY\_CD8\_TCELL\_UP | MSigDB lists | GSE21360\_TERTIARY\_VS\_QUATERNARY\_MEMORY\_CD8\_TCELL\_UP | 46 | 7 | 4161 | 170 | app,sws,step,CG5493,Acbp2,dnc,LRP1 | | 2.334e-03 | -6.06 | GSE11924\_TH2\_VS\_TH17\_CD4\_TCELL\_UP | MSigDB lists | GSE11924\_TH2\_VS\_TH17\_CD4\_TCELL\_UP | 46 | 7 | 4161 | 170 | Pkc53E,mino,Ser,sdt,TfAP-2,dpp,Nha1 | | 2.334e-03 | -6.06 | PID\_PDGFRB\_PATHWAY | MSigDB lists | PID\_PDGFRB\_PATHWAY | 46 | 7 | 4161 | 170 | LRP1,Pkc53E,Pkc98E,csw,Pak,Ptp61F,Raf | | 2.335e-03 | -6.06 | specification of segmental identity, antennal segment | biological process | GO:0007383 | 2 | 2 | 12345 | 597 | Antp,hth | | 2.335e-03 | -6.06 | negative regulation of oxidative stress-induced cell death | biological process | GO:1903202 | 2 | 2 | 12345 | 597 | Pde8,Fer2 | | 2.335e-03 | -6.06 | Rac protein signal transduction | biological process | GO:0016601 | 2 | 2 | 12345 | 597 | Pak,Cdep | | 2.335e-03 | -6.06 | nuclear speck organization | biological process | GO:0035063 | 2 | 2 | 12345 | 597 | lncRNA:Hsromega,Iswi | | 2.335e-03 | -6.06 | dense core granule exocytosis | biological process | GO:1990504 | 2 | 2 | 12345 | 597 | Cadps,unc-13 | | 2.335e-03 | -6.06 | regulation of cAMP-dependent protein kinase activity | biological process | GO:2000479 | 2 | 2 | 12345 | 597 | step,sws | | 2.386e-03 | -6.04 | GO\_REGULATION\_OF\_IMMUNE\_SYSTEM\_PROCESS | MSigDB lists | GO\_REGULATION\_OF\_IMMUNE\_SYSTEM\_PROCESS | 302 | 23 | 4161 | 170 | sxc,Sod1,Pkc53E,RhoGAP1A,Flo2,EcR,Pkc98E,Ser,Act87E,Tsp96F,Pak,csw,Ca-beta,Tor,IP3K2,fz2,mino,Syn2,REG,dnc,scramb1,Nha1,Ptp61F | | 2.401e-03 | -6.03 | GO\_REGULATION\_OF\_CELL\_DEVELOPMENT | MSigDB lists | GO\_REGULATION\_OF\_CELL\_DEVELOPMENT | 163 | 15 | 4161 | 170 | LRP1,Ser,dpp,Tif-IA,Raf,Nha1,unc-13,LpR2,Dys,chb,Frq2,CG42672,Tor,eIF4E7,Frq1 | | 2.410e-03 | -6.03 | poe (purity of essence) | protein interactions | 46243 | 19 | 5 | 9204 | 481 | flw,ine,rictor,Cam,sgg | | 2.412e-03 | -6.03 | negative regulation of transcription by RNA polymerase II | biological process | GO:0000122 | 221 | 21 | 12345 | 597 | B-H2,cic,TfAP-2,Su(H),CG12071,polybromo,foxo,toe,bi,CG43689,CG12605,CG31224,CG7368,mid,E5,Antp,scrib,Ubx,retn,EcR,esg | | 2.423e-03 | -6.02 | GO\_PHOSPHORIC\_DIESTER\_HYDROLASE\_ACTIVITY | MSigDB lists | GO\_PHOSPHORIC\_DIESTER\_HYDROLASE\_ACTIVITY | 24 | 5 | 4161 | 170 | Pde8,CG3376,Pde1c,Pde6,dnc | | 2.423e-03 | -6.02 | EPPERT\_HSC\_R | MSigDB lists | EPPERT\_HSC\_R | 24 | 5 | 4161 | 170 | Iswi,sxc,Rbp6,Mctp,Ank | | 2.452e-03 | -6.01 | positive regulation of intracellular signal transduction | biological process | GO:1902533 | 178 | 18 | 12345 | 597 | ex,Raf,bnl,MESK2,Tor,lncRNA:Hsromega,Lpin,Src42A,olf186-F,Pde8,siz,step,rictor,Mos,Exn,melt,RhoGAP1A,CG10188 | | 2.469e-03 | -6.00 | digestive system development | biological process | GO:0055123 | 110 | 13 | 12345 | 597 | Inx2,GATAe,shg,rib,tin,opa,dpp,shd,Antp,raw,EcR,tnc,Ubx | | 2.469e-03 | -6.00 | digestive tract development | biological process | GO:0048565 | 110 | 13 | 12345 | 597 | rib,shg,Inx2,GATAe,tin,opa,dpp,shd,raw,Antp,tnc,EcR,Ubx | | 2.477e-03 | -6.00 | GO\_REGULATION\_OF\_ION\_TRANSPORT | MSigDB lists | GO\_REGULATION\_OF\_ION\_TRANSPORT | 116 | 12 | 4161 | 170 | Sh,dnc,Pkc98E,Dys,Octbeta2R,Khc,AstA-R1,CCAP-R,Ca-beta,Tor,Ank,ZnT63C | | 2.513e-03 | -5.99 | CTTTGA\_LEF1\_Q2 | MSigDB lists | CTTTGA\_LEF1\_Q2 | 285 | 22 | 4161 | 170 | Ser,CG6966,CG15093,CG14480,ABCD,fus,eIF4E7,polybromo,sdt,wake,RhoGAP1A,Dys,Pde1c,CCAP-R,scramb1,Syx1A,Rbp6,Pde6,Cadps,Syn2,Ank,CG3961 | | 2.520e-03 | -5.98 | yuri (yuri gagarin) | protein interactions | 34894 | 6 | 3 | 9204 | 481 | IP3K2,Ubx,Rbp6 | | 2.520e-03 | -5.98 | Ire1 (Inositol-requiring enzyme-1) | protein interactions | 42358 | 6 | 3 | 9204 | 481 | CG4622,nemy,Vha36-3 | | 2.564e-03 | -5.97 | PE/DAG-bd | interpro domains | IPR002219 | 28 | 6 | 11383 | 585 | Pkc98E,Raf,Stacl,rdgA,Pkc53E,unc-13 | | 2.569e-03 | -5.96 | GO\_REGULATION\_OF\_CELLULAR\_COMPONENT\_MOVEMENT | MSigDB lists | GO\_REGULATION\_OF\_CELLULAR\_COMPONENT\_MOVEMENT | 132 | 13 | 4161 | 170 | CG17387,Elp3,RhoGAP1A,Pkc53E,Pak,chb,Raf,Tsp96F,dnc,LRP1,Ser,dpp,Pkc98E | | 2.575e-03 | -5.96 | GSE43863\_DAY6\_EFF\_VS\_DAY150\_MEM\_TH1\_CD4\_TCELL\_UP | MSigDB lists | GSE43863\_DAY6\_EFF\_VS\_DAY150\_MEM\_TH1\_CD4\_TCELL\_UP | 35 | 6 | 4161 | 170 | pod1,Frq1,Nha1,Frq2,Pkc53E,Su(H) | | 2.575e-03 | -5.96 | GSE14415\_ACT\_TCONV\_VS\_ACT\_NATURAL\_TREG\_UP | MSigDB lists | GSE14415\_ACT\_TCONV\_VS\_ACT\_NATURAL\_TREG\_UP | 35 | 6 | 4161 | 170 | Sh,Stt3B,dsd,CG5339,sno,CNT1 | | 2.577e-03 | -5.96 | BOYLAN\_MULTIPLE\_MYELOMA\_PCA3\_DN | MSigDB lists | BOYLAN\_MULTIPLE\_MYELOMA\_PCA3\_DN | 15 | 4 | 4161 | 170 | Meltrin,Cdep,CG10126,Flo2 | | 2.578e-03 | -5.96 | crb (crumbs) | protein interactions | 42896 | 57 | 9 | 9204 | 481 | scrib,rib,Mbs,Pak,shg,Ser,sdt,ex,flw | | 2.637e-03 | -5.94 | RPEL\_repeat | interpro domains | IPR004018 | 2 | 2 | 11383 | 585 | Mrtf,CG32264 | | 2.637e-03 | -5.94 | NDRG | interpro domains | IPR004142 | 2 | 2 | 11383 | 585 | MESK2,CG2082 | | 2.637e-03 | -5.94 | Ankyrin\_UPA | interpro domains | IPR040745 | 2 | 2 | 11383 | 585 | Ank,Ank2 | | 2.637e-03 | -5.94 | CAPS\_dom | interpro domains | IPR010439 | 2 | 2 | 11383 | 585 | unc-13,Cadps | | 2.637e-03 | -5.94 | MSI\_RRM2 | interpro domains | IPR034126 | 2 | 2 | 11383 | 585 | Rbp6,msi | | 2.638e-03 | -5.94 | ovarian follicle cell migration | biological process | GO:0007297 | 124 | 14 | 12345 | 597 | ex,egh,rols,bbg,RhoL,cindr,Raf,kay,dsh,Mrtf,shg,bun,EcR,shd | | 2.649e-03 | -5.93 | GSE19941\_IL10\_KO\_VS\_IL10\_KO\_AND\_NFKBP50\_KO\_UNSTIM\_MACROPHAGE\_DN | MSigDB lists | GSE19941\_IL10\_KO\_VS\_IL10\_KO\_AND\_NFKBP50\_KO\_UNSTIM\_MACROPHAGE\_DN | 47 | 7 | 4161 | 170 | KrT95D,Frq1,Frq2,sdt,CG17883,AGO3,LRP1 | | 2.653e-03 | -5.93 | axon part | cellular component | GO:0033267 | 105 | 12 | 11940 | 538 | sky,Syx1A,pod1,cpx,g,unc-13,gukh,mnb,chb,Khc,Ser,Ank2 | | 2.674e-03 | -5.92 | EGFR interacts with phospholipase C-gamma | REACTOME pathways | R-DME-212718 | 29 | 6 | 3923 | 197 | Cam,Pkc53E,Pkc98E,Gprk1,Pde1c,ACXD | | 2.687e-03 | -5.92 | GO\_SYNAPTIC\_SIGNALING | MSigDB lists | GO\_SYNAPTIC\_SIGNALING | 102 | 11 | 4161 | 170 | kcc,Flo2,KaiR1D,Ca-beta,mGluR,Octbeta2R,Nmdar2,unc-13,AstA-R1,Syx1A,Cadps | | 2.712e-03 | -5.91 | negative regulation of macromolecule metabolic process | biological process | GO:0010605 | 777 | 55 | 12345 | 597 | toe,chinmo,foxo,HPS4,AGO3,TfAP-2,CG12071,tin,Sox102F,GATAe,lncRNA:bxd,REG,Ptp61F,EcR,AGO2,Antp,scrib,dpp,CG7368,Achl,Maf1,mid,Marf1,CG31224,CG43689,bi,msi,raw,et,tld,kek1,Iswi,cic,sgg,numb,B-H2,polybromo,Su(H),Tor,hdc,crol,mnb,retn,Ubx,flw,Khc,esg,CG1894,E5,bru1,bol,sws,Spn88Eb,CG12605,Acp62F | | 2.726e-03 | -5.91 | CG11110 (uncharacterized protein) | protein interactions | 37337 | 2 | 2 | 9204 | 481 | Ptp99A,E5 | | 2.726e-03 | -5.91 | CG17344 (uncharacterized protein) | protein interactions | 35185 | 2 | 2 | 9204 | 481 | Dmtn,CG9650 | | 2.726e-03 | -5.91 | wry (weary) | protein interactions | 33361 | 2 | 2 | 9204 | 481 | Ser,Poxm | | 2.726e-03 | -5.91 | CG17278 (uncharacterized protein) | protein interactions | 50050 | 2 | 2 | 9204 | 481 | sgg,dsh | | 2.726e-03 | -5.91 | CG10343 (uncharacterized protein) | protein interactions | 35127 | 2 | 2 | 9204 | 481 | rictor,Stacl | | 2.728e-03 | -5.90 | cardioblast differentiation | biological process | GO:0010002 | 13 | 4 | 12345 | 597 | dpp,mid,EcR,tin | | 2.728e-03 | -5.90 | wing and notum subfield formation | biological process | GO:0035309 | 13 | 4 | 12345 | 597 | dsh,Raf,sgg,dpp | | 2.750e-03 | -5.90 | DUF1041 | pfam domains | PF06292 | 2 | 2 | 10480 | 550 | Cadps,unc-13 | | 2.750e-03 | -5.90 | RPEL | pfam domains | PF02755 | 2 | 2 | 10480 | 550 | Mrtf,CG32264 | | 2.750e-03 | -5.90 | UPA\_2 | pfam domains | PF17809 | 2 | 2 | 10480 | 550 | Ank,Ank2 | | 2.750e-03 | -5.90 | Ndr | pfam domains | PF03096 | 2 | 2 | 10480 | 550 | CG2082,MESK2 | | 2.774e-03 | -5.89 | negative regulation of synaptic transmission | biological process | GO:0050805 | 21 | 5 | 12345 | 597 | qvr,cpx,Ank2,sky,Nlg2 | | 2.838e-03 | -5.86 | cellular response to chemical stimulus | biological process | GO:0070887 | 378 | 31 | 12345 | 597 | rictor,Sh,Atf6,gce,Der-1,dpp,opa,csw,Syt12,Ptp61F,Cat,Gpb5,eIF2Bgamma,EcR,CG7362,Mrtf,Cam,Set2,ex,Tor,ths,bnl,Raf,foxo,et,St4,Iswi,msi,bun,Lpin,Sod1 | | 2.843e-03 | -5.86 | regulation of MAPK cascade | biological process | GO:0043408 | 125 | 14 | 12345 | 597 | Cln3,lncRNA:Hsromega,Pak,step,bnl,Raf,dpp,sgg,Pde8,Src42A,raw,flw,Mos,Ptp61F | | 2.846e-03 | -5.86 | homeostatic process | biological process | GO:0042592 | 475 | 37 | 12345 | 597 | Pdfr,Vha100-5,upd3,Src42A,ZnT63C,Mondo,Fhos,Ank2,esg,ine,KaiR1D,Dys,CG40178,woc,nrv2,sws,shakB,l(2)01289,CG33310,Cln3,Glut1,Lpin,bun,foxo,Raf,rdgA,GATAe,Cam,EcR,Acbp2,Cat,ddbt,Qsox2,MCU,Hr96,shg,mag | | 2.862e-03 | -5.86 | GO\_AXON | MSigDB lists | GO\_AXON | 118 | 12 | 4161 | 170 | Sh,unc-13,Octbeta2R,mGluR,Pak,Frq2,Sod1,Maf1,sky,KaiR1D,Ank,Frq1 | | 2.916e-03 | -5.84 | regulation of protein modification process | biological process | GO:0031399 | 347 | 29 | 12345 | 597 | sws,dpp,Pak,Cln3,rictor,step,flw,Ptp61F,CycT,Mos,Tor,bnl,Raf,Yeti,sgg,SteXh:CG42398,mnb,lncRNA:Hsromega,kay,chinmo,Stlk,CG6966,Pde8,Ste:CG33247,kek1,CG41520,et,raw,Src42A | | 2.923e-03 | -5.84 | mam (mastermind) | protein interactions | 36555 | 69 | 10 | 9204 | 481 | fz2,sgg,Su(H),Raf,sno,EcR,Antp,dsh,Ser,RpL38 | | 2.931e-03 | -5.83 | GO\_INOSITOL\_LIPID\_MEDIATED\_SIGNALING | MSigDB lists | GO\_INOSITOL\_LIPID\_MEDIATED\_SIGNALING | 25 | 5 | 4161 | 170 | csw,Frq2,sxc,Tor,Frq1 | | 2.931e-03 | -5.83 | GO\_ACTIVATION\_OF\_MAPK\_ACTIVITY | MSigDB lists | GO\_ACTIVATION\_OF\_MAPK\_ACTIVITY | 25 | 5 | 4161 | 170 | Tsp96F,dpp,wbl,Sod1,csw | | 2.931e-03 | -5.83 | WNT\_UP.V1\_DN | MSigDB lists | WNT\_UP.V1\_DN | 25 | 5 | 4161 | 170 | Meltrin,step,Cln3,CG5493,Sh | | 2.931e-03 | -5.83 | GSE16450\_IMMATURE\_VS\_MATURE\_NEURON\_CELL\_LINE\_6H\_IFNA\_STIM\_UP | MSigDB lists | GSE16450\_IMMATURE\_VS\_MATURE\_NEURON\_CELL\_LINE\_6H\_IFNA\_STIM\_UP | 25 | 5 | 4161 | 170 | CG17387,Src42A,CG4341,Sh,Rbp6 | | 2.931e-03 | -5.83 | GO\_EAR\_MORPHOGENESIS | MSigDB lists | GO\_EAR\_MORPHOGENESIS | 25 | 5 | 4161 | 170 | RpL38,TfAP-2,RhoGAP1A,Sod1,dsh | | 2.931e-03 | -5.83 | GO\_LYMPHOCYTE\_DIFFERENTIATION | MSigDB lists | GO\_LYMPHOCYTE\_DIFFERENTIATION | 25 | 5 | 4161 | 170 | IP3K2,fz2,Su(H),Ptp61F,Raf | | 2.931e-03 | -5.83 | GSE9988\_LPS\_VS\_LOW\_LPS\_MONOCYTE\_DN | MSigDB lists | GSE9988\_LPS\_VS\_LOW\_LPS\_MONOCYTE\_DN | 25 | 5 | 4161 | 170 | CG4341,Syn2,melt,Nlg2,PCB | | 2.931e-03 | -5.83 | GO\_TRANSCRIPTIONAL\_ACTIVATOR\_ACTIVITY\_RNA\_POLYMERASE\_II\_TRANSCRIPTION\_REGULATORY\_REGION\_SEQUENCE\_SPECIFIC\_BINDING | MSigDB lists | GO\_TRANSCRIPTIONAL\_ACTIVATOR\_ACTIVITY\_RNA\_POLYMERASE\_II\_TRANSCRIPTION\_REGULATORY\_REGION\_SEQUENCE\_SPECIFIC\_BINDING | 25 | 5 | 4161 | 170 | scramb1,Hr96,TfAP-2,Su(H),EcR | | 2.933e-03 | -5.83 | GO\_REGULATION\_OF\_NERVOUS\_SYSTEM\_DEVELOPMENT | MSigDB lists | GO\_REGULATION\_OF\_NERVOUS\_SYSTEM\_DEVELOPMENT | 150 | 14 | 4161 | 170 | LpR2,unc-13,Raf,dsh,Dys,Ser,Tif-IA,dpp,LRP1,eIF4E7,Frq1,Frq2,Tor,CG42672 | | 2.950e-03 | -5.83 | GO\_CELL\_JUNCTION | MSigDB lists | GO\_CELL\_JUNCTION | 307 | 23 | 4161 | 170 | RpL38,Frq2,Ank,Cadps,fz2,Mlp84B,chb,Cat,Dys,Nmdar2,Syx1A,sdt,Frq1,Nlg2,Flo2,KaiR1D,Ser,Act87E,LRP1,flw,Pak,unc-13,Tsp96F | | 2.954e-03 | -5.82 | negative regulation of RNA biosynthetic process | biological process | GO:1902679 | 379 | 31 | 12345 | 597 | retn,Ubx,CG1894,AGO2,Khc,esg,EcR,E5,Antp,scrib,CG7368,Maf1,mid,CG12605,CG31224,toe,bi,CG43689,foxo,HPS4,Iswi,B-H2,TfAP-2,cic,CG12071,Su(H),polybromo,tin,lncRNA:bxd,Sox102F,crol | | 2.954e-03 | -5.82 | negative regulation of nucleic acid-templated transcription | biological process | GO:1903507 | 379 | 31 | 12345 | 597 | HPS4,Iswi,foxo,toe,bi,CG43689,lncRNA:bxd,Sox102F,crol,tin,Su(H),CG12071,polybromo,B-H2,cic,TfAP-2,scrib,Antp,E5,CG1894,AGO2,esg,EcR,Khc,retn,Ubx,CG12605,CG31224,mid,Maf1,CG7368 | | 2.987e-03 | -5.81 | CAGCCTC\_MIR4855P | MSigDB lists | CAGCCTC\_MIR4855P | 36 | 6 | 4161 | 170 | dsd,dsh,Pak,CG33090,wake,Rbp6 | | 2.997e-03 | -5.81 | GSE7460\_CTRL\_VS\_TGFB\_TREATED\_ACT\_FOXP3\_MUT\_TCONV\_UP | MSigDB lists | GSE7460\_CTRL\_VS\_TGFB\_TREATED\_ACT\_FOXP3\_MUT\_TCONV\_UP | 48 | 7 | 4161 | 170 | CG4341,dnc,Su(H),Src42A,Glut1,MCU,Tsp96F | | 3.043e-03 | -5.79 | GO\_REGULATION\_OF\_HEMOPOIESIS | MSigDB lists | GO\_REGULATION\_OF\_HEMOPOIESIS | 61 | 8 | 4161 | 170 | Ser,Ptp61F,Nha1,IP3K2,Tor,Pkc53E,Sod1,sxc | | 3.058e-03 | -5.79 | EF\_hand\_dom | interpro domains | IPR002048 | 94 | 12 | 11383 | 585 | TpnC25D,TpnC41C,CG10126,Scp1,CG14362,TpnC4,Cam,Frq1,Actn,Frq2,CG13526,CG13898 | | 3.125e-03 | -5.77 | axon guidance | biological process | GO:0007411 | 271 | 24 | 12345 | 597 | CadN2,shg,pod1,Pak,klg,Ptp99A,grn,egh,mid,fru,retn,Ptp61F,RhoGAP100F,Khc,fz2,dsh,Cam,haf,msps,beat-IIa,Src42A,trol,gukh,chb | | 3.140e-03 | -5.76 | DNA-binding transcription repressor activity, RNA polymerase II-specific | molecular function | GO:0001227 | 64 | 9 | 12176 | 580 | mid,cic,Su(H),esg,CG43689,E5,CG31224,CG12071,CG12605 | | 3.140e-03 | -5.76 | DNA-binding transcription repressor activity | molecular function | GO:0001217 | 64 | 9 | 12176 | 580 | CG43689,E5,CG12605,CG12071,CG31224,esg,cic,Su(H),mid | | 3.197e-03 | -5.75 | regulation of cell division | biological process | GO:0051302 | 41 | 7 | 12345 | 597 | Mrtf,numb,flw,nuf,chinmo,cindr,Spn88Eb | | 3.216e-03 | -5.74 | C2 | pfam domains | PF00168 | 38 | 7 | 10480 | 550 | unc-13,Pkc98E,Mctp,Pkc53E,RhoGAP100F,Syt12,Fife | | 3.224e-03 | -5.74 | GO\_REGULATION\_OF\_GLUCONEOGENESIS | MSigDB lists | GO\_REGULATION\_OF\_GLUCONEOGENESIS | 8 | 3 | 4161 | 170 | Doa,Ptp61F,sxc | | 3.224e-03 | -5.74 | MODULE\_121 | MSigDB lists | MODULE\_121 | 8 | 3 | 4161 | 170 | Tsp96F,Mlp84B,LRP1 | | 3.224e-03 | -5.74 | GO\_ARTERY\_MORPHOGENESIS | MSigDB lists | GO\_ARTERY\_MORPHOGENESIS | 8 | 3 | 4161 | 170 | Ser,LRP1,Su(H) | | 3.224e-03 | -5.74 | GO\_POSITIVE\_REGULATION\_OF\_SODIUM\_ION\_TRANSPORT | MSigDB lists | GO\_POSITIVE\_REGULATION\_OF\_SODIUM\_ION\_TRANSPORT | 8 | 3 | 4161 | 170 | Octbeta2R,Ank,Dys | | 3.224e-03 | -5.74 | GO\_ENDOCARDIAL\_CUSHION\_DEVELOPMENT | MSigDB lists | GO\_ENDOCARDIAL\_CUSHION\_DEVELOPMENT | 8 | 3 | 4161 | 170 | Ser,dpp,Su(H) | | 3.234e-03 | -5.73 | PH\_domain | interpro domains | IPR001849 | 71 | 10 | 11383 | 585 | CG10188,Cdep,cert,Gprk1,step,Cadps,RhoGAP1A,CG5522,CG5004,melt | | 3.259e-03 | -5.73 | transcription cofactor binding | molecular function | GO:0001221 | 7 | 3 | 12176 | 580 | hth,EcR,mid | | 3.259e-03 | -5.73 | mannose-ethanolamine phosphotransferase activity | molecular function | GO:0051377 | 7 | 3 | 12176 | 580 | PIG-G,CG6790,PIG-O | | 3.260e-03 | -5.73 | regulation of ERK1 and ERK2 cascade | biological process | GO:0070372 | 31 | 6 | 12345 | 597 | Ptp61F,Raf,bnl,Src42A,step,Pde8 | | 3.267e-03 | -5.72 | compound eye photoreceptor fate commitment | biological process | GO:0001752 | 52 | 8 | 12345 | 597 | hth,kay,csw,Raf,klg,scrib,melt,msi | | 3.267e-03 | -5.72 | eye photoreceptor cell fate commitment | biological process | GO:0042706 | 52 | 8 | 12345 | 597 | melt,msi,scrib,csw,Raf,klg,hth,kay | | 3.289e-03 | -5.72 | response to endogenous stimulus | biological process | GO:0009719 | 212 | 20 | 12345 | 597 | bnl,ths,Tor,Raf,ex,Set2,CG7362,Lpin,bun,Iswi,foxo,csw,opa,dpp,gce,Sh,Pak,rictor,EcR,Gpb5 | | 3.313e-03 | -5.71 | mating behavior | biological process | GO:0007617 | 183 | 18 | 12345 | 597 | Obp56h,egh,fru,Sh,retn,Fad2,bol,Ebp,lncRNA:flam,dnc,TfAP-2,Gr22c,mGluR,Gr39a,shep,JhI-26,Nlg2,Pde1c | | 3.319e-03 | -5.71 | negative regulation of cellular metabolic process | biological process | GO:0031324 | 681 | 49 | 12345 | 597 | mnb,crol,sgg,B-H2,cic,numb,Tor,Su(H),polybromo,kek1,et,raw,Iswi,msi,bi,CG43689,Acp62F,CG12605,sws,Spn88Eb,bru1,E5,bol,Ubx,retn,CG1894,Khc,esg,flw,tin,lncRNA:bxd,Sox102F,TfAP-2,CG12071,Raf,foxo,HPS4,toe,chinmo,CG31224,CG7368,mid,Maf1,Antp,scrib,REG,Ptp61F,AGO2,EcR | | 3.319e-03 | -5.71 | negative regulation of RNA metabolic process | biological process | GO:0051253 | 398 | 32 | 12345 | 597 | CG12605,CG1894,Khc,esg,Ubx,retn,bol,E5,Su(H),polybromo,B-H2,cic,crol,bi,CG43689,Iswi,Maf1,mid,CG7368,CG31224,AGO2,EcR,scrib,Antp,CG12071,TfAP-2,lncRNA:bxd,Sox102F,tin,toe,HPS4,foxo | | 3.328e-03 | -5.71 | GO\_REGULATION\_OF\_CALCIUM\_ION\_TRANSPORT\_INTO\_CYTOSOL | MSigDB lists | GO\_REGULATION\_OF\_CALCIUM\_ION\_TRANSPORT\_INTO\_CYTOSOL | 16 | 4 | 4161 | 170 | CCAP-R,Dys,Pkc98E,dnc | | 3.328e-03 | -5.71 | GO\_REGULATION\_OF\_RELEASE\_OF\_SEQUESTERED\_CALCIUM\_ION\_INTO\_CYTOSOL | MSigDB lists | GO\_REGULATION\_OF\_RELEASE\_OF\_SEQUESTERED\_CALCIUM\_ION\_INTO\_CYTOSOL | 16 | 4 | 4161 | 170 | dnc,Pkc98E,Dys,CCAP-R | | 3.328e-03 | -5.71 | GO\_POSITIVE\_REGULATION\_OF\_ERK1\_AND\_ERK2\_CASCADE | MSigDB lists | GO\_POSITIVE\_REGULATION\_OF\_ERK1\_AND\_ERK2\_CASCADE | 16 | 4 | 4161 | 170 | Raf,dpp,Pkc53E,csw | | 3.328e-03 | -5.71 | actin cytoskeleton organization | biological process | GO:0030036 | 242 | 22 | 12345 | 597 | Src42A,RhoGAP18B,gukh,Mrtf,nuf,sick,cindr,p130CAS,Msp300,Act88F,bru1,Khc,Actn,RhoGEF3,CG34417,pod1,rictor,CG32264,Prm,rols,Mlp84B,siz | | 3.344e-03 | -5.70 | fly chr26C1-26C1\|2-19 cM | chromosome location | fly chr26C1-26C1\|2-19 cM | 3 | 2 | 25010 | 845 | lncRNA:CR42735,AANATL2 | | 3.344e-03 | -5.70 | fly chr59F5-59F5\|2-104 | chromosome location | fly chr59F5-59F5\|2-104 | 3 | 2 | 25010 | 845 | Pde8,retn | | 3.344e-03 | -5.70 | fly chr66A11-66A11\|3-21 | chromosome location | fly chr66A11-66A11\|3-21 | 3 | 2 | 25010 | 845 | CG13685,CG8492 | | 3.344e-03 | -5.70 | fly chr102E5-102E5\|4-0 cM | chromosome location | fly chr102E5-102E5\|4-0 cM | 3 | 2 | 25010 | 845 | asRNA:CR44029,asRNA:CR43425 | | 3.344e-03 | -5.70 | fly chr59F5-59F5\|2-104 cM | chromosome location | fly chr59F5-59F5\|2-104 cM | 3 | 2 | 25010 | 845 | retn,Pde8 | | 3.344e-03 | -5.70 | fly chr53A4-53A4\|2-79 | chromosome location | fly chr53A4-53A4\|2-79 | 3 | 2 | 25010 | 845 | cato,prim | | 3.344e-03 | -5.70 | fly chr88A1-88A1\|3-54 | chromosome location | fly chr88A1-88A1\|3-54 | 3 | 2 | 25010 | 845 | CG14362,E5 | | 3.344e-03 | -5.70 | fly chr88A1-88A1\|3-54 cM | chromosome location | fly chr88A1-88A1\|3-54 cM | 3 | 2 | 25010 | 845 | CG14362,E5 | | 3.344e-03 | -5.70 | fly chr53A4-53A4\|2-79 cM | chromosome location | fly chr53A4-53A4\|2-79 cM | 3 | 2 | 25010 | 845 | cato,prim | | 3.344e-03 | -5.70 | fly chr36D3-36D3\|2-53 cM | chromosome location | fly chr36D3-36D3\|2-53 cM | 3 | 2 | 25010 | 845 | CG42830,CadN2 | | 3.344e-03 | -5.70 | fly chr26C1-26C1\|2-19 | chromosome location | fly chr26C1-26C1\|2-19 | 3 | 2 | 25010 | 845 | lncRNA:CR42735,AANATL2 | | 3.344e-03 | -5.70 | fly chr66A11-66A11\|3-21 cM | chromosome location | fly chr66A11-66A11\|3-21 cM | 3 | 2 | 25010 | 845 | CG13685,CG8492 | | 3.344e-03 | -5.70 | fly chr59A4-59A4\|2-101 cM | chromosome location | fly chr59A4-59A4\|2-101 cM | 3 | 2 | 25010 | 845 | CG13526,CG13527 | | 3.344e-03 | -5.70 | fly chr102E5-102E5\|4-0 | chromosome location | fly chr102E5-102E5\|4-0 | 3 | 2 | 25010 | 845 | asRNA:CR44029,asRNA:CR43425 | | 3.344e-03 | -5.70 | fly chr59A4-59A4\|2-101 | chromosome location | fly chr59A4-59A4\|2-101 | 3 | 2 | 25010 | 845 | CG13526,CG13527 | | 3.344e-03 | -5.70 | fly chr36D3-36D3\|2-53 | chromosome location | fly chr36D3-36D3\|2-53 | 3 | 2 | 25010 | 845 | CG42830,CadN2 | | 3.347e-03 | -5.70 | regulation of circadian rhythm | biological process | GO:0042752 | 88 | 11 | 12345 | 597 | CrebB,Iswi,AANAT1,EcR,Pdfr,qvr,Sh,Mbs,wake,TfAP-2,sgg | | 3.350e-03 | -5.70 | GO\_EMBRYONIC\_ORGAN\_DEVELOPMENT | MSigDB lists | GO\_EMBRYONIC\_ORGAN\_DEVELOPMENT | 90 | 10 | 4161 | 170 | dpp,fz2,dsh,RpL38,TfAP-2,RhoGAP1A,Su(H),Sod1,CG7483,CG17387 | | 3.379e-03 | -5.69 | GSE16697\_CD4\_TCELL\_VS\_TFH\_CD4\_TCELL\_DN | MSigDB lists | GSE16697\_CD4\_TCELL\_VS\_TFH\_CD4\_TCELL\_DN | 49 | 7 | 4161 | 170 | Flo2,Syx1A,MESK2,CG5339,Cln3,Su(H),rdgA | | 3.379e-03 | -5.69 | GSE37301\_PRO\_BCELL\_VS\_RAG2\_KO\_NK\_CELL\_DN | MSigDB lists | GSE37301\_PRO\_BCELL\_VS\_RAG2\_KO\_NK\_CELL\_DN | 49 | 7 | 4161 | 170 | Tsp96F,Mctp,CG17698,Pkc53E,CG12547,Sh,CG5522 | | 3.383e-03 | -5.69 | enhancer binding | molecular function | GO:0035326 | 42 | 7 | 12176 | 580 | tin,ems,Su(H),Antp,hth,Ubx,opa | | 3.403e-03 | -5.68 | lamellocyte differentiation | biological process | GO:0035171 | 7 | 3 | 12345 | 597 | Tor,ths,Raf | | 3.403e-03 | -5.68 | muscle cell fate specification | biological process | GO:0042694 | 7 | 3 | 12345 | 597 | Ubx,numb,Antp | | 3.414e-03 | -5.68 | heart development | biological process | GO:0007507 | 101 | 12 | 12345 | 597 | mid,sgg,numb,nrv2,dpp,shg,dsh,tin,EcR,Cat,Ubx,Antp | | 3.414e-03 | -5.68 | second-messenger-mediated signaling | biological process | GO:0019932 | 101 | 12 | 12345 | 597 | ACXD,CG13526,Ca-beta,Cam,TpnC25D,Pde9,CG13898,Octbeta2R,TpnC41C,dnc,CG11318,Pdfr | | 3.414e-03 | -5.68 | GO\_LYTIC\_VACUOLE | MSigDB lists | GO\_LYTIC\_VACUOLE | 169 | 15 | 4161 | 170 | Sod1,Cln3,Tor,CG41099,kcc,DNaseII,CG3842,Ank,Sk1,LRP1,sws,CG3860,Octbeta2R,CG3376,Cat | | 3.426e-03 | -5.68 | GO\_CELLULAR\_RESPONSE\_TO\_ENDOGENOUS\_STIMULUS | MSigDB lists | GO\_CELLULAR\_RESPONSE\_TO\_ENDOGENOUS\_STIMULUS | 238 | 19 | 4161 | 170 | LpR2,Tsp96F,Pak,EcR,Pkc98E,dpp,CG7483,fus,sxc,Sod1,TfIIFbeta,Ptp61F,mino,Ugt301D1,dnc,Hr96,Uros1,csw,CG42672 | | 3.447e-03 | -5.67 | GOZGIT\_ESR1\_TARGETS\_UP | MSigDB lists | GOZGIT\_ESR1\_TARGETS\_UP | 37 | 6 | 4161 | 170 | Ptp61F,CG3961,CG3860,PCB,Aldh-III,TfAP-2 | | 3.447e-03 | -5.67 | GSE1460\_INTRATHYMIC\_T\_PROGENITOR\_VS\_DP\_THYMOCYTE\_DN | MSigDB lists | GSE1460\_INTRATHYMIC\_T\_PROGENITOR\_VS\_DP\_THYMOCYTE\_DN | 37 | 6 | 4161 | 170 | IP3K2,dnc,nompB,EcR,Argl,kcc | | 3.447e-03 | -5.67 | GSE28726\_NAIVE\_VS\_ACTIVATED\_CD4\_TCELL\_UP | MSigDB lists | GSE28726\_NAIVE\_VS\_ACTIVATED\_CD4\_TCELL\_UP | 37 | 6 | 4161 | 170 | Flo2,Glut1,RhoGAP1A,PhKgamma,rdgA,Cadps | | 3.449e-03 | -5.67 | cardiocyte differentiation | biological process | GO:0035051 | 22 | 5 | 12345 | 597 | tin,mid,EcR,dpp,Ubx | | 3.480e-03 | -5.66 | GO\_POSITIVE\_REGULATION\_OF\_CELLULAR\_COMPONENT\_ORGANIZATION | MSigDB lists | GO\_POSITIVE\_REGULATION\_OF\_CELLULAR\_COMPONENT\_ORGANIZATION | 311 | 23 | 4161 | 170 | Pkc98E,EcR,dpp,CG8184,AGO3,LpR2,unc-13,Raf,Pak,Khc,sxc,RhoGAP1A,fz2,Tif-IA,Fhos,TfIIFbeta,chb,Dys,Octbeta2R,Tor,CG42672,CG41099,Ank | | 3.495e-03 | -5.66 | DNA-binding transcription factor binding | molecular function | GO:0140297 | 65 | 9 | 12176 | 580 | Mrtf,ex,bi,kay,EcR,mid,retn,Su(H),cic | | 3.510e-03 | -5.65 | GSE39110\_UNTREATED\_VS\_IL2\_TREATED\_CD8\_TCELL\_DAY6\_POST\_IMMUNIZATION\_UP | MSigDB lists | GSE39110\_UNTREATED\_VS\_IL2\_TREATED\_CD8\_TCELL\_DAY6\_POST\_IMMUNIZATION\_UP | 26 | 5 | 4161 | 170 | Cln3,Meltrin,Pgant6,Glut1,Ptp61F | | 3.529e-03 | -5.65 | GO\_NEURON\_PART | MSigDB lists | GO\_NEURON\_PART | 330 | 24 | 4161 | 170 | Tor,Maf1,Frq2,Cln3,Ank,Pde6,Cadps,Sh,Root,mGluR,Dys,Octbeta2R,Syx1A,sxc,Sod1,Frq1,CG7483,sky,KaiR1D,LRP1,Pak,Khc,unc-13,Raf | | 3.537e-03 | -5.64 | morphogenesis of embryonic epithelium | biological process | GO:0016331 | 128 | 14 | 12345 | 597 | sdt,dpp,rib,Inx2,Pak,Mbs,kay,step,tin,dsh,shd,scrib,raw,Src42A | | 3.608e-03 | -5.62 | Zig-8 | interpro domains | IPR037448 | 21 | 5 | 11383 | 585 | dpr12,dpr5,dpr15,dpr18,dpr21 | | 3.609e-03 | -5.62 | GO\_INTRINSIC\_COMPONENT\_OF\_PLASMA\_MEMBRANE | MSigDB lists | GO\_INTRINSIC\_COMPONENT\_OF\_PLASMA\_MEMBRANE | 257 | 20 | 4161 | 170 | Nlg2,Glut1,KaiR1D,kcc,Tsp96F,CNT1,LRP1,dpp,Ser,dsd,Octbeta2R,Treh,mGluR,Syx1A,AstA-R1,Nmdar2,CCAP-R,scramb1,Sh,Ugt301D1 | | 3.609e-03 | -5.62 | kirre (kin of irre) | protein interactions | 31292 | 13 | 4 | 9204 | 481 | rols,siz,rst,Actn | | 3.618e-03 | -5.62 | cell projection part | cellular component | GO:0044463 | 447 | 33 | 11940 | 538 | Or65c,dpr12,chb,Or42b,Khc,chp,dpr5,sky,dpr18,CG17387,Root,Syx1A,AGO2,CG14182,Gr22c,hdc,Ser,nompB,g,path,EcR,cpx,pod1,unc-13,Cln3,mnb,kl-5,Gr39a,Or85b,dpr15,CG5921,gukh,Ank2 | | 3.618e-03 | -5.62 | plasma membrane bounded cell projection part | cellular component | GO:0120038 | 447 | 33 | 11940 | 538 | gukh,Ank2,CG5921,Cln3,unc-13,mnb,kl-5,Gr39a,Or85b,dpr15,EcR,cpx,pod1,hdc,Gr22c,Ser,nompB,g,path,dpr5,sky,dpr18,Root,CG17387,AGO2,Syx1A,CG14182,chb,Khc,Or42b,chp,Or65c,dpr12 | | 3.639e-03 | -5.62 | GO\_RESPONSE\_TO\_PEPTIDE | MSigDB lists | GO\_RESPONSE\_TO\_PEPTIDE | 106 | 11 | 4161 | 170 | Tor,sxc,Pak,Ptp61F,Cat,LpR2,Raf,Ser,CG5493,eIF2Bgamma,mino | | 3.641e-03 | -5.62 | contractile fiber part | cellular component | GO:0044449 | 69 | 9 | 11940 | 538 | zormin,rols,Mlp84B,TpnC25D,CG3078,Msp300,Actn,Act88F,Prm | | 3.649e-03 | -5.61 | Kr (Kruppel) | protein interactions | 38012 | 39 | 7 | 9204 | 481 | dpp,chinmo,bun,ex,Antp,crol,fru | | 3.657e-03 | -5.61 | programmed cell death involved in cell development | biological process | GO:0010623 | 89 | 11 | 12345 | 597 | EcR,IP3K2,Eip74EF,rst,DNaseII,dpp,Tor,Fhos,hdc,Pak,Cam | | 3.675e-03 | -5.61 | synaptic transmission, glutamatergic | biological process | GO:0035249 | 14 | 4 | 12345 | 597 | unc-13,GluRIIC,Cadps,KaiR1D | | 3.689e-03 | -5.60 | development of primary sexual characteristics | biological process | GO:0045137 | 53 | 8 | 12345 | 597 | raw,Tor,mid,shg,rib,fz2,tin,dsh | | 3.689e-03 | -5.60 | gonad development | biological process | GO:0008406 | 53 | 8 | 12345 | 597 | mid,Tor,shg,rib,tin,dsh,fz2,raw | | 3.704e-03 | -5.60 | cell-cell adhesion | biological process | GO:0098609 | 102 | 12 | 12345 | 597 | scrib,Src42A,Nlg2,fw,CadN2,shg,chp,beat-IIa,p130CAS,klg,rst,egh | | 3.733e-03 | -5.59 | cell cortex | cellular component | GO:0005938 | 183 | 17 | 11940 | 538 | step,scrib,spri,numb,sdt,unc-13,Khc,dsh,shg,Fife,app,cindr,nuf,Hil,Mbs,Ser,Dys | | 3.734e-03 | -5.59 | GO\_REGULATION\_OF\_DEVELOPMENTAL\_GROWTH | MSigDB lists | GO\_REGULATION\_OF\_DEVELOPMENTAL\_GROWTH | 63 | 8 | 4161 | 170 | mino,Octbeta2R,unc-13,Tor,csw,Sod1,Su(H),dsd | | 3.738e-03 | -5.59 | GO\_KINASE\_BINDING | MSigDB lists | GO\_KINASE\_BINDING | 154 | 14 | 4161 | 170 | Ca-beta,Tor,Frq2,csw,Stlk,Frq1,kcc,Cadps,fz2,flw,Ptp61F,Pak,Raf,Syx1A | | 3.796e-03 | -5.57 | GSE15735\_CTRL\_VS\_HDAC\_INHIBITOR\_TREATED\_CD4\_TCELL\_12H\_DN | MSigDB lists | GSE15735\_CTRL\_VS\_HDAC\_INHIBITOR\_TREATED\_CD4\_TCELL\_12H\_DN | 50 | 7 | 4161 | 170 | CG5339,Rat1,rdgA,IP3K2,Pgant6,CG17698,Cln3 | | 3.796e-03 | -5.57 | AP4\_Q6\_01 | MSigDB lists | AP4\_Q6\_01 | 50 | 7 | 4161 | 170 | Nmdar2,Dys,dsh,Iswi,RhoGAP1A,Act87E,Cadps | | 3.808e-03 | -5.57 | cell adhesion | biological process | GO:0007155 | 171 | 17 | 12345 | 597 | CadN2,shg,Pak,Flo2,nrv2,egh,klg,rst,scrib,EcR,fw,crol,chp,beat-IIa,p130CAS,Src42A,Nlg2 | | 3.812e-03 | -5.57 | Opioid Signalling | REACTOME pathways | R-DME-111885 | 31 | 6 | 3923 | 197 | Cam,dnc,Pkc53E,Gprk1,ACXD,Pde1c | | 3.814e-03 | -5.57 | RPEL | prosite domains | PS51073 | 2 | 2 | 6243 | 386 | CG32264,Mrtf | | 3.824e-03 | -5.57 | DUF1041 | smart domains | SM01145 | 2 | 2 | 5086 | 315 | Cadps,unc-13 | | 3.824e-03 | -5.57 | RPEL | smart domains | SM00707 | 2 | 2 | 5086 | 315 | CG32264,Mrtf | | 3.832e-03 | -5.56 | cellular component organization or biogenesis | biological process | GO:0071840 | 2975 | 172 | 12345 | 597 | CG12782,DIP-delta,Yeti,nuf,aust,Stacl,Mrtf,dpr18,Cam,lncRNA:Hsromega,luna,CG17387,trol,HPS4,shg,CG34417,AGO2,mino,RhoGAP1A,g,sgg,Fife,app,polybromo,msps,bnl,fz2,kay,mGluR,CG4301,Mob2,Set2,dpr21,Syn2,Nnf1b,Src42A,raw,CG33158,Art2,cert,Dlic,spri,shakB,Mbs,jim,RhoGAP100F,Frq2,Pld,IFT46,cindr,His1:CG33837,fs(1)N,rib,CG30036,chp,sqz,bun,Nlg2,klg,grn,siz,egh,rols,csw,ems,Ptp61F,EcR,Ank,E(Pc),ddbt,B-H2,scro,ths,Pomp,Syx1A,Ank2,Strn-Mlck,path,crol,mamo,dpr15,sws,woc,fru,CG32264,Prm,CG41562,by,retn,CG6839,Actn,dpr12,CG18109,bru1,CG7483,CG41099,Raf,Msp300,dsh,Root,CG40813,Meltrin,chb,Act87E,chinmo,CG41520,foxo,CG14105,dpp,CG5059,CG5921,HDAC4,dnc,spd-2,Snoo,Tor,cpx,Tusp,Frq1,sky,Stlk,Spt20,nrv2,RhoGEF3,Cln3,nompB,Pak,Rcd6,Khc,CG1894,Pkc98E,Act88F,Cep97,beat-IIa,ham,sick,CG42355,RhoGAP18B,CG12200,Epg5,CG34172,Mlp84B,wake,Ptp99A,mid,scramb1,Sh,dpr5,eRF3,CrebB,scrib,DNaseII,cic,haf,p130CAS,Poxm,hdc,Elp3,Ca-beta,unc-13,gukh,can,CG9981,Iswi,sdt,Doa,rictor,CadN2,pod1,fw,flw,CG3594 | | 3.852e-03 | -5.56 | homeostasis of number of cells | biological process | GO:0048872 | 32 | 6 | 12345 | 597 | upd3,Src42A,GATAe,bun,Cat,esg | | 3.853e-03 | -5.56 | wg (wingless) | protein interactions | 34009 | 120 | 14 | 9204 | 481 | sgg,bi,CG46385,eRF3,fz2,sno,Raf,dpp,dsh,mid,shg,Ser,ex,Dys | | 3.873e-03 | -5.55 | C2\_dom | interpro domains | IPR000008 | 40 | 7 | 11383 | 585 | Pkc98E,RhoGAP100F,Pkc53E,Syt12,Mctp,unc-13,Fife | | 3.875e-03 | -5.55 | embryonic pattern specification | biological process | GO:0009880 | 230 | 21 | 12345 | 597 | CG7483,scrib,Antp,AGO2,Khc,Ubx,retn,hth,shg,ems,Doa,csw,dpp,tld,dsh,fz2,fs(1)N,Raf,msps,cic,sgg | | 3.915e-03 | -5.54 | GO\_REGULATION\_OF\_SYSTEM\_PROCESS | MSigDB lists | GO\_REGULATION\_OF\_SYSTEM\_PROCESS | 107 | 11 | 4161 | 170 | dnc,EcR,Octbeta2R,Dys,Nmdar2,Ca-beta,Tor,Sod1,csw,Pkc53E,bru1 | | 3.915e-03 | -5.54 | GO\_SEQUENCE\_SPECIFIC\_DNA\_BINDING | MSigDB lists | GO\_SEQUENCE\_SPECIFIC\_DNA\_BINDING | 107 | 11 | 4161 | 170 | Rat1,Hr96,scro,Awh,Maf1,Tor,TfAP-2,Su(H),Tif-IA,Act87E,EcR | | 3.957e-03 | -5.53 | GSE30083\_SP3\_VS\_SP4\_THYMOCYTE\_UP | MSigDB lists | GSE30083\_SP3\_VS\_SP4\_THYMOCYTE\_UP | 38 | 6 | 4161 | 170 | Ank,scro,CG3860,app,CG17698,Root | | 3.957e-03 | -5.53 | GSE1791\_CTRL\_VS\_NEUROMEDINU\_IN\_T\_CELL\_LINE\_0.8H\_DN | MSigDB lists | GSE1791\_CTRL\_VS\_NEUROMEDINU\_IN\_T\_CELL\_LINE\_0.8H\_DN | 38 | 6 | 4161 | 170 | IP3K2,TfAP-2,Glut1,Pak,bru1,Syx1A | | 3.957e-03 | -5.53 | GO\_TISSUE\_HOMEOSTASIS | MSigDB lists | GO\_TISSUE\_HOMEOSTASIS | 38 | 6 | 4161 | 170 | Act88F,Sod1,Root,csw,Act87E,Octbeta2R | | 3.957e-03 | -5.53 | GO\_CELLULAR\_RESPONSE\_TO\_INORGANIC\_SUBSTANCE | MSigDB lists | GO\_CELLULAR\_RESPONSE\_TO\_INORGANIC\_SUBSTANCE | 38 | 6 | 4161 | 170 | Raf,CG7483,Ank,Sod1,IP3K2,TfAP-2 | | 3.979e-03 | -5.53 | system process | biological process | GO:0003008 | 636 | 46 | 12345 | 597 | Ank2,mGluR,Fhos,Gr39a,Mob2,mnb,sgg,Su(H),upd3,Obp56e,Tk,dpr15,drd,nompB,sws,nrv2,Or65c,Pkc98E,CG5541,ine,dpr12,Gr22c,dpr18,Cam,Root,rdgA,Or85b,geko,CG5921,Sh,CG31750,Obp56h,klg,Gld2,MCU,Nmdar2,Or42b,mid,scrib,dnc,Nha1,CrebB,dpr5,EcR,nemy,HDAC4 | | 4.008e-03 | -5.52 | positive regulation of cell communication | biological process | GO:0010647 | 403 | 32 | 12345 | 597 | lncRNA:Hsromega,Raf,MESK2,olf186-F,Pde8,trol,foxo,Lpin,Gprk1,Glut1,sno,step,dpp,siz,RhoGAP1A,melt,bnl,Tor,ex,Fife,Src42A,tld,hth,sxc,rictor,Doa,Flo2,CG10188,KaiR1D,flw,Exn,Mos | | 4.008e-03 | -5.52 | positive regulation of signaling | biological process | GO:0023056 | 403 | 32 | 12345 | 597 | KaiR1D,CG10188,Exn,Mos,flw,rictor,sxc,hth,Flo2,Doa,tld,Src42A,ex,Fife,Tor,bnl,RhoGAP1A,melt,step,sno,siz,dpp,trol,foxo,olf186-F,Pde8,Glut1,Lpin,Gprk1,lncRNA:Hsromega,Raf,MESK2 | | 4.071e-03 | -5.50 | positive regulation of protein phosphorylation | biological process | GO:0001934 | 130 | 14 | 12345 | 597 | rictor,step,Pak,lncRNA:Hsromega,dpp,Tor,Raf,bnl,CG41520,Src42A,Pde8,Stlk,CycT,Mos | | 4.168e-03 | -5.48 | MORF\_LCAT | MSigDB lists | MORF\_LCAT | 27 | 5 | 4161 | 170 | Src42A,Sh,nompB,PhKgamma,Spf45 | | 4.202e-03 | -5.47 | actin filament-based process | biological process | GO:0030029 | 262 | 23 | 12345 | 597 | Actn,Khc,bru1,Mlp84B,siz,rols,rictor,CG32264,Prm,RhoGEF3,CG34417,pod1,gukh,RhoGAP18B,Src42A,Act88F,nuf,cindr,sick,p130CAS,Msp300,Cam,Mrtf | | 4.203e-03 | -5.47 | renal system development | biological process | GO:0072001 | 78 | 10 | 12345 | 597 | numb,dpp,rols,tin,hth,rib,Ubx,EcR,raw,scrib | | 4.203e-03 | -5.47 | urogenital system development | biological process | GO:0001655 | 78 | 10 | 12345 | 597 | rib,hth,tin,rols,numb,dpp,scrib,raw,EcR,Ubx | | 4.214e-03 | -5.47 | regionalization | biological process | GO:0003002 | 503 | 38 | 12345 | 597 | scrib,Antp,EcR,AGO2,Ser,sno,shg,step,ems,csw,mid,dpp,wbl,Lpin,fs(1)N,dsh,nuf,Raf,TfAP-2,CG7483,Pkc98E,Dys,Khc,flw,retn,Ubx,hth,Doa,tld,boi,bi,fz2,msps,Su(H),numb,sgg,cic,B-H2 | | 4.215e-03 | -5.47 | GO\_REGULATION\_OF\_NEUROTRANSMITTER\_TRANSPORT | MSigDB lists | GO\_REGULATION\_OF\_NEUROTRANSMITTER\_TRANSPORT | 17 | 4 | 4161 | 170 | ZnT63C,Syx1A,Raf,unc-13 | | 4.215e-03 | -5.47 | GO\_CYCLIC\_NUCLEOTIDE\_METABOLIC\_PROCESS | MSigDB lists | GO\_CYCLIC\_NUCLEOTIDE\_METABOLIC\_PROCESS | 17 | 4 | 4161 | 170 | Ca-beta,Pde6,dnc,Pde8 | | 4.215e-03 | -5.47 | MORF\_IL9 | MSigDB lists | MORF\_IL9 | 17 | 4 | 4161 | 170 | Src42A,Sh,Spf45,Nlg2 | | 4.215e-03 | -5.47 | GO\_REGULATION\_OF\_WOUND\_HEALING | MSigDB lists | GO\_REGULATION\_OF\_WOUND\_HEALING | 17 | 4 | 4161 | 170 | Pkc53E,Pkc98E,Octbeta2R,chb | | 4.215e-03 | -5.47 | GO\_VOLTAGE\_GATED\_CATION\_CHANNEL\_ACTIVITY | MSigDB lists | GO\_VOLTAGE\_GATED\_CATION\_CHANNEL\_ACTIVITY | 17 | 4 | 4161 | 170 | Sh,Ca-beta,Frq2,Frq1 | | 4.231e-03 | -5.47 | Signaling by Rho GTPases | REACTOME pathways | R-DME-194315 | 114 | 13 | 3923 | 197 | RhoGAP1A,dsh,CG10188,Strn-Mlck,CG6424,RhoGAP100F,flw,Pak,Mbs,Mrtf,Exn,Cam,RhoGAP18B | | 4.234e-03 | -5.46 | positive regulation of ERK1 and ERK2 cascade | biological process | GO:0070374 | 23 | 5 | 12345 | 597 | Src42A,step,Pde8,Raf,bnl | | 4.239e-03 | -5.46 | wun (wunen) | protein interactions | 35966 | 7 | 3 | 9204 | 481 | Iswi,rdgA,Dys | | 4.239e-03 | -5.46 | apical junction complex | cellular component | GO:0043296 | 58 | 8 | 11940 | 538 | Mbs,fw,sdt,nrv2,scrib,ex,shg,bbg | | 4.252e-03 | -5.46 | GSE24671\_CTRL\_VS\_BAKIMULC\_INFECTED\_MOUSE\_SPLENOCYTES\_UP | MSigDB lists | GSE24671\_CTRL\_VS\_BAKIMULC\_INFECTED\_MOUSE\_SPLENOCYTES\_UP | 51 | 7 | 4161 | 170 | D2hgdh,wbl,flw,step,CG12547,Meltrin,bru1 | | 4.252e-03 | -5.46 | MEIS1\_01 | MSigDB lists | MEIS1\_01 | 51 | 7 | 4161 | 170 | Cln3,Epg5,CG14480,Stt3B,REG,DNaseII,polybromo | | 4.252e-03 | -5.46 | GSE42021\_CD24HI\_VS\_CD24INT\_TCONV\_THYMUS\_UP | MSigDB lists | GSE42021\_CD24HI\_VS\_CD24INT\_TCONV\_THYMUS\_UP | 51 | 7 | 4161 | 170 | Pgant6,Der-1,step,RhoGAP1A,Odc1,MESK2,CG5339 | | 4.257e-03 | -5.46 | GO\_PLASMA\_MEMBRANE\_PROTEIN\_COMPLEX | MSigDB lists | GO\_PLASMA\_MEMBRANE\_PROTEIN\_COMPLEX | 93 | 10 | 4161 | 170 | Syx1A,Nmdar2,Dys,Syn2,Sh,dnc,dpp,KaiR1D,Flo2,Ca-beta | | 4.266e-03 | -5.46 | Signaling by GPCR | REACTOME pathways | R-DME-372790 | 227 | 21 | 3923 | 197 | Prosbeta5R1,Pkc53E,Pde1c,CG41520,CG30054,dnc,CrebB,ACXD,Gprk1,Pkc98E,REG,RhoGAP1A,Octbeta2R,Pde8,Cam,Exn,rdgA,bnl,CG10188,mGluR,Raf | | 4.284e-03 | -5.45 | establishment or maintenance of cell polarity | biological process | GO:0007163 | 217 | 20 | 12345 | 597 | msps,Raf,p130CAS,sgg,rib,kay,dsh,chb,trol,csw,sdt,egh,wake,shg,CadN2,rictor,Khc,fw,scrib,Dys | | 4.293e-03 | -5.45 | bZIP\_1 | pfam domains | PF00170 | 7 | 3 | 10480 | 550 | Atf6,CrebB,kay | | 4.317e-03 | -5.45 | TGTTTGY\_HNF3\_Q6 | MSigDB lists | TGTTTGY\_HNF3\_Q6 | 124 | 12 | 4161 | 170 | Dys,CG8272,CCAP-R,CG9132,dnc,Ser,Rbp6,fus,KaiR1D,app,IP3K2,sxc | | 4.317e-03 | -5.45 | GO\_DENDRITE | MSigDB lists | GO\_DENDRITE | 124 | 12 | 4161 | 170 | Frq1,Ank,CG7483,KaiR1D,Maf1,Tor,Sod1,Frq2,Octbeta2R,mGluR,Pak,LRP1 | | 4.344e-03 | -5.44 | circulatory system development | biological process | GO:0072359 | 104 | 12 | 12345 | 597 | Antp,EcR,Ubx,Cat,shg,dsh,tin,mid,sgg,nrv2,numb,dpp | | 4.344e-03 | -5.44 | striated muscle cell differentiation | biological process | GO:0051146 | 104 | 12 | 12345 | 597 | Actn,Khc,bru1,foxo,Act88F,siz,rst,Mlp84B,Tor,rols,Prm,tin | | 4.345e-03 | -5.44 | circadian rhythm | biological process | GO:0007623 | 91 | 11 | 12345 | 597 | CrebB,dnc,foxo,ITP,Fer2,Pdfr,mnb,sxc,kay,sgg,tyf | | 4.348e-03 | -5.44 | negative regulation of response to stimulus | biological process | GO:0048585 | 389 | 31 | 12345 | 597 | Fer2,dnc,Ser,PGRP-LA,Ptp61F,HDAC4,kek5,flw,kek1,raw,et,foxo,Src42A,trol,Pde8,chb,CG14182,Gprk1,Lpin,qvr,mnb,crol,Cam,CG9380,numb,sgg,GlyP,Snoo,pirk,Su(H),Raf | | 4.362e-03 | -5.43 | regulation of protein kinase activity | biological process | GO:0045859 | 131 | 14 | 12345 | 597 | step,kay,rictor,Pak,mnb,sgg,SteXh:CG42398,sws,Raf,CG41520,kek1,Ste:CG33247,CycT,Stlk | | 4.386e-03 | -5.43 | EGF-like\_dom | interpro domains | IPR000742 | 74 | 10 | 11383 | 585 | LRP1,CG13654,dsd,CadN2,Meltrin,tld,shg,trol,LpR2,Ser | | 4.419e-03 | -5.42 | actin cytoskeleton | cellular component | GO:0015629 | 126 | 13 | 11940 | 538 | Msp300,Act88F,Myo81F,pod1,CG34417,Prm,CG34435,Cam,Mlp84B,TpnC25D,Pak,Khc,Act87E | | 4.420e-03 | -5.42 | contractile fiber | cellular component | GO:0043292 | 71 | 9 | 11940 | 538 | zormin,rols,CG3078,Actn,Msp300,Act88F,Prm,TpnC25D,Mlp84B | | 4.420e-03 | -5.42 | apical plasma membrane | cellular component | GO:0016324 | 71 | 9 | 11940 | 538 | Inx2,ZnT63C,ex,rst,kek1,Ser,Cln3,sdt,Pde6 | | 4.446e-03 | -5.42 | GO\_MOLECULAR\_FUNCTION\_REGULATOR | MSigDB lists | GO\_MOLECULAR\_FUNCTION\_REGULATOR | 280 | 21 | 4161 | 170 | Ugt301D1,REG,Octbeta2R,mGluR,Syx1A,Nmdar2,CG42672,step,ZnT63C,Stlk,CG34408,dpp,CG5522,Pkc98E,eIF2Bgamma,Elp3,CG17883,RhoGAP1A,sxc,Cdep,sky | | 4.471e-03 | -5.41 | GO\_REGULATION\_OF\_PROTEIN\_MODIFICATION\_PROCESS | MSigDB lists | GO\_REGULATION\_OF\_PROTEIN\_MODIFICATION\_PROCESS | 413 | 28 | 4161 | 170 | Ptp61F,dsh,Dys,CG3376,Octbeta2R,REG,dnc,fz2,Stlk,Tor,IP3K2,CG42672,Der-1,csw,CG17698,Pak,LpR2,Raf,Tsp96F,dpp,AGO3,Pkc98E,KaiR1D,Elp3,Sod1,sxc,Pkc53E,wbl | | 4.482e-03 | -5.41 | mating | biological process | GO:0007618 | 203 | 19 | 12345 | 597 | Pde1c,Nlg2,shep,JhI-26,Gr39a,mGluR,Gr22c,TfAP-2,dnc,lncRNA:flam,Ebp,bol,Fad2,retn,Sh,Acp62F,fru,Obp56h,egh | | 4.496e-03 | -5.40 | ZHOU\_INFLAMMATORY\_RESPONSE\_FIMA\_UP | MSigDB lists | ZHOU\_INFLAMMATORY\_RESPONSE\_FIMA\_UP | 79 | 9 | 4161 | 170 | Cadps,Ser,Sk1,CG12121,CNT1,CG32320,CG6418,eIF4E7,KaiR1D | | 4.500e-03 | -5.40 | microtubule organizing center | cellular component | GO:0005815 | 171 | 16 | 11940 | 538 | chb,Cep97,CG18109,wake,spd-2,Rcd6,IFT46,nompB,nuf,Cam,CG13526,Dlic,CG34417,CG13251,msps,sgg | | 4.519e-03 | -5.40 | ecdysteroid metabolic process | biological process | GO:0045455 | 33 | 6 | 12345 | 597 | foxo,CG9519,CG9514,shd,spok,woc | | 4.522e-03 | -5.40 | GO\_LEARNING | MSigDB lists | GO\_LEARNING | 39 | 6 | 4161 | 170 | Tor,Cln3,Nlg2,Octbeta2R,CG7483,Raf | | 4.522e-03 | -5.40 | RUTELLA\_RESPONSE\_TO\_HGF\_DN | MSigDB lists | RUTELLA\_RESPONSE\_TO\_HGF\_DN | 39 | 6 | 4161 | 170 | step,Meltrin,bru1,Glut1,CG5339,scramb1 | | 4.522e-03 | -5.40 | GSE37416\_CTRL\_VS\_48H\_F\_TULARENSIS\_LVS\_NEUTROPHIL\_UP | MSigDB lists | GSE37416\_CTRL\_VS\_48H\_F\_TULARENSIS\_LVS\_NEUTROPHIL\_UP | 39 | 6 | 4161 | 170 | CG5522,CG30036,TfAP-2,Syn2,mino,CG32196 | | 4.522e-03 | -5.40 | GSE40274\_SATB1\_VS\_FOXP3\_AND\_SATB1\_TRANSDUCED\_ACTIVATED\_CD4\_TCELL\_UP | MSigDB lists | GSE40274\_SATB1\_VS\_FOXP3\_AND\_SATB1\_TRANSDUCED\_ACTIVATED\_CD4\_TCELL\_UP | 39 | 6 | 4161 | 170 | Mlp84B,Pak,pod1,CNT1,CCAP-R,sws | | 4.522e-03 | -5.40 | GO\_LEUKOCYTE\_DIFFERENTIATION | MSigDB lists | GO\_LEUKOCYTE\_DIFFERENTIATION | 39 | 6 | 4161 | 170 | Raf,Cdep,Ptp61F,Su(H),fz2,IP3K2 | | 4.522e-03 | -5.40 | GSE24210\_CTRL\_VS\_IL35\_TREATED\_TCONV\_CD4\_TCELL\_UP | MSigDB lists | GSE24210\_CTRL\_VS\_IL35\_TREATED\_TCONV\_CD4\_TCELL\_UP | 39 | 6 | 4161 | 170 | Raf,Pak,bru1,Uros1,step,CG42672 | | 4.546e-03 | -5.39 | biological adhesion | biological process | GO:0022610 | 174 | 17 | 12345 | 597 | scrib,fw,EcR,Pak,shg,CadN2,klg,Flo2,rst,nrv2,egh,Src42A,Nlg2,chp,crol,p130CAS,beat-IIa | | 4.592e-03 | -5.38 | GO\_LIPID\_METABOLIC\_PROCESS | MSigDB lists | GO\_LIPID\_METABOLIC\_PROCESS | 375 | 26 | 4161 | 170 | CG3842,Acbp2,LpR2,Aldh-III,EcR,ABCD,PIG-G,Sk1,rdgA,LRP1,CG3961,Hr96,CG3530,csw,PIG-O,Cln3,sws,scramb1,CG3860,AstA-R1,PCB,Cat,CG3376,CG33090,mino,Ugt301D1 | | 4.593e-03 | -5.38 | scrib (scribble) | protein interactions | 44448 | 51 | 8 | 9204 | 481 | hth,scrib,Raf,gukh,kay,sdt,Pak,Pino | | 4.617e-03 | -5.38 | aop (anterior open) | protein interactions | 33392 | 62 | 9 | 9204 | 481 | Su(H),raw,Raf,bun,chb,Syx1A,retn,shg,dsh | | 4.676e-03 | -5.37 | EF\_HAND\_1 | prosite domains | PS00018 | 83 | 12 | 6243 | 386 | TpnC25D,TpnC41C,TpnC4,Cam,CG10126,CG14362,Frq2,Actn,Scp1,Frq1,CG13898,CG13526 | | 4.692e-03 | -5.36 | GO\_IONOTROPIC\_GLUTAMATE\_RECEPTOR\_SIGNALING\_PATHWAY | MSigDB lists | GO\_IONOTROPIC\_GLUTAMATE\_RECEPTOR\_SIGNALING\_PATHWAY | 9 | 3 | 4161 | 170 | Cln3,KaiR1D,Nmdar2 | | 4.692e-03 | -5.36 | GO\_NEGATIVE\_REGULATION\_OF\_CYCLIC\_NUCLEOTIDE\_METABOLIC\_PROCESS | MSigDB lists | GO\_NEGATIVE\_REGULATION\_OF\_CYCLIC\_NUCLEOTIDE\_METABOLIC\_PROCESS | 9 | 3 | 4161 | 170 | Pkc53E,mGluR,AstA-R1 | | 4.692e-03 | -5.36 | GO\_HEART\_VALVE\_DEVELOPMENT | MSigDB lists | GO\_HEART\_VALVE\_DEVELOPMENT | 9 | 3 | 4161 | 170 | Ser,Tor,dpp | | 4.692e-03 | -5.36 | GO\_POSITIVE\_REGULATION\_OF\_POTASSIUM\_ION\_TRANSPORT | MSigDB lists | GO\_POSITIVE\_REGULATION\_OF\_POTASSIUM\_ION\_TRANSPORT | 9 | 3 | 4161 | 170 | AstA-R1,Octbeta2R,Khc | | 4.692e-03 | -5.36 | MODULE\_418 | MSigDB lists | MODULE\_418 | 9 | 3 | 4161 | 170 | CG34408,KrT95D,unc-13 | | 4.692e-03 | -5.36 | GO\_GLUTAMATE\_RECEPTOR\_ACTIVITY | MSigDB lists | GO\_GLUTAMATE\_RECEPTOR\_ACTIVITY | 9 | 3 | 4161 | 170 | KaiR1D,Nmdar2,mGluR | | 4.748e-03 | -5.35 | KEGG\_CALCIUM\_SIGNALING\_PATHWAY | MSigDB lists | KEGG\_CALCIUM\_SIGNALING\_PATHWAY | 52 | 7 | 4161 | 170 | Pkc53E,PhKgamma,Sk1,IP3K2,Pde1c,Nmdar2,Octbeta2R | | 4.748e-03 | -5.35 | KEGG\_INSULIN\_SIGNALING\_PATHWAY | MSigDB lists | KEGG\_INSULIN\_SIGNALING\_PATHWAY | 52 | 7 | 4161 | 170 | flw,PhKgamma,Tor,Raf,Flo2,eIF4E7,GlyP | | 4.748e-03 | -5.35 | VECCHI\_GASTRIC\_CANCER\_EARLY\_DN | MSigDB lists | VECCHI\_GASTRIC\_CANCER\_EARLY\_DN | 52 | 7 | 4161 | 170 | melt,Iswi,Mctp,LpR2,CNT1,Octbeta2R,Dys | | 4.748e-03 | -5.35 | CACTTTG\_MIR520G\_MIR520H | MSigDB lists | CACTTTG\_MIR520G\_MIR520H | 52 | 7 | 4161 | 170 | eIF4E7,LpR2,Dys,Glut1,CG4341,Awh,Maf1 | | 4.794e-03 | -5.34 | eve (even skipped) | protein interactions | 36039 | 22 | 5 | 9204 | 481 | trol,Antp,Ubx,fru,grn | | 4.836e-03 | -5.33 | integral component of membrane | cellular component | GO:0016021 | 1438 | 85 | 11940 | 538 | Ir10a,MCU,chk,dpr5,PIG-G,CG33090,Nha1,CG13229,beat-IIa,Syx1A,Nlg2,CG14606,fz4,AstA-R1,CG33934,Sh,CG4301,Mctp,Der-1,CG11318,Qsox2,Ir56a,CG9981,CG10960,nrv2,CG6790,Meltrin,ine,Octbeta2R,olf186-F,Gr39a,fz2,Glut1,CG5059,CG8468,CG12121,rst,CCAP-R,kek5,CadN2,Gr22c,Irk3,CG13654,Ser,CG1572,CG3168,egh,Nmdar2,Ir41a,ABCD,PGRP-LA,CG3285,Msp300,dpr18,MsR2,kek1,Or42b,CG14024,CG13794,CG32320,ACXD,GluRIIC,Or65c,scrib,CG33310,CG7882,CG42826,Pdfr,CG41128,Syt12,CNT1,CG17600,kcc,Pzl,DIP-lambda,shg,Tsp96F,Inx2,ZnT63C,CG14439,DIP-delta,Cyt-b5-r,Or85b,Ca-beta,mGluR | | 4.845e-03 | -5.33 | GO\_CELLULAR\_RESPONSE\_TO\_ETHANOL | MSigDB lists | GO\_CELLULAR\_RESPONSE\_TO\_ETHANOL | 3 | 2 | 4161 | 170 | Pkc98E,Ugt301D1 | | 4.845e-03 | -5.33 | MODULE\_156 | MSigDB lists | MODULE\_156 | 3 | 2 | 4161 | 170 | Frq1,Frq2 | | 4.845e-03 | -5.33 | GO\_POSITIVE\_REGULATION\_OF\_SODIUM\_ION\_TRANSMEMBRANE\_TRANSPORTER\_ACTIVITY | MSigDB lists | GO\_POSITIVE\_REGULATION\_OF\_SODIUM\_ION\_TRANSMEMBRANE\_TRANSPORTER\_ACTIVITY | 3 | 2 | 4161 | 170 | Dys,Ank | | 4.845e-03 | -5.33 | GO\_NEGATIVE\_REGULATION\_OF\_EPITHELIAL\_CELL\_APOPTOTIC\_PROCESS | MSigDB lists | GO\_NEGATIVE\_REGULATION\_OF\_EPITHELIAL\_CELL\_APOPTOTIC\_PROCESS | 3 | 2 | 4161 | 170 | Tor,Raf | | 4.845e-03 | -5.33 | GO\_REGULATION\_OF\_FATTY\_ACID\_BETA\_OXIDATION | MSigDB lists | GO\_REGULATION\_OF\_FATTY\_ACID\_BETA\_OXIDATION | 3 | 2 | 4161 | 170 | Tor,ABCD | | 4.845e-03 | -5.33 | HALLMARK\_APICAL\_SURFACE | MSigDB lists | HALLMARK\_APICAL\_SURFACE | 3 | 2 | 4161 | 170 | Root,Flo2 | | 4.845e-03 | -5.33 | BRUNEAU\_HEART\_GREAT\_VESSELS\_AND\_VALVULOGENESIS | MSigDB lists | BRUNEAU\_HEART\_GREAT\_VESSELS\_AND\_VALVULOGENESIS | 3 | 2 | 4161 | 170 | csw,Ser | | 4.845e-03 | -5.33 | GO\_3\_5\_CYCLIC\_GMP\_PHOSPHODIESTERASE\_ACTIVITY | MSigDB lists | GO\_3\_5\_CYCLIC\_GMP\_PHOSPHODIESTERASE\_ACTIVITY | 3 | 2 | 4161 | 170 | Pde1c,Pde6 | | 4.845e-03 | -5.33 | GO\_REGULATION\_OF\_CELL\_COMMUNICATION\_BY\_ELECTRICAL\_COUPLING | MSigDB lists | GO\_REGULATION\_OF\_CELL\_COMMUNICATION\_BY\_ELECTRICAL\_COUPLING | 3 | 2 | 4161 | 170 | dnc,Ank | | 4.845e-03 | -5.33 | GO\_REGULATION\_OF\_MACROPHAGE\_DIFFERENTIATION | MSigDB lists | GO\_REGULATION\_OF\_MACROPHAGE\_DIFFERENTIATION | 3 | 2 | 4161 | 170 | Pkc53E,Ptp61F | | 4.845e-03 | -5.33 | GO\_COSTAMERE | MSigDB lists | GO\_COSTAMERE | 3 | 2 | 4161 | 170 | Dys,Ank | | 4.845e-03 | -5.33 | GO\_NEGATIVE\_REGULATION\_OF\_LEUKOCYTE\_APOPTOTIC\_PROCESS | MSigDB lists | GO\_NEGATIVE\_REGULATION\_OF\_LEUKOCYTE\_APOPTOTIC\_PROCESS | 3 | 2 | 4161 | 170 | IP3K2,mino | | 4.845e-03 | -5.33 | GO\_POSITIVE\_T\_CELL\_SELECTION | MSigDB lists | GO\_POSITIVE\_T\_CELL\_SELECTION | 3 | 2 | 4161 | 170 | Raf,IP3K2 | | 4.845e-03 | -5.33 | GO\_ACUTE\_PHASE\_RESPONSE | MSigDB lists | GO\_ACUTE\_PHASE\_RESPONSE | 3 | 2 | 4161 | 170 | scramb1,Ugt301D1 | | 4.847e-03 | -5.33 | unc-13 (unc-13) | protein interactions | 43841 | 14 | 4 | 9204 | 481 | RhoGAP100F,CG15198,Cam,flw | | 4.875e-03 | -5.32 | Abl (Abl tyrosine kinase) | protein interactions | 45821 | 41 | 7 | 9204 | 481 | msps,flw,cindr,Ptp99A,Ptp61F,dsh,chb | | 4.875e-03 | -5.32 | hop (hopscotch) | protein interactions | 32080 | 41 | 7 | 9204 | 481 | et,Raf,Ser,Ptp61F,CG1572,kay,csw | | 4.893e-03 | -5.32 | GTGCCAA\_MIR96 | MSigDB lists | GTGCCAA\_MIR96 | 80 | 9 | 4161 | 170 | sxc,Ca-beta,Tor,CG40045,Flo2,bru1,Pkc98E,CG5522,Pak | | 4.909e-03 | -5.32 | BROWNE\_HCMV\_INFECTION\_24HR\_DN | MSigDB lists | BROWNE\_HCMV\_INFECTION\_24HR\_DN | 28 | 5 | 4161 | 170 | Act87E,Pkc53E,Root,Uros1,KaiR1D | | 4.909e-03 | -5.32 | GSE9960\_HEALTHY\_VS\_SEPSIS\_PBMC\_DN | MSigDB lists | GSE9960\_HEALTHY\_VS\_SEPSIS\_PBMC\_DN | 28 | 5 | 4161 | 170 | Rat1,Nlg2,AGO3,Elp3,Pkc98E | | 4.909e-03 | -5.32 | GO\_RESPONSE\_TO\_ORGANOPHOSPHORUS | MSigDB lists | GO\_RESPONSE\_TO\_ORGANOPHOSPHORUS | 28 | 5 | 4161 | 170 | GlyP,Raf,dnc,CG5493,Sod1 | | 4.909e-03 | -5.32 | GO\_REGULATION\_OF\_CAMP\_METABOLIC\_PROCESS | MSigDB lists | GO\_REGULATION\_OF\_CAMP\_METABOLIC\_PROCESS | 28 | 5 | 4161 | 170 | dnc,Pkc53E,mGluR,Octbeta2R,AstA-R1 | | 4.909e-03 | -5.32 | GO\_REGULATION\_OF\_CALCIUM\_ION\_TRANSMEMBRANE\_TRANSPORT | MSigDB lists | GO\_REGULATION\_OF\_CALCIUM\_ION\_TRANSMEMBRANE\_TRANSPORT | 28 | 5 | 4161 | 170 | dnc,Ca-beta,Pkc98E,Dys,CCAP-R | | 4.988e-03 | -5.30 | PTPc\_motif | smart domains | SM00404 | 19 | 5 | 5086 | 315 | Ptp99A,Ptp36E,CG3530,Ptp61F,csw | | 4.991e-03 | -5.30 | GO\_EMBRYONIC\_ORGAN\_MORPHOGENESIS | MSigDB lists | GO\_EMBRYONIC\_ORGAN\_MORPHOGENESIS | 66 | 8 | 4161 | 170 | TfAP-2,RpL38,Sod1,RhoGAP1A,CG17387,CG7483,fz2,dsh | | 5.014e-03 | -5.30 | Nucleotide-bd\_a/b\_plait\_sf | interpro domains | IPR012677 | 152 | 16 | 11383 | 585 | bol,trv,barc,CG41562,msi,Achl,Rbp6,CG3594,CG42458,CG40813,fus,bru1,Spf45,CG44249,shep,Marf1 | | 5.040e-03 | -5.29 | intrinsic component of membrane | cellular component | GO:0031224 | 1460 | 86 | 11940 | 538 | MCU,Ir10a,chk,beat-IIa,Syx1A,dpr5,CG33090,Nha1,PIG-G,CG13229,CG14606,chp,Nlg2,Sh,CG4301,Der-1,Mctp,fz4,AstA-R1,CG33934,CG10960,nrv2,CG6790,CG11318,Qsox2,Ir56a,CG9981,Octbeta2R,olf186-F,Meltrin,ine,fz2,Glut1,CG5059,Gr39a,rst,CCAP-R,kek5,CG8468,CG12121,CG3168,egh,Gr22c,CadN2,Irk3,CG13654,Ser,CG1572,Msp300,CG3285,dpr18,Nmdar2,Ir41a,PGRP-LA,ABCD,CG14024,MsR2,Or42b,kek1,Or65c,GluRIIC,scrib,CG13794,ACXD,CG32320,CG41128,Syt12,CNT1,CG33310,CG7882,CG42826,Pdfr,shg,Tsp96F,Inx2,ZnT63C,CG14439,CG17600,kcc,Pzl,DIP-lambda,mGluR,DIP-delta,Cyt-b5-r,Or85b,Ca-beta | | 5.047e-03 | -5.29 | hemopoiesis | biological process | GO:0030097 | 80 | 10 | 12345 | 597 | Ser,chb,shg,Elp3,Tor,ths,Su(H),Raf,RhoL,dpp | | 5.052e-03 | -5.29 | positive regulation of multicellular organismal process | biological process | GO:0051240 | 266 | 23 | 12345 | 597 | shg,hth,Pak,Sh,step,rictor,spri,dpp,wake,Octbeta2R,EcR,Ubx,Sox102F,mnb,kay,Ank2,dsh,qvr,Tor,Raf,Tk,Src42A,bun | | 5.070e-03 | -5.28 | Wnt signaling | KEGG pathways | dme\_M00677 | 14 | 4 | 3198 | 170 | fz2,dsh,sgg,fz4 | | 5.070e-03 | -5.28 | Wnt signaling | KEGG pathways | M00677 | 14 | 4 | 3198 | 170 | fz2,dsh,fz4,sgg | | 5.139e-03 | -5.27 | striated muscle tissue development | biological process | GO:0014706 | 24 | 5 | 12345 | 597 | Msp300,Khc,flw,EcR,tin | | 5.143e-03 | -5.27 | MODULE\_27 | MSigDB lists | MODULE\_27 | 40 | 6 | 4161 | 170 | fz2,Pkc53E,LRP1,KaiR1D,mGluR,Octbeta2R | | 5.143e-03 | -5.27 | GO\_REGULATION\_OF\_RAS\_PROTEIN\_SIGNAL\_TRANSDUCTION | MSigDB lists | GO\_REGULATION\_OF\_RAS\_PROTEIN\_SIGNAL\_TRANSDUCTION | 40 | 6 | 4161 | 170 | IP3K2,CG5522,sxc,step,RhoGAP1A,Cdep | | 5.143e-03 | -5.27 | GSE29949\_MICROGLIA\_BRAIN\_VS\_CD8\_NEG\_DC\_SPLEEN\_DN | MSigDB lists | GSE29949\_MICROGLIA\_BRAIN\_VS\_CD8\_NEG\_DC\_SPLEEN\_DN | 40 | 6 | 4161 | 170 | Dys,Raf,kcc,Hr96,Tif-IA,ABCD | | 5.143e-03 | -5.27 | GSE37301\_CD4\_TCELL\_VS\_GRANULOCYTE\_MONOCYTE\_PROGENITOR\_DN | MSigDB lists | GSE37301\_CD4\_TCELL\_VS\_GRANULOCYTE\_MONOCYTE\_PROGENITOR\_DN | 40 | 6 | 4161 | 170 | IP3K2,Pgant6,Pde6,scramb1,eIF4E7,Mctp | | 5.143e-03 | -5.27 | GO\_INFLAMMATORY\_RESPONSE | MSigDB lists | GO\_INFLAMMATORY\_RESPONSE | 40 | 6 | 4161 | 170 | Su(H),CG5493,Ugt301D1,dpp,scramb1,dsd | | 5.186e-03 | -5.26 | dlg1 (discs large 1) | protein interactions | 32083 | 52 | 8 | 9204 | 481 | wake,dsh,CG42672,shg,Pak,gukh,Sh,scrib | | 5.186e-03 | -5.26 | S (Star) | protein interactions | 33281 | 52 | 8 | 9204 | 481 | Raf,Vha36-3,bun,step,bru1,csw,Src42A,shg | | 5.202e-03 | -5.26 | positive regulation of cellular biosynthetic process | biological process | GO:0031328 | 594 | 43 | 12345 | 597 | bol,CG1894,esg,retn,hth,rictor,CG42830,gce,tyf,Eip74EF,Iswi,bi,CG43689,mnb,kay,Su(H),Tor,polybromo,scro,TfIIFbeta,B-H2,CrebB,Antp,Awh,EcR,CycT,CG14252,Hr96,Atf6,sno,ems,mid,opa,CG4374,foxo,Lpin,GATAe,Sox102F,Mrtf,tin,CG12071,CG10462,TfAP-2 | | 5.202e-03 | -5.26 | regulation of multicellular organismal development | biological process | GO:2000026 | 594 | 43 | 12345 | 597 | Octbeta2R,Khc,Ubx,Mbs,Pak,hth,spri,rictor,Dlic,Src42A,raw,Syn2,mnb,Mob2,Elp3,hdc,kay,Syx1A,Ank2,bnl,Tor,Su(H),barc,sgg,ex,numb,CG9380,scrib,Antp,EcR,Ser,PGRP-LA,shg,mid,csw,rst,dpp,Mlp84B,bun,CG42797,dsh,Raf,ham | | 5.202e-03 | -5.26 | positive regulation of biosynthetic process | biological process | GO:0009891 | 594 | 43 | 12345 | 597 | mnb,kay,scro,polybromo,Tor,Su(H),TfIIFbeta,B-H2,Iswi,Eip74EF,CG43689,bi,hth,rictor,CG42830,tyf,gce,bol,esg,CG1894,retn,Mrtf,Sox102F,GATAe,tin,CG12071,CG10462,TfAP-2,CG4374,foxo,Lpin,Atf6,sno,Hr96,CG14252,ems,opa,mid,CrebB,Antp,Awh,EcR,CycT | | 5.209e-03 | -5.26 | regulation of endocytosis | biological process | GO:0030100 | 56 | 8 | 12345 | 597 | scrib,numb,cindr,sdt,step,Pkc53E,spri,mnb | | 5.240e-03 | -5.25 | regulation of cellular response to growth factor stimulus | biological process | GO:0090287 | 68 | 9 | 12345 | 597 | Lpin,kek5,Ptp61F,trol,GluRIIC,tld,Snoo,hth,sxc | | 5.240e-03 | -5.25 | negative regulation of protein phosphorylation | biological process | GO:0001933 | 68 | 9 | 12345 | 597 | flw,chinmo,Ptp61F,kek1,et,raw,sws,sgg,mnb | | 5.249e-03 | -5.25 | ELVIDGE\_HIF1A\_TARGETS\_DN | MSigDB lists | ELVIDGE\_HIF1A\_TARGETS\_DN | 18 | 4 | 4161 | 170 | Mlp84B,Su(H),CG30036,LpR2 | | 5.249e-03 | -5.25 | RPS14\_DN.V1\_UP | MSigDB lists | RPS14\_DN.V1\_UP | 18 | 4 | 4161 | 170 | dnc,Ser,Ca-beta,Syn2 | | 5.249e-03 | -5.25 | GSE7218\_UNSTIM\_VS\_ANTIGEN\_STIM\_THROUGH\_IGM\_BCELL\_UP | MSigDB lists | GSE7218\_UNSTIM\_VS\_ANTIGEN\_STIM\_THROUGH\_IGM\_BCELL\_UP | 18 | 4 | 4161 | 170 | Syx1A,Gas8,Stt3B,fz2 | | 5.250e-03 | -5.25 | anterior head segmentation | biological process | GO:0035288 | 8 | 3 | 12345 | 597 | hth,ems,Antp | | 5.250e-03 | -5.25 | anterior head development | biological process | GO:0097065 | 8 | 3 | 12345 | 597 | ems,Antp,hth | | 5.250e-03 | -5.25 | positive regulation of feeding behavior | biological process | GO:2000253 | 8 | 3 | 12345 | 597 | mnb,CrebB,foxo | | 5.252e-03 | -5.25 | C1 | smart domains | SM00109 | 27 | 6 | 5086 | 315 | rdgA,Stacl,unc-13,Pkc53E,Pkc98E,Raf | | 5.287e-03 | -5.24 | GCM\_MAP4K4 | MSigDB lists | GCM\_MAP4K4 | 53 | 7 | 4161 | 170 | dsd,KrT95D,Dlic,CG41099,CG12121,CG42672,CG33090 | | 5.287e-03 | -5.24 | CACCAGC\_MIR138 | MSigDB lists | CACCAGC\_MIR138 | 53 | 7 | 4161 | 170 | Ptp61F,Doa,MCU,csw,CG5522,CG6966,Rbp6 | | 5.315e-03 | -5.24 | GO\_RESPONSE\_TO\_OXYGEN\_LEVELS | MSigDB lists | GO\_RESPONSE\_TO\_OXYGEN\_LEVELS | 81 | 9 | 4161 | 170 | Su(H),Tor,Pkc98E,dpp,LpR2,Cat,Octbeta2R,GlyP,Pak | | 5.335e-03 | -5.23 | tetrapyrrole binding | molecular function | GO:0046906 | 136 | 14 | 12176 | 580 | Cyp6u1,Cyp4p2,Cyp12a4,Cyt-b5-r,Cyp304a1,spok,Cyp28d1,CG3556,glob3,shd,Cyp28d2,Cat,Cyp6a13,Cyp309a1 | | 5.338e-03 | -5.23 | localization | biological process | GO:0051179 | 2206 | 131 | 12345 | 597 | Gas8,shakB,hth,Flo2,cert,Dlic,ninaD,KaiR1D,Syt12,RhoGAP100F,ine,tamo,CG8184,Fhos,fz2,kay,CG4301,Fife,bnl,msps,app,CG3285,RhoL,Src42A,raw,Nnf1b,shg,Obp56h,CG7720,CG17883,g,shd,AGO2,Rab9Fa,lncRNA:Hsromega,Mrtf,ABCD,CG12782,nuf,CG17387,by,CG18659,sws,CG8468,CG7483,Actn,path,Syx1A,numb,ths,CG9132,Vha100-5,egh,CG33934,csw,Nmdar2,rols,CG13794,Ank,EcR,Inx2,rib,Frq2,IFT46,cindr,olf186-F,bun,Glut1,Nlg2,Cadps,Pak,nompB,CG33310,Cln3,nrv2,KrT95D,Vha36-3,CG14606,kcc,Khc,Tusp,cpx,ex,Tor,bbg,CG42826,Tk,sky,Frq1,boi,CG14439,chk,dpp,CG5059,opa,Irk3,CG30503,CG3168,dsh,tin,CG42235,Raf,Msp300,CG41099,wbl,chb,chinmo,rictor,pod1,CG7882,Doa,sdt,CG10960,CG3556,esg,flw,unc-13,Ca-beta,ZnT63C,p130CAS,gukh,CG9981,scramb1,Sh,CNT1,wake,MCU,scrib,Nha1,Ucp4A,GluRIIC | | 5.420e-03 | -5.22 | guanyl-nucleotide exchange factor activity | molecular function | GO:0005085 | 82 | 10 | 12176 | 580 | CG5522,Cdep,RhoGAP1A,Exn,step,RhoGEF3,siz,CG18659,CG10188,spri | | 5.425e-03 | -5.22 | shg (shotgun) | protein interactions | 37386 | 87 | 11 | 9204 | 481 | raw,RhoL,cindr,sgg,step,Inx2,jus,Src42A,Pak,sdt,dsh | | 5.434e-03 | -5.22 | signaling receptor binding | molecular function | GO:0005102 | 256 | 22 | 12176 | 580 | spri,scrib,numb,CG30054,wake,Src42A,bnl,Ser,upd3,kek1,Gpb5,Tk,Neb-cGP,RYa,Nlg2,Gprk1,pirk,dpp,Idgf6,dsh,ITP,ths | | 5.460e-03 | -5.21 | intercellular bridge | cellular component | GO:0045171 | 26 | 5 | 11940 | 538 | Lpin,sgg,chb,mino,cindr | | 5.601e-03 | -5.18 | svp (seven up) | protein interactions | 41491 | 32 | 6 | 9204 | 481 | Raf,csw,Antp,EcR,Ubx,kay | | 5.712e-03 | -5.17 | negative regulation of metabolic process | biological process | GO:0009892 | 823 | 56 | 12345 | 597 | E5,bru1,bol,retn,Ubx,esg,flw,Khc,CG1894,CG12605,Acp62F,sws,Spn88Eb,raw,et,tld,kek1,Iswi,CG43689,msi,bi,hdc,crol,mnb,numb,sgg,B-H2,cic,polybromo,Su(H),Tor,Antp,scrib,REG,Ptp61F,EcR,AGO2,Marf1,CG31224,dpp,CG7368,Achl,Maf1,mid,foxo,HPS4,toe,chinmo,tin,Sox102F,GATAe,lncRNA:bxd,AGO3,TfAP-2,CG12071,Raf | | 5.740e-03 | -5.16 | BMI1\_DN.V1\_UP | MSigDB lists | BMI1\_DN.V1\_UP | 29 | 5 | 4161 | 170 | LpR2,Syx1A,Octbeta2R,fz2,Ser | | 5.740e-03 | -5.16 | GO\_REGULATION\_OF\_ERK1\_AND\_ERK2\_CASCADE | MSigDB lists | GO\_REGULATION\_OF\_ERK1\_AND\_ERK2\_CASCADE | 29 | 5 | 4161 | 170 | Raf,Ptp61F,csw,Pkc53E,dpp | | 5.740e-03 | -5.16 | MORF\_MAP2K7 | MSigDB lists | MORF\_MAP2K7 | 29 | 5 | 4161 | 170 | dnc,fz2,CG40178,Dys,Hr96 | | 5.740e-03 | -5.16 | GO\_REGULATION\_OF\_LIPID\_BIOSYNTHETIC\_PROCESS | MSigDB lists | GO\_REGULATION\_OF\_LIPID\_BIOSYNTHETIC\_PROCESS | 29 | 5 | 4161 | 170 | dpp,Tor,mino,EcR,Sod1 | | 5.740e-03 | -5.16 | MORF\_BCL2L11 | MSigDB lists | MORF\_BCL2L11 | 29 | 5 | 4161 | 170 | Hr96,Dys,CG40178,fz2,dnc | | 5.775e-03 | -5.15 | Malpighian tubule development | biological process | GO:0072002 | 69 | 9 | 12345 | 597 | dpp,numb,rols,rib,hth,Ubx,EcR,raw,scrib | | 5.775e-03 | -5.15 | renal tubule development | biological process | GO:0061326 | 69 | 9 | 12345 | 597 | numb,dpp,rols,rib,hth,Ubx,EcR,raw,scrib | | 5.825e-03 | -5.15 | AML1\_Q6 | MSigDB lists | AML1\_Q6 | 41 | 6 | 4161 | 170 | eIF2Bgamma,CG12547,CG3961,DNaseII,Glut1,Ank | | 5.825e-03 | -5.15 | AML1\_01 | MSigDB lists | AML1\_01 | 41 | 6 | 4161 | 170 | eIF2Bgamma,CG12547,CG3961,DNaseII,Glut1,Ank | | 5.861e-03 | -5.14 | nervous system process | biological process | GO:0050877 | 564 | 41 | 12345 | 597 | Tk,dpr15,Obp56e,mnb,Mob2,Gr39a,mGluR,Ank2,Su(H),sgg,Pkc98E,dpr12,ine,CG5541,nompB,Or65c,sws,nrv2,geko,rdgA,dpr18,Root,Cam,Gr22c,Or85b,CrebB,scrib,dnc,HDAC4,nemy,EcR,dpr5,Sh,CG5921,Or42b,MCU,Gld2,Nmdar2,Obp56h,klg,CG31750 | | 5.864e-03 | -5.14 | structural constituent of muscle | molecular function | GO:0008307 | 16 | 4 | 12176 | 580 | rols,Syn2,Mlp84B,Dys | | 5.870e-03 | -5.14 | GO\_RESPONSE\_TO\_INSULIN | MSigDB lists | GO\_RESPONSE\_TO\_INSULIN | 54 | 7 | 4161 | 170 | mino,sxc,Tor,LpR2,Cat,Pak,Ptp61F | | 5.870e-03 | -5.14 | AP4\_01 | MSigDB lists | AP4\_01 | 54 | 7 | 4161 | 170 | EcR,Iswi,LRP1,eIF4E7,Nmdar2,fus,Dys | | 5.901e-03 | -5.13 | Facilitative Na+-independent glucose transporters | REACTOME pathways | R-DME-428790 | 24 | 5 | 3923 | 197 | CG10960,CG3285,CG14606,CG7882,Glut1 | | 5.995e-03 | -5.12 | FORTSCHEGGER\_PHF8\_TARGETS\_UP | MSigDB lists | FORTSCHEGGER\_PHF8\_TARGETS\_UP | 68 | 8 | 4161 | 170 | Cln3,Glut1,dnc,dpp,fz2,Ptp61F,Pde1c,LpR2 | | 5.995e-03 | -5.12 | WGGAATGY\_TEF1\_Q6 | MSigDB lists | WGGAATGY\_TEF1\_Q6 | 68 | 8 | 4161 | 170 | Pde1c,Dys,dsh,Fhos,kcc,fus,dsd,csw | | 6.081e-03 | -5.10 | regulation of kinase activity | biological process | GO:0043549 | 136 | 14 | 12345 | 597 | Ste:CG33247,CG41520,kek1,CycT,Stlk,Pak,mnb,kay,step,rictor,Raf,sgg,sws,SteXh:CG42398 | | 6.104e-03 | -5.10 | calcium-mediated signaling | biological process | GO:0019722 | 35 | 6 | 12345 | 597 | TpnC25D,Ca-beta,Cam,CG13898,CG13526,TpnC41C | | 6.104e-03 | -5.10 | positive regulation of neuromuscular junction development | biological process | GO:1904398 | 35 | 6 | 12345 | 597 | Tor,rictor,Ank2,Octbeta2R,Pak,mnb | | 6.104e-03 | -5.10 | cell fate commitment involved in pattern specification | biological process | GO:0060581 | 35 | 6 | 12345 | 597 | Su(H),nuf,EcR,numb,Ubx,scrib | | 6.104e-03 | -5.10 | positive regulation of synaptic growth at neuromuscular junction | biological process | GO:0045887 | 35 | 6 | 12345 | 597 | Tor,Pak,mnb,Octbeta2R,rictor,Ank2 | | 6.153e-03 | -5.09 | GO\_PLASMA\_MEMBRANE\_REGION | MSigDB lists | GO\_PLASMA\_MEMBRANE\_REGION | 215 | 17 | 4161 | 170 | Syn2,dnc,Nmdar2,mGluR,Dys,Octbeta2R,Frq2,Cln3,Ank,Ser,unc-13,Tsp96F,Pak,Flo2,KaiR1D,Frq1,Nlg2 | | 6.182e-03 | -5.09 | animal organ formation | biological process | GO:0048645 | 46 | 7 | 12345 | 597 | dsh,hth,B-H2,numb,Ubx,Su(H),nuf | | 6.182e-03 | -5.09 | cellular hormone metabolic process | biological process | GO:0034754 | 46 | 7 | 12345 | 597 | CG9519,foxo,shd,CG9514,spok,CG3842,woc | | 6.186e-03 | -5.09 | positive regulation of circadian sleep/wake cycle, sleep | biological process | GO:0045938 | 16 | 4 | 12345 | 597 | Sh,qvr,EcR,wake | | 6.186e-03 | -5.09 | canonical Wnt signaling pathway | biological process | GO:0060070 | 16 | 4 | 12345 | 597 | flw,fz2,fz4,dsh | | 6.186e-03 | -5.09 | larval lymph gland hemocyte differentiation | biological process | GO:0035168 | 16 | 4 | 12345 | 597 | dpp,Raf,Tor,ths | | 6.191e-03 | -5.08 | KOYAMA\_SEMA3B\_TARGETS\_DN | MSigDB lists | KOYAMA\_SEMA3B\_TARGETS\_DN | 98 | 10 | 4161 | 170 | app,dsd,Cdep,pod1,csw,CG40045,Raf,sno,CG33090,Ser | | 6.216e-03 | -5.08 | GO\_SOMATODENDRITIC\_COMPARTMENT | MSigDB lists | GO\_SOMATODENDRITIC\_COMPARTMENT | 163 | 14 | 4161 | 170 | Maf1,Tor,Frq2,sxc,Sod1,CG7483,Frq1,Ank,KaiR1D,LRP1,Pde6,Octbeta2R,mGluR,Pak | | 6.340e-03 | -5.06 | CadN (Cadherin-N) | protein interactions | 35070 | 15 | 4 | 9204 | 481 | shg,jus,sgg,siz | | 6.351e-03 | -5.06 | negative regulation of phosphorylation | biological process | GO:0042326 | 70 | 9 | 12345 | 597 | mnb,sgg,sws,et,raw,kek1,chinmo,flw,Ptp61F | | 6.356e-03 | -5.06 | GO\_POSITIVE\_REGULATION\_OF\_HYDROLASE\_ACTIVITY | MSigDB lists | GO\_POSITIVE\_REGULATION\_OF\_HYDROLASE\_ACTIVITY | 198 | 16 | 4161 | 170 | CG34408,step,Octbeta2R,dsh,scramb1,Nmdar2,REG,Cdep,sky,CG17883,RhoGAP1A,dpp,CG5522,Pkc98E,EcR,eIF2Bgamma | | 6.372e-03 | -5.06 | regulation of neuron differentiation | biological process | GO:0045664 | 180 | 17 | 12345 | 597 | Tor,Raf,ham,sgg,CG42797,Elp3,mnb,dsh,hdc,Src42A,csw,Dlic,hth,Mbs,Pak,rictor,EcR | | 6.387e-03 | -5.05 | up (upheld) | protein interactions | 32314 | 43 | 7 | 9204 | 481 | TpnC4,Scp1,TpnC41C,Actn,sgg,Act88F,Prm | | 6.394e-03 | -5.05 | sarcomere | cellular component | GO:0030017 | 62 | 8 | 11940 | 538 | rols,zormin,Prm,Act88F,Msp300,Actn,TpnC25D,Mlp84B | | 6.396e-03 | -5.05 | regulation of transferase activity | biological process | GO:0051338 | 151 | 15 | 12345 | 597 | Raf,SteXh:CG42398,sws,sgg,Pak,mnb,step,kay,rictor,CG6966,CycT,Stlk,Ste:CG33247,CG41520,kek1 | | 6.440e-03 | -5.05 | SESTO\_RESPONSE\_TO\_UV\_C2 | MSigDB lists | SESTO\_RESPONSE\_TO\_UV\_C2 | 19 | 4 | 4161 | 170 | Ser,TfAP-2,scramb1,Khc | | 6.440e-03 | -5.05 | GSE9946\_MATURE\_STIMULATORY\_VS\_PROSTAGLANDINE2\_TREATED\_MATURE\_DC\_DN | MSigDB lists | GSE9946\_MATURE\_STIMULATORY\_VS\_PROSTAGLANDINE2\_TREATED\_MATURE\_DC\_DN | 19 | 4 | 4161 | 170 | LpR2,CG14105,Ser,wake | | 6.440e-03 | -5.05 | GO\_INNER\_EAR\_MORPHOGENESIS | MSigDB lists | GO\_INNER\_EAR\_MORPHOGENESIS | 19 | 4 | 4161 | 170 | TfAP-2,RhoGAP1A,Sod1,dsh | | 6.440e-03 | -5.05 | GO\_POSITIVE\_REGULATION\_OF\_CATION\_TRANSMEMBRANE\_TRANSPORT | MSigDB lists | GO\_POSITIVE\_REGULATION\_OF\_CATION\_TRANSMEMBRANE\_TRANSPORT | 19 | 4 | 4161 | 170 | Ank,Dys,AstA-R1,CCAP-R | | 6.443e-03 | -5.04 | GO\_REGULATION\_OF\_CELLULAR\_COMPONENT\_BIOGENESIS | MSigDB lists | GO\_REGULATION\_OF\_CELLULAR\_COMPONENT\_BIOGENESIS | 216 | 17 | 4161 | 170 | Cdep,sky,CG17883,Pkc53E,Pak,Raf,nompB,Pkc98E,EcR,Tor,csw,chb,TfIIFbeta,Syx1A,Tif-IA,Fhos,fz2 | | 6.452e-03 | -5.04 | GTPase binding | molecular function | GO:0051020 | 198 | 18 | 12176 | 580 | CG17883,CG10188,tamo,Fife,Pak,CG18659,Gas8,CG42390,Cdep,nuf,Exn,step,RhoGEF3,siz,spri,RhoGAP1A,CG5522,RhoGAP18B | | 6.500e-03 | -5.04 | GSE28237\_EARLY\_VS\_LATE\_GC\_BCELL\_DN | MSigDB lists | GSE28237\_EARLY\_VS\_LATE\_GC\_BCELL\_DN | 55 | 7 | 4161 | 170 | Rat1,CG3842,kcc,rdgA,Root,RhoGAP1A,PIG-O | | 6.500e-03 | -5.04 | GSE1432\_CTRL\_VS\_IFNG\_1H\_MICROGLIA\_DN | MSigDB lists | GSE1432\_CTRL\_VS\_IFNG\_1H\_MICROGLIA\_DN | 55 | 7 | 4161 | 170 | Ca-beta,CG6966,fz2,ABCD,Pak,GlyP,CNT1 | | 6.503e-03 | -5.04 | BREDEMEYER\_RAG\_SIGNALING\_VIA\_ATM\_NOT\_VIA\_NFKB\_DN | MSigDB lists | BREDEMEYER\_RAG\_SIGNALING\_VIA\_ATM\_NOT\_VIA\_NFKB\_DN | 10 | 3 | 4161 | 170 | D2hgdh,Khc,Ank | | 6.503e-03 | -5.04 | GO\_POSITIVE\_REGULATION\_OF\_ENDOTHELIAL\_CELL\_PROLIFERATION | MSigDB lists | GO\_POSITIVE\_REGULATION\_OF\_ENDOTHELIAL\_CELL\_PROLIFERATION | 10 | 3 | 4161 | 170 | Pkc53E,dpp,Tor | | 6.503e-03 | -5.04 | PID\_IFNG\_PATHWAY | MSigDB lists | PID\_IFNG\_PATHWAY | 10 | 3 | 4161 | 170 | Ptp61F,Tor,csw | | 6.503e-03 | -5.04 | AMIT\_EGF\_RESPONSE\_60\_HELA | MSigDB lists | AMIT\_EGF\_RESPONSE\_60\_HELA | 10 | 3 | 4161 | 170 | csw,Glut1,scramb1 | | 6.503e-03 | -5.04 | GO\_REGULATION\_OF\_T\_CELL\_DIFFERENTIATION | MSigDB lists | GO\_REGULATION\_OF\_T\_CELL\_DIFFERENTIATION | 10 | 3 | 4161 | 170 | IP3K2,Sod1,Ptp61F | | 6.503e-03 | -5.04 | MODULE\_455 | MSigDB lists | MODULE\_455 | 10 | 3 | 4161 | 170 | CG34408,KrT95D,unc-13 | | 6.503e-03 | -5.04 | GO\_REGULATION\_OF\_NEUROLOGICAL\_SYSTEM\_PROCESS | MSigDB lists | GO\_REGULATION\_OF\_NEUROLOGICAL\_SYSTEM\_PROCESS | 10 | 3 | 4161 | 170 | Nmdar2,Octbeta2R,Tor | | 6.503e-03 | -5.04 | GO\_AORTA\_DEVELOPMENT | MSigDB lists | GO\_AORTA\_DEVELOPMENT | 10 | 3 | 4161 | 170 | Su(H),Ser,LRP1 | | 6.503e-03 | -5.04 | GO\_CALCIUM\_CHANNEL\_REGULATOR\_ACTIVITY | MSigDB lists | GO\_CALCIUM\_CHANNEL\_REGULATOR\_ACTIVITY | 10 | 3 | 4161 | 170 | Syx1A,ZnT63C,mGluR | | 6.503e-03 | -5.04 | GO\_RIBONUCLEOTIDE\_CATABOLIC\_PROCESS | MSigDB lists | GO\_RIBONUCLEOTIDE\_CATABOLIC\_PROCESS | 10 | 3 | 4161 | 170 | Pde8,Pde6,dnc | | 6.507e-03 | -5.03 | protein binding | molecular function | GO:0005515 | 2410 | 139 | 12176 | 580 | melt,AGO2,rols,Ank2,siz,wbl,dsh,Yeti,pod1,CycT,Fife,eIF4E7,zormin,Pde6,unc-13,et,mid,IP3K2,PhKgamma,CG5445,CG11970,bun,Dlic,fz2,Gld2,CG13251,shg,Actn,sdt,Gas8,Atf6,Ank,csw,kek1,Raf,TfIIFbeta,Sod1,msps,scrib,bi,wake,chb,Su(H),Idgf6,Cdep,Vha100-5,CG12782,Cam,CG10188,Gpb5,CG7483,Tk,Tor,Ser,upd3,sgg,cindr,nompB,CG42673,GlyP,CG1894,nuf,dpp,Meltrin,pirk,ex,tyf,Neb-cGP,CG17883,RYa,cic,Der-1,Khc,Hil,Mlp84B,spri,Msp300,Mbs,Dys,CG30054,cato,gce,Fhos,Exn,Syt12,Set2,CG18109,Pak,CG42390,gukh,CG18135,RhoGAP18B,Ptp61F,Mob2,Hr96,sws,numb,Maf1,CG34417,Src42A,CG32264,Snoo,hth,Cpsf160,ITP,Mondo,CG18659,rst,Mrtf,RhoGAP1A,retn,kay,RhoGEF3,Ubx,Gprk1,step,boi,EcR,CadN2,ths,CG5921,tamo,bnl,Fer2,CG5522,Tif-IA,Syn2,by,woc,bru1,foxo,Iswi,fz4,Pfdn4,Pde1c,Pdfr,Syx1A,cpx,Nlg2 | | 6.521e-03 | -5.03 | Wnk (Wnk kinase) | protein interactions | 40391 | 8 | 3 | 9204 | 481 | dsh,fz2,Awh | | 6.523e-03 | -5.03 | regulation of hormone levels | biological process | GO:0010817 | 96 | 11 | 12345 | 597 | spok,Glut1,CG3842,Fad2,foxo,CG9514,shd,Obp56h,CG9650,woc,CG9519 | | 6.538e-03 | -5.03 | fly chr24C8-24C8\|2-11 | chromosome location | fly chr24C8-24C8\|2-11 | 4 | 2 | 25010 | 845 | TTLL4B,Cep97 | | 6.538e-03 | -5.03 | fly chr24C8-24C8\|2-11 cM | chromosome location | fly chr24C8-24C8\|2-11 cM | 4 | 2 | 25010 | 845 | TTLL4B,Cep97 | | 6.538e-03 | -5.03 | fly chr76B5-76B5\|3-45 | chromosome location | fly chr76B5-76B5\|3-45 | 4 | 2 | 25010 | 845 | CG9449,CG9451 | | 6.538e-03 | -5.03 | fly chr34A1-34A1\|2-47 cM | chromosome location | fly chr34A1-34A1\|2-47 cM | 4 | 2 | 25010 | 845 | lncRNA:CR44971,kek1 | | 6.538e-03 | -5.03 | fly chr98B7-98B7\|3-94 | chromosome location | fly chr98B7-98B7\|3-94 | 4 | 2 | 25010 | 845 | lncRNA:CR46087,trv | | 6.538e-03 | -5.03 | fly chr34A1-34A1\|2-47 | chromosome location | fly chr34A1-34A1\|2-47 | 4 | 2 | 25010 | 845 | lncRNA:CR44971,kek1 | | 6.538e-03 | -5.03 | fly chr76B5-76B5\|3-45 cM | chromosome location | fly chr76B5-76B5\|3-45 cM | 4 | 2 | 25010 | 845 | CG9449,CG9451 | | 6.538e-03 | -5.03 | fly chr98B7-98B7\|3-94 cM | chromosome location | fly chr98B7-98B7\|3-94 cM | 4 | 2 | 25010 | 845 | lncRNA:CR46087,trv | | 6.550e-03 | -5.03 | GO\_REGULATION\_OF\_ACTIN\_FILAMENT\_BASED\_PROCESS | MSigDB lists | GO\_REGULATION\_OF\_ACTIN\_FILAMENT\_BASED\_PROCESS | 69 | 8 | 4161 | 170 | Tor,Pkc98E,dnc,Fhos,LRP1,Raf,Pak,chb | | 6.571e-03 | -5.03 | CHIANG\_LIVER\_CANCER\_SUBCLASS\_CTNNB1\_UP | MSigDB lists | CHIANG\_LIVER\_CANCER\_SUBCLASS\_CTNNB1\_UP | 42 | 6 | 4161 | 170 | LpR2,CG3961,Octbeta2R,mino,wake,CG15093 | | 6.571e-03 | -5.03 | GSE20727\_CTRL\_VS\_H2O2\_TREATED\_DC\_DN | MSigDB lists | GSE20727\_CTRL\_VS\_H2O2\_TREATED\_DC\_DN | 42 | 6 | 4161 | 170 | Elp3,Act87E,bru1,GlyP,Dys,Ank | | 6.571e-03 | -5.03 | CAIRO\_HEPATOBLASTOMA\_CLASSES\_DN | MSigDB lists | CAIRO\_HEPATOBLASTOMA\_CLASSES\_DN | 42 | 6 | 4161 | 170 | Cat,ZnT63C,Tsp96F,CG3961,TfAP-2,Mlp84B | | 6.571e-03 | -5.03 | TCCATTKW\_UNKNOWN | MSigDB lists | TCCATTKW\_UNKNOWN | 42 | 6 | 4161 | 170 | CG4341,dpp,mRpL24,CG12547,chb,DNaseII | | 6.614e-03 | -5.02 | negative regulation of nucleobase-containing compound metabolic process | biological process | GO:0045934 | 417 | 32 | 12345 | 597 | Iswi,CG43689,bi,crol,B-H2,cic,polybromo,Su(H),E5,bol,Ubx,retn,Khc,esg,CG1894,CG12605,foxo,HPS4,toe,tin,Sox102F,lncRNA:bxd,TfAP-2,CG12071,Antp,scrib,EcR,AGO2,CG31224,CG7368,mid,Maf1 | | 6.635e-03 | -5.02 | DBL\_dom\_sf | interpro domains | IPR035899 | 24 | 5 | 11383 | 585 | Exn,RhoGAP1A,RhoGEF3,Cdep,CG10188 | | 6.635e-03 | -5.02 | DH-domain | interpro domains | IPR000219 | 24 | 5 | 11383 | 585 | Exn,RhoGAP1A,Cdep,RhoGEF3,CG10188 | | 6.665e-03 | -5.01 | GO\_CATION\_CHANNEL\_COMPLEX | MSigDB lists | GO\_CATION\_CHANNEL\_COMPLEX | 30 | 5 | 4161 | 170 | Ca-beta,dnc,Sh,Syx1A,MCU | | 6.665e-03 | -5.01 | BMI1\_DN\_MEL18\_DN.V1\_UP | MSigDB lists | BMI1\_DN\_MEL18\_DN.V1\_UP | 30 | 5 | 4161 | 170 | Meltrin,Ser,LpR2,Syx1A,Octbeta2R | | 6.665e-03 | -5.01 | GSE7768\_OVA\_ALONE\_VS\_OVA\_WITH\_LPS\_IMMUNIZED\_MOUSE\_WHOLE\_SPLEEN\_6H\_DN | MSigDB lists | GSE7768\_OVA\_ALONE\_VS\_OVA\_WITH\_LPS\_IMMUNIZED\_MOUSE\_WHOLE\_SPLEEN\_6H\_DN | 30 | 5 | 4161 | 170 | Sh,sdt,RhoGAP1A,Treh,scramb1 | | 6.665e-03 | -5.01 | PID\_MTOR\_4PATHWAY | MSigDB lists | PID\_MTOR\_4PATHWAY | 30 | 5 | 4161 | 170 | Raf,eIF4E7,Pkc53E,Tif-IA,Tor | | 6.755e-03 | -5.00 | GO\_POSITIVE\_REGULATION\_OF\_CELL\_DEVELOPMENT | MSigDB lists | GO\_POSITIVE\_REGULATION\_OF\_CELL\_DEVELOPMENT | 84 | 9 | 4161 | 170 | Dys,LpR2,unc-13,Nha1,Raf,dpp,Tif-IA,Tor,CG42672 | | 6.767e-03 | -5.00 | Response to elevated platelet cytosolic Ca2+ | REACTOME pathways | R-DME-76005 | 107 | 12 | 3923 | 197 | Pkc53E,CG12547,kek5,ninaD,fw,kek1,Tsf1,CG41520,Cam,Actn,Qsox2,CG32264 | | 6.776e-03 | -4.99 | reproductive behavior | biological process | GO:0019098 | 196 | 18 | 12345 | 597 | Nlg2,JhI-26,shep,Pde1c,TfAP-2,Gr39a,Gr22c,mGluR,Fad2,retn,Ebp,lncRNA:flam,dnc,bol,fru,Obp56h,egh,Sh | | 6.780e-03 | -4.99 | embryonic heart tube anterior/posterior pattern specification | biological process | GO:0035054 | 3 | 2 | 12345 | 597 | EcR,Ubx | | 6.780e-03 | -4.99 | positive regulation of ARF protein signal transduction | biological process | GO:0032014 | 3 | 2 | 12345 | 597 | step,siz | | 6.780e-03 | -4.99 | regulation of calcineurin-NFAT signaling cascade | biological process | GO:0070884 | 3 | 2 | 12345 | 597 | sgg,olf186-F | | 6.780e-03 | -4.99 | regulation of calcineurin-mediated signaling | biological process | GO:0106056 | 3 | 2 | 12345 | 597 | olf186-F,sgg | | 6.780e-03 | -4.99 | negative regulation of response to oxidative stress | biological process | GO:1902883 | 3 | 2 | 12345 | 597 | Fer2,Pde8 | | 6.780e-03 | -4.99 | negative regulation of cellular response to oxidative stress | biological process | GO:1900408 | 3 | 2 | 12345 | 597 | Pde8,Fer2 | | 6.780e-03 | -4.99 | specification of segmental identity, thorax | biological process | GO:0007384 | 3 | 2 | 12345 | 597 | Antp,Ubx | | 6.780e-03 | -4.99 | regulation of oxidative stress-induced cell death | biological process | GO:1903201 | 3 | 2 | 12345 | 597 | Pde8,Fer2 | | 6.808e-03 | -4.99 | border follicle cell migration | biological process | GO:0007298 | 110 | 12 | 12345 | 597 | shg,Mrtf,dsh,Raf,cindr,bbg,rols,RhoL,egh,ex,shd,EcR | | 6.808e-03 | -4.99 | PH\_DOMAIN | prosite domains | PS50003 | 66 | 10 | 6243 | 386 | cert,CG10188,Gprk1,RhoGAP1A,CG5004,Cdep,step,Cadps,CG5522,melt | | 6.832e-03 | -4.99 | RBD\_domain\_sf | interpro domains | IPR035979 | 157 | 16 | 11383 | 585 | CG44249,Spf45,Marf1,shep,CG40813,bru1,fus,CG42458,CG3594,Rbp6,Achl,CG41562,bol,trv,barc,msi | | 6.886e-03 | -4.98 | TGTTTAC\_MIR30A5P\_MIR30C\_MIR30D\_MIR30B\_MIR30E5P | MSigDB lists | TGTTTAC\_MIR30A5P\_MIR30C\_MIR30D\_MIR30B\_MIR30E5P | 148 | 13 | 4161 | 170 | Awh,Src42A,Meltrin,bru1,dnc,rdgA,CG5522,REG,Cadps,Dys,mGluR,Ptp61F,CG9132 | | 6.892e-03 | -4.98 | positive regulation of signal transduction | biological process | GO:0009967 | 369 | 29 | 12345 | 597 | Lpin,Gprk1,tld,foxo,trol,Src42A,olf186-F,Pde8,ex,Raf,MESK2,bnl,Tor,lncRNA:Hsromega,Exn,Mos,melt,flw,RhoGAP1A,CG10188,Flo2,siz,dpp,Doa,rictor,step,sxc,sno,hth | | 6.969e-03 | -4.97 | steroid metabolic process | biological process | GO:0008202 | 47 | 7 | 12345 | 597 | woc,spok,CG9514,shd,CG33090,CG9519,foxo | | 6.969e-03 | -4.97 | VEGFR2 mediated vascular permeability | REACTOME pathways | R-DME-5218920 | 16 | 4 | 3923 | 197 | rictor,Pak,Cam,Tor | | 7.032e-03 | -4.96 | adherens junction organization | biological process | GO:0034332 | 36 | 6 | 12345 | 597 | sdt,cindr,scrib,CadN2,shg,Src42A | | 7.036e-03 | -4.96 | synapse part | cellular component | GO:0044456 | 211 | 18 | 11940 | 538 | Ank2,Pak,gukh,Frq2,KaiR1D,g,Exn,Cadps,Syx1A,sky,Fife,Nlg2,mnb,unc-13,RhoGAP100F,cpx,scrib,GluRIIC | | 7.081e-03 | -4.95 | CaM pathway | REACTOME pathways | R-DME-111997 | 25 | 5 | 3923 | 197 | Gprk1,Pde1c,ACXD,Cam,Pkc53E | | 7.081e-03 | -4.95 | Ca-dependent events | REACTOME pathways | R-DME-111996 | 25 | 5 | 3923 | 197 | Pkc53E,Cam,Pde1c,ACXD,Gprk1 | | 7.081e-03 | -4.95 | Calmodulin induced events | REACTOME pathways | R-DME-111933 | 25 | 5 | 3923 | 197 | Pkc53E,Cam,ACXD,Pde1c,Gprk1 | | 7.128e-03 | -4.94 | reproductive structure development | biological process | GO:0048608 | 84 | 10 | 12345 | 597 | tin,dsh,fz2,shg,rib,TfAP-2,mid,Tor,raw,tnc | | 7.128e-03 | -4.94 | reproductive system development | biological process | GO:0061458 | 84 | 10 | 12345 | 597 | rib,shg,dsh,tin,fz2,Tor,mid,TfAP-2,raw,tnc | | 7.179e-03 | -4.94 | GO\_REGULATION\_OF\_SMALL\_GTPASE\_MEDIATED\_SIGNAL\_TRANSDUCTION | MSigDB lists | GO\_REGULATION\_OF\_SMALL\_GTPASE\_MEDIATED\_SIGNAL\_TRANSDUCTION | 56 | 7 | 4161 | 170 | Cdep,CG34408,CG5522,IP3K2,step,sxc,RhoGAP1A | | 7.179e-03 | -4.94 | STAT\_01 | MSigDB lists | STAT\_01 | 56 | 7 | 4161 | 170 | Ank,scramb1,eIF4E7,CG8272,rdgA,Sod1,mRpL24 | | 7.229e-03 | -4.93 | GO\_SENSORY\_ORGAN\_DEVELOPMENT | MSigDB lists | GO\_SENSORY\_ORGAN\_DEVELOPMENT | 116 | 11 | 4161 | 170 | dsh,Ser,dpp,fz2,RpL38,TfAP-2,CG17883,csw,Sod1,Su(H),RhoGAP1A | | 7.236e-03 | -4.93 | RhoGEF | pfam domains | PF00621 | 24 | 5 | 10480 | 550 | RhoGAP1A,RhoGEF3,Exn,CG10188,Cdep | | 7.247e-03 | -4.93 | Signaling by EGFR | REACTOME pathways | R-DME-177929 | 178 | 17 | 3923 | 197 | foxo,Pkc53E,Tor,Prosbeta5R1,CG41520,Pde1c,Cam,Raf,rictor,bnl,CrebB,cindr,REG,Pkc98E,Gprk1,ACXD,csw | | 7.262e-03 | -4.93 | tsr (twinstar) | protein interactions | 37841 | 44 | 7 | 9204 | 481 | Pak,scrib,Act87E,tobi,sick,Act88F,cindr | | 7.279e-03 | -4.92 | Neurofascin interactions | REACTOME pathways | R-DME-447043 | 3 | 2 | 3923 | 197 | Ank2,Ank | | 7.279e-03 | -4.92 | MAPK6/MAPK4 signaling | REACTOME pathways | R-DME-5687128 | 3 | 2 | 3923 | 197 | Pak,foxo | | 7.279e-03 | -4.92 | Regulation of PLK1 Activity at G2/M Transition | REACTOME pathways | R-DME-2565942 | 3 | 2 | 3923 | 197 | Mbs,flw | | 7.321e-03 | -4.92 | gcm (glial cells missing) | protein interactions | 34277 | 55 | 8 | 9204 | 481 | Su(H),fru,crol,upd3,Pkc53E,esg,Dys,mid | | 7.348e-03 | -4.91 | positive regulation of phosphorylation | biological process | GO:0042327 | 139 | 14 | 12345 | 597 | Stlk,CycT,Mos,Pde8,CG41520,Src42A,Tor,bnl,Raf,dpp,Pak,lncRNA:Hsromega,rictor,step | | 7.348e-03 | -4.91 | regulation of cellular localization | biological process | GO:0060341 | 139 | 14 | 12345 | 597 | sdt,Fife,Frq2,sgg,mnb,Mctp,Syx1A,cpx,Khc,nemy,Frq1,Exn,sky,Dys | | 7.384e-03 | -4.91 | GSE43955\_10H\_VS\_60H\_ACT\_CD4\_TCELL\_WITH\_TGFB\_IL6\_DN | MSigDB lists | GSE43955\_10H\_VS\_60H\_ACT\_CD4\_TCELL\_WITH\_TGFB\_IL6\_DN | 43 | 6 | 4161 | 170 | Stt3B,Cln3,Doa,Odc1,fus,Cat | | 7.384e-03 | -4.91 | GO\_REGULATION\_OF\_TRANSPORTER\_ACTIVITY | MSigDB lists | GO\_REGULATION\_OF\_TRANSPORTER\_ACTIVITY | 43 | 6 | 4161 | 170 | AstA-R1,Ank,Dys,Pkc98E,dnc,Ca-beta | | 7.384e-03 | -4.91 | GSE40277\_EOS\_AND\_LEF1\_TRANSDUCED\_VS\_GATA1\_AND\_SATB1\_TRANSDUCED\_CD4\_TCELL\_UP | MSigDB lists | GSE40277\_EOS\_AND\_LEF1\_TRANSDUCED\_VS\_GATA1\_AND\_SATB1\_TRANSDUCED\_CD4\_TCELL\_UP | 43 | 6 | 4161 | 170 | CG5522,Mlp84B,Su(H),CG10126,scramb1,kcc | | 7.384e-03 | -4.91 | GSE2770\_IL12\_ACT\_VS\_ACT\_CD4\_TCELL\_2H\_UP | MSigDB lists | GSE2770\_IL12\_ACT\_VS\_ACT\_CD4\_TCELL\_2H\_UP | 43 | 6 | 4161 | 170 | D2hgdh,CG5339,Glut1,Syx1A,Mctp,sws | | 7.421e-03 | -4.90 | Notch Signaling Pathway | WikiPathways | WP208 | 11 | 3 | 499 | 20 | dsh,numb,Ser | | 7.448e-03 | -4.90 | Downstream signal transduction | REACTOME pathways | R-DME-186763 | 164 | 16 | 3923 | 197 | CrebB,Pkc98E,Gprk1,REG,ACXD,csw,Cam,Raf,rictor,bnl,Pkc53E,Prosbeta5R1,Tor,CG41520,Pde1c,foxo | | 7.448e-03 | -4.90 | DAP12 signaling | REACTOME pathways | R-DME-2424491 | 164 | 16 | 3923 | 197 | Prosbeta5R1,Tor,Pkc53E,Pde1c,CG41520,foxo,CrebB,csw,ACXD,Pkc98E,Gprk1,REG,Cam,bnl,rictor,Raf | | 7.448e-03 | -4.90 | Signaling by PDGF | REACTOME pathways | R-DME-186797 | 164 | 16 | 3923 | 197 | Cam,bnl,rictor,Raf,CrebB,csw,ACXD,Gprk1,Pkc98E,REG,foxo,Prosbeta5R1,Tor,Pkc53E,Pde1c,CG41520 | | 7.448e-03 | -4.90 | DAP12 interactions | REACTOME pathways | R-DME-2172127 | 164 | 16 | 3923 | 197 | Cam,bnl,Raf,rictor,CrebB,ACXD,csw,REG,Pkc98E,Gprk1,foxo,Pkc53E,Prosbeta5R1,Tor,Pde1c,CG41520 | | 7.491e-03 | -4.89 | cellular response to organic substance | biological process | GO:0071310 | 275 | 23 | 12345 | 597 | Lpin,bun,Iswi,foxo,et,Tor,Raf,ths,bnl,ex,Set2,Cam,CG7362,Gpb5,EcR,csw,opa,gce,Der-1,dpp,Sh,Atf6,rictor | | 7.559e-03 | -4.88 | GO\_ION\_HOMEOSTASIS | MSigDB lists | GO\_ION\_HOMEOSTASIS | 133 | 12 | 4161 | 170 | Pkc98E,mino,AstA-R1,MCU,GlyP,Sod1,Cln3,Ca-beta,CG4341,KaiR1D,ZnT63C,Ank | | 7.588e-03 | -4.88 | PH | smart domains | SM00233 | 67 | 10 | 5086 | 315 | step,cert,RhoGAP1A,CG5004,CG5522,Gprk1,Cdep,Cadps,melt,CG10188 | | 7.594e-03 | -4.88 | zonula adherens assembly | biological process | GO:0045186 | 9 | 3 | 12345 | 597 | shg,scrib,sdt | | 7.607e-03 | -4.88 | Rho1 (Rho1) | protein interactions | 36775 | 91 | 11 | 9204 | 481 | RhoGAP18B,flw,shg,Pak,dsh,RhoGAP100F,Mbs,Fhos,cindr,sgg,step | | 7.635e-03 | -4.87 | neuron fate commitment | biological process | GO:0048663 | 72 | 9 | 12345 | 597 | msi,melt,scrib,klg,csw,Raf,mid,kay,hth | | 7.638e-03 | -4.87 | enzyme linked receptor protein signaling pathway | biological process | GO:0007167 | 154 | 15 | 12345 | 597 | dpp,cic,ex,csw,opa,Raf,bnl,ths,Tor,fs(1)N,rictor,bun,Src42A,foxo,CG41520 | | 7.641e-03 | -4.87 | Cadherin\_cytoplasmic-dom | interpro domains | IPR000233 | 3 | 2 | 11383 | 585 | shg,CadN2 | | 7.641e-03 | -4.87 | Fibroblast\_GF\_fam | interpro domains | IPR002209 | 3 | 2 | 11383 | 585 | ths,bnl | | 7.641e-03 | -4.87 | IL1/FGF | interpro domains | IPR008996 | 3 | 2 | 11383 | 585 | bnl,ths | | 7.641e-03 | -4.87 | CLASP\_N\_dom | interpro domains | IPR024395 | 3 | 2 | 11383 | 585 | msps,chb | | 7.643e-03 | -4.87 | Cyt\_P450\_CS | interpro domains | IPR017972 | 80 | 10 | 11383 | 585 | Cyp28d1,Cyp309a1,Cyp304a1,spok,shd,Cyp12a4,Cyp6a13,Cyp28d2,Cyp4p2,Cyp6u1 | | 7.660e-03 | -4.87 | PTPase\_domain | interpro domains | IPR000242 | 16 | 4 | 11383 | 585 | csw,Ptp36E,Ptp61F,Ptp99A | | 7.675e-03 | -4.87 | negative regulation of cellular macromolecule biosynthetic process | biological process | GO:2000113 | 438 | 33 | 12345 | 597 | scrib,Antp,EcR,AGO2,CG31224,mid,Maf1,CG7368,HPS4,foxo,toe,Sox102F,lncRNA:bxd,tin,CG12071,TfAP-2,E5,bru1,esg,Khc,CG1894,retn,Ubx,CG12605,Iswi,CG43689,msi,bi,crol,polybromo,Su(H),B-H2,cic | | 7.689e-03 | -4.87 | ELVIDGE\_HYPOXIA\_UP | MSigDB lists | ELVIDGE\_HYPOXIA\_UP | 31 | 5 | 4161 | 170 | CG30036,Pkc53E,Mlp84B,Su(H),LpR2 | | 7.689e-03 | -4.87 | GSE15324\_ELF4\_KO\_VS\_WT\_NAIVE\_CD8\_TCELL\_DN | MSigDB lists | GSE15324\_ELF4\_KO\_VS\_WT\_NAIVE\_CD8\_TCELL\_DN | 31 | 5 | 4161 | 170 | Argl,DNaseII,scro,Ank,CG17387 | | 7.689e-03 | -4.87 | AREB6\_01 | MSigDB lists | AREB6\_01 | 31 | 5 | 4161 | 170 | PCB,Syx1A,fus,CG17698,TfAP-2 | | 7.689e-03 | -4.87 | GO\_CALCIUM\_ION\_TRANSMEMBRANE\_TRANSPORT | MSigDB lists | GO\_CALCIUM\_ION\_TRANSMEMBRANE\_TRANSPORT | 31 | 5 | 4161 | 170 | MCU,Frq1,Frq2,Pkc98E,Ca-beta | | 7.785e-03 | -4.86 | terminal region determination | biological process | GO:0007362 | 17 | 4 | 12345 | 597 | fs(1)N,Raf,csw,cic | | 7.785e-03 | -4.86 | anesthesia-resistant memory | biological process | GO:0007615 | 17 | 4 | 12345 | 597 | klg,MCU,CrebB,dnc | | 7.798e-03 | -4.85 | GO\_RESPONSE\_TO\_CADMIUM\_ION | MSigDB lists | GO\_RESPONSE\_TO\_CADMIUM\_ION | 20 | 4 | 4161 | 170 | ZnT63C,Cat,Sod1,mino | | 7.798e-03 | -4.85 | GO\_T\_CELL\_DIFFERENTIATION | MSigDB lists | GO\_T\_CELL\_DIFFERENTIATION | 20 | 4 | 4161 | 170 | IP3K2,fz2,Ptp61F,Raf | | 7.798e-03 | -4.85 | GO\_TRANSCRIPTIONAL\_ACTIVATOR\_ACTIVITY\_RNA\_POLYMERASE\_II\_CORE\_PROMOTER\_PROXIMAL\_REGION\_SEQUENCE\_SPECIFIC\_BINDING | MSigDB lists | GO\_TRANSCRIPTIONAL\_ACTIVATOR\_ACTIVITY\_RNA\_POLYMERASE\_II\_CORE\_PROMOTER\_PROXIMAL\_REGION\_SEQUENCE\_SPECIFIC\_BINDING | 20 | 4 | 4161 | 170 | EcR,Su(H),TfAP-2,scramb1 | | 7.798e-03 | -4.85 | KEGG\_NATURAL\_KILLER\_CELL\_MEDIATED\_CYTOTOXICITY | MSigDB lists | KEGG\_NATURAL\_KILLER\_CELL\_MEDIATED\_CYTOTOXICITY | 20 | 4 | 4161 | 170 | Pkc53E,csw,Raf,Pak | | 7.798e-03 | -4.85 | GO\_NEUROEPITHELIAL\_CELL\_DIFFERENTIATION | MSigDB lists | GO\_NEUROEPITHELIAL\_CELL\_DIFFERENTIATION | 20 | 4 | 4161 | 170 | Su(H),Sod1,dpp,Ser | | 7.798e-03 | -4.85 | GO\_REGULATION\_OF\_SEQUESTERING\_OF\_CALCIUM\_ION | MSigDB lists | GO\_REGULATION\_OF\_SEQUESTERING\_OF\_CALCIUM\_ION | 20 | 4 | 4161 | 170 | Pkc98E,dnc,CCAP-R,Dys | | 7.798e-03 | -4.85 | GO\_PURINE\_CONTAINING\_COMPOUND\_CATABOLIC\_PROCESS | MSigDB lists | GO\_PURINE\_CONTAINING\_COMPOUND\_CATABOLIC\_PROCESS | 20 | 4 | 4161 | 170 | Cat,Pde8,dnc,Pde6 | | 7.798e-03 | -4.85 | GO\_RESPONSE\_TO\_CAMP | MSigDB lists | GO\_RESPONSE\_TO\_CAMP | 20 | 4 | 4161 | 170 | dnc,CG5493,GlyP,Raf | | 7.798e-03 | -4.85 | GSE9946\_LISTERIA\_INF\_MATURE\_VS\_PROSTAGLANDINE2\_TREATED\_MATURE\_DC\_DN | MSigDB lists | GSE9946\_LISTERIA\_INF\_MATURE\_VS\_PROSTAGLANDINE2\_TREATED\_MATURE\_DC\_DN | 20 | 4 | 4161 | 170 | Uros1,Aldh-III,LpR2,dpp | | 7.798e-03 | -4.85 | GSE41176\_UNSTIM\_VS\_ANTI\_IGM\_STIM\_TAK1\_KO\_BCELL\_3H\_DN | MSigDB lists | GSE41176\_UNSTIM\_VS\_ANTI\_IGM\_STIM\_TAK1\_KO\_BCELL\_3H\_DN | 20 | 4 | 4161 | 170 | GlyP,Dys,CG4341,melt | | 7.798e-03 | -4.85 | GO\_POSITIVE\_REGULATION\_OF\_OSSIFICATION | MSigDB lists | GO\_POSITIVE\_REGULATION\_OF\_OSSIFICATION | 20 | 4 | 4161 | 170 | Octbeta2R,TfAP-2,dpp,Ser | | 7.832e-03 | -4.85 | blastoderm segmentation | biological process | GO:0007350 | 214 | 19 | 12345 | 597 | hth,shg,ems,Doa,csw,dpp,scrib,CG7483,Antp,AGO2,Khc,Ubx,dsh,fs(1)N,fz2,Raf,msps,sgg,cic | | 7.893e-03 | -4.84 | laza (lazaro) | protein interactions | 40468 | 3 | 2 | 9204 | 481 | Pld,rdgA | | 7.893e-03 | -4.84 | CG6873 (uncharacterized protein) | protein interactions | 32861 | 3 | 2 | 9204 | 481 | Act88F,Act87E | | 7.893e-03 | -4.84 | bc10 (bc10) | protein interactions | 38628 | 3 | 2 | 9204 | 481 | CG46385,rdgA | | 7.893e-03 | -4.84 | Cyp310a1 (Cyp310a1) | protein interactions | 35115 | 3 | 2 | 9204 | 481 | dsh,sgg | | 7.893e-03 | -4.84 | Apoltp (Apolipoprotein lipid transfer particle) | protein interactions | 34283 | 3 | 2 | 9204 | 481 | LpR2,sgg | | 7.893e-03 | -4.84 | Inx3 (Innexin 3) | protein interactions | 44266 | 3 | 2 | 9204 | 481 | Inx2,shg | | 7.893e-03 | -4.84 | NKAIN (Na,K-ATPase Interacting) | protein interactions | 37979 | 3 | 2 | 9204 | 481 | esn,nrv2 | | 7.893e-03 | -4.84 | CG10014 (uncharacterized protein) | protein interactions | 41481 | 3 | 2 | 9204 | 481 | Poxm,Cam | | 7.893e-03 | -4.84 | ClC-a (Chloride channel-a) | protein interactions | 41428 | 3 | 2 | 9204 | 481 | Doa,Pak | | 7.893e-03 | -4.84 | CG11426 (uncharacterized protein) | protein interactions | 40471 | 3 | 2 | 9204 | 481 | rdgA,Pld | | 7.930e-03 | -4.84 | photoreceptor cell fate commitment | biological process | GO:0046552 | 60 | 8 | 12345 | 597 | scrib,msi,melt,kay,hth,klg,csw,Raf | | 7.961e-03 | -4.83 | FGF | pfam domains | PF00167 | 3 | 2 | 10480 | 550 | bnl,ths | | 7.961e-03 | -4.83 | CLASP\_N | pfam domains | PF12348 | 3 | 2 | 10480 | 550 | chb,msps | | 7.961e-03 | -4.83 | Cadherin\_C | pfam domains | PF01049 | 3 | 2 | 10480 | 550 | shg,CadN2 | | 8.014e-03 | -4.83 | receptor regulator activity | molecular function | GO:0030545 | 114 | 12 | 12176 | 580 | Neb-cGP,Tk,Gpb5,RYa,CG41520,dpp,upd3,Ser,bnl,ths,ITP,kek1 | | 8.020e-03 | -4.83 | Ras guanyl-nucleotide exchange factor activity | molecular function | GO:0005088 | 61 | 8 | 12176 | 580 | RhoGEF3,spri,CG10188,CG18659,Cdep,RhoGAP1A,CG5522,Exn | | 8.091e-03 | -4.82 | Rho GTPase binding | molecular function | GO:0017048 | 49 | 7 | 12176 | 580 | RhoGEF3,Cdep,RhoGAP1A,CG10188,Pak,Exn,RhoGAP18B | | 8.110e-03 | -4.81 | ms(3)76Cc (male sterile (3) 76Cc) | protein interactions | 40142 | 16 | 4 | 9204 | 481 | sgg,Cam,Ubx,Aldh-III | | 8.110e-03 | -4.81 | fmt (fiery mountain) | protein interactions | 38714 | 16 | 4 | 9204 | 481 | melt,Raf,Ptp61F,scrib | | 8.110e-03 | -4.81 | Cln3 (Cln3) | protein interactions | 39981 | 16 | 4 | 9204 | 481 | Cat,Sod1,foxo,bol | | 8.110e-03 | -4.81 | Sh (Shaker) | protein interactions | 32780 | 16 | 4 | 9204 | 481 | jus,qvr,Khc,LpR2 | | 8.239e-03 | -4.80 | Y\_phosphatase | pfam domains | PF00102 | 16 | 4 | 10480 | 550 | Ptp36E,Ptp99A,csw,Ptp61F | | 8.246e-03 | -4.80 | enzyme activator activity | molecular function | GO:0008047 | 143 | 14 | 12176 | 580 | CG42390,sky,CG17883,REG,RhoGAP100F,CG41520,nrv2,rictor,RhoGAP18B,CG34408,CycT,CG33310,RhoGAP1A,Frq1 | | 8.268e-03 | -4.80 | GINESTIER\_BREAST\_CANCER\_20Q13\_AMPLIFICATION\_DN | MSigDB lists | GINESTIER\_BREAST\_CANCER\_20Q13\_AMPLIFICATION\_DN | 44 | 6 | 4161 | 170 | CG3961,Dys,step,CG33090,rdgA,Qsox2 | | 8.316e-03 | -4.79 | GO\_RECEPTOR\_ACTIVITY | MSigDB lists | GO\_RECEPTOR\_ACTIVITY | 186 | 15 | 4161 | 170 | fz2,EcR,Sk1,LRP1,LpR2,CCAP-R,Nmdar2,AstA-R1,mGluR,Octbeta2R,Der-1,Hr96,KaiR1D,dsd,Nlg2 | | 8.367e-03 | -4.78 | CD28 co-stimulation | REACTOME pathways | R-DME-389356 | 9 | 3 | 3923 | 197 | Tor,rictor,Pak | | 8.391e-03 | -4.78 | eye photoreceptor cell development | biological process | GO:0042462 | 86 | 10 | 12345 | 597 | Syn2,RhoGAP100F,CG13251,Doa,Raf,CG5921,chp,Mob2,CadN2,Cam | | 8.430e-03 | -4.78 | PH | pfam domains | PF00169 | 45 | 7 | 10480 | 550 | CG5522,Cdep,step,Gprk1,CG5004,Cadps,cert | | 8.454e-03 | -4.77 | GSE16450\_CTRL\_VS\_IFNA\_6H\_STIM\_MATURE\_NEURON\_CELL\_LINE\_DN | MSigDB lists | GSE16450\_CTRL\_VS\_IFNA\_6H\_STIM\_MATURE\_NEURON\_CELL\_LINE\_DN | 72 | 8 | 4161 | 170 | Sh,Tsp96F,sws,Spf45,Octbeta2R,IP3K2,CG4849,Glut1 | | 8.471e-03 | -4.77 | extrinsic component of plasma membrane | cellular component | GO:0019897 | 40 | 6 | 11940 | 538 | Src42A,D2hgdh,shg,klg,CG30054,CadN2 | | 8.500e-03 | -4.77 | negative regulation of signal transduction | biological process | GO:0009968 | 326 | 26 | 12345 | 597 | kek1,Src42A,raw,trol,et,foxo,CG14182,Lpin,Gprk1,chb,mnb,Cam,crol,qvr,Raf,pirk,Snoo,Su(H),sgg,numb,dnc,HDAC4,kek5,flw,Ser,Ptp61F | | 8.532e-03 | -4.76 | regulation of growth | biological process | GO:0040008 | 310 | 25 | 12345 | 597 | EcR,Neb-cGP,Ubx,scrib,Octbeta2R,dpp,Pak,shg,step,rictor,Syn2,bun,wbl,foxo,Src42A,Tor,Raf,ex,sgg,cic,mnb,Mob2,dsh,path,Ank2 | | 8.574e-03 | -4.76 | CAGGTA\_AREB6\_01 | MSigDB lists | CAGGTA\_AREB6\_01 | 152 | 13 | 4161 | 170 | Iswi,TfAP-2,Acbp2,IP3K2,CG3961,fus,MESK2,Ser,dnc,Aldh-III,Ptp61F,Octbeta2R,Dys | | 8.606e-03 | -4.76 | cytoskeletal part | cellular component | GO:0044430 | 456 | 32 | 11940 | 538 | Dlic,CG34417,msps,Gas8,CG34435,Cam,CG13526,pod1,Act88F,IFT46,Asator,TpnC25D,spd-2,CG18109,kl-5,Cep97,Myo81F,CG13251,Root,Msp300,sgg,CG17387,nuf,nompB,aust,Pak,Prm,Rcd6,wake,chb,Khc,Act87E | | 8.676e-03 | -4.75 | GO\_REGULATION\_OF\_SODIUM\_ION\_TRANSMEMBRANE\_TRANSPORT | MSigDB lists | GO\_REGULATION\_OF\_SODIUM\_ION\_TRANSMEMBRANE\_TRANSPORT | 11 | 3 | 4161 | 170 | Pkc98E,Ank,Dys | | 8.676e-03 | -4.75 | GO\_SCHWANN\_CELL\_DIFFERENTIATION | MSigDB lists | GO\_SCHWANN\_CELL\_DIFFERENTIATION | 11 | 3 | 4161 | 170 | sdt,TfAP-2,Sod1 | | 8.676e-03 | -4.75 | PID\_IL2\_PI3K\_PATHWAY | MSigDB lists | PID\_IL2\_PI3K\_PATHWAY | 11 | 3 | 4161 | 170 | CG3376,csw,Tor | | 8.676e-03 | -4.75 | BROWNE\_HCMV\_INFECTION\_4HR\_UP | MSigDB lists | BROWNE\_HCMV\_INFECTION\_4HR\_UP | 11 | 3 | 4161 | 170 | KaiR1D,csw,dnc | | 8.676e-03 | -4.75 | GO\_SCHWANN\_CELL\_DEVELOPMENT | MSigDB lists | GO\_SCHWANN\_CELL\_DEVELOPMENT | 11 | 3 | 4161 | 170 | sdt,TfAP-2,Sod1 | | 8.676e-03 | -4.75 | GO\_POSITIVE\_REGULATION\_OF\_JUN\_KINASE\_ACTIVITY | MSigDB lists | GO\_POSITIVE\_REGULATION\_OF\_JUN\_KINASE\_ACTIVITY | 11 | 3 | 4161 | 170 | dsh,Pak,fz2 | | 8.676e-03 | -4.75 | GO\_REGULATION\_OF\_STEROID\_METABOLIC\_PROCESS | MSigDB lists | GO\_REGULATION\_OF\_STEROID\_METABOLIC\_PROCESS | 11 | 3 | 4161 | 170 | Ugt301D1,dpp,Sod1 | | 8.676e-03 | -4.75 | GO\_REGULATION\_OF\_CELLULAR\_RESPONSE\_TO\_INSULIN\_STIMULUS | MSigDB lists | GO\_REGULATION\_OF\_CELLULAR\_RESPONSE\_TO\_INSULIN\_STIMULUS | 11 | 3 | 4161 | 170 | csw,sxc,Ptp61F | | 8.676e-03 | -4.75 | chr7p15 | MSigDB lists | chr7p15 | 11 | 3 | 4161 | 170 | CG15093,Act87E,CG32196 | | 8.676e-03 | -4.75 | GCM\_AQP4 | MSigDB lists | GCM\_AQP4 | 11 | 3 | 4161 | 170 | CG41099,Pde8,Dlic | | 8.689e-03 | -4.75 | GO\_RESPONSE\_TO\_OXYGEN\_CONTAINING\_COMPOUND | MSigDB lists | GO\_RESPONSE\_TO\_OXYGEN\_CONTAINING\_COMPOUND | 316 | 22 | 4161 | 170 | Cat,CG3376,Octbeta2R,Ptp61F,dnc,CG5493,Ugt301D1,mino,Doa,Tor,CG4849,TfAP-2,GlyP,Pak,Raf,LpR2,Ser,Pkc98E,eIF2Bgamma,RhoGAP1A,sxc,Sod1 | | 8.751e-03 | -4.74 | GO\_IMMUNE\_SYSTEM\_DEVELOPMENT | MSigDB lists | GO\_IMMUNE\_SYSTEM\_DEVELOPMENT | 103 | 10 | 4161 | 170 | IP3K2,Ca-beta,Su(H),csw,Sod1,Cdep,Ser,fz2,Ptp61F,Raf | | 8.752e-03 | -4.74 | cellular response to growth factor stimulus | biological process | GO:0071363 | 61 | 8 | 12345 | 597 | bun,ex,dpp,bnl,ths,Raf,opa,csw | | 8.752e-03 | -4.74 | anatomical structure regression | biological process | GO:0060033 | 61 | 8 | 12345 | 597 | EcR,IP3K2,Eip74EF,Tor,Fhos,hdc,Cam,Pak | | 8.752e-03 | -4.74 | cell junction assembly | biological process | GO:0034329 | 61 | 8 | 12345 | 597 | cindr,sdt,nrv2,shg,CadN2,shakB,Pak,scrib | | 8.791e-03 | -4.73 | Exo70 (Exocyst 70) | protein interactions | 38959 | 35 | 6 | 9204 | 481 | CG6424,Actn,CG41520,Khc,Prm,Dys | | 8.816e-03 | -4.73 | GO\_SYNAPSE | MSigDB lists | GO\_SYNAPSE | 205 | 16 | 4161 | 170 | unc-13,Nlg2,Frq1,sky,KaiR1D,Octbeta2R,Dys,mGluR,Syx1A,Nmdar2,Cadps,Ank,Maf1,Ca-beta,Cln3,Frq2 | | 8.818e-03 | -4.73 | HFH3\_01 | MSigDB lists | HFH3\_01 | 32 | 5 | 4161 | 170 | Rbp6,flw,Dys,mGluR,KaiR1D | | 8.818e-03 | -4.73 | GO\_CELLULAR\_RESPONSE\_TO\_BIOTIC\_STIMULUS | MSigDB lists | GO\_CELLULAR\_RESPONSE\_TO\_BIOTIC\_STIMULUS | 32 | 5 | 4161 | 170 | LpR2,dnc,fz2,sxc,Pkc98E | | 8.818e-03 | -4.73 | GSE36392\_EOSINOPHIL\_VS\_NEUTROPHIL\_IL25\_TREATED\_LUNG\_UP | MSigDB lists | GSE36392\_EOSINOPHIL\_VS\_NEUTROPHIL\_IL25\_TREATED\_LUNG\_UP | 32 | 5 | 4161 | 170 | MCU,sws,Fhos,CG5522,CG17698 | | 8.818e-03 | -4.73 | GO\_RESPONSE\_TO\_PURINE\_CONTAINING\_COMPOUND | MSigDB lists | GO\_RESPONSE\_TO\_PURINE\_CONTAINING\_COMPOUND | 32 | 5 | 4161 | 170 | dnc,CG5493,Sod1,GlyP,Raf | | 8.818e-03 | -4.73 | BROCKE\_APOPTOSIS\_REVERSED\_BY\_IL6 | MSigDB lists | BROCKE\_APOPTOSIS\_REVERSED\_BY\_IL6 | 32 | 5 | 4161 | 170 | Glut1,scramb1,eIF4E7,Cadps,csw | | 8.818e-03 | -4.73 | GO\_REGULATION\_OF\_CYCLIC\_NUCLEOTIDE\_METABOLIC\_PROCESS | MSigDB lists | GO\_REGULATION\_OF\_CYCLIC\_NUCLEOTIDE\_METABOLIC\_PROCESS | 32 | 5 | 4161 | 170 | AstA-R1,Octbeta2R,mGluR,Pkc53E,dnc | | 8.870e-03 | -4.73 | cell-cell signaling | biological process | GO:0007267 | 311 | 25 | 12345 | 597 | Frq1,sky,Nlg2,upd3,GluRIIC,Fife,Frq2,fz2,cpx,Syx1A,dsh,Tusp,Ca-beta,unc-13,CG5541,Cat,Ser,flw,Dys,KaiR1D,dnc,Obp56h,Cadps,fz4,scramb1 | | 8.908e-03 | -4.72 | Ras GTPase binding | molecular function | GO:0017016 | 174 | 16 | 12176 | 580 | CG18659,Fife,Pak,Gas8,CG42390,CG10188,tamo,CG17883,Exn,nuf,Cdep,spri,RhoGEF3,RhoGAP18B,CG5522,RhoGAP1A | | 8.987e-03 | -4.71 | Actinin\_actin-bd\_CS | interpro domains | IPR001589 | 9 | 3 | 11383 | 585 | Actn,Msp300,Dys | | 9.022e-03 | -4.71 | C2 | smart domains | SM00239 | 39 | 7 | 5086 | 315 | Mctp,Pkc53E,Pkc98E,Fife,unc-13,RhoGAP100F,Syt12 | | 9.083e-03 | -4.70 | regulation of cell development | biological process | GO:0060284 | 344 | 27 | 12345 | 597 | Dlic,csw,spri,rictor,Pak,Mbs,hth,shg,EcR,Khc,Antp,numb,sgg,Raf,Tor,ham,barc,hdc,dsh,Fhos,Syx1A,kay,mnb,CG42797,Elp3,bun,Src42A | | 9.083e-03 | -4.70 | columnar/cuboidal epithelial cell differentiation | biological process | GO:0002065 | 344 | 27 | 12345 | 597 | bun,RhoL,polybromo,bbg,cindr,Raf,numb,sgg,cic,ex,Mrtf,GATAe,fs(1)N,kay,tin,dsh,EcR,Khc,shd,lncRNA:flam,scrib,rols,csw,mid,dpp,egh,shg | | 9.121e-03 | -4.70 | GO\_RESPONSE\_TO\_BACTERIUM | MSigDB lists | GO\_RESPONSE\_TO\_BACTERIUM | 88 | 9 | 4161 | 170 | Pkc98E,fz2,Ugt301D1,dnc,LpR2,sxc,RhoGAP1A,Su(H),TfAP-2 | | 9.172e-03 | -4.69 | GO\_POSITIVE\_REGULATION\_OF\_NERVOUS\_SYSTEM\_DEVELOPMENT | MSigDB lists | GO\_POSITIVE\_REGULATION\_OF\_NERVOUS\_SYSTEM\_DEVELOPMENT | 73 | 8 | 4161 | 170 | CG42672,Tor,Tif-IA,dpp,Dys,Raf,unc-13,LpR2 | | 9.187e-03 | -4.69 | oocyte dorsal/ventral axis specification | biological process | GO:0007310 | 38 | 6 | 12345 | 597 | Khc,flw,cic,dpp,wbl,tld | | 9.187e-03 | -4.69 | positive regulation of endocytosis | biological process | GO:0045807 | 38 | 6 | 12345 | 597 | mnb,Pkc53E,spri,sdt,cindr,numb | | 9.225e-03 | -4.69 | OCT1\_02 | MSigDB lists | OCT1\_02 | 45 | 6 | 4161 | 170 | Dys,Ank,CG9132,Pde1c,dnc,Ser | | 9.225e-03 | -4.69 | YAUCH\_HEDGEHOG\_SIGNALING\_PARACRINE\_DN | MSigDB lists | YAUCH\_HEDGEHOG\_SIGNALING\_PARACRINE\_DN | 45 | 6 | 4161 | 170 | Syn2,Pkc53E,Pkc98E,Ucp4A,Treh,GlyP | | 9.225e-03 | -4.69 | LEF1\_Q6 | MSigDB lists | LEF1\_Q6 | 45 | 6 | 4161 | 170 | dnc,ABCD,RhoGAP1A,Meltrin,Pak,chb | | 9.225e-03 | -4.69 | HALLMARK\_HYPOXIA | MSigDB lists | HALLMARK\_HYPOXIA | 45 | 6 | 4161 | 170 | Mlp84B,Su(H),Pkc53E,GlyP,Glut1,LpR2 | | 9.225e-03 | -4.69 | HNF3ALPHA\_Q6 | MSigDB lists | HNF3ALPHA\_Q6 | 45 | 6 | 4161 | 170 | DNaseII,Dys,fus,flw,dnc,IP3K2 | | 9.225e-03 | -4.69 | GSE46606\_IRF4MID\_VS\_WT\_CD40L\_IL2\_IL5\_DAY1\_STIMULATED\_BCELL\_UP | MSigDB lists | GSE46606\_IRF4MID\_VS\_WT\_CD40L\_IL2\_IL5\_DAY1\_STIMULATED\_BCELL\_UP | 45 | 6 | 4161 | 170 | CG5339,Ucp4A,ABCD,CG33090,CG12547,rdgA | | 9.292e-03 | -4.68 | myofibril | cellular component | GO:0030016 | 66 | 8 | 11940 | 538 | rols,zormin,Prm,Act88F,Actn,Msp300,Mlp84B,TpnC25D | | 9.332e-03 | -4.67 | KEGG\_AMYOTROPHIC\_LATERAL\_SCLEROSIS\_ALS | MSigDB lists | KEGG\_AMYOTROPHIC\_LATERAL\_SCLEROSIS\_ALS | 21 | 4 | 4161 | 170 | Nmdar2,Cat,Sod1,Der-1 | | 9.332e-03 | -4.67 | GO\_REGULATION\_OF\_ADENYLATE\_CYCLASE\_ACTIVITY | MSigDB lists | GO\_REGULATION\_OF\_ADENYLATE\_CYCLASE\_ACTIVITY | 21 | 4 | 4161 | 170 | mGluR,Octbeta2R,AstA-R1,Pkc53E | | 9.332e-03 | -4.67 | KEGG\_MTOR\_SIGNALING\_PATHWAY | MSigDB lists | KEGG\_MTOR\_SIGNALING\_PATHWAY | 21 | 4 | 4161 | 170 | Raf,eIF4E7,Stlk,Tor | | 9.376e-03 | -4.67 | MAPK signaling pathway - fly | KEGG pathways | ko04013 | 93 | 11 | 3198 | 170 | RhoL,dpp,kay,kek1,csw,fs(1)N,Raf,Src42A,cic,raw,bi | | 9.376e-03 | -4.67 | MAPK signaling pathway - fly | KEGG pathways | dme04013 | 93 | 11 | 3198 | 170 | raw,cic,bi,Src42A,Raf,csw,fs(1)N,kay,kek1,dpp,RhoL | | 9.398e-03 | -4.67 | GO\_SYNAPSE\_PART | MSigDB lists | GO\_SYNAPSE\_PART | 171 | 14 | 4161 | 170 | mGluR,Dys,Octbeta2R,Nmdar2,unc-13,Syx1A,Cadps,Ank,Nlg2,Frq1,KaiR1D,sky,Frq2,Cln3 | | 9.405e-03 | -4.67 | dpr17 (defective proboscis extension response 17) | protein interactions | 41470 | 9 | 3 | 9204 | 481 | CG5386,CG18304,PIG-G | | 9.430e-03 | -4.66 | GO\_REGULATION\_OF\_SHORT\_TERM\_NEURONAL\_SYNAPTIC\_PLASTICITY | MSigDB lists | GO\_REGULATION\_OF\_SHORT\_TERM\_NEURONAL\_SYNAPTIC\_PLASTICITY | 4 | 2 | 4161 | 170 | KaiR1D,unc-13 | | 9.430e-03 | -4.66 | GO\_T\_CELL\_SELECTION | MSigDB lists | GO\_T\_CELL\_SELECTION | 4 | 2 | 4161 | 170 | IP3K2,Raf | | 9.430e-03 | -4.66 | GO\_NEGATIVE\_REGULATION\_OF\_LIPID\_STORAGE | MSigDB lists | GO\_NEGATIVE\_REGULATION\_OF\_LIPID\_STORAGE | 4 | 2 | 4161 | 170 | EcR,Ptp61F | | 9.430e-03 | -4.66 | GO\_MUSCLE\_CELL\_CELLULAR\_HOMEOSTASIS | MSigDB lists | GO\_MUSCLE\_CELL\_CELLULAR\_HOMEOSTASIS | 4 | 2 | 4161 | 170 | Sod1,Dys | | 9.430e-03 | -4.66 | GO\_REGULATION\_OF\_PLATELET\_DERIVED\_GROWTH\_FACTOR\_RECEPTOR\_SIGNALING\_PATHWAY | MSigDB lists | GO\_REGULATION\_OF\_PLATELET\_DERIVED\_GROWTH\_FACTOR\_RECEPTOR\_SIGNALING\_PATHWAY | 4 | 2 | 4161 | 170 | LRP1,Ptp61F | | 9.430e-03 | -4.66 | MODULE\_433 | MSigDB lists | MODULE\_433 | 4 | 2 | 4161 | 170 | Ser,dpp | | 9.430e-03 | -4.66 | GO\_REGULATION\_OF\_ODONTOGENESIS | MSigDB lists | GO\_REGULATION\_OF\_ODONTOGENESIS | 4 | 2 | 4161 | 170 | dpp,TfAP-2 | | 9.430e-03 | -4.66 | GO\_MICROTUBULE\_PLUS\_END | MSigDB lists | GO\_MICROTUBULE\_PLUS\_END | 4 | 2 | 4161 | 170 | msps,chb | | 9.430e-03 | -4.66 | HALLMARK\_IL6\_JAK\_STAT3\_SIGNALING | MSigDB lists | HALLMARK\_IL6\_JAK\_STAT3\_SIGNALING | 4 | 2 | 4161 | 170 | csw,Ptp61F | | 9.430e-03 | -4.66 | GO\_NEURONAL\_ACTION\_POTENTIAL | MSigDB lists | GO\_NEURONAL\_ACTION\_POTENTIAL | 4 | 2 | 4161 | 170 | Ank,KaiR1D | | 9.430e-03 | -4.66 | GO\_POSITIVE\_REGULATION\_OF\_CHOLESTEROL\_EFFLUX | MSigDB lists | GO\_POSITIVE\_REGULATION\_OF\_CHOLESTEROL\_EFFLUX | 4 | 2 | 4161 | 170 | EcR,LRP1 | | 9.430e-03 | -4.66 | CROONQUIST\_IL6\_DEPRIVATION\_UP | MSigDB lists | CROONQUIST\_IL6\_DEPRIVATION\_UP | 4 | 2 | 4161 | 170 | Pde1c,dpp | | 9.493e-03 | -4.66 | RRM\_dom | interpro domains | IPR000504 | 135 | 14 | 11383 | 585 | Rbp6,CG42458,CG3594,CG40813,fus,bru1,trv,barc,bol,CG44249,CG41562,Spf45,shep,msi | | 9.520e-03 | -4.65 | Ig\_2 | pfam domains | PF13895 | 9 | 3 | 10480 | 550 | rst,Strn-Mlck,trol | | 9.533e-03 | -4.65 | GO\_MUSCLE\_CELL\_DIFFERENTIATION | MSigDB lists | GO\_MUSCLE\_CELL\_DIFFERENTIATION | 59 | 7 | 4161 | 170 | Dys,Ca-beta,dpp,Ser,Tor,Mlp84B,Su(H) | | 9.533e-03 | -4.65 | ONDER\_CDH1\_TARGETS\_2\_DN | MSigDB lists | ONDER\_CDH1\_TARGETS\_2\_DN | 59 | 7 | 4161 | 170 | fz2,dpp,Dys,Octbeta2R,Ank,fus,Glut1 | | 9.533e-03 | -4.65 | GSE369\_PRE\_VS\_POST\_IL6\_INJECTION\_SOCS3\_KO\_LIVER\_UP | MSigDB lists | GSE369\_PRE\_VS\_POST\_IL6\_INJECTION\_SOCS3\_KO\_LIVER\_UP | 59 | 7 | 4161 | 170 | Meltrin,Su(H),Tif-IA,CG6966,Pfdn4,Glut1,Dlic | | 9.623e-03 | -4.64 | GO\_RESPONSE\_TO\_ABIOTIC\_STIMULUS | MSigDB lists | GO\_RESPONSE\_TO\_ABIOTIC\_STIMULUS | 262 | 19 | 4161 | 170 | Octbeta2R,Dys,Cat,Syx1A,TfAP-2,Ca-beta,Tor,Doa,dpp,eIF2Bgamma,flw,IntS3,Pkc98E,GlyP,Pak,LpR2,Raf,Sod1,Su(H) | | 9.633e-03 | -4.64 | regulation of transmembrane transporter activity | biological process | GO:0022898 | 18 | 4 | 12345 | 597 | qvr,Ca-beta,CG42797,Cam | | 9.633e-03 | -4.64 | regulation of ion transmembrane transporter activity | biological process | GO:0032412 | 18 | 4 | 12345 | 597 | Ca-beta,CG42797,Cam,qvr | | 9.633e-03 | -4.64 | aggressive behavior | biological process | GO:0002118 | 18 | 4 | 12345 | 597 | fru,Nlg2,TfAP-2,Tk | | 9.633e-03 | -4.64 | medium-term memory | biological process | GO:0072375 | 18 | 4 | 12345 | 597 | MCU,Nmdar2,CrebB,dnc | | 9.633e-03 | -4.64 | regulation of transporter activity | biological process | GO:0032409 | 18 | 4 | 12345 | 597 | qvr,Cam,Ca-beta,CG42797 | | 9.633e-03 | -4.64 | regulation of feeding behavior | biological process | GO:0060259 | 18 | 4 | 12345 | 597 | RYa,CrebB,mnb,foxo | | 9.633e-03 | -4.64 | adherens junction assembly | biological process | GO:0034333 | 18 | 4 | 12345 | 597 | scrib,shg,cindr,sdt | | 9.636e-03 | -4.64 | response to growth factor | biological process | GO:0070848 | 62 | 8 | 12345 | 597 | ex,dpp,ths,Raf,bnl,csw,opa,bun | | 9.659e-03 | -4.64 | ZF\_DAG\_PE\_1 | prosite domains | PS00479 | 22 | 5 | 6243 | 386 | unc-13,Raf,Pkc53E,Stacl,Pkc98E | | 9.774e-03 | -4.63 | regulation of ion transport | biological process | GO:0043269 | 50 | 7 | 12345 | 597 | CG3078,qvr,Syt12,olf186-F,Cam,CG42797,Ca-beta | | 9.780e-03 | -4.63 | pnr (pannier) | protein interactions | 44849 | 107 | 12 | 9204 | 481 | Ubx,Antp,dpp,mid,esg,mir-278,ps,CG46385,bun,sgg,tin,Iswi | | 9.800e-03 | -4.63 | GO\_GLAND\_DEVELOPMENT | MSigDB lists | GO\_GLAND\_DEVELOPMENT | 89 | 9 | 4161 | 170 | Ca-beta,Su(H),Sod1,CG17387,dpp,Ugt301D1,CG5493,Octbeta2R,Raf | | 9.802e-03 | -4.63 | synapse | cellular component | GO:0045202 | 285 | 22 | 11940 | 538 | Dys,Ank2,gukh,Pak,KaiR1D,Frq2,g,Cadps,Exn,Syx1A,sky,Fife,sgg,Nlg2,unc-13,Elp3,mnb,RhoGAP100F,cpx,pod1,scrib,GluRIIC | | 9.806e-03 | -4.62 | negative regulation of gene expression | biological process | GO:0010629 | 582 | 41 | 12345 | 597 | retn,Ubx,CG1894,esg,Khc,E5,bru1,CG12605,msi,bi,CG43689,tld,Iswi,cic,numb,B-H2,Su(H),polybromo,hdc,crol,AGO2,EcR,Antp,scrib,CG7368,dpp,mid,Achl,Maf1,CG31224,Marf1,toe,foxo,HPS4,TfAP-2,AGO3,CG12071,tin,GATAe,lncRNA:bxd,Sox102F | | 9.819e-03 | -4.62 | negative regulation of cell population proliferation | biological process | GO:0008285 | 88 | 10 | 12345 | 597 | numb,ex,dpp,Doa,ham,foxo,Pkc98E,scrib,Ptp61F,EcR | | 9.926e-03 | -4.61 | negative regulation of macromolecule biosynthetic process | biological process | GO:0010558 | 446 | 33 | 12345 | 597 | B-H2,cic,polybromo,Su(H),crol,CG43689,bi,msi,Iswi,CG12605,Ubx,retn,Khc,esg,CG1894,E5,bru1,TfAP-2,CG12071,tin,Sox102F,lncRNA:bxd,toe,foxo,HPS4,CG7368,Maf1,mid,CG31224,EcR,AGO2,Antp,scrib | | 1.006e-02 | -4.60 | GSE34156\_UNTREATED\_VS\_24H\_TLR1\_TLR2\_LIGAND\_TREATED\_MONOCYTE\_UP | MSigDB lists | GSE34156\_UNTREATED\_VS\_24H\_TLR1\_TLR2\_LIGAND\_TREATED\_MONOCYTE\_UP | 33 | 5 | 4161 | 170 | Maf1,Frq2,csw,melt,Frq1 | | 1.006e-02 | -4.60 | HALLMARK\_UV\_RESPONSE\_DN | MSigDB lists | HALLMARK\_UV\_RESPONSE\_DN | 33 | 5 | 4161 | 170 | LpR2,dsd,bru1,Pkc98E,Pkc53E | | 1.006e-02 | -4.60 | GSE23114\_PERITONEAL\_CAVITY\_B1A\_BCELL\_VS\_SPLEEN\_BCELL\_IN\_SLE2C1\_MOUSE\_UP | MSigDB lists | GSE23114\_PERITONEAL\_CAVITY\_B1A\_BCELL\_VS\_SPLEEN\_BCELL\_IN\_SLE2C1\_MOUSE\_UP | 33 | 5 | 4161 | 170 | Glut1,GlyP,Cadps,Pkc98E,Cln3 | | 1.006e-02 | -4.60 | IZADPANAH\_STEM\_CELL\_ADIPOSE\_VS\_BONE\_UP | MSigDB lists | IZADPANAH\_STEM\_CELL\_ADIPOSE\_VS\_BONE\_UP | 33 | 5 | 4161 | 170 | CG3961,Odc1,Ser,TfAP-2,Fhos | | 1.006e-02 | -4.60 | GSE29949\_CD8\_NEG\_DC\_SPLEEN\_VS\_DC\_BRAIN\_UP | MSigDB lists | GSE29949\_CD8\_NEG\_DC\_SPLEEN\_VS\_DC\_BRAIN\_UP | 33 | 5 | 4161 | 170 | kcc,CNT1,Octbeta2R,CG7483,ABCD | | 1.008e-02 | -4.60 | GO\_NEURON\_DIFFERENTIATION | MSigDB lists | GO\_NEURON\_DIFFERENTIATION | 208 | 16 | 4161 | 170 | Ank,CG42672,Awh,csw,AstA-R1,CG33090,fz2,Cdep,sky,Su(H),Iswi,Sod1,Khc,Pak,LpR2,Ser | | 1.009e-02 | -4.60 | cytoplasmic region | cellular component | GO:0099568 | 252 | 20 | 11940 | 538 | spri,scrib,step,Khc,dsh,kl-5,unc-13,sdt,numb,Hil,cindr,nuf,app,AGO2,shg,CG17387,Fife,Ser,Dys,Mbs | | 1.010e-02 | -4.60 | term (terminus) | protein interactions | 47208 | 36 | 6 | 9204 | 481 | Ubx,Glut1,REG,CG14147,CG3081,Marf1 | | 1.014e-02 | -4.59 | regulation of cell fate specification | biological process | GO:0042659 | 28 | 5 | 12345 | 597 | mid,Su(H),Ubx,bun,hth | | 1.014e-02 | -4.59 | zygotic determination of anterior/posterior axis, embryo | biological process | GO:0007354 | 28 | 5 | 12345 | 597 | fs(1)N,dpp,cic,csw,Raf | | 1.017e-02 | -4.59 | Pak3 (Pak3) | protein interactions | 41995 | 17 | 4 | 9204 | 481 | Pak,scrib,rictor,foxo | | 1.017e-02 | -4.59 | Lasp (lasp) | protein interactions | 39864 | 17 | 4 | 9204 | 481 | Strn-Mlck,Act88F,Actn,Act87E | | 1.023e-02 | -4.58 | apical cortex | cellular component | GO:0045179 | 54 | 7 | 11940 | 538 | sdt,app,Mbs,Ser,dsh,cindr,nuf | | 1.024e-02 | -4.58 | presynapse | cellular component | GO:0098793 | 155 | 14 | 11940 | 538 | sky,Fife,Syx1A,Exn,cpx,Cadps,g,RhoGAP100F,Frq2,KaiR1D,mnb,unc-13,gukh,Ank2 | | 1.026e-02 | -4.58 | GSE28726\_ACT\_CD4\_TCELL\_VS\_ACT\_VA24NEG\_NKTCELL\_DN | MSigDB lists | GSE28726\_ACT\_CD4\_TCELL\_VS\_ACT\_VA24NEG\_NKTCELL\_DN | 46 | 6 | 4161 | 170 | Cln3,step,dnc,IP3K2,RpL38,Flo2 | | 1.030e-02 | -4.58 | intrinsic component of plasma membrane | cellular component | GO:0031226 | 480 | 33 | 11940 | 538 | CadN2,beat-IIa,dpr18,CG3285,PGRP-LA,dpr5,Nmdar2,chp,CG14606,Nlg2,kek1,MsR2,scrib,GluRIIC,Sh,ACXD,AstA-R1,CNT1,nrv2,CG10960,Pdfr,CG33310,CG11318,olf186-F,Octbeta2R,shg,DIP-lambda,kcc,mGluR,fz2,DIP-delta,CCAP-R,kek5 | | 1.039e-02 | -4.57 | plasma membrane protein complex | cellular component | GO:0098797 | 171 | 15 | 11940 | 538 | CadN2,CG33310,Dys,CG30054,Ca-beta,nrv2,sdt,Syn2,mGluR,Nmdar2,Sh,GluRIIC,shg,Flo2,scrib | | 1.040e-02 | -4.57 | ACTININ\_2 | prosite domains | PS00020 | 8 | 3 | 6243 | 386 | Dys,Msp300,Actn | | 1.041e-02 | -4.57 | small GTPase binding | molecular function | GO:0031267 | 177 | 16 | 12176 | 580 | RhoGEF3,spri,CG5522,RhoGAP1A,RhoGAP18B,CG17883,Gas8,CG42390,CG18659,Fife,Pak,CG10188,tamo,Cdep,Exn,nuf | | 1.042e-02 | -4.56 | Mlc1 (Myosin alkali light chain 1) | protein interactions | 43323 | 47 | 7 | 9204 | 481 | CG34172,Scp1,TpnC41C,Actn,Prm,Act88F,CG5522 | | 1.043e-02 | -4.56 | positive regulation of synapse assembly | biological process | GO:0051965 | 39 | 6 | 12345 | 597 | Ank2,rictor,Octbeta2R,Pak,mnb,Tor | | 1.043e-02 | -4.56 | GSE33425\_CD161\_HIGH\_VS\_NEG\_CD8\_TCELL\_UP | MSigDB lists | GSE33425\_CD161\_HIGH\_VS\_NEG\_CD8\_TCELL\_UP | 60 | 7 | 4161 | 170 | IP3K2,Stlk,CG3376,Octbeta2R,CG41099,Nha1,Syx1A | | 1.043e-02 | -4.56 | GO\_REGULATION\_OF\_NEURON\_DEATH | MSigDB lists | GO\_REGULATION\_OF\_NEURON\_DEATH | 60 | 7 | 4161 | 170 | KaiR1D,Raf,TfAP-2,Tor,LRP1,Sod1,Cln3 | | 1.043e-02 | -4.56 | GO\_REGULATION\_OF\_RESPONSE\_TO\_WOUNDING | MSigDB lists | GO\_REGULATION\_OF\_RESPONSE\_TO\_WOUNDING | 60 | 7 | 4161 | 170 | RhoGAP1A,Pkc53E,Pkc98E,Raf,Octbeta2R,Ptp61F,chb | | 1.046e-02 | -4.56 | pericardial nephrocyte differentiation | biological process | GO:0061320 | 10 | 3 | 12345 | 597 | dpp,numb,tin | | 1.046e-02 | -4.56 | intracellular steroid hormone receptor signaling pathway | biological process | GO:0030518 | 10 | 3 | 12345 | 597 | EcR,Set2,Iswi | | 1.046e-02 | -4.56 | ecdysone receptor-mediated signaling pathway | biological process | GO:0035076 | 10 | 3 | 12345 | 597 | EcR,Iswi,Set2 | | 1.046e-02 | -4.56 | GO\_REGULATION\_OF\_TRANSFERASE\_ACTIVITY | MSigDB lists | GO\_REGULATION\_OF\_TRANSFERASE\_ACTIVITY | 283 | 20 | 4161 | 170 | Pak,LpR2,Tsp96F,Raf,dpp,Elp3,Sod1,wbl,dsh,Ptp61F,CG3376,Octbeta2R,REG,Ugt301D1,fz2,Stlk,Tor,CG42672,csw,CG17698 | | 1.052e-02 | -4.55 | GO\_REGULATORY\_REGION\_NUCLEIC\_ACID\_BINDING | MSigDB lists | GO\_REGULATORY\_REGION\_NUCLEIC\_ACID\_BINDING | 90 | 9 | 4161 | 170 | Su(H),Maf1,Tor,TfAP-2,Hr96,Rat1,EcR,Act87E,Tif-IA | | 1.052e-02 | -4.55 | GO\_POSITIVE\_REGULATION\_OF\_TRANSFERASE\_ACTIVITY | MSigDB lists | GO\_POSITIVE\_REGULATION\_OF\_TRANSFERASE\_ACTIVITY | 191 | 15 | 4161 | 170 | CG42672,CG17698,wbl,Sod1,csw,Stlk,dpp,REG,fz2,Octbeta2R,dsh,Pak,Tsp96F,Raf,LpR2 | | 1.062e-02 | -4.55 | GO\_CELL\_CELL\_SIGNALING | MSigDB lists | GO\_CELL\_CELL\_SIGNALING | 139 | 12 | 4161 | 170 | Ca-beta,KaiR1D,kcc,Flo2,Cadps,dpp,Octbeta2R,mGluR,AstA-R1,Syx1A,unc-13,Nmdar2 | | 1.065e-02 | -4.54 | fly chr64B17-64B17\|3-12 | chromosome location | fly chr64B17-64B17\|3-12 | 5 | 2 | 25010 | 845 | CG18808,lncRNA:CR45821 | | 1.065e-02 | -4.54 | fly chr70D2-70D2\|3-42 cM | chromosome location | fly chr70D2-70D2\|3-42 cM | 5 | 2 | 25010 | 845 | lncRNA:CR45887,Fbp1 | | 1.065e-02 | -4.54 | fly chr70D2-70D2\|3-42 | chromosome location | fly chr70D2-70D2\|3-42 | 5 | 2 | 25010 | 845 | lncRNA:CR45887,Fbp1 | | 1.065e-02 | -4.54 | fly chr25C10-25C10\|2-15 cM | chromosome location | fly chr25C10-25C10\|2-15 cM | 5 | 2 | 25010 | 845 | Cyp28d2,Cyp28d1 | | 1.065e-02 | -4.54 | fly chr64B17-64B17\|3-12 cM | chromosome location | fly chr64B17-64B17\|3-12 cM | 5 | 2 | 25010 | 845 | CG18808,lncRNA:CR45821 | | 1.065e-02 | -4.54 | fly chr25C10-25C10\|2-15 | chromosome location | fly chr25C10-25C10\|2-15 | 5 | 2 | 25010 | 845 | Cyp28d1,Cyp28d2 | | 1.069e-02 | -4.54 | reproductive process | biological process | GO:0022414 | 1436 | 88 | 12345 | 597 | tnc,tld,upd3,kek1,raw,Tor,polybromo,msps,RhoL,bbg,spd-2,ex,sgg,Gr39a,mGluR,fz2,kay,Khc,Mos,Dlic,Mbs,lncRNA:CR43753,Gas8,chinmo,shep,chb,wbl,Pde1c,Raf,Msp300,Yeti,TfAP-2,lncRNA:Hsromega,Mrtf,dsh,aust,tin,Gr22c,AGO2,dnc,shd,opa,dany,Obp56h,dpp,shg,msi,can,Eip74EF,Iswi,mamo,cic,DNaseII,CG18109,Fad2,flw,retn,CG7483,lncRNA:flam,Ebp,bol,bru1,Mst87F,Octbeta2R,Doa,fru,Acp62F,drd,lncRNA:CR43484,CG12200,Nlg2,JhI-26,bun,cindr,lncRNA:CR44585,CG42355,Inx2,lncRNA:CR44420,rib,fs(1)N,EcR,scrib,lncRNA:CR43802,mid,rols,csw,egh,Sh | | 1.073e-02 | -4.53 | actin filament | cellular component | GO:0005884 | 42 | 6 | 11940 | 538 | CG34417,pod1,Act88F,Msp300,Act87E,Pak | | 1.079e-02 | -4.53 | dorsal/ventral axis specification | biological process | GO:0009950 | 76 | 9 | 12345 | 597 | wbl,Pkc98E,tld,Khc,flw,csw,Raf,dpp,cic | | 1.081e-02 | -4.53 | supramolecular fiber | cellular component | GO:0099512 | 204 | 17 | 11940 | 538 | Mlp84B,CG34417,Gas8,Actn,Msp300,msps,CG3078,Pak,zormin,TpnC25D,Prm,Act88F,pod1,Khc,Act87E,chb,rols | | 1.083e-02 | -4.53 | G alpha (q) signalling events | REACTOME pathways | R-DME-416476 | 18 | 4 | 3923 | 197 | Pkc98E,Gprk1,rdgA,CG30054 | | 1.105e-02 | -4.51 | MORF\_CD8A | MSigDB lists | MORF\_CD8A | 22 | 4 | 4161 | 170 | Dys,Hr96,dnc,fz2 | | 1.105e-02 | -4.51 | GO\_NEGATIVE\_REGULATION\_OF\_SYNAPTIC\_TRANSMISSION | MSigDB lists | GO\_NEGATIVE\_REGULATION\_OF\_SYNAPTIC\_TRANSMISSION | 22 | 4 | 4161 | 170 | Raf,KaiR1D,CG7483,ZnT63C | | 1.105e-02 | -4.51 | GO\_REGULATION\_OF\_POTASSIUM\_ION\_TRANSPORT | MSigDB lists | GO\_REGULATION\_OF\_POTASSIUM\_ION\_TRANSPORT | 22 | 4 | 4161 | 170 | Octbeta2R,Khc,Ank,AstA-R1 | | 1.105e-02 | -4.51 | GSE40274\_CTRL\_VS\_FOXP3\_AND\_PBX1\_TRANSDUCED\_ACTIVATED\_CD4\_TCELL\_UP | MSigDB lists | GSE40274\_CTRL\_VS\_FOXP3\_AND\_PBX1\_TRANSDUCED\_ACTIVATED\_CD4\_TCELL\_UP | 22 | 4 | 4161 | 170 | CNT1,Glut1,Pgant6,CG42672 | | 1.105e-02 | -4.51 | GO\_POSITIVE\_REGULATION\_OF\_EXOCYTOSIS | MSigDB lists | GO\_POSITIVE\_REGULATION\_OF\_EXOCYTOSIS | 22 | 4 | 4161 | 170 | Frq2,chb,Frq1,Syx1A | | 1.105e-02 | -4.51 | GO\_MULTICELLULAR\_ORGANISMAL\_SIGNALING | MSigDB lists | GO\_MULTICELLULAR\_ORGANISMAL\_SIGNALING | 22 | 4 | 4161 | 170 | Ank,KaiR1D,Ca-beta,Sod1 | | 1.105e-02 | -4.51 | GO\_NEGATIVE\_REGULATION\_OF\_TRANSMEMBRANE\_TRANSPORT | MSigDB lists | GO\_NEGATIVE\_REGULATION\_OF\_TRANSMEMBRANE\_TRANSPORT | 22 | 4 | 4161 | 170 | ZnT63C,Ank,Pkc98E,Tor | | 1.106e-02 | -4.50 | terminal bouton | cellular component | GO:0043195 | 68 | 8 | 11940 | 538 | g,Ank2,gukh,unc-13,mnb,Syx1A,sky,cpx | | 1.107e-02 | -4.50 | Rho guanyl-nucleotide exchange factor activity | molecular function | GO:0005089 | 29 | 5 | 12176 | 580 | Exn,CG10188,Cdep,RhoGEF3,RhoGAP1A | | 1.107e-02 | -4.50 | GO\_RESPONSE\_TO\_WOUNDING | MSigDB lists | GO\_RESPONSE\_TO\_WOUNDING | 123 | 11 | 4161 | 170 | Pkc98E,rdgA,Act87E,scramb1,Tsp96F,Pak,Octbeta2R,csw,Sod1,Pkc53E,Tor | | 1.112e-02 | -4.50 | Mlc2 (Myosin light chain 2) | protein interactions | 43587 | 59 | 8 | 9204 | 481 | Prm,zormin,Scp1,CG34172,Act88F,TpnC4,TpnC41C,Actn | | 1.112e-02 | -4.50 | Hrs (Hepatocyte growth factor regulated tyrosine kinase substrate) | protein interactions | 33458 | 59 | 8 | 9204 | 481 | dsh,rictor,Msp300,Cdep,hth,Actn,polybromo,sky | | 1.122e-02 | -4.49 | GO\_REGULATION\_OF\_CARDIAC\_MUSCLE\_TISSUE\_DEVELOPMENT | MSigDB lists | GO\_REGULATION\_OF\_CARDIAC\_MUSCLE\_TISSUE\_DEVELOPMENT | 12 | 3 | 4161 | 170 | Su(H),Tor,dpp | | 1.122e-02 | -4.49 | GO\_POSITIVE\_REGULATION\_OF\_BIOMINERAL\_TISSUE\_DEVELOPMENT | MSigDB lists | GO\_POSITIVE\_REGULATION\_OF\_BIOMINERAL\_TISSUE\_DEVELOPMENT | 12 | 3 | 4161 | 170 | Octbeta2R,dpp,TfAP-2 | | 1.122e-02 | -4.49 | PID\_CD8\_TCR\_DOWNSTREAM\_PATHWAY | MSigDB lists | PID\_CD8\_TCR\_DOWNSTREAM\_PATHWAY | 12 | 3 | 4161 | 170 | Pkc98E,Pkc53E,Raf | | 1.122e-02 | -4.49 | GO\_OUTFLOW\_TRACT\_MORPHOGENESIS | MSigDB lists | GO\_OUTFLOW\_TRACT\_MORPHOGENESIS | 12 | 3 | 4161 | 170 | dsh,Su(H),Ser | | 1.122e-02 | -4.49 | GO\_NEUROMUSCULAR\_JUNCTION\_DEVELOPMENT | MSigDB lists | GO\_NEUROMUSCULAR\_JUNCTION\_DEVELOPMENT | 12 | 3 | 4161 | 170 | Ca-beta,Ank,unc-13 | | 1.122e-02 | -4.49 | GO\_REGULATION\_OF\_CARBOHYDRATE\_CATABOLIC\_PROCESS | MSigDB lists | GO\_REGULATION\_OF\_CARBOHYDRATE\_CATABOLIC\_PROCESS | 12 | 3 | 4161 | 170 | PhKgamma,sxc,flw | | 1.122e-02 | -4.49 | REACTOME\_POST\_NMDA\_RECEPTOR\_ACTIVATION\_EVENTS | MSigDB lists | REACTOME\_POST\_NMDA\_RECEPTOR\_ACTIVATION\_EVENTS | 12 | 3 | 4161 | 170 | Nmdar2,Raf,CG17698 | | 1.122e-02 | -4.49 | GO\_THYMUS\_DEVELOPMENT | MSigDB lists | GO\_THYMUS\_DEVELOPMENT | 12 | 3 | 4161 | 170 | Raf,Ca-beta,Sod1 | | 1.122e-02 | -4.49 | GO\_RESPONSE\_TO\_COCAINE | MSigDB lists | GO\_RESPONSE\_TO\_COCAINE | 12 | 3 | 4161 | 170 | Tor,CG4849,CG3376 | | 1.137e-02 | -4.48 | GO\_REGULATION\_OF\_PEPTIDE\_SECRETION | MSigDB lists | GO\_REGULATION\_OF\_PEPTIDE\_SECRETION | 47 | 6 | 4161 | 170 | Pkc53E,csw,Pkc98E,MCU,Pde8,Syx1A | | 1.137e-02 | -4.48 | GO\_REGULATION\_OF\_NEURON\_APOPTOTIC\_PROCESS | MSigDB lists | GO\_REGULATION\_OF\_NEURON\_APOPTOTIC\_PROCESS | 47 | 6 | 4161 | 170 | Sod1,Cln3,TfAP-2,LRP1,Raf,KaiR1D | | 1.137e-02 | -4.48 | EVI1\_04 | MSigDB lists | EVI1\_04 | 47 | 6 | 4161 | 170 | CG9132,CG12121,chb,Dys,dnc,LRP1 | | 1.137e-02 | -4.48 | GO\_POSITIVE\_REGULATION\_OF\_ION\_TRANSPORT | MSigDB lists | GO\_POSITIVE\_REGULATION\_OF\_ION\_TRANSPORT | 47 | 6 | 4161 | 170 | Khc,Octbeta2R,Ank,Dys,CCAP-R,AstA-R1 | | 1.137e-02 | -4.48 | GSE15624\_CTRL\_VS\_3H\_HALOFUGINONE\_TREATED\_CD4\_TCELL\_UP | MSigDB lists | GSE15624\_CTRL\_VS\_3H\_HALOFUGINONE\_TREATED\_CD4\_TCELL\_UP | 47 | 6 | 4161 | 170 | dsh,CG4301,polybromo,Rbp6,Pde6,CG42672 | | 1.137e-02 | -4.48 | GSE11961\_UNSTIM\_VS\_ANTI\_IGM\_AND\_CD40\_STIM\_6H\_FOLLICULAR\_BCELL\_DN | MSigDB lists | GSE11961\_UNSTIM\_VS\_ANTI\_IGM\_AND\_CD40\_STIM\_6H\_FOLLICULAR\_BCELL\_DN | 47 | 6 | 4161 | 170 | CG10126,Doa,Nha1,Syx1A,Sh,Pgant6 | | 1.137e-02 | -4.48 | GSE6269\_E\_COLI\_VS\_STAPH\_AUREUS\_INF\_PBMC\_DN | MSigDB lists | GSE6269\_E\_COLI\_VS\_STAPH\_AUREUS\_INF\_PBMC\_DN | 47 | 6 | 4161 | 170 | Flo2,Cln3,Argl,EcR,Fhos,Ser | | 1.137e-02 | -4.48 | CREBP1\_01 | MSigDB lists | CREBP1\_01 | 47 | 6 | 4161 | 170 | sdt,Mctp,chb,bru1,Octbeta2R,Dys | | 1.139e-02 | -4.48 | ATF4\_Q2 | MSigDB lists | ATF4\_Q2 | 61 | 7 | 4161 | 170 | Iswi,Der-1,dnc,Ser,Raf,Pak,mGluR | | 1.139e-02 | -4.48 | GO\_POSITIVE\_REGULATION\_OF\_CELL\_PROJECTION\_ORGANIZATION | MSigDB lists | GO\_POSITIVE\_REGULATION\_OF\_CELL\_PROJECTION\_ORGANIZATION | 61 | 7 | 4161 | 170 | Dys,LpR2,unc-13,Raf,Tor,Tif-IA,CG42672 | | 1.141e-02 | -4.47 | GSE36476\_CTRL\_VS\_TSST\_ACT\_16H\_MEMORY\_CD4\_TCELL\_OLD\_UP | MSigDB lists | GSE36476\_CTRL\_VS\_TSST\_ACT\_16H\_MEMORY\_CD4\_TCELL\_OLD\_UP | 34 | 5 | 4161 | 170 | CNT1,MESK2,Glut1,flw,dnc | | 1.141e-02 | -4.47 | GSE2770\_TGFB\_AND\_IL4\_VS\_IL12\_TREATED\_ACT\_CD4\_TCELL\_2H\_DN | MSigDB lists | GSE2770\_TGFB\_AND\_IL4\_VS\_IL12\_TREATED\_ACT\_CD4\_TCELL\_2H\_DN | 34 | 5 | 4161 | 170 | Ank,Syx1A,Der-1,dnc,PIG-G | | 1.141e-02 | -4.47 | GSE8685\_IL15\_ACT\_IL2\_STARVED\_VS\_IL21\_ACT\_IL2\_STARVED\_CD4\_TCELL\_UP | MSigDB lists | GSE8685\_IL15\_ACT\_IL2\_STARVED\_VS\_IL21\_ACT\_IL2\_STARVED\_CD4\_TCELL\_UP | 34 | 5 | 4161 | 170 | CG12121,CG12547,Sh,Rbp6,rdgA | | 1.141e-02 | -4.47 | anatomical structure formation involved in morphogenesis | biological process | GO:0048646 | 553 | 39 | 12345 | 597 | EcR,AGO2,dpp,Mlp84B,egh,rst,siz,rols,mid,shg,bun,chb,foxo,Pld,Act88F,nuf,Raf,fs(1)N,dsh,tin,Inx2,Ubx,Actn,Khc,flw,bru1,Dlic,Prm,hth,tld,numb,B-H2,cic,polybromo,Tor,Su(H),bnl,ths,Syx1A | | 1.144e-02 | -4.47 | catenin complex | cellular component | GO:0016342 | 4 | 2 | 11940 | 538 | shg,CadN2 | | 1.149e-02 | -4.47 | GO\_RESPONSE\_TO\_ORGANIC\_CYCLIC\_COMPOUND | MSigDB lists | GO\_RESPONSE\_TO\_ORGANIC\_CYCLIC\_COMPOUND | 211 | 16 | 4161 | 170 | CG5493,Ugt301D1,dnc,Octbeta2R,Cat,CG3376,CG4849,Tor,Hr96,dpp,EcR,Pkc98E,GlyP,Raf,Sod1,CG7483 | | 1.153e-02 | -4.46 | dock (dreadlocks) | protein interactions | 33262 | 37 | 6 | 9204 | 481 | sick,Pak,Src42A,Ptp61F,fru,rst | | 1.159e-02 | -4.46 | NGF signalling via TRKA from the plasma membrane | REACTOME pathways | R-DME-187037 | 172 | 16 | 3923 | 197 | Pde1c,CG41520,Prosbeta5R1,Tor,Pkc53E,foxo,csw,ACXD,Pkc98E,Gprk1,REG,CrebB,bnl,rictor,Raf,Cam | | 1.160e-02 | -4.46 | integral component of plasma membrane | cellular component | GO:0005887 | 466 | 32 | 11940 | 538 | CadN2,dpr5,Nmdar2,PGRP-LA,dpr18,CG3285,beat-IIa,MsR2,Nlg2,kek1,CG14606,AstA-R1,ACXD,Sh,GluRIIC,scrib,CG33310,CG11318,Pdfr,nrv2,CG10960,CNT1,DIP-lambda,kcc,Octbeta2R,shg,olf186-F,DIP-delta,fz2,mGluR,kek5,CCAP-R | | 1.160e-02 | -4.46 | hormone metabolic process | biological process | GO:0042445 | 64 | 8 | 12345 | 597 | Fad2,CG3842,spok,shd,CG9514,foxo,woc,CG9519 | | 1.160e-02 | -4.46 | GO\_REGULATION\_OF\_CELLULAR\_KETONE\_METABOLIC\_PROCESS | MSigDB lists | GO\_REGULATION\_OF\_CELLULAR\_KETONE\_METABOLIC\_PROCESS | 76 | 8 | 4161 | 170 | Tor,Odc1,Pkc98E,EcR,ABCD,dpp,REG,Ugt301D1 | | 1.165e-02 | -4.45 | solute:cation symporter activity | molecular function | GO:0015294 | 65 | 8 | 12176 | 580 | CG14606,ine,CG45782,kcc,CNT1,CG3285,CG13794,CG10960 | | 1.167e-02 | -4.45 | CG34168 (uncharacterized protein) | protein interactions | 50457 | 48 | 7 | 9204 | 481 | tyf,esg,fru,opa,CG15564,CG17777,Mob2 | | 1.174e-02 | -4.44 | steroid hormone biosynthetic process | biological process | GO:0120178 | 19 | 4 | 12345 | 597 | spok,woc,foxo,shd | | 1.174e-02 | -4.44 | ecdysteroid biosynthetic process | biological process | GO:0045456 | 19 | 4 | 12345 | 597 | woc,spok,shd,foxo | | 1.177e-02 | -4.44 | female germ-line stem cell population maintenance | biological process | GO:0036099 | 29 | 5 | 12345 | 597 | shg,foxo,Tif-IA,bru1,dpp | | 1.177e-02 | -4.44 | RhoGEF | smart domains | SM00325 | 23 | 5 | 5086 | 315 | Cdep,RhoGEF3,CG10188,RhoGAP1A,Exn | | 1.177e-02 | -4.44 | CH | smart domains | SM00033 | 23 | 5 | 5086 | 315 | Msp300,Dys,CG34417,sick,Actn | | 1.178e-02 | -4.44 | regulation of cell fate commitment | biological process | GO:0010453 | 40 | 6 | 12345 | 597 | Su(H),mid,Ubx,bun,hth,Src42A | | 1.181e-02 | -4.44 | mTOR signaling pathway | KEGG pathways | ko04150 | 96 | 11 | 3198 | 170 | eIF4E7,Stlk,rictor,Tor,Vha36-3,dsh,fz2,fz4,sgg,Pkc53E,Raf | | 1.181e-02 | -4.44 | mTOR signaling pathway | KEGG pathways | dme04150 | 96 | 11 | 3198 | 170 | dsh,fz2,Vha36-3,Tor,rictor,eIF4E7,Stlk,Raf,Pkc53E,sgg,fz4 | | 1.184e-02 | -4.44 | supramolecular polymer | cellular component | GO:0099081 | 206 | 17 | 11940 | 538 | Mlp84B,CG34417,Actn,Gas8,Msp300,msps,CG3078,Pak,zormin,TpnC25D,Prm,Act88F,pod1,Khc,Act87E,chb,rols | | 1.186e-02 | -4.43 | cDIP (Common Dpr-interacting protein) | protein interactions | 42512 | 27 | 5 | 9204 | 481 | dpr5,DIP-delta,dpr12,dpr21,klg | | 1.191e-02 | -4.43 | regulation of cell cycle process | biological process | GO:0010564 | 177 | 16 | 12345 | 597 | esg,flw,Mos,REG,bru1,CG12200,chb,foxo,nuf,Cep97,cindr,Tor,Su(H),sgg,ex,kay | | 1.196e-02 | -4.43 | GO\_CELLULAR\_HOMEOSTASIS | MSigDB lists | GO\_CELLULAR\_HOMEOSTASIS | 176 | 14 | 4161 | 170 | Cln3,Sod1,sxc,Ca-beta,Qsox2,KaiR1D,kcc,ZnT63C,Pkc98E,AstA-R1,Dys,MCU,GlyP,CG40178 | | 1.196e-02 | -4.43 | GCANCTGNY\_MYOD\_Q6 | MSigDB lists | GCANCTGNY\_MYOD\_Q6 | 176 | 14 | 4161 | 170 | eIF2Bgamma,Act87E,Cadps,Ser,Pde1c,Nmdar2,Dys,dsh,RhoGAP1A,CG17698,Iswi,CG4341,polybromo,Odc1 | | 1.199e-02 | -4.42 | C2\_domain\_sf | interpro domains | IPR035892 | 49 | 7 | 11383 | 585 | RhoGAP100F,Pkc98E,Fife,Mctp,unc-13,Syt12,Pkc53E | | 1.206e-02 | -4.42 | GO\_G\_PROTEIN\_COUPLED\_RECEPTOR\_SIGNALING\_PATHWAY | MSigDB lists | GO\_G\_PROTEIN\_COUPLED\_RECEPTOR\_SIGNALING\_PATHWAY | 108 | 10 | 4161 | 170 | Ca-beta,fz2,dnc,LRP1,rdgA,Sk1,AstA-R1,CCAP-R,Octbeta2R,mGluR | | 1.210e-02 | -4.41 | PIP3 activates AKT signaling | REACTOME pathways | R-DME-1257604 | 39 | 6 | 3923 | 197 | bnl,csw,rictor,foxo,Tor,CrebB | | 1.217e-02 | -4.41 | heme binding | molecular function | GO:0020037 | 135 | 13 | 12176 | 580 | spok,Cyp304a1,Cyt-b5-r,Cyp6u1,Cyp12a4,Cyp4p2,Cyp6a13,Cyp309a1,shd,Cyp28d2,Cat,glob3,Cyp28d1 | | 1.223e-02 | -4.40 | epithelial cell differentiation | biological process | GO:0030855 | 419 | 31 | 12345 | 597 | bun,fs(1)N,kay,dsh,tin,Mrtf,GATAe,ex,numb,cic,sgg,bbg,app,polybromo,CG41099,RhoL,Raf,cindr,scrib,lncRNA:flam,shd,fw,EcR,Khc,shg,Mbs,dpp,egh,csw,rols,mid | | 1.230e-02 | -4.40 | GO\_CENTRAL\_NERVOUS\_SYSTEM\_DEVELOPMENT | MSigDB lists | GO\_CENTRAL\_NERVOUS\_SYSTEM\_DEVELOPMENT | 231 | 17 | 4161 | 170 | csw,Tor,Awh,TfAP-2,dsd,CG33090,Iswi,Su(H),RhoGAP1A,sxc,Elp3,eIF2Bgamma,LRP1,Act87E,Sk1,dpp,LpR2 | | 1.241e-02 | -4.39 | ER\_Q6\_02 | MSigDB lists | ER\_Q6\_02 | 62 | 7 | 4161 | 170 | Raf,Ucp4A,CG3376,Epg5,Iswi,Syn2,CG15093 | | 1.241e-02 | -4.39 | T3R\_Q6 | MSigDB lists | T3R\_Q6 | 62 | 7 | 4161 | 170 | CG3961,eIF4E7,Raf,fus,sxc,RhoGAP1A,Iswi | | 1.242e-02 | -4.39 | GO\_PHOSPHOLIPID\_METABOLIC\_PROCESS | MSigDB lists | GO\_PHOSPHOLIPID\_METABOLIC\_PROCESS | 125 | 11 | 4161 | 170 | CG3376,scramb1,AstA-R1,Sk1,mino,PIG-G,CG3530,Acbp2,csw,Cln3,PIG-O | | 1.242e-02 | -4.39 | GO\_ALCOHOL\_METABOLIC\_PROCESS | MSigDB lists | GO\_ALCOHOL\_METABOLIC\_PROCESS | 125 | 11 | 4161 | 170 | CG3860,Aldh-III,AstA-R1,LpR2,CG3376,Cat,Sk1,CG3842,Cln3,IP3K2,Acbp2 | | 1.251e-02 | -4.38 | NUYTTEN\_EZH2\_TARGETS\_UP | MSigDB lists | NUYTTEN\_EZH2\_TARGETS\_UP | 213 | 16 | 4161 | 170 | flw,Pkc98E,MCU,Su(H),sxc,Stt3B,sdt,Frq1,fz2,scramb1,Octbeta2R,CG40178,step,Frq2,CG4341,CG41099 | | 1.252e-02 | -4.38 | EF\_HAND\_2 | prosite domains | PS50222 | 94 | 12 | 6243 | 386 | TpnC25D,TpnC41C,TpnC4,Cam,Frq2,CG14362,Frq1,Scp1,Actn,CG10126,CG13898,CG13526 | | 1.255e-02 | -4.38 | mtSSB (mitochondrial single stranded DNA-binding protein) | protein interactions | 41968 | 18 | 4 | 9204 | 481 | CG15564,TfAP-2,Sod1,CG14907 | | 1.257e-02 | -4.38 | WHITFIELD\_CELL\_CYCLE\_M\_G1 | MSigDB lists | WHITFIELD\_CELL\_CYCLE\_M\_G1 | 48 | 6 | 4161 | 170 | eIF4E7,Khc,Frq1,Frq2,Elp3,eRF3 | | 1.257e-02 | -4.38 | NERF\_Q2 | MSigDB lists | NERF\_Q2 | 48 | 6 | 4161 | 170 | Frq1,mGluR,CG12121,Ser,IP3K2,Frq2 | | 1.257e-02 | -4.38 | GSE22140\_HEALTHY\_VS\_ARTHRITIC\_GERMFREE\_MOUSE\_CD4\_TCELL\_DN | MSigDB lists | GSE22140\_HEALTHY\_VS\_ARTHRITIC\_GERMFREE\_MOUSE\_CD4\_TCELL\_DN | 48 | 6 | 4161 | 170 | CG12121,CG7483,dsd,Glut1,Cln3,Meltrin | | 1.257e-02 | -4.38 | GO\_REGULATION\_OF\_MUSCLE\_SYSTEM\_PROCESS | MSigDB lists | GO\_REGULATION\_OF\_MUSCLE\_SYSTEM\_PROCESS | 48 | 6 | 4161 | 170 | Tor,dnc,Sod1,Pkc53E,Dys,Octbeta2R | | 1.257e-02 | -4.38 | ZHOU\_INFLAMMATORY\_RESPONSE\_LPS\_UP | MSigDB lists | ZHOU\_INFLAMMATORY\_RESPONSE\_LPS\_UP | 48 | 6 | 4161 | 170 | CG6418,Syn2,Cadps,Ser,eIF4E7,CNT1 | | 1.257e-02 | -4.38 | GO\_REGULATION\_OF\_EXOCYTOSIS | MSigDB lists | GO\_REGULATION\_OF\_EXOCYTOSIS | 48 | 6 | 4161 | 170 | RhoGAP1A,Frq2,Frq1,chb,Syx1A,Raf | | 1.257e-02 | -4.38 | ELF1\_Q6 | MSigDB lists | ELF1\_Q6 | 48 | 6 | 4161 | 170 | Der-1,rdgA,mRpL24,Raf,Nha1,CG3961 | | 1.265e-02 | -4.37 | GO\_POSITIVE\_REGULATION\_OF\_PROTEIN\_METABOLIC\_PROCESS | MSigDB lists | GO\_POSITIVE\_REGULATION\_OF\_PROTEIN\_METABOLIC\_PROCESS | 366 | 24 | 4161 | 170 | fz2,REG,Octbeta2R,CG3376,dsh,CG17698,csw,CG42672,Der-1,Tor,Stlk,Pkc98E,EcR,AGO3,dpp,Tsp96F,Raf,LpR2,Pak,Pkc53E,wbl,sxc,Sod1,CG7483 | | 1.266e-02 | -4.37 | segmentation | biological process | GO:0035282 | 256 | 21 | 12345 | 597 | Doa,csw,dpp,shg,hth,ems,AGO2,Khc,Ser,Ubx,scrib,CG7483,Antp,Raf,msps,cic,TfAP-2,sgg,dsh,fz2,fs(1)N | | 1.273e-02 | -4.36 | symporter activity | molecular function | GO:0015293 | 66 | 8 | 12176 | 580 | CG14606,ine,CG45782,CNT1,kcc,CG13794,CG3285,CG10960 | | 1.288e-02 | -4.35 | GO\_POSITIVE\_REGULATION\_OF\_DEVELOPMENTAL\_GROWTH | MSigDB lists | GO\_POSITIVE\_REGULATION\_OF\_DEVELOPMENTAL\_GROWTH | 35 | 5 | 4161 | 170 | unc-13,Octbeta2R,Su(H),mino,Tor | | 1.288e-02 | -4.35 | PID\_MET\_PATHWAY | MSigDB lists | PID\_MET\_PATHWAY | 35 | 5 | 4161 | 170 | eIF4E7,Pak,Ptp61F,csw,Tor | | 1.288e-02 | -4.35 | GSE25088\_CTRL\_VS\_IL4\_STIM\_STAT6\_KO\_MACROPHAGE\_DN | MSigDB lists | GSE25088\_CTRL\_VS\_IL4\_STIM\_STAT6\_KO\_MACROPHAGE\_DN | 35 | 5 | 4161 | 170 | kcc,AstA-R1,Pak,Src42A,Su(H) | | 1.288e-02 | -4.35 | GSE22886\_NAIVE\_BCELL\_VS\_BLOOD\_PLASMA\_CELL\_UP | MSigDB lists | GSE22886\_NAIVE\_BCELL\_VS\_BLOOD\_PLASMA\_CELL\_UP | 35 | 5 | 4161 | 170 | CG5522,IP3K2,CG12547,CG40178,CG41099 | | 1.288e-02 | -4.35 | GSE22935\_UNSTIM\_VS\_48H\_MBOVIS\_BCG\_STIM\_MYD88\_KO\_MACROPHAGE\_UP | MSigDB lists | GSE22935\_UNSTIM\_VS\_48H\_MBOVIS\_BCG\_STIM\_MYD88\_KO\_MACROPHAGE\_UP | 35 | 5 | 4161 | 170 | CG5493,rdgA,Iswi,PIG-O,PCB | | 1.288e-02 | -4.35 | GO\_CELLULAR\_RESPONSE\_TO\_OXYGEN\_LEVELS | MSigDB lists | GO\_CELLULAR\_RESPONSE\_TO\_OXYGEN\_LEVELS | 35 | 5 | 4161 | 170 | LpR2,Octbeta2R,Su(H),Pkc98E,Tor | | 1.292e-02 | -4.35 | lab (labial) | protein interactions | 40817 | 10 | 3 | 9204 | 481 | hth,Antp,Ubx | | 1.292e-02 | -4.35 | ec (echinus) | protein interactions | 31339 | 10 | 3 | 9204 | 481 | flw,sgg,shg | | 1.292e-02 | -4.35 | CG14315 (uncharacterized protein) | protein interactions | 42164 | 10 | 3 | 9204 | 481 | Actn,Prm,l(2)01289 | | 1.294e-02 | -4.35 | supramolecular complex | cellular component | GO:0099080 | 208 | 17 | 11940 | 538 | TpnC25D,pod1,Act88F,Prm,chb,Act87E,Khc,rols,Mlp84B,msps,CG3078,Msp300,Gas8,Actn,CG34417,Pak,zormin | | 1.296e-02 | -4.35 | GO\_CAMERA\_TYPE\_EYE\_MORPHOGENESIS | MSigDB lists | GO\_CAMERA\_TYPE\_EYE\_MORPHOGENESIS | 23 | 4 | 4161 | 170 | TfAP-2,Ser,CG17883,fz2 | | 1.296e-02 | -4.35 | GO\_HEART\_PROCESS | MSigDB lists | GO\_HEART\_PROCESS | 23 | 4 | 4161 | 170 | dnc,Tor,Sod1,Dys | | 1.296e-02 | -4.35 | GO\_REGULATION\_OF\_PEPTIDYL\_TYROSINE\_PHOSPHORYLATION | MSigDB lists | GO\_REGULATION\_OF\_PEPTIDYL\_TYROSINE\_PHOSPHORYLATION | 23 | 4 | 4161 | 170 | Pkc98E,Tor,Tsp96F,Ptp61F | | 1.296e-02 | -4.35 | KEGG\_ERBB\_SIGNALING\_PATHWAY | MSigDB lists | KEGG\_ERBB\_SIGNALING\_PATHWAY | 23 | 4 | 4161 | 170 | Tor,Pkc53E,Pak,Raf | | 1.296e-02 | -4.35 | PID\_VEGFR1\_2\_PATHWAY | MSigDB lists | PID\_VEGFR1\_2\_PATHWAY | 23 | 4 | 4161 | 170 | Raf,Ptp61F,csw,Pkc53E | | 1.296e-02 | -4.35 | CHESLER\_BRAIN\_HIGHEST\_EXPRESSION | MSigDB lists | CHESLER\_BRAIN\_HIGHEST\_EXPRESSION | 23 | 4 | 4161 | 170 | Pkc98E,flw,Act87E,Cadps | | 1.296e-02 | -4.35 | GSE9509\_LPS\_VS\_LPS\_AND\_IL10\_STIM\_IL10\_KO\_MACROPHAGE\_10MIN\_UP | MSigDB lists | GSE9509\_LPS\_VS\_LPS\_AND\_IL10\_STIM\_IL10\_KO\_MACROPHAGE\_10MIN\_UP | 23 | 4 | 4161 | 170 | scramb1,step,Pkc98E,Cln3 | | 1.296e-02 | -4.35 | GO\_NEURON\_NEURON\_SYNAPTIC\_TRANSMISSION | MSigDB lists | GO\_NEURON\_NEURON\_SYNAPTIC\_TRANSMISSION | 23 | 4 | 4161 | 170 | Ca-beta,Octbeta2R,unc-13,KaiR1D | | 1.296e-02 | -4.35 | LDLR\_class-A\_CS | interpro domains | IPR023415 | 28 | 5 | 11383 | 585 | LpR2,trol,CG31221,LRP1,CG32432 | | 1.302e-02 | -4.34 | mAcon1 (Mitochondrial aconitase 1) | protein interactions | 44149 | 49 | 7 | 9204 | 481 | TpnC4,Actn,Scp1,TpnC41C,CG34172,Act88F,Prm | | 1.313e-02 | -4.33 | positive regulation of RNA splicing | biological process | GO:0033120 | 4 | 2 | 12345 | 597 | barc,ps | | 1.313e-02 | -4.33 | phosphatidic acid biosynthetic process | biological process | GO:0006654 | 4 | 2 | 12345 | 597 | rdgA,Pld | | 1.313e-02 | -4.33 | specification of animal organ identity | biological process | GO:0010092 | 4 | 2 | 12345 | 597 | Ubx,hth | | 1.314e-02 | -4.33 | GO\_SIGNALING\_RECEPTOR\_ACTIVITY | MSigDB lists | GO\_SIGNALING\_RECEPTOR\_ACTIVITY | 126 | 11 | 4161 | 170 | Nmdar2,LpR2,CCAP-R,AstA-R1,mGluR,Octbeta2R,fz2,EcR,Sk1,Hr96,KaiR1D | | 1.315e-02 | -4.33 | positive regulation of phosphorus metabolic process | biological process | GO:0010562 | 149 | 14 | 12345 | 597 | dpp,bnl,Tor,Raf,step,rictor,Pak,lncRNA:Hsromega,Stlk,Mos,CycT,CG41520,Src42A,Pde8 | | 1.315e-02 | -4.33 | positive regulation of phosphate metabolic process | biological process | GO:0045937 | 149 | 14 | 12345 | 597 | bnl,Raf,Tor,dpp,lncRNA:Hsromega,Pak,rictor,step,CycT,Mos,Stlk,Pde8,Src42A,CG41520 | | 1.324e-02 | -4.32 | apicolateral plasma membrane | cellular component | GO:0016327 | 21 | 4 | 11940 | 538 | Inx2,sdt,scrib,ex | | 1.325e-02 | -4.32 | regulation of hemocyte proliferation | biological process | GO:0035206 | 41 | 6 | 12345 | 597 | Cat,EcR,foxo,Pkc53E,fz2,CG1572 | | 1.333e-02 | -4.32 | regulation of tube size | biological process | GO:0035150 | 53 | 7 | 12345 | 597 | raw,dsh,rib,dpp,nrv2,tnc,EcR | | 1.345e-02 | -4.31 | PLC beta mediated events | REACTOME pathways | R-DME-112043 | 29 | 5 | 3923 | 197 | Cam,Pkc53E,Gprk1,Pde1c,ACXD | | 1.347e-02 | -4.31 | BROWNE\_HCMV\_INFECTION\_16HR\_UP | MSigDB lists | BROWNE\_HCMV\_INFECTION\_16HR\_UP | 78 | 8 | 4161 | 170 | Raf,LpR2,CG9132,scramb1,dnc,dpp,dsd,csw | | 1.349e-02 | -4.31 | GSE39022\_LN\_VS\_SPLEEN\_DC\_DN | MSigDB lists | GSE39022\_LN\_VS\_SPLEEN\_DC\_DN | 63 | 7 | 4161 | 170 | MESK2,Syx1A,Nha1,Cpsf160,sws,CG42672,D2hgdh | | 1.349e-02 | -4.31 | YYCATTCAWW\_UNKNOWN | MSigDB lists | YYCATTCAWW\_UNKNOWN | 63 | 7 | 4161 | 170 | Su(H),csw,Frq2,eIF4E7,Ank,Frq1,Dys | | 1.349e-02 | -4.31 | TTCYNRGAA\_STAT5B\_01 | MSigDB lists | TTCYNRGAA\_STAT5B\_01 | 63 | 7 | 4161 | 170 | rdgA,mRpL24,Ank,CG3376,scramb1,eIF4E7,CG8272 | | 1.349e-02 | -4.31 | Adh (Alcohol dehydrogenase) | protein interactions | 3771877 | 61 | 8 | 9204 | 481 | Scp1,CG34172,Prm,TpnC41C,Actn,GlyP,Tsf1,Act88F | | 1.359e-02 | -4.30 | GO\_CELL\_PROJECTION | MSigDB lists | GO\_CELL\_PROJECTION | 469 | 29 | 4161 | 170 | sky,KaiR1D,Flo2,CG7483,Frq1,CG17387,sxc,Sod1,Raf,unc-13,Gas8,Khc,Pak,nompB,LRP1,rdgA,Ank,Cln3,Frq2,Maf1,Tor,Syx1A,Octbeta2R,Dys,mGluR,Syn2,Root,Sh,Fhos | | 1.360e-02 | -4.30 | EGF\_1 | prosite domains | PS00022 | 52 | 8 | 6243 | 386 | CG13654,shg,trol,LRP1,CadN2,Ser,dsd,Meltrin | | 1.366e-02 | -4.29 | PI3K/AKT activation | REACTOME pathways | R-DME-198203 | 40 | 6 | 3923 | 197 | CrebB,foxo,Tor,rictor,bnl,csw | | 1.367e-02 | -4.29 | courtship behavior | biological process | GO:0007619 | 135 | 13 | 12345 | 597 | Gr39a,Sh,Gr22c,mGluR,fru,egh,TfAP-2,dnc,lncRNA:flam,Fad2,Nlg2,retn,shep | | 1.378e-02 | -4.28 | GO\_DOUBLE\_STRANDED\_DNA\_BINDING | MSigDB lists | GO\_DOUBLE\_STRANDED\_DNA\_BINDING | 94 | 9 | 4161 | 170 | Hr96,Rat1,Su(H),Tor,TfAP-2,Maf1,EcR,Tif-IA,Act87E | | 1.383e-02 | -4.28 | Nplp2 (Neuropeptide-like precursor 2) | protein interactions | 50286 | 28 | 5 | 9204 | 481 | TpnC41C,Scp1,Cpr64Ac,Prm,Tsf1 | | 1.383e-02 | -4.28 | ptc (patched) | protein interactions | 35851 | 153 | 15 | 9204 | 481 | boi,CG8184,dpp,trol,EcR,bnl,zormin,Mlp84B,crol,rdgA,Fhos,LRP1,Pgant6,CG46385,fz2 | | 1.384e-02 | -4.28 | regulation of transmembrane receptor protein serine/threonine kinase signaling pathway | biological process | GO:0090092 | 66 | 8 | 12345 | 597 | Snoo,dpp,hth,kek5,Lpin,tld,GluRIIC,foxo | | 1.385e-02 | -4.28 | GSE25677\_MPL\_VS\_MPL\_AND\_R848\_STIM\_BCELL\_UP | MSigDB lists | GSE25677\_MPL\_VS\_MPL\_AND\_R848\_STIM\_BCELL\_UP | 49 | 6 | 4161 | 170 | Frq2,Meltrin,Frq1,MESK2,CG3961,Raf | | 1.385e-02 | -4.28 | GO\_HEART\_MORPHOGENESIS | MSigDB lists | GO\_HEART\_MORPHOGENESIS | 49 | 6 | 4161 | 170 | CG17387,dsh,Tor,dpp,Ser,Su(H) | | 1.386e-02 | -4.28 | zip (zipper) | protein interactions | 38001 | 86 | 10 | 9204 | 481 | chb,sgg,step,rib,AANAT1,raw,Mbs,dsh,Uros1,flw | | 1.388e-02 | -4.28 | negative regulation of biosynthetic process | biological process | GO:0009890 | 457 | 33 | 12345 | 597 | Su(H),polybromo,B-H2,cic,crol,bi,msi,CG43689,Iswi,CG12605,CG1894,esg,Khc,Ubx,retn,bru1,E5,CG12071,TfAP-2,lncRNA:bxd,Sox102F,tin,toe,HPS4,foxo,Maf1,mid,CG7368,CG31224,AGO2,EcR,scrib,Antp | | 1.388e-02 | -4.28 | negative regulation of cellular biosynthetic process | biological process | GO:0031327 | 457 | 33 | 12345 | 597 | crol,cic,B-H2,polybromo,Su(H),Iswi,CG43689,msi,bi,CG12605,E5,bru1,retn,Ubx,esg,Khc,CG1894,tin,Sox102F,lncRNA:bxd,TfAP-2,CG12071,foxo,HPS4,toe,CG31224,CG7368,Maf1,mid,Antp,scrib,EcR,AGO2 | | 1.395e-02 | -4.27 | RRM\_1 | pfam domains | PF00076 | 125 | 13 | 10480 | 550 | CG42458,shep,trv,CG3594,Spf45,CG44249,CG40813,Rbp6,barc,msi,CG41562,bru1,bol | | 1.401e-02 | -4.27 | GO\_RECEPTOR\_BINDING | MSigDB lists | GO\_RECEPTOR\_BINDING | 253 | 18 | 4161 | 170 | Tsp96F,EcR,LRP1,Ser,dpp,Flo2,Nlg2,Src42A,Acbp2,scramb1,Octbeta2R,Cat,dsh,Ptp61F,dnc,csw,Der-1,Maf1 | | 1.408e-02 | -4.26 | GO\_REGULATION\_OF\_CELL\_DEATH | MSigDB lists | GO\_REGULATION\_OF\_CELL\_DEATH | 330 | 22 | 4161 | 170 | LRP1,dpp,Sk1,Raf,RhoGAP1A,wbl,Pkc53E,Sod1,sxc,KaiR1D,mino,REG,Tif-IA,CG3376,Octbeta2R,Cat,Ptp61F,Cln3,TfAP-2,IP3K2,Tor,CG3961 | | 1.408e-02 | -4.26 | DH\_2 | prosite domains | PS50010 | 24 | 5 | 6243 | 386 | RhoGAP1A,CG10188,RhoGEF3,Exn,Cdep | | 1.413e-02 | -4.26 | somatic stem cell population maintenance | biological process | GO:0035019 | 20 | 4 | 12345 | 597 | dsh,shg,sgg,esg | | 1.413e-02 | -4.26 | regulation of cation transmembrane transport | biological process | GO:1904062 | 20 | 4 | 12345 | 597 | Ca-beta,CG42797,Cam,qvr | | 1.416e-02 | -4.26 | HU\_GENOTOXIN\_ACTION\_DIRECT\_VS\_INDIRECT\_4HR | MSigDB lists | HU\_GENOTOXIN\_ACTION\_DIRECT\_VS\_INDIRECT\_4HR | 13 | 3 | 4161 | 170 | Act87E,EcR,Doa | | 1.416e-02 | -4.26 | MARSON\_FOXP3\_TARGETS\_UP | MSigDB lists | MARSON\_FOXP3\_TARGETS\_UP | 13 | 3 | 4161 | 170 | Glut1,bru1,Tsp96F | | 1.416e-02 | -4.26 | GO\_AUDITORY\_RECEPTOR\_CELL\_DIFFERENTIATION | MSigDB lists | GO\_AUDITORY\_RECEPTOR\_CELL\_DIFFERENTIATION | 13 | 3 | 4161 | 170 | Ser,Sod1,Su(H) | | 1.416e-02 | -4.26 | SNIJDERS\_AMPLIFIED\_IN\_HEAD\_AND\_NECK\_TUMORS | MSigDB lists | SNIJDERS\_AMPLIFIED\_IN\_HEAD\_AND\_NECK\_TUMORS | 13 | 3 | 4161 | 170 | Pak,Su(H),Ser | | 1.416e-02 | -4.26 | GO\_REGULATION\_OF\_JUN\_KINASE\_ACTIVITY | MSigDB lists | GO\_REGULATION\_OF\_JUN\_KINASE\_ACTIVITY | 13 | 3 | 4161 | 170 | fz2,dsh,Pak | | 1.423e-02 | -4.25 | Phosphatidylinositol signaling system | KEGG pathways | ko04070 | 61 | 8 | 3198 | 170 | CG13526,IP3K2,Pkc53E,rdgA,Ipk1,Cam,CG13898,CG3530 | | 1.423e-02 | -4.25 | Phosphatidylinositol signaling system | KEGG pathways | dme04070 | 61 | 8 | 3198 | 170 | CG3530,CG13898,Cam,Ipk1,rdgA,Pkc53E,IP3K2,CG13526 | | 1.423e-02 | -4.25 | Hippo signaling pathway - fly | KEGG pathways | ko04391 | 61 | 8 | 3198 | 170 | Act87E,upd3,sdt,Act88F,hth,scrib,app,ex | | 1.423e-02 | -4.25 | Hippo signaling pathway - fly | KEGG pathways | dme04391 | 61 | 8 | 3198 | 170 | hth,Act88F,scrib,app,ex,Act87E,upd3,sdt | | 1.429e-02 | -4.25 | Dl (Delta) | protein interactions | 42313 | 126 | 13 | 9204 | 481 | sno,Su(H),htk,CG9650,csw,boi,HDAC4,Dys,rst,Ser,dsh,Ubx,cic | | 1.429e-02 | -4.25 | positive regulation of developmental process | biological process | GO:0051094 | 259 | 21 | 12345 | 597 | Ubx,EcR,Octbeta2R,bru1,dpp,step,rictor,spri,shg,hth,Pak,bun,Src42A,foxo,Raf,Tor,kay,Ank2,dsh,Fhos,mnb | | 1.432e-02 | -4.25 | C2 | prosite domains | PS50004 | 33 | 6 | 6243 | 386 | Pkc98E,Pkc53E,Fife,unc-13,Syt12,Mctp | | 1.435e-02 | -4.24 | TGACAGNY\_MEIS1\_01 | MSigDB lists | TGACAGNY\_MEIS1\_01 | 198 | 15 | 4161 | 170 | Syn2,CG14480,Rbp6,REG,Raf,Aldh-III,LpR2,Pde1c,CG5339,Dys,CG6418,Stt3B,CG4341,polybromo,eIF4E7 | | 1.441e-02 | -4.24 | Homeobox-like\_sf | interpro domains | IPR009057 | 142 | 14 | 11383 | 585 | scro,tin,Antp,Poxm,Awh,dany,rib,toe,ems,hth,Ubx,Iswi,B-H2,E5 | | 1.445e-02 | -4.24 | TYR\_PHOSPHATASE\_PTP | prosite domains | PS50055 | 16 | 4 | 6243 | 386 | Ptp36E,Ptp61F,Ptp99A,csw | | 1.448e-02 | -4.23 | CEBP\_C | MSigDB lists | CEBP\_C | 36 | 5 | 4161 | 170 | dnc,Dlic,Dys,Octbeta2R,CG3961 | | 1.448e-02 | -4.23 | GO\_NEGATIVE\_REGULATION\_OF\_NEURON\_APOPTOTIC\_PROCESS | MSigDB lists | GO\_NEGATIVE\_REGULATION\_OF\_NEURON\_APOPTOTIC\_PROCESS | 36 | 5 | 4161 | 170 | KaiR1D,Raf,LRP1,Cln3,Sod1 | | 1.448e-02 | -4.23 | GSE360\_HIGH\_DOSE\_B\_MALAYI\_VS\_M\_TUBERCULOSIS\_DC\_DN | MSigDB lists | GSE360\_HIGH\_DOSE\_B\_MALAYI\_VS\_M\_TUBERCULOSIS\_DC\_DN | 36 | 5 | 4161 | 170 | kcc,scramb1,Acbp2,CG5493,Meltrin | | 1.448e-02 | -4.23 | GSE22886\_CD4\_TCELL\_VS\_BCELL\_NAIVE\_UP | MSigDB lists | GSE22886\_CD4\_TCELL\_VS\_BCELL\_NAIVE\_UP | 36 | 5 | 4161 | 170 | Pkc53E,rdgA,IP3K2,KrT95D,CG3376 | | 1.448e-02 | -4.23 | REACTOME\_SIGNALING\_BY\_FGFR | MSigDB lists | REACTOME\_SIGNALING\_BY\_FGFR | 36 | 5 | 4161 | 170 | Pkc98E,Pkc53E,Tor,Pde1c,Raf | | 1.448e-02 | -4.23 | GSE30083\_SP1\_VS\_SP4\_THYMOCYTE\_DN | MSigDB lists | GSE30083\_SP1\_VS\_SP4\_THYMOCYTE\_DN | 36 | 5 | 4161 | 170 | Meltrin,sws,fus,CG4301,MESK2 | | 1.448e-02 | -4.23 | AML\_Q6 | MSigDB lists | AML\_Q6 | 36 | 5 | 4161 | 170 | Rbp6,eIF2Bgamma,CG12547,Glut1,Nmdar2 | | 1.450e-02 | -4.23 | distal axon | cellular component | GO:0150034 | 100 | 10 | 11940 | 538 | mnb,unc-13,gukh,Ank2,chb,g,cpx,sky,Syx1A,pod1 | | 1.450e-02 | -4.23 | PTPc | smart domains | SM00194 | 16 | 4 | 5086 | 315 | Ptp61F,csw,Ptp36E,Ptp99A | | 1.456e-02 | -4.23 | Cyt\_P450\_sf | interpro domains | IPR036396 | 88 | 10 | 11383 | 585 | Cyp309a1,Cyp28d1,Cyp304a1,shd,Cyp12a4,Cyp6a13,spok,Cyp6u1,Cyp28d2,Cyp4p2 | | 1.456e-02 | -4.23 | Cyt\_P450 | interpro domains | IPR001128 | 88 | 10 | 11383 | 585 | Cyp304a1,Cyp28d1,Cyp309a1,Cyp4p2,Cyp28d2,Cyp6u1,spok,shd,Cyp6a13,Cyp12a4 | | 1.464e-02 | -4.22 | GSE22886\_NAIVE\_BCELL\_VS\_BM\_PLASMA\_CELL\_UP | MSigDB lists | GSE22886\_NAIVE\_BCELL\_VS\_BM\_PLASMA\_CELL\_UP | 64 | 7 | 4161 | 170 | CG12547,IP3K2,rdgA,RpL38,Cat,sno,CG40178 | | 1.469e-02 | -4.22 | transcription factor binding | molecular function | GO:0008134 | 184 | 16 | 12176 | 580 | Hr96,kay,TfIIFbeta,mid,retn,cic,gce,bi,EcR,Su(H),hth,Iswi,foxo,Mrtf,Mondo,ex | | 1.469e-02 | -4.22 | cellular ketone metabolic process | biological process | GO:0042180 | 54 | 7 | 12345 | 597 | woc,spok,Kyat,CG9514,shd,CG9519,foxo | | 1.471e-02 | -4.22 | Ras85D (Ras oncogene at 85D) | protein interactions | 41140 | 241 | 21 | 9204 | 481 | scrib,chb,step,IntS3,CrebB,Raf,spri,cindr,dpp,bun,Tor,csw,upd3,Src42A,egh,Ptp61F,kay,cic,retn,MESK2,CG7483 | | 1.476e-02 | -4.22 | regulation of developmental growth | biological process | GO:0048638 | 228 | 19 | 12345 | 597 | Syn2,wbl,Src42A,foxo,Raf,Tor,ex,sgg,Mob2,mnb,Ank2,dsh,Ubx,scrib,Octbeta2R,dpp,Pak,rictor,step | | 1.476e-02 | -4.22 | ZU5\_dom | interpro domains | IPR000906 | 4 | 2 | 11383 | 585 | Ank2,Ank | | 1.476e-02 | -4.22 | Phosphodiest/P\_Trfase | interpro domains | IPR002591 | 4 | 2 | 11383 | 585 | PIG-G,PIG-O | | 1.476e-02 | -4.22 | Api\_m\_3 | interpro domains | IPR029630 | 4 | 2 | 11383 | 585 | CG9449,CG9451 | | 1.476e-02 | -4.22 | Catenin\_binding\_dom\_sf | interpro domains | IPR027397 | 4 | 2 | 11383 | 585 | CadN2,shg | | 1.476e-02 | -4.22 | Munc13\_1 | interpro domains | IPR014770 | 4 | 2 | 11383 | 585 | unc-13,Cadps | | 1.476e-02 | -4.22 | Tubby-like\_C | interpro domains | IPR025659 | 4 | 2 | 11383 | 585 | scramb1,Tusp | | 1.480e-02 | -4.21 | GO\_CELLULAR\_LIPID\_METABOLIC\_PROCESS | MSigDB lists | GO\_CELLULAR\_LIPID\_METABOLIC\_PROCESS | 312 | 21 | 4161 | 170 | CG3842,Acbp2,Aldh-III,Sk1,rdgA,LRP1,EcR,ABCD,PIG-G,CG3530,CG3961,csw,PIG-O,Cln3,Cat,CG3376,scramb1,AstA-R1,Ugt301D1,CG33090,mino | | 1.483e-02 | -4.21 | msi (musashi) | protein interactions | 43087 | 39 | 6 | 9204 | 481 | CG13157,ps,Marf1,CG5386,bru1,msi | | 1.489e-02 | -4.21 | ACTININ\_1 | prosite domains | PS00019 | 9 | 3 | 6243 | 386 | Actn,Dys,Msp300 | | 1.495e-02 | -4.20 | protein serine/threonine kinase activity | molecular function | GO:0004674 | 200 | 17 | 12176 | 580 | Strn-Mlck,Tor,Pkc53E,Ste:CG33247,mnb,Pak,Pkc98E,Nuak1,Gprk1,Stlk,Asator,Doa,PhKgamma,Raf,CG4839,CG17698,sgg | | 1.501e-02 | -4.20 | Tyr\_Pase\_AS | interpro domains | IPR016130 | 29 | 5 | 11383 | 585 | csw,Ptp36E,CG3530,Ptp61F,Ptp99A | | 1.505e-02 | -4.20 | GO\_REGULATION\_OF\_IMMUNE\_RESPONSE | MSigDB lists | GO\_REGULATION\_OF\_IMMUNE\_RESPONSE | 181 | 14 | 4161 | 170 | Flo2,RhoGAP1A,csw,Ca-beta,Tsp96F,scramb1,Pak,Ptp61F,Pkc98E,EcR,fz2,Act87E,dnc,REG | | 1.505e-02 | -4.20 | GO\_POSITIVE\_REGULATION\_OF\_TRANSCRIPTION\_FROM\_RNA\_POLYMERASE\_II\_PROMOTER | MSigDB lists | GO\_POSITIVE\_REGULATION\_OF\_TRANSCRIPTION\_FROM\_RNA\_POLYMERASE\_II\_PROMOTER | 181 | 14 | 4161 | 170 | Hr96,Su(H),sxc,TfAP-2,AstA-R1,Tsp96F,TfIIFbeta,scramb1,Octbeta2R,fz2,EcR,Fhos,Ser,dpp | | 1.506e-02 | -4.20 | GO\_REGULATION\_OF\_GENERATION\_OF\_PRECURSOR\_METABOLITES\_AND\_ENERGY | MSigDB lists | GO\_REGULATION\_OF\_GENERATION\_OF\_PRECURSOR\_METABOLITES\_AND\_ENERGY | 24 | 4 | 4161 | 170 | PhKgamma,Tor,flw,sxc | | 1.506e-02 | -4.20 | GO\_CARDIOCYTE\_DIFFERENTIATION | MSigDB lists | GO\_CARDIOCYTE\_DIFFERENTIATION | 24 | 4 | 4161 | 170 | Ser,Tor,dpp,Su(H) | | 1.506e-02 | -4.20 | GSE45365\_WT\_VS\_IFNAR\_KO\_CD11B\_DC\_MCMV\_INFECTION\_DN | MSigDB lists | GSE45365\_WT\_VS\_IFNAR\_KO\_CD11B\_DC\_MCMV\_INFECTION\_DN | 24 | 4 | 4161 | 170 | MCU,Pak,Src42A,Ser | | 1.506e-02 | -4.20 | GO\_POSITIVE\_REGULATION\_OF\_TRANSMEMBRANE\_TRANSPORT | MSigDB lists | GO\_POSITIVE\_REGULATION\_OF\_TRANSMEMBRANE\_TRANSPORT | 24 | 4 | 4161 | 170 | Ank,Dys,CCAP-R,AstA-R1 | | 1.522e-02 | -4.19 | HEB\_Q6 | MSigDB lists | HEB\_Q6 | 50 | 6 | 4161 | 170 | ZnT63C,dsh,Nmdar2,Cadps,REG,Act87E | | 1.522e-02 | -4.19 | BROWN\_MYELOID\_CELL\_DEVELOPMENT\_DN | MSigDB lists | BROWN\_MYELOID\_CELL\_DEVELOPMENT\_DN | 50 | 6 | 4161 | 170 | Sod1,Rbp6,CG4341,eIF4E7,PCB,Odc1 | | 1.522e-02 | -4.19 | GSE5589\_UNSTIM\_VS\_180MIN\_LPS\_AND\_IL10\_STIM\_MACROPHAGE\_DN | MSigDB lists | GSE5589\_UNSTIM\_VS\_180MIN\_LPS\_AND\_IL10\_STIM\_MACROPHAGE\_DN | 50 | 6 | 4161 | 170 | polybromo,scramb1,MCU,Stlk,Sod1,Pfdn4 | | 1.522e-02 | -4.19 | GO\_POSITIVE\_REGULATION\_OF\_CYTOSKELETON\_ORGANIZATION | MSigDB lists | GO\_POSITIVE\_REGULATION\_OF\_CYTOSKELETON\_ORGANIZATION | 50 | 6 | 4161 | 170 | Raf,chb,Pak,Pkc98E,Tor,Fhos | | 1.524e-02 | -4.18 | Cht6 (Chitinase 6) | protein interactions | 31935 | 4 | 2 | 9204 | 481 | flw,sgg | | 1.524e-02 | -4.18 | yata (yata) | protein interactions | 43508 | 4 | 2 | 9204 | 481 | sgg,Raf | | 1.524e-02 | -4.18 | Rim (Rab3 interacting molecule) | protein interactions | 42150 | 4 | 2 | 9204 | 481 | sgg,Fife | | 1.524e-02 | -4.18 | sano (serrano) | protein interactions | 37185 | 4 | 2 | 9204 | 481 | dsh,chb | | 1.524e-02 | -4.18 | bigmax (bigmax) | protein interactions | 43293 | 4 | 2 | 9204 | 481 | Mondo,Aldh-III | | 1.524e-02 | -4.18 | RhoL (Rho-like) | protein interactions | 41136 | 4 | 2 | 9204 | 481 | shg,RhoGAP100F | | 1.524e-02 | -4.18 | CG6227 (uncharacterized protein) | protein interactions | 32464 | 4 | 2 | 9204 | 481 | sgg,rictor | | 1.524e-02 | -4.18 | Asap (ArfGAP with SH3 domain, ankyrin repeat and PH domain) | protein interactions | 35783 | 4 | 2 | 9204 | 481 | cindr,sgg | | 1.524e-02 | -4.18 | Rsf1 (Repressor splicing factor 1) | protein interactions | 34370 | 4 | 2 | 9204 | 481 | Iswi,ps | | 1.526e-02 | -4.18 | CG5890 (uncharacterized protein) | protein interactions | 43126 | 19 | 4 | 9204 | 481 | cert,IP3K2,Spn88Eb,klg | | 1.526e-02 | -4.18 | Ras64B (Ras oncogene at 64B) | protein interactions | 38494 | 19 | 4 | 9204 | 481 | spri,foxo,Raf,dpp | | 1.526e-02 | -4.18 | fln (flightin) | protein interactions | 40185 | 19 | 4 | 9204 | 481 | Strn-Mlck,Act88F,TpnC4,CG34172 | | 1.526e-02 | -4.18 | Cp110 (Centriolar coiled coil protein 110kDa) | protein interactions | 33136 | 19 | 4 | 9204 | 481 | CG8117,Cam,Cep97,spd-2 | | 1.526e-02 | -4.18 | Platelet degranulation | REACTOME pathways | R-DME-114608 | 105 | 11 | 3923 | 197 | Actn,Qsox2,CG32264,Cam,CG41520,Tsf1,fw,kek1,CG12547,ninaD,kek5 | | 1.530e-02 | -4.18 | GO\_POSITIVE\_REGULATION\_OF\_CARDIAC\_MUSCLE\_CELL\_DIFFERENTIATION | MSigDB lists | GO\_POSITIVE\_REGULATION\_OF\_CARDIAC\_MUSCLE\_CELL\_DIFFERENTIATION | 5 | 2 | 4161 | 170 | Tor,Su(H) | | 1.530e-02 | -4.18 | GO\_NEGATIVE\_REGULATION\_OF\_LIPID\_TRANSPORT | MSigDB lists | GO\_NEGATIVE\_REGULATION\_OF\_LIPID\_TRANSPORT | 5 | 2 | 4161 | 170 | EcR,csw | | 1.530e-02 | -4.18 | GO\_POSITIVE\_REGULATION\_OF\_SYNAPTIC\_TRANSMISSION\_GABAERGIC | MSigDB lists | GO\_POSITIVE\_REGULATION\_OF\_SYNAPTIC\_TRANSMISSION\_GABAERGIC | 5 | 2 | 4161 | 170 | Khc,Pkc98E | | 1.530e-02 | -4.18 | GO\_REGULATION\_OF\_EPITHELIAL\_CELL\_APOPTOTIC\_PROCESS | MSigDB lists | GO\_REGULATION\_OF\_EPITHELIAL\_CELL\_APOPTOTIC\_PROCESS | 5 | 2 | 4161 | 170 | Raf,Tor | | 1.530e-02 | -4.18 | GO\_EMBRYONIC\_CAMERA\_TYPE\_EYE\_MORPHOGENESIS | MSigDB lists | GO\_EMBRYONIC\_CAMERA\_TYPE\_EYE\_MORPHOGENESIS | 5 | 2 | 4161 | 170 | fz2,TfAP-2 | | 1.530e-02 | -4.18 | GO\_REGULATION\_OF\_CHOLESTEROL\_EFFLUX | MSigDB lists | GO\_REGULATION\_OF\_CHOLESTEROL\_EFFLUX | 5 | 2 | 4161 | 170 | EcR,LRP1 | | 1.530e-02 | -4.18 | REACTOME\_CGMP\_EFFECTS | MSigDB lists | REACTOME\_CGMP\_EFFECTS | 5 | 2 | 4161 | 170 | Pde6,Pde1c | | 1.530e-02 | -4.18 | GO\_LABYRINTHINE\_LAYER\_BLOOD\_VESSEL\_DEVELOPMENT | MSigDB lists | GO\_LABYRINTHINE\_LAYER\_BLOOD\_VESSEL\_DEVELOPMENT | 5 | 2 | 4161 | 170 | fz2,Su(H) | | 1.530e-02 | -4.18 | GO\_POSITIVE\_REGULATION\_OF\_NEUROTRANSMITTER\_TRANSPORT | MSigDB lists | GO\_POSITIVE\_REGULATION\_OF\_NEUROTRANSMITTER\_TRANSPORT | 5 | 2 | 4161 | 170 | unc-13,Syx1A | | 1.530e-02 | -4.18 | GO\_NEUROTROPHIN\_SIGNALING\_PATHWAY | MSigDB lists | GO\_NEUROTROPHIN\_SIGNALING\_PATHWAY | 5 | 2 | 4161 | 170 | csw,CG42672 | | 1.530e-02 | -4.18 | GO\_EMBRYONIC\_EYE\_MORPHOGENESIS | MSigDB lists | GO\_EMBRYONIC\_EYE\_MORPHOGENESIS | 5 | 2 | 4161 | 170 | TfAP-2,fz2 | | 1.530e-02 | -4.18 | LEE\_EARLY\_T\_LYMPHOCYTE\_DN | MSigDB lists | LEE\_EARLY\_T\_LYMPHOCYTE\_DN | 5 | 2 | 4161 | 170 | Ank,Glut1 | | 1.530e-02 | -4.18 | GO\_POSITIVE\_REGULATION\_OF\_NEUROTRANSMITTER\_SECRETION | MSigDB lists | GO\_POSITIVE\_REGULATION\_OF\_NEUROTRANSMITTER\_SECRETION | 5 | 2 | 4161 | 170 | Syx1A,unc-13 | | 1.530e-02 | -4.18 | BOYLAN\_MULTIPLE\_MYELOMA\_C\_DN | MSigDB lists | BOYLAN\_MULTIPLE\_MYELOMA\_C\_DN | 5 | 2 | 4161 | 170 | dnc,ZnT63C | | 1.530e-02 | -4.18 | GO\_CARDIAC\_EPITHELIAL\_TO\_MESENCHYMAL\_TRANSITION | MSigDB lists | GO\_CARDIAC\_EPITHELIAL\_TO\_MESENCHYMAL\_TRANSITION | 5 | 2 | 4161 | 170 | dpp,Su(H) | | 1.530e-02 | -4.18 | GO\_LOW\_DENSITY\_LIPOPROTEIN\_RECEPTOR\_ACTIVITY | MSigDB lists | GO\_LOW\_DENSITY\_LIPOPROTEIN\_RECEPTOR\_ACTIVITY | 5 | 2 | 4161 | 170 | LpR2,LRP1 | | 1.530e-02 | -4.18 | GO\_REGULATION\_OF\_THYMOCYTE\_AGGREGATION | MSigDB lists | GO\_REGULATION\_OF\_THYMOCYTE\_AGGREGATION | 5 | 2 | 4161 | 170 | Sod1,Ptp61F | | 1.530e-02 | -4.18 | GO\_NEGATIVE\_REGULATION\_OF\_SMOOTH\_MUSCLE\_CONTRACTION | MSigDB lists | GO\_NEGATIVE\_REGULATION\_OF\_SMOOTH\_MUSCLE\_CONTRACTION | 5 | 2 | 4161 | 170 | Sod1,Octbeta2R | | 1.530e-02 | -4.18 | GO\_MYELIN\_MAINTENANCE | MSigDB lists | GO\_MYELIN\_MAINTENANCE | 5 | 2 | 4161 | 170 | sdt,Sod1 | | 1.530e-02 | -4.18 | PID\_TCR\_RAS\_PATHWAY | MSigDB lists | PID\_TCR\_RAS\_PATHWAY | 5 | 2 | 4161 | 170 | Raf,Pkc53E | | 1.530e-02 | -4.18 | GO\_REGULATION\_OF\_TYPE\_I\_INTERFERON\_MEDIATED\_SIGNALING\_PATHWAY | MSigDB lists | GO\_REGULATION\_OF\_TYPE\_I\_INTERFERON\_MEDIATED\_SIGNALING\_PATHWAY | 5 | 2 | 4161 | 170 | Ptp61F,csw | | 1.530e-02 | -4.18 | GO\_REGULATION\_OF\_VOLTAGE\_GATED\_CALCIUM\_CHANNEL\_ACTIVITY | MSigDB lists | GO\_REGULATION\_OF\_VOLTAGE\_GATED\_CALCIUM\_CHANNEL\_ACTIVITY | 5 | 2 | 4161 | 170 | Dys,Ca-beta | | 1.530e-02 | -4.18 | GO\_NEGATIVE\_REGULATION\_OF\_ALCOHOL\_BIOSYNTHETIC\_PROCESS | MSigDB lists | GO\_NEGATIVE\_REGULATION\_OF\_ALCOHOL\_BIOSYNTHETIC\_PROCESS | 5 | 2 | 4161 | 170 | Sod1,dpp | | 1.530e-02 | -4.18 | GO\_GLYCOPROTEIN\_COMPLEX | MSigDB lists | GO\_GLYCOPROTEIN\_COMPLEX | 5 | 2 | 4161 | 170 | Syn2,Dys | | 1.530e-02 | -4.18 | GO\_REGULATION\_OF\_FATTY\_ACID\_OXIDATION | MSigDB lists | GO\_REGULATION\_OF\_FATTY\_ACID\_OXIDATION | 5 | 2 | 4161 | 170 | Tor,ABCD | | 1.530e-02 | -4.18 | GO\_RESPONSE\_TO\_MAGNESIUM\_ION | MSigDB lists | GO\_RESPONSE\_TO\_MAGNESIUM\_ION | 5 | 2 | 4161 | 170 | D2hgdh,Ank | | 1.530e-02 | -4.18 | GO\_NEGATIVE\_REGULATION\_OF\_NEUROTRANSMITTER\_SECRETION | MSigDB lists | GO\_NEGATIVE\_REGULATION\_OF\_NEUROTRANSMITTER\_SECRETION | 5 | 2 | 4161 | 170 | ZnT63C,Raf | | 1.530e-02 | -4.18 | GO\_POSITIVE\_REGULATION\_OF\_INTERFERON\_GAMMA\_PRODUCTION | MSigDB lists | GO\_POSITIVE\_REGULATION\_OF\_INTERFERON\_GAMMA\_PRODUCTION | 5 | 2 | 4161 | 170 | fz2,dnc | | 1.530e-02 | -4.18 | GO\_MICROTUBULE\_END | MSigDB lists | GO\_MICROTUBULE\_END | 5 | 2 | 4161 | 170 | msps,chb | | 1.530e-02 | -4.18 | HERNANDEZ\_MITOTIC\_ARREST\_BY\_DOCETAXEL\_2\_DN | MSigDB lists | HERNANDEZ\_MITOTIC\_ARREST\_BY\_DOCETAXEL\_2\_DN | 5 | 2 | 4161 | 170 | Frq2,Frq1 | | 1.530e-02 | -4.18 | GO\_REGULATION\_BY\_VIRUS\_OF\_VIRAL\_PROTEIN\_LEVELS\_IN\_HOST\_CELL | MSigDB lists | GO\_REGULATION\_BY\_VIRUS\_OF\_VIRAL\_PROTEIN\_LEVELS\_IN\_HOST\_CELL | 5 | 2 | 4161 | 170 | CG17883,Der-1 | | 1.530e-02 | -4.18 | LI\_ADIPOGENESIS\_BY\_ACTIVATED\_PPARG | MSigDB lists | LI\_ADIPOGENESIS\_BY\_ACTIVATED\_PPARG | 5 | 2 | 4161 | 170 | PCB,EcR | | 1.530e-02 | -4.18 | GO\_CELL\_SURFACE\_RECEPTOR\_SIGNALING\_PATHWAY\_INVOLVED\_IN\_HEART\_DEVELOPMENT | MSigDB lists | GO\_CELL\_SURFACE\_RECEPTOR\_SIGNALING\_PATHWAY\_INVOLVED\_IN\_HEART\_DEVELOPMENT | 5 | 2 | 4161 | 170 | Ser,Su(H) | | 1.530e-02 | -4.18 | BIOCARTA\_LONGEVITY\_PATHWAY | MSigDB lists | BIOCARTA\_LONGEVITY\_PATHWAY | 5 | 2 | 4161 | 170 | Cat,Sod1 | | 1.530e-02 | -4.18 | GO\_POSITIVE\_REGULATION\_BY\_HOST\_OF\_VIRAL\_PROCESS | MSigDB lists | GO\_POSITIVE\_REGULATION\_BY\_HOST\_OF\_VIRAL\_PROCESS | 5 | 2 | 4161 | 170 | CG17883,PCB | | 1.530e-02 | -4.18 | FUKUSHIMA\_TNFSF11\_TARGETS | MSigDB lists | FUKUSHIMA\_TNFSF11\_TARGETS | 5 | 2 | 4161 | 170 | Su(H),Ser | | 1.530e-02 | -4.18 | GO\_CAMP\_BINDING | MSigDB lists | GO\_CAMP\_BINDING | 5 | 2 | 4161 | 170 | dnc,Pde6 | | 1.532e-02 | -4.18 | GO\_REGULATION\_OF\_GTPASE\_ACTIVITY | MSigDB lists | GO\_REGULATION\_OF\_GTPASE\_ACTIVITY | 146 | 12 | 4161 | 170 | eIF2Bgamma,CG5522,Nmdar2,dsh,Sod1,step,RhoGAP1A,CG17883,Tor,sky,Cdep,CG34408 | | 1.532e-02 | -4.18 | GO\_CELL\_MORPHOGENESIS\_INVOLVED\_IN\_DIFFERENTIATION | MSigDB lists | GO\_CELL\_MORPHOGENESIS\_INVOLVED\_IN\_DIFFERENTIATION | 112 | 10 | 4161 | 170 | dpp,LpR2,Khc,Pak,Su(H),Sod1,csw,CG42672,CG17883,Ank | | 1.535e-02 | -4.18 | Role of LAT2/NTAL/LAB on calcium mobilization | REACTOME pathways | R-DME-2730905 | 41 | 6 | 3923 | 197 | csw,bnl,rictor,foxo,Tor,CrebB | | 1.537e-02 | -4.18 | Phosphodiest | pfam domains | PF01663 | 4 | 2 | 10480 | 550 | PIG-G,PIG-O | | 1.537e-02 | -4.18 | ZU5 | pfam domains | PF00791 | 4 | 2 | 10480 | 550 | Ank,Ank2 | | 1.540e-02 | -4.17 | regulation of cell population proliferation | biological process | GO:0042127 | 229 | 19 | 12345 | 597 | dpp,Doa,Pkc53E,hth,Cat,Ptp61F,EcR,esg,Pkc98E,CG1572,scrib,ex,numb,Raf,ham,fz2,bun,foxo,upd3 | | 1.542e-02 | -4.17 | cell cortex part | cellular component | GO:0044448 | 116 | 11 | 11940 | 538 | unc-13,Ser,Khc,dsh,sdt,Mbs,cindr,nuf,step,Fife,app | | 1.544e-02 | -4.17 | WH-like\_DNA-bd\_sf | interpro domains | IPR036388 | 102 | 11 | 11383 | 585 | foxo,Poxm,Achl,dsh,Eip74EF,toe,CG32006,TfIIFbeta,CG1894,dany,FoxL1 | | 1.547e-02 | -4.17 | regulation of neurogenesis | biological process | GO:0050767 | 261 | 21 | 12345 | 597 | Antp,EcR,rictor,hth,Pak,Mbs,csw,Dlic,Src42A,bun,hdc,dsh,CG42797,Elp3,mnb,numb,sgg,barc,Tor,Raf,ham | | 1.548e-02 | -4.17 | LEE\_BMP2\_TARGETS\_UP | MSigDB lists | LEE\_BMP2\_TARGETS\_UP | 129 | 11 | 4161 | 170 | Ser,LRP1,LpR2,scramb1,sws,Maf1,CG4341,Iswi,Cln3,bru1,Ank | | 1.549e-02 | -4.17 | V-set | pfam domains | PF07686 | 19 | 4 | 10480 | 550 | side-VIII,dpr15,dpr5,nrm | | 1.550e-02 | -4.17 | G-protein mediated events | REACTOME pathways | R-DME-112040 | 30 | 5 | 3923 | 197 | ACXD,Pde1c,Gprk1,Pkc53E,Cam | | 1.556e-02 | -4.16 | intracellular receptor signaling pathway | biological process | GO:0030522 | 31 | 5 | 12345 | 597 | Hr96,Set2,Iswi,EcR,gce | | 1.556e-02 | -4.16 | cyclic nucleotide metabolic process | biological process | GO:0009187 | 31 | 5 | 12345 | 597 | Pde6,Pde9,Pde1c,ACXD,Pde8 | | 1.556e-02 | -4.16 | establishment or maintenance of epithelial cell apical/basal polarity | biological process | GO:0045197 | 31 | 5 | 12345 | 597 | rib,scrib,trol,dsh,sdt | | 1.562e-02 | -4.16 | fly chr77C4-77C4\|3-47 | chromosome location | fly chr77C4-77C4\|3-47 | 6 | 2 | 25010 | 845 | CG5955,mag | | 1.562e-02 | -4.16 | fly chr96F8-96F8\|3-89 cM | chromosome location | fly chr96F8-96F8\|3-89 cM | 6 | 2 | 25010 | 845 | CG42235,CCAP-R | | 1.562e-02 | -4.16 | fly chr77C4-77C4\|3-47 cM | chromosome location | fly chr77C4-77C4\|3-47 cM | 6 | 2 | 25010 | 845 | CG5955,mag | | 1.562e-02 | -4.16 | fly chr3A6-3A6\|1-1 | chromosome location | fly chr3A6-3A6\|1-1 | 6 | 2 | 25010 | 845 | Vha36-3,egh | | 1.562e-02 | -4.16 | fly chr3A6-3A6\|1-1 cM | chromosome location | fly chr3A6-3A6\|1-1 cM | 6 | 2 | 25010 | 845 | egh,Vha36-3 | | 1.562e-02 | -4.16 | fly chr96F8-96F8\|3-89 | chromosome location | fly chr96F8-96F8\|3-89 | 6 | 2 | 25010 | 845 | CG42235,CCAP-R | | 1.565e-02 | -4.16 | GO\_CIRCULATORY\_SYSTEM\_DEVELOPMENT | MSigDB lists | GO\_CIRCULATORY\_SYSTEM\_DEVELOPMENT | 164 | 13 | 4161 | 170 | Pkc53E,Su(H),csw,Meltrin,Tor,CG17387,fz2,LRP1,dpp,Sk1,Ser,LpR2,dsh | | 1.586e-02 | -4.14 | TAGCTTT\_MIR9 | MSigDB lists | TAGCTTT\_MIR9 | 65 | 7 | 4161 | 170 | CG40178,bru1,msps,eRF3,sdt,Pkc98E,Iswi | | 1.586e-02 | -4.14 | GO\_RESPONSE\_TO\_ACID\_CHEMICAL | MSigDB lists | GO\_RESPONSE\_TO\_ACID\_CHEMICAL | 65 | 7 | 4161 | 170 | Tor,CG5493,TfAP-2,sxc,Pkc98E,Doa,Cat | | 1.596e-02 | -4.14 | Neuronal System | REACTOME pathways | R-DME-112316 | 134 | 13 | 3923 | 197 | Nmdar2,Actn,Raf,Sh,KaiR1D,GluRIIC,Cam,ACXD,CG10000,Flo2,CG17698,Pkc53E,CrebB | | 1.597e-02 | -4.14 | repressing transcription factor binding | molecular function | GO:0070491 | 21 | 4 | 12176 | 580 | cic,retn,EcR,kay | | 1.597e-02 | -4.14 | carbohydrate:cation symporter activity | molecular function | GO:0005402 | 21 | 4 | 12176 | 580 | CG14606,CG10960,CG45782,CG3285 | | 1.597e-02 | -4.14 | carbohydrate:proton symporter activity | molecular function | GO:0005351 | 21 | 4 | 12176 | 580 | CG3285,CG14606,CG45782,CG10960 | | 1.599e-02 | -4.14 | cell junction organization | biological process | GO:0034330 | 81 | 9 | 12345 | 597 | shakB,Pak,shg,CadN2,cindr,sdt,nrv2,scrib,Src42A | | 1.601e-02 | -4.13 | sog (short gastrulation) | protein interactions | 32498 | 29 | 5 | 9204 | 481 | tld,dpp,sgg,kek5,fw | | 1.601e-02 | -4.13 | if (inflated) | protein interactions | 32661 | 29 | 5 | 9204 | 481 | crol,Raf,csw,EcR,by | | 1.620e-02 | -4.12 | GSE14415\_FOXP3\_KO\_NATURAL\_TREG\_VS\_TCONV\_DN | MSigDB lists | GSE14415\_FOXP3\_KO\_NATURAL\_TREG\_VS\_TCONV\_DN | 37 | 5 | 4161 | 170 | Stt3B,Pgant6,Elp3,fus,CG5339 | | 1.620e-02 | -4.12 | GO\_SYNAPSE\_ORGANIZATION | MSigDB lists | GO\_SYNAPSE\_ORGANIZATION | 37 | 5 | 4161 | 170 | Ca-beta,fz2,Ank,Nlg2,unc-13 | | 1.620e-02 | -4.12 | GO\_DRUG\_BINDING | MSigDB lists | GO\_DRUG\_BINDING | 37 | 5 | 4161 | 170 | Hr96,GlyP,Octbeta2R,Tor,dnc | | 1.620e-02 | -4.12 | GSE24142\_EARLY\_THYMIC\_PROGENITOR\_VS\_DN3\_THYMOCYTE\_ADULT\_UP | MSigDB lists | GSE24142\_EARLY\_THYMIC\_PROGENITOR\_VS\_DN3\_THYMOCYTE\_ADULT\_UP | 37 | 5 | 4161 | 170 | Pkc98E,Pak,Tsp96F,CNT1,CG3860 | | 1.620e-02 | -4.12 | GSE41867\_NAIVE\_VS\_DAY6\_LCMV\_EFFECTOR\_CD8\_TCELL\_DN | MSigDB lists | GSE41867\_NAIVE\_VS\_DAY6\_LCMV\_EFFECTOR\_CD8\_TCELL\_DN | 37 | 5 | 4161 | 170 | step,D2hgdh,CG17883,CG9132,Ucp4A | | 1.620e-02 | -4.12 | GO\_GLIAL\_CELL\_DIFFERENTIATION | MSigDB lists | GO\_GLIAL\_CELL\_DIFFERENTIATION | 37 | 5 | 4161 | 170 | TfAP-2,sdt,csw,eIF2Bgamma,Sod1 | | 1.620e-02 | -4.12 | ROZANOV\_MMP14\_TARGETS\_UP | MSigDB lists | ROZANOV\_MMP14\_TARGETS\_UP | 37 | 5 | 4161 | 170 | Rat1,sws,Ser,CG7878,Iswi | | 1.620e-02 | -4.12 | GSE40274\_CTRL\_VS\_LEF1\_TRANSDUCED\_ACTIVATED\_CD4\_TCELL\_UP | MSigDB lists | GSE40274\_CTRL\_VS\_LEF1\_TRANSDUCED\_ACTIVATED\_CD4\_TCELL\_UP | 37 | 5 | 4161 | 170 | Ser,Ca-beta,Glut1,Tsp96F,CNT1 | | 1.620e-02 | -4.12 | HP1SITEFACTOR\_Q6 | MSigDB lists | HP1SITEFACTOR\_Q6 | 37 | 5 | 4161 | 170 | Dys,Dlic,LpR2,Ser,flw | | 1.620e-02 | -4.12 | GSE3982\_EOSINOPHIL\_VS\_BASOPHIL\_UP | MSigDB lists | GSE3982\_EOSINOPHIL\_VS\_BASOPHIL\_UP | 37 | 5 | 4161 | 170 | EcR,flw,Ser,Cadps,sno | | 1.620e-02 | -4.12 | MORF\_IFNA1 | MSigDB lists | MORF\_IFNA1 | 37 | 5 | 4161 | 170 | fz2,Pkc53E,dnc,Hr96,Dys | | 1.620e-02 | -4.12 | GSE17974\_CTRL\_VS\_ACT\_IL4\_AND\_ANTI\_IL12\_4H\_CD4\_TCELL\_UP | MSigDB lists | GSE17974\_CTRL\_VS\_ACT\_IL4\_AND\_ANTI\_IL12\_4H\_CD4\_TCELL\_UP | 37 | 5 | 4161 | 170 | Glut1,sws,Sh,dnc,Pkc53E | | 1.623e-02 | -4.12 | PDZ | interpro domains | IPR001478 | 64 | 8 | 11383 | 585 | Fife,sdt,CG5921,scrib,Syn2,dsh,bbg,RhoGAP100F | | 1.630e-02 | -4.12 | Ldl\_recept\_a | pfam domains | PF00057 | 29 | 5 | 10480 | 550 | LpR2,LRP1,CG32432,CG31221,trol | | 1.633e-02 | -4.11 | GO\_ENDOCYTOSIS | MSigDB lists | GO\_ENDOCYTOSIS | 130 | 11 | 4161 | 170 | CG12121,Tsp96F,LpR2,CG9132,Octbeta2R,Pak,Pkc98E,Act87E,LRP1,CG41099,Cln3 | | 1.635e-02 | -4.11 | embryonic development via the syncytial blastoderm | biological process | GO:0001700 | 199 | 17 | 12345 | 597 | scrib,shd,AGO2,EcR,ems,step,Mbs,Pak,shg,dpp,upd3,Src42A,raw,dsh,kay,rib,sgg | | 1.636e-02 | -4.11 | cell leading edge | cellular component | GO:0031252 | 117 | 11 | 11940 | 538 | dpr5,dpp,dpr12,dpr18,step,Or65c,Or85b,Or42b,Ser,Pak,dpr15 | | 1.643e-02 | -4.11 | PDZ | pfam domains | PF00595 | 51 | 7 | 10480 | 550 | bbg,Syn2,dsh,CG5921,RhoGAP100F,scrib,sdt | | 1.659e-02 | -4.10 | signaling receptor activator activity | molecular function | GO:0030546 | 111 | 11 | 12176 | 580 | CG41520,dpp,bnl,upd3,Ser,ITP,ths,Gpb5,Tk,Neb-cGP,RYa | | 1.660e-02 | -4.10 | embryo development ending in birth or egg hatching | biological process | GO:0009792 | 215 | 18 | 12345 | 597 | wbl,raw,Src42A,upd3,rib,kay,dsh,sgg,shd,scrib,EcR,AGO2,shg,Mbs,Pak,step,ems,dpp | | 1.661e-02 | -4.10 | p450 | pfam domains | PF00067 | 88 | 10 | 10480 | 550 | spok,Cyp6u1,Cyp28d2,Cyp309a1,shd,Cyp4p2,Cyp12a4,Cyp304a1,Cyp28d1,Cyp6a13 | | 1.665e-02 | -4.10 | GO\_RESPONSE\_TO\_HORMONE | MSigDB lists | GO\_RESPONSE\_TO\_HORMONE | 220 | 16 | 4161 | 170 | sxc,Pak,Raf,LpR2,EcR,Pkc98E,eIF2Bgamma,Hr96,Tor,csw,Cat,Octbeta2R,Ptp61F,Ugt301D1,CG5493,mino | | 1.666e-02 | -4.09 | GO\_POSITIVE\_REGULATION\_OF\_GENE\_EXPRESSION | MSigDB lists | GO\_POSITIVE\_REGULATION\_OF\_GENE\_EXPRESSION | 395 | 25 | 4161 | 170 | EcR,AGO3,Act87E,Ser,dpp,Tsp96F,Raf,Su(H),wbl,Iswi,sxc,CG7483,fz2,Fhos,Tif-IA,AstA-R1,scramb1,TfIIFbeta,Octbeta2R,Cat,dsh,TfAP-2,Tor,Hr96,Ank | | 1.668e-02 | -4.09 | GSE6875\_WT\_VS\_FOXP3\_KO\_TREG\_DN | MSigDB lists | GSE6875\_WT\_VS\_FOXP3\_KO\_TREG\_DN | 51 | 6 | 4161 | 170 | Tsp96F,CNT1,Dys,Frq1,Frq2,nompB | | 1.668e-02 | -4.09 | GSE27786\_LSK\_VS\_CD4\_TCELL\_DN | MSigDB lists | GSE27786\_LSK\_VS\_CD4\_TCELL\_DN | 51 | 6 | 4161 | 170 | Sk1,CG42672,ZnT63C,CG34408,Rat1,MESK2 | | 1.668e-02 | -4.09 | GO\_REGULATION\_OF\_NUCLEOTIDE\_METABOLIC\_PROCESS | MSigDB lists | GO\_REGULATION\_OF\_NUCLEOTIDE\_METABOLIC\_PROCESS | 51 | 6 | 4161 | 170 | AstA-R1,mGluR,Octbeta2R,sxc,Pkc53E,dnc | | 1.668e-02 | -4.09 | REACTOME\_NGF\_SIGNALLING\_VIA\_TRKA\_FROM\_THE\_PLASMA\_MEMBRANE | MSigDB lists | REACTOME\_NGF\_SIGNALLING\_VIA\_TRKA\_FROM\_THE\_PLASMA\_MEMBRANE | 51 | 6 | 4161 | 170 | Pkc53E,Pkc98E,CG42672,Tor,Raf,Pde1c | | 1.668e-02 | -4.09 | GSE360\_L\_DONOVANI\_VS\_M\_TUBERCULOSIS\_MAC\_UP | MSigDB lists | GSE360\_L\_DONOVANI\_VS\_M\_TUBERCULOSIS\_MAC\_UP | 51 | 6 | 4161 | 170 | PCB,KaiR1D,Sh,RhoGAP1A,Iswi,Cln3 | | 1.668e-02 | -4.09 | GSE17580\_TREG\_VS\_TEFF\_DN | MSigDB lists | GSE17580\_TREG\_VS\_TEFF\_DN | 51 | 6 | 4161 | 170 | mRpS5,sws,CG4301,CG3376,fz2,Elp3 | | 1.673e-02 | -4.09 | SUGAR\_TRANSPORT\_2 | prosite domains | PS00217 | 25 | 5 | 6243 | 386 | CG10960,Glut1,CG3285,CG3168,CG14606 | | 1.680e-02 | -4.09 | BMP signaling pathway | biological process | GO:0030509 | 21 | 4 | 12345 | 597 | dpp,bun,ex,opa | | 1.680e-02 | -4.09 | regulation of ion transmembrane transport | biological process | GO:0034765 | 21 | 4 | 12345 | 597 | qvr,CG42797,Ca-beta,Cam | | 1.680e-02 | -4.09 | cellular response to BMP stimulus | biological process | GO:0071773 | 21 | 4 | 12345 | 597 | ex,dpp,bun,opa | | 1.680e-02 | -4.09 | regulation of transmembrane transport | biological process | GO:0034762 | 21 | 4 | 12345 | 597 | qvr,Cam,Ca-beta,CG42797 | | 1.680e-02 | -4.09 | ecdysis, chitin-based cuticle | biological process | GO:0018990 | 21 | 4 | 12345 | 597 | Idgf6,Cht6,EcR,CCAP-R | | 1.683e-02 | -4.08 | Cyt\_P450\_E\_grp-I | interpro domains | IPR002401 | 77 | 9 | 11383 | 585 | shd,Cyp6a13,Cyp12a4,spok,Cyp6u1,Cyp4p2,Cyp309a1,Cyp28d1,Cyp304a1 | | 1.684e-02 | -4.08 | PRTYPHPHTASE | prints domains | PR00700 | 16 | 4 | 2185 | 142 | Ptp36E,Ptp61F,csw,Ptp99A | | 1.684e-02 | -4.08 | LDLRECEPTOR | prints domains | PR00261 | 16 | 4 | 2185 | 142 | LRP1,LpR2,trol,CG32432 | | 1.696e-02 | -4.08 | signaling receptor activity | molecular function | GO:0038023 | 419 | 30 | 12176 | 580 | PGRP-LA,CCAP-R,klg,ine,KaiR1D,Or42b,Gr22c,CG9380,Hr96,CG11318,Nmdar2,shakB,et,AstA-R1,Gr39a,Ptp99A,Nlg2,Pdfr,MsR2,CG31750,Or85b,EcR,CG13229,fz4,Octbeta2R,GluRIIC,boi,fz2,mGluR,Or65c | | 1.707e-02 | -4.07 | multi-organism process | biological process | GO:0051704 | 1653 | 98 | 12345 | 597 | Mrtf,lncRNA:Hsromega,Gr22c,tin,dsh,aust,Raf,Msp300,GlyP,TfAP-2,wbl,foxo,Pde1c,chb,shep,shg,opa,dpp,dany,Obp56h,shd,LysP,dnc,AGO2,Gr39a,kay,mGluR,bbg,polybromo,msps,RhoL,Tor,pirk,ex,CG9380,spd-2,sgg,Tk,Src42A,et,kek1,upd3,tld,RhoGEF3,Cyp6a13,lncRNA:CR43753,Dlic,Spn88Eb,tyf,Flo2,Khc,Mos,Exn,lncRNA:CR44420,CG42355,Inx2,GATAe,fs(1)N,cindr,sick,Drsl2,lncRNA:CR44585,Nlg2,bun,JhI-26,Sh,csw,rols,mid,egh,scrib,lncRNA:CR43802,EcR,PGRP-LA,Tsf1,DNaseII,cic,Iswi,mamo,Eip74EF,can,msi,Acp62F,drd,lncRNA:CR43484,fru,Doa,CG8492,Ebp,CG7483,lncRNA:flam,Mst87F,Octbeta2R,bru1,bol,flw,Fad2,retn | | 1.709e-02 | -4.07 | CG8319 (uncharacterized protein) | protein interactions | 41165 | 11 | 3 | 9204 | 481 | esg,CG15073,opa | | 1.709e-02 | -4.07 | seq (sequoia) | protein interactions | 3772396 | 11 | 3 | 9204 | 481 | ham,dsh,bnl | | 1.709e-02 | -4.07 | Strn-Mlck (Stretchin-Mlck) | protein interactions | 36753 | 11 | 3 | 9204 | 481 | Act88F,CG34172,Cam | | 1.709e-02 | -4.07 | kek1 (kekkon 1) | protein interactions | 34688 | 11 | 3 | 9204 | 481 | Dys,kek1,kek5 | | 1.709e-02 | -4.07 | CG8677 (uncharacterized protein) | protein interactions | 35397 | 11 | 3 | 9204 | 481 | CG3078,Iswi,E(Pc) | | 1.709e-02 | -4.07 | Aldh (Aldehyde dehydrogenase) | protein interactions | 34256 | 11 | 3 | 9204 | 481 | rictor,foxo,Raf | | 1.714e-02 | -4.07 | cellular component assembly involved in morphogenesis | biological process | GO:0010927 | 154 | 14 | 12345 | 597 | bun,Actn,Khc,bru1,dpp,Mlp84B,cic,Act88F,rols,polybromo,bnl,Raf,fs(1)N,Prm | | 1.715e-02 | -4.07 | GSE37605\_C57BL6\_VS\_NOD\_FOXP3\_FUSION\_GFP\_TCONV\_UP | MSigDB lists | GSE37605\_C57BL6\_VS\_NOD\_FOXP3\_FUSION\_GFP\_TCONV\_UP | 66 | 7 | 4161 | 170 | Pkc53E,Su(H),nompB,REG,DNaseII,Tsp96F,Mctp | | 1.718e-02 | -4.06 | GAB1 signalosome | REACTOME pathways | R-DME-180292 | 42 | 6 | 3923 | 197 | bnl,csw,rictor,foxo,Tor,CrebB | | 1.727e-02 | -4.06 | TYR\_PHOSPHATASE\_dom | interpro domains | IPR000387 | 30 | 5 | 11383 | 585 | CG3530,csw,Ptp36E,Ptp61F,Ptp99A | | 1.733e-02 | -4.06 | bZIP | interpro domains | IPR004827 | 20 | 4 | 11383 | 585 | CrebB,Octbeta2R,kay,Atf6 | | 1.738e-02 | -4.05 | GSE17974\_0H\_VS\_6H\_IN\_VITRO\_ACT\_CD4\_TCELL\_UP | MSigDB lists | GSE17974\_0H\_VS\_6H\_IN\_VITRO\_ACT\_CD4\_TCELL\_UP | 25 | 4 | 4161 | 170 | Sh,dnc,Pkc53E,Glut1 | | 1.738e-02 | -4.05 | GO\_GLIAL\_CELL\_DEVELOPMENT | MSigDB lists | GO\_GLIAL\_CELL\_DEVELOPMENT | 25 | 4 | 4161 | 170 | Sod1,eIF2Bgamma,sdt,TfAP-2 | | 1.738e-02 | -4.05 | GGTGAAG\_MIR412 | MSigDB lists | GGTGAAG\_MIR412 | 25 | 4 | 4161 | 170 | CG5522,flw,Doa,eIF4E7 | | 1.738e-02 | -4.05 | GO\_NUCLEOSIDE\_PHOSPHATE\_CATABOLIC\_PROCESS | MSigDB lists | GO\_NUCLEOSIDE\_PHOSPHATE\_CATABOLIC\_PROCESS | 25 | 4 | 4161 | 170 | Pde8,Cat,Pde6,dnc | | 1.738e-02 | -4.05 | MEL18\_DN.V1\_UP | MSigDB lists | MEL18\_DN.V1\_UP | 25 | 4 | 4161 | 170 | Syx1A,LpR2,Ser,Meltrin | | 1.738e-02 | -4.05 | GSE16450\_CTRL\_VS\_IFNA\_12H\_STIM\_MATURE\_NEURON\_CELL\_LINE\_UP | MSigDB lists | GSE16450\_CTRL\_VS\_IFNA\_12H\_STIM\_MATURE\_NEURON\_CELL\_LINE\_UP | 25 | 4 | 4161 | 170 | CG17387,Ank,EcR,CG4849 | | 1.738e-02 | -4.05 | GO\_GLYCOPROTEIN\_BINDING | MSigDB lists | GO\_GLYCOPROTEIN\_BINDING | 25 | 4 | 4161 | 170 | chb,Dys,LpR2,Syx1A | | 1.738e-02 | -4.05 | GO\_SYSTEM\_PROCESS | MSigDB lists | GO\_SYSTEM\_PROCESS | 278 | 19 | 4161 | 170 | KaiR1D,Nlg2,CG7483,Sod1,RhoGAP1A,LpR2,Raf,Ank,Cln3,TfAP-2,Tor,Ca-beta,Awh,RpL38,Nmdar2,AstA-R1,Octbeta2R,Dys,dnc | | 1.748e-02 | -4.05 | GGGTGGRR\_PAX4\_03 | MSigDB lists | GGGTGGRR\_PAX4\_03 | 240 | 17 | 4161 | 170 | Nmdar2,mGluR,chb,CG12547,Fhos,app,IP3K2,PhKgamma,eRF3,Khc,GlyP,Pkc98E,LRP1,kcc,eIF4E7,Iswi,RhoGAP1A | | 1.749e-02 | -4.05 | GO\_CELL\_CELL\_CONTACT\_ZONE | MSigDB lists | GO\_CELL\_CELL\_CONTACT\_ZONE | 14 | 3 | 4161 | 170 | Ank,Pak,Flo2 | | 1.749e-02 | -4.05 | GO\_ARTERY\_DEVELOPMENT | MSigDB lists | GO\_ARTERY\_DEVELOPMENT | 14 | 3 | 4161 | 170 | Su(H),LRP1,Ser | | 1.749e-02 | -4.05 | GO\_CALCIUM\_CHANNEL\_COMPLEX | MSigDB lists | GO\_CALCIUM\_CHANNEL\_COMPLEX | 14 | 3 | 4161 | 170 | Ca-beta,dnc,MCU | | 1.749e-02 | -4.05 | MIKKELSEN\_MEF\_ICP\_WITH\_H3K27ME3 | MSigDB lists | MIKKELSEN\_MEF\_ICP\_WITH\_H3K27ME3 | 14 | 3 | 4161 | 170 | mGluR,Pde8,Pde6 | | 1.749e-02 | -4.05 | REACTOME\_ACTIVATION\_OF\_NMDA\_RECEPTOR\_UPON\_GLUTAMATE\_BINDING\_AND\_POSTSYNAPTIC\_EVENTS | MSigDB lists | REACTOME\_ACTIVATION\_OF\_NMDA\_RECEPTOR\_UPON\_GLUTAMATE\_BINDING\_AND\_POSTSYNAPTIC\_EVENTS | 14 | 3 | 4161 | 170 | Nmdar2,Raf,CG17698 | | 1.749e-02 | -4.05 | GO\_REGULATION\_OF\_ENDOTHELIAL\_CELL\_PROLIFERATION | MSigDB lists | GO\_REGULATION\_OF\_ENDOTHELIAL\_CELL\_PROLIFERATION | 14 | 3 | 4161 | 170 | dpp,Tor,Pkc53E | | 1.749e-02 | -4.05 | ULE\_SPLICING\_VIA\_NOVA2 | MSigDB lists | ULE\_SPLICING\_VIA\_NOVA2 | 14 | 3 | 4161 | 170 | KaiR1D,Ank,chb | | 1.749e-02 | -4.05 | GO\_REGULATION\_OF\_BIOMINERAL\_TISSUE\_DEVELOPMENT | MSigDB lists | GO\_REGULATION\_OF\_BIOMINERAL\_TISSUE\_DEVELOPMENT | 14 | 3 | 4161 | 170 | dpp,TfAP-2,Octbeta2R | | 1.749e-02 | -4.05 | GO\_LYMPHOCYTE\_COSTIMULATION | MSigDB lists | GO\_LYMPHOCYTE\_COSTIMULATION | 14 | 3 | 4161 | 170 | Pak,csw,Tor | | 1.749e-02 | -4.05 | WINZEN\_DEGRADED\_VIA\_KHSRP | MSigDB lists | WINZEN\_DEGRADED\_VIA\_KHSRP | 14 | 3 | 4161 | 170 | Ser,dpp,Octbeta2R | | 1.749e-02 | -4.05 | REACTOME\_DAG\_AND\_IP3\_SIGNALING | MSigDB lists | REACTOME\_DAG\_AND\_IP3\_SIGNALING | 14 | 3 | 4161 | 170 | Pkc53E,Pkc98E,Pde1c | | 1.749e-02 | -4.05 | KEGG\_BASAL\_CELL\_CARCINOMA | MSigDB lists | KEGG\_BASAL\_CELL\_CARCINOMA | 14 | 3 | 4161 | 170 | dsh,fz2,dpp | | 1.749e-02 | -4.05 | GO\_RNA\_POLYMERASE\_BINDING | MSigDB lists | GO\_RNA\_POLYMERASE\_BINDING | 14 | 3 | 4161 | 170 | Elp3,Tif-IA,Maf1 | | 1.749e-02 | -4.05 | HALLMARK\_NOTCH\_SIGNALING | MSigDB lists | HALLMARK\_NOTCH\_SIGNALING | 14 | 3 | 4161 | 170 | Ser,Pkc53E,fz2 | | 1.749e-02 | -4.05 | MODULE\_223 | MSigDB lists | MODULE\_223 | 14 | 3 | 4161 | 170 | Tsp96F,Mlp84B,LRP1 | | 1.767e-02 | -4.04 | Developmental Biology | REACTOME pathways | R-DME-1266738 | 211 | 18 | 3923 | 197 | CrebB,Ank2,REG,Cdep,boi,Syx1A,Exn,Raf,CG10188,bnl,Nmdar2,Pak,Ank,Prosbeta5R1,CG41520,numb,foxo,toe | | 1.773e-02 | -4.03 | maintenance of protein location in cell | biological process | GO:0032507 | 32 | 5 | 12345 | 597 | by,p130CAS,Msp300,Ank,Actn | | 1.773e-02 | -4.03 | regulation of Rho protein signal transduction | biological process | GO:0035023 | 32 | 5 | 12345 | 597 | RhoGEF3,CG10188,RhoGAP1A,Cdep,Exn | | 1.774e-02 | -4.03 | ey (eyeless) | protein interactions | 43812 | 52 | 7 | 9204 | 481 | foxo,Mctp,shg,dpp,dsh,hth,Antp | | 1.775e-02 | -4.03 | GO\_REGULATION\_OF\_HOMEOSTATIC\_PROCESS | MSigDB lists | GO\_REGULATION\_OF\_HOMEOSTATIC\_PROCESS | 98 | 9 | 4161 | 170 | IP3K2,Ank,ZnT63C,dnc,mino,Pkc98E,EcR,Dys,CCAP-R | | 1.775e-02 | -4.03 | GO\_RESPONSE\_TO\_METAL\_ION | MSigDB lists | GO\_RESPONSE\_TO\_METAL\_ION | 98 | 9 | 4161 | 170 | Sod1,TfAP-2,IP3K2,D2hgdh,Ank,ZnT63C,mino,Raf,Cat | | 1.777e-02 | -4.03 | ZWANG\_TRANSIENTLY\_UP\_BY\_1ST\_EGF\_PULSE\_ONLY | MSigDB lists | ZWANG\_TRANSIENTLY\_UP\_BY\_1ST\_EGF\_PULSE\_ONLY | 298 | 20 | 4161 | 170 | Cat,CG4301,CG3860,Ugt301D1,dnc,mino,Ank,TfAP-2,CG4341,Pak,Aldh-III,dpp,LRP1,Pkc98E,IntS3,Glut1,Nlg2,DNaseII,Pfdn4,Src42A | | 1.784e-02 | -4.03 | REACTOME\_NEURONAL\_SYSTEM | MSigDB lists | REACTOME\_NEURONAL\_SYSTEM | 82 | 8 | 4161 | 170 | Ca-beta,Pkc53E,CG17698,KaiR1D,Sh,Raf,Syx1A,Nmdar2 | | 1.784e-02 | -4.03 | ARNT\_01 | MSigDB lists | ARNT\_01 | 82 | 8 | 4161 | 170 | LpR2,CG14480,REG,Ser,polybromo,eIF4E7,Odc1,Cln3 | | 1.785e-02 | -4.03 | regulation of cation channel activity | biological process | GO:2001257 | 12 | 3 | 12345 | 597 | qvr,Cam,Ca-beta | | 1.785e-02 | -4.03 | mesodermal cell fate specification | biological process | GO:0007501 | 12 | 3 | 12345 | 597 | tin,Ubx,ths | | 1.785e-02 | -4.03 | anterior Malpighian tubule development | biological process | GO:0061327 | 12 | 3 | 12345 | 597 | rols,dpp,Ubx | | 1.785e-02 | -4.03 | axonal defasciculation | biological process | GO:0007414 | 12 | 3 | 12345 | 597 | trol,p130CAS,Ptp99A | | 1.785e-02 | -4.03 | phospholipid translocation | biological process | GO:0045332 | 12 | 3 | 12345 | 597 | CG4301,scramb1,CG9981 | | 1.785e-02 | -4.03 | determination of left/right symmetry | biological process | GO:0007368 | 12 | 3 | 12345 | 597 | shg,tin,CG17387 | | 1.787e-02 | -4.02 | cell surface | cellular component | GO:0009986 | 74 | 8 | 11940 | 538 | boi,fs(1)N,rols,Ser,Nlg2,CadN2,Alp12,qvr | | 1.788e-02 | -4.02 | Sphingolipid metabolism | REACTOME pathways | R-DME-428157 | 67 | 8 | 3923 | 197 | Sk1,cert,Aldh-III,CG33090,CG3376,Ugt301D1,Ugt302K1,Ugt304A1 | | 1.804e-02 | -4.01 | regulation of neurotransmitter levels | biological process | GO:0001505 | 140 | 13 | 12345 | 597 | nemy,sky,Frq1,Exn,AANAT1,Dys,Fife,Frq2,Mctp,unc-13,Cadps,cpx,Syx1A | | 1.806e-02 | -4.01 | GSE7852\_TREG\_VS\_TCONV\_THYMUS\_UP | MSigDB lists | GSE7852\_TREG\_VS\_TCONV\_THYMUS\_UP | 38 | 5 | 4161 | 170 | CG10126,Odc1,scramb1,CG8184,Mlp84B | | 1.806e-02 | -4.01 | GO\_HOMEOSTASIS\_OF\_NUMBER\_OF\_CELLS | MSigDB lists | GO\_HOMEOSTASIS\_OF\_NUMBER\_OF\_CELLS | 38 | 5 | 4161 | 170 | Tif-IA,IP3K2,Sod1,csw,Ptp61F | | 1.806e-02 | -4.01 | GO\_DEVELOPMENTAL\_MATURATION | MSigDB lists | GO\_DEVELOPMENTAL\_MATURATION | 38 | 5 | 4161 | 170 | Su(H),fz2,dpp,unc-13,Cdep | | 1.806e-02 | -4.01 | GSE3982\_MAST\_CELL\_VS\_EFF\_MEMORY\_CD4\_TCELL\_DN | MSigDB lists | GSE3982\_MAST\_CELL\_VS\_EFF\_MEMORY\_CD4\_TCELL\_DN | 38 | 5 | 4161 | 170 | step,Sh,AGO3,Flo2,GlyP | | 1.806e-02 | -4.01 | REACTOME\_SIGNALING\_BY\_FGFR\_IN\_DISEASE | MSigDB lists | REACTOME\_SIGNALING\_BY\_FGFR\_IN\_DISEASE | 38 | 5 | 4161 | 170 | Raf,Pde1c,Tor,Pkc53E,Pkc98E | | 1.806e-02 | -4.01 | HNF3\_Q6 | MSigDB lists | HNF3\_Q6 | 38 | 5 | 4161 | 170 | Dys,KaiR1D,DNaseII,IP3K2,flw | | 1.806e-02 | -4.01 | GSE19941\_UNSTIM\_VS\_LPS\_STIM\_IL10\_KO\_MACROPHAGE\_UP | MSigDB lists | GSE19941\_UNSTIM\_VS\_LPS\_STIM\_IL10\_KO\_MACROPHAGE\_UP | 38 | 5 | 4161 | 170 | KrT95D,melt,CG17883,Der-1,LRP1 | | 1.806e-02 | -4.01 | GSE20727\_CTRL\_VS\_H2O2\_TREATED\_DC\_UP | MSigDB lists | GSE20727\_CTRL\_VS\_H2O2\_TREATED\_DC\_UP | 38 | 5 | 4161 | 170 | Epg5,Act87E,Raf,Tsp96F,Dys | | 1.806e-02 | -4.01 | IGv | smart domains | SM00406 | 17 | 4 | 5086 | 315 | dpr21,DIP-lambda,dpr15,Strn-Mlck | | 1.810e-02 | -4.01 | Egfr (Epidermal growth factor receptor) | protein interactions | 37455 | 158 | 15 | 9204 | 481 | csw,step,kek1,scrib,bun,Mbs,upd3,Raf,Ubx,kay,dpp,Ptp61F,trol,Src42A,shg | | 1.813e-02 | -4.01 | GO\_REGULATION\_OF\_CELL\_PROLIFERATION | MSigDB lists | GO\_REGULATION\_OF\_CELL\_PROLIFERATION | 260 | 18 | 4161 | 170 | polybromo,Su(H),Pkc53E,Src42A,Pak,Raf,Tsp96F,Gas8,Sk1,dpp,Ser,Odc1,TfAP-2,Tor,Octbeta2R,Ptp61F,mino,fz2 | | 1.817e-02 | -4.01 | JOHNSTONE\_PARVB\_TARGETS\_3\_UP | MSigDB lists | JOHNSTONE\_PARVB\_TARGETS\_3\_UP | 115 | 10 | 4161 | 170 | Flo2,CG3961,ZnT63C,Ser,CG8184,Act87E,LRP1,CG12121,Aldh-III,Cat | | 1.824e-02 | -4.00 | GO\_CALCIUM\_ION\_TRANSPORT | MSigDB lists | GO\_CALCIUM\_ION\_TRANSPORT | 52 | 6 | 4161 | 170 | Pkc98E,Frq2,Ca-beta,Frq1,MCU,ZnT63C | | 1.824e-02 | -4.00 | MORF\_CAMK4 | MSigDB lists | MORF\_CAMK4 | 52 | 6 | 4161 | 170 | Dys,CG40178,Hr96,dnc,Pkc53E,fz2 | | 1.824e-02 | -4.00 | GSE29617\_CTRL\_VS\_DAY3\_TIV\_FLU\_VACCINE\_PBMC\_2008\_UP | MSigDB lists | GSE29617\_CTRL\_VS\_DAY3\_TIV\_FLU\_VACCINE\_PBMC\_2008\_UP | 52 | 6 | 4161 | 170 | CG9132,Raf,Glut1,Dlic,Qsox2,Rbp6 | | 1.824e-02 | -4.00 | GO\_REGULATION\_OF\_NEUROTRANSMITTER\_LEVELS | MSigDB lists | GO\_REGULATION\_OF\_NEUROTRANSMITTER\_LEVELS | 52 | 6 | 4161 | 170 | Syx1A,Raf,unc-13,ZnT63C,Cln3,Cadps | | 1.824e-02 | -4.00 | GTGCCAT\_MIR183 | MSigDB lists | GTGCCAT\_MIR183 | 52 | 6 | 4161 | 170 | Frq1,bru1,Pkc53E,Frq2,sxc,dnc | | 1.824e-02 | -4.00 | GO\_NEGATIVE\_REGULATION\_OF\_RESPONSE\_TO\_EXTERNAL\_STIMULUS | MSigDB lists | GO\_NEGATIVE\_REGULATION\_OF\_RESPONSE\_TO\_EXTERNAL\_STIMULUS | 52 | 6 | 4161 | 170 | Cln3,RhoGAP1A,Tor,Ptp61F,chb,Octbeta2R | | 1.824e-02 | -4.00 | GSE5142\_CTRL\_VS\_HTERT\_TRANSDUCED\_CD8\_TCELL\_EARLY\_PASSAGE\_CLONE\_UP | MSigDB lists | GSE5142\_CTRL\_VS\_HTERT\_TRANSDUCED\_CD8\_TCELL\_EARLY\_PASSAGE\_CLONE\_UP | 52 | 6 | 4161 | 170 | scramb1,Tsp96F,Syx1A,Doa,fus,Pkc53E | | 1.825e-02 | -4.00 | germline ring canal | cellular component | GO:0045172 | 23 | 4 | 11940 | 538 | sgg,Lpin,chb,mino | | 1.830e-02 | -4.00 | Med (Medea) | protein interactions | 43725 | 20 | 4 | 9204 | 481 | dpp,Snoo,CG32264,CG14634 | | 1.830e-02 | -4.00 | DAAM (Dishevelled Associated Activator of Morphogenesis) | protein interactions | 31075 | 20 | 4 | 9204 | 481 | Src42A,CG10164,dsh,Act88F | | 1.836e-02 | -4.00 | dorsal closure | biological process | GO:0007391 | 111 | 11 | 12345 | 597 | raw,Src42A,scrib,shd,dsh,step,kay,Mbs,Pak,rib,dpp | | 1.841e-02 | -3.99 | tsh (teashirt) | protein interactions | 35430 | 30 | 5 | 9204 | 481 | cindr,hth,Ser,Antp,Ubx | | 1.842e-02 | -3.99 | homophilic cell adhesion via plasma membrane adhesion molecules | biological process | GO:0007156 | 44 | 6 | 12345 | 597 | fw,klg,rst,shg,CadN2,chp | | 1.842e-02 | -3.99 | maintenance of protein location | biological process | GO:0045185 | 44 | 6 | 12345 | 597 | by,p130CAS,Msp300,Ank,boi,Actn | | 1.851e-02 | -3.99 | GO\_MEMBRANE\_LIPID\_METABOLIC\_PROCESS | MSigDB lists | GO\_MEMBRANE\_LIPID\_METABOLIC\_PROCESS | 67 | 7 | 4161 | 170 | Sk1,Cln3,PIG-G,PIG-O,CG33090,CG3376,Aldh-III | | 1.851e-02 | -3.99 | SATO\_SILENCED\_BY\_METHYLATION\_IN\_PANCREATIC\_CANCER\_1 | MSigDB lists | SATO\_SILENCED\_BY\_METHYLATION\_IN\_PANCREATIC\_CANCER\_1 | 67 | 7 | 4161 | 170 | kcc,CG12121,dsh,CG3376,Iswi,CG5493,CG7878 | | 1.855e-02 | -3.99 | myofilament | cellular component | GO:0036379 | 13 | 3 | 11940 | 538 | TpnC25D,Prm,Act88F | | 1.855e-02 | -3.99 | pericentriolar material | cellular component | GO:0000242 | 13 | 3 | 11940 | 538 | Rcd6,spd-2,Cep97 | | 1.855e-02 | -3.99 | microtubule plus-end | cellular component | GO:0035371 | 13 | 3 | 11940 | 538 | msps,chb,Khc | | 1.855e-02 | -3.99 | female germline ring canal | cellular component | GO:0035324 | 13 | 3 | 11940 | 538 | mino,sgg,Lpin | | 1.862e-02 | -3.98 | 14-3-3zeta (14-3-3zeta) | protein interactions | 36059 | 90 | 10 | 9204 | 481 | csw,Raf,melt,rictor,CG16854,Src42A,Elp3,foxo,HDAC4,Syx1A | | 1.873e-02 | -3.98 | pnt (pointed) | protein interactions | 42757 | 41 | 6 | 9204 | 481 | Raf,foxo,Ubx,cic,csw,Antp | | 1.874e-02 | -3.98 | CH | pfam domains | PF00307 | 30 | 5 | 10480 | 550 | sick,Msp300,CG34417,Dys,Actn | | 1.883e-02 | -3.97 | Ion transport by P-type ATPases | REACTOME pathways | R-DME-936837 | 21 | 4 | 3923 | 197 | Cam,CG4301,CG9981,nrv2 | | 1.883e-02 | -3.97 | Antimicrobial peptides | REACTOME pathways | R-DME-6803157 | 21 | 4 | 3923 | 197 | CG8492,PGRP-LA,LysP,Tsf1 | | 1.886e-02 | -3.97 | GO\_MORPHOGENESIS\_OF\_AN\_EPITHELIUM | MSigDB lists | GO\_MORPHOGENESIS\_OF\_AN\_EPITHELIUM | 99 | 9 | 4161 | 170 | Pak,dsh,dpp,Ser,fz2,CG17387,TfAP-2,sdt,Su(H) | | 1.899e-02 | -3.96 | Ser/Thr\_kinase\_AS | interpro domains | IPR008271 | 147 | 14 | 11383 | 585 | CG17698,Pak,CG4839,Pkc98E,Gprk1,Mos,PhKgamma,Nuak1,Pkc53E,mnb,Raf,sgg,Strn-Mlck,Doa | | 1.912e-02 | -3.96 | Signalling by NGF | REACTOME pathways | R-DME-166520 | 182 | 16 | 3923 | 197 | Cam,bnl,rictor,Raf,CrebB,csw,ACXD,Pkc98E,Gprk1,REG,foxo,Prosbeta5R1,Tor,Pkc53E,Pde1c,CG41520 | | 1.924e-02 | -3.95 | axon terminus | cellular component | GO:0043679 | 75 | 8 | 11940 | 538 | Ank2,mnb,unc-13,gukh,g,cpx,Syx1A,sky | | 1.925e-02 | -3.95 | monovalent inorganic cation transmembrane transporter activity | molecular function | GO:0015077 | 238 | 19 | 12176 | 580 | CG45782,CNT1,Vha100-5,Nha1,CG3285,GluRIIC,CG34172,Irk3,CG10960,Vha36-3,Sh,CG14606,ine,KaiR1D,kcc,CG33310,CG13794,nrv2,ND-MLRQ | | 1.939e-02 | -3.94 | cell-cell junction assembly | biological process | GO:0007043 | 57 | 7 | 12345 | 597 | shg,CadN2,scrib,shakB,Pak,sdt,nrv2 | | 1.939e-02 | -3.94 | detection of light stimulus | biological process | GO:0009583 | 57 | 7 | 12345 | 597 | Pld,ninaD,Gprk1,Sh,shakB,rdgA,Cam | | 1.946e-02 | -3.94 | Innate Immune System | REACTOME pathways | R-DME-168249 | 651 | 44 | 3923 | 197 | CG8184,CG8492,Prosbeta5R1,CG9981,Tsf1,Pde1c,Vha36-3,CG9451,Aldh-III,Pld,Dlic,Cat,Qsox2,CG7362,CrebB,CG40470,Src42A,Pkc98E,PGRP-LA,REG,Glut1,csw,CG7882,RhoGAP18B,Raf,Pak,ninaD,Tor,Pkc53E,Vha100-5,CG41520,kay,fw,CG4301,foxo,CG9449,LysP,Gprk1,Meltrin,ACXD,GlyP,Cam,rictor,bnl | | 1.950e-02 | -3.94 | imaginal disc pattern formation | biological process | GO:0007447 | 112 | 11 | 12345 | 597 | nuf,Su(H),mid,dpp,B-H2,cic,hth,sno,boi,bi,Ser | | 1.950e-02 | -3.94 | transmembrane receptor protein tyrosine kinase signaling pathway | biological process | GO:0007169 | 112 | 11 | 12345 | 597 | foxo,CG41520,Src42A,Raf,bnl,ths,Tor,csw,cic,fs(1)N,rictor | | 1.952e-02 | -3.94 | GO\_HEAD\_DEVELOPMENT | MSigDB lists | GO\_HEAD\_DEVELOPMENT | 187 | 14 | 4161 | 170 | Raf,LpR2,Act87E,LRP1,dpp,Sk1,dsd,Tor,Awh,Su(H),Iswi,RhoGAP1A,sxc,csw | | 1.953e-02 | -3.94 | GO\_REGULATION\_OF\_PROTEIN\_LOCALIZATION | MSigDB lists | GO\_REGULATION\_OF\_PROTEIN\_LOCALIZATION | 243 | 17 | 4161 | 170 | mino,fz2,Syx1A,dsh,csw,Tor,Ank,EcR,Pkc98E,LRP1,CG8184,Khc,MCU,wbl,Pkc53E,sxc,Pde8 | | 1.960e-02 | -3.93 | Costimulation by the CD28 family | REACTOME pathways | R-DME-388841 | 12 | 3 | 3923 | 197 | Tor,Pak,rictor | | 1.960e-02 | -3.93 | Non-integrin membrane-ECM interactions | REACTOME pathways | R-DME-3000171 | 12 | 3 | 3923 | 197 | Pkc53E,bnl,trol | | 1.961e-02 | -3.93 | Mad (Mothers against dpp) | protein interactions | 33529 | 104 | 11 | 9204 | 481 | tamo,rib,csw,scrib,Aldh-III,Snoo,Ubx,Antp,shg,numb,dpp | | 1.964e-02 | -3.93 | programmed cell death | biological process | GO:0012501 | 172 | 15 | 12345 | 597 | Eip74EF,IP3K2,EcR,CG6839,Cam,Pak,hdc,Fhos,Damm,CG5059,Tor,dpp,DNaseII,rst,sws | | 1.977e-02 | -3.92 | leg disc pattern formation | biological process | GO:0035223 | 22 | 4 | 12345 | 597 | hth,dpp,B-H2,mid | | 1.977e-02 | -3.92 | response to fibroblast growth factor | biological process | GO:0071774 | 22 | 4 | 12345 | 597 | csw,bnl,ths,Raf | | 1.977e-02 | -3.92 | response to BMP | biological process | GO:0071772 | 22 | 4 | 12345 | 597 | opa,dpp,bun,ex | | 1.977e-02 | -3.92 | cellular response to fibroblast growth factor stimulus | biological process | GO:0044344 | 22 | 4 | 12345 | 597 | bnl,ths,Raf,csw | | 1.977e-02 | -3.92 | fibroblast growth factor receptor signaling pathway | biological process | GO:0008543 | 22 | 4 | 12345 | 597 | csw,bnl,ths,Raf | | 1.977e-02 | -3.92 | torso signaling pathway | biological process | GO:0008293 | 22 | 4 | 12345 | 597 | cic,csw,Raf,fs(1)N | | 1.980e-02 | -3.92 | Ank\_2 | pfam domains | PF12796 | 65 | 8 | 10480 | 550 | CG41099,rdgA,CG6966,Ank2,rols,Ank,Mbs,CG42672 | | 1.983e-02 | -3.92 | cell projection membrane | cellular component | GO:0031253 | 105 | 10 | 11940 | 538 | dpr5,CG14182,dpr12,dpr18,Or65c,Ser,Or42b,Or85b,Cln3,dpr15 | | 1.989e-02 | -3.92 | SOX9\_B1 | MSigDB lists | SOX9\_B1 | 53 | 6 | 4161 | 170 | Cadps,Pak,Dys,Octbeta2R,CG9132,LpR2 | | 1.989e-02 | -3.92 | GO\_G\_PROTEIN\_COUPLED\_RECEPTOR\_ACTIVITY | MSigDB lists | GO\_G\_PROTEIN\_COUPLED\_RECEPTOR\_ACTIVITY | 53 | 6 | 4161 | 170 | fz2,Sk1,CCAP-R,AstA-R1,mGluR,Octbeta2R | | 1.990e-02 | -3.92 | GO\_REGULATION\_OF\_LYASE\_ACTIVITY | MSigDB lists | GO\_REGULATION\_OF\_LYASE\_ACTIVITY | 26 | 4 | 4161 | 170 | Pkc53E,AstA-R1,mGluR,Octbeta2R | | 1.990e-02 | -3.92 | ATAAGCT\_MIR21 | MSigDB lists | ATAAGCT\_MIR21 | 26 | 4 | 4161 | 170 | polybromo,CG40178,Ser,Sh | | 1.990e-02 | -3.92 | GSE46606\_UNSTIM\_VS\_CD40L\_IL2\_IL5\_1DAY\_STIMULATED\_IRF4\_KO\_BCELL\_DN | MSigDB lists | GSE46606\_UNSTIM\_VS\_CD40L\_IL2\_IL5\_1DAY\_STIMULATED\_IRF4\_KO\_BCELL\_DN | 26 | 4 | 4161 | 170 | dnc,step,Pkc98E,kcc | | 1.990e-02 | -3.92 | GO\_POSITIVE\_REGULATION\_OF\_STRESS\_ACTIVATED\_PROTEIN\_KINASE\_SIGNALING\_CASCADE | MSigDB lists | GO\_POSITIVE\_REGULATION\_OF\_STRESS\_ACTIVATED\_PROTEIN\_KINASE\_SIGNALING\_CASCADE | 26 | 4 | 4161 | 170 | Pak,dsh,dpp,fz2 | | 1.990e-02 | -3.92 | BROWNE\_HCMV\_INFECTION\_12HR\_UP | MSigDB lists | BROWNE\_HCMV\_INFECTION\_12HR\_UP | 26 | 4 | 4161 | 170 | dpp,step,Raf,scramb1 | | 1.990e-02 | -3.92 | KEGG\_LONG\_TERM\_POTENTIATION | MSigDB lists | KEGG\_LONG\_TERM\_POTENTIATION | 26 | 4 | 4161 | 170 | Pkc53E,flw,Raf,Nmdar2 | | 1.990e-02 | -3.92 | GSE30962\_ACUTE\_VS\_CHRONIC\_LCMV\_SECONDARY\_INF\_CD8\_TCELL\_UP | MSigDB lists | GSE30962\_ACUTE\_VS\_CHRONIC\_LCMV\_SECONDARY\_INF\_CD8\_TCELL\_UP | 26 | 4 | 4161 | 170 | CG17698,mRpL24,Pgant6,Syx1A | | 1.990e-02 | -3.92 | GO\_ENERGY\_RESERVE\_METABOLIC\_PROCESS | MSigDB lists | GO\_ENERGY\_RESERVE\_METABOLIC\_PROCESS | 26 | 4 | 4161 | 170 | GlyP,Tor,PhKgamma,flw | | 1.990e-02 | -3.92 | GO\_ION\_CHANNEL\_BINDING | MSigDB lists | GO\_ION\_CHANNEL\_BINDING | 26 | 4 | 4161 | 170 | KrT95D,Ank,Syx1A,dnc | | 1.990e-02 | -3.92 | RTTTNNNYTGGM\_UNKNOWN | MSigDB lists | RTTTNNNYTGGM\_UNKNOWN | 26 | 4 | 4161 | 170 | sxc,LRP1,Dys,fus | | 1.990e-02 | -3.92 | DACOSTA\_UV\_RESPONSE\_VIA\_ERCC3\_TTD\_DN | MSigDB lists | DACOSTA\_UV\_RESPONSE\_VIA\_ERCC3\_TTD\_DN | 26 | 4 | 4161 | 170 | Dys,CG34408,dsd,IntS3 | | 2.002e-02 | -3.91 | GO\_POSTSYNAPSE | MSigDB lists | GO\_POSTSYNAPSE | 100 | 9 | 4161 | 170 | Frq2,Frq1,Nlg2,Ank,KaiR1D,Octbeta2R,Dys,mGluR,Nmdar2 | | 2.005e-02 | -3.91 | GO\_REGULATION\_OF\_STRESS\_ACTIVATED\_PROTEIN\_KINASE\_SIGNALING\_CASCADE | MSigDB lists | GO\_REGULATION\_OF\_STRESS\_ACTIVATED\_PROTEIN\_KINASE\_SIGNALING\_CASCADE | 39 | 5 | 4161 | 170 | dpp,fz2,dsh,Pak,KaiR1D | | 2.005e-02 | -3.91 | GSE7852\_TREG\_VS\_TCONV\_LN\_UP | MSigDB lists | GSE7852\_TREG\_VS\_TCONV\_LN\_UP | 39 | 5 | 4161 | 170 | Tsp96F,Nha1,Glut1,Odc1,Dys | | 2.005e-02 | -3.91 | GSE14769\_UNSTIM\_VS\_60MIN\_LPS\_BMDM\_DN | MSigDB lists | GSE14769\_UNSTIM\_VS\_60MIN\_LPS\_BMDM\_DN | 39 | 5 | 4161 | 170 | CG6966,dpp,fz2,Odc1,scramb1 | | 2.005e-02 | -3.91 | MORF\_PRKCA | MSigDB lists | MORF\_PRKCA | 39 | 5 | 4161 | 170 | Pkc53E,fz2,dnc,Hr96,Dys | | 2.005e-02 | -3.91 | TGTATGA\_MIR4853P | MSigDB lists | TGTATGA\_MIR4853P | 39 | 5 | 4161 | 170 | msps,Ptp61F,Ank,Ser,dnc | | 2.005e-02 | -3.91 | GSE13411\_NAIVE\_BCELL\_VS\_PLASMA\_CELL\_UP | MSigDB lists | GSE13411\_NAIVE\_BCELL\_VS\_PLASMA\_CELL\_UP | 39 | 5 | 4161 | 170 | Glut1,Doa,Pgant6,RhoGAP1A,CG14480 | | 2.005e-02 | -3.91 | ATGCAGT\_MIR217 | MSigDB lists | ATGCAGT\_MIR217 | 39 | 5 | 4161 | 170 | Syx1A,KaiR1D,Dlic,sxc,Sh | | 2.005e-02 | -3.91 | GSE25123\_WT\_VS\_PPARG\_KO\_MACROPHAGE\_ROSIGLITAZONE\_STIM\_UP | MSigDB lists | GSE25123\_WT\_VS\_PPARG\_KO\_MACROPHAGE\_ROSIGLITAZONE\_STIM\_UP | 39 | 5 | 4161 | 170 | sdt,Dys,Octbeta2R,CG41099,app | | 2.005e-02 | -3.91 | GSE369\_SOCS3\_KO\_VS\_WT\_LIVER\_POST\_IL6\_INJECTION\_DN | MSigDB lists | GSE369\_SOCS3\_KO\_VS\_WT\_LIVER\_POST\_IL6\_INJECTION\_DN | 39 | 5 | 4161 | 170 | step,Pkc98E,Meltrin,Rbp6,Sh | | 2.005e-02 | -3.91 | HALLMARK\_BILE\_ACID\_METABOLISM | MSigDB lists | HALLMARK\_BILE\_ACID\_METABOLISM | 39 | 5 | 4161 | 170 | Sod1,ABCD,Hr96,CG3961,Cat | | 2.005e-02 | -3.91 | GO\_REGULATION\_OF\_OSSIFICATION | MSigDB lists | GO\_REGULATION\_OF\_OSSIFICATION | 39 | 5 | 4161 | 170 | Octbeta2R,Su(H),dpp,TfAP-2,Ser | | 2.009e-02 | -3.91 | supramolecular fiber organization | biological process | GO:0097435 | 188 | 16 | 12345 | 597 | rols,Mlp84B,pod1,Prm,CG18109,Khc,Actn,bru1,p130CAS,Msp300,sick,cindr,msps,spd-2,Act88F,Src42A | | 2.020e-02 | -3.90 | G alpha (s) signalling events | REACTOME pathways | R-DME-418555 | 32 | 5 | 3923 | 197 | ACXD,Pde1c,Octbeta2R,Pde8,dnc | | 2.028e-02 | -3.90 | CHICAS\_RB1\_TARGETS\_CONFLUENT | MSigDB lists | CHICAS\_RB1\_TARGETS\_CONFLUENT | 117 | 10 | 4161 | 170 | Tsp96F,CG3860,CG3376,Pak,Act87E,Ser,Meltrin,Epg5,CG4341,Pgant6 | | 2.036e-02 | -3.89 | regulation of cellular catabolic process | biological process | GO:0031329 | 204 | 17 | 12345 | 597 | CG18304,Tor,Pomp,Raf,GlyP,AGO3,sgg,foxo,Cpsf160,Sod1,Marf1,CG3530,Doa,Fer2,bol,EcR,REG | | 2.037e-02 | -3.89 | GO\_POSITIVE\_REGULATION\_OF\_RESPONSE\_TO\_STIMULUS | MSigDB lists | GO\_POSITIVE\_REGULATION\_OF\_RESPONSE\_TO\_STIMULUS | 402 | 25 | 4161 | 170 | Ca-beta,Tor,IP3K2,CG42672,csw,REG,dnc,fz2,Ptp61F,dsh,Cat,Octbeta2R,scramb1,Syx1A,Sod1,Pkc53E,Su(H),wbl,Ser,dpp,Act87E,Pkc98E,Pak,Raf,Tsp96F | | 2.042e-02 | -3.89 | neuroblast differentiation | biological process | GO:0014016 | 45 | 6 | 12345 | 597 | ham,mid,numb,sqz,ems,Antp | | 2.058e-02 | -3.88 | sphingolipid binding | molecular function | GO:0046625 | 5 | 2 | 12176 | 580 | cert,Ser | | 2.058e-02 | -3.88 | protein kinase C activity | molecular function | GO:0004697 | 5 | 2 | 12176 | 580 | Pkc53E,Pkc98E | | 2.058e-02 | -3.88 | transcription coactivator binding | molecular function | GO:0001223 | 5 | 2 | 12176 | 580 | hth,EcR | | 2.058e-02 | -3.88 | fibroblast growth factor receptor binding | molecular function | GO:0005104 | 5 | 2 | 12176 | 580 | bnl,ths | | 2.059e-02 | -3.88 | Transport of glucose and other sugars, bile salts and organic acids, metal ions and amine compounds | REACTOME pathways | R-DME-425366 | 82 | 9 | 3923 | 197 | ZnT63C,CG33934,CG42235,CG3285,CG10960,Glut1,CG14606,CG7882,CG7720 | | 2.062e-02 | -3.88 | GO\_REGULATION\_OF\_HYDROLASE\_ACTIVITY | MSigDB lists | GO\_REGULATION\_OF\_HYDROLASE\_ACTIVITY | 283 | 19 | 4161 | 170 | Nmdar2,scramb1,Octbeta2R,dsh,REG,CG34408,step,Tor,eIF2Bgamma,EcR,Pkc98E,LRP1,dpp,CG5522,sky,Cdep,RhoGAP1A,Sod1,CG17883 | | 2.067e-02 | -3.88 | - | gene3d domains | 4.10.900.10 | 4 | 2 | 4929 | 302 | CadN2,shg | | 2.069e-02 | -3.88 | neuron projection terminus | cellular component | GO:0044306 | 76 | 8 | 11940 | 538 | cpx,sky,Syx1A,unc-13,gukh,mnb,Ank2,g | | 2.090e-02 | -3.87 | GGGAGGRR\_MAZ\_Q6 | MSigDB lists | GGGAGGRR\_MAZ\_Q6 | 506 | 30 | 4161 | 170 | Frq1,Rat1,eIF4E7,kcc,Iswi,Su(H),Meltrin,CG8272,LpR2,Act87E,LRP1,dpp,rdgA,Ser,flw,Pkc98E,pod1,app,PhKgamma,eRF3,PIG-O,Frq2,Dys,CG10126,Nmdar2,Sh,Pde6,Rbp6,Cadps,fz2 | | 2.098e-02 | -3.86 | EFTu-like\_2 | interpro domains | IPR004161 | 12 | 3 | 11383 | 585 | CG33158,eRF3,CG4849 | | 2.104e-02 | -3.86 | PyK (Pyruvate kinase) | protein interactions | 42620 | 31 | 5 | 9204 | 481 | csw,rictor,Raf,melt,Aldh-III | | 2.104e-02 | -3.86 | vg (vestigial) | protein interactions | 36421 | 31 | 5 | 9204 | 481 | sno,RpL38,Su(H),Ser,dpp | | 2.105e-02 | -3.86 | ZU5 | prosite domains | PS51145 | 4 | 2 | 6243 | 386 | Ank,Ank2 | | 2.105e-02 | -3.86 | MHD1 | prosite domains | PS51258 | 4 | 2 | 6243 | 386 | Cadps,unc-13 | | 2.111e-02 | -3.86 | ZU5 | smart domains | SM00218 | 4 | 2 | 5086 | 315 | Ank,Ank2 | | 2.116e-02 | -3.86 | neuromuscular synaptic transmission | biological process | GO:0007274 | 58 | 7 | 12345 | 597 | cpx,Syx1A,Dys,GluRIIC,Nlg2,CG5541,sky | | 2.118e-02 | -3.85 | synaptic vesicle membrane organization | biological process | GO:0048499 | 5 | 2 | 12345 | 597 | g,Nlg2 | | 2.118e-02 | -3.85 | negative regulation of signaling receptor activity | biological process | GO:2000272 | 5 | 2 | 12345 | 597 | kek1,qvr | | 2.118e-02 | -3.85 | calcium ion-regulated exocytosis of neurotransmitter | biological process | GO:0048791 | 5 | 2 | 12345 | 597 | Fife,unc-13 | | 2.118e-02 | -3.85 | phosphatidic acid metabolic process | biological process | GO:0046473 | 5 | 2 | 12345 | 597 | rdgA,Pld | | 2.118e-02 | -3.85 | establishment of epithelial cell planar polarity | biological process | GO:0090163 | 5 | 2 | 12345 | 597 | dsh,sgg | | 2.118e-02 | -3.85 | positive regulation of phosphatidylinositol 3-kinase signaling | biological process | GO:0014068 | 5 | 2 | 12345 | 597 | step,Lpin | | 2.118e-02 | -3.85 | metarhodopsin inactivation | biological process | GO:0016060 | 5 | 2 | 12345 | 597 | Cam,Gprk1 | | 2.118e-02 | -3.85 | negative regulation of neurotransmitter transport | biological process | GO:0051589 | 5 | 2 | 12345 | 597 | sky,cpx | | 2.118e-02 | -3.85 | head morphogenesis | biological process | GO:0060323 | 5 | 2 | 12345 | 597 | hth,dpp | | 2.118e-02 | -3.85 | negative regulation of neurotransmitter secretion | biological process | GO:0046929 | 5 | 2 | 12345 | 597 | sky,cpx | | 2.118e-02 | -3.85 | ornithine metabolic process | biological process | GO:0006591 | 5 | 2 | 12345 | 597 | Odc1,Argl | | 2.122e-02 | -3.85 | ST\_FAS\_SIGNALING\_PATHWAY | MSigDB lists | ST\_FAS\_SIGNALING\_PATHWAY | 15 | 3 | 4161 | 170 | Frq2,CG3376,Frq1 | | 2.122e-02 | -3.85 | RUAN\_RESPONSE\_TO\_TNF\_TROGLITAZONE\_DN | MSigDB lists | RUAN\_RESPONSE\_TO\_TNF\_TROGLITAZONE\_DN | 15 | 3 | 4161 | 170 | PCB,mino,ABCD | | 2.122e-02 | -3.85 | GO\_NEGATIVE\_REGULATION\_OF\_NUCLEOTIDE\_METABOLIC\_PROCESS | MSigDB lists | GO\_NEGATIVE\_REGULATION\_OF\_NUCLEOTIDE\_METABOLIC\_PROCESS | 15 | 3 | 4161 | 170 | Pkc53E,AstA-R1,mGluR | | 2.122e-02 | -3.85 | GO\_POSITIVE\_REGULATION\_OF\_EPITHELIAL\_CELL\_MIGRATION | MSigDB lists | GO\_POSITIVE\_REGULATION\_OF\_EPITHELIAL\_CELL\_MIGRATION | 15 | 3 | 4161 | 170 | Pkc98E,Pkc53E,chb | | 2.122e-02 | -3.85 | GO\_REGULATION\_OF\_SYNAPTIC\_TRANSMISSION\_GLUTAMATERGIC | MSigDB lists | GO\_REGULATION\_OF\_SYNAPTIC\_TRANSMISSION\_GLUTAMATERGIC | 15 | 3 | 4161 | 170 | mGluR,KaiR1D,unc-13 | | 2.122e-02 | -3.85 | GO\_PRESYNAPTIC\_MEMBRANE | MSigDB lists | GO\_PRESYNAPTIC\_MEMBRANE | 15 | 3 | 4161 | 170 | KaiR1D,unc-13,mGluR | | 2.122e-02 | -3.85 | YAMASHITA\_LIVER\_CANCER\_STEM\_CELL\_DN | MSigDB lists | YAMASHITA\_LIVER\_CANCER\_STEM\_CELL\_DN | 15 | 3 | 4161 | 170 | CG3961,Hr96,Sod1 | | 2.122e-02 | -3.85 | KEGG\_NOTCH\_SIGNALING\_PATHWAY | MSigDB lists | KEGG\_NOTCH\_SIGNALING\_PATHWAY | 15 | 3 | 4161 | 170 | dsh,Su(H),Ser | | 2.122e-02 | -3.85 | KRASNOSELSKAYA\_ILF3\_TARGETS\_DN | MSigDB lists | KRASNOSELSKAYA\_ILF3\_TARGETS\_DN | 15 | 3 | 4161 | 170 | Dys,bru1,PCB | | 2.122e-02 | -3.85 | GO\_POSITIVE\_REGULATION\_OF\_NEURON\_DEATH | MSigDB lists | GO\_POSITIVE\_REGULATION\_OF\_NEURON\_DEATH | 15 | 3 | 4161 | 170 | TfAP-2,Tor,KaiR1D | | 2.122e-02 | -3.85 | KEGG\_ARRHYTHMOGENIC\_RIGHT\_VENTRICULAR\_CARDIOMYOPATHY\_ARVC | MSigDB lists | KEGG\_ARRHYTHMOGENIC\_RIGHT\_VENTRICULAR\_CARDIOMYOPATHY\_ARVC | 15 | 3 | 4161 | 170 | Dys,Act87E,Ca-beta | | 2.122e-02 | -3.85 | BIOCARTA\_EIF4\_PATHWAY | MSigDB lists | BIOCARTA\_EIF4\_PATHWAY | 15 | 3 | 4161 | 170 | Pkc53E,Tor,eIF4E7 | | 2.122e-02 | -3.85 | CCAGGTT\_MIR490 | MSigDB lists | CCAGGTT\_MIR490 | 15 | 3 | 4161 | 170 | sxc,REG,polybromo | | 2.122e-02 | -3.85 | GO\_MYOBLAST\_DIFFERENTIATION | MSigDB lists | GO\_MYOBLAST\_DIFFERENTIATION | 15 | 3 | 4161 | 170 | Su(H),Mlp84B,Ser | | 2.122e-02 | -3.85 | GO\_CALCIUM\_ION\_IMPORT | MSigDB lists | GO\_CALCIUM\_ION\_IMPORT | 15 | 3 | 4161 | 170 | MCU,ZnT63C,Pkc98E | | 2.139e-02 | -3.84 | fly chr37E1-37E1\|2-54 | chromosome location | fly chr37E1-37E1\|2-54 | 7 | 2 | 25010 | 845 | CG10237,CG10026 | | 2.139e-02 | -3.84 | fly chr32D2-32D2\|2-44 cM | chromosome location | fly chr32D2-32D2\|2-44 cM | 7 | 2 | 25010 | 845 | CG16854,CG43725 | | 2.139e-02 | -3.84 | fly chr37E1-37E1\|2-54 cM | chromosome location | fly chr37E1-37E1\|2-54 cM | 7 | 2 | 25010 | 845 | CG10237,CG10026 | | 2.139e-02 | -3.84 | fly chr11E3-11E3\|1-42 cM | chromosome location | fly chr11E3-11E3\|1-42 cM | 7 | 2 | 25010 | 845 | sno,REG | | 2.139e-02 | -3.84 | fly chr11E3-11E3\|1-42 | chromosome location | fly chr11E3-11E3\|1-42 | 7 | 2 | 25010 | 845 | sno,REG | | 2.139e-02 | -3.84 | fly chr2F2-2F2\|1-0.9 cM | chromosome location | fly chr2F2-2F2\|1-0.9 cM | 7 | 2 | 25010 | 845 | CG3091,CG3078 | | 2.139e-02 | -3.84 | fly chr2F2-2F2\|1-0.9 | chromosome location | fly chr2F2-2F2\|1-0.9 | 7 | 2 | 25010 | 845 | CG3078,CG3091 | | 2.139e-02 | -3.84 | fly chr32D2-32D2\|2-44 | chromosome location | fly chr32D2-32D2\|2-44 | 7 | 2 | 25010 | 845 | CG43725,CG16854 | | 2.146e-02 | -3.84 | E2F1\_Q6\_01 | MSigDB lists | E2F1\_Q6\_01 | 69 | 7 | 4161 | 170 | Stt3B,Iswi,Sk1,eRF3,dpp,LRP1,Fhos | | 2.146e-02 | -3.84 | GCM\_RAN | MSigDB lists | GCM\_RAN | 69 | 7 | 4161 | 170 | MESK2,CG12121,CG42672,eRF3,CG5522,Su(H),CG33090 | | 2.149e-02 | -3.84 | abd-A (abdominal A) | protein interactions | 42037 | 54 | 7 | 9204 | 481 | tin,Su(H),ems,Poxm,hth,mid,Ubx | | 2.164e-02 | -3.83 | Protein\_kinase\_ATP\_BS | interpro domains | IPR017441 | 164 | 15 | 11383 | 585 | Asator,Pkc53E,mnb,Raf,sgg,Doa,Strn-Mlck,CG17698,Pak,Gprk1,Mos,Src42A,Pkc98E,PhKgamma,Nuak1 | | 2.165e-02 | -3.83 | GSE21546\_WT\_VS\_SAP1A\_KO\_AND\_ELK1\_KO\_ANTI\_CD3\_STIM\_DP\_THYMOCYTES\_DN | MSigDB lists | GSE21546\_WT\_VS\_SAP1A\_KO\_AND\_ELK1\_KO\_ANTI\_CD3\_STIM\_DP\_THYMOCYTES\_DN | 54 | 6 | 4161 | 170 | CG41099,CG10126,Pkc53E,mino,Meltrin,Tor | | 2.165e-02 | -3.83 | GSE17721\_CTRL\_VS\_LPS\_0.5H\_BMDC\_UP | MSigDB lists | GSE17721\_CTRL\_VS\_LPS\_0.5H\_BMDC\_UP | 54 | 6 | 4161 | 170 | nompB,Der-1,CG5493,CG4301,bru1,Flo2 | | 2.165e-02 | -3.83 | ATGAAGG\_MIR205 | MSigDB lists | ATGAAGG\_MIR205 | 54 | 6 | 4161 | 170 | LRP1,CG40045,Act87E,Pkc98E,Src42A,CG9132 | | 2.165e-02 | -3.83 | GO\_REGULATION\_OF\_PEPTIDE\_TRANSPORT | MSigDB lists | GO\_REGULATION\_OF\_PEPTIDE\_TRANSPORT | 54 | 6 | 4161 | 170 | Pkc98E,csw,Pkc53E,MCU,Pde8,Syx1A | | 2.165e-02 | -3.83 | GO\_RECEPTOR\_MEDIATED\_ENDOCYTOSIS | MSigDB lists | GO\_RECEPTOR\_MEDIATED\_ENDOCYTOSIS | 54 | 6 | 4161 | 170 | LRP1,Cln3,Octbeta2R,LpR2,Tsp96F,CG12121 | | 2.165e-02 | -3.83 | MORF\_SUPT3H | MSigDB lists | MORF\_SUPT3H | 54 | 6 | 4161 | 170 | Pkc53E,fz2,dnc,Hr96,Dys,CG40178 | | 2.170e-02 | -3.83 | Liprin-alpha (Liprin-alpha) | protein interactions | 249072 | 21 | 4 | 9204 | 481 | RhoGAP100F,sgg,Khc,unc-13 | | 2.170e-02 | -3.83 | coro (coro) | protein interactions | 35596 | 21 | 4 | 9204 | 481 | Syx1A,dpp,rictor,PhKgamma | | 2.170e-02 | -3.83 | Hmg-2 (High mobility group protein 2) | protein interactions | 37407 | 21 | 4 | 9204 | 481 | JhI-26,CG13476,REG,kay | | 2.170e-02 | -3.83 | Plp (Pericentrin-like protein) | protein interactions | 3772382 | 21 | 4 | 9204 | 481 | Cam,sgg,Cep97,spd-2 | | 2.191e-02 | -3.82 | CG18476 (uncharacterized protein) | protein interactions | 41404 | 12 | 3 | 9204 | 481 | Doa,sgg,bru1 | | 2.191e-02 | -3.82 | msps (mini spindles) | protein interactions | 41952 | 12 | 3 | 9204 | 481 | chb,sgg,CG10164 | | 2.191e-02 | -3.82 | mid (midline) | protein interactions | 33770 | 12 | 3 | 9204 | 481 | Ubx,Antp,tin | | 2.191e-02 | -3.82 | Nrx-1 (Neurexin 1) | protein interactions | 42646 | 12 | 3 | 9204 | 481 | RhoGAP100F,scrib,Nlg2 | | 2.193e-02 | -3.82 | carbohydrate transmembrane transporter activity | molecular function | GO:0015144 | 23 | 4 | 12176 | 580 | CG3285,CG10960,CG45782,CG14606 | | 2.195e-02 | -3.82 | neuron recognition | biological process | GO:0008038 | 114 | 11 | 12345 | 597 | dsh,fz2,fru,p130CAS,mid,esn,Ptp99A,nrm,trol,dpr12,dsd | | 2.202e-02 | -3.82 | reproduction | biological process | GO:0000003 | 1651 | 97 | 12345 | 597 | Dlic,Mbs,Gas8,lncRNA:CR43753,Khc,Mos,polybromo,bbg,RhoL,msps,Tor,ex,sgg,spd-2,Gr39a,CG42823,fz2,kay,mGluR,tnc,raw,tld,kek1,upd3,opa,dpp,Obp56h,dany,shg,AGO2,shd,ND-51L2,dnc,CG34435,Yeti,Raf,Msp300,TfAP-2,Mrtf,lncRNA:Hsromega,Gr22c,tin,dsh,aust,chinmo,chb,shep,wbl,Pde1c,fru,Doa,Acp62F,lncRNA:CR43484,drd,flw,Fad2,CG18109,retn,Ebp,CG7483,lncRNA:flam,Octbeta2R,bru1,bol,Mst87F,CG18135,DNaseII,CG10000,cic,CG16854,can,msi,Iswi,mamo,Obp56e,Eip74EF,csw,rols,mid,egh,Sh,EcR,scrib,lncRNA:CR43802,Qsox2,cindr,lncRNA:CR44585,lncRNA:CR44420,rib,CG42355,Inx2,fs(1)N,CG1428,CG12200,Nlg2,bun,JhI-26 | | 2.211e-02 | -3.81 | BUYTAERT\_PHOTODYNAMIC\_THERAPY\_STRESS\_DN | MSigDB lists | BUYTAERT\_PHOTODYNAMIC\_THERAPY\_STRESS\_DN | 190 | 14 | 4161 | 170 | Pgant6,Acbp2,PIG-O,mRpL24,dsd,Uros1,CG3961,CG41099,Fhos,Ser,REG,CG14480,CG12547,Octbeta2R | | 2.216e-02 | -3.81 | developmental maturation | biological process | GO:0021700 | 206 | 17 | 12345 | 597 | egh,dpp,shg,Khc,flw,EcR,CG7483,scrib,cic,msps,unc-13,Ca-beta,shep,chb,Nlg2,tld,wbl | | 2.217e-02 | -3.81 | GTP\_EFTU\_D2 | pfam domains | PF03144 | 12 | 3 | 10480 | 550 | CG33158,CG4849,eRF3 | | 2.217e-02 | -3.81 | RA | pfam domains | PF00788 | 12 | 3 | 10480 | 550 | spri,wake,Myo81F | | 2.218e-02 | -3.81 | GSE360\_L\_MAJOR\_VS\_T\_GONDII\_MAC\_UP | MSigDB lists | GSE360\_L\_MAJOR\_VS\_T\_GONDII\_MAC\_UP | 40 | 5 | 4161 | 170 | Tif-IA,PIG-O,Syx1A,Flo2,Pde1c | | 2.218e-02 | -3.81 | MEISSNER\_NPC\_HCP\_WITH\_H3K4ME2\_AND\_H3K27ME3 | MSigDB lists | MEISSNER\_NPC\_HCP\_WITH\_H3K4ME2\_AND\_H3K27ME3 | 40 | 5 | 4161 | 170 | Awh,Cadps,Sh,Syn2,scro | | 2.218e-02 | -3.81 | GSE37534\_UNTREATED\_VS\_ROSIGLITAZONE\_TREATED\_CD4\_TCELL\_PPARG1\_AND\_FOXP3\_TRASDUCED\_UP | MSigDB lists | GSE37534\_UNTREATED\_VS\_ROSIGLITAZONE\_TREATED\_CD4\_TCELL\_PPARG1\_AND\_FOXP3\_TRASDUCED\_UP | 40 | 5 | 4161 | 170 | mRpL24,flw,Sk1,LRP1,KaiR1D | | 2.218e-02 | -3.81 | STAT5B\_01 | MSigDB lists | STAT5B\_01 | 40 | 5 | 4161 | 170 | Sod1,mRpL24,Ank,CG3376,scramb1 | | 2.218e-02 | -3.81 | AAAGGAT\_MIR501 | MSigDB lists | AAAGGAT\_MIR501 | 40 | 5 | 4161 | 170 | Doa,bru1,MCU,flw,Pkc98E | | 2.218e-02 | -3.81 | GSE15659\_TREG\_VS\_TCONV\_DN | MSigDB lists | GSE15659\_TREG\_VS\_TCONV\_DN | 40 | 5 | 4161 | 170 | CG41099,Ank,Uros1,dnc,CG8184 | | 2.218e-02 | -3.81 | CAGCAGG\_MIR370 | MSigDB lists | CAGCAGG\_MIR370 | 40 | 5 | 4161 | 170 | sxc,Src42A,RhoGAP1A,polybromo,Ank | | 2.233e-02 | -3.80 | GO\_CYTOKINE\_SECRETION | MSigDB lists | GO\_CYTOKINE\_SECRETION | 6 | 2 | 4161 | 170 | Su(H),mino | | 2.233e-02 | -3.80 | KORKOLA\_YOLK\_SAC\_TUMOR | MSigDB lists | KORKOLA\_YOLK\_SAC\_TUMOR | 6 | 2 | 4161 | 170 | fz2,dpp | | 2.233e-02 | -3.80 | MODULE\_357 | MSigDB lists | MODULE\_357 | 6 | 2 | 4161 | 170 | Octbeta2R,eIF2Bgamma | | 2.233e-02 | -3.80 | GO\_GLYCOSYLCERAMIDE\_METABOLIC\_PROCESS | MSigDB lists | GO\_GLYCOSYLCERAMIDE\_METABOLIC\_PROCESS | 6 | 2 | 4161 | 170 | CG33090,Cln3 | | 2.233e-02 | -3.80 | GO\_POSITIVE\_REGULATION\_OF\_CATION\_CHANNEL\_ACTIVITY | MSigDB lists | GO\_POSITIVE\_REGULATION\_OF\_CATION\_CHANNEL\_ACTIVITY | 6 | 2 | 4161 | 170 | Ank,AstA-R1 | | 2.233e-02 | -3.80 | GO\_DEVELOPMENTAL\_INDUCTION | MSigDB lists | GO\_DEVELOPMENTAL\_INDUCTION | 6 | 2 | 4161 | 170 | dpp,fz2 | | 2.233e-02 | -3.80 | MODULE\_154 | MSigDB lists | MODULE\_154 | 6 | 2 | 4161 | 170 | Octbeta2R,eIF2Bgamma | | 2.233e-02 | -3.80 | GO\_ENDOCARDIAL\_CUSHION\_MORPHOGENESIS | MSigDB lists | GO\_ENDOCARDIAL\_CUSHION\_MORPHOGENESIS | 6 | 2 | 4161 | 170 | Su(H),dpp | | 2.233e-02 | -3.80 | GO\_MODULATION\_BY\_HOST\_OF\_VIRAL\_PROCESS | MSigDB lists | GO\_MODULATION\_BY\_HOST\_OF\_VIRAL\_PROCESS | 6 | 2 | 4161 | 170 | PCB,CG17883 | | 2.233e-02 | -3.80 | MODULE\_292 | MSigDB lists | MODULE\_292 | 6 | 2 | 4161 | 170 | scramb1,Meltrin | | 2.233e-02 | -3.80 | GO\_REGULATION\_OF\_PLATELET\_AGGREGATION | MSigDB lists | GO\_REGULATION\_OF\_PLATELET\_AGGREGATION | 6 | 2 | 4161 | 170 | Pkc53E,Octbeta2R | | 2.233e-02 | -3.80 | ROZANOV\_MMP14\_CORRELATED | MSigDB lists | ROZANOV\_MMP14\_CORRELATED | 6 | 2 | 4161 | 170 | Ser,Iswi | | 2.233e-02 | -3.80 | GO\_REGULATION\_OF\_TRANSCRIPTION\_FROM\_RNA\_POLYMERASE\_III\_PROMOTER | MSigDB lists | GO\_REGULATION\_OF\_TRANSCRIPTION\_FROM\_RNA\_POLYMERASE\_III\_PROMOTER | 6 | 2 | 4161 | 170 | Maf1,Tor | | 2.233e-02 | -3.80 | GO\_REGULATION\_OF\_PLATELET\_ACTIVATION | MSigDB lists | GO\_REGULATION\_OF\_PLATELET\_ACTIVATION | 6 | 2 | 4161 | 170 | Octbeta2R,Pkc53E | | 2.233e-02 | -3.80 | GO\_POSITIVE\_REGULATION\_OF\_FATTY\_ACID\_METABOLIC\_PROCESS | MSigDB lists | GO\_POSITIVE\_REGULATION\_OF\_FATTY\_ACID\_METABOLIC\_PROCESS | 6 | 2 | 4161 | 170 | EcR,ABCD | | 2.233e-02 | -3.80 | GHO\_ATF5\_TARGETS\_UP | MSigDB lists | GHO\_ATF5\_TARGETS\_UP | 6 | 2 | 4161 | 170 | nompB,CG4301 | | 2.233e-02 | -3.80 | GO\_B\_CELL\_DIFFERENTIATION | MSigDB lists | GO\_B\_CELL\_DIFFERENTIATION | 6 | 2 | 4161 | 170 | Su(H),Ptp61F | | 2.233e-02 | -3.80 | MODULE\_366 | MSigDB lists | MODULE\_366 | 6 | 2 | 4161 | 170 | Uros1,kcc | | 2.233e-02 | -3.80 | GO\_PROTEIN\_KINASE\_C\_ACTIVITY | MSigDB lists | GO\_PROTEIN\_KINASE\_C\_ACTIVITY | 6 | 2 | 4161 | 170 | Pkc53E,Pkc98E | | 2.233e-02 | -3.80 | GO\_RESPONSE\_TO\_ISOQUINOLINE\_ALKALOID | MSigDB lists | GO\_RESPONSE\_TO\_ISOQUINOLINE\_ALKALOID | 6 | 2 | 4161 | 170 | Tor,Pkc98E | | 2.233e-02 | -3.80 | GO\_POSITIVE\_REGULATION\_OF\_SODIUM\_ION\_TRANSMEMBRANE\_TRANSPORT | MSigDB lists | GO\_POSITIVE\_REGULATION\_OF\_SODIUM\_ION\_TRANSMEMBRANE\_TRANSPORT | 6 | 2 | 4161 | 170 | Ank,Dys | | 2.233e-02 | -3.80 | GO\_NITRIC\_OXIDE\_SYNTHASE\_BINDING | MSigDB lists | GO\_NITRIC\_OXIDE\_SYNTHASE\_BINDING | 6 | 2 | 4161 | 170 | Dys,Act87E | | 2.233e-02 | -3.80 | PID\_WNT\_CANONICAL\_PATHWAY | MSigDB lists | PID\_WNT\_CANONICAL\_PATHWAY | 6 | 2 | 4161 | 170 | fz2,dsh | | 2.233e-02 | -3.80 | EHLERS\_ANEUPLOIDY\_DN | MSigDB lists | EHLERS\_ANEUPLOIDY\_DN | 6 | 2 | 4161 | 170 | Frq1,Frq2 | | 2.233e-02 | -3.80 | GO\_POSITIVE\_REGULATION\_OF\_TRANSCRIPTION\_FROM\_RNA\_POLYMERASE\_II\_PROMOTER\_INVOLVED\_IN\_CELLULAR\_RESPONSE\_TO\_CHEMICAL\_STIMULUS | MSigDB lists | GO\_POSITIVE\_REGULATION\_OF\_TRANSCRIPTION\_FROM\_RNA\_POLYMERASE\_II\_PROMOTER\_INVOLVED\_IN\_CELLULAR\_RESPONSE\_TO\_CHEMICAL\_STIMULUS | 6 | 2 | 4161 | 170 | Su(H),dpp | | 2.233e-02 | -3.80 | GO\_LIPOPROTEIN\_PARTICLE\_RECEPTOR\_ACTIVITY | MSigDB lists | GO\_LIPOPROTEIN\_PARTICLE\_RECEPTOR\_ACTIVITY | 6 | 2 | 4161 | 170 | LpR2,LRP1 | | 2.233e-02 | -3.80 | GO\_REGULATION\_OF\_RESPONSE\_TO\_INTERFERON\_GAMMA | MSigDB lists | GO\_REGULATION\_OF\_RESPONSE\_TO\_INTERFERON\_GAMMA | 6 | 2 | 4161 | 170 | EcR,Ptp61F | | 2.233e-02 | -3.80 | MODULE\_297 | MSigDB lists | MODULE\_297 | 6 | 2 | 4161 | 170 | eIF2Bgamma,Octbeta2R | | 2.238e-02 | -3.80 | potassium ion import across plasma membrane | biological process | GO:1990573 | 13 | 3 | 12345 | 597 | nrv2,Irk3,CG33310 | | 2.238e-02 | -3.80 | renal filtration cell differentiation | biological process | GO:0061318 | 13 | 3 | 12345 | 597 | tin,numb,dpp | | 2.238e-02 | -3.80 | haltere development | biological process | GO:0007482 | 13 | 3 | 12345 | 597 | Syx1A,hth,Ubx | | 2.238e-02 | -3.80 | regulation of response to drug | biological process | GO:2001023 | 13 | 3 | 12345 | 597 | Tor,Pde8,Syt12 | | 2.238e-02 | -3.80 | ecdysone biosynthetic process | biological process | GO:0006697 | 13 | 3 | 12345 | 597 | woc,spok,foxo | | 2.238e-02 | -3.80 | lipid translocation | biological process | GO:0034204 | 13 | 3 | 12345 | 597 | CG9981,CG4301,scramb1 | | 2.238e-02 | -3.80 | cholesterol homeostasis | biological process | GO:0042632 | 13 | 3 | 12345 | 597 | mag,Hr96,EcR | | 2.238e-02 | -3.80 | nephrocyte differentiation | biological process | GO:0061319 | 13 | 3 | 12345 | 597 | dpp,numb,tin | | 2.245e-02 | -3.80 | CH-domain | interpro domains | IPR001715 | 32 | 5 | 11383 | 585 | Msp300,Actn,CG34417,Dys,sick | | 2.247e-02 | -3.80 | Transmission across Chemical Synapses | REACTOME pathways | R-DME-112315 | 97 | 10 | 3923 | 197 | Pkc53E,CrebB,ACXD,CG17698,KaiR1D,Cam,GluRIIC,Nmdar2,Actn,Raf | | 2.255e-02 | -3.79 | myofibril assembly | biological process | GO:0030239 | 46 | 6 | 12345 | 597 | rols,Act88F,Actn,Mlp84B,Prm,bru1 | | 2.255e-02 | -3.79 | positive regulation of Ras protein signal transduction | biological process | GO:0046579 | 46 | 6 | 12345 | 597 | step,RhoGAP1A,CG10188,Exn,siz,MESK2 | | 2.255e-02 | -3.79 | positive regulation of small GTPase mediated signal transduction | biological process | GO:0051057 | 46 | 6 | 12345 | 597 | MESK2,Exn,siz,CG10188,step,RhoGAP1A | | 2.263e-02 | -3.79 | lipid binding | molecular function | GO:0008289 | 162 | 14 | 12176 | 580 | CG18659,Pld,cert,gce,qvr,Gprk1,Syt12,step,Ser,unc-13,melt,mGluR,dsh,EcR | | 2.263e-02 | -3.79 | DAIRKEE\_TERT\_TARGETS\_DN | MSigDB lists | DAIRKEE\_TERT\_TARGETS\_DN | 27 | 4 | 4161 | 170 | bru1,Dlic,polybromo,CG5522 | | 2.263e-02 | -3.79 | VILLANUEVA\_LIVER\_CANCER\_KRT19\_DN | MSigDB lists | VILLANUEVA\_LIVER\_CANCER\_KRT19\_DN | 27 | 4 | 4161 | 170 | LRP1,CG15093,Sod1,Hr96 | | 2.263e-02 | -3.79 | CRX\_NRL\_DN.V1\_DN | MSigDB lists | CRX\_NRL\_DN.V1\_DN | 27 | 4 | 4161 | 170 | fz2,Octbeta2R,GlyP,KaiR1D | | 2.263e-02 | -3.79 | GO\_REGULATION\_OF\_JNK\_CASCADE | MSigDB lists | GO\_REGULATION\_OF\_JNK\_CASCADE | 27 | 4 | 4161 | 170 | fz2,KaiR1D,dsh,Pak | | 2.263e-02 | -3.79 | GSE36078\_UNTREATED\_VS\_AD5\_INF\_MOUSE\_LUNG\_DC\_UP | MSigDB lists | GSE36078\_UNTREATED\_VS\_AD5\_INF\_MOUSE\_LUNG\_DC\_UP | 27 | 4 | 4161 | 170 | dsh,sky,Nha1,Tsp96F | | 2.263e-02 | -3.79 | PKCA\_DN.V1\_DN | MSigDB lists | PKCA\_DN.V1\_DN | 27 | 4 | 4161 | 170 | ZnT63C,Pkc53E,PIG-O,CG14105 | | 2.265e-02 | -3.79 | regulation of metal ion transport | biological process | GO:0010959 | 34 | 5 | 12345 | 597 | qvr,CG3078,Ca-beta,CG42797,olf186-F | | 2.269e-02 | -3.79 | Cam-PDE 1 activation | REACTOME pathways | R-DME-111957 | 5 | 2 | 3923 | 197 | Cam,Pde1c | | 2.269e-02 | -3.79 | Interaction between L1 and Ankyrins | REACTOME pathways | R-DME-445095 | 5 | 2 | 3923 | 197 | Ank,Ank2 | | 2.269e-02 | -3.79 | AKT phosphorylates targets in the nucleus | REACTOME pathways | R-DME-198693 | 5 | 2 | 3923 | 197 | foxo,CrebB | | 2.269e-02 | -3.79 | Role of phospholipids in phagocytosis | REACTOME pathways | R-DME-2029485 | 5 | 2 | 3923 | 197 | Pld,Pkc98E | | 2.271e-02 | -3.79 | SH3-like\_dom\_sf | interpro domains | IPR036028 | 68 | 8 | 11383 | 585 | p130CAS,Ca-beta,Stacl,RhoGEF3,sdt,cindr,Src42A,Exn | | 2.271e-02 | -3.79 | PDZ\_sf | interpro domains | IPR036034 | 68 | 8 | 11383 | 585 | dsh,RhoGAP100F,bbg,Syn2,sdt,CG5921,scrib,Fife | | 2.277e-02 | -3.78 | - | gene3d domains | 3.30.70.330 | 152 | 16 | 4929 | 302 | CG44249,fus,barc,trv,Marf1,msi,shep,Rbp6,bru1,CG40813,CG3594,CG41562,bol,CG42458,Achl,Spf45 | | 2.284e-02 | -3.78 | microtubule end | cellular component | GO:1990752 | 14 | 3 | 11940 | 538 | Khc,chb,msps | | 2.297e-02 | -3.77 | molecular transducer activity | molecular function | GO:0060089 | 447 | 31 | 12176 | 580 | CG31750,MsR2,Pdfr,Nlg2,Ptp99A,Or65c,mGluR,fz2,boi,GluRIIC,CG13229,Octbeta2R,EcR,Or85b,fz4,Or42b,KaiR1D,ine,klg,CCAP-R,PGRP-LA,AstA-R1,Gr39a,et,shakB,Nmdar2,CG11318,CG4839,Hr96,CG9380,Gr22c | | 2.299e-02 | -3.77 | ZF\_DAG\_PE\_2 | prosite domains | PS50081 | 27 | 5 | 6243 | 386 | Raf,Pkc98E,Stacl,Pkc53E,unc-13 | | 2.304e-02 | -3.77 | phototransduction, visible light | biological process | GO:0007603 | 23 | 4 | 12345 | 597 | Gprk1,ninaD,Cam,rdgA | | 2.304e-02 | -3.77 | negative regulation of cellular response to insulin stimulus | biological process | GO:1900077 | 23 | 4 | 12345 | 597 | foxo,sgg,Ptp61F,HDAC4 | | 2.304e-02 | -3.77 | positive regulation of circadian rhythm | biological process | GO:0042753 | 23 | 4 | 12345 | 597 | qvr,Sh,wake,EcR | | 2.304e-02 | -3.77 | negative regulation of insulin receptor signaling pathway | biological process | GO:0046627 | 23 | 4 | 12345 | 597 | HDAC4,sgg,Ptp61F,foxo | | 2.304e-02 | -3.77 | phospholipid transport | biological process | GO:0015914 | 23 | 4 | 12345 | 597 | scramb1,CG4301,cert,CG9981 | | 2.305e-02 | -3.77 | REACTOME\_TRANSMISSION\_ACROSS\_CHEMICAL\_SYNAPSES | MSigDB lists | REACTOME\_TRANSMISSION\_ACROSS\_CHEMICAL\_SYNAPSES | 70 | 7 | 4161 | 170 | Nmdar2,KaiR1D,Syx1A,Raf,Ca-beta,CG17698,Pkc53E | | 2.305e-02 | -3.77 | GSE17721\_0.5H\_VS\_24H\_PAM3CSK4\_BMDC\_DN | MSigDB lists | GSE17721\_0.5H\_VS\_24H\_PAM3CSK4\_BMDC\_DN | 70 | 7 | 4161 | 170 | EcR,Sh,Acbp2,Ser,PCB,CG3961,CG3842 | | 2.316e-02 | -3.77 | FOSTER\_TOLERANT\_MACROPHAGE\_DN | MSigDB lists | FOSTER\_TOLERANT\_MACROPHAGE\_DN | 86 | 8 | 4161 | 170 | IP3K2,eRF3,CG4341,kcc,ZnT63C,Odc1,fz2,CG4301 | | 2.316e-02 | -3.77 | GO\_METAL\_ION\_TRANSMEMBRANE\_TRANSPORTER\_ACTIVITY | MSigDB lists | GO\_METAL\_ION\_TRANSMEMBRANE\_TRANSPORTER\_ACTIVITY | 86 | 8 | 4161 | 170 | Ca-beta,Frq2,Frq1,ZnT63C,kcc,Sh,MCU,CNT1 | | 2.317e-02 | -3.76 | orb2 (orb2) | protein interactions | 39018 | 80 | 9 | 9204 | 481 | Nlg2,bnl,Flo2,ps,Root,Rbp6,path,bol,sky | | 2.327e-02 | -3.76 | Scr (Sex combs reduced) | protein interactions | 40833 | 43 | 6 | 9204 | 481 | ems,Su(H),tin,Antp,hth,Poxm | | 2.335e-02 | -3.76 | GO\_NEGATIVE\_REGULATION\_OF\_CELL\_DEATH | MSigDB lists | GO\_NEGATIVE\_REGULATION\_OF\_CELL\_DEATH | 210 | 15 | 4161 | 170 | KaiR1D,Cln3,Pkc53E,Sod1,sxc,IP3K2,Tor,TfAP-2,Raf,Cat,mino,LRP1,REG,Tif-IA,Sk1 | | 2.335e-02 | -3.76 | GO\_REGULATION\_OF\_ANATOMICAL\_STRUCTURE\_MORPHOGENESIS | MSigDB lists | GO\_REGULATION\_OF\_ANATOMICAL\_STRUCTURE\_MORPHOGENESIS | 210 | 15 | 4161 | 170 | sky,Su(H),RhoGAP1A,Pkc53E,TfAP-2,CG17883,unc-13,Raf,chb,dsh,Octbeta2R,fz2,REG,dpp,nompB | | 2.350e-02 | -3.75 | GSE5589\_IL6\_KO\_VS\_IL10\_KO\_LPS\_STIM\_MACROPHAGE\_180MIN\_UP | MSigDB lists | GSE5589\_IL6\_KO\_VS\_IL10\_KO\_LPS\_STIM\_MACROPHAGE\_180MIN\_UP | 55 | 6 | 4161 | 170 | Frq2,Frq1,CG4301,mGluR,Odc1,CG12121 | | 2.350e-02 | -3.75 | GTTTGTT\_MIR495 | MSigDB lists | GTTTGTT\_MIR495 | 55 | 6 | 4161 | 170 | bru1,dsd,Dlic,REG,dnc,sxc | | 2.377e-02 | -3.74 | EFG\_II | interpro domains | IPR041095 | 5 | 2 | 11383 | 585 | CG4849,CG33158 | | 2.377e-02 | -3.74 | Transcription\_factor\_GATA | interpro domains | IPR039355 | 5 | 2 | 11383 | 585 | GATAe,grn | | 2.377e-02 | -3.74 | Homeobox\_Antennapedia\_CS | interpro domains | IPR001827 | 5 | 2 | 11383 | 585 | Ubx,Antp | | 2.377e-02 | -3.74 | Frizzled/Smoothened\_TM | interpro domains | IPR000539 | 5 | 2 | 11383 | 585 | fz2,fz4 | | 2.377e-02 | -3.74 | 7TM\_GPCR\_serpentine\_rcpt\_Srw | interpro domains | IPR019427 | 5 | 2 | 11383 | 585 | MsR2,CG13229 | | 2.377e-02 | -3.74 | Piwi | interpro domains | IPR003165 | 5 | 2 | 11383 | 585 | AGO3,AGO2 | | 2.377e-02 | -3.74 | Sec7\_dom\_sf | interpro domains | IPR035999 | 5 | 2 | 11383 | 585 | siz,step | | 2.377e-02 | -3.74 | Sec7\_C\_sf | interpro domains | IPR023394 | 5 | 2 | 11383 | 585 | siz,step | | 2.377e-02 | -3.74 | TOG | interpro domains | IPR034085 | 5 | 2 | 11383 | 585 | chb,msps | | 2.377e-02 | -3.74 | Frizzled/SFRP | interpro domains | IPR015526 | 5 | 2 | 11383 | 585 | fz4,fz2 | | 2.378e-02 | -3.74 | regulation of nervous system development | biological process | GO:0051960 | 356 | 26 | 12345 | 597 | sgg,numb,barc,Raf,Tor,ham,Ank2,hdc,dsh,Elp3,CG42797,Mob2,mnb,bun,Syn2,Src42A,Dlic,csw,mid,rictor,hth,Pak,Mbs,EcR,Antp,Octbeta2R | | 2.381e-02 | -3.74 | GO\_WOUND\_HEALING | MSigDB lists | GO\_WOUND\_HEALING | 103 | 9 | 4161 | 170 | Pkc98E,rdgA,Act87E,scramb1,Pak,Octbeta2R,csw,Pkc53E,Tor | | 2.390e-02 | -3.73 | sip2 (septin interacting protein 2) | protein interactions | 33979 | 32 | 5 | 9204 | 481 | foxo,spd-2,Raf,rictor,chb | | 2.390e-02 | -3.73 | CG13838 (uncharacterized protein) | protein interactions | 42719 | 32 | 5 | 9204 | 481 | CG15468,Vha36-3,Dys,Fhos,htk | | 2.405e-02 | -3.73 | Alkaline\_phosphatase\_core\_sf | interpro domains | IPR017850 | 44 | 6 | 11383 | 585 | PIG-G,Alp12,CG16854,PIG-O,CG1428,CG6790 | | 2.415e-02 | -3.72 | fusome | cellular component | GO:0045169 | 37 | 5 | 11940 | 538 | mino,chb,scrib,Ank,sgg | | 2.420e-02 | -3.72 | positive regulation of cell cycle | biological process | GO:0045787 | 73 | 8 | 12345 | 597 | CG12200,CycT,foxo,bru1,Tor,Su(H),sgg,crol | | 2.431e-02 | -3.72 | zf-C2H2 | pfam domains | PF00096 | 207 | 18 | 10480 | 550 | CG31224,CG4374,luna,crol,mamo,CG15073,CG12605,esg,chinmo,ham,CG10462,CG7368,sqz,CG9650,opa,l(3)neo38,jim,CG12071 | | 2.441e-02 | -3.71 | GRYDER\_PAX3FOXO1\_ENHANCERS\_KO\_DOWN | MSigDB lists | GRYDER\_PAX3FOXO1\_ENHANCERS\_KO\_DOWN | 138 | 11 | 4161 | 170 | Act87E,Rbp6,Tsp96F,CG12121,Khc,Su(H),Sod1,CG40045,IP3K2,bru1,dsd | | 2.445e-02 | -3.71 | GSE21927\_SPLEEN\_C57BL6\_VS\_4T1\_TUMOR\_BALBC\_MONOCYTES\_DN | MSigDB lists | GSE21927\_SPLEEN\_C57BL6\_VS\_4T1\_TUMOR\_BALBC\_MONOCYTES\_DN | 41 | 5 | 4161 | 170 | EcR,Sh,CG41099,scramb1,CG3860 | | 2.445e-02 | -3.71 | GSE19941\_UNSTIM\_VS\_LPS\_AND\_IL10\_STIM\_IL10\_KO\_MACROPHAGE\_UP | MSigDB lists | GSE19941\_UNSTIM\_VS\_LPS\_AND\_IL10\_STIM\_IL10\_KO\_MACROPHAGE\_UP | 41 | 5 | 4161 | 170 | CG40178,KrT95D,CG3842,melt,Src42A | | 2.445e-02 | -3.71 | GSE24634\_TREG\_VS\_TCONV\_POST\_DAY3\_IL4\_CONVERSION\_DN | MSigDB lists | GSE24634\_TREG\_VS\_TCONV\_POST\_DAY3\_IL4\_CONVERSION\_DN | 41 | 5 | 4161 | 170 | Ser,dsd,Cat,Mctp,CG3860 | | 2.445e-02 | -3.71 | BILD\_HRAS\_ONCOGENIC\_SIGNATURE | MSigDB lists | BILD\_HRAS\_ONCOGENIC\_SIGNATURE | 41 | 5 | 4161 | 170 | dpp,Octbeta2R,Odc1,Glut1,Syx1A | | 2.445e-02 | -3.71 | GSE21927\_EL4\_VS\_MCA203\_TUMOR\_MONOCYTES\_UP | MSigDB lists | GSE21927\_EL4\_VS\_MCA203\_TUMOR\_MONOCYTES\_UP | 41 | 5 | 4161 | 170 | Ca-beta,dnc,Fhos,Act87E,CG40045 | | 2.445e-02 | -3.71 | TCF4\_Q5 | MSigDB lists | TCF4\_Q5 | 41 | 5 | 4161 | 170 | ABCD,CG6966,Syx1A,eIF4E7,scramb1 | | 2.445e-02 | -3.71 | RREB1\_01 | MSigDB lists | RREB1\_01 | 41 | 5 | 4161 | 170 | Nmdar2,flw,Pkc98E,CG15093,LRP1 | | 2.445e-02 | -3.71 | GSE44955\_MCSF\_VS\_MCSF\_AND\_IL27\_STIM\_MACROPHAGE\_DN | MSigDB lists | GSE44955\_MCSF\_VS\_MCSF\_AND\_IL27\_STIM\_MACROPHAGE\_DN | 41 | 5 | 4161 | 170 | Flo2,CG4301,IntS3,Fhos,rdgA | | 2.445e-02 | -3.71 | LHX3\_01 | MSigDB lists | LHX3\_01 | 41 | 5 | 4161 | 170 | Dys,Octbeta2R,bru1,TfAP-2,Syn2 | | 2.445e-02 | -3.71 | GSE3039\_CD4\_TCELL\_VS\_ALPHABETA\_CD8\_TCELL\_UP | MSigDB lists | GSE3039\_CD4\_TCELL\_VS\_ALPHABETA\_CD8\_TCELL\_UP | 41 | 5 | 4161 | 170 | Mctp,MCU,Glut1,Meltrin,dnc | | 2.445e-02 | -3.71 | GSE17721\_POLYIC\_VS\_GARDIQUIMOD\_16H\_BMDC\_UP | MSigDB lists | GSE17721\_POLYIC\_VS\_GARDIQUIMOD\_16H\_BMDC\_UP | 41 | 5 | 4161 | 170 | KrT95D,Spf45,scramb1,Cadps,EcR | | 2.445e-02 | -3.71 | chr16q22 | MSigDB lists | chr16q22 | 41 | 5 | 4161 | 170 | CG8272,kcc,Dlic,fus,Fhos | | 2.445e-02 | -3.71 | GSE40666\_NAIVE\_VS\_EFFECTOR\_CD8\_TCELL\_UP | MSigDB lists | GSE40666\_NAIVE\_VS\_EFFECTOR\_CD8\_TCELL\_UP | 41 | 5 | 4161 | 170 | CG5522,Ucp4A,MCU,KrT95D,pod1 | | 2.453e-02 | -3.71 | CG34353 (uncharacterized protein) | protein interactions | 5740590 | 5 | 2 | 9204 | 481 | CG32264,CG14634 | | 2.453e-02 | -3.71 | x16 (x16 splicing factor) | protein interactions | 33967 | 5 | 2 | 9204 | 481 | rictor,Doa | | 2.453e-02 | -3.71 | RpII18 (RNA polymerase II 18kD subunit) | protein interactions | 48312 | 5 | 2 | 9204 | 481 | CG8117,CG10164 | | 2.453e-02 | -3.71 | CG1785 (uncharacterized protein) | protein interactions | 31810 | 5 | 2 | 9204 | 481 | foxo,rictor | | 2.453e-02 | -3.71 | Tom (Twin of m4) | protein interactions | 39619 | 5 | 2 | 9204 | 481 | Su(H),tin | | 2.453e-02 | -3.71 | bap (bagpipe) | protein interactions | 42537 | 5 | 2 | 9204 | 481 | Antp,Ubx | | 2.453e-02 | -3.71 | Irc (Immune-regulated catalase) | protein interactions | 42049 | 5 | 2 | 9204 | 481 | Cat,Treh | | 2.453e-02 | -3.71 | CG9915 (uncharacterized protein) | protein interactions | 32593 | 5 | 2 | 9204 | 481 | rictor,Doa | | 2.453e-02 | -3.71 | Dnah3 (dynein, axonemal, heavy chain 3) | protein interactions | 38586 | 5 | 2 | 9204 | 481 | flw,sgg | | 2.453e-02 | -3.71 | Etl1 (Etl1) | protein interactions | 34311 | 5 | 2 | 9204 | 481 | Syx1A,sgg | | 2.453e-02 | -3.71 | GstE3 (Glutathione S transferase E3) | protein interactions | 37108 | 5 | 2 | 9204 | 481 | tamo,GATAe | | 2.453e-02 | -3.71 | CG3071 (uncharacterized protein) | protein interactions | 31213 | 5 | 2 | 9204 | 481 | Mbs,ZnT63C | | 2.453e-02 | -3.71 | Nos (Nitric oxide synthase) | protein interactions | 34495 | 5 | 2 | 9204 | 481 | Spf45,foxo | | 2.453e-02 | -3.71 | Claspin (claspin) | protein interactions | 326205 | 5 | 2 | 9204 | 481 | rictor,foxo | | 2.453e-02 | -3.71 | sNPF-R (short neuropeptide F receptor) | protein interactions | 40195 | 5 | 2 | 9204 | 481 | mnb,CrebB | | 2.456e-02 | -3.71 | Glycogen breakdown (glycogenolysis) | REACTOME pathways | R-DME-70221 | 13 | 3 | 3923 | 197 | PhKgamma,Cam,GlyP | | 2.457e-02 | -3.71 | SH3\_domain | interpro domains | IPR001452 | 69 | 8 | 11383 | 585 | p130CAS,Ca-beta,Stacl,RhoGEF3,sdt,cindr,Src42A,Exn | | 2.473e-02 | -3.70 | EF-hand\_1 | pfam domains | PF00036 | 5 | 2 | 10480 | 550 | Frq1,Frq2 | | 2.473e-02 | -3.70 | Frizzled | pfam domains | PF01534 | 5 | 2 | 10480 | 550 | fz2,fz4 | | 2.473e-02 | -3.70 | EFG\_II | pfam domains | PF14492 | 5 | 2 | 10480 | 550 | CG4849,CG33158 | | 2.473e-02 | -3.70 | Sec7 | pfam domains | PF01369 | 5 | 2 | 10480 | 550 | step,siz | | 2.473e-02 | -3.70 | 7TM\_GPCR\_Srw | pfam domains | PF10324 | 5 | 2 | 10480 | 550 | CG13229,MsR2 | | 2.473e-02 | -3.70 | Piwi | pfam domains | PF02171 | 5 | 2 | 10480 | 550 | AGO3,AGO2 | | 2.474e-02 | -3.70 | CYTOCHROME\_P450 | prosite domains | PS00086 | 80 | 10 | 6243 | 386 | Cyp4p2,Cyp304a1,Cyp28d1,Cyp6a13,Cyp309a1,Cyp6u1,Cyp28d2,spok,Cyp12a4,shd | | 2.483e-02 | -3.70 | positive regulation of cell cycle process | biological process | GO:0090068 | 47 | 6 | 12345 | 597 | bru1,foxo,Su(H),Tor,CG12200,sgg | | 2.494e-02 | -3.69 | GO\_EPITHELIUM\_DEVELOPMENT | MSigDB lists | GO\_EPITHELIUM\_DEVELOPMENT | 193 | 14 | 4161 | 170 | Su(H),Sod1,TfAP-2,sdt,CG17883,Flo2,CG17387,fz2,dnc,Ser,dpp,Cat,Pak,dsh | | 2.504e-02 | -3.69 | lymph gland development | biological process | GO:0048542 | 60 | 7 | 12345 | 597 | ths,Tor,Raf,dpp,Ser,Antp,tin | | 2.521e-02 | -3.68 | sexual reproduction | biological process | GO:0019953 | 1300 | 78 | 12345 | 597 | Eip74EF,mamo,Iswi,msi,can,cic,DNaseII,CG7483,lncRNA:flam,Ebp,Mst87F,Octbeta2R,bol,bru1,Fad2,flw,retn,Acp62F,drd,lncRNA:CR43484,fru,Doa,Nlg2,JhI-26,bun,Inx2,CG42355,lncRNA:CR44420,fs(1)N,cindr,lncRNA:CR44585,scrib,lncRNA:CR43802,EcR,Sh,mid,rols,csw,egh,tld,kek1,upd3,Gr39a,mGluR,kay,Tor,RhoL,bbg,msps,polybromo,sgg,spd-2,ex,Khc,Mos,lncRNA:CR43753,Dlic,wbl,Pde1c,shep,chb,lncRNA:Hsromega,Mrtf,dsh,aust,tin,Gr22c,Msp300,Raf,TfAP-2,shd,dnc,AGO2,shg,opa,Obp56h,dany,dpp | | 2.521e-02 | -3.68 | multi-organism reproductive process | biological process | GO:0044703 | 1300 | 78 | 12345 | 597 | Acp62F,drd,lncRNA:CR43484,fru,Doa,lncRNA:flam,CG7483,Ebp,Mst87F,bru1,bol,Octbeta2R,Fad2,flw,retn,cic,DNaseII,Eip74EF,Iswi,mamo,msi,can,Sh,mid,csw,rols,egh,scrib,lncRNA:CR43802,EcR,CG42355,Inx2,lncRNA:CR44420,fs(1)N,cindr,lncRNA:CR44585,Nlg2,JhI-26,bun,lncRNA:CR43753,Dlic,Khc,Mos,Gr39a,mGluR,kay,Tor,polybromo,bbg,msps,RhoL,sgg,ex,spd-2,kek1,tld,upd3,shg,opa,Obp56h,dany,dpp,shd,dnc,AGO2,lncRNA:Hsromega,Mrtf,tin,dsh,aust,Gr22c,Raf,Msp300,TfAP-2,wbl,Pde1c,shep,chb | | 2.533e-02 | -3.68 | developmental process involved in reproduction | biological process | GO:0003006 | 928 | 58 | 12345 | 597 | bun,chb,chinmo,wbl,TfAP-2,Raf,Msp300,cindr,fs(1)N,dsh,tin,rib,Mrtf,Inx2,CG42355,lncRNA:Hsromega,EcR,AGO2,shd,scrib,dpp,egh,rols,csw,mid,shg,can,msi,tnc,raw,kek1,tld,upd3,Iswi,Eip74EF,DNaseII,cic,sgg,ex,polybromo,msps,bbg,RhoL,Tor,fz2,kay,retn,Mos,Khc,flw,bru1,bol,CG7483,lncRNA:flam,Dlic,Doa,fru,drd | | 2.535e-02 | -3.68 | REACTOME\_PHOSPHOLIPASE\_C\_MEDIATED\_CASCADE | MSigDB lists | REACTOME\_PHOSPHOLIPASE\_C\_MEDIATED\_CASCADE | 16 | 3 | 4161 | 170 | Pkc98E,Pkc53E,Pde1c | | 2.535e-02 | -3.68 | GO\_PERIPHERAL\_NERVOUS\_SYSTEM\_DEVELOPMENT | MSigDB lists | GO\_PERIPHERAL\_NERVOUS\_SYSTEM\_DEVELOPMENT | 16 | 3 | 4161 | 170 | Sod1,sdt,TfAP-2 | | 2.535e-02 | -3.68 | GSE17974\_IL4\_AND\_ANTI\_IL12\_VS\_UNTREATED\_24H\_ACT\_CD4\_TCELL\_DN | MSigDB lists | GSE17974\_IL4\_AND\_ANTI\_IL12\_VS\_UNTREATED\_24H\_ACT\_CD4\_TCELL\_DN | 16 | 3 | 4161 | 170 | fz2,Iswi,scramb1 | | 2.535e-02 | -3.68 | GO\_REGULATION\_OF\_STRIATED\_MUSCLE\_CELL\_DIFFERENTIATION | MSigDB lists | GO\_REGULATION\_OF\_STRIATED\_MUSCLE\_CELL\_DIFFERENTIATION | 16 | 3 | 4161 | 170 | Tor,dpp,Su(H) | | 2.535e-02 | -3.68 | GO\_NEUROMUSCULAR\_JUNCTION | MSigDB lists | GO\_NEUROMUSCULAR\_JUNCTION | 16 | 3 | 4161 | 170 | unc-13,sky,Ank | | 2.535e-02 | -3.68 | GO\_TISSUE\_REMODELING | MSigDB lists | GO\_TISSUE\_REMODELING | 16 | 3 | 4161 | 170 | Octbeta2R,Ser,Su(H) | | 2.535e-02 | -3.68 | GO\_POSITIVE\_REGULATION\_OF\_CARBOHYDRATE\_METABOLIC\_PROCESS | MSigDB lists | GO\_POSITIVE\_REGULATION\_OF\_CARBOHYDRATE\_METABOLIC\_PROCESS | 16 | 3 | 4161 | 170 | Ptp61F,PhKgamma,Pkc98E | | 2.535e-02 | -3.68 | KIM\_LIVER\_CANCER\_POOR\_SURVIVAL\_DN | MSigDB lists | KIM\_LIVER\_CANCER\_POOR\_SURVIVAL\_DN | 16 | 3 | 4161 | 170 | CG3961,ZnT63C,CG5493 | | 2.535e-02 | -3.68 | FRIDMAN\_SENESCENCE\_UP | MSigDB lists | FRIDMAN\_SENESCENCE\_UP | 16 | 3 | 4161 | 170 | CG3376,TfAP-2,Sod1 | | 2.535e-02 | -3.68 | GO\_REGULATION\_OF\_LIPASE\_ACTIVITY | MSigDB lists | GO\_REGULATION\_OF\_LIPASE\_ACTIVITY | 16 | 3 | 4161 | 170 | EcR,Pkc98E,LRP1 | | 2.535e-02 | -3.68 | KEGG\_GLIOMA | MSigDB lists | KEGG\_GLIOMA | 16 | 3 | 4161 | 170 | Raf,Tor,Pkc53E | | 2.535e-02 | -3.68 | REACTOME\_PKB\_MEDIATED\_EVENTS | MSigDB lists | REACTOME\_PKB\_MEDIATED\_EVENTS | 16 | 3 | 4161 | 170 | Tor,eIF4E7,Stlk | | 2.535e-02 | -3.68 | GO\_NEGATIVE\_REGULATION\_OF\_CATION\_TRANSMEMBRANE\_TRANSPORT | MSigDB lists | GO\_NEGATIVE\_REGULATION\_OF\_CATION\_TRANSMEMBRANE\_TRANSPORT | 16 | 3 | 4161 | 170 | Ank,ZnT63C,Pkc98E | | 2.535e-02 | -3.68 | MORF\_RAB3A | MSigDB lists | MORF\_RAB3A | 16 | 3 | 4161 | 170 | Spf45,Nlg2,Sh | | 2.535e-02 | -3.68 | GO\_POSITIVE\_REGULATION\_OF\_TRANSPORTER\_ACTIVITY | MSigDB lists | GO\_POSITIVE\_REGULATION\_OF\_TRANSPORTER\_ACTIVITY | 16 | 3 | 4161 | 170 | AstA-R1,Dys,Ank | | 2.535e-02 | -3.68 | snRNA binding | molecular function | GO:0017069 | 24 | 4 | 12176 | 580 | CG4849,scaRNA:PsiU2-35.45,CG1239,scaRNA:MeU2-C41 | | 2.535e-02 | -3.68 | growth factor receptor binding | molecular function | GO:0070851 | 24 | 4 | 12176 | 580 | ths,kek1,Idgf6,bnl | | 2.537e-02 | -3.67 | HMG\_box\_dom\_sf | interpro domains | IPR036910 | 33 | 5 | 11383 | 585 | polybromo,Sox102F,CG42355,cic,bbx | | 2.540e-02 | -3.67 | detection of visible light | biological process | GO:0009584 | 35 | 5 | 12345 | 597 | Gprk1,ninaD,Cam,rdgA,Sh | | 2.540e-02 | -3.67 | synaptic growth at neuromuscular junction | biological process | GO:0051124 | 35 | 5 | 12345 | 597 | Nlg2,sgg,Elp3,cpx,kay | | 2.545e-02 | -3.67 | Exo84 (Exocyst 84) | protein interactions | 43163 | 22 | 4 | 9204 | 481 | unc-13,htk,cindr,Msp300 | | 2.546e-02 | -3.67 | GSE43955\_TH0\_VS\_TGFB\_IL6\_TH17\_ACT\_CD4\_TCELL\_60H\_UP | MSigDB lists | GSE43955\_TH0\_VS\_TGFB\_IL6\_TH17\_ACT\_CD4\_TCELL\_60H\_UP | 56 | 6 | 4161 | 170 | Sh,Tor,Octbeta2R,CG9132,Flo2,Nmdar2 | | 2.546e-02 | -3.67 | GSE17721\_CTRL\_VS\_LPS\_0.5H\_BMDC\_DN | MSigDB lists | GSE17721\_CTRL\_VS\_LPS\_0.5H\_BMDC\_DN | 56 | 6 | 4161 | 170 | Nha1,Ptp61F,Stlk,Uros1,CG17883,PhKgamma | | 2.546e-02 | -3.67 | GSE8921\_UNSTIM\_VS\_TLR1\_2\_STIM\_MONOCYTE\_6H\_UP | MSigDB lists | GSE8921\_UNSTIM\_VS\_TLR1\_2\_STIM\_MONOCYTE\_6H\_UP | 56 | 6 | 4161 | 170 | CG3842,Ucp4A,Raf,Syx1A,CG15093,Stt3B | | 2.546e-02 | -3.67 | GO\_MUSCLE\_SYSTEM\_PROCESS | MSigDB lists | GO\_MUSCLE\_SYSTEM\_PROCESS | 56 | 6 | 4161 | 170 | Sod1,dnc,Tor,AstA-R1,Octbeta2R,Dys | | 2.546e-02 | -3.67 | GO\_REGULATION\_OF\_HORMONE\_SECRETION | MSigDB lists | GO\_REGULATION\_OF\_HORMONE\_SECRETION | 56 | 6 | 4161 | 170 | Pkc53E,Pkc98E,csw,MCU,Pde8,Syx1A | | 2.546e-02 | -3.67 | GSE20715\_0H\_VS\_24H\_OZONE\_TLR4\_KO\_LUNG\_DN | MSigDB lists | GSE20715\_0H\_VS\_24H\_OZONE\_TLR4\_KO\_LUNG\_DN | 56 | 6 | 4161 | 170 | CG5493,eRF3,dpp,csw,fus,eIF4E7 | | 2.559e-02 | -3.67 | GO\_CALCIUM\_ION\_TRANSMEMBRANE\_TRANSPORTER\_ACTIVITY | MSigDB lists | GO\_CALCIUM\_ION\_TRANSMEMBRANE\_TRANSPORTER\_ACTIVITY | 28 | 4 | 4161 | 170 | Ca-beta,Frq2,Frq1,MCU | | 2.559e-02 | -3.67 | CHIARADONNA\_NEOPLASTIC\_TRANSFORMATION\_CDC25\_UP | MSigDB lists | CHIARADONNA\_NEOPLASTIC\_TRANSFORMATION\_CDC25\_UP | 28 | 4 | 4161 | 170 | CG5493,Meltrin,Ank,scramb1 | | 2.559e-02 | -3.67 | LABBE\_TARGETS\_OF\_TGFB1\_AND\_WNT3A\_UP | MSigDB lists | LABBE\_TARGETS\_OF\_TGFB1\_AND\_WNT3A\_UP | 28 | 4 | 4161 | 170 | GlyP,Tsp96F,Ser,Act87E | | 2.559e-02 | -3.67 | GSE7509\_FCGRIIB\_VS\_TNFA\_IL1B\_IL6\_PGE\_STIM\_DC\_DN | MSigDB lists | GSE7509\_FCGRIIB\_VS\_TNFA\_IL1B\_IL6\_PGE\_STIM\_DC\_DN | 28 | 4 | 4161 | 170 | Root,Awh,rdgA,Doa | | 2.559e-02 | -3.67 | GO\_REGULATION\_OF\_MUSCLE\_TISSUE\_DEVELOPMENT | MSigDB lists | GO\_REGULATION\_OF\_MUSCLE\_TISSUE\_DEVELOPMENT | 28 | 4 | 4161 | 170 | dpp,Tor,Su(H),Octbeta2R | | 2.559e-02 | -3.67 | GO\_CHANNEL\_REGULATOR\_ACTIVITY | MSigDB lists | GO\_CHANNEL\_REGULATOR\_ACTIVITY | 28 | 4 | 4161 | 170 | Syx1A,mGluR,ZnT63C,Octbeta2R | | 2.559e-02 | -3.67 | chr17q23 | MSigDB lists | chr17q23 | 28 | 4 | 4161 | 170 | CG6418,wake,Rbp6,Stlk | | 2.583e-02 | -3.66 | HMG\_box | pfam domains | PF00505 | 22 | 4 | 10480 | 550 | Sox102F,bbx,cic,polybromo | | 2.583e-02 | -3.66 | EGF | pfam domains | PF00008 | 22 | 4 | 10480 | 550 | Ser,trol,LRP1,CadN2 | | 2.620e-02 | -3.64 | CATTTCA\_MIR203 | MSigDB lists | CATTTCA\_MIR203 | 88 | 8 | 4161 | 170 | dnc,rdgA,flw,CG4341,Pkc53E,MESK2,Rat1,eIF4E7 | | 2.621e-02 | -3.64 | Ankyrin\_rpt | interpro domains | IPR002110 | 83 | 9 | 11383 | 585 | wake,CG6966,CG42672,Ank2,Ank,rdgA,CG41099,Mbs,rols | | 2.625e-02 | -3.64 | Iono\_rcpt\_met | interpro domains | IPR001508 | 13 | 3 | 11383 | 585 | Nmdar2,GluRIIC,KaiR1D | | 2.625e-02 | -3.64 | RA\_dom | interpro domains | IPR000159 | 13 | 3 | 11383 | 585 | Myo81F,spri,wake | | 2.648e-02 | -3.63 | GO\_SECRETORY\_GRANULE | MSigDB lists | GO\_SECRETORY\_GRANULE | 72 | 7 | 4161 | 170 | CG3376,Frq1,Syx1A,Flo2,sxc,Frq2,Sod1 | | 2.656e-02 | -3.63 | organic hydroxy compound metabolic process | biological process | GO:1901615 | 163 | 14 | 12345 | 597 | RhoGEF3,Ipk1,Flo2,RhoL,woc,foxo,CG33090,AOX1,D2hgdh,IP3K2,AANAT1,spok,Exn,CG3842 | | 2.659e-02 | -3.63 | Notch signaling pathway | KEGG pathways | dme04330 | 22 | 4 | 3198 | 170 | Ser,numb,dsh,Su(H) | | 2.659e-02 | -3.63 | Notch signaling pathway | KEGG pathways | ko04330 | 22 | 4 | 3198 | 170 | numb,Ser,dsh,Su(H) | | 2.661e-02 | -3.63 | LDLRA\_1 | prosite domains | PS01209 | 28 | 5 | 6243 | 386 | trol,LpR2,CG32432,CG31221,LRP1 | | 2.661e-02 | -3.63 | GO\_ENZYME\_ACTIVATOR\_ACTIVITY | MSigDB lists | GO\_ENZYME\_ACTIVATOR\_ACTIVITY | 105 | 9 | 4161 | 170 | CG34408,Stlk,sky,CG17883,RhoGAP1A,sxc,dpp,REG,Pkc98E | | 2.662e-02 | -3.63 | multi-organism behavior | biological process | GO:0051705 | 24 | 4 | 12345 | 597 | Tk,TfAP-2,fru,Nlg2 | | 2.662e-02 | -3.63 | inorganic cation import across plasma membrane | biological process | GO:0098659 | 24 | 4 | 12345 | 597 | nrv2,CG33310,KaiR1D,Irk3 | | 2.662e-02 | -3.63 | inorganic ion import across plasma membrane | biological process | GO:0099587 | 24 | 4 | 12345 | 597 | CG33310,Irk3,KaiR1D,nrv2 | | 2.674e-02 | -3.62 | DACOSTA\_UV\_RESPONSE\_VIA\_ERCC3\_DN | MSigDB lists | DACOSTA\_UV\_RESPONSE\_VIA\_ERCC3\_DN | 291 | 19 | 4161 | 170 | Su(H),Iswi,Pkc53E,sxc,eIF4E7,msps,IntS3,LpR2,CG6418,step,CG42672,CG40045,dsd,scramb1,TfIIFbeta,CG4301,Dys,chb,Ptp61F | | 2.676e-02 | -3.62 | RA | smart domains | SM00314 | 11 | 3 | 5086 | 315 | wake,spri,Myo81F | | 2.679e-02 | -3.62 | multicellular organismal reproductive process | biological process | GO:0048609 | 1285 | 77 | 12345 | 597 | AGO2,dnc,shd,opa,dany,Obp56h,dpp,shg,shep,chb,wbl,Pde1c,Raf,Msp300,TfAP-2,lncRNA:Hsromega,Mrtf,dsh,aust,tin,Gr22c,Khc,Mos,Dlic,lncRNA:CR43753,tld,upd3,kek1,Tor,msps,bbg,RhoL,polybromo,sgg,ex,Gr39a,mGluR,kay,EcR,scrib,lncRNA:CR43802,mid,csw,rols,egh,Sh,Nlg2,JhI-26,bun,cindr,lncRNA:CR44585,Inx2,CG42355,lncRNA:CR44420,fs(1)N,Fad2,flw,retn,lncRNA:flam,CG7483,Ebp,bol,bru1,Mst87F,Octbeta2R,fru,Doa,Acp62F,lncRNA:CR43484,drd,msi,can,Eip74EF,mamo,Iswi,cic,DNaseII | | 2.687e-02 | -3.62 | GSE17721\_LPS\_VS\_GARDIQUIMOD\_2H\_BMDC\_UP | MSigDB lists | GSE17721\_LPS\_VS\_GARDIQUIMOD\_2H\_BMDC\_UP | 42 | 5 | 4161 | 170 | AGO3,Sh,Ser,kcc,scramb1 | | 2.687e-02 | -3.62 | GO\_LIPOSACCHARIDE\_METABOLIC\_PROCESS | MSigDB lists | GO\_LIPOSACCHARIDE\_METABOLIC\_PROCESS | 42 | 5 | 4161 | 170 | CG3376,CG33090,PIG-O,Cln3,PIG-G | | 2.687e-02 | -3.62 | GSE5589\_WT\_VS\_IL6\_KO\_LPS\_STIM\_MACROPHAGE\_45MIN\_DN | MSigDB lists | GSE5589\_WT\_VS\_IL6\_KO\_LPS\_STIM\_MACROPHAGE\_45MIN\_DN | 42 | 5 | 4161 | 170 | Pfdn4,Frq2,Frq1,CG3860,kcc | | 2.687e-02 | -3.62 | GO\_NEGATIVE\_REGULATION\_OF\_NEURON\_DEATH | MSigDB lists | GO\_NEGATIVE\_REGULATION\_OF\_NEURON\_DEATH | 42 | 5 | 4161 | 170 | KaiR1D,Raf,Sod1,Cln3,LRP1 | | 2.687e-02 | -3.62 | GSE13306\_RA\_VS\_UNTREATED\_TCONV\_DN | MSigDB lists | GSE13306\_RA\_VS\_UNTREATED\_TCONV\_DN | 42 | 5 | 4161 | 170 | Hr96,TfIIFbeta,Odc1,CG4849,TfAP-2 | | 2.687e-02 | -3.62 | GO\_DIVALENT\_INORGANIC\_CATION\_TRANSMEMBRANE\_TRANSPORTER\_ACTIVITY | MSigDB lists | GO\_DIVALENT\_INORGANIC\_CATION\_TRANSMEMBRANE\_TRANSPORTER\_ACTIVITY | 42 | 5 | 4161 | 170 | Frq2,Ca-beta,ZnT63C,MCU,Frq1 | | 2.687e-02 | -3.62 | COREBINDINGFACTOR\_Q6 | MSigDB lists | COREBINDINGFACTOR\_Q6 | 42 | 5 | 4161 | 170 | Ank,Glut1,CG12547,eIF2Bgamma,Rbp6 | | 2.687e-02 | -3.62 | GSE12366\_NAIVE\_VS\_MEMORY\_BCELL\_DN | MSigDB lists | GSE12366\_NAIVE\_VS\_MEMORY\_BCELL\_DN | 42 | 5 | 4161 | 170 | CG5522,dnc,CG40045,Tsp96F,Nha1 | | 2.694e-02 | -3.61 | TORC2 complex | cellular component | GO:0031932 | 6 | 2 | 11940 | 538 | Tor,rictor | | 2.699e-02 | -3.61 | regulation of synapse structure or activity | biological process | GO:0050803 | 179 | 15 | 12345 | 597 | sgg,Tor,rictor,Ank2,Mob2,Sh,Pak,mnb,Nlg2,Sod1,Syn2,Src42A,KaiR1D,Octbeta2R,scrib | | 2.699e-02 | -3.61 | kis (kismet) | protein interactions | 33185 | 33 | 5 | 9204 | 481 | Dys,foxo,Raf,Syx1A,rictor | | 2.704e-02 | -3.61 | regulation of catabolic process | biological process | GO:0009894 | 227 | 18 | 12345 | 597 | REG,EcR,bol,Fer2,Doa,CG3530,Marf1,Sod1,Cpsf160,foxo,sgg,AGO3,GlyP,Raf,Tor,Pomp,CG42797,CG18304 | | 2.710e-02 | -3.61 | leg disc development | biological process | GO:0035218 | 103 | 10 | 12345 | 597 | hth,Pak,polybromo,mid,dpp,TfAP-2,B-H2,Exn,Ubx,Ser | | 2.715e-02 | -3.61 | striated muscle cell development | biological process | GO:0055002 | 61 | 7 | 12345 | 597 | Khc,rols,Actn,Act88F,Mlp84B,bru1,Prm | | 2.735e-02 | -3.60 | potassium ion transmembrane transporter activity | molecular function | GO:0015079 | 62 | 7 | 12176 | 580 | GluRIIC,Irk3,nrv2,KaiR1D,kcc,CG33310,Sh | | 2.740e-02 | -3.60 | CG3630 (uncharacterized protein) | protein interactions | 31189 | 13 | 3 | 9204 | 481 | Sod1,CG7017,Cam | | 2.740e-02 | -3.60 | Doc2 (Dorsocross2) | protein interactions | 39038 | 13 | 3 | 9204 | 481 | Antp,Ubx,tin | | 2.740e-02 | -3.60 | mir-14 (mir-14 stem loop) | protein interactions | 12798384 | 13 | 3 | 9204 | 481 | IP3K2,EcR,esg | | 2.740e-02 | -3.60 | Ir54a (Ionotropic receptor 54a) | protein interactions | 37022 | 13 | 3 | 9204 | 481 | Fhos,ana,CG15894 | | 2.740e-02 | -3.60 | POSH (Plenty of SH3s) | protein interactions | 36990 | 13 | 3 | 9204 | 481 | ex,nompB,Dys | | 2.748e-02 | -3.59 | sterol homeostasis | biological process | GO:0055092 | 14 | 3 | 12345 | 597 | EcR,mag,Hr96 | | 2.748e-02 | -3.59 | regulation of membrane lipid distribution | biological process | GO:0097035 | 14 | 3 | 12345 | 597 | CG4301,scramb1,CG9981 | | 2.748e-02 | -3.59 | haltere disc development | biological process | GO:0035216 | 14 | 3 | 12345 | 597 | Ubx,hth,Syx1A | | 2.748e-02 | -3.59 | muscle cell fate commitment | biological process | GO:0042693 | 14 | 3 | 12345 | 597 | numb,Ubx,Antp | | 2.748e-02 | -3.59 | regulation of phosphatidylinositol 3-kinase signaling | biological process | GO:0014066 | 14 | 3 | 12345 | 597 | Lpin,Ptp61F,step | | 2.753e-02 | -3.59 | GSE40274\_CTRL\_VS\_FOXP3\_AND\_HELIOS\_TRANSDUCED\_ACTIVATED\_CD4\_TCELL\_UP | MSigDB lists | GSE40274\_CTRL\_VS\_FOXP3\_AND\_HELIOS\_TRANSDUCED\_ACTIVATED\_CD4\_TCELL\_UP | 57 | 6 | 4161 | 170 | CNT1,Syx1A,sws,CG41099,Glut1,CG17387 | | 2.753e-02 | -3.59 | GSE3982\_CTRL\_VS\_LPS\_48H\_DC\_UP | MSigDB lists | GSE3982\_CTRL\_VS\_LPS\_48H\_DC\_UP | 57 | 6 | 4161 | 170 | DNaseII,CG3860,LpR2,CG3530,IP3K2,CG5522 | | 2.753e-02 | -3.59 | HNF4\_01\_B | MSigDB lists | HNF4\_01\_B | 57 | 6 | 4161 | 170 | CG3961,fus,CG14480,sdt,LRP1,Act87E | | 2.753e-02 | -3.59 | HLF\_01 | MSigDB lists | HLF\_01 | 57 | 6 | 4161 | 170 | dnc,Dys,Octbeta2R,bru1,chb,app | | 2.753e-02 | -3.59 | GSE14308\_TH1\_VS\_INDUCED\_TREG\_UP | MSigDB lists | GSE14308\_TH1\_VS\_INDUCED\_TREG\_UP | 57 | 6 | 4161 | 170 | Ca-beta,RpL38,CG42672,Der-1,Rat1,Cat | | 2.753e-02 | -3.59 | GSE17721\_CTRL\_VS\_POLYIC\_0.5H\_BMDC\_DN | MSigDB lists | GSE17721\_CTRL\_VS\_POLYIC\_0.5H\_BMDC\_DN | 57 | 6 | 4161 | 170 | PhKgamma,CG17883,Stlk,Uros1,Nha1,Syx1A | | 2.753e-02 | -3.59 | SP1\_01 | MSigDB lists | SP1\_01 | 57 | 6 | 4161 | 170 | Pak,LpR2,LRP1,sdt,Iswi,CG17698 | | 2.753e-02 | -3.59 | GO\_RECEPTOR\_COMPLEX | MSigDB lists | GO\_RECEPTOR\_COMPLEX | 57 | 6 | 4161 | 170 | dpp,LRP1,LpR2,Nmdar2,KaiR1D,Octbeta2R | | 2.762e-02 | -3.59 | ovarian follicle cell development | biological process | GO:0030707 | 327 | 24 | 12345 | 597 | dsh,fs(1)N,kay,Mrtf,ex,sgg,cic,Raf,cindr,bbg,polybromo,RhoL,bun,shg,egh,dpp,mid,csw,rols,lncRNA:flam,shd,scrib,Khc,EcR | | 2.763e-02 | -3.59 | GO\_REGULATION\_OF\_INTRACELLULAR\_TRANSPORT | MSigDB lists | GO\_REGULATION\_OF\_INTRACELLULAR\_TRANSPORT | 177 | 13 | 4161 | 170 | Ank,CG17883,Tor,csw,sxc,Dys,CCAP-R,Raf,Syx1A,CG8184,dnc,fz2,Pkc98E | | 2.782e-02 | -3.58 | negative regulation of phosphate metabolic process | biological process | GO:0045936 | 89 | 9 | 12345 | 597 | chinmo,flw,Ptp61F,kek1,raw,et,sws,sgg,mnb | | 2.782e-02 | -3.58 | negative regulation of phosphorus metabolic process | biological process | GO:0010563 | 89 | 9 | 12345 | 597 | sws,sgg,mnb,Ptp61F,flw,chinmo,kek1,et,raw | | 2.788e-02 | -3.58 | fly chr50E4-50E4\|2-69 cM | chromosome location | fly chr50E4-50E4\|2-69 cM | 8 | 2 | 25010 | 845 | CG8468,mir-1016 | | 2.788e-02 | -3.58 | fly chr99E3-99E3\|3-100 | chromosome location | fly chr99E3-99E3\|3-100 | 8 | 2 | 25010 | 845 | CG9682,CG9737 | | 2.788e-02 | -3.58 | fly chr50E4-50E4\|2-69 | chromosome location | fly chr50E4-50E4\|2-69 | 8 | 2 | 25010 | 845 | CG8468,mir-1016 | | 2.788e-02 | -3.58 | fly chr100A6-100A6\|3-101 cM | chromosome location | fly chr100A6-100A6\|3-101 cM | 8 | 2 | 25010 | 845 | cindr,CG12071 | | 2.788e-02 | -3.58 | fly chr68A1-68A1\|3-34 cM | chromosome location | fly chr68A1-68A1\|3-34 cM | 8 | 2 | 25010 | 845 | CG14147,Fad2 | | 2.788e-02 | -3.58 | fly chr100A6-100A6\|3-101 | chromosome location | fly chr100A6-100A6\|3-101 | 8 | 2 | 25010 | 845 | CG12071,cindr | | 2.788e-02 | -3.58 | fly chr99E3-99E3\|3-100 cM | chromosome location | fly chr99E3-99E3\|3-100 cM | 8 | 2 | 25010 | 845 | CG9737,CG9682 | | 2.788e-02 | -3.58 | fly chr68A1-68A1\|3-34 | chromosome location | fly chr68A1-68A1\|3-34 | 8 | 2 | 25010 | 845 | CG14147,Fad2 | | 2.804e-02 | -3.57 | GO\_SIGNAL\_TRANSDUCER\_ACTIVITY | MSigDB lists | GO\_SIGNAL\_TRANSDUCER\_ACTIVITY | 196 | 14 | 4161 | 170 | Sk1,fz2,Pkc98E,EcR,Octbeta2R,Pak,mGluR,Raf,AstA-R1,Nmdar2,CCAP-R,LpR2,KaiR1D,Hr96 | | 2.806e-02 | -3.57 | HMG\_box\_dom | interpro domains | IPR009071 | 23 | 4 | 11383 | 585 | bbx,cic,polybromo,Sox102F | | 2.807e-02 | -3.57 | Ankyrin\_rpt-contain\_sf | interpro domains | IPR036770 | 84 | 9 | 11383 | 585 | CG6966,CG42672,wake,Mbs,CG41099,rols,Ank2,Ank,rdgA | | 2.807e-02 | -3.57 | Ankyrin\_rpt-contain\_dom | interpro domains | IPR020683 | 84 | 9 | 11383 | 585 | wake,CG42672,CG6966,rdgA,Ank,Ank2,rols,Mbs,CG41099 | | 2.809e-02 | -3.57 | YOSHIMURA\_MAPK8\_TARGETS\_DN | MSigDB lists | YOSHIMURA\_MAPK8\_TARGETS\_DN | 141 | 11 | 4161 | 170 | mino,Ser,Cat,Src42A,sxc,CG40045,Acbp2,CG4849,kcc,Dlic,Odc1 | | 2.819e-02 | -3.57 | cell death | biological process | GO:0008219 | 180 | 15 | 12345 | 597 | CG6839,EcR,Eip74EF,IP3K2,sws,rst,dpp,DNaseII,Tor,Damm,CG5059,hdc,Fhos,Pak,Cam | | 2.824e-02 | -3.57 | - | gene3d domains | 2.60.40.150 | 49 | 7 | 4929 | 302 | Pkc53E,Syt12,Fife,RhoGAP100F,Mctp,unc-13,Pkc98E | | 2.837e-02 | -3.56 | maintenance of location in cell | biological process | GO:0051651 | 36 | 5 | 12345 | 597 | Ank,Msp300,p130CAS,Actn,by | | 2.837e-02 | -3.56 | epithelial cell migration, open tracheal system | biological process | GO:0007427 | 36 | 5 | 12345 | 597 | dpp,bnl,csw,esg,rib | | 2.837e-02 | -3.56 | larval locomotory behavior | biological process | GO:0008345 | 36 | 5 | 12345 | 597 | EcR,Khc,Nlg2,Sh,MsR2 | | 2.837e-02 | -3.56 | midgut development | biological process | GO:0007494 | 36 | 5 | 12345 | 597 | Antp,GATAe,shd,Ubx,opa | | 2.837e-02 | -3.56 | cellular monovalent inorganic cation homeostasis | biological process | GO:0030004 | 36 | 5 | 12345 | 597 | nrv2,Vha100-5,Raf,CG33310,Cln3 | | 2.846e-02 | -3.56 | Rac1 (Rac1) | protein interactions | 38146 | 124 | 12 | 9204 | 481 | RhoGAP18B,RhoGAP100F,Pak,shg,numb,dsh,spri,Mbs,Fhos,Raf,scrib,RhoGEF3 | | 2.850e-02 | -3.56 | columnar/cuboidal epithelial cell development | biological process | GO:0002066 | 328 | 24 | 12345 | 597 | Raf,cindr,polybromo,RhoL,bbg,sgg,ex,cic,Mrtf,dsh,kay,fs(1)N,bun,mid,rols,csw,egh,dpp,shg,EcR,Khc,shd,lncRNA:flam,scrib | | 2.853e-02 | -3.56 | LDL\_receptor-like\_sf | interpro domains | IPR036055 | 34 | 5 | 11383 | 585 | LpR2,trol,CG31221,LRP1,CG32432 | | 2.876e-02 | -3.55 | GO\_VOLTAGE\_GATED\_ION\_CHANNEL\_ACTIVITY | MSigDB lists | GO\_VOLTAGE\_GATED\_ION\_CHANNEL\_ACTIVITY | 29 | 4 | 4161 | 170 | Frq2,Ca-beta,Sh,Frq1 | | 2.876e-02 | -3.55 | GO\_REGULATION\_OF\_FATTY\_ACID\_METABOLIC\_PROCESS | MSigDB lists | GO\_REGULATION\_OF\_FATTY\_ACID\_METABOLIC\_PROCESS | 29 | 4 | 4161 | 170 | Tor,Ugt301D1,EcR,ABCD | | 2.876e-02 | -3.55 | GO\_TRANSCRIPTION\_FACTOR\_ACTIVITY\_RNA\_POLYMERASE\_II\_CORE\_PROMOTER\_PROXIMAL\_REGION\_SEQUENCE\_SPECIFIC\_BINDING | MSigDB lists | GO\_TRANSCRIPTION\_FACTOR\_ACTIVITY\_RNA\_POLYMERASE\_II\_CORE\_PROMOTER\_PROXIMAL\_REGION\_SEQUENCE\_SPECIFIC\_BINDING | 29 | 4 | 4161 | 170 | scramb1,EcR,Su(H),TfAP-2 | | 2.876e-02 | -3.55 | GSE18791\_CTRL\_VS\_NEWCASTLE\_VIRUS\_DC\_10H\_DN | MSigDB lists | GSE18791\_CTRL\_VS\_NEWCASTLE\_VIRUS\_DC\_10H\_DN | 29 | 4 | 4161 | 170 | scramb1,Der-1,CG6966,Mlp84B | | 2.876e-02 | -3.55 | REACTOME\_DOWNSTREAM\_SIGNALING\_OF\_ACTIVATED\_FGFR | MSigDB lists | REACTOME\_DOWNSTREAM\_SIGNALING\_OF\_ACTIVATED\_FGFR | 29 | 4 | 4161 | 170 | Pde1c,Tor,Pkc98E,Pkc53E | | 2.876e-02 | -3.55 | GSE18791\_CTRL\_VS\_NEWCASTLE\_VIRUS\_DC\_12H\_DN | MSigDB lists | GSE18791\_CTRL\_VS\_NEWCASTLE\_VIRUS\_DC\_12H\_DN | 29 | 4 | 4161 | 170 | CG6966,Mlp84B,scramb1,unc-13 | | 2.876e-02 | -3.55 | GO\_NEGATIVE\_REGULATION\_OF\_RESPONSE\_TO\_WOUNDING | MSigDB lists | GO\_NEGATIVE\_REGULATION\_OF\_RESPONSE\_TO\_WOUNDING | 29 | 4 | 4161 | 170 | Ptp61F,chb,Octbeta2R,RhoGAP1A | | 2.876e-02 | -3.55 | GSE21360\_PRIMARY\_VS\_TERTIARY\_MEMORY\_CD8\_TCELL\_DN | MSigDB lists | GSE21360\_PRIMARY\_VS\_TERTIARY\_MEMORY\_CD8\_TCELL\_DN | 29 | 4 | 4161 | 170 | CG12121,Ptp61F,CG3376,CG6966 | | 2.876e-02 | -3.55 | GSE36476\_CTRL\_VS\_TSST\_ACT\_72H\_MEMORY\_CD4\_TCELL\_YOUNG\_UP | MSigDB lists | GSE36476\_CTRL\_VS\_TSST\_ACT\_72H\_MEMORY\_CD4\_TCELL\_YOUNG\_UP | 29 | 4 | 4161 | 170 | flw,RpL38,CG5522,Glut1 | | 2.876e-02 | -3.55 | GSE23114\_PERITONEAL\_CAVITY\_B1A\_BCELL\_VS\_SPLEEN\_BCELL\_UP | MSigDB lists | GSE23114\_PERITONEAL\_CAVITY\_B1A\_BCELL\_VS\_SPLEEN\_BCELL\_UP | 29 | 4 | 4161 | 170 | AGO3,Frq2,Frq1,DNaseII | | 2.876e-02 | -3.55 | GO\_PROTEASE\_BINDING | MSigDB lists | GO\_PROTEASE\_BINDING | 29 | 4 | 4161 | 170 | Der-1,LRP1,dsh,Flo2 | | 2.876e-02 | -3.55 | GSE36826\_WT\_VS\_IL1R\_KO\_SKIN\_STAPH\_AUREUS\_INF\_DN | MSigDB lists | GSE36826\_WT\_VS\_IL1R\_KO\_SKIN\_STAPH\_AUREUS\_INF\_DN | 29 | 4 | 4161 | 170 | Meltrin,fz2,scramb1,Octbeta2R | | 2.905e-02 | -3.54 | cell recognition | biological process | GO:0008037 | 119 | 11 | 12345 | 597 | trol,nrm,dsd,dpr12,dsh,fz2,Ptp99A,p130CAS,fru,mid,esn | | 2.907e-02 | -3.54 | WONG\_ADULT\_TISSUE\_STEM\_MODULE | MSigDB lists | WONG\_ADULT\_TISSUE\_STEM\_MODULE | 124 | 10 | 4161 | 170 | Pkc98E,Rbp6,LpR2,CG3860,Nha1,GlyP,Octbeta2R,Dys,eRF3,Ank | | 2.914e-02 | -3.54 | PEREZ\_TP53\_TARGETS | MSigDB lists | PEREZ\_TP53\_TARGETS | 197 | 14 | 4161 | 170 | bru1,Stlk,fus,Ank,RhoGAP1A,Src42A,IP3K2,Pgant6,AstA-R1,dsh,Octbeta2R,Root,rdgA,dpp | | 2.925e-02 | -3.53 | GGGYGTGNY\_UNKNOWN | MSigDB lists | GGGYGTGNY\_UNKNOWN | 160 | 12 | 4161 | 170 | Acbp2,CG17698,RhoGAP1A,sxc,pod1,Act87E,rdgA,Ser,Pde6,MCU,mGluR,Syx1A | | 2.933e-02 | -3.53 | Pkinase | pfam domains | PF00069 | 167 | 15 | 10480 | 550 | Pkc98E,Pkc53E,mnb,Pak,sgg,CG17698,PhKgamma,Mos,Nuak1,Doa,CG4839,Asator,Stlk,Strn-Mlck,Gprk1 | | 2.938e-02 | -3.53 | muscle cell development | biological process | GO:0055001 | 62 | 7 | 12345 | 597 | Prm,bru1,rols,Khc,Mlp84B,Act88F,Actn | | 2.943e-02 | -3.53 | GSE3982\_MAST\_CELL\_VS\_BASOPHIL\_DN | MSigDB lists | GSE3982\_MAST\_CELL\_VS\_BASOPHIL\_DN | 43 | 5 | 4161 | 170 | dnc,Act87E,CG9132,KaiR1D,Ptp61F | | 2.943e-02 | -3.53 | GSE2770\_IL4\_ACT\_VS\_ACT\_CD4\_TCELL\_2H\_DN | MSigDB lists | GSE2770\_IL4\_ACT\_VS\_ACT\_CD4\_TCELL\_2H\_DN | 43 | 5 | 4161 | 170 | CG4301,CG9132,scramb1,CG4341,CG8184 | | 2.943e-02 | -3.53 | SHEN\_SMARCA2\_TARGETS\_DN | MSigDB lists | SHEN\_SMARCA2\_TARGETS\_DN | 43 | 5 | 4161 | 170 | CG30036,Tor,CG3530,Cpsf160,Aldh-III | | 2.943e-02 | -3.53 | ATACTGT\_MIR144 | MSigDB lists | ATACTGT\_MIR144 | 43 | 5 | 4161 | 170 | bru1,Dys,eRF3,dnc,Iswi | | 2.943e-02 | -3.53 | GSE21546\_SAP1A\_KO\_VS\_SAP1A\_KO\_AND\_ELK1\_KO\_DP\_THYMOCYTES\_DN | MSigDB lists | GSE21546\_SAP1A\_KO\_VS\_SAP1A\_KO\_AND\_ELK1\_KO\_DP\_THYMOCYTES\_DN | 43 | 5 | 4161 | 170 | mino,Mlp84B,Pkc53E,step,IP3K2 | | 2.943e-02 | -3.53 | GSE4142\_NAIVE\_VS\_GC\_BCELL\_UP | MSigDB lists | GSE4142\_NAIVE\_VS\_GC\_BCELL\_UP | 43 | 5 | 4161 | 170 | Awh,PhKgamma,Cdep,CG3860,mRpS5 | | 2.943e-02 | -3.53 | GSE28726\_NAIVE\_VS\_ACTIVATED\_VA24NEG\_NKTCELL\_DN | MSigDB lists | GSE28726\_NAIVE\_VS\_ACTIVATED\_VA24NEG\_NKTCELL\_DN | 43 | 5 | 4161 | 170 | PhKgamma,Cadps,Argl,GlyP,Flo2 | | 2.943e-02 | -3.53 | GSE12963\_ENV\_NEF\_VS\_ENV\_NEF\_AND\_VPR\_DEFICIENT\_HIV1\_INF\_CD4\_TCELL\_UP | MSigDB lists | GSE12963\_ENV\_NEF\_VS\_ENV\_NEF\_AND\_VPR\_DEFICIENT\_HIV1\_INF\_CD4\_TCELL\_UP | 43 | 5 | 4161 | 170 | CG32196,GlyP,fus,CG41099,Pkc98E | | 2.943e-02 | -3.53 | IK2\_01 | MSigDB lists | IK2\_01 | 43 | 5 | 4161 | 170 | Ank,Act87E,LRP1,TfAP-2,CG33090 | | 2.943e-02 | -3.53 | GSE22886\_NAIVE\_CD4\_TCELL\_VS\_12H\_ACT\_TH1\_UP | MSigDB lists | GSE22886\_NAIVE\_CD4\_TCELL\_VS\_12H\_ACT\_TH1\_UP | 43 | 5 | 4161 | 170 | CG3376,Flo2,rdgA,Cln3,wbl | | 2.943e-02 | -3.53 | GO\_RESPONSE\_TO\_ETHANOL | MSigDB lists | GO\_RESPONSE\_TO\_ETHANOL | 43 | 5 | 4161 | 170 | Cat,Ugt301D1,CG5493,Pkc98E,Sod1 | | 2.943e-02 | -3.53 | GO\_GLIOGENESIS | MSigDB lists | GO\_GLIOGENESIS | 43 | 5 | 4161 | 170 | eIF2Bgamma,Sod1,csw,TfAP-2,sdt | | 2.943e-02 | -3.53 | GSE7460\_TCONV\_VS\_TREG\_THYMUS\_DN | MSigDB lists | GSE7460\_TCONV\_VS\_TREG\_THYMUS\_DN | 43 | 5 | 4161 | 170 | CG10126,Odc1,scramb1,Mlp84B,Meltrin | | 2.943e-02 | -3.53 | GSE360\_L\_DONOVANI\_VS\_L\_MAJOR\_MAC\_DN | MSigDB lists | GSE360\_L\_DONOVANI\_VS\_L\_MAJOR\_MAC\_DN | 43 | 5 | 4161 | 170 | GlyP,Pde1c,Ser,step,EcR | | 2.948e-02 | -3.52 | receptor ligand activity | molecular function | GO:0048018 | 106 | 10 | 12176 | 580 | ths,ITP,dpp,Ser,upd3,bnl,RYa,Neb-cGP,Tk,Gpb5 | | 2.948e-02 | -3.52 | phospholipid binding | molecular function | GO:0005543 | 106 | 10 | 12176 | 580 | cert,Pld,CG18659,dsh,melt,unc-13,Syt12,step,qvr,Gprk1 | | 2.950e-02 | -3.52 | SANSOM\_APC\_TARGETS\_DN | MSigDB lists | SANSOM\_APC\_TARGETS\_DN | 90 | 8 | 4161 | 170 | Meltrin,RhoGAP1A,Pgant6,sky,Ank,Sk1,LRP1,PCB | | 2.958e-02 | -3.52 | Socs36E (Suppressor of cytokine signaling at 36E) | protein interactions | 35085 | 23 | 4 | 9204 | 481 | upd3,CG18135,Tor,scrib | | 2.958e-02 | -3.52 | cora (coracle) | protein interactions | 37205 | 23 | 4 | 9204 | 481 | rictor,GluRIIC,nrv2,foxo | | 2.963e-02 | -3.52 | GO\_LYTIC\_VACUOLE\_MEMBRANE | MSigDB lists | GO\_LYTIC\_VACUOLE\_MEMBRANE | 107 | 9 | 4161 | 170 | CG3842,CG41099,kcc,Tor,Cln3,CG3860,sws,LRP1,Sk1 | | 2.964e-02 | -3.52 | gastrulation | biological process | GO:0007369 | 90 | 9 | 12345 | 597 | Pld,dpp,ths,Raf,tin,shg,Ptp61F,Ubx,tld | | 2.970e-02 | -3.52 | E4BP4\_01 | MSigDB lists | E4BP4\_01 | 58 | 6 | 4161 | 170 | sxc,sdt,app,chb,Octbeta2R,Dys | | 2.970e-02 | -3.52 | MORF\_MLLT10 | MSigDB lists | MORF\_MLLT10 | 58 | 6 | 4161 | 170 | fz2,Pkc53E,dnc,Hr96,CG40178,Dys | | 2.970e-02 | -3.52 | CEBPB\_02 | MSigDB lists | CEBPB\_02 | 58 | 6 | 4161 | 170 | Dys,Octbeta2R,CG5339,dnc,Iswi,flw | | 2.970e-02 | -3.52 | CTATGCA\_MIR153 | MSigDB lists | CTATGCA\_MIR153 | 58 | 6 | 4161 | 170 | CG6966,Ser,Sk1,Meltrin,bru1,Dys | | 2.970e-02 | -3.52 | GO\_MULTICELLULAR\_ORGANISMAL\_HOMEOSTASIS | MSigDB lists | GO\_MULTICELLULAR\_ORGANISMAL\_HOMEOSTASIS | 58 | 6 | 4161 | 170 | Octbeta2R,Root,Sod1,csw,Act88F,Act87E | | 2.970e-02 | -3.52 | MORF\_ERCC4 | MSigDB lists | MORF\_ERCC4 | 58 | 6 | 4161 | 170 | Hr96,CG40178,Dys,fz2,Pkc53E,dnc | | 2.970e-02 | -3.52 | GO\_GUANYL\_NUCLEOTIDE\_EXCHANGE\_FACTOR\_ACTIVITY | MSigDB lists | GO\_GUANYL\_NUCLEOTIDE\_EXCHANGE\_FACTOR\_ACTIVITY | 58 | 6 | 4161 | 170 | eIF2Bgamma,step,RhoGAP1A,CG5522,Nmdar2,Cdep | | 2.974e-02 | -3.52 | Hexose transport | REACTOME pathways | R-DME-189200 | 24 | 4 | 3923 | 197 | CG7720,CG7882,Glut1,CG42235 | | 2.984e-02 | -3.51 | regulation of epidermal growth factor receptor signaling pathway | biological process | GO:0042058 | 49 | 6 | 12345 | 597 | kek5,Ptp61F,scrib,kek1,Src42A,step | | 2.984e-02 | -3.51 | regulation of ERBB signaling pathway | biological process | GO:1901184 | 49 | 6 | 12345 | 597 | scrib,kek1,step,Src42A,kek5,Ptp61F | | 2.984e-02 | -3.51 | myotube differentiation | biological process | GO:0014902 | 49 | 6 | 12345 | 597 | Khc,rols,Tor,rst,siz,foxo | | 2.984e-02 | -3.51 | negative regulation of transport | biological process | GO:0051051 | 49 | 6 | 12345 | 597 | sky,CG9650,Pde6,qvr,cpx,CG42797 | | 2.987e-02 | -3.51 | GCM\_MAPK10 | MSigDB lists | GCM\_MAPK10 | 17 | 3 | 4161 | 170 | Dlic,Frq1,Frq2 | | 2.987e-02 | -3.51 | KEGG\_HYPERTROPHIC\_CARDIOMYOPATHY\_HCM | MSigDB lists | KEGG\_HYPERTROPHIC\_CARDIOMYOPATHY\_HCM | 17 | 3 | 4161 | 170 | Dys,Ca-beta,Act87E | | 2.987e-02 | -3.51 | NADLER\_OBESITY\_DN | MSigDB lists | NADLER\_OBESITY\_DN | 17 | 3 | 4161 | 170 | PCB,Sod1,Acbp2 | | 2.987e-02 | -3.51 | MARTORIATI\_MDM4\_TARGETS\_NEUROEPITHELIUM\_DN | MSigDB lists | MARTORIATI\_MDM4\_TARGETS\_NEUROEPITHELIUM\_DN | 17 | 3 | 4161 | 170 | Pak,Cadps,ABCD | | 2.987e-02 | -3.51 | GO\_GLYCOSPHINGOLIPID\_METABOLIC\_PROCESS | MSigDB lists | GO\_GLYCOSPHINGOLIPID\_METABOLIC\_PROCESS | 17 | 3 | 4161 | 170 | CG3376,CG33090,Cln3 | | 2.987e-02 | -3.51 | GO\_HAIR\_CELL\_DIFFERENTIATION | MSigDB lists | GO\_HAIR\_CELL\_DIFFERENTIATION | 17 | 3 | 4161 | 170 | Su(H),Sod1,Ser | | 2.987e-02 | -3.51 | GO\_MECHANORECEPTOR\_DIFFERENTIATION | MSigDB lists | GO\_MECHANORECEPTOR\_DIFFERENTIATION | 17 | 3 | 4161 | 170 | Su(H),Sod1,Ser | | 2.987e-02 | -3.51 | GO\_REGULATION\_OF\_LYMPHOCYTE\_DIFFERENTIATION | MSigDB lists | GO\_REGULATION\_OF\_LYMPHOCYTE\_DIFFERENTIATION | 17 | 3 | 4161 | 170 | IP3K2,Sod1,Ptp61F | | 2.987e-02 | -3.51 | GSE21546\_UNSTIM\_VS\_ANTI\_CD3\_STIM\_ELK1\_KO\_DP\_THYMOCYTES\_UP | MSigDB lists | GSE21546\_UNSTIM\_VS\_ANTI\_CD3\_STIM\_ELK1\_KO\_DP\_THYMOCYTES\_UP | 17 | 3 | 4161 | 170 | CG14105,scramb1,DNaseII | | 2.987e-02 | -3.51 | MODULE\_46 | MSigDB lists | MODULE\_46 | 17 | 3 | 4161 | 170 | Tsp96F,LRP1,Mlp84B | | 2.987e-02 | -3.51 | GSE19888\_ADENOSINE\_A3R\_INH\_VS\_TCELL\_MEMBRANES\_ACT\_MAST\_CELL\_UP | MSigDB lists | GSE19888\_ADENOSINE\_A3R\_INH\_VS\_TCELL\_MEMBRANES\_ACT\_MAST\_CELL\_UP | 17 | 3 | 4161 | 170 | Sh,CG4341,step | | 2.987e-02 | -3.51 | MORF\_KDR | MSigDB lists | MORF\_KDR | 17 | 3 | 4161 | 170 | fz2,Hr96,Dys | | 2.987e-02 | -3.51 | GO\_CALCIUM\_ION\_REGULATED\_EXOCYTOSIS | MSigDB lists | GO\_CALCIUM\_ION\_REGULATED\_EXOCYTOSIS | 17 | 3 | 4161 | 170 | Cadps,unc-13,Syx1A | | 2.988e-02 | -3.51 | sgg (shaggy) | protein interactions | 31248 | 183 | 16 | 9204 | 481 | crol,trol,Doa,dsh,Ank2,zormin,msps,CG15185,tyf,htk,Prm,CG42784,shg,kay,Cht6,Msp300 | | 2.990e-02 | -3.51 | glutamate-gated calcium ion channel activity | molecular function | GO:0022849 | 6 | 2 | 12176 | 580 | GluRIIC,KaiR1D | | 2.990e-02 | -3.51 | Wnt-activated receptor activity | molecular function | GO:0042813 | 6 | 2 | 12176 | 580 | fz4,fz2 | | 2.990e-02 | -3.51 | glycolipid binding | molecular function | GO:0051861 | 6 | 2 | 12176 | 580 | qvr,Ser | | 2.990e-02 | -3.51 | gap junction channel activity | molecular function | GO:0005243 | 6 | 2 | 12176 | 580 | shakB,Inx2 | | 2.995e-02 | -3.51 | GO\_RESPONSE\_TO\_LIPID | MSigDB lists | GO\_RESPONSE\_TO\_LIPID | 179 | 13 | 4161 | 170 | Doa,Hr96,TfAP-2,RhoGAP1A,sxc,Octbeta2R,Cat,LpR2,dnc,CG5493,Ugt301D1,EcR,Pkc98E | | 3.000e-02 | -3.51 | EGF\_CA | pfam domains | PF07645 | 23 | 4 | 10480 | 550 | LpR2,Ser,tld,LRP1 | | 3.014e-02 | -3.50 | C2DOMAIN | prints domains | PR00360 | 11 | 3 | 2185 | 142 | unc-13,Pkc53E,Mctp | | 3.023e-02 | -3.50 | GSE360\_DC\_VS\_MAC\_B\_MALAYI\_LOW\_DOSE\_UP | MSigDB lists | GSE360\_DC\_VS\_MAC\_B\_MALAYI\_LOW\_DOSE\_UP | 74 | 7 | 4161 | 170 | dsd,Cpsf160,msps,CG8184,CG42672,Sh,flw | | 3.023e-02 | -3.50 | CEBP\_Q2\_01 | MSigDB lists | CEBP\_Q2\_01 | 74 | 7 | 4161 | 170 | sdt,dnc,flw,chb,Octbeta2R,Dys,CG3961 | | 3.027e-02 | -3.50 | cofactor binding | molecular function | GO:0048037 | 387 | 27 | 12176 | 580 | Cyt-b5-r,Acbp2,Cyp4p2,PCB,CG9519,Cyp304a1,CG5955,AOX1,glob3,ND-51L2,CG15093,Cyp309a1,shd,Cat,Kyat,Cyp6u1,Cyp12a4,spok,CG1434,CG3556,Cyp28d1,GlyP,D2hgdh,Elp3,Cyp6a13,CG9514,Cyp28d2 | | 3.034e-02 | -3.50 | vn (vein) | protein interactions | 38657 | 34 | 5 | 9204 | 481 | bun,Ubx,csw,boi,Su(H) | | 3.034e-02 | -3.50 | ap (apterous) | protein interactions | 35509 | 34 | 5 | 9204 | 481 | fru,Ubx,sqz,Ser,bi | | 3.044e-02 | -3.49 | GO\_POSITIVE\_REGULATION\_OF\_TRIGLYCERIDE\_BIOSYNTHETIC\_PROCESS | MSigDB lists | GO\_POSITIVE\_REGULATION\_OF\_TRIGLYCERIDE\_BIOSYNTHETIC\_PROCESS | 7 | 2 | 4161 | 170 | mino,EcR | | 3.044e-02 | -3.49 | GO\_POSITIVE\_REGULATION\_OF\_STEROL\_TRANSPORT | MSigDB lists | GO\_POSITIVE\_REGULATION\_OF\_STEROL\_TRANSPORT | 7 | 2 | 4161 | 170 | EcR,LRP1 | | 3.044e-02 | -3.49 | GO\_PROTEIN\_TARGETING\_TO\_PLASMA\_MEMBRANE | MSigDB lists | GO\_PROTEIN\_TARGETING\_TO\_PLASMA\_MEMBRANE | 7 | 2 | 4161 | 170 | KrT95D,Ank | | 3.044e-02 | -3.49 | chr12q21 | MSigDB lists | chr12q21 | 7 | 2 | 4161 | 170 | CG4341,Mlp84B | | 3.044e-02 | -3.49 | GO\_REGULATION\_OF\_RELEASE\_OF\_SEQUESTERED\_CALCIUM\_ION\_INTO\_CYTOSOL\_BY\_SARCOPLASMIC\_RETICULUM | MSigDB lists | GO\_REGULATION\_OF\_RELEASE\_OF\_SEQUESTERED\_CALCIUM\_ION\_INTO\_CYTOSOL\_BY\_SARCOPLASMIC\_RETICULUM | 7 | 2 | 4161 | 170 | Dys,dnc | | 3.044e-02 | -3.49 | GO\_MUSCLE\_FIBER\_DEVELOPMENT | MSigDB lists | GO\_MUSCLE\_FIBER\_DEVELOPMENT | 7 | 2 | 4161 | 170 | Ca-beta,Dys | | 3.044e-02 | -3.49 | CLIMENT\_BREAST\_CANCER\_COPY\_NUMBER\_UP | MSigDB lists | CLIMENT\_BREAST\_CANCER\_COPY\_NUMBER\_UP | 7 | 2 | 4161 | 170 | Pak,Ser | | 3.044e-02 | -3.49 | PARENT\_MTOR\_SIGNALING\_DN | MSigDB lists | PARENT\_MTOR\_SIGNALING\_DN | 7 | 2 | 4161 | 170 | Meltrin,dpp | | 3.044e-02 | -3.49 | GO\_BETA\_CATENIN\_DESTRUCTION\_COMPLEX\_DISASSEMBLY | MSigDB lists | GO\_BETA\_CATENIN\_DESTRUCTION\_COMPLEX\_DISASSEMBLY | 7 | 2 | 4161 | 170 | dsh,fz2 | | 3.044e-02 | -3.49 | GO\_PLACENTA\_BLOOD\_VESSEL\_DEVELOPMENT | MSigDB lists | GO\_PLACENTA\_BLOOD\_VESSEL\_DEVELOPMENT | 7 | 2 | 4161 | 170 | fz2,Su(H) | | 3.044e-02 | -3.49 | GO\_NEGATIVE\_REGULATION\_OF\_LIPID\_BIOSYNTHETIC\_PROCESS | MSigDB lists | GO\_NEGATIVE\_REGULATION\_OF\_LIPID\_BIOSYNTHETIC\_PROCESS | 7 | 2 | 4161 | 170 | dpp,Sod1 | | 3.044e-02 | -3.49 | GO\_TRANSCRIPTION\_FACTOR\_ACTIVITY\_DIRECT\_LIGAND\_REGULATED\_SEQUENCE\_SPECIFIC\_DNA\_BINDING | MSigDB lists | GO\_TRANSCRIPTION\_FACTOR\_ACTIVITY\_DIRECT\_LIGAND\_REGULATED\_SEQUENCE\_SPECIFIC\_DNA\_BINDING | 7 | 2 | 4161 | 170 | Hr96,EcR | | 3.044e-02 | -3.49 | GO\_REGULATION\_OF\_SYNAPTIC\_VESICLE\_EXOCYTOSIS | MSigDB lists | GO\_REGULATION\_OF\_SYNAPTIC\_VESICLE\_EXOCYTOSIS | 7 | 2 | 4161 | 170 | Raf,Syx1A | | 3.044e-02 | -3.49 | GO\_ADRENERGIC\_RECEPTOR\_SIGNALING\_PATHWAY | MSigDB lists | GO\_ADRENERGIC\_RECEPTOR\_SIGNALING\_PATHWAY | 7 | 2 | 4161 | 170 | dnc,Octbeta2R | | 3.044e-02 | -3.49 | GO\_VOLTAGE\_GATED\_CALCIUM\_CHANNEL\_COMPLEX | MSigDB lists | GO\_VOLTAGE\_GATED\_CALCIUM\_CHANNEL\_COMPLEX | 7 | 2 | 4161 | 170 | dnc,Ca-beta | | 3.044e-02 | -3.49 | GO\_POSITIVE\_REGULATION\_OF\_TRIGLYCERIDE\_METABOLIC\_PROCESS | MSigDB lists | GO\_POSITIVE\_REGULATION\_OF\_TRIGLYCERIDE\_METABOLIC\_PROCESS | 7 | 2 | 4161 | 170 | mino,EcR | | 3.044e-02 | -3.49 | REACTOME\_CREB\_PHOSPHORYLATION\_THROUGH\_THE\_ACTIVATION\_OF\_RAS | MSigDB lists | REACTOME\_CREB\_PHOSPHORYLATION\_THROUGH\_THE\_ACTIVATION\_OF\_RAS | 7 | 2 | 4161 | 170 | Raf,Nmdar2 | | 3.044e-02 | -3.49 | GO\_MODULATION\_OF\_EXCITATORY\_POSTSYNAPTIC\_POTENTIAL | MSigDB lists | GO\_MODULATION\_OF\_EXCITATORY\_POSTSYNAPTIC\_POTENTIAL | 7 | 2 | 4161 | 170 | CG7483,Syx1A | | 3.044e-02 | -3.49 | GO\_NEGATIVE\_REGULATION\_OF\_NEUROTRANSMITTER\_TRANSPORT | MSigDB lists | GO\_NEGATIVE\_REGULATION\_OF\_NEUROTRANSMITTER\_TRANSPORT | 7 | 2 | 4161 | 170 | Raf,ZnT63C | | 3.044e-02 | -3.49 | GHANDHI\_DIRECT\_IRRADIATION\_UP | MSigDB lists | GHANDHI\_DIRECT\_IRRADIATION\_UP | 7 | 2 | 4161 | 170 | ZnT63C,dpp | | 3.044e-02 | -3.49 | REACTOME\_NITRIC\_OXIDE\_STIMULATES\_GUANYLATE\_CYCLASE | MSigDB lists | REACTOME\_NITRIC\_OXIDE\_STIMULATES\_GUANYLATE\_CYCLASE | 7 | 2 | 4161 | 170 | Pde6,Pde1c | | 3.044e-02 | -3.49 | LIU\_TARGETS\_OF\_VMYB\_VS\_CMYB\_DN | MSigDB lists | LIU\_TARGETS\_OF\_VMYB\_VS\_CMYB\_DN | 7 | 2 | 4161 | 170 | Octbeta2R,Sh | | 3.044e-02 | -3.49 | GO\_REGULATION\_OF\_RYANODINE\_SENSITIVE\_CALCIUM\_RELEASE\_CHANNEL\_ACTIVITY | MSigDB lists | GO\_REGULATION\_OF\_RYANODINE\_SENSITIVE\_CALCIUM\_RELEASE\_CHANNEL\_ACTIVITY | 7 | 2 | 4161 | 170 | Dys,dnc | | 3.044e-02 | -3.49 | GO\_ESTABLISHMENT\_OF\_EPITHELIAL\_CELL\_POLARITY | MSigDB lists | GO\_ESTABLISHMENT\_OF\_EPITHELIAL\_CELL\_POLARITY | 7 | 2 | 4161 | 170 | chb,sdt | | 3.044e-02 | -3.49 | GO\_ACUTE\_INFLAMMATORY\_RESPONSE | MSigDB lists | GO\_ACUTE\_INFLAMMATORY\_RESPONSE | 7 | 2 | 4161 | 170 | Ugt301D1,scramb1 | | 3.044e-02 | -3.49 | PID\_VEGFR1\_PATHWAY | MSigDB lists | PID\_VEGFR1\_PATHWAY | 7 | 2 | 4161 | 170 | csw,Pkc53E | | 3.044e-02 | -3.49 | GO\_NEGATIVE\_REGULATION\_OF\_MUSCLE\_CONTRACTION | MSigDB lists | GO\_NEGATIVE\_REGULATION\_OF\_MUSCLE\_CONTRACTION | 7 | 2 | 4161 | 170 | Sod1,Octbeta2R | | 3.044e-02 | -3.49 | GO\_POSITIVE\_REGULATION\_OF\_GLIAL\_CELL\_DIFFERENTIATION | MSigDB lists | GO\_POSITIVE\_REGULATION\_OF\_GLIAL\_CELL\_DIFFERENTIATION | 7 | 2 | 4161 | 170 | Tor,dpp | | 3.044e-02 | -3.49 | GO\_RESPONSE\_TO\_MONOAMINE | MSigDB lists | GO\_RESPONSE\_TO\_MONOAMINE | 7 | 2 | 4161 | 170 | Octbeta2R,dnc | | 3.044e-02 | -3.49 | WU\_ALZHEIMER\_DISEASE\_DN | MSigDB lists | WU\_ALZHEIMER\_DISEASE\_DN | 7 | 2 | 4161 | 170 | Pde1c,Ank | | 3.044e-02 | -3.49 | BOYLAN\_MULTIPLE\_MYELOMA\_D\_CLUSTER\_DN | MSigDB lists | BOYLAN\_MULTIPLE\_MYELOMA\_D\_CLUSTER\_DN | 7 | 2 | 4161 | 170 | Cdep,Meltrin | | 3.044e-02 | -3.49 | GO\_EXTRACELLULAR\_GLUTAMATE\_GATED\_ION\_CHANNEL\_ACTIVITY | MSigDB lists | GO\_EXTRACELLULAR\_GLUTAMATE\_GATED\_ION\_CHANNEL\_ACTIVITY | 7 | 2 | 4161 | 170 | Nmdar2,KaiR1D | | 3.044e-02 | -3.49 | GO\_NEGATIVE\_REGULATION\_OF\_RESPONSE\_TO\_CYTOKINE\_STIMULUS | MSigDB lists | GO\_NEGATIVE\_REGULATION\_OF\_RESPONSE\_TO\_CYTOKINE\_STIMULUS | 7 | 2 | 4161 | 170 | EcR,Ptp61F | | 3.044e-02 | -3.49 | GO\_BLOOD\_VESSEL\_REMODELING | MSigDB lists | GO\_BLOOD\_VESSEL\_REMODELING | 7 | 2 | 4161 | 170 | Ser,Su(H) | | 3.044e-02 | -3.49 | BROWNE\_INTERFERON\_RESPONSIVE\_GENES | MSigDB lists | BROWNE\_INTERFERON\_RESPONSIVE\_GENES | 7 | 2 | 4161 | 170 | scramb1,step | | 3.044e-02 | -3.49 | PID\_IL8\_CXCR1\_PATHWAY | MSigDB lists | PID\_IL8\_CXCR1\_PATHWAY | 7 | 2 | 4161 | 170 | Pkc53E,Pkc98E | | 3.044e-02 | -3.49 | GO\_REGULATION\_OF\_STEROID\_BIOSYNTHETIC\_PROCESS | MSigDB lists | GO\_REGULATION\_OF\_STEROID\_BIOSYNTHETIC\_PROCESS | 7 | 2 | 4161 | 170 | Sod1,dpp | | 3.045e-02 | -3.49 | cellular response to hormone stimulus | biological process | GO:0032870 | 105 | 10 | 12345 | 597 | gce,Tor,CG7362,rictor,Set2,Gpb5,Lpin,EcR,foxo,Iswi | | 3.048e-02 | -3.49 | oogenesis | biological process | GO:0048477 | 682 | 44 | 12345 | 597 | drd,Doa,Dlic,CG7483,lncRNA:flam,bru1,flw,Khc,Mos,retn,kay,Tor,msps,polybromo,RhoL,bbg,sgg,cic,ex,DNaseII,Eip74EF,upd3,tld,kek1,shg,mid,csw,rols,egh,dpp,scrib,shd,EcR,Inx2,lncRNA:Hsromega,Mrtf,dsh,fs(1)N,Raf,cindr,Msp300,wbl,chb,bun | | 3.052e-02 | -3.49 | synaptic vesicle localization | biological process | GO:0097479 | 25 | 4 | 12345 | 597 | g,Khc,RhoGAP100F,Fife | | 3.052e-02 | -3.49 | sensory organ boundary specification | biological process | GO:0008052 | 25 | 4 | 12345 | 597 | numb,B-H2,Su(H),nuf | | 3.052e-02 | -3.49 | regulation of exocytosis | biological process | GO:0017157 | 25 | 4 | 12345 | 597 | Fife,Cadps,Syt12,Syx1A | | 3.052e-02 | -3.49 | cell fate commitment involved in formation of primary germ layer | biological process | GO:0060795 | 25 | 4 | 12345 | 597 | tin,ths,dpp,Ubx | | 3.057e-02 | -3.49 | Stat92E (Signal-transducer and activator of transcription protein at 92E) | protein interactions | 42428 | 58 | 7 | 9204 | 481 | csw,Src42A,scrib,dpp,Ptp61F,et,upd3 | | 3.057e-02 | -3.49 | TYR\_PHOSPHATASE\_1 | prosite domains | PS00383 | 29 | 5 | 6243 | 386 | CG3530,csw,Ptp99A,Ptp61F,Ptp36E | | 3.076e-02 | -3.48 | GO\_REGULATION\_OF\_CYTOPLASMIC\_TRANSPORT | MSigDB lists | GO\_REGULATION\_OF\_CYTOPLASMIC\_TRANSPORT | 143 | 11 | 4161 | 170 | csw,CG17883,Tor,Pkc98E,fz2,dnc,CG8184,Raf,Syx1A,CCAP-R,Dys | | 3.076e-02 | -3.48 | rRNA 5'-end processing | biological process | GO:0000967 | 6 | 2 | 12345 | 597 | CG41562,CG40813 | | 3.076e-02 | -3.48 | positive regulation of microtubule polymerization | biological process | GO:0031116 | 6 | 2 | 12345 | 597 | chb,msps | | 3.076e-02 | -3.48 | positive regulation of cation channel activity | biological process | GO:2001259 | 6 | 2 | 12345 | 597 | Cam,qvr | | 3.076e-02 | -3.48 | endonucleolytic cleavage to generate mature 5'-end of SSU-rRNA from (SSU-rRNA, 5.8S rRNA, LSU-rRNA) | biological process | GO:0000472 | 6 | 2 | 12345 | 597 | CG41562,CG40813 | | 3.076e-02 | -3.48 | endonucleolytic cleavage in 5'-ETS of tricistronic rRNA transcript (SSU-rRNA, 5.8S rRNA, LSU-rRNA) | biological process | GO:0000480 | 6 | 2 | 12345 | 597 | CG41562,CG40813 | | 3.076e-02 | -3.48 | amnioserosa formation | biological process | GO:0007378 | 6 | 2 | 12345 | 597 | tld,dpp | | 3.076e-02 | -3.48 | determination of digestive tract left/right asymmetry | biological process | GO:0071907 | 6 | 2 | 12345 | 597 | shg,tin | | 3.076e-02 | -3.48 | small RNA loading onto RISC | biological process | GO:0070922 | 6 | 2 | 12345 | 597 | HPS4,AGO2 | | 3.076e-02 | -3.48 | small-subunit processome assembly | biological process | GO:0034462 | 6 | 2 | 12345 | 597 | CG41562,CG40813 | | 3.077e-02 | -3.48 | MFS\_sugar\_transport-like | interpro domains | IPR005828 | 59 | 7 | 11383 | 585 | CG3168,CG10960,CG42826,Glut1,CG14606,CG3285,CG7882 | | 3.092e-02 | -3.48 | RRM | prosite domains | PS50102 | 131 | 14 | 6243 | 386 | Spf45,CG44249,msi,barc,CG41562,CG42458,shep,CG3594,fus,trv,bol,bru1,CG40813,Rbp6 | | 3.116e-02 | -3.47 | GO\_RESPONSE\_TO\_BIOTIC\_STIMULUS | MSigDB lists | GO\_RESPONSE\_TO\_BIOTIC\_STIMULUS | 180 | 13 | 4161 | 170 | TfAP-2,RhoGAP1A,Su(H),sxc,Odc1,dnc,Ugt301D1,Syn2,mino,fz2,Pkc98E,scramb1,LpR2 | | 3.123e-02 | -3.47 | GO\_POSITIVE\_REGULATION\_OF\_CELLULAR\_COMPONENT\_BIOGENESIS | MSigDB lists | GO\_POSITIVE\_REGULATION\_OF\_CELLULAR\_COMPONENT\_BIOGENESIS | 108 | 9 | 4161 | 170 | Pkc53E,Tor,Pkc98E,EcR,Fhos,Raf,TfIIFbeta,Pak,chb | | 3.141e-02 | -3.46 | myosin complex | cellular component | GO:0016459 | 27 | 4 | 11940 | 538 | Prm,Myo81F,Cam,CG34435 | | 3.154e-02 | -3.46 | formation of animal organ boundary | biological process | GO:0010160 | 37 | 5 | 12345 | 597 | Su(H),nuf,B-H2,numb,hth | | 3.173e-02 | -3.45 | cell-cell junction organization | biological process | GO:0045216 | 63 | 7 | 12345 | 597 | CadN2,shg,Pak,scrib,shakB,nrv2,sdt | | 3.182e-02 | -3.45 | GO\_NEURON\_DEVELOPMENT | MSigDB lists | GO\_NEURON\_DEVELOPMENT | 162 | 12 | 4161 | 170 | CG33090,LpR2,AstA-R1,Pak,Khc,csw,Sod1,Awh,CG42672,sky,Cdep,Ank | | 3.183e-02 | -3.45 | central nervous system development | biological process | GO:0007417 | 281 | 21 | 12345 | 597 | shd,Antp,E5,EcR,esg,CycT,Dmtn,hth,shg,ems,fru,mid,csw,opa,siz,raw,chinmo,bun,rib,tin,numb | | 3.193e-02 | -3.44 | LDrepeatLR\_classA\_rpt | interpro domains | IPR002172 | 35 | 5 | 11383 | 585 | LpR2,trol,CG32432,LRP1,CG31221 | | 3.193e-02 | -3.44 | CH\_dom\_sf | interpro domains | IPR036872 | 35 | 5 | 11383 | 585 | sick,Dys,CG34417,Msp300,Actn | | 3.194e-02 | -3.44 | regulation of molecular function | biological process | GO:0065009 | 523 | 35 | 12345 | 597 | step,CG17883,RhoGAP1A,REG,CycT,melt,HDAC4,qvr,Cam,CG42797,SteXh:CG42398,CG34408,CG42390,Raf,CG41520,RhoGAP18B,rictor,CG33310,Acp62F,Pak,Mbs,sws,nrv2,Spn88Eb,RhoGAP100F,kay,Strn-Mlck,Ca-beta,mnb,sgg,kek1,Ste:CG33247,CG6966,sky,Stlk | | 3.198e-02 | -3.44 | GSE27859\_DC\_VS\_CD11C\_INT\_F480\_INT\_DC\_UP | MSigDB lists | GSE27859\_DC\_VS\_CD11C\_INT\_F480\_INT\_DC\_UP | 59 | 6 | 4161 | 170 | Maf1,RpL38,Pgant6,Der-1,Ptp61F,CG3376 | | 3.198e-02 | -3.44 | USF\_Q6\_01 | MSigDB lists | USF\_Q6\_01 | 59 | 6 | 4161 | 170 | eIF4E7,LpR2,Stt3B,Cln3,RhoGAP1A,CG4849 | | 3.202e-02 | -3.44 | GO\_REGULATION\_OF\_GROWTH | MSigDB lists | GO\_REGULATION\_OF\_GROWTH | 126 | 10 | 4161 | 170 | Tor,csw,Sod1,Su(H),dsd,MESK2,mino,Octbeta2R,unc-13,Tsp96F | | 3.214e-02 | -3.44 | GTATTAT\_MIR3693P | MSigDB lists | GTATTAT\_MIR3693P | 44 | 5 | 4161 | 170 | Dys,Pkc53E,Iswi,dnc,LRP1 | | 3.214e-02 | -3.44 | GSE5589\_LPS\_VS\_LPS\_AND\_IL10\_STIM\_IL10\_KO\_MACROPHAGE\_45MIN\_UP | MSigDB lists | GSE5589\_LPS\_VS\_LPS\_AND\_IL10\_STIM\_IL10\_KO\_MACROPHAGE\_45MIN\_UP | 44 | 5 | 4161 | 170 | Octbeta2R,Syx1A,LRP1,CG42672,mino | | 3.214e-02 | -3.44 | GSE46242\_CTRL\_VS\_EGR2\_DELETED\_ANERGIC\_TH1\_CD4\_TCELL\_UP | MSigDB lists | GSE46242\_CTRL\_VS\_EGR2\_DELETED\_ANERGIC\_TH1\_CD4\_TCELL\_UP | 44 | 5 | 4161 | 170 | rdgA,Maf1,LpR2,AstA-R1,CG4301 | | 3.214e-02 | -3.44 | GSE29949\_CD8\_POS\_DC\_SPLEEN\_VS\_MONOCYTE\_BONE\_MARROW\_UP | MSigDB lists | GSE29949\_CD8\_POS\_DC\_SPLEEN\_VS\_MONOCYTE\_BONE\_MARROW\_UP | 44 | 5 | 4161 | 170 | KaiR1D,Pde8,Odc1,KrT95D,Cadps | | 3.214e-02 | -3.44 | GSE34156\_TLR1\_TLR2\_LIGAND\_VS\_NOD2\_AND\_TLR1\_TLR2\_LIGAND\_24H\_TREATED\_MONOCYTE\_UP | MSigDB lists | GSE34156\_TLR1\_TLR2\_LIGAND\_VS\_NOD2\_AND\_TLR1\_TLR2\_LIGAND\_24H\_TREATED\_MONOCYTE\_UP | 44 | 5 | 4161 | 170 | LRP1,Act87E,Pkc98E,Frq2,Frq1 | | 3.214e-02 | -3.44 | GO\_RNA\_POLYMERASE\_II\_TRANSCRIPTION\_FACTOR\_ACTIVITY\_SEQUENCE\_SPECIFIC\_DNA\_BINDING | MSigDB lists | GO\_RNA\_POLYMERASE\_II\_TRANSCRIPTION\_FACTOR\_ACTIVITY\_SEQUENCE\_SPECIFIC\_DNA\_BINDING | 44 | 5 | 4161 | 170 | TfAP-2,Su(H),EcR,scramb1,Hr96 | | 3.214e-02 | -3.44 | GSE15750\_DAY6\_VS\_DAY10\_EFF\_CD8\_TCELL\_DN | MSigDB lists | GSE15750\_DAY6\_VS\_DAY10\_EFF\_CD8\_TCELL\_DN | 44 | 5 | 4161 | 170 | Pkc98E,Pkc53E,CG17698,Cdep,ZnT63C | | 3.214e-02 | -3.44 | GO\_MEMBRANE\_LIPID\_BIOSYNTHETIC\_PROCESS | MSigDB lists | GO\_MEMBRANE\_LIPID\_BIOSYNTHETIC\_PROCESS | 44 | 5 | 4161 | 170 | Sk1,PIG-O,PIG-G,CG3376,Aldh-III | | 3.214e-02 | -3.44 | GSE27786\_LIN\_NEG\_VS\_MONO\_MAC\_DN | MSigDB lists | GSE27786\_LIN\_NEG\_VS\_MONO\_MAC\_DN | 44 | 5 | 4161 | 170 | Khc,pod1,MCU,CG12121,Maf1 | | 3.214e-02 | -3.44 | SMID\_BREAST\_CANCER\_NORMAL\_LIKE\_UP | MSigDB lists | SMID\_BREAST\_CANCER\_NORMAL\_LIKE\_UP | 44 | 5 | 4161 | 170 | CG5493,Dys,sno,bru1,kcc | | 3.214e-02 | -3.44 | GO\_NEGATIVE\_REGULATION\_OF\_LOCOMOTION | MSigDB lists | GO\_NEGATIVE\_REGULATION\_OF\_LOCOMOTION | 44 | 5 | 4161 | 170 | chb,Ptp61F,Raf,LRP1,RhoGAP1A | | 3.214e-02 | -3.44 | GO\_SPHINGOLIPID\_METABOLIC\_PROCESS | MSigDB lists | GO\_SPHINGOLIPID\_METABOLIC\_PROCESS | 44 | 5 | 4161 | 170 | Aldh-III,CG3376,CG33090,Cln3,Sk1 | | 3.214e-02 | -3.44 | GSE3982\_EOSINOPHIL\_VS\_NKCELL\_UP | MSigDB lists | GSE3982\_EOSINOPHIL\_VS\_NKCELL\_UP | 44 | 5 | 4161 | 170 | IP3K2,Ser,Su(H),fus,CG7483 | | 3.216e-02 | -3.44 | GO\_ORGANELLE\_LOCALIZATION | MSigDB lists | GO\_ORGANELLE\_LOCALIZATION | 144 | 11 | 4161 | 170 | Dlic,CG41099,CG17883,sxc,Cln3,chb,Khc,unc-13,Syx1A,Cadps,Root | | 3.216e-02 | -3.44 | SERVITJA\_ISLET\_HNF1A\_TARGETS\_DN | MSigDB lists | SERVITJA\_ISLET\_HNF1A\_TARGETS\_DN | 30 | 4 | 4161 | 170 | Src42A,Ugt301D1,CG5493,LpR2 | | 3.216e-02 | -3.44 | HALLMARK\_SPERMATOGENESIS | MSigDB lists | HALLMARK\_SPERMATOGENESIS | 30 | 4 | 4161 | 170 | Tor,PhKgamma,nompB,Raf | | 3.216e-02 | -3.44 | GO\_RESPONSE\_TO\_ALKALOID | MSigDB lists | GO\_RESPONSE\_TO\_ALKALOID | 30 | 4 | 4161 | 170 | Pkc98E,Tor,CG4849,CG3376 | | 3.216e-02 | -3.44 | PID\_CDC42\_PATHWAY | MSigDB lists | PID\_CDC42\_PATHWAY | 30 | 4 | 4161 | 170 | Raf,Pak,Pkc98E,Tor | | 3.216e-02 | -3.44 | GO\_SARCOLEMMA | MSigDB lists | GO\_SARCOLEMMA | 30 | 4 | 4161 | 170 | Dys,Ank,Octbeta2R,ZnT63C | | 3.216e-02 | -3.44 | ACEVEDO\_LIVER\_CANCER\_WITH\_H3K27ME3\_UP | MSigDB lists | ACEVEDO\_LIVER\_CANCER\_WITH\_H3K27ME3\_UP | 30 | 4 | 4161 | 170 | CG14480,MCU,Pde1c,DNaseII | | 3.216e-02 | -3.44 | GO\_EYE\_MORPHOGENESIS | MSigDB lists | GO\_EYE\_MORPHOGENESIS | 30 | 4 | 4161 | 170 | fz2,TfAP-2,CG17883,Ser | | 3.216e-02 | -3.44 | GSE12003\_MIR223\_KO\_VS\_WT\_BM\_PROGENITOR\_8D\_CULTURE\_DN | MSigDB lists | GSE12003\_MIR223\_KO\_VS\_WT\_BM\_PROGENITOR\_8D\_CULTURE\_DN | 30 | 4 | 4161 | 170 | Acbp2,CCAP-R,CG12121,CG3376 | | 3.216e-02 | -3.44 | GSE3982\_DC\_VS\_NEUTROPHIL\_DN | MSigDB lists | GSE3982\_DC\_VS\_NEUTROPHIL\_DN | 30 | 4 | 4161 | 170 | Glut1,bru1,CG9132,Pgant6 | | 3.216e-02 | -3.44 | GSE7852\_TREG\_VS\_TCONV\_DN | MSigDB lists | GSE7852\_TREG\_VS\_TCONV\_DN | 30 | 4 | 4161 | 170 | CG5339,csw,Cln3,Pgant6 | | 3.216e-02 | -3.44 | Sugar/inositol\_transpt | interpro domains | IPR003663 | 14 | 3 | 11383 | 585 | CG7882,CG10960,Glut1 | | 3.223e-02 | -3.43 | regulation of autophagy | biological process | GO:0010506 | 106 | 10 | 12345 | 597 | CG18304,CG3530,Raf,Tor,Doa,foxo,Fer2,Cpsf160,EcR,Sod1 | | 3.224e-02 | -3.43 | REACTOME\_SIGNALLING\_BY\_NGF | MSigDB lists | REACTOME\_SIGNALLING\_BY\_NGF | 75 | 7 | 4161 | 170 | Pde1c,Raf,Tor,CG42672,Pkc98E,RhoGAP1A,Pkc53E | | 3.224e-02 | -3.43 | GSE27670\_BLIMP1\_VS\_LMP1\_TRANSDUCED\_GC\_BCELL\_DN | MSigDB lists | GSE27670\_BLIMP1\_VS\_LMP1\_TRANSDUCED\_GC\_BCELL\_DN | 75 | 7 | 4161 | 170 | kcc,Flo2,CG41099,MESK2,Stlk,mino,Pgant6 | | 3.234e-02 | -3.43 | iron ion binding | molecular function | GO:0005506 | 154 | 13 | 12176 | 580 | Cyp28d1,AOX1,shd,Cyp28d2,Cyp6a13,Cyp309a1,Cyp6u1,Cyp4p2,Cyp12a4,CG5493,Cyp304a1,spok,Fad2 | | 3.245e-02 | -3.43 | nucleus | cellular component | GO:0005634 | 2809 | 145 | 11940 | 538 | REG,CG7483,grn,trv,kay,bol,CG1894,Mrtf,toe,FoxL1,shep,CG42458,can,Eip74EF,CG32971,IntS3,CG14480,Tif-IA,Rbp6,Spt20,Asator,Awh,Art2,sno,rib,Marf1,bru1,CG12071,bi,opa,CG40813,CG15073,Fhos,gce,hth,TfIIFbeta,CG31224,TfAP-2,CG10462,CG9449,Elp3,chb,CG17698,Sod1,dany,fru,CG12200,lncRNA:Hsromega,Pomp,Set2,Iswi,CG12782,barc,PhKgamma,CG42390,woc,retn,Doa,CG4374,ps,Cpsf160,Poxm,Cam,esg,Stlk,E(Pc),CG4849,Dlic,Usp1,luna,CG11970,mamo,scaRNA:MeU2-C41,polybromo,scaRNA:PsiU2-35.45,E5,ems,Msp300,Ubx,sgg,fus,CG9650,htk,AGO2,GATAe,CG8184,ddbt,Tor,CG46385,EcR,bbg,Rat1,ham,jigr1,mnb,chinmo,CG32006,CG44249,cic,Mlp84B,Snoo,CG14252,His1:CG33837,Atf6,crol,scro,Ste:CG33247,Spf45,CrebB,Ptp61F,CG11023,Su(H),foxo,Act87E,Yeti,Nuak1,CG41562,HDAC4,CG9451,CG32264,dnc,Lpin,CycT,sqz,CG3594,CG12605,Antp,Maf1,msi,Prosbeta5R1,Hr96,tin,flw,CG42830,fz2,lncRNA:bxd,CG5059,mid,CG6418,CG43689,Mondo,CG7368,Ipk1,Gld2,B-H2 | | 3.288e-02 | -3.41 | GO\_REGULATION\_OF\_BODY\_FLUID\_LEVELS | MSigDB lists | GO\_REGULATION\_OF\_BODY\_FLUID\_LEVELS | 109 | 9 | 4161 | 170 | Act87E,rdgA,CG5493,EcR,Pkc98E,Octbeta2R,scramb1,Pkc53E,csw | | 3.307e-02 | -3.41 | - | gene3d domains | 1.10.1000.11 | 5 | 2 | 4929 | 302 | siz,step | | 3.311e-02 | -3.41 | solute:proton symporter activity | molecular function | GO:0015295 | 26 | 4 | 12176 | 580 | CG14606,CG45782,CG10960,CG3285 | | 3.311e-02 | -3.41 | activating transcription factor binding | molecular function | GO:0033613 | 26 | 4 | 12176 | 580 | Mrtf,mid,ex,bi | | 3.315e-02 | -3.41 | regulation of muscle system process | biological process | GO:0090257 | 15 | 3 | 12345 | 597 | Tk,CG3078,bru1 | | 3.315e-02 | -3.41 | ecdysone metabolic process | biological process | GO:0008205 | 15 | 3 | 12345 | 597 | woc,spok,foxo | | 3.315e-02 | -3.41 | plasmatocyte differentiation | biological process | GO:0042387 | 15 | 3 | 12345 | 597 | Tor,ths,Raf | | 3.315e-02 | -3.41 | regulation of regulated secretory pathway | biological process | GO:1903305 | 15 | 3 | 12345 | 597 | Syt12,Cadps,Fife | | 3.315e-02 | -3.41 | presynaptic active zone organization | biological process | GO:1990709 | 15 | 3 | 12345 | 597 | Nlg2,Fife,RhoGAP100F | | 3.315e-02 | -3.41 | positive regulation of ion transport | biological process | GO:0043270 | 15 | 3 | 12345 | 597 | qvr,Cam,olf186-F | | 3.315e-02 | -3.41 | crystal cell differentiation | biological process | GO:0042688 | 15 | 3 | 12345 | 597 | Su(H),ths,Ser | | 3.315e-02 | -3.41 | inter-male aggressive behavior | biological process | GO:0002121 | 15 | 3 | 12345 | 597 | Tk,Nlg2,TfAP-2 | | 3.319e-02 | -3.41 | Hsp60A (Heat shock protein 60A) | protein interactions | 32045 | 59 | 7 | 9204 | 481 | rictor,csw,ex,zormin,Raf,Cyp6a13,D2hgdh | | 3.329e-02 | -3.40 | anterior/posterior pattern specification | biological process | GO:0009952 | 200 | 16 | 12345 | 597 | Khc,EcR,Ubx,retn,Ser,CG7483,scrib,Antp,csw,dpp,shg,bi,msps,Raf,cic,fs(1)N | | 3.337e-02 | -3.40 | cell maturation | biological process | GO:0048469 | 184 | 15 | 12345 | 597 | tld,CG7483,scrib,wbl,shep,chb,EcR,Khc,flw,shg,Ca-beta,egh,cic,dpp,msps | | 3.356e-02 | -3.39 | SMC1 (Structural maintenance of chromosomes 1) | protein interactions | 42853 | 14 | 3 | 9204 | 481 | sgg,EcR,CG3078 | | 3.356e-02 | -3.39 | MED26 (Mediator complex subunit 26) | protein interactions | 43816 | 14 | 3 | 9204 | 481 | Prosbeta5R1,EcR,CG46385 | | 3.356e-02 | -3.39 | CG10948 (uncharacterized protein) | protein interactions | 39459 | 14 | 3 | 9204 | 481 | CG3081,Dys,Mob2 | | 3.356e-02 | -3.39 | caz (cabeza) | protein interactions | 32587 | 14 | 3 | 9204 | 481 | lncRNA:Hsromega,flw,csw | | 3.356e-02 | -3.39 | CG15293 (uncharacterized protein) | protein interactions | 34828 | 14 | 3 | 9204 | 481 | CG40045,tamo,shg | | 3.356e-02 | -3.39 | kto (kohtalo) | protein interactions | 44830 | 14 | 3 | 9204 | 481 | EcR,flw,Su(H) | | 3.366e-02 | -3.39 | PIWI | prosite domains | PS50822 | 5 | 2 | 6243 | 386 | AGO3,AGO2 | | 3.366e-02 | -3.39 | SEC7 | prosite domains | PS50190 | 5 | 2 | 6243 | 386 | siz,step | | 3.366e-02 | -3.39 | ANTENNAPEDIA | prosite domains | PS00032 | 5 | 2 | 6243 | 386 | Ubx,Antp | | 3.366e-02 | -3.39 | GATA\_ZN\_FINGER\_1 | prosite domains | PS00344 | 5 | 2 | 6243 | 386 | grn,GATAe | | 3.366e-02 | -3.39 | GATA\_ZN\_FINGER\_2 | prosite domains | PS50114 | 5 | 2 | 6243 | 386 | GATAe,grn | | 3.375e-02 | -3.39 | Piwi | smart domains | SM00950 | 5 | 2 | 5086 | 315 | AGO3,AGO2 | | 3.375e-02 | -3.39 | Frizzled | smart domains | SM01330 | 5 | 2 | 5086 | 315 | fz2,fz4 | | 3.375e-02 | -3.39 | TOG | smart domains | SM01349 | 5 | 2 | 5086 | 315 | chb,msps | | 3.375e-02 | -3.39 | EFG\_C | smart domains | SM00838 | 5 | 2 | 5086 | 315 | CG33158,CG4849 | | 3.379e-02 | -3.39 | fly chr48C3-48C4\|2-65 | chromosome location | fly chr48C3-48C4\|2-65 | 1 | 1 | 25010 | 845 | ths | | 3.379e-02 | -3.39 | fly chr29E4-29E6\|2-33 cM | chromosome location | fly chr29E4-29E6\|2-33 cM | 1 | 1 | 25010 | 845 | raw | | 3.379e-02 | -3.39 | fly chr8B6-8B6\|1-25 | chromosome location | fly chr8B6-8B6\|1-25 | 1 | 1 | 25010 | 845 | Uros1 | | 3.379e-02 | -3.39 | fly chr80A-80B\|2-55 cM | chromosome location | fly chr80A-80B\|2-55 cM | 1 | 1 | 25010 | 845 | Scp1 | | 3.379e-02 | -3.39 | fly chr91E2-91E3\|3-65 | chromosome location | fly chr91E2-91E3\|3-65 | 1 | 1 | 25010 | 845 | gukh | | 3.379e-02 | -3.39 | fly chr84F12-84F12\|3-48 | chromosome location | fly chr84F12-84F12\|3-48 | 1 | 1 | 25010 | 845 | CG7443 | | 3.379e-02 | -3.39 | fly chr96E10-96F1\|3-88 | chromosome location | fly chr96E10-96F1\|3-88 | 1 | 1 | 25010 | 845 | LpR2 | | 3.379e-02 | -3.39 | fly chr85D19-85D22\|3-48 | chromosome location | fly chr85D19-85D22\|3-48 | 1 | 1 | 25010 | 845 | by | | 3.379e-02 | -3.39 | fly chr64C7-64C11\|3-13 cM | chromosome location | fly chr64C7-64C11\|3-13 cM | 1 | 1 | 25010 | 845 | shep | | 3.379e-02 | -3.39 | fly chr89E2-89E4\|3-59 | chromosome location | fly chr89E2-89E4\|3-59 | 1 | 1 | 25010 | 845 | lncRNA:iab8 | | 3.379e-02 | -3.39 | fly chr33E7-33E9\|2-46 | chromosome location | fly chr33E7-33E9\|2-46 | 1 | 1 | 25010 | 845 | bun | | 3.379e-02 | -3.39 | fly chr75E3-75E4\|3-45 cM | chromosome location | fly chr75E3-75E4\|3-45 cM | 1 | 1 | 25010 | 845 | CG3961 | | 3.379e-02 | -3.39 | fly chr27C1-27C4\|2-23 | chromosome location | fly chr27C1-27C4\|2-23 | 1 | 1 | 25010 | 845 | Nlg2 | | 3.379e-02 | -3.39 | fly chr3C3-3C4\|1-2 | chromosome location | fly chr3C3-3C4\|1-2 | 1 | 1 | 25010 | 845 | rst | | 3.379e-02 | -3.39 | fly chr41B3-41B3\|2-55 cM | chromosome location | fly chr41B3-41B3\|2-55 cM | 1 | 1 | 25010 | 845 | CG17883 | | 3.379e-02 | -3.39 | fly chr56B3-56B3\|2-88 | chromosome location | fly chr56B3-56B3\|2-88 | 1 | 1 | 25010 | 845 | Ir56a | | 3.379e-02 | -3.39 | fly chr16F3-16F6\|1-59 | chromosome location | fly chr16F3-16F6\|1-59 | 1 | 1 | 25010 | 845 | Sh | | 3.379e-02 | -3.39 | fly chr91C6-91D1\|3-64 | chromosome location | fly chr91C6-91D1\|3-64 | 1 | 1 | 25010 | 845 | CG7720 | | 3.379e-02 | -3.39 | fly chr68C12-68C13\|3-36 cM | chromosome location | fly chr68C12-68C13\|3-36 cM | 1 | 1 | 25010 | 845 | Mob2 | | 3.379e-02 | -3.39 | fly chr40F2-40F3\|2-54 cM | chromosome location | fly chr40F2-40F3\|2-54 cM | 1 | 1 | 25010 | 845 | CG3635 | | 3.379e-02 | -3.39 | fly chr9B14-9C1\|1-31 cM | chromosome location | fly chr9B14-9C1\|1-31 cM | 1 | 1 | 25010 | 845 | flw | | 3.379e-02 | -3.39 | fly chr99E4-99F2\|3-100 | chromosome location | fly chr99E4-99F2\|3-100 | 1 | 1 | 25010 | 845 | hdc | | 3.379e-02 | -3.39 | fly chr10C5-10C5\|1-35 | chromosome location | fly chr10C5-10C5\|1-35 | 1 | 1 | 25010 | 845 | CG1572 | | 3.379e-02 | -3.39 | fly chr42F1-42F1\|2-55 | chromosome location | fly chr42F1-42F1\|2-55 | 1 | 1 | 25010 | 845 | esn | | 3.379e-02 | -3.39 | fly chr22C3-22D1\|2-4 | chromosome location | fly chr22C3-22D1\|2-4 | 1 | 1 | 25010 | 845 | GlyP | | 3.379e-02 | -3.39 | fly chr41F1-41F1\|2-55 | chromosome location | fly chr41F1-41F1\|2-55 | 1 | 1 | 25010 | 845 | stw | | 3.379e-02 | -3.39 | fly chr96A19-96A19\|3-85 cM | chromosome location | fly chr96A19-96A19\|3-85 cM | 1 | 1 | 25010 | 845 | tld | | 3.379e-02 | -3.39 | fly chr11E8-11E9\|1-42 | chromosome location | fly chr11E8-11E9\|1-42 | 1 | 1 | 25010 | 845 | HDAC4 | | 3.379e-02 | -3.39 | fly chr8B6-8B6\|1-25 cM | chromosome location | fly chr8B6-8B6\|1-25 cM | 1 | 1 | 25010 | 845 | Uros1 | | 3.379e-02 | -3.39 | fly chr57B3-57B4\|2-92 | chromosome location | fly chr57B3-57B4\|2-92 | 1 | 1 | 25010 | 845 | Ipk1 | | 3.379e-02 | -3.39 | fly chr60B12-60C1\|2-106 | chromosome location | fly chr60B12-60C1\|2-106 | 1 | 1 | 25010 | 845 | AANAT1 | | 3.379e-02 | -3.39 | fly chr15E3-15E5\|1-57 cM | chromosome location | fly chr15E3-15E5\|1-57 cM | 1 | 1 | 25010 | 845 | CG5004 | | 3.379e-02 | -3.39 | fly chr86D1-86D1\|3-50 | chromosome location | fly chr86D1-86D1\|3-50 | 1 | 1 | 25010 | 845 | lncRNA:CR46007 | | 3.379e-02 | -3.39 | fly chr88A5-88A8\|3-54 | chromosome location | fly chr88A5-88A8\|3-54 | 1 | 1 | 25010 | 845 | foxo | | 3.379e-02 | -3.39 | fly chr5E8-5F1\|1-16 | chromosome location | fly chr5E8-5F1\|1-16 | 1 | 1 | 25010 | 845 | CG15894 | | 3.379e-02 | -3.39 | fly chr51F3-51F4\|2-74 | chromosome location | fly chr51F3-51F4\|2-74 | 1 | 1 | 25010 | 845 | ND-51L2 | | 3.379e-02 | -3.39 | fly chr57A3-57A3\|2-91 cM | chromosome location | fly chr57A3-57A3\|2-91 cM | 1 | 1 | 25010 | 845 | side-VIII | | 3.379e-02 | -3.39 | fly chr19E6-19E6\|1-65 | chromosome location | fly chr19E6-19E6\|1-65 | 1 | 1 | 25010 | 845 | CG11666 | | 3.379e-02 | -3.39 | fly chr41D4-41E1\|2-55 cM | chromosome location | fly chr41D4-41E1\|2-55 cM | 1 | 1 | 25010 | 845 | Atf6 | | 3.379e-02 | -3.39 | fly chr33A3-33A5\|2-45 | chromosome location | fly chr33A3-33A5\|2-45 | 1 | 1 | 25010 | 845 | Pde1c | | 3.379e-02 | -3.39 | fly chr99B10-99C1\|3-99 | chromosome location | fly chr99B10-99C1\|3-99 | 1 | 1 | 25010 | 845 | kay | | 3.379e-02 | -3.39 | fly chr29F3-29F4\|2-34 | chromosome location | fly chr29F3-29F4\|2-34 | 1 | 1 | 25010 | 845 | CG34398 | | 3.379e-02 | -3.39 | fly chr22F4-23A1\|2-6 cM | chromosome location | fly chr22F4-23A1\|2-6 cM | 1 | 1 | 25010 | 845 | Cyp309a1 | | 3.379e-02 | -3.39 | fly chr68C14-68C15\|3-36 cM | chromosome location | fly chr68C14-68C15\|3-36 cM | 1 | 1 | 25010 | 845 | CG7368 | | 3.379e-02 | -3.39 | fly chr89F1-89F3\|3-60 cM | chromosome location | fly chr89F1-89F3\|3-60 cM | 1 | 1 | 25010 | 845 | Mur89F | | 3.379e-02 | -3.39 | fly chr9B14-9C1\|1-31 | chromosome location | fly chr9B14-9C1\|1-31 | 1 | 1 | 25010 | 845 | flw | | 3.379e-02 | -3.39 | fly chr78D8-78E1\|3-47 cM | chromosome location | fly chr78D8-78E1\|3-47 cM | 1 | 1 | 25010 | 845 | TfAP-2 | | 3.379e-02 | -3.39 | fly chr13B5-13B6\|1-50 cM | chromosome location | fly chr13B5-13B6\|1-50 cM | 1 | 1 | 25010 | 845 | gce | | 3.379e-02 | -3.39 | fly chr16F7-17A1\|1-59 | chromosome location | fly chr16F7-17A1\|1-59 | 1 | 1 | 25010 | 845 | Frq1 | | 3.379e-02 | -3.39 | fly chr42A1-42A1\|2-55 cM | chromosome location | fly chr42A1-42A1\|2-55 cM | 1 | 1 | 25010 | 845 | dpr12 | | 3.379e-02 | -3.39 | fly chr96B16-96B17\|3-86 | chromosome location | fly chr96B16-96B17\|3-86 | 1 | 1 | 25010 | 845 | Hr96 | | 3.379e-02 | -3.39 | fly chr29E4-29E6\|2-33 | chromosome location | fly chr29E4-29E6\|2-33 | 1 | 1 | 25010 | 845 | raw | | 3.379e-02 | -3.39 | fly chr47F13-47F13\|2-64 cM | chromosome location | fly chr47F13-47F13\|2-64 cM | 1 | 1 | 25010 | 845 | qvr | | 3.379e-02 | -3.39 | fly chr5E8-5F1\|1-16 cM | chromosome location | fly chr5E8-5F1\|1-16 cM | 1 | 1 | 25010 | 845 | CG15894 | | 3.379e-02 | -3.39 | fly chr91D3-91D3\|3-65 cM | chromosome location | fly chr91D3-91D3\|3-65 cM | 1 | 1 | 25010 | 845 | CG14298 | | 3.379e-02 | -3.39 | fly chr41B3-41C2\|2-55 cM | chromosome location | fly chr41B3-41C2\|2-55 cM | 1 | 1 | 25010 | 845 | Gprk1 | | 3.379e-02 | -3.39 | fly chr66A17-66A18\|3-21 | chromosome location | fly chr66A17-66A18\|3-21 | 1 | 1 | 25010 | 845 | CG32365 | | 3.379e-02 | -3.39 | fly chr52E2-52E2\|2-77 cM | chromosome location | fly chr52E2-52E2\|2-77 cM | 1 | 1 | 25010 | 845 | lncRNA:CR43415 | | 3.379e-02 | -3.39 | fly chr26A2-26A2\|2-18 | chromosome location | fly chr26A2-26A2\|2-18 | 1 | 1 | 25010 | 845 | lncRNA:CR45288 | | 3.379e-02 | -3.39 | fly chr91C6-91C6\|3-64 cM | chromosome location | fly chr91C6-91C6\|3-64 cM | 1 | 1 | 25010 | 845 | Epg5 | | 3.379e-02 | -3.39 | fly chr52D4-52D9\|2-76 | chromosome location | fly chr52D4-52D9\|2-76 | 1 | 1 | 25010 | 845 | Strn-Mlck | | 3.379e-02 | -3.39 | fly chr64D1-64D1\|3-14 | chromosome location | fly chr64D1-64D1\|3-14 | 1 | 1 | 25010 | 845 | DIP-delta | | 3.379e-02 | -3.39 | fly chr98A10-98A10\|3-94 | chromosome location | fly chr98A10-98A10\|3-94 | 1 | 1 | 25010 | 845 | dsd | | 3.379e-02 | -3.39 | fly chr52B3-52B5\|2-75 cM | chromosome location | fly chr52B3-52B5\|2-75 cM | 1 | 1 | 25010 | 845 | fus | | 3.379e-02 | -3.39 | fly chr85D16-85D17\|3-48 | chromosome location | fly chr85D16-85D17\|3-48 | 1 | 1 | 25010 | 845 | ps | | 3.379e-02 | -3.39 | fly chr66A1-66A1\|3-20 | chromosome location | fly chr66A1-66A1\|3-20 | 1 | 1 | 25010 | 845 | CG42458 | | 3.379e-02 | -3.39 | fly chr62F2-62F3\|3-4 | chromosome location | fly chr62F2-62F3\|3-4 | 1 | 1 | 25010 | 845 | Mrtf | | 3.379e-02 | -3.39 | fly chr83E5-83E6\|3-47.5 cM | chromosome location | fly chr83E5-83E6\|3-47.5 cM | 1 | 1 | 25010 | 845 | Dmtn | | 3.379e-02 | -3.39 | fly chr93D4-93D5\|3-71 cM | chromosome location | fly chr93D4-93D5\|3-71 cM | 1 | 1 | 25010 | 845 | lncRNA:Hsromega | | 3.379e-02 | -3.39 | fly chr6C11-6C12\|1-18 cM | chromosome location | fly chr6C11-6C12\|1-18 cM | 1 | 1 | 25010 | 845 | CG14439 | | 3.379e-02 | -3.39 | fly chr40F6-40F7\|2-54 | chromosome location | fly chr40F6-40F7\|2-54 | 1 | 1 | 25010 | 845 | Tif-IA | | 3.379e-02 | -3.39 | fly chr67C1-67C1\|3-29 | chromosome location | fly chr67C1-67C1\|3-29 | 1 | 1 | 25010 | 845 | CG42673 | | 3.379e-02 | -3.39 | fly chr49B11-49C1\|2-67 | chromosome location | fly chr49B11-49C1\|2-67 | 1 | 1 | 25010 | 845 | nemy | | 3.379e-02 | -3.39 | fly chr92A5-92A10\|3-66 | chromosome location | fly chr92A5-92A10\|3-66 | 1 | 1 | 25010 | 845 | Dys | | 3.379e-02 | -3.39 | fly chr98D2-98D2\|3-96 cM | chromosome location | fly chr98D2-98D2\|3-96 cM | 1 | 1 | 25010 | 845 | CG1894 | | 3.379e-02 | -3.39 | fly chr102E1-102E3\|4-0 | chromosome location | fly chr102E1-102E3\|4-0 | 1 | 1 | 25010 | 845 | Sox102F | | 3.379e-02 | -3.39 | fly chr47D1-47D4\|2-63 cM | chromosome location | fly chr47D1-47D4\|2-63 cM | 1 | 1 | 25010 | 845 | luna | | 3.379e-02 | -3.39 | fly chr55F1-55F2\|2-86 cM | chromosome location | fly chr55F1-55F2\|2-86 cM | 1 | 1 | 25010 | 845 | Mctp | | 3.379e-02 | -3.39 | fly chr52B3-52B5\|2-75 | chromosome location | fly chr52B3-52B5\|2-75 | 1 | 1 | 25010 | 845 | fus | | 3.379e-02 | -3.39 | fly chr66A17-66A18\|3-21 cM | chromosome location | fly chr66A17-66A18\|3-21 cM | 1 | 1 | 25010 | 845 | CG32365 | | 3.379e-02 | -3.39 | fly chr41C5-41C5\|2-55 cM | chromosome location | fly chr41C5-41C5\|2-55 cM | 1 | 1 | 25010 | 845 | CG44102 | | 3.379e-02 | -3.39 | fly chr100D2-100D3\|3-102 | chromosome location | fly chr100D2-100D3\|3-102 | 1 | 1 | 25010 | 845 | RhoGAP100F | | 3.379e-02 | -3.39 | fly chr92B2-92B3\|3-67 cM | chromosome location | fly chr92B2-92B3\|3-67 cM | 1 | 1 | 25010 | 845 | bnl | | 3.379e-02 | -3.39 | fly chr68C14-68C15\|3-36 | chromosome location | fly chr68C14-68C15\|3-36 | 1 | 1 | 25010 | 845 | CG7368 | | 3.379e-02 | -3.39 | fly chr69F7-69F7\|3-39 cM | chromosome location | fly chr69F7-69F7\|3-39 cM | 1 | 1 | 25010 | 845 | lncRNA:CR43652 | | 3.379e-02 | -3.39 | fly chr41E5-41E6\|2-55 cM | chromosome location | fly chr41E5-41E6\|2-55 cM | 1 | 1 | 25010 | 845 | lncRNA:CR42646 | | 3.379e-02 | -3.39 | fly chr73E5-74A1\|3-45 cM | chromosome location | fly chr73E5-74A1\|3-45 cM | 1 | 1 | 25010 | 845 | Rbp6 | | 3.379e-02 | -3.39 | fly chr61F7-62A2\|3-0 | chromosome location | fly chr61F7-62A2\|3-0 | 1 | 1 | 25010 | 845 | Ptp61F | | 3.379e-02 | -3.39 | fly chr55C10-55D1\|2-85 | chromosome location | fly chr55C10-55D1\|2-85 | 1 | 1 | 25010 | 845 | CG30116 | | 3.379e-02 | -3.39 | fly chr42A6-42A7\|2-55 cM | chromosome location | fly chr42A6-42A7\|2-55 cM | 1 | 1 | 25010 | 845 | Src42A | | 3.379e-02 | -3.39 | fly chr77F1-77F3\|3-47 | chromosome location | fly chr77F1-77F3\|3-47 | 1 | 1 | 25010 | 845 | CG32432 | | 3.379e-02 | -3.39 | fly chr62C4-62D1\|3-1.5 cM | chromosome location | fly chr62C4-62D1\|3-1.5 cM | 1 | 1 | 25010 | 845 | zormin | | 3.379e-02 | -3.39 | fly chr42A1-42A1\|2-55 | chromosome location | fly chr42A1-42A1\|2-55 | 1 | 1 | 25010 | 845 | dpr12 | | 3.379e-02 | -3.39 | fly chr44B4-44B5\|2-58 cM | chromosome location | fly chr44B4-44B5\|2-58 cM | 1 | 1 | 25010 | 845 | Lpin | | 3.379e-02 | -3.39 | fly chr33F4-33F4\|2-47 | chromosome location | fly chr33F4-33F4\|2-47 | 1 | 1 | 25010 | 845 | mir-9382 | | 3.379e-02 | -3.39 | fly chr86C1-86C3\|3-50 cM | chromosome location | fly chr86C1-86C3\|3-50 cM | 1 | 1 | 25010 | 845 | hth | | 3.379e-02 | -3.39 | fly chr47D5-47D6\|2-63 | chromosome location | fly chr47D5-47D6\|2-63 | 1 | 1 | 25010 | 845 | CG13229 | | 3.379e-02 | -3.39 | fly chr11F4-11F5\|1-43 cM | chromosome location | fly chr11F4-11F5\|1-43 cM | 1 | 1 | 25010 | 845 | Set2 | | 3.379e-02 | -3.39 | fly chr41C6-41D1\|2-55 | chromosome location | fly chr41C6-41D1\|2-55 | 1 | 1 | 25010 | 845 | Ir41a | | 3.379e-02 | -3.39 | fly chr87A10-87B1\|3-51 | chromosome location | fly chr87A10-87B1\|3-51 | 1 | 1 | 25010 | 845 | dpr15 | | 3.379e-02 | -3.39 | fly chr75E5-75E6\|3-45 cM | chromosome location | fly chr75E5-75E6\|3-45 cM | 1 | 1 | 25010 | 845 | CG6839 | | 3.379e-02 | -3.39 | fly chr52A3-52A1\|2-74 cM | chromosome location | fly chr52A3-52A1\|2-74 cM | 1 | 1 | 25010 | 845 | Stacl | | 3.379e-02 | -3.39 | fly chr87C1-87C2\|3-52 cM | chromosome location | fly chr87C1-87C2\|3-52 cM | 1 | 1 | 25010 | 845 | Octbeta2R | | 3.379e-02 | -3.39 | fly chr66B8-66B10\|3-22 | chromosome location | fly chr66B8-66B10\|3-22 | 1 | 1 | 25010 | 845 | CG13676 | | 3.379e-02 | -3.39 | fly chr49B9-49B9\|2-67 | chromosome location | fly chr49B9-49B9\|2-67 | 1 | 1 | 25010 | 845 | CG30054 | | 3.379e-02 | -3.39 | fly chr10B4-10B5\|1-34 cM | chromosome location | fly chr10B4-10B5\|1-34 cM | 1 | 1 | 25010 | 845 | dsh | | 3.379e-02 | -3.39 | fly chr44F2-44F3\|2-59 cM | chromosome location | fly chr44F2-44F3\|2-59 cM | 1 | 1 | 25010 | 845 | CG8272 | | 3.379e-02 | -3.39 | fly chr75F9-76A3\|3-45 cM | chromosome location | fly chr75F9-76A3\|3-45 cM | 1 | 1 | 25010 | 845 | fz2 | | 3.379e-02 | -3.39 | fly chr82D8-82E1\|3-47.1 cM | chromosome location | fly chr82D8-82E1\|3-47.1 cM | 1 | 1 | 25010 | 845 | opa | | 3.379e-02 | -3.39 | fly chr2C5-2C5\|1-0.5 cM | chromosome location | fly chr2C5-2C5\|1-0.5 cM | 1 | 1 | 25010 | 845 | mir-4954 | | 3.379e-02 | -3.39 | fly chr13A12-13B1\|1-49 cM | chromosome location | fly chr13A12-13B1\|1-49 cM | 1 | 1 | 25010 | 845 | drd | | 3.379e-02 | -3.39 | fly chr70D3-70D4\|3-42 | chromosome location | fly chr70D3-70D4\|3-42 | 1 | 1 | 25010 | 845 | nuf | | 3.379e-02 | -3.39 | fly chr89B1-89B2\|3-58 cM | chromosome location | fly chr89B1-89B2\|3-58 cM | 1 | 1 | 25010 | 845 | msps | | 3.379e-02 | -3.39 | fly chr6F10-7A5\|1-19 | chromosome location | fly chr6F10-7A5\|1-19 | 1 | 1 | 25010 | 845 | CG9650 | | 3.379e-02 | -3.39 | fly chr27B1-27B2\|2-22 | chromosome location | fly chr27B1-27B2\|2-22 | 1 | 1 | 25010 | 845 | nrv2 | | 3.379e-02 | -3.39 | fly chr33D2-33D5\|2-46 | chromosome location | fly chr33D2-33D5\|2-46 | 1 | 1 | 25010 | 845 | bru1 | | 3.379e-02 | -3.39 | fly chr20A1-20A2\|1-66 | chromosome location | fly chr20A1-20A2\|1-66 | 1 | 1 | 25010 | 845 | lncRNA:flam | | 3.379e-02 | -3.39 | fly chr17A1-17A2\|1-60 | chromosome location | fly chr17A1-17A2\|1-60 | 1 | 1 | 25010 | 845 | Frq2 | | 3.379e-02 | -3.39 | fly chr2D1-2D2\|1-0.6 | chromosome location | fly chr2D1-2D2\|1-0.6 | 1 | 1 | 25010 | 845 | csw | | 3.379e-02 | -3.39 | fly chr18A3-18A4\|1-62 cM | chromosome location | fly chr18A3-18A4\|1-62 cM | 1 | 1 | 25010 | 845 | RhoGAP18B | | 3.379e-02 | -3.39 | fly chr84A6-84B2\|3-48 | chromosome location | fly chr84A6-84B2\|3-48 | 1 | 1 | 25010 | 845 | Antp | | 3.379e-02 | -3.39 | fly chr4C8-4C8\|1-8 cM | chromosome location | fly chr4C8-4C8\|1-8 cM | 1 | 1 | 25010 | 845 | CG12684 | | 3.379e-02 | -3.39 | fly chr33E4-33E5\|2-46 cM | chromosome location | fly chr33E4-33E5\|2-46 cM | 1 | 1 | 25010 | 845 | eRF3 | | 3.379e-02 | -3.39 | fly chr70B3-70B4\|3-40 | chromosome location | fly chr70B3-70B4\|3-40 | 1 | 1 | 25010 | 845 | Spt20 | | 3.379e-02 | -3.39 | fly chr15A10-15A10\|1-55 | chromosome location | fly chr15A10-15A10\|1-55 | 1 | 1 | 25010 | 845 | lncRNA:CR44129 | | 3.379e-02 | -3.39 | fly chr57B16-57B19\|2-92 cM | chromosome location | fly chr57B16-57B19\|2-92 cM | 1 | 1 | 25010 | 845 | Treh | | 3.379e-02 | -3.39 | fly chr80A2-80A4\|3-47 cM | chromosome location | fly chr80A2-80A4\|3-47 cM | 1 | 1 | 25010 | 845 | jim | | 3.379e-02 | -3.39 | fly chr36E1-36E1\|2-53 | chromosome location | fly chr36E1-36E1\|2-53 | 1 | 1 | 25010 | 845 | lncRNA:CR43420 | | 3.379e-02 | -3.39 | fly chr96A22-96A23\|3-85 cM | chromosome location | fly chr96A22-96A23\|3-85 cM | 1 | 1 | 25010 | 845 | polybromo | | 3.379e-02 | -3.39 | fly chr57F6-57F7\|2-96 cM | chromosome location | fly chr57F6-57F7\|2-96 cM | 1 | 1 | 25010 | 845 | pirk | | 3.379e-02 | -3.39 | fly chr13B5-13B6\|1-50 | chromosome location | fly chr13B5-13B6\|1-50 | 1 | 1 | 25010 | 845 | gce | | 3.379e-02 | -3.39 | fly chr40F6-40F7\|2-54 cM | chromosome location | fly chr40F6-40F7\|2-54 cM | 1 | 1 | 25010 | 845 | Tif-IA | | 3.379e-02 | -3.39 | fly chr97E6-97E8\|3-92 cM | chromosome location | fly chr97E6-97E8\|3-92 cM | 1 | 1 | 25010 | 845 | Ser | | 3.379e-02 | -3.39 | fly chr2C4-2C7\|1-0.5 | chromosome location | fly chr2C4-2C7\|1-0.5 | 1 | 1 | 25010 | 845 | Actn | | 3.379e-02 | -3.39 | fly chr15F7-16A1\|1-57 | chromosome location | fly chr15F7-16A1\|1-57 | 1 | 1 | 25010 | 845 | CG5445 | | 3.379e-02 | -3.39 | fly chr70E4-70E4\|3-42 cM | chromosome location | fly chr70E4-70E4\|3-42 cM | 1 | 1 | 25010 | 845 | shd | | 3.379e-02 | -3.39 | fly chr6F10-7A5\|1-19 cM | chromosome location | fly chr6F10-7A5\|1-19 cM | 1 | 1 | 25010 | 845 | CG9650 | | 3.379e-02 | -3.39 | fly chr96C4-96C4\|3-87 cM | chromosome location | fly chr96C4-96C4\|3-87 cM | 1 | 1 | 25010 | 845 | CG13653 | | 3.379e-02 | -3.39 | fly chr75B11-75B12\|3-45 | chromosome location | fly chr75B11-75B12\|3-45 | 1 | 1 | 25010 | 845 | geko | | 3.379e-02 | -3.39 | fly chr70E4-70E4\|3-42 | chromosome location | fly chr70E4-70E4\|3-42 | 1 | 1 | 25010 | 845 | shd | | 3.379e-02 | -3.39 | fly chr11C4-11D1\|1-40 | chromosome location | fly chr11C4-11D1\|1-40 | 1 | 1 | 25010 | 845 | Pde9 | | 3.379e-02 | -3.39 | fly chr57D8-57D11\|2-94 | chromosome location | fly chr57D8-57D11\|2-94 | 1 | 1 | 25010 | 845 | CG42672 | | 3.379e-02 | -3.39 | fly chr12C8-12C7\|1-46 cM | chromosome location | fly chr12C8-12C7\|1-46 cM | 1 | 1 | 25010 | 845 | mamo | | 3.379e-02 | -3.39 | fly chr75B11-75B12\|3-45 cM | chromosome location | fly chr75B11-75B12\|3-45 cM | 1 | 1 | 25010 | 845 | geko | | 3.379e-02 | -3.39 | fly chr94A4-94A5\|3-74 cM | chromosome location | fly chr94A4-94A5\|3-74 cM | 1 | 1 | 25010 | 845 | asRNA:CR46261 | | 3.379e-02 | -3.39 | fly chr53D2-53D2\|2-80 | chromosome location | fly chr53D2-53D2\|2-80 | 1 | 1 | 25010 | 845 | Idgf6 | | 3.379e-02 | -3.39 | fly chr42A6-42A7\|2-55 | chromosome location | fly chr42A6-42A7\|2-55 | 1 | 1 | 25010 | 845 | Src42A | | 3.379e-02 | -3.39 | fly chr95D5-95D8\|3-82 | chromosome location | fly chr95D5-95D8\|3-82 | 1 | 1 | 25010 | 845 | KrT95D | | 3.379e-02 | -3.39 | fly chr86D1-86D1\|3-50 cM | chromosome location | fly chr86D1-86D1\|3-50 cM | 1 | 1 | 25010 | 845 | lncRNA:CR46007 | | 3.379e-02 | -3.39 | fly chr72A4-72A4\|3-43 | chromosome location | fly chr72A4-72A4\|3-43 | 1 | 1 | 25010 | 845 | lncRNA:CR44050 | | 3.379e-02 | -3.39 | fly chrh2-h3 | chromosome location | fly chrh2-h3 | 1 | 1 | 25010 | 845 | kl-5 | | 3.379e-02 | -3.39 | fly chr44B4-44B5\|2-58 | chromosome location | fly chr44B4-44B5\|2-58 | 1 | 1 | 25010 | 845 | Lpin | | 3.379e-02 | -3.39 | fly chr4C8-4C8\|1-8 | chromosome location | fly chr4C8-4C8\|1-8 | 1 | 1 | 25010 | 845 | CG12684 | | 3.379e-02 | -3.39 | fly chr19E2-19E3\|1-65 | chromosome location | fly chr19E2-19E3\|1-65 | 1 | 1 | 25010 | 845 | shakB | | 3.379e-02 | -3.39 | fly chr33F4-33F4\|2-47 cM | chromosome location | fly chr33F4-33F4\|2-47 cM | 1 | 1 | 25010 | 845 | mir-9382 | | 3.379e-02 | -3.39 | fly chr33A1-33A2\|2-45 cM | chromosome location | fly chr33A1-33A2\|2-45 cM | 1 | 1 | 25010 | 845 | crol | | 3.379e-02 | -3.39 | fly chr2C5-2C5\|1-0.5 | chromosome location | fly chr2C5-2C5\|1-0.5 | 1 | 1 | 25010 | 845 | mir-4954 | | 3.379e-02 | -3.39 | fly chr82A1-82A3\|3-47.1 cM | chromosome location | fly chr82A1-82A3\|3-47.1 cM | 1 | 1 | 25010 | 845 | cpx | | 3.379e-02 | -3.39 | fly chr5A7-5A8\|1-12 cM | chromosome location | fly chr5A7-5A8\|1-12 cM | 1 | 1 | 25010 | 845 | CG34435 | | 3.379e-02 | -3.39 | fly chr49A3-49A4\|2-67 | chromosome location | fly chr49A3-49A4\|2-67 | 1 | 1 | 25010 | 845 | CG13157 | | 3.379e-02 | -3.39 | fly chr43E13-43E16\|2-57 | chromosome location | fly chr43E13-43E16\|2-57 | 1 | 1 | 25010 | 845 | CG46385 | | 3.379e-02 | -3.39 | fly chr33E4-33E5\|2-46 | chromosome location | fly chr33E4-33E5\|2-46 | 1 | 1 | 25010 | 845 | eRF3 | | 3.379e-02 | -3.39 | fly chr80C3-80C3\|3-47 | chromosome location | fly chr80C3-80C3\|3-47 | 1 | 1 | 25010 | 845 | nrm | | 3.379e-02 | -3.39 | fly chr70D3-70D4\|3-42 cM | chromosome location | fly chr70D3-70D4\|3-42 cM | 1 | 1 | 25010 | 845 | nuf | | 3.379e-02 | -3.39 | fly chr62F2-62F3\|3-4 cM | chromosome location | fly chr62F2-62F3\|3-4 cM | 1 | 1 | 25010 | 845 | Mrtf | | 3.379e-02 | -3.39 | fly chr47D1-47D4\|2-63 | chromosome location | fly chr47D1-47D4\|2-63 | 1 | 1 | 25010 | 845 | luna | | 3.379e-02 | -3.39 | fly chr91A7-91B3\|3-63 | chromosome location | fly chr91A7-91B3\|3-63 | 1 | 1 | 25010 | 845 | fru | | 3.379e-02 | -3.39 | fly chr62F2-62F2\|3-4 | chromosome location | fly chr62F2-62F2\|3-4 | 1 | 1 | 25010 | 845 | Acp62F | | 3.379e-02 | -3.39 | fly chr100B2-100B3\|3-102 cM | chromosome location | fly chr100B2-100B3\|3-102 cM | 1 | 1 | 25010 | 845 | CG11318 | | 3.379e-02 | -3.39 | fly chr75E5-75E6\|3-45 | chromosome location | fly chr75E5-75E6\|3-45 | 1 | 1 | 25010 | 845 | CG6839 | | 3.379e-02 | -3.39 | fly chr16F3-16F6\|1-59 cM | chromosome location | fly chr16F3-16F6\|1-59 cM | 1 | 1 | 25010 | 845 | Sh | | 3.379e-02 | -3.39 | fly chr70D7-70E1\|3-42 | chromosome location | fly chr70D7-70E1\|3-42 | 1 | 1 | 25010 | 845 | bbg | | 3.379e-02 | -3.39 | fly chr94B8-94B11\|3-75 cM | chromosome location | fly chr94B8-94B11\|3-75 cM | 1 | 1 | 25010 | 845 | wake | | 3.379e-02 | -3.39 | fly chr44B8-44C2\|2-58 | chromosome location | fly chr44B8-44C2\|2-58 | 1 | 1 | 25010 | 845 | LRP1 | | 3.379e-02 | -3.39 | fly chr13A12-13B1\|1-49 | chromosome location | fly chr13A12-13B1\|1-49 | 1 | 1 | 25010 | 845 | drd | | 3.379e-02 | -3.39 | fly chr69F7-69F7\|3-39 | chromosome location | fly chr69F7-69F7\|3-39 | 1 | 1 | 25010 | 845 | lncRNA:CR43652 | | 3.379e-02 | -3.39 | fly chr45A9-45A10\|2-60 cM | chromosome location | fly chr45A9-45A10\|2-60 cM | 1 | 1 | 25010 | 845 | CG18659 | | 3.379e-02 | -3.39 | fly chr33A3-33A5\|2-45 cM | chromosome location | fly chr33A3-33A5\|2-45 cM | 1 | 1 | 25010 | 845 | Pde1c | | 3.379e-02 | -3.39 | fly chr27D2-27D2\|2-24 | chromosome location | fly chr27D2-27D2\|2-24 | 1 | 1 | 25010 | 845 | lncRNA:CR44326 | | 3.379e-02 | -3.39 | fly chr17A1-17A2\|1-60 cM | chromosome location | fly chr17A1-17A2\|1-60 cM | 1 | 1 | 25010 | 845 | Frq2 | | 3.379e-02 | -3.39 | fly chr91E5-91E5\|3-65 cM | chromosome location | fly chr91E5-91E5\|3-65 cM | 1 | 1 | 25010 | 845 | lncRNA:CR46041 | | 3.379e-02 | -3.39 | fly chr51B4-51B4\|2-72 | chromosome location | fly chr51B4-51B4\|2-72 | 1 | 1 | 25010 | 845 | CG12869 | | 3.379e-02 | -3.39 | fly chr96E2-96E4\|3-88 cM | chromosome location | fly chr96E2-96E4\|3-88 cM | 1 | 1 | 25010 | 845 | msi | | 3.379e-02 | -3.39 | fly chr40A1-40A2\|2-54 | chromosome location | fly chr40A1-40A2\|2-54 | 1 | 1 | 25010 | 845 | step | | 3.379e-02 | -3.39 | fly chr99E5-99E5\|3-100 cM | chromosome location | fly chr99E5-99E5\|3-100 cM | 1 | 1 | 25010 | 845 | CG34300 | | 3.379e-02 | -3.39 | fly chr97B9-97C1\|3-91 | chromosome location | fly chr97B9-97C1\|3-91 | 1 | 1 | 25010 | 845 | scrib | | 3.379e-02 | -3.39 | fly chr96B19-96B20\|3-86 | chromosome location | fly chr96B19-96B20\|3-86 | 1 | 1 | 25010 | 845 | Stt3B | | 3.379e-02 | -3.39 | fly chr11E8-11E9\|1-42 cM | chromosome location | fly chr11E8-11E9\|1-42 cM | 1 | 1 | 25010 | 845 | HDAC4 | | 3.379e-02 | -3.39 | fly chr64D1-64D1\|3-14 cM | chromosome location | fly chr64D1-64D1\|3-14 cM | 1 | 1 | 25010 | 845 | DIP-delta | | 3.379e-02 | -3.39 | fly chr94F1-94F4\|3-79 cM | chromosome location | fly chr94F1-94F4\|3-79 cM | 1 | 1 | 25010 | 845 | CG4374 | | 3.379e-02 | -3.39 | fly chr13-13\|1-49 cM | chromosome location | fly chr13-13\|1-49 cM | 1 | 1 | 25010 | 845 | Flo2 | | 3.379e-02 | -3.39 | fly chr57B3-57B4\|2-92 cM | chromosome location | fly chr57B3-57B4\|2-92 cM | 1 | 1 | 25010 | 845 | Ipk1 | | 3.379e-02 | -3.39 | fly chr16E4-16E4\|1-59 | chromosome location | fly chr16E4-16E4\|1-59 | 1 | 1 | 25010 | 845 | mnb | | 3.379e-02 | -3.39 | fly chr25D5-25D5\|2-16 | chromosome location | fly chr25D5-25D5\|2-16 | 1 | 1 | 25010 | 845 | CG14024 | | 3.379e-02 | -3.39 | fly chr69C1-69C1\|3-38 cM | chromosome location | fly chr69C1-69C1\|3-38 cM | 1 | 1 | 25010 | 845 | toe | | 3.379e-02 | -3.39 | fly chr96B16-96B17\|3-86 cM | chromosome location | fly chr96B16-96B17\|3-86 cM | 1 | 1 | 25010 | 845 | Hr96 | | 3.379e-02 | -3.39 | fly chr73A8-73A8\|3-44 cM | chromosome location | fly chr73A8-73A8\|3-44 cM | 1 | 1 | 25010 | 845 | CG43206 | | 3.379e-02 | -3.39 | fly chr6B3-6C1\|1-17 cM | chromosome location | fly chr6B3-6C1\|1-17 cM | 1 | 1 | 25010 | 845 | CG34417 | | 3.379e-02 | -3.39 | fly chr72F1-73A3\|3-44 | chromosome location | fly chr72F1-73A3\|3-44 | 1 | 1 | 25010 | 845 | CG33158 | | 3.379e-02 | -3.39 | fly chr64C7-64C11\|3-13 | chromosome location | fly chr64C7-64C11\|3-13 | 1 | 1 | 25010 | 845 | shep | | 3.379e-02 | -3.39 | fly chr18A3-18A4\|1-62 | chromosome location | fly chr18A3-18A4\|1-62 | 1 | 1 | 25010 | 845 | RhoGAP18B | | 3.379e-02 | -3.39 | fly chr65E4-65E5\|3-19 | chromosome location | fly chr65E4-65E5\|3-19 | 1 | 1 | 25010 | 845 | melt | | 3.379e-02 | -3.39 | fly chr33E7-33E9\|2-46 cM | chromosome location | fly chr33E7-33E9\|2-46 cM | 1 | 1 | 25010 | 845 | bun | | 3.379e-02 | -3.39 | fly chr17B-17B\|1-60 | chromosome location | fly chr17B-17B\|1-60 | 1 | 1 | 25010 | 845 | CrebB | | 3.379e-02 | -3.39 | fly chr37D5-37D5\|2-54 cM | chromosome location | fly chr37D5-37D5\|2-54 cM | 1 | 1 | 25010 | 845 | snmRNA:359 | | 3.379e-02 | -3.39 | fly chr87C1-87C2\|3-52 | chromosome location | fly chr87C1-87C2\|3-52 | 1 | 1 | 25010 | 845 | Octbeta2R | | 3.379e-02 | -3.39 | fly chr33D2-33D3\|2-46 | chromosome location | fly chr33D2-33D3\|2-46 | 1 | 1 | 25010 | 845 | atilla | | 3.379e-02 | -3.39 | fly chr33F2-33F2\|2-47 cM | chromosome location | fly chr33F2-33F2\|2-47 cM | 1 | 1 | 25010 | 845 | Nepl7 | | 3.379e-02 | -3.39 | fly chr10B4-10B5\|1-34 | chromosome location | fly chr10B4-10B5\|1-34 | 1 | 1 | 25010 | 845 | dsh | | 3.379e-02 | -3.39 | fly chr37D5-37D5\|2-54 | chromosome location | fly chr37D5-37D5\|2-54 | 1 | 1 | 25010 | 845 | snmRNA:359 | | 3.379e-02 | -3.39 | fly chr60A9-60A11\|2-105 cM | chromosome location | fly chr60A9-60A11\|2-105 cM | 1 | 1 | 25010 | 845 | kcc | | 3.379e-02 | -3.39 | fly chr41E4-41E5\|2-55 cM | chromosome location | fly chr41E4-41E5\|2-55 cM | 1 | 1 | 25010 | 845 | sxc | | 3.379e-02 | -3.39 | fly chr37E4-37E5\|2-54 cM | chromosome location | fly chr37E4-37E5\|2-54 cM | 1 | 1 | 25010 | 845 | CG10188 | | 3.379e-02 | -3.39 | fly chr10B13-10B13\|1-34 | chromosome location | fly chr10B13-10B13\|1-34 | 1 | 1 | 25010 | 845 | Sk1 | | 3.379e-02 | -3.39 | fly chr3A4-3A6\|1-1 cM | chromosome location | fly chr3A4-3A6\|1-1 cM | 1 | 1 | 25010 | 845 | Pdfr | | 3.379e-02 | -3.39 | fly chr52B3-52B3\|2-75 cM | chromosome location | fly chr52B3-52B3\|2-75 cM | 1 | 1 | 25010 | 845 | mir-278 | | 3.379e-02 | -3.39 | fly chr53D8-53D9\|2-80 cM | chromosome location | fly chr53D8-53D9\|2-80 cM | 1 | 1 | 25010 | 845 | CG5522 | | 3.379e-02 | -3.39 | fly chr93D4-93D5\|3-71 | chromosome location | fly chr93D4-93D5\|3-71 | 1 | 1 | 25010 | 845 | lncRNA:Hsromega | | 3.379e-02 | -3.39 | fly chr98A10-98A12\|3-94 | chromosome location | fly chr98A10-98A12\|3-94 | 1 | 1 | 25010 | 845 | Tusp | | 3.379e-02 | -3.39 | fly chr73A10-73A11\|3-44 | chromosome location | fly chr73A10-73A11\|3-44 | 1 | 1 | 25010 | 845 | spd-2 | | 3.379e-02 | -3.39 | fly chr73C3-73C3\|3-44 | chromosome location | fly chr73C3-73C3\|3-44 | 1 | 1 | 25010 | 845 | lncRNA:CR46214 | | 3.379e-02 | -3.39 | fly chr6D7-6D8\|1-18 cM | chromosome location | fly chr6D7-6D8\|1-18 cM | 1 | 1 | 25010 | 845 | lncRNA:CR45531 | | 3.379e-02 | -3.39 | fly chr53D8-53D9\|2-80 | chromosome location | fly chr53D8-53D9\|2-80 | 1 | 1 | 25010 | 845 | CG5522 | | 3.379e-02 | -3.39 | fly chr3A1-3A4\|1-1 | chromosome location | fly chr3A1-3A4\|1-1 | 1 | 1 | 25010 | 845 | trol | | 3.379e-02 | -3.39 | fly chr61B3-61C1\|3-1 cM | chromosome location | fly chr61B3-61C1\|3-1 cM | 1 | 1 | 25010 | 845 | RhoGEF3 | | 3.379e-02 | -3.39 | fly chr54F1-54F3\|2-84 cM | chromosome location | fly chr54F1-54F3\|2-84 cM | 1 | 1 | 25010 | 845 | olf186-F | | 3.379e-02 | -3.39 | fly chr84F11-84F12\|3-48 cM | chromosome location | fly chr84F11-84F12\|3-48 cM | 1 | 1 | 25010 | 845 | Poxm | | 3.379e-02 | -3.39 | fly chr11D9-11D10\|1-42 | chromosome location | fly chr11D9-11D10\|1-42 | 1 | 1 | 25010 | 845 | IP3K2 | | 3.379e-02 | -3.39 | fly chr3A8-3B1\|1-1 cM | chromosome location | fly chr3A8-3B1\|1-1 cM | 1 | 1 | 25010 | 845 | sgg | | 3.379e-02 | -3.39 | fly chr31A3-31A3\|2-39 | chromosome location | fly chr31A3-31A3\|2-39 | 1 | 1 | 25010 | 845 | CG4839 | | 3.379e-02 | -3.39 | fly chr99F1-99F1\|3-100 | chromosome location | fly chr99F1-99F1\|3-100 | 1 | 1 | 25010 | 845 | lncRNA:CR46114 | | 3.379e-02 | -3.39 | fly chr66A1-66A1\|3-20 cM | chromosome location | fly chr66A1-66A1\|3-20 cM | 1 | 1 | 25010 | 845 | CG42458 | | 3.379e-02 | -3.39 | fly chr33F2-33F2\|2-47 | chromosome location | fly chr33F2-33F2\|2-47 | 1 | 1 | 25010 | 845 | Nepl7 | | 3.379e-02 | -3.39 | fly chr30B3-30B5\|2-35 cM | chromosome location | fly chr30B3-30B5\|2-35 cM | 1 | 1 | 25010 | 845 | numb | | 3.379e-02 | -3.39 | fly chr57B15-57B16\|2-92 cM | chromosome location | fly chr57B15-57B16\|2-92 cM | 1 | 1 | 25010 | 845 | shg | | 3.379e-02 | -3.39 | fly chr51F3-51F4\|2-74 cM | chromosome location | fly chr51F3-51F4\|2-74 cM | 1 | 1 | 25010 | 845 | ND-51L2 | | 3.379e-02 | -3.39 | fly chr8F2-8F2\|1-29 | chromosome location | fly chr8F2-8F2\|1-29 | 1 | 1 | 25010 | 845 | CG42797 | | 3.379e-02 | -3.39 | fly chr40F2-40F3\|2-54 | chromosome location | fly chr40F2-40F3\|2-54 | 1 | 1 | 25010 | 845 | CG3635 | | 3.379e-02 | -3.39 | fly chr92A3-92A5\|3-66 cM | chromosome location | fly chr92A3-92A5\|3-66 cM | 1 | 1 | 25010 | 845 | CG31221 | | 3.379e-02 | -3.39 | fly chr3A4-3A6\|1-1 | chromosome location | fly chr3A4-3A6\|1-1 | 1 | 1 | 25010 | 845 | Pdfr | | 3.379e-02 | -3.39 | fly chr102E5-102F1\|4-0 cM | chromosome location | fly chr102E5-102F1\|4-0 cM | 1 | 1 | 25010 | 845 | unc-13 | | 3.379e-02 | -3.39 | fly chr73E1-73E2\|3-45 cM | chromosome location | fly chr73E1-73E2\|3-45 cM | 1 | 1 | 25010 | 845 | Exn | | 3.379e-02 | -3.39 | fly chr27C4-27C6\|2-23 cM | chromosome location | fly chr27C4-27C6\|2-23 cM | 1 | 1 | 25010 | 845 | CG18304 | | 3.379e-02 | -3.39 | fly chr27D2-27D2\|2-24 cM | chromosome location | fly chr27D2-27D2\|2-24 cM | 1 | 1 | 25010 | 845 | lncRNA:CR44326 | | 3.379e-02 | -3.39 | fly chr83E8-83F1\|3-47.5 cM | chromosome location | fly chr83E8-83F1\|3-47.5 cM | 1 | 1 | 25010 | 845 | CG15186 | | 3.379e-02 | -3.39 | fly chr86D5-86D6\|3-50 cM | chromosome location | fly chr86D5-86D6\|3-50 cM | 1 | 1 | 25010 | 845 | Ugt304A1 | | 3.379e-02 | -3.39 | fly chr43E13-43E16\|2-57 cM | chromosome location | fly chr43E13-43E16\|2-57 cM | 1 | 1 | 25010 | 845 | CG46385 | | 3.379e-02 | -3.39 | fly chr13E8-13E12\|1-51 cM | chromosome location | fly chr13E8-13E12\|1-51 cM | 1 | 1 | 25010 | 845 | CG8184 | | 3.379e-02 | -3.39 | fly chr44F2-44F3\|2-59 | chromosome location | fly chr44F2-44F3\|2-59 | 1 | 1 | 25010 | 845 | CG8272 | | 3.379e-02 | -3.39 | fly chr2C4-2C7\|1-0.5 cM | chromosome location | fly chr2C4-2C7\|1-0.5 cM | 1 | 1 | 25010 | 845 | Actn | | 3.379e-02 | -3.39 | fly chr94C9-94C9\|3-76 | chromosome location | fly chr94C9-94C9\|3-76 | 1 | 1 | 25010 | 845 | lncRNA:CR44350 | | 3.379e-02 | -3.39 | fly chr53C7-53C7\|2-80 | chromosome location | fly chr53C7-53C7\|2-80 | 1 | 1 | 25010 | 845 | Syn2 | | 3.379e-02 | -3.39 | fly chr90A5-90A6\|3-60 cM | chromosome location | fly chr90A5-90A6\|3-60 cM | 1 | 1 | 25010 | 845 | CG42823 | | 3.379e-02 | -3.39 | fly chr2B2-2B4\|1-0.2 | chromosome location | fly chr2B2-2B4\|1-0.2 | 1 | 1 | 25010 | 845 | Mur2B | | 3.379e-02 | -3.39 | fly chr17B-17B\|1-60 cM | chromosome location | fly chr17B-17B\|1-60 cM | 1 | 1 | 25010 | 845 | CrebB | | 3.379e-02 | -3.39 | fly chr62E3-62E4\|3-3 cM | chromosome location | fly chr62E3-62E4\|3-3 cM | 1 | 1 | 25010 | 845 | CG1146 | | 3.379e-02 | -3.39 | fly chr41B3-41B3\|2-55 | chromosome location | fly chr41B3-41B3\|2-55 | 1 | 1 | 25010 | 845 | CG17883 | | 3.379e-02 | -3.39 | fly chr6C9-6C10\|1-18 | chromosome location | fly chr6C9-6C10\|1-18 | 1 | 1 | 25010 | 845 | CG3168 | | 3.379e-02 | -3.39 | fly chr92A5-92A10\|3-66 cM | chromosome location | fly chr92A5-92A10\|3-66 cM | 1 | 1 | 25010 | 845 | Dys | | 3.379e-02 | -3.39 | fly chr41F1-41F1\|2-55 cM | chromosome location | fly chr41F1-41F1\|2-55 cM | 1 | 1 | 25010 | 845 | stw | | 3.379e-02 | -3.39 | fly chr31A3-31A3\|2-39 cM | chromosome location | fly chr31A3-31A3\|2-39 cM | 1 | 1 | 25010 | 845 | CG4839 | | 3.379e-02 | -3.39 | fly chr88B1-88B3\|3-55 cM | chromosome location | fly chr88B1-88B3\|3-55 cM | 1 | 1 | 25010 | 845 | red | | 3.379e-02 | -3.39 | fly chr3C9-3D1\|1-2 cM | chromosome location | fly chr3C9-3D1\|1-2 cM | 1 | 1 | 25010 | 845 | dnc | | 3.379e-02 | -3.39 | fly chr12E10-12E10\|1-48 cM | chromosome location | fly chr12E10-12E10\|1-48 cM | 1 | 1 | 25010 | 845 | CG9411 | | 3.379e-02 | -3.39 | fly chr4C3-4C4\|1-8 | chromosome location | fly chr4C3-4C4\|1-8 | 1 | 1 | 25010 | 845 | bi | | 3.379e-02 | -3.39 | fly chr98F1-98F3\|3-97 | chromosome location | fly chr98F1-98F3\|3-97 | 1 | 1 | 25010 | 845 | Doa | | 3.379e-02 | -3.39 | fly chr5F1-5F2\|1-16 | chromosome location | fly chr5F1-5F2\|1-16 | 1 | 1 | 25010 | 845 | CG42240 | | 3.379e-02 | -3.39 | fly chr76D7-76D7\|3-46 cM | chromosome location | fly chr76D7-76D7\|3-46 cM | 1 | 1 | 25010 | 845 | CG14182 | | 3.379e-02 | -3.39 | fly chr69C1-69C1\|3-38 | chromosome location | fly chr69C1-69C1\|3-38 | 1 | 1 | 25010 | 845 | toe | | 3.379e-02 | -3.39 | fly chr6D7-6D8\|1-18 | chromosome location | fly chr6D7-6D8\|1-18 | 1 | 1 | 25010 | 845 | lncRNA:CR45531 | | 3.379e-02 | -3.39 | fly chr15A10-15A10\|1-55 cM | chromosome location | fly chr15A10-15A10\|1-55 cM | 1 | 1 | 25010 | 845 | lncRNA:CR44129 | | 3.379e-02 | -3.39 | fly chr18B6-18B6\|1-62 cM | chromosome location | fly chr18B6-18B6\|1-62 cM | 1 | 1 | 25010 | 845 | rictor | | 3.379e-02 | -3.39 | fly chr25D5-25D5\|2-16 cM | chromosome location | fly chr25D5-25D5\|2-16 cM | 1 | 1 | 25010 | 845 | CG14024 | | 3.379e-02 | -3.39 | fly chr57B15-57B16\|2-92 | chromosome location | fly chr57B15-57B16\|2-92 | 1 | 1 | 25010 | 845 | shg | | 3.379e-02 | -3.39 | fly chr88B1-88B3\|3-55 | chromosome location | fly chr88B1-88B3\|3-55 | 1 | 1 | 25010 | 845 | red | | 3.379e-02 | -3.39 | fly chr57F6-57F7\|2-96 | chromosome location | fly chr57F6-57F7\|2-96 | 1 | 1 | 25010 | 845 | pirk | | 3.379e-02 | -3.39 | fly chr22A5-22B1\|2-3 cM | chromosome location | fly chr22A5-22B1\|2-3 cM | 1 | 1 | 25010 | 845 | chinmo | | 3.379e-02 | -3.39 | fly chr100B2-100B3\|3-102 | chromosome location | fly chr100B2-100B3\|3-102 | 1 | 1 | 25010 | 845 | CG11318 | | 3.379e-02 | -3.39 | fly chr62C4-62D1\|3-1.5 | chromosome location | fly chr62C4-62D1\|3-1.5 | 1 | 1 | 25010 | 845 | zormin | | 3.379e-02 | -3.39 | fly chr10B13-10B13\|1-34 cM | chromosome location | fly chr10B13-10B13\|1-34 cM | 1 | 1 | 25010 | 845 | Sk1 | | 3.379e-02 | -3.39 | fly chr90A5-90A6\|3-60 | chromosome location | fly chr90A5-90A6\|3-60 | 1 | 1 | 25010 | 845 | CG42823 | | 3.379e-02 | -3.39 | fly chr80C3-80C3\|3-47 cM | chromosome location | fly chr80C3-80C3\|3-47 cM | 1 | 1 | 25010 | 845 | nrm | | 3.379e-02 | -3.39 | fly chr37A2-37A4\|2-54 cM | chromosome location | fly chr37A2-37A4\|2-54 cM | 1 | 1 | 25010 | 845 | ham | | 3.379e-02 | -3.39 | fly chr95D5-95D8\|3-82 cM | chromosome location | fly chr95D5-95D8\|3-82 cM | 1 | 1 | 25010 | 845 | KrT95D | | 3.379e-02 | -3.39 | fly chr57D8-57D11\|2-94 cM | chromosome location | fly chr57D8-57D11\|2-94 cM | 1 | 1 | 25010 | 845 | CG42672 | | 3.379e-02 | -3.39 | fly chr84D11-84D11\|3-48 cM | chromosome location | fly chr84D11-84D11\|3-48 cM | 1 | 1 | 25010 | 845 | lncRNA:CR44332 | | 3.379e-02 | -3.39 | fly chr61B1-61B2\|3-1 | chromosome location | fly chr61B1-61B2\|3-1 | 1 | 1 | 25010 | 845 | p130CAS | | 3.379e-02 | -3.39 | fly chr42C6-42C7\|2-55 | chromosome location | fly chr42C6-42C7\|2-55 | 1 | 1 | 25010 | 845 | l(2)01289 | | 3.379e-02 | -3.39 | fly chr98D2-98D2\|3-96 | chromosome location | fly chr98D2-98D2\|3-96 | 1 | 1 | 25010 | 845 | CG1894 | | 3.379e-02 | -3.39 | fly chr78D1-78D2\|3-47 | chromosome location | fly chr78D1-78D2\|3-47 | 1 | 1 | 25010 | 845 | barc | | 3.379e-02 | -3.39 | fly chr6D2-6D3\|1-18 | chromosome location | fly chr6D2-6D3\|1-18 | 1 | 1 | 25010 | 845 | pod1 | | 3.379e-02 | -3.39 | fly chr11C4-11D1\|1-40 cM | chromosome location | fly chr11C4-11D1\|1-40 cM | 1 | 1 | 25010 | 845 | Pde9 | | 3.379e-02 | -3.39 | fly chr18B6-18B6\|1-62 | chromosome location | fly chr18B6-18B6\|1-62 | 1 | 1 | 25010 | 845 | rictor | | 3.379e-02 | -3.39 | fly chr62F2-62F2\|3-4 cM | chromosome location | fly chr62F2-62F2\|3-4 cM | 1 | 1 | 25010 | 845 | Acp62F | | 3.379e-02 | -3.39 | fly chr49D4-49D6\|2-67 | chromosome location | fly chr49D4-49D6\|2-67 | 1 | 1 | 25010 | 845 | CG17574 | | 3.379e-02 | -3.39 | fly chr32F1-32F1\|2-44 | chromosome location | fly chr32F1-32F1\|2-44 | 1 | 1 | 25010 | 845 | lncRNA:CR43681 | | 3.379e-02 | -3.39 | fly chr78D8-78E1\|3-47 | chromosome location | fly chr78D8-78E1\|3-47 | 1 | 1 | 25010 | 845 | TfAP-2 | | 3.379e-02 | -3.39 | fly chr91F3-91F3\|3-66 cM | chromosome location | fly chr91F3-91F3\|3-66 cM | 1 | 1 | 25010 | 845 | Cyp12a4 | | 3.379e-02 | -3.39 | fly chr66A10-66A9\|3-21 | chromosome location | fly chr66A10-66A9\|3-21 | 1 | 1 | 25010 | 845 | Ank2 | | 3.379e-02 | -3.39 | fly chr8C1-8C4\|1-26 | chromosome location | fly chr8C1-8C4\|1-26 | 1 | 1 | 25010 | 845 | lncRNA:CR43836 | | 3.379e-02 | -3.39 | fly chr91D3-91D3\|3-65 | chromosome location | fly chr91D3-91D3\|3-65 | 1 | 1 | 25010 | 845 | CG14298 | | 3.379e-02 | -3.39 | fly chr91F3-91F3\|3-66 | chromosome location | fly chr91F3-91F3\|3-66 | 1 | 1 | 25010 | 845 | Cyp12a4 | | 3.379e-02 | -3.39 | fly chr53A3-53A4\|2-79 | chromosome location | fly chr53A3-53A4\|2-79 | 1 | 1 | 25010 | 845 | Khc | | 3.379e-02 | -3.39 | fly chr29F3-29F4\|2-34 cM | chromosome location | fly chr29F3-29F4\|2-34 cM | 1 | 1 | 25010 | 845 | CG34398 | | 3.379e-02 | -3.39 | fly chr27B1-27B2\|2-22 cM | chromosome location | fly chr27B1-27B2\|2-22 cM | 1 | 1 | 25010 | 845 | nrv2 | | 3.379e-02 | -3.39 | fly chr41C6-41D1\|2-55 cM | chromosome location | fly chr41C6-41D1\|2-55 cM | 1 | 1 | 25010 | 845 | Ir41a | | 3.379e-02 | -3.39 | fly chr15A8-15A9\|1-55 cM | chromosome location | fly chr15A8-15A9\|1-55 cM | 1 | 1 | 25010 | 845 | CG9132 | | 3.379e-02 | -3.39 | fly chr62E3-62E4\|3-3 | chromosome location | fly chr62E3-62E4\|3-3 | 1 | 1 | 25010 | 845 | CG1146 | | 3.379e-02 | -3.39 | fly chr89B1-89B2\|3-58 | chromosome location | fly chr89B1-89B2\|3-58 | 1 | 1 | 25010 | 845 | msps | | 3.379e-02 | -3.39 | fly chr41A3-41A3\|2-55 cM | chromosome location | fly chr41A3-41A3\|2-55 cM | 1 | 1 | 25010 | 845 | Stlk | | 3.379e-02 | -3.39 | fly chr51B4-51B4\|2-72 cM | chromosome location | fly chr51B4-51B4\|2-72 cM | 1 | 1 | 25010 | 845 | CG12869 | | 3.379e-02 | -3.39 | fly chr69E5-69E6\|3-39 | chromosome location | fly chr69E5-69E6\|3-39 | 1 | 1 | 25010 | 845 | CG10960 | | 3.379e-02 | -3.39 | fly chr22F4-23A1\|2-6 | chromosome location | fly chr22F4-23A1\|2-6 | 1 | 1 | 25010 | 845 | Cyp309a1 | | 3.379e-02 | -3.39 | fly chr86D5-86D6\|3-50 | chromosome location | fly chr86D5-86D6\|3-50 | 1 | 1 | 25010 | 845 | Ugt304A1 | | 3.379e-02 | -3.39 | fly chr48C3-48C4\|2-65 cM | chromosome location | fly chr48C3-48C4\|2-65 cM | 1 | 1 | 25010 | 845 | ths | | 3.379e-02 | -3.39 | fly chr3D6-3E2\|1-4 cM | chromosome location | fly chr3D6-3E2\|1-4 cM | 1 | 1 | 25010 | 845 | AstA-R1 | | 3.379e-02 | -3.39 | fly chr15E3-15E5\|1-57 | chromosome location | fly chr15E3-15E5\|1-57 | 1 | 1 | 25010 | 845 | CG5004 | | 3.379e-02 | -3.39 | fly chr94E3-94E4\|3-78 cM | chromosome location | fly chr94E3-94E4\|3-78 cM | 1 | 1 | 25010 | 845 | CG46310 | | 3.379e-02 | -3.39 | fly chr100B5-100B6\|3-102 | chromosome location | fly chr100B5-100B6\|3-102 | 1 | 1 | 25010 | 845 | chp | | 3.379e-02 | -3.39 | fly chr66A10-66A9\|3-21 cM | chromosome location | fly chr66A10-66A9\|3-21 cM | 1 | 1 | 25010 | 845 | Ank2 | | 3.379e-02 | -3.39 | fly chr42F1-42F1\|2-55 cM | chromosome location | fly chr42F1-42F1\|2-55 cM | 1 | 1 | 25010 | 845 | esn | | 3.379e-02 | -3.39 | fly chr22A5-22B1\|2-3 | chromosome location | fly chr22A5-22B1\|2-3 | 1 | 1 | 25010 | 845 | chinmo | | 3.379e-02 | -3.39 | fly chr63E8-63F1\|3-8 cM | chromosome location | fly chr63E8-63F1\|3-8 cM | 1 | 1 | 25010 | 845 | CG32264 | | 3.379e-02 | -3.39 | fly chr92B2-92B3\|3-67 | chromosome location | fly chr92B2-92B3\|3-67 | 1 | 1 | 25010 | 845 | bnl | | 3.379e-02 | -3.39 | fly chr30B3-30B5\|2-35 | chromosome location | fly chr30B3-30B5\|2-35 | 1 | 1 | 25010 | 845 | numb | | 3.379e-02 | -3.39 | fly chr94F1-94F4\|3-79 | chromosome location | fly chr94F1-94F4\|3-79 | 1 | 1 | 25010 | 845 | CG4374 | | 3.379e-02 | -3.39 | fly chr93B11-93B11\|3-70 | chromosome location | fly chr93B11-93B11\|3-70 | 1 | 1 | 25010 | 845 | lncRNA:CR44178 | | 3.379e-02 | -3.39 | fly chr47F13-47F14\|2-64 cM | chromosome location | fly chr47F13-47F14\|2-64 cM | 1 | 1 | 25010 | 845 | E(Pc) | | 3.379e-02 | -3.39 | fly chr4C3-4C4\|1-8 cM | chromosome location | fly chr4C3-4C4\|1-8 cM | 1 | 1 | 25010 | 845 | bi | | 3.379e-02 | -3.39 | fly chr102F1-102F3\|4-0 cM | chromosome location | fly chr102F1-102F3\|4-0 cM | 1 | 1 | 25010 | 845 | mGluR | | 3.379e-02 | -3.39 | fly chr88F5-88F5\|3-57 cM | chromosome location | fly chr88F5-88F5\|3-57 cM | 1 | 1 | 25010 | 845 | Act88F | | 3.379e-02 | -3.39 | fly chr55F1-55F2\|2-86 | chromosome location | fly chr55F1-55F2\|2-86 | 1 | 1 | 25010 | 845 | Mctp | | 3.379e-02 | -3.39 | fly chr43C4-43C4\|2-56 | chromosome location | fly chr43C4-43C4\|2-56 | 1 | 1 | 25010 | 845 | Aldh-III | | 3.379e-02 | -3.39 | fly chr3A1-3A1\|1-0.9 cM | chromosome location | fly chr3A1-3A1\|1-0.9 cM | 1 | 1 | 25010 | 845 | Raf | | 3.379e-02 | -3.39 | fly chr89A1-89A2\|3-57 cM | chromosome location | fly chr89A1-89A2\|3-57 cM | 1 | 1 | 25010 | 845 | AOX1 | | 3.379e-02 | -3.39 | fly chr7D17-7D22\|1-22 cM | chromosome location | fly chr7D17-7D22\|1-22 cM | 1 | 1 | 25010 | 845 | sdt | | 3.379e-02 | -3.39 | fly chr99A9-99B1\|3-98 cM | chromosome location | fly chr99A9-99B1\|3-98 cM | 1 | 1 | 25010 | 845 | Ptp99A | | 3.379e-02 | -3.39 | fly chr12E10-12E10\|1-48 | chromosome location | fly chr12E10-12E10\|1-48 | 1 | 1 | 25010 | 845 | CG9411 | | 3.379e-02 | -3.39 | fly chr60B12-60C1\|2-106 cM | chromosome location | fly chr60B12-60C1\|2-106 cM | 1 | 1 | 25010 | 845 | AANAT1 | | 3.379e-02 | -3.39 | fly chr94B8-94B11\|3-75 | chromosome location | fly chr94B8-94B11\|3-75 | 1 | 1 | 25010 | 845 | wake | | 3.379e-02 | -3.39 | fly chr63C6-63D1\|3-7 cM | chromosome location | fly chr63C6-63D1\|3-7 cM | 1 | 1 | 25010 | 845 | ZnT63C | | 3.379e-02 | -3.39 | fly chr85D16-85D17\|3-48 cM | chromosome location | fly chr85D16-85D17\|3-48 cM | 1 | 1 | 25010 | 845 | ps | | 3.379e-02 | -3.39 | fly chr8B8-8C3\|1-25 cM | chromosome location | fly chr8B8-8C3\|1-25 cM | 1 | 1 | 25010 | 845 | rdgA | | 3.379e-02 | -3.39 | fly chr8C1-8C4\|1-26 cM | chromosome location | fly chr8C1-8C4\|1-26 cM | 1 | 1 | 25010 | 845 | lncRNA:CR43836 | | 3.379e-02 | -3.39 | fly chr89B19-89B19\|3-58 cM | chromosome location | fly chr89B19-89B19\|3-58 cM | 1 | 1 | 25010 | 845 | lncRNA:CR46030 | | 3.379e-02 | -3.39 | fly chr66D12-66D15\|3-26 | chromosome location | fly chr66D12-66D15\|3-26 | 1 | 1 | 25010 | 845 | Fhos | | 3.379e-02 | -3.39 | fly chr6C9-6C10\|1-18 cM | chromosome location | fly chr6C9-6C10\|1-18 cM | 1 | 1 | 25010 | 845 | CG3168 | | 3.379e-02 | -3.39 | fly chr41C5-41C5\|2-55 | chromosome location | fly chr41C5-41C5\|2-55 | 1 | 1 | 25010 | 845 | CG44102 | | 3.379e-02 | -3.39 | fly chr75F9-76A3\|3-45 | chromosome location | fly chr75F9-76A3\|3-45 | 1 | 1 | 25010 | 845 | fz2 | | 3.379e-02 | -3.39 | fly chr34C3-34C3\|2-48 cM | chromosome location | fly chr34C3-34C3\|2-48 cM | 1 | 1 | 25010 | 845 | CG42784 | | 3.379e-02 | -3.39 | fly chr99A9-99B1\|3-98 | chromosome location | fly chr99A9-99B1\|3-98 | 1 | 1 | 25010 | 845 | Ptp99A | | 3.379e-02 | -3.39 | fly chr10C5-10C5\|1-35 cM | chromosome location | fly chr10C5-10C5\|1-35 cM | 1 | 1 | 25010 | 845 | CG1572 | | 3.379e-02 | -3.39 | fly chr98F13-99A1\|3-98 cM | chromosome location | fly chr98F13-99A1\|3-98 cM | 1 | 1 | 25010 | 845 | jus | | 3.379e-02 | -3.39 | fly chr78D1-78D2\|3-47 cM | chromosome location | fly chr78D1-78D2\|3-47 cM | 1 | 1 | 25010 | 845 | barc | | 3.379e-02 | -3.39 | fly chr66D12-66D15\|3-26 cM | chromosome location | fly chr66D12-66D15\|3-26 cM | 1 | 1 | 25010 | 845 | Fhos | | 3.379e-02 | -3.39 | fly chr69E5-69E6\|3-39 cM | chromosome location | fly chr69E5-69E6\|3-39 cM | 1 | 1 | 25010 | 845 | CG10960 | | 3.379e-02 | -3.39 | fly chr61F7-62A2\|3-0 cM | chromosome location | fly chr61F7-62A2\|3-0 cM | 1 | 1 | 25010 | 845 | Ptp61F | | 3.379e-02 | -3.39 | fly chr13E8-13E12\|1-51 | chromosome location | fly chr13E8-13E12\|1-51 | 1 | 1 | 25010 | 845 | CG8184 | | 3.379e-02 | -3.39 | fly chr84D11-84D11\|3-48 | chromosome location | fly chr84D11-84D11\|3-48 | 1 | 1 | 25010 | 845 | lncRNA:CR44332 | | 3.379e-02 | -3.39 | fly chr42C6-42C7\|2-55 cM | chromosome location | fly chr42C6-42C7\|2-55 cM | 1 | 1 | 25010 | 845 | l(2)01289 | | 3.379e-02 | -3.39 | fly chr98A10-98A12\|3-94 cM | chromosome location | fly chr98A10-98A12\|3-94 cM | 1 | 1 | 25010 | 845 | Tusp | | 3.379e-02 | -3.39 | fly chr96B19-96B20\|3-86 cM | chromosome location | fly chr96B19-96B20\|3-86 cM | 1 | 1 | 25010 | 845 | Stt3B | | 3.379e-02 | -3.39 | fly chr82A1-82A3\|3-47.1 | chromosome location | fly chr82A1-82A3\|3-47.1 | 1 | 1 | 25010 | 845 | cpx | | 3.379e-02 | -3.39 | fly chr83C2-83C3\|3-47.5 | chromosome location | fly chr83C2-83C3\|3-47.5 | 1 | 1 | 25010 | 845 | CG2082 | | 3.379e-02 | -3.39 | fly chr8F2-8F2\|1-29 cM | chromosome location | fly chr8F2-8F2\|1-29 cM | 1 | 1 | 25010 | 845 | CG42797 | | 3.379e-02 | -3.39 | fly chr15F7-16A1\|1-57 cM | chromosome location | fly chr15F7-16A1\|1-57 cM | 1 | 1 | 25010 | 845 | CG5445 | | 3.379e-02 | -3.39 | fly chr91C6-91C6\|3-64 | chromosome location | fly chr91C6-91C6\|3-64 | 1 | 1 | 25010 | 845 | Epg5 | | 3.379e-02 | -3.39 | fly chr83E5-83E6\|3-47.5 | chromosome location | fly chr83E5-83E6\|3-47.5 | 1 | 1 | 25010 | 845 | Dmtn | | 3.379e-02 | -3.39 | fly chr89F1-89F3\|3-60 | chromosome location | fly chr89F1-89F3\|3-60 | 1 | 1 | 25010 | 845 | Mur89F | | 3.379e-02 | -3.39 | fly chr96A22-96A23\|3-85 | chromosome location | fly chr96A22-96A23\|3-85 | 1 | 1 | 25010 | 845 | polybromo | | 3.379e-02 | -3.39 | fly chr16B12-16B12\|1-57 cM | chromosome location | fly chr16B12-16B12\|1-57 cM | 1 | 1 | 25010 | 845 | CG12990 | | 3.379e-02 | -3.39 | fly chr89B19-89B19\|3-58 | chromosome location | fly chr89B19-89B19\|3-58 | 1 | 1 | 25010 | 845 | lncRNA:CR46030 | | 3.379e-02 | -3.39 | fly chr34C3-34C3\|2-48 | chromosome location | fly chr34C3-34C3\|2-48 | 1 | 1 | 25010 | 845 | CG42784 | | 3.379e-02 | -3.39 | fly chr2D1-2D1\|1-0.6 | chromosome location | fly chr2D1-2D1\|1-0.6 | 1 | 1 | 25010 | 845 | snoRNA:Me18S-C1024 | | 3.379e-02 | -3.39 | fly chr72F1-73A3\|3-44 cM | chromosome location | fly chr72F1-73A3\|3-44 cM | 1 | 1 | 25010 | 845 | CG33158 | | 3.379e-02 | -3.39 | fly chr96B15-96B15\|3-86 | chromosome location | fly chr96B15-96B15\|3-86 | 1 | 1 | 25010 | 845 | tnc | | 3.379e-02 | -3.39 | fly chr16B12-16B12\|1-57 | chromosome location | fly chr16B12-16B12\|1-57 | 1 | 1 | 25010 | 845 | CG12990 | | 3.379e-02 | -3.39 | fly chr86E7-86E8\|3-51 cM | chromosome location | fly chr86E7-86E8\|3-51 cM | 1 | 1 | 25010 | 845 | dpr5 | | 3.379e-02 | -3.39 | fly chr37A2-37A4\|2-54 | chromosome location | fly chr37A2-37A4\|2-54 | 1 | 1 | 25010 | 845 | ham | | 3.379e-02 | -3.39 | fly chr52A3-52A1\|2-74 | chromosome location | fly chr52A3-52A1\|2-74 | 1 | 1 | 25010 | 845 | Stacl | | 3.379e-02 | -3.39 | fly chr97B9-97C1\|3-91 cM | chromosome location | fly chr97B9-97C1\|3-91 cM | 1 | 1 | 25010 | 845 | scrib | | 3.379e-02 | -3.39 | fly chr82D8-82E1\|3-47.1 | chromosome location | fly chr82D8-82E1\|3-47.1 | 1 | 1 | 25010 | 845 | opa | | 3.379e-02 | -3.39 | fly chr84A6-84B2\|3-48 cM | chromosome location | fly chr84A6-84B2\|3-48 cM | 1 | 1 | 25010 | 845 | Antp | | 3.379e-02 | -3.39 | fly chr50B6-50C3\|2-68 | chromosome location | fly chr50B6-50C3\|2-68 | 1 | 1 | 25010 | 845 | CG17716 | | 3.379e-02 | -3.39 | fly chr84F11-84F12\|3-48 | chromosome location | fly chr84F11-84F12\|3-48 | 1 | 1 | 25010 | 845 | Poxm | | 3.379e-02 | -3.39 | fly chr52B3-52B3\|2-75 | chromosome location | fly chr52B3-52B3\|2-75 | 1 | 1 | 25010 | 845 | mir-278 | | 3.379e-02 | -3.39 | fly chr99E4-99F2\|3-100 cM | chromosome location | fly chr99E4-99F2\|3-100 cM | 1 | 1 | 25010 | 845 | hdc | | 3.379e-02 | -3.39 | fly chr3C9-3D1\|1-2 | chromosome location | fly chr3C9-3D1\|1-2 | 1 | 1 | 25010 | 845 | dnc | | 3.379e-02 | -3.39 | fly chr96A19-96A19\|3-85 | chromosome location | fly chr96A19-96A19\|3-85 | 1 | 1 | 25010 | 845 | tld | | 3.379e-02 | -3.39 | fly chr15A8-15A9\|1-55 | chromosome location | fly chr15A8-15A9\|1-55 | 1 | 1 | 25010 | 845 | CG9132 | | 3.379e-02 | -3.39 | fly chr2D1-2D2\|1-0.6 cM | chromosome location | fly chr2D1-2D2\|1-0.6 cM | 1 | 1 | 25010 | 845 | csw | | 3.379e-02 | -3.39 | fly chr87A10-87B1\|3-51 cM | chromosome location | fly chr87A10-87B1\|3-51 cM | 1 | 1 | 25010 | 845 | dpr15 | | 3.379e-02 | -3.39 | fly chr45A9-45A10\|2-60 | chromosome location | fly chr45A9-45A10\|2-60 | 1 | 1 | 25010 | 845 | CG18659 | | 3.379e-02 | -3.39 | fly chr65B4-65B5\|3-17 cM | chromosome location | fly chr65B4-65B5\|3-17 cM | 1 | 1 | 25010 | 845 | MCU | | 3.379e-02 | -3.39 | fly chr75B1-75B1\|3-45 cM | chromosome location | fly chr75B1-75B1\|3-45 cM | 1 | 1 | 25010 | 845 | lncRNA:CR45921 | | 3.379e-02 | -3.39 | fly chr74D4-74E2\|3-45 cM | chromosome location | fly chr74D4-74E2\|3-45 cM | 1 | 1 | 25010 | 845 | Eip74EF | | 3.379e-02 | -3.39 | fly chr90A2-90A3\|3-60 | chromosome location | fly chr90A2-90A3\|3-60 | 1 | 1 | 25010 | 845 | beat-IIa | | 3.379e-02 | -3.39 | fly chr22F1-22F3\|2-6 cM | chromosome location | fly chr22F1-22F3\|2-6 cM | 1 | 1 | 25010 | 845 | dpp | | 3.379e-02 | -3.39 | fly chr56B3-56B3\|2-88 cM | chromosome location | fly chr56B3-56B3\|2-88 cM | 1 | 1 | 25010 | 845 | Ir56a | | 3.379e-02 | -3.39 | fly chr33D2-33D5\|2-46 cM | chromosome location | fly chr33D2-33D5\|2-46 cM | 1 | 1 | 25010 | 845 | bru1 | | 3.379e-02 | -3.39 | fly chr41D4-41E1\|2-55 | chromosome location | fly chr41D4-41E1\|2-55 | 1 | 1 | 25010 | 845 | Atf6 | | 3.379e-02 | -3.39 | fly chr14C3-14C4\|1-53 cM | chromosome location | fly chr14C3-14C4\|1-53 cM | 1 | 1 | 25010 | 845 | CG9981 | | 3.379e-02 | -3.39 | fly chr57A3-57A3\|2-91 | chromosome location | fly chr57A3-57A3\|2-91 | 1 | 1 | 25010 | 845 | side-VIII | | 3.379e-02 | -3.39 | fly chr53C7-53C7\|2-80 cM | chromosome location | fly chr53C7-53C7\|2-80 cM | 1 | 1 | 25010 | 845 | Syn2 | | 3.379e-02 | -3.39 | fly chr88C10-88D1\|3-55 | chromosome location | fly chr88C10-88D1\|3-55 | 1 | 1 | 25010 | 845 | Meltrin | | 3.379e-02 | -3.39 | fly chr88F5-88F5\|3-57 | chromosome location | fly chr88F5-88F5\|3-57 | 1 | 1 | 25010 | 845 | Act88F | | 3.379e-02 | -3.39 | fly chr98F13-99A1\|3-98 | chromosome location | fly chr98F13-99A1\|3-98 | 1 | 1 | 25010 | 845 | jus | | 3.379e-02 | -3.39 | fly chr73A10-73A11\|3-44 cM | chromosome location | fly chr73A10-73A11\|3-44 cM | 1 | 1 | 25010 | 845 | spd-2 | | 3.379e-02 | -3.39 | fly chr94E3-94E4\|3-78 | chromosome location | fly chr94E3-94E4\|3-78 | 1 | 1 | 25010 | 845 | CG46310 | | 3.379e-02 | -3.39 | fly chr94A4-94A5\|3-74 | chromosome location | fly chr94A4-94A5\|3-74 | 1 | 1 | 25010 | 845 | asRNA:CR46261 | | 3.379e-02 | -3.39 | fly chr19E2-19E3\|1-65 cM | chromosome location | fly chr19E2-19E3\|1-65 cM | 1 | 1 | 25010 | 845 | shakB | | 3.379e-02 | -3.39 | fly chr102F1-102F3\|4-0 | chromosome location | fly chr102F1-102F3\|4-0 | 1 | 1 | 25010 | 845 | mGluR | | 3.379e-02 | -3.39 | fly chr96E10-96F1\|3-88 cM | chromosome location | fly chr96E10-96F1\|3-88 cM | 1 | 1 | 25010 | 845 | LpR2 | | 3.379e-02 | -3.39 | fly chr83C2-83C3\|3-47.5 cM | chromosome location | fly chr83C2-83C3\|3-47.5 cM | 1 | 1 | 25010 | 845 | CG2082 | | 3.379e-02 | -3.39 | fly chr100D2-100D3\|3-102 cM | chromosome location | fly chr100D2-100D3\|3-102 cM | 1 | 1 | 25010 | 845 | RhoGAP100F | | 3.379e-02 | -3.39 | fly chr66C4-66C5\|3-24 cM | chromosome location | fly chr66C4-66C5\|3-24 cM | 1 | 1 | 25010 | 845 | cert | | 3.379e-02 | -3.39 | fly chr91A7-91B3\|3-63 cM | chromosome location | fly chr91A7-91B3\|3-63 cM | 1 | 1 | 25010 | 845 | fru | | 3.379e-02 | -3.39 | fly chr16F7-17A1\|1-59 cM | chromosome location | fly chr16F7-17A1\|1-59 cM | 1 | 1 | 25010 | 845 | Frq1 | | 3.379e-02 | -3.39 | fly chr92A3-92A5\|3-66 | chromosome location | fly chr92A3-92A5\|3-66 | 1 | 1 | 25010 | 845 | CG31221 | | 3.379e-02 | -3.39 | fly chr100B5-100B6\|3-102 cM | chromosome location | fly chr100B5-100B6\|3-102 cM | 1 | 1 | 25010 | 845 | chp | | 3.379e-02 | -3.39 | fly chr88C10-88D1\|3-55 cM | chromosome location | fly chr88C10-88D1\|3-55 cM | 1 | 1 | 25010 | 845 | Meltrin | | 3.379e-02 | -3.39 | fly chr75E3-75E4\|3-45 | chromosome location | fly chr75E3-75E4\|3-45 | 1 | 1 | 25010 | 845 | CG3961 | | 3.379e-02 | -3.39 | fly chr3A1-3A1\|1-0.9 | chromosome location | fly chr3A1-3A1\|1-0.9 | 1 | 1 | 25010 | 845 | Raf | | 3.379e-02 | -3.39 | fly chr57B16-57B19\|2-92 | chromosome location | fly chr57B16-57B19\|2-92 | 1 | 1 | 25010 | 845 | Treh | | 3.379e-02 | -3.39 | fly chr16E4-16E4\|1-59 cM | chromosome location | fly chr16E4-16E4\|1-59 cM | 1 | 1 | 25010 | 845 | mnb | | 3.379e-02 | -3.39 | fly chr96E2-96E4\|3-88 | chromosome location | fly chr96E2-96E4\|3-88 | 1 | 1 | 25010 | 845 | msi | | 3.379e-02 | -3.39 | fly chr99B10-99C1\|3-99 cM | chromosome location | fly chr99B10-99C1\|3-99 cM | 1 | 1 | 25010 | 845 | kay | | 3.379e-02 | -3.39 | fly chr78A5-78B1\|3-47 cM | chromosome location | fly chr78A5-78B1\|3-47 cM | 1 | 1 | 25010 | 845 | siz | | 3.379e-02 | -3.39 | fly chr70D7-70E1\|3-42 cM | chromosome location | fly chr70D7-70E1\|3-42 cM | 1 | 1 | 25010 | 845 | bbg | | 3.379e-02 | -3.39 | fly chr49B11-49C1\|2-67 cM | chromosome location | fly chr49B11-49C1\|2-67 cM | 1 | 1 | 25010 | 845 | nemy | | 3.379e-02 | -3.39 | fly chr3C3-3C4\|1-2 cM | chromosome location | fly chr3C3-3C4\|1-2 cM | 1 | 1 | 25010 | 845 | rst | | 3.379e-02 | -3.39 | fly chr78A5-78B1\|3-47 | chromosome location | fly chr78A5-78B1\|3-47 | 1 | 1 | 25010 | 845 | siz | | 3.379e-02 | -3.39 | fly chr55C10-55D1\|2-85 cM | chromosome location | fly chr55C10-55D1\|2-85 cM | 1 | 1 | 25010 | 845 | CG30116 | | 3.379e-02 | -3.39 | fly chr27C1-27C4\|2-23 cM | chromosome location | fly chr27C1-27C4\|2-23 cM | 1 | 1 | 25010 | 845 | Nlg2 | | 3.379e-02 | -3.39 | fly chr91C6-91D1\|3-64 cM | chromosome location | fly chr91C6-91D1\|3-64 cM | 1 | 1 | 25010 | 845 | CG7720 | | 3.379e-02 | -3.39 | fly chr41A3-41A3\|2-55 | chromosome location | fly chr41A3-41A3\|2-55 | 1 | 1 | 25010 | 845 | Stlk | | 3.379e-02 | -3.39 | fly chr75B1-75B1\|3-45 | chromosome location | fly chr75B1-75B1\|3-45 | 1 | 1 | 25010 | 845 | lncRNA:CR45921 | | 3.379e-02 | -3.39 | fly chr6D2-6D3\|1-18 cM | chromosome location | fly chr6D2-6D3\|1-18 cM | 1 | 1 | 25010 | 845 | pod1 | | 3.379e-02 | -3.39 | fly chr75C4-75C4 | chromosome location | fly chr75C4-75C4 | 1 | 1 | 25010 | 845 | lncRNA:CR46353 | | 3.379e-02 | -3.39 | fly chr50B6-50C3\|2-68 cM | chromosome location | fly chr50B6-50C3\|2-68 cM | 1 | 1 | 25010 | 845 | CG17716 | | 3.379e-02 | -3.39 | fly chr43C4-43C4\|2-56 cM | chromosome location | fly chr43C4-43C4\|2-56 cM | 1 | 1 | 25010 | 845 | Aldh-III | | 3.379e-02 | -3.39 | fly chr2D1-2D1\|1-0.6 cM | chromosome location | fly chr2D1-2D1\|1-0.6 cM | 1 | 1 | 25010 | 845 | snoRNA:Me18S-C1024 | | 3.379e-02 | -3.39 | fly chr98F1-98F3\|3-97 cM | chromosome location | fly chr98F1-98F3\|3-97 cM | 1 | 1 | 25010 | 845 | Doa | | 3.379e-02 | -3.39 | fly chr65E4-65E5\|3-19 cM | chromosome location | fly chr65E4-65E5\|3-19 cM | 1 | 1 | 25010 | 845 | melt | | 3.379e-02 | -3.39 | fly chr85D19-85D22\|3-48 cM | chromosome location | fly chr85D19-85D22\|3-48 cM | 1 | 1 | 25010 | 845 | by | | 3.379e-02 | -3.39 | fly chr37E4-37E5\|2-54 | chromosome location | fly chr37E4-37E5\|2-54 | 1 | 1 | 25010 | 845 | CG10188 | | 3.379e-02 | -3.39 | fly chr33D2-33D3\|2-46 cM | chromosome location | fly chr33D2-33D3\|2-46 cM | 1 | 1 | 25010 | 845 | atilla | | 3.379e-02 | -3.39 | fly chr96B15-96B15\|3-86 cM | chromosome location | fly chr96B15-96B15\|3-86 cM | 1 | 1 | 25010 | 845 | tnc | | 3.379e-02 | -3.39 | fly chr73C3-73C3\|3-44 cM | chromosome location | fly chr73C3-73C3\|3-44 cM | 1 | 1 | 25010 | 845 | lncRNA:CR46214 | | 3.379e-02 | -3.39 | fly chr47D5-47D6\|2-63 cM | chromosome location | fly chr47D5-47D6\|2-63 cM | 1 | 1 | 25010 | 845 | CG13229 | | 3.379e-02 | -3.39 | fly chr61B3-61C1\|3-1 | chromosome location | fly chr61B3-61C1\|3-1 | 1 | 1 | 25010 | 845 | RhoGEF3 | | 3.379e-02 | -3.39 | fly chr35B4-35B5\|2-50 | chromosome location | fly chr35B4-35B5\|2-50 | 1 | 1 | 25010 | 845 | CG42685 | | 3.379e-02 | -3.39 | fly chr16A1-16A1\|1-57 cM | chromosome location | fly chr16A1-16A1\|1-57 cM | 1 | 1 | 25010 | 845 | B-H2 | | 3.379e-02 | -3.39 | fly chr74D4-74E2\|3-45 | chromosome location | fly chr74D4-74E2\|3-45 | 1 | 1 | 25010 | 845 | Eip74EF | | 3.379e-02 | -3.39 | fly chr102E1-102E3\|4-0 cM | chromosome location | fly chr102E1-102E3\|4-0 cM | 1 | 1 | 25010 | 845 | Sox102F | | 3.379e-02 | -3.39 | fly chr32F1-32F1\|2-44 cM | chromosome location | fly chr32F1-32F1\|2-44 cM | 1 | 1 | 25010 | 845 | lncRNA:CR43681 | | 3.379e-02 | -3.39 | fly chr99E5-99E5\|3-100 | chromosome location | fly chr99E5-99E5\|3-100 | 1 | 1 | 25010 | 845 | CG34300 | | 3.379e-02 | -3.39 | fly chr3A1-3A4\|1-1 cM | chromosome location | fly chr3A1-3A4\|1-1 cM | 1 | 1 | 25010 | 845 | trol | | 3.379e-02 | -3.39 | fly chr94C9-94C9\|3-76 cM | chromosome location | fly chr94C9-94C9\|3-76 cM | 1 | 1 | 25010 | 845 | lncRNA:CR44350 | | 3.379e-02 | -3.39 | fly chr88C3-88C6\|3-55 cM | chromosome location | fly chr88C3-88C6\|3-55 cM | 1 | 1 | 25010 | 845 | Pde6 | | 3.379e-02 | -3.39 | fly chr49B9-49B9\|2-67 cM | chromosome location | fly chr49B9-49B9\|2-67 cM | 1 | 1 | 25010 | 845 | CG30054 | | 3.379e-02 | -3.39 | fly chr97E6-97E8\|3-92 | chromosome location | fly chr97E6-97E8\|3-92 | 1 | 1 | 25010 | 845 | Ser | | 3.379e-02 | -3.39 | fly chr89D6-89D9\|3-59 | chromosome location | fly chr89D6-89D9\|3-59 | 1 | 1 | 25010 | 845 | Ubx | | 3.379e-02 | -3.39 | fly chr89A1-89A2\|3-57 | chromosome location | fly chr89A1-89A2\|3-57 | 1 | 1 | 25010 | 845 | AOX1 | | 3.379e-02 | -3.39 | fly chr80A-80B\|2-55 | chromosome location | fly chr80A-80B\|2-55 | 1 | 1 | 25010 | 845 | Scp1 | | 3.379e-02 | -3.39 | fly chr33A1-33A2\|2-45 | chromosome location | fly chr33A1-33A2\|2-45 | 1 | 1 | 25010 | 845 | crol | | 3.379e-02 | -3.39 | fly chr57E8-57E9\|2-95 cM | chromosome location | fly chr57E8-57E9\|2-95 cM | 1 | 1 | 25010 | 845 | MESK2 | | 3.379e-02 | -3.39 | fly chr47F13-47F14\|2-64 | chromosome location | fly chr47F13-47F14\|2-64 | 1 | 1 | 25010 | 845 | E(Pc) | | 3.379e-02 | -3.39 | fly chr44B8-44C2\|2-58 cM | chromosome location | fly chr44B8-44C2\|2-58 cM | 1 | 1 | 25010 | 845 | LRP1 | | 3.379e-02 | -3.39 | fly chr9C4-9C5\|1-31 cM | chromosome location | fly chr9C4-9C5\|1-31 cM | 1 | 1 | 25010 | 845 | Neb-cGP | | 3.379e-02 | -3.39 | fly chr93B11-93B11\|3-70 cM | chromosome location | fly chr93B11-93B11\|3-70 cM | 1 | 1 | 25010 | 845 | lncRNA:CR44178 | | 3.379e-02 | -3.39 | fly chr41E5-41E6\|2-55 | chromosome location | fly chr41E5-41E6\|2-55 | 1 | 1 | 25010 | 845 | lncRNA:CR42646 | | 3.379e-02 | -3.39 | fly chr98A10-98A10\|3-94 cM | chromosome location | fly chr98A10-98A10\|3-94 cM | 1 | 1 | 25010 | 845 | dsd | | 3.379e-02 | -3.39 | fly chr49A3-49A4\|2-67 cM | chromosome location | fly chr49A3-49A4\|2-67 cM | 1 | 1 | 25010 | 845 | CG13157 | | 3.379e-02 | -3.39 | fly chr54F1-54F3\|2-84 | chromosome location | fly chr54F1-54F3\|2-84 | 1 | 1 | 25010 | 845 | olf186-F | | 3.379e-02 | -3.39 | fly chr66C4-66C5\|3-24 | chromosome location | fly chr66C4-66C5\|3-24 | 1 | 1 | 25010 | 845 | cert | | 3.379e-02 | -3.39 | fly chr40A1-40A2\|2-54 cM | chromosome location | fly chr40A1-40A2\|2-54 cM | 1 | 1 | 25010 | 845 | step | | 3.379e-02 | -3.39 | fly chr102E5-102F1\|4-0 | chromosome location | fly chr102E5-102F1\|4-0 | 1 | 1 | 25010 | 845 | unc-13 | | 3.379e-02 | -3.39 | fly chr26A2-26A2\|2-18 cM | chromosome location | fly chr26A2-26A2\|2-18 cM | 1 | 1 | 25010 | 845 | lncRNA:CR45288 | | 3.379e-02 | -3.39 | fly chr70B3-70B4\|3-40 cM | chromosome location | fly chr70B3-70B4\|3-40 cM | 1 | 1 | 25010 | 845 | Spt20 | | 3.379e-02 | -3.39 | fly chr35B4-35B5\|2-50 cM | chromosome location | fly chr35B4-35B5\|2-50 cM | 1 | 1 | 25010 | 845 | CG42685 | | 3.379e-02 | -3.39 | fly chr66B8-66B10\|3-22 cM | chromosome location | fly chr66B8-66B10\|3-22 cM | 1 | 1 | 25010 | 845 | CG13676 | | 3.379e-02 | -3.39 | fly chr47F13-47F13\|2-64 | chromosome location | fly chr47F13-47F13\|2-64 | 1 | 1 | 25010 | 845 | qvr | | 3.379e-02 | -3.39 | fly chr91E2-91E3\|3-65 cM | chromosome location | fly chr91E2-91E3\|3-65 cM | 1 | 1 | 25010 | 845 | gukh | | 3.379e-02 | -3.39 | fly chr18C3-18C7\|1-62 | chromosome location | fly chr18C3-18C7\|1-62 | 1 | 1 | 25010 | 845 | kek5 | | 3.379e-02 | -3.39 | fly chr61D4-61E1\|3-0.5 cM | chromosome location | fly chr61D4-61E1\|3-0.5 cM | 1 | 1 | 25010 | 845 | Glut1 | | 3.379e-02 | -3.39 | fly chr89D6-89D9\|3-59 cM | chromosome location | fly chr89D6-89D9\|3-59 cM | 1 | 1 | 25010 | 845 | Ubx | | 3.379e-02 | -3.39 | fly chr16A1-16A1\|1-57 | chromosome location | fly chr16A1-16A1\|1-57 | 1 | 1 | 25010 | 845 | B-H2 | | 3.379e-02 | -3.39 | fly chr27C4-27C6\|2-23 | chromosome location | fly chr27C4-27C6\|2-23 | 1 | 1 | 25010 | 845 | CG18304 | | 3.379e-02 | -3.39 | fly chr2B2-2B4\|1-0.2 cM | chromosome location | fly chr2B2-2B4\|1-0.2 cM | 1 | 1 | 25010 | 845 | Mur2B | | 3.379e-02 | -3.39 | fly chr53D2-53D2\|2-80 cM | chromosome location | fly chr53D2-53D2\|2-80 cM | 1 | 1 | 25010 | 845 | Idgf6 | | 3.379e-02 | -3.39 | fly chr60A9-60A11\|2-105 | chromosome location | fly chr60A9-60A11\|2-105 | 1 | 1 | 25010 | 845 | kcc | | 3.379e-02 | -3.39 | fly chr8B8-8C3\|1-25 | chromosome location | fly chr8B8-8C3\|1-25 | 1 | 1 | 25010 | 845 | rdgA | | 3.379e-02 | -3.39 | fly chr5F1-5F2\|1-16 cM | chromosome location | fly chr5F1-5F2\|1-16 cM | 1 | 1 | 25010 | 845 | CG42240 | | 3.379e-02 | -3.39 | fly chr20A1-20A2\|1-66 cM | chromosome location | fly chr20A1-20A2\|1-66 cM | 1 | 1 | 25010 | 845 | lncRNA:flam | | 3.379e-02 | -3.39 | fly chr73A8-73A8\|3-44 | chromosome location | fly chr73A8-73A8\|3-44 | 1 | 1 | 25010 | 845 | CG43206 | | 3.379e-02 | -3.39 | fly chr76D7-76D7\|3-46 | chromosome location | fly chr76D7-76D7\|3-46 | 1 | 1 | 25010 | 845 | CG14182 | | 3.379e-02 | -3.39 | fly chr88C3-88C6\|3-55 | chromosome location | fly chr88C3-88C6\|3-55 | 1 | 1 | 25010 | 845 | Pde6 | | 3.379e-02 | -3.39 | fly chr52E2-52E2\|2-77 | chromosome location | fly chr52E2-52E2\|2-77 | 1 | 1 | 25010 | 845 | lncRNA:CR43415 | | 3.379e-02 | -3.39 | fly chr90A2-90A3\|3-60 cM | chromosome location | fly chr90A2-90A3\|3-60 cM | 1 | 1 | 25010 | 845 | beat-IIa | | 3.379e-02 | -3.39 | fly chr86C1-86C3\|3-50 | chromosome location | fly chr86C1-86C3\|3-50 | 1 | 1 | 25010 | 845 | hth | | 3.379e-02 | -3.39 | fly chr99F1-99F1\|3-100 cM | chromosome location | fly chr99F1-99F1\|3-100 cM | 1 | 1 | 25010 | 845 | lncRNA:CR46114 | | 3.379e-02 | -3.39 | fly chr83E8-83F1\|3-47.5 | chromosome location | fly chr83E8-83F1\|3-47.5 | 1 | 1 | 25010 | 845 | CG15186 | | 3.379e-02 | -3.39 | fly chr14C3-14C4\|1-53 | chromosome location | fly chr14C3-14C4\|1-53 | 1 | 1 | 25010 | 845 | CG9981 | | 3.379e-02 | -3.39 | fly chr53A3-53A4\|2-79 cM | chromosome location | fly chr53A3-53A4\|2-79 cM | 1 | 1 | 25010 | 845 | Khc | | 3.379e-02 | -3.39 | fly chr63C6-63D1\|3-7 | chromosome location | fly chr63C6-63D1\|3-7 | 1 | 1 | 25010 | 845 | ZnT63C | | 3.379e-02 | -3.39 | fly chr11D9-11D10\|1-42 cM | chromosome location | fly chr11D9-11D10\|1-42 cM | 1 | 1 | 25010 | 845 | IP3K2 | | 3.379e-02 | -3.39 | fly chr77F1-77F3\|3-47 cM | chromosome location | fly chr77F1-77F3\|3-47 cM | 1 | 1 | 25010 | 845 | CG32432 | | 3.379e-02 | -3.39 | fly chr5A7-5A8\|1-12 | chromosome location | fly chr5A7-5A8\|1-12 | 1 | 1 | 25010 | 845 | CG34435 | | 3.379e-02 | -3.39 | fly chr80A2-80A4\|3-47 | chromosome location | fly chr80A2-80A4\|3-47 | 1 | 1 | 25010 | 845 | jim | | 3.379e-02 | -3.39 | fly chr66F5-67A1\|3-28 | chromosome location | fly chr66F5-67A1\|3-28 | 1 | 1 | 25010 | 845 | bol | | 3.379e-02 | -3.39 | fly chr68C12-68C13\|3-36 | chromosome location | fly chr68C12-68C13\|3-36 | 1 | 1 | 25010 | 845 | Mob2 | | 3.379e-02 | -3.39 | fly chr18C3-18C7\|1-62 cM | chromosome location | fly chr18C3-18C7\|1-62 cM | 1 | 1 | 25010 | 845 | kek5 | | 3.379e-02 | -3.39 | fly chr96C4-96C4\|3-87 | chromosome location | fly chr96C4-96C4\|3-87 | 1 | 1 | 25010 | 845 | CG13653 | | 3.379e-02 | -3.39 | fly chr36E1-36E1\|2-53 cM | chromosome location | fly chr36E1-36E1\|2-53 cM | 1 | 1 | 25010 | 845 | lncRNA:CR43420 | | 3.379e-02 | -3.39 | fly chr96F5-96F5\|3-89 | chromosome location | fly chr96F5-96F5\|3-89 | 1 | 1 | 25010 | 845 | Tsp96F | | 3.379e-02 | -3.39 | fly chr41B3-41C2\|2-55 | chromosome location | fly chr41B3-41C2\|2-55 | 1 | 1 | 25010 | 845 | Gprk1 | | 3.379e-02 | -3.39 | fly chr73E5-74A1\|3-45 | chromosome location | fly chr73E5-74A1\|3-45 | 1 | 1 | 25010 | 845 | Rbp6 | | 3.379e-02 | -3.39 | fly chr86E7-86E8\|3-51 | chromosome location | fly chr86E7-86E8\|3-51 | 1 | 1 | 25010 | 845 | dpr5 | | 3.379e-02 | -3.39 | fly chr49D4-49D6\|2-67 cM | chromosome location | fly chr49D4-49D6\|2-67 cM | 1 | 1 | 25010 | 845 | CG17574 | | 3.379e-02 | -3.39 | fly chr88A5-88A8\|3-54 cM | chromosome location | fly chr88A5-88A8\|3-54 cM | 1 | 1 | 25010 | 845 | foxo | | 3.379e-02 | -3.39 | fly chr7D17-7D22\|1-22 | chromosome location | fly chr7D17-7D22\|1-22 | 1 | 1 | 25010 | 845 | sdt | | 3.379e-02 | -3.39 | fly chr6B3-6C1\|1-17 | chromosome location | fly chr6B3-6C1\|1-17 | 1 | 1 | 25010 | 845 | CG34417 | | 3.379e-02 | -3.39 | fly chr3D6-3E2\|1-4 | chromosome location | fly chr3D6-3E2\|1-4 | 1 | 1 | 25010 | 845 | AstA-R1 | | 3.379e-02 | -3.39 | fly chr19E6-19E6\|1-65 cM | chromosome location | fly chr19E6-19E6\|1-65 cM | 1 | 1 | 25010 | 845 | CG11666 | | 3.379e-02 | -3.39 | fly chr73E1-73E2\|3-45 | chromosome location | fly chr73E1-73E2\|3-45 | 1 | 1 | 25010 | 845 | Exn | | 3.379e-02 | -3.39 | fly chr22C3-22D1\|2-4 cM | chromosome location | fly chr22C3-22D1\|2-4 cM | 1 | 1 | 25010 | 845 | GlyP | | 3.379e-02 | -3.39 | fly chr22F1-22F3\|2-6 | chromosome location | fly chr22F1-22F3\|2-6 | 1 | 1 | 25010 | 845 | dpp | | 3.379e-02 | -3.39 | fly chr61B1-61B2\|3-1 cM | chromosome location | fly chr61B1-61B2\|3-1 cM | 1 | 1 | 25010 | 845 | p130CAS | | 3.379e-02 | -3.39 | fly chr14A6-14A7\|1-52 | chromosome location | fly chr14A6-14A7\|1-52 | 1 | 1 | 25010 | 845 | dpr18 | | 3.379e-02 | -3.39 | fly chr11F4-11F5\|1-43 | chromosome location | fly chr11F4-11F5\|1-43 | 1 | 1 | 25010 | 845 | Set2 | | 3.379e-02 | -3.39 | fly chr3A8-3B1\|1-1 | chromosome location | fly chr3A8-3B1\|1-1 | 1 | 1 | 25010 | 845 | sgg | | 3.379e-02 | -3.39 | fly chr96F5-96F5\|3-89 cM | chromosome location | fly chr96F5-96F5\|3-89 cM | 1 | 1 | 25010 | 845 | Tsp96F | | 3.379e-02 | -3.39 | fly chr13-13\|1-49 | chromosome location | fly chr13-13\|1-49 | 1 | 1 | 25010 | 845 | Flo2 | | 3.379e-02 | -3.39 | fly chr12C8-12C7\|1-46 | chromosome location | fly chr12C8-12C7\|1-46 | 1 | 1 | 25010 | 845 | mamo | | 3.379e-02 | -3.39 | fly chr61D4-61E1\|3-0.5 | chromosome location | fly chr61D4-61E1\|3-0.5 | 1 | 1 | 25010 | 845 | Glut1 | | 3.379e-02 | -3.39 | fly chr67C1-67C1\|3-29 cM | chromosome location | fly chr67C1-67C1\|3-29 cM | 1 | 1 | 25010 | 845 | CG42673 | | 3.379e-02 | -3.39 | fly chr6C11-6C12\|1-18 | chromosome location | fly chr6C11-6C12\|1-18 | 1 | 1 | 25010 | 845 | CG14439 | | 3.379e-02 | -3.39 | fly chr89E2-89E4\|3-59 cM | chromosome location | fly chr89E2-89E4\|3-59 cM | 1 | 1 | 25010 | 845 | lncRNA:iab8 | | 3.379e-02 | -3.39 | fly chr57E8-57E9\|2-95 | chromosome location | fly chr57E8-57E9\|2-95 | 1 | 1 | 25010 | 845 | MESK2 | | 3.379e-02 | -3.39 | fly chr91E5-91E5\|3-65 | chromosome location | fly chr91E5-91E5\|3-65 | 1 | 1 | 25010 | 845 | lncRNA:CR46041 | | 3.379e-02 | -3.39 | fly chr52D4-52D9\|2-76 cM | chromosome location | fly chr52D4-52D9\|2-76 cM | 1 | 1 | 25010 | 845 | Strn-Mlck | | 3.379e-02 | -3.39 | fly chr66F5-67A1\|3-28 cM | chromosome location | fly chr66F5-67A1\|3-28 cM | 1 | 1 | 25010 | 845 | bol | | 3.379e-02 | -3.39 | fly chr14A6-14A7\|1-52 cM | chromosome location | fly chr14A6-14A7\|1-52 cM | 1 | 1 | 25010 | 845 | dpr18 | | 3.379e-02 | -3.39 | fly chr41E4-41E5\|2-55 | chromosome location | fly chr41E4-41E5\|2-55 | 1 | 1 | 25010 | 845 | sxc | | 3.379e-02 | -3.39 | fly chr9C4-9C5\|1-31 | chromosome location | fly chr9C4-9C5\|1-31 | 1 | 1 | 25010 | 845 | Neb-cGP | | 3.379e-02 | -3.39 | fly chr63E8-63F1\|3-8 | chromosome location | fly chr63E8-63F1\|3-8 | 1 | 1 | 25010 | 845 | CG32264 | | 3.379e-02 | -3.39 | fly chr84F12-84F12\|3-48 cM | chromosome location | fly chr84F12-84F12\|3-48 cM | 1 | 1 | 25010 | 845 | CG7443 | | 3.379e-02 | -3.39 | fly chr65B4-65B5\|3-17 | chromosome location | fly chr65B4-65B5\|3-17 | 1 | 1 | 25010 | 845 | MCU | | 3.379e-02 | -3.39 | fly chr72A4-72A4\|3-43 cM | chromosome location | fly chr72A4-72A4\|3-43 cM | 1 | 1 | 25010 | 845 | lncRNA:CR44050 | | 3.391e-02 | -3.38 | Sugar\_tr | pfam domains | PF00083 | 59 | 7 | 10480 | 550 | CG3285,CG14606,Glut1,CG7882,CG10960,CG3168,CG42826 | | 3.393e-02 | -3.38 | hth (homothorax) | protein interactions | 41273 | 35 | 5 | 9204 | 481 | Ubx,Pkc98E,Antp,scrib,tamo | | 3.393e-02 | -3.38 | eIF4G2 (eukaryotic translation initiation factor 4G2) | protein interactions | 42819 | 35 | 5 | 9204 | 481 | CG3961,CG40160,Marf1,CG13157,eIF4E7 | | 3.396e-02 | -3.38 | BZIP\_BASIC | prosite domains | PS00036 | 12 | 3 | 6243 | 386 | Octbeta2R,CrebB,kay | | 3.404e-02 | -3.38 | GO\_RESPONSE\_TO\_EXTERNAL\_STIMULUS | MSigDB lists | GO\_RESPONSE\_TO\_EXTERNAL\_STIMULUS | 380 | 23 | 4161 | 170 | Pkc98E,flw,LpR2,Khc,RhoGAP1A,Su(H),sxc,Sod1,Syn2,mino,fz2,dnc,Ugt301D1,Nmdar2,scramb1,Dys,Octbeta2R,Cat,Ca-beta,TfAP-2,Tor,Ank,Odc1 | | 3.407e-02 | -3.38 | tim (timeless) | protein interactions | 33571 | 24 | 4 | 9204 | 481 | Sox102F,bi,sgg,tyf | | 3.407e-02 | -3.38 | CG13032 (uncharacterized protein) | protein interactions | 39863 | 24 | 4 | 9204 | 481 | Tk,Hr96,Prm,Marf1 | | 3.407e-02 | -3.38 | E(bx) (Enhancer of bithorax) | protein interactions | 44811 | 24 | 4 | 9204 | 481 | Ubx,EcR,chb,Iswi | | 3.407e-02 | -3.38 | CG5177 (uncharacterized protein) | protein interactions | 34017 | 24 | 4 | 9204 | 481 | Prm,TpnC41C,Actn,Scp1 | | 3.407e-02 | -3.38 | TpnC41C (Troponin C at 41C) | protein interactions | 35473 | 24 | 4 | 9204 | 481 | TpnC4,Actn,Scp1,Prm | | 3.407e-02 | -3.38 | Rac2 (Rac2) | protein interactions | 38831 | 24 | 4 | 9204 | 481 | RhoGAP100F,CG17777,Pak,spri | | 3.414e-02 | -3.38 | inorganic cation transmembrane transporter activity | molecular function | GO:0022890 | 339 | 24 | 12176 | 580 | Nmdar2,nrv2,ZnT63C,ND-MLRQ,kcc,CG33310,CG13794,CG14606,KaiR1D,ine,Ca-beta,Sh,Vha36-3,GluRIIC,Irk3,CG34172,MCU,olf186-F,CG10960,Vha100-5,CNT1,Nha1,CG3285,CG45782 | | 3.420e-02 | -3.38 | establishment of cell polarity | biological process | GO:0030010 | 64 | 7 | 12345 | 597 | dsh,rictor,Dys,scrib,sgg,wake,msps | | 3.431e-02 | -3.37 | germ-line stem cell population maintenance | biological process | GO:0030718 | 78 | 8 | 12345 | 597 | Tif-IA,shg,Inx2,dpp,foxo,bru1,Ser,esg | | 3.432e-02 | -3.37 | phosphatidylinositol binding | molecular function | GO:0035091 | 65 | 7 | 12176 | 580 | cert,melt,Pld,step,Gprk1,qvr,CG18659 | | 3.438e-02 | -3.37 | GSE17812\_WT\_VS\_THPOK\_KO\_MEMORY\_CD8\_TCELL\_UP | MSigDB lists | GSE17812\_WT\_VS\_THPOK\_KO\_MEMORY\_CD8\_TCELL\_UP | 60 | 6 | 4161 | 170 | CG5339,MESK2,Pgant6,Elp3,Sh,step | | 3.438e-02 | -3.37 | CREBP1\_Q2 | MSigDB lists | CREBP1\_Q2 | 60 | 6 | 4161 | 170 | Sk1,dnc,RhoGAP1A,mGluR,Dys,Raf | | 3.438e-02 | -3.37 | NKX25\_02 | MSigDB lists | NKX25\_02 | 60 | 6 | 4161 | 170 | eIF4E7,LpR2,Ank,Dys,LRP1,Rbp6 | | 3.438e-02 | -3.37 | GO\_PLASMA\_MEMBRANE\_ORGANIZATION | MSigDB lists | GO\_PLASMA\_MEMBRANE\_ORGANIZATION | 60 | 6 | 4161 | 170 | scramb1,Flo2,Ank,KrT95D,Sod1,sdt | | 3.438e-02 | -3.37 | CEBPB\_01 | MSigDB lists | CEBPB\_01 | 60 | 6 | 4161 | 170 | flw,LRP1,Flo2,CG3961,CG5339,Octbeta2R | | 3.439e-02 | -3.37 | GO\_POSITIVE\_REGULATION\_OF\_BIOSYNTHETIC\_PROCESS | MSigDB lists | GO\_POSITIVE\_REGULATION\_OF\_BIOSYNTHETIC\_PROCESS | 401 | 24 | 4161 | 170 | Hr96,TfAP-2,Tor,Octbeta2R,Cat,dsh,Ptp61F,AstA-R1,TfIIFbeta,scramb1,Fhos,Tif-IA,mino,fz2,CG7483,Su(H),Iswi,Pkc53E,sxc,Tsp96F,AGO3,Ser,dpp,EcR | | 3.445e-02 | -3.37 | ABT1/ESF2\_RRM | interpro domains | IPR034353 | 6 | 2 | 11383 | 585 | CG41562,CG40813 | | 3.445e-02 | -3.37 | Cellulose/chitin-bd\_N | interpro domains | IPR004302 | 6 | 2 | 11383 | 585 | CG42749,CG42598 | | 3.445e-02 | -3.37 | ARID\_dom\_sf | interpro domains | IPR036431 | 6 | 2 | 11383 | 585 | htk,retn | | 3.445e-02 | -3.37 | P\_typ\_ATPase\_c | interpro domains | IPR032630 | 6 | 2 | 11383 | 585 | CG4301,CG9981 | | 3.445e-02 | -3.37 | Sec7\_dom | interpro domains | IPR000904 | 6 | 2 | 11383 | 585 | siz,step | | 3.445e-02 | -3.37 | Na/K\_ATPase\_sub\_beta | interpro domains | IPR000402 | 6 | 2 | 11383 | 585 | nrv2,CG33310 | | 3.445e-02 | -3.37 | P-type\_ATPase\_IV | interpro domains | IPR006539 | 6 | 2 | 11383 | 585 | CG9981,CG4301 | | 3.445e-02 | -3.37 | P-type\_ATPase\_N | interpro domains | IPR032631 | 6 | 2 | 11383 | 585 | CG9981,CG4301 | | 3.445e-02 | -3.37 | Acid\_Pase\_AS | interpro domains | IPR033379 | 6 | 2 | 11383 | 585 | CG9451,CG9449 | | 3.445e-02 | -3.37 | Znf\_GATA | interpro domains | IPR000679 | 6 | 2 | 11383 | 585 | grn,GATAe | | 3.445e-02 | -3.37 | Na/K\_ATPase\_sub\_beta\_sf | interpro domains | IPR038702 | 6 | 2 | 11383 | 585 | CG33310,nrv2 | | 3.445e-02 | -3.37 | PAZ\_dom\_sf | interpro domains | IPR036085 | 6 | 2 | 11383 | 585 | AGO2,AGO3 | | 3.445e-02 | -3.37 | TIL\_dom | interpro domains | IPR002919 | 6 | 2 | 11383 | 585 | Acp62F,CG45546 | | 3.445e-02 | -3.37 | EFG\_V-like | interpro domains | IPR000640 | 6 | 2 | 11383 | 585 | CG4849,CG33158 | | 3.445e-02 | -3.37 | ABT1/Esf2 | interpro domains | IPR039119 | 6 | 2 | 11383 | 585 | CG41562,CG40813 | | 3.445e-02 | -3.37 | EFG\_III/V | interpro domains | IPR035647 | 6 | 2 | 11383 | 585 | CG4849,CG33158 | | 3.445e-02 | -3.37 | ARID\_dom | interpro domains | IPR001606 | 6 | 2 | 11383 | 585 | retn,htk | | 3.445e-02 | -3.37 | mys (myospheroid) | protein interactions | 44885 | 47 | 6 | 9204 | 481 | EcR,sws,Ptp99A,p130CAS,tnc,crol | | 3.460e-02 | -3.36 | GO\_HEART\_DEVELOPMENT | MSigDB lists | GO\_HEART\_DEVELOPMENT | 110 | 9 | 4161 | 170 | CG17387,Tor,Su(H),csw,Meltrin,dsh,LpR2,dpp,Ser | | 3.473e-02 | -3.36 | negative regulation of protein kinase activity | biological process | GO:0006469 | 26 | 4 | 12345 | 597 | kek1,mnb,sws,sgg | | 3.473e-02 | -3.36 | mechanosensory behavior | biological process | GO:0007638 | 26 | 4 | 12345 | 597 | jus,kcc,Pzl,Root | | 3.480e-02 | -3.36 | chr10q25 | MSigDB lists | chr10q25 | 18 | 3 | 4161 | 170 | mino,CG3961,Uros1 | | 3.480e-02 | -3.36 | GCM\_IL6ST | MSigDB lists | GCM\_IL6ST | 18 | 3 | 4161 | 170 | Dlic,KrT95D,CG41099 | | 3.480e-02 | -3.36 | ELVIDGE\_HIF1A\_AND\_HIF2A\_TARGETS\_DN | MSigDB lists | ELVIDGE\_HIF1A\_AND\_HIF2A\_TARGETS\_DN | 18 | 3 | 4161 | 170 | LpR2,Mlp84B,Su(H) | | 3.480e-02 | -3.36 | GO\_NEGATIVE\_REGULATION\_OF\_PROTEIN\_SECRETION | MSigDB lists | GO\_NEGATIVE\_REGULATION\_OF\_PROTEIN\_SECRETION | 18 | 3 | 4161 | 170 | csw,wbl,Pde8 | | 3.480e-02 | -3.36 | GO\_REGULATION\_OF\_LIPID\_TRANSPORT | MSigDB lists | GO\_REGULATION\_OF\_LIPID\_TRANSPORT | 18 | 3 | 4161 | 170 | csw,EcR,LRP1 | | 3.480e-02 | -3.36 | GO\_ENDOTHELIAL\_CELL\_DIFFERENTIATION | MSigDB lists | GO\_ENDOTHELIAL\_CELL\_DIFFERENTIATION | 18 | 3 | 4161 | 170 | Su(H),dnc,Ser | | 3.480e-02 | -3.36 | GO\_APICAL\_JUNCTION\_COMPLEX | MSigDB lists | GO\_APICAL\_JUNCTION\_COMPLEX | 18 | 3 | 4161 | 170 | Ank,fz2,sdt | | 3.480e-02 | -3.36 | GO\_POSITIVE\_REGULATION\_OF\_LIPID\_BIOSYNTHETIC\_PROCESS | MSigDB lists | GO\_POSITIVE\_REGULATION\_OF\_LIPID\_BIOSYNTHETIC\_PROCESS | 18 | 3 | 4161 | 170 | EcR,mino,Tor | | 3.480e-02 | -3.36 | GGCKCATGS\_UNKNOWN | MSigDB lists | GGCKCATGS\_UNKNOWN | 18 | 3 | 4161 | 170 | mGluR,Ptp61F,Ser | | 3.480e-02 | -3.36 | RASHI\_RESPONSE\_TO\_IONIZING\_RADIATION\_2 | MSigDB lists | RASHI\_RESPONSE\_TO\_IONIZING\_RADIATION\_2 | 18 | 3 | 4161 | 170 | Mlp84B,Su(H),scramb1 | | 3.480e-02 | -3.36 | GO\_CELLULAR\_CARBOHYDRATE\_CATABOLIC\_PROCESS | MSigDB lists | GO\_CELLULAR\_CARBOHYDRATE\_CATABOLIC\_PROCESS | 18 | 3 | 4161 | 170 | PhKgamma,Treh,GlyP | | 3.480e-02 | -3.36 | PID\_TCR\_PATHWAY | MSigDB lists | PID\_TCR\_PATHWAY | 18 | 3 | 4161 | 170 | csw,Pkc98E,Pkc53E | | 3.480e-02 | -3.36 | MORF\_DCC | MSigDB lists | MORF\_DCC | 18 | 3 | 4161 | 170 | Src42A,nompB,Spf45 | | 3.480e-02 | -3.36 | MODULE\_75 | MSigDB lists | MODULE\_75 | 18 | 3 | 4161 | 170 | LRP1,Mlp84B,Tsp96F | | 3.489e-02 | -3.36 | TYR\_PHOSPHATASE\_2 | prosite domains | PS50056 | 30 | 5 | 6243 | 386 | Ptp61F,Ptp36E,CG3530,csw,Ptp99A | | 3.501e-02 | -3.35 | GSE14415\_INDUCED\_TREG\_VS\_FOXP3\_KO\_INDUCED\_TREG\_UP | MSigDB lists | GSE14415\_INDUCED\_TREG\_VS\_FOXP3\_KO\_INDUCED\_TREG\_UP | 45 | 5 | 4161 | 170 | Elp3,Pgant6,sdt,Syx1A,CG5339 | | 3.501e-02 | -3.35 | GSE21546\_WT\_VS\_ELK1\_KO\_DP\_THYMOCYTES\_DN | MSigDB lists | GSE21546\_WT\_VS\_ELK1\_KO\_DP\_THYMOCYTES\_DN | 45 | 5 | 4161 | 170 | CG40045,rdgA,CG17698,MESK2,CG3376 | | 3.501e-02 | -3.35 | GSE7596\_AKT\_TRANSD\_VS\_CTRL\_CD4\_TCONV\_WITH\_TGFB\_UP | MSigDB lists | GSE7596\_AKT\_TRANSD\_VS\_CTRL\_CD4\_TCONV\_WITH\_TGFB\_UP | 45 | 5 | 4161 | 170 | Rat1,CG12121,CNT1,CG42672,CG6418 | | 3.501e-02 | -3.35 | GSE360\_L\_DONOVANI\_VS\_M\_TUBERCULOSIS\_DC\_DN | MSigDB lists | GSE360\_L\_DONOVANI\_VS\_M\_TUBERCULOSIS\_DC\_DN | 45 | 5 | 4161 | 170 | Meltrin,CG40178,GlyP,Aldh-III,Spf45 | | 3.501e-02 | -3.35 | GO\_NEGATIVE\_REGULATION\_OF\_ESTABLISHMENT\_OF\_PROTEIN\_LOCALIZATION | MSigDB lists | GO\_NEGATIVE\_REGULATION\_OF\_ESTABLISHMENT\_OF\_PROTEIN\_LOCALIZATION | 45 | 5 | 4161 | 170 | Pde8,Tor,wbl,sxc,csw | | 3.501e-02 | -3.35 | GSE36392\_TYPE\_2\_MYELOID\_VS\_NEUTROPHIL\_IL25\_TREATED\_LUNG\_UP | MSigDB lists | GSE36392\_TYPE\_2\_MYELOID\_VS\_NEUTROPHIL\_IL25\_TREATED\_LUNG\_UP | 45 | 5 | 4161 | 170 | rdgA,CG5493,sxc,CG17698,Glut1 | | 3.501e-02 | -3.35 | GO\_ORGANOPHOSPHATE\_CATABOLIC\_PROCESS | MSigDB lists | GO\_ORGANOPHOSPHATE\_CATABOLIC\_PROCESS | 45 | 5 | 4161 | 170 | Pde6,dnc,Pde8,Cat,CG3376 | | 3.501e-02 | -3.35 | GO\_CELLULAR\_RESPONSE\_TO\_INSULIN\_STIMULUS | MSigDB lists | GO\_CELLULAR\_RESPONSE\_TO\_INSULIN\_STIMULUS | 45 | 5 | 4161 | 170 | sxc,mino,LpR2,Ptp61F,Pak | | 3.520e-02 | -3.35 | phosphoric ester hydrolase activity | molecular function | GO:0042578 | 221 | 17 | 12176 | 580 | Alp12,Pld,CG3530,Ptp36E,Ptp61F,csw,dnc,Lpin,CG9449,Pde1c,Pde6,Pde9,Pde8,Ptp99A,CG18135,CG9451,flw | | 3.520e-02 | -3.35 | oocyte construction | biological process | GO:0007308 | 138 | 12 | 12345 | 597 | msps,dpp,cic,egh,shg,flw,Khc,chb,wbl,scrib,CG7483,tld | | 3.538e-02 | -3.34 | actin-based cell projection | cellular component | GO:0098858 | 28 | 4 | 11940 | 538 | shg,chp,CG5921,Cln3 | | 3.538e-02 | -3.34 | focal adhesion | cellular component | GO:0005925 | 28 | 4 | 11940 | 538 | Pak,by,p130CAS,Actn | | 3.549e-02 | -3.34 | ci (cubitus interruptus) | protein interactions | 43767 | 187 | 16 | 9204 | 481 | boi,dpp,mid,trol,CG7017,hth,bru1,Cdep,D2hgdh,Antp,Msp300,LRP1,Su(H),bi,sgg,CG33127 | | 3.553e-02 | -3.34 | CG14671 (uncharacterized protein) | protein interactions | 40671 | 6 | 2 | 9204 | 481 | raw,CG13526 | | 3.553e-02 | -3.34 | RpI1 (RNA polymerase I subunit) | protein interactions | 36617 | 6 | 2 | 9204 | 481 | rictor,foxo | | 3.553e-02 | -3.34 | CG42748 (uncharacterized protein) | protein interactions | 35442 | 6 | 2 | 9204 | 481 | flw,scrib | | 3.553e-02 | -3.34 | CG3817 (uncharacterized protein) | protein interactions | 41805 | 6 | 2 | 9204 | 481 | foxo,rictor | | 3.553e-02 | -3.34 | CG13728 (uncharacterized protein) | protein interactions | 39937 | 6 | 2 | 9204 | 481 | CG17777,opa | | 3.553e-02 | -3.34 | kek2 (kekkon 2) | protein interactions | 34582 | 6 | 2 | 9204 | 481 | kek1,kek5 | | 3.553e-02 | -3.34 | Sox15 (Sox box protein 15) | protein interactions | 36575 | 6 | 2 | 9204 | 481 | CG14634,by | | 3.553e-02 | -3.34 | mir-1000 (mir-1000 stem loop) | protein interactions | 12798261 | 6 | 2 | 9204 | 481 | mGluR,AGO2 | | 3.553e-02 | -3.34 | CG17221 (uncharacterized protein) | protein interactions | 33521 | 6 | 2 | 9204 | 481 | nompB,CG14634 | | 3.553e-02 | -3.34 | sktl (skittles) | protein interactions | 37356 | 6 | 2 | 9204 | 481 | Pld,rdgA | | 3.553e-02 | -3.34 | dbo (diablo) | protein interactions | 53556 | 6 | 2 | 9204 | 481 | dsh,Pak | | 3.553e-02 | -3.34 | shf (shifted) | protein interactions | 31617 | 6 | 2 | 9204 | 481 | fz2,boi | | 3.557e-02 | -3.34 | EGF-like\_Ca-bd\_dom | interpro domains | IPR001881 | 36 | 5 | 11383 | 585 | LpR2,Ser,CadN2,tld,LRP1 | | 3.578e-02 | -3.33 | GSE2585\_CD80\_HIGH\_VS\_LOW\_AIRE\_KO\_MTEC\_UP | MSigDB lists | GSE2585\_CD80\_HIGH\_VS\_LOW\_AIRE\_KO\_MTEC\_UP | 31 | 4 | 4161 | 170 | dsh,Ser,RhoGAP1A,Meltrin | | 3.578e-02 | -3.33 | GO\_ESTABLISHMENT\_OF\_PROTEIN\_LOCALIZATION\_TO\_PLASMA\_MEMBRANE | MSigDB lists | GO\_ESTABLISHMENT\_OF\_PROTEIN\_LOCALIZATION\_TO\_PLASMA\_MEMBRANE | 31 | 4 | 4161 | 170 | KrT95D,Ank,Flo2,sdt | | 3.578e-02 | -3.33 | GSE20484\_MCSG\_VS\_CXCL4\_MONOCYTE\_DERIVED\_MACROPHAGE\_DN | MSigDB lists | GSE20484\_MCSG\_VS\_CXCL4\_MONOCYTE\_DERIVED\_MACROPHAGE\_DN | 31 | 4 | 4161 | 170 | CG3961,chb,CG42672,sdt | | 3.578e-02 | -3.33 | GSE3982\_MEMORY\_CD4\_TCELL\_VS\_TH2\_UP | MSigDB lists | GSE3982\_MEMORY\_CD4\_TCELL\_VS\_TH2\_UP | 31 | 4 | 4161 | 170 | CG8184,CG5522,CG9132,CG34408 | | 3.578e-02 | -3.33 | REACTOME\_DOWNSTREAM\_SIGNAL\_TRANSDUCTION | MSigDB lists | REACTOME\_DOWNSTREAM\_SIGNAL\_TRANSDUCTION | 31 | 4 | 4161 | 170 | Pkc98E,Pkc53E,Tor,Pde1c | | 3.578e-02 | -3.33 | MORF\_CTSB | MSigDB lists | MORF\_CTSB | 31 | 4 | 4161 | 170 | Dys,Hr96,fz2,Pkc53E | | 3.578e-02 | -3.33 | KEGG\_FC\_GAMMA\_R\_MEDIATED\_PHAGOCYTOSIS | MSigDB lists | KEGG\_FC\_GAMMA\_R\_MEDIATED\_PHAGOCYTOSIS | 31 | 4 | 4161 | 170 | Pak,Sk1,Pkc98E,Pkc53E | | 3.578e-02 | -3.33 | GO\_NEGATIVE\_REGULATION\_OF\_ION\_TRANSPORT | MSigDB lists | GO\_NEGATIVE\_REGULATION\_OF\_ION\_TRANSPORT | 31 | 4 | 4161 | 170 | Ank,ZnT63C,Tor,Pkc98E | | 3.578e-02 | -3.33 | PICCALUGA\_ANGIOIMMUNOBLASTIC\_LYMPHOMA\_DN | MSigDB lists | PICCALUGA\_ANGIOIMMUNOBLASTIC\_LYMPHOMA\_DN | 31 | 4 | 4161 | 170 | dnc,ZnT63C,Glut1,eIF4E7 | | 3.578e-02 | -3.33 | GO\_POLYSACCHARIDE\_METABOLIC\_PROCESS | MSigDB lists | GO\_POLYSACCHARIDE\_METABOLIC\_PROCESS | 31 | 4 | 4161 | 170 | GlyP,flw,PhKgamma,CG30036 | | 3.582e-02 | -3.33 | PhoLip\_ATPase\_C | pfam domains | PF16212 | 6 | 2 | 10480 | 550 | CG4301,CG9981 | | 3.582e-02 | -3.33 | Na\_K-ATPase | pfam domains | PF00287 | 6 | 2 | 10480 | 550 | CG33310,nrv2 | | 3.582e-02 | -3.33 | ARID | pfam domains | PF01388 | 6 | 2 | 10480 | 550 | htk,retn | | 3.582e-02 | -3.33 | GATA | pfam domains | PF00320 | 6 | 2 | 10480 | 550 | grn,GATAe | | 3.582e-02 | -3.33 | TIL | pfam domains | PF01826 | 6 | 2 | 10480 | 550 | Acp62F,CG45546 | | 3.582e-02 | -3.33 | EFG\_C | pfam domains | PF00679 | 6 | 2 | 10480 | 550 | CG4849,CG33158 | | 3.582e-02 | -3.33 | LPMO\_10 | pfam domains | PF03067 | 6 | 2 | 10480 | 550 | CG42749,CG42598 | | 3.582e-02 | -3.33 | PhoLip\_ATPase\_N | pfam domains | PF16209 | 6 | 2 | 10480 | 550 | CG4301,CG9981 | | 3.587e-02 | -3.33 | GO\_VESICLE\_MEDIATED\_TRANSPORT | MSigDB lists | GO\_VESICLE\_MEDIATED\_TRANSPORT | 382 | 23 | 4161 | 170 | Sod1,Epg5,CG17883,unc-13,LpR2,Tsp96F,Pak,Pkc98E,LRP1,Act87E,CG41099,pod1,Dlic,Ank,step,Cln3,CG9132,CG12121,Syx1A,CG3860,chb,Octbeta2R,Cadps | | 3.589e-02 | -3.33 | Z disc | cellular component | GO:0030018 | 41 | 5 | 11940 | 538 | rols,Actn,Msp300,zormin,Mlp84B | | 3.592e-02 | -3.33 | polo (polo) | protein interactions | 40232 | 73 | 8 | 9204 | 481 | numb,zormin,fs(1)N,by,bi,chb,nuf,CG5004 | | 3.595e-02 | -3.33 | Abd-B (Abdominal B) | protein interactions | 47763 | 60 | 7 | 9204 | 481 | Poxm,hth,mid,Ubx,tin,ems,Su(H) | | 3.597e-02 | -3.33 | GO\_ORGANIC\_HYDROXY\_COMPOUND\_METABOLIC\_PROCESS | MSigDB lists | GO\_ORGANIC\_HYDROXY\_COMPOUND\_METABOLIC\_PROCESS | 165 | 12 | 4161 | 170 | CG3842,Cln3,IP3K2,Acbp2,LpR2,Aldh-III,AstA-R1,CG3860,CG3376,Cat,CG33090,Sk1 | | 3.638e-02 | -3.31 | GO\_NEGATIVE\_REGULATION\_OF\_PHOSPHORUS\_METABOLIC\_PROCESS | MSigDB lists | GO\_NEGATIVE\_REGULATION\_OF\_PHOSPHORUS\_METABOLIC\_PROCESS | 111 | 9 | 4161 | 170 | CG3376,Dys,mGluR,Ptp61F,AstA-R1,dnc,Tor,Pkc53E,sxc | | 3.653e-02 | -3.31 | Homeobox\_dom | interpro domains | IPR001356 | 102 | 10 | 11383 | 585 | scro,Awh,Antp,tin,E5,B-H2,ems,Ubx,hth,toe | | 3.660e-02 | -3.31 | sodium:potassium-exchanging ATPase complex | cellular component | GO:0005890 | 7 | 2 | 11940 | 538 | nrv2,CG33310 | | 3.660e-02 | -3.31 | cation-transporting ATPase complex | cellular component | GO:0090533 | 7 | 2 | 11940 | 538 | CG33310,nrv2 | | 3.683e-02 | -3.30 | CAGTATT\_MIR200B\_MIR200C\_MIR429 | MSigDB lists | CAGTATT\_MIR200B\_MIR200C\_MIR429 | 129 | 10 | 4161 | 170 | LpR2,Syx1A,chb,LRP1,Fhos,Ank,Frq1,sxc,Frq2,sdt | | 3.683e-02 | -3.30 | GATAZNFINGER | prints domains | PR00619 | 5 | 2 | 2185 | 142 | grn,GATAe | | 3.683e-02 | -3.30 | FRIZZLED | prints domains | PR00489 | 5 | 2 | 2185 | 142 | fz4,fz2 | | 3.688e-02 | -3.30 | GO\_LEUKOCYTE\_ACTIVATION | MSigDB lists | GO\_LEUKOCYTE\_ACTIVATION | 61 | 6 | 4161 | 170 | Raf,Ptp61F,Su(H),Pkc98E,fz2,IP3K2 | | 3.688e-02 | -3.30 | GSE17580\_TREG\_VS\_TEFF\_S\_MANSONI\_INF\_UP | MSigDB lists | GSE17580\_TREG\_VS\_TEFF\_S\_MANSONI\_INF\_UP | 61 | 6 | 4161 | 170 | Glut1,Odc1,Nha1,Tsp96F,scramb1,Mlp84B | | 3.688e-02 | -3.30 | GO\_GATED\_CHANNEL\_ACTIVITY | MSigDB lists | GO\_GATED\_CHANNEL\_ACTIVITY | 61 | 6 | 4161 | 170 | Frq2,Sh,Ca-beta,KaiR1D,Nmdar2,Frq1 | | 3.688e-02 | -3.30 | STK33\_NOMO\_DN | MSigDB lists | STK33\_NOMO\_DN | 61 | 6 | 4161 | 170 | CG4341,rdgA,Argl,CG12547,CG3860,Nha1 | | 3.688e-02 | -3.30 | GSE40655\_FOXO1\_KO\_VS\_WT\_NTREG\_UP | MSigDB lists | GSE40655\_FOXO1\_KO\_VS\_WT\_NTREG\_UP | 61 | 6 | 4161 | 170 | Cln3,RpL38,CG8184,Cpsf160,sws,CG3376 | | 3.688e-02 | -3.30 | RRAGTTGT\_UNKNOWN | MSigDB lists | RRAGTTGT\_UNKNOWN | 61 | 6 | 4161 | 170 | CG4341,TfAP-2,Rbp6,app,Dys,dsd | | 3.691e-02 | -3.30 | RHO GTPase Effectors | REACTOME pathways | R-DME-195258 | 63 | 7 | 3923 | 197 | Strn-Mlck,Pak,dsh,flw,Mbs,Mrtf,Cam | | 3.691e-02 | -3.30 | - | gene3d domains | 3.40.720.10 | 41 | 6 | 4929 | 302 | PIG-O,CG1428,CG6790,CG16854,PIG-G,Alp12 | | 3.699e-02 | -3.30 | ANF\_lig-bd\_rcpt | interpro domains | IPR001828 | 25 | 4 | 11383 | 585 | Nmdar2,mGluR,GluRIIC,KaiR1D | | 3.743e-02 | -3.29 | TAATTA\_CHX10\_01 | MSigDB lists | TAATTA\_CHX10\_01 | 166 | 12 | 4161 | 170 | DNaseII,Doa,bru1,sxc,TfAP-2,sdt,CG4341,LpR2,Pde1c,Dys,Syn2,CG5493 | | 3.752e-02 | -3.28 | oocyte development | biological process | GO:0048599 | 155 | 13 | 12345 | 597 | shg,dpp,egh,cic,msps,tld,CG7483,wbl,scrib,Mos,chb,Khc,flw | | 3.771e-02 | -3.28 | kdn (knockdown) | protein interactions | 31579 | 48 | 6 | 9204 | 481 | Prm,Act88F,CG34172,TpnC41C,Actn,Scp1 | | 3.776e-02 | -3.28 | positive regulation of developmental growth | biological process | GO:0048639 | 94 | 9 | 12345 | 597 | Octbeta2R,Tor,dpp,Pak,mnb,dsh,step,Ank2,rictor | | 3.777e-02 | -3.28 | RpII215 (RNA polymerase II 215kD subunit) | protein interactions | 32100 | 36 | 5 | 9204 | 481 | CG8117,Set2,TfIIFbeta,AGO2,Iswi | | 3.781e-02 | -3.28 | tkv (thickveins) | protein interactions | 33753 | 101 | 10 | 9204 | 481 | Khc,Snoo,Ubx,shg,dpp,dsh,Raf,kek5,csw,scrib | | 3.789e-02 | -3.27 | phosphotransferase activity, for other substituted phosphate groups | molecular function | GO:0016780 | 16 | 3 | 12176 | 580 | PIG-G,CG6790,PIG-O | | 3.801e-02 | -3.27 | regulation of synaptic growth at neuromuscular junction | biological process | GO:0008582 | 109 | 10 | 12345 | 597 | Mob2,Pak,mnb,rictor,Ank2,Tor,sgg,Src42A,Octbeta2R,Syn2 | | 3.803e-02 | -3.27 | REACTOME\_NEUROTRANSMITTER\_RECEPTOR\_BINDING\_AND\_DOWNSTREAM\_TRANSMISSION\_IN\_THE\_POSTSYNAPTIC\_CELL | MSigDB lists | REACTOME\_NEUROTRANSMITTER\_RECEPTOR\_BINDING\_AND\_DOWNSTREAM\_TRANSMISSION\_IN\_THE\_POSTSYNAPTIC\_CELL | 46 | 5 | 4161 | 170 | KaiR1D,Raf,Nmdar2,CG17698,Pkc53E | | 3.803e-02 | -3.27 | GSE31082\_DN\_VS\_DP\_THYMOCYTE\_DN | MSigDB lists | GSE31082\_DN\_VS\_DP\_THYMOCYTE\_DN | 46 | 5 | 4161 | 170 | Khc,Raf,PhKgamma,Maf1,EcR | | 3.803e-02 | -3.27 | LEF1\_UP.V1\_DN | MSigDB lists | LEF1\_UP.V1\_DN | 46 | 5 | 4161 | 170 | Cat,Ank,Pak,CG3961,CG41099 | | 3.803e-02 | -3.27 | GO\_MUSCLE\_CONTRACTION | MSigDB lists | GO\_MUSCLE\_CONTRACTION | 46 | 5 | 4161 | 170 | AstA-R1,Octbeta2R,Dys,dnc,Tor | | 3.803e-02 | -3.27 | GSE31082\_DN\_VS\_CD8\_SP\_THYMOCYTE\_DN | MSigDB lists | GSE31082\_DN\_VS\_CD8\_SP\_THYMOCYTE\_DN | 46 | 5 | 4161 | 170 | step,CG17883,Pgant6,rdgA,CNT1 | | 3.803e-02 | -3.27 | RUTELLA\_RESPONSE\_TO\_HGF\_VS\_CSF2RB\_AND\_IL4\_DN | MSigDB lists | RUTELLA\_RESPONSE\_TO\_HGF\_VS\_CSF2RB\_AND\_IL4\_DN | 46 | 5 | 4161 | 170 | Meltrin,fz2,step,Ser,bru1 | | 3.803e-02 | -3.27 | GSE21360\_NAIVE\_VS\_PRIMARY\_MEMORY\_CD8\_TCELL\_DN | MSigDB lists | GSE21360\_NAIVE\_VS\_PRIMARY\_MEMORY\_CD8\_TCELL\_DN | 46 | 5 | 4161 | 170 | step,LRP1,CG5493,app,KrT95D | | 3.803e-02 | -3.27 | GSE11961\_FOLLICULAR\_BCELL\_VS\_MARGINAL\_ZONE\_BCELL\_DN | MSigDB lists | GSE11961\_FOLLICULAR\_BCELL\_VS\_MARGINAL\_ZONE\_BCELL\_DN | 46 | 5 | 4161 | 170 | Odc1,Cat,Tsp96F,Sod1,Argl | | 3.803e-02 | -3.27 | GSE29618\_MONOCYTE\_VS\_PDC\_DAY7\_FLU\_VACCINE\_DN | MSigDB lists | GSE29618\_MONOCYTE\_VS\_PDC\_DAY7\_FLU\_VACCINE\_DN | 46 | 5 | 4161 | 170 | Khc,Odc1,KrT95D,Meltrin,sxc | | 3.803e-02 | -3.27 | GSE43955\_1H\_VS\_20H\_ACT\_CD4\_TCELL\_DN | MSigDB lists | GSE43955\_1H\_VS\_20H\_ACT\_CD4\_TCELL\_DN | 46 | 5 | 4161 | 170 | Stt3B,Pkc98E,nompB,sky,CG41099 | | 3.803e-02 | -3.27 | GSE360\_HIGH\_DOSE\_B\_MALAYI\_VS\_M\_TUBERCULOSIS\_MAC\_DN | MSigDB lists | GSE360\_HIGH\_DOSE\_B\_MALAYI\_VS\_M\_TUBERCULOSIS\_MAC\_DN | 46 | 5 | 4161 | 170 | Ptp61F,dsh,GlyP,Glut1,dnc | | 3.803e-02 | -3.27 | GO\_EXCITATORY\_SYNAPSE | MSigDB lists | GO\_EXCITATORY\_SYNAPSE | 46 | 5 | 4161 | 170 | KaiR1D,mGluR,Frq1,Nlg2,Frq2 | | 3.803e-02 | -3.27 | GSE8921\_UNSTIM\_0H\_VS\_TLR1\_2\_STIM\_MONOCYTE\_12H\_DN | MSigDB lists | GSE8921\_UNSTIM\_0H\_VS\_TLR1\_2\_STIM\_MONOCYTE\_12H\_DN | 46 | 5 | 4161 | 170 | CG42672,TfAP-2,Cat,dsd,mRpS5 | | 3.803e-02 | -3.27 | TGTGTGA\_MIR377 | MSigDB lists | TGTGTGA\_MIR377 | 46 | 5 | 4161 | 170 | eIF4E7,app,bru1,Sod1,Ser | | 3.803e-02 | -3.27 | OCT1\_03 | MSigDB lists | OCT1\_03 | 46 | 5 | 4161 | 170 | Ser,sdt,CG4341,Dys,Mctp | | 3.803e-02 | -3.27 | GSE360\_L\_MAJOR\_VS\_M\_TUBERCULOSIS\_MAC\_DN | MSigDB lists | GSE360\_L\_MAJOR\_VS\_M\_TUBERCULOSIS\_MAC\_DN | 46 | 5 | 4161 | 170 | GlyP,bru1,CG34408,chb,msps | | 3.853e-02 | -3.26 | phototransduction | biological process | GO:0007602 | 52 | 6 | 12345 | 597 | rdgA,shakB,Cam,Gprk1,ninaD,Pld | | 3.866e-02 | -3.25 | positive regulation of molecular function | biological process | GO:0044093 | 287 | 21 | 12345 | 597 | CG34408,CG42390,Raf,qvr,Strn-Mlck,Cam,sky,Stlk,RhoGAP18B,CG41520,nrv2,CG17883,step,rictor,CG33310,Pak,RhoGAP100F,REG,CycT,HDAC4,RhoGAP1A | | 3.875e-02 | -3.25 | - | gene3d domains | 1.10.630.10 | 87 | 10 | 4929 | 302 | Cyp4p2,shd,spok,Cyp6u1,Cyp28d2,Cyp309a1,Cyp6a13,Cyp12a4,Cyp28d1,Cyp304a1 | | 3.879e-02 | -3.25 | GO\_VESICLE\_LOCALIZATION | MSigDB lists | GO\_VESICLE\_LOCALIZATION | 78 | 7 | 4161 | 170 | Cadps,CG17883,Cln3,Khc,chb,Syx1A,unc-13 | | 3.895e-02 | -3.25 | Gdh (Glutamate dehydrogenase) | protein interactions | 42832 | 25 | 4 | 9204 | 481 | Actn,Scp1,CG15564,Prm | | 3.895e-02 | -3.25 | GlyP (Glycogen phosphorylase) | protein interactions | 33386 | 25 | 4 | 9204 | 481 | Ubx,Scp1,CG46385,CG1894 | | 3.896e-02 | -3.25 | PDZ | prosite domains | PS50106 | 63 | 8 | 6243 | 386 | sdt,bbg,Syn2,scrib,dsh,CG5921,Fife,RhoGAP100F | | 3.901e-02 | -3.24 | bsk (basket) | protein interactions | 44801 | 130 | 12 | 9204 | 481 | Cln3,foxo,flw,dsh,Pak,Src42A,kay,raw,cindr,sgg,by,scrib | | 3.906e-02 | -3.24 | sensory perception of mechanical stimulus | biological process | GO:0050954 | 80 | 8 | 12345 | 597 | CG5921,rdgA,nompB,Root,Cam,Ank2,Nmdar2,nrv2 | | 3.906e-02 | -3.24 | wing disc pattern formation | biological process | GO:0035222 | 80 | 8 | 12345 | 597 | sno,nuf,Su(H),dpp,cic,boi,bi,Ser | | 3.926e-02 | -3.24 | negative regulation of kinase activity | biological process | GO:0033673 | 27 | 4 | 12345 | 597 | mnb,kek1,sgg,sws | | 3.926e-02 | -3.24 | body morphogenesis | biological process | GO:0010171 | 27 | 4 | 12345 | 597 | hth,TwdlG,dpp,TwdlY | | 3.926e-02 | -3.24 | negative regulation of JNK cascade | biological process | GO:0046329 | 27 | 4 | 12345 | 597 | raw,flw,sgg,Ptp61F | | 3.926e-02 | -3.24 | negative regulation of gene silencing | biological process | GO:0060969 | 27 | 4 | 12345 | 597 | HPS4,g,polybromo,Ubx | | 3.926e-02 | -3.24 | positive regulation of cell cycle phase transition | biological process | GO:1901989 | 27 | 4 | 12345 | 597 | Tor,Su(H),sgg,bru1 | | 3.929e-02 | -3.24 | I band | cellular component | GO:0031674 | 42 | 5 | 11940 | 538 | rols,zormin,Msp300,Actn,Mlp84B | | 3.937e-02 | -3.23 | wing disc anterior/posterior pattern formation | biological process | GO:0048100 | 16 | 3 | 12345 | 597 | dpp,Ser,bi | | 3.937e-02 | -3.23 | secondary alcohol biosynthetic process | biological process | GO:1902653 | 16 | 3 | 12345 | 597 | spok,woc,foxo | | 3.937e-02 | -3.23 | regulation of muscle organ development | biological process | GO:0048634 | 16 | 3 | 12345 | 597 | rst,dpp,Ubx | | 3.937e-02 | -3.23 | establishment of epithelial cell polarity | biological process | GO:0090162 | 16 | 3 | 12345 | 597 | sgg,dsh,scrib | | 3.937e-02 | -3.23 | regulation of synaptic vesicle cycle | biological process | GO:0098693 | 16 | 3 | 12345 | 597 | mnb,Fife,sky | | 3.946e-02 | -3.23 | Sugar\_transporter\_CS | interpro domains | IPR005829 | 37 | 5 | 11383 | 585 | CG3168,CG10960,Glut1,CG3285,CG14606 | | 3.949e-02 | -3.23 | ANF\_receptor | pfam domains | PF01094 | 25 | 4 | 10480 | 550 | Nmdar2,KaiR1D,GluRIIC,mGluR | | 3.951e-02 | -3.23 | GO\_KINESIN\_BINDING | MSigDB lists | GO\_KINESIN\_BINDING | 8 | 2 | 4161 | 170 | Root,Act87E | | 3.951e-02 | -3.23 | WHITESIDE\_CISPLATIN\_RESISTANCE\_DN | MSigDB lists | WHITESIDE\_CISPLATIN\_RESISTANCE\_DN | 8 | 2 | 4161 | 170 | Act87E,Sod1 | | 3.951e-02 | -3.23 | GO\_HEAD\_MORPHOGENESIS | MSigDB lists | GO\_HEAD\_MORPHOGENESIS | 8 | 2 | 4161 | 170 | Raf,csw | | 3.951e-02 | -3.23 | GO\_EMBRYONIC\_CAMERA\_TYPE\_EYE\_DEVELOPMENT | MSigDB lists | GO\_EMBRYONIC\_CAMERA\_TYPE\_EYE\_DEVELOPMENT | 8 | 2 | 4161 | 170 | fz2,TfAP-2 | | 3.951e-02 | -3.23 | GO\_REGULATION\_OF\_LIPID\_STORAGE | MSigDB lists | GO\_REGULATION\_OF\_LIPID\_STORAGE | 8 | 2 | 4161 | 170 | EcR,Ptp61F | | 3.951e-02 | -3.23 | YAP1\_DN | MSigDB lists | YAP1\_DN | 8 | 2 | 4161 | 170 | Ca-beta,PIG-O | | 3.951e-02 | -3.23 | GO\_REGULATION\_OF\_TRIGLYCERIDE\_BIOSYNTHETIC\_PROCESS | MSigDB lists | GO\_REGULATION\_OF\_TRIGLYCERIDE\_BIOSYNTHETIC\_PROCESS | 8 | 2 | 4161 | 170 | EcR,mino | | 3.951e-02 | -3.23 | GO\_NEGATIVE\_REGULATION\_OF\_REGULATED\_SECRETORY\_PATHWAY | MSigDB lists | GO\_NEGATIVE\_REGULATION\_OF\_REGULATED\_SECRETORY\_PATHWAY | 8 | 2 | 4161 | 170 | RhoGAP1A,Raf | | 3.951e-02 | -3.23 | VERRECCHIA\_EARLY\_RESPONSE\_TO\_TGFB1 | MSigDB lists | VERRECCHIA\_EARLY\_RESPONSE\_TO\_TGFB1 | 8 | 2 | 4161 | 170 | Pak,LRP1 | | 3.951e-02 | -3.23 | GO\_IONOTROPIC\_GLUTAMATE\_RECEPTOR\_COMPLEX | MSigDB lists | GO\_IONOTROPIC\_GLUTAMATE\_RECEPTOR\_COMPLEX | 8 | 2 | 4161 | 170 | Nmdar2,KaiR1D | | 3.951e-02 | -3.23 | BIOCARTA\_FEEDER\_PATHWAY | MSigDB lists | BIOCARTA\_FEEDER\_PATHWAY | 8 | 2 | 4161 | 170 | Treh,GlyP | | 3.951e-02 | -3.23 | GO\_REGULATION\_OF\_HEART\_MORPHOGENESIS | MSigDB lists | GO\_REGULATION\_OF\_HEART\_MORPHOGENESIS | 8 | 2 | 4161 | 170 | dpp,Su(H) | | 3.951e-02 | -3.23 | GO\_PERIPHERAL\_NERVOUS\_SYSTEM\_AXON\_ENSHEATHMENT | MSigDB lists | GO\_PERIPHERAL\_NERVOUS\_SYSTEM\_AXON\_ENSHEATHMENT | 8 | 2 | 4161 | 170 | Sod1,sdt | | 3.951e-02 | -3.23 | GO\_MOTILE\_CILIUM\_ASSEMBLY | MSigDB lists | GO\_MOTILE\_CILIUM\_ASSEMBLY | 8 | 2 | 4161 | 170 | Dys,CG17387 | | 3.951e-02 | -3.23 | PID\_BETA\_CATENIN\_DEG\_PATHWAY | MSigDB lists | PID\_BETA\_CATENIN\_DEG\_PATHWAY | 8 | 2 | 4161 | 170 | dsh,fz2 | | 3.951e-02 | -3.23 | GO\_SPHINGOMYELIN\_METABOLIC\_PROCESS | MSigDB lists | GO\_SPHINGOMYELIN\_METABOLIC\_PROCESS | 8 | 2 | 4161 | 170 | CG3376,Cln3 | | 3.951e-02 | -3.23 | GO\_CALMODULIN\_DEPENDENT\_PROTEIN\_KINASE\_ACTIVITY | MSigDB lists | GO\_CALMODULIN\_DEPENDENT\_PROTEIN\_KINASE\_ACTIVITY | 8 | 2 | 4161 | 170 | CG17698,PhKgamma | | 3.951e-02 | -3.23 | GO\_NEGATIVE\_REGULATION\_OF\_EXOCYTOSIS | MSigDB lists | GO\_NEGATIVE\_REGULATION\_OF\_EXOCYTOSIS | 8 | 2 | 4161 | 170 | RhoGAP1A,Raf | | 3.951e-02 | -3.23 | GO\_REGULATION\_OF\_LEUKOCYTE\_APOPTOTIC\_PROCESS | MSigDB lists | GO\_REGULATION\_OF\_LEUKOCYTE\_APOPTOTIC\_PROCESS | 8 | 2 | 4161 | 170 | IP3K2,mino | | 3.951e-02 | -3.23 | GO\_PHOSPHOTRANSFERASE\_ACTIVITY\_FOR\_OTHER\_SUBSTITUTED\_PHOSPHATE\_GROUPS | MSigDB lists | GO\_PHOSPHOTRANSFERASE\_ACTIVITY\_FOR\_OTHER\_SUBSTITUTED\_PHOSPHATE\_GROUPS | 8 | 2 | 4161 | 170 | PIG-G,PIG-O | | 3.951e-02 | -3.23 | HOEGERKORP\_CD44\_TARGETS\_DIRECT\_UP | MSigDB lists | HOEGERKORP\_CD44\_TARGETS\_DIRECT\_UP | 8 | 2 | 4161 | 170 | Syx1A,Octbeta2R | | 3.951e-02 | -3.23 | GO\_MULTICELLULAR\_ORGANISMAL\_MOVEMENT | MSigDB lists | GO\_MULTICELLULAR\_ORGANISMAL\_MOVEMENT | 8 | 2 | 4161 | 170 | Tor,Octbeta2R | | 3.951e-02 | -3.23 | MYOGNF1\_01 | MSigDB lists | MYOGNF1\_01 | 8 | 2 | 4161 | 170 | CG33090,TfAP-2 | | 3.951e-02 | -3.23 | PID\_RAS\_PATHWAY | MSigDB lists | PID\_RAS\_PATHWAY | 8 | 2 | 4161 | 170 | Pkc53E,Pkc98E | | 3.951e-02 | -3.23 | GO\_POSITIVE\_REGULATION\_OF\_HEART\_CONTRACTION | MSigDB lists | GO\_POSITIVE\_REGULATION\_OF\_HEART\_CONTRACTION | 8 | 2 | 4161 | 170 | Octbeta2R,dnc | | 3.951e-02 | -3.23 | MODULE\_563 | MSigDB lists | MODULE\_563 | 8 | 2 | 4161 | 170 | Syx1A,Cadps | | 3.951e-02 | -3.23 | ZHENG\_GLIOBLASTOMA\_PLASTICITY\_DN | MSigDB lists | ZHENG\_GLIOBLASTOMA\_PLASTICITY\_DN | 8 | 2 | 4161 | 170 | Cadps,app | | 3.951e-02 | -3.23 | GO\_POSITIVE\_REGULATION\_OF\_BLOOD\_PRESSURE | MSigDB lists | GO\_POSITIVE\_REGULATION\_OF\_BLOOD\_PRESSURE | 8 | 2 | 4161 | 170 | dnc,Octbeta2R | | 3.951e-02 | -3.23 | GO\_VASODILATION | MSigDB lists | GO\_VASODILATION | 8 | 2 | 4161 | 170 | Octbeta2R,Sod1 | | 3.951e-02 | -3.23 | REACTOME\_ARMS\_MEDIATED\_ACTIVATION | MSigDB lists | REACTOME\_ARMS\_MEDIATED\_ACTIVATION | 8 | 2 | 4161 | 170 | CG42672,Raf | | 3.951e-02 | -3.23 | GO\_RHO\_GUANYL\_NUCLEOTIDE\_EXCHANGE\_FACTOR\_ACTIVITY | MSigDB lists | GO\_RHO\_GUANYL\_NUCLEOTIDE\_EXCHANGE\_FACTOR\_ACTIVITY | 8 | 2 | 4161 | 170 | Cdep,RhoGAP1A | | 3.951e-02 | -3.23 | CHYLA\_CBFA2T3\_TARGETS\_UP | MSigDB lists | CHYLA\_CBFA2T3\_TARGETS\_UP | 62 | 6 | 4161 | 170 | IP3K2,ABCD,bru1,Ank,Mctp,CNT1 | | 3.951e-02 | -3.23 | MORF\_ATF2 | MSigDB lists | MORF\_ATF2 | 62 | 6 | 4161 | 170 | dnc,Pkc53E,fz2,Dys,CG40178,Hr96 | | 3.951e-02 | -3.23 | PTF1BETA\_Q6 | MSigDB lists | PTF1BETA\_Q6 | 62 | 6 | 4161 | 170 | TfAP-2,eRF3,CG4341,Ptp61F,Dys,Ank | | 3.951e-02 | -3.23 | GSE17721\_LPS\_VS\_CPG\_2H\_BMDC\_DN | MSigDB lists | GSE17721\_LPS\_VS\_CPG\_2H\_BMDC\_DN | 62 | 6 | 4161 | 170 | sxc,CG6418,CG9132,unc-13,KrT95D,Uros1 | | 3.951e-02 | -3.23 | GSE24142\_DN2\_VS\_DN3\_THYMOCYTE\_ADULT\_DN | MSigDB lists | GSE24142\_DN2\_VS\_DN3\_THYMOCYTE\_ADULT\_DN | 62 | 6 | 4161 | 170 | Doa,CG17698,Stt3B,CG15093,eRF3,CG8184 | | 3.951e-02 | -3.23 | GO\_POSITIVE\_REGULATION\_OF\_LOCOMOTION | MSigDB lists | GO\_POSITIVE\_REGULATION\_OF\_LOCOMOTION | 62 | 6 | 4161 | 170 | Pkc53E,Pkc98E,Elp3,dpp,chb,Pak | | 3.951e-02 | -3.23 | LIM\_MAMMARY\_STEM\_CELL\_UP | MSigDB lists | LIM\_MAMMARY\_STEM\_CELL\_UP | 62 | 6 | 4161 | 170 | Mlp84B,Ca-beta,Ser,LRP1,kcc,Glut1 | | 3.953e-02 | -3.23 | sensory perception of sound | biological process | GO:0007605 | 66 | 7 | 12345 | 597 | Root,Cam,rdgA,CG5921,nompB,Ank2,nrv2 | | 3.964e-02 | -3.23 | GO\_BINDING\_BRIDGING | MSigDB lists | GO\_BINDING\_BRIDGING | 32 | 4 | 4161 | 170 | Syx1A,Ank,dsh,csw | | 3.964e-02 | -3.23 | GSE22432\_MULTIPOTENT\_PROGENITOR\_VS\_CDC\_UP | MSigDB lists | GSE22432\_MULTIPOTENT\_PROGENITOR\_VS\_CDC\_UP | 32 | 4 | 4161 | 170 | CNT1,AstA-R1,Pde6,Sk1 | | 3.964e-02 | -3.23 | MODULE\_182 | MSigDB lists | MODULE\_182 | 32 | 4 | 4161 | 170 | csw,sno,Dys,Raf | | 3.964e-02 | -3.23 | GSE21927\_SPLEEN\_VS\_BONE\_MARROW\_MONOCYTE\_BALBC\_UP | MSigDB lists | GSE21927\_SPLEEN\_VS\_BONE\_MARROW\_MONOCYTE\_BALBC\_UP | 32 | 4 | 4161 | 170 | scramb1,Cdep,chb,CG4341 | | 3.964e-02 | -3.23 | GSE30962\_ACUTE\_VS\_CHRONIC\_LCMV\_PRIMARY\_INF\_CD8\_TCELL\_UP | MSigDB lists | GSE30962\_ACUTE\_VS\_CHRONIC\_LCMV\_PRIMARY\_INF\_CD8\_TCELL\_UP | 32 | 4 | 4161 | 170 | Pgant6,sws,Glut1,Octbeta2R | | 3.964e-02 | -3.23 | GSE15733\_BM\_VS\_SPLEEN\_MEMORY\_CD4\_TCELL\_UP | MSigDB lists | GSE15733\_BM\_VS\_SPLEEN\_MEMORY\_CD4\_TCELL\_UP | 32 | 4 | 4161 | 170 | Iswi,Mlp84B,Src42A,sdt | | 3.964e-02 | -3.23 | LTE2\_UP.V1\_DN | MSigDB lists | LTE2\_UP.V1\_DN | 32 | 4 | 4161 | 170 | scramb1,Ser,TfAP-2,Tif-IA | | 3.964e-02 | -3.23 | GSE13522\_WT\_VS\_IFNAR\_KO\_SKIN\_DN | MSigDB lists | GSE13522\_WT\_VS\_IFNAR\_KO\_SKIN\_DN | 32 | 4 | 4161 | 170 | scramb1,Odc1,Meltrin,Su(H) | | 3.964e-02 | -3.23 | GSE14769\_UNSTIM\_VS\_120MIN\_LPS\_BMDM\_DN | MSigDB lists | GSE14769\_UNSTIM\_VS\_120MIN\_LPS\_BMDM\_DN | 32 | 4 | 4161 | 170 | dpp,CG6966,fz2,Cln3 | | 3.964e-02 | -3.23 | REACTOME\_SIGNALING\_BY\_PDGF | MSigDB lists | REACTOME\_SIGNALING\_BY\_PDGF | 32 | 4 | 4161 | 170 | Pde1c,Tor,Pkc98E,Pkc53E | | 3.964e-02 | -3.23 | GSE41978\_WT\_VS\_ID2\_KO\_KLRG1\_LOW\_EFFECTOR\_CD8\_TCELL\_UP | MSigDB lists | GSE41978\_WT\_VS\_ID2\_KO\_KLRG1\_LOW\_EFFECTOR\_CD8\_TCELL\_UP | 32 | 4 | 4161 | 170 | dsd,Khc,CG17698,AGO3 | | 3.964e-02 | -3.23 | GSE8921\_UNSTIM\_VS\_TLR1\_2\_STIM\_MONOCYTE\_6H\_DN | MSigDB lists | GSE8921\_UNSTIM\_VS\_TLR1\_2\_STIM\_MONOCYTE\_6H\_DN | 32 | 4 | 4161 | 170 | Acbp2,Ca-beta,Root,CG3961 | | 3.964e-02 | -3.23 | HUMMERICH\_SKIN\_CANCER\_PROGRESSION\_DN | MSigDB lists | HUMMERICH\_SKIN\_CANCER\_PROGRESSION\_DN | 32 | 4 | 4161 | 170 | CG3961,PCB,GlyP,flw | | 3.999e-02 | -3.22 | GO\_REGULATION\_OF\_RESPONSE\_TO\_STRESS | MSigDB lists | GO\_REGULATION\_OF\_RESPONSE\_TO\_STRESS | 407 | 24 | 4161 | 170 | Pak,Raf,nompB,dpp,EcR,Pkc98E,KaiR1D,Flo2,RhoGAP1A,wbl,Pkc53E,sxc,Sod1,Octbeta2R,dsh,Ptp61F,chb,scramb1,REG,Syn2,fz2,Tor,Cln3,csw | | 4.012e-02 | -3.22 | GO\_NEUTRAL\_LIPID\_METABOLIC\_PROCESS | MSigDB lists | GO\_NEUTRAL\_LIPID\_METABOLIC\_PROCESS | 19 | 3 | 4161 | 170 | Cat,csw,mino | | 4.012e-02 | -3.22 | MODULE\_180 | MSigDB lists | MODULE\_180 | 19 | 3 | 4161 | 170 | Rbp6,Ank,fus | | 4.012e-02 | -3.22 | GO\_CORE\_PROMOTER\_SEQUENCE\_SPECIFIC\_DNA\_BINDING | MSigDB lists | GO\_CORE\_PROMOTER\_SEQUENCE\_SPECIFIC\_DNA\_BINDING | 19 | 3 | 4161 | 170 | Tif-IA,TfAP-2,Su(H) | | 4.012e-02 | -3.22 | GO\_REGULATION\_OF\_CILIUM\_ASSEMBLY | MSigDB lists | GO\_REGULATION\_OF\_CILIUM\_ASSEMBLY | 19 | 3 | 4161 | 170 | CG17883,nompB,sky | | 4.012e-02 | -3.22 | MORF\_FRK | MSigDB lists | MORF\_FRK | 19 | 3 | 4161 | 170 | Src42A,nompB,Spf45 | | 4.012e-02 | -3.22 | GO\_MYELOID\_CELL\_HOMEOSTASIS | MSigDB lists | GO\_MYELOID\_CELL\_HOMEOSTASIS | 19 | 3 | 4161 | 170 | Ptp61F,Sod1,IP3K2 | | 4.012e-02 | -3.22 | CCATCCA\_MIR432 | MSigDB lists | CCATCCA\_MIR432 | 19 | 3 | 4161 | 170 | wbl,flw,Doa | | 4.012e-02 | -3.22 | GO\_POSITIVE\_REGULATION\_OF\_MUSCLE\_TISSUE\_DEVELOPMENT | MSigDB lists | GO\_POSITIVE\_REGULATION\_OF\_MUSCLE\_TISSUE\_DEVELOPMENT | 19 | 3 | 4161 | 170 | Tor,Su(H),Octbeta2R | | 4.012e-02 | -3.22 | GO\_GLYCEROLIPID\_METABOLIC\_PROCESS | MSigDB lists | GO\_GLYCEROLIPID\_METABOLIC\_PROCESS | 113 | 9 | 4161 | 170 | PIG-O,csw,Acbp2,CG3530,PIG-G,mino,AstA-R1,scramb1,Cat | | 4.036e-02 | -3.21 | Mst87F (Male-specific RNA 87F) | protein interactions | 41693 | 15 | 3 | 9204 | 481 | opa,Mob2,Pkc98E | | 4.036e-02 | -3.21 | B-H1 (BarH1) | protein interactions | 32724 | 15 | 3 | 9204 | 481 | Raf,B-H2,Pkc98E | | 4.036e-02 | -3.21 | slou (slouch) | protein interactions | 42547 | 15 | 3 | 9204 | 481 | Mst87F,Antp,Ubx | | 4.036e-02 | -3.21 | Met (Methoprene-tolerant) | protein interactions | 32114 | 15 | 3 | 9204 | 481 | EcR,gce,Khc | | 4.036e-02 | -3.21 | beta'COP (Coat Protein (coatomer) beta') | protein interactions | 45757 | 15 | 3 | 9204 | 481 | rictor,CG7878,foxo | | 4.036e-02 | -3.21 | NAT1 (NAT1) | protein interactions | 46020 | 15 | 3 | 9204 | 481 | foxo,Raf,rictor | | 4.056e-02 | -3.21 | PDZ domain binding | molecular function | GO:0030165 | 7 | 2 | 12176 | 580 | rst,fz2 | | 4.056e-02 | -3.21 | ARF guanyl-nucleotide exchange factor activity | molecular function | GO:0005086 | 7 | 2 | 12176 | 580 | siz,step | | 4.056e-02 | -3.21 | diacylglycerol kinase activity | molecular function | GO:0004143 | 7 | 2 | 12176 | 580 | rdgA,Sk1 | | 4.056e-02 | -3.21 | polynucleotide adenylyltransferase activity | molecular function | GO:0004652 | 7 | 2 | 12176 | 580 | Gld2,CG46385 | | 4.056e-02 | -3.21 | wide pore channel activity | molecular function | GO:0022829 | 7 | 2 | 12176 | 580 | shakB,Inx2 | | 4.057e-02 | -3.20 | multicellular organism reproduction | biological process | GO:0032504 | 1503 | 87 | 12345 | 597 | rols,csw,mid,egh,Sh,EcR,scrib,lncRNA:CR43802,Qsox2,cindr,lncRNA:CR44585,lncRNA:CR44420,CG42355,Inx2,fs(1)N,CG1428,Nlg2,bun,JhI-26,fru,Doa,Acp62F,drd,lncRNA:CR43484,flw,Fad2,retn,Ebp,CG7483,lncRNA:flam,bru1,bol,Mst87F,Octbeta2R,CG18135,DNaseII,CG10000,cic,CG16854,can,msi,Iswi,Obp56e,mamo,Eip74EF,opa,dpp,Obp56h,dany,shg,AGO2,CG34435,shd,dnc,ND-51L2,Raf,Msp300,TfAP-2,Mrtf,lncRNA:Hsromega,Gr22c,tin,aust,dsh,chb,shep,wbl,Pde1c,Dlic,Mbs,lncRNA:CR43753,Khc,Mos,bbg,msps,RhoL,polybromo,Tor,sgg,ex,Gr39a,kay,CG42823,mGluR,upd3,tld,kek1 | | 4.086e-02 | -3.20 | GO\_POSITIVE\_REGULATION\_OF\_CELL\_FATE\_COMMITMENT | MSigDB lists | GO\_POSITIVE\_REGULATION\_OF\_CELL\_FATE\_COMMITMENT | 1 | 1 | 4161 | 170 | Su(H) | | 4.086e-02 | -3.20 | GO\_NEGATIVE\_REGULATION\_OF\_ACTIN\_FILAMENT\_BUNDLE\_ASSEMBLY | MSigDB lists | GO\_NEGATIVE\_REGULATION\_OF\_ACTIN\_FILAMENT\_BUNDLE\_ASSEMBLY | 1 | 1 | 4161 | 170 | chb | | 4.086e-02 | -3.20 | KYNG\_ENVIRONMENTAL\_STRESS\_RESPONSE\_NOT\_BY\_UV\_IN\_WS | MSigDB lists | KYNG\_ENVIRONMENTAL\_STRESS\_RESPONSE\_NOT\_BY\_UV\_IN\_WS | 1 | 1 | 4161 | 170 | Sod1 | | 4.086e-02 | -3.20 | MIZUKAMI\_HYPOXIA\_DN | MSigDB lists | MIZUKAMI\_HYPOXIA\_DN | 1 | 1 | 4161 | 170 | LRP1 | | 4.086e-02 | -3.20 | chr12q11 | MSigDB lists | chr12q11 | 1 | 1 | 4161 | 170 | ABCD | | 4.086e-02 | -3.20 | GO\_TRIPARTITE\_REGIONAL\_SUBDIVISION | MSigDB lists | GO\_TRIPARTITE\_REGIONAL\_SUBDIVISION | 1 | 1 | 4161 | 170 | fz2 | | 4.086e-02 | -3.20 | FUNG\_IL2\_SIGNALING\_2 | MSigDB lists | FUNG\_IL2\_SIGNALING\_2 | 1 | 1 | 4161 | 170 | ZnT63C | | 4.086e-02 | -3.20 | MIKKELSEN\_PARTIALLY\_REPROGRAMMED\_TO\_PLURIPOTENCY | MSigDB lists | MIKKELSEN\_PARTIALLY\_REPROGRAMMED\_TO\_PLURIPOTENCY | 1 | 1 | 4161 | 170 | TfAP-2 | | 4.086e-02 | -3.20 | GO\_REGULATION\_OF\_GRANULOCYTE\_DIFFERENTIATION | MSigDB lists | GO\_REGULATION\_OF\_GRANULOCYTE\_DIFFERENTIATION | 1 | 1 | 4161 | 170 | sxc | | 4.086e-02 | -3.20 | JI\_CARCINOGENESIS\_BY\_KRAS\_AND\_STK11\_UP | MSigDB lists | JI\_CARCINOGENESIS\_BY\_KRAS\_AND\_STK11\_UP | 1 | 1 | 4161 | 170 | TfAP-2 | | 4.086e-02 | -3.20 | GO\_POSITIVE\_REGULATION\_OF\_MACROPHAGE\_DIFFERENTIATION | MSigDB lists | GO\_POSITIVE\_REGULATION\_OF\_MACROPHAGE\_DIFFERENTIATION | 1 | 1 | 4161 | 170 | Pkc53E | | 4.086e-02 | -3.20 | BENNETT\_SYSTEMIC\_LUPUS\_ERYTHEMATOSUS | MSigDB lists | BENNETT\_SYSTEMIC\_LUPUS\_ERYTHEMATOSUS | 1 | 1 | 4161 | 170 | scramb1 | | 4.086e-02 | -3.20 | GO\_MEGAKARYOCYTE\_DEVELOPMENT | MSigDB lists | GO\_MEGAKARYOCYTE\_DEVELOPMENT | 1 | 1 | 4161 | 170 | csw | | 4.086e-02 | -3.20 | GO\_NEGATIVE\_REGULATION\_OF\_STRESS\_FIBER\_ASSEMBLY | MSigDB lists | GO\_NEGATIVE\_REGULATION\_OF\_STRESS\_FIBER\_ASSEMBLY | 1 | 1 | 4161 | 170 | chb | | 4.086e-02 | -3.20 | GO\_ACROSOME\_ASSEMBLY | MSigDB lists | GO\_ACROSOME\_ASSEMBLY | 1 | 1 | 4161 | 170 | CG17883 | | 4.086e-02 | -3.20 | GO\_NEGATIVE\_REGULATION\_OF\_ENDOTHELIAL\_CELL\_APOPTOTIC\_PROCESS | MSigDB lists | GO\_NEGATIVE\_REGULATION\_OF\_ENDOTHELIAL\_CELL\_APOPTOTIC\_PROCESS | 1 | 1 | 4161 | 170 | Raf | | 4.086e-02 | -3.20 | GO\_POSITIVE\_REGULATION\_OF\_OSTEOBLAST\_PROLIFERATION | MSigDB lists | GO\_POSITIVE\_REGULATION\_OF\_OSTEOBLAST\_PROLIFERATION | 1 | 1 | 4161 | 170 | dpp | | 4.086e-02 | -3.20 | GO\_MEGAKARYOCYTE\_DIFFERENTIATION | MSigDB lists | GO\_MEGAKARYOCYTE\_DIFFERENTIATION | 1 | 1 | 4161 | 170 | csw | | 4.086e-02 | -3.20 | BEGUM\_TARGETS\_OF\_PAX3\_FOXO1\_FUSION\_AND\_PAX3 | MSigDB lists | BEGUM\_TARGETS\_OF\_PAX3\_FOXO1\_FUSION\_AND\_PAX3 | 1 | 1 | 4161 | 170 | Ser | | 4.086e-02 | -3.20 | GO\_NEGATIVE\_REGULATION\_OF\_T\_CELL\_APOPTOTIC\_PROCESS | MSigDB lists | GO\_NEGATIVE\_REGULATION\_OF\_T\_CELL\_APOPTOTIC\_PROCESS | 1 | 1 | 4161 | 170 | mino | | 4.086e-02 | -3.20 | GO\_NEGATIVE\_REGULATION\_OF\_CARDIAC\_MUSCLE\_TISSUE\_DEVELOPMENT | MSigDB lists | GO\_NEGATIVE\_REGULATION\_OF\_CARDIAC\_MUSCLE\_TISSUE\_DEVELOPMENT | 1 | 1 | 4161 | 170 | dpp | | 4.086e-02 | -3.20 | GO\_EMBRYONIC\_CAMERA\_TYPE\_EYE\_FORMATION | MSigDB lists | GO\_EMBRYONIC\_CAMERA\_TYPE\_EYE\_FORMATION | 1 | 1 | 4161 | 170 | TfAP-2 | | 4.086e-02 | -3.20 | MYLLYKANGAS\_AMPLIFICATION\_HOT\_SPOT\_30 | MSigDB lists | MYLLYKANGAS\_AMPLIFICATION\_HOT\_SPOT\_30 | 1 | 1 | 4161 | 170 | csw | | 4.086e-02 | -3.20 | GO\_RETINOID\_X\_RECEPTOR\_BINDING | MSigDB lists | GO\_RETINOID\_X\_RECEPTOR\_BINDING | 1 | 1 | 4161 | 170 | EcR | | 4.086e-02 | -3.20 | LEE\_SP4\_THYMOCYTE | MSigDB lists | LEE\_SP4\_THYMOCYTE | 1 | 1 | 4161 | 170 | dnc | | 4.086e-02 | -3.20 | GO\_REGULATION\_OF\_INTERLEUKIN\_5\_PRODUCTION | MSigDB lists | GO\_REGULATION\_OF\_INTERLEUKIN\_5\_PRODUCTION | 1 | 1 | 4161 | 170 | dnc | | 4.086e-02 | -3.20 | GO\_ESTROUS\_CYCLE | MSigDB lists | GO\_ESTROUS\_CYCLE | 1 | 1 | 4161 | 170 | Octbeta2R | | 4.086e-02 | -3.20 | GO\_REGULATION\_OF\_ODONTOGENESIS\_OF\_DENTIN\_CONTAINING\_TOOTH | MSigDB lists | GO\_REGULATION\_OF\_ODONTOGENESIS\_OF\_DENTIN\_CONTAINING\_TOOTH | 1 | 1 | 4161 | 170 | dpp | | 4.086e-02 | -3.20 | VANDESLUIS\_COMMD1\_TARGETS\_GROUP\_4\_UP | MSigDB lists | VANDESLUIS\_COMMD1\_TARGETS\_GROUP\_4\_UP | 1 | 1 | 4161 | 170 | Glut1 | | 4.086e-02 | -3.20 | ST\_PAC1\_RECEPTOR\_PATHWAY | MSigDB lists | ST\_PAC1\_RECEPTOR\_PATHWAY | 1 | 1 | 4161 | 170 | IP3K2 | | 4.086e-02 | -3.20 | MCCOLLUM\_GELDANAMYCIN\_RESISTANCE\_UP | MSigDB lists | MCCOLLUM\_GELDANAMYCIN\_RESISTANCE\_UP | 1 | 1 | 4161 | 170 | CG3376 | | 4.086e-02 | -3.20 | GO\_NEGATIVE\_REGULATION\_OF\_TUMOR\_NECROSIS\_FACTOR\_MEDIATED\_SIGNALING\_PATHWAY | MSigDB lists | GO\_NEGATIVE\_REGULATION\_OF\_TUMOR\_NECROSIS\_FACTOR\_MEDIATED\_SIGNALING\_PATHWAY | 1 | 1 | 4161 | 170 | Ptp61F | | 4.086e-02 | -3.20 | MASRI\_RESISTANCE\_TO\_TAMOXIFEN\_AND\_AROMATASE\_INHIBITORS\_UP | MSigDB lists | MASRI\_RESISTANCE\_TO\_TAMOXIFEN\_AND\_AROMATASE\_INHIBITORS\_UP | 1 | 1 | 4161 | 170 | Dys | | 4.086e-02 | -3.20 | TESAR\_JAK\_TARGETS\_MOUSE\_ES\_D3\_DN | MSigDB lists | TESAR\_JAK\_TARGETS\_MOUSE\_ES\_D3\_DN | 1 | 1 | 4161 | 170 | AGO3 | | 4.086e-02 | -3.20 | GO\_REGULATION\_OF\_LIPOPROTEIN\_LIPASE\_ACTIVITY | MSigDB lists | GO\_REGULATION\_OF\_LIPOPROTEIN\_LIPASE\_ACTIVITY | 1 | 1 | 4161 | 170 | EcR | | 4.086e-02 | -3.20 | NIKOLSKY\_BREAST\_CANCER\_20P13\_AMPLICON | MSigDB lists | NIKOLSKY\_BREAST\_CANCER\_20P13\_AMPLICON | 1 | 1 | 4161 | 170 | dsd | | 4.086e-02 | -3.20 | NIKOLSKY\_BREAST\_CANCER\_19Q13.1\_AMPLICON | MSigDB lists | NIKOLSKY\_BREAST\_CANCER\_19Q13.1\_AMPLICON | 1 | 1 | 4161 | 170 | crol | | 4.086e-02 | -3.20 | GO\_POSITIVE\_REGULATION\_OF\_ACTIVATED\_T\_CELL\_PROLIFERATION | MSigDB lists | GO\_POSITIVE\_REGULATION\_OF\_ACTIVATED\_T\_CELL\_PROLIFERATION | 1 | 1 | 4161 | 170 | mino | | 4.115e-02 | -3.19 | GO\_TRANSPORTER\_COMPLEX | MSigDB lists | GO\_TRANSPORTER\_COMPLEX | 79 | 7 | 4161 | 170 | Sh,dnc,Ca-beta,MCU,Syx1A,KaiR1D,Nmdar2 | | 4.117e-02 | -3.19 | fly chr67F1-67F1\|3-33 | chromosome location | fly chr67F1-67F1\|3-33 | 23 | 3 | 25010 | 845 | CG6418,Cpr67Fa1,lncRNA:CR45247 | | 4.117e-02 | -3.19 | fly chr67F1-67F1\|3-33 cM | chromosome location | fly chr67F1-67F1\|3-33 cM | 23 | 3 | 25010 | 845 | CG6418,Cpr67Fa1,lncRNA:CR45247 | | 4.118e-02 | -3.19 | negative regulation of developmental process | biological process | GO:0051093 | 222 | 17 | 12345 | 597 | numb,sgg,ex,CG9380,Su(H),Snoo,hdc,Mob2,bun,Src42A,foxo,dpp,PGRP-LA,Ser,esg,EcR,scrib | | 4.120e-02 | -3.19 | GSE14699\_NAIVE\_VS\_ACT\_CD8\_TCELL\_UP | MSigDB lists | GSE14699\_NAIVE\_VS\_ACT\_CD8\_TCELL\_UP | 47 | 5 | 4161 | 170 | scramb1,Glut1,Odc1,Mlp84B,mRpL24 | | 4.120e-02 | -3.19 | MORF\_MAGEA8 | MSigDB lists | MORF\_MAGEA8 | 47 | 5 | 4161 | 170 | Dys,Hr96,dnc,fz2,Pkc53E | | 4.120e-02 | -3.19 | GSE22935\_UNSTIM\_VS\_24H\_MBOVIS\_BCG\_STIM\_MACROPHAGE\_DN | MSigDB lists | GSE22935\_UNSTIM\_VS\_24H\_MBOVIS\_BCG\_STIM\_MACROPHAGE\_DN | 47 | 5 | 4161 | 170 | Cln3,Rat1,Raf,eIF4E7,TfIIFbeta | | 4.120e-02 | -3.19 | GSE26928\_CENTR\_MEMORY\_VS\_CXCR5\_POS\_CD4\_TCELL\_UP | MSigDB lists | GSE26928\_CENTR\_MEMORY\_VS\_CXCR5\_POS\_CD4\_TCELL\_UP | 47 | 5 | 4161 | 170 | mRpL24,Argl,Syn2,dnc,Cpsf160 | | 4.120e-02 | -3.19 | GSE360\_L\_MAJOR\_VS\_B\_MALAYI\_LOW\_DOSE\_MAC\_UP | MSigDB lists | GSE360\_L\_MAJOR\_VS\_B\_MALAYI\_LOW\_DOSE\_MAC\_UP | 47 | 5 | 4161 | 170 | Pfdn4,flw,EcR,Pde1c,kcc | | 4.152e-02 | -3.18 | transporter activity | molecular function | GO:0005215 | 745 | 46 | 12176 | 580 | ND-MLRQ,chk,ZnT63C,nrv2,Nmdar2,CG33310,CG13794,Ir10a,KaiR1D,Ca-beta,ine,CG14606,Sh,CG7720,CG42826,CG10960,CG34172,Nha1,Ir56a,Ir41a,CG42235,CG8468,CG45782,CG7882,Pzl,cpx,shakB,CG33934,kcc,CG3168,Glut1,Vha36-3,MCU,olf186-F,Irk3,scramb1,GluRIIC,CG3285,Ucp4A,Vha100-5,CNT1,cert,path,Tk,Inx2,ABCD | | 4.165e-02 | -3.18 | transmembrane transporter activity | molecular function | GO:0022857 | 708 | 44 | 12176 | 580 | GluRIIC,olf186-F,Irk3,MCU,Ucp4A,CG3285,CNT1,Vha100-5,path,Tk,Inx2,ABCD,CG33934,shakB,kcc,CG3168,Glut1,Vha36-3,CG10960,CG34172,Ir56a,Nha1,Ir41a,CG42235,CG8468,CG45782,CG7882,Pzl,cpx,ND-MLRQ,Nmdar2,chk,ZnT63C,nrv2,CG33310,CG13794,Ir10a,ine,Ca-beta,KaiR1D,CG14606,CG7720,Sh,CG42826 | | 4.170e-02 | -3.18 | axon extension involved in axon guidance | biological process | GO:0048846 | 7 | 2 | 12345 | 597 | CadN2,RhoGAP100F | | 4.170e-02 | -3.18 | regulation of cellular response to oxidative stress | biological process | GO:1900407 | 7 | 2 | 12345 | 597 | Fer2,Pde8 | | 4.170e-02 | -3.18 | negative regulation of transporter activity | biological process | GO:0032410 | 7 | 2 | 12345 | 597 | qvr,CG42797 | | 4.170e-02 | -3.18 | regulation of transcription by RNA polymerase III | biological process | GO:0006359 | 7 | 2 | 12345 | 597 | Maf1,Tor | | 4.170e-02 | -3.18 | gonad morphogenesis | biological process | GO:0035262 | 7 | 2 | 12345 | 597 | shg,raw | | 4.170e-02 | -3.18 | paracrine signaling | biological process | GO:0038001 | 7 | 2 | 12345 | 597 | upd3,Cat | | 4.170e-02 | -3.18 | positive regulation of muscle organ development | biological process | GO:0048636 | 7 | 2 | 12345 | 597 | dpp,Ubx | | 4.170e-02 | -3.18 | cardioblast cell fate commitment | biological process | GO:0042684 | 7 | 2 | 12345 | 597 | mid,tin | | 4.170e-02 | -3.18 | regulation of microtubule polymerization | biological process | GO:0031113 | 7 | 2 | 12345 | 597 | chb,msps | | 4.170e-02 | -3.18 | regulation of ARF protein signal transduction | biological process | GO:0032012 | 7 | 2 | 12345 | 597 | step,siz | | 4.170e-02 | -3.18 | cytoskeletal matrix organization at active zone | biological process | GO:0048789 | 7 | 2 | 12345 | 597 | RhoGAP100F,Elp3 | | 4.170e-02 | -3.18 | regulation of signaling receptor activity | biological process | GO:0010469 | 7 | 2 | 12345 | 597 | qvr,kek1 | | 4.170e-02 | -3.18 | negative regulation of ion transmembrane transport | biological process | GO:0034766 | 7 | 2 | 12345 | 597 | CG42797,qvr | | 4.170e-02 | -3.18 | specification of segmental identity, trunk | biological process | GO:0035292 | 7 | 2 | 12345 | 597 | Ubx,Antp | | 4.170e-02 | -3.18 | negative regulation of cation transmembrane transport | biological process | GO:1904063 | 7 | 2 | 12345 | 597 | CG42797,qvr | | 4.170e-02 | -3.18 | negative regulation of ion transmembrane transporter activity | biological process | GO:0032413 | 7 | 2 | 12345 | 597 | CG42797,qvr | | 4.170e-02 | -3.18 | hemocyte development | biological process | GO:0007516 | 7 | 2 | 12345 | 597 | RhoL,dpp | | 4.170e-02 | -3.18 | second mitotic wave involved in compound eye morphogenesis | biological process | GO:0016330 | 7 | 2 | 12345 | 597 | kay,Ser | | 4.170e-02 | -3.18 | receptor localization to synapse | biological process | GO:0097120 | 7 | 2 | 12345 | 597 | Khc,scrib | | 4.170e-02 | -3.18 | neuron projection extension involved in neuron projection guidance | biological process | GO:1902284 | 7 | 2 | 12345 | 597 | CadN2,RhoGAP100F | | 4.170e-02 | -3.18 | regulation of compound eye pigmentation | biological process | GO:0048076 | 7 | 2 | 12345 | 597 | dsh,esg | | 4.170e-02 | -3.18 | nuclear body organization | biological process | GO:0030575 | 7 | 2 | 12345 | 597 | Iswi,lncRNA:Hsromega | | 4.170e-02 | -3.18 | negative regulation of transmembrane transport | biological process | GO:0034763 | 7 | 2 | 12345 | 597 | qvr,CG42797 | | 4.170e-02 | -3.18 | regulation of eye pigmentation | biological process | GO:0048073 | 7 | 2 | 12345 | 597 | dsh,esg | | 4.173e-02 | -3.18 | phosphotransferase activity, alcohol group as acceptor | molecular function | GO:0016773 | 346 | 24 | 12176 | 580 | IP3K2,PhKgamma,rdgA,Src42A,Asator,CG7362,Doa,CG17698,sgg,Raf,CG4839,Sk1,Ste:CG33247,mnb,Pak,Ipk1,Tor,Strn-Mlck,Mos,Pkc53E,Nuak1,Stlk,Gprk1,Pkc98E | | 4.175e-02 | -3.18 | regulation of BMP signaling pathway | biological process | GO:0030510 | 53 | 6 | 12345 | 597 | Snoo,kek5,Lpin,GluRIIC,tld,hth | | 4.175e-02 | -3.18 | carbohydrate biosynthetic process | biological process | GO:0016051 | 53 | 6 | 12345 | 597 | CG30036,PhKgamma,CG14024,CG10000,PCB,Pgant6 | | 4.187e-02 | -3.17 | brp (bruchpilot) | protein interactions | 35977 | 37 | 5 | 9204 | 481 | unc-13,Elp3,RhoGAP100F,Mst87F,CG17777 | | 4.187e-02 | -3.17 | ana2 (anastral spindle 2) | protein interactions | 35878 | 37 | 5 | 9204 | 481 | rictor,Cep97,foxo,woc,spd-2 | | 4.187e-02 | -3.17 | CCT1 (Chaperonin containing TCP1 subunit 1) | protein interactions | 42649 | 37 | 5 | 9204 | 481 | foxo,Raf,melt,csw,rictor | | 4.187e-02 | -3.17 | skd (skuld) | protein interactions | 43906 | 37 | 5 | 9204 | 481 | Su(H),sxc,dsh,step,EcR | | 4.191e-02 | -3.17 | metal ion binding | molecular function | GO:0046872 | 1047 | 62 | 12176 | 580 | Sod1,grn,Dys,CG14362,CG40470,CG41099,Cyp28d1,PGRP-LA,CG7362,CG17698,TpnC25D,TpnC4,CG4301,Cyp6u1,loopin-1,stw,Cyp12a4,esn,Cadps,CG43689,shd,CG13898,CG10126,tld,LRP1,Pkc53E,PCB,Fad2,CG15073,Cyp304a1,CG4622,Syt12,EcR,CadN2,Cam,Mctp,Cyp6a13,Frq2,Cyp28d2,CG13526,CG12200,spok,unc-13,Ser,CG3530,CG8117,Frq1,LpR2,Hr96,CG5493,CG9981,Scp1,shg,Cyp309a1,Actn,CAH13,AOX1,woc,CG6839,TpnC41C,GATAe,Cyp4p2 | | 4.200e-02 | -3.17 | AGC-kinase\_C | interpro domains | IPR000961 | 26 | 4 | 11383 | 585 | Gprk1,Pkc98E,Pkc53E,CG4839 | | 4.200e-02 | -3.17 | Peripla\_BP\_I | interpro domains | IPR028082 | 26 | 4 | 11383 | 585 | KaiR1D,GluRIIC,mGluR,Nmdar2 | | 4.200e-02 | -3.17 | Znf\_NHR/GATA | interpro domains | IPR013088 | 26 | 4 | 11383 | 585 | grn,EcR,Hr96,GATAe | | 4.212e-02 | -3.17 | GO\_RESPONSE\_TO\_INORGANIC\_SUBSTANCE | MSigDB lists | GO\_RESPONSE\_TO\_INORGANIC\_SUBSTANCE | 132 | 10 | 4161 | 170 | mino,Raf,Cat,Sod1,TfAP-2,D2hgdh,IP3K2,CG7483,Ank,ZnT63C | | 4.218e-02 | -3.17 | RA | prosite domains | PS50200 | 13 | 3 | 6243 | 386 | wake,Myo81F,spri | | 4.225e-02 | -3.16 | BACH2\_01 | MSigDB lists | BACH2\_01 | 63 | 6 | 4161 | 170 | Frq1,Dys,eIF4E7,dpp,CG17698,Frq2 | | 4.225e-02 | -3.16 | TARTE\_PLASMA\_CELL\_VS\_PLASMABLAST\_UP | MSigDB lists | TARTE\_PLASMA\_CELL\_VS\_PLASMABLAST\_UP | 63 | 6 | 4161 | 170 | Syx1A,Dys,CG3376,IP3K2,LRP1,Act87E | | 4.235e-02 | -3.16 | monovalent inorganic cation homeostasis | biological process | GO:0055067 | 40 | 5 | 12345 | 597 | Cln3,CG33310,Vha100-5,nrv2,Raf | | 4.236e-02 | -3.16 | microtubule organizing center part | cellular component | GO:0044450 | 57 | 6 | 11940 | 538 | nompB,CG18109,Rcd6,Cam,Cep97,spd-2 | | 4.238e-02 | -3.16 | positive regulation of protein kinase activity | biological process | GO:0045860 | 67 | 7 | 12345 | 597 | Raf,Stlk,CycT,Pak,CG41520,step,rictor | | 4.239e-02 | -3.16 | CG17666 (uncharacterized protein) | protein interactions | 39452 | 103 | 10 | 9204 | 481 | CG5011,opa,PIG-G,esg,gce,rols,fru,CG3078,Mbs,CG15564 | | 4.240e-02 | -3.16 | PDZ | smart domains | SM00228 | 64 | 8 | 5086 | 315 | dsh,bbg,RhoGAP100F,CG5921,Fife,Syn2,scrib,sdt | | 4.286e-02 | -3.15 | fly chr15A3-15A3\|1-55 | chromosome location | fly chr15A3-15A3\|1-55 | 10 | 2 | 25010 | 845 | CG4653,TwdlY | | 4.286e-02 | -3.15 | fly chr95E1-95E1\|3-83 | chromosome location | fly chr95E1-95E1\|3-83 | 10 | 2 | 25010 | 845 | Syx1A,Root | | 4.286e-02 | -3.15 | fly chr15A3-15A3\|1-55 cM | chromosome location | fly chr15A3-15A3\|1-55 cM | 10 | 2 | 25010 | 845 | CG4653,TwdlY | | 4.286e-02 | -3.15 | fly chr95E1-95E1\|3-83 cM | chromosome location | fly chr95E1-95E1\|3-83 cM | 10 | 2 | 25010 | 845 | Syx1A,Root | | 4.287e-02 | -3.15 | postsynapse | cellular component | GO:0098794 | 72 | 7 | 11940 | 538 | Nlg2,Pak,unc-13,scrib,KaiR1D,GluRIIC,sky | | 4.305e-02 | -3.15 | - | gene3d domains | 4.10.400.10 | 32 | 5 | 4929 | 302 | CG32432,trol,CG31221,LRP1,LpR2 | | 4.306e-02 | -3.15 | Activation of Ca-permeable Kainate Receptor | REACTOME pathways | R-DME-451308 | 16 | 3 | 3923 | 197 | Cam,GluRIIC,KaiR1D | | 4.306e-02 | -3.15 | Activation of Kainate Receptors upon glutamate binding | REACTOME pathways | R-DME-451326 | 16 | 3 | 3923 | 197 | KaiR1D,GluRIIC,Cam | | 4.306e-02 | -3.15 | Ionotropic activity of Kainate Receptors | REACTOME pathways | R-DME-451306 | 16 | 3 | 3923 | 197 | GluRIIC,Cam,KaiR1D | | 4.355e-02 | -3.13 | HMG | smart domains | SM00398 | 22 | 4 | 5086 | 315 | polybromo,cic,Sox102F,bbx | | 4.359e-02 | -3.13 | Prot-tyrosine\_phosphatase-like | interpro domains | IPR029021 | 38 | 5 | 11383 | 585 | Ptp99A,Ptp61F,csw,Ptp36E,CG3530 | | 4.371e-02 | -3.13 | GO\_NEGATIVE\_REGULATION\_OF\_MULTICELLULAR\_ORGANISMAL\_PROCESS | MSigDB lists | GO\_NEGATIVE\_REGULATION\_OF\_MULTICELLULAR\_ORGANISMAL\_PROCESS | 189 | 13 | 4161 | 170 | Tor,IP3K2,Su(H),RhoGAP1A,Sod1,csw,eIF4E7,dnc,LRP1,dpp,Ser,Octbeta2R,Ptp61F | | 4.372e-02 | -3.13 | GSE21774\_CD62L\_POS\_CD56\_DIM\_VS\_CD62L\_NEG\_CD56\_DIM\_NK\_CELL\_UP | MSigDB lists | GSE21774\_CD62L\_POS\_CD56\_DIM\_VS\_CD62L\_NEG\_CD56\_DIM\_NK\_CELL\_UP | 33 | 4 | 4161 | 170 | Pak,Mlp84B,CG17698,nompB | | 4.372e-02 | -3.13 | GSE30971\_CTRL\_VS\_LPS\_STIM\_MACROPHAGE\_WBP7\_KO\_2H\_UP | MSigDB lists | GSE30971\_CTRL\_VS\_LPS\_STIM\_MACROPHAGE\_WBP7\_KO\_2H\_UP | 33 | 4 | 4161 | 170 | Meltrin,Glut1,CG3961,Spf45 | | 4.372e-02 | -3.13 | GSE13946\_CTRL\_VS\_DSS\_COLITIS\_GD\_TCELL\_FROM\_COLON\_DN | MSigDB lists | GSE13946\_CTRL\_VS\_DSS\_COLITIS\_GD\_TCELL\_FROM\_COLON\_DN | 33 | 4 | 4161 | 170 | Elp3,Mlp84B,CG5339,CNT1 | | 4.372e-02 | -3.13 | GSE36476\_CTRL\_VS\_TSST\_ACT\_16H\_MEMORY\_CD4\_TCELL\_YOUNG\_UP | MSigDB lists | GSE36476\_CTRL\_VS\_TSST\_ACT\_16H\_MEMORY\_CD4\_TCELL\_YOUNG\_UP | 33 | 4 | 4161 | 170 | flw,dnc,MESK2,Glut1 | | 4.372e-02 | -3.13 | GSE3982\_MAC\_VS\_NEUTROPHIL\_DN | MSigDB lists | GSE3982\_MAC\_VS\_NEUTROPHIL\_DN | 33 | 4 | 4161 | 170 | Treh,Pak,Act87E,Pgant6 | | 4.372e-02 | -3.13 | GO\_COLUMNAR\_CUBOIDAL\_EPITHELIAL\_CELL\_DIFFERENTIATION | MSigDB lists | GO\_COLUMNAR\_CUBOIDAL\_EPITHELIAL\_CELL\_DIFFERENTIATION | 33 | 4 | 4161 | 170 | dpp,Ser,Sod1,Su(H) | | 4.372e-02 | -3.13 | DACOSTA\_UV\_RESPONSE\_VIA\_ERCC3\_XPCS\_DN | MSigDB lists | DACOSTA\_UV\_RESPONSE\_VIA\_ERCC3\_XPCS\_DN | 33 | 4 | 4161 | 170 | LpR2,CG34408,dsd,IntS3 | | 4.372e-02 | -3.13 | KEGG\_TIGHT\_JUNCTION | MSigDB lists | KEGG\_TIGHT\_JUNCTION | 33 | 4 | 4161 | 170 | Pkc53E,Pkc98E,Act87E,sdt | | 4.372e-02 | -3.13 | GSE29618\_BCELL\_VS\_MDC\_DAY7\_FLU\_VACCINE\_UP | MSigDB lists | GSE29618\_BCELL\_VS\_MDC\_DAY7\_FLU\_VACCINE\_UP | 33 | 4 | 4161 | 170 | Sh,Meltrin,Odc1,Tsp96F | | 4.372e-02 | -3.13 | GSE20715\_0H\_VS\_48H\_OZONE\_TLR4\_KO\_LUNG\_UP | MSigDB lists | GSE20715\_0H\_VS\_48H\_OZONE\_TLR4\_KO\_LUNG\_UP | 33 | 4 | 4161 | 170 | Glut1,CG3842,CG6418,CG17698 | | 4.372e-02 | -3.13 | MODULE\_97 | MSigDB lists | MODULE\_97 | 33 | 4 | 4161 | 170 | csw,Raf,Dys,sno | | 4.372e-02 | -3.13 | GO\_NEGATIVE\_REGULATION\_OF\_DEFENSE\_RESPONSE | MSigDB lists | GO\_NEGATIVE\_REGULATION\_OF\_DEFENSE\_RESPONSE | 33 | 4 | 4161 | 170 | Ptp61F,Octbeta2R,EcR,RhoGAP1A | | 4.372e-02 | -3.13 | GO\_STEM\_CELL\_DIFFERENTIATION | MSigDB lists | GO\_STEM\_CELL\_DIFFERENTIATION | 33 | 4 | 4161 | 170 | Rbp6,dpp,Ser,Su(H) | | 4.372e-02 | -3.13 | GO\_REGULATION\_OF\_MUSCLE\_CELL\_DIFFERENTIATION | MSigDB lists | GO\_REGULATION\_OF\_MUSCLE\_CELL\_DIFFERENTIATION | 33 | 4 | 4161 | 170 | Flo2,dpp,Tor,Su(H) | | 4.399e-02 | -3.12 | GO\_CELLULAR\_CHEMICAL\_HOMEOSTASIS | MSigDB lists | GO\_CELLULAR\_CHEMICAL\_HOMEOSTASIS | 133 | 10 | 4161 | 170 | sxc,Sod1,Cln3,Ca-beta,KaiR1D,ZnT63C,Pkc98E,AstA-R1,MCU,GlyP | | 4.408e-02 | -3.12 | drk (downstream of receptor kinase) | protein interactions | 36497 | 76 | 8 | 9204 | 481 | RhoGAP18B,ex,Src42A,cindr,Raf,CG5004,csw,CG18659 | | 4.412e-02 | -3.12 | terminal branching, open tracheal system | biological process | GO:0007430 | 28 | 4 | 12345 | 597 | hdc,Raf,bnl,Tor | | 4.412e-02 | -3.12 | regulation of generation of precursor metabolites and energy | biological process | GO:0043467 | 28 | 4 | 12345 | 597 | Hr96,CG34172,EcR,GlyP | | 4.412e-02 | -3.12 | endocytic recycling | biological process | GO:0032456 | 28 | 4 | 12345 | 597 | CG18659,ex,nuf,Tor | | 4.412e-02 | -3.12 | cellular response to alcohol | biological process | GO:0097306 | 28 | 4 | 12345 | 597 | Set2,Iswi,Cam,EcR | | 4.412e-02 | -3.12 | regulation of G protein-coupled receptor signaling pathway | biological process | GO:0008277 | 28 | 4 | 12345 | 597 | Gprk1,Cam,rdgA,foxo | | 4.412e-02 | -3.12 | hormone biosynthetic process | biological process | GO:0042446 | 28 | 4 | 12345 | 597 | woc,spok,shd,foxo | | 4.418e-02 | -3.12 | Signaling by FGFR2 | REACTOME pathways | R-DME-5654738 | 52 | 6 | 3923 | 197 | bnl,csw,TfIIFbeta,Raf,trv,fus | | 4.419e-02 | -3.12 | CG31343 (uncharacterized protein) | protein interactions | 326133 | 26 | 4 | 9204 | 481 | Cat,Scp1,GlyP,Tsf1 | | 4.419e-02 | -3.12 | Ppn (Papilin) | protein interactions | 43872 | 26 | 4 | 9204 | 481 | GlyP,CG18135,bru1,rols | | 4.419e-02 | -3.12 | RhoGEF2 (Rho guanine nucleotide exchange factor 2) | protein interactions | 36915 | 26 | 4 | 9204 | 481 | dsh,flw,Raf,foxo | | 4.419e-02 | -3.12 | Vang (Van Gogh) | protein interactions | 35922 | 26 | 4 | 9204 | 481 | dsh,scrib,shg,fw | | 4.419e-02 | -3.12 | pyd (polychaetoid) | protein interactions | 41062 | 26 | 4 | 9204 | 481 | rst,rols,ex,shg | | 4.419e-02 | -3.12 | Cpn (Calphotin) | protein interactions | 41474 | 26 | 4 | 9204 | 481 | pod1,CG34056,Act87E,TwdlY | | 4.422e-02 | -3.12 | compound eye photoreceptor development | biological process | GO:0042051 | 82 | 8 | 12345 | 597 | RhoGAP100F,Syn2,Cam,Mob2,CadN2,chp,Raf,Doa | | 4.453e-02 | -3.11 | GO\_LIPOPROTEIN\_METABOLIC\_PROCESS | MSigDB lists | GO\_LIPOPROTEIN\_METABOLIC\_PROCESS | 48 | 5 | 4161 | 170 | app,PIG-G,PIG-O,Acbp2,LRP1 | | 4.453e-02 | -3.11 | TITF1\_Q3 | MSigDB lists | TITF1\_Q3 | 48 | 5 | 4161 | 170 | Dys,CG10126,Odc1,app,IP3K2 | | 4.453e-02 | -3.11 | GSE32164\_RESTING\_DIFFERENTIATED\_VS\_CMYC\_INHIBITED\_MACROPHAGE\_UP | MSigDB lists | GSE32164\_RESTING\_DIFFERENTIATED\_VS\_CMYC\_INHIBITED\_MACROPHAGE\_UP | 48 | 5 | 4161 | 170 | Tsp96F,Nmdar2,app,Octbeta2R,TfAP-2 | | 4.453e-02 | -3.11 | GSE21033\_CTRL\_VS\_POLYIC\_STIM\_DC\_6H\_UP | MSigDB lists | GSE21033\_CTRL\_VS\_POLYIC\_STIM\_DC\_6H\_UP | 48 | 5 | 4161 | 170 | Frq2,Der-1,CG3961,CG12121,Frq1 | | 4.453e-02 | -3.11 | GSE21927\_SPLEEN\_C57BL6\_VS\_EL4\_TUMOR\_BALBC\_MONOCYTES\_UP | MSigDB lists | GSE21927\_SPLEEN\_C57BL6\_VS\_EL4\_TUMOR\_BALBC\_MONOCYTES\_UP | 48 | 5 | 4161 | 170 | sxc,CG12547,RhoGAP1A,chb,Octbeta2R | | 4.453e-02 | -3.11 | GSE19923\_E2A\_KO\_VS\_HEB\_AND\_E2A\_KO\_DP\_THYMOCYTE\_UP | MSigDB lists | GSE19923\_E2A\_KO\_VS\_HEB\_AND\_E2A\_KO\_DP\_THYMOCYTE\_UP | 48 | 5 | 4161 | 170 | Glut1,kcc,Sh,Pkc98E,Stt3B | | 4.453e-02 | -3.11 | HNF6\_Q6 | MSigDB lists | HNF6\_Q6 | 48 | 5 | 4161 | 170 | flw,TfIIFbeta,LpR2,KaiR1D,mGluR | | 4.453e-02 | -3.11 | GSE21360\_NAIVE\_VS\_TERTIARY\_MEMORY\_CD8\_TCELL\_UP | MSigDB lists | GSE21360\_NAIVE\_VS\_TERTIARY\_MEMORY\_CD8\_TCELL\_UP | 48 | 5 | 4161 | 170 | MESK2,CG3961,Nmdar2,PCB,Pkc53E | | 4.453e-02 | -3.11 | GO\_CATION\_CHANNEL\_ACTIVITY | MSigDB lists | GO\_CATION\_CHANNEL\_ACTIVITY | 48 | 5 | 4161 | 170 | MCU,Frq1,Frq2,Ca-beta,Sh | | 4.453e-02 | -3.11 | GSE36009\_WT\_VS\_NLRP10\_KO\_DC\_DN | MSigDB lists | GSE36009\_WT\_VS\_NLRP10\_KO\_DC\_DN | 48 | 5 | 4161 | 170 | sxc,Fhos,CG3961,scramb1,CG9132 | | 4.453e-02 | -3.11 | GO\_REGULATION\_OF\_HEART\_CONTRACTION | MSigDB lists | GO\_REGULATION\_OF\_HEART\_CONTRACTION | 48 | 5 | 4161 | 170 | dnc,Ca-beta,Octbeta2R,Dys,bru1 | | 4.453e-02 | -3.11 | PEA3\_Q6 | MSigDB lists | PEA3\_Q6 | 48 | 5 | 4161 | 170 | CG3961,Ank,Glut1,rdgA,IP3K2 | | 4.453e-02 | -3.11 | REACTOME\_GPCR\_LIGAND\_BINDING | MSigDB lists | REACTOME\_GPCR\_LIGAND\_BINDING | 48 | 5 | 4161 | 170 | fz2,CCAP-R,AstA-R1,mGluR,Octbeta2R | | 4.453e-02 | -3.11 | GO\_RESPONSE\_TO\_NUTRIENT | MSigDB lists | GO\_RESPONSE\_TO\_NUTRIENT | 48 | 5 | 4161 | 170 | sxc,Tor,Ugt301D1,LpR2,Cat | | 4.453e-02 | -3.11 | GSE36891\_UNSTIM\_VS\_POLYIC\_TLR3\_STIM\_PERITONEAL\_MACROPHAGE\_DN | MSigDB lists | GSE36891\_UNSTIM\_VS\_POLYIC\_TLR3\_STIM\_PERITONEAL\_MACROPHAGE\_DN | 48 | 5 | 4161 | 170 | ZnT63C,Rat1,CNT1,CG9132,step | | 4.459e-02 | -3.11 | CREB phosphorylation through the activation of Ras | REACTOME pathways | R-DME-442742 | 7 | 2 | 3923 | 197 | Raf,CrebB | | 4.459e-02 | -3.11 | CD28 dependent PI3K/Akt signaling | REACTOME pathways | R-DME-389357 | 7 | 2 | 3923 | 197 | Tor,rictor | | 4.459e-02 | -3.11 | Unblocking of NMDA receptor, glutamate binding and activation | REACTOME pathways | R-DME-438066 | 7 | 2 | 3923 | 197 | Nmdar2,Actn | | 4.459e-02 | -3.11 | larval development | biological process | GO:0002164 | 159 | 13 | 12345 | 597 | sgg,dpp,Raf,Poxm,Msp300,Tor,ths,fz2,Elp3,Set2,Ser,EcR,scrib | | 4.461e-02 | -3.11 | CH | prosite domains | PS50021 | 32 | 5 | 6243 | 386 | Actn,sick,CG34417,Msp300,Dys | | 4.479e-02 | -3.11 | CkIIalpha (casein kinase IIalpha) | protein interactions | 48448 | 50 | 6 | 9204 | 481 | Raf,dsh,Pdfr,Rbp6,Pkc98E,Ank2 | | 4.506e-02 | -3.10 | proteasome activator complex | cellular component | GO:0008537 | 1 | 1 | 11940 | 538 | REG | | 4.506e-02 | -3.10 | microtubule organizing center attachment site | cellular component | GO:0034992 | 1 | 1 | 11940 | 538 | Msp300 | | 4.506e-02 | -3.10 | nuclear membrane protein complex | cellular component | GO:0106083 | 1 | 1 | 11940 | 538 | Msp300 | | 4.506e-02 | -3.10 | dense core granule lumen | cellular component | GO:0098898 | 1 | 1 | 11940 | 538 | dpp | | 4.506e-02 | -3.10 | axolemma | cellular component | GO:0030673 | 1 | 1 | 11940 | 538 | Ser | | 4.506e-02 | -3.10 | G protein-coupled receptor homodimeric complex | cellular component | GO:0038038 | 1 | 1 | 11940 | 538 | mGluR | | 4.506e-02 | -3.10 | vesicle lumen | cellular component | GO:0031983 | 1 | 1 | 11940 | 538 | dpp | | 4.506e-02 | -3.10 | cell leading edge cell cortex | cellular component | GO:1904269 | 1 | 1 | 11940 | 538 | step | | 4.506e-02 | -3.10 | presynaptic endosome | cellular component | GO:0098830 | 1 | 1 | 11940 | 538 | g | | 4.506e-02 | -3.10 | cytoplasmic vesicle lumen | cellular component | GO:0060205 | 1 | 1 | 11940 | 538 | dpp | | 4.506e-02 | -3.10 | secretory granule lumen | cellular component | GO:0034774 | 1 | 1 | 11940 | 538 | dpp | | 4.506e-02 | -3.10 | nuclear membrane microtubule tethering complex | cellular component | GO:0106094 | 1 | 1 | 11940 | 538 | Msp300 | | 4.506e-02 | -3.10 | meiotic nuclear membrane microtubule tethering complex | cellular component | GO:0034993 | 1 | 1 | 11940 | 538 | Msp300 | | 4.506e-02 | -3.10 | U11/U12 snRNP | cellular component | GO:0034693 | 1 | 1 | 11940 | 538 | CG44249 | | 4.510e-02 | -3.10 | GSE27786\_NKCELL\_VS\_ERYTHROBLAST\_UP | MSigDB lists | GSE27786\_NKCELL\_VS\_ERYTHROBLAST\_UP | 64 | 6 | 4161 | 170 | sno,crol,scramb1,Spf45,Flo2,Argl | | 4.510e-02 | -3.10 | HATADA\_METHYLATED\_IN\_LUNG\_CANCER\_UP | MSigDB lists | HATADA\_METHYLATED\_IN\_LUNG\_CANCER\_UP | 64 | 6 | 4161 | 170 | Cadps,IP3K2,TfAP-2,Act87E,Pde8,Tsp96F | | 4.515e-02 | -3.10 | GO\_GOLGI\_APPARATUS | MSigDB lists | GO\_GOLGI\_APPARATUS | 433 | 25 | 4161 | 170 | Frq1,Qsox2,CG17883,Pgant6,Acbp2,Meltrin,Pak,Gas8,Pkc98E,ZnT63C,pod1,Ank,TfAP-2,Tor,step,Frq2,Cln3,chb,Cat,CG4301,scramb1,CG12121,CG30036,CG33090,fz2 | | 4.519e-02 | -3.10 | epithelial cell development | biological process | GO:0002064 | 361 | 25 | 12345 | 597 | ex,sgg,cic,Raf,cindr,CG41099,bbg,polybromo,RhoL,dsh,kay,fs(1)N,Mrtf,bun,egh,dpp,mid,rols,csw,shg,Khc,EcR,shd,scrib,lncRNA:flam | | 4.536e-02 | -3.09 | glycerolipid biosynthetic process | biological process | GO:0045017 | 68 | 7 | 12345 | 597 | rdgA,CG6790,mino,PIG-O,Lpin,Pld,PIG-G | | 4.540e-02 | -3.09 | RODRIGUES\_THYROID\_CARCINOMA\_POORLY\_DIFFERENTIATED\_DN | MSigDB lists | RODRIGUES\_THYROID\_CARCINOMA\_POORLY\_DIFFERENTIATED\_DN | 171 | 12 | 4161 | 170 | sky,step,Stt3B,Pgant6,eRF3,Pak,Dys,CG5339,Cat,flw,fz2,Rbp6 | | 4.556e-02 | -3.09 | GO\_VACUOLE | MSigDB lists | GO\_VACUOLE | 350 | 21 | 4161 | 170 | CG4301,CG3376,Octbeta2R,Cat,sws,CG3860,CG12121,fz2,Ank,CG41099,Tor,CG42672,Der-1,Cln3,Sk1,LRP1,CG3842,kcc,Flo2,DNaseII,Sod1 | | 4.582e-02 | -3.08 | GO\_NEGATIVE\_REGULATION\_OF\_INFLAMMATORY\_RESPONSE | MSigDB lists | GO\_NEGATIVE\_REGULATION\_OF\_INFLAMMATORY\_RESPONSE | 20 | 3 | 4161 | 170 | Octbeta2R,Ptp61F,RhoGAP1A | | 4.582e-02 | -3.08 | GO\_ADENYLATE\_CYCLASE\_ACTIVATING\_G\_PROTEIN\_COUPLED\_RECEPTOR\_SIGNALING\_PATHWAY | MSigDB lists | GO\_ADENYLATE\_CYCLASE\_ACTIVATING\_G\_PROTEIN\_COUPLED\_RECEPTOR\_SIGNALING\_PATHWAY | 20 | 3 | 4161 | 170 | dnc,AstA-R1,Octbeta2R | | 4.582e-02 | -3.08 | GO\_REGULATION\_OF\_ORGAN\_GROWTH | MSigDB lists | GO\_REGULATION\_OF\_ORGAN\_GROWTH | 20 | 3 | 4161 | 170 | Tor,Sod1,Su(H) | | 4.582e-02 | -3.08 | GO\_CARDIAC\_CHAMBER\_MORPHOGENESIS | MSigDB lists | GO\_CARDIAC\_CHAMBER\_MORPHOGENESIS | 20 | 3 | 4161 | 170 | Ser,Su(H),dsh | | 4.582e-02 | -3.08 | GO\_REGULATION\_OF\_CALCIUM\_ION\_TRANSMEMBRANE\_TRANSPORTER\_ACTIVITY | MSigDB lists | GO\_REGULATION\_OF\_CALCIUM\_ION\_TRANSMEMBRANE\_TRANSPORTER\_ACTIVITY | 20 | 3 | 4161 | 170 | Ca-beta,dnc,Dys | | 4.582e-02 | -3.08 | KEGG\_RENAL\_CELL\_CARCINOMA | MSigDB lists | KEGG\_RENAL\_CELL\_CARCINOMA | 20 | 3 | 4161 | 170 | csw,Pak,Raf | | 4.582e-02 | -3.08 | GERHOLD\_ADIPOGENESIS\_UP | MSigDB lists | GERHOLD\_ADIPOGENESIS\_UP | 20 | 3 | 4161 | 170 | mino,PCB,LpR2 | | 4.582e-02 | -3.08 | GO\_ENDOTHELIUM\_DEVELOPMENT | MSigDB lists | GO\_ENDOTHELIUM\_DEVELOPMENT | 20 | 3 | 4161 | 170 | Su(H),Ser,dnc | | 4.582e-02 | -3.08 | GSE18791\_CTRL\_VS\_NEWCASTLE\_VIRUS\_DC\_8H\_DN | MSigDB lists | GSE18791\_CTRL\_VS\_NEWCASTLE\_VIRUS\_DC\_8H\_DN | 20 | 3 | 4161 | 170 | CG41099,scramb1,unc-13 | | 4.582e-02 | -3.08 | GO\_SYNAPTIC\_VESICLE\_CYCLE | MSigDB lists | GO\_SYNAPTIC\_VESICLE\_CYCLE | 20 | 3 | 4161 | 170 | unc-13,Syx1A,Cadps | | 4.582e-02 | -3.08 | GO\_NEUROMUSCULAR\_PROCESS | MSigDB lists | GO\_NEUROMUSCULAR\_PROCESS | 20 | 3 | 4161 | 170 | Nmdar2,RhoGAP1A,Cln3 | | 4.582e-02 | -3.08 | GO\_ATPASE\_BINDING | MSigDB lists | GO\_ATPASE\_BINDING | 20 | 3 | 4161 | 170 | EcR,dnc,Der-1 | | 4.582e-02 | -3.08 | GO\_POSITIVE\_REGULATION\_OF\_EPITHELIAL\_CELL\_PROLIFERATION | MSigDB lists | GO\_POSITIVE\_REGULATION\_OF\_EPITHELIAL\_CELL\_PROLIFERATION | 20 | 3 | 4161 | 170 | dpp,Tor,Pkc53E | | 4.582e-02 | -3.08 | GSE42021\_TREG\_PLN\_VS\_TREG\_PRECURSORS\_THYMUS\_DN | MSigDB lists | GSE42021\_TREG\_PLN\_VS\_TREG\_PRECURSORS\_THYMUS\_DN | 20 | 3 | 4161 | 170 | scramb1,CG41099,step | | 4.582e-02 | -3.08 | GO\_NEGATIVE\_REGULATION\_OF\_LIPID\_METABOLIC\_PROCESS | MSigDB lists | GO\_NEGATIVE\_REGULATION\_OF\_LIPID\_METABOLIC\_PROCESS | 20 | 3 | 4161 | 170 | Sod1,dpp,Ugt301D1 | | 4.582e-02 | -3.08 | HEDENFALK\_BREAST\_CANCER\_HEREDITARY\_VS\_SPORADIC | MSigDB lists | HEDENFALK\_BREAST\_CANCER\_HEREDITARY\_VS\_SPORADIC | 20 | 3 | 4161 | 170 | LRP1,flw,LpR2 | | 4.587e-02 | -3.08 | Ribosomal\_S5\_D2-typ\_fold\_subgr | interpro domains | IPR014721 | 16 | 3 | 11383 | 585 | CG4849,CG33158,mRpS5 | | 4.597e-02 | -3.08 | TGACCTY\_ERR1\_Q2 | MSigDB lists | TGACCTY\_ERR1\_Q2 | 269 | 17 | 4161 | 170 | CG3376,Dys,Ucp4A,Pde1c,Rbp6,Syn2,pod1,Ank,Hr96,GlyP,Khc,Raf,CG15093,flw,eIF4E7,Iswi,RhoGAP1A | | 4.614e-02 | -3.08 | regulation of actomyosin structure organization | biological process | GO:0110020 | 17 | 3 | 12345 | 597 | flw,Fhos,step | | 4.614e-02 | -3.08 | negative regulation of secretion by cell | biological process | GO:1903531 | 17 | 3 | 12345 | 597 | cpx,sky,CG9650 | | 4.614e-02 | -3.08 | cGMP metabolic process | biological process | GO:0046068 | 17 | 3 | 12345 | 597 | Pde1c,Pde6,Pde9 | | 4.614e-02 | -3.08 | apical protein localization | biological process | GO:0045176 | 17 | 3 | 12345 | 597 | shg,Tor,sdt | | 4.614e-02 | -3.08 | deactivation of rhodopsin mediated signaling | biological process | GO:0016059 | 17 | 3 | 12345 | 597 | Gprk1,Cam,rdgA | | 4.614e-02 | -3.08 | intercellular bridge organization | biological process | GO:0043063 | 17 | 3 | 12345 | 597 | cindr,flw,Mbs | | 4.614e-02 | -3.08 | positive regulation of Rho protein signal transduction | biological process | GO:0035025 | 17 | 3 | 12345 | 597 | Exn,CG10188,RhoGAP1A | | 4.616e-02 | -3.08 | CHARAFE\_BREAST\_CANCER\_LUMINAL\_VS\_MESENCHYMAL\_UP | MSigDB lists | CHARAFE\_BREAST\_CANCER\_LUMINAL\_VS\_MESENCHYMAL\_UP | 81 | 7 | 4161 | 170 | Ank,fus,CG32196,eRF3,CG5522,wbl,Cln3 | | 4.623e-02 | -3.07 | exd (extradenticle) | protein interactions | 32567 | 38 | 5 | 9204 | 481 | tamo,Antp,Ubx,hth,CG7443 | | 4.627e-02 | -3.07 | positive regulation of protein modification process | biological process | GO:0031401 | 176 | 14 | 12345 | 597 | lncRNA:Hsromega,Pak,rictor,step,Tor,bnl,Raf,dpp,Pde8,CG41520,Src42A,Stlk,CycT,Mos | | 4.634e-02 | -3.07 | - | gene3d domains | 1.10.10.10 | 102 | 11 | 4929 | 302 | FoxL1,CG32006,Eip74EF,toe,dany,CG1894,foxo,Poxm,Achl,dsh,TfIIFbeta | | 4.639e-02 | -3.07 | establishment or maintenance of bipolar cell polarity | biological process | GO:0061245 | 41 | 5 | 12345 | 597 | rib,scrib,trol,dsh,sdt | | 4.639e-02 | -3.07 | establishment or maintenance of apical/basal cell polarity | biological process | GO:0035088 | 41 | 5 | 12345 | 597 | sdt,trol,dsh,rib,scrib | | 4.645e-02 | -3.07 | arm (armadillo) | protein interactions | 31151 | 119 | 11 | 9204 | 481 | Iswi,crol,Su(H),sgg,foxo,shg,Src42A,dsh,dpp,sdt,Ubx | | 4.649e-02 | -3.07 | chemical homeostasis | biological process | GO:0048878 | 259 | 19 | 12345 | 597 | EcR,Acbp2,ine,KaiR1D,MCU,nrv2,Cln3,shakB,Hr96,CG33310,mag,Pdfr,Lpin,Glut1,Vha100-5,foxo,Raf,ZnT63C,Mondo | | 4.662e-02 | -3.07 | FAD-bd\_PCMH | interpro domains | IPR016166 | 7 | 2 | 11383 | 585 | AOX1,D2hgdh | | 4.662e-02 | -3.07 | Ser\_inhib-like\_sf | interpro domains | IPR036084 | 7 | 2 | 11383 | 585 | CG45546,Acp62F | | 4.662e-02 | -3.07 | HEAT\_type\_2 | interpro domains | IPR021133 | 7 | 2 | 11383 | 585 | chb,msps | | 4.662e-02 | -3.07 | FAD-bd\_PCMH-like\_sf | interpro domains | IPR036318 | 7 | 2 | 11383 | 585 | AOX1,D2hgdh | | 4.662e-02 | -3.07 | PAZ\_dom | interpro domains | IPR003100 | 7 | 2 | 11383 | 585 | AGO3,AGO2 | | 4.662e-02 | -3.07 | Actin\_CS | interpro domains | IPR004001 | 7 | 2 | 11383 | 585 | Act88F,Act87E | | 4.666e-02 | -3.06 | cell division site part | cellular component | GO:0032155 | 44 | 5 | 11940 | 538 | CG18109,step,nuf,cindr,Hil | | 4.677e-02 | -3.06 | regulation of neuromuscular junction development | biological process | GO:1904396 | 113 | 10 | 12345 | 597 | Octbeta2R,Src42A,Syn2,Ank2,rictor,Pak,mnb,Mob2,sgg,Tor | | 4.694e-02 | -3.06 | Immune System | REACTOME pathways | R-DME-168256 | 744 | 47 | 3923 | 197 | fw,kay,CG41520,Tor,Pkc53E,Vha100-5,ninaD,CG9449,foxo,CG4301,olf186-F,ACXD,LysP,Gprk1,Meltrin,bnl,rictor,Cam,GlyP,Syx1A,Vha36-3,Pde1c,Ptp61F,Tsf1,Prosbeta5R1,CG8492,CG9981,CG8184,Cat,Dlic,CG7362,Qsox2,Pld,Aldh-III,CG9451,Glut1,csw,CG7882,PGRP-LA,Pkc98E,REG,Src42A,CG40470,CrebB,Pak,Raf,RhoGAP18B | | 4.699e-02 | -3.06 | neuron projection membrane | cellular component | GO:0032589 | 89 | 8 | 11940 | 538 | dpr18,Or65c,dpr12,dpr5,dpr15,Or85b,Ser,Or42b | | 4.707e-02 | -3.06 | voltage-gated cation channel activity | molecular function | GO:0022843 | 29 | 4 | 12176 | 580 | Nmdar2,Irk3,Ca-beta,Sh | | 4.728e-02 | -3.05 | negative regulation of protein modification process | biological process | GO:0031400 | 98 | 9 | 12345 | 597 | mnb,sws,sgg,raw,et,kek1,Ptp61F,flw,chinmo | | 4.733e-02 | -3.05 | sty (sprouty) | protein interactions | 38424 | 105 | 10 | 9204 | 481 | LRP1,Raf,csw,CG13492,Mob2,rst,Msp300,kay,trol,Ser | | 4.737e-02 | -3.05 | striated muscle thin filament | cellular component | GO:0005865 | 8 | 2 | 11940 | 538 | TpnC25D,Act88F | | 4.737e-02 | -3.05 | gap junction | cellular component | GO:0005921 | 8 | 2 | 11940 | 538 | shakB,Inx2 | | 4.750e-02 | -3.05 | NMDARECEPTOR | prints domains | PR00177 | 13 | 3 | 2185 | 142 | GluRIIC,KaiR1D,Nmdar2 | | 4.757e-02 | -3.05 | Glycerophospholipid metabolism | KEGG pathways | dme00564 | 63 | 7 | 3198 | 170 | CG30503,Gpo3,mino,Lpin,sws,rdgA,Pld | | 4.757e-02 | -3.05 | Glycerophospholipid metabolism | KEGG pathways | ko00564 | 63 | 7 | 3198 | 170 | Gpo3,mino,CG30503,Lpin,sws,Pld,rdgA | | 4.762e-02 | -3.04 | - | gene3d domains | 2.60.40.1660 | 6 | 2 | 4929 | 302 | CG33310,nrv2 | | 4.762e-02 | -3.04 | - | gene3d domains | 1.10.150.60 | 6 | 2 | 4929 | 302 | htk,retn | | 4.763e-02 | -3.04 | juvenile hormone binding | molecular function | GO:0005500 | 1 | 1 | 12176 | 580 | gce | | 4.763e-02 | -3.04 | RNA binding involved in posttranscriptional gene silencing | molecular function | GO:0150100 | 1 | 1 | 12176 | 580 | Marf1 | | 4.763e-02 | -3.04 | beta-1,4-mannosyltransferase activity | molecular function | GO:0019187 | 1 | 1 | 12176 | 580 | egh | | 4.763e-02 | -3.04 | vasopressin receptor activity | molecular function | GO:0005000 | 1 | 1 | 12176 | 580 | CCAP-R | | 4.763e-02 | -3.04 | phospholipase D activity | molecular function | GO:0004630 | 1 | 1 | 12176 | 580 | Pld | | 4.763e-02 | -3.04 | sterol esterase activity | molecular function | GO:0004771 | 1 | 1 | 12176 | 580 | mag | | 4.763e-02 | -3.04 | D-lactate dehydrogenase (cytochrome) activity | molecular function | GO:0004458 | 1 | 1 | 12176 | 580 | D2hgdh | | 4.763e-02 | -3.04 | RNA polymerase I general transcription initiation factor activity | molecular function | GO:0001181 | 1 | 1 | 12176 | 580 | Tif-IA | | 4.763e-02 | -3.04 | biotin binding | molecular function | GO:0009374 | 1 | 1 | 12176 | 580 | PCB | | 4.763e-02 | -3.04 | glycogen phosphorylase activity | molecular function | GO:0008184 | 1 | 1 | 12176 | 580 | GlyP | | 4.763e-02 | -3.04 | RNA polymerase II transcription corepressor binding | molecular function | GO:0001226 | 1 | 1 | 12176 | 580 | mid | | 4.763e-02 | -3.04 | protein-lipid complex binding | molecular function | GO:0071814 | 1 | 1 | 12176 | 580 | LpR2 | | 4.763e-02 | -3.04 | U2 snRNP binding | molecular function | GO:1990447 | 1 | 1 | 12176 | 580 | barc | | 4.763e-02 | -3.04 | glycerol-3-phosphate O-acyltransferase activity | molecular function | GO:0004366 | 1 | 1 | 12176 | 580 | mino | | 4.763e-02 | -3.04 | nitric-oxide synthase binding | molecular function | GO:0050998 | 1 | 1 | 12176 | 580 | CG42673 | | 4.763e-02 | -3.04 | kynurenine aminotransferase activity | molecular function | GO:0036137 | 1 | 1 | 12176 | 580 | Kyat | | 4.763e-02 | -3.04 | arylamine N-acetyltransferase activity | molecular function | GO:0004060 | 1 | 1 | 12176 | 580 | AANAT1 | | 4.763e-02 | -3.04 | osmosensor activity | molecular function | GO:0005034 | 1 | 1 | 12176 | 580 | ine | | 4.763e-02 | -3.04 | kynurenine-oxoglutarate transaminase activity | molecular function | GO:0016212 | 1 | 1 | 12176 | 580 | Kyat | | 4.763e-02 | -3.04 | diacylglycerol binding | molecular function | GO:0019992 | 1 | 1 | 12176 | 580 | unc-13 | | 4.763e-02 | -3.04 | G protein-coupled glutamate receptor activity | molecular function | GO:0098988 | 1 | 1 | 12176 | 580 | mGluR | | 4.763e-02 | -3.04 | ecdysone 20-monooxygenase activity | molecular function | GO:0004501 | 1 | 1 | 12176 | 580 | shd | | 4.763e-02 | -3.04 | calcium sensitive guanylate cyclase activator activity | molecular function | GO:0008048 | 1 | 1 | 12176 | 580 | Frq1 | | 4.763e-02 | -3.04 | phosphatidylinositol-3,5-bisphosphate 3-phosphatase activity | molecular function | GO:0052629 | 1 | 1 | 12176 | 580 | CG3530 | | 4.763e-02 | -3.04 | cysteine dioxygenase activity | molecular function | GO:0017172 | 1 | 1 | 12176 | 580 | CG5493 | | 4.763e-02 | -3.04 | CP2 mannose-ethanolamine phosphotransferase activity | molecular function | GO:0051267 | 1 | 1 | 12176 | 580 | PIG-G | | 4.763e-02 | -3.04 | RNA polymerase I core binding | molecular function | GO:0001042 | 1 | 1 | 12176 | 580 | Tif-IA | | 4.763e-02 | -3.04 | myosin V binding | molecular function | GO:0031489 | 1 | 1 | 12176 | 580 | Cam | | 4.763e-02 | -3.04 | cholesterol binding | molecular function | GO:0015485 | 1 | 1 | 12176 | 580 | mGluR | | 4.763e-02 | -3.04 | syntaxin-1 binding | molecular function | GO:0017075 | 1 | 1 | 12176 | 580 | unc-13 | | 4.763e-02 | -3.04 | protein kinase A catalytic subunit binding | molecular function | GO:0034236 | 1 | 1 | 12176 | 580 | sws | | 4.763e-02 | -3.04 | neuropeptide binding | molecular function | GO:0042923 | 1 | 1 | 12176 | 580 | AstA-R1 | | 4.763e-02 | -3.04 | pyridoxal oxidase activity | molecular function | GO:0004732 | 1 | 1 | 12176 | 580 | AOX1 | | 4.763e-02 | -3.04 | 5'-3' exoribonuclease activity | molecular function | GO:0004534 | 1 | 1 | 12176 | 580 | Rat1 | | 4.763e-02 | -3.04 | calcium-dependent protein binding | molecular function | GO:0048306 | 1 | 1 | 12176 | 580 | IP3K2 | | 4.763e-02 | -3.04 | inositol pentakisphosphate 2-kinase activity | molecular function | GO:0035299 | 1 | 1 | 12176 | 580 | Ipk1 | | 4.763e-02 | -3.04 | 3-hydroxyisobutyrate dehydrogenase activity | molecular function | GO:0008442 | 1 | 1 | 12176 | 580 | CG15093 | | 4.763e-02 | -3.04 | cysteine-S-conjugate beta-lyase activity | molecular function | GO:0047804 | 1 | 1 | 12176 | 580 | Kyat | | 4.763e-02 | -3.04 | oxidoreductase activity, acting on the CH-OH group of donors, cytochrome as acceptor | molecular function | GO:0016898 | 1 | 1 | 12176 | 580 | D2hgdh | | 4.763e-02 | -3.04 | acyl-CoA N-acyltransferase activity | molecular function | GO:0019186 | 1 | 1 | 12176 | 580 | AANATL2 | | 4.763e-02 | -3.04 | argininosuccinate lyase activity | molecular function | GO:0004056 | 1 | 1 | 12176 | 580 | Argl | | 4.763e-02 | -3.04 | RNA polymerase II transcription coactivator binding | molecular function | GO:0001225 | 1 | 1 | 12176 | 580 | EcR | | 4.763e-02 | -3.04 | (R)-2-hydroxyglutarate dehydrogenase activity | molecular function | GO:0051990 | 1 | 1 | 12176 | 580 | D2hgdh | | 4.763e-02 | -3.04 | CCR4-NOT complex binding | molecular function | GO:1905762 | 1 | 1 | 12176 | 580 | Marf1 | | 4.763e-02 | -3.04 | adenylate cyclase inhibiting G protein-coupled glutamate receptor activity | molecular function | GO:0001640 | 1 | 1 | 12176 | 580 | mGluR | | 4.763e-02 | -3.04 | RNA polymerase III core binding | molecular function | GO:0000994 | 1 | 1 | 12176 | 580 | Maf1 | | 4.763e-02 | -3.04 | mRNA binding involved in posttranscriptional gene silencing | molecular function | GO:1903231 | 1 | 1 | 12176 | 580 | Marf1 | | 4.763e-02 | -3.04 | pyruvate carboxylase activity | molecular function | GO:0004736 | 1 | 1 | 12176 | 580 | PCB | | 4.763e-02 | -3.04 | lipoprotein particle binding | molecular function | GO:0071813 | 1 | 1 | 12176 | 580 | LpR2 | | 4.763e-02 | -3.04 | cobalamin binding | molecular function | GO:0031419 | 1 | 1 | 12176 | 580 | CG3556 | | 4.763e-02 | -3.04 | lysophospholipase activity | molecular function | GO:0004622 | 1 | 1 | 12176 | 580 | sws | | 4.763e-02 | -3.04 | group II metabotropic glutamate receptor activity | molecular function | GO:0001641 | 1 | 1 | 12176 | 580 | mGluR | | 4.763e-02 | -3.04 | uniporter activity | molecular function | GO:0015292 | 1 | 1 | 12176 | 580 | MCU | | 4.763e-02 | -3.04 | calcium- and calmodulin-regulated 3',5'-cyclic-GMP phosphodiesterase activity | molecular function | GO:0048101 | 1 | 1 | 12176 | 580 | Pde1c | | 4.763e-02 | -3.04 | rhodopsin kinase activity | molecular function | GO:0050254 | 1 | 1 | 12176 | 580 | Gprk1 | | 4.763e-02 | -3.04 | phosphatidylinositol-3,5-bisphosphate phosphatase activity | molecular function | GO:0106018 | 1 | 1 | 12176 | 580 | CG3530 | | 4.763e-02 | -3.04 | L-threonine 3-dehydrogenase activity | molecular function | GO:0008743 | 1 | 1 | 12176 | 580 | CG5955 | | 4.763e-02 | -3.04 | tachykinin receptor binding | molecular function | GO:0071861 | 1 | 1 | 12176 | 580 | Tk | | 4.763e-02 | -3.04 | deoxyribonuclease II activity | molecular function | GO:0004531 | 1 | 1 | 12176 | 580 | DNaseII | | 4.780e-02 | -3.04 | CG2691 (uncharacterized protein) | protein interactions | 32320 | 16 | 3 | 9204 | 481 | nuf,rictor,foxo | | 4.780e-02 | -3.04 | TrpA1 (Transient receptor potential cation channel A1) | protein interactions | 39015 | 16 | 3 | 9204 | 481 | Tk,Cam,shakB | | 4.780e-02 | -3.04 | CG9248 (uncharacterized protein) | protein interactions | 35384 | 16 | 3 | 9204 | 481 | EcR,Mst87F,Rbp6 | | 4.780e-02 | -3.04 | CG10581 (uncharacterized protein) | protein interactions | 40326 | 16 | 3 | 9204 | 481 | Prm,TpnC41C,Scp1 | | 4.780e-02 | -3.04 | cac (cacophony) | protein interactions | 32158 | 16 | 3 | 9204 | 481 | GluRIIC,Frq2,jus | | 4.780e-02 | -3.04 | CG8929 (uncharacterized protein) | protein interactions | 37306 | 16 | 3 | 9204 | 481 | Doa,E5,IP3K2 | | 4.780e-02 | -3.04 | Su(var)3-3 (Suppressor of variegation 3-3) | protein interactions | 40217 | 16 | 3 | 9204 | 481 | Su(H),polybromo,dpp | | 4.780e-02 | -3.04 | TfIIB (Transcription factor IIB) | protein interactions | 34430 | 16 | 3 | 9204 | 481 | Ubx,Mlp84B,Antp | | 4.780e-02 | -3.04 | hb (hunchback) | protein interactions | 41032 | 16 | 3 | 9204 | 481 | bnl,fru,cindr | | 4.780e-02 | -3.04 | Ect4 (Ectoderm-expressed 4) | protein interactions | 38895 | 16 | 3 | 9204 | 481 | foxo,mRpL24,rictor | | 4.780e-02 | -3.04 | Mpi (Mannose phosphate isomerase) | protein interactions | 41192 | 16 | 3 | 9204 | 481 | egh,Ptp61F,rols | | 4.780e-02 | -3.04 | nkd (naked cuticle) | protein interactions | 44843 | 16 | 3 | 9204 | 481 | dsh,sgg,shg | | 4.783e-02 | -3.04 | Rho GTPase cycle | REACTOME pathways | R-DME-194840 | 53 | 6 | 3923 | 197 | RhoGAP100F,RhoGAP1A,CG6424,CG10188,RhoGAP18B,Exn | | 4.802e-02 | -3.04 | GSE41867\_NAIVE\_VS\_DAY8\_LCMV\_EFFECTOR\_CD8\_TCELL\_DN | MSigDB lists | GSE41867\_NAIVE\_VS\_DAY8\_LCMV\_EFFECTOR\_CD8\_TCELL\_DN | 49 | 5 | 4161 | 170 | Der-1,Meltrin,step,pod1,Octbeta2R | | 4.802e-02 | -3.04 | MORF\_PTPRB | MSigDB lists | MORF\_PTPRB | 49 | 5 | 4161 | 170 | dnc,fz2,Pkc53E,Dys,Hr96 | | 4.802e-02 | -3.04 | GSE27859\_CD11C\_INT\_F480\_HI\_MACROPHAGE\_VS\_CD11C\_ING\_F480\_INT\_DC\_DN | MSigDB lists | GSE27859\_CD11C\_INT\_F480\_HI\_MACROPHAGE\_VS\_CD11C\_ING\_F480\_INT\_DC\_DN | 49 | 5 | 4161 | 170 | Pgant6,LRP1,AGO3,Glut1,Hr96 | | 4.802e-02 | -3.04 | GSE360\_T\_GONDII\_VS\_B\_MALAYI\_HIGH\_DOSE\_MAC\_DN | MSigDB lists | GSE360\_T\_GONDII\_VS\_B\_MALAYI\_HIGH\_DOSE\_MAC\_DN | 49 | 5 | 4161 | 170 | GlyP,Aldh-III,Flo2,Sk1,PIG-O | | 4.802e-02 | -3.04 | ZHONG\_RESPONSE\_TO\_AZACITIDINE\_AND\_TSA\_UP | MSigDB lists | ZHONG\_RESPONSE\_TO\_AZACITIDINE\_AND\_TSA\_UP | 49 | 5 | 4161 | 170 | Mlp84B,CG5493,rdgA,sno,Odc1 | | 4.802e-02 | -3.04 | MODULE\_195 | MSigDB lists | MODULE\_195 | 49 | 5 | 4161 | 170 | Pfdn4,nompB,Ser,CG4849,dpp | | 4.802e-02 | -3.04 | GSE360\_L\_DONOVANI\_VS\_B\_MALAYI\_HIGH\_DOSE\_MAC\_UP | MSigDB lists | GSE360\_L\_DONOVANI\_VS\_B\_MALAYI\_HIGH\_DOSE\_MAC\_UP | 49 | 5 | 4161 | 170 | ZnT63C,dsh,Glut1,kcc,Su(H) | | 4.802e-02 | -3.04 | GSE13493\_DP\_VS\_CD8POS\_THYMOCYTE\_UP | MSigDB lists | GSE13493\_DP\_VS\_CD8POS\_THYMOCYTE\_UP | 49 | 5 | 4161 | 170 | Tsp96F,KaiR1D,kcc,CG42672,Su(H) | | 4.802e-02 | -3.04 | MORF\_MDM2 | MSigDB lists | MORF\_MDM2 | 49 | 5 | 4161 | 170 | fz2,Pkc53E,CG40178,Dys,Hr96 | | 4.802e-02 | -3.04 | HALLMARK\_PEROXISOME | MSigDB lists | HALLMARK\_PEROXISOME | 49 | 5 | 4161 | 170 | ABCD,Sod1,Cat,Hr96,CG3961 | | 4.802e-02 | -3.04 | GSE23114\_WT\_VS\_SLE2C1\_MOUSE\_PERITONEAL\_CAVITY\_B1A\_BCELL\_UP | MSigDB lists | GSE23114\_WT\_VS\_SLE2C1\_MOUSE\_PERITONEAL\_CAVITY\_B1A\_BCELL\_UP | 49 | 5 | 4161 | 170 | scramb1,DNaseII,CG10126,CG12547,Tor | | 4.802e-02 | -3.04 | GSE37416\_12H\_VS\_48H\_F\_TULARENSIS\_LVS\_NEUTROPHIL\_DN | MSigDB lists | GSE37416\_12H\_VS\_48H\_F\_TULARENSIS\_LVS\_NEUTROPHIL\_DN | 49 | 5 | 4161 | 170 | sno,Sh,Acbp2,TfAP-2,RpL38 | | 4.802e-02 | -3.04 | GSE22196\_HEALTHY\_VS\_OBESE\_MOUSE\_SKIN\_GAMMADELTA\_TCELL\_UP | MSigDB lists | GSE22196\_HEALTHY\_VS\_OBESE\_MOUSE\_SKIN\_GAMMADELTA\_TCELL\_UP | 49 | 5 | 4161 | 170 | sxc,PhKgamma,CG3376,Ptp61F,CG32196 | | 4.802e-02 | -3.04 | GSE360\_L\_MAJOR\_VS\_B\_MALAYI\_LOW\_DOSE\_DC\_UP | MSigDB lists | GSE360\_L\_MAJOR\_VS\_B\_MALAYI\_LOW\_DOSE\_DC\_UP | 49 | 5 | 4161 | 170 | step,Syx1A,Pde1c,Glut1,bru1 | | 4.802e-02 | -3.04 | GSE45365\_WT\_VS\_IFNAR\_KO\_CD8A\_DC\_MCMV\_INFECTION\_UP | MSigDB lists | GSE45365\_WT\_VS\_IFNAR\_KO\_CD8A\_DC\_MCMV\_INFECTION\_UP | 49 | 5 | 4161 | 170 | chb,Doa,dsh,REG,Fhos | | 4.802e-02 | -3.04 | GSE24142\_EARLY\_THYMIC\_PROGENITOR\_VS\_DN2\_THYMOCYTE\_FETAL\_UP | MSigDB lists | GSE24142\_EARLY\_THYMIC\_PROGENITOR\_VS\_DN2\_THYMOCYTE\_FETAL\_UP | 34 | 4 | 4161 | 170 | GlyP,Pak,Tsp96F,CG3860 | | 4.802e-02 | -3.04 | GSE26495\_NAIVE\_VS\_PD1HIGH\_CD8\_TCELL\_DN | MSigDB lists | GSE26495\_NAIVE\_VS\_PD1HIGH\_CD8\_TCELL\_DN | 34 | 4 | 4161 | 170 | Der-1,Elp3,Octbeta2R,app | | 4.802e-02 | -3.04 | GSE46242\_TH1\_VS\_ANERGIC\_TH1\_CD4\_TCELL\_WITH\_EGR2\_DELETED\_DN | MSigDB lists | GSE46242\_TH1\_VS\_ANERGIC\_TH1\_CD4\_TCELL\_WITH\_EGR2\_DELETED\_DN | 34 | 4 | 4161 | 170 | CG4301,CG5339,Octbeta2R,rdgA | | 4.802e-02 | -3.04 | GSE46242\_CTRL\_VS\_EGR2\_DELETED\_TH1\_CD4\_TCELL\_DN | MSigDB lists | GSE46242\_CTRL\_VS\_EGR2\_DELETED\_TH1\_CD4\_TCELL\_DN | 34 | 4 | 4161 | 170 | LpR2,CG12121,Dys,Acbp2 | | 4.802e-02 | -3.04 | GO\_REGULATION\_OF\_MUSCLE\_CONTRACTION | MSigDB lists | GO\_REGULATION\_OF\_MUSCLE\_CONTRACTION | 34 | 4 | 4161 | 170 | Dys,Octbeta2R,dnc,Sod1 | | 4.802e-02 | -3.04 | GSE19923\_WT\_VS\_HEB\_KO\_DP\_THYMOCYTE\_UP | MSigDB lists | GSE19923\_WT\_VS\_HEB\_KO\_DP\_THYMOCYTE\_UP | 34 | 4 | 4161 | 170 | Gas8,Tsp96F,Meltrin,Mlp84B | | 4.802e-02 | -3.04 | GSE28726\_NAIVE\_CD4\_TCELL\_VS\_NAIVE\_VA24NEG\_NKTCELL\_DN | MSigDB lists | GSE28726\_NAIVE\_CD4\_TCELL\_VS\_NAIVE\_VA24NEG\_NKTCELL\_DN | 34 | 4 | 4161 | 170 | Flo2,Glut1,GlyP,RhoGAP1A | | 4.805e-02 | -3.04 | CG14480 (uncharacterized protein) | protein interactions | 36988 | 7 | 2 | 9204 | 481 | Pomp,REG | | 4.805e-02 | -3.04 | Lsm12 (LSM12 homolog) | protein interactions | 32167 | 7 | 2 | 9204 | 481 | tyf,CG11883 | | 4.805e-02 | -3.04 | CG14829 (uncharacterized protein) | protein interactions | 38792 | 7 | 2 | 9204 | 481 | Mondo,Khc | | 4.805e-02 | -3.04 | Synj (Synaptojanin) | protein interactions | 37517 | 7 | 2 | 9204 | 481 | flw,mnb | | 4.805e-02 | -3.04 | CG9775 (uncharacterized protein) | protein interactions | 40576 | 7 | 2 | 9204 | 481 | eIF4E7,Raf | | 4.805e-02 | -3.04 | tap (target of Poxn) | protein interactions | 39935 | 7 | 2 | 9204 | 481 | dsh,Act87E | | 4.805e-02 | -3.04 | LpR2 (Lipophorin receptor 2) | protein interactions | 43105 | 7 | 2 | 9204 | 481 | Sh,Hil | | 4.805e-02 | -3.04 | Tps1 (Trehalose-6-phosphate synthase 1) | protein interactions | 33642 | 7 | 2 | 9204 | 481 | Stacl,zormin | | 4.805e-02 | -3.04 | miple1 (midkine and pleiotrophin 1) | protein interactions | 38047 | 7 | 2 | 9204 | 481 | Ptp99A,shg | | 4.805e-02 | -3.04 | TfIIFalpha (Transcription factor IIFalpha) | protein interactions | 40790 | 7 | 2 | 9204 | 481 | CG8117,TfIIFbeta | | 4.805e-02 | -3.04 | ry (rosy) | protein interactions | 41605 | 7 | 2 | 9204 | 481 | flw,ZnT63C | | 4.805e-02 | -3.04 | Rdl (Resistant to dieldrin) | protein interactions | 39054 | 7 | 2 | 9204 | 481 | kcc,wake | | 4.807e-02 | -3.04 | GO\_STEROID\_METABOLIC\_PROCESS | MSigDB lists | GO\_STEROID\_METABOLIC\_PROCESS | 65 | 6 | 4161 | 170 | CG33090,Ugt301D1,CG3860,Hr96,LpR2,Cat | | 4.807e-02 | -3.04 | MODULE\_2 | MSigDB lists | MODULE\_2 | 65 | 6 | 4161 | 170 | Glut1,Dys,Ank,CG6418,Cadps,LRP1 | | 4.830e-02 | -3.03 | - | gene3d domains | 1.10.418.10 | 33 | 5 | 4929 | 302 | CG34417,Actn,sick,Msp300,Dys | | 4.833e-02 | -3.03 | Laminin\_G\_2 | pfam domains | PF02210 | 16 | 3 | 10480 | 550 | shg,trol,CadN2 | | 4.836e-02 | -3.03 | ER to Golgi ceramide transport | biological process | GO:0035621 | 1 | 1 | 12345 | 597 | cert | | 4.836e-02 | -3.03 | neurotransmitter receptor transport, endosome to plasma membrane | biological process | GO:0099639 | 1 | 1 | 12345 | 597 | scrib | | 4.836e-02 | -3.03 | regulation of octopamine signaling pathway | biological process | GO:2000128 | 1 | 1 | 12345 | 597 | foxo | | 4.836e-02 | -3.03 | regulation of octopamine or tyramine signaling pathway | biological process | GO:2000125 | 1 | 1 | 12345 | 597 | foxo | | 4.836e-02 | -3.03 | hypotonic salinity response | biological process | GO:0042539 | 1 | 1 | 12345 | 597 | kcc | | 4.836e-02 | -3.03 | regulation of cell cycle arrest | biological process | GO:0071156 | 1 | 1 | 12345 | 597 | foxo | | 4.836e-02 | -3.03 | negative regulation of response to reactive oxygen species | biological process | GO:1901032 | 1 | 1 | 12345 | 597 | Pde8 | | 4.836e-02 | -3.03 | presynaptic dense core vesicle exocytosis | biological process | GO:0099525 | 1 | 1 | 12345 | 597 | unc-13 | | 4.836e-02 | -3.03 | positive regulation of semaphorin-plexin signaling pathway | biological process | GO:2001262 | 1 | 1 | 12345 | 597 | trol | | 4.836e-02 | -3.03 | synaptic vesicle tethering involved in synaptic vesicle exocytosis | biological process | GO:0099069 | 1 | 1 | 12345 | 597 | unc-13 | | 4.836e-02 | -3.03 | regulation of histone H3-K9 acetylation | biological process | GO:2000615 | 1 | 1 | 12345 | 597 | Yeti | | 4.836e-02 | -3.03 | oligosaccharide catabolic process | biological process | GO:0009313 | 1 | 1 | 12345 | 597 | Treh | | 4.836e-02 | -3.03 | regulation of Ral protein signal transduction | biological process | GO:0032485 | 1 | 1 | 12345 | 597 | CG5522 | | 4.836e-02 | -3.03 | positive regulation of digestive system process | biological process | GO:0060456 | 1 | 1 | 12345 | 597 | Tk | | 4.836e-02 | -3.03 | L-cysteine metabolic process | biological process | GO:0046439 | 1 | 1 | 12345 | 597 | CG5493 | | 4.836e-02 | -3.03 | establishment of protein localization to postsynaptic membrane | biological process | GO:1903540 | 1 | 1 | 12345 | 597 | scrib | | 4.836e-02 | -3.03 | brush border assembly | biological process | GO:1904970 | 1 | 1 | 12345 | 597 | CG5921 | | 4.836e-02 | -3.03 | regulation of hindgut contraction | biological process | GO:0043134 | 1 | 1 | 12345 | 597 | Tk | | 4.836e-02 | -3.03 | regulation of ubiquitin-dependent endocytosis | biological process | GO:2000395 | 1 | 1 | 12345 | 597 | sdt | | 4.836e-02 | -3.03 | hormone transport | biological process | GO:0009914 | 1 | 1 | 12345 | 597 | Obp56h | | 4.836e-02 | -3.03 | negative regulation of synaptic vesicle recycling | biological process | GO:1903422 | 1 | 1 | 12345 | 597 | sky | | 4.836e-02 | -3.03 | cellular hypotonic salinity response | biological process | GO:0071477 | 1 | 1 | 12345 | 597 | kcc | | 4.836e-02 | -3.03 | trehalose catabolic process | biological process | GO:0005993 | 1 | 1 | 12345 | 597 | Treh | | 4.836e-02 | -3.03 | tRNA wobble base 5-methoxycarbonylmethyl-2-thiouridinylation | biological process | GO:0002926 | 1 | 1 | 12345 | 597 | Elp3 | | 4.836e-02 | -3.03 | primary amino compound catabolic process | biological process | GO:1901161 | 1 | 1 | 12345 | 597 | AANAT1 | | 4.836e-02 | -3.03 | presynaptic active zone assembly | biological process | GO:1904071 | 1 | 1 | 12345 | 597 | Fife | | 4.836e-02 | -3.03 | protein localization to early endosome | biological process | GO:1902946 | 1 | 1 | 12345 | 597 | scrib | | 4.836e-02 | -3.03 | positive regulation of calcineurin-mediated signaling | biological process | GO:0106058 | 1 | 1 | 12345 | 597 | olf186-F | | 4.836e-02 | -3.03 | negative regulation of nucleobase-containing compound transport | biological process | GO:0032240 | 1 | 1 | 12345 | 597 | Pde6 | | 4.836e-02 | -3.03 | mating pheromone secretion | biological process | GO:0071834 | 1 | 1 | 12345 | 597 | Obp56h | | 4.836e-02 | -3.03 | negative regulation of male germ cell proliferation | biological process | GO:2000255 | 1 | 1 | 12345 | 597 | Doa | | 4.836e-02 | -3.03 | negative regulation of oxidative stress-induced neuron death | biological process | GO:1903204 | 1 | 1 | 12345 | 597 | Fer2 | | 4.836e-02 | -3.03 | positive regulation of calcineurin-NFAT signaling cascade | biological process | GO:0070886 | 1 | 1 | 12345 | 597 | olf186-F | | 4.836e-02 | -3.03 | anterograde axonal transport of mitochondrion | biological process | GO:0098957 | 1 | 1 | 12345 | 597 | Khc | | 4.836e-02 | -3.03 | cellular anion homeostasis | biological process | GO:0030002 | 1 | 1 | 12345 | 597 | Acbp2 | | 4.836e-02 | -3.03 | hypotonic response | biological process | GO:0006971 | 1 | 1 | 12345 | 597 | kcc | | 4.836e-02 | -3.03 | bile acid metabolic process | biological process | GO:0008206 | 1 | 1 | 12345 | 597 | CG33090 | | 4.836e-02 | -3.03 | stress granule disassembly | biological process | GO:0035617 | 1 | 1 | 12345 | 597 | Khc | | 4.836e-02 | -3.03 | positive regulation of octopamine signaling pathway | biological process | GO:2000130 | 1 | 1 | 12345 | 597 | foxo | | 4.836e-02 | -3.03 | regulation of cellular response to drug | biological process | GO:2001038 | 1 | 1 | 12345 | 597 | Pde8 | | 4.836e-02 | -3.03 | cell communication by electrical coupling | biological process | GO:0010644 | 1 | 1 | 12345 | 597 | shakB | | 4.836e-02 | -3.03 | ectodermal cell fate specification | biological process | GO:0001715 | 1 | 1 | 12345 | 597 | dpp | | 4.836e-02 | -3.03 | positive regulation of hindgut contraction | biological process | GO:0060450 | 1 | 1 | 12345 | 597 | Tk | | 4.836e-02 | -3.03 | dorsal vessel aortic cell fate commitment | biological process | GO:0035052 | 1 | 1 | 12345 | 597 | Ubx | | 4.836e-02 | -3.03 | negative regulation of germ cell proliferation | biological process | GO:1905937 | 1 | 1 | 12345 | 597 | Doa | | 4.836e-02 | -3.03 | exon-exon junction complex assembly | biological process | GO:1903040 | 1 | 1 | 12345 | 597 | CG7483 | | 4.836e-02 | -3.03 | retinal cone cell development | biological process | GO:0046549 | 1 | 1 | 12345 | 597 | CG5921 | | 4.836e-02 | -3.03 | positive regulation of synaptic vesicle clustering | biological process | GO:2000809 | 1 | 1 | 12345 | 597 | Fife | | 4.836e-02 | -3.03 | neurotransmitter receptor transport postsynaptic membrane to endosome | biological process | GO:0098968 | 1 | 1 | 12345 | 597 | scrib | | 4.836e-02 | -3.03 | hormone secretion | biological process | GO:0046879 | 1 | 1 | 12345 | 597 | Obp56h | | 4.836e-02 | -3.03 | inositol phosphorylation | biological process | GO:0052746 | 1 | 1 | 12345 | 597 | Ipk1 | | 4.836e-02 | -3.03 | protein localization to endosome | biological process | GO:0036010 | 1 | 1 | 12345 | 597 | scrib | | 4.836e-02 | -3.03 | peptide hormone secretion | biological process | GO:0030072 | 1 | 1 | 12345 | 597 | Obp56h | | 4.836e-02 | -3.03 | negative regulation of excitatory postsynaptic potential | biological process | GO:0090394 | 1 | 1 | 12345 | 597 | qvr | | 4.836e-02 | -3.03 | serotonin catabolic process | biological process | GO:0042429 | 1 | 1 | 12345 | 597 | AANAT1 | | 4.836e-02 | -3.03 | cuticle chitin catabolic process | biological process | GO:0006036 | 1 | 1 | 12345 | 597 | Idgf6 | | 4.836e-02 | -3.03 | cellular hypotonic response | biological process | GO:0071476 | 1 | 1 | 12345 | 597 | kcc | | 4.836e-02 | -3.03 | cysteine catabolic process | biological process | GO:0009093 | 1 | 1 | 12345 | 597 | CG5493 | | 4.836e-02 | -3.03 | epithelial cilium movement involved in determination of left/right asymmetry | biological process | GO:0060287 | 1 | 1 | 12345 | 597 | CG17387 | | 4.836e-02 | -3.03 | cellular acyl-CoA homeostasis | biological process | GO:0042049 | 1 | 1 | 12345 | 597 | Acbp2 | | 4.836e-02 | -3.03 | regulation of light-activated channel activity | biological process | GO:0016061 | 1 | 1 | 12345 | 597 | Cam | | 4.836e-02 | -3.03 | regulation of response to osmotic stress | biological process | GO:0047484 | 1 | 1 | 12345 | 597 | ine | | 4.836e-02 | -3.03 | dense core granule priming | biological process | GO:0061789 | 1 | 1 | 12345 | 597 | unc-13 | | 4.836e-02 | -3.03 | negative regulation of hydrogen peroxide-induced cell death | biological process | GO:1903206 | 1 | 1 | 12345 | 597 | Pde8 | | 4.836e-02 | -3.03 | negative regulation of fat cell proliferation | biological process | GO:0070345 | 1 | 1 | 12345 | 597 | foxo | | 4.836e-02 | -3.03 | neurotransmitter receptor transport to plasma membrane | biological process | GO:0098877 | 1 | 1 | 12345 | 597 | scrib | | 4.836e-02 | -3.03 | lactate oxidation | biological process | GO:0019516 | 1 | 1 | 12345 | 597 | D2hgdh | | 4.836e-02 | -3.03 | maintenance of imaginal histoblast diploidy | biological process | GO:0007489 | 1 | 1 | 12345 | 597 | esg | | 4.836e-02 | -3.03 | neurotransmitter receptor transport to postsynaptic membrane | biological process | GO:0098969 | 1 | 1 | 12345 | 597 | scrib | | 4.836e-02 | -3.03 | positive regulation of microtubule nucleation | biological process | GO:0090063 | 1 | 1 | 12345 | 597 | msps | | 4.836e-02 | -3.03 | disaccharide catabolic process | biological process | GO:0046352 | 1 | 1 | 12345 | 597 | Treh | | 4.836e-02 | -3.03 | regulation of synaptic vesicle clustering | biological process | GO:2000807 | 1 | 1 | 12345 | 597 | Fife | | 4.836e-02 | -3.03 | neuronal dense core vesicle exocytosis | biological process | GO:0099011 | 1 | 1 | 12345 | 597 | unc-13 | | 4.836e-02 | -3.03 | gap junction assembly | biological process | GO:0016264 | 1 | 1 | 12345 | 597 | shakB | | 4.836e-02 | -3.03 | positive regulation of octopamine or tyramine signaling pathway | biological process | GO:2000127 | 1 | 1 | 12345 | 597 | foxo | | 4.836e-02 | -3.03 | positive regulation of cAMP-dependent protein kinase activity | biological process | GO:2000481 | 1 | 1 | 12345 | 597 | step | | 4.836e-02 | -3.03 | regulation of cyclin-dependent protein serine/threonine kinase activity involved in G2/M transition of mitotic cell cycle | biological process | GO:0031660 | 1 | 1 | 12345 | 597 | kay | | 4.836e-02 | -3.03 | regulation of hydrogen peroxide-induced cell death | biological process | GO:1903205 | 1 | 1 | 12345 | 597 | Pde8 | | 4.836e-02 | -3.03 | synaptic vesicle budding from endosome | biological process | GO:0016182 | 1 | 1 | 12345 | 597 | g | | 4.836e-02 | -3.03 | miRNA loading onto RISC involved in gene silencing by miRNA | biological process | GO:0035280 | 1 | 1 | 12345 | 597 | HPS4 | | 4.836e-02 | -3.03 | L-cysteine catabolic process | biological process | GO:0019448 | 1 | 1 | 12345 | 597 | CG5493 | | 4.836e-02 | -3.03 | regulation of acetylcholine-gated cation channel activity | biological process | GO:1903048 | 1 | 1 | 12345 | 597 | qvr | | 4.836e-02 | -3.03 | cytoskeletal anchoring at nuclear membrane | biological process | GO:0090286 | 1 | 1 | 12345 | 597 | Msp300 | | 4.836e-02 | -3.03 | regulation of male germ cell proliferation | biological process | GO:2000254 | 1 | 1 | 12345 | 597 | Doa | | 4.836e-02 | -3.03 | regulation of postsynaptic neurotransmitter receptor activity | biological process | GO:0098962 | 1 | 1 | 12345 | 597 | qvr | | 4.836e-02 | -3.03 | positive regulation of cell cycle arrest | biological process | GO:0071158 | 1 | 1 | 12345 | 597 | foxo | | 4.836e-02 | -3.03 | cyclic nucleotide catabolic process | biological process | GO:0009214 | 1 | 1 | 12345 | 597 | Pde9 | | 4.836e-02 | -3.03 | negative regulation of response to drug | biological process | GO:2001024 | 1 | 1 | 12345 | 597 | Pde8 | | 4.836e-02 | -3.03 | carotenoid transport | biological process | GO:0046867 | 1 | 1 | 12345 | 597 | ninaD | | 4.836e-02 | -3.03 | pyridoxal metabolic process | biological process | GO:0042817 | 1 | 1 | 12345 | 597 | AOX1 | | 4.836e-02 | -3.03 | tetraterpenoid transport | biological process | GO:0046866 | 1 | 1 | 12345 | 597 | ninaD | | 4.836e-02 | -3.03 | cGMP catabolic process | biological process | GO:0046069 | 1 | 1 | 12345 | 597 | Pde9 | | 4.836e-02 | -3.03 | regulation of microtubule nucleation | biological process | GO:0010968 | 1 | 1 | 12345 | 597 | msps | | 4.836e-02 | -3.03 | regulation of actomyosin contractile ring contraction | biological process | GO:0031991 | 1 | 1 | 12345 | 597 | flw | | 4.836e-02 | -3.03 | octopamine catabolic process | biological process | GO:0046334 | 1 | 1 | 12345 | 597 | AANAT1 | | 4.836e-02 | -3.03 | positive regulation of ubiquitin-dependent endocytosis | biological process | GO:2000397 | 1 | 1 | 12345 | 597 | sdt | | 4.836e-02 | -3.03 | neurotransmitter receptor transport, endosome to postsynaptic membrane | biological process | GO:0098887 | 1 | 1 | 12345 | 597 | scrib | | 4.836e-02 | -3.03 | cobalamin transport | biological process | GO:0015889 | 1 | 1 | 12345 | 597 | CG3556 | | 4.836e-02 | -3.03 | negative regulation of acetylcholine-gated cation channel activity | biological process | GO:1903049 | 1 | 1 | 12345 | 597 | qvr | | 4.836e-02 | -3.03 | arginine biosynthetic process via ornithine | biological process | GO:0042450 | 1 | 1 | 12345 | 597 | Argl | | 4.836e-02 | -3.03 | retinal rod cell development | biological process | GO:0046548 | 1 | 1 | 12345 | 597 | CG13251 | | 4.836e-02 | -3.03 | negative regulation of cellular response to drug | biological process | GO:2001039 | 1 | 1 | 12345 | 597 | Pde8 | | 4.836e-02 | -3.03 | defecation | biological process | GO:0030421 | 1 | 1 | 12345 | 597 | drd | | 4.836e-02 | -3.03 | positive regulation of calcium ion-dependent exocytosis | biological process | GO:0045956 | 1 | 1 | 12345 | 597 | Cadps | | 4.836e-02 | -3.03 | negative regulation of cAMP-dependent protein kinase activity | biological process | GO:2000480 | 1 | 1 | 12345 | 597 | sws | | 4.836e-02 | -3.03 | regulation of establishment of bipolar cell polarity | biological process | GO:0061172 | 1 | 1 | 12345 | 597 | wake | | 4.837e-02 | -3.03 | MODULE\_66 | MSigDB lists | MODULE\_66 | 117 | 9 | 4161 | 170 | Ank,bru1,Glut1,Pde8,Mlp84B,Sk1,CG5522,Syx1A,CG9132 | | 4.837e-02 | -3.03 | MODULE\_100 | MSigDB lists | MODULE\_100 | 117 | 9 | 4161 | 170 | Glut1,bru1,Pde8,Ank,Sk1,CG5522,Mlp84B,CG9132,Syx1A | | 4.837e-02 | -3.03 | GO\_PHOSPHORIC\_ESTER\_HYDROLASE\_ACTIVITY | MSigDB lists | GO\_PHOSPHORIC\_ESTER\_HYDROLASE\_ACTIVITY | 117 | 9 | 4161 | 170 | Pde1c,CG3376,Ptp61F,flw,dnc,Pde6,CG3530,Pde8,csw | | 4.843e-02 | -3.03 | Death | pfam domains | PF00531 | 7 | 2 | 10480 | 550 | Ank2,Ank | | 4.843e-02 | -3.03 | EF-hand\_6 | pfam domains | PF13405 | 7 | 2 | 10480 | 550 | CG13526,Actn | | 4.843e-02 | -3.03 | PAZ | pfam domains | PF02170 | 7 | 2 | 10480 | 550 | AGO3,AGO2 | | 4.844e-02 | -3.03 | ARID | prosite domains | PS51011 | 6 | 2 | 6243 | 386 | retn,htk | | 4.844e-02 | -3.03 | ACTINS\_1 | prosite domains | PS00406 | 6 | 2 | 6243 | 386 | Act88F,Act87E | | 4.844e-02 | -3.03 | HIS\_ACID\_PHOSPHAT\_1 | prosite domains | PS00616 | 6 | 2 | 6243 | 386 | CG9449,CG9451 | | 4.847e-02 | -3.03 | positive regulation of kinase activity | biological process | GO:0033674 | 69 | 7 | 12345 | 597 | step,CG41520,rictor,Pak,Stlk,CycT,Raf | | 4.850e-02 | -3.03 | rho (rhomboid) | protein interactions | 38168 | 64 | 7 | 9204 | 481 | Mbs,hth,dpp,Src42A,Ser,shg,csw | | 4.858e-02 | -3.02 | ZnF\_GATA | smart domains | SM00401 | 6 | 2 | 5086 | 315 | grn,GATAe | | 4.858e-02 | -3.02 | BRIGHT | smart domains | SM00501 | 6 | 2 | 5086 | 315 | retn,htk | | 4.858e-02 | -3.02 | Sec7 | smart domains | SM00222 | 6 | 2 | 5086 | 315 | step,siz | | 4.859e-02 | -3.02 | - | gene3d domains | 1.20.900.10 | 23 | 4 | 4929 | 302 | RhoGAP1A,CG10188,Cdep,Exn | | 4.868e-02 | -3.02 | somatic stem cell division | biological process | GO:0048103 | 55 | 6 | 12345 | 597 | sdt,msps,esg,wake,numb,shg | | 4.868e-02 | -3.02 | embryonic organ development | biological process | GO:0048568 | 55 | 6 | 12345 | 597 | shg,tin,mid,RhoL,numb,chb | | 4.880e-02 | -3.02 | GSE11961\_FOLLICULAR\_BCELL\_VS\_MEMORY\_BCELL\_DAY7\_DN | MSigDB lists | GSE11961\_FOLLICULAR\_BCELL\_VS\_MEMORY\_BCELL\_DAY7\_DN | 82 | 7 | 4161 | 170 | Cdep,CG9132,eIF4E7,Raf,CG3860,Stt3B,mino | | 4.880e-02 | -3.02 | KYNG\_WERNER\_SYNDROM\_AND\_NORMAL\_AGING\_DN | MSigDB lists | KYNG\_WERNER\_SYNDROM\_AND\_NORMAL\_AGING\_DN | 82 | 7 | 4161 | 170 | Khc,Doa,msps,Pfdn4,RhoGAP1A,Iswi,step | | 4.929e-02 | -3.01 | negative regulation of stress-activated protein kinase signaling cascade | biological process | GO:0070303 | 29 | 4 | 12345 | 597 | raw,Ptp61F,sgg,flw | | 4.929e-02 | -3.01 | steroid biosynthetic process | biological process | GO:0006694 | 29 | 4 | 12345 | 597 | spok,woc,foxo,shd | | 4.929e-02 | -3.01 | cellular response to hypoxia | biological process | GO:0071456 | 29 | 4 | 12345 | 597 | Ptp61F,msi,eIF2Bgamma,Mrtf | | 4.929e-02 | -3.01 | establishment of mitotic spindle localization | biological process | GO:0040001 | 29 | 4 | 12345 | 597 | msps,wake,chb,dsh | | 4.929e-02 | -3.01 | negative regulation of stress-activated MAPK cascade | biological process | GO:0032873 | 29 | 4 | 12345 | 597 | flw,sgg,Ptp61F,raw | | 4.929e-02 | -3.01 | negative regulation of transferase activity | biological process | GO:0051348 | 29 | 4 | 12345 | 597 | kek1,mnb,sws,sgg | | 4.929e-02 | -3.01 | establishment or maintenance of polarity of follicular epithelium | biological process | GO:0016334 | 29 | 4 | 12345 | 597 | scrib,Raf,csw,egh | | 4.945e-02 | -3.01 | GO\_PITUITARY\_GLAND\_DEVELOPMENT | MSigDB lists | GO\_PITUITARY\_GLAND\_DEVELOPMENT | 9 | 2 | 4161 | 170 | Su(H),dpp | | 4.945e-02 | -3.01 | REACTOME\_NUCLEAR\_RECEPTOR\_TRANSCRIPTION\_PATHWAY | MSigDB lists | REACTOME\_NUCLEAR\_RECEPTOR\_TRANSCRIPTION\_PATHWAY | 9 | 2 | 4161 | 170 | EcR,Hr96 | | 4.945e-02 | -3.01 | DAUER\_STAT3\_TARGETS\_UP | MSigDB lists | DAUER\_STAT3\_TARGETS\_UP | 9 | 2 | 4161 | 170 | Glut1,CG6966 | | 4.945e-02 | -3.01 | GO\_PROTEIN\_TYROSINE\_KINASE\_BINDING | MSigDB lists | GO\_PROTEIN\_TYROSINE\_KINASE\_BINDING | 9 | 2 | 4161 | 170 | csw,Ptp61F | | 4.945e-02 | -3.01 | GO\_REGULATION\_OF\_PROTEIN\_TARGETING\_TO\_MEMBRANE | MSigDB lists | GO\_REGULATION\_OF\_PROTEIN\_TARGETING\_TO\_MEMBRANE | 9 | 2 | 4161 | 170 | sxc,Ank | | 4.945e-02 | -3.01 | PID\_AURORA\_A\_PATHWAY | MSigDB lists | PID\_AURORA\_A\_PATHWAY | 9 | 2 | 4161 | 170 | Pak,msps | | 4.945e-02 | -3.01 | ALONSO\_METASTASIS\_EMT\_UP | MSigDB lists | ALONSO\_METASTASIS\_EMT\_UP | 9 | 2 | 4161 | 170 | Iswi,Pkc53E | | 4.945e-02 | -3.01 | GO\_PROTEIN\_PALMITOYLATION | MSigDB lists | GO\_PROTEIN\_PALMITOYLATION | 9 | 2 | 4161 | 170 | Acbp2,app | | 4.945e-02 | -3.01 | HUANG\_FOXA2\_TARGETS\_UP | MSigDB lists | HUANG\_FOXA2\_TARGETS\_UP | 9 | 2 | 4161 | 170 | Mctp,Ser | | 4.945e-02 | -3.01 | GO\_ACTIVATION\_OF\_MAPKK\_ACTIVITY | MSigDB lists | GO\_ACTIVATION\_OF\_MAPKK\_ACTIVITY | 9 | 2 | 4161 | 170 | Raf,CG42672 | | 4.945e-02 | -3.01 | GO\_MEMBRANE\_LIPID\_CATABOLIC\_PROCESS | MSigDB lists | GO\_MEMBRANE\_LIPID\_CATABOLIC\_PROCESS | 9 | 2 | 4161 | 170 | CG33090,CG3376 | | 4.945e-02 | -3.01 | GO\_POSITIVE\_REGULATION\_OF\_RESPONSE\_TO\_WOUNDING | MSigDB lists | GO\_POSITIVE\_REGULATION\_OF\_RESPONSE\_TO\_WOUNDING | 9 | 2 | 4161 | 170 | Pkc98E,Raf | | 4.945e-02 | -3.01 | REACTOME\_O\_LINKED\_GLYCOSYLATION\_OF\_MUCINS | MSigDB lists | REACTOME\_O\_LINKED\_GLYCOSYLATION\_OF\_MUCINS | 9 | 2 | 4161 | 170 | CG30036,Pgant6 | | 4.945e-02 | -3.01 | YRCCAKNNGNCGC\_UNKNOWN | MSigDB lists | YRCCAKNNGNCGC\_UNKNOWN | 9 | 2 | 4161 | 170 | PIG-O,fz2 | | 4.945e-02 | -3.01 | GO\_RNA\_POLYMERASE\_CORE\_ENZYME\_BINDING | MSigDB lists | GO\_RNA\_POLYMERASE\_CORE\_ENZYME\_BINDING | 9 | 2 | 4161 | 170 | Maf1,Elp3 | | 4.945e-02 | -3.01 | REACTOME\_GLYCOGEN\_BREAKDOWN\_GLYCOGENOLYSIS | MSigDB lists | REACTOME\_GLYCOGEN\_BREAKDOWN\_GLYCOGENOLYSIS | 9 | 2 | 4161 | 170 | GlyP,PhKgamma | | 4.945e-02 | -3.01 | PID\_UPA\_UPAR\_PATHWAY | MSigDB lists | PID\_UPA\_UPAR\_PATHWAY | 9 | 2 | 4161 | 170 | LRP1,LpR2 | | 4.945e-02 | -3.01 | GO\_CHANNEL\_INHIBITOR\_ACTIVITY | MSigDB lists | GO\_CHANNEL\_INHIBITOR\_ACTIVITY | 9 | 2 | 4161 | 170 | Syx1A,ZnT63C | | 4.945e-02 | -3.01 | DORSAM\_HOXA9\_TARGETS\_DN | MSigDB lists | DORSAM\_HOXA9\_TARGETS\_DN | 9 | 2 | 4161 | 170 | sno,Dlic | | 4.945e-02 | -3.01 | GO\_REGULATION\_OF\_INTERFERON\_GAMMA\_PRODUCTION | MSigDB lists | GO\_REGULATION\_OF\_INTERFERON\_GAMMA\_PRODUCTION | 9 | 2 | 4161 | 170 | dnc,fz2 | | 4.945e-02 | -3.01 | GO\_CYCLIC\_NUCLEOTIDE\_BINDING | MSigDB lists | GO\_CYCLIC\_NUCLEOTIDE\_BINDING | 9 | 2 | 4161 | 170 | Pde6,dnc | | 4.945e-02 | -3.01 | GO\_REGULATION\_OF\_ALCOHOL\_BIOSYNTHETIC\_PROCESS | MSigDB lists | GO\_REGULATION\_OF\_ALCOHOL\_BIOSYNTHETIC\_PROCESS | 9 | 2 | 4161 | 170 | dpp,Sod1 | | 4.945e-02 | -3.01 | HOQUE\_METHYLATED\_IN\_CANCER | MSigDB lists | HOQUE\_METHYLATED\_IN\_CANCER | 9 | 2 | 4161 | 170 | TfAP-2,dpp | | 4.945e-02 | -3.01 | REACTOME\_PROLONGED\_ERK\_ACTIVATION\_EVENTS | MSigDB lists | REACTOME\_PROLONGED\_ERK\_ACTIVATION\_EVENTS | 9 | 2 | 4161 | 170 | Raf,CG42672 | | 4.945e-02 | -3.01 | GO\_H4\_HISTONE\_ACETYLTRANSFERASE\_ACTIVITY | MSigDB lists | GO\_H4\_HISTONE\_ACETYLTRANSFERASE\_ACTIVITY | 9 | 2 | 4161 | 170 | Elp3,sxc | | 4.945e-02 | -3.01 | LANDIS\_ERBB2\_BREAST\_TUMORS\_65\_DN | MSigDB lists | LANDIS\_ERBB2\_BREAST\_TUMORS\_65\_DN | 9 | 2 | 4161 | 170 | mino,Der-1 | | 4.945e-02 | -3.01 | GO\_FACE\_DEVELOPMENT | MSigDB lists | GO\_FACE\_DEVELOPMENT | 9 | 2 | 4161 | 170 | Raf,csw | | 4.945e-02 | -3.01 | MODULE\_196 | MSigDB lists | MODULE\_196 | 9 | 2 | 4161 | 170 | Dlic,Octbeta2R | | 4.945e-02 | -3.01 | GO\_APOLIPOPROTEIN\_BINDING | MSigDB lists | GO\_APOLIPOPROTEIN\_BINDING | 9 | 2 | 4161 | 170 | LRP1,LpR2 | | 4.945e-02 | -3.01 | GO\_POSITIVE\_REGULATION\_OF\_GLUCOSE\_METABOLIC\_PROCESS | MSigDB lists | GO\_POSITIVE\_REGULATION\_OF\_GLUCOSE\_METABOLIC\_PROCESS | 9 | 2 | 4161 | 170 | Ptp61F,PhKgamma | | 4.945e-02 | -3.01 | MODULE\_533 | MSigDB lists | MODULE\_533 | 9 | 2 | 4161 | 170 | step,Syx1A | | 4.945e-02 | -3.01 | GO\_STEROID\_HORMONE\_RECEPTOR\_ACTIVITY | MSigDB lists | GO\_STEROID\_HORMONE\_RECEPTOR\_ACTIVITY | 9 | 2 | 4161 | 170 | EcR,Hr96 | | 4.945e-02 | -3.01 | GO\_NEGATIVE\_REGULATION\_OF\_MACROAUTOPHAGY | MSigDB lists | GO\_NEGATIVE\_REGULATION\_OF\_MACROAUTOPHAGY | 9 | 2 | 4161 | 170 | Cln3,Tor | | 4.945e-02 | -3.01 | JAZAERI\_BREAST\_CANCER\_BRCA1\_VS\_BRCA2\_DN | MSigDB lists | JAZAERI\_BREAST\_CANCER\_BRCA1\_VS\_BRCA2\_DN | 9 | 2 | 4161 | 170 | sno,Mlp84B | | 4.945e-02 | -3.01 | GO\_EXPLORATION\_BEHAVIOR | MSigDB lists | GO\_EXPLORATION\_BEHAVIOR | 9 | 2 | 4161 | 170 | Pkc98E,CG7483 | | 4.945e-02 | -3.01 | GO\_REGULATION\_OF\_OSTEOCLAST\_DIFFERENTIATION | MSigDB lists | GO\_REGULATION\_OF\_OSTEOCLAST\_DIFFERENTIATION | 9 | 2 | 4161 | 170 | Nha1,Tor | | 4.945e-02 | -3.01 | GO\_MONOSACCHARIDE\_TRANSPORT | MSigDB lists | GO\_MONOSACCHARIDE\_TRANSPORT | 9 | 2 | 4161 | 170 | Raf,Glut1 | | 4.945e-02 | -3.01 | GO\_REGULATION\_OF\_INSULIN\_RECEPTOR\_SIGNALING\_PATHWAY | MSigDB lists | GO\_REGULATION\_OF\_INSULIN\_RECEPTOR\_SIGNALING\_PATHWAY | 9 | 2 | 4161 | 170 | sxc,Ptp61F | | 4.945e-02 | -3.01 | GO\_REGULATION\_OF\_COAGULATION | MSigDB lists | GO\_REGULATION\_OF\_COAGULATION | 9 | 2 | 4161 | 170 | Pkc53E,Octbeta2R | | 4.945e-02 | -3.01 | FOURNIER\_ACINAR\_DEVELOPMENT\_LATE\_DN | MSigDB lists | FOURNIER\_ACINAR\_DEVELOPMENT\_LATE\_DN | 9 | 2 | 4161 | 170 | Odc1,Act87E | | 4.964e-02 | -3.00 | leading edge membrane | cellular component | GO:0031256 | 90 | 8 | 11940 | 538 | Or85b,Ser,Or42b,dpr15,dpr5,Or65c,dpr18,dpr12 | | 4.982e-02 | -3.00 | Lmpt (Limpet) | protein interactions | 39889 | 27 | 4 | 9204 | 481 | dsh,Mlp84B,CG7368,rdgA | | 4.982e-02 | -3.00 | spen (split ends) | protein interactions | 44205 | 27 | 4 | 9204 | 481 | Antp,dpp,Raf,MESK2 |
