## Supplementary material for "A single DPE core promoter motif contributes to *in vivo* transcriptional regulation and affects cardiac function": SupFile2: knownResults.html

./cluster.allRlog.c300.t8//1365.cluster.txt.motifs/ - Homer Known Motif Enrichment Results


### Homer Known Motif Enrichment Results (./cluster.allRlog.c300.t8//1365.cluster.txt.motifs/)

Homer *de novo* Motif Results  
Gene Ontology Enrichment Results  
Known Motif Enrichment Results (txt file)  
Total Target Sequences = 939, Total Background Sequences = 28699

|  |  |  |  |  |  |  |  |  |  |  |  |
| --- | --- | --- | --- | --- | --- | --- | --- | --- | --- | --- | --- |
| Rank | Motif | Name | P-value | log P-pvalue | q-value (Benjamini) | # Target Sequences with Motif | % of Targets Sequences with Motif | # Background Sequences with Motif | % of Background Sequences with Motif | Motif File | SVG |
