## Supplementary material for "A single DPE core promoter motif contributes to *in vivo* transcriptional regulation and affects cardiac function": SupFile2: geneOntology.html

Gene Ontology Results

### Gene Ontology Enrichment Results

Homer *de novo* Motif Enrichment Results  
Known Motif Enrichment Results  

###### Text file version of complete results (i.e. open with Excel) - biological process: Functional groupings of proteins (Gene Ontology) - molecular function: Mechanistic actions of proteins (Gene Ontology) - cellular component: Protein localization (Gene Ontology) - chromosome location: Genes with similar chromosome localization (NCBI Gene) - KEGG pathways: Groups of proteins in the same pathways (KEGG) - protein interactions: "Proteins interacting with a common protein (BIND, EcoCyc, HPRD)" (NCBI Gene) - interpro domains: Proteins with similar domains and features (Interpro) - pfam domains: Proteins with similar domains and features (Pfam) - smart domains: Proteins with similar domains and features (SMART) - gene3d domains: Proteins with similar domains and features (Gene3D) - prosite domains: Proteins with similar domains and features (Prosite) - prints domains: Proteins with similar domains and features (PRINTS) - MSigDB lists: "Genes sets for pathways, factor/miRNA target predictions, expression patterns, etc." (MSigDB) - BIOCYC pathways: Groups of proteins in the same pathways (BIOCYC) - COSMIC cancer mutations: Genes mutated in similar cancers (COSMIC) - GWAS genes: Genes mutated in similar diseases (GWAS Catalog) - Lipid Maps pathways: Groups of proteins in the same lipid pathways (Lipid Maps/Biosystems) - Pathway Interaction DB: Groups of proteins in the same pathways (Pathway Interaction Database) - REACTOME pathways: Groups of proteins in the same pathways (REACTOME) - SMPDB pathways: Groups of proteins in the same pathways (SMPDB) - WikiPathways: Groups of proteins in the same pathways (Wikipathways) Enriched Categories | | | | | | | | | | | | --- | --- | --- | --- | --- | --- | --- | --- | --- | --- | | P-value | ln(P) | Term | GO Tree | GO ID | # of Genes in Term | # of Target Genes in Term | # of Total Genes | # of Target Genes | Common Genes | | 1.401e-28 | -64.13 | fly chr3-47 | chromosome location | fly chr3-47 | 46 | 27 | 25010 | 823 | asRNA:CR45182,eIF4B,CG41284,lncRNA:CR46252,CG40472,ND-AGGG,lncRNA:CR43942,CG42598,lncRNA:CR40053,CG40045,lncRNA:CR45180,Snap25,lncRNA:CR45159,mRpS5,FASN3,CG40160,CG40228,lncRNA:CR46250,UQCR-11,CG45782,Set1,lncRNA:CR45177,nvd,CG17514,CG40178,lncRNA:CR43941,scro | | 1.401e-28 | -64.13 | fly chr3-47 cM | chromosome location | fly chr3-47 cM | 46 | 27 | 25010 | 823 | CG40228,CG40160,FASN3,mRpS5,lncRNA:CR45159,Snap25,lncRNA:CR45180,lncRNA:CR43941,scro,CG40178,CG17514,nvd,lncRNA:CR45177,Set1,UQCR-11,CG45782,lncRNA:CR46250,CG40472,ND-AGGG,CG41284,lncRNA:CR46252,eIF4B,asRNA:CR45182,CG40045,CG42598,lncRNA:CR40053,lncRNA:CR43942 | | 6.470e-15 | -32.67 | fly chr2-55 cM | chromosome location | fly chr2-55 cM | 36 | 16 | 25010 | 823 | MFS17,RYa,CG40498,Maf1,RNASEK,CG17683,asRNA:CR45139,DIP-lambda,mir-2279,CG41378,CG41242,CG17684,dpr21,CG40191,CG17691,CG41520 | | 6.470e-15 | -32.67 | fly chr2-55 | chromosome location | fly chr2-55 | 36 | 16 | 25010 | 823 | CG41242,dpr21,CG17684,CG41378,CG41520,CG40191,CG17691,MFS17,RYa,asRNA:CR45139,CG17683,mir-2279,DIP-lambda,Maf1,RNASEK,CG40498 | | 7.927e-15 | -32.47 | fly chr3-47.1 cM | chromosome location | fly chr3-47.1 cM | 17 | 12 | 25010 | 823 | CG42402,Alg-2,lncRNA:CR46122,Myo81F,CG41128,spok,CG41099,lncRNA:CR46123,Pzl,CG40198,Tim17b,lncRNA:CR45220 | | 7.927e-15 | -32.47 | fly chr3-47.1 | chromosome location | fly chr3-47.1 | 17 | 12 | 25010 | 823 | Tim17b,lncRNA:CR45220,CG40198,Pzl,lncRNA:CR46123,CG41128,CG41099,spok,Myo81F,CG42402,Alg-2,lncRNA:CR46122 | | 2.821e-07 | -15.08 | fly chr2-54 | chromosome location | fly chr2-54 | 22 | 8 | 25010 | 823 | CG17715,lncRNA:CR43242,Cht10,cta,lncRNA:CR43927,lncRNA:CR46253,Tim23,CG40006 | | 2.821e-07 | -15.08 | fly chr2-54 cM | chromosome location | fly chr2-54 cM | 22 | 8 | 25010 | 823 | Tim23,CG40006,lncRNA:CR46253,Cht10,lncRNA:CR43927,cta,CG17715,lncRNA:CR43242 | | 3.396e-07 | -14.90 | regulation of cellular process | biological process | GO:0050794 | 3560 | 219 | 12345 | 571 | mbc,mlt,Ulp1,hth,Trxr-2,Ir40a,Src42A,Ppm1,Parp,Rh3,CG12147,CG9389,CG12200,dgt5,CG41106,tay,par-1,RYa,TrissinR,capu,Diap1,CG34334,rl,sdk,Octbeta1R,sif,Slmap,pk,Glut1,fru,step,CG12769,Pi3K68D,CG34408,SPR,Mical,Tak1,Nipped-A,CG41520,d4,Spn42De,lola,en,dnt,cic,cpx,Trl,mrt,Ca-alpha1D,fuss,Mnt,sna,cv-2,CycB,dimm,GABA-B-R3,Gprk1,Coq7,heph,Hipk,Sema1a,CG40178,rno,CG9650,Clk,cta,fd96Ca,Thor,rau,SteXh:CG42398,tej,caup,hdc,Git,shakB,salm,CG30440,CG13284,Atf6,shot,CG42637,RhoGAP100F,CG10188,Mef2,Marf1,ewg,TpnC41C,gus,fbl,Pld,Dys,PNPase,br,norpA,mthl8,mthl11,tou,Toll-7,neo,Tgi,Hsc70-3,tj,CG14184,SCAP,Fer3,Catsup,cpo,Myo81F,mxt,Mp,Nlg3,kni,Sod1,dnr1,CG5270,Wdr59,Wdr62,Octbeta2R,orb,svp,CG32944,Spn,dnc,scw,fray,boi,AGO3,bru1,lz,CG42541,pho,5-HT2A,Rgk1,Pkg21D,Bruce,Cip4,zen,CG32006,Ptip,rdgC,5-HT2B,pan,Magi,mav,lqf,pico,Setd3,E5,conu,alpha4GT1,jing,HP6,Psc,tinc,bi,Parg,CG12206,CG40228,Camta,AP-2alpha,lncRNA:bxd,HisCl1,SoxN,zld,kirre,CG5160,Ten-a,ru,CG33557,CG6888,rib,babos,Syt7,Su(z)2,Alg-2,mim,grim,bru3,Ac78C,Stlk,Vsx2,Nlg2,Snap25,Spn75F,CG5862,PEK,Der-1,CdGAPr,CG17514,ct,Sdc,spel1,ap,vap,CG34357,CG1620,Myc,nAChRalpha3,CG9932,CG9380,ArfGAP1,CG31750,RSG7,CG12772,Gr2a,Slob,pum,toy,dsx,bab2,fz2,Maf1,nAChRalpha4,scro,nullo,HPS4,tld,sky,RabX2 | | 3.752e-07 | -14.80 | signaling | biological process | GO:0023052 | 1169 | 91 | 12345 | 571 | Octbeta1R,mav,rl,Magi,pan,pico,lqf,SPR,Pi3K68D,HisCl1,CG41520,Tak1,cpx,cic,dnt,sna,Ca-alpha1D,CG32944,mbc,Spn,Syx18,Src42A,Ir40a,unc-13-4A,scw,fray,dnc,boi,Rh3,CG9389,5-HT2A,Rgk1,CG12147,CG42541,Pkg21D,par-1,TrissinR,5-HT2B,RYa,rdgC,gus,vlc,TpnC41C,Der-1,Pld,Dys,CdGAPr,nAChRalpha3,Myc,mthl11,mthl8,CG34357,spel1,norpA,SCAP,RSG7,CG31750,Hsc70-3,Toll-7,pum,Slob,Gr2a,cpo,Myo81F,nAChRalpha4,fz2,RabX2,sky,svp,Octbeta2R,Gprk1,CG5160,GABA-B-R3,Ten-a,Atpalpha,be,Sema1a,Syt7,cta,grim,Thor,shakB,Ac78C,Atf6,Nlg2,Stlk,RhoGAP100F,Snap25,shot,CG42637,PEK | | 7.919e-07 | -14.05 | wing disc morphogenesis | biological process | GO:0007472 | 308 | 35 | 12345 | 571 | heph,Hipk,Src42A,cv-2,sn,hth,caup,salm,par-1,Cip4,tay,shot,fbl,Dys,pk,ct,pan,rl,gus,bi,fru,ap,cmb,step,Myc,ck,jing,zld,lola,en,Tak1,fuss,tld,cic,Trl | | 8.044e-07 | -14.03 | biological regulation | biological process | GO:0065007 | 4350 | 256 | 12345 | 571 | dnc,fray,scw,Acox57D-d,Npc2b,Spn,rdgC,spo,zen,SmydA-8,CG32006,5-HT2A,Rgk1,bi,tinc,CG12206,Parg,E5,conu,Psc,HP6,alpha4GT1,Setd3,pan,mav,CG30495,Camta,AP-2alpha,lncRNA:bxd,Su(z)2,grim,rib,ru,CG5160,CG5862,PEK,IA-2,Snap25,Stlk,CG31750,ArfGAP1,vap,spel1,CG9932,Myc,THADA,Der-1,sky,RabX2,PMCA,scro,nAChRalpha4,Tbc1d15-17,inc,toy,dsx,Gr2a,Rh3,bond,unc-13-4A,Trxr-2,Ir40a,Ppm1,Khc-73,mlt,DNAlig4,RYa,TrissinR,capu,par-1,CG41106,dgt5,spok,CG12200,Mical,CG34408,CG12769,Glut1,fru,Slmap,CG34334,Octbeta1R,sdk,drosha,dnt,cic,CG9981,mrt,en,CG41520,rau,fd96Ca,Thor,SteXh:CG42398,Hipk,Gprk1,Marf1,CG10077,SmydA-9,CG30440,Atf6,caup,shakB,salm,CG10184,Hsc70-3,tj,Tgi,Toll-7,SCAP,CG14184,mthl8,mthl11,Ars2,Pld,PNPase,kune,gus,ABCB7,TpnC41C,jigr1,Wdr62,orb,Wdr59,Octbeta2R,svp,Mp,kni,Myo81F,mxt,Catsup,lz,boi,bru1,AGO3,CG32944,5-HT2B,Cip4,Ptip,Pkg21D,Bruce,pho,CG42541,knk,jing,lqf,pico,Magi,kirre,zld,CG40228,Teh2,SoxN,HisCl1,CG9514,Alg-2,MCU,mim,babos,Syt7,CG6888,CG33557,Ten-a,Spn75F,Nlg2,Vsx2,bru3,Ac78C,RSG7,CG1620,CG34357,ap,nAChRalpha3,CG9380,CdGAPr,Sdc,ct,CG17514,nullo,HPS4,tld,fz2,Maf1,dati,bab2,Slob,CG12772,Acp36DE,pum,Parp,SK,Src42A,mbc,hth,Ulp1,tay,Lsd-2,CG9389,CG12147,SPR,Pi3K68D,vtd,step,sif,Msp300,pk,rl,Diap1,sna,fuss,Ca-alpha1D,Mnt,cpx,KCNQ,Trl,Spn42De,lola,Tak1,d4,Nipped-A,tej,rno,CG40178,cta,CG9650,Clk,Sema1a,heph,Coq7,CycB,cv-2,GABA-B-R3,Atpalpha,dimm,Mef2,ewg,RhoGAP100F,CG42637,CG10188,shot,CG13284,Git,hdc,neo,norpA,tou,Dys,fbl,br,CG5270,nvd,Sod1,dnr1,Nlg3,cpo,Fer3 | | 8.464e-07 | -13.98 | regulation of biological process | biological process | GO:0050789 | 3862 | 232 | 12345 | 571 | lz,dnc,scw,boi,fray,bru1,AGO3,Spn,CG32944,rdgC,5-HT2B,zen,Cip4,SmydA-8,Ptip,CG32006,Pkg21D,Bruce,CG42541,pho,5-HT2A,Rgk1,bi,tinc,CG12206,Parg,conu,E5,HP6,jing,Psc,alpha4GT1,lqf,Setd3,pico,Magi,pan,mav,kirre,zld,Camta,AP-2alpha,CG40228,Teh2,lncRNA:bxd,HisCl1,SoxN,Alg-2,Su(z)2,grim,mim,rib,babos,Syt7,CG6888,ru,CG33557,Ten-a,CG5160,CG5862,PEK,IA-2,Spn75F,Snap25,Stlk,Vsx2,Nlg2,bru3,Ac78C,CG31750,ArfGAP1,RSG7,CG34357,CG1620,ap,spel1,vap,CG9380,CG9932,nAChRalpha3,Myc,CdGAPr,Sdc,CG17514,THADA,ct,Der-1,nullo,RabX2,sky,tld,HPS4,Maf1,fz2,bab2,scro,nAChRalpha4,inc,toy,dsx,Slob,Gr2a,CG12772,pum,Parp,Rh3,Src42A,Ir40a,Trxr-2,Ppm1,mlt,mbc,hth,Ulp1,RYa,capu,TrissinR,par-1,tay,Lsd-2,CG41106,dgt5,CG9389,CG12200,CG12147,SPR,Mical,Pi3K68D,CG34408,vtd,CG12769,Glut1,fru,step,sif,Slmap,pk,Diap1,CG34334,rl,Octbeta1R,sdk,sna,Mnt,fuss,Ca-alpha1D,drosha,cpx,KCNQ,cic,dnt,mrt,Trl,Spn42De,lola,en,Tak1,d4,CG41520,Nipped-A,rau,fd96Ca,Thor,SteXh:CG42398,tej,rno,CG40178,cta,Clk,CG9650,Sema1a,Hipk,heph,Coq7,CycB,cv-2,Gprk1,GABA-B-R3,dimm,ewg,Mef2,Marf1,CG10077,RhoGAP100F,shot,CG42637,CG10188,SmydA-9,CG30440,Atf6,CG13284,caup,Git,shakB,salm,hdc,Hsc70-3,Tgi,tj,neo,Toll-7,SCAP,CG14184,mthl11,mthl8,norpA,Ars2,tou,Dys,Pld,fbl,PNPase,br,gus,jigr1,TpnC41C,CG5270,Wdr59,Wdr62,orb,Octbeta2R,svp,Mp,dnr1,Sod1,kni,Nlg3,Myo81F,cpo,mxt,Catsup,Fer3 | | 9.390e-07 | -13.88 | nervous system development | biological process | GO:0007399 | 1108 | 86 | 12345 | 571 | Cip4,par-1,spo,pho,dnc,fray,boi,fdl,lz,sn,Ulp1,hth,mbc,Spn,Src42A,dnt,cpx,drosha,Ca-alpha1D,fuss,sna,SoxN,d4,lola,en,zld,ck,Psc,jing,fru,E5,vtd,CadN2,tinc,Mical,mav,sdk,wb,rl,pk,sif,shot,RhoGAP100F,ewg,shep,Ptp99A,hdc,Git,salm,Nlg2,klg,Stlk,babos,rib,rno,Dscam2,grim,rau,dimm,Atpalpha,Ten-a,ru,Sema1a,rg,Nlg3,scro,Mp,dati,fz2,svp,sky,pum,nrm,toy,dsx,Myc,tou,ap,cmb,vap,Toll-7,kune,ct,Sdc,br,CdGAPr | | 1.078e-06 | -13.74 | tube development | biological process | GO:0035295 | 747 | 64 | 12345 | 571 | Src42A,sn,hth,mbc,lz,scw,boi,fray,CG17211,knk,wge,spo,lama,Ptip,ect,par-1,Cip4,tay,pk,pico,wb,pan,rl,Diap1,bi,ck,jing,step,fru,lola,en,zld,Nipped-A,Tak1,fuss,sna,Trl,cic,heph,ru,Hipk,cv-2,Idgf4,rib,hdc,salm,caup,Mef2,ewg,shot,IA-2,ImpE2,ct,fbl,Dys,gus,Myc,ap,cmb,toy,dsx,tld,svp,bab2 | | 1.280e-06 | -13.57 | system development | biological process | GO:0048731 | 1808 | 125 | 12345 | 571 | rib,rno,babos,rau,Dscam2,grim,Idgf4,cv-2,Atpalpha,dimm,Ten-a,ru,heph,Hipk,Sema1a,Nna1,ImpE2,shot,RhoGAP100F,IA-2,shep,Ptp99A,ewg,Mef2,caup,hdc,salm,Git,klg,Stlk,Nlg2,cmb,vap,ap,Myc,CAP,tou,Toll-7,tj,gus,fbl,Dys,CdGAPr,kune,ct,br,Sdc,Mp,bab2,fz2,dati,Nlg3,rg,kni,scro,tld,sky,svp,nrm,pum,dsx,toy,dnc,boi,scw,fray,lz,cato,fdl,mbc,Ulp1,sn,NAT1,hth,Src42A,Spn,Cip4,tay,par-1,zen,Ptip,spo,lama,wge,knk,pho,Pkg21D,step,fru,E5,vtd,ck,Psc,jing,tinc,Mical,bi,CadN2,pan,rl,Diap1,mav,sdk,wb,sif,Msp300,pk,pico,dnt,cic,drosha,cpx,Trl,fuss,kirre,Ca-alpha1D,sna,Tak1,Nipped-A,SoxN,fred,d4,zld,en,lola | | 1.381e-06 | -13.49 | cell communication | biological process | GO:0007154 | 1239 | 93 | 12345 | 571 | CG41520,HisCl1,Tak1,cpx,cic,dnt,sna,Ca-alpha1D,Octbeta1R,mav,rl,Magi,pan,pico,lqf,SPR,Pi3K68D,CG9389,Rgk1,5-HT2A,CG12147,CG42541,Bruce,Pkg21D,par-1,TrissinR,rdgC,RYa,5-HT2B,mbc,CG32944,Syx18,Spn,Src42A,Ir40a,unc-13-4A,scw,fray,dnc,boi,Rh3,pum,Slob,Gr2a,cpo,Myo81F,nAChRalpha4,fz2,svp,sky,RabX2,Octbeta2R,Wdr59,gus,Der-1,TpnC41C,CG7536,CdGAPr,Pld,Dys,nAChRalpha3,Myc,CG34357,mthl8,mthl11,norpA,spel1,SCAP,RSG7,Hsc70-3,CG31750,Toll-7,shakB,Ac78C,Atf6,Nlg2,Stlk,shot,Snap25,RhoGAP100F,CG42637,PEK,Gprk1,GABA-B-R3,Ten-a,CG5160,Atpalpha,be,Sema1a,Syt7,cta,grim,Thor | | 1.440e-06 | -13.45 | post-embryonic development | biological process | GO:0009791 | 642 | 57 | 12345 | 571 | Trl,bab2,cic,drosha,fz2,tld,fuss,kirre,nvd,SoxN,Tak1,lola,en,zld,ck,Myc,jing,step,cmb,fru,ap,bi,gus,pan,rl,Diap1,pk,ct,br,fbl,Msp300,Dys,shot,ImpE2,tay,par-1,Cip4,wge,ewg,spo,hdc,salm,spok,caup,Pkg21D,grim,lz,CG9514,sn,hth,cv-2,mbc,Nna1,Npc2b,heph,Hipk,Src42A | | 2.066e-06 | -13.09 | epithelial tube morphogenesis | biological process | GO:0060562 | 495 | 47 | 12345 | 571 | wge,ewg,par-1,Cip4,ImpE2,tay,shot,knk,caup,hdc,salm,CG17211,rib,heph,Src42A,Hipk,mbc,cv-2,sn,hth,fuss,sna,tld,svp,cic,bab2,Trl,zld,en,lola,Tak1,bi,ap,cmb,fru,step,ck,Myc,jing,fbl,Dys,pk,ct,pan,rl,Diap1,gus | | 2.311e-06 | -12.98 | metamorphosis | biological process | GO:0007552 | 497 | 47 | 12345 | 571 | lola,en,zld,Tak1,tld,fuss,nvd,Trl,bab2,cic,drosha,ct,pk,br,fbl,Dys,gus,pan,rl,Diap1,bi,ck,Myc,jing,step,cmb,fru,ap,hdc,salm,caup,wge,ewg,shep,shot,Cip4,ImpE2,tay,par-1,heph,Hipk,Src42A,sn,hth,mbc,cv-2,Parp | | 2.631e-06 | -12.85 | instar larval or pupal development | biological process | GO:0002165 | 576 | 52 | 12345 | 571 | ct,pk,br,fbl,Dys,gus,pan,Diap1,rl,bi,Myc,ck,jing,step,fru,cmb,ap,lola,en,zld,SoxN,Tak1,tld,fuss,nvd,Trl,bab2,cic,drosha,fz2,Npc2b,heph,Hipk,Src42A,sn,hth,cv-2,mbc,grim,CG9514,hdc,salm,spok,caup,wge,ewg,spo,shot,tay,par-1,ImpE2,Cip4 | | 2.905e-06 | -12.75 | imaginal disc-derived wing morphogenesis | biological process | GO:0007476 | 298 | 33 | 12345 | 571 | jing,ck,Myc,fru,cmb,ap,step,bi,rl,pan,ct,pk,Dys,fbl,Trl,cic,tld,fuss,en,lola,zld,hth,sn,cv-2,Hipk,Src42A,heph,shot,tay,Cip4,par-1,salm,caup | | 3.563e-06 | -12.55 | negative regulation of gene expression | biological process | GO:0010629 | 582 | 52 | 12345 | 571 | HP6,jing,CG9932,Psc,Ars2,CG1620,E5,Parg,bi,Hsc70-3,Tgi,pan,Diap1,ct,br,PNPase,kni,mrt,Trl,cic,drosha,Maf1,HPS4,tld,svp,Mnt,sna,orb,lncRNA:bxd,d4,pum,Fer3,en,dsx,zld,bru1,AGO3,Clk,scw,Thor,Su(z)2,lz,heph,SmydA-9,SmydA-8,CG10077,zen,Marf1,hdc,bru3,pho,Vsx2 | | 4.687e-06 | -12.27 | anatomical structure morphogenesis | biological process | GO:0009653 | 1579 | 110 | 12345 | 571 | pk,wb,sdk,rl,Diap1,pan,CadN2,Mical,bi,Psc,jing,ck,fru,step,lola,en,zld,fred,SoxN,Tak1,sna,fuss,kirre,Trl,drosha,dnt,cic,Src42A,hth,fan,Ulp1,sn,mlt,mbc,Parp,lz,bru1,scw,boi,dnc,CG17211,5-HT2A,pho,knk,capu,wge,lama,spo,CG32006,zen,Cip4,tay,par-1,br,Sdc,ct,Dys,Pld,CdGAPr,fbl,gus,tj,neo,Toll-7,Myc,cmb,ap,dsx,toy,pum,svp,tld,nullo,orb,scro,bt,rg,fz2,TwdlG,bab2,Mp,Hipk,Sema1a,heph,ru,Ten-a,Atpalpha,cv-2,grim,Dscam2,rau,fd96Ca,Thor,cta,rno,rib,klg,Stlk,salm,hdc,jvl,caup,Irp-1A,ewg,Mef2,Ptp99A,CG41099,RhoGAP100F,shot,ImpE2 | | 4.787e-06 | -12.25 | imaginal disc morphogenesis | biological process | GO:0007560 | 420 | 41 | 12345 | 571 | Hipk,Src42A,heph,mbc,cv-2,hth,sn,ewg,wge,Cip4,ImpE2,tay,par-1,shot,caup,salm,bi,cmb,fru,step,ap,jing,Myc,ck,Dys,fbl,ct,pk,Diap1,rl,pan,gus,fuss,tld,cic,bab2,Trl,zld,en,lola,Tak1 | | 4.787e-06 | -12.25 | post-embryonic animal organ morphogenesis | biological process | GO:0048563 | 420 | 41 | 12345 | 571 | bi,Myc,ck,jing,ap,cmb,step,fru,pk,ct,fbl,Dys,gus,pan,rl,Diap1,tld,fuss,Trl,cic,bab2,lola,en,zld,Tak1,heph,Src42A,Hipk,sn,hth,mbc,cv-2,wge,ewg,shot,par-1,Cip4,ImpE2,tay,salm,caup | | 5.032e-06 | -12.20 | animal organ morphogenesis | biological process | GO:0009887 | 782 | 64 | 12345 | 571 | tj,Myc,ap,cmb,br,ct,Dys,fbl,gus,svp,tld,rg,fz2,bab2,toy,rau,rno,rib,Hipk,ru,heph,Atpalpha,cv-2,ewg,RhoGAP100F,shot,ImpE2,klg,salm,hdc,caup,CadN2,bi,jing,ck,step,fru,pk,wb,sdk,Diap1,rl,pan,fuss,kirre,Trl,drosha,dnt,cic,en,lola,zld,fred,Tak1,lz,boi,Src42A,hth,sn,mbc,wge,Cip4,par-1,tay | | 5.074e-06 | -12.19 | post-embryonic animal organ development | biological process | GO:0048569 | 481 | 45 | 12345 | 571 | par-1,Cip4,tay,ImpE2,shot,wge,ewg,caup,salm,Pkg21D,lz,cv-2,mbc,sn,hth,heph,Src42A,Hipk,bab2,cic,Trl,fuss,kirre,tld,Tak1,zld,en,lola,ap,cmb,step,fru,Myc,ck,jing,bi,pan,Diap1,rl,gus,fbl,Dys,Msp300,pk,ct | | 5.548e-06 | -12.10 | imaginal disc development | biological process | GO:0007444 | 591 | 52 | 12345 | 571 | par-1,tay,ImpE2,Cip4,Ptip,shot,lama,wge,ewg,caup,salm,fray,boi,lz,Idgf4,mbc,cv-2,sn,hth,ru,heph,Hipk,Src42A,bab2,cic,Trl,fuss,tld,Tak1,Nipped-A,zld,lola,en,toy,dsx,ap,step,fru,cmb,ck,Myc,jing,bi,pan,rl,Diap1,gus,fbl,Dys,pk,pico,ct | | 5.665e-06 | -12.08 | epithelium development | biological process | GO:0060429 | 1020 | 78 | 12345 | 571 | lz,Parp,boi,fray,Src42A,sn,hth,mbc,wge,capu,spo,lama,Ptip,Cip4,par-1,tay,CG17211,knk,CadN2,bi,ck,jing,step,fru,pk,pico,pan,Dlg5,Diap1,rl,fuss,sna,Trl,cic,lola,en,zld,fred,Nipped-A,Tak1,AP-2alpha,mim,Idgf4,cta,rib,ru,lncRNA:flam,heph,Hipk,dimm,cv-2,ewg,Mef2,shot,CG41099,ImpE2,toc,hdc,salm,caup,jvl,neo,tj,Myc,cmb,ap,ct,br,fbl,Dys,gus,tld,svp,nullo,orb,bab2,toy,dsx | | 5.970e-06 | -12.03 | post-embryonic animal morphogenesis | biological process | GO:0009886 | 484 | 45 | 12345 | 571 | heph,Src42A,Hipk,mbc,cv-2,sn,hth,grim,caup,hdc,salm,wge,ewg,ImpE2,tay,Cip4,par-1,shot,fbl,Dys,pk,ct,br,pan,Diap1,rl,gus,bi,step,fru,cmb,ap,Myc,ck,jing,zld,en,lola,Tak1,fuss,tld,bab2,cic,drosha,Trl | | 6.208e-06 | -11.99 | morphogenesis of an epithelium | biological process | GO:0002009 | 625 | 54 | 12345 | 571 | hdc,salm,CG17211,knk,caup,wge,ewg,spo,shot,par-1,Cip4,ImpE2,tay,heph,Hipk,Src42A,sn,hth,mbc,cv-2,Parp,cta,rib,lola,en,zld,fred,Tak1,tld,svp,fuss,nullo,orb,sna,Trl,bab2,cic,pk,ct,fbl,Dys,gus,pan,rl,Diap1,CadN2,bi,ck,Myc,jing,ap,cmb,step,fru | | 6.371e-06 | -11.96 | instar larval or pupal morphogenesis | biological process | GO:0048707 | 470 | 44 | 12345 | 571 | bi,jing,ck,Myc,fru,ap,cmb,step,br,pk,ct,Dys,fbl,gus,Diap1,rl,pan,tld,fuss,Trl,drosha,bab2,cic,en,lola,zld,Tak1,Src42A,Hipk,heph,hth,sn,mbc,cv-2,ewg,wge,shot,ImpE2,tay,Cip4,par-1,salm,hdc,caup | | 6.403e-06 | -11.96 | animal organ development | biological process | GO:0048513 | 1267 | 92 | 12345 | 571 | Ptip,zen,tay,par-1,Cip4,wge,lama,spo,Pkg21D,fray,scw,boi,fdl,cato,lz,hth,NAT1,sn,mbc,Src42A,Trl,drosha,cic,dnt,sna,fuss,kirre,Nipped-A,fred,Tak1,lola,en,zld,jing,Psc,ck,E5,step,fru,CadN2,bi,wb,sdk,Diap1,rl,pan,pk,pico,sif,Msp300,shot,RhoGAP100F,ImpE2,Mef2,ewg,salm,Git,hdc,caup,klg,rno,rib,Idgf4,rau,Ten-a,Atpalpha,dimm,cv-2,Nna1,Sema1a,Hipk,ru,heph,rg,fz2,dati,bab2,svp,tld,toy,dsx,CAP,Myc,vap,ap,cmb,tj,gus,br,ct,Dys,fbl | | 7.208e-06 | -11.84 | tissue development | biological process | GO:0009888 | 1113 | 83 | 12345 | 571 | nullo,orb,tld,svp,bab2,bt,toy,dsx,neo,tj,ap,cmb,Myc,fbl,Dys,ct,br,gus,Mef2,ewg,ImpE2,shot,CG41099,toc,caup,jvl,hdc,salm,mim,Idgf4,rib,cta,lncRNA:flam,heph,ru,Hipk,Sema1a,cv-2,dimm,fuss,sna,cic,Trl,zld,lola,en,Tak1,AP-2alpha,Nipped-A,fred,bi,CadN2,step,fru,ck,jing,Msp300,pico,pk,pan,Dlg5,Diap1,rl,wb,spo,lama,wge,capu,Cip4,tay,par-1,Ptip,knk,CG17211,lz,Parp,scw,fray,boi,Src42A,mbc,sn,hth | | 7.618e-06 | -11.78 | neuron differentiation | biological process | GO:0030182 | 710 | 59 | 12345 | 571 | dnc,rno,boi,Dscam2,rau,lz,hth,Ten-a,Ulp1,sn,Spn,Src42A,Sema1a,ru,RhoGAP100F,shot,Cip4,par-1,Ptp99A,shep,salm,hdc,pho,Stlk,klg,jing,Psc,ck,vtd,E5,cmb,ap,fru,CadN2,Mical,Toll-7,wb,rl,Sdc,br,ct,pk,CdGAPr,scro,rg,fz2,dati,Mp,dnt,svp,sky,fuss,Ca-alpha1D,pum,SoxN,nrm,dsx,lola,en | | 1.163e-05 | -11.36 | multicellular organism development | biological process | GO:0007275 | 2496 | 157 | 12345 | 571 | boi,fray,scw,dnc,lz,cato,NAT1,fan,CG17387,Npc2b,Spn,zen,Cip4,ect,Ptip,lama,spo,wge,pho,knk,CG17211,5-HT2A,Pkg21D,E5,jing,Psc,ck,bi,tinc,CadN2,pan,mav,pico,kirre,fred,SoxN,karr,zld,rib,babos,CG9514,grim,Ten-a,ru,Nna1,ImpE2,CG41099,IA-2,Stlk,toc,Nlg2,vap,ap,cmb,Myc,CdGAPr,Sdc,ct,dati,stw,fz2,bab2,scro,rg,nullo,sky,tld,nrm,pum,dsx,toy,Parp,fdl,mlt,mbc,hth,Ulp1,sn,Src42A,par-1,tay,capu,spok,vtd,step,fru,Mical,Diap1,rl,wb,sdk,sif,Msp300,pk,KCNQ,drosha,cpx,cic,dnt,Trl,sna,fuss,Ca-alpha1D,Tak1,d4,Nipped-A,en,lola,rno,cta,Clk,rau,Thor,Idgf4,Dscam2,Cpr49Ac,CycB,cv-2,Atpalpha,dimm,Hipk,Sema1a,Coq7,heph,RhoGAP100F,shot,Ptp99A,shep,Mef2,ewg,jvl,caup,Irp-1A,salm,Git,hdc,klg,mthl11,mthl8,tou,CAP,tj,Toll-7,gus,Dys,Pld,fbl,br,kune,Mp,TwdlG,Sod1,Nlg3,kni,nvd,orb,svp | | 1.469e-05 | -11.13 | tissue morphogenesis | biological process | GO:0048729 | 644 | 54 | 12345 | 571 | Trl,cic,bab2,tld,svp,fuss,sna,nullo,orb,fred,Tak1,lola,en,zld,Myc,ck,jing,fru,ap,cmb,step,CadN2,bi,gus,pan,Diap1,rl,ct,pk,fbl,Dys,shot,par-1,Cip4,tay,ImpE2,wge,ewg,spo,hdc,CG17211,salm,caup,knk,cta,rib,Parp,sn,hth,cv-2,mbc,heph,Hipk,Src42A | | 1.734e-05 | -10.96 | cell differentiation | biological process | GO:0030154 | 1773 | 118 | 12345 | 571 | pho,Bruce,Cip4,par-1,CG32006,lama,spo,capu,wge,mlt,mbc,fan,hth,sn,Tpl94D,Ulp1,Src42A,Spn,dnc,boi,scw,bru1,Parp,lz,AP-2alpha,SoxN,zld,lola,en,drosha,dnt,cic,Trl,sna,Ca-alpha1D,kirre,fuss,DNaseII,rl,Diap1,Dlg5,pan,wb,mav,sdk,sif,Msp300,pk,vtd,E5,fru,Psc,jing,ck,Mical,bi,tinc,CadN2,jvl,caup,salm,hdc,klg,Stlk,toc,CG41099,RhoGAP100F,shot,Ptp99A,shep,Mef2,CycB,Ten-a,dimm,Sema1a,heph,ru,lncRNA:flam,rno,cta,babos,rau,fd96Ca,Thor,tej,Rbp9,Dscam2,grim,mim,Catsup,nrm,pum,mei-P26,dsx,fz2,dati,Mp,scro,bt,rg,orb,svp,sky,tld,gus,CdGAPr,fbl,Sdc,br,kune,ct,ap,cmb,Myc,fzo,tj,Toll-7,neo | | 1.741e-05 | -10.96 | tube morphogenesis | biological process | GO:0035239 | 536 | 47 | 12345 | 571 | Src42A,Hipk,heph,cv-2,mbc,hth,sn,rib,caup,knk,CG17211,salm,hdc,ewg,wge,par-1,tay,Cip4,ImpE2,shot,Dys,fbl,pk,ct,Diap1,rl,pan,gus,bi,ap,fru,step,cmb,jing,ck,Myc,zld,lola,en,Tak1,sna,fuss,svp,tld,cic,bab2,Trl | | 1.887e-05 | -10.88 | signal transduction | biological process | GO:0007165 | 968 | 73 | 12345 | 571 | Myo81F,Gr2a,RabX2,svp,Octbeta2R,nAChRalpha4,fz2,Pld,CdGAPr,gus,Der-1,TpnC41C,SCAP,RSG7,CG31750,Hsc70-3,Toll-7,nAChRalpha3,Myc,CG34357,mthl11,mthl8,spel1,norpA,Atf6,Stlk,shakB,Ac78C,PEK,RhoGAP100F,CG42637,Sema1a,Gprk1,CG5160,GABA-B-R3,grim,Thor,cta,CG41520,HisCl1,Tak1,Ca-alpha1D,cic,dnt,pico,Octbeta1R,mav,Magi,rl,pan,SPR,Pi3K68D,Pkg21D,5-HT2A,Rgk1,CG9389,CG12147,CG42541,TrissinR,RYa,5-HT2B,rdgC,par-1,Spn,Src42A,Ir40a,CG32944,mbc,Rh3,scw,dnc,fray,boi | | 1.936e-05 | -10.85 | imaginal disc-derived appendage morphogenesis | biological process | GO:0035114 | 355 | 35 | 12345 | 571 | caup,salm,tay,Cip4,par-1,shot,cv-2,sn,hth,heph,Hipk,Src42A,lz,zld,en,lola,cic,Trl,fuss,tld,pan,rl,Diap1,fbl,Dys,ct,pk,ap,fru,step,cmb,ck,Myc,jing,bi | | 2.316e-05 | -10.67 | appendage morphogenesis | biological process | GO:0035107 | 358 | 35 | 12345 | 571 | zld,en,lola,fuss,tld,cic,Trl,Dys,fbl,pk,ct,rl,Diap1,pan,bi,cmb,fru,step,ap,jing,Myc,ck,caup,salm,Cip4,tay,par-1,shot,Hipk,Src42A,heph,cv-2,hth,sn,lz | | 2.428e-05 | -10.63 | Ig-like\_dom\_sf | interpro domains | IPR036179 | 133 | 19 | 11383 | 557 | beat-VII,bt,DIP-delta,sdk,zormin,CG14964,Ptp99A,Dscam2,fred,CG45263,side-II,DIP-lambda,nrm,side-III,kirre,klg,boi,dpr21,babos | | 2.592e-05 | -10.56 | Ig\_sub | interpro domains | IPR003599 | 111 | 17 | 11383 | 557 | dpr21,boi,klg,babos,DIP-lambda,side-II,CG45263,fred,kirre,side-III,nrm,CG14964,Dscam2,zormin,sdk,DIP-delta,bt | | 2.696e-05 | -10.52 | movement of cell or subcellular component | biological process | GO:0006928 | 724 | 58 | 12345 | 571 | par-1,Klp59C,RhoGAP100F,shot,Ptp99A,jvl,salm,klg,Lsd-2,rib,cta,CG1486,mim,Dscam2,mbc,CycB,CG9492,CG17387,sn,Ten-a,ru,Src42A,Sema1a,Khc-73,Mp,dnt,fz2,kni,SoxN,pum,CG5359,lola,en,dsx,l(2)41Ab,ap,fru,ck,Psc,jing,Pi3K68D,Toll-7,neo,tj,Mical,CadN2,Diap1,Dlg5,gus,wb,Msp300,CdGAPr,pk,ct,Dhc16F,Sdc,CG31275,kl-3 | | 2.870e-05 | -10.46 | CD80\_C2-set | interpro domains | IPR013162 | 20 | 7 | 11383 | 557 | side-II,fred,CG45263,beat-VII,nrm,kirre,side-III | | 2.921e-05 | -10.44 | wing disc development | biological process | GO:0035220 | 407 | 38 | 12345 | 571 | tld,fuss,Trl,cic,en,lola,zld,Nipped-A,Tak1,bi,ck,Myc,jing,fru,cmb,step,ap,ct,pk,fbl,Dys,gus,pan,rl,shot,par-1,Cip4,tay,salm,caup,boi,fray,heph,Src42A,Hipk,sn,hth,cv-2 | | 2.927e-05 | -10.44 | imaginal disc-derived appendage development | biological process | GO:0048737 | 362 | 35 | 12345 | 571 | ct,pk,Dys,fbl,Diap1,rl,pan,bi,jing,Myc,ck,cmb,ap,fru,step,en,lola,zld,tld,fuss,Trl,cic,Src42A,Hipk,heph,hth,sn,cv-2,lz,salm,caup,shot,Cip4,tay,par-1 | | 3.188e-05 | -10.35 | generation of neurons | biological process | GO:0048699 | 880 | 67 | 12345 | 571 | par-1,Cip4,pho,lz,boi,dnc,Src42A,Spn,hth,Ulp1,sn,sna,fuss,Ca-alpha1D,dnt,zld,lola,en,SoxN,Mical,tinc,CadN2,E5,vtd,fru,Psc,jing,ck,sif,pk,rl,wb,mav,sdk,Ptp99A,shep,RhoGAP100F,shot,Stlk,klg,salm,hdc,rau,Dscam2,rno,babos,Sema1a,ru,Ten-a,svp,sky,fz2,dati,Mp,scro,rg,dsx,nrm,pum,Toll-7,cmb,ap,Myc,CdGAPr,br,Sdc,ct | | 3.292e-05 | -10.32 | negative regulation of biological process | biological process | GO:0048519 | 1502 | 102 | 12345 | 571 | Maf1,kni,dnr1,orb,tld,HPS4,sky,svp,CG12772,Fer3,Slob,pum,dsx,Ars2,norpA,spel1,vap,CG1620,CG9380,CG9932,Tgi,Hsc70-3,ct,THADA,PNPase,br,CG10077,SmydA-9,shot,Spn75F,ewg,Marf1,caup,bru3,hdc,salm,CG13284,Vsx2,Nlg2,rno,Clk,CG9650,Thor,Su(z)2,mim,grim,tej,Gprk1,heph,Sema1a,Hipk,cic,drosha,cpx,mrt,Trl,Mnt,fuss,sna,lncRNA:bxd,CG41520,SoxN,d4,zld,Spn42De,en,conu,E5,alpha4GT1,Psc,jing,HP6,Pi3K68D,bi,Parg,pan,rl,Diap1,sdk,Octbeta1R,lqf,Slmap,tay,par-1,Cip4,zen,SmydA-8,RYa,pho,CG12147,Rgk1,Bruce,Lsd-2,scw,dnc,AGO3,bru1,lz,Ulp1,Src42A,Spn,Ppm1 | | 3.441e-05 | -10.28 | locomotion | biological process | GO:0040011 | 664 | 54 | 12345 | 571 | svp,Mp,dnt,fz2,kni,en,lola,dsx,SoxN,HisCl1,pum,Toll-7,Pi3K68D,tj,Mical,CadN2,norpA,fru,ap,Psc,jing,fbl,CdGAPr,Msp300,Dhc16F,ct,br,Sdc,Diap1,Dlg5,gus,wb,shep,rdgC,Ptp99A,par-1,shot,RhoGAP100F,klg,l(3)L1231,salm,CG1486,mim,Dscam2,rib,cta,ru,Hipk,Src42A,Sema1a,mbc,sn,CG17387,Ten-a | | 3.456e-05 | -10.27 | C2-set\_2 | pfam domains | PF08205 | 20 | 7 | 10480 | 528 | nrm,side-II,fred,beat-VII,kirre,CG45263,side-III | | 3.551e-05 | -10.25 | fly chr41D3-41D3\|2-55 cM | chromosome location | fly chr41D3-41D3\|2-55 cM | 3 | 3 | 25010 | 823 | gus,CG3107,lncRNA:CR43256 | | 3.551e-05 | -10.25 | fly chr41D3-41D3\|2-55 | chromosome location | fly chr41D3-41D3\|2-55 | 3 | 3 | 25010 | 823 | gus,lncRNA:CR43256,CG3107 | | 3.640e-05 | -10.22 | cell periphery | cellular component | GO:0071944 | 1409 | 94 | 11940 | 534 | Or67d,Octbeta2R,Ten-a,TrissinR,Cip4,CG9981,capu,5-HT2A,CG1213,Teh2,norpA,conu,step,fdl,Dys,Dlg5,Rh3,Ca-alpha1D,Nlg3,Ac78C,CG7536,mbc,Syt7,Git,PMCA,CG42541,Mical,Slob,SK,Octbeta1R,Oatp33Ea,kirre,CG33970,Dscam2,Rift,par-1,Or98a,CG8249,ck,CG34357,Or69a,Toll-7,ru,CG15449,Gprk1,DIP-delta,fz2,Atpalpha,HisCl1,KCNQ,SdhA,CG31342,Pi3K68D,GABA-B-R3,CG42637,Pzl,fbl,nAChRalpha4,Snap25,CG2061,cta,klg,AP-2alpha,CG15570,Ptp99A,dnt,CG15408,nAChRalpha3,Tsp5D,Magi,Syx18,igl,Or83c,Nlg2,pk,Rgk1,sdk,Ca-Ma2d,CG31689,sbm,CG8888,Sdc,DIP-lambda,Src42A,CG2003,shakB,nrm,gus,Spn,toc,CadN2,SPR,fan,5-HT2B | | 3.681e-05 | -10.21 | appendage development | biological process | GO:0048736 | 366 | 35 | 12345 | 571 | shot,par-1,tay,Cip4,salm,caup,lz,heph,Hipk,Src42A,sn,hth,cv-2,tld,fuss,Trl,cic,lola,en,zld,bi,ck,Myc,jing,ap,cmb,step,fru,pk,ct,fbl,Dys,pan,Diap1,rl | | 4.277e-05 | -10.06 | developmental process | biological process | GO:0032502 | 2988 | 179 | 12345 | 571 | lz,cato,fray,boi,scw,dnc,bru1,Npc2b,Spn,CG17387,NAT1,fan,spo,lama,wge,Cip4,zen,Ptip,CG32006,ect,Pkg21D,Bruce,knk,pho,CG17211,5-HT2A,tinc,bi,Parg,CadN2,E5,ck,jing,Psc,pico,pan,Dlg5,mav,kirre,DNaseII,zld,AP-2alpha,fred,karr,SoxN,CG9514,mim,grim,rib,babos,ru,Nna1,Ten-a,ImpE2,IA-2,CG41099,toc,Stlk,Nlg2,Vsx2,cmb,ap,vap,Myc,CdGAPr,ct,Sdc,nullo,tld,sky,bab2,stw,fz2,dati,bt,rg,scro,dsx,toy,nrm,pum,Parp,fdl,Src42A,mbc,mlt,Ulp1,sn,Tpl94D,hth,capu,tay,par-1,spok,Mical,fru,step,vtd,Msp300,sif,pk,rl,Diap1,sdk,wb,fuss,Ca-alpha1D,sna,cic,dnt,drosha,cpx,KCNQ,Trl,en,lola,Tak1,Yp2,Nipped-A,d4,fd96Ca,Thor,rau,Cpr49Ac,Idgf4,Dscam2,tej,Rbp9,rno,Clk,cta,lncRNA:flam,Coq7,heph,Sema1a,Hipk,cv-2,CycB,Atpalpha,dimm,shep,Ptp99A,Mef2,ewg,shot,RhoGAP100F,klg,Irp-1A,caup,jvl,hdc,Git,salm,neo,Toll-7,fzo,tj,mthl8,mthl11,tou,CAP,fbl,Dys,Pld,kune,br,gus,orb,nvd,svp,Mp,TwdlG,Nlg3,kni,Sod1,cpo,mei-P26,mxt,Catsup | | 5.109e-05 | -9.88 | multicellular organismal process | biological process | GO:0032501 | 3844 | 221 | 12345 | 571 | SPR,Mical,vtd,step,fru,twz,sif,Msp300,pk,Diap1,rl,Obp85a,wb,sdk,CheB42a,sna,Ca-alpha1D,fuss,drosha,KCNQ,cpx,dnt,cic,Trl,en,lola,Tak1,d4,Nipped-A,Parp,Rh3,fdl,bond,Gdh,Src42A,Ir40a,mlt,mbc,hth,sn,Tpl94D,Ulp1,capu,par-1,tay,spok,Or98a,fzo,Hsc70-3,tj,Toll-7,CG11630,mthl8,mthl11,norpA,tou,CAP,Pld,Dys,fbl,br,kune,fest,gus,Octbeta2R,orb,nvd,svp,TwdlG,Mp,dnr1,Sod1,kni,Nlg3,mei-P26,Mst87F,mxt,Catsup,CG4546,rau,Thor,Cpr49Ac,tej,Dscam2,Rbp9,Idgf4,rno,cta,Clk,Or83c,Hipk,Sema1a,heph,Coq7,lncRNA:flam,lncRNA:CR44344,CycB,CG9492,be,cv-2,GABA-B-R3,dimm,Atpalpha,Ptp99A,shep,Mef2,ewg,shot,RhoGAP100F,klg,jvl,Irp-1A,caup,SKIP,Git,salm,shakB,hdc,bi,tinc,Pgant9,CadN2,2mit,Parg,E5,jing,HP6,Psc,ck,pico,Dlg5,pan,mav,kirre,DNaseII,zld,AP-2alpha,fred,HisCl1,karr,SoxN,lz,cato,fray,boi,scw,dnc,bru1,Gr93d,Npc2b,Spn,NAT1,fan,CG17387,rdgC,lama,spo,Sfp77F,wge,zen,Cip4,ect,Ptip,Pkg21D,Bruce,pho,knk,CG17211,5-HT2A,Rgk1,CG31750,vap,ap,cmb,nAChRalpha3,Or69a,Myc,CdGAPr,Sdc,ct,nullo,sky,tld,fz2,dati,stw,bab2,scro,bt,nAChRalpha4,rg,snky,toy,dsx,inc,Acp36DE,CG30486,nrm,Gr2a,pum,CG9514,MCU,grim,mim,rib,babos,lncRNA:CR44412,Tdc2,Gr39a,ru,Nna1,Ten-a,Or67d,ImpE2,Spn75F,CG41099,IA-2,toc,Stlk,Nlg2,Ac78C | | 5.224e-05 | -9.86 | sex differentiation | biological process | GO:0007548 | 100 | 15 | 12345 | 571 | fz2,scw,rib,bab2,fru,tj,lz,salm,wb,Yp2,dsx,br,ct,lola,en | | 5.383e-05 | -9.83 | cellular developmental process | biological process | GO:0048869 | 1818 | 118 | 12345 | 571 | scro,rg,bt,dati,fz2,Mp,sky,svp,tld,orb,pum,Catsup,nrm,dsx,mei-P26,Myc,ap,cmb,tj,fzo,neo,Toll-7,gus,Sdc,br,ct,kune,CdGAPr,fbl,CG41099,shot,RhoGAP100F,Mef2,Ptp99A,shep,salm,hdc,jvl,caup,klg,toc,Stlk,cta,babos,rno,Dscam2,tej,grim,Rbp9,mim,rau,Thor,fd96Ca,Ten-a,dimm,CycB,Sema1a,heph,ru,lncRNA:flam,Trl,drosha,cic,dnt,DNaseII,sna,Ca-alpha1D,kirre,fuss,SoxN,AP-2alpha,lola,en,zld,jing,Psc,ck,E5,vtd,fru,CadN2,Mical,bi,tinc,wb,sdk,mav,Dlg5,rl,Diap1,pan,pk,sif,Msp300,CG32006,par-1,Cip4,capu,wge,lama,spo,pho,Bruce,bru1,dnc,scw,boi,lz,Parp,fan,hth,sn,Ulp1,Tpl94D,mlt,mbc,Spn,Src42A | | 5.796e-05 | -9.76 | anatomical structure development | biological process | GO:0048856 | 2924 | 175 | 12345 | 571 | Catsup,mei-P26,Mp,TwdlG,kni,Nlg3,Sod1,nvd,orb,svp,gus,fbl,Dys,Pld,kune,br,mthl8,mthl11,CAP,tou,Toll-7,neo,tj,fzo,Irp-1A,caup,jvl,hdc,Git,salm,klg,shot,RhoGAP100F,shep,Ptp99A,Mef2,ewg,cv-2,CycB,dimm,Atpalpha,Coq7,lncRNA:flam,heph,Sema1a,Hipk,rno,Clk,cta,Thor,fd96Ca,rau,tej,Cpr49Ac,Dscam2,Rbp9,Idgf4,Tak1,Nipped-A,d4,lola,en,dnt,cic,KCNQ,cpx,drosha,Trl,fuss,Ca-alpha1D,sna,Diap1,rl,sdk,wb,Msp300,sif,pk,step,fru,vtd,Mical,spok,tay,par-1,capu,mbc,mlt,Tpl94D,Ulp1,sn,hth,Src42A,Parp,fdl,nrm,pum,toy,dsx,bab2,dati,stw,fz2,rg,bt,scro,nullo,tld,sky,CdGAPr,ct,Sdc,vap,cmb,ap,Myc,toc,Stlk,Nlg2,Vsx2,ImpE2,IA-2,CG41099,Ten-a,ru,Nna1,rib,babos,CG9514,mim,grim,AP-2alpha,SoxN,fred,karr,zld,kirre,DNaseII,pan,Dlg5,mav,pico,E5,ck,Psc,jing,tinc,bi,CadN2,knk,pho,CG17211,5-HT2A,Pkg21D,Bruce,Cip4,zen,Ptip,CG32006,ect,spo,lama,wge,CG17387,NAT1,fan,Npc2b,Spn,boi,dnc,scw,fray,bru1,lz,cato | | 5.975e-05 | -9.73 | binding | molecular function | GO:0005488 | 5743 | 306 | 12176 | 554 | Tdc2,CG42637,Hmt-1,Takl2,kni,rl,dimm,spok,Ranbp16,CG5862,Ubr1,Npc2b,Ten-a,Trxr-2,CG17211,RpL15,CG32006,CG5897,CG34334,Cyp6a8,Su(z)2,CG30094,CG3588,CG40813,scro,mav,Slmap,Parp,heph,ABCB7,capu,CadN2,Trl,ck,CG42813,CG10077,Atf6,CG14301,CG2256,thw,rib,Myc,THADA,sif,CG9044,cato,cpx,CG30440,CG9747,Or83c,pico,Or69a,Clk,Toll-7,Nadsyn,Hsc70-3,d4,Nna1,CG12200,vtd,CG9492,Src42A,par-1,CG8219,Cip4,cbs,CG1620,dsx,AP-2alpha,Meltrin,Acp36DE,CG9650,lz,Tbc1d15-17,Sema1a,igl,CG7029,CG9514,kl-3,Dscam2,CG5160,CG1486,Mef2,SmydA-9,Syx18,CG43783,alpha-Man-IIb,fray,su(w[a]),Pld,CG3556,spo,CG31703,RabX2,Klp59C,br,dnr1,CG33557,NAT1,Marf1,mRpS5,ewg,svp,lola,Psc,Scp1,fz2,mim,Rbp9,cpo,TpnC41C,Glut4EF,Obp85a,CG12147,CG33970,inc,Or98a,CG8539,AGO3,CG10565,CG12769,mlt,shep,shot,mrt,CG32944,CheB42a,tou,CG17683,Sod1,Magi,CG10188,Thor,Acbp5,scw,Rgk1,bab2,step,Maf1,Nlg2,RpL22-like,CG34357,CG15725,Diap1,prage,CG17361,CG41562,5-HT2A,CG3107,CG41099,Slob,wge,Pkg21D,Mp,rg,CG42402,Ars2,cic,CG30059,ct,CG32772,zen,Acox57D-d,Mur89F,Set1,godzilla,bru1,E5,Syt7,rdgC,Or67d,Der-1,sna,Stlk,Cht10,CG31689,bru3,orb,Myo81F,Spn,mIF2,Rpb12,conu,sn,CG17514,Cyp6g2,FASN3,CG14184,fzo,pan,SoxN,zormin,spel1,CG40228,Khc-73,Nlg3,Lsd-2,CG5789,Snap25,salm,pk,CG31275,snky,CG9380,grim,hth,boi,Atpalpha,CG34347,RpL6,SK,vap,CG14891,cv-2,nullo,CG9107,bma,Svil,fd96Ca,jing,HP6,SmydA-8,fbl,Msp300,Mical,PNPase,cmb,Hipk,Tak1,cert,arg,pum,jigr1,Fer3,CG44002,pho,mxt,zld,CycB,Vsx2,CG5270,CG7536,CG14304,CG42255,en,Idgf4,ap,Cyp316a1,cta,Amyrel,Tgi,bi,norpA,caup,Gprk1,CG14820,tld,CG9743,Git,Nrd1,toy,Gfat1,mei-P26,dati,CG43672,stw,Camta,CG14721,bt,CG10939,dnt,eIF4B,PMCA,nvd,Trs23,Dys,mbc,gus,Dhc16F,PEK,Pi3K68D,Scp2,KCNQ,mus308,SCAP,CG9932,fru,fuss,RYa,GCS2alpha,Cyp309a1,drosha,SdhA,Ppm1,DNAlig4,tj,CG9981,fan,rau,Mnt,CG42541,Alg-2,CG5359,kirre,Gdh,Irp-1A | | 6.098e-05 | -9.71 | post-embryonic appendage morphogenesis | biological process | GO:0035120 | 345 | 33 | 12345 | 571 | heph,Src42A,Hipk,sn,hth,cv-2,salm,caup,shot,par-1,tay,Cip4,pk,ct,fbl,Dys,pan,rl,bi,Myc,ck,jing,step,fru,cmb,ap,en,lola,zld,tld,fuss,Trl,cic | | 7.158e-05 | -9.54 | DNA-binding transcription factor activity | molecular function | GO:0003700 | 463 | 40 | 12176 | 554 | fd96Ca,ewg,svp,CG32006,bab2,toy,CG9932,fru,sna,CG33557,caup,bi,Octbeta2R,Mnt,SoxN,scro,pan,CG9650,lola,tj,dsx,Trl,kni,Mef2,Vsx2,zld,cic,Atf6,zen,Camta,Fer3,lz,E5,ap,Clk,CG12769,hth,rib,Myc,en | | 7.389e-05 | -9.51 | Ig-like\_dom | interpro domains | IPR007110 | 132 | 18 | 11383 | 557 | side-III,kirre,nrm,CG45263,fred,DIP-lambda,side-II,babos,klg,dpr21,boi,bt,sdk,zormin,DIP-delta,beat-VII,Dscam2,CG14964 | | 7.549e-05 | -9.49 | Ig\_sub2 | interpro domains | IPR003598 | 109 | 16 | 11383 | 557 | side-II,DIP-lambda,fred,CG45263,nrm,kirre,side-III,boi,dpr21,klg,DIP-delta,zormin,sdk,bt,CG14964,Dscam2 | | 7.719e-05 | -9.47 | neurogenesis | biological process | GO:0022008 | 923 | 68 | 12345 | 571 | br,Sdc,ct,kune,CdGAPr,Myc,cmb,ap,Toll-7,pum,nrm,dsx,scro,rg,fz2,dati,Mp,sky,svp,Ten-a,Sema1a,ru,babos,rno,Dscam2,rau,salm,hdc,Stlk,klg,RhoGAP100F,shot,Ptp99A,shep,wb,sdk,mav,rl,pk,sif,jing,Psc,ck,vtd,E5,fru,CadN2,Mical,tinc,SoxN,lola,en,zld,dnt,sna,Ca-alpha1D,fuss,hth,Ulp1,sn,Spn,Src42A,boi,dnc,lz,pho,par-1,Cip4 | | 7.759e-05 | -9.46 | synapse | cellular component | GO:0045202 | 285 | 28 | 11940 | 534 | Magi,rl,Syx18,Catsup,RhoGAP100F,Nlg2,Syt7,Rgk1,sif,Dys,Snap25,Spn,Nlg3,nAChRalpha3,Sdc,sky,nAChRalpha4,par-1,unc-13-4A,mav,cpx,Ten-a,Git,orb,Mical,Cip4,pum,HisCl1 | | 8.022e-05 | -9.43 | taxis | biological process | GO:0042330 | 365 | 34 | 12345 | 571 | l(3)L1231,klg,RhoGAP100F,shot,shep,rdgC,Ptp99A,Ten-a,Src42A,Hipk,Sema1a,Dscam2,HisCl1,SoxN,lola,en,dsx,dnt,Mp,fz2,svp,wb,ct,Sdc,br,CdGAPr,Psc,jing,ap,norpA,fru,CadN2,Toll-7,Mical | | 1.012e-04 | -9.20 | female sex differentiation | biological process | GO:0046660 | 25 | 7 | 12345 | 571 | tj,lz,en,ct,dsx,bab2,fz2 | | 1.063e-04 | -9.15 | GSE3982\_DC\_VS\_BCELL\_DN | MSigDB lists | GSE3982\_DC\_VS\_BCELL\_DN | 39 | 8 | 4161 | 164 | Khc-73,Nipped-A,Git,step,Ars2,ABCB7,RpL6,Parp | | 1.100e-04 | -9.11 | identical protein binding | molecular function | GO:0042802 | 159 | 19 | 12176 | 554 | dsx,cbs,rau,mim,step,grim,Gdh,Meltrin,bru3,Sod1,Camta,Ten-a,Khc-73,Atf6,inc,svp,Dscam2,Trl,CadN2 | | 1.178e-04 | -9.05 | reproductive system development | biological process | GO:0061458 | 84 | 13 | 12345 | 571 | bab2,rib,scw,fz2,tj,lz,salm,wb,en,ct,lola,dsx,br | | 1.178e-04 | -9.05 | reproductive structure development | biological process | GO:0048608 | 84 | 13 | 12345 | 571 | bab2,fz2,scw,rib,tj,lz,salm,wb,lola,en,ct,br,dsx | | 1.335e-04 | -8.92 | development of primary sexual characteristics | biological process | GO:0045137 | 53 | 10 | 12345 | 571 | ct,en,lola,wb,salm,tj,rib,scw,fz2,bab2 | | 1.335e-04 | -8.92 | gonad development | biological process | GO:0008406 | 53 | 10 | 12345 | 571 | ct,en,lola,salm,wb,tj,bab2,rib,scw,fz2 | | 1.440e-04 | -8.85 | neuron development | biological process | GO:0048666 | 584 | 47 | 12345 | 571 | Psc,jing,vtd,fru,ap,CadN2,Mical,Toll-7,wb,br,Sdc,ct,CdGAPr,scro,rg,fz2,dati,Mp,dnt,sky,fuss,Ca-alpha1D,pum,SoxN,nrm,dsx,lola,en,rno,dnc,Dscam2,rau,lz,Ten-a,sn,Ulp1,Spn,Src42A,Sema1a,shot,RhoGAP100F,Ptp99A,shep,hdc,pho,Stlk,klg | | 1.566e-04 | -8.76 | negative regulation of metabolic process | biological process | GO:0009892 | 823 | 61 | 12345 | 571 | Hsc70-3,bi,Tgi,Parg,spel1,Ars2,CG1620,E5,Psc,CG9932,HP6,jing,ct,PNPase,br,pan,Diap1,rl,Mnt,orb,sna,HPS4,tld,svp,cic,Maf1,drosha,kni,dnr1,mrt,Trl,Spn42De,zld,en,dsx,Fer3,CG12772,lncRNA:bxd,pum,d4,Su(z)2,Thor,lz,scw,AGO3,bru1,Clk,heph,Ppm1,Marf1,zen,CG10077,SmydA-9,SmydA-8,Spn75F,Bruce,Vsx2,Lsd-2,bru3,pho,hdc | | 1.705e-04 | -8.68 | cell development | biological process | GO:0048468 | 1436 | 95 | 12345 | 571 | mav,wb,rl,Diap1,Dlg5,Msp300,jing,Psc,fru,vtd,CadN2,Mical,SoxN,AP-2alpha,lola,en,Trl,cic,dnt,drosha,DNaseII,Ca-alpha1D,fuss,Tpl94D,Ulp1,sn,fan,mbc,mlt,Spn,Src42A,bru1,scw,dnc,Parp,lz,pho,Bruce,Cip4,par-1,capu,spo,lama,gus,kune,ct,Sdc,br,fbl,CdGAPr,ap,neo,Toll-7,fzo,tj,pum,nrm,Catsup,dsx,mei-P26,rg,bt,scro,Mp,fz2,dati,tld,svp,sky,orb,Ten-a,CycB,heph,lncRNA:flam,Sema1a,cta,rno,mim,Rbp9,Dscam2,tej,grim,rau,hdc,salm,jvl,Stlk,toc,klg,RhoGAP100F,shot,CG41099,Mef2,shep,Ptp99A | | 1.755e-04 | -8.65 | CG15370 (uncharacterized protein) | protein interactions | 31864 | 18 | 6 | 9204 | 464 | CG13492,CG31626,CG3081,rau,Mst87F,fest | | 1.757e-04 | -8.65 | membrane | cellular component | GO:0016020 | 2722 | 157 | 11940 | 534 | rg,MFS16,fan,sdk,sbm,Sdc,Ir94e,Tim23,Src42A,kune,bru1,CG2061,Snap25,bond,ND-AGGG,Ptp99A,Tsp5D,Catsup,Nlg2,alpha-Man-IIb,pk,snky,SCAP,HisCl1,SdhA,CG42637,fbl,CG33233,nAChRalpha4,CG8249,CG34357,CG45076,THADA,ND-30,Meltrin,ru,neo,PMCA,CG7255,CG42541,Coq7,Dscam2,Rift,ABCB7,Ir41a,CG14184,par-1,CG12121,Dlg5,Rh3,Ca-alpha1D,Atf6,Nlg3,CG41128,mbc,Syt7,HPS4,Octbeta2R,Or67d,CG9747,CG9981,CG1213,step,fdl,CG9743,Gr93d,CG14160,CG14196,Ir60e,toc,CadN2,MCU,SPR,Mct1,5-HT2B,RabX2,cpx,Hmt-1,Ca-Ma2d,CG45782,CG31689,CG40472,CG8888,DIP-lambda,CG2003,Ulp1,shakB,nrm,cta,klg,AP-2alpha,CG15408,Pgant9,dnt,ImpE2,CG40006,nAChRalpha3,Magi,Syx18,igl,Gr39a,Or83c,CG15459,Msp300,Rgk1,UQCR-11,Gprk1,CG15449,DIP-delta,ct,Atpalpha,fz2,CG41106,Tim17b,KCNQ,GABA-B-R3,CG31342,Pi3K68D,Pzl,Sema1a,ArfGAP1,Or98a,Glut1,Or69a,Toll-7,CG10171,CG5789,ND-MLRQ,CG43672,Mical,Slob,SK,Octbeta1R,kirre,Oatp33Ea,CG33970,mthl8,Dys,shot,Ac78C,CG7536,CG14024,mthl11,Ten-a,TrissinR,CG10345,Cip4,CG14744,Hsc70-3,CG5160,5-HT2A,Teh2,norpA,Gr2a,Ir40a,fzo,Der-1 | | 1.829e-04 | -8.61 | signaling receptor activity | molecular function | GO:0038023 | 419 | 36 | 12176 | 554 | 5-HT2B,Or83c,SK,Or67d,Ptp99A,Toll-7,Or69a,shakB,Gr39a,Gr93d,CG9380,boi,HisCl1,Or98a,nAChRalpha4,GABA-B-R3,Nlg3,klg,CG42637,dnt,Gr2a,Ir40a,5-HT2A,nAChRalpha3,Octbeta2R,CG31750,Rh3,mthl11,Nlg2,fz2,CG34357,Octbeta1R,TrissinR,svp,mthl8,SPR | | 1.885e-04 | -8.58 | TCF1P\_Q6 | MSigDB lists | TCF1P\_Q6 | 65 | 10 | 4161 | 164 | Rbp9,CG10077,CG2061,PMCA,toy,mIF2,lqf,Mp,Nlg3,Ac78C | | 1.901e-04 | -8.57 | HFH3\_01 | MSigDB lists | HFH3\_01 | 32 | 7 | 4161 | 164 | Mef2,Rbp9,Dys,vtd,lqf,Pi3K68D,twz | | 1.912e-04 | -8.56 | cuticle pattern formation | biological process | GO:0035017 | 36 | 8 | 12345 | 571 | gus,SoxN,sn,fred,pan,ck,boi,fz2 | | 1.966e-04 | -8.53 | plasma membrane | cellular component | GO:0005886 | 1265 | 83 | 11940 | 534 | ru,Or69a,Toll-7,Or98a,CG8249,CG34357,Rift,Dscam2,CG33970,par-1,Octbeta1R,SK,Oatp33Ea,kirre,CG42541,Mical,Slob,PMCA,mbc,Syt7,Ac78C,CG7536,Nlg3,Dlg5,Dys,Rh3,Ca-alpha1D,CG1213,5-HT2A,norpA,step,Teh2,fdl,TrissinR,Cip4,CG9981,Or67d,Octbeta2R,Ten-a,5-HT2B,SPR,fan,CadN2,CG2003,Src42A,shakB,nrm,CG8888,Sdc,DIP-lambda,Ca-Ma2d,CG31689,sdk,Nlg2,Or83c,pk,Rgk1,Tsp5D,Magi,Ptp99A,dnt,CG15408,nAChRalpha3,Snap25,CG2061,cta,klg,AP-2alpha,nAChRalpha4,GABA-B-R3,CG42637,Pi3K68D,CG31342,Pzl,fbl,DIP-delta,fz2,Atpalpha,HisCl1,KCNQ,SdhA,CG15449,Gprk1 | | 2.121e-04 | -8.46 | neuron projection development | biological process | GO:0031175 | 447 | 38 | 12345 | 571 | pho,hdc,Stlk,klg,shot,RhoGAP100F,Ptp99A,Ten-a,Ulp1,sn,Src42A,Sema1a,Spn,dnc,Dscam2,pum,SoxN,dsx,en,lola,dati,fz2,dnt,Mp,scro,sky,wb,CdGAPr,Sdc,br,ct,fru,ap,Psc,jing,Mical,Toll-7,CadN2 | | 2.609e-04 | -8.25 | central nervous system development | biological process | GO:0007417 | 281 | 27 | 12345 | 571 | en,toy,SoxN,Git,spo,sna,svp,ewg,dati,drosha,rg,shot,ct,mbc,dimm,Ulp1,hth,Ten-a,grim,fdl,fru,vap,ap,rib,E5,jing,Psc | | 3.042e-04 | -8.10 | HNF3ALPHA\_Q6 | MSigDB lists | HNF3ALPHA\_Q6 | 45 | 8 | 4161 | 164 | DNaseII,twz,Rbp9,dnc,Pi3K68D,Mef2,Dys,Cyp6a8 | | 3.124e-04 | -8.07 | molecular transducer activity | molecular function | GO:0060089 | 447 | 37 | 12176 | 554 | shakB,Gr39a,Toll-7,Or69a,Or67d,SK,Ptp99A,5-HT2B,Or83c,boi,CG9380,Gr93d,Or98a,HisCl1,Ir40a,Gr2a,CG42637,dnt,Pkg21D,GABA-B-R3,Nlg3,klg,nAChRalpha4,mthl11,Rh3,Octbeta2R,CG31750,nAChRalpha3,5-HT2A,CG34357,fz2,Nlg2,svp,TrissinR,Octbeta1R,SPR,mthl8 | | 3.162e-04 | -8.06 | IG | smart domains | SM00409 | 111 | 17 | 5086 | 308 | Dscam2,nrm,babos,sdk,side-II,kirre,side-III,dpr21,CG45263,bt,fred,boi,zormin,klg,DIP-lambda,CG14964,DIP-delta | | 3.251e-04 | -8.03 | HLF\_01 | MSigDB lists | HLF\_01 | 57 | 9 | 4161 | 164 | dnc,Ranbp16,Mp,svp,Octbeta2R,bru1,Dys,CG10077,Rbp9 | | 3.621e-04 | -7.92 | receptor serine/threonine kinase binding | molecular function | GO:0033612 | 4 | 3 | 12176 | 554 | cv-2,mav,scw | | 3.621e-04 | -7.92 | transmembrane receptor protein serine/threonine kinase binding | molecular function | GO:0070696 | 4 | 3 | 12176 | 554 | mav,scw,cv-2 | | 3.621e-04 | -7.92 | BMP receptor binding | molecular function | GO:0070700 | 4 | 3 | 12176 | 554 | scw,mav,cv-2 | | 3.626e-04 | -7.92 | cellular response to stimulus | biological process | GO:0051716 | 1467 | 95 | 12345 | 571 | Pkg21D,Bruce,CG42541,CG12147,CG9389,Rgk1,5-HT2A,rdgC,5-HT2B,RYa,TrissinR,par-1,Ir40a,Src42A,Spn,Ppm1,CG32944,mbc,Ulp1,DNAlig4,Parp,Rh3,scw,dnc,fray,boi,Tak1,CG41520,Nipped-A,HisCl1,Ca-alpha1D,dnt,cic,pico,pan,rl,Magi,mav,Octbeta1R,Pi3K68D,SPR,vtd,Stlk,Atf6,Ac78C,shakB,shep,PEK,RhoGAP100F,CG42637,Sema1a,CG6888,GABA-B-R3,Gprk1,CG5160,Thor,grim,Syt7,cta,Myo81F,godzilla,Gr2a,Octbeta2R,CG5270,Wdr59,svp,RabX2,fz2,nAChRalpha4,Sod1,fbl,Cyp6a8,CdGAPr,Pld,br,mus308,CG7536,TpnC41C,Der-1,gus,Toll-7,CG31750,Hsc70-3,CG14184,RSG7,SCAP,norpA,spel1,mthl11,mthl8,St1,CG34357,Cyp6g2,Myc,nAChRalpha3 | | 3.720e-04 | -7.90 | MORF\_ERCC4 | MSigDB lists | MORF\_ERCC4 | 58 | 9 | 4161 | 164 | Coq7,fz2,mbc,RSG7,Pi3K68D,dnc,CG40178,Dys,toy | | 4.047e-04 | -7.81 | pattern specification process | biological process | GO:0007389 | 544 | 43 | 12345 | 571 | ap,step,Myc,ck,jing,bi,CadN2,pan,rl,gus,Dys,ct,pk,cic,fz2,kni,fuss,sna,orb,tld,Tak1,fred,Nipped-A,SoxN,pum,en,scw,boi,lz,Parp,cv-2,CG17387,sn,hth,heph,ru,par-1,zen,Ptip,capu,caup,jvl,salm | | 4.084e-04 | -7.80 | DACOSTA\_UV\_RESPONSE\_VIA\_ERCC3\_TTD\_DN | MSigDB lists | DACOSTA\_UV\_RESPONSE\_VIA\_ERCC3\_TTD\_DN | 26 | 6 | 4161 | 164 | Pi3K68D,CG34408,shot,Dys,CG10565,IntS3 | | 4.126e-04 | -7.79 | AAANWWTGC\_UNKNOWN | MSigDB lists | AAANWWTGC\_UNKNOWN | 36 | 7 | 4161 | 164 | GCS2alpha,Ca-alpha1D,svp,Dys,Mef2,Rbp9,toy | | 4.126e-04 | -7.79 | GSE23321\_CD8\_STEM\_CELL\_MEMORY\_VS\_CENTRAL\_MEMORY\_CD8\_TCELL\_UP | MSigDB lists | GSE23321\_CD8\_STEM\_CELL\_MEMORY\_VS\_CENTRAL\_MEMORY\_CD8\_TCELL\_UP | 36 | 7 | 4161 | 164 | Git,step,cv-2,IntS3,Tsp5D,Dys,svp | | 4.242e-04 | -7.77 | R7 cell development | biological process | GO:0045467 | 15 | 5 | 12345 | 571 | RhoGAP100F,lz,rau,lola,CadN2 | | 4.244e-04 | -7.76 | MORF\_ARL3 | MSigDB lists | MORF\_ARL3 | 59 | 9 | 4161 | 164 | dnc,Pi3K68D,CG40178,Octbeta1R,RSG7,toy,Coq7,fz2,mbc | | 4.923e-04 | -7.62 | GSE7219\_WT\_VS\_NIK\_NFKB2\_KO\_LPS\_AND\_ANTI\_CD40\_STIM\_DC\_DN | MSigDB lists | GSE7219\_WT\_VS\_NIK\_NFKB2\_KO\_LPS\_AND\_ANTI\_CD40\_STIM\_DC\_DN | 37 | 7 | 4161 | 164 | dnr1,CG12121,norpA,twz,cv-2,DNAlig4,Ca-alpha1D | | 5.229e-04 | -7.56 | regulation of biological quality | biological process | GO:0065008 | 1352 | 88 | 12345 | 571 | nvd,Octbeta2R,nullo,sky,svp,PMCA,dati,nAChRalpha4,Sod1,Acp36DE,Catsup,Slob,pum,neo,Hsc70-3,CG14184,norpA,Myc,nAChRalpha3,fbl,Dys,kune,THADA,Sdc,PNPase,ABCB7,Mef2,ewg,shot,Snap25,Nlg2,CG10184,shakB,MCU,Thor,CG9514,CG40178,rib,Clk,CG9650,Syt7,cta,Hipk,Sema1a,CG6888,dimm,Atpalpha,Ten-a,KCNQ,cpx,CG9981,zld,CG30495,lola,HisCl1,SPR,Parg,CG12206,Glut1,fru,conu,HP6,lqf,sif,Msp300,pico,pk,rl,Octbeta1R,spo,capu,Cip4,Lsd-2,knk,spok,Parp,SK,bond,Acox57D-d,boi,bru1,unc-13-4A,Trxr-2,Npc2b,Src42A,Khc-73,Spn,DNAlig4 | | 5.329e-04 | -7.54 | embryo development | biological process | GO:0009790 | 568 | 44 | 12345 | 571 | capu,spo,Ptip,shot,CG41099,par-1,zen,toc,5-HT2A,Irp-1A,knk,lz,cta,scw,rib,Src42A,sn,hth,mbc,CycB,tld,svp,kirre,orb,nullo,sna,kni,Trl,cic,KCNQ,fz2,en,SoxN,pum,Tak1,Myc,jing,Psc,step,ct,Pld,gus,pan,rl | | 5.659e-04 | -7.48 | Ig-like\_fold | interpro domains | IPR013783 | 195 | 21 | 11383 | 557 | DIP-lambda,side-II,CG45263,Camta,fred,kirre,side-III,nrm,dpr21,boi,klg,fan,babos,beat-VII,zormin,sdk,DIP-delta,bt,Ptp99A,CG14964,Dscam2 | | 5.699e-04 | -7.47 | neuron projection guidance | biological process | GO:0097485 | 280 | 26 | 12345 | 571 | fru,ap,jing,Psc,Toll-7,Mical,Dscam2,CadN2,wb,Ten-a,Src42A,CdGAPr,Sema1a,ct,Sdc,dnt,Mp,fz2,RhoGAP100F,shot,Ptp99A,SoxN,klg,lola,en,dsx | | 5.840e-04 | -7.45 | negative regulation of cellular macromolecule biosynthetic process | biological process | GO:2000113 | 438 | 36 | 12345 | 571 | Trl,kni,Maf1,cic,svp,HPS4,orb,sna,Mnt,pum,d4,lncRNA:bxd,Fer3,dsx,en,zld,jing,CG9932,Psc,HP6,E5,CG1620,Parg,bi,Tgi,pan,ct,pho,bru3,Vsx2,bru1,Clk,lz,Su(z)2,Thor,heph | | 5.868e-04 | -7.44 | Spectrin/alpha-actinin | interpro domains | IPR018159 | 9 | 4 | 11383 | 557 | zormin,shot,Dys,Msp300 | | 5.868e-04 | -7.44 | Spectrin\_repeat | interpro domains | IPR002017 | 9 | 4 | 11383 | 557 | shot,Dys,zormin,Msp300 | | 5.870e-04 | -7.44 | negative regulation of macromolecule metabolic process | biological process | GO:0010605 | 777 | 56 | 12345 | 571 | scw,Clk,AGO3,bru1,lz,Su(z)2,Thor,heph,Ppm1,zen,CG10077,SmydA-8,Spn75F,SmydA-9,Marf1,pho,bru3,hdc,Vsx2,E5,CG1620,Ars2,spel1,jing,Psc,HP6,CG9932,Hsc70-3,Tgi,bi,Parg,Diap1,pan,PNPase,br,ct,Maf1,drosha,cic,Trl,mrt,kni,sna,orb,Mnt,svp,tld,HPS4,Fer3,pum,d4,lncRNA:bxd,Spn42De,zld,dsx,en | | 5.916e-04 | -7.43 | neuromuscular junction | cellular component | GO:0031594 | 77 | 11 | 11940 | 534 | Ten-a,cpx,pum,Cip4,sky,Syx18,rl,Magi,Sdc,par-1,Nlg2 | | 6.127e-04 | -7.40 | IG\_LIKE | prosite domains | PS50835 | 132 | 18 | 6243 | 365 | DIP-lambda,Dscam2,klg,beat-VII,dpr21,CG45263,babos,fred,side-II,sdk,DIP-delta,zormin,side-III,kirre,CG14964,nrm,boi,bt | | 6.210e-04 | -7.38 | protein homodimerization activity | molecular function | GO:0042803 | 140 | 16 | 12176 | 554 | Trl,CadN2,Atf6,svp,inc,Camta,Ten-a,Khc-73,bru3,Sod1,mim,step,grim,dsx,cbs,rau | | 6.214e-04 | -7.38 | GO\_CIRCULATORY\_SYSTEM\_DEVELOPMENT | MSigDB lists | GO\_CIRCULATORY\_SYSTEM\_DEVELOPMENT | 164 | 16 | 4161 | 164 | rl,Mp,CycB,svp,vap,fz2,CG17387,l(2)41Ab,Meltrin,cta,Mef2,toy,CG9492,cv-2,Alg-2,nAChRalpha4 | | 6.256e-04 | -7.38 | response to stimulus | biological process | GO:0050896 | 2655 | 155 | 12345 | 571 | Gr39a,Tdc2,CG6888,Ten-a,CG5160,MCU,grim,PPO2,Syt7,CG7255,Stlk,l(3)L1231,Ac78C,Ir41a,PEK,Or67d,Pzl,Rift,CdGAPr,ct,mus308,Sdc,CG7536,Der-1,CG31750,RSG7,vap,ap,spel1,CG34357,Myc,Cyp6g2,Or69a,CG9380,nAChRalpha3,dsx,Gr2a,Slob,pum,RabX2,fz2,nAChRalpha4,rg,Spn,CG32944,lz,fray,dnc,boi,scw,Gr93d,Pkg21D,Bruce,CG42541,5-HT2A,Rgk1,rdgC,5-HT2B,pico,pan,Magi,Dlg5,mav,Parg,CadN2,jing,Psc,Camta,HisCl1,SoxN,CG17691,Sema1a,Hipk,Atpalpha,GABA-B-R3,Gprk1,Thor,Dscam2,Idgf4,Clk,Or83c,cta,klg,Atf6,shakB,shep,Ptp99A,Mef2,RhoGAP100F,CG42637,shot,fbl,Cyp6a8,Pld,br,TpnC41C,gus,Toll-7,Hsc70-3,CG14184,SCAP,norpA,Ars2,mthl8,St1,mthl11,Myo81F,godzilla,Catsup,Wdr59,CG5270,Octbeta2R,svp,RNASEK,Mp,Sod1,CG3556,Ir40a,Src42A,Ppm1,mbc,Ulp1,sn,DNAlig4,Parp,Rh3,Eig71Ef,CG12147,Or98a,CG9389,RYa,Hmt-1,TrissinR,tay,par-1,Ir60e,rl,wb,Octbeta1R,Pi3K68D,SPR,Mical,fru,vtd,en,lola,Tak1,Ir94e,CG41520,Nipped-A,Ca-alpha1D,CheB42a,dnt,cic,Trl | | 6.526e-04 | -7.33 | negative regulation of biosynthetic process | biological process | GO:0009890 | 457 | 37 | 12345 | 571 | Vsx2,bru3,pho,Thor,Su(z)2,lz,bru1,Clk,heph,Mnt,orb,sna,HPS4,svp,cic,Maf1,kni,dnr1,Trl,zld,en,dsx,Fer3,lncRNA:bxd,pum,d4,Tgi,bi,Parg,CG1620,E5,jing,HP6,Psc,CG9932,ct,pan | | 6.526e-04 | -7.33 | negative regulation of cellular biosynthetic process | biological process | GO:0031327 | 457 | 37 | 12345 | 571 | Thor,Su(z)2,lz,Clk,bru1,heph,Vsx2,bru3,pho,Parg,bi,Tgi,HP6,CG9932,Psc,jing,CG1620,E5,ct,pan,HPS4,svp,Mnt,sna,orb,kni,dnr1,Trl,cic,Maf1,en,dsx,zld,lncRNA:bxd,d4,pum,Fer3 | | 6.552e-04 | -7.33 | Spectrin | pfam domains | PF00435 | 9 | 4 | 10480 | 528 | shot,zormin,Msp300,Dys | | 6.638e-04 | -7.32 | HADDAD\_T\_LYMPHOCYTE\_AND\_NK\_PROGENITOR\_UP | MSigDB lists | HADDAD\_T\_LYMPHOCYTE\_AND\_NK\_PROGENITOR\_UP | 19 | 5 | 4161 | 164 | arg,norpA,Hsc70-3,spel1,vap | | 6.696e-04 | -7.31 | neuron projection morphogenesis | biological process | GO:0048812 | 425 | 35 | 12345 | 571 | CdGAPr,ct,br,Sdc,wb,Toll-7,Mical,CadN2,fru,ap,jing,Psc,lola,en,dsx,SoxN,pum,dnt,Mp,fz2,scro,Src42A,Sema1a,sn,Ulp1,Ten-a,Dscam2,dnc,klg,Stlk,pho,hdc,Ptp99A,shot,RhoGAP100F | | 6.869e-04 | -7.28 | plasma membrane bounded cell projection organization | biological process | GO:0120036 | 626 | 47 | 12345 | 571 | pk,ct,Sdc,br,kl-3,CdGAPr,wb,CadN2,Toll-7,Mical,ck,Psc,jing,fru,cmb,ap,en,lola,dsx,l(2)41Ab,SoxN,pum,sky,scro,Mp,dnt,fz2,dati,Spn,Src42A,Sema1a,sn,Ulp1,CG17387,Ten-a,mim,Dscam2,dnc,klg,Stlk,hdc,pho,Ptp99A,shot,RhoGAP100F,Cip4,par-1 | | 6.889e-04 | -7.28 | gastrulation with mouth forming first | biological process | GO:0001703 | 43 | 8 | 12345 | 571 | Pld,5-HT2A,Irp-1A,tld,sna,cta,scw,zen | | 6.889e-04 | -7.28 | gastrulation involving germ band extension | biological process | GO:0010004 | 43 | 8 | 12345 | 571 | Pld,Irp-1A,5-HT2A,sna,tld,scw,zen,cta | | 6.984e-04 | -7.27 | plasma membrane bounded cell projection morphogenesis | biological process | GO:0120039 | 426 | 35 | 12345 | 571 | pum,SoxN,dsx,lola,en,fz2,dnt,Mp,scro,wb,CdGAPr,Sdc,br,ct,ap,fru,Psc,jing,Mical,Toll-7,CadN2,pho,hdc,klg,Stlk,RhoGAP100F,shot,Ptp99A,Ten-a,sn,Ulp1,Sema1a,Src42A,dnc,Dscam2 | | 7.051e-04 | -7.26 | muscle structure development | biological process | GO:0061061 | 224 | 22 | 12345 | 571 | Msp300,Dys,br,mbc,hth,Thor,Mical,ap,fru,bru1,CAP,Pkg21D,lola,caup,salm,Git,kirre,Mef2,ewg,dnt,shot,bt | | 7.104e-04 | -7.25 | GO\_EMBRYONIC\_ORGAN\_DEVELOPMENT | MSigDB lists | GO\_EMBRYONIC\_ORGAN\_DEVELOPMENT | 90 | 11 | 4161 | 164 | Bruce,CG17387,fz2,Sod1,Mef2,toy,svp,lqf,rl,ck,l(2)41Ab | | 7.595e-04 | -7.18 | cell projection morphogenesis | biological process | GO:0048858 | 428 | 35 | 12345 | 571 | Mical,Toll-7,CadN2,ap,fru,jing,Psc,CdGAPr,br,Sdc,ct,wb,fz2,Mp,dnt,scro,dsx,lola,en,pum,SoxN,Dscam2,dnc,Src42A,Sema1a,Ten-a,Ulp1,sn,Ptp99A,shot,RhoGAP100F,klg,Stlk,pho,hdc | | 7.637e-04 | -7.18 | MORF\_MAP2K7 | MSigDB lists | MORF\_MAP2K7 | 29 | 6 | 4161 | 164 | CG40178,fz2,dnc,RSG7,toy,Dys | | 7.658e-04 | -7.17 | IGc2 | smart domains | SM00408 | 109 | 16 | 5086 | 308 | klg,DIP-lambda,boi,zormin,DIP-delta,CG14964,kirre,side-II,sdk,Dscam2,nrm,fred,bt,CG45263,dpr21,side-III | | 7.946e-04 | -7.14 | axon guidance | biological process | GO:0007411 | 271 | 25 | 12345 | 571 | SoxN,klg,dsx,lola,en,fz2,dnt,Mp,RhoGAP100F,shot,Ptp99A,Ten-a,wb,CdGAPr,Sema1a,Src42A,Sdc,ap,fru,Psc,jing,Mical,Toll-7,CadN2,Dscam2 | | 8.128e-04 | -7.12 | negative regulation of macromolecule biosynthetic process | biological process | GO:0010558 | 446 | 36 | 12345 | 571 | Parg,Tgi,bi,jing,CG9932,HP6,Psc,E5,CG1620,ct,pan,svp,HPS4,orb,sna,Mnt,Trl,kni,Maf1,cic,dsx,en,zld,pum,d4,lncRNA:bxd,Fer3,lz,Thor,Su(z)2,Clk,bru1,heph,Vsx2,pho,bru3 | | 8.611e-04 | -7.06 | z (zeste) | protein interactions | 31230 | 41 | 8 | 9204 | 464 | Git,lz,ewg,sna,Psc,ap,ct,Su(z)2 | | 8.956e-04 | -7.02 | GO\_REGULATION\_OF\_SYSTEM\_PROCESS | MSigDB lists | GO\_REGULATION\_OF\_SYSTEM\_PROCESS | 107 | 12 | 4161 | 164 | Parp,PMCA,Ca-alpha1D,nAChRalpha4,Atpalpha,GLS,Nlg3,dnc,Octbeta2R,bru1,Sod1,Dys | | 8.987e-04 | -7.01 | GO\_EYE\_PHOTORECEPTOR\_CELL\_DIFFERENTIATION | MSigDB lists | GO\_EYE\_PHOTORECEPTOR\_CELL\_DIFFERENTIATION | 12 | 4 | 4161 | 164 | toy,sn,sdk,ck | | 9.134e-04 | -7.00 | positive regulation of nitrogen compound metabolic process | biological process | GO:0051173 | 827 | 58 | 12345 | 571 | Trl,scro,sna,orb,pum,d4,SoxN,CG41520,Camta,CG40228,Tak1,toy,dsx,en,mxt,lola,zld,tou,jing,Myc,CG12769,ap,step,SCAP,CG14184,tj,bi,gus,mav,Diap1,pan,br,Setd3,Ptip,zen,PEK,Mef2,Marf1,CG12200,CG12147,pho,caup,Atf6,CG41106,Stlk,Clk,AGO3,fray,scw,grim,Parp,Alg-2,lz,Su(z)2,hth,dimm,mbc,Src42A | | 9.373e-04 | -6.97 | regulation of gene expression | biological process | GO:0010468 | 1547 | 97 | 12345 | 571 | mrt,Trl,drosha,cic,sna,fuss,Mnt,d4,lncRNA:bxd,SoxN,Nipped-A,Camta,CG40228,Tak1,en,lola,zld,Psc,jing,HP6,E5,CG12769,vtd,fru,Parg,bi,mav,rl,Diap1,CG34334,pan,Setd3,SmydA-8,CG32006,zen,pho,bru1,AGO3,scw,lz,Parp,hth,scro,kni,Maf1,bab2,svp,tld,HPS4,Octbeta2R,orb,pum,Fer3,dsx,toy,mxt,CG9932,tou,Myc,CG1620,ap,Ars2,SCAP,tj,Tgi,Hsc70-3,jigr1,PNPase,Sdc,br,CG17514,ct,SmydA-9,CG10077,Mef2,Marf1,PEK,ewg,CG5862,salm,hdc,bru3,caup,Atf6,Vsx2,Clk,CG9650,rib,Thor,Su(z)2,fd96Ca,dimm,cv-2,Hipk,heph,Coq7,CG33557 | | 9.818e-04 | -6.93 | positive regulation of metabolic process | biological process | GO:0009893 | 937 | 64 | 12345 | 571 | hth,mbc,Src42A,AGO3,fray,scw,lz,Parp,CG12147,CG12200,pho,CG41106,Ptip,zen,mav,pan,Diap1,Setd3,jing,step,vtd,CG12769,bi,CG41520,SoxN,d4,CG40228,Tak1,Camta,AP-2alpha,en,lola,zld,Trl,sna,dimm,Hipk,Clk,grim,Su(z)2,Alg-2,caup,Atf6,Stlk,PEK,Marf1,Mef2,gus,CG17514,br,Myc,tou,ap,CG14184,SCAP,Toll-7,tj,pum,mxt,dsx,toy,scro,HPS4,orb | | 9.842e-04 | -6.92 | imaginal disc-derived wing vein specification | biological process | GO:0007474 | 56 | 9 | 12345 | 571 | en,Dys,cv-2,caup,rl,fuss,jing,step,cic | | 9.940e-04 | -6.91 | Dorso-ventral axis formation | KEGG pathways | ko04320 | 28 | 6 | 3198 | 138 | rl,bru1,capu,orb,br,AGO3 | | 9.940e-04 | -6.91 | Dorso-ventral axis formation | KEGG pathways | dme04320 | 28 | 6 | 3198 | 138 | orb,br,AGO3,rl,bru1,capu | | 9.945e-04 | -6.91 | cell projection organization | biological process | GO:0030030 | 637 | 47 | 12345 | 571 | Spn,Src42A,Sema1a,CG17387,Ulp1,sn,Ten-a,mim,Dscam2,dnc,klg,Stlk,hdc,pho,Ptp99A,shot,RhoGAP100F,par-1,Cip4,pk,ct,kl-3,Sdc,br,CdGAPr,wb,CadN2,Toll-7,Mical,ck,Psc,jing,ap,fru,cmb,lola,en,dsx,l(2)41Ab,SoxN,pum,sky,scro,dnt,Mp,fz2,dati | | 1.058e-03 | -6.85 | negative regulation of transcription, DNA-templated | biological process | GO:0045892 | 371 | 31 | 12345 | 571 | pan,ct,Clk,CG9932,Psc,HP6,jing,CG1620,E5,Parg,Su(z)2,bi,lz,Tgi,lncRNA:bxd,pum,d4,Fer3,pho,en,Vsx2,dsx,zld,kni,Trl,cic,Maf1,HPS4,svp,Mnt,sna | | 1.062e-03 | -6.85 | chemotaxis | biological process | GO:0006935 | 292 | 26 | 12345 | 571 | SoxN,klg,lola,en,dsx,dnt,Mp,fz2,shot,RhoGAP100F,Ptp99A,wb,Ten-a,Src42A,Sema1a,CdGAPr,ct,Sdc,fru,ap,Psc,jing,Toll-7,Mical,Dscam2,CadN2 | | 1.071e-03 | -6.84 | protein binding | molecular function | GO:0005515 | 2410 | 139 | 12176 | 554 | igl,CG7029,pho,kl-3,Sema1a,Tbc1d15-17,Fer3,zld,Syx18,CycB,Mef2,SmydA-9,Dscam2,mxt,Idgf4,cta,CG31703,bi,CG33557,Gprk1,Klp59C,Tgi,svp,Git,mim,fz2,lola,dnt,bt,Camta,Dys,Trs23,inc,Pi3K68D,shot,gus,mbc,Dhc16F,mlt,CheB42a,KCNQ,CG10188,Sod1,RYa,Magi,fuss,tou,bab2,scw,Thor,rau,Diap1,Nlg2,fan,step,DNAlig4,Maf1,Gdh,CG5359,kirre,Mnt,rg,Slob,Ranbp16,cic,dimm,Ars2,rl,kni,Der-1,rdgC,CG5862,Syt7,bru1,bru3,CG17211,sna,Ten-a,conu,sn,Spn,orb,CG17514,mav,pan,SoxN,zormin,CG14184,CG3588,Snap25,pk,Nlg3,Lsd-2,capu,Parp,Slmap,CG40228,Khc-73,Atf6,ck,CadN2,Trl,CG31275,cpx,CG30440,Myc,THADA,sif,boi,CG9044,cato,hth,grim,Clk,CG34347,vap,SK,pico,d4,nullo,cv-2,Hsc70-3,Src42A,HP6,Svil,dsx,CG1620,cbs,Cip4,Msp300,CG8219,SmydA-8,Tak1,Hipk,Acp36DE,cmb,Meltrin,Mical,AP-2alpha | | 1.078e-03 | -6.83 | leg disc proximal/distal pattern formation | biological process | GO:0007479 | 18 | 5 | 12345 | 571 | ru,hth,jing,ap,rl | | 1.082e-03 | -6.83 | fly chr71B8-71B8\|3-42 cM | chromosome location | fly chr71B8-71B8\|3-42 cM | 2 | 2 | 25010 | 823 | lncRNA:CR44562,CG6888 | | 1.082e-03 | -6.83 | fly chr68C1-68C1\|3-36 cM | chromosome location | fly chr68C1-68C1\|3-36 cM | 2 | 2 | 25010 | 823 | lncRNA:CR45404,lncRNA:CR44720 | | 1.082e-03 | -6.83 | fly chr6F1-6F1\|1-19 | chromosome location | fly chr6F1-6F1\|1-19 | 2 | 2 | 25010 | 823 | nullo,CG14427 | | 1.082e-03 | -6.83 | fly chr41D1-41D1\|2-55 | chromosome location | fly chr41D1-41D1\|2-55 | 2 | 2 | 25010 | 823 | lncRNA:CR44105,TpnC41C | | 1.082e-03 | -6.83 | fly chr71B8-71B8\|3-42 | chromosome location | fly chr71B8-71B8\|3-42 | 2 | 2 | 25010 | 823 | lncRNA:CR44562,CG6888 | | 1.082e-03 | -6.83 | fly chr81F-81F\|3-47.1 | chromosome location | fly chr81F-81F\|3-47.1 | 2 | 2 | 25010 | 823 | Gfat1,Parp | | 1.082e-03 | -6.83 | fly chr68C1-68C1\|3-36 | chromosome location | fly chr68C1-68C1\|3-36 | 2 | 2 | 25010 | 823 | lncRNA:CR45404,lncRNA:CR44720 | | 1.082e-03 | -6.83 | fly chr6F1-6F1\|1-19 cM | chromosome location | fly chr6F1-6F1\|1-19 cM | 2 | 2 | 25010 | 823 | CG14427,nullo | | 1.082e-03 | -6.83 | fly chr20F2-20F2\|1-66 cM | chromosome location | fly chr20F2-20F2\|1-66 cM | 2 | 2 | 25010 | 823 | lncRNA:CR41352,CG41106 | | 1.082e-03 | -6.83 | fly chr41D1-41D1\|2-55 cM | chromosome location | fly chr41D1-41D1\|2-55 cM | 2 | 2 | 25010 | 823 | TpnC41C,lncRNA:CR44105 | | 1.082e-03 | -6.83 | fly chr90C10-90C10\|3-61 | chromosome location | fly chr90C10-90C10\|3-61 | 2 | 2 | 25010 | 823 | CG12347,CG43445 | | 1.082e-03 | -6.83 | fly chr20F2-20F2\|1-66 | chromosome location | fly chr20F2-20F2\|1-66 | 2 | 2 | 25010 | 823 | CG41106,lncRNA:CR41352 | | 1.082e-03 | -6.83 | fly chr81F-81F\|3-47.1 cM | chromosome location | fly chr81F-81F\|3-47.1 cM | 2 | 2 | 25010 | 823 | Gfat1,Parp | | 1.082e-03 | -6.83 | fly chr90C10-90C10\|3-61 cM | chromosome location | fly chr90C10-90C10\|3-61 cM | 2 | 2 | 25010 | 823 | CG12347,CG43445 | | 1.085e-03 | -6.83 | SCHUETZ\_BREAST\_CANCER\_DUCTAL\_INVASIVE\_UP | MSigDB lists | SCHUETZ\_BREAST\_CANCER\_DUCTAL\_INVASIVE\_UP | 54 | 8 | 4161 | 164 | Glut1,Gfat1,bru1,Mef2,Meltrin,Mp,GLS,CG41378 | | 1.093e-03 | -6.82 | TCCATTKW\_UNKNOWN | MSigDB lists | TCCATTKW\_UNKNOWN | 42 | 7 | 4161 | 164 | Rbp9,CG12547,shot,Ca-alpha1D,CstF50,DNaseII,CG4341 | | 1.096e-03 | -6.82 | cell part morphogenesis | biological process | GO:0032990 | 437 | 35 | 12345 | 571 | Ptp99A,RhoGAP100F,shot,Stlk,klg,hdc,pho,Dscam2,dnc,Sema1a,Src42A,Ten-a,sn,Ulp1,scro,fz2,Mp,dnt,dsx,en,lola,pum,SoxN,CadN2,Mical,Toll-7,Psc,jing,fru,ap,br,Sdc,ct,CdGAPr,wb | | 1.097e-03 | -6.81 | R7 cell differentiation | biological process | GO:0045466 | 46 | 8 | 12345 | 571 | lola,klg,salm,rl,CadN2,rau,lz,RhoGAP100F | | 1.125e-03 | -6.79 | fly chr80F9-80F9\|3-47 | chromosome location | fly chr80F9-80F9\|3-47 | 7 | 3 | 25010 | 823 | CG40298,AGO3,lncRNA:CR45174 | | 1.125e-03 | -6.79 | fly chr80F9-80F9\|3-47 cM | chromosome location | fly chr80F9-80F9\|3-47 cM | 7 | 3 | 25010 | 823 | lncRNA:CR45174,AGO3,CG40298 | | 1.180e-03 | -6.74 | CG5986 (uncharacterized protein) | protein interactions | 192506 | 5 | 3 | 9204 | 464 | Diap1,Sema1a,Rtnl1 | | 1.203e-03 | -6.72 | cell junction organization | biological process | GO:0034330 | 81 | 11 | 12345 | 571 | kirre,nullo,sna,CadN2,CG15449,Magi,rl,Atpalpha,shakB,Src42A,kune | | 1.205e-03 | -6.72 | axonogenesis | biological process | GO:0007409 | 326 | 28 | 12345 | 571 | CdGAPr,Src42A,Sema1a,Sdc,wb,Ten-a,Ulp1,Mical,Toll-7,Dscam2,CadN2,dnc,ap,fru,jing,Psc,klg,dsx,en,lola,SoxN,hdc,Ptp99A,fz2,dnt,Mp,RhoGAP100F,shot | | 1.228e-03 | -6.70 | GSE37301\_MULTIPOTENT\_PROGENITOR\_VS\_RAG2\_KO\_NK\_CELL\_UP | MSigDB lists | GSE37301\_MULTIPOTENT\_PROGENITOR\_VS\_RAG2\_KO\_NK\_CELL\_UP | 68 | 9 | 4161 | 164 | GCS2alpha,SdhA,rl,Ranbp16,mlt,Odc2,CstF50,St1,CG10077 | | 1.228e-03 | -6.70 | antennal development | biological process | GO:0007469 | 27 | 6 | 12345 | 571 | ck,sn,salm,Diap1,ct,lz | | 1.232e-03 | -6.70 | ion binding | molecular function | GO:0043167 | 2058 | 121 | 12176 | 554 | Sod1,d4,CG12200,Nna1,Cyp309a1,Nadsyn,cv-2,Hsc70-3,Acbp5,Rgk1,Ppm1,CG9492,Src42A,bma,SdhA,Cip4,CG34357,Diap1,CG17361,dsx,par-1,step,DNAlig4,fbl,CG9981,CG41099,Tak1,Hipk,5-HT2A,CG3107,Mical,Alg-2,CG42541,dnt,CG5789,pk,spel1,Parp,Khc-73,CG14721,CG33970,ABCB7,bt,Dys,CG42813,CG10077,CG8539,CadN2,ck,PMCA,Pi3K68D,shot,Atpalpha,CG2256,Dhc16F,PEK,snky,CG32944,mus308,CG9747,Scp2,dnr1,Klp59C,Stlk,norpA,Gprk1,CG31689,CG14820,Amyrel,Trxr-2,svp,Rpb12,Nrd1,tld,Myo81F,CG9743,mIF2,Cyp6g2,FASN3,TpnC41C,Su(z)2,CG43672,CG30094,stw,Psc,Scp1,mim,Cyp6a8,mei-P26,CG12147,fzo,Hmt-1,CG42637,Sema1a,CG9514,kl-3,cert,arg,Tdc2,lz,CG44002,Pkg21D,spok,Acox57D-d,CG5160,CG1486,Takl2,kni,rl,fray,CG42255,Pld,godzilla,CG5270,CG7536,Cyp316a1,RabX2,Ubr1,Syt7,spo,rdgC | | 1.251e-03 | -6.68 | cell-cell signaling | biological process | GO:0007267 | 311 | 27 | 12345 | 571 | dnc,unc-13-4A,nAChRalpha3,Syt7,pan,be,Atpalpha,Ten-a,lqf,Dys,Syx18,dnt,fz2,cpx,nAChRalpha4,shot,Snap25,sna,sky,svp,Slob,HisCl1,CG12147,5-HT2A,pum,cpo,Nlg2 | | 1.263e-03 | -6.67 | IKEDA\_MIR30\_TARGETS\_UP | MSigDB lists | IKEDA\_MIR30\_TARGETS\_UP | 43 | 7 | 4161 | 164 | vap,Gfat1,bru1,norpA,cta,Tbc1d15-17,Dys | | 1.263e-03 | -6.67 | GSE45365\_CD8A\_DC\_VS\_CD11B\_DC\_IFNAR\_KO\_MCMV\_INFECTION\_UP | MSigDB lists | GSE45365\_CD8A\_DC\_VS\_CD11B\_DC\_IFNAR\_KO\_MCMV\_INFECTION\_UP | 43 | 7 | 4161 | 164 | CG41099,CG3107,par-1,CG14820,Glut1,CG34408,dnr1 | | 1.274e-03 | -6.67 | neuromuscular synaptic transmission | biological process | GO:0007274 | 58 | 9 | 12345 | 571 | Nlg2,Syx18,Dys,Ten-a,be,Slob,sky,Snap25,cpx | | 1.290e-03 | -6.65 | positive regulation of cellular metabolic process | biological process | GO:0031325 | 875 | 60 | 12345 | 571 | Toll-7,tj,bi,CG14184,SCAP,ap,step,CG12769,Myc,tou,jing,CG17514,Setd3,br,pan,Diap1,mav,orb,sna,scro,Trl,zld,lola,mxt,en,dsx,toy,Tak1,CG40228,Camta,AP-2alpha,SoxN,CG41520,pum,d4,Su(z)2,lz,Parp,Alg-2,grim,scw,fray,Clk,AGO3,Src42A,mbc,dimm,hth,PEK,Marf1,Mef2,zen,Ptip,Stlk,Atf6,CG41106,caup,pho,CG12147,CG12200 | | 1.294e-03 | -6.65 | Apoptosis - fly | KEGG pathways | dme04214 | 63 | 9 | 3198 | 138 | br,rl,PEK,Bruce,Parp,grim,fzo,Tak1,Takl2 | | 1.294e-03 | -6.65 | Apoptosis - fly | KEGG pathways | ko04214 | 63 | 9 | 3198 | 138 | grim,Parp,fzo,Tak1,Takl2,br,PEK,rl,Bruce | | 1.303e-03 | -6.64 | SPEC | smart domains | SM00150 | 9 | 4 | 5086 | 308 | shot,Dys,zormin,Msp300 | | 1.307e-03 | -6.64 | RNA polymerase II regulatory region sequence-specific DNA binding | molecular function | GO:0000977 | 254 | 23 | 12176 | 554 | ewg,Mef2,Atf6,zen,toy,ct,svp,cic,CG9932,sna,bi,pho,br,E5,scro,ap,Clk,pan,tj,CG12769,dsx,en,Myc | | 1.324e-03 | -6.63 | positive regulation of macromolecule metabolic process | biological process | GO:0010604 | 858 | 59 | 12345 | 571 | Stlk,CG41106,Atf6,caup,pho,CG12147,CG12200,Marf1,Mef2,PEK,zen,Src42A,Hipk,mbc,dimm,hth,Su(z)2,lz,Alg-2,grim,scw,fray,AGO3,Clk,zld,lola,en,mxt,dsx,toy,CG40228,Tak1,Camta,SoxN,CG41520,d4,pum,orb,sna,HPS4,Trl,scro,br,Setd3,pan,Diap1,mav,gus,bi,tj,CG14184,SCAP,step,ap,CG12769,vtd,Myc,tou,jing | | 1.366e-03 | -6.60 | PID\_P53\_DOWNSTREAM\_PATHWAY | MSigDB lists | PID\_P53\_DOWNSTREAM\_PATHWAY | 22 | 5 | 4161 | 164 | Nipped-A,CG10077,spel1,CycB,drosha | | 1.386e-03 | -6.58 | GO\_MUSCLE\_SYSTEM\_PROCESS | MSigDB lists | GO\_MUSCLE\_SYSTEM\_PROCESS | 56 | 8 | 4161 | 164 | Sod1,Mef2,Dys,arg,Octbeta2R,Pi3K68D,dnc,Atpalpha | | 1.398e-03 | -6.57 | muscle organ development | biological process | GO:0007517 | 95 | 12 | 12345 | 571 | dnt,fru,ap,shot,CAP,kirre,Mef2,ewg,mbc,salm,Dys,Msp300 | | 1.489e-03 | -6.51 | negative regulation of RNA biosynthetic process | biological process | GO:1902679 | 379 | 31 | 12345 | 571 | d4,pum,lncRNA:bxd,pho,Fer3,dsx,en,Vsx2,zld,Trl,kni,Maf1,cic,svp,HPS4,sna,Mnt,pan,ct,HP6,Psc,jing,CG9932,Clk,CG1620,E5,Parg,bi,lz,Tgi,Su(z)2 | | 1.489e-03 | -6.51 | negative regulation of nucleic acid-templated transcription | biological process | GO:1903507 | 379 | 31 | 12345 | 571 | pan,ct,Clk,jing,CG9932,HP6,Psc,CG1620,E5,Parg,Su(z)2,bi,Tgi,lz,lncRNA:bxd,pum,d4,Fer3,pho,en,Vsx2,dsx,zld,kni,Trl,cic,Maf1,HPS4,svp,Mnt,sna | | 1.515e-03 | -6.49 | GO\_REGULATION\_OF\_MEMBRANE\_POTENTIAL | MSigDB lists | GO\_REGULATION\_OF\_MEMBRANE\_POTENTIAL | 70 | 9 | 4161 | 164 | nAChRalpha4,Atpalpha,nAChRalpha3,Ca-alpha1D,Octbeta2R,Sod1,Mef2,Dys,Nlg3 | | 1.543e-03 | -6.47 | Sirt2 (Sirtuin 2) | protein interactions | 42414 | 26 | 6 | 9204 | 464 | Vm34Ca,dimm,gus,CG17508,E5,heph | | 1.543e-03 | -6.47 | membrane part | cellular component | GO:0044425 | 2068 | 119 | 11940 | 534 | step,norpA,Teh2,CG14744,5-HT2A,CG1213,Der-1,fzo,CG9743,Gr2a,Ir40a,Ten-a,mthl11,CG9747,Octbeta2R,Or67d,CG9981,TrissinR,Cip4,CG7536,Ac78C,CG41128,Syt7,CG14024,mbc,Ca-alpha1D,Atf6,Dys,Rh3,Nlg3,Octbeta1R,SK,Oatp33Ea,mthl8,CG14184,Ir41a,par-1,CG12121,Dscam2,CG33970,Rift,ABCB7,CG43672,Toll-7,ND-30,CG10171,Or69a,CG45076,THADA,neo,ND-MLRQ,Meltrin,CG5789,ru,CG8249,Or98a,Glut1,CG34357,Sema1a,CG42637,Pi3K68D,GABA-B-R3,Pzl,nAChRalpha4,CG33233,CG15449,UQCR-11,Tim17b,HisCl1,SCAP,KCNQ,SdhA,snky,Atpalpha,CG41106,DIP-delta,fz2,Syx18,Tsp5D,Magi,Rgk1,Or83c,Nlg2,Gr39a,CG15459,Msp300,klg,bond,ND-AGGG,AP-2alpha,Snap25,cta,CG15408,dnt,Pgant9,nAChRalpha3,ImpE2,Tim23,Ir94e,DIP-lambda,CG2003,Src42A,kune,sdk,Hmt-1,cpx,CG40472,Ca-Ma2d,fan,SPR,MCU,Mct1,MFS16,CG14196,Ir60e,Gr93d,rg,CG14160,CadN2 | | 1.544e-03 | -6.47 | GO\_ACTIVATION\_OF\_TRANSMEMBRANE\_RECEPTOR\_PROTEIN\_TYROSINE\_KINASE\_ACTIVITY | MSigDB lists | GO\_ACTIVATION\_OF\_TRANSMEMBRANE\_RECEPTOR\_PROTEIN\_TYROSINE\_KINASE\_ACTIVITY | 2 | 2 | 4161 | 164 | nAChRalpha4,Octbeta2R | | 1.544e-03 | -6.47 | DUNNE\_TARGETS\_OF\_AML1\_MTG8\_FUSION\_DN | MSigDB lists | DUNNE\_TARGETS\_OF\_AML1\_MTG8\_FUSION\_DN | 2 | 2 | 4161 | 164 | Mef2,Glut1 | | 1.544e-03 | -6.47 | GO\_HMG\_BOX\_DOMAIN\_BINDING | MSigDB lists | GO\_HMG\_BOX\_DOMAIN\_BINDING | 2 | 2 | 4161 | 164 | toy,Mef2 | | 1.559e-03 | -6.46 | MODULE\_84 | MSigDB lists | MODULE\_84 | 57 | 8 | 4161 | 164 | CG41099,CG41378,bru1,Gfat1,Cyp6a8,Octbeta1R,Meltrin,unc-13-4A | | 1.563e-03 | -6.46 | GSE13762\_CTRL\_VS\_125\_VITAMIND\_DAY12\_DC\_UP | MSigDB lists | GSE13762\_CTRL\_VS\_125\_VITAMIND\_DAY12\_DC\_UP | 33 | 6 | 4161 | 164 | CG2061,CG14721,Hsc70-3,norpA,rg,St1 | | 1.628e-03 | -6.42 | PH | pfam domains | PF00169 | 45 | 8 | 10480 | 528 | cert,Gprk1,vap,sif,osp,CG30440,step,pico | | 1.664e-03 | -6.40 | S8\_01 | MSigDB lists | S8\_01 | 45 | 7 | 4161 | 164 | CG4341,Nlg3,twz,Mef2,CG10077,Dys,bru1 | | 1.680e-03 | -6.39 | regulation of epidermal growth factor receptor signaling pathway | biological process | GO:0042058 | 49 | 8 | 12345 | 571 | rau,Pi3K68D,rno,tay,step,Src42A,ru,pum | | 1.680e-03 | -6.39 | regulation of ERBB signaling pathway | biological process | GO:1901184 | 49 | 8 | 12345 | 571 | pum,ru,Src42A,step,tay,rno,Pi3K68D,rau | | 1.687e-03 | -6.38 | negative regulation of cellular process | biological process | GO:0048523 | 1291 | 82 | 12345 | 571 | CG1620,norpA,vap,spel1,CG9932,CG9380,Tgi,ct,Maf1,dnr1,kni,orb,svp,sky,HPS4,Fer3,CG12772,pum,dsx,rno,CG9650,Clk,Thor,Su(z)2,tej,grim,mim,Gprk1,Sema1a,Hipk,heph,Spn75F,shot,ewg,bru3,salm,hdc,Nlg2,Vsx2,CG13284,E5,jing,Psc,HP6,alpha4GT1,bi,Pi3K68D,Parg,rl,Diap1,pan,Octbeta1R,sdk,lqf,Slmap,cpx,cic,Trl,sna,Mnt,fuss,d4,CG41520,lncRNA:bxd,SoxN,zld,Spn42De,en,dnc,bru1,AGO3,lz,Ulp1,Src42A,Ppm1,Spn,par-1,Cip4,tay,pho,CG12147,Bruce | | 1.690e-03 | -6.38 | ubiquitin-specific protease binding | molecular function | GO:1990381 | 6 | 3 | 12176 | 554 | Der-1,Diap1,Myc | | 1.708e-03 | -6.37 | GO\_CELLULAR\_IRON\_ION\_HOMEOSTASIS | MSigDB lists | GO\_CELLULAR\_IRON\_ION\_HOMEOSTASIS | 14 | 4 | 4161 | 164 | Sod1,Hmt-1,ABCB7,Irp-1A | | 1.708e-03 | -6.37 | GO\_EMBRYONIC\_PLACENTA\_DEVELOPMENT | MSigDB lists | GO\_EMBRYONIC\_PLACENTA\_DEVELOPMENT | 14 | 4 | 4161 | 164 | rl,fz2,Bruce,svp | | 1.713e-03 | -6.37 | cell morphogenesis | biological process | GO:0000902 | 533 | 40 | 12345 | 571 | pho,hdc,Stlk,klg,CG41099,RhoGAP100F,shot,Ptp99A,mbc,Ten-a,Ulp1,sn,Sema1a,Src42A,dnc,cta,Dscam2,pum,SoxN,dsx,lola,en,fz2,dnt,Mp,scro,pan,wb,CdGAPr,br,Sdc,ct,ap,fru,Psc,jing,Mical,Toll-7,neo,CadN2 | | 1.756e-03 | -6.34 | fly chr40F7-40F7\|2-54 | chromosome location | fly chr40F7-40F7\|2-54 | 8 | 3 | 25010 | 823 | Marf1,Slmap,snoRNA:Psi28S-2996 | | 1.756e-03 | -6.34 | fly chr40F7-40F7\|2-54 cM | chromosome location | fly chr40F7-40F7\|2-54 cM | 8 | 3 | 25010 | 823 | snoRNA:Psi28S-2996,Slmap,Marf1 | | 1.773e-03 | -6.34 | amnioserosa formation | biological process | GO:0007378 | 6 | 3 | 12345 | 571 | zen,scw,tld | | 1.794e-03 | -6.32 | positive regulation of cellular process | biological process | GO:0048522 | 1464 | 91 | 12345 | 571 | Trl,sna,Camta,AP-2alpha,Tak1,CG40228,d4,CG41520,SoxN,zld,en,lola,CG12769,conu,Glut1,step,jing,bi,Diap1,rl,pan,mav,sif,lqf,Setd3,Slmap,pico,pk,zen,Ptip,capu,pho,CG12200,CG12147,CG41106,fray,scw,AGO3,bru1,Parp,lz,mlt,mbc,hth,Src42A,Mp,scro,Wdr59,Octbeta2R,orb,svp,tld,Catsup,pum,toy,dsx,mxt,norpA,ap,tou,Myc,tj,Toll-7,SCAP,CG14184,Der-1,Pld,br,Sdc,CG17514,shot,CG10188,Marf1,Mef2,ewg,PEK,caup,Git,Stlk,CG30440,Atf6,Clk,rau,Alg-2,Su(z)2,grim,Gprk1,dimm,Hipk,Sema1a,ru | | 1.821e-03 | -6.31 | SH3\_domain | interpro domains | IPR001452 | 69 | 10 | 11383 | 557 | CAP,vap,CdGAPr,mbc,Src42A,shot,SKIP,l(3)05822,ck,Cip4 | | 1.836e-03 | -6.30 | GSE36392\_TYPE\_2\_MYELOID\_VS\_MAC\_IL25\_TREATED\_LUNG\_UP | MSigDB lists | GSE36392\_TYPE\_2\_MYELOID\_VS\_MAC\_IL25\_TREATED\_LUNG\_UP | 34 | 6 | 4161 | 164 | cta,CG30495,Glut1,Meltrin,Octbeta1R,RSG7 | | 1.836e-03 | -6.30 | GSE28737\_BCL6\_HET\_VS\_BCL6\_KO\_MARGINAL\_ZONE\_BCELL\_UP | MSigDB lists | GSE28737\_BCL6\_HET\_VS\_BCL6\_KO\_MARGINAL\_ZONE\_BCELL\_UP | 34 | 6 | 4161 | 164 | SdhA,sdk,CG9886,Atpalpha,CG9492,bru1 | | 1.869e-03 | -6.28 | RNA polymerase II regulatory region DNA binding | molecular function | GO:0001012 | 261 | 23 | 12176 | 554 | Myc,en,dsx,CG12769,tj,pan,ap,Clk,scro,E5,br,pho,bi,sna,CG9932,cic,svp,ct,toy,Atf6,zen,Mef2,ewg | | 1.872e-03 | -6.28 | GO\_PLACENTA\_BLOOD\_VESSEL\_DEVELOPMENT | MSigDB lists | GO\_PLACENTA\_BLOOD\_VESSEL\_DEVELOPMENT | 7 | 3 | 4161 | 164 | svp,fz2,rl | | 1.889e-03 | -6.27 | axon development | biological process | GO:0061564 | 336 | 28 | 12345 | 571 | Dscam2,CadN2,Toll-7,Mical,Psc,jing,ap,fru,dnc,Sdc,Src42A,Sema1a,CdGAPr,Ulp1,Ten-a,wb,Ptp99A,shot,RhoGAP100F,Mp,dnt,fz2,en,lola,dsx,klg,hdc,SoxN | | 1.894e-03 | -6.27 | cell morphogenesis involved in differentiation | biological process | GO:0000904 | 468 | 36 | 12345 | 571 | ct,Sdc,br,CdGAPr,wb,CadN2,neo,Toll-7,Mical,jing,Psc,fru,ap,en,lola,dsx,SoxN,pum,scro,Mp,dnt,fz2,Sema1a,Src42A,Ulp1,Ten-a,Dscam2,cta,dnc,klg,hdc,pho,Ptp99A,shot,RhoGAP100F,CG41099 | | 1.957e-03 | -6.24 | GSE22033\_UNTREATED\_VS\_ROSIGLITAZONE\_TREATED\_MEF\_DN | MSigDB lists | GSE22033\_UNTREATED\_VS\_ROSIGLITAZONE\_TREATED\_MEF\_DN | 59 | 8 | 4161 | 164 | Pdk,GLS,Coq7,Pgant5,CG12121,Khc-73,Cyp6a8,Syx18 | | 1.957e-03 | -6.24 | OCT1\_04 | MSigDB lists | OCT1\_04 | 59 | 8 | 4161 | 164 | toy,Mef2,Cyp6a8,Dys,CG4341,Ca-alpha1D,Pgant5,Rbp9 | | 1.959e-03 | -6.24 | chemical synaptic transmission | biological process | GO:0007268 | 227 | 21 | 12345 | 571 | dnc,Syt7,nAChRalpha3,unc-13-4A,lqf,Dys,Syx18,be,Atpalpha,Ten-a,svp,sky,cpx,nAChRalpha4,Snap25,cpo,Nlg2,Slob,HisCl1,5-HT2A,pum | | 1.959e-03 | -6.24 | anterograde trans-synaptic signaling | biological process | GO:0098916 | 227 | 21 | 12345 | 571 | nAChRalpha4,Snap25,cpx,sky,svp,pum,5-HT2A,HisCl1,Slob,Nlg2,cpo,nAChRalpha3,Syt7,unc-13-4A,dnc,Ten-a,Atpalpha,be,Syx18,Dys,lqf | | 1.974e-03 | -6.23 | regionalization | biological process | GO:0003002 | 503 | 38 | 12345 | 571 | cic,fz2,kni,fuss,orb,sna,tld,fred,SoxN,Nipped-A,pum,en,ap,step,ck,jing,bi,pan,rl,gus,Dys,ct,pk,par-1,zen,Ptip,capu,caup,jvl,salm,scw,boi,lz,Parp,cv-2,sn,hth,ru | | 1.987e-03 | -6.22 | regulation of cell communication | biological process | GO:0010646 | 855 | 58 | 12345 | 571 | Ppm1,ru,Hipk,Src42A,dimm,Ulp1,hth,Gprk1,cv-2,mbc,Alg-2,rau,CG9650,dnc,scw,rno,Nlg2,CG30440,CG12147,Git,ewg,PEK,rdgC,Snap25,CG10188,par-1,tay,Slmap,lqf,Dys,sif,mav,Diap1,rl,CG14184,RSG7,CG34408,Pi3K68D,ArfGAP1,vap,step,norpA,Glut1,conu,SoxN,pum,Tak1,Slob,Camta,tld,svp,sky,fuss,Wdr59,Nlg3,Sod1,Mp,cpx | | 1.987e-03 | -6.22 | regulation of signaling | biological process | GO:0023051 | 855 | 58 | 12345 | 571 | Git,CG12147,CG30440,Nlg2,par-1,tay,Snap25,CG10188,rdgC,ewg,PEK,cv-2,mbc,hth,Gprk1,Ulp1,dimm,Hipk,Src42A,ru,Ppm1,scw,dnc,rno,CG9650,Alg-2,rau,Camta,Slob,Tak1,pum,SoxN,cpx,Mp,Sod1,Nlg3,Wdr59,fuss,sky,svp,tld,rl,Diap1,mav,Dys,sif,lqf,Slmap,conu,Glut1,vap,norpA,step,ArfGAP1,Pi3K68D,CG34408,RSG7,CG14184 | | 2.012e-03 | -6.21 | Chi (Chip) | protein interactions | 37837 | 68 | 10 | 9204 | 464 | dimm,tou,lz,pan,Nipped-A,Scp2,lola,CG9932,ct,ap | | 2.056e-03 | -6.19 | Hipk (Homeodomain interacting protein kinase) | protein interactions | 38070 | 19 | 5 | 9204 | 464 | Dys,Hipk,toy,pan,fz2 | | 2.061e-03 | -6.18 | posttranscriptional gene silencing | biological process | GO:0016441 | 62 | 9 | 12345 | 571 | hdc,pum,AGO3,Ars2,drosha,CG10077,HPS4,Marf1,Hsc70-3 | | 2.067e-03 | -6.18 | glutaminase activity | molecular function | GO:0004359 | 2 | 2 | 12176 | 554 | Nadsyn,GLS | | 2.068e-03 | -6.18 | HIRSCH\_CELLULAR\_TRANSFORMATION\_SIGNATURE\_DN | MSigDB lists | HIRSCH\_CELLULAR\_TRANSFORMATION\_SIGNATURE\_DN | 24 | 5 | 4161 | 164 | Ars2,drosha,shot,svp,Dys | | 2.123e-03 | -6.15 | repressing transcription factor binding | molecular function | GO:0070491 | 21 | 5 | 12176 | 554 | CG1620,cic,pan,kni,sna | | 2.136e-03 | -6.15 | haltere morphogenesis | biological process | GO:0048735 | 2 | 2 | 12345 | 571 | ap,hth | | 2.145e-03 | -6.14 | GSE3982\_DC\_VS\_NKCELL\_DN | MSigDB lists | GSE3982\_DC\_VS\_NKCELL\_DN | 35 | 6 | 4161 | 164 | RpL6,dnc,step,CG32944,Nipped-A,Mef2 | | 2.180e-03 | -6.13 | trans-synaptic signaling | biological process | GO:0099537 | 229 | 21 | 12345 | 571 | cpo,Nlg2,Slob,pum,5-HT2A,HisCl1,sky,svp,cpx,Snap25,nAChRalpha4,Dys,lqf,Syx18,be,Ten-a,Atpalpha,dnc,nAChRalpha3,Syt7,unc-13-4A | | 2.221e-03 | -6.11 | cell morphogenesis involved in neuron differentiation | biological process | GO:0048667 | 422 | 33 | 12345 | 571 | Toll-7,Mical,CadN2,fru,ap,Psc,jing,CdGAPr,ct,Sdc,br,wb,dnt,Mp,fz2,scro,lola,en,dsx,SoxN,pum,Dscam2,dnc,Src42A,Sema1a,Ulp1,Ten-a,Ptp99A,shot,RhoGAP100F,klg,pho,hdc | | 2.230e-03 | -6.11 | sequence-specific DNA binding | molecular function | GO:0043565 | 515 | 38 | 12176 | 554 | ap,Clk,E5,Myc,en,CG12769,hth,Vsx2,zld,cic,zen,Atf6,ct,Trl,kni,Mef2,pho,Camta,lz,pan,scro,CG9650,Mnt,dsx,Maf1,Psc,tj,CG32006,svp,bab2,toy,fd96Ca,ewg,br,caup,bi,CG9932,sna | | 2.249e-03 | -6.10 | Atg1 (Autophagy-related 1) | protein interactions | 39454 | 69 | 10 | 9204 | 464 | Diap1,gus,E5,Nrd1,CG17508,orb,rl,norpA,Myc,vtd | | 2.258e-03 | -6.09 | GO\_PHOTORECEPTOR\_CELL\_DIFFERENTIATION | MSigDB lists | GO\_PHOTORECEPTOR\_CELL\_DIFFERENTIATION | 15 | 4 | 4161 | 164 | toy,sdk,sn,ck | | 2.271e-03 | -6.09 | sqz (squeeze) | protein interactions | 42300 | 6 | 3 | 9204 | 464 | ap,ct,dimm | | 2.299e-03 | -6.08 | synaptic signaling | biological process | GO:0099536 | 230 | 21 | 12345 | 571 | Nlg2,cpo,5-HT2A,pum,HisCl1,Slob,svp,sky,Snap25,nAChRalpha4,cpx,Syx18,Dys,lqf,Ten-a,Atpalpha,be,Syt7,unc-13-4A,nAChRalpha3,dnc | | 2.309e-03 | -6.07 | Mitochondrial ABC transporters | REACTOME pathways | R-DME-1369007 | 2 | 2 | 3923 | 189 | Hmt-1,ABCB7 | | 2.309e-03 | -6.07 | POLB-Dependent Long Patch Base Excision Repair | REACTOME pathways | R-DME-110362 | 2 | 2 | 3923 | 189 | Parg,Parp | | 2.318e-03 | -6.07 | positive regulation of ERBB signaling pathway | biological process | GO:1901186 | 13 | 4 | 12345 | 571 | step,Src42A,rau,ru | | 2.318e-03 | -6.07 | female gonad development | biological process | GO:0008585 | 13 | 4 | 12345 | 571 | ct,tj,fz2,bab2 | | 2.318e-03 | -6.07 | development of primary female sexual characteristics | biological process | GO:0046545 | 13 | 4 | 12345 | 571 | bab2,fz2,tj,ct | | 2.318e-03 | -6.07 | positive regulation of epidermal growth factor receptor signaling pathway | biological process | GO:0045742 | 13 | 4 | 12345 | 571 | step,rau,Src42A,ru | | 2.348e-03 | -6.05 | - | gene3d domains | 2.60.40.10 | 195 | 21 | 4929 | 273 | dpr21,Ptp99A,side-III,sdk,babos,side-II,kirre,nrm,zormin,fan,klg,bt,fred,boi,Dscam2,Camta,CG45263,CG14964,DIP-lambda,beat-VII,DIP-delta | | 2.368e-03 | -6.05 | GO\_VASCULATURE\_DEVELOPMENT | MSigDB lists | GO\_VASCULATURE\_DEVELOPMENT | 89 | 10 | 4161 | 164 | Alg-2,vap,fz2,cv-2,cta,toy,Mef2,svp,Mp,rl | | 2.390e-03 | -6.04 | MAPKKK7 | interpro domains | IPR017421 | 2 | 2 | 11383 | 557 | Takl2,Tak1 | | 2.399e-03 | -6.03 | transcription regulator activity | molecular function | GO:0140110 | 605 | 43 | 12176 | 554 | toy,bab2,CG32006,svp,ewg,fd96Ca,CG33557,caup,bi,Nipped-A,sna,CG9932,fru,CG9650,scro,pan,SoxN,Mnt,Octbeta2R,CG1620,Setd3,dsx,tj,lola,zen,Atf6,cic,zld,Vsx2,Mef2,kni,Trl,lz,Fer3,Camta,Clk,ap,E5,en,Myc,rib,hth,CG12769 | | 2.424e-03 | -6.02 | actin filament-based process | biological process | GO:0030029 | 262 | 23 | 12345 | 571 | Src42A,Msp300,sif,Setd3,pico,Spn,mbc,sn,CG34347,Mical,Pi3K68D,neo,bond,CG43901,CAP,cta,ck,bru1,jvl,CG10939,capu,bt,shot | | 2.445e-03 | -6.01 | TITF1\_Q3 | MSigDB lists | TITF1\_Q3 | 48 | 7 | 4161 | 164 | Nlg3,twz,Odc2,toy,Mef2,Dys,par-1 | | 2.445e-03 | -6.01 | GO\_REGULATION\_OF\_HEART\_CONTRACTION | MSigDB lists | GO\_REGULATION\_OF\_HEART\_CONTRACTION | 48 | 7 | 4161 | 164 | Ca-alpha1D,PMCA,dnc,Octbeta2R,bru1,Atpalpha,Dys | | 2.445e-03 | -6.01 | AAAYWAACM\_HFH4\_01 | MSigDB lists | AAAYWAACM\_HFH4\_01 | 48 | 7 | 4161 | 164 | Pi3K68D,sdk,Dys,Mef2,vtd,svp,dnr1 | | 2.520e-03 | -5.98 | Ig\_I-set | interpro domains | IPR013098 | 72 | 10 | 11383 | 557 | bt,DIP-delta,sdk,kirre,zormin,DIP-lambda,Dscam2,CG14964,klg,dpr21 | | 2.531e-03 | -5.98 | cellular component morphogenesis | biological process | GO:0032989 | 667 | 47 | 12345 | 571 | Dscam2,bru1,cta,dnc,Sema1a,Src42A,Ulp1,sn,Ten-a,mbc,lama,Ptp99A,shot,RhoGAP100F,CG41099,klg,Stlk,hdc,pho,jvl,CadN2,Toll-7,neo,Mical,Psc,jing,fru,ap,ct,Sdc,br,CdGAPr,gus,wb,pan,rl,bt,scro,dnt,Mp,cic,fz2,en,lola,dsx,SoxN,pum | | 2.536e-03 | -5.98 | Octbeta2R (Octopamine beta2 receptor) | protein interactions | 41549 | 2 | 2 | 9204 | 464 | Octbeta1R,Myc | | 2.536e-03 | -5.98 | CG15145 (uncharacterized protein) | protein interactions | 35064 | 2 | 2 | 9204 | 464 | Rtnl1,Sema1a | | 2.536e-03 | -5.98 | Reck (Reversion-inducing-cysteine-rich protein with kazal motifs) | protein interactions | 39629 | 2 | 2 | 9204 | 464 | Vsx2,CG14820 | | 2.536e-03 | -5.98 | rgr (regular) | protein interactions | 35843 | 2 | 2 | 9204 | 464 | rau,ct | | 2.536e-03 | -5.98 | ATP8B (ATPase 8B) | protein interactions | 41469 | 2 | 2 | 9204 | 464 | Or67d,Or83c | | 2.536e-03 | -5.98 | CG31740 (uncharacterized protein) | protein interactions | 261618 | 2 | 2 | 9204 | 464 | dsx,br | | 2.536e-03 | -5.98 | Ude (Uracil-DNA degrading factor) | protein interactions | 42953 | 2 | 2 | 9204 | 464 | cmb,Sema1a | | 2.536e-03 | -5.98 | CAH16 (Carbonic anhydrase 16) | protein interactions | 50101 | 2 | 2 | 9204 | 464 | Atpalpha,CG41099 | | 2.536e-03 | -5.98 | CG11110 (uncharacterized protein) | protein interactions | 37337 | 2 | 2 | 9204 | 464 | E5,Ptp99A | | 2.552e-03 | -5.97 | N (Notch) | protein interactions | 31293 | 301 | 27 | 9204 | 464 | Toll-7,vtd,ap,ct,Hipk,klg,jigr1,par-1,pk,ND-30,fred,Pi3K68D,bi,fru,PEK,Myc,AP-2alpha,caup,lola,Nipped-A,RpL15,lqf,pan,Catsup,CG41099,Mnt,alpha4GT1 | | 2.559e-03 | -5.97 | plasma membrane part | cellular component | GO:0044459 | 820 | 54 | 11940 | 534 | CG8249,Or98a,CG34357,CadN2,ru,Or69a,Toll-7,Ca-Ma2d,sdk,Rift,CG33970,Dscam2,Src42A,par-1,Octbeta1R,DIP-lambda,Oatp33Ea,Nlg3,dnt,CG15408,nAChRalpha3,Dys,Rh3,cta,klg,Ca-alpha1D,AP-2alpha,mbc,Nlg2,Or83c,Rgk1,Ac78C,Tsp5D,Magi,TrissinR,Cip4,Atpalpha,DIP-delta,fz2,KCNQ,HisCl1,SdhA,Octbeta2R,Or67d,Ten-a,CG15449,nAChRalpha4,CG42637,GABA-B-R3,5-HT2A,CG1213,Teh2,step,norpA | | 2.570e-03 | -5.96 | fly chr35B3-35B3\|2-50 | chromosome location | fly chr35B3-35B3\|2-50 | 9 | 3 | 25010 | 823 | lncRNA:CR44856,lncRNA:CR43411,CG42680 | | 2.570e-03 | -5.96 | fly chr35B3-35B3\|2-50 cM | chromosome location | fly chr35B3-35B3\|2-50 cM | 9 | 3 | 25010 | 823 | lncRNA:CR43411,lncRNA:CR44856,CG42680 | | 2.647e-03 | -5.93 | G protein-coupled neurotransmitter receptor activity | molecular function | GO:0099528 | 22 | 5 | 12176 | 554 | 5-HT2A,5-HT2B,Octbeta1R,Octbeta2R,GABA-B-R3 | | 2.659e-03 | -5.93 | regulation of metabolic process | biological process | GO:0019222 | 2165 | 126 | 12345 | 571 | Stlk,Vsx2,Atf6,caup,bru3,hdc,salm,CG5862,ewg,Marf1,PEK,Mef2,CG10077,SmydA-9,Spn75F,CG33557,heph,Coq7,Hipk,cv-2,CycB,dimm,Su(z)2,Thor,fd96Ca,Alg-2,SteXh:CG42398,grim,rib,Clk,CG9650,mxt,toy,dsx,CG12772,Fer3,Catsup,pum,orb,Octbeta2R,HPS4,tld,svp,bab2,Maf1,kni,Sod1,dnr1,scro,CG17514,ct,br,Sdc,PNPase,jigr1,gus,Toll-7,Tgi,tj,Hsc70-3,CG14184,SCAP,spel1,ap,Ars2,CG1620,Myc,CG9932,tou,Bruce,CG41106,Lsd-2,pho,CG12147,CG12200,par-1,zen,CG32006,Ptip,SmydA-8,Src42A,Ppm1,mbc,hth,lz,Parp,scw,fray,AGO3,bru1,Spn42De,zld,en,lola,CG40228,Tak1,Camta,AP-2alpha,lncRNA:bxd,SoxN,CG41520,Nipped-A,d4,Mnt,fuss,sna,cic,drosha,Trl,mrt,Setd3,pan,rl,CG34334,Diap1,mav,bi,Parg,step,fru,CG12769,E5,vtd,Psc,HP6,jing | | 2.699e-03 | -5.91 | MORF\_ATF2 | MSigDB lists | MORF\_ATF2 | 62 | 8 | 4161 | 164 | Coq7,fz2,CG40178,dnc,Pi3K68D,RSG7,toy,Dys | | 2.716e-03 | -5.91 | G protein-coupled receptor activity | molecular function | GO:0004930 | 118 | 13 | 12176 | 554 | GABA-B-R3,mthl8,Gr2a,SPR,Octbeta1R,5-HT2B,5-HT2A,Octbeta2R,Or67d,TrissinR,CG31750,Rh3,mthl11 | | 2.759e-03 | -5.89 | MORF\_PTPRB | MSigDB lists | MORF\_PTPRB | 49 | 7 | 4161 | 164 | toy,Dys,RSG7,fz2,Coq7,dnc,Pi3K68D | | 2.759e-03 | -5.89 | ODONNELL\_TFRC\_TARGETS\_UP | MSigDB lists | ODONNELL\_TFRC\_TARGETS\_UP | 49 | 7 | 4161 | 164 | dnr1,Pkg21D,Glut1,heph,Wdr59,ck,PMCA | | 2.775e-03 | -5.89 | SKP1/BTB/POZ\_sf | interpro domains | IPR011333 | 85 | 11 | 11383 | 557 | lola,rib,fru,bab2,inc,br,CG15725,CG41099,CG12857,twz,Trl | | 2.792e-03 | -5.88 | sensory organ development | biological process | GO:0007423 | 496 | 37 | 12345 | 571 | Src42A,Hipk,ru,Nna1,NAT1,hth,sn,rau,lz,cato,boi,rno,klg,caup,salm,zen,shot,RhoGAP100F,sif,br,pk,Diap1,rl,sdk,bi,CadN2,ck,dsx,toy,lola,Tak1,fred,sna,kirre,svp,cic,rg | | 2.837e-03 | -5.86 | drug binding | molecular function | GO:0008144 | 828 | 55 | 12176 | 554 | bma,Myo81F,Src42A,CG9492,Hsc70-3,Nadsyn,Cht10,Gprk1,CG31689,Klp59C,Stlk,Obp85a,Tak1,Hipk,CG12147,fbl,CG5897,CG9981,par-1,DNAlig4,CG43672,CG34357,ck,PMCA,Takl2,rl,CG1486,CG10077,CG14721,CG33970,lz,Pkg21D,ABCB7,bt,spel1,Tdc2,Khc-73,kl-3,Hmt-1,dnt,CG42637,CG5789,CG3556,mus308,CG32944,CG14304,CG7536,thw,CG14301,PEK,Mur89F,Dhc16F,Atpalpha,Idgf4,fray | | 2.843e-03 | -5.86 | leg disc pattern formation | biological process | GO:0035223 | 22 | 5 | 12345 | 571 | jing,hth,rl,ap,ru | | 2.843e-03 | -5.86 | proximal/distal pattern formation, imaginal disc | biological process | GO:0007449 | 22 | 5 | 12345 | 571 | ru,hth,jing,ap,rl | | 2.856e-03 | -5.86 | eIF4E1 (eukaryotic translation initiation factor 4E1) | protein interactions | 45525 | 49 | 8 | 9204 | 464 | Diap1,fest,pum,mxt,ewg,ct,Thor,bru1 | | 2.901e-03 | -5.84 | Abd-B (Abdominal B) | protein interactions | 47763 | 60 | 9 | 9204 | 464 | ap,Mef2,CG1486,hth,kni,svp,pan,salm,Trl | | 2.909e-03 | -5.84 | GO\_REACTIVE\_OXYGEN\_SPECIES\_BIOSYNTHETIC\_PROCESS | MSigDB lists | GO\_REACTIVE\_OXYGEN\_SPECIES\_BIOSYNTHETIC\_PROCESS | 8 | 3 | 4161 | 164 | Sod1,arg,CG7255 | | 2.909e-03 | -5.84 | GO\_MOTILE\_CILIUM\_ASSEMBLY | MSigDB lists | GO\_MOTILE\_CILIUM\_ASSEMBLY | 8 | 3 | 4161 | 164 | CG17387,Dys,l(2)41Ab | | 2.928e-03 | -5.83 | synapse part | cellular component | GO:0044456 | 211 | 19 | 11940 | 534 | Mical,pum,cpx,mav,unc-13-4A,Git,Ten-a,nAChRalpha4,par-1,sky,nAChRalpha3,Snap25,Spn,RhoGAP100F,Nlg2,Rgk1,Syt7,Magi,Catsup | | 2.949e-03 | -5.83 | regulatory region nucleic acid binding | molecular function | GO:0001067 | 369 | 29 | 12176 | 554 | mrt,E5,pan,ap,scro,Clk,tj,hth,CG12769,Maf1,dsx,en,Myc,ewg,Mef2,ct,toy,Atf6,zen,zld,cic,svp,jigr1,lz,sna,CG9932,pho,bi,br | | 2.949e-03 | -5.83 | transcription regulatory region DNA binding | molecular function | GO:0044212 | 369 | 29 | 12176 | 554 | svp,cic,zld,Atf6,zen,toy,ct,ewg,Mef2,br,pho,bi,CG9932,lz,jigr1,sna,ap,scro,Clk,pan,E5,mrt,Myc,en,dsx,Maf1,CG12769,hth,tj | | 2.978e-03 | -5.82 | RNA polymerase II proximal promoter sequence-specific DNA binding | molecular function | GO:0000978 | 92 | 11 | 12176 | 554 | CG12769,CG9932,tj,Myc,pho,bi,E5,ewg,scro,zen,Atf6 | | 3.003e-03 | -5.81 | GSE12839\_CTRL\_VS\_IL12\_TREATED\_PBMC\_DN | MSigDB lists | GSE12839\_CTRL\_VS\_IL12\_TREATED\_PBMC\_DN | 26 | 5 | 4161 | 164 | dnr1,CG10077,RSG7,Ac78C,CG41099 | | 3.003e-03 | -5.81 | GSE18791\_UNSTIM\_VS\_NEWCATSLE\_VIRUS\_DC\_2H\_DN | MSigDB lists | GSE18791\_UNSTIM\_VS\_NEWCATSLE\_VIRUS\_DC\_2H\_DN | 26 | 5 | 4161 | 164 | alpha4GT1,par-1,Ubr1,mIF2,CG1486 | | 3.039e-03 | -5.80 | embryonic morphogenesis | biological process | GO:0048598 | 251 | 22 | 12345 | 571 | cta,step,rib,scw,pan,mbc,ct,Src42A,Pld,shot,CG41099,zen,tld,svp,spo,sna,nullo,pum,5-HT2A,Irp-1A,knk,Tak1 | | 3.094e-03 | -5.78 | BTB/POZ\_dom | interpro domains | IPR000210 | 74 | 10 | 11383 | 557 | CG41099,Trl,twz,fru,rib,lola,CG15725,br,inc,bab2 | | 3.103e-03 | -5.78 | I-set | pfam domains | PF07679 | 72 | 10 | 10480 | 528 | DIP-lambda,bt,CG14964,kirre,zormin,klg,Dscam2,dpr21,sdk,DIP-delta | | 3.104e-03 | -5.77 | GSE20152\_SPHK1\_KO\_VS\_HTNFA\_OVEREXPRESS\_ANKLE\_UP | MSigDB lists | GSE20152\_SPHK1\_KO\_VS\_HTNFA\_OVEREXPRESS\_ANKLE\_UP | 50 | 7 | 4161 | 164 | CG14621,Aldh-III,twz,Tsp5D,Alg9,CG34408,CG30495 | | 3.167e-03 | -5.76 | negative regulation of RNA metabolic process | biological process | GO:0051253 | 398 | 31 | 12345 | 571 | zld,Vsx2,en,dsx,Fer3,pho,lncRNA:bxd,d4,pum,Mnt,sna,HPS4,svp,cic,Maf1,kni,Trl,ct,pan,Su(z)2,Tgi,lz,bi,Parg,CG1620,E5,Clk,Psc,jing,HP6,CG9932 | | 3.167e-03 | -5.75 | CG12116 (uncharacterized protein) | protein interactions | 31782 | 13 | 4 | 9204 | 464 | Odc2,CG7029,grim,CG8539 | | 3.174e-03 | -5.75 | fly chr21B4-21B4\|2-0.5 cM | chromosome location | fly chr21B4-21B4\|2-0.5 cM | 3 | 2 | 25010 | 823 | Tbc1d15-17,mRpL10 | | 3.174e-03 | -5.75 | fly chr80F-80F\|3-47 | chromosome location | fly chr80F-80F\|3-47 | 3 | 2 | 25010 | 823 | RpL15,vtd | | 3.174e-03 | -5.75 | fly chr87C6-87C6\|3-52 | chromosome location | fly chr87C6-87C6\|3-52 | 3 | 2 | 25010 | 823 | CG12279,mus308 | | 3.174e-03 | -5.75 | fly chr21B4-21B4\|2-0.5 | chromosome location | fly chr21B4-21B4\|2-0.5 | 3 | 2 | 25010 | 823 | Tbc1d15-17,mRpL10 | | 3.174e-03 | -5.75 | fly chr87C6-87C6\|3-52 cM | chromosome location | fly chr87C6-87C6\|3-52 cM | 3 | 2 | 25010 | 823 | CG12279,mus308 | | 3.174e-03 | -5.75 | fly chr80F-80F\|3-47 cM | chromosome location | fly chr80F-80F\|3-47 cM | 3 | 2 | 25010 | 823 | RpL15,vtd | | 3.174e-03 | -5.75 | fly chrh41-h41\|2-55 cM | chromosome location | fly chrh41-h41\|2-55 cM | 3 | 2 | 25010 | 823 | rl,l(2)41Ab | | 3.174e-03 | -5.75 | fly chrh41-h41\|2-55 | chromosome location | fly chrh41-h41\|2-55 | 3 | 2 | 25010 | 823 | l(2)41Ab,rl | | 3.190e-03 | -5.75 | GO\_EMBRYO\_DEVELOPMENT | MSigDB lists | GO\_EMBRYO\_DEVELOPMENT | 227 | 18 | 4161 | 164 | ck,l(2)41Ab,toy,Mef2,cta,lqf,fz2,ap,vap,CG9107,Bruce,CG17387,rl,DNAlig4,CycB,spel1,Sod1,svp | | 3.265e-03 | -5.72 | positive regulation of protein modification process | biological process | GO:0031401 | 176 | 17 | 12345 | 571 | fray,scw,step,CG14184,Alg-2,Su(z)2,mav,Diap1,mbc,Src42A,Trl,PEK,sna,CG41520,Tak1,CG41106,Stlk | | 3.286e-03 | -5.72 | GO\_PLASMA\_MEMBRANE\_PROTEIN\_COMPLEX | MSigDB lists | GO\_PLASMA\_MEMBRANE\_PROTEIN\_COMPLEX | 93 | 10 | 4161 | 164 | Ca-alpha1D,nAChRalpha3,Atpalpha,nAChRalpha4,AP-2alpha,dnc,RSG7,cta,Hmt-1,Dys | | 3.309e-03 | -5.71 | GSE39556\_UNTREATED\_VS\_3H\_POLYIC\_INJ\_MOUSE\_NK\_CELL\_DN | MSigDB lists | GSE39556\_UNTREATED\_VS\_3H\_POLYIC\_INJ\_MOUSE\_NK\_CELL\_DN | 38 | 6 | 4161 | 164 | CG30495,GLS,CG30059,Tsp5D,Mef2,DNAlig4 | | 3.309e-03 | -5.71 | HNF3\_Q6 | MSigDB lists | HNF3\_Q6 | 38 | 6 | 4161 | 164 | Mef2,Dys,Cyp6a8,Pi3K68D,DNaseII,twz | | 3.336e-03 | -5.70 | Serotonin receptors | REACTOME pathways | R-DME-390666 | 7 | 3 | 3923 | 189 | 5-HT2A,Octbeta1R,Octbeta2R | | 3.349e-03 | -5.70 | response to chemical | biological process | GO:0042221 | 1191 | 75 | 12345 | 571 | Ten-a,Gr39a,CG6888,Sema1a,Tdc2,Or83c,Clk,CG7255,Syt7,Thor,Dscam2,Ac78C,klg,Atf6,RhoGAP100F,shot,Rift,Ptp99A,Ir41a,PEK,Or67d,Der-1,CdGAPr,Cyp6a8,ct,br,Sdc,vap,ap,norpA,St1,Myc,Or69a,Cyp6g2,Toll-7,CG31750,Hsc70-3,CG14184,Gr2a,Catsup,pum,godzilla,dsx,Mp,fz2,rg,Sod1,CG3556,svp,Ulp1,Ir40a,Src42A,scw,dnc,Gr93d,Or98a,Ir60e,Hmt-1,rl,mav,wb,fru,Psc,jing,Mical,CadN2,Tak1,Ir94e,SoxN,lola,en,dnt,CG17691,Trl,CheB42a | | 3.355e-03 | -5.70 | ash2 (absent, small, or homeotic discs 2) | protein interactions | 42936 | 30 | 6 | 9204 | 464 | Set1,cv-2,Ptip,rl,kni,Myc | | 3.428e-03 | -5.68 | Rho GTPase cycle | REACTOME pathways | R-DME-194840 | 53 | 8 | 3923 | 189 | conu,CdGAPr,CG30440,RhoGAP100F,Cip4,sif,cta,CG10188 | | 3.438e-03 | -5.67 | negative regulation of nucleobase-containing compound metabolic process | biological process | GO:0045934 | 417 | 32 | 12345 | 571 | Maf1,cic,Trl,kni,sna,Mnt,svp,HPS4,Fer3,d4,pum,lncRNA:bxd,zld,dsx,en,E5,CG1620,spel1,Psc,jing,CG9932,HP6,bi,Tgi,Parg,pan,ct,pho,Vsx2,Clk,lz,Su(z)2 | | 3.453e-03 | -5.67 | Ig\_V-set | interpro domains | IPR013106 | 31 | 6 | 11383 | 557 | DIP-lambda,dpr21,nrm,side-III,kirre,sdk | | 3.479e-03 | -5.66 | AACTTT\_UNKNOWN | MSigDB lists | AACTTT\_UNKNOWN | 421 | 28 | 4161 | 164 | Ranbp16,dnc,CG9389,Dys,svp,CG32944,dnr1,Gfat1,ap,fz2,nAChRalpha3,twz,Tsp5D,Bruce,vtd,Ac78C,Mef2,CG2528,cta,toy,Cyp6a8,Octbeta2R,Ca-alpha1D,sbm,CG4341,Rbp9,GLS,MCU | | 3.481e-03 | -5.66 | GSE8868\_SPLEEN\_VS\_INTESTINE\_CD11B\_POS\_CD11C\_NEG\_DC\_DN | MSigDB lists | GSE8868\_SPLEEN\_VS\_INTESTINE\_CD11B\_POS\_CD11C\_NEG\_DC\_DN | 51 | 7 | 4161 | 164 | Pdk,Alg9,CG30059,MFS16,CG42813,Meltrin,Hsc70-3 | | 3.509e-03 | -5.65 | anion binding | molecular function | GO:0043168 | 1099 | 69 | 12176 | 554 | Acox57D-d,rl,Takl2,CG1486,CG5160,kl-3,CG9514,Sema1a,CG42637,Hmt-1,Pkg21D,lz,Tdc2,cert,RabX2,Syt7,Pld,fray,CG7536,CG5270,mIF2,Myo81F,Gprk1,CG31689,Stlk,Klp59C,Trxr-2,CG12147,fzo,CG43672,FASN3,mim,CG10077,PMCA,ck,CG5789,dnt,bt,ABCB7,CG14721,CG33970,Khc-73,spel1,mus308,CG32944,Atpalpha,Pi3K68D,Dhc16F,PEK,CG9492,Src42A,Acbp5,Rgk1,bma,SdhA,Hsc70-3,cv-2,Nadsyn,Tak1,Hipk,CG42541,Mical,CG34357,Cip4,CG9981,fbl,DNAlig4,par-1,step | | 3.569e-03 | -5.64 | GCANCTGNY\_MYOD\_Q6 | MSigDB lists | GCANCTGNY\_MYOD\_Q6 | 176 | 15 | 4161 | 164 | Ranbp16,Nlg3,ck,Hsc70-3,Dys,Khc-73,svp,Odc2,CG4341,sn,Atpalpha,Tsp5D,Rbp9,rg,shot | | 3.569e-03 | -5.64 | GSE21379\_WT\_VS\_SAP\_KO\_TFH\_CD4\_TCELL\_UP | MSigDB lists | GSE21379\_WT\_VS\_SAP\_KO\_TFH\_CD4\_TCELL\_UP | 27 | 5 | 4161 | 164 | CG30059,mbc,CG7255,Src42A,bru3 | | 3.570e-03 | -5.64 | MEISSNER\_BRAIN\_HCP\_WITH\_H3K4ME3\_AND\_H3K27ME3 | MSigDB lists | MEISSNER\_BRAIN\_HCP\_WITH\_H3K4ME3\_AND\_H3K27ME3 | 142 | 13 | 4161 | 164 | Mp,Mct1,toy,arg,Octbeta2R,CG5254,mbc,CG7255,fz2,ap,alpha4GT1,gus,CG6788 | | 3.594e-03 | -5.63 | positive regulation of transcription by RNA polymerase II | biological process | GO:0045944 | 368 | 29 | 12345 | 571 | zld,toy,Atf6,dsx,en,Camta,caup,d4,SoxN,sna,Mef2,zen,Trl,scro,Setd3,br,pan,hth,dimm,lz,bi,tj,SCAP,CG12769,ap,jing,tou,Myc,Clk | | 3.643e-03 | -5.61 | cup (cup) | protein interactions | 33934 | 62 | 9 | 9204 | 464 | Rbp9,CG40045,Cht10,pum,orb,gus,Pdk,Tak1,bru1 | | 3.702e-03 | -5.60 | MORF\_IL9 | MSigDB lists | MORF\_IL9 | 17 | 4 | 4161 | 164 | cv-2,Src42A,nAChRalpha4,Nlg2 | | 3.702e-03 | -5.60 | GO\_SYNAPSE\_ASSEMBLY | MSigDB lists | GO\_SYNAPSE\_ASSEMBLY | 17 | 4 | 4161 | 164 | Nlg2,fz2,Nlg3,sdk | | 3.767e-03 | -5.58 | positive regulation of phosphorus metabolic process | biological process | GO:0010562 | 149 | 15 | 12345 | 571 | CG17514,CG41106,Src42A,Stlk,mav,CG41520,mbc,Tak1,CG14184,PEK,Alg-2,Trl,step,fray,scw | | 3.767e-03 | -5.58 | positive regulation of phosphate metabolic process | biological process | GO:0045937 | 149 | 15 | 12345 | 571 | Stlk,Src42A,CG17514,CG41106,Tak1,mbc,CG41520,mav,Alg-2,CG14184,PEK,step,scw,fray,Trl | | 3.786e-03 | -5.58 | MORF\_PRKCA | MSigDB lists | MORF\_PRKCA | 39 | 6 | 4161 | 164 | dnc,Pi3K68D,fz2,Coq7,Dys,toy | | 3.786e-03 | -5.58 | RUTELLA\_RESPONSE\_TO\_HGF\_DN | MSigDB lists | RUTELLA\_RESPONSE\_TO\_HGF\_DN | 39 | 6 | 4161 | 164 | Mef2,CG9389,Glut1,bru1,step,Meltrin | | 3.826e-03 | -5.57 | puf (puffyeye) | protein interactions | 42935 | 7 | 3 | 9204 | 464 | Diap1,Myc,Sema1a | | 3.826e-03 | -5.57 | CG6996 (uncharacterized protein) | protein interactions | 40212 | 7 | 3 | 9204 | 464 | Vsx2,zld,dsx | | 3.840e-03 | -5.56 | GO\_IN\_UTERO\_EMBRYONIC\_DEVELOPMENT | MSigDB lists | GO\_IN\_UTERO\_EMBRYONIC\_DEVELOPMENT | 95 | 10 | 4161 | 164 | Bruce,CG9107,fz2,cta,svp,lqf,rl,spel1,DNAlig4,CycB | | 3.892e-03 | -5.55 | MORF\_CAMK4 | MSigDB lists | MORF\_CAMK4 | 52 | 7 | 4161 | 164 | toy,Dys,CG40178,fz2,dnc,Coq7,RSG7 | | 3.892e-03 | -5.55 | GSE28737\_WT\_VS\_BCL6\_HET\_FOLLICULAR\_BCELL\_UP | MSigDB lists | GSE28737\_WT\_VS\_BCL6\_HET\_FOLLICULAR\_BCELL\_UP | 52 | 7 | 4161 | 164 | unc-13-4A,mbc,CG2061,SCAP,CG42813,Khc-73,CG1486 | | 3.892e-03 | -5.55 | GO\_SENSORY\_ORGAN\_MORPHOGENESIS | MSigDB lists | GO\_SENSORY\_ORGAN\_MORPHOGENESIS | 52 | 7 | 4161 | 164 | Sod1,toy,rl,fz2,ck,sdk,sn | | 3.892e-03 | -5.55 | GSE2706\_UNSTIM\_VS\_2H\_R848\_DC\_UP | MSigDB lists | GSE2706\_UNSTIM\_VS\_2H\_R848\_DC\_UP | 52 | 7 | 4161 | 164 | dnr1,mRpS5,Ubr1,CG42813,drosha,SdhA,Parg | | 3.892e-03 | -5.55 | GFI1\_01 | MSigDB lists | GFI1\_01 | 52 | 7 | 4161 | 164 | dnc,CG4341,fbl,Rbp9,Mef2,CG2528,svp | | 3.902e-03 | -5.55 | GPCR downstream signaling | REACTOME pathways | R-DME-388396 | 91 | 11 | 3923 | 189 | CG30440,dnc,5-HT2A,Octbeta1R,norpA,Gprk1,Octbeta2R,sif,Ac78C,cta,CG10188 | | 3.920e-03 | -5.54 | TGACATY\_UNKNOWN | MSigDB lists | TGACATY\_UNKNOWN | 127 | 12 | 4161 | 164 | Mef2,Dys,Khc-73,svp,dnr1,AP-2alpha,dnc,ck,Rbp9,Bruce,PMCA,Ca-alpha1D | | 3.945e-03 | -5.54 | integral component of membrane | cellular component | GO:0016021 | 1438 | 85 | 11940 | 534 | CG43672,SK,Octbeta1R,Oatp33Ea,Rift,CG33970,Dscam2,ABCB7,Ir41a,mthl8,CG12121,CG8249,Or98a,Glut1,CG34357,Or69a,CG45076,CG10171,Meltrin,CG5789,ru,neo,Or67d,Octbeta2R,mthl11,TrissinR,CG9981,CG14744,5-HT2A,CG1213,Teh2,Gr2a,Ir40a,fzo,Der-1,Rh3,Ca-alpha1D,Nlg3,Ac78C,CG41128,CG7536,Syt7,CG14024,Hmt-1,sdk,Ca-Ma2d,DIP-lambda,Ir94e,Tim23,CG2003,kune,Gr93d,CG14160,rg,MFS16,CG14196,Ir60e,CadN2,MCU,SPR,Mct1,CG15449,snky,fz2,Atpalpha,DIP-delta,KCNQ,Tim17b,HisCl1,GABA-B-R3,CG42637,Pzl,Sema1a,CG33233,nAChRalpha4,bond,dnt,Pgant9,CG15408,nAChRalpha3,Tsp5D,Or83c,Gr39a,Nlg2,Msp300 | | 3.986e-03 | -5.52 | vg (vestigial) | protein interactions | 36421 | 31 | 6 | 9204 | 464 | Diap1,Nipped-A,Mef2,ap,Tgi,salm | | 4.001e-03 | -5.52 | sli (slit) | protein interactions | 36746 | 41 | 7 | 9204 | 464 | Sdc,lola,shot,jing,Mp,wb,Dys | | 4.012e-03 | -5.52 | GO\_EMBRYONIC\_ORGAN\_MORPHOGENESIS | MSigDB lists | GO\_EMBRYONIC\_ORGAN\_MORPHOGENESIS | 66 | 8 | 4161 | 164 | rl,ck,l(2)41Ab,toy,Mef2,Sod1,fz2,CG17387 | | 4.012e-03 | -5.52 | MORF\_RAD51L3 | MSigDB lists | MORF\_RAD51L3 | 66 | 8 | 4161 | 164 | Coq7,fz2,toy,Dys,CG40178,dnc,Pi3K68D,RSG7 | | 4.110e-03 | -5.49 | compound eye cone cell differentiation | biological process | GO:0042675 | 15 | 4 | 12345 | 571 | rg,sdk,ru,lz | | 4.139e-03 | -5.49 | positive regulation of biological process | biological process | GO:0048518 | 1642 | 98 | 12345 | 571 | CG14184,SCAP,Toll-7,tj,Myc,tou,ap,norpA,CG17514,br,Sdc,Pld,Der-1,gus,HPS4,tld,svp,Octbeta2R,orb,Wdr59,scro,Mp,mxt,toy,dsx,pum,Catsup,grim,Su(z)2,rau,Alg-2,Clk,ru,Sema1a,Hipk,dimm,Ten-a,Gprk1,ewg,Mef2,Marf1,PEK,CG5862,shot,CG10188,Atf6,Stlk,CG30440,Git,caup,bi,jing,step,Glut1,conu,vtd,CG12769,Slmap,pico,pk,Setd3,lqf,sif,mav,pan,Diap1,rl,sna,Trl,lola,en,zld,CG41520,SoxN,d4,CG40228,Tak1,Camta,AP-2alpha,lz,Parp,bru1,AGO3,fray,scw,Src42A,hth,mbc,mlt,capu,Ptip,zen,CG41106,Lsd-2,CG12147,5-HT2A,CG12200,pho | | 4.153e-03 | -5.48 | proximal promoter sequence-specific DNA binding | molecular function | GO:0000987 | 96 | 11 | 12176 | 554 | scro,Atf6,zen,E5,ewg,Myc,pho,bi,CG12769,tj,CG9932 | | 4.197e-03 | -5.47 | actin cytoskeleton organization | biological process | GO:0030036 | 242 | 21 | 12345 | 571 | jvl,CG10939,capu,bt,shot,sif,Msp300,Src42A,Spn,pico,Setd3,mbc,sn,CG34347,Pi3K68D,Mical,CG43901,bond,bru1,ck,CAP | | 4.209e-03 | -5.47 | GSE9960\_HEALTHY\_VS\_SEPSIS\_PBMC\_DN | MSigDB lists | GSE9960\_HEALTHY\_VS\_SEPSIS\_PBMC\_DN | 28 | 5 | 4161 | 164 | CG30059,Nlg2,AGO3,ck,mlt | | 4.209e-03 | -5.47 | DR3\_Q4 | MSigDB lists | DR3\_Q4 | 28 | 5 | 4161 | 164 | lqf,toy,rg,Dys,CG14621 | | 4.237e-03 | -5.46 | KEGG\_PROXIMAL\_TUBULE\_BICARBONATE\_RECLAMATION | MSigDB lists | KEGG\_PROXIMAL\_TUBULE\_BICARBONATE\_RECLAMATION | 9 | 3 | 4161 | 164 | Atpalpha,GLS,Gdh | | 4.237e-03 | -5.46 | GO\_SOMATIC\_DIVERSIFICATION\_OF\_IMMUNOGLOBULINS | MSigDB lists | GO\_SOMATIC\_DIVERSIFICATION\_OF\_IMMUNOGLOBULINS | 9 | 3 | 4161 | 164 | mus308,spel1,DNAlig4 | | 4.237e-03 | -5.46 | PID\_SYNDECAN\_2\_PATHWAY | MSigDB lists | PID\_SYNDECAN\_2\_PATHWAY | 9 | 3 | 4161 | 164 | Trs23,vap,rl | | 4.259e-03 | -5.46 | Fak (Focal adhesion kinase) | protein interactions | 37233 | 14 | 4 | 9204 | 464 | rl,dnc,Sema1a,CdGAPr | | 4.339e-03 | -5.44 | GSE3982\_MAST\_CELL\_VS\_DC\_UP | MSigDB lists | GSE3982\_MAST\_CELL\_VS\_DC\_UP | 53 | 7 | 4161 | 164 | Khc-73,Nipped-A,shot,rg,heph,RpL6,drosha | | 4.452e-03 | -5.41 | regulation of transcription by RNA polymerase II | biological process | GO:0006357 | 741 | 50 | 12345 | 571 | br,Setd3,ct,pan,mav,bi,tj,SCAP,CG1620,E5,CG12769,fru,ap,jing,tou,CG9932,Myc,zld,toy,dsx,en,lola,Camta,d4,SoxN,sna,Mnt,fuss,svp,tld,cic,bab2,scro,Trl,kni,cv-2,hth,dimm,lz,Su(z)2,fd96Ca,scw,Clk,Atf6,Vsx2,caup,ewg,Mef2,zen,CG32006 | | 4.482e-03 | -5.41 | GO\_POSITIVE\_REGULATION\_OF\_HYDROLASE\_ACTIVITY | MSigDB lists | GO\_POSITIVE\_REGULATION\_OF\_HYDROLASE\_ACTIVITY | 198 | 16 | 4161 | 164 | mbc,CG34408,Alg-2,Git,Octbeta2R,CG10565,cta,Mef2,vap,sky,Hsc70-3,spel1,RSG7,Trs23,step,Tbc1d15-17 | | 4.488e-03 | -5.41 | regulation of macromolecule metabolic process | biological process | GO:0060255 | 2019 | 117 | 12345 | 571 | Parp,lz,fray,scw,AGO3,bru1,Src42A,Ppm1,mbc,hth,par-1,zen,CG32006,SmydA-8,CG41106,pho,CG12147,CG12200,bi,Parg,step,fru,vtd,E5,CG12769,Psc,HP6,jing,Setd3,pan,rl,CG34334,Diap1,mav,fuss,Mnt,sna,cic,drosha,mrt,Trl,Spn42De,zld,en,lola,Tak1,CG40228,Camta,CG41520,SoxN,Nipped-A,lncRNA:bxd,d4,Su(z)2,fd96Ca,Thor,Alg-2,SteXh:CG42398,grim,rib,CG9650,Clk,CG33557,heph,Coq7,Hipk,cv-2,CycB,dimm,CG5862,ewg,PEK,Mef2,Marf1,CG10077,SmydA-9,Spn75F,Stlk,Vsx2,Atf6,bru3,caup,hdc,salm,tj,Tgi,Hsc70-3,CG14184,SCAP,Ars2,ap,spel1,CG1620,Myc,CG9932,tou,CG17514,ct,Sdc,br,PNPase,jigr1,gus,orb,Octbeta2R,HPS4,tld,svp,bab2,Maf1,kni,scro,mxt,dsx,toy,Fer3,pum | | 4.513e-03 | -5.40 | GO\_IRON\_COORDINATION\_ENTITY\_TRANSPORT | MSigDB lists | GO\_IRON\_COORDINATION\_ENTITY\_TRANSPORT | 3 | 2 | 4161 | 164 | Hmt-1,ABCB7 | | 4.513e-03 | -5.40 | chrxp21 | MSigDB lists | chrxp21 | 3 | 2 | 4161 | 164 | Dys,CG2061 | | 4.513e-03 | -5.40 | GO\_REGULATION\_OF\_ALPHA\_AMINO\_3\_HYDROXY\_5\_METHYL\_4\_ISOXAZOLE\_PROPIONATE\_SELECTIVE\_GLUTAMATE\_RECEPTOR\_ACTIVITY | MSigDB lists | GO\_REGULATION\_OF\_ALPHA\_AMINO\_3\_HYDROXY\_5\_METHYL\_4\_ISOXAZOLE\_PROPIONATE\_SELECTIVE\_GLUTAMATE\_RECEPTOR\_ACTIVITY | 3 | 2 | 4161 | 164 | Nlg3,Mef2 | | 4.513e-03 | -5.40 | GO\_NEURON\_CELL\_CELL\_ADHESION | MSigDB lists | GO\_NEURON\_CELL\_CELL\_ADHESION | 3 | 2 | 4161 | 164 | Nlg3,Nlg2 | | 4.513e-03 | -5.40 | GO\_MAINTENANCE\_OF\_CELL\_POLARITY | MSigDB lists | GO\_MAINTENANCE\_OF\_CELL\_POLARITY | 3 | 2 | 4161 | 164 | shot,ap | | 4.513e-03 | -5.40 | GO\_SIGNAL\_TRANSDUCTION\_INVOLVED\_IN\_REGULATION\_OF\_GENE\_EXPRESSION | MSigDB lists | GO\_SIGNAL\_TRANSDUCTION\_INVOLVED\_IN\_REGULATION\_OF\_GENE\_EXPRESSION | 3 | 2 | 4161 | 164 | Parp,toy | | 4.513e-03 | -5.40 | ZEMBUTSU\_SENSITIVITY\_TO\_DOXORUBICIN | MSigDB lists | ZEMBUTSU\_SENSITIVITY\_TO\_DOXORUBICIN | 3 | 2 | 4161 | 164 | Pi3K68D,CycB | | 4.513e-03 | -5.40 | GO\_B\_CELL\_RECEPTOR\_SIGNALING\_PATHWAY | MSigDB lists | GO\_B\_CELL\_RECEPTOR\_SIGNALING\_PATHWAY | 3 | 2 | 4161 | 164 | Mef2,rl | | 4.530e-03 | -5.40 | EPH-Ephrin signaling | REACTOME pathways | R-DME-2682334 | 44 | 7 | 3923 | 189 | Git,vap,Act42A,Sdc,CG10188,AP-2alpha,sif | | 4.598e-03 | -5.38 | Psc (Posterior sex combs) | protein interactions | 36431 | 42 | 7 | 9204 | 464 | vtd,ct,Su(z)2,CycB,Rtnl1,Ulp1,Psc | | 4.616e-03 | -5.38 | HOOI\_ST7\_TARGETS\_DN | MSigDB lists | HOOI\_ST7\_TARGETS\_DN | 18 | 4 | 4161 | 164 | twz,CG9492,Pdk,Ubr1 | | 4.629e-03 | -5.38 | imaginal disc-derived female genitalia development | biological process | GO:0007486 | 8 | 3 | 12345 | 571 | lz,en,dsx | | 4.681e-03 | -5.36 | GTPase activator activity | molecular function | GO:0005096 | 84 | 10 | 12176 | 554 | norpA,Tbc1d15-17,RhoGAP100F,sky,conu,CdGAPr,CG34408,ArfGAP1,vap,Git | | 4.708e-03 | -5.36 | nmo (nemo) | protein interactions | 38890 | 53 | 8 | 9204 | 464 | mthl8,Tak1,Clk,unc-13-4A,pk,Ptp99A,pan,fz2 | | 4.712e-03 | -5.36 | neurotransmitter receptor activity | molecular function | GO:0030594 | 71 | 9 | 12176 | 554 | Ir40a,GABA-B-R3,nAChRalpha4,HisCl1,Octbeta2R,5-HT2B,Octbeta1R,5-HT2A,nAChRalpha3 | | 4.824e-03 | -5.33 | MORF\_SUPT3H | MSigDB lists | MORF\_SUPT3H | 54 | 7 | 4161 | 164 | CG40178,fz2,dnc,Coq7,RSG7,toy,Dys | | 4.889e-03 | -5.32 | GO\_G\_PROTEIN\_COUPLED\_RECEPTOR\_SIGNALING\_PATHWAY\_COUPLED\_TO\_CYCLIC\_NUCLEOTIDE\_SECOND\_MESSENGER | MSigDB lists | GO\_G\_PROTEIN\_COUPLED\_RECEPTOR\_SIGNALING\_PATHWAY\_COUPLED\_TO\_CYCLIC\_NUCLEOTIDE\_SECOND\_MESSENGER | 41 | 6 | 4161 | 164 | Octbeta2R,cta,Octbeta1R,Ac78C,Ca-alpha1D,dnc | | 4.889e-03 | -5.32 | GSE8621\_UNSTIM\_VS\_LPS\_PRIMED\_AND\_LPS\_STIM\_MACROPHAGE\_DN | MSigDB lists | GSE8621\_UNSTIM\_VS\_LPS\_PRIMED\_AND\_LPS\_STIM\_MACROPHAGE\_DN | 41 | 6 | 4161 | 164 | SCAP,Parp,Ranbp16,AP-2alpha,Bruce,inc | | 4.889e-03 | -5.32 | AML1\_Q6 | MSigDB lists | AML1\_Q6 | 41 | 6 | 4161 | 164 | Glut1,CG12547,MFS16,toy,Tsp5D,DNaseII | | 4.889e-03 | -5.32 | CATGTAA\_MIR496 | MSigDB lists | CATGTAA\_MIR496 | 41 | 6 | 4161 | 164 | vtd,Mef2,rg,Dys,Mp,CG40045 | | 4.889e-03 | -5.32 | AML1\_01 | MSigDB lists | AML1\_01 | 41 | 6 | 4161 | 164 | DNaseII,CG12547,Glut1,MFS16,toy,Tsp5D | | 4.927e-03 | -5.31 | KMCATNNWGGA\_UNKNOWN | MSigDB lists | KMCATNNWGGA\_UNKNOWN | 29 | 5 | 4161 | 164 | svp,PNPase,Dys,toy,dnc | | 4.927e-03 | -5.31 | GO\_PLACENTA\_DEVELOPMENT | MSigDB lists | GO\_PLACENTA\_DEVELOPMENT | 29 | 5 | 4161 | 164 | Bruce,Sod1,svp,fz2,rl | | 5.017e-03 | -5.29 | positive regulation of phosphorylation | biological process | GO:0042327 | 139 | 14 | 12345 | 571 | Trl,step,fray,scw,CG14184,PEK,Alg-2,CG41520,mav,Tak1,mbc,CG17514,Src42A,Stlk | | 5.038e-03 | -5.29 | ttm50 (tiny tim 50) | protein interactions | 31266 | 23 | 5 | 9204 | 464 | Diap1,tobi,CG4691,jvl,CG7367 | | 5.038e-03 | -5.29 | RYBP (Ring and YY1 Binding Protein) | protein interactions | 37601 | 23 | 5 | 9204 | 464 | Diap1,pho,tay,Su(z)2,Psc | | 5.116e-03 | -5.28 | axon midline choice point recognition | biological process | GO:0016199 | 25 | 5 | 12345 | 571 | shot,fru,lola,dsx,Sema1a | | 5.390e-03 | -5.22 | Polyamine biosynthesis, arginine => ornithine => putrescine | KEGG pathways | dme\_M00134 | 3 | 2 | 3198 | 138 | Odc2,arg | | 5.390e-03 | -5.22 | D-Glutamine and D-glutamate metabolism | KEGG pathways | dme00471 | 3 | 2 | 3198 | 138 | Gdh,GLS | | 5.390e-03 | -5.22 | D-Glutamine and D-glutamate metabolism | KEGG pathways | ko00471 | 3 | 2 | 3198 | 138 | Gdh,GLS | | 5.390e-03 | -5.22 | Polyamine biosynthesis, arginine => ornithine => putrescine | KEGG pathways | M00134 | 3 | 2 | 3198 | 138 | Odc2,arg | | 5.467e-03 | -5.21 | plasma membrane bounded cell projection | cellular component | GO:0120025 | 626 | 42 | 11940 | 534 | par-1,Dscam2,sky,SK,Rtnl1,orb,Mical,CG9492,mlt,cpx,Dhc16F,Or69a,Or98a,ck,Spn,Gr93d,nAChRalpha4,Gr2a,norpA,CG17387,kl-3,5-HT2A,HisCl1,pum,Or67d,pico,Syt7,Or83c,Gr39a,mbc,hdc,Mst87F,Catsup,sn,boi,Magi,shot,nAChRalpha3,Ptp99A,klg,Snap25,cta | | 5.489e-03 | -5.21 | imaginal disc-derived wing vein morphogenesis | biological process | GO:0008586 | 47 | 7 | 12345 | 571 | ck,salm,tay,tld,heph,Hipk,Dys | | 5.535e-03 | -5.20 | GO\_ACTIVE\_TRANSMEMBRANE\_TRANSPORTER\_ACTIVITY | MSigDB lists | GO\_ACTIVE\_TRANSMEMBRANE\_TRANSPORTER\_ACTIVITY | 116 | 11 | 4161 | 164 | Tim17b,Tim23,PMCA,sbm,CG7255,MCU,MFS16,Atpalpha,ABCB7,Mct1,Hmt-1 | | 5.535e-03 | -5.20 | MODULE\_137 | MSigDB lists | MODULE\_137 | 116 | 11 | 4161 | 164 | Rbp9,Tim17b,ap,bru1,Gfat1,Glut1,Mef2,toy,Octbeta1R,RSG7,unc-13-4A | | 5.580e-03 | -5.19 | Rbfox1 (RNA-binding Fox protein 1) | protein interactions | 39198 | 15 | 4 | 9204 | 464 | ewg,bru1,pum,mir-980 | | 5.642e-03 | -5.18 | SH3-like\_dom\_sf | interpro domains | IPR036028 | 68 | 9 | 11383 | 557 | Cip4,ck,SKIP,l(3)05822,mbc,Src42A,CAP,vap,CdGAPr | | 5.669e-03 | -5.17 | GO\_MEIOSIS\_I | MSigDB lists | GO\_MEIOSIS\_I | 19 | 4 | 4161 | 164 | vtd,Ubr1,Rpn9,spel1 | | 5.680e-03 | -5.17 | adherens junction organization | biological process | GO:0034332 | 36 | 6 | 12345 | 571 | CadN2,kirre,Src42A,sna,Magi,rl | | 5.724e-03 | -5.16 | SH3 | prosite domains | PS50002 | 68 | 10 | 6243 | 365 | shot,ck,vap,l(3)05822,CdGAPr,Cip4,Src42A,CAP,SKIP,mbc | | 5.728e-03 | -5.16 | GO\_EYE\_MORPHOGENESIS | MSigDB lists | GO\_EYE\_MORPHOGENESIS | 30 | 5 | 4161 | 164 | fz2,sdk,sn,ck,toy | | 5.728e-03 | -5.16 | CROMER\_METASTASIS\_UP | MSigDB lists | CROMER\_METASTASIS\_UP | 30 | 5 | 4161 | 164 | Parp,GCS2alpha,CG30059,Glut1,CG12121 | | 5.728e-03 | -5.16 | IL2\_UP.V1\_UP | MSigDB lists | IL2\_UP.V1\_UP | 30 | 5 | 4161 | 164 | Meltrin,Octbeta1R,fz2,svp,Glut1 | | 5.822e-03 | -5.15 | GO\_SYSTEM\_PROCESS | MSigDB lists | GO\_SYSTEM\_PROCESS | 278 | 20 | 4161 | 164 | arg,Dys,Sod1,Pi3K68D,rdgC,dnc,rl,Atpalpha,toy,Mef2,Octbeta2R,Nlg3,ck,Ac78C,GLS,Nlg2,Irp-1A,nAChRalpha4,Ca-alpha1D,sn | | 5.850e-03 | -5.14 | Rho1 (Rho1) | protein interactions | 36775 | 91 | 11 | 9204 | 464 | cta,kirre,capu,RhoGAP100F,Parp,step,Sema1a,CG10939,br,conu,pk | | 5.880e-03 | -5.14 | GO\_INTRINSIC\_COMPONENT\_OF\_MITOCHONDRIAL\_INNER\_MEMBRANE | MSigDB lists | GO\_INTRINSIC\_COMPONENT\_OF\_MITOCHONDRIAL\_INNER\_MEMBRANE | 10 | 3 | 4161 | 164 | MCU,Tim17b,Tim23 | | 5.880e-03 | -5.14 | GO\_COFACTOR\_TRANSPORTER\_ACTIVITY | MSigDB lists | GO\_COFACTOR\_TRANSPORTER\_ACTIVITY | 10 | 3 | 4161 | 164 | Mct1,ABCB7,Hmt-1 | | 5.880e-03 | -5.14 | AMIT\_EGF\_RESPONSE\_480\_MCF10A | MSigDB lists | AMIT\_EGF\_RESPONSE\_480\_MCF10A | 10 | 3 | 4161 | 164 | Odc2,Git,rl | | 5.880e-03 | -5.14 | KEGG\_PRION\_DISEASES | MSigDB lists | KEGG\_PRION\_DISEASES | 10 | 3 | 4161 | 164 | Hsc70-3,rl,Sod1 | | 5.880e-03 | -5.14 | GO\_LABYRINTHINE\_LAYER\_DEVELOPMENT | MSigDB lists | GO\_LABYRINTHINE\_LAYER\_DEVELOPMENT | 10 | 3 | 4161 | 164 | rl,fz2,Bruce | | 5.880e-03 | -5.14 | GO\_RESPONSE\_TO\_ANTIBIOTIC | MSigDB lists | GO\_RESPONSE\_TO\_ANTIBIOTIC | 10 | 3 | 4161 | 164 | Hsc70-3,Mef2,Sod1 | | 5.880e-03 | -5.14 | MODULE\_375 | MSigDB lists | MODULE\_375 | 10 | 3 | 4161 | 164 | Git,cta,Octbeta2R | | 5.880e-03 | -5.14 | GO\_IMMUNOGLOBULIN\_PRODUCTION | MSigDB lists | GO\_IMMUNOGLOBULIN\_PRODUCTION | 10 | 3 | 4161 | 164 | DNAlig4,spel1,mus308 | | 5.880e-03 | -5.14 | GO\_EYE\_PHOTORECEPTOR\_CELL\_DEVELOPMENT | MSigDB lists | GO\_EYE\_PHOTORECEPTOR\_CELL\_DEVELOPMENT | 10 | 3 | 4161 | 164 | ck,sn,toy | | 5.894e-03 | -5.13 | qvr (quiver) | protein interactions | 2768718 | 8 | 3 | 9204 | 464 | nAChRalpha4,ewg,nAChRalpha3 | | 5.894e-03 | -5.13 | CG9609 (uncharacterized protein) | protein interactions | 32663 | 8 | 3 | 9204 | 464 | rau,toy,Scp2 | | 5.894e-03 | -5.13 | hang (hangover) | protein interactions | 32613 | 8 | 3 | 9204 | 464 | Myc,dnc,rau | | 5.905e-03 | -5.13 | MODULE\_100 | MSigDB lists | MODULE\_100 | 117 | 11 | 4161 | 164 | unc-13-4A,RSG7,Octbeta1R,toy,Mef2,Gfat1,bru1,Glut1,ap,Tim17b,Rbp9 | | 5.905e-03 | -5.13 | MODULE\_66 | MSigDB lists | MODULE\_66 | 117 | 11 | 4161 | 164 | Rbp9,Tim17b,ap,Gfat1,bru1,Glut1,toy,Mef2,RSG7,Octbeta1R,unc-13-4A | | 5.913e-03 | -5.13 | intrinsic component of membrane | cellular component | GO:0031224 | 1460 | 85 | 11940 | 534 | Or83c,Nlg2,Gr39a,Msp300,Tsp5D,dnt,Pgant9,CG15408,nAChRalpha3,bond,CG33233,nAChRalpha4,GABA-B-R3,CG42637,Pzl,Sema1a,snky,Atpalpha,fz2,DIP-delta,Tim17b,HisCl1,KCNQ,CG15449,Mct1,MCU,SPR,CadN2,Gr93d,CG14160,rg,MFS16,CG14196,Ir60e,CG2003,kune,Ir94e,DIP-lambda,Tim23,Ca-Ma2d,Hmt-1,sdk,Syt7,CG14024,CG41128,Ac78C,CG7536,Nlg3,Rh3,Ca-alpha1D,Ir40a,Gr2a,Der-1,fzo,CG14744,5-HT2A,CG1213,Teh2,TrissinR,CG9981,Or67d,Octbeta2R,mthl11,Meltrin,CG5789,ru,neo,Or69a,CG45076,CG10171,Or98a,CG8249,Glut1,CG34357,Dscam2,Rift,CG33970,ABCB7,mthl8,Ir41a,CG12121,Octbeta1R,SK,Oatp33Ea,CG43672 | | 5.924e-03 | -5.13 | apolpp (apolipophorin) | protein interactions | 43827 | 55 | 8 | 9204 | 464 | 2mit,Rtnl1,klg,PPO2,Yp2,Sema1a,Idgf4,Scp1 | | 5.933e-03 | -5.13 | cell projection | cellular component | GO:0042995 | 629 | 42 | 11940 | 534 | Or67d,HisCl1,pum,CG17387,norpA,5-HT2A,kl-3,nAChRalpha4,Gr2a,klg,Snap25,cta,nAChRalpha3,shot,Ptp99A,Catsup,Magi,sn,boi,Syt7,pico,hdc,Mst87F,mbc,Gr39a,Or83c,CG9492,cpx,mlt,orb,Mical,sky,Rtnl1,SK,par-1,Dscam2,Spn,Gr93d,ck,Or98a,Or69a,Dhc16F | | 5.970e-03 | -5.12 | regulation of GTPase activity | biological process | GO:0043087 | 113 | 12 | 12345 | 571 | ArfGAP1,CG34408,sky,vap,norpA,conu,RhoGAP100F,sif,Tbc1d15-17,CdGAPr,mbc,Git | | 5.981e-03 | -5.12 | Egfr (Epidermal growth factor receptor) | protein interactions | 37455 | 158 | 16 | 9204 | 464 | gus,pico,Pi3K68D,pum,svp,step,Ppm1,rg,vap,kni,SoxN,tay,CG33993,jing,Src42A,rl | | 6.012e-03 | -5.11 | heme transporter activity | molecular function | GO:0015232 | 3 | 2 | 12176 | 554 | ABCB7,Hmt-1 | | 6.014e-03 | -5.11 | adenyl ribonucleotide binding | molecular function | GO:0032559 | 655 | 44 | 12176 | 554 | CG10077,rl,ck,Takl2,PMCA,kl-3,CG5789,Hmt-1,CG42637,dnt,ABCB7,bt,Pkg21D,CG33970,CG14721,lz,Khc-73,spel1,mus308,CG32944,Atpalpha,fray,Dhc16F,PEK,CG9492,Src42A,Acbp5,bma,Myo81F,CG31689,Gprk1,Klp59C,Stlk,Hsc70-3,Nadsyn,CG12147,Hipk,Tak1,CG43672,CG34357,CG9981,fbl,DNAlig4,par-1 | | 6.098e-03 | -5.10 | regulation of presynapse organization | biological process | GO:0099174 | 26 | 5 | 12345 | 571 | Sod1,Ten-a,Nlg2,Spn,Thor | | 6.209e-03 | -5.08 | fly chr89E1-89E1\|3-59 cM | chromosome location | fly chr89E1-89E1\|3-59 cM | 4 | 2 | 25010 | 823 | CG31275,lncRNA:bxd | | 6.209e-03 | -5.08 | fly chr21E3-21E3\|2-1.5 | chromosome location | fly chr21E3-21E3\|2-1.5 | 4 | 2 | 25010 | 823 | IA-2,lncRNA:CR44801 | | 6.209e-03 | -5.08 | fly chr89E1-89E1\|3-59 | chromosome location | fly chr89E1-89E1\|3-59 | 4 | 2 | 25010 | 823 | lncRNA:bxd,CG31275 | | 6.209e-03 | -5.08 | fly chr85A1-85A1\|3-48 | chromosome location | fly chr85A1-85A1\|3-48 | 4 | 2 | 25010 | 823 | Obp85a,CG45263 | | 6.209e-03 | -5.08 | fly chr40E3-40E3\|2-54 cM | chromosome location | fly chr40E3-40E3\|2-54 cM | 4 | 2 | 25010 | 823 | CG31703,CG6675 | | 6.209e-03 | -5.08 | fly chr19D2-19D2\|1-64 cM | chromosome location | fly chr19D2-19D2\|1-64 cM | 4 | 2 | 25010 | 823 | Tak1,CG15461 | | 6.209e-03 | -5.08 | fly chr40E3-40E3\|2-54 | chromosome location | fly chr40E3-40E3\|2-54 | 4 | 2 | 25010 | 823 | CG6675,CG31703 | | 6.209e-03 | -5.08 | fly chr19D2-19D2\|1-64 | chromosome location | fly chr19D2-19D2\|1-64 | 4 | 2 | 25010 | 823 | Tak1,CG15461 | | 6.209e-03 | -5.08 | fly chr21E3-21E3\|2-1.5 cM | chromosome location | fly chr21E3-21E3\|2-1.5 cM | 4 | 2 | 25010 | 823 | IA-2,lncRNA:CR44801 | | 6.209e-03 | -5.08 | fly chr85A1-85A1\|3-48 cM | chromosome location | fly chr85A1-85A1\|3-48 cM | 4 | 2 | 25010 | 823 | Obp85a,CG45263 | | 6.211e-03 | -5.08 | haltere disc morphogenesis | biological process | GO:0007481 | 3 | 2 | 12345 | 571 | hth,ap | | 6.211e-03 | -5.08 | actin modification | biological process | GO:0030047 | 3 | 2 | 12345 | 571 | Setd3,Mical | | 6.211e-03 | -5.08 | iron coordination entity transport | biological process | GO:1901678 | 3 | 2 | 12345 | 571 | ABCB7,Hmt-1 | | 6.211e-03 | -5.08 | heme transport | biological process | GO:0015886 | 3 | 2 | 12345 | 571 | Hmt-1,ABCB7 | | 6.213e-03 | -5.08 | ATACTGT\_MIR144 | MSigDB lists | ATACTGT\_MIR144 | 43 | 6 | 4161 | 164 | fbl,dnc,ap,bru1,svp,Dys | | 6.213e-03 | -5.08 | GSE23321\_CENTRAL\_MEMORY\_VS\_NAIVE\_CD8\_TCELL\_DN | MSigDB lists | GSE23321\_CENTRAL\_MEMORY\_VS\_NAIVE\_CD8\_TCELL\_DN | 43 | 6 | 4161 | 164 | ck,Mp,PMCA,CG9389,norpA,Trxr-2 | | 6.213e-03 | -5.08 | GO\_REGULATION\_OF\_TRANSPORTER\_ACTIVITY | MSigDB lists | GO\_REGULATION\_OF\_TRANSPORTER\_ACTIVITY | 43 | 6 | 4161 | 164 | Ca-alpha1D,dnc,fray,Nlg3,Mef2,Dys | | 6.295e-03 | -5.07 | MODULE\_11 | MSigDB lists | MODULE\_11 | 118 | 11 | 4161 | 164 | RSG7,Octbeta1R,unc-13-4A,Glut1,bru1,Gfat1,Mef2,toy,Tim17b,ap,Rbp9 | | 6.303e-03 | -5.07 | GO\_MOVEMENT\_OF\_CELL\_OR\_SUBCELLULAR\_COMPONENT | MSigDB lists | GO\_MOVEMENT\_OF\_CELL\_OR\_SUBCELLULAR\_COMPONENT | 280 | 20 | 4161 | 164 | ck,l(2)41Ab,AP-2alpha,Dhc16F,toy,CG9492,cta,Mef2,sbm,mbc,shot,dnc,rl,svp,dnr1,Khc-73,Dys,Sod1,ap,CG17387 | | 6.386e-03 | -5.05 | ATP binding | molecular function | GO:0005524 | 639 | 43 | 12176 | 554 | Takl2,PMCA,ck,rl,CG10077,spel1,Khc-73,CG14721,lz,CG33970,Pkg21D,ABCB7,bt,CG42637,Hmt-1,dnt,CG5789,kl-3,mus308,CG32944,Dhc16F,PEK,fray,Atpalpha,bma,Myo81F,Src42A,CG9492,Nadsyn,Hsc70-3,Stlk,Klp59C,CG31689,Gprk1,Tak1,Hipk,CG12147,par-1,DNAlig4,fbl,CG9981,CG34357,CG43672 | | 6.396e-03 | -5.05 | neuron part | cellular component | GO:0097458 | 540 | 37 | 11940 | 534 | Or69a,Spn,Gr93d,rg,Or98a,d4,sky,SK,Rtnl1,par-1,Dscam2,Git,unc-13-4A,cpx,Mical,orb,Catsup,sn,Magi,Syt7,Rgk1,Or83c,RhoGAP100F,Gr39a,hdc,klg,Snap25,shot,nAChRalpha3,Ptp99A,5-HT2A,nAChRalpha4,Gr2a,Ten-a,Or67d,HisCl1,pum | | 6.397e-03 | -5.05 | positive regulation of GTPase activity | biological process | GO:0043547 | 100 | 11 | 12345 | 571 | RhoGAP100F,vap,norpA,conu,sky,CG34408,ArfGAP1,Git,mbc,CdGAPr,Tbc1d15-17 | | 6.406e-03 | -5.05 | neurexin family protein binding | molecular function | GO:0042043 | 9 | 3 | 12176 | 554 | Spn,Nlg3,Nlg2 | | 6.406e-03 | -5.05 | SH3 domain binding | molecular function | GO:0017124 | 9 | 3 | 12176 | 554 | Pi3K68D,mim,CG3588 | | 6.436e-03 | -5.05 | gene silencing | biological process | GO:0016458 | 143 | 14 | 12345 | 571 | pho,pum,hdc,CG10077,drosha,Ars2,mrt,Psc,AGO3,Hsc70-3,Su(z)2,Marf1,HPS4,Parg | | 6.484e-03 | -5.04 | sensory system development | biological process | GO:0048880 | 401 | 30 | 12345 | 571 | CadN2,bi,lz,rau,boi,rno,pk,Nna1,br,ru,sif,Src42A,Hipk,sdk,hth,rl,svp,kirre,sna,rg,RhoGAP100F,cic,zen,lola,toy,klg,fred,salm,Tak1,caup | | 6.484e-03 | -5.04 | visual system development | biological process | GO:0150063 | 401 | 30 | 12345 | 571 | svp,sna,kirre,RhoGAP100F,rg,zen,cic,toy,lola,klg,salm,fred,Tak1,caup,CadN2,rau,bi,lz,rno,boi,br,Nna1,pk,Src42A,Hipk,sif,ru,hth,sdk,rl | | 6.484e-03 | -5.04 | eye development | biological process | GO:0001654 | 401 | 30 | 12345 | 571 | ru,Hipk,sif,Src42A,pk,br,Nna1,rl,sdk,hth,lz,bi,rau,CadN2,boi,rno,klg,lola,toy,caup,Tak1,fred,salm,kirre,sna,svp,cic,zen,RhoGAP100F,rg | | 6.524e-03 | -5.03 | GSE35825\_UNTREATED\_VS\_IFNG\_STIM\_MACROPHAGE\_DN | MSigDB lists | GSE35825\_UNTREATED\_VS\_IFNG\_STIM\_MACROPHAGE\_DN | 57 | 7 | 4161 | 164 | AP-2alpha,fbl,sky,Mef2,CG30059,bru1,lqf | | 6.524e-03 | -5.03 | GO\_REGULATION\_OF\_BLOOD\_CIRCULATION | MSigDB lists | GO\_REGULATION\_OF\_BLOOD\_CIRCULATION | 57 | 7 | 4161 | 164 | Octbeta2R,bru1,Atpalpha,Dys,PMCA,Ca-alpha1D,dnc | | 6.616e-03 | -5.02 | GSE3982\_MEMORY\_CD4\_TCELL\_VS\_TH2\_UP | MSigDB lists | GSE3982\_MEMORY\_CD4\_TCELL\_VS\_TH2\_UP | 31 | 5 | 4161 | 164 | CG10077,CG34408,CG14721,CG8184,Wdr59 | | 6.616e-03 | -5.02 | GRE\_C | MSigDB lists | GRE\_C | 31 | 5 | 4161 | 164 | Atpalpha,Rbp9,Mef2,CG7255,twz | | 6.616e-03 | -5.02 | MORF\_RPA1 | MSigDB lists | MORF\_RPA1 | 31 | 5 | 4161 | 164 | Sod1,vtd,SdhA,Parg,spel1 | | 6.659e-03 | -5.01 | glutamine family amino acid catabolic process | biological process | GO:0009065 | 17 | 4 | 12345 | 571 | GLS,Ate1,Gdh,arg | | 6.659e-03 | -5.01 | anesthesia-resistant memory | biological process | GO:0007615 | 17 | 4 | 12345 | 571 | MCU,klg,dnc,Rgk1 | | 6.659e-03 | -5.01 | syncytial blastoderm mitotic cell cycle | biological process | GO:0035186 | 17 | 4 | 12345 | 571 | CycB,Trl,Psc,toc | | 6.659e-03 | -5.01 | deactivation of rhodopsin mediated signaling | biological process | GO:0016059 | 17 | 4 | 12345 | 571 | rdgC,Camta,norpA,Gprk1 | | 6.685e-03 | -5.01 | adenyl nucleotide binding | molecular function | GO:0030554 | 659 | 44 | 12176 | 554 | fray,Atpalpha,PEK,Dhc16F,mus308,CG32944,Hmt-1,CG42637,dnt,CG5789,kl-3,spel1,Khc-73,CG14721,lz,CG33970,Pkg21D,ABCB7,bt,CG10077,PMCA,ck,Takl2,rl,CG34357,CG43672,par-1,DNAlig4,fbl,CG9981,Tak1,Hipk,CG12147,Klp59C,Stlk,Gprk1,CG31689,Nadsyn,Hsc70-3,Acbp5,Src42A,CG9492,bma,Myo81F | | 6.707e-03 | -5.00 | female genitalia development | biological process | GO:0030540 | 9 | 3 | 12345 | 571 | lz,dsx,en | | 6.707e-03 | -5.00 | central nervous system neuron development | biological process | GO:0021954 | 9 | 3 | 12345 | 571 | en,Ulp1,fru | | 6.707e-03 | -5.00 | maintenance of synapse structure | biological process | GO:0099558 | 9 | 3 | 12345 | 571 | Nlg2,RhoGAP100F,Ten-a | | 6.707e-03 | -5.00 | maintenance of presynaptic active zone structure | biological process | GO:0048790 | 9 | 3 | 12345 | 571 | RhoGAP100F,Ten-a,Nlg2 | | 6.707e-03 | -5.00 | larval visceral muscle development | biological process | GO:0007523 | 9 | 3 | 12345 | 571 | mbc,kirre,Msp300 | | 6.741e-03 | -5.00 | GO\_PATTERN\_SPECIFICATION\_PROCESS | MSigDB lists | GO\_PATTERN\_SPECIFICATION\_PROCESS | 87 | 9 | 4161 | 164 | l(2)41Ab,svp,cta,Mef2,toy,CG9492,fz2,ap,CG17387 | | 6.794e-03 | -4.99 | cellular component organization | biological process | GO:0016043 | 2830 | 156 | 12345 | 571 | bru3,CG31703,Stlk,Nlg2,Meltrin,CG41099,Snap25,CG10939,Ten-a,Nna1,rib,Su(z)2,CG15449,mim,CG43901,snRNA:U1:82Eb,pum,snky,inc,dsx,fz2,dati,scro,bt,rg,nullo,sky,CG32087,HPS4,Rpn9,Set1,CdGAPr,Sdc,ct,CG1620,su(w[a]),ap,cmb,spel1,Myc,pho,Act42A,Tim17b,Cip4,SmydA-8,Ptip,lama,fan,CG34347,CG17387,Spn,Syx18,dnc,DIP-delta,bru1,AP-2alpha,CG40228,SoxN,CG40813,zld,l(2)41Ab,RpL6,kirre,DNaseII,Magi,pan,CG9107,Setd3,pico,conu,HP6,Psc,jing,ck,CadN2,Parg,jvl,shakB,hdc,klg,CG17683,CG13284,shot,RhoGAP100F,Svil,Ptp99A,CycB,Atpalpha,Sema1a,heph,eIF4B,cta,Thor,Dscam2,tej,cert,Mp,Nlg3,orb,Wdr62,ABCB7,gus,fest,cbs,Pld,fbl,br,kune,tou,CAP,fzo,Toll-7,neo,CG14184,CG12200,dgt5,Tim23,par-1,capu,mlt,mbc,DNAlig4,sn,Ulp1,CG41562,Tpl94D,Src42A,Khc-73,unc-13-4A,Parp,bond,dpr21,d4,CG41520,Nipped-A,en,lola,cpx,dnt,cic,CG9981,Trl,sna,rl,Diap1,wb,Msp300,sif,kl-3,pk,vtd,fru,twz,Mical,Pi3K68D,Rtnl1 | | 6.846e-03 | -4.98 | YKACATTT\_UNKNOWN | MSigDB lists | YKACATTT\_UNKNOWN | 72 | 8 | 4161 | 164 | PMCA,CG30495,Bruce,Pdk,svp,Khc-73,Dys,Mef2 | | 6.873e-03 | -4.98 | GO\_IRON\_ION\_HOMEOSTASIS | MSigDB lists | GO\_IRON\_ION\_HOMEOSTASIS | 20 | 4 | 4161 | 164 | Sod1,Hmt-1,ABCB7,Irp-1A | | 6.873e-03 | -4.98 | GO\_ADENYLATE\_CYCLASE\_ACTIVATING\_G\_PROTEIN\_COUPLED\_RECEPTOR\_SIGNALING\_PATHWAY | MSigDB lists | GO\_ADENYLATE\_CYCLASE\_ACTIVATING\_G\_PROTEIN\_COUPLED\_RECEPTOR\_SIGNALING\_PATHWAY | 20 | 4 | 4161 | 164 | Octbeta2R,cta,Ac78C,dnc | | 6.889e-03 | -4.98 | second-messenger-mediated signaling | biological process | GO:0019932 | 101 | 11 | 12345 | 571 | Ca-alpha1D,Octbeta2R,rdgC,CG42637,norpA,dnc,CG34357,Spn,TpnC41C,Ac78C,Octbeta1R | | 6.925e-03 | -4.97 | negative regulation of transport | biological process | GO:0051051 | 49 | 7 | 12345 | 571 | THADA,mim,sky,Slob,cpx,CG9650,Rgk1 | | 6.938e-03 | -4.97 | Glyco\_hydro\_31 | interpro domains | IPR000322 | 3 | 2 | 11383 | 557 | GCS2alpha,tobi | | 6.938e-03 | -4.97 | GDS\_CDC24\_CS | interpro domains | IPR001331 | 3 | 2 | 11383 | 557 | CG30440,sif | | 7.049e-03 | -4.95 | GTPase regulator activity | molecular function | GO:0030695 | 89 | 10 | 12176 | 554 | sky,RhoGAP100F,Tbc1d15-17,norpA,Git,vap,ArfGAP1,CG34408,CdGAPr,conu | | 7.131e-03 | -4.94 | positive regulation of protein phosphorylation | biological process | GO:0001934 | 130 | 13 | 12345 | 571 | scw,fray,step,Trl,Alg-2,PEK,CG14184,Tak1,mbc,CG41520,mav,Src42A,Stlk | | 7.163e-03 | -4.94 | enzyme binding | molecular function | GO:0019899 | 481 | 34 | 12176 | 554 | step,Maf1,SmydA-8,CG17514,CG8219,Msp300,cbs,rau,Diap1,CG1620,CG14184,pan,Tak1,Magi,CG10188,mbc,Pi3K68D,Myc,sif,CG9044,CG30440,pico,CG5862,CG31703,Der-1,Slmap,CG40228,Tbc1d15-17,CG7029,rl,SmydA-9,Trs23,CycB,Ranbp16 | | 7.179e-03 | -4.94 | GSE37301\_MULTIPOTENT\_PROGENITOR\_VS\_PRO\_BCELL\_DN | MSigDB lists | GSE37301\_MULTIPOTENT\_PROGENITOR\_VS\_PRO\_BCELL\_DN | 58 | 7 | 4161 | 164 | CG9389,MFS16,CG30495,Pgant5,Gdh,CG4341,DNAlig4 | | 7.179e-03 | -4.94 | GSE7768\_OVA\_ALONE\_VS\_OVA\_WITH\_MPL\_IMMUNIZED\_MOUSE\_WHOLE\_SPLEEN\_6H\_UP | MSigDB lists | GSE7768\_OVA\_ALONE\_VS\_OVA\_WITH\_MPL\_IMMUNIZED\_MOUSE\_WHOLE\_SPLEEN\_6H\_UP | 58 | 7 | 4161 | 164 | Mef2,Irp-1A,MFS16,Ubr1,CG34408,fbl,ck | | 7.179e-03 | -4.94 | MORF\_MLLT10 | MSigDB lists | MORF\_MLLT10 | 58 | 7 | 4161 | 164 | Dys,toy,Pi3K68D,Coq7,fz2,dnc,CG40178 | | 7.205e-03 | -4.93 | axon choice point recognition | biological process | GO:0016198 | 27 | 5 | 12345 | 571 | fru,shot,Sema1a,dsx,lola | | 7.223e-03 | -4.93 | sequence-specific double-stranded DNA binding | molecular function | GO:1990837 | 359 | 27 | 12176 | 554 | Mef2,ewg,toy,ct,bab2,Atf6,zen,zld,cic,svp,sna,CG9932,pho,bi,br,E5,Clk,scro,pan,ap,hth,tj,CG12769,Maf1,en,dsx,Myc | | 7.262e-03 | -4.93 | MORF\_FOSL1 | MSigDB lists | MORF\_FOSL1 | 88 | 9 | 4161 | 164 | toy,CG40178,dnc,Pi3K68D,Octbeta1R,Irp-1A,mbc,Coq7,fz2 | | 7.337e-03 | -4.91 | perivitelline space | cellular component | GO:0098595 | 18 | 4 | 11940 | 534 | Sdc,sdk,Meltrin,babos | | 7.347e-03 | -4.91 | Glyco\_hydro\_31 | pfam domains | PF01055 | 3 | 2 | 10480 | 528 | tobi,GCS2alpha | | 7.354e-03 | -4.91 | CG42795 (uncharacterized protein) | protein interactions | 41252 | 3 | 2 | 9204 | 464 | Sema1a,Rtnl1 | | 7.354e-03 | -4.91 | eIF4E4 (eukaryotic translation initiation factor 4E4) | protein interactions | 38743 | 3 | 2 | 9204 | 464 | mxt,Thor | | 7.354e-03 | -4.91 | Max (Max) | protein interactions | 40095 | 3 | 2 | 9204 | 464 | Myc,Mnt | | 7.354e-03 | -4.91 | CG32694 (uncharacterized protein) | protein interactions | 31933 | 3 | 2 | 9204 | 464 | CG32772,CG2061 | | 7.354e-03 | -4.91 | ClC-b (Chloride channel-b) | protein interactions | 36381 | 3 | 2 | 9204 | 464 | CG32772,par-1 | | 7.354e-03 | -4.91 | mRpL50 (mitochondrial ribosomal protein L50) | protein interactions | 38796 | 3 | 2 | 9204 | 464 | Rtnl1,Slmap | | 7.354e-03 | -4.91 | Roe1 (Roe1) | protein interactions | 36508 | 3 | 2 | 9204 | 464 | ewg,Nrd1 | | 7.354e-03 | -4.91 | hec (hector) | protein interactions | 32246 | 3 | 2 | 9204 | 464 | fru,en | | 7.354e-03 | -4.91 | Osi21 (Osiris 21) | protein interactions | 34566 | 3 | 2 | 9204 | 464 | norpA,rdgC | | 7.414e-03 | -4.90 | B52 (B52) | protein interactions | 41670 | 35 | 6 | 9204 | 464 | CG5953,CG32772,bru1,lola,hth,Sdc | | 7.494e-03 | -4.89 | PH\_domain | interpro domains | IPR001849 | 71 | 9 | 11383 | 557 | pico,CG30440,Gprk1,cert,sif,step,osp,vap,CG10188 | | 7.505e-03 | -4.89 | Nucleotide-bd\_a/b\_plait\_sf | interpro domains | IPR012677 | 152 | 15 | 11383 | 557 | bru3,CG40813,CG9107,eIF4B,CG14891,orb,heph,Set1,CG34334,cpo,bru1,Rbp9,Marf1,CG41562,shep | | 7.528e-03 | -4.89 | actin binding | molecular function | GO:0003779 | 133 | 13 | 12176 | 554 | Mical,ck,Svil,Spn,sn,Dys,zormin,mim,bt,capu,nullo,Msp300,shot | | 7.597e-03 | -4.88 | GO\_CELLULAR\_RESPONSE\_TO\_BIOTIC\_STIMULUS | MSigDB lists | GO\_CELLULAR\_RESPONSE\_TO\_BIOTIC\_STIMULUS | 32 | 5 | 4161 | 164 | rl,dnc,fz2,Hsc70-3,Mef2 | | 7.633e-03 | -4.88 | GO\_NEUROLOGICAL\_SYSTEM\_PROCESS | MSigDB lists | GO\_NEUROLOGICAL\_SYSTEM\_PROCESS | 173 | 14 | 4161 | 164 | ck,Nlg3,Ac78C,rl,rdgC,Octbeta2R,toy,Sod1,Mef2,sn,Ca-alpha1D,Atpalpha,Nlg2,GLS | | 7.780e-03 | -4.86 | OCT1\_02 | MSigDB lists | OCT1\_02 | 45 | 6 | 4161 | 164 | Ca-alpha1D,dnc,Rbp9,Tsp5D,toy,Dys | | 7.780e-03 | -4.86 | GSE46606\_IRF4MID\_VS\_WT\_CD40L\_IL2\_IL5\_DAY1\_STIMULATED\_BCELL\_UP | MSigDB lists | GSE46606\_IRF4MID\_VS\_WT\_CD40L\_IL2\_IL5\_DAY1\_STIMULATED\_BCELL\_UP | 45 | 6 | 4161 | 164 | spel1,fbl,CG10171,drosha,CG12547,MFS16 | | 7.819e-03 | -4.85 | GO\_ENZYME\_ACTIVATOR\_ACTIVITY | MSigDB lists | GO\_ENZYME\_ACTIVATOR\_ACTIVITY | 105 | 10 | 4161 | 164 | mbc,vap,sky,CG34408,Git,RSG7,CG10565,CG3107,Stlk,Tbc1d15-17 | | 7.844e-03 | -4.85 | Actinin\_actin-bd\_CS | interpro domains | IPR001589 | 9 | 3 | 11383 | 557 | Msp300,shot,Dys | | 7.853e-03 | -4.85 | GO\_EXCITATORY\_POSTSYNAPTIC\_POTENTIAL | MSigDB lists | GO\_EXCITATORY\_POSTSYNAPTIC\_POTENTIAL | 11 | 3 | 4161 | 164 | nAChRalpha4,Octbeta2R,Mef2 | | 7.853e-03 | -4.85 | GO\_PRODUCTION\_OF\_MOLECULAR\_MEDIATOR\_OF\_IMMUNE\_RESPONSE | MSigDB lists | GO\_PRODUCTION\_OF\_MOLECULAR\_MEDIATOR\_OF\_IMMUNE\_RESPONSE | 11 | 3 | 4161 | 164 | mus308,spel1,DNAlig4 | | 7.853e-03 | -4.85 | MODULE\_171 | MSigDB lists | MODULE\_171 | 11 | 3 | 4161 | 164 | MCU,PNPase,Meltrin | | 7.853e-03 | -4.85 | YNTTTNNNANGCARM\_UNKNOWN | MSigDB lists | YNTTTNNNANGCARM\_UNKNOWN | 11 | 3 | 4161 | 164 | Dys,CG9492,dnc | | 7.853e-03 | -4.85 | GO\_AXONEMAL\_DYNEIN\_COMPLEX\_ASSEMBLY | MSigDB lists | GO\_AXONEMAL\_DYNEIN\_COMPLEX\_ASSEMBLY | 11 | 3 | 4161 | 164 | l(2)41Ab,CG17387,CG9492 | | 7.882e-03 | -4.84 | ONDER\_CDH1\_TARGETS\_2\_DN | MSigDB lists | ONDER\_CDH1\_TARGETS\_2\_DN | 59 | 7 | 4161 | 164 | CG41378,fz2,Dys,arg,shot,Glut1,Octbeta2R | | 7.908e-03 | -4.84 | CTGCAGY\_UNKNOWN | MSigDB lists | CTGCAGY\_UNKNOWN | 156 | 13 | 4161 | 164 | gus,rg,Atpalpha,Rbp9,CG4341,CG42813,Dys,toy,cta,GCS2alpha,Nlg3,dnc,Git | | 7.915e-03 | -4.84 | posttranscriptional regulation of gene expression | biological process | GO:0010608 | 208 | 18 | 12345 | 571 | CG17514,PNPase,heph,Thor,Hsc70-3,bru1,AGO3,Ars2,mxt,hdc,pum,bru3,HPS4,PEK,Marf1,orb,CG10077,drosha | | 7.916e-03 | -4.84 | response to external stimulus | biological process | GO:0009605 | 987 | 62 | 12345 | 571 | Sdc,br,ct,Pld,CdGAPr,wb,rl,Dlg5,CG7536,CadN2,Mical,Toll-7,CG9380,Psc,jing,Myc,mthl8,mthl11,fru,Ars2,ap,norpA,dsx,lola,en,HisCl1,SoxN,Slob,Tak1,svp,RNASEK,Wdr59,fz2,CG17691,dnt,Mp,Src42A,Sema1a,Hipk,Gprk1,Ten-a,Atpalpha,Rh3,Dscam2,grim,Parp,Thor,Eig71Ef,PPO2,Clk,Bruce,klg,shakB,l(3)L1231,Mef2,Ptp99A,rdgC,shep,RhoGAP100F,Pzl,shot,par-1 | | 7.964e-03 | -4.83 | DNA-binding transcription factor activity, RNA polymerase II-specific | molecular function | GO:0000981 | 277 | 22 | 12176 | 554 | Mef2,kni,fd96Ca,toy,zen,cic,CG32006,svp,CG9932,Camta,caup,bi,E5,pan,ap,SoxN,tj,hth,CG12769,dsx,en,Myc | | 7.988e-03 | -4.83 | synapse organization | biological process | GO:0050808 | 177 | 16 | 12345 | 571 | rg,RhoGAP100F,Nlg3,par-1,cpx,fz2,Nlg2,klg,dpr21,Mical,DIP-delta,Syx18,Sema1a,Atpalpha,Ulp1,Ten-a | | 8.011e-03 | -4.83 | intracellular signal transduction | biological process | GO:0035556 | 373 | 28 | 12345 | 571 | cta,norpA,spel1,fray,dnc,CG34357,grim,RSG7,Pi3K68D,TpnC41C,gus,Octbeta1R,CG32944,rl,Spn,Pld,CdGAPr,RhoGAP100F,CG42637,par-1,RabX2,Ca-alpha1D,rdgC,Octbeta2R,Rgk1,Ac78C,Tak1,Stlk | | 8.178e-03 | -4.81 | extrinsic component of plasma membrane | cellular component | GO:0019897 | 40 | 6 | 11940 | 534 | mbc,Src42A,CadN2,cta,norpA,klg | | 8.253e-03 | -4.80 | regulation of rhodopsin mediated signaling pathway | biological process | GO:0022400 | 18 | 4 | 12345 | 571 | rdgC,Gprk1,norpA,Camta | | 8.355e-03 | -4.78 | EPHB-mediated forward signaling | REACTOME pathways | R-DME-3928662 | 27 | 5 | 3923 | 189 | sif,CG10188,Act42A,Sdc,vap | | 8.384e-03 | -4.78 | GO\_EPITHELIUM\_DEVELOPMENT | MSigDB lists | GO\_EPITHELIUM\_DEVELOPMENT | 193 | 15 | 4161 | 164 | l(2)41Ab,ck,dnc,rl,svp,arg,cta,Mef2,Sod1,toy,cv-2,ap,fz2,St1,CG17387 | | 8.412e-03 | -4.78 | transcription regulatory region sequence-specific DNA binding | molecular function | GO:0000976 | 329 | 25 | 12176 | 554 | zld,cic,svp,ct,toy,Atf6,zen,Mef2,ewg,br,bi,pho,sna,CG9932,pan,ap,Clk,scro,E5,Myc,dsx,en,CG12769,hth,tj | | 8.445e-03 | -4.77 | proximal/distal pattern formation | biological process | GO:0009954 | 28 | 5 | 12345 | 571 | ru,rl,ap,jing,hth | | 8.445e-03 | -4.77 | regulation of G protein-coupled receptor signaling pathway | biological process | GO:0008277 | 28 | 5 | 12345 | 571 | Gprk1,norpA,Camta,RSG7,rdgC | | 8.445e-03 | -4.77 | hormone biosynthetic process | biological process | GO:0042446 | 28 | 5 | 12345 | 571 | bond,spo,Npc2b,nvd,spok | | 8.466e-03 | -4.77 | response to xenobiotic stimulus | biological process | GO:0009410 | 39 | 6 | 12345 | 571 | St1,Ac78C,Cyp6g2,Clk,Tdc2,Cyp6a8 | | 8.513e-03 | -4.77 | cep290 (cep290) | protein interactions | 38099 | 9 | 3 | 9204 | 464 | sna,spel1,Rbp9 | | 8.513e-03 | -4.77 | uzip (unzipped) | protein interactions | 38002 | 9 | 3 | 9204 | 464 | Vsx2,dsx,CG3081 | | 8.513e-03 | -4.77 | Ten-m (Tenascin major) | protein interactions | 40464 | 9 | 3 | 9204 | 464 | Rtnl1,Sema1a,Ten-a | | 8.513e-03 | -4.77 | DUBAI (Deubiquitinating apoptotic inhibitor) | protein interactions | 36361 | 9 | 3 | 9204 | 464 | Diap1,Sod1,grim | | 8.574e-03 | -4.76 | regulation of gene expression, epigenetic | biological process | GO:0040029 | 163 | 15 | 12345 | 571 | Su(z)2,Hsc70-3,HPS4,Parg,Marf1,Ars2,drosha,CG10077,AGO3,Trl,Psc,CG40228,pho,hdc,pum | | 8.601e-03 | -4.76 | yki (yorkie) | protein interactions | 37851 | 109 | 12 | 9204 | 464 | Tgi,caup,hth,Hsc70-3,rl,Atpalpha,Hipk,Myc,Trl,CG2186,Diap1,par-1 | | 8.635e-03 | -4.75 | GSE24634\_IL4\_VS\_CTRL\_TREATED\_NAIVE\_CD4\_TCELL\_DAY3\_UP | MSigDB lists | GSE24634\_IL4\_VS\_CTRL\_TREATED\_NAIVE\_CD4\_TCELL\_DAY3\_UP | 60 | 7 | 4161 | 164 | ap,sbm,AP-2alpha,Mp,CG3107,CG30059,CG32944 | | 8.662e-03 | -4.75 | GSE17721\_CTRL\_VS\_LPS\_2H\_BMDC\_DN | MSigDB lists | GSE17721\_CTRL\_VS\_LPS\_2H\_BMDC\_DN | 46 | 6 | 4161 | 164 | Ca-alpha1D,fz2,cta,Tsp5D,arg,Ugt50B3 | | 8.662e-03 | -4.75 | GSE6269\_HEALTHY\_VS\_E\_COLI\_INF\_PBMC\_UP | MSigDB lists | GSE6269\_HEALTHY\_VS\_E\_COLI\_INF\_PBMC\_UP | 46 | 6 | 4161 | 164 | Octbeta2R,CG1486,GLS,SCAP,Wdr59,CG40178 | | 8.662e-03 | -4.75 | GO\_MUSCLE\_CONTRACTION | MSigDB lists | GO\_MUSCLE\_CONTRACTION | 46 | 6 | 4161 | 164 | dnc,Pi3K68D,Octbeta2R,Atpalpha,Dys,arg | | 8.674e-03 | -4.75 | DACOSTA\_UV\_RESPONSE\_VIA\_ERCC3\_XPCS\_DN | MSigDB lists | DACOSTA\_UV\_RESPONSE\_VIA\_ERCC3\_XPCS\_DN | 33 | 5 | 4161 | 164 | Pi3K68D,shot,CG34408,norpA,IntS3 | | 8.697e-03 | -4.74 | fs(1)h (female sterile (1) homeotic) | protein interactions | 31722 | 26 | 5 | 9204 | 464 | Trl,Psc,Rtnl1,Su(z)2,ct | | 8.792e-03 | -4.73 | GO\_REGULATION\_OF\_RESPIRATORY\_GASEOUS\_EXCHANGE\_BY\_NEUROLOGICAL\_SYSTEM\_PROCESS | MSigDB lists | GO\_REGULATION\_OF\_RESPIRATORY\_GASEOUS\_EXCHANGE\_BY\_NEUROLOGICAL\_SYSTEM\_PROCESS | 4 | 2 | 4161 | 164 | Nlg3,GLS | | 8.792e-03 | -4.73 | GO\_POST\_EMBRYONIC\_ORGAN\_DEVELOPMENT | MSigDB lists | GO\_POST\_EMBRYONIC\_ORGAN\_DEVELOPMENT | 4 | 2 | 4161 | 164 | fz2,ck | | 8.792e-03 | -4.73 | GO\_TELENCEPHALON\_REGIONALIZATION | MSigDB lists | GO\_TELENCEPHALON\_REGIONALIZATION | 4 | 2 | 4161 | 164 | toy,ap | | 8.792e-03 | -4.73 | GO\_SOMATIC\_DIVERSIFICATION\_OF\_IMMUNE\_RECEPTORS\_VIA\_SOMATIC\_MUTATION | MSigDB lists | GO\_SOMATIC\_DIVERSIFICATION\_OF\_IMMUNE\_RECEPTORS\_VIA\_SOMATIC\_MUTATION | 4 | 2 | 4161 | 164 | spel1,mus308 | | 8.792e-03 | -4.73 | VALK\_AML\_CLUSTER\_1 | MSigDB lists | VALK\_AML\_CLUSTER\_1 | 4 | 2 | 4161 | 164 | Mef2,Glut1 | | 8.792e-03 | -4.73 | GO\_REGULATION\_OF\_CILIUM\_MOVEMENT | MSigDB lists | GO\_REGULATION\_OF\_CILIUM\_MOVEMENT | 4 | 2 | 4161 | 164 | l(2)41Ab,CG17387 | | 8.792e-03 | -4.73 | CHESLER\_BRAIN\_D6MIT150\_QTL\_TRANS | MSigDB lists | CHESLER\_BRAIN\_D6MIT150\_QTL\_TRANS | 4 | 2 | 4161 | 164 | rl,Octbeta1R | | 8.792e-03 | -4.73 | GO\_REGULATION\_OF\_CHOLESTEROL\_BIOSYNTHETIC\_PROCESS | MSigDB lists | GO\_REGULATION\_OF\_CHOLESTEROL\_BIOSYNTHETIC\_PROCESS | 4 | 2 | 4161 | 164 | SCAP,Sod1 | | 8.792e-03 | -4.73 | GO\_GLYCOSPHINGOLIPID\_BIOSYNTHETIC\_PROCESS | MSigDB lists | GO\_GLYCOSPHINGOLIPID\_BIOSYNTHETIC\_PROCESS | 4 | 2 | 4161 | 164 | Ugt50B3,alpha4GT1 | | 8.792e-03 | -4.73 | PLASARI\_TGFB1\_SIGNALING\_VIA\_NFIC\_10HR\_DN | MSigDB lists | PLASARI\_TGFB1\_SIGNALING\_VIA\_NFIC\_10HR\_DN | 4 | 2 | 4161 | 164 | bru1,MFS16 | | 8.792e-03 | -4.73 | GO\_CELLULAR\_RESPONSE\_TO\_ANTIBIOTIC | MSigDB lists | GO\_CELLULAR\_RESPONSE\_TO\_ANTIBIOTIC | 4 | 2 | 4161 | 164 | Mef2,Hsc70-3 | | 8.792e-03 | -4.73 | GO\_DORSAL\_VENTRAL\_AXIS\_SPECIFICATION | MSigDB lists | GO\_DORSAL\_VENTRAL\_AXIS\_SPECIFICATION | 4 | 2 | 4161 | 164 | toy,fz2 | | 8.792e-03 | -4.73 | GO\_MIRNA\_BINDING | MSigDB lists | GO\_MIRNA\_BINDING | 4 | 2 | 4161 | 164 | Mef2,PNPase | | 8.792e-03 | -4.73 | SATO\_SILENCED\_EPIGENETICALLY\_IN\_PANCREATIC\_CANCER | MSigDB lists | SATO\_SILENCED\_EPIGENETICALLY\_IN\_PANCREATIC\_CANCER | 4 | 2 | 4161 | 164 | toy,ap | | 8.792e-03 | -4.73 | BIOCARTA\_ERK5\_PATHWAY | MSigDB lists | BIOCARTA\_ERK5\_PATHWAY | 4 | 2 | 4161 | 164 | Mef2,rl | | 8.792e-03 | -4.73 | GO\_MUSCLE\_CELL\_CELLULAR\_HOMEOSTASIS | MSigDB lists | GO\_MUSCLE\_CELL\_CELLULAR\_HOMEOSTASIS | 4 | 2 | 4161 | 164 | Dys,Sod1 | | 8.856e-03 | -4.73 | BTB | smart domains | SM00225 | 70 | 10 | 5086 | 308 | twz,rib,fru,inc,bab2,br,CG15725,CG41099,lola,Trl | | 8.857e-03 | -4.73 | regulation of transcription, DNA-templated | biological process | GO:0006355 | 1144 | 70 | 12345 | 571 | CG33557,dimm,cv-2,Su(z)2,fd96Ca,Clk,CG9650,rib,Atf6,Vsx2,salm,caup,Mef2,ewg,CG5862,br,ct,SCAP,Tgi,tj,tou,CG9932,Myc,CG1620,ap,dsx,toy,pum,Fer3,svp,HPS4,tld,Octbeta2R,scro,kni,Maf1,bab2,hth,lz,Parp,scw,pho,CG32006,zen,Setd3,mav,rl,pan,Parg,bi,HP6,jing,Psc,E5,CG12769,fru,lola,en,zld,d4,lncRNA:bxd,SoxN,Nipped-A,Camta,CG40228,sna,Mnt,fuss,Trl,cic | | 8.902e-03 | -4.72 | ACTININ\_2 | prosite domains | PS00020 | 8 | 3 | 6243 | 365 | Msp300,shot,Dys | | 8.981e-03 | -4.71 | upSET (upSET) | protein interactions | 39551 | 17 | 4 | 9204 | 464 | CG6621,Sema1a,Tak1,Rtnl1 | | 8.997e-03 | -4.71 | NUYTTEN\_EZH2\_TARGETS\_UP | MSigDB lists | NUYTTEN\_EZH2\_TARGETS\_UP | 213 | 16 | 4161 | 164 | fray,cta,Octbeta2R,PNPase,Pgant5,CG4341,MCU,step,CG40178,Hsc70-3,svp,Gfat1,fz2,CG41099,Tsp5D,MFS16 | | 9.009e-03 | -4.71 | synapse assembly | biological process | GO:0007416 | 64 | 8 | 12345 | 571 | Nlg3,par-1,cpx,Mical,Atpalpha,Ten-a,Nlg2,Sema1a | | 9.058e-03 | -4.70 | aos (argos) | protein interactions | 39833 | 59 | 8 | 9204 | 464 | rg,Dys,rl,Diap1,salm,Pi3K68D,rno,lz | | 9.203e-03 | -4.69 | carbohydrate derivative binding | molecular function | GO:0097367 | 1011 | 62 | 12176 | 554 | CG43672,CG34357,CG9981,CG5897,fbl,Gfat1,DNAlig4,par-1,CG12147,Tak1,Hipk,CG42541,fzo,Cht10,Gprk1,CG31689,Stlk,Klp59C,Hsc70-3,cv-2,Nadsyn,CG9492,Src42A,Acbp5,Rgk1,mIF2,bma,Myo81F,Idgf4,Atpalpha,fray,thw,CG14304,PEK,Dhc16F,Mur89F,CG9380,CG14301,RabX2,CG32944,mus308,kl-3,CG5789,CG42637,Hmt-1,Sema1a,dnt,ABCB7,bt,Pkg21D,CG33970,CG14721,lz,Khc-73,spel1,CG10077,CG30059,rl,PMCA,Takl2,ck,CG5160 | | 9.256e-03 | -4.68 | positive regulation of nuclear-transcribed mRNA poly(A) tail shortening | biological process | GO:0060213 | 10 | 3 | 12345 | 571 | Marf1,pum,AGO3 | | 9.256e-03 | -4.68 | central nervous system neuron differentiation | biological process | GO:0021953 | 10 | 3 | 12345 | 571 | Ulp1,fru,en | | 9.256e-03 | -4.68 | regulation of nuclear-transcribed mRNA poly(A) tail shortening | biological process | GO:0060211 | 10 | 3 | 12345 | 571 | AGO3,pum,Marf1 | | 9.305e-03 | -4.68 | C2\_domain\_sf | interpro domains | IPR035892 | 49 | 7 | 11383 | 557 | norpA,mbc,Pi3K68D,vap,RhoGAP100F,unc-13-4A,Syt7 | | 9.345e-03 | -4.67 | cytoplasmic region | cellular component | GO:0099568 | 252 | 20 | 11940 | 534 | CG9492,Ten-a,Git,mlt,sbm,pum,capu,CG17387,conu,step,kl-3,par-1,Spn,gus,Dys,ck,toc,Syx18,Dhc16F,igl | | 9.409e-03 | -4.67 | negative regulation of nitrogen compound metabolic process | biological process | GO:0051172 | 625 | 42 | 12345 | 571 | en,dsx,zld,Spn42De,lncRNA:bxd,pum,d4,Fer3,HPS4,svp,Mnt,orb,sna,kni,dnr1,Trl,cic,Maf1,ct,pan,Diap1,Parg,bi,Tgi,Psc,CG9932,jing,HP6,spel1,E5,CG1620,Vsx2,bru3,pho,Spn75F,Ppm1,heph,Su(z)2,Thor,lz,bru1,Clk | | 9.418e-03 | -4.67 | positive regulation of gene expression | biological process | GO:0010628 | 571 | 39 | 12345 | 571 | Trl,scro,HPS4,orb,sna,SoxN,d4,pum,Tak1,CG40228,Camta,en,lola,mxt,dsx,toy,zld,Myc,tou,jing,ap,CG12769,vtd,SCAP,tj,bi,pan,Setd3,br,zen,Mef2,caup,pho,Atf6,Clk,lz,dimm,hth,Hipk | | 9.439e-03 | -4.66 | GSE22611\_UNSTIM\_VS\_6H\_MDP\_STIM\_MUTANT\_NOD2\_TRANSDUCED\_HEK293T\_CELL\_UP | MSigDB lists | GSE22611\_UNSTIM\_VS\_6H\_MDP\_STIM\_MUTANT\_NOD2\_TRANSDUCED\_HEK293T\_CELL\_UP | 61 | 7 | 4161 | 164 | bru1,CG2528,Stlk,Nadsyn,Syx18,CstF50,mbc | | 9.485e-03 | -4.66 | GO\_G\_PROTEIN\_COUPLED\_RECEPTOR\_SIGNALING\_PATHWAY | MSigDB lists | GO\_G\_PROTEIN\_COUPLED\_RECEPTOR\_SIGNALING\_PATHWAY | 108 | 10 | 4161 | 164 | dnc,unc-13-4A,Ac78C,Octbeta1R,RSG7,cta,Octbeta2R,fz2,Ca-alpha1D,CG30495 | | 9.511e-03 | -4.66 | BONOME\_OVARIAN\_CANCER\_SURVIVAL\_SUBOPTIMAL\_DEBULKING | MSigDB lists | BONOME\_OVARIAN\_CANCER\_SURVIVAL\_SUBOPTIMAL\_DEBULKING | 142 | 12 | 4161 | 164 | Mef2,svp,lqf,CG12547,Ars2,Mp,l(2)41Ab,GLS,Alg-2,Irp-1A,shot,vap | | 9.545e-03 | -4.65 | eye photoreceptor cell differentiation | biological process | GO:0001754 | 165 | 15 | 12345 | 571 | klg,ru,br,lola,rl,hth,salm,lz,rau,svp,CadN2,boi,rno,rg,RhoGAP100F | | 9.569e-03 | -4.65 | DACOSTA\_UV\_RESPONSE\_VIA\_ERCC3\_DN | MSigDB lists | DACOSTA\_UV\_RESPONSE\_VIA\_ERCC3\_DN | 291 | 20 | 4161 | 164 | spel1,RSG7,Pi3K68D,step,Nipped-A,svp,IntS3,Dys,Parg,CG40045,vap,norpA,par-1,fray,mIF2,CG10565,mbc,Pgant5,Ubr1,shot | | 9.579e-03 | -4.65 | eye photoreceptor cell fate commitment | biological process | GO:0042706 | 52 | 7 | 12345 | 571 | lola,svp,br,ru,klg,hth,rl | | 9.579e-03 | -4.65 | compound eye photoreceptor fate commitment | biological process | GO:0001752 | 52 | 7 | 12345 | 571 | rl,hth,klg,ru,br,svp,lola | | 9.614e-03 | -4.64 | MORF\_MAGEA8 | MSigDB lists | MORF\_MAGEA8 | 47 | 6 | 4161 | 164 | Dys,toy,RSG7,Coq7,fz2,dnc | | 9.614e-03 | -4.64 | OCT1\_05 | MSigDB lists | OCT1\_05 | 47 | 6 | 4161 | 164 | DNAlig4,Pgant5,dnc,Cyp6a8,Dys,toy | | 9.635e-03 | -4.64 | presynapse | cellular component | GO:0098793 | 155 | 14 | 11940 | 534 | RhoGAP100F,Syt7,par-1,Rgk1,Magi,sky,Catsup,Mical,pum,unc-13-4A,cpx,Snap25,Ten-a,Git | | 9.686e-03 | -4.64 | Homeobox-like\_sf | interpro domains | IPR009057 | 142 | 14 | 11383 | 557 | zen,CG10565,hth,ap,ct,scro,caup,bab2,en,Vsx2,E5,toy,CG1620,rib | | 9.698e-03 | -4.64 | regulation of nitrogen compound metabolic process | biological process | GO:0051171 | 1870 | 107 | 12345 | 571 | pum,Fer3,Catsup,mxt,toy,dsx,kni,dnr1,scro,bab2,Maf1,tld,HPS4,svp,orb,Octbeta2R,gus,CG17514,ct,PNPase,br,Myc,CG9932,tou,spel1,ap,CG1620,CG14184,SCAP,tj,Tgi,salm,bru3,caup,Vsx2,Atf6,Stlk,Spn75F,Mef2,Marf1,PEK,ewg,CG5862,dimm,cv-2,CycB,heph,CG33557,Clk,CG9650,rib,SteXh:CG42398,grim,fd96Ca,Su(z)2,Thor,Alg-2,lncRNA:bxd,SoxN,Nipped-A,CG41520,d4,CG40228,Tak1,Camta,en,lola,Spn42De,zld,Trl,cic,fuss,Mnt,sna,mav,pan,CG34334,Diap1,rl,Setd3,jing,HP6,Psc,step,fru,CG12769,E5,Parg,bi,CG12147,CG12200,pho,CG41106,Ptip,CG32006,par-1,zen,hth,mbc,Ppm1,Src42A,bru1,AGO3,fray,scw,lz,Parp | | 9.713e-03 | -4.63 | GO\_EMBRYO\_DEVELOPMENT\_ENDING\_IN\_BIRTH\_OR\_EGG\_HATCHING | MSigDB lists | GO\_EMBRYO\_DEVELOPMENT\_ENDING\_IN\_BIRTH\_OR\_EGG\_HATCHING | 160 | 13 | 4161 | 164 | spel1,CycB,DNAlig4,rl,svp,lqf,Mef2,toy,cta,fz2,ap,Bruce,CG9107 | | 9.744e-03 | -4.63 | plasma membrane protein complex | cellular component | GO:0098797 | 171 | 15 | 11940 | 534 | AP-2alpha,Ca-alpha1D,cta,Dys,CadN2,nAChRalpha3,SdhA,KCNQ,HisCl1,Ca-Ma2d,Atpalpha,norpA,GABA-B-R3,nAChRalpha4,CG33970 | | 9.759e-03 | -4.63 | GO\_EXTRINSIC\_COMPONENT\_OF\_CYTOPLASMIC\_SIDE\_OF\_PLASMA\_MEMBRANE | MSigDB lists | GO\_EXTRINSIC\_COMPONENT\_OF\_CYTOPLASMIC\_SIDE\_OF\_PLASMA\_MEMBRANE | 22 | 4 | 4161 | 164 | RSG7,Src42A,step,cta | | 9.759e-03 | -4.63 | GO\_CELLULAR\_TRANSITION\_METAL\_ION\_HOMEOSTASIS | MSigDB lists | GO\_CELLULAR\_TRANSITION\_METAL\_ION\_HOMEOSTASIS | 22 | 4 | 4161 | 164 | Irp-1A,ABCB7,Sod1,Hmt-1 | | 9.759e-03 | -4.63 | MORF\_CDH4 | MSigDB lists | MORF\_CDH4 | 22 | 4 | 4161 | 164 | fz2,RSG7,toy,Dys | | 9.759e-03 | -4.63 | GO\_REGULATION\_OF\_POTASSIUM\_ION\_TRANSPORT | MSigDB lists | GO\_REGULATION\_OF\_POTASSIUM\_ION\_TRANSPORT | 22 | 4 | 4161 | 164 | fray,Ca-alpha1D,norpA,Octbeta2R | | 9.759e-03 | -4.63 | GO\_STRIATED\_MUSCLE\_CONTRACTION | MSigDB lists | GO\_STRIATED\_MUSCLE\_CONTRACTION | 22 | 4 | 4161 | 164 | Atpalpha,arg,Dys,Octbeta2R | | 9.759e-03 | -4.63 | MENSE\_HYPOXIA\_UP | MSigDB lists | MENSE\_HYPOXIA\_UP | 22 | 4 | 4161 | 164 | Glut1,Hmt-1,cta,Nadsyn | | 9.767e-03 | -4.63 | negative regulation of cellular metabolic process | biological process | GO:0031324 | 681 | 45 | 12345 | 571 | bru3,pho,Vsx2,Bruce,Spn75F,heph,Ppm1,bru1,Clk,Su(z)2,Thor,lz,Fer3,CG12772,lncRNA:bxd,pum,d4,zld,Spn42De,en,dsx,cic,Maf1,kni,Trl,dnr1,Mnt,sna,orb,HPS4,svp,pan,Diap1,rl,ct,spel1,CG1620,E5,Psc,jing,CG9932,HP6,bi,Tgi,Parg | | 9.832e-03 | -4.62 | DH\_1 | prosite domains | PS00741 | 3 | 2 | 6243 | 365 | sif,CG30440 | | 9.851e-03 | -4.62 | fly chr100D1-100D1\|3-102 cM | chromosome location | fly chr100D1-100D1\|3-102 cM | 14 | 3 | 25010 | 823 | CstF50,lncRNA:CR45565,CG42693 | | 9.851e-03 | -4.62 | fly chr100D1-100D1\|3-102 | chromosome location | fly chr100D1-100D1\|3-102 | 14 | 3 | 25010 | 823 | CG42693,CstF50,lncRNA:CR45565 | | 9.853e-03 | -4.62 | GSE33162\_HDAC3\_KO\_VS\_HDAC3\_KO\_4H\_LPS\_STIM\_MACROPHAGE\_DN | MSigDB lists | GSE33162\_HDAC3\_KO\_VS\_HDAC3\_KO\_4H\_LPS\_STIM\_MACROPHAGE\_DN | 34 | 5 | 4161 | 164 | Ugt50B3,MFS16,alpha4GT1,CG10171,Gdh | | 9.853e-03 | -4.62 | GO\_REGULATION\_OF\_MUSCLE\_CONTRACTION | MSigDB lists | GO\_REGULATION\_OF\_MUSCLE\_CONTRACTION | 34 | 5 | 4161 | 164 | dnc,Sod1,Dys,nAChRalpha4,Octbeta2R | | 9.996e-03 | -4.61 | RBD\_domain\_sf | interpro domains | IPR035979 | 157 | 15 | 11383 | 557 | eIF4B,CG40813,bru3,CG9107,orb,CG34334,heph,Set1,CG14891,Rbp9,Marf1,cpo,bru1,CG41562,shep | | 1.008e-02 | -4.60 | ecdysteroid biosynthetic process | biological process | GO:0045456 | 19 | 4 | 12345 | 571 | nvd,Npc2b,spo,spok | | 1.008e-02 | -4.60 | steroid hormone biosynthetic process | biological process | GO:0120178 | 19 | 4 | 12345 | 571 | nvd,Npc2b,spo,spok | | 1.013e-02 | -4.59 | rl (rolled) | protein interactions | 3354888 | 85 | 10 | 9204 | 464 | cic,pum,svp,tay,step,Svil,rl,Src42A,Ppm1,rg | | 1.017e-02 | -4.59 | CHESLER\_BRAIN\_HIGHEST\_GENETIC\_VARIANCE | MSigDB lists | CHESLER\_BRAIN\_HIGHEST\_GENETIC\_VARIANCE | 12 | 3 | 4161 | 164 | Mef2,CG1486,mIF2 | | 1.017e-02 | -4.59 | GO\_SOMATIC\_DIVERSIFICATION\_OF\_IMMUNE\_RECEPTORS | MSigDB lists | GO\_SOMATIC\_DIVERSIFICATION\_OF\_IMMUNE\_RECEPTORS | 12 | 3 | 4161 | 164 | DNAlig4,spel1,mus308 | | 1.017e-02 | -4.59 | GO\_CILIUM\_MOVEMENT | MSigDB lists | GO\_CILIUM\_MOVEMENT | 12 | 3 | 4161 | 164 | CG17387,CG9492,l(2)41Ab | | 1.030e-02 | -4.58 | GSE30971\_CTRL\_VS\_LPS\_STIM\_MACROPHAGE\_WBP7\_HET\_4H\_DN | MSigDB lists | GSE30971\_CTRL\_VS\_LPS\_STIM\_MACROPHAGE\_WBP7\_HET\_4H\_DN | 62 | 7 | 4161 | 164 | Ars2,Gdh,vap,AP-2alpha,MFS16,CG9389,Mef2 | | 1.030e-02 | -4.58 | MODULE\_49 | MSigDB lists | MODULE\_49 | 62 | 7 | 4161 | 164 | CG9492,Rbp9,CG32944,vtd,CG2256,PMCA,l(2)41Ab | | 1.038e-02 | -4.57 | SH3 | smart domains | SM00326 | 61 | 9 | 5086 | 308 | mbc,vap,Src42A,CdGAPr,SKIP,CAP,Cip4,ck,l(3)05822 | | 1.038e-02 | -4.57 | regulation of developmental process | biological process | GO:0050793 | 757 | 49 | 12345 | 571 | pum,SoxN,Catsup,cpo,zld,fz2,bab2,nullo,Octbeta2R,fuss,kirre,Octbeta1R,sdk,mav,Diap1,rl,Sdc,br,pico,pk,sif,lqf,CG9380,Myc,norpA,step,Mical,tinc,neo,Git,salm,hdc,Bruce,shot,par-1,Cip4,Mef2,ewg,hth,mbc,Spn,Hipk,Src42A,Sema1a,cta,babos,bru1,rib,lz | | 1.048e-02 | -4.56 | Fatty acid biosynthesis, elongation | KEGG pathways | dme\_M00083 | 4 | 2 | 3198 | 138 | CG12170,FASN3 | | 1.048e-02 | -4.56 | Fatty acid biosynthesis, elongation | KEGG pathways | M00083 | 4 | 2 | 3198 | 138 | CG12170,FASN3 | | 1.048e-02 | -4.56 | integral component of plasma membrane | cellular component | GO:0005887 | 466 | 32 | 11940 | 534 | CG8249,CadN2,CG34357,ru,sdk,Octbeta1R,Oatp33Ea,DIP-lambda,Rift,Dscam2,Rh3,dnt,CG15408,nAChRalpha3,Nlg3,Ac78C,Tsp5D,Nlg2,CG15449,Octbeta2R,HisCl1,KCNQ,TrissinR,fz2,DIP-delta,Atpalpha,Teh2,GABA-B-R3,CG42637,5-HT2A,CG1213,nAChRalpha4 | | 1.055e-02 | -4.55 | regulation of cellular macromolecule biosynthetic process | biological process | GO:2000112 | 1307 | 78 | 12345 | 571 | zld,lola,en,Camta,CG40228,d4,lncRNA:bxd,SoxN,Nipped-A,sna,Mnt,fuss,cic,Trl,Setd3,rl,pan,mav,bi,Parg,E5,CG12769,fru,Psc,HP6,jing,pho,zen,CG32006,hth,lz,Parp,scw,bru1,toy,dsx,mxt,Fer3,pum,Octbeta2R,orb,svp,tld,HPS4,Maf1,bab2,scro,kni,br,CG17514,ct,tj,Tgi,SCAP,CG1620,ap,CG9932,tou,Myc,Atf6,Vsx2,caup,bru3,salm,CG5862,PEK,ewg,Mef2,CG33557,heph,cv-2,dimm,Su(z)2,fd96Ca,Thor,rib,Clk,CG9650 | | 1.064e-02 | -4.54 | DURAND\_STROMA\_S\_UP | MSigDB lists | DURAND\_STROMA\_S\_UP | 48 | 6 | 4161 | 164 | CG42813,MFS16,dnc,cv-2,Meltrin,alpha4GT1 | | 1.064e-02 | -4.54 | GSE22140\_GERMFREE\_VS\_SPF\_ARTHRITIC\_MOUSE\_CD4\_TCELL\_UP | MSigDB lists | GSE22140\_GERMFREE\_VS\_SPF\_ARTHRITIC\_MOUSE\_CD4\_TCELL\_UP | 48 | 6 | 4161 | 164 | ck,DNAlig4,Mp,RpL15,CG1486,norpA | | 1.064e-02 | -4.54 | GO\_PEPTIDYL\_SERINE\_MODIFICATION | MSigDB lists | GO\_PEPTIDYL\_SERINE\_MODIFICATION | 48 | 6 | 4161 | 164 | CycB,rl,Pgant5,CG32944,Pkg21D,Pdk | | 1.064e-02 | -4.54 | GSE43955\_TH0\_VS\_TGFB\_IL6\_IL23\_TH17\_ACT\_CD4\_TCELL\_60H\_DN | MSigDB lists | GSE43955\_TH0\_VS\_TGFB\_IL6\_IL23\_TH17\_ACT\_CD4\_TCELL\_60H\_DN | 48 | 6 | 4161 | 164 | Coq7,Ars2,Gdh,vap,arg,Mef2 | | 1.064e-02 | -4.54 | GO\_REGULATION\_OF\_MUSCLE\_SYSTEM\_PROCESS | MSigDB lists | GO\_REGULATION\_OF\_MUSCLE\_SYSTEM\_PROCESS | 48 | 6 | 4161 | 164 | Dys,Sod1,Octbeta2R,nAChRalpha4,dnc,Parp | | 1.070e-02 | -4.54 | transmembrane signaling receptor activity | molecular function | GO:0004888 | 353 | 26 | 12176 | 554 | nAChRalpha4,GABA-B-R3,klg,mthl8,Gr2a,dnt,SPR,Ir40a,Octbeta1R,TrissinR,HisCl1,Or98a,Gr93d,fz2,5-HT2B,nAChRalpha3,5-HT2A,Or83c,Or67d,Octbeta2R,Ptp99A,CG31750,Rh3,Or69a,Gr39a,mthl11 | | 1.078e-02 | -4.53 | WTGAAAT\_UNKNOWN | MSigDB lists | WTGAAAT\_UNKNOWN | 127 | 11 | 4161 | 164 | Gdh,Dys,toy,Mef2,CG12547,vap,CG4341,Bruce,rg,Rbp9,CG2256 | | 1.080e-02 | -4.53 | brain development | biological process | GO:0007420 | 122 | 12 | 12345 | 571 | rg,shot,dati,E5,drosha,vap,fdl,Ten-a,hth,Git,toy,en | | 1.086e-02 | -4.52 | neuron projection | cellular component | GO:0043005 | 431 | 30 | 11940 | 534 | Syt7,Gr39a,Or83c,hdc,Catsup,sn,Or69a,Magi,Or98a,shot,nAChRalpha3,Ptp99A,klg,Spn,Gr93d,Snap25,nAChRalpha4,par-1,Dscam2,Gr2a,sky,SK,5-HT2A,Rtnl1,HisCl1,pum,orb,Mical,Or67d,cpx | | 1.086e-02 | -4.52 | sgg (shaggy) | protein interactions | 31248 | 183 | 17 | 9204 | 464 | pan,Nipped-A,cv-2,Dlg5,PEK,Or69a,cmb,zormin,Msp300,CG33288,Khc-73,Rtnl1,CG42637,ck,CG8219,CG1387,Bruce | | 1.101e-02 | -4.51 | positive regulation of nucleobase-containing compound metabolic process | biological process | GO:0045935 | 523 | 36 | 12345 | 571 | CG12200,caup,pho,Atf6,zen,Marf1,Mef2,dimm,hth,Clk,AGO3,lz,SoxN,d4,pum,CG40228,Camta,en,lola,dsx,toy,zld,Trl,scro,sna,pan,Setd3,br,Myc,jing,tou,ap,CG12769,SCAP,bi,tj | | 1.109e-02 | -4.50 | C2 | pfam domains | PF00168 | 38 | 6 | 10480 | 528 | unc-13-4A,norpA,RhoGAP100F,vap,Pi3K68D,Syt7 | | 1.110e-02 | -4.50 | elav (embryonic lethal abnormal vision) | protein interactions | 31000 | 18 | 4 | 9204 | 464 | Rbp9,heph,pum,ewg | | 1.111e-02 | -4.50 | CycJ (Cyclin J) | protein interactions | 38428 | 38 | 6 | 9204 | 464 | dimm,CG17508,E5,gus,CG7518,heph | | 1.114e-02 | -4.50 | GATA4\_Q3 | MSigDB lists | GATA4\_Q3 | 35 | 5 | 4161 | 164 | PMCA,Mp,dnc,Dys,svp | | 1.114e-02 | -4.50 | FLORIO\_NEOCORTEX\_BASAL\_RADIAL\_GLIA\_UP | MSigDB lists | FLORIO\_NEOCORTEX\_BASAL\_RADIAL\_GLIA\_UP | 35 | 5 | 4161 | 164 | St1,Octbeta1R,Mct1,Src42A,cv-2 | | 1.114e-02 | -4.50 | SWEET\_KRAS\_ONCOGENIC\_SIGNATURE | MSigDB lists | SWEET\_KRAS\_ONCOGENIC\_SIGNATURE | 35 | 5 | 4161 | 164 | RpL6,rl,mIF2,Pdk,CG30059 | | 1.121e-02 | -4.49 | MIKKELSEN\_MCV6\_HCP\_WITH\_H3K27ME3 | MSigDB lists | MIKKELSEN\_MCV6\_HCP\_WITH\_H3K27ME3 | 63 | 7 | 4161 | 164 | CG6788,nAChRalpha4,Mp,Ca-alpha1D,PMCA,sdk,scro | | 1.130e-02 | -4.48 | FN3\_dom | interpro domains | IPR003961 | 63 | 8 | 11383 | 557 | fred,bt,sdk,klg,CG14964,Ptp99A,boi,Dscam2 | | 1.130e-02 | -4.48 | FN3\_sf | interpro domains | IPR036116 | 63 | 8 | 11383 | 557 | Ptp99A,boi,klg,CG14964,Dscam2,fred,sdk,bt | | 1.135e-02 | -4.48 | male sex differentiation | biological process | GO:0046661 | 30 | 5 | 12345 | 571 | fru,salm,tj,br,dsx | | 1.146e-02 | -4.47 | GSE5589\_WT\_VS\_IL6\_KO\_LPS\_AND\_IL10\_STIM\_MACROPHAGE\_180MIN\_UP | MSigDB lists | GSE5589\_WT\_VS\_IL6\_KO\_LPS\_AND\_IL10\_STIM\_MACROPHAGE\_180MIN\_UP | 23 | 4 | 4161 | 164 | rg,nAChRalpha3,CG4341,fz2 | | 1.146e-02 | -4.47 | ACAGGGT\_MIR10A\_MIR10B | MSigDB lists | ACAGGGT\_MIR10A\_MIR10B | 23 | 4 | 4161 | 164 | CG41099,Pgant5,step,bru1 | | 1.146e-02 | -4.47 | KEGG\_LONG\_TERM\_DEPRESSION | MSigDB lists | KEGG\_LONG\_TERM\_DEPRESSION | 23 | 4 | 4161 | 164 | rl,Pkg21D,cta,norpA | | 1.146e-02 | -4.47 | WANG\_PROSTATE\_CANCER\_ANDROGEN\_INDEPENDENT | MSigDB lists | WANG\_PROSTATE\_CANCER\_ANDROGEN\_INDEPENDENT | 23 | 4 | 4161 | 164 | shot,Ugt50B3,Mp,AP-2alpha | | 1.146e-02 | -4.47 | GO\_HEART\_PROCESS | MSigDB lists | GO\_HEART\_PROCESS | 23 | 4 | 4161 | 164 | dnc,Atpalpha,Sod1,Dys | | 1.146e-02 | -4.47 | GO\_INTRINSIC\_COMPONENT\_OF\_MITOCHONDRIAL\_MEMBRANE | MSigDB lists | GO\_INTRINSIC\_COMPONENT\_OF\_MITOCHONDRIAL\_MEMBRANE | 23 | 4 | 4161 | 164 | Tim23,Tim17b,Hmt-1,MCU | | 1.156e-02 | -4.46 | regulation of nucleic acid-templated transcription | biological process | GO:1903506 | 1158 | 70 | 12345 | 571 | br,ct,Tgi,tj,SCAP,CG1620,ap,tou,CG9932,Myc,dsx,toy,Fer3,pum,Octbeta2R,svp,tld,HPS4,Maf1,bab2,scro,kni,CG33557,cv-2,dimm,Su(z)2,fd96Ca,rib,Clk,CG9650,Atf6,Vsx2,caup,salm,CG5862,ewg,Mef2,Setd3,rl,pan,mav,bi,Parg,CG12769,E5,fru,jing,HP6,Psc,zld,en,lola,Camta,CG40228,d4,Nipped-A,lncRNA:bxd,SoxN,sna,fuss,Mnt,cic,Trl,hth,Parp,lz,scw,pho,zen,CG32006 | | 1.156e-02 | -4.46 | regulation of RNA biosynthetic process | biological process | GO:2001141 | 1158 | 70 | 12345 | 571 | cic,Trl,sna,Mnt,fuss,Camta,CG40228,d4,SoxN,Nipped-A,lncRNA:bxd,zld,lola,en,E5,CG12769,fru,Psc,HP6,jing,bi,Parg,rl,pan,mav,Setd3,zen,CG32006,pho,scw,lz,Parp,hth,Maf1,bab2,scro,kni,Octbeta2R,svp,tld,HPS4,Fer3,pum,toy,dsx,CG1620,ap,tou,CG9932,Myc,tj,Tgi,SCAP,br,ct,CG5862,ewg,Mef2,caup,salm,Atf6,Vsx2,rib,CG9650,Clk,Su(z)2,fd96Ca,cv-2,dimm,CG33557 | | 1.166e-02 | -4.45 | ion channel binding | molecular function | GO:0044325 | 4 | 2 | 12176 | 554 | CheB42a,THADA | | 1.166e-02 | -4.45 | eukaryotic initiation factor 4E binding | molecular function | GO:0008190 | 4 | 2 | 12176 | 554 | Thor,mxt | | 1.166e-02 | -4.45 | calcium-dependent phospholipase A2 activity | molecular function | GO:0047498 | 4 | 2 | 12176 | 554 | CG3009,CG42237 | | 1.168e-02 | -4.45 | epithelial cell differentiation | biological process | GO:0030855 | 419 | 30 | 12345 | 571 | cmb,jing,cta,ck,tj,Parp,neo,mim,rl,Diap1,Dlg5,mbc,gus,dimm,sn,lncRNA:flam,br,ct,pk,par-1,Cip4,cic,CG41099,lama,orb,capu,Mef2,jvl,AP-2alpha,toc | | 1.169e-02 | -4.45 | regulation of cellular biosynthetic process | biological process | GO:0031326 | 1371 | 81 | 12345 | 571 | fru,E5,CG12769,jing,HP6,Psc,bi,Parg,pan,rl,mav,Setd3,cic,Trl,Mnt,fuss,sna,CG40228,Tak1,Camta,lncRNA:bxd,Nipped-A,SoxN,d4,zld,lola,en,scw,bru1,Parp,lz,hth,zen,CG32006,Ptip,pho,ap,CG1620,Myc,CG9932,tou,Tgi,tj,SCAP,ct,CG17514,br,bab2,Maf1,kni,scro,dnr1,Octbeta2R,orb,HPS4,tld,svp,Fer3,pum,mxt,dsx,toy,rib,CG9650,Clk,fd96Ca,Thor,Su(z)2,cv-2,dimm,CG33557,heph,CG5862,PEK,ewg,Mef2,caup,bru3,salm,Vsx2,Atf6 | | 1.171e-02 | -4.45 | CG9436 (uncharacterized protein) | protein interactions | 35586 | 10 | 3 | 9204 | 464 | Atpalpha,Sema1a,Rtnl1 | | 1.171e-02 | -4.45 | CG3021 (uncharacterized protein) | protein interactions | 53546 | 10 | 3 | 9204 | 464 | Nna1,CG14427,ArfGAP1 | | 1.171e-02 | -4.45 | prg (piragua) | protein interactions | 34177 | 10 | 3 | 9204 | 464 | Tak1,E5,rau | | 1.171e-02 | -4.45 | Idgf4 (Imaginal disc growth factor 4) | protein interactions | 31926 | 10 | 3 | 9204 | 464 | CG8219,Nrd1,Rtnl1 | | 1.171e-02 | -4.45 | TfIIA-S (Transcription-factor-IIA-S) | protein interactions | 42822 | 10 | 3 | 9204 | 464 | CG3081,svp,rl | | 1.171e-02 | -4.45 | su(r) (suppressor of rudimentary) | protein interactions | 31858 | 10 | 3 | 9204 | 464 | Vsx2,Syt7,ect | | 1.171e-02 | -4.45 | zda (zonda) | protein interactions | 37144 | 10 | 3 | 9204 | 464 | SKIP,CG33970,Myc | | 1.171e-02 | -4.45 | sv (shaven) | protein interactions | 43825 | 10 | 3 | 9204 | 464 | lz,Myc,rg | | 1.174e-02 | -4.45 | GSE21670\_UNTREATED\_VS\_TGFB\_TREATED\_STAT3\_KO\_CD4\_TCELL\_DN | MSigDB lists | GSE21670\_UNTREATED\_VS\_TGFB\_TREATED\_STAT3\_KO\_CD4\_TCELL\_DN | 49 | 6 | 4161 | 164 | rl,vap,Git,step,Meltrin,CG3107 | | 1.174e-02 | -4.45 | OCT1\_Q5\_01 | MSigDB lists | OCT1\_Q5\_01 | 49 | 6 | 4161 | 164 | dnc,Pgant5,toy,Cyp6a8,Dys,CG30495 | | 1.174e-02 | -4.45 | GSE25123\_WT\_VS\_PPARG\_KO\_MACROPHAGE\_IL4\_STIM\_UP | MSigDB lists | GSE25123\_WT\_VS\_PPARG\_KO\_MACROPHAGE\_IL4\_STIM\_UP | 49 | 6 | 4161 | 164 | mRpS5,CG9886,CG13284,Tsp5D,Parp,Mct1 | | 1.175e-02 | -4.44 | adult behavior | biological process | GO:0030534 | 169 | 15 | 12345 | 571 | Tdc2,klg,Nlg2,rl,pum,shakB,Atpalpha,shep,dati,dnc,vap,tay,norpA,twz,Clk | | 1.176e-02 | -4.44 | protease binding | molecular function | GO:0002020 | 11 | 3 | 12176 | 554 | Der-1,Diap1,Myc | | 1.183e-02 | -4.44 | GO\_NUCLEOSIDE\_TRIPHOSPHATASE\_REGULATOR\_ACTIVITY | MSigDB lists | GO\_NUCLEOSIDE\_TRIPHOSPHATASE\_REGULATOR\_ACTIVITY | 79 | 8 | 4161 | 164 | RSG7,Git,CG10565,Tbc1d15-17,vap,mbc,CG34408,sky | | 1.183e-02 | -4.44 | MORF\_MAGEA9 | MSigDB lists | MORF\_MAGEA9 | 79 | 8 | 4161 | 164 | fz2,Coq7,dnc,Pi3K68D,CG40178,RSG7,Dys,toy | | 1.183e-02 | -4.44 | double-stranded DNA binding | molecular function | GO:0003690 | 391 | 28 | 12176 | 554 | cic,zld,svp,toy,ct,bab2,zen,Atf6,ewg,Mef2,br,bi,pho,spel1,sna,CG9932,scro,pan,ap,Clk,E5,Myc,dsx,en,CG12769,Maf1,tj,hth | | 1.204e-02 | -4.42 | negative regulation of calcium ion transmembrane transport | biological process | GO:1903170 | 4 | 2 | 12345 | 571 | THADA,Rgk1 | | 1.204e-02 | -4.42 | negative regulation of calcium ion transmembrane transporter activity | biological process | GO:1901020 | 4 | 2 | 12345 | 571 | THADA,Rgk1 | | 1.205e-02 | -4.42 | CkIIalpha (casein kinase IIalpha) | protein interactions | 48448 | 50 | 7 | 9204 | 464 | Myc,par-1,pan,Clk,CG3907,orb,Rtnl1 | | 1.211e-02 | -4.41 | BTB | pfam domains | PF00651 | 62 | 8 | 10480 | 528 | Trl,br,bab2,CG15725,fru,lola,rib,CG41099 | | 1.214e-02 | -4.41 | regulation of nucleobase-containing compound metabolic process | biological process | GO:0019219 | 1315 | 78 | 12345 | 571 | d4,Nipped-A,SoxN,lncRNA:bxd,Camta,CG40228,lola,en,zld,Trl,cic,sna,fuss,Mnt,mav,CG34334,rl,pan,Setd3,jing,Psc,HP6,E5,CG12769,fru,Parg,bi,CG12200,pho,CG32006,zen,hth,AGO3,bru1,scw,lz,Parp,pum,Fer3,toy,dsx,scro,kni,Maf1,bab2,svp,HPS4,tld,Octbeta2R,br,PNPase,ct,CG9932,tou,Myc,CG1620,ap,spel1,SCAP,Tgi,tj,salm,bru3,caup,Atf6,Vsx2,Marf1,Mef2,ewg,CG5862,dimm,cv-2,CG33557,CG9650,Clk,rib,Su(z)2,fd96Ca | | 1.215e-02 | -4.41 | endoplasmic reticulum unfolded protein response | biological process | GO:0030968 | 20 | 4 | 12345 | 571 | Der-1,Hsc70-3,PEK,Atf6 | | 1.215e-02 | -4.41 | cell cycle comprising mitosis without cytokinesis | biological process | GO:0033301 | 20 | 4 | 12345 | 571 | Trl,Psc,CycB,toc | | 1.215e-02 | -4.41 | chitin-based larval cuticle pattern formation | biological process | GO:0035293 | 20 | 4 | 12345 | 571 | SoxN,ck,pan,fz2 | | 1.222e-02 | -4.40 | Glyco\_hydro\_b | interpro domains | IPR013780 | 19 | 4 | 11383 | 557 | alpha-Man-IIb,tobi,Amyrel,GCS2alpha | | 1.228e-02 | -4.40 | YOSHIMURA\_MAPK8\_TARGETS\_UP | MSigDB lists | YOSHIMURA\_MAPK8\_TARGETS\_UP | 298 | 20 | 4161 | 164 | Sod1,Gdh,dnc,spel1,CycB,Atpalpha,CG7255,ap,nAChRalpha3,Cyp6a8,Pkg21D,Octbeta2R,lqf,Dhc16F,Nlg3,Octbeta1R,Rbp9,Coq7,Ca-alpha1D,PMCA | | 1.229e-02 | -4.40 | presynaptic membrane organization | biological process | GO:0097090 | 11 | 3 | 12345 | 571 | Nlg2,Ten-a,Nlg3 | | 1.229e-02 | -4.40 | positive regulation of nuclear-transcribed mRNA catabolic process, deadenylation-dependent decay | biological process | GO:1900153 | 11 | 3 | 12345 | 571 | Marf1,AGO3,pum | | 1.229e-02 | -4.40 | regulation of nuclear-transcribed mRNA catabolic process, deadenylation-dependent decay | biological process | GO:1900151 | 11 | 3 | 12345 | 571 | pum,AGO3,Marf1 | | 1.230e-02 | -4.40 | regulation of biosynthetic process | biological process | GO:0009889 | 1374 | 81 | 12345 | 571 | Ptip,CG32006,zen,pho,bru1,scw,lz,Parp,hth,Trl,cic,sna,Mnt,fuss,d4,lncRNA:bxd,Nipped-A,SoxN,Camta,Tak1,CG40228,lola,en,zld,Psc,jing,HP6,CG12769,E5,fru,Parg,bi,mav,rl,pan,Setd3,ewg,Mef2,PEK,CG5862,salm,caup,bru3,Atf6,Vsx2,CG9650,Clk,rib,Thor,fd96Ca,Su(z)2,dimm,cv-2,heph,CG33557,scro,dnr1,kni,Maf1,bab2,svp,tld,HPS4,orb,Octbeta2R,pum,Fer3,dsx,toy,mxt,tou,CG9932,Myc,CG1620,ap,SCAP,tj,Tgi,br,CG17514,ct | | 1.234e-02 | -4.40 | FERM/acyl-CoA-bd\_prot\_sf | interpro domains | IPR014352 | 29 | 5 | 11383 | 557 | Acbp5,CG34347,Myo81F,dnr1,ck | | 1.239e-02 | -4.39 | regulation of transport | biological process | GO:0051049 | 267 | 21 | 12345 | 571 | AP-2alpha,Slob,Teh2,Rgk1,Lsd-2,cpx,IA-2,sky,dimm,Pld,Dys,ru,THADA,vap,step,Glut1,norpA,Syt7,CG9650,ArfGAP1,mim | | 1.242e-02 | -4.39 | C2\_dom | interpro domains | IPR000008 | 40 | 6 | 11383 | 557 | norpA,Pi3K68D,Syt7,unc-13-4A,vap,RhoGAP100F | | 1.253e-02 | -4.38 | GO\_NEGATIVE\_REGULATION\_OF\_NEURON\_APOPTOTIC\_PROCESS | MSigDB lists | GO\_NEGATIVE\_REGULATION\_OF\_NEURON\_APOPTOTIC\_PROCESS | 36 | 5 | 4161 | 164 | Mef2,Sod1,vap,DNAlig4,spel1 | | 1.253e-02 | -4.38 | KEGG\_VASCULAR\_SMOOTH\_MUSCLE\_CONTRACTION | MSigDB lists | KEGG\_VASCULAR\_SMOOTH\_MUSCLE\_CONTRACTION | 36 | 5 | 4161 | 164 | Ac78C,rl,Ca-alpha1D,cta,norpA | | 1.255e-02 | -4.38 | protein phosphorylation | biological process | GO:0006468 | 284 | 22 | 12345 | 571 | Src42A,Hipk,CG9886,Pdk,rl,CycB,CG32944,Gprk1,Takl2,fray,CG34357,Pkg21D,Stlk,Slob,Tak1,CG12147,PEK,par-1,dnt,bma,CG42637,bt | | 1.257e-02 | -4.38 | regulation of cellular metabolic process | biological process | GO:0031323 | 1987 | 112 | 12345 | 571 | Clk,CG9650,rib,grim,SteXh:CG42398,Alg-2,Su(z)2,Thor,fd96Ca,dimm,CycB,cv-2,Coq7,heph,CG33557,Spn75F,PEK,Mef2,Marf1,ewg,CG5862,salm,bru3,caup,Atf6,Vsx2,Stlk,CG9932,tou,Myc,CG1620,spel1,ap,SCAP,CG14184,Tgi,tj,Toll-7,PNPase,br,CG17514,ct,Sod1,scro,dnr1,kni,Maf1,bab2,svp,HPS4,tld,Octbeta2R,orb,pum,Catsup,Fer3,CG12772,dsx,toy,mxt,bru1,AGO3,fray,scw,lz,Parp,hth,mbc,Ppm1,Src42A,Ptip,CG32006,zen,par-1,CG12200,CG12147,pho,CG41106,Bruce,jing,HP6,Psc,CG12769,E5,fru,step,Parg,bi,mav,Diap1,CG34334,rl,pan,Setd3,Trl,cic,sna,Mnt,fuss,d4,lncRNA:bxd,Nipped-A,CG41520,SoxN,Camta,AP-2alpha,Tak1,CG40228,lola,en,Spn42De,zld | | 1.258e-02 | -4.38 | Signal Transduction | REACTOME pathways | R-DME-162582 | 717 | 47 | 3923 | 189 | Ac78C,conu,dnc,Pdk,Rpb12,norpA,Gprk1,Octbeta2R,sif,pico,eIF4B,mbc,CG41520,Parp,CG30440,Stlk,RhoGAP100F,ck,Rtnl1,Octbeta1R,5-HT2A,Cip4,Diap1,CG14184,rl,pk,fz2,cta,Act42A,par-1,rdgC,Rpn9,lqf,Tak1,vap,Rh3,CG10188,boi,SoxN,heph,CdGAPr,Nipped-A,pan,CG34357,CG12147,Prosalpha3T,Thor | | 1.258e-02 | -4.38 | plu (plutonium) | protein interactions | 44848 | 39 | 6 | 9204 | 464 | pan,pum,CycB,CG17508,dimm,unc-13-4A | | 1.278e-02 | -4.36 | ACTININ\_1 | prosite domains | PS00019 | 9 | 3 | 6243 | 365 | Msp300,shot,Dys | | 1.284e-02 | -4.35 | GO\_REGULATION\_OF\_SKELETAL\_MUSCLE\_TISSUE\_DEVELOPMENT | MSigDB lists | GO\_REGULATION\_OF\_SKELETAL\_MUSCLE\_TISSUE\_DEVELOPMENT | 13 | 3 | 4161 | 164 | Mef2,CG10077,Octbeta2R | | 1.284e-02 | -4.35 | PID\_AVB3\_INTEGRIN\_PATHWAY | MSigDB lists | PID\_AVB3\_INTEGRIN\_PATHWAY | 13 | 3 | 4161 | 164 | Pi3K68D,Mp,rl | | 1.284e-02 | -4.35 | GO\_PANCREAS\_DEVELOPMENT | MSigDB lists | GO\_PANCREAS\_DEVELOPMENT | 13 | 3 | 4161 | 164 | CG17387,toy,l(2)41Ab | | 1.284e-02 | -4.35 | GO\_PHOTORECEPTOR\_CELL\_DEVELOPMENT | MSigDB lists | GO\_PHOTORECEPTOR\_CELL\_DEVELOPMENT | 13 | 3 | 4161 | 164 | sn,ck,toy | | 1.291e-02 | -4.35 | IQ | smart domains | SM00015 | 24 | 5 | 5086 | 308 | ck,igl,rdgC,Myo81F,Camta | | 1.291e-02 | -4.35 | PEREZ\_TP63\_TARGETS | MSigDB lists | PEREZ\_TP63\_TARGETS | 50 | 6 | 4161 | 164 | arg,bru1,dnr1,Gfat1,step,CG14621 | | 1.292e-02 | -4.35 | compound eye development | biological process | GO:0048749 | 370 | 27 | 12345 | 571 | sdk,hth,rl,pk,br,ru,Hipk,Src42A,rno,boi,CadN2,bi,lz,rau,fred,salm,Tak1,caup,lola,toy,klg,RhoGAP100F,rg,zen,svp,kirre,sna | | 1.295e-02 | -4.35 | GO\_POSITIVE\_REGULATION\_OF\_CATALYTIC\_ACTIVITY | MSigDB lists | GO\_POSITIVE\_REGULATION\_OF\_CATALYTIC\_ACTIVITY | 401 | 25 | 4161 | 164 | sky,vap,fz2,Sod1,Tbc1d15-17,Stlk,CycB,spel1,RSG7,Trs23,Hsc70-3,rl,step,nAChRalpha4,CG34408,Alg-2,mbc,Octbeta2R,cta,Mef2,CG10565,CG3107,Ac78C,Rpn9,Git | | 1.303e-02 | -4.34 | cilium movement | biological process | GO:0003341 | 31 | 5 | 12345 | 571 | CG17387,CG9492,l(2)41Ab,kl-3,Dhc16F | | 1.303e-02 | -4.34 | ketone biosynthetic process | biological process | GO:0042181 | 31 | 5 | 12345 | 571 | Npc2b,spo,Coq7,nvd,spok | | 1.305e-02 | -4.34 | actin filament binding | molecular function | GO:0051015 | 56 | 7 | 12176 | 554 | ck,Svil,bt,Spn,Msp300,zormin,sn | | 1.314e-02 | -4.33 | regulation of hormone levels | biological process | GO:0010817 | 96 | 10 | 12345 | 571 | spok,dimm,CG30495,Npc2b,CG9650,Glut1,bond,nvd,CG9514,spo | | 1.319e-02 | -4.33 | regulation of RNA metabolic process | biological process | GO:0051252 | 1281 | 76 | 12345 | 571 | Myc,CG9932,tou,ap,CG1620,SCAP,tj,Tgi,ct,PNPase,br,kni,scro,bab2,Maf1,HPS4,tld,svp,Octbeta2R,pum,Fer3,toy,dsx,Clk,CG9650,rib,Su(z)2,fd96Ca,dimm,cv-2,CG33557,Mef2,Marf1,ewg,CG5862,salm,caup,bru3,Vsx2,Atf6,jing,Psc,HP6,fru,E5,CG12769,Parg,bi,mav,pan,CG34334,rl,Setd3,Trl,cic,Mnt,fuss,sna,lncRNA:bxd,SoxN,Nipped-A,d4,CG40228,Camta,lola,en,zld,AGO3,bru1,scw,lz,Parp,hth,CG32006,zen,pho | | 1.334e-02 | -4.32 | MORF\_IL4 | MSigDB lists | MORF\_IL4 | 24 | 4 | 4161 | 164 | Dys,toy,dnc,fz2 | | 1.334e-02 | -4.32 | GSE7852\_THYMUS\_VS\_FAT\_TCONV\_DN | MSigDB lists | GSE7852\_THYMUS\_VS\_FAT\_TCONV\_DN | 24 | 4 | 4161 | 164 | Hsc70-3,Odc2,PMCA,unc-13-4A | | 1.334e-02 | -4.32 | GSE45365\_WT\_VS\_IFNAR\_KO\_CD11B\_DC\_MCMV\_INFECTION\_DN | MSigDB lists | GSE45365\_WT\_VS\_IFNAR\_KO\_CD11B\_DC\_MCMV\_INFECTION\_DN | 24 | 4 | 4161 | 164 | MCU,norpA,Src42A,rdgC | | 1.334e-02 | -4.32 | AACTGAC\_MIR223 | MSigDB lists | AACTGAC\_MIR223 | 24 | 4 | 4161 | 164 | vap,dnc,Mef2,Der-1 | | 1.334e-02 | -4.32 | IL21\_UP.V1\_UP | MSigDB lists | IL21\_UP.V1\_UP | 24 | 4 | 4161 | 164 | Octbeta1R,unc-13-4A,Mp,Ugt50B3 | | 1.343e-02 | -4.31 | Catenin\_binding\_dom\_sf | interpro domains | IPR027397 | 4 | 2 | 11383 | 557 | pan,CadN2 | | 1.343e-02 | -4.31 | Ketoacyl\_synth\_N | interpro domains | IPR014030 | 4 | 2 | 11383 | 557 | FASN3,CG12170 | | 1.343e-02 | -4.31 | Ketoacyl\_synth\_C | interpro domains | IPR014031 | 4 | 2 | 11383 | 557 | CG12170,FASN3 | | 1.343e-02 | -4.31 | DUF1736 | interpro domains | IPR013618 | 4 | 2 | 11383 | 557 | CG4341,CG31690 | | 1.343e-02 | -4.31 | PKS\_Beta-ketoAc\_synthase\_dom | interpro domains | IPR020841 | 4 | 2 | 11383 | 557 | CG12170,FASN3 | | 1.343e-02 | -4.31 | BIR\_rpt | interpro domains | IPR001370 | 4 | 2 | 11383 | 557 | Bruce,Diap1 | | 1.343e-02 | -4.31 | RAWUL | interpro domains | IPR032443 | 4 | 2 | 11383 | 557 | Su(z)2,Psc | | 1.343e-02 | -4.31 | Ketoacyl\_synth\_AS | interpro domains | IPR018201 | 4 | 2 | 11383 | 557 | FASN3,CG12170 | | 1.343e-02 | -4.31 | OST-HTH/LOTUS\_dom | interpro domains | IPR025605 | 4 | 2 | 11383 | 557 | tej,Marf1 | | 1.351e-02 | -4.30 | Trim9 (Tripartite motif containing 9) | protein interactions | 34453 | 19 | 4 | 9204 | 464 | dsx,zld,pico,Vsx2 | | 1.351e-02 | -4.30 | slbo (slow border cells) | protein interactions | 37889 | 19 | 4 | 9204 | 464 | jing,AP-2alpha,mbc,tj | | 1.359e-02 | -4.30 | actin filament organization | biological process | GO:0007015 | 111 | 11 | 12345 | 571 | jvl,sn,Msp300,Src42A,pico,Spn,ck,bt,CAP,Mical,capu | | 1.365e-02 | -4.29 | SMTTTTGT\_UNKNOWN | MSigDB lists | SMTTTTGT\_UNKNOWN | 81 | 8 | 4161 | 164 | Gfat1,lqf,Dys,dnc,CG10077,Rbp9,twz,fz2 | | 1.368e-02 | -4.29 | protein kinase activity | molecular function | GO:0004672 | 274 | 21 | 12176 | 554 | Src42A,CycB,rl,Takl2,Pdk,bma,Gprk1,dnt,CG42637,Stlk,bt,Pkg21D,Slob,CG12147,Tak1,Hipk,CG32944,CG34357,fray,PEK,par-1 | | 1.368e-02 | -4.29 | carbohydrate:cation symporter activity | molecular function | GO:0005402 | 21 | 4 | 12176 | 554 | CG15408,CG8249,CG45782,CG1213 | | 1.368e-02 | -4.29 | carbohydrate:proton symporter activity | molecular function | GO:0005351 | 21 | 4 | 12176 | 554 | CG15408,CG8249,CG45782,CG1213 | | 1.368e-02 | -4.29 | channel regulator activity | molecular function | GO:0016247 | 21 | 4 | 12176 | 554 | Rgk1,Teh2,CheB42a,CG42541 | | 1.369e-02 | -4.29 | positive regulation of RNA metabolic process | biological process | GO:0051254 | 513 | 35 | 12345 | 571 | Atf6,caup,pho,Marf1,Mef2,zen,dimm,hth,lz,AGO3,Clk,lola,en,dsx,toy,zld,SoxN,d4,pum,CG40228,Camta,sna,Trl,scro,Setd3,br,pan,SCAP,bi,tj,Myc,jing,tou,ap,CG12769 | | 1.388e-02 | -4.28 | PH\_DOMAIN | prosite domains | PS50003 | 66 | 9 | 6243 | 365 | Gprk1,vap,step,cert,osp,pico,sif,CG30440,CG10188 | | 1.393e-02 | -4.27 | phospholipase activity | molecular function | GO:0004620 | 32 | 5 | 12176 | 554 | Yp2,norpA,Pld,CG3009,CG42237 | | 1.398e-02 | -4.27 | purine nucleotide binding | molecular function | GO:0017076 | 821 | 51 | 12176 | 554 | CG34357,CG43672,DNAlig4,par-1,CG9981,fbl,CG12147,Hipk,Tak1,CG42541,fzo,Klp59C,Stlk,Gprk1,CG31689,Nadsyn,Hsc70-3,Rgk1,Acbp5,Src42A,CG9492,Myo81F,bma,mIF2,fray,Atpalpha,Dhc16F,PEK,mus308,RabX2,CG32944,cta,CG5789,Hmt-1,CG42637,dnt,kl-3,Khc-73,spel1,ABCB7,Pkg21D,bt,lz,CG33970,CG14721,CG10077,CG5160,rl,ck,Takl2,PMCA | | 1.404e-02 | -4.27 | cell migration | biological process | GO:0016477 | 287 | 22 | 12345 | 571 | cta,jing,rib,mim,tj,CG1486,Pi3K68D,gus,sn,Dlg5,Diap1,mbc,Sdc,Msp300,Sema1a,Src42A,ru,kni,fz2,par-1,salm,pum | | 1.404e-02 | -4.27 | GSE28737\_WT\_VS\_BCL6\_KO\_MARGINAL\_ZONE\_BCELL\_UP | MSigDB lists | GSE28737\_WT\_VS\_BCL6\_KO\_MARGINAL\_ZONE\_BCELL\_UP | 37 | 5 | 4161 | 164 | CG9107,drosha,sdk,scro,AGO3 | | 1.404e-02 | -4.27 | MODULE\_176 | MSigDB lists | MODULE\_176 | 37 | 5 | 4161 | 164 | Mef2,Ugt50B3,Octbeta2R,Mp,RSG7 | | 1.404e-02 | -4.27 | HFH4\_01 | MSigDB lists | HFH4\_01 | 37 | 5 | 4161 | 164 | Mef2,Dys,Ca-alpha1D,Pi3K68D,twz | | 1.404e-02 | -4.27 | MORF\_IFNA1 | MSigDB lists | MORF\_IFNA1 | 37 | 5 | 4161 | 164 | toy,Dys,Coq7,dnc,fz2 | | 1.404e-02 | -4.27 | GO\_ADENYLATE\_CYCLASE\_MODULATING\_G\_PROTEIN\_COUPLED\_RECEPTOR\_SIGNALING\_PATHWAY | MSigDB lists | GO\_ADENYLATE\_CYCLASE\_MODULATING\_G\_PROTEIN\_COUPLED\_RECEPTOR\_SIGNALING\_PATHWAY | 37 | 5 | 4161 | 164 | Octbeta2R,cta,Ac78C,dnc,Ca-alpha1D | | 1.404e-02 | -4.27 | GSE19198\_CTRL\_VS\_IL21\_TREATED\_TCELL\_6H\_DN | MSigDB lists | GSE19198\_CTRL\_VS\_IL21\_TREATED\_TCELL\_6H\_DN | 37 | 5 | 4161 | 164 | Khc-73,Pi3K68D,vap,AP-2alpha,ck | | 1.408e-02 | -4.26 | compound eye photoreceptor cell differentiation | biological process | GO:0001751 | 157 | 14 | 12345 | 571 | lola,br,ru,klg,hth,salm,rl,svp,CadN2,lz,rau,RhoGAP100F,boi,rno | | 1.417e-02 | -4.26 | MORF\_NOS2A | MSigDB lists | MORF\_NOS2A | 51 | 6 | 4161 | 164 | toy,Octbeta1R,Pi3K68D,dnc,fz2,CG40178 | | 1.417e-02 | -4.26 | P53\_DECAMER\_Q2 | MSigDB lists | P53\_DECAMER\_Q2 | 51 | 6 | 4161 | 164 | Nlg3,sbm,Dhc16F,shot,inc,Dys | | 1.418e-02 | -4.26 | fn3 | pfam domains | PF00041 | 40 | 6 | 10480 | 528 | boi,Ptp99A,sdk,CG14964,Dscam2,bt | | 1.419e-02 | -4.26 | GO\_SYNAPSE | MSigDB lists | GO\_SYNAPSE | 205 | 15 | 4161 | 164 | nAChRalpha3,sdk,Atpalpha,Nlg2,norpA,sky,nAChRalpha4,Maf1,unc-13-4A,Trs23,Nlg3,ck,Mef2,Dys,Octbeta2R | | 1.420e-02 | -4.25 | GO\_ANATOMICAL\_STRUCTURE\_FORMATION\_INVOLVED\_IN\_MORPHOGENESIS | MSigDB lists | GO\_ANATOMICAL\_STRUCTURE\_FORMATION\_INVOLVED\_IN\_MORPHOGENESIS | 224 | 16 | 4161 | 164 | cv-2,Ugt50B3,Alg-2,l(2)41Ab,Mef2,toy,cta,CG9492,sdk,ap,fz2,CG17387,CG9107,Mp,rl,Dys | | 1.420e-02 | -4.25 | OST-HTH | pfam domains | PF12872 | 4 | 2 | 10480 | 528 | tej,Marf1 | | 1.420e-02 | -4.25 | Ketoacyl-synt\_C | pfam domains | PF02801 | 4 | 2 | 10480 | 528 | FASN3,CG12170 | | 1.420e-02 | -4.25 | BIR | pfam domains | PF00653 | 4 | 2 | 10480 | 528 | Bruce,Diap1 | | 1.420e-02 | -4.25 | RAWUL | pfam domains | PF16207 | 4 | 2 | 10480 | 528 | Psc,Su(z)2 | | 1.420e-02 | -4.25 | ketoacyl-synt | pfam domains | PF00109 | 4 | 2 | 10480 | 528 | FASN3,CG12170 | | 1.420e-02 | -4.25 | DUF1736 | pfam domains | PF08409 | 4 | 2 | 10480 | 528 | CG4341,CG31690 | | 1.422e-02 | -4.25 | Rgk1 (Rad, Gem/Kir family member 1) | protein interactions | 14462605 | 4 | 2 | 9204 | 464 | Rtnl1,dnc | | 1.422e-02 | -4.25 | Spn42Da (Serpin 42Da) | protein interactions | 49805 | 4 | 2 | 9204 | 464 | norpA,Nrd1 | | 1.422e-02 | -4.25 | Pcd (pterin-4a-carbinolamine dehydratase) | protein interactions | 43499 | 4 | 2 | 9204 | 464 | tobi,2mit | | 1.427e-02 | -4.25 | GO\_REGULATION\_OF\_RESPIRATORY\_SYSTEM\_PROCESS | MSigDB lists | GO\_REGULATION\_OF\_RESPIRATORY\_SYSTEM\_PROCESS | 5 | 2 | 4161 | 164 | GLS,Nlg3 | | 1.427e-02 | -4.25 | REACTOME\_PRESYNAPTIC\_NICOTINIC\_ACETYLCHOLINE\_RECEPTORS | MSigDB lists | REACTOME\_PRESYNAPTIC\_NICOTINIC\_ACETYLCHOLINE\_RECEPTORS | 5 | 2 | 4161 | 164 | nAChRalpha3,nAChRalpha4 | | 1.427e-02 | -4.25 | GO\_INHIBITORY\_SYNAPSE | MSigDB lists | GO\_INHIBITORY\_SYNAPSE | 5 | 2 | 4161 | 164 | Nlg3,Maf1 | | 1.427e-02 | -4.25 | GO\_REGULATION\_OF\_NEUROTRANSMITTER\_RECEPTOR\_ACTIVITY | MSigDB lists | GO\_REGULATION\_OF\_NEUROTRANSMITTER\_RECEPTOR\_ACTIVITY | 5 | 2 | 4161 | 164 | Nlg3,Mef2 | | 1.427e-02 | -4.25 | GO\_NEGATIVE\_REGULATION\_OF\_STEROID\_METABOLIC\_PROCESS | MSigDB lists | GO\_NEGATIVE\_REGULATION\_OF\_STEROID\_METABOLIC\_PROCESS | 5 | 2 | 4161 | 164 | SCAP,Sod1 | | 1.427e-02 | -4.25 | GO\_REGULATION\_OF\_SKELETAL\_MUSCLE\_CELL\_DIFFERENTIATION | MSigDB lists | GO\_REGULATION\_OF\_SKELETAL\_MUSCLE\_CELL\_DIFFERENTIATION | 5 | 2 | 4161 | 164 | CG10077,Mef2 | | 1.427e-02 | -4.25 | GO\_NEGATIVE\_REGULATION\_OF\_ALCOHOL\_BIOSYNTHETIC\_PROCESS | MSigDB lists | GO\_NEGATIVE\_REGULATION\_OF\_ALCOHOL\_BIOSYNTHETIC\_PROCESS | 5 | 2 | 4161 | 164 | Sod1,SCAP | | 1.427e-02 | -4.25 | GO\_INNER\_EAR\_RECEPTOR\_STEREOCILIUM\_ORGANIZATION | MSigDB lists | GO\_INNER\_EAR\_RECEPTOR\_STEREOCILIUM\_ORGANIZATION | 5 | 2 | 4161 | 164 | Sod1,ck | | 1.427e-02 | -4.25 | GO\_NEUREXIN\_FAMILY\_PROTEIN\_BINDING | MSigDB lists | GO\_NEUREXIN\_FAMILY\_PROTEIN\_BINDING | 5 | 2 | 4161 | 164 | Nlg3,Nlg2 | | 1.427e-02 | -4.25 | GO\_POSITIVE\_REGULATION\_OF\_INTERFERON\_GAMMA\_PRODUCTION | MSigDB lists | GO\_POSITIVE\_REGULATION\_OF\_INTERFERON\_GAMMA\_PRODUCTION | 5 | 2 | 4161 | 164 | dnc,fz2 | | 1.427e-02 | -4.25 | SCHLESINGER\_METHYLATED\_DE\_NOVO\_IN\_CANCER | MSigDB lists | SCHLESINGER\_METHYLATED\_DE\_NOVO\_IN\_CANCER | 5 | 2 | 4161 | 164 | toy,svp | | 1.427e-02 | -4.25 | GO\_CELL\_DIFFERENTIATION\_INVOLVED\_IN\_EMBRYONIC\_PLACENTA\_DEVELOPMENT | MSigDB lists | GO\_CELL\_DIFFERENTIATION\_INVOLVED\_IN\_EMBRYONIC\_PLACENTA\_DEVELOPMENT | 5 | 2 | 4161 | 164 | svp,fz2 | | 1.427e-02 | -4.25 | GO\_LABYRINTHINE\_LAYER\_BLOOD\_VESSEL\_DEVELOPMENT | MSigDB lists | GO\_LABYRINTHINE\_LAYER\_BLOOD\_VESSEL\_DEVELOPMENT | 5 | 2 | 4161 | 164 | rl,fz2 | | 1.427e-02 | -4.25 | GO\_DNA\_LIGATION\_INVOLVED\_IN\_DNA\_REPAIR | MSigDB lists | GO\_DNA\_LIGATION\_INVOLVED\_IN\_DNA\_REPAIR | 5 | 2 | 4161 | 164 | Parp,DNAlig4 | | 1.427e-02 | -4.25 | PID\_SYNDECAN\_1\_PATHWAY | MSigDB lists | PID\_SYNDECAN\_1\_PATHWAY | 5 | 2 | 4161 | 164 | rl,Mp | | 1.427e-02 | -4.25 | GO\_DOPAMINE\_RECEPTOR\_BINDING | MSigDB lists | GO\_DOPAMINE\_RECEPTOR\_BINDING | 5 | 2 | 4161 | 164 | Atpalpha,cta | | 1.427e-02 | -4.25 | GO\_REGULATION\_OF\_GOLGI\_ORGANIZATION | MSigDB lists | GO\_REGULATION\_OF\_GOLGI\_ORGANIZATION | 5 | 2 | 4161 | 164 | Syx18,rl | | 1.427e-02 | -4.25 | GO\_EMBRYONIC\_CAMERA\_TYPE\_EYE\_MORPHOGENESIS | MSigDB lists | GO\_EMBRYONIC\_CAMERA\_TYPE\_EYE\_MORPHOGENESIS | 5 | 2 | 4161 | 164 | fz2,toy | | 1.427e-02 | -4.25 | GO\_REGULATION\_OF\_CHOLESTEROL\_METABOLIC\_PROCESS | MSigDB lists | GO\_REGULATION\_OF\_CHOLESTEROL\_METABOLIC\_PROCESS | 5 | 2 | 4161 | 164 | SCAP,Sod1 | | 1.427e-02 | -4.25 | GO\_NEGATIVE\_REGULATION\_OF\_SMOOTH\_MUSCLE\_CONTRACTION | MSigDB lists | GO\_NEGATIVE\_REGULATION\_OF\_SMOOTH\_MUSCLE\_CONTRACTION | 5 | 2 | 4161 | 164 | Octbeta2R,Sod1 | | 1.427e-02 | -4.25 | GO\_REGULATION\_OF\_TRANSCRIPTION\_INVOLVED\_IN\_CELL\_FATE\_COMMITMENT | MSigDB lists | GO\_REGULATION\_OF\_TRANSCRIPTION\_INVOLVED\_IN\_CELL\_FATE\_COMMITMENT | 5 | 2 | 4161 | 164 | toy,svp | | 1.427e-02 | -4.25 | GO\_EMBRYONIC\_EYE\_MORPHOGENESIS | MSigDB lists | GO\_EMBRYONIC\_EYE\_MORPHOGENESIS | 5 | 2 | 4161 | 164 | fz2,toy | | 1.430e-02 | -4.25 | ERR1\_Q2 | MSigDB lists | ERR1\_Q2 | 66 | 7 | 4161 | 164 | fbl,ap,CG17691,Pkg21D,MFS16,Tbc1d15-17,Mef2 | | 1.431e-02 | -4.25 | GO\_CELL\_ACTIVATION | MSigDB lists | GO\_CELL\_ACTIVATION | 98 | 9 | 4161 | 164 | Khc-73,Mef2,cta,spel1,DNAlig4,rl,cv-2,CG7255,fz2 | | 1.431e-02 | -4.25 | TGCACTT\_MIR519C\_MIR519B\_MIR519A | MSigDB lists | TGCACTT\_MIR519C\_MIR519B\_MIR519A | 98 | 9 | 4161 | 164 | CG41099,mus308,Bruce,CG10077,rg,CG30059,Tsp5D,CG40178,bru1 | | 1.432e-02 | -4.25 | DHC\_fam | interpro domains | IPR026983 | 11 | 3 | 11383 | 557 | Dhc16F,CG9492,kl-3 | | 1.432e-02 | -4.25 | DNAH3\_AAA\_lid\_1 | interpro domains | IPR041589 | 11 | 3 | 11383 | 557 | kl-3,CG9492,Dhc16F | | 1.447e-02 | -4.24 | imaginal disc pattern formation | biological process | GO:0007447 | 112 | 11 | 12345 | 571 | en,ru,hth,Nipped-A,rl,bi,jing,Ptip,boi,cic,ap | | 1.452e-02 | -4.23 | GO\_REGULATION\_OF\_HYDROLASE\_ACTIVITY | MSigDB lists | GO\_REGULATION\_OF\_HYDROLASE\_ACTIVITY | 283 | 19 | 4161 | 164 | Tbc1d15-17,Sod1,step,Hsc70-3,spel1,RSG7,Trs23,sky,Bruce,vap,CG10565,Mef2,cta,Octbeta2R,Git,Alg-2,CG34408,Spn42De,mbc | | 1.460e-02 | -4.23 | GO\_POSITIVE\_REGULATION\_OF\_MOLECULAR\_FUNCTION | MSigDB lists | GO\_POSITIVE\_REGULATION\_OF\_MOLECULAR\_FUNCTION | 468 | 28 | 4161 | 164 | Parp,mbc,nAChRalpha4,CG34408,Alg-2,Rpn9,Ac78C,Git,Octbeta2R,CG10565,CG3107,Mef2,cta,fz2,vap,Der-1,sky,Hsc70-3,spel1,RSG7,Trs23,CycB,step,rl,Stlk,Dys,Tbc1d15-17,Sod1 | | 1.466e-02 | -4.22 | regulation of macromolecule biosynthetic process | biological process | GO:0010556 | 1326 | 78 | 12345 | 571 | Setd3,mav,pan,rl,Parg,bi,HP6,Psc,jing,fru,E5,CG12769,en,lola,zld,Nipped-A,SoxN,lncRNA:bxd,d4,CG40228,Camta,fuss,Mnt,sna,Trl,cic,hth,lz,Parp,bru1,scw,pho,CG32006,zen,CG17514,ct,br,SCAP,tj,Tgi,Myc,tou,CG9932,ap,CG1620,mxt,toy,dsx,pum,Fer3,tld,HPS4,svp,orb,Octbeta2R,kni,scro,bab2,Maf1,heph,CG33557,dimm,cv-2,fd96Ca,Su(z)2,Thor,Clk,CG9650,rib,Vsx2,Atf6,salm,bru3,caup,PEK,Mef2,ewg,CG5862 | | 1.482e-02 | -4.21 | UQCR-C1 (Ubiquinol-cytochrome c reductase core protein 1) | protein interactions | 41800 | 52 | 7 | 9204 | 464 | Nrd1,bab2,Sema1a,Tak1,Rtnl1,UQCR-11,Thor | | 1.483e-02 | -4.21 | 14-3-3zeta (14-3-3zeta) | protein interactions | 36059 | 90 | 10 | 9204 | 464 | Khc-73,par-1,Pi3K68D,Nrd1,ct,Src42A,Slob,RNASEK,Sema1a,jvl | | 1.485e-02 | -4.21 | fly chr7B6-7B6\|1-20 cM | chromosome location | fly chr7B6-7B6\|1-20 cM | 6 | 2 | 25010 | 823 | CG15478,lncRNA:CR45665 | | 1.485e-02 | -4.21 | fly chr7B6-7B6\|1-20 | chromosome location | fly chr7B6-7B6\|1-20 | 6 | 2 | 25010 | 823 | CG15478,lncRNA:CR45665 | | 1.493e-02 | -4.20 | CG4617 (uncharacterized protein) | protein interactions | 31657 | 77 | 9 | 9204 | 464 | CG13405,RpL15,fzo,Git,CG7518,ewg,en,zormin,Yp2 | | 1.515e-02 | -4.19 | translation repressor activity, mRNA regulatory element binding | molecular function | GO:0000900 | 12 | 3 | 12176 | 554 | heph,orb,pum | | 1.515e-02 | -4.19 | photoreceptor activity | molecular function | GO:0009881 | 12 | 3 | 12176 | 554 | SK,Rh3,shakB | | 1.517e-02 | -4.19 | phosphotransferase activity, alcohol group as acceptor | molecular function | GO:0016773 | 346 | 25 | 12176 | 554 | dnt,Stlk,CG42637,Gprk1,CG3544,Slob,Pkg21D,bt,CycB,Src42A,bma,Pdk,Takl2,rl,Pi3K68D,CG34357,fray,par-1,PEK,fbl,CG32944,Tak1,Hipk,CG12147,CG9886 | | 1.521e-02 | -4.19 | anatomical structure formation involved in morphogenesis | biological process | GO:0048646 | 553 | 37 | 12345 | 571 | kirre,nullo,orb,sna,tld,cic,drosha,bt,zld,Mical,Pld,ct,br,pan,rl,gus,lama,capu,Mef2,par-1,Cip4,zen,shot,Irp-1A,knk,caup,jvl,salm,Thor,lz,scw,bru1,heph,mbc,mlt,hth,fan | | 1.526e-02 | -4.18 | GO\_REGULATION\_OF\_ION\_TRANSPORT | MSigDB lists | GO\_REGULATION\_OF\_ION\_TRANSPORT | 116 | 10 | 4161 | 164 | nAChRalpha3,Ca-alpha1D,nAChRalpha4,norpA,Nlg3,fray,dnc,Octbeta2R,Mef2,Dys | | 1.534e-02 | -4.18 | regulation of cell differentiation | biological process | GO:0045595 | 428 | 30 | 12345 | 571 | hth,mav,sdk,Diap1,rl,mbc,pk,sif,Src42A,Sema1a,lqf,babos,CG9380,bru1,Myc,norpA,Mical,lz,tinc,salm,hdc,SoxN,Catsup,Bruce,zld,shot,bab2,par-1,kirre,fuss | | 1.540e-02 | -4.17 | REACTOME\_G\_ALPHA\_S\_SIGNALLING\_EVENTS | MSigDB lists | REACTOME\_G\_ALPHA\_S\_SIGNALLING\_EVENTS | 25 | 4 | 4161 | 164 | Octbeta2R,Octbeta1R,Ac78C,dnc | | 1.540e-02 | -4.17 | GO\_GLYCOPROTEIN\_BINDING | MSigDB lists | GO\_GLYCOPROTEIN\_BINDING | 25 | 4 | 4161 | 164 | vap,Hsc70-3,Dys,Atpalpha | | 1.540e-02 | -4.17 | GSE4748\_CYANOBACTERIUM\_LPSLIKE\_VS\_LPS\_AND\_CYANOBACTERIUM\_LPSLIKE\_STIM\_DC\_3H\_DN | MSigDB lists | GSE4748\_CYANOBACTERIUM\_LPSLIKE\_VS\_LPS\_AND\_CYANOBACTERIUM\_LPSLIKE\_STIM\_DC\_3H\_DN | 25 | 4 | 4161 | 164 | CG40228,Maf1,St1,par-1 | | 1.540e-02 | -4.17 | GO\_CELL\_SURFACE\_RECEPTOR\_SIGNALING\_PATHWAY\_INVOLVED\_IN\_CELL\_CELL\_SIGNALING | MSigDB lists | GO\_CELL\_SURFACE\_RECEPTOR\_SIGNALING\_PATHWAY\_INVOLVED\_IN\_CELL\_CELL\_SIGNALING | 25 | 4 | 4161 | 164 | nAChRalpha3,Octbeta2R,nAChRalpha4,Mef2 | | 1.540e-02 | -4.17 | LEF1\_UP.V1\_UP | MSigDB lists | LEF1\_UP.V1\_UP | 25 | 4 | 4161 | 164 | Meltrin,toy,arg,Glut1 | | 1.540e-02 | -4.17 | GSE10463\_CD40L\_AND\_VA347\_VS\_CD40L\_IN\_DC\_DN | MSigDB lists | GSE10463\_CD40L\_AND\_VA347\_VS\_CD40L\_IN\_DC\_DN | 25 | 4 | 4161 | 164 | Octbeta2R,lqf,Nlg3,twz | | 1.546e-02 | -4.17 | GO\_BLOOD\_VESSEL\_MORPHOGENESIS | MSigDB lists | GO\_BLOOD\_VESSEL\_MORPHOGENESIS | 67 | 7 | 4161 | 164 | Alg-2,cta,svp,fz2,Mp,vap,cv-2 | | 1.549e-02 | -4.17 | AAA\_lid\_1 | pfam domains | PF17857 | 11 | 3 | 10480 | 528 | Dhc16F,CG9492,kl-3 | | 1.550e-02 | -4.17 | molecular function regulator | molecular function | GO:0098772 | 563 | 37 | 12176 | 554 | CG14298,mbc,Catsup,CG30440,sif,vap,CheB42a,CG34276,RhoGAP100F,sky,Sema1a,Tbc1d15-17,Trs23,CycB,step,CG17514,CG41520,Diap1,CG42541,CG14184,mav,Acp36DE,CG34408,CG5639,Spn75F,norpA,SteXh:CG42398,CG10188,Spn42De,RYa,ArfGAP1,Git,CdGAPr,conu,scw,Teh2,Rgk1 | | 1.551e-02 | -4.17 | slo (slowpoke) | protein interactions | 42940 | 11 | 3 | 9204 | 464 | dnc,Slob,Ca-alpha1D | | 1.551e-02 | -4.17 | CG10147 (uncharacterized protein) | protein interactions | 38733 | 11 | 3 | 9204 | 464 | toy,rau,fest | | 1.551e-02 | -4.17 | CG4270 (uncharacterized protein) | protein interactions | 33408 | 11 | 3 | 9204 | 464 | CG10164,br,CG34347 | | 1.551e-02 | -4.17 | pcm (pacman) | protein interactions | 32935 | 11 | 3 | 9204 | 464 | Psc,Diap1,Myc | | 1.551e-02 | -4.17 | GSE27859\_CD11C\_INT\_F480\_HI\_MACROPHAGE\_VS\_CD11C\_ING\_F480\_INT\_DC\_UP | MSigDB lists | GSE27859\_CD11C\_INT\_F480\_HI\_MACROPHAGE\_VS\_CD11C\_ING\_F480\_INT\_DC\_UP | 52 | 6 | 4161 | 164 | Maf1,IntS3,Atpalpha,SCAP,CG10171,CG41099 | | 1.551e-02 | -4.17 | MORF\_FSHR | MSigDB lists | MORF\_FSHR | 52 | 6 | 4161 | 164 | RSG7,Coq7,fz2,dnc,Dys,toy | | 1.551e-02 | -4.17 | KEGG\_CALCIUM\_SIGNALING\_PATHWAY | MSigDB lists | KEGG\_CALCIUM\_SIGNALING\_PATHWAY | 52 | 6 | 4161 | 164 | Octbeta1R,Ac78C,PMCA,Ca-alpha1D,Octbeta2R,norpA | | 1.554e-02 | -4.16 | cell-cell junction assembly | biological process | GO:0007043 | 57 | 7 | 12345 | 571 | kune,CadN2,CG15449,kirre,nullo,Atpalpha,shakB | | 1.557e-02 | -4.16 | intrinsic component of plasma membrane | cellular component | GO:0031226 | 480 | 32 | 11940 | 534 | HisCl1,KCNQ,DIP-delta,fz2,Atpalpha,TrissinR,CG15449,Octbeta2R,nAChRalpha4,Teh2,CG1213,5-HT2A,GABA-B-R3,CG42637,nAChRalpha3,dnt,CG15408,Nlg3,Rh3,Nlg2,Tsp5D,Ac78C,sdk,Rift,Dscam2,DIP-lambda,Oatp33Ea,Octbeta1R,CG34357,CadN2,CG8249,ru | | 1.559e-02 | -4.16 | larval development | biological process | GO:0002164 | 159 | 14 | 12345 | 571 | lola,Nna1,Msp300,Pkg21D,SoxN,mbc,pan,rl,kirre,nvd,lz,ck,ap,fz2 | | 1.566e-02 | -4.16 | MODULE\_45 | MSigDB lists | MODULE\_45 | 83 | 8 | 4161 | 164 | Cyp6a8,CG9389,bru1,Gfat1,Glut1,Git,Meltrin,Octbeta1R | | 1.566e-02 | -4.16 | compound eye morphogenesis | biological process | GO:0001745 | 290 | 22 | 12345 | 571 | rg,RhoGAP100F,kirre,svp,Tak1,caup,fred,salm,klg,lola,boi,rno,rau,bi,lz,CadN2,rl,sdk,hth,ru,pk,br | | 1.567e-02 | -4.16 | MIYAGAWA\_TARGETS\_OF\_EWSR1\_ETS\_FUSIONS\_DN | MSigDB lists | MIYAGAWA\_TARGETS\_OF\_EWSR1\_ETS\_FUSIONS\_DN | 38 | 5 | 4161 | 164 | Meltrin,Glut1,svp,CG6788,Gfat1 | | 1.567e-02 | -4.16 | GO\_CELLULAR\_RESPONSE\_TO\_INORGANIC\_SUBSTANCE | MSigDB lists | GO\_CELLULAR\_RESPONSE\_TO\_INORGANIC\_SUBSTANCE | 38 | 5 | 4161 | 164 | Sod1,Mef2,CycB,DNAlig4,Hsc70-3 | | 1.567e-02 | -4.16 | GSE40273\_GATA1\_KO\_VS\_WT\_TREG\_DN | MSigDB lists | GSE40273\_GATA1\_KO\_VS\_WT\_TREG\_DN | 38 | 5 | 4161 | 164 | step,CG41099,Mef2,Glut1,Ate1 | | 1.567e-02 | -4.16 | MORF\_MSH2 | MSigDB lists | MORF\_MSH2 | 38 | 5 | 4161 | 164 | Sod1,vtd,Nipped-A,Parg,spel1 | | 1.567e-02 | -4.16 | GSE1791\_CTRL\_VS\_NEUROMEDINU\_IN\_T\_CELL\_LINE\_0.8H\_DN | MSigDB lists | GSE1791\_CTRL\_VS\_NEUROMEDINU\_IN\_T\_CELL\_LINE\_0.8H\_DN | 38 | 5 | 4161 | 164 | bru3,Dhc16F,bru1,svp,Glut1 | | 1.584e-02 | -4.15 | positive regulation of mRNA catabolic process | biological process | GO:0061014 | 12 | 3 | 12345 | 571 | Marf1,pum,AGO3 | | 1.584e-02 | -4.15 | mRNA splice site selection | biological process | GO:0006376 | 12 | 3 | 12345 | 571 | bru1,su(w[a]),bru3 | | 1.588e-02 | -4.14 | GO\_PHOTORECEPTOR\_INNER\_SEGMENT | MSigDB lists | GO\_PHOTORECEPTOR\_INNER\_SEGMENT | 14 | 3 | 4161 | 164 | CG30495,rdgC,ck | | 1.588e-02 | -4.14 | GO\_AXONEME\_ASSEMBLY | MSigDB lists | GO\_AXONEME\_ASSEMBLY | 14 | 3 | 4161 | 164 | l(2)41Ab,CG9492,CG17387 | | 1.588e-02 | -4.14 | GAANYNYGACNY\_UNKNOWN | MSigDB lists | GAANYNYGACNY\_UNKNOWN | 14 | 3 | 4161 | 164 | toy,svp,Mp | | 1.588e-02 | -4.14 | GO\_CALCIUM\_CHANNEL\_COMPLEX | MSigDB lists | GO\_CALCIUM\_CHANNEL\_COMPLEX | 14 | 3 | 4161 | 164 | dnc,Ca-alpha1D,MCU | | 1.588e-02 | -4.14 | GO\_SUBSTANTIA\_NIGRA\_DEVELOPMENT | MSigDB lists | GO\_SUBSTANTIA\_NIGRA\_DEVELOPMENT | 14 | 3 | 4161 | 164 | Gdh,Hsc70-3,ND-30 | | 1.588e-02 | -4.14 | YAO\_TEMPORAL\_RESPONSE\_TO\_PROGESTERONE\_CLUSTER\_8 | MSigDB lists | YAO\_TEMPORAL\_RESPONSE\_TO\_PROGESTERONE\_CLUSTER\_8 | 14 | 3 | 4161 | 164 | Syx18,Glut1,Pdk | | 1.588e-02 | -4.14 | GAZDA\_DIAMOND\_BLACKFAN\_ANEMIA\_MYELOID\_DN | MSigDB lists | GAZDA\_DIAMOND\_BLACKFAN\_ANEMIA\_MYELOID\_DN | 14 | 3 | 4161 | 164 | RpL15,rl,dnr1 | | 1.588e-02 | -4.14 | ODONNELL\_METASTASIS\_UP | MSigDB lists | ODONNELL\_METASTASIS\_UP | 14 | 3 | 4161 | 164 | Ac78C,ck,norpA | | 1.588e-02 | -4.14 | ZHANG\_PROLIFERATING\_VS\_QUIESCENT | MSigDB lists | ZHANG\_PROLIFERATING\_VS\_QUIESCENT | 14 | 3 | 4161 | 164 | Sod1,CycB,RpL6 | | 1.588e-02 | -4.14 | GO\_SPERM\_MOTILITY | MSigDB lists | GO\_SPERM\_MOTILITY | 14 | 3 | 4161 | 164 | l(2)41Ab,CG17387,CG9492 | | 1.589e-02 | -4.14 | lipid binding | molecular function | GO:0008289 | 162 | 14 | 12176 | 554 | Syt7,CheB42a,Acox57D-d,mim,CG5270,cert,step,Npc2b,CG43783,SCAP,Pld,Gprk1,Pi3K68D,Cip4 | | 1.598e-02 | -4.14 | Cyp1 (Cyclophilin 1) | protein interactions | 32595 | 30 | 5 | 9204 | 464 | gus,Sod1,Sema1a,Rtnl1,CG8219 | | 1.603e-02 | -4.13 | head development | biological process | GO:0060322 | 144 | 13 | 12345 | 571 | Ten-a,Git,hth,en,toy,vap,drosha,dati,E5,rg,shot,jing,fdl | | 1.614e-02 | -4.13 | G protein-coupled amine receptor activity | molecular function | GO:0008227 | 22 | 4 | 12176 | 554 | 5-HT2A,5-HT2B,Octbeta1R,Octbeta2R | | 1.618e-02 | -4.12 | GO\_POSTSYNAPSE | MSigDB lists | GO\_POSTSYNAPSE | 100 | 9 | 4161 | 164 | nAChRalpha3,nAChRalpha4,norpA,Nlg2,Atpalpha,Nlg3,Octbeta2R,Mef2,Dys | | 1.623e-02 | -4.12 | Mes2 (Mesoderm-expressed 2) | protein interactions | 40514 | 20 | 4 | 9204 | 464 | CG12768,eIF4B,CG3081,Scp2 | | 1.623e-02 | -4.12 | dia (diaphanous) | protein interactions | 35340 | 20 | 4 | 9204 | 464 | Myc,Cip4,CycB,Sema1a | | 1.634e-02 | -4.11 | HLH | pfam domains | PF00010 | 53 | 7 | 10480 | 528 | cato,Myc,Fer3,CG33557,Clk,dimm,Mnt | | 1.637e-02 | -4.11 | eye-antennal disc development | biological process | GO:0035214 | 71 | 8 | 12345 | 571 | bab2,ck,lz,Diap1,sn,salm,ct,toy | | 1.658e-02 | -4.10 | purine ribonucleoside triphosphate binding | molecular function | GO:0035639 | 791 | 49 | 12176 | 554 | CG10077,CG5160,PMCA,ck,Takl2,rl,Hmt-1,CG42637,dnt,CG5789,kl-3,spel1,Khc-73,lz,CG14721,CG33970,bt,Pkg21D,ABCB7,RabX2,mus308,CG32944,fray,Atpalpha,PEK,Dhc16F,Rgk1,CG9492,Src42A,bma,Myo81F,mIF2,Klp59C,Stlk,Gprk1,CG31689,Nadsyn,Hsc70-3,Tak1,Hipk,CG12147,CG42541,fzo,CG34357,CG43672,par-1,DNAlig4,fbl,CG9981 | | 1.668e-02 | -4.09 | GO\_MITOCHONDRIAL\_PROTEIN\_COMPLEX | MSigDB lists | GO\_MITOCHONDRIAL\_PROTEIN\_COMPLEX | 68 | 7 | 4161 | 164 | Tim23,CG17691,SdhA,Tim17b,ND-30,MCU,PNPase | | 1.668e-02 | -4.09 | GSE17580\_UNINFECTED\_VS\_S\_MANSONI\_INF\_TEFF\_UP | MSigDB lists | GSE17580\_UNINFECTED\_VS\_S\_MANSONI\_INF\_TEFF\_UP | 68 | 7 | 4161 | 164 | inc,Mef2,Octbeta2R,nAChRalpha4,AP-2alpha,CstF50,RpL6 | | 1.688e-02 | -4.08 | ecdysteroid metabolic process | biological process | GO:0045455 | 33 | 5 | 12345 | 571 | Npc2b,spo,nvd,CG9514,spok | | 1.691e-02 | -4.08 | purine ribonucleotide binding | molecular function | GO:0032555 | 811 | 50 | 12176 | 554 | Stlk,Klp59C,Gprk1,CG31689,Nadsyn,Hsc70-3,Acbp5,Rgk1,CG9492,Src42A,bma,Myo81F,mIF2,CG34357,CG43672,DNAlig4,par-1,CG9981,fbl,CG12147,Hipk,Tak1,fzo,CG42541,CG5789,Hmt-1,CG42637,dnt,kl-3,Khc-73,spel1,bt,Pkg21D,ABCB7,lz,CG33970,CG14721,CG10077,CG5160,rl,Takl2,ck,PMCA,fray,Atpalpha,PEK,Dhc16F,mus308,RabX2,CG32944 | | 1.694e-02 | -4.08 | GSE17721\_CTRL\_VS\_CPG\_24H\_BMDC\_UP | MSigDB lists | GSE17721\_CTRL\_VS\_CPG\_24H\_BMDC\_UP | 53 | 6 | 4161 | 164 | bru1,ABCB7,CG10077,Der-1,step,unc-13-4A | | 1.694e-02 | -4.08 | GSE30971\_CTRL\_VS\_LPS\_STIM\_MACROPHAGE\_WBP7\_HET\_2H\_DN | MSigDB lists | GSE30971\_CTRL\_VS\_LPS\_STIM\_MACROPHAGE\_WBP7\_HET\_2H\_DN | 53 | 6 | 4161 | 164 | MFS16,cta,CG9389,Mef2,Gdh,vap | | 1.700e-02 | -4.07 | posttranscriptional gene silencing by RNA | biological process | GO:0035194 | 58 | 7 | 12345 | 571 | Ars2,CG10077,drosha,hdc,AGO3,Hsc70-3,HPS4 | | 1.702e-02 | -4.07 | - | gene3d domains | 4.10.900.10 | 4 | 2 | 4929 | 273 | CadN2,pan | | 1.704e-02 | -4.07 | axon | cellular component | GO:0030424 | 233 | 18 | 11940 | 534 | sky,Rtnl1,par-1,Dscam2,Gr2a,cpx,pum,Mical,Catsup,Magi,Syt7,Gr39a,hdc,klg,Gr93d,Snap25,shot,Ptp99A | | 1.710e-02 | -4.07 | fly chr35D2-35D2\|2-51 | chromosome location | fly chr35D2-35D2\|2-51 | 17 | 3 | 25010 | 823 | CG15258,lncRNA:CR44859,sna | | 1.710e-02 | -4.07 | fly chr42E1-42E1\|2-55 cM | chromosome location | fly chr42E1-42E1\|2-55 cM | 17 | 3 | 25010 | 823 | mim,CheB42a,CG3358 | | 1.710e-02 | -4.07 | fly chr35D2-35D2\|2-51 cM | chromosome location | fly chr35D2-35D2\|2-51 cM | 17 | 3 | 25010 | 823 | lncRNA:CR44859,CG15258,sna | | 1.710e-02 | -4.07 | fly chr42E1-42E1\|2-55 | chromosome location | fly chr42E1-42E1\|2-55 | 17 | 3 | 25010 | 823 | CheB42a,mim,CG3358 | | 1.716e-02 | -4.07 | G alpha (s) signalling events | REACTOME pathways | R-DME-418555 | 32 | 5 | 3923 | 189 | 5-HT2A,Octbeta1R,Octbeta2R,dnc,Ac78C | | 1.716e-02 | -4.07 | cell surface | cellular component | GO:0009986 | 74 | 8 | 11940 | 534 | Nlg2,boi,Sdc,kirre,Toll-7,Nlg3,CadN2,cv-2 | | 1.721e-02 | -4.06 | - | gene3d domains | 2.60.40.150 | 49 | 7 | 4929 | 273 | RhoGAP100F,mbc,vap,norpA,Pi3K68D,Syt7,unc-13-4A | | 1.733e-02 | -4.06 | ameboidal-type cell migration | biological process | GO:0001667 | 177 | 15 | 12345 | 571 | mbc,Dlg5,Diap1,salm,ru,Sema1a,Sdc,par-1,rib,kni,cta,jing,CG1486,tj,mim | | 1.738e-02 | -4.05 | eye morphogenesis | biological process | GO:0048592 | 310 | 23 | 12345 | 571 | br,pk,ru,hth,sdk,rl,CadN2,bi,rau,lz,rno,boi,lola,klg,salm,fred,caup,Tak1,svp,kirre,rg,RhoGAP100F,cic | | 1.738e-02 | -4.05 | sensory organ morphogenesis | biological process | GO:0090596 | 310 | 23 | 12345 | 571 | Tak1,caup,salm,fred,klg,lola,cic,RhoGAP100F,rg,kirre,svp,rl,hth,sdk,ru,br,pk,boi,rno,bi,lz,rau,CadN2 | | 1.739e-02 | -4.05 | GO\_KINASE\_BINDING | MSigDB lists | GO\_KINASE\_BINDING | 154 | 12 | 4161 | 164 | toy,Stlk,Khc-73,AP-2alpha,rl,rdgC,CycB,spel1,norpA,rg,fz2,Parp | | 1.741e-02 | -4.05 | GSE22589\_SIV\_VS\_HIV\_AND\_SIV\_INFECTED\_DC\_DN | MSigDB lists | GSE22589\_SIV\_VS\_HIV\_AND\_SIV\_INFECTED\_DC\_DN | 39 | 5 | 4161 | 164 | vtd,Ubr1,rg,heph,Tim17b | | 1.741e-02 | -4.05 | GSE369\_SOCS3\_KO\_VS\_WT\_LIVER\_POST\_IL6\_INJECTION\_DN | MSigDB lists | GSE369\_SOCS3\_KO\_VS\_WT\_LIVER\_POST\_IL6\_INJECTION\_DN | 39 | 5 | 4161 | 164 | step,Meltrin,DNAlig4,Mef2,MFS16 | | 1.741e-02 | -4.05 | GO\_LEUKOCYTE\_DIFFERENTIATION | MSigDB lists | GO\_LEUKOCYTE\_DIFFERENTIATION | 39 | 5 | 4161 | 164 | Parp,DNAlig4,spel1,fz2,Mef2 | | 1.741e-02 | -4.05 | GSE25088\_CTRL\_VS\_IL4\_STIM\_MACROPHAGE\_DN | MSigDB lists | GSE25088\_CTRL\_VS\_IL4\_STIM\_MACROPHAGE\_DN | 39 | 5 | 4161 | 164 | lqf,MFS16,norpA,nAChRalpha3,Ranbp16 | | 1.741e-02 | -4.05 | FOXM1\_01 | MSigDB lists | FOXM1\_01 | 39 | 5 | 4161 | 164 | DNaseII,Dys,Mef2,toy,Rbp9 | | 1.747e-02 | -4.05 | FERM\_domain | interpro domains | IPR000299 | 21 | 4 | 11383 | 557 | Myo81F,dnr1,ck,CG34347 | | 1.754e-02 | -4.04 | Acyl chain remodelling of PE | REACTOME pathways | R-DME-1482839 | 12 | 3 | 3923 | 189 | CG42237,CG3009,lama | | 1.757e-02 | -4.04 | MAPK signaling pathway - fly | KEGG pathways | dme04013 | 93 | 9 | 3198 | 138 | Mef2,Src42A,vap,rl,Tak1,Takl2,bi,lz,cic | | 1.757e-02 | -4.04 | MAPK signaling pathway - fly | KEGG pathways | ko04013 | 93 | 9 | 3198 | 138 | cic,bi,lz,Takl2,Tak1,rl,Mef2,Src42A,vap | | 1.765e-02 | -4.04 | MODULE\_123 | MSigDB lists | MODULE\_123 | 26 | 4 | 4161 | 164 | ap,cta,toy,Octbeta2R | | 1.765e-02 | -4.04 | GSE40274\_CTRL\_VS\_FOXP3\_AND\_SATB1\_TRANSDUCED\_ACTIVATED\_CD4\_TCELL\_DN | MSigDB lists | GSE40274\_CTRL\_VS\_FOXP3\_AND\_SATB1\_TRANSDUCED\_ACTIVATED\_CD4\_TCELL\_DN | 26 | 4 | 4161 | 164 | twz,mlt,sdk,CG41378 | | 1.765e-02 | -4.04 | PID\_FAK\_PATHWAY | MSigDB lists | PID\_FAK\_PATHWAY | 26 | 4 | 4161 | 164 | vap,rl,Git,mbc | | 1.796e-02 | -4.02 | PHONG\_TNF\_RESPONSE\_NOT\_VIA\_P38 | MSigDB lists | PHONG\_TNF\_RESPONSE\_NOT\_VIA\_P38 | 69 | 7 | 4161 | 164 | Hsc70-3,Trs23,fz2,Gfat1,svp,norpA,Mef2 | | 1.800e-02 | -4.02 | shi (shibire) | protein interactions | 45928 | 54 | 7 | 9204 | 464 | Pld,lqf,rdgC,par-1,Cip4,Sema1a,pk | | 1.819e-02 | -4.01 | calcium channel complex | cellular component | GO:0034704 | 13 | 3 | 11940 | 534 | Ca-alpha1D,MCU,Ca-Ma2d | | 1.828e-02 | -4.00 | br (broad) | protein interactions | 44505 | 31 | 5 | 9204 | 464 | pan,Mef2,CG4270,br,CG15459 | | 1.840e-02 | -4.00 | regulation of primary metabolic process | biological process | GO:0080090 | 1894 | 106 | 12345 | 571 | Trl,cic,sna,fuss,Mnt,d4,SoxN,CG41520,Nipped-A,lncRNA:bxd,Camta,Tak1,CG40228,lola,en,zld,Spn42De,Psc,jing,HP6,E5,CG12769,step,fru,Parg,bi,mav,rl,CG34334,Diap1,pan,Setd3,CG32006,zen,par-1,CG12200,CG12147,pho,Lsd-2,CG41106,AGO3,bru1,scw,fray,Parp,lz,hth,mbc,Ppm1,Src42A,scro,kni,Maf1,bab2,svp,tld,HPS4,orb,Octbeta2R,pum,Catsup,Fer3,dsx,toy,mxt,tou,CG9932,Myc,CG1620,spel1,ap,SCAP,CG14184,Tgi,tj,gus,br,PNPase,CG17514,ct,Spn75F,ewg,Mef2,Marf1,PEK,CG5862,salm,bru3,caup,Atf6,Vsx2,Stlk,CG9650,Clk,rib,SteXh:CG42398,grim,Alg-2,fd96Ca,Thor,Su(z)2,dimm,CycB,cv-2,heph,CG33557 | | 1.841e-02 | -3.99 | Dynein\_AAA5\_ext | interpro domains | IPR041466 | 12 | 3 | 11383 | 557 | Dhc16F,kl-3,CG9492 | | 1.841e-02 | -3.99 | Dynein\_C | interpro domains | IPR041228 | 12 | 3 | 11383 | 557 | Dhc16F,kl-3,CG9492 | | 1.844e-02 | -3.99 | - | gene3d domains | 2.60.40.1180 | 19 | 4 | 4929 | 273 | alpha-Man-IIb,GCS2alpha,Amyrel,tobi | | 1.845e-02 | -3.99 | GSE40685\_NAIVE\_CD4\_TCELL\_VS\_TREG\_UP | MSigDB lists | GSE40685\_NAIVE\_CD4\_TCELL\_VS\_TREG\_UP | 54 | 6 | 4161 | 164 | CG12170,GLS,CG3107,RpL6,SCAP,CG14194 | | 1.854e-02 | -3.99 | synaptic vesicle recycling | biological process | GO:0036465 | 46 | 6 | 12345 | 571 | Nlg3,Git,AP-2alpha,Nlg2,sky,lqf | | 1.854e-02 | -3.99 | cellular hormone metabolic process | biological process | GO:0034754 | 46 | 6 | 12345 | 571 | spo,Npc2b,nvd,CG30495,CG9514,spok | | 1.860e-02 | -3.98 | PH-like\_dom\_sf | interpro domains | IPR011993 | 139 | 13 | 11383 | 557 | CG10188,vap,CG34347,osp,Myo81F,Pld,step,sif,cert,ck,CG30440,Gprk1,pico | | 1.865e-02 | -3.98 | PH | smart domains | SM00233 | 67 | 9 | 5086 | 308 | vap,Gprk1,cert,CG30440,step,sif,osp,CG10188,pico | | 1.885e-02 | -3.97 | carbohydrate transmembrane transporter activity | molecular function | GO:0015144 | 23 | 4 | 12176 | 554 | CG15408,CG1213,CG45782,CG8249 | | 1.885e-02 | -3.97 | calcium channel regulator activity | molecular function | GO:0005246 | 5 | 2 | 12176 | 554 | Rgk1,CG42541 | | 1.885e-02 | -3.97 | transcription coactivator binding | molecular function | GO:0001223 | 5 | 2 | 12176 | 554 | hth,Tgi | | 1.890e-02 | -3.97 | BIR\_REPEAT\_2 | prosite domains | PS50143 | 4 | 2 | 6243 | 365 | Diap1,Bruce | | 1.890e-02 | -3.97 | B\_KETOACYL\_SYNTHASE | prosite domains | PS00606 | 4 | 2 | 6243 | 365 | FASN3,CG12170 | | 1.890e-02 | -3.97 | HTH\_OST | prosite domains | PS51644 | 4 | 2 | 6243 | 365 | tej,Marf1 | | 1.896e-02 | -3.97 | cytoskeleton organization | biological process | GO:0007010 | 580 | 38 | 12345 | 571 | capu,Svil,CG10939,RhoGAP100F,shot,par-1,dgt5,Act42A,jvl,CG43901,bond,bru1,cta,rib,Spn,Khc-73,heph,Src42A,sn,CG34347,CG17387,mbc,mlt,Wdr62,bt,l(2)41Ab,Pi3K68D,Mical,ck,CAP,pk,pico,kl-3,Setd3,cbs,sif,Msp300,fest | | 1.906e-02 | -3.96 | negative regulation of ERBB signaling pathway | biological process | GO:1901185 | 34 | 5 | 12345 | 571 | pum,rno,tay,Src42A,Pi3K68D | | 1.906e-02 | -3.96 | negative regulation of epidermal growth factor receptor signaling pathway | biological process | GO:0042059 | 34 | 5 | 12345 | 571 | Src42A,Pi3K68D,pum,rno,tay | | 1.906e-02 | -3.96 | genitalia development | biological process | GO:0048806 | 34 | 5 | 12345 | 571 | lz,br,dsx,en,salm | | 1.924e-02 | -3.95 | FERM\_M | pfam domains | PF00373 | 21 | 4 | 10480 | 528 | Myo81F,dnr1,CG34347,ck | | 1.926e-02 | -3.95 | lig (lingerer) | protein interactions | 35771 | 21 | 4 | 9204 | 464 | orb,Tak1,CG12521,Diap1 | | 1.926e-02 | -3.95 | twin (twin) | protein interactions | 42880 | 21 | 4 | 9204 | 464 | mei-P26,orb,AGO3,pum | | 1.929e-02 | -3.95 | GSE3982\_EOSINOPHIL\_VS\_MAST\_CELL\_UP | MSigDB lists | GSE3982\_EOSINOPHIL\_VS\_MAST\_CELL\_UP | 40 | 5 | 4161 | 164 | Pdk,Tsp5D,CG30495,step,Git | | 1.929e-02 | -3.95 | GSE15659\_TREG\_VS\_TCONV\_DN | MSigDB lists | GSE15659\_TREG\_VS\_TCONV\_DN | 40 | 5 | 4161 | 164 | svp,CG41099,fbl,dnc,CG8184 | | 1.929e-02 | -3.95 | GSE3982\_MEMORY\_CD4\_TCELL\_VS\_TH1\_UP | MSigDB lists | GSE3982\_MEMORY\_CD4\_TCELL\_VS\_TH1\_UP | 40 | 5 | 4161 | 164 | CG34408,rl,vap,RpL6,Nadsyn | | 1.929e-02 | -3.95 | AAAGGAT\_MIR501 | MSigDB lists | AAAGGAT\_MIR501 | 40 | 5 | 4161 | 164 | Rbp9,Mef2,MCU,bru1,svp | | 1.929e-02 | -3.95 | BIOCARTA\_MET\_PATHWAY | MSigDB lists | BIOCARTA\_MET\_PATHWAY | 15 | 3 | 4161 | 164 | mbc,rl,vap | | 1.929e-02 | -3.95 | KRAS.PROSTATE\_UP.V1\_UP | MSigDB lists | KRAS.PROSTATE\_UP.V1\_UP | 15 | 3 | 4161 | 164 | nAChRalpha4,rg,CG14721 | | 1.929e-02 | -3.95 | GO\_REGULATION\_OF\_POSTSYNAPTIC\_MEMBRANE\_POTENTIAL | MSigDB lists | GO\_REGULATION\_OF\_POSTSYNAPTIC\_MEMBRANE\_POTENTIAL | 15 | 3 | 4161 | 164 | Octbeta2R,nAChRalpha4,Mef2 | | 1.929e-02 | -3.95 | KRASNOSELSKAYA\_ILF3\_TARGETS\_DN | MSigDB lists | KRASNOSELSKAYA\_ILF3\_TARGETS\_DN | 15 | 3 | 4161 | 164 | Dys,bru1,Ugt50B3 | | 1.929e-02 | -3.95 | CCAGGGG\_MIR331 | MSigDB lists | CCAGGGG\_MIR331 | 15 | 3 | 4161 | 164 | rg,twz,Ranbp16 | | 1.929e-02 | -3.95 | ACEVEDO\_LIVER\_CANCER\_WITH\_H3K9ME3\_UP | MSigDB lists | ACEVEDO\_LIVER\_CANCER\_WITH\_H3K9ME3\_UP | 15 | 3 | 4161 | 164 | DNaseII,cv-2,Spn42De | | 1.929e-02 | -3.95 | GO\_REGULATION\_OF\_HEART\_RATE | MSigDB lists | GO\_REGULATION\_OF\_HEART\_RATE | 15 | 3 | 4161 | 164 | dnc,Ca-alpha1D,Dys | | 1.932e-02 | -3.95 | GSE15330\_WT\_VS\_IKAROS\_KO\_GRANULOCYTE\_MONOCYTE\_PROGENITOR\_UP | MSigDB lists | GSE15330\_WT\_VS\_IKAROS\_KO\_GRANULOCYTE\_MONOCYTE\_PROGENITOR\_UP | 70 | 7 | 4161 | 164 | CG42813,PNPase,bru1,Dys,par-1,CG17691,CG41378 | | 1.935e-02 | -3.95 | Ras85D (Ras oncogene at 85D) | protein interactions | 41140 | 241 | 20 | 9204 | 464 | RpL15,rau,tay,vap,rno,lz,IntS3,Src42A,Myc,rl,jing,svp,pum,cic,RpL6,pico,rg,Ppm1,Tak1,step | | 1.944e-02 | -3.94 | cell-cell adhesion | biological process | GO:0098609 | 102 | 10 | 12345 | 571 | Nlg3,CadN2,Dscam2,CG32087,kirre,beat-VII,sdk,Nlg2,Src42A,klg | | 1.946e-02 | -3.94 | antennal joint development | biological process | GO:0048098 | 5 | 2 | 12345 | 571 | ct,salm | | 1.946e-02 | -3.94 | ornithine metabolic process | biological process | GO:0006591 | 5 | 2 | 12345 | 571 | Odc2,arg | | 1.946e-02 | -3.94 | nucleolus organization | biological process | GO:0007000 | 5 | 2 | 12345 | 571 | Myc,Parp | | 1.946e-02 | -3.94 | regulation of cysteine-type endopeptidase activity involved in execution phase of apoptosis | biological process | GO:2001270 | 5 | 2 | 12345 | 571 | grim,Diap1 | | 1.946e-02 | -3.94 | positive regulation of cytosolic calcium ion concentration involved in phospholipase C-activating G protein-coupled signaling pathway | biological process | GO:0051482 | 5 | 2 | 12345 | 571 | norpA,Octbeta1R | | 1.946e-02 | -3.94 | maintenance of protein location in nucleus | biological process | GO:0051457 | 5 | 2 | 12345 | 571 | Parg,Parp | | 1.946e-02 | -3.94 | synaptic vesicle membrane organization | biological process | GO:0048499 | 5 | 2 | 12345 | 571 | Nlg2,Ten-a | | 1.946e-02 | -3.94 | mesodermal cell fate determination | biological process | GO:0007500 | 5 | 2 | 12345 | 571 | pan,sna | | 1.946e-02 | -3.94 | Cajal body organization | biological process | GO:0030576 | 5 | 2 | 12345 | 571 | Parp,Parg | | 1.946e-02 | -3.94 | negative regulation of neurotransmitter secretion | biological process | GO:0046929 | 5 | 2 | 12345 | 571 | cpx,sky | | 1.946e-02 | -3.94 | peptidyl-lysine dimethylation | biological process | GO:0018027 | 5 | 2 | 12345 | 571 | Set1,Setd3 | | 1.946e-02 | -3.94 | digestive tract mesoderm development | biological process | GO:0007502 | 5 | 2 | 12345 | 571 | scw,wb | | 1.946e-02 | -3.94 | negative regulation of neurotransmitter transport | biological process | GO:0051589 | 5 | 2 | 12345 | 571 | sky,cpx | | 1.969e-02 | -3.93 | reproductive behavior | biological process | GO:0019098 | 196 | 16 | 12345 | 571 | lncRNA:flam,Gr39a,Tdc2,fru,dnc,Gr93d,twz,SPR,Acp36DE,Gr2a,Nlg2,dsx,dati,shep,CheB42a,Or67d | | 1.988e-02 | -3.92 | extrinsic component of membrane | cellular component | GO:0019898 | 76 | 8 | 11940 | 534 | cta,klg,ImpE2,CadN2,Pi3K68D,norpA,mbc,Src42A | | 1.989e-02 | -3.92 | EF-hand\_5 | pfam domains | PF13202 | 12 | 3 | 10480 | 528 | Scp2,Scp1,rdgC | | 1.989e-02 | -3.92 | Dynein\_AAA\_lid | pfam domains | PF17852 | 12 | 3 | 10480 | 528 | CG9492,Dhc16F,kl-3 | | 1.989e-02 | -3.92 | RA | pfam domains | PF00788 | 12 | 3 | 10480 | 528 | Myo81F,pico,rau | | 1.989e-02 | -3.92 | Dynein\_C | pfam domains | PF18199 | 12 | 3 | 10480 | 528 | Dhc16F,CG9492,kl-3 | | 1.990e-02 | -3.92 | spiracle morphogenesis, open tracheal system | biological process | GO:0035277 | 23 | 4 | 12345 | 571 | salm,pan,ct,en | | 1.992e-02 | -3.92 | sov (small ovary) | protein interactions | 31615 | 12 | 3 | 9204 | 464 | Sema1a,Rtnl1,toy | | 1.992e-02 | -3.92 | SA (Stromalin) | protein interactions | 33974 | 12 | 3 | 9204 | 464 | Sema1a,ct,vtd | | 1.992e-02 | -3.92 | Nrx-1 (Neurexin 1) | protein interactions | 42646 | 12 | 3 | 9204 | 464 | RhoGAP100F,Nlg2,Spn | | 1.992e-02 | -3.92 | wde (windei) | protein interactions | 36133 | 12 | 3 | 9204 | 464 | ct,CG13405,jvl | | 1.992e-02 | -3.92 | jus (julius seizure) | protein interactions | 43453 | 12 | 3 | 9204 | 464 | shakB,mei-P26,cpo | | 1.992e-02 | -3.92 | Mtl (Mig-2-like) | protein interactions | 43319 | 12 | 3 | 9204 | 464 | pk,RhoGAP100F,CG15136 | | 1.994e-02 | -3.91 | positive regulation of transcription, DNA-templated | biological process | GO:0045893 | 473 | 32 | 12345 | 571 | Mef2,zen,Atf6,caup,pho,lz,Clk,dimm,hth,sna,scro,Trl,zld,lola,en,dsx,toy,CG40228,Camta,SoxN,d4,bi,tj,SCAP,ap,CG12769,Myc,jing,tou,br,Setd3,pan | | 1.996e-02 | -3.91 | - | gene3d domains | 1.20.80.10 | 29 | 5 | 4929 | 273 | Acbp5,dnr1,CG34347,Myo81F,ck | | 2.001e-02 | -3.91 | GO\_TRANSMEMBRANE\_RECEPTOR\_PROTEIN\_TYROSINE\_KINASE\_SIGNALING\_PATHWAY | MSigDB lists | GO\_TRANSMEMBRANE\_RECEPTOR\_PROTEIN\_TYROSINE\_KINASE\_SIGNALING\_PATHWAY | 121 | 10 | 4161 | 164 | AP-2alpha,rl,Pi3K68D,Src42A,Octbeta2R,vap,mbc,Alg-2,heph,nAChRalpha4 | | 2.006e-02 | -3.91 | TGATTTRY\_GFI1\_01 | MSigDB lists | TGATTTRY\_GFI1\_01 | 55 | 6 | 4161 | 164 | Ac78C,Mef2,Rbp9,Tsp5D,CG14820,Dys | | 2.006e-02 | -3.91 | GSE5589\_IL6\_KO\_VS\_IL10\_KO\_LPS\_STIM\_MACROPHAGE\_180MIN\_UP | MSigDB lists | GSE5589\_IL6\_KO\_VS\_IL10\_KO\_LPS\_STIM\_MACROPHAGE\_180MIN\_UP | 55 | 6 | 4161 | 164 | Hsc70-3,Odc2,Pi3K68D,CG12121,Mef2,rg | | 2.010e-02 | -3.91 | KEGG\_MELANOGENESIS | MSigDB lists | KEGG\_MELANOGENESIS | 27 | 4 | 4161 | 164 | norpA,rl,fz2,Ac78C | | 2.010e-02 | -3.91 | WNT\_UP.V1\_UP | MSigDB lists | WNT\_UP.V1\_UP | 27 | 4 | 4161 | 164 | Ugt50B3,CG7255,CG14721,alpha4GT1 | | 2.010e-02 | -3.91 | GO\_REGULATION\_OF\_RECEPTOR\_ACTIVITY | MSigDB lists | GO\_REGULATION\_OF\_RECEPTOR\_ACTIVITY | 27 | 4 | 4161 | 164 | Mef2,Dys,Nlg3,dnc | | 2.018e-02 | -3.90 | photoreceptor cell fate commitment | biological process | GO:0046552 | 60 | 7 | 12345 | 571 | ru,klg,lola,svp,br,rl,hth | | 2.022e-02 | -3.90 | PKS\_KS | smart domains | SM00825 | 4 | 2 | 5086 | 308 | FASN3,CG12170 | | 2.022e-02 | -3.90 | BIR | smart domains | SM00238 | 4 | 2 | 5086 | 308 | Diap1,Bruce | | 2.025e-02 | -3.90 | - | gene3d domains | 3.30.70.330 | 152 | 15 | 4929 | 273 | CG9107,Marf1,CG14891,CG40813,Set1,CG34334,shep,bru3,Rbp9,heph,eIF4B,bru1,CG41562,cpo,orb | | 2.027e-02 | -3.90 | nucleoside phosphate binding | molecular function | GO:1901265 | 955 | 57 | 12176 | 554 | Nadsyn,Trxr-2,Hsc70-3,Klp59C,Stlk,Gprk1,CG31689,Myo81F,SdhA,bma,mIF2,Acbp5,Rgk1,Src42A,CG9492,par-1,DNAlig4,fbl,CG9981,CG34357,CG43672,Mical,CG42541,fzo,Tak1,Hipk,CG12147,spel1,Parp,Khc-73,CG14721,lz,CG33970,bt,ABCB7,Pkg21D,Hmt-1,dnt,CG42637,CG5789,CG9514,kl-3,CG5160,PMCA,ck,Takl2,rl,CG10077,Acox57D-d,Dhc16F,PEK,fray,Atpalpha,cta,RabX2,CG32944,mus308 | | 2.027e-02 | -3.90 | nucleotide binding | molecular function | GO:0000166 | 955 | 57 | 12176 | 554 | Acox57D-d,CG10077,Takl2,PMCA,ck,rl,CG5160,kl-3,CG9514,dnt,CG42637,Hmt-1,CG5789,CG33970,CG14721,lz,Pkg21D,bt,ABCB7,Parp,spel1,Khc-73,CG32944,cta,RabX2,mus308,Atpalpha,fray,Dhc16F,PEK,CG9492,Src42A,Acbp5,Rgk1,mIF2,Myo81F,bma,SdhA,Gprk1,CG31689,Klp59C,Stlk,Trxr-2,Hsc70-3,Nadsyn,Hipk,Tak1,CG12147,Mical,CG42541,fzo,CG43672,CG34357,fbl,CG9981,par-1,DNAlig4 | | 2.035e-02 | -3.89 | fly chr16F6-16F6\|1-59 | chromosome location | fly chr16F6-16F6\|1-59 | 7 | 2 | 25010 | 823 | Dhc16F,CG7536 | | 2.035e-02 | -3.89 | fly chr22F4-22F4\|2-6 | chromosome location | fly chr22F4-22F4\|2-6 | 7 | 2 | 25010 | 823 | lncRNA:CR44112,lncRNA:CR44113 | | 2.035e-02 | -3.89 | fly chr16F6-16F6\|1-59 cM | chromosome location | fly chr16F6-16F6\|1-59 cM | 7 | 2 | 25010 | 823 | CG7536,Dhc16F | | 2.035e-02 | -3.89 | fly chr22F4-22F4\|2-6 cM | chromosome location | fly chr22F4-22F4\|2-6 cM | 7 | 2 | 25010 | 823 | lncRNA:CR44113,lncRNA:CR44112 | | 2.035e-02 | -3.89 | fly chr89A2-89A2\|3-58 cM | chromosome location | fly chr89A2-89A2\|3-58 cM | 7 | 2 | 25010 | 823 | CG34274,lncRNA:CR45635 | | 2.035e-02 | -3.89 | fly chr89A2-89A2\|3-58 | chromosome location | fly chr89A2-89A2\|3-58 | 7 | 2 | 25010 | 823 | lncRNA:CR45635,CG34274 | | 2.055e-02 | -3.89 | embryonic pattern specification | biological process | GO:0009880 | 230 | 18 | 12345 | 571 | rl,pan,gus,hth,scw,jing,Myc,pum,SoxN,en,fz2,par-1,cic,kni,Ptip,orb,capu,tld | | 2.055e-02 | -3.89 | FERM\_central | interpro domains | IPR019748 | 22 | 4 | 11383 | 557 | CG34347,Myo81F,dnr1,ck | | 2.055e-02 | -3.89 | FERM\_2 | interpro domains | IPR035963 | 22 | 4 | 11383 | 557 | ck,dnr1,Myo81F,CG34347 | | 2.080e-02 | -3.87 | CG31999 (uncharacterized protein) | protein interactions | 43777 | 32 | 5 | 9204 | 464 | Vsx2,fru,eIF4B,zld,dsx | | 2.080e-02 | -3.87 | CG13838 (uncharacterized protein) | protein interactions | 42719 | 32 | 5 | 9204 | 464 | zen,fdl,Dys,CG14071,CG15468 | | 2.080e-02 | -3.87 | GO\_MEMBRANE\_PROTEIN\_COMPLEX | MSigDB lists | GO\_MEMBRANE\_PROTEIN\_COMPLEX | 314 | 20 | 4161 | 164 | nAChRalpha3,CG41099,Der-1,Atpalpha,dnc,Pi3K68D,SdhA,RSG7,fbl,Dys,Hmt-1,Ca-alpha1D,Tim23,Tim17b,ND-30,MCU,nAChRalpha4,AP-2alpha,Syx18,cta | | 2.086e-02 | -3.87 | KEGG\_NITROGEN\_METABOLISM | MSigDB lists | KEGG\_NITROGEN\_METABOLISM | 6 | 2 | 4161 | 164 | Gdh,GLS | | 2.086e-02 | -3.87 | ZWANG\_CLASS\_2\_TRANSIENTLY\_INDUCED\_BY\_EGF | MSigDB lists | ZWANG\_CLASS\_2\_TRANSIENTLY\_INDUCED\_BY\_EGF | 6 | 2 | 4161 | 164 | Gfat1,GLS | | 2.086e-02 | -3.87 | MODULE\_297 | MSigDB lists | MODULE\_297 | 6 | 2 | 4161 | 164 | Octbeta2R,shot | | 2.086e-02 | -3.87 | GO\_R\_SMAD\_BINDING | MSigDB lists | GO\_R\_SMAD\_BINDING | 6 | 2 | 4161 | 164 | Parp,toy | | 2.086e-02 | -3.87 | PID\_S1P\_S1P4\_PATHWAY | MSigDB lists | PID\_S1P\_S1P4\_PATHWAY | 6 | 2 | 4161 | 164 | rl,cta | | 2.086e-02 | -3.87 | MEF2\_04 | MSigDB lists | MEF2\_04 | 6 | 2 | 4161 | 164 | Rbp9,CG2528 | | 2.086e-02 | -3.87 | GO\_CARBOHYDRATE\_TRANSMEMBRANE\_TRANSPORT | MSigDB lists | GO\_CARBOHYDRATE\_TRANSMEMBRANE\_TRANSPORT | 6 | 2 | 4161 | 164 | CG14621,Glut1 | | 2.086e-02 | -3.87 | GO\_REGULATION\_OF\_MICROTUBULE\_BASED\_MOVEMENT | MSigDB lists | GO\_REGULATION\_OF\_MICROTUBULE\_BASED\_MOVEMENT | 6 | 2 | 4161 | 164 | l(2)41Ab,CG17387 | | 2.086e-02 | -3.87 | REACTOME\_ACETYLCHOLINE\_BINDING\_AND\_DOWNSTREAM\_EVENTS | MSigDB lists | REACTOME\_ACETYLCHOLINE\_BINDING\_AND\_DOWNSTREAM\_EVENTS | 6 | 2 | 4161 | 164 | nAChRalpha3,nAChRalpha4 | | 2.086e-02 | -3.87 | GO\_REGULATION\_OF\_RESPIRATORY\_GASEOUS\_EXCHANGE | MSigDB lists | GO\_REGULATION\_OF\_RESPIRATORY\_GASEOUS\_EXCHANGE | 6 | 2 | 4161 | 164 | GLS,Nlg3 | | 2.086e-02 | -3.87 | PID\_HNF3A\_PATHWAY | MSigDB lists | PID\_HNF3A\_PATHWAY | 6 | 2 | 4161 | 164 | Sod1,svp | | 2.086e-02 | -3.87 | GO\_EPITHELIAL\_CILIUM\_MOVEMENT | MSigDB lists | GO\_EPITHELIAL\_CILIUM\_MOVEMENT | 6 | 2 | 4161 | 164 | l(2)41Ab,CG17387 | | 2.086e-02 | -3.87 | GO\_RESPONSE\_TO\_X\_RAY | MSigDB lists | GO\_RESPONSE\_TO\_X\_RAY | 6 | 2 | 4161 | 164 | spel1,DNAlig4 | | 2.086e-02 | -3.87 | GO\_CELL\_COMMUNICATION\_INVOLVED\_IN\_CARDIAC\_CONDUCTION | MSigDB lists | GO\_CELL\_COMMUNICATION\_INVOLVED\_IN\_CARDIAC\_CONDUCTION | 6 | 2 | 4161 | 164 | Ca-alpha1D,Atpalpha | | 2.086e-02 | -3.87 | GO\_NEGATIVE\_REGULATION\_OF\_DNA\_RECOMBINATION | MSigDB lists | GO\_NEGATIVE\_REGULATION\_OF\_DNA\_RECOMBINATION | 6 | 2 | 4161 | 164 | spel1,mus308 | | 2.086e-02 | -3.87 | GO\_NITRIC\_OXIDE\_METABOLIC\_PROCESS | MSigDB lists | GO\_NITRIC\_OXIDE\_METABOLIC\_PROCESS | 6 | 2 | 4161 | 164 | CG7255,arg | | 2.086e-02 | -3.87 | GO\_ACETYLCHOLINE\_BINDING | MSigDB lists | GO\_ACETYLCHOLINE\_BINDING | 6 | 2 | 4161 | 164 | nAChRalpha4,nAChRalpha3 | | 2.086e-02 | -3.87 | GO\_INNER\_DYNEIN\_ARM\_ASSEMBLY | MSigDB lists | GO\_INNER\_DYNEIN\_ARM\_ASSEMBLY | 6 | 2 | 4161 | 164 | CG17387,l(2)41Ab | | 2.086e-02 | -3.87 | GO\_AUDITORY\_RECEPTOR\_CELL\_MORPHOGENESIS | MSigDB lists | GO\_AUDITORY\_RECEPTOR\_CELL\_MORPHOGENESIS | 6 | 2 | 4161 | 164 | ck,Sod1 | | 2.086e-02 | -3.87 | POS\_RESPONSE\_TO\_HISTAMINE\_DN | MSigDB lists | POS\_RESPONSE\_TO\_HISTAMINE\_DN | 6 | 2 | 4161 | 164 | CG3358,CG9886 | | 2.086e-02 | -3.87 | REACTOME\_THROMBIN\_SIGNALLING\_THROUGH\_PROTEINASE\_ACTIVATED\_RECEPTORS\_PARS | MSigDB lists | REACTOME\_THROMBIN\_SIGNALLING\_THROUGH\_PROTEINASE\_ACTIVATED\_RECEPTORS\_PARS | 6 | 2 | 4161 | 164 | cta,rl | | 2.086e-02 | -3.87 | MODULE\_357 | MSigDB lists | MODULE\_357 | 6 | 2 | 4161 | 164 | Octbeta2R,shot | | 2.086e-02 | -3.87 | REACTOME\_HIGHLY\_CALCIUM\_PERMEABLE\_POSTSYNAPTIC\_NICOTINIC\_ACETYLCHOLINE\_RECEPTORS | MSigDB lists | REACTOME\_HIGHLY\_CALCIUM\_PERMEABLE\_POSTSYNAPTIC\_NICOTINIC\_ACETYLCHOLINE\_RECEPTORS | 6 | 2 | 4161 | 164 | nAChRalpha4,nAChRalpha3 | | 2.086e-02 | -3.87 | MODULE\_154 | MSigDB lists | MODULE\_154 | 6 | 2 | 4161 | 164 | shot,Octbeta2R | | 2.096e-02 | -3.87 | transporter activity | molecular function | GO:0005215 | 745 | 46 | 12176 | 554 | Tim17b,CG8219,Ir60e,CG43672,Ir94e,Ca-alpha1D,nAChRalpha3,CG33233,CG14196,Oatp33Ea,sbm,CG14160,Ir41a,MCU,ND-MLRQ,CG31689,CG7255,CG7536,Tim23,Glut1,Catsup,cpx,Atpalpha,SK,KCNQ,shakB,Rift,ABCB7,nAChRalpha4,CG15408,CG5254,cert,Ir40a,MFS16,CG5789,CG45782,Pzl,Hmt-1,PMCA,Ca-Ma2d,CG14621,CG1213,Ranbp16,Mct1,CG8249,HisCl1 | | 2.102e-02 | -3.86 | HMG\_box\_dom\_sf | interpro domains | IPR036910 | 33 | 5 | 11383 | 557 | SoxN,cic,Tpl94D,CG12104,pan | | 2.116e-02 | -3.86 | ribonucleotide binding | molecular function | GO:0032553 | 822 | 50 | 12176 | 554 | CG12147,Hipk,Tak1,CG42541,fzo,CG43672,CG34357,CG9981,fbl,DNAlig4,par-1,Src42A,CG9492,Acbp5,Rgk1,mIF2,Myo81F,bma,Gprk1,CG31689,Klp59C,Stlk,Hsc70-3,Nadsyn,CG32944,mus308,RabX2,Atpalpha,fray,Dhc16F,PEK,CG10077,rl,Takl2,PMCA,ck,CG5160,kl-3,CG5789,Hmt-1,dnt,CG42637,Pkg21D,ABCB7,bt,lz,CG14721,CG33970,Khc-73,spel1 | | 2.129e-02 | -3.85 | MORF\_PAX7 | MSigDB lists | MORF\_PAX7 | 41 | 5 | 4161 | 164 | toy,fz2,dnc,CG40178,Octbeta1R | | 2.129e-02 | -3.85 | chr8q24 | MSigDB lists | chr8q24 | 41 | 5 | 4161 | 164 | Der-1,CG3358,Maf1,vtd,Ac78C | | 2.129e-02 | -3.85 | LHX3\_01 | MSigDB lists | LHX3\_01 | 41 | 5 | 4161 | 164 | twz,Nlg3,Dys,bru1,Octbeta2R | | 2.129e-02 | -3.85 | PRC2\_EZH2\_UP.V1\_UP | MSigDB lists | PRC2\_EZH2\_UP.V1\_UP | 41 | 5 | 4161 | 164 | Dys,Cyp6a8,Pkg21D,Gfat1,Glut1 | | 2.129e-02 | -3.85 | GSE28130\_ACTIVATED\_VS\_INDUCEED\_TREG\_DN | MSigDB lists | GSE28130\_ACTIVATED\_VS\_INDUCEED\_TREG\_DN | 41 | 5 | 4161 | 164 | Aldh-III,ap,Coq7,Ca-alpha1D,CG41378 | | 2.129e-02 | -3.85 | GSE3039\_CD4\_TCELL\_VS\_ALPHABETA\_CD8\_TCELL\_UP | MSigDB lists | GSE3039\_CD4\_TCELL\_VS\_ALPHABETA\_CD8\_TCELL\_UP | 41 | 5 | 4161 | 164 | MCU,Glut1,Tim17b,Meltrin,dnc | | 2.129e-02 | -3.85 | BILD\_HRAS\_ONCOGENIC\_SIGNATURE | MSigDB lists | BILD\_HRAS\_ONCOGENIC\_SIGNATURE | 41 | 5 | 4161 | 164 | arg,Glut1,Octbeta2R,Odc2,fray | | 2.129e-02 | -3.85 | MORF\_DMPK | MSigDB lists | MORF\_DMPK | 41 | 5 | 4161 | 164 | Dys,toy,fz2,Pi3K68D,Coq7 | | 2.129e-02 | -3.85 | GSE37301\_HEMATOPOIETIC\_STEM\_CELL\_VS\_GRAN\_MONO\_PROGENITOR\_DN | MSigDB lists | GSE37301\_HEMATOPOIETIC\_STEM\_CELL\_VS\_GRAN\_MONO\_PROGENITOR\_DN | 41 | 5 | 4161 | 164 | CG9389,MCU,Pgant5,Gdh,CG4341 | | 2.137e-02 | -3.85 | localization of cell | biological process | GO:0051674 | 351 | 25 | 12345 | 571 | Sdc,Dhc16F,Src42A,Msp300,Sema1a,ru,gus,sn,CG17387,Diap1,Dlg5,mbc,mim,tj,CG1486,Pi3K68D,cta,jing,rib,salm,pum,jvl,kni,fz2,par-1 | | 2.142e-02 | -3.84 | axon extension | biological process | GO:0048675 | 35 | 5 | 12345 | 571 | RhoGAP100F,hdc,dnc,fz2,CadN2 | | 2.142e-02 | -3.84 | calcium-mediated signaling | biological process | GO:0019722 | 35 | 5 | 12345 | 571 | Spn,Ca-alpha1D,rdgC,TpnC41C,norpA | | 2.142e-02 | -3.84 | synaptic growth at neuromuscular junction | biological process | GO:0051124 | 35 | 5 | 12345 | 571 | Nlg2,Ten-a,Atpalpha,cpx,par-1 | | 2.157e-02 | -3.84 | kinase activity | molecular function | GO:0016301 | 375 | 26 | 12176 | 554 | Gprk1,Stlk,dnt,CG42637,bt,Pkg21D,CG3544,Slob,Src42A,CycB,Takl2,rl,bma,Pdk,fray,Pi3K68D,CG34357,fbl,par-1,PEK,Hipk,Tak1,CG12147,CG4546,CG32944,CG9886 | | 2.162e-02 | -3.83 | regulation of secretion | biological process | GO:0051046 | 89 | 9 | 12345 | 571 | cpx,Glut1,IA-2,Syt7,CG9650,sky,dimm,Dys,ru | | 2.164e-02 | -3.83 | REACTOME\_SIGNALING\_BY\_GPCR | MSigDB lists | REACTOME\_SIGNALING\_BY\_GPCR | 105 | 9 | 4161 | 164 | norpA,fz2,cta,Octbeta2R,dnc,rl,Ac78C,RSG7,Octbeta1R | | 2.166e-02 | -3.83 | TGF-b\_propeptide | interpro domains | IPR001111 | 5 | 2 | 11383 | 557 | mav,scw | | 2.166e-02 | -3.83 | LOTUS-like | interpro domains | IPR041966 | 5 | 2 | 11383 | 557 | Marf1,tej | | 2.167e-02 | -3.83 | DODD\_NASOPHARYNGEAL\_CARCINOMA\_UP | MSigDB lists | DODD\_NASOPHARYNGEAL\_CARCINOMA\_UP | 295 | 19 | 4161 | 164 | CG17387,CG2256,alpha4GT1,Aldh-III,arg,CG9389,Khc-73,CG12547,rg,MCU,nAChRalpha4,CG8184,Mef2,CG9492,Dhc16F,CG42813,unc-13-4A,Syx18,l(2)41Ab | | 2.176e-02 | -3.83 | GSE29949\_MICROGLIA\_VS\_DC\_BRAIN\_UP | MSigDB lists | GSE29949\_MICROGLIA\_VS\_DC\_BRAIN\_UP | 56 | 6 | 4161 | 164 | CG34408,Meltrin,Parp,DNAlig4,Git,CG8184 | | 2.176e-02 | -3.83 | GSE37416\_0H\_VS\_6H\_F\_TULARENSIS\_LVS\_NEUTROPHIL\_UP | MSigDB lists | GSE37416\_0H\_VS\_6H\_F\_TULARENSIS\_LVS\_NEUTROPHIL\_UP | 56 | 6 | 4161 | 164 | Meltrin,vap,Gdh,St1,CG9389,IntS3 | | 2.183e-02 | -3.82 | cell recognition | biological process | GO:0008037 | 119 | 11 | 12345 | 571 | fz2,fru,shot,Ptp99A,Dscam2,igl,nrm,Ten-a,Sema1a,dsx,lola | | 2.192e-02 | -3.82 | negative regulation of cellular amide metabolic process | biological process | GO:0034249 | 61 | 7 | 12345 | 571 | bru3,dnr1,pum,bru1,orb,heph,Thor | | 2.192e-02 | -3.82 | cell junction assembly | biological process | GO:0034329 | 61 | 7 | 12345 | 571 | Atpalpha,shakB,kune,CG15449,CadN2,kirre,nullo | | 2.196e-02 | -3.82 | regulation of small GTPase mediated signal transduction | biological process | GO:0051056 | 104 | 10 | 12345 | 571 | CG34408,ArfGAP1,CG10188,vap,step,conu,Slmap,sif,CG30440,mbc | | 2.206e-02 | -3.81 | tsr (twinstar) | protein interactions | 37841 | 44 | 6 | 9204 | 464 | fbl,inc,tobi,Rtnl1,Tak1,Sema1a | | 2.219e-02 | -3.81 | GO\_CELLULAR\_RESPONSE\_TO\_NITROGEN\_COMPOUND | MSigDB lists | GO\_CELLULAR\_RESPONSE\_TO\_NITROGEN\_COMPOUND | 123 | 10 | 4161 | 164 | Ubr1,drosha,Parp,Sod1,Mef2,Pi3K68D,dnc,Ars2,rl,Ac78C | | 2.224e-02 | -3.81 | ACTGAAA\_MIR30A3P\_MIR30E3P | MSigDB lists | ACTGAAA\_MIR30A3P\_MIR30E3P | 72 | 7 | 4161 | 164 | CG40045,Pgant5,step,RSG7,Mef2,Bruce,bru1 | | 2.227e-02 | -3.80 | Rac1 (Rac1) | protein interactions | 38146 | 124 | 12 | 9204 | 464 | mbc,Diap1,cpo,Rgk1,conu,pk,CG30440,tou,CdGAPr,RhoGAP100F,ct,ck | | 2.238e-02 | -3.80 | bHLH\_dom | interpro domains | IPR011598 | 58 | 7 | 11383 | 557 | CG33557,Mnt,dimm,Clk,cato,Myc,Fer3 | | 2.238e-02 | -3.80 | HLH\_DNA-bd\_sf | interpro domains | IPR036638 | 58 | 7 | 11383 | 557 | cato,Myc,Fer3,CG33557,Mnt,dimm,Clk | | 2.248e-02 | -3.80 | BTB | prosite domains | PS50097 | 60 | 8 | 6243 | 365 | rib,CG15725,lola,CG41099,bab2,br,fru,Trl | | 2.250e-02 | -3.79 | Sphingolipid de novo biosynthesis | REACTOME pathways | R-DME-1660661 | 23 | 4 | 3923 | 189 | Aldh-III,scw,fan,cert | | 2.251e-02 | -3.79 | courtship behavior | biological process | GO:0007619 | 135 | 12 | 12345 | 571 | Gr39a,lncRNA:flam,Nlg2,dsx,Gr2a,shep,CheB42a,Or67d,fru,dnc,Gr93d,twz | | 2.260e-02 | -3.79 | HMG\_box | pfam domains | PF00505 | 22 | 4 | 10480 | 528 | SoxN,cic,pan,CG12104 | | 2.263e-02 | -3.79 | ph-d (polyhomeotic distal) | protein interactions | 44889 | 22 | 4 | 9204 | 464 | pho,Su(z)2,salm,Psc | | 2.269e-02 | -3.79 | positive regulation of protein metabolic process | biological process | GO:0051247 | 318 | 23 | 12345 | 571 | Trl,sna,orb,PEK,Tak1,pum,CG41520,CG12147,Stlk,CG41106,mxt,scw,fray,step,Alg-2,Su(z)2,grim,CG14184,Diap1,mbc,gus,mav,Src42A | | 2.270e-02 | -3.79 | mating behavior | biological process | GO:0007617 | 183 | 15 | 12345 | 571 | Or67d,CheB42a,SPR,shep,twz,Gr93d,dati,dnc,fru,dsx,Nlg2,lncRNA:flam,Gr39a,Acp36DE,Gr2a | | 2.270e-02 | -3.79 | photoreceptor cell differentiation | biological process | GO:0046530 | 183 | 15 | 12345 | 571 | CadN2,svp,lz,rau,rg,RhoGAP100F,boi,rno,br,lola,klg,ru,salm,hth,rl | | 2.275e-02 | -3.78 | GSE22611\_UNSTIM\_VS\_6H\_MDP\_STIM\_MUTANT\_NOD2\_TRANSDUCED\_HEK293T\_CELL\_DN | MSigDB lists | GSE22611\_UNSTIM\_VS\_6H\_MDP\_STIM\_MUTANT\_NOD2\_TRANSDUCED\_HEK293T\_CELL\_DN | 28 | 4 | 4161 | 164 | Mef2,rl,SCAP,Ac78C | | 2.275e-02 | -3.78 | GSE9946\_IMMATURE\_VS\_LISTERIA\_INF\_MATURE\_DC\_UP | MSigDB lists | GSE9946\_IMMATURE\_VS\_LISTERIA\_INF\_MATURE\_DC\_UP | 28 | 4 | 4161 | 164 | CG4341,CG14621,Pkg21D,rg | | 2.275e-02 | -3.78 | GTTGNYNNRGNAAC\_UNKNOWN | MSigDB lists | GTTGNYNNRGNAAC\_UNKNOWN | 28 | 4 | 4161 | 164 | Syx18,Nlg3,CG4341,Rbp9 | | 2.275e-02 | -3.78 | GSE19888\_ADENOSINE\_A3R\_INH\_VS\_INH\_PRETREAT\_AND\_ACT\_WITH\_TCELL\_MEMBRANES\_MAST\_CELL\_DN | MSigDB lists | GSE19888\_ADENOSINE\_A3R\_INH\_VS\_INH\_PRETREAT\_AND\_ACT\_WITH\_TCELL\_MEMBRANES\_MAST\_CELL\_DN | 28 | 4 | 4161 | 164 | nAChRalpha3,CstF50,CG40178,Rbp9 | | 2.275e-02 | -3.78 | ABRAMSON\_INTERACT\_WITH\_AIRE | MSigDB lists | ABRAMSON\_INTERACT\_WITH\_AIRE | 28 | 4 | 4161 | 164 | Parp,spel1,CG10077,vtd | | 2.275e-02 | -3.78 | GO\_REGULATION\_OF\_MUSCLE\_TISSUE\_DEVELOPMENT | MSigDB lists | GO\_REGULATION\_OF\_MUSCLE\_TISSUE\_DEVELOPMENT | 28 | 4 | 4161 | 164 | CG10077,Mef2,Octbeta2R,CycB | | 2.275e-02 | -3.78 | FOXD3\_01 | MSigDB lists | FOXD3\_01 | 28 | 4 | 4161 | 164 | twz,Pi3K68D,Mef2,Dys | | 2.289e-02 | -3.78 | Phospholip\_A2\_2 | pfam domains | PF05826 | 5 | 2 | 10480 | 528 | CG42237,CG3009 | | 2.289e-02 | -3.78 | TGFb\_propeptide | pfam domains | PF00688 | 5 | 2 | 10480 | 528 | mav,scw | | 2.291e-02 | -3.78 | Usp15-31 (Ubiquitin specific protease 15/31) | protein interactions | 37954 | 5 | 2 | 9204 | 464 | Diap1,grim | | 2.291e-02 | -3.78 | CG15170 (uncharacterized protein) | protein interactions | 35160 | 5 | 2 | 9204 | 464 | Vsx2,dsx | | 2.291e-02 | -3.78 | CG9674 (uncharacterized protein) | protein interactions | 39878 | 5 | 2 | 9204 | 464 | Sema1a,Rtnl1 | | 2.291e-02 | -3.78 | CG13337 (uncharacterized protein) | protein interactions | 36518 | 5 | 2 | 9204 | 464 | CG12857,CG30480 | | 2.291e-02 | -3.78 | CG14062 (uncharacterized protein) | protein interactions | 43389 | 5 | 2 | 9204 | 464 | l(3)L1231,CycB | | 2.291e-02 | -3.78 | robl22E (robl22E) | protein interactions | 33412 | 5 | 2 | 9204 | 464 | CG31275,Idgf4 | | 2.291e-02 | -3.78 | CG4078 (uncharacterized protein) | protein interactions | 31497 | 5 | 2 | 9204 | 464 | SoxN,Sema1a | | 2.291e-02 | -3.78 | tapas (tapas) | protein interactions | 37304 | 5 | 2 | 9204 | 464 | tej,AGO3 | | 2.291e-02 | -3.78 | CG4042 (uncharacterized protein) | protein interactions | 40291 | 5 | 2 | 9204 | 464 | Rtnl1,Sema1a | | 2.291e-02 | -3.78 | qua (quail) | protein interactions | 35058 | 5 | 2 | 9204 | 464 | sn,CycB | | 2.291e-02 | -3.78 | CG5180 (uncharacterized protein) | protein interactions | 192508 | 5 | 2 | 9204 | 464 | dsx,Git | | 2.303e-02 | -3.77 | postsynapse organization | biological process | GO:0099173 | 24 | 4 | 12345 | 571 | Nlg3,Ulp1,Ten-a,Nlg2 | | 2.303e-02 | -3.77 | muscle system process | biological process | GO:0003012 | 24 | 4 | 12345 | 571 | bt,Mp,KCNQ,Ca-alpha1D | | 2.306e-02 | -3.77 | MODULE\_440 | MSigDB lists | MODULE\_440 | 16 | 3 | 4161 | 164 | arg,Gdh,Odc2 | | 2.306e-02 | -3.77 | GO\_REGULATION\_OF\_CARDIAC\_CONDUCTION | MSigDB lists | GO\_REGULATION\_OF\_CARDIAC\_CONDUCTION | 16 | 3 | 4161 | 164 | Atpalpha,PMCA,dnc | | 2.306e-02 | -3.77 | GO\_ADAPTIVE\_IMMUNE\_RESPONSE\_BASED\_ON\_SOMATIC\_RECOMBINATION\_OF\_IMMUNE\_RECEPTORS\_BUILT\_FROM\_IMMUNOGLOBULIN\_SUPERFAMILY\_DOMAINS | MSigDB lists | GO\_ADAPTIVE\_IMMUNE\_RESPONSE\_BASED\_ON\_SOMATIC\_RECOMBINATION\_OF\_IMMUNE\_RECEPTORS\_BUILT\_FROM\_IMMUNOGLOBULIN\_SUPERFAMILY\_DOMAINS | 16 | 3 | 4161 | 164 | Mef2,DNAlig4,spel1 | | 2.306e-02 | -3.77 | HOEBEKE\_LYMPHOID\_STEM\_CELL\_UP | MSigDB lists | HOEBEKE\_LYMPHOID\_STEM\_CELL\_UP | 16 | 3 | 4161 | 164 | mbc,Ubr1,CG32944 | | 2.306e-02 | -3.77 | GO\_REGULATION\_OF\_SMOOTH\_MUSCLE\_CONTRACTION | MSigDB lists | GO\_REGULATION\_OF\_SMOOTH\_MUSCLE\_CONTRACTION | 16 | 3 | 4161 | 164 | Sod1,nAChRalpha4,Octbeta2R | | 2.306e-02 | -3.77 | BIOCARTA\_AGR\_PATHWAY | MSigDB lists | BIOCARTA\_AGR\_PATHWAY | 16 | 3 | 4161 | 164 | Dys,rl,Git | | 2.306e-02 | -3.77 | MODULE\_289 | MSigDB lists | MODULE\_289 | 16 | 3 | 4161 | 164 | Git,cta,Octbeta2R | | 2.306e-02 | -3.77 | GO\_POSITIVE\_REGULATION\_OF\_INSULIN\_SECRETION | MSigDB lists | GO\_POSITIVE\_REGULATION\_OF\_INSULIN\_SECRETION | 16 | 3 | 4161 | 164 | Gdh,Nlg3,MCU | | 2.306e-02 | -3.77 | GO\_RESPONSE\_TO\_DSRNA | MSigDB lists | GO\_RESPONSE\_TO\_DSRNA | 16 | 3 | 4161 | 164 | drosha,Ars2,rl | | 2.307e-02 | -3.77 | gastrulation | biological process | GO:0007369 | 90 | 9 | 12345 | 571 | scw,zen,cta,sna,tld,Irp-1A,pan,5-HT2A,Pld | | 2.308e-02 | -3.77 | Dynein\_HC\_stalk | interpro domains | IPR024743 | 13 | 3 | 11383 | 557 | Dhc16F,CG9492,kl-3 | | 2.308e-02 | -3.77 | Dynein\_2\_C | interpro domains | IPR042228 | 13 | 3 | 11383 | 557 | CG9492,kl-3,Dhc16F | | 2.308e-02 | -3.77 | RA\_dom | interpro domains | IPR000159 | 13 | 3 | 11383 | 557 | rau,Myo81F,pico | | 2.308e-02 | -3.77 | Dynein\_heavy\_D6\_P-loop | interpro domains | IPR004273 | 13 | 3 | 11383 | 557 | CG9492,kl-3,Dhc16F | | 2.308e-02 | -3.77 | AAA\_lid\_11 | interpro domains | IPR041658 | 13 | 3 | 11383 | 557 | Dhc16F,CG9492,kl-3 | | 2.308e-02 | -3.77 | Dynein\_2\_N | interpro domains | IPR042222 | 13 | 3 | 11383 | 557 | CG9492,kl-3,Dhc16F | | 2.308e-02 | -3.77 | Dynein\_heavy\_chain\_D4\_dom | interpro domains | IPR024317 | 13 | 3 | 11383 | 557 | Dhc16F,kl-3,CG9492 | | 2.308e-02 | -3.77 | AAA\_6 | interpro domains | IPR035699 | 13 | 3 | 11383 | 557 | Dhc16F,kl-3,CG9492 | | 2.308e-02 | -3.77 | AAA\_9 | interpro domains | IPR035706 | 13 | 3 | 11383 | 557 | CG9492,kl-3,Dhc16F | | 2.308e-02 | -3.77 | AAA\_lid\_11\_sf | interpro domains | IPR042219 | 13 | 3 | 11383 | 557 | kl-3,CG9492,Dhc16F | | 2.322e-02 | -3.76 | fly chr1A1-1A1\|1-0 cM | chromosome location | fly chr1A1-1A1\|1-0 cM | 19 | 3 | 25010 | 823 | lncRNA:CR40469,mir-980,ewg | | 2.322e-02 | -3.76 | fly chr1A1-1A1\|1-0 | chromosome location | fly chr1A1-1A1\|1-0 | 19 | 3 | 25010 | 823 | ewg,mir-980,lncRNA:CR40469 | | 2.342e-02 | -3.75 | GSE22935\_UNSTIM\_VS\_48H\_MBOVIS\_BCG\_STIM\_MACROPHAGE\_UP | MSigDB lists | GSE22935\_UNSTIM\_VS\_48H\_MBOVIS\_BCG\_STIM\_MACROPHAGE\_UP | 42 | 5 | 4161 | 164 | bru1,dnc,Meltrin,alpha4GT1,nAChRalpha3 | | 2.342e-02 | -3.75 | MODULE\_112 | MSigDB lists | MODULE\_112 | 42 | 5 | 4161 | 164 | RSG7,Mp,Octbeta2R,Ugt50B3,Mef2 | | 2.342e-02 | -3.75 | GO\_NEGATIVE\_REGULATION\_OF\_NEURON\_DEATH | MSigDB lists | GO\_NEGATIVE\_REGULATION\_OF\_NEURON\_DEATH | 42 | 5 | 4161 | 164 | Mef2,Sod1,DNAlig4,spel1,vap | | 2.342e-02 | -3.75 | COREBINDINGFACTOR\_Q6 | MSigDB lists | COREBINDINGFACTOR\_Q6 | 42 | 5 | 4161 | 164 | MFS16,CG12547,Glut1,toy,twz | | 2.342e-02 | -3.75 | GSE32901\_NAIVE\_VS\_TH17\_NEG\_CD4\_TCELL\_UP | MSigDB lists | GSE32901\_NAIVE\_VS\_TH17\_NEG\_CD4\_TCELL\_UP | 42 | 5 | 4161 | 164 | CG41378,Odc2,MFS16,CG9389,Mef2 | | 2.343e-02 | -3.75 | polypeptide N-acetylgalactosaminyltransferase activity | molecular function | GO:0004653 | 14 | 3 | 12176 | 554 | Pgant5,Pgant1,Pgant9 | | 2.350e-02 | -3.75 | SH3\_1 | pfam domains | PF00018 | 33 | 5 | 10480 | 528 | Cip4,Src42A,CAP,l(3)05822,vap | | 2.355e-02 | -3.75 | GSE18281\_CORTICAL\_THYMOCYTE\_VS\_WHOLE\_CORTEX\_THYMUS\_DN | MSigDB lists | GSE18281\_CORTICAL\_THYMOCYTE\_VS\_WHOLE\_CORTEX\_THYMUS\_DN | 57 | 6 | 4161 | 164 | fray,vap,DNAlig4,par-1,Tbc1d15-17,Sod1 | | 2.355e-02 | -3.75 | OCT1\_01 | MSigDB lists | OCT1\_01 | 57 | 6 | 4161 | 164 | CG9492,toy,Cyp6a8,Dys,svp,dnc | | 2.362e-02 | -3.75 | Signaling by GPCR | REACTOME pathways | R-DME-372790 | 227 | 18 | 3923 | 189 | rl,cta,Ac78C,par-1,dnc,sif,norpA,Rpn9,Octbeta2R,Gprk1,CG10188,CG41520,vap,Rh3,Octbeta1R,5-HT2A,CG30440,Prosalpha3T | | 2.372e-02 | -3.74 | establishment or maintenance of cell polarity | biological process | GO:0007163 | 217 | 17 | 12345 | 571 | rib,cta,Parp,CadN2,sn,Dys,heph,Khc-73,pk,par-1,orb,svp,jvl,caup,Tak1,fred,lola | | 2.376e-02 | -3.74 | establishment of ommatidial planar polarity | biological process | GO:0042067 | 62 | 7 | 12345 | 571 | svp,CadN2,lola,pk,Tak1,caup,fred | | 2.381e-02 | -3.74 | REACTOME\_GPCR\_DOWNSTREAM\_SIGNALING | MSigDB lists | REACTOME\_GPCR\_DOWNSTREAM\_SIGNALING | 73 | 7 | 4161 | 164 | Octbeta2R,cta,norpA,Octbeta1R,RSG7,Ac78C,dnc | | 2.390e-02 | -3.73 | Fas2 (Fasciclin 2) | protein interactions | 31364 | 294 | 23 | 9204 | 464 | par-1,Hsc70-3,CG9044,CG5270,CG9932,Sema1a,shot,sif,toc,CAP,mbc,Nipped-A,Mical,CG7518,tou,Dys,CG31342,rl,jvl,Dhc16F,CG10188,Msp300,Psc | | 2.391e-02 | -3.73 | gro (groucho) | protein interactions | 43162 | 70 | 8 | 9204 | 464 | rl,Myc,en,Hipk,CG3199,cic,pan,kni | | 2.393e-02 | -3.73 | Band\_41\_domain | interpro domains | IPR019749 | 23 | 4 | 11383 | 557 | CG34347,Myo81F,ck,dnr1 | | 2.393e-02 | -3.73 | HMG\_box\_dom | interpro domains | IPR009071 | 23 | 4 | 11383 | 557 | SoxN,cic,pan,CG12104 | | 2.396e-02 | -3.73 | epithelial cell migration, open tracheal system | biological process | GO:0007427 | 36 | 5 | 12345 | 571 | kni,salm,rib,Sdc,ru | | 2.410e-02 | -3.73 | FN3 | smart domains | SM00060 | 48 | 7 | 5086 | 308 | Dscam2,CG14964,sdk,Ptp99A,boi,fred,bt | | 2.431e-02 | -3.72 | cellular component organization or biogenesis | biological process | GO:0071840 | 2975 | 158 | 12345 | 571 | dpr21,d4,CG41520,Nipped-A,en,lola,Trl,CG9981,cpx,cic,dnt,sna,wb,rl,Diap1,kl-3,pk,Msp300,sif,twz,vtd,fru,Rtnl1,Mical,Pi3K68D,CG12200,dgt5,Tim23,par-1,capu,DNAlig4,sn,Ulp1,Tpl94D,CG41562,mlt,mbc,Khc-73,Src42A,unc-13-4A,bond,Parp,Nlg3,cert,Mp,Wdr62,orb,gus,ABCB7,fest,br,kune,Pld,cbs,fbl,CAP,tou,CG14184,fzo,Toll-7,neo,shakB,hdc,jvl,CG17683,CG13284,klg,shot,RhoGAP100F,Ptp99A,Svil,Atpalpha,CycB,eIF4B,Sema1a,heph,cta,Dscam2,tej,Thor,SoxN,AP-2alpha,CG40228,l(2)41Ab,RpL6,CG40813,zld,DNaseII,kirre,Magi,pan,Setd3,CG9107,pico,Psc,jing,HP6,ck,conu,CadN2,Parg,pho,Act42A,CG33217,SmydA-8,Ptip,Tim17b,Cip4,lama,fan,CG17387,CG34347,Spn,Syx18,bru1,dnc,DIP-delta,snRNA:U1:82Eb,pum,inc,dsx,snky,scro,rg,bt,fz2,dati,sky,CG32087,HPS4,nullo,Set1,Rpn9,Sdc,ct,CdGAPr,Myc,CG1620,su(w[a]),spel1,ap,cmb,CG31703,mRpL10,bru3,Nlg2,Stlk,CG41099,Snap25,Meltrin,CG10939,Ten-a,Nna1,rib,CG15449,mim,CG43901,Su(z)2 | | 2.443e-02 | -3.71 | Arginine biosynthesis | KEGG pathways | ko00220 | 15 | 3 | 3198 | 138 | arg,Gdh,GLS | | 2.443e-02 | -3.71 | Arginine biosynthesis | KEGG pathways | dme00220 | 15 | 3 | 3198 | 138 | GLS,arg,Gdh | | 2.447e-02 | -3.71 | positive regulation of mRNA metabolic process | biological process | GO:1903313 | 14 | 3 | 12345 | 571 | Marf1,pum,AGO3 | | 2.447e-02 | -3.71 | negative regulation of ion transport | biological process | GO:0043271 | 14 | 3 | 12345 | 571 | Rgk1,Slob,THADA | | 2.447e-02 | -3.71 | postsynaptic membrane organization | biological process | GO:0001941 | 14 | 3 | 12345 | 571 | Ten-a,Nlg3,Nlg2 | | 2.450e-02 | -3.71 | positive regulation of cellular protein metabolic process | biological process | GO:0032270 | 303 | 22 | 12345 | 571 | Stlk,CG41106,mxt,Tak1,pum,CG41520,CG12147,sna,orb,PEK,Trl,Src42A,Diap1,mbc,mav,Alg-2,Su(z)2,grim,CG14184,fray,scw,step | | 2.466e-02 | -3.70 | imaginal disc-derived wing hair organization | biological process | GO:0035317 | 49 | 6 | 12345 | 571 | ck,sn,cmb,Cip4,par-1,pk | | 2.491e-02 | -3.69 | MT | pfam domains | PF12777 | 13 | 3 | 10480 | 528 | Dhc16F,CG9492,kl-3 | | 2.491e-02 | -3.69 | AAA\_9 | pfam domains | PF12781 | 13 | 3 | 10480 | 528 | Dhc16F,CG9492,kl-3 | | 2.491e-02 | -3.69 | AAA\_8 | pfam domains | PF12780 | 13 | 3 | 10480 | 528 | Dhc16F,CG9492,kl-3 | | 2.491e-02 | -3.69 | AAA\_lid\_11 | pfam domains | PF18198 | 13 | 3 | 10480 | 528 | kl-3,Dhc16F,CG9492 | | 2.491e-02 | -3.69 | AAA\_6 | pfam domains | PF12774 | 13 | 3 | 10480 | 528 | kl-3,Dhc16F,CG9492 | | 2.491e-02 | -3.69 | Dynein\_heavy | pfam domains | PF03028 | 13 | 3 | 10480 | 528 | kl-3,CG9492,Dhc16F | | 2.494e-02 | -3.69 | Atg18a (Autophagy-related 18a) | protein interactions | 38913 | 13 | 3 | 9204 | 464 | orb,vtd,Myc | | 2.494e-02 | -3.69 | POSH (Plenty of SH3s) | protein interactions | 36990 | 13 | 3 | 9204 | 464 | Alg-2,Dys,Tak1 | | 2.494e-02 | -3.69 | tsg (twisted gastrulation) | protein interactions | 32160 | 13 | 3 | 9204 | 464 | dsx,Vsx2,scw | | 2.506e-02 | -3.69 | calmodulin binding | molecular function | GO:0005516 | 37 | 5 | 12176 | 554 | Camta,SK,rdgC,KCNQ,igl | | 2.507e-02 | -3.69 | protein domain specific binding | molecular function | GO:0019904 | 78 | 8 | 12176 | 554 | Pi3K68D,mim,fz2,fan,Dys,CG3588,Trl,pico | | 2.522e-02 | -3.68 | RA | smart domains | SM00314 | 11 | 3 | 5086 | 308 | Myo81F,pico,rau | | 2.522e-02 | -3.68 | BRCT | smart domains | SM00292 | 11 | 3 | 5086 | 308 | Parp,DNAlig4,Ptip | | 2.544e-02 | -3.67 | CTATGCA\_MIR153 | MSigDB lists | CTATGCA\_MIR153 | 58 | 6 | 4161 | 164 | Meltrin,vap,Gfat1,bru1,Dys,inc | | 2.544e-02 | -3.67 | E4BP4\_01 | MSigDB lists | E4BP4\_01 | 58 | 6 | 4161 | 164 | Dys,CG10077,Rbp9,Octbeta2R,Mp,Ranbp16 | | 2.544e-02 | -3.67 | MODULE\_12 | MSigDB lists | MODULE\_12 | 58 | 6 | 4161 | 164 | Mp,toy,Gfat1,Ugt50B3,Glut1,shot | | 2.544e-02 | -3.67 | GSE360\_L\_DONOVANI\_VS\_L\_MAJOR\_DC\_DN | MSigDB lists | GSE360\_L\_DONOVANI\_VS\_L\_MAJOR\_DC\_DN | 58 | 6 | 4161 | 164 | mbc,CG40178,Aldh-III,DNAlig4,Atpalpha,Trxr-2 | | 2.560e-02 | -3.67 | TGFB\_UP.V1\_DN | MSigDB lists | TGFB\_UP.V1\_DN | 29 | 4 | 4161 | 164 | Octbeta2R,svp,Ugt50B3,Git | | 2.560e-02 | -3.67 | GSE21379\_WT\_VS\_SAP\_KO\_TFH\_CD4\_TCELL\_DN | MSigDB lists | GSE21379\_WT\_VS\_SAP\_KO\_TFH\_CD4\_TCELL\_DN | 29 | 4 | 4161 | 164 | Pi3K68D,Git,Nlg3,rg | | 2.560e-02 | -3.67 | REACTOME\_OPIOID\_SIGNALLING | MSigDB lists | REACTOME\_OPIOID\_SIGNALLING | 29 | 4 | 4161 | 164 | dnc,rl,Ac78C,norpA | | 2.560e-02 | -3.67 | MORF\_BCL2L11 | MSigDB lists | MORF\_BCL2L11 | 29 | 4 | 4161 | 164 | fz2,dnc,CG40178,Dys | | 2.560e-02 | -3.67 | GO\_TRANSITION\_METAL\_ION\_HOMEOSTASIS | MSigDB lists | GO\_TRANSITION\_METAL\_ION\_HOMEOSTASIS | 29 | 4 | 4161 | 164 | Irp-1A,Sod1,Hmt-1,ABCB7 | | 2.560e-02 | -3.67 | GSE11884\_WT\_VS\_FURIN\_KO\_NAIVE\_CD4\_TCELL\_DN | MSigDB lists | GSE11884\_WT\_VS\_FURIN\_KO\_NAIVE\_CD4\_TCELL\_DN | 29 | 4 | 4161 | 164 | Aldh-III,Tim23,shot,Alg9 | | 2.560e-02 | -3.67 | ATF2\_S\_UP.V1\_UP | MSigDB lists | ATF2\_S\_UP.V1\_UP | 29 | 4 | 4161 | 164 | norpA,Octbeta2R,ap,ck | | 2.560e-02 | -3.67 | GSE17974\_CTRL\_VS\_ACT\_IL4\_AND\_ANTI\_IL12\_12H\_CD4\_TCELL\_UP | MSigDB lists | GSE17974\_CTRL\_VS\_ACT\_IL4\_AND\_ANTI\_IL12\_12H\_CD4\_TCELL\_UP | 29 | 4 | 4161 | 164 | dnc,St1,dnr1,Glut1 | | 2.560e-02 | -3.67 | GSE36826\_WT\_VS\_IL1R\_KO\_SKIN\_STAPH\_AUREUS\_INF\_DN | MSigDB lists | GSE36826\_WT\_VS\_IL1R\_KO\_SKIN\_STAPH\_AUREUS\_INF\_DN | 29 | 4 | 4161 | 164 | mbc,fz2,Meltrin,Octbeta2R | | 2.566e-02 | -3.66 | DNA-binding transcription repressor activity, RNA polymerase II-specific | molecular function | GO:0001227 | 64 | 7 | 12176 | 554 | cic,svp,en,dsx,E5,kni,CG9932 | | 2.566e-02 | -3.66 | DNA-binding transcription repressor activity | molecular function | GO:0001217 | 64 | 7 | 12176 | 554 | CG9932,kni,E5,dsx,en,cic,svp | | 2.568e-02 | -3.66 | GO\_CYTOPLASMIC\_SIDE\_OF\_MEMBRANE | MSigDB lists | GO\_CYTOPLASMIC\_SIDE\_OF\_MEMBRANE | 43 | 5 | 4161 | 164 | cta,RSG7,Src42A,vap,step | | 2.568e-02 | -3.66 | GSE22611\_NOD2\_TRANSD\_VS\_CTRL\_TRANSD\_HEK293\_MDP\_STIM\_2H\_UP | MSigDB lists | GSE22611\_NOD2\_TRANSD\_VS\_CTRL\_TRANSD\_HEK293\_MDP\_STIM\_2H\_UP | 43 | 5 | 4161 | 164 | Spn42De,Parg,unc-13-4A,nAChRalpha4,CG12121 | | 2.568e-02 | -3.66 | TEF\_Q6 | MSigDB lists | TEF\_Q6 | 43 | 5 | 4161 | 164 | Ca-alpha1D,bru1,toy,arg,ABCB7 | | 2.568e-02 | -3.66 | GSE43863\_DAY6\_EFF\_VS\_DAY150\_MEM\_TFH\_CD4\_TCELL\_DN | MSigDB lists | GSE43863\_DAY6\_EFF\_VS\_DAY150\_MEM\_TFH\_CD4\_TCELL\_DN | 43 | 5 | 4161 | 164 | rg,Spn42De,CG14721,Ac78C,Meltrin | | 2.571e-02 | -3.66 | cell-cell junction organization | biological process | GO:0045216 | 63 | 7 | 12345 | 571 | shakB,Atpalpha,CadN2,CG15449,kune,nullo,kirre | | 2.601e-02 | -3.65 | Facilitative Na+-independent glucose transporters | REACTOME pathways | R-DME-428790 | 24 | 4 | 3923 | 189 | CG15408,CG1213,CG8249,Glut1 | | 2.634e-02 | -3.64 | lolal (lola like) | protein interactions | 44703 | 23 | 4 | 9204 | 464 | CG7518,rib,Trl,Rpb12 | | 2.634e-02 | -3.64 | mating | biological process | GO:0007618 | 203 | 16 | 12345 | 571 | Gr2a,Acp36DE,Nlg2,dsx,dati,Or67d,shep,CheB42a,lncRNA:flam,Gr39a,Gr93d,twz,fru,Gdh,dnc,SPR | | 2.636e-02 | -3.64 | positive regulation of biosynthetic process | biological process | GO:0009891 | 594 | 38 | 12345 | 571 | dsx,toy,mxt,en,lola,zld,d4,pum,SoxN,Camta,CG40228,Tak1,sna,orb,Trl,scro,Setd3,br,pan,SCAP,bi,tj,jing,tou,Myc,CG12769,ap,Atf6,pho,caup,Mef2,Ptip,zen,hth,dimm,lz,Parp,Clk | | 2.636e-02 | -3.64 | positive regulation of cellular biosynthetic process | biological process | GO:0031328 | 594 | 38 | 12345 | 571 | en,lola,mxt,dsx,toy,zld,SoxN,d4,pum,Tak1,CG40228,Camta,sna,orb,Trl,scro,Setd3,br,pan,SCAP,tj,bi,Myc,tou,jing,ap,CG12769,Atf6,caup,pho,Mef2,Ptip,zen,dimm,hth,Parp,lz,Clk | | 2.645e-02 | -3.63 | regulation of mRNA catabolic process | biological process | GO:0061013 | 25 | 4 | 12345 | 571 | PNPase,Marf1,AGO3,pum | | 2.645e-02 | -3.63 | regulation of terminal button organization | biological process | GO:2000331 | 25 | 4 | 12345 | 571 | Nlg2,Thor,Ten-a,Sod1 | | 2.654e-02 | -3.63 | fly chr94A5-94A5\|3-74 | chromosome location | fly chr94A5-94A5\|3-74 | 8 | 2 | 25010 | 823 | CG5388,asRNA:CR46054 | | 2.654e-02 | -3.63 | fly chr86E16-86E16\|3-51 | chromosome location | fly chr86E16-86E16\|3-51 | 8 | 2 | 25010 | 823 | CG14720,Fer3 | | 2.654e-02 | -3.63 | fly chr94A5-94A5\|3-74 cM | chromosome location | fly chr94A5-94A5\|3-74 cM | 8 | 2 | 25010 | 823 | CG5388,asRNA:CR46054 | | 2.654e-02 | -3.63 | fly chr86E16-86E16\|3-51 cM | chromosome location | fly chr86E16-86E16\|3-51 cM | 8 | 2 | 25010 | 823 | Fer3,CG14720 | | 2.656e-02 | -3.63 | astral microtubule | cellular component | GO:0000235 | 6 | 2 | 11940 | 534 | Khc-73,shot | | 2.665e-02 | -3.62 | membrane organization | biological process | GO:0061024 | 220 | 17 | 12345 | 571 | Rtnl1,mim,rib,fbl,Syx18,CycB,Ten-a,fan,nullo,cert,Cip4,CG9981,Snap25,Nlg3,snky,Nlg2,AP-2alpha | | 2.668e-02 | -3.62 | neuron projection extension | biological process | GO:1990138 | 37 | 5 | 12345 | 571 | fz2,dnc,hdc,RhoGAP100F,CadN2 | | 2.695e-02 | -3.61 | Wnt signaling pathway | KEGG pathways | dme04310 | 100 | 9 | 3198 | 138 | CG12147,fz2,Tak1,SteXh:CG42398,pan,norpA,Takl2,Myc,pk | | 2.695e-02 | -3.61 | Wnt signaling pathway | KEGG pathways | ko04310 | 100 | 9 | 3198 | 138 | CG12147,Tak1,fz2,pan,SteXh:CG42398,norpA,Takl2,Myc,pk | | 2.696e-02 | -3.61 | non-sensory hair organization | biological process | GO:0035316 | 50 | 6 | 12345 | 571 | par-1,Cip4,cmb,sn,ck,pk | | 2.696e-02 | -3.61 | hair cell differentiation | biological process | GO:0035315 | 50 | 6 | 12345 | 571 | sn,ck,Cip4,cmb,par-1,pk | | 2.704e-02 | -3.61 | Acyl chain remodelling of PC | REACTOME pathways | R-DME-1482788 | 14 | 3 | 3923 | 189 | CG42237,lama,CG3009 | | 2.705e-02 | -3.61 | regulation of signal transduction | biological process | GO:0009966 | 745 | 46 | 12345 | 571 | CG14184,RSG7,ArfGAP1,Pi3K68D,CG34408,vap,step,norpA,conu,Slmap,lqf,sif,mav,Diap1,tld,svp,fuss,Wdr59,Mp,SoxN,pum,Tak1,Camta,rau,Alg-2,dnc,rno,scw,Ppm1,ru,Src42A,Hipk,Ulp1,hth,Gprk1,cv-2,mbc,ewg,PEK,rdgC,CG10188,tay,par-1,CG30440,CG12147,Git | | 2.709e-02 | -3.61 | SWEET\_LUNG\_CANCER\_KRAS\_UP | MSigDB lists | SWEET\_LUNG\_CANCER\_KRAS\_UP | 164 | 12 | 4161 | 164 | RpL6,CG41378,CG30495,CG30059,sky,Pdk,Meltrin,Hsc70-3,rl,Mp,mIF2,arg | | 2.718e-02 | -3.61 | GSE27670\_BLIMP1\_VS\_LMP1\_TRANSDUCED\_GC\_BCELL\_DN | MSigDB lists | GSE27670\_BLIMP1\_VS\_LMP1\_TRANSDUCED\_GC\_BCELL\_DN | 75 | 7 | 4161 | 164 | Tsp5D,Pdk,Stlk,CG41099,vap,rl,CG17691 | | 2.721e-02 | -3.60 | ROSS\_ACUTE\_MYELOID\_LEUKEMIA\_CBF | MSigDB lists | ROSS\_ACUTE\_MYELOID\_LEUKEMIA\_CBF | 17 | 3 | 4161 | 164 | CG32944,unc-13-4A,CG14721 | | 2.721e-02 | -3.60 | GO\_NEURAL\_NUCLEUS\_DEVELOPMENT | MSigDB lists | GO\_NEURAL\_NUCLEUS\_DEVELOPMENT | 17 | 3 | 4161 | 164 | Gdh,Hsc70-3,ND-30 | | 2.721e-02 | -3.60 | CCAATNNSNNNGCG\_UNKNOWN | MSigDB lists | CCAATNNSNNNGCG\_UNKNOWN | 17 | 3 | 4161 | 164 | ap,toy,Mef2 | | 2.721e-02 | -3.60 | NKX25\_01 | MSigDB lists | NKX25\_01 | 17 | 3 | 4161 | 164 | Octbeta1R,Mef2,toy | | 2.721e-02 | -3.60 | GO\_COLUMNAR\_CUBOIDAL\_EPITHELIAL\_CELL\_DEVELOPMENT | MSigDB lists | GO\_COLUMNAR\_CUBOIDAL\_EPITHELIAL\_CELL\_DEVELOPMENT | 17 | 3 | 4161 | 164 | Sod1,toy,ck | | 2.721e-02 | -3.60 | BROWNE\_HCMV\_INFECTION\_8HR\_UP | MSigDB lists | BROWNE\_HCMV\_INFECTION\_8HR\_UP | 17 | 3 | 4161 | 164 | cta,CG30059,CG40045 | | 2.721e-02 | -3.60 | GO\_ADP\_BINDING | MSigDB lists | GO\_ADP\_BINDING | 17 | 3 | 4161 | 164 | spel1,ck,Gdh | | 2.721e-02 | -3.60 | MODULE\_456 | MSigDB lists | MODULE\_456 | 17 | 3 | 4161 | 164 | Hsc70-3,alpha4GT1,Glut1 | | 2.721e-02 | -3.60 | KRAS.AMP.LUNG\_UP.V1\_UP | MSigDB lists | KRAS.AMP.LUNG\_UP.V1\_UP | 17 | 3 | 4161 | 164 | Pkg21D,sbm,cv-2 | | 2.721e-02 | -3.60 | GO\_GLUTAMINE\_METABOLIC\_PROCESS | MSigDB lists | GO\_GLUTAMINE\_METABOLIC\_PROCESS | 17 | 3 | 4161 | 164 | Gfat1,GLS,Gdh | | 2.721e-02 | -3.60 | VANTVEER\_BREAST\_CANCER\_POOR\_PROGNOSIS | MSigDB lists | VANTVEER\_BREAST\_CANCER\_POOR\_PROGNOSIS | 17 | 3 | 4161 | 164 | CG3107,CG32944,Glut1 | | 2.721e-02 | -3.60 | REACTOME\_PLC\_BETA\_MEDIATED\_EVENTS | MSigDB lists | REACTOME\_PLC\_BETA\_MEDIATED\_EVENTS | 17 | 3 | 4161 | 164 | norpA,Ac78C,rl | | 2.734e-02 | -3.60 | - | gene3d domains | 1.10.10.1880 | 5 | 2 | 4929 | 273 | tej,Marf1 | | 2.736e-02 | -3.60 | Homeobox\_dom | interpro domains | IPR001356 | 102 | 10 | 11383 | 557 | zen,ap,hth,scro,caup,ct,en,E5,Vsx2,toy | | 2.742e-02 | -3.60 | HOLLMANN\_APOPTOSIS\_VIA\_CD40\_UP | MSigDB lists | HOLLMANN\_APOPTOSIS\_VIA\_CD40\_UP | 59 | 6 | 4161 | 164 | Mef2,shot,CG41378,Ranbp16,Parp,Meltrin | | 2.742e-02 | -3.60 | GSE22886\_IGG\_IGA\_MEMORY\_BCELL\_VS\_BM\_PLASMA\_CELL\_UP | MSigDB lists | GSE22886\_IGG\_IGA\_MEMORY\_BCELL\_VS\_BM\_PLASMA\_CELL\_UP | 59 | 6 | 4161 | 164 | CG10077,dnr1,rl,CG40178,CG8184,Git | | 2.744e-02 | -3.60 | octopamine receptor activity | molecular function | GO:0004989 | 6 | 2 | 12176 | 554 | Octbeta2R,Octbeta1R | | 2.744e-02 | -3.60 | aldehyde-lyase activity | molecular function | GO:0016832 | 6 | 2 | 12176 | 554 | CG1486,CG10184 | | 2.747e-02 | -3.59 | MOOTHA\_HUMAN\_MITODB\_6\_2002 | MSigDB lists | MOOTHA\_HUMAN\_MITODB\_6\_2002 | 242 | 16 | 4161 | 164 | arg,Hmt-1,Gdh,SdhA,Pdk,Odc2,CG5254,Trxr-2,mIF2,GLS,ABCB7,ND-30,Coq7,CG17691,Tim23,Tim17b | | 2.756e-02 | -3.59 | fly chr21E2-21E2\|2-1.5 | chromosome location | fly chr21E2-21E2\|2-1.5 | 70 | 6 | 25010 | 823 | lncRNA:CR44988,GABA-B-R3,Pkg21D,lncRNA:CR45261,CG4341,CG3544 | | 2.756e-02 | -3.59 | fly chr21E2-21E2\|2-1.5 cM | chromosome location | fly chr21E2-21E2\|2-1.5 cM | 70 | 6 | 25010 | 823 | lncRNA:CR45261,lncRNA:CR44988,GABA-B-R3,Pkg21D,CG3544,CG4341 | | 2.769e-02 | -3.59 | negative regulation of transcription by RNA polymerase II | biological process | GO:0000122 | 221 | 17 | 12345 | 571 | kni,cic,svp,Mnt,sna,en,Vsx2,dsx,Clk,CG9932,jing,CG1620,E5,Su(z)2,bi,lz,ct | | 2.776e-02 | -3.58 | hormone metabolic process | biological process | GO:0042445 | 64 | 7 | 12345 | 571 | spok,bond,CG9514,nvd,CG30495,Npc2b,spo | | 2.782e-02 | -3.58 | enhancer sequence-specific DNA binding | molecular function | GO:0001158 | 38 | 5 | 12176 | 554 | Mef2,hth,en,pho,pan | | 2.782e-02 | -3.58 | mRNA 3'-UTR binding | molecular function | GO:0003730 | 38 | 5 | 12176 | 554 | orb,heph,pum,bru1,shep | | 2.801e-02 | -3.58 | sav (salvador) | protein interactions | 252554 | 59 | 7 | 9204 | 464 | CG9411,par-1,Pdk,Magi,CG3081,kune,Tgi | | 2.807e-02 | -3.57 | nucleoside-triphosphatase regulator activity | molecular function | GO:0060589 | 110 | 10 | 12176 | 554 | sky,RhoGAP100F,Tbc1d15-17,norpA,Git,vap,ArfGAP1,CG34408,conu,CdGAPr | | 2.808e-02 | -3.57 | GSE27786\_LIN\_NEG\_VS\_MONO\_MAC\_DN | MSigDB lists | GSE27786\_LIN\_NEG\_VS\_MONO\_MAC\_DN | 44 | 5 | 4161 | 164 | ck,AP-2alpha,Maf1,MCU,CG12121 | | 2.808e-02 | -3.57 | GTATTAT\_MIR3693P | MSigDB lists | GTATTAT\_MIR3693P | 44 | 5 | 4161 | 164 | dnc,Git,Dys,toy,svp | | 2.808e-02 | -3.57 | GSE5099\_DAY3\_VS\_DAY7\_MCSF\_TREATED\_MACROPHAGE\_DN | MSigDB lists | GSE5099\_DAY3\_VS\_DAY7\_MCSF\_TREATED\_MACROPHAGE\_DN | 44 | 5 | 4161 | 164 | cta,Alg-2,nAChRalpha4,shot,mbc | | 2.808e-02 | -3.57 | GSE37416\_0H\_VS\_48H\_F\_TULARENSIS\_LVS\_NEUTROPHIL\_UP | MSigDB lists | GSE37416\_0H\_VS\_48H\_F\_TULARENSIS\_LVS\_NEUTROPHIL\_UP | 44 | 5 | 4161 | 164 | Ubr1,IntS3,SCAP,Gdh,vap | | 2.808e-02 | -3.57 | SMID\_BREAST\_CANCER\_NORMAL\_LIKE\_UP | MSigDB lists | SMID\_BREAST\_CANCER\_NORMAL\_LIKE\_UP | 44 | 5 | 4161 | 164 | Git,Dys,Mef2,bru1,shot | | 2.821e-02 | -3.57 | C2 | smart domains | SM00239 | 39 | 6 | 5086 | 308 | norpA,unc-13-4A,vap,RhoGAP100F,Syt7,Pi3K68D | | 2.830e-02 | -3.56 | maternal specification of dorsal/ventral axis, oocyte, soma encoded | biological process | GO:0007313 | 6 | 2 | 12345 | 571 | tld,cic | | 2.830e-02 | -3.56 | analia development | biological process | GO:0007487 | 6 | 2 | 12345 | 571 | en,dsx | | 2.830e-02 | -3.56 | central nervous system projection neuron axonogenesis | biological process | GO:0021952 | 6 | 2 | 12345 | 571 | Ulp1,en | | 2.830e-02 | -3.56 | rRNA 5'-end processing | biological process | GO:0000967 | 6 | 2 | 12345 | 571 | CG40813,CG41562 | | 2.830e-02 | -3.56 | endonucleolytic cleavage to generate mature 5'-end of SSU-rRNA from (SSU-rRNA, 5.8S rRNA, LSU-rRNA) | biological process | GO:0000472 | 6 | 2 | 12345 | 571 | CG41562,CG40813 | | 2.830e-02 | -3.56 | endonucleolytic cleavage in 5'-ETS of tricistronic rRNA transcript (SSU-rRNA, 5.8S rRNA, LSU-rRNA) | biological process | GO:0000480 | 6 | 2 | 12345 | 571 | CG41562,CG40813 | | 2.830e-02 | -3.56 | small-subunit processome assembly | biological process | GO:0034462 | 6 | 2 | 12345 | 571 | CG41562,CG40813 | | 2.833e-02 | -3.56 | Sugar/inositol\_transpt | interpro domains | IPR003663 | 14 | 3 | 11383 | 557 | Glut1,CG1213,CG8249 | | 2.833e-02 | -3.56 | Dynein\_heavy\_dom-2 | interpro domains | IPR013602 | 14 | 3 | 11383 | 557 | CG9492,kl-3,Dhc16F | | 2.833e-02 | -3.56 | ATPase\_P-typ\_P\_site | interpro domains | IPR018303 | 14 | 3 | 11383 | 557 | Atpalpha,PMCA,CG9981 | | 2.833e-02 | -3.56 | Ricin\_B-like\_lectins | interpro domains | IPR035992 | 14 | 3 | 11383 | 557 | Pgant9,Pgant1,Pgant5 | | 2.833e-02 | -3.56 | P\_typ\_ATPase | interpro domains | IPR001757 | 14 | 3 | 11383 | 557 | Atpalpha,CG9981,PMCA | | 2.833e-02 | -3.56 | BRCT\_dom | interpro domains | IPR001357 | 14 | 3 | 11383 | 557 | Ptip,DNAlig4,Parp | | 2.846e-02 | -3.56 | CHEN\_LVAD\_SUPPORT\_OF\_FAILING\_HEART\_DN | MSigDB lists | CHEN\_LVAD\_SUPPORT\_OF\_FAILING\_HEART\_DN | 7 | 2 | 4161 | 164 | Atpalpha,dnc | | 2.846e-02 | -3.56 | BIOCARTA\_AKAP95\_PATHWAY | MSigDB lists | BIOCARTA\_AKAP95\_PATHWAY | 7 | 2 | 4161 | 164 | CG10077,CycB | | 2.846e-02 | -3.56 | RODRIGUES\_NTN1\_AND\_DCC\_TARGETS | MSigDB lists | RODRIGUES\_NTN1\_AND\_DCC\_TARGETS | 7 | 2 | 4161 | 164 | Khc-73,Pi3K68D | | 2.846e-02 | -3.56 | GO\_NEGATIVE\_REGULATION\_OF\_PEPTIDYL\_SERINE\_PHOSPHORYLATION | MSigDB lists | GO\_NEGATIVE\_REGULATION\_OF\_PEPTIDYL\_SERINE\_PHOSPHORYLATION | 7 | 2 | 4161 | 164 | dnc,Dys | | 2.846e-02 | -3.56 | ST\_GRANULE\_CELL\_SURVIVAL\_PATHWAY | MSigDB lists | ST\_GRANULE\_CELL\_SURVIVAL\_PATHWAY | 7 | 2 | 4161 | 164 | rl,Parp | | 2.846e-02 | -3.56 | chr2q34 | MSigDB lists | chr2q34 | 7 | 2 | 4161 | 164 | fz2,CG14820 | | 2.846e-02 | -3.56 | GO\_DNA\_LIGATION | MSigDB lists | GO\_DNA\_LIGATION | 7 | 2 | 4161 | 164 | DNAlig4,Parp | | 2.846e-02 | -3.56 | MODULE\_200 | MSigDB lists | MODULE\_200 | 7 | 2 | 4161 | 164 | toy,Octbeta2R | | 2.846e-02 | -3.56 | BROWNE\_INTERFERON\_RESPONSIVE\_GENES | MSigDB lists | BROWNE\_INTERFERON\_RESPONSIVE\_GENES | 7 | 2 | 4161 | 164 | step,CG41378 | | 2.846e-02 | -3.56 | GO\_POLY\_PYRIMIDINE\_TRACT\_BINDING | MSigDB lists | GO\_POLY\_PYRIMIDINE\_TRACT\_BINDING | 7 | 2 | 4161 | 164 | PNPase,heph | | 2.846e-02 | -3.56 | SA\_CASPASE\_CASCADE | MSigDB lists | SA\_CASPASE\_CASCADE | 7 | 2 | 4161 | 164 | Parp,SCAP | | 2.846e-02 | -3.56 | GO\_CELLULAR\_RESPONSE\_TO\_TOXIC\_SUBSTANCE | MSigDB lists | GO\_CELLULAR\_RESPONSE\_TO\_TOXIC\_SUBSTANCE | 7 | 2 | 4161 | 164 | Hsc70-3,Mef2 | | 2.846e-02 | -3.56 | GNF2\_PTPRC | MSigDB lists | GNF2\_PTPRC | 7 | 2 | 4161 | 164 | CG10077,step | | 2.846e-02 | -3.56 | GO\_HETEROTRIMERIC\_G\_PROTEIN\_COMPLEX | MSigDB lists | GO\_HETEROTRIMERIC\_G\_PROTEIN\_COMPLEX | 7 | 2 | 4161 | 164 | RSG7,cta | | 2.846e-02 | -3.56 | GO\_AUDITORY\_RECEPTOR\_CELL\_DEVELOPMENT | MSigDB lists | GO\_AUDITORY\_RECEPTOR\_CELL\_DEVELOPMENT | 7 | 2 | 4161 | 164 | ck,Sod1 | | 2.846e-02 | -3.56 | GO\_SOMATIC\_RECOMBINATION\_OF\_IMMUNOGLOBULIN\_GENE\_SEGMENTS | MSigDB lists | GO\_SOMATIC\_RECOMBINATION\_OF\_IMMUNOGLOBULIN\_GENE\_SEGMENTS | 7 | 2 | 4161 | 164 | DNAlig4,spel1 | | 2.846e-02 | -3.56 | BIOCARTA\_SPRY\_PATHWAY | MSigDB lists | BIOCARTA\_SPRY\_PATHWAY | 7 | 2 | 4161 | 164 | vap,rl | | 2.846e-02 | -3.56 | PID\_S1P\_S1P3\_PATHWAY | MSigDB lists | PID\_S1P\_S1P3\_PATHWAY | 7 | 2 | 4161 | 164 | cta,rl | | 2.846e-02 | -3.56 | GO\_IMMUNOGLOBULIN\_PRODUCTION\_INVOLVED\_IN\_IMMUNOGLOBULIN\_MEDIATED\_IMMUNE\_RESPONSE | MSigDB lists | GO\_IMMUNOGLOBULIN\_PRODUCTION\_INVOLVED\_IN\_IMMUNOGLOBULIN\_MEDIATED\_IMMUNE\_RESPONSE | 7 | 2 | 4161 | 164 | spel1,DNAlig4 | | 2.846e-02 | -3.56 | GO\_REGULATION\_OF\_RELEASE\_OF\_SEQUESTERED\_CALCIUM\_ION\_INTO\_CYTOSOL\_BY\_SARCOPLASMIC\_RETICULUM | MSigDB lists | GO\_REGULATION\_OF\_RELEASE\_OF\_SEQUESTERED\_CALCIUM\_ION\_INTO\_CYTOSOL\_BY\_SARCOPLASMIC\_RETICULUM | 7 | 2 | 4161 | 164 | dnc,Dys | | 2.846e-02 | -3.56 | HALLMARK\_INTERFERON\_ALPHA\_RESPONSE | MSigDB lists | HALLMARK\_INTERFERON\_ALPHA\_RESPONSE | 7 | 2 | 4161 | 164 | PNPase,CG41378 | | 2.846e-02 | -3.56 | GO\_NEGATIVE\_REGULATION\_OF\_LIPID\_BIOSYNTHETIC\_PROCESS | MSigDB lists | GO\_NEGATIVE\_REGULATION\_OF\_LIPID\_BIOSYNTHETIC\_PROCESS | 7 | 2 | 4161 | 164 | SCAP,Sod1 | | 2.846e-02 | -3.56 | PID\_VEGFR1\_PATHWAY | MSigDB lists | PID\_VEGFR1\_PATHWAY | 7 | 2 | 4161 | 164 | vap,rl | | 2.846e-02 | -3.56 | GO\_PROTEIN\_CHANNEL\_ACTIVITY | MSigDB lists | GO\_PROTEIN\_CHANNEL\_ACTIVITY | 7 | 2 | 4161 | 164 | Tim23,Tim17b | | 2.846e-02 | -3.56 | GO\_ADRENERGIC\_RECEPTOR\_SIGNALING\_PATHWAY | MSigDB lists | GO\_ADRENERGIC\_RECEPTOR\_SIGNALING\_PATHWAY | 7 | 2 | 4161 | 164 | Octbeta2R,dnc | | 2.846e-02 | -3.56 | GO\_MITOGEN\_ACTIVATED\_PROTEIN\_KINASE\_KINASE\_KINASE\_BINDING | MSigDB lists | GO\_MITOGEN\_ACTIVATED\_PROTEIN\_KINASE\_KINASE\_KINASE\_BINDING | 7 | 2 | 4161 | 164 | rl,rdgC | | 2.846e-02 | -3.56 | YU\_MYC\_TARGETS\_DN | MSigDB lists | YU\_MYC\_TARGETS\_DN | 7 | 2 | 4161 | 164 | CG30059,ck | | 2.846e-02 | -3.56 | GO\_REGULATION\_OF\_STEROID\_BIOSYNTHETIC\_PROCESS | MSigDB lists | GO\_REGULATION\_OF\_STEROID\_BIOSYNTHETIC\_PROCESS | 7 | 2 | 4161 | 164 | Sod1,SCAP | | 2.846e-02 | -3.56 | GO\_VOLTAGE\_GATED\_CALCIUM\_CHANNEL\_COMPLEX | MSigDB lists | GO\_VOLTAGE\_GATED\_CALCIUM\_CHANNEL\_COMPLEX | 7 | 2 | 4161 | 164 | dnc,Ca-alpha1D | | 2.846e-02 | -3.56 | GO\_REGULATION\_OF\_RYANODINE\_SENSITIVE\_CALCIUM\_RELEASE\_CHANNEL\_ACTIVITY | MSigDB lists | GO\_REGULATION\_OF\_RYANODINE\_SENSITIVE\_CALCIUM\_RELEASE\_CHANNEL\_ACTIVITY | 7 | 2 | 4161 | 164 | dnc,Dys | | 2.846e-02 | -3.56 | GO\_ACETYLCHOLINE\_RECEPTOR\_ACTIVITY | MSigDB lists | GO\_ACETYLCHOLINE\_RECEPTOR\_ACTIVITY | 7 | 2 | 4161 | 164 | nAChRalpha3,nAChRalpha4 | | 2.846e-02 | -3.56 | GO\_RESPONSE\_TO\_MONOAMINE | MSigDB lists | GO\_RESPONSE\_TO\_MONOAMINE | 7 | 2 | 4161 | 164 | dnc,Octbeta2R | | 2.846e-02 | -3.56 | GO\_DICARBOXYLIC\_ACID\_BIOSYNTHETIC\_PROCESS | MSigDB lists | GO\_DICARBOXYLIC\_ACID\_BIOSYNTHETIC\_PROCESS | 7 | 2 | 4161 | 164 | Gdh,GLS | | 2.846e-02 | -3.56 | GO\_NEGATIVE\_REGULATION\_OF\_MUSCLE\_CONTRACTION | MSigDB lists | GO\_NEGATIVE\_REGULATION\_OF\_MUSCLE\_CONTRACTION | 7 | 2 | 4161 | 164 | Octbeta2R,Sod1 | | 2.846e-02 | -3.56 | GO\_ISOTYPE\_SWITCHING | MSigDB lists | GO\_ISOTYPE\_SWITCHING | 7 | 2 | 4161 | 164 | DNAlig4,spel1 | | 2.846e-02 | -3.56 | chr12q15 | MSigDB lists | chr12q15 | 7 | 2 | 4161 | 164 | CG10171,Tbc1d15-17 | | 2.846e-02 | -3.56 | GO\_NEGATIVE\_REGULATION\_OF\_POTASSIUM\_ION\_TRANSPORT | MSigDB lists | GO\_NEGATIVE\_REGULATION\_OF\_POTASSIUM\_ION\_TRANSPORT | 7 | 2 | 4161 | 164 | norpA,fray | | 2.846e-02 | -3.56 | GO\_INTERMEDIATE\_FILAMENT\_BASED\_PROCESS | MSigDB lists | GO\_INTERMEDIATE\_FILAMENT\_BASED\_PROCESS | 7 | 2 | 4161 | 164 | shot,Sod1 | | 2.846e-02 | -3.56 | GO\_CILIUM\_OR\_FLAGELLUM\_DEPENDENT\_CELL\_MOTILITY | MSigDB lists | GO\_CILIUM\_OR\_FLAGELLUM\_DEPENDENT\_CELL\_MOTILITY | 7 | 2 | 4161 | 164 | CG17387,Dhc16F | | 2.846e-02 | -3.56 | ABE\_VEGFA\_TARGETS\_2HR | MSigDB lists | ABE\_VEGFA\_TARGETS\_2HR | 7 | 2 | 4161 | 164 | Ranbp16,Rbp9 | | 2.846e-02 | -3.56 | GO\_ATPASE\_COMPLEX | MSigDB lists | GO\_ATPASE\_COMPLEX | 7 | 2 | 4161 | 164 | Hmt-1,Atpalpha | | 2.855e-02 | -3.56 | regulation of trans-synaptic signaling | biological process | GO:0099177 | 124 | 11 | 12345 | 571 | rl,Slob,pum,Dys,Src42A,Nlg2,cpx,Sod1,Nlg3,Snap25,sky | | 2.855e-02 | -3.56 | modulation of chemical synaptic transmission | biological process | GO:0050804 | 124 | 11 | 12345 | 571 | pum,Slob,rl,Nlg2,Dys,Src42A,Nlg3,Snap25,Sod1,cpx,sky | | 2.860e-02 | -3.55 | solute:proton symporter activity | molecular function | GO:0015295 | 26 | 4 | 12176 | 554 | CG15408,CG1213,CG45782,CG8249 | | 2.866e-02 | -3.55 | XU\_GH1\_EXOGENOUS\_TARGETS\_DN | MSigDB lists | XU\_GH1\_EXOGENOUS\_TARGETS\_DN | 30 | 4 | 4161 | 164 | Rbp9,ND-30,rdgC,Pi3K68D | | 2.866e-02 | -3.55 | KEGG\_ARGININE\_AND\_PROLINE\_METABOLISM | MSigDB lists | KEGG\_ARGININE\_AND\_PROLINE\_METABOLISM | 30 | 4 | 4161 | 164 | GLS,arg,Odc2,Gdh | | 2.866e-02 | -3.55 | GSE3982\_DC\_VS\_NEUTROPHIL\_DN | MSigDB lists | GSE3982\_DC\_VS\_NEUTROPHIL\_DN | 30 | 4 | 4161 | 164 | CG9389,Glut1,bru1,Git | | 2.866e-02 | -3.55 | GO\_SARCOLEMMA | MSigDB lists | GO\_SARCOLEMMA | 30 | 4 | 4161 | 164 | Ca-alpha1D,Octbeta2R,Dys,Atpalpha | | 2.866e-02 | -3.55 | GSE9037\_WT\_VS\_IRAK4\_KO\_LPS\_1H\_STIM\_BMDM\_DN | MSigDB lists | GSE9037\_WT\_VS\_IRAK4\_KO\_LPS\_1H\_STIM\_BMDM\_DN | 30 | 4 | 4161 | 164 | CG3358,CG42813,CG2256,Pkg21D | | 2.884e-02 | -3.55 | epithelial cell development | biological process | GO:0002064 | 361 | 25 | 12345 | 571 | jing,cta,mim,tj,Parp,neo,gus,rl,Dlg5,Diap1,mbc,br,ct,lncRNA:flam,CG41099,Cip4,cic,par-1,capu,Mef2,orb,lama,jvl,AP-2alpha,toc | | 2.929e-02 | -3.53 | system process | biological process | GO:0003008 | 636 | 40 | 12345 | 571 | MCU,Rh3,dnc,Gr93d,Or83c,Ir40a,ru,Gr39a,be,CG9492,Atpalpha,rdgC,Or67d,klg,SKIP,Or98a,salm,Ac78C,Rgk1,CG31750,2mit,norpA,Or69a,ck,nAChRalpha3,CAP,sif,ct,Obp85a,Ca-alpha1D,kirre,CheB42a,Mp,KCNQ,nAChRalpha4,rg,bt,Gr2a,HisCl1,pum | | 2.938e-02 | -3.53 | BRCT | prosite domains | PS50172 | 12 | 3 | 6243 | 365 | DNAlig4,Parp,Ptip | | 2.948e-02 | -3.52 | cell motility | biological process | GO:0048870 | 344 | 24 | 12345 | 571 | kni,par-1,fz2,pum,salm,mim,Pi3K68D,CG1486,tj,cta,jing,rib,Dhc16F,Sdc,ru,Msp300,Src42A,Sema1a,sn,CG17387,gus,mbc,Dlg5,Diap1 | | 2.951e-02 | -3.52 | GSE17721\_LPS\_VS\_GARDIQUIMOD\_8H\_BMDC\_UP | MSigDB lists | GSE17721\_LPS\_VS\_GARDIQUIMOD\_8H\_BMDC\_UP | 60 | 6 | 4161 | 164 | Tbc1d15-17,Pdk,cta,nAChRalpha4,Hsc70-3,Trs23 | | 2.951e-02 | -3.52 | CEBP\_Q2 | MSigDB lists | CEBP\_Q2 | 60 | 6 | 4161 | 164 | toy,Cyp6a8,svp,Octbeta2R,dnc,Syx18 | | 2.956e-02 | -3.52 | cellular response to xenobiotic stimulus | biological process | GO:0071466 | 15 | 3 | 12345 | 571 | Cyp6a8,St1,Cyp6g2 | | 2.956e-02 | -3.52 | antennal morphogenesis | biological process | GO:0048800 | 15 | 3 | 12345 | 571 | Diap1,ck,sn | | 2.956e-02 | -3.52 | regulation of tube diameter | biological process | GO:0035296 | 15 | 3 | 12345 | 571 | Atpalpha,rib,knk | | 2.956e-02 | -3.52 | xenobiotic metabolic process | biological process | GO:0006805 | 15 | 3 | 12345 | 571 | Cyp6a8,Cyp6g2,St1 | | 2.956e-02 | -3.52 | regulation of oskar mRNA translation | biological process | GO:0046011 | 15 | 3 | 12345 | 571 | bru1,heph,orb | | 2.956e-02 | -3.52 | male courtship behavior, veined wing generated song production | biological process | GO:0045433 | 15 | 3 | 12345 | 571 | fru,Nlg2,dsx | | 2.956e-02 | -3.52 | visceral muscle development | biological process | GO:0007522 | 15 | 3 | 12345 | 571 | mbc,Msp300,kirre | | 2.956e-02 | -3.52 | presynaptic active zone organization | biological process | GO:1990709 | 15 | 3 | 12345 | 571 | Ten-a,RhoGAP100F,Nlg2 | | 2.959e-02 | -3.52 | Vha68-2 (Vacuolar H[+] ATPase 68 kDa subunit 2) | protein interactions | 45012 | 47 | 6 | 9204 | 464 | ct,Atpalpha,CG8888,Sema1a,Rtnl1,CG14820 | | 2.959e-02 | -3.52 | RPA2 (Replication protein A2) | protein interactions | 35364 | 47 | 6 | 9204 | 464 | Parp,CG12521,CG14113,CycB,CG30440,CG17508 | | 2.959e-02 | -3.52 | glial cell development | biological process | GO:0021782 | 38 | 5 | 12345 | 571 | rau,kune,svp,Sdc,salm | | 2.963e-02 | -3.52 | IQ\_motif\_EF-hand-BS | interpro domains | IPR000048 | 36 | 5 | 11383 | 557 | rdgC,ck,igl,Camta,Myo81F | | 3.015e-02 | -3.50 | regulation of imaginal disc-derived wing size | biological process | GO:0044719 | 26 | 4 | 12345 | 571 | Myc,fru,lola,zld | | 3.046e-02 | -3.49 | TGTTTAC\_MIR30A5P\_MIR30C\_MIR30D\_MIR30B\_MIR30E5P | MSigDB lists | TGTTTAC\_MIR30A5P\_MIR30C\_MIR30D\_MIR30B\_MIR30E5P | 148 | 11 | 4161 | 164 | Dys,inc,Tbc1d15-17,Gfat1,bru1,dnc,Meltrin,Hsc70-3,Src42A,Pgant5,vap | | 3.055e-02 | -3.49 | DHC\_N2 | pfam domains | PF08393 | 14 | 3 | 10480 | 528 | kl-3,CG9492,Dhc16F | | 3.055e-02 | -3.49 | Ricin\_B\_lectin | pfam domains | PF00652 | 14 | 3 | 10480 | 528 | Pgant1,Pgant5,Pgant9 | | 3.058e-02 | -3.49 | shtd (shattered) | protein interactions | 32473 | 14 | 3 | 9204 | 464 | Sema1a,Diap1,CG5359 | | 3.058e-02 | -3.49 | tra (transformer) | protein interactions | 39849 | 14 | 3 | 9204 | 464 | CheB42a,dsx,fru | | 3.062e-02 | -3.49 | GSE22935\_UNSTIM\_VS\_48H\_MBOVIS\_BCG\_STIM\_MYD88\_KO\_MACROPHAGE\_DN | MSigDB lists | GSE22935\_UNSTIM\_VS\_48H\_MBOVIS\_BCG\_STIM\_MYD88\_KO\_MACROPHAGE\_DN | 45 | 5 | 4161 | 164 | Meltrin,ck,CG2256,Gfat1,Dys | | 3.062e-02 | -3.49 | GSE36392\_TYPE\_2\_MYELOID\_VS\_NEUTROPHIL\_IL25\_TREATED\_LUNG\_UP | MSigDB lists | GSE36392\_TYPE\_2\_MYELOID\_VS\_NEUTROPHIL\_IL25\_TREATED\_LUNG\_UP | 45 | 5 | 4161 | 164 | Tsp5D,Glut1,Git,Octbeta1R,nAChRalpha3 | | 3.062e-02 | -3.49 | MORF\_TFDP2 | MSigDB lists | MORF\_TFDP2 | 45 | 5 | 4161 | 164 | mbc,CG40178,Coq7,dnc,Pi3K68D | | 3.062e-02 | -3.49 | HNF1\_C | MSigDB lists | HNF1\_C | 45 | 5 | 4161 | 164 | Dys,Cyp6a8,Nlg3,sbm,PMCA | | 3.062e-02 | -3.49 | LYF1\_01 | MSigDB lists | LYF1\_01 | 45 | 5 | 4161 | 164 | fbl,PMCA,Mef2,CG9389,GLS | | 3.062e-02 | -3.49 | GSE36888\_STAT5\_AB\_KNOCKIN\_VS\_WT\_TCELL\_IL2\_TREATED\_2H\_UP | MSigDB lists | GSE36888\_STAT5\_AB\_KNOCKIN\_VS\_WT\_TCELL\_IL2\_TREATED\_2H\_UP | 45 | 5 | 4161 | 164 | CG14721,vap,Pdk,cta,Mef2 | | 3.062e-02 | -3.49 | IPF1\_Q4 | MSigDB lists | IPF1\_Q4 | 45 | 5 | 4161 | 164 | mbc,svp,Rbp9,Mef2,Dys | | 3.062e-02 | -3.49 | GSE21546\_WT\_VS\_ELK1\_KO\_DP\_THYMOCYTES\_DN | MSigDB lists | GSE21546\_WT\_VS\_ELK1\_KO\_DP\_THYMOCYTES\_DN | 45 | 5 | 4161 | 164 | nAChRalpha3,CG40045,SdhA,nAChRalpha4,Alg-2 | | 3.075e-02 | -3.48 | GO\_EPITHELIAL\_CELL\_DIFFERENTIATION | MSigDB lists | GO\_EPITHELIAL\_CELL\_DIFFERENTIATION | 94 | 8 | 4161 | 164 | svp,Mef2,toy,Sod1,ck,dnc,St1,cv-2 | | 3.082e-02 | -3.48 | axis specification | biological process | GO:0009798 | 207 | 16 | 12345 | 571 | rl,gus,heph,pk,scw,Parp,jvl,pum,zen,par-1,cic,kni,sna,orb,capu,tld | | 3.117e-02 | -3.47 | FERM\_3 | prosite domains | PS50057 | 21 | 4 | 6243 | 365 | Myo81F,ck,dnr1,CG34347 | | 3.132e-02 | -3.46 | small molecule binding | molecular function | GO:0036094 | 1078 | 62 | 12176 | 554 | CG43672,CG34357,FASN3,fbl,CG9981,par-1,DNAlig4,Hipk,Tak1,CG12147,Mical,CG42541,fzo,CG31689,Gprk1,Klp59C,Stlk,Trxr-2,Hsc70-3,Nadsyn,Src42A,CG9492,Rgk1,Acbp5,mIF2,SdhA,bma,Myo81F,Atpalpha,fray,CG7536,Dhc16F,PEK,RabX2,cta,CG32944,mus308,CG3556,CG9514,kl-3,Hmt-1,dnt,CG42637,CG5789,lz,CG14721,CG33970,bt,ABCB7,Pkg21D,Parp,spel1,Tdc2,Khc-73,Acox57D-d,CG10077,PMCA,ck,Takl2,rl,CG5160,CG1486 | | 3.144e-02 | -3.46 | ABT1/ESF2\_RRM | interpro domains | IPR034353 | 6 | 2 | 11383 | 557 | CG40813,CG41562 | | 3.144e-02 | -3.46 | ABT1/Esf2 | interpro domains | IPR039119 | 6 | 2 | 11383 | 557 | CG41562,CG40813 | | 3.144e-02 | -3.46 | ATPase\_P-typ\_cation-transptr\_C | interpro domains | IPR006068 | 6 | 2 | 11383 | 557 | Atpalpha,PMCA | | 3.144e-02 | -3.46 | Cellulose/chitin-bd\_N | interpro domains | IPR004302 | 6 | 2 | 11383 | 557 | CG42598,CG41284 | | 3.144e-02 | -3.46 | MyTH4\_dom\_sf | interpro domains | IPR038185 | 6 | 2 | 11383 | 557 | Myo81F,ck | | 3.144e-02 | -3.46 | ATPase\_P-typ\_cation-transptr\_N | interpro domains | IPR004014 | 6 | 2 | 11383 | 557 | Atpalpha,PMCA | | 3.144e-02 | -3.46 | MyTH4\_dom | interpro domains | IPR000857 | 6 | 2 | 11383 | 557 | Myo81F,ck | | 3.144e-02 | -3.46 | PLipase\_A2\_dom | interpro domains | IPR016090 | 6 | 2 | 11383 | 557 | CG3009,CG42237 | | 3.155e-02 | -3.46 | SUGRTRNSPORT | prints domains | PR00171 | 14 | 3 | 2185 | 112 | CG8249,Glut1,CG1213 | | 3.170e-02 | -3.45 | GO\_LEUKOCYTE\_ACTIVATION | MSigDB lists | GO\_LEUKOCYTE\_ACTIVATION | 61 | 6 | 4161 | 164 | DNAlig4,spel1,fz2,CG7255,Khc-73,Mef2 | | 3.170e-02 | -3.45 | GSE32255\_WT\_UNSTIM\_VS\_JMJD2D\_KNOCKDOWN\_4H\_LPS\_STIM\_DC\_DN | MSigDB lists | GSE32255\_WT\_UNSTIM\_VS\_JMJD2D\_KNOCKDOWN\_4H\_LPS\_STIM\_DC\_DN | 61 | 6 | 4161 | 164 | Gdh,CG4341,norpA,shot,MFS16,Gfat1 | | 3.170e-02 | -3.45 | terminal bouton | cellular component | GO:0043195 | 68 | 7 | 11940 | 534 | par-1,pum,Mical,sky,Magi,Snap25,cpx | | 3.173e-02 | -3.45 | GO\_POSITIVE\_REGULATION\_OF\_PEPTIDE\_SECRETION | MSigDB lists | GO\_POSITIVE\_REGULATION\_OF\_PEPTIDE\_SECRETION | 18 | 3 | 4161 | 164 | MCU,Nlg3,Gdh | | 3.173e-02 | -3.45 | MORF\_DCC | MSigDB lists | MORF\_DCC | 18 | 3 | 4161 | 164 | cv-2,Src42A,nAChRalpha4 | | 3.173e-02 | -3.45 | GO\_MICROTUBULE\_BUNDLE\_FORMATION | MSigDB lists | GO\_MICROTUBULE\_BUNDLE\_FORMATION | 18 | 3 | 4161 | 164 | l(2)41Ab,CG9492,CG17387 | | 3.173e-02 | -3.45 | GSE7218\_UNSTIM\_VS\_ANTIGEN\_STIM\_THROUGH\_IGM\_BCELL\_UP | MSigDB lists | GSE7218\_UNSTIM\_VS\_ANTIGEN\_STIM\_THROUGH\_IGM\_BCELL\_UP | 18 | 3 | 4161 | 164 | fz2,AP-2alpha,Wdr59 | | 3.173e-02 | -3.45 | RICKMAN\_TUMOR\_DIFFERENTIATED\_WELL\_VS\_MODERATELY\_DN | MSigDB lists | RICKMAN\_TUMOR\_DIFFERENTIATED\_WELL\_VS\_MODERATELY\_DN | 18 | 3 | 4161 | 164 | CG9389,inc,fray | | 3.173e-02 | -3.45 | GO\_ENDOTHELIAL\_CELL\_DIFFERENTIATION | MSigDB lists | GO\_ENDOTHELIAL\_CELL\_DIFFERENTIATION | 18 | 3 | 4161 | 164 | svp,cv-2,dnc | | 3.173e-02 | -3.45 | GO\_AMMONIUM\_ION\_BINDING | MSigDB lists | GO\_AMMONIUM\_ION\_BINDING | 18 | 3 | 4161 | 164 | nAChRalpha3,nAChRalpha4,Octbeta2R | | 3.173e-02 | -3.45 | LEE\_NEURAL\_CREST\_STEM\_CELL\_DN | MSigDB lists | LEE\_NEURAL\_CREST\_STEM\_CELL\_DN | 18 | 3 | 4161 | 164 | Dys,ap,fz2 | | 3.173e-02 | -3.45 | MCLACHLAN\_DENTAL\_CARIES\_UP | MSigDB lists | MCLACHLAN\_DENTAL\_CARIES\_UP | 18 | 3 | 4161 | 164 | CG41378,Mp,Glut1 | | 3.173e-02 | -3.45 | GGGCATT\_MIR365 | MSigDB lists | GGGCATT\_MIR365 | 18 | 3 | 4161 | 164 | dnr1,norpA,toy | | 3.177e-02 | -3.45 | - | gene3d domains | 1.20.140.100 | 13 | 3 | 4929 | 273 | CG9492,Dhc16F,kl-3 | | 3.177e-02 | -3.45 | - | gene3d domains | 3.20.180.20 | 13 | 3 | 4929 | 273 | Dhc16F,CG9492,kl-3 | | 3.177e-02 | -3.45 | - | gene3d domains | 3.40.50.10190 | 13 | 3 | 4929 | 273 | Parp,Ptip,DNAlig4 | | 3.177e-02 | -3.45 | - | gene3d domains | 1.10.8.720 | 13 | 3 | 4929 | 273 | Dhc16F,CG9492,kl-3 | | 3.182e-02 | -3.45 | transmembrane transporter activity | molecular function | GO:0022857 | 708 | 43 | 12176 | 554 | ND-MLRQ,CG31689,MCU,CG7255,Ir94e,Ca-alpha1D,CG43672,Ir60e,Tim17b,Ir41a,CG14160,Oatp33Ea,sbm,CG14196,nAChRalpha3,CG33233,Hmt-1,Pzl,CG5789,CG45782,Ir40a,MFS16,nAChRalpha4,CG5254,CG15408,ABCB7,HisCl1,CG8249,Mct1,CG1213,CG14621,Ca-Ma2d,PMCA,cpx,Atpalpha,Catsup,Glut1,Tim23,CG7536,Rift,shakB,KCNQ,SK | | 3.193e-02 | -3.44 | GO\_CILIARY\_PLASM | MSigDB lists | GO\_CILIARY\_PLASM | 31 | 4 | 4161 | 164 | CG17387,Dhc16F,CG9492,l(2)41Ab | | 3.193e-02 | -3.44 | GSE21546\_ELK1\_KO\_VS\_SAP1A\_KO\_AND\_ELK1\_KO\_DP\_THYMOCYTES\_UP | MSigDB lists | GSE21546\_ELK1\_KO\_VS\_SAP1A\_KO\_AND\_ELK1\_KO\_DP\_THYMOCYTES\_UP | 31 | 4 | 4161 | 164 | CG10077,PNPase,CG41378,CG5254 | | 3.193e-02 | -3.44 | GO\_REACTIVE\_OXYGEN\_SPECIES\_METABOLIC\_PROCESS | MSigDB lists | GO\_REACTIVE\_OXYGEN\_SPECIES\_METABOLIC\_PROCESS | 31 | 4 | 4161 | 164 | ND-30,Sod1,arg,CG7255 | | 3.193e-02 | -3.44 | GSE15324\_ELF4\_KO\_VS\_WT\_NAIVE\_CD8\_TCELL\_DN | MSigDB lists | GSE15324\_ELF4\_KO\_VS\_WT\_NAIVE\_CD8\_TCELL\_DN | 31 | 4 | 4161 | 164 | CG17387,Ca-alpha1D,DNaseII,scro | | 3.193e-02 | -3.44 | ATF2\_UP.V1\_UP | MSigDB lists | ATF2\_UP.V1\_UP | 31 | 4 | 4161 | 164 | DNaseII,unc-13-4A,St1,norpA | | 3.200e-02 | -3.44 | phototransduction | biological process | GO:0007602 | 52 | 6 | 12345 | 571 | Pld,rdgC,Rh3,norpA,shakB,Gprk1 | | 3.211e-02 | -3.44 | dl (dorsal) | protein interactions | 35047 | 74 | 8 | 9204 | 464 | rau,RpL15,dsx,Vsx2,RNASEK,CG3081,vtd,Ulp1 | | 3.280e-02 | -3.42 | RRM\_dom | interpro domains | IPR000504 | 135 | 12 | 11383 | 557 | CG41562,shep,cpo,bru1,Rbp9,orb,Set1,CG34334,heph,CG40813,bru3,eIF4B | | 3.281e-02 | -3.42 | GO\_MICROTUBULE\_BASED\_MOVEMENT | MSigDB lists | GO\_MICROTUBULE\_BASED\_MOVEMENT | 78 | 7 | 4161 | 164 | l(2)41Ab,AP-2alpha,Khc-73,Dhc16F,CG17387,Sod1,CG9492 | | 3.286e-02 | -3.42 | transmembrane transporter complex | cellular component | GO:1902495 | 99 | 9 | 11940 | 534 | Ca-alpha1D,nAChRalpha3,HisCl1,KCNQ,Atpalpha,Ca-Ma2d,MCU,nAChRalpha4,CG33970 | | 3.291e-02 | -3.41 | fly chr63B9-63B9\|3-6 | chromosome location | fly chr63B9-63B9\|3-6 | 1 | 1 | 25010 | 823 | CG14964 | | 3.291e-02 | -3.41 | fly chr51F3-51F3\|2-74 cM | chromosome location | fly chr51F3-51F3\|2-74 cM | 1 | 1 | 25010 | 823 | lncRNA:CR44241 | | 3.291e-02 | -3.41 | fly chr43E5-43E7\|2-57 | chromosome location | fly chr43E5-43E7\|2-57 | 1 | 1 | 25010 | 823 | CG43340 | | 3.291e-02 | -3.41 | fly chr75C4-75C4 | chromosome location | fly chr75C4-75C4 | 1 | 1 | 25010 | 823 | lncRNA:CR46353 | | 3.291e-02 | -3.41 | fly chr99A9-99B1\|3-98 | chromosome location | fly chr99A9-99B1\|3-98 | 1 | 1 | 25010 | 823 | Ptp99A | | 3.291e-02 | -3.41 | fly chr61F2-61F3\|3-0 | chromosome location | fly chr61F2-61F3\|3-0 | 1 | 1 | 25010 | 823 | bab2 | | 3.291e-02 | -3.41 | fly chr41E3-41E4\|2-55 | chromosome location | fly chr41E3-41E4\|2-55 | 1 | 1 | 25010 | 823 | d4 | | 3.291e-02 | -3.41 | fly chr4F2-4F3\|1-10 cM | chromosome location | fly chr4F2-4F3\|1-10 cM | 1 | 1 | 25010 | 823 | rg | | 3.291e-02 | -3.41 | fly chr13E8-13E12\|1-51 cM | chromosome location | fly chr13E8-13E12\|1-51 cM | 1 | 1 | 25010 | 823 | CG8184 | | 3.291e-02 | -3.41 | fly chr78B3-78C1\|3-47 | chromosome location | fly chr78B3-78C1\|3-47 | 1 | 1 | 25010 | 823 | Ac78C | | 3.291e-02 | -3.41 | fly chr89F1-89F3\|3-60 cM | chromosome location | fly chr89F1-89F3\|3-60 cM | 1 | 1 | 25010 | 823 | Mur89F | | 3.291e-02 | -3.41 | fly chr20C2-20C2\|1-66 cM | chromosome location | fly chr20C2-20C2\|1-66 cM | 1 | 1 | 25010 | 823 | CG17601 | | 3.291e-02 | -3.41 | fly chr97A8-97A10\|3-90 | chromosome location | fly chr97A8-97A10\|3-90 | 1 | 1 | 25010 | 823 | CG33970 | | 3.291e-02 | -3.41 | fly chr24C8-24C9\|2-11 | chromosome location | fly chr24C8-24C9\|2-11 | 1 | 1 | 25010 | 823 | capu | | 3.291e-02 | -3.41 | fly chr3C9-3D1\|1-2 | chromosome location | fly chr3C9-3D1\|1-2 | 1 | 1 | 25010 | 823 | dnc | | 3.291e-02 | -3.41 | fly chr47A-47B\|2-62 | chromosome location | fly chr47A-47B\|2-62 | 1 | 1 | 25010 | 823 | lola | | 3.291e-02 | -3.41 | fly chr61C3-61C5\|3-0.5 | chromosome location | fly chr61C3-61C5\|3-0.5 | 1 | 1 | 25010 | 823 | Hipk | | 3.291e-02 | -3.41 | fly chr46F3-46F5\|2-62 | chromosome location | fly chr46F3-46F5\|2-62 | 1 | 1 | 25010 | 823 | KCNQ | | 3.291e-02 | -3.41 | fly chr91D3-91D3\|3-65 | chromosome location | fly chr91D3-91D3\|3-65 | 1 | 1 | 25010 | 823 | CG14298 | | 3.291e-02 | -3.41 | fly chr25F5-26A1\|2-17 | chromosome location | fly chr25F5-26A1\|2-17 | 1 | 1 | 25010 | 823 | TrissinR | | 3.291e-02 | -3.41 | fly chr48A2-48A3\|2-64 cM | chromosome location | fly chr48A2-48A3\|2-64 cM | 1 | 1 | 25010 | 823 | tou | | 3.291e-02 | -3.41 | fly chr82B4-82C2\|3-47.1 | chromosome location | fly chr82B4-82C2\|3-47.1 | 1 | 1 | 25010 | 823 | CG34357 | | 3.291e-02 | -3.41 | fly chr50F1-50F1\|2-70 cM | chromosome location | fly chr50F1-50F1\|2-70 cM | 1 | 1 | 25010 | 823 | tej | | 3.291e-02 | -3.41 | fly chr85E2-85E3\|3-49 | chromosome location | fly chr85E2-85E3\|3-49 | 1 | 1 | 25010 | 823 | CG9492 | | 3.291e-02 | -3.41 | fly chr64C12-64C13\|3-13 cM | chromosome location | fly chr64C12-64C13\|3-13 cM | 1 | 1 | 25010 | 823 | lama | | 3.291e-02 | -3.41 | fly chr102D5-102E1\|4-0 cM | chromosome location | fly chr102D5-102E1\|4-0 cM | 1 | 1 | 25010 | 823 | bt | | 3.291e-02 | -3.41 | fly chr94C2-94C3\|3-76 cM | chromosome location | fly chr94C2-94C3\|3-76 cM | 1 | 1 | 25010 | 823 | wge | | 3.291e-02 | -3.41 | fly chr9C1-9C1\|1-31 cM | chromosome location | fly chr9C1-9C1\|1-31 cM | 1 | 1 | 25010 | 823 | RabX2 | | 3.291e-02 | -3.41 | fly chr50C3-50C3\|2-68 cM | chromosome location | fly chr50C3-50C3\|2-68 cM | 1 | 1 | 25010 | 823 | CG6280 | | 3.291e-02 | -3.41 | fly chr67D1-67D2\|3-32 cM | chromosome location | fly chr67D1-67D2\|3-32 cM | 1 | 1 | 25010 | 823 | bma | | 3.291e-02 | -3.41 | fly chr46C4-46C7\|2-61 | chromosome location | fly chr46C4-46C7\|2-61 | 1 | 1 | 25010 | 823 | Mef2 | | 3.291e-02 | -3.41 | fly chr66E4-66E5\|3-27 | chromosome location | fly chr66E4-66E5\|3-27 | 1 | 1 | 25010 | 823 | CG43783 | | 3.291e-02 | -3.41 | fly chr86A7-86A8\|3-49 | chromosome location | fly chr86A7-86A8\|3-49 | 1 | 1 | 25010 | 823 | Bruce | | 3.291e-02 | -3.41 | fly chr52A11-52A12\|2-74 | chromosome location | fly chr52A11-52A12\|2-74 | 1 | 1 | 25010 | 823 | CG12963 | | 3.291e-02 | -3.41 | fly chr75F9-76A3\|3-45 | chromosome location | fly chr75F9-76A3\|3-45 | 1 | 1 | 25010 | 823 | fz2 | | 3.291e-02 | -3.41 | fly chr86C1-86C3\|3-50 cM | chromosome location | fly chr86C1-86C3\|3-50 cM | 1 | 1 | 25010 | 823 | hth | | 3.291e-02 | -3.41 | fly chr80C3-80C3\|3-47 | chromosome location | fly chr80C3-80C3\|3-47 | 1 | 1 | 25010 | 823 | nrm | | 3.291e-02 | -3.41 | fly chr87C1-87C2\|3-52 cM | chromosome location | fly chr87C1-87C2\|3-52 cM | 1 | 1 | 25010 | 823 | Octbeta2R | | 3.291e-02 | -3.41 | fly chr91A7-91B3\|3-63 cM | chromosome location | fly chr91A7-91B3\|3-63 cM | 1 | 1 | 25010 | 823 | fru | | 3.291e-02 | -3.41 | fly chr61C3-61C5\|3-0.5 cM | chromosome location | fly chr61C3-61C5\|3-0.5 cM | 1 | 1 | 25010 | 823 | Hipk | | 3.291e-02 | -3.41 | fly chr40A4-40A4\|2-54 cM | chromosome location | fly chr40A4-40A4\|2-54 cM | 1 | 1 | 25010 | 823 | lncRNA:CR44786 | | 3.291e-02 | -3.41 | fly chr70C12-70C14\|3-41 | chromosome location | fly chr70C12-70C14\|3-41 | 1 | 1 | 25010 | 823 | Ptip | | 3.291e-02 | -3.41 | fly chr2E1-2E2\|1-0.8 cM | chromosome location | fly chr2E1-2E2\|1-0.8 cM | 1 | 1 | 25010 | 823 | Mct1 | | 3.291e-02 | -3.41 | fly chr95B7-95C1\|3-80 | chromosome location | fly chr95B7-95C1\|3-80 | 1 | 1 | 25010 | 823 | CG33111 | | 3.291e-02 | -3.41 | fly chr74F1-74F1\|3-45 cM | chromosome location | fly chr74F1-74F1\|3-45 cM | 1 | 1 | 25010 | 823 | lncRNA:CR45976 | | 3.291e-02 | -3.41 | fly chr92C5-92C5\|3-67 cM | chromosome location | fly chr92C5-92C5\|3-67 cM | 1 | 1 | 25010 | 823 | Rh3 | | 3.291e-02 | -3.41 | fly chr15A11-15B1\|1-55 | chromosome location | fly chr15A11-15B1\|1-55 | 1 | 1 | 25010 | 823 | RSG7 | | 3.291e-02 | -3.41 | fly chr65B4-65B5\|3-17 cM | chromosome location | fly chr65B4-65B5\|3-17 cM | 1 | 1 | 25010 | 823 | MCU | | 3.291e-02 | -3.41 | fly chr10E3-10E4\|1-36 cM | chromosome location | fly chr10E3-10E4\|1-36 cM | 1 | 1 | 25010 | 823 | Hsc70-3 | | 3.291e-02 | -3.41 | fly chr43E7-43E8\|2-57 | chromosome location | fly chr43E7-43E8\|2-57 | 1 | 1 | 25010 | 823 | CG30495 | | 3.291e-02 | -3.41 | fly chr4F5-4F9\|1-11 cM | chromosome location | fly chr4F5-4F9\|1-11 cM | 1 | 1 | 25010 | 823 | SK | | 3.291e-02 | -3.41 | fly chr60E10-60E10\|2-107 | chromosome location | fly chr60E10-60E10\|2-107 | 1 | 1 | 25010 | 823 | lncRNA:CR44830 | | 3.291e-02 | -3.41 | fly chr41E1-41E3\|2-55 cM | chromosome location | fly chr41E1-41E3\|2-55 cM | 1 | 1 | 25010 | 823 | Nipped-A | | 3.291e-02 | -3.41 | fly chr41C6-41D1\|2-55 | chromosome location | fly chr41C6-41D1\|2-55 | 1 | 1 | 25010 | 823 | Ir41a | | 3.291e-02 | -3.41 | fly chr62C4-62D1\|3-1.5 | chromosome location | fly chr62C4-62D1\|3-1.5 | 1 | 1 | 25010 | 823 | zormin | | 3.291e-02 | -3.41 | fly chr18F2-18F4\|1-64 | chromosome location | fly chr18F2-18F4\|1-64 | 1 | 1 | 25010 | 823 | pico | | 3.291e-02 | -3.41 | fly chr90D1-90E1\|3-62 cM | chromosome location | fly chr90D1-90E1\|3-62 cM | 1 | 1 | 25010 | 823 | cpo | | 3.291e-02 | -3.41 | fly chr11B12-11B12\|1-39 cM | chromosome location | fly chr11B12-11B12\|1-39 cM | 1 | 1 | 25010 | 823 | CG15725 | | 3.291e-02 | -3.41 | fly chr88C10-88D1\|3-55 cM | chromosome location | fly chr88C10-88D1\|3-55 cM | 1 | 1 | 25010 | 823 | Meltrin | | 3.291e-02 | -3.41 | fly chr65D3-65D4\|3-18 | chromosome location | fly chr65D3-65D4\|3-18 | 1 | 1 | 25010 | 823 | CG10077 | | 3.291e-02 | -3.41 | fly chr94C4-94C8\|3-76 | chromosome location | fly chr94C4-94C8\|3-76 | 1 | 1 | 25010 | 823 | CG7029 | | 3.291e-02 | -3.41 | fly chr85A3-85A5\|3-48 cM | chromosome location | fly chr85A3-85A5\|3-48 cM | 1 | 1 | 25010 | 823 | 5-HT2B | | 3.291e-02 | -3.41 | fly chr73F3-73F3\|3-45 | chromosome location | fly chr73F3-73F3\|3-45 | 1 | 1 | 25010 | 823 | asRNA:CR43433 | | 3.291e-02 | -3.41 | fly chr62E4-62E5\|3-3 cM | chromosome location | fly chr62E4-62E5\|3-3 cM | 1 | 1 | 25010 | 823 | Spn | | 3.291e-02 | -3.41 | fly chr82A1-82A3\|3-47.1 cM | chromosome location | fly chr82A1-82A3\|3-47.1 cM | 1 | 1 | 25010 | 823 | cpx | | 3.291e-02 | -3.41 | fly chr67D7-67D8\|3-32 | chromosome location | fly chr67D7-67D8\|3-32 | 1 | 1 | 25010 | 823 | CG14160 | | 3.291e-02 | -3.41 | fly chr25D5-25D5\|2-16 cM | chromosome location | fly chr25D5-25D5\|2-16 cM | 1 | 1 | 25010 | 823 | CG14024 | | 3.291e-02 | -3.41 | fly chr6C13-6C13\|1-18 cM | chromosome location | fly chr6C13-6C13\|1-18 cM | 1 | 1 | 25010 | 823 | Coq7 | | 3.291e-02 | -3.41 | fly chr47F17-48A1\|2-64 | chromosome location | fly chr47F17-48A1\|2-64 | 1 | 1 | 25010 | 823 | en | | 3.291e-02 | -3.41 | fly chr33F4-33F4\|2-47 cM | chromosome location | fly chr33F4-33F4\|2-47 cM | 1 | 1 | 25010 | 823 | mir-9382 | | 3.291e-02 | -3.41 | fly chr41F9-41F8\|2-55 | chromosome location | fly chr41F9-41F8\|2-55 | 1 | 1 | 25010 | 823 | vlc | | 3.291e-02 | -3.41 | fly chr41F2-41F2\|2-55 cM | chromosome location | fly chr41F2-41F2\|2-55 cM | 1 | 1 | 25010 | 823 | CG30440 | | 3.291e-02 | -3.41 | fly chr4F2-4F3\|1-10 | chromosome location | fly chr4F2-4F3\|1-10 | 1 | 1 | 25010 | 823 | rg | | 3.291e-02 | -3.41 | fly chr80B2-80B3\|3-47 | chromosome location | fly chr80B2-80B3\|3-47 | 1 | 1 | 25010 | 823 | CG12768 | | 3.291e-02 | -3.41 | fly chr43E7-43E8\|2-57 cM | chromosome location | fly chr43E7-43E8\|2-57 cM | 1 | 1 | 25010 | 823 | CG30495 | | 3.291e-02 | -3.41 | fly chr57D12-57E1\|2-95 | chromosome location | fly chr57D12-57E1\|2-95 | 1 | 1 | 25010 | 823 | cv-2 | | 3.291e-02 | -3.41 | fly chr61A5-61A5\|3-1 | chromosome location | fly chr61A5-61A5\|3-1 | 1 | 1 | 25010 | 823 | mthl8 | | 3.291e-02 | -3.41 | fly chr68D4-68D6\|3-36 | chromosome location | fly chr68D4-68D6\|3-36 | 1 | 1 | 25010 | 823 | Pi3K68D | | 3.291e-02 | -3.41 | fly chr35A4-35B1\|2-50 cM | chromosome location | fly chr35A4-35B1\|2-50 cM | 1 | 1 | 25010 | 823 | spel1 | | 3.291e-02 | -3.41 | fly chr61A5-61A5\|3-1 cM | chromosome location | fly chr61A5-61A5\|3-1 cM | 1 | 1 | 25010 | 823 | mthl8 | | 3.291e-02 | -3.41 | fly chr87B6-87B7\|3-52 | chromosome location | fly chr87B6-87B7\|3-52 | 1 | 1 | 25010 | 823 | CG10013 | | 3.291e-02 | -3.41 | fly chr29F3-29F4\|2-34 | chromosome location | fly chr29F3-29F4\|2-34 | 1 | 1 | 25010 | 823 | CG34398 | | 3.291e-02 | -3.41 | fly chr99A9-99B1\|3-98 cM | chromosome location | fly chr99A9-99B1\|3-98 cM | 1 | 1 | 25010 | 823 | Ptp99A | | 3.291e-02 | -3.41 | fly chr9F8-9F11\|1-33 cM | chromosome location | fly chr9F8-9F11\|1-33 cM | 1 | 1 | 25010 | 823 | CG43901 | | 3.291e-02 | -3.41 | fly chr40F3-40F3\|2-54 cM | chromosome location | fly chr40F3-40F3\|2-54 cM | 1 | 1 | 25010 | 823 | Ir40a | | 3.291e-02 | -3.41 | fly chr80B2-80B3\|3-47 cM | chromosome location | fly chr80B2-80B3\|3-47 cM | 1 | 1 | 25010 | 823 | CG12768 | | 3.291e-02 | -3.41 | fly chr46F5-46F5\|2-62 cM | chromosome location | fly chr46F5-46F5\|2-62 cM | 1 | 1 | 25010 | 823 | CG34221 | | 3.291e-02 | -3.41 | fly chr70C12-70C14\|3-41 cM | chromosome location | fly chr70C12-70C14\|3-41 cM | 1 | 1 | 25010 | 823 | Ptip | | 3.291e-02 | -3.41 | fly chr50F1-50F1\|2-70 | chromosome location | fly chr50F1-50F1\|2-70 | 1 | 1 | 25010 | 823 | tej | | 3.291e-02 | -3.41 | fly chr64C7-64C11\|3-13 | chromosome location | fly chr64C7-64C11\|3-13 | 1 | 1 | 25010 | 823 | shep | | 3.291e-02 | -3.41 | fly chr32D2-32D3\|2-44 | chromosome location | fly chr32D2-32D3\|2-44 | 1 | 1 | 25010 | 823 | Wdr59 | | 3.291e-02 | -3.41 | fly chr8C15-8C17\|1-26 cM | chromosome location | fly chr8C15-8C17\|1-26 cM | 1 | 1 | 25010 | 823 | mei-P26 | | 3.291e-02 | -3.41 | fly chr96D2-96D3\|3-87 cM | chromosome location | fly chr96D2-96D3\|3-87 cM | 1 | 1 | 25010 | 823 | alrm | | 3.291e-02 | -3.41 | fly chr41F1-41F1\|2-55 cM | chromosome location | fly chr41F1-41F1\|2-55 cM | 1 | 1 | 25010 | 823 | stw | | 3.291e-02 | -3.41 | fly chr3C9-3D1\|1-2 cM | chromosome location | fly chr3C9-3D1\|1-2 cM | 1 | 1 | 25010 | 823 | dnc | | 3.291e-02 | -3.41 | fly chr70A7-70A8\|3-40 cM | chromosome location | fly chr70A7-70A8\|3-40 cM | 1 | 1 | 25010 | 823 | CG10171 | | 3.291e-02 | -3.41 | fly chr90C6-90C7\|3-61 | chromosome location | fly chr90C6-90C7\|3-61 | 1 | 1 | 25010 | 823 | tinc | | 3.291e-02 | -3.41 | fly chr25A8-25B1\|2-13 | chromosome location | fly chr25A8-25B1\|2-13 | 1 | 1 | 25010 | 823 | mxt | | 3.291e-02 | -3.41 | fly chr14C3-14C4\|1-53 cM | chromosome location | fly chr14C3-14C4\|1-53 cM | 1 | 1 | 25010 | 823 | CG9981 | | 3.291e-02 | -3.41 | fly chr75F9-75F9\|3-45 cM | chromosome location | fly chr75F9-75F9\|3-45 cM | 1 | 1 | 25010 | 823 | Spn75F | | 3.291e-02 | -3.41 | fly chr80A-80B\|2-55 cM | chromosome location | fly chr80A-80B\|2-55 cM | 1 | 1 | 25010 | 823 | Scp1 | | 3.291e-02 | -3.41 | fly chr41F1-41F1\|2-55 | chromosome location | fly chr41F1-41F1\|2-55 | 1 | 1 | 25010 | 823 | stw | | 3.291e-02 | -3.41 | fly chr100E1-100E1\|3-103 cM | chromosome location | fly chr100E1-100E1\|3-103 cM | 1 | 1 | 25010 | 823 | CG2003 | | 3.291e-02 | -3.41 | fly chr22B4-22B6\|2-3 cM | chromosome location | fly chr22B4-22B6\|2-3 cM | 1 | 1 | 25010 | 823 | Wdr62 | | 3.291e-02 | -3.41 | fly chr4F11-5A1\|1-11 | chromosome location | fly chr4F11-5A1\|1-11 | 1 | 1 | 25010 | 823 | SPR | | 3.291e-02 | -3.41 | fly chr25C4-25C6\|2-15 cM | chromosome location | fly chr25C4-25C6\|2-15 cM | 1 | 1 | 25010 | 823 | Pgant5 | | 3.291e-02 | -3.41 | fly chr4C3-4C4\|1-8 | chromosome location | fly chr4C3-4C4\|1-8 | 1 | 1 | 25010 | 823 | bi | | 3.291e-02 | -3.41 | fly chr102B5-102B5\|4-0 cM | chromosome location | fly chr102B5-102B5\|4-0 cM | 1 | 1 | 25010 | 823 | PMCA | | 3.291e-02 | -3.41 | fly chr6F10-7A5\|1-19 | chromosome location | fly chr6F10-7A5\|1-19 | 1 | 1 | 25010 | 823 | CG9650 | | 3.291e-02 | -3.41 | fly chr49E6-49E6\|2-67 | chromosome location | fly chr49E6-49E6\|2-67 | 1 | 1 | 25010 | 823 | Psc | | 3.291e-02 | -3.41 | fly chr22A5-22A5\|2-3 | chromosome location | fly chr22A5-22A5\|2-3 | 1 | 1 | 25010 | 823 | lncRNA:CR45695 | | 3.291e-02 | -3.41 | fly chr66C4-66C5\|3-24 | chromosome location | fly chr66C4-66C5\|3-24 | 1 | 1 | 25010 | 823 | cert | | 3.291e-02 | -3.41 | fly chr87D7-87D8\|3-53 | chromosome location | fly chr87D7-87D8\|3-53 | 1 | 1 | 25010 | 823 | CG7518 | | 3.291e-02 | -3.41 | fly chr65B4-65B5\|3-17 | chromosome location | fly chr65B4-65B5\|3-17 | 1 | 1 | 25010 | 823 | MCU | | 3.291e-02 | -3.41 | fly chr96E4-96E5\|3-88 cM | chromosome location | fly chr96E4-96E5\|3-88 cM | 1 | 1 | 25010 | 823 | CG5079 | | 3.291e-02 | -3.41 | fly chr24C8-24C9\|2-11 cM | chromosome location | fly chr24C8-24C9\|2-11 cM | 1 | 1 | 25010 | 823 | capu | | 3.291e-02 | -3.41 | fly chr80A-80B\|2-55 | chromosome location | fly chr80A-80B\|2-55 | 1 | 1 | 25010 | 823 | Scp1 | | 3.291e-02 | -3.41 | fly chr82B4-82C2\|3-47.1 cM | chromosome location | fly chr82B4-82C2\|3-47.1 cM | 1 | 1 | 25010 | 823 | CG34357 | | 3.291e-02 | -3.41 | fly chr50B9-50B9\|2-68 | chromosome location | fly chr50B9-50B9\|2-68 | 1 | 1 | 25010 | 823 | CG6209 | | 3.291e-02 | -3.41 | fly chr2A3-2A4\|1-0.2 cM | chromosome location | fly chr2A3-2A4\|1-0.2 cM | 1 | 1 | 25010 | 823 | Gr2a | | 3.291e-02 | -3.41 | fly chr41F2-41F2\|2-55 | chromosome location | fly chr41F2-41F2\|2-55 | 1 | 1 | 25010 | 823 | CG30440 | | 3.291e-02 | -3.41 | fly chr40F3-40F3\|2-54 | chromosome location | fly chr40F3-40F3\|2-54 | 1 | 1 | 25010 | 823 | Ir40a | | 3.291e-02 | -3.41 | fly chr3E4-3E5\|1-4 cM | chromosome location | fly chr3E4-3E5\|1-4 cM | 1 | 1 | 25010 | 823 | Mnt | | 3.291e-02 | -3.41 | fly chr10E3-10E4\|1-36 | chromosome location | fly chr10E3-10E4\|1-36 | 1 | 1 | 25010 | 823 | Hsc70-3 | | 3.291e-02 | -3.41 | fly chr85E14-85F2\|3-49 | chromosome location | fly chr85E14-85F2\|3-49 | 1 | 1 | 25010 | 823 | Glut4EF | | 3.291e-02 | -3.41 | fly chr7D1-7D2\|1-22 cM | chromosome location | fly chr7D1-7D2\|1-22 cM | 1 | 1 | 25010 | 823 | sn | | 3.291e-02 | -3.41 | fly chr18F-18F\|1-64 cM | chromosome location | fly chr18F-18F\|1-64 cM | 1 | 1 | 25010 | 823 | zld | | 3.291e-02 | -3.41 | fly chr80D5-80E3\|3-47 cM | chromosome location | fly chr80D5-80E3\|3-47 cM | 1 | 1 | 25010 | 823 | nAChRalpha4 | | 3.291e-02 | -3.41 | fly chr37E4-37E5\|2-54 cM | chromosome location | fly chr37E4-37E5\|2-54 cM | 1 | 1 | 25010 | 823 | CG10188 | | 3.291e-02 | -3.41 | fly chr36A2-36A3\|2-52 | chromosome location | fly chr36A2-36A3\|2-52 | 1 | 1 | 25010 | 823 | CG5953 | | 3.291e-02 | -3.41 | fly chr41A3-41A3\|2-55 | chromosome location | fly chr41A3-41A3\|2-55 | 1 | 1 | 25010 | 823 | Stlk | | 3.291e-02 | -3.41 | fly chr36A2-36A3\|2-52 cM | chromosome location | fly chr36A2-36A3\|2-52 cM | 1 | 1 | 25010 | 823 | CG5953 | | 3.291e-02 | -3.41 | fly chr91E5-91E5\|3-65 | chromosome location | fly chr91E5-91E5\|3-65 | 1 | 1 | 25010 | 823 | lncRNA:CR46041 | | 3.291e-02 | -3.41 | fly chr22B4-22B6\|2-3 | chromosome location | fly chr22B4-22B6\|2-3 | 1 | 1 | 25010 | 823 | Wdr62 | | 3.291e-02 | -3.41 | fly chr22F4-23A1\|2-6 | chromosome location | fly chr22F4-23A1\|2-6 | 1 | 1 | 25010 | 823 | Cyp309a1 | | 3.291e-02 | -3.41 | fly chr96A19-96A19\|3-85 cM | chromosome location | fly chr96A19-96A19\|3-85 cM | 1 | 1 | 25010 | 823 | tld | | 3.291e-02 | -3.41 | fly chr100B-100B\|3-102 cM | chromosome location | fly chr100B-100B\|3-102 cM | 1 | 1 | 25010 | 823 | Prosalpha3T | | 3.291e-02 | -3.41 | fly chr72D1-72D5\|3-44 cM | chromosome location | fly chr72D1-72D5\|3-44 cM | 1 | 1 | 25010 | 823 | Diap1 | | 3.291e-02 | -3.41 | fly chr25F5-26A1\|2-17 cM | chromosome location | fly chr25F5-26A1\|2-17 cM | 1 | 1 | 25010 | 823 | TrissinR | | 3.291e-02 | -3.41 | fly chr40D1-40D2\|2-54 cM | chromosome location | fly chr40D1-40D2\|2-54 cM | 1 | 1 | 25010 | 823 | CG2528 | | 3.291e-02 | -3.41 | fly chr11C2-11C2\|1-40 | chromosome location | fly chr11C2-11C2\|1-40 | 1 | 1 | 25010 | 823 | Nep6 | | 3.291e-02 | -3.41 | fly chr84D2-84D3\|3-48 cM | chromosome location | fly chr84D2-84D3\|3-48 cM | 1 | 1 | 25010 | 823 | thw | | 3.291e-02 | -3.41 | fly chr7D1-7D2\|1-22 | chromosome location | fly chr7D1-7D2\|1-22 | 1 | 1 | 25010 | 823 | sn | | 3.291e-02 | -3.41 | fly chr46F3-46F5\|2-62 cM | chromosome location | fly chr46F3-46F5\|2-62 cM | 1 | 1 | 25010 | 823 | KCNQ | | 3.291e-02 | -3.41 | fly chr85E14-85F2\|3-49 cM | chromosome location | fly chr85E14-85F2\|3-49 cM | 1 | 1 | 25010 | 823 | Glut4EF | | 3.291e-02 | -3.41 | fly chr84E5-84E6\|3-48 cM | chromosome location | fly chr84E5-84E6\|3-48 cM | 1 | 1 | 25010 | 823 | dsx | | 3.291e-02 | -3.41 | fly chr80D2-80D2\|3-47 | chromosome location | fly chr80D2-80D2\|3-47 | 1 | 1 | 25010 | 823 | CG33217 | | 3.291e-02 | -3.41 | fly chr41F-41F8\|2-55 cM | chromosome location | fly chr41F-41F8\|2-55 cM | 1 | 1 | 25010 | 823 | ap | | 3.291e-02 | -3.41 | fly chr32F1-32F2\|2-44 | chromosome location | fly chr32F1-32F2\|2-44 | 1 | 1 | 25010 | 823 | salm | | 3.291e-02 | -3.41 | fly chr56C11-56D1\|2-88 cM | chromosome location | fly chr56C11-56D1\|2-88 cM | 1 | 1 | 25010 | 823 | Rgk1 | | 3.291e-02 | -3.41 | fly chr43C5-43C6\|2-56 | chromosome location | fly chr43C5-43C6\|2-56 | 1 | 1 | 25010 | 823 | CG1620 | | 3.291e-02 | -3.41 | fly chr6A1-6A1\|1-17 cM | chromosome location | fly chr6A1-6A1\|1-17 cM | 1 | 1 | 25010 | 823 | CG33668 | | 3.291e-02 | -3.41 | fly chr5F6-5F6\|1-16 | chromosome location | fly chr5F6-5F6\|1-16 | 1 | 1 | 25010 | 823 | CG33669 | | 3.291e-02 | -3.41 | fly chr46F5-46F5\|2-62 | chromosome location | fly chr46F5-46F5\|2-62 | 1 | 1 | 25010 | 823 | CG34221 | | 3.291e-02 | -3.41 | fly chr91A1-91A2\|3-63 | chromosome location | fly chr91A1-91A2\|3-63 | 1 | 1 | 25010 | 823 | l(3)05822 | | 3.291e-02 | -3.41 | fly chr7E5-7E6\|1-23 | chromosome location | fly chr7E5-7E6\|1-23 | 1 | 1 | 25010 | 823 | CG1387 | | 3.291e-02 | -3.41 | fly chr50C6-50C9\|2-68 | chromosome location | fly chr50C6-50C9\|2-68 | 1 | 1 | 25010 | 823 | shot | | 3.291e-02 | -3.41 | fly chr25B9-25C1\|2-14 cM | chromosome location | fly chr25B9-25C1\|2-14 cM | 1 | 1 | 25010 | 823 | Rtnl1 | | 3.291e-02 | -3.41 | fly chr41E3-41E4\|2-55 cM | chromosome location | fly chr41E3-41E4\|2-55 cM | 1 | 1 | 25010 | 823 | d4 | | 3.291e-02 | -3.41 | fly chr86F7-86F8\|3-51 cM | chromosome location | fly chr86F7-86F8\|3-51 cM | 1 | 1 | 25010 | 823 | HisCl1 | | 3.291e-02 | -3.41 | fly chr3C3-3C7\|1-1.5 cM | chromosome location | fly chr3C3-3C7\|1-1.5 cM | 1 | 1 | 25010 | 823 | kirre | | 3.291e-02 | -3.41 | fly chr13E16-13E17\|1-51 | chromosome location | fly chr13E16-13E17\|1-51 | 1 | 1 | 25010 | 823 | CG43672 | | 3.291e-02 | -3.41 | fly chr89E2-89E4\|3-59 | chromosome location | fly chr89E2-89E4\|3-59 | 1 | 1 | 25010 | 823 | lncRNA:iab8 | | 3.291e-02 | -3.41 | fly chr66C4-66C5\|3-24 cM | chromosome location | fly chr66C4-66C5\|3-24 cM | 1 | 1 | 25010 | 823 | cert | | 3.291e-02 | -3.41 | fly chr80C3-80C3\|3-47 cM | chromosome location | fly chr80C3-80C3\|3-47 cM | 1 | 1 | 25010 | 823 | nrm | | 3.291e-02 | -3.41 | fly chr7B4-7B6\|1-20 | chromosome location | fly chr7B4-7B6\|1-20 | 1 | 1 | 25010 | 823 | ct | | 3.291e-02 | -3.41 | fly chr29E3-29E4\|2-33 | chromosome location | fly chr29E3-29E4\|2-33 | 1 | 1 | 25010 | 823 | Trs23 | | 3.291e-02 | -3.41 | fly chr52E2-52E2\|2-77 | chromosome location | fly chr52E2-52E2\|2-77 | 1 | 1 | 25010 | 823 | lncRNA:CR43415 | | 3.291e-02 | -3.41 | fly chr75B5-75B5\|3-45 cM | chromosome location | fly chr75B5-75B5\|3-45 cM | 1 | 1 | 25010 | 823 | CG42393 | | 3.291e-02 | -3.41 | fly chr72D1-72D5\|3-44 | chromosome location | fly chr72D1-72D5\|3-44 | 1 | 1 | 25010 | 823 | Diap1 | | 3.291e-02 | -3.41 | fly chr11D9-11D9\|1-41 | chromosome location | fly chr11D9-11D9\|1-41 | 1 | 1 | 25010 | 823 | mir-971 | | 3.291e-02 | -3.41 | fly chr69B5-69B5\|3-38 | chromosome location | fly chr69B5-69B5\|3-38 | 1 | 1 | 25010 | 823 | ArfGAP1 | | 3.291e-02 | -3.41 | fly chr99E4-99F2\|3-100 | chromosome location | fly chr99E4-99F2\|3-100 | 1 | 1 | 25010 | 823 | hdc | | 3.291e-02 | -3.41 | fly chr50C3-50C3\|2-68 | chromosome location | fly chr50C3-50C3\|2-68 | 1 | 1 | 25010 | 823 | CG6280 | | 3.291e-02 | -3.41 | fly chr48D8-48E1\|2-66 | chromosome location | fly chr48D8-48E1\|2-66 | 1 | 1 | 25010 | 823 | CG8888 | | 3.291e-02 | -3.41 | fly chr67D1-67D2\|3-32 | chromosome location | fly chr67D1-67D2\|3-32 | 1 | 1 | 25010 | 823 | bma | | 3.291e-02 | -3.41 | fly chr77B9-77C1\|3-47 cM | chromosome location | fly chr77B9-77C1\|3-47 cM | 1 | 1 | 25010 | 823 | fbl | | 3.291e-02 | -3.41 | fly chr85F5-85F7\|3-49 | chromosome location | fly chr85F5-85F7\|3-49 | 1 | 1 | 25010 | 823 | Mical | | 3.291e-02 | -3.41 | fly chr74F1-74F1\|3-45 | chromosome location | fly chr74F1-74F1\|3-45 | 1 | 1 | 25010 | 823 | lncRNA:CR45976 | | 3.291e-02 | -3.41 | fly chr78B3-78C1\|3-47 cM | chromosome location | fly chr78B3-78C1\|3-47 cM | 1 | 1 | 25010 | 823 | Ac78C | | 3.291e-02 | -3.41 | fly chr67D7-67D8\|3-32 cM | chromosome location | fly chr67D7-67D8\|3-32 cM | 1 | 1 | 25010 | 823 | CG14160 | | 3.291e-02 | -3.41 | fly chr96A12-96A13\|3-85 cM | chromosome location | fly chr96A12-96A13\|3-85 cM | 1 | 1 | 25010 | 823 | Syx18 | | 3.291e-02 | -3.41 | fly chr73F3-73F3\|3-45 cM | chromosome location | fly chr73F3-73F3\|3-45 cM | 1 | 1 | 25010 | 823 | asRNA:CR43433 | | 3.291e-02 | -3.41 | fly chr102B6-102B7\|4-0 | chromosome location | fly chr102B6-102B7\|4-0 | 1 | 1 | 25010 | 823 | dati | | 3.291e-02 | -3.41 | fly chr75F9-75F9\|3-45 | chromosome location | fly chr75F9-75F9\|3-45 | 1 | 1 | 25010 | 823 | Spn75F | | 3.291e-02 | -3.41 | fly chr40A1-40A2\|2-54 cM | chromosome location | fly chr40A1-40A2\|2-54 cM | 1 | 1 | 25010 | 823 | step | | 3.291e-02 | -3.41 | fly chr88C10-88D1\|3-55 | chromosome location | fly chr88C10-88D1\|3-55 | 1 | 1 | 25010 | 823 | Meltrin | | 3.291e-02 | -3.41 | fly chr8D5-8D6\|1-27 | chromosome location | fly chr8D5-8D6\|1-27 | 1 | 1 | 25010 | 823 | lz | | 3.291e-02 | -3.41 | fly chr7E5-7E6\|1-23 cM | chromosome location | fly chr7E5-7E6\|1-23 cM | 1 | 1 | 25010 | 823 | CG1387 | | 3.291e-02 | -3.41 | fly chr8D5-8D6\|1-27 cM | chromosome location | fly chr8D5-8D6\|1-27 cM | 1 | 1 | 25010 | 823 | lz | | 3.291e-02 | -3.41 | fly chr46F9-47A1\|2-62 | chromosome location | fly chr46F9-47A1\|2-62 | 1 | 1 | 25010 | 823 | CAP | | 3.291e-02 | -3.41 | fly chr11D9-11D9\|1-41 cM | chromosome location | fly chr11D9-11D9\|1-41 cM | 1 | 1 | 25010 | 823 | mir-971 | | 3.291e-02 | -3.41 | fly chr69B5-69B5\|3-38 cM | chromosome location | fly chr69B5-69B5\|3-38 cM | 1 | 1 | 25010 | 823 | ArfGAP1 | | 3.291e-02 | -3.41 | fly chr37C7-37D1\|2-54 | chromosome location | fly chr37C7-37D1\|2-54 | 1 | 1 | 25010 | 823 | lncRNA:CR44189 | | 3.291e-02 | -3.41 | fly chr70B5-70C4\|3-40 | chromosome location | fly chr70B5-70C4\|3-40 | 1 | 1 | 25010 | 823 | bru3 | | 3.291e-02 | -3.41 | fly chr95B7-95C1\|3-80 cM | chromosome location | fly chr95B7-95C1\|3-80 cM | 1 | 1 | 25010 | 823 | CG33111 | | 3.291e-02 | -3.41 | fly chr3C3-3C7\|1-1.5 | chromosome location | fly chr3C3-3C7\|1-1.5 | 1 | 1 | 25010 | 823 | kirre | | 3.291e-02 | -3.41 | fly chr57F1-57F2\|2-96 cM | chromosome location | fly chr57F1-57F2\|2-96 cM | 1 | 1 | 25010 | 823 | twz | | 3.291e-02 | -3.41 | fly chr3D2-3D4\|1-3 cM | chromosome location | fly chr3D2-3D4\|1-3 cM | 1 | 1 | 25010 | 823 | Myc | | 3.291e-02 | -3.41 | fly chr15C4-15C6\|1-56 | chromosome location | fly chr15C4-15C6\|1-56 | 1 | 1 | 25010 | 823 | Ubr1 | | 3.291e-02 | -3.41 | fly chr90C6-90C7\|3-61 cM | chromosome location | fly chr90C6-90C7\|3-61 cM | 1 | 1 | 25010 | 823 | tinc | | 3.291e-02 | -3.41 | fly chr56D9-56D11\|2-89 cM | chromosome location | fly chr56D9-56D11\|2-89 cM | 1 | 1 | 25010 | 823 | par-1 | | 3.291e-02 | -3.41 | fly chr60B3-60B4\|2-106 cM | chromosome location | fly chr60B3-60B4\|2-106 cM | 1 | 1 | 25010 | 823 | CG3907 | | 3.291e-02 | -3.41 | fly chr4B6-4C1\|1-7 | chromosome location | fly chr4B6-4C1\|1-7 | 1 | 1 | 25010 | 823 | norpA | | 3.291e-02 | -3.41 | fly chr92C5-92C5\|3-67 | chromosome location | fly chr92C5-92C5\|3-67 | 1 | 1 | 25010 | 823 | Rh3 | | 3.291e-02 | -3.41 | fly chr52E10-52E11\|2-77 cM | chromosome location | fly chr52E10-52E11\|2-77 cM | 1 | 1 | 25010 | 823 | CG30094 | | 3.291e-02 | -3.41 | fly chr41B3-41C2\|2-55 cM | chromosome location | fly chr41B3-41C2\|2-55 cM | 1 | 1 | 25010 | 823 | Gprk1 | | 3.291e-02 | -3.41 | fly chr57C2-57C3\|2-93 | chromosome location | fly chr57C2-57C3\|2-93 | 1 | 1 | 25010 | 823 | Magi | | 3.291e-02 | -3.41 | fly chr78B2-78B3\|3-47 cM | chromosome location | fly chr78B2-78B3\|3-47 cM | 1 | 1 | 25010 | 823 | CG10565 | | 3.291e-02 | -3.41 | fly chr35B8-35B9\|2-50 cM | chromosome location | fly chr35B8-35B9\|2-50 cM | 1 | 1 | 25010 | 823 | ck | | 3.291e-02 | -3.41 | fly chr15C4-15C6\|1-56 cM | chromosome location | fly chr15C4-15C6\|1-56 cM | 1 | 1 | 25010 | 823 | Ubr1 | | 3.291e-02 | -3.41 | fly chr95B5-95B7\|3-80 cM | chromosome location | fly chr95B5-95B7\|3-80 cM | 1 | 1 | 25010 | 823 | mbc | | 3.291e-02 | -3.41 | fly chr20C2-20C2\|1-66 | chromosome location | fly chr20C2-20C2\|1-66 | 1 | 1 | 25010 | 823 | CG17601 | | 3.291e-02 | -3.41 | fly chr57E1-57E6\|2-95 | chromosome location | fly chr57E1-57E6\|2-95 | 1 | 1 | 25010 | 823 | Sdc | | 3.291e-02 | -3.41 | fly chr82A6-82B1\|3-47.1 cM | chromosome location | fly chr82A6-82B1\|3-47.1 cM | 1 | 1 | 25010 | 823 | CG32944 | | 3.291e-02 | -3.41 | fly chr62E4-62E5\|3-3 | chromosome location | fly chr62E4-62E5\|3-3 | 1 | 1 | 25010 | 823 | Spn | | 3.291e-02 | -3.41 | fly chr93A4-93A4\|3-69 | chromosome location | fly chr93A4-93A4\|3-69 | 1 | 1 | 25010 | 823 | Atpalpha | | 3.291e-02 | -3.41 | fly chr37C7-37D1\|2-54 cM | chromosome location | fly chr37C7-37D1\|2-54 cM | 1 | 1 | 25010 | 823 | lncRNA:CR44189 | | 3.291e-02 | -3.41 | fly chr80C2-80C2\|3-47 | chromosome location | fly chr80C2-80C2\|3-47 | 1 | 1 | 25010 | 823 | CG32457 | | 3.291e-02 | -3.41 | fly chr33D2-33D5\|2-46 | chromosome location | fly chr33D2-33D5\|2-46 | 1 | 1 | 25010 | 823 | bru1 | | 3.291e-02 | -3.41 | fly chr60E10-60E10\|2-107 cM | chromosome location | fly chr60E10-60E10\|2-107 cM | 1 | 1 | 25010 | 823 | lncRNA:CR44830 | | 3.291e-02 | -3.41 | fly chr92A5-92A10\|3-66 cM | chromosome location | fly chr92A5-92A10\|3-66 cM | 1 | 1 | 25010 | 823 | Dys | | 3.291e-02 | -3.41 | fly chr75B5-75B5\|3-45 | chromosome location | fly chr75B5-75B5\|3-45 | 1 | 1 | 25010 | 823 | CG42393 | | 3.291e-02 | -3.41 | fly chr41E1-41E3\|2-55 | chromosome location | fly chr41E1-41E3\|2-55 | 1 | 1 | 25010 | 823 | Nipped-A | | 3.291e-02 | -3.41 | fly chr57C2-57C3\|2-93 cM | chromosome location | fly chr57C2-57C3\|2-93 cM | 1 | 1 | 25010 | 823 | Magi | | 3.291e-02 | -3.41 | fly chr54B6-54B7\|2-82 cM | chromosome location | fly chr54B6-54B7\|2-82 cM | 1 | 1 | 25010 | 823 | CG10939 | | 3.291e-02 | -3.41 | fly chr14A5-14A6\|1-52 | chromosome location | fly chr14A5-14A6\|1-52 | 1 | 1 | 25010 | 823 | vap | | 3.291e-02 | -3.41 | fly chr13A8-13A9\|1-49 | chromosome location | fly chr13A8-13A9\|1-49 | 1 | 1 | 25010 | 823 | Lsd-2 | | 3.291e-02 | -3.41 | fly chr53D1-53D4\|2-80 | chromosome location | fly chr53D1-53D4\|2-80 | 1 | 1 | 25010 | 823 | Pgant9 | | 3.291e-02 | -3.41 | fly chr96C6-96C7\|3-87 cM | chromosome location | fly chr96C6-96C7\|3-87 cM | 1 | 1 | 25010 | 823 | Alg9 | | 3.291e-02 | -3.41 | fly chr48A2-48A3\|2-64 | chromosome location | fly chr48A2-48A3\|2-64 | 1 | 1 | 25010 | 823 | tou | | 3.291e-02 | -3.41 | fly chr87B6-87B7\|3-52 cM | chromosome location | fly chr87B6-87B7\|3-52 cM | 1 | 1 | 25010 | 823 | CG10013 | | 3.291e-02 | -3.41 | fly chr83C6-83C8\|3-47.5 | chromosome location | fly chr83C6-83C8\|3-47.5 | 1 | 1 | 25010 | 823 | side-III | | 3.291e-02 | -3.41 | fly chr8A1-8A1\|1-24 | chromosome location | fly chr8A1-8A1\|1-24 | 1 | 1 | 25010 | 823 | CG12772 | | 3.291e-02 | -3.41 | fly chr14C3-14C4\|1-53 | chromosome location | fly chr14C3-14C4\|1-53 | 1 | 1 | 25010 | 823 | CG9981 | | 3.291e-02 | -3.41 | fly chr57E1-57E6\|2-95 cM | chromosome location | fly chr57E1-57E6\|2-95 cM | 1 | 1 | 25010 | 823 | Sdc | | 3.291e-02 | -3.41 | fly chr49E6-49E6\|2-67 cM | chromosome location | fly chr49E6-49E6\|2-67 cM | 1 | 1 | 25010 | 823 | Psc | | 3.291e-02 | -3.41 | fly chr20A1-20A2\|1-66 | chromosome location | fly chr20A1-20A2\|1-66 | 1 | 1 | 25010 | 823 | lncRNA:flam | | 3.291e-02 | -3.41 | fly chr9C1-9C1\|1-31 | chromosome location | fly chr9C1-9C1\|1-31 | 1 | 1 | 25010 | 823 | RabX2 | | 3.291e-02 | -3.41 | fly chr12E10-12E10\|1-48 | chromosome location | fly chr12E10-12E10\|1-48 | 1 | 1 | 25010 | 823 | CG9411 | | 3.291e-02 | -3.41 | fly chr49A1-49A2\|2-67 cM | chromosome location | fly chr49A1-49A2\|2-67 cM | 1 | 1 | 25010 | 823 | Cpr49Ac | | 3.291e-02 | -3.41 | fly chr35B3-35B4\|2-50 cM | chromosome location | fly chr35B3-35B4\|2-50 cM | 1 | 1 | 25010 | 823 | osp | | 3.291e-02 | -3.41 | fly chr52F3-52F3\|2-78 | chromosome location | fly chr52F3-52F3\|2-78 | 1 | 1 | 25010 | 823 | CG8435 | | 3.291e-02 | -3.41 | fly chr94B1-94B2\|3-75 cM | chromosome location | fly chr94B1-94B2\|3-75 cM | 1 | 1 | 25010 | 823 | Octbeta1R | | 3.291e-02 | -3.41 | fly chr70B2-70B3\|3-40 | chromosome location | fly chr70B2-70B3\|3-40 | 1 | 1 | 25010 | 823 | Tgi | | 3.291e-02 | -3.41 | fly chr70C10-70C11\|3-41 cM | chromosome location | fly chr70C10-70C11\|3-41 cM | 1 | 1 | 25010 | 823 | CG17361 | | 3.291e-02 | -3.41 | fly chr80D2-80D2\|3-47 cM | chromosome location | fly chr80D2-80D2\|3-47 cM | 1 | 1 | 25010 | 823 | CG33217 | | 3.291e-02 | -3.41 | fly chr32D2-32D3\|2-44 cM | chromosome location | fly chr32D2-32D3\|2-44 cM | 1 | 1 | 25010 | 823 | Wdr59 | | 3.291e-02 | -3.41 | fly chr15B1-15B3\|1-56 cM | chromosome location | fly chr15B1-15B3\|1-56 cM | 1 | 1 | 25010 | 823 | CG13003 | | 3.291e-02 | -3.41 | fly chr99E5-99E5\|3-100 cM | chromosome location | fly chr99E5-99E5\|3-100 cM | 1 | 1 | 25010 | 823 | CG34300 | | 3.291e-02 | -3.41 | fly chr19C6-19D1\|1-64 cM | chromosome location | fly chr19C6-19D1\|1-64 cM | 1 | 1 | 25010 | 823 | CG15459 | | 3.291e-02 | -3.41 | fly chr86F7-86F8\|3-51 | chromosome location | fly chr86F7-86F8\|3-51 | 1 | 1 | 25010 | 823 | HisCl1 | | 3.291e-02 | -3.41 | fly chr84D2-84D3\|3-48 | chromosome location | fly chr84D2-84D3\|3-48 | 1 | 1 | 25010 | 823 | thw | | 3.291e-02 | -3.41 | fly chr99E5-99E5\|3-100 | chromosome location | fly chr99E5-99E5\|3-100 | 1 | 1 | 25010 | 823 | CG34300 | | 3.291e-02 | -3.41 | fly chr99E1-99E2\|3-100 | chromosome location | fly chr99E1-99E2\|3-100 | 1 | 1 | 25010 | 823 | CG9747 | | 3.291e-02 | -3.41 | fly chr4F11-5A1\|1-11 cM | chromosome location | fly chr4F11-5A1\|1-11 cM | 1 | 1 | 25010 | 823 | SPR | | 3.291e-02 | -3.41 | fly chr77A4-77B1\|3-47 cM | chromosome location | fly chr77A4-77B1\|3-47 cM | 1 | 1 | 25010 | 823 | rdgC | | 3.291e-02 | -3.41 | fly chr94C9-94C9\|3-76 | chromosome location | fly chr94C9-94C9\|3-76 | 1 | 1 | 25010 | 823 | lncRNA:CR44350 | | 3.291e-02 | -3.41 | fly chr33F4-33F4\|2-47 | chromosome location | fly chr33F4-33F4\|2-47 | 1 | 1 | 25010 | 823 | mir-9382 | | 3.291e-02 | -3.41 | fly chr95B5-95B7\|3-80 | chromosome location | fly chr95B5-95B7\|3-80 | 1 | 1 | 25010 | 823 | mbc | | 3.291e-02 | -3.41 | fly chr65E10-65F4\|3-19 | chromosome location | fly chr65E10-65F4\|3-19 | 1 | 1 | 25010 | 823 | unc-13-4A | | 3.291e-02 | -3.41 | fly chr45D5-45D7\|2-60 cM | chromosome location | fly chr45D5-45D7\|2-60 cM | 1 | 1 | 25010 | 823 | Pdk | | 3.291e-02 | -3.41 | fly chr25D5-25D5\|2-16 | chromosome location | fly chr25D5-25D5\|2-16 | 1 | 1 | 25010 | 823 | CG14024 | | 3.291e-02 | -3.41 | fly chr6E3-6E4\|1-18 | chromosome location | fly chr6E3-6E4\|1-18 | 1 | 1 | 25010 | 823 | Setd3 | | 3.291e-02 | -3.41 | fly chr85F5-85F7\|3-49 cM | chromosome location | fly chr85F5-85F7\|3-49 cM | 1 | 1 | 25010 | 823 | Mical | | 3.291e-02 | -3.41 | fly chr66D14-66D14\|3-26 cM | chromosome location | fly chr66D14-66D14\|3-26 cM | 1 | 1 | 25010 | 823 | Acbp5 | | 3.291e-02 | -3.41 | fly chr99E1-99E2\|3-100 cM | chromosome location | fly chr99E1-99E2\|3-100 cM | 1 | 1 | 25010 | 823 | CG9747 | | 3.291e-02 | -3.41 | fly chr43C4-43C4\|2-56 | chromosome location | fly chr43C4-43C4\|2-56 | 1 | 1 | 25010 | 823 | Aldh-III | | 3.291e-02 | -3.41 | fly chr11C2-11C2\|1-40 cM | chromosome location | fly chr11C2-11C2\|1-40 cM | 1 | 1 | 25010 | 823 | Nep6 | | 3.291e-02 | -3.41 | fly chr68D4-68D6\|3-36 cM | chromosome location | fly chr68D4-68D6\|3-36 cM | 1 | 1 | 25010 | 823 | Pi3K68D | | 3.291e-02 | -3.41 | fly chr29F3-29F4\|2-34 cM | chromosome location | fly chr29F3-29F4\|2-34 cM | 1 | 1 | 25010 | 823 | CG34398 | | 3.291e-02 | -3.41 | fly chrh8-h8 | chromosome location | fly chrh8-h8 | 1 | 1 | 25010 | 823 | kl-3 | | 3.291e-02 | -3.41 | fly chr94B1-94B2\|3-75 | chromosome location | fly chr94B1-94B2\|3-75 | 1 | 1 | 25010 | 823 | Octbeta1R | | 3.291e-02 | -3.41 | fly chr94A2-94A4\|3-74 | chromosome location | fly chr94A2-94A4\|3-74 | 1 | 1 | 25010 | 823 | SKIP | | 3.291e-02 | -3.41 | fly chr42B1-42B2\|2-55 | chromosome location | fly chr42B1-42B2\|2-55 | 1 | 1 | 25010 | 823 | jing | | 3.291e-02 | -3.41 | fly chr43E5-43E7\|2-57 cM | chromosome location | fly chr43E5-43E7\|2-57 cM | 1 | 1 | 25010 | 823 | CG43340 | | 3.291e-02 | -3.41 | fly chr15B1-15B3\|1-56 | chromosome location | fly chr15B1-15B3\|1-56 | 1 | 1 | 25010 | 823 | CG13003 | | 3.291e-02 | -3.41 | fly chr28B4-28C1\|2-27 | chromosome location | fly chr28B4-28C1\|2-27 | 1 | 1 | 25010 | 823 | Slob | | 3.291e-02 | -3.41 | fly chr1C3-1C4\|1-0 cM | chromosome location | fly chr1C3-1C4\|1-0 cM | 1 | 1 | 25010 | 823 | CG5254 | | 3.291e-02 | -3.41 | fly chr60B3-60B4\|2-106 | chromosome location | fly chr60B3-60B4\|2-106 | 1 | 1 | 25010 | 823 | CG3907 | | 3.291e-02 | -3.41 | fly chr27C1-27C4\|2-23 | chromosome location | fly chr27C1-27C4\|2-23 | 1 | 1 | 25010 | 823 | Nlg2 | | 3.291e-02 | -3.41 | fly chr42F3-43A1\|2-55 cM | chromosome location | fly chr42F3-43A1\|2-55 cM | 1 | 1 | 25010 | 823 | pk | | 3.291e-02 | -3.41 | fly chr102B6-102B7\|4-0 cM | chromosome location | fly chr102B6-102B7\|4-0 cM | 1 | 1 | 25010 | 823 | dati | | 3.291e-02 | -3.41 | fly chr101F1-102A4\|4-0 cM | chromosome location | fly chr101F1-102A4\|4-0 cM | 1 | 1 | 25010 | 823 | pan | | 3.291e-02 | -3.41 | fly chr83C6-83C7\|3-47.5 cM | chromosome location | fly chr83C6-83C7\|3-47.5 cM | 1 | 1 | 25010 | 823 | lncRNA:CR45761 | | 3.291e-02 | -3.41 | fly chr7D5-7D6\|1-22 | chromosome location | fly chr7D5-7D6\|1-22 | 1 | 1 | 25010 | 823 | CG2256 | | 3.291e-02 | -3.41 | fly chr64C12-64C13\|3-13 | chromosome location | fly chr64C12-64C13\|3-13 | 1 | 1 | 25010 | 823 | lama | | 3.291e-02 | -3.41 | fly chr42A6-42A7\|2-55 cM | chromosome location | fly chr42A6-42A7\|2-55 cM | 1 | 1 | 25010 | 823 | Src42A | | 3.291e-02 | -3.41 | fly chr91D3-91D3\|3-65 cM | chromosome location | fly chr91D3-91D3\|3-65 cM | 1 | 1 | 25010 | 823 | CG14298 | | 3.291e-02 | -3.41 | fly chr12B2-12B4\|1-45 | chromosome location | fly chr12B2-12B4\|1-45 | 1 | 1 | 25010 | 823 | Nadsyn | | 3.291e-02 | -3.41 | fly chr41C5-41C5\|2-55 cM | chromosome location | fly chr41C5-41C5\|2-55 cM | 1 | 1 | 25010 | 823 | CG44102 | | 3.291e-02 | -3.41 | fly chr80C2-80C2\|3-47 cM | chromosome location | fly chr80C2-80C2\|3-47 cM | 1 | 1 | 25010 | 823 | CG32457 | | 3.291e-02 | -3.41 | fly chr41A3-41A3\|2-55 cM | chromosome location | fly chr41A3-41A3\|2-55 cM | 1 | 1 | 25010 | 823 | Stlk | | 3.291e-02 | -3.41 | fly chr62A8-62A9\|3-0.5 | chromosome location | fly chr62A8-62A9\|3-0.5 | 1 | 1 | 25010 | 823 | ABCB7 | | 3.291e-02 | -3.41 | fly chr64B1-64B2\|3-12 cM | chromosome location | fly chr64B1-64B2\|3-12 cM | 1 | 1 | 25010 | 823 | Cip4 | | 3.291e-02 | -3.41 | fly chr41C6-41D1\|2-55 cM | chromosome location | fly chr41C6-41D1\|2-55 cM | 1 | 1 | 25010 | 823 | Ir41a | | 3.291e-02 | -3.41 | fly chr4B6-4C1\|1-7 cM | chromosome location | fly chr4B6-4C1\|1-7 cM | 1 | 1 | 25010 | 823 | norpA | | 3.291e-02 | -3.41 | fly chr35E5-35E6\|2-51 cM | chromosome location | fly chr35E5-35E6\|2-51 cM | 1 | 1 | 25010 | 823 | Ca-alpha1D | | 3.291e-02 | -3.41 | fly chr25A8-25B1\|2-13 cM | chromosome location | fly chr25A8-25B1\|2-13 cM | 1 | 1 | 25010 | 823 | mxt | | 3.291e-02 | -3.41 | fly chr75B1-75B1\|3-45 cM | chromosome location | fly chr75B1-75B1\|3-45 cM | 1 | 1 | 25010 | 823 | lncRNA:CR45921 | | 3.291e-02 | -3.41 | fly chr41D4-41E1\|2-55 | chromosome location | fly chr41D4-41E1\|2-55 | 1 | 1 | 25010 | 823 | Atf6 | | 3.291e-02 | -3.41 | fly chr89E2-89E4\|3-59 cM | chromosome location | fly chr89E2-89E4\|3-59 cM | 1 | 1 | 25010 | 823 | lncRNA:iab8 | | 3.291e-02 | -3.41 | fly chr19E2-19E3\|1-65 | chromosome location | fly chr19E2-19E3\|1-65 | 1 | 1 | 25010 | 823 | shakB | | 3.291e-02 | -3.41 | fly chr70C10-70C11\|3-41 | chromosome location | fly chr70C10-70C11\|3-41 | 1 | 1 | 25010 | 823 | CG17361 | | 3.291e-02 | -3.41 | fly chr84E5-84E6\|3-48 | chromosome location | fly chr84E5-84E6\|3-48 | 1 | 1 | 25010 | 823 | dsx | | 3.291e-02 | -3.41 | fly chr15A11-15B1\|1-55 cM | chromosome location | fly chr15A11-15B1\|1-55 cM | 1 | 1 | 25010 | 823 | RSG7 | | 3.291e-02 | -3.41 | fly chr46F9-47A1\|2-62 cM | chromosome location | fly chr46F9-47A1\|2-62 cM | 1 | 1 | 25010 | 823 | CAP | | 3.291e-02 | -3.41 | fly chr23D1-23D2\|2-8 cM | chromosome location | fly chr23D1-23D2\|2-8 cM | 1 | 1 | 25010 | 823 | toc | | 3.291e-02 | -3.41 | fly chr57D7-57D7\|2-94 | chromosome location | fly chr57D7-57D7\|2-94 | 1 | 1 | 25010 | 823 | MFS16 | | 3.291e-02 | -3.41 | fly chr82E5-82E5\|3-47.1 cM | chromosome location | fly chr82E5-82E5\|3-47.1 cM | 1 | 1 | 25010 | 823 | lncRNA:CR45585 | | 3.291e-02 | -3.41 | fly chr94C2-94C3\|3-76 | chromosome location | fly chr94C2-94C3\|3-76 | 1 | 1 | 25010 | 823 | wge | | 3.291e-02 | -3.41 | fly chr93A4-93A4\|3-69 cM | chromosome location | fly chr93A4-93A4\|3-69 cM | 1 | 1 | 25010 | 823 | Atpalpha | | 3.291e-02 | -3.41 | fly chr52A11-52A12\|2-74 cM | chromosome location | fly chr52A11-52A12\|2-74 cM | 1 | 1 | 25010 | 823 | CG12963 | | 3.291e-02 | -3.41 | fly chr90D1-90E1\|3-62 | chromosome location | fly chr90D1-90E1\|3-62 | 1 | 1 | 25010 | 823 | cpo | | 3.291e-02 | -3.41 | fly chr62D7-62E1\|3-3 cM | chromosome location | fly chr62D7-62E1\|3-3 cM | 1 | 1 | 25010 | 823 | Svil | | 3.291e-02 | -3.41 | fly chr4C8-4C8\|1-8 | chromosome location | fly chr4C8-4C8\|1-8 | 1 | 1 | 25010 | 823 | CG12684 | | 3.291e-02 | -3.41 | fly chr65E10-65F4\|3-19 cM | chromosome location | fly chr65E10-65F4\|3-19 cM | 1 | 1 | 25010 | 823 | unc-13-4A | | 3.291e-02 | -3.41 | fly chr94A2-94A4\|3-74 cM | chromosome location | fly chr94A2-94A4\|3-74 cM | 1 | 1 | 25010 | 823 | SKIP | | 3.291e-02 | -3.41 | fly chr33C4-33D2\|2-46 | chromosome location | fly chr33C4-33D2\|2-46 | 1 | 1 | 25010 | 823 | CG17211 | | 3.291e-02 | -3.41 | fly chr29E1-29E3\|2-33 | chromosome location | fly chr29E1-29E3\|2-33 | 1 | 1 | 25010 | 823 | Sema1a | | 3.291e-02 | -3.41 | fly chr75B13-75B13\|3-45 cM | chromosome location | fly chr75B13-75B13\|3-45 cM | 1 | 1 | 25010 | 823 | CG7330 | | 3.291e-02 | -3.41 | fly chr94E6-94E6\|3-78 | chromosome location | fly chr94E6-94E6\|3-78 | 1 | 1 | 25010 | 823 | fzo | | 3.291e-02 | -3.41 | fly chr94C4-94C8\|3-76 cM | chromosome location | fly chr94C4-94C8\|3-76 cM | 1 | 1 | 25010 | 823 | CG7029 | | 3.291e-02 | -3.41 | fly chr2A3-2A4\|1-0.2 | chromosome location | fly chr2A3-2A4\|1-0.2 | 1 | 1 | 25010 | 823 | Gr2a | | 3.291e-02 | -3.41 | fly chr40D1-40D2\|2-54 | chromosome location | fly chr40D1-40D2\|2-54 | 1 | 1 | 25010 | 823 | CG2528 | | 3.291e-02 | -3.41 | fly chr40A1-40A2\|2-54 | chromosome location | fly chr40A1-40A2\|2-54 | 1 | 1 | 25010 | 823 | step | | 3.291e-02 | -3.41 | fly chr70A7-70A8\|3-40 | chromosome location | fly chr70A7-70A8\|3-40 | 1 | 1 | 25010 | 823 | CG10171 | | 3.291e-02 | -3.41 | fly chr98D4-98D4\|3-96 cM | chromosome location | fly chr98D4-98D4\|3-96 cM | 1 | 1 | 25010 | 823 | mIF2 | | 3.291e-02 | -3.41 | fly chr87D13-87D13\|3-53 cM | chromosome location | fly chr87D13-87D13\|3-53 cM | 1 | 1 | 25010 | 823 | 2mit | | 3.291e-02 | -3.41 | fly chr95C13-95D1\|3-81 | chromosome location | fly chr95C13-95D1\|3-81 | 1 | 1 | 25010 | 823 | Gdh | | 3.291e-02 | -3.41 | fly chr56D9-56D11\|2-89 | chromosome location | fly chr56D9-56D11\|2-89 | 1 | 1 | 25010 | 823 | par-1 | | 3.291e-02 | -3.41 | fly chr35B3-35B4\|2-50 | chromosome location | fly chr35B3-35B4\|2-50 | 1 | 1 | 25010 | 823 | osp | | 3.291e-02 | -3.41 | fly chr41B3-41C2\|2-55 | chromosome location | fly chr41B3-41C2\|2-55 | 1 | 1 | 25010 | 823 | Gprk1 | | 3.291e-02 | -3.41 | fly chr102E4-102E4\|4-0 | chromosome location | fly chr102E4-102E4\|4-0 | 1 | 1 | 25010 | 823 | lncRNA:CR45123 | | 3.291e-02 | -3.41 | fly chr47C3-47C5\|2-63 | chromosome location | fly chr47C3-47C5\|2-63 | 1 | 1 | 25010 | 823 | CG30015 | | 3.291e-02 | -3.41 | fly chr91A1-91A2\|3-63 cM | chromosome location | fly chr91A1-91A2\|3-63 cM | 1 | 1 | 25010 | 823 | l(3)05822 | | 3.291e-02 | -3.41 | fly chr8C15-8C17\|1-26 | chromosome location | fly chr8C15-8C17\|1-26 | 1 | 1 | 25010 | 823 | mei-P26 | | 3.291e-02 | -3.41 | fly chr12E10-12E10\|1-48 cM | chromosome location | fly chr12E10-12E10\|1-48 cM | 1 | 1 | 25010 | 823 | CG9411 | | 3.291e-02 | -3.41 | fly chr98D4-98D4\|3-96 | chromosome location | fly chr98D4-98D4\|3-96 | 1 | 1 | 25010 | 823 | mIF2 | | 3.291e-02 | -3.41 | fly chr100D2-100D3\|3-102 | chromosome location | fly chr100D2-100D3\|3-102 | 1 | 1 | 25010 | 823 | RhoGAP100F | | 3.291e-02 | -3.41 | fly chr75B13-75B13\|3-45 | chromosome location | fly chr75B13-75B13\|3-45 | 1 | 1 | 25010 | 823 | CG7330 | | 3.291e-02 | -3.41 | fly chr6C13-6C13\|1-18 | chromosome location | fly chr6C13-6C13\|1-18 | 1 | 1 | 25010 | 823 | Coq7 | | 3.291e-02 | -3.41 | fly chr6A1-6A1\|1-17 | chromosome location | fly chr6A1-6A1\|1-17 | 1 | 1 | 25010 | 823 | CG33668 | | 3.291e-02 | -3.41 | fly chr85C4-85D1\|3-48 cM | chromosome location | fly chr85C4-85D1\|3-48 cM | 1 | 1 | 25010 | 823 | pum | | 3.291e-02 | -3.41 | fly chr8A1-8A1\|1-24 cM | chromosome location | fly chr8A1-8A1\|1-24 cM | 1 | 1 | 25010 | 823 | CG12772 | | 3.291e-02 | -3.41 | fly chr64D1-64D1\|3-14 | chromosome location | fly chr64D1-64D1\|3-14 | 1 | 1 | 25010 | 823 | DIP-delta | | 3.291e-02 | -3.41 | fly chr88D2-88D3\|3-56 | chromosome location | fly chr88D2-88D3\|3-56 | 1 | 1 | 25010 | 823 | jvl | | 3.291e-02 | -3.41 | fly chr100D3-100E2\|3-103 cM | chromosome location | fly chr100D3-100E2\|3-103 cM | 1 | 1 | 25010 | 823 | heph | | 3.291e-02 | -3.41 | fly chr62D7-62E1\|3-3 | chromosome location | fly chr62D7-62E1\|3-3 | 1 | 1 | 25010 | 823 | Svil | | 3.291e-02 | -3.41 | fly chr3F4-3F7\|1-5 cM | chromosome location | fly chr3F4-3F7\|1-5 cM | 1 | 1 | 25010 | 823 | CG42541 | | 3.291e-02 | -3.41 | fly chr102B5-102B5\|4-0 | chromosome location | fly chr102B5-102B5\|4-0 | 1 | 1 | 25010 | 823 | PMCA | | 3.291e-02 | -3.41 | fly chr51E3-51E5\|2-74 cM | chromosome location | fly chr51E3-51E5\|2-74 cM | 1 | 1 | 25010 | 823 | igl | | 3.291e-02 | -3.41 | fly chr56C11-56D1\|2-88 | chromosome location | fly chr56C11-56D1\|2-88 | 1 | 1 | 25010 | 823 | Rgk1 | | 3.291e-02 | -3.41 | fly chr57D7-57D7\|2-94 cM | chromosome location | fly chr57D7-57D7\|2-94 cM | 1 | 1 | 25010 | 823 | MFS16 | | 3.291e-02 | -3.41 | fly chr87D13-87D13\|3-53 | chromosome location | fly chr87D13-87D13\|3-53 | 1 | 1 | 25010 | 823 | 2mit | | 3.291e-02 | -3.41 | fly chr47C3-47C5\|2-63 cM | chromosome location | fly chr47C3-47C5\|2-63 cM | 1 | 1 | 25010 | 823 | CG30015 | | 3.291e-02 | -3.41 | fly chr47A-47B\|2-62 cM | chromosome location | fly chr47A-47B\|2-62 cM | 1 | 1 | 25010 | 823 | lola | | 3.291e-02 | -3.41 | fly chr25B9-25C1\|2-14 | chromosome location | fly chr25B9-25C1\|2-14 | 1 | 1 | 25010 | 823 | Rtnl1 | | 3.291e-02 | -3.41 | fly chr20A1-20A2\|1-66 cM | chromosome location | fly chr20A1-20A2\|1-66 cM | 1 | 1 | 25010 | 823 | lncRNA:flam | | 3.291e-02 | -3.41 | fly chr7B4-7B6\|1-20 cM | chromosome location | fly chr7B4-7B6\|1-20 cM | 1 | 1 | 25010 | 823 | ct | | 3.291e-02 | -3.41 | fly chr82A6-82B1\|3-47.1 | chromosome location | fly chr82A6-82B1\|3-47.1 | 1 | 1 | 25010 | 823 | CG32944 | | 3.291e-02 | -3.41 | fly chr6E3-6E4\|1-18 cM | chromosome location | fly chr6E3-6E4\|1-18 cM | 1 | 1 | 25010 | 823 | Setd3 | | 3.291e-02 | -3.41 | fly chr58E9-58F1\|2-100 cM | chromosome location | fly chr58E9-58F1\|2-100 cM | 1 | 1 | 25010 | 823 | dnr1 | | 3.291e-02 | -3.41 | fly chr61D4-61E1\|3-0.5 | chromosome location | fly chr61D4-61E1\|3-0.5 | 1 | 1 | 25010 | 823 | Glut1 | | 3.291e-02 | -3.41 | fly chr61F2-61F3\|3-0 cM | chromosome location | fly chr61F2-61F3\|3-0 cM | 1 | 1 | 25010 | 823 | bab2 | | 3.291e-02 | -3.41 | fly chr89B20-89D4\|3-58 | chromosome location | fly chr89B20-89D4\|3-58 | 1 | 1 | 25010 | 823 | Scp2 | | 3.291e-02 | -3.41 | fly chr82A1-82A3\|3-47.1 | chromosome location | fly chr82A1-82A3\|3-47.1 | 1 | 1 | 25010 | 823 | cpx | | 3.291e-02 | -3.41 | fly chr29E1-29E3\|2-33 cM | chromosome location | fly chr29E1-29E3\|2-33 cM | 1 | 1 | 25010 | 823 | Sema1a | | 3.291e-02 | -3.41 | fly chr37E4-37E5\|2-54 | chromosome location | fly chr37E4-37E5\|2-54 | 1 | 1 | 25010 | 823 | CG10188 | | 3.291e-02 | -3.41 | fly chr43F2-43F3\|2-57 cM | chromosome location | fly chr43F2-43F3\|2-57 cM | 1 | 1 | 25010 | 823 | drosha | | 3.291e-02 | -3.41 | fly chr42B1-42B2\|2-55 cM | chromosome location | fly chr42B1-42B2\|2-55 cM | 1 | 1 | 25010 | 823 | jing | | 3.291e-02 | -3.41 | fly chr52F3-52F3\|2-78 cM | chromosome location | fly chr52F3-52F3\|2-78 cM | 1 | 1 | 25010 | 823 | CG8435 | | 3.291e-02 | -3.41 | fly chr1C2-1C3\|1-0 | chromosome location | fly chr1C2-1C3\|1-0 | 1 | 1 | 25010 | 823 | sdk | | 3.291e-02 | -3.41 | fly chr96A6-96A7\|3-85 | chromosome location | fly chr96A6-96A7\|3-85 | 1 | 1 | 25010 | 823 | CG5789 | | 3.291e-02 | -3.41 | fly chr43C5-43C6\|2-56 cM | chromosome location | fly chr43C5-43C6\|2-56 cM | 1 | 1 | 25010 | 823 | CG1620 | | 3.291e-02 | -3.41 | fly chr43C4-43C4\|2-56 cM | chromosome location | fly chr43C4-43C4\|2-56 cM | 1 | 1 | 25010 | 823 | Aldh-III | | 3.291e-02 | -3.41 | fly chr38A1-38A1\|2-54 | chromosome location | fly chr38A1-38A1\|2-54 | 1 | 1 | 25010 | 823 | scw | | 3.291e-02 | -3.41 | fly chr89B20-89D4\|3-58 cM | chromosome location | fly chr89B20-89D4\|3-58 cM | 1 | 1 | 25010 | 823 | Scp2 | | 3.291e-02 | -3.41 | fly chr19C6-19D1\|1-64 | chromosome location | fly chr19C6-19D1\|1-64 | 1 | 1 | 25010 | 823 | CG15459 | | 3.291e-02 | -3.41 | fly chr85A3-85A5\|3-48 | chromosome location | fly chr85A3-85A5\|3-48 | 1 | 1 | 25010 | 823 | 5-HT2B | | 3.291e-02 | -3.41 | fly chr97E3-97E3\|3-92 | chromosome location | fly chr97E3-97E3\|3-92 | 1 | 1 | 25010 | 823 | CG6059 | | 3.291e-02 | -3.41 | fly chr14A1-14B1\|1-52 cM | chromosome location | fly chr14A1-14B1\|1-52 cM | 1 | 1 | 25010 | 823 | tay | | 3.291e-02 | -3.41 | fly chr22A5-22A5\|2-3 cM | chromosome location | fly chr22A5-22A5\|2-3 cM | 1 | 1 | 25010 | 823 | lncRNA:CR45695 | | 3.291e-02 | -3.41 | fly chr85E2-85E3\|3-49 cM | chromosome location | fly chr85E2-85E3\|3-49 cM | 1 | 1 | 25010 | 823 | CG9492 | | 3.291e-02 | -3.41 | fly chr45D5-45D7\|2-60 | chromosome location | fly chr45D5-45D7\|2-60 | 1 | 1 | 25010 | 823 | Pdk | | 3.291e-02 | -3.41 | fly chr13E16-13E17\|1-51 cM | chromosome location | fly chr13E16-13E17\|1-51 cM | 1 | 1 | 25010 | 823 | CG43672 | | 3.291e-02 | -3.41 | fly chr35E5-35E6\|2-51 | chromosome location | fly chr35E5-35E6\|2-51 | 1 | 1 | 25010 | 823 | Ca-alpha1D | | 3.291e-02 | -3.41 | fly chr2B4-2B5\|1-0.2 cM | chromosome location | fly chr2B4-2B5\|1-0.2 cM | 1 | 1 | 25010 | 823 | br | | 3.291e-02 | -3.41 | fly chr35A4-35B1\|2-50 | chromosome location | fly chr35A4-35B1\|2-50 | 1 | 1 | 25010 | 823 | spel1 | | 3.291e-02 | -3.41 | fly chr91A7-91B3\|3-63 | chromosome location | fly chr91A7-91B3\|3-63 | 1 | 1 | 25010 | 823 | fru | | 3.291e-02 | -3.41 | fly chr19E2-19E3\|1-65 cM | chromosome location | fly chr19E2-19E3\|1-65 cM | 1 | 1 | 25010 | 823 | shakB | | 3.291e-02 | -3.41 | fly chr61C5-61C5\|3-0.5 cM | chromosome location | fly chr61C5-61C5\|3-0.5 cM | 1 | 1 | 25010 | 823 | Ppm1 | | 3.291e-02 | -3.41 | fly chr86D9-86E1\|3-51 | chromosome location | fly chr86D9-86E1\|3-51 | 1 | 1 | 25010 | 823 | lncRNA:CR31386 | | 3.291e-02 | -3.41 | fly chr98A8-98A10\|3-94 cM | chromosome location | fly chr98A8-98A10\|3-94 cM | 1 | 1 | 25010 | 823 | CG5639 | | 3.291e-02 | -3.41 | fly chr25C4-25C6\|2-15 | chromosome location | fly chr25C4-25C6\|2-15 | 1 | 1 | 25010 | 823 | Pgant5 | | 3.291e-02 | -3.41 | fly chr64C7-64C11\|3-13 cM | chromosome location | fly chr64C7-64C11\|3-13 cM | 1 | 1 | 25010 | 823 | shep | | 3.291e-02 | -3.41 | fly chr42A7-42A7\|2-55 cM | chromosome location | fly chr42A7-42A7\|2-55 cM | 1 | 1 | 25010 | 823 | Act42A | | 3.291e-02 | -3.41 | fly chr97E3-97E3\|3-92 cM | chromosome location | fly chr97E3-97E3\|3-92 cM | 1 | 1 | 25010 | 823 | CG6059 | | 3.291e-02 | -3.41 | fly chr64D1-64D1\|3-14 cM | chromosome location | fly chr64D1-64D1\|3-14 cM | 1 | 1 | 25010 | 823 | DIP-delta | | 3.291e-02 | -3.41 | fly chr54B6-54B7\|2-82 | chromosome location | fly chr54B6-54B7\|2-82 | 1 | 1 | 25010 | 823 | CG10939 | | 3.291e-02 | -3.41 | fly chr82E5-82E5\|3-47.1 | chromosome location | fly chr82E5-82E5\|3-47.1 | 1 | 1 | 25010 | 823 | lncRNA:CR45585 | | 3.291e-02 | -3.41 | fly chr38A1-38A1\|2-54 cM | chromosome location | fly chr38A1-38A1\|2-54 cM | 1 | 1 | 25010 | 823 | scw | | 3.291e-02 | -3.41 | fly chr102D5-102E1\|4-0 | chromosome location | fly chr102D5-102E1\|4-0 | 1 | 1 | 25010 | 823 | bt | | 3.291e-02 | -3.41 | fly chr23D1-23D2\|2-8 | chromosome location | fly chr23D1-23D2\|2-8 | 1 | 1 | 25010 | 823 | toc | | 3.291e-02 | -3.41 | fly chr1C2-1C3\|1-0 cM | chromosome location | fly chr1C2-1C3\|1-0 cM | 1 | 1 | 25010 | 823 | sdk | | 3.291e-02 | -3.41 | fly chr83C6-83C7\|3-47.5 | chromosome location | fly chr83C6-83C7\|3-47.5 | 1 | 1 | 25010 | 823 | lncRNA:CR45761 | | 3.291e-02 | -3.41 | fly chr32F1-32F2\|2-44 cM | chromosome location | fly chr32F1-32F2\|2-44 cM | 1 | 1 | 25010 | 823 | salm | | 3.291e-02 | -3.41 | fly chr33C4-33D2\|2-46 cM | chromosome location | fly chr33C4-33D2\|2-46 cM | 1 | 1 | 25010 | 823 | CG17211 | | 3.291e-02 | -3.41 | fly chr12B2-12B4\|1-45 cM | chromosome location | fly chr12B2-12B4\|1-45 cM | 1 | 1 | 25010 | 823 | Nadsyn | | 3.291e-02 | -3.41 | fly chr100D3-100E2\|3-103 | chromosome location | fly chr100D3-100E2\|3-103 | 1 | 1 | 25010 | 823 | heph | | 3.291e-02 | -3.41 | fly chr84D10-84D11\|3-48 cM | chromosome location | fly chr84D10-84D11\|3-48 cM | 1 | 1 | 25010 | 823 | Nlg3 | | 3.291e-02 | -3.41 | fly chr63B9-63B9\|3-6 cM | chromosome location | fly chr63B9-63B9\|3-6 cM | 1 | 1 | 25010 | 823 | CG14964 | | 3.291e-02 | -3.41 | fly chr52E10-52E11\|2-77 | chromosome location | fly chr52E10-52E11\|2-77 | 1 | 1 | 25010 | 823 | CG30094 | | 3.291e-02 | -3.41 | fly chr65D3-65D4\|3-18 cM | chromosome location | fly chr65D3-65D4\|3-18 cM | 1 | 1 | 25010 | 823 | CG10077 | | 3.291e-02 | -3.41 | fly chr86B3-86B4\|3-49 cM | chromosome location | fly chr86B3-86B4\|3-49 cM | 1 | 1 | 25010 | 823 | mthl11 | | 3.291e-02 | -3.41 | fly chr96A6-96A7\|3-85 cM | chromosome location | fly chr96A6-96A7\|3-85 cM | 1 | 1 | 25010 | 823 | CG5789 | | 3.291e-02 | -3.41 | fly chr8E12-8E12\|1-28 | chromosome location | fly chr8E12-8E12\|1-28 | 1 | 1 | 25010 | 823 | lncRNA:CR45543 | | 3.291e-02 | -3.41 | fly chr7E1-7E4\|1-23 | chromosome location | fly chr7E1-7E4\|1-23 | 1 | 1 | 25010 | 823 | nAChRalpha3 | | 3.291e-02 | -3.41 | fly chr57F1-57F2\|2-96 | chromosome location | fly chr57F1-57F2\|2-96 | 1 | 1 | 25010 | 823 | twz | | 3.291e-02 | -3.41 | fly chr61D4-61E1\|3-0.5 cM | chromosome location | fly chr61D4-61E1\|3-0.5 cM | 1 | 1 | 25010 | 823 | Glut1 | | 3.291e-02 | -3.41 | fly chr102F4-102F5\|4-0 cM | chromosome location | fly chr102F4-102F5\|4-0 cM | 1 | 1 | 25010 | 823 | toy | | 3.291e-02 | -3.41 | fly chr35C4-35C4\|2-51 | chromosome location | fly chr35C4-35C4\|2-51 | 1 | 1 | 25010 | 823 | CG43230 | | 3.291e-02 | -3.41 | fly chr75B1-75B1\|3-45 | chromosome location | fly chr75B1-75B1\|3-45 | 1 | 1 | 25010 | 823 | lncRNA:CR45921 | | 3.291e-02 | -3.41 | fly chr29E3-29E4\|2-33 cM | chromosome location | fly chr29E3-29E4\|2-33 cM | 1 | 1 | 25010 | 823 | Trs23 | | 3.291e-02 | -3.41 | fly chr14A5-14A6\|1-52 cM | chromosome location | fly chr14A5-14A6\|1-52 cM | 1 | 1 | 25010 | 823 | vap | | 3.291e-02 | -3.41 | fly chr96A12-96A13\|3-85 | chromosome location | fly chr96A12-96A13\|3-85 | 1 | 1 | 25010 | 823 | Syx18 | | 3.291e-02 | -3.41 | fly chr100E1-100E1\|3-103 | chromosome location | fly chr100E1-100E1\|3-103 | 1 | 1 | 25010 | 823 | CG2003 | | 3.291e-02 | -3.41 | fly chr84D10-84D11\|3-48 | chromosome location | fly chr84D10-84D11\|3-48 | 1 | 1 | 25010 | 823 | Nlg3 | | 3.291e-02 | -3.41 | fly chr91E5-91E5\|3-65 cM | chromosome location | fly chr91E5-91E5\|3-65 cM | 1 | 1 | 25010 | 823 | lncRNA:CR46041 | | 3.291e-02 | -3.41 | fly chr97C3-97C4\|3-91 | chromosome location | fly chr97C3-97C4\|3-91 | 1 | 1 | 25010 | 823 | beat-VII | | 3.291e-02 | -3.41 | fly chr49A1-49A2\|2-67 | chromosome location | fly chr49A1-49A2\|2-67 | 1 | 1 | 25010 | 823 | Cpr49Ac | | 3.291e-02 | -3.41 | fly chr95C13-95D1\|3-81 cM | chromosome location | fly chr95C13-95D1\|3-81 cM | 1 | 1 | 25010 | 823 | Gdh | | 3.291e-02 | -3.41 | fly chr1C3-1C4\|1-0 | chromosome location | fly chr1C3-1C4\|1-0 | 1 | 1 | 25010 | 823 | CG5254 | | 3.291e-02 | -3.41 | fly chr61C5-61C5\|3-0.5 | chromosome location | fly chr61C5-61C5\|3-0.5 | 1 | 1 | 25010 | 823 | Ppm1 | | 3.291e-02 | -3.41 | fly chr99F1-99F1\|3-100 | chromosome location | fly chr99F1-99F1\|3-100 | 1 | 1 | 25010 | 823 | lncRNA:CR46114 | | 3.291e-02 | -3.41 | fly chr87D7-87D8\|3-53 cM | chromosome location | fly chr87D7-87D8\|3-53 cM | 1 | 1 | 25010 | 823 | CG7518 | | 3.291e-02 | -3.41 | fly chr96E4-96E5\|3-88 | chromosome location | fly chr96E4-96E5\|3-88 | 1 | 1 | 25010 | 823 | CG5079 | | 3.291e-02 | -3.41 | fly chr88D2-88D3\|3-56 cM | chromosome location | fly chr88D2-88D3\|3-56 cM | 1 | 1 | 25010 | 823 | jvl | | 3.291e-02 | -3.41 | fly chr2E1-2E2\|1-0.8 | chromosome location | fly chr2E1-2E2\|1-0.8 | 1 | 1 | 25010 | 823 | Mct1 | | 3.291e-02 | -3.41 | fly chr97C3-97C4\|3-91 cM | chromosome location | fly chr97C3-97C4\|3-91 cM | 1 | 1 | 25010 | 823 | beat-VII | | 3.291e-02 | -3.41 | fly chr41F-41F8\|2-55 | chromosome location | fly chr41F-41F8\|2-55 | 1 | 1 | 25010 | 823 | ap | | 3.291e-02 | -3.41 | fly chr11B12-11B12\|1-39 | chromosome location | fly chr11B12-11B12\|1-39 | 1 | 1 | 25010 | 823 | CG15725 | | 3.291e-02 | -3.41 | fly chr67E1-67E2\|3-33 | chromosome location | fly chr67E1-67E2\|3-33 | 1 | 1 | 25010 | 823 | lncRNA:CR46006 | | 3.291e-02 | -3.41 | fly chr75F9-76A3\|3-45 cM | chromosome location | fly chr75F9-76A3\|3-45 cM | 1 | 1 | 25010 | 823 | fz2 | | 3.291e-02 | -3.41 | fly chr51E3-51E5\|2-74 | chromosome location | fly chr51E3-51E5\|2-74 | 1 | 1 | 25010 | 823 | igl | | 3.291e-02 | -3.41 | fly chr101F1-102A4\|4-0 | chromosome location | fly chr101F1-102A4\|4-0 | 1 | 1 | 25010 | 823 | pan | | 3.291e-02 | -3.41 | fly chr51E8-51E9\|2-74 cM | chromosome location | fly chr51E8-51E9\|2-74 cM | 1 | 1 | 25010 | 823 | asRNA:CR44370 | | 3.291e-02 | -3.41 | fly chr96C6-96C7\|3-87 | chromosome location | fly chr96C6-96C7\|3-87 | 1 | 1 | 25010 | 823 | Alg9 | | 3.291e-02 | -3.41 | fly chr85C4-85D1\|3-48 | chromosome location | fly chr85C4-85D1\|3-48 | 1 | 1 | 25010 | 823 | pum | | 3.291e-02 | -3.41 | fly chr87C1-87C2\|3-52 | chromosome location | fly chr87C1-87C2\|3-52 | 1 | 1 | 25010 | 823 | Octbeta2R | | 3.291e-02 | -3.41 | fly chr92A5-92A10\|3-66 | chromosome location | fly chr92A5-92A10\|3-66 | 1 | 1 | 25010 | 823 | Dys | | 3.291e-02 | -3.41 | fly chr57D12-57E1\|2-95 cM | chromosome location | fly chr57D12-57E1\|2-95 cM | 1 | 1 | 25010 | 823 | cv-2 | | 3.291e-02 | -3.41 | fly chr43F2-43F3\|2-57 | chromosome location | fly chr43F2-43F3\|2-57 | 1 | 1 | 25010 | 823 | drosha | | 3.291e-02 | -3.41 | fly chr99E4-99F2\|3-100 cM | chromosome location | fly chr99E4-99F2\|3-100 cM | 1 | 1 | 25010 | 823 | hdc | | 3.291e-02 | -3.41 | fly chr62C4-62D1\|3-1.5 cM | chromosome location | fly chr62C4-62D1\|3-1.5 cM | 1 | 1 | 25010 | 823 | zormin | | 3.291e-02 | -3.41 | fly chr13E8-13E12\|1-51 | chromosome location | fly chr13E8-13E12\|1-51 | 1 | 1 | 25010 | 823 | CG8184 | | 3.291e-02 | -3.41 | fly chr4F5-4F9\|1-11 | chromosome location | fly chr4F5-4F9\|1-11 | 1 | 1 | 25010 | 823 | SK | | 3.291e-02 | -3.41 | fly chr77B9-77C1\|3-47 | chromosome location | fly chr77B9-77C1\|3-47 | 1 | 1 | 25010 | 823 | fbl | | 3.291e-02 | -3.41 | fly chr102E4-102E4\|4-0 cM | chromosome location | fly chr102E4-102E4\|4-0 cM | 1 | 1 | 25010 | 823 | lncRNA:CR45123 | | 3.291e-02 | -3.41 | fly chr9F8-9F11\|1-33 | chromosome location | fly chr9F8-9F11\|1-33 | 1 | 1 | 25010 | 823 | CG43901 | | 3.291e-02 | -3.41 | fly chr33D2-33D5\|2-46 cM | chromosome location | fly chr33D2-33D5\|2-46 cM | 1 | 1 | 25010 | 823 | bru1 | | 3.291e-02 | -3.41 | fly chr22F4-23A1\|2-6 cM | chromosome location | fly chr22F4-23A1\|2-6 cM | 1 | 1 | 25010 | 823 | Cyp309a1 | | 3.291e-02 | -3.41 | fly chr6F10-7A5\|1-19 cM | chromosome location | fly chr6F10-7A5\|1-19 cM | 1 | 1 | 25010 | 823 | CG9650 | | 3.291e-02 | -3.41 | fly chr50B9-50B9\|2-68 cM | chromosome location | fly chr50B9-50B9\|2-68 cM | 1 | 1 | 25010 | 823 | CG6209 | | 3.291e-02 | -3.41 | fly chr62A8-62A9\|3-0.5 cM | chromosome location | fly chr62A8-62A9\|3-0.5 cM | 1 | 1 | 25010 | 823 | ABCB7 | | 3.291e-02 | -3.41 | fly chr87B4-87B5\|3-52 | chromosome location | fly chr87B4-87B5\|3-52 | 1 | 1 | 25010 | 823 | svp | | 3.291e-02 | -3.41 | fly chr64B1-64B2\|3-12 | chromosome location | fly chr64B1-64B2\|3-12 | 1 | 1 | 25010 | 823 | Cip4 | | 3.291e-02 | -3.41 | fly chr96D2-96D3\|3-87 | chromosome location | fly chr96D2-96D3\|3-87 | 1 | 1 | 25010 | 823 | alrm | | 3.291e-02 | -3.41 | fly chr86C1-86C3\|3-50 | chromosome location | fly chr86C1-86C3\|3-50 | 1 | 1 | 25010 | 823 | hth | | 3.291e-02 | -3.41 | fly chr3E4-3E5\|1-4 | chromosome location | fly chr3E4-3E5\|1-4 | 1 | 1 | 25010 | 823 | Mnt | | 3.291e-02 | -3.41 | fly chr87B4-87B5\|3-52 cM | chromosome location | fly chr87B4-87B5\|3-52 cM | 1 | 1 | 25010 | 823 | svp | | 3.291e-02 | -3.41 | fly chr2B9-2B12\|1-0.2 | chromosome location | fly chr2B9-2B12\|1-0.2 | 1 | 1 | 25010 | 823 | prage | | 3.291e-02 | -3.41 | fly chr52E2-52E2\|2-77 cM | chromosome location | fly chr52E2-52E2\|2-77 cM | 1 | 1 | 25010 | 823 | lncRNA:CR43415 | | 3.291e-02 | -3.41 | fly chr18F-18F\|1-64 | chromosome location | fly chr18F-18F\|1-64 | 1 | 1 | 25010 | 823 | zld | | 3.291e-02 | -3.41 | fly chr99F1-99F1\|3-100 cM | chromosome location | fly chr99F1-99F1\|3-100 cM | 1 | 1 | 25010 | 823 | lncRNA:CR46114 | | 3.291e-02 | -3.41 | fly chr7E1-7E4\|1-23 cM | chromosome location | fly chr7E1-7E4\|1-23 cM | 1 | 1 | 25010 | 823 | nAChRalpha3 | | 3.291e-02 | -3.41 | fly chr82E6-82E6 | chromosome location | fly chr82E6-82E6 | 1 | 1 | 25010 | 823 | lncRNA:CR46352 | | 3.291e-02 | -3.41 | fly chr67E1-67E2\|3-33 cM | chromosome location | fly chr67E1-67E2\|3-33 cM | 1 | 1 | 25010 | 823 | lncRNA:CR46006 | | 3.291e-02 | -3.41 | fly chr42F3-43A1\|2-55 | chromosome location | fly chr42F3-43A1\|2-55 | 1 | 1 | 25010 | 823 | pk | | 3.291e-02 | -3.41 | fly chr3D2-3D4\|1-3 | chromosome location | fly chr3D2-3D4\|1-3 | 1 | 1 | 25010 | 823 | Myc | | 3.291e-02 | -3.41 | fly chr51F3-51F3\|2-74 | chromosome location | fly chr51F3-51F3\|2-74 | 1 | 1 | 25010 | 823 | lncRNA:CR44241 | | 3.291e-02 | -3.41 | fly chr86A7-86A8\|3-49 cM | chromosome location | fly chr86A7-86A8\|3-49 cM | 1 | 1 | 25010 | 823 | Bruce | | 3.291e-02 | -3.41 | fly chr42A7-42A7\|2-55 | chromosome location | fly chr42A7-42A7\|2-55 | 1 | 1 | 25010 | 823 | Act42A | | 3.291e-02 | -3.41 | fly chr47F17-48A1\|2-64 cM | chromosome location | fly chr47F17-48A1\|2-64 cM | 1 | 1 | 25010 | 823 | en | | 3.291e-02 | -3.41 | fly chr70B2-70B3\|3-40 cM | chromosome location | fly chr70B2-70B3\|3-40 cM | 1 | 1 | 25010 | 823 | Tgi | | 3.291e-02 | -3.41 | fly chr100B-100B\|3-102 | chromosome location | fly chr100B-100B\|3-102 | 1 | 1 | 25010 | 823 | Prosalpha3T | | 3.291e-02 | -3.41 | fly chr18F2-18F4\|1-64 cM | chromosome location | fly chr18F2-18F4\|1-64 cM | 1 | 1 | 25010 | 823 | pico | | 3.291e-02 | -3.41 | fly chr51E8-51E9\|2-74 | chromosome location | fly chr51E8-51E9\|2-74 | 1 | 1 | 25010 | 823 | asRNA:CR44370 | | 3.291e-02 | -3.41 | fly chr58E9-58F1\|2-100 | chromosome location | fly chr58E9-58F1\|2-100 | 1 | 1 | 25010 | 823 | dnr1 | | 3.291e-02 | -3.41 | fly chr76C2-76C5\|3-46 cM | chromosome location | fly chr76C2-76C5\|3-46 cM | 1 | 1 | 25010 | 823 | CG42637 | | 3.291e-02 | -3.41 | fly chr52A6-52A10\|2-74 | chromosome location | fly chr52A6-52A10\|2-74 | 1 | 1 | 25010 | 823 | Khc-73 | | 3.291e-02 | -3.41 | fly chr28B4-28C1\|2-27 cM | chromosome location | fly chr28B4-28C1\|2-27 cM | 1 | 1 | 25010 | 823 | Slob | | 3.291e-02 | -3.41 | fly chr86D9-86E1\|3-51 cM | chromosome location | fly chr86D9-86E1\|3-51 cM | 1 | 1 | 25010 | 823 | lncRNA:CR31386 | | 3.291e-02 | -3.41 | fly chr48D8-48E1\|2-66 cM | chromosome location | fly chr48D8-48E1\|2-66 cM | 1 | 1 | 25010 | 823 | CG8888 | | 3.291e-02 | -3.41 | fly chr14A1-14B1\|1-52 | chromosome location | fly chr14A1-14B1\|1-52 | 1 | 1 | 25010 | 823 | tay | | 3.291e-02 | -3.41 | fly chr96A19-96A19\|3-85 | chromosome location | fly chr96A19-96A19\|3-85 | 1 | 1 | 25010 | 823 | tld | | 3.291e-02 | -3.41 | fly chr5F6-5F6\|1-16 cM | chromosome location | fly chr5F6-5F6\|1-16 cM | 1 | 1 | 25010 | 823 | CG33669 | | 3.291e-02 | -3.41 | fly chr100D2-100D3\|3-102 cM | chromosome location | fly chr100D2-100D3\|3-102 cM | 1 | 1 | 25010 | 823 | RhoGAP100F | | 3.291e-02 | -3.41 | fly chr91B5-91B6\|3-64 | chromosome location | fly chr91B5-91B6\|3-64 | 1 | 1 | 25010 | 823 | CG14304 | | 3.291e-02 | -3.41 | fly chr80D5-80E3\|3-47 | chromosome location | fly chr80D5-80E3\|3-47 | 1 | 1 | 25010 | 823 | nAChRalpha4 | | 3.291e-02 | -3.41 | fly chr79C1-79C1\|3-47 cM | chromosome location | fly chr79C1-79C1\|3-47 cM | 1 | 1 | 25010 | 823 | lncRNA:CR45236 | | 3.291e-02 | -3.41 | fly chr7D5-7D6\|1-22 cM | chromosome location | fly chr7D5-7D6\|1-22 cM | 1 | 1 | 25010 | 823 | CG2256 | | 3.291e-02 | -3.41 | fly chr40A4-40A4\|2-54 | chromosome location | fly chr40A4-40A4\|2-54 | 1 | 1 | 25010 | 823 | lncRNA:CR44786 | | 3.291e-02 | -3.41 | fly chr76C2-76C5\|3-46 | chromosome location | fly chr76C2-76C5\|3-46 | 1 | 1 | 25010 | 823 | CG42637 | | 3.291e-02 | -3.41 | fly chr97A8-97A10\|3-90 cM | chromosome location | fly chr97A8-97A10\|3-90 cM | 1 | 1 | 25010 | 823 | CG33970 | | 3.291e-02 | -3.41 | fly chr4C8-4C8\|1-8 cM | chromosome location | fly chr4C8-4C8\|1-8 cM | 1 | 1 | 25010 | 823 | CG12684 | | 3.291e-02 | -3.41 | fly chr66E4-66E5\|3-27 cM | chromosome location | fly chr66E4-66E5\|3-27 cM | 1 | 1 | 25010 | 823 | CG43783 | | 3.291e-02 | -3.41 | fly chr52A6-52A10\|2-74 cM | chromosome location | fly chr52A6-52A10\|2-74 cM | 1 | 1 | 25010 | 823 | Khc-73 | | 3.291e-02 | -3.41 | fly chr42A6-42A7\|2-55 | chromosome location | fly chr42A6-42A7\|2-55 | 1 | 1 | 25010 | 823 | Src42A | | 3.291e-02 | -3.41 | fly chr89F1-89F3\|3-60 | chromosome location | fly chr89F1-89F3\|3-60 | 1 | 1 | 25010 | 823 | Mur89F | | 3.291e-02 | -3.41 | fly chr70B5-70C4\|3-40 cM | chromosome location | fly chr70B5-70C4\|3-40 cM | 1 | 1 | 25010 | 823 | bru3 | | 3.291e-02 | -3.41 | fly chr46C4-46C7\|2-61 cM | chromosome location | fly chr46C4-46C7\|2-61 cM | 1 | 1 | 25010 | 823 | Mef2 | | 3.291e-02 | -3.41 | fly chr79C1-79C1\|3-47 | chromosome location | fly chr79C1-79C1\|3-47 | 1 | 1 | 25010 | 823 | lncRNA:CR45236 | | 3.291e-02 | -3.41 | fly chr2B9-2B12\|1-0.2 cM | chromosome location | fly chr2B9-2B12\|1-0.2 cM | 1 | 1 | 25010 | 823 | prage | | 3.291e-02 | -3.41 | fly chr98A8-98A10\|3-94 | chromosome location | fly chr98A8-98A10\|3-94 | 1 | 1 | 25010 | 823 | CG5639 | | 3.291e-02 | -3.41 | fly chr35C4-35C4\|2-51 cM | chromosome location | fly chr35C4-35C4\|2-51 cM | 1 | 1 | 25010 | 823 | CG43230 | | 3.291e-02 | -3.41 | fly chr94E6-94E6\|3-78 cM | chromosome location | fly chr94E6-94E6\|3-78 cM | 1 | 1 | 25010 | 823 | fzo | | 3.291e-02 | -3.41 | fly chr66D14-66D14\|3-26 | chromosome location | fly chr66D14-66D14\|3-26 | 1 | 1 | 25010 | 823 | Acbp5 | | 3.291e-02 | -3.41 | fly chr94C9-94C9\|3-76 cM | chromosome location | fly chr94C9-94C9\|3-76 cM | 1 | 1 | 25010 | 823 | lncRNA:CR44350 | | 3.291e-02 | -3.41 | fly chr41C5-41C5\|2-55 | chromosome location | fly chr41C5-41C5\|2-55 | 1 | 1 | 25010 | 823 | CG44102 | | 3.291e-02 | -3.41 | fly chr66A4-66A5\|3-21 cM | chromosome location | fly chr66A4-66A5\|3-21 cM | 1 | 1 | 25010 | 823 | lqf | | 3.291e-02 | -3.41 | fly chr41D4-41E1\|2-55 cM | chromosome location | fly chr41D4-41E1\|2-55 cM | 1 | 1 | 25010 | 823 | Atf6 | | 3.291e-02 | -3.41 | fly chr91B5-91B6\|3-64 cM | chromosome location | fly chr91B5-91B6\|3-64 cM | 1 | 1 | 25010 | 823 | CG14304 | | 3.291e-02 | -3.41 | fly chr41F9-41F8\|2-55 cM | chromosome location | fly chr41F9-41F8\|2-55 cM | 1 | 1 | 25010 | 823 | vlc | | 3.291e-02 | -3.41 | fly chr4C3-4C4\|1-8 cM | chromosome location | fly chr4C3-4C4\|1-8 cM | 1 | 1 | 25010 | 823 | bi | | 3.291e-02 | -3.41 | fly chr2B4-2B5\|1-0.2 | chromosome location | fly chr2B4-2B5\|1-0.2 | 1 | 1 | 25010 | 823 | br | | 3.291e-02 | -3.41 | fly chr27C1-27C4\|2-23 cM | chromosome location | fly chr27C1-27C4\|2-23 cM | 1 | 1 | 25010 | 823 | Nlg2 | | 3.291e-02 | -3.41 | fly chr8E12-8E12\|1-28 cM | chromosome location | fly chr8E12-8E12\|1-28 cM | 1 | 1 | 25010 | 823 | lncRNA:CR45543 | | 3.291e-02 | -3.41 | fly chr50C6-50C9\|2-68 cM | chromosome location | fly chr50C6-50C9\|2-68 cM | 1 | 1 | 25010 | 823 | shot | | 3.291e-02 | -3.41 | fly chr53D1-53D4\|2-80 cM | chromosome location | fly chr53D1-53D4\|2-80 cM | 1 | 1 | 25010 | 823 | Pgant9 | | 3.291e-02 | -3.41 | fly chr66A4-66A5\|3-21 | chromosome location | fly chr66A4-66A5\|3-21 | 1 | 1 | 25010 | 823 | lqf | | 3.291e-02 | -3.41 | fly chr77A4-77B1\|3-47 | chromosome location | fly chr77A4-77B1\|3-47 | 1 | 1 | 25010 | 823 | rdgC | | 3.291e-02 | -3.41 | fly chr102F4-102F5\|4-0 | chromosome location | fly chr102F4-102F5\|4-0 | 1 | 1 | 25010 | 823 | toy | | 3.291e-02 | -3.41 | fly chr86B3-86B4\|3-49 | chromosome location | fly chr86B3-86B4\|3-49 | 1 | 1 | 25010 | 823 | mthl11 | | 3.291e-02 | -3.41 | fly chr78B2-78B3\|3-47 | chromosome location | fly chr78B2-78B3\|3-47 | 1 | 1 | 25010 | 823 | CG10565 | | 3.291e-02 | -3.41 | fly chr35B8-35B9\|2-50 | chromosome location | fly chr35B8-35B9\|2-50 | 1 | 1 | 25010 | 823 | ck | | 3.291e-02 | -3.41 | fly chr3F4-3F7\|1-5 | chromosome location | fly chr3F4-3F7\|1-5 | 1 | 1 | 25010 | 823 | CG42541 | | 3.291e-02 | -3.41 | fly chr83C6-83C8\|3-47.5 cM | chromosome location | fly chr83C6-83C8\|3-47.5 cM | 1 | 1 | 25010 | 823 | side-III | | 3.291e-02 | -3.41 | fly chr13A8-13A9\|1-49 cM | chromosome location | fly chr13A8-13A9\|1-49 cM | 1 | 1 | 25010 | 823 | Lsd-2 | | 3.293e-02 | -3.41 | Sugar\_transporter\_CS | interpro domains | IPR005829 | 37 | 5 | 11383 | 557 | CG33233,CG1213,Glut1,CG8249,CG15408 | | 3.297e-02 | -3.41 | protein dimerization activity | molecular function | GO:0046983 | 390 | 26 | 12176 | 554 | Khc-73,Ten-a,Camta,Fer3,dnt,Sod1,bru3,CG33557,Trl,CadN2,Mef2,svp,scw,inc,dimm,Atf6,grim,step,mim,rau,cato,Myc,cbs,dsx,Mnt,Clk | | 3.320e-02 | -3.41 | MyTH4 | pfam domains | PF00784 | 6 | 2 | 10480 | 528 | Myo81F,ck | | 3.320e-02 | -3.41 | Cation\_ATPase\_N | pfam domains | PF00690 | 6 | 2 | 10480 | 528 | Atpalpha,PMCA | | 3.320e-02 | -3.41 | LPMO\_10 | pfam domains | PF03067 | 6 | 2 | 10480 | 528 | CG42598,CG41284 | | 3.320e-02 | -3.41 | Cation\_ATPase\_C | pfam domains | PF00689 | 6 | 2 | 10480 | 528 | Atpalpha,PMCA | | 3.323e-02 | -3.40 | CG5027 (uncharacterized protein) | protein interactions | 39775 | 6 | 2 | 9204 | 464 | Parg,CG9107 | | 3.323e-02 | -3.40 | shf (shifted) | protein interactions | 31617 | 6 | 2 | 9204 | 464 | fz2,boi | | 3.323e-02 | -3.40 | Svil (Supervillin) | protein interactions | 38306 | 6 | 2 | 9204 | 464 | Mef2,rl | | 3.323e-02 | -3.40 | Plzf (Promyelocytic leukemia zinc finger) | protein interactions | 34622 | 6 | 2 | 9204 | 464 | CG3199,rl | | 3.323e-02 | -3.40 | Adat1 (Adenosine deaminase, tRNA-specific 1) | protein interactions | 34787 | 6 | 2 | 9204 | 464 | CG10077,RNASEK | | 3.323e-02 | -3.40 | Irp-1A (Iron regulatory protein 1A) | protein interactions | 42689 | 6 | 2 | 9204 | 464 | CG17999,Nrd1 | | 3.323e-02 | -3.40 | bmm (brummer) | protein interactions | 39611 | 6 | 2 | 9204 | 464 | Lsd-2,eIF4B | | 3.323e-02 | -3.40 | rumi (rumi) | protein interactions | 326122 | 6 | 2 | 9204 | 464 | CG15136,Hsc70-3 | | 3.323e-02 | -3.40 | cv (crossveinless) | protein interactions | 44510 | 6 | 2 | 9204 | 464 | Vsx2,dsx | | 3.323e-02 | -3.40 | Mnt (Mnt) | protein interactions | 31331 | 6 | 2 | 9204 | 464 | Myc,toy | | 3.323e-02 | -3.40 | sktl (skittles) | protein interactions | 37356 | 6 | 2 | 9204 | 464 | Pld,CG14820 | | 3.326e-02 | -3.40 | developmental process involved in reproduction | biological process | GO:0003006 | 928 | 55 | 12345 | 571 | mei-P26,en,lola,dsx,Yp2,AP-2alpha,Catsup,pum,orb,tld,DNaseII,cic,bab2,drosha,fz2,Trl,fbl,Msp300,ct,br,Dlg5,Diap1,rl,wb,gus,tj,fzo,fru,jing,toc,Bruce,jvl,salm,spo,lama,capu,Mef2,par-1,Cip4,lncRNA:flam,heph,mbc,mlt,CycB,Tpl94D,fan,lz,Parp,mim,grim,Rbp9,tej,scw,rib,bru1 | | 3.329e-02 | -3.40 | REACTOME\_NEUROTRANSMITTER\_RECEPTOR\_BINDING\_AND\_DOWNSTREAM\_TRANSMISSION\_IN\_THE\_POSTSYNAPTIC\_CELL | MSigDB lists | REACTOME\_NEUROTRANSMITTER\_RECEPTOR\_BINDING\_AND\_DOWNSTREAM\_TRANSMISSION\_IN\_THE\_POSTSYNAPTIC\_CELL | 46 | 5 | 4161 | 164 | AP-2alpha,rl,Ac78C,nAChRalpha3,nAChRalpha4 | | 3.329e-02 | -3.40 | GSE11961\_FOLLICULAR\_BCELL\_VS\_MARGINAL\_ZONE\_BCELL\_UP | MSigDB lists | GSE11961\_FOLLICULAR\_BCELL\_VS\_MARGINAL\_ZONE\_BCELL\_UP | 46 | 5 | 4161 | 164 | CG9492,Alg-2,Mef2,ABCB7,Glut1 | | 3.329e-02 | -3.40 | MORF\_PCNA | MSigDB lists | MORF\_PCNA | 46 | 5 | 4161 | 164 | Sod1,ND-30,vtd,Parp,spel1 | | 3.329e-02 | -3.40 | RUTELLA\_RESPONSE\_TO\_HGF\_VS\_CSF2RB\_AND\_IL4\_DN | MSigDB lists | RUTELLA\_RESPONSE\_TO\_HGF\_VS\_CSF2RB\_AND\_IL4\_DN | 46 | 5 | 4161 | 164 | bru1,GLS,Meltrin,step,fz2 | | 3.329e-02 | -3.40 | LEF1\_Q2 | MSigDB lists | LEF1\_Q2 | 46 | 5 | 4161 | 164 | Gdh,fbl,toy,ND-30,Rbp9 | | 3.329e-02 | -3.40 | GSE21360\_TERTIARY\_VS\_QUATERNARY\_MEMORY\_CD8\_TCELL\_UP | MSigDB lists | GSE21360\_TERTIARY\_VS\_QUATERNARY\_MEMORY\_CD8\_TCELL\_UP | 46 | 5 | 4161 | 164 | Cyp6a8,Tsp5D,MFS16,step,dnc | | 3.332e-02 | -3.40 | FN3 | prosite domains | PS50853 | 53 | 7 | 6243 | 365 | CG14964,Ptp99A,fred,bt,Dscam2,sdk,boi | | 3.339e-02 | -3.40 | fly chr95A7-95A7\|3-80 cM | chromosome location | fly chr95A7-95A7\|3-80 cM | 9 | 2 | 25010 | 823 | asRNA:CR43903,CG18754 | | 3.339e-02 | -3.40 | fly chr86E5-86E5\|3-51 cM | chromosome location | fly chr86E5-86E5\|3-51 cM | 9 | 2 | 25010 | 823 | CG5270,mir-4945 | | 3.339e-02 | -3.40 | fly chr86E5-86E5\|3-51 | chromosome location | fly chr86E5-86E5\|3-51 | 9 | 2 | 25010 | 823 | mir-4945,CG5270 | | 3.339e-02 | -3.40 | fly chr56E4-56E4\|2-90 cM | chromosome location | fly chr56E4-56E4\|2-90 cM | 9 | 2 | 25010 | 823 | lncRNA:CR44570,Toll-7 | | 3.339e-02 | -3.40 | fly chr56E4-56E4\|2-90 | chromosome location | fly chr56E4-56E4\|2-90 | 9 | 2 | 25010 | 823 | Toll-7,lncRNA:CR44570 | | 3.339e-02 | -3.40 | fly chr95A7-95A7\|3-80 | chromosome location | fly chr95A7-95A7\|3-80 | 9 | 2 | 25010 | 823 | asRNA:CR43903,CG18754 | | 3.366e-02 | -3.39 | detection of stimulus | biological process | GO:0051606 | 278 | 20 | 12345 | 571 | Gprk1,Ir40a,Gr39a,Pld,norpA,Or83c,Gr93d,Or69a,CG31750,Rh3,Gr2a,Ir94e,Or98a,shakB,Ac78C,Ir60e,rdgC,CheB42a,Ir41a,Or67d | | 3.371e-02 | -3.39 | translation repressor activity | molecular function | GO:0030371 | 16 | 3 | 12176 | 554 | heph,pum,orb | | 3.395e-02 | -3.38 | positive regulation of nucleic acid-templated transcription | biological process | GO:1903508 | 494 | 32 | 12345 | 571 | Mef2,zen,Atf6,pho,caup,lz,Clk,hth,dimm,sna,Trl,scro,zld,dsx,toy,lola,en,Camta,CG40228,d4,SoxN,bi,tj,SCAP,CG12769,ap,tou,jing,Myc,br,Setd3,pan | | 3.395e-02 | -3.38 | positive regulation of RNA biosynthetic process | biological process | GO:1902680 | 494 | 32 | 12345 | 571 | Trl,scro,sna,d4,SoxN,Camta,CG40228,toy,dsx,lola,en,zld,jing,tou,Myc,CG12769,ap,SCAP,tj,bi,pan,br,Setd3,zen,Mef2,pho,caup,Atf6,Clk,lz,hth,dimm | | 3.399e-02 | -3.38 | GO\_ACTIVATION\_OF\_PROTEIN\_KINASE\_ACTIVITY | MSigDB lists | GO\_ACTIVATION\_OF\_PROTEIN\_KINASE\_ACTIVITY | 62 | 6 | 4161 | 164 | Ac78C,rl,nAChRalpha4,Octbeta2R,Sod1,Stlk | | 3.399e-02 | -3.38 | LIM\_MAMMARY\_STEM\_CELL\_UP | MSigDB lists | LIM\_MAMMARY\_STEM\_CELL\_UP | 62 | 6 | 4161 | 164 | CG2256,shot,Glut1,Mef2,sdk,Nlg3 | | 3.399e-02 | -3.38 | ER\_Q6\_02 | MSigDB lists | ER\_Q6\_02 | 62 | 6 | 4161 | 164 | Gfat1,toy,gus,RSG7,Tim23,Gdh | | 3.414e-02 | -3.38 | gravitaxis | biological process | GO:0042332 | 27 | 4 | 12345 | 571 | l(3)L1231,Hipk,shep,br | | 3.414e-02 | -3.38 | response to gravity | biological process | GO:0009629 | 27 | 4 | 12345 | 571 | br,shep,Hipk,l(3)L1231 | | 3.414e-02 | -3.38 | response to alkaloid | biological process | GO:0043279 | 27 | 4 | 12345 | 571 | Cyp6a8,Tdc2,Clk,Ac78C | | 3.414e-02 | -3.38 | activation of GTPase activity | biological process | GO:0090630 | 27 | 4 | 12345 | 571 | RhoGAP100F,mbc,Tbc1d15-17,CG34408 | | 3.414e-02 | -3.38 | GO\_REGULATION\_OF\_CELLULAR\_COMPONENT\_MOVEMENT | MSigDB lists | GO\_REGULATION\_OF\_CELLULAR\_COMPONENT\_MOVEMENT | 132 | 10 | 4161 | 164 | rl,dnc,l(2)41Ab,Hsc70-3,cta,toy,svp,cv-2,Alg-2,CG17387 | | 3.416e-02 | -3.38 | ATPase\_P-typ\_cyto\_dom\_N | interpro domains | IPR023299 | 15 | 3 | 11383 | 557 | CG9981,PMCA,Atpalpha | | 3.416e-02 | -3.38 | BRCT\_dom\_sf | interpro domains | IPR036420 | 15 | 3 | 11383 | 557 | Parp,DNAlig4,Ptip | | 3.416e-02 | -3.38 | ATPase\_P-typ\_transduc\_dom\_A\_sf | interpro domains | IPR008250 | 15 | 3 | 11383 | 557 | Atpalpha,CG9981,PMCA | | 3.416e-02 | -3.38 | Ricin\_B\_lectin | interpro domains | IPR000772 | 15 | 3 | 11383 | 557 | Pgant1,Pgant9,Pgant5 | | 3.418e-02 | -3.38 | positive regulation of cell communication | biological process | GO:0010647 | 403 | 27 | 12345 | 571 | hth,Gprk1,mav,dimm,Diap1,mbc,Slmap,Hipk,Src42A,lqf,ru,scw,step,Glut1,CG14184,rau,Alg-2,Git,Tak1,CG30440,CG10188,Mp,svp,PEK,ewg,tld,Wdr59 | | 3.418e-02 | -3.38 | positive regulation of signaling | biological process | GO:0023056 | 403 | 27 | 12345 | 571 | mbc,Diap1,dimm,mav,hth,Gprk1,ru,lqf,Hipk,Src42A,Slmap,Glut1,step,scw,Alg-2,rau,CG14184,Tak1,Git,CG30440,Mp,CG10188,Wdr59,tld,svp,ewg,PEK | | 3.429e-02 | -3.37 | GO\_INTRINSIC\_COMPONENT\_OF\_ORGANELLE\_MEMBRANE | MSigDB lists | GO\_INTRINSIC\_COMPONENT\_OF\_ORGANELLE\_MEMBRANE | 96 | 8 | 4161 | 164 | Hsc70-3,Hmt-1,Tim17b,alpha4GT1,Tim23,MCU,MFS16,Der-1 | | 3.443e-02 | -3.37 | slmb (supernumerary limbs) | protein interactions | 42504 | 75 | 8 | 9204 | 464 | Dys,Hipk,norpA,en,Mical,tou,IA-2,par-1 | | 3.446e-02 | -3.37 | vesicle-mediated transport in synapse | biological process | GO:0099003 | 112 | 10 | 12345 | 571 | Snap25,Nlg3,unc-13-4A,Syt7,cpx,sky,Git,AP-2alpha,Nlg2,lqf | | 3.446e-02 | -3.37 | synaptic vesicle cycle | biological process | GO:0099504 | 112 | 10 | 12345 | 571 | Nlg3,Snap25,unc-13-4A,Syt7,cpx,sky,Git,AP-2alpha,Nlg2,lqf | | 3.456e-02 | -3.37 | presynaptic active zone | cellular component | GO:0048786 | 28 | 4 | 11940 | 534 | RhoGAP100F,cpx,Git,Ten-a | | 3.456e-02 | -3.37 | PcG protein complex | cellular component | GO:0031519 | 28 | 4 | 11940 | 534 | Su(z)2,jing,pho,Psc | | 3.456e-02 | -3.37 | euchromatin | cellular component | GO:0000791 | 28 | 4 | 11940 | 534 | Parp,Set1,Trl,HP6 | | 3.457e-02 | -3.36 | gene silencing by RNA | biological process | GO:0031047 | 67 | 7 | 12345 | 571 | HPS4,Hsc70-3,AGO3,hdc,Ars2,CG10077,drosha | | 3.457e-02 | -3.36 | maintenance of location | biological process | GO:0051235 | 67 | 7 | 12345 | 571 | boi,Mef2,Lsd-2,Parg,pk,Parp,Msp300 | | 3.458e-02 | -3.36 | GO\_CHEMICAL\_HOMEOSTASIS | MSigDB lists | GO\_CHEMICAL\_HOMEOSTASIS | 209 | 14 | 4161 | 164 | Ac78C,dnr1,cta,Sod1,toy,Hmt-1,CG4341,PMCA,Irp-1A,MCU,ABCB7,Der-1,Pdk,Atpalpha | | 3.473e-02 | -3.36 | epidermal cell differentiation | biological process | GO:0009913 | 53 | 6 | 12345 | 571 | pk,cmb,par-1,Cip4,ck,sn | | 3.473e-02 | -3.36 | skeletal muscle organ development | biological process | GO:0060538 | 53 | 6 | 12345 | 571 | Msp300,Mef2,dnt,mbc,shot,CAP | | 3.475e-02 | -3.36 | distal axon | cellular component | GO:0150034 | 100 | 9 | 11940 | 534 | cpx,Snap25,shot,pum,Mical,sky,Magi,Rtnl1,par-1 | | 3.477e-02 | -3.36 | CG13423 (uncharacterized protein) | protein interactions | 37315 | 25 | 4 | 9204 | 464 | kni,Atf6,CycB,CstF50 | | 3.485e-02 | -3.36 | GO\_TRANSPORTER\_COMPLEX | MSigDB lists | GO\_TRANSPORTER\_COMPLEX | 79 | 7 | 4161 | 164 | dnc,Ca-alpha1D,nAChRalpha3,Hmt-1,Atpalpha,nAChRalpha4,MCU | | 3.485e-02 | -3.36 | GSE12198\_CTRL\_VS\_LOW\_IL2\_STIM\_NK\_CELL\_DN | MSigDB lists | GSE12198\_CTRL\_VS\_LOW\_IL2\_STIM\_NK\_CELL\_DN | 79 | 7 | 4161 | 164 | Rpn9,Syx18,AGO3,CG9107,Trxr-2,IntS3,mIF2 | | 3.485e-02 | -3.36 | MODULE\_532 | MSigDB lists | MODULE\_532 | 79 | 7 | 4161 | 164 | Nadsyn,CG41099,unc-13-4A,dnc,bru1,Alg-2,gus | | 3.516e-02 | -3.35 | male courtship behavior, veined wing vibration | biological process | GO:0016545 | 16 | 3 | 12345 | 571 | fru,Nlg2,dsx | | 3.516e-02 | -3.35 | cellular response to reactive oxygen species | biological process | GO:0034614 | 16 | 3 | 12345 | 571 | Sod1,rl,CG6888 | | 3.516e-02 | -3.35 | nurse cell apoptotic process | biological process | GO:0045476 | 16 | 3 | 12345 | 571 | lola,grim,DNaseII | | 3.521e-02 | -3.35 | cell cortex | cellular component | GO:0005938 | 183 | 14 | 11940 | 534 | igl,par-1,Syx18,step,conu,toc,capu,ck,sbm,gus,Dys,Git,Spn,Ten-a | | 3.534e-02 | -3.34 | SWI/SNF superfamily-type complex | cellular component | GO:0070603 | 55 | 6 | 11940 | 534 | en,tou,Yp2,Act42A,pho,d4 | | 3.540e-02 | -3.34 | GO\_EXTRINSIC\_COMPONENT\_OF\_PLASMA\_MEMBRANE | MSigDB lists | GO\_EXTRINSIC\_COMPONENT\_OF\_PLASMA\_MEMBRANE | 32 | 4 | 4161 | 164 | cta,RSG7,Src42A,step | | 3.540e-02 | -3.34 | KEGG\_GNRH\_SIGNALING\_PATHWAY | MSigDB lists | KEGG\_GNRH\_SIGNALING\_PATHWAY | 32 | 4 | 4161 | 164 | Ac78C,rl,Ca-alpha1D,norpA | | 3.540e-02 | -3.34 | GSE29618\_PRE\_VS\_DAY7\_POST\_LAIV\_FLU\_VACCINE\_PDC\_UP | MSigDB lists | GSE29618\_PRE\_VS\_DAY7\_POST\_LAIV\_FLU\_VACCINE\_PDC\_UP | 32 | 4 | 4161 | 164 | sn,DNaseII,Spn42De,CG2528 | | 3.540e-02 | -3.34 | KEGG\_GAP\_JUNCTION | MSigDB lists | KEGG\_GAP\_JUNCTION | 32 | 4 | 4161 | 164 | rl,Ac78C,norpA,Pkg21D | | 3.540e-02 | -3.34 | GSE41978\_WT\_VS\_ID2\_KO\_KLRG1\_LOW\_EFFECTOR\_CD8\_TCELL\_UP | MSigDB lists | GSE41978\_WT\_VS\_ID2\_KO\_KLRG1\_LOW\_EFFECTOR\_CD8\_TCELL\_UP | 32 | 4 | 4161 | 164 | gus,Ugt50B3,Ca-alpha1D,AGO3 | | 3.540e-02 | -3.34 | GSE14769\_UNSTIM\_VS\_120MIN\_LPS\_BMDM\_DN | MSigDB lists | GSE14769\_UNSTIM\_VS\_120MIN\_LPS\_BMDM\_DN | 32 | 4 | 4161 | 164 | CG7255,fz2,arg,cta | | 3.558e-02 | -3.34 | regulation of protein phosphorylation | biological process | GO:0001932 | 245 | 18 | 12345 | 571 | Stlk,CG41520,Tak1,PEK,Trl,par-1,Ppm1,CG17514,Src42A,mav,CycB,mbc,SteXh:CG42398,CG14184,Alg-2,fray,scw,step | | 3.570e-02 | -3.33 | GO\_BEHAVIOR | MSigDB lists | GO\_BEHAVIOR | 133 | 10 | 4161 | 164 | nAChRalpha4,GLS,Nlg2,Atpalpha,Nlg3,Ac78C,Git,Octbeta2R,Mef2,Sod1 | | 3.571e-02 | -3.33 | Homeobox\_CS | interpro domains | IPR017970 | 92 | 9 | 11383 | 557 | zen,ap,ct,scro,caup,en,E5,Vsx2,toy | | 3.592e-02 | -3.33 | GO\_SYNAPSE\_PART | MSigDB lists | GO\_SYNAPSE\_PART | 171 | 12 | 4161 | 164 | Nlg3,Trs23,unc-13-4A,Octbeta2R,Mef2,Dys,nAChRalpha3,nAChRalpha4,Atpalpha,Nlg2,norpA,sky | | 3.593e-02 | -3.33 | positive regulation of macromolecule biosynthetic process | biological process | GO:0010557 | 552 | 35 | 12345 | 571 | SoxN,d4,pum,CG40228,Camta,lola,mxt,en,dsx,toy,zld,Trl,scro,sna,orb,pan,br,Setd3,Myc,tou,jing,ap,CG12769,SCAP,bi,tj,caup,pho,Atf6,zen,Mef2,dimm,hth,Clk,lz | | 3.600e-02 | -3.32 | RNA interference | biological process | GO:0016246 | 40 | 5 | 12345 | 571 | hdc,CG10077,drosha,Ars2,Hsc70-3 | | 3.607e-02 | -3.32 | NAKAMURA\_TUMOR\_ZONE\_PERIPHERAL\_VS\_CENTRAL\_DN | MSigDB lists | NAKAMURA\_TUMOR\_ZONE\_PERIPHERAL\_VS\_CENTRAL\_DN | 152 | 11 | 4161 | 164 | Dys,arg,Glut1,dnc,RpL15,Hsc70-3,mRpL10,CG14621,ABCB7,CG7255,Spn42De | | 3.610e-02 | -3.32 | intercalary heterochromatin | cellular component | GO:0005725 | 7 | 2 | 11940 | 534 | Psc,Su(z)2 | | 3.610e-02 | -3.32 | voltage-gated calcium channel complex | cellular component | GO:0005891 | 7 | 2 | 11940 | 534 | Ca-Ma2d,Ca-alpha1D | | 3.611e-02 | -3.32 | HNF1\_01 | MSigDB lists | HNF1\_01 | 47 | 5 | 4161 | 164 | sbm,PMCA,heph,Cyp6a8,Dys | | 3.611e-02 | -3.32 | GO\_REGULATION\_OF\_NEURON\_APOPTOTIC\_PROCESS | MSigDB lists | GO\_REGULATION\_OF\_NEURON\_APOPTOTIC\_PROCESS | 47 | 5 | 4161 | 164 | vap,DNAlig4,spel1,Mef2,Sod1 | | 3.611e-02 | -3.32 | CREBP1\_01 | MSigDB lists | CREBP1\_01 | 47 | 5 | 4161 | 164 | Octbeta2R,bru1,Rbp9,Dys,Mp | | 3.611e-02 | -3.32 | HFH1\_01 | MSigDB lists | HFH1\_01 | 47 | 5 | 4161 | 164 | Dys,vtd,Pi3K68D,Ca-alpha1D,mbc | | 3.611e-02 | -3.32 | NFAT\_Q6 | MSigDB lists | NFAT\_Q6 | 47 | 5 | 4161 | 164 | mIF2,svp,rg,CG14820,dnc | | 3.611e-02 | -3.32 | EVI1\_04 | MSigDB lists | EVI1\_04 | 47 | 5 | 4161 | 164 | Rbp9,Dys,CG12121,svp,dnc | | 3.611e-02 | -3.32 | NKX62\_Q2 | MSigDB lists | NKX62\_Q2 | 47 | 5 | 4161 | 164 | Dys,Mef2,Rbp9,Nlg3,sbm | | 3.611e-02 | -3.32 | GSE7568\_IL4\_TGFB\_DEXAMETHASONE\_VS\_IL4\_TGFB\_TREATED\_MACROPHAGE\_DN | MSigDB lists | GSE7568\_IL4\_TGFB\_DEXAMETHASONE\_VS\_IL4\_TGFB\_TREATED\_MACROPHAGE\_DN | 47 | 5 | 4161 | 164 | Glut1,Ubr1,Maf1,Trs23,mlt | | 3.611e-02 | -3.32 | GAUSSMANN\_MLL\_AF4\_FUSION\_TARGETS\_A\_UP | MSigDB lists | GAUSSMANN\_MLL\_AF4\_FUSION\_TARGETS\_A\_UP | 47 | 5 | 4161 | 164 | Irp-1A,CG3107,Git,CG8184,RpL15 | | 3.614e-02 | -3.32 | positive regulation of signal transduction | biological process | GO:0009967 | 369 | 25 | 12345 | 571 | Mp,CG10188,Wdr59,svp,PEK,ewg,tld,Tak1,Git,CG30440,scw,step,rau,Alg-2,CG14184,Diap1,mbc,Gprk1,hth,mav,Hipk,Src42A,lqf,ru,Slmap | | 3.624e-02 | -3.32 | membrane protein complex | cellular component | GO:0098796 | 533 | 33 | 11940 | 534 | nAChRalpha4,Der-1,Pi3K68D,GABA-B-R3,norpA,CG41106,Atpalpha,SdhA,Tim17b,KCNQ,HisCl1,UQCR-11,CG15459,Msp300,CG41128,Syx18,nAChRalpha3,Snap25,cta,Dys,AP-2alpha,Ca-alpha1D,ND-AGGG,CG33970,CG14184,Tim23,Ca-Ma2d,CG40472,cpx,ND-MLRQ,MCU,ND-30,CadN2 | | 3.639e-02 | -3.31 | GO\_REGULATION\_OF\_METAL\_ION\_TRANSPORT | MSigDB lists | GO\_REGULATION\_OF\_METAL\_ION\_TRANSPORT | 63 | 6 | 4161 | 164 | Dys,norpA,Octbeta2R,dnc,fray,Ca-alpha1D | | 3.639e-02 | -3.31 | LINDGREN\_BLADDER\_CANCER\_CLUSTER\_2B | MSigDB lists | LINDGREN\_BLADDER\_CANCER\_CLUSTER\_2B | 63 | 6 | 4161 | 164 | DNAlig4,Meltrin,CG41378,Mp,CG7255,bru1 | | 3.646e-02 | -3.31 | - | gene3d domains | 4.10.280.10 | 57 | 7 | 4929 | 273 | Myc,dimm,Mnt,cato,CG33557,Fer3,Clk | | 3.649e-02 | -3.31 | dynein intermediate chain binding | molecular function | GO:0045505 | 28 | 4 | 12176 | 554 | CG5359,kl-3,Dhc16F,CG31275 | | 3.650e-02 | -3.31 | dendrite development | biological process | GO:0016358 | 145 | 12 | 12345 | 571 | br,lola,ct,Sema1a,pum,sn,Ulp1,pho,Mical,scro,shot,fru | | 3.657e-02 | -3.31 | RA | prosite domains | PS50200 | 13 | 3 | 6243 | 365 | pico,rau,Myo81F | | 3.662e-02 | -3.31 | TAKEDA\_TARGETS\_OF\_NUP98\_HOXA9\_FUSION\_10D\_UP | MSigDB lists | TAKEDA\_TARGETS\_OF\_NUP98\_HOXA9\_FUSION\_10D\_UP | 19 | 3 | 4161 | 164 | nAChRalpha3,Dys,arg | | 3.662e-02 | -3.31 | GO\_POSITIVE\_REGULATION\_OF\_MUSCLE\_TISSUE\_DEVELOPMENT | MSigDB lists | GO\_POSITIVE\_REGULATION\_OF\_MUSCLE\_TISSUE\_DEVELOPMENT | 19 | 3 | 4161 | 164 | CycB,Mef2,Octbeta2R | | 3.662e-02 | -3.31 | MIKKELSEN\_MEF\_HCP\_WITH\_H3\_UNMETHYLATED | MSigDB lists | MIKKELSEN\_MEF\_HCP\_WITH\_H3\_UNMETHYLATED | 19 | 3 | 4161 | 164 | Nlg3,sn,drosha | | 3.662e-02 | -3.31 | GO\_REGULATION\_OF\_FIBROBLAST\_PROLIFERATION | MSigDB lists | GO\_REGULATION\_OF\_FIBROBLAST\_PROLIFERATION | 19 | 3 | 4161 | 164 | CG41378,CycB,DNAlig4 | | 3.662e-02 | -3.31 | MORF\_FRK | MSigDB lists | MORF\_FRK | 19 | 3 | 4161 | 164 | nAChRalpha4,Src42A,cv-2 | | 3.662e-02 | -3.31 | GO\_B\_CELL\_ACTIVATION | MSigDB lists | GO\_B\_CELL\_ACTIVATION | 19 | 3 | 4161 | 164 | Mef2,DNAlig4,spel1 | | 3.671e-02 | -3.30 | vret (vreteno) | protein interactions | 42695 | 37 | 5 | 9204 | 464 | cbs,tinc,jigr1,AGO3,Tak1 | | 3.683e-02 | -3.30 | knrl (knirps-like) | protein interactions | 40285 | 15 | 3 | 9204 | 464 | Vsx2,CG5953,kni | | 3.683e-02 | -3.30 | Cpr100A (Cuticular protein 100A) | protein interactions | 43657 | 15 | 3 | 9204 | 464 | CG9411,hth,dnt | | 3.683e-02 | -3.30 | Idh3b (Isocitrate dehydrogenase 3b) | protein interactions | 42586 | 15 | 3 | 9204 | 464 | Nrd1,CG8888,Tak1 | | 3.683e-02 | -3.30 | PH4alphaEFB (prolyl-4-hydroxylase-alpha EFB) | protein interactions | 43620 | 15 | 3 | 9204 | 464 | Rtnl1,Sema1a,CG11060 | | 3.690e-02 | -3.30 | GO\_TISSUE\_DEVELOPMENT | MSigDB lists | GO\_TISSUE\_DEVELOPMENT | 292 | 18 | 4161 | 164 | Bruce,CG17387,fz2,ap,sdk,arg,Sod1,svp,dnc,rl,CycB,St1,cv-2,toy,Mef2,cta,ck,l(2)41Ab | | 3.697e-02 | -3.30 | GO\_REGULATION\_OF\_CARDIAC\_MUSCLE\_CELL\_PROLIFERATION | MSigDB lists | GO\_REGULATION\_OF\_CARDIAC\_MUSCLE\_CELL\_PROLIFERATION | 8 | 2 | 4161 | 164 | Mef2,CycB | | 3.697e-02 | -3.30 | KUNINGER\_IGF1\_VS\_PDGFB\_TARGETS\_DN | MSigDB lists | KUNINGER\_IGF1\_VS\_PDGFB\_TARGETS\_DN | 8 | 2 | 4161 | 164 | Atpalpha,Pkg21D | | 3.697e-02 | -3.30 | GO\_POSITIVE\_REGULATION\_OF\_SODIUM\_ION\_TRANSPORT | MSigDB lists | GO\_POSITIVE\_REGULATION\_OF\_SODIUM\_ION\_TRANSPORT | 8 | 2 | 4161 | 164 | Dys,Octbeta2R | | 3.697e-02 | -3.30 | ABE\_VEGFA\_TARGETS\_30MIN | MSigDB lists | ABE\_VEGFA\_TARGETS\_30MIN | 8 | 2 | 4161 | 164 | ck,Rbp9 | | 3.697e-02 | -3.30 | GO\_POSITIVE\_REGULATION\_OF\_CARDIAC\_MUSCLE\_CELL\_PROLIFERATION | MSigDB lists | GO\_POSITIVE\_REGULATION\_OF\_CARDIAC\_MUSCLE\_CELL\_PROLIFERATION | 8 | 2 | 4161 | 164 | CycB,Mef2 | | 3.697e-02 | -3.30 | PARK\_HSC\_VS\_MULTIPOTENT\_PROGENITORS\_DN | MSigDB lists | PARK\_HSC\_VS\_MULTIPOTENT\_PROGENITORS\_DN | 8 | 2 | 4161 | 164 | RpL6,vtd | | 3.697e-02 | -3.30 | VALK\_AML\_CLUSTER\_13 | MSigDB lists | VALK\_AML\_CLUSTER\_13 | 8 | 2 | 4161 | 164 | CG32944,unc-13-4A | | 3.697e-02 | -3.30 | GO\_INTRINSIC\_APOPTOTIC\_SIGNALING\_PATHWAY\_BY\_P53\_CLASS\_MEDIATOR | MSigDB lists | GO\_INTRINSIC\_APOPTOTIC\_SIGNALING\_PATHWAY\_BY\_P53\_CLASS\_MEDIATOR | 8 | 2 | 4161 | 164 | spel1,CG10077 | | 3.697e-02 | -3.30 | TENEDINI\_MEGAKARYOCYTE\_MARKERS | MSigDB lists | TENEDINI\_MEGAKARYOCYTE\_MARKERS | 8 | 2 | 4161 | 164 | vap,CycB | | 3.697e-02 | -3.30 | GO\_REGULATION\_OF\_PROTEIN\_HOMODIMERIZATION\_ACTIVITY | MSigDB lists | GO\_REGULATION\_OF\_PROTEIN\_HOMODIMERIZATION\_ACTIVITY | 8 | 2 | 4161 | 164 | Mef2,Hsc70-3 | | 3.697e-02 | -3.30 | GO\_VASODILATION | MSigDB lists | GO\_VASODILATION | 8 | 2 | 4161 | 164 | Sod1,Octbeta2R | | 3.697e-02 | -3.30 | GO\_CARDIAC\_MUSCLE\_CELL\_ACTION\_POTENTIAL | MSigDB lists | GO\_CARDIAC\_MUSCLE\_CELL\_ACTION\_POTENTIAL | 8 | 2 | 4161 | 164 | Ca-alpha1D,Dys | | 3.697e-02 | -3.30 | KYNG\_ENVIRONMENTAL\_STRESS\_RESPONSE\_NOT\_BY\_GAMMA\_IN\_WS | MSigDB lists | KYNG\_ENVIRONMENTAL\_STRESS\_RESPONSE\_NOT\_BY\_GAMMA\_IN\_WS | 8 | 2 | 4161 | 164 | Sod1,par-1 | | 3.697e-02 | -3.30 | GO\_B\_CELL\_ACTIVATION\_INVOLVED\_IN\_IMMUNE\_RESPONSE | MSigDB lists | GO\_B\_CELL\_ACTIVATION\_INVOLVED\_IN\_IMMUNE\_RESPONSE | 8 | 2 | 4161 | 164 | spel1,DNAlig4 | | 3.697e-02 | -3.30 | GO\_POSITIVE\_REGULATION\_OF\_HEART\_CONTRACTION | MSigDB lists | GO\_POSITIVE\_REGULATION\_OF\_HEART\_CONTRACTION | 8 | 2 | 4161 | 164 | Octbeta2R,dnc | | 3.697e-02 | -3.30 | GO\_REACTIVE\_NITROGEN\_SPECIES\_METABOLIC\_PROCESS | MSigDB lists | GO\_REACTIVE\_NITROGEN\_SPECIES\_METABOLIC\_PROCESS | 8 | 2 | 4161 | 164 | arg,CG7255 | | 3.697e-02 | -3.30 | GO\_FOREBRAIN\_REGIONALIZATION | MSigDB lists | GO\_FOREBRAIN\_REGIONALIZATION | 8 | 2 | 4161 | 164 | ap,toy | | 3.697e-02 | -3.30 | PID\_ARF6\_PATHWAY | MSigDB lists | PID\_ARF6\_PATHWAY | 8 | 2 | 4161 | 164 | Octbeta2R,Khc-73 | | 3.697e-02 | -3.30 | GO\_ORGAN\_FORMATION | MSigDB lists | GO\_ORGAN\_FORMATION | 8 | 2 | 4161 | 164 | Mef2,rl | | 3.697e-02 | -3.30 | GO\_POSITIVE\_REGULATION\_OF\_BLOOD\_PRESSURE | MSigDB lists | GO\_POSITIVE\_REGULATION\_OF\_BLOOD\_PRESSURE | 8 | 2 | 4161 | 164 | dnc,Octbeta2R | | 3.697e-02 | -3.30 | BOGNI\_TREATMENT\_RELATED\_MYELOID\_LEUKEMIA\_DN | MSigDB lists | BOGNI\_TREATMENT\_RELATED\_MYELOID\_LEUKEMIA\_DN | 8 | 2 | 4161 | 164 | sn,CG2528 | | 3.697e-02 | -3.30 | GO\_VENTRICULAR\_CARDIAC\_MUSCLE\_CELL\_DIFFERENTIATION | MSigDB lists | GO\_VENTRICULAR\_CARDIAC\_MUSCLE\_CELL\_DIFFERENTIATION | 8 | 2 | 4161 | 164 | CycB,Mef2 | | 3.697e-02 | -3.30 | GO\_MODIFIED\_AMINO\_ACID\_TRANSMEMBRANE\_TRANSPORTER\_ACTIVITY | MSigDB lists | GO\_MODIFIED\_AMINO\_ACID\_TRANSMEMBRANE\_TRANSPORTER\_ACTIVITY | 8 | 2 | 4161 | 164 | Mct1,sbm | | 3.697e-02 | -3.30 | GO\_NEUROTRANSMITTER\_BINDING | MSigDB lists | GO\_NEUROTRANSMITTER\_BINDING | 8 | 2 | 4161 | 164 | nAChRalpha4,nAChRalpha3 | | 3.697e-02 | -3.30 | GO\_EMBRYONIC\_CAMERA\_TYPE\_EYE\_DEVELOPMENT | MSigDB lists | GO\_EMBRYONIC\_CAMERA\_TYPE\_EYE\_DEVELOPMENT | 8 | 2 | 4161 | 164 | fz2,toy | | 3.697e-02 | -3.30 | MODULE\_186 | MSigDB lists | MODULE\_186 | 8 | 2 | 4161 | 164 | sbm,CG7255 | | 3.697e-02 | -3.30 | GAUSSMANN\_MLL\_AF4\_FUSION\_TARGETS\_D\_UP | MSigDB lists | GAUSSMANN\_MLL\_AF4\_FUSION\_TARGETS\_D\_UP | 8 | 2 | 4161 | 164 | Ranbp16,Mct1 | | 3.697e-02 | -3.30 | YIH\_RESPONSE\_TO\_ARSENITE\_C1 | MSigDB lists | YIH\_RESPONSE\_TO\_ARSENITE\_C1 | 8 | 2 | 4161 | 164 | CycB,vap | | 3.727e-02 | -3.29 | transforming growth factor beta receptor binding | molecular function | GO:0005160 | 7 | 2 | 12176 | 554 | scw,mav | | 3.727e-02 | -3.29 | stearoyl-CoA 9-desaturase activity | molecular function | GO:0004768 | 7 | 2 | 12176 | 554 | CG9747,CG9743 | | 3.727e-02 | -3.29 | phospholipase A2 activity | molecular function | GO:0004623 | 7 | 2 | 12176 | 554 | CG42237,CG3009 | | 3.727e-02 | -3.29 | acyl-CoA desaturase activity | molecular function | GO:0016215 | 7 | 2 | 12176 | 554 | CG9747,CG9743 | | 3.727e-02 | -3.29 | transcription cofactor binding | molecular function | GO:0001221 | 7 | 2 | 12176 | 554 | hth,Tgi | | 3.727e-02 | -3.29 | Rac guanyl-nucleotide exchange factor activity | molecular function | GO:0030676 | 7 | 2 | 12176 | 554 | mbc,sif | | 3.727e-02 | -3.29 | acetylcholine receptor activity | molecular function | GO:0015464 | 7 | 2 | 12176 | 554 | nAChRalpha3,nAChRalpha4 | | 3.727e-02 | -3.29 | oxidoreductase activity, acting on paired donors, with oxidation of a pair of donors resulting in the reduction of molecular oxygen to two molecules of water | molecular function | GO:0016717 | 7 | 2 | 12176 | 554 | CG9743,CG9747 | | 3.731e-02 | -3.29 | GO\_TUBE\_DEVELOPMENT | MSigDB lists | GO\_TUBE\_DEVELOPMENT | 134 | 10 | 4161 | 164 | rl,l(2)41Ab,CycB,arg,cta,Mef2,toy,ap,cv-2,CG17387 | | 3.760e-02 | -3.28 | cellular ketone metabolic process | biological process | GO:0042180 | 54 | 6 | 12345 | 571 | Npc2b,Coq7,spo,CG9514,nvd,spok | | 3.760e-02 | -3.28 | regulation of multicellular organismal process | biological process | GO:0051239 | 781 | 47 | 12345 | 571 | ewg,Mef2,Cip4,par-1,shot,Bruce,salm,Git,hdc,lz,cta,babos,Clk,bru1,Sema1a,Src42A,Hipk,Spn,mbc,hth,Octbeta2R,kirre,KCNQ,fz2,zld,cpo,inc,Catsup,pum,SoxN,SPR,Mical,tinc,neo,norpA,step,CG9380,Myc,sif,lqf,Sdc,pk,Diap1,rl,Octbeta1R,sdk,mav | | 3.771e-02 | -3.28 | lipid localization | biological process | GO:0010876 | 83 | 8 | 12345 | 571 | CG43672,Mef2,cert,CG9981,Npc2b,Lsd-2,CG42237,CG3009 | | 3.771e-02 | -3.28 | neuromuscular junction development | biological process | GO:0007528 | 83 | 8 | 12345 | 571 | Nlg2,Syx18,Atpalpha,Ten-a,Nlg3,rg,par-1,cpx | | 3.790e-02 | -3.27 | extracellular exosome | cellular component | GO:0070062 | 17 | 3 | 11940 | 534 | CG3544,CG6888,CG9449 | | 3.808e-02 | -3.27 | GO\_SENSORY\_ORGAN\_DEVELOPMENT | MSigDB lists | GO\_SENSORY\_ORGAN\_DEVELOPMENT | 116 | 9 | 4161 | 164 | fz2,ap,sdk,cv-2,sn,rl,ck,toy,Sod1 | | 3.820e-02 | -3.26 | regulation of synapse structure or activity | biological process | GO:0050803 | 179 | 14 | 12345 | 571 | Cip4,Sod1,Thor,Octbeta2R,ewg,Octbeta1R,pum,Ten-a,sif,Src42A,Khc-73,Nlg2,Spn,Sdc | | 3.823e-02 | -3.26 | neuron recognition | biological process | GO:0008038 | 114 | 10 | 12345 | 571 | shot,fz2,fru,Dscam2,Ptp99A,Ten-a,nrm,dsx,lola,Sema1a | | 3.842e-02 | -3.26 | G alpha (12/13) signalling events | REACTOME pathways | R-DME-416482 | 27 | 4 | 3923 | 189 | sif,CG30440,cta,CG10188 | | 3.842e-02 | -3.26 | zygotic determination of anterior/posterior axis, embryo | biological process | GO:0007354 | 28 | 4 | 12345 | 571 | kni,rl,scw,cic | | 3.842e-02 | -3.26 | positive regulation of transmembrane receptor protein serine/threonine kinase signaling pathway | biological process | GO:0090100 | 28 | 4 | 12345 | 571 | scw,mav,hth,tld | | 3.842e-02 | -3.26 | response to oxygen radical | biological process | GO:0000305 | 7 | 2 | 12345 | 571 | CG6888,Sod1 | | 3.842e-02 | -3.26 | SMAD protein signal transduction | biological process | GO:0060395 | 7 | 2 | 12345 | 571 | mav,scw | | 3.842e-02 | -3.26 | icosanoid secretion | biological process | GO:0032309 | 7 | 2 | 12345 | 571 | CG42237,CG3009 | | 3.842e-02 | -3.26 | regulation of calcium ion transmembrane transport | biological process | GO:1903169 | 7 | 2 | 12345 | 571 | Rgk1,THADA | | 3.842e-02 | -3.26 | maintenance of protein localization in organelle | biological process | GO:0072595 | 7 | 2 | 12345 | 571 | Parp,Parg | | 3.842e-02 | -3.26 | arachidonic acid secretion | biological process | GO:0050482 | 7 | 2 | 12345 | 571 | CG42237,CG3009 | | 3.842e-02 | -3.26 | nuclear body organization | biological process | GO:0030575 | 7 | 2 | 12345 | 571 | Parp,Parg | | 3.842e-02 | -3.26 | negative regulation of cation transmembrane transport | biological process | GO:1904063 | 7 | 2 | 12345 | 571 | THADA,Rgk1 | | 3.842e-02 | -3.26 | double-strand break repair via nonhomologous end joining | biological process | GO:0006303 | 7 | 2 | 12345 | 571 | mus308,DNAlig4 | | 3.842e-02 | -3.26 | primary miRNA processing | biological process | GO:0031053 | 7 | 2 | 12345 | 571 | drosha,Ars2 | | 3.842e-02 | -3.26 | gonad morphogenesis | biological process | GO:0035262 | 7 | 2 | 12345 | 571 | tj,bab2 | | 3.842e-02 | -3.26 | postsynaptic membrane assembly | biological process | GO:0097104 | 7 | 2 | 12345 | 571 | Nlg2,Nlg3 | | 3.842e-02 | -3.26 | negative regulation of transporter activity | biological process | GO:0032410 | 7 | 2 | 12345 | 571 | Rgk1,THADA | | 3.842e-02 | -3.26 | fatty acid derivative transport | biological process | GO:1901571 | 7 | 2 | 12345 | 571 | CG42237,CG3009 | | 3.842e-02 | -3.26 | axon extension involved in axon guidance | biological process | GO:0048846 | 7 | 2 | 12345 | 571 | CadN2,RhoGAP100F | | 3.842e-02 | -3.26 | insecticide metabolic process | biological process | GO:0017143 | 7 | 2 | 12345 | 571 | Cyp6a8,Cyp6g2 | | 3.842e-02 | -3.26 | regulation of transcription by RNA polymerase III | biological process | GO:0006359 | 7 | 2 | 12345 | 571 | Maf1,Myc | | 3.842e-02 | -3.26 | negative regulation of retinal cell programmed cell death | biological process | GO:0046671 | 7 | 2 | 12345 | 571 | norpA,Diap1 | | 3.842e-02 | -3.26 | microvillus organization | biological process | GO:0032528 | 7 | 2 | 12345 | 571 | sn,ck | | 3.842e-02 | -3.26 | removal of superoxide radicals | biological process | GO:0019430 | 7 | 2 | 12345 | 571 | CG6888,Sod1 | | 3.842e-02 | -3.26 | negative regulation of compound eye retinal cell programmed cell death | biological process | GO:0046673 | 7 | 2 | 12345 | 571 | Diap1,norpA | | 3.842e-02 | -3.26 | icosanoid transport | biological process | GO:0071715 | 7 | 2 | 12345 | 571 | CG42237,CG3009 | | 3.842e-02 | -3.26 | cellular response to oxygen radical | biological process | GO:0071450 | 7 | 2 | 12345 | 571 | Sod1,CG6888 | | 3.842e-02 | -3.26 | response to superoxide | biological process | GO:0000303 | 7 | 2 | 12345 | 571 | Sod1,CG6888 | | 3.842e-02 | -3.26 | negative regulation of ion transmembrane transport | biological process | GO:0034766 | 7 | 2 | 12345 | 571 | Rgk1,THADA | | 3.842e-02 | -3.26 | regulation of ARF protein signal transduction | biological process | GO:0032012 | 7 | 2 | 12345 | 571 | step,ArfGAP1 | | 3.842e-02 | -3.26 | arachidonate transport | biological process | GO:1903963 | 7 | 2 | 12345 | 571 | CG42237,CG3009 | | 3.842e-02 | -3.26 | cellular response to osmotic stress | biological process | GO:0071470 | 7 | 2 | 12345 | 571 | grim,Ppm1 | | 3.842e-02 | -3.26 | negative regulation of calcium ion transport | biological process | GO:0051926 | 7 | 2 | 12345 | 571 | Rgk1,THADA | | 3.842e-02 | -3.26 | regulation of calcium ion transmembrane transporter activity | biological process | GO:1901019 | 7 | 2 | 12345 | 571 | Rgk1,THADA | | 3.842e-02 | -3.26 | cellular response to superoxide | biological process | GO:0071451 | 7 | 2 | 12345 | 571 | Sod1,CG6888 | | 3.842e-02 | -3.26 | endoplasmic reticulum localization | biological process | GO:0051643 | 7 | 2 | 12345 | 571 | fan,Msp300 | | 3.842e-02 | -3.26 | negative regulation of transmembrane transport | biological process | GO:0034763 | 7 | 2 | 12345 | 571 | THADA,Rgk1 | | 3.842e-02 | -3.26 | neuron projection extension involved in neuron projection guidance | biological process | GO:1902284 | 7 | 2 | 12345 | 571 | RhoGAP100F,CadN2 | | 3.842e-02 | -3.26 | central nervous system neuron axonogenesis | biological process | GO:0021955 | 7 | 2 | 12345 | 571 | Ulp1,en | | 3.842e-02 | -3.26 | negative regulation of ion transmembrane transporter activity | biological process | GO:0032413 | 7 | 2 | 12345 | 571 | THADA,Rgk1 | | 3.842e-02 | -3.26 | regulation of short-term neuronal synaptic plasticity | biological process | GO:0048172 | 7 | 2 | 12345 | 571 | Dys,Snap25 | | 3.864e-02 | -3.25 | grp (grapes) | protein interactions | 34993 | 50 | 6 | 9204 | 464 | PEK,pan,heph,CG17508,E5,dimm | | 3.877e-02 | -3.25 | Amine ligand-binding receptors | REACTOME pathways | R-DME-375280 | 16 | 3 | 3923 | 189 | Octbeta2R,5-HT2A,Octbeta1R | | 3.890e-02 | -3.25 | AR\_Q6 | MSigDB lists | AR\_Q6 | 64 | 6 | 4161 | 164 | Octbeta1R,Mef2,Dys,rg,shot,lqf | | 3.890e-02 | -3.25 | GSE20715\_WT\_VS\_TLR4\_KO\_6H\_OZONE\_LUNG\_DN | MSigDB lists | GSE20715\_WT\_VS\_TLR4\_KO\_6H\_OZONE\_LUNG\_DN | 64 | 6 | 4161 | 164 | fray,AP-2alpha,mlt,Meltrin,CG40228,Hmt-1 | | 3.907e-02 | -3.24 | GSE20727\_ROS\_INH\_VS\_ROS\_INH\_AND\_DNFB\_ALLERGEN\_TREATED\_DC\_UP | MSigDB lists | GSE20727\_ROS\_INH\_VS\_ROS\_INH\_AND\_DNFB\_ALLERGEN\_TREATED\_DC\_UP | 48 | 5 | 4161 | 164 | Glut1,cta,arg,ck,AP-2alpha | | 3.907e-02 | -3.24 | GSE17721\_LPS\_VS\_PAM3CSK4\_8H\_BMDC\_UP | MSigDB lists | GSE17721\_LPS\_VS\_PAM3CSK4\_8H\_BMDC\_UP | 48 | 5 | 4161 | 164 | vtd,PNPase,nAChRalpha4,CG10565,ND-30 | | 3.907e-02 | -3.24 | GSE10325\_BCELL\_VS\_MYELOID\_UP | MSigDB lists | GSE10325\_BCELL\_VS\_MYELOID\_UP | 48 | 5 | 4161 | 164 | Parp,Meltrin,RpL6,Odc2,GLS | | 3.907e-02 | -3.24 | GSE7460\_CTRL\_VS\_TGFB\_TREATED\_ACT\_FOXP3\_MUT\_TCONV\_UP | MSigDB lists | GSE7460\_CTRL\_VS\_TGFB\_TREATED\_ACT\_FOXP3\_MUT\_TCONV\_UP | 48 | 5 | 4161 | 164 | MCU,Glut1,dnc,CG4341,Src42A | | 3.907e-02 | -3.24 | OCT\_Q6 | MSigDB lists | OCT\_Q6 | 48 | 5 | 4161 | 164 | dnc,toy,CG9492,Dys,Cyp6a8 | | 3.907e-02 | -3.24 | OCT1\_06 | MSigDB lists | OCT1\_06 | 48 | 5 | 4161 | 164 | dnc,Ca-alpha1D,Octbeta1R,inc,bru1 | | 3.909e-02 | -3.24 | GSE34156\_UNTREATED\_VS\_24H\_TLR1\_TLR2\_LIGAND\_TREATED\_MONOCYTE\_UP | MSigDB lists | GSE34156\_UNTREATED\_VS\_24H\_TLR1\_TLR2\_LIGAND\_TREATED\_MONOCYTE\_UP | 33 | 4 | 4161 | 164 | CG41378,Tsp5D,inc,Maf1 | | 3.909e-02 | -3.24 | MORF\_NF1 | MSigDB lists | MORF\_NF1 | 33 | 4 | 4161 | 164 | Octbeta1R,dnc,mbc,CG40178 | | 3.909e-02 | -3.24 | NKX22\_01 | MSigDB lists | NKX22\_01 | 33 | 4 | 4161 | 164 | Dys,shot,Coq7,Nlg3 | | 3.909e-02 | -3.24 | GO\_COLUMNAR\_CUBOIDAL\_EPITHELIAL\_CELL\_DIFFERENTIATION | MSigDB lists | GO\_COLUMNAR\_CUBOIDAL\_EPITHELIAL\_CELL\_DIFFERENTIATION | 33 | 4 | 4161 | 164 | ck,Sod1,Mef2,toy | | 3.909e-02 | -3.24 | REACTOME\_METABOLISM\_OF\_VITAMINS\_AND\_COFACTORS | MSigDB lists | REACTOME\_METABOLISM\_OF\_VITAMINS\_AND\_COFACTORS | 33 | 4 | 4161 | 164 | Glut1,CG14721,fbl,Nadsyn | | 3.909e-02 | -3.24 | GSE29614\_CTRL\_VS\_TIV\_FLU\_VACCINE\_PBMC\_2007\_DN | MSigDB lists | GSE29614\_CTRL\_VS\_TIV\_FLU\_VACCINE\_PBMC\_2007\_DN | 33 | 4 | 4161 | 164 | SdhA,CG10171,Hsc70-3,CycB | | 3.909e-02 | -3.24 | GSE20715\_0H\_VS\_48H\_OZONE\_TLR4\_KO\_LUNG\_UP | MSigDB lists | GSE20715\_0H\_VS\_48H\_OZONE\_TLR4\_KO\_LUNG\_UP | 33 | 4 | 4161 | 164 | Ac78C,Glut1,dnr1,Mef2 | | 3.909e-02 | -3.24 | GSE17974\_IL4\_AND\_ANTI\_IL12\_VS\_UNTREATED\_6H\_ACT\_CD4\_TCELL\_DN | MSigDB lists | GSE17974\_IL4\_AND\_ANTI\_IL12\_VS\_UNTREATED\_6H\_ACT\_CD4\_TCELL\_DN | 33 | 4 | 4161 | 164 | Pgant5,dnc,Src42A,nAChRalpha3 | | 3.910e-02 | -3.24 | Fatty acid metabolism | KEGG pathways | dme01212 | 44 | 5 | 3198 | 138 | CG9747,Acox57D-d,CG9743,CG12170,FASN3 | | 3.910e-02 | -3.24 | Fatty acid metabolism | KEGG pathways | ko01212 | 44 | 5 | 3198 | 138 | Acox57D-d,CG9747,CG12170,FASN3,CG9743 | | 3.917e-02 | -3.24 | microtubule-based movement | biological process | GO:0007018 | 163 | 13 | 12345 | 571 | Dhc16F,Khc-73,pk,l(2)41Ab,CG31275,Lsd-2,kl-3,CG5359,CG17387,CG9492,jvl,CycB,Klp59C | | 3.918e-02 | -3.24 | CTACCTC\_LET7A\_LET7B\_LET7C\_LET7D\_LET7E\_LET7F\_MIR98\_LET7G\_LET7I | MSigDB lists | CTACCTC\_LET7A\_LET7B\_LET7C\_LET7D\_LET7E\_LET7F\_MIR98\_LET7G\_LET7I | 81 | 7 | 4161 | 164 | Pgant5,PMCA,CG41099,Tim17b,Octbeta1R,Dys,Octbeta2R | | 3.918e-02 | -3.24 | GSE25123\_ROSIGLITAZONE\_VS\_IL4\_AND\_ROSIGLITAZONE\_STIM\_MACROPHAGE\_DAY10\_UP | MSigDB lists | GSE25123\_ROSIGLITAZONE\_VS\_IL4\_AND\_ROSIGLITAZONE\_STIM\_MACROPHAGE\_DAY10\_UP | 81 | 7 | 4161 | 164 | Trs23,Parp,Pgant5,mRpS5,CG30495,CG9886,CG3107 | | 3.941e-02 | -3.23 | TURJANSKI\_MAPK7\_TARGETS | MSigDB lists | TURJANSKI\_MAPK7\_TARGETS | 1 | 1 | 4161 | 164 | Mef2 | | 3.941e-02 | -3.23 | KYNG\_ENVIRONMENTAL\_STRESS\_RESPONSE\_NOT\_BY\_UV\_IN\_WS | MSigDB lists | KYNG\_ENVIRONMENTAL\_STRESS\_RESPONSE\_NOT\_BY\_UV\_IN\_WS | 1 | 1 | 4161 | 164 | Sod1 | | 3.941e-02 | -3.23 | TESAR\_ALK\_TARGETS\_HUMAN\_ES\_5D\_UP | MSigDB lists | TESAR\_ALK\_TARGETS\_HUMAN\_ES\_5D\_UP | 1 | 1 | 4161 | 164 | toy | | 3.941e-02 | -3.23 | CGGTGTG\_MIR220 | MSigDB lists | CGGTGTG\_MIR220 | 1 | 1 | 4161 | 164 | ap | | 3.941e-02 | -3.23 | GO\_TRANSCRIPTIONAL\_ACTIVATOR\_ACTIVITY\_RNA\_POLYMERASE\_II\_DISTAL\_ENHANCER\_SEQUENCE\_SPECIFIC\_BINDING | MSigDB lists | GO\_TRANSCRIPTIONAL\_ACTIVATOR\_ACTIVITY\_RNA\_POLYMERASE\_II\_DISTAL\_ENHANCER\_SEQUENCE\_SPECIFIC\_BINDING | 1 | 1 | 4161 | 164 | Mef2 | | 3.941e-02 | -3.23 | GO\_PHAGOLYSOSOME\_ASSEMBLY | MSigDB lists | GO\_PHAGOLYSOSOME\_ASSEMBLY | 1 | 1 | 4161 | 164 | ck | | 3.941e-02 | -3.23 | MARIADASON\_RESPONSE\_TO\_BUTYRATE\_CURCUMIN\_SULINDAC\_TSA\_1 | MSigDB lists | MARIADASON\_RESPONSE\_TO\_BUTYRATE\_CURCUMIN\_SULINDAC\_TSA\_1 | 1 | 1 | 4161 | 164 | norpA | | 3.941e-02 | -3.23 | VANDESLUIS\_COMMD1\_TARGETS\_GROUP\_4\_UP | MSigDB lists | VANDESLUIS\_COMMD1\_TARGETS\_GROUP\_4\_UP | 1 | 1 | 4161 | 164 | Glut1 | | 3.941e-02 | -3.23 | CHOI\_ATL\_ACUTE\_STAGE | MSigDB lists | CHOI\_ATL\_ACUTE\_STAGE | 1 | 1 | 4161 | 164 | ap | | 3.941e-02 | -3.23 | GRANDVAUX\_IRF3\_TARGETS\_UP | MSigDB lists | GRANDVAUX\_IRF3\_TARGETS\_UP | 1 | 1 | 4161 | 164 | arg | | 3.941e-02 | -3.23 | UROSEVIC\_RESPONSE\_TO\_IMIQUIMOD | MSigDB lists | UROSEVIC\_RESPONSE\_TO\_IMIQUIMOD | 1 | 1 | 4161 | 164 | CG41378 | | 3.941e-02 | -3.23 | GO\_JAK\_STAT\_CASCADE\_INVOLVED\_IN\_GROWTH\_HORMONE\_SIGNALING\_PATHWAY | MSigDB lists | GO\_JAK\_STAT\_CASCADE\_INVOLVED\_IN\_GROWTH\_HORMONE\_SIGNALING\_PATHWAY | 1 | 1 | 4161 | 164 | rl | | 3.941e-02 | -3.23 | CLIMENT\_BREAST\_CANCER\_COPY\_NUMBER\_DN | MSigDB lists | CLIMENT\_BREAST\_CANCER\_COPY\_NUMBER\_DN | 1 | 1 | 4161 | 164 | Khc-73 | | 3.941e-02 | -3.23 | RIZ\_ERYTHROID\_DIFFERENTIATION\_APOBEC2 | MSigDB lists | RIZ\_ERYTHROID\_DIFFERENTIATION\_APOBEC2 | 1 | 1 | 4161 | 164 | Mef2 | | 3.941e-02 | -3.23 | SASAI\_TARGETS\_OF\_CXCR6\_AND\_PTCH1\_UP | MSigDB lists | SASAI\_TARGETS\_OF\_CXCR6\_AND\_PTCH1\_UP | 1 | 1 | 4161 | 164 | norpA | | 3.941e-02 | -3.23 | GO\_REGULATION\_OF\_INTERLEUKIN\_5\_PRODUCTION | MSigDB lists | GO\_REGULATION\_OF\_INTERLEUKIN\_5\_PRODUCTION | 1 | 1 | 4161 | 164 | dnc | | 3.941e-02 | -3.23 | TURJANSKI\_MAPK11\_TARGETS | MSigDB lists | TURJANSKI\_MAPK11\_TARGETS | 1 | 1 | 4161 | 164 | Mef2 | | 3.941e-02 | -3.23 | TESAR\_JAK\_TARGETS\_MOUSE\_ES\_D3\_DN | MSigDB lists | TESAR\_JAK\_TARGETS\_MOUSE\_ES\_D3\_DN | 1 | 1 | 4161 | 164 | AGO3 | | 3.941e-02 | -3.23 | KEGG\_MATURITY\_ONSET\_DIABETES\_OF\_THE\_YOUNG | MSigDB lists | KEGG\_MATURITY\_ONSET\_DIABETES\_OF\_THE\_YOUNG | 1 | 1 | 4161 | 164 | toy | | 3.941e-02 | -3.23 | GO\_SPONGIOTROPHOBLAST\_LAYER\_DEVELOPMENT | MSigDB lists | GO\_SPONGIOTROPHOBLAST\_LAYER\_DEVELOPMENT | 1 | 1 | 4161 | 164 | Bruce | | 3.941e-02 | -3.23 | GO\_TRIPARTITE\_REGIONAL\_SUBDIVISION | MSigDB lists | GO\_TRIPARTITE\_REGIONAL\_SUBDIVISION | 1 | 1 | 4161 | 164 | fz2 | | 3.941e-02 | -3.23 | GO\_REGULATION\_OF\_VASCULAR\_ENDOTHELIAL\_GROWTH\_FACTOR\_RECEPTOR\_SIGNALING\_PATHWAY | MSigDB lists | GO\_REGULATION\_OF\_VASCULAR\_ENDOTHELIAL\_GROWTH\_FACTOR\_RECEPTOR\_SIGNALING\_PATHWAY | 1 | 1 | 4161 | 164 | Alg-2 | | 3.941e-02 | -3.23 | GO\_ESTROUS\_CYCLE | MSigDB lists | GO\_ESTROUS\_CYCLE | 1 | 1 | 4161 | 164 | Octbeta2R | | 3.941e-02 | -3.23 | TURJANSKI\_MAPK14\_TARGETS | MSigDB lists | TURJANSKI\_MAPK14\_TARGETS | 1 | 1 | 4161 | 164 | Mef2 | | 3.941e-02 | -3.23 | GO\_VASCULAR\_SMOOTH\_MUSCLE\_CONTRACTION | MSigDB lists | GO\_VASCULAR\_SMOOTH\_MUSCLE\_CONTRACTION | 1 | 1 | 4161 | 164 | Pi3K68D | | 3.941e-02 | -3.23 | NIKOLSKY\_OVERCONNECTED\_IN\_BREAST\_CANCER | MSigDB lists | NIKOLSKY\_OVERCONNECTED\_IN\_BREAST\_CANCER | 1 | 1 | 4161 | 164 | Mef2 | | 3.941e-02 | -3.23 | MEISSNER\_BRAIN\_HCP\_WITH\_H3K4ME2 | MSigDB lists | MEISSNER\_BRAIN\_HCP\_WITH\_H3K4ME2 | 1 | 1 | 4161 | 164 | CG2061 | | 3.941e-02 | -3.23 | TESAR\_ALK\_AND\_JAK\_TARGETS\_MOUSE\_ES\_D4\_UP | MSigDB lists | TESAR\_ALK\_AND\_JAK\_TARGETS\_MOUSE\_ES\_D4\_UP | 1 | 1 | 4161 | 164 | toy | | 3.941e-02 | -3.23 | MASRI\_RESISTANCE\_TO\_TAMOXIFEN\_AND\_AROMATASE\_INHIBITORS\_UP | MSigDB lists | MASRI\_RESISTANCE\_TO\_TAMOXIFEN\_AND\_AROMATASE\_INHIBITORS\_UP | 1 | 1 | 4161 | 164 | Dys | | 3.941e-02 | -3.23 | TCGATGG\_MIR213 | MSigDB lists | TCGATGG\_MIR213 | 1 | 1 | 4161 | 164 | GCS2alpha | | 3.941e-02 | -3.23 | LEE\_SP4\_THYMOCYTE | MSigDB lists | LEE\_SP4\_THYMOCYTE | 1 | 1 | 4161 | 164 | dnc | | 3.947e-02 | -3.23 | SET | pfam domains | PF00856 | 26 | 4 | 10480 | 528 | SmydA-9,SmydA-8,Set1,Setd3 | | 3.952e-02 | -3.23 | dally (division abnormally delayed) | protein interactions | 39013 | 26 | 4 | 9204 | 464 | fz2,Sema1a,cv-2,Sdc | | 3.952e-02 | -3.23 | Mef2 (Myocyte enhancer factor 2) | protein interactions | 36032 | 26 | 4 | 9204 | 464 | br,Mef2,Myc,Svil | | 3.953e-02 | -3.23 | - | gene3d domains | 1.25.40.530 | 6 | 2 | 4929 | 273 | ck,Myo81F | | 3.953e-02 | -3.23 | - | gene3d domains | 1.20.90.10 | 6 | 2 | 4929 | 273 | CG3009,CG42237 | | 3.959e-02 | -3.23 | ATP-dependent microtubule motor activity, minus-end-directed | molecular function | GO:0008569 | 17 | 3 | 12176 | 554 | CG9492,kl-3,Dhc16F | | 3.967e-02 | -3.23 | lipid transport | biological process | GO:0006869 | 69 | 7 | 12345 | 571 | cert,CG42237,CG9981,CG3009,Npc2b,CG43672,Lsd-2 | | 3.984e-02 | -3.22 | cell fate commitment | biological process | GO:0045165 | 301 | 21 | 12345 | 571 | caup,salm,klg,lola,en,par-1,drosha,shot,sna,orb,svp,pan,rl,hth,ru,br,ap,scw,lz,bi,tj | | 3.987e-02 | -3.22 | regulation of response to stimulus | biological process | GO:0048583 | 958 | 56 | 12345 | 571 | rno,dnc,scw,Syt7,rau,Alg-2,Parp,mbc,cv-2,Ulp1,Gprk1,hth,ru,Sema1a,Hipk,Src42A,Ppm1,tay,par-1,Ptip,CG10188,rdgC,ewg,PEK,CG12147,Git,CG30440,step,vap,norpA,conu,CG9380,Toll-7,Pi3K68D,CG34408,ArfGAP1,CG14184,Parg,RSG7,Diap1,rl,mav,lqf,sif,pk,Slmap,Mp,dnr1,fuss,Wdr59,tld,svp,Tak1,Camta,SoxN,pum | | 3.997e-02 | -3.22 | RICIN | smart domains | SM00458 | 13 | 3 | 5086 | 308 | Pgant1,Pgant5,Pgant9 | | 4.007e-02 | -3.22 | somatic muscle development | biological process | GO:0007525 | 84 | 8 | 12345 | 571 | hth,Git,salm,Pkg21D,lola,ap,Thor,ewg | | 4.012e-02 | -3.22 | GO\_SENSORY\_PERCEPTION | MSigDB lists | GO\_SENSORY\_PERCEPTION | 99 | 8 | 4161 | 164 | Ca-alpha1D,sn,Sod1,toy,rl,rdgC,ck,Nlg3 | | 4.059e-02 | -3.20 | HMG | smart domains | SM00398 | 22 | 4 | 5086 | 308 | CG12104,SoxN,cic,pan | | 4.069e-02 | -3.20 | regulation of anatomical structure size | biological process | GO:0090066 | 249 | 18 | 12345 | 571 | shot,Cip4,capu,knk,lola,zld,Myc,bru1,fru,rib,conu,Thor,Atpalpha,rl,pico,pk,kune,Sema1a | | 4.071e-02 | -3.20 | enhancer binding | molecular function | GO:0035326 | 42 | 5 | 12176 | 554 | pho,en,pan,hth,Mef2 | | 4.072e-02 | -3.20 | SET\_dom | interpro domains | IPR001214 | 27 | 4 | 11383 | 557 | Set1,Setd3,SmydA-9,SmydA-8 | | 4.085e-02 | -3.20 | fly chr66A2-66A2\|3-21 | chromosome location | fly chr66A2-66A2\|3-21 | 10 | 2 | 25010 | 823 | Cyp316a1,CG8539 | | 4.085e-02 | -3.20 | fly chr66A2-66A2\|3-21 cM | chromosome location | fly chr66A2-66A2\|3-21 cM | 10 | 2 | 25010 | 823 | CG8539,Cyp316a1 | | 4.086e-02 | -3.20 | acetylgalactosaminyltransferase activity | molecular function | GO:0008376 | 29 | 4 | 12176 | 554 | Pgant1,Pgant5,alpha4GT1,Pgant9 | | 4.086e-02 | -3.20 | Rho guanyl-nucleotide exchange factor activity | molecular function | GO:0005089 | 29 | 4 | 12176 | 554 | mbc,CG30440,CG10188,sif | | 4.086e-02 | -3.20 | C2 | prosite domains | PS50004 | 33 | 5 | 6243 | 365 | unc-13-4A,norpA,Syt7,vap,Pi3K68D | | 4.104e-02 | -3.19 | GO\_RESPONSE\_TO\_NITROGEN\_COMPOUND | MSigDB lists | GO\_RESPONSE\_TO\_NITROGEN\_COMPOUND | 214 | 14 | 4161 | 164 | dnc,Ars2,Pi3K68D,rl,Ac78C,Sod1,Mef2,Octbeta2R,drosha,nAChRalpha3,Parp,SCAP,Ubr1,nAChRalpha4 | | 4.126e-02 | -3.19 | male courtship behavior, veined wing extension | biological process | GO:0048065 | 17 | 3 | 12345 | 571 | fru,Nlg2,dsx | | 4.126e-02 | -3.19 | negative regulation of secretion by cell | biological process | GO:1903531 | 17 | 3 | 12345 | 571 | cpx,CG9650,sky | | 4.126e-02 | -3.19 | chitin catabolic process | biological process | GO:0006032 | 17 | 3 | 12345 | 571 | Cht10,Idgf4,fdl | | 4.126e-02 | -3.19 | cofactor transport | biological process | GO:0051181 | 17 | 3 | 12345 | 571 | CG3556,ABCB7,Hmt-1 | | 4.126e-02 | -3.19 | muscle contraction | biological process | GO:0006936 | 17 | 3 | 12345 | 571 | Ca-alpha1D,KCNQ,bt | | 4.132e-02 | -3.19 | Keratan sulfate degradation | REACTOME pathways | R-DME-2022857 | 7 | 2 | 3923 | 189 | fdl,CG30059 | | 4.132e-02 | -3.19 | Transcriptional regulation by the AP-2 (TFAP2) family of transcription factors | REACTOME pathways | R-DME-8864260 | 7 | 2 | 3923 | 189 | Myc,twz | | 4.132e-02 | -3.19 | Activation of SMO | REACTOME pathways | R-DME-5635838 | 7 | 2 | 3923 | 189 | Gprk1,CG12147 | | 4.145e-02 | -3.18 | postsynapse | cellular component | GO:0098794 | 72 | 7 | 11940 | 534 | Spn,sky,Magi,nAChRalpha4,nAChRalpha3,pum,Nlg2 | | 4.147e-02 | -3.18 | MODULE\_459 | MSigDB lists | MODULE\_459 | 82 | 7 | 4161 | 164 | Alg-2,gus,bru1,unc-13-4A,dnc,Nadsyn,CG41099 | | 4.151e-02 | -3.18 | JISON\_SICKLE\_CELL\_DISEASE\_DN | MSigDB lists | JISON\_SICKLE\_CELL\_DISEASE\_DN | 65 | 6 | 4161 | 164 | Ars2,RpL6,spel1,GCS2alpha,CG10077,GLS | | 4.151e-02 | -3.18 | GO\_EXTRINSIC\_COMPONENT\_OF\_MEMBRANE | MSigDB lists | GO\_EXTRINSIC\_COMPONENT\_OF\_MEMBRANE | 65 | 6 | 4161 | 164 | cta,dnr1,Pi3K68D,step,RSG7,Src42A | | 4.151e-02 | -3.18 | RFX1\_02 | MSigDB lists | RFX1\_02 | 65 | 6 | 4161 | 164 | Nlg3,Syx18,unc-13-4A,Atpalpha,Dys,CG13284 | | 4.166e-02 | -3.18 | mesoderm development | biological process | GO:0007498 | 100 | 9 | 12345 | 571 | Msp300,wb,pan,Mef2,sna,bt,cta,Cip4,scw | | 4.174e-02 | -3.18 | intrinsic component of mitochondrial membrane | cellular component | GO:0098573 | 43 | 5 | 11940 | 534 | Tim17b,fzo,Tim23,MCU,CG41128 | | 4.187e-02 | -3.17 | GO\_EMBRYONIC\_HEART\_TUBE\_MORPHOGENESIS | MSigDB lists | GO\_EMBRYONIC\_HEART\_TUBE\_MORPHOGENESIS | 20 | 3 | 4161 | 164 | l(2)41Ab,CG17387,Mef2 | | 4.187e-02 | -3.17 | GSE42021\_TREG\_PLN\_VS\_TREG\_PRECURSORS\_THYMUS\_DN | MSigDB lists | GSE42021\_TREG\_PLN\_VS\_TREG\_PRECURSORS\_THYMUS\_DN | 20 | 3 | 4161 | 164 | step,CG41378,CG41099 | | 4.187e-02 | -3.17 | GO\_ENDOTHELIUM\_DEVELOPMENT | MSigDB lists | GO\_ENDOTHELIUM\_DEVELOPMENT | 20 | 3 | 4161 | 164 | dnc,cv-2,svp | | 4.187e-02 | -3.17 | GO\_REGULATION\_OF\_ORGAN\_GROWTH | MSigDB lists | GO\_REGULATION\_OF\_ORGAN\_GROWTH | 20 | 3 | 4161 | 164 | CycB,Sod1,Mef2 | | 4.187e-02 | -3.17 | chr10q26 | MSigDB lists | chr10q26 | 20 | 3 | 4161 | 164 | mbc,Ate1,CG12547 | | 4.187e-02 | -3.17 | GSE18791\_CTRL\_VS\_NEWCASTLE\_VIRUS\_DC\_8H\_DN | MSigDB lists | GSE18791\_CTRL\_VS\_NEWCASTLE\_VIRUS\_DC\_8H\_DN | 20 | 3 | 4161 | 164 | PNPase,CG41099,PMCA | | 4.187e-02 | -3.17 | GO\_REGULATION\_OF\_ENDOTHELIAL\_CELL\_MIGRATION | MSigDB lists | GO\_REGULATION\_OF\_ENDOTHELIAL\_CELL\_MIGRATION | 20 | 3 | 4161 | 164 | cv-2,Alg-2,svp | | 4.187e-02 | -3.17 | GO\_ADAPTIVE\_IMMUNE\_RESPONSE | MSigDB lists | GO\_ADAPTIVE\_IMMUNE\_RESPONSE | 20 | 3 | 4161 | 164 | Mef2,DNAlig4,spel1 | | 4.187e-02 | -3.17 | MORF\_PTPRR | MSigDB lists | MORF\_PTPRR | 20 | 3 | 4161 | 164 | fz2,Dys,toy | | 4.187e-02 | -3.17 | DACOSTA\_UV\_RESPONSE\_VIA\_ERCC3\_COMMON\_UP | MSigDB lists | DACOSTA\_UV\_RESPONSE\_VIA\_ERCC3\_COMMON\_UP | 20 | 3 | 4161 | 164 | ap,Hmt-1,CG30059 | | 4.187e-02 | -3.17 | GO\_NEUROEPITHELIAL\_CELL\_DIFFERENTIATION | MSigDB lists | GO\_NEUROEPITHELIAL\_CELL\_DIFFERENTIATION | 20 | 3 | 4161 | 164 | Mef2,Sod1,ck | | 4.205e-02 | -3.17 | regulation of anatomical structure morphogenesis | biological process | GO:0022603 | 250 | 18 | 12345 | 571 | kirre,nullo,par-1,shot,neo,tinc,lz,norpA,rib,cta,babos,Sema1a,sif,Src42A,pk,rl,Diap1,hth | | 4.209e-02 | -3.17 | HMG\_BOX\_2 | prosite domains | PS50118 | 23 | 4 | 6243 | 365 | SoxN,cic,pan,CG12104 | | 4.218e-02 | -3.17 | GSE20715\_WT\_VS\_TLR4\_KO\_LUNG\_DN | MSigDB lists | GSE20715\_WT\_VS\_TLR4\_KO\_LUNG\_DN | 49 | 5 | 4161 | 164 | CG41099,Nlg3,Pkg21D,CG1486,Atpalpha | | 4.218e-02 | -3.17 | GSE41867\_NAIVE\_VS\_DAY8\_LCMV\_EFFECTOR\_CD8\_TCELL\_DN | MSigDB lists | GSE41867\_NAIVE\_VS\_DAY8\_LCMV\_EFFECTOR\_CD8\_TCELL\_DN | 49 | 5 | 4161 | 164 | Meltrin,step,Octbeta2R,GLS,Der-1 | | 4.218e-02 | -3.17 | GSE17721\_POLYIC\_VS\_GARDIQUIMOD\_12H\_BMDC\_UP | MSigDB lists | GSE17721\_POLYIC\_VS\_GARDIQUIMOD\_12H\_BMDC\_UP | 49 | 5 | 4161 | 164 | nAChRalpha3,Nlg3,Maf1,PNPase,gus | | 4.218e-02 | -3.17 | GSE8384\_CTRL\_VS\_B\_ABORTUS\_4H\_MAC\_CELL\_LINE\_UP | MSigDB lists | GSE8384\_CTRL\_VS\_B\_ABORTUS\_4H\_MAC\_CELL\_LINE\_UP | 49 | 5 | 4161 | 164 | Mp,Parg,toy,Mef2,sky | | 4.218e-02 | -3.17 | KARLSSON\_TGFB1\_TARGETS\_DN | MSigDB lists | KARLSSON\_TGFB1\_TARGETS\_DN | 49 | 5 | 4161 | 164 | bru1,Maf1,CG42813,mlt,Nlg3 | | 4.218e-02 | -3.17 | GSE30971\_CTRL\_VS\_LPS\_STIM\_MACROPHAGE\_WBP7\_KO\_2H\_DN | MSigDB lists | GSE30971\_CTRL\_VS\_LPS\_STIM\_MACROPHAGE\_WBP7\_KO\_2H\_DN | 49 | 5 | 4161 | 164 | Gdh,Ars2,vap,CG9389,MFS16 | | 4.218e-02 | -3.17 | GSE25088\_ROSIGLITAZONE\_VS\_IL4\_AND\_ROSIGLITAZONE\_STIM\_STAT6\_KO\_MACROPHAGE\_DAY10\_UP | MSigDB lists | GSE25088\_ROSIGLITAZONE\_VS\_IL4\_AND\_ROSIGLITAZONE\_STIM\_STAT6\_KO\_MACROPHAGE\_DAY10\_UP | 49 | 5 | 4161 | 164 | SCAP,rl,step,bru1,Cyp6a8 | | 4.218e-02 | -3.17 | MORF\_MDM2 | MSigDB lists | MORF\_MDM2 | 49 | 5 | 4161 | 164 | Coq7,fz2,CG40178,Dys,toy | | 4.218e-02 | -3.17 | FOXJ2\_01 | MSigDB lists | FOXJ2\_01 | 49 | 5 | 4161 | 164 | Pi3K68D,twz,Mef2,Dys,Cyp6a8 | | 4.218e-02 | -3.17 | ZHONG\_RESPONSE\_TO\_AZACITIDINE\_AND\_TSA\_UP | MSigDB lists | ZHONG\_RESPONSE\_TO\_AZACITIDINE\_AND\_TSA\_UP | 49 | 5 | 4161 | 164 | GLS,CG3107,Odc2,CG41378,unc-13-4A | | 4.245e-02 | -3.16 | GOZGIT\_ESR1\_TARGETS\_DN | MSigDB lists | GOZGIT\_ESR1\_TARGETS\_DN | 137 | 10 | 4161 | 164 | CG14721,svp,CG42813,arg,sdk,twz,CG4341,CG7255,Pdk,rg | | 4.252e-02 | -3.16 | regulation of secretion by cell | biological process | GO:1903530 | 85 | 8 | 12345 | 571 | sky,cpx,Glut1,Syt7,CG9650,Dys,ru,dimm | | 4.259e-02 | -3.16 | regulation of intracellular signal transduction | biological process | GO:1902531 | 339 | 23 | 12345 | 571 | dnc,scw,conu,vap,step,Alg-2,ArfGAP1,CG34408,CG14184,mbc,mav,Hipk,Src42A,sif,Ppm1,Slmap,par-1,CG10188,Wdr59,PEK,Tak1,Git,CG30440 | | 4.261e-02 | -3.16 | Metalloenz\_LuxS/M16 | interpro domains | IPR011249 | 7 | 2 | 11383 | 557 | Nrd1,CG3107 | | 4.261e-02 | -3.16 | TGF-beta-rel | interpro domains | IPR015615 | 7 | 2 | 11383 | 557 | mav,scw | | 4.261e-02 | -3.16 | Peptidase\_M16\_C | interpro domains | IPR007863 | 7 | 2 | 11383 | 557 | Nrd1,CG3107 | | 4.261e-02 | -3.16 | PLipase\_A2\_dom\_sf | interpro domains | IPR036444 | 7 | 2 | 11383 | 557 | CG3009,CG42237 | | 4.261e-02 | -3.16 | Pept\_M16\_N | interpro domains | IPR011765 | 7 | 2 | 11383 | 557 | CG3107,Nrd1 | | 4.261e-02 | -3.16 | PLipase\_A2\_His\_AS | interpro domains | IPR033113 | 7 | 2 | 11383 | 557 | CG3009,CG42237 | | 4.261e-02 | -3.16 | Hud\_Sxl\_RNA | interpro domains | IPR002343 | 7 | 2 | 11383 | 557 | Rbp9,shep | | 4.261e-02 | -3.16 | TGFb\_CS | interpro domains | IPR017948 | 7 | 2 | 11383 | 557 | mav,scw | | 4.261e-02 | -3.16 | TGF-b\_C | interpro domains | IPR001839 | 7 | 2 | 11383 | 557 | mav,scw | | 4.261e-02 | -3.16 | NHL\_repeat\_subgr | interpro domains | IPR013017 | 7 | 2 | 11383 | 557 | CG12547,mei-P26 | | 4.261e-02 | -3.16 | Homeobox\_KN\_domain | interpro domains | IPR008422 | 7 | 2 | 11383 | 557 | hth,caup | | 4.261e-02 | -3.16 | Acyl-CoA\_DS | interpro domains | IPR015876 | 7 | 2 | 11383 | 557 | CG9743,CG9747 | | 4.261e-02 | -3.16 | NHL\_repeat | interpro domains | IPR001258 | 7 | 2 | 11383 | 557 | CG12547,mei-P26 | | 4.277e-02 | -3.15 | reproductive process | biological process | GO:0022414 | 1436 | 80 | 12345 | 571 | pum,Catsup,Acp36DE,Gr2a,igl,dsx,Mst87F,mei-P26,snky,dati,fz2,bab2,tld,Octbeta2R,orb,gus,fest,br,ct,fbl,tj,fzo,salm,jvl,Nlg2,toc,Mef2,Or67d,shep,CycB,prage,lncRNA:CR44344,Tdc2,lncRNA:flam,Gr39a,heph,lncRNA:CR44412,rib,grim,Rbp9,tej,mim,AP-2alpha,Yp2,lola,en,Trl,drosha,cic,DNaseII,CheB42a,wb,Diap1,rl,Dlg5,Msp300,jing,HP6,twz,fru,SPR,CG12200,Bruce,Cip4,par-1,capu,lama,spo,fan,Tpl94D,mlt,mbc,bru1,Gr93d,scw,dnc,Gdh,bond,Parp,lz | | 4.299e-02 | -3.15 | GSE46242\_CTRL\_VS\_EGR2\_DELETED\_TH1\_CD4\_TCELL\_DN | MSigDB lists | GSE46242\_CTRL\_VS\_EGR2\_DELETED\_TH1\_CD4\_TCELL\_DN | 34 | 4 | 4161 | 164 | Dys,Alg9,CG32944,CG12121 | | 4.299e-02 | -3.15 | GSE5099\_CLASSICAL\_M1\_VS\_ALTERNATIVE\_M2\_MACROPHAGE\_DN | MSigDB lists | GSE5099\_CLASSICAL\_M1\_VS\_ALTERNATIVE\_M2\_MACROPHAGE\_DN | 34 | 4 | 4161 | 164 | Rbp9,CG10077,shot,nAChRalpha4 | | 4.299e-02 | -3.15 | MORF\_STK17A | MSigDB lists | MORF\_STK17A | 34 | 4 | 4161 | 164 | CG40178,mbc,Pi3K68D,Octbeta1R | | 4.299e-02 | -3.15 | GSE2770\_TGFB\_AND\_IL4\_VS\_IL12\_TREATED\_ACT\_CD4\_TCELL\_2H\_DN | MSigDB lists | GSE2770\_TGFB\_AND\_IL4\_VS\_IL12\_TREATED\_ACT\_CD4\_TCELL\_2H\_DN | 34 | 4 | 4161 | 164 | Khc-73,GLS,Der-1,dnc | | 4.299e-02 | -3.15 | MODULE\_480 | MSigDB lists | MODULE\_480 | 34 | 4 | 4161 | 164 | step,CycB,Dys,Ate1 | | 4.299e-02 | -3.15 | EVI1\_05 | MSigDB lists | EVI1\_05 | 34 | 4 | 4161 | 164 | Rbp9,Tsp5D,Dys,Ca-alpha1D | | 4.299e-02 | -3.15 | GSE8685\_IL15\_ACT\_IL2\_STARVED\_VS\_IL21\_ACT\_IL2\_STARVED\_CD4\_TCELL\_UP | MSigDB lists | GSE8685\_IL15\_ACT\_IL2\_STARVED\_VS\_IL21\_ACT\_IL2\_STARVED\_CD4\_TCELL\_UP | 34 | 4 | 4161 | 164 | CG12547,CG12121,Gfat1,Nlg3 | | 4.299e-02 | -3.15 | GSE37301\_MULTIPOTENT\_PROGENITOR\_VS\_LYMPHOID\_PRIMED\_MPP\_DN | MSigDB lists | GSE37301\_MULTIPOTENT\_PROGENITOR\_VS\_LYMPHOID\_PRIMED\_MPP\_DN | 34 | 4 | 4161 | 164 | Odc2,Ac78C,CG9389,shot | | 4.299e-02 | -3.15 | GSE40277\_EOS\_AND\_LEF1\_TRANSDUCED\_VS\_CTRL\_CD4\_TCELL\_UP | MSigDB lists | GSE40277\_EOS\_AND\_LEF1\_TRANSDUCED\_VS\_CTRL\_CD4\_TCELL\_UP | 34 | 4 | 4161 | 164 | bru3,DNAlig4,Maf1,MCU | | 4.299e-02 | -3.15 | steroid biosynthetic process | biological process | GO:0006694 | 29 | 4 | 12345 | 571 | spok,spo,Npc2b,nvd | | 4.303e-02 | -3.15 | - | gene3d domains | 2.30.29.30 | 139 | 13 | 4929 | 273 | CG30440,osp,Myo81F,Gprk1,cert,CG34347,sif,Pld,ck,pico,vap,step,CG10188 | | 4.316e-02 | -3.14 | integral component of mitochondrial inner membrane | cellular component | GO:0031305 | 30 | 4 | 11940 | 534 | Tim17b,Tim23,CG41128,MCU | | 4.319e-02 | -3.14 | Class A/1 (Rhodopsin-like receptors) | REACTOME pathways | R-DME-373076 | 28 | 4 | 3923 | 189 | Rh3,Octbeta2R,5-HT2A,Octbeta1R | | 4.336e-02 | -3.14 | GO\_CELLULAR\_HOMEOSTASIS | MSigDB lists | GO\_CELLULAR\_HOMEOSTASIS | 176 | 12 | 4161 | 164 | Irp-1A,MCU,Pdk,ABCB7,Atpalpha,PMCA,Dys,Trxr-2,Hmt-1,cta,Sod1,CG40178 | | 4.345e-02 | -3.14 | Prot\_kinase\_dom | interpro domains | IPR000719 | 221 | 17 | 11383 | 557 | Pkg21D,Gprk1,CG12147,bt,CG32944,CG34357,PEK,dnt,fray,Stlk,Hipk,bma,Tak1,Takl2,rl,par-1,Src42A | | 4.364e-02 | -3.13 | Glycos\_transf\_2 | pfam domains | PF00535 | 16 | 3 | 10480 | 528 | Pgant9,Pgant5,Pgant1 | | 4.369e-02 | -3.13 | rut (rutabaga) | protein interactions | 32406 | 16 | 3 | 9204 | 464 | dnc,Rgk1,Sema1a | | 4.369e-02 | -3.13 | spz (spatzle) | protein interactions | 43256 | 16 | 3 | 9204 | 464 | Toll-7,CG3081,cic | | 4.369e-02 | -3.13 | Zpr1 (Zpr1) | protein interactions | 31851 | 16 | 3 | 9204 | 464 | Atpalpha,CG14926,Sod1 | | 4.369e-02 | -3.13 | TrpA1 (Transient receptor potential cation channel A1) | protein interactions | 39015 | 16 | 3 | 9204 | 464 | Or67d,shakB,norpA | | 4.369e-02 | -3.13 | hb (hunchback) | protein interactions | 41032 | 16 | 3 | 9204 | 464 | pum,fru,kni | | 4.369e-02 | -3.13 | Apc (APC-like) | protein interactions | 44642 | 16 | 3 | 9204 | 464 | ewg,pan,Src42A | | 4.371e-02 | -3.13 | MYTH4 | prosite domains | PS51016 | 6 | 2 | 6243 | 365 | Myo81F,ck | | 4.377e-02 | -3.13 | ZWANG\_TRANSIENTLY\_UP\_BY\_1ST\_EGF\_PULSE\_ONLY | MSigDB lists | ZWANG\_TRANSIENTLY\_UP\_BY\_1ST\_EGF\_PULSE\_ONLY | 298 | 18 | 4161 | 164 | Pkg21D,Nlg2,rg,St1,DNaseII,CG4341,CG10171,cv-2,IntS3,arg,Glut1,Khc-73,dnc,rdgC,DNAlig4,Src42A,CG5254,Aldh-III | | 4.380e-02 | -3.13 | dorsal appendage formation | biological process | GO:0046843 | 56 | 6 | 12345 | 571 | gus,cic,rl,jvl,br,lama | | 4.380e-02 | -3.13 | regulation of endocytosis | biological process | GO:0030100 | 56 | 6 | 12345 | 571 | ArfGAP1,mim,vap,norpA,step,AP-2alpha | | 4.380e-02 | -3.13 | negative regulation of translation | biological process | GO:0017148 | 56 | 6 | 12345 | 571 | Thor,heph,orb,bru3,bru1,pum | | 4.385e-02 | -3.13 | MODULE\_378 | MSigDB lists | MODULE\_378 | 83 | 7 | 4161 | 164 | bru1,Alg-2,gus,Nadsyn,CG41099,unc-13-4A,dnc | | 4.406e-02 | -3.12 | GO\_ENZYME\_LINKED\_RECEPTOR\_PROTEIN\_SIGNALING\_PATHWAY | MSigDB lists | GO\_ENZYME\_LINKED\_RECEPTOR\_PROTEIN\_SIGNALING\_PATHWAY | 157 | 11 | 4161 | 164 | mbc,vap,Parp,heph,Alg-2,nAChRalpha4,Pi3K68D,AP-2alpha,rl,Src42A,Octbeta2R | | 4.423e-02 | -3.12 | GSE17721\_LPS\_VS\_CPG\_6H\_BMDC\_UP | MSigDB lists | GSE17721\_LPS\_VS\_CPG\_6H\_BMDC\_UP | 66 | 6 | 4161 | 164 | vap,ap,cta,norpA,arg,CG30495 | | 4.423e-02 | -3.12 | MYB\_Q5\_01 | MSigDB lists | MYB\_Q5\_01 | 66 | 6 | 4161 | 164 | CG30495,Rbp9,Dys,DNAlig4,mbc,Ranbp16 | | 4.423e-02 | -3.12 | MEISSNER\_NPC\_HCP\_WITH\_H3K4ME2 | MSigDB lists | MEISSNER\_NPC\_HCP\_WITH\_H3K4ME2 | 66 | 6 | 4161 | 164 | Octbeta2R,MFS16,Pkg21D,Atpalpha,Mct1,twz | | 4.423e-02 | -3.12 | GSE17721\_POLYIC\_VS\_CPG\_24H\_BMDC\_DN | MSigDB lists | GSE17721\_POLYIC\_VS\_CPG\_24H\_BMDC\_DN | 66 | 6 | 4161 | 164 | CG3107,CG9389,nAChRalpha3,AGO3,Coq7,fray | | 4.426e-02 | -3.12 | GRYDER\_PAX3FOXO1\_ENHANCERS\_KO\_DOWN | MSigDB lists | GRYDER\_PAX3FOXO1\_ENHANCERS\_KO\_DOWN | 138 | 10 | 4161 | 164 | CG40045,CG7255,Parp,Atpalpha,Tsp5D,Git,Sod1,CG12121,svp,bru1 | | 4.454e-02 | -3.11 | positive regulation of molecular function | biological process | GO:0044093 | 287 | 20 | 12345 | 571 | rl,mbc,CdGAPr,CG17514,fray,conu,step,norpA,vap,ArfGAP1,CG34408,grim,Tak1,Git,CG41520,Tbc1d15-17,Stlk,RhoGAP100F,CG5862,sky | | 4.458e-02 | -3.11 | RICIN\_B\_LECTIN | prosite domains | PS50231 | 14 | 3 | 6243 | 365 | Pgant5,Pgant1,Pgant9 | | 4.458e-02 | -3.11 | ATPASE\_E1\_E2 | prosite domains | PS00154 | 14 | 3 | 6243 | 365 | Atpalpha,PMCA,CG9981 | | 4.461e-02 | -3.11 | btl (breathless) | protein interactions | 39564 | 27 | 4 | 9204 | 464 | jing,Catsup,vap,kni | | 4.468e-02 | -3.11 | anterior/posterior pattern specification | biological process | GO:0009952 | 200 | 15 | 12345 | 571 | jvl,rl,gus,pum,pk,ct,en,par-1,cic,scw,kni,Ptip,orb,bi,capu | | 4.472e-02 | -3.11 | hemidesmosome | cellular component | GO:0030056 | 1 | 1 | 11940 | 534 | shot | | 4.472e-02 | -3.11 | MutSbeta complex | cellular component | GO:0032302 | 1 | 1 | 11940 | 534 | spel1 | | 4.472e-02 | -3.11 | meiotic nuclear membrane microtubule tethering complex | cellular component | GO:0034993 | 1 | 1 | 11940 | 534 | Msp300 | | 4.472e-02 | -3.11 | DNA-dependent protein kinase-DNA ligase 4 complex | cellular component | GO:0005958 | 1 | 1 | 11940 | 534 | DNAlig4 | | 4.472e-02 | -3.11 | cell leading edge cell cortex | cellular component | GO:1904269 | 1 | 1 | 11940 | 534 | step | | 4.472e-02 | -3.11 | microtubule organizing center attachment site | cellular component | GO:0034992 | 1 | 1 | 11940 | 534 | Msp300 | | 4.472e-02 | -3.11 | nuclear membrane protein complex | cellular component | GO:0106083 | 1 | 1 | 11940 | 534 | Msp300 | | 4.472e-02 | -3.11 | nuclear membrane microtubule tethering complex | cellular component | GO:0106094 | 1 | 1 | 11940 | 534 | Msp300 | | 4.472e-02 | -3.11 | external encapsulating structure part | cellular component | GO:0044462 | 1 | 1 | 11940 | 534 | ck | | 4.472e-02 | -3.11 | nonhomologous end joining complex | cellular component | GO:0070419 | 1 | 1 | 11940 | 534 | DNAlig4 | | 4.472e-02 | -3.11 | DNA ligase IV complex | cellular component | GO:0032807 | 1 | 1 | 11940 | 534 | DNAlig4 | | 4.472e-02 | -3.11 | CLOCK-BMAL transcription complex | cellular component | GO:1990513 | 1 | 1 | 11940 | 534 | Clk | | 4.472e-02 | -3.11 | micropyle | cellular component | GO:0070825 | 1 | 1 | 11940 | 534 | ck | | 4.486e-02 | -3.10 | positive regulation of catalytic activity | biological process | GO:0043085 | 252 | 18 | 12345 | 571 | ArfGAP1,CG34408,grim,norpA,step,vap,conu,fray,CdGAPr,CG17514,mbc,sky,RhoGAP100F,Tbc1d15-17,Stlk,Tak1,CG41520,Git | | 4.495e-02 | -3.10 | Peptidase\_M16 | pfam domains | PF00675 | 7 | 2 | 10480 | 528 | Nrd1,CG3107 | | 4.495e-02 | -3.10 | Peptidase\_M16\_C | pfam domains | PF05193 | 7 | 2 | 10480 | 528 | CG3107,Nrd1 | | 4.495e-02 | -3.10 | Homeobox\_KN | pfam domains | PF05920 | 7 | 2 | 10480 | 528 | hth,caup | | 4.495e-02 | -3.10 | TGF\_beta | pfam domains | PF00019 | 7 | 2 | 10480 | 528 | mav,scw | | 4.495e-02 | -3.10 | NHL | pfam domains | PF01436 | 7 | 2 | 10480 | 528 | mei-P26,CG12547 | | 4.499e-02 | -3.10 | atms (antimeros) | protein interactions | 40593 | 7 | 2 | 9204 | 464 | CG12769,Myc | | 4.499e-02 | -3.10 | whe (what else) | protein interactions | 33121 | 7 | 2 | 9204 | 464 | ewg,rl | | 4.499e-02 | -3.10 | lmgA (lemming A) | protein interactions | 5740853 | 7 | 2 | 9204 | 464 | Psc,CycB | | 4.499e-02 | -3.10 | Nost (Nostrin) | protein interactions | 8673992 | 7 | 2 | 9204 | 464 | Cip4,Khc-73 | | 4.499e-02 | -3.10 | stai (stathmin) | protein interactions | 33863 | 7 | 2 | 9204 | 464 | Tak1,ct | | 4.499e-02 | -3.10 | CG13312 (uncharacterized protein) | protein interactions | 39010 | 7 | 2 | 9204 | 464 | Vsx2,zld | | 4.499e-02 | -3.10 | del (deadlock) | protein interactions | 35380 | 7 | 2 | 9204 | 464 | Dys,Git | | 4.528e-02 | -3.09 | axon part | cellular component | GO:0033267 | 105 | 9 | 11940 | 534 | cpx,Snap25,pum,shot,Mical,sky,Rtnl1,Magi,par-1 | | 4.543e-02 | -3.09 | GSE37301\_PRO\_BCELL\_VS\_GRANULOCYTE\_MONOCYTE\_PROGENITOR\_DN | MSigDB lists | GSE37301\_PRO\_BCELL\_VS\_GRANULOCYTE\_MONOCYTE\_PROGENITOR\_DN | 50 | 5 | 4161 | 164 | CG4341,Odc2,MFS16,MCU,Khc-73 | | 4.543e-02 | -3.09 | GO\_MEIOTIC\_CELL\_CYCLE | MSigDB lists | GO\_MEIOTIC\_CELL\_CYCLE | 50 | 5 | 4161 | 164 | vtd,Ubr1,Rpn9,AGO3,spel1 | | 4.543e-02 | -3.09 | CTTGTAT\_MIR381 | MSigDB lists | CTTGTAT\_MIR381 | 50 | 5 | 4161 | 164 | Ranbp16,twz,Bruce,Tbc1d15-17,rg | | 4.543e-02 | -3.09 | BROWN\_MYELOID\_CELL\_DEVELOPMENT\_DN | MSigDB lists | BROWN\_MYELOID\_CELL\_DEVELOPMENT\_DN | 50 | 5 | 4161 | 164 | spel1,CG4341,Odc2,rdgC,Sod1 | | 4.543e-02 | -3.09 | GSE26928\_NAIVE\_VS\_EFF\_MEMORY\_CD4\_TCELL\_DN | MSigDB lists | GSE26928\_NAIVE\_VS\_EFF\_MEMORY\_CD4\_TCELL\_DN | 50 | 5 | 4161 | 164 | CG30059,Octbeta2R,Glut1,Ars2,Odc2 | | 4.543e-02 | -3.09 | GSE5542\_UNTREATED\_VS\_IFNA\_TREATED\_EPITHELIAL\_CELLS\_6H\_DN | MSigDB lists | GSE5542\_UNTREATED\_VS\_IFNA\_TREATED\_EPITHELIAL\_CELLS\_6H\_DN | 50 | 5 | 4161 | 164 | rg,Gdh,AP-2alpha,CG17691,sdk | | 4.543e-02 | -3.09 | GSE5503\_MLN\_DC\_VS\_PLN\_DC\_ACTIVATED\_ALLOGENIC\_TCELL\_DN | MSigDB lists | GSE5503\_MLN\_DC\_VS\_PLN\_DC\_ACTIVATED\_ALLOGENIC\_TCELL\_DN | 50 | 5 | 4161 | 164 | Parp,Ranbp16,AP-2alpha,inc,Bruce | | 4.543e-02 | -3.09 | GSE19941\_UNSTIM\_VS\_LPS\_AND\_IL10\_STIM\_IL10\_KO\_NFKBP50\_KO\_MACROPHAGE\_DN | MSigDB lists | GSE19941\_UNSTIM\_VS\_LPS\_AND\_IL10\_STIM\_IL10\_KO\_NFKBP50\_KO\_MACROPHAGE\_DN | 50 | 5 | 4161 | 164 | spel1,rdgC,drosha,sbm,vtd | | 4.543e-02 | -3.09 | POU1F1\_Q6 | MSigDB lists | POU1F1\_Q6 | 50 | 5 | 4161 | 164 | Rbp9,Dys,MFS16,Ca-alpha1D,CG4341 | | 4.550e-02 | -3.09 | thiamine diphosphokinase activity | molecular function | GO:0004788 | 1 | 1 | 12176 | 554 | CG14721 | | 4.550e-02 | -3.09 | dinucleotide repeat insertion binding | molecular function | GO:0032181 | 1 | 1 | 12176 | 554 | spel1 | | 4.550e-02 | -3.09 | uniporter activity | molecular function | GO:0015292 | 1 | 1 | 12176 | 554 | MCU | | 4.550e-02 | -3.09 | beta-N-acetylgalactosaminidase activity | molecular function | GO:0032428 | 1 | 1 | 12176 | 554 | fdl | | 4.550e-02 | -3.09 | double-strand/single-strand DNA junction binding | molecular function | GO:0000406 | 1 | 1 | 12176 | 554 | spel1 | | 4.550e-02 | -3.09 | polyribonucleotide nucleotidyltransferase activity | molecular function | GO:0004654 | 1 | 1 | 12176 | 554 | PNPase | | 4.550e-02 | -3.09 | arginase activity | molecular function | GO:0004053 | 1 | 1 | 12176 | 554 | arg | | 4.550e-02 | -3.09 | alkane 1-monooxygenase activity | molecular function | GO:0018685 | 1 | 1 | 12176 | 554 | Cyp6a8 | | 4.550e-02 | -3.09 | pyruvate dehydrogenase (acetyl-transferring) kinase activity | molecular function | GO:0004740 | 1 | 1 | 12176 | 554 | Pdk | | 4.550e-02 | -3.09 | threonine aldolase activity | molecular function | GO:0004793 | 1 | 1 | 12176 | 554 | CG10184 | | 4.550e-02 | -3.09 | poly(ADP-ribose) glycohydrolase activity | molecular function | GO:0004649 | 1 | 1 | 12176 | 554 | Parg | | 4.550e-02 | -3.09 | dinucleotide insertion or deletion binding | molecular function | GO:0032139 | 1 | 1 | 12176 | 554 | spel1 | | 4.550e-02 | -3.09 | CCR4-NOT complex binding | molecular function | GO:1905762 | 1 | 1 | 12176 | 554 | Marf1 | | 4.550e-02 | -3.09 | riboflavin transmembrane transporter activity | molecular function | GO:0032217 | 1 | 1 | 12176 | 554 | Rift | | 4.550e-02 | -3.09 | single guanine insertion binding | molecular function | GO:0032142 | 1 | 1 | 12176 | 554 | spel1 | | 4.550e-02 | -3.09 | pheromone receptor activity | molecular function | GO:0016503 | 1 | 1 | 12176 | 554 | Or67d | | 4.550e-02 | -3.09 | deoxyribonuclease II activity | molecular function | GO:0004531 | 1 | 1 | 12176 | 554 | DNaseII | | 4.550e-02 | -3.09 | Gq/11-coupled serotonin receptor activity | molecular function | GO:0001587 | 1 | 1 | 12176 | 554 | 5-HT2A | | 4.550e-02 | -3.09 | RNA binding involved in posttranscriptional gene silencing | molecular function | GO:0150100 | 1 | 1 | 12176 | 554 | Marf1 | | 4.550e-02 | -3.09 | guanine/thymine mispair binding | molecular function | GO:0032137 | 1 | 1 | 12176 | 554 | spel1 | | 4.550e-02 | -3.09 | phospholipase D activity | molecular function | GO:0004630 | 1 | 1 | 12176 | 554 | Pld | | 4.550e-02 | -3.09 | glutamate dehydrogenase (NAD+) activity | molecular function | GO:0004352 | 1 | 1 | 12176 | 554 | Gdh | | 4.550e-02 | -3.09 | 3-oxoacyl-[acyl-carrier-protein] synthase activity | molecular function | GO:0004315 | 1 | 1 | 12176 | 554 | CG12170 | | 4.550e-02 | -3.09 | L-allo-threonine aldolase activity | molecular function | GO:0008732 | 1 | 1 | 12176 | 554 | CG10184 | | 4.550e-02 | -3.09 | NEDD8 ligase activity | molecular function | GO:0061663 | 1 | 1 | 12176 | 554 | Diap1 | | 4.550e-02 | -3.09 | mannosyl-oligosaccharide 1,6-alpha-mannosidase activity | molecular function | GO:0052767 | 1 | 1 | 12176 | 554 | alpha-Man-IIb | | 4.550e-02 | -3.09 | dopamine receptor binding | molecular function | GO:0050780 | 1 | 1 | 12176 | 554 | cta | | 4.550e-02 | -3.09 | RNA polymerase III core binding | molecular function | GO:0000994 | 1 | 1 | 12176 | 554 | Maf1 | | 4.550e-02 | -3.09 | small conductance calcium-activated potassium channel activity | molecular function | GO:0016286 | 1 | 1 | 12176 | 554 | SK | | 4.550e-02 | -3.09 | NAD+ synthase (glutamine-hydrolyzing) activity | molecular function | GO:0003952 | 1 | 1 | 12176 | 554 | Nadsyn | | 4.550e-02 | -3.09 | protein-histidine N-methyltransferase activity | molecular function | GO:0018064 | 1 | 1 | 12176 | 554 | Setd3 | | 4.550e-02 | -3.09 | 2-octoprenyl-3-methyl-6-methoxy-1,4-benzoquinone hydroxylase activity | molecular function | GO:0008682 | 1 | 1 | 12176 | 554 | Coq7 | | 4.550e-02 | -3.09 | glycerate kinase activity | molecular function | GO:0008887 | 1 | 1 | 12176 | 554 | CG9886 | | 4.550e-02 | -3.09 | oxidoreductase activity, acting on paired donors, with incorporation or reduction of molecular oxygen, reduced iron-sulfur protein as one donor, and incorporation of one atom of oxygen | molecular function | GO:0016713 | 1 | 1 | 12176 | 554 | Cyp6a8 | | 4.550e-02 | -3.09 | arginyltransferase activity | molecular function | GO:0004057 | 1 | 1 | 12176 | 554 | Ate1 | | 4.550e-02 | -3.09 | mRNA binding involved in posttranscriptional gene silencing | molecular function | GO:1903231 | 1 | 1 | 12176 | 554 | Marf1 | | 4.550e-02 | -3.09 | co-SMAD binding | molecular function | GO:0070410 | 1 | 1 | 12176 | 554 | fuss | | 4.550e-02 | -3.09 | protein ADP-ribosylase activity | molecular function | GO:1990404 | 1 | 1 | 12176 | 554 | Parp | | 4.550e-02 | -3.09 | virion binding | molecular function | GO:0046790 | 1 | 1 | 12176 | 554 | Toll-7 | | 4.550e-02 | -3.09 | cobalamin binding | molecular function | GO:0031419 | 1 | 1 | 12176 | 554 | CG3556 | | 4.550e-02 | -3.09 | thiamine binding | molecular function | GO:0030975 | 1 | 1 | 12176 | 554 | CG14721 | | 4.550e-02 | -3.09 | UDP-sugar diphosphatase activity | molecular function | GO:0008768 | 1 | 1 | 12176 | 554 | CG42813 | | 4.550e-02 | -3.09 | rhodopsin kinase activity | molecular function | GO:0050254 | 1 | 1 | 12176 | 554 | Gprk1 | | 4.550e-02 | -3.09 | D5 dopamine receptor binding | molecular function | GO:0031752 | 1 | 1 | 12176 | 554 | cta | | 4.550e-02 | -3.09 | elongation factor-2 kinase activity | molecular function | GO:0004686 | 1 | 1 | 12176 | 554 | PEK | | 4.591e-02 | -3.08 | Biosynthesis of unsaturated fatty acids | KEGG pathways | ko01040 | 19 | 3 | 3198 | 138 | CG9743,CG9747,Acox57D-d | | 4.591e-02 | -3.08 | Biosynthesis of unsaturated fatty acids | KEGG pathways | dme01040 | 19 | 3 | 3198 | 138 | Acox57D-d,CG9747,CG9743 | | 4.601e-02 | -3.08 | phosphorylation | biological process | GO:0016310 | 470 | 30 | 12345 | 571 | Pi3K68D,Takl2,fray,CG34357,fbl,CG9886,Hipk,Src42A,Pdk,CG32944,CycB,rl,Gprk1,CG3544,ND-MLRQ,PEK,dnt,par-1,bt,SdhA,CG42637,bma,Stlk,Pkg21D,UQCR-11,Tak1,Slob,CG4546,CG12147,CG9389 | | 4.618e-02 | -3.08 | HUDSXLRNA | prints domains | PR00961 | 7 | 2 | 2185 | 112 | Rbp9,shep | | 4.618e-02 | -3.08 | FACDDSATRASE | prints domains | PR00075 | 7 | 2 | 2185 | 112 | CG9747,CG9743 | | 4.625e-02 | -3.07 | cytoskeletal anchoring at nuclear membrane | biological process | GO:0090286 | 1 | 1 | 12345 | 571 | Msp300 | | 4.625e-02 | -3.07 | microvillus assembly | biological process | GO:0030033 | 1 | 1 | 12345 | 571 | sn | | 4.625e-02 | -3.07 | diacylglycerol biosynthetic process | biological process | GO:0006651 | 1 | 1 | 12345 | 571 | norpA | | 4.625e-02 | -3.07 | ATP generation from poly-ADP-D-ribose | biological process | GO:1990966 | 1 | 1 | 12345 | 571 | Parg | | 4.625e-02 | -3.07 | positive regulation of female receptivity | biological process | GO:0045925 | 1 | 1 | 12345 | 571 | dati | | 4.625e-02 | -3.07 | gap junction assembly | biological process | GO:0016264 | 1 | 1 | 12345 | 571 | shakB | | 4.625e-02 | -3.07 | adult chitin-based cuticle pattern formation | biological process | GO:0035018 | 1 | 1 | 12345 | 571 | fred | | 4.625e-02 | -3.07 | conversion of ds siRNA to ss siRNA | biological process | GO:0036404 | 1 | 1 | 12345 | 571 | Ars2 | | 4.625e-02 | -3.07 | arginine catabolic process to ornithine | biological process | GO:0019547 | 1 | 1 | 12345 | 571 | arg | | 4.625e-02 | -3.07 | mitochondrial mRNA catabolic process | biological process | GO:0000958 | 1 | 1 | 12345 | 571 | PNPase | | 4.625e-02 | -3.07 | negative regulation of synaptic vesicle recycling | biological process | GO:1903422 | 1 | 1 | 12345 | 571 | sky | | 4.625e-02 | -3.07 | regulation of presynaptic active zone assembly | biological process | GO:1905518 | 1 | 1 | 12345 | 571 | Spn | | 4.625e-02 | -3.07 | cardiac muscle adaptation | biological process | GO:0014887 | 1 | 1 | 12345 | 571 | Mp | | 4.625e-02 | -3.07 | protein arginylation | biological process | GO:0016598 | 1 | 1 | 12345 | 571 | Ate1 | | 4.625e-02 | -3.07 | muscle atrophy | biological process | GO:0014889 | 1 | 1 | 12345 | 571 | Mp | | 4.625e-02 | -3.07 | carbohydrate storage | biological process | GO:0052576 | 1 | 1 | 12345 | 571 | Mef2 | | 4.625e-02 | -3.07 | histone H3-K4 dimethylation | biological process | GO:0044648 | 1 | 1 | 12345 | 571 | Set1 | | 4.625e-02 | -3.07 | epithelial cilium movement involved in determination of left/right asymmetry | biological process | GO:0060287 | 1 | 1 | 12345 | 571 | CG17387 | | 4.625e-02 | -3.07 | muscle adaptation | biological process | GO:0043500 | 1 | 1 | 12345 | 571 | Mp | | 4.625e-02 | -3.07 | microvillar actin bundle assembly | biological process | GO:0030034 | 1 | 1 | 12345 | 571 | sn | | 4.625e-02 | -3.07 | innate immune response activating cell surface receptor signaling pathway | biological process | GO:0002220 | 1 | 1 | 12345 | 571 | Toll-7 | | 4.625e-02 | -3.07 | regulation of formation of translation initiation ternary complex | biological process | GO:1901190 | 1 | 1 | 12345 | 571 | mxt | | 4.625e-02 | -3.07 | skeletal muscle adaptation | biological process | GO:0043501 | 1 | 1 | 12345 | 571 | Mp | | 4.625e-02 | -3.07 | positive regulation of phosphorylation of RNA polymerase II C-terminal domain | biological process | GO:1901409 | 1 | 1 | 12345 | 571 | Trl | | 4.625e-02 | -3.07 | ER to Golgi ceramide transport | biological process | GO:0035621 | 1 | 1 | 12345 | 571 | cert | | 4.625e-02 | -3.07 | carbohydrate localization | biological process | GO:0052575 | 1 | 1 | 12345 | 571 | Mef2 | | 4.625e-02 | -3.07 | cobalamin transport | biological process | GO:0015889 | 1 | 1 | 12345 | 571 | CG3556 | | 4.625e-02 | -3.07 | absorption of UV light | biological process | GO:0016039 | 1 | 1 | 12345 | 571 | Rh3 | | 4.625e-02 | -3.07 | riboflavin transport | biological process | GO:0032218 | 1 | 1 | 12345 | 571 | Rift | | 4.625e-02 | -3.07 | lauric acid metabolic process | biological process | GO:0048252 | 1 | 1 | 12345 | 571 | Cyp6a8 | | 4.625e-02 | -3.07 | skeletal muscle atrophy | biological process | GO:0014732 | 1 | 1 | 12345 | 571 | Mp | | 4.625e-02 | -3.07 | negative regulation of presynaptic active zone assembly | biological process | GO:1905519 | 1 | 1 | 12345 | 571 | Spn | | 4.625e-02 | -3.07 | negative regulation of cysteine-type endopeptidase activity involved in execution phase of apoptosis | biological process | GO:2001271 | 1 | 1 | 12345 | 571 | Diap1 | | 4.625e-02 | -3.07 | pattern orientation | biological process | GO:0007633 | 1 | 1 | 12345 | 571 | tay | | 4.625e-02 | -3.07 | conversion of ds siRNA to ss siRNA involved in RNA interference | biological process | GO:0033168 | 1 | 1 | 12345 | 571 | Ars2 | | 4.625e-02 | -3.07 | cardiac muscle atrophy | biological process | GO:0014899 | 1 | 1 | 12345 | 571 | Mp | | 4.625e-02 | -3.07 | nucleologenesis | biological process | GO:0017126 | 1 | 1 | 12345 | 571 | Myc | | 4.625e-02 | -3.07 | striated muscle adaptation | biological process | GO:0014888 | 1 | 1 | 12345 | 571 | Mp | | 4.625e-02 | -3.07 | glutamine catabolic process | biological process | GO:0006543 | 1 | 1 | 12345 | 571 | GLS | | 4.625e-02 | -3.07 | posterior compartment specification | biological process | GO:0007388 | 1 | 1 | 12345 | 571 | en | | 4.625e-02 | -3.07 | thiamine-containing compound metabolic process | biological process | GO:0042723 | 1 | 1 | 12345 | 571 | CG14721 | | 4.625e-02 | -3.07 | double-strand break repair via alternative nonhomologous end joining | biological process | GO:0097681 | 1 | 1 | 12345 | 571 | mus308 | | 4.625e-02 | -3.07 | thiamine diphosphate metabolic process | biological process | GO:0042357 | 1 | 1 | 12345 | 571 | CG14721 | | 4.625e-02 | -3.07 | cell communication by electrical coupling | biological process | GO:0010644 | 1 | 1 | 12345 | 571 | shakB | | 4.625e-02 | -3.07 | thiamine metabolic process | biological process | GO:0006772 | 1 | 1 | 12345 | 571 | CG14721 | | 4.625e-02 | -3.07 | negative regulation of endoplasmic reticulum calcium ion concentration | biological process | GO:0032471 | 1 | 1 | 12345 | 571 | THADA | | 4.625e-02 | -3.07 | positive regulation of cAMP-dependent protein kinase activity | biological process | GO:2000481 | 1 | 1 | 12345 | 571 | step | | 4.625e-02 | -3.07 | cell surface pattern recognition receptor signaling pathway | biological process | GO:0002752 | 1 | 1 | 12345 | 571 | Toll-7 | | 4.625e-02 | -3.07 | female gonad morphogenesis | biological process | GO:0061040 | 1 | 1 | 12345 | 571 | bab2 | | 4.625e-02 | -3.07 | miRNA loading onto RISC involved in gene silencing by miRNA | biological process | GO:0035280 | 1 | 1 | 12345 | 571 | HPS4 | | 4.625e-02 | -3.07 | protein localization to presynapse | biological process | GO:1905383 | 1 | 1 | 12345 | 571 | Git | | 4.625e-02 | -3.07 | immune response-activating cell surface receptor signaling pathway | biological process | GO:0002429 | 1 | 1 | 12345 | 571 | Toll-7 | | 4.625e-02 | -3.07 | thiamine diphosphate biosynthetic process | biological process | GO:0009229 | 1 | 1 | 12345 | 571 | CG14721 | | 4.625e-02 | -3.07 | negative regulation of presynapse assembly | biological process | GO:1905607 | 1 | 1 | 12345 | 571 | Spn | | 4.625e-02 | -3.07 | regulation of presynapse assembly | biological process | GO:1905606 | 1 | 1 | 12345 | 571 | Spn | | 4.625e-02 | -3.07 | negative regulation of ATPase-coupled calcium transmembrane transporter activity | biological process | GO:1901895 | 1 | 1 | 12345 | 571 | THADA | | 4.625e-02 | -3.07 | peptidyl-histidine methylation | biological process | GO:0018021 | 1 | 1 | 12345 | 571 | Setd3 | | 4.625e-02 | -3.07 | striated muscle atrophy | biological process | GO:0014891 | 1 | 1 | 12345 | 571 | Mp | | 4.625e-02 | -3.07 | mitochondrial RNA catabolic process | biological process | GO:0000957 | 1 | 1 | 12345 | 571 | PNPase | | 4.625e-02 | -3.07 | anterior mRNA localization involved in anterior/posterior axis specification | biological process | GO:0060813 | 1 | 1 | 12345 | 571 | jvl | | 4.632e-02 | -3.07 | MODULE\_317 | MSigDB lists | MODULE\_317 | 84 | 7 | 4161 | 164 | gus,Alg-2,bru1,dnc,unc-13-4A,CG41099,Nadsyn | | 4.633e-02 | -3.07 | AMIT\_SERUM\_RESPONSE\_480\_MCF10A | MSigDB lists | AMIT\_SERUM\_RESPONSE\_480\_MCF10A | 9 | 2 | 4161 | 164 | Parg,Tim23 | | 4.633e-02 | -3.07 | GO\_REGULATION\_OF\_TRANSCRIPTION\_REGULATORY\_REGION\_DNA\_BINDING | MSigDB lists | GO\_REGULATION\_OF\_TRANSCRIPTION\_REGULATORY\_REGION\_DNA\_BINDING | 9 | 2 | 4161 | 164 | ap,Parp | | 4.633e-02 | -3.07 | COLLIS\_PRKDC\_REGULATORS | MSigDB lists | COLLIS\_PRKDC\_REGULATORS | 9 | 2 | 4161 | 164 | Parp,DNAlig4 | | 4.633e-02 | -3.07 | DAUER\_STAT3\_TARGETS\_UP | MSigDB lists | DAUER\_STAT3\_TARGETS\_UP | 9 | 2 | 4161 | 164 | Gfat1,Glut1 | | 4.633e-02 | -3.07 | KEGG\_ALDOSTERONE\_REGULATED\_SODIUM\_REABSORPTION | MSigDB lists | KEGG\_ALDOSTERONE\_REGULATED\_SODIUM\_REABSORPTION | 9 | 2 | 4161 | 164 | Atpalpha,rl | | 4.633e-02 | -3.07 | GO\_LAGGING\_STRAND\_ELONGATION | MSigDB lists | GO\_LAGGING\_STRAND\_ELONGATION | 9 | 2 | 4161 | 164 | Parp,DNAlig4 | | 4.633e-02 | -3.07 | MODULE\_208 | MSigDB lists | MODULE\_208 | 9 | 2 | 4161 | 164 | Meltrin,MCU | | 4.633e-02 | -3.07 | GO\_REGULATION\_OF\_INTERFERON\_GAMMA\_PRODUCTION | MSigDB lists | GO\_REGULATION\_OF\_INTERFERON\_GAMMA\_PRODUCTION | 9 | 2 | 4161 | 164 | fz2,dnc | | 4.633e-02 | -3.07 | SCHRAMM\_INHBA\_TARGETS\_DN | MSigDB lists | SCHRAMM\_INHBA\_TARGETS\_DN | 9 | 2 | 4161 | 164 | vap,cta | | 4.633e-02 | -3.07 | GO\_ADAPTATION\_OF\_SIGNALING\_PATHWAY | MSigDB lists | GO\_ADAPTATION\_OF\_SIGNALING\_PATHWAY | 9 | 2 | 4161 | 164 | Octbeta2R,CG30495 | | 4.633e-02 | -3.07 | GO\_POSITIVE\_REGULATION\_OF\_SKELETAL\_MUSCLE\_TISSUE\_DEVELOPMENT | MSigDB lists | GO\_POSITIVE\_REGULATION\_OF\_SKELETAL\_MUSCLE\_TISSUE\_DEVELOPMENT | 9 | 2 | 4161 | 164 | Mef2,Octbeta2R | | 4.633e-02 | -3.07 | MORI\_IMMATURE\_B\_LYMPHOCYTE\_UP | MSigDB lists | MORI\_IMMATURE\_B\_LYMPHOCYTE\_UP | 9 | 2 | 4161 | 164 | CG41378,CG30059 | | 4.633e-02 | -3.07 | GO\_B\_CELL\_MEDIATED\_IMMUNITY | MSigDB lists | GO\_B\_CELL\_MEDIATED\_IMMUNITY | 9 | 2 | 4161 | 164 | DNAlig4,spel1 | | 4.633e-02 | -3.07 | GO\_PROTEIN\_IMPORT\_INTO\_MITOCHONDRIAL\_MATRIX | MSigDB lists | GO\_PROTEIN\_IMPORT\_INTO\_MITOCHONDRIAL\_MATRIX | 9 | 2 | 4161 | 164 | Tim17b,Tim23 | | 4.633e-02 | -3.07 | GO\_CAMP\_MEDIATED\_SIGNALING | MSigDB lists | GO\_CAMP\_MEDIATED\_SIGNALING | 9 | 2 | 4161 | 164 | Ac78C,dnc | | 4.633e-02 | -3.07 | MORF\_PDCD1 | MSigDB lists | MORF\_PDCD1 | 9 | 2 | 4161 | 164 | toy,fz2 | | 4.633e-02 | -3.07 | NIKOLSKY\_BREAST\_CANCER\_5P15\_AMPLICON | MSigDB lists | NIKOLSKY\_BREAST\_CANCER\_5P15\_AMPLICON | 9 | 2 | 4161 | 164 | Alg-2,SdhA | | 4.633e-02 | -3.07 | LIAN\_LIPA\_TARGETS\_6M | MSigDB lists | LIAN\_LIPA\_TARGETS\_6M | 9 | 2 | 4161 | 164 | MFS16,shot | | 4.633e-02 | -3.07 | GO\_STEREOCILIUM\_BUNDLE | MSigDB lists | GO\_STEREOCILIUM\_BUNDLE | 9 | 2 | 4161 | 164 | ck,sn | | 4.633e-02 | -3.07 | GO\_PRE\_MRNA\_BINDING | MSigDB lists | GO\_PRE\_MRNA\_BINDING | 9 | 2 | 4161 | 164 | CG10077,heph | | 4.633e-02 | -3.07 | GO\_REGULATION\_OF\_ALCOHOL\_BIOSYNTHETIC\_PROCESS | MSigDB lists | GO\_REGULATION\_OF\_ALCOHOL\_BIOSYNTHETIC\_PROCESS | 9 | 2 | 4161 | 164 | SCAP,Sod1 | | 4.633e-02 | -3.07 | GO\_AXONEMAL\_DYNEIN\_COMPLEX | MSigDB lists | GO\_AXONEMAL\_DYNEIN\_COMPLEX | 9 | 2 | 4161 | 164 | Dhc16F,CG9492 | | 4.633e-02 | -3.07 | GO\_CYCLIC\_NUCLEOTIDE\_BINDING | MSigDB lists | GO\_CYCLIC\_NUCLEOTIDE\_BINDING | 9 | 2 | 4161 | 164 | Pkg21D,dnc | | 4.633e-02 | -3.07 | LANDIS\_ERBB2\_BREAST\_TUMORS\_65\_DN | MSigDB lists | LANDIS\_ERBB2\_BREAST\_TUMORS\_65\_DN | 9 | 2 | 4161 | 164 | CG17691,Der-1 | | 4.633e-02 | -3.07 | GYORFFY\_MITOXANTRONE\_RESISTANCE | MSigDB lists | GYORFFY\_MITOXANTRONE\_RESISTANCE | 9 | 2 | 4161 | 164 | vap,Rbp9 | | 4.633e-02 | -3.07 | HUNSBERGER\_EXERCISE\_REGULATED\_GENES | MSigDB lists | HUNSBERGER\_EXERCISE\_REGULATED\_GENES | 9 | 2 | 4161 | 164 | Odc2,rl | | 4.633e-02 | -3.07 | MODULE\_418 | MSigDB lists | MODULE\_418 | 9 | 2 | 4161 | 164 | CG34408,Mef2 | | 4.633e-02 | -3.07 | SHEPARD\_BMYB\_TARGETS | MSigDB lists | SHEPARD\_BMYB\_TARGETS | 9 | 2 | 4161 | 164 | Rbp9,CycB | | 4.633e-02 | -3.07 | YRCCAKNNGNCGC\_UNKNOWN | MSigDB lists | YRCCAKNNGNCGC\_UNKNOWN | 9 | 2 | 4161 | 164 | fz2,ap | | 4.633e-02 | -3.07 | GO\_NEUROMUSCULAR\_SYNAPTIC\_TRANSMISSION | MSigDB lists | GO\_NEUROMUSCULAR\_SYNAPTIC\_TRANSMISSION | 9 | 2 | 4161 | 164 | nAChRalpha4,nAChRalpha3 | | 4.633e-02 | -3.07 | GO\_RESPONSE\_TO\_OXYGEN\_RADICAL | MSigDB lists | GO\_RESPONSE\_TO\_OXYGEN\_RADICAL | 9 | 2 | 4161 | 164 | Trxr-2,Sod1 | | 4.633e-02 | -3.07 | GO\_REGULATION\_OF\_ALTERNATIVE\_MRNA\_SPLICING\_VIA\_SPLICEOSOME | MSigDB lists | GO\_REGULATION\_OF\_ALTERNATIVE\_MRNA\_SPLICING\_VIA\_SPLICEOSOME | 9 | 2 | 4161 | 164 | heph,CG10077 | | 4.633e-02 | -3.07 | KEGG\_DORSO\_VENTRAL\_AXIS\_FORMATION | MSigDB lists | KEGG\_DORSO\_VENTRAL\_AXIS\_FORMATION | 9 | 2 | 4161 | 164 | rl,AGO3 | | 4.633e-02 | -3.07 | GO\_SYNAPTIC\_TRANSMISSION\_CHOLINERGIC | MSigDB lists | GO\_SYNAPTIC\_TRANSMISSION\_CHOLINERGIC | 9 | 2 | 4161 | 164 | nAChRalpha4,nAChRalpha3 | | 4.639e-02 | -3.07 | Signaling by Rho GTPases | REACTOME pathways | R-DME-194315 | 114 | 10 | 3923 | 189 | Act42A,CG10188,cta,rl,sif,Cip4,CdGAPr,RhoGAP100F,conu,CG30440 | | 4.658e-02 | -3.07 | GO\_SYNAPTIC\_SIGNALING | MSigDB lists | GO\_SYNAPTIC\_SIGNALING | 102 | 8 | 4161 | 164 | nAChRalpha4,GLS,nAChRalpha3,Octbeta2R,Mef2,Octbeta1R,unc-13-4A,rl | | 4.660e-02 | -3.07 | - | gene3d domains | 3.40.1110.10 | 15 | 3 | 4929 | 273 | CG9981,PMCA,Atpalpha | | 4.662e-02 | -3.07 | MyTH4 | smart domains | SM00139 | 6 | 2 | 5086 | 308 | Myo81F,ck | | 4.662e-02 | -3.07 | Cation\_ATPase\_N | smart domains | SM00831 | 6 | 2 | 5086 | 308 | PMCA,Atpalpha | | 4.673e-02 | -3.06 | synaptonemal complex | cellular component | GO:0000795 | 8 | 2 | 11940 | 534 | CG12200,vtd | | 4.673e-02 | -3.06 | cytoskeleton of presynaptic active zone | cellular component | GO:0048788 | 8 | 2 | 11940 | 534 | Git,Ten-a | | 4.673e-02 | -3.06 | synaptonemal structure | cellular component | GO:0099086 | 8 | 2 | 11940 | 534 | vtd,CG12200 | | 4.673e-02 | -3.06 | presynaptic cytoskeleton | cellular component | GO:0099569 | 8 | 2 | 11940 | 534 | Ten-a,Git | | 4.688e-02 | -3.06 | B41 | smart domains | SM00295 | 23 | 4 | 5086 | 308 | CG34347,Myo81F,dnr1,ck | | 4.706e-02 | -3.06 | GSE17721\_CPG\_VS\_GARDIQUIMOD\_4H\_BMDC\_DN | MSigDB lists | GSE17721\_CPG\_VS\_GARDIQUIMOD\_4H\_BMDC\_DN | 67 | 6 | 4161 | 164 | dnr1,lqf,CG12170,IntS3,CG7255,Pi3K68D | | 4.710e-02 | -3.06 | GSE17974\_0H\_VS\_0.5H\_IN\_VITRO\_ACT\_CD4\_TCELL\_UP | MSigDB lists | GSE17974\_0H\_VS\_0.5H\_IN\_VITRO\_ACT\_CD4\_TCELL\_UP | 35 | 4 | 4161 | 164 | CG41378,nAChRalpha4,CG9886,CG9492 | | 4.710e-02 | -3.06 | GSE6259\_33D1\_POS\_DC\_VS\_CD4\_TCELL\_DN | MSigDB lists | GSE6259\_33D1\_POS\_DC\_VS\_CD4\_TCELL\_DN | 35 | 4 | 4161 | 164 | shot,CG30059,DNAlig4,ck | | 4.710e-02 | -3.06 | GSE16450\_IMMATURE\_VS\_MATURE\_NEURON\_CELL\_LINE\_12H\_IFNA\_STIM\_UP | MSigDB lists | GSE16450\_IMMATURE\_VS\_MATURE\_NEURON\_CELL\_LINE\_12H\_IFNA\_STIM\_UP | 35 | 4 | 4161 | 164 | step,norpA,CG13284,dnr1 | | 4.710e-02 | -3.06 | RB\_DN.V1\_DN | MSigDB lists | RB\_DN.V1\_DN | 35 | 4 | 4161 | 164 | Alg-2,step,Hsc70-3,CG41099 | | 4.710e-02 | -3.06 | GSE22886\_IGG\_IGA\_MEMORY\_BCELL\_VS\_BLOOD\_PLASMA\_CELL\_UP | MSigDB lists | GSE22886\_IGG\_IGA\_MEMORY\_BCELL\_VS\_BLOOD\_PLASMA\_CELL\_UP | 35 | 4 | 4161 | 164 | DNaseII,dnc,unc-13-4A,CG8184 | | 4.710e-02 | -3.06 | GSE22443\_NAIVE\_VS\_ACT\_AND\_IL2\_TREATED\_CD8\_TCELL\_UP | MSigDB lists | GSE22443\_NAIVE\_VS\_ACT\_AND\_IL2\_TREATED\_CD8\_TCELL\_UP | 35 | 4 | 4161 | 164 | CG10077,Gfat1,mRpS5,Ac78C | | 4.710e-02 | -3.06 | FREAC7\_01 | MSigDB lists | FREAC7\_01 | 35 | 4 | 4161 | 164 | lqf,Pgant5,Pi3K68D,twz | | 4.710e-02 | -3.06 | GSE37533\_PPARG1\_FOXP3\_VS\_PPARG2\_FOXP3\_TRANSDUCED\_CD4\_TCELL\_PIOGLITAZONE\_TREATED\_DN | MSigDB lists | GSE37533\_PPARG1\_FOXP3\_VS\_PPARG2\_FOXP3\_TRANSDUCED\_CD4\_TCELL\_PIOGLITAZONE\_TREATED\_DN | 35 | 4 | 4161 | 164 | Ubr1,CG41378,step,fz2 | | 4.712e-02 | -3.05 | detection of light stimulus | biological process | GO:0009583 | 57 | 6 | 12345 | 571 | shakB,Gprk1,norpA,Rh3,rdgC,Pld | | 4.712e-02 | -3.05 | cellular amino acid catabolic process | biological process | GO:0009063 | 57 | 6 | 12345 | 571 | GLS,arg,CG10184,Ate1,CG17691,Gdh | | 4.730e-02 | -3.05 | Glycoside\_hydrolase\_SF | interpro domains | IPR017853 | 54 | 6 | 11383 | 557 | fdl,Cht10,Amyrel,Idgf4,tobi,GCS2alpha | | 4.748e-02 | -3.05 | FXR\_IR1\_Q6 | MSigDB lists | FXR\_IR1\_Q6 | 21 | 3 | 4161 | 164 | Dys,inc,Ac78C | | 4.748e-02 | -3.05 | GCCNNNWTAAR\_UNKNOWN | MSigDB lists | GCCNNNWTAAR\_UNKNOWN | 21 | 3 | 4161 | 164 | Ranbp16,twz,Rbp9 | | 4.748e-02 | -3.05 | HALLMARK\_REACTIVE\_OXIGEN\_SPECIES\_PATHWAY | MSigDB lists | HALLMARK\_REACTIVE\_OXIGEN\_SPECIES\_PATHWAY | 21 | 3 | 4161 | 164 | fray,Sod1,Trxr-2 | | 4.748e-02 | -3.05 | GO\_PEPTIDYL\_THREONINE\_MODIFICATION | MSigDB lists | GO\_PEPTIDYL\_THREONINE\_MODIFICATION | 21 | 3 | 4161 | 164 | rl,Pgant5,fray | | 4.748e-02 | -3.05 | GO\_CARBOHYDRATE\_TRANSPORT | MSigDB lists | GO\_CARBOHYDRATE\_TRANSPORT | 21 | 3 | 4161 | 164 | Glut1,MFS16,CG14621 | | 4.750e-02 | -3.05 | Glyco\_trans\_2-like | interpro domains | IPR001173 | 17 | 3 | 11383 | 557 | Pgant5,Pgant9,Pgant1 | | 4.772e-02 | -3.04 | w (white) | protein interactions | 31271 | 80 | 8 | 9204 | 464 | drosha,Psc,su(w[a]),Su(z)2,Trl,fru,pho,Diap1 | | 4.772e-02 | -3.04 | cellular metal ion homeostasis | biological process | GO:0006875 | 87 | 8 | 12345 | 571 | MCU,norpA,PMCA,THADA,Catsup,Atpalpha,Octbeta1R,ABCB7 | | 4.786e-02 | -3.04 | positive regulation of JNK cascade | biological process | GO:0046330 | 30 | 4 | 12345 | 571 | PEK,Alg-2,mbc,Tak1 | | 4.786e-02 | -3.04 | mitotic cell cycle, embryonic | biological process | GO:0045448 | 30 | 4 | 12345 | 571 | toc,CycB,Psc,Trl | | 4.786e-02 | -3.04 | cellular component maintenance | biological process | GO:0043954 | 30 | 4 | 12345 | 571 | Nlg2,RhoGAP100F,Ten-a,rl | | 4.786e-02 | -3.04 | intrinsic component of mitochondrial inner membrane | cellular component | GO:0031304 | 31 | 4 | 11940 | 534 | Tim23,CG41128,MCU,Tim17b | | 4.787e-02 | -3.04 | aggressive behavior | biological process | GO:0002118 | 18 | 3 | 12345 | 571 | Nlg2,twz,fru | | 4.787e-02 | -3.04 | regulation of compound eye retinal cell programmed cell death | biological process | GO:0046669 | 18 | 3 | 12345 | 571 | Diap1,norpA,lz | | 4.787e-02 | -3.04 | medium-term memory | biological process | GO:0072375 | 18 | 3 | 12345 | 571 | dnc,Rgk1,MCU | | 4.787e-02 | -3.04 | cyclic purine nucleotide metabolic process | biological process | GO:0052652 | 18 | 3 | 12345 | 571 | CG34357,CG42637,Ac78C | | 4.787e-02 | -3.04 | negative regulation of secretion | biological process | GO:0051048 | 18 | 3 | 12345 | 571 | sky,CG9650,cpx | | 4.818e-02 | -3.03 | Ca2+ activated K+ channels | REACTOME pathways | R-DME-1296052 | 1 | 1 | 3923 | 189 | SK | | 4.818e-02 | -3.03 | Hydrolysis of LPC | REACTOME pathways | R-DME-1483115 | 1 | 1 | 3923 | 189 | lama | | 4.822e-02 | -3.03 | neuron fate commitment | biological process | GO:0048663 | 72 | 7 | 12345 | 571 | hth,rl,br,svp,lola,klg,ru | | 4.827e-02 | -3.03 | SET | prosite domains | PS50280 | 24 | 4 | 6243 | 365 | SmydA-8,Set1,SmydA-9,Setd3 | | 4.828e-02 | -3.03 | Regulation of cholesterol biosynthesis by SREBP (SREBF) | REACTOME pathways | R-DME-1655829 | 29 | 4 | 3923 | 189 | FASN3,CG9747,CG9743,SCAP | | 4.880e-02 | -3.02 | regulation of cell development | biological process | GO:0060284 | 344 | 23 | 12345 | 571 | shot,par-1,Bruce,zld,SoxN,hdc,Catsup,tinc,Mical,lz,Myc,babos,norpA,pk,sif,Src42A,Sema1a,mav,sdk,hth,mbc,rl,Diap1 | | 4.880e-02 | -3.02 | columnar/cuboidal epithelial cell differentiation | biological process | GO:0002065 | 344 | 23 | 12345 | 571 | toc,AP-2alpha,jvl,capu,Mef2,orb,lama,cic,par-1,Cip4,ct,br,lncRNA:flam,dimm,gus,mbc,rl,Dlg5,Diap1,mim,tj,Parp,jing | | 4.883e-02 | -3.02 | GSE20366\_TREG\_VS\_NAIVE\_CD4\_TCELL\_DEC205\_CONVERSION\_UP | MSigDB lists | GSE20366\_TREG\_VS\_NAIVE\_CD4\_TCELL\_DEC205\_CONVERSION\_UP | 51 | 5 | 4161 | 164 | Meltrin,Ca-alpha1D,CG40178,step,Glut1 | | 4.883e-02 | -3.02 | GSE2770\_IL12\_VS\_TGFB\_AND\_IL12\_TREATED\_ACT\_CD4\_TCELL\_48H\_UP | MSigDB lists | GSE2770\_IL12\_VS\_TGFB\_AND\_IL12\_TREATED\_ACT\_CD4\_TCELL\_48H\_UP | 51 | 5 | 4161 | 164 | Ubr1,Irp-1A,Mp,Aldh-III,ck | | 4.883e-02 | -3.02 | GO\_INNER\_MITOCHONDRIAL\_MEMBRANE\_PROTEIN\_COMPLEX | MSigDB lists | GO\_INNER\_MITOCHONDRIAL\_MEMBRANE\_PROTEIN\_COMPLEX | 51 | 5 | 4161 | 164 | Tim17b,Tim23,SdhA,MCU,ND-30 | | 4.883e-02 | -3.02 | MORF\_IL16 | MSigDB lists | MORF\_IL16 | 51 | 5 | 4161 | 164 | CG40178,Coq7,Pi3K68D,fz2,toy | | 4.883e-02 | -3.02 | GSE17721\_CTRL\_VS\_GARDIQUIMOD\_1H\_BMDC\_UP | MSigDB lists | GSE17721\_CTRL\_VS\_GARDIQUIMOD\_1H\_BMDC\_UP | 51 | 5 | 4161 | 164 | CG40228,RpL6,Gdh,dnr1,CG9107 | | 4.883e-02 | -3.02 | GSE40274\_CTRL\_VS\_FOXP3\_AND\_IRF4\_TRANSDUCED\_ACTIVATED\_CD4\_TCELL\_DN | MSigDB lists | GSE40274\_CTRL\_VS\_FOXP3\_AND\_IRF4\_TRANSDUCED\_ACTIVATED\_CD4\_TCELL\_DN | 51 | 5 | 4161 | 164 | nAChRalpha3,AP-2alpha,MFS16,Hmt-1,CG9886 | | 4.885e-02 | -3.02 | fly chr60F5-60F5\|2-107 | chromosome location | fly chr60F5-60F5\|2-107 | 11 | 2 | 25010 | 823 | CG9380,CG30430 | | 4.885e-02 | -3.02 | fly chr60F5-60F5\|2-107 cM | chromosome location | fly chr60F5-60F5\|2-107 cM | 11 | 2 | 25010 | 823 | CG30430,CG9380 | | 4.887e-02 | -3.02 | DANG\_MYC\_TARGETS\_UP | MSigDB lists | DANG\_MYC\_TARGETS\_UP | 85 | 7 | 4161 | 164 | spel1,mRpL10,CycB,Odc2,CG10077,CG2528,CG9389 | | 4.887e-02 | -3.02 | GSE7218\_IGM\_VS\_IGG\_SIGNAL\_THGOUGH\_ANTIGEN\_BCELL\_UP | MSigDB lists | GSE7218\_IGM\_VS\_IGG\_SIGNAL\_THGOUGH\_ANTIGEN\_BCELL\_UP | 85 | 7 | 4161 | 164 | mbc,CG17691,Parp,ck,Nadsyn,rg,inc | | 4.887e-02 | -3.02 | MODULE\_255 | MSigDB lists | MODULE\_255 | 85 | 7 | 4161 | 164 | Nadsyn,CG41099,unc-13-4A,dnc,bru1,Alg-2,gus | | 4.888e-02 | -3.02 | GO\_IMMUNE\_SYSTEM\_DEVELOPMENT | MSigDB lists | GO\_IMMUNE\_SYSTEM\_DEVELOPMENT | 103 | 8 | 4161 | 164 | Mef2,Sod1,rl,DNAlig4,spel1,mus308,fz2,Parp | | 4.893e-02 | -3.02 | ArAA\_b-elim\_lyase/Thr\_aldolase | interpro domains | IPR001597 | 1 | 1 | 11383 | 557 | CG10184 | | 4.893e-02 | -3.02 | Headcase\_mid | interpro domains | IPR031947 | 1 | 1 | 11383 | 557 | hdc | | 4.893e-02 | -3.02 | CD141 | interpro domains | IPR016316 | 1 | 1 | 11383 | 557 | knk | | 4.893e-02 | -3.02 | Ubq\_synth\_Coq7 | interpro domains | IPR011566 | 1 | 1 | 11383 | 557 | Coq7 | | 4.893e-02 | -3.02 | RBM33 | interpro domains | IPR039878 | 1 | 1 | 11383 | 557 | CG2186 | | 4.893e-02 | -3.02 | FAM193 | interpro domains | IPR029717 | 1 | 1 | 11383 | 557 | CG7518 | | 4.893e-02 | -3.02 | MCU\_C | interpro domains | IPR006769 | 1 | 1 | 11383 | 557 | MCU | | 4.893e-02 | -3.02 | ATP2B1 | interpro domains | IPR030320 | 1 | 1 | 11383 | 557 | PMCA | | 4.893e-02 | -3.02 | Rrp7A\_RRM | interpro domains | IPR034890 | 1 | 1 | 11383 | 557 | CG9107 | | 4.893e-02 | -3.02 | C2B\_Synaptotagmin-7 | interpro domains | IPR037741 | 1 | 1 | 11383 | 557 | Syt7 | | 4.893e-02 | -3.02 | E3\_Ub\_ligase\_DUF913 | interpro domains | IPR010314 | 1 | 1 | 11383 | 557 | CG8184 | | 4.893e-02 | -3.02 | HPS4 | interpro domains | IPR026091 | 1 | 1 | 11383 | 557 | HPS4 | | 4.893e-02 | -3.02 | RNAP\_P/RPABC4 | interpro domains | IPR006591 | 1 | 1 | 11383 | 557 | Rpb12 | | 4.893e-02 | -3.02 | Gprotein\_alpha\_12/13 | interpro domains | IPR000469 | 1 | 1 | 11383 | 557 | cta | | 4.893e-02 | -3.02 | ALG9-like | interpro domains | IPR039484 | 1 | 1 | 11383 | 557 | Alg9 | | 4.893e-02 | -3.02 | Receptor\_IA-2\_dom | interpro domains | IPR021613 | 1 | 1 | 11383 | 557 | IA-2 | | 4.893e-02 | -3.02 | NADH\_UbQ\_OxRdtase\_CS | interpro domains | IPR020396 | 1 | 1 | 11383 | 557 | ND-30 | | 4.893e-02 | -3.02 | Int\_cplx\_su3 | interpro domains | IPR019333 | 1 | 1 | 11383 | 557 | IntS3 | | 4.893e-02 | -3.02 | Arginase | interpro domains | IPR014033 | 1 | 1 | 11383 | 557 | arg | | 4.893e-02 | -3.02 | Fascin\_metazoans | interpro domains | IPR024703 | 1 | 1 | 11383 | 557 | sn | | 4.893e-02 | -3.02 | Poly(ADP-ribose)pol\_reg\_dom | interpro domains | IPR004102 | 1 | 1 | 11383 | 557 | Parp | | 4.893e-02 | -3.02 | Fascin | interpro domains | IPR010431 | 1 | 1 | 11383 | 557 | sn | | 4.893e-02 | -3.02 | Headcase | interpro domains | IPR026066 | 1 | 1 | 11383 | 557 | hdc | | 4.893e-02 | -3.02 | RasGAP\_C\_SH2 | interpro domains | IPR035842 | 1 | 1 | 11383 | 557 | vap | | 4.893e-02 | -3.02 | Ribosomal\_L15e | interpro domains | IPR000439 | 1 | 1 | 11383 | 557 | RpL15 | | 4.893e-02 | -3.02 | DUF1180 | interpro domains | IPR009565 | 1 | 1 | 11383 | 557 | CG12159 | | 4.893e-02 | -3.02 | CSTF1\_dimer\_sf | interpro domains | IPR038184 | 1 | 1 | 11383 | 557 | CstF50 | | 4.893e-02 | -3.02 | PLipase\_D\_fam | interpro domains | IPR015679 | 1 | 1 | 11383 | 557 | Pld | | 4.893e-02 | -3.02 | Threonine\_aldolase | interpro domains | IPR023603 | 1 | 1 | 11383 | 557 | CG10184 | | 4.893e-02 | -3.02 | Znf\_GAGA-bd\_fac | interpro domains | IPR015318 | 1 | 1 | 11383 | 557 | Trl | | 4.893e-02 | -3.02 | Rad21/Rec8-like | interpro domains | IPR039781 | 1 | 1 | 11383 | 557 | vtd | | 4.893e-02 | -3.02 | PTBP1/2/3\_RRM2 | interpro domains | IPR034798 | 1 | 1 | 11383 | 557 | heph | | 4.893e-02 | -3.02 | MCU\_fam | interpro domains | IPR039055 | 1 | 1 | 11383 | 557 | MCU | | 4.893e-02 | -3.02 | PNPase\_PH\_RNA-bd\_sf | interpro domains | IPR036456 | 1 | 1 | 11383 | 557 | PNPase | | 4.893e-02 | -3.02 | Sugar\_P\_transporter | interpro domains | IPR000849 | 1 | 1 | 11383 | 557 | MFS16 | | 4.893e-02 | -3.02 | Thi\_PPkinase | interpro domains | IPR006282 | 1 | 1 | 11383 | 557 | CG14721 | | 4.893e-02 | -3.02 | Ribosomal\_L6\_N | interpro domains | IPR005568 | 1 | 1 | 11383 | 557 | RpL6 | | 4.893e-02 | -3.02 | SoHo\_dom | interpro domains | IPR003127 | 1 | 1 | 11383 | 557 | CAP | | 4.893e-02 | -3.02 | DUF4787 | interpro domains | IPR031985 | 1 | 1 | 11383 | 557 | CG14321 | | 4.893e-02 | -3.02 | KIF13A | interpro domains | IPR032923 | 1 | 1 | 11383 | 557 | Khc-73 | | 4.893e-02 | -3.02 | ClpS\_core | interpro domains | IPR003769 | 1 | 1 | 11383 | 557 | Ubr1 | | 4.893e-02 | -3.02 | WDR59\_mRING-H2-C3H3C2 | interpro domains | IPR039456 | 1 | 1 | 11383 | 557 | Wdr59 | | 4.893e-02 | -3.02 | NDUFB2 | interpro domains | IPR026627 | 1 | 1 | 11383 | 557 | CG40472 | | 4.893e-02 | -3.02 | MADS\_MEF2-like | interpro domains | IPR033896 | 1 | 1 | 11383 | 557 | Mef2 | | 4.893e-02 | -3.02 | Rad21/Rec8\_C\_eu | interpro domains | IPR006909 | 1 | 1 | 11383 | 557 | vtd | | 4.893e-02 | -3.02 | Poly(ADP-ribose)pol\_reg\_dom\_sf | interpro domains | IPR036616 | 1 | 1 | 11383 | 557 | Parp | | 4.893e-02 | -3.02 | GIT\_SHD | interpro domains | IPR013724 | 1 | 1 | 11383 | 557 | Git | | 4.893e-02 | -3.02 | DDRGK\_dom-contain | interpro domains | IPR019153 | 1 | 1 | 11383 | 557 | CG5862 | | 4.893e-02 | -3.02 | DUF4739 | interpro domains | IPR031764 | 1 | 1 | 11383 | 557 | CG30015 | | 4.893e-02 | -3.02 | DNA\_polymerase\_A | interpro domains | IPR002298 | 1 | 1 | 11383 | 557 | mus308 | | 4.893e-02 | -3.02 | N-end\_Aminoacyl\_Trfase\_C | interpro domains | IPR007472 | 1 | 1 | 11383 | 557 | Ate1 | | 4.893e-02 | -3.02 | Ribosomal\_L15e\_CS | interpro domains | IPR020925 | 1 | 1 | 11383 | 557 | RpL15 | | 4.893e-02 | -3.02 | LIP1\_N | interpro domains | IPR031782 | 1 | 1 | 11383 | 557 | CG9044 | | 4.893e-02 | -3.02 | Thiosulphate\_STrfase\_CS | interpro domains | IPR001307 | 1 | 1 | 11383 | 557 | CG12279 | | 4.893e-02 | -3.02 | TPK\_catalytic\_sf | interpro domains | IPR036759 | 1 | 1 | 11383 | 557 | CG14721 | | 4.893e-02 | -3.02 | DNA\_mismatch\_repair\_Msh2 | interpro domains | IPR011184 | 1 | 1 | 11383 | 557 | spel1 | | 4.893e-02 | -3.02 | PLipase\_D\_euk | interpro domains | IPR016555 | 1 | 1 | 11383 | 557 | Pld | | 4.893e-02 | -3.02 | Beta-lactam/transpept-like | interpro domains | IPR012338 | 1 | 1 | 11383 | 557 | GLS | | 4.893e-02 | -3.02 | Fascin-domain | interpro domains | IPR022768 | 1 | 1 | 11383 | 557 | sn | | 4.893e-02 | -3.02 | DSX\_dimer | interpro domains | IPR014932 | 1 | 1 | 11383 | 557 | dsx | | 4.893e-02 | -3.02 | CAC1F\_C | interpro domains | IPR031688 | 1 | 1 | 11383 | 557 | Ca-alpha1D | | 4.893e-02 | -3.02 | His\_kinase\_dom | interpro domains | IPR005467 | 1 | 1 | 11383 | 557 | Pdk | | 4.893e-02 | -3.02 | HJURP\_C | interpro domains | IPR022102 | 1 | 1 | 11383 | 557 | Mef2 | | 4.893e-02 | -3.02 | PADR1 | interpro domains | IPR012982 | 1 | 1 | 11383 | 557 | Parp | | 4.893e-02 | -3.02 | PKNOX/Meis\_N | interpro domains | IPR032453 | 1 | 1 | 11383 | 557 | hth | | 4.893e-02 | -3.02 | CaM-bd\_dom | interpro domains | IPR004178 | 1 | 1 | 11383 | 557 | SK | | 4.893e-02 | -3.02 | PLC-beta\_CS | interpro domains | IPR009535 | 1 | 1 | 11383 | 557 | norpA | | 4.893e-02 | -3.02 | Glutaminase\_EF-hand | interpro domains | IPR041541 | 1 | 1 | 11383 | 557 | GLS | | 4.893e-02 | -3.02 | Glutaminase | interpro domains | IPR015868 | 1 | 1 | 11383 | 557 | GLS | | 4.893e-02 | -3.02 | MOFRL | interpro domains | IPR007835 | 1 | 1 | 11383 | 557 | CG9886 | | 4.893e-02 | -3.02 | DLG5\_SH3 | interpro domains | IPR035537 | 1 | 1 | 11383 | 557 | Dlg5 | | 4.893e-02 | -3.02 | Syndecan/Neurexin\_dom | interpro domains | IPR027789 | 1 | 1 | 11383 | 557 | Sdc | | 4.893e-02 | -3.02 | CCDC40 | interpro domains | IPR037386 | 1 | 1 | 11383 | 557 | l(2)41Ab | | 4.893e-02 | -3.02 | DUF4772 | interpro domains | IPR031940 | 1 | 1 | 11383 | 557 | Glut4EF | | 4.893e-02 | -3.02 | SERRATE/Ars2\_N | interpro domains | IPR021933 | 1 | 1 | 11383 | 557 | Ars2 | | 4.893e-02 | -3.02 | K\_chnl\_volt-dep\_KCNQ | interpro domains | IPR003937 | 1 | 1 | 11383 | 557 | KCNQ | | 4.893e-02 | -3.02 | Apc13 | interpro domains | IPR008401 | 1 | 1 | 11383 | 557 | CG33981 | | 4.893e-02 | -3.02 | CSTF1\_dimer | interpro domains | IPR032028 | 1 | 1 | 11383 | 557 | CstF50 | | 4.893e-02 | -3.02 | Receptor\_IA-2\_ectodomain\_sf | interpro domains | IPR038112 | 1 | 1 | 11383 | 557 | IA-2 | | 4.893e-02 | -3.02 | Antistasin-like | interpro domains | IPR004094 | 1 | 1 | 11383 | 557 | CG5639 | | 4.893e-02 | -3.02 | RAC\_head | interpro domains | IPR032003 | 1 | 1 | 11383 | 557 | CG10565 | | 4.893e-02 | -3.02 | AK/P\_DHK\_N\_sf | interpro domains | IPR036784 | 1 | 1 | 11383 | 557 | Pdk | | 4.893e-02 | -3.02 | Desmoplakin\_SH3 | interpro domains | IPR041615 | 1 | 1 | 11383 | 557 | shot | | 4.893e-02 | -3.02 | RasGAP\_N\_SH2 | interpro domains | IPR035841 | 1 | 1 | 11383 | 557 | vap | | 4.893e-02 | -3.02 | DUF2967 | interpro domains | IPR021349 | 1 | 1 | 11383 | 557 | Rgk1 | | 4.893e-02 | -3.02 | RNase\_III | interpro domains | IPR011907 | 1 | 1 | 11383 | 557 | drosha | | 4.893e-02 | -3.02 | TRA1/TRRAP | interpro domains | IPR033317 | 1 | 1 | 11383 | 557 | Nipped-A | | 4.893e-02 | -3.02 | NAD\_synthase | interpro domains | IPR003694 | 1 | 1 | 11383 | 557 | Nadsyn | | 4.893e-02 | -3.02 | Syntaxin-18\_N | interpro domains | IPR019529 | 1 | 1 | 11383 | 557 | Syx18 | | 4.893e-02 | -3.02 | 3-oxoacyl-ACP\_synth-2 | interpro domains | IPR017568 | 1 | 1 | 11383 | 557 | CG12170 | | 4.893e-02 | -3.02 | Rbsml\_L15e\_core\_dom\_sf | interpro domains | IPR024794 | 1 | 1 | 11383 | 557 | RpL15 | | 4.893e-02 | -3.02 | Dystrophin/utrophin | interpro domains | IPR035436 | 1 | 1 | 11383 | 557 | Dys | | 4.893e-02 | -3.02 | Type\_II\_PanK | interpro domains | IPR004567 | 1 | 1 | 11383 | 557 | fbl | | 4.893e-02 | -3.02 | Mxt | interpro domains | IPR040160 | 1 | 1 | 11383 | 557 | mxt | | 4.893e-02 | -3.02 | PNPase | interpro domains | IPR012162 | 1 | 1 | 11383 | 557 | PNPase | | 4.893e-02 | -3.02 | RALGAPB | interpro domains | IPR039930 | 1 | 1 | 11383 | 557 | CG34408 | | 4.893e-02 | -3.02 | DUF92\_TMEM19 | interpro domains | IPR002794 | 1 | 1 | 11383 | 557 | CG10171 | | 4.893e-02 | -3.02 | CTNNB1-bd\_N | interpro domains | IPR013558 | 1 | 1 | 11383 | 557 | pan | | 4.893e-02 | -3.02 | Plakin\_repeat\_sf | interpro domains | IPR035915 | 1 | 1 | 11383 | 557 | shot | | 4.893e-02 | -3.02 | LANC-like | interpro domains | IPR007822 | 1 | 1 | 11383 | 557 | CG2061 | | 4.893e-02 | -3.02 | Rubisco\_LSMT\_subst-bd | interpro domains | IPR015353 | 1 | 1 | 11383 | 557 | Setd3 | | 4.893e-02 | -3.02 | BIP\_NBD | interpro domains | IPR042050 | 1 | 1 | 11383 | 557 | Hsc70-3 | | 4.893e-02 | -3.02 | Synaptotagmin7 | interpro domains | IPR015427 | 1 | 1 | 11383 | 557 | Syt7 | | 4.893e-02 | -3.02 | K\_chnl\_volt-dep\_KCNQ\_C | interpro domains | IPR013821 | 1 | 1 | 11383 | 557 | KCNQ | | 4.893e-02 | -3.02 | GRB/APBB1IP | interpro domains | IPR039664 | 1 | 1 | 11383 | 557 | pico | | 4.893e-02 | -3.02 | DNase\_TatD-rel\_CS | interpro domains | IPR018228 | 1 | 1 | 11383 | 557 | CG3358 | | 4.893e-02 | -3.02 | MyoVII\_FERM\_C1 | interpro domains | IPR041793 | 1 | 1 | 11383 | 557 | ck | | 4.893e-02 | -3.02 | p120-RasGAP | interpro domains | IPR028554 | 1 | 1 | 11383 | 557 | vap | | 4.893e-02 | -3.02 | STARD11\_START | interpro domains | IPR041952 | 1 | 1 | 11383 | 557 | cert | | 4.893e-02 | -3.02 | PDE4\_UCR | interpro domains | IPR040844 | 1 | 1 | 11383 | 557 | dnc | | 4.893e-02 | -3.02 | WGR\_dom\_sf | interpro domains | IPR036930 | 1 | 1 | 11383 | 557 | Parp | | 4.893e-02 | -3.02 | Fe\_hydrogenase | interpro domains | IPR009016 | 1 | 1 | 11383 | 557 | CG17683 | | 4.893e-02 | -3.02 | RRM\_Rrp7 | interpro domains | IPR040447 | 1 | 1 | 11383 | 557 | CG9107 | | 4.893e-02 | -3.02 | Epsin-2\_metazoa | interpro domains | IPR027319 | 1 | 1 | 11383 | 557 | lqf | | 4.893e-02 | -3.02 | MTABC\_N | interpro domains | IPR032410 | 1 | 1 | 11383 | 557 | Hmt-1 | | 4.893e-02 | -3.02 | XPO7 | interpro domains | IPR040021 | 1 | 1 | 11383 | 557 | Ranbp16 | | 4.893e-02 | -3.02 | Arg-tRNA-P\_Trfase\_1\_euk | interpro domains | IPR017137 | 1 | 1 | 11383 | 557 | Ate1 | | 4.893e-02 | -3.02 | DUF4588 | interpro domains | IPR029196 | 1 | 1 | 11383 | 557 | CG4768 | | 4.893e-02 | -3.02 | KOW\_RPL6 | interpro domains | IPR041997 | 1 | 1 | 11383 | 557 | RpL6 | | 4.893e-02 | -3.02 | GIT1\_C | interpro domains | IPR022018 | 1 | 1 | 11383 | 557 | Git | | 4.893e-02 | -3.02 | Fe\_hydrogenase\_lsu\_C | interpro domains | IPR004108 | 1 | 1 | 11383 | 557 | CG17683 | | 4.893e-02 | -3.02 | Band4 | interpro domains | IPR030696 | 1 | 1 | 11383 | 557 | CG34347 | | 4.893e-02 | -3.02 | LanC-like\_prot\_euk | interpro domains | IPR020464 | 1 | 1 | 11383 | 557 | CG2061 | | 4.893e-02 | -3.02 | RasGAP\_SH3 | interpro domains | IPR035652 | 1 | 1 | 11383 | 557 | vap | | 4.893e-02 | -3.02 | Fe\_hydrogenase\_ssu | interpro domains | IPR003149 | 1 | 1 | 11383 | 557 | CG17683 | | 4.893e-02 | -3.02 | ZFYVE26 | interpro domains | IPR028730 | 1 | 1 | 11383 | 557 | CG5270 | | 4.893e-02 | -3.02 | Synaphin | interpro domains | IPR008849 | 1 | 1 | 11383 | 557 | cpx | | 4.893e-02 | -3.02 | Maf1\_sf | interpro domains | IPR038564 | 1 | 1 | 11383 | 557 | Maf1 | | 4.893e-02 | -3.02 | Fibrinogen\_a/b/g\_C\_2 | interpro domains | IPR014715 | 1 | 1 | 11383 | 557 | CG41520 | | 4.893e-02 | -3.02 | P-type\_ATPase\_IIB | interpro domains | IPR006408 | 1 | 1 | 11383 | 557 | PMCA | | 4.893e-02 | -3.02 | ATP\_Ca\_trans\_C | interpro domains | IPR022141 | 1 | 1 | 11383 | 557 | PMCA | | 4.893e-02 | -3.02 | MMM1\_dom | interpro domains | IPR019411 | 1 | 1 | 11383 | 557 | CG43783 | | 4.893e-02 | -3.02 | PLipase\_B-like | interpro domains | IPR007000 | 1 | 1 | 11383 | 557 | lama | | 4.893e-02 | -3.02 | Aldehyde\_DH\_NAD(P) | interpro domains | IPR012394 | 1 | 1 | 11383 | 557 | Aldh-III | | 4.893e-02 | -3.02 | MYLIP\_FERM\_C | interpro domains | IPR041790 | 1 | 1 | 11383 | 557 | dnr1 | | 4.893e-02 | -3.02 | 60S\_ribosomal\_L6E | interpro domains | IPR000915 | 1 | 1 | 11383 | 557 | RpL6 | | 4.893e-02 | -3.02 | TMEM65 | interpro domains | IPR019537 | 1 | 1 | 11383 | 557 | CG17715 | | 4.893e-02 | -3.02 | DUF4812 | interpro domains | IPR032084 | 1 | 1 | 11383 | 557 | CG3081 | | 4.893e-02 | -3.02 | MAPK\_ERK1/2 | interpro domains | IPR008349 | 1 | 1 | 11383 | 557 | rl | | 4.893e-02 | -3.02 | BIRC6 | interpro domains | IPR022103 | 1 | 1 | 11383 | 557 | Bruce | | 4.893e-02 | -3.02 | Syndecan\_inverte | interpro domains | IPR031198 | 1 | 1 | 11383 | 557 | Sdc | | 4.893e-02 | -3.02 | RRP7 | interpro domains | IPR040446 | 1 | 1 | 11383 | 557 | CG9107 | | 4.893e-02 | -3.02 | PARP | interpro domains | IPR008288 | 1 | 1 | 11383 | 557 | Parp | | 4.893e-02 | -3.02 | Beta-ketoacyl\_synthase | interpro domains | IPR000794 | 1 | 1 | 11383 | 557 | CG12170 | | 4.893e-02 | -3.02 | Formin\_Cappuccino\_subfam | interpro domains | IPR001265 | 1 | 1 | 11383 | 557 | capu | | 4.893e-02 | -3.02 | Rubisco\_LSMT\_subst-bd\_sf | interpro domains | IPR036464 | 1 | 1 | 11383 | 557 | Setd3 | | 4.893e-02 | -3.02 | Gcn1 | interpro domains | IPR033173 | 1 | 1 | 11383 | 557 | CG17514 | | 4.893e-02 | -3.02 | Neurabin-1/2\_PDZ | interpro domains | IPR040645 | 1 | 1 | 11383 | 557 | Spn | | 4.893e-02 | -3.02 | RRP7\_C | interpro domains | IPR024326 | 1 | 1 | 11383 | 557 | CG9107 | | 4.893e-02 | -3.02 | Optomoror-blind\_N | interpro domains | IPR021101 | 1 | 1 | 11383 | 557 | bi | | 4.893e-02 | -3.02 | VGLL4 | interpro domains | IPR028184 | 1 | 1 | 11383 | 557 | Tgi | | 4.893e-02 | -3.02 | C2A\_Synaptotagmin-7 | interpro domains | IPR037732 | 1 | 1 | 11383 | 557 | Syt7 | | 4.893e-02 | -3.02 | Marf1 | interpro domains | IPR024768 | 1 | 1 | 11383 | 557 | Marf1 | | 4.893e-02 | -3.02 | SCAP | interpro domains | IPR030225 | 1 | 1 | 11383 | 557 | SCAP | | 4.893e-02 | -3.02 | Poly\_GlycHdrlase | interpro domains | IPR007724 | 1 | 1 | 11383 | 557 | Parg | | 4.893e-02 | -3.02 | Lustrin\_cystein | interpro domains | IPR028150 | 1 | 1 | 11383 | 557 | CG5639 | | 4.893e-02 | -3.02 | MOFRL\_assoc\_dom | interpro domains | IPR025286 | 1 | 1 | 11383 | 557 | CG9886 | | 4.893e-02 | -3.02 | Syndecan | interpro domains | IPR001050 | 1 | 1 | 11383 | 557 | Sdc | | 4.893e-02 | -3.02 | COMPASS\_Set1\_N-SET | interpro domains | IPR024657 | 1 | 1 | 11383 | 557 | Set1 | | 4.893e-02 | -3.02 | Riboflavin\_transptr | interpro domains | IPR009357 | 1 | 1 | 11383 | 557 | Rift | | 4.893e-02 | -3.02 | C2orf42 | interpro domains | IPR026049 | 1 | 1 | 11383 | 557 | CG2321 | | 4.893e-02 | -3.02 | Peptidase\_M16C\_assoc | interpro domains | IPR013578 | 1 | 1 | 11383 | 557 | CG3107 | | 4.893e-02 | -3.02 | DNase\_II | interpro domains | IPR004947 | 1 | 1 | 11383 | 557 | DNaseII | | 4.893e-02 | -3.02 | NADH\_DH\_suC | interpro domains | IPR010218 | 1 | 1 | 11383 | 557 | ND-30 | | 4.893e-02 | -3.02 | Shootin-1 | interpro domains | IPR024849 | 1 | 1 | 11383 | 557 | CG33111 | | 4.893e-02 | -3.02 | LAMTOR1/MEH1 | interpro domains | IPR028209 | 1 | 1 | 11383 | 557 | CG14184 | | 4.893e-02 | -3.02 | C16orf72-like | interpro domains | IPR040308 | 1 | 1 | 11383 | 557 | CG4768 | | 4.893e-02 | -3.02 | BAZ2A/B\_Bromo | interpro domains | IPR037374 | 1 | 1 | 11383 | 557 | tou | | 4.893e-02 | -3.02 | K\_chnl\_Ca-activ\_SK | interpro domains | IPR015449 | 1 | 1 | 11383 | 557 | SK | | 4.893e-02 | -3.02 | Clathrin\_a-adaptin\_app\_sub\_C | interpro domains | IPR003164 | 1 | 1 | 11383 | 557 | AP-2alpha | | 4.893e-02 | -3.02 | TIAM1\_CC-Ex | interpro domains | IPR040655 | 1 | 1 | 11383 | 557 | sif | | 4.893e-02 | -3.02 | BCDHK/PDK\_N | interpro domains | IPR018955 | 1 | 1 | 11383 | 557 | Pdk | | 4.893e-02 | -3.02 | Cbl-bd\_prot | interpro domains | IPR002157 | 1 | 1 | 11383 | 557 | CG3556 | | 4.893e-02 | -3.02 | EIF4EBP | interpro domains | IPR008606 | 1 | 1 | 11383 | 557 | Thor | | 4.893e-02 | -3.02 | Znf\_PARP\_sf | interpro domains | IPR036957 | 1 | 1 | 11383 | 557 | Parp | | 4.893e-02 | -3.02 | COL15A1 | interpro domains | IPR039070 | 1 | 1 | 11383 | 557 | Mp | | 4.893e-02 | -3.02 | Msh2 | interpro domains | IPR032642 | 1 | 1 | 11383 | 557 | spel1 | | 4.893e-02 | -3.02 | N-end\_Aminoacyl\_Trfase\_N | interpro domains | IPR007471 | 1 | 1 | 11383 | 557 | Ate1 | | 4.893e-02 | -3.02 | B12D | interpro domains | IPR010530 | 1 | 1 | 11383 | 557 | ND-MLRQ | | 4.893e-02 | -3.02 | DUF1989 | interpro domains | IPR018959 | 1 | 1 | 11383 | 557 | CG10208 | | 4.893e-02 | -3.02 | Hirudin/antistatin | interpro domains | IPR011061 | 1 | 1 | 11383 | 557 | CG5639 | | 4.893e-02 | -3.02 | LIG4 | interpro domains | IPR029710 | 1 | 1 | 11383 | 557 | DNAlig4 | | 4.893e-02 | -3.02 | Quinoprot\_gluc/sorb\_DH | interpro domains | IPR011041 | 1 | 1 | 11383 | 557 | Ten-a | | 4.893e-02 | -3.02 | PNPase\_PH\_RNA-bd\_bac/org-type | interpro domains | IPR015848 | 1 | 1 | 11383 | 557 | PNPase | | 4.893e-02 | -3.02 | Ser/Thr-Pase\_EF-hand\_contain | interpro domains | IPR012008 | 1 | 1 | 11383 | 557 | rdgC | | 4.893e-02 | -3.02 | DOCK1\_homologue | interpro domains | IPR031057 | 1 | 1 | 11383 | 557 | mbc | | 4.893e-02 | -3.02 | Fibrinogen\_a/b/g\_C\_1 | interpro domains | IPR014716 | 1 | 1 | 11383 | 557 | CG41520 | | 4.893e-02 | -3.02 | PADR1\_dom\_sf | interpro domains | IPR038650 | 1 | 1 | 11383 | 557 | Parp | | 4.893e-02 | -3.02 | CPEB1\_RRM1 | interpro domains | IPR034977 | 1 | 1 | 11383 | 557 | orb | | 4.893e-02 | -3.02 | Maf1 | interpro domains | IPR015257 | 1 | 1 | 11383 | 557 | Maf1 | | 4.893e-02 | -3.02 | AUTS2 | interpro domains | IPR023246 | 1 | 1 | 11383 | 557 | tay | | 4.893e-02 | -3.02 | N-end\_Aminoacyl\_Trfase | interpro domains | IPR030700 | 1 | 1 | 11383 | 557 | Ate1 | | 4.893e-02 | -3.02 | Glyco\_hydro\_31\_N\_dom | interpro domains | IPR025887 | 1 | 1 | 11383 | 557 | GCS2alpha | | 4.893e-02 | -3.02 | Plectin\_repeat | interpro domains | IPR001101 | 1 | 1 | 11383 | 557 | shot | | 4.893e-02 | -3.02 | NADH\_UbQ\_OxRdtase\_30kDa\_su | interpro domains | IPR001268 | 1 | 1 | 11383 | 557 | ND-30 | | 4.893e-02 | -3.02 | DUF4819 | interpro domains | IPR032147 | 1 | 1 | 11383 | 557 | cic | | 4.893e-02 | -3.02 | MOFRL\_protein | interpro domains | IPR039760 | 1 | 1 | 11383 | 557 | CG9886 | | 4.893e-02 | -3.02 | AP2\_complex\_asu | interpro domains | IPR017104 | 1 | 1 | 11383 | 557 | AP-2alpha | | 4.893e-02 | -3.02 | RPABC4 | interpro domains | IPR039747 | 1 | 1 | 11383 | 557 | Rpb12 | | 4.893e-02 | -3.02 | M-RIP\_PH | interpro domains | IPR039597 | 1 | 1 | 11383 | 557 | osp | | 4.893e-02 | -3.02 | Egl-10 | interpro domains | IPR034483 | 1 | 1 | 11383 | 557 | RSG7 | | 4.893e-02 | -3.02 | E3\_Ub\_ligase\_DUF908 | interpro domains | IPR010309 | 1 | 1 | 11383 | 557 | CG8184 | | 4.893e-02 | -3.02 | PSMD13 | interpro domains | IPR035298 | 1 | 1 | 11383 | 557 | Rpn9 | | 4.893e-02 | -3.02 | DUF1088 | interpro domains | IPR010508 | 1 | 1 | 11383 | 557 | rg | | 4.893e-02 | -3.02 | PH\_APBB1IP | interpro domains | IPR039665 | 1 | 1 | 11383 | 557 | pico | | 4.893e-02 | -3.02 | HUWE1/Rev1\_UBM | interpro domains | IPR025527 | 1 | 1 | 11383 | 557 | CG8184 | | 4.893e-02 | -3.02 | SERRATE/Ars2\_C | interpro domains | IPR007042 | 1 | 1 | 11383 | 557 | Ars2 | | 4.893e-02 | -3.02 | Znf\_PARP | interpro domains | IPR001510 | 1 | 1 | 11383 | 557 | Parp | | 4.893e-02 | -3.02 | Gln-dep\_NAD\_synthase | interpro domains | IPR014445 | 1 | 1 | 11383 | 557 | Nadsyn | | 4.893e-02 | -3.02 | TPK\_B1-bd\_sf | interpro domains | IPR036371 | 1 | 1 | 11383 | 557 | CG14721 | | 4.893e-02 | -3.02 | Cpo\_RRM | interpro domains | IPR034788 | 1 | 1 | 11383 | 557 | cpo | | 4.893e-02 | -3.02 | BCKD/PDK | interpro domains | IPR039028 | 1 | 1 | 11383 | 557 | Pdk | | 4.893e-02 | -3.02 | WGR\_domain | interpro domains | IPR008893 | 1 | 1 | 11383 | 557 | Parp | | 4.893e-02 | -3.02 | Collagenase\_NC10/endostatin | interpro domains | IPR010515 | 1 | 1 | 11383 | 557 | Mp | | 4.893e-02 | -3.02 | SOX\_fam | interpro domains | IPR022097 | 1 | 1 | 11383 | 557 | SoxN | | 4.893e-02 | -3.02 | Pumilio\_RNA-bd | interpro domains | IPR033712 | 1 | 1 | 11383 | 557 | pum | | 4.893e-02 | -3.02 | Nrf1\_activation-bd | interpro domains | IPR019526 | 1 | 1 | 11383 | 557 | ewg | | 4.893e-02 | -3.02 | SE/Ars2 | interpro domains | IPR039727 | 1 | 1 | 11383 | 557 | Ars2 | | 4.893e-02 | -3.02 | TPK\_catalytic | interpro domains | IPR007371 | 1 | 1 | 11383 | 557 | CG14721 | | 4.893e-02 | -3.02 | Oxsr1 | interpro domains | IPR028749 | 1 | 1 | 11383 | 557 | fray | | 4.893e-02 | -3.02 | TCF/LEF | interpro domains | IPR024940 | 1 | 1 | 11383 | 557 | pan | | 4.893e-02 | -3.02 | Ureohydrolase | interpro domains | IPR006035 | 1 | 1 | 11383 | 557 | arg | | 4.893e-02 | -3.02 | Zf-tcix | interpro domains | IPR029269 | 1 | 1 | 11383 | 557 | CG2321 | | 4.893e-02 | -3.02 | CaM-bd\_dom\_sf | interpro domains | IPR036122 | 1 | 1 | 11383 | 557 | SK | | 4.893e-02 | -3.02 | TTC14 | interpro domains | IPR039190 | 1 | 1 | 11383 | 557 | CG6621 | | 4.893e-02 | -3.02 | Wge | interpro domains | IPR029616 | 1 | 1 | 11383 | 557 | wge | | 4.893e-02 | -3.02 | Syndecan\_CS | interpro domains | IPR030479 | 1 | 1 | 11383 | 557 | Sdc | | 4.893e-02 | -3.02 | Thiamin\_PyroPKinase\_B1-bd | interpro domains | IPR007373 | 1 | 1 | 11383 | 557 | CG14721 | | 4.893e-02 | -3.02 | CG-1\_dom | interpro domains | IPR005559 | 1 | 1 | 11383 | 557 | Camta | | 4.893e-02 | -3.02 | NADH\_quin\_OxRdtase\_su\_C/D-like | interpro domains | IPR037232 | 1 | 1 | 11383 | 557 | ND-30 | | 4.893e-02 | -3.02 | CCDC39 | interpro domains | IPR033290 | 1 | 1 | 11383 | 557 | CG17387 | | 4.940e-02 | -3.01 | GO\_CELL\_MOTILITY | MSigDB lists | GO\_CELL\_MOTILITY | 160 | 11 | 4161 | 164 | sbm,mbc,CG17387,shot,l(2)41Ab,dnc,Dhc16F,svp,Mef2,CG9492,toy | | 4.968e-02 | -3.00 | positive regulation of pathway-restricted SMAD protein phosphorylation | biological process | GO:0010862 | 8 | 2 | 12345 | 571 | scw,mav | | 4.968e-02 | -3.00 | regulation of muscle tissue development | biological process | GO:1901861 | 8 | 2 | 12345 | 571 | Mef2,kirre | | 4.968e-02 | -3.00 | negative regulation of oskar mRNA translation | biological process | GO:0007319 | 8 | 2 | 12345 | 571 | heph,bru1 | | 4.968e-02 | -3.00 | anterior head development | biological process | GO:0097065 | 8 | 2 | 12345 | 571 | en,hth | | 4.968e-02 | -3.00 | regulation of execution phase of apoptosis | biological process | GO:1900117 | 8 | 2 | 12345 | 571 | Diap1,grim | | 4.968e-02 | -3.00 | regulation of pathway-restricted SMAD protein phosphorylation | biological process | GO:0060393 | 8 | 2 | 12345 | 571 | mav,scw | | 4.968e-02 | -3.00 | neuron cell-cell adhesion | biological process | GO:0007158 | 8 | 2 | 12345 | 571 | Nlg3,Nlg2 | | 4.968e-02 | -3.00 | membrane assembly | biological process | GO:0071709 | 8 | 2 | 12345 | 571 | Nlg2,Nlg3 | | 4.968e-02 | -3.00 | ER-nucleus signaling pathway | biological process | GO:0006984 | 8 | 2 | 12345 | 571 | SCAP,PEK | | 4.968e-02 | -3.00 | anterior head segmentation | biological process | GO:0035288 | 8 | 2 | 12345 | 571 | en,hth | | 4.968e-02 | -3.00 | filopodium assembly | biological process | GO:0046847 | 8 | 2 | 12345 | 571 | mim,sn | | 4.968e-02 | -3.00 | male gonad development | biological process | GO:0008584 | 8 | 2 | 12345 | 571 | salm,tj | | 4.968e-02 | -3.00 | presynaptic membrane assembly | biological process | GO:0097105 | 8 | 2 | 12345 | 571 | Nlg3,Nlg2 | | 4.968e-02 | -3.00 | regulation of striated muscle tissue development | biological process | GO:0016202 | 8 | 2 | 12345 | 571 | kirre,Mef2 | | 4.968e-02 | -3.00 | acid secretion | biological process | GO:0046717 | 8 | 2 | 12345 | 571 | CG3009,CG42237 | | 4.968e-02 | -3.00 | postsynapse assembly | biological process | GO:0099068 | 8 | 2 | 12345 | 571 | Nlg2,Nlg3 | | 4.968e-02 | -3.00 | development of primary male sexual characteristics | biological process | GO:0046546 | 8 | 2 | 12345 | 571 | tj,salm | | 4.986e-02 | -3.00 | GO\_TRANSMEMBRANE\_TRANSPORTER\_ACTIVITY | MSigDB lists | GO\_TRANSMEMBRANE\_TRANSPORTER\_ACTIVITY | 261 | 16 | 4161 | 164 | Mct1,Ca-alpha1D,PMCA,Tim23,sbm,Tim17b,ABCB7,nAChRalpha4,MCU,Hmt-1,Glut1,CG5254,CG7255,nAChRalpha3,Atpalpha,MFS16 | | 4.991e-02 | -3.00 | axon terminus | cellular component | GO:0043679 | 75 | 7 | 11940 | 534 | sky,Magi,Snap25,cpx,par-1,pum,Mical | | 5.000e-02 | -3.00 | GSE360\_L\_DONOVANI\_VS\_B\_MALAYI\_LOW\_DOSE\_DC\_DN | MSigDB lists | GSE360\_L\_DONOVANI\_VS\_B\_MALAYI\_LOW\_DOSE\_DC\_DN | 68 | 6 | 4161 | 164 | CG9389,CG30495,shot,AP-2alpha,RpL6,ck |
