## Supplementary material for "A single DPE core promoter motif contributes to *in vivo* transcriptional regulation and affects cardiac function": SupFile2: geneOntology.html

Gene Ontology Results

### Gene Ontology Enrichment Results

Homer *de novo* Motif Enrichment Results  
Known Motif Enrichment Results  

###### Text file version of complete results (i.e. open with Excel) - biological process: Functional groupings of proteins (Gene Ontology) - molecular function: Mechanistic actions of proteins (Gene Ontology) - cellular component: Protein localization (Gene Ontology) - chromosome location: Genes with similar chromosome localization (NCBI Gene) - KEGG pathways: Groups of proteins in the same pathways (KEGG) - protein interactions: "Proteins interacting with a common protein (BIND, EcoCyc, HPRD)" (NCBI Gene) - interpro domains: Proteins with similar domains and features (Interpro) - pfam domains: Proteins with similar domains and features (Pfam) - smart domains: Proteins with similar domains and features (SMART) - gene3d domains: Proteins with similar domains and features (Gene3D) - prosite domains: Proteins with similar domains and features (Prosite) - prints domains: Proteins with similar domains and features (PRINTS) - MSigDB lists: "Genes sets for pathways, factor/miRNA target predictions, expression patterns, etc." (MSigDB) - BIOCYC pathways: Groups of proteins in the same pathways (BIOCYC) - COSMIC cancer mutations: Genes mutated in similar cancers (COSMIC) - GWAS genes: Genes mutated in similar diseases (GWAS Catalog) - Lipid Maps pathways: Groups of proteins in the same lipid pathways (Lipid Maps/Biosystems) - Pathway Interaction DB: Groups of proteins in the same pathways (Pathway Interaction Database) - REACTOME pathways: Groups of proteins in the same pathways (REACTOME) - SMPDB pathways: Groups of proteins in the same pathways (SMPDB) - WikiPathways: Groups of proteins in the same pathways (Wikipathways) Enriched Categories | | | | | | | | | | | | --- | --- | --- | --- | --- | --- | --- | --- | --- | --- | | P-value | ln(P) | Term | GO Tree | GO ID | # of Genes in Term | # of Target Genes in Term | # of Total Genes | # of Target Genes | Common Genes | | 9.644e-85 | -193.45 | intracellular | cellular component | GO:0005622 | 6762 | 917 | 11940 | 1105 | Atg6,insv,Cbl,Blos2,Ady43A,CG9186,CG10561,CG9436,ATPsynF,AGO3,CG2938,SP1029,Tak1,dco,l(1)G0196,CG7488,CG9643,milt,RpS14a,SerRS-m,Maf1,CG4880,ND-30,dob,nej,Spx,Smox,Droj2,RpS27A,Marf,Cul2,GCC88,Argk,Orc6,mRpS23,Ugt36E1,Arpc1,Trf2,Pym,stil,YL-1,Naa20A,isoQC,spn-E,CG14464,TfIIFalpha,CG3437,Ts,RpS5b,levy,Exo70,CG1344,Prosbeta3,Acyp2,mael,Sec22,CG7920,PPYR1,CG9393,mtTFB1,Adi1,Elba2,Fer1HCH,Gasz,tws,Ntan1,gus,CG33156,Cp1,CG5447,apt,RpS14b,Snx6,CG40472,Sry-alpha,CG8726,CG8311,CG4390,CG16865,CG17187,Xpac,Jafrac1,Tlk,alphaTub84B,CG6276,RpS16,Cchl,mus81,RpL23,Snm1,krimp,Clp,pen-2,Grip75,SNF4Agamma,RpS29,Girdin,CG10932,glob1,Fdh,Aladin,Ubc4,Atf3,GstT1,Cul5,CG6813,mute,Topors,pgc,Pex10,ACC,Bx,RpL17,CG1640,MED24,Moe,l(3)73Ah,Arpc2,SelD,P5CDh1,S2P,shu,Uch-L5,COX4,Nup62,Hr78,RfC38,Fkbp12,O-fut1,Nmt,Pfdn2,aay,CG4238,Chd64,Iml1,CG17768,Ankle2,CG1827,PIG-P,HP1c,E2f1,Echs1,GstT2,Tom40,Rel,CG32485,Afti,Cdk5,CG32473,cathD,CG32640,Efr,Wdfy2,CG42376,CG5026,Pfdn5,CG6967,alph,CG31759,Atg7,Psf3,pygo,bun,Cse1,larp,Seipin,ND-B18,CanA-14F,Kyat,not,Irp-1B,esc,CG2911,Prosbeta5,PpV,STUB1,lin-52,del,RpL24,PEK,Abi,Pdxk,grp,wal,yin,CG31633,Sod1,mbo,CG1602,EndoGI,egg,ATP8A,bys,cu,CG8611,Grip128,Ublcp1,Pex23,Rpt2,mop,mRpL16,Idh,velo,trr,squ,sea,RpL23A,mRpL48,Acox57D-p,mRpS5,BigH1,EMC8-9,Kap-alpha3,hb,Cen,BoYb,Smyd4-4,CG17068,EMRE,CG7414,l(1)G0004,APC7,RpS15,RpL34a,CG42526,CG3226,GluRS-m,mre11,Sgf11,Trs33,CG14671,CG4872,Unr,CycJ,bam,CG8009,CG2129,Pgk,Rbcn-3A,Gk2,Got1,CG32221,CHMP2B,mts,COX5B,Mf,CG8112,CG15141,CG14814,ovo,CG8475,Blos4,CG4882,eIF4A,RpL9,Ubc6,snRNA:U12,l(2)k01209,DppIII,muskelin,wash,CG31875,otu,Nc73EF,GlyP,mRpS21,stai,Mes2,AOX3,RpL30,blw,Ugt35A1,CG17691,Taf7,e(y)1,Pfrx,sbb,NHP2,CG12093,yip2,Idh3b,REPTOR,CkIIalpha,Med,stet,Pi4KIIalpha,CG14407,RYBP,fray,Mat89Ba,CG4586,CG11077,CG32536,RpL37a,CG31729,alphaTub67C,UQCR-11,PDCD-5,brwl,Atg4b,Bin1,CG4266,cype,CSN6,alt,Mybbp1A,tko,G9a,MAGE,Synj,Sec23,Fkbp59,ERp60,Roe1,PHGPx,CG4730,Thiolase,RpLP1,RecQ5,Actn,Edc3,Cbp80,dl,Utx,shep,COX5A,rept,Stat92E,CG8974,HmgD,CG8199,ATPsynbeta,CysRS,CG3309,Rnmt,CG17593,Dip-B,PpD3,Clc,Nup58,CycE,CG9316,cnn,opm,asf1,kl-2,Ndc1,Nap1,Atu,CG10038,Caf1-105,ATPsynE,Pgm1,Ssdp,tral,sol,Kap3,ari-1,RpL34b,CG11486,twe,Pka-R1,Pfas,Plap,jigr1,PIG-H,RpLP0,Cul1,mRpL17,AP-1-2beta,CG12018,UK114,Mitofilin,mxc,Crag,Mtpalpha,CG10863,Usp39,CG17202,Jabba,Uxt,IntS11,CG31342,Pax,CG12375,Sec15,Pex13,Aven,CG10344,ND-MWFE,Sod3,Gapdh1,ste14,bif,CG6707,CG4293,Pp2B-14D,bor,thoc6,chrb,Su(H),CG46385,smg,RpL38,lid,snRNA:U4:38AB,CanB2,CG6094,Pglym78,twin,UQCR-Q,Rtnl1,CG3587,Mctp,CG5734,CG5292,wapl,TBCC,CG12301,Eno,Aldh,RpS30,Pdk1,Larp7,CG11902,Mo25,Zwilch,cort,Ptpmeg2,spd-2,ago,NP15.6,Scsalpha1,spen,Klp10A,sau,CG4594,mino,ABCD,SelR,RpII140,RhoGAP54D,TrpRS-m,rig,Desat1,CycC,lt,Tip60,Rpn2,CG13551,whd,RpS24,Rpn3,CG14516,raptor,CG5003,RpL13,galectin,cyc,Lkb1,Pmi,mus304,Shmt,Usp10,Fkbp39,CG33977,CG4598,CG17454,CG2201,Pi3K92E,mRpL18,Top1,Vamp7,Hsp26,Rga,CG9143,Pp2A-29B,up,gw,RpL31,Ptpmeg,Herc4,COX8,Adh,CG14817,mEFTu1,CG5116,scny,CG10055,Ugalt,alc,cana,CG13344,eEF1beta,Wbp2,yps,Atf6,Naxe,CG10959,ATPsynD,NFAT,CG8204,Gdi,crm,Mcad,Gclc,Sgf29,dgt4,Arts,dnc,CG8004,CG17270,CG14408,DNApol-epsilon255,Cht11,mRpL35,Dph5,CG2199,Vha68-2,CG17294,Tango10,CG3939,Hnf4,CG9981,zf30C,GatA,CHOp24,fs(1)Ya,Prosalpha4,CG1890,CG31674,psq,Sccpdh1,LSm3,MED27,RpS7,mus205,Tao,Nf-YA,Non1,sws,PrBP,CG2162,mRpL10,mad2,CG11089,CG11070,CG6767,ATPsynO,CG2767,Marf1,vers,lin,Vha36-1,ND-75,EloC,RpS28b,CG9391,Coq7,Nadsyn,Bap170,Rpn13,CG11134,rl,bcd,Fur2,CG10418,CG11110,Rep,GstE11,Uba5,LysRS,Aos1,ben,tef,Coq6,Dhfr,CG1622,ND-24,Etf-QO,PyK,PAPLA1,Rbf,CG10638,fwe,ATPsynCF6,pps,Cdk7,tos,Pld,Nurf-38,Jafrac2,Mekk1,SdhD,CG5028,elg1,yl,CG10189,ND-49,p38a,Prosalpha5,RpS23,CG3223,Atpalpha,Hat1,colt,dx,Not10,HP1e,msps,ATPsyndelta,CG7728,MED20,yem,wdn,CG4860,Parp16,CG7716,Rbfox1,CG9773,Spt20,ix,Gk1,CG7655,Rrp40,RpL26,Pex12,Agpat2,Cyp1,CG11710,CG32441,ATPsynB,RpL29,CG2678,Nxf3,ND-23,Scox,csw,CG3631,sw,Cf2,nmdyn-D7,CG31053,pn,ScsbetaA,CycB,Cisd2,Neb-cGP,capt,foxo,tkv,cold,osk,AsnS,Axs,RpS2,alpha-Man-Ia,CG5969,mRpS10,Edem1,ND-51,SdhA,CG3511,CG11164,ATPsynG,Rpn12,Grip71,Archease,Tspo,obe,CG17328,Neos,CG8001,Gga,HDAC6,CG12288,CCT6,vret,CG7115,CG17931,DnaJ-H,MED4,mEFTu2,CG30496,Chrac-14,CCT3,r,Grip84,Rpb8,CG8507,CG4404,Surf4,Rbf2,CG10654,CG5880,SmydA-1,scu,Tfb4,mol,Tim8,pr,CG31523,Rab5,Lsm12,26-29-p,CG7997,CG5412,cN-IIIB,Sdhaf3,mRpL13,trbd,PIG-C,CG2034,Dd,Pomp,RpS4,Hip1,ND-18,Ppox,Tm1,SmE,Prosbeta6,CG11807,heph,GAPsec,sip3,exu,mRpL36,RhoGDI,CG10254,ND-B22,CG7328,CG17746,Gabat,Arpc4,r2d2,CG6610,AgmNAT,CG7009,eIF3m,CG9132,gammaTub37C,Ptip,Hmg-2,CG10336,RpL11,Cyt-c-p,X11L,Tomosyn,wisp,cbs,Vha13,SkpA,CG6178,d4,Coq8,Lnpk,RpS18,Trxr-1,lwr,cib,Pka-R2,GalT1,EcR,nmdyn-D6,Dcr-1,Imp,Cka,SRPK,IntS6,qkr58E-2,RpL10,grsm,Nprl2,pont,CG11562,CG9425,CG9399,AMPdeam,ND-B15,AIF,Gyg,CG13366,cup,CCT5,piwi,Vps16B,CG5510,Arf51F,APP-BP1,Xpc,Rcd5,CG3021,Sply,Pfk,Pus7,Ranbp21,RpL7A,Mtl,CG9941,CG11267,CG9265,SERCA,SuUR,shrb,CG4557,CG15525,Hsc70-3,CG8314,ND-39,Pen,CG7840,Tengl4,Trf4-2,Rab32,Fbw5,CG10470,mRpS11,Ufm1,armi,tho2,park,tapas,Pink1,Arl5,ph-d,Acph-1,fdl,shop,lqfR,tun,CG17698,toc,Karybeta3,ND-SGDH,cad,ASPP,Tom70,faf,Debcl,Orc1,CG10631,CG8728,RpL40,Mos,CG10425,CG9684,Mst89B,Uev1A,escl,GatB,Pdk,Uch,CG42375,mtrm,Traf6,cl,Prosalpha3,Prosbeta4,RpL13A,CG13601,nos,CG5903,parvin,Sirt7,enc,nbs,UGP,trsn,ste24a,Rab10,TyrRS,jvl,bru1,ND-MNLL,eIF4E1,CG14183,CG15715,SREBP,Exd2,tub,l(3)mbt,Phb2,spag,Cdk1,RpS27,CG6766,CG5986,CG1774,Syx13,cnc,RpL12,hng3,sbr,Iru,Vsp37A,DCAF12,crok,CG4842,CG14894,EMC3,awd,cact,Stam,CG4021,fal,RpS13,CG3803,Sec10,Ptp61F,snRNA:U4atac,Cenp-C,Sfxn2,CG3198,Gmap,Sld5,eIF3i | | 9.644e-85 | -193.45 | intracellular part | cellular component | GO:0044424 | 6762 | 917 | 11940 | 1105 | Cul1,RpLP0,PIG-H,Pfas,Plap,jigr1,Pka-R1,twe,CG11486,RpL34b,ari-1,Kap3,sol,Ssdp,tral,Pgm1,Jabba,CG17202,Usp39,CG10863,Mtpalpha,Crag,mxc,UK114,Mitofilin,AP-1-2beta,CG12018,mRpL17,Nup58,Clc,PpD3,CG17593,Dip-B,Rnmt,CysRS,CG3309,ATPsynbeta,CG8199,HmgD,CG8974,Stat92E,ATPsynE,CG10038,Caf1-105,Atu,Ndc1,Nap1,kl-2,asf1,opm,cnn,CG9316,CycE,Larp7,Pdk1,RpS30,Eno,Aldh,CG12301,wapl,TBCC,CG5292,CG5734,Mctp,CG3587,CG4594,sau,Klp10A,spen,Scsalpha1,NP15.6,ago,Ptpmeg2,spd-2,Zwilch,cort,Mo25,CG11902,CG4293,CG6707,bif,Gapdh1,Sod3,ste14,ND-MWFE,CG10344,Aven,Sec15,Pex13,Pax,CG12375,CG31342,IntS11,Uxt,Rtnl1,UQCR-Q,twin,Pglym78,CanB2,CG6094,snRNA:U4:38AB,lid,RpL38,smg,Su(H),CG46385,chrb,thoc6,bor,Pp2B-14D,Ugt35A1,blw,RpL30,AOX3,Mes2,stai,mRpS21,GlyP,Nc73EF,CG31875,otu,wash,muskelin,DppIII,l(2)k01209,CG12093,yip2,NHP2,sbb,Pfrx,e(y)1,CG17691,Taf7,CG2129,Pgk,CG8009,bam,CycJ,Unr,snRNA:U12,Ubc6,eIF4A,RpL9,CG4882,Blos4,CG8475,ovo,CG14814,CG15141,CG8112,COX5B,Mf,CHMP2B,mts,CG32221,Got1,Gk2,Rbcn-3A,Roe1,ERp60,Fkbp59,Sec23,Synj,MAGE,G9a,Mybbp1A,tko,alt,rept,Utx,COX5A,shep,dl,Cbp80,Edc3,Actn,Thiolase,RpLP1,RecQ5,CG4730,PHGPx,CG4586,Mat89Ba,fray,RYBP,CG14407,Pi4KIIalpha,stet,CkIIalpha,Med,REPTOR,Idh3b,CSN6,cype,CG4266,Bin1,Atg4b,brwl,PDCD-5,UQCR-11,alphaTub67C,RpL37a,CG32536,CG31729,CG11077,larp,bun,Cse1,pygo,Psf3,CG31759,Atg7,alph,CG6967,Pfdn5,CG5026,Efr,Wdfy2,CG42376,Cdk5,CG32640,cathD,CG32473,Afti,PpV,Prosbeta5,CG2911,not,Irp-1B,esc,Kyat,CanA-14F,ND-B18,Seipin,Pfdn2,Nmt,O-fut1,Fkbp12,RfC38,Hr78,COX4,Uch-L5,Nup62,shu,S2P,CG32485,Rel,Tom40,GstT2,Echs1,E2f1,HP1c,CG1827,Ankle2,PIG-P,CG17768,Iml1,Chd64,CG4238,aay,Cen,hb,Kap-alpha3,EMC8-9,BigH1,mRpS5,Acox57D-p,RpL23A,mRpL48,sea,squ,trr,Trs33,CG4872,CG14671,Sgf11,mre11,GluRS-m,CG42526,CG3226,RpL34a,RpS15,APC7,l(1)G0004,CG7414,EMRE,Smyd4-4,CG17068,BoYb,CG31633,Sod1,yin,grp,wal,Pdxk,RpL24,Abi,PEK,del,lin-52,STUB1,velo,Idh,mop,Rpt2,mRpL16,Pex23,CG8611,Grip128,Ublcp1,cu,bys,ATP8A,EndoGI,egg,CG1602,mbo,Naa20A,isoQC,spn-E,stil,Trf2,YL-1,Pym,Arpc1,Ugt36E1,mRpS23,Orc6,Argk,GCC88,Cul2,mael,Acyp2,Prosbeta3,CG1344,Exo70,levy,RpS5b,CG3437,Ts,CG14464,TfIIFalpha,CG9643,CG7488,l(1)G0196,dco,SP1029,Tak1,CG2938,AGO3,CG9436,ATPsynF,CG10561,CG9186,Ady43A,Blos2,Cbl,insv,Atg6,Marf,Droj2,RpS27A,Smox,Spx,nej,dob,ND-30,CG4880,Maf1,SerRS-m,RpS14a,milt,Atf3,Ubc4,Fdh,Aladin,glob1,Girdin,CG10932,RpS29,SNF4Agamma,Grip75,pen-2,Clp,krimp,Snm1,RpL23,mus81,Cchl,RpS16,CG6276,SelD,Arpc2,P5CDh1,l(3)73Ah,Moe,CG1640,MED24,RpL17,ACC,Bx,Pex10,Topors,pgc,mute,CG6813,Cul5,GstT1,CG5447,apt,Cp1,CG33156,gus,Ntan1,Gasz,tws,Fer1HCH,Elba2,Adi1,mtTFB1,CG9393,PPYR1,CG7920,Sec22,alphaTub84B,Tlk,Jafrac1,Xpac,CG17187,CG16865,CG4390,CG8311,Sry-alpha,CG8726,CG40472,Snx6,RpS14b,tun,shop,lqfR,fdl,ph-d,Acph-1,Arl5,Pink1,tapas,park,tho2,armi,Ufm1,Uev1A,Mst89B,CG9684,CG10425,Mos,RpL40,CG8728,Orc1,CG10631,Debcl,Tom70,faf,ASPP,ND-SGDH,cad,Karybeta3,CG17698,toc,CG4557,shrb,SuUR,SERCA,CG11267,CG9941,CG9265,Mtl,RpL7A,Pfk,Ranbp21,Pus7,Sply,CG3021,Rcd5,mRpS11,CG10470,Fbw5,Rab32,Trf4-2,Tengl4,CG7840,CG8314,Pen,ND-39,Hsc70-3,CG15525,cact,awd,EMC3,CG14894,CG4842,crok,DCAF12,Vsp37A,Iru,hng3,sbr,RpL12,Syx13,cnc,CG1774,RpS27,CG5986,CG6766,Sld5,eIF3i,Gmap,CG3198,Sfxn2,Cenp-C,snRNA:U4atac,Ptp61F,CG3803,Sec10,RpS13,fal,Stam,CG4021,parvin,CG5903,nos,CG13601,RpL13A,Prosbeta4,cl,Prosalpha3,Traf6,mtrm,Pdk,CG42375,Uch,GatB,escl,Cdk1,Phb2,l(3)mbt,spag,tub,Exd2,SREBP,CG15715,CG14183,eIF4E1,bru1,jvl,ND-MNLL,TyrRS,Rab10,ste24a,trsn,UGP,nbs,enc,Sirt7,Dd,CG2034,trbd,PIG-C,mRpL13,Sdhaf3,CG5412,cN-IIIB,CG7997,Rab5,Lsm12,26-29-p,CG31523,pr,Tim8,mol,Tfb4,ND-B22,CG7328,CG10254,exu,RhoGDI,mRpL36,sip3,GAPsec,heph,CG11807,Prosbeta6,Tm1,SmE,Ppox,ND-18,RpS4,Hip1,Pomp,Grip84,r,Chrac-14,CCT3,MED4,CG30496,mEFTu2,DnaJ-H,CG17931,CG7115,CCT6,vret,CG12288,Gga,HDAC6,CG8001,Neos,CG17328,obe,Tspo,scu,SmydA-1,CG5880,Rbf2,CG10654,Surf4,CG4404,CG8507,Rpb8,grsm,RpL10,qkr58E-2,IntS6,SRPK,Cka,Imp,Dcr-1,nmdyn-D6,EcR,Pka-R2,GalT1,cib,Xpc,Arf51F,APP-BP1,Vps16B,CG5510,piwi,CCT5,cup,CG13366,Gyg,AIF,ND-B15,AMPdeam,CG9399,CG9425,CG11562,Nprl2,pont,RpL11,CG10336,Hmg-2,CG9132,Ptip,gammaTub37C,CG7009,eIF3m,CG6610,r2d2,AgmNAT,Arpc4,Gabat,CG17746,lwr,Trxr-1,RpS18,Lnpk,Coq8,CG6178,d4,SkpA,Vha13,cbs,wisp,Tomosyn,X11L,Cyt-c-p,CG5028,SdhD,Mekk1,Nurf-38,Jafrac2,Pld,tos,Cdk7,pps,fwe,ATPsynCF6,CG10638,Rbf,PyK,PAPLA1,Etf-QO,yem,MED20,CG7728,msps,ATPsyndelta,Not10,HP1e,dx,Hat1,Atpalpha,colt,RpS23,CG3223,Prosalpha5,p38a,ND-49,CG10189,yl,elg1,Rep,CG11110,CG10418,Fur2,bcd,rl,CG11134,Bap170,Rpn13,Coq7,CG9391,Nadsyn,RpS28b,EloC,ND-75,CG1622,ND-24,Dhfr,Coq6,tef,ben,Aos1,LysRS,GstE11,Uba5,Axs,osk,AsnS,cold,tkv,foxo,capt,Neb-cGP,Cisd2,CycB,pn,ScsbetaA,CG31053,Cf2,nmdyn-D7,sw,Archease,Grip71,Rpn12,ATPsynG,CG3511,CG11164,SdhA,ND-51,Edem1,mRpS10,CG5969,alpha-Man-Ia,RpS2,Gk1,CG7655,ix,Spt20,CG9773,Rbfox1,CG7716,Parp16,CG4860,wdn,csw,CG3631,Scox,ND-23,CG2678,Nxf3,RpL29,ATPsynB,CG32441,CG11710,Cyp1,Agpat2,Pex12,RpL26,Rrp40,up,gw,Pp2A-29B,CG9143,Rga,Vamp7,Hsp26,Top1,mRpL18,Pi3K92E,CG2201,CG4598,CG17454,CG33977,Atf6,yps,Wbp2,eEF1beta,CG13344,cana,Ugalt,alc,CG10055,CG5116,scny,mEFTu1,CG14817,Adh,COX8,Ptpmeg,Herc4,RpL31,CG13551,Rpn2,lt,Tip60,Desat1,CycC,rig,TrpRS-m,RpII140,RhoGAP54D,SelR,ABCD,mino,Fkbp39,Shmt,Usp10,mus304,Pmi,cyc,Lkb1,galectin,RpL13,CG5003,raptor,CG14516,Rpn3,RpS24,whd,Tao,mus205,RpS7,MED27,LSm3,Sccpdh1,psq,CG31674,CG1890,fs(1)Ya,Prosalpha4,CHOp24,GatA,Vha36-1,lin,vers,Marf1,ATPsynO,CG2767,CG11070,CG6767,CG11089,mad2,mRpL10,CG2162,PrBP,Nf-YA,Non1,sws,DNApol-epsilon255,CG8004,CG17270,dnc,CG14408,dgt4,Sgf29,Arts,Gclc,Mcad,Gdi,crm,CG8204,ATPsynD,NFAT,CG10959,Naxe,CG9981,Hnf4,zf30C,Tango10,CG3939,CG17294,Vha68-2,CG2199,Dph5,mRpL35,Cht11 | | 2.297e-75 | -171.86 | cell | cellular component | GO:0005623 | 7811 | 978 | 11940 | 1105 | wal,grp,yin,Sod1,CG31633,lin-52,STUB1,Abi,RpL24,PEK,Pdxk,del,mop,mRpL16,Rpt2,Pex23,velo,Idh,CG1602,EndoGI,egg,mbo,cu,Ublcp1,CG8611,Grip128,ATP8A,bys,mtgo,mRpS5,Kap-alpha3,Cen,hb,BigH1,EMC8-9,GILT1,trr,squ,mRpL48,RpL23A,Acox57D-p,sea,dpr6,Sgf11,CG3226,CG42526,mre11,GluRS-m,CG4872,CG14671,Trs33,Smyd4-4,CG17068,EMRE,CG6927,BoYb,RpS15,RpL34a,CG7414,l(1)G0004,APC7,Fkbp12,DIP-lambda,Hr78,RfC38,O-fut1,Nmt,Pfdn2,CG42748,shu,S2P,Nup62,COX4,Uch-L5,Tom40,Rel,GstT2,CG32485,slif,Chd64,aay,CG4238,E2f1,HP1c,Echs1,Iml1,CG17768,CG1827,PIG-P,Ankle2,Psf3,CG31759,Atg7,larp,pygo,Cse1,bun,Efr,Wdfy2,CG42376,Afti,Cdk5,CG32473,CG32640,cathD,CG6967,alph,CG5026,Pfdn5,PpV,CG2911,Prosbeta5,ND-B18,Seipin,CanA-14F,not,esc,Kyat,Irp-1B,Nlg1,Adi1,Elba2,Fer1HCH,Cp1,CG33156,CG5447,apt,tws,Ntan1,Gasz,gus,Hexo1,Sec22,CG7920,PPYR1,CG9393,mtTFB1,CG4390,CG17187,CG16865,CG8311,Jafrac1,Tlk,alphaTub84B,Xpac,fs(1)N,CG16868,RpS14b,Snx6,CG8726,Sry-alpha,CG40472,CG31689,glob1,Aladin,Fdh,Grip75,CG10932,Girdin,RpS29,SNF4Agamma,Ubc4,Atf3,Ork1,Cchl,RpS16,mus81,CG6276,Clp,pen-2,RpL23,krimp,Snm1,RpL17,MED24,CG1640,Bx,ACC,l(3)73Ah,SelD,P5CDh1,Arpc2,Moe,CG6813,GstT1,Cul5,Pex10,pgc,mute,Topors,blot,SP1029,Tak1,AGO3,CG2938,l(1)G0196,CG9643,CG7488,dco,Blos2,Cbl,Ady43A,CG9186,Atg6,insv,CAH2,CG9436,ATPsynF,CG10561,nej,dob,RpS27A,Droj2,Marf,Spx,Smox,Maf1,SerRS-m,milt,RpS14a,ND-30,CG4880,mRpS23,Ugt36E1,CG14715,spn-E,Naa20A,isoQC,Arpc1,stil,Trf2,YL-1,Pym,Cul2,Argk,Orc6,GCC88,CG9911,Acyp2,mael,Prosbeta3,RpS5b,CG3437,Ts,CG14464,TfIIFalpha,CG1344,levy,Exo70,bif,CG6707,ste14,Sod3,Gapdh1,CG4293,fs(1)M3,CG31342,Uxt,IntS11,CG10344,Aven,ND-MWFE,CG12375,Pax,Pex13,Sec15,Pglym78,twin,lid,smg,RpL38,CanB2,CG6094,snRNA:U4:38AB,Rtnl1,UQCR-Q,bor,thoc6,Pp2B-14D,zyd,chrb,CG46385,Su(H),RpS30,CG12301,Aldh,Eno,Larp7,Pdk1,mAChR-A,Ac3,TMEM216,CG5734,CG3587,Mctp,wapl,TBCC,CG5292,NP15.6,Scsalpha1,spen,ago,CG4594,sau,SelT,Klp10A,Mo25,CG11902,Ptpmeg2,spd-2,Zwilch,cort,rpk,PpD3,CG17593,Dip-B,Clc,Nup58,HmgD,CG8974,CG8199,Stat92E,Rnmt,ATPsynbeta,CG3309,CysRS,Ndc1,Nap1,ATPsynE,Atu,Caf1-105,CG42258,CG10038,CG9316,CycE,asf1,kl-2,cnn,opm,Pka-R1,CG11486,twe,Cul1,RpLP0,Pfas,jigr1,Plap,kcc,PIG-H,Pgm1,Kap3,RpL34b,ari-1,Ssdp,tral,Ho,sol,Mtpalpha,CG10863,Crag,CG17202,Jabba,Usp39,mRpL17,mxc,CG12018,AP-1-2beta,UK114,Mitofilin,fray,RYBP,CG31195,CG4586,Mat89Ba,Idh3b,REPTOR,CG14407,Med,CkIIalpha,stet,Pi4KIIalpha,brwl,PDCD-5,Atg4b,UQCR-11,CSN6,cype,CG4266,Bin1,CG11077,alphaTub67C,CG32536,RpL37a,CG31729,Fkbp59,Sec23,Roe1,ERp60,G9a,tko,Mybbp1A,alt,Synj,MAGE,Edc3,Cbp80,CG16979,Actn,shep,rept,Utx,COX5A,dl,PHGPx,Thiolase,RecQ5,RpLP1,CG4730,Pgk,CG2129,CG8009,CycJ,bam,Unr,CG8475,Blos4,CG15141,CG14814,ovo,Ubc6,snRNA:U12,CG4882,Myd88,eIF4A,RpL9,CHMP2B,mts,Rbcn-3A,Gk2,Got1,Tsp42Ef,CG32221,CG8112,Mf,COX5B,Mes2,AOX3,stai,RpL30,blw,Ugt35A1,DppIII,l(2)k01209,GCS2beta,GlyP,Nc73EF,ZnT35C,mRpS21,muskelin,wash,otu,CG31875,sbb,yip2,NHP2,CG12093,Taf7,CG17691,Pfrx,e(y)1,Spt20,CG9773,CG7655,Gk1,ix,wdn,boi,CG4860,Parp16,CG7716,Rbfox1,RpL29,Ptp69D,ATPsynB,CG32441,Scox,CG3631,csw,Nxf3,CG2678,ND-23,Rrp40,Cyp1,CG11710,RpL26,Pex12,Agpat2,capt,foxo,tkv,cold,Neb-cGP,Axs,osk,AsnS,sw,Cf2,nmdyn-D7,CG31053,pn,ScsbetaA,CycB,Cisd2,Tsp66E,CG11164,CG3511,SdhA,ND-51,Archease,Grip71,ATPsynG,Rpn12,RpS2,mRpS10,CG5969,Edem1,alpha-Man-Ia,CG10418,Rep,CG11110,Coq7,CG9391,Nadsyn,Bap170,Rpn13,EloC,ND-75,RpS28b,bcd,rl,LpR2,Fur2,CG11134,ben,tef,ND-24,CG1622,Coq6,Dhfr,Ect4,LysRS,GstE11,Uba5,Aos1,tos,Pld,CG10960,SdhD,CG5028,Jafrac2,Nurf-38,Mekk1,Rbf,CG10638,Etf-QO,PyK,PAPLA1,pps,Cdk7,fwe,ATPsynCF6,HP1e,Not10,msps,ATPsyndelta,CG3223,RpS23,Atpalpha,dx,colt,Hat1,MED20,yem,CG12945,CG7728,elg1,yl,p38a,Prosalpha5,CG10189,ND-49,dgt4,Sgf29,Arts,CG33181,Mcad,Gclc,dnc,CG17270,CG14408,CG8004,DNApol-epsilon255,CG10959,Naxe,CG8204,Gdi,crm,NFAT,ATPsynD,Vha68-2,CG3939,Tango10,zf30C,CG9981,Hnf4,CG17294,nrv1,Cht11,Dph5,CG2199,mRpL35,LSm3,MED27,psq,Sccpdh1,mus205,RpS7,Tao,GatA,CHOp24,CG1890,CG31674,Prosalpha4,fs(1)Ya,CG2767,ATPsynO,mad2,CG11089,CG11070,CG6767,lin,Vha36-1,Marf1,vers,PrBP,sws,Non1,Nf-YA,mRpL10,CG2162,Desat1,CycC,lt,Tip60,TrpRS-m,rig,CG13551,Rpn2,mino,RpII140,RhoGAP54D,ABCD,SelR,mus304,Usp10,Shmt,fz4,Pmi,Fkbp39,galectin,RpL13,whd,RpS24,Rpn3,raptor,CG14516,CG5003,cyc,Lkb1,Vamp7,Hsp26,CG9143,Rga,Top1,Pp2A-29B,up,gw,CG33977,Pi3K92E,mRpL18,CG17454,CG4598,CG2201,cana,Ugalt,CG10055,alc,yps,eEF1beta,Wbp2,Atf6,CG13344,Herc4,COX8,Ptpmeg,RpL31,CG1309,Adh,CG14817,scny,CG5116,mEFTu1,Prosbeta4,Traf6,mtrm,Prosalpha3,cl,CG5903,parvin,RpL13A,nos,CG13601,escl,GatB,Pdk,CG42375,Uch,Exd2,tub,CG15715,SREBP,Cdk1,l(3)mbt,Phb2,spag,UGP,ste24a,trsn,enc,Sirt7,nbs,eIF4E1,CG14183,Rab10,TyrRS,jvl,bru1,ND-MNLL,crok,Vsp37A,Iru,DCAF12,awd,cact,CG14894,CG4842,EMC3,CG30291,RpS27,CG6766,CG5986,CG1774,RpL12,hng3,sbr,cnc,Syx13,Cenp-C,snRNA:U4atac,CG3198,Gmap,eIF3i,Sld5,Sfxn2,fal,Stam,CG4021,CG3803,Sec10,Ptp61F,RpS13,SuUR,CG9941,CG9265,CG11267,SERCA,CG4476,shrb,CG4557,CG3021,Rcd5,RpL7A,Mtl,Sply,Pus7,Pfk,Ranbp21,Rab32,Trf4-2,CG10470,Fbw5,mRpS11,CG15525,Hsc70-3,CG4332,Tengl4,Pen,ND-39,CG8314,CG7840,fdl,nSMase,CG2061,tapas,Pink1,Acph-1,Arl5,ph-d,tun,shop,lqfR,tho2,park,Ufm1,armi,CG10425,RpL40,Mos,Mst89B,CG9684,Uev1A,faf,Tom70,Debcl,Karybeta3,CG17698,toc,cad,ASPP,ND-SGDH,CG8728,CG10631,Orc1,Hmg-2,CG10336,RpL11,CG17746,slam,CG6610,r2d2,AgmNAT,CG9132,gammaTub37C,Ptip,eIF3m,CG7009,Gabat,Arpc4,Oseg1,Trxr-1,Lnpk,RpS18,lwr,wisp,Vha13,cbs,Cyt-c-p,Tomosyn,X11L,Coq8,SkpA,d4,CG6178,qkr58E-2,RpL10,SRPK,E23,Cka,IntS6,tor,grsm,GalT1,EcR,Pka-R2,cib,Dcr-1,nmdyn-D6,Imp,Tsp74F,CCT5,piwi,nAChRalpha4,CG13366,cup,Gyg,Arf51F,APP-BP1,Xpc,Vps16B,CG5510,Nprl2,pont,AMPdeam,CG9399,AIF,ND-B15,Drep2,CG11562,CG9425,CG7115,CG17931,CG12288,vret,CCT6,CCT3,Chrac-14,r,Grip84,ssh,DnaJ-H,mEFTu2,MED4,CG30496,Neos,CG8001,Tspo,obe,CG17328,HDAC6,Gga,CG5880,scu,SmydA-1,CG8507,Rpb8,Rbf2,CG10654,CG4404,Ptp10D,Surf4,Sdhaf3,CG2034,Dd,mRpL13,trbd,PIG-C,pr,CG31523,Tim8,NaPi-III,Lsm12,Rab5,26-29-p,mol,Ggt-1,Tfb4,cN-IIIB,CG5412,CG7997,sip3,RhoGDI,exu,mRpL36,GAPsec,CG7328,ND-B22,CG10254,Ppox,Tm1,SmE,Hip1,RpS4,Pomp,ND-18,heph,Prosbeta6,CG11807,zpg | | 2.297e-75 | -171.86 | cell part | cellular component | GO:0044464 | 7811 | 978 | 11940 | 1105 | Bx,ACC,CG1640,MED24,RpL17,Moe,Arpc2,SelD,P5CDh1,l(3)73Ah,Cul5,GstT1,CG6813,Topors,mute,pgc,blot,Pex10,SNF4Agamma,RpS29,CG10932,Girdin,Grip75,Fdh,Aladin,glob1,CG31689,Ork1,Atf3,Ubc4,CG6276,mus81,RpS16,Cchl,Snm1,krimp,RpL23,pen-2,Clp,CG8311,CG16865,CG17187,CG4390,fs(1)N,Xpac,alphaTub84B,Jafrac1,Tlk,Snx6,RpS14b,CG16868,CG40472,Sry-alpha,CG8726,Fer1HCH,Adi1,Elba2,Nlg1,gus,tws,Ntan1,Gasz,CG5447,apt,CG33156,Cp1,CG7920,Sec22,Hexo1,CG9393,mtTFB1,PPYR1,Prosbeta3,mael,Acyp2,TfIIFalpha,CG14464,RpS5b,Ts,CG3437,Exo70,levy,CG1344,Ugt36E1,mRpS23,stil,Pym,Trf2,YL-1,Arpc1,spn-E,CG14715,Naa20A,isoQC,Cul2,GCC88,CG9911,Orc6,Argk,dob,nej,Spx,Smox,Marf,Droj2,RpS27A,RpS14a,milt,SerRS-m,Maf1,CG4880,ND-30,CG2938,AGO3,SP1029,Tak1,dco,CG9643,CG7488,l(1)G0196,CAH2,Atg6,insv,CG9186,Ady43A,Cbl,Blos2,CG10561,CG9436,ATPsynF,GluRS-m,mre11,CG3226,CG42526,Sgf11,Trs33,CG4872,CG14671,BoYb,CG6927,EMRE,CG17068,Smyd4-4,APC7,CG7414,l(1)G0004,RpL34a,RpS15,mRpS5,mtgo,EMC8-9,BigH1,Cen,hb,Kap-alpha3,squ,trr,GILT1,sea,dpr6,Acox57D-p,mRpL48,RpL23A,Pex23,Rpt2,mop,mRpL16,Idh,velo,mbo,EndoGI,egg,CG1602,bys,ATP8A,Grip128,Ublcp1,CG8611,cu,grp,wal,Sod1,CG31633,yin,STUB1,lin-52,del,Pdxk,Abi,PEK,RpL24,Prosbeta5,CG2911,PpV,Seipin,ND-B18,Kyat,not,Irp-1B,esc,CanA-14F,CG31759,Atg7,Psf3,Cse1,bun,pygo,larp,CG32473,Cdk5,cathD,CG32640,Afti,Wdfy2,CG42376,Efr,Pfdn5,CG5026,alph,CG6967,GstT2,Rel,Tom40,CG32485,CG4238,aay,Chd64,slif,PIG-P,CG1827,Ankle2,CG17768,Iml1,Echs1,HP1c,E2f1,RfC38,Hr78,DIP-lambda,Fkbp12,CG42748,Pfdn2,Nmt,O-fut1,S2P,shu,Uch-L5,Nup62,COX4,Actn,CG16979,Cbp80,Edc3,dl,Utx,rept,shep,COX5A,PHGPx,CG4730,Thiolase,RecQ5,RpLP1,Sec23,Fkbp59,ERp60,Roe1,tko,alt,Mybbp1A,G9a,MAGE,Synj,UQCR-11,Atg4b,PDCD-5,brwl,cype,CG4266,Bin1,CSN6,CG11077,RpL37a,CG31729,CG32536,alphaTub67C,CG31195,RYBP,fray,Mat89Ba,CG4586,REPTOR,Idh3b,stet,Pi4KIIalpha,CkIIalpha,Med,CG14407,sbb,yip2,CG12093,NHP2,CG17691,Taf7,e(y)1,Pfrx,stai,AOX3,Mes2,Ugt35A1,blw,RpL30,l(2)k01209,DppIII,CG31875,otu,wash,muskelin,ZnT35C,mRpS21,GlyP,GCS2beta,Nc73EF,CG14814,ovo,CG15141,Blos4,CG8475,Myd88,eIF4A,RpL9,CG4882,snRNA:U12,Ubc6,CG32221,Tsp42Ef,Got1,Rbcn-3A,Gk2,CHMP2B,mts,COX5B,Mf,CG8112,CG8009,CG2129,Pgk,Unr,bam,CycJ,ago,Scsalpha1,spen,NP15.6,Klp10A,SelT,CG4594,sau,CG11902,Mo25,Zwilch,cort,spd-2,Ptpmeg2,Aldh,Eno,CG12301,RpS30,Ac3,Pdk1,Larp7,mAChR-A,Mctp,CG3587,CG5734,TMEM216,CG5292,wapl,TBCC,CG6094,snRNA:U4:38AB,CanB2,RpL38,smg,lid,twin,Pglym78,UQCR-Q,Rtnl1,zyd,Pp2B-14D,thoc6,bor,CG46385,Su(H),chrb,ste14,Sod3,Gapdh1,CG6707,bif,CG4293,fs(1)M3,Uxt,IntS11,CG31342,Sec15,Pex13,CG12375,Pax,ND-MWFE,Aven,CG10344,Crag,CG10863,Mtpalpha,Usp39,Jabba,CG17202,mRpL17,UK114,Mitofilin,CG12018,AP-1-2beta,mxc,twe,CG11486,Pka-R1,PIG-H,kcc,Pfas,jigr1,Plap,RpLP0,Cul1,Pgm1,sol,Ho,Ssdp,tral,RpL34b,ari-1,Kap3,Nap1,Ndc1,CG10038,CG42258,Caf1-105,Atu,ATPsynE,CycE,CG9316,opm,cnn,kl-2,asf1,CG17593,Dip-B,PpD3,rpk,Nup58,Clc,Stat92E,CG8199,CG8974,HmgD,CysRS,CG3309,ATPsynbeta,Rnmt,CG11070,CG6767,mad2,CG11089,CG2767,ATPsynO,vers,Marf1,Vha36-1,lin,Nf-YA,sws,Non1,PrBP,CG2162,mRpL10,Sccpdh1,psq,MED27,LSm3,Tao,mus205,RpS7,CHOp24,GatA,Prosalpha4,fs(1)Ya,CG31674,CG1890,Vha68-2,CG17294,zf30C,Hnf4,CG9981,Tango10,CG3939,nrv1,Cht11,mRpL35,CG2199,Dph5,Gclc,Mcad,CG33181,Sgf29,Arts,dgt4,DNApol-epsilon255,CG8004,CG14408,dnc,CG17270,Naxe,CG10959,ATPsynD,NFAT,Gdi,crm,CG8204,Ugalt,alc,CG10055,cana,CG13344,Atf6,Wbp2,eEF1beta,yps,RpL31,Ptpmeg,Herc4,COX8,mEFTu1,CG5116,scny,CG14817,Adh,CG1309,Top1,Rga,CG9143,Hsp26,Vamp7,gw,up,Pp2A-29B,CG33977,CG2201,CG4598,CG17454,mRpL18,Pi3K92E,Pmi,Usp10,Shmt,fz4,mus304,Fkbp39,CG14516,raptor,CG5003,Rpn3,RpS24,whd,galectin,RpL13,cyc,Lkb1,TrpRS-m,rig,lt,Tip60,CycC,Desat1,Rpn2,CG13551,mino,SelR,ABCD,RpII140,RhoGAP54D,SdhA,ND-51,CG3511,CG11164,Rpn12,Grip71,ATPsynG,Archease,RpS2,alpha-Man-Ia,Edem1,mRpS10,CG5969,Neb-cGP,cold,tkv,capt,foxo,osk,AsnS,Axs,CG31053,pn,ScsbetaA,nmdyn-D7,sw,Cf2,Tsp66E,Cisd2,CycB,CG32441,ATPsynB,Ptp69D,RpL29,ND-23,CG2678,Nxf3,csw,CG3631,Scox,Rrp40,Agpat2,Pex12,RpL26,CG11710,Cyp1,CG9773,Spt20,ix,Gk1,CG7655,wdn,CG7716,Rbfox1,CG4860,Parp16,boi,colt,Hat1,Atpalpha,dx,CG3223,RpS23,ATPsyndelta,msps,HP1e,Not10,CG7728,CG12945,yem,MED20,yl,elg1,ND-49,CG10189,Prosalpha5,p38a,CG10960,Pld,tos,Mekk1,Nurf-38,Jafrac2,CG5028,SdhD,PyK,PAPLA1,Etf-QO,CG10638,Rbf,fwe,ATPsynCF6,Cdk7,pps,tef,ben,Dhfr,Coq6,CG1622,ND-24,Uba5,GstE11,Ect4,LysRS,Aos1,CG10418,CG11110,Rep,RpS28b,ND-75,EloC,Rpn13,Bap170,CG9391,Nadsyn,Coq7,CG11134,Fur2,LpR2,bcd,rl,Gyg,nAChRalpha4,CG13366,cup,piwi,Tsp74F,CCT5,CG5510,Vps16B,Xpc,Arf51F,APP-BP1,Nprl2,pont,CG9425,CG11562,Drep2,AIF,ND-B15,CG9399,AMPdeam,IntS6,E23,Cka,SRPK,qkr58E-2,RpL10,grsm,tor,cib,EcR,Pka-R2,GalT1,Imp,nmdyn-D6,Dcr-1,RpS18,Lnpk,Trxr-1,Oseg1,lwr,X11L,Tomosyn,Cyt-c-p,Vha13,cbs,wisp,CG6178,d4,SkpA,Coq8,CG10336,Hmg-2,RpL11,slam,CG17746,Arpc4,Gabat,Ptip,CG7009,CG9132,gammaTub37C,eIF3m,AgmNAT,r2d2,CG6610,GAPsec,RhoGDI,mRpL36,exu,sip3,CG10254,CG7328,ND-B22,ND-18,RpS4,Hip1,Pomp,Tm1,SmE,Ppox,zpg,CG11807,Prosbeta6,heph,Sdhaf3,trbd,PIG-C,mRpL13,Dd,CG2034,mol,Ggt-1,Tfb4,26-29-p,Lsm12,Rab5,pr,CG31523,Tim8,NaPi-III,CG7997,cN-IIIB,CG5412,scu,SmydA-1,CG5880,Rpb8,CG8507,Surf4,Ptp10D,CG4404,CG10654,Rbf2,vret,CCT6,CG12288,CG17931,CG7115,CG30496,MED4,mEFTu2,DnaJ-H,ssh,Grip84,r,CCT3,Chrac-14,obe,Tspo,CG17328,CG8001,Neos,Gga,HDAC6,Cenp-C,snRNA:U4atac,Sfxn2,Sld5,eIF3i,CG3198,Gmap,CG4021,Stam,fal,RpS13,Ptp61F,Sec10,CG3803,DCAF12,Vsp37A,Iru,crok,EMC3,CG14894,CG4842,cact,awd,CG1774,CG6766,RpS27,CG5986,CG30291,Syx13,cnc,sbr,hng3,RpL12,SREBP,CG15715,tub,Exd2,l(3)mbt,Phb2,spag,Cdk1,nbs,Sirt7,enc,trsn,ste24a,UGP,ND-MNLL,bru1,jvl,TyrRS,Rab10,CG14183,eIF4E1,Prosalpha3,cl,Traf6,mtrm,Prosbeta4,nos,CG13601,RpL13A,parvin,CG5903,escl,Uch,Pdk,CG42375,GatB,Mos,RpL40,CG10425,Uev1A,Mst89B,CG9684,ND-SGDH,cad,ASPP,toc,Karybeta3,CG17698,Debcl,Tom70,faf,CG10631,Orc1,CG8728,Acph-1,Arl5,ph-d,Pink1,CG2061,tapas,nSMase,fdl,lqfR,shop,tun,armi,Ufm1,park,tho2,Trf4-2,Rab32,mRpS11,Fbw5,CG10470,Hsc70-3,CG15525,CG7840,ND-39,Pen,CG8314,CG4332,Tengl4,SERCA,CG9265,CG11267,CG9941,SuUR,CG4557,shrb,CG4476,Rcd5,CG3021,Ranbp21,Pus7,Pfk,Sply,Mtl,RpL7A | | 1.378e-71 | -163.16 | cytoplasm | cellular component | GO:0005737 | 4500 | 696 | 11940 | 1105 | CG10425,RpL40,Mos,Mst89B,faf,Tom70,Debcl,Karybeta3,CG17698,toc,ND-SGDH,cad,CG8728,fdl,tapas,Pink1,Arl5,tun,shop,lqfR,park,Ufm1,armi,Rab32,CG10470,mRpS11,Hsc70-3,Tengl4,ND-39,CG8314,Pen,CG7840,CG11267,CG9941,SERCA,shrb,CG4557,CG3021,RpL7A,Mtl,Sply,Ranbp21,Pfk,Gmap,eIF3i,Sfxn2,Stam,CG3803,Sec10,Ptp61F,RpS13,crok,Iru,Vsp37A,cact,CG4842,CG14894,EMC3,CG6766,RpS27,CG1774,RpL12,sbr,Syx13,Exd2,tub,CG15715,SREBP,Cdk1,spag,Phb2,l(3)mbt,UGP,trsn,ste24a,enc,eIF4E1,CG14183,Rab10,TyrRS,bru1,ND-MNLL,Prosbeta4,Traf6,Prosalpha3,cl,CG5903,parvin,RpL13A,nos,CG13601,GatB,CG42375,Pdk,Uch,sip3,RhoGDI,exu,mRpL36,GAPsec,CG7328,ND-B22,CG10254,Ppox,Tm1,Pomp,Hip1,RpS4,ND-18,heph,CG11807,Prosbeta6,Sdhaf3,Dd,mRpL13,trbd,PIG-C,Tim8,CG31523,pr,Rab5,26-29-p,mol,CG5412,cN-IIIB,CG7997,CG5880,scu,SmydA-1,CG8507,Surf4,CG7115,CG17931,CCT6,vret,CCT3,Grip84,r,DnaJ-H,CG30496,mEFTu2,CG8001,Tspo,HDAC6,Gga,CCT5,piwi,cup,Gyg,Arf51F,APP-BP1,Xpc,Vps16B,CG5510,Nprl2,AMPdeam,CG9399,AIF,ND-B15,CG11562,CG9425,RpL10,Cka,SRPK,grsm,EcR,GalT1,Pka-R2,cib,Dcr-1,nmdyn-D6,Imp,Trxr-1,Lnpk,RpS18,wisp,Vha13,cbs,Cyt-c-p,Tomosyn,X11L,Coq8,SkpA,CG6178,RpL11,CG17746,r2d2,AgmNAT,CG9132,CG7009,eIF3m,gammaTub37C,Gabat,ATPsyndelta,RpS23,CG3223,colt,dx,Atpalpha,yl,p38a,Prosalpha5,CG10189,ND-49,Pld,SdhD,CG5028,Nurf-38,Jafrac2,Mekk1,CG10638,Etf-QO,PyK,PAPLA1,Cdk7,ATPsynCF6,fwe,ben,ND-24,Coq6,Dhfr,LysRS,GstE11,Uba5,Aos1,CG10418,Rep,CG11110,Nadsyn,CG9391,Coq7,Rpn13,ND-75,RpS28b,rl,Fur2,CG11134,SdhA,ND-51,ATPsynG,Rpn12,Grip71,RpS2,mRpS10,CG5969,Edem1,alpha-Man-Ia,tkv,capt,foxo,cold,Neb-cGP,Axs,AsnS,osk,nmdyn-D7,Cf2,ScsbetaA,pn,CycB,RpL29,CG32441,ATPsynB,Scox,CG3631,csw,CG2678,ND-23,Rrp40,Cyp1,RpL26,Pex12,Agpat2,CG9773,Gk1,CG4860,Parp16,Rbfox1,CG7716,CG10055,alc,Ugalt,Wbp2,eEF1beta,yps,Atf6,COX8,Herc4,Ptpmeg,RpL31,Adh,CG14817,CG5116,mEFTu1,Vamp7,Hsp26,Rga,Top1,Pp2A-29B,gw,up,CG33977,Pi3K92E,mRpL18,CG17454,CG4598,CG2201,mus304,Shmt,Usp10,Pmi,galectin,RpL13,RpS24,whd,CG14516,raptor,CG5003,cyc,Lkb1,Desat1,lt,TrpRS-m,rig,CG13551,Rpn2,mino,RhoGAP54D,ABCD,SelR,CG2767,ATPsynO,CG11089,mad2,CG11070,CG6767,lin,Vha36-1,Marf1,PrBP,sws,mRpL10,LSm3,Sccpdh1,RpS7,Tao,GatA,CHOp24,CG1890,CG31674,Prosalpha4,fs(1)Ya,Vha68-2,CG3939,Tango10,CG9981,CG17294,Cht11,Dph5,mRpL35,Arts,Mcad,Gclc,CG8004,CG14408,CG17270,dnc,Naxe,Gdi,ATPsynD,NFAT,Mtpalpha,CG10863,Crag,mRpL17,AP-1-2beta,Mitofilin,UK114,Pka-R1,CG11486,twe,RpLP0,Plap,Pfas,PIG-H,Pgm1,ari-1,RpL34b,tral,sol,Nap1,ATPsynE,CycE,kl-2,opm,cnn,PpD3,CG17593,Dip-B,Clc,CG8974,CG8199,Stat92E,ATPsynbeta,CysRS,CG3309,NP15.6,Scsalpha1,CG4594,sau,Klp10A,Mo25,Ptpmeg2,spd-2,Zwilch,cort,RpS30,Eno,Aldh,Pdk1,CG5734,Mctp,TBCC,twin,Pglym78,smg,RpL38,CanB2,CG6094,Rtnl1,UQCR-Q,bor,Pp2B-14D,chrb,Su(H),CG46385,bif,CG6707,Sod3,Gapdh1,ste14,CG4293,CG31342,Uxt,IntS11,CG10344,Aven,ND-MWFE,CG12375,Pax,Sec15,Pex13,CG12093,yip2,CG17691,Pfrx,AOX3,stai,blw,RpL30,DppIII,l(2)k01209,GlyP,Nc73EF,mRpS21,muskelin,wash,otu,CG8475,Blos4,CG15141,Ubc6,CG4882,eIF4A,RpL9,CHMP2B,mts,Rbcn-3A,Gk2,Got1,CG8112,Mf,COX5B,Pgk,CG8009,bam,CycJ,Unr,Edc3,Actn,COX5A,shep,dl,PHGPx,RpLP1,Thiolase,RecQ5,Fkbp59,Sec23,Roe1,ERp60,alt,tko,Synj,PDCD-5,Atg4b,UQCR-11,cype,CG11077,alphaTub67C,CG32536,RpL37a,CG31729,fray,CG4586,Idh3b,REPTOR,CG14407,CkIIalpha,Med,stet,Pi4KIIalpha,PpV,CG2911,Prosbeta5,ND-B18,Seipin,CanA-14F,Kyat,Irp-1B,CG31759,Atg7,larp,Cse1,bun,Wdfy2,CG42376,Efr,Afti,cathD,CG32640,Cdk5,CG32473,CG6967,alph,CG5026,Pfdn5,Tom40,Rel,GstT2,CG32485,Chd64,aay,CG4238,Echs1,CG17768,Iml1,Ankle2,CG1827,PIG-P,Fkbp12,Nmt,O-fut1,Pfdn2,shu,S2P,COX4,Uch-L5,GluRS-m,CG14671,CG4872,Trs33,Smyd4-4,CG17068,EMRE,RpS15,RpL34a,CG7414,mRpS5,Kap-alpha3,EMC8-9,squ,RpL23A,mRpL48,Acox57D-p,sea,mRpL16,Rpt2,mop,Pex23,velo,Idh,egg,cu,Grip128,ATP8A,bys,wal,grp,yin,Sod1,STUB1,Abi,RpL24,PEK,Pdxk,Acyp2,mael,Prosbeta3,RpS5b,Ts,CG1344,levy,Exo70,mRpS23,Ugt36E1,Naa20A,isoQC,spn-E,Pym,Cul2,Argk,GCC88,dob,Droj2,RpS27A,Marf,Smox,SerRS-m,Maf1,milt,RpS14a,ND-30,CG4880,Tak1,SP1029,AGO3,CG2938,l(1)G0196,CG9643,CG7488,dco,Blos2,Cbl,Ady43A,CG9186,Atg6,ATPsynF,CG9436,CG10561,RpL17,CG1640,ACC,P5CDh1,SelD,Arpc2,Moe,GstT1,Pex10,pgc,Topors,glob1,Fdh,Grip75,Girdin,CG10932,SNF4Agamma,RpS29,Cchl,RpS16,pen-2,RpL23,krimp,CG4390,CG8311,Jafrac1,alphaTub84B,RpS14b,Snx6,Sry-alpha,CG8726,CG40472,Adi1,Fer1HCH,Cp1,CG33156,apt,CG5447,Gasz,tws,gus,Sec22,CG7920,PPYR1,mtTFB1,CG9393 | | 1.571e-57 | -130.80 | intracellular organelle | cellular component | GO:0043229 | 5391 | 750 | 11940 | 1105 | HP1c,E2f1,Echs1,CG17768,Iml1,Ankle2,PIG-P,CG1827,Chd64,CG32485,Tom40,Rel,Nup62,Uch-L5,COX4,shu,O-fut1,Hr78,RfC38,Irp-1B,Kyat,not,esc,ND-B18,Seipin,CG2911,Prosbeta5,CG6967,alph,CG42376,Efr,Wdfy2,Afti,CG32640,cathD,Cdk5,larp,pygo,Cse1,Psf3,CG31759,cu,CG8611,Grip128,Ublcp1,ATP8A,bys,CG1602,egg,EndoGI,mbo,velo,Idh,mRpL16,mop,Pex23,PEK,RpL24,del,lin-52,STUB1,yin,Sod1,wal,grp,RpS15,RpL34a,CG7414,l(1)G0004,APC7,Smyd4-4,CG17068,EMRE,Trs33,CG14671,Sgf11,CG42526,CG3226,mre11,GluRS-m,RpL23A,mRpL48,Acox57D-p,sea,trr,squ,Kap-alpha3,Cen,hb,BigH1,EMC8-9,mRpS5,ND-30,CG4880,Maf1,SerRS-m,milt,RpS14a,RpS27A,Marf,Spx,Smox,nej,dob,ATPsynF,Blos2,CG9186,Ady43A,insv,CG7488,dco,AGO3,CG2938,levy,RpS5b,Ts,CG3437,CG14464,TfIIFalpha,mael,Prosbeta3,Orc6,GCC88,spn-E,isoQC,Arpc1,stil,Pym,YL-1,Trf2,mRpS23,Ugt36E1,CG40472,RpS14b,Snx6,Tlk,alphaTub84B,Xpac,CG16865,CG17187,CG8311,mtTFB1,CG9393,Sec22,CG7920,Cp1,CG33156,CG5447,apt,tws,Gasz,Ntan1,gus,Elba2,Adi1,Fer1HCH,Pex10,pgc,mute,Topors,CG6813,l(3)73Ah,P5CDh1,Arpc2,Moe,RpL17,CG1640,MED24,Bx,Clp,pen-2,RpL23,Snm1,krimp,Cchl,RpS16,mus81,CG6276,Ubc4,Atf3,Aladin,Grip75,SNF4Agamma,RpS29,CG10932,Girdin,asf1,kl-2,opm,cnn,CycE,ATPsynE,Atu,CG10038,Caf1-105,Nap1,Ndc1,Rnmt,ATPsynbeta,CG3309,CG8974,HmgD,CG8199,Stat92E,Clc,Nup58,PpD3,CG17593,mxc,AP-1-2beta,CG12018,UK114,Mitofilin,mRpL17,CG17202,Jabba,Usp39,Mtpalpha,Crag,Kap3,ari-1,RpL34b,Ssdp,tral,Cul1,RpLP0,Plap,jigr1,PIG-H,CG11486,twe,Su(H),CG46385,bor,thoc6,Rtnl1,UQCR-Q,twin,Pglym78,RpL38,smg,lid,CG6094,snRNA:U4:38AB,CanB2,CG10344,Aven,ND-MWFE,CG12375,Pax,Sec15,Pex13,IntS11,Uxt,CG4293,bif,CG6707,Sod3,Gapdh1,ste14,spd-2,Zwilch,Mo25,CG11902,CG4594,sau,Klp10A,NP15.6,Scsalpha1,spen,wapl,CG5734,Mctp,Larp7,Pdk1,RpS30,CG12301,Aldh,CG8112,COX5B,Mf,CHMP2B,mts,Gk2,Got1,Ubc6,snRNA:U12,CG4882,eIF4A,RpL9,ovo,CG14814,CycJ,bam,CG2129,Pgk,CG8009,e(y)1,Taf7,CG17691,yip2,NHP2,sbb,Nc73EF,mRpS21,wash,CG31875,l(2)k01209,RpL30,blw,Ugt35A1,Mes2,stai,alphaTub67C,CG31729,RpL37a,CG32536,CG11077,CSN6,Bin1,CG4266,cype,PDCD-5,brwl,UQCR-11,CG14407,CkIIalpha,Med,Pi4KIIalpha,stet,Idh3b,REPTOR,CG4586,Mat89Ba,RYBP,RecQ5,Thiolase,RpLP1,CG4730,PHGPx,rept,shep,Utx,COX5A,dl,Edc3,Cbp80,Actn,Synj,MAGE,G9a,alt,tko,Mybbp1A,Roe1,ERp60,Sec23,Aos1,LysRS,ND-24,CG1622,Dhfr,Coq6,ben,tef,rl,bcd,Fur2,Coq7,Bap170,Rpn13,ND-75,EloC,RpS28b,Rep,CG11110,CG10418,p38a,Prosalpha5,ND-49,elg1,yl,MED20,yem,CG7728,HP1e,ATPsyndelta,msps,RpS23,colt,Atpalpha,Hat1,pps,Cdk7,fwe,ATPsynCF6,Rbf,Etf-QO,PyK,PAPLA1,CG5028,SdhD,Jafrac2,Nurf-38,tos,Pld,CG11710,Cyp1,RpL26,Pex12,Agpat2,Rrp40,Scox,CG3631,CG2678,Nxf3,ND-23,RpL29,ATPsynB,CG32441,Parp16,CG4860,Rbfox1,CG7716,wdn,CG7655,Gk1,ix,Spt20,CG9773,CG5969,mRpS10,Edem1,alpha-Man-Ia,RpS2,ATPsynG,Grip71,CG11164,CG3511,SdhA,ND-51,CycB,Cisd2,nmdyn-D7,sw,Cf2,ScsbetaA,pn,CG31053,Axs,osk,capt,tkv,foxo,Neb-cGP,Lkb1,cyc,RpL13,RpS24,whd,CG5003,Fkbp39,mus304,Shmt,Pmi,RpII140,SelR,ABCD,mino,CG13551,Rpn2,Desat1,CycC,Tip60,lt,rig,TrpRS-m,CG14817,scny,mEFTu1,Ptpmeg,Herc4,COX8,RpL31,yps,Wbp2,Atf6,CG13344,cana,CG10055,Ugalt,alc,mRpL18,CG4598,CG17454,CG33977,Pp2A-29B,gw,up,Vamp7,CG9143,Rga,Top1,CG2199,mRpL35,CG9981,Hnf4,zf30C,CG17294,Vha68-2,CG8204,Gdi,crm,ATPsynD,NFAT,CG10959,Naxe,CG8004,CG17270,dnc,DNApol-epsilon255,Arts,Sgf29,dgt4,Mcad,Gclc,mRpL10,CG2162,PrBP,sws,Nf-YA,Non1,lin,Vha36-1,Marf1,vers,ATPsynO,mad2,CG11070,CG1890,Prosalpha4,fs(1)Ya,GatA,CHOp24,RpS7,mus205,Tao,LSm3,MED27,psq,Sccpdh1,Tengl4,Pen,ND-39,CG8314,CG7840,CG15525,Hsc70-3,CG10470,mRpS11,Rab32,Trf4-2,RpL7A,Mtl,Sply,Pus7,Ranbp21,CG3021,Rcd5,shrb,CG4557,SuUR,CG9265,CG11267,SERCA,CG8728,Orc1,CG10631,Tom70,Debcl,CG17698,toc,Karybeta3,cad,ND-SGDH,ASPP,CG9684,Uev1A,CG10425,RpL40,tho2,park,Ufm1,armi,tun,shop,lqfR,fdl,Pink1,tapas,ph-d,Arl5,Acph-1,eIF4E1,CG14183,Rab10,bru1,ND-MNLL,jvl,trsn,ste24a,Sirt7,nbs,Cdk1,Phb2,l(3)mbt,Exd2,tub,SREBP,GatB,Pdk,CG42375,escl,CG5903,parvin,RpL13A,CG13601,nos,Prosbeta4,mtrm,Prosalpha3,CG3803,Ptp61F,RpS13,fal,Stam,CG4021,CG3198,Gmap,Sld5,Sfxn2,snRNA:U4atac,Cenp-C,RpL12,hng3,sbr,cnc,RpS27,CG6766,CG5986,CG1774,awd,CG14894,EMC3,Vsp37A,Rbf2,CG10654,CG4404,CG8507,Rpb8,CG5880,scu,Gga,CG8001,Neos,obe,Tspo,CG17328,Chrac-14,Grip84,mEFTu2,MED4,CG7115,CG12288,vret,heph,Prosbeta6,Ppox,Tm1,SmE,RpS4,Pomp,Hip1,ND-18,ND-B22,sip3,mRpL36,exu,CG5412,CG31523,pr,Tim8,26-29-p,Rab5,Lsm12,Tfb4,mol,Dd,mRpL13,PIG-C,trbd,Sdhaf3,Coq8,SkpA,d4,CG6178,Vha13,cbs,Cyt-c-p,Tomosyn,lwr,Trxr-1,Lnpk,RpS18,r2d2,CG6610,eIF3m,CG9132,gammaTub37C,Ptip,Gabat,Arpc4,CG17746,RpL11,Hmg-2,CG10336,CG9399,ND-B15,AIF,CG11562,CG9425,Nprl2,pont,Arf51F,Xpc,CG5510,Vps16B,piwi,CG13366,cup,nmdyn-D6,Dcr-1,Imp,GalT1,EcR,qkr58E-2,RpL10,Cka,SRPK,IntS6 | | 1.135e-55 | -126.52 | organelle | cellular component | GO:0043226 | 5470 | 753 | 11940 | 1105 | krimp,Snm1,RpL23,pen-2,Clp,CG6276,mus81,Cchl,RpS16,Atf3,Ubc4,RpS29,Girdin,CG10932,SNF4Agamma,Grip75,Aladin,mute,Topors,pgc,Pex10,CG6813,Moe,Arpc2,P5CDh1,l(3)73Ah,Bx,CG1640,MED24,RpL17,CG9393,mtTFB1,CG7920,Sec22,gus,Ntan1,Gasz,tws,apt,CG5447,Cp1,CG33156,Fer1HCH,Elba2,Adi1,CG40472,Snx6,RpS14b,Xpac,alphaTub84B,Tlk,CG8311,CG17187,CG16865,GCC88,Orc6,YL-1,Trf2,Pym,stil,Arpc1,spn-E,isoQC,Ugt36E1,mRpS23,levy,TfIIFalpha,CG14464,RpS5b,CG3437,Ts,Prosbeta3,mael,ATPsynF,insv,Ady43A,CG9186,Blos2,dco,CG7488,CG2938,AGO3,CG4880,ND-30,RpS14a,milt,SerRS-m,Maf1,Spx,Smox,Marf,RpS27A,dob,nej,sea,Acox57D-p,RpL23A,mRpL48,squ,trr,EMC8-9,BigH1,Cen,hb,Kap-alpha3,mRpS5,APC7,l(1)G0004,CG7414,RpL34a,RpS15,EMRE,CG17068,Smyd4-4,Trs33,CG14671,GluRS-m,mre11,CG3226,CG42526,Sgf11,del,RpL24,PEK,STUB1,lin-52,Sod1,yin,grp,wal,bys,ATP8A,Ublcp1,CG8611,Grip128,cu,mbo,EndoGI,egg,CG1602,Idh,velo,Pex23,mop,mRpL16,alph,CG6967,CG32640,Cdk5,cathD,Afti,Efr,Wdfy2,CG42376,Cse1,pygo,larp,CG31759,Psf3,esc,Kyat,not,Irp-1B,Seipin,ND-B18,Prosbeta5,CG2911,COX4,Nup62,Uch-L5,shu,O-fut1,RfC38,Hr78,CG1827,PIG-P,Ankle2,Iml1,CG17768,Echs1,HP1c,E2f1,Chd64,CG32485,Rel,Tom40,MAGE,Synj,tko,Mybbp1A,alt,G9a,ERp60,Roe1,Sec23,CG4730,RpLP1,RecQ5,Thiolase,PHGPx,dl,COX5A,rept,shep,Utx,Actn,Cbp80,Edc3,Pi4KIIalpha,stet,Med,CkIIalpha,CG14407,REPTOR,Idh3b,Mat89Ba,CG4586,RYBP,CG31729,RpL37a,CG32536,alphaTub67C,CG11077,CG4266,cype,Bin1,CSN6,UQCR-11,brwl,PDCD-5,CG31875,wash,mRpS21,Nc73EF,l(2)k01209,Ugt35A1,RpL30,blw,stai,Mes2,e(y)1,CG17691,Taf7,NHP2,yip2,sbb,bam,CycJ,CG8009,Pgk,CG2129,COX5B,Mf,CG8112,Got1,Gk2,mts,CHMP2B,eIF4A,RpL9,CG4882,snRNA:U12,Ubc6,CG14814,ovo,wapl,Mctp,CG5734,TMEM216,Larp7,Pdk1,Aldh,CG12301,RpS30,Zwilch,spd-2,CG11902,Mo25,Klp10A,CG4594,sau,spen,Scsalpha1,NP15.6,Sec15,Pex13,CG12375,Pax,ND-MWFE,CG10344,Aven,IntS11,Uxt,CG4293,ste14,Gapdh1,Sod3,CG6707,bif,Su(H),CG46385,thoc6,bor,UQCR-Q,Rtnl1,Arc2,CanB2,CG6094,snRNA:U4:38AB,RpL38,lid,smg,Pglym78,twin,tral,Ssdp,RpL34b,ari-1,Kap3,PIG-H,jigr1,Plap,RpLP0,Cul1,twe,CG11486,UK114,Mitofilin,AP-1-2beta,CG12018,mxc,mRpL17,Usp39,Jabba,CG17202,Crag,Mtpalpha,CG3309,ATPsynbeta,Rnmt,Stat92E,CG8199,CG8974,HmgD,Nup58,Clc,CG17593,PpD3,opm,cnn,kl-2,asf1,CycE,Caf1-105,CG10038,Atu,ATPsynE,Ndc1,Nap1,fs(1)Ya,Prosalpha4,CG1890,CHOp24,GatA,Tao,mus205,RpS7,Sccpdh1,psq,MED27,LSm3,CG2162,mRpL10,sws,Non1,Nf-YA,PrBP,vers,Marf1,Vha36-1,lin,CG11070,mad2,ATPsynO,ATPsynD,NFAT,Gdi,crm,CG8204,Naxe,CG10959,DNApol-epsilon255,dnc,CG8004,CG17270,Gclc,Mcad,Arts,Sgf29,dgt4,mRpL35,CG2199,CG17294,CG9981,zf30C,Hnf4,Vha68-2,CG4598,CG17454,mRpL18,CG33977,gw,up,Pp2A-29B,Top1,Rga,CG9143,Vamp7,scny,mEFTu1,CG14817,RpL31,Ptpmeg,Herc4,COX8,CG13344,Atf6,Wbp2,yps,Ugalt,CG10055,alc,cana,SelR,ABCD,RpII140,mino,Rpn2,CG13551,rig,TrpRS-m,lt,Tip60,CycC,Desat1,Lkb1,cyc,CG5003,whd,RpS24,RpL13,Fkbp39,Pmi,Shmt,mus304,Cisd2,CycB,ScsbetaA,CG31053,pn,nmdyn-D7,Cf2,sw,osk,Axs,Neb-cGP,capt,tkv,foxo,alpha-Man-Ia,Edem1,CG5969,mRpS10,RpS2,ATPsynG,Grip71,ND-51,SdhA,CG11164,CG3511,CG7716,Rbfox1,CG4860,Parp16,wdn,ix,Gk1,CG7655,CG9773,Spt20,Pex12,Agpat2,RpL26,Cyp1,CG11710,Rrp40,ND-23,CG2678,Nxf3,CG3631,Scox,ATPsynB,CG32441,RpL29,fwe,ATPsynCF6,Cdk7,pps,PAPLA1,PyK,Etf-QO,Rbf,Jafrac2,Nurf-38,CG5028,SdhD,Pld,tos,ND-49,Prosalpha5,p38a,yl,elg1,CG7728,yem,MED20,colt,Atpalpha,Hat1,RpS23,ATPsyndelta,msps,HP1e,Fur2,bcd,rl,RpS28b,EloC,ND-75,Bap170,Rpn13,Coq7,CG11110,Rep,CG10418,Aos1,LysRS,Dhfr,Coq6,ND-24,CG1622,tef,ben,Imp,Dcr-1,nmdyn-D6,EcR,GalT1,IntS6,SRPK,Cka,qkr58E-2,RpL10,CG9425,CG11562,ND-B15,AIF,CG9399,Nprl2,pont,Vps16B,CG5510,Xpc,Arf51F,cup,CG13366,piwi,Arpc4,Gabat,gammaTub37C,CG9132,eIF3m,Ptip,r2d2,CG6610,CG17746,RpL11,CG10336,Hmg-2,d4,CG6178,SkpA,Coq8,Tomosyn,Cyt-c-p,cbs,Vha13,lwr,RpS18,Lnpk,Trxr-1,Oseg1,CG5412,mol,Tfb4,26-29-p,Rab5,Lsm12,Tim8,pr,CG31523,PIG-C,trbd,mRpL13,Dd,Sdhaf3,Prosbeta6,heph,ND-18,Pomp,Hip1,RpS4,SmE,Tm1,Ppox,ND-B22,exu,mRpL36,sip3,Gga,Tspo,obe,CG17328,CG8001,Neos,mEFTu2,MED4,Grip84,Chrac-14,vret,CG12288,CG7115,CG4404,CG10654,Rbf2,Rpb8,CG8507,scu,CG5880,cnc,hng3,sbr,RpL12,CG1774,CG5986,RpS27,CG6766,EMC3,CG14894,awd,Vsp37A,RpS13,Ptp61F,CG3803,CG4021,Stam,fal,Sfxn2,Sld5,Gmap,CG3198,snRNA:U4atac,Cenp-C,CG42375,Pdk,GatB,escl,CG13601,nos,RpL13A,parvin,CG5903,Prosalpha3,mtrm,Prosbeta4,jvl,ND-MNLL,bru1,Rab10,CG14183,eIF4E1,nbs,Sirt7,trsn,ste24a,Phb2,l(3)mbt,Cdk1,SREBP,tub,Exd2,armi,Ufm1,park,tho2,shop,lqfR,tun,Acph-1,Arl5,ph-d,Pink1,tapas,fdl,CG10631,Orc1,CG8728,ASPP,ND-SGDH,cad,Karybeta3,CG17698,toc,Debcl,Tom70,Uev1A,CG9684,RpL40,CG10425,Pus7,Ranbp21,Sply,Mtl,RpL7A,Rcd5,CG3021,CG4557,shrb,SERCA,CG9265,CG11267,SuUR,CG7840,ND-39,Pen,CG8314,Tengl4,Hsc70-3,CG15525,mRpS11,CG10470,Trf4-2,Rab32 | | 8.599e-46 | -103.77 | membrane-bounded organelle | cellular component | GO:0043227 | 4716 | 659 | 11940 | 1105 | CG8009,CG2129,CycJ,eIF4A,CG4882,snRNA:U12,Ubc6,CG14814,ovo,COX5B,CG8112,Got1,Gk2,mts,CHMP2B,Ugt35A1,blw,Mes2,CG31875,wash,mRpS21,Nc73EF,l(2)k01209,yip2,NHP2,sbb,e(y)1,CG17691,Taf7,Mat89Ba,CG4586,RYBP,stet,Pi4KIIalpha,Med,CkIIalpha,CG14407,REPTOR,Idh3b,cype,Bin1,CG4266,CSN6,UQCR-11,brwl,PDCD-5,CG32536,CG31729,alphaTub67C,CG11077,ERp60,Roe1,Sec23,MAGE,Synj,Mybbp1A,tko,alt,G9a,dl,COX5A,Utx,shep,rept,Cbp80,CG4730,RecQ5,Thiolase,PHGPx,Nup58,Clc,CG17593,PpD3,CG3309,ATPsynbeta,Rnmt,Stat92E,CG8199,CG8974,HmgD,CG10038,Caf1-105,Atu,ATPsynE,Nap1,Ndc1,opm,asf1,CycE,PIG-H,Plap,jigr1,Cul1,twe,Ssdp,tral,ari-1,Usp39,CG17202,Crag,Mtpalpha,UK114,Mitofilin,AP-1-2beta,CG12018,mxc,mRpL17,CG4293,Sod3,ste14,CG6707,bif,Pex13,Sec15,CG12375,ND-MWFE,CG10344,Aven,Uxt,IntS11,UQCR-Q,Rtnl1,Arc2,CanB2,CG6094,snRNA:U4:38AB,lid,twin,CG46385,Su(H),thoc6,bor,Pdk1,Larp7,Aldh,CG12301,wapl,Mctp,CG5734,sau,CG4594,spen,Scsalpha1,NP15.6,spd-2,CG11902,Mo25,dco,CG7488,CG2938,AGO3,ATPsynF,insv,CG9186,Ady43A,Spx,Smox,Marf,RpS27A,nej,CG4880,ND-30,milt,Maf1,SerRS-m,stil,YL-1,Trf2,Pym,isoQC,spn-E,Ugt36E1,mRpS23,GCC88,Orc6,Prosbeta3,mael,levy,TfIIFalpha,CG14464,Ts,CG3437,gus,Gasz,Ntan1,tws,CG5447,apt,CG33156,Cp1,Fer1HCH,Elba2,Adi1,CG9393,mtTFB1,CG7920,Sec22,Xpac,alphaTub84B,Tlk,CG8311,CG17187,CG16865,CG40472,Snx6,Atf3,Ubc4,SNF4Agamma,CG10932,Aladin,Snm1,pen-2,Clp,CG6276,mus81,Cchl,Moe,P5CDh1,l(3)73Ah,Bx,CG1640,MED24,Topors,mute,Pex10,CG6813,O-fut1,RfC38,Hr78,Nup62,COX4,Uch-L5,CG32485,Rel,Tom40,Ankle2,PIG-P,CG1827,CG17768,Iml1,Echs1,E2f1,HP1c,Cse1,pygo,larp,CG31759,Psf3,alph,CG32640,Cdk5,cathD,Afti,CG42376,Wdfy2,Efr,Prosbeta5,CG2911,esc,not,Kyat,Irp-1B,Seipin,ND-B18,Sod1,yin,wal,grp,del,lin-52,Idh,velo,Pex23,mop,mRpL16,bys,ATP8A,CG8611,Ublcp1,cu,mbo,EndoGI,egg,CG1602,EMC8-9,BigH1,hb,Kap-alpha3,mRpS5,sea,Acox57D-p,mRpL48,trr,CG14671,Trs33,GluRS-m,mre11,CG3226,CG42526,Sgf11,APC7,l(1)G0004,RpS15,EMRE,Smyd4-4,mEFTu2,MED4,Chrac-14,CG12288,CG7115,Gga,Tspo,obe,CG17328,CG8001,Neos,scu,CG5880,CG4404,CG10654,Rbf2,Rpb8,CG8507,PIG-C,trbd,mRpL13,Dd,Sdhaf3,CG5412,Tfb4,mol,26-29-p,Lsm12,Rab5,pr,CG31523,Tim8,ND-B22,mRpL36,exu,sip3,Prosbeta6,heph,ND-18,Hip1,Pomp,SmE,Tm1,Ppox,RpL11,CG10336,Hmg-2,Gabat,Ptip,eIF3m,gammaTub37C,CG9132,r2d2,CG6610,CG17746,lwr,RpS18,Lnpk,Trxr-1,CG6178,d4,SkpA,Coq8,Tomosyn,Cyt-c-p,Vha13,cbs,IntS6,SRPK,Cka,qkr58E-2,Imp,nmdyn-D6,Dcr-1,GalT1,EcR,Vps16B,CG5510,Xpc,Arf51F,cup,piwi,CG11562,ND-B15,AIF,CG9399,Nprl2,pont,CG4557,shrb,SERCA,CG11267,SuUR,Pus7,Ranbp21,Sply,Rcd5,CG3021,mRpS11,CG10470,Trf4-2,Rab32,CG7840,CG8314,Pen,ND-39,Tengl4,Hsc70-3,CG15525,shop,lqfR,tun,Arl5,ph-d,Acph-1,Pink1,fdl,armi,Ufm1,park,tho2,Uev1A,CG9684,RpL40,CG10425,CG10631,Orc1,CG8728,ND-SGDH,cad,ASPP,Karybeta3,CG17698,toc,Debcl,Tom70,CG13601,CG5903,Prosalpha3,mtrm,Prosbeta4,Pdk,CG42375,GatB,escl,l(3)mbt,Phb2,Cdk1,SREBP,tub,Exd2,ND-MNLL,bru1,Rab10,nbs,Sirt7,ste24a,trsn,EMC3,CG14894,awd,Vsp37A,cnc,sbr,hng3,CG1774,CG6766,CG5986,Sfxn2,Sld5,CG3198,Gmap,snRNA:U4atac,Cenp-C,RpS13,Ptp61F,CG3803,Stam,CG4021,fal,Rpn2,CG13551,TrpRS-m,rig,Tip60,lt,Desat1,CycC,ABCD,SelR,RpII140,mino,Pmi,Shmt,mus304,cyc,Lkb1,CG5003,whd,gw,Top1,CG9143,Rga,Vamp7,CG4598,CG17454,mRpL18,CG33977,CG13344,Atf6,Wbp2,yps,alc,Ugalt,CG10055,mEFTu1,scny,CG14817,COX8,Herc4,Ptpmeg,DNApol-epsilon255,CG17270,dnc,CG8004,Gclc,Mcad,Sgf29,Arts,ATPsynD,NFAT,crm,Gdi,CG8204,Naxe,CG10959,CG17294,CG9981,Hnf4,zf30C,Vha68-2,mRpL35,CG2199,Tao,mus205,RpS7,Sccpdh1,psq,MED27,LSm3,fs(1)Ya,Prosalpha4,CHOp24,GatA,vers,Marf1,Vha36-1,lin,CG11070,mad2,ATPsynO,CG2162,mRpL10,sws,Non1,Nf-YA,PrBP,CG11110,Rep,CG10418,Fur2,rl,bcd,ND-75,EloC,Bap170,Rpn13,Coq7,Coq6,Dhfr,CG1622,ND-24,ben,Aos1,LysRS,Jafrac2,Nurf-38,CG5028,SdhD,Pld,tos,ATPsynCF6,fwe,Cdk7,pps,PAPLA1,Etf-QO,Rbf,CG7728,yem,MED20,colt,Hat1,Atpalpha,ATPsyndelta,HP1e,ND-49,Prosalpha5,p38a,yl,elg1,ix,CG7655,Gk1,CG9773,Spt20,Rbfox1,Parp16,CG4860,wdn,ND-23,Nxf3,CG2678,CG3631,Scox,ATPsynB,CG32441,Pex12,Agpat2,Cyp1,CG11710,Rrp40,osk,Axs,Neb-cGP,foxo,tkv,Cisd2,CycB,ScsbetaA,pn,CG31053,Cf2,nmdyn-D7,ATPsynG,ND-51,SdhA,CG3511,CG11164,alpha-Man-Ia,Edem1,CG5969,mRpS10,RpS2 | | 1.397e-45 | -103.28 | cellular metabolic process | biological process | GO:0044237 | 4466 | 631 | 12345 | 1129 | Rpb8,AGBE,ND-B15,fdl,Xpc,grp,GAPsec,CG1774,Pomp,Treh,S2P,CG5292,CG2201,CG18135,Plap,Not10,CG11164,CG7728,Usp16-45,Sod3,Mal-A5,mRpL17,Ts,DNApol-epsilon255,ari-1,Sod1,park,CG6767,CG32221,CG32473,CG7789,Sgf11,trsn,CG10425,Son,CG8475,CG8204,mal,Desat1,GCS2beta,CG9316,AMPdeam,SNF4Agamma,Trxr-1,whd,Eno,G9a,Ppox,CG5880,mre11,Mo25,CAH2,otu,Gk2,Pglym78,RpS13,CG10463,Dd,GstT1,eEF1beta,COX4,GluRS-m,Prosalpha4,EMC3,CG4572,Grx1,isoQC,CG17454,Drep2,Atu,Agpat2,dob,yip2,Cdk7,CSN6,CG4407,CG17333,CG13344,YL-1,CG3511,c(2)M,ND-23,Thiolase,faf,CG12301,RpL40,Sccpdh1,twe,PyK,Debcl,RpS27A,wisp,pn,ste24a,RpL26,Nadsyn,agt,nbs,SREBP,CG10932,ago,ND-24,cort,CG11089,Sp7,awd,Aldh,Atg7,Idh,RpS14b,CG11771,SelR,armi,Ptp69D,Idgf4,r2d2,Sld5,Edc3,Rab32,RpL7A,Pfas,nmdyn-D7,CG5026,shu,CG2034,levy,CG14407,RfC38,CG42554,ATPsynE,shop,CG3223,CG40045,Ntan1,CG17187,eIF3i,RpS15,O-fut1,Lkb1,RpL23A,d4,CycE,sgll,Chrac-14,PIG-P,kri,Sirt7,Mtpalpha,Gclc,Pp2A-29B,Tfb4,CG6094,Cht11,CG11486,CG3021,AIF,CG7328,Cyp1,CG17270,CG8726,CG4041,CG31633,SelD,RpS7,CG5028,CG1578,CG4586,CG6610,Tak1,pygo,RpS27,Gabat,nos,GatA,Iru,RpL9,Uch,PIG-C,fray,RpS28b,l(3)73Ah,mRpS10,AcCoAS,sws,Atg4b,velo,CG6766,CanB2,RpL11,Dhfr,shrb,SdhA,ND-30,mEFTu1,CG17691,Pcd,APP-BP1,scu,ND-51,E2f1,Npc2a,RpL10,Rpn3,Larp7,CG16865,Prosbeta3,mRpL13,ATPsynD,Coq7,Coq8,Gk1,RpS2,Sgf29,CG8311,CG17768,IntS6,RpL30,CG4598,Pfrx,eIF3m,Ptip,CG33156,Cchl,CG14516,tko,Ugt35A1,CG9143,smg,Cdk5,CG8188,mino,CG9485,CG3198,rl,ND-B18,tkv,Idgf2,RpL34b,CG5285,CG7414,Taf7,Rep,Dph5,Cyt-c-p,Ptp61F,Pgm1,Usp39,Tip60,RpS16,esc,Jafrac2,CG9391,Orc1,exu,oxt,RpL37a,snRNA:U12,Rga,scny,ND-SGDH,CG31053,l(2)k01209,Gapdh1,QC,RpS24,tun,Pex10,Ady43A,RpL23,CG7997,Ptpmeg2,Ho,Ptp10D,CkIIalpha,lwr,egg,Efr,CycB,mEFTu2,ScsbetaA,twin,CG46338,DhpD,dco,CG14100,Rpn2,Mekk1,Ubc4,CG1622,Pdxk,Mos,CG11070,Top1,tho2,Argk,RpL31,cype,Pdk,ACC,Pex12,CanA-14F,Pi3K92E,Pp2B-14D,bys,Gyg,RpL29,CG14997,ATPsynO,AsnS,Spx,mRpL36,CG14435,CG2100,regucalcin,Ugt36E1,Nmt,snRNA:U4:38AB,CG11134,CycJ,UQCR-Q,Echs1,Ubc6,IntS11,mus205,SerRS-m,CG4594,r,CG2911,mRpS21,Utx,eIF4E1,HDAC6,Vamp7,RpL17,vret,MED24,Prosbeta4,CG3631,Cul1,Aos1,Gmer,RpS23,Cbp80,TyrRS,Rpn12,CG7115,aay,AOX3,Tengl4,Pld,Rbf,LSm3,RpS29,alph,bru1,Arf51F,Nc73EF,ATPsynG,UQCR-11,GatB,COX5A,ATPsyndelta,Irp-1B,Rab5,cN-IIIB,CG17698,nej,Adi1,Edem1,SuUR,spen,Drep1,Fdh,Naa20A,CG5986,Prosalpha5,SRPK,Cp1,SkpA,CG7840,Hnf4,Acph-1,Ac3,Rrp40,Bin1,CG8728,dx,MED4,Exd2,eIF4A,Usp10,Clp,fal,Gip,PIG-H,Mat89Ba,sip3,Sply,STUB1,Fkbp12,Prosbeta6,P5CDh1,Cbl,Nap1,CG7920,CysRS,Nurf-38,yps,UbcE2H,Shmt,PEK,CG33977,fs(1)Ya,nSMase,Snm1,RpL12,EloC,CG7718,PpV,Fkbp59,CG31759,CG8199,Rcd5,RpS5b,png,tapas,RpLP0,Uev1A,CG13551,mRpL48,dl,Cul5,foxo,l(1)G0196,not,nmdyn-D6,piwi,pr,Tlk,Cka,GstE11,PpD3,Sptr,CG9643,SdhD,CG3587,Rpn13,NP15.6,Neos,Rpt2,CG5003,Kyat,Pus7,CG1344,NHP2,Scsalpha1,RpS4,Adh,rept,PIG-L,CG31523,e(y)1,mtTFB1,Fbw5,SERCA,sbr,ND-MNLL,CG10344,mRpS5,obe,Pmi,Acyp2,mRpL35,CG10418,trbd,CG11110,ND-39,CG9941,lid,CG8974,GstT2,ATPsynCF6,Prosbeta5,Pgk,tor,RpL13,26-29-p,Cdk1,Sdhaf3,Gasz,tos,CG3842,escl,CG8314,CG6178,Got1,mus304,Uba5,Seipin,RpS30,RecQ5,pen-2,CG12018,r-l,Idgf6,p38a,Spt20,CG10561,Prosalpha3,lt,ben,CG17294,blw,mRpL10,mRpS23,CG7009,RYBP,Tao,Coq6,Cul2,RpL24,Ufm1,CG2924,CG10336,Traf6,mRpS11,ND-75,Acox57D-p,RpII140,Trf4-2,pps,rig,CG3803,ATPsynB,CG4238,P5cr,mRpL18,snRNA:U4atac,Herc4,RpLP1,heph,raptor,UGP,pont,Jafrac1,Dcr-1,TfIIFalpha,Archease,EcR,gw,del,mRpL16,CG14817,gus,Orc6,Etf-QO,mus81,mts,Lsm12,CG11858,RpL34a,ND-18,RpL13A,Uch-L5,CG8112,csw,Hat1,RpS18,ND-49,ATPsynbeta,Atg6,RpS14a,cu,alpha-Man-Ia,CG1827,Vsp37A,CG9684,CG12375,CG6707,SmE,Mcad,RpL38,Marf,CycC,CG32640,ssh,CG17746,Hsc70-3,tws,trr,COX5B,PAPLA1,ste14,Imp,Parp16,CG10189,Pdk1,Topors,Ggt-1,Ptpmeg,Hexo1,MAGE,CG10254,Synj,Pi4KIIalpha,LysRS,GlyP,mop,Pxt,CG4860,CG4390,Ppcdc,Mtl,CG11899,SP1029,Ublcp1,Rnmt,mael,wal,Trf2,GalT1,tzn,Pfk,Idh3b,Cyp6v1,Xpac,CG30343,Pym,ND-B22,TrpRS-m,Pink1 | | 1.322e-44 | -101.03 | intracellular membrane-bounded organelle | cellular component | GO:0043231 | 4479 | 633 | 11940 | 1105 | e(y)1,Taf7,CG17691,yip2,NHP2,sbb,Nc73EF,mRpS21,wash,CG31875,l(2)k01209,blw,Ugt35A1,Mes2,CG8112,COX5B,mts,Gk2,Got1,Ubc6,snRNA:U12,CG4882,eIF4A,ovo,CG14814,CycJ,CG2129,CG8009,RecQ5,Thiolase,CG4730,PHGPx,Utx,shep,rept,COX5A,dl,Cbp80,Synj,MAGE,G9a,alt,Mybbp1A,tko,Roe1,ERp60,Sec23,alphaTub67C,CG31729,CG11077,CSN6,CG4266,cype,Bin1,brwl,PDCD-5,UQCR-11,CG14407,Med,CkIIalpha,stet,Pi4KIIalpha,Idh3b,REPTOR,CG4586,Mat89Ba,RYBP,mxc,CG12018,AP-1-2beta,Mitofilin,UK114,mRpL17,CG17202,Usp39,Mtpalpha,ari-1,tral,Ssdp,Cul1,jigr1,Plap,PIG-H,twe,asf1,opm,CycE,ATPsynE,Atu,Caf1-105,CG10038,Nap1,Ndc1,Rnmt,ATPsynbeta,HmgD,CG8974,CG8199,Stat92E,Clc,Nup58,PpD3,CG17593,spd-2,Mo25,CG11902,sau,CG4594,NP15.6,spen,Scsalpha1,wapl,Mctp,Larp7,Pdk1,CG12301,Aldh,Su(H),CG46385,bor,thoc6,Rtnl1,UQCR-Q,twin,lid,snRNA:U4:38AB,CG6094,CG10344,Aven,ND-MWFE,CG12375,Pex13,IntS11,Uxt,CG4293,bif,CG6707,ste14,Sod3,levy,CG3437,Ts,TfIIFalpha,CG14464,mael,Prosbeta3,Orc6,GCC88,isoQC,spn-E,Trf2,Pym,stil,YL-1,mRpS23,Ugt36E1,ND-30,CG4880,Maf1,SerRS-m,milt,RpS27A,Marf,Smox,Spx,nej,ATPsynF,Ady43A,CG9186,insv,CG7488,dco,AGO3,CG2938,Pex10,Topors,mute,CG6813,l(3)73Ah,P5CDh1,Moe,CG1640,MED24,Bx,Clp,Snm1,Cchl,mus81,CG6276,Ubc4,Atf3,Aladin,CG10932,SNF4Agamma,CG40472,Tlk,alphaTub84B,Xpac,CG16865,CG17187,CG8311,CG9393,mtTFB1,Sec22,CG7920,Cp1,CG33156,apt,CG5447,Ntan1,tws,Gasz,gus,Elba2,Adi1,Fer1HCH,Kyat,not,Irp-1B,esc,ND-B18,Seipin,CG2911,Prosbeta5,alph,CG42376,Efr,Afti,CG32640,cathD,Cdk5,larp,pygo,Cse1,Psf3,CG31759,HP1c,E2f1,Echs1,Iml1,CG17768,Ankle2,CG1827,PIG-P,CG32485,Tom40,Rel,Uch-L5,Nup62,COX4,O-fut1,Hr78,RfC38,RpS15,l(1)G0004,APC7,Smyd4-4,EMRE,Trs33,CG14671,Sgf11,CG42526,CG3226,GluRS-m,mre11,mRpL48,Acox57D-p,sea,trr,Kap-alpha3,hb,BigH1,EMC8-9,mRpS5,cu,CG8611,Ublcp1,ATP8A,bys,CG1602,egg,EndoGI,mbo,velo,Idh,mRpL16,Pex23,del,lin-52,Sod1,grp,wal,heph,Prosbeta6,Ppox,Tm1,SmE,Hip1,Pomp,ND-18,ND-B22,sip3,mRpL36,exu,CG5412,Tim8,pr,CG31523,26-29-p,Lsm12,Tfb4,mol,Dd,mRpL13,PIG-C,trbd,Sdhaf3,Rbf2,CG10654,CG4404,CG8507,Rpb8,CG5880,scu,Gga,Neos,CG8001,obe,Tspo,CG17328,Chrac-14,MED4,mEFTu2,CG7115,CG12288,CG9399,ND-B15,AIF,CG11562,pont,Nprl2,Xpc,CG5510,piwi,cup,nmdyn-D6,Dcr-1,Imp,GalT1,EcR,qkr58E-2,SRPK,Cka,IntS6,Coq8,SkpA,d4,CG6178,Vha13,cbs,Cyt-c-p,lwr,Trxr-1,Lnpk,RpS18,CG6610,r2d2,eIF3m,gammaTub37C,Ptip,Gabat,CG17746,RpL11,Hmg-2,CG10336,CG8728,CG10631,Orc1,Tom70,Debcl,CG17698,toc,Karybeta3,ASPP,cad,ND-SGDH,CG9684,Uev1A,CG10425,RpL40,tho2,park,Ufm1,armi,tun,lqfR,shop,fdl,Pink1,Acph-1,Arl5,ph-d,Tengl4,ND-39,CG8314,Pen,CG7840,CG15525,Hsc70-3,CG10470,mRpS11,Rab32,Trf4-2,Sply,Ranbp21,Pus7,CG3021,Rcd5,CG4557,SuUR,CG11267,SERCA,CG3803,Ptp61F,RpS13,fal,CG4021,Gmap,CG3198,Sld5,Sfxn2,snRNA:U4atac,Cenp-C,hng3,sbr,cnc,CG5986,CG6766,CG1774,awd,CG14894,EMC3,Vsp37A,Rab10,bru1,ND-MNLL,ste24a,trsn,Sirt7,nbs,Cdk1,l(3)mbt,Phb2,Exd2,tub,SREBP,GatB,Pdk,CG42375,escl,CG5903,CG13601,Prosbeta4,mtrm,Prosalpha3,CG14817,mEFTu1,scny,Herc4,COX8,Ptpmeg,yps,Wbp2,Atf6,CG13344,CG10055,Ugalt,alc,mRpL18,CG4598,CG17454,CG33977,gw,Vamp7,Rga,CG9143,Top1,Lkb1,cyc,whd,CG5003,mus304,Shmt,Pmi,RpII140,ABCD,SelR,mino,CG13551,Rpn2,CycC,Desat1,Tip60,TrpRS-m,rig,mRpL10,CG2162,PrBP,sws,Nf-YA,Non1,lin,Vha36-1,Marf1,vers,ATPsynO,mad2,CG11070,fs(1)Ya,Prosalpha4,GatA,CHOp24,mus205,RpS7,Tao,LSm3,MED27,psq,Sccpdh1,CG2199,mRpL35,CG9981,zf30C,Hnf4,CG17294,Vha68-2,CG8204,crm,ATPsynD,NFAT,CG10959,Naxe,CG17270,dnc,CG8004,DNApol-epsilon255,Sgf29,Arts,Mcad,Gclc,p38a,Prosalpha5,ND-49,elg1,MED20,yem,CG7728,HP1e,ATPsyndelta,Atpalpha,Hat1,colt,pps,Cdk7,ATPsynCF6,Rbf,Etf-QO,PAPLA1,CG5028,SdhD,Jafrac2,Nurf-38,tos,Aos1,LysRS,CG1622,ND-24,Coq6,Dhfr,ben,bcd,rl,Fur2,Coq7,Rpn13,Bap170,EloC,ND-75,Rep,CG11110,CG10418,mRpS10,CG5969,Edem1,alpha-Man-Ia,RpS2,ATPsynG,CG3511,CG11164,SdhA,ND-51,Cisd2,CycB,Cf2,nmdyn-D7,pn,CG31053,ScsbetaA,Axs,osk,foxo,Neb-cGP,CG11710,Cyp1,Pex12,Agpat2,Rrp40,Scox,CG3631,CG2678,Nxf3,ND-23,CG32441,ATPsynB,CG4860,Parp16,Rbfox1,wdn,Gk1,CG7655,ix,Spt20,CG9773 | | 1.587e-42 | -96.25 | cytoplasmic part | cellular component | GO:0044444 | 3495 | 528 | 11940 | 1105 | Edc3,Actn,shep,COX5A,dl,PHGPx,Thiolase,RpLP1,Sec23,Roe1,ERp60,alt,tko,Synj,Atg4b,PDCD-5,UQCR-11,cype,CG11077,alphaTub67C,RpL37a,CG31729,CG32536,CG4586,REPTOR,Idh3b,CG14407,stet,Pi4KIIalpha,CkIIalpha,yip2,CG17691,Pfrx,AOX3,stai,blw,RpL30,DppIII,mRpS21,Nc73EF,wash,Blos4,CG8475,eIF4A,RpL9,CG4882,CHMP2B,mts,Got1,Gk2,Rbcn-3A,CG8112,Mf,COX5B,Pgk,bam,Scsalpha1,NP15.6,sau,CG4594,Klp10A,Ptpmeg2,spd-2,RpS30,Eno,Aldh,Pdk1,CG5734,Mctp,twin,Pglym78,CanB2,CG6094,smg,RpL38,Rtnl1,UQCR-Q,bor,CG46385,CG6707,bif,Gapdh1,ste14,Sod3,CG4293,Uxt,ND-MWFE,Aven,CG10344,Pex13,Sec15,Pax,CG12375,CG10863,Mtpalpha,Crag,mRpL17,Mitofilin,UK114,AP-1-2beta,Pka-R1,CG11486,RpLP0,PIG-H,Pgm1,RpL34b,tral,Nap1,ATPsynE,kl-2,cnn,opm,PpD3,CG17593,Clc,CG8199,CG8974,Stat92E,CG3309,ATPsynbeta,CG1640,RpL17,P5CDh1,Arpc2,Moe,Pex10,Topors,pgc,Fdh,RpS29,CG10932,Grip75,RpS16,Cchl,pen-2,krimp,RpL23,CG4390,CG8311,alphaTub84B,Jafrac1,Snx6,RpS14b,Sry-alpha,CG40472,Fer1HCH,CG5447,Cp1,gus,tws,Gasz,CG7920,Sec22,mtTFB1,CG9393,mael,RpS5b,Ts,Exo70,levy,Ugt36E1,mRpS23,spn-E,Naa20A,isoQC,Cul2,Argk,GCC88,Marf,Droj2,RpS27A,Smox,SerRS-m,RpS14a,milt,ND-30,CG4880,CG2938,AGO3,CG7488,l(1)G0196,CG9186,Ady43A,Blos2,Cbl,Atg6,ATPsynF,CG9436,GluRS-m,CG14671,Trs33,EMRE,CG17068,RpL34a,RpS15,CG7414,mRpS5,EMC8-9,squ,Acox57D-p,RpL23A,mRpL48,sea,mop,mRpL16,Pex23,Idh,Grip128,ATP8A,grp,wal,Sod1,yin,STUB1,Pdxk,PEK,RpL24,Abi,Prosbeta5,CG2911,ND-B18,Seipin,Irp-1B,Kyat,CG31759,Atg7,larp,bun,Cse1,Wdfy2,CG42376,Efr,CG32640,Cdk5,cathD,Afti,alph,CG6967,Rel,Tom40,CG32485,Chd64,CG4238,Echs1,CG1827,PIG-P,Ankle2,CG17768,Iml1,O-fut1,shu,COX4,piwi,CCT5,Gyg,cup,Xpc,Arf51F,Vps16B,CG5510,Nprl2,ND-B15,AIF,CG9399,AMPdeam,CG9425,CG11562,RpL10,GalT1,Pka-R2,cib,nmdyn-D6,Trxr-1,RpS18,Lnpk,cbs,Vha13,Tomosyn,Cyt-c-p,Coq8,CG6178,RpL11,CG17746,gammaTub37C,eIF3m,CG9132,Gabat,exu,mRpL36,RhoGDI,sip3,ND-B22,Tm1,Ppox,ND-18,RpS4,Hip1,heph,Sdhaf3,Dd,PIG-C,mRpL13,26-29-p,Rab5,pr,CG31523,Tim8,mol,cN-IIIB,scu,CG5880,CG8507,Surf4,CG17931,CCT6,vret,Grip84,r,CCT3,mEFTu2,DnaJ-H,Tspo,Gga,eIF3i,Gmap,Sfxn2,Stam,Ptp61F,Sec10,CG3803,RpS13,Vsp37A,cact,EMC3,CG14894,CG1774,CG6766,RpS27,RpL12,Syx13,tub,Exd2,SREBP,Phb2,ste24a,trsn,UGP,CG14183,eIF4E1,ND-MNLL,bru1,Rab10,cl,CG5903,CG13601,nos,RpL13A,CG42375,Pdk,GatB,CG10425,RpL40,Debcl,Tom70,ND-SGDH,Karybeta3,toc,CG8728,fdl,Arl5,tapas,Pink1,tun,lqfR,shop,park,armi,Rab32,mRpS11,CG10470,Hsc70-3,Tengl4,CG7840,ND-39,CG8314,SERCA,CG11267,CG4557,shrb,CG3021,RpL7A,Pfk,Sply,CG2767,ATPsynO,CG11089,Vha36-1,Marf1,sws,mRpL10,LSm3,Sccpdh1,RpS7,CHOp24,GatA,CG31674,CG1890,Vha68-2,CG9981,CG3939,Tango10,mRpL35,Gclc,Mcad,CG8004,dnc,CG17270,Naxe,Gdi,ATPsynD,NFAT,Ugalt,Atf6,yps,eEF1beta,Wbp2,COX8,Ptpmeg,RpL31,mEFTu1,Adh,CG14817,Rga,Vamp7,Top1,gw,up,Pp2A-29B,CG33977,mRpL18,CG4598,Usp10,Shmt,Pmi,RpL13,galectin,CG5003,whd,RpS24,Lkb1,lt,Desat1,TrpRS-m,rig,CG13551,Rpn2,mino,ABCD,SelR,SdhA,ND-51,ATPsynG,Rpn12,Grip71,RpS2,Edem1,CG5969,mRpS10,alpha-Man-Ia,foxo,tkv,capt,Neb-cGP,Axs,osk,AsnS,ScsbetaA,pn,Cf2,nmdyn-D7,RpL29,CG32441,ATPsynB,CG3631,Scox,ND-23,CG2678,Rrp40,Cyp1,Agpat2,Pex12,RpL26,CG9773,Gk1,CG7716,CG4860,Parp16,ATPsyndelta,colt,RpS23,CG3223,yl,ND-49,CG10189,Pld,CG5028,SdhD,Jafrac2,CG10638,PyK,PAPLA1,Etf-QO,fwe,ATPsynCF6,ben,ND-24,Coq6,Dhfr,LysRS,Uba5,CG10418,Rep,CG11110,Coq7,RpS28b,ND-75,Fur2,rl | | 4.362e-42 | -95.24 | cellular process | biological process | GO:0009987 | 7485 | 889 | 12345 | 1129 | RhoGDI,RpS24,mute,QC,Gapdh1,Tom70,CG7997,tun,Pex10,Ady43A,RpL23,CkIIalpha,Ptpmeg2,opm,Ho,Ptp10D,Jafrac2,esc,Nf-YA,exu,oxt,RpL37a,CG9391,Orc1,Rga,CG17068,ND-SGDH,scny,snRNA:U12,Ankle2,CG31053,l(2)k01209,slam,Mekk1,Rpn2,Pdxk,Ubc4,CG1622,Mos,CG11070,lwr,shep,egg,Efr,CG9186,CG17272,parvin,ScsbetaA,mEFTu2,CycB,wash,twin,DhpD,CG46338,BicC,dco,CG14100,Hsp26,RpL31,cype,Med,Mitofilin,Pdk,Pex12,fs(1)M3,ACC,disp,CanA-14F,tho2,Top1,AGO3,Argk,anne,DnaJ-H,nAChRalpha4,RhoGAP54D,CG10055,CG14997,Spx,crq,ATPsynO,AsnS,fwe,mRpL36,CG14435,regucalcin,CG2100,Rtnl1,Nmt,Ugt36E1,Pi3K92E,Pp2B-14D,galectin,stil,bys,plu,RpL29,Gyg,RpS28b,yl,bor,fray,Abi,velo,Atg4b,CG6766,sws,l(3)73Ah,AcCoAS,mRpS10,CanB2,psq,RpL11,Klp10A,Dhfr,CG1890,CG1578,nos,GatA,pygo,Tak1,CG6610,CG4586,RpS27,Gabat,RpL9,PIG-C,Uch,Iru,CG14408,CG31729,Exo70,RpL10,colt,Grip71,CG16865,Rpn3,Larp7,stet,ATPsynD,alc,Coq7,Prosbeta3,mRpL13,SdhA,cnn,ND-30,shrb,mEFTu1,CG10465,milt,CG17691,ND-51,E2f1,Npc2a,Pcd,APP-BP1,scu,Clc,Ptip,eIF3m,Pfrx,Grip128,CG14516,Vha68-2,CG33156,Sec22,Cchl,Myd88,tko,Kap3,cup,Ugt35A1,Coq8,RpS2,Gk1,Sgf29,CG8311,mAChR-A,CG4598,IntS6,CG17768,RpL30,pgc,Ptp61F,enc,Cyt-c-p,Dph5,mtrm,Usp39,Pgm1,Tip60,Uxt,CG42748,CG10038,alphaTub67C,RpS16,Cdk5,CG8188,cathD,CG9143,smg,CG9485,rl,mino,CG3198,ND-B18,RpL34b,CG5285,CG7414,Idgf2,Pka-R1,tkv,Taf7,l(3)mbt,Rep,awd,Sp7,Aldh,Atg7,Sap47,Idh,Ptp69D,CG11771,SelR,RpS14b,armi,Sld5,cana,r2d2,Idgf4,CG10932,SREBP,hb,Pex13,ago,CG11089,cort,ND-24,zyd,CG5026,shu,osk,Scox,levy,CG14407,CG2034,ATPsynE,RfC38,CG42554,Grip84,RpL7A,Rab32,Edc3,Tomosyn,nmdyn-D7,Pfas,PIG-P,Chrac-14,sgll,CycE,Sec23,TMEM216,kri,Gclc,Pp2A-29B,Mtpalpha,Sirt7,Tfb4,Svil,CG6094,Ntan1,shop,CG3223,CG40045,RpS15,CG17187,eIF3i,RpL23A,O-fut1,Lkb1,d4,krimp,up,CG3894,CG17270,CG8726,bcd,CG4041,CG5903,CHOp24,Axs,Liprin-beta,RpS7,CG31633,SelD,spz,CG5028,DCAF12,Cht11,CG11486,AIF,CG7328,CG3021,Cyp1,UK114,ovo,S2P,Treh,CG5292,Plap,CG2201,CG18135,CG11164,Not10,CG9005,Sod3,bam,CG13366,CG7728,Usp16-45,AGBE,Rpb8,Lnpk,ND-B15,fdl,kcc,Xpc,grp,Tim8,Pomp,CG1774,Afti,GAPsec,mal,CG8204,jvl,CG9316,Desat1,GCS2beta,Son,CG8475,dnc,Rab10,Moe,ATP8A,SNF4Agamma,whd,Trxr-1,AMPdeam,ZnT35C,G9a,CCT5,Eno,Ts,ari-1,DNApol-epsilon255,zpg,Mal-A5,mRpL17,park,CG32221,CG6767,Sod1,CG7789,Sgf11,Stat92E,Actn,CG32473,Hr78,CG11077,CG10425,trsn,Grx1,Drep2,Crag,tral,Atu,isoQC,CG17454,trk,yip2,CG9981,Cdk7,Pka-R2,Agpat2,dob,CG13344,CG4407,CSN6,CG17333,Wnt10,Ppox,squ,CG5880,mre11,Mo25,RpS13,CG10463,GstT1,Dd,COX4,eEF1beta,otu,Grip75,CAH2,Gk2,Pglym78,CG4572,Zwilch,Prosalpha4,GluRS-m,EMC3,chrb,PyK,CG13933,twe,fs(1)N,Debcl,Rbfox1,RpS27A,Non1,wisp,X11L,ste24a,pn,RpL26,nbs,Nadsyn,agt,Cen,faf,Nup62,CG3511,YL-1,c(2)M,ND-23,Thiolase,cad,Sccpdh1,CG12301,RpL40,Uch-L5,RpL13A,Hat1,lqfR,CG8112,csw,RpS18,Atg6,RpS14a,ND-49,Ndc1,ATPsynbeta,Lsm12,CG11858,mts,mus81,mad2,Tango10,CG10470,ND-18,Vps16B,RpL34a,Marf,RpL38,SmE,Mcad,CycC,CG17746,CG32640,ssh,tws,Hsc70-3,cu,Pax,Girdin,alpha-Man-Ia,CG1827,Vsp37A,CG6707,CG12375,CG9684,fz4,apt,CG10254,Pen,Synj,Arpc2,MAGE,Nlg1,REPTOR,Gmap,CG14894,Pi4KIIalpha,bun,GlyP,mop,Pxt,nrv1,LysRS,gammaTub37C,CG4390,Ppcdc,HP1e,Cf2,CG4860,ste14,PAPLA1,trr,COX5B,crok,Pdk1,Imp,CG10189,Parp16,cold,Hexo1,Ggt-1,Ptpmeg,Topors,tzn,Pfk,cact,Pym,ND-B22,Idh3b,Xpac,Cyp6v1,CG30343,Pink1,TrpRS-m,Mtl,boi,CG11899,CG5510,Calx,stai,Fkbp39,mael,wal,SP1029,Ublcp1,Rnmt,GalT1,Trf2,CG8314,CG5844,msps,escl,Arpc4,Atf6,Su(H),Seipin,CG6178,Got1,mus304,Uba5,Prosbeta5,BoYb,GstT2,ATPsynCF6,26-29-p,RpL13,tor,wapl,Pgk,Cdk1,Sdhaf3,tos,Caf1-105,cnc,CG3842,Gasz,lt,Prosalpha3,blw,Elba2,mRpL10,mRpS23,Iml1,dgt4,ben,spd-2,CG17294,Tao,Sry-alpha,CG7009,RYBP,RpL24,Ufm1,Coq6,Cul2,Sox102F,RpS30,RecQ5,pen-2,rost,CG12018,r-l,HmgD,Rbf2,p38a,CG14671,Spt20,CG10561,Idgf6,Pfdn5,ATPsynB,P5cr,CG4238,mRpL18,snRNA:U4atac,TBCC,Chd64,RpLP1,Herc4,UGP,mbo,raptor,heph,CG10336,sau,Traf6,CG2924,CG11562,Acox57D-p,PHGPx,mRpS11,ND-75,rig,CG3803,RpII140,Trf4-2,Smyd4-4,pps,bif,gw,del,Archease,EcR,mRpL16,Marf1,CG14817,ph-d,gus,Orc6,Sec15,Etf-QO,cib,pont,Dcr-1,Jafrac1,elg1,TfIIFalpha,Oseg1,capt,PGRP-LE,STUB1,Sply,Fkbp12,scramb1,P5CDh1,CG42837,Prosbeta6,dpr6,Exd2,alphaTub84B,MED4,CG9393,Arl6IP1,fal,eIF4A,Clp,Usp10,ea,sip3,muskelin,PIG-H,Gip,Mat89Ba,EloC,RpL12,PpV,Fkbp59,CG7718,png,CG8199,Rcd5,Smox,CG31759,RpS5b,CG8004,CG13551,Uev1A,mRpL48,tapas,RpLP0,Cbl,Nurf-38,CysRS,thoc6,Mitf,ERp60,CG7920,Nap1,Stam,Shmt,yps,UbcE2H,CG33977,fs(1)Ya,nmd,nSMase,Snm1,PEK,CG7716,Rpn13,CG3587,SdhD,Hip1,spn-E,sbb,Neos,Pfdn2,Tom40,NP15.6,CG12773,CG5003,Rel,Kyat,Pus7,Rpt2,tef,Adh,CG1344,Scsalpha1,NHP2,RpS4,Cul5,foxo,Pura,dl,l(1)G0196,not,nmdyn-D6,pr,Tlk,Cka,Atpalpha,piwi,CG9643,Sptr,GstE11,PpD3,mRpL35,CG11267,tub,CCT6,CG10418,CG11110,trbd,ND-39,CG9941,Droj2,lid,CG8974,rept,CG9425,Sec10,mtTFB1,Fbw5,CG31523,e(y)1,PIG-L,RhoGAP5A,Tm1,cbs,SERCA,ND-MNLL,sbr,Rbcn-3A,CG10344,mRpS5,obe,Acyp2,Pmi,Roe1,SerRS-m,Arpc1,r,CG4594,CG2911,mRpS21,CG6927,CG7488,Vamp7,Utx,eIF4E1,HDAC6,shv,snRNA:U4:38AB,Ubc6,CG11134,Blos4,CycJ,UQCR-Q,Echs1,Unr,IntS11,mus205,yem,Nup358,TyrRS,Rpn12,CG7115,Fer1HCH,Cenp-C,Snx6,aay,AOX3,Tengl4,CCT3,MED24,vret,RpL17,Cul1,Aos1,toc,Prosbeta4,CG3631,CG9911,Cbp80,swa,Gmer,RpS23,COX5A,Arf51F,ATPsynG,Nc73EF,UQCR-11,GatB,ATPsyndelta,Irp-1B,cN-IIIB,kl-2,sw,Rab5,CG17698,mxc,Pld,larp,Rbf,RpS29,LSm3,CG3770,bru1,alph,EMRE,Hnf4,Acph-1,MED27,Ac3,Syx13,CG8728,dx,Rrp40,Bin1,Adi1,Edem1,CG5969,nej,spen,Drep1,SuUR,Nprl2,CG6967,Prosalpha5,CG3437,BigH1,SRPK,Naa20A,Fdh,CG5986,asf1,CG7840,SkpA,Cp1,Gdi | | 1.112e-36 | -82.79 | metabolic process | biological process | GO:0008152 | 5257 | 682 | 12345 | 1129 | bru1,alph,RpS29,LSm3,Pld,Rbf,CG17698,cN-IIIB,Rab5,Irp-1B,ATPsyndelta,COX5A,ATPsynG,Nc73EF,Arf51F,UQCR-11,GatB,CG7840,CG4842,Cp1,SkpA,Prosalpha5,SRPK,Fdh,Naa20A,CG5986,spen,SuUR,Drep1,Edem1,Adi1,nej,CG8728,CG2493,dx,Bin1,Rrp40,Ac3,Acph-1,Hnf4,CG16979,IntS11,mus205,Ubc6,CG11134,CycJ,UQCR-Q,Echs1,snRNA:U4:38AB,Vamp7,Utx,CG6144,eIF4E1,CG5966,HDAC6,r,CG4594,CG2911,mRpS21,CG6650,SerRS-m,Nxf3,Cbp80,RpS23,Gmer,Cul1,Aos1,Prosbeta4,CG3631,MED24,RpL17,vret,CG3609,Tengl4,aay,AOX3,TyrRS,Rpn12,CG7115,CG9643,Sptr,CG1275,GstE11,PpD3,pr,Tlk,Cka,piwi,l(1)G0196,not,nmdyn-D6,Cul5,foxo,dl,Adh,NHP2,Scsalpha1,CG1344,CG4267,RpS4,CG5003,Kyat,Pus7,Rpt2,Neos,NP15.6,Rpn13,CG3587,SdhD,Acyp2,Pmi,ND-MNLL,SERCA,sbr,CG10344,obe,mRpS5,mtTFB1,Fbw5,CG31523,e(y)1,PIG-L,rept,CG8974,lid,trbd,CG10418,CG11110,CG9941,ND-39,mRpL35,sip3,PIG-H,Gip,Mat89Ba,ea,fal,eIF4A,Usp10,Clp,Exd2,MED4,Prosbeta6,P5CDh1,Fkbp12,Sply,STUB1,CG33977,fs(1)Ya,Snm1,nSMase,PEK,Shmt,yps,UbcE2H,CysRS,thoc6,Nurf-38,Mitf,CG7920,Nap1,Cbl,CG13551,Uev1A,mRpL48,tapas,RpLP0,png,CG8199,Rcd5,CG31759,RpS5b,PpV,Fkbp59,stw,CG7718,EloC,RpL12,rig,CG3803,RpII140,Trf4-2,pps,Cyp6a22,PHGPx,Acox57D-p,mRpS11,ND-75,CG10336,Traf6,CG2924,UGP,mbo,heph,raptor,Herc4,RpLP1,mRpL18,CG4238,P5cr,snRNA:U4atac,ATPsynB,TfIIFalpha,PGRP-LE,Dcr-1,Jafrac1,pont,Etf-QO,CG14817,Orc6,gus,mRpL16,gw,del,Archease,EcR,tos,CG3842,Gasz,Cdk1,Sdhaf3,CG9953,tor,RpL13,26-29-p,Pgk,Prosbeta5,GstT2,ATPsynCF6,Seipin,mus304,Got1,CG6178,Uba5,CG8314,escl,p38a,Spt20,CG10561,Idgf6,sol,r-l,pen-2,CG12018,RecQ5,RpS30,CG2200,RpL24,Ufm1,Coq6,Cul2,Tao,RYBP,CG7009,blw,mRpS23,mRpL10,ben,CG17294,lt,Prosalpha3,Ggt-1,Hexo1,Ptpmeg,Topors,Pdk1,Imp,CG10189,Parp16,ste14,PAPLA1,trr,COX5B,CG4390,Ppcdc,CG4860,GlyP,mop,Pxt,CG8067,LysRS,CG13630,Pi4KIIalpha,CG10254,Synj,MAGE,Trf2,GalT1,wal,mael,SP1029,Ublcp1,Rnmt,CG11899,Mtl,Pink1,TrpRS-m,Pym,ND-B22,Cyp6v1,Idh3b,Xpac,CG30343,Pfk,tzn,ND-18,RpL34a,Nepl3,CG11858,Lsm12,mts,mus81,Atg6,RpS14a,ND-49,ATPsynbeta,RpS18,Hat1,csw,CG8112,Uch-L5,RpL13A,CG6707,CG12375,CG9684,CG1827,Vsp37A,alpha-Man-Ia,cu,Fur2,Hsc70-3,tws,CG17746,CG32640,ssh,CycC,RpL38,Marf,SmE,Mcad,CG4572,Prosalpha4,GluRS-m,EMC3,Ser7,CG10463,GstT1,CG10638,RpS13,Dd,COX4,eEF1beta,Gk2,otu,CAH2,Pglym78,CG5880,Mo25,mre11,Ppox,CG13344,CG4407,CSN6,CG17333,yip2,Cdk7,dob,Agpat2,Drep2,Atu,isoQC,CG17454,Grx1,Sccpdh1,CG12301,RpL40,CG31674,faf,YL-1,c(2)M,CG3511,Thiolase,ND-23,nbs,agt,CG1640,Nadsyn,Cyp6u1,ste24a,pn,RpL26,Debcl,wisp,RpS27A,PyK,twe,Pomp,CG1774,GAPsec,grp,ND-B15,fdl,grsm,Xpc,GILT1,AGBE,Rpb8,Sod3,CG7728,Usp16-45,CG11164,Not10,Plap,CG2201,CG18135,Treh,S2P,CG2767,CG5292,CG10425,trsn,CG7789,Sgf11,CG32473,park,CG32221,CG6767,Sod1,Ts,cl,ari-1,DNApol-epsilon255,Mal-A5,mRpL17,G9a,Eno,SNF4Agamma,whd,Trxr-1,AMPdeam,mal,CG8204,CG5390,CG9316,GCS2beta,Desat1,CG9436,Son,CG8475,d4,RpL23A,O-fut1,Lkb1,RpS15,CG17187,eIF3i,Ntan1,shop,CG3223,CG40045,Tfb4,CG6094,Gclc,Pp2A-29B,Sirt7,Mtpalpha,kri,sgll,Chrac-14,PIG-P,CycE,Cyp1,UK114,CG7328,AIF,CG3021,Cht11,adp,CG11486,CG5028,RpS7,CG31633,SelD,CG4041,CG17270,CG8726,CG11089,cort,ND-24,ago,SREBP,CG10932,Sld5,CG7550,r2d2,Idgf4,CG11771,Ptp69D,armi,RpS14b,SelR,Aldh,Atg7,Idh,awd,Sp7,nmdyn-D7,Pfas,RpL7A,Edc3,Rab32,CG7322,ATPsynE,RfC38,CG42554,levy,CG14407,CG2034,Dip-B,shu,CG5026,CG8311,CG4598,CG17768,RpL30,IntS6,Sgf29,RpS2,Gk1,Coq8,DppIII,Ugt35A1,tko,CG14516,CG33156,Vha68-2,Cchl,eIF3m,Ptip,Pfrx,Taf7,Rep,RpL34b,CG7414,CG5285,tkv,Idgf2,CG9485,mino,rl,CG3198,ND-B18,CG8188,Cdk5,cathD,CG9143,smg,RpS16,Tip60,Usp39,Pgm1,Ptp61F,Cyt-c-p,Dph5,RpL9,Uch,PIG-C,Iru,nos,sni,GatA,Tak1,CG6610,CG4586,pygo,Gabat,RpS27,CG1578,Dhfr,CanB2,RpL11,velo,Atg4b,sws,CG6766,l(3)73Ah,AcCoAS,mRpS10,RpS28b,fray,ND-51,Npc2a,E2f1,Pcd,APP-BP1,scu,CG17691,mEFTu1,SdhA,ND-30,shrb,ATPsynD,Coq7,Prosbeta3,mRpL13,CG16865,Rpn3,stet,Larp7,RpL10,CG2938,Argk,Top1,tho2,CanA-14F,Pex12,ACC,Pdk,cype,RpL31,RpL29,Gyg,bys,Pi3K92E,Pp2B-14D,snk,Nmt,Ugt36E1,mRpL36,CG14435,regucalcin,CG2100,Spx,ATPsynO,AsnS,CG14997,CG31053,l(2)k01209,Rga,ND-SGDH,scny,Cyp6a19,snRNA:U12,oxt,RpL37a,exu,Orc1,CG9391,Jafrac2,esc,CkIIalpha,Ho,Ptpmeg2,Ptp10D,CG7997,CG10863,tun,Pex10,Ady43A,RpL23,RpS24,QC,Gapdh1,dco,CG14100,twin,DhpD,CG46338,mEFTu2,ScsbetaA,CycB,lwr,egg,Efr,CG11070,Mos,gd,Pdxk,Ubc4,CG1622,Mekk1,Rpn2 | | 2.744e-33 | -74.98 | protein-containing complex | cellular component | GO:0032991 | 3016 | 452 | 11940 | 1105 | Scsalpha1,NP15.6,Cenp-C,ago,snRNA:U4atac,Sld5,eIF3i,CG3198,Klp10A,Stam,Sec10,Zwilch,RpS13,RpS30,Eno,DCAF12,Vsp37A,CG12301,Larp7,EMC3,CG3587,RpS27,CG5986,RpL12,Syx13,CG5292,twin,CG6094,snRNA:U4:38AB,CanB2,RpL38,smg,lid,UQCR-Q,l(3)mbt,thoc6,nbs,Pp2B-14D,eIF4E1,ND-MNLL,Su(H),bru1,TyrRS,Prosbeta4,Prosalpha3,Traf6,CG5903,nos,RpL13A,Uxt,IntS11,escl,ND-MWFE,Sec15,Pex13,GatB,Mtpalpha,png,RpL40,Usp39,Uev1A,CG9684,Tom70,mRpL17,ND-SGDH,CG8728,CG12018,Orc1,AP-1-2beta,Pka-R1,ph-d,tapas,CG11486,RpLP0,Cul1,PIG-H,jigr1,RpL34b,park,ari-1,tho2,Kap3,armi,tral,Ndc1,Nap1,Trf4-2,ATPsynE,mRpS11,Caf1-105,Fbw5,Atu,Hsc70-3,CG15525,CG9316,CycE,kl-2,asf1,cnn,ND-39,Nup58,Clc,shrb,CG8199,CG8974,Rcd5,RpL7A,Ranbp21,Pfk,ATPsynbeta,piwi,Cbp80,Edc3,CCT5,CG13366,nAChRalpha4,cup,Xpc,COX5A,shep,rept,APP-BP1,Utx,Vps16B,Nprl2,pont,ND-B15,Thiolase,RpLP1,CG9425,Fkbp59,RpL10,IntS6,Sec23,Cka,Roe1,tor,EcR,Pka-R2,tko,Imp,Dcr-1,MAGE,Oseg1,brwl,RpS18,UQCR-11,CSN6,Bin1,cype,lwr,Vha13,Tomosyn,d4,RpL37a,SkpA,CG10336,RpL11,Mat89Ba,eIF3m,CG9132,gammaTub37C,Ptip,r2d2,CG6610,Arpc4,Med,CkIIalpha,mRpL36,exu,sip3,ND-B22,NHP2,CG10254,CG17691,SmE,Taf7,Tm1,ND-18,Pomp,RpS4,Hip1,heph,Prosbeta6,e(y)1,Mes2,stai,Dd,RpL30,blw,CG2034,PIG-C,mRpL13,Tim8,Tfb4,mRpS21,GlyP,Nc73EF,CG31875,wash,scu,Blos4,CG8475,snRNA:U12,Ubc6,eIF4A,RpL9,mts,CHMP2B,CG32221,Rpb8,Rbcn-3A,Rbf2,COX5B,Mf,CG4404,spz,vret,CCT6,CG12288,CG8009,Grip84,Chrac-14,CCT3,MED4,CG30496,Neos,obe,CycJ,CG11164,CG3511,Sgf11,mre11,CG42526,SdhA,ND-51,Trs33,Archease,Grip71,ATPsynG,Rpn12,EMRE,CG17068,RpS2,RpL34a,mRpS10,RpS15,APC7,CG7414,mRpS5,tkv,Neb-cGP,EMC8-9,BigH1,osk,ScsbetaA,squ,trr,sw,mRpL48,RpL23A,CycB,Rpt2,RpL29,mRpL16,CG32441,ATPsynB,ND-23,Rrp40,mbo,Grip128,cu,bys,Pex12,RpL26,Spt20,wal,CG31633,ix,lin-52,Abi,PEK,RpL24,CG7716,msps,ATPsyndelta,Not10,Prosbeta5,Atpalpha,RpS23,yem,MED20,CG7728,ND-B18,elg1,Liprin-beta,Prosalpha5,not,esc,ND-49,CanA-14F,Psf3,SdhD,pygo,Nurf-38,Rbf,Cdk5,PyK,Afti,Cdk7,CG6967,pps,Pfdn5,ATPsynCF6,Tom40,ben,CG1622,ND-24,LysRS,Uba5,Aos1,E2f1,HP1c,Iml1,CG17768,RfC38,CG10418,Pfdn2,Rep,CG11110,Rpn13,Bap170,shu,RpS28b,EloC,ND-75,Uch-L5,COX4,Nup62,MED24,ATPsynO,RpL17,CG6767,CG11070,mad2,Arpc2,Vha36-1,l(3)73Ah,Moe,Cul5,Nf-YA,mRpL10,plu,pgc,CG2162,Aladin,MED27,LSm3,RpS29,psq,Grip75,RpS7,mus205,Ork1,mus81,RpS16,GatA,CG6276,pen-2,Clp,krimp,Prosalpha4,RpL23,CG17187,CG16865,Vha68-2,Xpac,CG16868,Snx6,nrv1,RpS14b,CG40472,mRpL35,Fer1HCH,Elba2,Nlg1,Sgf29,Gclc,DNApol-epsilon255,CG8004,gus,tws,Sec22,CG9393,CG8204,ATPsynD,NFAT,cana,alc,mael,yps,eEF1beta,Prosbeta3,RpS5b,COX8,RpL31,TfIIFalpha,Exo70,levy,CG14817,Adh,Rga,Vamp7,Top1,mRpS23,Naa20A,spn-E,Trf2,YL-1,Pym,up,gw,Pp2A-29B,Arpc1,CG33977,Cul2,Orc6,mRpL18,Pi3K92E,CG17454,nej,RpS27A,Smox,Spx,RpL13,CG5003,RpS14a,raptor,Rpn3,RpS24,cyc,ND-30,lt,Tip60,CycC,rig,AGO3,Rpn2,Blos2,Atg6,RpII140,ATPsynF | | 2.933e-29 | -65.70 | organelle part | cellular component | GO:0044422 | 3156 | 455 | 11940 | 1105 | tko,alt,Mybbp1A,G9a,EcR,MAGE,Synj,Imp,Sec23,IntS6,RpL10,Roe1,pont,CG9399,ND-B15,AIF,RpLP1,Thiolase,CG13366,cup,Actn,piwi,Cbp80,CG5510,dl,Vps16B,rept,Utx,COX5A,Xpc,Idh3b,Gabat,stet,Pi4KIIalpha,Arpc4,CG14407,CG6610,gammaTub37C,eIF3m,CG9132,Ptip,RYBP,CG10336,Mat89Ba,RpL11,Cyt-c-p,cbs,Vha13,CG11077,RpL37a,d4,CG31729,Coq8,alphaTub67C,Lnpk,UQCR-11,RpS18,cype,Bin1,lwr,CSN6,mol,Tfb4,Tim8,CG31523,wash,Nc73EF,mRpS21,stai,Sdhaf3,mRpL13,PIG-C,RpL30,blw,Dd,Hip1,RpS4,ND-18,Ppox,CG17691,SmE,Taf7,Tm1,e(y)1,sip3,exu,mRpL36,NHP2,ND-B22,Tspo,CG17328,obe,Neos,Gga,CG8009,CG12288,CG2129,Pgk,MED4,Chrac-14,Grip84,Rpb8,Got1,mts,CHMP2B,COX5B,Mf,CG8112,Rbf2,scu,RpL9,Ubc6,snRNA:U12,CG5986,RpS27,CG6766,Mctp,TMEM216,cnc,RpL12,sbr,CG12301,Vsp37A,Aldh,RpS30,EMC3,awd,Stam,CG4021,RpS13,Zwilch,CG3803,Ptp61F,spd-2,snRNA:U4atac,Cenp-C,NP15.6,Scsalpha1,Klp10A,sau,CG3198,Gmap,Sld5,IntS11,escl,Uxt,CG12375,Pdk,Pex13,ND-MWFE,Gapdh1,bif,CG6707,RpL13A,CG4293,CG5903,Sirt7,nbs,thoc6,ste24a,bru1,CG46385,ND-MNLL,Su(H),CG14183,RpL38,lid,snRNA:U4:38AB,SREBP,CG6094,Pglym78,UQCR-Q,l(3)mbt,Cdk1,tho2,Kap3,RpL34b,Arl5,twe,ph-d,PIG-H,lqfR,shop,RpLP0,toc,ND-SGDH,mRpL17,Tom70,Debcl,Orc1,CG12018,AP-1-2beta,Mitofilin,mxc,CG8728,RpL40,Mtpalpha,CG10425,CG9684,Usp39,Rcd5,HmgD,CG8974,CG8199,ATPsynbeta,RpL7A,CG11267,SERCA,SuUR,shrb,Clc,Nup58,CG15525,CycE,Hsc70-3,ND-39,opm,cnn,asf1,kl-2,Tengl4,Trf4-2,Rab32,Nap1,Ndc1,Atu,CG10470,mRpS11,ATPsynE,Cchl,RpS16,RpL23,fs(1)Ya,Snm1,Clp,psq,Grip75,RpS29,Girdin,LSm3,MED27,Aladin,RpS7,mus205,Tao,Nf-YA,sws,Non1,CG2162,Topors,mute,Pex10,mRpL10,mad2,RpL17,ATPsynO,MED24,Moe,vers,l(3)73Ah,Vha36-1,P5CDh1,Arpc2,Sec22,ATPsynD,NFAT,mtTFB1,CG9393,Elba2,Sgf29,dgt4,gus,CG17270,CG8004,DNApol-epsilon255,CG5447,RpS14b,mRpL35,CG40472,Vha68-2,CG8311,CG16865,CG17187,Xpac,Tlk,alphaTub84B,CG9981,CG33977,CG17454,GCC88,mRpL18,Orc6,mRpS23,Top1,Vamp7,CG9143,Pp2A-29B,Arpc1,up,Pym,gw,YL-1,stil,Trf2,RpL31,TfIIFalpha,RpS5b,CG3437,levy,CG14817,scny,CG10055,Ugalt,cana,yps,Atf6,mino,Blos2,ABCD,SelR,ATPsynF,RpII140,TrpRS-m,Desat1,CycC,Tip60,CG7488,RpS24,RpS14a,RpL13,CG4880,ND-30,Pmi,nej,Fkbp39,Spx,RpS27A,Marf,sw,trr,CG31053,pn,CycB,mRpL48,RpL23A,Neb-cGP,capt,mRpS5,BigH1,EMC8-9,Axs,hb,Cen,RpS2,EMRE,l(1)G0004,CG7414,alpha-Man-Ia,APC7,RpS15,mRpS10,CG5969,RpL34a,ND-51,SdhA,mre11,Sgf11,CG3511,CG11164,ATPsynG,Grip71,CG14671,Trs33,STUB1,lin-52,del,Parp16,Rbfox1,CG7716,RpL24,CG9773,wal,Spt20,ix,yin,CG7655,mbo,Rrp40,CG1602,egg,RpL26,ATP8A,bys,Pex12,Grip128,CG32441,Pex23,ATPsynB,RpL29,mRpL16,ND-23,Scox,Afti,Etf-QO,PAPLA1,CG32640,Rbf,Efr,ATPsynCF6,fwe,pps,Cdk7,Psf3,pygo,Nurf-38,Cse1,SdhD,elg1,Seipin,ND-B18,esc,ND-49,not,RpS23,Hat1,colt,Prosbeta5,HP1e,msps,ATPsyndelta,MED20,yem,EloC,ND-75,RpS28b,Coq7,Bap170,Nup62,COX4,Fur2,CG10418,RfC38,CG11110,Rep,Chd64,CG17768,Iml1,HP1c,E2f1,Aos1,Echs1,Tom40,Rel,ND-24,CG1622 | | 3.200e-29 | -65.61 | intracellular organelle part | cellular component | GO:0044446 | 3091 | 448 | 11940 | 1105 | Kap3,tho2,RpL34b,ph-d,Arl5,twe,shop,lqfR,PIG-H,RpLP0,toc,ND-SGDH,mRpL17,Tom70,Debcl,AP-1-2beta,CG12018,Orc1,Mitofilin,CG8728,mxc,RpL40,Mtpalpha,CG10425,CG9684,Usp39,Rcd5,HmgD,CG8974,CG8199,ATPsynbeta,RpL7A,CG11267,SERCA,SuUR,shrb,Clc,Nup58,CycE,CG15525,Hsc70-3,ND-39,opm,cnn,asf1,Tengl4,kl-2,Trf4-2,Rab32,Nap1,Ndc1,Atu,CG10470,mRpS11,ATPsynE,CG6766,CG5986,RpS27,Mctp,cnc,RpL12,sbr,CG12301,Vsp37A,Aldh,RpS30,EMC3,awd,Stam,CG4021,RpS13,Zwilch,CG3803,spd-2,Ptp61F,snRNA:U4atac,Cenp-C,NP15.6,Scsalpha1,Klp10A,Gmap,sau,CG3198,Sld5,Uxt,escl,IntS11,CG12375,Pdk,Pex13,ND-MWFE,bif,CG6707,RpL13A,CG5903,CG4293,Sirt7,nbs,thoc6,ste24a,ND-MNLL,Su(H),bru1,CG46385,CG14183,RpL38,lid,SREBP,CG6094,snRNA:U4:38AB,UQCR-Q,l(3)mbt,Cdk1,Tfb4,mol,Tim8,CG31523,wash,Nc73EF,mRpS21,stai,Sdhaf3,mRpL13,PIG-C,blw,RpL30,Dd,Hip1,RpS4,ND-18,Ppox,Tm1,CG17691,Taf7,SmE,e(y)1,sip3,mRpL36,exu,NHP2,ND-B22,obe,Tspo,CG17328,Neos,Gga,CG8009,CG12288,CG2129,MED4,Chrac-14,Grip84,Rpb8,Got1,mts,CHMP2B,Mf,COX5B,Rbf2,CG8112,scu,RpL9,Ubc6,snRNA:U12,alt,tko,Mybbp1A,EcR,G9a,MAGE,Imp,Sec23,IntS6,RpL10,Roe1,pont,CG9399,ND-B15,Thiolase,RpLP1,AIF,CG13366,cup,piwi,Cbp80,CG5510,Vps16B,dl,rept,COX5A,Utx,Xpc,Idh3b,Gabat,Pi4KIIalpha,stet,Arpc4,CG6610,CG14407,gammaTub37C,Ptip,eIF3m,CG9132,RYBP,CG10336,RpL11,Mat89Ba,Cyt-c-p,CG11077,cbs,Vha13,RpL37a,d4,CG31729,Coq8,alphaTub67C,UQCR-11,Lnpk,RpS18,Bin1,lwr,cype,CSN6,Afti,Etf-QO,CG32640,PAPLA1,Rbf,Efr,ATPsynCF6,fwe,pps,Cdk7,Psf3,pygo,Nurf-38,Cse1,SdhD,elg1,Seipin,ND-B18,esc,ND-49,not,RpS23,colt,Hat1,Prosbeta5,HP1e,msps,ATPsyndelta,MED20,yem,EloC,ND-75,RpS28b,Coq7,Bap170,COX4,Nup62,Fur2,CG10418,RfC38,CG11110,Rep,Chd64,Iml1,CG17768,HP1c,E2f1,Aos1,Echs1,Tom40,Rel,CG1622,ND-24,sw,trr,CG31053,pn,CycB,RpL23A,mRpL48,Neb-cGP,capt,mRpS5,BigH1,EMC8-9,Axs,Cen,hb,RpS2,EMRE,CG7414,l(1)G0004,APC7,alpha-Man-Ia,mRpS10,RpS15,CG5969,RpL34a,ND-51,SdhA,mre11,Sgf11,CG11164,CG3511,Grip71,ATPsynG,CG14671,Trs33,lin-52,del,Parp16,Rbfox1,CG7716,RpL24,CG9773,wal,Spt20,ix,yin,CG7655,mbo,Rrp40,CG1602,egg,ATP8A,RpL26,bys,Pex12,Grip128,ATPsynB,Pex23,CG32441,mRpL16,RpL29,ND-23,Scox,CG33977,GCC88,CG17454,Orc6,mRpL18,mRpS23,Top1,Vamp7,CG9143,Pp2A-29B,Arpc1,YL-1,Trf2,stil,Pym,up,gw,RpL31,TfIIFalpha,CG3437,RpS5b,CG14817,levy,scny,CG10055,Ugalt,cana,yps,Atf6,mino,Blos2,ABCD,SelR,ATPsynF,RpII140,TrpRS-m,CycC,Desat1,Tip60,CG7488,RpS24,RpS14a,RpL13,CG4880,ND-30,Pmi,nej,Fkbp39,Spx,RpS27A,Marf,RpS16,Cchl,fs(1)Ya,RpL23,Snm1,Clp,psq,Grip75,RpS29,Girdin,LSm3,MED27,Aladin,RpS7,mus205,Tao,Nf-YA,Non1,sws,Topors,mute,CG2162,Pex10,mRpL10,mad2,RpL17,MED24,ATPsynO,Moe,vers,l(3)73Ah,P5CDh1,Vha36-1,Arpc2,Sec22,ATPsynD,NFAT,CG9393,mtTFB1,Sgf29,dgt4,Elba2,gus,CG8004,CG17270,CG5447,DNApol-epsilon255,RpS14b,mRpL35,CG40472,Vha68-2,CG8311,CG17187,CG16865,Xpac,Tlk,alphaTub84B,CG9981 | | 7.904e-29 | -64.71 | catalytic activity | molecular function | GO:0003824 | 4067 | 545 | 12176 | 1119 | CycE,snk,Dcr-1,PGRP-LE,CG31053,COX8,CSN6,Sply,nmdyn-D7,Sirt7,nSMase,CG10254,Cyt-c-p,Gyg,CG5292,CG6178,ND-B18,Pfas,UK114,Rab5,UQCR-Q,escl,Iru,CG46385,DhpD,CG30343,Cht11,CG14100,Jheh2,CG31523,ScsbetaA,SdhD,CG8728,isoQC,CG9471,COX5B,CG13630,CG12375,Prosbeta6,GstE11,cu,Coq7,Exd2,AGBE,Pex10,CAH7,Gk2,CG8726,fal,RecQ5,Nadsyn,PHGPx,gammaTub37C,cathD,l(1)G0196,CG17746,l(2)k01209,nbs,AgmNAT,CG9485,Gs1l,CG11771,CG9391,kl-2,Adi1,whd,Acox57D-p,Agpat2,Pi3K92E,ND-23,CG8188,Mtl,Pfk,ari-1,UbcE2H,ND-SGDH,Trf4-2,CG7328,shop,Usp39,sip3,exu,G9a,Prosbeta4,mts,tor,UGP,CG5880,Prosalpha4,CG10344,CG7322,fdl,CG4390,CG2200,Ppcdc,CG10189,PpV,GluRS-m,awd,Traf6,blw,CG9143,ND-49,CG17691,Ublcp1,AMPdeam,Kap3,Rpt2,CG5390,CG32473,pr,Eno,twe,Uev1A,CG9316,CG9581,Utx,CG7718,CG8475,yip2,stet,CG32221,alpha-Man-Ia,Aldh,Ptp61F,faf,CG8974,GstT2,CG2493,regucalcin,Pink1,CG14435,Nmt,CG5026,ea,levy,CG11070,otu,dco,CanA-14F,Ac3,CG14407,Ugt36E1,CG4267,wal,ND-B22,cN-IIIB,Prosalpha3,AIF,e(y)1,S2P,CG6028,CG4586,nej,ND-75,aay,Klp10A,obe,CG2767,dx,CG7728,Rga,twin,ATPsynbeta,Cdk5,Mekk1,tos,CG7920,CG8112,tws,CG4502,mre11,Scox,Pld,armi,sgll,spd-2,Cchl,alc,Uba5,Idgf6,HDAC6,CG5844,mtTFB1,mop,sol,tun,Mcad,CG14516,PyK,Irp-1B,ben,CG4860,CG10638,r,Ntan1,CG14204,Got1,grp,ste24a,Ubc6,CG2201,Atg4b,glob1,grsm,Idh3b,ND-51,kri,CG1774,Sp7,ND-24,SelR,Cdk1,dnc,Top1,STUB1,CG3631,Uch-L5,CG7115,CG14997,CG2924,Ugt35A1,Uch,ERp60,mino,CG11089,Naa20A,CG1640,Hsc70-3,COX4,CG46338,CG12018,mus81,egg,nmdyn-D6,Dd,EndoGI,Kyat,CG6707,Cdk7,Arf51F,Tak1,Pex12,Gabat,Pdk1,CycB,Pp2B-14D,up,CG3511,Pgm1,CysRS,Nepl3,eIF4A,Parp16,Hexo1,UQCR-11,Mtpalpha,cl,lid,PEK,LysRS,CG11486,sw,pps,Psf3,Snm1,CG3894,esc,YL-1,GlyP,Tfb4,cana,alphaTub84B,ND-B15,Lkb1,CG4880,Cyp6u1,RpII140,CG13344,Scsalpha1,Aos1,Etf-QO,Marf,Desat1,Pi4KIIalpha,Fkbp12,Ptp10D,mal,Nurf-38,Cyp6a22,trbd,mEFTu1,Nc73EF,trr,GatB,CG7840,CG15141,Sld5,Pglym78,Usp16-45,sws,CkIIalpha,Rab10,Tao,CG9248,CG9953,ssh,Mal-A5,Pxt,trsn,CG17294,Ptp69D,ND-39,Edem1,SkpA,CG9941,PpD3,Rpn2,Cyp1,CG1827,Pka-R1,CG8679,Dip-B,spn-E,gd,Treh,MAGE,not,CG33156,CG17333,Pcd,Naxe,CG16979,pn,PAPLA1,CG12171,P5cr,CG10863,Mos,CG9643,Cp1,Sgf29,CG11134,Cbl,Acph-1,park,PIG-P,alpha-Est5,dob,CG2061,RpLP0,wisp,CG7997,pont,p38a,Ser7,Argk,APP-BP1,CG10425,CG4238,Prosalpha5,fray,CG10932,GstT1,Sptr,CG1344,GalT1,Chrac-14,ste14,CG8199,PIG-L,sni,milt,Ptpmeg2,CG9436,EloC,Pdxk,CG31548,ACC,Topors,Rnmt,Dph5,CG4842,CG4594,CG8314,PIG-C,Fkbp59,CG31674,tub,CG5966,RfC38,Tip60,AOX3,CG8611,Gk1,GatA,Tlk,DNApol-epsilon255,Pdk,Coq6,alph,Pgk,APC7,CG3021,Acyp2,tkv,Coq8,CG8311,CG5285,CG1578,Drep2,CG4407,CG7009,Atg7,CG8607,CAH2,GILT1,Trxr-1,Thiolase,CG6144,Usp10,CG5028,Adh,CG40045,Ts,CG5003,SelT,Gapdh1,Ho,agt,CG2100,Jafrac2,Ubc4,stw,CG10463,CCT3,SelD,TyrRS,scny,CG4572,Pfrx,ATPsynF,Marc,CG2938,cype,Cyp6a19,Prosbeta5,Shmt,ND-30,csw,Gmer,Rpb8,tzn,Ggt-1,Clp,CG18135,alphaTub67C,Idh,AcCoAS,CG7550,CG11858,CG3939,26-29-p,ND-18,Hat1,Fur2,SP1029,mEFTu2,TrpRS-m,Grx1,COX5A,AsnS,Fbw5,Gclc,Cyp6v1,Gip,QC,CG32549,clt,CG4041,velo,Pus7,CG31549,CG6767,Fdh,lwr,CG33977,P5CDh1,Ptpmeg,SerRS-m,Prosbeta3,CG1275,rl,CG3609,Sod1,CG11164,Rab32,rept,CycJ,CG17698,Mo25,CG3842,anne,png,CG3803,CG9265,CG8067,oxt,SRPK,Ady43A,DppIII,CG31633,SdhA,Pp2A-29B,ago,O-fut1,mus205,Pka-R2,CG10561,Herc4,Ppox,CG6650,ND-MNLL,Arl5,d4,Jafrac1,Echs1,CG11110,Dhfr,Sccpdh1,ATPsynCF6,Synj,CG6967,CG7789,r-l,Sod3,CG4598,Tengl4,CG31759,Rcd5,scu,Kcmf1,PrBP,CG14615,CG11899 | | 1.724e-28 | -63.93 | organic substance metabolic process | biological process | GO:0071704 | 4824 | 616 | 12345 | 1129 | Pdk1,Imp,Parp16,CG10189,PAPLA1,ste14,COX5B,trr,Hexo1,Ptpmeg,Ggt-1,Topors,CG13630,Pi4KIIalpha,CG10254,Synj,MAGE,CG4390,Ppcdc,CG4860,GlyP,mop,Pxt,LysRS,CG11899,Mtl,Trf2,GalT1,mael,wal,SP1029,Ublcp1,Rnmt,tzn,Pfk,Pink1,TrpRS-m,Pym,Xpac,Idh3b,CG30343,Lsm12,CG11858,Nepl3,mus81,mts,RpL34a,Hat1,CG8112,csw,Uch-L5,RpL13A,Atg6,RpS14a,ATPsynbeta,RpS18,alpha-Man-Ia,Fur2,cu,CG12375,CG6707,CG9684,CG1827,Vsp37A,CycC,RpL38,Marf,SmE,Mcad,tws,Hsc70-3,CG17746,CG32640,ssh,CG10336,Traf6,CG2924,rig,CG3803,RpII140,Trf4-2,pps,Acox57D-p,mRpS11,mRpL18,CG4238,P5cr,snRNA:U4atac,ATPsynB,mbo,UGP,heph,RpLP1,Herc4,Dcr-1,pont,TfIIFalpha,PGRP-LE,mRpL16,gw,del,Archease,EcR,Etf-QO,CG14817,gus,Orc6,26-29-p,RpL13,tor,Pgk,Prosbeta5,GstT2,ATPsynCF6,tos,CG3842,Gasz,Cdk1,CG9953,Sdhaf3,CG8314,escl,Seipin,CG6178,Got1,mus304,Uba5,pen-2,CG12018,RpS30,RecQ5,p38a,CG10561,Spt20,sol,Idgf6,r-l,blw,mRpS23,mRpL10,ben,lt,Prosalpha3,CG2200,Ufm1,RpL24,Coq6,Cul2,Tao,RYBP,CG7009,not,l(1)G0196,nmdyn-D6,Cul5,foxo,dl,Sptr,CG9643,GstE11,PpD3,pr,Tlk,Cka,piwi,Neos,CG3587,Rpn13,Adh,Scsalpha1,CG4267,NHP2,RpS4,CG5003,Pus7,Kyat,Rpt2,Fbw5,mtTFB1,PIG-L,CG31523,e(y)1,rept,sbr,SERCA,CG10344,obe,mRpS5,mRpL35,CG8974,lid,CG11110,CG10418,trbd,CG9941,ND-39,fal,eIF4A,Clp,Usp10,Exd2,MED4,sip3,PIG-H,Gip,Mat89Ba,ea,Prosbeta6,P5CDh1,STUB1,Sply,Fkbp12,Nurf-38,thoc6,CysRS,Mitf,CG7920,Nap1,Cbl,CG33977,fs(1)Ya,nSMase,Snm1,PEK,Shmt,yps,UbcE2H,PpV,Fkbp59,CG7718,EloC,RpL12,CG13551,Uev1A,mRpL48,tapas,RpLP0,png,CG8199,CG31759,Rcd5,RpS5b,RpS29,LSm3,Pld,Rbf,bru1,alph,ATPsyndelta,Irp-1B,ATPsynG,Arf51F,GatB,CG17698,cN-IIIB,Rab5,SuUR,Drep1,spen,Adi1,Edem1,nej,CG7840,Cp1,SkpA,Prosalpha5,SRPK,Naa20A,Fdh,CG5986,Hnf4,CG8728,CG2493,dx,Rrp40,Bin1,Ac3,Ubc6,CycJ,CG11134,Echs1,snRNA:U4:38AB,CG16979,mus205,IntS11,CG2911,CG4594,r,mRpS21,CG6650,SerRS-m,Nxf3,Utx,HDAC6,CG5966,eIF4E1,MED24,RpL17,vret,Cbp80,RpS23,Gmer,Cul1,Aos1,Prosbeta4,CG3631,TyrRS,Rpn12,CG7115,Tengl4,aay,AOX3,Top1,tho2,CG2938,Pdk,RpL31,CanA-14F,Pex12,ACC,Pi3K92E,Pp2B-14D,RpL29,Gyg,bys,Spx,ATPsynO,AsnS,snk,Nmt,Ugt36E1,mRpL36,CG14435,CG2100,regucalcin,RpL37a,exu,oxt,Orc1,CG9391,esc,CG31053,l(2)k01209,Rga,scny,snRNA:U12,RpS24,Gapdh1,QC,CkIIalpha,Ho,Ptpmeg2,Ptp10D,CG7997,tun,Pex10,Ady43A,RpL23,mEFTu2,ScsbetaA,CycB,lwr,egg,Efr,dco,CG14100,twin,DhpD,CG46338,Pdxk,CG1622,Ubc4,Mekk1,Rpn2,CG11070,Mos,gd,RpS2,Gk1,Coq8,CG8311,IntS6,RpL30,CG17768,CG4598,Sgf29,CG14516,Cchl,CG33156,Ptip,eIF3m,Pfrx,Ugt35A1,DppIII,tko,mino,CG3198,rl,CG9485,Cdk5,CG8188,cathD,smg,CG9143,Taf7,Rep,RpL34b,CG5285,CG7414,Idgf2,tkv,Usp39,Pgm1,Ptp61F,Dph5,RpS16,Tip60,nos,GatA,Tak1,CG4586,CG6610,pygo,Gabat,RpS27,CG1578,RpL9,PIG-C,Uch,Iru,sws,CG6766,velo,l(3)73Ah,mRpS10,AcCoAS,RpS28b,fray,Dhfr,CanB2,RpL11,mEFTu1,shrb,E2f1,Npc2a,APP-BP1,Pcd,scu,CG17691,CG16865,Rpn3,stet,Larp7,RpL10,ATPsynD,Coq7,Prosbeta3,mRpL13,RpS15,CG17187,eIF3i,Ntan1,shop,CG40045,CG3223,d4,RpL23A,O-fut1,Lkb1,kri,PIG-P,sgll,Chrac-14,CycE,Tfb4,CG6094,Gclc,Pp2A-29B,Mtpalpha,Sirt7,CG7328,CG3021,Cht11,adp,CG11486,Cyp1,UK114,CG4041,CG17270,CG8726,CG5028,RpS7,CG31633,SelD,ago,CG10932,SREBP,cort,CG11089,Aldh,Idh,Atg7,awd,Sp7,Sld5,Idgf4,r2d2,armi,RpS14b,SelR,CG11771,Ptp69D,CG7322,nmdyn-D7,Pfas,RpL7A,Edc3,Rab32,shu,CG5026,ATPsynE,RfC38,CG2034,Dip-B,CG5880,Mo25,mre11,Ppox,CG4572,GluRS-m,Prosalpha4,EMC3,RpS13,CG10463,Ser7,Dd,GstT1,eEF1beta,Gk2,otu,Pglym78,Drep2,Atu,isoQC,CG17454,CG13344,CSN6,CG4407,CG17333,yip2,Cdk7,dob,Agpat2,faf,YL-1,c(2)M,CG3511,Thiolase,Sccpdh1,CG12301,RpL40,RpS27A,wisp,PyK,twe,nbs,agt,Nadsyn,ste24a,pn,RpL26,fdl,grsm,Xpc,AGBE,Rpb8,Pomp,CG1774,GAPsec,grp,Plap,CG2201,CG18135,S2P,Treh,CG5292,CG7728,Usp16-45,CG11164,Not10,park,CG32221,CG6767,Ts,ari-1,DNApol-epsilon255,mRpL17,Mal-A5,trsn,CG10425,CG7789,Sgf11,CG32473,mal,CG8204,CG5390,GCS2beta,CG9316,Desat1,Son,CG8475,G9a,Eno,SNF4Agamma,whd,AMPdeam | | 1.943e-28 | -63.81 | primary metabolic process | biological process | GO:0044238 | 4472 | 582 | 12345 | 1129 | RpL34a,Lsm12,CG11858,Nepl3,mts,mus81,RpS18,RpS14a,Atg6,ATPsynbeta,Uch-L5,RpL13A,Hat1,CG8112,csw,Vsp37A,CG1827,CG6707,CG12375,CG9684,cu,Fur2,alpha-Man-Ia,CG17746,ssh,CG32640,tws,Hsc70-3,RpL38,Marf,Mcad,SmE,CycC,Ptpmeg,Hexo1,Ggt-1,Topors,ste14,PAPLA1,COX5B,trr,Pdk1,CG10189,Parp16,Imp,Pxt,GlyP,mop,LysRS,Ppcdc,CG4860,Synj,CG10254,MAGE,CG13630,Pi4KIIalpha,wal,mael,Ublcp1,Rnmt,SP1029,GalT1,Trf2,Mtl,CG11899,Pym,CG30343,Idh3b,Xpac,Pink1,TrpRS-m,Pfk,CG9953,Cdk1,CG3842,tos,Gasz,Prosbeta5,ATPsynCF6,26-29-p,RpL13,tor,Pgk,Seipin,Uba5,CG6178,Got1,mus304,CG8314,escl,r-l,Spt20,p38a,Idgf6,sol,RpS30,RecQ5,CG12018,pen-2,Tao,RYBP,CG7009,RpL24,Ufm1,CG2200,Cul2,lt,Prosalpha3,mRpS23,mRpL10,blw,ben,Acox57D-p,mRpS11,rig,pps,Trf4-2,RpII140,Traf6,CG10336,CG2924,Herc4,RpLP1,UGP,heph,ATPsynB,snRNA:U4atac,mRpL18,CG4238,P5cr,TfIIFalpha,pont,Dcr-1,gus,Orc6,CG14817,Etf-QO,del,gw,EcR,Archease,mRpL16,ea,sip3,Mat89Ba,PIG-H,Exd2,MED4,fal,Usp10,Clp,eIF4A,Sply,Fkbp12,STUB1,P5CDh1,Prosbeta6,Shmt,UbcE2H,yps,Snm1,nSMase,CG33977,PEK,Cbl,Mitf,Nurf-38,CysRS,CG7920,Nap1,png,RpS5b,Rcd5,CG8199,CG31759,mRpL48,Uev1A,RpLP0,tapas,EloC,RpL12,Fkbp59,PpV,CG7718,Tlk,Cka,piwi,CG9643,PpD3,Cul5,foxo,nmdyn-D6,l(1)G0196,not,Kyat,Pus7,CG5003,Rpt2,RpS4,CG4267,Scsalpha1,NHP2,CG3587,Rpn13,SdhD,Neos,CG10344,obe,mRpS5,sbr,SERCA,rept,Fbw5,mtTFB1,CG31523,e(y)1,PIG-L,CG9941,CG10418,trbd,CG11110,CG8974,lid,mRpL35,IntS11,mus205,CG16979,snRNA:U4:38AB,Ubc6,Echs1,CG11134,CycJ,HDAC6,eIF4E1,CG5966,Utx,CG6650,SerRS-m,mRpS21,CG2911,CG4594,r,Aos1,Cul1,CG3631,Prosbeta4,Cbp80,RpS23,Gmer,MED24,RpL17,vret,AOX3,aay,Tengl4,Rpn12,CG7115,TyrRS,bru1,alph,Rbf,Pld,RpS29,LSm3,cN-IIIB,Rab5,CG17698,GatB,Arf51F,ATPsynG,Nc73EF,Irp-1B,ATPsyndelta,SRPK,Prosalpha5,CG5986,Naa20A,CG7840,Cp1,SkpA,Adi1,Edem1,nej,spen,Drep1,Ac3,dx,CG2493,CG8728,Bin1,Rrp40,Hnf4,scny,Rga,snRNA:U12,l(2)k01209,CG31053,esc,exu,oxt,RpL37a,CG9391,Orc1,CG7997,RpL23,Ady43A,tun,Pex10,CkIIalpha,Ptp10D,Ptpmeg2,RpS24,Gapdh1,QC,CG46338,DhpD,twin,CG14100,dco,Efr,lwr,egg,mEFTu2,ScsbetaA,CycB,Mos,gd,CG11070,Mekk1,Rpn2,Ubc4,CG1622,CG2938,tho2,Top1,Pex12,ACC,CanA-14F,RpL31,Pdk,bys,RpL29,Gyg,Pp2B-14D,Pi3K92E,CG2100,regucalcin,mRpL36,CG14435,Nmt,snk,Ugt36E1,Spx,AsnS,ATPsynO,PIG-C,Uch,RpL9,Iru,CG1578,GatA,nos,Gabat,RpS27,CG6610,Tak1,pygo,CG4586,RpL11,CanB2,Dhfr,RpS28b,fray,CG6766,velo,sws,mRpS10,AcCoAS,l(3)73Ah,CG17691,E2f1,Npc2a,scu,APP-BP1,SdhA,shrb,mEFTu1,ATPsynD,mRpL13,Prosbeta3,RpL10,CG16865,Larp7,stet,Rpn3,Sgf29,CG4598,RpL30,CG17768,IntS6,CG8311,Coq8,RpS2,Gk1,tko,DppIII,Ugt35A1,Ptip,eIF3m,Pfrx,CG14516,Cchl,CG33156,CG5285,CG7414,RpL34b,tkv,Idgf2,Rep,Taf7,Cdk5,CG8188,smg,CG9143,cathD,CG3198,mino,rl,CG9485,Tip60,RpS16,Ptp61F,Dph5,Usp39,Pgm1,cort,CG11089,SREBP,CG10932,ago,SelR,Ptp69D,armi,RpS14b,CG11771,Sld5,Idgf4,r2d2,awd,Sp7,Atg7,RpL7A,Edc3,nmdyn-D7,Pfas,CG7322,Dip-B,CG2034,ATPsynE,RfC38,CG5026,shu,RpL23A,Lkb1,O-fut1,d4,Ntan1,CG40045,CG3223,RpS15,eIF3i,CG17187,Pp2A-29B,Sirt7,Mtpalpha,CG6094,Tfb4,PIG-P,Chrac-14,CycE,kri,Cyp1,CG11486,adp,Cht11,CG7328,CG3021,RpS7,SelD,CG31633,CG5028,CG8726,CG17270,CG4041,grp,CG1774,Pomp,GAPsec,AGBE,Rpb8,grsm,Xpc,fdl,CG11164,Not10,CG7728,Usp16-45,CG5292,Treh,S2P,Plap,CG18135,CG2201,Sgf11,CG7789,CG32473,CG10425,trsn,ari-1,DNApol-epsilon255,Ts,Mal-A5,mRpL17,CG32221,CG6767,park,SNF4Agamma,AMPdeam,G9a,Eno,Desat1,GCS2beta,CG5390,CG9316,CG8204,mal,CG8475,Son,eEF1beta,Ser7,RpS13,Dd,CG10463,Pglym78,otu,Gk2,CG4572,EMC3,GluRS-m,Prosalpha4,mre11,Mo25,CG5880,Cdk7,yip2,dob,Agpat2,CG13344,CG17333,CG4407,CSN6,Atu,Drep2,CG17454,isoQC,Sccpdh1,RpL40,CG12301,faf,Thiolase,CG3511,YL-1,c(2)M,RpL26,ste24a,nbs,agt,Nadsyn,PyK,twe,wisp,RpS27A | | 2.759e-26 | -58.85 | mitochondrion | cellular component | GO:0005739 | 800 | 169 | 11940 | 1105 | CG11077,Cyt-c-p,Coq8,CG40472,mRpL35,Trxr-1,UQCR-11,cype,CG7920,Idh3b,Naxe,mtTFB1,CG9393,CG14407,Gabat,ATPsynD,Mcad,CG8004,Gasz,PHGPx,ND-B15,Thiolase,mRpL10,AIF,CG9399,CG11562,CG1640,ATPsynO,P5CDh1,COX5A,Cchl,GatA,tko,nmdyn-D6,Sccpdh1,CG10932,Roe1,SerRS-m,Got1,whd,milt,Gk2,ND-30,COX5B,scu,Shmt,Pmi,Marf,CG4882,mino,Tspo,ATPsynF,TrpRS-m,AGO3,CG7488,CG13551,mEFTu2,Ts,CG17691,COX8,Ppox,Ptpmeg,ND-18,mEFTu1,levy,CG14817,mRpL36,ND-B22,yip2,pr,Tim8,mRpL18,mRpS21,Nc73EF,CG4598,Sdhaf3,mRpS23,isoQC,blw,mRpL13,bor,ND-MNLL,mRpL16,Exd2,ATPsynB,CG6094,Scox,Phb2,ND-23,UQCR-Q,Idh,ND-MWFE,CG42375,Pdk,CG12375,GatB,CG4860,wal,Gk1,CG5903,CG13601,EMRE,Ptp61F,spd-2,mRpS10,CG3803,Scsalpha1,NP15.6,GluRS-m,ND-51,SdhA,CG4594,Sfxn2,ATPsynG,CG1774,ScsbetaA,pn,nmdyn-D7,mRpL48,sea,mRpS5,Aldh,Neb-cGP,CG14894,LysRS,Tengl4,Echs1,ND-39,Rab32,Nap1,Tom40,ATPsynE,ND-24,Dhfr,Coq6,mRpS11,CG8199,Coq7,CG3021,ND-75,COX4,ATPsynbeta,CG11267,CG11110,Debcl,Tom70,ND-B18,mRpL17,ND-SGDH,CG8728,Kyat,ND-49,UK114,Mitofilin,Irp-1B,ATPsyndelta,Mtpalpha,colt,CG42376,Etf-QO,park,armi,ATPsynCF6,CG31759,Pink1,SdhD,CG5028,larp,shop | | 6.530e-21 | -46.48 | nitrogen compound metabolic process | biological process | GO:0006807 | 4233 | 532 | 12345 | 1129 | RpL23A,Lkb1,O-fut1,d4,Ntan1,CG3223,CG40045,shop,RpS15,eIF3i,CG17187,Pp2A-29B,Gclc,Sirt7,CG6094,Tfb4,PIG-P,sgll,Chrac-14,CycE,kri,UK114,Cyp1,CG11486,Cht11,CG3021,RpS7,SelD,CG31633,CG8726,CG4041,CG11089,cort,ago,CG11771,Ptp69D,RpS14b,SelR,armi,Sld5,Idgf4,r2d2,awd,Sp7,Atg7,RpL7A,Edc3,nmdyn-D7,Pfas,Dip-B,CG2034,ATPsynE,RfC38,CG5026,shu,eEF1beta,RpS13,Ser7,GstT1,Dd,CG10463,Pglym78,otu,CG4572,EMC3,Prosalpha4,GluRS-m,Ppox,Mo25,mre11,CG5880,Cdk7,CG13344,CSN6,CG4407,Atu,Drep2,CG17454,isoQC,RpL40,CG12301,faf,YL-1,CG3511,c(2)M,RpL26,ste24a,pn,nbs,agt,Nadsyn,PyK,twe,RpS27A,wisp,grp,Pomp,CG1774,GAPsec,Rpb8,grsm,Xpc,fdl,CG11164,Not10,Usp16-45,CG7728,CG5292,S2P,Plap,CG2201,Sgf11,CG32473,CG10425,trsn,ari-1,DNApol-epsilon255,Ts,mRpL17,CG32221,CG6767,park,whd,AMPdeam,G9a,Eno,CG9316,CG5390,GCS2beta,mal,CG8204,Son,tho2,Top1,Pex12,CanA-14F,RpL31,Pdk,bys,RpL29,Pp2B-14D,CG2100,mRpL36,CG14435,Nmt,snk,Ugt36E1,Spx,AsnS,ATPsynO,scny,Rga,snRNA:U12,l(2)k01209,CG31053,esc,RpL37a,oxt,exu,Orc1,CG7997,RpL23,Ady43A,Pex10,tun,CkIIalpha,Ptp10D,Ho,Ptpmeg2,RpS24,QC,Gapdh1,DhpD,CG46338,twin,CG14100,dco,Efr,egg,lwr,mEFTu2,ScsbetaA,CycB,Mos,gd,CG11070,Mekk1,Rpn2,Pdxk,Ubc4,CG1622,Sgf29,RpL30,IntS6,CG17768,Coq8,RpS2,tko,Ugt35A1,DppIII,eIF3m,Ptip,Pfrx,CG14516,Cchl,CG33156,CG7414,CG5285,RpL34b,Idgf2,tkv,Rep,Taf7,Cdk5,CG8188,CG9143,smg,cathD,CG3198,mino,rl,Tip60,RpS16,Ptp61F,Dph5,Usp39,Uch,PIG-C,RpL9,Iru,CG1578,GatA,nos,Gabat,RpS27,pygo,CG6610,Tak1,RpL11,CanB2,Dhfr,RpS28b,fray,CG6766,velo,sws,mRpS10,AcCoAS,l(3)73Ah,CG17691,E2f1,scu,Pcd,APP-BP1,shrb,mEFTu1,ATPsynD,mRpL13,Prosbeta3,RpL10,CG16865,Larp7,stet,Rpn3,Tlk,Cka,pr,piwi,CG9643,Sptr,PpD3,GstE11,Cul5,nmdyn-D6,not,Kyat,Pus7,CG5003,Rpt2,RpS4,Scsalpha1,NHP2,CG3587,Rpn13,Neos,CG10344,mRpS5,obe,sbr,rept,mtTFB1,Fbw5,PIG-L,e(y)1,CG31523,CG9941,CG11110,trbd,CG10418,lid,CG8974,mRpL35,ea,sip3,Mat89Ba,PIG-H,Exd2,MED4,fal,Usp10,Clp,eIF4A,Fkbp12,STUB1,Sply,P5CDh1,Prosbeta6,Shmt,UbcE2H,yps,Snm1,nSMase,CG33977,PEK,Cbl,CysRS,Nurf-38,Nap1,CG7920,png,RpS5b,CG8199,Rcd5,CG31759,CG13551,mRpL48,Uev1A,RpLP0,tapas,EloC,RpL12,PpV,Fkbp59,bru1,alph,Rbf,RpS29,LSm3,cN-IIIB,Rab5,CG17698,GatB,Arf51F,ATPsynG,ATPsyndelta,SRPK,Prosalpha5,CG5986,Naa20A,CG7840,SkpA,Cp1,Adi1,Edem1,nej,spen,Drep1,Ac3,dx,CG8728,CG2493,Bin1,Rrp40,IntS11,mus205,CG16979,snRNA:U4:38AB,Ubc6,CycJ,CG11134,HDAC6,eIF4E1,Utx,SerRS-m,mRpS21,CG2911,r,Aos1,Cul1,CG3631,Prosbeta4,Cbp80,Gmer,RpS23,MED24,vret,RpL17,AOX3,aay,Tengl4,CG7115,Rpn12,TyrRS,Ptpmeg,Hexo1,Ggt-1,Topors,ste14,PAPLA1,COX5B,trr,Pdk1,CG10189,Parp16,Imp,mop,LysRS,Ppcdc,CG10254,MAGE,CG13630,mael,Rnmt,Ublcp1,SP1029,Trf2,GalT1,Mtl,CG11899,Pym,CG30343,Xpac,Pink1,TrpRS-m,Pfk,RpL34a,Nepl3,Lsm12,CG11858,mts,mus81,RpS18,RpS14a,ATPsynbeta,Uch-L5,RpL13A,Hat1,csw,Vsp37A,CG1827,CG12375,CG9684,cu,Fur2,alpha-Man-Ia,CG17746,ssh,CG32640,tws,Hsc70-3,RpL38,SmE,CycC,mRpS11,CG3803,rig,pps,Trf4-2,RpII140,Traf6,CG10336,CG2924,Herc4,RpLP1,UGP,heph,ATPsynB,snRNA:U4atac,P5cr,CG4238,mRpL18,TfIIFalpha,PGRP-LE,pont,Dcr-1,Orc6,gus,CG14817,del,gw,Archease,mRpL16,CG9953,Cdk1,tos,Gasz,Prosbeta5,ATPsynCF6,GstT2,tor,26-29-p,RpL13,Pgk,Uba5,CG6178,Got1,mus304,CG8314,escl,r-l,CG10561,Spt20,p38a,sol,Idgf6,RecQ5,RpS30,CG12018,pen-2,Tao,RYBP,CG7009,Ufm1,RpL24,CG2200,Cul2,lt,Prosalpha3,mRpL10,mRpS23,blw,ben | | 2.707e-20 | -45.06 | phosphorus metabolic process | biological process | GO:0006793 | 920 | 170 | 12345 | 1129 | PIG-C,grp,Cdk1,CG1774,ATPsynCF6,tor,ND-B15,Pgk,UQCR-Q,Tak1,CycJ,CG11164,CanB2,Seipin,CG6178,fray,sws,AcCoAS,CG18135,CG2201,r,r-l,CG7789,CG3631,CG17691,p38a,ND-51,scu,Gmer,Ts,SdhA,CG6767,Tao,aay,AMPdeam,ATPsynD,Eno,mal,CG7115,blw,CG17294,COX4,Dd,Pglym78,ND-75,Gk2,CG8311,alph,Coq8,Pld,Mo25,Gk1,Cdk7,cN-IIIB,Agpat2,CG17698,UGP,Ugt35A1,CG17333,CG4407,COX5A,UQCR-11,ATPsynB,ATPsynG,Pfrx,ATPsyndelta,CG33156,SRPK,tkv,Cdk5,ND-23,ND-B18,mino,rl,Ac3,pn,Nadsyn,Etf-QO,Cyt-c-p,PyK,Ptp61F,twe,Acph-1,ND-SGDH,CG11089,l(2)k01209,ND-24,PIG-H,mts,fal,CG9391,Ptp69D,Ady43A,CkIIalpha,Ptp10D,ATPsynbeta,Ptpmeg2,ND-49,awd,csw,Gapdh1,nSMase,nmdyn-D7,CG33977,CG6707,dco,PEK,Pfas,ScsbetaA,Nurf-38,CG7920,CycB,png,Mos,levy,CG17746,ssh,CG8199,ATPsynE,tws,Mekk1,CG5026,Pdxk,PpV,CG7718,Tlk,Cka,Lkb1,Ptpmeg,PpD3,COX5B,Pdk1,nmdyn-D6,l(1)G0196,Argk,Pp2A-29B,mop,Ppcdc,Tfb4,CG1344,CanA-14F,Scsalpha1,Synj,sgll,PIG-P,SdhD,cype,CycE,Pi4KIIalpha,Pdk,kri,wal,Ublcp1,ND-MNLL,Pmi,Mtl,CG11486,Pp2B-14D,Pi3K92E,CG7328,AIF,PIG-L,ND-B22,ND-39,Pink1,Ugt36E1,CG8726,Pfk,CG4041,ATPsynO | | 1.116e-19 | -43.64 | catabolic process | biological process | GO:0009056 | 901 | 166 | 12345 | 1129 | CG32221,park,ari-1,shrb,CG10561,Cbp80,Idgf6,Cul1,CG32473,Prosbeta4,CG17691,Rpn3,lt,CG9316,Rpn12,Prosalpha3,Cul2,Eno,Prosbeta3,Tengl4,SNF4Agamma,AOX3,RYBP,Ubc6,fdl,26-29-p,Echs1,Pgk,Gabat,CG4586,Prosbeta5,Pomp,Uch,Plap,CG6766,Atg4b,CG18135,CG4594,Treh,Vamp7,Seipin,Usp16-45,CG5966,CG11164,Not10,Dcr-1,Drep1,Jafrac1,CG9485,Edem1,CG8188,faf,Thiolase,smg,cathD,CG7840,RpL40,Cp1,PGRP-LE,SkpA,Prosalpha5,Fdh,Idgf2,RpS27A,Debcl,Pgm1,PyK,gw,EcR,Hnf4,Etf-QO,Rrp40,gus,pn,LSm3,CG4598,CG17768,CG4572,Trf4-2,Prosalpha4,Acox57D-p,Pglym78,CG14516,CG4238,Pfrx,raptor,Herc4,yip2,cN-IIIB,dob,Mitf,Cbl,cu,Shmt,Vsp37A,DhpD,UbcE2H,Edc3,twin,EloC,Rpn2,Mcad,CG42554,CG11070,Hsc70-3,CG8199,CG32640,CG31759,fal,Usp10,CG9391,ago,Jafrac2,CG10932,mts,sip3,cort,scny,Rga,Atg7,Prosbeta6,P5CDh1,Gapdh1,Sply,Uch-L5,Atg6,Idgf4,Ho,CG7997,armi,Pex10,Fbw5,CG11486,Cht11,Pi3K92E,wal,UK114,SERCA,SP1029,tzn,Pfk,Pink1,CG8974,Pym,Cyp6v1,CG31633,trbd,not,Ntan1,Cul5,CG40045,CG3223,Ggt-1,Topors,Rpn13,Synj,CG10254,Adh,CG4390,CG4860,CG4267,Pp2A-29B,GlyP,CG5003,Rpt2,Mtpalpha | | 1.237e-19 | -43.54 | cellular catabolic process | biological process | GO:0044248 | 764 | 148 | 12345 | 1129 | Cul5,Ntan1,CG3223,CG40045,not,Ggt-1,Topors,CG10254,Rpn13,CG5003,GlyP,Pp2A-29B,Rpt2,Mtpalpha,CG4390,CG4860,Pi3K92E,Cht11,CG11486,Fbw5,wal,SERCA,SP1029,tzn,Pym,Cyp6v1,CG31633,trbd,Pink1,CG8974,CG10932,Jafrac2,mts,fal,ago,Usp10,Rga,scny,sip3,cort,Uch-L5,Sply,Atg7,P5CDh1,Prosbeta6,armi,CG7997,Pex10,Atg6,Ho,Idgf4,Cbl,cu,Shmt,twin,Edc3,DhpD,Vsp37A,UbcE2H,EloC,Mcad,Rpn2,CG32640,CG31759,CG8199,CG11070,Hsc70-3,CG42554,LSm3,Acox57D-p,CG4572,CG4598,CG17768,Prosalpha4,Trf4-2,CG14516,CG4238,cN-IIIB,yip2,Herc4,dob,raptor,faf,CG8188,Edem1,Thiolase,smg,Drep1,Dcr-1,CG9485,Jafrac1,Prosalpha5,Idgf2,Fdh,CG7840,SkpA,Cp1,RpL40,gw,EcR,Debcl,RpS27A,pn,gus,Rrp40,Etf-QO,Prosbeta5,fdl,26-29-p,Ubc6,CG4586,Gabat,Echs1,Uch,Pomp,Treh,Atg4b,Plap,CG6766,CG4594,CG18135,CG11164,Not10,Seipin,Vamp7,Usp16-45,ari-1,shrb,park,CG32221,Cul1,Prosbeta4,CG17691,CG32473,Cbp80,CG10561,Idgf6,CG9316,lt,Rpn12,Prosalpha3,Rpn3,SNF4Agamma,RYBP,AOX3,Tengl4,Prosbeta3,Cul2 | | 2.453e-19 | -42.85 | phosphate-containing compound metabolic process | biological process | GO:0006796 | 900 | 165 | 12345 | 1129 | scu,p38a,ND-51,CG3631,CG17691,r-l,CG7789,CG6767,Ts,SdhA,Eno,ATPsynD,AMPdeam,aay,Tao,CG17294,blw,CG7115,mal,CG1774,Cdk1,PIG-C,grp,Pgk,UQCR-Q,CycJ,Tak1,ND-B15,tor,ATPsynCF6,CG6178,Seipin,CG11164,CanB2,AcCoAS,CG18135,r,CG2201,sws,fray,tkv,SRPK,ND-B18,mino,rl,ND-23,Cdk5,Etf-QO,Nadsyn,Ac3,Acph-1,twe,Cyt-c-p,PyK,Ptp61F,alph,CG8311,ND-75,Pglym78,Gk2,COX4,Dd,Gk1,Mo25,Pld,Coq8,CG17333,CG4407,CG17698,Agpat2,Cdk7,cN-IIIB,CG33156,ATPsyndelta,UQCR-11,ATPsynB,Pfrx,ATPsynG,COX5A,PEK,Pfas,nmdyn-D7,nSMase,CG6707,dco,CG33977,CG7920,CycB,ScsbetaA,Nurf-38,tws,ATPsynE,ssh,CG8199,png,Mos,CG17746,levy,CG7718,Pdxk,PpV,CG5026,Mekk1,l(2)k01209,PIG-H,ND-24,CG11089,ND-SGDH,CG9391,fal,mts,Ptp10D,ATPsynbeta,Ptpmeg2,ND-49,CkIIalpha,Ady43A,Ptp69D,csw,Gapdh1,awd,Pmi,Ublcp1,ND-MNLL,wal,PIG-L,CG7328,AIF,CG11486,Pp2B-14D,Pi3K92E,Mtl,Pink1,ND-39,ND-B22,ATPsynO,Pfk,CG4041,CG8726,PpD3,Ptpmeg,Lkb1,Tlk,Cka,Argk,Pdk1,nmdyn-D6,l(1)G0196,COX5B,CG1344,Scsalpha1,CanA-14F,Ppcdc,Tfb4,Pp2A-29B,mop,Pi4KIIalpha,Pdk,kri,SdhD,cype,CycE,sgll,PIG-P,Synj | | 1.495e-18 | -41.04 | organic substance catabolic process | biological process | GO:1901575 | 807 | 151 | 12345 | 1129 | CG10932,fal,Usp10,CG9391,ago,scny,Rga,sip3,cort,Sply,Uch-L5,Prosbeta6,P5CDh1,Gapdh1,CG7997,armi,Pex10,Atg6,Idgf4,Ho,Cbl,cu,Mitf,Shmt,Vsp37A,DhpD,UbcE2H,Edc3,twin,EloC,Mcad,Rpn2,CG8199,CG32640,CG31759,Hsc70-3,CG11070,Ntan1,Cul5,CG40045,CG3223,not,Ggt-1,Topors,Synj,Rpn13,CG10254,CG5003,GlyP,Rpt2,Mtpalpha,Adh,CG4390,CG4860,CG4267,CG11486,Cht11,Fbw5,wal,UK114,SP1029,SERCA,tzn,Pfk,Pym,trbd,CG31633,Pink1,CG8974,Prosbeta5,Ubc6,fdl,26-29-p,Pgk,Echs1,Gabat,CG4586,Uch,Pomp,Treh,CG6766,Plap,CG18135,CG4594,CG11164,Not10,Seipin,CG5966,Usp16-45,ari-1,shrb,CG32221,park,Cul1,CG32473,CG17691,Prosbeta4,CG10561,Cbp80,Idgf6,CG9316,Prosalpha3,Rpn12,Rpn3,AOX3,RYBP,Cul2,Eno,Prosbeta3,Tengl4,LSm3,Acox57D-p,Pglym78,CG17768,CG4598,CG4572,Trf4-2,Prosalpha4,Pfrx,CG14516,CG4238,Herc4,cN-IIIB,yip2,dob,Edem1,CG8188,faf,smg,Thiolase,cathD,Dcr-1,Drep1,CG9485,Prosalpha5,Idgf2,Fdh,CG7840,RpL40,Cp1,SkpA,PGRP-LE,PyK,gw,Hnf4,RpS27A,Pgm1,gus,pn,Etf-QO,Rrp40 | | 2.052e-18 | -40.73 | cellular macromolecule metabolic process | biological process | GO:0044260 | 2688 | 366 | 12345 | 1129 | MAGE,RpL31,CG10254,Pdk,ACC,LysRS,mop,Pex12,GlyP,CanA-14F,trr,PAPLA1,Top1,ste14,Parp16,Pdk1,Topors,Hexo1,Ptpmeg,Ggt-1,Xpac,Pym,mRpL36,CG14435,TrpRS-m,Nmt,Pink1,Pp2B-14D,Mtl,Ublcp1,Rnmt,Gyg,Trf2,GalT1,RpL29,RpL13A,Uch-L5,Gapdh1,QC,csw,Hat1,RpS24,Pex10,tun,RpL23,RpS18,Ptpmeg2,Ptp10D,Atg6,RpS14a,CkIIalpha,esc,mts,mus81,CG11858,Orc1,oxt,RpL37a,Rga,scny,CG31053,RpL34a,Rpn2,RpL38,Mekk1,Ubc4,CG32640,ssh,Mos,CG17746,Hsc70-3,CG11070,tws,cu,egg,lwr,Efr,CycB,alpha-Man-Ia,mEFTu2,CG1827,twin,CG46338,Vsp37A,dco,CG14100,eIF3m,Ptip,mRpL18,Cchl,CG4238,tko,Herc4,RpLP1,UGP,CG2924,CG10336,Coq8,Traf6,RpS2,Sgf29,mRpS11,RpII140,pps,Trf4-2,RpL30,CG17768,Dph5,gw,Ptp61F,Pgm1,mRpL16,Usp39,CG14817,gus,Orc6,Tip60,RpS16,pont,smg,Cdk5,CG8188,CG9485,rl,Dcr-1,tkv,RpL34b,CG5285,CG7414,Taf7,Rep,TfIIFalpha,fray,RpS28b,l(3)73Ah,mRpS10,escl,CG8314,velo,CG6766,CanB2,RpL11,mus304,Uba5,CG1578,Prosbeta5,pygo,Tak1,RpS27,tor,RpL13,26-29-p,GatA,Cdk1,Iru,RpL9,Uch,PIG-C,tos,Prosalpha3,RpL10,lt,ben,Rpn3,mRpL10,mRpS23,CG7009,RYBP,Tao,Prosbeta3,mRpL13,Cul2,RpL24,Ufm1,RecQ5,RpS30,shrb,pen-2,CG12018,mEFTu1,APP-BP1,p38a,E2f1,Spt20,CycE,Rpn13,PIG-P,Chrac-14,Sirt7,Rpt2,CG5003,Pp2A-29B,RpS4,Tfb4,CG6094,CG40045,CG3223,Cul5,foxo,Ntan1,eIF3i,RpS15,not,O-fut1,Lkb1,Cka,RpL23A,Tlk,PpD3,d4,CG9643,CG8726,mRpL35,CG4041,trbd,CG11110,CG31633,CG9941,RpS7,CG8974,lid,CG11486,rept,e(y)1,PIG-L,mtTFB1,Fbw5,Cyp1,mRpS5,obe,CG10344,awd,Fkbp12,STUB1,Prosbeta6,Atg7,SelR,RpS14b,armi,Ptp69D,Sld5,MED4,Exd2,ago,eIF4A,Usp10,PIG-H,sip3,cort,CG5026,RpL12,EloC,Fkbp59,PpV,CG31759,Rcd5,RpS5b,png,RfC38,RpLP0,mRpL48,Uev1A,Cbl,Nap1,CysRS,Nurf-38,Edc3,UbcE2H,RpL7A,PEK,CG33977,fs(1)Ya,Snm1,Arf51F,GatB,isoQC,Atu,Rab5,Cdk7,CSN6,CG13344,CG17698,Rbf,CG5880,LSm3,Mo25,mre11,RpS29,otu,Dd,RpS13,eEF1beta,Prosalpha4,alph,GluRS-m,EMC3,CG4572,twe,RpS27A,ste24a,RpL26,Rrp40,agt,CG8728,nbs,dx,c(2)M,CG3511,YL-1,nej,faf,Edem1,spen,Drep1,SuUR,Naa20A,Prosalpha5,SRPK,SkpA,Cp1,RpL40,CG7840,SerRS-m,S2P,CG2911,mRpS21,Plap,Not10,CG11164,Utx,eIF4E1,Usp16-45,HDAC6,Rpb8,AGBE,CycJ,fdl,Xpc,Ubc6,grp,mus205,GAPsec,Pomp,TyrRS,CG8475,CG7115,Rpn12,mal,CG9316,GCS2beta,Tengl4,G9a,mRpL17,ari-1,DNApol-epsilon255,RpL17,MED24,park,CG32221,Prosbeta4,CG3631,Cul1,Sgf11,Aos1,RpS23,Cbp80 | | 4.859e-18 | -39.87 | mitochondrial part | cellular component | GO:0044429 | 480 | 106 | 11940 | 1105 | UQCR-Q,ND-23,cype,Scox,ATPsynB,CG6094,UQCR-11,mRpL16,CG40472,ND-MNLL,mRpL35,Coq8,Cyt-c-p,CG11077,CG8004,CG5903,wal,Pdk,Gabat,ATPsynD,CG12375,ND-MWFE,CG9393,mtTFB1,CG14407,Idh3b,ATPsynG,P5CDh1,COX5A,ND-51,SdhA,Scsalpha1,ATPsynO,NP15.6,mRpL10,Thiolase,ND-B15,Ptp61F,AIF,CG3803,mRpS10,CG9399,EMRE,Roe1,Aldh,Neb-cGP,mRpS5,mRpL48,pn,tko,Cchl,mRpS11,ATPsynE,Marf,ND-24,Pmi,scu,Tom40,ND-39,COX5B,Tengl4,Echs1,ND-30,Got1,CG11110,CG7488,CG11267,TrpRS-m,ATPsynbeta,COX4,ATPsynF,ND-75,Tspo,CG8199,mino,Coq7,ND-B22,colt,ATPsyndelta,mRpL36,Mtpalpha,Mitofilin,ND-49,CG14817,levy,CG8728,ND-SGDH,ND-18,CG17691,Debcl,Tom70,ND-B18,Ppox,mRpL17,shop,mRpL13,SdhD,blw,mRpS23,Sdhaf3,ATPsynCF6,mRpL18,mRpS21,Nc73EF,Etf-QO,Tim8 | | 1.250e-17 | -38.92 | binding | molecular function | GO:0005488 | 5743 | 663 | 12176 | 1119 | Arf51F,Tak1,MICAL-like,Cdk7,Kyat,Cbp80,cact,RpL26,RpL9,CycB,RpL7A,dl,spz,Pdk1,Gabat,stai,mRpL16,CG10654,Pex23,CG17260,RpS15,Moe,Gdi,Grip128,Bin1,boi,Pex12,Trf2,CG14997,CG7115,Debcl,Nxf3,RpL12,yl,STUB1,CG31729,Top1,SelR,Zwilch,Cdk1,Sry-alpha,Rbfox1,psq,insv,egg,mus81,CG12018,CG17454,Hsc70-3,CG1640,pre-lola-G,CG6610,spen,bun,glob1,mRpL13,CG9925,RpLP1,Ubc6,rig,CG15514,jigr1,sbr,CG9981,CG7716,RYBP,ND-51,grsm,PyK,CG14516,Crag,Hip1,Mcad,CG5734,CG31875,Nlg1,CG11710,grp,mael,brwl,Orc6,Got1,Pfdn2,r,Rbf2,CG4860,ben,CG30496,Abi,Pura,Irp-1B,Grip84,Scox,mre11,CG17187,EcR,Stip1,CG31689,tos,RpL24,eIF3m,HDAC6,Afti,Idgf6,SREBP,SERCA,CG2162,alc,mRpL18,sgll,armi,Pld,cnc,aay,ND-75,CG4586,nej,e(y)1,AIF,Exo70,nmd,Mekk1,CG8507,Npc2a,CG11267,CG10188,Cdk5,ATPsynbeta,CG44422,twin,RpL29,CG7728,Rga,dx,obe,Klp10A,dco,Nup58,eEF1beta,bam,CG14435,Pink1,regucalcin,E23,Scp1,CG8974,pygo,CG6094,cN-IIIB,crok,wal,shrb,piwi,CG34116,CanA-14F,cib,CG9581,elg1,Cul1,bcd,Uev1A,Eno,CG1890,Atf6,CG32473,CG10631,faf,crm,Ptp61F,SNF4Agamma,alpha-Man-Ia,Neos,snz,stet,CG8475,Utx,nrv1,Arpc4,Rpn13,CG9143,ND-49,Traf6,blw,RpS2,Myd88,Idgf4,awd,AP-1-2beta,Cf2,GluRS-m,CG12288,CG10189,Larp4B,X11L,Snx6,CG11807,Rpt2,wash,Nprl2,Kap3,zf30C,l(3)73Ah,Nup62,G9a,exu,sip3,Usp39,enc,CG7414,CG10492,shop,CG7806,hwt,Ranbp21,Smox,DnaJ-H,Nf-YA,HP1c,Gmap,AnxB10,tor,ari-1,Pfk,Mtl,Sec22,ND-23,tef,Acox57D-p,MED27,kl-2,yem,CG10055,l(3)mbt,SmydA-1,Sec15,CG3223,RpS5b,l(1)G0196,PPYR1,Blos4,CG7488,Pfdn5,CG4404,gammaTub37C,Nadsyn,Hnf4,RpL30,CG8726,RecQ5,Gs1l,Ork1,CG11902,CG5986,nbs,l(2)k01209,CG17746,CG12375,LSm3,Cka,Wbp2,CG17272,swa,CAH7,E2f1,Pex10,AGBE,CG2678,krimp,Exd2,CG13551,ScsbetaA,CG10336,CG1416,l(1)G0004,CG3077,CG14100,Cht11,RpS16,DhpD,Elba2,Rbf,tral,Iru,escl,Grip75,isoQC,CG8728,SdhD,Mat89Ba,Mctp,Arpc2,msps,mol,Liprin-beta,RpS14a,Rpn3,Imp,UQCR-Q,AGO3,Fer1HCH,Cen,Rab5,CG5116,nmdyn-D7,MED20,cnn,Sply,CSN6,CG31053,CG9684,PGRP-LE,Dcr-1,cyc,CycE,Nup358,ASPP,ovo,Rrp40,Cyt-c-p,bif,BigH1,HP1e,NFAT,CG10254,Sirt7,PDCD-5,CG17768,Edc3,shep,Rcd5,Tengl4,Vamp7,Sod3,RpS29,Synj,CG6967,Bap170,hng3,Dhfr,Arpc1,cup,PrBP,fs(1)N,Caf1-105,Kcmf1,Mitf,Grip71,CG4562,Taf7,CG32640,Pka-R2,RpL34a,mus205,Ect4,O-fut1,ago,SdhA,CG11590,CG17270,CCT5,TBCC,CCT6,d4,mRpS11,Arl5,Syx13,CG4021,capt,Ssdp,mRpS5,CG5335,CG17698,CycJ,Larp7,SRPK,Bx,CG14671,Cse1,ATP8A,png,anne,CG2258,SerRS-m,CG5789,Ptpmeg,CG10038,Roe1,rept,sbb,ix,Rab32,Sod1,Smyd4-4,rl,MtnA,QC,Karybeta3,Spx,asf1,NHP2,Cyp6v1,CG6276,Arts,Gclc,Fbw5,Gga,Nap1,mbo,r2d2,lwr,Fdh,CG6767,Calx,parvin,Pus7,Atf3,Mitofilin,CG4041,CG4853,CG5510,RpL13A,AcCoAS,Idh,TfIIFalpha,CG18135,alphaTub67C,Clp,SuUR,Ankle2,Gmer,csw,Idgf2,mEFTu2,TrpRS-m,Marf1,vret,ph-d,Girdin,SP1029,Cul2,Tm1,Actn,Hat1,CG4557,Non1,Son,LpR2,Pfrx,cort,scny,TyrRS,SelD,CCT3,CG3309,CG10463,RpS27A,Ubc4,stw,Neb-cGP,Shmt,Sgf11,Wdfy2,olf186-M,CG17328,Cyp6a19,shv,Marc,Adh,eIF4E1,RpS4,Stam,Trxr-1,CAH2,Xpc,CanB2,Jafrac2,nos,yps,Vps16B,RpS18,CG2100,CG32485,Mybbp1A,bys,Ho,Cul5,Rel,Gapdh1,lt,CG8963,Pgk,alph,Coq6,CG6813,DNApol-epsilon255,hb,spag,Tlk,MED24,CycC,CG4872,AOX3,Tip60,cad,CG8611,Sec23,RfC38,RpL11,RpS14b,CG3226,CG3198,tkv,Unr,tho2,raptor,Rnmt,Topors,ACC,Med,mad2,tub,CG31674,Fkbp59,CG2199,Uxt,cbs,CG4730,RpS13,eIF3i,gw,Chrac-14,Stat92E,Sec10,fray,CG10418,CG2129,Svil,ABCD,milt,fz4,bru1,CG17593,RpL40,Cbl,CG42748,Sgf29,CG11134,Mos,Maf1,del,Nca,CG4709,tyf,PAPLA1,APP-BP1,CG1602,p38a,pont,Chd64,galectin,smg,RpLP0,wisp,sau,park,Acph-1,Blos2,Atpalpha,Atu,Cyp1,Rpn2,CG9941,Cisd2,SkpA,Edem1,ND-39,foxo,REPTOR,trsn,Pxt,Hsp26,wdn,Rep,GAPsec,RpL23,RhoGDI,not,qkr58E-2,Reps,spn-E,Dip-B,trk,trr,Nc73EF,apt,mEFTu1,trbd,CG9425,Cyp6a22,Nurf-38,mal,HmgD,ssh,Tao,Rab10,CkIIalpha,sws,Fkbp39,Usp16-45,BicC,Vha68-2,CG15141,gus,Cyp6u1,Pym,CG4880,fs(1)Ya,CG14183,MED4,CG10959,Lkb1,Marf,Desat1,Clc,osk,heph,Aos1,Scsalpha1,lin,CG4266,CG13344,RpII140,CG11486,Wnt10,Kap-alpha3,Xpac,Plap,LysRS,Hr78,SLIRP1,PEK,Su(H),lid,Tomosyn,blot,Parp16,eIF4A,CysRS,Pgm1,up,Pp2B-14D,alphaTub84B,CG6923,cana,GCC88,GlyP,Droj2,esc,bor,Snm1,pps,Orc1,sw | | 1.267e-16 | -36.60 | cytoplasmic translation | biological process | GO:0002181 | 120 | 44 | 12345 | 1129 | RpL23A,RpS13,RpL9,RpL30,RpL34a,RpL37a,RpS2,RpS15,RpS29,RpL13,RpS27,eIF4A,RpLP1,RpL11,RpS14b,RpS18,RpL23,RpS14a,eIF4E1,RpS4,eIF3m,RpS28b,RpL31,RpL13A,RpS24,RpL7A,RpL34b,RpL29,RpL40,RpS23,RpS30,RpL17,RpS7,Pym,RpL26,RpS5b,CG7009,RpL24,RpS16,RpLP0,RpL38,RpL10,RpL12,RpS27A | | 3.911e-16 | -35.48 | ribosomal subunit | cellular component | GO:0044391 | 180 | 55 | 11940 | 1105 | RpL7A,RpL24,RpS28b,RpL11,RpL13A,mRpL35,RpL26,RpL37a,RpL13,RpS24,RpS14b,RpS14a,RpS27A,mRpS11,RpL9,mRpL16,RpL29,RpL38,RpS18,CG6094,RpL12,RpL23A,mRpL48,mRpS21,RpL34b,mRpL18,RpL23,RpS16,RpS27,tko,RpL30,RpS7,RpLP0,mRpL13,RpL10,mRpS5,RpS30,mRpS23,RpS29,RpS15,mRpS10,RpLP1,RpL34a,mRpL10,RpS13,CG7414,mRpL17,RpS5b,RpS4,RpL31,RpS2,RpL17,mRpL36,RpL40,RpS23 | | 4.529e-16 | -35.33 | generation of precursor metabolites and energy | biological process | GO:0006091 | 217 | 61 | 12345 | 1129 | Fdh,ND-MNLL,wal,Gyg,Pmi,ND-51,ND-23,SdhA,CG9485,ND-B18,AIF,ScsbetaA,Idh3b,ND-39,AOX3,CG14407,levy,ND-B22,Trxr-1,Eno,Etf-QO,CG5028,CG8475,PyK,Cyt-c-p,ATPsynO,Pgm1,blw,Pfk,Pglym78,ND-75,ND-SGDH,COX4,ND-24,ND-18,COX5B,foxo,AGBE,Coq8,UQCR-Q,Pgk,ND-B15,ACC,GlyP,Scsalpha1,ND-49,CG17333,ATPsynbeta,Atg6,UGP,Adh,Pfrx,Nc73EF,SdhD,UQCR-11,cype,Grx1,COX5A,Gapdh1,NP15.6,Irp-1B | | 5.687e-16 | -35.10 | mitochondrial protein complex | cellular component | GO:0098798 | 270 | 70 | 11940 | 1105 | ATPsynD,CG9393,ND-MWFE,CG5903,CG8004,wal,mRpL35,CG40472,ND-MNLL,cype,ND-23,UQCR-Q,UQCR-11,ATPsynB,CG6094,mRpL16,mRpL48,tko,Roe1,Neb-cGP,mRpS5,mRpS10,mRpL10,Thiolase,ND-B15,EMRE,ATPsynG,COX5A,ND-51,SdhA,NP15.6,Scsalpha1,ATPsynO,ATPsynbeta,ATPsynF,COX4,ND-75,CG8199,CG11110,ND-39,COX5B,ND-30,mRpS11,ND-24,ATPsynE,Tom40,scu,ATPsynCF6,Nc73EF,mRpS21,mRpL18,Tim8,mRpL13,blw,SdhD,mRpS23,levy,CG14817,ND-49,CG8728,ND-SGDH,ND-18,ND-B18,mRpL17,Tom70,CG17691,ND-B22,Mtpalpha,ATPsyndelta,mRpL36 | | 9.909e-16 | -34.55 | cellular protein metabolic process | biological process | GO:0044267 | 2099 | 293 | 12345 | 1129 | tko,Herc4,RpLP1,CG4238,mRpL18,Cchl,eIF3m,Ptip,RpL30,mRpS11,Sgf29,RpS2,CG2924,Traf6,Coq8,RpS16,gus,CG14817,Tip60,Usp39,mRpL16,Dph5,Ptp61F,Rep,tkv,CG7414,RpL34b,rl,pont,CG8188,Cdk5,Uba5,RpL11,CanB2,escl,mRpS10,l(3)73Ah,CG8314,velo,CG6766,fray,RpS28b,Iru,Cdk1,PIG-C,Uch,RpL9,RpS27,Tak1,GatA,26-29-p,RpL13,tor,CG1578,Prosbeta5,Cul2,mRpL13,Prosbeta3,RpL24,Ufm1,RYBP,CG7009,Tao,Rpn3,ben,mRpL10,mRpS23,Prosalpha3,RpL10,lt,APP-BP1,Spt20,p38a,pen-2,mEFTu1,shrb,RpS30,CanA-14F,LysRS,Pex12,mop,Pdk,RpL31,MAGE,CG10254,Topors,Ptpmeg,Ggt-1,Hexo1,Parp16,Pdk1,trr,ste14,PAPLA1,TrpRS-m,Pink1,Nmt,Pym,mRpL36,CG14435,GalT1,RpL29,Ublcp1,Pp2B-14D,Mtl,Ptp10D,Ptpmeg2,RpS14a,CkIIalpha,RpL23,tun,Pex10,RpS18,QC,Gapdh1,csw,RpS24,Hat1,RpL13A,Uch-L5,RpL34a,CG31053,scny,RpL37a,mts,esc,CG11858,tws,CG11070,Hsc70-3,ssh,CG32640,Mos,CG17746,Ubc4,Rpn2,RpL38,Mekk1,dco,Vsp37A,CG46338,CG1827,CycB,mEFTu2,Efr,egg,lwr,CSN6,CG17698,CG13344,Rab5,Cdk7,isoQC,Atu,GatB,Arf51F,EMC3,alph,GluRS-m,Prosalpha4,CG4572,otu,eEF1beta,RpS13,Dd,Mo25,CG5880,RpS29,dx,CG8728,RpL26,ste24a,RpS27A,twe,RpL40,SkpA,Cp1,CG7840,Naa20A,SRPK,Prosalpha5,nej,YL-1,CG3511,Edem1,faf,Usp16-45,HDAC6,eIF4E1,Utx,mRpS21,CG2911,Plap,SerRS-m,S2P,GAPsec,Pomp,grp,CycJ,Ubc6,fdl,G9a,CG7115,Rpn12,TyrRS,CG9316,mal,RpS23,CG3631,Prosbeta4,Sgf11,Aos1,Cul1,RpL17,CG32221,park,mRpL17,ari-1,RpS4,CG6094,Tfb4,Rpt2,Sirt7,Pp2A-29B,CG5003,CycE,PIG-P,Rpn13,Chrac-14,PpD3,CG9643,d4,Lkb1,O-fut1,RpL23A,Tlk,Cka,eIF3i,not,RpS15,CG3223,CG40045,Ntan1,Cul5,lid,CG8974,CG9941,trbd,CG11110,CG31633,RpS7,CG4041,CG8726,mRpL35,CG10344,mRpS5,Cyp1,e(y)1,PIG-L,Fbw5,CG11486,rept,RpS14b,Ptp69D,SelR,Prosbeta6,Atg7,STUB1,Fkbp12,awd,PIG-H,sip3,cort,Usp10,eIF4A,ago,RpLP0,Uev1A,mRpL48,RpS5b,Rcd5,png,Fkbp59,PpV,RpL12,CG5026,EloC,PEK,CG33977,UbcE2H,RpL7A,CysRS,Cbl | | 1.247e-15 | -34.32 | protein binding | molecular function | GO:0005515 | 2410 | 327 | 12176 | 1119 | CanA-14F,shv,shrb,piwi,Shmt,Sgf11,Wdfy2,olf186-M,CCT3,CG3309,RpS27A,CG8974,Ubc4,pygo,Neb-cGP,eEF1beta,scny,Pink1,TyrRS,dco,CG8475,Utx,nrv1,Mybbp1A,Cul5,Rel,faf,Ptp61F,CG32485,SNF4Agamma,Jafrac2,yps,nos,Vps16B,Atf6,CanB2,Uev1A,Trxr-1,CG1890,Cul1,bcd,Stam,eIF4E1,Adh,cib,CG4853,armi,SERCA,alc,Atf3,Mitofilin,CG4041,parvin,Afti,Idgf6,SREBP,eIF3m,lwr,Gga,Fbw5,mbo,Nap1,Gclc,Stip1,asf1,Arts,EcR,Grip84,CG4557,Klp10A,Actn,Tm1,dx,Hat1,twin,Cul2,Rga,CG8507,Mekk1,vret,ph-d,Girdin,CG11267,CG10188,Exo70,csw,Idgf2,SuUR,AIF,e(y)1,Ankle2,TfIIFalpha,CG18135,nej,cnc,Cse1,CG7716,Bx,CG14671,sbr,CycJ,Ubc6,rig,capt,Abi,Pura,Smyd4-4,rl,Rbf2,Sod1,ben,CG30496,Orc6,Rab32,ix,Pfdn2,rept,CG31875,Roe1,Nlg1,CG2258,Ptpmeg,Crag,Hip1,Gdi,Bin1,Grip128,boi,Pex12,Mitf,Caf1-105,Moe,stai,PrBP,CycB,dl,Arpc1,spz,cup,hng3,Dhfr,cact,Bap170,Vamp7,Rcd5,Tak1,MICAL-like,bun,d4,Syx13,insv,CG17270,egg,CCT6,Hsc70-3,CCT5,TBCC,Sry-alpha,Rbfox1,psq,ago,Zwilch,Cdk1,Taf7,CG32640,Pka-R2,RpL34a,STUB1,Ect4,O-fut1,Trf2,Grip71,Debcl,Rab5,lin,Aos1,Cen,Clc,Imp,mol,Pym,Liprin-beta,fs(1)Ya,MED4,Lkb1,Arpc2,gus,msps,Edc3,pps,sw,bif,NFAT,CG10254,ASPP,GlyP,Droj2,esc,GCC88,cana,CycE,Nup358,Pp2B-14D,up,CSN6,PGRP-LE,cyc,Dcr-1,Parp16,Su(H),lid,MED20,Tomosyn,cnn,Wnt10,CG11486,Kap-alpha3,Plap,CG13551,Reps,RpL23,RhoGDI,not,E2f1,GAPsec,Rep,Hsp26,Cka,trsn,REPTOR,foxo,SkpA,Blos2,Atu,Cyp1,BicC,sws,Fkbp39,Tao,CkIIalpha,Grip75,ssh,Rbf,tral,HmgD,apt,trbd,trk,trr,CG1416,CG3223,bru1,fz4,SmydA-1,Svil,milt,Sec15,CG10055,l(3)mbt,MED27,kl-2,Stat92E,Sec10,gw,Chrac-14,ari-1,Mtl,Pfk,Sec22,park,Acph-1,smg,wisp,sau,Ork1,Chd64,APP-BP1,pont,tyf,CG8726,Maf1,Pfdn5,RpL40,Cbl,CG42748,Sgf29,Blos4,raptor,tkv,Unr,wash,Nprl2,Kap3,CG3226,Rpt2,RpL11,Tip60,X11L,CG11807,Tlk,spag,MED24,CycC,Cf2,AP-1-2beta,Rpn13,Arpc4,Traf6,Myd88,awd,Idgf4,AnxB10,HP1c,Gmap,Fkbp59,Uxt,cbs,hwt,mad2,Ranbp21,tub,Smox,DnaJ-H,Med,sip3,exu,Topors | | 5.413e-15 | -32.85 | small molecule metabolic process | biological process | GO:0044281 | 794 | 140 | 12345 | 1129 | mino,Adi1,Thiolase,CG7840,Fdh,Pgm1,PyK,Etf-QO,Nadsyn,pn,Ac3,Gk1,Coq8,CG4598,GluRS-m,Acox57D-p,Pglym78,CAH2,Gk2,ATPsyndelta,Irp-1B,CG33156,P5cr,ATPsynB,GatB,ATPsynG,Pfrx,UGP,Ugt35A1,CG4407,cN-IIIB,yip2,CG6767,Ts,Npc2a,scu,CG10425,Pcd,Gmer,r-l,CG17691,blw,Desat1,TyrRS,Coq7,ATPsynD,Coq6,Eno,AOX3,AMPdeam,aay,GatA,Echs1,Gabat,Pgk,CG4586,CG11134,ATPsynCF6,CG3842,CG1774,Sdhaf3,AcCoAS,r,CG4594,SerRS-m,Seipin,Dhfr,Got1,CG6178,CG11164,CG7328,CG11899,CG31523,wal,SERCA,Pfk,tzn,AsnS,ATPsynO,CG17270,CG5028,TrpRS-m,Ugt36E1,regucalcin,SelD,ND-39,Idh3b,nmdyn-D6,l(1)G0196,Ntan1,foxo,shop,COX5B,Sptr,pr,Pdk,kri,Synj,sgll,Adh,Ppcdc,CG4390,CG4860,Scsalpha1,Pxt,Kyat,LysRS,Mtpalpha,ACC,ScsbetaA,CysRS,CG7920,CG7322,nmdyn-D7,Pfas,Shmt,DhpD,Pdxk,Marf,Mcad,ATPsynE,CG8199,oxt,fal,CG9391,CG10932,SREBP,CG11089,l(2)k01209,Gip,Idh,Aldh,P5CDh1,Gapdh1,CG8112,Sply,awd,ATPsynbeta,Ady43A,Pex10 | | 1.034e-14 | -32.20 | organonitrogen compound metabolic process | biological process | GO:1901564 | 3232 | 407 | 12345 | 1129 | RpL26,ste24a,pn,Ac3,dx,CG2493,CG8728,Nadsyn,PyK,twe,RpS27A,SRPK,Prosalpha5,Naa20A,CG7840,RpL40,SkpA,Cp1,Edem1,Adi1,faf,nej,CG3511,YL-1,Cdk7,cN-IIIB,Rab5,CG17698,CG13344,CSN6,CG4407,GatB,ATPsynG,Arf51F,Atu,ATPsyndelta,isoQC,eEF1beta,RpS13,GstT1,Dd,Ser7,Pglym78,otu,CG4572,EMC3,Prosalpha4,alph,GluRS-m,Ppox,RpS29,Mo25,CG5880,whd,AOX3,AMPdeam,aay,G9a,Eno,CG9316,CG5390,GCS2beta,mal,CG7115,Rpn12,TyrRS,Sgf11,Aos1,Cul1,CG32473,CG3631,Prosbeta4,RpS23,CG10425,ari-1,Ts,mRpL17,CG32221,CG6767,park,RpL17,eIF4E1,Usp16-45,HDAC6,Utx,SerRS-m,S2P,Plap,mRpS21,r,CG2201,CG2911,grp,Pomp,CG1774,GAPsec,CG16979,grsm,Ubc6,fdl,CG11134,CycJ,RpS7,SelD,CG9941,CG11110,trbd,CG31633,CG8974,lid,mRpL35,CG8726,CG4041,mRpS5,UK114,CG10344,Cyp1,rept,CG11486,Cht11,Fbw5,e(y)1,PIG-L,CG31523,Pp2A-29B,Kyat,CG5003,Gclc,Sirt7,Rpt2,CG6094,Tfb4,RpS4,Scsalpha1,Rpn13,sgll,PIG-P,Chrac-14,CycE,kri,RpL23A,Tlk,Cka,pr,Lkb1,O-fut1,CG9643,Sptr,d4,PpD3,GstE11,Ntan1,Cul5,CG3223,CG40045,nmdyn-D6,RpS15,not,eIF3i,png,RpS5b,Dip-B,Rcd5,CG8199,Uev1A,CG13551,mRpL48,ATPsynE,RpLP0,EloC,RpL12,CG5026,Fkbp59,PpV,RpL7A,Shmt,UbcE2H,nSMase,nmdyn-D7,CG33977,Pfas,PEK,Cbl,CysRS,CG7920,CG11771,SelR,RpS14b,Ptp69D,Idgf4,STUB1,Fkbp12,Sply,awd,Sp7,Atg7,P5CDh1,Prosbeta6,ea,cort,sip3,CG11089,PIG-H,fal,Usp10,eIF4A,ago,Tip60,gus,CG14817,RpS16,Ptp61F,Dph5,Usp39,mRpL16,CG7414,RpL34b,tkv,Idgf2,Rep,PGRP-LE,Cdk5,CG8188,pont,cathD,rl,Herc4,RpLP1,tko,DppIII,eIF3m,Ptip,ATPsynB,Pfrx,CG14516,CG33156,P5cr,Cchl,CG4238,mRpL18,mRpS11,Sgf29,RpL30,CG3803,Traf6,Coq8,CG2924,RpS2,Tao,CG7009,RYBP,RpL24,Ufm1,CG2200,ATPsynD,Cul2,mRpL13,Prosbeta3,lt,RpL10,Prosalpha3,mRpS23,mRpL10,blw,stet,Rpn3,ben,r-l,CG17691,CG10561,Spt20,p38a,scu,sol,Idgf6,Pcd,APP-BP1,shrb,RpS30,mEFTu1,pen-2,RpL11,CanB2,Uba5,Dhfr,CG6178,Got1,RpS28b,fray,velo,sws,CG6766,CG8314,escl,mRpS10,AcCoAS,l(3)73Ah,Uch,PIG-C,RpL9,CG9953,Cdk1,Iru,Prosbeta5,ATPsynCF6,CG1578,GstT2,GatA,RpL13,tor,26-29-p,Gabat,RpS27,Pgk,Tak1,mRpL36,CG14435,Pym,Pink1,Nmt,snk,TrpRS-m,Pfk,AsnS,ATPsynO,Ublcp1,SP1029,RpL29,GalT1,Mtl,Pp2B-14D,CG11899,mop,Pex12,LysRS,Ppcdc,CanA-14F,CG10254,RpL31,MAGE,CG13630,Pdk,Ggt-1,Hexo1,Ptpmeg,Topors,PAPLA1,ste14,trr,COX5B,Pdk1,Parp16,Mos,CG17746,gd,ssh,CG32640,Hsc70-3,CG11070,tws,Mekk1,RpL38,Rpn2,Pdxk,Ubc4,Vsp37A,DhpD,CG46338,CG1827,dco,Efr,egg,lwr,Fur2,mEFTu2,ScsbetaA,CycB,alpha-Man-Ia,CG7997,RpS18,RpL23,Ady43A,Pex10,tun,CkIIalpha,RpS14a,ATPsynbeta,Ptp10D,Ho,Ptpmeg2,Uch-L5,RpL13A,RpS24,Hat1,csw,Gapdh1,QC,scny,RpL34a,l(2)k01209,CG31053,Nepl3,CG11858,mts,esc,RpL37a,oxt | | 1.122e-14 | -32.12 | macromolecule metabolic process | biological process | GO:0043170 | 3775 | 461 | 12345 | 1129 | CG17746,Mos,CG32640,gd,ssh,CG11070,tws,Hsc70-3,Mekk1,RpL38,SmE,Rpn2,CycC,Ubc4,CG1622,twin,CG1827,CG46338,Vsp37A,dco,CG12375,CG14100,CG9684,lwr,egg,Efr,cu,Fur2,mEFTu2,alpha-Man-Ia,CycB,RpS18,Pex10,tun,RpL23,Atg6,RpS14a,CkIIalpha,Ptpmeg2,Ptp10D,Uch-L5,RpL13A,Hat1,RpS24,QC,Gapdh1,csw,Rga,scny,snRNA:U12,CG31053,RpL34a,Nepl3,CG11858,Lsm12,esc,mts,mus81,exu,RpL37a,oxt,Orc1,mRpL36,CG14435,Pym,CG2100,Xpac,CG30343,snk,Pink1,Nmt,TrpRS-m,Spx,mael,SP1029,Ublcp1,bys,Rnmt,RpL29,Gyg,GalT1,Trf2,Mtl,Pp2B-14D,Pex12,GlyP,mop,ACC,LysRS,CanA-14F,CG10254,RpL31,MAGE,CG13630,Pdk,Ptpmeg,Hexo1,Ggt-1,Topors,Top1,ste14,PAPLA1,tho2,trr,Pdk1,Imp,CG10189,Parp16,Tao,RYBP,CG7009,CG2200,RpL24,Ufm1,Prosbeta3,Cul2,mRpL13,lt,RpL10,Prosalpha3,CG16865,mRpL10,mRpS23,Rpn3,ben,Larp7,stet,p38a,E2f1,Spt20,APP-BP1,sol,scu,Idgf6,RecQ5,RpS30,shrb,mEFTu1,pen-2,CG12018,CanB2,RpL11,mus304,Uba5,RpS28b,fray,velo,CG8314,CG6766,l(3)73Ah,escl,mRpS10,RpL9,Uch,PIG-C,Cdk1,Iru,CG9953,tos,Gasz,Prosbeta5,CG1578,tor,26-29-p,nos,RpL13,GatA,pygo,CG6610,Tak1,RpS27,Tip60,CG14817,gus,Orc6,RpS16,gw,del,Ptp61F,Archease,Dph5,mRpL16,Usp39,Pgm1,RpL34b,CG7414,CG5285,tkv,Idgf2,TfIIFalpha,Taf7,PGRP-LE,Rep,CG8188,Cdk5,cathD,CG9143,smg,pont,Dcr-1,CG3198,CG9485,mino,rl,RpLP1,Herc4,tko,DppIII,mbo,UGP,heph,Ptip,eIF3m,CG14516,Cchl,CG4238,mRpL18,snRNA:U4atac,mRpS11,Sgf29,rig,IntS6,RpL30,CG17768,RpII140,Trf4-2,pps,Coq8,CG10336,Traf6,CG2924,RpS2,png,CG31759,Rcd5,CG2034,Dip-B,RpS5b,Uev1A,mRpL48,tapas,RfC38,RpLP0,EloC,CG5026,RpL12,PpV,shu,Fkbp59,RpL7A,Edc3,yps,UbcE2H,fs(1)Ya,CG33977,Snm1,PEK,Cbl,Nurf-38,thoc6,CysRS,Nap1,armi,RpS14b,SelR,CG11771,Ptp69D,Sld5,Idgf4,r2d2,awd,Fkbp12,STUB1,Sp7,Atg7,Prosbeta6,ea,sip3,cort,PIG-H,Mat89Ba,Exd2,MED4,eIF4A,ago,Clp,Usp10,RpS7,CG11110,CG10418,trbd,CG31633,CG9941,SelD,CG8974,lid,mRpL35,CG8726,CG4041,Cyp1,sbr,obe,CG10344,mRpS5,rept,Cht11,CG11486,mtTFB1,Fbw5,CG3021,PIG-L,e(y)1,CG5003,Pp2A-29B,Pus7,Sirt7,Rpt2,Tfb4,CG6094,NHP2,RpS4,Chrac-14,PIG-P,Rpn13,CG3587,CycE,Neos,Cka,RpL23A,Tlk,O-fut1,piwi,Lkb1,d4,CG9643,PpD3,Cul5,foxo,Ntan1,CG3223,CG40045,not,RpS15,CG17187,eIF3i,G9a,Tengl4,CG8204,mal,CG9316,CG5390,GCS2beta,Son,TyrRS,Rpn12,CG8475,CG7115,Cul1,Sgf11,Aos1,Prosbeta4,CG32473,CG3631,Cbp80,RpS23,trsn,ari-1,DNApol-epsilon255,mRpL17,MED24,park,CG32221,RpL17,vret,CG11164,Not10,Utx,Usp16-45,eIF4E1,HDAC6,CG7728,SerRS-m,S2P,Nxf3,CG5292,Plap,CG2911,mRpS21,grp,IntS11,mus205,Pomp,CG16979,GAPsec,Rpb8,AGBE,snRNA:U4:38AB,fdl,Ubc6,grsm,Xpc,CycJ,ste24a,RpL26,CG2493,CG8728,nbs,dx,Rrp40,Bin1,agt,twe,wisp,RpS27A,Prosalpha5,SRPK,Naa20A,CG5986,CG7840,SkpA,CG12301,Cp1,RpL40,faf,Edem1,CG3511,c(2)M,YL-1,nej,SuUR,Drep1,spen,Cdk7,Rab5,CG13344,CG17698,CSN6,Arf51F,GatB,Drep2,Atu,isoQC,CG17454,Dd,Ser7,CG10463,RpS13,eEF1beta,otu,CG4572,bru1,Prosalpha4,GluRS-m,alph,EMC3,Rbf,RpS29,LSm3,CG5880,Mo25,mre11 | | 1.439e-14 | -31.87 | Mdh2 (Malate dehydrogenase 2) | protein interactions | 42185 | 90 | 40 | 9204 | 1112 | Uch,Pglym78,Rtnl1,CG4598,capt,scu,Moe,Gdi,Argk,Idh,CG10863,HmgD,RhoGDI,ATPsynCF6,awd,Trxr-1,up,Mf,Adh,Fkbp12,Actn,Scp1,Eno,Gapdh1,CG3226,AnxB10,Phb2,regucalcin,Prosbeta5,Stip1,CG11267,Chd64,yip2,EndoGI,cib,PHGPx,Pgk,ATPsynbeta,CG10638,CG14715 | | 1.587e-14 | -31.77 | RpS21 (Ribosomal protein S21) | protein interactions | 33487 | 136 | 51 | 9204 | 1112 | RpL13A,RpS28b,RpL11,HP1e,RpS4,regucalcin,RpL30,RpS23,RpL40,RpL29,RpS16,RpS27,Pi3K92E,RpS14b,RpS7,RpS24,RpS2,CG14715,cib,RpL13,RpL34a,RpL37a,RpS18,RpL12,RpS27A,RpL26,RpLP0,Got1,RpL31,RpS13,RpS30,RpL17,Bx,RpS15,CG14036,Treh,RpS29,CG1532,RpL7A,RpL24,RpLP1,RpL23,RpL23A,RpL10,RpL38,Fkbp12,Sod1,RpL9,Rel,awd,RpS14a | | 3.303e-14 | -31.04 | heterocycle metabolic process | biological process | GO:0046483 | 1498 | 221 | 12345 | 1129 | CG8204,mal,lt,TyrRS,Son,blw,CG16865,ben,Larp7,AMPdeam,CG7009,AOX3,ATPsynD,Eno,Tengl4,Ts,DNApol-epsilon255,RecQ5,MED24,CG6767,vret,CG12018,r-l,CG17691,Cbp80,E2f1,Gmer,trsn,Pcd,scu,RpS28b,SerRS-m,CG5292,l(3)73Ah,CG2911,r,AcCoAS,CG11164,Not10,CG6178,mus304,CG7728,eIF4E1,Dhfr,Rpb8,snRNA:U4:38AB,ATPsynCF6,nos,GatA,Xpc,Ubc6,pygo,CG6610,Pgk,mus205,IntS11,tos,CG1774,Gasz,gw,del,PyK,Archease,wisp,Usp39,pn,Ac3,Orc6,RpS16,nbs,Bin1,Rrp40,Nadsyn,agt,c(2)M,CG9143,smg,Drep1,spen,Dcr-1,mino,CG3198,CG5285,CG5986,TfIIFalpha,Taf7,CG12301,ATPsynG,Pfrx,GatB,ATPsynB,Drep2,ATPsyndelta,P5cr,CG33156,snRNA:U4atac,CG17454,cN-IIIB,Cdk7,UGP,Ugt35A1,CG4407,heph,CG10336,Ppox,Rbf,LSm3,mre11,CG10463,mRpS11,Sgf29,Pglym78,rig,bru1,IntS6,CG17768,CG3803,GluRS-m,RpII140,Trf4-2,pps,SmE,Pdxk,CycC,shu,CG1622,CG2034,CG8199,CG31759,ATPsynE,CG13551,Uev1A,tapas,RfC38,cu,Nurf-38,CysRS,ScsbetaA,CG7920,Nap1,Shmt,RpL7A,twin,Edc3,yps,Rab32,DhpD,CG12375,CG14100,Snm1,nmdyn-D7,Pfas,CG9684,awd,Gapdh1,P5CDh1,armi,RpS14b,Ady43A,Sld5,RpS14a,Ho,ATPsynbeta,r2d2,Lsm12,Exd2,mus81,MED4,fal,exu,Orc1,Clp,Rga,snRNA:U12,CG11089,Mat89Ba,l(2)k01209,Spx,Pfk,ATPsynO,Pym,CG2100,RpS7,CG10418,Xpac,CG30343,SelD,TrpRS-m,Ugt36E1,CG11486,mtTFB1,CG3021,e(y)1,mael,sbr,bys,Rnmt,obe,Trf2,sgll,CG3587,CycE,MAGE,Neos,kri,Pus7,LysRS,Tfb4,Ppcdc,Scsalpha1,NHP2,tho2,Top1,COX5B,nmdyn-D6,Imp,CG17187,CG10189,pr,piwi,Sptr | | 4.805e-14 | -30.67 | RpS5a (Ribosomal protein S5a) | protein interactions | 32700 | 109 | 44 | 9204 | 1112 | RpL13A,foxo,ago,RpS28b,RpL11,Lkb1,RpS4,RpL30,RpL40,RpS23,RpL29,ScsbetaA,Pi3K92E,RpS27,RpS16,RpS14b,RpS7,RpS24,RpS2,RpL34a,RpS18,RpL37a,RpL13,RpL12,RpLP0,RpS27A,RpL26,RpL17,RpS13,RpS30,RpL31,RpS15,RpS29,RpLP1,RpL7A,RpL24,RpL23,RpL10,RpL23A,RpL38,slam,RpL9,PGRP-LE,RpS14a | | 1.263e-13 | -29.70 | 5.8SrRNA:CR40454 (5.8S ribosomal RNA) | protein interactions | 5740577 | 83 | 37 | 9204 | 1112 | RpL29,RpS27,RpS16,RpL30,RpS23,RpL40,RpL11,RpS4,RpL13A,RpS28b,RpL12,RpLP0,RpL26,RpS27A,RpS2,RpL37a,RpS18,RpL34a,RpL13,RpS7,RpS24,RpS14b,RpS29,RpS15,RpS13,RpS30,RpL17,RpL31,RpL9,RpS14a,RpL38,RpL23,RpL23A,RpL10,RpL7A,RpLP1,RpL24 | | 1.263e-13 | -29.70 | 18SrRNA:CR41548 (18S ribosomal RNA:CR41548) | protein interactions | 5740812 | 83 | 37 | 9204 | 1112 | RpS27,RpS16,RpL29,RpL40,RpS23,RpL30,RpS4,RpL11,RpS28b,RpL13A,RpLP0,RpL26,RpS27A,RpL12,RpL34a,RpS18,RpL37a,RpL13,RpS2,RpS24,RpS7,RpS14b,RpS29,RpS15,RpS13,RpS30,RpL17,RpL31,RpS14a,RpL9,RpL38,RpL23A,RpL10,RpL23,RpLP1,RpL24,RpL7A | | 1.266e-13 | -29.70 | RpL28 (Ribosomal protein L28) | protein interactions | 38397 | 99 | 41 | 9204 | 1112 | RpS24,RpS7,RpS14b,RpL26,RpS27A,RpLP0,Pdk1,RpL12,RpL13,RpL34a,RpS18,RpL37a,RpS2,Lkb1,RpS4,RpL11,RpS28b,RpL13A,RpS16,RpS27,Pi3K92E,RpL29,RpS23,RpL40,RpL30,RpL10,RpL23A,RpL23,RpL7A,RpL24,RpLP1,RpS14a,RpL9,RpL38,RpS15,RpL31,RpL17,RpS13,RpS30,RpS29,capt | | 1.330e-13 | -29.65 | nucleus | cellular component | GO:0005634 | 2809 | 362 | 11940 | 1105 | Rbf2,CG10654,CG4404,mts,Rpb8,snRNA:U12,Ubc6,eIF4A,CG14814,ovo,CycJ,Neos,CG8001,CG17328,obe,Chrac-14,MED4,CG2129,CG7115,CG8009,CG12288,heph,e(y)1,Prosbeta6,Taf7,SmE,Pomp,NHP2,exu,sbb,CG5412,CG31875,wash,Lsm12,Tfb4,l(2)k01209,Dd,trbd,Mes2,alphaTub67C,d4,SkpA,CSN6,CG4266,lwr,Bin1,brwl,PDCD-5,RpS18,gammaTub37C,Ptip,CG6610,r2d2,CkIIalpha,Med,CG17746,REPTOR,RpL11,Mat89Ba,CG10336,Hmg-2,RYBP,RecQ5,CG4730,pont,Nprl2,Xpc,Utx,rept,shep,dl,Cbp80,piwi,cup,Imp,Dcr-1,MAGE,G9a,EcR,Mybbp1A,qkr58E-2,IntS6,Cka,SRPK,Tengl4,asf1,Pen,Hsc70-3,CG15525,CycE,CG10038,Caf1-105,Atu,Nap1,Ndc1,Trf4-2,Rnmt,Pus7,Ranbp21,HmgD,Stat92E,Rcd5,Nup58,PpD3,SuUR,mxc,Orc1,CG12018,CG10631,cad,ASPP,toc,CG17698,Karybeta3,CG17202,Uev1A,Usp39,CG9684,RpL40,ari-1,tho2,Ufm1,Ssdp,Cul1,tun,lqfR,jigr1,Plap,ph-d,Acph-1,twe,CG46385,bru1,Su(H),thoc6,trsn,nbs,Sirt7,Cdk1,l(3)mbt,twin,tub,snRNA:U4:38AB,SREBP,lid,Aven,Uxt,escl,IntS11,Prosbeta4,bif,Prosalpha3,Sod3,mtrm,Ptp61F,RpS13,Mo25,fal,CG4021,CG11902,Sld5,CG3198,spen,snRNA:U4atac,Cenp-C,hng3,sbr,wapl,cnc,CG5986,Larp7,awd,Pdk1,CG12301,cyc,Lkb1,CG4880,Maf1,CG5003,RpS27A,Spx,Smox,Shmt,nej,mus304,RpII140,SelR,Ady43A,insv,dco,Rpn2,Tip60,CycC,rig,scny,CG3437,Herc4,TfIIFalpha,Atf6,mael,yps,Wbp2,CG13344,Prosbeta3,CG10055,alc,Orc6,CG17454,spn-E,YL-1,Trf2,stil,gw,Pym,CG9143,Rga,Top1,CG2199,alphaTub84B,zf30C,Hnf4,Tlk,CG17294,Xpac,CG17187,CG16865,crm,CG8204,NFAT,CG10959,apt,DNApol-epsilon255,dnc,CG33156,gus,Ntan1,tws,Arts,Elba2,Adi1,Sgf29,Gclc,mute,Topors,CG2162,CG6813,PrBP,Non1,Nf-YA,l(3)73Ah,lin,vers,Marf1,Moe,MED24,CG11070,Bx,mad2,Clp,Snm1,fs(1)Ya,Prosalpha4,mus81,CG6276,Tao,Atf3,mus205,Ubc4,RpS7,Aladin,MED27,LSm3,SNF4Agamma,psq,Aos1,E2f1,HP1c,CG17768,CG1622,Rel,ben,rl,Uch-L5,bcd,Nup62,Bap170,Rpn13,Coq7,EloC,Rep,RfC38,Hr78,CG10418,Prosalpha5,p38a,esc,not,elg1,yem,MED20,CG7728,HP1e,Atpalpha,Hat1,Prosbeta5,CG2911,alph,Cdk7,pps,Rbf,Cdk5,PAPLA1,Cse1,Nurf-38,pygo,tos,Psf3,Ublcp1,Cyp1,CG8611,CG11710,cu,bys,EndoGI,egg,Rrp40,CG1602,mbo,velo,CG2678,Nxf3,Rbfox1,Parp16,del,lin-52,wdn,CG7655,Sod1,ix,grp,Spt20,RpS15,APC7,l(1)G0004,Smyd4-4,RpS2,CG11164,CG3511,Sgf11,mre11,CG42526,CG3226,CycB,CG31053,trr,Cf2,Axs,hb,Kap-alpha3,osk,BigH1,foxo | | 1.437e-13 | -29.57 | organic cyclic compound metabolic process | biological process | GO:1901360 | 1599 | 230 | 12345 | 1129 | armi,RpS14b,Ady43A,RpS14a,Sld5,r2d2,ATPsynbeta,Ho,awd,Sp7,P5CDh1,Gapdh1,CG8112,Rga,snRNA:U12,CG11089,l(2)k01209,Mat89Ba,Exd2,Lsm12,MED4,mus81,fal,exu,Clp,Orc1,CG2034,CG8199,CG31759,Uev1A,CG13551,ATPsynE,tapas,RfC38,Marf,SmE,CycC,shu,Pdxk,CG1622,RpL7A,Shmt,DhpD,Edc3,yps,twin,Rab32,Snm1,nmdyn-D7,CG14100,CG12375,CG9684,Pfas,cu,ScsbetaA,CysRS,Nurf-38,CG7920,Nap1,Kyat,Pus7,LysRS,Ppcdc,Tfb4,Scsalpha1,NHP2,CG3587,sgll,MAGE,CycE,Neos,kri,pr,piwi,Sptr,Top1,foxo,tho2,COX5B,dl,nmdyn-D6,CG10189,Imp,CG17187,CG2100,RpS7,Pym,CG30343,SelD,CG10418,Xpac,TrpRS-m,Ugt36E1,CG17270,Pfk,Spx,ATPsynO,mael,obe,Rnmt,bys,sbr,Trf2,CG11486,mtTFB1,e(y)1,CG3021,CG11164,Not10,CG7728,eIF4E1,Dhfr,CG6178,mus304,RpS28b,CG5292,SerRS-m,AcCoAS,CG2911,l(3)73Ah,r,IntS11,mus205,CG1774,tos,Gasz,snRNA:U4:38AB,Rpb8,ATPsynCF6,GatA,Xpc,Ubc6,nos,Pgk,pygo,CG6610,AOX3,CG7009,AMPdeam,ATPsynD,Tengl4,Eno,lt,CG8204,mal,Son,TyrRS,CG16865,blw,Larp7,ben,r-l,CG17691,E2f1,Npc2a,Cbp80,scu,Pcd,trsn,Gmer,DNApol-epsilon255,Ts,RecQ5,CG6767,MED24,CG12018,vret,Cdk7,cN-IIIB,Ugt35A1,UGP,heph,CG4407,GatB,ATPsynB,Pfrx,ATPsynG,ATPsyndelta,Drep2,snRNA:U4atac,CG17454,CG33156,P5cr,CG10463,Pglym78,mRpS11,Sgf29,CG3803,bru1,CG17768,IntS6,rig,Trf4-2,pps,RpII140,GluRS-m,CG10336,Ppox,Rbf,mre11,LSm3,Orc6,Ac3,pn,nbs,RpS16,agt,Nadsyn,Bin1,Rrp40,del,PyK,gw,Archease,Usp39,wisp,CG5285,CG5986,TfIIFalpha,Taf7,CG12301,Cdk5,CG9143,smg,c(2)M,Dcr-1,spen,Drep1,mino,CG3198 | | 1.659e-13 | -29.43 | RpLP0 (Ribosomal protein LP0) | protein interactions | 40451 | 108 | 43 | 9204 | 1112 | RpLP1,RpL24,RpL7A,CG14818,RpL10,RpL23A,RpL23,RpL38,RpS14a,RpL9,fs(1)Ya,RpL31,RpL17,RpS30,RpS13,RpS15,RpS29,CCT3,RpS14b,RpS24,Trs33,RpS7,RpL13,RpL34a,RpL37a,RpS18,RpS2,RpS27A,RpL26,RpL12,ago,RpS28b,RpL13A,RpS4,RpL11,GILT1,RpL40,RpS23,RpL30,RpS16,RpS27,Nca,RpL29 | | 1.951e-13 | -29.27 | oxidation-reduction process | biological process | GO:0055114 | 608 | 112 | 12345 | 1129 | Sod3,Atg6,CG7550,Seipin,Dhfr,Ho,ND-49,CG6144,CG10863,SelR,Idh,Aldh,P5CDh1,CG4594,Gapdh1,CG2767,ND-18,ND-24,ND-SGDH,Cyp6a19,ND-B15,sni,Echs1,UQCR-Q,CG4586,GILT1,Jafrac2,AGBE,CG10932,Coq7,Coq6,Trxr-1,levy,CG14407,AOX3,blw,stw,Desat1,CG9436,CG8475,Mcad,CG10561,ND-51,ScsbetaA,Sod1,CG3609,ND-30,SdhA,Adh,UGP,CG17333,CG4860,Scsalpha1,Pxt,GlyP,yip2,Mtpalpha,ACC,Irp-1B,NP15.6,P5cr,COX5A,sgll,Grx1,SdhD,cype,UQCR-11,Nc73EF,CG3803,Sptr,CG1275,CG4598,Cyp6a22,COX4,PHGPx,Acox57D-p,CG10638,CG10463,ND-75,Ppox,Coq8,foxo,shop,COX5B,CG5028,Etf-QO,ND-B22,Cyp6u1,ND-39,Cyp6v1,Idh3b,Pfk,Pgm1,Cyt-c-p,CG14997,Hnf4,CG4842,Sccpdh1,CG31674,Gyg,Pmi,wal,SERCA,Fdh,ND-MNLL,AIF,Jafrac1,ND-B18,CG9485,Adi1,Thiolase,ND-23 | | 2.000e-13 | -29.24 | 5SrRNA:CR33353 (5S ribosomal RNA) | protein interactions | 3771903 | 84 | 37 | 9204 | 1112 | RpL30,RpL40,RpS23,RpL29,RpS16,RpS27,RpL13A,RpS28b,RpL11,RpS4,RpS2,RpL37a,RpL34a,RpS18,RpL13,RpL12,RpLP0,RpL26,RpS27A,RpS14b,RpS7,RpS24,RpS29,RpS13,RpL17,RpS30,RpL31,RpS15,RpL38,RpL9,RpS14a,RpL7A,RpLP1,RpL24,RpL23,RpL23A,RpL10 | | 2.034e-13 | -29.22 | biological regulation | biological process | GO:0065007 | 4350 | 511 | 12345 | 1129 | not,dl,Pura,foxo,Cul5,Ntan1,krimp,up,d4,CG2678,Atpalpha,piwi,O-fut1,Lkb1,Tlk,Cka,CG10188,sbb,Hip1,Sec23,spn-E,CG4872,CycE,Chrac-14,CG3587,Rpn13,Tfb4,CG4853,Sirt7,Rpt2,CG12773,CG5003,Rel,Pp2A-29B,e(y)1,CG7328,mtTFB1,Fbw5,adp,Sec10,CG11486,hwt,rept,CG9425,ovo,Rbcn-3A,SERCA,sbr,obe,RhoGAP5A,Maf1,Tm1,CHOp24,bcd,CG4041,tub,CG8726,CG3894,APC7,lid,DCAF12,spz,CG10418,trbd,AP-1-2beta,SelD,ago,eIF4A,CG2129,Usp10,MtnA,MED4,qkr58E-2,Exd2,SREBP,hb,Ssdp,muskelin,cort,cyc,sip3,ea,Aldh,Ranbp21,Sap47,Atg7,Sp7,scramb1,awd,STUB1,r2d2,Sld5,cana,armi,ERp60,Nap1,CG1416,Stam,Nurf-38,Mitf,Grip84,Cbl,PEK,tyf,Snm1,Edc3,Rab32,Tomosyn,Shmt,Mctp,CG17328,Ect4,PpV,Fkbp59,HP1c,osk,zyd,EloC,tapas,CG13551,Uev1A,Rcd5,CG31759,Smox,levy,CG14407,Scox,png,LSm3,mre11,Mo25,Pld,Rbf,squ,Wnt10,alph,spag,Zwilch,bru1,CG10631,otu,Dd,COX4,CG17454,trk,Crag,Drep2,tral,Atu,Arf51F,Grx1,COX5A,CSN6,CG4021,CG13344,Tspo,CG17698,CG6813,Pka-R2,dob,Rab5,CG9981,ix,Cdk7,Nprl2,crm,CG6967,spen,Drep1,SuUR,YL-1,nej,faf,Adi1,Nup62,Edem1,SkpA,Gdi,asf1,Naa20A,Fdh,CG4730,insv,cad,BigH1,Debcl,Rbfox1,wisp,CG13933,Hnf4,twe,fs(1)N,PyK,EMRE,chrb,Bin1,jigr1,Rrp40,CG11710,dx,nbs,Ac3,MED27,pn,Syx13,X11L,CycJ,GILT1,Ubc6,shv,GAPsec,Pomp,Nup358,Unr,grp,CG2911,Plap,Arpc1,S2P,Utx,eIF4E1,HDAC6,bam,Not10,CG6927,vret,Sod1,MED24,park,CCT3,Tsp86D,ari-1,trsn,CG9911,Cbp80,Hr78,swa,CG11077,Stat92E,Actn,CG32473,stas,Cul1,Sgf11,Aos1,Moe,Rab10,ATP8A,dnc,Son,TyrRS,Fer1HCH,Cenp-C,CG7115,Ork1,Desat1,Eno,G9a,ZnT35C,Snx6,SNF4Agamma,Trxr-1,Imp,Parp16,Pdk1,AGO3,trr,crok,Top1,Topors,RhoGAP54D,Ptpmeg,nAChRalpha4,cold,CG30496,anne,Nlg1,Med,REPTOR,Arpc2,MAGE,Synj,HP1e,Cf2,CanA-14F,disp,CG2199,mop,GlyP,fs(1)M3,bun,CG9925,nrv1,Calx,stai,Fkbp39,boi,Pi3K92E,Pp2B-14D,Mtl,Trf2,plu,mael,cact,Pfk,crq,snk,Pink1,Idh3b,fwe,Pym,CG13601,CG9391,CG8009,Nf-YA,exu,esc,mts,mus81,mad2,Jafrac2,CG31053,Vps16B,Ankle2,Rga,scny,CG10470,mute,Atf3,csw,CG8112,Mybbp1A,CG8963,lqfR,Hat1,RpL13A,RhoGDI,CG17202,opm,Ho,Ptp10D,Atg6,CkIIalpha,wash,CycB,CG14036,ScsbetaA,Girdin,Pax,CG17272,Fur2,cu,egg,E23,lwr,CG9186,apt,LpR2,fz4,dco,BicC,twin,CycC,NFAT,vers,Rpn2,Marf,Mekk1,Hsc70-3,tws,hng3,gd,ssh,CG17746,Mos,CG10336,Irk1,Traf6,Trf4-2,pps,Smyd4-4,slif,CG17768,mAChR-A,Bx,PHGPx,Acox57D-p,CG4238,Vha68-2,CG30291,eIF3m,Ptip,raptor,heph,mbo,Myd88,cup,CG11902,RpLP1,Chd64,Herc4,CG3198,rl,Jafrac1,Dcr-1,lin-52,pont,smg,Cdk5,Taf7,capt,PGRP-LE,l(3)mbt,Rep,TfIIFalpha,Pka-R1,Idgf2,tkv,CG7414,CG1602,mtrm,Archease,Dph5,EcR,gw,bif,pgc,Ptp61F,enc,Neb-cGP,Sec15,ph-d,gus,CG10038,Orc6,Tip60,Uxt,PDCD-5,Marf1,CG4586,pygo,Tak1,wapl,Pgk,Gabat,nos,tor,BoYb,CG1578,Prosbeta5,CG31729,CG14408,CG3842,Wbp2,Trs33,cnc,Cdk1,CG31875,msps,l(3)73Ah,ASPP,escl,velo,sws,Abi,fray,mus304,Dhfr,Seipin,Su(H),psq,Arpc4,CanB2,Atf6,pen-2,Aven,CG10959,RecQ5,shrb,lin,cnn,Sox102F,scu,p38a,MED20,Npc2a,Spt20,E2f1,HmgD,Rbf2,EndoGI,dgt4,Rpn3,ben,CG10654,stet,Grip71,blw,Iml1,Elba2,Exo70,lt,Bap170,mRpL13,Cul2,alc,ATPsynD,Coq7,SmydA-1,RYBP,Tao | | 2.458e-13 | -29.03 | cellular aromatic compound metabolic process | biological process | GO:0006725 | 1544 | 223 | 12345 | 1129 | E2f1,Cbp80,scu,trsn,Gmer,Pcd,r-l,CG17691,CG6767,MED24,CG12018,vret,DNApol-epsilon255,Ts,RecQ5,ATPsynD,Tengl4,Eno,AOX3,AMPdeam,CG7009,CG16865,blw,Larp7,ben,lt,mal,CG8204,Son,TyrRS,CG1774,tos,Gasz,IntS11,mus205,Xpc,Ubc6,GatA,nos,Pgk,pygo,CG6610,snRNA:U4:38AB,Rpb8,ATPsynCF6,Dhfr,eIF4E1,CG7728,mus304,CG6178,CG11164,Not10,AcCoAS,CG2911,r,l(3)73Ah,RpS28b,CG5292,SerRS-m,TfIIFalpha,CG12301,Taf7,CG5285,CG5986,Dcr-1,spen,Drep1,CG3198,mino,Cdk5,CG9143,smg,c(2)M,nbs,RpS16,Nadsyn,agt,Rrp40,Bin1,Orc6,pn,Ac3,wisp,Usp39,del,PyK,gw,Archease,CG3803,bru1,IntS6,CG17768,rig,pps,Trf4-2,GluRS-m,RpII140,CG10463,Pglym78,Sgf29,mRpS11,mre11,LSm3,Ppox,CG10336,Rbf,Ugt35A1,UGP,heph,CG4407,Cdk7,cN-IIIB,Drep2,ATPsyndelta,CG17454,snRNA:U4atac,CG33156,GatB,ATPsynB,Pfrx,ATPsynG,Snm1,nmdyn-D7,CG14100,CG12375,CG9684,Pfas,RpL7A,Shmt,DhpD,yps,Rab32,twin,Edc3,ScsbetaA,Nurf-38,CysRS,CG7920,Nap1,cu,CG13551,Uev1A,ATPsynE,tapas,RfC38,CG31759,CG8199,CG2034,CycC,shu,Pdxk,CG1622,SmE,CG11089,l(2)k01209,Mat89Ba,Rga,snRNA:U12,fal,exu,Clp,Orc1,Exd2,Lsm12,MED4,mus81,RpS14a,Sld5,r2d2,ATPsynbeta,Ho,RpS14b,armi,Ady43A,Gapdh1,awd,Sp7,Trf2,mael,obe,Rnmt,bys,sbr,mtTFB1,e(y)1,CG3021,CG11486,Ugt36E1,TrpRS-m,CG2100,RpS7,Pym,CG30343,SelD,CG10418,Xpac,Pfk,Spx,ATPsynO,Sptr,pr,piwi,nmdyn-D6,CG10189,CG17187,Imp,Top1,tho2,COX5B,dl,Ppcdc,Tfb4,NHP2,Scsalpha1,Pus7,Kyat,LysRS,Neos,kri,sgll,CG3587,MAGE,CycE | | 3.137e-13 | -28.79 | RpS10a (Ribosomal protein S10a) | protein interactions | 43321 | 85 | 37 | 9204 | 1112 | RpS14b,RpS24,RpS7,RpL13,RpS18,RpL34a,RpL37a,RpS2,RpS27A,RpL26,RpLP0,RpL12,RpS28b,RpL13A,RpS4,RpL11,RpL40,RpS23,RpL30,RpS27,RpS16,RpL29,RpL24,RpLP1,RpL7A,RpL23A,RpL10,RpL23,RpL38,RpS14a,RpL9,RpL31,RpL17,RpS13,RpS30,RpS15,RpS29 | | 3.530e-13 | -28.67 | RpS11 (Ribosomal protein S11) | protein interactions | 36321 | 110 | 43 | 9204 | 1112 | RpS4,RpL11,RpS28b,foxo,RpL13A,RpS27,RpS16,Pi3K92E,RpL29,CG11164,RpL40,RpS23,RpL30,RpS24,RpS7,RpS14b,RpL26,RpS27A,RpLP0,Pdk1,RpL12,RpL13,RpL34a,RpL37a,RpS18,RpS2,RpS15,RpL31,RpS30,RpL17,RpS13,RpS29,RpL10,RpL23A,RpL23,RpLP1,RpL24,RpL7A,RpS14a,PGRP-LE,RpL9,RpL38,CG9902 | | 3.633e-13 | -28.64 | organelle membrane | cellular component | GO:0031090 | 787 | 135 | 11940 | 1105 | Ugalt,sip3,Atf6,ND-B22,ND-18,Hip1,Ppox,levy,Vamp7,PIG-C,Dd,blw,CG33977,mol,CG31523,Tim8,Pmi,Marf,CHMP2B,COX5B,CG8112,ND-30,CG8009,Desat1,Tspo,mino,ABCD,ATPsynF,ATPsynO,CG5510,Vha36-1,COX5A,sws,Topors,ND-B15,CG9399,Pex10,Sec23,Roe1,alt,Cchl,Synj,CG8311,UQCR-11,Vha68-2,cype,Cyt-c-p,Vha13,CG11077,CG40472,Coq8,CG5447,CG17270,CG8004,Sec22,stet,Pi4KIIalpha,ATPsynD,CG9132,CG9393,colt,ATPsyndelta,CG10425,ND-SGDH,Seipin,toc,Debcl,ND-B18,Tom70,ND-49,Mitofilin,AP-1-2beta,CG8728,PIG-H,SdhD,CG32640,Afti,Etf-QO,Efr,ATPsynCF6,fwe,Tom40,CG10470,ATPsynE,ND-24,ND-39,Iml1,Tengl4,SERCA,CG11110,shrb,Clc,ND-75,CG8974,Coq7,ATPsynbeta,Fur2,COX4,ND-51,SdhA,NP15.6,ATPsynG,CG14671,Gmap,EMRE,alpha-Man-Ia,CG3803,CG5969,Neb-cGP,Vsp37A,EMC3,EMC8-9,Axs,Mctp,sbr,Pex23,ATPsynB,CG32441,SREBP,UQCR-Q,ND-23,Scox,ste24a,bru1,Pex12,ND-MNLL,CG9773,CG6707,bif,CG7655,CG5903,CG4293,yin,Pex13,ND-MWFE | | 3.728e-13 | -28.62 | ATP metabolic process | biological process | GO:0046034 | 140 | 43 | 12345 | 1129 | ND-SGDH,COX4,Pglym78,ND-75,ND-24,Coq8,ATPsynCF6,COX5B,ND-B15,Pgk,UQCR-Q,ATPsynbeta,ND-49,COX5A,cype,UQCR-11,SdhD,ATPsynB,ATPsynG,Pfrx,ATPsyndelta,Gapdh1,Vha68-2,wal,ND-MNLL,ND-51,Pmi,SdhA,ND-23,AIF,ND-B18,ND-B22,levy,ND-39,ATPsynE,ATPsynD,Etf-QO,Eno,Cyt-c-p,PyK,Pfk,blw,ATPsynO | | 4.196e-13 | -28.50 | RpL18 (Ribosomal protein L18) | protein interactions | 38794 | 102 | 41 | 9204 | 1112 | RpLP1,RpL7A,RpL24,RpL10,RpL23A,RpL23,RpL38,Not10,RpS14a,RpL9,RpL17,RpS13,RpS30,RpL31,RpS15,RpS29,RpS14b,RpS24,RpS7,RpL34a,RpL37a,RpS18,RpL13,RpS2,RpLP0,RpL26,RpS27A,RpL12,RpS28b,RpL13A,foxo,Lkb1,RpS4,RpL11,RpL40,RpS23,RpL30,Pi3K92E,RpS27,RpS16,RpL29 | | 4.477e-13 | -28.43 | RpS15 (Ribosomal protein S15) | protein interactions | 36851 | 98 | 40 | 9204 | 1112 | RpS29,RpL31,RpL17,RpS30,RpS13,RpL9,RpS14a,RpL38,RpL23,RpL23A,RpL10,RpL7A,RpL24,RpLP1,RpL29,RpS16,RpS27,Pi3K92E,RpL30,RpS23,RpL40,RpL11,HP1e,RpS4,foxo,RpL13A,RpS28b,Pdk1,RpL12,RpL26,RpS27A,RpLP0,RpS2,RpL13,RpL37a,RpL34a,RpS18,RpS7,RpS24,RpS14b | | 4.838e-13 | -28.36 | RpL37A (Ribosomal protein L37A) | protein interactions | 44783 | 90 | 38 | 9204 | 1112 | RpS29,RpS13,RpS30,RpL17,RpL31,RpS15,RpL38,RpS14a,RpL9,RpLP1,RpL7A,RpL24,RpL10,RpL23A,RpL23,csw,RpS23,RpL40,RpL30,RpS27,RpS16,RpL29,RpS28b,RpL13A,RpS4,RpL11,RpS18,RpL34a,RpL37a,RpL13,RpS2,RpLP0,RpL26,RpS27A,RpL12,RpS14b,RpS24,RpS7 | | 5.105e-13 | -28.30 | RpS23 (Ribosomal protein S23) | protein interactions | 36576 | 111 | 43 | 9204 | 1112 | RpS7,RpS24,RpS14b,Pdk1,RpL12,RpL26,RpS27A,RpLP0,RpS2,RpL13,RpL37a,RpL34a,RpS18,RpL11,HP1e,RpS4,Lkb1,foxo,RpL13A,RpS28b,RpL29,RpS27,RpS16,Pi3K92E,RpL30,RpL40,RpL23,RpL10,RpL23A,RpLP1,RpL7A,RpL24,RpL9,slam,RpS14a,CG15141,RpL38,RpS15,RpL31,RpL17,RpS30,RpS13,RpS29 | | 5.504e-13 | -28.23 | organelle envelope | cellular component | GO:0031967 | 432 | 88 | 11940 | 1105 | CG11077,Cyt-c-p,mbo,Coq8,ND-MNLL,CG40472,bru1,UQCR-11,SREBP,ATPsynB,Scox,cype,ND-23,UQCR-Q,ND-MWFE,CG9393,ATPsynD,Parp16,bif,CG5903,CG8004,CG7655,EMRE,CG9399,CG3803,Ptp61F,AIF,ND-B15,NP15.6,ATPsynO,SdhA,mad2,ND-51,COX5A,ATPsynG,Cchl,sbr,fs(1)Ya,Aladin,Neb-cGP,Roe1,Axs,ND-30,Tengl4,ND-39,COX5B,CG4880,Tom40,Ndc1,Pmi,ND-24,Marf,ATPsynE,Coq7,mino,ND-75,Tspo,ATPsynF,Nup62,COX4,ATPsynbeta,CG8009,Nup58,CG11110,Tom70,ND-B18,Ppox,Debcl,toc,ND-SGDH,ND-18,CG8728,levy,ND-49,Mitofilin,ATPsyndelta,colt,ND-B22,Tim8,Etf-QO,ATPsynCF6,wash,Sdhaf3,blw,Dd,SdhD,shop,Cse1 | | 5.504e-13 | -28.23 | envelope | cellular component | GO:0031975 | 432 | 88 | 11940 | 1105 | CG8004,CG5903,CG7655,bif,ATPsynD,Parp16,CG9393,ND-MWFE,cype,ND-23,UQCR-Q,Scox,UQCR-11,ATPsynB,SREBP,bru1,CG40472,ND-MNLL,Coq8,Cyt-c-p,mbo,CG11077,Roe1,Axs,Neb-cGP,Aladin,fs(1)Ya,sbr,Cchl,ATPsynG,COX5A,mad2,SdhA,ND-51,NP15.6,ATPsynO,CG9399,CG3803,ND-B15,Ptp61F,AIF,EMRE,CG11110,Nup58,CG8009,ATPsynbeta,COX4,Nup62,ATPsynF,Tspo,ND-75,mino,Coq7,ND-24,ATPsynE,Marf,Pmi,Tom40,Ndc1,COX5B,ND-39,CG4880,ND-30,Tengl4,Cse1,shop,blw,SdhD,Dd,Sdhaf3,wash,ATPsynCF6,Etf-QO,Tim8,ND-B22,colt,ATPsyndelta,levy,Mitofilin,ND-49,CG8728,toc,ND-18,ND-SGDH,ND-B18,Tom70,Ppox,Debcl | | 5.954e-13 | -28.15 | regulation of metabolic process | biological process | GO:0019222 | 2165 | 289 | 12345 | 1129 | sws,l(3)73Ah,ASPP,escl,fray,Abi,Seipin,psq,Atf6,Su(H),nos,tor,pygo,Tak1,Gabat,Pgk,CG1578,CG14408,Wbp2,cnc,CG31875,Cdk1,Elba2,CG10654,Rpn3,ben,Bap170,ATPsynD,Coq7,mRpL13,Cul2,Tao,SmydA-1,RYBP,Sox102F,CG10959,lin,p38a,MED20,Spt20,E2f1,Rbf2,HmgD,EndoGI,CG4238,CG30291,Ptip,eIF3m,heph,raptor,RpLP1,Myd88,cup,CG11902,CG10336,Traf6,CG17768,pps,Smyd4-4,Trf4-2,Bx,CG1602,pgc,gw,Ptp61F,Dph5,EcR,Tip60,Uxt,Marf1,ph-d,gus,CG10038,Dcr-1,CG3198,rl,lin-52,smg,pont,TfIIFalpha,l(3)mbt,Taf7,CG7414,Pka-R1,tkv,Hat1,Atf3,mute,Mybbp1A,CG8963,CG17202,RpL13A,CkIIalpha,Nf-YA,CG8009,Jafrac2,esc,mts,Ankle2,CG31053,Rga,scny,CycC,NFAT,Mekk1,vers,Rpn2,tws,Hsc70-3,Mos,CG17746,hng3,ScsbetaA,Pax,wash,CycB,egg,lwr,cu,dco,BicC,apt,twin,REPTOR,Med,Synj,CG2199,HP1e,Cf2,GlyP,bun,Pdk1,Parp16,trr,AGO3,Topors,CG30496,cact,Pink1,Pym,CG13601,Fkbp39,Pi3K92E,plu,Trf2,mael,CG2911,S2P,bam,Utx,HDAC6,Not10,Ubc6,CycJ,Pomp,Nup358,GAPsec,Unr,Desat1,Son,G9a,SNF4Agamma,MED24,park,vret,Sod1,ari-1,Hr78,Cbp80,trsn,Cul1,Sgf11,Aos1,Stat92E,Atu,CG17454,CG13344,Tspo,CG4021,Cdk7,ix,Pka-R2,CG6813,LSm3,squ,Rbf,CG10631,bru1,alph,Dd,otu,Debcl,Rbfox1,wisp,Hnf4,CG11710,Rrp40,Bin1,jigr1,MED27,pn,Drep1,SuUR,spen,Nprl2,CG6967,crm,faf,YL-1,nej,asf1,SkpA,insv,BigH1,cad,Fdh,CG4730,Atg7,STUB1,Sp7,r2d2,armi,eIF4A,ago,CG2129,SREBP,qkr58E-2,Exd2,hb,MED4,cort,cyc,sip3,Ssdp,PpV,osk,HP1c,CG17328,EloC,Scox,Rcd5,CG31759,Smox,Mitf,Nap1,Cbl,PEK,tyf,Rab32,Edc3,spn-E,sbb,CG4872,Chrac-14,CG3587,Rpn13,CycE,Tfb4,Rel,Pp2A-29B,Rpt2,Sirt7,not,foxo,dl,d4,CG2678,krimp,Cka,Tlk,piwi,Lkb1,bcd,spz,lid,AP-1-2beta,CG10418,SelD,CG7328,Fbw5,mtTFB1,e(y)1,rept,CG9425,CG11486,ovo,RhoGAP5A,Maf1,obe | | 7.339e-13 | -27.94 | RpS6 (Ribosomal protein S6) | protein interactions | 31700 | 112 | 43 | 9204 | 1112 | RpS23,RpL40,RpL30,RpS16,RpS27,Pi3K92E,RpL29,RpS28b,foxo,RpL13A,RpS4,RpL11,RpL13,RpL37a,RpL34a,RpS18,RpS2,RpS27A,RpL26,CycE,RpLP0,Pdk1,RpL12,RpS14b,RpS24,RpS7,RpS29,RpL31,RpL17,RpS13,RpS30,RpS15,RpL38,RpS14a,RpL9,Pink1,RpLP1,RpL7A,RpL24,Rcd5,RpL10,RpL23A,RpL23 | | 7.339e-13 | -27.94 | RpL38 (Ribosomal protein L38) | protein interactions | 3355144 | 112 | 43 | 9204 | 1112 | PGRP-LE,RpS14a,RpL9,RpL23A,RpL10,RpL23,Sld5,RpL7A,RpLP1,RpL24,RpS29,Gdi,RpS15,RpL31,RpS13,RpS30,RpL17,RpL26,RpS27A,RpLP0,CycE,RpL12,RpL13,RpL37a,RpL34a,RpS18,gus,RpS2,RpS24,RpS7,RpS14b,RpS27,RpS16,Pi3K92E,RpL29,RpS23,RpL40,RpL30,RpS4,RpL11,RpS28b,CG3021,RpL13A | | 7.358e-13 | -27.94 | RpL36A (Ribosomal protein L36A) | protein interactions | 34098 | 91 | 38 | 9204 | 1112 | RpL13,RpL34a,RpS18,RpL37a,RpS2,RpL26,RpS27A,RpLP0,Pdk1,RpL12,RpS14b,RpS24,RpS7,RpS23,RpL40,RpL30,RpS16,RpS27,RpL29,RpS28b,RpL13A,RpS4,RpL11,RpL38,RpS14a,RpL9,RpLP1,RpL7A,RpL24,RpL10,RpL23A,RpL23,RpS29,RpL31,RpS30,RpS13,RpL17,RpS15 | | 7.358e-13 | -27.94 | RpL35 (Ribosomal protein L35) | protein interactions | 31483 | 91 | 38 | 9204 | 1112 | RpS7,RpS24,RpS14b,RpL12,RpLP0,RpS27A,RpL26,RpS2,CG3939,RpL34a,RpL37a,RpS18,RpL13,RpL11,RpS4,RpL13A,RpS28b,RpL29,RpS16,RpS27,RpL30,RpS23,RpL40,RpL23,RpL10,RpL23A,RpL7A,RpL24,RpLP1,RpL9,RpS14a,RpL38,RpS15,RpL17,RpS30,RpS13,RpL31,RpS29 | | 7.383e-13 | -27.93 | macromolecule modification | biological process | GO:0043412 | 1320 | 196 | 12345 | 1129 | ari-1,CG32221,park,Aos1,Sgf11,Cul1,CG3631,Spt20,p38a,APP-BP1,CG9316,CG7115,ben,Tao,CG7009,G9a,Ufm1,Cul2,CG1578,Ubc6,fdl,tor,CycJ,Tak1,PIG-C,Uch,grp,Iru,Cdk1,CG5292,fray,CG8314,velo,escl,l(3)73Ah,CG2911,CanB2,HDAC6,Uba5,Usp16-45,Utx,CG8188,Cdk5,faf,pont,nej,CG3511,YL-1,rl,SRPK,tkv,Naa20A,CG7840,RpL40,Rep,SkpA,Ptp61F,Dph5,twe,RpS27A,Usp39,Tip60,dx,agt,Traf6,Coq8,CG2924,Mo25,CG5880,CG10463,Dd,Sgf29,otu,alph,Ptip,Arf51F,Atu,Cchl,CG4238,isoQC,Herc4,Cdk7,CG17698,CG13344,CSN6,Efr,Cbl,lwr,egg,CycB,UbcE2H,CG46338,CG1827,CG14100,dco,CG33977,PEK,EloC,Mekk1,CG5026,PpV,Fkbp59,Ubc4,png,Mos,CG17746,ssh,CG2034,Rcd5,Uev1A,tws,CG11070,CG11858,mts,esc,Usp10,ago,scny,sip3,CG31053,PIG-H,STUB1,Fkbp12,Uch-L5,awd,Atg7,Hat1,csw,Gapdh1,QC,Ptp69D,Pex10,tun,CkIIalpha,Ptp10D,Ptpmeg2,rept,Mtl,CG11486,Pp2B-14D,mtTFB1,Fbw5,PIG-L,e(y)1,CG3021,CG10344,obe,Ublcp1,Rnmt,Cyp1,GalT1,CG8726,CG4041,CG14435,SelD,CG9941,CG31633,trbd,Nmt,Pink1,lid,CG8974,ste14,Cul5,CG40045,trr,Pdk1,not,Parp16,CG10189,Cka,Tlk,Lkb1,O-fut1,Ptpmeg,Hexo1,CG9643,d4,PpD3,Topors,Chrac-14,CG3587,PIG-P,CG10254,MAGE,CycE,Pdk,Pp2A-29B,Pus7,CG5003,mop,Pex12,Sirt7,Tfb4,NHP2,CanA-14F | | 1.038e-12 | -27.59 | cytosol | cellular component | GO:0005829 | 1181 | 181 | 11940 | 1105 | Adh,Pfrx,heph,RpS4,RpL31,Ptpmeg,RpS5b,Ts,Tm1,Wbp2,eEF1beta,RhoGDI,cN-IIIB,Argk,DppIII,Pp2A-29B,RpL30,Top1,stai,AOX3,RpS24,CG5003,Got1,RpS14a,galectin,RpL13,RpL9,Smox,eIF4A,Droj2,RpS27A,Blos4,Usp10,Shmt,scu,SelR,CG9436,Gga,Blos2,Ady43A,Rpn2,DnaJ-H,l(1)G0196,CCT3,r,AGO3,CCT6,Pgk,CG17931,CG9425,AMPdeam,RpLP1,Moe,dl,Arf51F,shep,Xpc,CG11089,cup,Gyg,CCT5,RpL17,CG2767,CG1640,piwi,RpL23,CG1890,CG31674,cib,RpS16,Pka-R2,RpS7,RpS29,Fdh,RpL10,RpL37a,Sry-alpha,Cyt-c-p,RpS14b,Tango10,CG3939,Jafrac1,RpS18,CG4390,PDCD-5,Trxr-1,Atg4b,CkIIalpha,NFAT,Gdi,eIF3m,CG7920,REPTOR,CG17746,tws,RpL11,dnc,CG10189,UK114,Irp-1B,Karybeta3,RpL40,CG2911,RpS23,CG3223,Prosbeta5,CG10863,tral,armi,CG6967,RpL34b,alph,park,PyK,Cdk5,CG10638,Pgm1,Jafrac2,Cse1,bun,tun,larp,RpLP0,Pink1,Pka-R1,Uba5,CG4238,LysRS,Rel,Pfk,rl,RpL7A,RpS28b,Stat92E,Rep,PpD3,CG7414,RpS13,RpS15,RpL34a,Ptpmeg2,RpS2,CG17068,Rpn12,sau,Scsalpha1,RpL23A,RpL12,Cf2,RpS27,AsnS,cact,Pdk1,Eno,tkv,foxo,RpS30,RpL26,CG46385,eIF4E1,Cyp1,UGP,trsn,CG2678,Idh,Rtnl1,RpL38,RpL29,twin,tub,Pax,PEK,RpL24,Aven,Pdxk,RpL13A,Sod1,Sod3,Gapdh1,cl,grp | | 1.691e-12 | -27.11 | RpL9 (Ribosomal protein L9) | protein interactions | 34526 | 110 | 42 | 9204 | 1112 | RpS14a,RpL38,RpL23,RpL10,RpL23A,RpL24,RpL7A,RpLP1,RpS29,Neos,RpS15,RpL31,RpS13,RpS30,RpL17,Pdk1,RpL12,RpS27A,RpL26,RpLP0,RpS2,RpL13,RpL34a,RpL37a,RpS18,RpS7,COX5A,RpS24,RpS14b,RpL29,RpS27,RpS16,Pi3K92E,RpL30,RpS23,RpL40,RpL11,RpS4,Lkb1,foxo,RpL13A,RpS28b | | 1.774e-12 | -27.06 | macromolecule catabolic process | biological process | GO:0009057 | 490 | 94 | 12345 | 1129 | Prosalpha5,Idgf2,SkpA,Cp1,PGRP-LE,RpL40,faf,Edem1,CG8188,Cht11,cathD,CG11486,smg,Drep1,Dcr-1,Fbw5,CG9485,Pym,CG31633,trbd,gus,Pink1,Rrp40,CG8974,gw,RpS27A,CG4572,CG17768,Prosalpha4,Topors,Trf4-2,Cul5,Ntan1,CG3223,CG40045,not,LSm3,CG5003,GlyP,Herc4,Rpt2,CG10254,Rpn13,CG4238,Cul1,Prosbeta4,twin,Edc3,UbcE2H,Vsp37A,Cbp80,Idgf6,Cbl,ari-1,shrb,cu,park,CG32221,CG31759,CG32640,RYBP,Tengl4,Prosbeta3,CG11070,Hsc70-3,Cul2,CG9316,EloC,Rpn12,Rpn2,Prosalpha3,Rpn3,Rga,scny,Uch,sip3,cort,Pomp,Prosbeta5,26-29-p,fdl,Ubc6,ago,Usp10,armi,CG11164,Not10,Atg6,Usp16-45,Idgf4,Uch-L5,CG6766,Plap,Prosbeta6 | | 1.834e-12 | -27.02 | RpS17 (Ribosomal protein S17) | protein interactions | 39088 | 119 | 44 | 9204 | 1112 | RpL11,RpS4,foxo,RpL13A,CG4476,RpS28b,RpL29,RpS16,RpS27,Pi3K92E,RpL30,RpL40,RpS23,RpS7,RpS24,RpS14b,Pdk1,RpL12,RpL26,RpS27A,RpLP0,RpS2,RpL13,RpL34a,RpS18,RpL37a,RpS15,RpL31,RpS13,RpL17,RpS30,RpS29,RpL23,RpL10,RpL23A,RpLP1,Tak1,RpL7A,RpL24,RpL9,Rel,RpS14a,PGRP-LE,RpL38 | | 1.866e-12 | -27.01 | RpL22 (Ribosomal protein L22) | protein interactions | 31022 | 128 | 46 | 9204 | 1112 | RpS29,Moe,RpS15,RpS13,jigr1,RpS30,RpL17,RpL31,RpL9,PGRP-LE,RpS14a,Rel,RpL38,Fkbp12,RpL23,RpL23A,RpL10,Tak1,RpL24,RpL7A,RpLP1,RpL29,RpS16,RpS27,RpL30,RpL40,RpS23,RpL11,mus304,CkIIalpha,RpS4,RpL13A,RpS28b,RpL12,RpLP0,RpL26,RpS27A,cup,RpS2,RpL34a,RpS18,RpL37a,RpL13,RpS7,RpS24,RpS14b | | 2.353e-12 | -26.78 | cell cycle | biological process | GO:0007049 | 678 | 118 | 12345 | 1129 | mtrm,wisp,Non1,Axs,bif,Ptp61F,enc,del,twe,alphaTub67C,nbs,lid,CCT6,ph-d,Orc6,Syx13,SuUR,Dcr-1,rl,Nprl2,CG8188,Nup62,c(2)M,pont,Pp2B-14D,SkpA,mael,CG3437,cbs,SRPK,spn-E,CG7716,Arf51F,CycE,Grip128,tef,HP1e,mop,CG5003,sw,Cdk7,Pp2A-29B,gammaTub37C,cup,Rab5,mre11,CG10336,sau,larp,Rbf,Zwilch,krimp,alph,PpD3,Topors,Trf4-2,COX4,Tlk,Grip75,Lkb1,Grip71,PpV,Iml1,dgt4,spd-2,Cenp-C,tws,RfC38,Mos,Rcd5,ssh,mEFTu1,CycB,egg,lwr,cnn,DNApol-epsilon255,RecQ5,Grip84,shrb,BicC,fs(1)Ya,E2f1,toc,Arpc1,csw,msps,awd,Sld5,CkIIalpha,cana,bam,mus304,Klp10A,eIF4E1,CanB2,Pex10,nos,Ubc6,GatA,Xpc,eIF4A,CycJ,ago,wapl,Orc1,alphaTub84B,Prosbeta5,mts,mus81,mad2,cort,Ankle2,CG31053,slam,grp,Cdk1,yem | | 2.410e-12 | -26.75 | RpL10 (Ribosomal protein L10) | protein interactions | 43864 | 111 | 42 | 9204 | 1112 | RpS14b,RpS7,Dip-B,RpS24,RpS2,RpL13,RpS18,RpL37a,RpL34a,Pdk1,RpL12,RpL26,RpS27A,RpLP0,foxo,RpL13A,RpS28b,RpL11,Lkb1,RpS4,RpL30,RpL40,RpS23,RpL29,RpS27,RpS16,Pi3K92E,RpL24,RpLP1,RpL7A,RpL23,RpL23A,RpL38,RpL9,RpS14a,Rtnl1,RpL31,RpL17,RpS13,RpS30,RpS15,RpS29 | | 2.410e-12 | -26.75 | RpS27 (Ribosomal protein S27) | protein interactions | 43039 | 111 | 42 | 9204 | 1112 | RpL38,Fkbp12,RpL9,RpS14a,PGRP-LE,Rel,RpL7A,RpLP1,Tak1,RpL24,RpL23,RpL10,RpL23A,RpS29,RpS30,RpL17,RpS13,RpL31,RpS15,RpS2,RpL37a,RpL34a,RpS18,RpL13,RpL12,RpLP0,RpS27A,RpL26,RpS14b,RpS7,RpS24,RpL30,csw,RpL40,RpS23,RpL29,RpS16,RpL13A,CG4476,RpS28b,RpL11,RpS4 | | 2.473e-12 | -26.73 | RpL26 (Ribosomal protein L26) | protein interactions | 40060 | 94 | 38 | 9204 | 1112 | RpS14b,RpS24,RpS7,RpL13,RpS18,RpL37a,RpL34a,RpS2,RpS27A,RpLP0,Pdk1,RpL12,RpS28b,foxo,RpL13A,RpS4,RpL11,RpL40,RpS23,RpL30,RpS16,RpS27,RpL29,RpLP1,RpL7A,RpL24,RpL10,RpL23A,RpL23,RpL38,RpS14a,RpL9,RpL31,RpS30,RpS13,RpL17,RpS15,RpS29 | | 2.538e-12 | -26.70 | nucleobase-containing compound metabolic process | biological process | GO:0006139 | 1417 | 205 | 12345 | 1129 | CG5986,CG5285,CG12301,Taf7,TfIIFalpha,CG9143,smg,c(2)M,mino,CG3198,Dcr-1,Drep1,spen,Orc6,Ac3,Nadsyn,agt,Rrp40,Bin1,RpS16,nbs,Archease,PyK,del,gw,Usp39,wisp,Pglym78,Sgf29,mRpS11,CG10463,Trf4-2,pps,GluRS-m,RpII140,bru1,IntS6,CG17768,rig,Rbf,CG10336,mre11,LSm3,Cdk7,cN-IIIB,heph,CG4407,UGP,Ugt35A1,GatB,ATPsynB,ATPsynG,Pfrx,snRNA:U4atac,CG17454,CG33156,Drep2,ATPsyndelta,CG17691,r-l,scu,Gmer,trsn,E2f1,Cbp80,RecQ5,DNApol-epsilon255,Ts,CG12018,vret,CG6767,MED24,AOX3,AMPdeam,CG7009,Tengl4,Eno,ATPsynD,TyrRS,Son,CG8204,Larp7,ben,CG16865,blw,IntS11,mus205,Gasz,tos,CG1774,ATPsynCF6,snRNA:U4:38AB,Rpb8,Pgk,CG6610,pygo,Ubc6,GatA,Xpc,nos,Not10,CG11164,CG7728,eIF4E1,mus304,CG6178,CG5292,SerRS-m,RpS28b,AcCoAS,CG2911,l(3)73Ah,r,Rnmt,obe,bys,sbr,mael,Trf2,CG11486,e(y)1,CG3021,mtTFB1,SelD,CG30343,CG10418,Xpac,CG2100,RpS7,Pym,Ugt36E1,TrpRS-m,ATPsynO,Pfk,Spx,piwi,COX5B,Top1,tho2,CG10189,Imp,CG17187,nmdyn-D6,LysRS,Pus7,NHP2,Scsalpha1,Ppcdc,Tfb4,MAGE,CycE,CG3587,kri,Neos,DhpD,twin,yps,Edc3,RpL7A,Shmt,CG9684,Pfas,Snm1,CG14100,nmdyn-D7,CG12375,cu,Nap1,CG7920,ScsbetaA,CysRS,Nurf-38,CG31759,CG2034,CG8199,tapas,RfC38,Uev1A,ATPsynE,SmE,CG1622,CycC,shu,snRNA:U12,Rga,l(2)k01209,Mat89Ba,CG11089,MED4,mus81,Exd2,Lsm12,Clp,Orc1,exu,fal,Ady43A,armi,RpS14b,ATPsynbeta,r2d2,RpS14a,Sld5,awd,Gapdh1 | | 2.565e-12 | -26.69 | RpS14a (Ribosomal protein S14a) | protein interactions | 47218 | 120 | 44 | 9204 | 1112 | RpL13,RpL37a,RpS18,RpL34a,Pura,RpS2,RpS27A,RpL26,RpLP0,Pdk1,RpL12,RpS14b,RpS24,RpS7,RpS23,RpL40,RpL30,RpS16,RpS27,RpL29,RpS28b,foxo,RpL13A,RpS4,Lkb1,RpL11,RpL38,CG9117,Rel,PGRP-LE,RpL9,Tak1,RpLP1,RpL24,RpL7A,RpL10,RpL23A,RpL23,RpS29,RpL31,RpS30,RpL17,RpS13,RpS15 | | 2.620e-12 | -26.67 | RpL27A (Ribosomal protein L27A) | protein interactions | 33654 | 90 | 37 | 9204 | 1112 | RpL12,RpLP0,RpL26,RpS27A,RpS2,RpL34a,RpL37a,RpS18,RpL13,RpS7,RpS24,RpS14b,RpL29,RpS27,RpS16,RpL30,RpL40,RpS23,RpL11,RpS4,RpL13A,RpS28b,RpL9,RpS14a,RpL38,RpL23,RpL10,RpL23A,RpL7A,RpL24,RpLP1,RpS29,RpS15,RpS13,RpS30,RpL17,RpL31 | | 2.722e-12 | -26.63 | RpS28b (Ribosomal protein S28b) | protein interactions | 31897 | 107 | 41 | 9204 | 1112 | RpL24,Tak1,RpLP1,RpL7A,RpL23,RpL10,RpL23A,RpL38,Sry-alpha,RpL9,Rel,RpS14a,PGRP-LE,RpL31,RpL17,RpS30,RpS13,RpS15,RpS29,RpS14b,RpS7,COX5A,RpS24,RpS2,RpL13,RpL37a,RpS18,RpL34a,RpL12,RpL26,RpS27A,RpLP0,RpL13A,RpL11,RpS4,RpL30,RpS23,RpL40,RpL29,RpS16,RpS27 | | 3.299e-12 | -26.44 | cytoplasmic ribonucleoprotein granule | cellular component | GO:0036464 | 88 | 32 | 11940 | 1105 | spn-E,osk,gw,LSm3,Rga,tapas,CG11486,CG6967,tral,armi,krimp,squ,PyK,yps,mael,Edc3,exu,cup,pgc,CG9425,CG17068,Tm1,nos,AGO3,CG10418,vret,shu,eIF4A,smg,eIF4E1,CG17768,bru1 | | 3.651e-12 | -26.34 | RpS14b (Ribosomal protein S14b) | protein interactions | 47219 | 95 | 38 | 9204 | 1112 | RpLP1,RpL7A,RpL24,RpL23,RpL23A,RpL10,RpL38,RpL9,RpS14a,CG9117,RpL17,RpS30,RpS13,RpL31,RpS15,RpS29,RpS7,RpS24,RpS2,Pura,RpL37a,RpL34a,RpS18,RpL13,RpL12,RpLP0,RpS27A,RpL26,RpL13A,RpS28b,RpL11,RpS4,RpL30,RpS23,RpL40,RpL29,RpS27,RpS16 | | 3.651e-12 | -26.34 | RpL39 (Ribosomal protein L39) | protein interactions | 37849 | 95 | 38 | 9204 | 1112 | RpS13,RpS30,RpL17,RpL31,RpS15,RpS29,RpL7A,RpLP1,RpL24,RpL23,RpL10,RpL23A,RpL38,RpL9,RpS14a,RpL13A,RpS28b,RpL11,RpS4,RpL30,RpS23,RpL40,RpL29,RpS27,RpS16,RpS14b,Traf6,RpS7,RpS24,RpS2,RpL37a,RpS18,RpL34a,RpL13,RpL12,RpLP0,RpL26,RpS27A | | 3.651e-12 | -26.34 | RpS4 (Ribosomal protein S4) | protein interactions | 39484 | 95 | 38 | 9204 | 1112 | RpS14b,Kyat,RpS7,RpS24,Trs33,RpS2,RpL13,RpL34a,RpS18,RpL37a,RpL12,RpL26,RpS27A,RpLP0,RpL13A,RpS28b,RpL11,RpL30,RpS23,RpL40,RpL29,RpS16,RpS27,RpL24,RpLP1,RpL7A,RpL23,RpL23A,RpL10,RpL38,RpL9,RpS14a,RpL31,RpS30,RpL17,RpS13,RpS15,RpS29 | | 3.911e-12 | -26.27 | RpS19a (Ribosomal protein S19a) | protein interactions | 32635 | 91 | 37 | 9204 | 1112 | RpS29,RpS15,RpS13,RpL17,RpS30,RpL31,RpS14a,RpL9,RpL38,RpL23A,RpL10,RpL23,RpL24,RpLP1,RpL7A,RpS16,RpS27,RpL29,RpS23,RpL40,RpL30,RpS4,RpL11,RpS28b,RpL13A,RpLP0,RpS27A,RpL26,RpL12,RpS18,RpL37a,RpL34a,RpL13,RpS2,RpS24,RpS7,RpS14b | | 4.118e-12 | -26.22 | RpL34a (Ribosomal protein L34a) | protein interactions | 43169 | 87 | 36 | 9204 | 1112 | RpL12,RpS27A,RpL26,RpLP0,RpS2,RpL13,RpL37a,RpS18,RpS7,RpS24,RpS14b,RpL29,RpS27,RpS16,RpL30,RpL40,RpS23,RpL11,RpS4,RpL13A,RpS28b,RpL9,RpS14a,RpL38,RpL23,RpL10,RpL23A,RpLP1,RpL7A,RpL24,RpS29,RpS15,RpL31,RpL17,RpS30,RpS13 | | 4.162e-12 | -26.20 | RpL17 (Ribosomal protein L17) | protein interactions | 31613 | 117 | 43 | 9204 | 1112 | RpL11,Lkb1,RpS4,foxo,RpL13A,RpS28b,RpL29,RpS27,RpS16,Pi3K92E,RpL30,RpL40,RpS23,RpS7,RpS24,RpS14b,RpL12,RpS27A,RpL26,RpLP0,RpS2,RpL13,RpS18,RpL34a,RpL37a,RpS15,CG4730,RpL31,RpS30,RpS13,RpS29,RpL23,RpL23A,RpL10,RpLP1,RpL7A,Spx,RpL24,RpL9,CG9117,RpS14a,RpL38,dl | | 4.701e-12 | -26.08 | ribonucleoprotein granule | cellular component | GO:0035770 | 89 | 32 | 11940 | 1105 | yps,mael,Edc3,exu,cup,pgc,CG9425,CG17068,Tm1,spn-E,osk,gw,LSm3,Rga,CG11486,tapas,CG6967,tral,armi,krimp,PyK,squ,eIF4A,smg,eIF4E1,CG17768,bru1,nos,CG10418,AGO3,vret,shu | | 5.124e-12 | -26.00 | protein modification process | biological process | GO:0036211 | 1215 | 181 | 12345 | 1129 | Ublcp1,CG10344,Cyp1,GalT1,Pp2B-14D,CG11486,Mtl,rept,PIG-L,e(y)1,Fbw5,CG9941,trbd,CG31633,CG14435,CG8974,lid,Nmt,Pink1,CG8726,CG4041,Lkb1,O-fut1,Cka,Tlk,PpD3,Topors,CG9643,Ptpmeg,Hexo1,d4,CG40045,trr,Cul5,ste14,Parp16,Pdk1,not,Sirt7,Pp2A-29B,CG5003,Pex12,mop,CanA-14F,Tfb4,MAGE,CycE,Chrac-14,PIG-P,CG10254,Pdk,UbcE2H,CG46338,CG1827,PEK,dco,CG33977,Efr,egg,lwr,Cbl,CycB,ssh,Rcd5,png,Mos,CG17746,CG11070,tws,Uev1A,CG5026,EloC,Mekk1,Ubc4,Fkbp59,PpV,scny,CG31053,PIG-H,sip3,esc,mts,CG11858,Usp10,ago,Pex10,tun,Ptp69D,Ptp10D,Ptpmeg2,CkIIalpha,Fkbp12,STUB1,Uch-L5,awd,csw,Gapdh1,QC,Atg7,Hat1,Naa20A,tkv,SRPK,RpL40,Rep,SkpA,CG7840,pont,nej,CG3511,YL-1,Cdk5,CG8188,faf,rl,Tip60,dx,Dph5,twe,Ptp61F,Usp39,RpS27A,otu,Sgf29,Dd,alph,CG2924,Traf6,Coq8,Mo25,CG5880,Cdk7,Herc4,CSN6,CG17698,CG13344,Arf51F,Ptip,CG4238,isoQC,Cchl,Atu,CG3631,Sgf11,Aos1,Cul1,APP-BP1,Spt20,p38a,ari-1,CG32221,park,Tao,Cul2,Ufm1,G9a,CG7115,CG9316,ben,Cdk1,Iru,PIG-C,Uch,grp,CG1578,CycJ,Tak1,Ubc6,fdl,tor,CanB2,Usp16-45,HDAC6,Uba5,Utx,fray,escl,l(3)73Ah,CG2911,velo,CG8314 | | 5.124e-12 | -26.00 | cellular protein modification process | biological process | GO:0006464 | 1215 | 181 | 12345 | 1129 | CG4041,CG8726,CG8974,lid,Nmt,Pink1,CG31633,trbd,CG9941,CG14435,PIG-L,e(y)1,Fbw5,CG11486,Pp2B-14D,rept,Mtl,GalT1,Cyp1,CG10344,Ublcp1,Pdk,CycE,MAGE,CG10254,PIG-P,Chrac-14,CanA-14F,Tfb4,Sirt7,mop,CG5003,Pex12,Pp2A-29B,Parp16,not,Pdk1,trr,CG40045,ste14,Cul5,PpD3,Topors,d4,Ptpmeg,Hexo1,CG9643,O-fut1,Lkb1,Cka,Tlk,Ubc4,Fkbp59,PpV,CG5026,Mekk1,EloC,CG11070,tws,Uev1A,Rcd5,ssh,CG17746,Mos,png,CycB,Cbl,lwr,egg,Efr,PEK,dco,CG33977,CG1827,UbcE2H,CG46338,QC,csw,Gapdh1,Hat1,Atg7,awd,Uch-L5,STUB1,Fkbp12,Ptpmeg2,Ptp10D,CkIIalpha,Pex10,tun,Ptp69D,ago,Usp10,mts,esc,CG11858,PIG-H,CG31053,sip3,scny,RpS27A,Usp39,Dph5,twe,Ptp61F,dx,Tip60,rl,YL-1,CG3511,nej,pont,faf,Cdk5,CG8188,SkpA,RpL40,Rep,CG7840,tkv,Naa20A,SRPK,CG4238,isoQC,Cchl,Atu,Arf51F,Ptip,CSN6,CG13344,CG17698,Herc4,Cdk7,CG5880,Mo25,CG2924,Coq8,Traf6,alph,otu,Sgf29,Dd,ben,CG7115,CG9316,Cul2,G9a,Ufm1,Tao,park,CG32221,ari-1,APP-BP1,p38a,Spt20,CG3631,Cul1,Sgf11,Aos1,CG2911,l(3)73Ah,escl,CG8314,velo,fray,Utx,HDAC6,Usp16-45,Uba5,CanB2,CycJ,Tak1,tor,fdl,Ubc6,CG1578,Cdk1,Iru,grp,Uch,PIG-C | | 5.353e-12 | -25.95 | RpS30 (Ribosomal protein S30) | protein interactions | 42470 | 96 | 38 | 9204 | 1112 | RpS14b,RpS24,RpS7,RpL13,RpL37a,RpL34a,RpS18,RpS2,RpL26,RpS27A,RpLP0,RpL12,RpS28b,RpL13A,RpS4,RpL11,RpS23,RpL40,RpL30,RpS16,RpS27,RpL29,RpLP1,RpL7A,RpL24,RpL23A,RpL10,RpL23,RpL38,PGRP-LE,RpS14a,RpL9,RpL31,RpS13,RpL17,CG10959,RpS15,RpS29 | | 5.518e-12 | -25.92 | RpL35A (Ribosomal protein L35A) | protein interactions | 40654 | 109 | 41 | 9204 | 1112 | RpS29,RpL31,RpL17,RpS13,RpS30,RpS15,RpL38,RpL9,RpS14a,Archease,RpLP1,RpL7A,RpL24,RpL23,RpL10,RpL23A,RpL30,RpS23,RpL40,RpL29,RpS16,RpS27,Pi3K92E,foxo,RpL13A,RpS28b,RpL11,RpS4,Lkb1,RpS2,RpL13,RpS18,RpL37a,RpL34a,RpL12,RpS27A,RpL26,RpLP0,RpS14b,RpS7,RpS24 | | 5.518e-12 | -25.92 | RpL10Ab (Ribosomal protein L10Ab) | protein interactions | 39338 | 109 | 41 | 9204 | 1112 | RpS15,RpL31,RpS13,RpL17,RpS30,RpS29,RpL10,gw,RpL23A,RpL23,RpLP1,RpL7A,RpL24,PGRP-LE,RpS14a,RpL9,RpL38,RpS4,RpL11,eIF4E1,RpS28b,RpL13A,Rga,RpS16,RpS27,RpL29,RpS23,RpL40,RpL30,RpS24,RpS7,RpS14b,RpS27A,RpL26,RpLP0,RpL12,RpL13,RpL37a,RpS18,RpL34a,RpS2 | | 5.657e-12 | -25.90 | cellular macromolecule catabolic process | biological process | GO:0044265 | 425 | 84 | 12345 | 1129 | Cul2,Hsc70-3,CG11070,Prosbeta3,Tengl4,CG31759,CG32640,RYBP,Rpn3,EloC,CG9316,Prosalpha3,Rpn2,Rpn12,Cbp80,Cul1,UbcE2H,Vsp37A,Prosbeta4,Edc3,twin,CG32221,park,ari-1,Cbl,cu,shrb,Usp16-45,CG11164,armi,Not10,Plap,CG6766,Prosbeta6,Uch-L5,cort,Pomp,sip3,scny,Uch,Rga,Ubc6,26-29-p,Usp10,ago,Prosbeta5,Pink1,CG8974,Rrp40,Pym,gus,trbd,CG31633,RpS27A,gw,RpL40,SkpA,Cp1,Prosalpha5,Fbw5,Dcr-1,Drep1,CG8188,Edem1,faf,smg,CG11486,Herc4,CG5003,Rpt2,CG4238,Rpn13,CG10254,CG17768,CG4572,Trf4-2,Topors,Prosalpha4,not,LSm3,Ntan1,Cul5,CG40045,CG3223 | | 5.796e-12 | -25.87 | RpL32 (Ribosomal protein L32) | protein interactions | 43573 | 92 | 37 | 9204 | 1112 | RpL9,RpS14a,RpL38,RpL23,RpL23A,RpL10,RpLP1,RpL7A,RpL24,RpS29,RpS15,RpS30,RpL17,RpS13,RpL31,RpL12,RpLP0,RpS27A,RpL26,RpS2,RpL34a,RpL37a,RpS18,RpL13,RpS7,RpS24,RpS14b,RpL29,RpS16,RpS27,RpL30,RpL40,RpS23,RpL11,RpS4,RpL13A,RpS28b | | 6.263e-12 | -25.80 | vig2 (vig2) | protein interactions | 43016 | 105 | 40 | 9204 | 1112 | RpS29,RpL31,RpS13,RpL17,RpS30,RpS15,RpL38,RpS14a,RpL9,RpL24,RpLP1,RpL7A,RpL10,RpL23A,RpL23,RpL40,RpS23,RpL30,RpS27,RpS16,Pi3K92E,RpL29,RpS28b,foxo,RpL13A,RpS4,RpL11,RpL13,RpS18,RpL37a,RpL34a,RpS2,RpL26,RpS27A,RpLP0,Pdk1,RpL12,RpS14b,RpS24,RpS7 | | 7.030e-12 | -25.68 | RpL41 (Ribosomal protein L41) | protein interactions | 251466 | 101 | 39 | 9204 | 1112 | RpS30,RpL17,RpS13,RpL31,RpS15,RpS29,RpL24,RpLP1,RpL7A,RpL23,CG1275,RpL10,RpL23A,RpL38,RpL9,RpS14a,RpL13A,RpS28b,RpL11,RpS4,RpL30,RpS23,RpL40,RpL29,RpS27,RpS16,RpS14b,hwt,RpS7,RpS24,RpS2,RpL37a,RpL34a,RpS18,RpL13,RpL12,RpLP0,RpS27A,RpL26 | | 7.048e-12 | -25.68 | regulation of biological process | biological process | GO:0050789 | 3862 | 456 | 12345 | 1129 | krimp,up,d4,CG2678,piwi,O-fut1,Lkb1,Tlk,Cka,not,dl,foxo,Cul5,Pura,Ntan1,CG4853,Tfb4,Sirt7,Rpt2,CG5003,Pp2A-29B,Rel,CG10188,Hip1,Sec23,spn-E,sbb,CG4872,CycE,CG3587,Rpn13,Chrac-14,ovo,SERCA,sbr,obe,Maf1,RhoGAP5A,Tm1,e(y)1,CG7328,Fbw5,mtTFB1,adp,hwt,CG11486,CG9425,rept,APC7,lid,DCAF12,spz,CG10418,trbd,AP-1-2beta,SelD,CHOp24,bcd,tub,CG8726,CG3894,Ssdp,muskelin,cyc,cort,sip3,ea,eIF4A,CG2129,ago,Usp10,MED4,Exd2,SREBP,qkr58E-2,hb,r2d2,Sld5,cana,armi,Aldh,Sap47,Ranbp21,Atg7,Sp7,STUB1,PEK,tyf,Rab32,Edc3,Tomosyn,Shmt,Mctp,ERp60,Stam,Nap1,Nurf-38,Mitf,Grip84,Cbl,tapas,Uev1A,Smox,Rcd5,CG31759,CG14407,Scox,png,CG17328,Fkbp59,Ect4,PpV,osk,HP1c,EloC,zyd,alph,spag,CG10631,Zwilch,bru1,otu,Dd,COX4,LSm3,Mo25,mre11,Pld,Rbf,squ,Wnt10,CG4021,CG13344,Tspo,CG17698,Pka-R2,CG6813,Rab5,ix,Cdk7,CG17454,trk,Drep2,Crag,tral,Atu,Grx1,COX5A,SkpA,Gdi,asf1,Naa20A,Fdh,CG4730,insv,BigH1,cad,Nprl2,CG6967,crm,Drep1,spen,SuUR,YL-1,nej,faf,Adi1,Edem1,Rrp40,Bin1,jigr1,CG11710,nbs,dx,MED27,Ac3,pn,Debcl,Rbfox1,wisp,CG13933,twe,fs(1)N,Hnf4,chrb,EMRE,GAPsec,Pomp,Nup358,grp,Unr,CycJ,GILT1,Ubc6,shv,Utx,HDAC6,eIF4E1,bam,Not10,CG6927,CG2911,Arpc1,S2P,trsn,Cbp80,CG9911,Hr78,swa,Stat92E,Cul1,stas,Aos1,Sgf11,Sod1,vret,park,MED24,Tsp86D,ari-1,G9a,Snx6,ZnT35C,SNF4Agamma,Trxr-1,Rab10,Moe,dnc,TyrRS,Son,CG7115,Cenp-C,Ork1,Desat1,Topors,RhoGAP54D,Ptpmeg,nAChRalpha4,CG30496,Imp,Parp16,Pdk1,trr,crok,AGO3,Top1,Cf2,HP1e,CanA-14F,disp,CG2199,bun,GlyP,CG9925,fs(1)M3,mop,Nlg1,REPTOR,Med,Arpc2,MAGE,Synj,Trf2,plu,mael,Calx,stai,Fkbp39,boi,Pi3K92E,Pp2B-14D,Mtl,snk,Pink1,Idh3b,Pym,fwe,CG13601,cact,crq,CG31053,Vps16B,Ankle2,Rga,scny,CG9391,Nf-YA,CG8009,exu,mus81,mts,esc,mad2,Jafrac2,opm,Ptp10D,Atg6,CkIIalpha,mute,csw,Atf3,CG8963,Mybbp1A,Hat1,RpL13A,RhoGDI,CG17202,LpR2,fz4,apt,BicC,dco,twin,wash,CycB,ScsbetaA,Pax,CG17272,Fur2,cu,lwr,egg,E23,Hsc70-3,tws,hng3,ssh,gd,Mos,CG17746,CycC,NFAT,vers,Rpn2,Mekk1,Marf,pps,Trf4-2,Smyd4-4,slif,CG17768,mAChR-A,Bx,CG10336,Traf6,heph,raptor,mbo,Myd88,cup,CG11902,Chd64,RpLP1,Herc4,CG4238,Ptip,CG30291,eIF3m,capt,l(3)mbt,Taf7,PGRP-LE,Rep,TfIIFalpha,Idgf2,Pka-R1,tkv,CG7414,CG3198,rl,Jafrac1,Dcr-1,lin-52,smg,pont,Cdk5,Neb-cGP,ph-d,CG10038,gus,Uxt,Tip60,Marf1,PDCD-5,CG1602,mtrm,Archease,EcR,Dph5,gw,pgc,bif,Ptp61F,enc,CG14408,cnc,Wbp2,Cdk1,CG31875,wapl,Tak1,pygo,Pgk,Gabat,tor,nos,BoYb,CG1578,Prosbeta5,mus304,Seipin,Su(H),psq,Arpc4,Atf6,ASPP,l(3)73Ah,msps,escl,velo,sws,Abi,fray,MED20,p38a,E2f1,Spt20,Npc2a,HmgD,Rbf2,EndoGI,pen-2,Aven,RecQ5,CG10959,shrb,lin,Sox102F,cnn,mRpL13,Cul2,ATPsynD,Coq7,SmydA-1,RYBP,Tao,dgt4,ben,CG10654,Rpn3,stet,Grip71,Iml1,Elba2,Bap170,lt | | 7.792e-12 | -25.58 | RpL12 (Ribosomal protein L12) | protein interactions | 45329 | 110 | 41 | 9204 | 1112 | RpS7,RpS24,RpS14b,RpL26,RpS27A,RpLP0,RpS2,RpL13,RpS18,RpL37a,RpL34a,RpL11,RpS4,RpL13A,RpS28b,RpL29,RpS27,RpS16,RpL30,RpL40,RpS23,RpL23,CG4853,RpL23A,RpL10,RpL24,Tak1,RpLP1,RpL7A,RpL9,slam,Rel,RpS14a,PGRP-LE,RpL38,RpS15,RpL31,RpS30,RpS13,RpL17,RpS29 | | 7.792e-12 | -25.58 | RpS12 (Ribosomal protein S12) | protein interactions | 39480 | 110 | 41 | 9204 | 1112 | RpS24,RpS7,RpS14b,RpS27A,RpL26,CycE,RpLP0,RpL12,RpL13,RpL37a,RpS18,RpL34a,RpS2,RpS4,CG46385,RpL11,RpS28b,RpL13A,RpS16,RpS27,RpL29,RpL40,RpS23,RpL30,RpL23A,RpL10,RpL23,RpL7A,RpLP1,RpL24,RpS14a,RpL9,Not10,RpL38,RpS15,cnn,RpL31,RpL17,RpS13,RpS30,RpS29 | | 7.792e-12 | -25.58 | RpS15Aa (Ribosomal protein S15Aa) | protein interactions | 44150 | 110 | 41 | 9204 | 1112 | bcd,RpS15,RpL31,RpL17,RpS30,RpS13,RpS29,RpL23,RpL10,RpL23A,RpL7A,RpLP1,RpL24,RpL9,RpS14a,RpL38,RpL11,Lkb1,RpS4,RpL13A,RpS28b,RpL29,RpS16,RpS27,Pi3K92E,RpL30,RpS23,RpL40,RpS7,RpS24,RpS14b,Pdk1,RpL12,RpS27A,RpL26,RpLP0,RpS2,RpL13,RpL34a,RpL37a,RpS18 | | 9.388e-12 | -25.39 | mitochondrial membrane part | cellular component | GO:0044455 | 193 | 50 | 11940 | 1105 | CG5903,CG8004,CG11110,ND-MWFE,CG9393,ATPsynF,COX4,ATPsynD,ATPsynbeta,ND-75,ATPsynE,ND-24,Marf,UQCR-Q,ND-23,cype,Tom40,ATPsynB,Pmi,UQCR-11,ND-30,Coq8,ND-MNLL,CG40472,ND-39,COX5B,SdhD,blw,Roe1,Neb-cGP,ATPsynCF6,Tim8,ND-B22,COX5A,ATPsynG,ATPsyndelta,ATPsynO,NP15.6,ND-51,SdhA,ND-B15,CG9399,ND-49,levy,EMRE,Ppox,ND-B18,Tom70,ND-SGDH,ND-18 | | 1.095e-11 | -25.24 | RpS9 (Ribosomal protein S9) | protein interactions | 39108 | 111 | 41 | 9204 | 1112 | RpS23,RpL40,RpL30,Pi3K92E,RpS27,RpS16,RpL29,RpS28b,RpL13A,foxo,RpS4,RpL11,RpL37a,RpL34a,RpS18,RpL13,RpS2,RpLP0,RpS27A,RpL26,RpL12,Pdk1,RpS14b,RpS24,RpS7,RpS29,RpS13,RpS30,RpL17,RpL31,RpS15,RpL38,RpS14a,RpL9,Pink1,RpL24,RpL7A,RpLP1,RpL10,RpL23A,RpL23 | | 1.128e-11 | -25.21 | RpL6 (Ribosomal protein L6) | protein interactions | 43723 | 98 | 38 | 9204 | 1112 | RpS4,RpL11,RpS28b,RpL13A,RpS27,RpS16,Pi3K92E,RpL29,RpL40,RpS23,RpL30,RpS24,RpS7,RpS14b,RpL26,RpS27A,RpLP0,RpL12,RpL13,RpS18,RpL37a,RpL34a,RpS2,RpS15,RpL31,RpS13,RpL17,RpS30,RpS29,RpL10,RpL23A,RpL23,RpL7A,RpLP1,RpL24,RpS14a,RpL9,RpL38 | | 1.137e-11 | -25.20 | regulation of macromolecule metabolic process | biological process | GO:0060255 | 2019 | 268 | 12345 | 1129 | Atu,CG17454,CG13344,CG4021,Cdk7,ix,Pka-R2,CG6813,LSm3,Rbf,squ,CG10631,bru1,alph,Dd,otu,Rbfox1,Hnf4,CG11710,Bin1,jigr1,Rrp40,pn,MED27,Drep1,spen,SuUR,CG6967,crm,faf,YL-1,nej,asf1,SkpA,insv,cad,BigH1,Fdh,CG4730,CG2911,S2P,bam,Utx,HDAC6,Not10,Ubc6,CycJ,Pomp,Nup358,GAPsec,Unr,Son,G9a,SNF4Agamma,park,MED24,vret,ari-1,Cbp80,Hr78,trsn,Cul1,Sgf11,Aos1,Stat92E,spn-E,sbb,CG4872,Chrac-14,CG3587,Rpn13,CycE,Tfb4,Pp2A-29B,Rel,Rpt2,Sirt7,not,foxo,dl,d4,CG2678,krimp,Tlk,Cka,piwi,Lkb1,bcd,lid,CG10418,CG7328,Fbw5,mtTFB1,e(y)1,CG9425,rept,CG11486,ovo,Maf1,RhoGAP5A,obe,STUB1,r2d2,armi,eIF4A,ago,CG2129,Exd2,SREBP,qkr58E-2,hb,MED4,cort,sip3,cyc,Ssdp,PpV,HP1c,osk,CG17328,EloC,Smox,Rcd5,CG31759,Mitf,Nap1,Cbl,PEK,tyf,Edc3,CG4238,eIF3m,CG30291,raptor,heph,RpLP1,Myd88,cup,CG11902,CG10336,Traf6,CG17768,pps,Trf4-2,Smyd4-4,Bx,CG1602,pgc,gw,Ptp61F,EcR,Dph5,Tip60,Uxt,Marf1,ph-d,CG10038,gus,Dcr-1,rl,CG3198,lin-52,smg,pont,TfIIFalpha,l(3)mbt,Taf7,CG7414,tkv,Pka-R1,sws,ASPP,l(3)73Ah,escl,Abi,fray,psq,Atf6,Su(H),tor,nos,pygo,Tak1,CG1578,CG14408,cnc,Wbp2,CG31875,Cdk1,Elba2,Rpn3,ben,CG10654,Bap170,ATPsynD,Coq7,mRpL13,Cul2,Tao,RYBP,SmydA-1,Sox102F,CG10959,lin,MED20,p38a,Spt20,E2f1,HmgD,Rbf2,EndoGI,Med,REPTOR,CG2199,HP1e,Cf2,bun,GlyP,Pdk1,Parp16,AGO3,trr,Topors,CG30496,cact,Pym,Pi3K92E,plu,Trf2,mael,Hat1,mute,Atf3,Mybbp1A,CG8963,CG17202,RpL13A,CkIIalpha,CG8009,Nf-YA,Jafrac2,esc,mts,Ankle2,CG31053,Rga,scny,CycC,NFAT,Mekk1,vers,Rpn2,tws,Hsc70-3,CG17746,Mos,hng3,Pax,ScsbetaA,CycB,wash,lwr,egg,cu,BicC,dco,apt,twin | | 1.260e-11 | -25.10 | RpLP2 (Ribosomal protein LP2) | protein interactions | 36855 | 107 | 40 | 9204 | 1112 | Sgf11,RpL11,RpS4,RpL13A,foxo,RpS28b,RpL29,RpS27,RpS16,RpL30,RpS23,RpL40,RpS7,RpS24,RpS14b,RpL12,RpLP0,RpS27A,RpL26,RpS2,RpL37a,RpL34a,RpS18,RpL13,RpS15,RpS30,RpS13,RpL17,RpL31,RpS29,RpL23,RpL10,RpL23A,RpLP1,RpL7A,RpL24,RpL9,RpS14a,RpL38,dl | | 1.280e-11 | -25.08 | RpL7A (Ribosomal protein L7A) | protein interactions | 31588 | 125 | 44 | 9204 | 1112 | RpS7,RpS24,RpS14b,RpL12,RpLP0,RpS27A,RpL26,RpS2,RpS18,RpL37a,RpL34a,RpL13,RpL11,Lkb1,RpS4,RpL13A,foxo,RpS28b,ago,RpL29,Pi3K92E,RpS27,RpS16,RpL30,RpL40,RpS23,RpL23,RpL10,RpL23A,Tak1,RpLP1,RpL24,slam,RpL9,RpS14a,PGRP-LE,Rel,RpL38,RpS15,RpL17,RpS30,RpS13,RpL31,RpS29 | | 1.282e-11 | -25.08 | eEF2 (eukaryotic translation elongation factor 2) | protein interactions | 35422 | 158 | 51 | 9204 | 1112 | Prosbeta3,RpS14b,RpS24,RpS7,RpL13,RpS18,RpL34a,RpL37a,RpS2,CG10638,RpL26,RpS27A,Chd64,RpLP0,RpL12,RpS28b,Eno,RpL13A,RpS4,NHP2,RpL11,RpL40,RpS23,RpL30,regucalcin,CG10265,RpS27,RpS16,AnxB10,RpL29,RpL24,Tak1,RpL7A,RpLP1,RpL10,RpL23A,RpL23,RpL38,awd,Rel,PGRP-LE,RpS14a,RpL9,RpL31,RpS30,RpS13,RpL17,RpS15,RpS29,Moe,capt | | 1.316e-11 | -25.05 | mitochondrial envelope | cellular component | GO:0005740 | 313 | 68 | 11940 | 1105 | CG40472,ND-MNLL,Coq8,Cyt-c-p,CG11077,ND-23,UQCR-Q,cype,Scox,ATPsynB,UQCR-11,ATPsynD,ND-MWFE,CG9393,CG8004,CG5903,ND-B15,Ptp61F,AIF,CG3803,CG9399,EMRE,ATPsynG,COX5A,SdhA,ND-51,ATPsynO,NP15.6,Cchl,Roe1,Neb-cGP,ND-39,COX5B,Tengl4,ND-30,Marf,ND-24,ATPsynE,Pmi,Tom40,ATPsynbeta,COX4,ATPsynF,ND-75,Tspo,mino,Coq7,CG11110,Mitofilin,ND-49,levy,CG8728,ND-18,ND-SGDH,Debcl,ND-B18,Ppox,Tom70,ND-B22,colt,ATPsyndelta,ATPsynCF6,Etf-QO,Tim8,shop,SdhD,blw,Sdhaf3 | | 1.437e-11 | -24.97 | RpL40 (Ribosomal protein L40) | protein interactions | 33629 | 103 | 39 | 9204 | 1112 | RpS29,trbd,RpS15,RpL31,RpL17,RpS13,RpS30,RpS14a,RpL9,RpL38,RpL23A,RpL10,RpL23,RpL24,RpLP1,RpL7A,RpS27,RpS16,RpL29,Sap47,RpS23,RpL30,RpS4,RpL11,RpS28b,RpL13A,RpS27A,RpL26,RpLP0,RpL12,RpL13,RpL37a,RpL34a,RpS18,RpS2,Wbp2,RpS24,RpS7,RpS14b | | 1.496e-11 | -24.93 | phosphorylation | biological process | GO:0016310 | 470 | 89 | 12345 | 1129 | Cdk5,Mtl,CG11486,ND-23,Pi3K92E,CG7328,AIF,ND-B18,rl,SRPK,wal,tkv,ND-MNLL,Pmi,Cyt-c-p,PyK,Ptp61F,CG8726,Pfk,CG4041,ATPsynO,ND-B22,ND-39,Pink1,Etf-QO,Coq8,COX5B,nmdyn-D6,Pdk1,Mo25,Gk1,Argk,Tlk,Cka,COX4,ND-75,Pglym78,Lkb1,Gk2,Ptpmeg,COX5A,cype,SdhD,UQCR-11,CycE,Pfrx,Pi4KIIalpha,Pdk,kri,Cdk7,CG17698,Tfb4,CG1344,Scsalpha1,SdhA,CycB,CG7789,CG3631,nmdyn-D7,dco,p38a,ND-51,PEK,Mekk1,Tao,png,levy,Mos,Eno,ND-B15,tor,UQCR-Q,Pgk,CycJ,Tak1,CG9391,ND-SGDH,grp,Cdk1,l(2)k01209,ND-24,awd,fray,CG2201,Gapdh1,CkIIalpha,ATPsynbeta,ND-49,Ptpmeg2 | | 1.529e-11 | -24.90 | RpL18A (Ribosomal protein L18A) | protein interactions | 36985 | 112 | 41 | 9204 | 1112 | RpL38,RpS14a,RpL9,RpLP1,RpL7A,RpL24,RpL10,RpL23A,RpL23,RpS29,RpS30,RpS13,RpL17,RpL31,RpS15,RpL37a,RpL34a,RpS18,RpL13,RpS2,RpLP0,RpS27A,RpL26,RpL12,Pdk1,RpS14b,CG8974,RpS24,RpS7,RpL40,RpS23,RpL30,RpS27,RpS16,RpL29,RpS28b,RpL13A,foxo,RpS4,Lkb1,RpL11 | | 1.529e-11 | -24.90 | RpS2 (Ribosomal protein S2) | protein interactions | 34309 | 112 | 41 | 9204 | 1112 | RpS7,RpS24,RpS14b,RpL12,RpS27A,RpL26,RpLP0,RpL13,RpL37a,RpS18,RpL34a,RpL11,RpS4,RpL13A,ago,RpS28b,RpL29,RpS27,RpS16,RpL30,RpL40,RpS23,RpL23,RpL23A,RpL10,RpL7A,RpL24,RpLP1,RpL9,CG9117,RpS14a,RpL38,dl,CG18135,RpS15,RpL31,RpS13,RpS30,RpL17,Smox,RpS29 | | 1.742e-11 | -24.77 | RpS20 (Ribosomal protein S20) | protein interactions | 42464 | 126 | 44 | 9204 | 1112 | RpL9,Rel,PGRP-LE,RpS14a,RpL38,RpL23,RpL10,RpL23A,RpL24,RpL7A,Tak1,RpLP1,PPYR1,RpS29,RpS15,RpL31,RpS13,RpL17,RpS30,RpL12,RpS27A,RpL26,RpLP0,RpS2,RpL13,RpL34a,RpL37a,RpS18,RpS7,RpS24,RpS14b,Kyat,RpL29,RpS16,RpS27,RpL30,Drep1,RpL40,RpS23,RpL11,HP1e,RpS4,RpL13A,RpS28b | | 1.751e-11 | -24.77 | RpLP1 (Ribosomal protein LP1) | protein interactions | 33214 | 140 | 47 | 9204 | 1112 | RpL13A,foxo,RpS28b,RpL11,RpS4,RpL30,RpS23,RpL40,RpL29,Pi3K92E,RpS27,RpS16,Mes2,RpS14b,CCT3,squ,Kyat,RpS7,COX5A,RpS24,RpS2,CCT6,RpL34a,RpS18,RpL37a,RpL13,RpL12,grsm,Pdk1,RpLP0,grp,RpL26,RpS27A,RpS13,RpL17,RpS30,RpL31,RpS15,RpS29,RpL24,RpL7A,RpL23,RpL23A,RpL10,RpL38,RpL9,RpS14a | | 1.773e-11 | -24.76 | RpL19 (Ribosomal protein L19) | protein interactions | 37995 | 108 | 40 | 9204 | 1112 | RpS14a,RpL9,RpL38,RpL10,RpL23A,RpL23,RpLP1,RpL7A,RpL24,RpS29,RpS15,RpL31,RpL17,RpS13,RpS30,RpS27A,RpL26,RpLP0,RpL12,RpL13,RpS18,RpL37a,RpL34a,RpS2,RpS24,RpS7,TyrRS,RpS14b,RpS16,RpS27,Pi3K92E,RpL29,RpS23,RpL40,RpL30,RpS4,RpL11,RpS28b,foxo,RpL13A | | 2.317e-11 | -24.49 | RpS18 (Ribosomal protein S18) | protein interactions | 37292 | 100 | 38 | 9204 | 1112 | RpL11,RpS4,RpL13A,RpS28b,RpL29,RpS16,RpS27,RpL30,RpS23,RpL40,RpS7,RpS24,RpS14b,RpL12,RpS27A,RpL26,RpLP0,RpS2,RpL13,RpL37a,RpL34a,RpS15,CG15525,RpL31,RpS13,RpS30,RpL17,RpS29,Pen,RpL23,RpL10,RpL23A,RpL24,RpL7A,RpLP1,RpL9,RpS14a,RpL38 | | 2.825e-11 | -24.29 | transferase activity | molecular function | GO:0016740 | 1384 | 198 | 12176 | 1119 | CG31633,Cdk1,EloC,Pdxk,ago,mus205,O-fut1,STUB1,CG3631,sip3,Topors,Herc4,Rnmt,G9a,CG2924,Ugt35A1,tor,Dph5,CG6650,mino,UGP,d4,CG11089,Naa20A,CG8314,PIG-C,CG1640,CG5880,tub,CG46338,CG12018,egg,CG10344,Gk1,Tip60,nmdyn-D6,CG10189,r-l,Tlk,DNApol-epsilon255,Pdk,Kyat,Pgk,awd,APC7,Cdk7,Arf51F,Traf6,CG3021,Tak1,Rcd5,Pex12,Kcmf1,tkv,CG8311,Coq8,CG14615,CG11899,Gabat,Pdk1,CG4407,CycB,CG8726,Mos,CG9643,l(1)G0196,Sgf29,PyK,Cbl,rl,park,PIG-P,l(2)k01209,wisp,ben,r,CG14204,Got1,AgmNAT,p38a,pont,CG9485,rept,Argk,CG4238,grp,fray,CG10932,GstT1,Ubc6,CycJ,whd,Agpat2,CG2201,CG1344,Chrac-14,GalT1,CG17698,Mo25,Pi3K92E,ste14,CG8188,Pfk,ari-1,png,CG8067,oxt,Ady43A,kri,SRPK,UbcE2H,CG7328,Trf4-2,escl,Pi4KIIalpha,Iru,Rpb8,mal,Ggt-1,CG46385,e(y)1,nej,CG14100,trr,CG31523,CG15141,Hat1,dx,CkIIalpha,isoQC,Cdk5,Tao,Mekk1,Fbw5,CG8112,CG4502,SkpA,CG9941,GstE11,QC,MAGE,CG4041,alc,CG33156,AGBE,Pex10,CG6767,lwr,Gk2,CG33977,mtTFB1,CG7009,CycE,Atg7,Parp16,CG31053,CG9316,Uev1A,lid,Thiolase,nmdyn-D7,PEK,CG11486,CG40045,Ts,CG5003,CG7718,CG8475,Gapdh1,yip2,CG3894,CG32221,CG10254,esc,CG2100,YL-1,agt,GlyP,Gyg,Tfb4,Ubc4,CG8974,CG6178,GstT2,SelD,Lkb1,Pink1,CG14435,Nmt,Pfrx,CG11070,dco,RpII140,CG13344,CG2938,Ugt36E1,Scsalpha1,Shmt | | 2.871e-11 | -24.27 | RpL7 (Ribosomal protein L7) | protein interactions | 34352 | 105 | 39 | 9204 | 1112 | RpL31,RpS30,RpS13,RpL17,RpS15,CG2061,RpS29,RpL7A,RpL24,RpLP1,RpL23,RpL10,RpL23A,RpL38,RpL9,RpS14a,RpL13A,RpS28b,RpL11,RpS4,RpL30,RpL40,RpS23,RpL29,RpS27,RpS16,RpS14b,RpS7,RpS24,RpS2,RpL13,RpL37a,RpS18,RpL34a,RpL12,RpS27A,RpL26,MED4,RpLP0 | | 2.871e-11 | -24.27 | RpL13 (Ribosomal protein L13) | protein interactions | 34329 | 105 | 39 | 9204 | 1112 | RpL17,RpS30,RpS13,RpL31,RpS15,RpS29,Rcd5,RpL24,RpL7A,RpLP1,RpL23,RpL10,RpL23A,RpL38,RpL9,RpS14a,RpL13A,RpS28b,CG14204,RpL11,Lkb1,RpS4,RpL30,RpL40,RpS23,RpL29,RpS16,RpS27,RpS14b,RpS7,RpS24,RpS2,RpL37a,RpL34a,RpS18,RpL12,RpLP0,RpL26,RpS27A | | 2.937e-11 | -24.25 | cellular nitrogen compound metabolic process | biological process | GO:0034641 | 2226 | 288 | 12345 | 1129 | mRpL35,CG11110,CG10418,SelD,RpS7,CG3021,e(y)1,mtTFB1,CG11486,sbr,obe,mRpS5,kri,Neos,CycE,CG3587,sgll,Scsalpha1,NHP2,RpS4,Tfb4,CG6094,Gclc,Pus7,CG17187,eIF3i,RpS15,nmdyn-D6,Ntan1,GstE11,Sptr,piwi,pr,RpL23A,shu,RpL12,tapas,RfC38,RpLP0,ATPsynE,mRpL48,CG13551,Uev1A,CG31759,CG8199,CG2034,RpS5b,CG7920,Nap1,CysRS,Nurf-38,Pfas,nSMase,nmdyn-D7,Snm1,yps,Edc3,Shmt,RpL7A,awd,r2d2,Sld5,armi,RpS14b,CG11771,eIF4A,Clp,fal,MED4,Exd2,Mat89Ba,CG11089,wisp,RpS27A,PyK,Bin1,Rrp40,agt,Nadsyn,nbs,Ac3,RpL26,spen,Drep1,c(2)M,CG12301,RpL40,CG5986,CG17454,Drep2,ATPsyndelta,ATPsynG,GatB,CG4407,cN-IIIB,Cdk7,LSm3,mre11,RpS29,Rbf,Ppox,GluRS-m,bru1,Pglym78,RpS13,GstT1,CG10463,eEF1beta,TyrRS,Son,CG8204,Eno,Tengl4,AMPdeam,AOX3,whd,vret,RpL17,MED24,CG6767,mRpL17,Ts,DNApol-epsilon255,trsn,RpS23,Gmer,Cbp80,CG32473,r,CG2911,mRpS21,SerRS-m,CG5292,CG7728,eIF4E1,Not10,CG11164,Xpc,Ubc6,Rpb8,snRNA:U4:38AB,GAPsec,CG1774,IntS11,mus205,ATPsynO,AsnS,Spx,Pfk,Ugt36E1,TrpRS-m,Xpac,CG30343,Pym,mRpL36,CG2100,Trf2,RpL29,SP1029,bys,Rnmt,mael,MAGE,RpL31,Ppcdc,LysRS,Imp,CG10189,COX5B,tho2,Top1,Ggt-1,CG1622,CycC,Pdxk,SmE,RpL38,mEFTu2,ScsbetaA,cu,CG9684,CG12375,CG14100,twin,DhpD,Gapdh1,RpS24,RpL13A,Ho,ATPsynbeta,RpS14a,Ady43A,RpL23,RpS18,CG7997,Orc1,RpL37a,exu,mus81,Lsm12,l(2)k01209,RpL34a,snRNA:U12,Rga,mRpL16,Usp39,Archease,gw,del,RpS16,CG14817,Orc6,mino,CG3198,Dcr-1,CG9143,smg,Taf7,TfIIFalpha,RpL34b,CG5285,CG7414,CG33156,mRpL18,snRNA:U4atac,CG14516,Pfrx,ATPsynB,eIF3m,heph,Ugt35A1,UGP,tko,RpLP1,RpS2,CG10336,RpII140,Trf4-2,pps,rig,CG17768,IntS6,CG3803,RpL30,Sgf29,mRpS11,ben,Larp7,blw,CG16865,mRpS23,mRpL10,RpL10,mRpL13,ATPsynD,RpL24,CG7009,pen-2,CG12018,mEFTu1,RpS30,RecQ5,Pcd,scu,CG10561,E2f1,CG17691,r-l,l(3)73Ah,AcCoAS,mRpS10,RpS28b,CG6178,mus304,Dhfr,RpL11,CG6610,pygo,RpS27,Pgk,nos,RpL13,GatA,GstT2,ATPsynCF6,Gasz,tos,RpL9 | | 2.941e-11 | -24.25 | RpS16 (Ribosomal protein S16) | protein interactions | 37580 | 114 | 41 | 9204 | 1112 | RpS7,RpS24,RpS14b,RpL12,Pdk1,RpLP0,RpS27A,RpL26,RpS2,RpL34a,RpL37a,RpS18,RpL13,RpL11,RpS4,RpL13A,RpS28b,RpL29,RpS27,RpL30,RpL40,RpS23,RpL23,RpL10,RpL23A,Tak1,RpL7A,RpL24,RpLP1,RpL9,RpS14a,PGRP-LE,Rel,RpL38,dl,RpS15,RpS13,RpL17,RpS30,RpL31,RpS29 | | 2.961e-11 | -24.24 | female gamete generation | biological process | GO:0007292 | 761 | 125 | 12345 | 1129 | Abi,yl,csw,msps,Atg7,lqfR,psq,armi,Klp10A,eIF4E1,r2d2,Ptpmeg2,mus304,bam,CkIIalpha,mts,eIF4A,wapl,CycJ,Orc1,GatA,exu,nos,tor,yem,ea,Cdk1,grp,muskelin,cnc,cort,Cenp-C,jvl,Moe,Rab10,osk,Iml1,shu,Grip71,Tao,Mos,Cul2,shrb,cnn,ari-1,lwr,egg,Cbl,wash,vret,Pax,milt,park,twin,Stat92E,yps,toc,apt,E2f1,swa,BicC,Grip128,CycE,Pen,Med,spn-E,Rab5,gammaTub37C,cup,Pxt,fs(1)M3,mop,bun,heph,raptor,Cf2,squ,Rbf,Cul5,Top1,tho2,Pdk1,Lkb1,otu,Grip75,piwi,krimp,bru1,fs(1)N,EcR,enc,del,bif,cact,Axs,wisp,Rbfox1,bcd,mtrm,gus,Syx13,lid,spz,Sec15,nej,Pp2B-14D,cathD,Pi3K92E,Nup62,e(y)1,elg1,stai,rl,Nprl2,Dcr-1,Drep1,stil,tkv,Pka-R1,SERCA,Tm1,SRPK,mael,capt,ovo | | 3.209e-11 | -24.16 | cellular process involved in reproduction in multicellular organism | biological process | GO:0022412 | 997 | 153 | 12345 | 1129 | bif,pgc,del,enc,EcR,twe,fs(1)N,mtrm,Rbfox1,wisp,Syx13,gus,Sec15,faf,Nup62,cathD,nej,Drep1,Dcr-1,rl,Nprl2,elg1,Jafrac1,SRPK,Pka-R1,tkv,capt,l(3)mbt,eIF3m,Arf51F,Grip128,sw,Rab5,cup,Kap3,CSN6,raptor,heph,sau,larp,Rbf,squ,otu,Grip75,bru1,jvl,Cenp-C,Grip71,Iml1,Moe,Rab10,Tao,Cul2,ari-1,cnn,zpg,shrb,park,milt,vret,toc,Stat92E,swa,E2f1,yl,Abi,msps,psq,bam,mus304,Klp10A,eIF4E1,BoYb,tor,nos,GatA,wapl,CycJ,grp,Cdk1,yem,cnc,Nup358,bcd,Axs,cact,spz,lid,Pi3K92E,Pp2B-14D,stai,e(y)1,mael,Tm1,SERCA,stil,ovo,Pen,CycE,spn-E,Med,fs(1)M3,bun,mop,Pex12,Pxt,gammaTub37C,tef,disp,HP1e,Cf2,Top1,Cul5,tho2,piwi,Lkb1,krimp,Topors,shu,osk,Mos,egg,Cbl,lwr,Pax,ScsbetaA,wash,CycB,Nap1,twin,yps,CG46338,BicC,nmd,apt,lqfR,Atg7,csw,armi,Pex10,CkIIalpha,Ptpmeg2,r2d2,mts,mad2,exu,eIF4A,Pex13,Orc1,ea,cort,muskelin,slam | | 3.290e-11 | -24.14 | RpL36 (Ribosomal protein L36) | protein interactions | 31009 | 101 | 38 | 9204 | 1112 | RpS4,RpL11,RpS28b,RpL13A,Atpalpha,RpS16,RpS27,RpL29,RpL40,RpS23,RpL30,RpS24,RpS7,RpS14b,RpS27A,RpL26,RpLP0,RpL12,RpL13,RpL37a,RpS18,RpL34a,RpS2,RpS15,RpL31,RpS30,RpS13,RpL17,RpS29,RpL10,RpL23A,RpL23,RpL24,RpL7A,RpLP1,RpS14a,RpL9,RpL38 | | 3.305e-11 | -24.13 | ribonucleoprotein complex | cellular component | GO:1990904 | 754 | 125 | 11940 | 1105 | CG7728,Usp39,CG9684,mael,NHP2,yps,RpS23,RpL40,exu,mRpL36,CG14817,heph,RpS4,RpL31,SmE,RpS5b,Tm1,mRpL17,gw,mRpL13,spn-E,RpLP0,RpL30,CG11486,mRpS23,tapas,Rga,armi,tral,CG17454,mRpS21,mRpL18,RpL34b,pps,CG6967,PyK,eIF4A,mRpS11,RpL9,Spx,CG1622,snRNA:U12,RpS27A,scu,CG17768,RpS14a,RpS24,RpL13,CG15525,vret,CG10418,AGO3,rig,CG12288,RpL7A,RpS28b,obe,Neos,shu,CG3198,shep,snRNA:U4atac,cup,CG3511,Cbp80,Edc3,RpL17,CG9425,RpS13,CG7414,pgc,RpL34a,RpLP1,mRpL10,mRpS10,RpS15,RpS2,CG17068,osk,Larp7,RpS7,RpS29,CG12301,RpL10,mRpS5,RpS30,LSm3,krimp,RpL23,Imp,RpL23A,Dcr-1,mRpL48,RpL12,squ,CG5986,tko,RpS27,RpS16,Bin1,snRNA:U4:38AB,CG6094,RpS18,smg,RpL38,RpL29,CG17187,mRpL16,CG16865,bys,bru1,RpL37a,RpL26,mRpL35,eIF4E1,RpS14b,nos,Mat89Ba,RpL13A,RpL11,CG8204,CG6610,r2d2,PEK,RpL24 | | 4.023e-11 | -23.94 | RpL8 (Ribosomal protein L8) | protein interactions | 44251 | 106 | 39 | 9204 | 1112 | RpS2,RpL37a,RpL34a,RpS18,RpL13,RpL12,Pdk1,RpLP0,RpL26,RpS27A,RpS14b,RpS7,RpS24,RpL30,RpL40,RpS23,RpL29,Pi3K92E,RpS16,RpS27,RpL13A,RpS28b,RpL11,RpS4,RpL38,RpL9,RpS14a,RpL7A,RpLP1,RpL24,RpL23,RpL23A,RpL10,RpS29,RpL17,RpS30,RpS13,RpL31,RpS15 | | 4.023e-11 | -23.94 | sta (stubarista) | protein interactions | 31106 | 106 | 39 | 9204 | 1112 | RpL10,RpL23A,RpL23,RpL7A,RpL24,RpLP1,RpS14a,RpL9,RpL38,RpS15,RpL31,RpL17,RpS13,RpS30,CG4511,RpS29,RpS24,RpS7,RpS14b,RpL26,RpS27A,RpLP0,RpL12,RpL13,RpL34a,RpL37a,RpS18,RpS2,RpS4,RpL11,RpS28b,RpL13A,RpS27,RpS16,Pi3K92E,RpL29,RpS23,RpL40,RpL30 | | 4.023e-11 | -23.94 | RpL30 (Ribosomal protein L30) | protein interactions | 44059 | 106 | 39 | 9204 | 1112 | RpLP1,RpL24,RpL7A,RpL10,RpL23A,RpL23,RpL38,RpS14a,RpL9,RpS13,RpS30,RpL17,RpL31,RpS15,RpS29,RpS14b,RpS24,RpS7,RpL37a,RpS18,RpL34a,RpL13,RpS2,RpLP0,RpL26,RpS27A,RpL12,RpS28b,RpL13A,RpS4,Lkb1,RpL11,E2f1,RpS23,RpL40,tos,RpS16,RpS27,RpL29 | | 4.049e-11 | -23.93 | RpL14 (Ribosomal protein L14) | protein interactions | 38983 | 115 | 41 | 9204 | 1112 | RpL30,RpS23,RpL40,RpL29,Pi3K92E,RpS27,RpS16,RpL13A,foxo,RpS28b,RpL11,RpS4,asf1,Lkb1,RpS2,RpL37a,RpL34a,RpS18,RpL13,RpL12,RpLP0,RpS27A,RpL26,RpS14b,RpS7,RpS24,RpS29,RpS30,RpS13,RpL17,RpL31,RpS15,RpL38,RpL9,RpS14a,RpL7A,RpLP1,RpL24,RpL23,RpL23A,RpL10 | | 4.180e-11 | -23.90 | RpL29 (Ribosomal protein L29) | protein interactions | 37430 | 93 | 36 | 9204 | 1112 | RpL11,RpS4,RpL13A,RpS28b,RpS16,RpS27,RpL30,RpL40,RpS23,RpS7,RpS24,RpS14b,RpL12,RpS27A,RpL26,RpLP0,RpS2,RpL13,RpL34a,RpS18,RpL37a,RpS15,RpL31,RpL17,RpS13,RpS30,RpS29,RpL23,RpL23A,RpL10,RpL7A,RpLP1,RpL24,RpL9,RpS14a,RpL38 | | 4.339e-11 | -23.86 | Nonsense Mediated Decay (NMD) independent of the Exon Junction Complex (EJC) | REACTOME pathways | R-DME-975956 | 95 | 40 | 3923 | 574 | RpS15,RpL31,RpS5b,RpL10,RpS4,RpLP0,RpL23A,RpLP1,RpL34a,RpL13,RpS13,RpS2,RpS27,RpS30,RpL29,RpS14b,RpL40,RpS16,RpS18,RpL23,RpL17,RpL9,RpL13A,RpS14a,RpL7A,RpL12,RpL38,RpL37a,RpL34b,RpS29,RpL24,RpS23,RpL26,RpL30,RpL11,RpS27A,RpS24,RpS28b,Cbp80,RpS7 | | 4.645e-11 | -23.79 | RpL21 (Ribosomal protein L21) | protein interactions | 35453 | 102 | 38 | 9204 | 1112 | RpS24,RpS7,RpS14b,RpLP0,RpL26,RpS27A,RpL12,RpS18,RpL34a,RpL37a,RpL13,RpS2,RpS4,RpL11,RpS28b,RpL13A,Pi3K92E,RpS27,RpS16,RpL29,RpS23,RpL40,RpL30,RpL10,RpL23A,RpL23,RpLP1,RpL24,RpL7A,RpS14a,RpL9,RpL38,RpS15,RpS30,RpL17,RpS13,RpL31,RpS29 | | 5.205e-11 | -23.68 | RpS26 (Ribosomal protein S26) | protein interactions | 35098 | 125 | 43 | 9204 | 1112 | RpS27A,RpL26,RpLP0,Pdk1,RpL12,RpL13,RpL34a,RpS18,RpL37a,RpS2,cup,RpS24,RpS7,RpS14b,RpS16,RpS27,Pi3K92E,RpL29,RpS23,RpL40,RpL30,RpS4,RpL11,RpS28b,foxo,RpL13A,RpS14a,RpL9,dl,CG9902,RpL38,RpL10,RpL23A,RpL23,RpLP1,RpL7A,RpL24,RpS29,RpS15,RpL31,RpS13,RpS30,RpL17 | | 5.737e-11 | -23.58 | RpS7 (Ribosomal protein S7) | protein interactions | 43594 | 130 | 44 | 9204 | 1112 | RpS18,RpL37a,RpL34a,RpL13,RpS2,RpLP0,RpL26,RpS27A,RpL12,cad,Pdk1,RpS14b,RpS24,RpS23,RpL40,RpL30,Pi3K92E,RpS27,RpS16,RpL29,RpS28b,RpL13A,foxo,RpS4,HP1e,RpL11,RpL38,Archease,RpS14a,RpL9,Sld5,RpL24,RpLP1,Snm1,RpL7A,RpL10,RpL23A,RpL23,RpS29,RpS30,RpS13,RpL17,RpL31,RpS15 | | 6.001e-11 | -23.54 | RpL11 (Ribosomal protein L11) | protein interactions | 37235 | 94 | 36 | 9204 | 1112 | RpL30,RpL40,RpS23,RpL29,RpS27,RpS16,RpL13A,RpS28b,RpS4,RpS2,RpL34a,RpS18,RpL37a,RpL13,RpL12,RpLP0,RpS27A,RpL26,RpS14b,RpS7,RpS24,RpS29,RpS30,RpL17,RpS13,RpL31,RpS15,RpL38,RpL9,RpS14a,RpLP1,RpL7A,RpL24,RpL23,RpL23A,RpL10 | | 6.520e-11 | -23.45 | RpS25 (Ribosomal protein S25) | protein interactions | 41343 | 103 | 38 | 9204 | 1112 | RpS15,RpL31,RpS30,RpS13,RpL17,RpS29,RpL10,RpL23A,RpL23,RpL24,RpLP1,RpL7A,RpS14a,RpL9,RpL38,RpS4,RpL11,RpS28b,RpL13A,RpS16,RpS27,RpL29,RpS23,RpL40,Cyp6v1,RpL30,RpS24,RpS7,RpS14b,RpL26,RpS27A,RpLP0,RpL12,RpL13,RpS18,RpL34a,RpL37a,RpS2 | | 6.591e-11 | -23.44 | RpS13 (Ribosomal protein S13) | protein interactions | 34149 | 112 | 40 | 9204 | 1112 | RpL13A,RpS28b,RpL11,Sgf11,CG46385,RpS4,RpL30,RpS23,RpL40,RpL29,RpS16,RpS27,Pi3K92E,RpS14b,RpS7,RpS24,RpS2,RpL13,RpS18,RpL37a,RpL34a,RpL12,RpS27A,RpL26,RpLP0,RpL31,RpS30,RpL17,RpS15,RpS29,RpL7A,RpLP1,RpL24,RpL23,CG4853,RpL23A,RpL10,RpL38,RpL9,RpS14a | | 7.154e-11 | -23.36 | non-membrane-bounded organelle | cellular component | GO:0043228 | 1752 | 238 | 11940 | 1105 | Mybbp1A,tko,EcR,RpS16,G9a,RpL23,MAGE,krimp,Snm1,CG1890,Grip75,Sccpdh1,Girdin,RpS29,LSm3,RpL10,RpS7,Tao,pont,Non1,pgc,Topors,CG9425,RecQ5,mRpL10,RpLP1,mad2,CG13366,cup,Actn,RpL17,Edc3,piwi,Moe,rept,Utx,Arpc2,Arpc4,eIF3m,gammaTub37C,Elba2,dgt4,CG10336,Mat89Ba,RpL11,DNApol-epsilon255,RpS14b,cbs,mRpL35,RpL37a,d4,alphaTub67C,RpS18,Bin1,lwr,Tlk,alphaTub84B,Orc6,mRpL18,mRpS21,mRpS23,Top1,stai,CG9143,Rga,Arpc1,Pp2A-29B,mRpL13,stil,Trf2,gw,up,RpL30,spn-E,Hip1,RpL31,RpS4,Ptpmeg,CG3437,Tm1,RpS5b,CG14817,scny,cana,mRpL36,exu,NHP2,yps,mael,Blos2,CG9186,bam,CG12288,AGO3,vret,Pgk,CG2129,Tip60,Chrac-14,Grip84,Rpb8,RpS24,RpS14a,RpL13,mts,Mf,Rbf2,dob,nej,eIF4A,RpL9,Fkbp39,RpS27A,Ubc6,RpS27,trr,sw,squ,CG31053,CycB,cnc,RpL12,RpL23A,mRpL48,wapl,CG12301,capt,mRpS5,RpS30,osk,BigH1,awd,Cen,Axs,RpS2,CG17068,RpS13,CG7414,l(1)G0004,Zwilch,mRpS10,RpS15,RpL34a,spd-2,Cenp-C,mre11,Grip71,Klp10A,Sld5,Uxt,STUB1,lin-52,Pax,del,CG7716,RpL24,PEK,Gapdh1,RpL13A,nos,parvin,Sirt7,nbs,mbo,egg,RpL26,jvl,Su(H),bys,bru1,eIF4E1,CG14183,Grip128,RpL38,lid,smg,CG6094,Pglym78,RpL29,mRpL16,l(3)mbt,Cdk1,PyK,Rbf,tral,armi,pps,CG6967,Kap3,RpL34b,Cdk7,CG11486,tapas,twe,ph-d,Psf3,Nurf-38,RpLP0,elg1,toc,mRpL17,CG12018,Orc1,esc,RpL40,RpS23,Hat1,HP1e,msps,Jabba,yem,Rcd5,EloC,RpS28b,HmgD,shu,Bap170,Nup62,RpL7A,Mtl,CG9265,CG10418,RfC38,SuUR,Clc,Rep,CycE,Chd64,CG17768,cnn,HP1c,asf1,kl-2,tef,Trf4-2,Rel,Nap1,mRpS11 | | 7.154e-11 | -23.36 | intracellular non-membrane-bounded organelle | cellular component | GO:0043232 | 1752 | 238 | 11940 | 1105 | egg,nbs,mbo,Sirt7,CG14183,Grip128,eIF4E1,Su(H),bru1,jvl,bys,RpL26,mRpL16,RpL29,Pglym78,CG6094,lid,RpL38,smg,Cdk1,l(3)mbt,lin-52,STUB1,Uxt,RpL24,PEK,CG7716,Pax,del,Gapdh1,parvin,nos,RpL13A,CG17068,RpS2,RpL34a,spd-2,RpS15,mRpS10,Zwilch,RpS13,CG7414,l(1)G0004,mre11,Cenp-C,Sld5,Klp10A,Grip71,CG31053,squ,trr,RpS27,sw,RpL12,RpL23A,wapl,mRpL48,cnc,CycB,mRpS5,RpS30,capt,CG12301,Axs,Cen,awd,osk,BigH1,Chd64,CycE,kl-2,HP1c,asf1,cnn,CG17768,Rel,Nap1,tef,Trf4-2,mRpS11,Bap170,shu,HmgD,RpS28b,Rcd5,EloC,Mtl,Nup62,RpL7A,SuUR,RfC38,CG9265,CG10418,Clc,Rep,mRpL17,elg1,toc,esc,CG12018,Orc1,msps,HP1e,Hat1,RpS23,RpL40,Jabba,yem,Rbf,PyK,RpL34b,Cdk7,pps,Kap3,CG6967,armi,tral,Psf3,twe,ph-d,CG11486,tapas,RpLP0,Nurf-38,cbs,RpS14b,alphaTub67C,d4,RpL37a,mRpL35,RpS18,alphaTub84B,Tlk,Bin1,lwr,gammaTub37C,eIF3m,Arpc4,CG10336,dgt4,Elba2,DNApol-epsilon255,RpL11,Mat89Ba,Non1,pont,mRpL10,RecQ5,RpLP1,CG9425,pgc,Topors,piwi,Edc3,RpL17,Actn,CG13366,cup,mad2,Arpc2,Utx,rept,Moe,G9a,EcR,RpS16,Mybbp1A,tko,CG1890,Snm1,krimp,MAGE,RpL23,RpL10,LSm3,RpS29,Girdin,Sccpdh1,Grip75,Tao,RpS7,mts,RpL13,RpS14a,Rpb8,RpS24,Rbf2,Mf,nej,dob,Ubc6,RpS27A,RpL9,eIF4A,Fkbp39,CG9186,Blos2,bam,Tip60,CG2129,Pgk,vret,CG12288,AGO3,Grip84,Chrac-14,Tm1,CG3437,RpS5b,Ptpmeg,RpL31,Hip1,RpS4,scny,CG14817,exu,mRpL36,cana,mael,yps,NHP2,mRpL18,mRpS21,Orc6,CG9143,Rga,stai,Top1,mRpS23,spn-E,RpL30,stil,Trf2,up,gw,Arpc1,mRpL13,Pp2A-29B | | 8.500e-11 | -23.19 | RpS3A (Ribosomal protein S3A) | protein interactions | 43768 | 122 | 42 | 9204 | 1112 | RpS29,RpS15,RpS30,RpS13,RpL17,RpL31,RpL9,RpS14a,RpL38,RpL23,RpL23A,RpL10,RpL24,RpL7A,RpLP1,RpL29,Pi3K92E,RpS27,RpS16,RpL30,RpL40,RpS23,RpL11,mus304,RpS4,RpL13A,dgt4,RpS28b,RpL12,RpLP0,Dcr-1,RpL26,RpS27A,RpS2,RpL34a,RpL37a,RpS18,RpL13,RpS7,RpS24,RpS14b,CG8974 | | 8.984e-11 | -23.13 | mitochondrial membrane | cellular component | GO:0031966 | 298 | 64 | 11940 | 1105 | ATPsynD,ND-MWFE,CG9393,CG5903,CG8004,Cyt-c-p,CG11077,ND-MNLL,CG40472,Coq8,UQCR-11,ATPsynB,cype,ND-23,UQCR-Q,Scox,Cchl,Neb-cGP,Roe1,EMRE,CG9399,CG3803,ND-B15,ND-51,SdhA,NP15.6,ATPsynO,ATPsynG,COX5A,Tspo,ND-75,mino,Coq7,ATPsynbeta,COX4,ATPsynF,CG11110,COX5B,ND-39,ND-30,Tengl4,Pmi,Tom40,ND-24,ATPsynE,Marf,Etf-QO,Tim8,ATPsynCF6,blw,SdhD,ND-18,ND-SGDH,Tom70,ND-B18,Ppox,Debcl,levy,Mitofilin,ND-49,CG8728,colt,ATPsyndelta,ND-B22 | | 9.605e-11 | -23.07 | mitochondrial inner membrane | cellular component | GO:0005743 | 244 | 56 | 11940 | 1105 | cype,UQCR-Q,ND-23,Scox,ATPsynE,ND-24,UQCR-11,Pmi,ATPsynB,COX5B,ND-39,CG40472,ND-MNLL,Coq8,ND-30,Tengl4,Cyt-c-p,CG11077,CG11110,CG5903,ATPsynbeta,ATPsynD,COX4,ATPsynF,ND-MWFE,ND-75,Coq7,ATPsynG,COX5A,ND-B22,ND-51,SdhA,colt,NP15.6,ATPsynO,ATPsyndelta,levy,ND-49,Mitofilin,CG3803,CG9399,ND-B15,CG8728,ND-18,ND-SGDH,ND-B18,Ppox,EMRE,blw,Roe1,SdhD,Neb-cGP,ATPsynCF6,Etf-QO,Tim8,Cchl | | 1.030e-10 | -23.00 | energy derivation by oxidation of organic compounds | biological process | GO:0015980 | 151 | 41 | 12345 | 1129 | CG8475,Pgm1,Idh3b,ND-39,levy,ND-B22,Etf-QO,CG5028,ND-23,SdhA,CG9485,ND-B18,AIF,ScsbetaA,ND-MNLL,Gyg,Pmi,ND-51,Nc73EF,SdhD,UQCR-11,cype,COX5A,NP15.6,Irp-1B,ACC,GlyP,Scsalpha1,ND-49,Atg6,UGP,COX5B,AGBE,Coq8,foxo,UQCR-Q,ND-B15,ND-75,ND-SGDH,COX4,ND-24 | | 1.056e-10 | -22.97 | RpS24 (Ribosomal protein S24) | protein interactions | 37589 | 100 | 37 | 9204 | 1112 | RpS29,RpS15,RpS30,RpS13,RpL17,RpL31,RpL9,RpS14a,RpL38,RpL23,RpL10,RpL23A,RpLP1,RpL24,RpL7A,RpL29,RpS27,RpS16,RpL30,RpL40,RpS23,RpL11,RpS4,RpL13A,RpS28b,RpL12,CycE,RpLP0,RpL26,RpS27A,RpS2,RpS18,RpL37a,RpL34a,RpL13,RpS7,RpS14b | | 1.073e-10 | -22.96 | AnxB10 (Annexin B10) | protein interactions | 33019 | 109 | 39 | 9204 | 1112 | yip2,Chd64,CG11267,CG14715,CG10638,ATPsynbeta,Pgk,PHGPx,Cyp1,EndoGI,cib,Stip1,Prosbeta5,Prosbeta3,CG5844,ERp60,Naa20A,regucalcin,Eno,Gapdh1,SelT,Trxr-1,awd,Sod1,Fkbp12,Aos1,RhoGDI,HmgD,ATPsynCF6,Usp39,CG4598,Gdi,capt,scu,Moe,Prosalpha4,Uch,Prosbeta6,Pglym78 | | 1.103e-10 | -22.93 | cellular component organization or biogenesis | biological process | GO:0071840 | 2975 | 362 | 12345 | 1129 | Nap1,Mitf,Nurf-38,Grip84,Snm1,fs(1)Ya,Tomosyn,Edc3,Rab32,RpL7A,Shmt,osk,RpL12,RpLP0,RfC38,CG8004,RpS5b,Smox,Rcd5,Scox,Pex13,ago,CG9393,Arl6IP1,alphaTub84B,SREBP,Mat89Ba,cort,P5CDh1,Atg7,scramb1,awd,r2d2,cana,dpr6,armi,Ptp69D,RpS14b,SelR,e(y)1,mtTFB1,Sec10,rept,Pmi,Roe1,ovo,obe,cbs,Tm1,RhoGAP5A,Axs,CHOp24,CG5903,CG4041,lid,spz,SelD,CG11110,trbd,RpS7,CCT6,Liprin-beta,not,RpS15,Cul5,Pura,foxo,up,krimp,d4,Lkb1,piwi,Atpalpha,Tlk,RpL23A,Tom40,spn-E,Hip1,Sec23,TMEM216,sbb,CycE,Rpn13,Chrac-14,CG7716,NHP2,Svil,CG6094,tef,Tfb4,Sirt7,Rel,Pp2A-29B,CG12773,Gclc,park,ari-1,swa,CG11077,Actn,Stat92E,Sgf11,ATP8A,Moe,Rab10,dnc,Cenp-C,Desat1,CG8204,jvl,Tengl4,G9a,Snx6,Blos4,Xpc,Ubc6,Lnpk,Pomp,yem,Tim8,Unr,grp,Arpc1,CG12288,eIF4E1,HDAC6,CG7728,Utx,CG13366,Vamp7,bam,CG7488,Nprl2,SuUR,Drep1,spen,nej,ND-23,c(2)M,CG5969,YL-1,Nup62,CG12301,SkpA,asf1,Naa20A,SRPK,BigH1,CG3437,wisp,Non1,Debcl,Hnf4,twe,fs(1)N,Rrp40,Bin1,nbs,CG8728,Syx13,RpL26,mre11,LSm3,squ,Rbf,larp,Pld,EMC3,bru1,Grip75,COX4,Dd,tral,Atu,Crag,Arf51F,Rab5,Pka-R2,CG9981,sw,wash,CycB,Pax,Girdin,parvin,CG9186,lwr,egg,dco,BicC,SmE,Marf,Mekk1,tws,ssh,Orc1,exu,mad2,esc,mts,mus81,Vps16B,slam,RpL34a,CG31053,ND-18,Ankle2,scny,Tom70,csw,Hat1,Ptp10D,Ndc1,opm,RpS14a,CkIIalpha,Atg6,Pex10,stai,CG5510,Pp2B-14D,Pi3K92E,Mtl,Trf2,CG8611,bys,mael,cact,crq,Pfk,Rtnl1,Pink1,Xpac,Pym,Imp,Pdk1,crok,trr,Top1,Topors,CG10055,Ptpmeg,cold,Pi4KIIalpha,Mitofilin,CG14894,Med,Gmap,Nlg1,Arpc2,Synj,Pen,HP1e,Cf2,gammaTub37C,ACC,Pxt,fs(1)M3,mop,Pex12,bun,mEFTu1,milt,CG10465,shrb,RecQ5,cnn,E2f1,Spt20,p38a,CG14671,HmgD,spd-2,dgt4,ben,Iml1,mRpL10,Elba2,Grip71,RpL10,colt,lt,RpL24,Sry-alpha,RpS27,wapl,GatA,nos,Prosbeta5,CG1890,CG31729,cnc,Caf1-105,Sdhaf3,Cdk1,PIG-C,escl,msps,l(3)73Ah,velo,sws,Abi,bor,RpS28b,Klp10A,mus304,Seipin,RpL11,Arpc4,CanB2,psq,rl,CG3198,mino,Dcr-1,CG9143,pont,Cdk5,CG8188,cib,capt,Taf7,Oseg1,Pka-R1,tkv,RpL34b,Usp39,mtrm,EcR,del,enc,Ptp61F,bif,alphaTub67C,RpS16,Sec15,CG10038,gus,Orc6,ph-d,CG42748,Tip60,CG11562,sau,Trf4-2,CG17768,rig,mRpS11,Sgf29,Sec22,Cchl,Vha68-2,Grip128,eIF3m,Ptip,Clc,heph,raptor,mbo,Kap3,cup,TBCC | | 1.104e-10 | -22.93 | negative regulation of biological process | biological process | GO:0048519 | 1502 | 208 | 12345 | 1129 | Hnf4,EcR,pgc,gw,bif,Ptp61F,chrb,CG1602,Rbfox1,mtrm,Debcl,ph-d,pn,MED27,CG10038,Tip60,Uxt,Marf1,Rrp40,Bin1,nbs,dx,YL-1,lin-52,nej,smg,faf,Nprl2,rl,CG6967,Drep1,SuUR,Dcr-1,Idgf2,tkv,Naa20A,insv,BigH1,PGRP-LE,l(3)mbt,SkpA,asf1,COX5A,CG4238,Drep2,tral,cup,Rab5,ix,Herc4,heph,raptor,mbo,Rbf,squ,CG10336,Traf6,LSm3,mre11,Dd,COX4,alph,Trf4-2,Smyd4-4,slif,spag,CG10631,Zwilch,bru1,CG17768,Bap170,dnc,Iml1,Elba2,RYBP,SmydA-1,Tao,Cul2,mRpL13,ATPsynD,CG10959,RecQ5,shrb,Sox102F,vret,park,Aven,Rbf2,Stat92E,EndoGI,Cul1,trsn,p38a,Cbp80,E2f1,escl,sws,velo,Not10,Su(H),psq,Utx,mus304,bam,BoYb,Prosbeta5,wapl,pygo,nos,tor,Ubc6,Cdk1,grp,Unr,CG31875,CG14408,CG8726,cact,bcd,SelD,Pym,lid,Pink1,Pi3K92E,adp,CG11486,CG9425,rept,Fkbp39,mael,Maf1,plu,ovo,CycE,MAGE,Chrac-14,spn-E,sbb,Med,CG4872,Sirt7,bun,GlyP,Rel,Pp2A-29B,CG2199,AGO3,dl,foxo,Cul5,Ntan1,Pdk1,piwi,Lkb1,CG30496,Tlk,Cka,krimp,d4,Ptpmeg,Mekk1,CG17328,PpV,Ect4,NFAT,Smox,CG31759,ssh,CG17746,Mos,tapas,Hsc70-3,tws,cu,Fur2,Cbl,lwr,egg,wash,twin,Edc3,apt,BicC,dco,RpL13A,STUB1,csw,Mybbp1A,Aldh,Hat1,Atg7,armi,Ptp10D,r2d2,Atg6,CkIIalpha,mus81,mts,esc,mad2,MED4,SREBP,hb,ago,Rga,scny,cyc,Ankle2 | | 1.280e-10 | -22.78 | organophosphate metabolic process | biological process | GO:0019637 | 383 | 75 | 12345 | 1129 | Pld,COX5B,nmdyn-D6,l(1)G0196,Gk1,Pglym78,Gk2,CG8311,sgll,PIG-P,Synj,ATPsynB,ATPsynG,Pfrx,ATPsyndelta,Pi4KIIalpha,CG33156,cN-IIIB,Agpat2,Ppcdc,CG17333,Scsalpha1,CG4407,Pi3K92E,PIG-L,mino,PyK,Pfk,ATPsynO,Ac3,Nadsyn,ATPsynCF6,fal,Pgk,CG9391,PIG-C,CG11089,CG1774,l(2)k01209,PIG-H,awd,sws,CG18135,AcCoAS,Gapdh1,r,CG2201,CG11164,Ady43A,Seipin,ATPsynbeta,CG6178,Ts,ScsbetaA,CG6767,CG7920,CG7789,r-l,CG17691,nSMase,nmdyn-D7,CG6707,CG33977,scu,Pfas,mal,CG5026,Pdxk,blw,CG7718,AMPdeam,CG8199,ATPsynE,ATPsynD,Eno | | 1.317e-10 | -22.75 | SRP-dependent cotranslational protein targeting to membrane | REACTOME pathways | R-DME-1799339 | 94 | 39 | 3923 | 574 | RpL38,RpL34b,RpL37a,RpL17,RpL9,RpS14a,RpL13A,RpL12,RpL7A,RpL11,RpL30,RpS24,RpS27A,RpS7,RpS28b,RpS29,RpL26,RpL24,RpS23,RpL13,RpS13,RpS2,RpL10,RpS5b,RpL31,RpS15,RpLP0,RpS4,RpL23A,RpL34a,RpLP1,RpS16,RpL23,RpS18,RpS27,RpS14b,RpL29,RpS30,RpL40 | | 1.359e-10 | -22.72 | modification-dependent macromolecule catabolic process | biological process | GO:0043632 | 297 | 63 | 12345 | 1129 | Rpt2,Herc4,CG5003,Usp16-45,Rpn13,Uch-L5,CG10254,Prosbeta6,CG4238,Plap,CG6766,Uch,scny,Topors,Trf4-2,Prosalpha4,cort,sip3,Pomp,CG3223,CG40045,Prosbeta5,Ntan1,Cul5,Usp10,ago,Ubc6,not,gus,CG31633,trbd,CG32640,RYBP,Cul2,Prosbeta3,CG8974,CG11070,Hsc70-3,Rrp40,Pink1,Prosalpha3,Rpn12,Rpn2,CG9316,EloC,Rpn3,RpS27A,Vsp37A,UbcE2H,Prosbeta4,Prosalpha5,Cul1,RpL40,SkpA,shrb,CG8188,ari-1,Edem1,faf,Cbl,Fbw5,CG32221,park | | 1.469e-10 | -22.64 | inner mitochondrial membrane protein complex | cellular component | GO:0098800 | 151 | 41 | 11940 | 1105 | Tim8,ATPsynCF6,Neb-cGP,Roe1,blw,SdhD,ND-B18,EMRE,ND-18,ND-SGDH,ND-B15,levy,ND-49,NP15.6,ATPsynO,ATPsyndelta,ND-51,SdhA,COX5A,ND-B22,ATPsynG,ND-75,ATPsynF,COX4,ND-MWFE,ATPsynbeta,ATPsynD,CG5903,CG11110,ND-30,ND-39,COX5B,ND-MNLL,CG40472,UQCR-11,ATPsynB,ND-24,ATPsynE,cype,UQCR-Q,ND-23 | | 1.745e-10 | -22.47 | RpL37a (Ribosomal protein L37a) | protein interactions | 32446 | 106 | 38 | 9204 | 1112 | RpS29,RpS15,RpL31,RpS13,RpS30,RpL17,RpL9,RpS14a,RpL38,RpL23,RpL10,RpL23A,RpLP1,RpL7A,RpL24,RpL29,RpS16,RpS27,RpL30,RpL40,RpS23,csw,RpL11,CG12301,RpS4,RpL13A,RpS28b,RpL12,RpL26,RpS27A,RpLP0,RpS2,RpL13,RpL34a,RpS18,RpS7,RpS24,RpS14b | | 1.959e-10 | -22.35 | Eno (Enolase) | protein interactions | 33351 | 93 | 35 | 9204 | 1112 | CG4598,Argk,Gdi,Moe,capt,scu,Uch,Pglym78,Adh,Trxr-1,awd,Fkbp12,Sod1,RhoGDI,CG4853,ATPsynCF6,AnxB10,regucalcin,Eno,Scp1,yip2,RpS27A,CG11267,Chd64,CG6707,Pgk,ATPsynbeta,CG14715,CG10638,cib,EndoGI,Cyp1,PHGPx,Stip1,Prosbeta5 | | 2.044e-10 | -22.31 | regulation of cellular metabolic process | biological process | GO:0031323 | 1987 | 259 | 12345 | 1129 | Su(H),psq,Atf6,Seipin,Abi,fray,l(3)73Ah,ASPP,escl,sws,Cdk1,CG31875,CG14408,cnc,Wbp2,CG1578,pygo,Tak1,Pgk,Gabat,nos,tor,RYBP,Tao,Cul2,mRpL13,ATPsynD,Coq7,Bap170,CG10654,ben,Elba2,HmgD,Rbf2,EndoGI,MED20,Spt20,E2f1,CG10959,lin,Sox102F,Myd88,CG11902,cup,RpLP1,heph,raptor,CG30291,Ptip,eIF3m,Bx,pps,CG10336,Traf6,ph-d,CG10038,Uxt,Tip60,Marf1,Dph5,EcR,gw,pgc,Ptp61F,CG1602,Pka-R1,tkv,CG7414,Taf7,l(3)mbt,TfIIFalpha,lin-52,pont,smg,rl,CkIIalpha,RpL13A,CG17202,mute,Atf3,Mybbp1A,CG8963,Hat1,Rga,scny,CG31053,Ankle2,mts,esc,Jafrac2,Nf-YA,CG8009,hng3,Mos,CG17746,tws,vers,Mekk1,CycC,NFAT,twin,apt,dco,BicC,egg,lwr,CycB,wash,Pax,ScsbetaA,GlyP,bun,HP1e,Cf2,CG2199,Synj,Med,REPTOR,CG30496,Topors,AGO3,trr,Parp16,Pdk1,Pink1,cact,mael,Trf2,plu,Pi3K92E,Fkbp39,Not10,Utx,HDAC6,bam,S2P,CG2911,Unr,GAPsec,Pomp,CycJ,Ubc6,SNF4Agamma,G9a,Son,Desat1,Stat92E,Cul1,Aos1,Sgf11,Hr78,ari-1,Sod1,park,MED24,CG6813,Pka-R2,Cdk7,ix,CG4021,CG13344,Tspo,CG17454,Atu,otu,Dd,alph,bru1,CG10631,Rbf,pn,MED27,Bin1,CG11710,Hnf4,Debcl,Rbfox1,wisp,Fdh,CG4730,insv,BigH1,cad,SkpA,asf1,YL-1,nej,faf,Nprl2,crm,SuUR,spen,Drep1,r2d2,Sp7,STUB1,Atg7,Ssdp,cyc,sip3,cort,MED4,qkr58E-2,Exd2,SREBP,hb,eIF4A,ago,CG2129,Rcd5,Smox,Scox,EloC,CG17328,PpV,osk,HP1c,Rab32,PEK,tyf,Cbl,Nap1,Mitf,Rpt2,Sirt7,Rel,Pp2A-29B,Tfb4,CycE,CG3587,Chrac-14,Rpn13,spn-E,sbb,CG4872,piwi,Lkb1,Tlk,Cka,krimp,d4,CG2678,dl,foxo,not,AP-1-2beta,CG10418,SelD,lid,spz,bcd,obe,Maf1,RhoGAP5A,ovo,rept,e(y)1,CG7328,mtTFB1 | | 2.501e-10 | -22.11 | RpL23A (Ribosomal protein L23A) | protein interactions | 38208 | 121 | 41 | 9204 | 1112 | RpL38,PGRP-LE,RpS14a,Rel,RpL9,RpLP1,Tak1,RpL7A,RpL24,RpL10,RpL23,RpS29,Utx,RpS30,RpS13,RpL17,RpL31,RpS15,RpS18,RpL37a,RpL34a,RpL13,RpS2,RpLP0,RpL26,RpS27A,RpL12,RpS14b,RpS24,RpS7,RpS23,RpL40,RpL30,RpS27,RpS16,RpL29,RpS28b,RpL13A,foxo,RpS4,RpL11 | | 2.702e-10 | -22.03 | germ cell development | biological process | GO:0007281 | 820 | 129 | 12345 | 1129 | eIF4A,Orc1,exu,mad2,mts,muskelin,ea,csw,Atg7,lqfR,r2d2,Ptpmeg2,CkIIalpha,Pex10,armi,Nap1,CycB,wash,Pax,ScsbetaA,egg,Cbl,apt,nmd,BicC,CG46338,yps,twin,osk,shu,Mos,Top1,Cul5,tho2,krimp,Lkb1,piwi,Med,spn-E,CycE,Pen,Cf2,HP1e,gammaTub37C,Pxt,fs(1)M3,Pex12,bun,mop,e(y)1,stai,Pi3K92E,ovo,stil,SERCA,Tm1,mael,cact,bcd,lid,spz,CycJ,GatA,nos,tor,BoYb,Nup358,cnc,Cdk1,msps,Abi,yl,eIF4E1,bam,psq,vret,milt,park,shrb,zpg,ari-1,swa,E2f1,Stat92E,toc,Rab10,Moe,Iml1,Grip71,jvl,Cul2,Tao,squ,Rbf,bru1,Grip75,otu,Grip128,eIF3m,heph,raptor,CSN6,Rab5,Kap3,cup,sw,elg1,Jafrac1,rl,Nprl2,Dcr-1,Drep1,nej,cathD,Nup62,capt,l(3)mbt,tkv,Pka-R1,SRPK,wisp,Rbfox1,EcR,fs(1)N,enc,del,pgc,Sec15,gus | | 2.732e-10 | -22.02 | RpL23 (Ribosomal protein L23) | protein interactions | 37628 | 112 | 39 | 9204 | 1112 | RpL31,RpS30,RpL17,RpS13,RpS15,RpS29,RpL7A,RpLP1,RpL24,RpL10,RpL23A,RpL38,RpL9,RpS14a,PGRP-LE,foxo,RpL13A,RpS28b,RpL11,RpS4,Lkb1,RpL30,RpL40,RpS23,RpL29,RpS27,RpS16,RpS14b,RpS7,RpS24,RpS2,RpL13,RpL37a,RpS18,RpL34a,RpL12,RpL26,RpS27A,RpLP0 | | 2.732e-10 | -22.02 | RpL13A (Ribosomal protein L13A) | protein interactions | 40687 | 112 | 39 | 9204 | 1112 | RpS29,RpL31,RpL17,RpS30,RpS13,RpS15,RpL38,RpL9,RpS14a,RpLP1,RpL24,RpL7A,RpL23,RpL10,RpL23A,RpL30,RpL40,RpS23,RpL29,RpS27,RpS16,Pi3K92E,foxo,RpS28b,RpL11,RpS4,RpS2,RpL13,RpL37a,RpS18,RpL34a,Pdk1,RpL12,RpS27A,RpL26,RpLP0,RpS14b,RpS7,RpS24 | | 2.837e-10 | -21.98 | RpL24 (Ribosomal protein L24) | protein interactions | 34754 | 103 | 37 | 9204 | 1112 | RpS4,RpL11,RpS28b,RpL13A,RpS27,RpS16,RpL29,RpS23,RpL40,RpL30,RpS24,RpS7,RpS14b,RpS27A,RpL26,RpLP0,RpL12,RpL13,RpL34a,RpL37a,RpS18,RpS2,RpS15,RpL31,RpS13,RpL17,RpS30,RpS29,RpL10,RpL23A,RpL23,RpLP1,RpL7A,RpS14a,PGRP-LE,RpL9,RpL38 | | 2.859e-10 | -21.98 | organophosphate biosynthetic process | biological process | GO:0090407 | 213 | 50 | 12345 | 1129 | PIG-L,mino,CG6767,Pi3K92E,Ts,Pfas,nmdyn-D7,CG33977,CG17691,r-l,CG7718,ATPsynO,blw,Pdxk,mal,Nadsyn,ATPsynE,ATPsynD,CG8199,Ac3,AMPdeam,Gk1,nmdyn-D6,l(1)G0196,ATPsynCF6,Pld,COX5B,l(2)k01209,PIG-H,CG8311,CG11089,Gk2,PIG-C,AcCoAS,r,CG33156,CG2201,ATPsyndelta,ATPsynB,ATPsynG,sgll,PIG-P,awd,ATPsynbeta,CG4407,CG6178,Scsalpha1,Ppcdc,Agpat2,Ady43A | | 3.048e-10 | -21.91 | RpL27 (Ribosomal protein L27) | protein interactions | 43103 | 117 | 40 | 9204 | 1112 | RpL10,RpL23A,RpL23,RpL24,RpL7A,RpLP1,RpS14a,RpL9,RpL38,RpS15,cnn,RpL31,RpL17,RpS30,RpS13,RpS29,RpS24,RpS7,RpS14b,RpS27A,RpL26,RpLP0,Pdk1,RpL12,RpL13,RpL34a,RpL37a,RpS18,RpS2,Lkb1,RpS4,RpL11,RpS28b,RpL13A,RpS16,RpS27,RpL29,RpL40,RpS23,RpL30 | | 3.144e-10 | -21.88 | fabp (fatty acid binding protein) | protein interactions | 3772232 | 73 | 30 | 9204 | 1112 | ERp60,Stip1,RhoGDI,HmgD,CG14715,CG10638,cib,Cyp1,EndoGI,PHGPx,Fkbp59,Fkbp12,Sod1,yip2,CG11267,awd,Chd64,Eno,Pglym78,CG11771,Uch,regucalcin,capt,Moe,scu,CG4598,CG9186,AnxB10,CG3226,CG10265 | | 3.194e-10 | -21.86 | Ribosome, eukaryotes | KEGG pathways | M00177 | 92 | 39 | 3198 | 496 | RpS15,RpL9,RpL34b,RpL7A,RpL13,RpL29,RpS18,RpL30,RpS2,RpS14a,RpS24,RpL17,RpL37a,RpL26,RpL31,RpS27,RpL38,RpS13,RpS5b,RpS16,RpL10,RpL12,RpS28b,RpLP0,RpL40,RpS30,RpL13A,RpS7,RpLP1,RpL34a,RpS23,RpS29,RpL23,RpS27A,RpL11,RpS4,RpL24,RpL23A,RpS14b | | 3.194e-10 | -21.86 | Ribosome, eukaryotes | KEGG pathways | dme\_M00177 | 92 | 39 | 3198 | 496 | RpLP0,RpL40,RpS30,RpL13A,RpS7,RpS16,RpS5b,RpL10,RpS28b,RpL12,RpL23,RpS27A,RpL11,RpS4,RpL23A,RpL24,RpS14b,RpLP1,RpL34a,RpS23,RpS29,RpL30,RpS2,RpS15,RpL9,RpL34b,RpL7A,RpL13,RpL29,RpS18,RpS13,RpS14a,RpS24,RpL37a,RpL17,RpL31,RpL26,RpL38,RpS27 | | 3.831e-10 | -21.68 | modification-dependent protein catabolic process | biological process | GO:0019941 | 290 | 61 | 12345 | 1129 | CG40045,CG3223,Ntan1,Prosbeta5,Cul5,Usp10,ago,Ubc6,not,scny,Uch,Topors,Prosalpha4,sip3,cort,Pomp,Rpn13,Uch-L5,CG10254,Prosbeta6,CG4238,CG6766,Plap,Rpt2,Herc4,CG5003,Usp16-45,shrb,ari-1,CG8188,Edem1,faf,Cbl,Fbw5,CG32221,park,UbcE2H,Vsp37A,Prosbeta4,Prosalpha5,Cul1,RpL40,SkpA,Rpn2,Rpn12,Prosalpha3,EloC,CG9316,Rpn3,RpS27A,gus,RYBP,CG31633,trbd,CG32640,Cul2,Hsc70-3,CG8974,CG11070,Prosbeta3,Pink1 | | 4.158e-10 | -21.60 | Formation of a pool of free 40S subunits | REACTOME pathways | R-DME-72689 | 105 | 41 | 3923 | 574 | RpL13,RpS13,RpS2,RpS5b,RpL10,RpL31,RpS15,RpLP0,RpS4,RpL34a,RpL23A,RpLP1,RpS16,RpL23,RpS18,eIF3i,RpS27,RpS14b,RpL29,RpS30,RpL40,RpL38,RpL34b,RpL37a,RpL17,RpL9,RpL13A,RpS14a,RpL12,RpL7A,RpL11,RpL30,eIF3m,RpS24,RpS27A,RpS7,RpS28b,RpS29,RpL26,RpL24,RpS23 | | 4.873e-10 | -21.44 | oxidoreductase activity | molecular function | GO:0016491 | 658 | 109 | 12176 | 1119 | CG9471,Grx1,CG31674,COX4,ERp60,CG7840,Ppox,ND-18,Jafrac1,SdhD,ND-MNLL,CG4842,CG2767,Idh,CG4586,CG31548,ND-75,CG14997,CG7550,CG3939,Nc73EF,CG10561,Gmer,shop,CG9436,Cyp6a22,SdhA,AIF,SelR,ND-24,CG31549,Fdh,sgll,scu,CG17691,Coq6,Cyp6v1,Sod3,ND-39,ND-49,Coq7,Scox,Pxt,Dhfr,CG7322,COX5A,AOX3,Sccpdh1,COX5B,Cyt-c-p,CG10425,Jafrac2,CG3609,CG4860,Ho,CG10638,Aldh,Sod1,Gapdh1,SelT,cl,Thiolase,Mtpalpha,CG5028,Adh,CG1275,CG6144,P5cr,P5CDh1,CG12171,COX8,CG10863,PHGPx,GILT1,Trxr-1,Mcad,UQCR-11,ND-B22,Etf-QO,Desat1,ND-SGDH,UQCR-Q,ND-30,Idh3b,CG8199,CG14407,cype,Marc,sni,wal,CG9265,ND-51,CG3803,Cyp6a19,ND-23,CG3842,glob1,Cyp6u1,levy,CG10463,Adi1,GstT1,ND-B18,ND-B15,GstT2,stw,CG6178,Sptr,Acox57D-p | | 4.892e-10 | -21.44 | organelle inner membrane | cellular component | GO:0019866 | 254 | 56 | 11940 | 1105 | Neb-cGP,blw,Roe1,SdhD,Tim8,Cchl,Etf-QO,ATPsynCF6,NP15.6,ATPsyndelta,ATPsynO,ND-51,SdhA,colt,COX5A,ND-B22,ATPsynG,ND-B18,Ppox,EMRE,ND-SGDH,ND-18,CG9399,CG3803,ND-B15,CG8728,levy,Mitofilin,ND-49,CG5903,CG11110,Coq7,ND-75,COX4,ATPsynF,ND-MWFE,ATPsynbeta,ATPsynD,Pmi,UQCR-11,ATPsynB,Scox,ATPsynE,ND-24,cype,UQCR-Q,ND-23,CG11077,Cyt-c-p,Coq8,ND-30,Tengl4,ND-39,COX5B,ND-MNLL,CG40472 | | 4.984e-10 | -21.42 | cellular component organization | biological process | GO:0016043 | 2830 | 344 | 12345 | 1129 | scramb1,awd,P5CDh1,Atg7,dpr6,Ptp69D,SelR,RpS14b,armi,r2d2,cana,Arl6IP1,CG9393,alphaTub84B,SREBP,ago,Pex13,cort,RpL12,osk,RpS5b,Smox,Rcd5,Scox,RpLP0,RfC38,CG8004,Grip84,Nap1,Mitf,Nurf-38,Tomosyn,Rab32,Edc3,Shmt,Snm1,fs(1)Ya,CycE,Chrac-14,Rpn13,CG7716,Tom40,Hip1,TMEM216,sbb,Sec23,spn-E,Sirt7,Pp2A-29B,Rel,CG12773,Gclc,Svil,CG6094,tef,Tfb4,foxo,Cul5,Pura,not,RpS15,Lkb1,piwi,Atpalpha,Tlk,RpL23A,up,krimp,d4,CHOp24,Axs,CG5903,CG4041,SelD,trbd,CG11110,CCT6,Liprin-beta,lid,spz,Sec10,rept,e(y)1,obe,Tm1,cbs,RhoGAP5A,Roe1,Pmi,ovo,Arpc1,eIF4E1,HDAC6,CG13366,Utx,Vamp7,bam,CG7488,Lnpk,Blos4,Xpc,Ubc6,yem,Tim8,Unr,grp,Pomp,Cenp-C,Desat1,jvl,CG8204,ATP8A,Rab10,Moe,dnc,Snx6,Tengl4,G9a,ari-1,park,Actn,Stat92E,Sgf11,CG11077,swa,Arf51F,tral,Atu,Crag,Rab5,Pka-R2,CG9981,sw,Rbf,squ,Pld,larp,mre11,LSm3,Grip75,COX4,Dd,EMC3,bru1,fs(1)N,twe,Hnf4,wisp,Non1,Debcl,Syx13,Bin1,nbs,CG8728,ND-23,nej,c(2)M,YL-1,CG5969,Nup62,Nprl2,SuUR,spen,Drep1,Naa20A,SRPK,BigH1,CG3437,SkpA,asf1,Tom70,csw,Hat1,Pex10,Ptp10D,opm,Ndc1,CkIIalpha,RpS14a,Atg6,mad2,esc,mus81,mts,Orc1,exu,scny,Vps16B,slam,CG31053,ND-18,Ankle2,SmE,Marf,Mekk1,ssh,tws,CG9186,egg,lwr,CycB,wash,Girdin,Pax,parvin,BicC,dco,Arpc2,Synj,Pen,Pi4KIIalpha,CG14894,Mitofilin,Med,Gmap,Nlg1,gammaTub37C,ACC,Pxt,bun,fs(1)M3,mop,Pex12,HP1e,Cf2,crok,trr,Top1,Imp,Pdk1,cold,Topors,CG10055,Ptpmeg,cact,crq,Pfk,Xpac,Pym,Rtnl1,Pink1,Pp2B-14D,Pi3K92E,Mtl,stai,CG5510,mael,Trf2,bor,Abi,RpS28b,escl,l(3)73Ah,msps,sws,velo,RpL11,Arpc4,CanB2,psq,Klp10A,mus304,Seipin,Prosbeta5,CG1890,RpS27,wapl,GatA,nos,Sdhaf3,Cdk1,PIG-C,CG31729,cnc,Caf1-105,colt,RpL10,lt,spd-2,dgt4,ben,Elba2,Iml1,Grip71,Sry-alpha,RpL24,shrb,RecQ5,cnn,milt,CG10465,mEFTu1,HmgD,Spt20,E2f1,p38a,CG14671,Grip128,eIF3m,Ptip,Clc,Vha68-2,Cchl,Sec22,cup,Kap3,TBCC,heph,raptor,sau,CG11562,mRpS11,Sgf29,Trf4-2,CG17768,rig,EcR,Ptp61F,enc,del,bif,Usp39,mtrm,gus,CG10038,Orc6,ph-d,CG42748,Tip60,alphaTub67C,Sec15,pont,CG8188,Cdk5,cib,CG3198,rl,mino,Dcr-1,Pka-R1,tkv,capt,Taf7,Oseg1 | | 4.986e-10 | -21.42 | Rack1 (Receptor of activated protein kinase C 1) | protein interactions | 34070 | 133 | 43 | 9204 | 1112 | Sry-alpha,RpL38,CG11486,Rel,RpS14a,RpL9,RpL24,RpLP1,RpL7A,RpL10,RpL23A,RpL23,RpS29,RpL31,RpS30,RpS13,RpL17,RpS15,RpL13,RpL34a,RpL37a,RpS18,RpS2,RpL26,RpS27A,RpLP0,Pdk1,RpL12,Kyat,RpS14b,RpS24,RpS7,RpS23,RpL40,RpL30,RpS16,RpS27,RpL29,RpS28b,RpL13A,RpS4,RpL11,Sgf11 | | 5.135e-10 | -21.39 | Akt1 (Akt1) | protein interactions | 41957 | 199 | 56 | 9204 | 1112 | RpL23,CycC,Cul1,r-l,RpLP1,Sply,Pp2A-29B,Hsc70-3,Kap-alpha3,RpL9,RpS14a,Desat1,Ptp61F,Shmt,heph,CCT5,Rpt2,RpL17,wal,RpL31,Mcad,cathD,yps,Nc73EF,RpS7,Scox,Girdin,CG5844,CG32425,blw,Kcmf1,Rpn13,Pdk1,26-29-p,gus,ATPsynbeta,CCT6,SERCA,Vha68-2,GCS2beta,CG9316,foxo,eIF4A,ScsbetaA,Pi3K92E,CG17508,ND-75,Atpalpha,Ntan1,RpS16,Nap1,Droj2,EcR,PyK,CG1416,ben | | 5.450e-10 | -21.33 | RpS29 (Ribosomal protein S29) | protein interactions | 41200 | 119 | 40 | 9204 | 1112 | RpL38,RpL9,PGRP-LE,RpS14a,RpL7A,RpLP1,RpL24,RpL23,l(2)k01209,RpL23A,RpL10,RpL31,RpS13,RpL17,RpS30,RpS15,RpS2,gus,RpL13,RpS18,RpL37a,RpL34a,RpL12,RpS27A,RpL26,RpLP0,RpS14b,RpS7,RpS24,RpL30,RpS23,RpL40,RpL29,RpS27,RpS16,RpL13A,RpS28b,RpL11,Sgf11,RpS4 | | 5.765e-10 | -21.27 | Nonsense Mediated Decay (NMD) enhanced by the Exon Junction Complex (EJC) | REACTOME pathways | R-DME-975957 | 110 | 42 | 3923 | 574 | RpS27,RpL29,RpS14b,RpS30,mts,RpL40,RpS16,RpL23,RpS18,RpS5b,RpL10,RpL31,RpS15,RpLP0,RpS4,RpL23A,RpLP1,RpL34a,RpL13,RpS13,RpS2,RpS29,RpL26,RpL24,RpS23,RpL11,RpL30,RpS27A,RpS24,RpS7,Cbp80,RpS28b,Pp2A-29B,RpL9,RpL17,RpS14a,RpL13A,RpL12,RpL7A,RpL38,RpL34b,RpL37a | | 5.765e-10 | -21.27 | Nonsense-Mediated Decay (NMD) | REACTOME pathways | R-DME-927802 | 110 | 42 | 3923 | 574 | RpS2,RpS13,RpL13,RpL23A,RpL34a,RpLP1,RpS4,RpLP0,RpS15,RpL31,RpS5b,RpL10,RpS18,RpL23,RpS16,RpL40,mts,RpS30,RpL29,RpS14b,RpS27,RpL37a,RpL34b,RpL38,RpL7A,RpL12,RpL13A,RpS14a,RpL17,RpL9,Pp2A-29B,Cbp80,RpS28b,RpS7,RpS24,RpS27A,RpL30,RpL11,RpL24,RpS23,RpL26,RpS29 | | 5.969e-10 | -21.24 | ubiquitin-dependent protein catabolic process | biological process | GO:0006511 | 286 | 60 | 12345 | 1129 | Cul1,Prosalpha5,Prosbeta4,UbcE2H,Vsp37A,SkpA,RpL40,Cbl,faf,CG8188,Edem1,ari-1,shrb,park,Fbw5,CG32221,RYBP,CG32640,trbd,CG31633,gus,Pink1,Hsc70-3,CG11070,Prosbeta3,CG8974,Cul2,CG9316,EloC,Prosalpha3,Rpn2,Rpn12,Rpn3,Uch,scny,sip3,cort,Pomp,Prosalpha4,Topors,Cul5,Prosbeta5,Ntan1,CG3223,CG40045,not,Ubc6,ago,Usp10,CG5003,Herc4,Rpt2,Usp16-45,Uch-L5,CG10254,Rpn13,Plap,CG6766,CG4238,Prosbeta6 | | 6.029e-10 | -21.23 | RpL31 (Ribosomal protein L31) | protein interactions | 35988 | 110 | 38 | 9204 | 1112 | RpS14a,RpL9,RpL38,RpL10,RpL23A,RpL23,RpLP1,RpL7A,RpL24,RpS29,RpS15,RpS30,RpS13,RpL17,RpL26,RpS27A,RpLP0,Pdk1,RpL12,RpL13,RpL37a,RpS18,RpL34a,RpS2,RpS24,RpS7,RpS14b,RpS16,RpS27,RpL29,RpS23,RpL40,RpL30,Lkb1,RpS4,RpL11,RpS28b,RpL13A | | 6.372e-10 | -21.17 | proteolysis involved in cellular protein catabolic process | biological process | GO:0051603 | 315 | 64 | 12345 | 1129 | CG4572,Prosalpha4,Topors,Cul5,Ntan1,CG40045,CG3223,not,CG5003,Herc4,Rpt2,CG10254,Rpn13,CG4238,Prosalpha5,SkpA,Cp1,RpL40,faf,Edem1,CG8188,Fbw5,CG31633,trbd,gus,Pink1,CG8974,RpS27A,scny,Uch,Pomp,sip3,cort,Prosbeta5,26-29-p,Ubc6,ago,Usp10,Usp16-45,Uch-L5,CG6766,Plap,Prosbeta6,Cul1,Prosbeta4,Vsp37A,UbcE2H,Cbl,ari-1,shrb,park,CG32221,CG32640,RYBP,Prosbeta3,Hsc70-3,CG11070,Cul2,CG9316,EloC,Prosalpha3,Rpn12,Rpn2,Rpn3 | | 6.889e-10 | -21.10 | GO\_CELLULAR\_CATABOLIC\_PROCESS | MSigDB lists | GO\_CELLULAR\_CATABOLIC\_PROCESS | 581 | 142 | 4161 | 644 | Prosbeta6,CG17593,mus81,Ubc4,APC7,ABCD,RpS24,CG8974,Rpn3,RpL12,Mtpalpha,Cdk5,CG32549,RpL26,RpS27,CG4872,Dcr-1,Fdh,dnc,CG11070,SkpA,CG8475,scu,Vsp37A,Rpn2,CG8188,ste24a,CG10465,eIF4A,Mcad,Cdk1,wal,Prosalpha5,CG14516,Unr,Herc4,Got1,shrb,Sply,CG15111,CG6766,CG9485,faf,RpS2,P5CDh1,CG8963,CG5285,Pfk,CG7322,RpL24,LSm3,Hsc70-3,RpL37a,Gk2,CG7550,tzn,Not10,eIF4E1,Rrp40,RpL13A,RpS13,trbd,Prosalpha4,Rpt2,CG11164,Cul2,CG9471,STUB1,Pcd,RpS27A,Faf2,RpS28b,Nc73EF,RpL40,RpL38,RpS4,RpS7,CG1640,Pex13,RpS16,CG4390,RpL23,Cul1,Thiolase,RpS18,CycB,Prosbeta4,Uch,CG32473,Gyg,RpL11,CG10932,Tip60,CG4572,Jafrac1,CG17691,HDAC6,UbcE2H,Pgm1,Prosbeta3,mad2,Treh,Shmt,Fbw5,Uch-L5,CG8199,RpS23,CG10418,Cbp80,not,CG40045,Etf-QO,DhpD,Edc3,Atg6,ari-1,RpL30,Edem1,lwr,CG3376,Prosbeta5,Gabat,Pp2A-29B,Iru,Prosalpha3,cyc,CG6610,RpS30,RpLP0,Rpn12,mts,RpL13,AIF,Cul5,Echs1,GlyP,CG7997,Jheh2,RpL9,twin,CG7840,snz | | 7.732e-10 | -20.98 | aub (aubergine) | protein interactions | 34524 | 67 | 28 | 9204 | 1112 | bru1,BicC,pgc,nos,PyK,bam,bif,dnc,CycB,l(3)mbt,Cbl,scny,eIF4E1,eIF4A,Eno,tapas,osk,cnc,cib,tkv,AGO3,Pgk,vret,wisp,squ,twin,CG4991,smg | | 8.357e-10 | -20.90 | cellular protein catabolic process | biological process | GO:0044257 | 317 | 64 | 12345 | 1129 | Plap,CG6766,Prosbeta6,Uch-L5,Usp16-45,26-29-p,Ubc6,ago,Usp10,Prosbeta5,sip3,cort,Pomp,scny,Uch,Rpn3,CG9316,EloC,Rpn12,Rpn2,Prosalpha3,Prosbeta3,Hsc70-3,CG11070,Cul2,CG32640,RYBP,park,CG32221,Cbl,ari-1,shrb,Cul1,Prosbeta4,Vsp37A,UbcE2H,CG4238,CG10254,Rpn13,CG5003,Herc4,Rpt2,not,Cul5,Ntan1,CG40045,CG3223,CG4572,Prosalpha4,Topors,RpS27A,Pink1,CG8974,trbd,CG31633,gus,Fbw5,faf,Edem1,CG8188,SkpA,Cp1,RpL40,Prosalpha5 | | 8.357e-10 | -20.90 | regulation of cell cycle | biological process | GO:0051726 | 317 | 64 | 12345 | 1129 | APC7,nbs,mtrm,twe,l(3)mbt,SkpA,plu,Nprl2,SuUR,nej,Pi3K92E,raptor,Cdk7,CG5003,CycE,CG30291,COX5A,bru1,Zwilch,otu,Tlk,mre11,Rbf,Ntan1,foxo,CG10336,tws,Smox,png,SNF4Agamma,Mos,CG17746,ben,Moe,dgt4,Iml1,Grip71,CycC,Son,PEK,E2f1,Rbf2,Aos1,Stam,CycB,Aven,Grip84,RecQ5,cnn,mus304,CkIIalpha,cana,Sld5,Su(H),armi,CG31053,Cdk1,grp,wapl,CycJ,ago,mad2,mus81,Prosbeta5 | | 8.770e-10 | -20.85 | Calr (Calreticulin) | protein interactions | 41166 | 93 | 34 | 9204 | 1112 | CG4598,Arpc2,scu,capt,Moe,Uch,Prosalpha4,Pglym78,Rtnl1,awd,Trxr-1,CG1092,Sod1,Fkbp12,HmgD,RhoGDI,CG10265,AnxB10,regucalcin,CG14204,Eno,CG11267,Chd64,yip2,PHGPx,Cyp1,EndoGI,cib,ATPsynbeta,CG14715,CG10638,Stip1,Nepl3,ERp60 | | 9.185e-10 | -20.81 | Prx2540-2 (Peroxiredoxin 2540-2) | protein interactions | 36098 | 80 | 31 | 9204 | 1112 | CG2767,Eno,Pglym78,Uch,GstT1,Gdi,regucalcin,capt,Moe,scu,CG3226,AnxB10,CG4598,CG5844,Stip1,RhoGDI,CG9391,Prosbeta5,CG10638,CG14715,Pgk,Fkbp59,Fkbp12,PHGPx,cib,EndoGI,yip2,Chd64,Trxr-1,CG11267,awd | | 9.583e-10 | -20.77 | RpL5 (Ribosomal protein L5) | protein interactions | 3355124 | 121 | 40 | 9204 | 1112 | RpS14b,CG2678,squ,RpS7,RpS24,RpS2,RpL13,RpL37a,RpS18,RpL34a,RpL12,RpL26,RpS27A,RpLP0,RpL13A,RpS28b,RpL11,RpS4,RpL30,RpL40,RpS23,RpL29,RpS27,RpS16,Pi3K92E,RpL24,RpLP1,RpL7A,RpL23,RpL23A,RpL10,RpL38,RpL9,RpS14a,RpL31,RpL17,RpS13,RpS30,RpS15,RpS29 | | 1.013e-09 | -20.71 | CG6084 (uncharacterized protein) | protein interactions | 39304 | 76 | 30 | 9204 | 1112 | CG2767,Pglym78,Rtnl1,Eno,Neos,Uch,Prosalpha4,scu,Moe,capt,Gdi,regucalcin,AnxB10,ATPsynCF6,Tak1,Stip1,RhoGDI,CG10863,Fkbp12,PHGPx,EndoGI,cib,CG10638,CG14715,PGRP-LE,CG11267,awd,Trxr-1,Rel,Chd64 | | 1.054e-09 | -20.67 | protein metabolic process | biological process | GO:0019538 | 2600 | 319 | 12345 | 1129 | TyrRS,CG7115,Rpn12,mal,CG9316,CG5390,GCS2beta,G9a,mRpL17,ari-1,RpL17,park,CG32221,Prosbeta4,CG32473,CG3631,Cul1,Aos1,Sgf11,RpS23,S2P,SerRS-m,CG2911,mRpS21,Plap,Utx,Usp16-45,eIF4E1,HDAC6,CycJ,fdl,Ubc6,grsm,grp,CG16979,GAPsec,Pomp,twe,RpS27A,ste24a,RpL26,CG8728,CG2493,dx,CG3511,YL-1,nej,faf,Edem1,Naa20A,Prosalpha5,SRPK,Cp1,SkpA,RpL40,CG7840,Arf51F,GatB,isoQC,Atu,Rab5,Cdk7,CSN6,CG13344,CG17698,CG5880,Mo25,RpS29,otu,Ser7,Dd,RpS13,eEF1beta,alph,GluRS-m,Prosalpha4,EMC3,CG4572,CG5026,RpL12,EloC,Fkbp59,PpV,Rcd5,Dip-B,RpS5b,png,RpLP0,Uev1A,mRpL48,Cbl,CysRS,UbcE2H,RpL7A,PEK,CG33977,Sp7,awd,Fkbp12,STUB1,Prosbeta6,Atg7,SelR,Ptp69D,RpS14b,CG11771,ago,eIF4A,Usp10,ea,PIG-H,cort,sip3,CG8726,mRpL35,CG4041,CG11110,CG31633,trbd,CG9941,RpS7,lid,CG8974,CG11486,rept,e(y)1,PIG-L,Fbw5,Cyp1,mRpS5,CG10344,CycE,Chrac-14,PIG-P,Rpn13,Sirt7,Rpt2,CG5003,Pp2A-29B,RpS4,Tfb4,CG6094,CG3223,CG40045,Cul5,Ntan1,eIF3i,not,RpS15,O-fut1,Lkb1,RpL23A,Tlk,Cka,PpD3,d4,CG9643,RpL10,Prosalpha3,lt,ben,Rpn3,stet,mRpL10,mRpS23,RYBP,CG7009,Tao,Prosbeta3,Cul2,mRpL13,CG2200,Ufm1,RpL24,RpS30,shrb,pen-2,mEFTu1,APP-BP1,sol,p38a,Spt20,fray,RpS28b,l(3)73Ah,escl,mRpS10,CG6766,CG8314,velo,CanB2,RpL11,Uba5,CG1578,Prosbeta5,Tak1,RpS27,RpL13,26-29-p,tor,GatA,Iru,Cdk1,CG9953,RpL9,PIG-C,Uch,Dph5,Ptp61F,mRpL16,Usp39,CG14817,gus,Tip60,RpS16,cathD,pont,Cdk5,CG8188,rl,tkv,RpL34b,CG7414,Rep,eIF3m,Ptip,mRpL18,Cchl,CG4238,CG14516,tko,RpLP1,Herc4,DppIII,CG2924,Coq8,Traf6,RpS2,mRpS11,Sgf29,RpL30,Rpn2,RpL38,Mekk1,Ubc4,CG32640,ssh,gd,CG17746,Mos,tws,CG11070,Hsc70-3,Fur2,lwr,egg,Efr,alpha-Man-Ia,CycB,mEFTu2,CG1827,Vsp37A,CG46338,dco,RpL13A,Uch-L5,Gapdh1,QC,csw,Hat1,RpS24,tun,Pex10,RpL23,RpS18,Ptpmeg2,Ptp10D,RpS14a,CkIIalpha,mts,esc,CG11858,Nepl3,RpL37a,oxt,scny,CG31053,RpL34a,mRpL36,Pym,CG14435,TrpRS-m,snk,Nmt,Pink1,Pp2B-14D,Mtl,SP1029,Ublcp1,GalT1,RpL29,MAGE,RpL31,CG10254,Pdk,CG13630,LysRS,mop,Pex12,CanA-14F,trr,ste14,PAPLA1,Parp16,Pdk1,Topors,Ptpmeg,Hexo1,Ggt-1 | | 1.056e-09 | -20.67 | positive regulation of biological process | biological process | GO:0048518 | 1642 | 219 | 12345 | 1129 | CG11710,dx,Tip60,Marf1,PDCD-5,pn,gus,Rbfox1,Debcl,wisp,CG1602,gw,pgc,EMRE,Archease,twe,EcR,TfIIFalpha,SkpA,PGRP-LE,Taf7,BigH1,cad,Idgf2,tkv,Pka-R1,spen,SuUR,CG3198,Nprl2,rl,Adi1,nej,pont,smg,CG13344,Tspo,raptor,Cdk7,RpLP1,Herc4,Myd88,Crag,tral,Atu,CG4238,eIF3m,COX5A,Ptip,bru1,Dd,otu,Bx,CG10336,Traf6,Pld,G9a,SNF4Agamma,Tao,RYBP,Snx6,Iml1,ben,Moe,stet,Bap170,Desat1,MED20,Hr78,E2f1,Cul1,stas,Aos1,Sgf11,Stat92E,EndoGI,park,MED24,ari-1,Sox102F,lin,Tsp86D,Seipin,Utx,eIF4E1,Arpc4,Atf6,Su(H),Arpc1,msps,ASPP,fray,Abi,S2P,Pomp,Nup358,Wbp2,cnc,tor,Ubc6,pygo,GILT1,Tak1,Pgk,shv,snk,spz,Pink1,APC7,lid,fwe,AP-1-2beta,trbd,Idh3b,SelD,bcd,tub,ovo,Trf2,SERCA,sbr,Fbw5,e(y)1,Mtl,rept,Pi3K92E,CG11486,CG4853,CG2199,disp,Cf2,HP1e,CG9925,mop,GlyP,bun,Rel,Rpt2,spn-E,Sec23,Hip1,Med,REPTOR,CG10188,Rpn13,Synj,CycE,Arpc2,d4,Cka,O-fut1,piwi,Lkb1,not,Pdk1,Imp,Parp16,Pura,foxo,crok,trr,AGO3,dl,tws,Scox,Mos,png,Rcd5,gd,hng3,CycC,Ect4,osk,HP1c,NFAT,CG17328,Mekk1,Marf,EloC,vers,dco,PEK,apt,LpR2,tyf,Mitf,ScsbetaA,wash,egg,Cbl,lwr,Sld5,CkIIalpha,armi,Hat1,Atg7,Mybbp1A,CG8963,CG17202,STUB1,Sp7,cort,cyc,Ankle2,Ssdp,CG31053,scny,ea,Nf-YA,CG8009,CG2129,Usp10,Exd2,SREBP,Jafrac2,hb,mts,esc,MED4 | | 1.239e-09 | -20.51 | organonitrogen compound catabolic process | biological process | GO:1901565 | 510 | 89 | 12345 | 1129 | shrb,Cbl,ari-1,park,CG32221,Prosbeta4,CG17691,DhpD,CG32473,Vsp37A,UbcE2H,Cul1,Shmt,Idgf6,CG10561,Prosalpha3,Rpn12,Rpn2,CG9316,EloC,Rpn3,RYBP,CG32640,CG8199,AOX3,Hsc70-3,CG11070,Prosbeta3,Cul2,Prosbeta5,ago,Gabat,Usp10,26-29-p,fdl,fal,Ubc6,scny,Uch,Pomp,sip3,cort,Uch-L5,Sply,Prosbeta6,P5CDh1,CG6766,Plap,CG7997,Ho,Usp16-45,Idgf4,cathD,Cht11,faf,Edem1,CG8188,Fbw5,SP1029,Idgf2,UK114,Prosalpha5,PGRP-LE,Cp1,SkpA,RpL40,RpS27A,trbd,CG31633,gus,CG8974,Pink1,CG3223,CG40045,Cul5,Ntan1,not,Prosalpha4,Topors,CG4572,Ggt-1,CG10254,Rpn13,CG4238,CG14516,Rpt2,cN-IIIB,CG5003,Herc4 | | 1.263e-09 | -20.49 | RpS8 (Ribosomal protein S8) | protein interactions | 43532 | 122 | 40 | 9204 | 1112 | RpL9,RpS14a,X11L,RpL38,RpL23,RpL23A,RpL10,RpL7A,RpL24,RpLP1,RpS29,Trf4-2,RpS15,RpL31,RpL17,RpS13,RpS30,RpL12,RpL26,RpS27A,RpLP0,RpS2,RpL13,RpL37a,RpL34a,RpS18,RpS7,RpS24,RpS14b,RpL29,RpS27,RpS16,RpL30,RpS23,RpL40,RpL11,CG46385,RpS4,RpL13A,RpS28b | | 1.263e-09 | -20.49 | RpL15 (Ribosomal protein L15) | protein interactions | 3354918 | 122 | 40 | 9204 | 1112 | dl,RpL38,RpS14a,RpL9,RpLP1,RpL7A,RpL24,RpL23A,RpL10,RpL23,RpS29,RpL31,RpS13,RpL17,RpS30,RpS15,RpL13,RpS18,RpL34a,RpL37a,RpS2,RpL26,RpS27A,RpLP0,RpL12,RpS14b,RpS24,RpS7,RpS23,RpL40,RpL30,RpS16,RpS27,RpL29,RpS28b,foxo,RpL13A,RpS4,osk,RpL11 | | 1.329e-09 | -20.44 | GO\_CATABOLIC\_PROCESS | MSigDB lists | GO\_CATABOLIC\_PROCESS | 705 | 164 | 4161 | 644 | CG9248,Dcr-1,CG4872,RpL26,CG32549,RpS27,Mtpalpha,Cdk5,Eno,Rpn3,CG8974,RpL12,RpS24,APC7,ABCD,Prosbeta6,Ubc4,mus81,CG17593,shrb,Herc4,Got1,Unr,CG15111,Sply,Prosalpha5,CG14516,Cdk1,wal,CG9391,eIF4A,CG10465,Mcad,Cisd2,Rpn2,scu,Vsp37A,ste24a,CG8188,SkpA,CG8475,Fdh,CG11070,dnc,LSm3,Hsc70-3,RpL37a,RpL24,Pfk,CG7322,CG8963,P5CDh1,CG5285,faf,RpS2,CG6766,CG9485,CG5966,Nc73EF,RpS27A,PAPLA1,Pcd,Pi4KIIalpha,Faf2,RpS28b,CG9471,CG3609,mRpS10,STUB1,Rpt2,Prosalpha4,trbd,Cul2,CG11164,Not10,RpS13,Rrp40,RpL13A,eIF4E1,tzn,CG7550,Gk2,Prosbeta4,Uch,RpL11,Gyg,CG32473,RpS18,CycB,Cul1,Pglym78,Thiolase,RpL23,Pex13,CG4390,RpS16,RpL38,RpS4,CG1640,RpS7,Synj,RpL40,RpS23,Aldh,CG8199,Shmt,mad2,Treh,Fbw5,Uch-L5,Pgk,UbcE2H,Gapdh1,Prosbeta3,Pgm1,HDAC6,ND-B22,CG17691,Tip60,CG10932,Jafrac1,CG5734,CG4572,Prosalpha3,Pp2A-29B,Gabat,Iru,Prosbeta5,Edem1,CG3376,lwr,RpL30,Edc3,ari-1,Atg6,sgll,CG40045,Etf-QO,DhpD,cathD,CG10418,Atg4b,not,Atg7,Cbp80,snz,CG7840,twin,RpL9,Jheh2,Caf1-105,AIF,crq,GlyP,CG7997,Cul5,Echs1,RpL13,mts,RpS30,Rpn12,RpLP0,CG6610,cyc | | 1.453e-09 | -20.35 | RpL4 (Ribosomal protein L4) | protein interactions | 43349 | 113 | 38 | 9204 | 1112 | RpL38,RpS14a,RpL9,RpL7A,RpLP1,RpL24,RpL10,RpL23A,RpL23,RpS29,RpL31,RpL17,RpS13,RpS30,RpS15,RpL13,RpL34a,RpL37a,RpS18,RpS2,RpL26,RpS27A,RpLP0,RpL12,RpS14b,RpS24,RpS7,RpS23,RpL40,RpL30,RpS16,RpS27,RpL29,RpS28b,RpL13A,RpS4,RpL11,mus304 | | 1.875e-09 | -20.09 | gamete generation | biological process | GO:0007276 | 1097 | 158 | 12345 | 1129 | wash,CycB,Nap1,ScsbetaA,Pax,Grip84,egg,Cbl,lwr,apt,BicC,nmd,yps,twin,CG46338,shu,osk,Mos,Pex13,eIF4A,Orc1,exu,mts,mad2,muskelin,slam,cort,ea,csw,lqfR,Atg7,Ptpmeg2,r2d2,CkIIalpha,Pex10,armi,stai,e(y)1,Pi3K92E,Pp2B-14D,ovo,SERCA,stil,mael,Tm1,cact,Axs,bcd,lid,spz,Imp,Pdk1,Cul5,Top1,PAPLA1,tho2,krimp,Topors,piwi,Lkb1,spn-E,Med,CycE,Pen,Cf2,HP1e,disp,tef,gammaTub37C,bun,fs(1)M3,Pex12,mop,Pxt,vret,park,milt,zpg,shrb,cnn,ari-1,E2f1,swa,Stat92E,toc,Rab10,Moe,Grip71,Iml1,Cenp-C,jvl,Cul2,Mst89B,Tao,CycJ,wapl,tor,nos,GatA,BoYb,cnc,Nup358,Cdk1,yem,grp,msps,Abi,yl,mus304,Klp10A,eIF4E1,bam,psq,Nprl2,rl,elg1,Jafrac1,Drep1,Dcr-1,cathD,nej,faf,Nup62,l(3)mbt,capt,tkv,Pka-R1,SRPK,mtrm,Rbfox1,wisp,twe,EcR,fs(1)N,bif,pgc,enc,del,Sec15,gus,Syx13,larp,squ,Rbf,sau,bru1,Grip75,otu,Arf51F,Grip128,eIF3m,CSN6,heph,raptor,Kap3,Rab5,cup,sw | | 1.880e-09 | -20.09 | electron transport chain | biological process | GO:0022900 | 116 | 33 | 12345 | 1129 | Etf-QO,ND-39,AOX3,Trxr-1,ND-B22,levy,CG14407,blw,Cyt-c-p,Pmi,ND-51,ND-MNLL,wal,ND-B18,AIF,ND-23,SdhA,ND-49,NP15.6,SdhD,cype,UQCR-11,COX5A,Grx1,ND-24,ND-18,ND-75,COX4,ND-SGDH,UQCR-Q,ND-B15,COX5B,Coq8 | | 2.017e-09 | -20.02 | catalytic complex | cellular component | GO:1902494 | 999 | 148 | 11940 | 1105 | Gclc,Sgf29,tws,gus,DNApol-epsilon255,r2d2,Ptip,UQCR-11,CG16865,Bin1,SkpA,CG40472,d4,LSm3,mus205,mus81,Prosalpha4,MAGE,Dcr-1,Imp,CG11070,CG6767,Cbp80,Utx,l(3)73Ah,rept,pont,Cul5,plu,ND-B15,CG8009,Tip60,Rpn2,CG30496,Chrac-14,Atg6,CycJ,RpII140,CG8475,nej,scu,Ubc6,Rpn3,Rpb8,CG5003,CG32221,mts,ND-30,Pp2A-29B,Trf2,YL-1,PIG-C,Dd,Naa20A,CG33977,Tfb4,Cul2,Pi3K92E,Nc73EF,alc,sip3,CG10254,Prosbeta3,yps,NHP2,ND-B22,TfIIFalpha,Pomp,ND-18,Taf7,SmE,CG17691,e(y)1,Prosbeta6,Prosalpha3,wal,Spt20,Prosbeta4,CG31633,escl,ND-MWFE,lid,CanB2,Rpt2,ND-23,UQCR-Q,Pp2B-14D,Rrp40,ND-MNLL,DCAF12,Eno,sw,trr,ScsbetaA,CG5292,CycB,ND-51,SdhA,ago,NP15.6,Sgf11,Scsalpha1,CG3511,Rpn12,APC7,CG10418,RfC38,Rep,Rcd5,ND-75,EloC,Bap170,Rpn13,CG8199,Pfk,Uch-L5,ben,Nap1,Fbw5,Atu,ND-24,CycE,CG9316,ND-39,Aos1,kl-2,ph-d,Nurf-38,PIG-H,Cul1,SdhD,Cdk5,park,Cdk7,ari-1,Prosbeta5,png,CG9684,Uev1A,elg1,ND-SGDH,ND-B18,CanA-14F,CG12018,esc,ND-49,not,Prosalpha5 | | 2.105e-09 | -19.98 | sima (similar) | protein interactions | 43580 | 206 | 56 | 9204 | 1112 | RpL23,CG2199,RpL23A,RpL10,RpL7A,piwi,RpLP1,Spx,Hsc70-3,ATPsynCF6,Rbcn-3A,RpL9,Kap-alpha3,RpS14a,Shmt,Imp,park,RpS15,spd-2,RpL31,wal,RpL17,Rpt2,Mcad,cathD,Nc73EF,yps,RpS7,eIF3i,Mybbp1A,blw,Usp10,ATPsynbeta,26-29-p,RpL13,eIF3m,SERCA,CCT6,Vha68-2,shrb,eIF4A,shv,RpL13A,Hnf4,ago,ScsbetaA,Nap1,RpS16,RpS27,whd,Droj2,RpL30,RpS23,TfIIFalpha,CG1416,PyK | | 2.603e-09 | -19.77 | regulation of primary metabolic process | biological process | GO:0080090 | 1894 | 244 | 12345 | 1129 | Dd,Bx,CG10631,bru1,alph,pps,Traf6,Rbf,RpLP1,ix,Cdk7,CG6813,Myd88,Pka-R2,cup,CG11902,CG13344,CG4021,raptor,heph,eIF3m,CG30291,Atu,CG4238,CG17454,insv,CG7414,BigH1,cad,Fdh,Pka-R1,tkv,CG4730,asf1,TfIIFalpha,l(3)mbt,Taf7,SkpA,faf,lin-52,YL-1,nej,pont,smg,Drep1,spen,rl,Nprl2,crm,Uxt,Tip60,Marf1,MED27,ph-d,CG10038,gus,CG11710,Bin1,gw,pgc,Ptp61F,Dph5,EcR,Hnf4,Rbfox1,CG1602,Unr,CG31875,Cdk1,Pomp,CG14408,cnc,Wbp2,GAPsec,CG1578,nos,tor,Ubc6,pygo,CycJ,Tak1,Gabat,psq,Atf6,Not10,Su(H),Seipin,bam,Utx,HDAC6,Abi,fray,S2P,sws,CG2911,ASPP,l(3)73Ah,escl,Cul1,Aos1,Sgf11,HmgD,Rbf2,Stat92E,EndoGI,MED20,Hr78,E2f1,Spt20,ari-1,Sox102F,CG10959,lin,MED24,park,SNF4Agamma,Tao,RYBP,ATPsynD,G9a,Cul2,mRpL13,Bap170,Desat1,Son,Elba2,Rpn3,CG10654,ben,CG30496,Cka,Tlk,piwi,Lkb1,d4,CG2678,krimp,Topors,foxo,trr,dl,AGO3,not,Pdk1,Parp16,bun,GlyP,Pp2A-29B,Rel,Sirt7,Rpt2,CG2199,Tfb4,HP1e,Cf2,CG3587,Rpn13,Chrac-14,CycE,sbb,spn-E,CG4872,Med,REPTOR,RhoGAP5A,mael,Maf1,obe,ovo,Trf2,rept,Pi3K92E,CG7328,Fbw5,mtTFB1,e(y)1,CG13601,CG10418,lid,bcd,cact,Rga,scny,Ankle2,cyc,sip3,cort,Ssdp,CG31053,SREBP,Exd2,qkr58E-2,hb,Jafrac2,mts,esc,MED4,CG8009,Nf-YA,eIF4A,ago,CG2129,CkIIalpha,r2d2,CG17202,STUB1,RpL13A,Hat1,mute,Atf3,CG8963,Mybbp1A,twin,BicC,dco,PEK,apt,tyf,egg,Cbl,lwr,Mitf,ScsbetaA,Pax,CycB,wash,Nap1,CG17746,Scox,Mos,Smox,Rcd5,hng3,tws,Mekk1,EloC,vers,Rpn2,CycC,PpV,osk,NFAT,HP1c,CG17328 | | 2.635e-09 | -19.75 | positive regulation of cellular process | biological process | GO:0048522 | 1464 | 198 | 12345 | 1129 | Pld,CG10336,Traf6,bru1,otu,Bx,Dd,Crag,Atu,eIF3m,Ptip,COX5A,raptor,Tspo,Myd88,Cdk7,RpLP1,Herc4,Nprl2,rl,SuUR,spen,smg,pont,nej,SkpA,Taf7,TfIIFalpha,Idgf2,tkv,BigH1,cad,CG1602,Rbfox1,Debcl,wisp,Archease,twe,EcR,pgc,gw,CG11710,dx,pn,Tip60,Marf1,PDCD-5,pygo,Tak1,Pgk,tor,Ubc6,shv,Pomp,cnc,Nup358,Wbp2,msps,ASPP,Arpc1,S2P,fray,Abi,Utx,eIF4E1,Seipin,Su(H),Arpc4,Atf6,MED24,park,Tsp86D,lin,Sox102F,ari-1,MED20,E2f1,Stat92E,EndoGI,Cul1,stas,Aos1,Sgf11,Moe,ben,stet,Iml1,Bap170,G9a,Snx6,RYBP,SNF4Agamma,Tao,Imp,Parp16,not,Pdk1,AGO3,dl,trr,foxo,Pura,d4,O-fut1,piwi,Lkb1,Cka,CG10188,Sec23,Hip1,spn-E,REPTOR,Med,CycE,Arpc2,Synj,Rpn13,Cf2,HP1e,CG4853,CG2199,disp,Rpt2,GlyP,bun,mop,Rel,e(y)1,Pi3K92E,CG11486,rept,Mtl,Trf2,ovo,bcd,APC7,lid,snk,spz,Pink1,trbd,AP-1-2beta,Idh3b,SelD,fwe,CG2129,Usp10,CG8009,Nf-YA,esc,mts,MED4,SREBP,Exd2,Jafrac2,hb,Ssdp,CG31053,cyc,Ankle2,cort,ea,scny,CG8963,Mybbp1A,Atg7,Sp7,STUB1,CG17202,Sld5,CkIIalpha,armi,wash,ScsbetaA,Mitf,egg,Cbl,PEK,tyf,apt,dco,CG17328,Ect4,CycC,NFAT,HP1c,osk,vers,Marf,Mekk1,EloC,tws,Rcd5,gd,hng3,Scox,Mos,png | | 3.057e-09 | -19.61 | oogenesis | biological process | GO:0048477 | 682 | 109 | 12345 | 1129 | CkIIalpha,bam,Ptpmeg2,r2d2,psq,armi,lqfR,Atg7,csw,msps,yl,Abi,cnc,muskelin,Cdk1,ea,tor,nos,exu,GatA,Orc1,eIF4A,CycJ,mts,Cul2,Mos,Tao,shu,Grip71,Iml1,osk,Rab10,Moe,jvl,BicC,swa,E2f1,apt,toc,twin,yps,Stat92E,park,Pax,milt,vret,wash,egg,Cbl,ari-1,shrb,Cf2,heph,raptor,bun,mop,fs(1)M3,Pxt,Rab5,gammaTub37C,cup,spn-E,Med,Pen,CycE,Grip128,bru1,krimp,otu,piwi,Grip75,Lkb1,Cul5,tho2,Top1,squ,Rbf,Sec15,spz,lid,gus,bcd,Rbfox1,wisp,cact,enc,del,EcR,fs(1)N,ovo,capt,mael,Tm1,SRPK,tkv,SERCA,Pka-R1,stil,Drep1,Dcr-1,rl,stai,Nprl2,e(y)1,elg1,Nup62,Pi3K92E,cathD,nej | | 3.222e-09 | -19.55 | cytosolic large ribosomal subunit | cellular component | GO:0022625 | 58 | 22 | 11940 | 1105 | RpL10,RpL11,RpL13A,RpL30,RpLP0,RpL23,RpL12,RpL23A,RpL24,RpL7A,RpL34b,RpL40,RpL38,RpL17,RpL29,RpL9,RpL31,RpL13,RpL26,RpL37a,RpLP1,RpL34a | | 3.262e-09 | -19.54 | regulation of nitrogen compound metabolic process | biological process | GO:0051171 | 1870 | 241 | 12345 | 1129 | CG10959,lin,ari-1,Sox102F,park,MED24,Stat92E,Rbf2,HmgD,EndoGI,Cul1,Sgf11,Aos1,MED20,Hr78,Spt20,E2f1,Son,Bap170,CG10654,ben,Rpn3,Elba2,RYBP,SNF4Agamma,Tao,mRpL13,Cul2,ATPsynD,G9a,CG1578,Tak1,CycJ,pygo,Gabat,nos,tor,Ubc6,Cdk1,Unr,CG31875,GAPsec,Pomp,CG14408,cnc,Wbp2,fray,S2P,Abi,l(3)73Ah,CG2911,ASPP,escl,sws,Not10,Su(H),psq,Atf6,Utx,HDAC6,bam,YL-1,lin-52,nej,pont,smg,faf,rl,crm,Drep1,spen,tkv,Pka-R1,Fdh,CG4730,insv,BigH1,CG7414,cad,Taf7,l(3)mbt,SkpA,asf1,TfIIFalpha,EcR,Hnf4,Dph5,gw,pgc,Ptp61F,CG1602,Rbfox1,wisp,ph-d,MED27,CG10038,gus,Uxt,Tip60,Marf1,Bin1,CG11710,Rbf,Traf6,Bx,Dd,alph,pps,CG10631,bru1,Ptip,eIF3m,CG30291,CG4238,CG17454,Atu,Pka-R2,CG6813,Myd88,CG11902,cup,Cdk7,ix,RpLP1,raptor,heph,CG4021,CG13344,Cbl,lwr,egg,CycB,wash,Nap1,Pax,ScsbetaA,Mitf,twin,PEK,apt,tyf,BicC,dco,vers,Rpn2,Mekk1,EloC,CG17328,CycC,PpV,HP1c,osk,NFAT,Rcd5,Smox,hng3,CG17746,Mos,Scox,tws,esc,mts,MED4,Exd2,SREBP,qkr58E-2,hb,Jafrac2,CG2129,ago,eIF4A,CG8009,Nf-YA,Rga,scny,CG31053,Ssdp,cort,Ankle2,cyc,sip3,RpL13A,CG17202,STUB1,Atf3,mute,Mybbp1A,CG8963,Hat1,r2d2,CkIIalpha,Pi3K92E,rept,e(y)1,Fbw5,mtTFB1,obe,Maf1,RhoGAP5A,mael,Trf2,ovo,cact,bcd,CG10418,lid,spz,AGO3,trr,dl,foxo,Parp16,not,Pdk1,piwi,Lkb1,CG30496,Cka,Tlk,krimp,Topors,d4,CG2678,CycE,Rpn13,Chrac-14,CG3587,sbb,spn-E,REPTOR,CG4872,Med,Sirt7,Rpt2,bun,Pp2A-29B,Rel,HP1e,Cf2,Tfb4,CG2199 | | 3.411e-09 | -19.50 | P granule | cellular component | GO:0043186 | 40 | 18 | 11940 | 1105 | eIF4A,mael,Tm1,pgc,bru1,CG9425,tapas,AGO3,vret,osk,spn-E,PyK,squ,shu,tral,krimp,armi,CG6967 | | 3.533e-09 | -19.46 | nucleoside phosphate metabolic process | biological process | GO:0006753 | 201 | 46 | 12345 | 1129 | Eno,Nadsyn,ATPsynD,ATPsynE,Ac3,CG8199,AMPdeam,ATPsynO,blw,Pfk,PyK,Pfas,scu,nmdyn-D7,CG17691,r-l,CG7920,CG6767,ScsbetaA,Ts,Scsalpha1,CG6178,CG4407,ATPsynbeta,Ppcdc,Ady43A,cN-IIIB,CG11164,CG33156,Gapdh1,r,AcCoAS,ATPsyndelta,ATPsynG,Pfrx,ATPsynB,awd,l(2)k01209,CG11089,CG1774,Pglym78,Pgk,nmdyn-D6,fal,COX5B,ATPsynCF6 | | 3.614e-09 | -19.44 | L13a-mediated translational silencing of Ceruloplasmin expression | REACTOME pathways | R-DME-156827 | 120 | 43 | 3923 | 574 | RpS29,RpL26,RpS23,RpL24,RpS7,RpS28b,RpL11,RpL30,RpS24,RpS27A,eIF3m,RpS14a,RpL13A,RpL12,RpL7A,RpL17,RpL9,RpL34b,RpL37a,RpL38,RpL40,eIF4A,RpS27,RpL29,RpS14b,RpS30,eIF3i,RpS16,RpL23,RpS18,eIF4E1,RpLP1,RpL23A,RpL34a,RpS5b,RpL10,RpS15,RpL31,RpLP0,RpS4,RpS13,RpS2,RpL13 | | 3.723e-09 | -19.41 | pole plasm | cellular component | GO:0045495 | 49 | 20 | 11940 | 1105 | Tm1,pgc,bru1,CG9425,eIF4A,mael,squ,PyK,shu,tral,armi,krimp,CG6967,AGO3,tapas,vret,osk,gus,nos,spn-E | | 4.299e-09 | -19.26 | organelle organization | biological process | GO:0006996 | 1885 | 242 | 12345 | 1129 | bif,del,enc,Hnf4,twe,Debcl,mtrm,Non1,wisp,Tip60,ph-d,Orc6,CG10038,Syx13,CG8728,alphaTub67C,nbs,Bin1,cib,Nup62,CG8188,YL-1,c(2)M,nej,pont,ND-23,SuUR,CG3437,BigH1,SRPK,tkv,Naa20A,Pka-R1,asf1,Oseg1,capt,SkpA,eIF3m,Ptip,Arf51F,Grip128,tral,Atu,Sec22,Vha68-2,sw,Rab5,Kap3,cup,heph,sau,larp,Pld,Rbf,CG11562,LSm3,mre11,Dd,COX4,mRpS11,Sgf29,Grip75,bru1,CG17768,EMC3,Trf4-2,jvl,lt,RpL10,Cenp-C,Grip71,Elba2,dgt4,Moe,spd-2,Sry-alpha,Snx6,G9a,RpL24,ari-1,cnn,RecQ5,park,milt,mEFTu1,Sgf11,HmgD,Actn,Spt20,swa,CG11077,RpS28b,Abi,bor,Arpc1,l(3)73Ah,msps,escl,CanB2,Arpc4,psq,RpL11,CG7488,Seipin,bam,Vamp7,mus304,Utx,CG13366,HDAC6,eIF4E1,Klp10A,Prosbeta5,Lnpk,nos,Ubc6,GatA,Xpc,Blos4,wapl,RpS27,grp,PIG-C,Tim8,Cdk1,Sdhaf3,yem,Caf1-105,cnc,CG4041,crq,CG5903,CHOp24,Axs,CCT6,trbd,CG11110,Xpac,SelD,Pink1,Rtnl1,lid,rept,Mtl,Pp2B-14D,CG5510,stai,e(y)1,mael,cbs,Tm1,ovo,Roe1,Pmi,CG7716,Chrac-14,Arpc2,TMEM216,spn-E,Hip1,Sec23,Gmap,Tom40,Mitofilin,CG14894,Pi4KIIalpha,mop,Pex12,Pp2A-29B,Sirt7,gammaTub37C,tef,Svil,HP1e,Top1,foxo,trr,RpS15,not,RpL23A,Tlk,piwi,Lkb1,d4,krimp,up,Topors,Marf,RpL12,osk,Scox,Rcd5,RpS5b,CG8004,RfC38,tws,RpLP0,lwr,egg,CG9186,Grip84,parvin,Nurf-38,Girdin,Pax,Mitf,wash,CycB,Nap1,Rab32,Edc3,fs(1)Ya,BicC,Snm1,Hat1,Atg7,Tom70,P5CDh1,RpS14b,armi,SelR,Pex10,Atg6,cana,RpS14a,opm,Ndc1,r2d2,alphaTub84B,mts,esc,mus81,Arl6IP1,CG9393,mad2,Pex13,scny,cort,Ankle2,ND-18,CG31053,Vps16B | | 4.455e-09 | -19.23 | CG2862 (uncharacterized protein) | protein interactions | 33471 | 59 | 25 | 9204 | 1112 | Arf51F,RhoGDI,Stip1,CG10638,CG14715,PHGPx,EndoGI,cib,Fkbp12,Sod1,mts,yip2,awd,CG11267,Chd64,Trxr-1,Eno,Uch,regucalcin,Moe,capt,scu,CG4598,CG3226,AnxB10 | | 4.479e-09 | -19.22 | ribose phosphate metabolic process | biological process | GO:0019693 | 163 | 40 | 12345 | 1129 | blw,Pfk,ATPsynO,PyK,ATPsynD,ATPsynE,Eno,AMPdeam,Ac3,CG8199,ScsbetaA,CG6767,CG7920,nmdyn-D7,Pfas,scu,r-l,CG17691,ATPsyndelta,Gapdh1,r,AcCoAS,awd,ATPsynG,Pfrx,ATPsynB,Ppcdc,Scsalpha1,CG6178,ATPsynbeta,CG11164,Ady43A,nmdyn-D6,Pgk,COX5B,ATPsynCF6,CG11089,CG1774,l(2)k01209,Pglym78 | | 4.618e-09 | -19.19 | oxidative phosphorylation | biological process | GO:0006119 | 91 | 28 | 12345 | 1129 | ND-51,ND-24,Pmi,COX4,ND-SGDH,wal,ND-75,ND-MNLL,AIF,ND-B15,ND-B18,UQCR-Q,Coq8,SdhA,ND-23,COX5B,ATPsynbeta,Etf-QO,ND-49,ND-B22,levy,ND-39,ATPsynO,Cyt-c-p,COX5A,UQCR-11,SdhD,cype | | 4.642e-09 | -19.19 | cytosolic small ribosomal subunit | cellular component | GO:0022627 | 45 | 19 | 11940 | 1105 | RpS7,RpS30,RpS29,RpS16,RpS27,RpS28b,RpS27A,RpL40,RpS23,RpS18,RpS15,CG7414,RpS13,RpS5b,RpS24,RpS4,RpS14b,RpS14a,RpS2 | | 4.769e-09 | -19.16 | PGRP-LE (Peptidoglycan recognition protein LE) | protein interactions | 32534 | 147 | 44 | 9204 | 1112 | CCT5,Cp1,Jafrac1,RpS30,bor,RpS29,scu,Idh3b,RpS5b,RpL7A,RpL24,Mitofilin,Tak1,Hsc70-3,RpL23,CG2199,RpL23A,alphaTub84B,CG4853,RpL38,Sod1,PGRP-LE,RpS14a,Sgf29,Rel,GCS2beta,eIF4A,RpS28b,rept,Nup358,pps,RpS27,RpS16,CCT3,blw,CG9391,pont,cup,SERCA,CCT6,SkpA,RpL12,Echs1,vret | | 4.852e-09 | -19.14 | protein catabolic process | biological process | GO:0030163 | 338 | 65 | 12345 | 1129 | Rpn12,Rpn2,Prosalpha3,EloC,CG9316,Rpn3,CG32640,RYBP,Cul2,CG11070,Prosbeta3,Hsc70-3,shrb,ari-1,Cbl,CG32221,park,Vsp37A,UbcE2H,Prosbeta4,Cul1,Uch-L5,Prosbeta6,Plap,CG6766,Usp16-45,Prosbeta5,Usp10,ago,Ubc6,26-29-p,scny,Uch,cort,Pomp,sip3,RpS27A,gus,trbd,CG31633,CG8974,Pink1,cathD,Edem1,CG8188,faf,Fbw5,Prosalpha5,RpL40,SkpA,Cp1,Rpn13,CG10254,CG4238,Rpt2,Herc4,CG5003,CG3223,CG40045,Ntan1,Cul5,not,Topors,Prosalpha4,CG4572 | | 5.121e-09 | -19.09 | Sh3beta (Sh3beta) | protein interactions | 38820 | 94 | 33 | 9204 | 1112 | awd,Fkbp59,Fkbp12,Sod1,RhoGDI,CG10863,CG4598,scu,capt,Moe,Uch,Prosalpha4,Pglym78,CG11267,Chd64,yip2,PHGPx,Cyp1,cib,EndoGI,CG14715,CG10638,ATPsynbeta,Pgk,Prosbeta5,Stip1,CG9391,ERp60,CG10265,AnxB10,regucalcin,CG2767,Eno | | 5.205e-09 | -19.07 | Pdi (Protein disulfide isomerase) | protein interactions | 39651 | 85 | 31 | 9204 | 1112 | CG10265,CG4598,Usp39,AnxB10,Moe,scu,capt,regucalcin,Uch,Prosalpha4,Idgf6,Pglym78,Eno,Chd64,CG11267,Trxr-1,awd,CG1092,PHGPx,cib,Fkbp12,Sod1,ATPsynbeta,CG14715,Nf-YA,CG10638,Fer1HCH,RhoGDI,HmgD,Stip1,ERp60 | | 5.987e-09 | -18.93 | Fkbp14 (FK506-binding protein 14) | protein interactions | 37449 | 68 | 27 | 9204 | 1112 | scu,Moe,capt,GstT1,regucalcin,AnxB10,CG4598,CG2767,Eno,Pglym78,Uch,Prosalpha4,Fkbp12,PHGPx,cib,CG9436,EndoGI,CG10638,CG14715,Pgk,awd,Trxr-1,CG11267,Chd64,yip2,RhoGDI,Nurf-38 | | 6.040e-09 | -18.92 | large ribosomal subunit | cellular component | GO:0015934 | 108 | 31 | 11940 | 1105 | RpL12,mRpL48,RpL24,RpL23A,RpL7A,mRpL18,RpL34b,RpL23,RpL30,RpLP0,RpL13A,mRpL13,RpL11,RpL10,mRpL10,RpLP1,RpL34a,RpL26,mRpL35,RpL37a,mRpL17,RpL13,RpL31,RpL9,RpL17,mRpL16,RpL29,mRpL36,RpL40,RpL38,CG6094 | | 6.267e-09 | -18.89 | Cpr78E (Cuticular protein 78E) | protein interactions | 40408 | 11 | 10 | 9204 | 1112 | ATPsynF,ATPsynB,ATPsynE,ATPsyndelta,blw,ATPsynD,ATPsynCF6,ATPsynO,ATPsynG,ATPsynbeta | | 6.474e-09 | -18.86 | GTP hydrolysis and joining of the 60S ribosomal subunit | REACTOME pathways | R-DME-72706 | 122 | 43 | 3923 | 574 | RpS4,RpLP0,RpL31,RpS15,RpS5b,RpL10,RpLP1,RpL23A,RpL34a,eIF4E1,RpL13,RpS2,RpS13,RpS30,RpS14b,RpL29,RpS27,eIF4A,RpL40,RpS18,RpL23,RpS16,eIF3i,RpL9,RpL17,RpL7A,RpL12,RpS14a,RpL13A,RpL38,RpL37a,RpL34b,RpS23,RpL24,RpL26,RpS29,RpS24,RpS27A,eIF3m,RpL30,RpL11,RpS28b,RpS7 | | 6.578e-09 | -18.84 | nucleotide metabolic process | biological process | GO:0009117 | 198 | 45 | 12345 | 1129 | Ts,CG6767,ScsbetaA,CG7920,r-l,CG17691,nmdyn-D7,Pfas,scu,PyK,blw,Pfk,ATPsynO,AMPdeam,Ac3,CG8199,ATPsynD,ATPsynE,Eno,Nadsyn,COX5B,ATPsynCF6,fal,nmdyn-D6,Pgk,Pglym78,CG11089,CG1774,l(2)k01209,awd,ATPsynG,Pfrx,ATPsynB,ATPsyndelta,CG33156,r,Gapdh1,AcCoAS,CG11164,Ady43A,Ppcdc,Scsalpha1,CG6178,CG4407,ATPsynbeta | | 7.213e-09 | -18.75 | CG10638 (uncharacterized protein) | protein interactions | 39424 | 86 | 31 | 9204 | 1112 | CG3226,AnxB10,CG4598,regucalcin,Gdi,Moe,capt,scu,Prosalpha4,Uch,Gapdh1,Pglym78,Eno,yip2,CG11267,Chd64,awd,Trxr-1,CG14715,Pgk,ATPsynbeta,Fkbp12,Sod1,cib,PHGPx,CG9436,EndoGI,RhoGDI,Stip1,Idh,ERp60 | | 8.555e-09 | -18.58 | ribonucleotide metabolic process | biological process | GO:0009259 | 160 | 39 | 12345 | 1129 | l(2)k01209,CG11089,CG1774,Pglym78,Pgk,nmdyn-D6,COX5B,ATPsynCF6,CG6178,Scsalpha1,ATPsynbeta,Ppcdc,Ady43A,CG11164,r,Gapdh1,AcCoAS,ATPsyndelta,ATPsynG,Pfrx,ATPsynB,awd,Pfas,scu,nmdyn-D7,CG17691,r-l,CG7920,ScsbetaA,Eno,ATPsynD,ATPsynE,Ac3,CG8199,AMPdeam,ATPsynO,blw,Pfk,PyK | | 8.890e-09 | -18.54 | germ plasm | cellular component | GO:0060293 | 42 | 18 | 11940 | 1105 | eIF4A,mael,Tm1,pgc,CG9425,bru1,AGO3,tapas,vret,osk,spn-E,PyK,squ,shu,tral,armi,krimp,CG6967 | | 9.477e-09 | -18.47 | axis specification | biological process | GO:0009798 | 207 | 46 | 12345 | 1129 | Grip128,spn-E,Med,csw,msps,trk,psq,fs(1)M3,armi,Myd88,gammaTub37C,CkIIalpha,heph,hb,dl,tor,nos,exu,Grip75,ea,Lkb1,cnc,krimp,jvl,enc,fs(1)N,bcd,Grip71,osk,Moe,cact,gus,gd,snk,spz,faf,Cbl,lin,milt,rl,mael,cad,Tm1,Pka-R1,swa,capt | | 1.006e-08 | -18.41 | cell cycle process | biological process | GO:0022402 | 564 | 93 | 12345 | 1129 | msps,Arpc1,Pex10,eIF4E1,Klp10A,mus304,cana,bam,mad2,mts,mus81,Prosbeta5,ago,wapl,Orc1,CycJ,GatA,Ubc6,Xpc,nos,yem,Cdk1,grp,slam,CG31053,Ankle2,cort,Cenp-C,spd-2,dgt4,Iml1,Grip71,PpV,Rcd5,Mos,tws,RfC38,Grip84,shrb,RecQ5,cnn,lwr,CycB,E2f1,Grip128,CycE,Arf51F,CG7716,spn-E,Rab5,cup,gammaTub37C,Pp2A-29B,sw,CG5003,mop,tef,Rbf,larp,sau,CG10336,mre11,Lkb1,Grip75,Trf4-2,Topors,krimp,Zwilch,twe,del,enc,bif,Axs,Non1,wisp,mtrm,Syx13,CCT6,lid,alphaTub67C,nbs,pont,Pp2B-14D,c(2)M,Nup62,CG8188,Nprl2,SuUR,cbs,SRPK,CG3437,mael,SkpA | | 1.030e-08 | -18.39 | regulation of cellular process | biological process | GO:0050794 | 3560 | 409 | 12345 | 1129 | Not10,CG6927,Utx,HDAC6,eIF4E1,bam,S2P,CG2911,Arpc1,grp,Unr,GAPsec,Pomp,Nup358,shv,CycJ,Ubc6,Snx6,ZnT35C,SNF4Agamma,Trxr-1,G9a,TyrRS,Son,CG7115,Cenp-C,Desat1,Rab10,Moe,dnc,Stat92E,stas,Cul1,Aos1,Sgf11,CG9911,Hr78,swa,Tsp86D,ari-1,Sod1,vret,park,MED24,CG6813,Pka-R2,Rab5,ix,Cdk7,CG4021,CG13344,Tspo,CG17698,Grx1,COX5A,trk,CG17454,Crag,Drep2,Atu,tral,otu,Dd,COX4,alph,spag,Zwilch,CG10631,bru1,Pld,Rbf,Wnt10,mre11,Mo25,MED27,Ac3,pn,Bin1,CG11710,nbs,dx,CG13933,Hnf4,fs(1)N,twe,chrb,Rbfox1,Debcl,wisp,Fdh,Naa20A,CG4730,insv,BigH1,cad,SkpA,Gdi,asf1,YL-1,nej,faf,Edem1,Nprl2,crm,SuUR,spen,Drep1,armi,r2d2,Sld5,cana,Sp7,STUB1,Aldh,Atg7,Sap47,ea,Ssdp,cort,cyc,sip3,MED4,qkr58E-2,SREBP,Exd2,hb,CG2129,eIF4A,ago,Usp10,Rcd5,Smox,Scox,CG14407,png,tapas,Uev1A,zyd,EloC,CG17328,Ect4,PpV,osk,HP1c,Rab32,Tomosyn,Mctp,PEK,tyf,Grip84,Cbl,ERp60,Nap1,Stam,Nurf-38,Mitf,Rpt2,Sirt7,CG5003,Pp2A-29B,Rel,CG4853,Tfb4,CycE,CG3587,Rpn13,Chrac-14,CG10188,Hip1,spn-E,Sec23,sbb,CG4872,O-fut1,piwi,Lkb1,Tlk,Cka,krimp,d4,CG2678,dl,Pura,foxo,Cul5,Ntan1,not,CG10418,AP-1-2beta,trbd,SelD,APC7,lid,DCAF12,spz,CG3894,bcd,tub,SERCA,obe,Maf1,RhoGAP5A,Tm1,ovo,CG11486,rept,e(y)1,CG7328,mtTFB1,Su(H),Arpc4,psq,Atf6,mus304,Seipin,Abi,fray,ASPP,l(3)73Ah,msps,escl,sws,velo,Cdk1,CG31875,CG14408,Wbp2,cnc,BoYb,CG1578,Prosbeta5,pygo,Tak1,wapl,Pgk,Gabat,tor,nos,RYBP,Tao,mRpL13,Cul2,ATPsynD,Coq7,lt,Bap170,ben,CG10654,dgt4,stet,Grip71,Iml1,Elba2,HmgD,Rbf2,EndoGI,MED20,p38a,Spt20,Npc2a,E2f1,RecQ5,CG10959,lin,cnn,Sox102F,pen-2,Aven,Myd88,cup,CG11902,Herc4,Chd64,RpLP1,heph,raptor,CG30291,Ptip,eIF3m,CG4238,Bx,pps,mAChR-A,CG10336,Traf6,ph-d,CG10038,gus,Tip60,Uxt,Marf1,PDCD-5,Archease,Dph5,EcR,bif,gw,pgc,Ptp61F,enc,CG1602,mtrm,tkv,Pka-R1,Idgf2,CG7414,capt,l(3)mbt,Taf7,PGRP-LE,Rep,TfIIFalpha,lin-52,smg,pont,Cdk5,rl,Jafrac1,Ptp10D,CkIIalpha,RpL13A,RhoGDI,CG17202,csw,mute,Atf3,Mybbp1A,CG8963,Hat1,Rga,scny,CG31053,Vps16B,Ankle2,mus81,mts,esc,mad2,Jafrac2,CG9391,Nf-YA,CG8009,exu,ssh,gd,hng3,CG17746,Mos,tws,Hsc70-3,vers,Mekk1,Marf,CycC,NFAT,twin,apt,fz4,BicC,dco,CG17272,egg,lwr,CycB,wash,ScsbetaA,Pax,GlyP,bun,mop,fs(1)M3,CanA-14F,HP1e,Cf2,disp,CG2199,Arpc2,MAGE,Synj,Nlg1,REPTOR,Med,CG30496,Topors,RhoGAP54D,Ptpmeg,nAChRalpha4,trr,AGO3,crok,Imp,Parp16,Pdk1,Idh3b,fwe,snk,Pink1,cact,crq,mael,Trf2,plu,Pi3K92E,Pp2B-14D,Mtl,Calx,stai,Fkbp39,boi | | 1.098e-08 | -18.33 | ion binding | molecular function | GO:0043167 | 2058 | 259 | 12176 | 1119 | Sply,PEK,nmdyn-D7,Hr78,LysRS,up,Pgm1,CysRS,eIF4A,PGRP-LE,CG31053,CG9684,CSN6,Droj2,GlyP,cana,CG6923,alphaTub84B,Orc1,Sirt7,bor,Cyp6u1,Rpn3,Lkb1,CG14183,CG4880,Fer1HCH,Marf,Desat1,CG5116,CG13344,Rab5,CG3077,mEFTu1,Nc73EF,ScsbetaA,Iru,mal,DhpD,CG9425,Nurf-38,Cyp6a22,CkIIalpha,isoQC,Rab10,Tao,Vha68-2,CG15141,Usp16-45,Mctp,CG8728,Edem1,CG9941,Rpn2,Atpalpha,CG2678,AGBE,Pex10,CAH7,CG17272,Dip-B,spn-E,Reps,not,krimp,Mos,CG7488,l(1)G0196,CG11134,Cbl,RecQ5,PAPLA1,CG8726,Hnf4,Nca,CG4709,Nadsyn,gammaTub37C,CG11902,pont,p38a,Gs1l,CG17746,l(2)k01209,sau,Mtl,Pfk,ari-1,kl-2,fray,Acox57D-p,ABCD,SmydA-1,l(3)mbt,CG17593,Usp39,ACC,sip3,Topors,G9a,CG7806,shop,CG10492,CG4730,tor,AnxB10,CG6813,DNApol-epsilon255,GluRS-m,alph,Coq6,Pgk,awd,blw,lt,Myd88,Traf6,CG9143,Sec23,CG11807,Snx6,AOX3,CG8611,Tlk,Rpt2,Nup62,l(3)73Ah,tkv,CG9581,CanB2,CG32473,CAH2,Eno,Trxr-1,nos,Utx,stet,snz,alpha-Man-Ia,Pink1,TyrRS,CG14435,Pfrx,dco,LpR2,stw,CG8974,Scp1,CG10463,E23,CCT3,SelD,regucalcin,CG17328,crok,cN-IIIB,Wdfy2,Shmt,Marc,Cyp6a19,wal,nej,CG4586,Clp,alphaTub67C,aay,Idh,AcCoAS,nmd,Exo70,AIF,CG7728,CG44422,ATPsynbeta,Cdk5,SP1029,CG11267,ph-d,mEFTu2,TrpRS-m,Mekk1,CG8507,Non1,Klp10A,obe,dx,Actn,EcR,Cyp6v1,mre11,Scox,QC,CG31689,HDAC6,Calx,CG6767,Fdh,Pld,sgll,CG4041,SERCA,Hip1,CG14516,PyK,CG11710,CG5789,SerRS-m,CG5734,Mcad,Rab32,Got1,rept,grp,MtnA,rl,CG4860,Sod1,r,CG17698,glob1,CG9925,CG9981,SRPK,png,grsm,ND-51,ATP8A,yl,CG7115,CG4562,CG14997,CG11590,SdhA,Cdk1,SelR,CG31729,CG1640,CCT5,Hsc70-3,CCT6,egg,Arl5,d4,RpS29,Sod3,Tengl4,Kyat,Cdk7,Arf51F,Tak1,fs(1)N,Gabat,Pdk1,Pex12,Kcmf1,Moe,CG17260,Pex23,CG10654 | | 1.108e-08 | -18.32 | nucleobase-containing small molecule metabolic process | biological process | GO:0055086 | 264 | 54 | 12345 | 1129 | PyK,ATPsynO,Pfk,blw,AOX3,CG8199,Ac3,AMPdeam,Nadsyn,Ugt36E1,Eno,ATPsynE,ATPsynD,Ts,CG7920,CG6767,ScsbetaA,DhpD,CG17691,Shmt,r-l,scu,Pfas,Gmer,nmdyn-D7,ATPsynB,ATPsynG,Pfrx,awd,AcCoAS,Gapdh1,CG33156,kri,r,ATPsyndelta,Ady43A,CG11164,cN-IIIB,ATPsynbeta,Scsalpha1,CG4407,CG6178,Ppcdc,UGP,Ugt35A1,ATPsynCF6,COX5B,Pgk,nmdyn-D6,fal,Pglym78,l(2)k01209,CG11089,CG1774 | | 1.117e-08 | -18.31 | PHGPx (PHGPx) | protein interactions | 38413 | 74 | 28 | 9204 | 1112 | capt,Moe,scu,regucalcin,CG10265,CG11899,Usp39,CG4598,CG3226,AnxB10,Eno,Pglym78,Uch,cib,EndoGI,Fkbp12,Pgk,CG14715,CG10638,awd,CG11267,Chd64,yip2,ATPsynCF6,Cisd2,HmgD,Stip1,RhoGDI | | 1.150e-08 | -18.28 | small ribosomal subunit | cellular component | GO:0015935 | 77 | 25 | 11940 | 1105 | RpL40,RpS23,RpS18,mRpS11,RpS27A,RpS24,RpS14b,RpS4,RpS14a,RpS2,RpS5b,CG7414,RpS13,RpS15,mRpS10,mRpS23,RpS29,RpS30,mRpS5,RpS7,RpS27,tko,RpS28b,RpS16,mRpS21 | | 1.222e-08 | -18.22 | ribose phosphate biosynthetic process | biological process | GO:0046390 | 89 | 27 | 12345 | 1129 | l(2)k01209,Pfas,nmdyn-D7,CG11089,CG17691,r-l,nmdyn-D6,CG6767,ATPsynCF6,COX5B,ATPsynbeta,CG6178,Ppcdc,ATPsynE,ATPsynD,Ady43A,Ac3,CG8199,AMPdeam,AcCoAS,ATPsynO,r,ATPsyndelta,blw,ATPsynB,ATPsynG,awd | | 1.271e-08 | -18.18 | protein modification by small protein conjugation or removal | biological process | GO:0070647 | 331 | 63 | 12345 | 1129 | CG9316,EloC,Ubc4,ben,RpS27A,Usp39,CG9941,CG31633,trbd,CG14435,Cul2,CG11070,CG8974,Uev1A,Ufm1,dx,ari-1,CG8188,lwr,faf,Cbl,Fbw5,CG32221,park,UbcE2H,CG46338,Aos1,Sgf11,Cul1,RpL40,SkpA,APP-BP1,MAGE,STUB1,CG10254,Uch-L5,l(3)73Ah,CG4238,Atg7,velo,Pex10,Herc4,Pex12,CG5003,Usp16-45,Uba5,CSN6,CG13344,CG40045,CG2924,Traf6,Cul5,Usp10,ago,Ubc6,not,Iru,otu,scny,Uch,Topors,CG31053,sip3 | | 1.498e-08 | -18.02 | nucleoside phosphate biosynthetic process | biological process | GO:1901293 | 119 | 32 | 12345 | 1129 | ATPsynbeta,Scsalpha1,CG6178,CG4407,Ppcdc,Ady43A,AcCoAS,CG33156,r,ATPsyndelta,ATPsynB,ATPsynG,awd,l(2)k01209,CG11089,nmdyn-D6,ATPsynCF6,COX5B,Nadsyn,ATPsynD,ATPsynE,CG8199,Ac3,AMPdeam,ATPsynO,blw,Pfas,nmdyn-D7,CG17691,r-l,CG6767,Ts | | 1.531e-08 | -17.99 | regucalcin (regucalcin) | protein interactions | 32165 | 93 | 32 | 9204 | 1112 | CG4598,CG1532,scu,Moe,capt,Gdi,Uch,Prosalpha4,Pglym78,Trxr-1,awd,Fkbp12,Sod1,RhoGDI,AnxB10,Seipin,Eno,Gapdh1,Chd64,CG11267,yip2,Cyp1,EndoGI,cib,PHGPx,CG14715,CG10638,Pgk,ATPsynbeta,Stip1,blw,ERp60 | | 1.688e-08 | -17.90 | RpS27A (Ribosomal protein S27A) | protein interactions | 34420 | 179 | 49 | 9204 | 1112 | RpLP1,RpL7A,RpL24,RpL23,alphaTub84B,RpL23A,RpL10,RpL38,dl,RpL9,RpS14a,RpL31,RpL17,RpS30,RpS13,bcd,RpS15,Argk,Moe,RpS29,bbx,Uch-L5,RpS14b,Traf6,RpS7,RpS24,RpS2,RpL13,RpL37a,RpS18,RpL34a,Rpn13,RpL12,RpL26,RpLP0,eIF4A,Eno,RpL13A,RpS28b,RpL11,RpS4,Hsp26,RpL30,RpL40,RpS23,RpL29,RpS27,RpS16,Atpalpha | | 1.688e-08 | -17.90 | RpS3 (Ribosomal protein S3) | protein interactions | 42761 | 179 | 49 | 9204 | 1112 | NaPi-III,RpS29,RpS15,RpL31,RpL17,RpS30,Usp16-45,RpS13,RpL9,Rel,RpS14a,PGRP-LE,RpL38,RpL23,RpL23A,RpL10,Tak1,RpL7A,RpL24,piwi,RpLP1,Mst89B,RpL29,RpS16,RpS27,Pi3K92E,RpL30,RpS23,RpL40,RpL11,RpS4,RpL13A,ago,RpS28b,RpL12,mts,RpS27A,RpL26,RpLP0,RpS2,Maf1,RpL13,RpL37a,RpS18,RpL34a,RpS7,RpS24,CG8974,RpS14b | | 1.760e-08 | -17.86 | fatty acid beta-oxidation | biological process | GO:0006635 | 31 | 15 | 12345 | 1129 | Seipin,CG4598,Etf-QO,CG4860,wal,yip2,Acox57D-p,SERCA,Mtpalpha,CG4586,CG4594,Echs1,CG10932,Mcad,Thiolase | | 1.804e-08 | -17.83 | developmental process involved in reproduction | biological process | GO:0003006 | 928 | 135 | 12345 | 1129 | Atg7,lqfR,csw,armi,Pex10,CkIIalpha,r2d2,Ptpmeg2,mad2,mts,exu,eIF4A,Orc1,ea,cort,muskelin,osk,shu,Mos,egg,Cbl,Pax,ScsbetaA,Nap1,CycB,wash,CG46338,yps,twin,nmd,BicC,apt,Pen,CycE,Med,spn-E,Pxt,bun,mop,Pex12,fs(1)M3,gammaTub37C,HP1e,Cf2,Top1,Cul5,tho2,Pdk1,Lkb1,piwi,krimp,bcd,cact,spz,Pink1,lid,Pi3K92E,e(y)1,stai,Tm1,mael,stil,SERCA,ovo,yl,Abi,msps,psq,bam,eIF4E1,BoYb,GatA,nos,tor,CycJ,Cdk1,cnc,Nup358,jvl,Iml1,Grip71,Rab10,Moe,Tao,Cul2,cnn,ari-1,shrb,zpg,milt,park,vret,toc,Stat92E,swa,E2f1,eIF3m,Grip128,sw,ix,Kap3,Rab5,cup,heph,raptor,CSN6,squ,Rbf,Grip75,otu,bru1,del,enc,pgc,EcR,fs(1)N,wisp,Debcl,Rbfox1,gus,Sec15,Nup62,nej,cathD,Dcr-1,Drep1,Jafrac1,elg1,rl,Nprl2,SRPK,Pka-R1,tkv,l(3)mbt,capt | | 1.986e-08 | -17.73 | ribosome | cellular component | GO:0005840 | 287 | 57 | 11940 | 1105 | RpL26,mRpL35,RpL37a,RpS14b,RpS24,RpS14a,RpL13,RpL9,mRpS11,RpS27A,RpL38,RpS18,CG6094,RpL29,mRpL16,RpL7A,PEK,RpL24,RpS28b,RpL11,RpL13A,CG7414,CG14817,RpS13,RpS15,mRpS10,RpL34a,RpLP1,mRpL10,RpL31,RpS4,RpS2,mRpL17,RpS5b,RpS23,RpL40,RpL17,mRpL36,RpL23,RpL23A,mRpL48,RpL12,mRpS21,mRpL18,RpL34b,tko,RpS27,RpS16,mRpL13,RpL30,RpS7,RpLP0,mRpS23,RpS29,mRpS5,RpL10,RpS30 | | 2.306e-08 | -17.59 | negative regulation of cellular process | biological process | GO:0048523 | 1291 | 175 | 12345 | 1129 | Pink1,lid,SelD,bcd,cact,plu,ovo,mael,Maf1,Fkbp39,rept,Pi3K92E,CG2199,bun,Rel,Pp2A-29B,Sirt7,spn-E,sbb,CG4872,Chrac-14,CycE,MAGE,d4,Ptpmeg,krimp,CG30496,Cka,Tlk,piwi,Lkb1,Pdk1,foxo,Cul5,Ntan1,dl,AGO3,tws,tapas,Mos,CG17746,Smox,ssh,Ect4,PpV,NFAT,CG17328,Mekk1,dco,BicC,apt,twin,wash,Cbl,egg,lwr,CkIIalpha,r2d2,Ptp10D,armi,Aldh,Hat1,Atg7,csw,Mybbp1A,STUB1,RpL13A,cyc,Ankle2,Rga,scny,ago,SREBP,hb,mts,esc,mus81,mad2,MED4,dx,nbs,Bin1,Tip60,Uxt,ph-d,MED27,pn,CG10038,Rbfox1,Debcl,mtrm,CG1602,gw,bif,pgc,Ptp61F,chrb,EcR,Hnf4,asf1,SkpA,PGRP-LE,l(3)mbt,insv,BigH1,Idgf2,Naa20A,SuUR,Drep1,rl,Nprl2,faf,lin-52,nej,smg,heph,raptor,ix,Herc4,cup,Rab5,Drep2,tral,CG4238,COX5A,spag,CG10631,Zwilch,bru1,alph,Dd,COX4,mre11,CG10336,Traf6,Rbf,ATPsynD,Cul2,mRpL13,RYBP,dnc,Elba2,Iml1,Bap170,E2f1,Cul1,Stat92E,Rbf2,EndoGI,park,Aven,vret,Sox102F,CG10959,RecQ5,bam,Utx,mus304,psq,Su(H),Not10,velo,sws,escl,CG14408,grp,Unr,CG31875,Cdk1,nos,Ubc6,pygo,wapl,Prosbeta5,BoYb | | 2.439e-08 | -17.53 | regulation of catabolic process | biological process | GO:0009894 | 227 | 48 | 12345 | 1129 | SkpA,PEK,dco,Rab32,Cul1,Fkbp39,Nprl2,Sod1,rl,Pax,Fbw5,Mitf,park,smg,Pi3K92E,ari-1,gus,Smox,AP-1-2beta,Marf1,Rpn3,osk,Debcl,CycC,EcR,Rpn2,Desat1,gw,Pomp,cort,Lkb1,piwi,scny,ago,Tak1,Ubc6,AGO3,Traf6,foxo,SREBP,Utx,CkIIalpha,Rpt2,GlyP,CG4238,Atg7,spn-E,Synj | | 2.590e-08 | -17.47 | REACTOME\_METABOLISM\_OF\_MRNA | MSigDB lists | REACTOME\_METABOLISM\_OF\_MRNA | 146 | 49 | 4161 | 644 | Edc3,RpS2,RpS16,RpL23,RpL30,CG8963,RpS24,CG10418,Cbp80,RpL40,RpS4,RpL38,Prosbeta6,RpS7,Pp2A-29B,RpL26,RpL24,RpS27,Prosbeta4,LSm3,RpL37a,Prosalpha3,RpL11,Rpn3,RpL12,RpS18,p38a,Prosbeta5,Rpn2,Not10,eIF4E1,Rrp40,CG6610,RpL13A,RpLP0,RpS13,Rpn12,RpS30,Prosalpha4,eIF4A,Rpt2,mts,Prosbeta3,Prosalpha5,RpS27A,RpL9,RpS23,RpS28b,RpL13 | | 2.702e-08 | -17.43 | lipid oxidation | biological process | GO:0034440 | 36 | 16 | 12345 | 1129 | CG4598,wal,Acox57D-p,SERCA,CG4586,Echs1,CG10932,Thiolase,Seipin,Etf-QO,CG4860,yip2,Mtpalpha,CG4594,Mcad,Hnf4 | | 3.207e-08 | -17.26 | ribonucleotide biosynthetic process | biological process | GO:0009260 | 87 | 26 | 12345 | 1129 | CG11089,nmdyn-D7,Pfas,l(2)k01209,r-l,CG17691,nmdyn-D6,COX5B,ATPsynCF6,ATPsynE,ATPsynD,Ppcdc,CG6178,ATPsynbeta,Ac3,CG8199,AMPdeam,Ady43A,blw,ATPsyndelta,r,ATPsynO,AcCoAS,awd,ATPsynG,ATPsynB | | 3.224e-08 | -17.25 | negative regulation of metabolic process | biological process | GO:0009892 | 823 | 122 | 12345 | 1129 | escl,Mybbp1A,sws,Hat1,RpL13A,r2d2,Utx,CkIIalpha,bam,Su(H),Not10,armi,psq,nos,mts,esc,hb,SREBP,CG14408,Ankle2,cyc,scny,CG31875,Unr,Rga,CG17328,Elba2,PpV,mRpL13,Hsc70-3,ATPsynD,CG31759,SmydA-1,Smox,CG17746,wash,park,cu,CG10959,Sox102F,Cbl,egg,apt,trsn,E2f1,Cbp80,BicC,dco,EndoGI,Edc3,twin,Rbf2,CG4872,spn-E,sbb,CycE,Chrac-14,heph,CG2199,cup,Sirt7,LSm3,squ,Rbf,dl,AGO3,CG10336,foxo,Smyd4-4,Trf4-2,alph,krimp,CG10631,bru1,CG17768,d4,Lkb1,piwi,Tlk,CG30496,CG1602,bcd,Rbfox1,EcR,Hnf4,Ptp61F,pgc,gw,Rrp40,Bin1,CG10038,SelD,ph-d,Marf1,Pym,Tip60,Uxt,Fkbp39,CG6967,rl,Nprl2,Dcr-1,SuUR,Drep1,CG11486,smg,YL-1,lin-52,Pi3K92E,faf,rept,CG9425,l(3)mbt,SkpA,ovo,plu,asf1,tkv,BigH1,mael,Maf1,insv | | 3.335e-08 | -17.22 | structural constituent of ribosome | molecular function | GO:0003735 | 271 | 54 | 12176 | 1119 | RpS13,mRpL17,mRpS11,RpLP0,mRpL36,RpS18,RpS28b,mRpL10,RpL29,RpL10,mRpS23,RpS16,RpL30,RpL40,mRpS10,RpL13A,RpS4,mRpL48,RpL23A,RpL31,RpL17,RpL12,RpL34a,RpL13,mRpL35,tko,CG14817,RpL23,RpS15,mRpL18,RpS5b,RpL7A,RpL37a,RpS30,RpL11,RpS14b,mRpL16,mRpS21,RpL38,RpS14a,RpLP1,RpL26,RpL9,RpL34b,RpS27A,RpS24,RpS23,RpL24,RpS2,mRpL13,RpS7,RpS29,RpS27,mRpS5 | | 3.356e-08 | -17.21 | Eukaryotic Translation Initiation | REACTOME pathways | R-DME-72613 | 128 | 43 | 3923 | 574 | RpL13A,RpS14a,RpL7A,RpL12,RpL17,RpL9,RpL37a,RpL34b,RpL38,RpS29,RpS23,RpL24,RpL26,RpS28b,RpS7,RpL30,RpL11,RpS27A,RpS24,eIF3m,RpLP1,RpL23A,RpL34a,eIF4E1,RpL31,RpS15,RpS5b,RpL10,RpS4,RpLP0,RpS13,RpS2,RpL13,RpL40,RpS27,eIF4A,RpS30,RpS14b,RpL29,eIF3i,RpS16,RpS18,RpL23 | | 3.356e-08 | -17.21 | Cap-dependent Translation Initiation | REACTOME pathways | R-DME-72737 | 128 | 43 | 3923 | 574 | RpS7,RpS28b,RpL11,RpL30,RpS24,eIF3m,RpS27A,RpS29,RpL26,RpS23,RpL24,RpL34b,RpL37a,RpL38,RpS14a,RpL13A,RpL12,RpL7A,RpL17,RpL9,eIF3i,RpS16,RpL23,RpS18,RpL40,eIF4A,RpS27,RpL29,RpS14b,RpS30,RpS13,RpS2,RpL13,eIF4E1,RpLP1,RpL34a,RpL23A,RpS5b,RpL10,RpS15,RpL31,RpLP0,RpS4 | | 3.408e-08 | -17.19 | purine-containing compound metabolic process | biological process | GO:0072521 | 181 | 41 | 12345 | 1129 | Shmt,CG17691,DhpD,nmdyn-D7,Pfas,scu,CG6767,ScsbetaA,CG7920,CG8199,AMPdeam,Ac3,AOX3,ATPsynE,ATPsynD,Eno,PyK,blw,Pfk,ATPsynO,Pglym78,CG1774,CG11089,COX5B,ATPsynCF6,fal,nmdyn-D6,Pgk,cN-IIIB,Ady43A,Ppcdc,Scsalpha1,CG6178,ATPsynbeta,awd,Pfrx,ATPsynG,ATPsynB,ATPsyndelta,Gapdh1,AcCoAS | | 3.658e-08 | -17.12 | nucleotide biosynthetic process | biological process | GO:0009165 | 117 | 31 | 12345 | 1129 | CG6178,CG4407,Nadsyn,ATPsynbeta,ATPsynE,ATPsynD,Ppcdc,CG8199,Ac3,AMPdeam,Ady43A,CG33156,r,ATPsynO,AcCoAS,ATPsyndelta,blw,ATPsynG,ATPsynB,awd,Pfas,l(2)k01209,CG11089,nmdyn-D7,CG17691,r-l,nmdyn-D6,CG6767,COX5B,ATPsynCF6,Ts | | 3.718e-08 | -17.11 | purine nucleotide metabolic process | biological process | GO:0006163 | 155 | 37 | 12345 | 1129 | CG11089,CG1774,Pglym78,Pgk,nmdyn-D6,fal,ATPsynCF6,COX5B,ATPsynbeta,CG6178,Scsalpha1,Ppcdc,Ady43A,AcCoAS,Gapdh1,ATPsyndelta,ATPsynB,Pfrx,ATPsynG,awd,scu,Pfas,nmdyn-D7,CG17691,CG7920,CG6767,ScsbetaA,Eno,ATPsynE,ATPsynD,CG8199,Ac3,AMPdeam,ATPsynO,Pfk,blw,PyK | | 3.775e-08 | -17.09 | Trx-2 (thioredoxin-2) | protein interactions | 34281 | 56 | 23 | 9204 | 1112 | cib,PHGPx,dl,Cyp1,AIF,Fkbp12,Sod1,CG14715,CG10638,Trxr-1,Chd64,CG11267,awd,CG9391,scu,Uev1A,regucalcin,Treh,CG1532,AnxB10,Got1,Eno,Uch | | 3.862e-08 | -17.07 | GO\_MACROMOLECULE\_CATABOLIC\_PROCESS | MSigDB lists | GO\_MACROMOLECULE\_CATABOLIC\_PROCESS | 407 | 103 | 4161 | 644 | cyc,Not10,CG6610,RpL13A,Rrp40,eIF4E1,RpS30,RpS13,Rpn12,RpLP0,trbd,Prosalpha4,Rpt2,mts,CG11164,Cul2,RpL13,AIF,Cul5,STUB1,GlyP,RpL9,RpS27A,RpS28b,Faf2,twin,snz,CG10418,cathD,Cbp80,not,CG6766,CG9485,CG40045,faf,Edc3,ari-1,Atg6,RpS2,CG8963,RpL30,Edem1,lwr,Pfk,Prosbeta5,Pp2A-29B,Iru,RpL24,Hsc70-3,LSm3,RpL37a,Prosalpha3,Tip60,CG4572,CG11070,CG8475,SkpA,Vsp37A,Rpn2,HDAC6,CG8188,ste24a,CG10465,eIF4A,UbcE2H,Prosbeta3,Pgm1,mad2,Uch-L5,Fbw5,Cdk1,Prosalpha5,RpS23,Herc4,Unr,shrb,RpL40,Prosbeta6,RpS4,RpL38,RpS7,CG17593,mus81,Ubc4,RpS16,APC7,RpL23,RpS24,CG8974,Cul1,Rpn3,RpL12,RpS18,Cdk5,CycB,RpL26,RpS27,Prosbeta4,Uch,CG32473,Dcr-1,RpL11,Gyg | | 4.278e-08 | -16.97 | cellular developmental process | biological process | GO:0048869 | 1818 | 230 | 12345 | 1129 | sw,TBCC,Rab5,cup,Kap3,Pka-R2,mxc,raptor,heph,CSN6,eIF3m,COX5A,Grip128,Arf51F,COX4,CG3770,Pglym78,otu,Grip75,bru1,EMC3,Smyd4-4,alph,Wnt10,squ,Rbf,gus,MED27,Sec15,Ptp61F,enc,PyK,del,bif,pgc,Hnf4,EcR,fs(1)N,Archease,wisp,Rbfox1,Debcl,SRPK,cad,Pka-R1,tkv,l(3)mbt,capt,SkpA,Nup62,Cdk5,faf,nej,cathD,Dcr-1,spen,Drep1,Jafrac1,elg1,Nprl2,rl,Arpc4,CanB2,psq,Su(H),bam,Seipin,eIF4E1,Klp10A,CG13366,Utx,yl,Abi,Arpc1,velo,msps,PIG-C,Cdk1,Nup358,cnc,Afti,BoYb,GatA,nos,tor,Pgk,CycJ,Tao,RYBP,G9a,Cul2,jvl,colt,Iml1,dnc,Grip71,ben,Moe,Rab10,toc,Cul1,Rbf2,Actn,Stat92E,Npc2a,swa,E2f1,Hr78,cnn,Sox102F,ari-1,shrb,zpg,milt,MED24,park,rost,vret,Pxt,Rel,bun,mop,fs(1)M3,Pex12,Gclc,gammaTub37C,ACC,Cf2,HP1e,Pen,MAGE,Arpc2,CycE,Med,Pfdn2,spn-E,Sec23,sbb,Mitofilin,Lkb1,cold,piwi,Ptpmeg,up,krimp,tho2,Top1,foxo,Pura,Cul5,dl,not,Imp,Liprin-beta,Idh3b,Pink1,spz,DCAF12,lid,Pfk,bcd,cact,Tm1,mael,stil,SERCA,ovo,Trf2,Pmi,Mtl,rept,Pi3K92E,galectin,AIF,boi,e(y)1,stai,armi,SelR,Ptp69D,Pex10,CkIIalpha,Atg6,r2d2,Ptp10D,Ptpmeg2,Atg7,lqfR,Gapdh1,csw,CG17068,ea,cort,Ankle2,muskelin,hb,SREBP,mad2,MED4,mts,exu,Nf-YA,eIF4A,ago,Orc1,Mos,ssh,Smox,tws,Marf,osk,shu,CG46338,twin,yps,nmd,BicC,dco,apt,shep,Cbl,egg,Girdin,Pax,ScsbetaA,parvin,Nap1,Stam,wash,CycB | | 4.283e-08 | -16.97 | awd (abnormal wing discs) | protein interactions | 43739 | 116 | 36 | 9204 | 1112 | CG11267,Chd64,yip2,cib,EndoGI,Cyp1,PHGPx,CG14715,CG10638,ATPsynbeta,Pgk,Stip1,ERp60,CG10265,AnxB10,Rab5,regucalcin,Sgf11,ago,Eno,CG15141,awd,Trxr-1,Fkbp12,Sod1,RhoGDI,HmgD,ATPsynCF6,CG4598,capt,Moe,scu,Uch,Prosalpha4,Pglym78,Rtnl1 | | 4.358e-08 | -16.95 | nucleoside triphosphate metabolic process | biological process | GO:0009141 | 37 | 16 | 12345 | 1129 | Scsalpha1,ATPsynbeta,ATPsynD,ATPsynE,ATPsynO,r,blw,ATPsyndelta,ATPsynG,ATPsynB,awd,nmdyn-D7,nmdyn-D6,fal,COX5B,ATPsynCF6 | | 4.478e-08 | -16.92 | oxidoreductase activity, acting on NAD(P)H | molecular function | GO:0016651 | 66 | 22 | 12176 | 1119 | ND-49,CG3939,cl,ND-23,ND-39,ND-75,AIF,Trxr-1,ND-24,ND-B18,ND-B15,CG7322,Grx1,ND-SGDH,ND-30,Fdh,ND-B22,ND-51,SelT,CG3803,ND-MNLL,ND-18 | | 4.854e-08 | -16.84 | GO\_PROTEIN\_TARGETING | MSigDB lists | GO\_PROTEIN\_TARGETING | 207 | 62 | 4161 | 644 | Karybeta3,Arl6IP1,CHOp24,GCC88,cyc,ste14,Arts,CG2100,RpS13,RpS30,RpLP0,Cse1,CG9393,RpL13A,rl,cact,Tom40,ERp60,Pen,RpL13,milt,shrb,RpS27A,RpL9,RpS23,RpS28b,RpL38,RpS4,RpS7,Rep,Tspo,Atg4b,RpL40,mbo,RpL23,RpL30,CG1847,Pex10,CG8728,RpS24,Phb2,lt,Pp2B-14D,Pex13,CG11110,RpS16,RpS2,Atg6,RpS18,Tim8,CG8004,Kap-alpha3,Marf,RpL12,CG8314,Pex12,CanA-14F,RpL11,RpL37a,RpL26,RpL24,RpS27 | | 5.144e-08 | -16.78 | RpL3 (Ribosomal protein L3) | protein interactions | 41347 | 137 | 40 | 9204 | 1112 | RpS15,RpL31,RpL17,RpS13,RpS30,RpS29,RpL23,RpL10,RpL23A,RpL24,RpLP1,RpL7A,RpL9,RpS14a,CG15141,RpL38,RpL11,RpS4,RpL13A,RpS28b,RpL29,RpS16,RpS27,RpL30,RpS23,RpL40,RpS7,RpS24,RpS14b,CG17328,RpL12,RpS27A,RpL26,RpLP0,RpS2,cup,RpL13,RpL37a,RpL34a,RpS18 | | 5.169e-08 | -16.78 | CG14715 (uncharacterized protein) | protein interactions | 41403 | 61 | 24 | 9204 | 1112 | RhoGDI,CG10638,Pgk,Fkbp12,Sod1,cib,PHGPx,EndoGI,yip2,Chd64,awd,CG11267,Trxr-1,Eno,Pglym78,Prosalpha4,l(1)G0004,Uch,regucalcin,capt,Moe,scu,AnxB10,CG4598 | | 5.269e-08 | -16.76 | respiratory chain complex | cellular component | GO:0098803 | 88 | 26 | 11940 | 1105 | ND-B15,ND-30,CG40472,ND-49,ND-MNLL,ND-39,levy,COX5B,COX8,ND-B18,ND-SGDH,ND-18,ND-24,ND-B22,COX5A,ND-23,UQCR-Q,cype,NP15.6,SdhA,ND-51,UQCR-11,ND-MWFE,COX4,ND-75,SdhD | | 5.725e-08 | -16.68 | GO\_ESTABLISHMENT\_OF\_PROTEIN\_LOCALIZATION\_TO\_ORGANELLE | MSigDB lists | GO\_ESTABLISHMENT\_OF\_PROTEIN\_LOCALIZATION\_TO\_ORGANELLE | 194 | 59 | 4161 | 644 | Tim8,CG8004,RpS18,Marf,RpL12,Kap-alpha3,CanA-14F,RpL11,RpL37a,Pex12,RpS27,RpL24,RpL26,RpS7,RpS4,RpL38,Tspo,RpL40,mbo,RpS24,CG1847,RpL30,RpL23,CG8728,Pex10,RpS16,Atg6,RpS2,lt,Phb2,Oseg1,CG11110,Pp2B-14D,Pex13,Tom40,rl,cact,RpL13,ERp60,Pen,shrb,Rab10,RpS28b,RpL9,RpS27A,RpS23,Karybeta3,rept,CG31729,Arts,GCC88,cyc,CG2100,RpS30,Cse1,RpLP0,RpS13,CG9393,RpL13A | | 6.159e-08 | -16.60 | cellular localization | biological process | GO:0051641 | 1000 | 141 | 12345 | 1129 | Rab32,Tomosyn,Vsp37A,Blos2,Efr,thoc6,Girdin,Pax,CycB,wash,Stam,ssh,CG8004,Marf,osk,sip3,Vps16B,Mat89Ba,MICAL-like,alphaTub84B,CG9393,exu,Pex13,Pex10,Atg6,CkIIalpha,cana,Ndc1,opm,ATPsynbeta,lqfR,Ranbp21,CG42837,Tom70,mael,Tm1,sbr,SERCA,obe,CG10344,Roe1,Sec10,stai,CG5510,Nup58,fwe,CG11110,AP-1-2beta,Gga,Pink1,CG4041,ATPsynO,CHOp24,Lkb1,Aladin,Topors,tho2,PAPLA1,COX5B,Pex12,mop,gammaTub37C,CHMP2B,Pen,Synj,CG5734,spn-E,Nlg1,Sec23,CG14894,Arts,Tom40,Pi4KIIalpha,Actn,swa,Npc2a,CG4293,shrb,Tsp86D,milt,Snx6,jvl,Exo70,lt,Cenp-C,Fer1HCH,Grip71,Moe,Rab10,Cse1,Tim8,cnc,Trs33,Nup358,GAPsec,Afti,Ubc6,CanB2,Vamp7,HDAC6,Klp10A,fray,Nxf3,sws,CG8314,CG6766,Arpc1,msps,Oseg1,l(3)mbt,Gdi,Rep,faf,Nup62,Cdk5,nej,SuUR,X11L,CG5447,Syx13,Sec15,CG8728,alphaTub67C,Rrp40,Arl5,wisp,Grip75,Kap-alpha3,sau,larp,sw,Karybeta3,Rab5,Kap3,cup,mbo,Clc,Arf51F,Grip128,Vha68-2,Sec22 | | 6.797e-08 | -16.50 | respirasome | cellular component | GO:0070469 | 89 | 26 | 11940 | 1105 | SdhA,ND-51,UQCR-11,NP15.6,cype,UQCR-Q,ND-23,COX5A,ND-24,ND-B22,ND-SGDH,ND-18,ND-B18,COX8,ND-39,COX5B,levy,CG40472,ND-49,ND-MNLL,ND-30,ND-B15,SdhD,ND-75,COX4,ND-MWFE | | 8.059e-08 | -16.33 | establishment of localization in cell | biological process | GO:0051649 | 784 | 116 | 12345 | 1129 | Tim8,Mat89Ba,GAPsec,Vps16B,Afti,sip3,Nup358,Trs33,CG9393,MICAL-like,Pex13,Ubc6,Pex10,CanB2,Ndc1,opm,ATPsynbeta,HDAC6,Klp10A,Atg6,cana,Vamp7,Nxf3,msps,Tom70,CG42837,CG6766,lqfR,CG8314,Ranbp21,Arpc1,Rab32,Vsp37A,Tomosyn,CG4293,Blos2,Npc2a,shrb,Tsp86D,Efr,CycB,wash,Stam,thoc6,Girdin,milt,Snx6,ssh,CG8004,Fer1HCH,Cenp-C,Exo70,jvl,Marf,lt,Moe,Rab10,Cse1,Kap-alpha3,Aladin,Topors,COX5B,PAPLA1,tho2,Karybeta3,Rab5,Kap3,gammaTub37C,Pex12,mop,sw,mbo,CHMP2B,Arf51F,Pen,Clc,CG5734,Synj,CG14894,Arts,Vha68-2,Sec22,Tom40,Sec23,Nlg1,sbr,SERCA,CG10344,Roe1,l(3)mbt,Rep,Gdi,Oseg1,Sec10,nej,faf,Cdk5,Nup62,CG5510,stai,CG11110,AP-1-2beta,Syx13,fwe,Nup58,X11L,Gga,CG8728,Sec15,alphaTub67C,Pink1,Arl5,ATPsynO,CHOp24,CG4041,wisp | | 8.221e-08 | -16.31 | mitochondrial matrix | cellular component | GO:0005759 | 171 | 39 | 11940 | 1105 | CG14407,mtTFB1,CG12375,Gabat,Pdk,CG8199,Idh3b,CG7488,wal,CG11267,TrpRS-m,Echs1,mRpL35,Got1,mRpS11,mRpL16,CG6094,Nc73EF,mRpL48,mRpS21,mRpL18,tko,pn,Roe1,mRpL13,mRpS5,mRpS23,Aldh,mRpS10,CG8728,mRpL10,Thiolase,CG14817,mRpL17,CG17691,P5CDh1,Mtpalpha,mRpL36,Scsalpha1 | | 8.347e-08 | -16.30 | nucleoside triphosphate biosynthetic process | biological process | GO:0009142 | 34 | 15 | 12345 | 1129 | Scsalpha1,ATPsynbeta,ATPsynD,ATPsynE,nmdyn-D7,r,ATPsynO,blw,ATPsyndelta,nmdyn-D6,COX5B,ATPsynG,ATPsynB,ATPsynCF6,awd | | 9.019e-08 | -16.22 | intracellular transport | biological process | GO:0046907 | 640 | 99 | 12345 | 1129 | Nxf3,Arpc1,Ranbp21,CG6766,CG8314,CG42837,Tom70,Pex10,Vamp7,Atg6,ATPsynbeta,Ndc1,opm,MICAL-like,CG9393,Ubc6,Pex13,Tim8,Nup358,Trs33,sip3,GAPsec,Afti,Vps16B,Mat89Ba,lt,jvl,Marf,Cse1,Rab10,Moe,ssh,Snx6,CG8004,Efr,Tsp86D,shrb,milt,Girdin,thoc6,Stam,wash,Vsp37A,Rab32,Npc2a,Blos2,CG4293,Synj,CG5734,Clc,Pen,Arf51F,Sec23,CG14894,Tom40,Vha68-2,Sec22,Arts,sw,Pex12,mop,gammaTub37C,Rab5,Kap3,Karybeta3,CHMP2B,mbo,PAPLA1,tho2,COX5B,Kap-alpha3,Topors,Aladin,Arl5,wisp,CG4041,CHOp24,ATPsynO,fwe,Nup58,Syx13,CG11110,AP-1-2beta,Pink1,alphaTub67C,Gga,CG8728,Sec15,Nup62,faf,stai,CG5510,CG10344,sbr,SERCA,Oseg1,Rep,l(3)mbt,Roe1 | | 9.475e-08 | -16.17 | purine nucleoside triphosphate metabolic process | biological process | GO:0009144 | 30 | 14 | 12345 | 1129 | awd,ATPsynG,COX5B,ATPsynCF6,ATPsynB,ATPsyndelta,blw,nmdyn-D6,fal,ATPsynO,ATPsynE,ATPsynD,nmdyn-D7,ATPsynbeta | | 9.506e-08 | -16.17 | oxoacid metabolic process | biological process | GO:0043436 | 419 | 72 | 12345 | 1129 | Desat1,TyrRS,Mcad,Eno,CG8199,aay,CysRS,scu,Shmt,CG17691,Aldh,Idh,CG4594,r,Gapdh1,P5CDh1,AcCoAS,Sply,SerRS-m,Seipin,CG6178,Got1,Dhfr,Pex10,oxt,GatA,CG11134,CG4586,Pgk,Echs1,Gabat,CG10932,SREBP,Gip,Sdhaf3,tzn,Pfk,AsnS,PyK,CG5028,TrpRS-m,Etf-QO,regucalcin,pn,Idh3b,SelD,CG11899,mino,CG31523,Adi1,Thiolase,wal,SERCA,Irp-1B,P5cr,Pdk,Pfrx,GatB,Scsalpha1,CG4860,yip2,Pxt,Kyat,ACC,Mtpalpha,LysRS,Ntan1,shop,CG4598,GluRS-m,Acox57D-p,Pglym78 | | 9.736e-08 | -16.14 | Nlp (Nucleoplasmin) | protein interactions | 43560 | 90 | 30 | 9204 | 1112 | ATPsynbeta,CG10638,CG14715,PHGPx,Cyp1,EndoGI,cib,Sod1,Fkbp12,yip2,CG11267,Trxr-1,awd,Chd64,ERp60,Stip1,HmgD,RhoGDI,regucalcin,capt,scu,Moe,CG4598,AnxB10,CG10265,Pglym78,Eno,Rtnl1,CG14204,Uch | | 9.757e-08 | -16.14 | cellular biosynthetic process | biological process | GO:0044249 | 1702 | 216 | 12345 | 1129 | aay,AMPdeam,CG7009,ATPsynD,Coq7,RpL24,Coq6,mRpL13,mal,lt,Desat1,TyrRS,RpL10,blw,mRpS23,mRpL10,r-l,CG17691,E2f1,Npc2a,CG10425,Pcd,Gmer,RpS23,Ts,DNApol-epsilon255,RpS30,RecQ5,mRpL17,MED24,mEFTu1,CG6767,RpL17,CG12018,RpL11,CG6178,Got1,Dhfr,eIF4E1,RpS28b,SerRS-m,CG8314,r,CG2911,CG2201,mRpS10,mRpS21,AcCoAS,RpL9,mus205,PIG-C,GAPsec,Rpb8,AGBE,CG1578,ATPsynCF6,RpL13,fdl,GatA,CG11134,pygo,RpS27,Ac3,CG14817,RpL26,Orc6,RpS16,Nadsyn,Dph5,mRpL16,RpS27A,Pgm1,RpL34b,CG7414,CG7840,TfIIFalpha,Sccpdh1,SkpA,Taf7,RpL40,Adi1,SuUR,spen,mino,CG9485,cN-IIIB,Cdk7,RpLP1,tko,Agpat2,CG4407,eIF3m,ATPsynG,GatB,ATPsynB,ATPsyndelta,mRpL18,CG33156,P5cr,RpS13,eEF1beta,mRpS11,Sgf29,CG8311,CG3803,RpL30,GluRS-m,RpII140,pps,EMC3,Ppox,CG10336,Coq8,Pld,Rbf,RpS29,RpS2,CG5880,CG8199,RpS5b,ATPsynE,CG13551,mRpL48,RfC38,RpLP0,Marf,RpL38,RpL12,Pdxk,CG7718,Shmt,RpL7A,Rab32,CG33977,fs(1)Ya,nmdyn-D7,Pfas,Efr,CysRS,Nurf-38,mEFTu2,RpS18,RpS14b,Ady43A,RpL23,Sld5,RpS14a,ATPsynbeta,awd,RpL13A,Atg7,RpS24,Rga,CG11089,PIG-H,RpL34a,l(2)k01209,CG10932,SREBP,MED4,fal,oxt,RpL37a,eIF4A,Orc1,ago,Pym,mRpL36,RpS7,regucalcin,ND-39,SelD,Nmt,TrpRS-m,mRpL35,CG17270,ATPsynO,AsnS,mRpS5,CG10344,RpL29,Gyg,Trf2,GalT1,Pi3K92E,CG11899,mtTFB1,CG31523,PIG-L,e(y)1,Gclc,Pxt,ACC,LysRS,Tfb4,CG6094,Ppcdc,Scsalpha1,RpS4,sgll,PIG-P,CycE,RpL31,pr,RpL23A,O-fut1,Ggt-1,Sptr,Hexo1,foxo,PAPLA1,Top1,COX5B,RpS15,nmdyn-D6,eIF3i | | 9.956e-08 | -16.12 | organic substance biosynthetic process | biological process | GO:1901576 | 1731 | 219 | 12345 | 1129 | Pi3K92E,CG11899,mtTFB1,PIG-L,e(y)1,CG31523,SERCA,CG10344,mRpS5,RpL29,Gyg,Trf2,GalT1,mRpL35,CG17270,ATPsynO,AsnS,Pym,mRpL36,RpS7,regucalcin,ND-39,SelD,Nmt,TrpRS-m,Top1,foxo,PAPLA1,COX5B,l(1)G0196,RpS15,nmdyn-D6,eIF3i,pr,RpL23A,O-fut1,Hexo1,Ggt-1,Sptr,PIG-P,sgll,CycE,RpL31,Gclc,Pxt,ACC,LysRS,Tfb4,CG6094,Ppcdc,Scsalpha1,RpS4,Efr,Nurf-38,CysRS,mEFTu2,Shmt,RpL7A,Rab32,CG33977,fs(1)Ya,nmdyn-D7,Pfas,Marf,RpL38,RpL12,Pdxk,CG7718,CG8199,RpS5b,ATPsynE,CG13551,mRpL48,RfC38,RpLP0,CG10932,SREBP,MED4,RpL37a,oxt,fal,ago,eIF4A,Orc1,Rga,CG11089,PIG-H,l(2)k01209,RpL34a,awd,RpL13A,RpS24,Atg7,RpS14b,RpS18,Ady43A,RpL23,Sld5,RpS14a,ATPsynbeta,Adi1,SuUR,spen,CG9485,mino,RpL34b,CG7414,CG7840,Sccpdh1,TfIIFalpha,SkpA,Taf7,RpL40,mRpL16,RpS27A,Pgm1,Ac3,CG14817,RpL26,Orc6,RpS16,Nadsyn,CG10336,Coq8,Ppox,Pld,Rbf,RpS29,RpS2,CG5880,Gk1,RpS13,eEF1beta,mRpS11,Gk2,Sgf29,CG8311,RpL30,CG3803,GluRS-m,RpII140,EMC3,pps,eIF3m,ATPsynG,ATPsynB,GatB,ATPsyndelta,P5cr,mRpL18,CG33156,cN-IIIB,RpLP1,Cdk7,tko,Agpat2,CG4407,Ts,DNApol-epsilon255,RecQ5,RpS30,mRpL17,MED24,mEFTu1,CG6767,RpL17,CG12018,r-l,CG17691,Npc2a,E2f1,Gmer,RpS23,Pcd,CG10425,mal,Desat1,lt,TyrRS,RpL10,blw,mRpL10,mRpS23,AMPdeam,CG7009,aay,ATPsynD,RpL24,Coq7,Coq6,mRpL13,AGBE,Rpb8,ATPsynCF6,RpL13,fdl,GatA,pygo,CG11134,Pgk,RpS27,RpL9,PIG-C,mus205,GAPsec,RpS28b,SerRS-m,CG8314,r,CG2201,AcCoAS,mRpS21,mRpS10,RpL11,Seipin,CG6178,Got1,Dhfr,eIF4E1 | | 9.959e-08 | -16.12 | IKKbeta (I-kappaB kinase beta) | protein interactions | 44432 | 172 | 46 | 9204 | 1112 | cact,pont,CG9391,blw,CCT3,CG15715,vret,RpL12,SERCA,CCT6,RpL13,cup,AMPdeam,RpS2,ATPsynbeta,rept,RpL11,RpS28b,GCS2beta,RpS27,Atpalpha,RpS16,PyK,Nup358,RpL23A,mtgo,alphaTub84B,CG4853,RpL23,CG8963,Hsc70-3,RpL24,RpL7A,Mitofilin,Sgf29,RpS14a,Rel,CG4557,RpL9,Sod1,dl,eEF1beta,Cp1,CCT5,Jafrac1,scu | | 1.029e-07 | -16.09 | positive regulation of cellular metabolic process | biological process | GO:0031325 | 875 | 126 | 12345 | 1129 | EcR,gw,CG1602,Debcl,Rbfox1,bcd,wisp,AP-1-2beta,pn,Tip60,Marf1,lid,CG11710,spz,Pi3K92E,pont,smg,nej,rept,Nprl2,e(y)1,SuUR,BigH1,cad,Taf7,SkpA,Trf2,TfIIFalpha,ovo,Rpn13,eIF3m,Ptip,Synj,spn-E,REPTOR,Atu,Rpt2,Myd88,GlyP,Cdk7,Rel,RpLP1,Cf2,raptor,CG2199,trr,AGO3,dl,foxo,Traf6,Parp16,not,Pdk1,piwi,otu,Bx,Lkb1,Dd,Cka,d4,vers,Mekk1,EloC,Bap170,ben,CG17328,CycC,NFAT,osk,RYBP,Rcd5,hng3,Scox,Mos,Tao,G9a,lin,Sox102F,ari-1,MED24,park,ScsbetaA,Mitf,Stat92E,Cul1,Sgf11,PEK,tyf,apt,MED20,dco,E2f1,Abi,fray,S2P,Sp7,CG17202,STUB1,ASPP,Mybbp1A,CG8963,Atg7,Su(H),Atf6,Utx,esc,MED4,Exd2,SREBP,Jafrac2,hb,CG2129,pygo,Tak1,Pgk,CG8009,tor,Nf-YA,Ubc6,CG31053,cyc,Pomp,cort,Ankle2,Wbp2,cnc | | 1.076e-07 | -16.04 | female meiotic nuclear division | biological process | GO:0007143 | 86 | 25 | 12345 | 1129 | Rab5,cup,gammaTub37C,Syx13,Klp10A,eIF4E1,mus304,Grip128,Cenp-C,bif,Axs,msps,wisp,mtrm,Grip71,yem,Grip75,SRPK,grp,cort,Pp2B-14D,cnn,lwr,wapl,nos | | 1.106e-07 | -16.02 | biosynthetic process | biological process | GO:0009058 | 1781 | 224 | 12345 | 1129 | COX5B,PAPLA1,Top1,foxo,eIF3i,nmdyn-D6,RpS15,l(1)G0196,O-fut1,RpL23A,pr,Hexo1,Sptr,Ggt-1,RpL31,CycE,sgll,PIG-P,LysRS,ACC,Kyat,Pxt,Gclc,RpS4,Scsalpha1,CG6094,Ppcdc,Tfb4,Pi3K92E,CG31523,PIG-L,e(y)1,mtTFB1,CG11899,mRpS5,CG10344,SERCA,Trf2,GalT1,Gyg,RpL29,CG17270,mRpL35,AsnS,ATPsynO,ND-39,SelD,RpS7,regucalcin,mRpL36,Pym,TrpRS-m,Nmt,MED4,SREBP,CG10932,eIF4A,Orc1,ago,oxt,RpL37a,fal,Rga,RpL34a,l(2)k01209,PIG-H,CG11089,RpL13A,awd,RpS24,Atg7,RpL23,Ady43A,RpS18,RpS14b,ATPsynbeta,RpS14a,Sld5,Efr,mEFTu2,CysRS,Nurf-38,Rab32,RpL7A,Shmt,Pfas,nmdyn-D7,fs(1)Ya,CG33977,RpL12,Marf,RpL38,CG7718,Pdxk,RpS5b,CG8199,RpLP0,RfC38,CG13551,mRpL48,ATPsynE,Rbf,Pld,Ppox,Coq8,CG10336,Gk1,CG5880,RpS2,RpS29,Sgf29,mRpS11,Gk2,eEF1beta,RpS13,EMC3,pps,RpII140,GluRS-m,CG3803,RpL30,CG8311,GatB,ATPsynB,ATPsynG,eIF3m,CG33156,P5cr,mRpL18,ATPsyndelta,Agpat2,tko,RpLP1,Cdk7,cN-IIIB,CG4407,Adi1,CG9485,mino,SuUR,spen,CG7414,RpL34b,RpL40,SkpA,Taf7,TfIIFalpha,Sccpdh1,CG7840,Dph5,Pgm1,RpS27A,mRpL16,RpL26,Orc6,Ac3,CG14817,CG1640,Nadsyn,RpS16,ATPsynCF6,CG1578,Rpb8,AGBE,Pgk,RpS27,pygo,CG11134,GatA,RpL13,fdl,PIG-C,mus205,RpL9,GAPsec,SerRS-m,RpS28b,AcCoAS,mRpS21,mRpS10,CG2911,CG2201,r,CG8314,RpL11,eIF4E1,Dhfr,Got1,CG6178,Seipin,mRpL17,RecQ5,RpS30,DNApol-epsilon255,Ts,CG12018,RpL17,CG6767,mEFTu1,MED24,CG17691,r-l,RpS23,CG10425,Gmer,Pcd,E2f1,Npc2a,RpL10,TyrRS,lt,Desat1,mal,mRpL10,mRpS23,blw,CG7009,aay,AMPdeam,mRpL13,Coq6,RpL24,Coq7,ATPsynD | | 1.155e-07 | -15.97 | organic acid metabolic process | biological process | GO:0006082 | 421 | 72 | 12345 | 1129 | Gip,Sdhaf3,oxt,GatA,CG11134,CG4586,Gabat,Pgk,Echs1,SREBP,CG10932,Seipin,CG6178,Got1,Dhfr,Pex10,Aldh,Idh,r,CG4594,Gapdh1,P5CDh1,AcCoAS,Sply,SerRS-m,scu,Shmt,CG17691,CysRS,Eno,aay,CG8199,Desat1,TyrRS,Mcad,CG4598,GluRS-m,Acox57D-p,Pglym78,Ntan1,shop,Scsalpha1,CG4860,yip2,Kyat,Pxt,ACC,Mtpalpha,LysRS,Irp-1B,P5cr,Pdk,Pfrx,GatB,wal,SERCA,CG11899,mino,CG31523,Adi1,Thiolase,CG5028,TrpRS-m,Etf-QO,regucalcin,Idh3b,pn,SelD,Pfk,tzn,AsnS,PyK | | 1.325e-07 | -15.84 | meiotic cell cycle | biological process | GO:0051321 | 261 | 51 | 12345 | 1129 | Cenp-C,twe,enc,del,bif,Axs,wisp,Grip71,mtrm,Syx13,lid,Pp2B-14D,Grip84,c(2)M,cnn,Nup62,egg,lwr,SRPK,mael,Grip128,Arf51F,CG7716,msps,spn-E,gammaTub37C,Rab5,cup,Pex10,CanB2,eIF4E1,Klp10A,mus304,bam,tef,larp,mus81,sau,mre11,wapl,GatA,Xpc,nos,yem,Grip75,Cdk1,grp,Topors,krimp,CG31053,cort | | 1.338e-07 | -15.83 | fatty acid oxidation | biological process | GO:0019395 | 35 | 15 | 12345 | 1129 | CG4586,CG4594,Echs1,CG10932,Mcad,Thiolase,Seipin,CG4598,Etf-QO,CG4860,yip2,wal,Acox57D-p,Mtpalpha,SERCA | | 1.355e-07 | -15.81 | Pgk (Phosphoglycerate kinase) | protein interactions | 33461 | 39 | 18 | 9204 | 1112 | CG32473,CG4598,AnxB10,scu,Moe,Argk,regucalcin,Uch,GlyP,Eno,Scp1,Pglym78,awd,Adh,PHGPx,CG14715,CG10638,Stip1 | | 1.432e-07 | -15.76 | regulation of cellular catabolic process | biological process | GO:0031329 | 204 | 43 | 12345 | 1129 | Tak1,ago,Ubc6,AGO3,foxo,SREBP,Traf6,Pomp,cort,piwi,Lkb1,scny,spn-E,Atg7,Synj,Utx,CkIIalpha,Rpt2,GlyP,Nprl2,Sod1,rl,Fkbp39,park,Pax,Mitf,Pi3K92E,smg,ari-1,SkpA,PEK,dco,Rab32,Cul1,CycC,Debcl,osk,EcR,gw,Desat1,AP-1-2beta,Smox,Marf1 | | 1.536e-07 | -15.69 | GO\_OXIDATION\_REDUCTION\_PROCESS | MSigDB lists | GO\_OXIDATION\_REDUCTION\_PROCESS | 381 | 96 | 4161 | 644 | CG4872,Gyg,Sccpdh1,Cyp6a22,CG14407,p38a,ND-B18,Mtpalpha,Gdi,Eno,SelR,Pglym78,Thiolase,ND-30,shop,CG10463,Dhfr,Pex13,ABCD,Rep,RpL40,cl,Scsalpha1,ND-39,Ppox,Irp-1B,Utx,CG8199,SdhA,Aldh,wal,CG3803,Pgk,Cyp6a19,Gapdh1,ND-49,Mcad,Pgm1,scu,Sod1,CG9265,ND-SGDH,lid,ND-B22,CG8475,CG17691,CG3842,Fdh,CG5028,Jafrac1,ND-23,ND-51,COX5A,Pfk,CG7322,Jafrac2,CG6665,Coq7,CG10425,ND-24,P5CDh1,P5cr,CG1275,CG11110,ScsbetaA,Cyt-c-p,sgll,Etf-QO,CG9485,AGBE,Adi1,Nc73EF,Desat1,Idh,CG7840,Sptr,Trxr-1,Pcd,RpS27A,Idh3b,AIF,AcCoAS,CG9471,CG3609,ND-75,GlyP,Echs1,UQCR-Q,ERp60,Gmer,CG6144,Coq6,levy,CG7550,CG4562,Grx1 | | 1.583e-07 | -15.66 | ribonucleoside triphosphate biosynthetic process | biological process | GO:0009201 | 31 | 14 | 12345 | 1129 | COX5B,ATPsynG,ATPsynCF6,ATPsynB,awd,ATPsynO,r,blw,ATPsyndelta,nmdyn-D6,ATPsynbeta,ATPsynE,ATPsynD,nmdyn-D7 | | 1.583e-07 | -15.66 | ribonucleoside triphosphate metabolic process | biological process | GO:0009199 | 31 | 14 | 12345 | 1129 | ATPsynbeta,ATPsynD,ATPsynE,nmdyn-D7,COX5B,ATPsynG,ATPsynCF6,ATPsynB,awd,r,ATPsynO,blw,ATPsyndelta,nmdyn-D6 | | 1.673e-07 | -15.60 | purine nucleotide biosynthetic process | biological process | GO:0006164 | 82 | 24 | 12345 | 1129 | AcCoAS,ATPsynO,ATPsyndelta,blw,ATPsynB,ATPsynG,awd,ATPsynbeta,CG6178,Ppcdc,ATPsynD,ATPsynE,Ady43A,CG8199,Ac3,AMPdeam,CG6767,nmdyn-D6,ATPsynCF6,COX5B,Pfas,nmdyn-D7,CG11089,CG17691 | | 1.807e-07 | -15.53 | positive regulation of metabolic process | biological process | GO:0009893 | 937 | 132 | 12345 | 1129 | lid,spz,AP-1-2beta,bcd,Trf2,ovo,e(y)1,Fbw5,Pi3K92E,rept,Cf2,CG2199,Rpt2,Rel,GlyP,REPTOR,spn-E,Synj,Rpn13,d4,Lkb1,piwi,Cka,Parp16,Pdk1,not,trr,AGO3,dl,foxo,hng3,Rcd5,Scox,Mos,CG17328,osk,NFAT,CycC,vers,EloC,Mekk1,tyf,apt,PEK,dco,ScsbetaA,Mitf,egg,Mybbp1A,CG8963,Atg7,Sp7,STUB1,CG17202,CG31053,cyc,Ankle2,cort,CG2129,CG8009,Nf-YA,MED4,esc,Jafrac2,hb,Exd2,SREBP,CG11710,gus,pn,Marf1,Tip60,CG1602,wisp,Debcl,Rbfox1,EcR,gw,SkpA,Taf7,TfIIFalpha,cad,BigH1,Nprl2,CG3198,SuUR,pont,smg,nej,raptor,Myd88,RpLP1,Cdk7,CG4238,Atu,Ptip,eIF3m,Bx,otu,Dd,Traf6,G9a,RYBP,Tao,ben,Bap170,E2f1,MED20,Stat92E,Sgf11,Cul1,MED24,park,lin,Sox102F,ari-1,Utx,Su(H),Atf6,ASPP,Abi,fray,S2P,Nup358,Wbp2,cnc,Pomp,Pgk,Tak1,pygo,Ubc6,tor | | 1.834e-07 | -15.51 | CG2852 (uncharacterized protein) | protein interactions | 37591 | 69 | 25 | 9204 | 1112 | p38a,CG4598,CG3226,AnxB10,CG10265,regucalcin,Uev1A,Moe,scu,capt,Uch,Eno,Trxr-1,awd,CG11267,Chd64,CG14715,CG10638,PHGPx,cib,EndoGI,Sod1,Fkbp12,RhoGDI,Stip1 | | 1.841e-07 | -15.51 | Chd64 (Chd64) | protein interactions | 38490 | 107 | 33 | 9204 | 1112 | regucalcin,EcR,AnxB10,CG10265,Eno,Fkbp39,Larp4B,ATPsynbeta,CG14715,CG10638,cib,PHGPx,Cyp1,CG11267,ERp60,Stip1,red,Moe,capt,scu,Usp39,CG4598,Pglym78,Uch,Fkbp12,Sod1,sol,Trxr-1,awd,CG15141,Tak1,HmgD,RhoGDI | | 1.878e-07 | -15.49 | purine ribonucleotide metabolic process | biological process | GO:0009150 | 151 | 35 | 12345 | 1129 | Ac3,CG8199,AMPdeam,ATPsynE,ATPsynD,Eno,PyK,Pfk,blw,ATPsynO,CG17691,nmdyn-D7,scu,Pfas,ScsbetaA,CG7920,Ady43A,Ppcdc,ATPsynbeta,CG6178,Scsalpha1,awd,ATPsynB,ATPsynG,Pfrx,ATPsyndelta,AcCoAS,Gapdh1,Pglym78,CG11089,CG1774,ATPsynCF6,COX5B,nmdyn-D6,Pgk | | 2.004e-07 | -15.42 | HALLMARK\_OXIDATIVE\_PHOSPHORYLATION | MSigDB lists | HALLMARK\_OXIDATIVE\_PHOSPHORYLATION | 124 | 42 | 4161 | 644 | Etf-QO,Roe1,tko,CG8728,Vha36-1,Phb2,colt,Cyt-c-p,ScsbetaA,ND-B18,Mtpalpha,Tim8,CG8004,ND-24,Marf,Thiolase,ND-30,Mitofilin,COX5A,yip2,levy,CG10932,CG5028,ND-51,ND-23,ND-49,Mcad,scu,ND-SGDH,SdhA,CG8199,AIF,ND-75,wal,Echs1,CG3803,UQCR-Q,Nc73EF,Scsalpha1,Idh,ND-39,Idh3b | | 2.105e-07 | -15.37 | MODULE\_83 | MSigDB lists | MODULE\_83 | 182 | 55 | 4161 | 644 | UQCR-Q,eEF1beta,Pen,RpL13,RpS23,RpS27A,RpL9,Nap1,RpS28b,Rab10,Idh,Fkbp59,RhoGDI,levy,CysRS,eIF3i,CHOp24,CG31715,Rpn2,Nurf-38,RpLP0,Rpn12,RpS13,RpS30,Sod1,CG6610,RpL13A,Prosbeta3,Mcad,Kap-alpha3,ND-24,RpL12,ND-B18,CCT3,RpS18,CG14715,Prosbeta5,RpL26,RpS27,RpL24,eIF3m,RpL11,Prosalpha3,RpL37a,Rab5,RpL40,Prosbeta6,RpL38,RpS4,Pfdn5,Phb2,RpS2,RpS16,RpL23,RpS24 | | 2.119e-07 | -15.37 | GSE24634\_NAIVE\_CD4\_TCELL\_VS\_DAY3\_IL4\_CONV\_TREG\_DN | MSigDB lists | GSE24634\_NAIVE\_CD4\_TCELL\_VS\_DAY3\_IL4\_CONV\_TREG\_DN | 95 | 35 | 4161 | 644 | mRpL17,Mitofilin,CycE,MED20,Tim8,Pfdn2,CCT5,CCT3,Pdxk,lwr,CG12018,Non1,RhoGAP54D,mus81,mtTFB1,CG17593,NHP2,CG1416,tko,heph,Idh,Prosalpha5,Pus7,CG7997,rl,mad2,Rpn12,Rpn2,scu,CG42542,DppIII,ste14,Su(H),Grx1,opm | | 2.204e-07 | -15.33 | mitochondrial respirasome | cellular component | GO:0005746 | 88 | 25 | 11940 | 1105 | ND-39,COX5B,levy,CG40472,ND-49,ND-MNLL,ND-30,ND-B15,ND-18,ND-SGDH,ND-B18,cype,UQCR-Q,ND-23,COX5A,ND-B22,ND-24,ND-51,SdhA,UQCR-11,NP15.6,COX4,ND-MWFE,ND-75,SdhD | | 2.247e-07 | -15.31 | carboxylic acid metabolic process | biological process | GO:0019752 | 404 | 69 | 12345 | 1129 | TyrRS,Mcad,Desat1,aay,CG8199,Eno,CysRS,CG17691,Shmt,scu,SerRS-m,Sply,r,Gapdh1,CG4594,AcCoAS,P5CDh1,Aldh,Idh,Pex10,Got1,CG6178,Dhfr,Seipin,SREBP,CG10932,CG4586,CG11134,Gabat,Echs1,Pgk,GatA,Sdhaf3,Gip,PyK,AsnS,tzn,Pfk,Idh3b,SelD,regucalcin,TrpRS-m,Etf-QO,CG5028,Thiolase,Adi1,mino,CG31523,CG11899,SERCA,wal,Pfrx,GatB,P5cr,Pdk,Irp-1B,ACC,Mtpalpha,LysRS,yip2,Pxt,Kyat,Scsalpha1,CG4860,Ntan1,Pglym78,Acox57D-p,GluRS-m,CG4598 | | 2.539e-07 | -15.19 | posttranscriptional regulation of gene expression | biological process | GO:0010608 | 208 | 43 | 12345 | 1129 | CG8963,CG2911,spn-E,RpL13A,heph,r2d2,bam,cup,Not10,armi,nos,CG1578,AGO3,Exd2,GAPsec,krimp,bru1,sip3,piwi,Rga,Unr,osk,Rbfox1,bcd,Dph5,gw,mRpL13,Hsc70-3,Marf1,CG6967,mtTFB1,Dcr-1,smg,Pi3K92E,CG9425,apt,tyf,PEK,trsn,Cbp80,BicC,twin,CG7414 | | 2.623e-07 | -15.15 | monocarboxylic acid catabolic process | biological process | GO:0072329 | 51 | 18 | 12345 | 1129 | Seipin,CG4860,Etf-QO,yip2,Mtpalpha,Pex10,CG4594,Mcad,CG4598,wal,Acox57D-p,SERCA,CG4586,Echs1,Gabat,CG10932,Ntan1,Thiolase | | 2.835e-07 | -15.08 | cell differentiation | biological process | GO:0030154 | 1773 | 221 | 12345 | 1129 | Rbf,squ,Wnt10,otu,Grip75,Pglym78,CG3770,COX4,alph,EMC3,Smyd4-4,bru1,Arf51F,Grip128,COX5A,eIF3m,Pka-R2,Rab5,Kap3,cup,sw,CSN6,mxc,heph,raptor,cathD,nej,faf,Nup62,Cdk5,rl,Nprl2,elg1,Jafrac1,spen,Drep1,Dcr-1,Pka-R1,tkv,cad,SRPK,capt,SkpA,l(3)mbt,Archease,Hnf4,EcR,fs(1)N,bif,pgc,PyK,del,Ptp61F,enc,Debcl,Rbfox1,wisp,MED27,gus,Sec15,BoYb,CycJ,Pgk,nos,tor,GatA,Cdk1,PIG-C,Afti,Nup358,cnc,Abi,yl,msps,velo,Arpc1,Su(H),CanB2,psq,Utx,CG13366,eIF4E1,Klp10A,Seipin,bam,zpg,shrb,cnn,ari-1,Sox102F,vret,rost,MED24,park,milt,Stat92E,Rbf2,Actn,Cul1,toc,Hr78,swa,Npc2a,E2f1,colt,jvl,ben,Rab10,Moe,Grip71,dnc,Iml1,RYBP,Tao,Cul2,G9a,dl,Pura,Cul5,Top1,foxo,tho2,Imp,not,piwi,cold,Lkb1,krimp,up,Ptpmeg,CycE,MAGE,Pen,spn-E,Sec23,sbb,Med,Pfdn2,gammaTub37C,mop,Gclc,Pex12,bun,fs(1)M3,Rel,Pxt,HP1e,Cf2,Pi3K92E,Mtl,rept,stai,e(y)1,AIF,boi,galectin,SERCA,stil,mael,Tm1,Trf2,ovo,cact,bcd,Pfk,Idh3b,Liprin-beta,lid,DCAF12,spz,Pink1,mts,MED4,mad2,hb,Orc1,eIF4A,ago,Nf-YA,exu,ea,CG17068,muskelin,cort,Gapdh1,csw,lqfR,Atg7,Pex10,armi,Ptp69D,SelR,Ptpmeg2,Ptp10D,r2d2,Atg6,CkIIalpha,Cbl,shep,egg,CycB,wash,Nap1,Stam,parvin,Pax,Girdin,ScsbetaA,yps,twin,CG46338,apt,dco,BicC,nmd,shu,osk,Smox,ssh,Mos,tws | | 2.838e-07 | -15.07 | respiratory electron transport chain | biological process | GO:0022904 | 90 | 25 | 12345 | 1129 | ND-23,COX5B,Coq8,SdhA,ND-B18,UQCR-Q,ND-B15,AIF,ND-75,ND-MNLL,ND-SGDH,COX4,ND-24,Pmi,ND-51,SdhD,UQCR-11,cype,COX5A,NP15.6,ND-39,ND-B22,levy,Etf-QO,ND-49 | | 2.905e-07 | -15.05 | yip2 (yippee interacting protein 2) | protein interactions | 34313 | 33 | 16 | 9204 | 1112 | Uch,CG2767,Eno,AnxB10,capt,scu,Moe,regucalcin,Gdi,Stip1,awd,cib,PHGPx,EndoGI,CG10638,CG14715 | | 2.992e-07 | -15.02 | negative regulation of macromolecule metabolic process | biological process | GO:0010605 | 777 | 113 | 12345 | 1129 | CG14408,cyc,Rga,Unr,CG31875,nos,mts,esc,SREBP,hb,Utx,r2d2,bam,CkIIalpha,Su(H),Not10,psq,armi,escl,Mybbp1A,sws,Hat1,RpL13A,trsn,apt,BicC,dco,Cbp80,E2f1,Edc3,twin,Rbf2,EndoGI,wash,park,CG10959,cu,Cbl,egg,Sox102F,Hsc70-3,mRpL13,ATPsynD,SmydA-1,CG31759,CG17746,CG17328,PpV,Elba2,alph,krimp,Trf4-2,Smyd4-4,d4,CG10631,bru1,CG17768,piwi,CG30496,Tlk,LSm3,AGO3,dl,Rbf,squ,foxo,CG10336,heph,CG2199,Sirt7,cup,spn-E,sbb,CG4872,CycE,Chrac-14,l(3)mbt,SkpA,plu,asf1,ovo,tkv,insv,Maf1,mael,BigH1,CG6967,SuUR,Drep1,Dcr-1,lin-52,YL-1,CG11486,smg,faf,CG9425,rept,Rrp40,Bin1,ph-d,CG10038,Pym,Tip60,Uxt,Marf1,CG1602,bcd,Rbfox1,Hnf4,EcR,pgc,gw,Ptp61F | | 3.145e-07 | -14.97 | regulation of protein metabolic process | biological process | GO:0051246 | 648 | 98 | 12345 | 1129 | Pi3K92E,smg,faf,Fbw5,mtTFB1,Fdh,Pka-R1,RhoGAP5A,CG7414,SkpA,Dph5,gw,Ptp61F,bcd,Rbfox1,gus,trr,Parp16,Pdk1,Lkb1,CG30496,Cka,krimp,alph,bru1,CycE,eIF3m,Rpn13,CG30291,CG4238,Rpt2,Pka-R2,cup,Pp2A-29B,RpLP1,Cdk7,heph,raptor,Cbl,ari-1,CycB,MED24,park,ScsbetaA,twin,Cul1,PEK,apt,tyf,dco,BicC,Rpn2,Mekk1,ben,Rpn3,CycC,PpV,osk,CG17746,SNF4Agamma,Mos,Tao,tws,mRpL13,ATPsynD,G9a,mts,esc,CG1578,Exd2,Jafrac2,ago,Tak1,CycJ,CG8009,nos,tor,Ubc6,Rga,Unr,GAPsec,CG14408,Ankle2,sip3,cort,Pomp,Abi,fray,RpL13A,STUB1,ASPP,CG2911,CG8963,sws,Not10,Utx,CkIIalpha,bam | | 3.239e-07 | -14.94 | regulation of gene expression | biological process | GO:0010468 | 1547 | 197 | 12345 | 1129 | scny,Rga,cyc,sip3,Ssdp,hb,Exd2,SREBP,qkr58E-2,MED4,esc,Nf-YA,eIF4A,CG2129,armi,r2d2,CG17202,RpL13A,Hat1,CG8963,Mybbp1A,Atf3,mute,twin,Edc3,BicC,apt,tyf,PEK,egg,lwr,cu,Mitf,Nap1,wash,hng3,Rcd5,CG31759,Smox,Hsc70-3,EloC,vers,NFAT,osk,HP1c,CycC,CG17328,Tlk,piwi,CG2678,d4,Topors,krimp,foxo,trr,dl,AGO3,not,Rel,bun,Sirt7,CG2199,Tfb4,HP1e,Cf2,CG3587,Chrac-14,REPTOR,CG4872,Med,spn-E,sbb,Maf1,mael,obe,ovo,Trf2,rept,CG9425,CG11486,Pi3K92E,mtTFB1,e(y)1,Pym,CG10418,lid,bcd,cact,CG31875,Unr,Nup358,Wbp2,cnc,GAPsec,CG1578,nos,Tak1,pygo,CycJ,Atf6,psq,Not10,Su(H),bam,HDAC6,Utx,S2P,escl,l(3)73Ah,CG2911,Aos1,Sgf11,HmgD,Stat92E,Rbf2,Spt20,E2f1,p38a,Cbp80,MED20,Hr78,trsn,Sox102F,lin,CG10959,MED24,vret,SmydA-1,RYBP,Coq7,G9a,Cul2,mRpL13,Bap170,Son,Elba2,CG10654,Dd,Bx,CG17768,CG10631,bru1,Trf4-2,pps,Smyd4-4,Traf6,squ,Rbf,LSm3,Cdk7,ix,CG11902,cup,CG6813,Myd88,CG13344,CG4021,heph,Atu,CG17454,cad,CG7414,BigH1,insv,CG4730,tkv,TfIIFalpha,asf1,Taf7,l(3)mbt,pont,nej,smg,lin-52,YL-1,Dcr-1,spen,crm,CG6967,CG3198,rl,Marf1,Tip60,CG10038,MED27,ph-d,CG11710,Rrp40,jigr1,Bin1,pgc,gw,Hnf4,Dph5,EcR,Rbfox1,CG1602 | | 3.259e-07 | -14.94 | mitotic cell cycle | biological process | GO:0000278 | 432 | 72 | 12345 | 1129 | mEFTu1,CycB,DNApol-epsilon255,RecQ5,Grip84,shrb,fs(1)Ya,BicC,E2f1,toc,Grip71,PpV,dgt4,Cenp-C,tws,ssh,eIF4A,wapl,CycJ,alphaTub84B,mts,mus81,mad2,Ankle2,grp,Cdk1,csw,msps,awd,Sld5,cana,CkIIalpha,mus304,Klp10A,eIF4E1,Dcr-1,rl,CG8188,pont,SkpA,Non1,wisp,del,Ptp61F,twe,nbs,alphaTub67C,CCT6,ph-d,Syx13,Orc6,mre11,CG10336,sau,larp,Rbf,Zwilch,alph,PpD3,COX4,Tlk,Grip75,Lkb1,spn-E,CG7716,CycE,Grip128,HP1e,CG5003,Cdk7,Rab5,gammaTub37C | | 3.326e-07 | -14.92 | cellular respiration | biological process | GO:0045333 | 128 | 31 | 12345 | 1129 | Etf-QO,Scsalpha1,ND-49,CG5028,ND-39,Idh3b,ND-B22,levy,NP15.6,Irp-1B,SdhD,cype,UQCR-11,Nc73EF,COX5A,Pmi,ND-24,ND-51,ND-75,ND-MNLL,ND-SGDH,COX4,ND-B18,UQCR-Q,ScsbetaA,ND-B15,AIF,ND-23,COX5B,Coq8,SdhA | | 3.352e-07 | -14.91 | purine ribonucleotide biosynthetic process | biological process | GO:0009152 | 79 | 23 | 12345 | 1129 | CG17691,CG11089,nmdyn-D7,Pfas,COX5B,ATPsynCF6,nmdyn-D6,Ac3,CG8199,AMPdeam,Ady43A,ATPsynD,ATPsynE,Ppcdc,CG6178,ATPsynbeta,awd,ATPsynG,ATPsynB,blw,ATPsyndelta,ATPsynO,AcCoAS | | 3.607e-07 | -14.84 | cell development | biological process | GO:0048468 | 1436 | 185 | 12345 | 1129 | Rbfox1,Debcl,wisp,fs(1)N,EcR,bif,pgc,Ptp61F,enc,del,Sec15,gus,Nprl2,rl,elg1,Jafrac1,Drep1,spen,Dcr-1,cathD,nej,Cdk5,Nup62,SkpA,l(3)mbt,capt,Pka-R1,tkv,SRPK,Grip128,eIF3m,CSN6,heph,raptor,Pka-R2,Rab5,Kap3,cup,sw,squ,Rbf,Smyd4-4,EMC3,bru1,otu,Grip75,CG3770,ben,Moe,Rab10,Grip71,dnc,Iml1,colt,jvl,Cul2,RYBP,Tao,vret,park,MED24,milt,zpg,shrb,ari-1,cnn,swa,Npc2a,E2f1,Actn,Stat92E,toc,msps,velo,Arpc1,Abi,yl,Utx,CG13366,eIF4E1,bam,CanB2,psq,CycJ,tor,nos,GatA,BoYb,Afti,cnc,Nup358,Cdk1,PIG-C,cact,bcd,lid,DCAF12,spz,Pink1,Idh3b,Liprin-beta,stai,e(y)1,galectin,AIF,Pi3K92E,rept,Trf2,ovo,SERCA,stil,mael,Tm1,spn-E,sbb,Med,CycE,Pen,HP1e,Cf2,gammaTub37C,mop,Gclc,bun,fs(1)M3,Pex12,Pxt,Rel,Imp,not,dl,Top1,Pura,foxo,tho2,Cul5,krimp,up,Ptpmeg,piwi,Lkb1,cold,shu,osk,Smox,ssh,Mos,CycB,wash,Nap1,Stam,parvin,Girdin,ScsbetaA,Pax,shep,egg,Cbl,apt,BicC,nmd,yps,twin,CG46338,csw,lqfR,Atg7,Ptpmeg2,r2d2,Ptp10D,Atg6,CkIIalpha,Pex10,armi,Ptp69D,SelR,eIF4A,Orc1,ago,exu,mts,mad2,muskelin,cort,ea | | 3.627e-07 | -14.83 | fatty acid catabolic process | biological process | GO:0009062 | 42 | 16 | 12345 | 1129 | CG10932,Thiolase,CG4586,Echs1,wal,Acox57D-p,SERCA,CG4598,Mcad,CG4594,yip2,Pex10,Mtpalpha,Seipin,CG4860,Etf-QO | | 3.628e-07 | -14.83 | cellular lipid catabolic process | biological process | GO:0044242 | 68 | 21 | 12345 | 1129 | CG10932,Thiolase,Echs1,CG4586,Acox57D-p,wal,SERCA,CG4598,CG7840,Sply,Mcad,CG18135,CG4594,CG7997,yip2,dob,Mtpalpha,Pex10,Seipin,Etf-QO,CG4860 | | 4.021e-07 | -14.73 | monocarboxylic acid metabolic process | biological process | GO:0032787 | 183 | 39 | 12345 | 1129 | Thiolase,CG31523,mino,wal,SERCA,scu,PyK,Desat1,Mcad,Pfk,tzn,AsnS,Etf-QO,Eno,Ntan1,SREBP,CG10932,Pgk,Gabat,Echs1,CG4586,Acox57D-p,Pglym78,CG4598,Gip,Pfrx,Aldh,AcCoAS,Pdk,CG4594,Gapdh1,Pxt,yip2,ACC,Pex10,Mtpalpha,Seipin,CG4860,CG6178 | | 4.037e-07 | -14.72 | ATPsynE (ATP synthase, subunit E) | protein interactions | 41745 | 14 | 10 | 9204 | 1112 | ATPsyndelta,ATPsynB,ATPsynF,Neos,ATPsynCF6,ATPsynD,blw,ATPsynO,ATPsynG,ATPsynbeta | | 4.162e-07 | -14.69 | Translation | REACTOME pathways | R-DME-72766 | 143 | 44 | 3923 | 574 | RpS29,RpL24,RpS23,RpL26,RpS28b,RpS7,RpL30,RpL11,RpS24,RpS27A,eIF3m,RpS14a,RpL13A,RpL7A,RpL12,RpL17,RpL9,RpL37a,RpL34b,RpL38,RpL40,RpS27,eIF4A,RpS30,RpL29,RpS14b,eIF3i,RpS16,RpS18,eEF1beta,RpL23,RpLP1,RpL34a,RpL23A,eIF4E1,RpS15,RpL31,RpS5b,RpL10,RpS4,RpLP0,RpS13,RpS2,RpL13 | | 4.493e-07 | -14.62 | purine-containing compound biosynthetic process | biological process | GO:0072522 | 92 | 25 | 12345 | 1129 | ATPsynbeta,CG6178,Ppcdc,ATPsynE,ATPsynD,Ady43A,CG8199,Ac3,AMPdeam,AcCoAS,ATPsynO,ATPsyndelta,blw,ATPsynB,ATPsynG,awd,Pfas,nmdyn-D7,CG11089,CG17691,Shmt,nmdyn-D6,CG6767,ATPsynCF6,COX5B | | 4.720e-07 | -14.57 | endomembrane system | cellular component | GO:0012505 | 1113 | 151 | 11940 | 1105 | Efr,Wdfy2,CG32640,Afti,park,Ho,fwe,tral,fdl,Arl5,lqfR,Cse1,PIG-H,Jafrac2,Seipin,toc,AP-1-2beta,CG10425,Crag,CG12945,CG8974,Fur2,Nup62,CG3309,Sply,SERCA,CG17593,Nup58,O-fut1,Clc,Rep,CG4557,shrb,Hsc70-3,CG4332,CG7840,Ankle2,PIG-P,opm,CG8314,Rab32,Ndc1,CG32485,CG10470,CG5734,CG30291,Mctp,CG6766,sbr,Syx13,tkv,Vsp37A,Axs,EMC8-9,EMC3,Stam,Edem1,CG5969,alpha-Man-Ia,CG14671,Trs33,Gmap,sau,SelT,CG10344,Sec15,Parp16,CG6707,bif,CG9773,ste14,CG7655,CG4293,ste24a,mbo,Agpat2,bru1,ATP8A,Rab10,mop,CG32441,CanB2,SREBP,Rtnl1,CG3631,Rab5,CG31523,mol,CG33977,GCS2beta,GCC88,CG9911,wash,Vamp7,Ugt36E1,Dd,CG14715,PIG-C,Tm1,Hip1,CG1309,sip3,Ugalt,Atf6,CG9186,mino,Gga,lt,Desat1,CG2938,CG8009,CHMP2B,CG8507,whd,Lkb1,CG8112,CG4880,Surf4,CG5880,CHOp24,GalT1,alt,pen-2,fs(1)Ya,Aladin,Sec23,ERp60,sws,PHGPx,mad2,CG16979,cup,Arf51F,CG5510,Vps16B,Sec22,Gdi,eIF3m,Pi4KIIalpha,stet,Fer1HCH,CG5447,CG17270,cbs,Snx6,Tomosyn,CG31729,CG32536,CG8311,Lnpk,CG9981 | | 4.920e-07 | -14.52 | purine nucleoside triphosphate biosynthetic process | biological process | GO:0009145 | 29 | 13 | 12345 | 1129 | ATPsynbeta,ATPsynD,ATPsynE,nmdyn-D7,COX5B,ATPsynG,ATPsynCF6,ATPsynB,awd,ATPsynO,blw,ATPsyndelta,nmdyn-D6 | | 4.920e-07 | -14.52 | purine ribonucleoside triphosphate metabolic process | biological process | GO:0009205 | 29 | 13 | 12345 | 1129 | ATPsynO,ATPsyndelta,blw,nmdyn-D6,COX5B,ATPsynG,ATPsynB,ATPsynCF6,awd,ATPsynbeta,ATPsynE,ATPsynD,nmdyn-D7 | | 4.920e-07 | -14.52 | purine ribonucleoside triphosphate biosynthetic process | biological process | GO:0009206 | 29 | 13 | 12345 | 1129 | ATPsynD,ATPsynE,nmdyn-D7,ATPsynbeta,ATPsyndelta,blw,nmdyn-D6,ATPsynO,awd,COX5B,ATPsynG,ATPsynB,ATPsynCF6 | | 5.069e-07 | -14.49 | REACTOME\_NONSENSE\_MEDIATED\_DECAY\_ENHANCED\_BY\_THE\_EXON\_JUNCTION\_COMPLEX | MSigDB lists | REACTOME\_NONSENSE\_MEDIATED\_DECAY\_ENHANCED\_BY\_THE\_EXON\_JUNCTION\_COMPLEX | 70 | 28 | 4161 | 644 | RpL24,RpS27,RpS28b,RpS27A,RpS23,RpL9,RpL26,Pp2A-29B,RpL11,RpL37a,RpL13,RpL12,RpS18,RpS13,RpLP0,RpS30,RpL13A,RpS16,RpS2,RpS24,mts,RpL23,RpL30,RpL40,Cbp80,RpS7,RpS4,RpL38 | | 5.107e-07 | -14.49 | REACTOME\_METABOLISM\_OF\_RNA | MSigDB lists | REACTOME\_METABOLISM\_OF\_RNA | 168 | 51 | 4161 | 644 | RpL13,RpL9,RpS27A,RpS23,Prosalpha5,RpS28b,eIF4A,Rpt2,Prosalpha4,Prosbeta3,mts,Not10,Rpn2,RpLP0,Rpn12,RpS13,RpS30,CG6610,RpL13A,eIF4E1,Rrp40,p38a,RpS18,Prosbeta5,Rpn3,RpL12,Prosbeta4,LSm3,RpL11,Prosalpha3,RpL37a,RpL26,Pp2A-29B,RpL24,RpS27,Prosbeta6,RpS4,RpL38,RpS7,CG10418,RpL40,mbo,Cbp80,RpL30,CG8963,RpL23,RpS24,Edc3,Aladin,RpS2,RpS16 | | 5.203e-07 | -14.47 | porin (porin) | protein interactions | 34500 | 77 | 26 | 9204 | 1112 | Eno,Surf4,wal,Tspo,Vha68-2,ATPsynB,regucalcin,Phb2,Mcad,AnxB10,ATPsynO,COX4,blw,ATPsynCF6,Cisd2,CG15098,ATPsynF,ATPsyndelta,ATPsynbeta,PHGPx,SERCA,CG10425,up,Chd64,Marf,Mf | | 5.513e-07 | -14.41 | hydrolase activity, acting on ester bonds | molecular function | GO:0016788 | 493 | 79 | 12176 | 1119 | regucalcin,CG5026,Pfrx,CanA-14F,Scsalpha1,CG4267,Ptpmeg2,cN-IIIB,fal,Pp2B-14D,PAPLA1,Ptpmeg,Dcr-1,twe,CG11486,Acph-1,alpha-Est5,CG17746,RpLP0,nSMase,dob,Snm1,Ptp61F,CG11164,CG9391,tos,CG17294,trsn,CG7789,Synj,tws,CG4390,Ptp69D,Dd,DNApol-epsilon255,PpV,CG12375,Tengl4,CG6707,alph,PpD3,mre11,CG31759,cu,clt,Pld,CG8679,Pka-R1,CG32549,scu,Exd2,Ublcp1,PrBP,CG17333,mop,Drep2,pn,CG16979,csw,Ptp10D,CG1774,tzn,dnc,CG30343,Pp2A-29B,CG18135,Clp,Pka-R2,mus205,aay,exu,CG7115,mts,sws,Rga,twin,ssh,CG5966,mus81 | | 5.549e-07 | -14.40 | coenzyme biosynthetic process | biological process | GO:0009108 | 64 | 20 | 12345 | 1129 | Coq8,Sptr,Pcd,pr,CG17691,Pdxk,AcCoAS,CG33156,sgll,mal,Coq7,Ppcdc,Dhfr,Nadsyn,CG6178,Coq6,CG4407,regucalcin,ND-39,CG8199 | | 6.282e-07 | -14.28 | cytosolic part | cellular component | GO:0044445 | 256 | 49 | 11940 | 1105 | RpL13,RpS14a,RpS24,RpS14b,RpL37a,RpL26,RpL29,Blos4,RpS18,RpL38,RpS27A,RpL9,Blos2,RpS28b,PEK,RpL7A,RpL24,Pfk,CCT6,CCT3,Rpn2,RpL11,RpL13A,RpS5b,RpS2,RpL31,RpS4,RpL34a,RpLP1,RpS15,RpS13,CG7414,CCT5,RpL17,RpL40,RpS23,RpS16,RpS27,RpL34b,RpL23A,RpL12,RpL23,RpL10,RpS30,Eno,RpS29,RpLP0,RpS7,RpL30 | | 6.322e-07 | -14.27 | positive regulation of catabolic process | biological process | GO:0009896 | 125 | 30 | 12345 | 1129 | Utx,AP-1-2beta,Rpt2,gus,GlyP,Marf1,CG4238,spn-E,Debcl,CycC,Atg7,gw,Synj,PEK,SkpA,cort,dco,Pomp,piwi,Cul1,Nprl2,park,Fbw5,Mitf,Ubc6,AGO3,smg,foxo,ari-1,Traf6 | | 6.498e-07 | -14.25 | membrane protein complex | cellular component | GO:0098796 | 533 | 84 | 11940 | 1105 | NP15.6,ATPsynO,ND-51,nAChRalpha4,SdhA,COX5A,Vha36-1,ATPsynG,Moe,Vps16B,EMRE,ND-B15,Fkbp59,Neb-cGP,Vsp37A,Sec23,Roe1,Ork1,EMC8-9,EMC3,pen-2,Syx13,UQCR-11,Vha68-2,CG32441,ATPsynB,cype,UQCR-Q,ND-23,CG16868,Vha13,nrv1,Tomosyn,Snx6,CG40472,ND-MNLL,CG8004,CG5903,Sec22,CG9393,CG9132,ND-MWFE,ATPsynD,sip3,ATPsyndelta,Atpalpha,ND-B22,Tom70,COX8,ND-B18,Hip1,ND-SGDH,ND-18,AP-1-2beta,levy,ND-49,Vamp7,blw,Dd,SdhD,PIG-C,PIG-H,Tim8,Afti,Pi3K92E,ATPsynCF6,Tom40,ATPsynE,ND-24,CHMP2B,ND-30,COX5B,ND-39,lt,CG8009,Clc,shrb,CG11110,CG8974,Atg6,ND-75,ATPsynF,COX4,ATPsynbeta | | 6.987e-07 | -14.17 | ATP synthesis coupled electron transport | biological process | GO:0042773 | 82 | 23 | 12345 | 1129 | UQCR-11,cype,SdhD,COX5A,ND-39,ND-B22,levy,ND-49,ND-23,COX5B,Coq8,SdhA,UQCR-Q,ND-B18,ND-B15,AIF,ND-75,ND-MNLL,ND-SGDH,COX4,Pmi,ND-24,ND-51 | | 7.711e-07 | -14.08 | Df31 (Decondensation factor 31) | protein interactions | 35418 | 108 | 32 | 9204 | 1112 | RhoGDI,CG32549,slam,Trxr-1,awd,Fkbp12,Sod1,Uch,Pglym78,CG11771,CG1532,CG4598,scu,capt,Moe,Stip1,ERp60,Chd64,CG11267,RpS27A,ATPsynbeta,CG14715,CG10638,PHGPx,cib,Cyp1,Eno,ago,AnxB10,CG1309,CG10265,regucalcin | | 7.850e-07 | -14.06 | Fadd (Fas-associated death domain) | protein interactions | 42594 | 103 | 31 | 9204 | 1112 | ScsbetaA,RpS16,RpS5b,Nup358,RpL30,scu,PyK,bor,rept,GCS2beta,Cp1,Jafrac1,RpS28b,RpL12,RpS14a,PGRP-LE,Myd88,Sod1,RYBP,SERCA,CCT6,CG9391,Sp7,RpL23A,alphaTub84B,pont,CCT3,Tak1,RpL7A,Mitofilin,blw | | 8.218e-07 | -14.01 | cytosolic ribosome | cellular component | GO:0022626 | 200 | 41 | 11940 | 1105 | RpS23,RpL40,RpL17,CG7414,RpS13,RpLP1,RpL34a,RpS15,RpS2,RpS4,RpL31,RpS5b,RpLP0,RpS7,RpL30,RpS29,RpS30,RpL10,RpL23,RpL34b,RpL23A,RpL12,RpS27,RpS16,RpL9,RpS27A,RpS18,RpL38,RpL29,RpL37a,RpL26,RpS14a,RpS14b,RpS24,RpL13,RpL11,RpL13A,PEK,RpL7A,RpL24,RpS28b | | 8.291e-07 | -14.00 | GO\_PHOSPHATE\_CONTAINING\_COMPOUND\_METABOLIC\_PROCESS | MSigDB lists | GO\_PHOSPHATE\_CONTAINING\_COMPOUND\_METABOLIC\_PROCESS | 671 | 147 | 4161 | 644 | CG6767,Traf6,CycC,CG5285,SelD,fray,nSMase,CanB2,r,CanA-14F,PIG-C,Pfk,CG7322,Jafrac2,CG7530,Rpt2,Prosalpha4,CG7115,CG11164,ACC,Gk2,CG11089,PIG-L,PAPLA1,RpS27A,Abi,Pi4KIIalpha,Nc73EF,Idh,UQCR-Q,Ptpmeg,Tlk,mal,alc,Ptp10D,Dd,Dhfr,Tao,Prosbeta6,CG32549,Acph-1,Rpn3,ND-30,ND-B18,Mtpalpha,Cdk5,Eno,alph,AMPdeam,Nurf-38,Rpn2,Prip,mino,ND-SGDH,ND-49,Cdk7,CG5028,dnc,ND-51,SkpA,CG8475,Stoml2,Prosalpha5,CG15111,Scsalpha1,ND-39,CG9391,l(1)G0196,Cdk1,rl,CG3803,Pp2B-14D,Cyt-c-p,CG7718,CG17294,CkIIalpha,CG2201,Pdk,CG17698,scramb1,Plap,CG8726,sgll,Nprl2,CG17333,Ts,PpV,DhpD,COX5A,Pp2A-29B,Prosalpha3,Ppcdc,CG3376,ND-24,r-l,Prosbeta5,nmdyn-D7,Rpn12,Nadsyn,LysRS,CG3631,csw,mts,CG4041,Ublcp1,levy,Mekk1,Idh3b,CG7840,Top1,ND-75,dco,aay,RpL40,Synj,Pi3K92E,Ptp61F,PpD3,CycE,CG7414,Prosbeta4,CG7789,Pfas,Cul1,nmdyn-D6,Pglym78,Thiolase,p38a,SRPK,Pdxk,Ac3,Pdk1,CycB,grp,CG33156,Gapdh1,Prosbeta3,Pgm1,CG4407,ND-23,ND-B22,Mo25,tkv,Shmt,Pgk,kri,Tfb4 | | 8.530e-07 | -13.97 | Pdk1 (Phosphoinositide-dependent kinase 1) | protein interactions | 38017 | 124 | 35 | 9204 | 1112 | RpS15,Rpt2,RpL31,cathD,Mcad,yps,RpL10,Pp2A-29B,Hsc70-3,CG9925,RpLP1,RpS14a,RpL9,Shmt,larp,levy,GCS2beta,RpL13A,eIF4A,shv,Pi3K92E,RpS16,ScsbetaA,CG1416,RpS23,Aldh,Droj2,Scox,eIF3i,RpS7,Kcmf1,RpL26,SERCA,eIF3m,CCT6 | | 9.132e-07 | -13.91 | KEGG\_RIBOSOME | MSigDB lists | KEGG\_RIBOSOME | 64 | 26 | 4161 | 644 | RpS18,RpL13,RpL12,RpL11,RpL37a,RpS27,RpL24,RpS28b,RpS23,RpL9,RpL26,RpS27A,RpS7,RpL38,RpS4,RpL40,RpS24,RpL30,RpL23,RpS30,RpLP0,mRpL13,RpS13,RpS2,RpL13A,RpS16 | | 9.202e-07 | -13.90 | Uch (Ubiquitin carboxy-terminal hydrolase) | protein interactions | 33397 | 65 | 23 | 9204 | 1112 | capt,scu,Moe,GstT1,Gdi,regucalcin,Ntan1,AnxB10,CG4598,Eno,Fkbp12,EndoGI,cib,PHGPx,CG14715,CG10638,Pgk,awd,Chd64,yip2,Stip1,RhoGDI,CG10863 | | 1.017e-06 | -13.80 | nucleic acid metabolic process | biological process | GO:0090304 | 1149 | 152 | 12345 | 1129 | armi,RpS14b,r2d2,RpS14a,Sld5,snRNA:U12,Rga,Mat89Ba,MED4,mus81,Exd2,Lsm12,Clp,Orc1,exu,CG2034,CG31759,tapas,RfC38,Uev1A,SmE,CG1622,CycC,shu,yps,Edc3,twin,RpL7A,CG9684,CG14100,Snm1,CG12375,cu,Nap1,CysRS,Nurf-38,LysRS,Pus7,NHP2,Tfb4,MAGE,CycE,CG3587,Neos,piwi,Top1,tho2,CG10189,Imp,CG17187,SelD,CG30343,Xpac,CG10418,RpS7,CG2100,Pym,TrpRS-m,Spx,obe,Rnmt,bys,sbr,mael,Trf2,CG11486,e(y)1,CG3021,mtTFB1,Not10,CG11164,eIF4E1,CG7728,mus304,CG5292,SerRS-m,RpS28b,CG2911,mus205,IntS11,Gasz,tos,snRNA:U4:38AB,Rpb8,CG6610,pygo,Xpc,GatA,Ubc6,nos,CG7009,Tengl4,Son,TyrRS,CG8204,Larp7,ben,CG16865,scu,trsn,E2f1,Cbp80,RecQ5,DNApol-epsilon255,CG12018,vret,MED24,Cdk7,heph,GatB,CG17454,snRNA:U4atac,Drep2,mRpS11,Sgf29,CG10463,pps,Trf4-2,RpII140,GluRS-m,CG17768,IntS6,bru1,rig,Rbf,CG10336,mre11,LSm3,Orc6,agt,Bin1,Rrp40,RpS16,nbs,Archease,del,gw,Usp39,wisp,CG5986,CG5285,CG12301,Taf7,TfIIFalpha,smg,CG9143,c(2)M,mino,CG3198,Dcr-1,Drep1,spen | | 1.074e-06 | -13.74 | protein modification by small protein conjugation | biological process | GO:0032446 | 271 | 50 | 12345 | 1129 | Cul5,Traf6,CG2924,CG40045,Ubc6,ago,Iru,sip3,CG31053,Topors,CG10254,STUB1,MAGE,Atg7,l(3)73Ah,CG4238,CG5003,Pex12,Herc4,Pex10,CG13344,Uba5,lwr,Cbl,ari-1,CG8188,park,Fbw5,CG32221,Cul1,Aos1,UbcE2H,CG46338,APP-BP1,SkpA,RpL40,CG9316,EloC,RpS27A,ben,Ubc4,CG14435,CG31633,CG9941,Ufm1,Uev1A,dx,CG11070,CG8974,Cul2 | | 1.078e-06 | -13.74 | regulation of biosynthetic process | biological process | GO:0009889 | 1374 | 176 | 12345 | 1129 | S2P,l(3)73Ah,CG2911,escl,psq,Atf6,Su(H),Not10,Seipin,bam,Utx,HDAC6,CG1578,nos,pygo,Tak1,Unr,CG31875,Cdk1,cnc,Wbp2,GAPsec,Bap170,Elba2,CG10654,RYBP,Cul2,mRpL13,Sox102F,CG10959,lin,MED24,Sgf11,Aos1,Rbf2,Stat92E,HmgD,Hr78,MED20,E2f1,Spt20,Ptip,Atu,Cdk7,ix,CG6813,Myd88,cup,CG11902,CG13344,heph,CG10336,Traf6,Rbf,Dd,otu,Bx,CG10631,bru1,pgc,gw,EcR,Hnf4,Dph5,Rbfox1,wisp,CG1602,Uxt,Tip60,pn,MED27,ph-d,CG10038,CG11710,Bin1,lin-52,YL-1,pont,nej,smg,SuUR,spen,rl,crm,insv,CG7414,BigH1,cad,tkv,CG4730,asf1,TfIIFalpha,l(3)mbt,Taf7,CG17202,RpL13A,Hat1,mute,Atf3,CG8963,Mybbp1A,r2d2,SREBP,Exd2,hb,esc,MED4,Nf-YA,CG8009,CG2129,Rga,scny,sip3,cyc,Ssdp,EloC,vers,CycC,NFAT,osk,HP1c,CG17328,Scox,Smox,Rcd5,hng3,egg,Cbl,lwr,Mitf,Pax,wash,Nap1,twin,BicC,PEK,tyf,apt,CG3587,Chrac-14,CycE,spn-E,sbb,Med,REPTOR,CG4872,bun,Rel,Sirt7,CG2199,Tfb4,Cf2,HP1e,foxo,trr,dl,not,Tlk,piwi,d4,CG2678,krimp,Topors,bcd,cact,spz,lid,rept,Pi3K92E,mtTFB1,e(y)1,mael,Maf1,plu,ovo,Trf2 | | 1.095e-06 | -13.72 | mitochondrial ATP synthesis coupled electron transport | biological process | GO:0042775 | 78 | 22 | 12345 | 1129 | ND-24,Pmi,ND-51,ND-MNLL,ND-SGDH,COX4,UQCR-Q,ND-B18,AIF,ND-B15,ND-23,COX5B,SdhA,Coq8,ND-49,ND-39,ND-B22,levy,UQCR-11,SdhD,cype,COX5A | | 1.137e-06 | -13.69 | multi-organism reproductive process | biological process | GO:0044703 | 1300 | 168 | 12345 | 1129 | CycE,Pen,spn-E,Med,gammaTub37C,Pex12,mop,fs(1)M3,bun,Pxt,HP1e,Cf2,disp,tef,Cul5,Top1,PAPLA1,tho2,Imp,Pdk1,piwi,Lkb1,krimp,Topors,Axs,cact,bcd,lid,spz,Pi3K92E,Pp2B-14D,stai,e(y)1,SERCA,stil,mael,Tm1,plu,ovo,csw,lqfR,Atg7,Pex10,armi,Ptpmeg2,r2d2,CkIIalpha,mts,mad2,Orc1,Pex13,eIF4A,exu,ea,muskelin,slam,cort,shu,osk,Mos,png,Grip84,lwr,Cbl,egg,shep,CycB,wash,Nap1,Pax,ScsbetaA,twin,yps,CG46338,apt,fs(1)Ya,BicC,nmd,Arf51F,Grip128,eIF3m,tko,Rab5,cup,Kap3,sw,CSN6,raptor,heph,larp,squ,Rbf,sau,otu,Grip75,bru1,twe,EcR,fs(1)N,bif,pgc,del,enc,Rbfox1,mtrm,wisp,gus,Syx13,Sec15,clt,alphaTub67C,cathD,nej,faf,Nup62,Nprl2,rl,Jafrac1,elg1,Drep1,Dcr-1,tkv,Pka-R1,SRPK,l(3)mbt,capt,Abi,yl,msps,psq,mus304,Klp10A,eIF4E1,bam,BoYb,CycJ,wapl,nos,tor,GatA,Cdk1,yem,grp,cnc,Nup358,Cenp-C,jvl,Desat1,Rab10,Moe,spd-2,Grip71,dnc,Iml1,Tao,Mst89B,Cul2,zpg,shrb,ari-1,cnn,vret,park,milt,Stat92E,toc,swa,E2f1 | | 1.137e-06 | -13.69 | sexual reproduction | biological process | GO:0019953 | 1300 | 168 | 12345 | 1129 | Pp2B-14D,Pi3K92E,e(y)1,stai,stil,SERCA,Tm1,mael,ovo,plu,cact,Axs,bcd,lid,spz,tho2,Top1,Cul5,PAPLA1,Imp,Pdk1,Lkb1,piwi,Topors,krimp,CycE,Pen,Med,spn-E,gammaTub37C,Pxt,mop,fs(1)M3,Pex12,bun,Cf2,HP1e,tef,disp,Grip84,egg,Cbl,lwr,shep,Nap1,CycB,wash,Pax,ScsbetaA,CG46338,yps,twin,apt,nmd,BicC,fs(1)Ya,osk,shu,png,Mos,mad2,mts,Orc1,Pex13,eIF4A,exu,ea,slam,muskelin,cort,csw,Atg7,lqfR,Pex10,armi,r2d2,Ptpmeg2,CkIIalpha,nej,cathD,Nup62,faf,elg1,Jafrac1,rl,Nprl2,Dcr-1,Drep1,tkv,Pka-R1,SRPK,capt,l(3)mbt,EcR,fs(1)N,twe,del,enc,pgc,bif,wisp,mtrm,Rbfox1,Syx13,gus,alphaTub67C,Sec15,clt,Rbf,squ,larp,sau,Grip75,otu,bru1,Grip128,Arf51F,eIF3m,Rab5,cup,Kap3,tko,sw,raptor,heph,CSN6,shrb,zpg,ari-1,cnn,vret,milt,park,Stat92E,toc,E2f1,swa,Cenp-C,Desat1,jvl,spd-2,Rab10,Moe,Iml1,dnc,Grip71,Mst89B,Tao,Cul2,BoYb,CycJ,wapl,GatA,nos,tor,yem,Cdk1,grp,cnc,Nup358,Abi,yl,msps,psq,Klp10A,eIF4E1,mus304,bam | | 1.253e-06 | -13.59 | cib (ciboulot) | protein interactions | 31359 | 66 | 23 | 9204 | 1112 | regucalcin,Moe,capt,scu,AnxB10,Eno,Pglym78,CG2767,Uch,CG10638,CG14715,Sod1,Fkbp12,EndoGI,PHGPx,yip2,Chd64,CG11267,awd,RhoGDI,HmgD,Stip1,Scox | | 1.276e-06 | -13.57 | CG8036 (uncharacterized protein) | protein interactions | 41027 | 57 | 21 | 9204 | 1112 | regucalcin,capt,scu,Moe,AnxB10,CG4598,Eno,Seipin,Uch,CG10638,CG14715,cib,PHGPx,grsm,yip2,Pdk1,awd,Chd64,Trxr-1,Stip1,RhoGDI | | 1.276e-06 | -13.57 | ATPsynF (ATP synthase, subunit F) | protein interactions | 37931 | 36 | 16 | 9204 | 1112 | ATPsyndelta,ATPsynE,smg,COX5A,ATPsynCF6,blw,COX4,Mf,up,CG10425,ATPsynG,ATPsynbeta,ATPsynB,ATPsynD,Rtnl1,ATPsynO | | 1.282e-06 | -13.57 | negative regulation of translation | biological process | GO:0017148 | 56 | 18 | 12345 | 1129 | RpL13A,gw,Rbfox1,bcd,cup,Not10,mRpL13,heph,bam,smg,nos,twin,Unr,Rga,apt,krimp,bru1,BicC | | 1.286e-06 | -13.56 | cell maturation | biological process | GO:0048469 | 184 | 38 | 12345 | 1129 | swa,capt,SkpA,mael,Tm1,Pka-R1,milt,Stam,shep,shrb,spz,gus,Grip71,wisp,osk,Moe,jvl,enc,EcR,cnc,krimp,muskelin,ea,Grip75,Lkb1,nos,exu,Imp,Atg6,CkIIalpha,heph,armi,mop,psq,gammaTub37C,spn-E,msps,Grip128 | | 1.303e-06 | -13.55 | meiotic cell cycle process | biological process | GO:1903046 | 250 | 47 | 12345 | 1129 | yem,Cdk1,Grip75,grp,Topors,krimp,CG31053,cort,mus81,larp,sau,mre11,wapl,Xpc,GatA,nos,Rab5,cup,gammaTub37C,Pex10,Klp10A,eIF4E1,mus304,bam,tef,Grip128,Arf51F,msps,spn-E,SRPK,mael,Pp2B-14D,c(2)M,cnn,Nup62,lwr,Syx13,lid,twe,Cenp-C,enc,del,bif,Axs,wisp,mtrm,Grip71 | | 1.303e-06 | -13.55 | regulation of multi-organism process | biological process | GO:0043900 | 250 | 47 | 12345 | 1129 | PDCD-5,spz,G9a,enc,EMRE,EcR,wisp,Ect4,tub,cact,Aos1,cad,mael,Stat92E,swa,E2f1,Hr78,Cbp80,p38a,PGRP-LE,Cbl,lwr,faf,shrb,Herc4,Rel,mop,CG9925,GlyP,cup,Myd88,Atg6,Ptip,Sp7,Atg7,Hat1,Crag,spn-E,scny,cnc,Ptpmeg,slif,Traf6,dl,exu,Tak1,GILT1 | | 1.346e-06 | -13.52 | positive regulation of nitrogen compound metabolic process | biological process | GO:0051173 | 827 | 116 | 12345 | 1129 | gw,EcR,wisp,bcd,Rbfox1,CG1602,Marf1,Tip60,gus,spz,CG11710,lid,rept,nej,pont,smg,Pi3K92E,Fbw5,e(y)1,cad,BigH1,TfIIFalpha,ovo,Trf2,Taf7,SkpA,Rpn13,eIF3m,Ptip,REPTOR,Atu,spn-E,CG4238,Rel,Cdk7,RpLP1,Rpt2,Myd88,CG2199,raptor,Cf2,foxo,AGO3,trr,dl,Pdk1,not,Parp16,Cka,Lkb1,Bx,piwi,d4,EloC,Bap170,Mekk1,vers,osk,NFAT,CycC,CG17328,ben,Tao,Scox,Mos,hng3,Rcd5,RYBP,G9a,Sox102F,ari-1,lin,ScsbetaA,Mitf,park,MED24,Sgf11,Cul1,Stat92E,E2f1,dco,MED20,tyf,apt,PEK,STUB1,CG17202,Abi,S2P,fray,CG8963,Mybbp1A,ASPP,Atf6,Su(H),Utx,Jafrac2,hb,SREBP,Exd2,MED4,esc,Ubc6,CG8009,tor,Nf-YA,pygo,Tak1,CG2129,Wbp2,cnc,cyc,Pomp,Ankle2,cort,CG31053 | | 1.426e-06 | -13.46 | positive regulation of cellular catabolic process | biological process | GO:0031331 | 110 | 27 | 12345 | 1129 | cort,Pomp,dco,SkpA,PEK,Cul1,piwi,park,Ubc6,Mitf,Nprl2,foxo,Traf6,ari-1,AGO3,smg,Utx,GlyP,Marf1,Rpt2,AP-1-2beta,spn-E,CycC,Debcl,Atg7,gw,Synj | | 1.462e-06 | -13.44 | regulation of cellular biosynthetic process | biological process | GO:0031326 | 1371 | 175 | 12345 | 1129 | lid,spz,cact,bcd,Maf1,mael,Trf2,ovo,plu,Pi3K92E,rept,e(y)1,mtTFB1,Sirt7,Rel,bun,HP1e,Cf2,CG2199,Tfb4,CycE,Chrac-14,CG3587,Med,REPTOR,CG4872,spn-E,sbb,piwi,Tlk,Topors,krimp,CG2678,d4,dl,trr,foxo,not,hng3,Rcd5,Smox,Scox,vers,EloC,CG17328,osk,HP1c,NFAT,CycC,twin,apt,tyf,PEK,BicC,lwr,Cbl,egg,Nap1,wash,Mitf,Pax,r2d2,RpL13A,CG17202,Mybbp1A,CG8963,mute,Atf3,Hat1,scny,Rga,Ssdp,cyc,sip3,MED4,esc,hb,SREBP,Exd2,CG2129,Nf-YA,CG8009,CG10038,pn,MED27,ph-d,Uxt,Tip60,Bin1,CG11710,Dph5,Hnf4,EcR,pgc,gw,CG1602,wisp,Rbfox1,CG4730,tkv,cad,CG7414,BigH1,insv,l(3)mbt,Taf7,TfIIFalpha,asf1,nej,pont,smg,YL-1,lin-52,crm,rl,spen,SuUR,CG11902,cup,CG6813,Myd88,Cdk7,ix,heph,CG13344,Ptip,Atu,Bx,otu,Dd,CG10631,bru1,Rbf,Traf6,CG10336,RYBP,mRpL13,Cul2,Bap170,CG10654,Elba2,Stat92E,HmgD,Rbf2,Aos1,Sgf11,Spt20,E2f1,MED20,Hr78,lin,CG10959,Sox102F,MED24,Not10,Su(H),Atf6,psq,HDAC6,Utx,bam,S2P,escl,l(3)73Ah,CG2911,Cdk1,CG31875,Unr,GAPsec,cnc,Wbp2,CG1578,pygo,Tak1,nos | | 1.642e-06 | -13.32 | Respiratory electron transport, ATP synthesis by chemiosmotic coupling, and heat production by uncoupling proteins. | REACTOME pathways | R-DME-163200 | 57 | 23 | 3923 | 574 | NP15.6,wal,ATPsynB,ATPsynCF6,ND-51,ND-30,ND-75,ND-24,ND-SGDH,blw,ATPsynD,ND-39,ND-49,ND-B18,CG8067,ATPsynF,ATPsyndelta,Etf-QO,ND-23,ATPsynG,ATPsynO,ATPsynbeta,ND-B22 | | 1.690e-06 | -13.29 | GNF2\_TPT1 | MSigDB lists | GNF2\_TPT1 | 27 | 15 | 4161 | 644 | RpL12,RpL40,RpS18,RpL38,RpS4,RpS23,RpL9,RpS2,RpL13A,RpS13,RpS27,RpL23,RpL30,RpL37a,RpS24 | | 1.699e-06 | -13.29 | regulation of mitotic cell cycle | biological process | GO:0007346 | 186 | 38 | 12345 | 1129 | Su(H),Sld5,cana,mus304,raptor,CG30291,CycE,grp,Cdk1,Zwilch,bru1,CG10336,mus81,Rbf,mad2,ago,mre11,png,Smox,nbs,APC7,tws,twe,Iml1,Moe,ben,dgt4,Aos1,Rbf2,plu,SkpA,l(3)mbt,PEK,RecQ5,Pi3K92E,nej,Aven,Nprl2 | | 1.729e-06 | -13.27 | positive regulation of macromolecule metabolic process | biological process | GO:0010604 | 858 | 119 | 12345 | 1129 | S2P,Abi,fray,CG17202,STUB1,ASPP,Mybbp1A,CG8963,Su(H),Atf6,Utx,esc,MED4,Exd2,SREBP,hb,Jafrac2,Tak1,CG2129,pygo,Nf-YA,CG8009,tor,Ubc6,CG31053,cort,cyc,Ankle2,Pomp,Wbp2,cnc,Nup358,vers,Mekk1,EloC,Bap170,ben,CG17328,CycC,osk,NFAT,Rcd5,RYBP,hng3,Mos,Tao,G9a,lin,egg,Sox102F,ari-1,MED24,park,ScsbetaA,Mitf,Stat92E,Cul1,Sgf11,PEK,apt,tyf,MED20,dco,E2f1,eIF3m,Rpn13,CG4238,spn-E,Atu,REPTOR,Myd88,Rpt2,GlyP,Cdk7,RpLP1,Rel,Cf2,raptor,CG2199,AGO3,dl,trr,foxo,Parp16,not,Pdk1,otu,piwi,Lkb1,Bx,Cka,d4,EcR,gw,CG1602,Rbfox1,bcd,pn,gus,Tip60,Marf1,lid,CG11710,Pi3K92E,nej,pont,smg,rept,CG3198,e(y)1,SuUR,Fbw5,BigH1,cad,SkpA,Taf7,Trf2,ovo,TfIIFalpha | | 1.737e-06 | -13.26 | negative regulation of cellular metabolic process | biological process | GO:0031324 | 681 | 99 | 12345 | 1129 | sbb,spn-E,CG4872,CycE,Chrac-14,heph,CG2199,Sirt7,cup,dl,Rbf,CG10336,foxo,alph,krimp,d4,bru1,CG10631,piwi,Lkb1,CG30496,Tlk,CG1602,Rbfox1,bcd,EcR,Hnf4,pgc,gw,Ptp61F,Bin1,ph-d,CG10038,SelD,Uxt,Tip60,rl,Nprl2,Fkbp39,SuUR,Drep1,Pi3K92E,lin-52,smg,rept,faf,l(3)mbt,SkpA,plu,asf1,ovo,mael,Maf1,insv,BigH1,escl,Mybbp1A,sws,Hat1,RpL13A,Utx,r2d2,CkIIalpha,bam,Su(H),Not10,psq,nos,mts,esc,SREBP,hb,Ankle2,cyc,CG14408,Rga,Unr,CG31875,scny,CG17328,PpV,Elba2,mRpL13,ATPsynD,Smox,CG17746,wash,park,CG10959,Cbl,egg,Sox102F,apt,dco,BicC,E2f1,Rbf2,twin,EndoGI | | 1.751e-06 | -13.26 | KEGG\_ALZHEIMERS\_DISEASE | MSigDB lists | KEGG\_ALZHEIMERS\_DISEASE | 62 | 25 | 4161 | 644 | ND-49,Gapdh1,Pp2B-14D,scu,Cyt-c-p,ND-SGDH,ND-B22,levy,CanB2,ND-23,ND-51,APP-BP1,SERCA,ND-39,CanA-14F,COX5A,rl,ND-B18,SdhA,ND-75,Cdk5,UQCR-Q,ND-30,ND-24,pen-2 | | 1.838e-06 | -13.21 | MORF\_ACTG1 | MSigDB lists | MORF\_ACTG1 | 86 | 31 | 4161 | 644 | RpL30,eIF4A,RpL23,RpS24,RpS16,RpL13A,RpS30,RpLP0,RpS13,Prosbeta6,RpL38,levy,RpS4,RpS7,Arpc2,RpL37a,RpL11,Nap1,RpS23,RpL9,RpS27A,RpS28b,RpL24,RpS27,RpS18,Eno,eEF1beta,RpL12,Pgk,RpL13,lwr | | 1.869e-06 | -13.19 | smt3 (smt3) | protein interactions | 33981 | 112 | 32 | 9204 | 1112 | velo,HmgD,Gip,lwr,awd,Sod1,Fkbp12,dl,Aos1,Uch,CG4598,tub,capt,Moe,scu,Stip1,CG14814,Prosbeta4,CG11267,Chd64,CG14715,CG10638,Myd88,CG9641,eIF3m,Cyp1,cib,PHGPx,Eno,eIF4E1,AnxB10,regucalcin | | 1.921e-06 | -13.16 | GO\_NUCLEAR\_TRANSCRIBED\_MRNA\_CATABOLIC\_PROCESS\_NONSENSE\_MEDIATED\_DECAY | MSigDB lists | GO\_NUCLEAR\_TRANSCRIBED\_MRNA\_CATABOLIC\_PROCESS\_NONSENSE\_MEDIATED\_DECAY | 74 | 28 | 4161 | 644 | RpL12,RpL13,RpS18,Pp2A-29B,RpS23,RpL26,RpS27A,RpL9,RpS28b,RpS27,RpL24,RpL37a,RpL11,Cbp80,RpL40,RpL38,RpS4,RpS7,RpL13A,RpS2,RpS16,RpS30,RpLP0,RpS13,RpL30,RpL23,mts,RpS24 | | 1.965e-06 | -13.14 | GO\_ENERGY\_DERIVATION\_BY\_OXIDATION\_OF\_ORGANIC\_COMPOUNDS | MSigDB lists | GO\_ENERGY\_DERIVATION\_BY\_OXIDATION\_OF\_ORGANIC\_COMPOUNDS | 103 | 35 | 4161 | 644 | RpL40,CG9485,Etf-QO,AGBE,CG11110,Cyt-c-p,ScsbetaA,ND-30,ND-24,ND-B18,Pfk,COX5A,Gyg,CG5028,ND-51,ND-23,ND-B22,CG8475,levy,ND-SGDH,ND-49,Pgm1,UQCR-Q,SdhA,CG3803,ND-75,wal,GlyP,Irp-1B,Idh3b,RpS27A,ND-39,Scsalpha1,Idh,Nc73EF | | 1.970e-06 | -13.14 | response to oxygen-containing compound | biological process | GO:1901700 | 339 | 58 | 12345 | 1129 | G9a,spz,Tao,whd,Idh3b,aay,MED27,dnc,Pdxk,Pfk,Rab10,Desat1,PyK,EcR,Hnf4,p38a,dco,Trf2,apt,Shmt,Fdh,Pka-R1,Stat92E,CG5412,MED24,Nurf-38,CG7328,Sod1,rl,Jafrac1,rept,Edem1,adp,Pi3K92E,Sod3,CG4390,Adh,Utx,raptor,Pka-R2,ACC,Agpat2,Aldh,AcCoAS,sgll,CG17734,Arf51F,rig,cyc,mAChR-A,Vps16B,ND-SGDH,PHGPx,Bx,Pdk1,Tak1,foxo,trr | | 2.049e-06 | -13.10 | cellular response to stress | biological process | GO:0033554 | 537 | 82 | 12345 | 1129 | CG9005,Atf6,Utx,mus304,Sld5,Atg6,Sod3,fray,S2P,CG5844,Idh,Atg7,CG6766,Cdk1,mus205,grp,cnc,tos,sip3,mus81,Jafrac2,SREBP,ago,Tak1,kcc,Ubc6,Xpc,CG32640,Tao,SNF4Agamma,CG17746,Hsc70-3,CG11070,Ufm1,Uev1A,lt,Marf,Mekk1,ben,Iml1,PEK,Snm1,p38a,RecQ5,DNApol-epsilon255,shep,Sod1,ERp60,Mitf,thoc6,Rel,raptor,Tfb4,MAGE,Chrac-14,Grx1,Tom40,PHGPx,Cka,Rbf,Traf6,CG10336,foxo,tho2,mre11,Parp16,Xpac,agt,CG8974,nbs,Ptp61F,Debcl,obe,CG5285,Pi3K92E,Cdk5,Edem1,Mtl,Jafrac1,Calx,Nprl2,rl | | 2.093e-06 | -13.08 | lipid modification | biological process | GO:0030258 | 93 | 24 | 12345 | 1129 | Echs1,CG4586,CG9391,CG10932,Thiolase,Pi3K92E,CG4598,CG6707,Acox57D-p,wal,CG7789,SERCA,Pi4KIIalpha,CG4594,CG8112,Synj,Mcad,Hnf4,CG5026,Seipin,Etf-QO,CG4860,yip2,Mtpalpha | | 2.102e-06 | -13.07 | regulation of cellular protein metabolic process | biological process | GO:0032268 | 623 | 92 | 12345 | 1129 | bcd,Rbfox1,Ptp61F,gw,Dph5,mtTFB1,faf,smg,Pi3K92E,SkpA,CG7414,RhoGAP5A,Fdh,Pka-R1,CG30291,Rpn13,eIF3m,CycE,raptor,heph,RpLP1,Pp2A-29B,Cdk7,cup,Pka-R2,Rpt2,Pdk1,Parp16,trr,bru1,alph,krimp,Cka,CG30496,Lkb1,osk,PpV,CycC,ben,Mekk1,G9a,ATPsynD,mRpL13,tws,Tao,SNF4Agamma,CG17746,Mos,ScsbetaA,park,MED24,CycB,ari-1,Cbl,dco,BicC,tyf,apt,PEK,Cul1,twin,sws,CG8963,ASPP,CG2911,STUB1,RpL13A,fray,Abi,bam,CkIIalpha,Utx,Not10,Ubc6,tor,CG8009,nos,Tak1,CycJ,Jafrac2,Exd2,CG1578,mts,esc,CG14408,cort,Pomp,Ankle2,sip3,GAPsec,Unr,Rga | | 2.243e-06 | -13.01 | developmental process | biological process | GO:0032502 | 2988 | 338 | 12345 | 1129 | CycE,Sec23,spn-E,sbb,Pfdn2,Mtpalpha,Gclc,Rel,CG4853,dl,Cul5,foxo,Pura,not,piwi,Atpalpha,Lkb1,Cka,Tlk,krimp,up,snz,d4,CHOp24,tub,bcd,AP-1-2beta,SelD,Liprin-beta,lid,DCAF12,spz,rept,e(y)1,AIF,Vha13,SERCA,Rbcn-3A,RhoGAP5A,Tm1,Mes2,Pmi,ovo,awd,Sply,Atg7,SelR,armi,Ptp69D,r2d2,Idgf4,MED4,SREBP,hb,ago,eIF4A,ea,muskelin,Ssdp,cyc,cort,EloC,stw,shu,osk,Smox,levy,Cbl,Stam,Nap1,Mitf,yps,nmd,Arf51F,COX5A,trk,Pka-R2,Rab5,ix,sw,CSN6,mxc,Tspo,larp,Pld,squ,Rbf,Wnt10,Grip75,otu,Pglym78,Dd,CG3770,COX4,alph,EMC3,bru1,twe,fs(1)N,Hnf4,chrb,PyK,Rbfox1,Debcl,wisp,MED27,pn,X11L,Rrp40,dx,Thiolase,nej,Cen,faf,Nup62,Edem1,Nprl2,crm,Drep1,spen,SRPK,cad,CG12301,SkpA,Treh,Arpc1,Utx,CG13366,eIF4E1,bam,AGBE,shv,CycJ,fdl,grp,Afti,Nup358,Fer1HCH,jvl,Desat1,Moe,Rab10,dnc,Trxr-1,G9a,CCT3,zpg,Tsp86D,ari-1,DNApol-epsilon255,vret,Sod1,MED24,park,Stat92E,Actn,Cul1,toc,Hr78,swa,Arpc2,MAGE,Pen,Mitofilin,Nlg1,Med,ACC,gammaTub37C,bun,fs(1)M3,mop,GlyP,Pex12,Pxt,Cf2,HP1e,disp,crok,trr,Top1,tho2,Imp,Pdk1,cold,Tsp42Ef,Ptpmeg,RhoGAP54D,cact,Pfk,Idh3b,fwe,snk,Pink1,Nmt,Pi3K92E,Pp2B-14D,Mtl,stai,boi,galectin,stil,wal,mael,Trf2,csw,Atf3,mute,Gapdh1,lqfR,Pex10,Ptpmeg2,Ptp10D,Atg6,CkIIalpha,mts,mad2,Orc1,Nf-YA,exu,CG17068,scny,ND-SGDH,slam,Ankle2,Marf,CycC,gd,ssh,Mos,tws,lwr,egg,shep,alpha-Man-Ia,CycB,wash,parvin,Girdin,Pax,ScsbetaA,twin,CG46338,Hsp26,apt,dco,BicC,Grip128,Ptip,eIF3m,Vha68-2,Myd88,CG3226,cup,Kap3,TBCC,PrBP,raptor,heph,Irk1,Traf6,Bx,Smyd4-4,Archease,EcR,bif,pgc,gw,enc,del,Ptp61F,ph-d,gus,Tip60,Sec15,alphaTub67C,cathD,lin-52,cib,Cdk5,rl,elg1,Jafrac1,Dcr-1,Idgf2,Pka-R1,tkv,PGRP-LE,capt,l(3)mbt,fray,Abi,yl,msps,velo,sws,Su(H),psq,Arpc4,CanB2,mus304,Klp10A,Seipin,BoYb,pygo,Tak1,Pgk,tor,nos,GatA,Cdk1,PIG-C,cnc,colt,lt,Bap170,ben,stet,Grip71,Iml1,RYBP,Sry-alpha,Tao,Cul2,ATPsynD,Coq7,lin,shrb,Sox102F,cnn,rost,milt,Rbf2,scu,Idgf6,sol,p38a,Npc2a,E2f1 | | 2.347e-06 | -12.96 | REACTOME\_TCA\_CYCLE\_AND\_RESPIRATORY\_ELECTRON\_TRANSPORT | MSigDB lists | REACTOME\_TCA\_CYCLE\_AND\_RESPIRATORY\_ELECTRON\_TRANSPORT | 59 | 24 | 4161 | 644 | ND-49,ND-SGDH,Cyt-c-p,ScsbetaA,ND-B22,Etf-QO,levy,ND-51,ND-23,CG5028,Pdk,ND-39,Nc73EF,Scsalpha1,Idh,Idh3b,COX5A,wal,ND-75,SdhA,ND-B18,ND-30,ND-24,UQCR-Q | | 2.376e-06 | -12.95 | Prosalpha1 (Proteasome alpha1 subunit) | protein interactions | 45780 | 50 | 19 | 9204 | 1112 | AnxB10,Moe,capt,regucalcin,Uch,Prosalpha4,Rpt2,Eno,Prosbeta6,awd,Rpn13,Pomp,Prosalpha5,Rpn2,CG10638,Prosbeta5,Prosbeta4,Prosalpha3,Prosbeta3 | | 2.376e-06 | -12.95 | ATPsynCF6 (ATP synthase, coupling factor 6) | protein interactions | 42759 | 50 | 19 | 9204 | 1112 | PHGPx,ATPsynbeta,Mf,awd,up,ATPsynG,blw,COX4,ATPsynE,ATPsyndelta,ATPsynF,CG15098,Argk,ATPsynO,AnxB10,ATPsynD,Scp1,Eno,ATPsynB | | 2.432e-06 | -12.93 | negative regulation of gene expression | biological process | GO:0010629 | 582 | 87 | 12345 | 1129 | SREBP,hb,esc,nos,Rga,Unr,cyc,RpL13A,Hat1,escl,armi,psq,Not10,Su(H),bam,r2d2,egg,Sox102F,CG10959,cu,wash,Edc3,twin,Rbf2,BicC,Cbp80,E2f1,trsn,apt,Elba2,CG17328,SmydA-1,CG31759,Hsc70-3,mRpL13,foxo,AGO3,dl,Rbf,squ,LSm3,Tlk,piwi,d4,CG10631,CG17768,bru1,krimp,Trf4-2,Smyd4-4,Chrac-14,sbb,spn-E,CG4872,Sirt7,cup,CG2199,heph,CG9425,rept,YL-1,lin-52,CG11486,smg,Dcr-1,CG6967,insv,Maf1,mael,tkv,asf1,ovo,l(3)mbt,gw,pgc,Hnf4,EcR,Rbfox1,bcd,CG1602,Tip60,Pym,Marf1,ph-d,CG10038,Bin1,Rrp40 | | 2.442e-06 | -12.92 | Fkbp12 (FK506-binding protein 12kD) | protein interactions | 37214 | 73 | 24 | 9204 | 1112 | Stip1,Prosalpha3,CG11267,CG11486,awd,Chd64,ATPsynbeta,CG10638,CG14715,Cyp1,cib,PHGPx,Sod1,Uch,Eno,CG4598,AnxB10,RpS27,CG10265,regucalcin,Moe,capt,scu,csw | | 2.447e-06 | -12.92 | Stip1 (Stress induced phosphoprotein 1) | protein interactions | 33202 | 59 | 21 | 9204 | 1112 | capt,scu,Moe,regucalcin,Gdi,AnxB10,Eno,Uch,PHGPx,EndoGI,cib,Fkbp12,Pgk,CG10638,CG11267,awd,Chd64,CG9117,yip2,piwi,HmgD | | 2.487e-06 | -12.90 | REACTOME\_PEPTIDE\_CHAIN\_ELONGATION | MSigDB lists | REACTOME\_PEPTIDE\_CHAIN\_ELONGATION | 63 | 25 | 4161 | 644 | RpS28b,RpS27,RpL24,RpL26,RpS27A,RpS23,RpL9,RpL37a,RpL11,RpL12,RpL13,RpS18,RpS16,RpS2,RpL13A,RpS13,RpS30,RpLP0,RpS24,RpL30,RpL23,RpL40,RpS7,RpL38,RpS4 | | 2.562e-06 | -12.87 | scu (scully) | protein interactions | 32789 | 78 | 25 | 9204 | 1112 | Uch,Eno,Pglym78,AnxB10,regucalcin,GstT1,Moe,capt,Stip1,RhoGDI,Prosbeta5,Tak1,yip2,CG11267,Rel,Chd64,awd,PGRP-LE,Pgk,CG14715,CG10638,PHGPx,EndoGI,cib,Fkbp12 | | 2.666e-06 | -12.83 | ATPsyngamma (ATP synthase, gamma subunit) | protein interactions | 43507 | 46 | 18 | 9204 | 1112 | SERCA,CCT6,ATPsynF,ATPsyndelta,CCT3,blw,ATPsynCF6,Mcad,ScsbetaA,ATPsynO,Rab5,CG1416,ATPsynB,cnn,dgt4,wal,mEFTu1,ATPsynD | | 2.911e-06 | -12.75 | cation binding | molecular function | GO:0043169 | 1072 | 142 | 12176 | 1119 | CG9581,Hr78,Eno,PGRP-LE,CSN6,CG9684,CG31053,Pgm1,CanB2,CG32473,up,CAH2,CG6923,nos,bor,stet,alpha-Man-Ia,Utx,Cyp6u1,LpR2,CG14435,regucalcin,CG8974,stw,Rpn3,Scp1,CG10463,Wdfy2,cN-IIIB,Shmt,Desat1,CG17328,Fer1HCH,Cyp6a19,CG13344,Marc,AcCoAS,Nc73EF,ScsbetaA,Clp,nej,Idh,aay,DhpD,Cyp6a22,CG9425,Nurf-38,Iru,mal,ph-d,CG11267,SP1029,CG44422,isoQC,Mctp,Usp16-45,Actn,CG8728,dx,CG15141,Scox,mre11,Rpn2,QC,Edem1,EcR,CG9941,Cyp6v1,CAH7,CG6767,Fdh,CG17272,AGBE,CG2678,Calx,HDAC6,Pex10,not,krimp,Reps,Dip-B,CG14516,Cbl,PyK,CG11134,Hnf4,CG11710,PAPLA1,Nca,CG4709,Gs1l,CG11902,Sod1,r,MtnA,CG17746,Pfk,glob1,ari-1,CG17698,CG9925,l(3)mbt,SmydA-1,CG9981,ATP8A,CG17593,grsm,Topors,CG7115,G9a,yl,Usp39,sip3,ACC,SelR,CG10492,CG31729,CG11590,shop,egg,CG4730,d4,AnxB10,Traf6,Tak1,lt,awd,GluRS-m,Sod3,CG6813,DNApol-epsilon255,RpS29,Tengl4,alph,AOX3,Sec23,CG17260,Kcmf1,CG10654,tkv,Pex12,l(3)73Ah | | 3.160e-06 | -12.67 | regulation of response to biotic stimulus | biological process | GO:0002831 | 162 | 34 | 12345 | 1129 | slif,scny,GILT1,Tak1,Traf6,dl,Atg6,Herc4,Rel,GlyP,CG9925,mop,Myd88,Atg7,Hat1,Crag,Ptip,Sp7,p38a,Hr78,PGRP-LE,Aos1,cad,Stat92E,lwr,faf,G9a,spz,PDCD-5,wisp,Ect4,tub,cact,EMRE | | 3.165e-06 | -12.66 | oocyte development | biological process | GO:0048599 | 155 | 33 | 12345 | 1129 | Grip128,spn-E,msps,psq,armi,cup,gammaTub37C,CkIIalpha,heph,nos,exu,Grip75,ea,Lkb1,cnc,muskelin,krimp,jvl,enc,del,Grip71,osk,wisp,Moe,Mos,gus,spz,milt,mael,Tm1,Pka-R1,swa,capt | | 3.285e-06 | -12.63 | GO\_GENERATION\_OF\_PRECURSOR\_METABOLITES\_AND\_ENERGY | MSigDB lists | GO\_GENERATION\_OF\_PRECURSOR\_METABOLITES\_AND\_ENERGY | 127 | 40 | 4161 | 644 | ND-B18,Pfk,Eno,Pglym78,ND-30,ND-24,Gyg,COX5A,Scox,CG9485,Etf-QO,AGBE,RpL40,CG11110,Cyt-c-p,ScsbetaA,SdhA,CG3803,wal,GlyP,ND-75,UQCR-Q,Pgk,ND-39,Scsalpha1,Idh,Nc73EF,Idh3b,Irp-1B,RpS27A,CG8475,ND-B22,levy,CG5028,ND-51,ND-23,Gapdh1,ND-49,Pgm1,ND-SGDH | | 3.322e-06 | -12.61 | transferase activity, transferring phosphorus-containing groups | molecular function | GO:0016772 | 482 | 75 | 12176 | 1119 | Trf4-2,CG7328,kri,Ady43A,SRPK,Scsalpha1,RpII140,png,Pfrx,Pfk,dco,CG17698,Pink1,Mo25,Pi3K92E,Lkb1,SelD,CG2201,CG1344,fray,CycJ,p38a,Tfb4,Argk,grp,CG2100,l(2)k01209,wisp,rl,CG8475,CG7718,l(1)G0196,CG11486,PyK,Mos,PEK,nmdyn-D7,CG8726,CycE,CG6767,Gk2,Pdk1,CG4407,CycB,CG4041,tkv,alc,Coq8,CG33156,CG8311,Cdk7,awd,Tak1,DNApol-epsilon255,Pdk,Pgk,Tlk,Gk1,nmdyn-D6,tub,CG12018,Mekk1,CkIIalpha,Tao,Cdk5,CG6650,UGP,tor,mus205,CG3631,Cdk1,Pdxk,Pi4KIIalpha,Rpb8,CG46385 | | 3.350e-06 | -12.61 | multicellular organismal reproductive process | biological process | GO:0048609 | 1285 | 164 | 12345 | 1129 | Mos,shu,osk,BicC,nmd,apt,twin,yps,CG46338,ScsbetaA,Pax,wash,CycB,Nap1,Cbl,egg,lwr,shep,Grip84,CkIIalpha,Ptpmeg2,r2d2,armi,Pex10,lqfR,Atg7,csw,cort,muskelin,slam,ea,exu,Pex13,Orc1,eIF4A,mts,mad2,spz,lid,bcd,cact,Axs,ovo,mael,Tm1,SERCA,stil,stai,e(y)1,Pi3K92E,Pp2B-14D,tef,disp,Cf2,HP1e,mop,bun,Pex12,fs(1)M3,Pxt,gammaTub37C,spn-E,Med,Pen,CycE,krimp,Topors,piwi,Lkb1,Pdk1,Imp,Cul5,PAPLA1,tho2,Top1,Cul2,Tao,Mst89B,Grip71,dnc,Iml1,Rab10,Moe,jvl,Desat1,Cenp-C,E2f1,swa,toc,Stat92E,park,milt,vret,ari-1,cnn,zpg,shrb,bam,mus304,Klp10A,eIF4E1,psq,msps,yl,Abi,cnc,Nup358,grp,Cdk1,yem,nos,tor,GatA,wapl,CycJ,BoYb,Sec15,clt,Syx13,gus,Rbfox1,mtrm,wisp,bif,pgc,del,enc,twe,fs(1)N,EcR,l(3)mbt,capt,SRPK,Pka-R1,tkv,Drep1,Dcr-1,Nprl2,rl,elg1,Jafrac1,faf,Nup62,cathD,nej,CSN6,raptor,heph,sw,ix,tko,Rab5,Kap3,cup,eIF3m,Arf51F,Grip128,bru1,otu,Grip75,sau,larp,squ,Rbf | | 3.484e-06 | -12.57 | developmental maturation | biological process | GO:0021700 | 206 | 40 | 12345 | 1129 | gus,spz,jvl,enc,EcR,Grip71,wisp,osk,ben,Moe,mael,Tm1,Pka-R1,swa,SkpA,capt,shep,Pi3K92E,shrb,milt,Stam,psq,mop,armi,gammaTub37C,Atg6,CkIIalpha,heph,Grip128,spn-E,msps,Grip75,ea,Lkb1,cnc,krimp,muskelin,nos,exu,Imp | | 3.529e-06 | -12.55 | oocyte differentiation | biological process | GO:0009994 | 177 | 36 | 12345 | 1129 | CkIIalpha,heph,psq,armi,gammaTub37C,cup,spn-E,msps,Grip128,cnc,krimp,muskelin,piwi,ea,Grip75,Lkb1,nos,exu,spz,Mos,gus,Rbfox1,Grip71,wisp,osk,Moe,jvl,del,enc,swa,capt,mael,Tm1,Pka-R1,Dcr-1,milt | | 3.560e-06 | -12.55 | REACTOME\_INFLUENZA\_VIRAL\_RNA\_TRANSCRIPTION\_AND\_REPLICATION | MSigDB lists | REACTOME\_INFLUENZA\_VIRAL\_RNA\_TRANSCRIPTION\_AND\_REPLICATION | 76 | 28 | 4161 | 644 | RpL12,RpL13,RpS18,RpS23,RpS27A,RpL9,RpL26,RpS28b,RpL24,RpS27,RpL37a,RpL11,RpL40,Rpb8,RpS4,Karybeta3,RpL38,RpS7,RpII140,RpS16,RpL13A,RpS2,RpS30,RpS13,RpLP0,RpL23,RpL30,RpS24 | | 3.878e-06 | -12.46 | regulation of cellular macromolecule biosynthetic process | biological process | GO:2000112 | 1307 | 166 | 12345 | 1129 | bun,Rel,Sirt7,Tfb4,CG2199,HP1e,Cf2,CG3587,Chrac-14,CycE,spn-E,sbb,Med,REPTOR,CG4872,Tlk,piwi,d4,CG2678,krimp,Topors,foxo,trr,dl,not,lid,bcd,cact,Maf1,mael,plu,ovo,Trf2,rept,Pi3K92E,mtTFB1,e(y)1,r2d2,CG17202,RpL13A,Hat1,Atf3,mute,CG8963,Mybbp1A,Rga,scny,sip3,cyc,Ssdp,Exd2,SREBP,hb,esc,MED4,Nf-YA,CG2129,Rcd5,Smox,hng3,EloC,vers,CycC,NFAT,osk,HP1c,CG17328,twin,BicC,PEK,apt,tyf,lwr,egg,Mitf,wash,Nap1,ix,Cdk7,Myd88,CG6813,cup,CG11902,CG13344,heph,Atu,Dd,otu,Bx,CG10631,bru1,CG10336,Traf6,Rbf,Tip60,ph-d,pn,MED27,CG10038,CG11710,Bin1,gw,pgc,EcR,Hnf4,Dph5,Rbfox1,CG1602,insv,BigH1,cad,CG7414,tkv,CG4730,asf1,TfIIFalpha,l(3)mbt,Taf7,lin-52,YL-1,nej,smg,pont,SuUR,spen,rl,crm,psq,Atf6,Su(H),Not10,bam,Utx,HDAC6,S2P,l(3)73Ah,CG2911,escl,Unr,CG31875,Cdk1,cnc,Wbp2,GAPsec,CG1578,nos,pygo,RYBP,Cul2,mRpL13,Bap170,Elba2,CG10654,Sgf11,Aos1,Rbf2,Stat92E,HmgD,MED20,Hr78,E2f1,Spt20,Sox102F,CG10959,lin,MED24 | | 3.898e-06 | -12.46 | regulation of macromolecule biosynthetic process | biological process | GO:0010556 | 1326 | 168 | 12345 | 1129 | CG1578,pygo,nos,Cdk1,Unr,CG31875,GAPsec,Wbp2,cnc,S2P,CG2911,l(3)73Ah,escl,Su(H),Not10,psq,Atf6,Utx,HDAC6,bam,CG10959,lin,Sox102F,MED24,HmgD,Rbf2,Stat92E,Sgf11,Aos1,MED20,Hr78,E2f1,Spt20,Bap170,CG10654,Elba2,RYBP,Cul2,mRpL13,Rbf,CG10336,Traf6,otu,Bx,Dd,CG10631,bru1,Atu,CG6813,Myd88,cup,CG11902,Cdk7,ix,heph,CG13344,lin-52,YL-1,pont,smg,nej,rl,crm,SuUR,spen,tkv,CG4730,insv,BigH1,CG7414,cad,l(3)mbt,Taf7,asf1,TfIIFalpha,Hnf4,EcR,Dph5,pgc,gw,CG1602,Rbfox1,MED27,ph-d,pn,CG10038,Tip60,Uxt,Bin1,CG11710,esc,MED4,SREBP,Exd2,hb,CG2129,Nf-YA,Rga,scny,Ssdp,sip3,cyc,RpL13A,CG17202,Atf3,mute,Mybbp1A,CG8963,Hat1,r2d2,egg,lwr,wash,Nap1,Mitf,Pax,twin,PEK,apt,tyf,BicC,vers,EloC,CG17328,CycC,NFAT,osk,HP1c,Smox,Rcd5,hng3,trr,dl,foxo,not,piwi,Tlk,krimp,Topors,d4,CG2678,CycE,CG3587,Chrac-14,spn-E,sbb,Med,REPTOR,CG4872,Sirt7,bun,Rel,Cf2,HP1e,CG2199,Tfb4,Pi3K92E,rept,e(y)1,mtTFB1,mael,Maf1,Trf2,plu,ovo,cact,bcd,lid | | 3.940e-06 | -12.44 | coenzyme metabolic process | biological process | GO:0006732 | 109 | 26 | 12345 | 1129 | regucalcin,CG8199,ND-39,Ppcdc,Coq7,Coq6,CG6178,CG4407,Scsalpha1,Nadsyn,Dhfr,mal,sgll,Pdxk,CG33156,AcCoAS,Shmt,pr,CG17691,CG1774,Sptr,Pcd,scu,Coq8,ScsbetaA,CG7920 | | 3.964e-06 | -12.44 | anatomical structure development | biological process | GO:0048856 | 2924 | 330 | 12345 | 1129 | Pmi,ovo,SERCA,Rbcn-3A,RhoGAP5A,Tm1,Mes2,e(y)1,AIF,Vha13,rept,lid,DCAF12,spz,AP-1-2beta,SelD,Liprin-beta,CHOp24,bcd,tub,krimp,up,snz,d4,piwi,Atpalpha,Lkb1,Cka,Tlk,not,dl,foxo,Cul5,Pura,CG4853,Mtpalpha,Gclc,Rel,sbb,spn-E,Sec23,Pfdn2,CycE,nmd,yps,Stam,Nap1,Mitf,Cbl,Smox,levy,stw,shu,osk,EloC,Ssdp,muskelin,cyc,cort,ea,eIF4A,ago,MED4,SREBP,hb,r2d2,Idgf4,Ptp69D,SelR,armi,Atg7,awd,Sply,CG12301,SkpA,SRPK,cad,Nprl2,crm,Drep1,spen,nej,Thiolase,faf,Cen,Nup62,Edem1,Rrp40,dx,pn,MED27,X11L,Rbfox1,Debcl,wisp,twe,fs(1)N,Hnf4,chrb,PyK,alph,EMC3,bru1,Grip75,otu,Pglym78,CG3770,Dd,COX4,larp,Pld,squ,Rbf,Wnt10,CSN6,mxc,Tspo,Pka-R2,Rab5,ix,sw,trk,Arf51F,COX5A,Hr78,swa,Stat92E,Actn,Cul1,toc,vret,Sod1,park,MED24,zpg,CCT3,Tsp86D,ari-1,G9a,Trxr-1,Rab10,Moe,dnc,Fer1HCH,jvl,Desat1,Afti,Nup358,grp,CycJ,fdl,AGBE,CG13366,Utx,eIF4E1,bam,Arpc1,Trf2,stil,wal,mael,stai,boi,galectin,Pi3K92E,Pp2B-14D,Mtl,snk,Nmt,Pink1,Idh3b,fwe,cact,Pfk,Tsp42Ef,Ptpmeg,RhoGAP54D,cold,Imp,Pdk1,crok,trr,tho2,Top1,HP1e,Cf2,disp,ACC,gammaTub37C,mop,Pex12,GlyP,fs(1)M3,bun,Pxt,Mitofilin,Nlg1,Med,MAGE,Arpc2,Pen,Hsp26,apt,BicC,dco,twin,CG46338,wash,alpha-Man-Ia,CycB,parvin,Pax,ScsbetaA,Girdin,shep,egg,lwr,tws,gd,ssh,Mos,CycC,Marf,slam,Ankle2,CG17068,ND-SGDH,Orc1,Nf-YA,exu,mts,mad2,Ptpmeg2,Ptp10D,Atg6,CkIIalpha,Pex10,csw,Gapdh1,mute,Atf3,lqfR,PGRP-LE,capt,l(3)mbt,Idgf2,Pka-R1,tkv,rl,elg1,Jafrac1,Dcr-1,lin-52,cathD,cib,Cdk5,Sec15,alphaTub67C,ph-d,gus,Tip60,Archease,EcR,gw,pgc,bif,enc,Ptp61F,del,Smyd4-4,Bx,Irk1,Traf6,PrBP,heph,raptor,CG3226,Myd88,Kap3,cup,TBCC,Vha68-2,Grip128,Ptip,eIF3m,scu,sol,Idgf6,p38a,Npc2a,E2f1,rost,milt,lin,shrb,cnn,Cul2,ATPsynD,Coq7,RYBP,Sry-alpha,Tao,ben,Grip71,Iml1,colt,lt,Bap170,cnc,Cdk1,PIG-C,Tak1,pygo,Pgk,nos,tor,GatA,BoYb,mus304,Seipin,Su(H),psq,Arpc4,CanB2,msps,velo,sws,fray,Abi,yl | | 4.161e-06 | -12.39 | MORF\_NPM1 | MSigDB lists | MORF\_NPM1 | 106 | 35 | 4161 | 644 | RpS27,RpL24,eIF3m,RpL11,RpL37a,lwr,RpL12,Eno,RpS18,RpS16,RpS2,Phb2,RpS24,RpL30,RpL23,not,RpS7,Prosbeta6,RpS4,RpL38,RpS28b,RpS23,RpS27A,RpL9,Nap1,CSN6,RpL13,Pgk,eEF1beta,RpLP0,RpS30,RpS13,RpL13A,eIF4A,levy | | 4.202e-06 | -12.38 | Electron Transport Chain | WikiPathways | WP542 | 48 | 25 | 499 | 116 | ND-B22,ATPsynbeta,SdhA,ND-SGDH,ND-75,ATPsynG,COX5A,ND-30,ATPsynD,levy,ND-24,UQCR-Q,ATPsynB,ATPsyndelta,ND-39,ND-49,Scox,ATPsynO,SdhD,ND-18,CG3803,ND-51,ND-B18,ND-23,blw | | 4.232e-06 | -12.37 | multi-organism process | biological process | GO:0051704 | 1653 | 202 | 12345 | 1129 | gammaTub37C,bun,CG9925,GlyP,Pex12,fs(1)M3,mop,Rel,Pxt,Cf2,HP1e,tef,disp,CycE,Pen,spn-E,Med,piwi,Lkb1,Tlk,krimp,Topors,dl,tho2,Cul5,PAPLA1,foxo,Top1,Imp,Pdk1,lid,spz,cact,Axs,bcd,tub,SERCA,stil,mael,Tm1,plu,ovo,Pi3K92E,Pp2B-14D,stai,e(y)1,AIF,Pex10,armi,Ptpmeg2,r2d2,Atg6,CkIIalpha,Sp7,csw,lqfR,Hat1,Atg7,ea,muskelin,Vps16B,slam,cort,mts,mad2,Jafrac2,alphaTub84B,eIF4A,Orc1,Pex13,exu,Mos,png,CG13551,Uev1A,shu,osk,twin,yps,CG46338,tyf,apt,fs(1)Ya,BicC,nmd,Grip84,lwr,Cbl,shep,egg,wash,CycB,Nap1,ScsbetaA,Pax,Myd88,tko,Kap3,Rab5,cup,Herc4,sw,CSN6,heph,raptor,mbo,Arf51F,Grip128,eIF3m,Crag,otu,Grip75,slif,bru1,larp,Rbf,squ,Traf6,sau,Syx13,gus,PDCD-5,clt,Sec15,alphaTub67C,fs(1)N,EcR,twe,bif,pgc,enc,del,EMRE,mtrm,Rbfox1,wisp,tkv,Pka-R1,SRPK,PGRP-LE,capt,l(3)mbt,cathD,nej,faf,Nup62,Cdk5,Nprl2,rl,elg1,Jafrac1,spen,Drep1,Dcr-1,psq,mus304,Klp10A,eIF4E1,bam,Abi,yl,msps,Cdk1,yem,grp,Nup358,cnc,BoYb,CycJ,wapl,Tak1,nos,tor,GatA,Mst89B,Tao,Cul2,Cenp-C,Fer1HCH,jvl,Desat1,Rab10,Moe,ben,spd-2,Grip71,dnc,Iml1,Stat92E,toc,Hr78,p38a,swa,Npc2a,E2f1,zpg,shrb,cnn,ari-1,vret,park,milt | | 4.259e-06 | -12.37 | membrane-enclosed lumen | cellular component | GO:0031974 | 1220 | 158 | 11940 | 1105 | mRpS23,Top1,Sdhaf3,CG9143,mRpL13,stil,YL-1,Trf2,gw,Tfb4,Tim8,Nc73EF,Orc6,mRpL18,mRpS21,CG10055,mRpL36,NHP2,TfIIFalpha,Taf7,CG17691,CG14817,scny,e(y)1,CG12288,TrpRS-m,CycC,Tip60,MED4,Chrac-14,CG7488,CG17328,SelR,RpII140,nej,Ubc6,Rpb8,Got1,mts,CG4880,MED27,Roe1,Mybbp1A,tko,fs(1)Ya,Snm1,Clp,mad2,MED24,piwi,dl,vers,rept,Utx,P5CDh1,pont,Non1,Topors,mute,AIF,Thiolase,mRpL10,RYBP,Elba2,Sgf29,CG10336,Mat89Ba,RpL11,gus,DNApol-epsilon255,Idh3b,Gabat,CG14407,mtTFB1,Ptip,RpS18,lwr,Bin1,Tlk,mRpL35,d4,ph-d,Psf3,Nurf-38,Cse1,shop,PAPLA1,pps,Cdk7,Prosbeta5,Hat1,Mtpalpha,MED20,mRpL17,CG12018,Orc1,esc,not,CG8728,mxc,CG11267,SuUR,EloC,Rcd5,HmgD,CG8199,Bap170,Trf4-2,Rel,Nap1,Atu,mRpS11,Hsc70-3,HP1c,Echs1,CG12301,Aldh,mRpS5,hb,CG6766,trr,pn,CG31053,mRpL48,sbr,Cenp-C,mre11,Sgf11,CG11164,Scsalpha1,Sld5,CG4021,RpS2,l(1)G0004,RpS13,mRpS10,RpS15,Spt20,wal,ix,escl,IntS11,lin-52,CG12375,Pdk,Rbfox1,lid,CG6094,mRpL16,Sirt7,mbo,nbs,Rrp40,CG1602,bru1,Su(H),CG46385,bys | | 4.259e-06 | -12.37 | intracellular organelle lumen | cellular component | GO:0070013 | 1220 | 158 | 11940 | 1105 | nbs,mbo,Sirt7,Rrp40,CG1602,Su(H),CG46385,bys,bru1,CG6094,lid,mRpL16,IntS11,escl,lin-52,Pdk,Rbfox1,CG12375,Spt20,wal,ix,RpS2,CG4021,l(1)G0004,RpS13,RpS15,mRpS10,mre11,Cenp-C,CG11164,Scsalpha1,Sgf11,Sld5,CG31053,pn,trr,CG6766,sbr,mRpL48,Aldh,CG12301,mRpS5,hb,Hsc70-3,Echs1,HP1c,Trf4-2,Rel,Nap1,mRpS11,Atu,Rcd5,EloC,Bap170,CG8199,HmgD,CG11267,SuUR,mRpL17,esc,not,Orc1,CG12018,CG8728,mxc,Prosbeta5,Hat1,Mtpalpha,MED20,PAPLA1,Cdk7,pps,ph-d,Psf3,Cse1,shop,Nurf-38,d4,mRpL35,RpS18,Bin1,lwr,Tlk,Idh3b,Gabat,Ptip,mtTFB1,CG14407,RYBP,CG10336,Sgf29,Elba2,gus,RpL11,Mat89Ba,DNApol-epsilon255,Non1,pont,Topors,mute,mRpL10,AIF,Thiolase,mad2,piwi,MED24,dl,vers,P5CDh1,Utx,rept,tko,Mybbp1A,Snm1,fs(1)Ya,Clp,MED27,Roe1,Got1,Rpb8,mts,CG4880,nej,Ubc6,CG17328,SelR,RpII140,TrpRS-m,CG12288,Tip60,CycC,MED4,CG7488,Chrac-14,TfIIFalpha,Taf7,CG17691,scny,e(y)1,CG14817,CG10055,mRpL36,NHP2,Tfb4,Tim8,mRpS21,Orc6,mRpL18,Nc73EF,Top1,mRpS23,CG9143,Sdhaf3,gw,Trf2,stil,YL-1,mRpL13 | | 4.259e-06 | -12.37 | organelle lumen | cellular component | GO:0043233 | 1220 | 158 | 11940 | 1105 | CG4880,mts,Rpb8,Got1,Ubc6,nej,RpII140,SelR,CG17328,Chrac-14,CG7488,MED4,CycC,Tip60,TrpRS-m,CG12288,CG14817,e(y)1,scny,CG17691,Taf7,TfIIFalpha,NHP2,mRpL36,CG10055,Nc73EF,Orc6,mRpL18,mRpS21,Tim8,Tfb4,mRpL13,stil,gw,Trf2,YL-1,CG9143,Sdhaf3,mRpS23,Top1,mRpL35,d4,Tlk,lwr,Bin1,RpS18,CG14407,mtTFB1,Ptip,Gabat,Idh3b,DNApol-epsilon255,RpL11,Mat89Ba,gus,Elba2,Sgf29,CG10336,RYBP,Thiolase,AIF,mRpL10,Topors,mute,pont,Non1,Utx,rept,P5CDh1,dl,vers,MED24,piwi,mad2,Clp,fs(1)Ya,Snm1,Mybbp1A,tko,Roe1,MED27,HP1c,Echs1,Hsc70-3,Atu,mRpS11,Nap1,Rel,Trf4-2,HmgD,CG8199,Bap170,Rcd5,EloC,SuUR,CG11267,CG8728,mxc,Orc1,CG12018,esc,not,mRpL17,MED20,Mtpalpha,Hat1,Prosbeta5,pps,Cdk7,PAPLA1,Nurf-38,Cse1,shop,Psf3,ph-d,CG46385,Su(H),bys,bru1,CG1602,Rrp40,Sirt7,nbs,mbo,mRpL16,lid,CG6094,CG12375,Rbfox1,Pdk,lin-52,IntS11,escl,ix,wal,Spt20,RpS15,mRpS10,RpS13,l(1)G0004,CG4021,RpS2,Sld5,Sgf11,CG11164,Scsalpha1,Cenp-C,mre11,mRpL48,sbr,CG6766,trr,CG31053,pn,hb,mRpS5,CG12301,Aldh | | 4.423e-06 | -12.33 | gamma-tubulin complex | cellular component | GO:0000930 | 13 | 8 | 11940 | 1105 | Grip75,Grip71,Grip84,Uxt,Blos2,CG7716,Grip128,gammaTub37C | | 4.428e-06 | -12.33 | negative regulation of macromolecule biosynthetic process | biological process | GO:0010558 | 446 | 70 | 12345 | 1129 | plu,asf1,ovo,l(3)mbt,insv,Maf1,mael,SuUR,rept,lin-52,smg,Bin1,Uxt,Tip60,ph-d,CG10038,bcd,Rbfox1,CG1602,gw,pgc,Hnf4,EcR,d4,bru1,CG10631,krimp,Tlk,piwi,foxo,CG10336,dl,Rbf,CG2199,heph,Sirt7,cup,sbb,spn-E,CG4872,Chrac-14,BicC,E2f1,apt,twin,Rbf2,wash,egg,Sox102F,CG10959,mRpL13,Elba2,CG17328,cyc,Rga,Unr,CG31875,nos,SREBP,hb,esc,bam,r2d2,psq,Not10,Su(H),Hat1,escl,Mybbp1A,RpL13A | | 4.478e-06 | -12.32 | regulation of cellular amide metabolic process | biological process | GO:0034248 | 186 | 37 | 12345 | 1129 | Dph5,gw,Rbfox1,bcd,wisp,osk,mRpL13,spz,Pi3K92E,smg,mtTFB1,twin,CG7414,cad,PEK,tyf,apt,BicC,RpL13A,Ptip,CG2911,CG8963,Not10,cup,Rel,heph,bam,CG1578,Exd2,Tak1,nos,Unr,Rga,krimp,GAPsec,sip3,bru1 | | 4.903e-06 | -12.23 | regulation of translation | biological process | GO:0006417 | 144 | 31 | 12345 | 1129 | cup,Not10,mRpL13,heph,bam,Dph5,RpL13A,gw,CG8963,CG2911,osk,bcd,Rbfox1,twin,CG7414,Rga,Unr,apt,tyf,GAPsec,PEK,krimp,bru1,sip3,BicC,CG1578,smg,Pi3K92E,Exd2,mtTFB1,nos | | 4.989e-06 | -12.21 | reproductive process | biological process | GO:0022414 | 1436 | 179 | 12345 | 1129 | Grip128,Arf51F,eIF3m,cup,Kap3,Rab5,tko,ix,sw,raptor,heph,CSN6,squ,Rbf,larp,sau,mre11,Bx,Grip75,otu,bru1,fs(1)N,twe,EcR,del,enc,bif,pgc,wisp,Rbfox1,mtrm,Debcl,Syx13,gus,alphaTub67C,clt,Sec15,nej,cathD,c(2)M,Nup62,faf,elg1,Jafrac1,rl,Nprl2,Dcr-1,Drep1,Pka-R1,tkv,SRPK,capt,l(3)mbt,Abi,yl,msps,psq,CanB2,eIF4E1,Klp10A,mus304,bam,BoYb,CycJ,wapl,Xpc,GatA,tor,nos,yem,Cdk1,grp,cnc,Nup358,Cenp-C,Desat1,jvl,spd-2,Rab10,Moe,Iml1,dnc,Grip71,Tao,Mst89B,Cul2,shrb,zpg,ari-1,cnn,vret,milt,park,Stat92E,toc,swa,E2f1,CycE,CG7716,Pen,Med,spn-E,gammaTub37C,Pxt,Pex12,mop,bun,fs(1)M3,HP1e,Cf2,tef,disp,tho2,Top1,PAPLA1,Cul5,Imp,Pdk1,Lkb1,piwi,Topors,krimp,cact,Axs,bcd,lid,spz,Pink1,Pp2B-14D,Pi3K92E,e(y)1,stai,stil,SERCA,Tm1,mael,ovo,plu,csw,Atg7,lqfR,Pex10,armi,r2d2,Ptpmeg2,CkIIalpha,mad2,mus81,mts,Pex13,Orc1,eIF4A,exu,ea,slam,CG31053,muskelin,cort,osk,shu,png,Mos,Grip84,egg,Cbl,lwr,shep,Nap1,CycB,wash,ScsbetaA,Pax,CG46338,twin,yps,apt,nmd,fs(1)Ya,BicC | | 5.074e-06 | -12.19 | GO\_RNA\_CATABOLIC\_PROCESS | MSigDB lists | GO\_RNA\_CATABOLIC\_PROCESS | 129 | 40 | 4161 | 644 | RpS13,RpS30,RpLP0,RpL13A,CG6610,eIF4E1,Rrp40,Not10,mts,CG11164,eIF4A,RpS28b,RpS23,RpS27A,RpL9,twin,Unr,RpL13,RpS16,RpS2,Edc3,RpS24,RpL23,RpL30,CG8963,RpL40,Cbp80,CG10418,RpS7,RpS4,RpL38,RpS27,RpL24,RpL26,Pp2A-29B,RpL11,RpL37a,LSm3,RpL12,RpS18 | | 5.095e-06 | -12.19 | negative regulation of cellular amide metabolic process | biological process | GO:0034249 | 61 | 18 | 12345 | 1129 | bam,heph,mRpL13,cup,Not10,bcd,Rbfox1,gw,RpL13A,bru1,BicC,apt,krimp,Rga,Unr,twin,nos,smg | | 5.341e-06 | -12.14 | proton-transporting ATP synthase complex | cellular component | GO:0045259 | 25 | 11 | 11940 | 1105 | ATPsynE,blw,ATPsynG,ATPsyndelta,ATPsynO,ATPsynB,Neb-cGP,ATPsynF,ATPsynbeta,ATPsynD,ATPsynCF6 | | 5.341e-06 | -12.14 | mitochondrial proton-transporting ATP synthase complex | cellular component | GO:0005753 | 25 | 11 | 11940 | 1105 | ATPsynF,ATPsynD,ATPsynbeta,ATPsynCF6,ATPsynO,ATPsyndelta,ATPsynB,Neb-cGP,ATPsynE,blw,ATPsynG | | 5.355e-06 | -12.14 | piRNA metabolic process | biological process | GO:0034587 | 17 | 9 | 12345 | 1129 | tapas,Gasz,piwi,armi,mael,vret,mino,shu,del | | 5.413e-06 | -12.13 | response to endoplasmic reticulum stress | biological process | GO:0034976 | 79 | 21 | 12345 | 1129 | Mekk1,Marf,S2P,CG6766,Atf6,CG32640,CG9005,Ufm1,CG8974,CG11070,Hsc70-3,shep,Edem1,Cdk5,Calx,ERp60,Parp16,PHGPx,sip3,cnc,PEK | | 5.455e-06 | -12.12 | negative regulation of cellular biosynthetic process | biological process | GO:0031327 | 457 | 71 | 12345 | 1129 | nos,esc,hb,SREBP,cyc,CG31875,Unr,Rga,escl,Mybbp1A,Hat1,RpL13A,r2d2,bam,Su(H),Not10,psq,wash,CG10959,Sox102F,egg,Cbl,apt,E2f1,BicC,Rbf2,twin,CG17328,Elba2,mRpL13,Rbf,dl,CG10336,foxo,krimp,bru1,CG10631,d4,piwi,Tlk,CG4872,spn-E,sbb,Chrac-14,heph,CG2199,cup,Sirt7,SuUR,smg,lin-52,rept,l(3)mbt,ovo,asf1,plu,Maf1,mael,insv,CG1602,Rbfox1,bcd,Hnf4,EcR,pgc,gw,Bin1,CG10038,ph-d,Uxt,Tip60 | | 5.455e-06 | -12.12 | negative regulation of biosynthetic process | biological process | GO:0009890 | 457 | 71 | 12345 | 1129 | Hat1,escl,Mybbp1A,RpL13A,bam,r2d2,psq,Su(H),Not10,nos,SREBP,hb,esc,cyc,Rga,Unr,CG31875,Elba2,CG17328,mRpL13,wash,egg,Cbl,Sox102F,CG10959,BicC,E2f1,apt,twin,Rbf2,sbb,spn-E,CG4872,Chrac-14,CG2199,heph,Sirt7,cup,foxo,CG10336,dl,Rbf,d4,CG10631,bru1,krimp,Tlk,piwi,bcd,Rbfox1,CG1602,pgc,gw,Hnf4,EcR,Bin1,Uxt,Tip60,ph-d,CG10038,SuUR,rept,lin-52,smg,asf1,plu,ovo,l(3)mbt,insv,mael,Maf1 | | 5.468e-06 | -12.12 | positive regulation of cell cycle | biological process | GO:0045787 | 73 | 20 | 12345 | 1129 | CG31053,SkpA,E2f1,bru1,otu,Nprl2,SuUR,foxo,APC7,Sld5,Su(H),armi,SNF4Agamma,png,ben,Moe,Iml1,CycE,twe,COX5A | | 5.481e-06 | -12.11 | MODULE\_151 | MSigDB lists | MODULE\_151 | 171 | 49 | 4161 | 644 | eEF1beta,UQCR-Q,RpL13,Idh3b,Nap1,RpS23,RpL9,RpS27A,RpS28b,Arpc2,Rab10,eIF3i,CG31715,Nurf-38,CG6610,RpL13A,Rpn12,Sod1,RpS13,ste24a,RpLP0,Mcad,Kap-alpha3,RpL12,ND-24,CG13630,CG15715,CCT3,Jafrac2,ND-B18,CG14715,RpL26,RpL24,RpS27,LSm3,Gyg,Prosalpha3,eIF3m,RpL11,Snx6,RpL40,Prosbeta6,RpL38,RpS4,Phb2,RpS16,RpS2,RpL23,RpS24 | | 5.484e-06 | -12.11 | response to stress | biological process | GO:0006950 | 1136 | 147 | 12345 | 1129 | Desat1,lt,Iml1,ben,SNF4Agamma,Trxr-1,Tao,whd,ZnT35C,ATPsynD,Ufm1,DNApol-epsilon255,RecQ5,park,Sod1,Stat92E,p38a,Hr78,Npc2a,trsn,Idgf6,fray,S2P,CG6766,CG5844,Arpc4,Atf6,CG9005,Seipin,Sod3,Utx,mus304,Xpc,kcc,Ubc6,Tak1,grp,mus205,Cdk1,Sdhaf3,tos,cnc,Ptp61F,EMRE,chrb,Debcl,PDCD-5,nbs,agt,faf,Cdk5,Edem1,cathD,spen,Dcr-1,Nprl2,rl,Jafrac1,mol,CG5285,PGRP-LE,Grx1,Crag,Herc4,tko,Myd88,UGP,raptor,CG10336,Traf6,Rbf,mre11,PHGPx,slif,alph,EMC3,Mekk1,Marf,Ect4,NFAT,CG30010,CG17746,CG32640,CG13551,Uev1A,CG11070,Hsc70-3,shep,thoc6,Mitf,ERp60,Stam,CG46338,Snm1,Hsp26,PEK,tyf,Sp7,Hat1,Atg7,Gs1l,Idh,Atg6,Sld5,Ho,r2d2,Idgf4,SREBP,Jafrac2,mus81,ago,ND-SGDH,sip3,CG11089,cyc,tub,tzn,Xpac,Droj2,spz,Pink1,CG8974,Mtl,adp,Pi3K92E,AIF,Calx,Tm1,obe,Chrac-14,MAGE,REPTOR,Tom40,CG9925,GlyP,bun,Pxt,Rel,Mtpalpha,Tfb4,foxo,tho2,dl,Imp,Parp16,Cka,cold | | 6.005e-06 | -12.02 | meiotic nuclear division | biological process | GO:0140013 | 203 | 39 | 12345 | 1129 | msps,Grip128,Arf51F,eIF4E1,Klp10A,mus304,bam,tef,gammaTub37C,cup,Rab5,Pex10,wapl,Xpc,nos,larp,mus81,sau,Topors,CG31053,cort,yem,Grip75,Cdk1,grp,Axs,wisp,mtrm,Grip71,Cenp-C,twe,bif,Syx13,Pp2B-14D,c(2)M,cnn,lwr,SRPK,mael | | 6.070e-06 | -12.01 | cup (cup) | protein interactions | 33934 | 62 | 21 | 9204 | 1112 | PGRP-LE,Rel,tral,gus,smg,exu,CG40045,twin,Tak1,nos,bru1,Tango10,BicC,tos,yps,cnn,osk,otu,Pdk,eIF4E1,Rga | | 6.070e-06 | -12.01 | Sod1 (Superoxide dismutase 1) | protein interactions | 39251 | 62 | 21 | 9204 | 1112 | Tak1,CG10638,CG14715,cib,Cyp1,park,Sod1,Fkbp12,Pink1,awd,Rel,Chd64,Trxr-1,Marf,PGRP-LE,Eno,Got1,regucalcin,Treh,CG1532,AnxB10 | | 6.070e-06 | -12.01 | Rpt4 (Regulatory particle triple-A ATPase 4) | protein interactions | 31567 | 62 | 21 | 9204 | 1112 | Droj2,CG1416,Pi3K92E,Rpn3,foxo,GCS2beta,Prosbeta6,Rpt2,Prosalpha4,Rpn2,CCT6,Pomp,Prosalpha5,Pdk1,mts,Rpn13,CCT3,Prosalpha3,Prosbeta3,Prosbeta4,Prosbeta5 | | 6.512e-06 | -11.94 | Pglym78 (Phosphoglyceromutase 78) | protein interactions | 43447 | 53 | 19 | 9204 | 1112 | RhoGDI,CG14715,CG10638,Pgk,PHGPx,cib,Adh,Chd64,awd,Eno,Scp1,GlyP,CG2493,Argk,regucalcin,scu,Moe,AnxB10,CG4598 | | 6.576e-06 | -11.93 | ribonucleoprotein complex assembly | biological process | GO:0022618 | 139 | 30 | 12345 | 1129 | Usp39,osk,spn-E,RpL10,RpL12,SmE,RpS28b,CG8204,r2d2,RpLP0,RpS14a,RpL24,CG7488,RpS5b,RpL11,RpS14b,RpS27,CG3198,LSm3,Dcr-1,RpS15,pont,rept,CG10055,CG17768,bru1,rig,Edc3,mRpS11,RpL23A | | 6.693e-06 | -11.91 | regulation of cell death | biological process | GO:0010941 | 257 | 46 | 12345 | 1129 | Pink1,DCAF12,fwe,Tip60,PDCD-5,RYBP,Idh3b,Debcl,Ect4,ben,EcR,dco,E2f1,SkpA,Cul1,Rab32,Naa20A,Idgf2,Drep1,MED24,park,ScsbetaA,rept,Cbl,lin-52,Tspo,Utx,bun,ix,Drep2,Aldh,velo,Atg7,csw,CG4238,eIF3m,CycE,MAGE,spag,scny,Lkb1,nos,Pdk1,Ubc6,Jafrac2,Rbf | | 6.830e-06 | -11.89 | Gpdh1 (Glycerol-3-phosphate dehydrogenase 1) | protein interactions | 33824 | 72 | 23 | 9204 | 1112 | Cyt-c-p,CG32473,ATPsynO,Argk,Nc73EF,GlyP,ATPsynB,Pglym78,Scp1,Gapdh1,Eno,ND-18,CG7920,Adh,Mf,up,Actn,Gk1,Pgk,ATPsynF,Fer1HCH,COX4,ATPsynCF6 | | 6.850e-06 | -11.89 | MODULE\_114 | MSigDB lists | MODULE\_114 | 177 | 50 | 4161 | 644 | RpS28b,RpS23,RpS27A,RpL9,Nap1,Rab10,Arpc2,RpL13,UQCR-Q,eEF1beta,Rpn12,RpS30,Sod1,RpS13,RpLP0,RpL13A,CG6610,Nurf-38,Mcad,RhoGDI,CG31715,eIF3i,RpS27,RpL24,RpL26,eIF3m,RpL11,Gyg,Prosalpha3,LSm3,CG15715,ND-24,CG13630,RpL12,Kap-alpha3,CG14715,ND-B18,CCT3,Jafrac2,RpS16,RpS2,Phb2,Pfdn5,RpS24,RpL23,RpL40,Snx6,RpL38,Prosbeta6,RpS4 | | 7.092e-06 | -11.86 | COX5A (Cytochrome c oxidase subunit 5A) | protein interactions | 41432 | 28 | 13 | 9204 | 1112 | ATPsynF,Cul1,ago,RpS28b,levy,Prosbeta6,COX4,cype,RpLP1,lwr,RpL9,SNF4Agamma,CG17163 | | 7.416e-06 | -11.81 | response to organic substance | biological process | GO:0010033 | 529 | 79 | 12345 | 1129 | aay,Smox,CG32640,Tao,whd,Hsc70-3,CG11070,G9a,Desat1,Rab10,dnc,Pdxk,Stat92E,Shmt,PEK,apt,Hr78,p38a,dco,lwr,MED24,Nurf-38,Atf6,Utx,CG17734,csw,AcCoAS,CG6766,Aldh,Vps16B,cyc,sip3,Nup358,SREBP,Tak1,MED27,Idh3b,Cyp6v1,CG8974,nbs,EcR,Hnf4,PyK,tub,Pfk,Pka-R1,Fdh,tkv,CG5412,Trf2,Pi3K92E,rept,Edem1,rl,Fkbp39,CG7328,ACC,Pka-R2,Agpat2,bun,Chd64,raptor,CG4390,Adh,Arf51F,MAGE,sgll,Med,REPTOR,Bx,PHGPx,alph,rig,mAChR-A,dl,trr,foxo,Parp16,Pdk1 | | 7.609e-06 | -11.79 | small molecule catabolic process | biological process | GO:0044282 | 154 | 32 | 12345 | 1129 | Thiolase,CG17691,Fdh,SERCA,Shmt,wal,CG7840,Mcad,Pgm1,tzn,Pfk,CG8199,Etf-QO,CG10932,Ntan1,CG4586,CG9391,Echs1,Gabat,Acox57D-p,CG4598,Synj,CG4594,P5CDh1,Mtpalpha,Pex10,cN-IIIB,yip2,CG4860,Seipin,CG4390,Adh | | 7.657e-06 | -11.78 | mAcon1 (Mitochondrial aconitase 1) | protein interactions | 44149 | 49 | 18 | 9204 | 1112 | Pink1,Adh,CG7920,up,Mf,Actn,ATPsynF,Trs33,ATPsynCF6,Cyt-c-p,ATPsynO,Argk,Nc73EF,ATPsynB,Eno,Scp1,ND-18,ATPsynD | | 8.082e-06 | -11.73 | GstO3 (Glutathione S transferase O3) | protein interactions | 38972 | 63 | 21 | 9204 | 1112 | PHGPx,EndoGI,cib,Fkbp12,CG10638,CG14715,Chd64,awd,Trxr-1,yip2,Stip1,RhoGDI,scu,Moe,capt,Gdi,regucalcin,CG4598,AnxB10,Eno,Uch | | 8.209e-06 | -11.71 | GO\_PHOSPHORYLATION | MSigDB lists | GO\_PHOSPHORYLATION | 376 | 89 | 4161 | 644 | Nc73EF,Pi4KIIalpha,Abi,RpS27A,Mekk1,ND-75,Top1,UQCR-Q,CG7115,Prosalpha4,csw,Rpt2,CG7530,CG3631,Rpn12,nmdyn-D7,levy,CG4041,Gk2,Prosalpha3,COX5A,Prosbeta5,Jafrac2,Pfk,ND-24,CG2201,CkIIalpha,CycC,Cyt-c-p,Traf6,CG6767,r,Nprl2,fray,SelD,CG8726,CG17698,Pdk,ND-39,tkv,Mo25,Prosalpha5,CG3803,Tfb4,rl,l(1)G0196,Cdk1,Pgk,CG9391,Cdk7,Pgm1,Prosbeta3,Gapdh1,ND-49,CG33156,ND-SGDH,Rpn2,grp,ND-B22,CG8475,SkpA,ND-51,ND-23,CG7789,CG7414,Prosbeta4,CycE,PpD3,CG32549,Eno,Cdk5,CycB,Pdk1,SRPK,Pdxk,p38a,ND-B18,ND-30,Pglym78,nmdyn-D6,Rpn3,Cul1,dco,alc,Tlk,Pi3K92E,Prosbeta6,RpL40,Tao | | 8.563e-06 | -11.67 | Bacc (Bacchus) | protein interactions | 33462 | 68 | 22 | 9204 | 1112 | HmgD,RhoGDI,Stip1,CG10638,CG14715,Fkbp12,PHGPx,cib,grsm,Tip60,Chd64,CG11267,awd,Ho,Pglym78,Eno,ago,Moe,capt,scu,AnxB10,pr | | 8.751e-06 | -11.65 | Tak1 (TGF-beta activated kinase 1) | protein interactions | 39659 | 153 | 38 | 9204 | 1112 | Chd64,vret,RpL12,SERCA,CCT6,cup,cact,pont,Traf6,blw,CCT3,RpS16,RpS27,ben,CG1416,Nup358,rept,larp,RpS28b,GCS2beta,CG4557,Sgf29,RpS14a,PGRP-LE,Sod1,CG4853,alphaTub84B,RpL23A,stai,Tak1,RpL7A,alph,scu,Idh3b,trbd,Jafrac1,Cp1,CCT5 | | 8.915e-06 | -11.63 | Gp93 (Glycoprotein 93) | protein interactions | 43354 | 83 | 25 | 9204 | 1112 | CCT3,blw,Pp2A-29B,spag,Fer1HCH,cup,CCT6,SERCA,Kap-alpha3,CG10425,CG1092,GCS2beta,Cp1,Rtnl1,mEFTu1,wal,Vha68-2,rl,Rab5,Droj2,CG1416,ScsbetaA,Mcad,CG9186,cathD | | 8.979e-06 | -11.62 | GO\_T\_CELL\_RECEPTOR\_SIGNALING\_PATHWAY | MSigDB lists | GO\_T\_CELL\_RECEPTOR\_SIGNALING\_PATHWAY | 59 | 23 | 4161 | 644 | Uev1A,Cul1,Rpn3,Prosbeta5,Pdk1,rl,cact,Prosalpha5,RpS27A,Prosalpha3,Prosbeta4,RpL40,dnc,Stoml2,SkpA,Prosbeta6,Pi3K92E,Traf6,Rpn12,Rpn2,Prosbeta3,Prosalpha4,Rpt2 | | 9.518e-06 | -11.56 | cellular component biogenesis | biological process | GO:0044085 | 1270 | 160 | 12345 | 1129 | crok,Imp,RpS15,Pdk1,Atpalpha,Lkb1,cold,RpL23A,CG10055,up,CycE,Arpc2,CG7716,Rpn13,Chrac-14,TMEM216,Nlg1,Hip1,spn-E,gammaTub37C,mop,fs(1)M3,bun,Pxt,Pp2A-29B,Cf2,NHP2,rept,Mtl,stai,mtTFB1,bys,mael,Tm1,Trf2,CG8611,cact,Pfk,CG11110,Liprin-beta,RpS7,CCT6,mts,Orc1,exu,Mat89Ba,Vps16B,RpL34a,ND-18,Hat1,Atg7,armi,SelR,RpS14b,Ndc1,r2d2,Atg6,RpS14a,Grip84,wash,Nap1,parvin,Pax,Edc3,Tomosyn,Shmt,RpL7A,SmE,RpL12,osk,Rcd5,RpS5b,Scox,tws,RpLP0,larp,Rbf,squ,sau,LSm3,CG11562,mRpS11,Sgf29,Grip75,rig,bru1,CG17768,Grip128,Clc,Cchl,Crag,Kap3,TBCC,mbo,CG5969,ND-23,pont,nej,CG9143,rl,CG3198,SuUR,Dcr-1,tkv,RpL34b,CG3437,BigH1,SRPK,Taf7,CG12301,asf1,Oseg1,fs(1)N,Usp39,wisp,Non1,RpL26,gus,Rrp40,alphaTub67C,RpS16,CG1890,RpS27,Sdhaf3,yem,grp,Unr,PIG-C,Caf1-105,Pomp,cnc,CG12288,Abi,RpS28b,msps,Arpc4,CanB2,psq,RpL11,CG13366,Klp10A,CG7728,CG7488,cnn,milt,CG10465,Actn,CG14671,E2f1,swa,RpL10,Cenp-C,CG8204,jvl,ben,Moe,Grip71,mRpL10,Snx6,RpL24 | | 9.631e-06 | -11.55 | phosphoric ester hydrolase activity | molecular function | GO:0042578 | 221 | 41 | 12176 | 1119 | CG9391,ssh,Ptp61F,nSMase,CG17746,mts,Acph-1,CG7115,aay,CG18135,Pka-R2,twe,Pp2A-29B,dnc,Ptpmeg,Ptp10D,csw,Pp2B-14D,fal,mop,pn,cN-IIIB,Ptpmeg2,PrBP,Ublcp1,CanA-14F,Pld,Pka-R1,CG32549,Pfrx,PpD3,CG5026,CG6707,alph,PpV,Ptp69D,Dd,Synj,CG7789,tws,CG17294 | | 1.000e-05 | -11.51 | oxidoreductase complex | cellular component | GO:1990204 | 100 | 24 | 11940 | 1105 | ND-18,ND-SGDH,ND-B18,CG17691,ND-39,ND-MNLL,CG40472,ND-49,ND-30,ND-B15,SdhA,ND-51,UQCR-11,NP15.6,ND-23,UQCR-Q,ND-B22,ND-24,ND-75,CG8199,Nc73EF,ND-MWFE,wal,SdhD | | 1.029e-05 | -11.48 | proteasome-mediated ubiquitin-dependent protein catabolic process | biological process | GO:0043161 | 200 | 38 | 12345 | 1129 | Rpn3,Prosalpha3,Rpn2,Rpn12,CG9316,Cul2,CG11070,Prosbeta3,Hsc70-3,gus,CG31633,RYBP,CG32640,Fbw5,CG32221,CG8188,Edem1,SkpA,Prosbeta4,Prosalpha5,Cul1,Prosbeta6,CG4238,Plap,CG6766,Rpn13,Uch-L5,Rpt2,CG5003,ago,Ubc6,CG40045,Prosbeta5,Cul5,Prosalpha4,sip3,Pomp,cort | | 1.031e-05 | -11.48 | REACTOME\_RESPIRATORY\_ELECTRON\_TRANSPORT\_ATP\_SYNTHESIS\_BY\_CHEMIOSMOTIC\_COUPLING\_AND\_HEAT\_PRODUCTION\_BY\_UNCOUPLING\_PROTEINS\_ | MSigDB lists | REACTOME\_RESPIRATORY\_ELECTRON\_TRANSPORT\_ATP\_SYNTHESIS\_BY\_CHEMIOSMOTIC\_COUPLING\_AND\_HEAT\_PRODUCTION\_BY\_UNCOUPLING\_PROTEINS\_ | 37 | 17 | 4161 | 644 | Etf-QO,levy,ND-B22,ND-51,ND-23,ND-49,Cyt-c-p,ND-SGDH,ND-75,wal,SdhA,ND-B18,ND-24,ND-30,UQCR-Q,ND-39,COX5A | | 1.031e-05 | -11.48 | REACTOME\_RESPIRATORY\_ELECTRON\_TRANSPORT | MSigDB lists | REACTOME\_RESPIRATORY\_ELECTRON\_TRANSPORT | 37 | 17 | 4161 | 644 | wal,ND-75,SdhA,ND-B18,ND-24,ND-30,UQCR-Q,ND-39,COX5A,Etf-QO,levy,ND-B22,ND-51,ND-23,ND-49,ND-SGDH,Cyt-c-p | | 1.047e-05 | -11.47 | Tab2 (TAK1-associated binding protein 2) | protein interactions | 37216 | 183 | 43 | 9204 | 1112 | RpS28b,sbb,GCS2beta,eIF4A,shv,rept,RpL11,bru1,Nup358,RpS27,RpS16,blw,Traf6,Kcmf1,CCT3,pont,CG9391,CCT6,RpS18,SERCA,RpL13,AMPdeam,RpS2,RpL12,RpL17,CCT5,Jafrac1,trbd,scu,zf30C,Hsc70-3,Pfdn5,Tak1,RpL24,RpL23A,alphaTub84B,RpL23,Sod1,Actn,RpL38,RpS14a,CG4557,RpL9 | | 1.049e-05 | -11.46 | Gdh (Glutamate dehydrogenase) | protein interactions | 42832 | 25 | 12 | 9204 | 1112 | ATPsynD,CG15564,Scp1,Rtnl1,ATPsynB,ATPsynF,Actn,Nc73EF,Argk,Mf,up,Adh | | 1.063e-05 | -11.45 | dorsal/ventral axis specification | biological process | GO:0009950 | 76 | 20 | 12345 | 1129 | snk,spz,CkIIalpha,Myd88,gd,gammaTub37C,csw,cact,spn-E,Med,jvl,enc,krimp,cnc,ea,mael,milt,dl,faf,Cbl | | 1.066e-05 | -11.45 | CG9117 (uncharacterized protein) | protein interactions | 33846 | 9 | 7 | 9204 | 1112 | RpL17,RpS14b,Mitofilin,RpS2,RpS14a,CycB,Stip1 | | 1.144e-05 | -11.38 | Hsc70Cb (Hsc70Cb) | protein interactions | 39557 | 29 | 13 | 9204 | 1112 | Eno,Stip1,Moe,cib,EndoGI,CG10638,CG14715,Gdi,regucalcin,awd,Chd64,AnxB10,yip2 | | 1.208e-05 | -11.32 | organonitrogen compound biosynthetic process | biological process | GO:1901566 | 1162 | 148 | 12345 | 1129 | mRpL13,RpL24,ATPsynD,aay,AMPdeam,CG7009,mRpL10,mRpS23,blw,RpL10,TyrRS,mal,RpS23,Pcd,CG10425,CG17691,r-l,RpL17,CG6767,mEFTu1,mRpL17,RpS30,Ts,Dhfr,eIF4E1,CG6178,Got1,RpL11,mRpS10,mRpS21,AcCoAS,r,CG2201,CG8314,SerRS-m,RpS28b,GAPsec,PIG-C,RpL9,RpS27,CG11134,GatA,RpL13,fdl,ATPsynCF6,Nadsyn,RpS16,RpL26,CG14817,Ac3,RpS27A,mRpL16,RpL40,CG7840,CG7414,RpL34b,Adi1,CG4407,tko,RpLP1,mRpL18,P5cr,CG33156,ATPsyndelta,GatB,ATPsynB,ATPsynG,eIF3m,EMC3,GluRS-m,RpL30,CG3803,mRpS11,eEF1beta,RpS13,CG5880,RpS2,RpS29,Ppox,RpLP0,CG13551,mRpL48,ATPsynE,RpS5b,CG8199,Pdxk,RpL12,RpL38,Pfas,nmdyn-D7,CG33977,RpL7A,Shmt,mEFTu2,CysRS,Efr,ATPsynbeta,RpS14a,RpL23,Ady43A,RpS18,RpS14b,Atg7,RpS24,RpL13A,awd,l(2)k01209,RpL34a,PIG-H,CG11089,eIF4A,RpL37a,oxt,TrpRS-m,Nmt,SelD,RpS7,mRpL36,Pym,AsnS,ATPsynO,mRpL35,GalT1,RpL29,CG10344,mRpS5,PIG-L,CG31523,CG11899,RpS4,CG6094,Ppcdc,LysRS,Gclc,RpL31,PIG-P,sgll,Ggt-1,Hexo1,Sptr,O-fut1,RpL23A,pr,eIF3i,nmdyn-D6,RpS15,COX5B,PAPLA1 | | 1.211e-05 | -11.32 | transferase complex | cellular component | GO:1990234 | 496 | 75 | 11940 | 1105 | Ptip,escl,gus,CG31633,DNApol-epsilon255,Spt20,Sgf29,SkpA,d4,lid,MAGE,CycB,trr,ScsbetaA,mus205,DCAF12,plu,APC7,pont,Cul5,rept,l(3)73Ah,Utx,CG6767,ago,CG11070,Sgf11,Pfk,CycJ,RpII140,EloC,Atg6,Rcd5,CG30496,Rep,Chrac-14,Tip60,Rpb8,CG32221,CG5003,CycE,CG9316,Atu,Fbw5,Ubc6,ben,CG8475,nej,Nc73EF,Pi3K92E,Cdk7,ari-1,park,CG33977,Tfb4,Cul2,Cdk5,PIG-C,YL-1,Trf2,PIG-H,Naa20A,Cul1,ph-d,CG12018,not,esc,e(y)1,TfIIFalpha,Taf7,Uev1A,CG10254,png,alc,sip3 | | 1.228e-05 | -11.31 | proteasomal protein catabolic process | biological process | GO:0010498 | 209 | 39 | 12345 | 1129 | Cul1,Prosalpha5,Prosbeta4,SkpA,Edem1,CG8188,CG32221,Fbw5,gus,CG32640,RYBP,CG31633,Cul2,CG8974,Hsc70-3,CG11070,Prosbeta3,CG9316,Rpn2,Prosalpha3,Rpn12,Rpn3,cort,Pomp,sip3,Prosalpha4,Prosbeta5,Cul5,CG40045,Ubc6,ago,CG5003,Rpt2,Rpn13,Uch-L5,Plap,CG6766,Prosbeta6,CG4238 | | 1.246e-05 | -11.29 | smg (smaug) | protein interactions | 39034 | 46 | 17 | 9204 | 1112 | nos,BicC,bru1,ND-75,Nap1,ND-24,eIF4E1,osk,cup,SdhA,ATPsynbeta,AGO3,SdhD,UQCR-11,twin,Unr,ATPsynF | | 1.248e-05 | -11.29 | GO\_ANTIGEN\_RECEPTOR\_MEDIATED\_SIGNALING\_PATHWAY | MSigDB lists | GO\_ANTIGEN\_RECEPTOR\_MEDIATED\_SIGNALING\_PATHWAY | 60 | 23 | 4161 | 644 | Uev1A,Cul1,Rpn3,Prosbeta5,rl,cact,Pdk1,RpS27A,Prosalpha5,Prosalpha3,Prosbeta4,dnc,RpL40,Pi3K92E,Prosbeta6,SkpA,Stoml2,Traf6,Rpn12,Rpn2,Prosbeta3,Rpt2,Prosalpha4 | | 1.277e-05 | -11.27 | oxidoreductase activity, acting on the CH-OH group of donors, NAD or NADP as acceptor | molecular function | GO:0016616 | 102 | 24 | 12176 | 1119 | scu,Idh3b,CG4842,CG2767,CG10638,sni,CG9265,CG31549,Fdh,CG31674,CG10425,CG7322,CG12171,Gmer,CG9436,Sptr,CG10863,CG31548,Mtpalpha,CG3842,Thiolase,Idh,CG5028,Adh | | 1.339e-05 | -11.22 | metal ion binding | molecular function | GO:0046872 | 1047 | 136 | 12176 | 1119 | QC,Scox,mre11,Rpn2,Cyp6v1,CG9941,Edem1,EcR,krimp,not,Reps,Dip-B,CG17272,CG6767,Fdh,CAH7,HDAC6,Calx,Pex10,CG2678,Nurf-38,CG9425,Cyp6a22,DhpD,mal,Iru,ScsbetaA,Idh,aay,Clp,nej,Actn,CG8728,dx,Mctp,Usp16-45,CG15141,ph-d,SP1029,CG11267,CG44422,isoQC,regucalcin,Rpn3,Scp1,CG10463,CG8974,stw,LpR2,Cyp6u1,CG14435,Cyp6a19,CG13344,Marc,Shmt,Desat1,Wdfy2,cN-IIIB,Fer1HCH,CG17328,CSN6,CG9684,CG31053,Eno,PGRP-LE,CAH2,Pgm1,up,CanB2,CG32473,CG9581,Hr78,alpha-Man-Ia,bor,stet,Utx,CG6923,nos,AOX3,Sec23,lt,Tak1,Traf6,awd,Tengl4,alph,GluRS-m,CG6813,DNApol-epsilon255,RpS29,Sod3,CG10654,tkv,CG17260,Kcmf1,l(3)73Ah,Pex12,CG31729,SelR,CG10492,shop,CG11590,G9a,Topors,sip3,ACC,yl,Usp39,d4,AnxB10,egg,CG4730,CG9925,glob1,ari-1,CG17698,ATP8A,grsm,CG17593,l(3)mbt,CG9981,SmydA-1,CG4709,Nca,Hnf4,CG11710,PAPLA1,PyK,Cbl,CG11134,CG14516,r,Sod1,MtnA,Gs1l,CG11902 | | 1.356e-05 | -11.21 | kinase activity | molecular function | GO:0016301 | 375 | 60 | 12176 | 1119 | CG11486,l(1)G0196,Mos,nmdyn-D7,PEK,CG3631,Pdxk,Cdk1,CycE,CG8726,Pi4KIIalpha,grp,Mekk1,Tfb4,p38a,Argk,tub,Cdk5,Tao,CkIIalpha,l(2)k01209,CG8475,tor,rl,dco,Pfrx,awd,Cdk7,Pfk,Tak1,Pi3K92E,Pgk,Pink1,CG17698,Pdk,Mo25,Tlk,CG1344,CG2201,SelD,Lkb1,nmdyn-D6,CycJ,Gk1,fray,CG7328,CycB,Pdk1,Gk2,SRPK,Ady43A,kri,tkv,alc,Scsalpha1,CG8311,CG33156,Coq8,CG4041,png | | 1.376e-05 | -11.19 | negative regulation of oskar mRNA translation | biological process | GO:0007319 | 8 | 6 | 12345 | 1129 | krimp,apt,heph,BicC,bru1,cup | | 1.392e-05 | -11.18 | macromolecule localization | biological process | GO:0033036 | 868 | 116 | 12345 | 1129 | CG6766,CG8314,sws,Arpc1,msps,Tom70,fray,scramb1,Nxf3,Atg6,Seipin,CkIIalpha,opm,Klp10A,armi,Pex10,exu,Tango10,Pex13,CG9393,sip3,cnc,Nup358,CG31729,Mat89Ba,GAPsec,slam,Vps16B,Tim8,Grip71,blw,osk,Rab10,Moe,Cse1,ATP8A,jvl,CG8204,lt,Cenp-C,CG8004,SNF4Agamma,thoc6,milt,Pax,Stam,CG9186,Tsp86D,BicC,Npc2a,swa,Gmer,Actn,Rab32,Vsp37A,spn-E,Nlg1,Crag,Sec23,Sec22,CG14894,Arts,Tom40,Pi4KIIalpha,Clc,Pen,CG5734,eIF3m,Arf51F,Grip128,mbo,CHMP2B,Pex12,sw,CG9981,Karybeta3,Rab5,cup,tho2,sau,CG4572,Topors,Grip75,Kap-alpha3,Lkb1,CG4041,wisp,cact,CHOp24,Arl5,Sec15,Gga,CG8728,alphaTub67C,Rrp40,X11L,AP-1-2beta,CG11110,gus,Syx13,CG5447,boi,SuUR,Dcr-1,Nup62,smg,Oseg1,Roe1,Rep,Gdi,CG3077,mol,mael,Tm1,sbr,CG10344 | | 1.425e-05 | -11.16 | NADH dehydrogenase complex | cellular component | GO:0030964 | 53 | 16 | 11940 | 1105 | ND-30,ND-B15,ND-39,ND-49,CG40472,ND-MNLL,ND-B18,ND-SGDH,ND-18,ND-B22,ND-24,ND-23,NP15.6,ND-51,ND-MWFE,ND-75 | | 1.425e-05 | -11.16 | respiratory chain complex I | cellular component | GO:0045271 | 53 | 16 | 11940 | 1105 | NP15.6,ND-51,ND-B22,ND-24,ND-23,ND-B18,ND-18,ND-SGDH,ND-30,ND-B15,ND-39,ND-49,CG40472,ND-MNLL,ND-75,ND-MWFE | | 1.425e-05 | -11.16 | mitochondrial respiratory chain complex I | cellular component | GO:0005747 | 53 | 16 | 11940 | 1105 | ND-B18,ND-18,ND-SGDH,ND-B15,ND-30,ND-49,CG40472,ND-MNLL,ND-39,NP15.6,ND-51,ND-B22,ND-24,ND-23,ND-75,ND-MWFE | | 1.444e-05 | -11.15 | Thor (thor) | protein interactions | 33569 | 70 | 22 | 9204 | 1112 | RpL13A,foxo,RpL17,eIF4E1,Ect4,yps,raptor,RpL30,RpS23,RpL7A,Hsc70-3,RpL23,RpS7,RpL10,Cul1,CG8728,park,RpL13,Pink1,RpL9,RpS14a,alt | | 1.458e-05 | -11.14 | Tm2 (Tropomyosin 2) | protein interactions | 41853 | 75 | 23 | 9204 | 1112 | Sld5,ATPsynCF6,Exo70,COX4,Tm1,ATPsynF,ATPsyndelta,SERCA,Actn,up,Pp2B-14D,Mf,Adh,CG7920,ND-18,Eno,Scp1,osk,ATPsynB,Nc73EF,Argk,ATPsynO,Cyt-c-p | | 1.463e-05 | -11.13 | P-body | cellular component | GO:0000932 | 37 | 13 | 11940 | 1105 | Edc3,LSm3,exu,Rga,CG11486,CG10418,cup,smg,gw,CG17068,eIF4E1,CG17768,tral | | 1.467e-05 | -11.13 | GO\_CYTOSOLIC\_PART | MSigDB lists | GO\_CYTOSOLIC\_PART | 107 | 34 | 4161 | 644 | RpL13A,RpS13,RpS30,RpLP0,Rpn2,Rpt2,RpL13,CCT6,Pi4KIIalpha,RpS28b,RpL9,RpS23,RpS27A,RpL40,RpS7,RpS4,RpL38,RpS16,RpS2,RpS24,RpL30,RpL23,RpL12,mRpS5,CCT5,Eno,CCT3,RpS18,Pfk,RpL24,RpS27,RpL26,RpL37a,RpL11 | | 1.472e-05 | -11.13 | UQCR-C2 (Ubiquinol-cytochrome c reductase core protein 2) | protein interactions | 39846 | 117 | 31 | 9204 | 1112 | GCS2beta,Rpt2,wal,mEFTu1,Tspo,Karybeta3,bor,Rab5,Droj2,CG1416,csw,PyK,ScsbetaA,Mcad,cathD,Atpalpha,UQCR-Q,COX4,CCT3,blw,Pp2A-29B,CG15098,ATPsynbeta,CCT6,SERCA,CG10425,Kap-alpha3,Pdk1,UQCR-11,PGRP-LE,Rel | | 1.482e-05 | -11.12 | cellular glucan metabolic process | biological process | GO:0006073 | 23 | 10 | 12345 | 1129 | Pgm1,CG9485,foxo,AGBE,CG8475,UGP,Atg6,Gyg,GlyP,ACC | | 1.482e-05 | -11.12 | glucan metabolic process | biological process | GO:0044042 | 23 | 10 | 12345 | 1129 | Gyg,UGP,Atg6,ACC,GlyP,CG9485,Pgm1,CG8475,AGBE,foxo | | 1.482e-05 | -11.12 | energy reserve metabolic process | biological process | GO:0006112 | 23 | 10 | 12345 | 1129 | AGBE,foxo,CG8475,CG9485,Pgm1,GlyP,ACC,UGP,Atg6,Gyg | | 1.482e-05 | -11.12 | glycogen metabolic process | biological process | GO:0005977 | 23 | 10 | 12345 | 1129 | Pgm1,CG9485,CG8475,AGBE,foxo,Gyg,UGP,Atg6,ACC,GlyP | | 1.487e-05 | -11.12 | lipid catabolic process | biological process | GO:0016042 | 110 | 25 | 12345 | 1129 | Sply,Hnf4,Mcad,CG18135,CG4594,CG7997,yip2,dob,Pex10,Mtpalpha,Seipin,CG5966,CG4860,Etf-QO,CG4267,CG10932,Thiolase,Mitf,Echs1,CG4586,Acox57D-p,wal,SERCA,CG4598,CG7840 | | 1.516e-05 | -11.10 | GO\_MITOCHONDRION | MSigDB lists | GO\_MITOCHONDRION | 715 | 149 | 4161 | 644 | CG8974,ND-30,mRpS5,Mtpalpha,ND-B18,CG8004,sws,CG14407,yip2,CG4872,Tspo,mtTFB1,mRpS23,colt,CG31759,AnxB10,rl,Cdk1,Tom40,CG3803,wal,Got1,CG1774,SERCA,Unr,ND-39,Scsalpha1,Fdh,CG5028,ND-51,Stoml2,GluRS-m,CG3842,scu,sau,mino,ND-SGDH,CG9393,ND-49,CG3021,Cisd2,mRpL18,Mcad,Cdk7,Coq7,Jafrac2,pr,Hsc70-3,CG7488,mRpL10,Mitofilin,CanA-14F,Mat89Ba,CanB2,Traf6,CG8728,P5CDh1,UQCR-Q,Ect4,Calx,SerRS-m,mRpS10,Rab32,Pi4KIIalpha,GatA,mRpS21,TrpRS-m,Idh,Nc73EF,ACC,CG11089,Gk2,CG7550,tzn,Coq6,CG13601,nmdyn-D6,Thiolase,shop,Marf,p38a,Pfdn2,SelR,Sccpdh1,Pmi,stet,CG9581,Roe1,CG1640,CG1532,Shmt,milt,SdhA,Aldh,CG8199,Ppox,Irp-1B,Fkbp59,CG10932,CG4407,ND-23,CG17691,ND-B22,Sod1,CG2100,Prosbeta3,ND-24,CHMP2B,Tim8,Pp2A-29B,Gabat,COX5A,Scox,mRpL17,Pdk,CG12375,Cbp80,tko,Cyp1,Atg4b,Ts,Etf-QO,nmd,Pp2B-14D,CG11110,Phb2,ScsbetaA,Cyt-c-p,bor,Atg6,CG11858,CG7718,P5cr,mRpL16,mEFTu1,AIF,Echs1,ND-75,Idh3b,Trxr-1,Nmt,Grx1,levy,LysRS,mRpL13,Sfxn2,CG10189,csw,mts | | 1.526e-05 | -11.09 | magnesium ion binding | molecular function | GO:0000287 | 48 | 15 | 12176 | 1119 | Pgm1,Eno,ATP8A,CG31729,Nurf-38,Idh,CG9981,aay,alph,Tak1,CG6767,Gs1l,awd,cN-IIIB,PyK | | 1.608e-05 | -11.04 | fly chr88F1-88F1\|3-57 cM | chromosome location | fly chr88F1-88F1\|3-57 cM | 23 | 8 | 25010 | 1343 | Surf4,ea,CG42727,Atg4b,Trs33,Fkbp39,SerRS-m,CG6276 | | 1.608e-05 | -11.04 | fly chr88F1-88F1\|3-57 | chromosome location | fly chr88F1-88F1\|3-57 | 23 | 8 | 25010 | 1343 | CG6276,Trs33,Atg4b,SerRS-m,Fkbp39,CG42727,ea,Surf4 | | 1.618e-05 | -11.03 | microtubule organizing center organization | biological process | GO:0031023 | 78 | 20 | 12345 | 1129 | cbs,CG3437,mael,grp,Topors,SkpA,Grip84,mts,cnn,Prosbeta5,Ubc6,Rcd5,Pp2A-29B,CCT6,armi,Klp10A,tws,spd-2,msps,Grip71 | | 1.657e-05 | -11.01 | RNA metabolic process | biological process | GO:0016070 | 908 | 120 | 12345 | 1129 | CG3021,mino,CG3198,e(y)1,spen,mtTFB1,Dcr-1,CG11486,CG9143,smg,Taf7,CG12301,Trf2,TfIIFalpha,sbr,obe,bys,CG5986,Rnmt,mael,Spx,wisp,Usp39,Archease,gw,del,TrpRS-m,Rrp40,Bin1,RpS16,CG10418,CG30343,SelD,Pym,RpS7,CG2100,LSm3,CG17187,Imp,CG10189,tho2,RpII140,GluRS-m,Trf4-2,pps,rig,bru1,IntS6,CG17768,piwi,mRpS11,Sgf29,CG10463,snRNA:U4atac,CG17454,Neos,GatB,CG3587,NHP2,heph,Tfb4,LysRS,Cdk7,Pus7,vret,Nurf-38,CysRS,MED24,cu,trsn,CG9684,scu,Cbp80,CG12375,CG14100,Edc3,twin,yps,RpL7A,CG1622,Larp7,shu,CycC,CG16865,SmE,TyrRS,Son,CG8204,Tengl4,tapas,CG7009,CG31759,CG2034,CG6610,pygo,Clp,nos,GatA,exu,MED4,Lsm12,Rpb8,snRNA:U4:38AB,Gasz,Mat89Ba,snRNA:U12,Rga,IntS11,CG2911,SerRS-m,CG5292,RpS28b,eIF4E1,CG7728,r2d2,RpS14a,Not10,RpS14b,armi,CG11164 | | 1.671e-05 | -11.00 | cofactor biosynthetic process | biological process | GO:0051188 | 104 | 24 | 12345 | 1129 | Nadsyn,Dhfr,CG4407,Coq6,CG6178,Ppcdc,CG13551,Coq7,ND-39,CG8199,regucalcin,Gclc,AcCoAS,CG33156,Pdxk,sgll,mal,Pcd,CG3803,Sptr,CG17691,pr,Ppox,Coq8 | | 1.681e-05 | -10.99 | GO\_PROTEIN\_LOCALIZATION\_TO\_ORGANELLE | MSigDB lists | GO\_PROTEIN\_LOCALIZATION\_TO\_ORGANELLE | 262 | 66 | 4161 | 644 | RpL40,Tspo,RpS7,RpS4,RpL38,RpS16,Pex13,lt,RpS24,Dd,CG1847,RpL23,RpL12,Marf,Kap-alpha3,CG8004,RpS18,RpS27,RpL26,Arf51F,RpL11,Arts,GCC88,Karybeta3,CG9393,sau,CG2100,Tom40,cact,rl,Cdk1,RpS23,Rab10,shrb,mbo,RpS2,Atg6,Pp2B-14D,CG11110,Phb2,Oseg1,Vha36-1,Pex10,CG8728,RpL30,Tim8,RpL24,RpL37a,CanA-14F,Hsc70-3,Pex12,CG31729,rept,cyc,RpL13A,RpLP0,RpS13,RpS30,Cse1,CG7530,RpL13,Pen,ERp60,RpS28b,RpL9,RpS27A | | 1.684e-05 | -10.99 | negative regulation of cellular macromolecule biosynthetic process | biological process | GO:2000113 | 438 | 67 | 12345 | 1129 | SuUR,rept,lin-52,smg,asf1,plu,ovo,l(3)mbt,mael,Maf1,insv,Rbfox1,bcd,CG1602,pgc,gw,EcR,Hnf4,Bin1,Tip60,ph-d,CG10038,foxo,CG10336,dl,Rbf,d4,bru1,CG10631,krimp,Tlk,piwi,sbb,spn-E,CG4872,Chrac-14,CG2199,heph,Sirt7,cup,wash,egg,Sox102F,CG10959,BicC,E2f1,apt,twin,Rbf2,Elba2,CG17328,mRpL13,nos,SREBP,hb,esc,cyc,Unr,Rga,Hat1,escl,RpL13A,bam,r2d2,psq,Not10,Su(H) | | 1.728e-05 | -10.97 | MORF\_RAN | MSigDB lists | MORF\_RAN | 168 | 47 | 4161 | 644 | Sod1,RpS30,Rpn12,RpS13,LysRS,Prosbeta3,Rpt2,rept,ND-51,CHOp24,Rpb8,levy,Stoml2,SkpA,RpS27A,Idh3b,Nap1,CSN6,RpL13,mEFTu1,Pgk,Chrac-14,SdhA,Cyt-c-p,Phb2,RpS24,RpL30,Dd,tko,heph,CG10418,Pka-R1,Prosbeta6,RpL24,Clc,COX5A,Pp2A-29B,RpL11,Prosalpha3,eIF3m,Prosbeta4,CG13630,lwr,ND-30,Thiolase,Pglym78,CCT3 | | 1.779e-05 | -10.94 | GO\_PROTEIN\_LOCALIZATION\_TO\_ENDOPLASMIC\_RETICULUM | MSigDB lists | GO\_PROTEIN\_LOCALIZATION\_TO\_ENDOPLASMIC\_RETICULUM | 86 | 29 | 4161 | 644 | RpS24,RpL23,RpL30,RpS13,RpLP0,RpS30,RpS16,RpL13A,RpS2,RpS7,RpL38,RpS4,RpL40,RpL11,RpL37a,shrb,Rab10,Hsc70-3,RpL24,RpS27,RpS28b,RpL9,RpS23,RpS27A,RpL26,RpS18,RpL13,RpL12,ERp60 | | 1.855e-05 | -10.89 | GO\_OXIDOREDUCTASE\_ACTIVITY | MSigDB lists | GO\_OXIDOREDUCTASE\_ACTIVITY | 304 | 74 | 4161 | 644 | ND-39,cl,Ppox,Utx,CG3803,wal,Aldh,SdhA,CG8199,Cyp6a19,Mcad,ND-49,Gapdh1,ND-SGDH,Sod1,CG9265,scu,CG3842,CG17691,ND-B22,lid,ND-51,ND-23,Jafrac1,Fdh,CG5028,CG4872,CG14407,Cyp6a22,Sccpdh1,SelR,Gdi,Mtpalpha,ND-B18,ND-30,shop,Thiolase,Dhfr,CG10463,Rep,CG7840,Sptr,Nc73EF,Desat1,Idh,Idh3b,Pcd,Trxr-1,ND-75,CG3609,CG9471,AIF,ERp60,UQCR-Q,CG6144,Gmer,Coq6,CG7550,CG4562,levy,Grx1,COX5A,CG10425,CG6665,Jafrac2,Coq7,CG7322,ND-24,P5cr,CG1275,P5CDh1,sgll,Etf-QO,Adi1 | | 1.867e-05 | -10.89 | small molecule biosynthetic process | biological process | GO:0044283 | 228 | 41 | 12345 | 1129 | pr,Sptr,Coq8,foxo,CG10932,SREBP,Pgk,CG11134,nmdyn-D6,fal,l(1)G0196,ACC,Pxt,cN-IIIB,Dhfr,Got1,CG6178,sgll,awd,AcCoAS,P5cr,r,Shmt,Pcd,nmdyn-D7,Npc2a,CG7840,Adi1,CG31523,CG11899,ND-39,SelD,aay,regucalcin,Coq6,Coq7,CG17270,Desat1,Marf,AsnS,Pdxk | | 1.924e-05 | -10.86 | REACTOME\_DARPP\_32\_EVENTS | MSigDB lists | REACTOME\_DARPP\_32\_EVENTS | 13 | 9 | 4161 | 644 | dnc,Cdk5,Pka-R1,CanB2,Pp2A-29B,Pp2B-14D,Pka-R2,CanA-14F,mts | | 1.954e-05 | -10.84 | catalytic activity, acting on a protein | molecular function | GO:0140096 | 1481 | 181 | 12176 | 1119 | Ubc6,CycJ,Prosalpha5,fray,ste24a,ari-1,ste14,CG8188,CG17698,Atg4b,Mo25,png,grsm,UbcE2H,oxt,SRPK,Ptpmeg2,Ptpmeg,sol,CG8726,Prosbeta3,Cbl,CG14516,Mos,cathD,CG9643,Cp1,ben,park,CG17746,rl,CG4238,CG11771,grp,Ser7,p38a,Dd,Tlk,CG2200,CG10344,Cdk7,APC7,Tak1,Traf6,alph,PpV,Pdk,tkv,Coq8,Kcmf1,Ublcp1,Pex12,CG5390,CycB,Pdk1,Pp2A-29B,ago,Cdk1,EloC,DppIII,CG31633,Sp7,G9a,Topors,CG7115,Herc4,sip3,Uch-L5,O-fut1,STUB1,CG3631,Usp39,ERp60,tor,Prosbeta4,Uch,CG2924,mts,egg,CG11110,CG5880,CG46338,tub,Prosalpha4,Fkbp59,CG8314,Lkb1,CG2493,Ubc4,CG8974,dco,CG11070,CG4572,otu,scny,ea,CG5026,Pink1,CG14435,CanA-14F,CG13344,Prosbeta5,CG31053,twe,Uev1A,CSN6,CG9316,CycE,Nepl3,snk,CG3511,CG32473,Pp2B-14D,CG11486,CG40045,Usp10,CG9581,lid,PEK,stet,Gapdh1,Sirt7,CG3894,CG32221,CG10254,CG8475,CG5003,Utx,Tfb4,faf,esc,Ptp61F,Ptp69D,tws,CG13630,Fbw5,QC,PpD3,Rpn2,Cyp1,CG9941,Prosbeta6,CG4502,SkpA,alc,CG4041,not,MAGE,Dip-B,gd,CG16979,mop,lwr,Pex10,HDAC6,velo,S2P,Ggt-1,Prosalpha3,escl,Fkbp12,Iru,Ptp10D,csw,CG11858,trr,trbd,dx,CG8728,Usp16-45,CG15141,26-29-p,Mekk1,SP1029,ssh,CG9953,isoQC,Tao,Fur2,Cdk5,CkIIalpha | | 1.983e-05 | -10.83 | Rpt5 (Regulatory particle triple-A ATPase 5) | protein interactions | 42805 | 52 | 18 | 9204 | 1112 | Pdk1,Rpn13,RpS27A,Rpn2,CCT6,Prosalpha5,Pomp,Prosbeta5,Prosalpha3,CCT3,Prosbeta3,Prosbeta4,Rpn3,CG1416,Prosalpha4,Prosbeta6,wal,Rpt2 | | 2.023e-05 | -10.81 | GO\_ESTABLISHMENT\_OF\_PROTEIN\_LOCALIZATION\_TO\_ENDOPLASMIC\_RETICULUM | MSigDB lists | GO\_ESTABLISHMENT\_OF\_PROTEIN\_LOCALIZATION\_TO\_ENDOPLASMIC\_RETICULUM | 78 | 27 | 4161 | 644 | RpL40,RpS7,RpL38,RpS4,RpS16,RpL13A,RpS2,RpS13,RpS30,RpLP0,RpS24,RpL30,RpL23,RpL12,RpL13,RpS18,RpS28b,RpL24,RpS27,RpS27A,RpL26,RpS23,RpL9,RpL37a,RpL11,shrb,Rab10 | | 2.029e-05 | -10.81 | RhoGDI (RhoGDI) | protein interactions | 40179 | 43 | 16 | 9204 | 1112 | cib,PHGPx,CG14715,CG10638,Chd64,awd,Pglym78,Eno,CG2767,Uch,scu,capt,Moe,GstT1,regucalcin,AnxB10 | | 2.029e-05 | -10.81 | EndoGI (Endonuclease G inhibitor) | protein interactions | 34953 | 43 | 16 | 9204 | 1112 | Stip1,CG10863,cib,PHGPx,CG10638,CG14715,awd,yip2,Eno,Uch,capt,Moe,scu,Gdi,regucalcin,AnxB10 | | 2.132e-05 | -10.76 | GO\_CYTOSOLIC\_RIBOSOME | MSigDB lists | GO\_CYTOSOLIC\_RIBOSOME | 74 | 26 | 4161 | 644 | RpS27,RpL24,RpS28b,RpS27A,RpL26,RpS23,RpL9,RpL11,RpL37a,RpL13,mRpS5,RpL12,RpS18,RpLP0,RpS13,RpS30,RpL13A,RpS2,RpS16,RpS24,RpL30,RpL23,RpL40,RpS7,RpL38,RpS4 | | 2.180e-05 | -10.73 | cellular component assembly | biological process | GO:0022607 | 1109 | 141 | 12345 | 1129 | mts,Orc1,exu,Vps16B,ND-18,Hat1,Atg7,RpS14b,SelR,armi,Ndc1,r2d2,Atg6,RpS14a,Grip84,wash,Nap1,parvin,Pax,Edc3,Tomosyn,Shmt,SmE,RpL12,osk,Rcd5,RpS5b,Scox,tws,RpLP0,crok,Imp,RpS15,Pdk1,Atpalpha,Lkb1,cold,RpL23A,CG10055,up,CycE,Arpc2,CG7716,Chrac-14,Rpn13,TMEM216,Nlg1,Hip1,spn-E,gammaTub37C,mop,bun,fs(1)M3,Pp2A-29B,Pxt,Cf2,rept,Mtl,stai,mael,Tm1,Trf2,cact,Pfk,CG11110,Liprin-beta,CCT6,CG1890,RpS27,yem,Sdhaf3,grp,Unr,Caf1-105,Pomp,cnc,Abi,RpS28b,msps,psq,CanB2,Arpc4,RpL11,CG13366,Klp10A,CG7488,cnn,milt,CG10465,Actn,CG14671,E2f1,swa,Cenp-C,RpL10,CG8204,jvl,Moe,ben,Grip71,Snx6,RpL24,larp,squ,Rbf,sau,LSm3,CG11562,mRpS11,Sgf29,Grip75,rig,CG17768,bru1,Grip128,Clc,Cchl,Crag,Kap3,TBCC,CG5969,pont,nej,ND-23,CG3198,rl,SuUR,Dcr-1,tkv,CG3437,SRPK,BigH1,Taf7,asf1,Oseg1,fs(1)N,Non1,wisp,Usp39,gus,alphaTub67C | | 2.182e-05 | -10.73 | protein ubiquitination | biological process | GO:0016567 | 245 | 43 | 12345 | 1129 | ago,Ubc6,CG2924,CG40045,Traf6,Cul5,Topors,sip3,Iru,l(3)73Ah,CG4238,Atg7,STUB1,CG10254,CG13344,Pex10,Herc4,Pex12,CG5003,CG32221,Fbw5,park,CG8188,ari-1,Cbl,RpL40,SkpA,CG46338,UbcE2H,Cul1,Ubc4,ben,RpS27A,CG9316,EloC,Cul2,CG11070,CG8974,dx,Uev1A,CG9941,CG31633,CG14435 | | 2.204e-05 | -10.72 | CG33303 (uncharacterized protein) | protein interactions | 2768916 | 87 | 25 | 9204 | 1112 | GCS2beta,Cp1,r,Rpt2,mEFTu1,wal,Vha68-2,Tspo,Rab5,PyK,CG10470,CG1416,ScsbetaA,Mcad,cathD,Atpalpha,CCT3,blw,Pp2A-29B,SERCA,CCT6,Kap-alpha3,CG10425,PGRP-LE,Rel | | 2.246e-05 | -10.70 | heterocycle biosynthetic process | biological process | GO:0018130 | 476 | 71 | 12345 | 1129 | nmdyn-D6,Ppox,Rbf,COX5B,Sptr,CG3803,pps,RpII140,pr,Sgf29,ATPsyndelta,CG33156,P5cr,sgll,ATPsynB,CycE,ATPsynG,Ppcdc,Tfb4,CG4407,Scsalpha1,Cdk7,cN-IIIB,mtTFB1,spen,e(y)1,TfIIFalpha,Trf2,Taf7,ATPsynO,Nadsyn,Ac3,fal,Orc1,pygo,Rpb8,ATPsynCF6,MED4,CG11089,l(2)k01209,mus205,Rga,AcCoAS,r,awd,Dhfr,ATPsynbeta,CG6178,Ady43A,CG6767,Nurf-38,MED24,Ts,nmdyn-D7,E2f1,Pcd,Gmer,Pfas,Shmt,r-l,CG17691,Rab32,Pdxk,blw,lt,mal,CG13551,ATPsynD,ATPsynE,AMPdeam,CG8199 | | 2.312e-05 | -10.68 | regulation of immune system process | biological process | GO:0002682 | 230 | 41 | 12345 | 1129 | scny,slif,dl,foxo,Traf6,Tak1,Myd88,mop,GlyP,CG9925,Herc4,Rel,Atg6,Sp7,Ptip,Hat1,Crag,Atg7,Atu,Stat92E,Aos1,cad,PGRP-LE,p38a,Hr78,E2f1,faf,lwr,Nurf-38,RYBP,PDCD-5,spz,G9a,TyrRS,EcR,EMRE,cact,tub,Ect4,crq,wisp | | 2.333e-05 | -10.67 | ATP synthesis coupled proton transport | biological process | GO:0015986 | 24 | 10 | 12345 | 1129 | ATPsynD,ATPsynE,ATPsynbeta,ATPsyndelta,blw,ATPsynO,ATPsynG,COX5B,ATPsynCF6,ATPsynB | | 2.333e-05 | -10.67 | regulation of transposition | biological process | GO:0010528 | 24 | 10 | 12345 | 1129 | BoYb,AGO3,vret,tral,spn-E,piwi,mael,tapas,Nup358,ovo | | 2.333e-05 | -10.67 | energy coupled proton transport, down electrochemical gradient | biological process | GO:0015985 | 24 | 10 | 12345 | 1129 | ATPsynE,ATPsynD,ATPsynbeta,COX5B,ATPsynG,ATPsynB,ATPsynCF6,ATPsyndelta,blw,ATPsynO | | 2.343e-05 | -10.66 | regulation of autophagy | biological process | GO:0010506 | 106 | 24 | 12345 | 1129 | foxo,SREBP,Traf6,Pi3K92E,Mitf,Pax,rl,ago,Nprl2,Tak1,Sod1,Fkbp39,scny,Rab32,PEK,Synj,Desat1,EcR,Debcl,CycC,Atg7,AP-1-2beta,Smox,Utx | | 2.434e-05 | -10.62 | HONMA\_DOCETAXEL\_RESISTANCE | MSigDB lists | HONMA\_DOCETAXEL\_RESISTANCE | 16 | 10 | 4161 | 644 | mRpL17,Gapdh1,Hsc70-3,APC7,PDCD-5,Ufm1,Eno,RpL38,CG4332,ND-30 | | 2.453e-05 | -10.62 | GO\_PROTEIN\_TARGETING\_TO\_MEMBRANE | MSigDB lists | GO\_PROTEIN\_TARGETING\_TO\_MEMBRANE | 96 | 31 | 4161 | 644 | RpL13A,RpS2,RpS16,RpLP0,RpS30,RpS13,RpL30,RpL23,RpS24,RpL40,Atg4b,ste14,RpS4,RpL38,RpS7,Rep,CHOp24,Arl6IP1,RpS27A,RpS23,RpL9,RpL26,RpS28b,RpL24,RpS27,shrb,RpL37a,RpL11,RpL12,RpL13,RpS18 | | 2.461e-05 | -10.61 | REACTOME\_3\_UTR\_MEDIATED\_TRANSLATIONAL\_REGULATION | MSigDB lists | REACTOME\_3\_UTR\_MEDIATED\_TRANSLATIONAL\_REGULATION | 83 | 28 | 4161 | 644 | RpL13,RpL12,RpS18,RpS27A,RpL26,RpS23,RpL9,RpS27,RpL24,RpS28b,RpL11,RpL37a,RpL40,RpS4,RpL38,eIF3i,RpS7,RpS30,RpS13,RpLP0,RpL13A,RpS2,RpS16,eIF4E1,eIF4A,RpL23,RpL30,RpS24 | | 2.497e-05 | -10.60 | organic cyclic compound biosynthetic process | biological process | GO:1901362 | 512 | 75 | 12345 | 1129 | spen,mtTFB1,e(y)1,TfIIFalpha,Taf7,Trf2,ATPsynO,CG17270,Nadsyn,Ac3,nmdyn-D6,Ppox,foxo,COX5B,Rbf,Sptr,CG3803,RpII140,pps,pr,Sgf29,ATPsyndelta,P5cr,CG33156,sgll,CycE,ATPsynG,ATPsynB,Tfb4,Ppcdc,Scsalpha1,CG4407,cN-IIIB,Cdk7,Nurf-38,MED24,CG6767,Ts,nmdyn-D7,Npc2a,E2f1,Pcd,Gmer,Pfas,Shmt,r-l,Rab32,CG17691,Pdxk,blw,mal,Marf,lt,ATPsynD,ATPsynE,CG13551,CG8199,AMPdeam,fal,pygo,Orc1,Rpb8,MED4,ATPsynCF6,CG11089,l(2)k01209,Rga,mus205,r,AcCoAS,awd,CG6178,ATPsynbeta,Dhfr,Ady43A | | 2.511e-05 | -10.59 | aromatic compound biosynthetic process | biological process | GO:0019438 | 469 | 70 | 12345 | 1129 | Rpb8,MED4,ATPsynCF6,fal,Orc1,pygo,mus205,Rga,CG11089,l(2)k01209,awd,AcCoAS,r,Ady43A,Dhfr,ATPsynbeta,CG6178,Ts,CG6767,MED24,Nurf-38,r-l,Shmt,Rab32,CG17691,E2f1,nmdyn-D7,Pfas,Gmer,Pcd,lt,mal,Pdxk,blw,AMPdeam,CG8199,CG13551,ATPsynE,ATPsynD,Ppox,Rbf,COX5B,nmdyn-D6,pr,Sgf29,Sptr,CG3803,pps,RpII140,sgll,ATPsynB,CycE,ATPsynG,ATPsyndelta,CG33156,Cdk7,cN-IIIB,Ppcdc,Tfb4,Scsalpha1,CG4407,mtTFB1,spen,e(y)1,TfIIFalpha,Trf2,Taf7,ATPsynO,Ac3,Nadsyn | | 2.699e-05 | -10.52 | GO\_NUCLEOBASE\_CONTAINING\_SMALL\_MOLECULE\_METABOLIC\_PROCESS | MSigDB lists | GO\_NUCLEOBASE\_CONTAINING\_SMALL\_MOLECULE\_METABOLIC\_PROCESS | 235 | 60 | 4161 | 644 | CG11134,Ppcdc,Pfas,CG7789,CG32549,COX5A,r-l,Ac3,CG7322,Pfk,ND-B18,Eno,Pglym78,nmdyn-D6,ND-30,ND-24,Dhfr,CG5285,mal,CG6767,Cyt-c-p,Ts,CG17333,CG4511,DhpD,r,ND-39,Scsalpha1,Nc73EF,Idh,Idh3b,kri,rl,CG3803,ND-75,UQCR-Q,Shmt,Pgk,Gapdh1,CG33156,ND-49,CG11164,Pgm1,Prip,AMPdeam,nmdyn-D7,LysRS,Nadsyn,ND-SGDH,ND-B22,Stoml2,levy,CG5028,CG4407,ACC,ND-23,ND-51,CG11089,dnc,ste14 | | 2.744e-05 | -10.50 | DIAZ\_CHRONIC\_MEYLOGENOUS\_LEUKEMIA\_UP | MSigDB lists | DIAZ\_CHRONIC\_MEYLOGENOUS\_LEUKEMIA\_UP | 665 | 139 | 4161 | 644 | CanA-14F,Mitofilin,CG10425,Pfk,Jafrac2,PIG-C,CG8963,P5CDh1,CG6767,faf,r,AGBE,fray,CG9485,SelD,Arl5,Nc73EF,Pcd,CG9471,Rpt2,Tsp66E,eIF4E1,ste14,CG11089,yip2,Hmg-2,Vamp7,Gclc,CG8004,alph,Mtpalpha,Hat1,Kap-alpha3,Rpn3,Moe,Kap3,alc,Sec10,Pka-R1,Ubc4,CG4511,Prosbeta6,RfC38,Med,Tm1,Scsalpha1,CG1774,Unr,Grip75,YL-1,wal,CG3803,rl,CG9391,Mcad,Cdk7,ste24a,ND-SGDH,sau,RpII140,Rpn2,CHOp24,CG42542,Karybeta3,Stoml2,CG8475,SkpA,CG11070,Fdh,Prosalpha3,Pp2A-29B,Prosbeta5,CCT3,ND-24,CG12018,CkIIalpha,TyrRS,MED4,Atg6,ScsbetaA,Cyt-c-p,Phb2,Pp2B-14D,CG40045,CG17454,Etf-QO,PpV,not,Cbp80,scramb1,Plap,Nmt,Idh3b,Echs1,Cul5,AIF,Top1,Chrac-14,mts,Rpn12,LysRS,CG6610,nmdyn-D7,CG4562,eIF3i,DppIII,eIF3m,SREBP,Prosbeta4,Cyp6a22,Arpc1,pont,SRPK,Thiolase,shop,Non1,lt,heph,Fkbp59,Mo25,Ppox,Irp-1B,Klp10A,Uch-L5,CCT6,Cyp6a19,Pgk,Shmt,mad2,CG3226,Sod1,HDAC6,isoQC,parvin,CG17691,CG4572,Su(H),l(1)G0004,CG10932 | | 2.749e-05 | -10.50 | Sod2 (Superoxide dismutase 2 (Mn)) | protein interactions | 36878 | 16 | 9 | 9204 | 1112 | Pink1,ND-75,tws,park,Sod1,Sdhaf3,Idh,Uch,Eno | | 2.754e-05 | -10.50 | ribonucleoprotein complex subunit organization | biological process | GO:0071826 | 149 | 30 | 12345 | 1129 | RpL11,RpS14b,RpS5b,RpL24,RpS14a,CG7488,RpLP0,r2d2,RpS28b,CG8204,RpL12,RpL10,SmE,Usp39,osk,spn-E,RpL23A,Edc3,mRpS11,CG17768,bru1,rig,CG10055,rept,pont,Dcr-1,RpS15,RpS27,LSm3,CG3198 | | 2.754e-05 | -10.50 | ubiquitin-like protein transferase activity | molecular function | GO:0019787 | 254 | 44 | 12176 | 1119 | Cbl,CG40045,Topors,Herc4,sip3,STUB1,CG31053,Uev1A,CG9316,ago,EloC,CG31633,Iru,CG4238,CG46338,dx,ben,CG3894,CG32221,CG10254,park,CG5003,CG15141,CG2924,ari-1,APC7,CG11070,Traf6,CG8188,CG9941,CG4502,SkpA,CG14435,Ubc6,Fbw5,Ubc4,CG8974,UbcE2H,lwr,Pex10,Kcmf1,MAGE,CG13344,Pex12 | | 2.755e-05 | -10.50 | Cyt-c1 (Cytochrome c1) | protein interactions | 38612 | 31 | 13 | 9204 | 1112 | COX5A,ATPsynF,Scp1,UQCR-Q,ATPsynCF6,Adh,up,UQCR-11,Mf,Nc73EF,Argk,regucalcin,Actn | | 2.799e-05 | -10.48 | GSE2405\_0H\_VS\_24H\_A\_PHAGOCYTOPHILUM\_STIM\_NEUTROPHIL\_UP | MSigDB lists | GSE2405\_0H\_VS\_24H\_A\_PHAGOCYTOPHILUM\_STIM\_NEUTROPHIL\_UP | 101 | 32 | 4161 | 644 | eIF3i,levy,RpL13A,Sod1,RpLP0,eEF1beta,RpL13,Arpc2,Npc2a,RpL9,RpS23,RpL38,RpS4,RpS7,RpL30,RpL23,RpS16,RpS2,CG1890,CG4390,Mtpalpha,RpS18,Ntan1,RpL12,Hsc70-3,Arf51F,RpL37a,RpL11,COX5A,RpL26,RpS27,RpL24 | | 2.857e-05 | -10.46 | Nacalpha (Nascent polypeptide associated complex protein alpha subunit) | protein interactions | 36409 | 83 | 24 | 9204 | 1112 | Fkbp12,PHGPx,cib,EndoGI,CG14715,CG10638,ATPsynbeta,RpLP0,awd,Chd64,CG11267,Echs1,CG32549,HmgD,Stip1,capt,Moe,scu,regucalcin,AnxB10,CG4598,Eno,Pglym78,Uch | | 2.873e-05 | -10.46 | oocyte axis specification | biological process | GO:0007309 | 135 | 28 | 12345 | 1129 | psq,armi,gammaTub37C,gus,spz,CkIIalpha,heph,enc,jvl,Grip128,osk,spn-E,Grip71,Moe,msps,Tm1,mael,Lkb1,Grip75,ea,Pka-R1,cnc,swa,capt,krimp,milt,exu,nos | | 2.896e-05 | -10.45 | response to alcohol | biological process | GO:0097305 | 128 | 27 | 12345 | 1129 | rept,trr,MED24,Nurf-38,Pdk1,rl,Shmt,Pka-R1,rig,Trf2,apt,CG17734,Arf51F,EcR,dnc,Aldh,AcCoAS,Tao,whd,Idh3b,Pka-R2,aay,MED27,Agpat2,G9a,Adh,Utx | | 2.956e-05 | -10.43 | PCNA (Proliferating cell nuclear antigen) | protein interactions | 37290 | 78 | 23 | 9204 | 1112 | Arf51F,RhoGDI,SuUR,squ,CG11267,lwr,Chd64,awd,grp,cib,PHGPx,Fkbp12,CG14715,CG10638,Cul2,Uch,Eno,CG4598,AnxB10,Moe,capt,scu,regucalcin | | 3.029e-05 | -10.40 | ATPsynbeta (ATP synthase, beta subunit) | protein interactions | 43829 | 68 | 21 | 9204 | 1112 | ATPsynCF6,blw,ATPsynE,ATPsyndelta,ATPsynF,smg,park,SERCA,Fkbp12,CG10638,Chd64,awd,ATPsynG,ATPsynD,Eno,ATPsynB,csw,regucalcin,Sod3,ATPsynO,AnxB10 | | 3.035e-05 | -10.40 | heterocyclic compound binding | molecular function | GO:1901363 | 3069 | 339 | 12176 | 1119 | Cdk5,ATPsynbeta,CG7728,TrpRS-m,mEFTu2,Marf1,Mekk1,Klp10A,Son,Non1,obe,TfIIFalpha,Idh,CG4586,Clp,alphaTub67C,cnc,RpL13A,AcCoAS,nmd,e(y)1,AIF,Pus7,SREBP,eIF3m,r2d2,CG6767,sgll,armi,Atf3,SERCA,CG2162,CG4041,mRpL18,Spx,NHP2,Cyp6v1,CG17187,EcR,CG6276,tos,RpL24,CG31689,CG2100,crm,RpS18,SNF4Agamma,CG10631,nos,yps,Utx,bys,Ho,Mybbp1A,Gapdh1,Rel,Neos,bcd,RpS4,eIF4E1,elg1,Xpc,Atf6,Trxr-1,CG17328,Shmt,CG6094,Marc,wal,CG34116,Cyp6a19,piwi,eEF1beta,TyrRS,Pink1,bam,dco,Pfrx,CG10463,E23,CCT3,SelD,CG1640,mus81,CG12018,egg,insv,Hsc70-3,CCT6,CCT5,CG17454,CG4021,pre-lola-G,CG6610,spen,mRpS11,Arl5,RpL12,Nxf3,Taf7,mus205,CG14997,Trf2,CG4562,Rbfox1,psq,Top1,CG31729,SdhA,Cdk1,mRpL16,dl,RpL7A,cup,Gabat,Pdk1,Mitf,CG10654,RpS15,Cbp80,Tengl4,Kyat,Rcd5,Cdk7,Tak1,Arf51F,hng3,RpL9,RpL26,Dhfr,Bap170,Synj,CG6967,Orc6,brwl,Rab32,Got1,mael,grp,sbb,rept,Irp-1B,rl,CG4860,r,Rbf2,Roe1,CG10038,CG31875,SerRS-m,CG5789,Mcad,CG9981,SRPK,RYBP,jigr1,Larp7,sbr,png,anne,ND-51,ATP8A,Ssdp,mRpS5,CG17698,glob1,mRpL13,CG15514,Rab10,Tao,CkIIalpha,BicC,Vha68-2,SdhD,Fkbp39,Mat89Ba,mEFTu1,l(1)G0004,apt,CG14100,ScsbetaA,Nc73EF,CG10336,Elba2,Rbf,mal,HmgD,Cyp6a22,CG9425,RpS16,E2f1,CG2678,wdn,swa,Exd2,spn-E,RpL23,qkr58E-2,CG12375,LSm3,Atpalpha,Pxt,trsn,Wbp2,foxo,GlyP,ovo,Rrp40,Cyt-c-p,Droj2,alphaTub84B,cana,Edc3,CG17768,shep,Orc1,NFAT,BigH1,Sirt7,PDCD-5,Snm1,bor,Su(H),lid,PEK,nmdyn-D7,Sply,CG11486,Hr78,SLIRP1,LysRS,Xpac,CysRS,eIF4A,cyc,Dcr-1,osk,AGO3,heph,Marf,CG5116,RpII140,Rab5,CG4266,Cyp6u1,Imp,Pym,CG4880,Lkb1,RpS14a,CG10959,CG14183,CG2199,Nf-YA,Fkbp59,CG4730,Smox,CG31674,tor,RpS13,ACC,Rnmt,exu,Med,shop,CG7806,enc,CG7414,CG10492,Rpt2,RpS14b,zf30C,tho2,l(3)73Ah,Nup62,tkv,CG3198,Unr,Coq6,Cf2,Pgk,CG6813,DNApol-epsilon255,CG12288,GluRS-m,ND-49,CG9143,awd,RpS2,blw,CG8963,RfC38,CG8611,cad,AOX3,CG10189,Tlk,hb,Larp4B,Chd64,CG11902,p38a,pont,CG1602,nbs,CG5986,l(2)k01209,smg,RpLP0,Mos,CG4404,CG7488,l(1)G0196,PPYR1,CG4709,del,Nadsyn,RecQ5,CG8726,RpL30,Hnf4,gammaTub37C,Maf1,CG2129,CG10418,ABCD,yem,l(3)mbt,RpS5b,bru1,gw,eIF3i,Pfk,Mtl,fray,kl-2,Stat92E,Acox57D-p,tef | | 3.066e-05 | -10.39 | MORF\_TPT1 | MSigDB lists | MORF\_TPT1 | 67 | 24 | 4161 | 644 | RpS7,RpS4,RpL38,RpS24,RpL23,eIF4A,RpL30,RpL13A,RpS16,RpS13,RpS30,RpLP0,RpS18,RpL12,Pgk,RpL13,eEF1beta,RpL37a,RpL11,RpL24,RpS27,RpS27A,RpS23,RpL9 | | 3.128e-05 | -10.37 | GO\_ELECTRON\_CARRIER\_ACTIVITY | MSigDB lists | GO\_ELECTRON\_CARRIER\_ACTIVITY | 51 | 20 | 4161 | 644 | CG14407,Idh3b,COX5A,CG3803,ND-75,wal,CG3609,AIF,Aldh,ND-30,ND-24,Mcad,P5CDh1,ND-49,Gmer,Cyt-c-p,levy,Etf-QO,Grx1,Fdh | | 3.148e-05 | -10.37 | hydrolase activity | molecular function | GO:0016787 | 1775 | 210 | 12176 | 1119 | CG2493,regucalcin,CG4880,CCT3,CG4572,Pfrx,UK114,ATPsynF,otu,scny,ea,CG5026,CG4267,Scsalpha1,CanA-14F,Rab5,cN-IIIB,Prosbeta5,Marf,Dcr-1,Hexo1,eIF4A,PGRP-LE,twe,CSN6,Pp2B-14D,CG32473,up,Nepl3,snk,Usp10,CG9581,CG11486,stet,Sirt7,Snm1,nSMase,alpha-Man-Ia,sw,pps,Psf3,cana,Tfb4,CG5292,alphaTub84B,Ptp61F,faf,tws,CG13630,Ptp69D,tos,Mal-A5,trsn,CG17294,PpD3,mre11,Rpn2,CG1827,cu,Edem1,CG12375,Prosbeta6,spd-2,not,CG8679,CG32549,Pka-R1,Pld,Dip-B,spn-E,gd,armi,clt,Exd2,Treh,CG16979,pn,mop,velo,CG17333,AGBE,Idgf6,HDAC6,DhpD,S2P,Nurf-38,CG30343,Cht11,Prosalpha3,Ptp10D,csw,tzn,Ggt-1,Jheh2,trbd,Clp,CG18135,alphaTub67C,aay,mEFTu1,Usp16-45,obe,Sld5,CG8728,sws,26-29-p,Klp10A,SP1029,ssh,CG9953,mEFTu2,Rga,CG7728,twin,Rab10,ATPsynbeta,Fur2,Prosalpha5,kl-2,ste24a,Mtl,Chrac-14,Atg4b,anne,PIG-L,grsm,milt,Ptpmeg2,tun,Ptpmeg,gammaTub37C,PAPLA1,RecQ5,fal,sol,Nadsyn,CG14516,Prosbeta3,Cp1,cathD,dob,RpLP0,nbs,r,Acph-1,alpha-Est5,CG17746,CG9485,Ser7,pont,rept,Gs1l,CG11771,CG9391,Ntan1,Rab32,CG11164,CG7997,CG4390,CG6967,Synj,CG7789,ATPsynCF6,CG2200,Dd,RfC38,fdl,CG8611,GatA,CG31759,blw,Arf51F,CG9143,DNApol-epsilon255,PpV,EndoGI,alph,Tengl4,CG6707,Acyp2,AMPdeam,Ublcp1,scu,Rpt2,CG5285,CG5390,Drep2,PrBP,Kap3,dnc,Pp2A-29B,Sp7,CG1774,DppIII,CG7115,exu,mus205,Pka-R2,Usp39,Uch-L5,Arl5,Uch,Prosbeta4,mts,Hsc70-3,Prosalpha4,mus81,CG5966,CG11110,CG11089 | | 3.239e-05 | -10.34 | gene silencing | biological process | GO:0016458 | 143 | 29 | 12345 | 1129 | mael,Tlk,piwi,Cbp80,trsn,rept,egg,CG9425,AGO3,esc,squ,Dcr-1,wash,CG6967,armi,psq,Marf1,ph-d,CG10038,Hsc70-3,Bin1,r2d2,gw,Chrac-14,Hnf4,spn-E,Hat1,Elba2,escl | | 3.321e-05 | -10.31 | Dsor1 (Downstream of raf1) | protein interactions | 31872 | 105 | 28 | 9204 | 1112 | vret,mts,RpL9,Rbcn-3A,SkpA,SERCA,Imp,RpS2,RpL10,Nepl3,fs(1)M3,RpL24,ScsbetaA,Mcad,csw,PyK,CG1416,rl,Droj2,tor,larp,RpS28b,RpS30,scny,eIF4E1,wal,cbs,GCS2beta | | 3.547e-05 | -10.25 | organelle fission | biological process | GO:0048285 | 306 | 50 | 12345 | 1129 | Grip128,Arf51F,spn-E,msps,cup,Rab5,gammaTub37C,Pex10,bam,cana,tef,Klp10A,eIF4E1,mus304,sau,Rbf,mus81,larp,Xpc,nos,wapl,grp,yem,Cdk1,Grip75,cort,Topors,CG31053,bif,twe,Cenp-C,wisp,Non1,mtrm,Grip71,Axs,Syx13,Pink1,alphaTub67C,cnn,CG8188,lwr,pont,Pp2B-14D,c(2)M,RecQ5,park,CycB,SRPK,mael | | 3.572e-05 | -10.24 | ATP biosynthetic process | biological process | GO:0006754 | 25 | 10 | 12345 | 1129 | ATPsynCF6,ATPsynB,ATPsynG,COX5B,ATPsynO,blw,ATPsyndelta,ATPsynbeta,ATPsynE,ATPsynD | | 3.640e-05 | -10.22 | GO\_CELLULAR\_RESPIRATION | MSigDB lists | GO\_CELLULAR\_RESPIRATION | 76 | 26 | 4161 | 644 | Irp-1B,Idh3b,COX5A,ND-39,Idh,Scsalpha1,Nc73EF,ND-30,ND-24,UQCR-Q,CG3803,wal,ND-75,ND-B18,SdhA,ND-SGDH,Cyt-c-p,ScsbetaA,CG11110,ND-49,ND-51,ND-23,CG5028,ND-B22,Etf-QO,levy | | 3.646e-05 | -10.22 | Oxidative phosphorylation | KEGG pathways | ko00190 | 144 | 41 | 3198 | 496 | cype,ND-49,ND-SGDH,ATPsynF,ND-MWFE,UQCR-11,ND-MNLL,blw,ATPsynE,COX8,ND-39,COX5B,ND-51,ND-75,ND-18,ATPsynO,levy,ND-B15,Nurf-38,ATPsynCF6,CG3803,COX5A,ATPsynG,SdhA,ATPsyndelta,Vha68-2,ND-23,ND-B22,Vha36-1,NP15.6,CG40472,UQCR-Q,ND-B18,Vha13,COX4,ND-30,ATPsynD,ND-24,SdhD,ATPsynB,ATPsynbeta | | 3.646e-05 | -10.22 | Oxidative phosphorylation | KEGG pathways | dme00190 | 144 | 41 | 3198 | 496 | ND-24,SdhD,ATPsynbeta,ATPsynB,COX4,Vha13,ATPsynD,ND-30,Vha36-1,NP15.6,ND-B22,ND-B18,CG40472,UQCR-Q,SdhA,ATPsynG,ND-23,Vha68-2,ATPsyndelta,ATPsynCF6,Nurf-38,COX5A,CG3803,levy,ND-B15,ND-18,ATPsynO,COX8,ATPsynE,ND-39,COX5B,UQCR-11,ND-MNLL,blw,ND-75,ND-51,cype,ATPsynF,ND-SGDH,ND-MWFE,ND-49 | | 3.682e-05 | -10.21 | organic cyclic compound binding | molecular function | GO:0097159 | 3086 | 340 | 12176 | 1119 | ACC,exu,Rnmt,CG7806,shop,Med,CG10492,CG7414,enc,CG4730,CG2199,Fkbp59,Nf-YA,Smox,CG31674,tor,RpS13,CG6813,CG12288,DNApol-epsilon255,GluRS-m,Coq6,Pgk,Cf2,awd,CG8963,blw,RpS2,ND-49,CG9143,RfC38,CG8611,cad,AOX3,Larp4B,CG10189,Tlk,hb,RpS14b,Rpt2,Nup62,zf30C,l(3)73Ah,tho2,Unr,tkv,CG3198,CG4404,Mos,CG7488,l(1)G0196,PPYR1,CG8726,RecQ5,RpL30,Hnf4,CG4709,Nadsyn,del,Maf1,gammaTub37C,CG11902,Chd64,pont,p38a,CG1602,l(2)k01209,smg,RpLP0,nbs,CG5986,gw,Mtl,Pfk,eIF3i,kl-2,fray,Acox57D-p,tef,Stat92E,ABCD,CG2129,CG10418,l(3)mbt,yem,RpS5b,bru1,CG14100,mEFTu1,apt,l(1)G0004,Nc73EF,CG10336,ScsbetaA,HmgD,mal,Rbf,Elba2,RpS16,Cyp6a22,CG9425,CkIIalpha,Rab10,Tao,Vha68-2,BicC,Mat89Ba,Fkbp39,SdhD,LSm3,CG12375,Atpalpha,Wbp2,Pxt,trsn,foxo,CG2678,E2f1,swa,wdn,spn-E,Exd2,qkr58E-2,RpL23,Sply,Su(H),lid,nmdyn-D7,PEK,SLIRP1,Hr78,Xpac,LysRS,CG11486,CysRS,Dcr-1,cyc,eIF4A,Cyt-c-p,Droj2,GlyP,ovo,Rrp40,cana,alphaTub84B,Orc1,shep,CG17768,Edc3,Snm1,PDCD-5,Sirt7,bor,NFAT,BigH1,Cyp6u1,Imp,CG10959,Lkb1,RpS14a,CG14183,CG4880,Pym,heph,AGO3,osk,Marf,RpII140,CG5116,Rab5,CG4266,mus205,Nxf3,RpL12,Taf7,CG4562,Trf2,CG14997,Rbfox1,psq,SdhA,Cdk1,Top1,CG31729,CG1640,Hsc70-3,CG17454,CCT6,CCT5,mus81,egg,CG12018,insv,CG6610,spen,CG4021,pre-lola-G,Arl5,mRpS11,Cbp80,Tengl4,Kyat,Cdk7,Tak1,Arf51F,Rcd5,Dhfr,hng3,RpL9,RpL26,Bap170,CG6967,Synj,mRpL16,cup,Gabat,Pdk1,RpL7A,dl,Mitf,RpS15,CG10654,CG10038,CG31875,Roe1,SerRS-m,CG5789,Mcad,Rab32,Got1,Orc6,brwl,sbb,rept,mael,grp,rl,Irp-1B,CG4860,Rbf2,r,mRpS5,CG17698,Ssdp,mRpL13,glob1,CG15514,RYBP,SRPK,CG9981,sbr,jigr1,Larp7,anne,png,ND-51,ATP8A,CG4586,Clp,alphaTub67C,TfIIFalpha,Idh,AcCoAS,cnc,RpL13A,nmd,AIF,e(y)1,CG7728,Cdk5,ATPsynbeta,Npc2a,Marf1,mEFTu2,TrpRS-m,Mekk1,Son,Non1,Klp10A,obe,EcR,CG6276,NHP2,CG17187,Cyp6v1,Spx,RpL24,tos,CG31689,Pus7,SREBP,CG6767,eIF3m,r2d2,armi,sgll,CG4041,mRpL18,Atf3,SERCA,CG2162,bcd,elg1,RpS4,eIF4E1,Atf6,Xpc,Trxr-1,SNF4Agamma,CG2100,crm,RpS18,nos,yps,CG10631,Utx,Rel,Gapdh1,Neos,Ho,bys,Mybbp1A,TyrRS,Pink1,bam,eEF1beta,Pfrx,dco,CG10463,E23,CCT3,SelD,CG17328,CG6094,Shmt,Marc,CG34116,Cyp6a19,piwi,wal | | 3.686e-05 | -10.21 | PPIase\_FKBP\_dom | interpro domains | IPR001179 | 8 | 6 | 11383 | 1233 | Fkbp12,CG1847,CG14715,Fkbp39,shu,Fkbp59 | | 3.716e-05 | -10.20 | enzyme binding | molecular function | GO:0019899 | 481 | 71 | 12176 | 1119 | Mekk1,tub,Ranbp21,CG10188,Cul2,cbs,Tm1,sws,Gmap,CG4557,Taf7,sip3,nej,ago,Ankle2,tral,CycB,eIF3m,lwr,GAPsec,Rep,wash,CG3226,alc,RhoGDI,Mitofilin,CG4041,Gdi,CG4853,CG13551,Atu,Tak1,MICAL-like,Arts,CG11807,GCC88,SNF4Agamma,CG32485,CG30496,ben,Cul5,CG10254,sau,park,Pura,Smyd4-4,rl,CG42748,RpL40,Cbl,Cul1,Tomosyn,Crag,Uev1A,Maf1,Parp16,CycE,Wdfy2,SmydA-1,Sec15,Aos1,Cse1,ari-1,Mtl,capt,eEF1beta,CycJ,CG3309,RpS27A,CG8974,Ubc4 | | 3.752e-05 | -10.19 | scf (supercoiling factor) | protein interactions | 38145 | 59 | 19 | 9204 | 1112 | Uch,Eno,AnxB10,regucalcin,capt,Moe,HmgD,Stip1,RpII140,yip2,RpLP0,CG11267,Chd64,awd,CG10638,CG14715,EndoGI,cib,PHGPx | | 3.755e-05 | -10.19 | CG4598 (uncharacterized protein) | protein interactions | 34315 | 36 | 14 | 9204 | 1112 | regucalcin,Pgk,CG10638,CG14715,PHGPx,Fkbp12,Moe,capt,AnxB10,Chd64,awd,Eno,Pglym78,Uch | | 3.813e-05 | -10.17 | FKBP\_C | pfam domains | PF00254 | 8 | 6 | 10480 | 1142 | shu,CG1847,Fkbp59,Fkbp39,CG14715,Fkbp12 | | 3.833e-05 | -10.17 | SERCA (Sarco/endoplasmic reticulum Ca(2+)-ATPase) | protein interactions | 49297 | 128 | 32 | 9204 | 1112 | CG31729,CG15098,CCT3,CG8974,blw,Pdk1,ATPsynbeta,CCT6,Tspo,Vha68-2,Seipin,GCS2beta,Surf4,ScsbetaA,Atpalpha,Droj2,Rab5,CG1416,Tak1,Pp2A-29B,CG10425,egg,Desat1,Rel,PGRP-LE,Bx,Karybeta3,Rtnl1,Cp1,wal,Mcad,cathD | | 3.866e-05 | -10.16 | Inos (myo-inositol-1-phosphate synthase) | protein interactions | 35671 | 69 | 21 | 9204 | 1112 | Moe,capt,scu,regucalcin,AnxB10,CG4598,CG11807,Eno,Uch,Fkbp12,cib,PHGPx,EndoGI,CG10638,CG14715,awd,Chd64,yip2,Echs1,CG32549,RhoGDI | | 3.915e-05 | -10.15 | carbon-nitrogen ligase activity, with glutamine as amido-N-donor | molecular function | GO:0016884 | 9 | 6 | 12176 | 1119 | Pfas,GatB,AsnS,GatA,Nadsyn,r | | 3.956e-05 | -10.14 | ribonucleoprotein complex biogenesis | biological process | GO:0022613 | 291 | 48 | 12345 | 1129 | RpS15,LSm3,RpS27,RpL23A,mRpS11,rig,CG17768,bru1,CG10055,Mat89Ba,RpL34a,RpS28b,CG12288,spn-E,RpS14b,RpL11,CG7488,mbo,RpS14a,NHP2,CG7728,r2d2,rept,pont,CG9143,mtTFB1,Dcr-1,CG3198,RpL34b,RpL7A,Edc3,bys,CG8611,CG12301,CG8204,SmE,RpL10,RpL12,mRpL10,osk,Usp39,RpS7,RpL26,RpS5b,RpS16,RpL24,Rrp40,RpLP0 | | 3.984e-05 | -10.13 | regulation of response to stress | biological process | GO:0080134 | 283 | 47 | 12345 | 1129 | Lkb1,Cka,scny,alph,slif,Traf6,Tak1,GILT1,Imp,ago,Ubc6,Myd88,Rel,GlyP,CG9925,armi,Seipin,Atg6,Sp7,Ptip,Atg7,Crag,Hat1,Stat92E,PGRP-LE,PEK,SkpA,p38a,Hr78,Mtl,faf,rl,park,PDCD-5,CG17746,G9a,spz,Archease,EMRE,Ptp61F,Mekk1,cact,ben,wisp,Ect4,PpV,tub | | 4.013e-05 | -10.12 | negative regulation of nitrogen compound metabolic process | biological process | GO:0051172 | 625 | 87 | 12345 | 1129 | CG14408,cyc,Unr,Rga,CG31875,nos,mts,esc,SREBP,hb,Utx,r2d2,CkIIalpha,bam,Su(H),Not10,psq,Mybbp1A,escl,Hat1,sws,RpL13A,apt,BicC,dco,E2f1,twin,Rbf2,EndoGI,wash,park,CG10959,egg,Cbl,Sox102F,mRpL13,ATPsynD,CG17746,CG17328,PpV,Elba2,krimp,alph,d4,bru1,CG10631,piwi,CG30496,Tlk,dl,Rbf,foxo,heph,CG2199,Sirt7,cup,sbb,spn-E,CG4872,CycE,Chrac-14,SkpA,l(3)mbt,asf1,ovo,Maf1,mael,insv,BigH1,Drep1,lin-52,smg,rept,faf,Bin1,ph-d,CG10038,Uxt,Tip60,CG1602,bcd,Rbfox1,Hnf4,EcR,pgc,gw,Ptp61F | | 4.043e-05 | -10.12 | Fatty acid degradation | KEGG pathways | dme00071 | 33 | 15 | 3198 | 496 | CG4586,CG10932,Thiolase,whd,Mtpalpha,CG4860,Aldh,CG4598,Acox57D-p,Adh,Mcad,yip2,Echs1,Fdh,CG4594 | | 4.043e-05 | -10.12 | Fatty acid degradation | KEGG pathways | ko00071 | 33 | 15 | 3198 | 496 | CG4598,Adh,Acox57D-p,Mcad,yip2,Echs1,Fdh,CG4594,CG4586,Thiolase,CG10932,Mtpalpha,whd,CG4860,Aldh | | 4.090e-05 | -10.10 | anion binding | molecular function | GO:0043168 | 1099 | 139 | 12176 | 1119 | GlyP,Droj2,alphaTub84B,cana,Orc1,Sirt7,snz,bor,nmdyn-D7,PEK,Sply,LysRS,CysRS,eIF4A,Trxr-1,crok,Marf,Shmt,Marc,CG5116,Rab5,wal,Pink1,TyrRS,dco,Pfrx,CG10463,CCT3,E23,CG4880,SelD,Lkb1,CG14183,Rab10,Cdk5,ATPsynbeta,Tao,CkIIalpha,CG7728,mEFTu2,TrpRS-m,Mekk1,CG8507,Klp10A,Vha68-2,Non1,obe,mEFTu1,CG4586,CG3077,alphaTub67C,ScsbetaA,Nc73EF,AcCoAS,mal,nmd,Exo70,AIF,CG6767,sgll,spn-E,Pld,SERCA,CG4041,Atpalpha,CG31689,Rab32,Got1,grp,rept,pont,p38a,rl,CG4860,r,l(2)k01209,sau,Mos,Hip1,CG7488,l(1)G0196,Nadsyn,CG8726,RecQ5,gammaTub37C,SerRS-m,CG5734,CG5789,Mcad,CG9981,SRPK,ABCD,png,ND-51,ATP8A,CG17698,Pfk,Mtl,fray,kl-2,Acox57D-p,CG1640,CCT5,CCT6,Hsc70-3,tor,AnxB10,Arl5,ACC,CG14997,CG4562,CG7806,CG31729,SdhA,Cdk1,fs(1)N,Rpt2,Gabat,Pdk1,Nup62,tkv,Moe,Pex23,Coq6,Kyat,Pgk,GluRS-m,CG9143,Cdk7,awd,Myd88,blw,Tak1,Arf51F,Snx6,AOX3,CG11807,CG8611,Tlk | | 4.130e-05 | -10.09 | ATPsynO (ATP synthase, oligomycin sensitivity conferring protein) | protein interactions | 41845 | 32 | 13 | 9204 | 1112 | ATPsynbeta,ATPsynG,Mf,up,COX4,Rtnl1,ATPsynD,blw,ATPsynCF6,ATPsynE,ATPsyndelta,ATPsynB,ATPsynF | | 4.221e-05 | -10.07 | The citric acid (TCA) cycle and respiratory electron transport | REACTOME pathways | R-DME-1428517 | 99 | 30 | 3923 | 574 | ND-B18,CG8067,Nc73EF,ND-49,Etf-QO,ATPsyndelta,ATPsynF,Pdk,ND-23,ND-B22,ATPsynbeta,ATPsynO,SdhA,ATPsynG,NP15.6,SdhD,ATPsynB,wal,tzn,ND-51,Idh3b,ATPsynCF6,ND-SGDH,blw,ND-39,ATPsynD,ND-75,ND-30,ND-24,CG5028 | | 4.234e-05 | -10.07 | determination of adult lifespan | biological process | GO:0008340 | 167 | 32 | 12345 | 1129 | AGBE,foxo,snz,alph,cnc,Tsp42Ef,Lkb1,Atpalpha,ND-SGDH,Atg7,raptor,Tspo,Mtpalpha,GlyP,bun,Jafrac1,rl,alpha-Man-Ia,Sod1,Thiolase,Cdk5,Edem1,Hsp26,PGRP-LE,p38a,EcR,lt,Coq7,ATPsynD,Smox,Trxr-1,levy | | 4.252e-05 | -10.07 | His3.3A (Histone H3.3A) | protein interactions | 33736 | 24 | 11 | 9204 | 1112 | Fkbp39,Cdk5,squ,asf1,Cdk7,trr,Caf1-105,p38a,CycJ,grp,Nap1 | | 4.363e-05 | -10.04 | oocyte construction | biological process | GO:0007308 | 138 | 28 | 12345 | 1129 | spz,CkIIalpha,heph,armi,psq,gammaTub37C,gus,osk,spn-E,Grip71,Moe,msps,enc,jvl,Grip128,cnc,swa,krimp,capt,Tm1,mael,Lkb1,ea,Grip75,Pka-R1,exu,milt,nos | | 4.507e-05 | -10.01 | positive regulation of response to external stimulus | biological process | GO:0032103 | 124 | 26 | 12345 | 1129 | Myd88,PDCD-5,Herc4,Rel,mop,CG9925,raptor,spz,Seipin,Sp7,Archease,EMRE,Ptip,wisp,Crag,tub,Hat1,cad,Aos1,PGRP-LE,Hr78,dl,lwr,Imp,Tak1,GILT1 | | 4.639e-05 | -9.98 | S6kII (Ribosomal protein S6 kinase II) | protein interactions | 33139 | 206 | 45 | 9204 | 1112 | rl,yps,bif,RpL31,RpL17,ACC,CCT5,RpS15,CG17593,Imp,alt,RpS14a,RpL9,Rbcn-3A,Kap-alpha3,Hsc70-3,RpL7A,RpLP1,CG4266,RpL10,RpL23A,TfIIFalpha,RpS23,Nup358,Aldh,Droj2,tos,RpS16,ScsbetaA,Ankle2,eIF4A,RpL13A,RpS4,rept,CkIIalpha,larp,RpL13,eIF3m,RpL26,Pp2B-14D,Pdk1,pont,CG42354,RpS7,eIF3i | | 4.727e-05 | -9.96 | nuclear division | biological process | GO:0000280 | 293 | 48 | 12345 | 1129 | alphaTub67C,Syx13,Axs,wisp,Non1,Grip71,mtrm,twe,Cenp-C,bif,SRPK,mael,CycB,pont,Pp2B-14D,c(2)M,RecQ5,cnn,CG8188,lwr,Klp10A,eIF4E1,mus304,cana,bam,tef,cup,Rab5,gammaTub37C,Pex10,msps,spn-E,Grip128,Arf51F,Topors,CG31053,cort,yem,Cdk1,Grip75,grp,wapl,Xpc,nos,Rbf,larp,mus81,sau | | 4.739e-05 | -9.96 | rRNA binding | molecular function | GO:0019843 | 41 | 13 | 12176 | 1119 | RpS14a,RpLP0,mRpL18,mRpS11,RpL23,RpS13,RpL9,RpS5b,RpS14b,CG7488,CG12288,RpL12,mRpL16 | | 4.782e-05 | -9.95 | regulation of immune response | biological process | GO:0050776 | 168 | 32 | 12345 | 1129 | slif,scny,Tak1,dl,Atg6,Myd88,GlyP,mop,CG9925,Rel,Herc4,Crag,Hat1,Sp7,Ptip,PGRP-LE,Hr78,p38a,Stat92E,Aos1,cad,faf,lwr,spz,RYBP,PDCD-5,cact,Ect4,tub,crq,wisp,EMRE | | 4.878e-05 | -9.93 | GO\_ORGANIC\_CYCLIC\_COMPOUND\_CATABOLIC\_PROCESS | MSigDB lists | GO\_ORGANIC\_CYCLIC\_COMPOUND\_CATABOLIC\_PROCESS | 199 | 52 | 4161 | 644 | mus81,DhpD,RpS7,RpL38,RpS4,Cbp80,RpL40,CG10418,RpS24,CG5285,CG8963,RpL30,RpL23,P5CDh1,RpS16,RpS2,Edc3,RpS18,RpL12,Dcr-1,RpL11,RpL37a,LSm3,RpL24,RpS27,RpL26,CG32549,Pp2A-29B,dnc,CG5734,mts,CG11164,eIF4A,RpS13,RpS30,RpLP0,eIF4E1,RpL13A,Rrp40,CG6610,Not10,AIF,crq,CG9471,RpL13,twin,Unr,RpS28b,RpS27A,RpL9,Pcd,RpS23 | | 4.951e-05 | -9.91 | GO\_ORGANONITROGEN\_COMPOUND\_METABOLIC\_PROCESS | MSigDB lists | GO\_ORGANONITROGEN\_COMPOUND\_METABOLIC\_PROCESS | 794 | 160 | 4161 | 644 | RpL23,aay,CG4390,RpS16,RpS4,RpL38,CG1640,RpS7,heph,RpL40,CG7414,eIF3m,Pfas,RpL11,CG7789,CG32473,RpS18,Pdxk,Ac3,nmdyn-D6,Pglym78,CG33156,Gapdh1,Pgm1,Sod1,CG17691,ND-B22,CG4407,CG10932,ND-23,RpS23,Ppox,CG8199,GalT1,kri,Tfb4,Shmt,Pgk,RpL30,P5cr,TyrRS,CG2201,CG11110,ScsbetaA,Atg6,Cyt-c-p,sgll,CG17333,Ts,DhpD,Adi1,scramb1,tko,mRpL17,Ppcdc,COX5A,Gabat,Pp2A-29B,r-l,CG3376,ND-24,mts,nmdyn-D7,RpLP0,mRpL13,RpS30,Nadsyn,LysRS,levy,Fur2,eIF3i,Grx1,Sptr,RpL9,Idh3b,CG7997,ND-75,mRpL16,eEF1beta,RpL13,mEFTu1,RpS24,Dhfr,colt,mal,mRpS23,CG4511,Tspo,CG11134,CG4872,RpL26,CG32549,RpS27,ND-B18,Gclc,Eno,mRpS5,ND-30,RpL12,eIF4A,ND-49,Mcad,CG11899,mRpL18,AMPdeam,Nurf-38,scu,Prip,ND-SGDH,CysRS,GluRS-m,Stoml2,CG5028,dnc,ND-51,Got1,CG15111,Sply,ND-39,CG14516,rl,CG3803,CG8963,P5CDh1,CG5285,CG6767,RpS2,nSMase,r,oxt,RpL37a,mRpL10,RpL24,Pfk,CG7322,Jafrac2,pr,CG10425,CG7115,RpS13,RpL13A,eIF4E1,tzn,CG31715,CG7550,ACC,ste14,CG11089,mRpS21,Idh,Nc73EF,TrpRS-m,RpS27A,Pcd,GatA,RpS28b,CG9471,SerRS-m,mRpS10,UQCR-Q | | 4.965e-05 | -9.91 | cofactor metabolic process | biological process | GO:0051186 | 214 | 38 | 12345 | 1129 | CG17691,Shmt,Pcd,scu,CG7920,Jafrac1,ScsbetaA,CG8199,ND-39,regucalcin,Coq6,Nadsyn,CG13551,Coq7,mal,Pdxk,GstT1,pr,GstE11,CG1774,CG3803,Ggt-1,Sptr,GstT2,Coq8,Ppox,Jafrac2,Gclc,Pxt,Scsalpha1,CG6178,CG4407,Ho,Dhfr,Ppcdc,sgll,CG33156,AcCoAS | | 4.969e-05 | -9.91 | GO\_ORGANONITROGEN\_COMPOUND\_BIOSYNTHETIC\_PROCESS | MSigDB lists | GO\_ORGANONITROGEN\_COMPOUND\_BIOSYNTHETIC\_PROCESS | 497 | 108 | 4161 | 644 | Ppox,RpS23,Got1,Sply,Shmt,GalT1,kri,CG3803,Tfb4,Prip,AMPdeam,Nurf-38,CG33156,eIF4A,mRpL18,CG11899,Mcad,CG4407,Stoml2,CysRS,GluRS-m,RpL26,RpS27,CG7414,CG11134,eIF3m,Pfas,RpL11,nmdyn-D6,RpL12,mRpS5,RpS18,Pdxk,Ac3,Gclc,mal,aay,colt,mRpS23,RpS16,CG4390,RpL23,Dhfr,RpS24,heph,RpL40,Tspo,RpL38,RpS4,CG4511,RpS7,CG1640,Pcd,RpL9,RpS27A,RpS28b,GatA,mRpS21,TrpRS-m,Sptr,eEF1beta,mRpL16,mEFTu1,RpL13,SerRS-m,mRpS10,nmdyn-D7,Nadsyn,RpL13A,eIF4E1,LysRS,RpLP0,RpS13,RpS30,mRpL13,CG7115,ACC,CG11089,eIF3i,Fur2,tzn,Gabat,RpL24,RpL37a,mRpL10,Ppcdc,mRpL17,CG3376,r-l,CG10425,pr,CG6767,RpS2,ScsbetaA,P5CDh1,CG8963,RpL30,TyrRS,CG2201,P5cr,scramb1,Adi1,tko,nSMase,Ts,sgll,oxt,r | | 5.078e-05 | -9.89 | regulation of programmed cell death | biological process | GO:0043067 | 222 | 39 | 12345 | 1129 | dco,E2f1,SkpA,Cul1,Naa20A,Idgf2,park,Drep1,MED24,ScsbetaA,rept,Cbl,lin-52,Pink1,DCAF12,fwe,Tip60,PDCD-5,RYBP,Idh3b,Debcl,EcR,spag,scny,Lkb1,nos,Ubc6,Pdk1,Jafrac2,Rbf,Tspo,Utx,bun,Atg7,csw,CG4238,eIF3m,CycE,MAGE | | 5.095e-05 | -9.88 | trans-Golgi network | cellular component | GO:0005802 | 41 | 13 | 11940 | 1105 | Afti,Gga,Fur2,Pi4KIIalpha,CG31729,AP-1-2beta,ATP8A,Rab32,Arl5,CG9981,Trs33,sau,lqfR | | 5.096e-05 | -9.88 | KEGG\_PROPANOATE\_METABOLISM | MSigDB lists | KEGG\_PROPANOATE\_METABOLISM | 17 | 10 | 4161 | 644 | Scsalpha1,Mcad,ScsbetaA,Gabat,Echs1,Mtpalpha,AcCoAS,Aldh,CG10932,ACC | | 5.282e-05 | -9.85 | ubiquitin-protein transferase activity | molecular function | GO:0004842 | 237 | 41 | 12176 | 1119 | CG8974,Ubc4,Fbw5,Ubc6,CG14435,SkpA,CG4502,CG8188,CG9941,Traf6,APC7,CG11070,ari-1,Pex12,CG13344,Kcmf1,Pex10,UbcE2H,Iru,CG31633,EloC,CG9316,Uev1A,ago,STUB1,sip3,Herc4,Topors,CG40045,Cbl,CG2924,CG15141,park,CG5003,CG32221,CG3894,CG10254,dx,ben,CG46338,CG4238 | | 5.342e-05 | -9.84 | MORF\_SOD1 | MSigDB lists | MORF\_SOD1 | 170 | 46 | 4161 | 644 | CG10418,CG1416,Prosbeta6,NHP2,Pka-R1,Phb2,CG1890,Cyt-c-p,Dd,ND-24,Hat1,CG13630,ND-30,Thiolase,CCT3,Jafrac2,CycB,CCT5,COX5A,Pp2A-29B,Prosbeta4,LSm3,Prosalpha3,rept,ND-23,CG11089,ND-51,levy,eIF3i,SkpA,Stoml2,CHOp24,Sod1,Rpn12,LysRS,csw,Rpt2,CG3631,ND-49,Chrac-14,Pen,mEFTu1,SdhA,Klp10A,ND-75,wal | | 5.366e-05 | -9.83 | GO\_TRANSLATIONAL\_INITIATION | MSigDB lists | GO\_TRANSLATIONAL\_INITIATION | 104 | 32 | 4161 | 644 | RpS27A,RpS23,RpL9,RpS28b,RpL13,RpL13A,eIF4E1,RpS13,RpS30,RpLP0,eIF4A,eIF3i,RpL26,RpS27,RpL24,CG7414,RpL37a,eIF3m,RpL11,RpL12,RpS18,RpS16,RpS2,CG8963,RpL23,RpL30,RpS24,heph,RpL40,RpS4,RpL38,RpS7 | | 5.366e-05 | -9.83 | GO\_RIBOSOMAL\_SUBUNIT | MSigDB lists | GO\_RIBOSOMAL\_SUBUNIT | 104 | 32 | 4161 | 644 | RpL12,mRpS5,RpS18,RpL26,RpL24,RpS27,RpL37a,mRpL10,RpL11,mRpL17,RpL40,tko,RpS4,RpL38,RpS7,RpS16,RpS2,RpL23,RpL30,RpS24,mRpL16,RpL13,RpS27A,RpS23,RpL9,RpS28b,mRpS21,RpL13A,mRpL13,RpLP0,RpS30,RpS13 | | 5.503e-05 | -9.81 | endoplasmic reticulum | cellular component | GO:0005783 | 475 | 70 | 11940 | 1105 | sws,Edem1,CG5969,alpha-Man-Ia,cup,CG14671,CG5510,CHOp24,GalT1,Mctp,alt,CG6766,Sec23,Axs,ERp60,EMC3,EMC8-9,ste24a,Agpat2,CG32441,CG8311,SREBP,Lnpk,CG9981,Rtnl1,CG3631,Sec22,CG10344,Parp16,bif,ste14,CG17270,CG4293,Tm1,Seipin,CG10425,sip3,Atf6,Efr,CG31523,CG32640,mol,CG33977,park,tral,fdl,Ugt36E1,Dd,PIG-H,PIG-C,Jafrac2,Hsc70-3,CG8507,CG8112,Ankle2,CG7840,opm,CG4880,PIG-P,CG8314,CG10470,CG9186,mino,CG8974,Sply,Desat1,SERCA,CG17593,O-fut1,CG4557 | | 5.634e-05 | -9.78 | small molecule binding | molecular function | GO:0036094 | 1078 | 136 | 12176 | 1119 | eIF4A,Trxr-1,CysRS,LysRS,nmdyn-D7,PEK,Sply,Gapdh1,Sirt7,bor,Orc1,alphaTub84B,cana,GlyP,SNF4Agamma,Droj2,CG4880,Lkb1,SelD,CG14183,CG10463,CCT3,E23,dco,Pfrx,TyrRS,Pink1,wal,Marc,RpII140,CG5116,Rab5,Marf,Shmt,AIF,mal,nmd,ScsbetaA,Nc73EF,AcCoAS,mEFTu1,Idh,CG4586,alphaTub67C,obe,Klp10A,Vha68-2,Non1,TrpRS-m,mEFTu2,Mekk1,Rab10,Cdk5,Tao,ATPsynbeta,CG7728,CkIIalpha,CG31689,Atpalpha,EcR,SERCA,CG4041,sgll,spn-E,CG5510,CG6767,gammaTub37C,CG5789,SerRS-m,Mcad,Roe1,Nadsyn,CG8726,RecQ5,CG7488,l(1)G0196,Mos,CG4860,r,l(2)k01209,rl,grp,pont,rept,p38a,Got1,Rab32,Acox57D-p,kl-2,fray,glob1,Pfk,Mtl,CG17698,ND-51,ATP8A,png,anne,SRPK,CG9981,ABCD,CG31729,SdhA,Cdk1,CG7806,CG14997,CG4562,ACC,mus205,Arl5,tor,CCT6,CCT5,Hsc70-3,CG31674,CG1640,Tlk,Dhfr,AOX3,CG8611,CG9143,ND-49,Cdk7,awd,blw,Tak1,Arf51F,Coq6,Pgk,Kyat,DNApol-epsilon255,GluRS-m,tkv,Rpt2,Gabat,Pdk1 | | 5.652e-05 | -9.78 | ubiquitin-like protein conjugating enzyme activity | molecular function | GO:0061650 | 31 | 11 | 12176 | 1119 | CG40045,UbcE2H,lwr,CG8188,CG4502,Uev1A,ben,CG10254,Ubc6,CG2924,Ubc4 | | 5.811e-05 | -9.75 | carbohydrate metabolic process | biological process | GO:0005975 | 247 | 42 | 12345 | 1129 | CG9485,CG7328,Mal-A5,Pi3K92E,CG7322,adp,Cht11,Idgf6,GalT1,Gyg,Idgf2,Pgm1,Pfk,CG8475,PyK,Eno,regucalcin,Pgk,Gk1,Imp,CG9391,oxt,l(1)G0196,fdl,AGBE,foxo,CG2938,Hexo1,Pglym78,Gk2,Gapdh1,Pfrx,Treh,Synj,CG6650,Idgf4,CG17333,UGP,Atg6,ACC,CG7997,GlyP | | 5.878e-05 | -9.74 | cellular response to oxygen-containing compound | biological process | GO:1901701 | 133 | 27 | 12345 | 1129 | Stat92E,Fdh,Shmt,Vps16B,mAChR-A,p38a,Pi3K92E,foxo,Jafrac1,Tak1,Sod1,rl,Pdk1,Nurf-38,MED27,Pka-R2,ACC,raptor,Utx,G9a,Sod3,CG4390,Hnf4,EcR,PyK,Rab10,Pdxk | | 5.878e-05 | -9.74 | response to oxidative stress | biological process | GO:0006979 | 133 | 27 | 12345 | 1129 | Pxt,whd,Ho,ATPsynD,Atg6,Sod3,spz,Pink1,Grx1,Mekk1,chrb,ben,Idh,Sdhaf3,mol,PHGPx,ND-SGDH,alph,p38a,cnc,Pi3K92E,foxo,Jafrac2,Sod1,rl,Jafrac1,park | | 5.982e-05 | -9.72 | GO\_COFACTOR\_BINDING | MSigDB lists | GO\_COFACTOR\_BINDING | 137 | 39 | 4161 | 644 | Dhfr,CG10463,mal,Sirt7,CG1640,Ts,sgll,Etf-QO,CG4872,Gabat,Gclc,Mtpalpha,Pdxk,shop,CG8112,Mcad,Gmer,ND-49,Gapdh1,scu,Coq6,ND-51,CG5028,CG10932,Sptr,ND-39,Sply,Scsalpha1,Nc73EF,Idh,Got1,Idh3b,Ppox,Trxr-1,wal,GlyP,SdhA,AIF,Shmt | | 6.010e-05 | -9.72 | transcription coregulator activity | molecular function | GO:0003712 | 147 | 29 | 12176 | 1119 | Wbp2,CG11710,pygo,CG31875,MED27,CG4872,hng3,MED4,CycC,MED24,Pax,nej,EcR,Taf7,CG17202,MED20,Spt20,trr,Bin1,Mybbp1A,ix,Uxt,rept,pont,sbb,mtTFB1,insv,Sgf11,CG10631 | | 6.075e-05 | -9.71 | multicellular organism aging | biological process | GO:0010259 | 170 | 32 | 12345 | 1129 | foxo,AGBE,ND-SGDH,Atpalpha,Lkb1,Tsp42Ef,cnc,alph,snz,Atg7,bun,GlyP,Mtpalpha,Tspo,raptor,Cdk5,Edem1,Thiolase,rl,alpha-Man-Ia,Sod1,Jafrac1,p38a,PGRP-LE,Hsp26,lt,EcR,levy,Trxr-1,Smox,ATPsynD,Coq7 | | 6.245e-05 | -9.68 | regulation of apoptotic process | biological process | GO:0042981 | 193 | 35 | 12345 | 1129 | Jafrac2,Rbf,nos,Pdk1,Ubc6,scny,Lkb1,spag,eIF3m,CycE,MAGE,Atg7,csw,CG4238,bun,Tspo,Cbl,lin-52,Drep1,park,MED24,ScsbetaA,Cul1,Idgf2,Naa20A,dco,E2f1,SkpA,EcR,Debcl,fwe,Tip60,PDCD-5,RYBP,Pink1 | | 6.291e-05 | -9.67 | NADH dehydrogenase activity | molecular function | GO:0003954 | 42 | 13 | 12176 | 1119 | ND-24,ND-MNLL,ND-51,ND-B15,ND-B18,ND-18,ND-30,ND-SGDH,ND-49,ND-23,ND-B22,ND-75,ND-39 | | 6.294e-05 | -9.67 | Adh (Alcohol dehydrogenase) | protein interactions | 3771877 | 61 | 19 | 9204 | 1112 | CG32473,Cyt-c-p,Nc73EF,Argk,GlyP,ovo,ND-18,Pglym78,Eno,Scp1,Rtnl1,up,Mf,CG7920,Pgk,Actn,Fer1HCH,ATPsyndelta,COX4 | | 6.343e-05 | -9.67 | Formation of ATP by chemiosmotic coupling | REACTOME pathways | R-DME-163210 | 15 | 9 | 3923 | 574 | ATPsyndelta,ATPsynB,ATPsynF,blw,ATPsynbeta,ATPsynO,ATPsynD,ATPsynG,ATPsynCF6 | | 6.349e-05 | -9.66 | BRUINS\_UVC\_RESPONSE\_LATE | MSigDB lists | BRUINS\_UVC\_RESPONSE\_LATE | 395 | 89 | 4161 | 644 | Cdk1,Nca,Uch-L5,Rab10,CG1774,Pka-R2,Prosalpha5,Utx,Mo25,spag,Stoml2,Fdh,Stam,Su(H),Gapdh1,Pgm1,CG11899,mRpL18,Pdxk,pont,CG8004,Kap-alpha3,Hat1,CG13630,ND-30,Tomosyn,Uch,Gyg,Vamp7,Hmg-2,CycE,gammaTub37C,Dph5,Vha13,RfC38,Ubc4,mus81,Pka-R1,CG17593,Ufm1,Tm1,Droj2,PrBP,CG1532,Rbf,AnxB10,CG5292,colt,aay,mal,CG4390,Kap3,AIF,Caf1-105,SerRS-m,AcCoAS,Trs33,CG16979,CG7997,CG5026,CG7840,TrpRS-m,Nmt,stas,GatA,Sld5,cyc,disp,ste14,CG11089,csw,Rpt2,Rrp40,Coq7,CG11710,CG10425,CG7488,Prosalpha3,mRpL17,Ppcdc,scramb1,Atg7,CG8963,CG17294,P5cr,Vha36-1,CkIIalpha,CG11110,CG11858 | | 6.381e-05 | -9.66 | ATPsyndelta (ATP synthase, delta subunit) | protein interactions | 31950 | 21 | 10 | 9204 | 1112 | ATPsynCF6,ATPsynD,blw,ATPsynF,ATPsynE,ATPsynB,ATPsynbeta,ATPsynG,Adh,ATPsynO | | 6.433e-05 | -9.65 | regulation of gene expression, epigenetic | biological process | GO:0040029 | 163 | 31 | 12345 | 1129 | Cbp80,trsn,Unr,Tlk,BigH1,piwi,Dcr-1,wash,CG6967,CG9425,rept,egg,AGO3,esc,Bin1,Hsc70-3,r2d2,psq,armi,Marf1,ph-d,CG10038,Hat1,spn-E,Elba2,l(3)73Ah,escl,gw,Chrac-14,Bap170,Hnf4 | | 6.623e-05 | -9.62 | Mec2 (Mec2) | protein interactions | 32905 | 87 | 24 | 9204 | 1112 | Rpt2,wal,Surf4,GCS2beta,Karybeta3,Vha68-2,Tspo,CG1416,Rab5,cathD,Pi3K92E,Atpalpha,ScsbetaA,Mcad,blw,Pp2A-29B,CCT3,CG15098,CG31729,CCT6,SERCA,CG10425,Kap-alpha3,Pdk1 | | 6.625e-05 | -9.62 | nuclear part | cellular component | GO:0044428 | 1416 | 172 | 11940 | 1105 | CSN6,Tlk,Bin1,Xpac,lwr,CG17187,CG16865,RpS18,d4,DNApol-epsilon255,gus,Mat89Ba,RpL11,CG10336,Sgf29,Elba2,RYBP,Ptip,CG6610,NFAT,Xpc,Utx,rept,l(3)73Ah,dl,vers,Cbp80,MED24,piwi,mad2,Topors,mute,CG2162,Non1,pont,Nf-YA,MED27,Aladin,LSm3,IntS6,psq,Imp,Clp,Snm1,fs(1)Ya,EcR,Mybbp1A,snRNA:U12,Ubc6,Spx,nej,Rbf2,CG4880,mts,Rpb8,Chrac-14,MED4,Tip60,CycC,CG8009,CG12288,RpII140,SelR,Neos,obe,CG17328,yps,NHP2,exu,CG10055,scny,e(y)1,Taf7,SmE,TfIIFalpha,Dd,Pym,YL-1,Trf2,stil,gw,CG9143,Top1,Orc6,CG17454,wash,Tfb4,SREBP,snRNA:U4:38AB,lid,bys,Su(H),bru1,CG46385,thoc6,CG1602,Rrp40,nbs,mbo,Sirt7,CG7655,ix,bif,Spt20,Rbfox1,Parp16,lin-52,escl,IntS11,Sld5,CG3198,CG11164,CG3511,Sgf11,Cenp-C,snRNA:U4atac,mre11,RpS15,APC7,RpS13,l(1)G0004,RpS2,CG4021,hb,Axs,awd,CG12301,sbr,CG31053,CG5986,trr,CG1622,Atu,Rel,Ndc1,Nap1,Trf4-2,Aos1,asf1,E2f1,HP1c,CG17768,CG15525,Nup58,SuUR,CG10418,Nup62,Bap170,HmgD,EloC,Rcd5,MED20,Usp39,CG9684,Prosbeta5,Hat1,mxc,not,esc,Orc1,CG12018,toc,Cse1,pygo,Nurf-38,Psf3,ph-d,Cdk7,pps,tho2,Rbf,PAPLA1 | | 6.868e-05 | -9.59 | ncRNA metabolic process | biological process | GO:0034660 | 322 | 51 | 12345 | 1129 | Dcr-1,mtTFB1,CysRS,vret,CG3021,mino,CG9143,Cbp80,scu,CG12301,RpL7A,mael,bys,shu,CycC,del,CG8204,Archease,TyrRS,RpS16,Rrp40,tapas,TrpRS-m,RpS7,SelD,CG30343,CG7009,CG2034,GatA,CG10189,IntS6,Trf4-2,Gasz,Mat89Ba,GluRS-m,IntS11,CG10463,mRpS11,piwi,CG2911,CG3587,RpS28b,CG5292,GatB,SerRS-m,RpS14a,CG7728,NHP2,armi,RpS14b,LysRS | | 6.874e-05 | -9.59 | Rheb (Ras homolog enriched in brain) | protein interactions | 117332 | 109 | 28 | 9204 | 1112 | eIF3m,Imp,RpL13,Actn,ATPsynbeta,RpS14a,alt,RpL9,Rbf,Hsc70-3,RpL7A,RpL10,eIF3i,RpS7,CG1416,RpS23,raptor,bif,Droj2,RpL30,Pi3K92E,tyf,Nap1,RpL17,CCT5,foxo,eIF4A,CG12301 | | 6.876e-05 | -9.58 | cell cycle phase transition | biological process | GO:0044770 | 37 | 12 | 12345 | 1129 | twe,CycE,CG8188,CG10336,CycJ,CycB,PpV,Cdk1,CG5003,tws,E2f1,cana | | 6.876e-05 | -9.58 | female meiosis chromosome segregation | biological process | GO:0016321 | 37 | 12 | 12345 | 1129 | Cenp-C,bif,lwr,cnn,wapl,Axs,nos,mtrm,Rab5,grp,mus304,eIF4E1 | | 6.962e-05 | -9.57 | UTP metabolic process | biological process | GO:0046051 | 4 | 4 | 12345 | 1129 | nmdyn-D6,nmdyn-D7,r,awd | | 6.962e-05 | -9.57 | UTP biosynthetic process | biological process | GO:0006228 | 4 | 4 | 12345 | 1129 | awd,nmdyn-D6,nmdyn-D7,r | | 7.055e-05 | -9.56 | UBQ-conjugat\_E2 | interpro domains | IPR000608 | 32 | 12 | 11383 | 1233 | ben,Uev1A,CG40045,CG8188,CG10254,Ubc6,CG2924,Ubc4,UbcE2H,lwr,CG46338,CG4502 | | 7.118e-05 | -9.55 | Diap2 (Death-associated inhibitor of apoptosis 2) | protein interactions | 36748 | 179 | 40 | 9204 | 1112 | Kcmf1,blw,CCT3,pont,cact,EndoGI,CCT6,SERCA,AMPdeam,cup,vret,Chd64,RpL12,RpS28b,GCS2beta,rept,shrb,larp,Nup358,RpS27,RpS16,CG7414,stai,RpL24,RpL7A,CG4853,RpL23A,alphaTub84B,Jafrac2,velo,AIF,Sod1,CG4557,RpS14a,Sgf29,Jafrac1,CCT5,Cp1,scu,CG9684 | | 7.371e-05 | -9.52 | GO\_FC\_EPSILON\_RECEPTOR\_SIGNALING\_PATHWAY | MSigDB lists | GO\_FC\_EPSILON\_RECEPTOR\_SIGNALING\_PATHWAY | 66 | 23 | 4161 | 644 | RpL40,CanB2,Prosbeta6,SkpA,Rpn12,Traf6,Rpn2,Pp2B-14D,Prosbeta3,Rpt2,Prosalpha4,Uev1A,Rpn3,Cul1,Prosbeta5,Pdk1,rl,cact,RpS27A,Prosalpha5,Prosalpha3,CanA-14F,Prosbeta4 | | 7.426e-05 | -9.51 | positive regulation of organelle organization | biological process | GO:0010638 | 142 | 28 | 12345 | 1129 | scny,Stat92E,piwi,bru1,dco,egg,Mtl,CG11486,esc,trr,Pld,Mitf,park,wash,Arpc4,G9a,Pink1,HP1e,APC7,Synj,pgc,Marf,Arpc2,Abi,Arpc1,HP1c,Moe,msps | | 7.464e-05 | -9.50 | UQ\_con | pfam domains | PF00179 | 32 | 12 | 10480 | 1142 | CG40045,UbcE2H,lwr,CG46338,ben,CG8188,CG4502,CG10254,Ubc6,CG2924,Uev1A,Ubc4 | | 7.490e-05 | -9.50 | Ald1 (Aldolase 1) | protein interactions | 43183 | 93 | 25 | 9204 | 1112 | Adh,Chd64,awd,up,Mf,Pgk,CG10638,Actn,CG14715,EndoGI,Fkbp12,RhoGDI,ATPsynCF6,AnxB10,Argk,regucalcin,Caf1-105,capt,scu,GlyP,Uch,CG15525,Pglym78,Eno,Scp1 | | 7.560e-05 | -9.49 | nucleoside diphosphate phosphorylation | biological process | GO:0006165 | 32 | 11 | 12345 | 1129 | Gapdh1,Pgk,nmdyn-D6,Pfk,Pfrx,awd,PyK,Scsalpha1,Eno,nmdyn-D7,Pglym78 | | 7.574e-05 | -9.49 | GstE6 (Glutathione S transferase E6) | protein interactions | 37111 | 38 | 14 | 9204 | 1112 | Stip1,RhoGDI,Uch,Eno,AnxB10,CG4598,Chd64,GstT1,CG10638,Gdi,regucalcin,Moe,scu,capt | | 7.675e-05 | -9.47 | programmed cell death | biological process | GO:0012501 | 172 | 32 | 12345 | 1129 | Atg7,sws,Drep2,raptor,Utx,Atg6,Cdk7,Tak1,Rbf,Jafrac2,crq,tub,Debcl,EcR,chrb,Mekk1,DCAF12,Tengl4,nbs,Pink1,Idh3b,RYBP,Tao,Dcr-1,MED24,AIF,Drep1,cathD,Cdk5,rept,Trf2,E2f1 | | 7.706e-05 | -9.47 | blw (bellwether) | protein interactions | 37617 | 52 | 17 | 9204 | 1112 | ATPsynB,ATPsynD,wal,ATPsynO,regucalcin,E2f1,CG15098,ATPsyndelta,ATPsynE,ATPsynF,Tak1,ATPsynCF6,ATPsynG,PGRP-LE,Rel,ATPsynbeta,SERCA | | 7.713e-05 | -9.47 | GO\_INTRACELLULAR\_PROTEIN\_TRANSPORT | MSigDB lists | GO\_INTRACELLULAR\_PROTEIN\_TRANSPORT | 359 | 82 | 4161 | 644 | Ranbp21,Snx6,Rab5,Atg4b,mbo,RpL30,Pex10,CG8728,Phb2,Oseg1,CG11110,Pp2B-14D,Atg6,RpS2,Tim8,Pex12,CanA-14F,RpL37a,RpL24,Syx13,Arl6IP1,cyc,ste14,CG4041,RpLP0,Cse1,RpS13,RpS30,RpL13A,ERp60,Pen,RpL13,RpS27A,RpL9,Rab32,RpS28b,Surf4,Faf2,RpS4,RpL38,RpS7,Rep,Tspo,RpL40,RpL23,CG1847,RpS24,lt,Pex13,GAPsec,RpS16,RpS18,Cdk5,CG8004,Kap-alpha3,Marf,CG8314,RpL12,Sec23,RpL11,Vamp7,Clc,RpL26,RpS27,Karybeta3,CHOp24,AP-1-2beta,GCC88,Stam,Arts,CG5734,CG2100,sau,HDAC6,CG9393,cact,rl,Tom40,milt,Rab10,shrb,RpS23 | | 7.727e-05 | -9.47 | GSE17721\_POLYIC\_VS\_PAM3CSK4\_16H\_BMDC\_DN | MSigDB lists | GSE17721\_POLYIC\_VS\_PAM3CSK4\_16H\_BMDC\_DN | 79 | 26 | 4161 | 644 | RpL38,Rpb8,PpV,CG8475,r,CG42542,Fdh,ND-23,CG13601,ND-49,CG17294,Mcad,scu,mRpS23,ABCD,RpS2,p38a,AIF,cact,Tim8,ND-24,Top1,shop,CSN6,RpL26,CG33181 | | 7.751e-05 | -9.47 | Rpt6 (Regulatory particle triple-A ATPase 6) | protein interactions | 33105 | 72 | 21 | 9204 | 1112 | Rpn3,Atg6,Droj2,CG1416,Prosalpha4,cnn,GCS2beta,Prosbeta6,Rpt2,wal,Rpn13,Pdk1,Rpn2,Prosalpha5,CCT6,Pomp,Prosbeta5,Prosbeta3,CCT3,Prosalpha3,Prosbeta4 | | 7.942e-05 | -9.44 | regulation of antimicrobial humoral response | biological process | GO:0002759 | 67 | 17 | 12345 | 1129 | Ptip,cact,tub,wisp,Myd88,mop,Herc4,Rel,spz,dl,lwr,Tak1,Stat92E,scny,Aos1,cad,p38a | | 8.046e-05 | -9.43 | cell death | biological process | GO:0008219 | 180 | 33 | 12345 | 1129 | Tak1,Rbf,Traf6,Jafrac2,sws,Drep2,Atg7,Utx,raptor,Atg6,Cdk7,Drep1,AIF,MED24,Dcr-1,cathD,rept,Cdk5,Trf2,E2f1,Debcl,tub,crq,EcR,Mekk1,chrb,Tengl4,DCAF12,nbs,Pink1,RYBP,Idh3b,Tao | | 8.068e-05 | -9.43 | UBQ-conjugating\_enzyme/RWD | interpro domains | IPR016135 | 42 | 14 | 11383 | 1233 | CG2924,CG13344,Ubc4,lwr,UbcE2H,CG17260,CG46338,CG4502,ben,Uev1A,CG40045,CG8188,CG10254,Ubc6 | | 8.155e-05 | -9.41 | nucleotide binding | molecular function | GO:0000166 | 955 | 122 | 12176 | 1119 | spn-E,sgll,CG4041,tkv,SERCA,Rpt2,CG6767,Pdk1,Dhfr,AOX3,CG8611,CG31689,Tlk,DNApol-epsilon255,GluRS-m,Coq6,Pgk,Cdk7,awd,Tak1,blw,Arf51F,Atpalpha,CG9143,ND-49,Vha68-2,Non1,tor,Klp10A,obe,Arl5,CG7728,CkIIalpha,Rab10,ATPsynbeta,Tao,Cdk5,CCT5,CCT6,Hsc70-3,CG31674,TrpRS-m,mEFTu2,Mekk1,CG7806,nmd,AIF,SdhA,Cdk1,CG31729,CG4586,mus205,alphaTub67C,mEFTu1,ACC,Idh,AcCoAS,CG4562,CG14997,ScsbetaA,CG5116,anne,Rab5,png,ND-51,ATP8A,wal,ABCD,CG9981,SRPK,Marf,fray,kl-2,CG10463,CCT3,E23,Lkb1,SelD,Acox57D-p,CG14183,CG4880,CG17698,TyrRS,Pink1,Pfrx,Pfk,Mtl,dco,rl,Orc1,Sirt7,l(2)k01209,Gapdh1,bor,CG4860,r,SNF4Agamma,Droj2,Rab32,cana,pont,rept,p38a,grp,alphaTub84B,RecQ5,CG8726,CysRS,Nadsyn,Roe1,CG5789,eIF4A,SerRS-m,Trxr-1,Mcad,gammaTub37C,Mos,PEK,nmdyn-D7,CG7488,LysRS,l(1)G0196 | | 8.155e-05 | -9.41 | nucleoside phosphate binding | molecular function | GO:1901265 | 955 | 122 | 12176 | 1119 | CG9143,ND-49,Atpalpha,awd,Cdk7,blw,Arf51F,Tak1,Coq6,Pgk,DNApol-epsilon255,GluRS-m,Tlk,CG31689,Dhfr,CG8611,AOX3,Rpt2,CG6767,Pdk1,tkv,SERCA,CG4041,sgll,spn-E,CG14997,ScsbetaA,AcCoAS,CG4562,mEFTu1,ACC,Idh,mus205,CG4586,alphaTub67C,CG31729,AIF,SdhA,Cdk1,CG7806,nmd,mEFTu2,TrpRS-m,Mekk1,CCT5,CCT6,Hsc70-3,CG31674,Rab10,Tao,ATPsynbeta,Cdk5,CG7728,CkIIalpha,obe,Arl5,tor,Klp10A,Vha68-2,Non1,dco,Pfrx,Pfk,Mtl,Pink1,CG17698,TyrRS,CG4880,Lkb1,SelD,Acox57D-p,CG14183,CG10463,E23,CCT3,kl-2,fray,Marf,SRPK,CG9981,ABCD,wal,ND-51,ATP8A,png,CG5116,anne,Rab5,CG7488,l(1)G0196,LysRS,Mos,nmdyn-D7,PEK,gammaTub37C,eIF4A,CG5789,SerRS-m,Trxr-1,Mcad,Roe1,Nadsyn,RecQ5,CG8726,CysRS,alphaTub84B,grp,pont,rept,p38a,cana,SNF4Agamma,Rab32,Droj2,CG4860,r,l(2)k01209,Sirt7,Gapdh1,bor,rl,Orc1 | | 8.191e-05 | -9.41 | GO\_POSITIVE\_REGULATION\_OF\_TRIGLYCERIDE\_BIOSYNTHETIC\_PROCESS | MSigDB lists | GO\_POSITIVE\_REGULATION\_OF\_TRIGLYCERIDE\_BIOSYNTHETIC\_PROCESS | 7 | 6 | 4161 | 644 | crq,Dd,SREBP,regucalcin,mino,EcR | | 8.191e-05 | -9.41 | GO\_POSITIVE\_REGULATION\_OF\_TRIGLYCERIDE\_METABOLIC\_PROCESS | MSigDB lists | GO\_POSITIVE\_REGULATION\_OF\_TRIGLYCERIDE\_METABOLIC\_PROCESS | 7 | 6 | 4161 | 644 | regucalcin,EcR,mino,Dd,crq,SREBP | | 8.210e-05 | -9.41 | GO\_ELECTRON\_TRANSPORT\_CHAIN | MSigDB lists | GO\_ELECTRON\_TRANSPORT\_CHAIN | 50 | 19 | 4161 | 644 | COX5A,ND-39,ND-24,ND-30,UQCR-Q,ND-75,wal,CG3803,ND-B18,SdhA,ND-SGDH,Cyt-c-p,CG11110,ND-49,ND-23,ND-51,Etf-QO,levy,ND-B22 | | 8.675e-05 | -9.35 | HSIAO\_HOUSEKEEPING\_GENES | MSigDB lists | HSIAO\_HOUSEKEEPING\_GENES | 198 | 51 | 4161 | 644 | Arl6IP1,Tsp74F,levy,CG11089,Prosbeta3,eIF4A,Rpt2,Gapdh1,Rpn12,RpLP0,Sod1,RpL13A,LysRS,Echs1,cact,RpL13,Pgk,eEF1beta,Fkbp59,Npc2a,CSN6,Arpc2,RpS28b,RpS23,RpL9,RpS27A,Nap1,YL-1,RpS7,RpL38,RpL40,CG2201,RpL23,CG4390,RpS2,RpS16,Atg6,Pfdn5,Phb2,Eno,Prosbeta5,RpS18,ND-24,capt,RpL12,Moe,Pglym78,RpL11,Prosbeta4,RpL24,Clc | | 8.746e-05 | -9.34 | Trxr-1 (Thioredoxin reductase-1) | protein interactions | 31760 | 34 | 13 | 9204 | 1112 | cl,Ho,Eno,awd,Chd64,cathD,AnxB10,CG42565,Sod1,Moe,regucalcin,CG10638,CG14715 | | 8.779e-05 | -9.34 | pyrimidine ribonucleotide biosynthetic process | biological process | GO:0009220 | 10 | 6 | 12345 | 1129 | l(2)k01209,r,nmdyn-D6,nmdyn-D7,r-l,awd | | 8.779e-05 | -9.34 | pyrimidine ribonucleotide metabolic process | biological process | GO:0009218 | 10 | 6 | 12345 | 1129 | r-l,awd,nmdyn-D6,nmdyn-D7,l(2)k01209,r | | 8.779e-05 | -9.34 | regulation of nuclear-transcribed mRNA poly(A) tail shortening | biological process | GO:0060211 | 10 | 6 | 12345 | 1129 | Marf1,gw,smg,AGO3,piwi,spn-E | | 8.779e-05 | -9.34 | positive regulation of nuclear-transcribed mRNA poly(A) tail shortening | biological process | GO:0060213 | 10 | 6 | 12345 | 1129 | gw,Marf1,piwi,AGO3,smg,spn-E | | 8.885e-05 | -9.33 | UBIQUITIN\_CONJUGAT\_2 | prosite domains | PS50127 | 32 | 12 | 6243 | 694 | CG46338,ben,Uev1A,Ubc6,lwr,CG10254,CG40045,CG2924,UbcE2H,CG4502,CG8188,Ubc4 | | 8.943e-05 | -9.32 | MODULE\_306 | MSigDB lists | MODULE\_306 | 21 | 11 | 4161 | 644 | SdhA,Pfk,Eno,CG5028,Pglym78,Pgk,Gapdh1,Idh,Scsalpha1,Irp-1B,Idh3b | | 8.984e-05 | -9.32 | Propanoate metabolism | KEGG pathways | dme00640 | 21 | 11 | 3198 | 496 | Mtpalpha,CG8199,CG10932,AcCoAS,ACC,Scsalpha1,Mcad,ScsbetaA,CG17691,Echs1,Gabat | | 8.984e-05 | -9.32 | Propanoate metabolism | KEGG pathways | ko00640 | 21 | 11 | 3198 | 496 | CG17691,Gabat,Echs1,Mcad,Scsalpha1,ACC,ScsbetaA,AcCoAS,Mtpalpha,CG10932,CG8199 | | 9.009e-05 | -9.31 | LI\_AMPLIFIED\_IN\_LUNG\_CANCER | MSigDB lists | LI\_AMPLIFIED\_IN\_LUNG\_CANCER | 71 | 24 | 4161 | 644 | lwr,RpL13,Top1,mEFTu1,Kap-alpha3,Eno,CCT5,SdhA,CCT3,Jafrac2,RpS27,RpL37a,Hsc70-3,Arl6IP1,RpS7,levy,RpS4,RpL38,Rpn12,RpS16,RpS2,Cyt-c-p,RpL30,Gmer | | 9.042e-05 | -9.31 | LOPEZ\_MBD\_TARGETS | MSigDB lists | LOPEZ\_MBD\_TARGETS | 339 | 78 | 4161 | 644 | CG9471,AcCoAS,Npc2a,CG8507,Nap1,eIF3i,RhoGDI,csw,Cse1,Rpn12,RpLP0,grsm,MED27,Pfk,CCT3,lwr,ND-24,Arpc4,CanA-14F,Prosalpha3,RpL37a,RpL24,Pp2A-29B,PDCD-5,tho2,sgll,PpV,Cyp1,Atg7,not,Rab5,CG17294,CG8728,RpS2,nrv1,CG1890,Cyt-c-p,Pp2B-14D,Tom40,Cdk1,cact,Ssdp,kcc,cl,Rab10,Got1,CG12929,Prosalpha5,Xpac,Rpb8,Jafrac1,ND-51,ND-23,CG5028,Prosbeta3,CG8188,Sod1,pont,Cdk5,Eno,CCT5,ND-B18,Stip1,ND-30,RpL11,mtgo,CG7414,gammaTub37C,mus81,RpS7,RfC38,RpL40,RpS24,CG30291,Dhfr,CG1847,Archease,GCS2beta | | 9.251e-05 | -9.29 | regulation of response to external stimulus | biological process | GO:0032101 | 228 | 39 | 12345 | 1129 | spz,G9a,PDCD-5,cact,wisp,tub,Ect4,TyrRS,Archease,EMRE,PGRP-LE,p38a,Hr78,Stat92E,Aos1,cad,faf,lwr,raptor,Seipin,Atg6,Myd88,Rel,Herc4,mop,GlyP,CG9925,Atg7,Hat1,Crag,Sp7,Ptip,slif,scny,Tak1,GILT1,Imp,dl,Traf6 | | 9.317e-05 | -9.28 | ND-PDSW (NADH dehydrogenase (ubiquinone) PDSW subunit) | protein interactions | 44228 | 18 | 9 | 9204 | 1112 | Adh,Mf,ND-75,up,Actn,Argk,ND-49,Nc73EF,Scp1 | | 9.389e-05 | -9.27 | positive regulation of response to biotic stimulus | biological process | GO:0002833 | 101 | 22 | 12345 | 1129 | GILT1,Tak1,lwr,dl,Hr78,PGRP-LE,cad,Aos1,wisp,Hat1,Crag,tub,Ptip,EMRE,Sp7,spz,PDCD-5,Herc4,Rel,mop,CG9925,Myd88 | | 9.490e-05 | -9.26 | organelle assembly | biological process | GO:0070925 | 385 | 58 | 12345 | 1129 | bru1,CG17768,up,Vps16B,RpL23A,grp,yem,Grip75,mRpS11,RpS15,RpS27,LSm3,sau,mts,larp,RpS14a,CG7488,Atg6,Klp10A,CG13366,RpL11,Pp2A-29B,SelR,RpS14b,mop,CanB2,gammaTub37C,Kap3,Atg7,TMEM216,RpS28b,CG7716,Grip128,Oseg1,SRPK,CG3437,Actn,Edc3,milt,Nap1,cnn,Grip84,pont,RpL24,alphaTub67C,RpLP0,tws,CCT6,RpS5b,Rcd5,Snx6,osk,Non1,wisp,Grip71,RpL12,Cenp-C,RpL10 | | 9.587e-05 | -9.25 | Hel25E (Helicase at 25E) | protein interactions | 33781 | 48 | 16 | 9204 | 1112 | Cp1,wal,mEFTu1,Vha68-2,Droj2,Rab5,CG1416,Mcad,ScsbetaA,Atpalpha,cathD,CCT3,CG2199,CCT6,SERCA,CG10425 | | 9.587e-05 | -9.25 | kdn (knockdown) | protein interactions | 31579 | 48 | 16 | 9204 | 1112 | Mf,Marf,awd,up,CG7920,yip2,Adh,cib,Actn,ATPsyndelta,ATPsynCF6,Cyt-c-p,Argk,ND-18,Eno,Scp1 | | 9.784e-05 | -9.23 | Cyp1 (Cyclophilin 1) | protein interactions | 32595 | 30 | 12 | 9204 | 1112 | Eno,Rtnl1,CG8204,CG14814,regucalcin,gus,Sod1,Fkbp12,AnxB10,awd,Chd64,CG6707 | | 9.925e-05 | -9.22 | MOOTHA\_PGC | MSigDB lists | MOOTHA\_PGC | 204 | 52 | 4161 | 644 | Mo25,stas,Prosalpha5,Scsalpha1,Pgk,mEFTu1,Shmt,GlyP,Tom40,AIF,SdhA,mRpL13,CG3860,ND-SGDH,CG9393,nmdyn-D7,sau,scu,Pgm1,Mcad,mRpL18,eIF4A,Gapdh1,Gk2,Jafrac1,rept,CG5028,levy,CG14407,Hsc70-3,CG3376,ND-24,Thiolase,ND-30,Pglym78,Tim8,Eno,Prosbeta5,CG8004,Pfk,Pdxk,ari-1,ScsbetaA,Cyt-c-p,Phb2,colt,Vha36-1,Non1,tko,CG14894,Roe1,CG17333 | | 9.941e-05 | -9.22 | F-type ATPase, eukaryotes | KEGG pathways | M00158 | 18 | 10 | 3198 | 496 | ATPsynO,ATPsynG,ATPsyndelta,ATPsynD,ATPsynF,blw,ATPsynCF6,ATPsynE,ATPsynB,ATPsynbeta | | 9.941e-05 | -9.22 | F-type ATPase, eukaryotes | KEGG pathways | dme\_M00158 | 18 | 10 | 3198 | 496 | ATPsynbeta,ATPsynB,ATPsynE,ATPsynCF6,blw,ATPsynF,ATPsynD,ATPsyndelta,ATPsynO,ATPsynG | | 1.004e-04 | -9.21 | MODULE\_42 | MSigDB lists | MODULE\_42 | 15 | 9 | 4161 | 644 | ND-51,ND-23,ND-30,ND-24,ND-B18,ND-75,ND-SGDH,ND-49,ND-39 | | 1.004e-04 | -9.21 | GO\_INTERSPECIES\_INTERACTION\_BETWEEN\_ORGANISMS | MSigDB lists | GO\_INTERSPECIES\_INTERACTION\_BETWEEN\_ORGANISMS | 271 | 65 | 4161 | 644 | RpL30,RpS2,Atg6,tho2,Ts,Atg7,mbo,Prosalpha3,RpL37a,RpL24,Smox,Prosbeta5,CHMP2B,lwr,Cul2,RpS13,RpLP0,RpS30,eIF4E1,RpL13A,LysRS,Fur2,Clp,RpS28b,RpL9,RpS27A,Abi,Cul5,crq,Top1,RpL13,Pen,RpS24,RpL23,RpS16,Aladin,RpS7,RpS4,RpL38,Prosbeta6,RpL40,heph,RpL11,Prosbeta4,RpS27,RpL26,thoc6,RpS18,SRPK,sbr,RpL12,Kap-alpha3,Cul1,Prosbeta3,eIF4A,Prip,Vsp37A,Karybeta3,Tip60,shrb,CSN6,CG14516,RpS23,cact,rl | | 1.004e-04 | -9.21 | GO\_MITOCHONDRIAL\_PART | MSigDB lists | GO\_MITOCHONDRIAL\_PART | 477 | 103 | 4161 | 644 | CG8199,Aldh,SdhA,wal,Tom40,CG3803,Ppox,CG1774,Unr,Scsalpha1,ND-39,CG4407,CG10932,CG5028,ND-51,ND-23,GluRS-m,ND-B22,CG17691,Stoml2,sau,scu,Sod1,CG9393,mino,ND-SGDH,ND-49,CG2100,Mcad,Cisd2,mRpL18,CG8974,mRpS5,Marf,ND-30,shop,Thiolase,ND-B18,Mtpalpha,CG8004,sws,yip2,CG14407,Pmi,stet,Tspo,Roe1,CG1640,mtTFB1,colt,mRpS23,CG31759,mRpL16,UQCR-Q,Calx,Ect4,mEFTu1,AIF,SerRS-m,ND-75,mRpS10,Echs1,Trxr-1,Idh3b,Rab32,mRpS21,Nc73EF,TrpRS-m,Gk2,Grx1,levy,tzn,Coq6,mRpL13,LysRS,Sfxn2,ND-24,Coq7,Tim8,COX5A,Gabat,Scox,CG7488,mRpL17,mRpL10,Mitofilin,Pdk,Cyp1,Atg4b,tko,Etf-QO,Ts,Phb2,CG11110,CG11858,Atg6,ScsbetaA,Cyt-c-p,bor,CG7718,CG8728,P5CDh1,P5cr | | 1.013e-04 | -9.20 | spindle assembly involved in female meiosis | biological process | GO:0007056 | 14 | 7 | 12345 | 1129 | Klp10A,Grip71,wisp,Grip75,gammaTub37C,Grip128,SRPK | | 1.019e-04 | -9.19 | regulation of cell population proliferation | biological process | GO:0042127 | 229 | 39 | 12345 | 1129 | Rab5,Rel,bun,Utx,MAGE,CycE,COX5A,Atu,COX4,Nup358,Rbf,MED4,mts,dl,hb,foxo,CG10336,Cul5,ago,tor,SelD,MED27,Smox,tws,EcR,Ptp61F,Moe,Stat92E,Idgf2,cad,l(3)mbt,E2f1,pont,Pi3K92E,rept,lwr,Cbl,rl,Nurf-38 | | 1.030e-04 | -9.18 | electron transfer activity | molecular function | GO:0009055 | 68 | 17 | 12176 | 1119 | levy,ND-75,COX8,COX5B,SdhA,UQCR-11,Trxr-1,COX5A,AOX3,UQCR-Q,Grx1,COX4,Etf-QO,Cyt-c-p,wal,CG14407,cype | | 1.038e-04 | -9.17 | regulation of immune effector process | biological process | GO:0002697 | 88 | 20 | 12345 | 1129 | PGRP-LE,Stat92E,scny,Aos1,cad,Tak1,dl,lwr,Traf6,spz,G9a,Myd88,mop,Rel,Herc4,cact,tub,wisp,Atg7,Ptip | | 1.038e-04 | -9.17 | meiotic chromosome segregation | biological process | GO:0045132 | 88 | 20 | 12345 | 1129 | c(2)M,mus81,lwr,cnn,wapl,nos,grp,CG31053,Topors,cort,Cenp-C,bif,Axs,mtrm,Rab5,gammaTub37C,cup,mus304,eIF4E1,tef | | 1.041e-04 | -9.17 | nucleotide phosphorylation | biological process | GO:0046939 | 33 | 11 | 12345 | 1129 | Pglym78,nmdyn-D7,Eno,Scsalpha1,awd,PyK,Pfrx,nmdyn-D6,Pfk,Gapdh1,Pgk | | 1.068e-04 | -9.14 | fatty acid metabolic process | biological process | GO:0006631 | 123 | 25 | 12345 | 1129 | mino,CG4586,Echs1,CG31523,Thiolase,SREBP,CG10932,scu,CG4598,SERCA,wal,Acox57D-p,CG4594,AcCoAS,Mcad,Desat1,CG6178,Etf-QO,CG4860,Seipin,Mtpalpha,Pex10,ACC,yip2,Pxt | | 1.079e-04 | -9.13 | aging | biological process | GO:0007568 | 175 | 32 | 12345 | 1129 | PGRP-LE,Hsp26,p38a,Thiolase,Edem1,Cdk5,rl,alpha-Man-Ia,Sod1,Jafrac1,Smox,levy,Trxr-1,ATPsynD,Coq7,EcR,lt,Atpalpha,Lkb1,ND-SGDH,alph,snz,Tsp42Ef,cnc,AGBE,foxo,Mtpalpha,GlyP,bun,raptor,Tspo,Atg7 | | 1.081e-04 | -9.13 | spindle assembly | biological process | GO:0051225 | 75 | 18 | 12345 | 1129 | Grip71,wisp,Non1,Grip128,CG7716,Klp10A,gammaTub37C,mop,Pp2A-29B,mts,larp,pont,Grip84,sau,Grip75,CG3437,grp,SRPK | | 1.085e-04 | -9.13 | spindle assembly involved in meiosis | biological process | GO:0090306 | 23 | 9 | 12345 | 1129 | Grip75,gammaTub37C,SRPK,Klp10A,larp,Grip128,sau,Grip71,wisp | | 1.085e-04 | -9.13 | cellular aldehyde metabolic process | biological process | GO:0006081 | 23 | 9 | 12345 | 1129 | Fdh,AOX3,CG4390,Adh,Gip,sgll,Pdxk,Aldh,tzn | | 1.097e-04 | -9.12 | pattern specification process | biological process | GO:0007389 | 544 | 76 | 12345 | 1129 | crm,rl,Dcr-1,spen,boi,faf,capt,ovo,tkv,Pka-R1,Tm1,cad,mael,cact,tub,Rbfox1,bcd,EcR,fs(1)N,twe,enc,spz,snk,gus,dl,Cul5,krimp,Bx,Lkb1,Grip75,Dd,trk,Med,spn-E,Sec23,sbb,Grip128,Ptip,heph,Cf2,disp,cup,gammaTub37C,Myd88,fs(1)M3,milt,lin,Cbl,apt,swa,nmd,Stat92E,Moe,osk,Grip71,CycC,EloC,jvl,tws,gd,pygo,Tak1,exu,tor,nos,mts,hb,Ssdp,cnc,ea,msps,csw,CkIIalpha,Su(H),psq,armi | | 1.099e-04 | -9.12 | protein dephosphorylation | biological process | GO:0006470 | 109 | 23 | 12345 | 1129 | ssh,mop,Ptp69D,CanB2,CG17746,Pp2A-29B,tws,Ptpmeg2,CanA-14F,Ptp10D,CG5026,twe,CG7115,Ptp61F,csw,PpV,Ublcp1,Dd,alph,PpD3,Ptpmeg,mts,Pp2B-14D | | 1.100e-04 | -9.11 | cellular nitrogen compound biosynthetic process | biological process | GO:0044271 | 1175 | 144 | 12345 | 1129 | Pdxk,RpL12,RpL38,RpLP0,mRpL48,CG13551,ATPsynE,RpS5b,CG8199,mEFTu2,Nurf-38,CysRS,Pfas,nmdyn-D7,RpL7A,Shmt,RpS24,RpL13A,awd,ATPsynbeta,RpS14a,Ady43A,RpL23,RpS14b,RpS18,Orc1,eIF4A,RpL37a,fal,MED4,RpL34a,l(2)k01209,CG11089,Rga,AsnS,ATPsynO,mRpL35,TrpRS-m,RpS7,Pym,mRpL36,e(y)1,mtTFB1,Trf2,RpL29,mRpS5,RpL31,CycE,sgll,RpS4,Scsalpha1,Ppcdc,CG6094,Tfb4,LysRS,Gclc,eIF3i,nmdyn-D6,RpS15,COX5B,Ggt-1,Sptr,RpL23A,pr,mRpL10,mRpS23,blw,RpL10,TyrRS,mRpL13,RpL24,ATPsynD,CG7009,AMPdeam,RpL17,CG6767,mEFTu1,MED24,mRpL17,RpS30,Ts,Pcd,RpS23,Gmer,E2f1,CG17691,r-l,AcCoAS,mRpS21,mRpS10,r,SerRS-m,RpS28b,eIF4E1,Dhfr,CG6178,RpL11,RpS27,pygo,GatA,RpL13,ATPsynCF6,Rpb8,GAPsec,mus205,RpL9,RpS27A,mRpL16,Nadsyn,RpS16,RpL26,CG14817,Ac3,spen,RpL40,Taf7,TfIIFalpha,CG7414,RpL34b,mRpL18,CG33156,ATPsyndelta,GatB,ATPsynB,ATPsynG,eIF3m,CG4407,tko,RpLP1,Cdk7,cN-IIIB,RpS2,RpS29,Rbf,Ppox,pps,GluRS-m,RpII140,CG3803,RpL30,mRpS11,Sgf29,eEF1beta,RpS13 | | 1.138e-04 | -9.08 | fly chr88A3-88A3\|3-54 cM | chromosome location | fly chr88A3-88A3\|3-54 cM | 6 | 4 | 25010 | 1343 | CG9925,asRNA:CR43460,CG9926,CG31517 | | 1.138e-04 | -9.08 | fly chr88A3-88A3\|3-54 | chromosome location | fly chr88A3-88A3\|3-54 | 6 | 4 | 25010 | 1343 | CG31517,CG9926,asRNA:CR43460,CG9925 | | 1.148e-04 | -9.07 | Chc (Clathrin heavy chain) | protein interactions | 32537 | 112 | 28 | 9204 | 1112 | Rab5,Nup358,Aldh,CG1416,Mcad,Clc,Atpalpha,r,foxo,Rpt2,mEFTu1,wal,faf,Vha68-2,Tspo,Karybeta3,bor,SERCA,CCT6,Kap-alpha3,Pdk1,RpLP0,CCT3,blw,Cdk5,Pp2A-29B,sau,Gga | | 1.176e-04 | -9.05 | phosphoprotein phosphatase activity | molecular function | GO:0004721 | 102 | 22 | 12176 | 1119 | Pp2B-14D,csw,Ptp10D,Dd,Pp2A-29B,twe,Ptp69D,Ptpmeg,tws,alph,CG5026,PpV,PpD3,CG7115,Ublcp1,CG17746,CanA-14F,mts,Ptpmeg2,Ptp61F,mop,ssh | | 1.205e-04 | -9.02 | Yb body | cellular component | GO:0070725 | 7 | 5 | 11940 | 1105 | armi,shu,piwi,vret,BoYb | | 1.239e-04 | -9.00 | me31B (maternal expression at 31B) | protein interactions | 34364 | 35 | 13 | 9204 | 1112 | cup,yps,bru1,BicC,nos,Lsm12,tral,tyf,eIF4E1,exu,GILT1,png,Edc3 | | 1.260e-04 | -8.98 | regulation of biological quality | biological process | GO:0065008 | 1352 | 162 | 12345 | 1129 | Cbl,CG9186,ERp60,wash,Nap1,CG14036,Girdin,Pax,Rab32,Tomosyn,Shmt,Mctp,dco,Snm1,Mekk1,Marf,zyd,osk,NFAT,ssh,CG14407,Scox,tws,Hsc70-3,MtnA,SREBP,Jafrac2,CG10470,sip3,cyc,scramb1,awd,mute,CG8112,Atf3,lqfR,Sap47,Ho,CkIIalpha,Sec10,adp,Pi3K92E,Mtl,boi,Rbcn-3A,SERCA,ovo,CG8726,cact,Pfk,fwe,Pink1,crok,foxo,Cul5,Pdk1,Atpalpha,cold,Lkb1,anne,up,nAChRalpha4,CycE,Arpc2,Synj,Sec23,Med,bun,CG12773,nrv1,Rel,CCT3,Sod1,park,Actn,CG32473,trsn,scu,p38a,CG9911,CG11077,Npc2a,Fer1HCH,Ork1,Exo70,Desat1,Bap170,Moe,ATP8A,blw,Iml1,ZnT35C,SNF4Agamma,Trxr-1,Tao,Eno,Cul2,ATPsynD,alc,shv,CG4586,Pgk,Gabat,nos,CG31729,cnc,CG3842,Abi,sws,Plap,Arpc1,Arpc4,CanB2,CG6927,mus304,HDAC6,Dhfr,Seipin,smg,nej,Cdk5,Nup62,Nprl2,rl,Jafrac1,SuUR,Fdh,tkv,BigH1,SkpA,capt,Rep,Gdi,Hnf4,EcR,bif,PyK,EMRE,Ptp61F,Debcl,Syx13,Tip60,X11L,Sec15,nbs,Rbf,Irk1,mre11,PHGPx,Acox57D-p,slif,bru1,Arf51F,Grx1,Vha68-2,dob,Rab5,cup,CG9981,CSN6,raptor,mbo | | 1.296e-04 | -8.95 | Kr-h2 (Kruppel homolog 2) | protein interactions | 33859 | 64 | 19 | 9204 | 1112 | ScsbetaA,Mcad,Tspo,wal,GCS2beta,Surf4,Kap-alpha3,CG10425,Echs1,SERCA,CCT6,Dip-B,CG15098,CG31729,Cisd2,ATPsynCF6,nSMase,Pp2A-29B,CCT3 | | 1.307e-04 | -8.94 | Mitochondrial Fatty Acid Beta-Oxidation | REACTOME pathways | R-DME-77289 | 13 | 8 | 3923 | 574 | CG4598,CG4594,Mcad,Mtpalpha,yip2,CG4860,Thiolase,Echs1 | | 1.312e-04 | -8.94 | dorsal/ventral pattern formation | biological process | GO:0009953 | 154 | 29 | 12345 | 1129 | spn-E,sbb,tub,Med,csw,cact,jvl,enc,snk,CkIIalpha,spz,Cf2,Su(H),Myd88,gammaTub37C,cup,gd,milt,Cbl,faf,dl,mts,cnc,nmd,krimp,capt,mael,tkv,ea | | 1.316e-04 | -8.94 | Myo10A (Myosin 10A) | protein interactions | 32028 | 59 | 18 | 9204 | 1112 | mts,Kap-alpha3,up,awd,grp,AGBE,CG18135,RpL34a,RpL23,smg,CG2258,squ,RpS5b,Hsp26,milt,BicC,osk,RpS13 | | 1.316e-04 | -8.94 | Mlc2 (Myosin light chain 2) | protein interactions | 43587 | 59 | 18 | 9204 | 1112 | Cyt-c-p,CG17260,ATPsynO,Nc73EF,Argk,APP-BP1,ATPsynB,Eno,Scp1,ND-18,nmd,Adh,up,Mf,Actn,ATPsynF,COX4,ATPsynCF6 | | 1.320e-04 | -8.93 | germ-line stem cell division | biological process | GO:0042078 | 45 | 13 | 12345 | 1129 | piwi,twin,Stat92E,tkv,bam,egg,shrb,mad2,BoYb,Dcr-1,Med,nos,eIF4A | | 1.332e-04 | -8.92 | localization | biological process | GO:0051179 | 2206 | 248 | 12345 | 1129 | Tsp86D,shrb,zpg,Prip,milt,Stat92E,Actn,CG4293,glob1,Gmer,rpk,Npc2a,swa,Hr78,Cenp-C,colt,Fer1HCH,CG31689,Ugalt,lt,jvl,Exo70,Ork1,CG8204,ATP8A,Cse1,Rab10,Moe,Grip71,blw,Snx6,ZnT35C,SNF4Agamma,ATPsynD,ATPsynCF6,kcc,Ubc6,nos,tor,Tim8,NaPi-III,Vha36-1,Afti,GAPsec,CG31729,Nup358,Trs33,cnc,Nxf3,fray,Abi,msps,CG9582,Arpc1,CG6766,CG8314,sws,CG6927,CG4476,CanB2,Klp10A,HDAC6,Vamp7,Seipin,nej,smg,Nup62,Cdk5,faf,Jafrac1,Dcr-1,SuUR,tkv,mol,CG7458,Rep,CG3077,Gdi,l(3)mbt,Oseg1,Sfxn2,EcR,Arl5,del,EMRE,pgc,wisp,Debcl,CG5447,gus,Syx13,X11L,CG6293,CG10960,Rrp40,alphaTub67C,dx,CG8728,Sec15,CG9132,larp,sau,Irk1,CG7091,sea,Kap-alpha3,Grip75,CG4572,slif,ATPsynB,Grip128,ATPsynG,Arf51F,eIF3m,CG7806,Clc,Sec22,Vha68-2,ATPsyndelta,Crag,cup,Kap3,Rab5,Karybeta3,CG9981,sw,mbo,Efr,CG9186,E23,lwr,Stam,wash,CycB,Girdin,Pax,thoc6,Tomosyn,Vsp37A,Rab32,apt,Blos2,yin,ATPsynF,BicC,dco,zyd,Marf,Mekk1,osk,ABCD,ssh,Scox,ATPsynE,CG8004,CG9393,alphaTub84B,MICAL-like,Pex13,exu,Tango10,Vps16B,slam,Mat89Ba,sip3,scramb1,Sply,CG5789,awd,CG42837,Tom70,csw,Ranbp21,lqfR,Pex10,armi,ATPsynbeta,Ptpmeg2,Ndc1,opm,cana,CG4991,CkIIalpha,Atg6,Sec10,Pi3K92E,Mtl,CG31262,stai,CG5510,Calx,Vha13,boi,CG10344,obe,sbr,SERCA,cbs,Tm1,mael,Roe1,CG9399,cact,CHOp24,ATPsynO,crq,CG4041,trbd,CG11110,AP-1-2beta,fwe,Nup58,GCC88,Pink1,Gga,COX5B,hoe1,CG33181,PAPLA1,tho2,not,Lkb1,Atpalpha,anne,Topors,Aladin,nAChRalpha4,CG5734,Synj,Pen,Pi4KIIalpha,Arts,Tom40,CG14894,Gmap,Nlg1,Sec23,CG12106,spn-E,gammaTub37C,blot,nrv1,mop,Pex12,bun,CG12773,CHMP2B,disp | | 1.339e-04 | -8.92 | regionalization | biological process | GO:0003002 | 503 | 71 | 12345 | 1129 | cact,tub,bcd,Rbfox1,fs(1)N,EcR,enc,snk,spz,gus,rl,crm,spen,boi,Dcr-1,faf,capt,ovo,tkv,mael,cad,Tm1,trk,Sec23,spn-E,sbb,Med,Grip128,Ptip,Cf2,disp,Myd88,gammaTub37C,cup,fs(1)M3,dl,Cul5,krimp,Grip75,Lkb1,Bx,Dd,Moe,Grip71,CycC,osk,jvl,EloC,tws,gd,milt,lin,Cbl,apt,swa,nmd,Stat92E,csw,msps,CkIIalpha,Su(H),armi,psq,pygo,nos,tor,exu,mts,hb,cnc,ea | | 1.359e-04 | -8.90 | GO\_NUCLEOSIDE\_TRIPHOSPHATE\_METABOLIC\_PROCESS | MSigDB lists | GO\_NUCLEOSIDE\_TRIPHOSPHATE\_METABOLIC\_PROCESS | 104 | 31 | 4161 | 644 | ND-30,Pgk,ND-24,Pglym78,UQCR-Q,nmdyn-D6,Eno,CG3803,ND-75,ND-B18,Pfk,COX5A,ND-39,Scsalpha1,Nc73EF,ND-23,ND-51,r,ND-B22,Stoml2,Ts,levy,Cyt-c-p,ND-SGDH,mal,AMPdeam,nmdyn-D7,Pgm1,ND-49,CG33156,Gapdh1 | | 1.366e-04 | -8.90 | establishment of localization | biological process | GO:0051234 | 1717 | 199 | 12345 | 1129 | CG9393,MICAL-like,Pex13,Tango10,exu,Mat89Ba,Vps16B,sip3,scramb1,CG5789,csw,Tom70,CG42837,lqfR,Ranbp21,Pex10,opm,Ndc1,ATPsynbeta,Atg6,cana,CG4991,E23,Efr,wash,CycB,Stam,thoc6,Girdin,Rab32,Vsp37A,Tomosyn,yin,Blos2,dco,ATPsynF,Marf,Mekk1,zyd,ABCD,ssh,Scox,CG8004,ATPsynE,COX5B,PAPLA1,tho2,hoe1,CG33181,Atpalpha,anne,Aladin,Topors,nAChRalpha4,Pen,Synj,CG5734,Tom40,Arts,CG14894,CG12106,Nlg1,Sec23,Gmap,gammaTub37C,CG12773,mop,Pex12,nrv1,blot,disp,CHMP2B,Sec10,Mtl,stai,CG5510,Calx,CG31262,Vha13,SERCA,sbr,CG10344,cbs,Roe1,CG9399,ATPsynO,CHOp24,CG4041,crq,AP-1-2beta,CG11110,Nup58,fwe,GCC88,Gga,Pink1,ATPsynCF6,kcc,Ubc6,NaPi-III,Tim8,Vha36-1,CG31729,Afti,GAPsec,Trs33,Nup358,Nxf3,CG9582,msps,CG8314,CG6766,Arpc1,CanB2,CG4476,CG6927,HDAC6,Klp10A,Vamp7,zpg,shrb,Tsp86D,Prip,milt,glob1,Gmer,CG4293,Hr78,Npc2a,rpk,Ugalt,colt,CG31689,Fer1HCH,Cenp-C,Ork1,jvl,Exo70,lt,Rab10,Moe,Cse1,ATP8A,blw,Snx6,ZnT35C,ATPsynD,CG9132,Irk1,sau,CG7091,Kap-alpha3,sea,slif,CG4572,ATPsynG,Arf51F,ATPsynB,CG7806,Clc,eIF3m,Vha68-2,Sec22,ATPsyndelta,Crag,Karybeta3,Rab5,Kap3,sw,CG9981,mbo,nej,smg,faf,Cdk5,Nup62,Dcr-1,mol,l(3)mbt,CG7458,Gdi,Rep,CG3077,Oseg1,Sfxn2,Arl5,EcR,EMRE,Debcl,wisp,CG5447,Syx13,CG6293,X11L,CG10960,Sec15,CG8728,alphaTub67C,dx | | 1.404e-04 | -8.87 | KIM\_BIPOLAR\_DISORDER\_OLIGODENDROCYTE\_DENSITY\_CORR\_UP | MSigDB lists | KIM\_BIPOLAR\_DISORDER\_OLIGODENDROCYTE\_DENSITY\_CORR\_UP | 311 | 72 | 4161 | 644 | Scsalpha1,Got1,YL-1,SdhA,Aldh,Pgk,Nca,Prosbeta3,Mcad,ND-SGDH,Sod1,scu,ND-51,ND-23,Gyg,RpL11,CG4872,RpS27,Clc,Eno,Cdk5,Gdi,Stip1,RpS18,ND-30,RpL12,CG13630,Cul1,RpS24,AnxB10,Kap3,CG14894,Pka-R1,RpS4,CG1416,Atpalpha,Npc2a,Arpc2,Jheh2,Idh3b,RpS27A,CG8507,PAPLA1,Echs1,mEFTu1,ERp60,UQCR-Q,Rpt2,Rpn12,RpS13,RpS30,tzn,levy,RpL37a,Prosalpha3,APP-BP1,CG11807,RpL24,Pp2A-29B,COX5A,pr,Prosbeta5,CCT3,CG7322,Pfk,capt,TyrRS,CG2493,CG9132,Phb2,Pfdn5,PpV | | 1.413e-04 | -8.86 | mitotic cell cycle phase transition | biological process | GO:0044772 | 34 | 11 | 12345 | 1129 | CG8188,twe,CycE,PpV,CycB,CycJ,CG5003,Cdk1,cana,E2f1,tws | | 1.427e-04 | -8.85 | response to lipid | biological process | GO:0033993 | 70 | 17 | 12345 | 1129 | Hnf4,EcR,Idh3b,MED27,G9a,Utx,rept,SREBP,trr,MED24,Nurf-38,Tak1,CG5412,Stat92E,Hr78,rig,Trf2 | | 1.427e-04 | -8.85 | transport | biological process | GO:0006810 | 1659 | 193 | 12345 | 1129 | zyd,Marf,Mekk1,ABCD,Scox,ssh,CG8004,ATPsynE,Efr,E23,Girdin,thoc6,Stam,wash,Vsp37A,Tomosyn,Rab32,dco,ATPsynF,Blos2,yin,CG5789,scramb1,Ranbp21,lqfR,CG42837,Tom70,csw,Pex10,CG4991,Atg6,ATPsynbeta,Ndc1,opm,MICAL-like,CG9393,Tango10,exu,Pex13,sip3,Vps16B,Mat89Ba,crq,CG4041,CHOp24,ATPsynO,Nup58,fwe,CG11110,AP-1-2beta,Pink1,Gga,GCC88,Mtl,Sec10,Vha13,CG31262,stai,Calx,CG5510,cbs,CG10344,sbr,SERCA,CG9399,Roe1,CG5734,Synj,Pen,Gmap,Nlg1,CG12106,Sec23,Tom40,Arts,CG14894,nrv1,blot,mop,CG12773,Pex12,gammaTub37C,CHMP2B,disp,CG33181,hoe1,tho2,PAPLA1,COX5B,anne,Atpalpha,nAChRalpha4,Topors,Aladin,lt,Ork1,jvl,Exo70,Fer1HCH,CG31689,colt,Ugalt,blw,ATP8A,Cse1,Moe,Rab10,Snx6,ZnT35C,ATPsynD,Prip,Tsp86D,shrb,zpg,milt,rpk,Npc2a,Hr78,CG4293,Gmer,glob1,Nxf3,Arpc1,CG6766,CG8314,CG9582,CG4476,CG6927,CanB2,Vamp7,HDAC6,ATPsynCF6,kcc,Ubc6,Vha36-1,NaPi-III,Tim8,Trs33,Nup358,GAPsec,Afti,CG31729,EMRE,EcR,Arl5,wisp,Debcl,X11L,CG6293,Syx13,CG5447,alphaTub67C,dx,CG8728,Sec15,CG10960,Cdk5,Nup62,faf,nej,Dcr-1,mol,Sfxn2,Oseg1,CG3077,CG7458,Rep,Gdi,l(3)mbt,eIF3m,Clc,CG7806,ATPsynB,Arf51F,ATPsynG,ATPsyndelta,Crag,Vha68-2,Sec22,CG9981,sw,Rab5,Kap3,Karybeta3,mbo,sau,Irk1,CG9132,CG7091,sea,Kap-alpha3,slif,CG4572 | | 1.440e-04 | -8.85 | CG11267 (uncharacterized protein) | protein interactions | 39476 | 40 | 14 | 9204 | 1112 | HmgD,Stip1,Eno,awd,Chd64,AnxB10,cib,PHGPx,scu,Fkbp12,Moe,regucalcin,CG10638,CG14715 | | 1.477e-04 | -8.82 | GO\_OXIDOREDUCTASE\_ACTIVITY\_ACTING\_ON\_NAD\_P\_H | MSigDB lists | GO\_OXIDOREDUCTASE\_ACTIVITY\_ACTING\_ON\_NAD\_P\_H | 40 | 16 | 4161 | 644 | ND-24,ND-30,ND-75,CG3803,CG7322,ND-B18,AIF,CG3609,Trxr-1,cl,ND-39,ND-23,ND-51,ND-B22,ND-SGDH,ND-49 | | 1.490e-04 | -8.81 | proton-transporting ATP synthase activity, rotational mechanism | molecular function | GO:0046933 | 19 | 8 | 12176 | 1119 | ATPsynO,ATPsyndelta,ATPsynD,ATPsynE,blw,ATPsynB,ATPsynbeta,ATPsynG | | 1.534e-04 | -8.78 | spindle organization | biological process | GO:0007051 | 133 | 26 | 12345 | 1129 | alphaTub67C,Klp10A,sw,Pp2A-29B,mop,gammaTub37C,wisp,Non1,Grip71,spd-2,Axs,dgt4,msps,del,CG7716,Grip128,SRPK,CG3437,grp,Lkb1,Grip75,sau,pont,Grip84,mts,larp | | 1.541e-04 | -8.78 | GO\_OXIDOREDUCTASE\_ACTIVITY\_ACTING\_ON\_NAD\_P\_H\_QUINONE\_OR\_SIMILAR\_COMPOUND\_AS\_ACCEPTOR | MSigDB lists | GO\_OXIDOREDUCTASE\_ACTIVITY\_ACTING\_ON\_NAD\_P\_H\_QUINONE\_OR\_SIMILAR\_COMPOUND\_AS\_ACCEPTOR | 22 | 11 | 4161 | 644 | ND-SGDH,ND-49,ND-39,ND-30,ND-51,ND-23,ND-24,ND-B22,CG7322,ND-B18,ND-75 | | 1.545e-04 | -8.78 | fly chr13A8-13A8\|1-49 | chromosome location | fly chr13A8-13A8\|1-49 | 3 | 3 | 25010 | 1343 | opm,ND-B18,dob | | 1.545e-04 | -8.78 | fly chr13A8-13A8\|1-49 cM | chromosome location | fly chr13A8-13A8\|1-49 cM | 3 | 3 | 25010 | 1343 | dob,opm,ND-B18 | | 1.553e-04 | -8.77 | GSE29617\_CTRL\_VS\_DAY3\_TIV\_FLU\_VACCINE\_PBMC\_2008\_DN | MSigDB lists | GSE29617\_CTRL\_VS\_DAY3\_TIV\_FLU\_VACCINE\_PBMC\_2008\_DN | 82 | 26 | 4161 | 644 | tho2,ste14,Naa20A,Mcad,CG17294,Prosbeta3,mRpL18,CG10470,Sec10,ND-SGDH,CG1890,nrv1,CG5510,p38a,Jafrac2,wal,Tim8,Eno,CCT5,Arpc4,Arpc2,Prosbeta4,Prosalpha3,Scsalpha1,Arf51F,yip2 | | 1.561e-04 | -8.77 | chic (chickadee) | protein interactions | 33834 | 70 | 20 | 9204 | 1112 | Uch,Eno,AnxB10,CycB,capt,scu,Moe,Stip1,HmgD,CG8204,yip2,Chd64,CG11267,awd,Pgk,CG10638,CG14715,PHGPx,Imp,cib | | 1.561e-04 | -8.76 | CCT6 (Chaperonin containing TCP1 subunit 6) | protein interactions | 32518 | 86 | 23 | 9204 | 1112 | Vha68-2,Tspo,Rpt2,mEFTu1,wal,CCT5,GCS2beta,foxo,cathD,ScsbetaA,Mcad,CG1416,Rab5,Droj2,Pp2A-29B,Tak1,RpLP1,CCT3,PGRP-LE,Rel,Kap-alpha3,Pdk1,SERCA | | 1.574e-04 | -8.76 | Tl (Toll) | protein interactions | 43222 | 27 | 11 | 9204 | 1112 | cact,ea,Stat92E,wisp,tub,Rel,spz,Myd88,mad2,Nmt,mxc | | 1.577e-04 | -8.75 | CG5174 (uncharacterized protein) | protein interactions | 37117 | 45 | 15 | 9204 | 1112 | capt,scu,Phb2,Drep1,EndoGI,CG10638,regucalcin,PGRP-LE,Rel,AnxB10,RpL13A,Tak1,Uch,RhoGDI,COX5A | | 1.580e-04 | -8.75 | microtubule cytoskeleton organization | biological process | GO:0000226 | 341 | 52 | 12345 | 1129 | Ubc6,foxo,Prosbeta5,alphaTub84B,sau,mts,larp,cnc,Topors,grp,Grip75,Lkb1,tral,msps,CG7716,Grip128,heph,Klp10A,mop,armi,sw,Pp2A-29B,gammaTub37C,Kap3,Girdin,wash,stai,cnn,pont,Grip84,BicC,capt,SkpA,mael,CG3437,cbs,SRPK,Pka-R1,Grip71,osk,wisp,Non1,Moe,dgt4,spd-2,Axs,jvl,del,alphaTub67C,tws,CCT6,Rcd5 | | 1.581e-04 | -8.75 | ATPsynG (ATP synthase, subunit G) | protein interactions | 46069 | 19 | 9 | 9204 | 1112 | ATPsynE,ATPsynB,ATPsyndelta,ATPsynF,ATPsynCF6,ATPsynD,blw,ATPsynO,ATPsynbeta | | 1.612e-04 | -8.73 | TUDOR | smart domains | SM00333 | 18 | 8 | 5086 | 506 | CG9925,otu,vret,tapas,spn-E,CG17454,krimp,CG9684 | | 1.613e-04 | -8.73 | MARTIN\_INTERACT\_WITH\_HDAC | MSigDB lists | MARTIN\_INTERACT\_WITH\_HDAC | 10 | 7 | 4161 | 644 | lwr,Pen,Nurf-38,Phb2,Tip60,rl,cact | | 1.613e-04 | -8.73 | GO\_REGULATION\_OF\_TRIGLYCERIDE\_METABOLIC\_PROCESS | MSigDB lists | GO\_REGULATION\_OF\_TRIGLYCERIDE\_METABOLIC\_PROCESS | 10 | 7 | 4161 | 644 | regucalcin,mino,EcR,CycC,crq,Dd,SREBP | | 1.638e-04 | -8.72 | cellular protein-containing complex assembly | biological process | GO:0034622 | 471 | 67 | 12345 | 1129 | rig,bru1,CG17768,CG10055,RpL23A,Grip75,mRpS11,Sgf29,RpS15,CG11562,LSm3,TBCC,gammaTub37C,spn-E,Hip1,Cchl,CG7716,Clc,Chrac-14,Rpn13,Grip128,asf1,Taf7,Trf2,BigH1,tkv,Dcr-1,CG3198,rept,CG5969,pont,ND-23,CG11110,Usp39,Pomp,Caf1-105,ND-18,Unr,Sdhaf3,RpS27,CG1890,CG7488,RpS14a,Ndc1,r2d2,CanB2,RpS14b,SelR,RpL11,Hat1,msps,RpS28b,CG14671,Edc3,Tomosyn,Nap1,Grip84,RpL24,RpLP0,Scox,RpS5b,osk,CG8204,SmE,RpL12,RpL10,Cenp-C | | 1.671e-04 | -8.70 | Nrd1 (Nardilysin) | protein interactions | 32143 | 60 | 18 | 9204 | 1112 | Roe1,Gdi,Nc73EF,AcCoAS,Idh3b,Moe,bor,CG7322,r,mEFTu1,RpS27A,tun,SRPK,Pgk,Shmt,CG3609,Fkbp12,Idgf4 | | 1.681e-04 | -8.69 | BLALOCK\_ALZHEIMERS\_DISEASE\_DN | MSigDB lists | BLALOCK\_ALZHEIMERS\_DISEASE\_DN | 545 | 114 | 4161 | 644 | CG10932,Fdh,Jafrac1,dnc,Karybeta3,CG2938,CHOp24,isoQC,Nurf-38,RpII140,Rpn2,ND-SGDH,Gapdh1,ND-49,mRpL18,CG11899,Prosbeta3,Pgm1,Cdk7,CG3226,Pgk,RhoGAP5A,rl,Aldh,SdhA,Klp10A,Rbcn-3A,wal,Mo25,Prosalpha5,asf1,Got1,Synj,CG1416,CG4511,Roe1,RpL38,Vha13,CG14894,Pka-R1,lt,Kap3,CG4390,CG1532,Pglym78,Cul1,ND-30,CG13630,p38a,Pfdn2,CCT5,CG8004,Cdk5,Vamp7,Arpc1,Sccpdh1,Hmg-2,Prosbeta4,Uch,CG4872,CG11089,levy,Crag,CG4562,CG6610,Cse1,mRpL13,mts,Cul2,Pen,UQCR-Q,mRpL16,CG5026,mEFTu1,Top1,LpR2,mRpS10,Bin1,Idh3b,Arpc2,Adi1,CG8204,CG40045,CanB2,Pp2B-14D,faf,CG6767,Cyt-c-p,Atg6,ScsbetaA,CG1890,CG8963,CG7718,CG9132,Vha36-1,CkIIalpha,capt,ND-24,CG15715,Jafrac2,CCT3,Pfk,Prosbeta5,pr,CG46338,PDCD-5,Rnmt,Gabat,COX5A,APP-BP1,LSm3,Hsc70-3,CanA-14F,CG4293 | | 1.683e-04 | -8.69 | AGO3 (Argonaute 3) | protein interactions | 3355150 | 12 | 7 | 9204 | 1112 | twin,tapas,vret,krimp,armi,tral,smg | | 1.684e-04 | -8.69 | Ntf-2 (Nuclear transport factor-2) | protein interactions | 33078 | 55 | 17 | 9204 | 1112 | Ho,Eno,Larp4B,AnxB10,regucalcin,scu,Moe,capt,CG2199,RhoGDI,awd,Trxr-1,CG10638,CG14715,cib,Cyp1,Sod1 | | 1.689e-04 | -8.69 | Lkb1 (Lkb1 kinase) | protein interactions | 41673 | 81 | 22 | 9204 | 1112 | Ankle2,tyf,Wdfy2,Nup358,RpL30,RpS23,Mo25,Ect4,Fkbp39,faf,RpL31,RpL17,RpL9,Rbcn-3A,Mat89Ba,alt,RpS14a,RpL13,RpL23,RpL10,RpL7A,Hsc70-3 | | 1.715e-04 | -8.67 | chromosome segregation | biological process | GO:0007059 | 187 | 33 | 12345 | 1129 | Axs,mtrm,Cenp-C,bif,RfC38,CycB,c(2)M,RecQ5,cnn,lwr,msps,spn-E,Klp10A,eIF4E1,mus304,cana,tef,gammaTub37C,Rab5,cup,Pp2A-29B,wapl,nos,Rbf,mus81,mts,Top1,Topors,Trf4-2,CG31053,cort,Tlk,grp | | 1.719e-04 | -8.67 | regulation of humoral immune response | biological process | GO:0002920 | 71 | 17 | 12345 | 1129 | Rel,Herc4,mop,Myd88,spz,Ptip,wisp,tub,cact,cad,Aos1,scny,Stat92E,p38a,lwr,dl,Tak1 | | 1.727e-04 | -8.66 | Gdi (GDP dissociation inhibitor) | protein interactions | 34264 | 36 | 13 | 9204 | 1112 | AnxB10,yip2,regucalcin,Rab5,CG10638,RpL38,EndoGI,capt,Rab10,Moe,Stip1,Uch,Eno | | 1.727e-04 | -8.66 | Rpn9 (Regulatory particle non-ATPase 9) | protein interactions | 42802 | 36 | 13 | 9204 | 1112 | Rpn2,Rpn12,Prosalpha5,Rpn13,Vamp7,Rpn3,Prosalpha3,Prosbeta3,Prosbeta6,Prosbeta4,Rpt2,Prosalpha4,Prosbeta5 | | 1.727e-04 | -8.66 | Pgd (Phosphogluconate dehydrogenase) | protein interactions | 31185 | 36 | 13 | 9204 | 1112 | Eno,Uch,regucalcin,CG14715,CG10638,PHGPx,EndoGI,Moe,capt,AnxB10,Chd64,Trxr-1,awd | | 1.745e-04 | -8.65 | GSE43863\_TH1\_VS\_LY6C\_INT\_CXCR5POS\_EFFECTOR\_CD4\_TCELL\_UP | MSigDB lists | GSE43863\_TH1\_VS\_LY6C\_INT\_CXCR5POS\_EFFECTOR\_CD4\_TCELL\_UP | 87 | 27 | 4161 | 644 | Prosbeta6,CG15141,r,Aos1,CG10418,Pdk,CG1532,CG11164,mRpL18,colt,Aladin,Rpn2,crq,Klp10A,CG7655,CG16979,Liprin-beta,mEFTu1,Uev1A,Uch-L5,f-cup,CG11134,Prosalpha3,cl,Idh3b,Nap1,DNApol-epsilon255 | | 1.750e-04 | -8.65 | regulation of oskar mRNA translation | biological process | GO:0046011 | 15 | 7 | 12345 | 1129 | BicC,osk,bru1,krimp,heph,apt,cup | | 1.782e-04 | -8.63 | regulation of nuclear-transcribed mRNA catabolic process, deadenylation-dependent decay | biological process | GO:1900151 | 11 | 6 | 12345 | 1129 | gw,Marf1,piwi,AGO3,smg,spn-E | | 1.782e-04 | -8.63 | positive regulation of nuclear-transcribed mRNA catabolic process, deadenylation-dependent decay | biological process | GO:1900153 | 11 | 6 | 12345 | 1129 | spn-E,smg,AGO3,piwi,Marf1,gw | | 1.820e-04 | -8.61 | negative regulation of cell death | biological process | GO:0060548 | 127 | 25 | 12345 | 1129 | Naa20A,Idgf2,Cul1,SkpA,spag,dco,Rbf,lin-52,Pdk1,Ubc6,nos,Drep1,RYBP,ix,bun,Tip60,Pink1,MAGE,CG4238,csw,Atg7,velo,Debcl,Drep2,Aldh | | 1.831e-04 | -8.61 | gamma-tubulin binding | molecular function | GO:0043015 | 11 | 6 | 12176 | 1119 | Grip128,CG7716,Grip75,Grip84,Blos2,Grip71 | | 1.884e-04 | -8.58 | PGRP-LC (Peptidoglycan recognition protein LC) | protein interactions | 39063 | 87 | 23 | 9204 | 1112 | blw,RpL7A,CCT3,CG4266,alphaTub84B,RpL23A,Sod1,SERCA,CCT6,cup,PGRP-LE,RpS14a,RpL12,CG17985,RpS28b,Jafrac1,GCS2beta,Cp1,CG1416,scu,Nup358,RpS27,RpS16 | | 1.912e-04 | -8.56 | TUDOR | prosite domains | PS50304 | 13 | 7 | 6243 | 694 | CG9925,CG9684,spn-E,krimp,otu,vret,CG17454 | | 1.935e-04 | -8.55 | cellular macromolecule localization | biological process | GO:0070727 | 635 | 85 | 12345 | 1129 | Cenp-C,jvl,lt,Rab10,Cse1,Grip71,osk,CG8004,Tsp86D,Stam,thoc6,Pax,milt,Rab32,Actn,Vsp37A,swa,fray,Nxf3,msps,Tom70,sws,CG8314,CG6766,Arpc1,Pex10,opm,Atg6,CkIIalpha,CG9393,Pex13,exu,Tim8,Mat89Ba,Vps16B,GAPsec,sip3,Nup358,cnc,Arl5,CHOp24,CG4041,wisp,AP-1-2beta,CG11110,Syx13,CG5447,X11L,Rrp40,CG8728,Sec15,Gga,Nup62,SuUR,sbr,CG10344,mael,Tm1,Roe1,Rep,Oseg1,Arf51F,Grip128,Clc,Pen,CG5734,Tom40,CG14894,Arts,Pi4KIIalpha,spn-E,Nlg1,Sec23,Karybeta3,Rab5,cup,Pex12,sw,mbo,tho2,sau,Grip75,Lkb1,Kap-alpha3,Topors | | 1.950e-04 | -8.54 | cellular response to chemical stimulus | biological process | GO:0070887 | 378 | 56 | 12345 | 1129 | CG5844,Tom40,csw,Idh,Med,Grx1,raptor,Utx,Sod3,CG4390,ACC,Pka-R2,Atf6,Chd64,bun,Parp16,ago,Tak1,Pdk1,foxo,Vps16B,mAChR-A,Nup358,sip3,Rab10,tub,Pdxk,EcR,Hnf4,Ptp61F,PyK,Mekk1,Ugt36E1,Hsc70-3,CG8974,G9a,CG32640,MED27,Cyp6v1,Smox,Trxr-1,Fkbp39,Jafrac1,rl,Sod1,Nurf-38,Pi3K92E,Edem1,lwr,PEK,p38a,Hr78,Stat92E,tkv,Fdh,Shmt | | 1.980e-04 | -8.53 | organic acid catabolic process | biological process | GO:0016054 | 106 | 22 | 12345 | 1129 | SERCA,CG17691,Acox57D-p,wal,Shmt,CG4598,Thiolase,Ntan1,CG10932,Gabat,Echs1,CG4586,Mtpalpha,CG8199,Pex10,yip2,CG4860,Etf-QO,Seipin,Mcad,P5CDh1,CG4594 | | 1.980e-04 | -8.53 | carboxylic acid catabolic process | biological process | GO:0046395 | 106 | 22 | 12345 | 1129 | Thiolase,CG10932,Ntan1,CG4586,Echs1,Gabat,CG17691,SERCA,wal,Shmt,Acox57D-p,CG4598,Mcad,CG4594,P5CDh1,Mtpalpha,Pex10,CG8199,yip2,CG4860,Etf-QO,Seipin | | 1.994e-04 | -8.52 | regulation of phosphate metabolic process | biological process | GO:0019220 | 302 | 47 | 12345 | 1129 | Pka-R2,Pp2A-29B,RpLP1,Cdk7,raptor,Tspo,CycE,Abi,fray,CG30291,ASPP,sws,Lkb1,Cka,alph,CG14408,Ankle2,mts,Pgk,Parp16,CycJ,Tak1,Pdk1,tor,CG8009,Tao,CG17746,Mos,Scox,SNF4Agamma,tws,Pink1,ATPsynD,Ptp61F,Mekk1,ben,CycC,PpV,Pka-R1,RhoGAP5A,SkpA,PEK,dco,Pi3K92E,Cbl,CycB,park | | 1.994e-04 | -8.52 | regulation of phosphorus metabolic process | biological process | GO:0051174 | 302 | 47 | 12345 | 1129 | CG14408,Ankle2,alph,Cka,Lkb1,CG8009,tor,Pdk1,Tak1,CycJ,Pgk,Parp16,mts,Tspo,raptor,RpLP1,Cdk7,Pp2A-29B,Pka-R2,sws,ASPP,CG30291,Abi,fray,CycE,dco,PEK,SkpA,RhoGAP5A,Pka-R1,park,CycB,Cbl,Pi3K92E,ATPsynD,Pink1,tws,SNF4Agamma,CG17746,Mos,Scox,Tao,PpV,CycC,ben,Mekk1,Ptp61F | | 1.994e-04 | -8.52 | negative regulation of apoptotic process | biological process | GO:0043066 | 99 | 21 | 12345 | 1129 | MAGE,CG4238,csw,Debcl,Atg7,RYBP,Tip60,bun,Pink1,lin-52,Rbf,nos,Drep1,Pdk1,Ubc6,Naa20A,Idgf2,Cul1,SkpA,spag,dco | | 1.994e-04 | -8.52 | positive regulation of immune response | biological process | GO:0050778 | 99 | 21 | 12345 | 1129 | Hr78,PGRP-LE,cad,Aos1,Tak1,lwr,dl,spz,Rel,PDCD-5,Herc4,mop,CG9925,Myd88,wisp,tub,Hat1,Crag,EMRE,Ptip,Sp7 | | 2.004e-04 | -8.52 | cellular lipid metabolic process | biological process | GO:0044255 | 387 | 57 | 12345 | 1129 | Thiolase,Pi3K92E,PIG-L,CG31523,mino,CG7789,wal,SERCA,nSMase,Sccpdh1,CG33977,CG6707,CG7840,scu,CG10425,Desat1,Mcad,Hnf4,CG17270,CG5026,CG7718,Etf-QO,CG10932,SREBP,Pld,Gk1,Echs1,CG4586,CG9391,Acox57D-p,PIG-C,Gk2,CG4598,CG3842,CG8311,PIG-H,Synj,PIG-P,Sply,sws,AcCoAS,CG18135,Pi4KIIalpha,CG2201,CG8112,CG4594,Pxt,CG7997,yip2,Agpat2,dob,ACC,Mtpalpha,Pex10,Seipin,CG4860,CG6178 | | 2.047e-04 | -8.49 | GSE22886\_NAIVE\_TCELL\_VS\_DC\_DN | MSigDB lists | GSE22886\_NAIVE\_TCELL\_VS\_DC\_DN | 70 | 23 | 4161 | 644 | Pcd,Rab32,CG14516,Pi4KIIalpha,Aldh,CG9471,CG7997,Cdk5,Gclc,Prosbeta5,ND-24,Marf,Top1,shop,Rpn2,isoQC,Irk1,CG17593,Stam,Cyp1,Atg7,Grx1,Synj | | 2.050e-04 | -8.49 | capt (capulet) | protein interactions | 45233 | 82 | 22 | 9204 | 1112 | Stip1,RhoGDI,Gip,CG10638,CG14715,cib,EndoGI,Pka-R1,PHGPx,Fkbp12,Echs1,yip2,Chd64,awd,Eno,Uch,Gdi,regucalcin,scu,Moe,CG4598,AnxB10 | | 2.050e-04 | -8.49 | Pp2A-29B (Protein phosphatase 2A at 29B) | protein interactions | 2768940 | 82 | 22 | 9204 | 1112 | ScsbetaA,Mcad,CG1416,Cka,Karybeta3,bor,Tspo,wal,GCS2beta,r,Ubc4,mts,Kap-alpha3,CG10425,Pdk1,SERCA,CCT6,Nf-YA,Cenp-C,REPTOR,CCT3,CG6175 | | 2.050e-04 | -8.49 | GO\_SINGLE\_ORGANISM\_CATABOLIC\_PROCESS | MSigDB lists | GO\_SINGLE\_ORGANISM\_CATABOLIC\_PROCESS | 347 | 78 | 4161 | 644 | CG9471,crq,CG3609,Caf1-105,mRpS10,STUB1,Echs1,GlyP,CG7997,CG7840,Nc73EF,PAPLA1,Pcd,Faf2,Pi4KIIalpha,CG7550,tzn,Gk2,Rpt2,Pfk,CG7322,Edem1,CG3376,Hsc70-3,Gabat,Iru,CG9485,sgll,Etf-QO,CG6766,CG5966,DhpD,cathD,Atg7,Atg4b,P5CDh1,CG5285,Atg6,Aldh,CG8199,wal,CG9391,Treh,Shmt,Pgk,Got1,Sply,CG15111,CG17691,ND-B22,CG8475,CG10932,Fdh,CG5734,Jafrac1,dnc,Gapdh1,Cisd2,Pgm1,Mcad,scu,HDAC6,Mtpalpha,Eno,Pglym78,CG8974,Thiolase,CG9248,CG32473,CG4872,Gyg,CG32549,CG17593,CG1640,Synj,Pex13,ABCD,CG4390 | | 2.061e-04 | -8.49 | centrosome cycle | biological process | GO:0007098 | 72 | 17 | 12345 | 1129 | Grip71,spd-2,msps,Pp2A-29B,CCT6,Rcd5,tws,Prosbeta5,cnn,Grip84,mts,Ubc6,cbs,grp,CG3437,Topors,SkpA | | 2.081e-04 | -8.48 | REACTOME\_ACTIVATION\_OF\_NF\_KAPPAB\_IN\_B\_CELLS | MSigDB lists | REACTOME\_ACTIVATION\_OF\_NF\_KAPPAB\_IN\_B\_CELLS | 41 | 16 | 4161 | 644 | Prosalpha3,Prosbeta4,Prosalpha5,RpS27A,Prosbeta5,cact,Cul1,Rpn3,Prosbeta3,Prosalpha4,Rpt2,Rpn12,Rpn2,SkpA,Prosbeta6,RpL40 | | 2.083e-04 | -8.48 | Tctp (Translationally controlled tumor protein) | protein interactions | 41341 | 46 | 15 | 9204 | 1112 | mre11,Sap47,cib,PHGPx,CG10638,CG14715,regucalcin,CycE,Chd64,awd,AnxB10,Snx6,Rtnl1,Eno,HP1c | | 2.107e-04 | -8.46 | nos (nanos) | protein interactions | 42297 | 61 | 18 | 9204 | 1112 | hb,Rga,eIF4E1,bcd,Tao,osk,bam,l(3)mbt,sw,CycB,twin,wisp,smg,cup,tkv,CG7655,tral,CG11486 | | 2.145e-04 | -8.45 | GSE36476\_CTRL\_VS\_TSST\_ACT\_16H\_MEMORY\_CD4\_TCELL\_YOUNG\_DN | MSigDB lists | GSE36476\_CTRL\_VS\_TSST\_ACT\_16H\_MEMORY\_CD4\_TCELL\_YOUNG\_DN | 79 | 25 | 4161 | 644 | DppIII,Karybeta3,CysRS,CG7550,Cbp80,Arts,Stam,rept,Gk2,alphaTub84B,Cyt-c-p,Jafrac2,r-l,Aldh,Eno,Pus7,Pglym78,bys,Marf,CG7728,Mitofilin,Ppcdc,Idh,Pfas,gammaTub37C | | 2.146e-04 | -8.45 | eEF5 (eukaryotic translation elongation factor 5) | protein interactions | 37846 | 51 | 16 | 9204 | 1112 | Bx,png,Eno,AnxB10,regucalcin,scu,Moe,capt,Stip1,Ptp61F,Chd64,awd,CG14715,CG10638,PHGPx,cib | | 2.150e-04 | -8.44 | cpb (capping protein beta) | protein interactions | 33346 | 56 | 17 | 9204 | 1112 | GlyP,Arpc1,osk,kri,Hsp26,mRpS5,Moe,Arpc2,RpLP1,CG1847,spag,RpL23,Arpc4,LysRS,RpS18,RecQ5,RpL12 | | 2.150e-04 | -8.44 | Rpn1 (Regulatory particle non-ATPase 1) | protein interactions | 40174 | 56 | 17 | 9204 | 1112 | Prosalpha4,Prosbeta6,Rpt2,wal,Rpn3,Rpn12,Prosbeta5,Prosbeta3,CCT3,Prosalpha3,Prosbeta4,Pp2A-29B,Rpn13,Pdk1,Rpn2,Pomp,Prosalpha5 | | 2.219e-04 | -8.41 | BASSO\_B\_LYMPHOCYTE\_NETWORK | MSigDB lists | BASSO\_B\_LYMPHOCYTE\_NETWORK | 66 | 22 | 4161 | 644 | mad2,bys,Tom40,Prosbeta5,CycB,Jafrac2,Cdk1,gammaTub37C,CG11807,LSm3,Prosbeta4,tko,CG1416,rept,Rpb8,Cyt-c-p,CG8188,Cse1,isoQC,Rpn2,CG12018,AnxB10 | | 2.235e-04 | -8.41 | ubiquitin conjugating enzyme activity | molecular function | GO:0061631 | 30 | 10 | 12176 | 1119 | UbcE2H,CG40045,CG8188,CG4502,ben,Uev1A,CG10254,Ubc6,CG2924,Ubc4 | | 2.254e-04 | -8.40 | imd (immune deficiency) | protein interactions | 44339 | 122 | 29 | 9204 | 1112 | RpL12,vret,Sgf29,PGRP-LE,RpS14a,Sod1,SERCA,CCT6,eIF3i,pont,alphaTub84B,RpL7A,Mitofilin,Tak1,CCT3,Hsc70-3,blw,RpS27,RpS16,Nup358,scu,bor,Jafrac1,GCS2beta,Cp1,scny,faf,RpS28b,RpL17 | | 2.281e-04 | -8.39 | response to organic cyclic compound | biological process | GO:0014070 | 107 | 22 | 12345 | 1129 | nbs,G9a,Utx,Tao,Cyp6v1,Pka-R2,Idh3b,MED27,EcR,MAGE,Hnf4,dco,rig,Hr78,cyc,Trf2,Shmt,Bx,MED24,Nurf-38,rept,trr | | 2.312e-04 | -8.37 | Rpn8 (Regulatory particle non-ATPase 8) | protein interactions | 37894 | 28 | 11 | 9204 | 1112 | Prosbeta5,Prosalpha4,Prosbeta4,Rpt2,Prosalpha3,Prosbeta6,Prosbeta3,Rpn3,Rpn13,Prosalpha5,Rpn2 | | 2.336e-04 | -8.36 | GO\_NUCLEOSIDE\_MONOPHOSPHATE\_METABOLIC\_PROCESS | MSigDB lists | GO\_NUCLEOSIDE\_MONOPHOSPHATE\_METABOLIC\_PROCESS | 121 | 34 | 4161 | 644 | kri,CG3803,ND-75,UQCR-Q,Shmt,Pgk,ND-39,Nc73EF,ND-B22,Stoml2,levy,ND-23,CG11089,ND-51,Gapdh1,ND-49,CG33156,Pgm1,AMPdeam,ND-SGDH,r-l,ND-B18,Pfk,Eno,Pglym78,ND-30,ND-24,Pfas,CG32549,COX5A,Ts,r,CG6767,Cyt-c-p | | 2.376e-04 | -8.34 | anterior/posterior axis specification | biological process | GO:0009948 | 144 | 27 | 12345 | 1129 | swa,cnc,mael,cad,Tm1,Grip75,Lkb1,tor,nos,exu,rl,hb,psq,armi,fs(1)M3,gus,Grip71,bcd,spn-E,osk,csw,Moe,msps,trk,jvl,Grip128,fs(1)N | | 2.389e-04 | -8.34 | regulation of phosphorylation | biological process | GO:0042325 | 271 | 43 | 12345 | 1129 | ASPP,sws,fray,CycE,Abi,CG30291,raptor,Tspo,Pka-R2,RpLP1,Cdk7,CycJ,Tak1,Pgk,Parp16,tor,Pdk1,mts,alph,Ankle2,CG14408,Lkb1,Cka,ben,CycC,PpV,Mekk1,Ptp61F,ATPsynD,Pink1,CG17746,SNF4Agamma,Mos,Tao,CycB,park,Pi3K92E,Cbl,PEK,SkpA,dco,Pka-R1,RhoGAP5A | | 2.410e-04 | -8.33 | WONG\_PROTEASOME\_GENE\_MODULE | MSigDB lists | WONG\_PROTEASOME\_GENE\_MODULE | 30 | 13 | 4161 | 644 | CG40045,CG15141,Usp39,Rpn3,Prosbeta4,Uch,Prosalpha3,Prosbeta3,Prosalpha5,CG2924,Rpn2,ari-1,Rpn12 | | 2.462e-04 | -8.31 | chromatin silencing | biological process | GO:0006342 | 73 | 17 | 12345 | 1129 | wash,esc,egg,rept,piwi,Tlk,escl,Elba2,Hat1,spn-E,Hnf4,Chrac-14,r2d2,Bin1,CG10038,ph-d,psq | | 2.462e-04 | -8.31 | negative regulation of gene expression, epigenetic | biological process | GO:0045814 | 73 | 17 | 12345 | 1129 | wash,esc,egg,rept,piwi,Tlk,escl,Elba2,Hat1,spn-E,Hnf4,Chrac-14,r2d2,Bin1,CG10038,ph-d,psq | | 2.462e-04 | -8.31 | chromatin organization involved in negative regulation of transcription | biological process | GO:0097549 | 73 | 17 | 12345 | 1129 | Elba2,spn-E,Hat1,escl,Chrac-14,Hnf4,r2d2,Bin1,psq,CG10038,ph-d,wash,rept,egg,esc,Tlk,piwi | | 2.547e-04 | -8.28 | MODULE\_43 | MSigDB lists | MODULE\_43 | 58 | 20 | 4161 | 644 | Mcad,ND-49,Cyt-c-p,AGBE,Etf-QO,ND-23,ND-51,Scsalpha1,Idh,Cyp6a22,Idh3b,Irp-1B,GlyP,wal,Echs1,ND-24,ND-30,Cyp6a19,Pglym78,Treh | | 2.596e-04 | -8.26 | GSE22886\_NAIVE\_BCELL\_VS\_NEUTROPHIL\_UP | MSigDB lists | GSE22886\_NAIVE\_BCELL\_VS\_NEUTROPHIL\_UP | 71 | 23 | 4161 | 644 | RpS18,Chrac-14,RpL11,RpL37a,RpS27A,RpL26,Mekk1,PAPLA1,RpL9,Nap1,RpS27,Smox,RpL38,RpS7,CG10932,eIF4A,RpL30,RpL23,RpS24,CG6767,RpS13,CG4390,RpS16 | | 2.616e-04 | -8.25 | Rel (Relish) | protein interactions | 41087 | 123 | 29 | 9204 | 1112 | CG9391,pont,cact,RpL23A,alphaTub84B,RpL7A,CCT3,Mitofilin,blw,RpL12,Rel,spz,RpS14a,PGRP-LE,cup,dl,CCT6,Sod1,SERCA,Atf3,bor,Jafrac1,Cp1,GCS2beta,RpS28b,RpS16,RpS27,Nup358,scu | | 2.671e-04 | -8.23 | response to nitrogen compound | biological process | GO:1901698 | 145 | 27 | 12345 | 1129 | Rab10,CG6766,EcR,Hnf4,MAGE,PyK,Hsc70-3,CG11070,CG8974,raptor,nbs,Pka-R2,CG32640,Tao,rl,CG7328,Pdk1,Pi3K92E,foxo,Edem1,Vps16B,sip3,cyc,dco,mAChR-A,Bx,Shmt | | 2.674e-04 | -8.23 | REACTOME\_GLUCOSE\_METABOLISM | MSigDB lists | REACTOME\_GLUCOSE\_METABOLISM | 34 | 14 | 4161 | 644 | Pp2A-29B,mts,Gyg,Pgm1,Got1,Gapdh1,Pgk,Pglym78,AGBE,Eno,GlyP,CG8475,CG9485,Pfk | | 2.693e-04 | -8.22 | Ost48 (Oligosaccharyltransferase 48kD subunit) | protein interactions | 31849 | 78 | 21 | 9204 | 1112 | Tspo,Vha68-2,Karybeta3,Surf4,wal,Mcad,ScsbetaA,ATPsynO,Atpalpha,Rab5,CG31729,CG15098,ATPsynF,ATPsyndelta,COX4,Pp2A-29B,ATPsynCF6,Kap-alpha3,CG10425,SERCA,CCT6 | | 2.780e-04 | -8.19 | GO\_PROTEIN\_UBIQUITINATION | MSigDB lists | GO\_PROTEIN\_UBIQUITINATION | 217 | 53 | 4161 | 644 | MED27,CG46338,CycB,Prosbeta5,Rpn3,Cul1,CG8974,Pex12,Prosbeta4,Prosalpha3,CG2924,MED20,Iru,CG40045,Prosbeta6,CG4502,Ubc4,Aos1,gus,not,Atg7,RpL40,CycC,MED24,Traf6,APC7,ari-1,Cdk1,Cul5,STUB1,mad2,Fbw5,Uev1A,Herc4,RpS27A,Prosalpha5,CG15141,SkpA,CG11070,UbcE2H,Rpt2,CG10465,Prosalpha4,Prosbeta3,Cul2,Rpn2,DCAF12,CG3894,HDAC6,adp,Rpn12,CG8188,CG1812 | | 2.802e-04 | -8.18 | KEGG\_OXIDATIVE\_PHOSPHORYLATION | MSigDB lists | KEGG\_OXIDATIVE\_PHOSPHORYLATION | 50 | 18 | 4161 | 644 | Vha13,levy,ND-B22,ND-51,ND-23,ND-49,Vha36-1,Nurf-38,ND-SGDH,ND-B18,SdhA,ND-75,CG3803,UQCR-Q,ND-24,ND-30,ND-39,COX5A | | 2.802e-04 | -8.18 | BILANGES\_SERUM\_AND\_RAPAMYCIN\_SENSITIVE\_GENES | MSigDB lists | BILANGES\_SERUM\_AND\_RAPAMYCIN\_SENSITIVE\_GENES | 50 | 18 | 4161 | 644 | eEF1beta,Exd2,RpL13,eIF3m,RpL26,RpS27A,RpS23,RpL24,RpS4,RpS7,RpL40,RpL23,RpL30,RpS24,RpS16,RpL13A,RpS30,RpS13 | | 2.833e-04 | -8.17 | GSE25846\_IL10\_POS\_VS\_NEG\_CD8\_TCELL\_DAY7\_POST\_CORONAVIRUS\_BRAIN\_DN | MSigDB lists | GSE25846\_IL10\_POS\_VS\_NEG\_CD8\_TCELL\_DAY7\_POST\_CORONAVIRUS\_BRAIN\_DN | 67 | 22 | 4161 | 644 | Clp,Idh,SREBP,Grip75,p38a,Aldh,Mtpalpha,thoc6,pr,Chrac-14,Uch-L5,ND-30,Sod1,CG9485,Vha13,DppIII,eIF3i,CHOp24,mus81,CG3842,Grx1,Arl5 | | 2.847e-04 | -8.16 | BIOCARTA\_CK1\_PATHWAY | MSigDB lists | BIOCARTA\_CK1\_PATHWAY | 8 | 6 | 4161 | 644 | mts,CanA-14F,Pka-R2,Pka-R1,Cdk5,Pp2B-14D | | 2.847e-04 | -8.16 | GO\_REGULATION\_OF\_TRIGLYCERIDE\_BIOSYNTHETIC\_PROCESS | MSigDB lists | GO\_REGULATION\_OF\_TRIGLYCERIDE\_BIOSYNTHETIC\_PROCESS | 8 | 6 | 4161 | 644 | mino,EcR,regucalcin,SREBP,crq,Dd | | 2.874e-04 | -8.15 | REACTOME\_SIGNALING\_BY\_WNT | MSigDB lists | REACTOME\_SIGNALING\_BY\_WNT | 46 | 17 | 4161 | 644 | Rpn2,Rpn12,Prosalpha4,Rpt2,mts,Prosbeta3,RpL40,SkpA,Prosbeta6,Pp2A-29B,Prosalpha5,RpS27A,Prosbeta4,Prosalpha3,Cul1,Rpn3,Prosbeta5 | | 2.884e-04 | -8.15 | GSE26156\_DOUBLE\_POSITIVE\_VS\_CD4\_SINGLE\_POSITIVE\_THYMOCYTE\_DN | MSigDB lists | GSE26156\_DOUBLE\_POSITIVE\_VS\_CD4\_SINGLE\_POSITIVE\_THYMOCYTE\_DN | 94 | 28 | 4161 | 644 | rept,Karybeta3,levy,RfC38,Ts,NHP2,CG8188,Rpn12,ScsbetaA,grp,P5cr,CCT6,Pgk,msps,CG9391,CycB,Tom40,CCT5,Prosbeta5,Eno,AIF,Coq7,Pdxk,yip2,CycE,gammaTub37C,Mitofilin,Prosbeta4 | | 2.888e-04 | -8.15 | GO\_RESPIRATORY\_CHAIN | MSigDB lists | GO\_RESPIRATORY\_CHAIN | 42 | 16 | 4161 | 644 | ND-49,ND-SGDH,Cyt-c-p,ND-B22,levy,ND-51,ND-23,ND-39,COX5A,ND-B18,SdhA,CG3803,ND-75,UQCR-Q,ND-30,ND-24 | | 2.902e-04 | -8.14 | FKBP\_PPIASE | prosite domains | PS50059 | 7 | 5 | 6243 | 694 | Fkbp59,Fkbp12,CG14715,Fkbp39,shu | | 2.917e-04 | -8.14 | Rpt1 (Regulatory particle triple-A ATPase 1) | protein interactions | 35701 | 95 | 24 | 9204 | 1112 | Prosbeta5,Prosalpha3,CCT3,Prosbeta3,Pp2A-29B,Prosbeta4,Pdk1,Rpn13,Rpn2,CCT6,Prosalpha5,SERCA,Pomp,Prosalpha4,GCS2beta,Prosbeta6,wal,Rpt2,Mcad,ScsbetaA,Rpn3,Droj2,Rab5,CG1416 | | 2.995e-04 | -8.11 | - | gene3d domains | 3.10.110.10 | 41 | 14 | 4929 | 622 | CG8188,lwr,CG2924,CG40045,CG4502,CG46338,CG17260,Ubc6,Uev1A,CG10254,Ubc4,CG13344,ben,UbcE2H | | 2.999e-04 | -8.11 | oxidoreductase activity, acting on CH-OH group of donors | molecular function | GO:0016614 | 123 | 24 | 12176 | 1119 | CG31674,Fdh,CG10425,CG31549,CG4842,CG2767,sni,CG9265,CG10638,Idh3b,scu,Adh,CG5028,CG3842,Mtpalpha,CG31548,Idh,Thiolase,CG10863,Sptr,CG12171,CG7322,CG9436,Gmer | | 3.018e-04 | -8.11 | positive regulation of response to stimulus | biological process | GO:0048584 | 481 | 67 | 12345 | 1129 | PGRP-LE,cad,tkv,spen,rl,Mtl,Pi3K92E,nej,pont,snk,dx,spz,PDCD-5,trbd,tub,wisp,EMRE,Archease,Cka,O-fut1,Pdk1,Imp,foxo,Pura,Traf6,dl,disp,CG4853,raptor,mop,CG9925,Herc4,Rel,Myd88,Crag,CG10188,Ptip,CycE,dco,Hr78,E2f1,PEK,Aos1,EndoGI,lwr,Tsp86D,tws,Mos,Tao,gd,Ect4,ben,stet,Mekk1,Ssdp,ea,tor,Tak1,GILT1,Usp10,shv,Jafrac2,mts,Seipin,CkIIalpha,Hat1,Sp7 | | 3.042e-04 | -8.10 | ELK1\_02 | MSigDB lists | ELK1\_02 | 113 | 32 | 4161 | 644 | RpS18,CG8004,Tim8,Hat1,Mitofilin,RpL37a,TBCC,Acph-1,Scox,Hmg-2,CG31759,CG5285,CG2493,Sec10,Tom40,ND-75,eEF1beta,mEFTu1,Herc4,Grip75,Faf2,Stoml2,CG8475,AP-1-2beta,CG31715,Arts,Naa20A,Gmer,Rpt2,CG6610,EcR,mRpL13 | | 3.059e-04 | -8.09 | regulation of cell cycle process | biological process | GO:0010564 | 177 | 31 | 12345 | 1129 | dgt4,Moe,CycE,twe,APC7,tws,mus304,Sld5,nbs,cana,Su(H),Mos,wapl,ago,mre11,SuUR,Aven,nej,mad2,Rbf,foxo,cnn,CG31053,PEK,bru1,E2f1,Zwilch,Rbf2,otu,Cdk1,grp | | 3.062e-04 | -8.09 | phosphatase activity | molecular function | GO:0016791 | 184 | 32 | 12176 | 1119 | Synj,CG7789,tws,Ptp69D,Dd,CG17294,PpD3,Pfrx,PpV,CG6707,CG5026,alph,CanA-14F,CG32549,Ublcp1,cN-IIIB,mop,Ptpmeg2,Ptpmeg,Pp2A-29B,twe,Ptp10D,csw,Pp2B-14D,CG7115,aay,Acph-1,mts,CG17746,ssh,CG9391,Ptp61F | | 3.068e-04 | -8.09 | MODULE\_8 | MSigDB lists | MODULE\_8 | 187 | 47 | 4161 | 644 | aay,CG12018,Vha36-1,P5CDh1,CG8963,Rab5,Tm1,r,NHP2,Ts,CG4511,CG9485,Arpc1,Mitofilin,Prosalpha3,Gyg,LSm3,Sec23,PIG-C,Hat1,CG13630,Chd64,bys,msps,pr,anne,CycB,r-l,SRPK,ND-B18,ste24a,Cse1,scu,ND-49,opm,eIF3i,Karybeta3,Tsp74F,CG8507,ND-39,Got1,CCT6,mad2,Chrac-14,Echs1,wal,Cdk1 | | 3.107e-04 | -8.08 | Akap200 (A kinase anchor protein 200) | protein interactions | 34170 | 68 | 19 | 9204 | 1112 | Pka-R2,Eno,Uch,capt,scu,Moe,Gdi,regucalcin,CG4598,AnxB10,Stip1,RhoGDI,cib,PHGPx,CG10638,RpS27A,awd,Chd64,Pdk1 | | 3.107e-04 | -8.08 | SHIPP\_DLBCL\_VS\_FOLLICULAR\_LYMPHOMA\_UP | MSigDB lists | SHIPP\_DLBCL\_VS\_FOLLICULAR\_LYMPHOMA\_UP | 20 | 10 | 4161 | 644 | Clc,Rpt2,cathD,Pglym78,Pen,CG11089,CCT3,CycB,CCT5,Eno | | 3.116e-04 | -8.07 | alcohol metabolic process | biological process | GO:0006066 | 95 | 20 | 12345 | 1129 | CG5028,Adh,Idh3b,Idh,CG8112,Synj,Marf,CG17270,CG3842,Sptr,CG7840,scu,Pcd,pr,Fdh,Gk2,l(1)G0196,Gk1,CG9391,foxo | | 3.181e-04 | -8.05 | GO\_ORGANOPHOSPHATE\_METABOLIC\_PROCESS | MSigDB lists | GO\_ORGANOPHOSPHATE\_METABOLIC\_PROCESS | 357 | 79 | 4161 | 644 | CG7718,CG2201,CG5285,CG6767,Cyt-c-p,nSMase,Ts,CG17333,sgll,r,DhpD,Plap,scramb1,Ppcdc,COX5A,r-l,CG7322,Pfk,PIG-C,CG3376,ND-24,csw,CG11164,nmdyn-D7,Nadsyn,LysRS,levy,PIG-L,ACC,CG11089,Gk2,CG7840,Nc73EF,Idh,Idh3b,PAPLA1,Pi4KIIalpha,ND-75,UQCR-Q,Dhfr,mal,Pi3K92E,Synj,CG7789,Pfas,CG32549,Pdxk,Mtpalpha,Ac3,ND-B18,Eno,Pglym78,nmdyn-D6,ND-30,Thiolase,Gapdh1,ND-49,CG33156,Pgm1,Prip,AMPdeam,mino,ND-SGDH,ND-B22,Stoml2,CG5028,CG4407,ND-23,ND-51,dnc,ND-39,Scsalpha1,CG15111,kri,l(1)G0196,CG3803,CG9391,Shmt,Pgk | | 3.181e-04 | -8.05 | peptide transport | biological process | GO:0015833 | 473 | 66 | 12345 | 1129 | CHMP2B,mbo,Rab5,Karybeta3,Pex12,CG14894,Arts,Sec22,Tom40,Crag,Sec23,Arf51F,CG5734,eIF3m,Pen,Clc,Topors,Kap-alpha3,sau,tho2,Gga,CG8728,Syx13,CG5447,CG11110,AP-1-2beta,CHOp24,CG4041,Arl5,CG3077,Rep,Gdi,Roe1,Oseg1,CG10344,sbr,mol,Nup62,opm,Atg6,Pex10,Tom70,Arpc1,CG8314,CG6766,Nxf3,GAPsec,Vps16B,Mat89Ba,Nup358,sip3,Tim8,Pex13,Tango10,CG9393,CG8004,Cse1,Rab10,lt,yin,Vsp37A,Rab32,Stam,milt,thoc6,Tsp86D | | 3.213e-04 | -8.04 | ribosome assembly | biological process | GO:0042255 | 55 | 14 | 12345 | 1129 | RpL10,RpL12,RpS28b,RpS27,RpS15,mRpS11,RpS5b,RpS14b,RpL11,RpL23A,RpLP0,CG7488,RpS14a,RpL24 | | 3.227e-04 | -8.04 | pyrimidine ribonucleoside triphosphate metabolic process | biological process | GO:0009208 | 5 | 4 | 12345 | 1129 | r,nmdyn-D7,nmdyn-D6,awd | | 3.227e-04 | -8.04 | pyrimidine ribonucleoside triphosphate biosynthetic process | biological process | GO:0009209 | 5 | 4 | 12345 | 1129 | awd,nmdyn-D6,nmdyn-D7,r | | 3.247e-04 | -8.03 | GSE22886\_DAY0\_VS\_DAY7\_MONOCYTE\_IN\_CULTURE\_DN | MSigDB lists | GSE22886\_DAY0\_VS\_DAY7\_MONOCYTE\_IN\_CULTURE\_DN | 90 | 27 | 4161 | 644 | Got1,Sptr,Irp-1B,Pcd,PAPLA1,Pi4KIIalpha,Acph-1,gammaTub37C,yip2,Jafrac2,Prosbeta5,CG8004,Eno,ND-75,Nca,UQCR-Q,Ntan1,Thiolase,Vha36-1,Etf-QO,parvin,Uba5,Fdh,CG10932,cathD,Grx1,ste14 | | 3.267e-04 | -8.03 | Vinc (Vinculin) | protein interactions | 31201 | 13 | 7 | 9204 | 1112 | Vamp7,ND-75,Su(H),Hip1,Pax,Fkbp12,Exo70 | | 3.269e-04 | -8.03 | nucleoside diphosphate metabolic process | biological process | GO:0009132 | 37 | 11 | 12345 | 1129 | Pfk,nmdyn-D6,Pgk,Gapdh1,PyK,awd,Pfrx,nmdyn-D7,Eno,Scsalpha1,Pglym78 | | 3.277e-04 | -8.02 | mRNA catabolic process | biological process | GO:0006402 | 68 | 16 | 12345 | 1129 | LSm3,smg,cu,CG11486,Trf4-2,CG17768,Cbp80,twin,Edc3,Rga,gw,Rrp40,Not10,CG31759,Pym,armi | | 3.282e-04 | -8.02 | nucleoside monophosphate biosynthetic process | biological process | GO:0009124 | 21 | 8 | 12345 | 1129 | CG6767,Ts,l(2)k01209,Pfas,CG11089,Ady43A,AMPdeam,r-l | | 3.288e-04 | -8.02 | positive regulation of mRNA catabolic process | biological process | GO:0061014 | 12 | 6 | 12345 | 1129 | spn-E,Marf1,gw,smg,AGO3,piwi | | 3.288e-04 | -8.02 | protein monoubiquitination | biological process | GO:0006513 | 12 | 6 | 12345 | 1129 | ari-1,CG10254,Cbl,Pex12,l(3)73Ah,park | | 3.316e-04 | -8.01 | REACTOME\_SRP\_DEPENDENT\_COTRANSLATIONAL\_PROTEIN\_TARGETING\_TO\_MEMBRANE | MSigDB lists | REACTOME\_SRP\_DEPENDENT\_COTRANSLATIONAL\_PROTEIN\_TARGETING\_TO\_MEMBRANE | 81 | 25 | 4161 | 644 | RpL40,RpS4,RpL38,RpS7,RpS2,RpL13A,RpS16,RpS30,RpS13,RpLP0,RpL30,RpL23,RpS24,RpL12,RpL13,RpS18,RpL26,RpL9,RpS23,RpS27A,RpS28b,RpL24,RpS27,RpL37a,RpL11 | | 3.321e-04 | -8.01 | sun (stunted) | protein interactions | 44046 | 29 | 11 | 9204 | 1112 | ATPsynbeta,ATPsynG,ATPsynO,Surf4,ATPsynCF6,ATPsynD,blw,ATPsynF,ATPsynB,ATPsyndelta,ATPsynE | | 3.321e-04 | -8.01 | CG11885 (uncharacterized protein) | protein interactions | 33215 | 29 | 11 | 9204 | 1112 | Pomp,Prosalpha5,dl,Rpn2,Rpn13,Prosbeta4,Prosbeta3,Prosbeta6,Prosalpha3,Prosbeta5,Prosalpha4 | | 3.324e-04 | -8.01 | dephosphorylation | biological process | GO:0016311 | 170 | 30 | 12345 | 1129 | Pp2A-29B,CG17746,cN-IIIB,mop,CanB2,Ptp69D,ssh,aay,Ptp10D,Ptpmeg2,tws,CanA-14F,Ptp61F,Synj,twe,CG7115,CG5026,Acph-1,PpV,CG17294,csw,Dd,Ublcp1,Ptpmeg,CG6707,PpD3,alph,Pp2B-14D,mts,CG9391 | | 3.360e-04 | -8.00 | regulation of defense response | biological process | GO:0031347 | 147 | 27 | 12345 | 1129 | Myd88,PDCD-5,Rel,GlyP,CG9925,spz,G9a,Atg6,Sp7,EMRE,Ptip,cact,Atg7,wisp,Crag,Hat1,Ect4,tub,Stat92E,scny,PGRP-LE,slif,Hr78,Traf6,faf,Tak1,GILT1 | | 3.388e-04 | -7.99 | RNA localization | biological process | GO:0006403 | 178 | 31 | 12345 | 1129 | mbo,alphaTub67C,CkIIalpha,Rrp40,cup,Grip71,spn-E,wisp,osk,msps,Moe,jvl,CG8204,lt,Grip128,Nxf3,CG4572,cnc,swa,Gmer,Mat89Ba,mael,Tm1,sbr,Grip75,Lkb1,thoc6,exu,Dcr-1,tho2,smg | | 3.444e-04 | -7.97 | spindle | cellular component | GO:0005819 | 116 | 23 | 11940 | 1105 | mad2,msps,Klp10A,Moe,alphaTub84B,Cdk1,toc,cbs,CG3437,Zwilch,cnn,Grip128,twe,Grip75,dgt4,Grip84,Axs,Tao,Clc,Rep,CG7716,CycB,gammaTub37C | | 3.453e-04 | -7.97 | ScsbetaG (Succinyl-coenzyme A synthetase beta subunit, GDP-forming) | protein interactions | 44001 | 96 | 24 | 9204 | 1112 | SERCA,CCT6,Pdk1,Kap-alpha3,CG11486,lwr,CCT3,Pp2A-29B,Pen,Droj2,Rab5,CG1416,Mcad,ScsbetaA,cathD,r,GCS2beta,Cp1,wal,mEFTu1,Rpt2,Tspo,bcd,Vha68-2 | | 3.467e-04 | -7.97 | REACTOME\_INFLUENZA\_LIFE\_CYCLE | MSigDB lists | REACTOME\_INFLUENZA\_LIFE\_CYCLE | 109 | 31 | 4161 | 644 | Rpb8,RpS4,Karybeta3,RpL38,RpS7,mbo,RpL40,RpL23,RpL30,RpS24,Aladin,RpII140,RpS16,RpL13A,RpS2,RpS13,RpLP0,RpS30,RpS18,RpL12,RpL13,RpL37a,RpL11,RpS27A,RpL9,RpS23,Clc,RpL26,RpS28b,RpS27,RpL24 | | 3.489e-04 | -7.96 | GO\_MITOCHONDRIAL\_MATRIX | MSigDB lists | GO\_MITOCHONDRIAL\_MATRIX | 245 | 58 | 4161 | 644 | Gabat,CG14407,CG7488,mRpL17,mRpL10,mRpS5,shop,ND-30,Thiolase,Mtpalpha,CG11858,bor,ScsbetaA,P5CDh1,CG8728,CG31759,P5cr,Pdk,Atg4b,Cyp1,tko,Roe1,Etf-QO,Ts,CG1640,mtTFB1,Trxr-1,Idh3b,mRpS21,CG1774,Nc73EF,Scsalpha1,ND-39,TrpRS-m,mRpL16,mEFTu1,CG8199,Aldh,SerRS-m,wal,ND-75,Echs1,scu,mRpL13,Sod1,LysRS,ND-49,CG2100,Mcad,mRpL18,CG4407,CG5028,CG10932,Grx1,ND-23,GluRS-m,CG17691,tzn | | 3.523e-04 | -7.95 | sstn (stepping stone) | protein interactions | 39643 | 48 | 15 | 9204 | 1112 | Sod1,SERCA,cup,aay,Adi1,CG9911,CycB,mts,Pax,faf,Marf1,cnn,smg,Pen,Pxt | | 3.523e-04 | -7.95 | Mf (Myofilin) | protein interactions | 41824 | 48 | 15 | 9204 | 1112 | ATPsynCF6,Scp1,ATPsynF,osk,ATPsynB,Bx,Ssdp,Nc73EF,Argk,Actn,up,ATPsynO,Cyt-c-p,Adh,CG7920 | | 3.526e-04 | -7.95 | GO\_INNATE\_IMMUNE\_RESPONSE\_ACTIVATING\_CELL\_SURFACE\_RECEPTOR\_SIGNALING\_PATHWAY | MSigDB lists | GO\_INNATE\_IMMUNE\_RESPONSE\_ACTIVATING\_CELL\_SURFACE\_RECEPTOR\_SIGNALING\_PATHWAY | 55 | 19 | 4161 | 644 | RpL40,SkpA,Prosbeta6,Rpn2,Traf6,Rpn12,Prosalpha4,Rpt2,Prosbeta3,Rpn3,Cul1,Uev1A,Pdk1,cact,Prosbeta5,Prosalpha5,RpS27A,Prosbeta4,Prosalpha3 | | 3.554e-04 | -7.94 | Moe (Moesin) | protein interactions | 31816 | 143 | 32 | 9204 | 1112 | Pglym78,Uch,Gdi,scu,Moe,capt,CG4598,REPTOR,RhoGDI,Fkbp12,Top1,Pld,Trxr-1,awd,CG2767,Eno,CG32512,otu,regucalcin,AnxB10,Stip1,Prosbeta5,CG14715,CG10638,Pgk,PHGPx,EndoGI,cib,yip2,Chd64,CG11267,RpS27A | | 3.583e-04 | -7.93 | homeostatic process | biological process | GO:0042592 | 475 | 66 | 12345 | 1129 | SERCA,Rbcn-3A,capt,Pi3K92E,Jafrac1,fwe,nbs,EcR,Hnf4,PyK,EMRE,Pfk,Debcl,Lkb1,Atpalpha,Acox57D-p,anne,PHGPx,up,slif,foxo,mre11,dob,nrv1,CG12773,bun,Grx1,Vha68-2,CG11077,Snm1,Npc2a,CG9911,CG14036,Nap1,ERp60,wash,park,ZnT35C,Tao,Trxr-1,CG14407,SNF4Agamma,Scox,Eno,Hsc70-3,alc,Fer1HCH,zyd,Desat1,CG10470,cnc,sip3,MtnA,shv,Jafrac2,SREBP,Pgk,CG4586,Ho,mus304,Seipin,mute,CG8112,Atf3,Plap,sws | | 3.594e-04 | -7.93 | response to organonitrogen compound | biological process | GO:0010243 | 140 | 26 | 12345 | 1129 | Pi3K92E,Edem1,foxo,rl,Pdk1,CG7328,Bx,Shmt,Vps16B,dco,sip3,cyc,Hnf4,EcR,MAGE,PyK,Rab10,CG6766,CG32640,Pka-R2,Tao,raptor,CG8974,Hsc70-3,CG11070,nbs | | 3.619e-04 | -7.92 | regulation of antimicrobial peptide production | biological process | GO:0002784 | 62 | 15 | 12345 | 1129 | Ptip,lwr,dl,wisp,tub,cact,Tak1,Rel,cad,Aos1,scny,Herc4,mop,Myd88,spz | | 3.627e-04 | -7.92 | Formation of the ternary complex, and subsequently, the 43S complex | REACTOME pathways | R-DME-72695 | 58 | 19 | 3923 | 574 | RpS14a,RpS4,RpS5b,RpS15,RpS2,RpS13,RpS14b,RpS23,RpS30,RpS27,RpS29,RpS7,RpS28b,eIF3i,RpS27A,RpS18,eIF3m,RpS24,RpS16 | | 3.718e-04 | -7.90 | GSE32255\_WT\_VS\_JMJD2D\_KNOCKDOWN\_4H\_LPS\_STIM\_DC\_DN | MSigDB lists | GSE32255\_WT\_VS\_JMJD2D\_KNOCKDOWN\_4H\_LPS\_STIM\_DC\_DN | 51 | 18 | 4161 | 644 | CG3226,P5CDh1,Nurf-38,Roe1,Stoml2,CG11089,Fdh,Rab5,Scsalpha1,eIF3m,Fkbp59,CG9143,RpS27A,CG5116,CG14715,Eno,Cdk1,ND-30 | | 3.751e-04 | -7.89 | Myc (Myc) | protein interactions | 31310 | 174 | 37 | 9204 | 1112 | Rab5,RpL30,RpS23,CG8188,RpS16,RpL13A,Larp4B,ago,cnc,ND-23,larp,lid,CkIIalpha,rept,CG11248,Usp10,Dcr-1,CG31523,RpS7,pont,yps,bif,BicC,fwe,RpL17,Ect4,Neos,tkv,Desat1,Atu,RpL7A,PEK,RpLP1,lqfR,Hsc70-3,RpL23A,gw | | 3.775e-04 | -7.88 | E2F mediated regulation of DNA replication | REACTOME pathways | R-DME-113510 | 25 | 11 | 3923 | 574 | Ts,Dhfr,E2f1,CycE,twe,Rbf2,Pp2A-29B,CycB,Rbf,Cdk1,Orc1 | | 3.846e-04 | -7.86 | MODULE\_4 | MSigDB lists | MODULE\_4 | 11 | 7 | 4161 | 644 | Uev1A,CG8188,CG11070,lwr,Ubc4,UbcE2H,CG40045 | | 3.851e-04 | -7.86 | nuclease activity | molecular function | GO:0004518 | 125 | 24 | 12176 | 1119 | CG8679,scu,Exd2,RpLP0,Snm1,CG11164,Rga,twin,mus81,Drep2,tos,trsn,Dcr-1,CG30343,Clp,DNApol-epsilon255,mus205,Tengl4,CG12375,exu,mre11,CG31759,cu,CG11486 | | 3.904e-04 | -7.85 | regulation of nucleobase-containing compound metabolic process | biological process | GO:0019219 | 1315 | 155 | 12345 | 1129 | CG6813,Myd88,CG11902,Cdk7,ix,CG4021,CG13344,CG17454,Atu,Bx,Dd,pps,bru1,CG10631,Rbf,Traf6,MED27,ph-d,CG10038,Uxt,Tip60,Marf1,Bin1,CG11710,EcR,Hnf4,pgc,gw,CG1602,Rbfox1,tkv,CG4730,insv,BigH1,cad,l(3)mbt,Taf7,asf1,TfIIFalpha,lin-52,YL-1,pont,smg,nej,rl,crm,spen,Drep1,Su(H),psq,Atf6,Utx,HDAC6,S2P,l(3)73Ah,escl,Cdk1,CG31875,Wbp2,cnc,pygo,nos,RYBP,Cul2,Son,Bap170,CG10654,Elba2,Rbf2,HmgD,Stat92E,EndoGI,Sgf11,Aos1,MED20,Hr78,Spt20,E2f1,CG10959,lin,Sox102F,MED24,Sirt7,bun,Rel,HP1e,Cf2,Tfb4,CG2199,CG3587,Chrac-14,spn-E,sbb,CG4872,REPTOR,Med,piwi,Tlk,Topors,d4,CG2678,trr,AGO3,dl,foxo,not,CG10418,lid,cact,bcd,obe,Maf1,mael,Trf2,ovo,rept,e(y)1,mtTFB1,r2d2,CG17202,Atf3,mute,Mybbp1A,Hat1,Rga,scny,CG31053,Ssdp,cyc,esc,MED4,qkr58E-2,SREBP,hb,CG2129,eIF4A,Nf-YA,Rcd5,Smox,hng3,Scox,vers,EloC,CG17328,CycC,HP1c,osk,NFAT,apt,lwr,egg,wash,Nap1,Pax,Mitf | | 3.917e-04 | -7.85 | PRTYPHPHTASE | prints domains | PR00700 | 16 | 7 | 2185 | 212 | Ptpmeg,Ptp10D,Ptpmeg2,Ptp69D,Ptp61F,csw,mop | | 3.968e-04 | -7.83 | fly chr69F5-69F5\|3-39 | chromosome location | fly chr69F5-69F5\|3-39 | 13 | 5 | 25010 | 1343 | snoRNA:Me28S-A774a,CG11267,MICAL-like,CG14117,ste14 | | 3.968e-04 | -7.83 | fly chr69F5-69F5\|3-39 cM | chromosome location | fly chr69F5-69F5\|3-39 cM | 13 | 5 | 25010 | 1343 | ste14,CG14117,MICAL-like,snoRNA:Me28S-A774a,CG11267 | | 4.148e-04 | -7.79 | Lactamase\_B | smart domains | SM00849 | 8 | 5 | 5086 | 506 | tzn,Snm1,CG12375,CG9117,IntS11 | | 4.158e-04 | -7.79 | stem cell division | biological process | GO:0017145 | 97 | 20 | 12345 | 1129 | bam,tws,Med,Pfdn2,msps,tkv,twin,piwi,Stat92E,Cdk1,Lkb1,nos,Dcr-1,eIF4A,egg,cnn,BoYb,mts,mad2,shrb | | 4.184e-04 | -7.78 | tsl (torso-like) | protein interactions | 42564 | 10 | 6 | 9204 | 1112 | bcd,tor,fs(1)N,trk,fs(1)M3,Stat92E | | 4.250e-04 | -7.76 | protein transport | biological process | GO:0015031 | 469 | 65 | 12345 | 1129 | Topors,Kap-alpha3,tho2,sau,mbo,CHMP2B,Karybeta3,Rab5,Pex12,Arts,CG14894,Sec22,Tom40,Sec23,Crag,Arf51F,Pen,Clc,eIF3m,CG5734,Roe1,Gdi,CG3077,Rep,Oseg1,sbr,CG10344,mol,Nup62,CG8728,Gga,CG11110,AP-1-2beta,Syx13,CG5447,CHOp24,CG4041,Arl5,Mat89Ba,Vps16B,GAPsec,sip3,Nup358,Tim8,Pex13,Tango10,CG9393,opm,Atg6,Pex10,Tom70,CG6766,CG8314,Arpc1,Nxf3,Rab32,Vsp37A,Stam,thoc6,milt,Tsp86D,CG8004,Rab10,Cse1,lt | | 4.261e-04 | -7.76 | S6k (Ribosomal protein S6 kinase) | protein interactions | 38654 | 103 | 25 | 9204 | 1112 | foxo,eIF4A,CCT5,levy,Scsalpha1,ago,Aldh,Droj2,raptor,bif,PyK,Mcad,ScsbetaA,cathD,Tak1,Hsc70-3,Kcmf1,Rbf,Cul1,bun,tws,Pink1,Pdk1,Kap-alpha3,Chd64 | | 4.276e-04 | -7.76 | regulation of protein phosphorylation | biological process | GO:0001932 | 245 | 39 | 12345 | 1129 | Mos,SNF4Agamma,CG17746,Tao,ATPsynD,Mekk1,Ptp61F,ben,CycC,PpV,Pka-R1,RhoGAP5A,SkpA,PEK,dco,Pi3K92E,Cbl,CycB,park,Pka-R2,Cdk7,RpLP1,raptor,Abi,fray,CycE,CG30291,ASPP,sws,Lkb1,Cka,alph,CG14408,mts,CycJ,Tak1,Parp16,tor,Pdk1 | | 4.290e-04 | -7.75 | Rpn2 (Regulatory particle non-ATPase 2) | protein interactions | 43449 | 39 | 13 | 9204 | 1112 | Rpn13,Rpn3,Uch-L5,Rpn12,Prosalpha5,Pomp,Prosalpha4,Prosbeta5,Prosbeta3,Prosbeta6,Prosalpha3,Rpt2,Prosbeta4 | | 4.305e-04 | -7.75 | positive regulation of nucleobase-containing compound metabolic process | biological process | GO:0045935 | 523 | 71 | 12345 | 1129 | d4,Bx,piwi,not,foxo,trr,AGO3,dl,CG2199,Cf2,Rel,Cdk7,Myd88,Atu,REPTOR,spn-E,TfIIFalpha,ovo,Trf2,Taf7,BigH1,cad,e(y)1,rept,pont,nej,smg,CG11710,lid,Marf1,Tip60,bcd,Rbfox1,CG1602,gw,EcR,Wbp2,cnc,cyc,CG31053,Nf-YA,pygo,CG2129,hb,SREBP,MED4,esc,Utx,Atf6,Su(H),Mybbp1A,CG17202,S2P,E2f1,MED20,apt,Sgf11,Stat92E,Mitf,Sox102F,lin,Scox,hng3,Rcd5,RYBP,NFAT,CycC,CG17328,EloC,Bap170,vers | | 4.306e-04 | -7.75 | anterior/posterior axis specification, embryo | biological process | GO:0008595 | 134 | 25 | 12345 | 1129 | gus,psq,armi,fs(1)M3,Moe,csw,msps,trk,spn-E,bcd,Grip71,osk,fs(1)N,Grip128,cnc,swa,Grip75,Lkb1,mael,Tm1,rl,tor,nos,exu,hb | | 4.306e-04 | -7.75 | tripartite regional subdivision | biological process | GO:0007351 | 134 | 25 | 12345 | 1129 | swa,cnc,Grip75,Lkb1,mael,Tm1,rl,nos,tor,exu,hb,gus,fs(1)M3,psq,armi,Moe,csw,msps,trk,spn-E,Grip71,bcd,osk,fs(1)N,Grip128 | | 4.343e-04 | -7.74 | dbe (dribble) | protein interactions | 33269 | 7 | 5 | 9204 | 1112 | alph,CG46385,Fkbp39,Droj2,alc | | 4.354e-04 | -7.74 | regulation of production of molecular mediator of immune response | biological process | GO:0002700 | 63 | 15 | 12345 | 1129 | spz,mop,Rel,Herc4,scny,Aos1,cad,Myd88,tub,wisp,Tak1,cact,lwr,Ptip,dl | | 4.354e-04 | -7.74 | locomotor rhythm | biological process | GO:0045475 | 63 | 15 | 12345 | 1129 | dnc,Irk1,Ork1,foxo,gw,cu,nej,CkIIalpha,cyc,dco,tyf,lid,Bx,Ac3,Pka-R2 | | 4.366e-04 | -7.74 | nitrogen compound transport | biological process | GO:0071705 | 605 | 80 | 12345 | 1129 | Efr,Tsp86D,milt,thoc6,Stam,Vsp37A,Rab32,yin,Gmer,lt,colt,Ugalt,Cse1,Rab10,Moe,CG8004,CG9393,kcc,Tango10,Pex13,Tim8,Nup358,sip3,Vps16B,Mat89Ba,GAPsec,Nxf3,Arpc1,CG6766,CG8314,Tom70,CG4476,Pex10,CG4991,Atg6,opm,Nup62,Dcr-1,mol,CG10344,sbr,Sfxn2,Oseg1,CG3077,Rep,Gdi,Roe1,Arl5,CG4041,CHOp24,CG5447,Syx13,CG11110,AP-1-2beta,alphaTub67C,CG8728,Gga,sau,tho2,Kap-alpha3,slif,CG4572,Topors,eIF3m,CG5734,Pen,Clc,Arf51F,Crag,Sec23,Sec22,Tom40,Arts,CG14894,Pex12,CG12773,Rab5,Karybeta3,CHMP2B,mbo | | 4.405e-04 | -7.73 | oxidoreductase activity, acting on the CH-CH group of donors | molecular function | GO:0016627 | 38 | 11 | 12176 | 1119 | CG9471,CG4586,CG3803,SdhA,Mcad,Acox57D-p,SdhD,CG4860,CG7840,Ppox,CG10463 | | 4.442e-04 | -7.72 | wupA (wings up A) | protein interactions | 32794 | 54 | 16 | 9204 | 1112 | Scp1,fwe,osk,ATPsynB,Argk,Nc73EF,Cyt-c-p,ATPsynO,ATPsynCF6,ATPsynF,Actn,SERCA,Adh,CG7920,up,Mf | | 4.460e-04 | -7.72 | GSE3982\_NEUTROPHIL\_VS\_BCELL\_DN | MSigDB lists | GSE3982\_NEUTROPHIL\_VS\_BCELL\_DN | 87 | 26 | 4161 | 644 | Pen,RpL13,RpS18,Rnmt,Mekk1,RpL9,RpS23,gammaTub37C,RpS27,Herc4,Clp,mRpL17,Adi1,rept,CG8475,DppIII,Stoml2,CG15141,RpS4,sau,RpS16,RpL13A,Pex10,Non1,Dhfr,TyrRS | | 4.481e-04 | -7.71 | eEF1delta (eukaryotic translation elongation factor 1 delta) | protein interactions | 34363 | 44 | 14 | 9204 | 1112 | PHGPx,Moe,capt,regucalcin,CG10638,awd,Chd64,alt,Echs1,AnxB10,ATPsynCF6,Eno,Uch,Stip1 | | 4.489e-04 | -7.71 | pyrimidine nucleotide biosynthetic process | biological process | GO:0006221 | 17 | 7 | 12345 | 1129 | r-l,Ts,awd,r,l(2)k01209,nmdyn-D7,nmdyn-D6 | | 4.489e-04 | -7.71 | pyrimidine nucleotide metabolic process | biological process | GO:0006220 | 17 | 7 | 12345 | 1129 | r-l,Ts,awd,nmdyn-D7,nmdyn-D6,l(2)k01209,r | | 4.509e-04 | -7.70 | MOOTHA\_HUMAN\_MITODB\_6\_2002 | MSigDB lists | MOOTHA\_HUMAN\_MITODB\_6\_2002 | 242 | 57 | 4161 | 644 | Echs1,CG3803,Tom40,mRpS10,ND-75,wal,CG8199,AIF,Aldh,SdhA,mEFTu1,Shmt,mRpL16,TrpRS-m,ND-39,Scsalpha1,Nc73EF,Got1,CG1774,Idh3b,Ppox,Trxr-1,CG17691,levy,ND-23,CG10932,CG5028,mRpL18,Mcad,ND-49,ND-SGDH,mRpL13,Tim8,Mtpalpha,Coq7,ND-B18,Thiolase,ND-30,ND-24,Marf,Mitofilin,mRpL17,yip2,Gabat,COX5A,mtTFB1,Etf-QO,tko,Tspo,Cyp1,Pdk,P5cr,P5CDh1,Cyt-c-p,ScsbetaA,CG11858,colt | | 4.519e-04 | -7.70 | key (kenny) | protein interactions | 37967 | 115 | 27 | 9204 | 1112 | blw,stai,RpL7A,Mitofilin,CCT3,pont,CG4853,RpL23A,alphaTub84B,CCT6,Sod1,cup,Rel,vret,Chd64,RpS14a,Sgf29,RpL12,RpS28b,Jafrac1,Cp1,GCS2beta,rept,scu,Nup358,RpS27,RpS16 | | 4.525e-04 | -7.70 | amide transport | biological process | GO:0042886 | 479 | 66 | 12345 | 1129 | sau,tho2,Kap-alpha3,Topors,CG5734,eIF3m,Clc,Pen,Arf51F,Crag,Sec23,Arts,CG14894,Sec22,Tom40,Pex12,Rab5,Karybeta3,CHMP2B,mbo,Nup62,mol,CG10344,sbr,Oseg1,CG3077,Rep,Gdi,Roe1,Arl5,CG4041,CHOp24,Syx13,CG5447,CG11110,AP-1-2beta,CG8728,Gga,CG9393,Tango10,Pex13,Tim8,Nup358,sip3,GAPsec,Vps16B,Mat89Ba,Nxf3,Arpc1,CG6766,CG8314,Tom70,Pex10,Atg6,opm,Tsp86D,milt,thoc6,Stam,Vsp37A,Rab32,yin,lt,Cse1,Rab10,CG8004 | | 4.542e-04 | -7.70 | Beta oxidation of octanoyl-CoA to hexanoyl-CoA | REACTOME pathways | R-DME-77348 | 4 | 4 | 3923 | 574 | Echs1,Thiolase,Mtpalpha,Mcad | | 4.542e-04 | -7.70 | Beta oxidation of decanoyl-CoA to octanoyl-CoA-CoA | REACTOME pathways | R-DME-77346 | 4 | 4 | 3923 | 574 | Mtpalpha,Mcad,Thiolase,Echs1 | | 4.645e-04 | -7.67 | regulation of protein serine/threonine kinase activity | biological process | GO:0071900 | 70 | 16 | 12345 | 1129 | CG17746,SNF4Agamma,Cdk7,Tao,raptor,Mekk1,CG30291,CycE,sws,CycC,alph,Pi3K92E,CycB,Tak1,CycJ,Parp16 | | 4.664e-04 | -7.67 | regulation of RNA metabolic process | biological process | GO:0051252 | 1281 | 151 | 12345 | 1129 | Su(H),Atf6,psq,HDAC6,Utx,S2P,escl,l(3)73Ah,CG31875,cnc,Wbp2,pygo,nos,RYBP,Cul2,Son,Bap170,CG10654,Elba2,EndoGI,Stat92E,Rbf2,HmgD,Aos1,Sgf11,E2f1,Spt20,Hr78,MED20,lin,CG10959,Sox102F,MED24,CG11902,Myd88,CG6813,Cdk7,ix,CG4021,CG13344,CG17454,Atu,Bx,Dd,pps,bru1,CG10631,Rbf,Traf6,CG10038,ph-d,MED27,Marf1,Tip60,Uxt,Bin1,CG11710,Hnf4,EcR,pgc,gw,CG1602,Rbfox1,CG4730,tkv,BigH1,cad,insv,Taf7,l(3)mbt,TfIIFalpha,asf1,nej,pont,smg,YL-1,lin-52,crm,rl,spen,r2d2,CG17202,Mybbp1A,Atf3,mute,Hat1,scny,Rga,Ssdp,cyc,MED4,esc,hb,SREBP,qkr58E-2,CG2129,eIF4A,Nf-YA,hng3,Rcd5,Smox,vers,EloC,CG17328,HP1c,osk,NFAT,CycC,apt,lwr,egg,Nap1,wash,Pax,Mitf,Sirt7,Rel,bun,Cf2,HP1e,CG2199,Tfb4,CG3587,Chrac-14,REPTOR,Med,CG4872,spn-E,sbb,piwi,Tlk,Topors,CG2678,d4,dl,AGO3,trr,foxo,not,CG10418,lid,cact,bcd,obe,Maf1,mael,Trf2,ovo,rept,e(y)1,mtTFB1 | | 4.677e-04 | -7.67 | phosphotransferase activity, alcohol group as acceptor | molecular function | GO:0016773 | 346 | 51 | 12176 | 1119 | Pi3K92E,Pdk,Mo25,CG17698,Pink1,dco,Tak1,Pfk,Cdk7,Pfrx,CycJ,Gk1,fray,Tlk,CG2201,Lkb1,SRPK,Ady43A,CycB,CG7328,Pdk1,Gk2,png,CG33156,Coq8,CG8311,alc,tkv,CG4041,PEK,Mos,CG3631,PyK,CG11486,CycE,CG8726,Pi4KIIalpha,Pdxk,Cdk1,Cdk5,Tao,CkIIalpha,grp,Mekk1,tub,Tfb4,p38a,tor,CG8475,rl,CG6650 | | 4.718e-04 | -7.66 | GSE22886\_NAIVE\_CD4\_TCELL\_VS\_48H\_ACT\_TH1\_DN | MSigDB lists | GSE22886\_NAIVE\_CD4\_TCELL\_VS\_48H\_ACT\_TH1\_DN | 106 | 30 | 4161 | 644 | Hat1,lwr,ND-30,Stip1,wal,ND-75,DNApol-epsilon255,asf1,CG7488,LSm3,Prosalpha3,Idh,mRpL17,Mitofilin,rept,Grx1,ND-51,levy,SelD,eIF3i,RfC38,CG10470,Edc3,Rpn2,Rpn12,CG8188,mRpL13,Prosbeta3,Cdk7,mRpL18 | | 4.747e-04 | -7.65 | negative regulation of transposition | biological process | GO:0010529 | 22 | 8 | 12345 | 1129 | tral,spn-E,vret,BoYb,AGO3,tapas,mael,piwi | | 4.756e-04 | -7.65 | PTPc | smart domains | SM00194 | 16 | 7 | 5086 | 506 | csw,mop,Ptpmeg2,Ptp69D,Ptp10D,Ptp61F,Ptpmeg | | 4.785e-04 | -7.64 | mitochondrial translation | biological process | GO:0032543 | 98 | 20 | 12345 | 1129 | mRpL13,TrpRS-m,mRpL48,CG6094,tko,mRpL36,mRpS21,mRpS10,mRpL18,mRpL10,mRpS23,mRpL16,GatB,SerRS-m,mRpL35,mRpS5,mRpS11,mEFTu2,mEFTu1,mRpL17 | | 4.812e-04 | -7.64 | protein-containing complex assembly | biological process | GO:0065003 | 516 | 70 | 12345 | 1129 | Pfk,Usp39,CG11110,CG3198,Dcr-1,pont,ND-23,CG5969,rept,Trf2,Taf7,asf1,tkv,BigH1,Cchl,spn-E,Hip1,Grip128,Rpn13,Chrac-14,CG7716,Clc,gammaTub37C,TBCC,LSm3,CG11562,RpS15,CG10055,CG17768,bru1,rig,mRpS11,Sgf29,Grip75,RpL23A,osk,Cenp-C,RpL12,RpL10,SmE,CG8204,RpLP0,RpL24,RpS5b,Scox,Nap1,CG10465,Grip84,CG14671,Tomosyn,Edc3,Shmt,msps,Hat1,RpS28b,r2d2,Ndc1,RpS14a,CG7488,RpL11,CanB2,RpS14b,SelR,RpS27,CG1890,ND-18,Caf1-105,Pomp,Sdhaf3,Unr | | 4.867e-04 | -7.63 | fly chr91E4-91E4\|3-65 cM | chromosome location | fly chr91E4-91E4\|3-65 cM | 8 | 4 | 25010 | 1343 | P5cr,NP15.6,GatA,CG14286 | | 4.867e-04 | -7.63 | fly chr91E4-91E4\|3-65 | chromosome location | fly chr91E4-91E4\|3-65 | 8 | 4 | 25010 | 1343 | CG14286,NP15.6,P5cr,GatA | | 4.869e-04 | -7.63 | peptidase complex | cellular component | GO:1905368 | 90 | 19 | 11940 | 1105 | Rpt2,Sgf11,Prosbeta5,Prosbeta3,Rpn12,Rpn3,Pomp,Prosalpha5,e(y)1,Prosbeta6,not,Prosbeta4,Spt20,Sgf29,Prosalpha3,Rpn2,Rpn13,Uch-L5,Prosalpha4 | | 4.869e-04 | -7.63 | endoplasmic reticulum part | cellular component | GO:0044432 | 252 | 40 | 11940 | 1105 | Dd,EMC3,PIG-C,EMC8-9,PIG-H,Sec23,Efr,CG31523,Mctp,mol,CG33977,CG32640,alt,CG6766,Atf6,CG14671,CG5510,CG10425,sip3,cup,CG5969,Seipin,sws,CG17270,CG4293,Desat1,bif,SERCA,Parp16,CG8974,Sec22,CG10470,CG32441,CG8311,SREBP,Lnpk,CG8112,CG4880,Hsc70-3,ste24a | | 4.915e-04 | -7.62 | GSE2405\_0H\_VS\_9H\_A\_PHAGOCYTOPHILUM\_STIM\_NEUTROPHIL\_DN | MSigDB lists | GSE2405\_0H\_VS\_9H\_A\_PHAGOCYTOPHILUM\_STIM\_NEUTROPHIL\_DN | 111 | 31 | 4161 | 644 | Arpc2,RpL11,RpL37a,RpL26,RpS23,COX5A,RpL9,RpS27,RpL24,RpS18,Mtpalpha,CCT5,eEF1beta,Pen,RpL13,RpL12,RpL30,RpL23,RpLP0,RpS16,RpS2,CG1890,RpL13A,levy,RpS4,RpL38,RfC38,eIF3i,Arl6IP1,Pka-R1,RpS7 | | 4.970e-04 | -7.61 | rictor (rapamycin-insensitive companion of Tor) | protein interactions | 32919 | 208 | 42 | 9204 | 1112 | eIF3i,RpS7,Mes2,mRpS23,blw,RpL13,larp,NHP2,rept,eIF4A,foxo,Ankle2,Pi3K92E,Nap1,Nup358,RpL30,Aldh,PyK,RpL23,shu,RpL10,RpL23A,CG4266,RpLP1,RpL7A,Spx,lqfR,Hsc70-3,Kap-alpha3,Pink1,Rbcn-3A,RpL9,RpS14a,Shmt,Imp,eEF1beta,Ect4,RpL17,faf,RpL31,yps,bif | | 4.997e-04 | -7.60 | CycA (Cyclin A) | protein interactions | 39340 | 65 | 18 | 9204 | 1112 | Cdk1,png,heph,otu,bcd,E2f1,EcR,Hsp26,bam,Ntan1,CG14881,CycB,CG17508,cort,twin,r-l,gus,CycE | | 5.017e-04 | -7.60 | GSE3982\_MAC\_VS\_NEUTROPHIL\_LPS\_STIM\_UP | MSigDB lists | GSE3982\_MAC\_VS\_NEUTROPHIL\_LPS\_STIM\_UP | 83 | 25 | 4161 | 644 | CG11710,Cul1,msps,CCT6,CG11134,mtgo,CG9773,Nap1,RpS28b,RpS27,CG8475,CG14894,Aos1,Fdh,mbo,CG11089,Tspo,tun,Cul2,RpII140,LysRS,CG6610,Rpn12,mRpL13,RpS13 | | 5.064e-04 | -7.59 | cytoskeletal protein binding | molecular function | GO:0008092 | 330 | 49 | 12176 | 1119 | Blos2,Grip84,Arpc2,Arpc4,msps,capt,MICAL-like,awd,RpL23,Moe,Grip128,Hsp26,Arpc1,stai,CG7716,wash,parvin,Svil,Kap3,milt,CG3226,CG2258,Ptpmeg,CG1890,Sry-alpha,CG8726,up,cib,Kap-alpha3,Grip71,RpL34a,CG18135,Tomosyn,cnn,Hip1,bif,Tm1,Actn,smg,AnxB10,Klp10A,sw,Grip75,ssh,Girdin,cana,TBCC,Uxt,Chd64 | | 5.105e-04 | -7.58 | GO\_PROTEIN\_MODIFICATION\_BY\_SMALL\_PROTEIN\_CONJUGATION\_OR\_REMOVAL | MSigDB lists | GO\_PROTEIN\_MODIFICATION\_BY\_SMALL\_PROTEIN\_CONJUGATION\_OR\_REMOVAL | 302 | 68 | 4161 | 644 | mad2,Uev1A,Fbw5,Uch-L5,Cdk1,Prosalpha5,Herc4,CSN6,Uba5,CG11070,SkpA,Rpn2,CG8188,CG1812,adp,HDAC6,CG10465,UbcE2H,Prosbeta3,CG8974,Cul1,Rpn3,CycB,MED20,CG2924,Uch,Prosbeta4,Usp39,RpL40,Prosbeta6,Ufm1,Ubc4,Aladin,APC7,CG30291,Top1,STUB1,Cul5,RpS27A,CG15141,CG3894,DCAF12,Rpn12,Prosalpha4,trbd,CG10189,Rpt2,Cul2,lwr,Prosbeta5,CG46338,MED27,Iru,APP-BP1,Pex12,Prosalpha3,Aos1,mbo,Atg7,gus,not,CG4502,CG40045,faf,ari-1,Traf6,CycC,MED24 | | 5.118e-04 | -7.58 | LANDIS\_ERBB2\_BREAST\_PRENEOPLASTIC\_DN | MSigDB lists | LANDIS\_ERBB2\_BREAST\_PRENEOPLASTIC\_DN | 21 | 10 | 4161 | 644 | CG17691,AGBE,Thiolase,ND-51,Gyg,Pgm1,Npc2a,yip2,Ptp10D,mino | | 5.132e-04 | -7.57 | polysaccharide metabolic process | biological process | GO:0005976 | 33 | 10 | 12345 | 1129 | Pgm1,CG9485,CG8475,AGBE,foxo,Gyg,UGP,Atg6,ACC,GlyP | | 5.132e-04 | -7.57 | cellular polysaccharide metabolic process | biological process | GO:0044264 | 33 | 10 | 12345 | 1129 | Pgm1,CG9485,CG8475,foxo,AGBE,Gyg,UGP,Atg6,ACC,GlyP | | 5.170e-04 | -7.57 | InR (Insulin-like receptor) | protein interactions | 42549 | 76 | 20 | 9204 | 1112 | foxo,RpS13,RpS28b,RpL30,Droj2,RpS23,Moe,Pi3K92E,Smox,STUB1,lqfR,RpLP1,RpS7,Cul1,RpL10,bun,Maf1,SkpA,Pdk1,Ptp61F | | 5.179e-04 | -7.57 | GO\_CYTOSOLIC\_SMALL\_RIBOSOMAL\_SUBUNIT | MSigDB lists | GO\_CYTOSOLIC\_SMALL\_RIBOSOMAL\_SUBUNIT | 32 | 13 | 4161 | 644 | RpS7,RpS18,RpS4,mRpS5,RpS24,RpS28b,RpS16,RpS2,RpS13,RpS27,RpS30,RpS23,RpS27A | | 5.308e-04 | -7.54 | Rtnl1 (Reticulon-like1) | protein interactions | 33721 | 177 | 37 | 9204 | 1112 | CycE,tral,Cyp1,mEFTu2,SERCA,ATPsynF,Idgf4,CG5986,Girdin,Mybbp1A,pps,dnc,tyf,ATPsynO,Phb2,PyK,ATP8A,CkIIalpha,Vha68-2,lid,awd,CG5390,Adh,CG10425,CG9485,CG9436,toc,Larp7,RpL10,CG13551,Bin1,CG12288,Argk,Karybeta3,GlyP,Rtnl1,Jafrac1 | | 5.372e-04 | -7.53 | lipid metabolic process | biological process | GO:0006629 | 482 | 66 | 12345 | 1129 | yip2,Pxt,Mtpalpha,ACC,Agpat2,dob,CG4267,CG4860,PIG-P,Synj,Pi4KIIalpha,Acox57D-p,Gk2,CG8311,CG4598,foxo,Pld,Gk1,Etf-QO,CG17270,EcR,Hnf4,wal,SERCA,CG7840,Sccpdh1,adp,Pi3K92E,Thiolase,mino,PIG-L,CG31523,CG7997,Pex10,Seipin,CG6178,CG5966,Sply,sws,CG4594,CG8112,CG2201,CG18135,AcCoAS,PIG-C,CG3842,PIG-H,SREBP,CG10932,CG9391,CG4586,Echs1,SNF4Agamma,Marf,Desat1,CG5026,Mcad,CG7718,CG7789,CG6707,CG33977,Npc2a,nSMase,CG10425,scu,Mitf | | 5.422e-04 | -7.52 | response to hormone | biological process | GO:0009725 | 136 | 25 | 12345 | 1129 | Rab10,PyK,EcR,Hnf4,G9a,Utx,raptor,Chd64,MED27,Idh3b,MED24,CG7328,Nurf-38,Pdk1,rl,Fkbp39,foxo,rept,trr,Pi3K92E,Hr78,rig,Nup358,mAChR-A,Trf2 | | 5.422e-04 | -7.52 | embryonic axis specification | biological process | GO:0000578 | 136 | 25 | 12345 | 1129 | hb,nos,tor,exu,rl,mael,Tm1,Grip75,Lkb1,swa,cnc,fs(1)N,Grip128,spn-E,Grip71,bcd,osk,msps,csw,Moe,trk,fs(1)M3,psq,armi,gus | | 5.433e-04 | -7.52 | GO\_STRUCTURAL\_CONSTITUENT\_OF\_RIBOSOME | MSigDB lists | GO\_STRUCTURAL\_CONSTITUENT\_OF\_RIBOSOME | 131 | 35 | 4161 | 644 | RpS27,RpL24,RpL26,mRpL17,RpL11,RpL37a,mRpL10,mRpS5,RpL12,RpS18,RpS2,RpS16,colt,mRpS23,RpS24,RpL23,RpL30,tko,RpL40,RpS7,RpL38,RpS4,RpS28b,RpL9,RpS27A,RpS23,mRpS21,RpL13,mRpL16,RpS13,RpS30,RpLP0,mRpL13,RpL13A,mRpL18 | | 5.573e-04 | -7.49 | MODULE\_3 | MSigDB lists | MODULE\_3 | 146 | 38 | 4161 | 644 | CG11070,CG11089,Tsp74F,eIF3i,Cse1,scu,ND-49,CCT6,Chrac-14,mad2,Echs1,Tom40,Cdk1,CG14516,Pcd,Nap1,ND-39,Tm1,Rab5,CG10418,r,AGBE,Ts,NHP2,CG12018,CG13630,CG8314,bys,Chd64,CycB,Smox,Arpc1,PpD3,Gyg,Prosalpha3,Mitofilin,Sec23,LSm3 | | 5.620e-04 | -7.48 | cellular carbohydrate metabolic process | biological process | GO:0044262 | 92 | 19 | 12345 | 1129 | ACC,GlyP,UGP,Atg6,CG8475,Pfrx,Treh,Synj,Pgm1,Gk2,Gyg,Mal-A5,foxo,AGBE,Gk1,CG9391,CG9485,CG7328,l(1)G0196 | | 5.635e-04 | -7.48 | cellular detoxification | biological process | GO:1990748 | 13 | 6 | 12345 | 1129 | CG4390,Sod3,Sod1,Jafrac1,Trxr-1,Fdh | | 5.635e-04 | -7.48 | response to cocaine | biological process | GO:0042220 | 13 | 6 | 12345 | 1129 | cyc,dco,Tao,Bx,EcR,Pka-R2 | | 5.644e-04 | -7.48 | germ-line stem cell population maintenance | biological process | GO:0030718 | 78 | 17 | 12345 | 1129 | Med,stet,bam,CSN6,lid,nos,eIF4A,shv,foxo,zpg,Pi3K92E,bru1,scny,mael,piwi,tkv,Stat92E | | 5.663e-04 | -7.48 | ATPsynB (ATP synthase, subunit B) | protein interactions | 39143 | 40 | 13 | 9204 | 1112 | ATPsynO,up,CG15141,Mf,ATPsynG,ATPsynbeta,ATPsynF,ATPsynE,ATPsyndelta,ATPsynCF6,blw,ATPsynD,Scp1 | | 5.693e-04 | -7.47 | REACTOME\_NFKB\_IS\_ACTIVATED\_AND\_SIGNALS\_SURVIVAL | MSigDB lists | REACTOME\_NFKB\_IS\_ACTIVATED\_AND\_SIGNALS\_SURVIVAL | 4 | 4 | 4161 | 644 | RpS27A,Traf6,RpL40,cact | | 5.772e-04 | -7.46 | meiosis I cell cycle process | biological process | GO:0061982 | 114 | 22 | 12345 | 1129 | GatA,Xpc,mre11,sau,Nup62,mus81,c(2)M,cort,CG31053,krimp,SRPK,mtrm,Grip71,spn-E,del,enc,twe,tef,Klp10A,mus304,lid,gammaTub37C | | 5.775e-04 | -7.46 | nuclear-transcribed mRNA catabolic process | biological process | GO:0000956 | 58 | 14 | 12345 | 1129 | CG11486,cu,smg,gw,LSm3,CG31759,Edc3,twin,armi,Pym,Rrp40,Trf4-2,Cbp80,CG17768 | | 5.792e-04 | -7.45 | Rpn13 (Regulatory particle non-ATPase 13) | protein interactions | 36545 | 45 | 14 | 9204 | 1112 | Uch-L5,RpS27A,Rpn3,Cen,Prosalpha5,Rpn2,Rpn12,Prosbeta5,Prosalpha4,Prosbeta4,Rpt2,Prosalpha3,Prosbeta3,Prosbeta6 | | 5.796e-04 | -7.45 | KAAB\_HEART\_ATRIUM\_VS\_VENTRICLE\_DN | MSigDB lists | KAAB\_HEART\_ATRIUM\_VS\_VENTRICLE\_DN | 93 | 27 | 4161 | 644 | yip2,Idh3b,COX5A,ND-39,Nc73EF,ND-30,ND-24,CG13630,Liprin-beta,UQCR-Q,Echs1,ND-75,Mtpalpha,RhoGAP5A,CG8199,AIF,SdhA,Pfk,colt,Pgm1,ND-49,UbcE2H,CG4572,CG10932,Fdh,AGBE,CG40045 | | 5.855e-04 | -7.44 | pll (pelle) | protein interactions | 43283 | 14 | 7 | 9204 | 1112 | cact,tub,dl,foxo,mad2,Cka,Myd88 | | 5.875e-04 | -7.44 | carbohydrate derivative biosynthetic process | biological process | GO:1901137 | 274 | 42 | 12345 | 1129 | nmdyn-D7,Sccpdh1,CG7840,CG33977,GalT1,Gmer,Pfas,r-l,CG17691,CG6767,PIG-L,Efr,Ts,ATPsynD,ATPsynE,CG8199,AMPdeam,Ac3,blw,ATPsynO,CG11089,l(2)k01209,PIG-H,PIG-C,Gk2,O-fut1,nmdyn-D6,oxt,Gk1,ATPsynCF6,COX5B,Ppcdc,ATPsynbeta,CG6178,Ady43A,ATPsyndelta,AcCoAS,r,PIG-P,awd,ATPsynB,ATPsynG | | 5.899e-04 | -7.44 | GSE36826\_NORMAL\_VS\_STAPH\_AUREUS\_INF\_SKIN\_DN | MSigDB lists | GSE36826\_NORMAL\_VS\_STAPH\_AUREUS\_INF\_SKIN\_DN | 57 | 19 | 4161 | 644 | CG10425,CG10344,Coq7,CG9471,crq,AIF,Pfk,Ntan1,UQCR-Q,Idh,Pex12,Irp-1B,Jheh2,Pcd,CG2938,Fur2,CG10932,cathD,CG6028 | | 5.899e-04 | -7.44 | GSE360\_T\_GONDII\_VS\_M\_TUBERCULOSIS\_DC\_UP | MSigDB lists | GSE360\_T\_GONDII\_VS\_M\_TUBERCULOSIS\_DC\_UP | 57 | 19 | 4161 | 644 | Exd2,Ect4,RpL12,Caf1-105,Aldh,SdhA,Ac3,r-l,Ppox,Rab32,obe,CG4872,SREBP,bru1,not,Nprl2,dco,ABCD,Gmer | | 5.932e-04 | -7.43 | fly chr89B9-89B9\|3-58 | chromosome location | fly chr89B9-89B9\|3-58 | 4 | 3 | 25010 | 1343 | bor,Mat89Ba,Mst89B | | 5.932e-04 | -7.43 | fly chr37B12-37B12\|2-54 cM | chromosome location | fly chr37B12-37B12\|2-54 cM | 4 | 3 | 25010 | 1343 | CG10492,Rpn3,CG10470 | | 5.932e-04 | -7.43 | fly chr89B9-89B9\|3-58 cM | chromosome location | fly chr89B9-89B9\|3-58 cM | 4 | 3 | 25010 | 1343 | Mst89B,Mat89Ba,bor | | 5.932e-04 | -7.43 | fly chr37B12-37B12\|2-54 | chromosome location | fly chr37B12-37B12\|2-54 | 4 | 3 | 25010 | 1343 | CG10470,Rpn3,CG10492 | | 5.972e-04 | -7.42 | enoyl-CoA hydratase activity | molecular function | GO:0004300 | 9 | 5 | 12176 | 1119 | Mtpalpha,Thiolase,CG4598,Echs1,CG4594 | | 5.994e-04 | -7.42 | GO\_PHOSPHORIC\_ESTER\_HYDROLASE\_ACTIVITY | MSigDB lists | GO\_PHOSPHORIC\_ESTER\_HYDROLASE\_ACTIVITY | 117 | 32 | 4161 | 644 | alph,mAChR-A,CG3376,Pglym78,CG7789,CanA-14F,CG9248,Acph-1,Pp2A-29B,PpD3,CG32549,CanB2,nSMase,Ptp61F,PpV,Synj,PrBP,CG17294,Dd,Ptp10D,Cyt-c-p,Pp2B-14D,aay,l(1)G0196,Ptpmeg,ERp60,CG9391,Ublcp1,dnc,CG7115,mts,csw | | 6.026e-04 | -7.41 | endoplasmic reticulum organization | biological process | GO:0007029 | 28 | 9 | 12345 | 1129 | CG5510,msps,Sec23,tral,Lnpk,Arl6IP1,Marf,ari-1,Rtnl1 | | 6.026e-04 | -7.41 | zygotic determination of anterior/posterior axis, embryo | biological process | GO:0007354 | 28 | 9 | 12345 | 1129 | fs(1)M3,csw,rl,trk,tor,bcd,osk,fs(1)N,hb | | 6.028e-04 | -7.41 | nuclear chromosome segregation | biological process | GO:0098813 | 168 | 29 | 12345 | 1129 | Rab5,gammaTub37C,cup,eIF4E1,Klp10A,RfC38,mus304,cana,tef,Cenp-C,bif,Axs,msps,spn-E,mtrm,grp,Trf4-2,Topors,CG31053,cort,Rbf,mus81,RecQ5,c(2)M,cnn,lwr,CycB,wapl,nos | | 6.048e-04 | -7.41 | negative regulation of programmed cell death | biological process | GO:0043069 | 107 | 21 | 12345 | 1129 | SkpA,spag,dco,Naa20A,Idgf2,Cul1,Drep1,nos,Pdk1,Ubc6,lin-52,Rbf,Pink1,RYBP,Tip60,bun,CG4238,csw,Debcl,Atg7,MAGE | | 6.068e-04 | -7.41 | nucleobase-containing compound biosynthetic process | biological process | GO:0034654 | 404 | 57 | 12345 | 1129 | TfIIFalpha,E2f1,nmdyn-D7,Trf2,Gmer,Taf7,Pfas,r-l,CG17691,mtTFB1,CG6767,spen,Nurf-38,MED24,e(y)1,Ts,ATPsynE,ATPsynD,Nadsyn,AMPdeam,CG8199,Ac3,blw,ATPsynO,CG11089,l(2)k01209,pps,RpII140,mus205,Rga,Sgf29,nmdyn-D6,fal,Orc1,pygo,Rpb8,ATPsynCF6,Rbf,MED4,COX5B,Ppcdc,Tfb4,ATPsynbeta,CG4407,CG6178,Scsalpha1,Cdk7,cN-IIIB,Ady43A,ATPsyndelta,AcCoAS,CG33156,r,awd,ATPsynB,ATPsynG,CycE | | 6.125e-04 | -7.40 | GSE22886\_NAIVE\_TCELL\_VS\_NEUTROPHIL\_UP | MSigDB lists | GSE22886\_NAIVE\_TCELL\_VS\_NEUTROPHIL\_UP | 98 | 28 | 4161 | 644 | LysRS,CG4390,CG6767,Phb2,Non1,CG1532,csw,CG8963,Fdh,Aos1,CG10932,Roe1,Karybeta3,CG14407,Idh3b,Nap1,RpL26,Nmt,RpL11,eIF3m,ND-30,mEFTu1,CG15715,ND-24,ERp60,Echs1,CG8199,AIF | | 6.125e-04 | -7.40 | GO\_MULTI\_ORGANISM\_METABOLIC\_PROCESS | MSigDB lists | GO\_MULTI\_ORGANISM\_METABOLIC\_PROCESS | 98 | 28 | 4161 | 644 | RpL12,RpL13,RpS18,RpL26,RpS27A,RpL9,RpS23,RpS28b,RpS27,RpL24,RpL37a,RpL11,heph,RpL40,mbo,RpS4,RpL38,RpS7,Aladin,RpS2,RpS16,RpL13A,RpLP0,RpS30,RpS13,RpL23,RpL30,RpS24 | | 6.170e-04 | -7.39 | mitochondrial proton-transporting ATP synthase complex, catalytic core F(1) | cellular component | GO:0000275 | 9 | 5 | 11940 | 1105 | ATPsyndelta,ATPsynO,blw,ATPsynbeta,ATPsynCF6 | | 6.170e-04 | -7.39 | proton-transporting ATP synthase complex, catalytic core F(1) | cellular component | GO:0045261 | 9 | 5 | 11940 | 1105 | ATPsynO,ATPsyndelta,ATPsynbeta,ATPsynCF6,blw | | 6.216e-04 | -7.38 | aerobic respiration | biological process | GO:0009060 | 65 | 15 | 12345 | 1129 | COX5B,Nc73EF,ND-23,UQCR-11,cype,SdhD,SdhA,COX5A,Irp-1B,ScsbetaA,Idh3b,levy,COX4,Scsalpha1,CG5028 | | 6.261e-04 | -7.38 | GCP\_C\_sf | interpro domains | IPR042241 | 5 | 4 | 11383 | 1233 | Grip128,CG7716,Grip75,Grip84 | | 6.279e-04 | -7.37 | GO\_MITOCHONDRION\_ORGANIZATION | MSigDB lists | GO\_MITOCHONDRION\_ORGANIZATION | 245 | 57 | 4161 | 644 | Pi4KIIalpha,GatA,Nmt,ND-39,SERCA,mRpS21,mEFTu1,mRpL16,mRpS10,Tom40,ND-75,Caf1-105,AIF,CG9393,ND-SGDH,HDAC6,mRpL13,scu,Cisd2,mRpL18,CG2100,ND-49,CG3021,CG13601,ND-51,ND-23,Stoml2,ND-B22,GluRS-m,yip2,Mitofilin,mRpL10,mRpL17,Pmi,ND-30,mRpS5,Marf,ND-24,CG8004,Tim8,Gclc,ND-B18,Atg6,Cyt-c-p,CG11110,Pex13,mRpS23,Phb2,CG8728,CG1847,tko,Tspo,Atg4b,Cyp1,mtTFB1,CanB2,sgll | | 6.303e-04 | -7.37 | GO\_MITOCHONDRIAL\_MEMBRANE\_PART | MSigDB lists | GO\_MITOCHONDRIAL\_MEMBRANE\_PART | 75 | 23 | 4161 | 644 | ND-49,ND-SGDH,CG11110,ND-B22,levy,Etf-QO,ND-23,ND-51,Cyp1,ND-39,Mitofilin,Pmi,Ppox,COX5A,Tom40,CG3803,ND-75,ND-B18,SdhA,Ect4,ND-30,Marf,ND-24 | | 6.330e-04 | -7.37 | positive regulation of biosynthetic process | biological process | GO:0009891 | 594 | 78 | 12345 | 1129 | Sox102F,lin,Mitf,Sgf11,Stat92E,MED20,E2f1,tyf,apt,EloC,Bap170,vers,CycC,osk,NFAT,CG17328,Scox,RYBP,Rcd5,hng3,SREBP,Exd2,hb,esc,MED4,CG8009,Nf-YA,Tak1,pygo,CG2129,cyc,cnc,Wbp2,CG17202,S2P,CG8963,Mybbp1A,Atf6,Su(H),Utx,rept,Pi3K92E,pont,nej,SuUR,e(y)1,cad,BigH1,TfIIFalpha,ovo,Taf7,Trf2,EcR,Rbfox1,bcd,wisp,CG1602,Tip60,pn,CG11710,spz,lid,foxo,dl,trr,not,Dd,otu,Bx,d4,Ptip,REPTOR,Atu,Rel,Cdk7,Myd88,CG2199,Cf2 | | 6.330e-04 | -7.37 | positive regulation of cellular biosynthetic process | biological process | GO:0031328 | 594 | 78 | 12345 | 1129 | Su(H),Atf6,Utx,S2P,CG17202,CG8963,Mybbp1A,cnc,Wbp2,cyc,MED4,esc,hb,Exd2,SREBP,Tak1,pygo,CG2129,CG8009,Nf-YA,hng3,RYBP,Rcd5,Scox,vers,Bap170,EloC,CG17328,osk,NFAT,CycC,Stat92E,Sgf11,apt,tyf,E2f1,MED20,lin,Sox102F,Mitf,Myd88,Rel,Cdk7,Cf2,CG2199,Ptip,REPTOR,Atu,Bx,otu,Dd,d4,trr,dl,foxo,not,pn,Tip60,lid,spz,CG11710,EcR,CG1602,wisp,bcd,Rbfox1,BigH1,cad,Trf2,Taf7,TfIIFalpha,ovo,pont,nej,Pi3K92E,rept,e(y)1,SuUR | | 6.336e-04 | -7.36 | CG5355 (uncharacterized protein) | protein interactions | 34421 | 18 | 8 | 9204 | 1112 | CG2767,cib,PHGPx,scu,CG14715,CG10638,awd,AnxB10 | | 6.350e-04 | -7.36 | MORF\_DAP | MSigDB lists | MORF\_DAP | 53 | 18 | 4161 | 644 | ND-23,CG11070,CG5028,CHOp24,SkpA,LysRS,Rpn12,Sod1,Thiolase,Pen,ND-75,Mtpalpha,SdhA,Pp2A-29B,Clc,SREBP,Prosbeta4,LSm3 | | 6.414e-04 | -7.35 | GO\_IMMUNE\_RESPONSE\_REGULATING\_CELL\_SURFACE\_RECEPTOR\_SIGNALING\_PATHWAY | MSigDB lists | GO\_IMMUNE\_RESPONSE\_REGULATING\_CELL\_SURFACE\_RECEPTOR\_SIGNALING\_PATHWAY | 103 | 29 | 4161 | 644 | Prosbeta3,Prosalpha4,Rpt2,Traf6,Rpn12,Pp2B-14D,Rpn2,CanB2,Stoml2,SkpA,Pi3K92E,Prosbeta6,RpL40,dnc,CanA-14F,Prosalpha3,Prosbeta4,Arpc2,Prosalpha5,RpS27A,Arpc1,Prosbeta5,Pdk1,rl,cact,Uev1A,Arpc4,Rpn3,Cul1 | | 6.414e-04 | -7.35 | MORF\_NME2 | MSigDB lists | MORF\_NME2 | 103 | 29 | 4161 | 644 | RpL13A,RpS16,Rpn12,RpS13,RpS30,RpLP0,Rpt2,RpL30,RpS24,Stoml2,RpS4,Rpb8,Prosbeta6,RpL38,levy,RpS7,RpL9,RpS27A,RpS23,RpL24,RpS27,CSN6,RpL11,RpL12,Pgk,lwr,RpL13,Eno,Bin1 | | 6.456e-04 | -7.35 | peptidyl-lysine modification | biological process | GO:0018205 | 153 | 27 | 12345 | 1129 | Ptip,Chrac-14,MAGE,velo,Hat1,escl,Tip60,Rcd5,G9a,lid,Utx,lwr,rept,egg,nej,pont,YL-1,trr,esc,e(y)1,Aos1,Sgf29,Naa20A,Spt20,CG9643,d4,CG31053 | | 6.468e-04 | -7.34 | CG13405 (uncharacterized protein) | protein interactions | 38016 | 31 | 11 | 9204 | 1112 | Droj2,sol,zf30C,RpS27A,ND-B22,CG13679,YL-1,RpS7,Marc,RpL23A,ATPsynF | | 6.468e-04 | -7.34 | ox (oxen) | protein interactions | 45401 | 31 | 11 | 9204 | 1112 | Tspo,eEF1beta,GCS2beta,UQCR-Q,wal,Mcad,UQCR-11,Droj2,Rab5,SERCA,CG1416 | | 6.468e-04 | -7.34 | enc (encore) | protein interactions | 44543 | 31 | 11 | 9204 | 1112 | otu,Cul1,Rbfox1,Ptip,nej,CG17002,Usp10,MED27,MED20,CycE,ix | | 6.600e-04 | -7.32 | ligase activity | molecular function | GO:0016874 | 122 | 23 | 12176 | 1119 | GatA,Nadsyn,Atg7,CysRS,CG6178,AsnS,Gclc,SerRS-m,ACC,GluRS-m,TyrRS,Pfas,ScsbetaA,AcCoAS,LysRS,GatB,Aos1,Scsalpha1,r,Uba5,APP-BP1,TrpRS-m,CG1578 | | 6.609e-04 | -7.32 | GO\_MITOCHONDRIAL\_ENVELOPE | MSigDB lists | GO\_MITOCHONDRIAL\_ENVELOPE | 321 | 71 | 4161 | 644 | shop,Thiolase,ND-30,Marf,mRpS5,CG8974,CG8004,Mtpalpha,ND-B18,yip2,sws,stet,Pmi,Tspo,colt,mRpS23,CG3803,Tom40,SdhA,Ppox,ND-39,Scsalpha1,Unr,ND-51,ND-23,CG10932,Stoml2,ND-B22,CG9393,mino,ND-SGDH,Sod1,sau,Cisd2,mRpL18,ND-49,ND-24,Tim8,Coq7,Scox,COX5A,mRpL10,Mitofilin,mRpL17,CG7488,tko,Cyp1,Ts,Etf-QO,Cyt-c-p,Atg6,bor,CG11110,Phb2,CG8728,CG7718,Ect4,Calx,UQCR-Q,mRpL16,mRpS10,ND-75,AIF,Rab32,Nc73EF,mRpS21,Gk2,levy,mRpL13,Coq6,Sfxn2 | | 6.689e-04 | -7.31 | positive regulation of RNA metabolic process | biological process | GO:0051254 | 513 | 69 | 12345 | 1129 | Atf6,Su(H),Utx,CG17202,S2P,Mybbp1A,cnc,Wbp2,cyc,hb,SREBP,MED4,esc,Nf-YA,CG2129,pygo,hng3,RYBP,Rcd5,EloC,Bap170,vers,NFAT,CycC,CG17328,Sgf11,Stat92E,E2f1,MED20,apt,Sox102F,lin,Mitf,Cdk7,Rel,Myd88,CG2199,Cf2,REPTOR,Atu,spn-E,Bx,piwi,d4,foxo,AGO3,trr,dl,not,Marf1,Tip60,CG11710,lid,gw,EcR,Rbfox1,bcd,CG1602,cad,BigH1,TfIIFalpha,ovo,Trf2,Taf7,rept,nej,smg,pont,e(y)1 | | 6.728e-04 | -7.30 | G1/S Transition | REACTOME pathways | R-DME-69206 | 114 | 30 | 3923 | 574 | Prosbeta3,Orc6,Cdk7,Rpn12,Orc1,Cul1,Rpn3,Cdk1,DNApol-epsilon255,CycB,Prosbeta6,Pp2A-29B,Rbf2,Rpn2,Prosbeta5,Prosalpha4,CycE,twe,Chrac-14,RpS27A,Prosalpha5,Dhfr,E2f1,Prosbeta4,Ts,Prosalpha3,RpL40,Rpt2,SkpA,Rbf | | 6.742e-04 | -7.30 | TER94 (TER94) | protein interactions | 36040 | 106 | 25 | 9204 | 1112 | cbs,Uch,bor,regucalcin,capt,Moe,AnxB10,sip3,CCT3,Ptpmeg,Stip1,Cul1,AGO3,Pgk,CG10638,PHGPx,cib,park,Echs1,Pink1,RpS27A,Chd64,awd,Marf,RpLP0 | | 6.753e-04 | -7.30 | chemical homeostasis | biological process | GO:0048878 | 259 | 40 | 12345 | 1129 | CG14036,wash,park,Pi3K92E,Npc2a,Rbcn-3A,SERCA,Pfk,EcR,Fer1HCH,Hnf4,Desat1,EMRE,zyd,PyK,Eno,ZnT35C,SNF4Agamma,Scox,fwe,CG4586,MtnA,SREBP,foxo,up,slif,Lkb1,Atpalpha,Acox57D-p,CG10470,anne,Atf3,Vha68-2,CG8112,Grx1,Ho,Seipin,dob,nrv1,CG12773 | | 6.767e-04 | -7.30 | REACTOME\_DOWNSTREAM\_SIGNALING\_EVENTS\_OF\_B\_CELL\_RECEPTOR\_BCR | MSigDB lists | REACTOME\_DOWNSTREAM\_SIGNALING\_EVENTS\_OF\_B\_CELL\_RECEPTOR\_BCR | 49 | 17 | 4161 | 644 | cact,Pdk1,Prosbeta5,Rpn3,Cul1,Prosbeta4,Prosalpha3,Prosalpha5,RpS27A,SkpA,Prosbeta6,RpL40,Prosalpha4,Rpt2,Prosbeta3,Rpn2,Rpn12 | | 6.769e-04 | -7.30 | ribonucleoside monophosphate biosynthetic process | biological process | GO:0009156 | 18 | 7 | 12345 | 1129 | r-l,AMPdeam,Ady43A,CG11089,CG6767,Pfas,l(2)k01209 | | 6.847e-04 | -7.29 | alpha-amino acid biosynthetic process | biological process | GO:1901607 | 40 | 11 | 12345 | 1129 | CG11899,AsnS,P5cr,CG11134,r,Adi1,Dhfr,Got1,Shmt,SelD,aay | | 6.867e-04 | -7.28 | Pi3K21B (Pi3K21B) | protein interactions | 33203 | 112 | 26 | 9204 | 1112 | SkpA,tkv,SdhA,Arpc4,Pp2B-14D,alt,RpS27A,Pink1,Usp10,Pp2A-29B,Hsc70-3,RpL7A,RpL23A,Cul1,CG2199,csw,CG1416,yps,Droj2,Pi3K92E,wal,Gapdh1,CG10418,bor,qkr58E-2,larp | | 6.873e-04 | -7.28 | GSE22886\_NEUTROPHIL\_VS\_MONOCYTE\_DN | MSigDB lists | GSE22886\_NEUTROPHIL\_VS\_MONOCYTE\_DN | 113 | 31 | 4161 | 644 | CG15715,Rpn3,ND-75,Eno,Cul5,CCT3,RpS27,COX5A,Nmt,RpS27A,RpL9,RpL26,Idh3b,Nap1,RpL11,eIF3m,ND-39,Mitofilin,RpL37a,CG1774,rept,Fdh,RpS7,Roe1,Ptp61F,RpS13,CG4390,LysRS,RpS24,TyrRS,RpL23 | | 6.915e-04 | -7.28 | Translation initiation complex formation | REACTOME pathways | R-DME-72649 | 70 | 21 | 3923 | 574 | RpS13,RpS2,RpS15,RpS5b,RpS4,RpS14a,eIF4E1,RpS16,RpS18,eIF3m,RpS27A,RpS24,eIF3i,RpS28b,RpS7,RpS29,RpS27,eIF4A,RpS23,RpS30,RpS14b | | 7.036e-04 | -7.26 | negative regulation of cellular protein metabolic process | biological process | GO:0032269 | 218 | 35 | 12345 | 1129 | RpL13A,CycE,sws,cup,Not10,heph,Utx,bam,CkIIalpha,mts,nos,Unr,Rga,CG30496,alph,krimp,bru1,CG14408,Ptp61F,gw,bcd,Rbfox1,PpV,CG17746,mRpL13,ATPsynD,smg,Cbl,faf,park,twin,apt,SkpA,dco,BicC | | 7.097e-04 | -7.25 | Rpn7 (Regulatory particle non-ATPase 7) | protein interactions | 42641 | 36 | 12 | 9204 | 1112 | Prosbeta5,Prosalpha4,Prosbeta4,Rpt2,Prosalpha3,Prosbeta6,Prosbeta3,Rpn3,Rpn13,Prosalpha5,Rpn12,Rpn2 | | 7.294e-04 | -7.22 | regulation of protein kinase activity | biological process | GO:0045859 | 131 | 24 | 12345 | 1129 | Parp16,CycJ,Tak1,CycB,Pdk1,Pi3K92E,Cbl,alph,CG14408,Lkb1,ASPP,CycC,sws,fray,CycE,Abi,CG30291,Mekk1,raptor,RpLP1,Cdk7,Tao,SNF4Agamma,CG17746 | | 7.369e-04 | -7.21 | REACTOME\_SCF\_BETA\_TRCP\_MEDIATED\_DEGRADATION\_OF\_EMI1 | MSigDB lists | REACTOME\_SCF\_BETA\_TRCP\_MEDIATED\_DEGRADATION\_OF\_EMI1 | 41 | 15 | 4161 | 644 | Cul1,Rpn3,RpL40,Prosbeta6,SkpA,Prosbeta5,RpS27A,Rpn2,Prosalpha5,Rpn12,Rpt2,Prosbeta4,Prosalpha4,Prosbeta3,Prosalpha3 | | 7.471e-04 | -7.20 | protein-containing complex subunit organization | biological process | GO:0043933 | 625 | 81 | 12345 | 1129 | Nap1,Nurf-38,CG10465,Grip84,CG14671,Edc3,Tomosyn,Shmt,osk,SmE,RpL12,RpL10,Cenp-C,CG8204,lt,RpLP0,RpL24,RpS5b,Scox,RpS27,CG1890,Caf1-105,Pomp,ND-18,yem,Sdhaf3,Unr,msps,Hat1,RpS28b,Ndc1,Klp10A,r2d2,CG7488,RpS14a,Vamp7,RpS14b,SelR,CanB2,RpL11,stai,CG3198,Dcr-1,YL-1,CG5969,pont,ND-23,rept,Taf7,Trf2,asf1,tkv,BigH1,Pfk,Usp39,CG11110,Tip60,Pym,LSm3,RpS15,CG11562,CG10055,rig,bru1,CG17768,mRpS11,Sgf29,Grip75,RpL23A,Cchl,Hip1,spn-E,Grip128,CG7716,Clc,Rpn13,Chrac-14,Tfb4,CG6094,gammaTub37C,TBCC | | 7.501e-04 | -7.20 | regulation of RNA biosynthetic process | biological process | GO:2001141 | 1158 | 137 | 12345 | 1129 | cnc,Wbp2,CG31875,pygo,nos,HDAC6,Utx,Su(H),Atf6,psq,escl,l(3)73Ah,S2P,Spt20,E2f1,MED20,Hr78,HmgD,Rbf2,Stat92E,Aos1,Sgf11,MED24,lin,CG10959,Sox102F,Cul2,RYBP,CG10654,Elba2,Bap170,CG10631,Bx,Dd,Rbf,Traf6,CG13344,CG11902,CG6813,Myd88,Cdk7,ix,Atu,l(3)mbt,Taf7,TfIIFalpha,asf1,CG4730,tkv,BigH1,cad,insv,crm,rl,spen,smg,nej,pont,lin-52,YL-1,Bin1,CG11710,CG10038,MED27,ph-d,Uxt,Tip60,CG1602,Rbfox1,EcR,Hnf4,pgc,Ssdp,cyc,scny,Rga,CG2129,Nf-YA,MED4,esc,hb,SREBP,r2d2,Mybbp1A,Atf3,mute,Hat1,CG17202,apt,Nap1,wash,Pax,Mitf,egg,lwr,hng3,Smox,Rcd5,CG17328,NFAT,HP1c,CycC,vers,EloC,Topors,CG2678,d4,piwi,Tlk,not,dl,trr,foxo,Cf2,HP1e,CG2199,Tfb4,Sirt7,Rel,bun,REPTOR,CG4872,Med,spn-E,sbb,Chrac-14,CG3587,Trf2,ovo,mael,Maf1,e(y)1,mtTFB1,rept,lid,cact,bcd | | 7.501e-04 | -7.20 | regulation of nucleic acid-templated transcription | biological process | GO:1903506 | 1158 | 137 | 12345 | 1129 | S2P,l(3)73Ah,escl,Su(H),psq,Atf6,Utx,HDAC6,pygo,nos,CG31875,Wbp2,cnc,Bap170,CG10654,Elba2,RYBP,Cul2,CG10959,lin,Sox102F,MED24,Rbf2,Stat92E,HmgD,Aos1,Sgf11,MED20,Hr78,Spt20,E2f1,Atu,CG6813,Myd88,CG11902,Cdk7,ix,CG13344,Rbf,Traf6,Bx,Dd,CG10631,EcR,Hnf4,pgc,CG1602,Rbfox1,MED27,ph-d,CG10038,Tip60,Uxt,Bin1,CG11710,YL-1,lin-52,pont,smg,nej,rl,crm,spen,tkv,CG4730,insv,cad,BigH1,Taf7,l(3)mbt,asf1,TfIIFalpha,CG17202,mute,Atf3,Mybbp1A,Hat1,r2d2,esc,MED4,SREBP,hb,CG2129,Nf-YA,Rga,scny,Ssdp,cyc,vers,EloC,CG17328,CycC,NFAT,HP1c,Rcd5,Smox,hng3,lwr,egg,wash,Nap1,Pax,Mitf,apt,Chrac-14,CG3587,sbb,spn-E,Med,REPTOR,CG4872,Sirt7,bun,Rel,HP1e,Cf2,Tfb4,CG2199,trr,dl,foxo,not,piwi,Tlk,Topors,d4,CG2678,cact,bcd,lid,rept,e(y)1,mtTFB1,mael,Maf1,Trf2,ovo | | 7.532e-04 | -7.19 | GSE360\_L\_DONOVANI\_VS\_M\_TUBERCULOSIS\_DC\_UP | MSigDB lists | GSE360\_L\_DONOVANI\_VS\_M\_TUBERCULOSIS\_DC\_UP | 58 | 19 | 4161 | 644 | Rbf,TyrRS,Pgm1,Stoml2,AP-1-2beta,Jafrac1,rept,Clp,SREBP,CG7806,YL-1,Rab32,Cyp6a22,crq,Aldh,eEF1beta,Exd2,mEFTu1,Cyp6a19 | | 7.621e-04 | -7.18 | GO\_UBIQUITIN\_LIKE\_PROTEIN\_LIGASE\_BINDING | MSigDB lists | GO\_UBIQUITIN\_LIKE\_PROTEIN\_LIGASE\_BINDING | 104 | 29 | 4161 | 644 | dnc,CG40045,Pka-R1,Ubc4,Traf6,CG8188,HDAC6,ari-1,Atg6,UbcE2H,ND-49,CG30291,Cul2,Cul1,lwr,Marf,Uev1A,cact,CG11710,CG46338,Cul5,STUB1,Abi,Smox,Faf2,Hsc70-3,APP-BP1,Pka-R2,CG30496 | | 7.629e-04 | -7.18 | ubiquitin protein ligase activity | molecular function | GO:0061630 | 101 | 20 | 12176 | 1119 | STUB1,SkpA,CG9941,sip3,Topors,CG11070,APC7,Herc4,Cbl,ari-1,Iru,CG8974,CG4238,CG15141,CG13344,CG5003,park,Kcmf1,CG3894,dx | | 7.631e-04 | -7.18 | endoplasmic reticulum tubular network organization | biological process | GO:0071786 | 3 | 3 | 12345 | 1129 | Lnpk,Arl6IP1,Rtnl1 | | 7.631e-04 | -7.18 | L-serine biosynthetic process | biological process | GO:0006564 | 3 | 3 | 12345 | 1129 | CG11899,aay,Shmt | | 7.639e-04 | -7.18 | positive regulation of antimicrobial peptide production | biological process | GO:0002225 | 53 | 13 | 12345 | 1129 | Myd88,cad,Herc4,Rel,Aos1,mop,spz,dl,Ptip,lwr,Tak1,wisp,tub | | 7.639e-04 | -7.18 | positive regulation of production of molecular mediator of immune response | biological process | GO:0002702 | 53 | 13 | 12345 | 1129 | mop,Aos1,Rel,cad,Herc4,Myd88,spz,lwr,Ptip,dl,tub,wisp,Tak1 | | 7.639e-04 | -7.18 | positive regulation of antimicrobial humoral response | biological process | GO:0002760 | 53 | 13 | 12345 | 1129 | lwr,Ptip,dl,tub,wisp,Tak1,mop,cad,Herc4,Aos1,Rel,Myd88,spz | | 7.639e-04 | -7.18 | positive regulation of immune effector process | biological process | GO:0002699 | 53 | 13 | 12345 | 1129 | Tak1,wisp,tub,dl,Ptip,lwr,spz,Myd88,cad,Herc4,Aos1,Rel,mop | | 7.657e-04 | -7.17 | negative regulation of protein metabolic process | biological process | GO:0051248 | 219 | 35 | 12345 | 1129 | twin,dco,BicC,apt,SkpA,Cbl,faf,smg,park,CG17746,ATPsynD,mRpL13,Ptp61F,gw,PpV,bcd,Rbfox1,Unr,Rga,CG30496,bru1,CG14408,krimp,alph,mts,nos,cup,Not10,bam,CkIIalpha,heph,Utx,RpL13A,CycE,sws | | 7.670e-04 | -7.17 | chromatin organization involved in regulation of transcription | biological process | GO:0034401 | 80 | 17 | 12345 | 1129 | Tlk,piwi,wash,rept,egg,esc,r2d2,Bin1,psq,CG10038,ph-d,Elba2,Hat1,spn-E,escl,Chrac-14,Hnf4 | | 7.743e-04 | -7.16 | molybdenum ion binding | molecular function | GO:0030151 | 3 | 3 | 12176 | 1119 | mal,shop,Marc | | 7.751e-04 | -7.16 | MODULE\_105 | MSigDB lists | MODULE\_105 | 76 | 23 | 4161 | 644 | Stip1,r-l,Pdxk,Mtpalpha,ND-B18,Cdk1,Pfk,pr,CycB,mad2,Chd64,PIG-C,Pcd,Arpc1,gammaTub37C,CysRS,CG40045,tzn,AGBE,CG5734,Jafrac1,CG12018,scu | | 7.799e-04 | -7.16 | positive regulation of immune system process | biological process | GO:0002684 | 109 | 21 | 12345 | 1129 | spz,Herc4,PDCD-5,Rel,CG9925,mop,Myd88,wisp,Crag,Hat1,tub,EMRE,Ptip,Sp7,Hr78,PGRP-LE,Aos1,cad,Tak1,lwr,dl | | 7.876e-04 | -7.15 | HALLMARK\_MTORC1\_SIGNALING | MSigDB lists | HALLMARK\_MTORC1\_SIGNALING | 109 | 30 | 4161 | 644 | CCT6,Uch-L5,Pgk,CG7997,Gclc,CG14715,Eno,Prosbeta5,Tom40,LpR2,Stip1,gammaTub37C,Nmt,Idh,Prosalpha3,Mitofilin,Got1,Hsc70-3,l(1)G0004,ACC,Ufm1,AGBE,Nurf-38,aay,Pgm1,CG3226,CG11899,Dhfr,Vha36-1,Gapdh1 | | 7.907e-04 | -7.14 | ubiquitin conjugating enzyme complex | cellular component | GO:0031371 | 3 | 3 | 11940 | 1105 | ben,Ubc6,Uev1A | | 7.959e-04 | -7.14 | Carbon metabolism | KEGG pathways | ko01200 | 114 | 31 | 3198 | 496 | ScsbetaA,Shmt,Scsalpha1,SdhA,Pglym78,Idh3b,CG6767,CG4390,CG10932,Irp-1B,Pfk,CG5028,CG17333,Got1,CG1640,PyK,Pgk,Mcad,Gapdh1,Fdh,Echs1,CG11899,Mtpalpha,Eno,SdhD,CG6650,AcCoAS,Idh,Nc73EF,aay,CG4860 | | 7.959e-04 | -7.14 | Carbon metabolism | KEGG pathways | dme01200 | 114 | 31 | 3198 | 496 | Echs1,Fdh,Gapdh1,CG1640,PyK,Pgk,Mcad,aay,Nc73EF,CG4860,CG6650,AcCoAS,Idh,Mtpalpha,SdhD,CG11899,Eno,CG4390,Idh3b,CG6767,Scsalpha1,SdhA,Pglym78,ScsbetaA,Shmt,Got1,Pfk,CG17333,CG5028,Irp-1B,CG10932 | | 8.003e-04 | -7.13 | REACTOME\_GLYCOLYSIS | MSigDB lists | REACTOME\_GLYCOLYSIS | 12 | 7 | 4161 | 644 | Pgk,Pp2A-29B,Pglym78,mts,Eno,Gapdh1,Pfk | | 8.003e-04 | -7.13 | REACTOME\_CIRCADIAN\_CLOCK | MSigDB lists | REACTOME\_CIRCADIAN\_CLOCK | 12 | 7 | 4161 | 644 | Cul1,RpS27A,cyc,dco,RpL40,SkpA,SREBP | | 8.009e-04 | -7.13 | regulation of protein modification process | biological process | GO:0031399 | 347 | 50 | 12345 | 1129 | raptor,Utx,Pka-R2,Cdk7,RpLP1,Pp2A-29B,ASPP,sws,fray,Abi,CycE,CG30291,alph,Ankle2,cort,CG14408,Lkb1,Cka,CG30496,Parp16,CycJ,Tak1,Pdk1,CG8009,tor,trr,esc,mts,tws,G9a,ATPsynD,Tao,CG17746,Mos,SNF4Agamma,ben,CycC,PpV,Ptp61F,Mekk1,PEK,SkpA,dco,Fdh,Pka-R1,RhoGAP5A,CycB,park,Pi3K92E,Cbl | | 8.029e-04 | -7.13 | dap (dacapo) | protein interactions | 36001 | 62 | 17 | 9204 | 1112 | CycC,Rbf,CG5390,Dcr-1,CycE,vret,lwr,RpS27A,CG11486,Cul2,Kap3,bcd,png,CG17508,Stam,Sod3,EcR | | 8.058e-04 | -7.12 | beta-Oxidation | KEGG pathways | M00087 | 12 | 7 | 3198 | 496 | Echs1,Mtpalpha,Mcad,yip2,CG4586,Thiolase,Acox57D-p | | 8.058e-04 | -7.12 | beta-Oxidation | KEGG pathways | dme\_M00087 | 12 | 7 | 3198 | 496 | Echs1,yip2,Mtpalpha,Mcad,Acox57D-p,CG4586,Thiolase | | 8.075e-04 | -7.12 | anatomical structure formation involved in morphogenesis | biological process | GO:0048646 | 553 | 73 | 12345 | 1129 | tor,Pgk,Orc1,eIF4A,CycJ,grp,ea,Cdk1,cort,slam,Gapdh1,armi,SelR,CanB2,Su(H),bam,CG13366,cnn,CCT3,ScsbetaA,Pax,rost,vret,CG46338,Actn,Stat92E,E2f1,jvl,osk,Iml1,Moe,Sry-alpha,gd,Cul5,foxo,squ,Rbf,Pld,Pglym78,piwi,bru1,up,eIF3m,Pen,CycE,Arf51F,spn-E,Sec23,Vha68-2,Pxt,sw,fs(1)M3,bun,Rab5,heph,Cf2,faf,Dcr-1,Nprl2,rl,mael,Rbcn-3A,tkv,del,PyK,enc,EcR,fs(1)N,Pfk,Rbfox1,cact,gus,spz | | 8.134e-04 | -7.11 | PX\_dom\_sf | interpro domains | IPR036871 | 16 | 7 | 11383 | 1233 | Snx6,CG8726,CG3077,CG5734,CG11807,Pld,snz | | 8.134e-04 | -7.11 | PTPase\_domain | interpro domains | IPR000242 | 16 | 7 | 11383 | 1233 | Ptp61F,mop,csw,Ptpmeg,Ptpmeg2,Ptp10D,Ptp69D | | 8.134e-04 | -7.11 | Phox | interpro domains | IPR001683 | 16 | 7 | 11383 | 1233 | Snx6,CG8726,snz,CG5734,CG3077,CG11807,Pld | | 8.179e-04 | -7.11 | regulation of cell cycle phase transition | biological process | GO:1901987 | 102 | 20 | 12345 | 1129 | Cdk1,Rbf2,grp,PEK,bru1,Zwilch,Rbf,mad2,mre11,ago,Aven,Su(H),Mos,mus304,APC7,tws,nbs,Sld5,twe,CycE | | 8.188e-04 | -7.11 | Dredd (Death related ced-3/Nedd2-like caspase) | protein interactions | 31011 | 73 | 19 | 9204 | 1112 | RpL12,CG11486,Rel,SERCA,RYBP,Sod1,CCT6,RpL23,alphaTub84B,CCT3,Tak1,Hsc70-3,ScsbetaA,RpS16,Nup358,scu,rept,GCS2beta,RpS28b | | 8.225e-04 | -7.10 | positive regulation of cell cycle process | biological process | GO:0090068 | 47 | 12 | 12345 | 1129 | CycE,twe,foxo,Moe,SuUR,Su(H),otu,APC7,CG31053,Sld5,bru1,E2f1 | | 8.255e-04 | -7.10 | MORF\_DAP3 | MSigDB lists | MORF\_DAP3 | 119 | 32 | 4161 | 644 | Pp2A-29B,eIF3m,Prosbeta4,Thiolase,PIG-C,ND-30,lwr,CG13630,CCT3,Phb2,Dd,CG10418,Pka-R1,Prosbeta6,CG8204,CG17454,Idh3b,CSN6,mEFTu1,Pgk,SdhA,ND-SGDH,LysRS,Rpn12,Prosbeta3,eIF4A,ND-51,rept,eIF3i,Stoml2,levy,Rpb8 | | 8.265e-04 | -7.10 | bgcn (benign gonial cell neoplasm) | protein interactions | 47873 | 11 | 6 | 9204 | 1112 | Pym,egg,zpg,nos,bam,twin | | 8.326e-04 | -7.09 | mitochondrial gene expression | biological process | GO:0140053 | 117 | 22 | 12345 | 1129 | tko,mRpL36,mRpL13,TrpRS-m,CG6094,mRpL48,GatB,SerRS-m,mRpL35,mRpS10,mRpS21,mRpL18,mRpL10,mRpS23,mRpL16,mRpS5,mRpS11,scu,mRpL17,mEFTu1,mEFTu2,mtTFB1 | | 8.329e-04 | -7.09 | CCT3 (Chaperonin containing TCP1 subunit 3) | protein interactions | 42029 | 90 | 22 | 9204 | 1112 | bor,DCAF12,wal,mEFTu1,r,CCT5,cathD,Mcad,ScsbetaA,CG1416,Droj2,Rab5,Pp2A-29B,RpLP1,Tak1,Rel,RpLP0,PGRP-LE,Kap-alpha3,mxc,CCT6,SERCA | | 8.345e-04 | -7.09 | GRUETZMANN\_PANCREATIC\_CANCER\_UP | MSigDB lists | GRUETZMANN\_PANCREATIC\_CANCER\_UP | 95 | 27 | 4161 | 644 | CG10425,Eno,Gdi,ND-B18,Pgk,Top1,Moe,ND-39,CanA-14F,Sec23,Smox,Pp2A-29B,Prosalpha5,Nmt,Arl6IP1,tko,Tspo,Gk2,RhoGDI,CG4407,bru1,Cdk7,Prosbeta3,trsn,Sod1,Cse1,Pp2B-14D | | 8.377e-04 | -7.08 | GO\_SMALL\_MOLECULE\_METABOLIC\_PROCESS | MSigDB lists | GO\_SMALL\_MOLECULE\_METABOLIC\_PROCESS | 736 | 143 | 4161 | 644 | Pcd,GatA,TrpRS-m,Idh,Nc73EF,UQCR-Q,AcCoAS,SerRS-m,CG3609,Coq6,CG11164,regucalcin,ACC,CG11089,ste14,Gk2,CG7550,tzn,O-fut1,CG11807,CG8112,Jafrac2,Coq7,CG7322,Pfk,pr,CG10425,CG6767,P5CDh1,CG5285,nSMase,oxt,r,Got1,Sply,ND-39,Scsalpha1,CG15111,CG9391,rl,l(1)G0196,CG3803,wal,scu,Prip,Nurf-38,AMPdeam,S2P,mino,ND-SGDH,ND-49,CG11899,Mcad,Fdh,CG5028,ND-51,dnc,Stoml2,GluRS-m,CysRS,CG3842,CG32549,yip2,CG11134,CG4872,ND-30,Mtpalpha,ND-B18,Eno,Gclc,mal,alc,ABCD,Dhfr,CG4511,Idh3b,CG7840,Npc2a,Desat1,crq,LpR2,Echs1,ND-75,nmdyn-D7,LysRS,Nadsyn,levy,Gabat,COX5A,Ppcdc,ND-24,CG3376,r-l,Cyt-c-p,ScsbetaA,TyrRS,CG2201,P5cr,Pdk,Adi1,Ts,CG17333,Etf-QO,sgll,DhpD,Irp-1B,Ugt35A1,Shmt,Cyp6a19,Pgk,GalT1,kri,CG8199,Aldh,SdhA,CG3860,CG9265,adp,Gapdh1,CG33156,Pgm1,CG10932,CG4407,CG5734,ND-23,ND-B22,CG17691,Cyp6a22,SREBP,Pfas,CG7789,Pglym78,nmdyn-D6,shop,Thiolase,Pdxk,Ac3,p38a,aay,Pex13,CG4390,Synj,CG1640 | | 8.393e-04 | -7.08 | GO\_REGULATION\_OF\_PROTEIN\_CATABOLIC\_PROCESS | MSigDB lists | GO\_REGULATION\_OF\_PROTEIN\_CATABOLIC\_PROCESS | 159 | 40 | 4161 | 644 | CG30496,RpS27A,Sec22,Prosalpha5,STUB1,Cdk1,Uch-L5,mad2,Prosbeta3,Rpt2,Prosalpha4,CG1812,Rpn12,Rpn2,SkpA,regucalcin,cyc,Prosalpha3,Prosbeta4,Wnt10,Smox,Gclc,CycB,Prosbeta5,Moe,Rpn3,Cul1,Edem1,CG30291,CkIIalpha,APC7,ari-1,dco,Tlk,Ubc4,Prosbeta6,CG6766,Med,Atg7,RpL40 | | 8.405e-04 | -7.08 | GSE5542\_UNTREATED\_VS\_IFNA\_AND\_IFNG\_TREATED\_EPITHELIAL\_CELLS\_24H\_UP | MSigDB lists | GSE5542\_UNTREATED\_VS\_IFNA\_AND\_IFNG\_TREATED\_EPITHELIAL\_CELLS\_24H\_UP | 81 | 24 | 4161 | 644 | CG17698,Su(H),CG6766,Karybeta3,CG17593,nmdyn-D7,grp,Phb2,RhoGAP54D,CG8188,CG4390,CG1532,CG12018,CG5292,UQCR-Q,milt,CCT6,Cyp6a19,alph,Nmt,Cyp6a22,Scox,eIF3m,Mitofilin | | 8.428e-04 | -7.08 | PX | pfam domains | PF00787 | 16 | 7 | 10480 | 1142 | snz,CG3077,Pld,CG11807,Snx6,CG8726,CG5734 | | 8.428e-04 | -7.08 | Y\_phosphatase | pfam domains | PF00102 | 16 | 7 | 10480 | 1142 | Ptpmeg2,Ptp10D,Ptp61F,Ptpmeg,csw,mop,Ptp69D | | 8.505e-04 | -7.07 | Activation of the mRNA upon binding of the cap-binding complex and eIFs, and subsequent binding to 43S | REACTOME pathways | R-DME-72662 | 71 | 21 | 3923 | 574 | eIF4E1,RpS14a,RpS5b,RpS15,RpS4,RpS13,RpS2,eIF4A,RpS27,RpS29,RpS14b,RpS30,RpS23,eIF3i,RpS7,RpS28b,RpS16,RpS27A,RpS18,eIF3m,RpS24 | | 8.505e-04 | -7.07 | Ribosomal scanning and start codon recognition | REACTOME pathways | R-DME-72702 | 71 | 21 | 3923 | 574 | RpS14a,eIF4E1,RpS15,RpS5b,RpS4,RpS13,RpS2,RpS27,RpS29,eIF4A,RpS23,RpS30,RpS14b,eIF3i,RpS28b,RpS7,RpS16,RpS27A,eIF3m,RpS24,RpS18 | | 8.530e-04 | -7.07 | GO\_ESTABLISHMENT\_OF\_PROTEIN\_LOCALIZATION\_TO\_MEMBRANE | MSigDB lists | GO\_ESTABLISHMENT\_OF\_PROTEIN\_LOCALIZATION\_TO\_MEMBRANE | 134 | 35 | 4161 | 644 | ste14,CHOp24,Arl6IP1,RpLP0,RpS30,RpS13,RpL13A,sau,Tsp86D,RpL13,RpS28b,RpL9,RpS23,RpS27A,Rab10,shrb,Atg4b,RpL40,RpS7,Rep,RpL38,RpS4,RpS16,RpS2,RpS24,RpL30,RpL23,RpL12,RpS18,RpS27,RpL24,RpL26,Vamp7,RpL11,RpL37a | | 8.589e-04 | -7.06 | Nfs1 (Nfs1 cysteine desulfurase) | protein interactions | 34613 | 57 | 16 | 9204 | 1112 | Rab5,cathD,Mcad,ScsbetaA,mEFTu1,wal,Cp1,r,bor,CCT6,SERCA,Pdk1,Kap-alpha3,Pp2A-29B,CCT3,Arf51F | | 8.594e-04 | -7.06 | coenzyme binding | molecular function | GO:0050662 | 203 | 33 | 12176 | 1119 | wal,ND-51,sgll,Marc,Shmt,Gabat,Acox57D-p,CG10463,AOX3,Dhfr,ND-49,Kyat,Coq6,ND-39,CG4860,Gapdh1,Sirt7,CG31674,CG1640,GlyP,Got1,Mcad,Trxr-1,SdhA,AIF,mal,shop,Gmer,CG14997,Nc73EF,Idh,CG4586,Sply | | 8.596e-04 | -7.06 | regulation of neuron death | biological process | GO:1901214 | 41 | 11 | 12345 | 1129 | Pink1,spag,Tip60,fwe,Naa20A,Rab32,Debcl,Drep2,Aldh,Ect4,MAGE | | 8.688e-04 | -7.05 | GSE360\_DC\_VS\_MAC\_B\_MALAYI\_HIGH\_DOSE\_UP | MSigDB lists | GSE360\_DC\_VS\_MAC\_B\_MALAYI\_HIGH\_DOSE\_UP | 72 | 22 | 4161 | 644 | Iru,CG14516,yip2,Sptr,mEFTu1,alph,Echs1,isoQC,Cyt-c-p,CG2493,mts,Tm1,Rab5,ste14,CG11089,Su(H),Fur2,Tsp74F,oxt,CG7550,CG4562,AP-1-2beta | | 8.712e-04 | -7.05 | GO\_COENZYME\_BINDING | MSigDB lists | GO\_COENZYME\_BINDING | 100 | 28 | 4161 | 644 | shop,CG8112,Mtpalpha,AIF,SdhA,Gclc,wal,Idh3b,Ppox,Trxr-1,ND-39,Sptr,CG4872,Idh,Nc73EF,CG10932,CG5028,ND-51,Etf-QO,Sirt7,scu,Coq6,Gapdh1,ND-49,Gmer,CG10463,Dhfr,Mcad | | 8.715e-04 | -7.05 | gene silencing by RNA | biological process | GO:0031047 | 67 | 15 | 12345 | 1129 | piwi,CG10038,mael,armi,trsn,Hsc70-3,r2d2,Cbp80,AGO3,squ,CG9425,gw,CG6967,spn-E,Dcr-1 | | 8.838e-04 | -7.03 | supramolecular fiber organization | biological process | GO:0097435 | 188 | 31 | 12345 | 1129 | gammaTub37C,CanB2,SelR,Arpc4,Sry-alpha,CG13366,Klp10A,Abi,Grip128,Arpc2,CG7716,jvl,msps,Moe,spd-2,Hip1,Arpc1,Pka-R1,Grip75,Actn,Tm1,capt,up,swa,bru1,Grip84,cib,Mtl,stai,wash,Girdin | | 8.896e-04 | -7.02 | RNA catabolic process | biological process | GO:0006401 | 81 | 17 | 12345 | 1129 | gw,Rrp40,CG11164,armi,Pym,Not10,CG31759,LSm3,CG11486,smg,cu,CG17768,Cbp80,Trf4-2,Rga,twin,Edc3 | | 8.931e-04 | -7.02 | exonuclease activity, active with either ribo- or deoxyribonucleic acids and producing 5'-phosphomonoesters | molecular function | GO:0016796 | 35 | 10 | 12176 | 1119 | cu,CG11486,mre11,CG31759,twin,Rga,DNApol-epsilon255,Snm1,Exd2,tos | | 8.977e-04 | -7.02 | pyrimidine nucleoside triphosphate biosynthetic process | biological process | GO:0009148 | 6 | 4 | 12345 | 1129 | awd,nmdyn-D6,nmdyn-D7,r | | 8.977e-04 | -7.02 | glucan biosynthetic process | biological process | GO:0009250 | 6 | 4 | 12345 | 1129 | AGBE,Gyg,CG9485,Pgm1 | | 8.977e-04 | -7.02 | glycogen biosynthetic process | biological process | GO:0005978 | 6 | 4 | 12345 | 1129 | Pgm1,Gyg,CG9485,AGBE | | 8.991e-04 | -7.01 | GSE17974\_CTRL\_VS\_ACT\_IL4\_AND\_ANTI\_IL12\_4H\_CD4\_TCELL\_DN | MSigDB lists | GSE17974\_CTRL\_VS\_ACT\_IL4\_AND\_ANTI\_IL12\_4H\_CD4\_TCELL\_DN | 86 | 25 | 4161 | 644 | bys,CG9911,Prosbeta5,esc,Gabat,Prosbeta4,Mitofilin,mRpL17,Idh,Prosalpha3,CG30496,escl,l(1)G0004,Cbp80,Naa20A,rept,Rpb8,CG15141,CysRS,CG7550,slif,Rpn12,CG12018,CG17187,CG3226 | | 9.017e-04 | -7.01 | NUCDPKINASE | prints domains | PR01243 | 3 | 3 | 2185 | 212 | nmdyn-D7,awd,nmdyn-D6 | | 9.023e-04 | -7.01 | REACTOME\_ANTIGEN\_PROCESSING\_UBIQUITINATION\_PROTEASOME\_DEGRADATION | MSigDB lists | REACTOME\_ANTIGEN\_PROCESSING\_UBIQUITINATION\_PROTEASOME\_DEGRADATION | 105 | 29 | 4161 | 644 | Cul2,Prosbeta3,Rpt2,UbcE2H,Prosalpha4,Rpn12,CG8188,APC7,Rpn2,Ubc4,Prosbeta6,CG40045,SkpA,CG11070,Atg7,gus,RpL40,Aos1,Uba5,Prosalpha3,Prosbeta4,CG2924,RpS27A,Prosalpha5,STUB1,Prosbeta5,Cul5,Cul1,Rpn3 | | 9.066e-04 | -7.01 | Prosalpha6 (Proteasome alpha6 subunit) | protein interactions | 34359 | 52 | 15 | 9204 | 1112 | Prosbeta5,Prosalpha4,Prosbeta4,Rpt2,Prosalpha3,Pglym78,Prosbeta6,Prosbeta3,Rpn13,AnxB10,cib,Pomp,Prosalpha5,Rpn2,CG10638 | | 9.066e-04 | -7.01 | cact (cactus) | protein interactions | 34969 | 52 | 15 | 9204 | 1112 | CG9925,STUB1,Tak1,cnn,ea,cact,CG14204,spag,CkIIalpha,dl,mad2,Rel,lwr,tub,spz | | 9.066e-04 | -7.01 | Prosalpha4 (Proteasome alpha4 subunit) | protein interactions | 32584 | 52 | 15 | 9204 | 1112 | Prosalpha3,Prosbeta3,Prosbeta6,Prosbeta4,Rpt2,Prosbeta5,Rpn2,regucalcin,CG14715,CG10638,Prosalpha5,Pomp,Rpn13,AnxB10,awd | | 9.102e-04 | -7.00 | positive regulation of mRNA metabolic process | biological process | GO:1903313 | 14 | 6 | 12345 | 1129 | spn-E,piwi,AGO3,smg,gw,Marf1 | | 9.147e-04 | -7.00 | nucleoside diphosphate kinase activity | molecular function | GO:0004550 | 6 | 4 | 12176 | 1119 | nmdyn-D6,nmdyn-D7,awd,Scsalpha1 | | 9.233e-04 | -6.99 | positive regulation of humoral immune response | biological process | GO:0002922 | 54 | 13 | 12345 | 1129 | wisp,tub,Tak1,Ptip,lwr,dl,spz,Herc4,Aos1,Rel,cad,mop,Myd88 | | 9.252e-04 | -6.99 | ribosomal small subunit assembly | biological process | GO:0000028 | 24 | 8 | 12345 | 1129 | RpS27,RpS15,RpS28b,CG7488,RpS14a,mRpS11,RpS5b,RpS14b | | 9.332e-04 | -6.98 | proton-transporting two-sector ATPase complex | cellular component | GO:0016469 | 60 | 14 | 11940 | 1105 | ATPsyndelta,ATPsynO,Vha68-2,Neb-cGP,ATPsynB,blw,Vha36-1,ATPsynE,ATPsynG,Vha13,ATPsynF,ATPsynD,ATPsynCF6,ATPsynbeta | | 9.383e-04 | -6.97 | KEGG\_PARKINSONS\_DISEASE | MSigDB lists | KEGG\_PARKINSONS\_DISEASE | 46 | 16 | 4161 | 644 | ND-B22,levy,CG40045,ND-23,ND-51,ND-49,ND-SGDH,Cyt-c-p,SdhA,ND-B18,ND-75,UQCR-Q,ND-30,ND-24,ND-39,COX5A | | 9.383e-04 | -6.97 | REACTOME\_CYCLIN\_E\_ASSOCIATED\_EVENTS\_DURING\_G1\_S\_TRANSITION\_ | MSigDB lists | REACTOME\_CYCLIN\_E\_ASSOCIATED\_EVENTS\_DURING\_G1\_S\_TRANSITION\_ | 46 | 16 | 4161 | 644 | Prosbeta5,Cul1,Rpn3,Prosbeta4,Prosalpha3,RpS27A,Prosalpha5,CycE,Prosbeta6,SkpA,RpL40,Rpt2,Cdk7,Prosbeta3,Rpn2,Rpn12 | | 9.440e-04 | -6.97 | TYR\_PHOSPHATASE\_PTP | prosite domains | PS50055 | 16 | 7 | 6243 | 694 | csw,Ptp61F,mop,Ptp69D,Ptpmeg2,Ptpmeg,Ptp10D | | 9.455e-04 | -6.96 | GO\_SMALL\_MOLECULE\_CATABOLIC\_PROCESS | MSigDB lists | GO\_SMALL\_MOLECULE\_CATABOLIC\_PROCESS | 165 | 41 | 4161 | 644 | CG1640,Etf-QO,Synj,P5CDh1,ABCD,CG4390,Pex13,Eno,Mtpalpha,CG7322,Pfk,Thiolase,CG3376,Pglym78,CG4872,Gabat,CG7550,tzn,CG17691,CG5734,Gk2,Fdh,CG10932,Mcad,Pgm1,Gapdh1,scu,Echs1,wal,crq,CG3609,CG8199,Aldh,Pgk,CG9391,Shmt,CG7840,Nc73EF,CG15111,Got1,Pcd | | 9.481e-04 | -6.96 | REACTOME\_G1\_S\_TRANSITION | MSigDB lists | REACTOME\_G1\_S\_TRANSITION | 77 | 23 | 4161 | 644 | RpL40,Prosbeta6,Ts,SkpA,Rpn2,Rpn12,Rpt2,Prosbeta3,Cdk7,mts,Dhfr,Rpn3,Cul1,Cdk1,CycB,Prosbeta5,RpS27A,DNApol-epsilon255,Pp2A-29B,Prosalpha5,CycE,Prosbeta4,Prosalpha3 | | 9.506e-04 | -6.96 | MODULE\_93 | MSigDB lists | MODULE\_93 | 91 | 26 | 4161 | 644 | ERp60,Cyp6a19,Thiolase,ND-30,ND-24,Mtpalpha,CG9471,Jafrac2,ND-B18,CG8199,wal,Idh3b,Cyp6a22,Sptr,Idh,ND-23,ND-51,CG17691,levy,scu,Cyt-c-p,Sod1,ND-49,P5CDh1,P5cr,Mcad | | 9.535e-04 | -6.96 | Rpt3 (Regulatory particle triple-A ATPase 3) | protein interactions | 32047 | 42 | 13 | 9204 | 1112 | Rpt2,Prosbeta4,Prosbeta6,Prosbeta3,Prosalpha3,Prosbeta5,Prosalpha4,Pomp,Prosalpha5,Rpn2,Rpn3,Rpn13,Pdk1 | | 9.581e-04 | -6.95 | CG16890 (uncharacterized protein) | protein interactions | 34781 | 5 | 4 | 9204 | 1112 | RpS14b,gus,RpL30,RpS14a | | 9.581e-04 | -6.95 | CG8187 (uncharacterized protein) | protein interactions | 36720 | 5 | 4 | 9204 | 1112 | RpL37a,RpL10,CG11267,CG1092 | | 9.643e-04 | -6.94 | REACTOME\_CLASS\_I\_MHC\_MEDIATED\_ANTIGEN\_PROCESSING\_PRESENTATION | MSigDB lists | REACTOME\_CLASS\_I\_MHC\_MEDIATED\_ANTIGEN\_PROCESSING\_PRESENTATION | 120 | 32 | 4161 | 644 | Prosalpha3,Sec23,Hsc70-3,Prosbeta4,CG2924,Prosbeta5,Cul1,Rpn3,APC7,Ubc4,Prosbeta6,CG40045,gus,Atg7,RpL40,Aos1,RpS27A,Prosalpha5,Cul5,STUB1,ERp60,Prosbeta3,Cul2,Rpt2,UbcE2H,Prosalpha4,Rpn12,CG8188,Rpn2,SkpA,CG11070,Uba5 | | 9.694e-04 | -6.94 | proton-transporting ATP synthase complex, coupling factor F(o) | cellular component | GO:0045263 | 14 | 6 | 11940 | 1105 | ATPsynB,ATPsynF,ATPsynE,ATPsynD,ATPsynCF6,ATPsynG | | 9.694e-04 | -6.94 | mitochondrial proton-transporting ATP synthase complex, coupling factor F(o) | cellular component | GO:0000276 | 14 | 6 | 11940 | 1105 | ATPsynG,ATPsynD,ATPsynCF6,ATPsynF,ATPsynE,ATPsynB | | 9.709e-04 | -6.94 | GO\_MITOCHONDRIAL\_PROTEIN\_COMPLEX | MSigDB lists | GO\_MITOCHONDRIAL\_PROTEIN\_COMPLEX | 68 | 21 | 4161 | 644 | ND-49,CG11110,ND-SGDH,ND-B22,CG17691,levy,ND-23,ND-51,Cyp1,Mitofilin,ND-39,COX5A,Mtpalpha,ND-B18,SdhA,CG8199,Tom40,ND-75,Tim8,ND-30,ND-24 | | 9.756e-04 | -6.93 | GO\_REGULATION\_OF\_CELLULAR\_PROTEIN\_CATABOLIC\_PROCESS | MSigDB lists | GO\_REGULATION\_OF\_CELLULAR\_PROTEIN\_CATABOLIC\_PROCESS | 125 | 33 | 4161 | 644 | Prosalpha5,RpS27A,CG30496,Uch-L5,mad2,STUB1,Cdk1,Rpn12,CG1812,Rpn2,Prosbeta3,Prosalpha4,Rpt2,regucalcin,SkpA,Prosalpha3,Wnt10,Prosbeta4,Rpn3,Cul1,Moe,Prosbeta5,Gclc,CycB,ari-1,APC7,dco,Tlk,CkIIalpha,CG30291,RpL40,Ubc4,Prosbeta6 | | 9.772e-04 | -6.93 | response to toxic substance | biological process | GO:0009636 | 165 | 28 | 12345 | 1129 | Pdk1,rl,Sod1,Jafrac1,Shmt,Pka-R1,Fdh,p38a,alph,apt,CG17734,Arf51F,Fer1HCH,Aldh,dnc,AcCoAS,Trxr-1,whd,Tao,ZnT35C,Pka-R2,Cyp6v1,aay,Agpat2,CG4390,Sod3,Adh,spz | | 9.783e-04 | -6.93 | Ldh (Lactate dehydrogenase) | protein interactions | 45880 | 63 | 17 | 9204 | 1112 | SERCA,CCT6,CG10425,CCT3,blw,CG15098,Rab5,SREBP,Droj2,PyK,CG1416,ScsbetaA,Mcad,cathD,GCS2beta,wal,Tspo | | 9.794e-04 | -6.93 | COX2 (mitochondrial Cytochrome c oxidase subunit II) | protein interactions | 19893535 | 19 | 8 | 9204 | 1112 | Rpt2,wal,COX4,Tspo,SERCA,CG1416,Rab5,Mcad | | 9.831e-04 | -6.92 | GO\_COENZYME\_METABOLIC\_PROCESS | MSigDB lists | GO\_COENZYME\_METABOLIC\_PROCESS | 145 | 37 | 4161 | 644 | Scsalpha1,Nc73EF,Idh,Desat1,Sptr,ND-39,CG1774,Pcd,Idh3b,AcCoAS,Pgk,Shmt,Pgm1,Gapdh1,CG33156,mino,Nadsyn,Coq6,CG11089,ACC,CG4407,CG5028,Ppcdc,CG11134,pr,Eno,CG7322,Pfk,Pdxk,Coq7,Pglym78,Dhfr,ScsbetaA,mal,sgll,CG17333,Ts | | 9.839e-04 | -6.92 | Mitofilin (mitofilin) | protein interactions | 42587 | 15 | 7 | 9204 | 1112 | Pink1,CG8004,PGRP-LE,CG9117,Rel,Mitofilin,Sap47 | | 9.839e-04 | -6.92 | NAT1 (NAT1) | protein interactions | 46020 | 15 | 7 | 9204 | 1112 | Sec10,Lkb1,mael,Drep2,foxo,eIF4A,mre11 | | 9.839e-04 | -6.92 | CG17776 (uncharacterized protein) | protein interactions | 50373 | 15 | 7 | 9204 | 1112 | Mf,CG32473,whd,Debcl,Bin1,CG11164,Hsp26 | | 9.858e-04 | -6.92 | REACTOME\_SCFSKP2\_MEDIATED\_DEGRADATION\_OF\_P27\_P21 | MSigDB lists | REACTOME\_SCFSKP2\_MEDIATED\_DEGRADATION\_OF\_P27\_P21 | 42 | 15 | 4161 | 644 | Rpn3,Cul1,RpL40,SkpA,Prosbeta6,Prosbeta5,Prosalpha5,Rpn2,RpS27A,CycE,Rpn12,Prosbeta4,Rpt2,Prosbeta3,Prosalpha3 | | 9.871e-04 | -6.92 | ubiquitin-like protein ligase activity | molecular function | GO:0061659 | 103 | 20 | 12176 | 1119 | CG5003,park,CG15141,CG13344,dx,Kcmf1,CG3894,CG4238,Iru,CG8974,sip3,CG9941,STUB1,SkpA,Cbl,ari-1,APC7,CG11070,Topors,Herc4 | | 9.878e-04 | -6.92 | response to ammonium ion | biological process | GO:0060359 | 19 | 7 | 12345 | 1129 | cyc,dco,mAChR-A,Pka-R2,Bx,EcR,Tao | | 9.970e-04 | -6.91 | negative regulation of cell cycle | biological process | GO:0045786 | 111 | 21 | 12345 | 1129 | Aven,wapl,Nprl2,mre11,CG10336,Ntan1,Prosbeta5,RecQ5,mus81,nej,mad2,Rbf,Zwilch,grp,Rbf2,Cdk1,mtrm,Iml1,nbs,mus304,Mos | | 9.973e-04 | -6.91 | KIM\_ALL\_DISORDERS\_OLIGODENDROCYTE\_NUMBER\_CORR\_UP | MSigDB lists | KIM\_ALL\_DISORDERS\_OLIGODENDROCYTE\_NUMBER\_CORR\_UP | 309 | 68 | 4161 | 644 | Gapdh1,Prosbeta3,AMPdeam,Sod1,ND-SGDH,CHOp24,ND-51,ND-23,Got1,Scsalpha1,YL-1,Aldh,SdhA,Nca,Pgk,Dd,AnxB10,Kap3,RpS16,RpS4,CG14894,CG1416,RpL11,CG4872,Clc,RpS27,Stip1,Mtpalpha,Gdi,Cdk5,CG8004,Eno,Thiolase,ND-30,Rpt2,Rpn12,RpS30,levy,tzn,rept,Arpc2,Npc2a,RpS27A,PAPLA1,CG8507,Idh3b,ND-75,Echs1,UQCR-Q,lqfR,CG5026,RpL13,mEFTu1,CG9132,TyrRS,Pfdn5,Atg6,Snx6,RpL37a,COX5A,Pp2A-29B,RpL24,CG5510,CG7322,CCT3,pr,Prosbeta5,capt | | 1.012e-03 | -6.90 | Gene Expression | REACTOME pathways | R-DME-74160 | 752 | 138 | 3923 | 574 | CG10418,RpL26,Smox,Ranbp21,Rpn2,RpL30,RpL11,Spx,eIF3m,MED27,RpS27A,Prosalpha5,CG6707,RpS14a,CG6610,Pp2A-29B,RpL9,RpL37a,l(1)G0004,trsn,RpS27,eIF3i,Prosalpha4,eEF1beta,CG3511,nrv1,RpLP1,eIF4E1,IntS11,RpS2,Prosbeta3,RpS29,Nurf-38,Rbf2,Prosbeta5,RpS7,Cwc25,RpS24,thoc6,cN-IIIB,CG6923,Ndc1,yps,MED20,MED4,Dcr-1,mts,MED24,RpL40,eIF4A,Rrp40,RpS14b,Taf7,RpL23,CG12301,Cdk1,Prosbeta6,CG30109,Rep,RpLP0,mbo,RpS13,Not10,faf,raptor,CG11486,twin,gw,EloC,Rbf,Aladin,RpS23,Nup358,RpS28b,SmE,Prosbeta4,alph,RpL7A,RpL12,RpL17,RpL34b,CG7728,Nup62,RpL38,Prosalpha3,Usp39,Rpt2,chrb,RpS16,Hr78,Rnmt,CG8963,RpL23A,bys,Rpn3,Jafrac1,RpL31,RpL10,RpS5b,CG17454,Nup58,RpS4,CycB,alc,Nxf3,tho2,IntS6,RpL13,lid,CG17768,heph,RpL24,SNF4Agamma,Cbp80,EcR,Rga,Med,RpL13A,TfIIFalpha,e(y)1,Edc3,Tfb4,RpS30,RpL29,dco,Rpb8,RpS18,Hnf4,LSm3,RpL34a,sbr,Atu,Rpn12,RpS15,Cdk7,Mat89Ba,CycC,Clp | | 1.019e-03 | -6.89 | GNF2\_EIF3S6 | MSigDB lists | GNF2\_EIF3S6 | 82 | 24 | 4161 | 644 | eEF1beta,RpL13,RpL12,RpS18,RpL9,RpS23,Nap1,RpL24,RpS28b,RpL11,RpL37a,RpL40,RpL38,RpS4,RpS7,RpS30,RpS13,RpS16,RpL13A,RpS2,LysRS,RpL23,RpL30,RpS24 | | 1.026e-03 | -6.88 | GSE24142\_EARLY\_THYMIC\_PROGENITOR\_VS\_DN2\_THYMOCYTE\_FETAL\_UP | MSigDB lists | GSE24142\_EARLY\_THYMIC\_PROGENITOR\_VS\_DN2\_THYMOCYTE\_FETAL\_UP | 34 | 13 | 4161 | 644 | Rab32,Pp2B-14D,CG3860,Tsp86D,Hsc70-3,CanA-14F,CG2493,PIG-C,Arl5,CG8475,CG2938,GlyP,Gclc | | 1.028e-03 | -6.88 | Respiratory electron transport | REACTOME pathways | R-DME-611105 | 40 | 14 | 3923 | 574 | ND-SGDH,ND-B22,ND-39,ND-75,ND-30,ND-24,ND-23,ND-51,wal,Etf-QO,ND-B18,CG8067,NP15.6,ND-49 | | 1.032e-03 | -6.88 | KEGG\_APOPTOSIS | MSigDB lists | KEGG\_APOPTOSIS | 19 | 9 | 4161 | 644 | Pka-R1,CanB2,cact,Pi3K92E,AIF,Cyt-c-p,Pp2B-14D,CanA-14F,Pka-R2 | | 1.033e-03 | -6.88 | Tudor | interpro domains | IPR002999 | 21 | 8 | 11383 | 1233 | CG9684,CG17454,otu,krimp,vret,tapas,CG9925,spn-E | | 1.041e-03 | -6.87 | establishment of protein localization | biological process | GO:0045184 | 485 | 65 | 12345 | 1129 | Tsp86D,Stam,milt,thoc6,Vsp37A,Rab32,lt,Cse1,Rab10,CG8004,CG9393,Pex13,Tango10,Tim8,Vps16B,Mat89Ba,GAPsec,Nup358,sip3,Nxf3,Tom70,Arpc1,CG8314,CG6766,Pex10,opm,Atg6,Nup62,CG10344,sbr,mol,Rep,CG3077,Gdi,Roe1,Oseg1,Arl5,CHOp24,CG4041,CG5447,Syx13,AP-1-2beta,CG11110,Gga,CG8728,sau,tho2,Kap-alpha3,Topors,Arf51F,eIF3m,CG5734,Clc,Pen,Arts,Sec22,Tom40,CG14894,Crag,Sec23,Rab5,Karybeta3,Pex12,CHMP2B,mbo | | 1.043e-03 | -6.87 | CARPB (Carbonic anhydrase-related protein B) | protein interactions | 31915 | 8 | 5 | 9204 | 1112 | Fer1HCH,RpS30,SERCA,CG4407,red | | 1.043e-03 | -6.87 | Grip163 (Grip163) | protein interactions | 39365 | 8 | 5 | 9204 | 1112 | Grip84,Grip128,Grip75,gammaTub37C,Rtnl1 | | 1.045e-03 | -6.86 | GSE5503\_LIVER\_DC\_VS\_PLN\_DC\_ACTIVATED\_ALLOGENIC\_TCELL\_UP | MSigDB lists | GSE5503\_LIVER\_DC\_VS\_PLN\_DC\_ACTIVATED\_ALLOGENIC\_TCELL\_UP | 55 | 18 | 4161 | 644 | Gdi,CG7655,Chrac-14,Cyp6a19,pen-2,Rab10,mRpS21,twin,CG8372,CG3587,Cyp6a22,cathD,Arts,CG4572,RpL30,Aladin,RpLP0,CG8188 | | 1.051e-03 | -6.86 | gig (gigas) | protein interactions | 40201 | 222 | 43 | 9204 | 1112 | Nup358,RpL30,TfIIFalpha,CG9205,Ankle2,ScsbetaA,RpS16,Atpalpha,Nap1,eIF4A,RpL13A,Mitf,CkIIalpha,larp,CG10418,eIF3m,RpL26,RpS27A,Mybbp1A,RpS7,eIF3i,bif,yps,Fbw5,faf,RpL31,RpL17,RpS15,Ect4,Imp,RpL9,Rbcn-3A,Mat89Ba,RpS14a,alt,RpL7A,RpLP1,Spx,Hsc70-3,Rbf,shu,RpL23,bun | | 1.056e-03 | -6.85 | cytoplasmic microtubule organization | biological process | GO:0031122 | 30 | 9 | 12345 | 1129 | Grip75,gammaTub37C,Girdin,Moe,msps,spd-2,CG7716,Grip84,Grip128 | | 1.056e-03 | -6.85 | nucleoside monophosphate metabolic process | biological process | GO:0009123 | 30 | 9 | 12345 | 1129 | Ts,CG6767,AMPdeam,Ady43A,cN-IIIB,r-l,Pfas,l(2)k01209,CG11089 | | 1.060e-03 | -6.85 | proton transmembrane transporter activity | molecular function | GO:0015078 | 111 | 21 | 12176 | 1119 | SERCA,cype,ATPsynD,Vha68-2,CG10960,COX4,ATPsynbeta,yin,COX8,ATPsynO,Vha36-1,ATPsynCF6,COX5B,ATPsyndelta,Vha13,COX5A,blw,levy,ATPsynE,ATPsynB,ATPsynG | | 1.069e-03 | -6.84 | carbohydrate derivative metabolic process | biological process | GO:1901135 | 577 | 75 | 12345 | 1129 | COX5B,Gk1,nmdyn-D6,Gk2,O-fut1,Pglym78,Hexo1,Pfrx,ATPsynG,ATPsynB,PIG-P,kri,ATPsyndelta,cN-IIIB,Scsalpha1,CG17333,UGP,Ugt35A1,Ppcdc,Cht11,Edem1,mino,PIG-L,Idgf2,PGRP-LE,GalT1,CG7840,Sccpdh1,PyK,ATPsynO,Pfk,Ac3,Ugt36E1,ATPsynCF6,Pgk,fdl,oxt,fal,PIG-C,PIG-H,l(2)k01209,CG1774,CG11089,awd,r,Gapdh1,AcCoAS,Ady43A,CG7997,CG11164,CG6178,Idgf4,ATPsynbeta,Ts,Efr,alpha-Man-Ia,CG7920,ScsbetaA,CG6767,CG1827,CG17691,r-l,Gmer,Pfas,scu,Idgf6,CG33977,nmdyn-D7,GCS2beta,blw,CG8199,AMPdeam,Eno,ATPsynE,ATPsynD | | 1.072e-03 | -6.84 | mitochondrial fatty acid beta-oxidation of saturated fatty acids | REACTOME pathways | R-DME-77286 | 7 | 5 | 3923 | 574 | Mcad,Mtpalpha,CG4860,Thiolase,Echs1 | | 1.080e-03 | -6.83 | serine family amino acid biosynthetic process | biological process | GO:0009070 | 10 | 5 | 12345 | 1129 | Dhfr,CG11899,aay,SelD,Shmt | | 1.090e-03 | -6.82 | MORF\_PSMC1 | MSigDB lists | MORF\_PSMC1 | 111 | 30 | 4161 | 644 | LSm3,Prosalpha3,COX5A,Clc,YL-1,Pp2A-29B,ND-75,wal,Bin1,Prosbeta5,Pen,ND-30,Rpt2,CG3631,Cdk7,scu,Sod1,Rpn12,LysRS,CG1890,Rpb8,levy,Prosbeta6,PpV,Stoml2,SkpA,CHOp24,RhoGDI,CG11070,CG11089 | | 1.104e-03 | -6.81 | cyclic-nucleotide phosphodiesterase activity | molecular function | GO:0004112 | 10 | 5 | 12176 | 1119 | Pka-R1,PrBP,Pka-R2,pn,dnc | | 1.104e-03 | -6.81 | oxidoreductase activity, acting on a sulfur group of donors, NAD(P) as acceptor | molecular function | GO:0016668 | 10 | 5 | 12176 | 1119 | CG3939,Grx1,SelT,Trxr-1,cl | | 1.107e-03 | -6.81 | regulation of mitotic cell cycle phase transition | biological process | GO:1901990 | 97 | 19 | 12345 | 1129 | Su(H),APC7,tws,mus304,Sld5,nbs,CycE,twe,Cdk1,Rbf2,grp,PEK,bru1,Zwilch,mad2,Rbf,ago,mre11,Aven | | 1.109e-03 | -6.80 | Toll Like Receptor 4 (TLR4) Cascade | REACTOME pathways | R-DME-166016 | 36 | 13 | 3923 | 574 | Pp2A-29B,p38a,dl,RpL40,RpS27A,rl,Myd88,Tak1,Ptpmeg,Uev1A,Rel,ben,Traf6 | | 1.126e-03 | -6.79 | pum (pumilio) | protein interactions | 41094 | 53 | 15 | 9204 | 1112 | bcd,png,plu,hb,Rbfox1,twin,eIF4E1,Rbf,Pink1,cad,CycB,bam,rl,cup,nos | | 1.126e-03 | -6.79 | Scp1 (Sarcoplasmic calcium-binding protein 1) | protein interactions | 3355102 | 53 | 15 | 9204 | 1112 | CG32473,Mf,up,Cyt-c-p,Adh,Actn,Pgk,Argk,Nc73EF,ATPsynB,Fer1HCH,GlyP,ATPsynCF6,Pglym78,Eno | | 1.131e-03 | -6.78 | - | gene3d domains | 1.20.120.1900 | 5 | 4 | 4929 | 622 | Grip75,Grip84,CG7716,Grip128 | | 1.135e-03 | -6.78 | GSE1925\_3H\_VS\_24H\_IFNG\_STIM\_IFNG\_PRIMED\_MACROPHAGE\_UP | MSigDB lists | GSE1925\_3H\_VS\_24H\_IFNG\_STIM\_IFNG\_PRIMED\_MACROPHAGE\_UP | 51 | 17 | 4161 | 644 | Uba5,Prosbeta6,CG10470,Kap3,ND-49,trbd,CkIIalpha,sbr,Jafrac2,cact,Pfk,PAPLA1,Grip75,CG11134,Wdfy2,Scsalpha1,Dcr-1 | | 1.135e-03 | -6.78 | GSE42021\_TCONV\_PLN\_VS\_CD24LO\_TCONV\_THYMUS\_UP | MSigDB lists | GSE42021\_TCONV\_PLN\_VS\_CD24LO\_TCONV\_THYMUS\_UP | 51 | 17 | 4161 | 644 | Rbf,ND-49,ari-1,Ptp10D,Traf6,CG10470,tzn,Xpac,CG40045,Gyg,cl,Dcr-1,Clp,mRpS21,mEFTu1,CG30010,Cul1 | | 1.136e-03 | -6.78 | GO\_FC\_RECEPTOR\_SIGNALING\_PATHWAY | MSigDB lists | GO\_FC\_RECEPTOR\_SIGNALING\_PATHWAY | 92 | 26 | 4161 | 644 | Prosbeta5,rl,cact,Pdk1,Uev1A,Arpc4,Rpn3,Cul1,CanA-14F,Prosalpha3,Prosbeta4,Arpc2,Prosalpha5,RpS27A,Arpc1,CanB2,SkpA,Prosbeta6,RpL40,Prosbeta3,Prosalpha4,Rpt2,Traf6,Rpn12,Pp2B-14D,Rpn2 | | 1.157e-03 | -6.76 | organelle subcompartment | cellular component | GO:0031984 | 111 | 21 | 11940 | 1105 | Pi4KIIalpha,Parp16,GCC88,Fur2,Gga,Afti,lqfR,Arl5,Sec23,opm,CG31729,ATP8A,AP-1-2beta,cbs,CG9981,Trs33,sau,Gmap,Ugalt,Lnpk,Rab32 | | 1.172e-03 | -6.75 | regulation of transferase activity | biological process | GO:0051338 | 151 | 26 | 12345 | 1129 | raptor,Tao,RpLP1,Cdk7,SNF4Agamma,CG17746,ASPP,sws,CycC,blw,Abi,CycE,fray,CG30291,Mekk1,alph,cort,CG14408,Lkb1,Parp16,CycJ,Tak1,CycB,Pdk1,Pi3K92E,Cbl | | 1.172e-03 | -6.75 | Synthesis of active ubiquitin: roles of E1 and E2 enzymes | REACTOME pathways | R-DME-8866652 | 24 | 10 | 3923 | 574 | faf,UbcE2H,RpS27A,Ubc4,RpL40,CG2924,Uch,CG8188,Ubc6,CG40045 | | 1.177e-03 | -6.74 | Rpn11 (Regulatory particle non-ATPase 11) | protein interactions | 33738 | 33 | 11 | 9204 | 1112 | Prosalpha4,Prosbeta5,Prosbeta6,Prosbeta3,Prosalpha3,Rpt2,Prosbeta4,Rpn13,Rpn3,Rpn2,Prosalpha5 | | 1.177e-03 | -6.74 | CG12321 (uncharacterized protein) | protein interactions | 42174 | 33 | 11 | 9204 | 1112 | Rpn13,Rpn2,Pomp,Prosalpha5,Prosalpha4,Prosbeta5,Prosalpha3,Prosbeta6,Prosbeta3,Prosbeta4,Rpt2 | | 1.178e-03 | -6.74 | cell population proliferation | biological process | GO:0008283 | 175 | 29 | 12345 | 1129 | CycB,Orc1,mEFTu1,Dcr-1,thoc6,mts,cnn,Top1,Cbl,l(3)mbt,Lkb1,Stat92E,Cdk1,COX4,msps,Iml1,Med,Pfdn2,tub,lqfR,Fer1HCH,CycE,enc,pgc,mxc,eIF4E1,tws,bam,SelD | | 1.183e-03 | -6.74 | Prosbeta6 (Proteasome beta6 subunit) | protein interactions | 39855 | 48 | 14 | 9204 | 1112 | AnxB10,Rpn13,Pomp,Prosalpha5,Rpn2,Prosbeta5,COX5A,Prosalpha4,Prosbeta4,faf,Rpt2,not,Prosalpha3,Prosbeta3 | | 1.183e-03 | -6.74 | cellular response to organic substance | biological process | GO:0071310 | 275 | 41 | 12345 | 1129 | MED27,CG32640,Smox,Hsc70-3,CG8974,G9a,EcR,Hnf4,PyK,Rab10,tub,Pdxk,Stat92E,tkv,Shmt,PEK,p38a,Hr78,Pi3K92E,Edem1,lwr,Fkbp39,rl,Nurf-38,ACC,Pka-R2,Atf6,Chd64,bun,raptor,Utx,csw,Med,Vps16B,mAChR-A,Nup358,sip3,foxo,Parp16,Tak1,Pdk1 | | 1.185e-03 | -6.74 | Cyp12d1-p (Cyp12d1-p) | protein interactions | 246648 | 64 | 17 | 9204 | 1112 | Rpt2,wal,mEFTu1,r,bor,Rab5,Rpn3,cathD,ScsbetaA,Mcad,Pp2A-29B,CCT3,Pen,SERCA,CCT6,Kap-alpha3,CG10425 | | 1.197e-03 | -6.73 | GO\_PURINE\_CONTAINING\_COMPOUND\_METABOLIC\_PROCESS | MSigDB lists | GO\_PURINE\_CONTAINING\_COMPOUND\_METABOLIC\_PROCESS | 172 | 42 | 4161 | 644 | levy,Stoml2,ND-B22,ACC,dnc,ste14,CG11089,ND-23,ND-51,ND-49,Gapdh1,CG33156,Pgm1,AMPdeam,nmdyn-D7,Prip,ND-SGDH,ND-75,CG3803,Shmt,UQCR-Q,Pgk,Nc73EF,ND-39,DhpD,mal,CG6767,Cyt-c-p,ND-B18,Pfk,Ac3,Eno,nmdyn-D6,Pglym78,ND-24,ND-30,CG11134,Pfas,CG7789,Ppcdc,COX5A,CG32549 | | 1.199e-03 | -6.73 | spindle pole | cellular component | GO:0000922 | 36 | 10 | 11940 | 1105 | CG7716,cnn,Grip128,toc,Klp10A,Rep,Grip84,Grip75,mad2,msps | | 1.204e-03 | -6.72 | MARTENS\_TRETINOIN\_RESPONSE\_DN | MSigDB lists | MARTENS\_TRETINOIN\_RESPONSE\_DN | 262 | 59 | 4161 | 644 | Phb2,Cyt-c-p,RpS16,CG4390,RpL30,RpL23,CG5292,CG12018,RpS24,Atg4b,Ts,CG17333,RpS4,RpL26,awd,RpS27,RpL24,Hsc70-3,Pex12,CG7488,RpL37a,TBCC,RpL11,CG30010,r-l,CG6665,CG7322,CCT5,CG14715,Tim8,CG3894,CG9393,RpL13A,CG6610,RpLP0,eIF4A,alphaTub84B,CG3226,CG2100,CG4407,Stoml2,CG7550,Idh3b,RpS27A,RpL9,RpS23,CG14516,CSN6,cl,mad2,eEF1beta,UQCR-Q,mEFTu1,Uch-L5,CCT6,pen-2,Cdk1,Bin1,CG7997 | | 1.218e-03 | -6.71 | Cp1 (Cysteine proteinase-1) | protein interactions | 36546 | 43 | 13 | 9204 | 1112 | Edc3,mEFTu1,Mitf,Tak1,Rel,Atpalpha,cathD,PGRP-LE,Mcad,ScsbetaA,SERCA,Droj2,Rab5 | | 1.218e-03 | -6.71 | up (upheld) | protein interactions | 32314 | 43 | 13 | 9204 | 1112 | Actn,Argk,Nc73EF,CG7920,Adh,Cyt-c-p,Mf,ATPsynO,Scp1,ATPsynCF6,ATPsynB,osk,ATPsynF | | 1.229e-03 | -6.70 | ATPase-coupled cation transmembrane transporter activity | molecular function | GO:0019829 | 62 | 14 | 12176 | 1119 | ATPsynG,ATPsynB,ATPsynbeta,ATPsynE,blw,Atpalpha,nrv1,Vha68-2,Vha13,ATPsyndelta,ATPsynD,Vha36-1,SERCA,ATPsynO | | 1.236e-03 | -6.70 | cytoskeleton organization | biological process | GO:0007010 | 580 | 75 | 12345 | 1129 | Abi,Arpc1,msps,CanB2,Arpc4,armi,SelR,Klp10A,CG13366,alphaTub84B,Prosbeta5,mts,Ubc6,grp,cnc,jvl,osk,Grip71,spd-2,Moe,dgt4,Sry-alpha,Rcd5,tws,cnn,Grip84,Girdin,Pax,parvin,wash,Actn,swa,BicC,CG7716,Grip128,Arpc2,Arf51F,tral,Hip1,Pp2A-29B,sw,mop,gammaTub37C,Kap3,Svil,heph,sau,foxo,larp,Lkb1,Grip75,bru1,up,Topors,del,wisp,Non1,Axs,CCT6,Orc6,trbd,alphaTub67C,Mtl,cib,pont,stai,cbs,Tm1,SRPK,CG3437,mael,Pka-R1,ovo,SkpA,capt | | 1.237e-03 | -6.70 | MODULE\_29 | MSigDB lists | MODULE\_29 | 23 | 10 | 4161 | 644 | RpS23,RpS28b,RpS16,RpS2,RpS13,RpS27,RpS24,RpS18,RpS4,RpS7 | | 1.253e-03 | -6.68 | antibiotic metabolic process | biological process | GO:0016999 | 25 | 8 | 12345 | 1129 | Jafrac1,AcCoAS,Irp-1B,Jafrac2,Adh,CG4390,Fdh,Pxt | | 1.253e-03 | -6.68 | GO\_ORGANELLE\_INNER\_MEMBRANE | MSigDB lists | GO\_ORGANELLE\_INNER\_MEMBRANE | 257 | 58 | 4161 | 644 | Ppox,mRpS21,Unr,Scsalpha1,ND-39,UQCR-Q,mRpL16,AIF,SdhA,CG7655,ND-75,mRpS10,CG3803,Coq6,ste24a,mRpL13,mino,ND-SGDH,Sfxn2,ND-49,mRpL18,CG10932,ND-51,ND-23,levy,Stoml2,ND-B22,COX5A,CG9773,Smox,yip2,Scox,Pmi,CG7488,mRpL17,Mitofilin,stet,mRpL10,ND-24,mRpS5,Thiolase,ND-30,ND-B18,Coq7,Mtpalpha,Tim8,Phb2,CG11110,colt,mRpS23,bor,Cyt-c-p,CG7718,CG8728,Cyp1,tko,Etf-QO,Ts | | 1.258e-03 | -6.68 | glycerolipid metabolic process | biological process | GO:0046486 | 136 | 24 | 12345 | 1129 | Seipin,ACC,dob,Agpat2,sws,CG2201,CG7718,Pi4KIIalpha,CG18135,Synj,PIG-P,CG5026,CG33977,CG6707,PIG-H,CG7789,PIG-C,Gk2,mino,CG9391,Gk1,PIG-L,Pi3K92E,Pld | | 1.258e-03 | -6.68 | regulation of kinase activity | biological process | GO:0043549 | 136 | 24 | 12345 | 1129 | SNF4Agamma,CG17746,RpLP1,Tao,Cdk7,raptor,CycE,fray,Abi,Mekk1,CG30291,ASPP,sws,CycC,Lkb1,alph,CG14408,Pi3K92E,Cbl,CycB,Tak1,CycJ,Parp16,Pdk1 | | 1.261e-03 | -6.68 | GO\_CELLULAR\_RESPONSE\_TO\_OXYGEN\_CONTAINING\_COMPOUND | MSigDB lists | GO\_CELLULAR\_RESPONSE\_TO\_OXYGEN\_CONTAINING\_COMPOUND | 188 | 45 | 4161 | 644 | Cyp1,Tspo,Tm1,Pdk,Pka-R1,Ptp61F,Vha13,Traf6,Pp2B-14D,Vha36-1,bys,Moe,CycB,mAChR-A,Ac3,Pdk1,p38a,PpD3,SREBP,CanA-14F,Wnt10,Grx1,rept,dnc,Jafrac1,ACC,Tip60,Karybeta3,mino,adp,HDAC6,Sod1,Prip,raptor,Calx,Shmt,cact,crq,rl,Cdk1,LpR2,AIF,Pka-R2,Got1,Rab10 | | 1.268e-03 | -6.67 | Nucleoside\_diP\_kinase | interpro domains | IPR001564 | 3 | 3 | 11383 | 1233 | nmdyn-D6,nmdyn-D7,awd | | 1.268e-03 | -6.67 | Ribosomal\_S11\_sf | interpro domains | IPR036967 | 3 | 3 | 11383 | 1233 | mRpS11,RpS14b,mRpL18 | | 1.275e-03 | -6.67 | regulation of transcription, DNA-templated | biological process | GO:0006355 | 1144 | 134 | 12345 | 1129 | Dd,Bx,CG10631,Traf6,Rbf,Cdk7,ix,CG6813,Myd88,CG11902,CG13344,Atu,insv,BigH1,cad,tkv,CG4730,asf1,TfIIFalpha,Taf7,l(3)mbt,YL-1,lin-52,pont,nej,smg,spen,rl,crm,Tip60,ph-d,MED27,CG10038,CG11710,Bin1,pgc,Hnf4,EcR,Rbfox1,CG1602,CG31875,cnc,Wbp2,nos,pygo,psq,Atf6,Su(H),Utx,HDAC6,S2P,l(3)73Ah,escl,Aos1,Sgf11,Stat92E,HmgD,Rbf2,Hr78,MED20,Spt20,E2f1,Sox102F,CG10959,lin,MED24,RYBP,Cul2,Bap170,Elba2,CG10654,Tlk,piwi,d4,CG2678,Topors,foxo,trr,dl,not,bun,Rel,Sirt7,CG2199,Tfb4,HP1e,Cf2,Chrac-14,CG3587,sbb,spn-E,CG4872,REPTOR,Med,Maf1,mael,ovo,Trf2,rept,e(y)1,lid,bcd,cact,Rga,scny,cyc,Ssdp,SREBP,hb,esc,MED4,Nf-YA,CG2129,r2d2,CG17202,Hat1,mute,Atf3,Mybbp1A,apt,lwr,egg,Mitf,wash,Nap1,Smox,Rcd5,hng3,EloC,vers,CycC,HP1c,NFAT,CG17328 | | 1.276e-03 | -6.66 | GEFmeso (Guanine nucleotide exchange factor in mesoderm) | protein interactions | 37134 | 24 | 9 | 9204 | 1112 | RpL23A,RpS14b,Fkbp39,spd-2,CG9911,RpS14a,CG6707,pgc,Tim8 | | 1.276e-03 | -6.66 | spn-F (spn-F) | protein interactions | 43700 | 24 | 9 | 9204 | 1112 | CG12107,TMEM216,CG11164,Ady43A,Rpn2,CG10654,CG31523,jvl,Utx | | 1.277e-03 | -6.66 | DER\_IFN\_ALPHA\_RESPONSE\_UP | MSigDB lists | DER\_IFN\_ALPHA\_RESPONSE\_UP | 16 | 8 | 4161 | 644 | Cyt-c-p,RpII140,Pp2B-14D,CanA-14F,Med,scramb1,SkpA,Pdxk | | 1.277e-03 | -6.66 | REACTOME\_G1\_PHASE | MSigDB lists | REACTOME\_G1\_PHASE | 16 | 8 | 4161 | 644 | RpS27A,Pp2A-29B,Rbf,Cdk7,mts,Cul1,RpL40,SkpA | | 1.277e-03 | -6.66 | GO\_PYRIMIDINE\_NUCLEOTIDE\_BIOSYNTHETIC\_PROCESS | MSigDB lists | GO\_PYRIMIDINE\_NUCLEOTIDE\_BIOSYNTHETIC\_PROCESS | 16 | 8 | 4161 | 644 | Ts,kri,r-l,r,nmdyn-D6,Shmt,nmdyn-D7,CG6767 | | 1.289e-03 | -6.65 | Rpt2 (Regulatory particle triple-A ATPase 2) | protein interactions | 42828 | 59 | 16 | 9204 | 1112 | Prosalpha4,wal,Prosbeta6,foxo,Rpn3,RpS5b,Prosbeta5,Prosbeta4,Prosbeta3,Prosalpha3,Rpn13,Pdk1,Prosalpha5,CCT6,Pomp,Rpn2 | | 1.298e-03 | -6.65 | protein localization | biological process | GO:0008104 | 693 | 87 | 12345 | 1129 | opm,Klp10A,Atg6,Pex10,armi,Tom70,CG8314,CG6766,sws,Arpc1,fray,Nxf3,slam,Vps16B,GAPsec,Mat89Ba,sip3,Nup358,Tim8,Pex13,Tango10,CG9393,CG8004,Rab10,Moe,Cse1,osk,Cenp-C,lt,BicC,Rab32,Actn,Vsp37A,Stam,thoc6,Pax,milt,Tsp86D,mbo,CHMP2B,Karybeta3,Rab5,Pex12,sw,Arts,CG14894,Tom40,Sec22,Pi4KIIalpha,Crag,Sec23,Nlg1,Arf51F,Pen,Clc,CG5734,eIF3m,Topors,Lkb1,Kap-alpha3,tho2,sau,CG8728,Sec15,Gga,AP-1-2beta,CG11110,Syx13,gus,CG5447,X11L,CHOp24,cact,CG4041,Arl5,Roe1,CG3077,Gdi,Rep,Oseg1,sbr,CG10344,mol,mael,SuUR,boi,Nup62 | | 1.306e-03 | -6.64 | nuclear outer membrane-endoplasmic reticulum membrane network | cellular component | GO:0042175 | 223 | 35 | 11940 | 1105 | EMC8-9,EMC3,PIG-H,PIG-C,Dd,Axs,alt,CG32640,Mctp,mol,CG33977,CG31523,Efr,CG5510,CG14671,Atf6,sip3,CG10425,CG5969,sws,Seipin,CG17270,CG4293,CG8009,SERCA,Desat1,bif,Sec22,CG8974,CG10470,SREBP,CG8311,CG32441,CG8112,ste24a | | 1.308e-03 | -6.64 | GO\_RIBOSOME | MSigDB lists | GO\_RIBOSOME | 137 | 35 | 4161 | 644 | RpL26,RpS27,RpL24,RpL11,mRpL17,mRpL10,RpL37a,mRpS5,RpL12,RpS18,mRpS23,RpS16,RpS2,RpL30,RpL23,RpS24,tko,RpL40,RpL38,RpS4,RpS7,RpS27A,RpS23,RpL9,RpS28b,mRpS21,mRpL16,RpL13,mRpS10,RpS30,mRpL13,RpS13,RpLP0,RpL13A,mRpL18 | | 1.314e-03 | -6.63 | endopeptidase complex | cellular component | GO:1905369 | 62 | 14 | 11940 | 1105 | Prosbeta3,Rpn12,Rpn2,Rpt2,Prosbeta4,Prosbeta5,Prosalpha3,Prosalpha5,Uch-L5,Prosbeta6,Prosalpha4,Rpn13,Rpn3,Pomp | | 1.314e-03 | -6.63 | proteasome complex | cellular component | GO:0000502 | 62 | 14 | 11940 | 1105 | Rpn13,Pomp,Rpn3,Prosalpha5,Uch-L5,Prosbeta6,Prosalpha4,Rpt2,Prosbeta4,Prosalpha3,Prosbeta5,Prosbeta3,Rpn12,Rpn2 | | 1.322e-03 | -6.63 | REACTOME\_SIGNALING\_BY\_THE\_B\_CELL\_RECEPTOR\_BCR | MSigDB lists | REACTOME\_SIGNALING\_BY\_THE\_B\_CELL\_RECEPTOR\_BCR | 56 | 18 | 4161 | 644 | SkpA,Prosbeta6,Pi3K92E,RpL40,Prosalpha4,Rpt2,Prosbeta3,Rpn2,Rpn12,Pdk1,cact,Prosbeta5,Cul1,Rpn3,Prosbeta4,Prosalpha3,Prosalpha5,RpS27A | | 1.333e-03 | -6.62 | reproduction | biological process | GO:0000003 | 1651 | 185 | 12345 | 1129 | Cul2,Mst89B,Tao,Iml1,dnc,Grip71,spd-2,Rab10,Moe,Desat1,jvl,Cenp-C,E2f1,swa,toc,Prosbeta4,Stat92E,milt,park,vret,ari-1,cnn,shrb,zpg,bam,eIF4E1,Klp10A,mus304,CanB2,psq,CG18135,msps,yl,Abi,Nup358,cnc,CG9471,grp,yem,Cdk1,GatA,Xpc,tor,nos,CycJ,wapl,BoYb,alphaTub67C,clt,Sec15,Syx13,gus,wisp,Debcl,Rbfox1,mtrm,enc,del,pgc,bif,twe,fs(1)N,EcR,capt,l(3)mbt,SRPK,Pka-R1,tkv,Dcr-1,Drep1,elg1,Jafrac1,Nprl2,rl,Nup62,faf,nej,cathD,c(2)M,heph,raptor,CSN6,ix,sw,Rab5,cup,Kap3,tko,eIF3m,Grip128,Arf51F,bru1,Bx,otu,Grip75,mre11,sau,squ,Rbf,larp,png,Mos,osk,shu,nmd,BicC,fs(1)Ya,apt,CG46338,twin,yps,ScsbetaA,Pax,Nap1,wash,CycB,lwr,Cbl,shep,egg,Grip84,CkIIalpha,r2d2,Ptpmeg2,armi,Pex10,Atg7,lqfR,PPYR1,csw,cort,slam,muskelin,CG31053,ea,exu,Orc1,Pex13,eIF4A,mad2,mts,mus81,spz,Pink1,lid,regucalcin,bcd,cact,Axs,ovo,plu,Tm1,mael,stil,SERCA,e(y)1,stai,Pp2B-14D,Pi3K92E,tef,disp,HP1e,Cf2,Pxt,Pex12,fs(1)M3,mop,bun,gammaTub37C,Med,spn-E,Pen,CG7716,CycE,Ggt-1,Topors,krimp,Lkb1,piwi,Pdk1,Imp,tho2,Cul5,Top1,PAPLA1 | | 1.341e-03 | -6.61 | JISON\_SICKLE\_CELL\_DISEASE\_DN | MSigDB lists | JISON\_SICKLE\_CELL\_DISEASE\_DN | 65 | 20 | 4161 | 644 | RpS28b,Iru,RpL24,Nap1,RpS27A,RpL9,RpL37a,RpL11,RpL13,eEF1beta,RpS13,RpLP0,dco,faf,RpS24,Rbf,eIF4A,RpL30,RpS7,Rpb8 | | 1.357e-03 | -6.60 | MORI\_MATURE\_B\_LYMPHOCYTE\_DN | MSigDB lists | MORI\_MATURE\_B\_LYMPHOCYTE\_DN | 27 | 11 | 4161 | 644 | CycB,Jafrac2,CG17333,Dhfr,CG11164,CG32473,Rpt2,Cyt-c-p,mino,Rpn12,CG6767 | | 1.357e-03 | -6.60 | WATANABE\_RECTAL\_CANCER\_RADIOTHERAPY\_RESPONSIVE\_DN | MSigDB lists | WATANABE\_RECTAL\_CANCER\_RADIOTHERAPY\_RESPONSIVE\_DN | 27 | 11 | 4161 | 644 | aay,ari-1,Sec23,Sptr,TBCC,mts,Cul1,CG1416,CCT6,crq,Prosbeta5 | | 1.366e-03 | -6.60 | REACTOME\_P53\_DEPENDENT\_G1\_DNA\_DAMAGE\_RESPONSE | MSigDB lists | REACTOME\_P53\_DEPENDENT\_G1\_DNA\_DAMAGE\_RESPONSE | 39 | 14 | 4161 | 644 | Rpn3,RpL40,Prosbeta6,Prosbeta5,RpS27A,Rpn2,Prosalpha5,CycE,Rpn12,Rpt2,Prosbeta4,Prosalpha4,Prosalpha3,Prosbeta3 | | 1.366e-03 | -6.60 | REACTOME\_P53\_INDEPENDENT\_G1\_S\_DNA\_DAMAGE\_CHECKPOINT | MSigDB lists | REACTOME\_P53\_INDEPENDENT\_G1\_S\_DNA\_DAMAGE\_CHECKPOINT | 39 | 14 | 4161 | 644 | Prosbeta6,Prosbeta5,Rpn3,RpL40,Rpt2,Prosalpha4,Prosbeta4,Prosbeta3,Prosalpha3,grp,RpS27A,Rpn2,Prosalpha5,Rpn12 | | 1.366e-03 | -6.60 | REACTOME\_VIF\_MEDIATED\_DEGRADATION\_OF\_APOBEC3G | MSigDB lists | REACTOME\_VIF\_MEDIATED\_DEGRADATION\_OF\_APOBEC3G | 39 | 14 | 4161 | 644 | Rpt2,Prosbeta4,Prosalpha4,Prosalpha3,Prosbeta3,RpS27A,Rpn2,Prosalpha5,Rpn12,Prosbeta6,Cul5,Prosbeta5,Rpn3,RpL40 | | 1.379e-03 | -6.59 | mitochondrial electron transport, NADH to ubiquinone | biological process | GO:0006120 | 37 | 10 | 12345 | 1129 | ND-51,ND-49,ND-24,ND-B22,ND-SGDH,ND-MNLL,ND-39,ND-B15,ND-B18,ND-23 | | 1.381e-03 | -6.58 | GO\_GLYCOSYL\_COMPOUND\_METABOLIC\_PROCESS | MSigDB lists | GO\_GLYCOSYL\_COMPOUND\_METABOLIC\_PROCESS | 168 | 41 | 4161 | 644 | Cyt-c-p,mal,CG6767,r,CG4511,Ts,CG7789,Ppcdc,CG11134,COX5A,CG32549,Eno,Pfk,ND-B18,r-l,ND-24,ND-30,nmdyn-D6,Pglym78,Pgm1,Gapdh1,CG33156,ND-49,ND-SGDH,nmdyn-D7,AMPdeam,levy,Stoml2,ND-B22,ste14,CG11089,ND-23,ND-51,ACC,Nc73EF,ND-39,ND-75,CG3803,kri,Pgk,UQCR-Q | | 1.383e-03 | -6.58 | Cnx99A (Calnexin 99A) | protein interactions | 44643 | 76 | 19 | 9204 | 1112 | Droj2,Rab5,cathD,Mcad,ScsbetaA,mEFTu1,wal,Cp1,GCS2beta,r,bor,CCT6,SERCA,CG1092,Kap-alpha3,CG10425,Pp2A-29B,CCT3,Pen | | 1.391e-03 | -6.58 | Mitotic G1-G1/S phases | REACTOME pathways | R-DME-453279 | 124 | 31 | 3923 | 574 | Orc6,Prosbeta3,Cdk7,DNApol-epsilon255,CycB,Pp2A-29B,Prosbeta6,Rpn12,Rpn3,Cul1,Orc1,Cdk1,RpS27A,Prosalpha5,Prosbeta4,Ts,E2f1,Dhfr,Rbf2,Rpn2,Prosbeta5,Prosalpha4,CycE,Chrac-14,twe,SkpA,Rbf,Prosalpha3,Rpt2,RpL40,lin-52 | | 1.400e-03 | -6.57 | REACTOME\_RECRUITMENT\_OF\_NUMA\_TO\_MITOTIC\_CENTROSOMES | MSigDB lists | REACTOME\_RECRUITMENT\_OF\_NUMA\_TO\_MITOTIC\_CENTROSOMES | 7 | 5 | 4161 | 644 | Cdk1,CycB,Grip128,gammaTub37C,Grip84 | | 1.400e-03 | -6.57 | PID\_SMAD2\_3PATHWAY | MSigDB lists | PID\_SMAD2\_3PATHWAY | 7 | 5 | 4161 | 644 | rl,Pen,lwr,Med,Smox | | 1.400e-03 | -6.57 | GO\_GAMMA\_TUBULIN\_COMPLEX | MSigDB lists | GO\_GAMMA\_TUBULIN\_COMPLEX | 7 | 5 | 4161 | 644 | msps,Grip128,Grip84,Grip75,gammaTub37C | | 1.400e-03 | -6.57 | regulation of triglyceride metabolic process | biological process | GO:0090207 | 15 | 6 | 12345 | 1129 | Dd,foxo,Lkb1,CG8009,Seipin,Nprl2 | | 1.415e-03 | -6.56 | multicellular organism reproduction | biological process | GO:0032504 | 1503 | 170 | 12345 | 1129 | psq,eIF4E1,Klp10A,mus304,bam,Abi,yl,CG18135,msps,yem,Cdk1,grp,CG9471,cnc,Nup358,BoYb,CycJ,wapl,GatA,nos,tor,Tao,Mst89B,Cul2,Cenp-C,Desat1,jvl,Rab10,Moe,Iml1,dnc,Grip71,Prosbeta4,Stat92E,toc,E2f1,swa,shrb,zpg,ari-1,cnn,vret,milt,park,cup,Kap3,Rab5,tko,sw,ix,heph,raptor,CSN6,Grip128,Arf51F,eIF3m,otu,Grip75,bru1,Rbf,squ,larp,sau,gus,Syx13,Sec15,clt,EcR,twe,fs(1)N,enc,del,bif,pgc,wisp,Rbfox1,mtrm,Pka-R1,tkv,SRPK,capt,l(3)mbt,nej,cathD,Nup62,faf,Jafrac1,elg1,Nprl2,rl,Dcr-1,Drep1,Pex10,armi,r2d2,Ptpmeg2,CkIIalpha,PPYR1,csw,Atg7,lqfR,ea,slam,muskelin,cort,mad2,mts,Orc1,Pex13,eIF4A,exu,Mos,osk,shu,CG46338,yps,twin,apt,nmd,BicC,Grip84,Cbl,egg,lwr,shep,Nap1,CycB,wash,Pax,ScsbetaA,gammaTub37C,Pxt,Pex12,bun,mop,fs(1)M3,Cf2,HP1e,tef,disp,CycE,Pen,Med,spn-E,Lkb1,piwi,Topors,krimp,Ggt-1,Top1,tho2,Cul5,PAPLA1,Imp,Pdk1,regucalcin,lid,spz,Axs,cact,bcd,stil,SERCA,Tm1,mael,ovo,Pp2B-14D,Pi3K92E,e(y)1,stai | | 1.424e-03 | -6.55 | fly chr5F4-5F4\|1-16 cM | chromosome location | fly chr5F4-5F4\|1-16 cM | 5 | 3 | 25010 | 1343 | PpV,swa,Marf | | 1.424e-03 | -6.55 | fly chr5F4-5F4\|1-16 | chromosome location | fly chr5F4-5F4\|1-16 | 5 | 3 | 25010 | 1343 | swa,Marf,PpV | | 1.424e-03 | -6.55 | fly chr7D6-7D6\|1-22 cM | chromosome location | fly chr7D6-7D6\|1-22 cM | 5 | 3 | 25010 | 1343 | Gclc,UbcE2H,CG2258 | | 1.424e-03 | -6.55 | fly chr7D6-7D6\|1-22 | chromosome location | fly chr7D6-7D6\|1-22 | 5 | 3 | 25010 | 1343 | CG2258,UbcE2H,Gclc | | 1.426e-03 | -6.55 | stem cell population maintenance | biological process | GO:0019827 | 99 | 19 | 12345 | 1129 | bru1,Stat92E,piwi,tkv,mael,scny,eIF4A,nos,Pi3K92E,zpg,foxo,shv,CSN6,lid,bam,stet,Med,Treh,CycE | | 1.426e-03 | -6.55 | maintenance of cell number | biological process | GO:0098727 | 99 | 19 | 12345 | 1129 | shv,foxo,zpg,Pi3K92E,nos,eIF4A,scny,mael,Stat92E,tkv,piwi,bru1,CycE,Treh,Med,stet,bam,lid,CSN6 | | 1.437e-03 | -6.55 | MORF\_PPP1CA | MSigDB lists | MORF\_PPP1CA | 103 | 28 | 4161 | 644 | scu,LysRS,CG1890,Sod1,Rpn12,Rpt2,ND-51,CG11089,CG11070,RhoGDI,PpV,Stoml2,SkpA,Prosbeta6,levy,AP-1-2beta,Pka-R1,CHOp24,Pp2A-29B,COX5A,Clc,LSm3,Prosbeta4,Chrac-14,ND-30,mEFTu1,ND-75,wal | | 1.437e-03 | -6.55 | nucleocytoplasmic carrier activity | molecular function | GO:0140142 | 15 | 6 | 12176 | 1119 | Arts,Karybeta3,Ranbp21,Pen,Kap-alpha3,Cse1 | | 1.450e-03 | -6.54 | GSE10239\_NAIVE\_VS\_DAY4.5\_EFF\_CD8\_TCELL\_UP | MSigDB lists | GSE10239\_NAIVE\_VS\_DAY4.5\_EFF\_CD8\_TCELL\_UP | 52 | 17 | 4161 | 644 | eEF1beta,sbr,RpL13,RpS18,RpL26,sws,Pex23,RpL11,Med,RpL38,tzn,RpS7,RpS30,RpLP0,UbcE2H,Dd,CG3021 | | 1.461e-03 | -6.53 | mst (misato) | protein interactions | 33119 | 20 | 8 | 9204 | 1112 | Smyd4-4,CCT5,STUB1,CCT3,Pfdn5,CCT6,PDCD-5,Pfdn2 | | 1.461e-03 | -6.53 | Pura (Puratrophin-1-like) | protein interactions | 2768948 | 20 | 8 | 9204 | 1112 | MED4,RpS14a,mad2,Cul2,galectin,CG13741,eIF4A,RpS14b | | 1.469e-03 | -6.52 | GSE21670\_IL6\_VS\_TGFB\_AND\_IL6\_TREATED\_STAT3\_KO\_CD4\_TCELL\_UP | MSigDB lists | GSE21670\_IL6\_VS\_TGFB\_AND\_IL6\_TREATED\_STAT3\_KO\_CD4\_TCELL\_UP | 70 | 21 | 4161 | 644 | Naa20A,Gdap2,Sap47,Arl6IP1,CG15141,Sld5,SkpA,shv,Rpn2,MED24,Hat1,mRpS5,lqfR,Pglym78,Caf1-105,Faf2,DNApol-epsilon255,Arpc1,stas,Tomosyn,Hsc70-3 | | 1.469e-03 | -6.52 | FLECHNER\_PBL\_KIDNEY\_TRANSPLANT\_OK\_VS\_DONOR\_UP | MSigDB lists | FLECHNER\_PBL\_KIDNEY\_TRANSPLANT\_OK\_VS\_DONOR\_UP | 70 | 21 | 4161 | 644 | ND-SGDH,RpLP0,CG6767,csw,Su(H),CG11070,CG10418,cathD,CHOp24,Pka-R1,r,CG9485,Grip84,Prosalpha5,RpS23,Mitofilin,Prosalpha3,RpL11,CG7728,lwr,eEF1beta | | 1.469e-03 | -6.52 | GO\_POSITIVE\_REGULATION\_OF\_WNT\_SIGNALING\_PATHWAY | MSigDB lists | GO\_POSITIVE\_REGULATION\_OF\_WNT\_SIGNALING\_PATHWAY | 70 | 21 | 4161 | 644 | Prosalpha5,RpS27A,Smox,Wnt10,Prosbeta4,Prosalpha3,Rpn3,Prosbeta5,dco,Rpn2,Rpn12,Prosalpha4,trbd,Rpt2,Dd,CkIIalpha,Prosbeta3,cyc,Su(H),RpL40,Prosbeta6 | | 1.476e-03 | -6.52 | GSE20727\_CTRL\_VS\_ROS\_INH\_AND\_DNFB\_ALLERGEN\_TREATED\_DC\_DN | MSigDB lists | GSE20727\_CTRL\_VS\_ROS\_INH\_AND\_DNFB\_ALLERGEN\_TREATED\_DC\_DN | 84 | 24 | 4161 | 644 | cathD,CG11070,ste14,CG4332,RhoGDI,Karybeta3,CysRS,Stoml2,Ufm1,sau,Aladin,ste24a,Sfxn2,Rpn3,lqfR,mRpL16,mad2,Pen,Cdk1,Prosbeta5,alpha-Man-Ia,Prosalpha5,gammaTub37C,Got1 | | 1.476e-03 | -6.52 | REACTOME\_MITOTIC\_G1\_G1\_S\_PHASES | MSigDB lists | REACTOME\_MITOTIC\_G1\_G1\_S\_PHASES | 84 | 24 | 4161 | 644 | Prosalpha5,Pp2A-29B,DNApol-epsilon255,RpS27A,CycE,Prosbeta4,Prosalpha3,Rpn3,Cul1,Cdk1,Prosbeta5,CycB,Rpn2,Rpn12,Rbf,Rpt2,Dhfr,mts,Prosbeta3,Cdk7,RpL40,SkpA,Ts,Prosbeta6 | | 1.485e-03 | -6.51 | CG15136 (uncharacterized protein) | protein interactions | 35033 | 12 | 6 | 9204 | 1112 | MAGE,Sec23,Arpc2,CG32267,Mtl,SdhA | | 1.485e-03 | -6.51 | CG7430 (uncharacterized protein) | protein interactions | 39988 | 12 | 6 | 9204 | 1112 | CG10638,Pgk,yip2,Uch,Atu,awd | | 1.488e-03 | -6.51 | GO\_POSITIVE\_REGULATION\_OF\_PROTEIN\_CATABOLIC\_PROCESS | MSigDB lists | GO\_POSITIVE\_REGULATION\_OF\_PROTEIN\_CATABOLIC\_PROCESS | 113 | 30 | 4161 | 644 | CkIIalpha,Prosbeta3,CG30291,Prosalpha4,Rpt2,ari-1,APC7,Rpn12,dco,Rpn2,SkpA,Prosbeta6,RpL40,Atg7,regucalcin,Prosalpha3,Prosbeta4,Prosalpha5,Sec22,RpS27A,STUB1,Prosbeta5,CycB,Gclc,Cdk1,mad2,Edem1,Cul1,Moe,Rpn3 | | 1.495e-03 | -6.51 | GO\_POSITIVE\_REGULATION\_OF\_CANONICAL\_WNT\_SIGNALING\_PATHWAY | MSigDB lists | GO\_POSITIVE\_REGULATION\_OF\_CANONICAL\_WNT\_SIGNALING\_PATHWAY | 61 | 19 | 4161 | 644 | Rpn12,Rpn2,dco,Prosbeta3,Rpt2,Dd,Prosalpha4,RpL40,Su(H),cyc,Prosbeta6,Smox,RpS27A,Prosalpha5,Prosalpha3,Wnt10,Prosbeta4,Rpn3,Prosbeta5 | | 1.504e-03 | -6.50 | MODULE\_303 | MSigDB lists | MODULE\_303 | 13 | 7 | 4161 | 644 | Echs1,Cdk5,CG33156,Cdk1,p38a,CG32549,grp | | 1.514e-03 | -6.49 | NADH dehydrogenase (ubiquinone) 1 beta subcomplex | KEGG pathways | M00147 | 13 | 7 | 3198 | 496 | NP15.6,ND-B15,ND-MNLL,ND-B22,ND-SGDH,ND-B18,CG40472 | | 1.514e-03 | -6.49 | NADH dehydrogenase (ubiquinone) 1 beta subcomplex | KEGG pathways | dme\_M00147 | 13 | 7 | 3198 | 496 | CG40472,ND-B18,ND-SGDH,ND-B22,ND-MNLL,ND-B15,NP15.6 | | 1.516e-03 | -6.49 | BERENJENO\_TRANSFORMED\_BY\_RHOA\_UP | MSigDB lists | BERENJENO\_TRANSFORMED\_BY\_RHOA\_UP | 232 | 53 | 4161 | 644 | Nmt,CG1774,Fkbp59,mad2,Pen,Shmt,mRpL16,Pgk,Uch-L5,CCT6,cact,Caf1-105,Cdk1,Tom40,mRpS10,Prip,grp,Nurf-38,Cse1,Gapdh1,mRpL18,CG11899,CG3226,Prosbeta3,escl,l(1)G0004,rept,Gk2,Karybeta3,Tsp74F,gammaTub37C,LSm3,Pglym78,bys,Moe,Hat1,CCT3,Stip1,Coq7,Prosbeta5,CycB,esc,CG14715,Cyt-c-p,CG4390,Dhfr,Aos1,CG10418,scramb1,Cbp80,NHP2,RfC38,AGBE | | 1.541e-03 | -6.48 | CG2046 (uncharacterized protein) | protein interactions | 40719 | 44 | 13 | 9204 | 1112 | Rpn13,Rpn2,Prosalpha5,Pomp,Prosalpha4,CG2199,Nurf-38,Prosbeta5,Prosbeta3,Prosbeta6,Prosalpha3,Rpt2,Prosbeta4 | | 1.541e-03 | -6.48 | Prosalpha2 (Proteasome alpha2 subunit) | protein interactions | 41531 | 44 | 13 | 9204 | 1112 | AnxB10,Rpn13,Rpn2,Pomp,Prosalpha5,Debcl,Prosalpha4,Prosbeta5,Prosalpha3,Prosbeta3,Prosbeta6,Prosbeta4,Rpt2 | | 1.541e-03 | -6.48 | GCS2beta (Glucosidase 2 beta subunit) | protein interactions | 35056 | 44 | 13 | 9204 | 1112 | Tspo,Pp2A-29B,wal,Tak1,Rel,PGRP-LE,Mcad,Pdk1,Kap-alpha3,CG1416,SERCA,CCT6,Droj2 | | 1.543e-03 | -6.47 | ubiquitin-like protein-specific protease activity | molecular function | GO:0019783 | 50 | 12 | 12176 | 1119 | Uch-L5,faf,scny,Usp39,velo,trbd,CG16979,otu,Usp10,Uch,not,Usp16-45 | | 1.543e-03 | -6.47 | histone acetyltransferase activity | molecular function | GO:0004402 | 50 | 12 | 12176 | 1119 | Chrac-14,nej,lid,YL-1,rept,pont,Sgf29,Rcd5,Tip60,Hat1,e(y)1,d4 | | 1.549e-03 | -6.47 | ribosome biogenesis | biological process | GO:0042254 | 211 | 33 | 12345 | 1129 | RpS27,RpS15,mRpS11,RpL23A,RpL34a,Mat89Ba,CG12288,RpS28b,RpL11,RpS14b,CG7728,NHP2,RpS14a,mbo,CG7488,CG9143,mtTFB1,bys,RpL7A,RpL34b,CG12301,CG8611,RpL10,RpL12,CG8204,mRpL10,RpS5b,RpL26,RpS7,RpLP0,Rrp40,RpS16,RpL24 | | 1.565e-03 | -6.46 | response to starvation | biological process | GO:0042594 | 146 | 25 | 12345 | 1129 | Desat1,lt,REPTOR,Iml1,Atg7,Debcl,Mtpalpha,whd,Rel,SNF4Agamma,raptor,Atg6,Seipin,Rbf,Pi3K92E,foxo,Jafrac1,Nprl2,Imp,rl,Dcr-1,Mitf,trsn,p38a,cyc | | 1.565e-03 | -6.46 | Argk (Arginine kinase) | protein interactions | 39041 | 60 | 16 | 9204 | 1112 | Actn,Pgk,Mf,up,RpS27A,CG7920,Adh,ATPsynCF6,Fer1HCH,Cyt-c-p,Fkbp39,Pglym78,Eno,Scp1,Rtnl1,GlyP | | 1.572e-03 | -6.46 | STARK\_PREFRONTAL\_CORTEX\_22Q11\_DELETION\_DN | MSigDB lists | STARK\_PREFRONTAL\_CORTEX\_22Q11\_DELETION\_DN | 227 | 52 | 4161 | 644 | Prosalpha5,Trxr-1,CG8507,Sec22,shrb,CSN6,ND-39,Idh,cl,Pen,CG5880,CG16979,ND-75,eIF4E1,CG3860,EcR,Rpt2,CG7115,Prosbeta3,CG3226,Fdh,CG5028,ND-23,Jafrac1,CG17691,Sld5,tzn,CG42542,PDCD-5,RpL26,gammaTub37C,mRpL17,ND-30,Ntan1,ND-24,Stip1,CCT3,CCT5,Cdk5,CG14715,Tim8,colt,mRpS23,Pfdn5,Cyt-c-p,Vha36-1,Usp39,mbo,PrBP,tko,CG1416,Nprl2 | | 1.573e-03 | -6.46 | cathD (cathD) | protein interactions | 45268 | 39 | 12 | 9204 | 1112 | CCT6,SERCA,Rab5,Droj2,Trxr-1,MED4,ScsbetaA,Pdk1,Mcad,wal,Cp1,CCT3 | | 1.588e-03 | -6.45 | REACTOME\_IMMUNE\_SYSTEM | MSigDB lists | REACTOME\_IMMUNE\_SYSTEM | 254 | 57 | 4161 | 644 | GCS2beta,Kap3,ari-1,APC7,Traf6,Aladin,Ubc4,Ptp61F,Pi3K92E,CG40045,Prosbeta6,mbo,RpL40,Atg7,gus,Aos1,cathD,Prosalpha3,Prosbeta4,Hsc70-3,Sec23,Pp2A-29B,Clc,CG2924,Prosbeta5,alph,Pdk1,p38a,Kap-alpha3,Rpn3,Cul1,mts,Prosbeta3,Cul2,Prosalpha4,UbcE2H,eIF4A,Rpt2,CG3860,eIF4E1,Rpn12,CG8188,Rpn2,AP-1-2beta,SkpA,CG11070,Uba5,Prosalpha5,RpS27A,STUB1,Cul5,rl,cact,Klp10A,Cdk1,Pen,ERp60 | | 1.614e-03 | -6.43 | MODULE\_25 | MSigDB lists | MODULE\_25 | 10 | 6 | 4161 | 644 | ND-SGDH,ND-51,ND-30,ND-23,ND-24,ND-49 | | 1.614e-03 | -6.43 | GO\_PROTEIN\_K63\_LINKED\_UBIQUITINATION | MSigDB lists | GO\_PROTEIN\_K63\_LINKED\_UBIQUITINATION | 10 | 6 | 4161 | 644 | STUB1,CG40045,Iru,Traf6,CG8188,Uev1A | | 1.618e-03 | -6.43 | positive regulation of autophagy | biological process | GO:0010508 | 44 | 11 | 12345 | 1129 | Nprl2,CycC,Debcl,Mitf,Atg7,foxo,Traf6,Synj,PEK,Utx,AP-1-2beta | | 1.618e-03 | -6.43 | cellular amino acid biosynthetic process | biological process | GO:0008652 | 44 | 11 | 12345 | 1129 | Dhfr,Got1,Shmt,SelD,aay,CG11899,AsnS,CG11134,P5cr,r,Adi1 | | 1.622e-03 | -6.42 | BIOCARTA\_MPR\_PATHWAY | MSigDB lists | BIOCARTA\_MPR\_PATHWAY | 20 | 9 | 4161 | 644 | Arpc1,Arpc2,Pka-R2,capt,Arpc4,Cdk1,rl,CycB,Pka-R1 | | 1.624e-03 | -6.42 | pigment biosynthetic process | biological process | GO:0046148 | 64 | 14 | 12345 | 1129 | Ppox,cl,mal,PAPLA1,lt,fdl,Rab32,DhpD,Shmt,pr,EMC3,CG13551,Hexo1,CG3803 | | 1.631e-03 | -6.42 | circadian behavior | biological process | GO:0048512 | 71 | 15 | 12345 | 1129 | Pka-R2,Ac3,Bx,lid,tyf,cyc,dco,CkIIalpha,cu,nej,Ork1,gw,foxo,Irk1,dnc | | 1.651e-03 | -6.41 | GSE27786\_LSK\_VS\_BCELL\_UP | MSigDB lists | GSE27786\_LSK\_VS\_BCELL\_UP | 66 | 20 | 4161 | 644 | Coq6,Pex13,dco,Mctp,HDAC6,CG1532,tun,CG17698,rept,Jafrac1,Vha13,CG15141,r,CG11807,Shmt,CG30010,p38a,SerRS-m,Pdxk,pont | | 1.660e-03 | -6.40 | GSE43955\_1H\_VS\_10H\_ACT\_CD4\_TCELL\_WITH\_TGFB\_IL6\_UP | MSigDB lists | GSE43955\_1H\_VS\_10H\_ACT\_CD4\_TCELL\_WITH\_TGFB\_IL6\_UP | 57 | 18 | 4161 | 644 | CG14715,Aldh,ND-B18,ND-30,Gyg,CG7789,Unr,Clc,Jheh2,CG17593,Snx6,Plap,Tsp66E,adp,ND-SGDH,bor,Phb2,alc | | 1.666e-03 | -6.40 | ribonucleoside monophosphate metabolic process | biological process | GO:0009161 | 26 | 8 | 12345 | 1129 | Pfas,l(2)k01209,CG11089,AMPdeam,Ady43A,cN-IIIB,r-l,CG6767 | | 1.666e-03 | -6.40 | reactive oxygen species metabolic process | biological process | GO:0072593 | 26 | 8 | 12345 | 1129 | ND-75,Pxt,p38a,Sod3,ND-30,Jafrac2,Jafrac1,Sod1 | | 1.678e-03 | -6.39 | response to endogenous stimulus | biological process | GO:0009719 | 212 | 33 | 12345 | 1129 | Hr78,Trf2,Shmt,tkv,MED24,CG7328,Nurf-38,Fkbp39,rl,lwr,rept,Pi3K92E,G9a,Idh3b,MED27,Smox,Rab10,PyK,Hnf4,EcR,Nup358,mAChR-A,rig,Vps16B,Pdk1,foxo,trr,raptor,Utx,Chd64,bun,Med,csw | | 1.700e-03 | -6.38 | GO\_OXIDATIVE\_PHOSPHORYLATION | MSigDB lists | GO\_OXIDATIVE\_PHOSPHORYLATION | 44 | 15 | 4161 | 644 | ND-39,ND-49,Cyt-c-p,ND-SGDH,COX5A,CG3803,ND-75,ND-B22,ND-B18,levy,ND-51,ND-30,ND-23,ND-24,UQCR-Q | | 1.704e-03 | -6.37 | response to antibiotic | biological process | GO:0046677 | 93 | 18 | 12345 | 1129 | rl,Pdk1,apt,p38a,Pka-R1,Shmt,AcCoAS,Aldh,dnc,Arf51F,CG17734,Adh,spz,aay,Pka-R2,Agpat2,whd,Tao | | 1.711e-03 | -6.37 | PHDP (Putative homeodomain protein) | protein interactions | 37788 | 66 | 17 | 9204 | 1112 | grp,CG13373,awd,squ,Cdk5,nej,PyK,Moe,CG1416,mad2,CG33156,AnxB10,CycJ,CG13742,png,CG11248,CkIIalpha | | 1.717e-03 | -6.37 | GCP\_C | interpro domains | IPR040457 | 6 | 4 | 11383 | 1233 | Grip84,Grip128,Grip75,CG7716 | | 1.717e-03 | -6.37 | Znf\_UBP | interpro domains | IPR001607 | 6 | 4 | 11383 | 1233 | HDAC6,Usp39,not,Usp16-45 | | 1.720e-03 | -6.37 | oocyte dorsal/ventral axis specification | biological process | GO:0007310 | 38 | 10 | 12345 | 1129 | enc,jvl,milt,gammaTub37C,ea,mael,krimp,spz,CkIIalpha,cnc | | 1.723e-03 | -6.36 | regulation of response to stimulus | biological process | GO:0048583 | 958 | 114 | 12345 | 1129 | spen,rl,Nprl2,rept,Mtl,faf,nej,pont,Pi3K92E,PGRP-LE,SkpA,cad,RhoGAP5A,insv,tkv,wisp,crq,tub,cact,chrb,Ptp61F,EMRE,Archease,spz,dx,snk,PDCD-5,pn,trbd,Pdk1,Imp,Traf6,Pura,foxo,Rbf,dl,RhoGAP54D,Ptpmeg,slif,alph,Cka,Dd,Lkb1,O-fut1,piwi,Crag,CG10188,Ptip,CycE,CG4853,disp,raptor,Pp2A-29B,Rel,Herc4,mop,GlyP,CG9925,Myd88,park,lwr,Cbl,Tsp86D,E2f1,p38a,Hr78,dco,apt,PEK,Aos1,Cul1,EndoGI,Stat92E,NFAT,Iml1,dnc,Ect4,PpV,stet,ben,Bap170,Mekk1,TyrRS,G9a,ATPsynD,Cul2,tws,Tao,CG17746,Mos,gd,RYBP,Ubc6,tor,Usp10,GILT1,ago,Tak1,shv,Jafrac2,mts,Ssdp,scny,ea,Atg7,Hat1,STUB1,Sp7,CkIIalpha,Seipin,Atg6,Ptp10D,armi,Su(H) | | 1.727e-03 | -6.36 | bounding membrane of organelle | cellular component | GO:0098588 | 329 | 47 | 11940 | 1105 | alpha-Man-Ia,AP-1-2beta,Topors,Pex10,Hip1,Debcl,Tom70,CG5510,Vha36-1,Gmap,Ugalt,fwe,Afti,Efr,Axs,Sec23,Vsp37A,Vamp7,Pex12,Iml1,CHMP2B,CG11077,Vha13,Marf,Pex23,Vha68-2,Tom40,Pex13,ABCD,Pi4KIIalpha,stet,CG9132,Fur2,CG9393,Tspo,Sec22,mino,shrb,CG5447,CG4293,Clc,yin,CG8004,SERCA,CG9773,CG6707,bif | | 1.738e-03 | -6.35 | Papain-like\_cys\_pep\_sf | interpro domains | IPR038765 | 67 | 16 | 11383 | 1233 | Usp16-45,Xpc,velo,CG16979,sol,Cp1,Usp39,Uch-L5,not,Usp10,faf,Uch,otu,Atg4b,scny,26-29-p | | 1.738e-03 | -6.35 | adenyl nucleotide binding | molecular function | GO:0030554 | 659 | 83 | 12176 | 1119 | obe,Klp10A,tor,Vha68-2,TrpRS-m,Mekk1,CCT6,Hsc70-3,CCT5,ATPsynbeta,Tao,Cdk5,CkIIalpha,CG31729,Cdk1,CG7806,nmd,ScsbetaA,AcCoAS,CG4562,ACC,tkv,SERCA,CG4041,spn-E,Rpt2,CG6767,Pdk1,Tlk,CG31689,CG8611,CG9143,Atpalpha,Cdk7,awd,Tak1,blw,Pgk,GluRS-m,r,l(2)k01209,bor,rl,Orc1,grp,p38a,pont,cana,rept,SNF4Agamma,Droj2,eIF4A,CG5789,SerRS-m,Roe1,Nadsyn,RecQ5,CG8726,CysRS,LysRS,l(1)G0196,Mos,PEK,nmdyn-D7,ATP8A,png,CG9981,SRPK,ABCD,CG4880,SelD,Lkb1,CG14183,CCT3,E23,kl-2,fray,dco,Pfrx,Pfk,TyrRS,Pink1,CG17698 | | 1.739e-03 | -6.35 | GO\_ENVELOPE | MSigDB lists | GO\_ENVELOPE | 456 | 93 | 4161 | 644 | ND-39,Scsalpha1,Unr,shrb,Ppox,Tom40,CG3803,SdhA,mad2,Cisd2,mRpL18,CG3226,ND-49,Gapdh1,ND-SGDH,mino,CG9393,Sod1,ste24a,Prip,sau,Stoml2,ND-B22,Karybeta3,ND-23,Arts,ND-51,CG10932,stet,SREBP,Pmi,yip2,CG9773,sws,CG8004,Mtpalpha,ND-B18,ND-30,shop,Thiolase,sbr,mRpS5,Marf,CG8974,Kap-alpha3,Dd,Non1,mRpS23,Aladin,colt,Tspo,Nc73EF,mRpS21,Rab32,mRpS10,ND-75,CG7655,AIF,Ect4,Calx,Pen,mRpL16,UQCR-Q,Sfxn2,Cse1,mRpL13,Coq6,levy,Gk2,mRpL10,Mitofilin,mRpL17,CG7488,Scox,Smox,COX5A,Tim8,Coq7,ND-24,lwr,CG8728,CG7718,Atg6,bor,Cyt-c-p,CG11110,Phb2,Ts,SelD,Etf-QO,mbo,tko,Cyp1 | | 1.752e-03 | -6.35 | regulation of macroautophagy | biological process | GO:0016241 | 32 | 9 | 12345 | 1129 | scny,rl,Nprl2,Fkbp39,Debcl,Atg7,Mitf,foxo,Synj | | 1.756e-03 | -6.34 | zf-UBP | pfam domains | PF02148 | 6 | 4 | 10480 | 1142 | Usp16-45,not,HDAC6,Usp39 | | 1.756e-03 | -6.34 | GCP\_C\_terminal | pfam domains | PF04130 | 6 | 4 | 10480 | 1142 | Grip84,CG7716,Grip75,Grip128 | | 1.759e-03 | -6.34 | CG4704 (uncharacterized protein) | protein interactions | 42706 | 3 | 3 | 9204 | 1112 | RpS24,RpL24,MtnA | | 1.759e-03 | -6.34 | Uch-L5 (Ubiquitin carboxy-terminal hydrolase L5) | protein interactions | 39102 | 3 | 3 | 9204 | 1112 | Rpn13,RpS27A,Rpn2 | | 1.784e-03 | -6.33 | heat shock protein binding | molecular function | GO:0031072 | 38 | 10 | 12176 | 1119 | Stip1,spag,Droj2,STUB1,Fkbp59,CG32640,lwr,Hsc70-3,CG1416,DnaJ-H | | 1.784e-03 | -6.33 | mRNA 3'-UTR binding | molecular function | GO:0003730 | 38 | 10 | 12176 | 1119 | bcd,heph,bam,apt,Unr,smg,bru1,Rbfox1,shep,Imp | | 1.790e-03 | -6.33 | GO\_POSITIVE\_REGULATION\_OF\_LIGASE\_ACTIVITY | MSigDB lists | GO\_POSITIVE\_REGULATION\_OF\_LIGASE\_ACTIVITY | 71 | 21 | 4161 | 644 | Prosalpha3,Prosbeta4,Prosalpha5,RpS27A,STUB1,Prosbeta5,CycB,Cdk1,mad2,Rpn3,Cul1,Prosbeta3,Prosalpha4,Rpt2,APC7,Rpn12,CG8188,Rpn2,SkpA,Prosbeta6,RpL40 | | 1.825e-03 | -6.31 | Gfat2 (Glutamine:fructose-6-phosphate aminotransferase 2) | protein interactions | 43373 | 50 | 14 | 9204 | 1112 | Vha68-2,CCT3,r,Cp1,Pp2A-29B,mEFTu1,wal,Pdk1,Mcad,ScsbetaA,cathD,Rab5,CCT6,SERCA | | 1.831e-03 | -6.30 | GO\_NADH\_DEHYDROGENASE\_COMPLEX | MSigDB lists | GO\_NADH\_DEHYDROGENASE\_COMPLEX | 24 | 10 | 4161 | 644 | ND-SGDH,ND-49,ND-39,ND-51,ND-30,ND-23,ND-24,ND-B22,ND-B18,ND-75 | | 1.834e-03 | -6.30 | perinuclear region of cytoplasm | cellular component | GO:0048471 | 71 | 15 | 11940 | 1105 | cbs,Cul2,Abi,alphaTub67C,Ptp61F,Gga,wash,krimp,tapas,ben,Gclc,Rep,alphaTub84B,mael,gus | | 1.837e-03 | -6.30 | positive regulation of cellular component organization | biological process | GO:0051130 | 290 | 42 | 12345 | 1129 | Mtl,egg,Cbl,Pi3K92E,CG11486,park,Mitf,wash,Stat92E,tkv,dco,Marf,pgc,Archease,HP1c,Moe,Tao,Pink1,G9a,spz,APC7,shv,esc,trr,Pld,Imp,scny,O-fut1,piwi,bru1,Synj,Abi,Arpc2,Sec23,Hip1,Arpc1,Med,msps,Arpc4,Seipin,HP1e,raptor | | 1.841e-03 | -6.30 | embryonic pattern specification | biological process | GO:0009880 | 230 | 35 | 12345 | 1129 | spn-E,csw,msps,trk,Ptip,Grip128,disp,psq,fs(1)M3,armi,nos,tor,exu,pygo,hb,cnc,Ssdp,Grip75,Lkb1,bcd,Grip71,osk,Moe,twe,fs(1)N,gus,Dcr-1,rl,lin,swa,apt,mael,cad,Tm1,Stat92E | | 1.849e-03 | -6.29 | response to ethanol | biological process | GO:0045471 | 79 | 16 | 12345 | 1129 | Arf51F,CG17734,AcCoAS,dnc,Aldh,aay,Pka-R2,Agpat2,Tao,whd,Adh,rl,Pdk1,Pka-R1,Shmt,apt | | 1.852e-03 | -6.29 | REACTOME\_REGULATION\_OF\_MITOTIC\_CELL\_CYCLE | MSigDB lists | REACTOME\_REGULATION\_OF\_MITOTIC\_CELL\_CYCLE | 62 | 19 | 4161 | 644 | Prosalpha5,RpS27A,Prosalpha3,Prosbeta4,mad2,Cul1,Rpn3,Prosbeta5,CycB,Cdk1,APC7,Rpn12,Rpn2,Prosbeta3,Prosalpha4,Rpt2,RpL40,SkpA,Prosbeta6 | | 1.852e-03 | -6.29 | antioxidant activity | molecular function | GO:0016209 | 51 | 12 | 12176 | 1119 | Sod3,Jafrac2,Grx1,glob1,Pxt,GstT1,GstT2,Jafrac1,Sod1,PHGPx,SelT,Trxr-1 | | 1.862e-03 | -6.29 | GO\_RESPONSE\_TO\_OXYGEN\_CONTAINING\_COMPOUND | MSigDB lists | GO\_RESPONSE\_TO\_OXYGEN\_CONTAINING\_COMPOUND | 316 | 68 | 4161 | 644 | GlyP,crq,LpR2,AIF,Ect4,Calx,Trxr-1,Grx1,Gk2,rept,ACC,mAChR-A,CG3376,Edem1,CanA-14F,Gabat,Cka,r,Ts,Cyp1,Pdk,Vha36-1,Atg6,Cyt-c-p,Traf6,Pp2B-14D,kri,rl,cact,Cdk1,Shmt,Pka-R2,Got1,Rab10,CG5412,Karybeta3,dnc,Jafrac1,Fdh,Tip60,raptor,mino,HDAC6,Sod1,adp,Prip,CycB,Cdk5,Ac3,Pdk1,Mtpalpha,p38a,bys,Moe,Kap-alpha3,SREBP,Wnt10,CycE,PpD3,Pka-R1,Ptp61F,Vha13,Gdap2,Tspo,Atpalpha,Tm1,Dhfr,aay | | 1.867e-03 | -6.28 | CLEC7A (Dectin-1) signaling | REACTOME pathways | R-DME-5607764 | 80 | 22 | 3923 | 574 | Rpn3,Cul1,Rpn12,dl,Prosbeta6,Rel,NFAT,Prosbeta3,CanB2,Rpt2,RpL40,Pp2B-14D,Prosalpha3,cact,SkpA,CanA-14F,Prosalpha4,Prosbeta5,Rpn2,Prosbeta4,Prosalpha5,RpS27A | | 1.870e-03 | -6.28 | p53 (p53) | protein interactions | 2768677 | 202 | 39 | 9204 | 1112 | CG42360,fdl,bcd,Svil,Utx,Tm1,mRpL16,Non1,cort,Rbf,CG6923,nbs,mtgo,Idh,CG8728,Nf-YA,Ady43A,Imp,lwr,CG11486,ago,asf1,Fdh,Hsp26,ScsbetaA,CG10932,Pi3K92E,sip3,Topors,COX5A,CG1602,Cyp6a22,Scox,gus,Ubc6,CycE,Dcr-1,Tom40,RpS27A | | 1.889e-03 | -6.27 | ZF\_UBP | prosite domains | PS50271 | 6 | 4 | 6243 | 694 | not,HDAC6,Usp39,Usp16-45 | | 1.899e-03 | -6.27 | positive regulation of macromolecule biosynthetic process | biological process | GO:0010557 | 552 | 71 | 12345 | 1129 | Atu,REPTOR,CG2199,Cf2,Rel,Cdk7,Myd88,not,foxo,dl,trr,d4,otu,Bx,bcd,Rbfox1,CG1602,EcR,CG11710,lid,Tip60,pn,SuUR,e(y)1,rept,Pi3K92E,nej,pont,ovo,TfIIFalpha,Taf7,Trf2,cad,BigH1,CG8963,Mybbp1A,CG17202,S2P,Utx,Atf6,Su(H),Nf-YA,CG2129,pygo,Exd2,SREBP,hb,esc,MED4,cyc,Wbp2,cnc,CycC,NFAT,osk,CG17328,Bap170,EloC,vers,Rcd5,RYBP,hng3,Mitf,Sox102F,lin,MED20,E2f1,apt,tyf,Sgf11,Stat92E | | 1.910e-03 | -6.26 | ribonucleotide binding | molecular function | GO:0032553 | 822 | 100 | 12176 | 1119 | Arf51F,blw,Tak1,Cdk7,awd,Atpalpha,CG9143,GluRS-m,Pgk,CG31689,Tlk,CG8611,Pdk1,Rpt2,CG6767,CG4041,SERCA,tkv,spn-E,sgll,AcCoAS,CG4562,ScsbetaA,alphaTub67C,ACC,mEFTu1,Cdk1,CG31729,CG7806,nmd,CCT5,CCT6,Hsc70-3,Mekk1,mEFTu2,TrpRS-m,CkIIalpha,CG7728,ATPsynbeta,Tao,Cdk5,Rab10,Arl5,obe,Non1,Vha68-2,Klp10A,tor,Mtl,Pfk,Pfrx,dco,TyrRS,CG17698,Pink1,CG14183,Lkb1,SelD,CG4880,fray,kl-2,E23,CCT3,Marf,ABCD,SRPK,CG9981,ATP8A,ND-51,Rab5,CG5116,png,l(1)G0196,LysRS,CG7488,nmdyn-D7,PEK,Mos,CG5789,SerRS-m,eIF4A,gammaTub37C,CysRS,RecQ5,CG8726,Nadsyn,cana,p38a,rept,pont,alphaTub84B,grp,Rab32,Droj2,SNF4Agamma,bor,l(2)k01209,r,Orc1,rl | | 1.928e-03 | -6.25 | DER\_IFN\_BETA\_RESPONSE\_UP | MSigDB lists | DER\_IFN\_BETA\_RESPONSE\_UP | 28 | 11 | 4161 | 644 | Pdxk,SkpA,Med,Thiolase,scramb1,CanA-14F,csw,Cyt-c-p,RpII140,PpD3,Pp2B-14D | | 1.928e-03 | -6.25 | KEGG\_VALINE\_LEUCINE\_AND\_ISOLEUCINE\_DEGRADATION | MSigDB lists | KEGG\_VALINE\_LEUCINE\_AND\_ISOLEUCINE\_DEGRADATION | 28 | 11 | 4161 | 644 | CG8199,Aldh,Mtpalpha,CG17691,Echs1,CG10932,Thiolase,Mcad,scu,Gabat,yip2 | | 1.937e-03 | -6.25 | pteridine-containing compound metabolic process | biological process | GO:0042558 | 21 | 7 | 12345 | 1129 | DhpD,pn,Shmt,pr,Dhfr,Pcd,Sptr | | 1.940e-03 | -6.24 | GO\_CELLULAR\_RESPONSE\_TO\_BIOTIC\_STIMULUS | MSigDB lists | GO\_CELLULAR\_RESPONSE\_TO\_BIOTIC\_STIMULUS | 32 | 12 | 4161 | 644 | Traf6,Hsc70-3,bys,dnc,Tspo,rl,crq,cact,p38a,LpR2,CG17593,mus81 | | 1.943e-03 | -6.24 | cellular response to biotic stimulus | biological process | GO:0071216 | 7 | 4 | 12345 | 1129 | EcR,Stat92E,Vps16B,Tak1 | | 1.943e-03 | -6.24 | cellular response to molecule of bacterial origin | biological process | GO:0071219 | 7 | 4 | 12345 | 1129 | Stat92E,EcR,Tak1,Vps16B | | 1.943e-03 | -6.24 | pyrimidine nucleoside triphosphate metabolic process | biological process | GO:0009147 | 7 | 4 | 12345 | 1129 | r,nmdyn-D6,nmdyn-D7,awd | | 1.943e-03 | -6.24 | positive regulation of triglyceride metabolic process | biological process | GO:0090208 | 7 | 4 | 12345 | 1129 | foxo,Dd,Nprl2,CG8009 | | 1.958e-03 | -6.24 | orb (oo18 RNA-binding protein) | protein interactions | 42752 | 30 | 10 | 9204 | 1112 | CkIIalpha,Atg7,osk,twin,wisp,RpL31,cup,yps,bru1,BicC | | 1.958e-03 | -6.24 | ncd (non-claret disjunctional) | protein interactions | 43517 | 30 | 10 | 9204 | 1112 | Pdk1,PGRP-LE,Pi3K92E,Rel,lwr,gammaTub37C,Cenp-C,Lkb1,alphaTub67C,Tak1 | | 1.979e-03 | -6.23 | mRNA 5'-UTR binding | molecular function | GO:0048027 | 7 | 4 | 12176 | 1119 | RpS14a,RpS14b,mRpS11,Shmt | | 1.988e-03 | -6.22 | phosphatidylinositol binding | molecular function | GO:0035091 | 65 | 14 | 12176 | 1119 | CG5734,Moe,Pex23,sau,snz,Pld,CG8726,CG11807,Exo70,Snx6,Myd88,Hip1,crok,CG3077 | | 1.993e-03 | -6.22 | MORF\_RAD21 | MSigDB lists | MORF\_RAD21 | 110 | 29 | 4161 | 644 | mts,Rbf,CG3631,Rpt2,LysRS,CG1890,Rpn12,Sod1,Pka-R1,CHOp24,Stoml2,SkpA,levy,CG17454,Prosalpha3,LSm3,Prosbeta4,COX5A,ND-75,wal,Klp10A,SdhA,ND-30,Hat1,CG13630,Pen,Shmt,Chrac-14,Cul1 | | 2.001e-03 | -6.21 | - | gene3d domains | 3.30.420.80 | 3 | 3 | 4929 | 622 | mRpS11,RpS14b,mRpL18 | | 2.007e-03 | -6.21 | Beta oxidation of hexanoyl-CoA to butanoyl-CoA | REACTOME pathways | R-DME-77350 | 5 | 4 | 3923 | 574 | CG4860,Mtpalpha,Thiolase,Echs1 | | 2.013e-03 | -6.21 | - | gene3d domains | 3.30.1520.10 | 16 | 7 | 4929 | 622 | CG5734,Pld,CG11807,snz,CG3077,CG8726,Snx6 | | 2.014e-03 | -6.21 | MORF\_PRDX3 | MSigDB lists | MORF\_PRDX3 | 49 | 16 | 4161 | 644 | LysRS,Cyt-c-p,Sod1,ND-51,CG1416,CG5028,Prosbeta4,COX5A,CCT5,ND-75,Mtpalpha,mEFTu1,Thiolase,Hat1,ND-24,Pen | | 2.021e-03 | -6.20 | YAMAZAKI\_TCEB3\_TARGETS\_DN | MSigDB lists | YAMAZAKI\_TCEB3\_TARGETS\_DN | 67 | 20 | 4161 | 644 | Pen,Hat1,Aldh,CCT3,esc,CG46338,CycE,escl,Fdh,rept,CG1416,l(1)G0004,Karybeta3,Stoml2,Arl6IP1,CG8188,CG3860,Cul2,alphaTub84B,Dhfr | | 2.032e-03 | -6.20 | equatorial microtubule organizing center | cellular component | GO:0000923 | 7 | 4 | 11940 | 1105 | Grip75,CG7716,Grip128,Grip84 | | 2.041e-03 | -6.19 | spidey (spidey) | protein interactions | 31703 | 67 | 17 | 9204 | 1112 | wal,Surf4,GCS2beta,Vha68-2,CG1416,CG10470,Rab5,Atpalpha,Pi3K92E,Mcad,Pp2A-29B,CCT3,CG15098,CCT6,SERCA,Kap-alpha3,CG10425 | | 2.043e-03 | -6.19 | Sc\_DH/Rdtase\_CS | interpro domains | IPR020904 | 23 | 8 | 11383 | 1233 | CG4842,CG31548,CG31549,CG10425,Adh,CG7322,scu,CG12171 | | 2.051e-03 | -6.19 | osk (oskar) | protein interactions | 41066 | 96 | 22 | 9204 | 1112 | bru1,nos,pgc,Nc73EF,yps,sw,swa,osk,Ork1,heph,larp,tkv,Imp,cup,Actn,up,Mf,slam,wisp,Tm1,apt,smg | | 2.067e-03 | -6.18 | MED17 (Mediator complex subunit 17) | protein interactions | 42175 | 56 | 15 | 9204 | 1112 | Usp10,Sec23,MED27,MED20,MED4,enc,ix,dl,Ssdp,CG2199,MED24,CG14036,Ptip,Larp4B,CG17002 | | 2.067e-03 | -6.18 | Eb1 (Eb1) | protein interactions | 35584 | 56 | 15 | 9204 | 1112 | Uch,Stip1,Tak1,CycE,Chd64,CG11267,awd,AnxB10,krimp,CG4598,scu,Moe,capt,CG10638,regucalcin | | 2.069e-03 | -6.18 | regulation of mitotic cell cycle, embryonic | biological process | GO:0009794 | 16 | 6 | 12345 | 1129 | grp,png,twe,mus304,ben,l(3)mbt | | 2.069e-03 | -6.18 | GSE360\_CTRL\_VS\_B\_MALAYI\_LOW\_DOSE\_MAC\_DN | MSigDB lists | GSE360\_CTRL\_VS\_B\_MALAYI\_LOW\_DOSE\_MAC\_DN | 58 | 18 | 4161 | 644 | Klp10A,Ac3,wal,Cul1,Chd64,Got1,Prosbeta4,Rab32,Iru,NHP2,parvin,tzn,Pex10,CG2201,Atg6,ari-1,CG1890,nrv1 | | 2.083e-03 | -6.17 | heterocycle catabolic process | biological process | GO:0046700 | 157 | 26 | 12345 | 1129 | smg,cu,CG11486,Dcr-1,fal,Drep1,LSm3,Rga,DhpD,twin,Edc3,CG17768,Cbp80,Trf4-2,gw,P5CDh1,CG11164,cN-IIIB,armi,Pym,AOX3,Not10,CG31759,Rrp40,Ho,Tengl4 | | 2.089e-03 | -6.17 | GO\_NEGATIVE\_REGULATION\_OF\_ERBB\_SIGNALING\_PATHWAY | MSigDB lists | GO\_NEGATIVE\_REGULATION\_OF\_ERBB\_SIGNALING\_PATHWAY | 17 | 8 | 4161 | 644 | Ptp61F,CG4709,Stam,RpL40,Snx6,Iru,RpS27A,Clc | | 2.113e-03 | -6.16 | CSN1a (COP9 signalosome subunit 1a) | protein interactions | 34857 | 21 | 8 | 9204 | 1112 | RpL23A,Stat92E,muskelin,RpL24,RpS29,dx,CG32267,RpL38 | | 2.113e-03 | -6.16 | TpnC25D (Troponin C at 25D) | protein interactions | 33752 | 9 | 5 | 9204 | 1112 | Fkbp12,CG40045,PHGPx,CG10265,awd | | 2.113e-03 | -6.16 | CG7148 (uncharacterized protein) | protein interactions | 40432 | 9 | 5 | 9204 | 1112 | SkpA,Cul2,Cul1,CG32221,Cyt-c-p | | 2.122e-03 | -6.16 | coated vesicle | cellular component | GO:0030135 | 65 | 14 | 11940 | 1105 | CHOp24,Sec22,Hip1,Afti,PAPLA1,yl,Gga,CG9132,AP-1-2beta,opm,Sec23,Clc,CG4293,CG5510 | | 2.124e-03 | -6.15 | TARTE\_PLASMA\_CELL\_VS\_PLASMABLAST\_DN | MSigDB lists | TARTE\_PLASMA\_CELL\_VS\_PLASMABLAST\_DN | 156 | 38 | 4161 | 644 | Karybeta3,lid,CG11089,escl,CG8188,Cse1,scu,ND-75,Klp10A,Cdk1,rl,Top1,CCT6,Pgk,Pen,mad2,ND-39,Prosalpha5,Ts,RfC38,Cyp1,CG12018,TyrRS,AnxB10,Cyt-c-p,CycB,esc,anne,CCT5,ND-B18,ND-24,bys,Moe,Prosbeta4,APP-BP1,Uch,CycE,gammaTub37C | | 2.127e-03 | -6.15 | positive regulation of mitotic cell cycle | biological process | GO:0045931 | 39 | 10 | 12345 | 1129 | Su(H),png,APC7,SkpA,bru1,Sld5,twe,CycE,ben,Moe | | 2.127e-03 | -6.15 | response to xenobiotic stimulus | biological process | GO:0009410 | 39 | 10 | 12345 | 1129 | Pka-R2,Cyp6v1,Bx,Tao,Ugt36E1,cyc,dco,nbs,MAGE,EcR | | 2.127e-03 | -6.15 | cellular response to stimulus | biological process | GO:0051716 | 1467 | 165 | 12345 | 1129 | PHGPx,mAChR-A,CG10336,Wnt10,Traf6,Pld,Rbf,mre11,Mo25,Chd64,Pka-R2,Myd88,Rab5,CG17698,raptor,Grx1,trk,CG5285,Fdh,Pka-R1,tkv,PGRP-LE,Rep,Gdi,Edem1,Cdk5,nej,spen,Nprl2,rl,Jafrac1,MED27,Ac3,gus,nbs,agt,bif,PyK,Ptp61F,CG13933,Hnf4,fs(1)N,EcR,Debcl,grp,mus205,Cdk1,CG14408,tos,Nup358,cnc,tor,Xpc,Ubc6,kcc,Tak1,pygo,Atf6,CG6927,Su(H),CG9005,Sod3,mus304,Utx,S2P,fray,CG6766,CG5844,Stat92E,p38a,Hr78,Npc2a,DNApol-epsilon255,RecQ5,Sod1,pen-2,SNF4Agamma,Trxr-1,Tao,Ufm1,G9a,lt,CG7115,dnc,Iml1,Rab10,ben,Cka,Tlk,O-fut1,Lkb1,RhoGAP54D,nAChRalpha4,tho2,foxo,dl,Pdk1,Parp16,bun,fs(1)M3,Rel,ACC,Tfb4,disp,CG4390,CanA-14F,Chrac-14,MAGE,Med,REPTOR,Tom40,obe,Mtl,Pi3K92E,Pp2B-14D,boi,Calx,Fkbp39,Xpac,Cyp6v1,spz,Ugt36E1,CG8974,CG3894,bcd,tub,crq,cact,sip3,Vps16B,SREBP,Jafrac2,mus81,CG9391,ago,Atg6,Sld5,RhoGDI,Idh,Atg7,csw,Shmt,Rab32,dco,Snm1,PEK,fz4,Cbl,lwr,shep,CG17272,Nurf-38,thoc6,Mitf,ERp60,CG17746,CG32640,Smox,Uev1A,Hsc70-3,CG11070,Mekk1,Marf,zyd,Pdxk | | 2.150e-03 | -6.14 | MyD88 cascade initiated on plasma membrane | REACTOME pathways | R-DME-975871 | 34 | 12 | 3923 | 574 | Rel,Uev1A,Tak1,ben,Traf6,rl,RpS27A,Myd88,RpL40,dl,Pp2A-29B,p38a | | 2.150e-03 | -6.14 | Toll Like Receptor TLR1:TLR2 Cascade | REACTOME pathways | R-DME-168179 | 34 | 12 | 3923 | 574 | p38a,Pp2A-29B,RpL40,dl,rl,RpS27A,Myd88,Rel,Uev1A,Tak1,Traf6,ben | | 2.150e-03 | -6.14 | Toll Like Receptor 5 (TLR5) Cascade | REACTOME pathways | R-DME-168176 | 34 | 12 | 3923 | 574 | dl,RpL40,Pp2A-29B,p38a,ben,Traf6,Tak1,Uev1A,Rel,Myd88,RpS27A,rl | | 2.150e-03 | -6.14 | MyD88:Mal cascade initiated on plasma membrane | REACTOME pathways | R-DME-166058 | 34 | 12 | 3923 | 574 | ben,Traf6,Tak1,Rel,Uev1A,Myd88,RpS27A,rl,dl,RpL40,p38a,Pp2A-29B | | 2.150e-03 | -6.14 | Toll Like Receptor TLR6:TLR2 Cascade | REACTOME pathways | R-DME-168188 | 34 | 12 | 3923 | 574 | RpL40,dl,p38a,Pp2A-29B,ben,Traf6,Rel,Uev1A,Tak1,Myd88,rl,RpS27A | | 2.150e-03 | -6.14 | Toll Like Receptor 2 (TLR2) Cascade | REACTOME pathways | R-DME-181438 | 34 | 12 | 3923 | 574 | Pp2A-29B,p38a,RpL40,dl,Myd88,rl,RpS27A,ben,Traf6,Rel,Uev1A,Tak1 | | 2.150e-03 | -6.14 | Activated TLR4 signalling | REACTOME pathways | R-DME-166054 | 34 | 12 | 3923 | 574 | dl,RpL40,Pp2A-29B,p38a,Tak1,Uev1A,Rel,ben,Traf6,RpS27A,rl,Myd88 | | 2.155e-03 | -6.14 | RNA processing | biological process | GO:0006396 | 564 | 72 | 12345 | 1129 | CG10463,mRpS11,bru1,IntS6,CG17768,rig,Trf4-2,tho2,CG10189,Imp,LSm3,CG17187,heph,NHP2,CG3587,Neos,CG17454,snRNA:U4atac,bys,obe,CG5986,Rnmt,sbr,CG12301,CG9143,CG11486,Dcr-1,mtTFB1,CG3198,CG3021,CG2100,RpS7,CG30343,SelD,CG10418,RpS16,Rrp40,Bin1,Archease,wisp,Usp39,Spx,IntS11,snRNA:U12,Mat89Ba,snRNA:U4:38AB,exu,nos,Clp,CG6610,RpS14b,RpS14a,CG7728,r2d2,RpS28b,CG5292,CG2911,RpL7A,yps,CG14100,Cbp80,CG9684,scu,CG2034,CG7009,CG8204,SmE,Son,CG16865,CycC,Larp7,CG1622 | | 2.158e-03 | -6.14 | regulation of organelle organization | biological process | GO:0033043 | 408 | 55 | 12345 | 1129 | msps,Arpc1,Sec23,Arpc2,CycE,Abi,Synj,Utx,HP1e,cana,psq,Arpc4,wapl,ago,Rbf,mad2,Pld,trr,esc,SREBP,Vps16B,bru1,Zwilch,piwi,Tlk,scny,dgt4,Moe,HP1c,Cenp-C,lt,Bap170,Mekk1,pgc,Marf,APC7,tws,G9a,Pink1,ssh,Mos,Sod1,stai,wash,Nprl2,Mitf,park,CG11486,nej,cnn,Mtl,egg,dco,Stat92E,BigH1 | | 2.160e-03 | -6.14 | p23 (p23) | protein interactions | 41173 | 73 | 18 | 9204 | 1112 | CG32549,HmgD,Stip1,RpLP0,CG11267,Chd64,awd,Echs1,Fkbp12,PHGPx,cib,CG10638,bor,AnxB10,capt,Moe,red,regucalcin | | 2.169e-03 | -6.13 | BLALOCK\_ALZHEIMERS\_DISEASE\_INCIPIENT\_DN | MSigDB lists | BLALOCK\_ALZHEIMERS\_DISEASE\_INCIPIENT\_DN | 72 | 21 | 4161 | 644 | CG1416,CG11089,Roe1,RpL38,CanB2,Crag,Nurf-38,Pp2B-14D,CG7718,Cul2,CG3226,Cdk7,CkIIalpha,Top1,SdhA,Aldh,CCT3,Pfdn2,CCT5,Hsc70-3,CanA-14F | | 2.169e-03 | -6.13 | GO\_UBIQUITIN\_LIKE\_PROTEIN\_LIGASE\_ACTIVITY | MSigDB lists | GO\_UBIQUITIN\_LIKE\_PROTEIN\_LIGASE\_ACTIVITY | 72 | 21 | 4161 | 644 | Cul5,STUB1,MED27,CG46338,Uev1A,lwr,CG8974,Cul1,Iru,MED20,Ubc4,SkpA,CG40045,CG11070,Cul2,MED24,CycC,UbcE2H,ari-1,CG8188,Traf6 | | 2.176e-03 | -6.13 | multicellular organism development | biological process | GO:0007275 | 2496 | 266 | 12345 | 1129 | Bx,Smyd4-4,Traf6,Irk1,cup,Myd88,CG3226,heph,raptor,PrBP,Ptip,eIF3m,Grip128,Vha68-2,Pka-R1,Idgf2,tkv,PGRP-LE,capt,l(3)mbt,Cdk5,cib,lin-52,Dcr-1,Jafrac1,rl,Tip60,gus,ph-d,alphaTub67C,Sec15,enc,Ptp61F,gw,bif,EcR,Archease,PIG-C,Cdk1,cnc,tor,nos,pygo,Tak1,psq,Su(H),Seipin,mus304,Abi,fray,sws,velo,msps,Npc2a,E2f1,p38a,Idgf6,sol,scu,cnn,shrb,lin,milt,Sry-alpha,Tao,Coq7,ATPsynD,Bap170,lt,colt,Grip71,ben,cold,RhoGAP54D,Ptpmeg,Tsp42Ef,Top1,crok,trr,Pdk1,Imp,bun,fs(1)M3,GlyP,mop,gammaTub37C,disp,HP1e,Cf2,Pen,MAGE,Med,Nlg1,wal,mael,stil,Trf2,Mtl,Pp2B-14D,Pi3K92E,galectin,boi,stai,fwe,Idh3b,Pink1,Nmt,snk,cact,ND-SGDH,CG17068,slam,mts,exu,Nf-YA,CkIIalpha,Atg6,Ptp10D,Atf3,mute,csw,CG46338,dco,apt,Hsp26,lwr,shep,Pax,Girdin,ScsbetaA,parvin,CycB,alpha-Man-Ia,wash,ssh,gd,tws,Marf,CycC,COX4,Dd,CG3770,otu,Grip75,EMC3,alph,Wnt10,larp,Pld,ix,sw,Rab5,Pka-R2,Tspo,mxc,COX5A,Arf51F,trk,cad,CG12301,SkpA,Edem1,faf,Cen,nej,Thiolase,spen,crm,X11L,pn,MED27,dx,Rrp40,chrb,fs(1)N,twe,Rbfox1,grp,Nup358,Afti,AGBE,fdl,CycJ,bam,CG13366,Utx,Arpc1,toc,Cul1,Stat92E,swa,Hr78,ari-1,Tsp86D,CCT3,MED24,Sod1,Trxr-1,G9a,Desat1,jvl,Fer1HCH,dnc,Moe,Cka,Tlk,Lkb1,piwi,Atpalpha,d4,up,snz,krimp,Pura,Cul5,foxo,dl,not,Rel,Gclc,Mtpalpha,CG4853,CycE,Pfdn2,sbb,spn-E,Sec23,Mes2,Tm1,RhoGAP5A,Rbcn-3A,ovo,rept,Vha13,Liprin-beta,SelD,AP-1-2beta,spz,DCAF12,lid,tub,bcd,CHOp24,ea,cyc,cort,Ssdp,hb,MED4,eIF4A,ago,Ptp69D,SelR,armi,Idgf4,awd,Atg7,nmd,Cbl,Mitf,Stam,levy,Smox,EloC,osk,stw | | 2.180e-03 | -6.13 | response to alkaloid | biological process | GO:0043279 | 27 | 8 | 12345 | 1129 | Tao,Bx,Pka-R2,nbs,cyc,dco,MAGE,EcR | | 2.182e-03 | -6.13 | negative regulation of mitotic cell cycle | biological process | GO:0045930 | 73 | 15 | 12345 | 1129 | Aven,Iml1,Nprl2,mre11,CG10336,mus81,RecQ5,Rbf,mad2,nbs,Zwilch,mus304,grp,Cdk1,Rbf2 | | 2.192e-03 | -6.12 | adenyl ribonucleotide binding | molecular function | GO:0032559 | 655 | 82 | 12176 | 1119 | CG9981,SRPK,ABCD,png,ATP8A,TyrRS,CG17698,Pink1,dco,Pfk,Pfrx,CCT3,E23,kl-2,fray,CG4880,CG14183,SelD,Lkb1,Droj2,SNF4Agamma,grp,rept,cana,pont,p38a,Orc1,rl,r,bor,l(2)k01209,PEK,nmdyn-D7,Mos,l(1)G0196,LysRS,Nadsyn,CysRS,CG8726,RecQ5,eIF4A,CG5789,SerRS-m,Pdk1,Rpt2,CG6767,spn-E,SERCA,tkv,CG4041,Pgk,GluRS-m,CG9143,Atpalpha,blw,Tak1,Cdk7,awd,CG8611,Tlk,CG31689,Tao,Cdk5,ATPsynbeta,CkIIalpha,Mekk1,TrpRS-m,CCT5,Hsc70-3,CCT6,Klp10A,tor,Vha68-2,obe,ACC,ScsbetaA,CG4562,AcCoAS,CG7806,nmd,CG31729,Cdk1 | | 2.199e-03 | -6.12 | respiratory chain complex IV | cellular component | GO:0045277 | 16 | 6 | 11940 | 1105 | cype,levy,COX5B,COX5A,COX4,COX8 | | 2.205e-03 | -6.12 | thiol-dependent ubiquitin-specific protease activity | molecular function | GO:0004843 | 39 | 10 | 12176 | 1119 | Uch,Usp16-45,not,Usp39,trbd,faf,Uch-L5,scny,otu,Usp10 | | 2.205e-03 | -6.12 | thiol-dependent ubiquitinyl hydrolase activity | molecular function | GO:0036459 | 39 | 10 | 12176 | 1119 | scny,Uch-L5,faf,trbd,Usp39,Usp10,otu,Uch,not,Usp16-45 | | 2.210e-03 | -6.11 | peptide-lysine-N-acetyltransferase activity | molecular function | GO:0061733 | 52 | 12 | 12176 | 1119 | YL-1,lid,Chrac-14,nej,Rcd5,Sgf29,pont,rept,Tip60,d4,e(y)1,Hat1 | | 2.210e-03 | -6.11 | protein serine/threonine phosphatase activity | molecular function | GO:0004722 | 52 | 12 | 12176 | 1119 | CanA-14F,mts,Pp2B-14D,CG17746,Ublcp1,tws,Dd,Pp2A-29B,PpV,alph,PpD3,CG7115 | | 2.217e-03 | -6.11 | germ-line cyst formation | biological process | GO:0048134 | 33 | 9 | 12345 | 1129 | CycB,Dcr-1,Iml1,enc,Cbl,pgc,eIF4E1,bam,Cdk1 | | 2.217e-03 | -6.11 | pole cell development | biological process | GO:0007277 | 33 | 9 | 12345 | 1129 | CycE,cnn,pgc,eIF4A,Dcr-1,osk,tor,piwi,cort | | 2.217e-03 | -6.11 | oocyte microtubule cytoskeleton organization | biological process | GO:0016325 | 33 | 9 | 12345 | 1129 | jvl,msps,osk,Lkb1,Pka-R1,mael,heph,capt,cnc | | 2.222e-03 | -6.11 | 6hp\_glycosidase-like\_sf | interpro domains | IPR012341 | 14 | 6 | 11383 | 1233 | Treh,CG2061,CG8475,CG9485,alpha-Man-Ia,Edem1 | | 2.222e-03 | -6.11 | ATPase\_P-typ\_P\_site | interpro domains | IPR018303 | 14 | 6 | 11383 | 1233 | anne,ATP8A,Atpalpha,CG9981,SERCA,CG31729 | | 2.222e-03 | -6.11 | P\_typ\_ATPase | interpro domains | IPR001757 | 14 | 6 | 11383 | 1233 | ATP8A,Atpalpha,CG9981,CG31729,SERCA,anne | | 2.239e-03 | -6.10 | zinc ion binding | molecular function | GO:0008270 | 516 | 67 | 12176 | 1119 | l(3)mbt,Shmt,CG17328,SmydA-1,CG13344,ari-1,CG14435,CG9925,CG8974,Rpn3,CG6923,nos,CG11902,bor,CG14516,Hr78,Cbl,CG11134,PGRP-LE,CSN6,CG9684,CG31053,Hnf4,CG11710,CG32473,CAH2,CAH7,Fdh,CG2678,HDAC6,Pex10,CG17260,Kcmf1,not,CG10654,Pex12,l(3)73Ah,lt,Traf6,Rpn2,QC,GluRS-m,EcR,RpS29,CG6813,DNApol-epsilon255,CG9941,Sec23,ph-d,SP1029,egg,CG4730,isoQC,Usp16-45,d4,dx,CG15141,Topors,G9a,Clp,Usp39,nej,sip3,SelR,DhpD,CG10492,Iru | | 2.257e-03 | -6.09 | whole membrane | cellular component | GO:0098805 | 247 | 37 | 11940 | 1105 | Vha36-1,Pex10,AP-1-2beta,Topors,Tom70,Debcl,Hip1,Axs,Vamp7,nSMase,Vsp37A,Sec23,fwe,Afti,Marf,Tom40,Vha68-2,Pex23,Iml1,Pex12,Vha13,CG11077,CHMP2B,CG8004,Clc,yin,shrb,CG6707,CG9393,CG9132,Pi4KIIalpha,ABCD,Pex13,Cbl,mino,Tspo,Sec22 | | 2.260e-03 | -6.09 | protein folding | biological process | GO:0006457 | 126 | 22 | 12345 | 1129 | CG32640,Droj2,Uxt,TBCC,CCT6,CCT5,Hsc70-3,CG11267,Pfdn5,Fkbp12,Fkbp59,Pfdn2,Cyp1,UK114,DnaJ-H,Roe1,Hsp26,EMC3,CCT3,CG1890,shv,ERp60 | | 2.266e-03 | -6.09 | GO\_PROTEASOMAL\_PROTEIN\_CATABOLIC\_PROCESS | MSigDB lists | GO\_PROTEASOMAL\_PROTEIN\_CATABOLIC\_PROCESS | 136 | 34 | 4161 | 644 | CG17593,Ubc4,Prosbeta6,CG6766,RpL40,APC7,Prosbeta5,CycB,CG8974,Edem1,Cul1,Rpn3,Prosalpha3,Prosbeta4,Hsc70-3,Iru,SkpA,cyc,Tip60,Prosbeta3,Prosalpha4,CG10465,Rpt2,UbcE2H,Rpn12,CG8188,Rpn2,STUB1,Cdk1,Fbw5,mad2,Faf2,Prosalpha5,RpS27A | | 2.267e-03 | -6.09 | GO\_PROTEIN\_CATABOLIC\_PROCESS | MSigDB lists | GO\_PROTEIN\_CATABOLIC\_PROCESS | 236 | 53 | 4161 | 644 | shrb,Herc4,RpS27A,Prosalpha5,Faf2,Cdk1,STUB1,Cul5,mad2,Fbw5,Uch-L5,UbcE2H,Rpt2,CG10465,Prosalpha4,trbd,Prosbeta3,Cul2,Rpn2,Vsp37A,HDAC6,CG8188,Rpn12,ste24a,SkpA,Tip60,cyc,CG11070,CG4572,Hsc70-3,Uch,Prosbeta4,Prosalpha3,CG32473,Iru,CycB,Prosbeta5,Rpn3,Cul1,CG8974,Edem1,lwr,faf,APC7,ari-1,Prosbeta6,CG40045,CG6766,Ubc4,CG17593,cathD,not,RpL40 | | 2.283e-03 | -6.08 | intracellular protein transport | biological process | GO:0006886 | 400 | 54 | 12345 | 1129 | CG9393,tho2,Pex13,Tim8,Kap-alpha3,Topors,Vps16B,GAPsec,Mat89Ba,sip3,Nup358,Arf51F,Nxf3,Clc,Pen,CG5734,Arts,CG14894,Tom40,Tom70,Sec23,CG8314,CG6766,Arpc1,Pex10,Karybeta3,Rab5,Pex12,opm,mbo,Tsp86D,Nup62,Stam,thoc6,milt,sbr,Rab32,Vsp37A,CG10344,Roe1,Rep,Oseg1,Arl5,lt,Rab10,Cse1,CHOp24,CG4041,AP-1-2beta,CG11110,Syx13,CG8004,Gga,CG8728 | | 2.291e-03 | -6.08 | TUDOR | pfam domains | PF00567 | 14 | 6 | 10480 | 1142 | CG9925,tapas,krimp,CG9684,vret,spn-E | | 2.292e-03 | -6.08 | REACTOME\_ADAPTIVE\_IMMUNE\_SYSTEM | MSigDB lists | REACTOME\_ADAPTIVE\_IMMUNE\_SYSTEM | 188 | 44 | 4161 | 644 | Prosalpha3,Hsc70-3,Prosbeta4,Sec23,Pp2A-29B,CG2924,Clc,Prosbeta5,Pdk1,Cul1,Rpn3,Kap3,Traf6,APC7,Ubc4,Pi3K92E,Prosbeta6,CG40045,RpL40,Atg7,gus,Aos1,cathD,Prosalpha5,RpS27A,Cul5,STUB1,cact,Klp10A,ERp60,mts,Prosbeta3,Cul2,Prosalpha4,UbcE2H,Rpt2,CG3860,CG8188,Rpn12,Rpn2,AP-1-2beta,SkpA,CG11070,Uba5 | | 2.295e-03 | -6.08 | GSE24210\_RESTING\_TREG\_VS\_TCONV\_DN | MSigDB lists | GSE24210\_RESTING\_TREG\_VS\_TCONV\_DN | 106 | 28 | 4161 | 644 | Cbp80,Grx1,CG1416,rept,Naa20A,heph,Adi1,CG4562,NHP2,Roe1,sgll,CG4390,mRpL13,Phb2,CG11164,mts,CG8314,Hat1,CCT6,msps,bys,Pus7,pont,CG8611,Irp-1B,Abi,PDCD-5,Prosalpha3 | | 2.296e-03 | -6.08 | GSE24634\_NAIVE\_CD4\_TCELL\_VS\_DAY10\_IL4\_CONV\_TREG\_DN | MSigDB lists | GSE24634\_NAIVE\_CD4\_TCELL\_VS\_DAY10\_IL4\_CONV\_TREG\_DN | 77 | 22 | 4161 | 644 | Mitofilin,Idh,mRpL17,Arpc2,Prosalpha5,CG11710,Pdxk,milt,Vha36-1,CG2493,RhoGAP54D,colt,CG17593,Ubc4,RfC38,Etf-QO,CG4511,Grx1,Su(H),opm,CG10418,CG10932 | | 2.301e-03 | -6.07 | REACTOME\_APC\_C\_CDC20\_MEDIATED\_DEGRADATION\_OF\_MITOTIC\_PROTEINS | MSigDB lists | REACTOME\_APC\_C\_CDC20\_MEDIATED\_DEGRADATION\_OF\_MITOTIC\_PROTEINS | 54 | 17 | 4161 | 644 | Rpn3,mad2,CycB,Prosbeta5,Cdk1,RpS27A,Prosalpha5,Prosalpha3,Prosbeta4,RpL40,Prosbeta6,APC7,Rpn12,Rpn2,Prosbeta3,Rpt2,Prosalpha4 | | 2.310e-03 | -6.07 | regulation of embryonic development | biological process | GO:0045995 | 88 | 17 | 12345 | 1129 | spn-E,ben,enc,twe,mus304,eIF4E1,png,cup,exu,Top1,Mtl,Pp2B-14D,cnc,swa,l(3)mbt,mael,grp | | 2.310e-03 | -6.07 | negative regulation of cell population proliferation | biological process | GO:0008285 | 88 | 17 | 12345 | 1129 | Rab5,MED27,Rel,tws,Utx,MAGE,EcR,Ptp61F,COX5A,COX4,l(3)mbt,MED4,mts,lwr,Cul5,foxo,ago | | 2.315e-03 | -6.07 | ATPase-coupled ion transmembrane transporter activity | molecular function | GO:0042625 | 66 | 14 | 12176 | 1119 | Atpalpha,ATPsynE,blw,ATPsynB,ATPsynbeta,ATPsynG,ATPsynO,SERCA,Vha36-1,ATPsyndelta,ATPsynD,Vha68-2,nrv1,Vha13 | | 2.322e-03 | -6.07 | GO\_SINGLE\_ORGANISM\_CELLULAR\_LOCALIZATION | MSigDB lists | GO\_SINGLE\_ORGANISM\_CELLULAR\_LOCALIZATION | 369 | 77 | 4161 | 644 | RpS28b,RpS27A,RpL9,CG44422,RpL13,Pen,ERp60,Trs33,RpL13A,Cse1,RpS13,RpLP0,RpS30,ste14,cyc,Arl6IP1,RpL24,RpL37a,CanA-14F,Pex12,Tim8,CHMP2B,RpS2,Atg6,CG11110,Pp2B-14D,Phb2,Oseg1,CG8728,Pex10,RpL30,mbo,Atg4b,PpV,Sec22,RpS23,SERCA,shrb,Rab10,Tom40,cact,rl,CG9393,sau,CG2100,Tsp86D,Arts,GCC88,CHOp24,Karybeta3,RpS27,Vamp7,RpL26,RpL11,Sec23,RpL12,Marf,Kap-alpha3,CG8004,CycB,Cdk5,RpS18,RpS16,Kap3,Pex13,lt,RpS24,RpL23,CG1847,Synj,RpL40,Tspo,RpS7,Rep,RpS4,RpL38,Pi3K92E | | 2.335e-03 | -6.06 | Metallo-B-lactamas | interpro domains | IPR001279 | 10 | 5 | 11383 | 1233 | tzn,IntS11,CG12375,Snm1,CG9117 | | 2.335e-03 | -6.06 | RibonucZ/Hydroxyglut\_hydro | interpro domains | IPR036866 | 10 | 5 | 11383 | 1233 | tzn,IntS11,CG12375,Snm1,CG9117 | | 2.355e-03 | -6.05 | CycG (Cyclin G) | protein interactions | 43724 | 103 | 23 | 9204 | 1112 | Stam,Ntan1,EcR,Nc73EF,dx,heph,plu,bcd,Cdk1,RpL13A,TBCC,Pglym78,CG5151,CG11486,mts,Cul2,26-29-p,gus,CycC,Aladin,Pp2A-29B,tzn,CG5003 | | 2.365e-03 | -6.05 | PID\_MYC\_ACTIV\_PATHWAY | MSigDB lists | PID\_MYC\_ACTIV\_PATHWAY | 41 | 14 | 4161 | 644 | Eno,r,CycB,pont,Pfk,Med,rept,Tip60,Shmt,RpL11,Gapdh1,eIF4E1,mino,Smox | | 2.365e-03 | -6.05 | MORF\_JUND | MSigDB lists | MORF\_JUND | 41 | 14 | 4161 | 644 | eEF1beta,sbr,RpL13,RpS7,RpL9,RpS27A,RpL13A,RpL24,RpS30,RpS27,eIF4A,RpL30,RpL11,RpS24 | | 2.376e-03 | -6.04 | response to ecdysone | biological process | GO:0035075 | 46 | 11 | 12345 | 1129 | trr,EcR,rept,MED24,Nurf-38,MED27,Idh3b,Utx,Trf2,rig,G9a | | 2.376e-03 | -6.04 | protein deubiquitination | biological process | GO:0016579 | 46 | 11 | 12345 | 1129 | faf,Uch-L5,Usp10,not,Usp39,otu,trbd,Sgf11,Uch,scny,Usp16-45 | | 2.376e-03 | -6.04 | response to sterol | biological process | GO:0036314 | 46 | 11 | 12345 | 1129 | Idh3b,MED27,Utx,Trf2,rig,G9a,trr,EcR,rept,MED24,Nurf-38 | | 2.390e-03 | -6.04 | Rga (Regena) | protein interactions | 40683 | 17 | 7 | 9204 | 1112 | nos,Larp4B,twin,bam,cup,gw,cu | | 2.391e-03 | -6.04 | ADH\_SHORT | prosite domains | PS00061 | 23 | 8 | 6243 | 694 | CG31549,CG10425,scu,Adh,CG7322,CG12171,CG4842,CG31548 | | 2.396e-03 | -6.03 | negative regulation of intracellular signal transduction | biological process | GO:1902532 | 111 | 20 | 12345 | 1129 | alph,SkpA,Cka,Cul1,Ubc6,park,Nprl2,Cbl,Rbf,mts,ATPsynD,Pp2A-29B,Herc4,CG17746,pn,NFAT,Iml1,PpV,dnc,Ptp61F | | 2.399e-03 | -6.03 | MOOTHA\_MITOCHONDRIA | MSigDB lists | MOOTHA\_MITOCHONDRIA | 242 | 54 | 4161 | 644 | Tspo,tko,Pdk,r,mtTFB1,Etf-QO,CG11858,Cyt-c-p,colt,P5cr,P5CDh1,ND-24,Marf,ND-30,Thiolase,Tim8,CG8004,ND-B18,Mtpalpha,Coq7,yip2,COX5A,Gabat,mRpL17,Mitofilin,ND-23,CG10932,levy,CG17691,mRpL13,CG9393,ND-SGDH,Nurf-38,Mcad,mRpL18,ND-49,mEFTu1,mRpL16,wal,ND-75,Tom40,mRpS10,CG3803,Echs1,SdhA,AIF,CG8199,Aldh,Trxr-1,Idh3b,Ppox,Scsalpha1,TrpRS-m,CG1774 | | 2.399e-03 | -6.03 | GO\_VIRAL\_LIFE\_CYCLE | MSigDB lists | GO\_VIRAL\_LIFE\_CYCLE | 157 | 38 | 4161 | 644 | RpL37a,RpL11,RpL26,RpL24,RpS27,RpS18,CHMP2B,thoc6,Kap-alpha3,RpL12,RpL23,RpL30,RpS24,Aladin,RpS2,RpS16,RpL38,RpS4,RpS7,tho2,heph,RpL40,mbo,shrb,RpL9,RpS27A,RpS23,RpS28b,CG14516,crq,Pen,RpL13,Vsp37A,RpL13A,RpLP0,RpS30,RpS13,Fur2 | | 2.400e-03 | -6.03 | Prosbeta2 (Proteasome beta2 subunit) | protein interactions | 39628 | 46 | 13 | 9204 | 1112 | Rpn13,HDAC6,Rpn2,Atg6,Prosalpha5,Pomp,Prosalpha4,Prosbeta5,Prosalpha3,Prosbeta6,Prosbeta3,Prosbeta4,Rpt2 | | 2.421e-03 | -6.02 | endomembrane system organization | biological process | GO:0010256 | 175 | 28 | 12345 | 1129 | COX4,Dd,Topors,Ankle2,Arl6IP1,Lnpk,Pld,sau,ari-1,mino,wash,CG5510,mEFTu1,opm,Rtnl1,scramb1,twe,Abi,eIF3m,Marf,Pi4KIIalpha,CHOp24,Vha68-2,msps,Gmap,tral,CG4041,Sec23 | | 2.438e-03 | -6.02 | dUTPase (Deoxyuridine triphosphatase) | protein interactions | 34529 | 26 | 9 | 9204 | 1112 | Eno,PHGPx,capt,Fkbp59,regucalcin,CG10638,RpS27A,Usp39,AnxB10 | | 2.438e-03 | -6.02 | Rbp6 (RNA-binding protein 6) | protein interactions | 39919 | 26 | 9 | 9204 | 1112 | CG13373,CycB,mad2,plu,png,smg,CkIIalpha,CG9248,Traf6 | | 2.455e-03 | -6.01 | GO\_PYRIMIDINE\_NUCLEOTIDE\_METABOLIC\_PROCESS | MSigDB lists | GO\_PYRIMIDINE\_NUCLEOTIDE\_METABOLIC\_PROCESS | 21 | 9 | 4161 | 644 | CG5285,nmdyn-D7,CG6767,r,kri,Ts,r-l,Shmt,nmdyn-D6 | | 2.459e-03 | -6.01 | MODULE\_152 | MSigDB lists | MODULE\_152 | 68 | 20 | 4161 | 644 | Cyt-c-p,ND-SGDH,ND-49,Vha36-1,ND-23,ND-51,levy,Idh3b,COX5A,yip2,ND-39,Scsalpha1,UQCR-Q,ND-30,ND-24,Mtpalpha,ND-B18,CG8199,SdhA,Echs1 | | 2.463e-03 | -6.01 | regulation of chromosome organization | biological process | GO:0033044 | 167 | 27 | 12345 | 1129 | piwi,Stat92E,Tlk,scny,BigH1,Zwilch,esc,trr,Rbf,mad2,egg,wapl,wash,psq,Mos,APC7,tws,HP1e,Utx,cana,G9a,Cenp-C,pgc,Mekk1,Bap170,Moe,HP1c | | 2.468e-03 | -6.00 | ATP binding | molecular function | GO:0005524 | 639 | 80 | 12176 | 1119 | spn-E,tkv,SERCA,CG4041,CG6767,Rpt2,Pdk1,CG8611,Tlk,CG31689,Pgk,GluRS-m,CG9143,Atpalpha,Cdk7,awd,blw,Tak1,tor,Klp10A,Vha68-2,obe,Cdk5,ATPsynbeta,Tao,CkIIalpha,TrpRS-m,Mekk1,CCT5,CCT6,Hsc70-3,nmd,CG7806,CG31729,Cdk1,ACC,ScsbetaA,CG4562,png,ATP8A,CG9981,SRPK,ABCD,E23,CCT3,fray,kl-2,CG4880,SelD,Lkb1,CG14183,Pink1,TyrRS,CG17698,dco,Pfrx,Pfk,rl,Orc1,r,l(2)k01209,bor,Droj2,grp,cana,p38a,rept,pont,Nadsyn,CG8726,RecQ5,CysRS,SerRS-m,eIF4A,CG5789,Mos,PEK,nmdyn-D7,l(1)G0196,LysRS | | 2.469e-03 | -6.00 | hydro-lyase activity | molecular function | GO:0016836 | 46 | 11 | 12176 | 1119 | Irp-1B,CAH2,CG4594,Eno,Thiolase,CG4598,Pcd,Mtpalpha,CG11134,CAH7,Echs1 | | 2.477e-03 | -6.00 | CG2767 (uncharacterized protein) | protein interactions | 40946 | 13 | 6 | 9204 | 1112 | Bx,RhoGDI,yip2,cib,Ssdp,Moe | | 2.496e-03 | -5.99 | BERENJENO\_ROCK\_SIGNALING\_NOT\_VIA\_RHOA\_UP | MSigDB lists | BERENJENO\_ROCK\_SIGNALING\_NOT\_VIA\_RHOA\_UP | 5 | 4 | 4161 | 644 | Aldh,Jheh2,yip2,Tspo | | 2.496e-03 | -5.99 | GO\_REGULATION\_OF\_INTERLEUKIN\_2\_PRODUCTION | MSigDB lists | GO\_REGULATION\_OF\_INTERLEUKIN\_2\_PRODUCTION | 5 | 4 | 4161 | 644 | CG30496,Tip60,dnc,Traf6 | | 2.496e-03 | -5.99 | REACTOME\_P75NTR\_SIGNALS\_VIA\_NFKB | MSigDB lists | REACTOME\_P75NTR\_SIGNALS\_VIA\_NFKB | 5 | 4 | 4161 | 644 | RpS27A,Traf6,RpL40,cact | | 2.496e-03 | -5.99 | XU\_HGF\_SIGNALING\_NOT\_VIA\_AKT1\_48HR\_DN | MSigDB lists | XU\_HGF\_SIGNALING\_NOT\_VIA\_AKT1\_48HR\_DN | 5 | 4 | 4161 | 644 | lt,mad2,Desat1,alph | | 2.511e-03 | -5.99 | rhythmic behavior | biological process | GO:0007622 | 74 | 15 | 12345 | 1129 | Bx,Ac3,Pka-R2,CkIIalpha,dco,cyc,tyf,lid,Irk1,Ork1,foxo,gw,nej,cu,dnc | | 2.511e-03 | -5.99 | proton transmembrane transport | biological process | GO:1902600 | 74 | 15 | 12345 | 1129 | ATPsynD,ATPsynE,ATPsynF,ATPsynbeta,Vha36-1,Atpalpha,ATPsyndelta,blw,Vha13,Vha68-2,ATPsynO,ATPsynG,COX5B,ATPsynB,ATPsynCF6 | | 2.511e-03 | -5.99 | Fatty acid biosynthesis, elongation, mitochondria | KEGG pathways | M00085 | 5 | 4 | 3198 | 496 | Echs1,Mtpalpha,yip2,Thiolase | | 2.511e-03 | -5.99 | Fatty acid biosynthesis, elongation, mitochondria | KEGG pathways | dme\_M00085 | 5 | 4 | 3198 | 496 | Thiolase,Mtpalpha,yip2,Echs1 | | 2.516e-03 | -5.99 | REACTOME\_REGULATION\_OF\_APOPTOSIS | MSigDB lists | REACTOME\_REGULATION\_OF\_APOPTOSIS | 37 | 13 | 4161 | 644 | Prosbeta3,Prosalpha3,Prosalpha4,Prosbeta4,Rpt2,Rpn12,Prosalpha5,Rpn2,RpS27A,Prosbeta5,Prosbeta6,RpL40,Rpn3 | | 2.516e-03 | -5.99 | LANDIS\_ERBB2\_BREAST\_TUMORS\_324\_DN | MSigDB lists | LANDIS\_ERBB2\_BREAST\_TUMORS\_324\_DN | 37 | 13 | 4161 | 644 | ND-51,Thiolase,AGBE,CG17691,Tsp74F,Ptp10D,mino,yip2,Npc2a,Idh,Pgm1,Gyg,Sec23 | | 2.516e-03 | -5.99 | REACTOME\_AUTODEGRADATION\_OF\_THE\_E3\_UBIQUITIN\_LIGASE\_COP1 | MSigDB lists | REACTOME\_AUTODEGRADATION\_OF\_THE\_E3\_UBIQUITIN\_LIGASE\_COP1 | 37 | 13 | 4161 | 644 | Prosbeta6,Prosbeta5,Rpn3,RpL40,Rpt2,Prosbeta4,Prosalpha4,Prosbeta3,Prosalpha3,RpS27A,Rpn2,Prosalpha5,Rpn12 | | 2.517e-03 | -5.98 | purine ribonucleoside triphosphate binding | molecular function | GO:0035639 | 791 | 96 | 12176 | 1119 | PEK,nmdyn-D7,Mos,LysRS,l(1)G0196,CG7488,Nadsyn,CysRS,CG8726,RecQ5,gammaTub37C,SerRS-m,CG5789,eIF4A,Rab32,Droj2,alphaTub84B,grp,rept,pont,cana,p38a,Orc1,rl,r,bor,l(2)k01209,CG17698,Pink1,TyrRS,dco,Mtl,Pfk,Pfrx,E23,CCT3,kl-2,fray,CG4880,CG14183,Lkb1,SelD,CG9981,SRPK,ABCD,Marf,png,Rab5,CG5116,ATP8A,ACC,mEFTu1,alphaTub67C,ScsbetaA,CG4562,nmd,CG7806,CG31729,Cdk1,ATPsynbeta,Cdk5,Tao,Rab10,CkIIalpha,CG7728,Mekk1,mEFTu2,TrpRS-m,CCT6,CCT5,Hsc70-3,Klp10A,tor,Non1,Vha68-2,Arl5,obe,Pgk,GluRS-m,Atpalpha,CG9143,Arf51F,blw,Tak1,Cdk7,awd,CG8611,Tlk,CG31689,Pdk1,CG6767,Rpt2,spn-E,SERCA,tkv,CG4041 | | 2.527e-03 | -5.98 | ATPASE\_E1\_E2 | prosite domains | PS00154 | 14 | 6 | 6243 | 694 | CG9981,CG31729,ATP8A,Atpalpha,SERCA,anne | | 2.527e-03 | -5.98 | Prosalpha3 (Proteasome alpha3 subunit) | protein interactions | 37378 | 41 | 12 | 9204 | 1112 | Rpt2,Prosbeta4,Prosbeta3,Prosbeta6,Prosbeta5,CG4853,Prosalpha4,Fkbp12,Pomp,Prosalpha5,Rpn2,Rpn13 | | 2.547e-03 | -5.97 | GSE27786\_NKTCELL\_VS\_NEUTROPHIL\_DN | MSigDB lists | GSE27786\_NKTCELL\_VS\_NEUTROPHIL\_DN | 50 | 16 | 4161 | 644 | RhoGAP54D,alc,EcR,galectin,CG10465,Mcad,CG42542,Rep,Arpc1,Cka,Ppox,ZnT35C,Tomosyn,Moe,Trs33,pr | | 2.547e-03 | -5.97 | GSE360\_CTRL\_VS\_L\_MAJOR\_MAC\_DN | MSigDB lists | GSE360\_CTRL\_VS\_L\_MAJOR\_MAC\_DN | 50 | 16 | 4161 | 644 | EcR,Rpn12,Sec15,tko,tho2,CG4562,CG17454,Iru,CycE,Vamp7,Top1,Pgk,kcc,lqfR,GlyP,pr | | 2.547e-03 | -5.97 | GO\_NIK\_NF\_KAPPAB\_SIGNALING | MSigDB lists | GO\_NIK\_NF\_KAPPAB\_SIGNALING | 50 | 16 | 4161 | 644 | Prosbeta5,Rpn3,Cul1,Prosalpha3,Prosbeta4,Prosalpha5,RpS27A,SkpA,Prosbeta6,RpL40,Prosbeta3,Prosalpha4,Rpt2,Traf6,Rpn12,Rpn2 | | 2.558e-03 | -5.97 | GSE27859\_DC\_VS\_CD11C\_INT\_F480\_INT\_DC\_UP | MSigDB lists | GSE27859\_DC\_VS\_CD11C\_INT\_F480\_INT\_DC\_UP | 59 | 18 | 4161 | 644 | CG7718,RpS16,colt,RpL38,Xpac,Ptp61F,cathD,Maf1,Scsalpha1,Idh,Cyp6a22,RpS27,Hmg-2,CG12929,alpha-Man-Ia,CG8199,CG3376,Cyp6a19 | | 2.567e-03 | -5.97 | asymmetric stem cell division | biological process | GO:0098722 | 67 | 14 | 12345 | 1129 | twin,piwi,Cdk1,Lkb1,tws,bam,mts,mad2,shrb,egg,cnn,msps,eIF4A,Pfdn2 | | 2.571e-03 | -5.96 | microtubule-based process | biological process | GO:0007017 | 466 | 61 | 12345 | 1129 | Prosbeta5,sau,alphaTub84B,foxo,mts,larp,Ubc6,grp,Lkb1,Grip75,cnc,Topors,awd,CG7716,Grip128,tral,CG42837,msps,Pp2A-29B,kl-2,sw,mop,armi,Kap3,gammaTub37C,cana,heph,Klp10A,cnn,pont,Grip84,Girdin,milt,wash,CycB,stai,cbs,SRPK,CG3437,mael,Pka-R1,Oseg1,BicC,capt,SkpA,del,jvl,Marf,osk,wisp,Non1,Grip71,spd-2,Axs,Moe,dgt4,CCT6,Rcd5,alphaTub67C,Pink1,tws | | 2.573e-03 | -5.96 | regulation of developmental process | biological process | GO:0050793 | 757 | 92 | 12345 | 1129 | COX4,Cka,Ptpmeg,bru1,alph,foxo,Cul5,Top1,dl,Pdk1,Imp,mop,bun,cup,raptor,COX5A,MAGE,Arpc2,CycE,Med,Atu,tral,spn-E,sbb,Sec23,mael,tkv,ovo,l(3)mbt,capt,PGRP-LE,SkpA,Cdk5,Mtl,Pp2B-14D,hwt,Pi3K92E,rl,MED27,Pink1,lid,enc,bif,twe,EcR,Archease,cact,scny,grp,cnc,Wbp2,hb,shv,MED4,mts,exu,tor,nos,ago,Arpc4,Su(H),Seipin,eIF4E1,mus304,Abi,Atg7,csw,Cul1,Stat92E,E2f1,swa,p38a,ari-1,lwr,Cbl,Pax,park,Nurf-38,Stam,Tao,png,ssh,Smox,G9a,Cul2,tws,Desat1,Marf,Cenp-C,vers,Moe,ben | | 2.583e-03 | -5.96 | tral (trailer hitch) | protein interactions | 39430 | 31 | 10 | 9204 | 1112 | yps,cup,AGO3,nos,BicC,Rtnl1,nSMase,spd-2,Trf2,png | | 2.599e-03 | -5.95 | CG12170 (uncharacterized protein) | protein interactions | 40692 | 6 | 4 | 9204 | 1112 | ATPsynF,ATPsynB,ATPsynCF6,ATPsynD | | 2.599e-03 | -5.95 | swa (swallow) | protein interactions | 31580 | 6 | 4 | 9204 | 1112 | gammaTub37C,alphaTub67C,osk,swa | | 2.599e-03 | -5.95 | levy (levy) | protein interactions | 37728 | 6 | 4 | 9204 | 1112 | cype,COX4,Pdk1,COX5A | | 2.599e-03 | -5.95 | CG4797 (uncharacterized protein) | protein interactions | 37770 | 6 | 4 | 9204 | 1112 | ATPsynCF6,ATPsynD,ATPsynF,ATPsynB | | 2.599e-03 | -5.95 | Act79B (Actin 79B) | protein interactions | 40444 | 36 | 11 | 9204 | 1112 | Actn,Argk,SERCA,glob1,Adh,Mf,PGRP-LE,up,Scp1,Cf2,osk | | 2.603e-03 | -5.95 | purine ribonucleotide binding | molecular function | GO:0032555 | 811 | 98 | 12176 | 1119 | E23,CCT3,fray,kl-2,CG4880,Lkb1,SelD,CG14183,Pink1,CG17698,TyrRS,dco,Pfrx,Mtl,Pfk,png,CG5116,Rab5,ATP8A,CG9981,SRPK,ABCD,Marf,Nadsyn,CG8726,RecQ5,CysRS,gammaTub37C,CG5789,eIF4A,SerRS-m,Mos,nmdyn-D7,PEK,CG7488,l(1)G0196,LysRS,rl,Orc1,r,l(2)k01209,bor,SNF4Agamma,Droj2,Rab32,alphaTub84B,grp,pont,rept,p38a,cana,CG8611,Tlk,CG31689,Pgk,GluRS-m,CG9143,Atpalpha,Cdk7,awd,Tak1,Arf51F,blw,spn-E,tkv,SERCA,CG4041,Rpt2,CG6767,Pdk1,CG7806,nmd,CG31729,Cdk1,mEFTu1,ACC,alphaTub67C,ScsbetaA,AcCoAS,CG4562,Klp10A,tor,Vha68-2,Non1,obe,Arl5,Rab10,Cdk5,Tao,ATPsynbeta,CG7728,CkIIalpha,TrpRS-m,mEFTu2,Mekk1,CCT5,CCT6,Hsc70-3 | | 2.607e-03 | -5.95 | GSE28726\_NAIVE\_CD4\_TCELL\_VS\_NAIVE\_VA24NEG\_NKTCELL\_UP | MSigDB lists | GSE28726\_NAIVE\_CD4\_TCELL\_VS\_NAIVE\_VA24NEG\_NKTCELL\_UP | 97 | 26 | 4161 | 644 | CG8188,S2P,RhoGAP54D,scu,P5cr,CG12018,Dhfr,muskelin,Gk2,rept,escl,AGBE,levy,RfC38,Ts,gammaTub37C,Pgk,esc,CycB,CG7997,Eno,CCT5,Tom40,LpR2,Cdk1,Pdxk | | 2.607e-03 | -5.95 | GO\_POSITIVE\_REGULATION\_OF\_CELLULAR\_PROTEIN\_CATABOLIC\_PROCESS | MSigDB lists | GO\_POSITIVE\_REGULATION\_OF\_CELLULAR\_PROTEIN\_CATABOLIC\_PROCESS | 97 | 26 | 4161 | 644 | Prosalpha3,Prosbeta4,RpS27A,Prosalpha5,Gclc,CycB,Prosbeta5,STUB1,Cdk1,Rpn3,Cul1,Moe,mad2,CG30291,Prosbeta3,Rpt2,Prosalpha4,APC7,Rpn12,ari-1,Rpn2,dco,Prosbeta6,SkpA,RpL40,regucalcin | | 2.607e-03 | -5.95 | regulation of cell maturation | biological process | GO:1903429 | 40 | 10 | 12345 | 1129 | cup,Cul1,mael,cnc,swa,EcR,Pi3K92E,enc,exu,spn-E | | 2.610e-03 | -5.95 | MODULE\_82 | MSigDB lists | MODULE\_82 | 14 | 7 | 4161 | 644 | CG40045,UbcE2H,Ubc4,lwr,CG11070,CG8188,Uev1A | | 2.610e-03 | -5.95 | PID\_CERAMIDE\_PATHWAY | MSigDB lists | PID\_CERAMIDE\_PATHWAY | 14 | 7 | 4161 | 644 | AIF,CG7414,rl,cact,CG3376,Cyt-c-p,cathD | | 2.611e-03 | -5.95 | C-type lectin receptors (CLRs) | REACTOME pathways | R-DME-5621481 | 87 | 23 | 3923 | 574 | Prosbeta6,Rpn3,Cul1,Rpn12,dl,Prosbeta3,NFAT,nej,Rel,cact,SkpA,CanA-14F,CanB2,RpL40,Rpt2,Pp2B-14D,Prosalpha3,Prosbeta4,Prosalpha5,RpS27A,Prosalpha4,Rpn2,Prosbeta5 | | 2.615e-03 | -5.95 | GSE13547\_CTRL\_VS\_ANTI\_IGM\_STIM\_ZFX\_KO\_BCELL\_2H\_UP | MSigDB lists | GSE13547\_CTRL\_VS\_ANTI\_IGM\_STIM\_ZFX\_KO\_BCELL\_2H\_UP | 73 | 21 | 4161 | 644 | SelD,Rab5,CG1416,grp,Rpn12,CG8188,CG5510,Caf1-105,Cdk1,Pfdn2,CG7655,CG14715,Edem1,mad2,Pen,ERp60,UQCR-Q,shop,Npc2a,Jheh2,DNApol-epsilon255 | | 2.615e-03 | -5.95 | WONG\_MITOCHONDRIA\_GENE\_MODULE | MSigDB lists | WONG\_MITOCHONDRIA\_GENE\_MODULE | 132 | 33 | 4161 | 644 | ScsbetaA,Vha36-1,ATP8A,Pdk,Tspo,Cyp1,COX5A,Scox,ND-30,ND-24,CG8004,anne,Tim8,scu,LysRS,ND-SGDH,mRpL13,Sfxn2,ND-49,mRpL18,CG10932,ND-23,ND-51,ND-B22,levy,ND-39,TrpRS-m,Scsalpha1,UQCR-Q,mEFTu1,AIF,Echs1,Tom40 | | 2.618e-03 | -5.95 | programmed cell death involved in cell development | biological process | GO:0010623 | 89 | 17 | 12345 | 1129 | Trf2,E2f1,AIF,MED24,Drep1,cathD,rept,DCAF12,raptor,Utx,Pink1,Atg6,RYBP,Idh3b,Atg7,Debcl,EcR | | 2.620e-03 | -5.94 | germline stem cell asymmetric division | biological process | GO:0098728 | 22 | 7 | 12345 | 1129 | eIF4A,bam,shrb,mad2,piwi,twin,egg | | 2.620e-03 | -5.94 | pyrimidine-containing compound biosynthetic process | biological process | GO:0072528 | 22 | 7 | 12345 | 1129 | nmdyn-D6,nmdyn-D7,r,l(2)k01209,r-l,awd,Ts | | 2.620e-03 | -5.94 | regulation of neuron apoptotic process | biological process | GO:0043523 | 22 | 7 | 12345 | 1129 | Naa20A,MAGE,Tip60,fwe,spag,Debcl,Pink1 | | 2.620e-03 | -5.94 | torso signaling pathway | biological process | GO:0008293 | 22 | 7 | 12345 | 1129 | fs(1)N,fs(1)M3,rl,csw,trk,tor,bcd | | 2.623e-03 | -5.94 | oxidoreductase activity, acting on a sulfur group of donors | molecular function | GO:0016667 | 53 | 12 | 12176 | 1119 | cl,Scox,Grx1,CG3939,CG14997,shop,CG14407,ERp60,Trxr-1,SelR,GILT1,SelT | | 2.629e-03 | -5.94 | TIEN\_INTESTINE\_PROBIOTICS\_6HR\_UP | MSigDB lists | TIEN\_INTESTINE\_PROBIOTICS\_6HR\_UP | 33 | 12 | 4161 | 644 | RpL13,Pgk,RpS7,Ac3,RpS27,RpLP0,RpS2,RpL9,RpS23,RpS24,RpL23,Gapdh1 | | 2.632e-03 | -5.94 | Golgi subcompartment | cellular component | GO:0098791 | 103 | 19 | 11940 | 1105 | Ugalt,Rab32,Gmap,sau,Trs33,CG9981,cbs,AP-1-2beta,ATP8A,opm,CG31729,Arl5,Sec23,lqfR,Afti,GCC88,Pi4KIIalpha,Fur2,Gga | | 2.640e-03 | -5.94 | FLECHNER\_BIOPSY\_KIDNEY\_TRANSPLANT\_REJECTED\_VS\_OK\_DN | MSigDB lists | FLECHNER\_BIOPSY\_KIDNEY\_TRANSPLANT\_REJECTED\_VS\_OK\_DN | 243 | 54 | 4161 | 644 | CG14715,alph,Thiolase,CG4872,mtgo,Mitofilin,CG32473,Sec23,yip2,Pp2A-29B,Cka,Gabat,Pka-R1,AGBE,CG9485,Med,CkIIalpha,CG1532,P5CDh1,CG4390,Cyt-c-p,ND-75,wal,Echs1,AIF,SdhA,Aldh,Pgk,Shmt,lqfR,UQCR-Q,Idh,Got1,CG14516,CG8507,Pcd,Irp-1B,tzn,CG4562,levy,CG8475,eIF3i,CG17691,Jafrac1,ND-23,regucalcin,CG10932,Fdh,opm,Mcad,Pgm1,Sod1,ND-SGDH,Prip | | 2.645e-03 | -5.94 | Valine, leucine and isoleucine degradation | KEGG pathways | dme00280 | 33 | 12 | 3198 | 496 | Aldh,CG4860,Mtpalpha,CG8199,CG10932,Thiolase,CG17691,Echs1,Gabat,yip2,scu,Mcad | | 2.645e-03 | -5.94 | Valine, leucine and isoleucine degradation | KEGG pathways | ko00280 | 33 | 12 | 3198 | 496 | Gabat,Echs1,CG17691,scu,Mcad,yip2,CG4860,Aldh,Thiolase,CG10932,CG8199,Mtpalpha | | 2.645e-03 | -5.94 | purine nucleotide binding | molecular function | GO:0017076 | 821 | 99 | 12176 | 1119 | dco,Mtl,Pfk,Pfrx,TyrRS,Pink1,CG17698,CG4880,CG14183,SelD,Lkb1,CCT3,E23,kl-2,fray,Marf,SRPK,CG9981,ABCD,ATP8A,png,Rab5,CG5116,l(1)G0196,LysRS,CG7488,PEK,nmdyn-D7,Mos,gammaTub37C,SerRS-m,CG5789,eIF4A,Roe1,Nadsyn,CysRS,CG8726,RecQ5,alphaTub84B,grp,p38a,cana,pont,rept,Rab32,Droj2,SNF4Agamma,r,bor,l(2)k01209,Orc1,rl,CG9143,Atpalpha,blw,Tak1,Arf51F,Cdk7,awd,Pgk,GluRS-m,Tlk,CG31689,CG8611,Pdk1,CG6767,Rpt2,SERCA,tkv,CG4041,spn-E,ScsbetaA,AcCoAS,CG4562,ACC,mEFTu1,alphaTub67C,CG31729,Cdk1,nmd,CG7806,Mekk1,mEFTu2,TrpRS-m,Hsc70-3,CCT5,CCT6,Tao,Cdk5,ATPsynbeta,Rab10,CG7728,CkIIalpha,Arl5,obe,tor,Klp10A,Non1,Vha68-2 | | 2.674e-03 | -5.92 | REACTOME\_TRANSLATION | MSigDB lists | REACTOME\_TRANSLATION | 112 | 29 | 4161 | 644 | RpLP0,RpS30,RpS13,eIF4E1,RpS16,RpS2,RpL13A,RpS24,RpL23,eIF4A,RpL30,RpL40,RpS7,RpL38,RpS4,eIF3i,RpS27,RpL24,RpS28b,RpS23,RpL26,RpS27A,RpL9,RpL11,RpL37a,RpL13,RpL12,eEF1beta,RpS18 | | 2.680e-03 | -5.92 | GO\_REGULATION\_OF\_LIPID\_BIOSYNTHETIC\_PROCESS | MSigDB lists | GO\_REGULATION\_OF\_LIPID\_BIOSYNTHETIC\_PROCESS | 29 | 11 | 4161 | 644 | Tspo,regucalcin,nSMase,crq,Sod1,adp,EcR,mino,Idh,SREBP,Dd | | 2.680e-03 | -5.92 | REACTOME\_OPIOID\_SIGNALLING | MSigDB lists | REACTOME\_OPIOID\_SIGNALLING | 29 | 11 | 4161 | 644 | CanA-14F,Pka-R2,mts,Pp2A-29B,Pp2B-14D,rl,Ac3,CanB2,Pka-R1,Cdk5,dnc | | 2.704e-03 | -5.91 | MED15 (Mediator complex subunit 15) | protein interactions | 33223 | 63 | 16 | 9204 | 1112 | CG17002,squ,nej,Ptip,Prosbeta4,ix,grp,CG13373,MED4,MED27,Usp10,Ssdp,EcR,enc,MED20,CycJ | | 2.728e-03 | -5.90 | organellar ribosome | cellular component | GO:0000313 | 81 | 16 | 11940 | 1105 | mRpL10,mRpS10,mRpL35,CG14817,mRpL17,mRpS11,mRpL16,mRpL36,CG6094,mRpS21,mRpL18,mRpL48,tko,mRpL13,mRpS5,mRpS23 | | 2.728e-03 | -5.90 | mitochondrial ribosome | cellular component | GO:0005761 | 81 | 16 | 11940 | 1105 | mRpL35,CG14817,mRpS10,mRpL10,mRpL17,mRpS11,CG6094,mRpL36,mRpL16,mRpL48,mRpL18,mRpS21,tko,mRpL13,mRpS23,mRpS5 | | 2.733e-03 | -5.90 | fly chr15B4-15B4\|1-56 | chromosome location | fly chr15B4-15B4\|1-56 | 6 | 3 | 25010 | 1343 | Nprl2,CG4872,CG4880 | | 2.733e-03 | -5.90 | fly chr15B4-15B4\|1-56 cM | chromosome location | fly chr15B4-15B4\|1-56 cM | 6 | 3 | 25010 | 1343 | CG4880,Nprl2,CG4872 | | 2.733e-03 | -5.90 | fly chr6B1-6B1\|1-17 | chromosome location | fly chr6B1-6B1\|1-17 | 6 | 3 | 25010 | 1343 | RpL7A,CG3342,snoRNA:Or-CD10 | | 2.733e-03 | -5.90 | fly chr6B1-6B1\|1-17 cM | chromosome location | fly chr6B1-6B1\|1-17 cM | 6 | 3 | 25010 | 1343 | snoRNA:Or-CD10,CG3342,RpL7A | | 2.740e-03 | -5.90 | Rossmann-like\_a/b/a\_fold | interpro domains | IPR014729 | 40 | 11 | 11383 | 1233 | CG10189,CG1578,wal,AsnS,CG3021,GluRS-m,CG4407,TrpRS-m,Nadsyn,CysRS,TyrRS | | 2.745e-03 | -5.90 | GSE7509\_UNSTIM\_VS\_FCGRIIB\_STIM\_MONOCYTE\_UP | MSigDB lists | GSE7509\_UNSTIM\_VS\_FCGRIIB\_STIM\_MONOCYTE\_UP | 78 | 22 | 4161 | 644 | SkpA,PpV,Tsp74F,tzn,Rab5,Jafrac1,Tspo,CG1847,CG11899,scu,isoQC,CG10470,LysRS,CCT3,CG7997,Pgk,mEFTu1,LSm3,CG7414,ND-39,PDCD-5,Nmt | | 2.745e-03 | -5.90 | WELCSH\_BRCA1\_TARGETS\_UP | MSigDB lists | WELCSH\_BRCA1\_TARGETS\_UP | 78 | 22 | 4161 | 644 | Hsc70-3,CG9248,Sec23,Npc2a,Arf51F,Scsalpha1,Idh,Arpc1,Pcd,Cdk1,Klp10A,ND-75,CG2493,isoQC,Sec10,Kap3,PpV,CG4511,tzn,CHOp24,Pka-R1,Tip60 | | 2.759e-03 | -5.89 | response to inorganic substance | biological process | GO:0010035 | 82 | 16 | 12345 | 1129 | Fer1HCH,Grx1,Mekk1,ZnT35C,Gclc,whd,Sod3,spz,adp,MtnA,CG11590,Sod1,rl,Jafrac1,CG7328,p38a | | 2.782e-03 | -5.88 | GO\_MITOCHONDRIAL\_TRANSPORT | MSigDB lists | GO\_MITOCHONDRIAL\_TRANSPORT | 64 | 19 | 4161 | 644 | colt,CG11110,CG9393,CG8728,CG1847,mRpL18,CG2100,ACC,Cyp1,Tspo,Stoml2,CanB2,Nmt,yip2,Marf,Calx,Tom40,CG8004,Tim8 | | 2.801e-03 | -5.88 | GO\_SMALL\_RIBOSOMAL\_SUBUNIT | MSigDB lists | GO\_SMALL\_RIBOSOMAL\_SUBUNIT | 46 | 15 | 4161 | 644 | tko,mRpS5,RpS18,RpS4,RpS7,RpS27A,RpS23,RpS2,RpS16,RpS28b,RpS27,RpS13,RpS30,mRpS21,RpS24 | | 2.803e-03 | -5.88 | proton-transporting two-sector ATPase complex, catalytic domain | cellular component | GO:0033178 | 22 | 7 | 11940 | 1105 | ATPsynO,ATPsyndelta,Vha68-2,Vha36-1,blw,ATPsynbeta,ATPsynCF6 | | 2.805e-03 | -5.88 | Beta Oxidation of Unsaturated Fatty Acids | WikiPathways | WP249 | 4 | 4 | 499 | 116 | Mtpalpha,Thiolase,Mcad,CG4598 | | 2.812e-03 | -5.87 | secondary alcohol metabolic process | biological process | GO:1902652 | 28 | 8 | 12345 | 1129 | CG8112,Idh,CG17270,foxo,Marf,scu,CG5028,Idh3b | | 2.843e-03 | -5.86 | CTP metabolic process | biological process | GO:0046036 | 4 | 3 | 12345 | 1129 | nmdyn-D6,nmdyn-D7,awd | | 2.843e-03 | -5.86 | CTP biosynthetic process | biological process | GO:0006241 | 4 | 3 | 12345 | 1129 | awd,nmdyn-D6,nmdyn-D7 | | 2.843e-03 | -5.86 | cellular response to lipopolysaccharide | biological process | GO:0071222 | 4 | 3 | 12345 | 1129 | Tak1,EcR,Stat92E | | 2.843e-03 | -5.86 | sperm aster formation | biological process | GO:0035044 | 4 | 3 | 12345 | 1129 | spd-2,wisp,alphaTub67C | | 2.843e-03 | -5.86 | regulation of cellular respiration | biological process | GO:0043457 | 4 | 3 | 12345 | 1129 | Pink1,MED27,EcR | | 2.843e-03 | -5.86 | aldehyde catabolic process | biological process | GO:0046185 | 4 | 3 | 12345 | 1129 | Fdh,tzn,CG4390 | | 2.843e-03 | -5.86 | GTP biosynthetic process | biological process | GO:0006183 | 4 | 3 | 12345 | 1129 | awd,nmdyn-D6,nmdyn-D7 | | 2.843e-03 | -5.86 | GTP metabolic process | biological process | GO:0046039 | 4 | 3 | 12345 | 1129 | awd,nmdyn-D6,nmdyn-D7 | | 2.843e-03 | -5.86 | oocyte maturation | biological process | GO:0001556 | 4 | 3 | 12345 | 1129 | muskelin,spn-E,wisp | | 2.846e-03 | -5.86 | Toll Like Receptor 10 (TLR10) Cascade | REACTOME pathways | R-DME-168142 | 35 | 12 | 3923 | 574 | Pp2A-29B,p38a,dl,RpL40,Myd88,RpS27A,rl,ben,Traf6,Tak1,Rel,Uev1A | | 2.848e-03 | -5.86 | GO\_CELLULAR\_AMIDE\_METABOLIC\_PROCESS | MSigDB lists | GO\_CELLULAR\_AMIDE\_METABOLIC\_PROCESS | 366 | 76 | 4161 | 644 | RpL12,mRpS5,RpS18,Gclc,RpL26,RpS27,CG7414,CG32473,RpL11,eIF3m,heph,RpL40,RpL38,RpS4,RpS7,mRpS23,colt,RpS16,RpL23,Dhfr,RpS24,Shmt,Tfb4,RpS23,CG14516,Sply,GluRS-m,CysRS,Nurf-38,Sod1,eIF4A,mRpL18,CG3376,Pp2A-29B,RpL24,RpL37a,mRpL10,mRpL17,tko,Ts,nSMase,r,CG11110,Atg6,RpS2,RpL30,CG8963,TyrRS,eEF1beta,mRpL16,mEFTu1,RpL13,SerRS-m,mRpS10,CG7997,RpL9,RpS27A,RpS28b,GatA,mRpS21,TrpRS-m,Idh,ACC,Grx1,CG11089,eIF3i,Fur2,tzn,LysRS,eIF4E1,RpL13A,RpS30,mRpL13,RpLP0,RpS13,mts | | 2.851e-03 | -5.86 | GSE12845\_IGD\_POS\_BLOOD\_VS\_DARKZONE\_GC\_TONSIL\_BCELL\_DN | MSigDB lists | GSE12845\_IGD\_POS\_BLOOD\_VS\_DARKZONE\_GC\_TONSIL\_BCELL\_DN | 83 | 23 | 4161 | 644 | Got1,Arpc2,Herc4,CG7488,Faf2,MED20,CCT5,Prosbeta5,Eno,Pus7,pont,r-l,AIF,Top1,Pglym78,Cse1,CG8188,Nurf-38,Crag,mtTFB1,Cbp80,Aos1,Pdk | | 2.852e-03 | -5.86 | PUJANA\_CHEK2\_PCC\_NETWORK | MSigDB lists | PUJANA\_CHEK2\_PCC\_NETWORK | 383 | 79 | 4161 | 644 | Prosalpha5,asf1,CG7806,mad2,Pgk,Cdk1,Tom40,wal,grp,RpII140,CG8188,escl,l(1)G0004,Rpb8,Karybeta3,MED20,Uch,eIF3m,Pfas,Rpn3,CG13630,Hat1,anne,CG8004,CCT5,pont,esc,CycB,RhoGAP54D,RpL23,Dhfr,CG1416,RfC38,Ptp61F,RpS7,Nap1,Idh3b,Nmt,RpS27A,Clp,CG30496,Pen,eEF1beta,Chrac-14,Top1,slif,eIF4E1,LysRS,CG6610,Cse1,CG11164,CG11089,DNApol-epsilon255,Iru,LSm3,APP-BP1,Mitofilin,Prosalpha3,lwr,ND-24,CCT3,CG6767,Cyt-c-p,CG1890,CycC,CG8728,CkIIalpha,MED24,mbo,Cbp80,Ts,NHP2,fray,SelD,CG8204,CG17454,CG9485,CG40045,r | | 2.868e-03 | -5.85 | pyridoxal phosphate binding | molecular function | GO:0030170 | 34 | 9 | 12176 | 1119 | mal,Marc,GlyP,Kyat,CG1640,Sply,Got1,Shmt,Gabat | | 2.868e-03 | -5.85 | vitamin B6 binding | molecular function | GO:0070279 | 34 | 9 | 12176 | 1119 | Got1,Sply,CG1640,Kyat,GlyP,Gabat,Shmt,Marc,mal | | 2.881e-03 | -5.85 | fly chr26B5-26B7\|2-18 | chromosome location | fly chr26B5-26B7\|2-18 | 2 | 2 | 25010 | 1343 | stai,Arpc4 | | 2.881e-03 | -5.85 | fly chr80D1-80D1\|3-47 cM | chromosome location | fly chr80D1-80D1\|3-47 cM | 2 | 2 | 25010 | 1343 | CkIIalpha,RpL10 | | 2.881e-03 | -5.85 | fly chr80D1-80D1\|3-47 | chromosome location | fly chr80D1-80D1\|3-47 | 2 | 2 | 25010 | 1343 | RpL10,CkIIalpha | | 2.881e-03 | -5.85 | fly chr43E17-43E17\|2-57 cM | chromosome location | fly chr43E17-43E17\|2-57 cM | 2 | 2 | 25010 | 1343 | mus205,Gapdh1 | | 2.881e-03 | -5.85 | fly chr43E17-43E17\|2-57 | chromosome location | fly chr43E17-43E17\|2-57 | 2 | 2 | 25010 | 1343 | Gapdh1,mus205 | | 2.881e-03 | -5.85 | fly chr83B1-83B2\|3-47.5 cM | chromosome location | fly chr83B1-83B2\|3-47.5 cM | 2 | 2 | 25010 | 1343 | Pi4KIIalpha,Snm1 | | 2.881e-03 | -5.85 | fly chr26B5-26B7\|2-18 cM | chromosome location | fly chr26B5-26B7\|2-18 cM | 2 | 2 | 25010 | 1343 | Arpc4,stai | | 2.881e-03 | -5.85 | fly chr86F9-86F9\|3-51 cM | chromosome location | fly chr86F9-86F9\|3-51 cM | 2 | 2 | 25010 | 1343 | COX5A,Kyat | | 2.881e-03 | -5.85 | fly chr86F9-86F9\|3-51 | chromosome location | fly chr86F9-86F9\|3-51 | 2 | 2 | 25010 | 1343 | COX5A,Kyat | | 2.881e-03 | -5.85 | fly chr83B1-83B2\|3-47.5 | chromosome location | fly chr83B1-83B2\|3-47.5 | 2 | 2 | 25010 | 1343 | Snm1,Pi4KIIalpha | | 2.884e-03 | -5.85 | acetyl-CoA C-acyltransferase activity | molecular function | GO:0003988 | 4 | 3 | 12176 | 1119 | yip2,Thiolase,CG10932 | | 2.894e-03 | -5.85 | fly chr94A1-94A1\|3-74 | chromosome location | fly chr94A1-94A1\|3-74 | 12 | 4 | 25010 | 1343 | CG6028,Cchl,Mitofilin,mRpL35 | | 2.894e-03 | -5.85 | fly chr94A1-94A1\|3-74 cM | chromosome location | fly chr94A1-94A1\|3-74 cM | 12 | 4 | 25010 | 1343 | mRpL35,Mitofilin,Cchl,CG6028 | | 2.897e-03 | -5.84 | cysteine-type peptidase activity | molecular function | GO:0008234 | 82 | 16 | 12176 | 1119 | Uch,26-29-p,not,Usp16-45,velo,faf,CG16979,sol,Usp39,trbd,Atg4b,Uch-L5,Cp1,scny,otu,Usp10 | | 2.904e-03 | -5.84 | water-soluble vitamin metabolic process | biological process | GO:0006767 | 12 | 5 | 12345 | 1129 | Shmt,sgll,regucalcin,Pdxk,Dhfr | | 2.904e-03 | -5.84 | regulation of antifungal peptide production | biological process | GO:0002788 | 12 | 5 | 12345 | 1129 | Rel,Myd88,spz,tub,cact | | 2.944e-03 | -5.83 | telomeric heterochromatin | cellular component | GO:0031933 | 4 | 3 | 11940 | 1105 | Sirt7,piwi,SuUR | | 2.949e-03 | -5.83 | Pi3K92E (Pi3K92E) | protein interactions | 42446 | 117 | 25 | 9204 | 1112 | RpL13A,foxo,eIF4A,RpS13,RpL17,larp,RpS15,yps,Droj2,PyK,RpS23,Pi3K92E,RpL7A,RpLP1,Hsc70-3,RpS7,RpL10,Cul1,RpL38,eIF3m,RpL9,Pdk1,CycE,alt,Ptp61F | | 2.952e-03 | -5.83 | terminal region determination | biological process | GO:0007362 | 17 | 6 | 12345 | 1129 | csw,rl,trk,tor,fs(1)N,fs(1)M3 | | 2.959e-03 | -5.82 | Mlc1 (Myosin alkali light chain 1) | protein interactions | 43323 | 47 | 13 | 9204 | 1112 | Argk,Actn,up,Mf,Cyt-c-p,Adh,mol,ND-18,ATPsynCF6,Scp1,Eno,COX4,ATPsynB | | 2.973e-03 | -5.82 | exu (exuperantia) | protein interactions | 37345 | 22 | 8 | 9204 | 1112 | CG4407,yps,cup,BicC,bcd,CG10880,eIF4E1,Cbp80 | | 2.973e-03 | -5.82 | CG13319 (uncharacterized protein) | protein interactions | 36423 | 22 | 8 | 9204 | 1112 | Prosalpha5,Pomp,Prosbeta4,Prosbeta3,Prosbeta6,Prosalpha3,Prosbeta5,Prosalpha4 | | 2.973e-03 | -5.82 | CG5506 (uncharacterized protein) | protein interactions | 39992 | 22 | 8 | 9204 | 1112 | Adh,CG32473,Pgk,Argk,GlyP,Fer1HCH,Scp1,Pglym78 | | 2.973e-03 | -5.82 | Opa1 (Optic atrophy 1) | protein interactions | 36578 | 22 | 8 | 9204 | 1112 | Mitofilin,RpLP1,cnn,park,ATPsynbeta,Marf,Tom40,Pink1 | | 2.976e-03 | -5.82 | endoplasmic reticulum membrane | cellular component | GO:0005789 | 217 | 33 | 11940 | 1105 | CG10425,sip3,CG5510,Atf6,CG14671,Seipin,sws,CG5969,EMC8-9,EMC3,PIG-H,PIG-C,Dd,Mctp,CG32640,CG33977,mol,alt,Efr,CG31523,CG32441,CG8311,SREBP,CG10470,ste24a,CG8112,SERCA,Desat1,bif,CG17270,CG4293,Sec22,CG8974 | | 2.980e-03 | -5.82 | MORF\_UBE2I | MSigDB lists | MORF\_UBE2I | 133 | 33 | 4161 | 644 | RpL11,eIF3m,RpL24,Pp2A-29B,ND-30,lwr,RpS24,Dd,RpL30,Phb2,RpS7,Prosbeta6,not,heph,CSN6,Idh3b,Nap1,RpL9,RpS27A,Pgk,mEFTu1,Top1,RpL13,Chrac-14,mts,Prosbeta3,eIF4A,LysRS,Rpn12,RpS13,RpS30,eIF3i,levy | | 2.998e-03 | -5.81 | CHIANG\_LIVER\_CANCER\_SUBCLASS\_UNANNOTATED\_DN | MSigDB lists | CHIANG\_LIVER\_CANCER\_SUBCLASS\_UNANNOTATED\_DN | 93 | 25 | 4161 | 644 | Arpc1,Prosalpha5,Mo25,Cdk1,Jafrac2,Stip1,Hat1,Pgk,Chrac-14,Chd64,mad2,mRpL13,Rpn12,RpS16,alc,CG6767,mRpS23,CG8204,Ts,RfC38,Stoml2,tko,CG11089,Fdh,CG10418 | | 3.002e-03 | -5.81 | Droj2 (DnaJ-like-2) | protein interactions | 41646 | 58 | 15 | 9204 | 1112 | CCT6,SERCA,Rab5,cathD,Pi3K92E,tub,ScsbetaA,CG17260,Mcad,Pdk1,wal,GCS2beta,Cp1,CCT3,Sec15 | | 3.004e-03 | -5.81 | protein modification by small protein removal | biological process | GO:0070646 | 61 | 13 | 12345 | 1129 | trbd,otu,Uch,scny,Sgf11,CSN6,Usp16-45,Uch-L5,faf,Usp10,not,velo,Usp39 | | 3.022e-03 | -5.80 | cellular nitrogen compound catabolic process | biological process | GO:0044270 | 153 | 25 | 12345 | 1129 | CG11486,cu,smg,Drep1,fal,Dcr-1,LSm3,Rga,Edc3,twin,DhpD,Cbp80,CG17768,Trf4-2,gw,cN-IIIB,Pym,armi,CG11164,CG31759,Not10,AOX3,Ho,Rrp40,Tengl4 | | 3.041e-03 | -5.80 | WANG\_TUMOR\_INVASIVENESS\_UP | MSigDB lists | WANG\_TUMOR\_INVASIVENESS\_UP | 180 | 42 | 4161 | 644 | CG8726,NHP2,Pi3K92E,CG4511,RpS7,CG17593,Ubc4,Pka-R1,Pex13,aay,Kap3,RpS16,ari-1,CG8728,RpL30,Rpn3,Stip1,CCT3,ND-B18,Eno,CCT5,Clc,CG11807,Hsc70-3,Prosalpha3,Fdh,DppIII,Karybeta3,RpL13A,RpS30,RpLP0,ND-49,Gmer,Pen,kcc,cact,SdhA,Bin1,Nap1,SERCA,Sply,Fkbp59 | | 3.054e-03 | -5.79 | GO\_INTRACELLULAR\_RECEPTOR\_SIGNALING\_PATHWAY | MSigDB lists | GO\_INTRACELLULAR\_RECEPTOR\_SIGNALING\_PATHWAY | 42 | 14 | 4161 | 644 | MED24,Cdk7,Fkbp59,EcR,MED4,CycE,Traf6,RpS27A,cact,CG11710,Uev1A,RpL40,Pdk,Tip60 | | 3.054e-03 | -5.79 | SCHLOSSER\_SERUM\_RESPONSE\_AUGMENTED\_BY\_MYC | MSigDB lists | SCHLOSSER\_SERUM\_RESPONSE\_AUGMENTED\_BY\_MYC | 42 | 14 | 4161 | 644 | CG9911,Tip60,UQCR-Q,RhoGDI,ND-51,Cul5,Lsm12,Idh3b,Pp2A-29B,PpD3,CG8188,CG1847,MED24,CkIIalpha | | 3.054e-03 | -5.79 | GO\_TUMOR\_NECROSIS\_FACTOR\_MEDIATED\_SIGNALING\_PATHWAY | MSigDB lists | GO\_TUMOR\_NECROSIS\_FACTOR\_MEDIATED\_SIGNALING\_PATHWAY | 42 | 14 | 4161 | 644 | Rpn3,RpL40,Prosbeta6,Prosbeta5,Rpn2,RpS27A,Prosalpha5,Rpn12,Rpt2,Prosbeta4,Prosalpha4,Prosbeta3,cl,Prosalpha3 | | 3.054e-03 | -5.79 | GSE3920\_IFNB\_VS\_IFNG\_TREATED\_ENDOTHELIAL\_CELL\_UP | MSigDB lists | GSE3920\_IFNB\_VS\_IFNG\_TREATED\_ENDOTHELIAL\_CELL\_UP | 42 | 14 | 4161 | 644 | RpL38,AGBE,CCT5,cyc,eEF1beta,RpL12,Pgk,mRpS5,RpL23,RpL37a,isoQC,Pi4KIIalpha,CG1812,RpL24 | | 3.054e-03 | -5.79 | GSE22886\_IGM\_MEMORY\_BCELL\_VS\_BM\_PLASMA\_CELL\_UP | MSigDB lists | GSE22886\_IGM\_MEMORY\_BCELL\_VS\_BM\_PLASMA\_CELL\_UP | 42 | 14 | 4161 | 644 | RpL12,CG9485,Pi3K92E,RfC38,CCT5,RpS7,RpL26,CG4390,GatA,RpS28b,RpS2,RpL23,Rbf,RpL11 | | 3.056e-03 | -5.79 | Spry regulation of FGF signaling | REACTOME pathways | R-DME-1295596 | 15 | 7 | 3923 | 574 | RpL40,mts,RpS27A,csw,Cbl,rl,Pp2A-29B | | 3.083e-03 | -5.78 | transition metal ion binding | molecular function | GO:0046914 | 720 | 88 | 12176 | 1119 | Hr78,CG14516,CG9581,CG11134,Cbl,CG32473,CG11710,Hnf4,CAH2,PGRP-LE,CG9684,CG31053,CSN6,CG11902,nos,CG6923,bor,Sod1,CG14435,Cyp6u1,glob1,ari-1,stw,CG8974,Rpn3,CG9925,CG17328,SmydA-1,Fer1HCH,l(3)mbt,Desat1,Shmt,Marc,CG13344,grsm,Cyp6a19,nej,Usp39,Clp,sip3,Topors,G9a,CG11590,Iru,shop,mal,CG10492,DhpD,SelR,Cyp6a22,CG4730,isoQC,SP1029,ph-d,egg,CG15141,Usp16-45,d4,dx,Sod3,EcR,DNApol-epsilon255,CG6813,RpS29,GluRS-m,alph,CG9941,Cyp6v1,mre11,Rpn2,Scox,Traf6,lt,QC,Sec23,AOX3,CG2678,Pex10,HDAC6,CAH7,Fdh,Dip-B,Pex12,l(3)73Ah,not,Kcmf1,CG17260,CG10654 | | 3.084e-03 | -5.78 | GO\_CELLULAR\_RESPONSE\_TO\_LIPID | MSigDB lists | GO\_CELLULAR\_RESPONSE\_TO\_LIPID | 108 | 28 | 4161 | 644 | Moe,bys,CycB,cact,crq,CG11710,rl,Ac3,LpR2,p38a,AIF,CycE,Fkbp59,SREBP,Wnt10,rept,dnc,Tspo,Atpalpha,Tip60,Pdk,ACC,Traf6,EcR,MED4,Prip,MED24,Cdk7 | | 3.089e-03 | -5.78 | MODULE\_50 | MSigDB lists | MODULE\_50 | 11 | 6 | 4161 | 644 | Prosalpha5,Prosalpha3,Prosbeta3,Prosbeta5,Prosbeta6,Prosbeta4 | | 3.089e-03 | -5.78 | GO\_POLYSACCHARIDE\_CATABOLIC\_PROCESS | MSigDB lists | GO\_POLYSACCHARIDE\_CATABOLIC\_PROCESS | 11 | 6 | 4161 | 644 | Pfk,CG9485,CG8475,GlyP,Pgm1,Gyg | | 3.089e-03 | -5.78 | MODULE\_249 | MSigDB lists | MODULE\_249 | 11 | 6 | 4161 | 644 | Mtpalpha,CG10932,isoQC,Thiolase,Hat1,yip2 | | 3.096e-03 | -5.78 | carbon-oxygen lyase activity | molecular function | GO:0016835 | 54 | 12 | 12176 | 1119 | CG11134,CAH7,Echs1,Thiolase,CG4598,Pcd,Mtpalpha,CG4594,Eno,Irp-1B,CAH2,pr | | 3.099e-03 | -5.78 | Mcad (Medium-chain acyl-CoA dehydrogenase) | protein interactions | 38864 | 93 | 21 | 9204 | 1112 | Tspo,Vha68-2,r,GCS2beta,Surf4,Cp1,mEFTu1,wal,ScsbetaA,Atpalpha,cathD,Droj2,Rab5,CG1416,CCT3,Pp2A-29B,Pdk1,Kap-alpha3,CG10425,CCT6,SERCA | | 3.114e-03 | -5.77 | maternal determination of anterior/posterior axis, embryo | biological process | GO:0008358 | 98 | 18 | 12345 | 1129 | armi,psq,gus,Grip128,osk,bcd,Grip71,spn-E,msps,Moe,Tm1,mael,Lkb1,Grip75,cnc,swa,exu,nos | | 3.118e-03 | -5.77 | Ethanol oxidation | REACTOME pathways | R-DME-71384 | 3 | 3 | 3923 | 574 | Aldh,Fdh,AcCoAS | | 3.118e-03 | -5.77 | Signaling by NODAL | REACTOME pathways | R-DME-1181150 | 3 | 3 | 3923 | 574 | Med,Smox,foxo | | 3.118e-03 | -5.77 | Beta oxidation of lauroyl-CoA to decanoyl-CoA-CoA | REACTOME pathways | R-DME-77310 | 3 | 3 | 3923 | 574 | Thiolase,Echs1,Mtpalpha | | 3.141e-03 | -5.76 | regulation of cellular component organization | biological process | GO:0051128 | 791 | 95 | 12345 | 1129 | Synj,CycE,Arpc2,spn-E,Sec23,Hip1,Crag,Med,bun,cup,Rab5,HP1e,raptor,foxo,Cul5,trr,Pld,Rbf,Pdk1,Imp,Tlk,O-fut1,piwi,Lkb1,Zwilch,bru1,pgc,bif,enc,Archease,crq,Tip60,Pink1,spz,APC7,Mtl,Cdk5,Pi3K92E,CG11486,nej,stai,Nprl2,mael,BigH1,Tm1,tkv,ovo,capt,SkpA,Abi,Arpc1,msps,psq,Arpc4,Seipin,cana,Utx,eIF4E1,SREBP,shv,esc,mad2,nos,exu,wapl,ago,scny,cnc,Vps16B,Mekk1,Marf,lt,Bap170,Cenp-C,HP1c,Rab10,Moe,dgt4,Mos,Tao,ssh,G9a,tws,Cul2,Cbl,egg,cnn,park,Mitf,Pax,wash,Sod1,Stat92E,dco,swa | | 3.157e-03 | -5.76 | Prosbeta7 (Proteasome beta7 subunit) | protein interactions | 40639 | 42 | 12 | 9204 | 1112 | Prosbeta5,Prosalpha4,Prosbeta4,Rpt2,Prosalpha3,Prosbeta6,Prosbeta3,AnxB10,Rpn13,Prosalpha5,Pomp,Rpn2 | | 3.171e-03 | -5.75 | establishment or maintenance of microtubule cytoskeleton polarity | biological process | GO:0030951 | 41 | 10 | 12345 | 1129 | msps,osk,jvl,heph,capt,cnc,Kap3,Lkb1,Pka-R1,mael | | 3.192e-03 | -5.75 | GSE14000\_TRANSLATED\_RNA\_VS\_MRNA\_4H\_LPS\_DC\_DN | MSigDB lists | GSE14000\_TRANSLATED\_RNA\_VS\_MRNA\_4H\_LPS\_DC\_DN | 51 | 16 | 4161 | 644 | CG4872,Wnt10,RpS27,RpS28b,Lsm12,CG2924,pr,CG6665,RpL12,UQCR-Q,eEF1beta,RpL30,l(3)73Ah,RpLP0,dco,CHOp24 | | 3.192e-03 | -5.75 | GO\_INNER\_MITOCHONDRIAL\_MEMBRANE\_PROTEIN\_COMPLEX | MSigDB lists | GO\_INNER\_MITOCHONDRIAL\_MEMBRANE\_PROTEIN\_COMPLEX | 51 | 16 | 4161 | 644 | ND-49,CG11110,ND-SGDH,levy,ND-B22,Cyp1,ND-23,ND-51,Mitofilin,ND-39,COX5A,ND-B18,SdhA,ND-75,ND-24,ND-30 | | 3.207e-03 | -5.74 | GO\_REGULATION\_OF\_CATABOLIC\_PROCESS | MSigDB lists | GO\_REGULATION\_OF\_CATABOLIC\_PROCESS | 267 | 58 | 4161 | 644 | Uch-L5,mad2,STUB1,AIF,Cdk1,crq,Sec22,RpS27A,Prosalpha5,Exo70,CG30496,Idh,shrb,regucalcin,cyc,SkpA,Rpn12,CG1812,HDAC6,Rpn2,Prosbeta3,Cisd2,Rpt2,Prosalpha4,Gapdh1,Cul1,Rpn3,Moe,Pglym78,Edem1,Gclc,pont,CycB,Cdk5,Prosbeta5,Smox,Prosalpha3,SREBP,Wnt10,Prosbeta4,Med,Tspo,Atg7,RpL40,Snx6,Ubc4,Prosbeta6,CG6766,Vha13,APC7,ABCD,ari-1,Atg6,dco,Tlk,CG30291,Vha36-1,CkIIalpha | | 3.257e-03 | -5.73 | MODULE\_77 | MSigDB lists | MODULE\_77 | 18 | 8 | 4161 | 644 | ND-39,ND-49,ND-SGDH,ND-B18,ND-24,ND-51,ND-23,ND-30 | | 3.257e-03 | -5.73 | GO\_NEGATIVE\_REGULATION\_OF\_TRANSFORMING\_GROWTH\_FACTOR\_BETA\_RECEPTOR\_SIGNALING\_PATHWAY | MSigDB lists | GO\_NEGATIVE\_REGULATION\_OF\_TRANSFORMING\_GROWTH\_FACTOR\_BETA\_RECEPTOR\_SIGNALING\_PATHWAY | 18 | 8 | 4161 | 644 | Snx6,RpL40,Pdk1,STUB1,RpS27A,Smox,Hsc70-3,snz | | 3.257e-03 | -5.73 | GO\_CELLULAR\_CARBOHYDRATE\_CATABOLIC\_PROCESS | MSigDB lists | GO\_CELLULAR\_CARBOHYDRATE\_CATABOLIC\_PROCESS | 18 | 8 | 4161 | 644 | Pgm1,Gyg,Gk2,Treh,GlyP,Pfk,CG9485,CG8475 | | 3.259e-03 | -5.73 | MODULE\_236 | MSigDB lists | MODULE\_236 | 8 | 5 | 4161 | 644 | SREBP,crq,LpR2,S2P,Nmt | | 3.259e-03 | -5.73 | REACTOME\_DOWNREGULATION\_OF\_SMAD2\_3\_SMAD4\_TRANSCRIPTIONAL\_ACTIVITY | MSigDB lists | REACTOME\_DOWNREGULATION\_OF\_SMAD2\_3\_SMAD4\_TRANSCRIPTIONAL\_ACTIVITY | 8 | 5 | 4161 | 644 | RpS27A,faf,Smox,Med,RpL40 | | 3.259e-03 | -5.73 | BIOCARTA\_VIP\_PATHWAY | MSigDB lists | BIOCARTA\_VIP\_PATHWAY | 8 | 5 | 4161 | 644 | Pka-R1,Pka-R2,CanA-14F,cact,Pp2B-14D | | 3.259e-03 | -5.73 | GO\_CYTOPLASMIC\_PATTERN\_RECOGNITION\_RECEPTOR\_SIGNALING\_PATHWAY | MSigDB lists | GO\_CYTOPLASMIC\_PATTERN\_RECOGNITION\_RECEPTOR\_SIGNALING\_PATHWAY | 8 | 5 | 4161 | 644 | RpL40,Uev1A,Traf6,RpS27A,cact | | 3.259e-03 | -5.73 | GO\_PROTEIN\_PHOSPHATASE\_TYPE\_2A\_COMPLEX | MSigDB lists | GO\_PROTEIN\_PHOSPHATASE\_TYPE\_2A\_COMPLEX | 8 | 5 | 4161 | 644 | tws,mts,Cyt-c-p,Cka,Pp2A-29B | | 3.259e-03 | -5.73 | GAZIN\_EPIGENETIC\_SILENCING\_BY\_KRAS | MSigDB lists | GAZIN\_EPIGENETIC\_SILENCING\_BY\_KRAS | 8 | 5 | 4161 | 644 | Pdk1,rl,esc,escl,asf1 | | 3.259e-03 | -5.73 | GO\_NUCLEOTIDE\_BINDING\_DOMAIN\_LEUCINE\_RICH\_REPEAT\_CONTAINING\_RECEPTOR\_SIGNALING\_PATHWAY | MSigDB lists | GO\_NUCLEOTIDE\_BINDING\_DOMAIN\_LEUCINE\_RICH\_REPEAT\_CONTAINING\_RECEPTOR\_SIGNALING\_PATHWAY | 8 | 5 | 4161 | 644 | RpS27A,RpL40,Uev1A,Traf6,cact | | 3.265e-03 | -5.72 | GSE13485\_DAY1\_VS\_DAY21\_YF17D\_VACCINE\_PBMC\_DN | MSigDB lists | GSE13485\_DAY1\_VS\_DAY21\_YF17D\_VACCINE\_PBMC\_DN | 79 | 22 | 4161 | 644 | CG3226,Mcad,nmdyn-D7,ND-SGDH,ScsbetaA,eIF4E1,Rrp40,SkpA,CG4502,CG4511,RpL38,CG14894,Fdh,CG7789,Prosalpha5,MED20,Sec22,r-l,CCT5,CG16979,CG9391,Hat1 | | 3.265e-03 | -5.72 | GO\_NUCLEOSIDE\_PHOSPHATE\_BIOSYNTHETIC\_PROCESS | MSigDB lists | GO\_NUCLEOSIDE\_PHOSPHATE\_BIOSYNTHETIC\_PROCESS | 79 | 22 | 4161 | 644 | Ppcdc,Scsalpha1,Pfas,r-l,kri,Ac3,Shmt,nmdyn-D6,CG33156,Dhfr,CG6767,Prip,AMPdeam,nmdyn-D7,LysRS,Nadsyn,Ts,Stoml2,r,ACC,CG4407,CG11089 | | 3.271e-03 | -5.72 | ATPsynD (ATP synthase, subunit D) | protein interactions | 42291 | 27 | 9 | 9204 | 1112 | ATPsynbeta,ATPsynO,ATPsynG,ATPsynCF6,blw,ATPsynF,ATPsyndelta,ATPsynE,ATPsynB | | 3.286e-03 | -5.72 | ubiquitinyl hydrolase activity | molecular function | GO:0101005 | 41 | 10 | 12176 | 1119 | Uch,Usp16-45,not,Uch-L5,faf,scny,Usp39,trbd,otu,Usp10 | | 3.295e-03 | -5.72 | GO\_CYTOSOLIC\_LARGE\_RIBOSOMAL\_SUBUNIT | MSigDB lists | GO\_CYTOSOLIC\_LARGE\_RIBOSOMAL\_SUBUNIT | 38 | 13 | 4161 | 644 | RpL37a,RpL11,RpL23,RpL30,RpL13A,RpL24,RpLP0,RpL26,RpL9,RpL38,RpL40,RpL12,RpL13 | | 3.295e-03 | -5.72 | MOOTHA\_VOXPHOS | MSigDB lists | MOOTHA\_VOXPHOS | 38 | 13 | 4161 | 644 | COX5A,Cyt-c-p,ND-SGDH,ND-49,ND-39,UQCR-Q,ND-30,ND-23,SdhA,levy,ND-B18,CG3803,ND-75 | | 3.295e-03 | -5.72 | GO\_REGENERATION | MSigDB lists | GO\_REGENERATION | 38 | 13 | 4161 | 644 | CycB,r,CG31715,p38a,Cdk1,Ts,Tspo,CG11089,regucalcin,Dhfr,Wnt10,CycE,CG6767 | | 3.302e-03 | -5.71 | Jupiter (jupiter) | protein interactions | 41392 | 37 | 11 | 9204 | 1112 | CG10638,CG14715,cib,dl,AnxB10,nmd,PGRP-LE,Rel,awd,Tak1,Bin1 | | 3.302e-03 | -5.71 | vret (vreteno) | protein interactions | 42695 | 37 | 11 | 9204 | 1112 | AGO3,CG3309,PGRP-LE,Tak1,piwi,cbs,Nepl3,jigr1,BoYb,armi,png | | 3.302e-03 | -5.71 | MED6 (Mediator complex subunit 6) | protein interactions | 44728 | 37 | 11 | 9204 | 1112 | MED27,Usp10,enc,MED4,MED20,ix,Ssdp,Imp,Larp4B,CG17002,Ptip | | 3.323e-03 | -5.71 | vesicle-mediated transport | biological process | GO:0016192 | 462 | 60 | 12345 | 1129 | CHOp24,Rab10,crq,EcR,Arl5,lt,Mekk1,Exo70,GCC88,dx,Gga,Sec15,CG5447,Syx13,Snx6,AP-1-2beta,X11L,fwe,Stam,CG5510,wash,shrb,Sec10,faf,Mtl,Rep,Blos2,Gdi,CG4293,dco,Vsp37A,Tomosyn,Rab32,cbs,Sec22,csw,Vha68-2,Gmap,Nlg1,lqfR,Arf51F,CG5734,Synj,Clc,opm,Vamp7,CHMP2B,Atg6,disp,Rab5,mop,CanB2,Ubc6,CG9132,sau,PAPLA1,MICAL-like,Vps16B,CG31729,Trs33 | | 3.329e-03 | -5.71 | MODULE\_18 | MSigDB lists | MODULE\_18 | 139 | 34 | 4161 | 644 | PpD3,Arpc1,Smox,Sec23,Gyg,Rcd5,Prosalpha3,Chd64,msps,Arpc4,PIG-C,CG13630,Hat1,pr,CycB,nrv1,CkIIalpha,Tm1,scramb1,RfC38,Ts,NHP2,AGBE,r,CG14516,mad2,Chrac-14,CCT6,Cdk1,isoQC,Cse1,ste14,Karybeta3,Tsp74F | | 3.357e-03 | -5.70 | Arl8 (ADP ribosylation factor-like 8) | protein interactions | 40961 | 32 | 10 | 9204 | 1112 | Rab5,CG1416,CCT6,SERCA,Mcad,ScsbetaA,CCT3,wal,Tspo,CG14818 | | 3.367e-03 | -5.69 | ATPase\_P-typ\_transduc\_dom\_A\_sf | interpro domains | IPR008250 | 15 | 6 | 11383 | 1233 | anne,CG31729,SERCA,Atpalpha,CG9981,ATP8A | | 3.367e-03 | -5.69 | ATPase\_P-typ\_cyto\_dom\_N | interpro domains | IPR023299 | 15 | 6 | 11383 | 1233 | anne,ATP8A,Atpalpha,CG9981,CG31729,SERCA | | 3.378e-03 | -5.69 | GSE5455\_EX\_VIVO\_VS\_POST\_24H\_INCUBATION\_MONOCYTES\_FROM\_TUMOR\_BEARING\_MOUSE\_UP | MSigDB lists | GSE5455\_EX\_VIVO\_VS\_POST\_24H\_INCUBATION\_MONOCYTES\_FROM\_TUMOR\_BEARING\_MOUSE\_UP | 65 | 19 | 4161 | 644 | CG4502,Ubc4,ND-23,CG1532,Psf3,CG11110,Rpn2,trsn,rl,Eno,Tim8,Pglym78,Liprin-beta,Chrac-14,nmdyn-D6,UQCR-Q,Pgk,milt,stas | | 3.378e-03 | -5.69 | DANG\_REGULATED\_BY\_MYC\_DN | MSigDB lists | DANG\_REGULATED\_BY\_MYC\_DN | 65 | 19 | 4161 | 644 | Npc2a,Unr,Rab10,RpS23,Nmt,Gdi,Caf1-105,LpR2,Arpc4,Uev1A,Moe,P5cr,AnxB10,ScsbetaA,Prip,Tsp74F,Synj,Tspo,Tm1 | | 3.399e-03 | -5.68 | response to ketone | biological process | GO:1901654 | 48 | 11 | 12345 | 1129 | G9a,rig,Trf2,Utx,Idh3b,MED27,Nurf-38,MED24,rept,EcR,trr | | 3.399e-03 | -5.68 | germ cell migration | biological process | GO:0008354 | 48 | 11 | 12345 | 1129 | disp,slam,gus,tkv,tor,nos,Jafrac1,stai,lwr,pgc,faf | | 3.402e-03 | -5.68 | Negative regulation of FGFR2 signaling | REACTOME pathways | R-DME-5654727 | 19 | 8 | 3923 | 574 | mts,RpL40,CG13398,Pp2A-29B,Cbl,csw,rl,RpS27A | | 3.402e-03 | -5.68 | Negative regulation of FGFR3 signaling | REACTOME pathways | R-DME-5654732 | 19 | 8 | 3923 | 574 | mts,CG13398,RpL40,Pp2A-29B,Cbl,csw,RpS27A,rl | | 3.402e-03 | -5.68 | Negative regulation of FGFR4 signaling | REACTOME pathways | R-DME-5654733 | 19 | 8 | 3923 | 574 | rl,RpS27A,Pp2A-29B,csw,Cbl,mts,RpL40,CG13398 | | 3.437e-03 | -5.67 | centrosome duplication | biological process | GO:0051298 | 35 | 9 | 12345 | 1129 | tws,SkpA,CG3437,Pp2A-29B,CCT6,Rcd5,msps,cnn,mts | | 3.463e-03 | -5.67 | GO\_AMIDE\_BIOSYNTHETIC\_PROCESS | MSigDB lists | GO\_AMIDE\_BIOSYNTHETIC\_PROCESS | 290 | 62 | 4161 | 644 | RpS7,r,nSMase,RpL38,RpS4,RpL40,tko,heph,TyrRS,Dhfr,RpS24,RpL23,CG8963,RpL30,RpS16,RpS2,mRpS23,colt,Gclc,RpS18,RpL12,CG3376,mRpS5,mRpL10,RpL37a,mRpL17,eIF3m,RpL11,CG7414,RpS27,RpL24,RpL26,tzn,eIF3i,Fur2,GluRS-m,CysRS,CG11089,mRpL18,eIF4A,LysRS,RpL13A,eIF4E1,mRpL13,RpLP0,RpS30,RpS13,Nurf-38,mRpS10,Tfb4,SerRS-m,mEFTu1,RpL13,eEF1beta,mRpL16,TrpRS-m,mRpS21,RpS28b,GatA,RpS27A,RpS23,RpL9 | | 3.473e-03 | -5.66 | negative regulation of insulin receptor signaling pathway | biological process | GO:0046627 | 23 | 7 | 12345 | 1129 | SkpA,foxo,Cul1,Ptp61F,Pp2A-29B,STUB1,mts | | 3.473e-03 | -5.66 | bicoid mRNA localization | biological process | GO:0045450 | 23 | 7 | 12345 | 1129 | Grip71,exu,swa,cnc,msps,Grip75,Grip128 | | 3.473e-03 | -5.66 | negative regulation of cellular response to insulin stimulus | biological process | GO:1900077 | 23 | 7 | 12345 | 1129 | SkpA,Pp2A-29B,Ptp61F,STUB1,foxo,Cul1,mts | | 3.490e-03 | -5.66 | posttranscriptional gene silencing | biological process | GO:0016441 | 62 | 13 | 12345 | 1129 | Cbp80,Hsc70-3,trsn,r2d2,armi,Marf1,piwi,spn-E,Dcr-1,CG6967,CG9425,gw,AGO3 | | 3.504e-03 | -5.65 | GO\_RRNA\_BINDING | MSigDB lists | GO\_RRNA\_BINDING | 34 | 12 | 4161 | 644 | RpLP0,RpS13,RpL9,RpL37a,mRpL18,RpL11,RpL23,CG7488,RpL12,mRpL16,RpS18,RpS4 | | 3.504e-03 | -5.65 | GO\_DEPHOSPHORYLATION | MSigDB lists | GO\_DEPHOSPHORYLATION | 94 | 25 | 4161 | 644 | Cyt-c-p,Ptp10D,Pp2B-14D,aay,CG7115,mts,CG17294,Dd,csw,Synj,Ublcp1,CanB2,PpV,Ptp61F,Acph-1,Pp2A-29B,CG32549,PpD3,CG7789,CanA-14F,Pglym78,CG9391,alph,l(1)G0196,Ptpmeg | | 3.508e-03 | -5.65 | eIF4B (eukaryotic translation initiation factor 4B) | protein interactions | 3355041 | 18 | 7 | 9204 | 1112 | cad,TyrRS,Chd64,Gip,Mes2,CG11771,capt | | 3.508e-03 | -5.65 | mael (maelstrom) | protein interactions | 40489 | 18 | 7 | 9204 | 1112 | rl,gammaTub37C,bam,CG9911,Dcr-1,cnn,msps | | 3.508e-03 | -5.65 | CG8117 (uncharacterized protein) | protein interactions | 32499 | 18 | 7 | 9204 | 1112 | awd,RpII140,TfIIFalpha,cib,Tim8,CG14715,regucalcin | | 3.519e-03 | -5.65 | wal (walrus) | protein interactions | 36252 | 100 | 22 | 9204 | 1112 | CG1416,csw,PyK,Rab5,Droj2,cathD,Atpalpha,ScsbetaA,Mcad,Rpt2,ago,mEFTu1,GCS2beta,Vha68-2,Tspo,CCT6,SERCA,Kap-alpha3,CG10425,blw,Pp2A-29B,CCT3 | | 3.525e-03 | -5.65 | Glucose metabolism | REACTOME pathways | R-DME-70326 | 59 | 17 | 3923 | 574 | Pgm1,CG8475,UGP,Pglym78,Gyg,Pgk,Gapdh1,Pfk,Pfrx,CG6650,GlyP,Got1,CG9485,PyK,sea,Eno,AGBE | | 3.527e-03 | -5.65 | Idh3a (Isocitrate dehydrogenase 3a) | protein interactions | 32940 | 82 | 19 | 9204 | 1112 | CG10425,Kap-alpha3,RpLP0,Actn,SERCA,CCT6,CG31729,CCT3,CG5028,Pp2A-29B,ScsbetaA,Mcad,Rab5,Argk,Karybeta3,bor,r,mEFTu1,wal | | 3.528e-03 | -5.65 | positive regulation of protein metabolic process | biological process | GO:0051247 | 318 | 44 | 12345 | 1129 | ben,osk,Mekk1,G9a,gus,Tao,Mos,Fbw5,ScsbetaA,park,MED24,Pi3K92E,ari-1,tyf,SkpA,PEK,dco,Cul1,CG8963,ASPP,CG4238,Abi,fray,Rpn13,eIF3m,STUB1,raptor,Rpt2,RpLP1,Parp16,Tak1,Pdk1,Ubc6,tor,CG8009,trr,esc,Jafrac2,Exd2,Pomp,Ankle2,cort,Lkb1,Cka | | 3.536e-03 | -5.64 | GO\_OXIDOREDUCTION\_COENZYME\_METABOLIC\_PROCESS | MSigDB lists | GO\_OXIDOREDUCTION\_COENZYME\_METABOLIC\_PROCESS | 56 | 17 | 4161 | 644 | Nadsyn,Coq6,Pgm1,Gapdh1,CG33156,CG5028,CG17333,Idh3b,Nc73EF,Idh,ND-39,Pgk,Pglym78,Eno,Pfk,CG7322,Coq7 | | 3.539e-03 | -5.64 | organic hydroxy compound metabolic process | biological process | GO:1901615 | 163 | 26 | 12345 | 1129 | CG3842,Sptr,CG7840,scu,Pcd,pr,Fdh,Gk2,l(1)G0196,Gk1,CG9391,Cdk5,foxo,dl,CG5028,Adh,Idh3b,Idh,tzn,Pdxk,CG8112,Synj,sgll,Marf,Sp7,CG17270 | | 3.539e-03 | -5.64 | GSE36476\_CTRL\_VS\_TSST\_ACT\_40H\_MEMORY\_CD4\_TCELL\_OLD\_DN | MSigDB lists | GSE36476\_CTRL\_VS\_TSST\_ACT\_40H\_MEMORY\_CD4\_TCELL\_OLD\_DN | 99 | 26 | 4161 | 644 | DppIII,RfC38,Ts,Rpb8,Cbp80,mRpL18,Psf3,alphaTub84B,Dhfr,Gapdh1,scu,grp,Prosbeta5,CycB,Stip1,CCT3,SRPK,Aldh,Pen,Pglym78,msps,Shmt,Scsalpha1,Idh,gammaTub37C,CG14516 | | 3.554e-03 | -5.64 | HAD\_sf | interpro domains | IPR023214 | 53 | 13 | 11383 | 1233 | CG9981,Dd,ATP8A,cN-IIIB,CG31729,SERCA,anne,CG17294,aay,Atpalpha,Gs1l,Ublcp1,CG32549 | | 3.576e-03 | -5.63 | GSE2770\_UNTREATED\_VS\_TGFB\_AND\_IL12\_TREATED\_ACT\_CD4\_TCELL\_48H\_DN | MSigDB lists | GSE2770\_UNTREATED\_VS\_TGFB\_AND\_IL12\_TREATED\_ACT\_CD4\_TCELL\_48H\_DN | 70 | 20 | 4161 | 644 | Wdfy2,TrpRS-m,Acph-1,RpL24,CG12929,pr,Tom40,Stip1,CG10344,p38a,CG9911,muskelin,eIF4A,Sec10,alc,RpS7,nSMase,cyc,Tip60,Pdk | | 3.576e-03 | -5.63 | GO\_REGULATION\_OF\_PROTEIN\_UBIQUITINATION\_INVOLVED\_IN\_UBIQUITIN\_DEPENDENT\_PROTEIN\_CATABOLIC\_PROCESS | MSigDB lists | GO\_REGULATION\_OF\_PROTEIN\_UBIQUITINATION\_INVOLVED\_IN\_UBIQUITIN\_DEPENDENT\_PROTEIN\_CATABOLIC\_PROCESS | 70 | 20 | 4161 | 644 | Prosalpha5,RpS27A,Prosalpha3,Prosbeta4,mad2,Cul1,Rpn3,Prosbeta5,CycB,Cdk1,APC7,Rpn12,Rpn2,Prosbeta3,CG30291,Prosalpha4,Rpt2,RpL40,SkpA,Prosbeta6 | | 3.577e-03 | -5.63 | autophagic cell death | biological process | GO:0048102 | 29 | 8 | 12345 | 1129 | rept,EcR,crq,MED24,Idh3b,DCAF12,Trf2,Utx | | 3.577e-03 | -5.63 | intestinal stem cell homeostasis | biological process | GO:0036335 | 29 | 8 | 12345 | 1129 | PHGPx,Tao,bun,SERCA,cnc,sip3,Hsc70-3,Pi3K92E | | 3.594e-03 | -5.63 | GO\_PHOSPHATASE\_COMPLEX | MSigDB lists | GO\_PHOSPHATASE\_COMPLEX | 22 | 9 | 4161 | 644 | CanB2,Pp2B-14D,Cka,Pp2A-29B,Cyt-c-p,Dd,mts,tws,CanA-14F | | 3.594e-03 | -5.63 | REACTOME\_NFKB\_AND\_MAP\_KINASES\_ACTIVATION\_MEDIATED\_BY\_TLR4\_SIGNALING\_REPERTOIRE | MSigDB lists | REACTOME\_NFKB\_AND\_MAP\_KINASES\_ACTIVATION\_MEDIATED\_BY\_TLR4\_SIGNALING\_REPERTOIRE | 22 | 9 | 4161 | 644 | mts,Pp2A-29B,RpS27A,Traf6,rl,cact,p38a,Cdk1,RpL40 | | 3.594e-03 | -5.63 | GO\_REGULATION\_OF\_ATPASE\_ACTIVITY | MSigDB lists | GO\_REGULATION\_OF\_ATPASE\_ACTIVITY | 22 | 9 | 4161 | 644 | nrv1,shv,Tm1,scramb1,regucalcin,Chd64,CG8726,Cyp1,CG1416 | | 3.594e-03 | -5.63 | GNF2\_GLTSCR2 | MSigDB lists | GNF2\_GLTSCR2 | 22 | 9 | 4161 | 644 | RpL38,eEF1beta,RpL12,RpL13,RpL30,RpS16,RpL13A,RpS30,RpS27 | | 3.594e-03 | -5.63 | GO\_GLUCAN\_METABOLIC\_PROCESS | MSigDB lists | GO\_GLUCAN\_METABOLIC\_PROCESS | 22 | 9 | 4161 | 644 | RpL40,Pfk,CG9485,CG8475,GlyP,AGBE,RpS27A,Gyg,Pgm1 | | 3.594e-03 | -5.63 | KYNG\_ENVIRONMENTAL\_STRESS\_RESPONSE\_UP | MSigDB lists | KYNG\_ENVIRONMENTAL\_STRESS\_RESPONSE\_UP | 22 | 9 | 4161 | 644 | Unr,Trxr-1,sau,PDCD-5,Sod1,CG1890,eIF4E1,Cyt-c-p,Jafrac2 | | 3.594e-03 | -5.63 | GO\_REGULATION\_OF\_ERBB\_SIGNALING\_PATHWAY | MSigDB lists | GO\_REGULATION\_OF\_ERBB\_SIGNALING\_PATHWAY | 22 | 9 | 4161 | 644 | Clc,RpS27A,Iru,Snx6,Su(H),RpL40,Stam,Ptp61F,CG4709 | | 3.605e-03 | -5.63 | small nucleolar ribonucleoprotein complex assembly | biological process | GO:0000491 | 8 | 4 | 12345 | 1129 | rept,CG8204,pont,CG10055 | | 3.605e-03 | -5.63 | protein import into peroxisome matrix | biological process | GO:0016558 | 8 | 4 | 12345 | 1129 | Pex13,Topors,Pex12,Pex10 | | 3.605e-03 | -5.63 | microtubule nucleation by interphase microtubule organizing center | biological process | GO:0051415 | 8 | 4 | 12345 | 1129 | Grip128,Grip84,Grip75,CG7716 | | 3.605e-03 | -5.63 | regulation of execution phase of apoptosis | biological process | GO:1900117 | 8 | 4 | 12345 | 1129 | Rbf,lin-52,eIF3m,E2f1 | | 3.605e-03 | -5.63 | female meiosis I | biological process | GO:0007144 | 8 | 4 | 12345 | 1129 | gammaTub37C,Grip71,cort,Klp10A | | 3.605e-03 | -5.63 | positive regulation of DNA-dependent DNA replication | biological process | GO:2000105 | 8 | 4 | 12345 | 1129 | pn,otu,E2f1,SuUR | | 3.605e-03 | -5.63 | negative regulation of ERK1 and ERK2 cascade | biological process | GO:0070373 | 8 | 4 | 12345 | 1129 | Ptp61F,Cbl,alph,ATPsynD | | 3.618e-03 | -5.62 | Cdk8 (Cyclin-dependent kinase 8) | protein interactions | 39157 | 48 | 13 | 9204 | 1112 | Usp10,MED27,MED4,MED20,enc,ix,dl,E2f1,EcR,Ssdp,CycC,Ptip,CG17002 | | 3.627e-03 | -5.62 | NDK | smart domains | SM00562 | 4 | 3 | 5086 | 506 | nmdyn-D6,nmdyn-D7,awd | | 3.627e-03 | -5.62 | PLDc | smart domains | SM00155 | 4 | 3 | 5086 | 506 | CG7718,CG9248,Pld | | 3.653e-03 | -5.61 | GO\_PROTEIN\_SERINE\_THREONINE\_PHOSPHATASE\_ACTIVITY | MSigDB lists | GO\_PROTEIN\_SERINE\_THREONINE\_PHOSPHATASE\_ACTIVITY | 30 | 11 | 4161 | 644 | CanB2,alph,PpV,Ublcp1,CanA-14F,CG7115,Dd,Cyt-c-p,Pp2A-29B,Pp2B-14D,PpD3 | | 3.662e-03 | -5.61 | HEAT\_type\_2 | interpro domains | IPR021133 | 7 | 4 | 11383 | 1233 | CG1344,Karybeta3,Pp2A-29B,msps | | 3.671e-03 | -5.61 | Hsp90 protein binding | molecular function | GO:0051879 | 8 | 4 | 12176 | 1119 | Stip1,CG1416,spag,STUB1 | | 3.671e-03 | -5.61 | poly(A)-specific ribonuclease activity | molecular function | GO:0004535 | 8 | 4 | 12176 | 1119 | CG11486,cu,Rga,twin | | 3.671e-03 | -5.61 | alditol:NADP+ 1-oxidoreductase activity | molecular function | GO:0004032 | 8 | 4 | 12176 | 1119 | CG9436,CG10863,CG10638,CG2767 | | 3.671e-03 | -5.61 | ubiquitin-like modifier activating enzyme activity | molecular function | GO:0008641 | 8 | 4 | 12176 | 1119 | Atg7,Aos1,Uba5,APP-BP1 | | 3.693e-03 | -5.60 | Cyclin E associated events during G1/S transition | REACTOME pathways | R-DME-69202 | 74 | 20 | 3923 | 574 | Rpn12,Rpn3,Cul1,Prosbeta6,Cdk7,Prosbeta3,Prosalpha3,RpL40,Rpt2,Rbf,SkpA,Prosalpha4,twe,CycE,Rbf2,Prosbeta5,Rpn2,Prosbeta4,RpS27A,Prosalpha5 | | 3.693e-03 | -5.60 | REACTOME\_IRAK2\_MEDIATED\_ACTIVATION\_OF\_TAK1\_COMPLEX\_UPON\_TLR7\_8\_OR\_9\_STIMULATION | MSigDB lists | REACTOME\_IRAK2\_MEDIATED\_ACTIVATION\_OF\_TAK1\_COMPLEX\_UPON\_TLR7\_8\_OR\_9\_STIMULATION | 3 | 3 | 4161 | 644 | RpL40,Traf6,RpS27A | | 3.693e-03 | -5.60 | MCGOWAN\_RSP6\_TARGETS\_UP | MSigDB lists | MCGOWAN\_RSP6\_TARGETS\_UP | 3 | 3 | 4161 | 644 | CG8314,Jheh2,Aladin | | 3.693e-03 | -5.60 | CHUNG\_BLISTER\_CYTOTOXICITY\_DN | MSigDB lists | CHUNG\_BLISTER\_CYTOTOXICITY\_DN | 3 | 3 | 4161 | 644 | Rbf,Tomosyn,Pdk | | 3.693e-03 | -5.60 | BIOCARTA\_AKT\_PATHWAY | MSigDB lists | BIOCARTA\_AKT\_PATHWAY | 3 | 3 | 4161 | 644 | Pdk1,cact,mts | | 3.701e-03 | -5.60 | GO\_NAD\_METABOLIC\_PROCESS | MSigDB lists | GO\_NAD\_METABOLIC\_PROCESS | 26 | 10 | 4161 | 644 | Pgk,CG5028,Pglym78,Eno,Pfk,Nadsyn,Idh3b,Nc73EF,Gapdh1,CG33156 | | 3.722e-03 | -5.59 | nuclear membrane | cellular component | GO:0031965 | 35 | 9 | 11940 | 1105 | CG7655,Axs,Dd,bif,CG8009,SREBP,sbr,bru1,toc | | 3.737e-03 | -5.59 | UBQ-conjugating\_AS | interpro domains | IPR023313 | 20 | 7 | 11383 | 1233 | CG8188,CG40045,ben,Ubc6,Ubc4,lwr,UbcE2H | | 3.753e-03 | -5.59 | intracellular mRNA localization | biological process | GO:0008298 | 92 | 17 | 12345 | 1129 | mael,Tm1,Grip75,Lkb1,cnc,swa,exu,cup,CkIIalpha,Rrp40,jvl,Grip128,spn-E,Grip71,osk,wisp,msps | | 3.765e-03 | -5.58 | organic substance transport | biological process | GO:0071702 | 719 | 87 | 12345 | 1129 | Roe1,CG3077,Gdi,Rep,CG9399,Sfxn2,Oseg1,sbr,CG10344,mol,Dcr-1,Nup62,Gga,CG8728,alphaTub67C,CG11110,AP-1-2beta,CG5447,Syx13,CHOp24,CG4041,Arl5,Topors,slif,CG4572,Kap-alpha3,sea,tho2,sau,mbo,CHMP2B,Karybeta3,Rab5,Pex12,CG12773,CG9981,Sec22,Arts,CG14894,Tom40,Sec23,Crag,Arf51F,Clc,Pen,eIF3m,CG5734,yin,Gmer,Npc2a,Rab32,Vsp37A,Stam,thoc6,milt,Tsp86D,Efr,CG8004,Moe,Rab10,ATP8A,Cse1,Ugalt,colt,lt,CG31729,GAPsec,Mat89Ba,Vps16B,sip3,Nup358,Tim8,Pex13,Tango10,kcc,CG9393,opm,Atg6,CG4991,Pex10,CG4476,Tom70,CG6766,CG8314,Arpc1,scramb1,Nxf3 | | 3.808e-03 | -5.57 | Lpin (Lipin) | protein interactions | 35790 | 10 | 5 | 9204 | 1112 | CG4853,Dd,Seipin,Rtnl1,rl | | 3.808e-03 | -5.57 | armi (armitage) | protein interactions | 38427 | 10 | 5 | 9204 | 1112 | piwi,AGO3,vret,armi,shu | | 3.808e-03 | -5.57 | cellular protein localization | biological process | GO:0034613 | 558 | 70 | 12345 | 1129 | tho2,sau,Lkb1,Kap-alpha3,Topors,Arf51F,Clc,Pen,CG5734,Tom40,CG14894,Arts,Pi4KIIalpha,Sec23,Nlg1,Karybeta3,Rab5,Pex12,sw,mbo,Nup62,SuUR,sbr,CG10344,Roe1,Rep,Oseg1,Arl5,CHOp24,CG4041,CG11110,AP-1-2beta,Syx13,CG5447,X11L,Gga,Sec15,CG8728,CG9393,Pex13,Tim8,Mat89Ba,GAPsec,Vps16B,sip3,Nup358,fray,Nxf3,Tom70,CG8314,sws,CG6766,Arpc1,Pex10,opm,Atg6,Tsp86D,Stam,thoc6,milt,Pax,Actn,Rab32,Vsp37A,Cenp-C,lt,Rab10,Cse1,osk,CG8004 | | 3.814e-03 | -5.57 | intracellular vesicle | cellular component | GO:0097708 | 378 | 51 | 11940 | 1105 | AP-1-2beta,Hip1,Stam,yl,Arf51F,Vps16B,CG5510,Crag,pen-2,wash,fwe,CHOp24,CG5734,Wdfy2,Rab5,Afti,PAPLA1,lqfR,Vamp7,tkv,fdl,Pld,Vsp37A,Sec23,Lkb1,Rab10,CG31729,CG32536,opm,CHMP2B,milt,Tomosyn,Snx6,mop,Rab32,CanB2,Gga,CG9132,Gdi,CG3309,Pi4KIIalpha,Sec15,Sec22,yin,Clc,CG4293,Rep,shrb,Desat1,lt,CG6707 | | 3.814e-03 | -5.57 | cytoplasmic vesicle | cellular component | GO:0031410 | 378 | 51 | 11940 | 1105 | Wdfy2,CHOp24,CG5734,Rab5,PAPLA1,Afti,pen-2,wash,fwe,Pld,fdl,Vamp7,tkv,Sec23,Vsp37A,lqfR,yl,Stam,Hip1,AP-1-2beta,Crag,Arf51F,CG5510,Vps16B,Sec22,Gga,Gdi,CG9132,CG3309,Pi4KIIalpha,Sec15,lt,CG6707,Desat1,Rep,yin,Clc,CG4293,shrb,CHMP2B,Tomosyn,Snx6,milt,Lkb1,opm,CG32536,CG31729,Rab10,mop,Rab32,CanB2 | | 3.820e-03 | -5.57 | PX | prosite domains | PS50195 | 15 | 6 | 6243 | 694 | Pld,Snx6,CG3077,CG8726,CG11807,CG5734 | | 3.828e-03 | -5.57 | meiotic spindle organization | biological process | GO:0000212 | 42 | 10 | 12345 | 1129 | Klp10A,Grip75,gammaTub37C,SRPK,Axs,Grip71,wisp,larp,Grip128,sau | | 3.829e-03 | -5.57 | GO\_ANAPHASE\_PROMOTING\_COMPLEX\_DEPENDENT\_CATABOLIC\_PROCESS | MSigDB lists | GO\_ANAPHASE\_PROMOTING\_COMPLEX\_DEPENDENT\_CATABOLIC\_PROCESS | 61 | 18 | 4161 | 644 | Rpn3,mad2,CycB,Prosbeta5,Cdk1,RpS27A,Prosalpha5,Prosalpha3,Prosbeta4,RpL40,Prosbeta6,APC7,Rpn12,CG8188,Rpn2,Prosbeta3,Rpt2,Prosalpha4 | | 3.829e-03 | -5.57 | GSE14000\_TRANSLATED\_RNA\_VS\_MRNA\_DC\_DN | MSigDB lists | GSE14000\_TRANSLATED\_RNA\_VS\_MRNA\_DC\_DN | 61 | 18 | 4161 | 644 | PDCD-5,CG5116,CG8372,RpL26,Lsm12,Arpc1,Clp,RpL11,eEF1beta,RpL12,RpS18,RpL13A,RpS13,RpLP0,RpL30,Dd,RpL23,Arl5 | | 3.834e-03 | -5.56 | REACTOME\_CELL\_CYCLE\_MITOTIC | MSigDB lists | REACTOME\_CELL\_CYCLE\_MITOTIC | 161 | 38 | 4161 | 644 | mad2,Klp10A,Cdk1,RpS27A,Prosalpha5,Grip84,Grip128,Sld5,SkpA,Rpn2,Rpn12,Rpt2,Prosalpha4,Prosbeta3,Cdk7,mts,Cul1,msps,Rpn3,CycB,Prosbeta5,DNApol-epsilon255,Pp2A-29B,CycE,RpS27,gammaTub37C,Prosbeta4,Prosalpha3,RpL40,Tao,Prosbeta6,Ts,RfC38,dco,APC7,Rbf,Dhfr,CG12018 | | 3.850e-03 | -5.56 | protein phosphorylation | biological process | GO:0006468 | 284 | 40 | 12345 | 1129 | Ptpmeg,Lkb1,Cdk1,Tlk,Cka,grp,Mo25,Tak1,CycJ,Pdk1,tor,Coq8,Ptpmeg2,CkIIalpha,CG17698,Tfb4,Cdk7,Pdk,fray,CycE,awd,PEK,dco,p38a,CG3631,tkv,SRPK,CycB,rl,CG11486,Cdk5,Mtl,Pink1,Tao,png,Mos,CG4041,CG8726,Ptp61F,Mekk1 | | 3.866e-03 | -5.56 | GSE2405\_HEAT\_KILLED\_VS\_LIVE\_A\_PHAGOCYTOPHILUM\_STIM\_NEUTROPHIL\_9H\_UP | MSigDB lists | GSE2405\_HEAT\_KILLED\_VS\_LIVE\_A\_PHAGOCYTOPHILUM\_STIM\_NEUTROPHIL\_9H\_UP | 80 | 22 | 4161 | 644 | Pglym78,pen-2,shrb,CG9143,Mat89Ba,Ppcdc,Arf51F,RpL37a,Nmt,COX5A,Axs,CG6766,levy,Arl6IP1,Adi1,cathD,Fdh,Jafrac1,Arl5,aay,shv,CG1890 | | 3.896e-03 | -5.55 | Prosbeta4R2 (Proteasome beta4 subunit-related 2) | protein interactions | 33451 | 14 | 6 | 9204 | 1112 | Prosbeta3,Prosbeta6,Prosalpha3,Pomp,Prosalpha5,Prosalpha4 | | 3.900e-03 | -5.55 | REACTOME\_ANTIGEN\_PROCESSING\_CROSS\_PRESENTATION | MSigDB lists | REACTOME\_ANTIGEN\_PROCESSING\_CROSS\_PRESENTATION | 43 | 14 | 4161 | 644 | Rpn12,Prosalpha5,Rpn2,RpS27A,Prosalpha3,Prosbeta3,Prosalpha4,Prosbeta4,Rpt2,RpL40,Rpn3,ERp60,Prosbeta5,Prosbeta6 | | 3.900e-03 | -5.55 | REACTOME\_ER\_PHAGOSOME\_PATHWAY | MSigDB lists | REACTOME\_ER\_PHAGOSOME\_PATHWAY | 43 | 14 | 4161 | 644 | Rpn12,RpS27A,Rpn2,Prosalpha5,Prosbeta3,Prosalpha3,Rpt2,Prosbeta4,Prosalpha4,RpL40,ERp60,Rpn3,Prosbeta5,Prosbeta6 | | 3.907e-03 | -5.54 | png (pan gu) | protein interactions | 31084 | 43 | 12 | 9204 | 1112 | mts,tral,CycE,Sod3,vret,CycB,BicC,Kap3,bcd,plu,png,CG4730 | | 3.938e-03 | -5.54 | GO\_ESTABLISHMENT\_OF\_PROTEIN\_LOCALIZATION | MSigDB lists | GO\_ESTABLISHMENT\_OF\_PROTEIN\_LOCALIZATION | 567 | 110 | 4161 | 644 | ste14,rept,CG4041,CG31729,opm,cyc,Arl6IP1,Fur2,RpS30,RpS13,Cse1,RpLP0,RpL13A,RpL13,ERp60,Pen,LpR2,Surf4,RpS28b,Faf2,RpL9,RpS27A,Rab32,Exo70,snz,Atg4b,Atg7,mbo,Snx6,Rab5,Ranbp21,RpS2,Atg6,Oseg1,Phb2,CG11110,Pp2B-14D,Vha36-1,CG9132,RpL30,CG8728,Pex10,CG3077,CG46338,Tim8,CG5510,CHMP2B,RpL24,Syx13,CanA-14F,RpL37a,Pex12,Stam,CG5734,Su(H),Arts,GCC88,CHOp24,AP-1-2beta,Karybeta3,HDAC6,CG9393,mino,sau,Vsp37A,CG2100,Tsp86D,mol,milt,Tom40,cact,rl,RpS23,Sec22,Rab10,shrb,Tspo,RpL40,RpS7,Rep,RpS4,RpL38,Vha13,RpS16,lt,Aladin,GAPsec,Sec10,Pex13,Sec15,RpS24,CG1847,RpL23,Marf,RpL12,CG8314,Kap-alpha3,Cdk5,Gdi,CG8004,RpS18,RpS27,Clc,RpL26,Vamp7,CG9773,RpL11,Arf51F,Tomosyn,Sec23 | | 3.964e-03 | -5.53 | GSE13547\_2H\_VS\_12\_H\_ANTI\_IGM\_STIM\_BCELL\_DN | MSigDB lists | GSE13547\_2H\_VS\_12\_H\_ANTI\_IGM\_STIM\_BCELL\_DN | 85 | 23 | 4161 | 644 | CG10418,l(1)G0004,RfC38,NHP2,Nurf-38,nmdyn-D7,CG9393,CG8188,mRpL18,CG11899,mad2,Shmt,Hat1,Cdk1,CG14715,CG7655,Prosalpha5,Lsm12,Surf4,gammaTub37C,Grip84,Prosbeta4,LSm3 | | 3.967e-03 | -5.53 | GSE32423\_MEMORY\_VS\_NAIVE\_CD8\_TCELL\_DN | MSigDB lists | GSE32423\_MEMORY\_VS\_NAIVE\_CD8\_TCELL\_DN | 52 | 16 | 4161 | 644 | Jheh2,obe,Chd64,Ssdp,Aldh,CG46338,dco,isoQC,ScsbetaA,CG8963,csw,CG7115,GCC88,ATP8A,CG17454,AGBE | | 3.967e-03 | -5.53 | GSE26030\_UNSTIM\_VS\_RESTIM\_TH1\_DAY15\_POST\_POLARIZATION\_UP | MSigDB lists | GSE26030\_UNSTIM\_VS\_RESTIM\_TH1\_DAY15\_POST\_POLARIZATION\_UP | 52 | 16 | 4161 | 644 | Arf51F,CG1774,LSm3,Acph-1,Smox,STUB1,alph,wal,SRPK,Kap-alpha3,Aladin,CG10470,Arl6IP1,Atg4b,Gk2,opm | | 3.984e-03 | -5.53 | CG1416 (uncharacterized protein) | protein interactions | 35426 | 54 | 14 | 9204 | 1112 | SdhD,CCT6,SERCA,Rab5,Pdk1,Mcad,Kap-alpha3,ScsbetaA,Pp2A-29B,wal,CCT3,Tak1,GCS2beta,Tspo | | 3.984e-03 | -5.53 | Prosbeta5 (Proteasome beta5 subunit) | protein interactions | 45269 | 54 | 14 | 9204 | 1112 | Prosbeta3,Prosbeta6,Eno,Prosalpha3,Rpt2,Prosbeta4,Prosalpha4,Rpn2,scu,Pomp,Prosalpha5,Moe,Rpn13,AnxB10 | | 4.004e-03 | -5.52 | fly chr88D6-88D6\|3-56 cM | chromosome location | fly chr88D6-88D6\|3-56 cM | 13 | 4 | 25010 | 1343 | CycC,CG3631,CG33332,RpS5b | | 4.004e-03 | -5.52 | fly chr88D6-88D6\|3-56 | chromosome location | fly chr88D6-88D6\|3-56 | 13 | 4 | 25010 | 1343 | CG33332,CycC,CG3631,RpS5b | | 4.022e-03 | -5.52 | HEAT\_REPEAT | prosite domains | PS50077 | 7 | 4 | 6243 | 694 | Karybeta3,Pp2A-29B,CG1344,msps | | 4.025e-03 | -5.52 | Hsp83 (Heat shock protein 83) | protein interactions | 38389 | 77 | 18 | 9204 | 1112 | spag,Stip1,smg,Sec10,piwi,blw,RpS27A,Chd64,CG12811,ATPsynbeta,cup,SERCA,tor,AnxB10,Nup358,regucalcin,nos,csw | | 4.061e-03 | -5.51 | positive regulation of nucleic acid-templated transcription | biological process | GO:1903508 | 494 | 63 | 12345 | 1129 | esc,dl,trr,MED4,SREBP,foxo,hb,CG2129,pygo,not,Nf-YA,Bx,cyc,d4,Wbp2,cnc,S2P,CG17202,Mybbp1A,Atu,REPTOR,Su(H),Myd88,Rel,Cdk7,Atf6,Cf2,Utx,CG2199,lin,pont,nej,rept,Sox102F,e(y)1,Mitf,Stat92E,BigH1,cad,Sgf11,Taf7,apt,Trf2,MED20,E2f1,ovo,TfIIFalpha,vers,EcR,EloC,Bap170,CG17328,CG1602,bcd,CycC,Rbfox1,NFAT,RYBP,Rcd5,hng3,Tip60,lid,CG11710 | | 4.061e-03 | -5.51 | positive regulation of RNA biosynthetic process | biological process | GO:1902680 | 494 | 63 | 12345 | 1129 | vers,EcR,EloC,Bap170,CG1602,CG17328,bcd,CycC,Rbfox1,NFAT,Rcd5,RYBP,hng3,Tip60,lid,CG11710,pont,nej,lin,rept,Sox102F,e(y)1,Mitf,Stat92E,cad,Sgf11,BigH1,Taf7,Trf2,apt,MED20,E2f1,TfIIFalpha,ovo,S2P,CG17202,Mybbp1A,Atu,REPTOR,Myd88,Su(H),Cdk7,Rel,Atf6,Utx,Cf2,CG2199,trr,esc,dl,MED4,foxo,SREBP,hb,pygo,CG2129,Nf-YA,not,Bx,cyc,d4,Wbp2,cnc | | 4.075e-03 | -5.50 | GSE3982\_MAC\_VS\_CENT\_MEMORY\_CD4\_TCELL\_UP | MSigDB lists | GSE3982\_MAC\_VS\_CENT\_MEMORY\_CD4\_TCELL\_UP | 66 | 19 | 4161 | 644 | Jheh2,Irp-1B,Acph-1,LSm3,CG9248,Idh,CG9911,CG8314,Ntan1,ND-24,Pdxk,Prosbeta5,Echs1,Tim8,isoQC,alphaTub84B,CG1275,CG4572,CG17593 | | 4.075e-03 | -5.50 | KYNG\_DNA\_DAMAGE\_DN | MSigDB lists | KYNG\_DNA\_DAMAGE\_DN | 66 | 19 | 4161 | 644 | shop,Jafrac2,Trxr-1,Nmt,Mo25,PDCD-5,Unr,Tm1,sau,Nurf-38,alc,scu,CG4390,Sod1,Cyt-c-p,CG1890,eIF4E1,CycC,mts | | 4.084e-03 | -5.50 | Not3 (CCR4-NOT transcription complex subunit 3) | protein interactions | 35501 | 23 | 8 | 9204 | 1112 | Rga,twin,cu,nos,CG11164,BicC,cup,bam | | 4.084e-03 | -5.50 | HmgD (High mobility group protein D) | protein interactions | 37481 | 23 | 8 | 9204 | 1112 | foxo,Stip1,PHGPx,cib,Chd64,awd,CG11267,AnxB10 | | 4.084e-03 | -5.50 | CG11672 (uncharacterized protein) | protein interactions | 40977 | 23 | 8 | 9204 | 1112 | EloC,esc,Sec10,Ady43A,ssh,Uev1A,RpL9,Gdap2 | | 4.084e-03 | -5.50 | CG7987 (uncharacterized protein) | protein interactions | 41768 | 23 | 8 | 9204 | 1112 | Rcd5,wal,RpL31,RpL23A,RpL10,RpS29,RpS27A,UbcE2H | | 4.085e-03 | -5.50 | carbohydrate catabolic process | biological process | GO:0016052 | 56 | 12 | 12345 | 1129 | PyK,Pfrx,Treh,Pfk,CG9485,Gapdh1,Pgm1,Pgk,GlyP,Pglym78,Atg6,Eno | | 4.090e-03 | -5.50 | female germ-line stem cell asymmetric division | biological process | GO:0048132 | 18 | 6 | 12345 | 1129 | bam,eIF4A,egg,shrb,twin,piwi | | 4.090e-03 | -5.50 | DNA endoreduplication | biological process | GO:0042023 | 18 | 6 | 12345 | 1129 | Rbf,SkpA,ago,Orc1,SuUR,E2f1 | | 4.102e-03 | -5.50 | MORF\_G22P1 | MSigDB lists | MORF\_G22P1 | 100 | 26 | 4161 | 644 | LysRS,RpS30,Rpn12,Phb2,Prosbeta3,Dd,ND-51,not,CG10418,heph,CG14894,AP-1-2beta,eIF3i,Prosbeta6,RpL24,Nap1,Idh3b,Prosalpha3,eIF3m,ND-30,mEFTu1,Pgk,lwr,ND-24,Mtpalpha,CCT3 | | 4.106e-03 | -5.50 | PENG\_LEUCINE\_DEPRIVATION\_DN | MSigDB lists | PENG\_LEUCINE\_DEPRIVATION\_DN | 105 | 27 | 4161 | 644 | Hsc70-3,Fkbp59,Prosalpha5,Nmt,Iru,r-l,Stip1,Pfk,CCT5,Tom40,CycB,CG14715,Pen,Pglym78,bys,Rpn3,Pgk,sbr,CCT6,Rpn2,Cyt-c-p,Cse1,CG8188,r,heph,tko,CG1416 | | 4.150e-03 | -5.48 | Pomp (pomp) | protein interactions | 35342 | 38 | 11 | 9204 | 1112 | Rpn2,Prosalpha5,Prosbeta3,Prosbeta6,Prosalpha3,Rpt2,Prosbeta4,Prosalpha4,Pen,Prosbeta5,CG46385 | | 4.150e-03 | -5.48 | Rpn6 (Regulatory particle non-ATPase 6) | protein interactions | 36638 | 38 | 11 | 9204 | 1112 | Prosbeta3,Prosbeta6,Prosalpha3,Rpt2,Prosbeta4,Prosalpha4,Prosbeta5,Rpn2,Prosalpha5,Rpn13,Rpn3 | | 4.180e-03 | -5.48 | PX | smart domains | SM00312 | 12 | 5 | 5086 | 506 | CG3077,Pld,CG8726,CG11807,CG5734 | | 4.181e-03 | -5.48 | ATPase activity, coupled to transmembrane movement of ions, rotational mechanism | molecular function | GO:0044769 | 49 | 11 | 12176 | 1119 | ATPsynG,ATPsynbeta,ATPsynB,ATPsynE,blw,Vha68-2,Vha13,ATPsyndelta,ATPsynD,Vha36-1,ATPsynO | | 4.185e-03 | -5.48 | response to nutrient levels | biological process | GO:0031667 | 165 | 26 | 12345 | 1129 | Imp,rl,Nprl2,Jafrac1,Mitf,Dcr-1,Pi3K92E,Rbf,foxo,trsn,p38a,cyc,CG17691,Debcl,Iml1,Atg7,REPTOR,Desat1,lt,raptor,Seipin,Atg6,Mtpalpha,SNF4Agamma,whd,Rel | | 4.191e-03 | -5.47 | transferase activity, transferring nitrogenous groups | molecular function | GO:0016769 | 18 | 6 | 12176 | 1119 | Got1,Kyat,CG1640,Gapdh1,Gabat,CG11899 | | 4.207e-03 | -5.47 | mRNA metabolic process | biological process | GO:0016071 | 357 | 48 | 12345 | 1129 | Son,SmE,gw,CG1622,Spx,Usp39,CG16865,wisp,CG10418,CG31759,Pym,Rrp40,Bin1,smg,CG11486,cu,CG3198,sbr,twin,Edc3,yps,obe,CG5986,Rnmt,CG9684,Cbp80,snRNA:U4atac,CG17454,Neos,Not10,armi,heph,tho2,snRNA:U4:38AB,CG6610,CG17187,Imp,LSm3,Clp,nos,exu,snRNA:U12,Rga,IntS11,Trf4-2,rig,CG17768,bru1 | | 4.218e-03 | -5.47 | regulation of DNA replication | biological process | GO:0006275 | 36 | 9 | 12345 | 1129 | pn,Cdk1,otu,plu,E2f1,CycE,Rbf,CG10336,SuUR | | 4.249e-03 | -5.46 | REACTOME\_NOTCH1\_INTRACELLULAR\_DOMAIN\_REGULATES\_TRANSCRIPTION | MSigDB lists | REACTOME\_NOTCH1\_INTRACELLULAR\_DOMAIN\_REGULATES\_TRANSCRIPTION | 15 | 7 | 4161 | 644 | SkpA,CycC,Cul1,RpS27A,HDAC6,Su(H),RpL40 | | 4.249e-03 | -5.46 | MOOTHA\_FFA\_OXYDATION | MSigDB lists | MOOTHA\_FFA\_OXYDATION | 15 | 7 | 4161 | 644 | Mtpalpha,Mcad,Echs1,CG10932,scu,yip2,Thiolase | | 4.259e-03 | -5.46 | REACTOME\_CDK\_MEDIATED\_PHOSPHORYLATION\_AND\_REMOVAL\_OF\_CDC6 | MSigDB lists | REACTOME\_CDK\_MEDIATED\_PHOSPHORYLATION\_AND\_REMOVAL\_OF\_CDC6 | 39 | 13 | 4161 | 644 | Prosbeta5,Prosbeta6,RpL40,Rpn3,Prosalpha3,Prosbeta3,Prosbeta4,Prosalpha4,Rpt2,Rpn12,Prosalpha5,Rpn2,RpS27A | | 4.260e-03 | -5.46 | Tapdelta (Translocon-associated protein delta) | protein interactions | 45680 | 60 | 15 | 9204 | 1112 | CCT3,Cp1,Surf4,Pp2A-29B,wal,blw,Tspo,Rab5,CG1416,SdhD,SERCA,CCT6,Mcad,ScsbetaA,cathD | | 4.276e-03 | -5.45 | GSE23308\_WT\_VS\_MINERALCORTICOID\_REC\_KO\_MACROPHAGE\_UP | MSigDB lists | GSE23308\_WT\_VS\_MINERALCORTICOID\_REC\_KO\_MACROPHAGE\_UP | 71 | 20 | 4161 | 644 | Herc4,Idh3b,PAPLA1,CG8507,yip2,Pdxk,SdhA,Aldh,Ssdp,Mcad,isoQC,CG6767,colt,Kap3,CG1812,CG6766,Etf-QO,Grx1,Naa20A,not | | 4.279e-03 | -5.45 | NADH dehydrogenase (ubiquinone) Fe-S protein/flavoprotein complex, mitochondria | KEGG pathways | M00143 | 15 | 7 | 3198 | 496 | ND-18,ND-24,ND-30,ND-49,ND-23,ND-75,ND-51 | | 4.279e-03 | -5.45 | NADH dehydrogenase (ubiquinone) Fe-S protein/flavoprotein complex, mitochondria | KEGG pathways | dme\_M00143 | 15 | 7 | 3198 | 496 | ND-18,ND-24,ND-30,ND-49,ND-75,ND-51,ND-23 | | 4.298e-03 | -5.45 | UBIQUITIN\_CONJUGAT\_1 | prosite domains | PS00183 | 20 | 7 | 6243 | 694 | Ubc4,Ubc6,CG8188,lwr,ben,UbcE2H,CG40045 | | 4.312e-03 | -5.45 | Myb (Myb oncogene-like) | protein interactions | 32543 | 28 | 9 | 9204 | 1112 | Rbf,nej,Cdk1,cnn,Rbf2,lin-52,Sry-alpha,Orc1,CycB | | 4.312e-03 | -5.45 | CG7352 (uncharacterized protein) | protein interactions | 40997 | 28 | 9 | 9204 | 1112 | ND-75,Actn,Pgm1,Grip71,RpS7,Hmg-2,cact,Eno,Kyat | | 4.331e-03 | -5.44 | GO\_NEGATIVE\_REGULATION\_OF\_MOLECULAR\_FUNCTION | MSigDB lists | GO\_NEGATIVE\_REGULATION\_OF\_MOLECULAR\_FUNCTION | 276 | 59 | 4161 | 644 | Uch-L5,mad2,CG7997,STUB1,Cdk1,cact,Faf2,CG8507,RpS27A,Prosalpha5,Mo25,Pka-R2,rept,regucalcin,CHOp24,Arl6IP1,Karybeta3,lid,Rpn12,adp,HDAC6,Rpn2,Prip,Prosbeta3,mts,Rpt2,eIF4A,CG7530,CG3376,Rpn3,Chd64,CycB,Prosbeta5,Pdk1,CG4709,Pp2A-29B,CanA-14F,Rcd5,Prosalpha3,Hsc70-3,Prosbeta4,Cyp1,CG8726,RpL40,PrBP,Snx6,Pka-R1,fray,Prosbeta6,Nprl2,Ptp61F,APC7,Phb2,Pp2B-14D,dco,CG30291,CkIIalpha,Non1,CG1847 | | 4.333e-03 | -5.44 | GSE19772\_CTRL\_VS\_HCMV\_INF\_MONOCYTES\_AND\_PI3K\_INHIBITION\_UP | MSigDB lists | GSE19772\_CTRL\_VS\_HCMV\_INF\_MONOCYTES\_AND\_PI3K\_INHIBITION\_UP | 57 | 17 | 4161 | 644 | Grx1,Atg4b,Pi3K92E,Cyt-c-p,CG8188,sau,mts,alphaTub84B,tws,CG13601,CG5026,Chrac-14,CG15525,stas,Sply,Mat89Ba,CG7728 | | 4.333e-03 | -5.44 | REACTOME\_CYTOKINE\_SIGNALING\_IN\_IMMUNE\_SYSTEM | MSigDB lists | REACTOME\_CYTOKINE\_SIGNALING\_IN\_IMMUNE\_SYSTEM | 57 | 17 | 4161 | 644 | RpS27A,alph,rl,Cdk1,Pen,Kap-alpha3,Cul1,eIF4A,eIF4E1,ari-1,Traf6,Aladin,SkpA,Ptp61F,Pi3K92E,RpL40,mbo | | 4.355e-03 | -5.44 | peptidyl-amino acid modification | biological process | GO:0018193 | 394 | 52 | 12345 | 1129 | CkIIalpha,Utx,Ptpmeg2,CG8314,Hat1,velo,escl,QC,CG2911,Gapdh1,isoQC,Fkbp12,Chrac-14,Ptip,awd,MAGE,CG9643,Ptpmeg,d4,CG31053,Tlk,Lkb1,Sgf29,Pdk1,tor,CG11858,CG1578,esc,trr,G9a,Nmt,lid,Tip60,Rcd5,Fkbp59,Ptp61F,Dph5,Spt20,dco,SRPK,Aos1,CG10344,Naa20A,Cyp1,e(y)1,rept,egg,lwr,pont,nej,CG3511,YL-1 | | 4.365e-03 | -5.43 | positive regulation of multi-organism process | biological process | GO:0043902 | 149 | 24 | 12345 | 1129 | Cbl,lwr,dl,Tak1,GILT1,cad,Aos1,E2f1,Hr78,PGRP-LE,EMRE,Ptip,Sp7,wisp,Hat1,tub,Crag,PDCD-5,Rel,Herc4,CG9925,mop,Myd88,spz | | 4.366e-03 | -5.43 | Cyclin A:Cdk2-associated events at S phase entry | REACTOME pathways | R-DME-69656 | 75 | 20 | 3923 | 574 | Prosalpha4,twe,CycE,Rbf2,Prosbeta5,Rpn2,Prosbeta4,RpS27A,Prosalpha5,Prosalpha3,RpL40,Rpt2,Rbf,SkpA,Cdk7,Prosbeta3,Rpn12,Cul1,Rpn3,Prosbeta6 | | 4.367e-03 | -5.43 | Toll signaling pathway | biological process | GO:0008063 | 13 | 5 | 12345 | 1129 | Myd88,dl,cact,tub,spz | | 4.367e-03 | -5.43 | regulation of reactive oxygen species metabolic process | biological process | GO:2000377 | 13 | 5 | 12345 | 1129 | Coq7,SelD,Fdh,Pi3K92E,Rel | | 4.376e-03 | -5.43 | Downstream TCR signaling | REACTOME pathways | R-DME-202424 | 70 | 19 | 3923 | 574 | cact,SkpA,Rpt2,RpL40,Prosalpha3,Prosbeta4,Prosalpha5,RpS27A,Pdk1,Prosalpha4,Rpn2,Prosbeta5,Prosbeta6,Rpn3,Cul1,dl,Rpn12,Prosbeta3,Rel | | 4.392e-03 | -5.43 | MED14 (Mediator complex subunit 14) | protein interactions | 38073 | 49 | 13 | 9204 | 1112 | MED24,CG2199,Larp4B,CG17002,Ptip,enc,MED20,MED4,MED27,Usp10,Ssdp,EcR,ix | | 4.392e-03 | -5.43 | CG4853 (uncharacterized protein) | protein interactions | 36974 | 49 | 13 | 9204 | 1112 | PGRP-LE,Roe1,RpL12,Utx,CG5903,RfC38,CG2938,CG11791,Bin1,RpS13,Eno,Tak1,Prosalpha3 | | 4.423e-03 | -5.42 | acetyltransferase activity | molecular function | GO:0016407 | 93 | 17 | 12176 | 1119 | CG10932,Tip60,e(y)1,lid,Chrac-14,nej,Rcd5,Sgf29,CG2938,d4,yip2,Hat1,YL-1,AgmNAT,Naa20A,pont,rept | | 4.423e-03 | -5.42 | unfolded protein binding | molecular function | GO:0051082 | 93 | 17 | 12176 | 1119 | Pfdn5,sip3,Cyp1,CCT3,Roe1,CG14671,Pfdn2,Droj2,Uxt,Hsc70-3,DnaJ-H,CG11267,CCT6,CCT5,CG10055,Hsp26,shv | | 4.433e-03 | -5.42 | GSE10325\_MYELOID\_VS\_LUPUS\_MYELOID\_DN | MSigDB lists | GSE10325\_MYELOID\_VS\_LUPUS\_MYELOID\_DN | 48 | 15 | 4161 | 644 | Acph-1,Sec22,Lsm12,CG7806,Cdk7,Top1,CG14194,Atg7,Gk2,Edem1,Cul1,cathD,scramb1,CG7550,Ubc4 | | 4.433e-03 | -5.42 | REACTOME\_ACTIVATION\_OF\_THE\_MRNA\_UPON\_BINDING\_OF\_THE\_CAP\_BINDING\_COMPLEX\_AND\_EIFS\_AND\_SUBSEQUENT\_BINDING\_TO\_43S | MSigDB lists | REACTOME\_ACTIVATION\_OF\_THE\_MRNA\_UPON\_BINDING\_OF\_THE\_CAP\_BINDING\_COMPLEX\_AND\_EIFS\_AND\_SUBSEQUENT\_BINDING\_TO\_43S | 48 | 15 | 4161 | 644 | RpS27,RpS13,RpS30,eIF4E1,RpS28b,RpS16,RpS2,RpS23,RpS27A,RpS24,eIF4A,RpS7,RpS4,eIF3i,RpS18 | | 4.433e-03 | -5.42 | GSE22140\_GERMFREE\_VS\_SPF\_ARTHRITIC\_MOUSE\_CD4\_TCELL\_DN | MSigDB lists | GSE22140\_GERMFREE\_VS\_SPF\_ARTHRITIC\_MOUSE\_CD4\_TCELL\_DN | 48 | 15 | 4161 | 644 | SdhA,Rpb8,Klp10A,Gk2,ND-24,Cul1,msps,Prosalpha3,CG2493,TyrRS,RpL37a,CG33156,ND-49,Phb2,RhoGAP54D | | 4.433e-03 | -5.42 | MODULE\_62 | MSigDB lists | MODULE\_62 | 48 | 15 | 4161 | 644 | ND-SGDH,Cyt-c-p,COX5A,Nurf-38,ND-39,Vha36-1,ND-49,ND-30,ND-51,ND-23,ND-24,UQCR-Q,ND-B18,SdhA,levy | | 4.433e-03 | -5.42 | GCM\_TPT1 | MSigDB lists | GCM\_TPT1 | 48 | 15 | 4161 | 644 | RpS18,RpL38,RpS7,RpL40,RpL23,RpL11,RpS24,RpL9,RpS27A,RpS23,RpS28b,RpL13A,RpS2,RpS16,RpLP0 | | 4.436e-03 | -5.42 | GSE19888\_CTRL\_VS\_T\_CELL\_MEMBRANES\_ACT\_MAST\_CELL\_UP | MSigDB lists | GSE19888\_CTRL\_VS\_T\_CELL\_MEMBRANES\_ACT\_MAST\_CELL\_UP | 76 | 21 | 4161 | 644 | Ufm1,Arl5,CG10418,Aos1,Uba5,shv,RpS30,Vsp37A,AMPdeam,Pdk1,Cul1,Liprin-beta,Shmt,Moe,Fkbp59,mRpL10,APP-BP1,CG11134,CG9773,Cka,COX5A | | 4.436e-03 | -5.42 | MORF\_AP3D1 | MSigDB lists | MORF\_AP3D1 | 76 | 21 | 4161 | 644 | mts,Dd,ND-49,LysRS,Rpn2,Pka-R1,CysRS,Prosbeta6,rept,CG10418,heph,Prosalpha3,Mitofilin,Prosbeta4,Idh3b,SdhA,capt,Thiolase,Pgk,Cul1,Pglym78 | | 4.441e-03 | -5.42 | glycerophospholipid metabolic process | biological process | GO:0006650 | 109 | 19 | 12345 | 1129 | PIG-L,CG9391,Pi3K92E,Pld,CG33977,CG6707,PIG-H,PIG-C,CG7789,sws,CG18135,Pi4KIIalpha,CG7718,CG2201,Synj,PIG-P,CG5026,Seipin,Agpat2 | | 4.493e-03 | -5.41 | negative regulation of neuron death | biological process | GO:1901215 | 30 | 8 | 12345 | 1129 | Pink1,spag,Tip60,Naa20A,Aldh,Drep2,Debcl,MAGE | | 4.493e-03 | -5.41 | mitotic cell cycle, embryonic | biological process | GO:0045448 | 30 | 8 | 12345 | 1129 | Dcr-1,CycB,CycJ,larp,HP1e,toc,Tlk,ph-d | | 4.522e-03 | -5.40 | mitotic DNA integrity checkpoint | biological process | GO:0044774 | 24 | 7 | 12345 | 1129 | mus81,Cdk1,grp,CG10336,mre11,mus304,nbs | | 4.523e-03 | -5.40 | positive regulation of defense response | biological process | GO:0031349 | 86 | 16 | 12345 | 1129 | GILT1,Tak1,Hr78,PGRP-LE,EMRE,Ptip,Sp7,tub,Crag,Hat1,wisp,CG9925,Rel,PDCD-5,Myd88,spz | | 4.588e-03 | -5.38 | pyruvate metabolic process | biological process | GO:0006090 | 43 | 10 | 12345 | 1129 | Eno,Pglym78,Pfk,Aldh,Pgk,Pdk,Gapdh1,PyK,CG10932,Pfrx | | 4.593e-03 | -5.38 | fly chr12F5-12F5\|1-49 | chromosome location | fly chr12F5-12F5\|1-49 | 7 | 3 | 25010 | 1343 | CG32590,CG14408,CG14407 | | 4.593e-03 | -5.38 | fly chr80F9-80F9\|3-47 | chromosome location | fly chr80F9-80F9\|3-47 | 7 | 3 | 25010 | 1343 | CG17454,CG17698,AGO3 | | 4.593e-03 | -5.38 | fly chr83B7-83B7\|3-47.5 | chromosome location | fly chr83B7-83B7\|3-47.5 | 7 | 3 | 25010 | 1343 | CG31548,CG12171,CG31549 | | 4.593e-03 | -5.38 | fly chr66C11-66C11\|3-24 cM | chromosome location | fly chr66C11-66C11\|3-24 cM | 7 | 3 | 25010 | 1343 | Oseg1,Exo70,mtrm | | 4.593e-03 | -5.38 | fly chr11D1-11D1\|1-41 | chromosome location | fly chr11D1-11D1\|1-41 | 7 | 3 | 25010 | 1343 | CG4404,Coq8,CG4407 | | 4.593e-03 | -5.38 | fly chr12F5-12F5\|1-49 cM | chromosome location | fly chr12F5-12F5\|1-49 cM | 7 | 3 | 25010 | 1343 | CG14408,CG14407,CG32590 | | 4.593e-03 | -5.38 | fly chr2B14-2B14\|1-0.2 | chromosome location | fly chr2B14-2B14\|1-0.2 | 7 | 3 | 25010 | 1343 | CG14818,trr,mRpL16 | | 4.593e-03 | -5.38 | fly chr11D1-11D1\|1-41 cM | chromosome location | fly chr11D1-11D1\|1-41 cM | 7 | 3 | 25010 | 1343 | CG4407,Coq8,CG4404 | | 4.593e-03 | -5.38 | fly chr83B7-83B7\|3-47.5 cM | chromosome location | fly chr83B7-83B7\|3-47.5 cM | 7 | 3 | 25010 | 1343 | CG12171,CG31548,CG31549 | | 4.593e-03 | -5.38 | fly chr17C1-17C1\|1-60 | chromosome location | fly chr17C1-17C1\|1-60 | 7 | 3 | 25010 | 1343 | Ggt-1,Bx,CG15047 | | 4.593e-03 | -5.38 | fly chr80F9-80F9\|3-47 cM | chromosome location | fly chr80F9-80F9\|3-47 cM | 7 | 3 | 25010 | 1343 | AGO3,CG17698,CG17454 | | 4.593e-03 | -5.38 | fly chr66C11-66C11\|3-24 | chromosome location | fly chr66C11-66C11\|3-24 | 7 | 3 | 25010 | 1343 | Exo70,Oseg1,mtrm | | 4.593e-03 | -5.38 | fly chr17C1-17C1\|1-60 cM | chromosome location | fly chr17C1-17C1\|1-60 cM | 7 | 3 | 25010 | 1343 | CG15047,Bx,Ggt-1 | | 4.593e-03 | -5.38 | fly chr2B14-2B14\|1-0.2 cM | chromosome location | fly chr2B14-2B14\|1-0.2 cM | 7 | 3 | 25010 | 1343 | trr,mRpL16,CG14818 | | 4.598e-03 | -5.38 | - | gene3d domains | 3.60.15.10 | 10 | 5 | 4929 | 622 | CG9117,tzn,CG12375,Snm1,IntS11 | | 4.598e-03 | -5.38 | GO\_CELLULAR\_RESPONSE\_TO\_OXYGEN\_LEVELS | MSigDB lists | GO\_CELLULAR\_RESPONSE\_TO\_OXYGEN\_LEVELS | 35 | 12 | 4161 | 644 | Cul2,yip2,RpS27A,Prip,CycB,AIF,LpR2,Atg7,Calx,RpL40,Su(H),Pdk | | 4.646e-03 | -5.37 | anterior/posterior pattern specification | biological process | GO:0009952 | 200 | 30 | 12345 | 1129 | osk,Grip71,bcd,spn-E,trk,msps,Moe,csw,Ptip,jvl,EcR,Grip128,fs(1)N,fs(1)M3,psq,armi,gus,exu,tor,nos,rl,hb,swa,cnc,Tm1,cad,mael,Lkb1,tkv,Grip75 | | 4.661e-03 | -5.37 | NDK-like\_dom | interpro domains | IPR034907 | 4 | 3 | 11383 | 1233 | awd,nmdyn-D7,nmdyn-D6 | | 4.661e-03 | -5.37 | RHD\_DNA\_bind\_dom | interpro domains | IPR011539 | 4 | 3 | 11383 | 1233 | Rel,dl,NFAT | | 4.661e-03 | -5.37 | PLipase\_D/transphosphatidylase | interpro domains | IPR001736 | 4 | 3 | 11383 | 1233 | CG7718,CG9248,Pld | | 4.661e-03 | -5.37 | RHD\_DNA\_bind\_dom\_sf | interpro domains | IPR037059 | 4 | 3 | 11383 | 1233 | Rel,NFAT,dl | | 4.661e-03 | -5.37 | OST-HTH/LOTUS\_dom | interpro domains | IPR025605 | 4 | 3 | 11383 | 1233 | Marf1,tapas,osk | | 4.661e-03 | -5.37 | NDK-like\_dom\_sf | interpro domains | IPR036850 | 4 | 3 | 11383 | 1233 | awd,nmdyn-D7,nmdyn-D6 | | 4.661e-03 | -5.37 | RHD\_dimer | interpro domains | IPR032397 | 4 | 3 | 11383 | 1233 | NFAT,dl,Rel | | 4.661e-03 | -5.37 | PPIase\_FKBP | interpro domains | IPR023566 | 4 | 3 | 11383 | 1233 | Fkbp39,Fkbp59,Fkbp12 | | 4.694e-03 | -5.36 | GO\_PROTEIN\_POLYUBIQUITINATION | MSigDB lists | GO\_PROTEIN\_POLYUBIQUITINATION | 111 | 28 | 4161 | 644 | STUB1,Prosbeta5,Cul1,Rpn3,Uev1A,Prosbeta4,Prosalpha3,CG2924,RpS27A,Prosalpha5,Iru,CG40045,Prosbeta6,SkpA,Ubc4,CG11070,RpL40,Rpt2,UbcE2H,Prosalpha4,Prosbeta3,Rpn2,Traf6,APC7,HDAC6,CG8188,Rpn12,ari-1 | | 4.702e-03 | -5.36 | TIEN\_INTESTINE\_PROBIOTICS\_24HR\_DN | MSigDB lists | TIEN\_INTESTINE\_PROBIOTICS\_24HR\_DN | 91 | 24 | 4161 | 644 | RpS2,ari-1,RpL13A,RpLP0,slif,RpS24,UbcE2H,RpL23,opm,RpS7,tho2,CHOp24,Faf2,RpS27,RpL9,RpS23,mtgo,Got1,Hsc70-3,RpL12,RpL13,ERp60,alpha-Man-Ia,RpS18 | | 4.743e-03 | -5.35 | OST-HTH | pfam domains | PF12872 | 4 | 3 | 10480 | 1142 | tapas,osk,Marf1 | | 4.743e-03 | -5.35 | NDK | pfam domains | PF00334 | 4 | 3 | 10480 | 1142 | nmdyn-D6,nmdyn-D7,awd | | 4.743e-03 | -5.35 | RHD\_DNA\_bind | pfam domains | PF00554 | 4 | 3 | 10480 | 1142 | dl,NFAT,Rel | | 4.743e-03 | -5.35 | RHD\_dimer | pfam domains | PF16179 | 4 | 3 | 10480 | 1142 | dl,NFAT,Rel | | 4.743e-03 | -5.35 | Lactamase\_B | pfam domains | PF00753 | 4 | 3 | 10480 | 1142 | CG9117,CG12375,tzn | | 4.747e-03 | -5.35 | response to metal ion | biological process | GO:0010038 | 50 | 11 | 12345 | 1129 | CG7328,rl,CG11590,Mekk1,Grx1,MtnA,Fer1HCH,p38a,whd,Gclc,ZnT35C | | 4.747e-03 | -5.35 | negative regulation of cell cycle phase transition | biological process | GO:1901988 | 50 | 11 | 12345 | 1129 | mre11,Aven,mad2,Rbf,mus304,Zwilch,nbs,Cdk1,Rbf2,grp,Mos | | 4.758e-03 | -5.35 | p75 NTR receptor-mediated signalling | REACTOME pathways | R-DME-193704 | 16 | 7 | 3923 | 574 | RpS27A,cact,Rel,RpL40,dl,Rtnl1,RhoGDI | | 4.785e-03 | -5.34 | Complex I biogenesis | REACTOME pathways | R-DME-6799198 | 37 | 12 | 3923 | 574 | CG8067,ND-B18,ND-49,NP15.6,ND-51,ND-23,ND-B22,ND-39,ND-SGDH,ND-24,ND-75,ND-30 | | 4.792e-03 | -5.34 | MORF\_DEK | MSigDB lists | MORF\_DEK | 142 | 34 | 4161 | 644 | Dd,CG6767,Phb2,CG17454,Prosbeta6,CG10418,Prosbeta4,eIF3m,Prosalpha3,yip2,Cul1,Thiolase,ND-30,CG13630,Hat1,lwr,eIF4A,mts,LysRS,Sod1,eIF3i,SkpA,Stoml2,ND-51,CSN6,Idh3b,Nap1,RpS27A,Klp10A,SdhA,Chrac-14,mEFTu1,Pgk,Top1 | | 4.794e-03 | -5.34 | Act57B (Actin 57B) | protein interactions | 37368 | 44 | 12 | 9204 | 1112 | PGRP-LE,CG10265,awd,glob1,Arpc2,Pdk1,capt,Arpc4,osk,Gip,Arpc1,Rpt2 | | 4.810e-03 | -5.34 | GO\_RESPONSE\_TO\_ENDOGENOUS\_STIMULUS | MSigDB lists | GO\_RESPONSE\_TO\_ENDOGENOUS\_STIMULUS | 356 | 73 | 4161 | 644 | Shmt,Cdk1,kri,rl,cact,tkv,Pka-R2,Fkbp59,Rab10,Got1,dnc,Tip60,CG10932,Karybeta3,Rpb8,Sod1,adp,mino,RpII140,Prip,Cdk7,raptor,Moe,Gclc,Cdk5,p38a,Pdk1,Mtpalpha,Ac3,CycE,nAChRalpha4,PpD3,SREBP,Prosbeta4,Wnt10,Tspo,Med,RpL40,Atpalpha,heph,Pka-R1,mus81,Vha13,Ptp61F,aay,Dhfr,Calx,GlyP,LpR2,AIF,RpS27A,Idh,rept,ACC,EcR,csw,CG3376,CG11710,Smox,Gabat,PDCD-5,Cka,CanA-14F,Cbp80,r,CG9485,Ts,MED4,Atg6,Pp2B-14D,faf,MED24,Vha36-1 | | 4.823e-03 | -5.33 | GO\_COFACTOR\_METABOLIC\_PROCESS | MSigDB lists | GO\_COFACTOR\_METABOLIC\_PROCESS | 179 | 41 | 4161 | 644 | Tspo,CG17333,sgll,Ts,mal,ScsbetaA,Dhfr,Pglym78,Pfk,CG7322,Coq7,Pdxk,pr,Eno,CG11134,Ppcdc,CG4407,ACC,CG5028,CG11089,Coq6,mino,Nadsyn,CG33156,Gapdh1,Pgm1,Shmt,Pgk,AcCoAS,CG9471,CG3803,Pcd,Idh3b,Ppox,CG1774,Desat1,Scsalpha1,Nc73EF,Idh,ND-39,Sptr | | 4.824e-03 | -5.33 | histone modification | biological process | GO:0016570 | 192 | 29 | 12345 | 1129 | pont,nej,trr,YL-1,esc,egg,rept,e(y)1,not,Sgf29,Naa20A,Tlk,Sgf11,scny,Spt20,d4,Ptip,Chrac-14,escl,l(3)73Ah,Atu,Hat1,Sirt7,Rcd5,Tip60,HDAC6,lid,Utx,G9a | | 4.824e-03 | -5.33 | covalent chromatin modification | biological process | GO:0016569 | 192 | 29 | 12345 | 1129 | Tlk,scny,Sgf11,Naa20A,Sgf29,Spt20,d4,egg,rept,nej,pont,trr,esc,YL-1,not,e(y)1,Tip60,Sirt7,Rcd5,G9a,HDAC6,lid,Utx,Chrac-14,Ptip,Atu,Hat1,escl,l(3)73Ah | | 4.826e-03 | -5.33 | drug binding | molecular function | GO:0008144 | 828 | 98 | 12176 | 1119 | bor,l(2)k01209,r,Orc1,rl,pont,rept,p38a,cana,grp,Got1,Droj2,GlyP,CG5789,SerRS-m,eIF4A,CysRS,CG8726,RecQ5,Nadsyn,l(1)G0196,LysRS,Sply,nmdyn-D7,PEK,Mos,ATP8A,Marc,png,Shmt,ABCD,SRPK,CG9981,CG14183,Lkb1,SelD,CG4880,fray,kl-2,E23,CCT3,Pfk,Pfrx,glob1,dco,Pink1,TyrRS,CG17698,obe,Fkbp39,Vha68-2,tor,Klp10A,CCT6,CCT5,Hsc70-3,Mekk1,TrpRS-m,CkIIalpha,Tao,ATPsynbeta,Cdk5,Fkbp59,CG1640,Cdk1,CG31729,Cht11,Idgf2,CG7806,nmd,mal,CG4562,AcCoAS,ScsbetaA,ACC,CG4041,SERCA,tkv,spn-E,Pdk1,Gabat,Rpt2,CG6767,Idgf6,CG31689,Tlk,CG8611,blw,Tak1,Cyp1,Cdk7,awd,Idgf4,Atpalpha,CG9143,GluRS-m,Pgk,Kyat | | 4.830e-03 | -5.33 | SMAD2/SMAD3:SMAD4 heterotrimer regulates transcription | REACTOME pathways | R-DME-2173796 | 24 | 9 | 3923 | 574 | RpL40,CG6923,Smox,Rbf,Rbf2,Med,CycC,nrv1,RpS27A | | 4.830e-03 | -5.33 | cytochrome complex | cellular component | GO:0070069 | 30 | 8 | 11940 | 1105 | COX4,levy,COX5B,COX8,COX5A,cype,UQCR-Q,UQCR-11 | | 4.830e-03 | -5.33 | - | gene3d domains | 1.50.10.10 | 14 | 6 | 4929 | 622 | CG9485,alpha-Man-Ia,Edem1,CG2061,Treh,CG8475 | | 4.849e-03 | -5.33 | peptide N-acetyltransferase activity | molecular function | GO:0034212 | 64 | 13 | 12176 | 1119 | d4,Hat1,e(y)1,Tip60,Rcd5,Sgf29,pont,rept,Naa20A,lid,YL-1,Chrac-14,nej | | 4.876e-03 | -5.32 | MORF\_ARAF1 | MSigDB lists | MORF\_ARAF1 | 19 | 8 | 4161 | 644 | Pdk1,kcc,CSN6,P5CDh1,RpS24,Grip84,CG9393,GCS2beta | | 4.876e-03 | -5.32 | GO\_NADH\_METABOLIC\_PROCESS | MSigDB lists | GO\_NADH\_METABOLIC\_PROCESS | 19 | 8 | 4161 | 644 | Idh3b,Nc73EF,Gapdh1,Pgk,Pglym78,CG5028,Eno,Pfk | | 4.876e-03 | -5.32 | KEGG\_FATTY\_ACID\_METABOLISM | MSigDB lists | KEGG\_FATTY\_ACID\_METABOLISM | 19 | 8 | 4161 | 644 | Echs1,Aldh,Mtpalpha,Thiolase,Fdh,CG10932,Mcad,yip2 | | 4.888e-03 | -5.32 | REACTOME\_SIGNALING\_BY\_NOTCH | MSigDB lists | REACTOME\_SIGNALING\_BY\_NOTCH | 31 | 11 | 4161 | 644 | RpS27A,HDAC6,SERCA,CycC,Cul1,RpL40,Su(H),pen-2,SkpA,O-fut1,CHOp24 | | 4.888e-03 | -5.32 | GO\_MITOCHONDRIAL\_RESPIRATORY\_CHAIN\_COMPLEX\_I\_BIOGENESIS | MSigDB lists | GO\_MITOCHONDRIAL\_RESPIRATORY\_CHAIN\_COMPLEX\_I\_BIOGENESIS | 31 | 11 | 4161 | 644 | ND-24,ND-23,ND-30,ND-51,AIF,ND-B18,ND-B22,ND-75,ND-SGDH,ND-49,ND-39 | | 4.888e-03 | -5.32 | GSE37534\_UNTREATED\_VS\_ROSIGLITAZONE\_TREATED\_CD4\_TCELL\_PPARG1\_AND\_FOXP3\_TRASDUCED\_DN | MSigDB lists | GSE37534\_UNTREATED\_VS\_ROSIGLITAZONE\_TREATED\_CD4\_TCELL\_PPARG1\_AND\_FOXP3\_TRASDUCED\_DN | 31 | 11 | 4161 | 644 | Ufm1,Rep,scramb1,PIG-C,CG12773,CG8112,Got1,Herc4,mal,S2P,MED4 | | 4.889e-03 | -5.32 | GSE360\_L\_MAJOR\_VS\_B\_MALAYI\_HIGH\_DOSE\_MAC\_DN | MSigDB lists | GSE360\_L\_MAJOR\_VS\_B\_MALAYI\_HIGH\_DOSE\_MAC\_DN | 67 | 19 | 4161 | 644 | CysRS,Etf-QO,Jafrac1,ND-51,CG4572,CG2493,Rpn12,trsn,ND-B18,Stip1,CycB,GlyP,Echs1,msps,Marf,CG3376,Idh3b,CG11807,Pi4KIIalpha | | 4.889e-03 | -5.32 | GSE10239\_NAIVE\_VS\_KLRG1HIGH\_EFF\_CD8\_TCELL\_UP | MSigDB lists | GSE10239\_NAIVE\_VS\_KLRG1HIGH\_EFF\_CD8\_TCELL\_UP | 53 | 16 | 4161 | 644 | RpL11,RpL26,RpL24,RpS27,CG8199,eEF1beta,RpL13,sbr,CG10189,CG17187,CG3894,RpL38,RpS7,tzn,Ublcp1,Med | | 4.908e-03 | -5.32 | PENG\_RAPAMYCIN\_RESPONSE\_DN | MSigDB lists | PENG\_RAPAMYCIN\_RESPONSE\_DN | 137 | 33 | 4161 | 644 | SREBP,Hsc70-3,Wnt10,gammaTub37C,Prosbeta5,CCT5,Stip1,Pfk,ND-30,sbr,Pglym78,bys,CG8728,P5CDh1,CycC,Cyt-c-p,tko,CG1416,Cyp1,heph,ND-39,Fkbp59,Idh3b,Nmt,Tom40,wal,Pgk,CCT6,Rpn12,Cse1,scu,RpII140,CHOp24 | | 4.908e-03 | -5.32 | nucleobase-containing compound catabolic process | biological process | GO:0034655 | 134 | 22 | 12345 | 1129 | Trf4-2,Cbp80,CG17768,twin,Edc3,Rga,LSm3,Drep1,Dcr-1,fal,smg,cu,CG11486,Rrp40,Tengl4,CG31759,Not10,Pym,armi,cN-IIIB,CG11164,gw | | 4.921e-03 | -5.31 | cellularization | biological process | GO:0007349 | 110 | 19 | 12345 | 1129 | Rab5,Sry-alpha,sw,heph,EcR,Pen,eIF3m,osk,piwi,CG46338,grp,slam,cort,Pld,faf,cnn,eIF4A,ScsbetaA,Dcr-1 | | 4.926e-03 | -5.31 | response to extracellular stimulus | biological process | GO:0009991 | 167 | 26 | 12345 | 1129 | trsn,p38a,cyc,CG17691,Imp,rl,Nprl2,Jafrac1,Dcr-1,Mitf,Pi3K92E,Rbf,foxo,raptor,Atg6,Seipin,Mtpalpha,SNF4Agamma,whd,Rel,Debcl,Atg7,Iml1,REPTOR,lt,Desat1 | | 4.939e-03 | -5.31 | organelle localization | biological process | GO:0051640 | 218 | 32 | 12345 | 1129 | cnc,larp,alphaTub84B,gammaTub37C,sw,Klp10A,mbo,cana,msps,Pi4KIIalpha,obe,mael,l(3)mbt,Blos2,Sec10,faf,Cdk5,Nup62,CycB,milt,Syx13,ssh,X11L,Sec15,alphaTub67C,Pink1,Cenp-C,jvl,Exo70,Marf,Rab10,wisp | | 4.956e-03 | -5.31 | Negative regulation of FGFR1 signaling | REACTOME pathways | R-DME-5654726 | 20 | 8 | 3923 | 574 | mts,CG13398,RpL40,Pp2A-29B,csw,Cbl,RpS27A,rl | | 4.977e-03 | -5.30 | transferase complex, transferring phosphorus-containing groups | cellular component | GO:0061695 | 157 | 25 | 11940 | 1105 | CG8475,png,CG6767,alc,Atu,CycE,Taf7,Rpb8,TfIIFalpha,plu,CG12018,e(y)1,Chrac-14,mus205,DNApol-epsilon255,Trf2,Atg6,Tfb4,Cdk5,Pi3K92E,CycJ,RpII140,Cdk7,CycB,Pfk | | 4.985e-03 | -5.30 | p75NTR signals via NF-kB | REACTOME pathways | R-DME-193639 | 9 | 5 | 3923 | 574 | RpS27A,cact,RpL40,Rel,dl | | 4.985e-03 | -5.30 | NF-kB is activated and signals survival | REACTOME pathways | R-DME-209560 | 9 | 5 | 3923 | 574 | Rel,RpL40,dl,RpS27A,cact | | 4.997e-03 | -5.30 | CG2064 (uncharacterized protein) | protein interactions | 35708 | 109 | 23 | 9204 | 1112 | CG5447,Rpt2,wal,Surf4,Karybeta3,bor,Tspo,Rab5,Nup358,Mcad,blw,Pp2A-29B,CCT3,CG15098,CG31729,Cisd2,SERCA,CCT6,ATPsynbeta,Desat1,Kap-alpha3,CG10425,Echs1 | | 4.998e-03 | -5.30 | Downstream signaling events of B Cell Receptor (BCR) | REACTOME pathways | R-DME-1168372 | 107 | 26 | 3923 | 574 | Prosbeta3,tor,Rel,csw,Prosbeta6,Pp2A-29B,CG13398,Rpn12,dl,Cul1,Rpn3,foxo,Pdk1,RpS27A,yl,Prosalpha5,Pi3K92E,Prosbeta4,Rpn2,Prosbeta5,Prosalpha4,SkpA,cact,Prosalpha3,RpL40,Rpt2 | | 5.007e-03 | -5.30 | anatomical structure morphogenesis | biological process | GO:0009653 | 1579 | 173 | 12345 | 1129 | Cbl,lwr,Pax,ScsbetaA,Girdin,Mitf,parvin,wash,CG46338,dco,EloC,Marf,osk,CycC,gd,ssh,Smox,SREBP,mts,Nf-YA,eIF4A,Orc1,ago,ea,cort,Ankle2,slam,awd,Atg7,Gapdh1,csw,SelR,Ptp69D,armi,CkIIalpha,Atg6,r2d2,Ptp10D,Mtl,rept,Pi3K92E,Vha13,boi,Tm1,RhoGAP5A,mael,wal,Rbcn-3A,ovo,Trf2,Pmi,Pfk,cact,Liprin-beta,Idh3b,Nmt,spz,DCAF12,Pura,Cul5,foxo,trr,not,Cka,Lkb1,Atpalpha,piwi,Ptpmeg,RhoGAP54D,up,Pen,Arpc2,CycE,Med,Sec23,spn-E,sbb,Mitofilin,Pxt,Rel,Gclc,bun,fs(1)M3,ACC,Cf2,cnn,shrb,lin,CCT3,MED24,rost,vret,Stat92E,Actn,E2f1,p38a,Bap170,jvl,colt,Iml1,dnc,ben,Moe,Sry-alpha,Cul2,nos,tor,Pgk,pygo,Tak1,CycJ,PIG-C,grp,Cdk1,cnc,Afti,Abi,Arpc1,velo,msps,psq,CanB2,Arpc4,Su(H),bam,CG13366,Utx,Cdk5,cib,faf,nej,Dcr-1,spen,Nprl2,rl,cad,tkv,capt,enc,Ptp61F,del,PyK,chrb,bif,EcR,fs(1)N,Debcl,Rbfox1,gus,pn,Sec15,Traf6,Rbf,squ,Pld,Dd,Bx,Pglym78,bru1,Smyd4-4,EMC3,alph,eIF3m,Arf51F,Vha68-2,ix,sw,TBCC,Rab5,Pka-R2,heph,raptor,CSN6 | | 5.019e-03 | -5.29 | REL\_2 | prosite domains | PS50254 | 4 | 3 | 6243 | 694 | dl,NFAT,Rel | | 5.019e-03 | -5.29 | PLD | prosite domains | PS50035 | 4 | 3 | 6243 | 694 | Pld,CG7718,CG9248 | | 5.019e-03 | -5.29 | HTH\_OST | prosite domains | PS51644 | 4 | 3 | 6243 | 694 | osk,Marf1,tapas | | 5.105e-03 | -5.28 | intracellular mRNA localization involved in pattern specification process | biological process | GO:0060810 | 72 | 14 | 12345 | 1129 | Grip75,Lkb1,mael,Tm1,CkIIalpha,cnc,swa,Grip128,jvl,msps,spn-E,Grip71,osk,exu | | 5.109e-03 | -5.28 | REACTOME\_TRIF\_MEDIATED\_TLR3\_SIGNALING | MSigDB lists | REACTOME\_TRIF\_MEDIATED\_TLR3\_SIGNALING | 23 | 9 | 4161 | 644 | Traf6,RpS27A,Pp2A-29B,mts,RpL40,Cdk1,p38a,rl,cact | | 5.109e-03 | -5.28 | GO\_REGULATION\_OF\_CELLULAR\_RESPONSE\_TO\_TRANSFORMING\_GROWTH\_FACTOR\_BETA\_STIMULUS | MSigDB lists | GO\_REGULATION\_OF\_CELLULAR\_RESPONSE\_TO\_TRANSFORMING\_GROWTH\_FACTOR\_BETA\_STIMULUS | 23 | 9 | 4161 | 644 | snz,Hsc70-3,Smox,RpS27A,STUB1,Pdk1,Med,RpL40,Snx6 | | 5.121e-03 | -5.27 | GO\_NEGATIVE\_REGULATION\_OF\_TRANSFERASE\_ACTIVITY | MSigDB lists | GO\_NEGATIVE\_REGULATION\_OF\_TRANSFERASE\_ACTIVITY | 127 | 31 | 4161 | 644 | RpL40,Snx6,regucalcin,Pka-R1,Ptp61F,Nprl2,Prosbeta6,Rpn12,APC7,Rpn2,mts,CG30291,Prosbeta3,eIF4A,Rpt2,CG7530,CG3376,mad2,Rpn3,Prosbeta5,CycB,Pdk1,CG4709,Cdk1,Pp2A-29B,Prosalpha5,RpS27A,Rcd5,Prosalpha3,Pka-R2,Prosbeta4 | | 5.131e-03 | -5.27 | cellular response to lipid | biological process | GO:0071396 | 37 | 9 | 12345 | 1129 | Hnf4,EcR,Nurf-38,Tak1,Stat92E,MED27,G9a,Hr78,Utx | | 5.152e-03 | -5.27 | negative regulation of nucleic acid-templated transcription | biological process | GO:1903507 | 379 | 50 | 12345 | 1129 | Chrac-14,CG4872,spn-E,sbb,Hat1,Mybbp1A,escl,psq,Su(H),Sirt7,CG2199,r2d2,hb,foxo,SREBP,Rbf,esc,dl,nos,Tlk,CG31875,piwi,CG10631,cyc,d4,pgc,EcR,Hnf4,Elba2,CG1602,CG17328,Uxt,Tip60,CG10038,ph-d,Bin1,Sox102F,egg,rept,CG10959,lin-52,wash,mael,Maf1,insv,Rbf2,E2f1,ovo,asf1,l(3)mbt | | 5.152e-03 | -5.27 | negative regulation of RNA biosynthetic process | biological process | GO:1902679 | 379 | 50 | 12345 | 1129 | wash,CG10959,lin-52,Sox102F,egg,rept,l(3)mbt,E2f1,ovo,asf1,Rbf2,Maf1,mael,insv,CG1602,CG17328,Elba2,EcR,Hnf4,pgc,Bin1,CG10038,ph-d,Uxt,Tip60,nos,Rbf,esc,dl,hb,foxo,SREBP,CG10631,cyc,d4,piwi,CG31875,Tlk,Mybbp1A,escl,CG4872,Hat1,sbb,spn-E,Chrac-14,r2d2,CG2199,Su(H),Sirt7,psq | | 5.182e-03 | -5.26 | TCR signaling | REACTOME pathways | R-DME-202403 | 71 | 19 | 3923 | 574 | Rpn3,Cul1,Rpn12,dl,Prosbeta6,Rel,Prosbeta3,RpL40,Rpt2,Prosalpha3,cact,SkpA,Prosalpha4,Prosbeta5,Rpn2,Prosbeta4,Prosalpha5,Pdk1,RpS27A | | 5.215e-03 | -5.26 | Jra (Jun-related antigen) | protein interactions | 36057 | 67 | 16 | 9204 | 1112 | CG14818,CG2199,nej,Stat92E,Tak1,Tm1,Chrac-14,yin,tkv,SkpA,Atf6,Atf3,faf,p38a,Cka,rl | | 5.225e-03 | -5.25 | microtubule organizing center part | cellular component | GO:0044450 | 57 | 12 | 11940 | 1105 | Grip71,Pp2A-29B,Grip84,Grip75,cnn,CG7716,gammaTub37C,Grip128,spd-2,Uxt,Blos2,mts | | 5.227e-03 | -5.25 | catalytic activity, acting on DNA | molecular function | GO:0140097 | 126 | 21 | 12176 | 1119 | Tengl4,mus205,DNApol-epsilon255,mre11,tos,RecQ5,Top1,Dcr-1,CG12018,mus81,Trf4-2,rept,pont,CG5285,Psf3,armi,nbs,obe,Sld5,Snm1,RpLP0 | | 5.233e-03 | -5.25 | dpp (decapentaplegic) | protein interactions | 33432 | 97 | 21 | 9204 | 1112 | nej,stet,Stat92E,Sec15,Dd,Med,tkv,mxc,ix,Cul2,spen,lwr,CycE,zpg,ago,eIF4A,cype,tor,csw,nos,trr | | 5.233e-03 | -5.25 | dom (domino) | protein interactions | 45655 | 97 | 21 | 9204 | 1112 | EcR,E2f1,Atg6,brwl,jigr1,YL-1,dco,rept,asf1,RYBP,mxc,Top1,tws,mts,Tip60,Rbf,Pp2A-29B,pont,Atg7,HP1c,Nurf-38 | | 5.266e-03 | -5.25 | oocyte anterior/posterior axis specification | biological process | GO:0007314 | 95 | 17 | 12345 | 1129 | nos,exu,Grip75,Lkb1,mael,Tm1,cnc,swa,Grip128,Moe,msps,Grip71,spn-E,osk,gus,armi,psq | | 5.271e-03 | -5.25 | GSE39820\_CTRL\_VS\_TGFBETA3\_IL6\_IL23A\_CD4\_TCELL\_DN | MSigDB lists | GSE39820\_CTRL\_VS\_TGFBETA3\_IL6\_IL23A\_CD4\_TCELL\_DN | 58 | 17 | 4161 | 644 | CG3376,Edem1,CG7997,Caf1-105,RpS18,CG2924,Scsalpha1,mtgo,AGO3,rept,Droj2,not,heph,EcR,ari-1,Pex13,TyrRS | | 5.271e-03 | -5.25 | GO\_LARGE\_RIBOSOMAL\_SUBUNIT | MSigDB lists | GO\_LARGE\_RIBOSOMAL\_SUBUNIT | 58 | 17 | 4161 | 644 | RpL38,RpL40,RpL23,RpL30,mRpL13,RpLP0,RpL13A,mRpL16,RpL13,RpL12,RpL11,mRpL17,mRpL10,RpL37a,RpL26,RpL9,RpL24 | | 5.324e-03 | -5.24 | Sensing of DNA Double Strand Breaks | REACTOME pathways | R-DME-5693548 | 6 | 4 | 3923 | 574 | nbs,mre11,Pen,Tip60 | | 5.348e-03 | -5.23 | MORF\_EI24 | MSigDB lists | MORF\_EI24 | 82 | 22 | 4161 | 644 | CG5028,CG10418,rept,CG1416,ND-51,CG4390,CG8188,Sod1,Cyt-c-p,ND-49,TyrRS,CG13630,Hat1,mEFTu1,SdhA,CCT3,crq,Jafrac2,CycB,CCT5,Prosbeta4,Mitofilin | | 5.348e-03 | -5.23 | GO\_REGULATION\_OF\_LIGASE\_ACTIVITY | MSigDB lists | GO\_REGULATION\_OF\_LIGASE\_ACTIVITY | 82 | 22 | 4161 | 644 | Rpt2,Prosalpha4,Prosbeta3,Rpn2,Rpn12,APC7,CG8188,Prosbeta6,SkpA,regucalcin,RpL40,Prosbeta4,Prosalpha3,RpS27A,Prosalpha5,Cdk1,CycB,Prosbeta5,STUB1,Rpn3,Cul1,mad2 | | 5.357e-03 | -5.23 | Ig\_E-set | interpro domains | IPR014756 | 68 | 15 | 11383 | 1233 | CG2993,shop,Irk1,Rel,AGBE,PrBP,CG4238,NFAT,dl,Su(H),Npc2a,alc,obe,RhoGDI,CG1105 | | 5.377e-03 | -5.23 | GO\_MODULATION\_BY\_VIRUS\_OF\_HOST\_MORPHOLOGY\_OR\_PHYSIOLOGY | MSigDB lists | GO\_MODULATION\_BY\_VIRUS\_OF\_HOST\_MORPHOLOGY\_OR\_PHYSIOLOGY | 12 | 6 | 4161 | 644 | Ts,Clp,Pen,Kap-alpha3,Smox,Atg7 | | 5.377e-03 | -5.23 | GO\_2\_OXOGLUTARATE\_METABOLIC\_PROCESS | MSigDB lists | GO\_2\_OXOGLUTARATE\_METABOLIC\_PROCESS | 12 | 6 | 4161 | 644 | CG5028,Idh3b,Nc73EF,Idh,CG1640,Got1 | | 5.377e-03 | -5.23 | GO\_GLUCAN\_BIOSYNTHETIC\_PROCESS | MSigDB lists | GO\_GLUCAN\_BIOSYNTHETIC\_PROCESS | 12 | 6 | 4161 | 644 | CG9485,Pgm1,Gyg,AGBE,RpS27A,RpL40 | | 5.381e-03 | -5.22 | negative regulation of response to stimulus | biological process | GO:0048585 | 389 | 51 | 12345 | 1129 | slif,Ptpmeg,alph,Dd,Cka,scny,piwi,Ubc6,ago,foxo,Traf6,mts,Rbf,Atg6,Ptp10D,GlyP,armi,Pp2A-29B,Herc4,Su(H),STUB1,dco,p38a,PGRP-LE,SkpA,apt,Cul1,insv,Stat92E,park,Nprl2,faf,lwr,Cbl,rept,nej,ATPsynD,dx,Cul2,CG17746,pn,RYBP,PpV,Ect4,dnc,Iml1,NFAT,cact,Ptp61F,chrb,Bap170 | | 5.383e-03 | -5.22 | apoptotic process | biological process | GO:0006915 | 103 | 18 | 12345 | 1129 | Tao,Tengl4,nbs,chrb,Mekk1,crq,tub,Debcl,Drep2,sws,E2f1,Rbf,cathD,Jafrac2,Cdk5,Tak1,Dcr-1,Drep1 | | 5.414e-03 | -5.22 | Toll-like receptor signaling | KEGG pathways | M00686 | 12 | 6 | 3198 | 496 | Traf6,Tak1,cact,tub,Rel,Myd88 | | 5.414e-03 | -5.22 | Toll-like receptor signaling | KEGG pathways | dme\_M00686 | 12 | 6 | 3198 | 496 | Myd88,Rel,cact,Tak1,tub,Traf6 | | 5.420e-03 | -5.22 | GSE22886\_DAY1\_VS\_DAY7\_MONOCYTE\_IN\_CULTURE\_DN | MSigDB lists | GSE22886\_DAY1\_VS\_DAY7\_MONOCYTE\_IN\_CULTURE\_DN | 87 | 23 | 4161 | 644 | CG9248,Idh,Scsalpha1,COX5A,CG14407,Aldh,AIF,Jafrac2,Cdk5,Echs1,UQCR-Q,ND-30,Thiolase,Mcad,Phb2,RpLP0,ND-SGDH,CG3860,CG9393,CG9485,Fdh,CG10932,CG5734 | | 5.440e-03 | -5.21 | transferase activity, transferring acyl groups | molecular function | GO:0016746 | 227 | 33 | 12176 | 1119 | nej,lid,Thiolase,CG31523,Sgf29,Ggt-1,e(y)1,CG14204,AgmNAT,isoQC,Naa20A,YL-1,CG8314,CG5880,rept,pont,yip2,mino,Hat1,d4,Chrac-14,Nmt,Rcd5,QC,CG10344,CG6178,CG10932,Tip60,CG8112,whd,Agpat2,CG14615,CG2938 | | 5.448e-03 | -5.21 | GO\_CELLULAR\_RESPONSE\_TO\_ORGANIC\_SUBSTANCE | MSigDB lists | GO\_CELLULAR\_RESPONSE\_TO\_ORGANIC\_SUBSTANCE | 415 | 83 | 4161 | 644 | Calx,STUB1,crq,AIF,LpR2,RpS27A,rept,ACC,shv,RpL13A,EcR,Rpn12,Prosalpha4,csw,Rpt2,Edem1,Prosbeta5,CG11710,Smox,Iru,PDCD-5,Prosalpha3,CanA-14F,Hsc70-3,Cbp80,Pdk,r,Atg6,Traf6,MED4,faf,Pp2B-14D,Vha36-1,MED24,Shmt,rl,cact,tkv,Prosalpha5,Fkbp59,Pka-R2,cl,Got1,Rab10,dnc,Tip60,Rpb8,Karybeta3,mino,S2P,adp,HDAC6,Sod1,Prip,RpII140,Rpn2,Cdk7,Prosbeta3,Gapdh1,raptor,Rpn3,bys,Moe,CycB,Ac3,Pdk1,p38a,CycE,SREBP,Dcr-1,Prosbeta4,Wnt10,RpL40,Med,Tspo,heph,Atpalpha,Pka-R1,Ubc4,Ptp61F,Vha13,Prosbeta6,CG30291 | | 5.450e-03 | -5.21 | regulation of protein catabolic process | biological process | GO:0042176 | 80 | 15 | 12345 | 1129 | park,Fbw5,Ubc6,CG4238,Rpn3,ari-1,Rpn2,Pomp,cort,dco,CkIIalpha,SkpA,Cul1,Rpt2,gus | | 5.452e-03 | -5.21 | CG5854 (uncharacterized protein) | protein interactions | 42831 | 34 | 10 | 9204 | 1112 | wal,mEFTu1,Cp1,CCT3,CCT6,SERCA,Rpn3,cathD,ScsbetaA,Mcad | | 5.459e-03 | -5.21 | GSE17974\_CTRL\_VS\_ACT\_IL4\_AND\_ANTI\_IL12\_48H\_CD4\_TCELL\_DN | MSigDB lists | GSE17974\_CTRL\_VS\_ACT\_IL4\_AND\_ANTI\_IL12\_48H\_CD4\_TCELL\_DN | 92 | 24 | 4161 | 644 | CG12018,CG3226,CG17187,Rpn12,colt,slif,CG42542,levy,Rpb8,SelD,Atg7,rept,CG10932,Arf51F,mRpL17,Prosbeta4,yip2,MED20,COX5A,Pcd,CG7655,Jafrac2,SerRS-m,Klp10A | | 5.461e-03 | -5.21 | single fertilization | biological process | GO:0007338 | 44 | 10 | 12345 | 1129 | spd-2,Nap1,msps,wisp,yem,png,alphaTub67C,cort,plu,fs(1)Ya | | 5.484e-03 | -5.21 | cype (cyclope) | protein interactions | 46040 | 7 | 4 | 9204 | 1112 | COX5A,ea,COX4,levy | | 5.484e-03 | -5.21 | CG9344 (uncharacterized protein) | protein interactions | 37372 | 7 | 4 | 9204 | 1112 | CG6610,LSm3,CG10418,CG17768 | | 5.484e-03 | -5.21 | CG2278 (uncharacterized protein) | protein interactions | 31751 | 7 | 4 | 9204 | 1112 | CG6276,CG3342,kri,Nca | | 5.484e-03 | -5.21 | CG32638 (uncharacterized protein) | protein interactions | 318133 | 7 | 4 | 9204 | 1112 | Tspo,Mcad,Clp,ago | | 5.484e-03 | -5.21 | Cht9 (Chitinase 9) | protein interactions | 37392 | 7 | 4 | 9204 | 1112 | crm,RpL29,CG12773,Rtnl1 | | 5.484e-03 | -5.21 | CG12126 (uncharacterized protein) | protein interactions | 35805 | 7 | 4 | 9204 | 1112 | CG11164,RpLP1,eIF4A,egg | | 5.484e-03 | -5.21 | ND-23 (NADH dehydrogenase (ubiquinone) 23 kDa subunit) | protein interactions | 44207 | 7 | 4 | 9204 | 1112 | ND-18,ND-75,ND-51,Pfk | | 5.488e-03 | -5.21 | Map60 (Microtubule-associated protein 60) | protein interactions | 35991 | 24 | 8 | 9204 | 1112 | PGRP-LE,Rel,lwr,CG14881,gammaTub37C,Pen,Tak1,jigr1 | | 5.488e-03 | -5.21 | CG42709 (uncharacterized protein) | protein interactions | 39453 | 24 | 8 | 9204 | 1112 | Cf2,CG8974,Rpt2,CycE,CycB,Nup358,scu,pgc | | 5.488e-03 | -5.21 | CG9951 (uncharacterized protein) | protein interactions | 39888 | 24 | 8 | 9204 | 1112 | Exo70,CG14818,Nup62,cnn,Abi,Blos2,CG5026,mbo | | 5.488e-03 | -5.21 | GO\_POSITIVE\_REGULATION\_OF\_TRANSFERASE\_ACTIVITY | MSigDB lists | GO\_POSITIVE\_REGULATION\_OF\_TRANSFERASE\_ACTIVITY | 191 | 43 | 4161 | 644 | mad2,STUB1,rl,LpR2,Cdk1,Prosalpha5,Mo25,Mekk1,RpS27A,Pka-R2,SkpA,Rpn12,CG8188,Sod1,Rpn2,mts,Prosbeta3,Prosalpha4,csw,Rpt2,raptor,lwr,Rpn3,Cul1,Prosbeta5,CycB,Cdk5,Pdk1,Ac3,p38a,nAChRalpha4,Prosalpha3,Prosbeta4,Hsc70-3,RpL40,CG17698,Pka-R1,RfC38,Prosbeta6,RpS2,Traf6,APC7,CycC | | 5.493e-03 | -5.20 | aromatic compound catabolic process | biological process | GO:0019439 | 160 | 25 | 12345 | 1129 | AOX3,Not10,CG31759,CG11164,armi,cN-IIIB,Pym,Rrp40,Tengl4,Ho,gw,DhpD,Edc3,twin,Rga,Trf4-2,CG17768,Cbp80,smg,cu,CG11486,LSm3,Dcr-1,fal,Drep1 | | 5.500e-03 | -5.20 | GSE41867\_LCMV\_ARMSTRONG\_VS\_CLONE13\_DAY8\_EFFECTOR\_CD8\_TCELL\_UP | MSigDB lists | GSE41867\_LCMV\_ARMSTRONG\_VS\_CLONE13\_DAY8\_EFFECTOR\_CD8\_TCELL\_UP | 49 | 15 | 4161 | 644 | lid,Jafrac2,CG7997,CG42542,Maf1,CG3376,Jafrac1,UbcE2H,Rbf,Gyg,l(3)mbt,CG10470,dco,RpLP0,ND-SGDH | | 5.500e-03 | -5.20 | GO\_CARBOHYDRATE\_CATABOLIC\_PROCESS | MSigDB lists | GO\_CARBOHYDRATE\_CATABOLIC\_PROCESS | 49 | 15 | 4161 | 644 | Eno,GlyP,CG8475,CG3609,CG9485,CG7322,Pfk,Pgk,Gk2,Treh,Pglym78,Nc73EF,Pgm1,Gyg,Gapdh1 | | 5.500e-03 | -5.20 | GSE30971\_CTRL\_VS\_LPS\_STIM\_MACROPHAGE\_WBP7\_KO\_2H\_DN | MSigDB lists | GSE30971\_CTRL\_VS\_LPS\_STIM\_MACROPHAGE\_WBP7\_KO\_2H\_DN | 49 | 15 | 4161 | 644 | scu,Lsm12,Wdfy2,CG7840,CG2493,Scsalpha1,Uch,ND-23,Arl5,Su(H),CG5028,disp,Tim8,CG9471,CG8199 | | 5.505e-03 | -5.20 | ubiquitin ligase complex | cellular component | GO:0000151 | 150 | 24 | 11940 | 1105 | ph-d,DCAF12,CG31633,Cul1,gus,CG30496,ScsbetaA,Cul2,EloC,park,ari-1,sip3,CG11070,ago,Ubc6,l(3)73Ah,CG10254,Fbw5,CG9316,CG5003,Cul5,CG32221,APC7,SkpA | | 5.508e-03 | -5.20 | posttranscriptional gene silencing by RNA | biological process | GO:0035194 | 58 | 12 | 12345 | 1129 | piwi,armi,r2d2,Hsc70-3,trsn,Cbp80,AGO3,CG9425,gw,CG6967,Dcr-1,spn-E | | 5.521e-03 | -5.20 | tRNA wobble base modification | biological process | GO:0002097 | 19 | 6 | 12345 | 1129 | CG10189,CG3021,CG2911,CG5292,CG2034,CG3587 | | 5.521e-03 | -5.20 | peptidoglycan recognition protein signaling pathway | biological process | GO:0061057 | 19 | 6 | 12345 | 1129 | Npc2a,Uev1A,Tak1,ben,PGRP-LE,Rel | | 5.521e-03 | -5.20 | oxidoreduction coenzyme metabolic process | biological process | GO:0006733 | 19 | 6 | 12345 | 1129 | Coq7,Coq6,CG33156,Nadsyn,Coq8,ND-39 | | 5.521e-03 | -5.20 | cell activation | biological process | GO:0001775 | 19 | 6 | 12345 | 1129 | cort,plu,wisp,crq,Jafrac1,png | | 5.521e-03 | -5.20 | negative regulation of neuron apoptotic process | biological process | GO:0043524 | 19 | 6 | 12345 | 1129 | Debcl,spag,Pink1,Tip60,Naa20A,MAGE | | 5.521e-03 | -5.20 | negative regulation of cell cycle G2/M phase transition | biological process | GO:1902750 | 19 | 6 | 12345 | 1129 | nbs,Aven,mus304,mre11,grp,Cdk1 | | 5.521e-03 | -5.20 | negative regulation of G2/M transition of mitotic cell cycle | biological process | GO:0010972 | 19 | 6 | 12345 | 1129 | grp,Cdk1,Aven,nbs,mre11,mus304 | | 5.545e-03 | -5.19 | fusome | cellular component | GO:0045169 | 37 | 9 | 11940 | 1105 | Fer1HCH,shrb,Moe,Cp1,Rtnl1,mino,tral,Surf4,bam | | 5.564e-03 | -5.19 | exonuclease activity | molecular function | GO:0004527 | 65 | 13 | 12176 | 1119 | CG11486,cu,CG31759,mre11,exu,twin,Tengl4,DNApol-epsilon255,mus205,Rga,Snm1,Exd2,tos | | 5.580e-03 | -5.19 | microtubule polymerization or depolymerization | biological process | GO:0031109 | 31 | 8 | 12345 | 1129 | Grip75,gammaTub37C,Klp10A,Grip84,Grip128,CG7716,msps,stai | | 5.580e-03 | -5.19 | intracellular receptor signaling pathway | biological process | GO:0030522 | 31 | 8 | 12345 | 1129 | EcR,Hnf4,Fkbp39,Nurf-38,Chd64,G9a,Nup358,Hr78 | | 5.580e-03 | -5.19 | positive regulation of proteasomal ubiquitin-dependent protein catabolic process | biological process | GO:0032436 | 31 | 8 | 12345 | 1129 | Ubc6,park,ari-1,dco,cort,Pomp,SkpA,Cul1 | | 5.595e-03 | -5.19 | Lsp2 (Larval serum protein 2) | protein interactions | 45326 | 29 | 9 | 9204 | 1112 | Fer1HCH,GlyP,Scp1,Pglym78,CG32473,Adh,Pgk,Argk,CG4390 | | 5.595e-03 | -5.19 | lace (lace) | protein interactions | 34910 | 29 | 9 | 9204 | 1112 | csw,Vha13,rl,RpS16,RpS27A,RpL31,Sply,GCS2beta,Su(H) | | 5.608e-03 | -5.18 | S Phase | REACTOME pathways | R-DME-69242 | 124 | 29 | 3923 | 574 | SkpA,Rbf,Sld5,Prosalpha3,RpL40,Rpt2,RpS27A,Prosalpha5,Prosbeta4,Rbf2,Prosbeta5,CG12018,Rpn2,Prosalpha4,twe,Chrac-14,CycE,DNApol-epsilon255,wapl,Prosbeta6,Rpn12,Orc1,Rpn3,Cul1,Psf3,Orc6,Prosbeta3,Cdk7,RfC38 | | 5.663e-03 | -5.17 | CCT7 (Chaperonin containing TCP1 subunit 7) | protein interactions | 41054 | 56 | 14 | 9204 | 1112 | Rel,MED4,PGRP-LE,Mcad,Kap-alpha3,CCT6,Karybeta3,Vha68-2,wal,Pp2A-29B,CCT3,Rtnl1,Tak1,CCT5 | | 5.667e-03 | -5.17 | membrane organization | biological process | GO:0061024 | 220 | 32 | 12345 | 1129 | Ankle2,slam,Topors,Vps16B,CG31729,Dd,sau,Arl6IP1,Vamp7,CG9981,Rab5,sws,Nlg1,Sec22,csw,scramb1,Pmi,wash,mino,CycB,Mtl,Rtnl1,Syx13,Snx6,crq,Debcl,ATP8A,CG5903,lt,Mekk1,EcR,twe | | 5.745e-03 | -5.16 | GO\_REGULATION\_OF\_MITOTIC\_CELL\_CYCLE | MSigDB lists | GO\_REGULATION\_OF\_MITOTIC\_CELL\_CYCLE | 170 | 39 | 4161 | 644 | mad2,Cdk1,Prosalpha5,RpS27A,shrb,twin,SkpA,Rpn2,Not10,grp,eIF4E1,CG1812,Rpn12,Prosalpha4,Rpt2,csw,CG11164,Prosbeta3,Rpn3,Cul1,nmdyn-D6,CHMP2B,Prosbeta5,CycB,Smox,APP-BP1,Prosbeta4,Wnt10,Prosalpha3,Aos1,RpL40,not,Prosbeta6,Phb2,Atg6,APC7,Rbf,CkIIalpha,CG30291 | | 5.793e-03 | -5.15 | regulation of mRNA catabolic process | biological process | GO:0061013 | 25 | 7 | 12345 | 1129 | spn-E,osk,gw,Marf1,AGO3,piwi,smg | | 5.793e-03 | -5.15 | glycolytic process | biological process | GO:0006096 | 25 | 7 | 12345 | 1129 | Pfk,Gapdh1,Eno,Pgk,PyK,Pfrx,Pglym78 | | 5.793e-03 | -5.15 | regulation of oocyte development | biological process | GO:0060281 | 25 | 7 | 12345 | 1129 | spn-E,exu,swa,cnc,mael,enc,cup | | 5.793e-03 | -5.15 | regulation of pole plasm oskar mRNA localization | biological process | GO:0007317 | 25 | 7 | 12345 | 1129 | cup,enc,mael,cnc,exu,swa,spn-E | | 5.793e-03 | -5.15 | ATP generation from ADP | biological process | GO:0006757 | 25 | 7 | 12345 | 1129 | Pgk,Eno,Gapdh1,Pfk,Pglym78,Pfrx,PyK | | 5.793e-03 | -5.15 | proteasomal ubiquitin-independent protein catabolic process | biological process | GO:0010499 | 25 | 7 | 12345 | 1129 | Prosalpha5,Prosbeta5,Prosbeta4,Prosalpha3,Prosalpha4,Prosbeta3,Prosbeta6 | | 5.831e-03 | -5.14 | GSE17974\_0H\_VS\_24H\_IN\_VITRO\_ACT\_CD4\_TCELL\_DN | MSigDB lists | GSE17974\_0H\_VS\_24H\_IN\_VITRO\_ACT\_CD4\_TCELL\_DN | 68 | 19 | 4161 | 644 | Psf3,CG17187,grp,Rrp40,CG42542,Cbp80,l(1)G0004,mRpL17,CG30496,Pfas,MED20,CycE,r-l,Stip1,Jafrac2,Pfk,Caf1-105,Cdk5,bys | | 5.837e-03 | -5.14 | Met (Methoprene-tolerant) | protein interactions | 32114 | 15 | 6 | 9204 | 1112 | Pfdn2,Chd64,Fkbp39,CG12106,Droj2,EcR | | 5.837e-03 | -5.14 | CG7054 (uncharacterized protein) | protein interactions | 42643 | 15 | 6 | 9204 | 1112 | AnxB10,CG11267,awd,CG10638,Surf4,CG11164 | | 5.879e-03 | -5.14 | HAD-like\_sf | interpro domains | IPR036412 | 56 | 13 | 11383 | 1233 | Atpalpha,Gs1l,Ublcp1,CG32549,CG31729,cN-IIIB,SERCA,ATP8A,Dd,CG9981,aay,CG17294,anne | | 5.917e-03 | -5.13 | pAbp (poly(A) binding protein) | protein interactions | 37070 | 62 | 15 | 9204 | 1112 | cup,BicC,Imp,Lsm12,tral,tyf,RpS27A,twin,Rga,eIF4E1,Unr,GILT1,larp,osk,gw | | 5.934e-03 | -5.13 | cell division site part | cellular component | GO:0032155 | 44 | 10 | 11940 | 1105 | Grip84,Cen,Arf51F,sau,Moe,Grip75,Grip128,Sry-alpha,CG7716,slam | | 5.950e-03 | -5.12 | oxidoreductase activity, acting on NAD(P)H, quinone or similar compound as acceptor | molecular function | GO:0016655 | 25 | 7 | 12176 | 1119 | CG7322,ND-18,ND-75,ND-B18,ND-B15,ND-51,ND-30 | | 5.951e-03 | -5.12 | TAKAO\_RESPONSE\_TO\_UVB\_RADIATION\_DN | MSigDB lists | TAKAO\_RESPONSE\_TO\_UVB\_RADIATION\_DN | 36 | 12 | 4161 | 644 | lqfR,Arpc4,PIG-C,CG11089,Echs1,Phb2,RpS23,CG4390,yip2,Cdk7,Prosalpha3,Nc73EF | | 5.951e-03 | -5.12 | GSE3920\_IFNA\_VS\_IFNB\_TREATED\_ENDOTHELIAL\_CELL\_UP | MSigDB lists | GSE3920\_IFNA\_VS\_IFNB\_TREATED\_ENDOTHELIAL\_CELL\_UP | 36 | 12 | 4161 | 644 | Chd64,disp,cyc,ND-30,PrBP,sws,Axs,adp,CG33181,RpS30,obe,Sec15 | | 5.979e-03 | -5.12 | GSE35543\_IN\_VITRO\_ITREG\_VS\_CONVERTED\_EX\_ITREG\_UP | MSigDB lists | GSE35543\_IN\_VITRO\_ITREG\_VS\_CONVERTED\_EX\_ITREG\_UP | 54 | 16 | 4161 | 644 | Uch,DNApol-epsilon255,Caf1-105,CG7322,Cdk1,CG7655,CG14715,mRpS10,CG15525,mad2,Ntan1,CG11164,CG6767,CG8188,HDAC6,ND-23 | | 5.979e-03 | -5.12 | SHETH\_LIVER\_CANCER\_VS\_TXNIP\_LOSS\_PAM1 | MSigDB lists | SHETH\_LIVER\_CANCER\_VS\_TXNIP\_LOSS\_PAM1 | 54 | 16 | 4161 | 644 | stas,Nap1,Axs,CG1774,Arf51F,nmdyn-D6,RpL12,PIG-C,crq,aay,CG8188,Tsp86D,Rpt2,P5cr,cyc,Tsp74F | | 6.020e-03 | -5.11 | GSE14415\_INDUCED\_TREG\_VS\_TCONV\_UP | MSigDB lists | GSE14415\_INDUCED\_TREG\_VS\_TCONV\_UP | 73 | 20 | 4161 | 644 | mRpL18,CG11899,CG9393,RfC38,NHP2,CG10418,LSm3,Prosbeta4,Surf4,Grip84,Prosalpha5,Lsm12,CG7655,crq,Cdk1,lwr,Hat1,mad2,Rpn3,Shmt | | 6.022e-03 | -5.11 | nuclear-transcribed mRNA poly(A) tail shortening | biological process | GO:0000289 | 9 | 4 | 12345 | 1129 | twin,smg,CG11486,cu | | 6.022e-03 | -5.11 | establishment of protein localization to peroxisome | biological process | GO:0072663 | 9 | 4 | 12345 | 1129 | Topors,Pex13,Pex10,Pex12 | | 6.022e-03 | -5.11 | positive regulation of meiotic cell cycle | biological process | GO:0051446 | 9 | 4 | 12345 | 1129 | twe,Iml1,Nprl2,CG31053 | | 6.022e-03 | -5.11 | protein localization to peroxisome | biological process | GO:0072662 | 9 | 4 | 12345 | 1129 | Pex13,Topors,Pex12,Pex10 | | 6.022e-03 | -5.11 | protein targeting to peroxisome | biological process | GO:0006625 | 9 | 4 | 12345 | 1129 | Topors,Pex13,Pex10,Pex12 | | 6.052e-03 | -5.11 | segmentation | biological process | GO:0035282 | 256 | 36 | 12345 | 1129 | fs(1)N,Grip71,CycC,bcd,osk,Moe,gus,spen,Dcr-1,rl,crm,mael,cad,Tm1,Stat92E,swa,ovo,apt,Grip128,spn-E,msps,csw,trk,fs(1)M3,armi,psq,disp,hb,tor,nos,exu,pygo,Grip75,Lkb1,Bx,cnc | | 6.087e-03 | -5.10 | TRAF6 mediated induction of NFkB and MAP kinases upon TLR7/8 or 9 activation | REACTOME pathways | R-DME-975138 | 38 | 12 | 3923 | 574 | Myd88,RpS27A,rl,ben,Traf6,Tak1,Uev1A,Rel,p38a,Pp2A-29B,dl,RpL40 | | 6.087e-03 | -5.10 | Toll Like Receptor 7/8 (TLR7/8) Cascade | REACTOME pathways | R-DME-168181 | 38 | 12 | 3923 | 574 | RpL40,dl,p38a,Pp2A-29B,Rel,Uev1A,Tak1,Traf6,ben,rl,RpS27A,Myd88 | | 6.087e-03 | -5.10 | MyD88 dependent cascade initiated on endosome | REACTOME pathways | R-DME-975155 | 38 | 12 | 3923 | 574 | RpL40,dl,Pp2A-29B,p38a,Uev1A,Rel,Tak1,ben,Traf6,rl,RpS27A,Myd88 | | 6.098e-03 | -5.10 | Endo/exonuclease/phosphatase | interpro domains | IPR005135 | 12 | 5 | 11383 | 1233 | cu,Synj,nSMase,CG31759,twin | | 6.098e-03 | -5.10 | Enoyl-CoA\_hydra/iso | interpro domains | IPR001753 | 12 | 5 | 11383 | 1233 | CG5844,CG4594,Mtpalpha,Echs1,CG4598 | | 6.098e-03 | -5.10 | Endo/exonu/phosph\_ase\_sf | interpro domains | IPR036691 | 12 | 5 | 11383 | 1233 | CG31759,twin,nSMase,Synj,cu | | 6.124e-03 | -5.10 | Hsc70-4 (Heat shock protein cognate 4) | protein interactions | 41840 | 86 | 19 | 9204 | 1112 | Zwilch,HDAC6,cathD,ph-d,EcR,dx,mad2,ago,RpS27A,Rel,CycE,PGRP-LE,cup,Cdk7,ND-MNLL,spag,Stip1,STUB1,Tak1 | | 6.129e-03 | -5.09 | microtubule minus-end binding | molecular function | GO:0051011 | 9 | 4 | 12176 | 1119 | Grip84,Grip75,CG7716,Grip128 | | 6.129e-03 | -5.09 | 3',5'-cyclic-nucleotide phosphodiesterase activity | molecular function | GO:0004114 | 9 | 4 | 12176 | 1119 | PrBP,Pka-R1,Pka-R2,dnc | | 6.129e-03 | -5.09 | small ribosomal subunit rRNA binding | molecular function | GO:0070181 | 9 | 4 | 12176 | 1119 | mRpS11,RpS13,RpS14a,RpS14b | | 6.129e-03 | -5.09 | C-acyltransferase activity | molecular function | GO:0016408 | 9 | 4 | 12176 | 1119 | yip2,CG6178,CG10932,Thiolase | | 6.165e-03 | -5.09 | bcd (bicoid) | protein interactions | 40830 | 80 | 18 | 9204 | 1112 | plu,png,tor,CG9248,eIF4E1,hb,CG11899,sw,CG31759,nos,trk,Bin1,exu,wisp,CG13373,RpS27A,cad,SkpA | | 6.189e-03 | -5.08 | negative regulation of stem cell proliferation | biological process | GO:2000647 | 38 | 9 | 12345 | 1129 | tws,MED27,COX4,Rel,mts,MED4,EcR,MAGE,COX5A | | 6.189e-03 | -5.08 | regulation of proteasomal ubiquitin-dependent protein catabolic process | biological process | GO:0032434 | 38 | 9 | 12345 | 1129 | Cul1,CkIIalpha,Pomp,cort,dco,SkpA,ari-1,Ubc6,park | | 6.189e-03 | -5.08 | detoxification | biological process | GO:0098754 | 38 | 9 | 12345 | 1129 | Jafrac1,Sod1,Fer1HCH,CG4390,Sod3,ZnT35C,Cyp6v1,Fdh,Trxr-1 | | 6.200e-03 | -5.08 | GO\_CELL\_CYCLE\_PHASE\_TRANSITION | MSigDB lists | GO\_CELL\_CYCLE\_PHASE\_TRANSITION | 108 | 27 | 4161 | 644 | Cul1,msps,Cdk1,Cul5,CycB,Pp2A-29B,RpS27A,DNApol-epsilon255,gammaTub37C,CycE,Wnt10,shrb,CanA-14F,RpL40,PpV,Ts,SkpA,Sirt7,dco,Pp2B-14D,grp,eIF4E1,CG8188,Dhfr,tws,Cdk7,Cul2 | | 6.206e-03 | -5.08 | nuclear envelope | cellular component | GO:0005635 | 119 | 20 | 11940 | 1105 | sbr,Nup62,fs(1)Ya,Parp16,wash,Aladin,bif,CG8009,Dd,CG7655,Nup58,Axs,Cse1,mbo,toc,CG4880,bru1,Ndc1,SREBP,mad2 | | 6.248e-03 | -5.08 | BIDUS\_METASTASIS\_UP | MSigDB lists | BIDUS\_METASTASIS\_UP | 83 | 22 | 4161 | 644 | eIF4E1,Cse1,faf,Tlk,grp,mts,Dhfr,Efr,CG8963,CG17593,Ts,Grip75,Nap1,Utx,Cka,stet,Sec23,mad2,msps,CycB,rl,Cdk1 | | 6.255e-03 | -5.07 | Exo\_endo\_phos | pfam domains | PF03372 | 12 | 5 | 10480 | 1142 | cu,Synj,CG31759,nSMase,twin | | 6.255e-03 | -5.07 | ECH\_1 | pfam domains | PF00378 | 12 | 5 | 10480 | 1142 | Echs1,CG5844,CG4594,Mtpalpha,CG4598 | | 6.286e-03 | -5.07 | mitochondrial tricarboxylic acid cycle enzyme complex | cellular component | GO:0030062 | 9 | 4 | 11940 | 1105 | Nc73EF,Scsalpha1,CG17691,CG8199 | | 6.287e-03 | -5.07 | regulation of defense response to fungus | biological process | GO:1900150 | 14 | 5 | 12345 | 1129 | cact,spz,tub,Myd88,Rel | | 6.287e-03 | -5.07 | cellular metabolic compound salvage | biological process | GO:0043094 | 14 | 5 | 12345 | 1129 | Pdxk,CG11134,l(2)k01209,Adi1,AMPdeam | | 6.287e-03 | -5.07 | nuclear envelope organization | biological process | GO:0006998 | 14 | 5 | 12345 | 1129 | Topors,wash,Ankle2,twe,Dd | | 6.292e-03 | -5.07 | Tudor-SN (Tudor staphylococcal nuclease) | protein interactions | 38045 | 11 | 5 | 9204 | 1112 | Moe,piwi,Rtnl1,Naa20A,Stip1 | | 6.292e-03 | -5.07 | CG13085 (uncharacterized protein) | protein interactions | 35219 | 11 | 5 | 9204 | 1112 | Idgf4,SkpA,CG30430,Prosbeta3,Nc73EF | | 6.292e-03 | -5.07 | PMCA (plasma membrane calcium ATPase) | protein interactions | 43787 | 11 | 5 | 9204 | 1112 | CG10425,mus304,CG31729,osk,ago | | 6.292e-03 | -5.07 | AnxB9 (Annexin B9) | protein interactions | 42492 | 11 | 5 | 9204 | 1112 | cyc,AnxB10,Uch,CG14715,PHGPx | | 6.369e-03 | -5.06 | GSE3982\_EOSINOPHIL\_VS\_EFF\_MEMORY\_CD4\_TCELL\_UP | MSigDB lists | GSE3982\_EOSINOPHIL\_VS\_EFF\_MEMORY\_CD4\_TCELL\_UP | 59 | 17 | 4161 | 644 | Crag,CG14894,Ubc4,Rab5,Adi1,scramb1,Synj,CG33156,SRPK,CG7997,CG8611,CG9911,PIG-C,shop,pen-2,Hat1,CG1774 | | 6.369e-03 | -5.06 | BROWNE\_HCMV\_INFECTION\_20HR\_UP | MSigDB lists | BROWNE\_HCMV\_INFECTION\_20HR\_UP | 59 | 17 | 4161 | 644 | Atpalpha,Pdk,escl,SelD,colt,CG8188,l(3)73Ah,Ptp10D,csw,CG3021,UbcE2H,CycC,CG9911,CG9391,esc,Gabat,Iru | | 6.369e-03 | -5.06 | GSE369\_PRE\_VS\_POST\_IL6\_INJECTION\_SOCS3\_KO\_LIVER\_UP | MSigDB lists | GSE369\_PRE\_VS\_POST\_IL6\_INJECTION\_SOCS3\_KO\_LIVER\_UP | 59 | 17 | 4161 | 644 | RpL23,muskelin,Gk2,Su(H),opm,CHOp24,CG7550,RpL24,RpL9,Cka,Arf51F,Arpc2,Pgk,Pglym78,CG7997,CHMP2B,Ac3 | | 6.369e-03 | -5.06 | GO\_PROTEIN\_DEPHOSPHORYLATION | MSigDB lists | GO\_PROTEIN\_DEPHOSPHORYLATION | 59 | 17 | 4161 | 644 | Ublcp1,CanB2,PpV,Ptp61F,Ptp10D,Cyt-c-p,Pp2B-14D,mts,CG7115,Dd,csw,alph,Ptpmeg,Acph-1,PpD3,Pp2A-29B,CanA-14F | | 6.378e-03 | -5.05 | MORF\_AATF | MSigDB lists | MORF\_AATF | 134 | 32 | 4161 | 644 | P5CDh1,Dd,Phb2,GCS2beta,CG8204,CG17454,r,CG10418,PrBP,CG1416,CG7728,APP-BP1,Mitofilin,eIF3m,pont,Cul1,Rpn3,Thiolase,CG13630,lwr,Marf,mts,Prosbeta3,CG9393,eIF3i,rept,CSN6,Idh3b,crq,SdhA,Chrac-14,mEFTu1 | | 6.396e-03 | -5.05 | regulation of molecular function | biological process | GO:0065009 | 523 | 65 | 12345 | 1129 | lwr,Cbl,CG1416,CycB,Pax,ScsbetaA,MED24,EndoGI,Tomosyn,Aos1,Rpn2,Mekk1,Rpn3,Iml1,blw,CycC,Tao,SNF4Agamma,CG17746,levy,tws,CG13551,esc,Jafrac2,wapl,CycJ,Tak1,GAPsec,Trs33,CG14408,cort,Abi,fray,S2P,ASPP,sws,Pi3K92E,rl,Drep1,SERCA,RhoGAP5A,CG8726,cact,CG4041,Orc6,Rbf,crok,Traf6,Parp16,Pdk1,Lkb1,alph,RhoGAP54D,CycE,Rpn13,CG30291,eIF3m,Crag,Sec23,Cdk7,Pp2A-29B,nrv1,RpLP1,raptor,CG13344 | | 6.401e-03 | -5.05 | CG13151 (uncharacterized protein) | protein interactions | 36374 | 4 | 3 | 9204 | 1112 | CG10959,CG2199,CG2129 | | 6.401e-03 | -5.05 | CG14260 (uncharacterized protein) | protein interactions | 43287 | 4 | 3 | 9204 | 1112 | Phb2,Rtnl1,Hsp26 | | 6.401e-03 | -5.05 | CG11839 (uncharacterized protein) | protein interactions | 43006 | 4 | 3 | 9204 | 1112 | SelD,CG14036,CG4860 | | 6.401e-03 | -5.05 | CG4278 (uncharacterized protein) | protein interactions | 34970 | 4 | 3 | 9204 | 1112 | isoQC,Arpc4,Sod1 | | 6.401e-03 | -5.05 | Blimp-1 (Blimp-1) | protein interactions | 38638 | 4 | 3 | 9204 | 1112 | RpL37a,SkpA,RpS29 | | 6.401e-03 | -5.05 | gd (gastrulation-defective) | protein interactions | 32159 | 4 | 3 | 9204 | 1112 | gd,ea,snk | | 6.401e-03 | -5.05 | CR31562 (pseudo) | protein interactions | 19835899 | 4 | 3 | 9204 | 1112 | CG11899,yl,EloC | | 6.401e-03 | -5.05 | CG7156 (uncharacterized protein) | protein interactions | 42187 | 4 | 3 | 9204 | 1112 | CG3309,Marf1,Prosalpha5 | | 6.401e-03 | -5.05 | CG14692 (uncharacterized protein) | protein interactions | 41298 | 4 | 3 | 9204 | 1112 | CG11164,CG2200,Smox | | 6.401e-03 | -5.05 | Spn88Eb (Serpin 88Eb) | protein interactions | 41829 | 4 | 3 | 9204 | 1112 | Ssdp,Rtnl1,Bx | | 6.404e-03 | -5.05 | GO\_NEGATIVE\_REGULATION\_OF\_PEPTIDYL\_TYROSINE\_PHOSPHORYLATION | MSigDB lists | GO\_NEGATIVE\_REGULATION\_OF\_PEPTIDYL\_TYROSINE\_PHOSPHORYLATION | 9 | 5 | 4161 | 644 | Pp2A-29B,Snx6,mts,CG4709,Ptp61F | | 6.404e-03 | -5.05 | REACTOME\_GLYCOGEN\_BREAKDOWN\_GLYCOGENOLYSIS | MSigDB lists | REACTOME\_GLYCOGEN\_BREAKDOWN\_GLYCOGENOLYSIS | 9 | 5 | 4161 | 644 | CG9485,CG8475,Gyg,GlyP,Pgm1 | | 6.404e-03 | -5.05 | KEGG\_ALDOSTERONE\_REGULATED\_SODIUM\_REABSORPTION | MSigDB lists | KEGG\_ALDOSTERONE\_REGULATED\_SODIUM\_REABSORPTION | 9 | 5 | 4161 | 644 | Pdk1,rl,Pi3K92E,nrv1,Atpalpha | | 6.404e-03 | -5.05 | BIOCARTA\_G1\_PATHWAY | MSigDB lists | BIOCARTA\_G1\_PATHWAY | 9 | 5 | 4161 | 644 | Cdk1,Dhfr,Med,CycE,Smox | | 6.404e-03 | -5.05 | LANDIS\_ERBB2\_BREAST\_TUMORS\_65\_DN | MSigDB lists | LANDIS\_ERBB2\_BREAST\_TUMORS\_65\_DN | 9 | 5 | 4161 | 644 | yip2,mino,Gyg,Pgm1,CG17691 | | 6.404e-03 | -5.05 | REACTOME\_SPRY\_REGULATION\_OF\_FGF\_SIGNALING | MSigDB lists | REACTOME\_SPRY\_REGULATION\_OF\_FGF\_SIGNALING | 9 | 5 | 4161 | 644 | RpL40,RpS27A,Pp2A-29B,mts,rl | | 6.404e-03 | -5.05 | LI\_WILMS\_TUMOR\_ANAPLASTIC\_UP | MSigDB lists | LI\_WILMS\_TUMOR\_ANAPLASTIC\_UP | 9 | 5 | 4161 | 644 | Cdk1,RfC38,Hat1,l(3)73Ah,Pen | | 6.459e-03 | -5.04 | eye pigmentation | biological process | GO:0048069 | 45 | 10 | 12345 | 1129 | DhpD,pn,Rab32,EMC3,Hexo1,lt,PAPLA1,mal,cl,fdl | | 6.527e-03 | -5.03 | process utilizing autophagic mechanism | biological process | GO:0061919 | 97 | 17 | 12345 | 1129 | Atg7,Atg4b,Debcl,EcR,lt,CG42554,raptor,Pink1,Vamp7,Atg6,Pp2A-29B,SNF4Agamma,Ubc6,park,shrb,mts,Pi3K92E | | 6.527e-03 | -5.03 | autophagy | biological process | GO:0006914 | 97 | 17 | 12345 | 1129 | Pink1,Vamp7,Atg6,raptor,CG42554,Pp2A-29B,SNF4Agamma,Atg7,Debcl,Atg4b,lt,EcR,Ubc6,park,shrb,mts,Pi3K92E | | 6.540e-03 | -5.03 | GO\_POSITIVE\_REGULATION\_OF\_PROTEIN\_METABOLIC\_PROCESS | MSigDB lists | GO\_POSITIVE\_REGULATION\_OF\_PROTEIN\_METABOLIC\_PROCESS | 366 | 74 | 4161 | 644 | STUB1,AIF,LpR2,RpS27A,Mekk1,CG30496,rept,regucalcin,CG31715,EcR,Rpn12,mts,Prosalpha4,csw,Rpt2,CG3376,CG8112,Edem1,Prosbeta5,CG46338,Smox,PDCD-5,Wdfy2,Prosalpha3,Hsc70-3,AGO3,Atg7,CG17698,ari-1,Cyt-c-p,Traf6,Phb2,CkIIalpha,CycC,CG8963,mad2,cact,rl,Cdk1,Mo25,Prosalpha5,Sec22,Pka-R2,Tip60,escl,SkpA,CG8188,Sod1,Rpn2,Prosbeta3,raptor,Rpn3,Cul1,Moe,Cdk5,esc,Gclc,CycB,Ac3,Pdk1,p38a,nAChRalpha4,SREBP,Prosbeta4,Tao,RpL40,Med,Ubc4,Pka-R1,Prosbeta6,RpS4,APC7,dco,CG30291 | | 6.560e-03 | -5.03 | GO\_REGULATION\_OF\_DNA\_TEMPLATED\_TRANSCRIPTION\_IN\_RESPONSE\_TO\_STRESS | MSigDB lists | GO\_REGULATION\_OF\_DNA\_TEMPLATED\_TRANSCRIPTION\_IN\_RESPONSE\_TO\_STRESS | 16 | 7 | 4161 | 644 | RpS27A,grp,S2P,Su(H),RpL40,Hsc70-3,Cul2 | | 6.560e-03 | -5.03 | GO\_ATP\_GENERATION\_FROM\_ADP | MSigDB lists | GO\_ATP\_GENERATION\_FROM\_ADP | 16 | 7 | 4161 | 644 | Pfk,Gapdh1,Nc73EF,Pgm1,Eno,Pglym78,Pgk | | 6.560e-03 | -5.03 | BIOCARTA\_CHREBP2\_PATHWAY | MSigDB lists | BIOCARTA\_CHREBP2\_PATHWAY | 16 | 7 | 4161 | 644 | Pp2A-29B,alc,ACC,mts,tws,Pka-R1,Pka-R2 | | 6.560e-03 | -5.03 | GO\_THREONINE\_TYPE\_PEPTIDASE\_ACTIVITY | MSigDB lists | GO\_THREONINE\_TYPE\_PEPTIDASE\_ACTIVITY | 16 | 7 | 4161 | 644 | Prosalpha5,Prosalpha3,Prosbeta3,Prosbeta5,Prosbeta6,Prosbeta4,Prosalpha4 | | 6.570e-03 | -5.03 | chr4q23 | MSigDB lists | chr4q23 | 6 | 4 | 4161 | 644 | Fdh,tkv,CG13630,CG1827 | | 6.570e-03 | -5.03 | GO\_DNA\_METHYLATION\_INVOLVED\_IN\_GAMETE\_GENERATION | MSigDB lists | GO\_DNA\_METHYLATION\_INVOLVED\_IN\_GAMETE\_GENERATION | 6 | 4 | 4161 | 644 | mael,AGO3,spn-E,shu | | 6.570e-03 | -5.03 | GO\_LIPOPOLYSACCHARIDE\_MEDIATED\_SIGNALING\_PATHWAY | MSigDB lists | GO\_LIPOPOLYSACCHARIDE\_MEDIATED\_SIGNALING\_PATHWAY | 6 | 4 | 4161 | 644 | p38a,rl,cact,crq | | 6.570e-03 | -5.03 | BIOCARTA\_STATHMIN\_PATHWAY | MSigDB lists | BIOCARTA\_STATHMIN\_PATHWAY | 6 | 4 | 4161 | 644 | Cdk1,CycB,Pka-R2,Pka-R1 | | 6.570e-03 | -5.03 | GO\_PIRNA\_METABOLIC\_PROCESS | MSigDB lists | GO\_PIRNA\_METABOLIC\_PROCESS | 6 | 4 | 4161 | 644 | mael,AGO3,spn-E,shu | | 6.570e-03 | -5.03 | GO\_CELLULAR\_RESPONSE\_TO\_OSMOTIC\_STRESS | MSigDB lists | GO\_CELLULAR\_RESPONSE\_TO\_OSMOTIC\_STRESS | 6 | 4 | 4161 | 644 | Tspo,Prip,Mo25,fray | | 6.570e-03 | -5.03 | REACTOME\_MITOCHONDRIAL\_FATTY\_ACID\_BETA\_OXIDATION | MSigDB lists | REACTOME\_MITOCHONDRIAL\_FATTY\_ACID\_BETA\_OXIDATION | 6 | 4 | 4161 | 644 | Mcad,Echs1,Mtpalpha,Thiolase | | 6.570e-03 | -5.03 | GO\_PYRIMIDINE\_NUCLEOSIDE\_TRIPHOSPHATE\_BIOSYNTHETIC\_PROCESS | MSigDB lists | GO\_PYRIMIDINE\_NUCLEOSIDE\_TRIPHOSPHATE\_BIOSYNTHETIC\_PROCESS | 6 | 4 | 4161 | 644 | Ts,r,nmdyn-D7,nmdyn-D6 | | 6.570e-03 | -5.03 | KEGG\_LEISHMANIA\_INFECTION | MSigDB lists | KEGG\_LEISHMANIA\_INFECTION | 6 | 4 | 4161 | 644 | cact,rl,p38a,Traf6 | | 6.605e-03 | -5.02 | cellular response to hormone stimulus | biological process | GO:0032870 | 105 | 18 | 12345 | 1129 | PyK,EcR,Hnf4,Rab10,Chd64,MED27,G9a,Utx,raptor,foxo,Pi3K92E,Nurf-38,Pdk1,rl,Fkbp39,Hr78,mAChR-A,Nup358 | | 6.614e-03 | -5.02 | neuronal ribonucleoprotein granule | cellular component | GO:0071598 | 14 | 5 | 11940 | 1105 | cup,yps,eIF4E1,nos,tral | | 6.614e-03 | -5.02 | clathrin vesicle coat | cellular component | GO:0030125 | 14 | 5 | 11940 | 1105 | Afti,Hip1,CG9132,Clc,AP-1-2beta | | 6.624e-03 | -5.02 | juvenile hormone mediated signaling pathway | biological process | GO:0035626 | 5 | 3 | 12345 | 1129 | Nup358,Fkbp39,Chd64 | | 6.624e-03 | -5.02 | phosphatidic acid metabolic process | biological process | GO:0046473 | 5 | 3 | 12345 | 1129 | Agpat2,Pld,Seipin | | 6.624e-03 | -5.02 | IMP biosynthetic process | biological process | GO:0006188 | 5 | 3 | 12345 | 1129 | AMPdeam,Pfas,CG11089 | | 6.624e-03 | -5.02 | positive regulation of DNA endoreduplication | biological process | GO:0032877 | 5 | 3 | 12345 | 1129 | otu,E2f1,SuUR | | 6.624e-03 | -5.02 | spindle assembly involved in female meiosis I | biological process | GO:0007057 | 5 | 3 | 12345 | 1129 | gammaTub37C,Grip71,Klp10A | | 6.624e-03 | -5.02 | L-serine metabolic process | biological process | GO:0006563 | 5 | 3 | 12345 | 1129 | CG11899,aay,Shmt | | 6.624e-03 | -5.02 | guanosine-containing compound biosynthetic process | biological process | GO:1901070 | 5 | 3 | 12345 | 1129 | awd,nmdyn-D7,nmdyn-D6 | | 6.624e-03 | -5.02 | fatty-acyl-CoA biosynthetic process | biological process | GO:0046949 | 5 | 3 | 12345 | 1129 | CG8199,CG17691,CG6178 | | 6.624e-03 | -5.02 | acetaldehyde metabolic process | biological process | GO:0006117 | 5 | 3 | 12345 | 1129 | Aldh,Adh,AOX3 | | 6.644e-03 | -5.01 | cellular component assembly involved in morphogenesis | biological process | GO:0010927 | 154 | 24 | 12345 | 1129 | bun,armi,SelR,fs(1)M3,CanB2,Pxt,gus,CG13366,Cf2,jvl,CycE,fs(1)N,spn-E,cact,mael,Actn,bru1,E2f1,up,Rbf,squ,Pax,Orc1,rl | | 6.679e-03 | -5.01 | GSE3920\_UNTREATED\_VS\_IFNA\_TREATED\_FIBROBLAST\_DN | MSigDB lists | GSE3920\_UNTREATED\_VS\_IFNA\_TREATED\_FIBROBLAST\_DN | 64 | 18 | 4161 | 644 | ACC,disp,CG1640,CG4511,Vha13,Ptp61F,Mctp,lt,CG2747,Sec15,Dd,CG8611,alpha-Man-Ia,r-l,CG8507,Scsalpha1,Pex12,CG9248 | | 6.682e-03 | -5.01 | protein tyrosine phosphatase activity | molecular function | GO:0004725 | 45 | 10 | 12176 | 1119 | Ptp10D,csw,Ptpmeg,twe,Ptp69D,Ptp61F,Ptpmeg2,CG5026,ssh,mop | | 6.699e-03 | -5.01 | GCP\_N | interpro domains | IPR041470 | 8 | 4 | 11383 | 1233 | Grip84,Grip128,CG7716,Grip75 | | 6.699e-03 | -5.01 | GCP | interpro domains | IPR007259 | 8 | 4 | 11383 | 1233 | Grip128,Grip75,CG7716,Grip84 | | 6.699e-03 | -5.01 | PRTase-like | interpro domains | IPR029057 | 8 | 4 | 11383 | 1233 | l(2)k01209,CG6767,r-l,kri | | 6.699e-03 | -5.01 | PRibTrfase\_dom | interpro domains | IPR000836 | 8 | 4 | 11383 | 1233 | kri,r-l,CG6767,l(2)k01209 | | 6.699e-03 | -5.01 | Ubiquitin-activating\_enz | interpro domains | IPR035985 | 8 | 4 | 11383 | 1233 | Aos1,APP-BP1,Uba5,Atg7 | | 6.699e-03 | -5.01 | ThiF\_NAD\_FAD-bd | interpro domains | IPR000594 | 8 | 4 | 11383 | 1233 | Atg7,Uba5,Aos1,APP-BP1 | | 6.699e-03 | -5.01 | Aldo/ket\_reductase\_CS | interpro domains | IPR018170 | 8 | 4 | 11383 | 1233 | CG10863,CG2767,CG10638,CG9436 | | 6.717e-03 | -5.00 | testosterone 17-beta-dehydrogenase (NADP+) activity | molecular function | GO:0047045 | 5 | 3 | 12176 | 1119 | CG31549,CG31548,CG12171 | | 6.717e-03 | -5.00 | large ribosomal subunit rRNA binding | molecular function | GO:0070180 | 5 | 3 | 12176 | 1119 | RpL12,RpLP0,RpL23 | | 6.726e-03 | -5.00 | ubiquitin protein ligase binding | molecular function | GO:0031625 | 52 | 11 | 12176 | 1119 | CG10254,ben,Cul5,CG30496,Ubc4,RpS27A,Tak1,RpL40,Cul2,Cul1,CG3226 | | 6.741e-03 | -5.00 | organic cyclic compound catabolic process | biological process | GO:1901361 | 171 | 26 | 12345 | 1129 | Rga,DhpD,twin,Edc3,CG17768,Cbp80,Trf4-2,smg,cu,CG11486,Dcr-1,fal,Drep1,LSm3,CG11164,cN-IIIB,Pym,armi,AOX3,CG31759,Not10,Ho,Tengl4,Rrp40,gw,P5CDh1 | | 6.761e-03 | -5.00 | Cell Cycle, Mitotic | REACTOME pathways | R-DME-69278 | 233 | 48 | 3923 | 574 | Prosbeta4,CG8009,E2f1,Prosalpha5,RpS27A,mad2,twe,Chrac-14,CycE,Prosbeta5,CG12018,Rpn2,Rbf2,Sld5,Dd,Nup358,Aladin,Rbf,lin-52,Nup62,rl,RfC38,Pp2A-29B,Ndc1,DNApol-epsilon255,Orc1,Dhfr,Ts,Prosalpha4,SkpA,RpL40,Rpt2,Ankle2,Prosalpha3,Cdk7,Prosbeta3,Orc6,Prosbeta6,APC7,wapl,mbo,CycB,Nup58,Cdk1,Psf3,Cul1,Rpn3,Rpn12 | | 6.793e-03 | -4.99 | blastoderm segmentation | biological process | GO:0007350 | 214 | 31 | 12345 | 1129 | disp,gus,psq,armi,fs(1)M3,trk,msps,Moe,csw,osk,bcd,Grip71,spn-E,Grip128,fs(1)N,apt,cnc,swa,Lkb1,Grip75,Stat92E,Tm1,cad,mael,rl,pygo,Dcr-1,exu,nos,tor,hb | | 6.803e-03 | -4.99 | GO\_NEGATIVE\_REGULATION\_OF\_CATALYTIC\_ACTIVITY | MSigDB lists | GO\_NEGATIVE\_REGULATION\_OF\_CATALYTIC\_ACTIVITY | 215 | 47 | 4161 | 644 | Arl6IP1,CHOp24,lid,Karybeta3,regucalcin,Prosbeta3,mts,CG7530,eIF4A,Rpt2,adp,HDAC6,Rpn12,Rpn2,Prip,CG7997,Cdk1,Uch-L5,mad2,Pka-R2,Faf2,RpS27A,Prosalpha5,Pka-R1,Prosbeta6,Nprl2,Ptp61F,CG8726,Cyp1,PrBP,RpL40,Snx6,CG30291,CkIIalpha,CG1847,APC7,CycB,Prosbeta5,Pdk1,CG4709,CG3376,Rpn3,Chd64,Prosalpha3,Rcd5,Prosbeta4,Pp2A-29B | | 6.824e-03 | -4.99 | usp (ultraspiracle) | protein interactions | 31165 | 35 | 10 | 9204 | 1112 | Utx,egg,Chd64,trr,rig,EcR,CycC,Fkbp39,foxo,ari-1 | | 6.844e-03 | -4.98 | zf-RanBP | pfam domains | PF00641 | 8 | 4 | 10480 | 1142 | sol,trbd,RYBP,Nup358 | | 6.844e-03 | -4.98 | ThiF | pfam domains | PF00899 | 8 | 4 | 10480 | 1142 | APP-BP1,Uba5,Atg7,Aos1 | | 6.844e-03 | -4.98 | GCP\_N\_terminal | pfam domains | PF17681 | 8 | 4 | 10480 | 1142 | Grip128,Grip84,CG7716,Grip75 | | 6.845e-03 | -4.98 | translation regulator activity | molecular function | GO:0045182 | 97 | 17 | 12176 | 1119 | bam,bcd,PEK,eEF1beta,mEFTu1,CG8963,eIF3i,eIF4E1,eIF4A,CG7414,heph,cup,CG6094,Shmt,eIF3m,mEFTu2,smg | | 6.852e-03 | -4.98 | Rb-E2F complex | cellular component | GO:0035189 | 5 | 3 | 11940 | 1105 | E2f1,Rbf2,Rbf | | 6.852e-03 | -4.98 | Lsm1-7-Pat1 complex | cellular component | GO:1990726 | 5 | 3 | 11940 | 1105 | CG6610,CG10418,LSm3 | | 6.856e-03 | -4.98 | oocyte karyosome formation | biological process | GO:0030717 | 32 | 8 | 12345 | 1129 | krimp,lid,SRPK,spn-E,GatA,del,Nup62,enc | | 6.856e-03 | -4.98 | karyosome formation | biological process | GO:0061988 | 32 | 8 | 12345 | 1129 | SRPK,krimp,lid,del,Nup62,enc,GatA,spn-E | | 6.856e-03 | -4.98 | homeostasis of number of cells | biological process | GO:0048872 | 32 | 8 | 12345 | 1129 | Hsc70-3,cnc,sip3,SERCA,PHGPx,Tao,bun,Pi3K92E | | 6.866e-03 | -4.98 | GSE21927\_SPLEEN\_C57BL6\_VS\_4T1\_TUMOR\_BALBC\_MONOCYTES\_DN | MSigDB lists | GSE21927\_SPLEEN\_C57BL6\_VS\_4T1\_TUMOR\_BALBC\_MONOCYTES\_DN | 41 | 13 | 4161 | 644 | Snx6,scramb1,PrBP,ND-24,muskelin,eIF4A,Ppcdc,mtgo,CG4293,Prosalpha5,CG3860,mRpL13,EcR | | 6.880e-03 | -4.98 | Clic (Chloride intracellular channel) | protein interactions | 32349 | 20 | 7 | 9204 | 1112 | AnxB10,CG10265,r-l,CG10638,regucalcin,PHGPx,SelD | | 6.880e-03 | -4.98 | Mes2 (Mesoderm-expressed 2) | protein interactions | 40514 | 20 | 7 | 9204 | 1112 | RpLP1,Mes2,CG15564,Echs1,CG6617,CG6707,Nca | | 6.886e-03 | -4.98 | ClpP/crotonase-like\_dom\_sf | interpro domains | IPR029045 | 17 | 6 | 11383 | 1233 | CG4598,ACC,Mtpalpha,Echs1,CG4594,CG5844 | | 6.916e-03 | -4.97 | GSE10239\_NAIVE\_VS\_KLRG1HIGH\_EFF\_CD8\_TCELL\_DN | MSigDB lists | GSE10239\_NAIVE\_VS\_KLRG1HIGH\_EFF\_CD8\_TCELL\_DN | 69 | 19 | 4161 | 644 | Grx1,Naa20A,Pdk,scramb1,CG42542,Ts,CG8204,RhoGAP54D,CG10470,grp,Tsp86D,ND-30,mad2,Pen,mRpS10,Coq7,Jafrac2,CycE,Uch | | 6.934e-03 | -4.97 | drug metabolic process | biological process | GO:0017144 | 276 | 38 | 12345 | 1129 | blw,Pdxk,ATPsynO,AMPdeam,ATPsynE,ATPsynD,Adi1,Cht11,Jafrac1,Shmt,Idgf2,Fdh,CG10561,Idgf6,Pcd,sgll,ATPsynB,ATPsynG,Irp-1B,ATPsyndelta,AcCoAS,P5cr,Pxt,Ady43A,Adh,CG4390,ATPsynbeta,Dhfr,Idgf4,Scsalpha1,Jafrac2,ATPsynCF6,COX5B,fdl,CG11134,pr,Sdhaf3,Sptr | | 6.973e-03 | -4.97 | positive regulation of cellular protein metabolic process | biological process | GO:0032270 | 303 | 41 | 12345 | 1129 | Pomp,cort,Ankle2,Cka,Lkb1,Pdk1,Ubc6,CG8009,tor,Parp16,Tak1,Jafrac2,Exd2,esc,trr,raptor,RpLP1,Rpt2,CG8963,ASPP,Rpn13,eIF3m,STUB1,Abi,fray,dco,tyf,SkpA,PEK,Cul1,ScsbetaA,park,MED24,ari-1,Pi3K92E,G9a,Tao,Mos,osk,ben,Mekk1 | | 7.011e-03 | -4.96 | cell division site | cellular component | GO:0032153 | 45 | 10 | 11940 | 1105 | CG7716,Grip128,Sry-alpha,slam,Moe,Grip84,Cen,Arf51F,sau,Grip75 | | 7.015e-03 | -4.96 | fly chr96C8-96C8\|3-87 cM | chromosome location | fly chr96C8-96C8\|3-87 cM | 15 | 4 | 25010 | 1343 | CG10425,RpS27,Nup358,CG11858 | | 7.015e-03 | -4.96 | fly chr96C8-96C8\|3-87 | chromosome location | fly chr96C8-96C8\|3-87 | 15 | 4 | 25010 | 1343 | CG10425,RpS27,CG11858,Nup358 | | 7.045e-03 | -4.96 | chr15q24 | MSigDB lists | chr15q24 | 20 | 8 | 4161 | 644 | Droj2,ari-1,Hmg-2,nAChRalpha4,Edc3,CG2924,Ppcdc,Prosalpha3 | | 7.045e-03 | -4.96 | GO\_PYRIMIDINE\_CONTAINING\_COMPOUND\_BIOSYNTHETIC\_PROCESS | MSigDB lists | GO\_PYRIMIDINE\_CONTAINING\_COMPOUND\_BIOSYNTHETIC\_PROCESS | 20 | 8 | 4161 | 644 | nmdyn-D6,Shmt,r,r-l,kri,Ts,nmdyn-D7,CG6767 | | 7.045e-03 | -4.96 | YOSHIOKA\_LIVER\_CANCER\_EARLY\_RECURRENCE\_DN | MSigDB lists | YOSHIOKA\_LIVER\_CANCER\_EARLY\_RECURRENCE\_DN | 20 | 8 | 4161 | 644 | CG10932,Kap-alpha3,sbr,Ufm1,alpha-Man-Ia,CG8611,CG12929,CG1847 | | 7.045e-03 | -4.96 | PID\_BCR\_5PATHWAY | MSigDB lists | PID\_BCR\_5PATHWAY | 20 | 8 | 4161 | 644 | Traf6,Pp2B-14D,CanA-14F,CkIIalpha,p38a,rl,Pdk1,cact | | 7.046e-03 | -4.96 | ScsbetaA (Succinyl-coenzyme A synthetase beta subunit, ADP-forming) | protein interactions | 41067 | 69 | 16 | 9204 | 1112 | Pdk1,CCT6,SkpA,SERCA,CCT3,Cdk5,Pp2A-29B,Mcad,cathD,Rab5,Droj2,CG1416,Tspo,Cp1,mEFTu1,wal | | 7.057e-03 | -4.95 | fly chr85E9-85E9\|3-49 | chromosome location | fly chr85E9-85E9\|3-49 | 8 | 3 | 25010 | 1343 | CG8507,CG12945,MtnA | | 7.057e-03 | -4.95 | fly chr84E8-84E8\|3-48 | chromosome location | fly chr84E8-84E8\|3-48 | 8 | 3 | 25010 | 1343 | CG2767,PIG-H,CG3223 | | 7.057e-03 | -4.95 | fly chr13B6-13B6\|1-50 cM | chromosome location | fly chr13B6-13B6\|1-50 cM | 8 | 3 | 25010 | 1343 | Top1,CG9114,HDAC6 | | 7.057e-03 | -4.95 | fly chr13B6-13B6\|1-50 | chromosome location | fly chr13B6-13B6\|1-50 | 8 | 3 | 25010 | 1343 | Top1,HDAC6,CG9114 | | 7.057e-03 | -4.95 | fly chr47A11-47A11\|2-62 cM | chromosome location | fly chr47A11-47A11\|2-62 cM | 8 | 3 | 25010 | 1343 | pre-lola-G,trsn,asRNA:CR45134 | | 7.057e-03 | -4.95 | fly chr85E9-85E9\|3-49 cM | chromosome location | fly chr85E9-85E9\|3-49 cM | 8 | 3 | 25010 | 1343 | MtnA,CG12945,CG8507 | | 7.057e-03 | -4.95 | fly chr73B1-73B1\|3-44 | chromosome location | fly chr73B1-73B1\|3-44 | 8 | 3 | 25010 | 1343 | Prosbeta6,Arts,Mo25 | | 7.057e-03 | -4.95 | fly chr84E8-84E8\|3-48 cM | chromosome location | fly chr84E8-84E8\|3-48 cM | 8 | 3 | 25010 | 1343 | PIG-H,CG3223,CG2767 | | 7.057e-03 | -4.95 | fly chr73B1-73B1\|3-44 cM | chromosome location | fly chr73B1-73B1\|3-44 cM | 8 | 3 | 25010 | 1343 | Arts,Prosbeta6,Mo25 | | 7.057e-03 | -4.95 | fly chr47A11-47A11\|2-62 | chromosome location | fly chr47A11-47A11\|2-62 | 8 | 3 | 25010 | 1343 | asRNA:CR45134,pre-lola-G,trsn | | 7.076e-03 | -4.95 | REACTOME\_TRAF6\_MEDIATED\_INDUCTION\_OF\_NFKB\_AND\_MAP\_KINASES\_UPON\_TLR7\_8\_OR\_9\_ACTIVATION | MSigDB lists | REACTOME\_TRAF6\_MEDIATED\_INDUCTION\_OF\_NFKB\_AND\_MAP\_KINASES\_UPON\_TLR7\_8\_OR\_9\_ACTIVATION | 24 | 9 | 4161 | 644 | RpL40,rl,cact,p38a,Cdk1,Pp2A-29B,RpS27A,Traf6,mts | | 7.076e-03 | -4.95 | KEGG\_TGF\_BETA\_SIGNALING\_PATHWAY | MSigDB lists | KEGG\_TGF\_BETA\_SIGNALING\_PATHWAY | 24 | 9 | 4161 | 644 | mts,Rbf,Smox,tkv,Pp2A-29B,rl,SkpA,Med,Cul1 | | 7.076e-03 | -4.95 | BURTON\_ADIPOGENESIS\_4 | MSigDB lists | BURTON\_ADIPOGENESIS\_4 | 24 | 9 | 4161 | 644 | CCT3,NHP2,tko,Uch-L5,CG15715,Fkbp59,SREBP,LSm3,Cse1 | | 7.076e-03 | -4.95 | REACTOME\_MYD88\_MAL\_CASCADE\_INITIATED\_ON\_PLASMA\_MEMBRANE | MSigDB lists | REACTOME\_MYD88\_MAL\_CASCADE\_INITIATED\_ON\_PLASMA\_MEMBRANE | 24 | 9 | 4161 | 644 | Cdk1,p38a,rl,cact,RpL40,mts,Traf6,RpS27A,Pp2A-29B | | 7.085e-03 | -4.95 | Cyclin D associated events in G1 | REACTOME pathways | R-DME-69231 | 17 | 7 | 3923 | 574 | Rbf2,RpL40,Cul1,Rbf,SkpA,RpS27A,Pp2A-29B | | 7.085e-03 | -4.95 | G1 Phase | REACTOME pathways | R-DME-69236 | 17 | 7 | 3923 | 574 | Pp2A-29B,Rbf,RpS27A,SkpA,RpL40,Cul1,Rbf2 | | 7.088e-03 | -4.95 | KEGG\_HUNTINGTONS\_DISEASE | MSigDB lists | KEGG\_HUNTINGTONS\_DISEASE | 74 | 20 | 4161 | 644 | Sod1,Cyt-c-p,ND-SGDH,RpII140,ND-49,ND-51,ND-23,Rpb8,kl-2,levy,ND-B22,COX5A,Clc,ND-39,ND-24,ND-30,UQCR-Q,ND-75,SdhA,ND-B18 | | 7.088e-03 | -4.95 | GSE14908\_ATOPIC\_VS\_NONATOPIC\_PATIENT\_RESTING\_CD4\_TCELL\_DN | MSigDB lists | GSE14908\_ATOPIC\_VS\_NONATOPIC\_PATIENT\_RESTING\_CD4\_TCELL\_DN | 74 | 20 | 4161 | 644 | eIF3i,Rpb8,levy,Ubc4,CG6767,aay,mRpS23,Kap3,shv,CycC,CG1532,CG9391,ND-30,f-cup,Irp-1B,Idh3b,Surf4,RpL24,CG7728,Fkbp59 | | 7.088e-03 | -4.95 | GSE17721\_0.5H\_VS\_12H\_PAM3CSK4\_BMDC\_DN | MSigDB lists | GSE17721\_0.5H\_VS\_12H\_PAM3CSK4\_BMDC\_DN | 74 | 20 | 4161 | 644 | Gdi,alpha-Man-Ia,STUB1,CG11710,AcCoAS,CG9391,Prosalpha3,CG4293,Unr,asf1,CG40045,Karybeta3,CG17691,Gk2,l(1)G0004,Imp,APC7,CG4390,Atg6,Tlk | | 7.115e-03 | -4.95 | nucleoplasm | cellular component | GO:0005654 | 557 | 69 | 11940 | 1105 | ix,gus,RYBP,Spt20,Sgf29,Elba2,Rbfox1,Ptip,escl,IntS11,lin-52,Bin1,Tlk,lid,Su(H),bru1,CG46385,d4,mbo,CG1602,hb,MED27,sbr,Clp,trr,vers,dl,Sld5,Utx,rept,mad2,MED24,piwi,CG11164,Sgf11,mute,CG4021,pont,MED4,Chrac-14,Tip60,CycC,SelR,RpII140,Rcd5,CG17328,EloC,Atu,nej,Rpb8,mts,Trf2,YL-1,Cse1,gw,Psf3,Cdk7,Tfb4,PAPLA1,MED20,CG10055,Prosbeta5,scny,not,esc,e(y)1,mxc,TfIIFalpha,Taf7 | | 7.153e-03 | -4.94 | Dp (DP transcription factor) | protein interactions | 36461 | 30 | 9 | 9204 | 1112 | Rbf,blw,eIF4A,Rbf2,lin-52,E2f1,RpL30,CycE,Tip60 | | 7.153e-03 | -4.94 | CG13887 (uncharacterized protein) | protein interactions | 38096 | 30 | 9 | 9204 | 1112 | Uch,Cisd2,ATPsynCF6,Surf4,COX4,ATPsynO,Chd64,CG4598,AnxB10 | | 7.163e-03 | -4.94 | GSE36826\_WT\_VS\_IL1R\_KO\_SKIN\_STAPH\_AUREUS\_INF\_UP | MSigDB lists | GSE36826\_WT\_VS\_IL1R\_KO\_SKIN\_STAPH\_AUREUS\_INF\_UP | 104 | 26 | 4161 | 644 | CG1774,Got1,Sptr,Gyg,Clc,Nmt,COX5A,CG11807,gammaTub37C,ND-75,CycB,Pen,Pglym78,UQCR-Q,Pex10,Gmer,Rpt2,Rpn12,DppIII,levy,Prosbeta6,CG14894,Arts,tko,CG11089,ste14 | | 7.203e-03 | -4.93 | protein kinase binding | molecular function | GO:0019901 | 82 | 15 | 12176 | 1119 | alc,sws,Tm1,Mitofilin,CycE,CycJ,rl,CG11807,Cbl,Mekk1,CycB,tub,Wdfy2,Mtl,SNF4Agamma | | 7.204e-03 | -4.93 | GSE32986\_UNSTIM\_VS\_CURDLAN\_HIGHDOSE\_STIM\_DC\_UP | MSigDB lists | GSE32986\_UNSTIM\_VS\_CURDLAN\_HIGHDOSE\_STIM\_DC\_UP | 79 | 21 | 4161 | 644 | msps,mad2,Pfdn2,Caf1-105,AcCoAS,SRPK,CycE,Prosalpha5,PpD3,mRpL17,Clp,Got1,Gk2,PrBP,AGBE,Rpb8,SelD,Oseg1,nmdyn-D7,RhoGAP54D,Aladin | | 7.204e-03 | -4.93 | BURTON\_ADIPOGENESIS\_6 | MSigDB lists | BURTON\_ADIPOGENESIS\_6 | 79 | 21 | 4161 | 644 | AGBE,CG17691,ND-B22,Tsp74F,Etf-QO,ND-SGDH,ABCD,CG11899,Pgm1,Thiolase,ND-30,mEFTu1,UQCR-Q,ND-75,crq,SdhA,ND-B18,yip2,ND-39,Nc73EF,Scsalpha1 | | 7.204e-03 | -4.93 | - | gene3d domains | 3.40.1110.10 | 15 | 6 | 4929 | 622 | SERCA,CG31729,CG9981,ATP8A,Atpalpha,anne | | 7.231e-03 | -4.93 | Ote (Otefin) | protein interactions | 37090 | 25 | 8 | 9204 | 1112 | CG2199,cnn,Tak1,PGRP-LE,CG4407,gammaTub37C,tkv,Med | | 7.231e-03 | -4.93 | GlyP (Glycogen phosphorylase) | protein interactions | 33386 | 25 | 8 | 9204 | 1112 | Scp1,Pglym78,Rtnl1,CG46385,Pgk,Argk,CG32473,Adh | | 7.231e-03 | -4.93 | Not1 (Not1) | protein interactions | 5740252 | 25 | 8 | 9204 | 1112 | cu,nos,cup,bam,Not10,gw,Rga,twin | | 7.231e-03 | -4.93 | yps (ypsilon schachtel) | protein interactions | 39377 | 25 | 8 | 9204 | 1112 | Pi3K92E,Pdk1,tral,cup,osk,eIF4E1,exu,foxo | | 7.231e-03 | -4.93 | Rbp4 (RNA-binding protein 4) | protein interactions | 41668 | 25 | 8 | 9204 | 1112 | 26-29-p,Sap47,CG11899,CycB,Dhfr,EloC,GlyP,CG2493 | | 7.250e-03 | -4.93 | - | gene3d domains | 3.30.70.141 | 4 | 3 | 4929 | 622 | nmdyn-D6,awd,nmdyn-D7 | | 7.250e-03 | -4.93 | - | gene3d domains | 2.60.40.340 | 4 | 3 | 4929 | 622 | NFAT,dl,Rel | | 7.257e-03 | -4.93 | GSE22886\_NAIVE\_VS\_IGG\_IGA\_MEMORY\_BCELL\_DN | MSigDB lists | GSE22886\_NAIVE\_VS\_IGG\_IGA\_MEMORY\_BCELL\_DN | 55 | 16 | 4161 | 644 | CG40045,Ptp61F,Iml1,CG10932,Tsp66E,Gapdh1,Nurf-38,sau,AMPdeam,Gclc,CCT5,ND-B18,AIF,CG9471,yip2,Mo25 | | 7.258e-03 | -4.93 | proteolysis | biological process | GO:0006508 | 844 | 98 | 12345 | 1129 | Usp39,RpS27A,Pink1,snk,CG8728,CG2493,CG8974,gus,CG11110,trbd,ste24a,CG31633,Fbw5,Edem1,CG8188,faf,cathD,RpL40,Cp1,SkpA,Prosalpha5,SP1029,CG14516,CG13630,CG4238,Rpn13,eIF3m,CG10254,DppIII,CSN6,Herc4,CG5003,Rpt2,not,Ntan1,Cul5,CG3223,CG40045,CG4572,Topors,Prosalpha4,Ser7,otu,stet,Rpn3,CG5390,EloC,CG9316,Rpn2,Rpn12,Prosalpha3,CG2200,Cul2,Prosbeta3,Hsc70-3,CG11070,gd,Dip-B,CG32640,RYBP,CG32221,park,pen-2,ari-1,Cbl,shrb,Fur2,sol,Sgf11,Cul1,CG32473,Vsp37A,UbcE2H,Prosbeta4,Plap,velo,CG6766,Prosbeta6,Uch-L5,Sp7,S2P,Usp16-45,CG11771,Ubc6,grsm,26-29-p,Usp10,ago,Prosbeta5,Nepl3,Pomp,sip3,cort,CG16979,scny,Uch,CG9953,ea | | 7.262e-03 | -4.93 | FCERI mediated NF-kB activation | REACTOME pathways | R-DME-2871837 | 68 | 18 | 3923 | 574 | cact,SkpA,Prosalpha3,Rpt2,RpL40,Prosbeta4,RpS27A,Prosalpha5,Prosalpha4,Rpn2,Prosbeta5,Prosbeta6,dl,Rpn12,Rpn3,Cul1,Prosbeta3,Rel | | 7.262e-03 | -4.93 | Activation of NF-kappaB in B cells | REACTOME pathways | R-DME-1169091 | 68 | 18 | 3923 | 574 | Prosbeta3,Rel,Prosbeta6,Rpn12,dl,Cul1,Rpn3,RpS27A,Prosalpha5,Prosbeta4,Prosbeta5,Rpn2,Prosalpha4,SkpA,cact,Prosalpha3,RpL40,Rpt2 | | 7.270e-03 | -4.92 | GSE17721\_POLYIC\_VS\_GARDIQUIMOD\_16H\_BMDC\_DN | MSigDB lists | GSE17721\_POLYIC\_VS\_GARDIQUIMOD\_16H\_BMDC\_DN | 84 | 22 | 4161 | 644 | RpL38,CG40045,CG17691,Stoml2,ACC,csw,CG17294,Mcad,scu,CG9393,ScsbetaA,cact,STUB1,bys,ERp60,Top1,ND-24,RpL12,CSN6,RpL26,CG33181,RpL24 | | 7.270e-03 | -4.92 | GSE27786\_CD8\_TCELL\_VS\_NEUTROPHIL\_UP | MSigDB lists | GSE27786\_CD8\_TCELL\_VS\_NEUTROPHIL\_UP | 84 | 22 | 4161 | 644 | Hmg-2,Grip84,yip2,CG9143,CG10038,ND-39,Arf51F,Shmt,Cul1,sbr,Stip1,Pus7,Coq6,mino,eIF4A,Dd,Gmer,Efr,Grip128,cyc,Med,Ptp61F | | 7.280e-03 | -4.92 | MORF\_HAT1 | MSigDB lists | MORF\_HAT1 | 94 | 24 | 4161 | 644 | csw,LysRS,Cyt-c-p,Sod1,SkpA,ND-51,CG1416,CG10418,Mitofilin,Prosbeta4,LSm3,Iru,COX5A,CCT5,ND-75,CycB,Klp10A,ND-30,mEFTu1,Hat1,ND-24,CG13630,Pen,Chrac-14 | | 7.287e-03 | -4.92 | response to sucrose | biological process | GO:0009744 | 20 | 6 | 12345 | 1129 | ACC,sgll,Desat1,PyK,Pfk,CG7328 | | 7.287e-03 | -4.92 | ribosomal large subunit assembly | biological process | GO:0000027 | 20 | 6 | 12345 | 1129 | RpL12,RpL10,RpL11,RpL23A,RpLP0,RpL24 | | 7.287e-03 | -4.92 | endoplasmic reticulum unfolded protein response | biological process | GO:0030968 | 20 | 6 | 12345 | 1129 | Parp16,PEK,Hsc70-3,sip3,Edem1,Atf6 | | 7.295e-03 | -4.92 | brm (brahma) | protein interactions | 39744 | 106 | 22 | 9204 | 1112 | asf1,rept,Lkb1,Bx,Bap170,Larp4B,foxo,enc,Utx,E2f1,pont,MED24,RpII140,Ptip,nej,Rbf,CycE,MED4,MED27,Usp10,mxc,tkv | | 7.296e-03 | -4.92 | asymmetric cell division | biological process | GO:0008356 | 106 | 18 | 12345 | 1129 | Lkb1,piwi,twin,Cdk1,cnn,egg,shrb,mad2,mts,eIF4A,Su(H),bam,tws,enc,CycE,Iml1,Pfdn2,msps | | 7.313e-03 | -4.92 | membrane invagination | biological process | GO:0010324 | 26 | 7 | 12345 | 1129 | slam,csw,crq,EcR,sau,Mekk1,Mtl | | 7.313e-03 | -4.92 | regulation of embryonic pattern specification | biological process | GO:1902875 | 26 | 7 | 12345 | 1129 | mael,enc,cup,spn-E,cnc,exu,swa | | 7.313e-03 | -4.92 | plasma membrane invagination | biological process | GO:0099024 | 26 | 7 | 12345 | 1129 | EcR,Mekk1,Mtl,sau,csw,slam,crq | | 7.322e-03 | -4.92 | GO\_NCRNA\_METABOLIC\_PROCESS | MSigDB lists | GO\_NCRNA\_METABOLIC\_PROCESS | 305 | 63 | 4161 | 644 | RpS24,TyrRS,CG5292,RpL30,Archease,RpL23,CG10463,CG11858,RpS16,RpS2,Sirt7,dco,mtTFB1,RpS7,RpL38,RpS4,NHP2,Cbp80,RpL40,RpL11,Mat89Ba,Dcr-1,RpL37a,AGO3,CG7728,RpS27,Smox,RpL24,RpL26,RpS18,RpL12,spn-E,bys,CG2100,Cdk7,CG3021,CG10189,RpS30,RpS13,RpLP0,LysRS,Rrp40,RpL13A,Nurf-38,RpII140,scu,CysRS,Rpb8,GluRS-m,l(1)G0004,Uba5,TrpRS-m,Clp,mael,shu,GatA,RpS28b,RpS27A,RpS23,RpL9,Pus7,SerRS-m,RpL13 | | 7.340e-03 | -4.91 | ALDOKETO\_REDUCTASE\_1 | prosite domains | PS00798 | 8 | 4 | 6243 | 694 | CG10863,CG9436,CG10638,CG2767 | | 7.340e-03 | -4.91 | ALDOKETO\_REDUCTASE\_2 | prosite domains | PS00062 | 8 | 4 | 6243 | 694 | CG2767,CG10638,CG9436,CG10863 | | 7.408e-03 | -4.91 | eye pigment biosynthetic process | biological process | GO:0006726 | 39 | 9 | 12345 | 1129 | fdl,lt,cl,PAPLA1,mal,Hexo1,EMC3,DhpD,Rab32 | | 7.461e-03 | -4.90 | 3'-5'-exoribonuclease activity | molecular function | GO:0000175 | 20 | 6 | 12176 | 1119 | twin,Exd2,Rga,cu,CG11486,CG31759 | | 7.528e-03 | -4.89 | anatomical structure maturation | biological process | GO:0071695 | 53 | 11 | 12345 | 1129 | mop,muskelin,SkpA,Atg6,shrb,EcR,shep,Stam,Imp,wisp,spn-E | | 7.594e-03 | -4.88 | fertilization | biological process | GO:0009566 | 46 | 10 | 12345 | 1129 | wisp,Nap1,spd-2,msps,png,yem,alphaTub67C,cort,fs(1)Ya,plu | | 7.599e-03 | -4.88 | GO\_DRUG\_BINDING | MSigDB lists | GO\_DRUG\_BINDING | 37 | 12 | 4161 | 644 | dnc,Cyp1,GlyP,CG14715,mAChR-A,Ts,shu,nAChRalpha4,Pp2B-14D,CanA-14F,Fkbp59,Dhfr | | 7.599e-03 | -4.88 | GO\_MITOCHONDRIAL\_RESPIRATORY\_CHAIN\_COMPLEX\_ASSEMBLY | MSigDB lists | GO\_MITOCHONDRIAL\_RESPIRATORY\_CHAIN\_COMPLEX\_ASSEMBLY | 37 | 12 | 4161 | 644 | ND-B22,ND-B18,AIF,ND-75,ND-51,ND-23,ND-30,ND-24,ND-49,ND-39,CG11110,ND-SGDH | | 7.599e-03 | -4.88 | GSE7509\_DC\_VS\_MONOCYTE\_WITH\_FCGRIIB\_STIM\_DN | MSigDB lists | GSE7509\_DC\_VS\_MONOCYTE\_WITH\_FCGRIIB\_STIM\_DN | 37 | 12 | 4161 | 644 | esc,Ptp61F,Su(H),Top1,Liprin-beta,scramb1,escl,Arf51F,Prosalpha3,Smox,CG8188,Tlk | | 7.631e-03 | -4.88 | GSE17721\_LPS\_VS\_CPG\_0.5H\_BMDC\_DN | MSigDB lists | GSE17721\_LPS\_VS\_CPG\_0.5H\_BMDC\_DN | 46 | 14 | 4161 | 644 | Sec22,CG3631,RpL11,CG2493,Chrac-14,Pglym78,pen-2,Stam,shop,Prosbeta6,sgll,Arl6IP1,Rep,CG42542 | | 7.631e-03 | -4.88 | GSE40685\_NAIVE\_CD4\_TCELL\_VS\_TREG\_DN | MSigDB lists | GSE40685\_NAIVE\_CD4\_TCELL\_VS\_TREG\_DN | 46 | 14 | 4161 | 644 | l(3)mbt,Scsalpha1,Wdfy2,RpL30,CG7718,mino,galectin,Lsm12,CG11110,Pfk,CG9485,sgll,CG8199,ATP8A | | 7.632e-03 | -4.88 | GO\_REGULATION\_OF\_TRANSFERASE\_ACTIVITY | MSigDB lists | GO\_REGULATION\_OF\_TRANSFERASE\_ACTIVITY | 283 | 59 | 4161 | 644 | CycE,Pp2A-29B,nAChRalpha4,Rcd5,Prosalpha3,Prosbeta4,Hsc70-3,lwr,CG3376,Cul1,Rpn3,Prosbeta5,Cdk5,CycB,CG4709,Pdk1,Ac3,p38a,RpS2,Traf6,APC7,alc,CG30291,Rbf,CycC,Non1,RpL40,Snx6,CG17698,Pka-R1,RfC38,Ptp61F,Prosbeta6,Nprl2,Prosalpha5,Mo25,Mekk1,RpS27A,Pka-R2,mad2,STUB1,rl,LpR2,Cdk1,Rpn12,CG8188,Sod1,Rpn2,mts,Cdk7,Prosbeta3,Prosalpha4,Rpt2,CG7530,csw,raptor,eIF4A,regucalcin,SkpA | | 7.643e-03 | -4.87 | disulfide oxidoreductase activity | molecular function | GO:0015036 | 39 | 9 | 12176 | 1119 | cl,Scox,Grx1,CG3939,CG14407,ERp60,Trxr-1,SelT,GILT1 | | 7.646e-03 | -4.87 | MORF\_PHB | MSigDB lists | MORF\_PHB | 60 | 17 | 4161 | 644 | crq,Marf,CG13630,Rpn3,kcc,Rcd5,CG7728,Clp,CG7488,CSN6,Grip84,Lsm12,r,Gmer,P5CDh1,CG9393,GCS2beta | | 7.766e-03 | -4.86 | Prosbeta1 (Proteasome beta1 subunit) | protein interactions | 46058 | 41 | 11 | 9204 | 1112 | Prosalpha4,Prosbeta5,Prosalpha3,Prosbeta6,Prosbeta3,Prosbeta4,Rpt2,Rpn13,Rpn2,Prosalpha5,Pomp | | 7.766e-03 | -4.86 | CG34159 (uncharacterized protein) | protein interactions | 34353 | 41 | 11 | 9204 | 1112 | png,Ptip,CG17002,Larp4B,Usp10,MED27,MED4,CG13373,MED20,ix,CG3939 | | 7.777e-03 | -4.86 | GO\_RESPONSE\_TO\_ORGANIC\_CYCLIC\_COMPOUND | MSigDB lists | GO\_RESPONSE\_TO\_ORGANIC\_CYCLIC\_COMPOUND | 211 | 46 | 4161 | 644 | bys,Moe,CG3376,CG11710,Ac3,CycB,Cdk5,Cka,PpD3,Gabat,nAChRalpha4,CycE,Smox,Prosbeta4,Wnt10,SREBP,Dcr-1,Atpalpha,Tspo,Ts,CG9485,r,aay,Atg6,MED4,Dhfr,MED24,Shmt,Calx,rl,cact,AIF,Cdk1,GlyP,Jheh2,Got1,Fkbp59,Idh,CG10932,Tip60,rept,dnc,Prip,EcR,Sod1,Cdk7 | | 7.796e-03 | -4.85 | proteasome core complex | cellular component | GO:0005839 | 26 | 7 | 11940 | 1105 | Prosbeta5,Prosalpha3,Prosbeta4,Prosalpha4,Prosbeta3,Prosbeta6,Prosalpha5 | | 7.867e-03 | -4.85 | Mhc (Myosin heavy chain) | protein interactions | 35007 | 58 | 14 | 9204 | 1112 | SERCA,bru1,Actn,ATPsynbeta,Argk,Mf,Ptp61F,up,Adh,CanB2,Scp1,ATPsynF,osk,ovo | | 7.881e-03 | -4.84 | Toll Like Receptor 3 (TLR3) Cascade | REACTOME pathways | R-DME-168164 | 30 | 10 | 3923 | 574 | Myd88,rl,RpS27A,Traf6,Rel,Tak1,Pp2A-29B,p38a,RpL40,dl | | 7.881e-03 | -4.84 | TRIF-mediated TLR3/TLR4 signaling | REACTOME pathways | R-DME-937061 | 30 | 10 | 3923 | 574 | Myd88,RpS27A,rl,Traf6,Tak1,Rel,Pp2A-29B,p38a,dl,RpL40 | | 7.881e-03 | -4.84 | MyD88-independent TLR3/TLR4 cascade | REACTOME pathways | R-DME-166166 | 30 | 10 | 3923 | 574 | Traf6,Tak1,Rel,Myd88,RpS27A,rl,dl,RpL40,Pp2A-29B,p38a | | 7.914e-03 | -4.84 | Sc2 (Sc2) | protein interactions | 38457 | 88 | 19 | 9204 | 1112 | Pex13,Sld5,Pp2A-29B,CG15098,CG31729,Cisd2,SERCA,CG10425,Kap-alpha3,CG15141,Surf4,CG5447,wal,Tspo,Karybeta3,Rab5,CG1416,Mcad,Atpalpha | | 7.944e-03 | -4.84 | GSE339\_CD4POS\_VS\_CD8POS\_DC\_IN\_CULTURE\_DN | MSigDB lists | GSE339\_CD4POS\_VS\_CD8POS\_DC\_IN\_CULTURE\_DN | 65 | 18 | 4161 | 644 | RpL37a,Npc2a,Pi4KIIalpha,Cdk1,Coq7,crq,RpS18,CG9911,CG10465,GCS2beta,RpL13A,CG1890,Kap3,colt,NHP2,Jafrac1,CG8726,CG5734 | | 7.944e-03 | -4.84 | GSE40274\_CTRL\_VS\_LEF1\_TRANSDUCED\_ACTIVATED\_CD4\_TCELL\_DN | MSigDB lists | GSE40274\_CTRL\_VS\_LEF1\_TRANSDUCED\_ACTIVATED\_CD4\_TCELL\_DN | 65 | 18 | 4161 | 644 | Grip84,Scox,PDCD-5,Prosalpha5,stas,Rab10,Pmi,CG11134,Hat1,mRpL16,SelR,r-l,CG4390,RpS2,LysRS,CG10189,ND-23,RfC38 | | 7.944e-03 | -4.84 | GNF2\_DAP3 | MSigDB lists | GNF2\_DAP3 | 65 | 18 | 4161 | 644 | RpL24,RpS28b,RpS23,RpL9,RpL37a,RpL12,LysRS,RpS2,RpS16,RpL13A,RpS24,RpL23,RpL40,heph,RpS7,RpL38,Prosbeta6,PpV | | 7.991e-03 | -4.83 | serine/threonine protein kinase complex | cellular component | GO:1902554 | 39 | 9 | 11940 | 1105 | png,CG8475,plu,CycB,CycJ,Cdk7,Tfb4,Cdk5,CycE | | 8.022e-03 | -4.83 | positive regulation of cell death | biological process | GO:0010942 | 99 | 17 | 12345 | 1129 | Lkb1,E2f1,rept,Cbl,Jafrac2,MED24,ScsbetaA,fwe,PDCD-5,Idh3b,Tspo,Utx,eIF3m,CycE,EcR,Debcl,ben | | 8.044e-03 | -4.82 | nucleocytoplasmic transport | biological process | GO:0006913 | 107 | 18 | 12345 | 1129 | Nup58,Karybeta3,mbo,Ndc1,Pen,Nxf3,Ranbp21,Cse1,Moe,Arts,Kap-alpha3,sbr,Nup358,Mat89Ba,Aladin,Nup62,tho2,thoc6 | | 8.044e-03 | -4.82 | nuclear transport | biological process | GO:0051169 | 107 | 18 | 12345 | 1129 | Nup62,tho2,thoc6,Kap-alpha3,sbr,Mat89Ba,Aladin,Nup358,Nxf3,Pen,Cse1,Arts,Moe,Ranbp21,Karybeta3,Nup58,Ndc1,mbo | | 8.057e-03 | -4.82 | Ranbp9 (Ranbp9) | protein interactions | 41367 | 64 | 15 | 9204 | 1112 | SERCA,CCT6,Rab5,Atpalpha,Pi3K92E,Pdk1,Mcad,Kap-alpha3,Pp2A-29B,wal,foxo,GCS2beta,Lkb1,Karybeta3,Tspo | | 8.067e-03 | -4.82 | N-acyltransferase activity | molecular function | GO:0016410 | 83 | 15 | 12176 | 1119 | Sgf29,Rcd5,pont,rept,lid,YL-1,Naa20A,AgmNAT,Nmt,nej,Chrac-14,d4,e(y)1,Hat1,Tip60 | | 8.087e-03 | -4.82 | GO\_PEPTIDE\_METABOLIC\_PROCESS | MSigDB lists | GO\_PEPTIDE\_METABOLIC\_PROCESS | 312 | 64 | 4161 | 644 | RpL24,RpL37a,mRpL10,mRpL17,tko,CG11110,RpS2,Atg6,CG8963,RpL30,TyrRS,eEF1beta,mRpL16,mEFTu1,RpL13,SerRS-m,mRpS10,RpL9,RpS27A,RpS28b,GatA,mRpS21,TrpRS-m,Idh,Grx1,eIF3i,Fur2,tzn,LysRS,RpL13A,eIF4E1,mRpL13,RpS13,RpLP0,RpS30,RpL12,mRpS5,RpS18,Gclc,RpL26,RpS27,CG7414,CG32473,eIF3m,RpL11,heph,RpL40,RpS4,RpL38,RpS7,mRpS23,colt,RpS16,RpL23,RpS24,Tfb4,RpS23,CG14516,GluRS-m,CysRS,Nurf-38,Sod1,eIF4A,mRpL18 | | 8.128e-03 | -4.81 | atl (atlastin) | protein interactions | 42934 | 76 | 17 | 9204 | 1112 | yip2,CG1105,SdhA,Mtpalpha,Hsc70-3,ERp60,snz,Rpn3,cathD,ND-75,Vha36-1,Pgm1,PyK,NHP2,Eno,Rtnl1,ACC | | 8.142e-03 | -4.81 | mitochondrion organization | biological process | GO:0007005 | 208 | 30 | 12345 | 1129 | bor,Marf,CG5903,Mitofilin,Tom40,CG14894,P5CDh1,Tom70,Debcl,Atg7,CG11110,SelD,Scox,CG8004,CG8728,Pink1,larp,CG9393,ND-23,park,CG11562,milt,Ubc6,Tim8,Sdhaf3,Pmi,Roe1,up,CG11077,ND-18 | | 8.160e-03 | -4.81 | GSE9006\_HEALTHY\_VS\_TYPE\_2\_DIABETES\_PBMC\_AT\_DX\_UP | MSigDB lists | GSE9006\_HEALTHY\_VS\_TYPE\_2\_DIABETES\_PBMC\_AT\_DX\_UP | 70 | 19 | 4161 | 644 | Pfdn2,thoc6,Ssdp,Arpc4,Edem1,Moe,ERp60,RpL37a,Iru,Lsm12,COX5A,CG3842,ND-23,heph,Rab5,AnxB10,Atg6,Nadsyn,CG8188 | | 8.253e-03 | -4.80 | GSE11961\_MARGINAL\_ZONE\_BCELL\_VS\_GERMINAL\_CENTER\_BCELL\_DAY40\_UP | MSigDB lists | GSE11961\_MARGINAL\_ZONE\_BCELL\_VS\_GERMINAL\_CENTER\_BCELL\_DAY40\_UP | 51 | 15 | 4161 | 644 | CG1812,yip2,Pp2B-14D,Jheh2,Gyg,CG10038,CanA-14F,CG10465,Arl5,cathD,CG7997,Trs33,Nprl2,Etf-QO,Tsp74F | | 8.255e-03 | -4.80 | PTPc\_motif | smart domains | SM00404 | 19 | 6 | 5086 | 506 | Ptpmeg,Ptp61F,Ptp10D,Ptp69D,Ptpmeg2,csw | | 8.279e-03 | -4.79 | GSE41978\_ID2\_KO\_VS\_ID2\_KO\_AND\_BIM\_KO\_KLRG1\_LOW\_EFFECTOR\_CD8\_TCELL\_DN | MSigDB lists | GSE41978\_ID2\_KO\_VS\_ID2\_KO\_AND\_BIM\_KO\_KLRG1\_LOW\_EFFECTOR\_CD8\_TCELL\_DN | 100 | 25 | 4161 | 644 | RpLP0,RpS30,RpS16,ND-SGDH,RpL23,Pka-R1,RpS7,levy,RpS4,RpL38,RpL24,RpS23,RpL9,COX5A,RpL26,RpL11,Arf51F,mRpL10,Arpc2,RpL13,RpL12,Pgk,Uev1A,Eno,RpS18 | | 8.281e-03 | -4.79 | Ubiquitin mediated proteolysis | KEGG pathways | ko04120 | 100 | 25 | 3198 | 496 | CG40045,Ubc6,APC7,Cul2,sip3,Cbl,ben,park,CG2924,lwr,Traf6,SkpA,STUB1,Ubc4,Herc4,CG4502,UbcE2H,CG8188,EloC,Cul5,Cul1,CG10254,Aos1,ago,CG11070 | | 8.281e-03 | -4.79 | Ubiquitin mediated proteolysis | KEGG pathways | dme04120 | 100 | 25 | 3198 | 496 | ben,park,Cbl,Cul2,APC7,sip3,CG40045,Ubc6,CG10254,Cul1,ago,CG11070,Aos1,CG4502,CG8188,EloC,UbcE2H,Cul5,Ubc4,STUB1,Herc4,CG2924,lwr,Traf6,SkpA | | 8.307e-03 | -4.79 | GSE26030\_TH1\_VS\_TH17\_RESTIMULATED\_DAY5\_POST\_POLARIZATION\_UP | MSigDB lists | GSE26030\_TH1\_VS\_TH17\_RESTIMULATED\_DAY5\_POST\_POLARIZATION\_UP | 75 | 20 | 4161 | 644 | Pfdn5,RpS30,AnxB10,CG6144,CG4572,ND-B22,eIF3i,RpL38,Prosbeta6,Irk1,CG7414,mRpL17,Prosalpha3,ND-30,Ntan1,lwr,CG3077,CG9471,Tom40,Tim8 | | 8.307e-03 | -4.79 | GSE17974\_0H\_VS\_72H\_IN\_VITRO\_ACT\_CD4\_TCELL\_DN | MSigDB lists | GSE17974\_0H\_VS\_72H\_IN\_VITRO\_ACT\_CD4\_TCELL\_DN | 75 | 20 | 4161 | 644 | Rpb8,fray,Arl6IP1,escl,rept,mbo,Tsp66E,CG17187,mRpL18,Klp10A,Caf1-105,AcCoAS,Jafrac2,esc,Shmt,Prosbeta4,mRpL17,Ppcdc,Mitofilin,COX5A | | 8.333e-03 | -4.79 | HAHTOLA\_SEZARY\_SYNDROM\_UP | MSigDB lists | HAHTOLA\_SEZARY\_SYNDROM\_UP | 33 | 11 | 4161 | 644 | Nmt,CG9393,Prosbeta4,Cdk7,Prosbeta3,Top1,Hat1,Ntan1,Pdxk,Ts,Jafrac2 | | 8.335e-03 | -4.79 | fly chr91C5-91C5\|3-64 | chromosome location | fly chr91C5-91C5\|3-64 | 3 | 2 | 25010 | 1343 | Mekk1,CG7718 | | 8.335e-03 | -4.79 | fly chr95D10-95D10\|3-82 cM | chromosome location | fly chr95D10-95D10\|3-82 cM | 3 | 2 | 25010 | 1343 | Atg6,CG5986 | | 8.335e-03 | -4.79 | fly chr8F1-8F1\|1-29 | chromosome location | fly chr8F1-8F1\|1-29 | 3 | 2 | 25010 | 1343 | Gga,LysRS | | 8.335e-03 | -4.79 | fly chr31F4-31F4\|2-43 cM | chromosome location | fly chr31F4-31F4\|2-43 cM | 3 | 2 | 25010 | 1343 | CG6094,CG7384 | | 8.335e-03 | -4.79 | fly chr63E1-63E1\|3-8 cM | chromosome location | fly chr63E1-63E1\|3-8 cM | 3 | 2 | 25010 | 1343 | armi,CycJ | | 8.335e-03 | -4.79 | fly chr70F6-70F6\|3-42 | chromosome location | fly chr70F6-70F6\|3-42 | 3 | 2 | 25010 | 1343 | NHP2,CG9425 | | 8.335e-03 | -4.79 | fly chr72F1-72F1\|3-44 cM | chromosome location | fly chr72F1-72F1\|3-44 cM | 3 | 2 | 25010 | 1343 | GluRS-m,TyrRS | | 8.335e-03 | -4.79 | fly chr31F4-31F4\|2-43 | chromosome location | fly chr31F4-31F4\|2-43 | 3 | 2 | 25010 | 1343 | CG6094,CG7384 | | 8.335e-03 | -4.79 | fly chr8F1-8F1\|1-29 cM | chromosome location | fly chr8F1-8F1\|1-29 cM | 3 | 2 | 25010 | 1343 | Gga,LysRS | | 8.335e-03 | -4.79 | fly chr91C5-91C5\|3-64 cM | chromosome location | fly chr91C5-91C5\|3-64 cM | 3 | 2 | 25010 | 1343 | Mekk1,CG7718 | | 8.335e-03 | -4.79 | fly chr72F1-72F1\|3-44 | chromosome location | fly chr72F1-72F1\|3-44 | 3 | 2 | 25010 | 1343 | GluRS-m,TyrRS | | 8.335e-03 | -4.79 | fly chr60A5-60A5\|2-105 cM | chromosome location | fly chr60A5-60A5\|2-105 cM | 3 | 2 | 25010 | 1343 | Thiolase,Cpes | | 8.335e-03 | -4.79 | fly chr95D10-95D10\|3-82 | chromosome location | fly chr95D10-95D10\|3-82 | 3 | 2 | 25010 | 1343 | CG5986,Atg6 | | 8.335e-03 | -4.79 | fly chr6C6-6C6\|1-18 | chromosome location | fly chr6C6-6C6\|1-18 | 3 | 2 | 25010 | 1343 | CG44303,Pink1 | | 8.335e-03 | -4.79 | fly chr45F5-45F5\|2-61 cM | chromosome location | fly chr45F5-45F5\|2-61 cM | 3 | 2 | 25010 | 1343 | CG1827,CG12929 | | 8.335e-03 | -4.79 | fly chr45F5-45F5\|2-61 | chromosome location | fly chr45F5-45F5\|2-61 | 3 | 2 | 25010 | 1343 | CG1827,CG12929 | | 8.335e-03 | -4.79 | fly chr3C9-3C9\|1-2 | chromosome location | fly chr3C9-3C9\|1-2 | 3 | 2 | 25010 | 1343 | CG3939,CG18508 | | 8.335e-03 | -4.79 | fly chr6C6-6C6\|1-18 cM | chromosome location | fly chr6C6-6C6\|1-18 cM | 3 | 2 | 25010 | 1343 | CG44303,Pink1 | | 8.335e-03 | -4.79 | fly chr63E1-63E1\|3-8 | chromosome location | fly chr63E1-63E1\|3-8 | 3 | 2 | 25010 | 1343 | CycJ,armi | | 8.335e-03 | -4.79 | fly chr70F6-70F6\|3-42 cM | chromosome location | fly chr70F6-70F6\|3-42 cM | 3 | 2 | 25010 | 1343 | NHP2,CG9425 | | 8.335e-03 | -4.79 | fly chr3C9-3C9\|1-2 cM | chromosome location | fly chr3C9-3C9\|1-2 cM | 3 | 2 | 25010 | 1343 | CG18508,CG3939 | | 8.335e-03 | -4.79 | fly chr60A5-60A5\|2-105 | chromosome location | fly chr60A5-60A5\|2-105 | 3 | 2 | 25010 | 1343 | Thiolase,Cpes | | 8.340e-03 | -4.79 | regulation of cell cycle G2/M phase transition | biological process | GO:1902749 | 33 | 8 | 12345 | 1129 | Cdk1,grp,PEK,mus304,nbs,twe,mre11,Aven | | 8.340e-03 | -4.79 | negative regulation of neural precursor cell proliferation | biological process | GO:2000178 | 33 | 8 | 12345 | 1129 | tws,COX4,MED27,COX5A,EcR,MED4,MAGE,mts | | 8.340e-03 | -4.79 | regulation of G2/M transition of mitotic cell cycle | biological process | GO:0010389 | 33 | 8 | 12345 | 1129 | nbs,mus304,PEK,grp,Cdk1,Aven,mre11,twe | | 8.340e-03 | -4.79 | negative regulation of neuroblast proliferation | biological process | GO:0007406 | 33 | 8 | 12345 | 1129 | COX4,MED27,tws,COX5A,MED4,EcR,MAGE,mts | | 8.349e-03 | -4.79 | positive regulation of chromosome organization | biological process | GO:2001252 | 61 | 12 | 12345 | 1129 | HP1c,Moe,pgc,egg,trr,esc,G9a,APC7,HP1e,scny,Stat92E,piwi | | 8.353e-03 | -4.79 | negative regulation of RNA metabolic process | biological process | GO:0051253 | 398 | 51 | 12345 | 1129 | lin-52,CG10959,egg,rept,Sox102F,wash,Rbf2,EndoGI,insv,mael,Maf1,l(3)mbt,asf1,E2f1,ovo,EcR,Hnf4,pgc,CG1602,CG17328,Elba2,ph-d,CG10038,Tip60,Uxt,Bin1,esc,dl,Rbf,foxo,SREBP,hb,nos,piwi,Tlk,CG31875,cyc,d4,CG10631,Chrac-14,Mybbp1A,escl,sbb,spn-E,Hat1,CG4872,Su(H),Sirt7,psq,r2d2,CG2199 | | 8.357e-03 | -4.78 | cytosol to endoplasmic reticulum transport | biological process | GO:0046967 | 2 | 2 | 12345 | 1129 | Efr,SERCA | | 8.357e-03 | -4.78 | endoplasmic reticulum to cytosol transport | biological process | GO:1903513 | 2 | 2 | 12345 | 1129 | CG6766,sip3 | | 8.357e-03 | -4.78 | formaldehyde catabolic process | biological process | GO:0046294 | 2 | 2 | 12345 | 1129 | Fdh,CG4390 | | 8.357e-03 | -4.78 | negative regulation of mitochondrial fusion | biological process | GO:0010637 | 2 | 2 | 12345 | 1129 | park,Pink1 | | 8.357e-03 | -4.78 | positive regulation of triglyceride biosynthetic process | biological process | GO:0010867 | 2 | 2 | 12345 | 1129 | CG8009,Dd | | 8.357e-03 | -4.78 | DNA dealkylation involved in DNA repair | biological process | GO:0006307 | 2 | 2 | 12345 | 1129 | obe,agt | | 8.357e-03 | -4.78 | positive regulation of locomotor rhythm | biological process | GO:1904061 | 2 | 2 | 12345 | 1129 | park,Pink1 | | 8.357e-03 | -4.78 | endoplasmic reticulum tubular network membrane organization | biological process | GO:1990809 | 2 | 2 | 12345 | 1129 | Arl6IP1,Rtnl1 | | 8.357e-03 | -4.78 | formaldehyde metabolic process | biological process | GO:0046292 | 2 | 2 | 12345 | 1129 | Fdh,CG4390 | | 8.357e-03 | -4.78 | negative regulation of cAMP-mediated signaling | biological process | GO:0043951 | 2 | 2 | 12345 | 1129 | pn,dnc | | 8.357e-03 | -4.78 | cellular response to aldehyde | biological process | GO:0110096 | 2 | 2 | 12345 | 1129 | CG4390,Fdh | | 8.357e-03 | -4.78 | neuron death | biological process | GO:0070997 | 2 | 2 | 12345 | 1129 | sws,Cdk5 | | 8.357e-03 | -4.78 | positive regulation of transmission of nerve impulse | biological process | GO:0051971 | 2 | 2 | 12345 | 1129 | park,Pink1 | | 8.357e-03 | -4.78 | protein exit from endoplasmic reticulum | biological process | GO:0032527 | 2 | 2 | 12345 | 1129 | sip3,CG6766 | | 8.357e-03 | -4.78 | positive regulation of action potential | biological process | GO:0045760 | 2 | 2 | 12345 | 1129 | park,Pink1 | | 8.357e-03 | -4.78 | neuron apoptotic process | biological process | GO:0051402 | 2 | 2 | 12345 | 1129 | Cdk5,sws | | 8.357e-03 | -4.78 | folic acid metabolic process | biological process | GO:0046655 | 2 | 2 | 12345 | 1129 | Dhfr,Shmt | | 8.357e-03 | -4.78 | positive regulation of eye pigmentation | biological process | GO:0048075 | 2 | 2 | 12345 | 1129 | Pink1,park | | 8.357e-03 | -4.78 | positive regulation of neuronal action potential | biological process | GO:1904457 | 2 | 2 | 12345 | 1129 | Pink1,park | | 8.357e-03 | -4.78 | cap-dependent translational initiation | biological process | GO:0002191 | 2 | 2 | 12345 | 1129 | Pym,eIF4E1 | | 8.357e-03 | -4.78 | positive regulation of chromatin silencing at telomere | biological process | GO:0031940 | 2 | 2 | 12345 | 1129 | HP1c,HP1e | | 8.357e-03 | -4.78 | cellular response to salt stress | biological process | GO:0071472 | 2 | 2 | 12345 | 1129 | kcc,Mekk1 | | 8.357e-03 | -4.78 | positive regulation of compound eye pigmentation | biological process | GO:0048078 | 2 | 2 | 12345 | 1129 | Pink1,park | | 8.357e-03 | -4.78 | retrograde protein transport, ER to cytosol | biological process | GO:0030970 | 2 | 2 | 12345 | 1129 | sip3,CG6766 | | 8.357e-03 | -4.78 | cellular detoxification of aldehyde | biological process | GO:0110095 | 2 | 2 | 12345 | 1129 | CG4390,Fdh | | 8.357e-03 | -4.78 | ethanol oxidation | biological process | GO:0006069 | 2 | 2 | 12345 | 1129 | Fdh,Adh | | 8.398e-03 | -4.78 | LA | smart domains | SM00715 | 5 | 3 | 5086 | 506 | Larp4B,Larp7,larp | | 8.398e-03 | -4.78 | GLECT | smart domains | SM00276 | 5 | 3 | 5086 | 506 | Pex23,galectin,CG5335 | | 8.399e-03 | -4.78 | Jwa (Jwa) | protein interactions | 35128 | 16 | 6 | 9204 | 1112 | ATPsynG,sol,ATPsynO,CG6707,grp,ATPsyndelta | | 8.399e-03 | -4.78 | Glo1 (Glyoxalase 1) | protein interactions | 35656 | 16 | 6 | 9204 | 1112 | EndoGI,nej,CG14715,CG10638,Uch,AnxB10 | | 8.399e-03 | -4.78 | Galphas (G protein alpha s subunit) | protein interactions | 37805 | 16 | 6 | 9204 | 1112 | RpL26,dnc,Vha36-1,Bx,RpS13,Ssdp | | 8.399e-03 | -4.78 | nocte (no circadian temperature entrainment) | protein interactions | 31976 | 16 | 6 | 9204 | 1112 | Rel,mael,PGRP-LE,gw,Tak1,Cse1 | | 8.439e-03 | -4.77 | protein-N-terminal asparagine amidohydrolase activity | molecular function | GO:0008418 | 2 | 2 | 12176 | 1119 | tun,Ntan1 | | 8.439e-03 | -4.77 | juvenile hormone response element binding | molecular function | GO:0070594 | 2 | 2 | 12176 | 1119 | Fkbp39,Chd64 | | 8.439e-03 | -4.77 | acetaldehyde dehydrogenase (acetylating) activity | molecular function | GO:0008774 | 2 | 2 | 12176 | 1119 | Aldh,Adh | | 8.439e-03 | -4.77 | inorganic diphosphatase activity | molecular function | GO:0004427 | 2 | 2 | 12176 | 1119 | Nurf-38,CG17294 | | 8.439e-03 | -4.77 | long-chain-3-hydroxyacyl-CoA dehydrogenase activity | molecular function | GO:0016509 | 2 | 2 | 12176 | 1119 | Thiolase,Mtpalpha | | 8.439e-03 | -4.77 | FK506 binding | molecular function | GO:0005528 | 2 | 2 | 12176 | 1119 | Fkbp39,Fkbp59 | | 8.439e-03 | -4.77 | macrolide binding | molecular function | GO:0005527 | 2 | 2 | 12176 | 1119 | Fkbp59,Fkbp39 | | 8.439e-03 | -4.77 | alpha-ketoacid dehydrogenase activity | molecular function | GO:0003826 | 2 | 2 | 12176 | 1119 | CG17691,CG8199 | | 8.439e-03 | -4.77 | ATP-dependent protein binding | molecular function | GO:0043008 | 2 | 2 | 12176 | 1119 | Mitofilin,Pink1 | | 8.439e-03 | -4.77 | cAMP-dependent protein kinase regulator activity | molecular function | GO:0008603 | 2 | 2 | 12176 | 1119 | Pka-R1,Pka-R2 | | 8.439e-03 | -4.77 | phosphofructokinase activity | molecular function | GO:0008443 | 2 | 2 | 12176 | 1119 | Pfrx,Pfk | | 8.449e-03 | -4.77 | Ptip (PAX transcription activation domain interacting protein) | protein interactions | 39552 | 36 | 10 | 9204 | 1112 | CG17002,ago,ix,Ssdp,MED27,Utx,Usp10,enc,MED4,MED20 | | 8.479e-03 | -4.77 | MILI\_PSEUDOPODIA\_HAPTOTAXIS\_UP | MSigDB lists | MILI\_PSEUDOPODIA\_HAPTOTAXIS\_UP | 201 | 44 | 4161 | 644 | Uch-L5,CG16865,CG16979,Klp10A,Grip75,Abi,Nap1,Herc4,l(1)G0004,Ublcp1,bru1,CG31715,SkpA,mRpL13,CG3511,sau,grp,Cisd2,CG3226,Cul2,CG10465,CG11710,CHMP2B,PDCD-5,Vamp7,mtgo,Arf51F,CG4293,CanA-14F,CG7414,APP-BP1,Arl5,Snx6,CG14894,Ptp61F,CG4511,ScsbetaA,Pp2B-14D,Tlk,lt,Vha36-1,CycC,CG8963,Non1 | | 8.516e-03 | -4.77 | negative regulation of nucleobase-containing compound metabolic process | biological process | GO:0045934 | 417 | 53 | 12345 | 1129 | Sox102F,egg,rept,CG10959,lin-52,Drep1,wash,BigH1,mael,Maf1,insv,EndoGI,Rbf2,E2f1,ovo,asf1,l(3)mbt,pgc,EcR,Hnf4,Elba2,CG1602,CG17328,Uxt,Tip60,CG10038,ph-d,Bin1,hb,SREBP,foxo,Rbf,esc,dl,nos,CG31875,Tlk,piwi,CG10631,cyc,d4,Chrac-14,CG4872,spn-E,Hat1,sbb,Mybbp1A,escl,psq,Su(H),Sirt7,CG2199,r2d2 | | 8.528e-03 | -4.76 | embryo development | biological process | GO:0009790 | 568 | 69 | 12345 | 1129 | l(3)mbt,cad,Tm1,wal,mael,tkv,Dcr-1,rl,faf,Mtl,Cen,Pi3K92E,Nmt,alphaTub67C,gus,ph-d,bcd,Ptp61F,chrb,gw,fs(1)N,EcR,twe,alph,Cka,Tlk,Lkb1,Grip75,Traf6,Pld,larp,disp,HP1e,fs(1)M3,Rab5,spn-E,trk,Ptip,Grip128,swa,apt,toc,Stat92E,CycB,cnn,lwr,lin,gd,osk,Grip71,Moe,cnc,slam,Ssdp,grp,ea,Cdk1,exu,nos,tor,pygo,CycJ,Tak1,hb,psq,armi,Arpc1,csw,msps | | 8.558e-03 | -4.76 | glutamyl-tRNA(Gln) amidotransferase complex | cellular component | GO:0030956 | 2 | 2 | 11940 | 1105 | GatB,GatA | | 8.558e-03 | -4.76 | UBC13-UEV1A complex | cellular component | GO:0035370 | 2 | 2 | 11940 | 1105 | ben,Uev1A | | 8.558e-03 | -4.76 | proton-transporting ATP synthase, central stalk | cellular component | GO:0045269 | 2 | 2 | 11940 | 1105 | ATPsynO,ATPsynCF6 | | 8.558e-03 | -4.76 | 3-methyl-2-oxobutanoate dehydrogenase (lipoamide) complex | cellular component | GO:0017086 | 2 | 2 | 11940 | 1105 | CG17691,CG8199 | | 8.558e-03 | -4.76 | mitochondrial fatty acid beta-oxidation multienzyme complex | cellular component | GO:0016507 | 2 | 2 | 11940 | 1105 | Thiolase,Mtpalpha | | 8.558e-03 | -4.76 | mitochondrial proton-transporting ATP synthase, central stalk | cellular component | GO:0005756 | 2 | 2 | 11940 | 1105 | ATPsynO,ATPsynCF6 | | 8.558e-03 | -4.76 | fatty acid beta-oxidation multienzyme complex | cellular component | GO:0036125 | 2 | 2 | 11940 | 1105 | Thiolase,Mtpalpha | | 8.575e-03 | -4.76 | REACTOME\_FORMATION\_OF\_THE\_TERNARY\_COMPLEX\_AND\_SUBSEQUENTLY\_THE\_43S\_COMPLEX | MSigDB lists | REACTOME\_FORMATION\_OF\_THE\_TERNARY\_COMPLEX\_AND\_SUBSEQUENTLY\_THE\_43S\_COMPLEX | 42 | 13 | 4161 | 644 | eIF3i,RpS18,RpS4,RpS7,RpS27A,RpS23,RpS16,RpS2,RpS28b,RpS30,RpS27,RpS13,RpS24 | | 8.575e-03 | -4.76 | REACTOME\_CDT1\_ASSOCIATION\_WITH\_THE\_CDC6\_ORC\_ORIGIN\_COMPLEX | MSigDB lists | REACTOME\_CDT1\_ASSOCIATION\_WITH\_THE\_CDC6\_ORC\_ORIGIN\_COMPLEX | 42 | 13 | 4161 | 644 | Prosbeta3,Prosalpha3,Rpt2,Prosbeta4,Prosalpha4,Rpn12,Rpn2,RpS27A,Prosalpha5,Prosbeta5,Prosbeta6,RpL40,Rpn3 | | 8.582e-03 | -4.76 | phosphoric diester hydrolase activity | molecular function | GO:0008081 | 33 | 8 | 12176 | 1119 | Pld,Pka-R1,nSMase,dnc,PrBP,Pka-R2,CG18135,pn | | 8.606e-03 | -4.76 | GO\_CELLULAR\_RESPONSE\_TO\_STRESS | MSigDB lists | GO\_CELLULAR\_RESPONSE\_TO\_STRESS | 498 | 96 | 4161 | 644 | SkpA,ND-B22,Stoml2,Xpac,Rpb8,Uba5,Tip60,Iml1,Su(H),Jafrac1,Cdk7,Prip,grp,RpII140,S2P,Sod1,HDAC6,rl,cact,Cdk1,Tfb4,Uev1A,Uch-L5,SERCA,CSN6,Mo25,asf1,RfC38,Ufm1,CG17593,Ubc4,mus81,Tm1,RpL40,Tao,Tspo,Rbf,Dhfr,CG30291,Tlk,Pex13,dco,p38a,CycB,pont,CG8974,Cul1,CG7414,Wnt10,obe,SREBP,PpD3,yip2,Sld5,cyc,Grx1,rept,csw,Rpt2,Cul2,Not10,shv,LpR2,Caf1-105,AIF,STUB1,mRpS10,CG9911,ERp60,Calx,mael,twin,RpS27A,Faf2,Pi4KIIalpha,fray,Nprl2,CG6766,sgll,Pdk,Imp,Cyp1,Atg7,CG12018,P5cr,CG5285,CG11110,Atg6,Traf6,Edem1,Exd2,lwr,Hsc70-3,Rcd5,DNApol-epsilon255,Iru | | 8.681e-03 | -4.75 | - | gene3d domains | 3.40.50.620 | 40 | 11 | 4929 | 622 | Nadsyn,CG4407,GluRS-m,CG1578,TrpRS-m,CG3021,CysRS,TyrRS,CG10189,wal,AsnS | | 8.693e-03 | -4.75 | cellular ketone metabolic process | biological process | GO:0042180 | 54 | 11 | 12345 | 1129 | ND-39,Kyat,Coq6,CG10425,scu,Coq7,Npc2a,Marf,foxo,Coq8,tzn | | 8.696e-03 | -4.74 | MARIADASON\_REGULATED\_BY\_HISTONE\_ACETYLATION\_DN | MSigDB lists | MARIADASON\_REGULATED\_BY\_HISTONE\_ACETYLATION\_DN | 13 | 6 | 4161 | 644 | Cul1,Sccpdh1,l(3)73Ah,Imp,Nc73EF,CG30496 | | 8.696e-03 | -4.74 | GO\_REGULATION\_OF\_TELOMERASE\_RNA\_LOCALIZATION\_TO\_CAJAL\_BODY | MSigDB lists | GO\_REGULATION\_OF\_TELOMERASE\_RNA\_LOCALIZATION\_TO\_CAJAL\_BODY | 13 | 6 | 4161 | 644 | CCT6,rept,pont,CCT5,CCT3,NHP2 | | 8.696e-03 | -4.74 | GO\_MODIFICATION\_BY\_SYMBIONT\_OF\_HOST\_MORPHOLOGY\_OR\_PHYSIOLOGY | MSigDB lists | GO\_MODIFICATION\_BY\_SYMBIONT\_OF\_HOST\_MORPHOLOGY\_OR\_PHYSIOLOGY | 13 | 6 | 4161 | 644 | Atg7,Smox,Pen,Kap-alpha3,Ts,Clp | | 8.696e-03 | -4.74 | REACTOME\_TRANSCRIPTIONAL\_ACTIVITY\_OF\_SMAD2\_SMAD3\_SMAD4\_HETEROTRIMER | MSigDB lists | REACTOME\_TRANSCRIPTIONAL\_ACTIVITY\_OF\_SMAD2\_SMAD3\_SMAD4\_HETEROTRIMER | 13 | 6 | 4161 | 644 | CycC,Smox,Med,RpL40,RpS27A,faf | | 8.696e-03 | -4.74 | MODULE\_295 | MSigDB lists | MODULE\_295 | 13 | 6 | 4161 | 644 | Thiolase,Pcd,Irp-1B,Echs1,pr,Mtpalpha | | 8.696e-03 | -4.74 | GO\_SOMITOGENESIS | MSigDB lists | GO\_SOMITOGENESIS | 13 | 6 | 4161 | 644 | Utx,Su(H),Smox,Med,O-fut1,CHOp24 | | 8.720e-03 | -4.74 | hpo (hippo) | protein interactions | 37247 | 95 | 20 | 9204 | 1112 | Pp2A-29B,CG2678,Herc4,bun,SERCA,RpS18,Ublcp1,CG6707,CycE,Pld,mts,mEFTu1,ago,Sox102F,Tao,bru1,CG13366,Cka,trr,Smox | | 8.729e-03 | -4.74 | protein homodimerization activity | molecular function | GO:0042803 | 140 | 22 | 12176 | 1119 | bun,Acph-1,park,Sod1,cbs,GlyP,Ork1,Orc6,GCC88,spz,Shmt,psq,Atf6,trsn,Roe1,Trxr-1,Liprin-beta,Debcl,exu,ari-1,Blos2,Adh | | 8.731e-03 | -4.74 | induction of programmed cell death | biological process | GO:0012502 | 15 | 5 | 12345 | 1129 | Utx,MED24,Idh3b,EcR,rept | | 8.731e-03 | -4.74 | mitochondrial electron transport, cytochrome c to oxygen | biological process | GO:0006123 | 15 | 5 | 12345 | 1129 | COX5A,COX4,levy,cype,COX5B | | 8.731e-03 | -4.74 | aerobic electron transport chain | biological process | GO:0019646 | 15 | 5 | 12345 | 1129 | levy,COX5A,COX4,COX5B,cype | | 8.731e-03 | -4.74 | vitamin metabolic process | biological process | GO:0006766 | 15 | 5 | 12345 | 1129 | Dhfr,Pdxk,Shmt,sgll,regucalcin | | 8.731e-03 | -4.74 | mitotic G2/M transition checkpoint | biological process | GO:0044818 | 15 | 5 | 12345 | 1129 | grp,Cdk1,nbs,mus304,mre11 | | 8.731e-03 | -4.74 | induction of programmed cell death by ecdysone | biological process | GO:0035078 | 15 | 5 | 12345 | 1129 | Utx,MED24,Idh3b,EcR,rept | | 8.731e-03 | -4.74 | induction of programmed cell death by hormones | biological process | GO:0035081 | 15 | 5 | 12345 | 1129 | rept,Idh3b,EcR,MED24,Utx | | 8.731e-03 | -4.74 | ecdysone-mediated induction of salivary gland cell autophagic cell death | biological process | GO:0035072 | 15 | 5 | 12345 | 1129 | MED24,Utx,rept,Idh3b,EcR | | 8.737e-03 | -4.74 | vesicle | cellular component | GO:0031982 | 422 | 54 | 11940 | 1105 | lqfR,Sec23,Acph-1,Vsp37A,Pld,fdl,tkv,Vamp7,wash,fwe,pen-2,Synj,PAPLA1,Afti,Wdfy2,CG5734,CHOp24,Rab5,CG5510,Vps16B,Arf51F,Crag,AP-1-2beta,yl,Hip1,Stam,shrb,CG4293,Clc,yin,Rep,CG6707,lt,Desat1,Pi4KIIalpha,CG3309,Sec15,Gdi,CG9132,Gga,Sec22,Arc2,CanB2,mop,Rab32,CG31729,CG32536,opm,Rab10,Lkb1,Tomosyn,Snx6,milt,CHMP2B | | 8.745e-03 | -4.74 | GSE37416\_0H\_VS\_3H\_F\_TULARENSIS\_LVS\_NEUTROPHIL\_DN | MSigDB lists | GSE37416\_0H\_VS\_3H\_F\_TULARENSIS\_LVS\_NEUTROPHIL\_DN | 56 | 16 | 4161 | 644 | Ppcdc,Rnmt,stas,Sec22,Stip1,cact,CHMP2B,anne,Eno,CG14715,mts,AMPdeam,ari-1,shv,fray,dnc | | 8.773e-03 | -4.74 | metal ion homeostasis | biological process | GO:0055065 | 116 | 19 | 12345 | 1129 | up,Atpalpha,SERCA,anne,CG10470,CG14036,park,MtnA,Ho,Seipin,ZnT35C,fwe,Scox,CG12773,nrv1,Fer1HCH,Grx1,zyd,EMRE | | 8.816e-03 | -4.73 | GO\_MRNA\_METABOLIC\_PROCESS | MSigDB lists | GO\_MRNA\_METABOLIC\_PROCESS | 268 | 56 | 4161 | 644 | RpL13,Bin1,Tfb4,RpS28b,RpL9,RpS23,RpS27A,twin,Clp,Unr,bru1,Rpb8,RpLP0,RpS13,RpS30,eIF4E1,CG6610,shv,Rrp40,RpL13A,Not10,RpII140,CG3511,Cdk7,mts,eIF4A,RpL12,thoc6,RpS18,SRPK,RpL24,RpS27,RpL26,Rnmt,Pp2A-29B,RpL11,RpL37a,LSm3,Cbp80,RpL40,Usp39,heph,CG10418,tho2,RpS7,RpS4,RpL38,CG17454,RpS16,RpS2,Edc3,RpS24,RpL30,RpL23,CG8963,CG31759 | | 8.853e-03 | -4.73 | endosomal transport | biological process | GO:0016197 | 108 | 18 | 12345 | 1129 | wash,Stam,Ubc6,shrb,MICAL-like,Blos2,Vps16B,Vsp37A,Vha68-2,Arl5,Synj,CG5734,Sec15,CHMP2B,Snx6,Rab5,fwe,mop | | 8.867e-03 | -4.73 | germarium-derived egg chamber formation | biological process | GO:0007293 | 100 | 17 | 12345 | 1129 | armi,bam,jvl,Pen,del,enc,Rbfox1,spn-E,Iml1,Cdk1,piwi,Stat92E,Cul5,Dcr-1,vret,Nprl2,CycJ | | 8.911e-03 | -4.72 | transaminase activity | molecular function | GO:0008483 | 15 | 5 | 12176 | 1119 | Got1,Kyat,CG1640,CG11899,Gabat | | 8.924e-03 | -4.72 | GO\_MITOTIC\_CELL\_CYCLE | MSigDB lists | GO\_MITOTIC\_CELL\_CYCLE | 302 | 62 | 4161 | 644 | mbo,RpL40,Ts,PpV,Prosbeta6,Atg6,APC7,dco,Aladin,Pp2B-14D,faf,Sirt7,CkIIalpha,Dhfr,CG30291,tws,Rbf,Dd,lwr,msps,Cul1,Rpn3,Prosbeta5,pont,CycB,CHMP2B,gammaTub37C,CycE,PpD3,Pp2A-29B,DNApol-epsilon255,CanA-14F,Prosalpha3,APP-BP1,Prosbeta4,Wnt10,Grip128,SkpA,eIF4E1,CG1812,Rpn12,CG8188,Not10,Rpn2,grp,mts,Cdk7,Prosbeta3,Cul2,Rpt2,csw,Fbw5,mad2,Cul5,Klp10A,Cdk1,Grip75,Grip84,Prosalpha5,RpS27A,twin,shrb | | 8.950e-03 | -4.72 | intracellular mRNA localization involved in anterior/posterior axis specification | biological process | GO:0060811 | 69 | 13 | 12345 | 1129 | jvl,Grip128,osk,exu,spn-E,Grip71,msps,Tm1,mael,Lkb1,Grip75,cnc,swa | | 8.953e-03 | -4.72 | PILON\_KLF1\_TARGETS\_DN | MSigDB lists | PILON\_KLF1\_TARGETS\_DN | 701 | 130 | 4161 | 644 | CG11089,ACC,CG7550,CG31715,CG15141,Rrp40,Cse1,CG11164,CG9911,CG16979,Rab32,Nap1,Idh,Aos1,Snx6,r,CG9485,CG9132,CG8112,CG15715,Chd64,CG10425,pr,Rnmt,DNApol-epsilon255,CanA-14F,CG10038,Hsc70-3,Fdh,CG42542,SkpA,Rpb8,Karybeta3,ste24a,scu,RpII140,Mcad,CG11899,Uev1A,CG9391,CG3803,rl,Grip75,Sec22,CG1774,Tspo,Med,Usp39,mRpS23,Dhfr,Rbf,Hat1,f-cup,Kap-alpha3,Moe,sws,Dcr-1,Naa20A,CG14194,cyc,bru1,CG4562,CG6610,Efr,CG10189,CG3631,CG5026,Bin1,Caf1-105,Npc2a,twin,Cbp80,CG8726,scramb1,Adi1,DhpD,Ranbp21,CG17454,CG40045,CG1890,bor,ScsbetaA,Cyt-c-p,Pp2B-14D,Phb2,CG10470,Psf3,CG17294,CG7718,r-l,CCT3,CHMP2B,Pp2A-29B,COX5A,Wdfy2,Su(H),CG10932,escl,grp,CG33156,CCT6,Uch-L5,mad2,Aldh,Mo25,spag,cl,Synj,Sap47,Ptp61F,Pi3K92E,Pex13,Thiolase,CG13630,nmdyn-D6,Cul1,pont,esc,Pdk1,SRPK,Pdxk,p38a,PpD3,MED20,Clc,Arf51F,Gyg,Pfas,CG7414,Tomosyn | | 9.005e-03 | -4.71 | ubiquitin-like protein ligase binding | molecular function | GO:0044389 | 54 | 11 | 12176 | 1119 | Ubc4,RpS27A,CG10254,ben,CG30496,Cul5,Cul2,Cul1,CG3226,Tak1,RpL40 | | 9.017e-03 | -4.71 | MORF\_CSNK1D | MSigDB lists | MORF\_CSNK1D | 29 | 10 | 4161 | 644 | Marf,CG13630,Arpc4,Gmer,P5CDh1,Grip84,Atg6,CG9393,GCS2beta,Lsm12 | | 9.022e-03 | -4.71 | CycD (Cyclin D) | protein interactions | 32551 | 31 | 9 | 9204 | 1112 | CycC,bcd,mute,Rbf,Cdk5,Stat92E,Cdk1,CG11486,EcR | | 9.022e-03 | -4.71 | Lrrk (Leucine-rich repeat kinase) | protein interactions | 42447 | 31 | 9 | 9204 | 1112 | Rbf,eIF4E1,foxo,Atg7,E2f1,park,Rab5,Dcr-1,Pink1 | | 9.036e-03 | -4.71 | Proteasome\_bsu\_CS | interpro domains | IPR016050 | 13 | 5 | 11383 | 1233 | Prosbeta5,Prosalpha3,Prosbeta6,Prosbeta4,Prosbeta3 | | 9.077e-03 | -4.70 | transcription corepressor activity | molecular function | GO:0003714 | 40 | 9 | 12176 | 1119 | Mybbp1A,CG4872,CG31875,Bin1,sbb,rept,insv,CG10631,Uxt | | 9.077e-03 | -4.70 | 3'-5' exonuclease activity | molecular function | GO:0008408 | 40 | 9 | 12176 | 1119 | twin,DNApol-epsilon255,mus205,Rga,CG11486,cu,CG31759,mre11,Exd2 | | 9.110e-03 | -4.70 | centriole assembly | biological process | GO:0098534 | 27 | 7 | 12345 | 1129 | mts,Rcd5,cnn,CCT6,Pp2A-29B,Klp10A,tws | | 9.110e-03 | -4.70 | regulation of intracellular mRNA localization | biological process | GO:1904580 | 27 | 7 | 12345 | 1129 | exu,cnc,swa,spn-E,enc,mael,cup | | 9.110e-03 | -4.70 | cellular response to toxic substance | biological process | GO:0097237 | 27 | 7 | 12345 | 1129 | Sod3,CG4390,Jafrac1,Sod1,Trxr-1,Pka-R2,Fdh | | 9.110e-03 | -4.70 | ADP metabolic process | biological process | GO:0046031 | 27 | 7 | 12345 | 1129 | Gapdh1,Eno,Pgk,Pfk,Pfrx,Pglym78,PyK | | 9.113e-03 | -4.70 | GSE21927\_SPLEEN\_C57BL6\_VS\_EL4\_TUMOR\_BALBC\_MONOCYTES\_DN | MSigDB lists | GSE21927\_SPLEEN\_C57BL6\_VS\_EL4\_TUMOR\_BALBC\_MONOCYTES\_DN | 111 | 27 | 4161 | 644 | CG8728,Non1,CkIIalpha,Psf3,CG12018,CG11899,CG8188,Rpn12,DppIII,Stoml2,NHP2,Rpb8,Karybeta3,Roe1,Uba5,tko,mbo,Pmi,Fkbp59,MED20,Clc,gammaTub37C,Tim8,ND-75,bys,UQCR-Q,CG15715 | | 9.113e-03 | -4.70 | RUTELLA\_RESPONSE\_TO\_HGF\_VS\_CSF2RB\_AND\_IL4\_UP | MSigDB lists | RUTELLA\_RESPONSE\_TO\_HGF\_VS\_CSF2RB\_AND\_IL4\_UP | 111 | 27 | 4161 | 644 | Cyp1,CG31729,cathD,scramb1,CG10932,Roe1,Mctp,CG3860,AMPdeam,CG7718,ND-49,CG15715,CG12773,CG16979,Rbcn-3A,CG7997,Prosbeta5,CG8004,anne,Mtpalpha,Pdxk,Smox,yip2,Sccpdh1,Sec22,Irp-1B,Fkbp59 | | 9.121e-03 | -4.70 | GSE20715\_WT\_VS\_TLR4\_KO\_6H\_OZONE\_LUNG\_UP | MSigDB lists | GSE20715\_WT\_VS\_TLR4\_KO\_6H\_OZONE\_LUNG\_UP | 61 | 17 | 4161 | 644 | Pka-R2,Hsc70-3,Cka,CG32549,COX5A,Gclc,p38a,AnxB10,eIF4E1,galectin,ABCD,HDAC6,Rpn12,CG10470,ste14,Atg7,Ublcp1 | | 9.175e-03 | -4.69 | Metabolic pathways | KEGG pathways | dme01100 | 999 | 178 | 3198 | 496 | Shmt,Ho,CG10863,CG9248,ND-49,Gabat,Sply,ND-MWFE,Thiolase,COX5B,Cyt-c-p,COX8,r-l,nmdyn-D6,ND-75,ND-B15,Pgk,CG3376,Echs1,Gapdh1,Coq7,AcCoAS,Gip,CG10638,Pglym78,P5cr,Vha68-2,CG9391,CG2201,Dip-B,shop,CG17333,Aldh,ND-B18,alpha-Man-Ia,CG4407,Pfas,CG14516,ATPsynD,Treh,Nadsyn,DNApol-epsilon255,CG5966,sgll,Pld,GatB,fdl,ATPsynE,Kyat,oxt,CG7718,Coq6,AGBE,CG12018,GalT1,CG9436,Chrac-14,Eno,aay,pr,GlyP,CG7322,Ppcdc,Pi4KIIalpha,CG6767,Gclc,SP1029,Ts,DhpD,CG10425,Ady43A,PIG-H,Dhfr,CG11089,Fdh,CG2767,CG7328,AsnS,cype,Adh,GluRS-m,PIG-L,Ppox,cN-IIIB,mino,ND-SGDH,ATPsynF,ND-MNLL,Irp-1B,P5CDh1,Pfk,Acph-1,CG5028,ND-51,Got1,CG8311,Pgm1,yip2,Synj,levy,Ugt35A1,UGP,Gk1,CG8199,ATPsynCF6,CG3803,PIG-C,Idh,CG9485,AOX3,ATPsynG,nmdyn-D7,ATPsyndelta,CG7550,Pdxk,ND-23,awd,Gk2,COX4,Sptr,ND-30,RpII140,SdhD,CG11899,kri,ND-24,Mtpalpha,CG6650,CG4860,Gmer,Acox57D-p,ScsbetaA,CG11134,Idh3b,CG17691,AMPdeam,blw,UQCR-11,CG10932,Ugt36E1,ND-39,CG32549,ATPsynO,ND-18,ACC,CG1640,l(2)k01209,Mcad,Cht11,Hexo1,r,Nc73EF,Adi1,COX5A,SelD,SdhA,Scsalpha1,scu,PIG-P,Rpb8,nSMase,GatA,CG4586,ND-B22,NP15.6,Vha36-1,UQCR-Q,CG40472,mus205,PyK,Vha13,CG33977,CG7789,Ggt-1,CG33156,Gyg,ATPsynB,ATPsynbeta | | 9.176e-03 | -4.69 | mitochondrial respiratory chain complex IV | cellular component | GO:0005751 | 15 | 5 | 11940 | 1105 | COX5A,COX4,cype,levy,COX5B | | 9.176e-03 | -4.69 | clathrin-coated vesicle membrane | cellular component | GO:0030665 | 15 | 5 | 11940 | 1105 | Hip1,Afti,AP-1-2beta,Clc,CG9132 | | 9.176e-03 | -4.69 | tricarboxylic acid cycle enzyme complex | cellular component | GO:0045239 | 15 | 5 | 11940 | 1105 | Nc73EF,ScsbetaA,Scsalpha1,CG8199,CG17691 | | 9.176e-03 | -4.69 | nuclear chromosome, telomeric region | cellular component | GO:0000784 | 15 | 5 | 11940 | 1105 | Snm1,SuUR,nbs,mre11,Sirt7 | | 9.176e-03 | -4.69 | clathrin coat | cellular component | GO:0030118 | 15 | 5 | 11940 | 1105 | Hip1,Afti,AP-1-2beta,Clc,CG9132 | | 9.199e-03 | -4.69 | Prosbeta3 (Proteasome beta3 subunit) | protein interactions | 41079 | 59 | 14 | 9204 | 1112 | Prosalpha4,Klp10A,Prosbeta5,Prosbeta6,CG9316,Prosalpha3,CG6175,Rpt2,Prosbeta4,Rpn13,AnxB10,Rpn2,Pomp,Prosalpha5 | | 9.199e-03 | -4.69 | ago (archipelago) | protein interactions | 38516 | 59 | 14 | 9204 | 1112 | COX5A,nrv1,mtgo,SP1029,RpL7A,Marf1,wal,Ptip,RpLP0,CycE,Atpalpha,awd,RpS2,SkpA | | 9.199e-03 | -4.69 | CR10032 (pseudo) | protein interactions | 40877 | 59 | 14 | 9204 | 1112 | sol,cad,tral,CG6707,tos,CG42565,CG14204,bcd,l(2)k01209,CG4404,Clp,alphaTub84B,CG2924,RpL17 | | 9.263e-03 | -4.68 | Stacl (Stac-like) | protein interactions | 5740318 | 21 | 7 | 9204 | 1112 | CG4557,CG10631,cnn,Hip1,Snx6,Kcmf1,Phb2 | | 9.263e-03 | -4.68 | Trf4-1 (Topoisomerase related function 4-1) | protein interactions | 31795 | 21 | 7 | 9204 | 1112 | SkpA,MtnA,RpL13A,ND-30,Naa20A,Cenp-C,CG11791 | | 9.263e-03 | -4.68 | Patr-1 (Protein associated with topo II related - 1) | protein interactions | 42058 | 21 | 7 | 9204 | 1112 | Lkb1,Sec10,gw,nos,twin,Rga,LSm3 | | 9.263e-03 | -4.68 | CG17508 (uncharacterized protein) | protein interactions | 49893 | 21 | 7 | 9204 | 1112 | Cdk5,squ,nej,Cul2,grp,plu,CycJ | | 9.263e-03 | -4.68 | Nf-YC (Nuclear factor Y-box C) | protein interactions | 31622 | 21 | 7 | 9204 | 1112 | Nf-YA,26-29-p,Marf1,Stat92E,Uev1A,cathD,CycB | | 9.263e-03 | -4.68 | twin (twin) | protein interactions | 42880 | 21 | 7 | 9204 | 1112 | smg,BicC,nos,bam,AGO3,cup,Rga | | 9.287e-03 | -4.68 | transferase activity, transferring acyl groups other than amino-acyl groups | molecular function | GO:0016747 | 209 | 30 | 12176 | 1119 | Agpat2,whd,e(y)1,Tip60,CG6178,CG10932,CG10344,CG8112,CG31523,Rcd5,Sgf29,Chrac-14,nej,Nmt,lid,Thiolase,Hat1,mino,yip2,d4,CG2938,CG5880,pont,rept,CG14204,CG14615,CG8314,YL-1,AgmNAT,Naa20A | | 9.306e-03 | -4.68 | transcription coactivator activity | molecular function | GO:0003713 | 69 | 13 | 12176 | 1119 | trr,pont,Sgf11,nej,EcR,Taf7,CG17202,MED20,MED4,Wbp2,CG11710,pygo,hng3 | | 9.317e-03 | -4.68 | regulation of locomotor rhythm | biological process | GO:1904059 | 10 | 4 | 12345 | 1129 | Pink1,spag,park,Pura | | 9.317e-03 | -4.68 | microtubule nucleation by microtubule organizing center | biological process | GO:0051418 | 10 | 4 | 12345 | 1129 | Grip128,Grip84,Grip75,CG7716 | | 9.317e-03 | -4.68 | G1/S transition of mitotic cell cycle | biological process | GO:0000082 | 10 | 4 | 12345 | 1129 | Cdk1,CycE,PpV,E2f1 | | 9.317e-03 | -4.68 | synaptic vesicle targeting | biological process | GO:0016080 | 10 | 4 | 12345 | 1129 | Sec15,X11L,Exo70,Sec10 | | 9.317e-03 | -4.68 | cell cycle G1/S phase transition | biological process | GO:0044843 | 10 | 4 | 12345 | 1129 | PpV,E2f1,CycE,Cdk1 | | 9.317e-03 | -4.68 | guanosine-containing compound metabolic process | biological process | GO:1901068 | 10 | 4 | 12345 | 1129 | awd,cN-IIIB,nmdyn-D7,nmdyn-D6 | | 9.317e-03 | -4.68 | peroxisomal membrane transport | biological process | GO:0015919 | 10 | 4 | 12345 | 1129 | Pex10,Pex12,Topors,Pex13 | | 9.317e-03 | -4.68 | negative regulation of macroautophagy | biological process | GO:0016242 | 10 | 4 | 12345 | 1129 | scny,Fkbp39,Nprl2,rl | | 9.317e-03 | -4.68 | peroxisomal transport | biological process | GO:0043574 | 10 | 4 | 12345 | 1129 | Topors,Pex13,Pex10,Pex12 | | 9.317e-03 | -4.68 | positive regulation of DNA replication | biological process | GO:0045740 | 10 | 4 | 12345 | 1129 | otu,pn,E2f1,SuUR | | 9.349e-03 | -4.67 | threonine-type peptidase activity | molecular function | GO:0070003 | 27 | 7 | 12176 | 1119 | Prosbeta6,Prosalpha3,Prosalpha5,Prosbeta4,Prosbeta3,Prosbeta5,Prosalpha4 | | 9.349e-03 | -4.67 | threonine-type endopeptidase activity | molecular function | GO:0004298 | 27 | 7 | 12176 | 1119 | Prosbeta5,Prosalpha4,Prosbeta3,Prosalpha5,Prosbeta4,Prosalpha3,Prosbeta6 | | 9.362e-03 | -4.67 | CG2070 (uncharacterized protein) | protein interactions | 35706 | 26 | 8 | 9204 | 1112 | SERCA,CG10425,CG31523,Surf4,SelT,CG15098,Tspo,CG31729 | | 9.362e-03 | -4.67 | CG5500 (uncharacterized protein) | protein interactions | 43233 | 26 | 8 | 9204 | 1112 | Adh,CG32473,Pgk,Argk,GlyP,Fer1HCH,Scp1,Pglym78 | | 9.362e-03 | -4.67 | CG15717 (uncharacterized protein) | protein interactions | 32262 | 26 | 8 | 9204 | 1112 | Drep2,AnxB10,Naa20A,Prosbeta3,Prosbeta6,SelT,Prosalpha4,Prosbeta5 | | 9.367e-03 | -4.67 | GSE10463\_CD40L\_AND\_VA347\_VS\_CD40L\_IN\_DC\_UP | MSigDB lists | GSE10463\_CD40L\_AND\_VA347\_VS\_CD40L\_IN\_DC\_UP | 47 | 14 | 4161 | 644 | Tomosyn,Pex12,muskelin,CG15111,MED24,DNApol-epsilon255,colt,CG3587,Faf2,RfC38,oxt,ATP8A,Med,dnc | | 9.367e-03 | -4.67 | MORF\_MBD4 | MSigDB lists | MORF\_MBD4 | 47 | 14 | 4161 | 644 | Rpt2,CG3631,Prosalpha3,COX5A,LysRS,CG1890,SkpA,PpV,Bin1,ND-75,CHOp24,Pka-R1,Thiolase,CG11070 | | 9.367e-03 | -4.67 | GSE6674\_UNSTIM\_VS\_PL2\_3\_STIM\_BCELL\_DN | MSigDB lists | GSE6674\_UNSTIM\_VS\_PL2\_3\_STIM\_BCELL\_DN | 47 | 14 | 4161 | 644 | PrBP,Top1,dnc,Pglym78,Chd64,Tip60,AIF,Scox,Surf4,asf1,CG6144,Tsp86D,Non1,Tsp66E | | 9.374e-03 | -4.67 | RIBOSOMALL34 | prints domains | PR01250 | 2 | 2 | 2185 | 212 | RpL34b,RpL34a | | 9.374e-03 | -4.67 | RABGDIREP | prints domains | PR00891 | 2 | 2 | 2185 | 212 | Gdi,Rep | | 9.376e-03 | -4.67 | GO\_CELLULAR\_PROTEIN\_COMPLEX\_ASSEMBLY | MSigDB lists | GO\_CELLULAR\_PROTEIN\_COMPLEX\_ASSEMBLY | 148 | 34 | 4161 | 644 | TBCC,nAChRalpha4,Vamp7,Smox,Scox,gammaTub37C,ND-B18,ND-24,ND-30,Arpc4,CG11110,CG1890,Ubc4,Maf1,Med,Synj,ND-39,Fkbp59,Grip84,Grip75,AIF,ND-75,CG3803,ND-49,CG8188,Rpn12,ND-SGDH,Karybeta3,ND-B22,CHOp24,Grip128,ND-51,Arts,ND-23 | | 9.394e-03 | -4.67 | GSE3720\_UNSTIM\_VS\_PMA\_STIM\_VD1\_GAMMADELTA\_TCELL\_UP | MSigDB lists | GSE3720\_UNSTIM\_VS\_PMA\_STIM\_VD1\_GAMMADELTA\_TCELL\_UP | 66 | 18 | 4161 | 644 | Archease,RpL23,CG11164,CG4332,ND-B22,MED20,RpS27A,RpL9,COX5A,shrb,Pex23,mad2,eEF1beta,mRpL16,UQCR-Q,RpL12,ND-B18,CCT5 | | 9.394e-03 | -4.67 | GSE42088\_UNINF\_VS\_LEISHMANIA\_INF\_DC\_2H\_DN | MSigDB lists | GSE42088\_UNINF\_VS\_LEISHMANIA\_INF\_DC\_2H\_DN | 66 | 18 | 4161 | 644 | RpS18,RpL13,RpL11,Ppcdc,RpL37a,RpL26,RpL24,RpS28b,RpL38,RpS7,Pdk,RpL40,RpL30,RpS24,Pfdn5,RpS30,RpS13,RpS16 | | 9.398e-03 | -4.67 | ATPase\_P-typ\_TM\_dom\_sf | interpro domains | IPR023298 | 18 | 6 | 11383 | 1233 | anne,CG31729,SERCA,Atpalpha,CG9981,ATP8A | | 9.427e-03 | -4.66 | nuclear-transcribed mRNA catabolic process, deadenylation-dependent decay | biological process | GO:0000288 | 21 | 6 | 12345 | 1129 | Rrp40,cu,smg,CG11486,twin,CG31759 | | 9.427e-03 | -4.66 | positive regulation of DNA-binding transcription factor activity | biological process | GO:0051091 | 21 | 6 | 12345 | 1129 | CG13344,rl,lwr,Aos1,Traf6,S2P | | 9.451e-03 | -4.66 | gene expression | biological process | GO:0010467 | 1554 | 168 | 12345 | 1129 | CG9143,spen,Dcr-1,CG3198,RpL34b,CG7414,Naa20A,CG5986,TfIIFalpha,Taf7,CG12301,RpL40,Archease,mRpL16,wisp,Usp39,RpS27A,ste24a,CG14817,RpL26,CG8728,RpS16,Rrp40,Bin1,RpS29,RpS2,LSm3,RpS13,CG10463,eEF1beta,mRpS11,Sgf29,rig,RpL30,IntS6,bru1,CG17768,RpII140,GluRS-m,Trf4-2,pps,eIF3m,GatB,mRpL18,snRNA:U4atac,CG17454,RpLP1,Cdk7,tko,mbo,heph,RpS30,mRpL17,MED24,mEFTu1,RpL17,CG32473,Cbp80,RpS23,scu,CG8204,TyrRS,Son,RpL10,mRpL10,CG16865,mRpS23,stet,Larp7,CG7009,RpL24,mRpL13,Rpb8,snRNA:U4:38AB,RpL13,nos,GatA,pygo,CG6610,RpS27,RpL9,IntS11,GAPsec,RpS28b,SerRS-m,S2P,Nxf3,CG5292,CG2911,mRpS10,mRpS21,RpL11,CG7728,eIF4E1,CG11486,mtTFB1,CG3021,e(y)1,sbr,obe,mRpS5,Rnmt,bys,RpL29,Trf2,mRpL35,Spx,Pym,mRpL36,RpS7,CG2100,CG11110,CG10418,CG30343,SelD,snk,TrpRS-m,tho2,RpS15,CG17187,Imp,eIF3i,CG10189,RpL23A,Ggt-1,CG3587,RpL31,Neos,LysRS,Tfb4,CG6094,NHP2,RpS4,Fur2,thoc6,Nurf-38,CysRS,mEFTu2,RpL7A,yps,CG14100,CG9684,RpL38,SmE,RpL12,CycC,CG1622,CG2034,gd,RpS5b,mRpL48,RpLP0,MED4,RpL37a,exu,eIF4A,Clp,Rga,snRNA:U12,Mat89Ba,RpL34a,RpL13A,RpS24,RpS18,RpS14b,RpL23,RpS14a,r2d2 | | 9.471e-03 | -4.66 | GO\_CELL\_CYCLE\_PROCESS | MSigDB lists | GO\_CELL\_CYCLE\_PROCESS | 383 | 76 | 4161 | 644 | Not10,Rpn12,eIF4E1,csw,Rpt2,Cul2,mts,RpS27A,Grip84,mael,twin,Cul5,Phb2,faf,Pp2B-14D,Atg6,CkIIalpha,mbo,PpV,Ts,DNApol-epsilon255,Pp2A-29B,Smox,APP-BP1,Prosalpha3,CanA-14F,spn-E,lwr,CHMP2B,Prosbeta5,Rpn2,grp,CG8188,CG1812,raptor,Prosbeta3,Cdk7,Grip128,SkpA,Mo25,Prosalpha5,shu,Grip75,shrb,mad2,Fbw5,Klp10A,Cdk1,dco,Sirt7,Aladin,alc,APC7,Dd,Rbf,CG30291,tws,Dhfr,RpL40,Tao,Prosbeta6,Pka-R1,PpD3,RpS27,CycE,gammaTub37C,Wnt10,Prosbeta4,Cul1,msps,Rpn3,Marf,p38a,Ac3,CycB,pont | | 9.490e-03 | -4.66 | cellular amide metabolic process | biological process | GO:0043603 | 872 | 100 | 12345 | 1129 | CG1774,RpL34a,GAPsec,RpL9,RpL37a,GatA,RpL13,RpS27,eIF4A,GstT2,RpS14a,Dhfr,eIF4E1,CG6178,RpL11,CG7997,RpS18,CG11771,RpS14b,RpL23,RpS24,mRpS10,AcCoAS,mRpS21,RpS28b,RpL13A,SerRS-m,nSMase,scu,RpS23,RpL7A,Shmt,CG32473,CG17691,mEFTu2,mEFTu1,ScsbetaA,CysRS,CG7920,RpL17,pen-2,mRpL17,RpS30,mRpL48,RpL24,RpLP0,mRpL13,RpS5b,CG8199,CG7009,mRpL10,mRpS23,RpL38,RpL10,RpL12,TyrRS,RpL30,Ggt-1,GstE11,GluRS-m,eEF1beta,RpL23A,RpS13,GstT1,mRpS11,RpS2,RpS29,RpS15,eIF3i,Ntan1,CG6094,RpS4,Scsalpha1,RpLP1,Gclc,LysRS,tko,CG14516,mRpL18,eIF3m,RpL31,GatB,RpL29,RpL40,CG7414,RpL34b,mRpS5,SP1029,RpS16,TrpRS-m,RpS7,Pym,mRpL36,RpL26,CG11110,CG14817,RpS27A,mRpL16,AsnS,mRpL35 | | 9.496e-03 | -4.66 | Cell Cycle | REACTOME pathways | R-DME-1640170 | 243 | 49 | 3923 | 574 | Ankle2,Rpt2,RpL40,Prosalpha3,SkpA,Prosalpha4,Ts,Dhfr,Cul1,Rpn3,Rpn12,Cdk1,Psf3,mbo,CycB,Nup58,Prosbeta6,wapl,APC7,Prosbeta3,Orc6,Cdk7,lin-52,Nup358,Rbf,Aladin,Sld5,Dd,CG12018,Prosbeta5,Rpn2,Rbf2,CycE,Chrac-14,twe,Prosalpha5,RpS27A,mad2,Prosbeta4,E2f1,CG8009,Orc1,grp,Ndc1,DNApol-epsilon255,Pp2A-29B,RfC38,rl,Nup62 | | 9.503e-03 | -4.66 | cellular component morphogenesis | biological process | GO:0032989 | 667 | 79 | 12345 | 1129 | spz,Sec15,Liprin-beta,gus,cact,Ptp61F,bif,EcR,fs(1)N,capt,Pmi,Tm1,mael,tkv,Dcr-1,spen,rl,Cdk5,Mtl,Pi3K92E,Cf2,Pxt,Rel,Gclc,bun,TBCC,fs(1)M3,Rab5,ACC,Pka-R2,Med,spn-E,sbb,Mitofilin,Arpc2,CycE,Ptpmeg,bru1,up,not,foxo,Pura,Rbf,squ,ssh,Smox,dnc,Moe,ben,jvl,Marf,colt,E2f1,Actn,Stat92E,Pax,Girdin,parvin,cnn,shrb,CkIIalpha,r2d2,Ptp10D,CG13366,SelR,armi,Ptp69D,CanB2,Arpc4,Arpc1,velo,msps,Abi,cnc,Ankle2,nos,Orc1,ago,SREBP | | 9.515e-03 | -4.65 | negative regulation of multi-organism process | biological process | GO:0043901 | 62 | 12 | 12345 | 1129 | cact,faf,EcR,shrb,p38a,slif,Atg6,Cbp80,Ptpmeg,GlyP,scny,Stat92E | | 9.535e-03 | -4.65 | GO\_POSITIVE\_REGULATION\_OF\_IMMUNE\_RESPONSE | MSigDB lists | GO\_POSITIVE\_REGULATION\_OF\_IMMUNE\_RESPONSE | 127 | 30 | 4161 | 644 | RpL40,dnc,scramb1,Ubc4,Stoml2,SkpA,Prosbeta6,Pi3K92E,Rrp40,Rpn12,Traf6,Rpn2,Prosbeta3,Prosalpha4,Rpt2,Arpc4,Uev1A,Cul1,Rpn3,Prosbeta5,Pdk1,cact,rl,Prosalpha5,Vamp7,Arpc1,RpS27A,Prosalpha3,Prosbeta4,Arpc2 | | 9.535e-03 | -4.65 | GSE24634\_NAIVE\_CD4\_TCELL\_VS\_DAY5\_IL4\_CONV\_TREG\_DN | MSigDB lists | GSE24634\_NAIVE\_CD4\_TCELL\_VS\_DAY5\_IL4\_CONV\_TREG\_DN | 127 | 30 | 4161 | 644 | Cyt-c-p,CG8188,scu,CkIIalpha,Vha36-1,CG17187,Non1,Su(H),CG1416,ste14,CG10932,CG42542,mus81,Ubc4,r,DppIII,Sld5,MED20,Mitofilin,Idh,Uch,mRpL16,Pfdn2,thoc6,Prosbeta5,Pus7,Cdk5,CCT3,CG9471,Coq7 | | 9.538e-03 | -4.65 | ovarian follicle cell development | biological process | GO:0030707 | 327 | 43 | 12345 | 1129 | CycE,Abi,csw,Med,spn-E,Rab5,cup,Pxt,bun,armi,mop,fs(1)M3,r2d2,Ptpmeg2,Cf2,Rbf,squ,Orc1,tor,Lkb1,Cdk1,EcR,fs(1)N,jvl,cact,gus,Tao,Sec15,nej,Pi3K92E,Cbl,e(y)1,rl,stai,Pax,SERCA,Stat92E,tkv,toc,mael,apt,E2f1,BicC | | 9.571e-03 | -4.65 | GSE24574\_BCL6\_HIGH\_VS\_LOW\_TFH\_CD4\_TCELL\_DN | MSigDB lists | GSE24574\_BCL6\_HIGH\_VS\_LOW\_TFH\_CD4\_TCELL\_DN | 96 | 24 | 4161 | 644 | RfC38,eIF3i,levy,Karybeta3,Gk2,CG12018,Dhfr,P5cr,ScsbetaA,scu,Rpn2,Echs1,Tom40,Eno,CG14715,CycB,Cdk1,CCT6,CG9391,msps,Pfas,CycE,Prosalpha5,Pcd | | 9.574e-03 | -4.65 | SASSON\_RESPONSE\_TO\_GONADOTROPHINS\_DN | MSigDB lists | SASSON\_RESPONSE\_TO\_GONADOTROPHINS\_DN | 25 | 9 | 4161 | 644 | Arpc4,Tm1,Rab5,CanB2,p38a,Pi3K92E,Pi4KIIalpha,Atg6,Npc2a | | 9.574e-03 | -4.65 | GO\_NUCLEOTIDE\_PHOSPHORYLATION | MSigDB lists | GO\_NUCLEOTIDE\_PHOSPHORYLATION | 25 | 9 | 4161 | 644 | Pgk,nmdyn-D6,Pglym78,Eno,Pfk,nmdyn-D7,Nc73EF,Pgm1,Gapdh1 | | 9.576e-03 | -4.65 | fly chr100D2-100D2\|3-102 cM | chromosome location | fly chr100D2-100D2\|3-102 cM | 25 | 5 | 25010 | 1343 | CG1896,sip3,awd,faf,CG1890 | | 9.576e-03 | -4.65 | fly chr100D2-100D2\|3-102 | chromosome location | fly chr100D2-100D2\|3-102 | 25 | 5 | 25010 | 1343 | awd,CG1890,faf,CG1896,sip3 | | 9.580e-03 | -4.65 | GSE360\_HIGH\_DOSE\_B\_MALAYI\_VS\_M\_TUBERCULOSIS\_MAC\_UP | MSigDB lists | GSE360\_HIGH\_DOSE\_B\_MALAYI\_VS\_M\_TUBERCULOSIS\_MAC\_UP | 71 | 19 | 4161 | 644 | Pi4KIIalpha,CG11807,Rab32,CSN6,RpL12,CG3376,Marf,eEF1beta,Echs1,CG9471,Ac3,TyrRS,MED24,CG2493,Gmer,CG8728,CG7550,Tsp74F,Etf-QO | | 9.580e-03 | -4.65 | RUTELLA\_RESPONSE\_TO\_CSF2RB\_AND\_IL4\_DN | MSigDB lists | RUTELLA\_RESPONSE\_TO\_CSF2RB\_AND\_IL4\_DN | 71 | 19 | 4161 | 644 | Sccpdh1,yip2,Smox,Fkbp59,CG15715,Mtpalpha,Pdxk,anne,Rbcn-3A,CG7997,AMPdeam,CG3860,Mctp,CG7718,CG10932,scramb1,cathD,Cyp1,Roe1 | | 9.580e-03 | -4.65 | GSE17721\_LPS\_VS\_POLYIC\_8H\_BMDC\_UP | MSigDB lists | GSE17721\_LPS\_VS\_POLYIC\_8H\_BMDC\_UP | 71 | 19 | 4161 | 644 | anne,CCT6,Moe,Cul1,Fkbp59,Unr,CG9143,CG33181,CG8372,eIF3i,SkpA,CG17691,Atg4b,tko,Cbp80,l(1)G0004,Grip128,mRpL13,AMPdeam | | 9.649e-03 | -4.64 | heme-copper terminal oxidase activity | molecular function | GO:0015002 | 21 | 6 | 12176 | 1119 | COX5A,cype,levy,COX5B,COX4,COX8 | | 9.649e-03 | -4.64 | exoribonuclease activity | molecular function | GO:0004532 | 21 | 6 | 12176 | 1119 | twin,Exd2,Rga,cu,CG11486,CG31759 | | 9.649e-03 | -4.64 | oxidoreductase activity, acting on a heme group of donors | molecular function | GO:0016675 | 21 | 6 | 12176 | 1119 | COX5A,cype,COX5B,levy,COX4,COX8 | | 9.649e-03 | -4.64 | cytochrome-c oxidase activity | molecular function | GO:0004129 | 21 | 6 | 12176 | 1119 | COX5A,cype,levy,COX5B,COX4,COX8 | | 9.649e-03 | -4.64 | oxidoreductase activity, acting on a heme group of donors, oxygen as acceptor | molecular function | GO:0016676 | 21 | 6 | 12176 | 1119 | levy,COX5B,COX4,COX8,COX5A,cype | | 9.649e-03 | -4.64 | exoribonuclease activity, producing 5'-phosphomonoesters | molecular function | GO:0016896 | 21 | 6 | 12176 | 1119 | CG31759,CG11486,cu,Rga,Exd2,twin | | 9.667e-03 | -4.64 | GSE14415\_INDUCED\_TREG\_VS\_FAILED\_INDUCED\_TREG\_UP | MSigDB lists | GSE14415\_INDUCED\_TREG\_VS\_FAILED\_INDUCED\_TREG\_UP | 91 | 23 | 4161 | 644 | Grip84,Lsm12,Uch,Prosbeta4,LSm3,CG8314,Pgk,Shmt,CG14715,Cdk1,r-l,Coq7,CG8188,Nurf-38,CG12018,mRpL18,CG1416,l(1)G0004,CG10418,r,CysRS,RfC38,NHP2 | | 9.667e-03 | -4.64 | HALLMARK\_ADIPOGENESIS | MSigDB lists | HALLMARK\_ADIPOGENESIS | 91 | 23 | 4161 | 644 | CG5028,levy,Tsp74F,Roe1,AGBE,mino,Sod1,Mcad,Pgm1,UQCR-Q,ND-30,crq,Aldh,ND-B18,CG8199,AIF,alph,Echs1,Nmt,yip2,Mitofilin,Scsalpha1,Idh | | 9.689e-03 | -4.64 | GSE27786\_LSK\_VS\_ERYTHROBLAST\_UP | MSigDB lists | GSE27786\_LSK\_VS\_ERYTHROBLAST\_UP | 76 | 20 | 4161 | 644 | r,RpS7,Rep,DppIII,CG11110,Sfxn2,CG10463,Rbf,ND-24,CG30010,mEFTu1,ND-30,Moe,pont,SdhA,crq,Hmg-2,Scox,Pfas,CG10038 | | 9.689e-03 | -4.64 | GSE29617\_CTRL\_VS\_TIV\_FLU\_VACCINE\_PBMC\_2008\_DN | MSigDB lists | GSE29617\_CTRL\_VS\_TIV\_FLU\_VACCINE\_PBMC\_2008\_DN | 76 | 20 | 4161 | 644 | tho2,Naa20A,mRpL18,CG17294,Mcad,Archease,RpS16,RhoGAP54D,Sec10,CG10470,Rpn2,Eno,CG14715,Tim8,Jafrac2,CG5510,UQCR-Q,Prosalpha3,Scsalpha1,Prosbeta4 | | 9.690e-03 | -4.64 | MODULE\_343 | MSigDB lists | MODULE\_343 | 17 | 7 | 4161 | 644 | Echs1,pr,Mcad,Mtpalpha,Thiolase,Irp-1B,Pcd | | 9.690e-03 | -4.64 | DER\_IFN\_GAMMA\_RESPONSE\_UP | MSigDB lists | DER\_IFN\_GAMMA\_RESPONSE\_UP | 17 | 7 | 4161 | 644 | Thiolase,Cyt-c-p,scramb1,Pp2B-14D,PpD3,CanA-14F,SkpA | | 9.690e-03 | -4.64 | GO\_POSITIVE\_REGULATION\_OF\_CELL\_CYCLE\_ARREST | MSigDB lists | GO\_POSITIVE\_REGULATION\_OF\_CELL\_CYCLE\_ARREST | 17 | 7 | 4161 | 644 | CycB,twin,Cdk1,Rbf,RpL40,Not10,RpS27A | | 9.716e-03 | -4.63 | U6 snRNP | cellular component | GO:0005688 | 10 | 4 | 11940 | 1105 | LSm3,CG10418,CG6610,CG17768 | | 9.716e-03 | -4.63 | DNA polymerase complex | cellular component | GO:0042575 | 10 | 4 | 11940 | 1105 | Chrac-14,mus205,DNApol-epsilon255,CG12018 | | 9.724e-03 | -4.63 | GSE25123\_WT\_VS\_PPARG\_KO\_MACROPHAGE\_DN | MSigDB lists | GSE25123\_WT\_VS\_PPARG\_KO\_MACROPHAGE\_DN | 86 | 22 | 4161 | 644 | Non1,CG2493,Nurf-38,CG10470,RpII140,LysRS,RpS16,RpS2,heph,obe,Prosbeta4,Prosalpha3,RpL37a,Fkbp59,RpS27A,Prosalpha5,RpL24,eEF1beta,CG15715,CG3376,CCT6,RpL12 | | 9.724e-03 | -4.63 | GO\_POSITIVE\_REGULATION\_OF\_DEFENSE\_RESPONSE | MSigDB lists | GO\_POSITIVE\_REGULATION\_OF\_DEFENSE\_RESPONSE | 86 | 22 | 4161 | 644 | Ubc4,Prosbeta6,SkpA,RpL40,scramb1,Prosbeta3,Rpt2,Prosalpha4,Traf6,Rpn12,Rpn2,Prosbeta5,cact,Pdk1,Uev1A,Rpn3,Cul1,Prosalpha3,Prosbeta4,RpS27A,Vamp7,Prosalpha5 | | 9.725e-03 | -4.63 | mitochondrial transport | biological process | GO:0006839 | 109 | 18 | 12345 | 1129 | CG9399,Sfxn2,Roe1,sea,Tim8,CG9393,COX5B,CG8004,CG8728,ATPsynbeta,CG11110,Debcl,Tom70,ATPsynO,CG14894,Tom40,EMRE,colt | | 9.726e-03 | -4.63 | gcl (germ cell-less) | protein interactions | 35864 | 12 | 5 | 9204 | 1112 | mre11,bru1,osk,CG2199,tor | | 9.726e-03 | -4.63 | Grip91 (Gamma-tubulin ring protein 91) | protein interactions | 48481 | 12 | 5 | 9204 | 1112 | Grip71,gammaTub37C,Grip75,Grip128,Grip84 | | 9.726e-03 | -4.63 | CG2224 (uncharacterized protein) | protein interactions | 43617 | 12 | 5 | 9204 | 1112 | shrb,CG8726,ND-23,mts,RpL13A | | 9.734e-03 | -4.63 | REACTOME\_S\_PHASE | MSigDB lists | REACTOME\_S\_PHASE | 81 | 21 | 4161 | 644 | Prosbeta4,Prosalpha3,RpS27A,DNApol-epsilon255,Prosalpha5,CycE,Prosbeta5,Cul1,Rpn3,Rpt2,Prosalpha4,Cdk7,Prosbeta3,CG12018,Rpn2,Rpn12,Sld5,Prosbeta6,RfC38,SkpA,RpL40 | | 9.734e-03 | -4.63 | GSE17721\_CTRL\_VS\_GARDIQUIMOD\_6H\_BMDC\_DN | MSigDB lists | GSE17721\_CTRL\_VS\_GARDIQUIMOD\_6H\_BMDC\_DN | 81 | 21 | 4161 | 644 | Rab10,APP-BP1,CG33181,Cka,Top1,CG13630,Hat1,Moe,nmdyn-D6,Chd64,CG17294,TyrRS,Tsp86D,CG4390,AGBE,CG14894,Imp,PrBP,Cbp80,Arts,scramb1 | | 9.753e-03 | -4.63 | eggshell chorion assembly | biological process | GO:0007306 | 93 | 16 | 12345 | 1129 | rl,Orc1,Pax,squ,Rbf,E2f1,mael,cact,spn-E,CycE,jvl,Cf2,gus,Pxt,bun,armi | | 9.851e-03 | -4.62 | GO\_CELL\_DEATH | MSigDB lists | GO\_CELL\_DEATH | 247 | 52 | 4161 | 644 | Sply,mael,SERCA,Top1,ND-75,Cul5,CG5880,AIF,Cdk1,rl,cact,Cse1,Prip,Cul2,mts,Gapdh1,Grx1,Arl6IP1,Xpac,Smox,yip2,CG11807,Acph-1,Pp2A-29B,PDCD-5,Dcr-1,Arf51F,CG11134,APP-BP1,Marf,Cul1,nmdyn-D6,Cdk5,Gclc,CG46338,thoc6,p38a,Pdk1,Atg6,Cyt-c-p,CG30291,TyrRS,CkIIalpha,AnxB10,Atg4b,Cyp1,Tspo,CG4332,Tao,scramb1,Ubc4,CG17454 | | 9.869e-03 | -4.62 | KENNY\_CTNNB1\_TARGETS\_DN | MSigDB lists | KENNY\_CTNNB1\_TARGETS\_DN | 21 | 8 | 4161 | 644 | Plap,SdhA,CG17593,CG42542,faf,Pp2B-14D,Faf2,CanA-14F | | 9.869e-03 | -4.62 | MODULE\_22 | MSigDB lists | MODULE\_22 | 21 | 8 | 4161 | 644 | ND-B18,ND-24,ND-51,ND-30,ND-23,ND-49,ND-39,ND-SGDH | | 9.869e-03 | -4.62 | GO\_REGULATION\_OF\_CELL\_CYCLE\_ARREST | MSigDB lists | GO\_REGULATION\_OF\_CELL\_CYCLE\_ARREST | 21 | 8 | 4161 | 644 | RpL40,Cdk1,CycB,Cdk5,RpS27A,Not10,Rbf,twin | | 9.869e-03 | -4.62 | HALLMARK\_REACTIVE\_OXIGEN\_SPECIES\_PATHWAY | MSigDB lists | HALLMARK\_REACTIVE\_OXIGEN\_SPECIES\_PATHWAY | 21 | 8 | 4161 | 644 | Jafrac2,fray,Gclc,Grx1,Jafrac1,ND-49,Trxr-1,Sod1 | | 9.879e-03 | -4.62 | PUJANA\_BRCA1\_PCC\_NETWORK | MSigDB lists | PUJANA\_BRCA1\_PCC\_NETWORK | 703 | 130 | 4161 | 644 | CG6610,LysRS,RpLP0,slif,eIF3i,rept,opm,Arpc2,Clp,Idh3b,Mekk1,Nmt,RpL9,alpha-Man-Ia,ND-75,AIF,Caf1-105,Top1,eEF1beta,Chrac-14,TyrRS,CkIIalpha,CG12018,P5cr,RpL30,CG1890,Cyt-c-p,Phb2,tho2,PpV,Ts,CG40045,CG17454,Cbp80,CG10418,Prosalpha3,CG7728,Iru,COX5A,CCT3,ND-24,lwr,Gapdh1,grp,l(1)G0004,escl,asf1,RpS23,Tfb4,Klp10A,Pgk,CCT6,mad2,RpL23,RhoGAP54D,RpS7,Ptp61F,Pi3K92E,heph,eIF3m,Pfas,RpL11,Prosbeta4,Uch,CycE,MED20,anne,CCT5,esc,pont,CycB,SRPK,CG13630,Cul1,CG11164,Cul2,Cse1,CG11089,CG30496,Nap1,RpS27A,Pen,lqfR,MED24,CG8728,CycC,CG6767,r,NHP2,fray,CG8204,mbo,RpL37a,Mitofilin,LSm3,APP-BP1,CG7488,RpL24,DNApol-epsilon255,Arpc4,PIG-C,Mcad,Cdk7,ste24a,CG8188,RpII140,Stoml2,Karybeta3,Rpb8,Prosalpha5,Tom40,wal,Cdk1,CG9391,Dhfr,RpS24,CG31759,Tlk,Ubc4,RfC38,Prosbeta6,CG1416,Vamp7,Eno,CG8004,RpL12,Hat1,bys,msps,Rpn3 | | 9.880e-03 | -4.62 | DUF3452 | smart domains | SM01367 | 2 | 2 | 5086 | 506 | Rbf2,Rbf | | 9.880e-03 | -4.62 | CARP | smart domains | SM00673 | 2 | 2 | 5086 | 506 | capt,TBCC | | 9.880e-03 | -4.62 | LSM14 | smart domains | SM01271 | 2 | 2 | 5086 | 506 | Edc3,tral | | 9.880e-03 | -4.62 | MGS | smart domains | SM00851 | 2 | 2 | 5086 | 506 | CG11089,r | | 9.880e-03 | -4.62 | FDF | smart domains | SM01199 | 2 | 2 | 5086 | 506 | Edc3,tral | | 9.880e-03 | -4.62 | DUF106 | smart domains | SM01415 | 2 | 2 | 5086 | 506 | EMC3,CG10470 | | 9.880e-03 | -4.62 | RB\_A | smart domains | SM01368 | 2 | 2 | 5086 | 506 | Rbf2,Rbf | | 9.926e-03 | -4.61 | Gas41 (Gas41) | protein interactions | 33973 | 8 | 4 | 9204 | 1112 | armi,Tip60,Dcr-1,pont | | 9.926e-03 | -4.61 | DNaseII (Deoxyribonuclease II) | protein interactions | 48228 | 8 | 4 | 9204 | 1112 | awd,CG12171,regucalcin,Drep1 | | 9.926e-03 | -4.61 | RpS5b (Ribosomal protein S5b) | protein interactions | 41807 | 8 | 4 | 9204 | 1112 | Rpt2,CG3939,PGRP-LE,vers | | 9.926e-03 | -4.61 | Grip75 (Grip75) | protein interactions | 34441 | 8 | 4 | 9204 | 1112 | gammaTub37C,Grip84,Grip128,cnn | | 9.938e-03 | -4.61 | PyrdxlP-dep\_Trfase\_dom1 | interpro domains | IPR015422 | 29 | 8 | 11383 | 1233 | Gabat,Kyat,Shmt,mal,CG1640,Got1,Sply,CG11899 | | 9.974e-03 | -4.61 | peroxisomal membrane | cellular component | GO:0005778 | 21 | 6 | 11940 | 1105 | Topors,Pex12,ABCD,Pex13,Pex10,CG11077 | | 9.974e-03 | -4.61 | microbody part | cellular component | GO:0044438 | 21 | 6 | 11940 | 1105 | CG11077,Topors,Pex12,ABCD,Pex13,Pex10 | | 9.974e-03 | -4.61 | microbody membrane | cellular component | GO:0031903 | 21 | 6 | 11940 | 1105 | Topors,ABCD,Pex13,Pex12,Pex10,CG11077 | | 9.974e-03 | -4.61 | peroxisomal part | cellular component | GO:0044439 | 21 | 6 | 11940 | 1105 | Pex10,Topors,Pex12,ABCD,Pex13,CG11077 | | 1.003e-02 | -4.60 | MODULE\_212 | MSigDB lists | MODULE\_212 | 138 | 32 | 4161 | 644 | sau,Sod1,Mcad,CG17691,tzn,Irp-1B,Pcd,Got1,Scsalpha1,Idh,Treh,ERp60,UQCR-Q,Cyp6a19,CG9471,Cdk1,Echs1,GlyP,CG11110,Phb2,P5CDh1,CkIIalpha,Rab5,CG9485,RpS27,Cyp6a22,CG32473,CG8112,Mtpalpha,Jafrac2,pr,Cdk5 | | 1.004e-02 | -4.60 | GO\_REGULATION\_OF\_IMMUNE\_RESPONSE | MSigDB lists | GO\_REGULATION\_OF\_IMMUNE\_RESPONSE | 181 | 40 | 4161 | 644 | rl,cact,Uev1A,Ect4,Arpc2,Prosalpha5,RpS27A,SkpA,Stoml2,dnc,Prosalpha4,csw,Rpt2,Prosbeta3,Rpn2,Rrp40,Rpn12,EcR,Pdk1,Prosbeta5,Cul1,Rpn3,Arpc4,Prosbeta4,CanA-14F,Prosalpha3,Vamp7,Arpc1,Smox,Ptp61F,Prosbeta6,Pi3K92E,CanB2,Ubc4,Snx6,scramb1,RpL40,Pp2B-14D,Phb2,Traf6 | | 1.004e-02 | -4.60 | columnar/cuboidal epithelial cell development | biological process | GO:0002066 | 328 | 43 | 12345 | 1129 | tor,Orc1,squ,Rbf,Lkb1,Cdk1,Med,spn-E,csw,Abi,CycE,r2d2,Cf2,Ptpmeg2,Pxt,fs(1)M3,bun,armi,mop,cup,Rab5,Pax,e(y)1,rl,stai,Cbl,nej,Pi3K92E,E2f1,BicC,apt,toc,mael,tkv,Stat92E,SERCA,cact,jvl,EcR,fs(1)N,Sec15,Tao,gus | | 1.010e-02 | -4.60 | negative regulation of phosphorylation | biological process | GO:0042326 | 70 | 13 | 12345 | 1129 | ATPsynD,CG14408,dco,Ankle2,alph,SkpA,CG17746,sws,PpV,park,Cbl,Ptp61F,mts | | 1.010e-02 | -4.59 | intrinsic component of organelle membrane | cellular component | GO:0031300 | 166 | 25 | 11940 | 1105 | EMRE,ste24a,Ppox,Seipin,CG9399,Pex10,Pex12,CG32441,CG8311,Ugalt,Pmi,Marf,CG10470,Efr,CG31523,alt,Fur2,stet,fwe,CG9773,CG5447,CG4293,CG5903,EMC8-9,EMC3 | | 1.015e-02 | -4.59 | RNA binding | molecular function | GO:0003723 | 929 | 106 | 12176 | 1119 | mEFTu2,Marf1,CG17454,CG7728,RpS13,mRpS11,Mat89Ba,BicC,CG4021,Son,CG6610,Rnmt,RpL13A,exu,l(1)G0004,apt,mEFTu1,RpL12,Nxf3,CG14100,CG9425,CG7414,RpS16,Rbfox1,r2d2,RpL7A,eIF3m,swa,cup,RpS14b,mRpL16,Pus7,CG3198,RpL23,mRpL18,RpS15,qkr58E-2,Unr,tho2,armi,spn-E,CG9143,Rcd5,RpS2,CG8963,CG12375,NHP2,Spx,Cbp80,LSm3,GluRS-m,CG12288,CG10189,CG6967,trsn,RpL26,RpL9,CG8611,RpL24,yps,nos,RpS18,Rrp40,CG2100,CG5986,bys,RpLP0,smg,Edc3,CG17768,Irp-1B,shep,eIF4E1,RpS4,CG11486,PPYR1,LysRS,SLIRP1,CG7488,bcd,Dcr-1,eIF4A,CG10038,CysRS,RpL30,Shmt,Larp7,CG6094,sbr,osk,AGO3,CG10418,heph,bru1,CG4266,piwi,RpS5b,eIF3i,mRpL13,eEF1beta,bam,mRpS5,TyrRS,Pym,RpS14a,Imp | | 1.016e-02 | -4.59 | yki (yorkie) | protein interactions | 37851 | 109 | 22 | 9204 | 1112 | tkv,Wbp2,CG9581,slam,Hsc70-3,piwi,cact,bun,Pen,EcR,CG11134,nos,CG31705,trr,rl,Atpalpha,Smox,l(3)mbt,Utx,mop,CG9425,Tao | | 1.017e-02 | -4.59 | fly chr62D4-62D4\|3-2 | chromosome location | fly chr62D4-62D4\|3-2 | 9 | 3 | 25010 | 1343 | CG2034,CG1275,oxt | | 1.017e-02 | -4.59 | fly chr85C2-85C2\|3-48 | chromosome location | fly chr85C2-85C2\|3-48 | 9 | 3 | 25010 | 1343 | Prosbeta3,Kcmf1,Iru | | 1.017e-02 | -4.59 | fly chr85C2-85C2\|3-48 cM | chromosome location | fly chr85C2-85C2\|3-48 cM | 9 | 3 | 25010 | 1343 | Prosbeta3,Iru,Kcmf1 | | 1.017e-02 | -4.59 | fly chr62D4-62D4\|3-2 cM | chromosome location | fly chr62D4-62D4\|3-2 cM | 9 | 3 | 25010 | 1343 | oxt,CG1275,CG2034 | | 1.028e-02 | -4.58 | organic acid biosynthetic process | biological process | GO:0016053 | 126 | 20 | 12345 | 1129 | CG11899,CG31523,CG11134,Adi1,SREBP,CG10932,Shmt,AcCoAS,AsnS,r,P5cr,Desat1,Dhfr,Got1,CG6178,regucalcin,Pxt,SelD,aay,ACC | | 1.028e-02 | -4.58 | carboxylic acid biosynthetic process | biological process | GO:0046394 | 126 | 20 | 12345 | 1129 | CG11134,CG31523,CG11899,CG10932,SREBP,Adi1,Shmt,r,P5cr,AsnS,AcCoAS,Desat1,Got1,CG6178,Dhfr,ACC,aay,SelD,Pxt,regucalcin | | 1.030e-02 | -4.58 | GO\_POSITIVE\_REGULATION\_OF\_CELLULAR\_PROTEIN\_LOCALIZATION | MSigDB lists | GO\_POSITIVE\_REGULATION\_OF\_CELLULAR\_PROTEIN\_LOCALIZATION | 112 | 27 | 4161 | 644 | CG30291,CG3631,Rpn12,Atg6,Pp2B-14D,CanB2,Karybeta3,Med,Aos1,CanA-14F,Rcd5,Arf51F,Smox,Nmt,PDCD-5,Cdk5,CycB,pont,CCT5,Cdk1,p38a,CCT3,mAChR-A,Pdk1,CCT6,Moe,Edem1 | | 1.036e-02 | -4.57 | Lam (Lamin) | protein interactions | 33782 | 37 | 10 | 9204 | 1112 | Rab5,rl,wash,EcR,fs(1)Ya,hb,Topors,Pen,Sec10,cnn | | 1.036e-02 | -4.57 | MED11 (Mediator complex subunit 11) | protein interactions | 40042 | 37 | 10 | 9204 | 1112 | MED27,Usp10,enc,MED20,MED4,ix,CG2199,Larp4B,CG17002,Ptip | | 1.037e-02 | -4.57 | MRP (Multidrug-Resistance like Protein 1) | protein interactions | 34686 | 84 | 18 | 9204 | 1112 | CG10425,Kap-alpha3,SERCA,CCT6,CG10132,Pen,CG31729,Pp2A-29B,Atpalpha,Mcad,mre11,Rab5,Karybeta3,bor,CG5447,wal,Surf4,foxo | | 1.039e-02 | -4.57 | pigment metabolic process involved in pigmentation | biological process | GO:0043474 | 41 | 9 | 12345 | 1129 | mal,PAPLA1,cl,lt,fdl,Rab32,DhpD,EMC3,Hexo1 | | 1.039e-02 | -4.57 | pigment metabolic process involved in developmental pigmentation | biological process | GO:0043324 | 41 | 9 | 12345 | 1129 | fdl,lt,mal,PAPLA1,cl,EMC3,Hexo1,DhpD,Rab32 | | 1.039e-02 | -4.57 | eye pigment metabolic process | biological process | GO:0042441 | 41 | 9 | 12345 | 1129 | Hexo1,EMC3,DhpD,Rab32,fdl,lt,mal,PAPLA1,cl | | 1.044e-02 | -4.56 | Ntn\_hydrolases\_N | interpro domains | IPR029055 | 41 | 10 | 11383 | 1233 | Prosalpha5,Prosbeta5,AsnS,Prosbeta4,Ggt-1,Prosalpha3,CG1827,Prosbeta6,Prosbeta3,Prosalpha4 | | 1.047e-02 | -4.56 | GSE13738\_TCR\_VS\_BYSTANDER\_ACTIVATED\_CD4\_TCELL\_UP | MSigDB lists | GSE13738\_TCR\_VS\_BYSTANDER\_ACTIVATED\_CD4\_TCELL\_UP | 57 | 16 | 4161 | 644 | Mcad,CG5292,CG11164,nmdyn-D7,Nurf-38,CG40045,ATP8A,Gk2,cl,CG32549,Hmg-2,Pi4KIIalpha,Cdk5,CG3803,milt,shop | | 1.047e-02 | -4.56 | GSE22886\_NAIVE\_TCELL\_VS\_DC\_UP | MSigDB lists | GSE22886\_NAIVE\_TCELL\_VS\_DC\_UP | 57 | 16 | 4161 | 644 | Iru,RpS27,RpL26,RpS23,RpL11,eEF1beta,RpS18,RpL13A,RpS30,Pfdn5,Rbf,RpL30,RpL23,RpL40,RpS7,RpL38 | | 1.047e-02 | -4.56 | Adaptive Immune System | REACTOME pathways | R-DME-1280218 | 186 | 39 | 3923 | 574 | Iru,Prosalpha4,Pi3K92E,Pdk1,Prosalpha3,Rpt2,RpL40,cact,SkpA,Rel,tor,cathD,Prosbeta3,dx,foxo,CG13398,Rpn12,dl,CG14435,Rpn3,Cul1,Sec23,Prosbeta6,csw,Cp1,Rpn2,Prosbeta5,Prosbeta4,ago,yl,RpS27A,Prosalpha5,Ubc6,crq,Herc4,gus,GILT1,Pp2A-29B,Fbw5 | | 1.053e-02 | -4.55 | positive regulation of signaling | biological process | GO:0023056 | 403 | 51 | 12345 | 1129 | Pi3K92E,nej,pont,Tsp86D,park,spen,tkv,EndoGI,stas,Aos1,PEK,dco,E2f1,Mekk1,ben,stet,Ect4,Snx6,trbd,gd,Mos,Tao,tws,snk,Pink1,spz,dx,mts,foxo,Pura,Traf6,shv,Jafrac2,Imp,Tak1,Usp10,tor,Pdk1,ea,O-fut1,Cka,Ssdp,CycE,CG10188,Myd88,mop,Rel,Herc4,CG4853,disp,CkIIalpha | | 1.053e-02 | -4.55 | positive regulation of cell communication | biological process | GO:0010647 | 403 | 51 | 12345 | 1129 | CG10188,CycE,disp,CG4853,CkIIalpha,mop,Herc4,Rel,Myd88,tor,Pdk1,Imp,Tak1,Usp10,foxo,Pura,shv,Jafrac2,Traf6,mts,Ssdp,Cka,ea,O-fut1,Ect4,ben,stet,Mekk1,snk,Pink1,spz,dx,tws,Mos,Tao,Snx6,trbd,gd,spen,park,Pi3K92E,nej,pont,Tsp86D,dco,E2f1,PEK,stas,Aos1,tkv,EndoGI | | 1.054e-02 | -4.55 | inorganic ion homeostasis | biological process | GO:0098771 | 143 | 22 | 12345 | 1129 | Vha68-2,EMRE,zyd,Grx1,Fer1HCH,Seipin,Ho,nrv1,Scox,fwe,CG12773,ZnT35C,park,CG14036,wash,MtnA,up,anne,CG10470,SERCA,Rbcn-3A,Atpalpha | | 1.056e-02 | -4.55 | Signaling by the B Cell Receptor (BCR) | REACTOME pathways | R-DME-983705 | 113 | 26 | 3923 | 574 | Prosbeta3,tor,Rel,csw,Prosbeta6,Pp2A-29B,Rpn12,CG13398,dl,Cul1,Rpn3,foxo,yl,RpS27A,Pdk1,Prosalpha5,Pi3K92E,Prosbeta4,Rpn2,Prosbeta5,Prosalpha4,SkpA,cact,Prosalpha3,RpL40,Rpt2 | | 1.058e-02 | -4.55 | GO\_POSITIVE\_REGULATION\_OF\_PROTEIN\_MODIFICATION\_PROCESS | MSigDB lists | GO\_POSITIVE\_REGULATION\_OF\_PROTEIN\_MODIFICATION\_PROCESS | 276 | 57 | 4161 | 644 | Traf6,APC7,Phb2,CG30291,CycC,Med,Atg7,Tao,RpL40,CG17698,Ubc4,Pka-R1,Prosbeta6,Smox,nAChRalpha4,Prosalpha3,SREBP,Wdfy2,AGO3,Hsc70-3,Prosbeta4,CG3376,Cul1,Rpn3,CG46338,Cdk5,esc,CycB,Prosbeta5,p38a,Pdk1,Ac3,Rpn12,CG8188,Sod1,Rpn2,Prosbeta3,mts,Rpt2,csw,raptor,Prosalpha4,rept,escl,Tip60,regucalcin,SkpA,Mekk1,RpS27A,Prosalpha5,Mo25,Pka-R2,mad2,STUB1,LpR2,Cdk1,rl | | 1.058e-02 | -4.55 | Pten (Phosphatase and tensin homolog) | protein interactions | 43991 | 103 | 21 | 9204 | 1112 | Rpt2,faf,wal,ACC,eIF4A,foxo,larp,EcR,Nup358,dx,Pi3K92E,Smox,slif,ScsbetaA,Hsc70-3,gw,Cul1,RpS7,eIF3m,alt,RpS14a | | 1.059e-02 | -4.55 | foxo (forkhead box, sub-group O) | protein interactions | 41709 | 182 | 33 | 9204 | 1112 | Aldh,Nup358,RpS23,Pi3K92E,ND-75,tyf,RpL13A,larp,qkr58E-2,CCT6,mts,RpL26,eIF3i,RpS7,yps,bif,Rpt2,RpL17,Ect4,RpS15,park,RpL9,Pink1,alt,RpS14a,RpL7A,RpLP1,Hsc70-3,HmgD,CG2199,RpL23,RpL10,RpL23A | | 1.060e-02 | -4.55 | GO\_POSITIVE\_REGULATION\_OF\_PROTEIN\_LOCALIZATION\_TO\_NUCLEUS | MSigDB lists | GO\_POSITIVE\_REGULATION\_OF\_PROTEIN\_LOCALIZATION\_TO\_NUCLEUS | 43 | 13 | 4161 | 644 | CanB2,CCT5,Karybeta3,p38a,Cdk1,CCT3,CCT6,Med,CanA-14F,CG30291,Rcd5,Smox,Pp2B-14D | | 1.064e-02 | -4.54 | KEGG\_PROTEASOME | MSigDB lists | KEGG\_PROTEASOME | 34 | 11 | 4161 | 644 | Rpn3,Prosbeta6,Prosbeta5,Rpn2,Prosalpha5,Rpn12,Rpt2,Prosalpha4,Prosbeta4,Prosbeta3,Prosalpha3 | | 1.064e-02 | -4.54 | GSE29618\_PDC\_VS\_MDC\_DN | MSigDB lists | GSE29618\_PDC\_VS\_MDC\_DN | 34 | 11 | 4161 | 644 | Nurf-38,Rab32,Gapdh1,Cyp1,Su(H),Cul1,Moe,AP-1-2beta,Eno,CHMP2B,CG9471 | | 1.064e-02 | -4.54 | GSE46242\_CTRL\_VS\_EGR2\_DELETED\_TH1\_CD4\_TCELL\_DN | MSigDB lists | GSE46242\_CTRL\_VS\_EGR2\_DELETED\_TH1\_CD4\_TCELL\_DN | 34 | 11 | 4161 | 644 | shv,Sccpdh1,Tsp66E,Pmi,Naa20A,disp,CG12375,CG9471,crq,LpR2,CG5510 | | 1.068e-02 | -4.54 | OSMAN\_BLADDER\_CANCER\_DN | MSigDB lists | OSMAN\_BLADDER\_CANCER\_DN | 128 | 30 | 4161 | 644 | eIF4A,RpS24,mts,RpS13,RpLP0,RpS30,RpL13A,RpS16,RpS4,RpL38,Pka-R1,RpS7,PIG-L,bru1,Cyp1,RpL40,Unr,CG30496,RpL37a,RpL9,stas,RpL24,RpS27,GatA,Dph5,CG11807,RpS18,eEF1beta,RpL13,RpL12 | | 1.070e-02 | -4.54 | protein polyubiquitination | biological process | GO:0000209 | 86 | 15 | 12345 | 1129 | park,Ubc6,ben,CG4238,Ubc4,Traf6,ari-1,CG40045,CG2924,sip3,Uev1A,CG11070,Topors,Herc4,UbcE2H | | 1.070e-02 | -4.54 | positive regulation of proteolysis | biological process | GO:0045862 | 86 | 15 | 12345 | 1129 | Cul1,Rpt2,dco,Pomp,cort,SkpA,eIF3m,STUB1,Jafrac2,Rpn13,ari-1,park,MED24,Ubc6,ScsbetaA | | 1.070e-02 | -4.54 | Rab1 (Rab1) | protein interactions | 42524 | 60 | 14 | 9204 | 1112 | cact,Cisd2,mus304,Tspo,sau,wal,COX4,Atpalpha,Mcad,ScsbetaA,SERCA,CG1416,Gdi,Rab5 | | 1.070e-02 | -4.54 | Cpsf6 (Cleavage and polyadenylation specific factor 6) | protein interactions | 38937 | 60 | 14 | 9204 | 1112 | ix,enc,MED20,MED4,MED27,Sec23,Usp10,CG17002,CG11755,Larp4B,Ptip,CG2199,MED24,l(1)G0004 | | 1.071e-02 | -4.54 | LOTUS-like | interpro domains | IPR041966 | 5 | 3 | 11383 | 1233 | tapas,osk,Marf1 | | 1.071e-02 | -4.54 | Thiolase | interpro domains | IPR002155 | 5 | 3 | 11383 | 1233 | CG10932,Thiolase,yip2 | | 1.071e-02 | -4.54 | La\_HTH | interpro domains | IPR006630 | 5 | 3 | 11383 | 1233 | larp,Larp4B,Larp7 | | 1.071e-02 | -4.54 | DUF4780 | interpro domains | IPR031961 | 5 | 3 | 11383 | 1233 | CG9641,CG7208,CG7824 | | 1.071e-02 | -4.54 | Cullin\_CS | interpro domains | IPR016157 | 5 | 3 | 11383 | 1233 | Cul5,Cul1,Cul2 | | 1.072e-02 | -4.54 | Mpcp2 (Mitochondrial phosphate carrier protein 2) | protein interactions | 39587 | 72 | 16 | 9204 | 1112 | Bx,bor,r,mEFTu1,wal,Mcad,colt,scu,nbs,Jafrac2,CCT3,Pp2A-29B,Kap-alpha3,Nf-YA,SERCA,CCT6 | | 1.074e-02 | -4.53 | GSE15930\_NAIVE\_VS\_24H\_IN\_VITRO\_STIM\_IL12\_CD8\_TCELL\_DN | MSigDB lists | GSE15930\_NAIVE\_VS\_24H\_IN\_VITRO\_STIM\_IL12\_CD8\_TCELL\_DN | 102 | 25 | 4161 | 644 | CysRS,r,scramb1,CG10418,Su(H),CG17294,CG11164,mRpL18,CG12018,Nurf-38,aay,Cdk1,CCT3,CG14715,CG8004,CG5880,mRpS10,msps,CG5026,mad2,Pglym78,CCT6,mRpL17,Sptr,gammaTub37C | | 1.076e-02 | -4.53 | phosphatase complex | cellular component | GO:1903293 | 34 | 8 | 11940 | 1105 | CG8009,CanB2,Pp2A-29B,tws,Dd,Pp2B-14D,mts,CanA-14F | | 1.076e-02 | -4.53 | protein serine/threonine phosphatase complex | cellular component | GO:0008287 | 34 | 8 | 11940 | 1105 | CanA-14F,mts,Pp2B-14D,Dd,Pp2A-29B,tws,CG8009,CanB2 | | 1.081e-02 | -4.53 | GSE17721\_POLYIC\_VS\_PAM3CSK4\_4H\_BMDC\_UP | MSigDB lists | GSE17721\_POLYIC\_VS\_PAM3CSK4\_4H\_BMDC\_UP | 62 | 17 | 4161 | 644 | mal,Mcad,CG2493,Vha36-1,Sfxn2,RpL38,Pi3K92E,Ptp61F,Pi4KIIalpha,YL-1,Prosalpha3,Gyg,ND-39,shrb,Hat1,pr,Echs1 | | 1.081e-02 | -4.53 | GSE40274\_CTRL\_VS\_FOXP3\_AND\_LEF1\_TRANSDUCED\_ACTIVATED\_CD4\_TCELL\_DN | MSigDB lists | GSE40274\_CTRL\_VS\_FOXP3\_AND\_LEF1\_TRANSDUCED\_ACTIVATED\_CD4\_TCELL\_DN | 62 | 17 | 4161 | 644 | PpD3,Irp-1B,CG14407,Grip84,Jafrac2,UQCR-Q,RpL23,Gip,Prip,Edc3,Oseg1,Nurf-38,nrv1,ND-B22,levy,ACC,Jafrac1 | | 1.086e-02 | -4.52 | Mitotic G2-G2/M phases | REACTOME pathways | R-DME-453274 | 81 | 20 | 3923 | 574 | Cdk7,Prosbeta3,Prosbeta6,CycB,Cdk1,Rpn3,Cul1,Rpn12,E2f1,Prosbeta4,Prosalpha5,RpS27A,twe,Prosalpha4,Rpn2,Prosbeta5,SkpA,RpL40,Rpt2,Prosalpha3 | | 1.088e-02 | -4.52 | tRNA metabolic process | biological process | GO:0006399 | 135 | 21 | 12345 | 1129 | CG10189,CG3021,GatA,CysRS,scu,GluRS-m,CG10463,CG2911,CG5292,GatB,SerRS-m,Archease,TyrRS,CG3587,Rrp40,TrpRS-m,CG30343,LysRS,SelD,CG2034,CG7009 | | 1.090e-02 | -4.52 | DUF4780 | pfam domains | PF16012 | 5 | 3 | 10480 | 1142 | CG7824,CG9641,CG7208 | | 1.090e-02 | -4.52 | La | pfam domains | PF05383 | 5 | 3 | 10480 | 1142 | Larp4B,Larp7,larp | | 1.090e-02 | -4.52 | GO\_PROTEIN\_LOCALIZATION\_TO\_MEMBRANE | MSigDB lists | GO\_PROTEIN\_LOCALIZATION\_TO\_MEMBRANE | 171 | 38 | 4161 | 644 | CG44422,shrb,Rab10,RpL9,RpS23,RpS27A,RpS28b,RpL13,Tsp86D,sau,RpL13A,RpS30,RpLP0,RpS13,Arl6IP1,CHOp24,ste14,RpL37a,RpL11,RpL26,Vamp7,RpL24,RpS27,RpS18,Cdk5,RpL12,RpL30,RpL23,RpS24,Oseg1,RpS16,RpS2,RpL38,RpS4,RpS7,Rep,RpL40,Atg4b | | 1.091e-02 | -4.52 | ribonuclease activity | molecular function | GO:0004540 | 78 | 14 | 12176 | 1119 | Tengl4,CG12375,twin,Clp,Rga,CG11164,cu,CG11486,CG31759,scu,trsn,Exd2,CG30343,Dcr-1 | | 1.092e-02 | -4.52 | GO\_CELLULAR\_RESPONSE\_TO\_CYTOKINE\_STIMULUS | MSigDB lists | GO\_CELLULAR\_RESPONSE\_TO\_CYTOKINE\_STIMULUS | 97 | 24 | 4161 | 644 | RpS27A,Prosalpha5,Prosalpha3,cl,Prosbeta4,Hsc70-3,bys,Rpn3,Prosbeta5,LpR2,cact,rl,Traf6,Rpn12,RpL13A,Rpn2,Prosbeta3,Rpt2,Prosalpha4,Gapdh1,Med,RpL40,Ubc4,Prosbeta6 | | 1.094e-02 | -4.52 | cell division | biological process | GO:0051301 | 266 | 36 | 12345 | 1129 | Iml1,enc,tws,alphaTub67C,CycB,Dcr-1,shrb,Cen,egg,cnn,tkv,Stat92E,twin,msps,Pfdn2,Med,CycE,Arf51F,Grip128,eIF4E1,bam,Su(H),Pex10,Pex12,Pex13,eIF4A,nos,mts,BoYb,mad2,sau,slam,Cdk1,Grip75,piwi,Lkb1 | | 1.095e-02 | -4.51 | monovalent inorganic cation transmembrane transporter activity | molecular function | GO:0015077 | 238 | 33 | 12176 | 1119 | ATPsynG,rpk,levy,blw,Irk1,Atpalpha,COX5A,Vha13,ATPsyndelta,COX5B,ATPsynCF6,Vha36-1,yin,Calx,CG4476,cype,ATPsynD,SERCA,kcc,ATPsynB,ATPsynE,NaPi-III,blot,COX8,ATPsynO,zyd,Ork1,ATPsynbeta,COX4,CG10960,Vha68-2,nrv1,CG12773 | | 1.099e-02 | -4.51 | Import of palmitoyl-CoA into the mitochondrial matrix | REACTOME pathways | R-DME-200425 | 7 | 4 | 3923 | 574 | colt,whd,SNF4Agamma,alc | | 1.100e-02 | -4.51 | ion homeostasis | biological process | GO:0050801 | 152 | 23 | 12345 | 1129 | Vha68-2,Fer1HCH,EMRE,zyd,Grx1,Ho,Seipin,ZnT35C,nrv1,fwe,Scox,CG12773,CG14036,wash,park,MtnA,SREBP,up,Rbcn-3A,Atpalpha,SERCA,CG10470,anne | | 1.100e-02 | -4.51 | MODULE\_32 | MSigDB lists | MODULE\_32 | 123 | 29 | 4161 | 644 | RpS24,eIF4A,LysRS,RpS16,RpS2,RpS13,RpLP0,RpS7,eIF3i,RpS4,CysRS,RpL38,Rab5,bru1,RpL37a,RpL11,Scsalpha1,Rab10,RpS28b,RpS27,RpL24,Rnmt,CG5116,RpL26,RpL9,RpS23,mEFTu1,RpL12,eEF1beta | | 1.105e-02 | -4.51 | GSE42088\_UNINF\_VS\_LEISHMANIA\_INF\_DC\_4H\_DN | MSigDB lists | GSE42088\_UNINF\_VS\_LEISHMANIA\_INF\_DC\_4H\_DN | 67 | 18 | 4161 | 644 | RpS18,RpL13,RpL11,Ppcdc,RpL37a,RpL26,RpS7,RpL38,RpS4,CG4572,RpL40,RpS24,RpL30,EcR,RpS13,RpS30,RpS16,RpL13A | | 1.105e-02 | -4.51 | GSE8515\_CTRL\_VS\_IL6\_4H\_STIM\_MAC\_UP | MSigDB lists | GSE8515\_CTRL\_VS\_IL6\_4H\_STIM\_MAC\_UP | 67 | 18 | 4161 | 644 | CG2938,CHOp24,lid,CG8204,PpV,Plap,muskelin,CG3860,Rpn2,Pus7,CG16979,CG13630,CG8112,mad2,GatA,RpL9,Mekk1,Utx | | 1.107e-02 | -4.50 | Raf (Raf oncogene) | protein interactions | 31221 | 149 | 28 | 9204 | 1112 | tkv,RpS18,SERCA,Nf-YA,spen,RpS27A,mts,Kcmf1,stet,PpV,PyK,csw,trk,Aldh,rl,alph,Mcad,ScsbetaA,wal,SelD,RpL17,Bap170,shv,eIF4A,GCS2beta,Cp1,CkIIalpha,tor | | 1.111e-02 | -4.50 | protein kinase activity | molecular function | GO:0004672 | 274 | 37 | 12176 | 1119 | CG11486,CG3631,Mos,PEK,Cdk1,CG8726,CycE,p38a,Tfb4,tub,Mekk1,grp,CkIIalpha,Tao,Cdk5,rl,CG8475,tor,Cdk7,Tak1,dco,Pink1,CG17698,Mo25,Pdk,Lkb1,Tlk,fray,CycJ,Pdk1,CycB,SRPK,CG4041,tkv,alc,Coq8,png | | 1.111e-02 | -4.50 | mitochondrial fatty acid beta-oxidation of unsaturated fatty acids | REACTOME pathways | R-DME-77288 | 4 | 3 | 3923 | 574 | Mcad,CG4598,CG4594 | | 1.111e-02 | -4.50 | Phosphorylation of proteins involved in G1/S transition by active Cyclin E:Cdk2 complexes | REACTOME pathways | R-DME-69200 | 4 | 3 | 3923 | 574 | Rbf,Rbf2,CycE | | 1.111e-02 | -4.50 | E2F-enabled inhibition of pre-replication complex formation | REACTOME pathways | R-DME-113507 | 4 | 3 | 3923 | 574 | CycB,Cdk1,twe | | 1.111e-02 | -4.50 | DARPP-32 events | REACTOME pathways | R-DME-180024 | 4 | 3 | 3923 | 574 | Pka-R1,Pka-R2,dnc | | 1.117e-02 | -4.49 | protein kinase complex | cellular component | GO:1902911 | 48 | 10 | 11940 | 1105 | CycE,Cdk5,Tfb4,CycJ,Cdk7,plu,CycB,CG8475,alc,png | | 1.118e-02 | -4.49 | GSE12845\_NAIVE\_VS\_DARKZONE\_GC\_TONSIL\_BCELL\_DN | MSigDB lists | GSE12845\_NAIVE\_VS\_DARKZONE\_GC\_TONSIL\_BCELL\_DN | 87 | 22 | 4161 | 644 | Herc4,Uch,Pmi,mRpL16,Top1,Prosbeta5,pont,aay,RpII140,Kap3,Cse1,Rpn12,Rpt2,alphaTub84B,Cdk7,Pgm1,Pdk,Iml1,CG11089,ste14,CG42542,Crag | | 1.119e-02 | -4.49 | GSE26290\_WT\_VS\_PDK1\_KO\_ANTI\_CD3\_AND\_IL2\_STIM\_CD8\_TCELL\_DN | MSigDB lists | GSE26290\_WT\_VS\_PDK1\_KO\_ANTI\_CD3\_AND\_IL2\_STIM\_CD8\_TCELL\_DN | 72 | 19 | 4161 | 644 | PAPLA1,Rab32,Acph-1,Aldh,Pdk1,thoc6,Chrac-14,Moe,ND-30,CycC,Mctp,ABCD,CG9485,eIF3i,nSMase,CG3842,Rep,Rab5,cyc | | 1.119e-02 | -4.49 | GO\_OXIDOREDUCTASE\_ACTIVITY\_ACTING\_ON\_THE\_CH\_NH\_GROUP\_OF\_DONORS\_NAD\_OR\_NADP\_AS\_ACCEPTOR | MSigDB lists | GO\_OXIDOREDUCTASE\_ACTIVITY\_ACTING\_ON\_THE\_CH\_NH\_GROUP\_OF\_DONORS\_NAD\_OR\_NADP\_AS\_ACCEPTOR | 10 | 5 | 4161 | 644 | P5CDh1,CG9471,CG4872,P5cr,Dhfr | | 1.119e-02 | -4.49 | SA\_PTEN\_PATHWAY | MSigDB lists | SA\_PTEN\_PATHWAY | 10 | 5 | 4161 | 644 | CG7789,rl,Pdk1,Rbf,Pi3K92E | | 1.119e-02 | -4.49 | GO\_PYRIDINE\_CONTAINING\_COMPOUND\_BIOSYNTHETIC\_PROCESS | MSigDB lists | GO\_PYRIDINE\_CONTAINING\_COMPOUND\_BIOSYNTHETIC\_PROCESS | 10 | 5 | 4161 | 644 | Nadsyn,sgll,Pdxk,CG33156,CG11899 | | 1.119e-02 | -4.49 | REACTOME\_SMAD2\_SMAD3\_SMAD4\_HETEROTRIMER\_REGULATES\_TRANSCRIPTION | MSigDB lists | REACTOME\_SMAD2\_SMAD3\_SMAD4\_HETEROTRIMER\_REGULATES\_TRANSCRIPTION | 10 | 5 | 4161 | 644 | CycC,Smox,Med,RpL40,RpS27A | | 1.121e-02 | -4.49 | regulation of generation of precursor metabolites and energy | biological process | GO:0043467 | 28 | 7 | 12345 | 1129 | Pgk,CG7328,Tspo,Pink1,MED27,EcR,GlyP | | 1.121e-02 | -4.49 | regulation of cyclin-dependent protein serine/threonine kinase activity | biological process | GO:0000079 | 28 | 7 | 12345 | 1129 | CycC,CycJ,CycB,CG30291,Cdk7,CG17746,CycE | | 1.121e-02 | -4.49 | salivary gland cell autophagic cell death | biological process | GO:0035071 | 28 | 7 | 12345 | 1129 | DCAF12,Trf2,Utx,MED24,EcR,Idh3b,rept | | 1.121e-02 | -4.49 | regulation of cyclin-dependent protein kinase activity | biological process | GO:1904029 | 28 | 7 | 12345 | 1129 | CycJ,CycB,CycC,CycE,Cdk7,CG30291,CG17746 | | 1.124e-02 | -4.49 | GstS1 (Glutathione S transferase S1) | protein interactions | 36927 | 32 | 9 | 9204 | 1112 | Eno,Tm1,COX4,ND-18,COX5A,ATPsyndelta,park,Adh,Pink1 | | 1.124e-02 | -4.49 | GO\_POSITIVE\_REGULATION\_OF\_INNATE\_IMMUNE\_RESPONSE | MSigDB lists | GO\_POSITIVE\_REGULATION\_OF\_INNATE\_IMMUNE\_RESPONSE | 82 | 21 | 4161 | 644 | Prosbeta6,SkpA,Ubc4,scramb1,RpL40,Rpt2,Prosalpha4,Prosbeta3,Rpn2,Rpn12,Traf6,cact,Pdk1,Prosbeta5,Cul1,Rpn3,Uev1A,Prosbeta4,Prosalpha3,RpS27A,Prosalpha5 | | 1.124e-02 | -4.49 | GSE22886\_NAIVE\_CD4\_TCELL\_VS\_DC\_DN | MSigDB lists | GSE22886\_NAIVE\_CD4\_TCELL\_VS\_DC\_DN | 82 | 21 | 4161 | 644 | Grx1,Stam,Pka-R1,Cyt-c-p,ste24a,CG10470,lt,Vha36-1,alphaTub84B,CkIIalpha,muskelin,Marf,Pglym78,Prosbeta5,Tim8,CG7997,wal,Gclc,Pi4KIIalpha,Clc,Idh | | 1.125e-02 | -4.49 | GSE14308\_TH1\_VS\_NAIVE\_CD4\_TCELL\_UP | MSigDB lists | GSE14308\_TH1\_VS\_NAIVE\_CD4\_TCELL\_UP | 77 | 20 | 4161 | 644 | Ptpmeg,AIF,CG3609,Axs,Arpc1,gammaTub37C,CG11134,Prosbeta4,CG10038,cl,ND-39,zyd,rept,Sld5,CG11110,EcR,GCS2beta,Kap3,CG1847,CycC | | 1.125e-02 | -4.49 | GSE14769\_20MIN\_VS\_360MIN\_LPS\_BMDM\_UP | MSigDB lists | GSE14769\_20MIN\_VS\_360MIN\_LPS\_BMDM\_UP | 77 | 20 | 4161 | 644 | ND-75,CG7997,SdhA,CG8314,shop,CG9391,eEF1beta,CG33181,Idh3b,PpD3,RpL9,CG17593,CG12713,Etf-QO,Maf1,CG10932,Psf3,RpS2,APC7,aay | | 1.131e-02 | -4.48 | ALDKETRDTASE | prints domains | PR00069 | 10 | 4 | 2185 | 212 | CG9436,CG10638,CG2767,CG10863 | | 1.136e-02 | -4.48 | WD40 | smart domains | SM00320 | 173 | 27 | 5086 | 506 | thoc6,cort,adp,Arpc1,Oseg1,Cka,esc,Plap,Rbcn-3A,eIF3i,CG3511,tws,sw,Fbw5,escl,Caf1-105,Tomosyn,CG10132,Grip71,ago,lt,DCAF12,raptor,Wdfy2,Aladin,rig,CG8001 | | 1.140e-02 | -4.47 | GSE22886\_DAY0\_VS\_DAY1\_MONOCYTE\_IN\_CULTURE\_UP | MSigDB lists | GSE22886\_DAY0\_VS\_DAY1\_MONOCYTE\_IN\_CULTURE\_UP | 48 | 14 | 4161 | 644 | RpS4,Pi3K92E,Nprl2,Aldh,p38a,RpS18,ATP8A,CG4572,RpL30,Rbf,Pgm1,RpL26,RpL13A,RpS2 | | 1.140e-02 | -4.47 | cation homeostasis | biological process | GO:0055080 | 144 | 22 | 12345 | 1129 | MtnA,CG14036,wash,park,Rbcn-3A,Atpalpha,SERCA,CG10470,anne,up,Fer1HCH,EMRE,zyd,Grx1,Vha68-2,ZnT35C,nrv1,CG12773,Scox,fwe,Ho,Seipin | | 1.144e-02 | -4.47 | dorsal appendage formation | biological process | GO:0046843 | 56 | 11 | 12345 | 1129 | squ,jvl,cact,rl,Pax,spn-E,gus,armi,bun,mael,Cf2 | | 1.144e-02 | -4.47 | chromosome organization involved in meiotic cell cycle | biological process | GO:0070192 | 56 | 11 | 12345 | 1129 | SRPK,eIF4E1,Topors,lid,CG31053,krimp,del,Nup62,enc,GatA,spn-E | | 1.145e-02 | -4.47 | regulation of cellular response to stress | biological process | GO:0080135 | 119 | 19 | 12345 | 1129 | Archease,Ptp61F,Mekk1,ben,PpV,CG17746,armi,Seipin,Imp,ago,Tak1,Ubc6,park,Lkb1,Cka,PEK,alph,SkpA,p38a | | 1.149e-02 | -4.47 | Fatty Acid Beta Oxidation 2 | WikiPathways | WP187 | 5 | 4 | 499 | 116 | Mtpalpha,Echs1,Thiolase,Mcad | | 1.151e-02 | -4.46 | HTH\_LA | prosite domains | PS50961 | 5 | 3 | 6243 | 694 | Larp7,Larp4B,larp | | 1.151e-02 | -4.46 | CULLIN\_1 | prosite domains | PS01256 | 5 | 3 | 6243 | 694 | Cul2,Cul1,Cul5 | | 1.151e-02 | -4.46 | ALDOKETO\_REDUCTASE\_3 | prosite domains | PS00063 | 5 | 3 | 6243 | 694 | CG10863,CG2767,CG10638 | | 1.157e-02 | -4.46 | GARY\_CD5\_TARGETS\_DN | MSigDB lists | GARY\_CD5\_TARGETS\_DN | 188 | 41 | 4161 | 644 | Non1,CG8728,APC7,mRpS23,Ubc4,Ufm1,SelD,CG4511,RfC38,tko,Cbp80,Adi1,Plap,obe,Hsc70-3,COX5A,PDCD-5,CycB,Prosbeta5,CG10425,ND-B18,Stip1,r-l,sbr,CG8112,Cul2,trsn,shv,Rpn2,CG3511,Karybeta3,l(1)G0004,CG4407,CG30496,GatA,Bin1,CG5880,Uch-L5,CCT6,Chrac-14,Pen | | 1.164e-02 | -4.45 | - | gene3d domains | 3.60.10.10 | 12 | 5 | 4929 | 622 | CG31759,nSMase,Synj,cu,twin | | 1.168e-02 | -4.45 | actin filament organization | biological process | GO:0007015 | 111 | 18 | 12345 | 1129 | wash,Girdin,Mtl,cib,capt,up,swa,Pka-R1,Actn,Tm1,Hip1,Arpc1,Abi,Arpc2,jvl,SelR,Arpc4,Sry-alpha | | 1.168e-02 | -4.45 | CG8173 (uncharacterized protein) | protein interactions | 32753 | 17 | 6 | 9204 | 1112 | tws,Droj2,capt,zf30C,CG3342,cu | | 1.168e-02 | -4.45 | CG8851 (uncharacterized protein) | protein interactions | 33575 | 17 | 6 | 9204 | 1112 | CG3342,cnn,CG11248,CG6276,Sec23,TBCC | | 1.170e-02 | -4.45 | histone acetylation | biological process | GO:0016573 | 79 | 14 | 12345 | 1129 | e(y)1,Hat1,YL-1,pont,nej,rept,Chrac-14,lid,d4,Spt20,Rcd5,Naa20A,Sgf29,Tip60 | | 1.170e-02 | -4.45 | COLDREN\_GEFITINIB\_RESISTANCE\_UP | MSigDB lists | COLDREN\_GEFITINIB\_RESISTANCE\_UP | 30 | 10 | 4161 | 644 | parvin,CG3803,Cyp1,CG7728,Got1,CG2100,Nap1,isoQC,slif,Dph5 | | 1.170e-02 | -4.45 | MODULE\_28 | MSigDB lists | MODULE\_28 | 30 | 10 | 4161 | 644 | Rpn3,Prosbeta6,Prosbeta5,Rpn2,Prosalpha5,Rpn12,Rpt2,Prosbeta4,Prosalpha3,Prosbeta3 | | 1.172e-02 | -4.45 | Nucleoside\_diP\_kinase\_AS | interpro domains | IPR023005 | 2 | 2 | 11383 | 1233 | nmdyn-D6,awd | | 1.172e-02 | -4.45 | Ribosomal\_L10e/L16\_sf | interpro domains | IPR036920 | 2 | 2 | 11383 | 1233 | RpL10,mRpL16 | | 1.172e-02 | -4.45 | CAP/MinC\_C | interpro domains | IPR016098 | 2 | 2 | 11383 | 1233 | TBCC,capt | | 1.172e-02 | -4.45 | RB\_N | interpro domains | IPR024599 | 2 | 2 | 11383 | 1233 | Rbf,Rbf2 | | 1.172e-02 | -4.45 | RB\_fam | interpro domains | IPR028309 | 2 | 2 | 11383 | 1233 | Rbf2,Rbf | | 1.172e-02 | -4.45 | Ribosomal\_S5\_N | interpro domains | IPR013810 | 2 | 2 | 11383 | 1233 | mRpS5,RpS2 | | 1.172e-02 | -4.45 | DUF953\_thioredox | interpro domains | IPR010357 | 2 | 2 | 11383 | 1233 | CG3939,cl | | 1.172e-02 | -4.45 | MGS-like\_dom | interpro domains | IPR011607 | 2 | 2 | 11383 | 1233 | r,CG11089 | | 1.172e-02 | -4.45 | RutC-like\_sf | interpro domains | IPR035959 | 2 | 2 | 11383 | 1233 | CG1578,UK114 | | 1.172e-02 | -4.45 | cAMP\_dep\_PK\_reg\_su | interpro domains | IPR012198 | 2 | 2 | 11383 | 1233 | Pka-R2,Pka-R1 | | 1.172e-02 | -4.45 | MGS-like\_dom\_sf | interpro domains | IPR036914 | 2 | 2 | 11383 | 1233 | CG11089,r | | 1.172e-02 | -4.45 | EF-Tu\_GTP-bd | interpro domains | IPR041709 | 2 | 2 | 11383 | 1233 | mEFTu2,mEFTu1 | | 1.172e-02 | -4.45 | EFTU\_2 | interpro domains | IPR033720 | 2 | 2 | 11383 | 1233 | mEFTu1,mEFTu2 | | 1.172e-02 | -4.45 | Ribosomal\_L34Ae | interpro domains | IPR008195 | 2 | 2 | 11383 | 1233 | RpL34b,RpL34a | | 1.172e-02 | -4.45 | PFD\_alpha\_archaea | interpro domains | IPR011599 | 2 | 2 | 11383 | 1233 | Pfdn5,Uxt | | 1.172e-02 | -4.45 | Ribosomal\_S5 | interpro domains | IPR000851 | 2 | 2 | 11383 | 1233 | RpS2,mRpS5 | | 1.172e-02 | -4.45 | Ribosomal\_S12/S23 | interpro domains | IPR006032 | 2 | 2 | 11383 | 1233 | RpS23,tko | | 1.172e-02 | -4.45 | CARP\_motif | interpro domains | IPR006599 | 2 | 2 | 11383 | 1233 | capt,TBCC | | 1.172e-02 | -4.45 | C-CAP\_CF\_C-like | interpro domains | IPR017901 | 2 | 2 | 11383 | 1233 | capt,TBCC | | 1.172e-02 | -4.45 | RB\_A | interpro domains | IPR002720 | 2 | 2 | 11383 | 1233 | Rbf2,Rbf | | 1.172e-02 | -4.45 | Ribosomal\_S11 | interpro domains | IPR001971 | 2 | 2 | 11383 | 1233 | mRpS11,RpS14b | | 1.172e-02 | -4.45 | RB\_B | interpro domains | IPR002719 | 2 | 2 | 11383 | 1233 | Rbf2,Rbf | | 1.172e-02 | -4.45 | ARPC2/4 | interpro domains | IPR034666 | 2 | 2 | 11383 | 1233 | Arpc4,Arpc2 | | 1.172e-02 | -4.45 | Ribosomal\_S5\_C | interpro domains | IPR005324 | 2 | 2 | 11383 | 1233 | mRpS5,RpS2 | | 1.172e-02 | -4.45 | Polycomb\_EED | interpro domains | IPR037352 | 2 | 2 | 11383 | 1233 | escl,esc | | 1.172e-02 | -4.45 | Lsm14-like\_N | interpro domains | IPR025609 | 2 | 2 | 11383 | 1233 | tral,Edc3 | | 1.172e-02 | -4.45 | YjgF/YER057c/UK114 | interpro domains | IPR006175 | 2 | 2 | 11383 | 1233 | UK114,CG1578 | | 1.172e-02 | -4.45 | TIP49\_P-loop | interpro domains | IPR010339 | 2 | 2 | 11383 | 1233 | pont,rept | | 1.172e-02 | -4.45 | Ribosomal\_L10e/L16 | interpro domains | IPR016180 | 2 | 2 | 11383 | 1233 | mRpL16,RpL10 | | 1.172e-02 | -4.45 | Ribosomal\_S5\_N\_CS | interpro domains | IPR018192 | 2 | 2 | 11383 | 1233 | RpS2,mRpS5 | | 1.172e-02 | -4.45 | RuvB-like\_C | interpro domains | IPR041048 | 2 | 2 | 11383 | 1233 | rept,pont | | 1.172e-02 | -4.45 | Glycerol\_kin | interpro domains | IPR005999 | 2 | 2 | 11383 | 1233 | Gk1,Gk2 | | 1.172e-02 | -4.45 | EMC3/TMCO1 | interpro domains | IPR002809 | 2 | 2 | 11383 | 1233 | EMC3,CG10470 | | 1.172e-02 | -4.45 | GDP\_dissociation\_inhibitor | interpro domains | IPR018203 | 2 | 2 | 11383 | 1233 | Rep,Gdi | | 1.172e-02 | -4.45 | MOSC\_N | interpro domains | IPR005303 | 2 | 2 | 11383 | 1233 | Marc,mal | | 1.172e-02 | -4.45 | MoCF\_Sase\_C | interpro domains | IPR005302 | 2 | 2 | 11383 | 1233 | Marc,mal | | 1.172e-02 | -4.45 | Ribosomal\_L34e\_CS | interpro domains | IPR018065 | 2 | 2 | 11383 | 1233 | RpL34b,RpL34a | | 1.172e-02 | -4.45 | DFDF | interpro domains | IPR025762 | 2 | 2 | 11383 | 1233 | Edc3,tral | | 1.172e-02 | -4.45 | Ribosomal\_L34Ae\_sf | interpro domains | IPR038562 | 2 | 2 | 11383 | 1233 | RpL34b,RpL34a | | 1.172e-02 | -4.45 | GK1-3\_metazoa | interpro domains | IPR042018 | 2 | 2 | 11383 | 1233 | Gk1,Gk2 | | 1.172e-02 | -4.45 | YjeF\_N\_dom | interpro domains | IPR004443 | 2 | 2 | 11383 | 1233 | Edc3,Naxe | | 1.172e-02 | -4.45 | RuvB-like | interpro domains | IPR027238 | 2 | 2 | 11383 | 1233 | rept,pont | | 1.172e-02 | -4.45 | FDF\_dom | interpro domains | IPR019050 | 2 | 2 | 11383 | 1233 | tral,Edc3 | | 1.173e-02 | -4.45 | GO\_CELLULAR\_RESPONSE\_TO\_ENDOGENOUS\_STIMULUS | MSigDB lists | GO\_CELLULAR\_RESPONSE\_TO\_ENDOGENOUS\_STIMULUS | 238 | 50 | 4161 | 644 | Moe,Pdk1,Ac3,CG11710,Smox,CycE,PDCD-5,SREBP,Wnt10,Med,RpL40,Cbp80,Atpalpha,heph,r,mus81,Pka-R1,Vha13,Ptp61F,MED4,Atg6,faf,MED24,Vha36-1,Calx,Shmt,AIF,LpR2,rl,tkv,RpS27A,Pka-R2,Fkbp59,Rab10,Got1,dnc,rept,ACC,Tip60,Karybeta3,Rpb8,adp,Sod1,EcR,mino,RpII140,Prip,Cdk7,csw,raptor | | 1.176e-02 | -4.44 | cellular response to reactive oxygen species | biological process | GO:0034614 | 16 | 5 | 12345 | 1129 | Sod3,p38a,Jafrac1,rl,Sod1 | | 1.176e-02 | -4.44 | serine family amino acid metabolic process | biological process | GO:0009069 | 16 | 5 | 12345 | 1129 | Shmt,aay,SelD,CG11899,Dhfr | | 1.187e-02 | -4.43 | Ribosom\_S12\_S23 | pfam domains | PF00164 | 2 | 2 | 10480 | 1142 | tko,RpS23 | | 1.187e-02 | -4.43 | MOSC\_N | pfam domains | PF03476 | 2 | 2 | 10480 | 1142 | Marc,mal | | 1.187e-02 | -4.43 | Ribosomal\_S11 | pfam domains | PF00411 | 2 | 2 | 10480 | 1142 | mRpS11,RpS14b | | 1.187e-02 | -4.43 | GDI | pfam domains | PF00996 | 2 | 2 | 10480 | 1142 | Gdi,Rep | | 1.187e-02 | -4.43 | RB\_A | pfam domains | PF01858 | 2 | 2 | 10480 | 1142 | Rbf,Rbf2 | | 1.187e-02 | -4.43 | Ribosomal\_L16 | pfam domains | PF00252 | 2 | 2 | 10480 | 1142 | mRpL16,RpL10 | | 1.187e-02 | -4.43 | EMC3\_TMCO1 | pfam domains | PF01956 | 2 | 2 | 10480 | 1142 | EMC3,CG10470 | | 1.187e-02 | -4.43 | DUF3452 | pfam domains | PF11934 | 2 | 2 | 10480 | 1142 | Rbf2,Rbf | | 1.187e-02 | -4.43 | Ribonuc\_L-PSP | pfam domains | PF01042 | 2 | 2 | 10480 | 1142 | CG1578,UK114 | | 1.187e-02 | -4.43 | Ribosomal\_S5 | pfam domains | PF00333 | 2 | 2 | 10480 | 1142 | mRpS5,RpS2 | | 1.187e-02 | -4.43 | RB\_B | pfam domains | PF01857 | 2 | 2 | 10480 | 1142 | Rbf,Rbf2 | | 1.187e-02 | -4.43 | TIP49\_C | pfam domains | PF17856 | 2 | 2 | 10480 | 1142 | rept,pont | | 1.187e-02 | -4.43 | TIP49 | pfam domains | PF06068 | 2 | 2 | 10480 | 1142 | rept,pont | | 1.187e-02 | -4.43 | DUF953 | pfam domains | PF06110 | 2 | 2 | 10480 | 1142 | cl,CG3939 | | 1.187e-02 | -4.43 | MGS | pfam domains | PF02142 | 2 | 2 | 10480 | 1142 | r,CG11089 | | 1.187e-02 | -4.43 | Ribosomal\_S5\_C | pfam domains | PF03719 | 2 | 2 | 10480 | 1142 | RpS2,mRpS5 | | 1.187e-02 | -4.43 | Ribosomal\_L34e | pfam domains | PF01199 | 2 | 2 | 10480 | 1142 | RpL34b,RpL34a | | 1.187e-02 | -4.43 | MOSC | pfam domains | PF03473 | 2 | 2 | 10480 | 1142 | Marc,mal | | 1.187e-02 | -4.43 | DNA replication | biological process | GO:0006260 | 87 | 15 | 12345 | 1129 | E2f1,fs(1)Ya,Sld5,SkpA,RfC38,Orc6,SuUR,CG12018,Orc1,ago,DNApol-epsilon255,Top1,CG10336,Rbf,RecQ5 | | 1.188e-02 | -4.43 | NIK-->noncanonical NF-kB signaling | REACTOME pathways | R-DME-5676590 | 66 | 17 | 3923 | 574 | Prosalpha4,Prosbeta5,Rpn2,Prosbeta4,RpS27A,Prosalpha5,Prosalpha3,Rpt2,RpL40,SkpA,Rel,Prosbeta3,Rpn12,dl,Cul1,Rpn3,Prosbeta6 | | 1.188e-02 | -4.43 | TNFR2 non-canonical NF-kB pathway | REACTOME pathways | R-DME-5668541 | 66 | 17 | 3923 | 574 | SkpA,Rpt2,RpL40,Prosalpha3,Prosalpha5,RpS27A,Prosbeta4,Rpn2,Prosbeta5,Prosalpha4,Prosbeta6,Cul1,Rpn3,dl,Rpn12,Prosbeta3,Rel | | 1.188e-02 | -4.43 | Dectin-1 mediated noncanonical NF-kB signaling | REACTOME pathways | R-DME-5607761 | 66 | 17 | 3923 | 574 | Rpt2,RpL40,Prosalpha3,SkpA,Prosbeta5,Rpn2,Prosalpha4,Prosalpha5,RpS27A,Prosbeta4,Rpn3,Cul1,dl,Rpn12,Prosbeta6,Rel,Prosbeta3 | | 1.190e-02 | -4.43 | Tor (Target of rapamycin) | protein interactions | 47396 | 49 | 12 | 9204 | 1112 | CycE,cathD,Pi3K92E,Pink1,Nprl2,raptor,pont,rept,Cul1,spag,REPTOR,lqfR | | 1.190e-02 | -4.43 | NRF2\_01 | MSigDB lists | NRF2\_01 | 108 | 26 | 4161 | 644 | Hmg-2,Acph-1,CG3587,TBCC,RpL37a,Herc4,pen-2,capt,mEFTu1,Tim8,Tom40,CG3803,RpS18,EcR,GCS2beta,Phb2,CG10470,Edc3,Sec10,Aladin,Vha36-1,Tip60,Uba5,CG31715,CG9485,CG8475 | | 1.190e-02 | -4.43 | GSE22886\_NAIVE\_CD4\_TCELL\_VS\_12H\_ACT\_TH1\_DN | MSigDB lists | GSE22886\_NAIVE\_CD4\_TCELL\_VS\_12H\_ACT\_TH1\_DN | 108 | 26 | 4161 | 644 | SelD,RfC38,Ts,Stoml2,CG42542,Plap,CG1416,Grx1,CG3226,CkIIalpha,CG10470,Rpn2,Caf1-105,r-l,MED27,CycB,mad2,ND-24,CCT6,ND-30,CG7488,Uch,Got1,CG7728,CG4293,CycE | | 1.190e-02 | -4.43 | regulation of innate immune response | biological process | GO:0045088 | 95 | 16 | 12345 | 1129 | faf,scny,PGRP-LE,slif,Hr78,Sp7,EMRE,Hat1,Crag,Ect4,Myd88,GlyP,CG9925,PDCD-5,Rel,Atg6 | | 1.191e-02 | -4.43 | Rab5 (Rab5) | protein interactions | 33418 | 130 | 25 | 9204 | 1112 | Nadsyn,CG1416,Droj2,Gdi,Atg6,cathD,Mcad,CG9205,ScsbetaA,wal,mop,Cp1,Tspo,Vha68-2,tkv,CCT6,SERCA,CG4041,awd,Rbcn-3A,CG10425,CCT3,sau,cDIP,CG15098 | | 1.193e-02 | -4.43 | Uba4 (Ubiquitin activating enzyme 4) | protein interactions | 34187 | 27 | 8 | 9204 | 1112 | Coq6,Pen,eIF4E1,Spx,CG11486,sol,zf30C,E2f1 | | 1.193e-02 | -4.43 | clu (clueless) | protein interactions | 36793 | 27 | 8 | 9204 | 1112 | RpL30,park,Pink1,Pdk1,Marf,RpL7A,RpS7,RpS4 | | 1.193e-02 | -4.43 | bsh (brain-specific homeobox) | protein interactions | 35266 | 27 | 8 | 9204 | 1112 | RpL37a,Sod3,Spx,RpL17,Bap170,Cbl,CG15098,galectin | | 1.193e-02 | -4.43 | negative regulation of mitotic cell cycle phase transition | biological process | GO:1901991 | 49 | 10 | 12345 | 1129 | mad2,Rbf,mre11,Aven,Rbf2,Cdk1,grp,mus304,Zwilch,nbs | | 1.193e-02 | -4.43 | positive regulation of chromatin organization | biological process | GO:1905269 | 49 | 10 | 12345 | 1129 | HP1c,esc,trr,pgc,egg,HP1e,G9a,Stat92E,piwi,scny | | 1.195e-02 | -4.43 | GSE4535\_BM\_DERIVED\_DC\_VS\_FOLLICULAR\_DC\_UP | MSigDB lists | GSE4535\_BM\_DERIVED\_DC\_VS\_FOLLICULAR\_DC\_UP | 39 | 12 | 4161 | 644 | Smox,snz,csw,CG1532,CG9143,LSm3,PrBP,scramb1,Liprin-beta,Pglym78,cyc,CG7322 | | 1.198e-02 | -4.42 | regulation of mitochondrion organization | biological process | GO:0010821 | 22 | 6 | 12345 | 1129 | park,Pink1,ago,Sod1,Marf,SREBP | | 1.198e-02 | -4.42 | response to disaccharide | biological process | GO:0034285 | 22 | 6 | 12345 | 1129 | ACC,Desat1,PyK,sgll,Pfk,CG7328 | | 1.198e-02 | -4.42 | regulation of meiotic cell cycle | biological process | GO:0051445 | 22 | 6 | 12345 | 1129 | Mos,twe,mtrm,Iml1,CG31053,Nprl2 | | 1.200e-02 | -4.42 | alcohol dehydrogenase (NADP+) activity | molecular function | GO:0008106 | 16 | 5 | 12176 | 1119 | CG3842,CG9436,CG2767,CG10638,CG10863 | | 1.200e-02 | -4.42 | thiolester hydrolase activity | molecular function | GO:0016790 | 16 | 5 | 12176 | 1119 | CG4390,CG16979,Scsalpha1,CG1774,tzn | | 1.200e-02 | -4.42 | translation repressor activity | molecular function | GO:0030371 | 16 | 5 | 12176 | 1119 | bam,heph,bcd,Shmt,smg | | 1.201e-02 | -4.42 | AACTGGA\_MIR145 | MSigDB lists | AACTGGA\_MIR145 | 53 | 15 | 4161 | 644 | CG2201,mtgo,CanA-14F,Herc4,eIF4A,shv,Smox,Nap1,Pp2B-14D,SkpA,CG16868,gus,lqfR,bru1,ACC | | 1.201e-02 | -4.42 | positive regulation of ubiquitin-dependent protein catabolic process | biological process | GO:2000060 | 35 | 8 | 12345 | 1129 | Ubc6,park,ari-1,cort,Pomp,dco,SkpA,Cul1 | | 1.209e-02 | -4.42 | FLECHNER\_BIOPSY\_KIDNEY\_TRANSPLANT\_OK\_VS\_DONOR\_UP | MSigDB lists | FLECHNER\_BIOPSY\_KIDNEY\_TRANSPLANT\_OK\_VS\_DONOR\_UP | 205 | 44 | 4161 | 644 | Bin1,wal,ND-75,ERp60,Pen,Pgk,Got1,Idh,Sec22,Nap1,asf1,CG8475,eIF3i,Fdh,opm,CG11070,Pgm1,ste24a,Jafrac2,alph,Moe,Hat1,sbr,Sec23,Hsc70-3,CG4872,Dcr-1,Prosalpha3,mtgo,Mitofilin,Arf51F,Pp2A-29B,Cka,Gabat,CG9485,CG4511,Pka-R1,AGBE,ATP8A,scramb1,Tspo,Rbf,CG1890,Cyt-c-p | | 1.217e-02 | -4.41 | positive regulation of proteasomal protein catabolic process | biological process | GO:1901800 | 42 | 9 | 12345 | 1129 | park,Ubc6,ari-1,dco,cort,Pomp,SkpA,Cul1,Rpt2 | | 1.217e-02 | -4.41 | nucleus localization | biological process | GO:0051647 | 42 | 9 | 12345 | 1129 | l(3)mbt,alphaTub67C,cnc,ssh,gammaTub37C,mael,wisp,jvl,faf | | 1.217e-02 | -4.41 | N-acetyltransferase activity | molecular function | GO:0008080 | 79 | 14 | 12176 | 1119 | Tip60,d4,Hat1,e(y)1,lid,Naa20A,YL-1,AgmNAT,Chrac-14,nej,Sgf29,Rcd5,pont,rept | | 1.220e-02 | -4.41 | fzr (fizzy-related) | protein interactions | 45922 | 22 | 7 | 9204 | 1112 | Cdk1,Grip71,spd-2,Grip84,cnn,CycE,CycB | | 1.220e-02 | -4.41 | CG2116 (uncharacterized protein) | protein interactions | 31735 | 22 | 7 | 9204 | 1112 | mad2,RpS23,wisp,RpL31,sol,RpL23A,RpL26 | | 1.220e-02 | -4.41 | jigr1 (jing interacting gene regulatory 1) | protein interactions | 43093 | 22 | 7 | 9204 | 1112 | Cenp-C,CG2199,Pen,vret,jigr1,Imp,Prosbeta4 | | 1.220e-02 | -4.41 | CG3847 (uncharacterized protein) | protein interactions | 31577 | 22 | 7 | 9204 | 1112 | sol,RpS7,ATPsynB,RpL30,RpS23,RpL40,RpL31 | | 1.220e-02 | -4.41 | DCP1 (Decapping protein 1) | protein interactions | 37790 | 22 | 7 | 9204 | 1112 | Nup58,Tak1,Edc3,cnn,osk,ND-30,Pink1 | | 1.222e-02 | -4.40 | positive regulation of protein catabolic process | biological process | GO:0045732 | 64 | 12 | 12345 | 1129 | SkpA,Pomp,cort,dco,Rpt2,gus,Cul1,CG4238,park,Ubc6,Fbw5,ari-1 | | 1.224e-02 | -4.40 | Signaling by TGF-beta Receptor Complex | REACTOME pathways | R-DME-170834 | 51 | 14 | 3923 | 574 | faf,Rbf2,STUB1,CycC,nrv1,Med,Fur2,RpS27A,RpL40,alph,CG10188,Smox,Rbf,CG6923 | | 1.224e-02 | -4.40 | positive regulation of signal transduction | biological process | GO:0009967 | 369 | 47 | 12345 | 1129 | Cka,ea,O-fut1,Ssdp,foxo,Pura,shv,Traf6,Jafrac2,mts,tor,Pdk1,Tak1,Imp,Usp10,mop,Rel,Herc4,Myd88,disp,CG4853,CkIIalpha,CycE,CG10188,Aos1,tkv,EndoGI,dco,E2f1,PEK,Pi3K92E,pont,nej,Tsp86D,spen,Mos,Tao,trbd,gd,snk,dx,spz,tws,Mekk1,Ect4,ben,stet | | 1.225e-02 | -4.40 | apolpp (apolipophorin) | protein interactions | 43827 | 55 | 13 | 9204 | 1112 | CG4390,Argk,Adh,e(y)1,CG32473,Rtnl1,Scp1,Pglym78,blw,Arf51F,Fer1HCH,Seipin,Idgf4 | | 1.234e-02 | -4.39 | S5\_DSRBD | prosite domains | PS50881 | 2 | 2 | 6243 | 694 | mRpS5,RpS2 | | 1.234e-02 | -4.39 | DFDF | prosite domains | PS51512 | 2 | 2 | 6243 | 694 | tral,Edc3 | | 1.234e-02 | -4.39 | FGGY\_KINASES\_1 | prosite domains | PS00933 | 2 | 2 | 6243 | 694 | Gk2,Gk1 | | 1.234e-02 | -4.39 | MOSC | prosite domains | PS51340 | 2 | 2 | 6243 | 694 | Marc,mal | | 1.234e-02 | -4.39 | C\_CAP\_COFACTOR\_C | prosite domains | PS51329 | 2 | 2 | 6243 | 694 | capt,TBCC | | 1.234e-02 | -4.39 | CPSASE\_1 | prosite domains | PS00866 | 2 | 2 | 6243 | 694 | ACC,r | | 1.234e-02 | -4.39 | RIBOSOMAL\_L34E | prosite domains | PS01145 | 2 | 2 | 6243 | 694 | RpL34a,RpL34b | | 1.234e-02 | -4.39 | RIBOSOMAL\_S12 | prosite domains | PS00055 | 2 | 2 | 6243 | 694 | tko,RpS23 | | 1.234e-02 | -4.39 | NDP\_KINASES | prosite domains | PS00469 | 2 | 2 | 6243 | 694 | awd,nmdyn-D6 | | 1.234e-02 | -4.39 | SOD\_CU\_ZN\_1 | prosite domains | PS00087 | 2 | 2 | 6243 | 694 | Sod1,Sod3 | | 1.234e-02 | -4.39 | CYTOSOL\_AP | prosite domains | PS00631 | 2 | 2 | 6243 | 694 | Dip-B,grsm | | 1.234e-02 | -4.39 | RIBOSOMAL\_S5 | prosite domains | PS00585 | 2 | 2 | 6243 | 694 | RpS2,mRpS5 | | 1.234e-02 | -4.39 | YJEF\_N | prosite domains | PS51385 | 2 | 2 | 6243 | 694 | Edc3,Naxe | | 1.235e-02 | -4.39 | tetrahydrobiopterin metabolic process | biological process | GO:0046146 | 6 | 3 | 12345 | 1129 | Sptr,Pcd,pr | | 1.235e-02 | -4.39 | box C/D snoRNP assembly | biological process | GO:0000492 | 6 | 3 | 12345 | 1129 | CG8204,rept,pont | | 1.235e-02 | -4.39 | cell cycle arrest | biological process | GO:0007050 | 6 | 3 | 12345 | 1129 | Mos,grp,mtrm | | 1.235e-02 | -4.39 | water-soluble vitamin biosynthetic process | biological process | GO:0042364 | 6 | 3 | 12345 | 1129 | Pdxk,sgll,regucalcin | | 1.235e-02 | -4.39 | cellular response to arsenic-containing substance | biological process | GO:0071243 | 6 | 3 | 12345 | 1129 | p38a,rl,Mekk1 | | 1.235e-02 | -4.39 | regulation of mitochondrial fusion | biological process | GO:0010635 | 6 | 3 | 12345 | 1129 | Marf,Pink1,park | | 1.235e-02 | -4.39 | tetrahydrobiopterin biosynthetic process | biological process | GO:0006729 | 6 | 3 | 12345 | 1129 | pr,Pcd,Sptr | | 1.235e-02 | -4.39 | cellular response to nitrogen levels | biological process | GO:0043562 | 6 | 3 | 12345 | 1129 | Atg6,Atg7,Nprl2 | | 1.235e-02 | -4.39 | IMP metabolic process | biological process | GO:0046040 | 6 | 3 | 12345 | 1129 | Pfas,CG11089,AMPdeam | | 1.235e-02 | -4.39 | cellular response to cadmium ion | biological process | GO:0071276 | 6 | 3 | 12345 | 1129 | Mekk1,rl,p38a | | 1.235e-02 | -4.39 | female meiosis II | biological process | GO:0007147 | 6 | 3 | 12345 | 1129 | Grip75,Grip128,cort | | 1.235e-02 | -4.39 | positive regulation of developmental pigmentation | biological process | GO:0048087 | 6 | 3 | 12345 | 1129 | park,Sec23,Pink1 | | 1.235e-02 | -4.39 | cellular response to nitrogen starvation | biological process | GO:0006995 | 6 | 3 | 12345 | 1129 | Nprl2,Atg6,Atg7 | | 1.236e-02 | -4.39 | Fatty acid, triacylglycerol, and ketone body metabolism | REACTOME pathways | R-DME-535734 | 98 | 23 | 3923 | 574 | colt,Echs1,CG10932,Mcad,Mtpalpha,dob,alc,whd,mino,Desat1,CG4598,CG31523,Gk1,Thiolase,ACC,SNF4Agamma,yip2,CG4860,Gk2,CG1774,sea,CG4594,Agpat2 | | 1.238e-02 | -4.39 | Sec61alpha (Sec61 alpha subunit) | protein interactions | 33905 | 67 | 15 | 9204 | 1112 | Pp2A-29B,wal,foxo,Surf4,Karybeta3,CG31729,CG8475,SERCA,Not10,Rab5,Atpalpha,Mcad,lt,Kap-alpha3,CG10425 | | 1.238e-02 | -4.39 | Rab2 (Rab2) | protein interactions | 35577 | 67 | 15 | 9204 | 1112 | CG1416,SERCA,CCT6,Gmap,Rab5,GCC88,ScsbetaA,Grip84,Mcad,lt,Echs1,Mitf,wal,CCT3,CG32485 | | 1.242e-02 | -4.39 | GSE24574\_BCL6\_LOW\_TFH\_VS\_TCONV\_CD4\_TCELL\_DN | MSigDB lists | GSE24574\_BCL6\_LOW\_TFH\_VS\_TCONV\_CD4\_TCELL\_DN | 98 | 24 | 4161 | 644 | levy,Karybeta3,Ts,eIF3i,CG5028,tko,Gapdh1,P5cr,Dhfr,RhoGAP54D,scu,AIF,r-l,CG14715,Tom40,Eno,Echs1,CCT6,Got1,Pfas,Mitofilin,Pcd,Prosalpha5,gammaTub37C | | 1.242e-02 | -4.39 | GSE17974\_CTRL\_VS\_ACT\_IL4\_AND\_ANTI\_IL12\_24H\_CD4\_TCELL\_DN | MSigDB lists | GSE17974\_CTRL\_VS\_ACT\_IL4\_AND\_ANTI\_IL12\_24H\_CD4\_TCELL\_DN | 98 | 24 | 4161 | 644 | CCT6,AIF,Caf1-105,Jafrac2,SRPK,esc,Cdk5,Sccpdh1,MED20,yip2,Prosbeta4,cl,CG4407,escl,CG10932,rept,Atg7,mbo,Stoml2,slif,colt,Rpn12,CG9393,mRpL18 | | 1.243e-02 | -4.39 | GDHRDH | prints domains | PR00081 | 47 | 10 | 2185 | 212 | CG3842,CG12171,CG31548,CG10425,CG9265,scu,Sptr,CG7322,sni,CG31549 | | 1.243e-02 | -4.39 | GO\_SINGLE\_ORGANISM\_BIOSYNTHETIC\_PROCESS | MSigDB lists | GO\_SINGLE\_ORGANISM\_BIOSYNTHETIC\_PROCESS | 546 | 103 | 4161 | 644 | SREBP,Pfas,Gyg,CG11134,Dph5,Cyp6a22,yip2,Eno,Pdxk,Ac3,mRpS5,Pglym78,nmdyn-D6,Dhfr,CG4390,mal,aay,mRpS23,alc,CG1640,RfC38,CG4511,Pi3K92E,RpL40,Synj,Tspo,ND-39,Sply,Scsalpha1,Got1,Ppox,CG3803,kri,GalT1,Cyp6a19,Pgk,CG9391,Shmt,mRpL18,CG11899,Pgm1,Mcad,Gapdh1,CG33156,mino,Prip,AMPdeam,Stoml2,GluRS-m,CG10932,CG4407,mRpL10,Ppcdc,mRpL17,Gabat,CG10425,r-l,Coq7,CG7322,PIG-C,CG3376,CG2201,CG12018,P5cr,P5CDh1,CG7718,ScsbetaA,CG6767,oxt,AGBE,r,Ts,nSMase,NHP2,CG9485,sgll,tko,scramb1,Adi1,CG7840,Desat1,mRpS21,Pi4KIIalpha,GatA,RpS27A,PAPLA1,mRpS10,crq,AcCoAS,mEFTu1,mRpL16,CG7115,csw,Nadsyn,LysRS,mRpL13,nmdyn-D7,Coq6,PIG-L,CG11089,Gk2,ACC,regucalcin | | 1.245e-02 | -4.39 | GSE360\_DC\_VS\_MAC\_DN | MSigDB lists | GSE360\_DC\_VS\_MAC\_DN | 58 | 16 | 4161 | 644 | RpS24,RpL23,AnxB10,nrv1,NHP2,Jafrac1,CG5734,Pex12,Prosbeta4,pont,CG7997,GlyP,Jafrac2,CG3376,ND-30,msps | | 1.245e-02 | -4.39 | GSE18804\_SPLEEN\_MACROPHAGE\_VS\_COLON\_TUMORAL\_MACROPHAGE\_DN | MSigDB lists | GSE18804\_SPLEEN\_MACROPHAGE\_VS\_COLON\_TUMORAL\_MACROPHAGE\_DN | 58 | 16 | 4161 | 644 | Karybeta3,cyc,CG4407,CG7115,Pgm1,CG8728,CG1847,Pp2B-14D,alph,GlyP,AIF,Mitofilin,CanA-14F,Unr,Smox,Lsm12 | | 1.245e-02 | -4.39 | AHR\_Q5 | MSigDB lists | AHR\_Q5 | 58 | 16 | 4161 | 644 | Su(H),escl,CG7550,AMPdeam,Sfxn2,eIF4A,esc,mAChR-A,Jafrac2,LpR2,gammaTub37C,Abi,Prosalpha5,Mitofilin,Pka-R2,Rab10 | | 1.245e-02 | -4.39 | GO\_CYTOKINE\_MEDIATED\_SIGNALING\_PATHWAY | MSigDB lists | GO\_CYTOKINE\_MEDIATED\_SIGNALING\_PATHWAY | 58 | 16 | 4161 | 644 | Med,RpL40,Prosbeta6,Rpn12,Traf6,Rpn2,Prosbeta3,Rpt2,Prosalpha4,Rpn3,Prosbeta5,RpS27A,Prosalpha5,Prosalpha3,cl,Prosbeta4 | | 1.247e-02 | -4.38 | - | gene3d domains | 3.40.50.1000 | 53 | 13 | 4929 | 622 | Ublcp1,Atpalpha,CG31729,CG32549,anne,ATP8A,Dd,CG17294,Gs1l,CG9981,aay,SERCA,cN-IIIB | | 1.248e-02 | -4.38 | GO\_HYDROLASE\_ACTIVITY\_ACTING\_ON\_ESTER\_BONDS | MSigDB lists | GO\_HYDROLASE\_ACTIVITY\_ACTING\_ON\_ESTER\_BONDS | 250 | 52 | 4161 | 644 | Rrp40,trsn,CG7115,mts,CG11164,csw,Ublcp1,dnc,regucalcin,tzn,twin,CG15111,CG1774,CG9391,ERp60,CG16979,Ptpmeg,l(1)G0196,Ptp10D,Cyt-c-p,CG4390,aay,Pp2B-14D,Edc3,CG17294,CG5285,CG31759,Dd,Synj,PrBP,CG5966,CanB2,mus81,PpV,Ptp61F,nSMase,CG17333,Acph-1,CG32549,Pp2A-29B,PpD3,sws,DNApol-epsilon255,CanA-14F,Dcr-1,CG7789,CG9248,Exd2,CG3376,Pglym78,alph,mAChR-A | | 1.250e-02 | -4.38 | chromosome | cellular component | GO:0005694 | 696 | 82 | 11940 | 1105 | d4,Su(H),cbs,egg,mbo,nbs,Sirt7,Tlk,l(3)mbt,lwr,Bin1,RpS18,lid,lin-52,Uxt,DNApol-epsilon255,RpL11,Mat89Ba,CG10336,Elba2,RecQ5,RpS15,Zwilch,RpS13,Topors,RpS2,pont,Sld5,rept,Utx,Klp10A,Moe,piwi,mre11,Cenp-C,mad2,wapl,cnc,Snm1,MAGE,CycB,EcR,G9a,CG31053,trr,BigH1,asf1,Rbf2,HP1c,Ubc6,Fkbp39,Rel,nej,Nap1,tef,Nup62,Bap170,HmgD,EloC,Rcd5,Chrac-14,Tip60,CG2129,SuUR,RfC38,esc,CG12018,Orc1,elg1,yem,msps,HP1e,Hat1,Orc6,pps,Rbf,Trf2,stil,Nurf-38,Psf3,ph-d,Top1 | | 1.250e-02 | -4.38 | Tyr\_Pase\_cat | interpro domains | IPR003595 | 19 | 6 | 11383 | 1233 | Ptp61F,csw,Ptpmeg,Ptp10D,Ptpmeg2,Ptp69D | | 1.250e-02 | -4.38 | LSM\_dom\_sf | interpro domains | IPR010920 | 19 | 6 | 11383 | 1233 | tral,LSm3,CG10418,CG17768,SmE,CG6610 | | 1.252e-02 | -4.38 | p53 binding | molecular function | GO:0002039 | 6 | 3 | 12176 | 1119 | Ubc6,ASPP,sip3 | | 1.252e-02 | -4.38 | imaginal disc growth factor receptor binding | molecular function | GO:0008084 | 6 | 3 | 12176 | 1119 | Idgf4,Idgf2,Idgf6 | | 1.254e-02 | -4.38 | Sxl (Sex lethal) | protein interactions | 3772180 | 111 | 22 | 9204 | 1112 | ND-49,bam,nos,CG11164,Mcad,pps,CycB,Eno,eIF4E1,Nup58,ovo,otu,CkIIalpha,NHP2,26-29-p,CG9485,tral,RpS14a,Stat92E,Unr,Kyat,hoe1 | | 1.257e-02 | -4.38 | Prosalpha5 (Proteasome alpha5 subunit) | protein interactions | 36951 | 38 | 10 | 9204 | 1112 | Rpn2,Pomp,Rpn13,Prosalpha3,Prosbeta3,Prosbeta6,Prosbeta4,Rpt2,Prosalpha4,Prosbeta5 | | 1.257e-02 | -4.38 | Ubi-p63E (Ubiquitin-63E) | protein interactions | 38456 | 38 | 10 | 9204 | 1112 | Sgf11,otu,Tak1,ago,not,Rbf,CG9186,Marf,lwr,bam | | 1.268e-02 | -4.37 | REACTOME\_DEADENYLATION\_DEPENDENT\_MRNA\_DECAY | MSigDB lists | REACTOME\_DEADENYLATION\_DEPENDENT\_MRNA\_DECAY | 26 | 9 | 4161 | 644 | CG10418,LSm3,CG8963,eIF4A,Rrp40,eIF4E1,CG6610,Edc3,Not10 | | 1.268e-02 | -4.37 | GO\_ENERGY\_RESERVE\_METABOLIC\_PROCESS | MSigDB lists | GO\_ENERGY\_RESERVE\_METABOLIC\_PROCESS | 26 | 9 | 4161 | 644 | Pfk,CG9485,CG8475,GlyP,AGBE,RpL40,Pgm1,Gyg,RpS27A | | 1.268e-02 | -4.37 | REACTOME\_ACTIVATED\_TLR4\_SIGNALLING | MSigDB lists | REACTOME\_ACTIVATED\_TLR4\_SIGNALLING | 26 | 9 | 4161 | 644 | mts,Pp2A-29B,RpS27A,Traf6,rl,cact,Cdk1,p38a,RpL40 | | 1.270e-02 | -4.37 | GO\_ESTABLISHMENT\_OF\_LOCALIZATION\_IN\_CELL | MSigDB lists | GO\_ESTABLISHMENT\_OF\_LOCALIZATION\_IN\_CELL | 642 | 119 | 4161 | 644 | Sec23,RpL11,RpL26,Clc,Vamp7,RpS27,yip2,RpS18,CG8004,thoc6,Cdk5,CycB,Kap-alpha3,CG8314,RpL12,sbr,Marf,CG1847,RpL23,RpS24,GAPsec,colt,Sec10,Pex13,Aladin,lt,Kap3,RpS16,RpL38,RpS4,Pi3K92E,Rep,RpS7,RpL40,Synj,Tspo,shrb,Rab10,Sec22,RpS23,rl,cact,Klp10A,Tom40,milt,alphaTub84B,mRpL18,CG2100,Vsp37A,sau,CG9393,Sod1,HDAC6,Stoml2,Karybeta3,AP-1-2beta,CG42542,CHOp24,GCC88,Arts,CG5734,Stam,Pex12,RpL37a,CG4293,CanA-14F,Mat89Ba,Syx13,Iru,RpL24,CG5510,CHMP2B,Tim8,CG46338,Pex10,CG8728,RpL30,Pp2B-14D,CG11110,Phb2,Oseg1,RpS2,Atg6,PpV,Ranbp21,tho2,CanB2,Rab5,ATP8A,Snx6,mbo,Cbp80,Cyp1,Atg4b,Npc2a,Rab32,RpS27A,RpL9,Nmt,RpS28b,Faf2,Surf4,Trs33,Pen,ERp60,RpL13,Calx,RpL13A,eIF4E1,Cse1,RpS30,RpS13,RpLP0,Arl6IP1,cyc,ACC,CG31729,CG4041,ste14 | | 1.275e-02 | -4.36 | GSE36009\_WT\_VS\_NLRP10\_KO\_DC\_LPS\_STIM\_DN | MSigDB lists | GSE36009\_WT\_VS\_NLRP10\_KO\_DC\_LPS\_STIM\_DN | 63 | 17 | 4161 | 644 | nmd,Ufm1,Etf-QO,cyc,CkIIalpha,eIF4A,Ptp10D,RpL13A,Sec10,ND-75,alpha-Man-Ia,RpL13,sbr,CCT6,Pka-R2,APP-BP1,Rnmt | | 1.275e-02 | -4.36 | response to stimulus | biological process | GO:0050896 | 2655 | 273 | 12345 | 1129 | rl,Jafrac1,Dcr-1,cathD,Cdk5,PGRP-LE,capt,Rep,Pka-R1,tkv,CG5285,EcR,bif,Ptp61F,Sec15,gus,PDCD-5,CG10336,Traf6,slif,rig,mAChR-A,Kap-alpha3,Bx,PHGPx,raptor,mbo,UGP,Myd88,tko,Herc4,Chd64,pen-2,RecQ5,Idgf6,p38a,Npc2a,E2f1,rpk,CG17691,ben,Iml1,lt,ATPsynD,Ufm1,Tao,pygo,Tak1,tor,tos,CG14408,cnc,Cdk1,Sdhaf3,CG5844,msps,AcCoAS,CG6766,velo,Abi,fray,mus304,Seipin,Su(H),Arpc4,CG4476,Atf6,Calx,Fkbp39,boi,Pi3K92E,Pp2B-14D,Mtl,Trf2,CG5412,cact,crq,tzn,Pfk,Ir41a,Ugt36E1,Pink1,Xpac,Cyp6v1,Idh3b,Imp,Parp16,Pdk1,trr,tho2,RhoGAP54D,nAChRalpha4,Ggt-1,Ptpmeg,cold,REPTOR,Med,MAGE,Pen,CanA-14F,CG4390,disp,ACC,bun,GlyP,fs(1)M3,CG9925,Pxt,alpha-Man-Ia,CG17272,cu,lwr,shep,Hsp26,fz4,apt,dco,CG46338,Pdxk,NFAT,Marf,Mekk1,CG11070,Hsc70-3,CG32640,CG17746,CG9391,mts,mus81,Jafrac2,Vps16B,ND-SGDH,csw,Atf3,Hat1,Gs1l,RhoGDI,Ho,Ptp10D,Atg6,CkIIalpha,Nprl2,spen,nej,faf,Edem1,Gdi,Fdh,mol,Debcl,CG13933,fs(1)N,Hnf4,PyK,chrb,EMRE,agt,nbs,MED27,Ac3,mre11,Mo25,Pld,Rbf,Wnt10,alph,EMC3,trk,Crag,Arf51F,Grx1,CG17698,Pka-R2,Agpat2,Rab5,Sod1,MED24,park,DNApol-epsilon255,trsn,glob1,Hr78,Stat92E,Rab10,Moe,dnc,TyrRS,CG7115,Fer1HCH,Desat1,G9a,aay,ZnT35C,SNF4Agamma,whd,Trxr-1,kcc,Ubc6,Xpc,Nup358,grp,mus205,S2P,CG17734,Utx,Sod3,CG9005,CG6927,CG7328,AIF,adp,rept,obe,Tm1,tub,bcd,CG3894,CG8974,spz,Droj2,not,dl,foxo,Atpalpha,O-fut1,Lkb1,Cka,Tlk,Tom40,sbb,sgll,Chrac-14,Tfb4,Adh,Mtpalpha,Gclc,Rel,ERp60,Stam,thoc6,Nurf-38,Mitf,Cbl,PEK,tyf,Snm1,Rab32,Shmt,Fkbp59,Ect4,CG30010,osk,zyd,Uev1A,CG13551,Smox,ago,CG11590,MtnA,SREBP,alphaTub84B,sip3,cyc,CG11089,Aldh,Atg7,Idh,Sp7,r2d2,Idgf4,Sld5,Ptp69D,SelR | | 1.277e-02 | -4.36 | gamma-tubulin ring complex | cellular component | GO:0008274 | 6 | 3 | 11940 | 1105 | Grip75,Grip128,Grip71 | | 1.277e-02 | -4.36 | gamma-tubulin large complex | cellular component | GO:0000931 | 6 | 3 | 11940 | 1105 | Grip75,Grip71,Grip128 | | 1.280e-02 | -4.36 | positive regulation of molecular function | biological process | GO:0044093 | 287 | 38 | 12345 | 1129 | Tak1,Parp16,Pdk1,esc,crok,Traf6,Jafrac2,GAPsec,cort,RhoGAP54D,Lkb1,ASPP,Sec23,Crag,fray,S2P,Abi,Rpn13,eIF3m,raptor,CG13344,nrv1,RpLP1,rl,CG1416,MED24,ScsbetaA,Pi3K92E,lwr,SERCA,Tomosyn,RhoGAP5A,Aos1,CG4041,Iml1,Mekk1,Orc6,Tao | | 1.282e-02 | -4.36 | ENTH\_VHS | interpro domains | IPR008942 | 14 | 5 | 11383 | 1233 | lqfR,Gga,Stam,Hip1,CG4266 | | 1.293e-02 | -4.35 | GSE17721\_CTRL\_VS\_PAM3CSK4\_12H\_BMDC\_DN | MSigDB lists | GSE17721\_CTRL\_VS\_PAM3CSK4\_12H\_BMDC\_DN | 68 | 18 | 4161 | 644 | TyrRS,Tsp66E,mino,CG4390,ste24a,AGBE,Imp,l(1)G0004,PrBP,Cyp1,Gk2,Prosalpha3,Prosbeta4,Pp2A-29B,CG3587,Rab32,Pus7,CG16979 | | 1.293e-02 | -4.35 | GSE13411\_NAIVE\_VS\_SWITCHED\_MEMORY\_BCELL\_DN | MSigDB lists | GSE13411\_NAIVE\_VS\_SWITCHED\_MEMORY\_BCELL\_DN | 68 | 18 | 4161 | 644 | CG8475,PpV,tzn,Iml1,CG8728,faf,Edc3,CG4709,CG10344,CG8199,Aldh,Pgk,mEFTu1,RpL13,ND-39,Sply,Utx,Hmg-2 | | 1.294e-02 | -4.35 | GSE37534\_UNTREATED\_VS\_GW1929\_TREATED\_CD4\_TCELL\_PPARG1\_AND\_FOXP3\_TRASDUCED\_DN | MSigDB lists | GSE37534\_UNTREATED\_VS\_GW1929\_TREATED\_CD4\_TCELL\_PPARG1\_AND\_FOXP3\_TRASDUCED\_DN | 83 | 21 | 4161 | 644 | Mctp,CG6610,Nurf-38,colt,Pp2B-14D,eIF4A,Rbf,not,Adi1,CG42542,SelD,RfC38,Hmg-2,PAPLA1,spag,Irp-1B,CanA-14F,CCT6,msps,Gclc,Cdk1 | | 1.294e-02 | -4.35 | GSE15930\_NAIVE\_VS\_72H\_IN\_VITRO\_STIM\_IFNAB\_CD8\_TCELL\_DN | MSigDB lists | GSE15930\_NAIVE\_VS\_72H\_IN\_VITRO\_STIM\_IFNAB\_CD8\_TCELL\_DN | 83 | 21 | 4161 | 644 | mRpL17,gammaTub37C,Cdk1,CG8004,CG5880,CG14715,mad2,Pglym78,msps,CG5026,CG3376,mRpL18,CG11164,CG11899,Pgm1,CG17294,aay,grp,CysRS,Su(H),ND-51 | | 1.301e-02 | -4.34 | PUJANA\_BRCA\_CENTERED\_NETWORK | MSigDB lists | PUJANA\_BRCA\_CENTERED\_NETWORK | 78 | 20 | 4161 | 644 | DNApol-epsilon255,Idh3b,MED20,asf1,Clp,APP-BP1,mad2,Top1,CG13630,Cdk1,RpII140,slif,CG6767,CG1890,Dhfr,CkIIalpha,CG11164,CG8204,RfC38,r | | 1.301e-02 | -4.34 | GSE2770\_IL12\_AND\_TGFB\_ACT\_VS\_ACT\_CD4\_TCELL\_6H\_DN | MSigDB lists | GSE2770\_IL12\_AND\_TGFB\_ACT\_VS\_ACT\_CD4\_TCELL\_6H\_DN | 78 | 20 | 4161 | 644 | Stip1,CCT3,Tom40,Pfdn2,CG10425,esc,bys,Chrac-14,Hsc70-3,Fkbp59,Prosalpha5,Roe1,Fur2,escl,ACC,l(1)G0004,CG1416,CG7718,CkIIalpha,LysRS | | 1.301e-02 | -4.34 | GO\_REGULATION\_OF\_LIPID\_METABOLIC\_PROCESS | MSigDB lists | GO\_REGULATION\_OF\_LIPID\_METABOLIC\_PROCESS | 78 | 20 | 4161 | 644 | Npc2a,SREBP,Idh,STUB1,crq,Rbf,CycC,Dd,CG13601,mino,EcR,Sod1,ABCD,adp,nSMase,Stoml2,Gk2,Tspo,regucalcin,Pdk | | 1.301e-02 | -4.34 | GSE3982\_MAST\_CELL\_VS\_BASOPHIL\_UP | MSigDB lists | GSE3982\_MAST\_CELL\_VS\_BASOPHIL\_UP | 73 | 19 | 4161 | 644 | anne,Eno,CCT3,CG8199,CG12773,Pka-R2,Dph5,Axs,CanB2,eIF3i,kl-2,CG17333,CG4511,Atg7,Uba5,Adi1,Cul2,Non1,CG4390 | | 1.301e-02 | -4.34 | GO\_ACTIVATION\_OF\_INNATE\_IMMUNE\_RESPONSE | MSigDB lists | GO\_ACTIVATION\_OF\_INNATE\_IMMUNE\_RESPONSE | 73 | 19 | 4161 | 644 | Prosbeta3,Prosalpha4,Rpt2,Rpn12,Traf6,Rpn2,SkpA,Prosbeta6,RpL40,Prosalpha3,Prosbeta4,Prosalpha5,RpS27A,Prosbeta5,Pdk1,cact,Uev1A,Rpn3,Cul1 | | 1.306e-02 | -4.34 | GO\_POSITIVE\_REGULATION\_OF\_INTERLEUKIN\_2\_PRODUCTION | MSigDB lists | GO\_POSITIVE\_REGULATION\_OF\_INTERLEUKIN\_2\_PRODUCTION | 4 | 3 | 4161 | 644 | dnc,Traf6,CG30496 | | 1.306e-02 | -4.34 | REACTOME\_TRAF6\_MEDIATED\_INDUCTION\_OF\_TAK1\_COMPLEX | MSigDB lists | REACTOME\_TRAF6\_MEDIATED\_INDUCTION\_OF\_TAK1\_COMPLEX | 4 | 3 | 4161 | 644 | RpS27A,RpL40,Traf6 | | 1.306e-02 | -4.34 | GO\_NEGATIVE\_REGULATION\_OF\_NITRIC\_OXIDE\_METABOLIC\_PROCESS | MSigDB lists | GO\_NEGATIVE\_REGULATION\_OF\_NITRIC\_OXIDE\_METABOLIC\_PROCESS | 4 | 3 | 4161 | 644 | regucalcin,Tspo,CG7997 | | 1.306e-02 | -4.34 | REACTOME\_P75NTR\_RECRUITS\_SIGNALLING\_COMPLEXES | MSigDB lists | REACTOME\_P75NTR\_RECRUITS\_SIGNALLING\_COMPLEXES | 4 | 3 | 4161 | 644 | RpS27A,Traf6,RpL40 | | 1.306e-02 | -4.34 | GO\_UTP\_METABOLIC\_PROCESS | MSigDB lists | GO\_UTP\_METABOLIC\_PROCESS | 4 | 3 | 4161 | 644 | nmdyn-D7,nmdyn-D6,r | | 1.306e-02 | -4.34 | GO\_ETHANOL\_METABOLIC\_PROCESS | MSigDB lists | GO\_ETHANOL\_METABOLIC\_PROCESS | 4 | 3 | 4161 | 644 | Fdh,AcCoAS,Aldh | | 1.306e-02 | -4.34 | SA\_G2\_AND\_M\_PHASES | MSigDB lists | SA\_G2\_AND\_M\_PHASES | 4 | 3 | 4161 | 644 | grp,Cdk1,Cdk7 | | 1.306e-02 | -4.34 | GO\_GABA\_RECEPTOR\_BINDING | MSigDB lists | GO\_GABA\_RECEPTOR\_BINDING | 4 | 3 | 4161 | 644 | milt,Maf1,mts | | 1.306e-02 | -4.34 | GO\_NEGATIVE\_REGULATION\_OF\_PROTEIN\_TYROSINE\_KINASE\_ACTIVITY | MSigDB lists | GO\_NEGATIVE\_REGULATION\_OF\_PROTEIN\_TYROSINE\_KINASE\_ACTIVITY | 4 | 3 | 4161 | 644 | Snx6,CG4709,Ptp61F | | 1.306e-02 | -4.34 | GO\_PSEUDOPODIUM | MSigDB lists | GO\_PSEUDOPODIUM | 4 | 3 | 4161 | 644 | Vamp7,Moe,rl | | 1.306e-02 | -4.34 | BANDRES\_RESPONSE\_TO\_CARMUSTIN\_WITHOUT\_MGMT\_48HR\_DN | MSigDB lists | BANDRES\_RESPONSE\_TO\_CARMUSTIN\_WITHOUT\_MGMT\_48HR\_DN | 4 | 3 | 4161 | 644 | Jheh2,Atpalpha,gammaTub37C | | 1.306e-02 | -4.34 | GO\_POSITIVE\_REGULATION\_OF\_TRANSCRIPTION\_FROM\_RNA\_POLYMERASE\_II\_PROMOTER\_IN\_RESPONSE\_TO\_STRESS | MSigDB lists | GO\_POSITIVE\_REGULATION\_OF\_TRANSCRIPTION\_FROM\_RNA\_POLYMERASE\_II\_PROMOTER\_IN\_RESPONSE\_TO\_STRESS | 4 | 3 | 4161 | 644 | Su(H),S2P,Hsc70-3 | | 1.306e-02 | -4.34 | GO\_NEGATIVE\_REGULATION\_OF\_LIPID\_STORAGE | MSigDB lists | GO\_NEGATIVE\_REGULATION\_OF\_LIPID\_STORAGE | 4 | 3 | 4161 | 644 | EcR,cact,Ptp61F | | 1.306e-02 | -4.34 | HALLMARK\_IL6\_JAK\_STAT3\_SIGNALING | MSigDB lists | HALLMARK\_IL6\_JAK\_STAT3\_SIGNALING | 4 | 3 | 4161 | 644 | Ptp61F,csw,Stam | | 1.310e-02 | -4.34 | Generic Transcription Pathway | REACTOME pathways | R-DME-212436 | 177 | 37 | 3923 | 574 | lid,raptor,Cdk7,twin,CycC,faf,alc,Not10,CycB,Rep,CG30109,Cdk1,Jafrac1,Rpb8,nrv1,Hr78,Hnf4,Tfb4,chrb,MED24,RpL40,MED4,MED20,CG6923,TfIIFalpha,alph,CG6707,EcR,Med,Rga,MED27,RpS27A,SNF4Agamma,Rbf2,Smox,Rbf,EloC | | 1.312e-02 | -4.33 | cnn (centrosomin) | protein interactions | 36491 | 105 | 21 | 9204 | 1112 | zf30C,Cen,CG4586,gammaTub37C,dx,mad2,mael,cnn,spd-2,Grip84,CG6927,MED4,alt,cup,Grip71,RhoGAP54D,apt,cact,bun,Grip75,wisp | | 1.326e-02 | -4.32 | GO\_OXIDOREDUCTASE\_ACTIVITY\_ACTING\_ON\_THE\_CH\_NH\_GROUP\_OF\_DONORS | MSigDB lists | GO\_OXIDOREDUCTASE\_ACTIVITY\_ACTING\_ON\_THE\_CH\_NH\_GROUP\_OF\_DONORS | 14 | 6 | 4161 | 644 | P5cr,CG4872,Dhfr,Etf-QO,CG9471,P5CDh1 | | 1.326e-02 | -4.32 | GO\_PYRIMIDINE\_NUCLEOSIDE\_BIOSYNTHETIC\_PROCESS | MSigDB lists | GO\_PYRIMIDINE\_NUCLEOSIDE\_BIOSYNTHETIC\_PROCESS | 14 | 6 | 4161 | 644 | r,Ts,r-l,kri,nmdyn-D6,nmdyn-D7 | | 1.326e-02 | -4.32 | GO\_NUCLEOSIDE\_TRIPHOSPHATE\_BIOSYNTHETIC\_PROCESS | MSigDB lists | GO\_NUCLEOSIDE\_TRIPHOSPHATE\_BIOSYNTHETIC\_PROCESS | 14 | 6 | 4161 | 644 | Ts,Stoml2,r,Scsalpha1,nmdyn-D7,nmdyn-D6 | | 1.326e-02 | -4.32 | GO\_POSITIVE\_REGULATION\_OF\_ATPASE\_ACTIVITY | MSigDB lists | GO\_POSITIVE\_REGULATION\_OF\_ATPASE\_ACTIVITY | 14 | 6 | 4161 | 644 | CG1416,shv,nrv1,scramb1,Tm1,regucalcin | | 1.326e-02 | -4.32 | MOOTHA\_TCA | MSigDB lists | MOOTHA\_TCA | 14 | 6 | 4161 | 644 | ScsbetaA,Idh3b,CG5028,Scsalpha1,Nc73EF,SdhA | | 1.326e-02 | -4.32 | GO\_UBIQUITIN\_LIKE\_PROTEIN\_CONJUGATING\_ENZYME\_ACTIVITY | MSigDB lists | GO\_UBIQUITIN\_LIKE\_PROTEIN\_CONJUGATING\_ENZYME\_ACTIVITY | 14 | 6 | 4161 | 644 | CG4502,CG40045,UbcE2H,lwr,CG8188,CG2924 | | 1.326e-02 | -4.32 | REACTOME\_TGF\_BETA\_RECEPTOR\_SIGNALING\_ACTIVATES\_SMADS | MSigDB lists | REACTOME\_TGF\_BETA\_RECEPTOR\_SIGNALING\_ACTIVATES\_SMADS | 14 | 6 | 4161 | 644 | RpS27A,Smox,Med,Uch-L5,RpL40,STUB1 | | 1.326e-02 | -4.32 | AGTCTTA\_MIR499 | MSigDB lists | AGTCTTA\_MIR499 | 14 | 6 | 4161 | 644 | CanA-14F,Uev1A,Top1,Rab5,Pp2B-14D,bru1 | | 1.329e-02 | -4.32 | DANG\_BOUND\_BY\_MYC | MSigDB lists | DANG\_BOUND\_BY\_MYC | 488 | 93 | 4161 | 644 | Gapdh1,CG3226,Mcad,Nurf-38,sau,AMPdeam,Sod1,HDAC6,ste24a,ND-SGDH,lid,CG17691,l(1)G0004,Unr,Scsalpha1,Sply,Prosalpha5,Aldh,kri,wal,CG3803,Shmt,mad2,alc,CG4390,GCS2beta,RpS16,Vha13,Prosbeta6,Ubc4,PrBP,RpL40,Wnt10,RpL26,Hmg-2,gammaTub37C,Pdk1,CG14715,CycB,Gclc,CCT5,Eno,Rpn3,Moe,Thiolase,muskelin,mts,slif,RpS13,Rpn12,RpL13A,eIF4E1,eIF3i,CG31715,CG4562,RpL9,RpS27A,Jheh2,Nap1,Idh3b,AIF,AcCoAS,ND-75,CG7997,Bin1,Cul5,STUB1,lqfR,Chrac-14,Top1,RpL13,CycC,P5cr,CG12018,Vha36-1,Phb2,Pp2B-14D,Nprl2,Ts,PpV,r,Pdk,Aos1,mbo,Cbp80,Rcd5,CanA-14F,mRpL10,RpL37a,Smox,Scox,Prosbeta5,Arpc4 | | 1.329e-02 | -4.32 | phospholipid metabolic process | biological process | GO:0006644 | 146 | 22 | 12345 | 1129 | Synj,PIG-P,CG5026,sws,CG2201,CG7718,Pi4KIIalpha,CG18135,Agpat2,Seipin,Pld,Pi3K92E,CG9391,mino,PIG-L,CG7789,PIG-C,CG33977,CG8311,CG6707,nSMase,PIG-H | | 1.341e-02 | -4.31 | SWEET\_KRAS\_ONCOGENIC\_SIGNATURE | MSigDB lists | SWEET\_KRAS\_ONCOGENIC\_SIGNATURE | 35 | 11 | 4161 | 644 | RpS2,Phb2,Nmt,Pcd,Npc2a,Gmer,mbo,eEF1beta,Pdk,CG14715,rl | | 1.341e-02 | -4.31 | EIF4E\_UP | MSigDB lists | EIF4E\_UP | 35 | 11 | 4161 | 644 | l(1)G0004,mEFTu1,CG1416,pr,CG8004,Pfdn2,YL-1,mRpL17,Idh,Uch,Pmi | | 1.341e-02 | -4.31 | REACTOME\_CROSS\_PRESENTATION\_OF\_SOLUBLE\_EXOGENOUS\_ANTIGENS\_ENDOSOMES | MSigDB lists | REACTOME\_CROSS\_PRESENTATION\_OF\_SOLUBLE\_EXOGENOUS\_ANTIGENS\_ENDOSOMES | 35 | 11 | 4161 | 644 | Prosalpha5,Rpn2,Rpn12,Prosalpha4,Prosbeta4,Rpt2,Prosalpha3,Prosbeta3,Rpn3,Prosbeta6,Prosbeta5 | | 1.341e-02 | -4.31 | CHEN\_HOXA5\_TARGETS\_9HR\_UP | MSigDB lists | CHEN\_HOXA5\_TARGETS\_9HR\_UP | 35 | 11 | 4161 | 644 | Ptp61F,cact,CysRS,cyc,Cdk7,CG7718,ari-1,Mo25,CG3511,Cka,Tlk | | 1.342e-02 | -4.31 | MORF\_RAD23B | MSigDB lists | MORF\_RAD23B | 109 | 26 | 4161 | 644 | SdhA,Klp10A,Prosbeta5,ND-75,wal,Pen,Shmt,Hat1,Prosbeta4,LSm3,Pp2A-29B,YL-1,COX5A,Stoml2,PpV,SkpA,Pka-R1,CHOp24,CG11070,Rpt2,CG3631,LysRS,Cyt-c-p,CG1890,Rpn12,Sod1 | | 1.342e-02 | -4.31 | GO\_SIGNAL\_TRANSDUCTION\_BY\_PROTEIN\_PHOSPHORYLATION | MSigDB lists | GO\_SIGNAL\_TRANSDUCTION\_BY\_PROTEIN\_PHOSPHORYLATION | 109 | 26 | 4161 | 644 | rl,p38a,Prosbeta5,Cul1,Rpn3,Prosbeta4,Prosalpha3,Mo25,PpD3,Prosalpha5,Mekk1,RpS27A,tkv,SkpA,Prosbeta6,fray,RpL40,Tao,Prosalpha4,CG7530,Rpt2,CG7115,Prosbeta3,Rpn2,Rpn12,Traf6 | | 1.343e-02 | -4.31 | negative regulation of transcription, DNA-templated | biological process | GO:0045892 | 371 | 47 | 12345 | 1129 | E2f1,ovo,asf1,l(3)mbt,insv,Maf1,mael,Rbf2,wash,Sox102F,egg,rept,CG10959,lin-52,Bin1,Tip60,CG10038,ph-d,Elba2,CG1602,CG17328,pgc,Hnf4,EcR,CG10631,cyc,d4,Tlk,piwi,nos,hb,SREBP,foxo,Rbf,dl,esc,CG2199,r2d2,psq,Sirt7,Su(H),CG4872,sbb,spn-E,Hat1,escl,Chrac-14 | | 1.346e-02 | -4.31 | YTTCCNNNGGAMR\_UNKNOWN | MSigDB lists | YTTCCNNNGGAMR\_UNKNOWN | 22 | 8 | 4161 | 644 | Smox,CG5285,CG2493,Ssdp,Tao,Arts,mEFTu1,AP-1-2beta | | 1.346e-02 | -4.31 | GO\_REGULATION\_OF\_INTRACELLULAR\_STEROID\_HORMONE\_RECEPTOR\_SIGNALING\_PATHWAY | MSigDB lists | GO\_REGULATION\_OF\_INTRACELLULAR\_STEROID\_HORMONE\_RECEPTOR\_SIGNALING\_PATHWAY | 22 | 8 | 4161 | 644 | HDAC6,Cka,PpD3,Phb2,Ufm1,CG16979,cyc,Uba5 | | 1.346e-02 | -4.31 | REACTOME\_SIGNALING\_BY\_TGF\_BETA\_RECEPTOR\_COMPLEX | MSigDB lists | REACTOME\_SIGNALING\_BY\_TGF\_BETA\_RECEPTOR\_COMPLEX | 22 | 8 | 4161 | 644 | CycC,faf,RpS27A,Smox,STUB1,RpL40,Med,Uch-L5 | | 1.346e-02 | -4.31 | Prosbeta4 (Proteasome beta4 subunit) | protein interactions | 34999 | 44 | 11 | 9204 | 1112 | Rpn2,Prosalpha5,Pomp,Rpn13,Prosalpha3,Prosbeta6,Prosbeta3,Rpt2,jigr1,Prosalpha4,Prosbeta5 | | 1.346e-02 | -4.31 | dgt6 (dim gamma-tubulin 6) | protein interactions | 43441 | 44 | 11 | 9204 | 1112 | gammaTub37C,RpL40,COX5B,Grip71,Atpalpha,msps,dgt4,ATPsynD,Pxt,cnc,vers | | 1.346e-02 | -4.31 | sqd (squid) | protein interactions | 41666 | 44 | 11 | 9204 | 1112 | cup,Imp,bru1,Pdk1,foxo,Rtnl1,Unr,Rbf,otu,bcd,osk | | 1.346e-02 | -4.31 | BIOCARTA\_NTHI\_PATHWAY | MSigDB lists | BIOCARTA\_NTHI\_PATHWAY | 7 | 4 | 4161 | 644 | cact,p38a,Med,Smox | | 1.346e-02 | -4.31 | CACGTTT\_MIR302A | MSigDB lists | CACGTTT\_MIR302A | 7 | 4 | 4161 | 644 | Cul2,Uch-L5,Imp,CG1309 | | 1.346e-02 | -4.31 | CHEN\_LVAD\_SUPPORT\_OF\_FAILING\_HEART\_DN | MSigDB lists | CHEN\_LVAD\_SUPPORT\_OF\_FAILING\_HEART\_DN | 7 | 4 | 4161 | 644 | Idh,CG4872,dnc,Atpalpha | | 1.346e-02 | -4.31 | WANG\_ADIPOGENIC\_GENES\_REPRESSED\_BY\_SIRT1 | MSigDB lists | WANG\_ADIPOGENIC\_GENES\_REPRESSED\_BY\_SIRT1 | 7 | 4 | 4161 | 644 | Pgk,Gk2,Gapdh1,Eno | | 1.346e-02 | -4.31 | GO\_RESPONSE\_TO\_EPIDERMAL\_GROWTH\_FACTOR | MSigDB lists | GO\_RESPONSE\_TO\_EPIDERMAL\_GROWTH\_FACTOR | 7 | 4 | 4161 | 644 | Atg6,rl,Pdk1,r | | 1.346e-02 | -4.31 | BIOCARTA\_CERAMIDE\_PATHWAY | MSigDB lists | BIOCARTA\_CERAMIDE\_PATHWAY | 7 | 4 | 4161 | 644 | AIF,rl,CG3376,Cyt-c-p | | 1.346e-02 | -4.31 | GO\_MITOCHONDRIAL\_ELECTRON\_TRANSPORT\_CYTOCHROME\_C\_TO\_OXYGEN | MSigDB lists | GO\_MITOCHONDRIAL\_ELECTRON\_TRANSPORT\_CYTOCHROME\_C\_TO\_OXYGEN | 7 | 4 | 4161 | 644 | COX5A,Cyt-c-p,levy,CG3803 | | 1.346e-02 | -4.31 | BIOCARTA\_AKAP95\_PATHWAY | MSigDB lists | BIOCARTA\_AKAP95\_PATHWAY | 7 | 4 | 4161 | 644 | mts,CycB,Pka-R2,Cdk1 | | 1.351e-02 | -4.30 | Ribosome | KEGG pathways | ko03010 | 240 | 50 | 3198 | 496 | mRpL35,RpL23A,RpL24,RpS14b,RpL23,RpS27A,RpL11,RpS4,RpS23,mRpS10,mRpS21,RpS29,mRpL17,RpLP1,RpL34a,RpL13A,RpS30,RpS7,RpLP0,RpL40,RpL10,RpS28b,RpL12,RpS5b,RpS16,RpS13,RpS27,RpL38,mRpS11,RpS14a,RpS24,mRpL10,RpL17,RpL37a,mRpL36,RpL26,RpL31,RpL30,tko,mRpL18,mRpL16,RpS2,RpL7A,mRpL13,RpL13,RpS18,RpL29,RpS15,RpL9,RpL34b | | 1.351e-02 | -4.30 | Ribosome | KEGG pathways | dme03010 | 240 | 50 | 3198 | 496 | RpS13,RpL38,RpS27,mRpS11,RpS14a,mRpL10,RpS24,RpL17,RpL37a,mRpL36,RpL26,RpL31,RpL30,tko,mRpL18,mRpL16,RpS2,RpL7A,RpL13,mRpL13,RpS18,RpL29,RpS15,RpL34b,RpL9,mRpL35,RpL24,RpL23A,RpS14b,RpL23,RpS27A,RpS4,RpL11,RpS23,mRpS10,mRpS21,RpS29,RpLP1,mRpL17,RpL34a,RpS30,RpL13A,RpS7,RpLP0,RpL40,RpL10,RpS28b,RpL12,RpS5b,RpS16 | | 1.354e-02 | -4.30 | Golgi apparatus part | cellular component | GO:0044431 | 196 | 28 | 11940 | 1105 | CG5447,CG4293,Clc,lqfR,Sec23,Arl5,CG9773,Gga,Fur2,Pi4KIIalpha,stet,GCC88,Efr,Sec22,Afti,Trs33,CG9981,Gmap,sau,CG5510,Rab32,Ugalt,CG31729,opm,alpha-Man-Ia,AP-1-2beta,ATP8A,cbs | | 1.354e-02 | -4.30 | Lsm 2-8 complex | KEGG pathways | M00396 | 7 | 4 | 3198 | 496 | LSm3,CG6610,CG10418,CG17768 | | 1.354e-02 | -4.30 | Lsm 1-7 complex | KEGG pathways | M00397 | 7 | 4 | 3198 | 496 | CG10418,CG17768,CG6610,LSm3 | | 1.354e-02 | -4.30 | Lsm 2-8 complex | KEGG pathways | dme\_M00396 | 7 | 4 | 3198 | 496 | LSm3,CG10418,CG17768,CG6610 | | 1.354e-02 | -4.30 | Lsm 1-7 complex | KEGG pathways | dme\_M00397 | 7 | 4 | 3198 | 496 | CG6610,CG17768,CG10418,LSm3 | | 1.358e-02 | -4.30 | regulation of reproductive process | biological process | GO:2000241 | 121 | 19 | 12345 | 1129 | mael,CG31053,Ptpmeg,E2f1,swa,cnc,Adi1,Cbl,Nprl2,exu,cup,Mos,opm,twe,enc,CHOp24,Iml1,mtrm,spn-E | | 1.360e-02 | -4.30 | thioester biosynthetic process | biological process | GO:0035384 | 11 | 4 | 12345 | 1129 | CG17691,CG8199,CG6178,AcCoAS | | 1.360e-02 | -4.30 | cellular oxidant detoxification | biological process | GO:0098869 | 11 | 4 | 12345 | 1129 | Trxr-1,Sod1,Jafrac1,Sod3 | | 1.360e-02 | -4.30 | tricarboxylic acid metabolic process | biological process | GO:0072350 | 11 | 4 | 12345 | 1129 | Irp-1B,Idh,CG5028,Idh3b | | 1.360e-02 | -4.30 | positive regulation of antifungal peptide production | biological process | GO:0002804 | 11 | 4 | 12345 | 1129 | spz,tub,Myd88,Rel | | 1.360e-02 | -4.30 | acyl-CoA biosynthetic process | biological process | GO:0071616 | 11 | 4 | 12345 | 1129 | CG6178,AcCoAS,CG17691,CG8199 | | 1.360e-02 | -4.30 | growth of a germarium-derived egg chamber | biological process | GO:0007295 | 11 | 4 | 12345 | 1129 | Cul5,cup,Rab10,Cul2 | | 1.360e-02 | -4.30 | Arp2/3 complex-mediated actin nucleation | biological process | GO:0034314 | 11 | 4 | 12345 | 1129 | Arpc2,Arpc4,wash,Arpc1 | | 1.360e-02 | -4.30 | oocyte nucleus localization involved in oocyte dorsal/ventral axis specification | biological process | GO:0051663 | 11 | 4 | 12345 | 1129 | gammaTub37C,mael,jvl,cnc | | 1.360e-02 | -4.30 | G2/M transition of mitotic cell cycle | biological process | GO:0000086 | 11 | 4 | 12345 | 1129 | CG5003,Cdk1,twe,CycB | | 1.364e-02 | -4.29 | pyrimidine-containing compound metabolic process | biological process | GO:0072527 | 29 | 7 | 12345 | 1129 | r-l,awd,Ts,nmdyn-D7,nmdyn-D6,r,l(2)k01209 | | 1.369e-02 | -4.29 | kinase binding | molecular function | GO:0019900 | 96 | 16 | 12176 | 1119 | rl,Mitofilin,alc,sws,Tm1,SNF4Agamma,Wdfy2,tub,Mekk1,CycB,CG11807,CycE,CycJ,Parp16,Mtl,Cbl | | 1.372e-02 | -4.29 | establishment or maintenance of cytoskeleton polarity | biological process | GO:0030952 | 50 | 10 | 12345 | 1129 | mael,Pka-R1,Lkb1,Kap3,cnc,capt,heph,jvl,osk,msps | | 1.376e-02 | -4.29 | GO\_REGULATION\_OF\_WNT\_SIGNALING\_PATHWAY | MSigDB lists | GO\_REGULATION\_OF\_WNT\_SIGNALING\_PATHWAY | 104 | 25 | 4161 | 644 | Smox,Prosalpha5,Pp2A-29B,RpS27A,Prosalpha3,Prosbeta4,Wnt10,Rpn3,Prosbeta5,p38a,Rpn12,dco,Rpn2,Pfdn5,CkIIalpha,mts,Prosbeta3,trbd,Prosalpha4,Rpt2,Dd,Su(H),RpL40,cyc,Prosbeta6 | | 1.376e-02 | -4.29 | GSE37416\_0H\_VS\_6H\_F\_TULARENSIS\_LVS\_NEUTROPHIL\_DN | MSigDB lists | GSE37416\_0H\_VS\_6H\_F\_TULARENSIS\_LVS\_NEUTROPHIL\_DN | 49 | 14 | 4161 | 644 | Su(H),CG4332,milt,Rab5,Eno,anne,CG14715,cact,fray,Sld5,stas,Ppcdc,Gapdh1,shrb | | 1.376e-02 | -4.29 | MORF\_UNG | MSigDB lists | MORF\_UNG | 49 | 14 | 4161 | 644 | Prosbeta4,Sod1,LysRS,Cyt-c-p,CCT3,NHP2,CCT5,Chrac-14,Pen,Hat1,CG1416,CG11089,mEFTu1,ND-30 | | 1.376e-02 | -4.29 | GO\_NUCLEAR\_IMPORT | MSigDB lists | GO\_NUCLEAR\_IMPORT | 49 | 14 | 4161 | 644 | Pen,cyc,ERp60,Kap-alpha3,Arts,mbo,rl,cact,Karybeta3,Pp2B-14D,Phb2,Cse1,RpL23,CanA-14F | | 1.376e-02 | -4.29 | KEGG\_WNT\_SIGNALING\_PATHWAY | MSigDB lists | KEGG\_WNT\_SIGNALING\_PATHWAY | 49 | 14 | 4161 | 644 | Wnt10,CkIIalpha,mts,CG3226,CanA-14F,Pp2A-29B,Pp2B-14D,dco,Smox,SkpA,pont,CanB2,Cul1,Med | | 1.377e-02 | -4.28 | Transcriptional activity of SMAD2/SMAD3:SMAD4 heterotrimer | REACTOME pathways | R-DME-2173793 | 37 | 11 | 3923 | 574 | CG6923,Rbf,Smox,alph,RpL40,RpS27A,nrv1,CycC,Med,Rbf2,faf | | 1.378e-02 | -4.28 | cellular response to endogenous stimulus | biological process | GO:0071495 | 181 | 26 | 12345 | 1129 | foxo,lwr,Pi3K92E,Pdk1,Nurf-38,Fkbp39,rl,Shmt,tkv,Nup358,mAChR-A,Hr78,Vps16B,PyK,EcR,Hnf4,Med,csw,Rab10,Chd64,bun,MED27,Smox,G9a,raptor,Utx | | 1.378e-02 | -4.28 | ribosomal small subunit biogenesis | biological process | GO:0042274 | 65 | 12 | 12345 | 1129 | RpS14b,RpS7,mRpS11,RpS5b,CG7488,RpS16,RpS14a,CG7728,RpS28b,CG12288,RpS15,RpS27 | | 1.378e-02 | -4.28 | ELVIDGE\_HIF1A\_TARGETS\_DN | MSigDB lists | ELVIDGE\_HIF1A\_TARGETS\_DN | 18 | 7 | 4161 | 644 | AGBE,Pgm1,GalT1,LpR2,Pgk,Su(H),GatA | | 1.378e-02 | -4.28 | NATSUME\_RESPONSE\_TO\_INTERFERON\_BETA\_DN | MSigDB lists | NATSUME\_RESPONSE\_TO\_INTERFERON\_BETA\_DN | 18 | 7 | 4161 | 644 | mts,Gapdh1,SkpA,cact,levy,RpS2,Sod1 | | 1.378e-02 | -4.28 | GO\_POSITIVE\_REGULATION\_OF\_LIPID\_BIOSYNTHETIC\_PROCESS | MSigDB lists | GO\_POSITIVE\_REGULATION\_OF\_LIPID\_BIOSYNTHETIC\_PROCESS | 18 | 7 | 4161 | 644 | EcR,mino,regucalcin,SREBP,Dd,crq,nSMase | | 1.378e-02 | -4.28 | MODULE\_350 | MSigDB lists | MODULE\_350 | 18 | 7 | 4161 | 644 | tko,ste24a,EcR,Iru,CG9248,CG6028,SREBP | | 1.383e-02 | -4.28 | nuclear import signal receptor activity | molecular function | GO:0061608 | 11 | 4 | 12176 | 1119 | Karybeta3,Kap-alpha3,Pen,Arts | | 1.383e-02 | -4.28 | fly chr46B4-46B4\|2-61 cM | chromosome location | fly chr46B4-46B4\|2-61 cM | 18 | 4 | 25010 | 1343 | Vamp7,Marc,Orc6,CG30010 | | 1.383e-02 | -4.28 | fly chr46B4-46B4\|2-61 | chromosome location | fly chr46B4-46B4\|2-61 | 18 | 4 | 25010 | 1343 | CG30010,Orc6,Marc,Vamp7 | | 1.384e-02 | -4.28 | ksr (kinase suppressor of ras) | protein interactions | 40660 | 33 | 9 | 9204 | 1112 | Droj2,PyK,SERCA,csw,sea,Mcad,GCS2beta,CkIIalpha,RpL10 | | 1.392e-02 | -4.27 | dl (dorsal) | protein interactions | 35047 | 74 | 16 | 9204 | 1112 | RpL17,mad2,CG10470,Cyt-c-p,RpS16,tub,mbo,nej,cact,RpS2,dl,Pp2B-14D,CG6707,lwr,Rel,RpS27A | | 1.392e-02 | -4.27 | fax (failed axon connections) | protein interactions | 39826 | 74 | 16 | 9204 | 1112 | Uch,CG5447,Surf4,Mcad,Moe,capt,Rab5,regucalcin,CG15098,blw,Chd64,RpS27A,CG10425,SERCA,PHGPx,CG10638 | | 1.392e-02 | -4.27 | pigment metabolic process | biological process | GO:0042440 | 113 | 18 | 12345 | 1129 | DhpD,Rab32,pr,Shmt,EMC3,Hexo1,CG3803,dl,Cdk5,cl,Ppox,PAPLA1,fdl,Ho,CG13551,Sp7,lt,mal | | 1.395e-02 | -4.27 | fly chr22B1-22B1\|2-3 cM | chromosome location | fly chr22B1-22B1\|2-3 cM | 10 | 3 | 25010 | 1343 | mRpL48,Rrp40,Eno | | 1.395e-02 | -4.27 | fly chr76C6-76C6\|3-46 cM | chromosome location | fly chr76C6-76C6\|3-46 cM | 10 | 3 | 25010 | 1343 | CG34116,TORIP,cyc | | 1.395e-02 | -4.27 | fly chr11A1-11A1\|1-38 cM | chromosome location | fly chr11A1-11A1\|1-38 cM | 10 | 3 | 25010 | 1343 | regucalcin,Lsm12,gd | | 1.395e-02 | -4.27 | fly chr11A1-11A1\|1-38 | chromosome location | fly chr11A1-11A1\|1-38 | 10 | 3 | 25010 | 1343 | gd,Lsm12,regucalcin | | 1.395e-02 | -4.27 | fly chr76C6-76C6\|3-46 | chromosome location | fly chr76C6-76C6\|3-46 | 10 | 3 | 25010 | 1343 | cyc,TORIP,CG34116 | | 1.395e-02 | -4.27 | fly chr22B1-22B1\|2-3 | chromosome location | fly chr22B1-22B1\|2-3 | 10 | 3 | 25010 | 1343 | Eno,Rrp40,mRpL48 | | 1.408e-02 | -4.26 | GO\_POSITIVE\_REGULATION\_OF\_PROTEIN\_MODIFICATION\_BY\_SMALL\_PROTEIN\_CONJUGATION\_OR\_REMOVAL | MSigDB lists | GO\_POSITIVE\_REGULATION\_OF\_PROTEIN\_MODIFICATION\_BY\_SMALL\_PROTEIN\_CONJUGATION\_OR\_REMOVAL | 99 | 24 | 4161 | 644 | Hsc70-3,Prosbeta4,Prosalpha3,Prosalpha5,RpS27A,Cdk1,STUB1,Prosbeta5,CycB,mad2,Cul1,Rpn3,Prosalpha4,Rpt2,Prosbeta3,CG30291,Rpn2,Rpn12,Traf6,CG8188,APC7,SkpA,Prosbeta6,RpL40 | | 1.414e-02 | -4.26 | import into cell | biological process | GO:0098657 | 164 | 24 | 12345 | 1129 | Mekk1,Clc,lt,Synj,Arf51F,EcR,Fer1HCH,lqfR,Nlg1,crq,csw,fwe,CG12773,nrv1,Rab5,Atg6,dx,faf,Mtl,CG9132,Atpalpha,dco,slif,CG31729 | | 1.418e-02 | -4.26 | acyl-CoA metabolic process | biological process | GO:0006637 | 43 | 9 | 12345 | 1129 | CG17691,CG8199,scu,CG6178,Scsalpha1,CG1774,AcCoAS,CG7920,ScsbetaA | | 1.418e-02 | -4.26 | thioester metabolic process | biological process | GO:0035383 | 43 | 9 | 12345 | 1129 | AcCoAS,CG7920,ScsbetaA,Scsalpha1,CG6178,scu,CG1774,CG8199,CG17691 | | 1.418e-02 | -4.26 | regulation of ubiquitin-dependent protein catabolic process | biological process | GO:2000058 | 43 | 9 | 12345 | 1129 | Cul1,dco,Pomp,cort,CkIIalpha,SkpA,ari-1,park,Ubc6 | | 1.418e-02 | -4.26 | regulation of sister chromatid segregation | biological process | GO:0033045 | 43 | 9 | 12345 | 1129 | Mos,Zwilch,cana,tws,APC7,Rbf,mad2,wapl,Moe | | 1.419e-02 | -4.26 | HDR through Homologous Recombination (HR) or Single Strand Annealing (SSA) | REACTOME pathways | R-DME-5693567 | 11 | 5 | 3923 | 574 | nbs,mus81,mre11,grp,Tip60 | | 1.424e-02 | -4.25 | DNA integrity checkpoint | biological process | GO:0031570 | 36 | 8 | 12345 | 1129 | Cdk1,grp,mus304,nbs,mus81,nej,CG10336,mre11 | | 1.426e-02 | -4.25 | CG15011 (uncharacterized protein) | protein interactions | 38518 | 13 | 5 | 9204 | 1112 | Tak1,CG11248,Rel,PGRP-LE,cnn | | 1.426e-02 | -4.25 | melt (melted) | protein interactions | 38785 | 13 | 5 | 9204 | 1112 | RpS4,RpS14a,Hsc70-3,PyK,Rpt2 | | 1.426e-02 | -4.25 | tub (tube) | protein interactions | 40554 | 13 | 5 | 9204 | 1112 | mxc,dl,Droj2,Myd88,cact | | 1.426e-02 | -4.25 | COX6B (Cytochrome c oxidase subunit 6B) | protein interactions | 32989 | 13 | 5 | 9204 | 1112 | levy,cype,COX4,COX5A,Cyt-c-p | | 1.426e-02 | -4.25 | bowl (brother of odd with entrails limited) | protein interactions | 33602 | 13 | 5 | 9204 | 1112 | lin,ND-18,bcd,tor,CG13373 | | 1.430e-02 | -4.25 | Toll Like Receptor 9 (TLR9) Cascade | REACTOME pathways | R-DME-168138 | 42 | 12 | 3923 | 574 | Myd88,RpS27A,rl,ben,Traf6,Tak1,Uev1A,Rel,p38a,Pp2A-29B,dl,RpL40 | | 1.431e-02 | -4.25 | STEIN\_ESRRA\_TARGETS\_UP | MSigDB lists | STEIN\_ESRRA\_TARGETS\_UP | 163 | 36 | 4161 | 644 | Roe1,Etf-QO,Cyp1,tko,AnxB10,tws,colt,ScsbetaA,Cyt-c-p,Pfk,CG8004,Marf,ND-30,Mitofilin,COX5A,yip2,CG10932,Gk2,ND-51,ND-23,CG6028,ND-49,Pgm1,Mcad,CG3860,ND-SGDH,AIF,SdhA,ND-75,CG9391,Got1,Nc73EF,Scsalpha1,ND-39,Idh3b,GatA | | 1.432e-02 | -4.25 | Proteasome\_sua/b | interpro domains | IPR001353 | 25 | 7 | 11383 | 1233 | Prosbeta3,Prosalpha4,Prosbeta6,Prosbeta4,Prosalpha3,Prosalpha5,Prosbeta5 | | 1.432e-02 | -4.25 | TF\_Adf1 | interpro domains | IPR039353 | 25 | 7 | 11383 | 1233 | hng3,CG6276,jigr1,CG42526,CG4404,Mes2,brwl | | 1.432e-02 | -4.25 | GSE7548\_DAY7\_VS\_DAY28\_PCC\_IMMUNIZATION\_CD4\_TCELL\_UP | MSigDB lists | GSE7548\_DAY7\_VS\_DAY28\_PCC\_IMMUNIZATION\_CD4\_TCELL\_UP | 54 | 15 | 4161 | 644 | CG11858,l(3)73Ah,Dcr-1,mts,CG2201,bru1,Liprin-beta,mad2,Thiolase,Su(H),Caf1-105,Ptp61F,Ac3,AGBE,anne | | 1.432e-02 | -4.25 | GSE19198\_CTRL\_VS\_IL21\_TREATED\_TCELL\_6H\_UP | MSigDB lists | GSE19198\_CTRL\_VS\_IL21\_TREATED\_TCELL\_6H\_UP | 54 | 15 | 4161 | 644 | CCT3,CG9471,crq,Jafrac2,Pfk,pont,CG5028,Snx6,Su(H),Arts,ND-51,CG12018,CG5292,Scsalpha1,Nc73EF | | 1.432e-02 | -4.25 | GSE17721\_12H\_VS\_24H\_POLYIC\_BMDC\_UP | MSigDB lists | GSE17721\_12H\_VS\_24H\_POLYIC\_BMDC\_UP | 54 | 15 | 4161 | 644 | ERp60,Stam,CG13630,PrBP,ND-23,CG40045,Mtpalpha,CG17691,alph,Ufm1,Lsm12,AnxB10,SERCA,CG1275,Prosalpha3 | | 1.441e-02 | -4.24 | signal release | biological process | GO:0023061 | 97 | 16 | 12345 | 1129 | Sec10,nej,Rep,Gdi,Tomosyn,Sec22,CHOp24,lqfR,Arf51F,Exo70,opm,HDAC6,Sec15,Syx13,CanB2,X11L | | 1.442e-02 | -4.24 | establishment or maintenance of cell polarity | biological process | GO:0007163 | 217 | 30 | 12345 | 1129 | nej,mts,Mtl,Cbl,Tak1,parvin,Lkb1,Tomosyn,Pka-R1,CG3770,mael,capt,cnc,dco,Abi,jvl,awd,csw,msps,Moe,Pfdn2,osk,Sec23,Kap3,Rab5,X11L,Klp10A,heph,tws,alc | | 1.443e-02 | -4.24 | MORF\_CSNK2B | MSigDB lists | MORF\_CSNK2B | 174 | 38 | 4161 | 644 | COX5A,Clc,Pp2A-29B,LSm3,Prosbeta4,ND-24,Hat1,ND-30,capt,Thiolase,CCT5,Mtpalpha,CCT3,Cyt-c-p,Phb2,Dd,CG10418,Prosbeta6,NHP2,mEFTu1,Pen,ND-75,SdhA,Klp10A,Rpn12,Sod1,LysRS,csw,ND-49,RhoGDI,rept,ND-23,ND-51,CG11089,CG5028,levy,SkpA,eIF3i | | 1.446e-02 | -4.24 | internal peptidyl-lysine acetylation | biological process | GO:0018393 | 81 | 14 | 12345 | 1129 | Rcd5,Sgf29,Naa20A,Tip60,lid,d4,Spt20,YL-1,pont,nej,rept,Chrac-14,e(y)1,Hat1 | | 1.446e-02 | -4.24 | peptidyl-lysine acetylation | biological process | GO:0018394 | 81 | 14 | 12345 | 1129 | lid,Spt20,d4,Naa20A,Sgf29,Rcd5,Tip60,e(y)1,Hat1,pont,nej,YL-1,Chrac-14,rept | | 1.446e-02 | -4.24 | internal protein amino acid acetylation | biological process | GO:0006475 | 81 | 14 | 12345 | 1129 | Hat1,e(y)1,rept,Chrac-14,YL-1,nej,pont,d4,Spt20,lid,Tip60,Sgf29,Naa20A,Rcd5 | | 1.450e-02 | -4.23 | COPII-coated ER to Golgi transport vesicle | cellular component | GO:0030134 | 29 | 7 | 11940 | 1105 | CG4293,CG5510,opm,CHOp24,PAPLA1,Sec23,Sec22 | | 1.451e-02 | -4.23 | positive regulation of programmed cell death | biological process | GO:0043068 | 89 | 15 | 12345 | 1129 | Idh3b,Lkb1,fwe,PDCD-5,Utx,Tspo,E2f1,CycE,EcR,rept,Cbl,Jafrac2,eIF3m,MED24,ScsbetaA | | 1.457e-02 | -4.23 | robl22E (robl22E) | protein interactions | 33412 | 5 | 3 | 9204 | 1112 | Prosbeta3,Idgf4,whd | | 1.457e-02 | -4.23 | CG11334 (uncharacterized protein) | protein interactions | 43713 | 5 | 3 | 9204 | 1112 | RpL23A,Bx,RpS13 | | 1.457e-02 | -4.23 | tapas (tapas) | protein interactions | 37304 | 5 | 3 | 9204 | 1112 | tapas,spn-E,AGO3 | | 1.457e-02 | -4.23 | s-cup (stanley-cup) | protein interactions | 36371 | 5 | 3 | 9204 | 1112 | Cyt-c-p,RpS27A,fwe | | 1.457e-02 | -4.23 | Mul1 (Mitochondrial E3 ubiquitin protein ligase 1) | protein interactions | 38472 | 5 | 3 | 9204 | 1112 | park,Marf,Pink1 | | 1.457e-02 | -4.23 | Sry-delta (Serendipity delta) | protein interactions | 43572 | 5 | 3 | 9204 | 1112 | bcd,piwi,CG10959 | | 1.457e-02 | -4.23 | Pdfr (Pigment-dispersing factor receptor) | protein interactions | 31234 | 5 | 3 | 9204 | 1112 | CkIIalpha,Ac3,dco | | 1.457e-02 | -4.23 | dpr5 (defective proboscis extension response 5) | protein interactions | 41381 | 5 | 3 | 9204 | 1112 | dpr6,cDIP,Echs1 | | 1.457e-02 | -4.23 | 128up (upstream of RpIII128) | protein interactions | 36288 | 5 | 3 | 9204 | 1112 | nos,smg,Utx | | 1.457e-02 | -4.23 | CG1354 (uncharacterized protein) | protein interactions | 31914 | 5 | 3 | 9204 | 1112 | GstT1,CG14036,Chd64 | | 1.457e-02 | -4.23 | CG7974 (uncharacterized protein) | protein interactions | 38207 | 5 | 3 | 9204 | 1112 | CG11164,Traf6,CG2199 | | 1.457e-02 | -4.23 | tej (tejas) | protein interactions | 36590 | 5 | 3 | 9204 | 1112 | spn-E,tapas,CG15141 | | 1.457e-02 | -4.23 | plx (pollux) | protein interactions | 40704 | 5 | 3 | 9204 | 1112 | Rtnl1,Arf51F,CycE | | 1.459e-02 | -4.23 | CG31548 (uncharacterized protein) | protein interactions | 318794 | 2 | 2 | 9204 | 1112 | CG12171,CG31549 | | 1.459e-02 | -4.23 | CG2063 (uncharacterized protein) | protein interactions | 35952 | 2 | 2 | 9204 | 1112 | sbr,mad2 | | 1.459e-02 | -4.23 | CG9416 (uncharacterized protein) | protein interactions | 37223 | 2 | 2 | 9204 | 1112 | RpS16,PyK | | 1.459e-02 | -4.23 | Mip (Myoinhibiting peptide precursor) | protein interactions | 39933 | 2 | 2 | 9204 | 1112 | awd,regucalcin | | 1.459e-02 | -4.23 | CG31522 (uncharacterized protein) | protein interactions | 40567 | 2 | 2 | 9204 | 1112 | ATPsynO,ATPsynB | | 1.459e-02 | -4.23 | sud1 (sudestada1) | protein interactions | 19834718 | 2 | 2 | 9204 | 1112 | PEK,CG5412 | | 1.459e-02 | -4.23 | CG14566 (uncharacterized protein) | protein interactions | 40421 | 2 | 2 | 9204 | 1112 | CG11164,exu | | 1.459e-02 | -4.23 | Cpr65Av (Cuticular protein 65Av) | protein interactions | 318014 | 2 | 2 | 9204 | 1112 | Drep2,slam | | 1.459e-02 | -4.23 | Ste:CG33237 (Ste:CG33237) | protein interactions | 2768872 | 2 | 2 | 9204 | 1112 | CkIIalpha,spn-E | | 1.459e-02 | -4.23 | CG13631 (uncharacterized protein) | protein interactions | 50073 | 2 | 2 | 9204 | 1112 | gus,EloC | | 1.459e-02 | -4.23 | CG14075 (uncharacterized protein) | protein interactions | 40071 | 2 | 2 | 9204 | 1112 | Nca,RpLP1 | | 1.459e-02 | -4.23 | fsd (fates-shifted) | protein interactions | 36463 | 2 | 2 | 9204 | 1112 | SkpA,bcd | | 1.459e-02 | -4.23 | SclA (Sarcolamban A) | protein interactions | 19835657 | 2 | 2 | 9204 | 1112 | CG11486,SERCA | | 1.459e-02 | -4.23 | Usp32 (Ubiquitin specific protease 32) | protein interactions | 40169 | 2 | 2 | 9204 | 1112 | mre11,park | | 1.459e-02 | -4.23 | CG31098 (uncharacterized protein) | protein interactions | 43057 | 2 | 2 | 9204 | 1112 | CG11771,CkIIalpha | | 1.459e-02 | -4.23 | CG2145 (uncharacterized protein) | protein interactions | 32027 | 2 | 2 | 9204 | 1112 | bun,RpL12 | | 1.459e-02 | -4.23 | CG15706 (uncharacterized protein) | protein interactions | 36799 | 2 | 2 | 9204 | 1112 | cad,ATPsynbeta | | 1.459e-02 | -4.23 | CG13901 (uncharacterized protein) | protein interactions | 38095 | 2 | 2 | 9204 | 1112 | pont,rept | | 1.459e-02 | -4.23 | CG5728 (uncharacterized protein) | protein interactions | 42899 | 2 | 2 | 9204 | 1112 | mre11,Fkbp39 | | 1.459e-02 | -4.23 | Agpat1 (1-Acylglycerol-3-phosphate O-acyltransferase 1) | protein interactions | 32230 | 2 | 2 | 9204 | 1112 | Prosalpha4,Prosbeta4 | | 1.459e-02 | -4.23 | Nmd3 (Nonsense-mediated mRNA 3) | protein interactions | 31195 | 2 | 2 | 9204 | 1112 | COX5A,CG9911 | | 1.459e-02 | -4.23 | Ucp4A (Ucp4A) | protein interactions | 32764 | 2 | 2 | 9204 | 1112 | Pink1,park | | 1.459e-02 | -4.23 | l(1)G0193 (lethal (1) G0193) | protein interactions | 31712 | 2 | 2 | 9204 | 1112 | Vamp7,Cyp6a22 | | 1.461e-02 | -4.23 | myosin binding | molecular function | GO:0017022 | 43 | 9 | 12176 | 1119 | Tomosyn,RpL34a,CG18135,milt,Hsp26,Kap-alpha3,smg,CG2258,RpL23 | | 1.471e-02 | -4.22 | REACTOME\_APOPTOSIS | MSigDB lists | REACTOME\_APOPTOSIS | 59 | 16 | 4161 | 644 | Prosalpha5,RpS27A,Nmt,Prosbeta4,Prosalpha3,Rpn3,Prosbeta5,Rpn2,Cyt-c-p,Rpn12,Prosalpha4,Rpt2,Prosbeta3,RpL40,Prosbeta6,CanB2 | | 1.471e-02 | -4.22 | GO\_CELL\_CYCLE\_CHECKPOINT | MSigDB lists | GO\_CELL\_CYCLE\_CHECKPOINT | 59 | 16 | 4161 | 644 | Not10,grp,CG1812,Rbf,csw,CkIIalpha,CG30291,Tao,RpL40,RpS27A,APP-BP1,twin,mad2,Cdk1,p38a,CycB | | 1.471e-02 | -4.22 | GSE37416\_CTRL\_VS\_24H\_F\_TULARENSIS\_LVS\_NEUTROPHIL\_UP | MSigDB lists | GSE37416\_CTRL\_VS\_24H\_F\_TULARENSIS\_LVS\_NEUTROPHIL\_UP | 59 | 16 | 4161 | 644 | CG11134,Pex12,Tfb4,mRpS5,Rbf,MED24,colt,CG6767,APC7,Pi3K92E,CG7550,oxt,Adi1,Naa20A,Jafrac1,CG5734 | | 1.471e-02 | -4.22 | GSE32901\_TH1\_VS\_TH17\_ENRICHED\_CD4\_TCELL\_DN | MSigDB lists | GSE32901\_TH1\_VS\_TH17\_ENRICHED\_CD4\_TCELL\_DN | 59 | 16 | 4161 | 644 | Sfxn2,Tsp66E,TyrRS,CG30291,CG2061,Phb2,CG9265,Karybeta3,ND-23,CG1416,Npc2a,Rcd5,Prosalpha5,PpD3,Tfb4,RpL13 | | 1.475e-02 | -4.22 | GSE17721\_CPG\_VS\_GARDIQUIMOD\_2H\_BMDC\_UP | MSigDB lists | GSE17721\_CPG\_VS\_GARDIQUIMOD\_2H\_BMDC\_UP | 40 | 12 | 4161 | 644 | colt,Rab32,RpLP0,mRpL13,RpS27,CG10465,mol,CG30010,gus,Etf-QO,CG7997,CCT5 | | 1.475e-02 | -4.22 | SAKAI\_CHRONIC\_HEPATITIS\_VS\_LIVER\_CANCER\_UP | MSigDB lists | SAKAI\_CHRONIC\_HEPATITIS\_VS\_LIVER\_CANCER\_UP | 40 | 12 | 4161 | 644 | CG17454,CanB2,Cul5,Cul1,Kap-alpha3,Hat1,Grx1,APP-BP1,Cul2,Rnmt,Traf6,Cse1 | | 1.475e-02 | -4.22 | CHAUHAN\_RESPONSE\_TO\_METHOXYESTRADIOL\_DN | MSigDB lists | CHAUHAN\_RESPONSE\_TO\_METHOXYESTRADIOL\_DN | 40 | 12 | 4161 | 644 | Dd,Npc2a,eIF3m,dco,Nap1,Phb2,CG8188,levy,Arl6IP1,RpL12,Pgk,lwr | | 1.475e-02 | -4.22 | SASSON\_RESPONSE\_TO\_FORSKOLIN\_UP | MSigDB lists | SASSON\_RESPONSE\_TO\_FORSKOLIN\_UP | 40 | 12 | 4161 | 644 | Atg6,Acph-1,Pp2A-29B,Rpn2,Arpc1,Pka-R1,Stip1,Pdxk,p38a,Arpc4,Pgk,Gk2 | | 1.476e-02 | -4.22 | G1/S-Specific Transcription | REACTOME pathways | R-DME-69205 | 15 | 6 | 3923 | 574 | Dhfr,Ts,E2f1,Orc1,CycE,Cdk1 | | 1.476e-02 | -4.22 | Proteasome | pfam domains | PF00227 | 25 | 7 | 10480 | 1142 | Prosalpha3,Prosbeta3,Prosalpha4,Prosbeta5,Prosbeta6,Prosalpha5,Prosbeta4 | | 1.480e-02 | -4.21 | behavioral response to ethanol | biological process | GO:0048149 | 58 | 11 | 12345 | 1129 | aay,Pka-R2,Agpat2,Shmt,apt,Adh,Arf51F,CG17734,rl,AcCoAS,dnc | | 1.481e-02 | -4.21 | VECCHI\_GASTRIC\_CANCER\_EARLY\_UP | MSigDB lists | VECCHI\_GASTRIC\_CANCER\_EARLY\_UP | 131 | 30 | 4161 | 644 | RfC38,DppIII,Ranbp21,Sld5,CG12375,Tm1,rept,Non1,Dhfr,CG5292,grp,mRpL13,Cse1,CCT3,Cdk1,mRpS10,Eno,Pfdn2,CycB,Pus7,mad2,bys,Arpc4,Uch,CG7728,CG15111,Idh3b,Pcd,yip2,CycE | | 1.486e-02 | -4.21 | chromosome organization | biological process | GO:0051276 | 680 | 79 | 12345 | 1129 | Rcd5,G9a,RfC38,Cenp-C,Elba2,Sgf11,HmgD,fs(1)Ya,Spt20,Snm1,egg,RecQ5,Nurf-38,CycB,wash,Nap1,armi,psq,cana,Utx,mus304,eIF4E1,Klp10A,r2d2,HDAC6,Hat1,msps,l(3)73Ah,escl,scny,yem,Caf1-105,CG31053,esc,mad2,GatA,wapl,Tip60,ph-d,Xpac,CG10038,nbs,Bin1,lid,del,enc,Hnf4,BigH1,SRPK,Naa20A,asf1,SkpA,rept,Nup62,YL-1,nej,pont,SuUR,e(y)1,Sirt7,gammaTub37C,Rab5,HP1e,Chrac-14,Ptip,spn-E,Atu,Tlk,piwi,Sgf29,d4,krimp,Topors,Trf4-2,Top1,trr,Rbf,not,mre11 | | 1.487e-02 | -4.21 | DNA Replication | REACTOME pathways | R-DME-69306 | 105 | 24 | 3923 | 574 | Psf3,Rpn3,Orc1,Rpn12,Prosbeta6,DNApol-epsilon255,RfC38,Prosbeta3,Orc6,Rpt2,RpL40,Prosalpha3,Sld5,Rbf,Chrac-14,Prosalpha4,Prosbeta5,CG12018,Rpn2,Rbf2,E2f1,Prosbeta4,Prosalpha5,RpS27A | | 1.494e-02 | -4.20 | GSE360\_L\_DONOVANI\_VS\_B\_MALAYI\_HIGH\_DOSE\_MAC\_DN | MSigDB lists | GSE360\_L\_DONOVANI\_VS\_B\_MALAYI\_HIGH\_DOSE\_MAC\_DN | 64 | 17 | 4161 | 644 | Nca,eEF1beta,Pen,CG3376,ND-30,RpL12,capt,AIF,CG8199,GlyP,Jheh2,Idh3b,Nc73EF,CG4572,AP-1-2beta,slif,Sod1 | | 1.494e-02 | -4.20 | PECE\_MAMMARY\_STEM\_CELL\_UP | MSigDB lists | PECE\_MAMMARY\_STEM\_CELL\_UP | 64 | 17 | 4161 | 644 | CG9143,RpL24,RpS28b,RpS18,Tim8,RpL13,Gapdh1,RpS13,RpS30,RpL13A,RpS16,LysRS,CG1890,RpS2,RpS7,Jafrac1,RpL40 | | 1.494e-02 | -4.20 | GSE15930\_NAIVE\_VS\_72H\_IN\_VITRO\_STIM\_IL12\_CD8\_TCELL\_DN | MSigDB lists | GSE15930\_NAIVE\_VS\_72H\_IN\_VITRO\_STIM\_IL12\_CD8\_TCELL\_DN | 64 | 17 | 4161 | 644 | mRpL17,gammaTub37C,CG14715,CG5880,Jafrac2,CG3376,msps,mad2,Pglym78,Pgm1,CG17294,mRpL18,CG11164,AGBE,CysRS,CG9485,Su(H) | | 1.494e-02 | -4.20 | GO\_NUCLEAR\_TRANSCRIBED\_MRNA\_CATABOLIC\_PROCESS\_DEADENYLATION\_DEPENDENT\_DECAY | MSigDB lists | GO\_NUCLEAR\_TRANSCRIBED\_MRNA\_CATABOLIC\_PROCESS\_DEADENYLATION\_DEPENDENT\_DECAY | 31 | 10 | 4161 | 644 | Edc3,Not10,CG6610,eIF4E1,Rrp40,LSm3,eIF4A,CG8963,twin,CG10418 | | 1.494e-02 | -4.20 | MODULE\_91 | MSigDB lists | MODULE\_91 | 31 | 10 | 4161 | 644 | Prosbeta3,Prosalpha3,Rpt2,Prosbeta4,Rpn12,Rpn2,Prosalpha5,Prosbeta5,Prosbeta6,Rpn3 | | 1.494e-02 | -4.20 | GO\_POLYSACCHARIDE\_METABOLIC\_PROCESS | MSigDB lists | GO\_POLYSACCHARIDE\_METABOLIC\_PROCESS | 31 | 10 | 4161 | 644 | Gyg,Pgm1,RpS27A,GalT1,CG8475,Pfk,CG9485,AGBE,GlyP,RpL40 | | 1.494e-02 | -4.20 | GSE21546\_ELK1\_KO\_VS\_SAP1A\_KO\_AND\_ELK1\_KO\_DP\_THYMOCYTES\_UP | MSigDB lists | GSE21546\_ELK1\_KO\_VS\_SAP1A\_KO\_AND\_ELK1\_KO\_DP\_THYMOCYTES\_UP | 31 | 10 | 4161 | 644 | Sply,CG17294,Tomosyn,CG1890,Tlk,alph,CG4041,Pglym78,CG9911,scramb1 | | 1.497e-02 | -4.20 | CG8993 (uncharacterized protein) | protein interactions | 38301 | 28 | 8 | 9204 | 1112 | awd,AnxB10,Sod1,Fkbp12,regucalcin,CG10638,Cdk5,Eno | | 1.497e-02 | -4.20 | pho (pleiohomeotic) | protein interactions | 43819 | 28 | 8 | 9204 | 1112 | rept,pont,esc,ph-d,E2f1,nos,mxc,RYBP | | 1.497e-02 | -4.20 | Nplp2 (Neuropeptide-like precursor 2) | protein interactions | 50286 | 28 | 8 | 9204 | 1112 | Adh,e(y)1,Mf,Argk,CG4390,Idgf4,Fer1HCH,Scp1 | | 1.497e-02 | -4.20 | Cat (Catalase) | protein interactions | 40048 | 28 | 8 | 9204 | 1112 | PGRP-LE,Trxr-1,Rel,Pink1,park,PHGPx,Idh,Tak1 | | 1.497e-02 | -4.20 | GSE43260\_BTLA\_POS\_VS\_NEG\_INTRATUMORAL\_CD8\_TCELL\_UP | MSigDB lists | GSE43260\_BTLA\_POS\_VS\_NEG\_INTRATUMORAL\_CD8\_TCELL\_UP | 79 | 20 | 4161 | 644 | stas,CG5116,mtgo,Sec23,Hsc70-3,ERp60,Gclc,CG14715,pr,ND-B18,CG9471,Rpn12,CG3631,UbcE2H,Cyp1,Tspo,cathD,opm,Fur2,NHP2 | | 1.497e-02 | -4.20 | GSE12392\_CD8A\_POS\_VS\_NEG\_SPLEEN\_IFNB\_KO\_DC\_UP | MSigDB lists | GSE12392\_CD8A\_POS\_VS\_NEG\_SPLEEN\_IFNB\_KO\_DC\_UP | 79 | 20 | 4161 | 644 | Rpn2,lt,DCAF12,Traf6,mRpL13,CG4390,CG1890,muskelin,mbo,PrBP,Xpac,Nmt,Grip75,eIF3m,Npc2a,Pglym78,ND-30,Echs1,Prosbeta5,Eno | | 1.498e-02 | -4.20 | regulation of DNA-dependent DNA replication | biological process | GO:0090329 | 23 | 6 | 12345 | 1129 | SuUR,E2f1,CG10336,pn,Cdk1,otu | | 1.498e-02 | -4.20 | pole cell formation | biological process | GO:0007279 | 23 | 6 | 12345 | 1129 | cnn,piwi,cort,osk,Dcr-1,eIF4A | | 1.505e-02 | -4.20 | GO\_REGULATION\_OF\_RNA\_STABILITY | MSigDB lists | GO\_REGULATION\_OF\_RNA\_STABILITY | 69 | 18 | 4161 | 644 | Rpn3,Prosbeta5,p38a,Prosalpha5,RpS27A,twin,Prosalpha3,Prosbeta4,RpL40,Imp,Prosbeta6,Rrp40,Rpn12,Rpn2,Prosbeta3,Prosalpha4,Rpt2,CG8963 | | 1.505e-02 | -4.20 | GO\_PROTEIN\_IMPORT | MSigDB lists | GO\_PROTEIN\_IMPORT | 69 | 18 | 4161 | 644 | cyc,Arts,mbo,Karybeta3,Pex13,Pp2B-14D,Phb2,Cse1,Pex10,RpL23,Pen,ERp60,Kap-alpha3,rl,cact,Tom40,Pex12,CanA-14F | | 1.505e-02 | -4.20 | GSE37301\_HEMATOPOIETIC\_STEM\_CELL\_VS\_LYMPHOID\_PRIMED\_MPP\_DN | MSigDB lists | GSE37301\_HEMATOPOIETIC\_STEM\_CELL\_VS\_LYMPHOID\_PRIMED\_MPP\_DN | 69 | 18 | 4161 | 644 | Rpn3,CG14715,Tfb4,Rnmt,gammaTub37C,RpL24,Rab10,CG7789,CG10418,CG17698,CG5734,CG8726,Tspo,Ptp61F,eIF4E1,Cyt-c-p,tun,csw | | 1.505e-02 | -4.20 | GSE14415\_TCONV\_VS\_FOXP3\_KO\_INDUCED\_TREG\_DN | MSigDB lists | GSE14415\_TCONV\_VS\_FOXP3\_KO\_INDUCED\_TREG\_DN | 74 | 19 | 4161 | 644 | yip2,Rnmt,Pen,mad2,ERp60,CG7655,CG14715,Cdk1,ScsbetaA,Rpn12,aay,grp,Nurf-38,Rpt2,Aos1,Pdk,Tm1,RfC38,CG15141 | | 1.505e-02 | -4.20 | GSE26351\_UNSTIM\_VS\_WNT\_PATHWAY\_STIM\_HEMATOPOIETIC\_PROGENITORS\_DN | MSigDB lists | GSE26351\_UNSTIM\_VS\_WNT\_PATHWAY\_STIM\_HEMATOPOIETIC\_PROGENITORS\_DN | 74 | 19 | 4161 | 644 | Cse1,lt,mts,CG9132,Pex10,Jafrac1,Gdap2,ATP8A,escl,PIG-L,CG17691,nSMase,CG8372,Pfas,CG8611,pont,esc,CG16865,Jafrac2 | | 1.513e-02 | -4.19 | egl (egalitarian) | protein interactions | 37757 | 39 | 10 | 9204 | 1112 | CG4949,RpL38,RpS29,pgc,nos,Sry-alpha,bcd,Xpc,osk,fwe | | 1.513e-02 | -4.19 | CG1910 (uncharacterized protein) | protein interactions | 43741 | 39 | 10 | 9204 | 1112 | SelD,r-l,RpS27A,awd,Chd64,CG4949,AnxB10,capt,Fkbp12,CG10638 | | 1.513e-02 | -4.19 | Lst8 (Lst8) | protein interactions | 31903 | 39 | 10 | 9204 | 1112 | raptor,CG4407,CCT6,Sec23,Pfdn2,mts,ND-75,Stip1,Fer1HCH,RpL10 | | 1.513e-02 | -4.19 | plu (plutonium) | protein interactions | 44848 | 39 | 10 | 9204 | 1112 | CycE,Stam,CG17508,CycB,CG11486,EcR,bcd,png,CycC,P5CDh1 | | 1.513e-02 | -4.19 | CG16826 (uncharacterized protein) | protein interactions | 34741 | 39 | 10 | 9204 | 1112 | GlyP,Fer1HCH,Scp1,Pglym78,Adh,Atpalpha,CG32473,Pgk,Argk,CG8490 | | 1.517e-02 | -4.19 | protein acylation | biological process | GO:0043543 | 114 | 18 | 12345 | 1129 | Hat1,CG8314,Chrac-14,Nmt,lid,Tip60,Rcd5,CG5880,e(y)1,rept,YL-1,nej,pont,d4,Spt20,Sgf29,Naa20A,CG10344 | | 1.532e-02 | -4.18 | NADH dehydrogenase (ubiquinone) activity | molecular function | GO:0008137 | 23 | 6 | 12176 | 1119 | ND-51,ND-30,ND-75,ND-18,ND-B15,ND-B18 | | 1.532e-02 | -4.18 | NADH dehydrogenase (quinone) activity | molecular function | GO:0050136 | 23 | 6 | 12176 | 1119 | ND-30,ND-51,ND-B15,ND-B18,ND-75,ND-18 | | 1.532e-02 | -4.18 | tRNA binding | molecular function | GO:0000049 | 23 | 6 | 12176 | 1119 | TyrRS,CysRS,GluRS-m,LysRS,CG7414,CG10189 | | 1.533e-02 | -4.18 | GO\_REGULATION\_OF\_CELL\_CYCLE\_PHASE\_TRANSITION | MSigDB lists | GO\_REGULATION\_OF\_CELL\_CYCLE\_PHASE\_TRANSITION | 126 | 29 | 4161 | 644 | Cdk1,Prosbeta5,CycB,mad2,Cul1,Rpn3,Wnt10,APP-BP1,Prosbeta4,twin,Prosalpha3,Prosalpha5,RpS27A,SkpA,Prosbeta6,RpL40,Prosalpha4,Rbf,Rpt2,CkIIalpha,CG11164,CG30291,Prosbeta3,Phb2,Not10,Rpn2,APC7,CG1812,Rpn12 | | 1.543e-02 | -4.17 | response to osmotic stress | biological process | GO:0006970 | 17 | 5 | 12345 | 1129 | NFAT,kcc,p38a,CG17746,Mekk1 | | 1.543e-02 | -4.17 | phagocytosis, engulfment | biological process | GO:0006911 | 17 | 5 | 12345 | 1129 | EcR,Mtl,Mekk1,csw,crq | | 1.543e-02 | -4.17 | tRNA wobble uridine modification | biological process | GO:0002098 | 17 | 5 | 12345 | 1129 | CG3021,CG2911,CG10189,CG2034,CG3587 | | 1.543e-02 | -4.17 | chitin catabolic process | biological process | GO:0006032 | 17 | 5 | 12345 | 1129 | Cht11,Idgf2,Idgf6,Idgf4,fdl | | 1.546e-02 | -4.17 | chorion-containing eggshell formation | biological process | GO:0007304 | 131 | 20 | 12345 | 1129 | cact,spn-E,fs(1)N,CycE,jvl,Cf2,gus,Pxt,armi,fs(1)M3,bun,Orc1,rl,Pax,tor,Rbf,squ,E2f1,tkv,mael | | 1.556e-02 | -4.16 | Gal-bind\_lectin | smart domains | SM00908 | 6 | 3 | 5086 | 506 | Pex23,CG5335,galectin | | 1.556e-02 | -4.16 | Cullin\_Nedd8 | smart domains | SM00884 | 6 | 3 | 5086 | 506 | Cul2,Cul5,Cul1 | | 1.556e-02 | -4.16 | PAZ | smart domains | SM00949 | 6 | 3 | 5086 | 506 | piwi,Dcr-1,AGO3 | | 1.571e-02 | -4.15 | regulation of proteasomal protein catabolic process | biological process | GO:0061136 | 51 | 10 | 12345 | 1129 | SkpA,Pomp,cort,dco,CkIIalpha,Rpt2,Cul1,park,Ubc6,ari-1 | | 1.572e-02 | -4.15 | Synthesis of DNA | REACTOME pathways | R-DME-69239 | 100 | 23 | 3923 | 574 | Prosbeta4,Prosalpha5,RpS27A,Chrac-14,Prosalpha4,Rpn2,Prosbeta5,CG12018,Rbf2,Sld5,Rbf,Rpt2,RpL40,Prosalpha3,Orc6,Prosbeta3,RfC38,Prosbeta6,DNApol-epsilon255,Psf3,Orc1,Rpn3,Rpn12 | | 1.574e-02 | -4.15 | aldo-keto reductase (NADP) activity | molecular function | GO:0004033 | 17 | 5 | 12176 | 1119 | CG3842,CG9436,CG2767,CG10638,CG10863 | | 1.575e-02 | -4.15 | SPE (Spatzle-Processing Enzyme) | protein interactions | 42791 | 23 | 7 | 9204 | 1112 | Hsp26,CG4407,Idh3b,exu,mts,Fer1HCH,ND-75 | | 1.575e-02 | -4.15 | PPO1 (Prophenoloxidase 1) | protein interactions | 37044 | 23 | 7 | 9204 | 1112 | CG32473,Fer1HCH,Adh,Sp7,Scp1,Jafrac1,Argk | | 1.575e-02 | -4.15 | corto (corto) | protein interactions | 40616 | 23 | 7 | 9204 | 1112 | esc,EloC,mxc,piwi,rl,ph-d,RpL12 | | 1.575e-02 | -4.15 | LFA1\_Q6 | MSigDB lists | LFA1\_Q6 | 45 | 13 | 4161 | 644 | CG14715,oxt,cact,Stoml2,sbr,gus,Su(H),Rab5,Rcd5,Npc2a,Mitofilin,ND-49,Smox | | 1.575e-02 | -4.15 | GSE10239\_NAIVE\_VS\_KLRG1INT\_EFF\_CD8\_TCELL\_UP | MSigDB lists | GSE10239\_NAIVE\_VS\_KLRG1INT\_EFF\_CD8\_TCELL\_UP | 45 | 13 | 4161 | 644 | Utx,Pi4KIIalpha,RpS27,CG10189,CG3021,Npc2a,RpL37a,RpL11,CG17187,Ublcp1,RpL13,RpL38,RpS7 | | 1.575e-02 | -4.15 | GO\_CELL\_CYCLE\_G2\_M\_PHASE\_TRANSITION | MSigDB lists | GO\_CELL\_CYCLE\_G2\_M\_PHASE\_TRANSITION | 45 | 13 | 4161 | 644 | gammaTub37C,dco,Pp2A-29B,RpS27A,grp,Cdk7,Wnt10,RpL40,msps,Cul1,CycB,SkpA,Cdk1 | | 1.575e-02 | -4.15 | GSE17974\_IL4\_AND\_ANTI\_IL12\_VS\_UNTREATED\_0.5H\_ACT\_CD4\_TCELL\_UP | MSigDB lists | GSE17974\_IL4\_AND\_ANTI\_IL12\_VS\_UNTREATED\_0.5H\_ACT\_CD4\_TCELL\_UP | 45 | 13 | 4161 | 644 | MED27,RpS7,nmdyn-D6,CG12375,CG4407,Snx6,Maf1,snz,TyrRS,Ppcdc,Rpt2,Lsm12,Pfdn5 | | 1.576e-02 | -4.15 | Tektin-C (Tektin C) | protein interactions | 38653 | 18 | 6 | 9204 | 1112 | CG13741,smg,CG11486,CG8679,Hsp26,Pgm1 | | 1.576e-02 | -4.15 | eIF2alpha (eukaryotic translation initiation factor 2 subunit alpha) | protein interactions | 32617 | 18 | 6 | 9204 | 1112 | Pdk1,PGRP-LE,Rel,Lkb1,Tak1,PEK | | 1.576e-02 | -4.15 | CSN5 (COP9 signalosome subunit 5) | protein interactions | 42000 | 18 | 6 | 9204 | 1112 | Trs33,CycE,EcR,CSN6,bam,mad2 | | 1.577e-02 | -4.15 | boca (boca) | protein interactions | 48986 | 75 | 16 | 9204 | 1112 | MED4,RpLP0,Usp10,MED27,Gk1,ix,Ser7,Ptip,CG17002,CCT3,MED20,Nup358,bor,CG13630,Larp4B,r | | 1.584e-02 | -4.15 | HALLMARK\_MYC\_TARGETS\_V1 | MSigDB lists | HALLMARK\_MYC\_TARGETS\_V1 | 137 | 31 | 4161 | 644 | Prosbeta3,CG12018,Rpn2,Phb2,RpLP0,Rpn12,LysRS,RpS2,eIF4E1,NHP2,Ts,r,CG10418,rept,Cbp80,Prosbeta4,Prosalpha3,COX5A,Nap1,SRPK,Jafrac2,CCT3,CCT5,Chrac-14,Cul1,Rpn3,Pen,eEF1beta,mad2,mEFTu1,Pgk | | 1.586e-02 | -4.14 | REACTOME\_METABOLISM\_OF\_AMINO\_ACIDS\_AND\_DERIVATIVES | MSigDB lists | REACTOME\_METABOLISM\_OF\_AMINO\_ACIDS\_AND\_DERIVATIVES | 121 | 28 | 4161 | 644 | Prosalpha3,Nc73EF,CG11134,Got1,Prosbeta4,Pcd,Prosalpha5,Gclc,Prosbeta5,CG8199,shop,Rpn3,Shmt,CG11899,P5cr,Prosbeta3,Rpt2,P5CDh1,Prosalpha4,Rpn12,Rpn2,scu,aay,CG1640,Prosbeta6,CG17691,Adi1,CG10932 | | 1.590e-02 | -4.14 | - | gene3d domains | 3.30.1460.20 | 2 | 2 | 4929 | 622 | Arpc4,Arpc2 | | 1.590e-02 | -4.14 | - | gene3d domains | 2.160.20.70 | 2 | 2 | 4929 | 622 | TBCC,capt | | 1.590e-02 | -4.14 | - | gene3d domains | 3.40.1800.40 | 2 | 2 | 4929 | 622 | RpL34b,RpL34a | | 1.590e-02 | -4.14 | - | gene3d domains | 3.30.1330.40 | 2 | 2 | 4929 | 622 | CG1578,UK114 | | 1.590e-02 | -4.14 | - | gene3d domains | 3.40.50.1380 | 2 | 2 | 4929 | 622 | CG11089,r | | 1.590e-02 | -4.14 | - | gene3d domains | 3.90.1170.10 | 2 | 2 | 4929 | 622 | RpL10,mRpL16 | | 1.591e-02 | -4.14 | GSE24634\_TEFF\_VS\_TCONV\_DAY5\_IN\_CULTURE\_UP | MSigDB lists | GSE24634\_TEFF\_VS\_TCONV\_DAY5\_IN\_CULTURE\_UP | 100 | 24 | 4161 | 644 | CG10932,Plap,Ts,eIF3i,Karybeta3,CG15141,Roe1,ste24a,alphaTub84B,Psf3,Dhfr,CG3226,mad2,Chrac-14,msps,Shmt,mRpL16,Stip1,Pfdn2,Tim8,Pus7,PDCD-5,Nmt,CG30496 | | 1.608e-02 | -4.13 | fly chr5D1-5D1\|1-15 cM | chromosome location | fly chr5D1-5D1\|1-15 cM | 4 | 2 | 25010 | 1343 | CG5966,fs(1)M3 | | 1.608e-02 | -4.13 | fly chr102F6-102F6\|4-0 cM | chromosome location | fly chr102F6-102F6\|4-0 cM | 4 | 2 | 25010 | 1343 | CG11077,ATPsynbeta | | 1.608e-02 | -4.13 | fly chr5D1-5D1\|1-15 | chromosome location | fly chr5D1-5D1\|1-15 | 4 | 2 | 25010 | 1343 | CG5966,fs(1)M3 | | 1.608e-02 | -4.13 | fly chr29B2-29B2\|2-31 cM | chromosome location | fly chr29B2-29B2\|2-31 cM | 4 | 2 | 25010 | 1343 | RpS13,Pp2A-29B | | 1.608e-02 | -4.13 | fly chr19D2-19D2\|1-64 | chromosome location | fly chr19D2-19D2\|1-64 | 4 | 2 | 25010 | 1343 | Tak1,CG1812 | | 1.608e-02 | -4.13 | fly chr76D5-76D5\|3-46 | chromosome location | fly chr76D5-76D5\|3-46 | 4 | 2 | 25010 | 1343 | RhoGDI,CG8004 | | 1.608e-02 | -4.13 | fly chr29B2-29B2\|2-31 | chromosome location | fly chr29B2-29B2\|2-31 | 4 | 2 | 25010 | 1343 | Pp2A-29B,RpS13 | | 1.608e-02 | -4.13 | fly chr3A3-3A3\|1-1 | chromosome location | fly chr3A3-3A3\|1-1 | 4 | 2 | 25010 | 1343 | boi,tko | | 1.608e-02 | -4.13 | fly chr30E4-30E4\|2-37 | chromosome location | fly chr30E4-30E4\|2-37 | 4 | 2 | 25010 | 1343 | yip2,CG4598 | | 1.608e-02 | -4.13 | fly chr67B6-67B6\|3-29 cM | chromosome location | fly chr67B6-67B6\|3-29 cM | 4 | 2 | 25010 | 1343 | Nf-YA,CG42526 | | 1.608e-02 | -4.13 | fly chr66B7-66B7\|3-22 | chromosome location | fly chr66B7-66B7\|3-22 | 4 | 2 | 25010 | 1343 | CG7550,mRpL36 | | 1.608e-02 | -4.13 | fly chr66B7-66B7\|3-22 cM | chromosome location | fly chr66B7-66B7\|3-22 cM | 4 | 2 | 25010 | 1343 | mRpL36,CG7550 | | 1.608e-02 | -4.13 | fly chr52A13-52A13\|2-74 | chromosome location | fly chr52A13-52A13\|2-74 | 4 | 2 | 25010 | 1343 | Cdk5,CG8204 | | 1.608e-02 | -4.13 | fly chr67B6-67B6\|3-29 | chromosome location | fly chr67B6-67B6\|3-29 | 4 | 2 | 25010 | 1343 | Nf-YA,CG42526 | | 1.608e-02 | -4.13 | fly chr52A13-52A13\|2-74 cM | chromosome location | fly chr52A13-52A13\|2-74 cM | 4 | 2 | 25010 | 1343 | Cdk5,CG8204 | | 1.608e-02 | -4.13 | fly chr27E4-27E4\|2-25 cM | chromosome location | fly chr27E4-27E4\|2-25 cM | 4 | 2 | 25010 | 1343 | CG4497,CG4502 | | 1.608e-02 | -4.13 | fly chr3A3-3A3\|1-1 cM | chromosome location | fly chr3A3-3A3\|1-1 cM | 4 | 2 | 25010 | 1343 | boi,tko | | 1.608e-02 | -4.13 | fly chr85B7-85B7\|3-48 | chromosome location | fly chr85B7-85B7\|3-48 | 4 | 2 | 25010 | 1343 | osk,ScsbetaA | | 1.608e-02 | -4.13 | fly chr19D2-19D2\|1-64 cM | chromosome location | fly chr19D2-19D2\|1-64 cM | 4 | 2 | 25010 | 1343 | CG1812,Tak1 | | 1.608e-02 | -4.13 | fly chr27E4-27E4\|2-25 | chromosome location | fly chr27E4-27E4\|2-25 | 4 | 2 | 25010 | 1343 | CG4497,CG4502 | | 1.608e-02 | -4.13 | fly chr102F6-102F6\|4-0 | chromosome location | fly chr102F6-102F6\|4-0 | 4 | 2 | 25010 | 1343 | ATPsynbeta,CG11077 | | 1.608e-02 | -4.13 | fly chr76D5-76D5\|3-46 cM | chromosome location | fly chr76D5-76D5\|3-46 cM | 4 | 2 | 25010 | 1343 | RhoGDI,CG8004 | | 1.608e-02 | -4.13 | fly chr56D7-56D7\|2-89 cM | chromosome location | fly chr56D7-56D7\|2-89 cM | 4 | 2 | 25010 | 1343 | RpL11,EloC | | 1.608e-02 | -4.13 | fly chr68A4-68A4\|3-34 cM | chromosome location | fly chr68A4-68A4\|3-34 cM | 4 | 2 | 25010 | 1343 | SuUR,Pfdn2 | | 1.608e-02 | -4.13 | fly chr56D7-56D7\|2-89 | chromosome location | fly chr56D7-56D7\|2-89 | 4 | 2 | 25010 | 1343 | RpL11,EloC | | 1.608e-02 | -4.13 | fly chr30E4-30E4\|2-37 cM | chromosome location | fly chr30E4-30E4\|2-37 cM | 4 | 2 | 25010 | 1343 | yip2,CG4598 | | 1.608e-02 | -4.13 | fly chr85B7-85B7\|3-48 cM | chromosome location | fly chr85B7-85B7\|3-48 cM | 4 | 2 | 25010 | 1343 | ScsbetaA,osk | | 1.608e-02 | -4.13 | fly chr68A4-68A4\|3-34 | chromosome location | fly chr68A4-68A4\|3-34 | 4 | 2 | 25010 | 1343 | Pfdn2,SuUR | | 1.610e-02 | -4.13 | GO\_RESPONSE\_TO\_NITROGEN\_COMPOUND | MSigDB lists | GO\_RESPONSE\_TO\_NITROGEN\_COMPOUND | 214 | 45 | 4161 | 644 | PpD3,Gabat,nAChRalpha4,Smox,CycE,Prosbeta4,Wnt10,SREBP,CanA-14F,Dcr-1,CG3376,Mtpalpha,Pdk1,Ac3,mAChR-A,p38a,Cdk5,Pp2B-14D,Dhfr,Vha36-1,Tspo,Ptp61F,Ts,Vha13,Pka-R1,r,Got1,Rab10,Pka-R2,Shmt,Calx,rl,cact,kri,Cdk1,LpR2,AIF,GlyP,Prip,mino,adp,Sod1,raptor,dnc,Karybeta3 | | 1.618e-02 | -4.12 | Ace (Acetylcholine esterase) | protein interactions | 41625 | 9 | 4 | 9204 | 1112 | CG14204,Kap3,dnc,Pka-R1 | | 1.618e-02 | -4.12 | CG10939 (uncharacterized protein) | protein interactions | 44155 | 9 | 4 | 9204 | 1112 | Sry-alpha,Moe,CG4853,Xpac | | 1.618e-02 | -4.12 | CG15177 (uncharacterized protein) | protein interactions | 40810 | 9 | 4 | 9204 | 1112 | Cyt-c-p,Pxt,ND-30,CG9917 | | 1.618e-02 | -4.12 | CG9925 (uncharacterized protein) | protein interactions | 41702 | 9 | 4 | 9204 | 1112 | pont,cact,Pdk1,CG3511 | | 1.618e-02 | -4.12 | CG11802 (uncharacterized protein) | protein interactions | 32145 | 9 | 4 | 9204 | 1112 | Orc6,Unr,mad2,Nup62 | | 1.618e-02 | -4.12 | slim (scruin like at the midline) | protein interactions | 37121 | 9 | 4 | 9204 | 1112 | EloC,p38a,RpS27A,Vha36-1 | | 1.618e-02 | -4.12 | CG8507 (uncharacterized protein) | protein interactions | 41205 | 9 | 4 | 9204 | 1112 | CG4511,lqfR,LpR2,Cwc25 | | 1.618e-02 | -4.12 | RpS19b (Ribosomal protein S19b) | protein interactions | 42830 | 9 | 4 | 9204 | 1112 | CycE,Gdap2,foxo,Tm1 | | 1.618e-02 | -4.12 | CG3517 (uncharacterized protein) | protein interactions | 42324 | 9 | 4 | 9204 | 1112 | Pfrx,MED4,CG6276,CG2199 | | 1.618e-02 | -4.12 | Spn27A (Serpin 27A) | protein interactions | 45815 | 9 | 4 | 9204 | 1112 | CG11164,ea,spz,Sp7 | | 1.618e-02 | -4.12 | messenger ribonucleoprotein complex | cellular component | GO:1990124 | 17 | 5 | 11940 | 1105 | cup,nos,tral,yps,eIF4E1 | | 1.628e-02 | -4.12 | ENK\_UV\_RESPONSE\_EPIDERMIS\_UP | MSigDB lists | ENK\_UV\_RESPONSE\_EPIDERMIS\_UP | 95 | 23 | 4161 | 644 | Prosbeta4,Nmt,DNApol-epsilon255,Prosalpha5,Grip84,gammaTub37C,CG11807,LpR2,AIF,pont,Rpn3,nmdyn-D6,Pglym78,CG2201,alc,Cse1,CG11858,DppIII,Ts,CG42542,scramb1,rept,CG11089 | | 1.634e-02 | -4.11 | Prosalpha7 (Proteasome alpha7 subunit) | protein interactions | 36018 | 51 | 12 | 9204 | 1112 | Rpn13,AnxB10,Rpn2,Prosalpha5,Pomp,Prosalpha4,Prosbeta5,Prosbeta6,Prosbeta3,Prosalpha3,Rpt2,Prosbeta4 | | 1.642e-02 | -4.11 | purine ribonucleoside diphosphate metabolic process | biological process | GO:0009179 | 30 | 7 | 12345 | 1129 | Pfk,Gapdh1,Eno,Pgk,PyK,Pfrx,Pglym78 | | 1.642e-02 | -4.11 | purine nucleoside diphosphate metabolic process | biological process | GO:0009135 | 30 | 7 | 12345 | 1129 | Pgk,Eno,Gapdh1,Pfk,Pglym78,Pfrx,PyK | | 1.642e-02 | -4.11 | ribonucleoside diphosphate metabolic process | biological process | GO:0009185 | 30 | 7 | 12345 | 1129 | Eno,Gapdh1,Pgk,Pfk,Pfrx,Pglym78,PyK | | 1.642e-02 | -4.11 | peptidyl-tyrosine dephosphorylation | biological process | GO:0035335 | 44 | 9 | 12345 | 1129 | Ptpmeg,Ptp10D,Ptpmeg2,mop,Ptp69D,ssh,Ptp61F,twe,CG5026 | | 1.643e-02 | -4.11 | - | gene3d domains | 1.10.10.1880 | 5 | 3 | 4929 | 622 | Marf1,tapas,osk | | 1.645e-02 | -4.11 | GO\_PROTEIN\_PHOSPHORYLATION | MSigDB lists | GO\_PROTEIN\_PHOSPHORYLATION | 248 | 51 | 4161 | 644 | Prosalpha5,Mo25,Abi,Mekk1,RpS27A,tkv,rl,Cdk1,Tfb4,Prosalpha4,CG7530,Rpt2,CG7115,Cdk7,Prosbeta3,grp,Rpn2,Rpn12,CG8475,SkpA,CG4041,Prosbeta4,CG7414,Prosalpha3,PpD3,CycE,SRPK,Pdk1,Jafrac2,p38a,Prosbeta5,Cdk5,CycB,Rpn3,Cul1,CycC,CkIIalpha,Tlk,alc,dco,Traf6,Prosbeta6,Pi3K92E,Nprl2,fray,r,CG17698,Pdk,Tao,RpL40,CG8726 | | 1.648e-02 | -4.11 | GO\_POSITIVE\_REGULATION\_OF\_CYTOSKELETON\_ORGANIZATION | MSigDB lists | GO\_POSITIVE\_REGULATION\_OF\_CYTOSKELETON\_ORGANIZATION | 50 | 14 | 4161 | 644 | Arpc4,Tm1,Grip128,Cdk5,CycB,Pfdn2,Smox,Grip84,Grip75,Atg6,Arpc1,Abi,Arf51F,Arpc2 | | 1.648e-02 | -4.11 | GO\_OXIDOREDUCTASE\_COMPLEX | MSigDB lists | GO\_OXIDOREDUCTASE\_COMPLEX | 50 | 14 | 4161 | 644 | ND-24,ND-30,ND-51,ND-23,CG8199,SdhA,ND-B18,ND-B22,CG17691,ND-75,ND-SGDH,ND-49,Nc73EF,ND-39 | | 1.648e-02 | -4.11 | GSE24726\_WT\_VS\_E2\_2\_KO\_PDC\_DAY4\_POST\_DELETION\_UP | MSigDB lists | GSE24726\_WT\_VS\_E2\_2\_KO\_PDC\_DAY4\_POST\_DELETION\_UP | 50 | 14 | 4161 | 644 | Cul2,Rab10,Nurf-38,Vsp37A,Tim8,Echs1,STUB1,Coq7,Tspo,CG13630,Gk2,UQCR-Q,scramb1,bys | | 1.648e-02 | -4.11 | GO\_CELLULAR\_CARBOHYDRATE\_BIOSYNTHETIC\_PROCESS | MSigDB lists | GO\_CELLULAR\_CARBOHYDRATE\_BIOSYNTHETIC\_PROCESS | 27 | 9 | 4161 | 644 | RpS27A,Got1,Gyg,Pgm1,CG9391,RpL40,GalT1,CG9485,AGBE | | 1.648e-02 | -4.11 | GO\_SIGNAL\_TRANSDUCTION\_IN\_RESPONSE\_TO\_DNA\_DAMAGE | MSigDB lists | GO\_SIGNAL\_TRANSDUCTION\_IN\_RESPONSE\_TO\_DNA\_DAMAGE | 27 | 9 | 4161 | 644 | RpS27A,Not10,Rbf,twin,Tip60,RpL40,Cdk1,p38a,CycB | | 1.648e-02 | -4.11 | CHIARADONNA\_NEOPLASTIC\_TRANSFORMATION\_CDC25\_DN | MSigDB lists | CHIARADONNA\_NEOPLASTIC\_TRANSFORMATION\_CDC25\_DN | 27 | 9 | 4161 | 644 | milt,Tspo,Aldh,Tsp74F,Acph-1,Prip,Nmt,CkIIalpha,Mitofilin | | 1.658e-02 | -4.10 | GO\_CELLULAR\_MACROMOLECULE\_LOCALIZATION | MSigDB lists | GO\_CELLULAR\_MACROMOLECULE\_LOCALIZATION | 498 | 94 | 4161 | 644 | RpL13,ERp60,Pen,Surf4,Faf2,RpS28b,RpS27A,RpL9,Rab32,CG44422,rept,CG4041,ste14,CG31729,cyc,Arl6IP1,Cse1,RpLP0,RpS13,RpS30,Rrp40,RpL13A,CG7530,Tim8,RpL24,Syx13,CanA-14F,RpL37a,Pex12,Hsc70-3,Atg4b,mbo,Snx6,Rab5,Ranbp21,Atg6,RpS2,Phb2,Oseg1,Pp2B-14D,CG11110,Vha36-1,RpL30,CG8728,Pex10,milt,Tom40,Cdk1,cact,rl,RpS23,Rab10,shrb,Stam,Arts,CG5734,GCC88,CHOp24,AP-1-2beta,Karybeta3,HDAC6,CG9393,sau,CG2100,Tsp86D,Marf,RpL12,CG8314,Kap-alpha3,Cdk5,CG8004,RpS18,RpS27,Vamp7,RpL26,Clc,RpL11,Arf51F,Sec23,Tspo,RpL40,RpS7,Rep,RpS4,RpL38,RpS16,lt,GAPsec,Pex13,dco,RpS24,CG1847,Dd,RpL23 | | 1.663e-02 | -4.10 | GSE22886\_IGM\_MEMORY\_BCELL\_VS\_BLOOD\_PLASMA\_CELL\_DN | MSigDB lists | GSE22886\_IGM\_MEMORY\_BCELL\_VS\_BLOOD\_PLASMA\_CELL\_DN | 90 | 22 | 4161 | 644 | CkIIalpha,CG2493,Aos1,DppIII,CysRS,Ufm1,CG42542,spag,PDCD-5,Faf2,CG1827,CSN6,mtgo,Top1,Hat1,Stip1,l(1)G0196,CCT5,alpha-Man-Ia,CG16979,Cdk5,pont | | 1.664e-02 | -4.10 | Rab GTPase binding | molecular function | GO:0017137 | 90 | 15 | 12176 | 1119 | MICAL-like,GCC88,Tomosyn,cbs,CG32485,Sec15,Crag,GAPsec,Rep,sau,CG4041,Gmap,CG4557,Gdi,CG3309 | | 1.675e-02 | -4.09 | positive regulation of canonical Wnt signaling pathway | biological process | GO:0090263 | 37 | 8 | 12345 | 1129 | tws,Ssdp,dco,trbd,spen,pont,nej,mts | | 1.675e-02 | -4.09 | negative regulation of MAPK cascade | biological process | GO:0043409 | 37 | 8 | 12345 | 1129 | ATPsynD,alph,SkpA,CG17746,PpV,park,Cbl,Ptp61F | | 1.683e-02 | -4.08 | Aldo/keto\_reductase | interpro domains | IPR020471 | 10 | 4 | 11383 | 1233 | CG10638,CG10863,CG2767,CG9436 | | 1.683e-02 | -4.08 | Cullin\_repeat-like\_dom\_sf | interpro domains | IPR016159 | 10 | 4 | 11383 | 1233 | Exo70,Cul5,Cul1,Cul2 | | 1.683e-02 | -4.08 | L30e-like | interpro domains | IPR029064 | 10 | 4 | 11383 | 1233 | RpL7A,CG14100,RpL30,NHP2 | | 1.688e-02 | -4.08 | GO\_CHEMICAL\_HOMEOSTASIS | MSigDB lists | GO\_CHEMICAL\_HOMEOSTASIS | 209 | 44 | 4161 | 644 | Mcad,csw,Sfxn2,EcR,Sod1,adp,mino,Prip,CG42542,Stoml2,Gk2,zyd,regucalcin,ACC,Pka-R2,Npc2a,SERCA,Got1,Irp-1B,GlyP,wal,crq,Calx,CG2493,CG1275,Vha36-1,AnxB10,nrv1,Pp2B-14D,Pka-R1,Irk1,CG17593,Vha13,Ptp61F,Med,Pdk,Atpalpha,CanA-14F,Mitofilin,Scox,anne,Pfk,Ac3,CG8112 | | 1.691e-02 | -4.08 | GO\_ACTIVATION\_OF\_IMMUNE\_RESPONSE | MSigDB lists | GO\_ACTIVATION\_OF\_IMMUNE\_RESPONSE | 111 | 26 | 4161 | 644 | RpS27A,Arpc1,Prosalpha5,Arpc2,Prosbeta4,Prosalpha3,Rpn3,Cul1,Arpc4,Uev1A,Pdk1,cact,rl,Prosbeta5,Rpn2,Rpn12,Traf6,Rpt2,Prosalpha4,Prosbeta3,dnc,RpL40,Prosbeta6,Pi3K92E,SkpA,Stoml2 | | 1.692e-02 | -4.08 | DANG\_MYC\_TARGETS\_UP | MSigDB lists | DANG\_MYC\_TARGETS\_UP | 85 | 21 | 4161 | 644 | mRpL10,Nap1,RpL9,RpL26,CCT5,Eno,CycB,kri,Top1,RpL13,Shmt,CG12018,P5cr,Gapdh1,CycC,eIF4E1,RpS16,RpS13,r,Ts,Prosbeta6 | | 1.692e-02 | -4.08 | GSE10239\_NAIVE\_VS\_DAY4.5\_EFF\_CD8\_TCELL\_DN | MSigDB lists | GSE10239\_NAIVE\_VS\_DAY4.5\_EFF\_CD8\_TCELL\_DN | 85 | 21 | 4161 | 644 | Pgk,Pen,mad2,Coq7,CycE,Prosalpha5,Uch,Naa20A,CG12375,scramb1,RfC38,Ts,SelD,fray,CG10470,grp,Dhfr,mRpL18,CG11899,CG13601,Tsp86D | | 1.693e-02 | -4.08 | EcR (Ecdysone receptor) | protein interactions | 35540 | 101 | 20 | 9204 | 1112 | rig,Cul2,MED27,Chd64,CG13373,Cdk5,Rbf,nej,G9a,Nurf-38,Sec10,CycC,trr,EcR,Utx,Fkbp39,Pax,ari-1,CG9248,plu | | 1.694e-02 | -4.08 | p53-Independent G1/S DNA damage checkpoint | REACTOME pathways | R-DME-69613 | 58 | 15 | 3923 | 574 | Prosbeta4,Prosbeta3,Prosalpha5,RpS27A,twe,Prosalpha4,Rpn2,Prosbeta5,Prosbeta6,grp,Rpn3,Rpt2,RpL40,Prosalpha3,Rpn12 | | 1.694e-02 | -4.08 | G1/S DNA Damage Checkpoints | REACTOME pathways | R-DME-69615 | 58 | 15 | 3923 | 574 | Prosbeta6,Rpn12,Prosalpha3,Rpt2,RpL40,Rpn3,grp,RpS27A,Prosbeta3,Prosalpha5,Prosbeta4,Prosbeta5,Rpn2,Prosalpha4,twe | | 1.694e-02 | -4.08 | p53-Independent DNA Damage Response | REACTOME pathways | R-DME-69610 | 58 | 15 | 3923 | 574 | Prosbeta6,RpL40,Rpn3,Rpt2,Prosalpha3,Rpn12,grp,Prosbeta3,Prosalpha5,RpS27A,Prosbeta4,Rpn2,Prosbeta5,twe,Prosalpha4 | | 1.694e-02 | -4.08 | Ubiquitin Mediated Degradation of Phosphorylated Cdc25A | REACTOME pathways | R-DME-69601 | 58 | 15 | 3923 | 574 | Prosalpha5,Prosbeta3,RpS27A,Prosbeta4,Rpn2,Prosbeta5,twe,Prosalpha4,Prosbeta6,RpL40,Rpn3,Rpt2,Rpn12,Prosalpha3,grp | | 1.697e-02 | -4.08 | GSE37416\_CTRL\_VS\_6H\_F\_TULARENSIS\_LVS\_NEUTROPHIL\_DN | MSigDB lists | GSE37416\_CTRL\_VS\_6H\_F\_TULARENSIS\_LVS\_NEUTROPHIL\_DN | 55 | 15 | 4161 | 644 | Rnmt,CG13601,Pgm1,Pglym78,Pgk,Su(H),RhoGDI,sbr,Pdxk,cact,Roe1,Eno,AGBE,CG16979,O-fut1 | | 1.697e-02 | -4.08 | GSE5542\_UNTREATED\_VS\_IFNA\_AND\_IFNG\_TREATED\_EPITHELIAL\_CELLS\_24H\_DN | MSigDB lists | GSE5542\_UNTREATED\_VS\_IFNA\_AND\_IFNG\_TREATED\_EPITHELIAL\_CELLS\_24H\_DN | 55 | 15 | 4161 | 644 | rl,p38a,CG15525,anne,Nca,cathD,Usp39,pen-2,eIF4A,snz,alc,Sirt7,Pp2A-29B,CG14407,Mctp | | 1.699e-02 | -4.08 | G2/M Transition | REACTOME pathways | R-DME-69275 | 79 | 19 | 3923 | 574 | Prosbeta5,Rpn2,twe,Prosalpha4,Prosalpha5,RpS27A,Prosbeta4,Rpt2,RpL40,Prosalpha3,SkpA,Prosbeta3,Cdk7,Rpn3,Cul1,Rpn12,Cdk1,CycB,Prosbeta6 | | 1.705e-02 | -4.07 | MORF\_AP2M1 | MSigDB lists | MORF\_AP2M1 | 127 | 29 | 4161 | 644 | SdhA,Bin1,Pglym78,lwr,Thiolase,Pgk,PIG-C,CSN6,Arpc2,Prosalpha3,Clc,Idh3b,Pp2A-29B,Prosbeta6,levy,Rpb8,Stoml2,SkpA,Pka-R1,CHOp24,AP-1-2beta,RhoGDI,tko,Rpt2,Dd,RpS30,Rpn12,Sod1,LysRS | | 1.717e-02 | -4.06 | GSE3982\_EOSINOPHIL\_VS\_MAST\_CELL\_DN | MSigDB lists | GSE3982\_EOSINOPHIL\_VS\_MAST\_CELL\_DN | 80 | 20 | 4161 | 644 | CG11134,CSN6,eIF3m,Pfas,ND-39,RpL26,Hmg-2,CG5510,r-l,CCT5,Eno,mRpL16,Rpn2,Cse1,RpS13,CG6610,RpS4,NHP2,Stoml2,CG14194 | | 1.717e-02 | -4.06 | lipid biosynthetic process | biological process | GO:0008610 | 211 | 29 | 12345 | 1129 | Pxt,Agpat2,ACC,Seipin,CG6178,PIG-P,Desat1,Marf,CG17270,AcCoAS,CG7718,CG2201,PIG-C,SERCA,Npc2a,Sccpdh1,CG7840,CG8311,CG33977,PIG-H,CG10425,SREBP,CG10932,foxo,Pld,Pi3K92E,PIG-L,CG31523,mino | | 1.721e-02 | -4.06 | ligase activity, forming carbon-sulfur bonds | molecular function | GO:0016877 | 37 | 8 | 12176 | 1119 | APP-BP1,ScsbetaA,AcCoAS,Atg7,CG6178,Uba5,Scsalpha1,Aos1 | | 1.728e-02 | -4.06 | GSE34156\_NOD2\_LIGAND\_VS\_NOD2\_AND\_TLR1\_TLR2\_LIGAND\_6H\_TREATED\_MONOCYTE\_UP | MSigDB lists | GSE34156\_NOD2\_LIGAND\_VS\_NOD2\_AND\_TLR1\_TLR2\_LIGAND\_6H\_TREATED\_MONOCYTE\_UP | 60 | 16 | 4161 | 644 | RpL11,Mat89Ba,Pmi,CG10344,RpS18,RpL13,eEF1beta,RpS13,CG3511,Pex13,CG2938,CG12713,RpL38,CG8204,RpL40,mbo | | 1.728e-02 | -4.06 | ZHONG\_SECRETOME\_OF\_LUNG\_CANCER\_AND\_FIBROBLAST | MSigDB lists | ZHONG\_SECRETOME\_OF\_LUNG\_CANCER\_AND\_FIBROBLAST | 60 | 16 | 4161 | 644 | Npc2a,Prosalpha3,Got1,Prosbeta4,Prosalpha5,Eno,Prosbeta5,Pgk,CG9391,Moe,Prosbeta3,CG1532,Prosbeta6,RpL40,RhoGDI,cathD | | 1.728e-02 | -4.06 | GSE22033\_WT\_VS\_PPARG\_KO\_MEF\_DN | MSigDB lists | GSE22033\_WT\_VS\_PPARG\_KO\_MEF\_DN | 60 | 16 | 4161 | 644 | Pp2B-14D,CG1890,Pi3K92E,tzn,CG4407,TBCC,Prosalpha3,Idh,CanA-14F,Mo25,RpS23,nAChRalpha4,Cdk1,CG16865,alph,Uch-L5 | | 1.728e-02 | -4.06 | GSE17721\_12H\_VS\_24H\_PAM3CSK4\_BMDC\_DN | MSigDB lists | GSE17721\_12H\_VS\_24H\_PAM3CSK4\_BMDC\_DN | 60 | 16 | 4161 | 644 | YL-1,CG11807,Pglym78,f-cup,ND-30,Pgk,shop,SdhA,CCT3,Cdk5,Gclc,EcR,Cse1,HDAC6,CG33156,Jafrac1 | | 1.734e-02 | -4.05 | GSE10239\_NAIVE\_VS\_KLRG1INT\_EFF\_CD8\_TCELL\_DN | MSigDB lists | GSE10239\_NAIVE\_VS\_KLRG1INT\_EFF\_CD8\_TCELL\_DN | 75 | 19 | 4161 | 644 | CycE,PDCD-5,Sccpdh1,DNApol-epsilon255,Uch,Prosbeta4,ND-30,mad2,mRpS10,r-l,RhoGAP54D,grp,CG10470,Dhfr,Tsp86D,Naa20A,scramb1,Ts,CG8204 | | 1.734e-02 | -4.05 | regulation of stress-activated MAPK cascade | biological process | GO:0032872 | 67 | 12 | 12345 | 1129 | PEK,SkpA,alph,CG17746,Cka,Lkb1,PpV,park,ben,Tak1,Mekk1,Ptp61F | | 1.734e-02 | -4.05 | positive regulation of protein kinase activity | biological process | GO:0045860 | 67 | 12 | 12345 | 1129 | raptor,Lkb1,RpLP1,Tao,Parp16,Tak1,ASPP,Pdk1,fray,Pi3K92E,Abi,Mekk1 | | 1.734e-02 | -4.05 | regulation of stress-activated protein kinase signaling cascade | biological process | GO:0070302 | 67 | 12 | 12345 | 1129 | Ptp61F,Mekk1,ben,Tak1,park,PpV,Lkb1,Cka,CG17746,alph,SkpA,PEK | | 1.737e-02 | -4.05 | SDR\_fam | interpro domains | IPR002347 | 57 | 12 | 11383 | 1233 | CG10425,CG31549,CG31548,CG4842,CG7322,Adh,sni,scu,CG9265,CG3842,Sptr,CG12171 | | 1.743e-02 | -4.05 | GSE12392\_CD8A\_POS\_VS\_NEG\_SPLEEN\_DC\_UP | MSigDB lists | GSE12392\_CD8A\_POS\_VS\_NEG\_SPLEEN\_DC\_UP | 70 | 18 | 4161 | 644 | mbo,Su(H),AGBE,CG15141,CG9485,CG4390,Pgm1,CG1532,CG16868,Prosbeta5,asf1,Surf4,Grip75,Pp2A-29B,cl,mtgo,Sply,Rab10 | | 1.743e-02 | -4.05 | GSE37301\_GRANULOCYTE\_MONOCYTE\_PROGENITOR\_VS\_RAG2\_KO\_NK\_CELL\_UP | MSigDB lists | GSE37301\_GRANULOCYTE\_MONOCYTE\_PROGENITOR\_VS\_RAG2\_KO\_NK\_CELL\_UP | 70 | 18 | 4161 | 644 | Ranbp21,Iml1,CG33156,tun,AnxB10,CG10189,Cdk7,Sec15,scu,AIF,Ptpmeg,Gdi,Cul1,Rab10,Uch,CG2924,Rnmt,Rab32 | | 1.743e-02 | -4.05 | GSE5542\_IFNG\_VS\_IFNA\_TREATED\_EPITHELIAL\_CELLS\_24H\_UP | MSigDB lists | GSE5542\_IFNG\_VS\_IFNA\_TREATED\_EPITHELIAL\_CELLS\_24H\_UP | 70 | 18 | 4161 | 644 | Prosbeta4,Gyg,Arpc1,CG32549,Ptpmeg,Coq7,thoc6,Chrac-14,Uch-L5,CG7530,Kap3,Rrp40,RfC38,Ubc4,CG42542,Uba5,Jafrac1,Cbp80 | | 1.743e-02 | -4.05 | GSE25087\_FETAL\_VS\_ADULT\_TREG\_UP | MSigDB lists | GSE25087\_FETAL\_VS\_ADULT\_TREG\_UP | 70 | 18 | 4161 | 644 | ND-39,Hsc70-3,LSm3,yip2,Faf2,Sccpdh1,Bin1,ND-B18,Stip1,CG9911,alphaTub84B,CG4390,mRpL13,ScsbetaA,colt,r,Cyp1,Uba5 | | 1.753e-02 | -4.04 | Biosynthesis of amino acids | KEGG pathways | ko01230 | 65 | 17 | 3198 | 496 | Eno,CG11899,Idh,aay,CG1640,PyK,Pgk,Gapdh1,Irp-1B,Pfk,CG5028,Got1,Shmt,Pglym78,P5cr,Idh3b,CG6767 | | 1.753e-02 | -4.04 | Biosynthesis of amino acids | KEGG pathways | dme01230 | 65 | 17 | 3198 | 496 | CG1640,PyK,Pgk,Gapdh1,CG11899,Eno,Idh,aay,Shmt,P5cr,Pglym78,Idh3b,CG6767,Irp-1B,Pfk,CG5028,Got1 | | 1.758e-02 | -4.04 | circadian rhythm | biological process | GO:0007623 | 91 | 15 | 12345 | 1129 | Ac3,Pka-R2,Bx,dco,cyc,CkIIalpha,lid,tyf,Ork1,Irk1,foxo,gw,cu,nej,dnc | | 1.772e-02 | -4.03 | GO\_CELL\_CYCLE | MSigDB lists | GO\_CELL\_CYCLE | 446 | 85 | 4161 | 644 | CkIIalpha,Pp2B-14D,faf,Phb2,Atg6,Ts,PpV,mbo,not,APP-BP1,AGO3,Prosalpha3,CanA-14F,Pp2A-29B,DNApol-epsilon255,Smox,CHMP2B,Prosbeta5,spn-E,lwr,piwi,Rpt2,csw,mts,Cul2,Not10,eIF4E1,Rpn12,mael,twin,RpS27A,Grip84,Caf1-105,Cul5,Rbf,Dd,Dhfr,CG30291,tws,dco,Aladin,Sirt7,alc,Tlk,APC7,Prosbeta6,Pka-R1,Usp39,Tao,RpL40,Wnt10,Prosbeta4,Arf51F,PpD3,gammaTub37C,RpS27,CycE,Ac3,p38a,pont,Cdk5,CycB,msps,Cul1,Rpn3,Marf,raptor,Prosbeta3,Cdk7,grp,Rpn2,CG1812,CG8188,SkpA,Grip128,shrb,Mo25,Prosalpha5,Grip75,shu,rl,Cdk1,Klp10A,mad2,Fbw5 | | 1.789e-02 | -4.02 | Prot-tyrosine\_phosphatase-like | interpro domains | IPR029021 | 38 | 9 | 11383 | 1233 | ssh,Ptp10D,Ptp69D,Ptpmeg2,Ptp61F,Ptpmeg,CG5026,mop,csw | | 1.791e-02 | -4.02 | REACTOME\_RECRUITMENT\_OF\_MITOTIC\_CENTROSOME\_PROTEINS\_AND\_COMPLEXES | MSigDB lists | REACTOME\_RECRUITMENT\_OF\_MITOTIC\_CENTROSOME\_PROTEINS\_AND\_COMPLEXES | 23 | 8 | 4161 | 644 | Grip128,msps,CycB,Cdk1,Grip84,gammaTub37C,Pp2A-29B,dco | | 1.791e-02 | -4.02 | LENAOUR\_DENDRITIC\_CELL\_MATURATION\_DN | MSigDB lists | LENAOUR\_DENDRITIC\_CELL\_MATURATION\_DN | 23 | 8 | 4161 | 644 | cact,CG7997,scramb1,Cyp1,Gk2,CG5734,scu,CG14516 | | 1.791e-02 | -4.02 | GCM\_BECN1 | MSigDB lists | GCM\_BECN1 | 23 | 8 | 4161 | 644 | RhoGDI,CG14894,SkpA,Atg6,CG4390,Tlk,YL-1,CG2201 | | 1.791e-02 | -4.02 | ribonucleoside bisphosphate metabolic process | biological process | GO:0033875 | 52 | 10 | 12345 | 1129 | CG7920,AcCoAS,ScsbetaA,scu,CG6178,Scsalpha1,Ppcdc,CG1774,CG8199,CG17691 | | 1.791e-02 | -4.02 | purine nucleoside bisphosphate metabolic process | biological process | GO:0034032 | 52 | 10 | 12345 | 1129 | CG6178,Scsalpha1,scu,CG1774,Ppcdc,CG17691,CG8199,CG7920,AcCoAS,ScsbetaA | | 1.791e-02 | -4.02 | nucleoside bisphosphate metabolic process | biological process | GO:0033865 | 52 | 10 | 12345 | 1129 | ScsbetaA,AcCoAS,CG7920,CG17691,CG8199,Ppcdc,CG1774,scu,Scsalpha1,CG6178 | | 1.794e-02 | -4.02 | fly chr19C1-19C1\|1-64 cM | chromosome location | fly chr19C1-19C1\|1-64 cM | 29 | 5 | 25010 | 1343 | Rab10,AnxB10,sw,CG9581,CG17068 | | 1.794e-02 | -4.02 | fly chr19C1-19C1\|1-64 | chromosome location | fly chr19C1-19C1\|1-64 | 29 | 5 | 25010 | 1343 | sw,CG9581,CG17068,Rab10,AnxB10 | | 1.795e-02 | -4.02 | REACTOME\_RIG\_I\_MDA5\_MEDIATED\_INDUCTION\_OF\_IFN\_ALPHA\_BETA\_PATHWAYS | MSigDB lists | REACTOME\_RIG\_I\_MDA5\_MEDIATED\_INDUCTION\_OF\_IFN\_ALPHA\_BETA\_PATHWAYS | 11 | 5 | 4161 | 644 | RpL40,Traf6,RpS27A,Ubc4,cact | | 1.795e-02 | -4.02 | REACTOME\_ION\_TRANSPORT\_BY\_P\_TYPE\_ATPASES | MSigDB lists | REACTOME\_ION\_TRANSPORT\_BY\_P\_TYPE\_ATPASES | 11 | 5 | 4161 | 644 | ATP8A,Atpalpha,nrv1,CG31729,SERCA | | 1.795e-02 | -4.02 | GO\_PYRIMIDINE\_NUCLEOSIDE\_MONOPHOSPHATE\_METABOLIC\_PROCESS | MSigDB lists | GO\_PYRIMIDINE\_NUCLEOSIDE\_MONOPHOSPHATE\_METABOLIC\_PROCESS | 11 | 5 | 4161 | 644 | r-l,Ts,kri,r,Shmt | | 1.795e-02 | -4.02 | GO\_MYELOID\_LEUKOCYTE\_ACTIVATION | MSigDB lists | GO\_MYELOID\_LEUKOCYTE\_ACTIVATION | 11 | 5 | 4161 | 644 | Pi3K92E,Vamp7,Traf6,Su(H),Pi4KIIalpha | | 1.795e-02 | -4.02 | GO\_INO80\_TYPE\_COMPLEX | MSigDB lists | GO\_INO80\_TYPE\_COMPLEX | 11 | 5 | 4161 | 644 | pont,Rcd5,rept,Uch-L5,Tip60 | | 1.795e-02 | -4.02 | LEE\_LIVER\_CANCER\_MYC\_TGFA\_UP | MSigDB lists | LEE\_LIVER\_CANCER\_MYC\_TGFA\_UP | 11 | 5 | 4161 | 644 | scramb1,aay,RpL12,Grip75,RpS24 | | 1.795e-02 | -4.02 | SIG\_CD40PATHWAYMAP | MSigDB lists | SIG\_CD40PATHWAYMAP | 11 | 5 | 4161 | 644 | Traf6,rl,cact,p38a,Pi3K92E | | 1.795e-02 | -4.02 | GO\_PYRIMIDINE\_RIBONUCLEOSIDE\_METABOLIC\_PROCESS | MSigDB lists | GO\_PYRIMIDINE\_RIBONUCLEOSIDE\_METABOLIC\_PROCESS | 11 | 5 | 4161 | 644 | r,kri,r-l,nmdyn-D7,nmdyn-D6 | | 1.795e-02 | -4.02 | GO\_PYRIMIDINE\_RIBONUCLEOTIDE\_METABOLIC\_PROCESS | MSigDB lists | GO\_PYRIMIDINE\_RIBONUCLEOTIDE\_METABOLIC\_PROCESS | 11 | 5 | 4161 | 644 | kri,r-l,r,nmdyn-D6,nmdyn-D7 | | 1.795e-02 | -4.02 | NGO\_MALIGNANT\_GLIOMA\_1P\_LOH | MSigDB lists | NGO\_MALIGNANT\_GLIOMA\_1P\_LOH | 11 | 5 | 4161 | 644 | Hsc70-3,Gapdh1,Eno,Echs1,Pgk | | 1.795e-02 | -4.02 | GROSS\_HIF1A\_TARGETS\_DN | MSigDB lists | GROSS\_HIF1A\_TARGETS\_DN | 11 | 5 | 4161 | 644 | Pgk,AGBE,Eno,Pgm1,Gapdh1 | | 1.795e-02 | -4.02 | REACTOME\_NEGATIVE\_REGULATION\_OF\_FGFR\_SIGNALING | MSigDB lists | REACTOME\_NEGATIVE\_REGULATION\_OF\_FGFR\_SIGNALING | 11 | 5 | 4161 | 644 | rl,mts,Pp2A-29B,RpS27A,RpL40 | | 1.805e-02 | -4.01 | Cyt-c-p (Cytochrome c proximal) | protein interactions | 34996 | 40 | 10 | 9204 | 1112 | Mf,up,slam,Adh,SkpA,dl,Sap47,Argk,CG46385,Scp1 | | 1.805e-02 | -4.01 | CG42369 (uncharacterized protein) | protein interactions | 33894 | 40 | 10 | 9204 | 1112 | RpS23,CG10470,Droj2,RpL9,wisp,ERp60,RpS13,CG4238,Sccpdh1,RpL23A | | 1.823e-02 | -4.00 | cellular homeostasis | biological process | GO:0019725 | 230 | 31 | 12345 | 1129 | ERp60,wash,Pgk,Jafrac1,park,Jafrac2,capt,up,CG9911,Atpalpha,SERCA,Rbcn-3A,CG10470,anne,Vha68-2,mute,Plap,Debcl,Fer1HCH,Grx1,EMRE,zyd,Seipin,alc,ZnT35C,CG12773,fwe,CG14407,Scox,nrv1,Trxr-1 | | 1.835e-02 | -4.00 | positive regulation of transcription, DNA-templated | biological process | GO:0045893 | 473 | 57 | 12345 | 1129 | S2P,Atu,REPTOR,Mybbp1A,Atf6,Rel,Cdk7,Su(H),Myd88,CG2199,Utx,Cf2,SREBP,foxo,hb,trr,dl,esc,not,Nf-YA,pygo,CG2129,Bx,d4,cyc,Wbp2,cnc,Bap170,EloC,vers,EcR,Rbfox1,CycC,bcd,NFAT,CG17328,CG1602,Tip60,Rcd5,RYBP,CG11710,lid,rept,Sox102F,nej,lin,Mitf,e(y)1,BigH1,Sgf11,cad,Stat92E,E2f1,TfIIFalpha,ovo,apt,Trf2 | | 1.840e-02 | -4.00 | MORF\_RPA2 | MSigDB lists | MORF\_RPA2 | 96 | 23 | 4161 | 644 | eIF3i,CG17454,Arl6IP1,r,PrBP,Cyp1,P5CDh1,Dd,CkIIalpha,Phb2,GCS2beta,CG9393,Tfb4,Pgk,Arpc4,Marf,CG13630,CSN6,Mitofilin,eIF3m,Rcd5,Nc73EF,Lsm12 | | 1.840e-02 | -4.00 | YAO\_TEMPORAL\_RESPONSE\_TO\_PROGESTERONE\_CLUSTER\_17 | MSigDB lists | YAO\_TEMPORAL\_RESPONSE\_TO\_PROGESTERONE\_CLUSTER\_17 | 96 | 23 | 4161 | 644 | Phb2,CG8188,Rpn12,mRpL18,Aos1,tko,DppIII,ND-B22,Roe1,Exo70,Nap1,CG8507,CSN6,UQCR-Q,mRpL16,ND-30,CG30010,ND-24,Coq7,ND-B18,CG15525,STUB1,pont | | 1.842e-02 | -3.99 | fly chr37A1-37A1\|2-54 | chromosome location | fly chr37A1-37A1\|2-54 | 11 | 3 | 25010 | 1343 | CG10336,Faf2,Grip71 | | 1.842e-02 | -3.99 | fly chr37A1-37A1\|2-54 cM | chromosome location | fly chr37A1-37A1\|2-54 cM | 11 | 3 | 25010 | 1343 | Grip71,Faf2,CG10336 | | 1.847e-02 | -3.99 | positive regulation of mitotic cell cycle phase transition | biological process | GO:1901992 | 24 | 6 | 12345 | 1129 | APC7,bru1,Sld5,twe,CycE,Su(H) | | 1.855e-02 | -3.99 | mask (multiple ankyrin repeats single KH domain) | protein interactions | 50070 | 29 | 8 | 9204 | 1112 | rl,csw,park,Pdk1,Pink1,PGRP-LE,Tak1,CG17328 | | 1.855e-02 | -3.99 | bru1 (bruno 1) | protein interactions | 34648 | 29 | 8 | 9204 | 1112 | osk,Cwc25,apt,smg,eIF4E1,Rbfox1,bru1,cup | | 1.855e-02 | -3.99 | gammaTub37C (gamma-Tubulin at 37C) | protein interactions | 35199 | 29 | 8 | 9204 | 1112 | swa,Grip128,Grip84,CG8475,Grip71,cnn,mael,Grip75 | | 1.855e-02 | -3.99 | CG11857 (uncharacterized protein) | protein interactions | 43042 | 29 | 8 | 9204 | 1112 | ATPsynO,ATPsynG,SERCA,CG31729,Tspo,CG15098,Tak1,Surf4 | | 1.855e-02 | -3.99 | calypso (calypso) | protein interactions | 36794 | 29 | 8 | 9204 | 1112 | RpL11,RpL23A,alphaTub84B,RpL7A,alphaTub67C,RpL12,RpS27A,RpL13 | | 1.855e-02 | -3.99 | Karybeta3 (Karyopherin beta 3) | protein interactions | 40581 | 29 | 8 | 9204 | 1112 | Gyg,SERCA,Kap-alpha3,Atpalpha,Rtnl1,Pp2A-29B,REPTOR,Vha68-2 | | 1.855e-02 | -3.99 | strat (stratum) | protein interactions | 34142 | 29 | 8 | 9204 | 1112 | Rab10,SdhA,Gdi,CG4949,CG32473,PrBP,Scox,Tspo | | 1.863e-02 | -3.98 | GO\_PROTEOLYSIS | MSigDB lists | GO\_PROTEOLYSIS | 394 | 76 | 4161 | 644 | Prosbeta3,CG10465,UbcE2H,S2P,HDAC6,CG8188,ste24a,Vsp37A,Rpn2,SkpA,Stoml2,CG4572,CG11070,Tip60,Herc4,shrb,CSN6,CG14516,Prosalpha5,Cdk1,Uch-L5,pen-2,Fbw5,mad2,APC7,CG17593,Ubc4,Prosbeta6,RpL40,Usp39,stet,CG32473,CG9581,Prosbeta4,Uch,CycB,CG13630,CG8974,Cul1,Rpn3,Cul2,Prosalpha4,trbd,Rpt2,Rpn12,grsm,DppIII,Fur2,cyc,Faf2,RpS27A,STUB1,Cul5,CG16979,AIF,ERp60,CG2493,CG8728,Cyt-c-p,ari-1,Traf6,CG11110,faf,CG40045,CG6766,Atg4b,not,cathD,Prosalpha3,CG1827,Hsc70-3,Iru,Smox,Prosbeta5,lwr,Edem1 | | 1.871e-02 | -3.98 | Past1 (Putative Achaete Scute Target 1) | protein interactions | 41569 | 46 | 11 | 9204 | 1112 | Pp2A-29B,wal,Rpt2,CCT3,bor,Vha68-2,CCT6,Atpalpha,RpLP0,Mcad,Kap-alpha3 | | 1.878e-02 | -3.97 | GO\_RRNA\_METABOLIC\_PROCESS | MSigDB lists | GO\_RRNA\_METABOLIC\_PROCESS | 155 | 34 | 4161 | 644 | RpL9,RpS27A,RpS23,RpS28b,RpL13,RpL13A,Rrp40,RpS30,RpLP0,RpS13,l(1)G0004,RpL26,RpL24,RpS27,CG7728,RpL37a,Mat89Ba,RpL11,bys,RpL12,RpS18,dco,RpS2,RpS16,CG11858,RpL30,RpL23,RpS24,RpL40,NHP2,RpL38,RpS4,RpS7,mtTFB1 | | 1.881e-02 | -3.97 | GO\_CARBON\_OXYGEN\_LYASE\_ACTIVITY | MSigDB lists | GO\_CARBON\_OXYGEN\_LYASE\_ACTIVITY | 32 | 10 | 4161 | 644 | Pcd,Irp-1B,CG5285,CG11134,Thiolase,Uba5,pr,Echs1,Eno,Mtpalpha | | 1.884e-02 | -3.97 | GO\_ANTIGEN\_PROCESSING\_AND\_PRESENTATION | MSigDB lists | GO\_ANTIGEN\_PROCESSING\_AND\_PRESENTATION | 91 | 22 | 4161 | 644 | Clc,Prosalpha5,Rab32,Prosalpha3,Rab10,Sec23,Prosbeta4,ERp60,Rpn3,Prosbeta5,Klp10A,Rpn12,Traf6,Kap3,CG3860,Rpn2,Prosbeta3,Rpt2,Prosalpha4,cathD,AP-1-2beta,Prosbeta6 | | 1.888e-02 | -3.97 | nucleobase-containing compound kinase activity | molecular function | GO:0019205 | 24 | 6 | 12176 | 1119 | Ady43A,nmdyn-D7,nmdyn-D6,l(2)k01209,awd,Scsalpha1 | | 1.888e-02 | -3.97 | ubiquitin-like protein conjugating enzyme binding | molecular function | GO:0044390 | 24 | 6 | 12176 | 1119 | Aos1,ari-1,Uev1A,ben,CG8974,park | | 1.891e-02 | -3.97 | GSE22886\_UNSTIM\_VS\_STIM\_MEMORY\_TCELL\_DN | MSigDB lists | GSE22886\_UNSTIM\_VS\_STIM\_MEMORY\_TCELL\_DN | 112 | 26 | 4161 | 644 | Top1,CCT6,CG8112,CG9391,esc,Tim8,anne,Echs1,COX5A,Clc,CG30496,Fkbp59,ste14,Arts,CG11089,Cbp80,escl,heph,CG10418,Plap,mRpL13,nmdyn-D7,Cdk7,CG11899,Dhfr,csw | | 1.891e-02 | -3.97 | GRAESSMANN\_APOPTOSIS\_BY\_SERUM\_DEPRIVATION\_UP | MSigDB lists | GRAESSMANN\_APOPTOSIS\_BY\_SERUM\_DEPRIVATION\_UP | 128 | 29 | 4161 | 644 | Pi4KIIalpha,Surf4,CG7789,mtgo,Shmt,lwr,CG30010,Ac3,Jafrac2,RpS18,CG11710,Cdk5,Pfdn5,Prip,Atg6,CG1890,Tsp86D,RpL30,Sfxn2,CG10463,CycC,Adi1,zyd,scramb1,opm,Cyp1,Stam,RpL40,CG2938 | | 1.892e-02 | -3.97 | regulation of chromosome segregation | biological process | GO:0051983 | 45 | 9 | 12345 | 1129 | Rbf,mad2,Moe,wapl,Mos,Zwilch,cana,APC7,tws | | 1.894e-02 | -3.97 | pyridine-containing compound biosynthetic process | biological process | GO:0072525 | 12 | 4 | 12345 | 1129 | Pdxk,Nadsyn,CG33156,sgll | | 1.894e-02 | -3.97 | purine nucleoside monophosphate biosynthetic process | biological process | GO:0009127 | 12 | 4 | 12345 | 1129 | AMPdeam,Ady43A,Pfas,CG11089 | | 1.894e-02 | -3.97 | ubiquinone metabolic process | biological process | GO:0006743 | 12 | 4 | 12345 | 1129 | ND-39,Coq8,Coq6,Coq7 | | 1.894e-02 | -3.97 | purine ribonucleoside monophosphate biosynthetic process | biological process | GO:0009168 | 12 | 4 | 12345 | 1129 | Ady43A,AMPdeam,Pfas,CG11089 | | 1.894e-02 | -3.97 | histone H3-K9 modification | biological process | GO:0061647 | 12 | 4 | 12345 | 1129 | G9a,lid,egg,esc | | 1.894e-02 | -3.97 | quinone metabolic process | biological process | GO:1901661 | 12 | 4 | 12345 | 1129 | ND-39,Coq8,Coq6,Coq7 | | 1.894e-02 | -3.97 | phospholipid translocation | biological process | GO:0045332 | 12 | 4 | 12345 | 1129 | CG31729,ATP8A,scramb1,CG9981 | | 1.894e-02 | -3.97 | quinone biosynthetic process | biological process | GO:1901663 | 12 | 4 | 12345 | 1129 | Coq8,ND-39,Coq7,Coq6 | | 1.894e-02 | -3.97 | ubiquinone biosynthetic process | biological process | GO:0006744 | 12 | 4 | 12345 | 1129 | Coq7,Coq6,Coq8,ND-39 | | 1.894e-02 | -3.97 | negative regulation of antimicrobial humoral response | biological process | GO:0008348 | 12 | 4 | 12345 | 1129 | cact,p38a,Stat92E,scny | | 1.895e-02 | -3.97 | REACTOME\_DESTABILIZATION\_OF\_MRNA\_BY\_AUF1\_HNRNP\_D0 | MSigDB lists | REACTOME\_DESTABILIZATION\_OF\_MRNA\_BY\_AUF1\_HNRNP\_D0 | 46 | 13 | 4161 | 644 | Prosbeta6,Prosbeta5,Rpn3,RpL40,Prosbeta4,Prosalpha4,Rpt2,Prosalpha3,Prosbeta3,Prosalpha5,RpS27A,Rpn2,Rpn12 | | 1.895e-02 | -3.97 | GSE4142\_NAIVE\_BCELL\_VS\_PLASMA\_CELL\_UP | MSigDB lists | GSE4142\_NAIVE\_BCELL\_VS\_PLASMA\_CELL\_UP | 46 | 13 | 4161 | 644 | Pi4KIIalpha,Ppox,faf,Pex13,CG5285,CG7115,tun,not,Tspo,milt,PrBP,Gdi,CG46338 | | 1.895e-02 | -3.97 | GO\_MRNA\_BINDING | MSigDB lists | GO\_MRNA\_BINDING | 46 | 13 | 4161 | 644 | Ts,Ranbp21,Imp,sbr,AGO3,Dhfr,Irp-1B,Edc3,trsn,RpS2,RpL13A,mRpL13,RpS13 | | 1.895e-02 | -3.97 | Atpalpha (Na pump alpha subunit) | protein interactions | 48971 | 58 | 13 | 9204 | 1112 | Mst89B,Kap-alpha3,Mcad,RpS27A,mxc,SERCA,CG9436,Sap47,Vha68-2,Karybeta3,Cp1,ago,wal | | 1.898e-02 | -3.96 | GO\_G1\_DNA\_DAMAGE\_CHECKPOINT | MSigDB lists | GO\_G1\_DNA\_DAMAGE\_CHECKPOINT | 19 | 7 | 4161 | 644 | Cdk1,Rbf,CycB,twin,Not10,RpS27A,RpL40 | | 1.898e-02 | -3.96 | GO\_PTERIDINE\_CONTAINING\_COMPOUND\_METABOLIC\_PROCESS | MSigDB lists | GO\_PTERIDINE\_CONTAINING\_COMPOUND\_METABOLIC\_PROCESS | 19 | 7 | 4161 | 644 | CG11089,Pcd,Shmt,Dhfr,Sptr,pr,Ts | | 1.898e-02 | -3.96 | MORF\_MYST2 | MSigDB lists | MORF\_MYST2 | 19 | 7 | 4161 | 644 | EcR,GCS2beta,RpS27A,kcc,Pex10,Pdk1,P5CDh1 | | 1.898e-02 | -3.96 | GO\_LIGASE\_ACTIVITY\_FORMING\_CARBON\_SULFUR\_BONDS | MSigDB lists | GO\_LIGASE\_ACTIVITY\_FORMING\_CARBON\_SULFUR\_BONDS | 19 | 7 | 4161 | 644 | APP-BP1,AcCoAS,Scsalpha1,Aos1,Uba5,ScsbetaA,Atg7 | | 1.899e-02 | -3.96 | structural molecule activity | molecular function | GO:0005198 | 558 | 66 | 12176 | 1119 | Clc,RpS30,RpL37a,RpS5b,CG14817,mRpS5,Nup58,RpS7,mRpL13,Arpc2,RpS24,RpS27A,RpLP1,RpL38,RpS14a,RpL10,RpS18,alphaTub84B,RpLP0,mRpL17,mRpL35,RpL23A,mRpL48,RpL31,RpL40,RpS4,RpL30,gammaTub37C,mRpL16,RpS14b,RpL11,Arpc1,RpL7A,Nup62,mRpL18,RpS15,tko,RpL23,RpS27,RpS29,RpS2,Arpc4,RpS23,RpL24,Ndc1,RpL26,mbo,RpL9,RpL34b,mRpS21,RpL29,RpS28b,mRpL10,mRpL36,Grip75,RpS13,mRpS11,RpL34a,alphaTub67C,RpL13,RpL12,RpL17,RpL13A,mRpS10,RpS16,mRpS23 | | 1.911e-02 | -3.96 | carbohydrate derivative binding | molecular function | GO:0097367 | 1011 | 112 | 12176 | 1119 | eIF4A,SerRS-m,CG5789,PGRP-LE,gammaTub37C,CG8726,RecQ5,CysRS,Nadsyn,CG7488,LysRS,l(1)G0196,Mos,PEK,nmdyn-D7,l(2)k01209,bor,galectin,r,rl,Orc1,p38a,cana,rept,pont,grp,alphaTub84B,SNF4Agamma,Rab32,Droj2,Lkb1,SelD,CG14183,CG4880,kl-2,fray,CCT3,E23,Pfrx,Pfk,Mtl,dco,CG17698,TyrRS,CG5335,Pink1,ND-51,ATP8A,CG5116,RpII140,Rab5,png,Marf,crok,ABCD,SRPK,CG9981,Cdk1,CG31729,Cht11,Idgf2,nmd,CG7806,CG4562,AcCoAS,ScsbetaA,alphaTub67C,mEFTu1,ACC,obe,Arl5,Vha68-2,Non1,Klp10A,tor,CCT6,Hsc70-3,CCT5,Npc2a,mEFTu2,TrpRS-m,Mekk1,CG8507,CkIIalpha,CG7728,Rab10,ATPsynbeta,Cdk5,Tao,CG31689,Tlk,CG8611,awd,Idgf4,Cdk7,blw,Tak1,Arf51F,Atpalpha,CG9143,GluRS-m,Pgk,CG4041,tkv,SERCA,spn-E,sgll,Rpt2,CG6767,Pdk1,fs(1)N,Idgf6 | | 1.917e-02 | -3.95 | GO\_MEMBRANE\_ORGANIZATION | MSigDB lists | GO\_MEMBRANE\_ORGANIZATION | 342 | 67 | 4161 | 644 | Arl6IP1,ste14,CG31729,mts,RpS30,RpLP0,RpS13,RpL13A,Trs33,RpL13,CG44422,Nmt,RpL9,RpS27A,RpS28b,PpV,CanB2,ATP8A,scramb1,Cyp1,Atg4b,Atg7,mbo,RpL30,Oseg1,RpS2,Atg6,CHMP2B,RpL37a,Mitofilin,Pp2A-29B,RpL24,Syx13,Stoml2,CHOp24,Tsp86D,sau,Vsp37A,Sod1,ste24a,Cdk1,shrb,Rab10,RpS23,Sec22,RpS4,RpL38,Rep,RpS7,RpL40,Dd,RpL23,RpS24,lt,Aladin,RpS16,RpS18,Cdk5,CycB,Marf,RpL12,Sec23,RpL11,RpL26,Vamp7,yip2,RpS27 | | 1.920e-02 | -3.95 | identical protein binding | molecular function | GO:0042802 | 159 | 23 | 12176 | 1119 | Sod1,park,bun,Acph-1,Shmt,GCC88,spz,GlyP,Orc6,Ork1,cbs,Liprin-beta,Trxr-1,Roe1,trsn,psq,Atf6,ari-1,Adh,Blos2,Debcl,Pfk,exu | | 1.925e-02 | -3.95 | GSE27859\_DC\_VS\_CD11C\_INT\_F480\_INT\_DC\_DN | MSigDB lists | GSE27859\_DC\_VS\_CD11C\_INT\_F480\_INT\_DC\_DN | 86 | 21 | 4161 | 644 | mRpL16,Chrac-14,bys,Rpn3,Bin1,CCT3,Cdk1,Dph5,Iru,ND-23,r,Ubc4,NHP2,CG9393,CG6767,RpII140,Coq6,CG2747,Prosbeta3,muskelin,Rpt2 | | 1.925e-02 | -3.95 | LOCKWOOD\_AMPLIFIED\_IN\_LUNG\_CANCER | MSigDB lists | LOCKWOOD\_AMPLIFIED\_IN\_LUNG\_CANCER | 86 | 21 | 4161 | 644 | ND-SGDH,aay,Cul2,CG1847,Gmer,ND-49,CG4332,Snx6,Maf1,Rpb8,DppIII,RpS27,Cka,CG11134,Sec23,CSN6,CCT6,Cdk5,CCT5,SdhA,cact | | 1.926e-02 | -3.95 | translation repressor activity, mRNA regulatory element binding | molecular function | GO:0000900 | 12 | 4 | 12176 | 1119 | bam,heph,Shmt,smg | | 1.927e-02 | -3.95 | REACTOME\_CITRIC\_ACID\_CYCLE\_TCA\_CYCLE | MSigDB lists | REACTOME\_CITRIC\_ACID\_CYCLE\_TCA\_CYCLE | 15 | 6 | 4161 | 644 | ScsbetaA,Idh3b,CG5028,Nc73EF,Scsalpha1,SdhA | | 1.927e-02 | -3.95 | GO\_REGULATION\_OF\_DENDRITE\_MORPHOGENESIS | MSigDB lists | GO\_REGULATION\_OF\_DENDRITE\_MORPHOGENESIS | 15 | 6 | 4161 | 644 | CanA-14F,Cdk5,Vamp7,nAChRalpha4,Pp2B-14D,Ect4 | | 1.927e-02 | -3.95 | GO\_SEGMENTATION | MSigDB lists | GO\_SEGMENTATION | 15 | 6 | 4161 | 644 | Utx,Su(H),Med,Smox,O-fut1,CHOp24 | | 1.927e-02 | -3.95 | MODULE\_184 | MSigDB lists | MODULE\_184 | 15 | 6 | 4161 | 644 | Aldh,CG9471,Scsalpha1,Mcad,Echs1,Gabat | | 1.931e-02 | -3.95 | CycK (Cyclin K) | protein interactions | 49816 | 83 | 17 | 9204 | 1112 | nej,CG2258,CG6175,CG11486,heph,plu,png,bcd,CG9248,mEFTu1,Cdk1,Sod3,CG17508,Ntan1,ph-d,CycJ,EcR | | 1.937e-02 | -3.94 | negative regulation of protein phosphorylation | biological process | GO:0001933 | 68 | 12 | 12345 | 1129 | Cbl,Ptp61F,mts,PpV,sws,park,CG17746,dco,ATPsynD,CG14408,alph,SkpA | | 1.937e-02 | -3.94 | negative regulation of cell cycle process | biological process | GO:0010948 | 68 | 12 | 12345 | 1129 | mus304,Zwilch,nbs,Cdk1,Rbf2,Mos,grp,mre11,wapl,Aven,mad2,Rbf | | 1.937e-02 | -3.94 | glycerolipid biosynthetic process | biological process | GO:0045017 | 68 | 12 | 12345 | 1129 | PIG-C,Agpat2,ACC,CG33977,PIG-H,PIG-P,Pld,Pi3K92E,PIG-L,CG7718,CG2201,mino | | 1.945e-02 | -3.94 | eggshell formation | biological process | GO:0030703 | 134 | 20 | 12345 | 1129 | jvl,fs(1)N,CycE,spn-E,cact,Pxt,armi,fs(1)M3,bun,gus,Cf2,squ,Rbf,Pax,tor,rl,Orc1,mael,tkv,E2f1 | | 1.947e-02 | -3.94 | Glycogen synthesis | REACTOME pathways | R-DME-3322077 | 8 | 4 | 3923 | 574 | UGP,Gyg,AGBE,Pgm1 | | 1.947e-02 | -3.94 | DNA Double Strand Break Response | REACTOME pathways | R-DME-5693606 | 8 | 4 | 3923 | 574 | Tip60,Pen,mre11,nbs | | 1.949e-02 | -3.94 | cofactor binding | molecular function | GO:0048037 | 387 | 48 | 12176 | 1119 | mal,shop,Gmer,Cyp6a22,Mcad,Trxr-1,SdhA,AIF,Idh,ND-75,Sply,CG4586,CG14997,Nc73EF,Irp-1B,SdhD,CG4860,Ho,Gapdh1,Sirt7,CG1640,GlyP,Got1,Cyt-c-p,CG31674,Pxt,CG10463,AOX3,Dhfr,Acox57D-p,Cyp6v1,Kyat,Coq6,ND-23,Cisd2,ND-39,glob1,ND-49,Cyp6u1,sgll,Marc,Scsalpha1,wal,Cyp6a19,ND-51,Shmt,UQCR-Q,Gabat | | 1.958e-02 | -3.93 | ketone biosynthetic process | biological process | GO:0042181 | 31 | 7 | 12345 | 1129 | Marf,Coq8,foxo,ND-39,Npc2a,Coq7,Coq6 | | 1.958e-02 | -3.93 | positive regulation of Toll signaling pathway | biological process | GO:0045752 | 31 | 7 | 12345 | 1129 | mop,Aos1,Herc4,ea,gd,snk,spz | | 1.958e-02 | -3.93 | response to carbohydrate | biological process | GO:0009743 | 31 | 7 | 12345 | 1129 | Desat1,sgll,PyK,ACC,Pdxk,CG7328,Pfk | | 1.958e-02 | -3.93 | DNA damage checkpoint | biological process | GO:0000077 | 31 | 7 | 12345 | 1129 | nbs,mre11,mus304,grp,CG10336,mus81,Cdk1 | | 1.959e-02 | -3.93 | GO\_REGULATION\_OF\_CELLULAR\_AMINE\_METABOLIC\_PROCESS | MSigDB lists | GO\_REGULATION\_OF\_CELLULAR\_AMINE\_METABOLIC\_PROCESS | 51 | 14 | 4161 | 644 | Prosbeta5,Prosbeta6,crq,mol,Rpn3,Prosbeta3,Prosalpha3,Rpt2,Prosalpha4,Prosbeta4,Rpn12,Rpn2,Prosalpha5,Gabat | | 1.959e-02 | -3.93 | GSE22103\_LPS\_VS\_GMCSF\_AND\_IFNG\_STIM\_NEUTROPHIL\_DN | MSigDB lists | GSE22103\_LPS\_VS\_GMCSF\_AND\_IFNG\_STIM\_NEUTROPHIL\_DN | 51 | 14 | 4161 | 644 | milt,Naa20A,Kap-alpha3,Liprin-beta,Edem1,anne,CG9471,PpV,DCAF12,CG5285,stet,Mitofilin,Arf51F,Pex12 | | 1.960e-02 | -3.93 | Signaling by Wnt | REACTOME pathways | R-DME-195721 | 147 | 31 | 3923 | 574 | Prosalpha5,Wnt10,Sox102F,RpS27A,Prosbeta4,Prosbeta5,Rpn2,Tak1,Prosalpha4,dco,CanA-14F,SkpA,opm,RpL40,Tip60,Rpt2,Pp2B-14D,Prosalpha3,CanB2,NFAT,Prosbeta3,pont,nej,Prosbeta6,Pp2A-29B,Cul1,Rpn3,Atu,trr,Rpn12,pygo | | 1.961e-02 | -3.93 | GRAHAM\_CML\_DIVIDING\_VS\_NORMAL\_QUIESCENT\_UP | MSigDB lists | GRAHAM\_CML\_DIVIDING\_VS\_NORMAL\_QUIESCENT\_UP | 81 | 20 | 4161 | 644 | CycE,CycB,Cdk1,Jafrac2,Hat1,Pen,mad2,Dhfr,CG8188,Cse1,ScsbetaA,CG6610,grp,RhoGAP54D,CG4562,Ts,RfC38,Jafrac1,Cyp1,Tm1 | | 1.961e-02 | -3.93 | MORF\_FBL | MSigDB lists | MORF\_FBL | 81 | 20 | 4161 | 644 | Chrac-14,lwr,mEFTu1,ND-30,CCT3,Nap1,Pp2A-29B,Idh3b,Prosbeta4,CG10418,rept,ND-51,Prosbeta6,eIF3i,Phb2,RpS30,Sod1,Rpn12,Cyt-c-p,LysRS | | 1.963e-02 | -3.93 | GO\_NEGATIVE\_REGULATION\_OF\_TRANSCRIPTION\_FROM\_RNA\_POLYMERASE\_II\_PROMOTER | MSigDB lists | GO\_NEGATIVE\_REGULATION\_OF\_TRANSCRIPTION\_FROM\_RNA\_POLYMERASE\_II\_PROMOTER | 123 | 28 | 4161 | 644 | Cka,YL-1,RpS27A,Smox,Hmg-2,mael,Wnt10,SREBP,CG4872,Dcr-1,lwr,CG4709,Coq7,LpR2,Eno,esc,Sirt7,faf,Traf6,EcR,adp,CG17187,Tip60,escl,Su(H),RpL40,Med,lid | | 1.971e-02 | -3.93 | ATP-grasp | interpro domains | IPR011761 | 6 | 3 | 11383 | 1233 | r,ACC,ScsbetaA | | 1.971e-02 | -3.93 | Thiolase\_C | interpro domains | IPR020617 | 6 | 3 | 11383 | 1233 | yip2,Thiolase,CG10932 | | 1.971e-02 | -3.93 | Pyrv\_Knase-like\_insert\_dom\_sf | interpro domains | IPR011037 | 6 | 3 | 11383 | 1233 | PyK,Marc,mal | | 1.971e-02 | -3.93 | Thiolase\_CS | interpro domains | IPR020613 | 6 | 3 | 11383 | 1233 | Thiolase,yip2,CG10932 | | 1.971e-02 | -3.93 | Thiolase\_N | interpro domains | IPR020616 | 6 | 3 | 11383 | 1233 | Thiolase,yip2,CG10932 | | 1.971e-02 | -3.93 | V\_SNARE\_CC | interpro domains | IPR042855 | 6 | 3 | 11383 | 1233 | Sec22,Vamp7,Tomosyn | | 1.971e-02 | -3.93 | Thiolase\_acyl\_enz\_int\_AS | interpro domains | IPR020615 | 6 | 3 | 11383 | 1233 | CG10932,yip2,Thiolase | | 1.971e-02 | -3.93 | PAZ\_dom\_sf | interpro domains | IPR036085 | 6 | 3 | 11383 | 1233 | Dcr-1,piwi,AGO3 | | 1.971e-02 | -3.93 | Cullin\_neddylation\_domain | interpro domains | IPR019559 | 6 | 3 | 11383 | 1233 | Cul5,Cul1,Cul2 | | 1.971e-02 | -3.93 | IDGF | interpro domains | IPR015520 | 6 | 3 | 11383 | 1233 | Idgf4,Idgf2,Idgf6 | | 1.971e-02 | -3.93 | P\_typ\_ATPase\_c | interpro domains | IPR032630 | 6 | 3 | 11383 | 1233 | CG31729,ATP8A,CG9981 | | 1.971e-02 | -3.93 | P-type\_ATPase\_N | interpro domains | IPR032631 | 6 | 3 | 11383 | 1233 | CG9981,ATP8A,CG31729 | | 1.971e-02 | -3.93 | P-type\_ATPase\_IV | interpro domains | IPR006539 | 6 | 3 | 11383 | 1233 | CG31729,ATP8A,CG9981 | | 1.971e-02 | -3.93 | ATPase\_F1/V1/A1\_a/bsu\_N\_sf | interpro domains | IPR036121 | 6 | 3 | 11383 | 1233 | blw,ATPsynbeta,Vha68-2 | | 1.971e-02 | -3.93 | GRIP\_dom | interpro domains | IPR000237 | 6 | 3 | 11383 | 1233 | GCC88,cbs,Gmap | | 1.971e-02 | -3.93 | Galectin\_CRD | interpro domains | IPR001079 | 6 | 3 | 11383 | 1233 | galectin,Pex23,CG5335 | | 1.972e-02 | -3.93 | Myb complex | cellular component | GO:0031523 | 12 | 4 | 11940 | 1105 | l(3)mbt,Rbf2,Rbf,lin-52 | | 1.979e-02 | -3.92 | purine nucleoside bisphosphate biosynthetic process | biological process | GO:0034033 | 18 | 5 | 12345 | 1129 | CG17691,CG8199,CG6178,AcCoAS,Ppcdc | | 1.979e-02 | -3.92 | ribonucleoside bisphosphate biosynthetic process | biological process | GO:0034030 | 18 | 5 | 12345 | 1129 | CG6178,AcCoAS,Ppcdc,CG17691,CG8199 | | 1.979e-02 | -3.92 | nucleoside bisphosphate biosynthetic process | biological process | GO:0033866 | 18 | 5 | 12345 | 1129 | CG6178,AcCoAS,Ppcdc,CG8199,CG17691 | | 1.979e-02 | -3.92 | mitotic DNA damage checkpoint | biological process | GO:0044773 | 18 | 5 | 12345 | 1129 | mre11,nbs,mus81,Cdk1,CG10336 | | 1.990e-02 | -3.92 | GO\_REGULATION\_OF\_CELLULAR\_KETONE\_METABOLIC\_PROCESS | MSigDB lists | GO\_REGULATION\_OF\_CELLULAR\_KETONE\_METABOLIC\_PROCESS | 76 | 19 | 4161 | 644 | Prosalpha4,Rpt2,Prosbeta3,Rpn2,EcR,Rpn12,adp,ABCD,Prosbeta6,Pdk,regucalcin,Gk2,Prosbeta4,SREBP,Prosalpha3,Prosalpha5,Prosbeta5,Rpn3,mol | | 1.990e-02 | -3.92 | YANG\_BCL3\_TARGETS\_UP | MSigDB lists | YANG\_BCL3\_TARGETS\_UP | 76 | 19 | 4161 | 644 | obe,ZnT35C,Pex12,CG8507,CycB,LpR2,Cdk1,Pfk,Mtpalpha,CG8112,ND-24,Mcad,ND-SGDH,mino,scu,tho2,Fur2,CG4502,Pdk | | 1.990e-02 | -3.92 | GSE9006\_TYPE\_1\_VS\_TYPE\_2\_DIABETES\_PBMC\_AT\_DX\_UP | MSigDB lists | GSE9006\_TYPE\_1\_VS\_TYPE\_2\_DIABETES\_PBMC\_AT\_DX\_UP | 76 | 19 | 4161 | 644 | CG1847,Gmer,AnxB10,CG17187,CG12018,CG8188,Nadsyn,CG17333,heph,RpL11,SREBP,RpL37a,Lsm12,Exo70,RpS27,thoc6,Pfdn2,ERp60,Arpc4 | | 1.990e-02 | -3.92 | GSE17721\_LPS\_VS\_PAM3CSK4\_1H\_BMDC\_UP | MSigDB lists | GSE17721\_LPS\_VS\_PAM3CSK4\_1H\_BMDC\_UP | 76 | 19 | 4161 | 644 | CG10470,ND-49,CG11899,CG12018,CG12375,escl,Jafrac1,Xpac,Prosalpha5,CG32549,CSN6,Uch,CG1309,capt,Thiolase,Pdk1,CCT3,esc,pont | | 1.991e-02 | -3.92 | REACTOME\_METABOLISM\_OF\_PROTEINS | MSigDB lists | REACTOME\_METABOLISM\_OF\_PROTEINS | 262 | 53 | 4161 | 644 | RpS28b,RpS27A,RpL9,RpS23,Fbw5,RpL13,CCT6,ERp60,eEF1beta,alpha-Man-Ia,Tom40,GalT1,RpS30,RpS13,RpLP0,eIF4E1,RpL13A,CG9393,alphaTub84B,eIF4A,CG5028,PIG-L,eIF3i,RpS27,RpL24,RpL26,RpL11,TBCC,RpL37a,Sec23,RpL12,PIG-C,Edem1,Tim8,CCT5,CG8004,Pfdn2,CCT3,RpS18,CG1890,RpS2,GCS2beta,RpS16,Pfdn5,RpS24,RpL30,RpL23,CG8728,RpL40,RpS7,RpL38,RpS4,Roe1 | | 1.992e-02 | -3.92 | regulation of cellular component biogenesis | biological process | GO:0044087 | 305 | 39 | 12345 | 1129 | Pi3K92E,CG11486,Mtl,Cdk5,wash,Nprl2,tkv,Stat92E,mael,Tm1,SkpA,dco,swa,Cenp-C,Mekk1,pgc,enc,lt,ssh,Cul2,spz,mad2,Cul5,nos,exu,Vps16B,cnc,Abi,Arpc2,Synj,msps,Mybbp1A,Sec23,spn-E,Med,Arpc1,cup,Arpc4,raptor | | 1.996e-02 | -3.91 | Formation of Incision Complex in GG-NER | REACTOME pathways | R-DME-5696395 | 34 | 10 | 3923 | 574 | Xpc,Tfb4,CG6923,RpL40,Xpac,Cdk7,RpS27A,Usp16-45,ben,Uev1A | | 1.997e-02 | -3.91 | spag (spaghetti) | protein interactions | 43958 | 24 | 7 | 9204 | 1112 | dco,lqfR,Hsp26,Hsc70-3,cact,rept,pont | | 1.997e-02 | -3.91 | HP1e (Heterochromatin Protein 1e) | protein interactions | 41108 | 24 | 7 | 9204 | 1112 | RpS15,Cen,CG7789,RpS7,Kcmf1,RpS23,CG4407 | | 1.997e-02 | -3.91 | CG15080 (uncharacterized protein) | protein interactions | 37167 | 24 | 7 | 9204 | 1112 | CG9205,CG7730,Kap3,glob1,RfC38,RpS30,PEK | | 1.997e-02 | -3.91 | AP-2mu (Adaptor Protein complex 2, mu subunit) | protein interactions | 42642 | 24 | 7 | 9204 | 1112 | Reps,BicC,PyK,Pdk1,lt,AP-1-2beta,Afti | | 1.997e-02 | -3.91 | gammaTub23C (gamma-Tubulin at 23C) | protein interactions | 33501 | 24 | 7 | 9204 | 1112 | Grip71,Grip75,gammaTub37C,cnn,Grip128,mael,Grip84 | | 1.997e-02 | -3.91 | CG5177 (uncharacterized protein) | protein interactions | 34017 | 24 | 7 | 9204 | 1112 | Actn,Argk,Scp1,Cyt-c-p,Adh,Mf,up | | 1.997e-02 | -3.91 | BicC (Bicaudal C) | protein interactions | 34946 | 24 | 7 | 9204 | 1112 | png,smg,tral,exu,BicC,cup,twin | | 1.998e-02 | -3.91 | GSE16385\_MONOCYTE\_VS\_12H\_ROSIGLITAZONE\_TREATED\_MACROPHAGE\_UP | MSigDB lists | GSE16385\_MONOCYTE\_VS\_12H\_ROSIGLITAZONE\_TREATED\_MACROPHAGE\_UP | 56 | 15 | 4161 | 644 | mRpL13,bor,CSN6,RpL23,LSm3,l(1)G0004,Pgk,RpL12,CG8204,Roe1,Trs33,Bin1,wal,CG16865,alph | | 1.998e-02 | -3.91 | CEBPA\_01 | MSigDB lists | CEBPA\_01 | 56 | 15 | 4161 | 644 | regucalcin,lqfR,Synj,dnc,rept,Rep,Irk1,dco,faf,Utx,Mctp,Wnt10,Unr,Archease,Arf51F | | 1.998e-02 | -3.91 | GSE19401\_PAM2CSK4\_VS\_RETINOIC\_ACID\_AND\_PAM2CSK4\_STIM\_FOLLICULAR\_DC\_DN | MSigDB lists | GSE19401\_PAM2CSK4\_VS\_RETINOIC\_ACID\_AND\_PAM2CSK4\_STIM\_FOLLICULAR\_DC\_DN | 56 | 15 | 4161 | 644 | CG5734,ERp60,Shmt,cathD,CG10932,Tsp74F,ND-SGDH,CG3860,PAPLA1,Trxr-1,Sccpdh1,CG6767,Mcad,mtgo,AnxB10 | | 1.998e-02 | -3.91 | GO\_NEGATIVE\_REGULATION\_OF\_MITOTIC\_CELL\_CYCLE | MSigDB lists | GO\_NEGATIVE\_REGULATION\_OF\_MITOTIC\_CELL\_CYCLE | 56 | 15 | 4161 | 644 | Smox,CG1812,Not10,grp,RpS27A,CG30291,CkIIalpha,twin,Rbf,APP-BP1,RpL40,nmdyn-D6,mad2,CycB,Cdk1 | | 2.001e-02 | -3.91 | dj-1beta (dj-1beta) | protein interactions | 43652 | 14 | 5 | 9204 | 1112 | cib,foxo,rl,PGRP-LE,Pink1 | | 2.001e-02 | -3.91 | psidin (phagocyte signaling impaired) | protein interactions | 42389 | 14 | 5 | 9204 | 1112 | Tm1,MED4,Neos,Naa20A,Sec10 | | 2.001e-02 | -3.91 | Arl2 (ADP ribosylation factor-like 2) | protein interactions | 40993 | 14 | 5 | 9204 | 1112 | CycJ,mts,alphaTub84B,msps,TBCC | | 2.001e-02 | -3.91 | Cnb (Centrobin) | protein interactions | 38258 | 14 | 5 | 9204 | 1112 | RpS13,Grip71,mRpL48,gammaTub37C,cnn | | 2.001e-02 | -3.91 | bel (belle) | protein interactions | 45826 | 14 | 5 | 9204 | 1112 | nos,eIF4E1,E2f1,cnn,CycB | | 2.001e-02 | -3.91 | ea (easter) | protein interactions | 41858 | 14 | 5 | 9204 | 1112 | gd,snk,spz,cact,cype | | 2.001e-02 | -3.91 | Zip99C (Zinc/iron regulated transporter-related protein 99C) | protein interactions | 43533 | 14 | 5 | 9204 | 1112 | PGRP-LE,Fer1HCH,Rel,piwi,Tak1 | | 2.001e-02 | -3.91 | CG1418 (uncharacterized protein) | protein interactions | 36035 | 14 | 5 | 9204 | 1112 | bcd,cad,Vha68-2,Rtnl1,SERCA | | 2.001e-02 | -3.91 | ncRNA processing | biological process | GO:0034470 | 259 | 34 | 12345 | 1129 | CycC,CG8204,Archease,RpS16,Rrp40,RpS7,CG7009,CG2034,SelD,CG30343,mtTFB1,Dcr-1,CG3021,CG9143,Cbp80,CG12301,scu,RpL7A,bys,CG2911,RpS28b,CG3587,CG5292,RpS14a,NHP2,CG7728,RpS14b,CG10189,IntS6,Mat89Ba,Trf4-2,CG10463,IntS11,mRpS11 | | 2.004e-02 | -3.91 | PhoLip\_ATPase\_C | pfam domains | PF16212 | 6 | 3 | 10480 | 1142 | CG31729,CG9981,ATP8A | | 2.004e-02 | -3.91 | Cullin\_Nedd8 | pfam domains | PF10557 | 6 | 3 | 10480 | 1142 | Cul5,Cul2,Cul1 | | 2.004e-02 | -3.91 | Thiolase\_C | pfam domains | PF02803 | 6 | 3 | 10480 | 1142 | yip2,Thiolase,CG10932 | | 2.004e-02 | -3.91 | PhoLip\_ATPase\_N | pfam domains | PF16209 | 6 | 3 | 10480 | 1142 | CG31729,ATP8A,CG9981 | | 2.004e-02 | -3.91 | Gal-bind\_lectin | pfam domains | PF00337 | 6 | 3 | 10480 | 1142 | galectin,CG5335,Pex23 | | 2.004e-02 | -3.91 | Thiolase\_N | pfam domains | PF00108 | 6 | 3 | 10480 | 1142 | CG10932,yip2,Thiolase | | 2.005e-02 | -3.91 | flavin adenine dinucleotide binding | molecular function | GO:0050660 | 68 | 12 | 12176 | 1119 | Coq6,CG4586,CG14997,CG10463,AOX3,CG4860,wal,AIF,SdhA,Trxr-1,Acox57D-p,Mcad | | 2.009e-02 | -3.91 | export from cell | biological process | GO:0140352 | 178 | 25 | 12345 | 1129 | CHOp24,Sec22,Rab10,lqfR,Crag,Arf51F,eIF3m,Exo70,HDAC6,opm,Vamp7,Atg6,Sec15,disp,Syx13,X11L,nrv1,CanB2,Tango10,nej,Sec10,Rep,Gdi,Tomosyn,Atpalpha | | 2.010e-02 | -3.91 | cysteine-type endopeptidase activity | molecular function | GO:0004197 | 38 | 8 | 12176 | 1119 | 26-29-p,sol,Usp16-45,Atg4b,velo,scny,Cp1,faf | | 2.010e-02 | -3.91 | ligase activity, forming carbon-nitrogen bonds | molecular function | GO:0016879 | 38 | 8 | 12176 | 1119 | Nadsyn,GatA,GatB,AsnS,Gclc,r,Pfas,CG1578 | | 2.010e-02 | -3.91 | GSE13547\_CTRL\_VS\_ANTI\_IGM\_STIM\_BCELL\_12H\_DN | MSigDB lists | GSE13547\_CTRL\_VS\_ANTI\_IGM\_STIM\_BCELL\_12H\_DN | 71 | 18 | 4161 | 644 | Scsalpha1,Fkbp59,yip2,asf1,Dph5,Axs,CG7655,pont,Pfdn2,RpL13,CCT6,mEFTu1,eEF1beta,Mcad,RpL13A,Pfdn5,isoQC,CG15141 | | 2.010e-02 | -3.91 | SASAKI\_ADULT\_T\_CELL\_LEUKEMIA | MSigDB lists | SASAKI\_ADULT\_T\_CELL\_LEUKEMIA | 71 | 18 | 4161 | 644 | Prosbeta4,Idh,Sccpdh1,SRPK,Jafrac2,CG8004,CycB,wal,mad2,AnxB10,MED24,Ts,RfC38,Rpb8,CG10418,ATP8A,tko,ste14 | | 2.010e-02 | -3.91 | MODULE\_60 | MSigDB lists | MODULE\_60 | 71 | 18 | 4161 | 644 | Gip,Prip,CG12018,P5CDh1,ste14,tzn,Tsp74F,Pcd,Jheh2,Gyg,Rcd5,CG4872,Got1,Echs1,ND-B18,CG8199,Pdxk,CG9471 | | 2.010e-02 | -3.91 | GO\_REGULATION\_OF\_PROTEOLYSIS | MSigDB lists | GO\_REGULATION\_OF\_PROTEOLYSIS | 200 | 42 | 4161 | 644 | regucalcin,SkpA,Arl6IP1,Prip,Rpn2,EcR,CG1812,Rpn12,HDAC6,Prosalpha4,Rpt2,Prosbeta3,mad2,Uch-L5,Cdk1,AIF,STUB1,Prosalpha5,RpS27A,CG30496,RpL40,Prosbeta6,Ubc4,Tlk,dco,Phb2,ari-1,Cyt-c-p,APC7,CkIIalpha,CG30291,Rpn3,Cul1,Prosbeta5,Cdk5,Gclc,CycB,PDCD-5,Smox,Wnt10,Prosbeta4,Prosalpha3 | | 2.016e-02 | -3.90 | G2 DNA damage checkpoint | biological process | GO:0031572 | 7 | 3 | 12345 | 1129 | Cdk1,nbs,mre11 | | 2.016e-02 | -3.90 | response to superoxide | biological process | GO:0000303 | 7 | 3 | 12345 | 1129 | Sod3,Jafrac1,Sod1 | | 2.016e-02 | -3.90 | isocitrate metabolic process | biological process | GO:0006102 | 7 | 3 | 12345 | 1129 | CG5028,Idh,Idh3b | | 2.016e-02 | -3.90 | response to lipopolysaccharide | biological process | GO:0032496 | 7 | 3 | 12345 | 1129 | EcR,Stat92E,Tak1 | | 2.016e-02 | -3.90 | circadian regulation of gene expression | biological process | GO:0032922 | 7 | 3 | 12345 | 1129 | lid,cyc,nej | | 2.016e-02 | -3.90 | piRNA biosynthetic process | biological process | GO:1990511 | 7 | 3 | 12345 | 1129 | tapas,armi,piwi | | 2.016e-02 | -3.90 | peptidyl-diphthamide metabolic process | biological process | GO:0017182 | 7 | 3 | 12345 | 1129 | CG2911,CG1578,Dph5 | | 2.016e-02 | -3.90 | sperm mitochondrion organization | biological process | GO:0030382 | 7 | 3 | 12345 | 1129 | park,milt,Pink1 | | 2.016e-02 | -3.90 | regulation of cytoplasmic translational elongation | biological process | GO:1900247 | 7 | 3 | 12345 | 1129 | CG1578,Dph5,CG2911 | | 2.016e-02 | -3.90 | cellular response to superoxide | biological process | GO:0071451 | 7 | 3 | 12345 | 1129 | Sod1,Jafrac1,Sod3 | | 2.016e-02 | -3.90 | pyrimidine nucleoside monophosphate biosynthetic process | biological process | GO:0009130 | 7 | 3 | 12345 | 1129 | r-l,Ts,l(2)k01209 | | 2.016e-02 | -3.90 | zymogen activation | biological process | GO:0031638 | 7 | 3 | 12345 | 1129 | eIF3m,gd,snk | | 2.016e-02 | -3.90 | positive regulation of NF-kappaB transcription factor activity | biological process | GO:0051092 | 7 | 3 | 12345 | 1129 | Traf6,Aos1,CG13344 | | 2.016e-02 | -3.90 | response to oxygen radical | biological process | GO:0000305 | 7 | 3 | 12345 | 1129 | Sod1,Jafrac1,Sod3 | | 2.016e-02 | -3.90 | cellular response to osmotic stress | biological process | GO:0071470 | 7 | 3 | 12345 | 1129 | CG17746,Mekk1,kcc | | 2.016e-02 | -3.90 | removal of superoxide radicals | biological process | GO:0019430 | 7 | 3 | 12345 | 1129 | Sod3,Sod1,Jafrac1 | | 2.016e-02 | -3.90 | mitotic G2 DNA damage checkpoint | biological process | GO:0007095 | 7 | 3 | 12345 | 1129 | Cdk1,nbs,mre11 | | 2.016e-02 | -3.90 | regulation of autophagy of mitochondrion | biological process | GO:1903146 | 7 | 3 | 12345 | 1129 | ago,Sod1,SREBP | | 2.016e-02 | -3.90 | TORC1 signaling | biological process | GO:0038202 | 7 | 3 | 12345 | 1129 | raptor,Nprl2,REPTOR | | 2.016e-02 | -3.90 | protein K63-linked ubiquitination | biological process | GO:0070534 | 7 | 3 | 12345 | 1129 | Traf6,ben,Uev1A | | 2.016e-02 | -3.90 | diol biosynthetic process | biological process | GO:0034312 | 7 | 3 | 12345 | 1129 | Sptr,Pcd,pr | | 2.016e-02 | -3.90 | peptidyl-diphthamide biosynthetic process from peptidyl-histidine | biological process | GO:0017183 | 7 | 3 | 12345 | 1129 | Dph5,CG1578,CG2911 | | 2.016e-02 | -3.90 | diol metabolic process | biological process | GO:0034311 | 7 | 3 | 12345 | 1129 | pr,Pcd,Sptr | | 2.016e-02 | -3.90 | phagosome maturation | biological process | GO:0090382 | 7 | 3 | 12345 | 1129 | Snx6,crq,Vps16B | | 2.016e-02 | -3.90 | cellular response to oxygen radical | biological process | GO:0071450 | 7 | 3 | 12345 | 1129 | Sod3,Sod1,Jafrac1 | | 2.016e-02 | -3.90 | pyrimidine nucleoside monophosphate metabolic process | biological process | GO:0009129 | 7 | 3 | 12345 | 1129 | Ts,r-l,l(2)k01209 | | 2.017e-02 | -3.90 | WINTER\_HYPOXIA\_METAGENE | MSigDB lists | WINTER\_HYPOXIA\_METAGENE | 61 | 16 | 4161 | 644 | Su(H),cathD,Pdk,CG10932,AGBE,Pgm1,Gapdh1,CCT6,Pgk,Pglym78,Eno,Cdk1,r-l,Fkbp59,CG32473,Hsc70-3 | | 2.017e-02 | -3.90 | GO\_PROTEIN\_LOCALIZATION\_TO\_NUCLEUS | MSigDB lists | GO\_PROTEIN\_LOCALIZATION\_TO\_NUCLEUS | 61 | 16 | 4161 | 644 | cyc,Arts,mbo,Karybeta3,Phb2,Pp2B-14D,Cse1,Dd,RpL23,ERp60,Kap-alpha3,Pen,cact,rl,CanA-14F,RpL11 | | 2.017e-02 | -3.90 | GSE11864\_CSF1\_VS\_CSF1\_IFNG\_IN\_MAC\_DN | MSigDB lists | GSE11864\_CSF1\_VS\_CSF1\_IFNG\_IN\_MAC\_DN | 61 | 16 | 4161 | 644 | isoQC,Rpn2,Nurf-38,LysRS,RpS2,Ranbp21,PIG-L,mtTFB1,Droj2,CG3587,stas,Prosalpha5,Prosbeta5,thoc6,pont,Rpn3 | | 2.017e-02 | -3.90 | ELVIDGE\_HYPOXIA\_DN | MSigDB lists | ELVIDGE\_HYPOXIA\_DN | 61 | 16 | 4161 | 644 | zyd,CG4407,tko,l(1)G0004,RhoGDI,Cyp1,P5CDh1,ste24a,rl,Stip1,Kap-alpha3,Arpc4,Fkbp59,Nc73EF,Gyg,GatA | | 2.018e-02 | -3.90 | clathrin binding | molecular function | GO:0030276 | 18 | 5 | 12176 | 1119 | Gga,Clc,AP-1-2beta,Afti,Hip1 | | 2.018e-02 | -3.90 | phosphotransferase activity, phosphate group as acceptor | molecular function | GO:0016776 | 18 | 5 | 12176 | 1119 | awd,l(1)G0196,Scsalpha1,nmdyn-D6,nmdyn-D7 | | 2.020e-02 | -3.90 | FOSTER\_KDM1A\_TARGETS\_DN | MSigDB lists | FOSTER\_KDM1A\_TARGETS\_DN | 66 | 17 | 4161 | 644 | grsm,CG33156,CG2201,CG5734,rept,CG15141,CG17333,AP-1-2beta,Rnmt,CG11807,shu,Grip84,mRpL10,f-cup,CG8199,CG7322,Echs1 | | 2.020e-02 | -3.90 | GSE17721\_PAM3CSK4\_VS\_GADIQUIMOD\_0.5H\_BMDC\_DN | MSigDB lists | GSE17721\_PAM3CSK4\_VS\_GADIQUIMOD\_0.5H\_BMDC\_DN | 66 | 17 | 4161 | 644 | rl,Arpc4,Uev1A,LSm3,Sply,CG12929,Lsm12,CG8372,lid,CG17691,CanB2,Adi1,Dhfr,faf,MED4,CG8188,CG9393 | | 2.020e-02 | -3.90 | GO\_CELLULAR\_CARBOHYDRATE\_METABOLIC\_PROCESS | MSigDB lists | GO\_CELLULAR\_CARBOHYDRATE\_METABOLIC\_PROCESS | 66 | 17 | 4161 | 644 | Gk2,RpL40,Synj,CG9485,CG8475,AGBE,Pgm1,CG9391,Treh,Pfk,l(1)G0196,GalT1,GlyP,RpS27A,Got1,Gyg,Idh | | 2.020e-02 | -3.90 | GSE3039\_ALPHABETA\_CD8\_TCELL\_VS\_B1\_BCELL\_DN | MSigDB lists | GSE3039\_ALPHABETA\_CD8\_TCELL\_VS\_B1\_BCELL\_DN | 66 | 17 | 4161 | 644 | Mitofilin,PAPLA1,PDCD-5,CG14516,Klp10A,pont,STUB1,Eno,Gmer,Pgm1,TyrRS,Rpn2,shv,NHP2,r,cathD,Su(H) | | 2.025e-02 | -3.90 | endocytosis | biological process | GO:0006897 | 126 | 19 | 12345 | 1129 | Mtl,faf,CG9132,dco,CG31729,Nlg1,lqfR,crq,csw,Clc,Mekk1,lt,Synj,Arf51F,EcR,Atg6,dx,fwe,Rab5 | | 2.033e-02 | -3.90 | positive regulation of innate immune response | biological process | GO:0045089 | 53 | 10 | 12345 | 1129 | Sp7,EMRE,Crag,Hat1,Myd88,CG9925,Rel,PDCD-5,PGRP-LE,Hr78 | | 2.033e-02 | -3.90 | gnu (giant nuclei) | protein interactions | 39615 | 35 | 9 | 9204 | 1112 | BicC,mxc,CycB,CG30430,wisp,png,plu,cnc,smg | | 2.033e-02 | -3.90 | MED7 (Mediator complex subunit 7) | protein interactions | 41288 | 35 | 9 | 9204 | 1112 | Ptip,CG17002,Larp4B,Usp10,MED27,MED4,MED20,enc,ix | | 2.033e-02 | -3.90 | eIF4G2 (eukaryotic translation initiation factor 4G2) | protein interactions | 42819 | 35 | 9 | 9204 | 1112 | Marf1,eIF4E1,Sld5,Chd64,CycB,cathD,26-29-p,Vsp37A,CG42565 | | 2.033e-02 | -3.90 | CycH (Cyclin H) | protein interactions | 40429 | 35 | 9 | 9204 | 1112 | Cdk1,CG5003,CycC,heph,gus,Cdk7,EcR,Ntan1,CG17508 | | 2.033e-02 | -3.90 | dimm (dimmed) | protein interactions | 35404 | 35 | 9 | 9204 | 1112 | Cul2,grp,CycJ,nej,RpL17,squ,STUB1,plu,RpS7 | | 2.036e-02 | -3.89 | GO\_POSITIVE\_REGULATION\_OF\_ORGANELLE\_ORGANIZATION | MSigDB lists | GO\_POSITIVE\_REGULATION\_OF\_ORGANELLE\_ORGANIZATION | 189 | 40 | 4161 | 644 | Phb2,Atg6,Tm1,Aos1,Cyp1,Med,Synj,CanB2,Arpc1,PDCD-5,Smox,AGO3,Arf51F,SREBP,Moe,Arpc4,CCT3,pont,Cdk5,esc,CycB,Pfdn2,CCT5,Rpn12,CG3631,Grip128,escl,rept,Stoml2,PAPLA1,Mekk1,Nmt,Abi,Grip84,Grip75,Surf4,Arpc2,twin,CCT6,rl | | 2.036e-02 | -3.89 | RUTELLA\_RESPONSE\_TO\_HGF\_UP | MSigDB lists | RUTELLA\_RESPONSE\_TO\_HGF\_UP | 118 | 27 | 4161 | 644 | SelD,CG31729,Cyp1,CG10932,escl,cathD,alphaTub84B,ND-49,CG3860,Mctp,GAPsec,anne,CG8004,Prosbeta5,Rbcn-3A,CG7997,esc,CG16979,Pdxk,CG12773,CG15715,Fkbp59,Sply,yip2,Irp-1B,Sec22,Sccpdh1 | | 2.038e-02 | -3.89 | response to abiotic stimulus | biological process | GO:0009628 | 447 | 54 | 12345 | 1129 | Tm1,Stat92E,Fdh,p38a,dco,rpk,Hsp26,shep,RecQ5,adp,nej,cu,thoc6,Calx,CG17746,Trxr-1,Xpac,Droj2,Sec15,Hsc70-3,Mekk1,Ptp61F,Debcl,dnc,Fkbp59,NFAT,osk,tzn,grp,Cka,Atpalpha,Kap-alpha3,Sdhaf3,Ggt-1,Ptpmeg,alph,tho2,Jafrac2,mts,Pld,tor,kcc,ago,CG4476,tko,Tfb4,Sod3,UGP,mus304,Utx,Pen,Chrac-14,CG5844,Tom40 | | 2.043e-02 | -3.89 | isocitrate dehydrogenase activity | molecular function | GO:0004448 | 7 | 3 | 12176 | 1119 | CG5028,Idh,Idh3b | | 2.043e-02 | -3.89 | cytokine binding | molecular function | GO:0019955 | 7 | 3 | 12176 | 1119 | tkv,dl,Imp | | 2.043e-02 | -3.89 | polynucleotide adenylyltransferase activity | molecular function | GO:0004652 | 7 | 3 | 12176 | 1119 | wisp,Trf4-2,CG46385 | | 2.043e-02 | -3.89 | GDP-dissociation inhibitor activity | molecular function | GO:0005092 | 7 | 3 | 12176 | 1119 | Rep,Gdi,RhoGDI | | 2.043e-02 | -3.89 | CASORELLI\_ACUTE\_PROMYELOCYTIC\_LEUKEMIA\_DN | MSigDB lists | CASORELLI\_ACUTE\_PROMYELOCYTIC\_LEUKEMIA\_DN | 291 | 58 | 4161 | 644 | Arpc4,Stip1,Jafrac2,Mtpalpha,CG7322,p38a,Pfk,Prosbeta5,Eno,CCT5,thoc6,CycB,esc,CG11134,Pdk,Tm1,Cbp80,Ts,RfC38,Roe1,RhoGAP54D,Edc3,Kap3,CG11858,CG1532,CG12018,Dhfr,CG17187,Pen,ERp60,Pgk,mEFTu1,AIF,Cdk1,SdhA,mRpS10,Tfb4,Idh3b,CG1774,Got1,Fkbp59,CG10932,escl,ACC,CG5734,Iml1,ND-51,CG11089,Gk2,Sld5,CG4562,Nurf-38,grp,Nadsyn,Cse1,Gapdh1,ND-49,CG11899 | | 2.054e-02 | -3.89 | Global Genome Nucleotide Excision Repair (GG-NER) | REACTOME pathways | R-DME-5696399 | 39 | 11 | 3923 | 574 | CG6923,Xpc,Tfb4,RpL40,Xpac,Cdk7,RpS27A,Usp16-45,ben,Uev1A,RfC38 | | 2.054e-02 | -3.89 | REACTOME\_REGULATION\_OF\_ORNITHINE\_DECARBOXYLASE\_ODC | MSigDB lists | REACTOME\_REGULATION\_OF\_ORNITHINE\_DECARBOXYLASE\_ODC | 37 | 11 | 4161 | 644 | Prosalpha4,Prosbeta4,Rpt2,Prosalpha3,Prosbeta3,Prosalpha5,Rpn2,Rpn12,Prosbeta6,Prosbeta5,Rpn3 | | 2.054e-02 | -3.89 | GO\_RESPONSE\_TO\_MECHANICAL\_STIMULUS | MSigDB lists | GO\_RESPONSE\_TO\_MECHANICAL\_STIMULUS | 37 | 11 | 4161 | 644 | Xpac,p38a,cact,nSMase,CycB,Irk1,CG31715,grp,aay,Prip,CG15111 | | 2.054e-02 | -3.89 | MORF\_GSPT1 | MSigDB lists | MORF\_GSPT1 | 37 | 11 | 4161 | 644 | CCT5,Tom40,CycB,NHP2,CG11089,CG1416,Hat1,Pen,Cyt-c-p,Sod1,Iru | | 2.054e-02 | -3.89 | ZHU\_CMV\_24\_HR\_UP | MSigDB lists | ZHU\_CMV\_24\_HR\_UP | 37 | 11 | 4161 | 644 | Cse1,CG8188,eIF4E1,Rpt2,Prosbeta4,Fkbp59,CG9391,Roe1,cact,Stip1,Eno | | 2.054e-02 | -3.89 | GO\_SPINDLE\_POLE | MSigDB lists | GO\_SPINDLE\_POLE | 37 | 11 | 4161 | 644 | mts,Grip84,CG1812,gammaTub37C,Grip75,Klp10A,p38a,CycB,msps,Grip128,mad2 | | 2.054e-02 | -3.89 | GO\_ANTIGEN\_PROCESSING\_AND\_PRESENTATION\_OF\_EXOGENOUS\_PEPTIDE\_ANTIGEN\_VIA\_MHC\_CLASS\_I | MSigDB lists | GO\_ANTIGEN\_PROCESSING\_AND\_PRESENTATION\_OF\_EXOGENOUS\_PEPTIDE\_ANTIGEN\_VIA\_MHC\_CLASS\_I | 37 | 11 | 4161 | 644 | Prosalpha3,Prosbeta3,Prosbeta4,Prosalpha4,Rpt2,Rpn12,Prosalpha5,Rpn2,Prosbeta5,Prosbeta6,Rpn3 | | 2.054e-02 | -3.89 | GO\_OXIDOREDUCTASE\_ACTIVITY\_ACTING\_ON\_THE\_CH\_CH\_GROUP\_OF\_DONORS | MSigDB lists | GO\_OXIDOREDUCTASE\_ACTIVITY\_ACTING\_ON\_THE\_CH\_CH\_GROUP\_OF\_DONORS | 37 | 11 | 4161 | 644 | wal,CG3803,SdhA,CG9471,CG3609,Cyp6a19,Mcad,CG7840,CG10463,Cyp6a22,Ppox | | 2.060e-02 | -3.88 | fly chr35F1-35F1\|2-52 cM | chromosome location | fly chr35F1-35F1\|2-52 cM | 30 | 5 | 25010 | 1343 | c(2)M,EndoGI,twe,cact,CG17328 | | 2.060e-02 | -3.88 | fly chr35F1-35F1\|2-52 | chromosome location | fly chr35F1-35F1\|2-52 | 30 | 5 | 25010 | 1343 | cact,CG17328,twe,c(2)M,EndoGI | | 2.072e-02 | -3.88 | Cyclin-like\_sf | interpro domains | IPR036915 | 21 | 6 | 11383 | 1233 | CycB,CycC,Rbf2,Rbf,CycJ,CycE | | 2.073e-02 | -3.88 | CG3726 (uncharacterized protein) | protein interactions | 31525 | 19 | 6 | 9204 | 1112 | Blos2,CG1532,CG14036,CG10265,Marf1,Hsp26 | | 2.073e-02 | -3.88 | mago (mago nashi) | protein interactions | 37402 | 19 | 6 | 9204 | 1112 | csw,nos,rl,Pym,osk,bcd | | 2.073e-02 | -3.88 | hoip (hoi-polloi) | protein interactions | 44173 | 19 | 6 | 9204 | 1112 | cnn,PGRP-LE,Rel,Fkbp39,Tak1,Sld5 | | 2.073e-02 | -3.88 | tyf (twenty-four) | protein interactions | 31363 | 19 | 6 | 9204 | 1112 | Lkb1,Lsm12,eIF4E1,Larp4B,Rtnl1,foxo | | 2.073e-02 | -3.88 | PAN2 (Poly(A)-binding protein-dependent poly(A) ribonuclease 2) | protein interactions | 35893 | 19 | 6 | 9204 | 1112 | gw,CG11486,Desat1,Arpc1,nos,RpL7A | | 2.073e-02 | -3.88 | Tengl3 (Testis EndoG-Like 3) | protein interactions | 33404 | 19 | 6 | 9204 | 1112 | grp,CG13373,CG9248,Cdk5,squ,nej | | 2.074e-02 | -3.88 | neuron projection cytoplasm | cellular component | GO:0120111 | 18 | 5 | 11940 | 1105 | cup,tral,nos,eIF4E1,yps | | 2.077e-02 | -3.87 | Negative regulation of MET activity | REACTOME pathways | R-DME-6807004 | 16 | 6 | 3923 | 574 | RpL40,Ptp61F,Ptp10D,tor,Cbl,RpS27A | | 2.077e-02 | -3.87 | Formation of the beta-catenin:TCF transactivating complex | REACTOME pathways | R-DME-201722 | 16 | 6 | 3923 | 574 | Atu,trr,Tip60,pygo,pont,nej | | 2.082e-02 | -3.87 | mitochondrial alpha-ketoglutarate dehydrogenase complex | cellular component | GO:0005947 | 7 | 3 | 11940 | 1105 | Nc73EF,CG17691,CG8199 | | 2.082e-02 | -3.87 | calcineurin complex | cellular component | GO:0005955 | 7 | 3 | 11940 | 1105 | CanA-14F,Pp2B-14D,CanB2 | | 2.083e-02 | -3.87 | GO\_REGULATION\_OF\_ORGANELLE\_ORGANIZATION | MSigDB lists | GO\_REGULATION\_OF\_ORGANELLE\_ORGANIZATION | 355 | 69 | 4161 | 644 | Arf51F,SREBP,Arpc1,yip2,gammaTub37C,Cdk5,esc,CycB,pont,Pfdn2,CCT5,nmdyn-D6,msps,Moe,lt,GAPsec,Tlk,Pka-R1,Tm1,Tspo,Med,Tao,Synj,shrb,Fkbp59,Sec22,Grip75,asf1,Klp10A,rl,mad2,CCT6,sau,grp,CG1812,HDAC6,lid,Stoml2,escl,Grip128,AGO3,Rcd5,PDCD-5,Pp2A-29B,Smox,CHMP2B,CCT3,Arpc4,CkIIalpha,Phb2,Atg6,CanB2,Aos1,Cyp1,Atg7,Arpc2,twin,Mekk1,Nmt,PAPLA1,Abi,Grip84,Surf4,CG3631,Rpn12,CG31715,CG4041,rept,RhoGDI | | 2.094e-02 | -3.87 | GO\_POSITIVE\_REGULATION\_OF\_PROTEOLYSIS | MSigDB lists | GO\_POSITIVE\_REGULATION\_OF\_PROTEOLYSIS | 129 | 29 | 4161 | 644 | Prosbeta6,SkpA,RpL40,regucalcin,Prosbeta3,CG30291,Rpt2,Prosalpha4,Rpn12,APC7,ari-1,Cyt-c-p,Rpn2,dco,Gclc,CycB,Prosbeta5,STUB1,Cdk1,AIF,Rpn3,Cul1,mad2,Prosalpha3,Prosbeta4,Smox,RpS27A,Prosalpha5,PDCD-5 | | 2.094e-02 | -3.87 | LEE\_BMP2\_TARGETS\_UP | MSigDB lists | LEE\_BMP2\_TARGETS\_UP | 129 | 29 | 4161 | 644 | CG14516,Pcd,sws,CG8507,Jheh2,SelR,CG8199,CG7322,LpR2,Pdxk,CG9471,crq,CG3077,CG2201,Tsp66E,AnxB10,Ptp10D,nrv1,Prip,isoQC,Gip,DhpD,gus,cathD,ATP8A,bru1,zyd,scramb1,Maf1 | | 2.106e-02 | -3.86 | MUNSHI\_MULTIPLE\_MYELOMA\_UP | MSigDB lists | MUNSHI\_MULTIPLE\_MYELOMA\_UP | 28 | 9 | 4161 | 644 | Kap-alpha3,not,cact,Prosbeta6,CG14715,faf,Pfdn5,Hsc70-3,Gapdh1 | | 2.106e-02 | -3.86 | GO\_RESPONSE\_TO\_ORGANOPHOSPHORUS | MSigDB lists | GO\_RESPONSE\_TO\_ORGANOPHOSPHORUS | 28 | 9 | 4161 | 644 | Ts,RfC38,GlyP,Calx,dnc,Wnt10,SREBP,Prip,Sod1 | | 2.106e-02 | -3.86 | KEGG\_GLYCOLYSIS\_GLUCONEOGENESIS | MSigDB lists | KEGG\_GLYCOLYSIS\_GLUCONEOGENESIS | 28 | 9 | 4161 | 644 | Pgm1,Gapdh1,Eno,AcCoAS,Pfk,Aldh,Pgk,Fdh,Pglym78 | | 2.110e-02 | -3.86 | GO\_NEGATIVE\_REGULATION\_OF\_PROTEIN\_METABOLIC\_PROCESS | MSigDB lists | GO\_NEGATIVE\_REGULATION\_OF\_PROTEIN\_METABOLIC\_PROCESS | 280 | 56 | 4161 | 644 | twin,snz,Pka-R2,CG30496,Prosalpha5,RpS27A,Cdk1,mad2,Uch-L5,Gapdh1,Rpt2,CG7530,mts,Prosbeta3,Prip,Rpn2,RpL13A,eIF4E1,Rpn12,HDAC6,CG1812,EcR,lid,Arl6IP1,regucalcin,dnc,Prosbeta4,Dcr-1,Prosalpha3,Pp2A-29B,PDCD-5,PpD3,Smox,CG4709,Pdk1,Prosbeta5,Gclc,CycB,Cdk5,Rpn3,CG3376,Non1,CkIIalpha,CG30291,Tlk,APC7,Ranbp21,Ptp61F,CG6766,Prosbeta6,Pka-R1,Snx6,RpL40,Imp,Med,Tspo | | 2.113e-02 | -3.86 | GRIP | prosite domains | PS50913 | 6 | 3 | 6243 | 694 | cbs,Gmap,GCC88 | | 2.113e-02 | -3.86 | THIOLASE\_1 | prosite domains | PS00098 | 6 | 3 | 6243 | 694 | yip2,Thiolase,CG10932 | | 2.113e-02 | -3.86 | GALECTIN | prosite domains | PS51304 | 6 | 3 | 6243 | 694 | CG5335,Pex23,galectin | | 2.113e-02 | -3.86 | ATP\_GRASP | prosite domains | PS50975 | 6 | 3 | 6243 | 694 | ACC,ScsbetaA,r | | 2.113e-02 | -3.86 | V\_SNARE | prosite domains | PS50892 | 6 | 3 | 6243 | 694 | Tomosyn,Vamp7,Sec22 | | 2.113e-02 | -3.86 | THIOLASE\_2 | prosite domains | PS00737 | 6 | 3 | 6243 | 694 | Thiolase,CG10932,yip2 | | 2.128e-02 | -3.85 | lwr (lesswright) | protein interactions | 33226 | 65 | 14 | 9204 | 1112 | cact,velo,Xpac,COX5A,squ,Axs,grp,lwr,CG13373,dl,Aos1,Med,Cul2,Hsp26 | | 2.137e-02 | -3.85 | ZnT63C (Zinc transporter 63C) | protein interactions | 38407 | 41 | 10 | 9204 | 1112 | Atpalpha,Mcad,CG10425,SERCA,Rab5,red,Karybeta3,CG31729,CG15098,Surf4 | | 2.138e-02 | -3.85 | Heme degradation | REACTOME pathways | R-DME-189483 | 2 | 2 | 3923 | 574 | CG9471,Ho | | 2.138e-02 | -3.85 | Interferon Signaling | REACTOME pathways | R-DME-913531 | 2 | 2 | 3923 | 574 | Ptp61F,Stat92E | | 2.138e-02 | -3.85 | Beta oxidation of myristoyl-CoA to lauroyl-CoA | REACTOME pathways | R-DME-77285 | 2 | 2 | 3923 | 574 | Thiolase,Mtpalpha | | 2.138e-02 | -3.85 | Regulation of IFNG signaling | REACTOME pathways | R-DME-877312 | 2 | 2 | 3923 | 574 | Stat92E,Ptp61F | | 2.138e-02 | -3.85 | Interferon gamma signaling | REACTOME pathways | R-DME-877300 | 2 | 2 | 3923 | 574 | Ptp61F,Stat92E | | 2.140e-02 | -3.84 | REACTOME\_CELL\_CYCLE | MSigDB lists | REACTOME\_CELL\_CYCLE | 212 | 44 | 4161 | 644 | Prosbeta6,RfC38,Ts,NHP2,RpL40,Tao,Rbf,Dhfr,CG12018,dco,APC7,CycB,pont,Prosbeta5,msps,Cul1,Rpn3,lwr,Prosbeta4,Prosalpha3,DNApol-epsilon255,Pp2A-29B,RpS27,CycE,gammaTub37C,Sld5,SkpA,Grip128,rept,Rpt2,Prosalpha4,Cdk7,Prosbeta3,mts,Rpn2,grp,Rpn12,Klp10A,Cdk1,mad2,RpS27A,Prosalpha5,shu,Grip84 | | 2.145e-02 | -3.84 | Mitophagy | REACTOME pathways | R-DME-5205647 | 12 | 5 | 3923 | 574 | RpL40,park,Pink1,RpS27A,Marf | | 2.145e-02 | -3.84 | Pink/Parkin Mediated Mitophagy | REACTOME pathways | R-DME-5205685 | 12 | 5 | 3923 | 574 | RpS27A,Marf,park,RpL40,Pink1 | | 2.146e-02 | -3.84 | transcription factor complex | cellular component | GO:0005667 | 177 | 25 | 11940 | 1105 | e(y)1,Su(H),CG4404,cyc,Rbf2,E2f1,Nf-YA,TfIIFalpha,Taf7,Smox,CG42526,brwl,CG31875,Med,NFAT,Cdk7,Tfb4,CG6276,Rbf,EcR,Trf2,MED4,jigr1,pygo,Mes2 | | 2.149e-02 | -3.84 | columnar/cuboidal epithelial cell differentiation | biological process | GO:0002065 | 344 | 43 | 12345 | 1129 | armi,fs(1)M3,bun,mop,Pxt,Rab5,cup,Cf2,Ptpmeg2,r2d2,Abi,CycE,spn-E,Med,csw,Cdk1,Lkb1,Rbf,squ,tor,Orc1,Tao,gus,Sec15,jvl,EcR,fs(1)N,cact,mael,toc,Stat92E,tkv,SERCA,BicC,E2f1,apt,Cbl,Pi3K92E,nej,Pax,stai,rl,e(y)1 | | 2.150e-02 | -3.84 | CYCLIN | smart domains | SM00385 | 17 | 5 | 5086 | 506 | CycC,CycB,CycE,CycJ,Rbf | | 2.150e-02 | -3.84 | Sm | smart domains | SM00651 | 17 | 5 | 5086 | 506 | CG17768,LSm3,CG10418,SmE,CG6610 | | 2.153e-02 | -3.84 | GO\_NCRNA\_PROCESSING | MSigDB lists | GO\_NCRNA\_PROCESSING | 229 | 47 | 4161 | 644 | l(1)G0004,Uba5,Rrp40,LysRS,RpL13A,RpLP0,RpS30,RpS13,scu,CG2100,CG10189,CG3021,RpL13,Pus7,RpS28b,RpS27A,RpS23,RpL9,Clp,RpL40,RpS7,mtTFB1,NHP2,RpS4,RpL38,RpS16,RpS2,CG11858,dco,CG5292,RpS24,CG10463,RpL23,Archease,RpL30,RpL12,bys,RpS18,RpS27,RpL24,Smox,RpL26,RpL37a,Dcr-1,Mat89Ba,RpL11,CG7728 | | 2.154e-02 | -3.84 | locomotory behavior | biological process | GO:0007626 | 188 | 26 | 12345 | 1129 | park,Cdk5,foxo,shep,Irk1,cu,nej,dco,cyc,rig,tyf,Bx,Atpalpha,SERCA,sbb,dnc,ben,gw,Ork1,Marf,EcR,CkIIalpha,G9a,lid,Pka-R2,Ac3 | | 2.156e-02 | -3.84 | positive regulation of apoptotic process | biological process | GO:0043065 | 69 | 12 | 12345 | 1129 | MED24,ScsbetaA,Cbl,eIF3m,Jafrac2,CycE,EcR,E2f1,Tspo,fwe,PDCD-5,Lkb1 | | 2.156e-02 | -3.84 | positive regulation of kinase activity | biological process | GO:0033674 | 69 | 12 | 12345 | 1129 | Pdk1,Tak1,ASPP,Parp16,Mekk1,fray,Abi,Pi3K92E,raptor,RpLP1,Tao,Lkb1 | | 2.163e-02 | -3.83 | DNA-dependent DNA replication | biological process | GO:0006261 | 77 | 13 | 12345 | 1129 | RecQ5,Rbf,CG10336,DNApol-epsilon255,ago,Orc1,CG12018,SuUR,Orc6,RfC38,SkpA,Sld5,E2f1 | | 2.163e-02 | -3.83 | cytoplasm organization | biological process | GO:0007028 | 77 | 13 | 12345 | 1129 | cnc,swa,mael,armi,psq,Tm1,gus,Lkb1,Rab5,spn-E,osk,exu,Moe | | 2.167e-02 | -3.83 | mitotic cell cycle checkpoint | biological process | GO:0007093 | 46 | 9 | 12345 | 1129 | Cdk1,grp,mus304,nbs,Zwilch,mad2,mus81,CG10336,mre11 | | 2.167e-02 | -3.83 | positive regulation of proteolysis involved in cellular protein catabolic process | biological process | GO:1903052 | 46 | 9 | 12345 | 1129 | ari-1,Ubc6,park,Cul1,Rpt2,cort,dco,Pomp,SkpA | | 2.167e-02 | -3.83 | Class I MHC mediated antigen processing & presentation | REACTOME pathways | R-DME-983169 | 114 | 25 | 3923 | 574 | Rpn3,Cul1,Rpn12,CG14435,dx,Herc4,gus,Fbw5,Prosbeta6,Sec23,Prosbeta3,crq,RpL40,Rpt2,Prosalpha3,SkpA,Ubc6,Prosbeta5,Rpn2,Prosalpha4,Iru,Prosalpha5,RpS27A,ago,Prosbeta4 | | 2.171e-02 | -3.83 | pan (pangolin) | protein interactions | 43769 | 59 | 13 | 9204 | 1112 | foxo,Rbf,squ,Cdk5,CkIIalpha,ovo,plu,tws,Cul2,E2f1,CG13373,grp,alph | | 2.178e-02 | -3.83 | REACTOME\_GENERIC\_TRANSCRIPTION\_PATHWAY | MSigDB lists | REACTOME\_GENERIC\_TRANSCRIPTION\_PATHWAY | 42 | 12 | 4161 | 644 | CycC,MED24,RpS27A,faf,MED20,Smox,MED4,EcR,MED27,Med,RpL40,Su(H) | | 2.178e-02 | -3.83 | TOOKER\_GEMCITABINE\_RESISTANCE\_DN | MSigDB lists | TOOKER\_GEMCITABINE\_RESISTANCE\_DN | 42 | 12 | 4161 | 644 | Grx1,Top1,CG1416,Nca,Karybeta3,CG3860,CG8188,Fkbp59,Sptr,TyrRS,CG11899,CG3226 | | 2.178e-02 | -3.83 | GSE7460\_TREG\_VS\_TCONV\_ACT\_WITH\_TGFB\_DN | MSigDB lists | GSE7460\_TREG\_VS\_TCONV\_ACT\_WITH\_TGFB\_DN | 42 | 12 | 4161 | 644 | Pex23,ND-39,CG9248,Surf4,GCS2beta,RpLP0,Sirt7,AcCoAS,CG8199,Arl5,Nca,cathD | | 2.178e-02 | -3.83 | GO\_ENDOPLASMIC\_RETICULUM\_GOLGI\_INTERMEDIATE\_COMPARTMENT | MSigDB lists | GO\_ENDOPLASMIC\_RETICULUM\_GOLGI\_INTERMEDIATE\_COMPARTMENT | 42 | 12 | 4161 | 644 | CG9911,opm,CHOp24,Ptp61F,CG14516,Surf4,Sec22,PAPLA1,CG4293,Dcr-1,SERCA,Hsc70-3 | | 2.182e-02 | -3.82 | GSE36392\_TYPE\_2\_MYELOID\_VS\_NEUTROPHIL\_IL25\_TREATED\_LUNG\_DN | MSigDB lists | GSE36392\_TYPE\_2\_MYELOID\_VS\_NEUTROPHIL\_IL25\_TREATED\_LUNG\_DN | 87 | 21 | 4161 | 644 | AP-1-2beta,Ubc4,Stoml2,CG8204,ste14,mRpL18,CG8728,ND-49,Tsp86D,Rpt2,GAPsec,Tom40,Bin1,CG8199,Pglym78,ND-39,RpL24,PpD3,Nap1,RpS23,RpL9 | | 2.182e-02 | -3.82 | Signaling by FGFR3 | REACTOME pathways | R-DME-5654741 | 25 | 8 | 3923 | 574 | rl,RpS27A,csw,Cbl,Pp2A-29B,RpL40,CG13398,mts | | 2.182e-02 | -3.82 | Signaling by FGFR4 | REACTOME pathways | R-DME-5654743 | 25 | 8 | 3923 | 574 | RpS27A,rl,Pp2A-29B,Cbl,csw,mts,CG13398,RpL40 | | 2.184e-02 | -3.82 | Vha68-2 (Vacuolar H[+] ATPase 68 kDa subunit 2) | protein interactions | 45012 | 47 | 11 | 9204 | 1112 | Mcad,Vha36-1,Atpalpha,Rab5,SERCA,CCT6,Coq6,Karybeta3,Rtnl1,Mitf,wal | | 2.184e-02 | -3.82 | SerRS (Seryl-tRNA synthetase) | protein interactions | 33518 | 47 | 11 | 9204 | 1112 | Pp2A-29B,wal,mEFTu1,r,CCT3,bor,Pen,CCT6,ScsbetaA,Kap-alpha3,Mcad | | 2.208e-02 | -3.81 | Cellular response to hypoxia | REACTOME pathways | R-DME-2262749 | 65 | 16 | 3923 | 574 | Rpn12,Rpn3,Prosbeta6,nej,Prosbeta3,EloC,Cul2,Prosalpha3,RpL40,Rpt2,Prosalpha4,Rpn2,Prosbeta5,Prosbeta4,RpS27A,Prosalpha5 | | 2.208e-02 | -3.81 | Regulation of Hypoxia-inducible Factor (HIF) by oxygen | REACTOME pathways | R-DME-1234174 | 65 | 16 | 3923 | 574 | nej,Prosbeta3,Rpn12,Rpn3,Prosbeta6,Prosbeta5,Rpn2,Prosalpha4,RpS27A,Prosalpha5,Prosbeta4,Prosalpha3,Rpt2,RpL40,EloC,Cul2 | | 2.217e-02 | -3.81 | regulation of localization | biological process | GO:0032879 | 478 | 57 | 12345 | 1129 | Rab32,sbr,SERCA,Tomosyn,mael,Mctp,PGRP-LE,LpR2,dco,swa,adp,Pi3K92E,Fur2,Cbl,wash,Stam,ZnT35C,Snx6,Tao,TyrRS,CG8726,gw,lt,Ptp61F,enc,Rab10,Moe,Fkbp59,crq,NFAT,O-fut1,Cdk1,Vps16B,Nup358,Ptpmeg,cnc,Pld,crok,Rbf,foxo,wapl,exu,Rab5,cup,mop,Rel,Herc4,raptor,mbo,Seipin,Atf3,Crag,Sec23,Hip1,spn-E,Ranbp21,tral | | 2.220e-02 | -3.81 | macromolecule biosynthetic process | biological process | GO:0009059 | 1224 | 132 | 12345 | 1129 | tko,Cdk7,RpLP1,GatB,eIF3m,mRpL18,mRpS11,Sgf29,eEF1beta,RpS13,EMC3,pps,RpII140,GluRS-m,RpL30,Rbf,CG10336,CG5880,RpS2,RpS29,RpL26,Orc6,CG14817,RpS16,Pgm1,RpS27A,mRpL16,CG7414,RpL34b,RpL40,Taf7,SkpA,TfIIFalpha,CG7840,CG9485,SuUR,spen,RpL11,eIF4E1,SerRS-m,RpS28b,mRpS10,mRpS21,CG8314,mus205,PIG-C,RpL9,GAPsec,AGBE,Rpb8,RpS27,pygo,GatA,fdl,RpL13,CG7009,mRpL13,RpL24,RpL10,TyrRS,mRpS23,mRpL10,RpS23,E2f1,mRpL17,RecQ5,RpS30,DNApol-epsilon255,CG12018,RpL17,mEFTu1,MED24,LysRS,RpS4,CG6094,Tfb4,RpL31,CycE,PIG-P,O-fut1,RpL23A,Ggt-1,Hexo1,PAPLA1,Top1,eIF3i,RpS15,RpS7,mRpL36,Pym,TrpRS-m,Nmt,mRpL35,CG10344,mRpS5,Trf2,GalT1,Gyg,RpL29,PIG-L,e(y)1,mtTFB1,RpL23,RpS14b,RpS18,RpS14a,Sld5,RpL13A,RpS24,Atg7,Rga,RpL34a,PIG-H,MED4,eIF4A,ago,Orc1,RpL37a,oxt,RpS5b,RpLP0,RfC38,mRpL48,RpL12,RpL38,RpL7A,fs(1)Ya,CG33977,Efr,mEFTu2,Nurf-38,CysRS | | 2.223e-02 | -3.81 | negative regulation of signal transduction | biological process | GO:0009968 | 326 | 41 | 12345 | 1129 | Ptp10D,Pp2A-29B,Herc4,Su(H),STUB1,Ptpmeg,alph,scny,Cka,Dd,piwi,Ubc6,Traf6,foxo,Rbf,mts,dx,ATPsynD,Cul2,CG17746,pn,NFAT,Iml1,Ect4,PpV,dnc,cact,chrb,Bap170,Ptp61F,dco,apt,SkpA,insv,Cul1,park,Nprl2,lwr,rept,Cbl,nej | | 2.230e-02 | -3.80 | SDRFAMILY | prints domains | PR00080 | 44 | 9 | 2185 | 212 | Adh,scu,CG9265,sni,CG31549,CG4842,CG7322,CG31548,CG12171 | | 2.230e-02 | -3.80 | GSE27786\_NKTCELL\_VS\_MONO\_MAC\_UP | MSigDB lists | GSE27786\_NKTCELL\_VS\_MONO\_MAC\_UP | 82 | 20 | 4161 | 644 | CG1827,CG9143,Rab10,Pfas,Idh3b,Stip1,ND-30,CCT6,RpL23,TyrRS,CG3226,nmdyn-D7,RpS16,GCS2beta,GluRS-m,Karybeta3,Pka-R1,Adi1,Arts,RhoGDI | | 2.230e-02 | -3.80 | HALLMARK\_GLYCOLYSIS | MSigDB lists | HALLMARK\_GLYCOLYSIS | 82 | 20 | 4161 | 644 | CG6767,Sod1,Gmer,MED24,Pdk,CG9485,CG17333,Got1,Hsc70-3,Idh,CG7789,CG7840,Fkbp59,Pglym78,Pgk,Cdk1,LpR2,Klp10A,Gclc,Eno | | 2.234e-02 | -3.80 | GO\_RESPONSE\_TO\_LIPID | MSigDB lists | GO\_RESPONSE\_TO\_LIPID | 179 | 38 | 4161 | 644 | Prip,EcR,Cdk7,Tip60,ACC,Fdh,rept,dnc,CG5412,Got1,Idh,Fkbp59,AIF,LpR2,crq,cact,rl,aay,Traf6,MED4,MED24,Atpalpha,Pdk,Tspo,Gdap2,CG9485,Ts,r,Cka,CycE,Wnt10,SREBP,Moe,bys,p38a,Ac3,CG11710,CycB | | 2.247e-02 | -3.80 | microtubule polymerization | biological process | GO:0046785 | 25 | 6 | 12345 | 1129 | msps,Grip75,Grip128,gammaTub37C,Grip84,CG7716 | | 2.249e-02 | -3.79 | Rab11 (Rab11) | protein interactions | 42501 | 78 | 16 | 9204 | 1112 | sau,Tm1,RpLP1,Sec15,Tak1,eIF3m,Vha68-2,Crag,Pglym78,ScsbetaA,CG9186,Rep,Gdi,scu,Moe,Phb2 | | 2.258e-02 | -3.79 | Beta Oxidation Meta MAPP | WikiPathways | WP443 | 23 | 10 | 499 | 116 | Thiolase,whd,Gk2,Mcad,CG10932,Echs1,AcCoAS,colt,Mtpalpha,Gk1 | | 2.259e-02 | -3.79 | MGGAAGTG\_GABP\_B | MSigDB lists | MGGAAGTG\_GABP\_B | 241 | 49 | 4161 | 644 | Tip60,CG11070,DppIII,CG8475,Rpb8,CG31715,CG2061,Not10,CG6610,CG13601,lqfR,kcc,ERp60,Ssdp,mEFTu1,pen-2,CG3803,Tom40,CG7997,Exo70,Herc4,Rab5,Synj,Ranbp21,Ptp61F,CG9485,Prosbeta6,Rep,Aladin,Pp2B-14D,Sec10,CG10470,Phb2,Edc3,GCS2beta,Vha36-1,capt,ND-30,Arpc4,PIG-C,Hat1,Marf,RpS18,Cdk5,Rnmt,Acph-1,TBCC,RpL37a,CanA-14F | | 2.261e-02 | -3.79 | GSE36888\_UNTREATED\_VS\_IL2\_TREATED\_TCELL\_17H\_UP | MSigDB lists | GSE36888\_UNTREATED\_VS\_IL2\_TREATED\_TCELL\_17H\_UP | 47 | 13 | 4161 | 644 | CG9471,Gclc,Chrac-14,Shmt,CG4572,CG9248,CG2493,Mcad,CG17294,CG7840,Pfdn5,CG3860,ND-SGDH | | 2.261e-02 | -3.79 | GO\_DNA\_INTEGRITY\_CHECKPOINT | MSigDB lists | GO\_DNA\_INTEGRITY\_CHECKPOINT | 47 | 13 | 4161 | 644 | Cdk1,p38a,CycB,RpL40,Tao,csw,Rbf,APP-BP1,CG30291,twin,RpS27A,Not10,grp | | 2.261e-02 | -3.79 | REACTOME\_AUTODEGRADATION\_OF\_CDH1\_BY\_CDH1\_APC\_C | MSigDB lists | REACTOME\_AUTODEGRADATION\_OF\_CDH1\_BY\_CDH1\_APC\_C | 47 | 13 | 4161 | 644 | Prosbeta6,Prosbeta5,Rpn3,RpL40,Rpt2,Prosbeta4,Prosalpha3,Prosbeta3,RpS27A,Rpn2,Prosalpha5,APC7,Rpn12 | | 2.261e-02 | -3.79 | GSE3982\_MAC\_VS\_NKCELL\_UP | MSigDB lists | GSE3982\_MAC\_VS\_NKCELL\_UP | 47 | 13 | 4161 | 644 | Irk1,Pdxk,Su(H),Gk2,Pglym78,CG9911,Idh,CG1275,muskelin,ND-SGDH,Acph-1,isoQC,Pcd | | 2.261e-02 | -3.79 | GSE19401\_UNSTIM\_VS\_PAM2CSK4\_STIM\_FOLLICULAR\_DC\_UP | MSigDB lists | GSE19401\_UNSTIM\_VS\_PAM2CSK4\_STIM\_FOLLICULAR\_DC\_UP | 47 | 13 | 4161 | 644 | RfC38,SRPK,Tm1,CG9911,TBCC,Arf51F,muskelin,CG7728,CG3631,obe,Clp,LysRS,dco | | 2.261e-02 | -3.79 | GSE21927\_UNTREATED\_VS\_GMCSF\_GCSF\_TREATED\_BONE\_MARROW\_DN | MSigDB lists | GSE21927\_UNTREATED\_VS\_GMCSF\_GCSF\_TREATED\_BONE\_MARROW\_DN | 47 | 13 | 4161 | 644 | CG4872,UbcE2H,Non1,CG14407,Ugt35A1,COX5A,anne,Ts,Pi3K92E,CG40045,Klp10A,Ssdp,Top1 | | 2.269e-02 | -3.79 | His2Av (Histone H2A variant) | protein interactions | 43229 | 30 | 8 | 9204 | 1112 | Tip60,Jabba,Imp,E2f1,nbs,pont,Fkbp39,ago | | 2.271e-02 | -3.79 | tricarboxylic acid cycle | biological process | GO:0006099 | 39 | 8 | 12345 | 1129 | Irp-1B,ScsbetaA,SdhA,Nc73EF,SdhD,CG5028,Scsalpha1,Idh3b | | 2.271e-02 | -3.79 | cellular response to oxidative stress | biological process | GO:0034599 | 39 | 8 | 12345 | 1129 | foxo,Grx1,Idh,Jafrac1,rl,Sod1,Sod3,p38a | | 2.273e-02 | -3.78 | GSE7568\_IL4\_VS\_IL4\_AND\_DEXAMETHASONE\_TREATED\_MACROPHAGE\_UP | MSigDB lists | GSE7568\_IL4\_VS\_IL4\_AND\_DEXAMETHASONE\_TREATED\_MACROPHAGE\_UP | 77 | 19 | 4161 | 644 | Psf3,CG11899,Archease,shv,RpS2,Rpn12,CG8188,Karybeta3,Sld5,ND-23,RhoGDI,CG10418,Hsc70-3,mRpS21,COX5A,Prosbeta5,CG7997,Pgk,mad2 | | 2.273e-02 | -3.78 | GO\_COENZYME\_BIOSYNTHETIC\_PROCESS | MSigDB lists | GO\_COENZYME\_BIOSYNTHETIC\_PROCESS | 77 | 19 | 4161 | 644 | mal,Coq6,Nadsyn,CG33156,Dhfr,CG4407,ACC,CG11089,sgll,Pcd,CG1774,ND-39,Ppcdc,Sptr,Desat1,Pdxk,Coq7,AcCoAS,pr | | 2.273e-02 | -3.78 | GCM\_NPM1 | MSigDB lists | GCM\_NPM1 | 77 | 19 | 4161 | 644 | CG4390,RpS2,RpL13A,RpS16,RpS24,RpL23,RpL40,RpS7,CG8204,RpS28b,RpL9,RpS27A,RpS23,Nap1,Prosalpha5,RpL11,eEF1beta,Bin1,RpS18 | | 2.273e-02 | -3.78 | GSE41176\_WT\_VS\_TAK1\_KO\_ANTI\_IGM\_STIM\_BCELL\_24H\_UP | MSigDB lists | GSE41176\_WT\_VS\_TAK1\_KO\_ANTI\_IGM\_STIM\_BCELL\_24H\_UP | 77 | 19 | 4161 | 644 | AGBE,Fur2,Tspo,Cyp1,CG2493,UbcE2H,ari-1,CG8188,Sod1,RpS30,CG7997,Gclc,CG9471,Cyp6a19,Nca,mtgo,Hsc70-3,Cyp6a22,stas | | 2.273e-02 | -3.78 | GSE8921\_UNSTIM\_0H\_VS\_TLR1\_2\_STIM\_MONOCYTE\_12H\_UP | MSigDB lists | GSE8921\_UNSTIM\_0H\_VS\_TLR1\_2\_STIM\_MONOCYTE\_12H\_UP | 77 | 19 | 4161 | 644 | ND-39,Nmt,pont,Tim8,wal,mRpS10,Pfdn2,Caf1-105,CCT6,Hat1,CG3226,Efr,Prosbeta3,CG12018,Dhfr,Non1,GCS2beta,ND-B22,Adi1 | | 2.275e-02 | -3.78 | regulation of vesicle-mediated transport | biological process | GO:0060627 | 119 | 18 | 12345 | 1129 | wash,Cbl,Pld,Vps16B,PGRP-LE,Tomosyn,O-fut1,crq,NFAT,Sec23,Hip1,Crag,Moe,Rab10,lt,Rel,Tao,Rab5 | | 2.288e-02 | -3.78 | regulation of catalytic activity | biological process | GO:0050790 | 450 | 54 | 12345 | 1129 | Sec23,Crag,sws,ASPP,Rpn13,eIF3m,CG30291,CycE,Abi,fray,raptor,nrv1,Pp2A-29B,RpLP1,Cdk7,Pdk1,Tak1,CycJ,Parp16,Jafrac2,esc,cort,CG14408,Trs33,RhoGAP54D,alph,GAPsec,Lkb1,CycC,blw,CG4041,Iml1,Rpn3,Mekk1,CG8726,Rpn2,CG13551,tws,CG17746,levy,SNF4Agamma,Tao,Orc6,MED24,Drep1,Pax,ScsbetaA,CycB,CG1416,Cbl,Pi3K92E,RhoGAP5A,Tomosyn,EndoGI | | 2.291e-02 | -3.78 | Toll-Like Receptors Cascades | REACTOME pathways | R-DME-168898 | 60 | 15 | 3923 | 574 | p38a,Pp2A-29B,dl,RpL40,Myd88,crq,RpS27A,rl,ben,Traf6,Ptpmeg,Cp1,Tak1,Rel,Uev1A | | 2.297e-02 | -3.77 | oxidoreductase activity, acting on the CH-NH group of donors | molecular function | GO:0016645 | 25 | 6 | 12176 | 1119 | P5cr,Dhfr,P5CDh1,Etf-QO,CG9471,CG10561 | | 2.297e-02 | -3.77 | ribosomal large subunit biogenesis | biological process | GO:0042273 | 54 | 10 | 12345 | 1129 | RpLP0,NHP2,RpL24,RpL26,RpL7A,RpL11,RpL23A,RpL10,RpL12,CG8204 | | 2.308e-02 | -3.77 | rhythmic process | biological process | GO:0048511 | 94 | 15 | 12345 | 1129 | tyf,lid,CkIIalpha,cyc,dco,Bx,Pka-R2,Ac3,dnc,cu,nej,gw,Ork1,foxo,Irk1 | | 2.308e-02 | -3.77 | Regulation of DNA replication | REACTOME pathways | R-DME-69304 | 76 | 18 | 3923 | 574 | Orc6,Prosbeta3,Rpn3,Orc1,Rpn12,Prosbeta6,Rpn2,Prosbeta5,Rbf2,Prosalpha4,Prosalpha5,RpS27A,E2f1,Prosbeta4,Rpt2,RpL40,Prosalpha3,Rbf | | 2.313e-02 | -3.77 | GSE360\_CTRL\_VS\_B\_MALAYI\_LOW\_DOSE\_DC\_UP | MSigDB lists | GSE360\_CTRL\_VS\_B\_MALAYI\_LOW\_DOSE\_DC\_UP | 52 | 14 | 4161 | 644 | Su(H),Pglym78,bru1,AGBE,CHOp24,gammaTub37C,ari-1,Idh3b,RpS27A,mts,Pgm1,RpS24,muskelin,CG44422 | | 2.313e-02 | -3.77 | DAVICIONI\_TARGETS\_OF\_PAX\_FOXO1\_FUSIONS\_UP | MSigDB lists | DAVICIONI\_TARGETS\_OF\_PAX\_FOXO1\_FUSIONS\_UP | 52 | 14 | 4161 | 644 | dco,Gabat,Rab32,CG8507,Rpn12,Dcr-1,CG1275,Nca,Moe,scramb1,capt,Ntan1,Fur2,MED27 | | 2.313e-02 | -3.77 | GSE14000\_TRANSLATED\_RNA\_VS\_MRNA\_16H\_LPS\_DC\_DN | MSigDB lists | GSE14000\_TRANSLATED\_RNA\_VS\_MRNA\_16H\_LPS\_DC\_DN | 52 | 14 | 4161 | 644 | CG15715,ste14,Arl5,RpL40,Tfb4,CHOp24,RpL38,shu,RpS2,Lsm12,CG5285,Cisd2,RpL23,RpL30 | | 2.313e-02 | -3.77 | MARTENS\_BOUND\_BY\_PML\_RARA\_FUSION | MSigDB lists | MARTENS\_BOUND\_BY\_PML\_RARA\_FUSION | 130 | 29 | 4161 | 644 | Tsp66E,CG33156,alphaTub84B,CG17333,PIG-L,AGBE,zyd,bru1,cathD,Pex12,Idh,Ppcdc,SREBP,RpL9,awd,stas,RpL24,Cyp6a22,Smox,Grip84,Aldh,CG11710,crq,Cdk5,pont,Tom40,Rpn3,Nca,Cyp6a19 | | 2.316e-02 | -3.77 | GO\_POSTTRANSCRIPTIONAL\_REGULATION\_OF\_GENE\_EXPRESSION | MSigDB lists | GO\_POSTTRANSCRIPTIONAL\_REGULATION\_OF\_GENE\_EXPRESSION | 185 | 39 | 4161 | 644 | rl,Irp-1B,Prosalpha5,RpS27A,twin,CG31715,eIF3i,eIF4E1,Rrp40,RpL13A,Rpn12,Not10,Rpn2,Prosbeta3,Gapdh1,Prosalpha4,piwi,eIF4A,Rpt2,CG13630,CG15715,Rpn3,Prosbeta5,p38a,Smox,eIF3m,Prosalpha3,Dcr-1,CG7414,Prosbeta4,AGO3,Imp,RpL40,Cbp80,Ranbp21,Prosbeta6,RpS4,RpL38,CG8963 | | 2.332e-02 | -3.76 | MORF\_EIF4A2 | MSigDB lists | MORF\_EIF4A2 | 67 | 17 | 4161 | 644 | eIF3m,RpL11,RpL24,RpS27A,RpL9,Idh3b,Nap1,lwr,RpL13,Pgk,eEF1beta,RpS24,RpL23,RpL30,eIF4A,RpS30,RpS7 | | 2.332e-02 | -3.76 | GO\_NEGATIVE\_REGULATION\_OF\_TRANSMEMBRANE\_RECEPTOR\_PROTEIN\_SERINE\_THREONINE\_KINASE\_SIGNALING\_PATHWAY | MSigDB lists | GO\_NEGATIVE\_REGULATION\_OF\_TRANSMEMBRANE\_RECEPTOR\_PROTEIN\_SERINE\_THREONINE\_KINASE\_SIGNALING\_PATHWAY | 24 | 8 | 4161 | 644 | Smox,RpS27A,snz,Hsc70-3,RpL40,Snx6,STUB1,Pdk1 | | 2.332e-02 | -3.76 | MARTINEZ\_RESPONSE\_TO\_TRABECTEDIN\_UP | MSigDB lists | MARTINEZ\_RESPONSE\_TO\_TRABECTEDIN\_UP | 24 | 8 | 4161 | 644 | Sod1,RpS27,Cyt-c-p,Jafrac1,Pgk,RpL40,ERp60,SkpA | | 2.332e-02 | -3.76 | PID\_P73PATHWAY | MSigDB lists | PID\_P73PATHWAY | 24 | 8 | 4161 | 644 | Pfdn5,grp,ND-49,Tip60,p38a,Cdk1,CycB,tzn | | 2.332e-02 | -3.76 | GSE13547\_WT\_VS\_ZFX\_KO\_BCELL\_UP | MSigDB lists | GSE13547\_WT\_VS\_ZFX\_KO\_BCELL\_UP | 24 | 8 | 4161 | 644 | Med,shop,eIF4A,Rbf,mino,sws,Pfdn5,Jheh2 | | 2.332e-02 | -3.76 | GO\_HYDRO\_LYASE\_ACTIVITY | MSigDB lists | GO\_HYDRO\_LYASE\_ACTIVITY | 24 | 8 | 4161 | 644 | Thiolase,Uba5,Echs1,Eno,Mtpalpha,Pcd,Irp-1B,CG11134 | | 2.332e-02 | -3.76 | KEGG\_T\_CELL\_RECEPTOR\_SIGNALING\_PATHWAY | MSigDB lists | KEGG\_T\_CELL\_RECEPTOR\_SIGNALING\_PATHWAY | 24 | 8 | 4161 | 644 | CanB2,Pdk1,rl,cact,p38a,Pi3K92E,Pp2B-14D,CanA-14F | | 2.332e-02 | -3.76 | GO\_NUCLEOBASE\_METABOLIC\_PROCESS | MSigDB lists | GO\_NUCLEOBASE\_METABOLIC\_PROCESS | 24 | 8 | 4161 | 644 | CG6767,Shmt,rl,kri,Ts,r-l,DhpD,r | | 2.337e-02 | -3.76 | GO\_PROTEIN\_TARGETING\_TO\_MITOCHONDRION | MSigDB lists | GO\_PROTEIN\_TARGETING\_TO\_MITOCHONDRION | 33 | 10 | 4161 | 644 | Marf,Tspo,Tom40,CG8004,Tim8,CG9393,CG11110,CG2100,CG8728,CG1847 | | 2.337e-02 | -3.76 | REACTOME\_ANTIVIRAL\_MECHANISM\_BY\_IFN\_STIMULATED\_GENES | MSigDB lists | REACTOME\_ANTIVIRAL\_MECHANISM\_BY\_IFN\_STIMULATED\_GENES | 33 | 10 | 4161 | 644 | eIF4A,RpS27A,Aladin,eIF4E1,ari-1,alph,Kap-alpha3,Pen,RpL40,mbo | | 2.337e-02 | -3.76 | GROSS\_HYPOXIA\_VIA\_ELK3\_DN | MSigDB lists | GROSS\_HYPOXIA\_VIA\_ELK3\_DN | 33 | 10 | 4161 | 644 | cact,CysRS,LpR2,Eno,AGBE,Su(H),Gmer,Hsc70-3,Traf6,CG14516 | | 2.337e-02 | -3.76 | BHATTACHARYA\_EMBRYONIC\_STEM\_CELL | MSigDB lists | BHATTACHARYA\_EMBRYONIC\_STEM\_CELL | 33 | 10 | 4161 | 644 | CycB,Cdk1,SelD,CG16868,Pen,RpS24,Idh,CycC,RpLP0,RpL24 | | 2.337e-02 | -3.76 | GSE34156\_NOD2\_LIGAND\_VS\_TLR1\_TLR2\_LIGAND\_6H\_TREATED\_MONOCYTE\_DN | MSigDB lists | GSE34156\_NOD2\_LIGAND\_VS\_TLR1\_TLR2\_LIGAND\_6H\_TREATED\_MONOCYTE\_DN | 33 | 10 | 4161 | 644 | Rbcn-3A,Irk1,anne,Tsp74F,CG9471,Gk2,cathD,CG1275,CG33156,CG14516 | | 2.337e-02 | -3.76 | GO\_CELLULAR\_RESPONSE\_TO\_REACTIVE\_OXYGEN\_SPECIES | MSigDB lists | GO\_CELLULAR\_RESPONSE\_TO\_REACTIVE\_OXYGEN\_SPECIES | 33 | 10 | 4161 | 644 | PpD3,Prip,HDAC6,Sod1,Tm1,Jafrac1,Cyp1,Grx1,AIF,Cdk1 | | 2.338e-02 | -3.76 | GSE19888\_NO\_PRETREAT\_VS\_ADENOSINE\_A3R\_INHIBITOR\_PRETREATED\_MAST\_CELL\_TCELL\_MEMBRANES\_ACT\_DN | MSigDB lists | GSE19888\_NO\_PRETREAT\_VS\_ADENOSINE\_A3R\_INHIBITOR\_PRETREATED\_MAST\_CELL\_TCELL\_MEMBRANES\_ACT\_DN | 57 | 15 | 4161 | 644 | Dhfr,Cisd2,mtgo,CG11858,CG1812,Nurf-38,Axs,nmdyn-D7,CG3842,CG16979,mRpS10,Karybeta3,Coq7,CG8726,Nca | | 2.338e-02 | -3.76 | GSE15750\_WT\_VS\_TRAF6KO\_DAY6\_EFF\_CD8\_TCELL\_UP | MSigDB lists | GSE15750\_WT\_VS\_TRAF6KO\_DAY6\_EFF\_CD8\_TCELL\_UP | 57 | 15 | 4161 | 644 | bys,ND-30,mEFTu1,CG13630,Tspo,CG9485,CG5510,CG3894,Sec22,CG9393,APC7,ND-49,CG10463,Sfxn2,CG15111 | | 2.338e-02 | -3.76 | GO\_OXIDOREDUCTASE\_ACTIVITY\_ACTING\_ON\_THE\_CH\_OH\_GROUP\_OF\_DONORS\_NAD\_OR\_NADP\_AS\_ACCEPTOR | MSigDB lists | GO\_OXIDOREDUCTASE\_ACTIVITY\_ACTING\_ON\_THE\_CH\_OH\_GROUP\_OF\_DONORS\_NAD\_OR\_NADP\_AS\_ACCEPTOR | 57 | 15 | 4161 | 644 | CG9265,Idh3b,scu,Sptr,Idh,Gmer,Thiolase,Fdh,CG5028,CG10425,CG4562,CG3842,Mtpalpha,CG3609,CG7322 | | 2.338e-02 | -3.76 | GSE36527\_CD62L\_HIGH\_VS\_CD62L\_LOW\_TREG\_CD69\_NEG\_KLRG1\_NEG\_UP | MSigDB lists | GSE36527\_CD62L\_HIGH\_VS\_CD62L\_LOW\_TREG\_CD69\_NEG\_KLRG1\_NEG\_UP | 57 | 15 | 4161 | 644 | regucalcin,Su(H),lwr,CG4332,CG15141,CG7997,Arl6IP1,wal,CG6767,Pi4KIIalpha,Hsc70-3,CG9143,shrb,Rab10,SREBP | | 2.338e-02 | -3.76 | YAMAZAKI\_TCEB3\_TARGETS\_UP | MSigDB lists | YAMAZAKI\_TCEB3\_TARGETS\_UP | 57 | 15 | 4161 | 644 | Cul1,CG9911,ERp60,parvin,CHOp24,alpha-Man-Ia,CG8507,CG2924,aay,slif,Surf4,CycC,Hsc70-3,mtgo,CG32473 | | 2.343e-02 | -3.75 | GSE14308\_INDUCED\_VS\_NATURAL\_TREG\_DN | MSigDB lists | GSE14308\_INDUCED\_VS\_NATURAL\_TREG\_DN | 62 | 16 | 4161 | 644 | RpL30,tun,P5CDh1,CG7115,CG6144,Sec10,RpS30,Ptp10D,nSMase,CG17593,Droj2,Stam,CG10038,l(1)G0196,sbr,Exd2 | | 2.343e-02 | -3.75 | GSE11961\_MEMORY\_BCELL\_DAY7\_VS\_MEMORY\_BCELL\_DAY40\_UP | MSigDB lists | GSE11961\_MEMORY\_BCELL\_DAY7\_VS\_MEMORY\_BCELL\_DAY40\_UP | 62 | 16 | 4161 | 644 | CG7414,Got1,Tomosyn,alpha-Man-Ia,Liprin-beta,Pgk,mRpS5,ND-49,Tsp86D,CG11899,isoQC,CG1890,Sod1,AP-1-2beta,ATP8A,Su(H) | | 2.343e-02 | -3.75 | TMTCGCGANR\_UNKNOWN | MSigDB lists | TMTCGCGANR\_UNKNOWN | 62 | 16 | 4161 | 644 | RpL12,Bin1,Jafrac2,Ptpmeg,p38a,RpL26,Prosbeta4,Unr,Su(H),Tao,ste14,CG5028,RpS7,CG9485,Tlk,CG10470 | | 2.343e-02 | -3.75 | GSE22601\_DOUBLE\_POSITIVE\_VS\_CD8\_SINGLE\_POSITIVE\_THYMOCYTE\_DN | MSigDB lists | GSE22601\_DOUBLE\_POSITIVE\_VS\_CD8\_SINGLE\_POSITIVE\_THYMOCYTE\_DN | 62 | 16 | 4161 | 644 | Pfas,r-l,SRPK,Pen,Dhfr,raptor,APC7,Aladin,mtTFB1,CG12713,Rpb8,CG15141,CG9485,not,CG10418,Aos1 | | 2.343e-02 | -3.75 | GO\_ACTIVATION\_OF\_PROTEIN\_KINASE\_ACTIVITY | MSigDB lists | GO\_ACTIVATION\_OF\_PROTEIN\_KINASE\_ACTIVITY | 62 | 16 | 4161 | 644 | Pdk1,rl,Ac3,p38a,Cdk1,Pka-R2,Mo25,Mekk1,nAChRalpha4,RpS27A,Pka-R1,RpL40,CG17698,csw,Sod1,Traf6 | | 2.343e-02 | -3.75 | GSE1925\_CTRL\_VS\_IFNG\_PRIMED\_MACROPHAGE\_24H\_IFNG\_STIM\_DN | MSigDB lists | GSE1925\_CTRL\_VS\_IFNG\_PRIMED\_MACROPHAGE\_24H\_IFNG\_STIM\_DN | 62 | 16 | 4161 | 644 | shrb,Idh3b,Ppox,CG7997,r-l,Hat1,mRpL16,Nca,Chd64,Pgm1,Vha36-1,Non1,Sod1,CG15141,Pi3K92E,Usp39 | | 2.343e-02 | -3.75 | GSE7460\_CTRL\_VS\_TGFB\_TREATED\_ACT\_CD8\_TCELL\_UP | MSigDB lists | GSE7460\_CTRL\_VS\_TGFB\_TREATED\_ACT\_CD8\_TCELL\_UP | 62 | 16 | 4161 | 644 | Caf1-105,Smox,CG33181,Mekk1,Axs,RpL26,spag,PpD3,Gyg,Clp,CG1105,Cse1,CG6610,Kap3,Aladin,CG17187 | | 2.343e-02 | -3.75 | MCBRYAN\_PUBERTAL\_BREAST\_4\_5WK\_DN | MSigDB lists | MCBRYAN\_PUBERTAL\_BREAST\_4\_5WK\_DN | 62 | 16 | 4161 | 644 | mino,Nurf-38,sau,P5cr,CkIIalpha,CG11070,Gk2,ACC,CG10932,Etf-QO,yip2,Idh3b,CG7789,Scsalpha1,Mitofilin,CG7488 | | 2.343e-02 | -3.75 | GSE19401\_RETINOIC\_ACID\_VS\_RETINOIC\_ACID\_AND\_PAM2CSK4\_STIM\_FOLLICULAR\_DC\_DN | MSigDB lists | GSE19401\_RETINOIC\_ACID\_VS\_RETINOIC\_ACID\_AND\_PAM2CSK4\_STIM\_FOLLICULAR\_DC\_DN | 62 | 16 | 4161 | 644 | mRpL16,CG8314,Mtpalpha,Bin1,Sccpdh1,Rab32,RpL24,shu,yip2,Prosbeta6,Pi3K92E,r,mtTFB1,O-fut1,RpS7,CG1532 | | 2.343e-02 | -3.75 | - | gene3d domains | 3.90.1150.10 | 29 | 8 | 4929 | 622 | Got1,CG11899,mal,Gabat,Kyat,CG1640,Sply,Shmt | | 2.355e-02 | -3.75 | regulation of basement membrane organization | biological process | GO:0110011 | 3 | 2 | 12345 | 1129 | Crag,Rab10 | | 2.355e-02 | -3.75 | maintenance of neuroblast polarity | biological process | GO:0045201 | 3 | 2 | 12345 | 1129 | Lkb1,tws | | 2.355e-02 | -3.75 | CMP-N-acetylneuraminate transmembrane transport | biological process | GO:0015782 | 3 | 2 | 12345 | 1129 | Efr,Ugalt | | 2.355e-02 | -3.75 | UMP biosynthetic process | biological process | GO:0006222 | 3 | 2 | 12345 | 1129 | r-l,l(2)k01209 | | 2.355e-02 | -3.75 | regulation of antifungal peptide biosynthetic process | biological process | GO:0002810 | 3 | 2 | 12345 | 1129 | spz,Rel | | 2.355e-02 | -3.75 | DNA dealkylation | biological process | GO:0035510 | 3 | 2 | 12345 | 1129 | agt,obe | | 2.355e-02 | -3.75 | succinate metabolic process | biological process | GO:0006105 | 3 | 2 | 12345 | 1129 | Scsalpha1,Sdhaf3 | | 2.355e-02 | -3.75 | histone H4-K8 acetylation | biological process | GO:0043982 | 3 | 2 | 12345 | 1129 | Rcd5,nej | | 2.355e-02 | -3.75 | ethanol metabolic process | biological process | GO:0006067 | 3 | 2 | 12345 | 1129 | Adh,Fdh | | 2.355e-02 | -3.75 | regulation of cAMP-mediated signaling | biological process | GO:0043949 | 3 | 2 | 12345 | 1129 | dnc,pn | | 2.355e-02 | -3.75 | glutaminyl-tRNAGln biosynthesis via transamidation | biological process | GO:0070681 | 3 | 2 | 12345 | 1129 | GatB,GatA | | 2.355e-02 | -3.75 | UMP metabolic process | biological process | GO:0046049 | 3 | 2 | 12345 | 1129 | r-l,l(2)k01209 | | 2.355e-02 | -3.75 | basolateral protein secretion | biological process | GO:0110010 | 3 | 2 | 12345 | 1129 | Crag,Rab10 | | 2.355e-02 | -3.75 | positive regulation of antifungal peptide biosynthetic process | biological process | GO:0006967 | 3 | 2 | 12345 | 1129 | Rel,spz | | 2.355e-02 | -3.75 | regulation of chromatin silencing at telomere | biological process | GO:0031938 | 3 | 2 | 12345 | 1129 | HP1c,HP1e | | 2.355e-02 | -3.75 | positive regulation of I-kappaB kinase/NF-kappaB signaling | biological process | GO:0043123 | 3 | 2 | 12345 | 1129 | Traf6,Myd88 | | 2.355e-02 | -3.75 | cleavage furrow formation | biological process | GO:0036089 | 3 | 2 | 12345 | 1129 | slam,sau | | 2.355e-02 | -3.75 | spectrosome organization | biological process | GO:0030721 | 3 | 2 | 12345 | 1129 | lwr,tkv | | 2.360e-02 | -3.75 | histone acetyltransferase complex | cellular component | GO:0000123 | 77 | 13 | 11940 | 1105 | Spt20,Sgf29,Sgf11,nej,Tip60,rept,Chrac-14,YL-1,Rcd5,pont,e(y)1,not,d4 | | 2.360e-02 | -3.75 | fly chr99D1-99D1\|3-100 | chromosome location | fly chr99D1-99D1\|3-100 | 12 | 3 | 25010 | 1343 | CG15525,Acph-1,CG7920 | | 2.360e-02 | -3.75 | fly chr99D1-99D1\|3-100 cM | chromosome location | fly chr99D1-99D1\|3-100 cM | 12 | 3 | 25010 | 1343 | CG7920,CG15525,Acph-1 | | 2.360e-02 | -3.75 | fly chr22D4-22D4\|2-5 | chromosome location | fly chr22D4-22D4\|2-5 | 12 | 3 | 25010 | 1343 | CG10880,Uch,sau | | 2.360e-02 | -3.75 | fly chr22D4-22D4\|2-5 cM | chromosome location | fly chr22D4-22D4\|2-5 cM | 12 | 3 | 25010 | 1343 | CG10880,Uch,sau | | 2.362e-02 | -3.75 | mitotic cell cycle process | biological process | GO:1903047 | 290 | 37 | 12345 | 1129 | del,twe,Cenp-C,PpV,wisp,Non1,dgt4,alphaTub67C,nbs,tws,CG8188,RecQ5,shrb,pont,CycB,E2f1,CycE,spn-E,msps,CG5003,gammaTub37C,cana,mus304,Klp10A,eIF4E1,CG10336,sau,mus81,Rbf,mad2,CycJ,mre11,grp,Cdk1,Lkb1,Ankle2,Zwilch | | 2.367e-02 | -3.74 | GO\_CELLULAR\_RESPONSE\_TO\_CAMP | MSigDB lists | GO\_CELLULAR\_RESPONSE\_TO\_CAMP | 8 | 4 | 4161 | 644 | Wnt10,Prip,dnc,Calx | | 2.367e-02 | -3.74 | GO\_CARBON\_NITROGEN\_LIGASE\_ACTIVITY\_WITH\_GLUTAMINE\_AS\_AMIDO\_N\_DONOR | MSigDB lists | GO\_CARBON\_NITROGEN\_LIGASE\_ACTIVITY\_WITH\_GLUTAMINE\_AS\_AMIDO\_N\_DONOR | 8 | 4 | 4161 | 644 | Nadsyn,GatA,r,Pfas | | 2.367e-02 | -3.74 | GO\_POSITIVE\_REGULATION\_OF\_CARDIAC\_MUSCLE\_CELL\_PROLIFERATION | MSigDB lists | GO\_POSITIVE\_REGULATION\_OF\_CARDIAC\_MUSCLE\_CELL\_PROLIFERATION | 8 | 4 | 4161 | 644 | p38a,Cdk1,CycB,Su(H) | | 2.367e-02 | -3.74 | GO\_PYRIMIDINE\_NUCLEOSIDE\_TRIPHOSPHATE\_METABOLIC\_PROCESS | MSigDB lists | GO\_PYRIMIDINE\_NUCLEOSIDE\_TRIPHOSPHATE\_METABOLIC\_PROCESS | 8 | 4 | 4161 | 644 | nmdyn-D7,nmdyn-D6,Ts,r | | 2.367e-02 | -3.74 | GO\_REGULATION\_OF\_NECROTIC\_CELL\_DEATH | MSigDB lists | GO\_REGULATION\_OF\_NECROTIC\_CELL\_DEATH | 8 | 4 | 4161 | 644 | RpS27A,RpL40,Tspo,Cyp1 | | 2.367e-02 | -3.74 | REACTOME\_NRIF\_SIGNALS\_CELL\_DEATH\_FROM\_THE\_NUCLEUS | MSigDB lists | REACTOME\_NRIF\_SIGNALS\_CELL\_DEATH\_FROM\_THE\_NUCLEUS | 8 | 4 | 4161 | 644 | pen-2,Traf6,RpL40,RpS27A | | 2.367e-02 | -3.74 | chr2p14 | MSigDB lists | chr2p14 | 8 | 4 | 4161 | 644 | Stam,l(1)G0004,Pex13,Sptr | | 2.367e-02 | -3.74 | GO\_NEGATIVE\_REGULATION\_OF\_ATPASE\_ACTIVITY | MSigDB lists | GO\_NEGATIVE\_REGULATION\_OF\_ATPASE\_ACTIVITY | 8 | 4 | 4161 | 644 | Chd64,regucalcin,CG8726,Cyp1 | | 2.367e-02 | -3.74 | REACTOME\_MRNA\_DECAY\_BY\_5\_TO\_3\_EXORIBONUCLEASE | MSigDB lists | REACTOME\_MRNA\_DECAY\_BY\_5\_TO\_3\_EXORIBONUCLEASE | 8 | 4 | 4161 | 644 | CG6610,CG10418,Edc3,LSm3 | | 2.367e-02 | -3.74 | BIOCARTA\_CELLCYCLE\_PATHWAY | MSigDB lists | BIOCARTA\_CELLCYCLE\_PATHWAY | 8 | 4 | 4161 | 644 | CycE,Cdk1,CycB,Cdk7 | | 2.367e-02 | -3.74 | TENEDINI\_MEGAKARYOCYTE\_MARKERS | MSigDB lists | TENEDINI\_MEGAKARYOCYTE\_MARKERS | 8 | 4 | 4161 | 644 | CycE,p38a,Cdk1,CycB | | 2.367e-02 | -3.74 | KYNG\_ENVIRONMENTAL\_STRESS\_RESPONSE\_NOT\_BY\_GAMMA\_IN\_WS | MSigDB lists | KYNG\_ENVIRONMENTAL\_STRESS\_RESPONSE\_NOT\_BY\_GAMMA\_IN\_WS | 8 | 4 | 4161 | 644 | Sod1,CG1890,PDCD-5,scu | | 2.367e-02 | -3.74 | GO\_RESPONSE\_TO\_PROGESTERONE | MSigDB lists | GO\_RESPONSE\_TO\_PROGESTERONE | 8 | 4 | 4161 | 644 | Tspo,CycE,SREBP,Ts | | 2.367e-02 | -3.74 | GO\_RIBONUCLEOSIDE\_TRIPHOSPHATE\_BIOSYNTHETIC\_PROCESS | MSigDB lists | GO\_RIBONUCLEOSIDE\_TRIPHOSPHATE\_BIOSYNTHETIC\_PROCESS | 8 | 4 | 4161 | 644 | nmdyn-D6,nmdyn-D7,Stoml2,r | | 2.367e-02 | -3.74 | GO\_DNA\_HELICASE\_COMPLEX | MSigDB lists | GO\_DNA\_HELICASE\_COMPLEX | 8 | 4 | 4161 | 644 | rept,Uch-L5,pont,Rcd5 | | 2.367e-02 | -3.74 | GO\_REGULATION\_OF\_MAST\_CELL\_ACTIVATION | MSigDB lists | GO\_REGULATION\_OF\_MAST\_CELL\_ACTIVATION | 8 | 4 | 4161 | 644 | Vamp7,Snx6,scramb1,Pdk1 | | 2.367e-02 | -3.74 | GO\_REGULATION\_OF\_CARDIAC\_MUSCLE\_CELL\_PROLIFERATION | MSigDB lists | GO\_REGULATION\_OF\_CARDIAC\_MUSCLE\_CELL\_PROLIFERATION | 8 | 4 | 4161 | 644 | Su(H),CycB,Cdk1,p38a | | 2.367e-02 | -3.74 | GO\_REGULATION\_OF\_LIPID\_STORAGE | MSigDB lists | GO\_REGULATION\_OF\_LIPID\_STORAGE | 8 | 4 | 4161 | 644 | cact,crq,Ptp61F,EcR | | 2.367e-02 | -3.74 | organelle fusion | biological process | GO:0048284 | 62 | 11 | 12345 | 1129 | fs(1)Ya,Vamp7,Vps16B,Snx6,Syx13,Rab5,wisp,Sec22,msps,Marf,lt | | 2.374e-02 | -3.74 | - | gene3d domains | 1.25.40.90 | 14 | 5 | 4929 | 622 | lqfR,Stam,Gga,Hip1,CG4266 | | 2.377e-02 | -3.74 | glutaminyl-tRNA synthase (glutamine-hydrolyzing) activity | molecular function | GO:0050567 | 3 | 2 | 12176 | 1119 | GatB,GatA | | 2.377e-02 | -3.74 | glutathione-disulfide reductase activity | molecular function | GO:0004362 | 3 | 2 | 12176 | 1119 | Trxr-1,Grx1 | | 2.377e-02 | -3.74 | CMP-N-acetylneuraminate transmembrane transporter activity | molecular function | GO:0005456 | 3 | 2 | 12176 | 1119 | Ugalt,Efr | | 2.377e-02 | -3.74 | thioredoxin-disulfide reductase activity | molecular function | GO:0004791 | 3 | 2 | 12176 | 1119 | Trxr-1,SelT | | 2.377e-02 | -3.74 | protein-disulfide reductase activity | molecular function | GO:0047134 | 3 | 2 | 12176 | 1119 | CG3939,cl | | 2.377e-02 | -3.74 | calmodulin-dependent protein phosphatase activity | molecular function | GO:0033192 | 3 | 2 | 12176 | 1119 | CanA-14F,Pp2B-14D | | 2.377e-02 | -3.74 | acetyl-CoA C-acetyltransferase activity | molecular function | GO:0003985 | 3 | 2 | 12176 | 1119 | CG10932,yip2 | | 2.377e-02 | -3.74 | succinate-CoA ligase (ADP-forming) activity | molecular function | GO:0004775 | 3 | 2 | 12176 | 1119 | ScsbetaA,Scsalpha1 | | 2.377e-02 | -3.74 | carbonyl reductase (NADPH) activity | molecular function | GO:0004090 | 3 | 2 | 12176 | 1119 | sni,CG7322 | | 2.377e-02 | -3.74 | AMP binding | molecular function | GO:0016208 | 3 | 2 | 12176 | 1119 | Pfk,AcCoAS | | 2.377e-02 | -3.74 | small protein activating enzyme binding | molecular function | GO:0044388 | 3 | 2 | 12176 | 1119 | lwr,Aos1 | | 2.377e-02 | -3.74 | ubiquitin activating enzyme binding | molecular function | GO:0033134 | 3 | 2 | 12176 | 1119 | lwr,Aos1 | | 2.382e-02 | -3.74 | fly chr32C1-32C1\|2-44 cM | chromosome location | fly chr32C1-32C1\|2-44 cM | 21 | 4 | 25010 | 1343 | RpL9,ATPsynG,YL-1,piwi | | 2.382e-02 | -3.74 | fly chr32C1-32C1\|2-44 | chromosome location | fly chr32C1-32C1\|2-44 | 21 | 4 | 25010 | 1343 | RpL9,ATPsynG,YL-1,piwi | | 2.387e-02 | -3.74 | positive regulation of transferase activity | biological process | GO:0051347 | 78 | 13 | 12345 | 1129 | raptor,cort,Lkb1,Tao,RpLP1,Parp16,Tak1,ASPP,Pdk1,Abi,Pi3K92E,fray,Mekk1 | | 2.392e-02 | -3.73 | GO\_MYELOID\_DENDRITIC\_CELL\_DIFFERENTIATION | MSigDB lists | GO\_MYELOID\_DENDRITIC\_CELL\_DIFFERENTIATION | 2 | 2 | 4161 | 644 | Traf6,Su(H) | | 2.392e-02 | -3.73 | GO\_NEGATIVE\_REGULATION\_OF\_MACROPHAGE\_DERIVED\_FOAM\_CELL\_DIFFERENTIATION | MSigDB lists | GO\_NEGATIVE\_REGULATION\_OF\_MACROPHAGE\_DERIVED\_FOAM\_CELL\_DIFFERENTIATION | 2 | 2 | 4161 | 644 | EcR,cact | | 2.392e-02 | -3.73 | GNF2\_CD7 | MSigDB lists | GNF2\_CD7 | 2 | 2 | 4161 | 644 | Ptpmeg,Nprl2 | | 2.392e-02 | -3.73 | GO\_POSITIVE\_REGULATION\_OF\_TRANSCRIPTION\_FROM\_RNA\_POLYMERASE\_II\_PROMOTER\_IN\_RESPONSE\_TO\_ENDOPLASMIC\_RETICULUM\_STRESS | MSigDB lists | GO\_POSITIVE\_REGULATION\_OF\_TRANSCRIPTION\_FROM\_RNA\_POLYMERASE\_II\_PROMOTER\_IN\_RESPONSE\_TO\_ENDOPLASMIC\_RETICULUM\_STRESS | 2 | 2 | 4161 | 644 | Hsc70-3,S2P | | 2.392e-02 | -3.73 | GO\_ENERGY\_HOMEOSTASIS | MSigDB lists | GO\_ENERGY\_HOMEOSTASIS | 2 | 2 | 4161 | 644 | AMPdeam,CG2493 | | 2.392e-02 | -3.73 | GO\_REGULATION\_OF\_MACROPHAGE\_DERIVED\_FOAM\_CELL\_DIFFERENTIATION | MSigDB lists | GO\_REGULATION\_OF\_MACROPHAGE\_DERIVED\_FOAM\_CELL\_DIFFERENTIATION | 2 | 2 | 4161 | 644 | cact,EcR | | 2.392e-02 | -3.73 | REACTOME\_ETHANOL\_OXIDATION | MSigDB lists | REACTOME\_ETHANOL\_OXIDATION | 2 | 2 | 4161 | 644 | Aldh,AcCoAS | | 2.392e-02 | -3.73 | MYLLYKANGAS\_AMPLIFICATION\_HOT\_SPOT\_18 | MSigDB lists | MYLLYKANGAS\_AMPLIFICATION\_HOT\_SPOT\_18 | 2 | 2 | 4161 | 644 | CG10425,Med | | 2.392e-02 | -3.73 | KEGG\_CIRCADIAN\_RHYTHM\_MAMMAL | MSigDB lists | KEGG\_CIRCADIAN\_RHYTHM\_MAMMAL | 2 | 2 | 4161 | 644 | cyc,dco | | 2.392e-02 | -3.73 | GO\_STEROL\_CATABOLIC\_PROCESS | MSigDB lists | GO\_STEROL\_CATABOLIC\_PROCESS | 2 | 2 | 4161 | 644 | CG5734,crq | | 2.392e-02 | -3.73 | GO\_PHOSPHOTYROSINE\_BINDING | MSigDB lists | GO\_PHOSPHOTYROSINE\_BINDING | 2 | 2 | 4161 | 644 | rl,Acph-1 | | 2.392e-02 | -3.73 | GNF2\_TTN | MSigDB lists | GNF2\_TTN | 2 | 2 | 4161 | 644 | GlyP,SERCA | | 2.392e-02 | -3.73 | GO\_DENDRITIC\_CELL\_DIFFERENTIATION | MSigDB lists | GO\_DENDRITIC\_CELL\_DIFFERENTIATION | 2 | 2 | 4161 | 644 | Su(H),Traf6 | | 2.392e-02 | -3.73 | GO\_POSITIVE\_REGULATION\_OF\_TRIGLYCERIDE\_LIPASE\_ACTIVITY | MSigDB lists | GO\_POSITIVE\_REGULATION\_OF\_TRIGLYCERIDE\_LIPASE\_ACTIVITY | 2 | 2 | 4161 | 644 | EcR,CG5966 | | 2.392e-02 | -3.73 | CHIARETTI\_T\_ALL\_REFRACTORY\_TO\_THERAPY | MSigDB lists | CHIARETTI\_T\_ALL\_REFRACTORY\_TO\_THERAPY | 2 | 2 | 4161 | 644 | aay,Jafrac1 | | 2.392e-02 | -3.73 | REACTOME\_TRAF6\_MEDIATED\_NFKB\_ACTIVATION | MSigDB lists | REACTOME\_TRAF6\_MEDIATED\_NFKB\_ACTIVATION | 2 | 2 | 4161 | 644 | Traf6,cact | | 2.392e-02 | -3.73 | GO\_POSITIVE\_REGULATION\_OF\_DIGESTIVE\_SYSTEM\_PROCESS | MSigDB lists | GO\_POSITIVE\_REGULATION\_OF\_DIGESTIVE\_SYSTEM\_PROCESS | 2 | 2 | 4161 | 644 | Prip,EcR | | 2.392e-02 | -3.73 | REACTOME\_TAK1\_ACTIVATES\_NFKB\_BY\_PHOSPHORYLATION\_AND\_ACTIVATION\_OF\_IKKS\_COMPLEX | MSigDB lists | REACTOME\_TAK1\_ACTIVATES\_NFKB\_BY\_PHOSPHORYLATION\_AND\_ACTIVATION\_OF\_IKKS\_COMPLEX | 2 | 2 | 4161 | 644 | cact,Traf6 | | 2.392e-02 | -3.73 | GO\_RESPIRATORY\_BURST | MSigDB lists | GO\_RESPIRATORY\_BURST | 2 | 2 | 4161 | 644 | Pglym78,Pi3K92E | | 2.392e-02 | -3.73 | GO\_MYELOID\_DENDRITIC\_CELL\_ACTIVATION | MSigDB lists | GO\_MYELOID\_DENDRITIC\_CELL\_ACTIVATION | 2 | 2 | 4161 | 644 | Traf6,Su(H) | | 2.392e-02 | -3.73 | GNF2\_ZAP70 | MSigDB lists | GNF2\_ZAP70 | 2 | 2 | 4161 | 644 | Nprl2,Ptpmeg | | 2.392e-02 | -3.73 | GO\_REGULATION\_OF\_ACTIVATED\_T\_CELL\_PROLIFERATION | MSigDB lists | GO\_REGULATION\_OF\_ACTIVATED\_T\_CELL\_PROLIFERATION | 2 | 2 | 4161 | 644 | Pka-R1,mino | | 2.392e-02 | -3.73 | ZERBINI\_RESPONSE\_TO\_SULINDAC\_DN | MSigDB lists | ZERBINI\_RESPONSE\_TO\_SULINDAC\_DN | 2 | 2 | 4161 | 644 | Cdk1,CycB | | 2.392e-02 | -3.73 | BIOCARTA\_NFKB\_PATHWAY | MSigDB lists | BIOCARTA\_NFKB\_PATHWAY | 2 | 2 | 4161 | 644 | cact,Traf6 | | 2.392e-02 | -3.73 | BIOCARTA\_CD40\_PATHWAY | MSigDB lists | BIOCARTA\_CD40\_PATHWAY | 2 | 2 | 4161 | 644 | Traf6,cact | | 2.392e-02 | -3.73 | MATZUK\_PREOVULATORY\_FOLLICLE | MSigDB lists | MATZUK\_PREOVULATORY\_FOLLICLE | 2 | 2 | 4161 | 644 | CG11110,Sod1 | | 2.392e-02 | -3.73 | GO\_TAU\_PROTEIN\_BINDING | MSigDB lists | GO\_TAU\_PROTEIN\_BINDING | 2 | 2 | 4161 | 644 | HDAC6,Fkbp59 | | 2.393e-02 | -3.73 | pole plasm assembly | biological process | GO:0007315 | 70 | 12 | 12345 | 1129 | cnc,swa,Tm1,psq,mael,armi,Lkb1,gus,exu,osk,spn-E,Moe | | 2.397e-02 | -3.73 | GSE36476\_CTRL\_VS\_TSST\_ACT\_16H\_MEMORY\_CD4\_TCELL\_OLD\_DN | MSigDB lists | GSE36476\_CTRL\_VS\_TSST\_ACT\_16H\_MEMORY\_CD4\_TCELL\_OLD\_DN | 93 | 22 | 4161 | 644 | alphaTub84B,CG11899,DppIII,Rpb8,Cbp80,tko,mRpL17,Got1,CG7728,Prosbeta4,gammaTub37C,Prosbeta5,Pus7,Tim8,CG7997,CCT3,SRPK,Stip1,Aldh,Pglym78,bys,msps | | 2.397e-02 | -3.73 | GSE36476\_CTRL\_VS\_TSST\_ACT\_40H\_MEMORY\_CD4\_TCELL\_YOUNG\_DN | MSigDB lists | GSE36476\_CTRL\_VS\_TSST\_ACT\_40H\_MEMORY\_CD4\_TCELL\_YOUNG\_DN | 93 | 22 | 4161 | 644 | Ts,Karybeta3,alphaTub84B,Dhfr,Psf3,thoc6,CycB,Pus7,Jafrac2,r-l,Stip1,SRPK,Aldh,Pglym78,bys,msps,Shmt,Ppcdc,Mitofilin,Pfas,gammaTub37C,CycE | | 2.409e-02 | -3.73 | GATOR1 complex | cellular component | GO:1990130 | 3 | 2 | 11940 | 1105 | Iml1,Nprl2 | | 2.409e-02 | -3.73 | cortical microtubule | cellular component | GO:0055028 | 3 | 2 | 11940 | 1105 | alphaTub67C,Klp10A | | 2.409e-02 | -3.73 | Nem1-Spo7 phosphatase complex | cellular component | GO:0071595 | 3 | 2 | 11940 | 1105 | Dd,CG8009 | | 2.409e-02 | -3.73 | plasma membrane respirasome | cellular component | GO:0070470 | 3 | 2 | 11940 | 1105 | SdhA,ND-75 | | 2.409e-02 | -3.73 | Mre11 complex | cellular component | GO:0030870 | 3 | 2 | 11940 | 1105 | mre11,nbs | | 2.409e-02 | -3.73 | peroxisomal importomer complex | cellular component | GO:1990429 | 3 | 2 | 11940 | 1105 | Pex12,Pex13 | | 2.409e-02 | -3.73 | Cul5-RING ubiquitin ligase complex | cellular component | GO:0031466 | 3 | 2 | 11940 | 1105 | gus,Cul5 | | 2.409e-02 | -3.73 | succinate-CoA ligase complex | cellular component | GO:0042709 | 3 | 2 | 11940 | 1105 | ScsbetaA,Scsalpha1 | | 2.409e-02 | -3.73 | nuclear telomeric heterochromatin | cellular component | GO:0005724 | 3 | 2 | 11940 | 1105 | SuUR,Sirt7 | | 2.409e-02 | -3.73 | R2TP complex | cellular component | GO:0097255 | 3 | 2 | 11940 | 1105 | pont,rept | | 2.409e-02 | -3.73 | Pex17p-Pex14p docking complex | cellular component | GO:1990415 | 3 | 2 | 11940 | 1105 | Pex12,Pex13 | | 2.411e-02 | -3.72 | Caf1-55 (Chromatin assembly factor 1, p55 subunit) | protein interactions | 41836 | 66 | 14 | 9204 | 1112 | foxo,Hnf4,Rbf,esc,Nurf-38,lin-52,Trf2,smg,asf1,Rbf2,Caf1-105,escl,Hat1,CG4557 | | 2.411e-02 | -3.72 | protein localization to organelle | biological process | GO:0033365 | 226 | 30 | 12345 | 1129 | CG9393,Nup62,Pex13,SuUR,Vsp37A,Kap-alpha3,Tim8,Topors,Roe1,Nup358,Oseg1,Cenp-C,Arl5,Pen,Tom70,Pi4KIIalpha,Cse1,Tom40,CG14894,Arts,osk,Karybeta3,Pex10,CG11110,sw,Pex12,mbo,CG8004,Atg6,CG8728 | | 2.427e-02 | -3.72 | WONG\_EMBRYONIC\_STEM\_CELL\_CORE | MSigDB lists | WONG\_EMBRYONIC\_STEM\_CELL\_CORE | 180 | 38 | 4161 | 644 | Stoml2,eIF3i,DppIII,Fdh,rept,ND-49,grp,mRpL13,Cse1,CG6610,Cdk1,wal,Echs1,mRpL16,mad2,Pen,RpL13,ND-39,RpS23,Prosalpha5,Nap1,CG40045,RfC38,NHP2,CG10418,CycC,CG6767,Cyt-c-p,RpS16,ND-B18,Stip1,pont,Tim8,Prosbeta5,CCT5,Eno,Hat1,RpS27 | | 2.429e-02 | -3.72 | REG (REG) | protein interactions | 32274 | 36 | 9 | 9204 | 1112 | Prosbeta3,Prosalpha3,Prosbeta4,Prosalpha4,CG11248,Hmg-2,Prosbeta5,Pomp,Prosalpha5 | | 2.429e-02 | -3.72 | Ufl1 (UFM1 specific ligase 1) | protein interactions | 40843 | 36 | 9 | 9204 | 1112 | RpL26,CycE,CG6707,Imp,alphaTub84B,RpS7,CG30291,Bin1,RpLP1 | | 2.432e-02 | -3.72 | MORF\_ANP32B | MSigDB lists | MORF\_ANP32B | 109 | 25 | 4161 | 644 | heph,ND-51,Prosbeta6,levy,SkpA,Phb2,Rpn12,RpS30,Sod1,LysRS,Rpt2,RpL30,RpS24,Chrac-14,lwr,mEFTu1,ND-30,SdhA,Mtpalpha,RpS27A,Pp2A-29B,Nap1,RpL24,CSN6,eIF3m | | 2.442e-02 | -3.71 | Dif (Dorsal-related immunity factor) | protein interactions | 35045 | 10 | 4 | 9204 | 1112 | cact,lwr,Rel,dl | | 2.442e-02 | -3.71 | CROT (Carnitine O-octanoyltransferase) | protein interactions | 43332 | 10 | 4 | 9204 | 1112 | CG10654,nbs,CG9917,kri | | 2.442e-02 | -3.71 | CG16908 (uncharacterized protein) | protein interactions | 41189 | 10 | 4 | 9204 | 1112 | spag,pont,rept,lqfR | | 2.442e-02 | -3.71 | CG12607 (uncharacterized protein) | protein interactions | 38547 | 10 | 4 | 9204 | 1112 | Fer1HCH,Sod3,Pxt,mts | | 2.442e-02 | -3.71 | CG11893 (uncharacterized protein) | protein interactions | 43056 | 10 | 4 | 9204 | 1112 | Bx,CG11164,Bin1,Ssdp | | 2.442e-02 | -3.71 | flr (flare) | protein interactions | 39505 | 10 | 4 | 9204 | 1112 | EndoGI,Moe,regucalcin,CG8204 | | 2.442e-02 | -3.71 | CG14556 (uncharacterized protein) | protein interactions | 43176 | 10 | 4 | 9204 | 1112 | Blos2,Blos4,CycE,Nup62 | | 2.442e-02 | -3.71 | MtnA (Metallothionein A) | protein interactions | 41202 | 10 | 4 | 9204 | 1112 | Drep1,PPYR1,Jafrac1,SdhA | | 2.442e-02 | -3.71 | sel (seele) | protein interactions | 36046 | 10 | 4 | 9204 | 1112 | Fer1HCH,Uch,PHGPx,EndoGI | | 2.442e-02 | -3.71 | CBP (sarcoplasmic calcium-binding protein) | protein interactions | 31669 | 10 | 4 | 9204 | 1112 | ago,trsn,Pka-R1,CG15525 | | 2.442e-02 | -3.71 | SoYb (Sister of Yb) | protein interactions | 34331 | 10 | 4 | 9204 | 1112 | Rtnl1,Tak1,PGRP-LE,Rel | | 2.450e-02 | -3.71 | Metabolism | REACTOME pathways | R-DME-1430728 | 1165 | 191 | 3923 | 574 | Prosalpha3,Aldh,Sptr,Agpat2,Rpt2,CG9265,Cyp6a19,ND-30,CG1774,l(1)G0196,Pi3K92E,ND-51,Npc2a,CG11089,CG14997,CG7328,Etf-QO,Rpn3,ND-49,CG17333,mal,ATPsynbeta,alc,CG9471,CG8475,Nadsyn,Gs1l,CG2201,wal,PyK,CG4860,Got1,Pi4KIIalpha,Jheh2,Prosbeta4,Idh3b,Rab5,CG4598,GlyP,Cyp6v1,Gapdh1,shop,nSMase,Pfas,SdhA,Sply,ND-B22,Ady43A,scu,CG7550,CG9205,AMPdeam,SdhD,Eno,CG9582,CG4594,ND-75,Shmt,yip2,Ts,CG17691,Pdk,Ac3,Rpn12,CG31262,Kyat,r-l,Acox57D-p,Pfk,ND-B18,Pka-R2,CG7322,Echs1,tzn,CG4586,CG33156,SNF4Agamma,CG31689,ATPsynD,EcR,GstT2,CG6650,sws,ATPsynF,mino,P5cr,CG8067,CG1847,P5CDh1,Pglym78,ATPsynG,Pgk,sgll,fdl,Pgm1,aay,CG7115,Ugt35A1,Pcd,Adh,CG9485,NP15.6,CG4390,CG31674,Prosalpha4,r,Cyp6a22,CG31523,Ppcdc,Dhfr,CG32549,CG10425,ACC,Thiolase,Desat1,UGP,Mtpalpha,CG10932,ATPsynO,CG8199,colt,Idh,oxt,CG11899,CG1640,Nf-YA,Hexo1,Prosbeta3,CG8112,S2P,CG6767,sea,SREBP,AGBE,ND-24,CG7997,ND-39,Rpn2,ND-SGDH,ATPsynCF6,CAH2,Prosalpha5,CG6707,ABCD,dob,CG6293,Pka-R1,Ho,CG4872,CG5028,Adi1,Pfrx,pr,CG4842,Nc73EF,Prosbeta6,whd,Gclc,Ggt-1,CG4407,Synj,Gyg,CG3803,CG3860,DhpD,Naxe,ATPsynB,CG4267,Ppox,Marc,Cyp6u1,awd,GstT1,Nurf-38,Fdh,CG7840,Gk2,Argk,blw,Prosbeta5,Gk1,CG3376,AcCoAS,Pld,ATPsyndelta,cN-IIIB,CG9248,Mcad,CG7789,Pdxk,ND-23 | | 2.453e-02 | -3.71 | intrinsic component of Golgi membrane | cellular component | GO:0031228 | 32 | 7 | 11940 | 1105 | CG4293,Fur2,CG5447,stet,Efr,CG9773,Ugalt | | 2.457e-02 | -3.71 | Antigen processing: Ubiquitination & Proteasome degradation | REACTOME pathways | R-DME-983168 | 104 | 23 | 3923 | 574 | SkpA,Ubc6,Prosalpha3,Rpt2,RpL40,RpS27A,Prosalpha5,ago,Prosbeta4,Prosbeta5,Rpn2,Iru,Prosalpha4,Fbw5,Prosbeta6,Rpn12,CG14435,Cul1,Rpn3,gus,Herc4,dx,Prosbeta3 | | 2.462e-02 | -3.70 | GO\_POSITIVE\_REGULATION\_OF\_CATABOLIC\_PROCESS | MSigDB lists | GO\_POSITIVE\_REGULATION\_OF\_CATABOLIC\_PROCESS | 147 | 32 | 4161 | 644 | Cul1,Moe,Rpn3,Edem1,Gclc,CycB,Prosbeta5,Prosalpha3,Prosbeta4,Atg7,RpL40,Prosbeta6,ABCD,APC7,Atg6,ari-1,dco,CG30291,CkIIalpha,mad2,STUB1,Cdk1,RpS27A,Sec22,Prosalpha5,regucalcin,SkpA,Rpn12,Rpn2,Prosbeta3,Rpt2,Prosalpha4 | | 2.464e-02 | -3.70 | GO\_NEGATIVE\_REGULATION\_OF\_PROTEIN\_MODIFICATION\_BY\_SMALL\_PROTEIN\_CONJUGATION\_OR\_REMOVAL | MSigDB lists | GO\_NEGATIVE\_REGULATION\_OF\_PROTEIN\_MODIFICATION\_BY\_SMALL\_PROTEIN\_CONJUGATION\_OR\_REMOVAL | 88 | 21 | 4161 | 644 | Cdk1,Gclc,Cdk5,CycB,Prosbeta5,Rpn3,mad2,Prosbeta4,CG30496,Prosalpha3,RpS27A,Prosalpha5,Prosbeta6,Tspo,RpL40,Rpt2,Non1,Prosbeta3,Rpn2,Rpn12,APC7 | | 2.464e-02 | -3.70 | GO\_REGULATION\_OF\_CANONICAL\_WNT\_SIGNALING\_PATHWAY | MSigDB lists | GO\_REGULATION\_OF\_CANONICAL\_WNT\_SIGNALING\_PATHWAY | 88 | 21 | 4161 | 644 | RpL40,Su(H),cyc,Prosbeta6,Rpn12,dco,Rpn2,Pfdn5,Prosbeta3,Prosalpha4,Dd,Rpt2,Rpn3,Prosbeta5,p38a,Smox,Prosalpha5,RpS27A,Prosalpha3,Wnt10,Prosbeta4 | | 2.464e-02 | -3.70 | GSE40225\_WT\_VS\_RIP\_B7X\_DIABETIC\_MOUSE\_PANCREATIC\_CD8\_TCELL\_DN | MSigDB lists | GSE40225\_WT\_VS\_RIP\_B7X\_DIABETIC\_MOUSE\_PANCREATIC\_CD8\_TCELL\_DN | 88 | 21 | 4161 | 644 | Gapdh1,CG8188,Aladin,SelD,RhoGDI,CG11070,CG5028,Rcd5,CG9248,gammaTub37C,Prosalpha5,CG3587,Eno,Echs1,ND-B18,AIF,SRPK,Top1,Uch-L5,kcc,mad2 | | 2.464e-02 | -3.70 | GSE43863\_TH1\_VS\_LY6C\_LOW\_CXCR5NEG\_EFFECTOR\_CD4\_TCELL\_UP | MSigDB lists | GSE43863\_TH1\_VS\_LY6C\_LOW\_CXCR5NEG\_EFFECTOR\_CD4\_TCELL\_UP | 88 | 21 | 4161 | 644 | DppIII,l(1)G0004,Aos1,mRpL18,CG11164,CG8188,grp,Rpn2,Tom40,Eno,CG7655,Caf1-105,Uev1A,Arpc4,Uch-L5,f-cup,UQCR-Q,Liprin-beta,CycE,yip2,PpD3 | | 2.464e-02 | -3.70 | GSE22611\_NOD2\_TRANSDUCED\_VS\_CTRL\_HEK293T\_STIMULATED\_WITH\_MDP\_2H\_DN | MSigDB lists | GSE22611\_NOD2\_TRANSDUCED\_VS\_CTRL\_HEK293T\_STIMULATED\_WITH\_MDP\_2H\_DN | 88 | 21 | 4161 | 644 | CG1890,slif,CG11110,CG12018,eIF4A,CG7718,CG1416,Arts,r,Karybeta3,levy,Ranbp21,eIF3i,Dph5,DNApol-epsilon255,COX5A,PpD3,Pfas,SREBP,Pus7,crq | | 2.464e-02 | -3.70 | Metabolism of proteins | REACTOME pathways | R-DME-392499 | 740 | 126 | 3923 | 574 | RpS13,faf,alpha-Man-Ia,cathD,CCT6,RpL34a,Rpn12,Atu,Rab32,Prosbeta6,RpS15,Sec23,RpLP0,Exo70,mbo,Rep,Rpn13,ERp60,Uch,Usp10,dco,Sec22,RpL23,RpS18,CG2924,RpL40,CG40045,eIF4A,RpS14b,RpL29,Uch-L5,RpS30,pont,not,CG32473,Trs33,ph-d,Atf6,Dph5,otu,Stam,RpL13A,CG9643,CHOp24,e(y)1,Ndc1,UbcE2H,CG7840,PIG-L,Prosbeta5,RpS7,CCT5,Med,RpS24,GCS2beta,CCT3,RpS29,Ubc6,RpL24,Atg6,nej,RpS2,RpL13,PEK,Prosbeta3,foxo,Gmer,eIF4E1,RpL23A,RpLP1,CG1578,Rpn3,trbd,RpL10,RpS5b,RpL31,Nup58,RpS4,eIF3i,Prosalpha4,RpS16,Ubc4,Rcd5,PAPLA1,Usp16-45,Fur2,lwr,eEF1beta,Rpt2,Prosalpha3,RpS27,RpL34b,RpL37a,Traf6,park,scny,Nup62,RpL38,RpS14a,RpL12,Sec15,RpL7A,RpL17,RpL9,twe,Rpn2,RpS28b,Prosbeta4,RpL11,RpL30,Edem1,Prosalpha5,Rab5,RpS27A,Ptip,ste14,eIF3m,Sec10,S2P,ZnT35C,CG8311,RpL26,Nup358,Aladin,CG8188,RpS23 | | 2.469e-02 | -3.70 | Fc epsilon receptor (FCERI) signaling | REACTOME pathways | R-DME-2454202 | 167 | 34 | 3923 | 574 | Rpn2,Prosbeta5,Prosbeta4,yl,RpS27A,Prosalpha5,CanB2,CG4853,rl,NFAT,Pp2A-29B,Prosalpha4,Pi3K92E,Pdk1,PpD3,CG7530,Pp2B-14D,Prosalpha3,RpL40,Rpt2,cact,SkpA,CanA-14F,Rel,tor,Prosbeta3,foxo,CG13398,Rpn12,dl,Rpn3,Cul1,Prosbeta6,csw | | 2.470e-02 | -3.70 | signal transduction by protein phosphorylation | biological process | GO:0023014 | 47 | 9 | 12345 | 1129 | p38a,Tao,Cka,Tak1,rl,Mo25,Mtl,Mekk1,fray | | 2.470e-02 | -3.70 | regulation of Toll signaling pathway | biological process | GO:0008592 | 47 | 9 | 12345 | 1129 | ea,gd,mop,Aos1,Herc4,snk,spz,lwr,cact | | 2.473e-02 | -3.70 | endonuclease activity | molecular function | GO:0004519 | 78 | 13 | 12176 | 1119 | mus81,mre11,Tengl4,CG12375,CG11164,Clp,CG30343,Dcr-1,RpLP0,trsn,scu,CG8679,tos | | 2.475e-02 | -3.70 | grk (gurken) | protein interactions | 34171 | 60 | 13 | 9204 | 1112 | cact,Pus7,heph,Cbl,exu,stet,sw,pr,bru1,Imp,BicC,obe,rig | | 2.475e-02 | -3.70 | 14-3-3epsilon (14-3-3epsilon) | protein interactions | 42186 | 60 | 13 | 9204 | 1112 | Cenp-C,tor,foxo,Eno,REPTOR,AnxB10,Chd64,awd,RpS27A,regucalcin,CG10638,EndoGI,EcR | | 2.477e-02 | -3.70 | ZBP1(DAI) mediated induction of type I IFNs | REACTOME pathways | R-DME-1606322 | 5 | 3 | 3923 | 574 | Rel,dl,Myd88 | | 2.477e-02 | -3.70 | RIG-I/MDA5 mediated induction of IFN-alpha/beta pathways | REACTOME pathways | R-DME-168928 | 5 | 3 | 3923 | 574 | Myd88,dl,Rel | | 2.477e-02 | -3.70 | TRAF6 mediated NF-kB activation | REACTOME pathways | R-DME-933542 | 5 | 3 | 3923 | 574 | Myd88,dl,Rel | | 2.477e-02 | -3.70 | RIP-mediated NFkB activation via ZBP1 | REACTOME pathways | R-DME-1810476 | 5 | 3 | 3923 | 574 | Rel,dl,Myd88 | | 2.477e-02 | -3.70 | TAK1 activates NFkB by phosphorylation and activation of IKKs complex | REACTOME pathways | R-DME-445989 | 5 | 3 | 3923 | 574 | Myd88,dl,Rel | | 2.477e-02 | -3.70 | DNA Damage/Telomere Stress Induced Senescence | REACTOME pathways | R-DME-2559586 | 5 | 3 | 3923 | 574 | mre11,nbs,Tip60 | | 2.479e-02 | -3.70 | cullin-RING ubiquitin ligase complex | cellular component | GO:0031461 | 102 | 16 | 11940 | 1105 | Fbw5,ago,APC7,SkpA,Cul5,CG5003,CG32221,CG9316,gus,CG30496,CG31633,Cul1,DCAF12,ScsbetaA,Cul2,EloC | | 2.489e-02 | -3.69 | glucosamine-containing compound catabolic process | biological process | GO:1901072 | 19 | 5 | 12345 | 1129 | Cht11,Idgf2,fdl,Idgf4,Idgf6 | | 2.489e-02 | -3.69 | cellular response to sterol | biological process | GO:0036315 | 19 | 5 | 12345 | 1129 | EcR,MED27,Utx,G9a,Nurf-38 | | 2.489e-02 | -3.69 | cellular response to ketone | biological process | GO:1901655 | 19 | 5 | 12345 | 1129 | Utx,G9a,Nurf-38,EcR,MED27 | | 2.489e-02 | -3.69 | microtubule nucleation | biological process | GO:0007020 | 19 | 5 | 12345 | 1129 | CG7716,gammaTub37C,Grip128,Grip84,Grip75 | | 2.489e-02 | -3.69 | amino sugar catabolic process | biological process | GO:0046348 | 19 | 5 | 12345 | 1129 | Idgf4,Idgf6,fdl,Cht11,Idgf2 | | 2.489e-02 | -3.69 | oocyte microtubule cytoskeleton polarization | biological process | GO:0008103 | 19 | 5 | 12345 | 1129 | capt,heph,osk,cnc,Pka-R1 | | 2.489e-02 | -3.69 | cellular response to ecdysone | biological process | GO:0071390 | 19 | 5 | 12345 | 1129 | Utx,G9a,Nurf-38,EcR,MED27 | | 2.501e-02 | -3.69 | MALONEY\_RESPONSE\_TO\_17AAG\_DN | MSigDB lists | MALONEY\_RESPONSE\_TO\_17AAG\_DN | 38 | 11 | 4161 | 644 | Pen,mad2,Hat1,Jafrac2,Prosalpha5,PDCD-5,Clc,Cyt-c-p,RpS16,RpL24,RpL30 | | 2.501e-02 | -3.69 | GSE41176\_WT\_VS\_TAK1\_KO\_ANTI\_IGM\_STIM\_BCELL\_6H\_UP | MSigDB lists | GSE41176\_WT\_VS\_TAK1\_KO\_ANTI\_IGM\_STIM\_BCELL\_6H\_UP | 38 | 11 | 4161 | 644 | Vamp7,colt,Atg6,Nadsyn,CycC,CG7789,scramb1,CG3077,CG14894,Echs1,Rep | | 2.501e-02 | -3.69 | GO\_ORGANELLE\_ENVELOPE\_LUMEN | MSigDB lists | GO\_ORGANELLE\_ENVELOPE\_LUMEN | 38 | 11 | 4161 | 644 | ND-75,Tim8,AIF,ND-B18,Stoml2,shop,CG3226,Sod1,Cyt-c-p,sau,Ppox | | 2.501e-02 | -3.69 | SAKAI\_TUMOR\_INFILTRATING\_MONOCYTES\_DN | MSigDB lists | SAKAI\_TUMOR\_INFILTRATING\_MONOCYTES\_DN | 38 | 11 | 4161 | 644 | Prosalpha5,ari-1,CycC,Hsc70-3,Cdk7,Jafrac1,Grx1,PpV,Pka-R1,Tfb4,Prosbeta5 | | 2.501e-02 | -3.69 | YAGI\_AML\_SURVIVAL | MSigDB lists | YAGI\_AML\_SURVIVAL | 38 | 11 | 4161 | 644 | Rbcn-3A,esc,Tfb4,Klp10A,Rpb8,cact,Tspo,escl,Npc2a,PAPLA1,Rab32 | | 2.501e-02 | -3.69 | REACTOME\_INTERFERON\_SIGNALING | MSigDB lists | REACTOME\_INTERFERON\_SIGNALING | 38 | 11 | 4161 | 644 | eIF4E1,ari-1,RpS27A,Aladin,eIF4A,mbo,RpL40,Kap-alpha3,Pen,alph,Ptp61F | | 2.504e-02 | -3.69 | Glycolysis | REACTOME pathways | R-DME-70171 | 21 | 7 | 3923 | 574 | Pfk,Eno,Gapdh1,PyK,Pgk,Pglym78,Pfrx | | 2.510e-02 | -3.68 | Psc (Posterior sex combs) | protein interactions | 36431 | 42 | 10 | 9204 | 1112 | ph-d,e(y)1,CycB,RYBP,mxc,psq,Cdk1,Rtnl1,esc,nej | | 2.510e-02 | -3.68 | Parp (Poly-(ADP-ribose) polymerase) | protein interactions | 3355109 | 42 | 10 | 9204 | 1112 | Pink1,RfC38,park,Argk,RpL30,RpS4,RpL23A,RpS13,CG15047,Gapdh1 | | 2.510e-02 | -3.68 | bam (bag of marbles) | protein interactions | 43038 | 42 | 10 | 9204 | 1112 | zpg,E2f1,tkv,nos,otu,mael,twin,eIF4A,Rga,eIF4E1 | | 2.510e-02 | -3.68 | Srpk79D (Serine-arginine protein kinase at 79D) | protein interactions | 40461 | 42 | 10 | 9204 | 1112 | Cka,cup,26-29-p,Plap,mts,Mst89B,Rcd5,mus205,CG6923,cnc | | 2.510e-02 | -3.68 | piwi (P-element induced wimpy testis) | protein interactions | 34521 | 42 | 10 | 9204 | 1112 | E2f1,l(3)mbt,Dcr-1,vret,squ,esc,Stip1,armi,shu,Klp10A | | 2.515e-02 | -3.68 | OstDelta (Oligosaccharide transferase Delta subunit) | protein interactions | 37029 | 79 | 16 | 9204 | 1112 | wal,mEFTu1,Surf4,Karybeta3,Rab5,Atpalpha,ScsbetaA,Mcad,Pp2A-29B,CCT3,CG15098,CG31729,SERCA,CCT6,CG10425,Kap-alpha3 | | 2.519e-02 | -3.68 | GSE28726\_NAIVE\_VS\_ACTIVATED\_VA24NEG\_NKTCELL\_UP | MSigDB lists | GSE28726\_NAIVE\_VS\_ACTIVATED\_VA24NEG\_NKTCELL\_UP | 104 | 24 | 4161 | 644 | CG5116,Pfas,Uch,Got1,CCT6,Pen,Cul1,Prosbeta5,Eno,Tom40,CG1890,ScsbetaA,CG8188,isoQC,Rpn2,mts,P5cr,Gmer,l(1)G0004,rept,CG7550,NHP2,Rpb8,levy | | 2.525e-02 | -3.68 | CULLIN | smart domains | SM00182 | 7 | 3 | 5086 | 506 | Cul2,Cul5,Cul1 | | 2.525e-02 | -3.68 | Iso\_dh | smart domains | SM01329 | 7 | 3 | 5086 | 506 | CG5028,Idh3b,Idh | | 2.525e-02 | -3.68 | IPT | smart domains | SM00429 | 7 | 3 | 5086 | 506 | Rel,NFAT,dl | | 2.528e-02 | -3.68 | MORF\_MAP2K2 | MSigDB lists | MORF\_MAP2K2 | 83 | 20 | 4161 | 644 | Prosbeta4,Idh3b,Pp2A-29B,Mtpalpha,SdhA,CCT5,ND-30,mEFTu1,Thiolase,Dd,LysRS,Cyt-c-p,Sod1,CG8188,Rpn12,eIF3i,CG5028,CG10418,ND-51,rept | | 2.528e-02 | -3.68 | PUJANA\_XPRSS\_INT\_NETWORK | MSigDB lists | PUJANA\_XPRSS\_INT\_NETWORK | 83 | 20 | 4161 | 644 | CG13630,Top1,mad2,Cdk1,asf1,DNApol-epsilon255,Idh3b,Clp,APP-BP1,r,CG8204,RfC38,CG8188,CG1890,RpII140,slif,CG6767,Dhfr,CkIIalpha,CG11164 | | 2.528e-02 | -3.68 | REACTOME\_CELL\_CYCLE\_CHECKPOINTS | MSigDB lists | REACTOME\_CELL\_CYCLE\_CHECKPOINTS | 83 | 20 | 4161 | 644 | CycE,RpS27A,Prosalpha5,Prosalpha3,Prosbeta4,Rpn3,mad2,CycB,Prosbeta5,Cdk1,APC7,Rpn12,grp,Rpn2,Prosbeta3,Rpt2,Prosalpha4,RpL40,Prosbeta6,RfC38 | | 2.528e-02 | -3.68 | GSE22886\_CD8\_VS\_CD4\_NAIVE\_TCELL\_UP | MSigDB lists | GSE22886\_CD8\_VS\_CD4\_NAIVE\_TCELL\_UP | 83 | 20 | 4161 | 644 | CG4293,Sccpdh1,Prosalpha5,Mo25,Uch-L5,nmdyn-D6,Pgm1,ste24a,CG10470,nmdyn-D7,Pex13,colt,Arl6IP1,Ubc4,CG4511,Nprl2,CysRS,Tm1,ATP8A,scramb1 | | 2.535e-02 | -3.68 | CG34168 (uncharacterized protein) | protein interactions | 50457 | 48 | 11 | 9204 | 1112 | cbs,CG15564,Larp4B,bcd,Clp,tos,Echs1,sol,cad,tyf,CG6707 | | 2.535e-02 | -3.68 | SCF(Skp2)-mediated degradation of p27/p21 | REACTOME pathways | R-DME-187577 | 66 | 16 | 3923 | 574 | Rpn12,Rpn3,Cul1,Prosbeta6,Prosbeta3,Prosalpha3,Rpt2,RpL40,SkpA,Rpn2,Prosbeta5,Prosalpha4,CycE,RpS27A,Prosalpha5,Prosbeta4 | | 2.541e-02 | -3.67 | GO\_DNA\_MODIFICATION | MSigDB lists | GO\_DNA\_MODIFICATION | 20 | 7 | 4161 | 644 | mael,obe,AGO3,CG5285,spn-E,Rrp40,shu | | 2.541e-02 | -3.67 | GO\_POLYSACCHARIDE\_BIOSYNTHETIC\_PROCESS | MSigDB lists | GO\_POLYSACCHARIDE\_BIOSYNTHETIC\_PROCESS | 20 | 7 | 4161 | 644 | GalT1,CG9485,AGBE,Pgm1,Gyg,RpS27A,RpL40 | | 2.541e-02 | -3.67 | GO\_STEROID\_BINDING | MSigDB lists | GO\_STEROID\_BINDING | 20 | 7 | 4161 | 644 | Atpalpha,scu,CG8112,Tspo,CG3860,AnxB10,Npc2a | | 2.541e-02 | -3.67 | GO\_ADP\_METABOLIC\_PROCESS | MSigDB lists | GO\_ADP\_METABOLIC\_PROCESS | 20 | 7 | 4161 | 644 | Pgk,Pglym78,Pgm1,Nc73EF,Eno,Pfk,Gapdh1 | | 2.542e-02 | -3.67 | pyridine-containing compound metabolic process | biological process | GO:0072524 | 13 | 4 | 12345 | 1129 | Nadsyn,CG33156,Pdxk,sgll | | 2.542e-02 | -3.67 | cell cycle G2/M phase transition | biological process | GO:0044839 | 13 | 4 | 12345 | 1129 | CG5003,Cdk1,twe,CycB | | 2.542e-02 | -3.67 | lipid translocation | biological process | GO:0034204 | 13 | 4 | 12345 | 1129 | scramb1,CG9981,ATP8A,CG31729 | | 2.542e-02 | -3.67 | axon target recognition | biological process | GO:0007412 | 13 | 4 | 12345 | 1129 | Liprin-beta,ben,not,sbb | | 2.542e-02 | -3.67 | vesicle targeting | biological process | GO:0006903 | 13 | 4 | 12345 | 1129 | Sec15,Sec10,Exo70,X11L | | 2.542e-02 | -3.67 | pteridine-containing compound biosynthetic process | biological process | GO:0042559 | 13 | 4 | 12345 | 1129 | Dhfr,Pcd,Sptr,pr | | 2.556e-02 | -3.67 | GO\_STRUCTURAL\_MOLECULE\_ACTIVITY | MSigDB lists | GO\_STRUCTURAL\_MOLECULE\_ACTIVITY | 220 | 45 | 4161 | 644 | RpL38,RpS4,RpS7,Tm1,tko,RpL40,RpL30,RpL23,RpS24,mRpS23,colt,RpS2,RpS16,RpS18,Moe,mRpS5,RpL12,Arpc4,mRpL17,RpL11,RpL37a,mRpL10,RpL26,Clc,RpL24,RpS27,gammaTub37C,Grip128,alphaTub84B,mRpL18,mRpL13,RpS30,RpS13,RpLP0,RpL13A,mRpL16,RpL13,mRpS21,Arpc2,RpS23,RpS27A,RpL9,Grip84,RpS28b,Grip75 | | 2.567e-02 | -3.66 | GO\_REGULATION\_OF\_CELLULAR\_PROTEIN\_LOCALIZATION | MSigDB lists | GO\_REGULATION\_OF\_CELLULAR\_PROTEIN\_LOCALIZATION | 164 | 35 | 4161 | 644 | CG3631,csw,Rpn12,Karybeta3,Nmt,cact,Cdk1,CCT6,CG30291,Atg6,dco,Pp2B-14D,CanB2,CG6766,Med,Aos1,Arf51F,SREBP,CanA-14F,Rcd5,Smox,PDCD-5,Vamp7,CCT5,alph,Gdi,Cdk5,pont,CycB,CCT3,Pdk1,mAChR-A,p38a,Edem1,Moe | | 2.573e-02 | -3.66 | SCHLOSSER\_SERUM\_RESPONSE\_DN | MSigDB lists | SCHLOSSER\_SERUM\_RESPONSE\_DN | 260 | 52 | 4161 | 644 | LpR2,l(1)G0196,Rbcn-3A,eEF1beta,CG5026,lqfR,Pgk,CG30496,Nap1,Prosalpha5,Nmt,SkpA,Xpac,CHOp24,opm,Fdh,escl,csw,Mcad,CG3226,Cdk7,isoQC,CG3511,RpII140,Rpn2,CHMP2B,alph,esc,Hat1,Hsc70-3,Sec23,TBCC,Prosalpha3,eIF3m,CanA-14F,CG32549,Gabat,PpV,Pi3K92E,CG17454,CG4511,CG40045,Pka-R1,ATP8A,Rbf,CG8963,Pp2B-14D,ScsbetaA,CG1890,Atg6,ari-1,Kap3 | | 2.574e-02 | -3.66 | GO\_CATALYTIC\_COMPLEX | MSigDB lists | GO\_CATALYTIC\_COMPLEX | 453 | 85 | 4161 | 644 | Cdk7,CG10465,ND-49,HDAC6,CG1812,CG8188,ND-SGDH,RpII140,Rpb8,lid,kl-2,CG8475,CG17691,SkpA,ND-B22,CG11070,ND-51,ND-23,escl,Tip60,Pka-R2,ND-39,Mo25,Utx,Tfb4,SdhA,CG8199,Fbw5,Uch-L5,Uev1A,tws,CG5292,Dd,APC7,alc,Pka-R1,Rep,Pi3K92E,Atpalpha,Dcr-1,CycE,MED20,esc,pont,Gclc,Eno,ND-B18,ND-30,Cul1,Cul2,mts,DCAF12,Naa20A,rept,CG30496,Nc73EF,Bin1,ND-75,Cul5,STUB1,Chrac-14,MED24,CkIIalpha,CycC,MED4,Cyt-c-p,Atg6,ari-1,nrv1,Pp2B-14D,CG6767,CanB2,CG9485,not,Aos1,Rcd5,CanA-14F,DNApol-epsilon255,Gabat,Pp2A-29B,Cka,MED27,Pfk,lwr,ND-24 | | 2.583e-02 | -3.66 | oxidoreductase activity, acting on the CH-NH group of donors, NAD or NADP as acceptor | molecular function | GO:0016646 | 13 | 4 | 12176 | 1119 | CG9471,P5cr,Dhfr,P5CDh1 | | 2.585e-02 | -3.66 | fly chr60A13-60A13\|2-105 cM | chromosome location | fly chr60A13-60A13\|2-105 cM | 5 | 2 | 25010 | 1343 | CG3803,Stoml2 | | 2.585e-02 | -3.66 | fly chr83B6-83B6\|3-47.5 cM | chromosome location | fly chr83B6-83B6\|3-47.5 cM | 5 | 2 | 25010 | 1343 | Atu,CG2911 | | 2.585e-02 | -3.66 | fly chr96E9-96E9\|3-88 cM | chromosome location | fly chr96E9-96E9\|3-88 cM | 5 | 2 | 25010 | 1343 | CG5028,CG4730 | | 2.585e-02 | -3.66 | fly chr69A1-69A1\|3-38 cM | chromosome location | fly chr69A1-69A1\|3-38 cM | 5 | 2 | 25010 | 1343 | CAH2,thoc6 | | 2.585e-02 | -3.66 | fly chr96E9-96E9\|3-88 | chromosome location | fly chr96E9-96E9\|3-88 | 5 | 2 | 25010 | 1343 | CG5028,CG4730 | | 2.585e-02 | -3.66 | fly chr1F4-1F4\|1-0.1 cM | chromosome location | fly chr1F4-1F4\|1-0.1 cM | 5 | 2 | 25010 | 1343 | png,CG12773 | | 2.585e-02 | -3.66 | fly chr69A1-69A1\|3-38 | chromosome location | fly chr69A1-69A1\|3-38 | 5 | 2 | 25010 | 1343 | thoc6,CAH2 | | 2.585e-02 | -3.66 | fly chr7D16-7D16\|1-22 cM | chromosome location | fly chr7D16-7D16\|1-22 cM | 5 | 2 | 25010 | 1343 | Traf6,CG10959 | | 2.585e-02 | -3.66 | fly chr94B5-94B5\|3-75 cM | chromosome location | fly chr94B5-94B5\|3-75 cM | 5 | 2 | 25010 | 1343 | CSN6,Dph5 | | 2.585e-02 | -3.66 | fly chr12E3-12E3\|1-48 | chromosome location | fly chr12E3-12E3\|1-48 | 5 | 2 | 25010 | 1343 | yl,CG11590 | | 2.585e-02 | -3.66 | fly chr57D8-57D8\|2-94 cM | chromosome location | fly chr57D8-57D8\|2-94 cM | 5 | 2 | 25010 | 1343 | Sgf29,RpL29 | | 2.585e-02 | -3.66 | fly chr57D8-57D8\|2-94 | chromosome location | fly chr57D8-57D8\|2-94 | 5 | 2 | 25010 | 1343 | Sgf29,RpL29 | | 2.585e-02 | -3.66 | fly chr92A11-92A11\|3-66 | chromosome location | fly chr92A11-92A11\|3-66 | 5 | 2 | 25010 | 1343 | Vha13,Nup58 | | 2.585e-02 | -3.66 | fly chr19F4-19F4\|1-65 cM | chromosome location | fly chr19F4-19F4\|1-65 cM | 5 | 2 | 25010 | 1343 | Dd,SLIRP1 | | 2.585e-02 | -3.66 | fly chr60A13-60A13\|2-105 | chromosome location | fly chr60A13-60A13\|2-105 | 5 | 2 | 25010 | 1343 | CG3803,Stoml2 | | 2.585e-02 | -3.66 | fly chr86E15-86E15\|3-51 cM | chromosome location | fly chr86E15-86E15\|3-51 cM | 5 | 2 | 25010 | 1343 | Ho,CG14715 | | 2.585e-02 | -3.66 | fly chr31F1-31F1\|2-43 | chromosome location | fly chr31F1-31F1\|2-43 | 5 | 2 | 25010 | 1343 | Grip75,CG6144 | | 2.585e-02 | -3.66 | fly chr92A11-92A11\|3-66 cM | chromosome location | fly chr92A11-92A11\|3-66 cM | 5 | 2 | 25010 | 1343 | Nup58,Vha13 | | 2.585e-02 | -3.66 | fly chr83B6-83B6\|3-47.5 | chromosome location | fly chr83B6-83B6\|3-47.5 | 5 | 2 | 25010 | 1343 | CG2911,Atu | | 2.585e-02 | -3.66 | fly chr7D16-7D16\|1-22 | chromosome location | fly chr7D16-7D16\|1-22 | 5 | 2 | 25010 | 1343 | CG10959,Traf6 | | 2.585e-02 | -3.66 | fly chr43D1-43D1\|2-56 cM | chromosome location | fly chr43D1-43D1\|2-56 cM | 5 | 2 | 25010 | 1343 | CG1602,mEFTu2 | | 2.585e-02 | -3.66 | fly chr31D1-31D1\|2-41 cM | chromosome location | fly chr31D1-31D1\|2-41 cM | 5 | 2 | 25010 | 1343 | nmd,CG5390 | | 2.585e-02 | -3.66 | fly chr94E13-94E13\|3-78 | chromosome location | fly chr94E13-94E13\|3-78 | 5 | 2 | 25010 | 1343 | ATPsynCF6,DNApol-epsilon255 | | 2.585e-02 | -3.66 | fly chr53C1-53C1\|2-80 cM | chromosome location | fly chr53C1-53C1\|2-80 cM | 5 | 2 | 25010 | 1343 | CG7997,Nup62 | | 2.585e-02 | -3.66 | fly chr19F4-19F4\|1-65 | chromosome location | fly chr19F4-19F4\|1-65 | 5 | 2 | 25010 | 1343 | SLIRP1,Dd | | 2.585e-02 | -3.66 | fly chr31D1-31D1\|2-41 | chromosome location | fly chr31D1-31D1\|2-41 | 5 | 2 | 25010 | 1343 | CG5390,nmd | | 2.585e-02 | -3.66 | fly chr86E15-86E15\|3-51 | chromosome location | fly chr86E15-86E15\|3-51 | 5 | 2 | 25010 | 1343 | Ho,CG14715 | | 2.585e-02 | -3.66 | fly chr94E13-94E13\|3-78 cM | chromosome location | fly chr94E13-94E13\|3-78 cM | 5 | 2 | 25010 | 1343 | DNApol-epsilon255,ATPsynCF6 | | 2.585e-02 | -3.66 | fly chr49C2-49C2\|2-67 cM | chromosome location | fly chr49C2-49C2\|2-67 cM | 5 | 2 | 25010 | 1343 | Mos,mRpL18 | | 2.585e-02 | -3.66 | fly chr94B5-94B5\|3-75 | chromosome location | fly chr94B5-94B5\|3-75 | 5 | 2 | 25010 | 1343 | Dph5,CSN6 | | 2.585e-02 | -3.66 | fly chr12E3-12E3\|1-48 cM | chromosome location | fly chr12E3-12E3\|1-48 cM | 5 | 2 | 25010 | 1343 | yl,CG11590 | | 2.585e-02 | -3.66 | fly chr1F4-1F4\|1-0.1 | chromosome location | fly chr1F4-1F4\|1-0.1 | 5 | 2 | 25010 | 1343 | CG12773,png | | 2.585e-02 | -3.66 | fly chr31F1-31F1\|2-43 cM | chromosome location | fly chr31F1-31F1\|2-43 cM | 5 | 2 | 25010 | 1343 | Grip75,CG6144 | | 2.585e-02 | -3.66 | fly chr49C2-49C2\|2-67 | chromosome location | fly chr49C2-49C2\|2-67 | 5 | 2 | 25010 | 1343 | mRpL18,Mos | | 2.585e-02 | -3.66 | fly chr43D1-43D1\|2-56 | chromosome location | fly chr43D1-43D1\|2-56 | 5 | 2 | 25010 | 1343 | mEFTu2,CG1602 | | 2.585e-02 | -3.66 | fly chr53C1-53C1\|2-80 | chromosome location | fly chr53C1-53C1\|2-80 | 5 | 2 | 25010 | 1343 | Nup62,CG7997 | | 2.586e-02 | -3.66 | GSE17721\_0.5H\_VS\_12H\_GARDIQUIMOD\_BMDC\_UP | MSigDB lists | GSE17721\_0.5H\_VS\_12H\_GARDIQUIMOD\_BMDC\_UP | 78 | 19 | 4161 | 644 | Sirt7,Dd,CG30291,Cbp80,sgll,CG15141,eIF3i,r,Trxr-1,CG12929,Idh3b,Dph5,Wdfy2,Chrac-14,CG15715,ND-30,RpS18,Echs1,CG8004 | | 2.586e-02 | -3.66 | GSE3982\_NEUTROPHIL\_VS\_CENT\_MEMORY\_CD4\_TCELL\_DN | MSigDB lists | GSE3982\_NEUTROPHIL\_VS\_CENT\_MEMORY\_CD4\_TCELL\_DN | 78 | 19 | 4161 | 644 | Ppox,Rnmt,RpS23,CG7806,CG7488,Gclc,esc,Pus7,RhoGAP5A,CG10344,RpL13A,RpS16,Sod1,Rpn2,CycC,Non1,CG11089,CG10418,escl | | 2.586e-02 | -3.66 | GSE20500\_CTRL\_VS\_RARA\_ANTAGONIST\_TREATED\_CD4\_TCELL\_DN | MSigDB lists | GSE20500\_CTRL\_VS\_RARA\_ANTAGONIST\_TREATED\_CD4\_TCELL\_DN | 78 | 19 | 4161 | 644 | tun,CG1532,CG9132,lt,RpII140,tko,Su(H),CG11089,CG1416,scramb1,CG4407,Idh,gammaTub37C,PpD3,Utx,Sccpdh1,pont,CG4709,Shmt | | 2.590e-02 | -3.65 | PyrdxlP-dep\_Trfase\_major | interpro domains | IPR015421 | 34 | 8 | 11383 | 1233 | CG11899,Sply,Got1,CG1640,mal,Shmt,Kyat,Gabat | | 2.590e-02 | -3.65 | PyrdxlP-dep\_Trfase | interpro domains | IPR015424 | 34 | 8 | 11383 | 1233 | CG1640,mal,Got1,Kyat,Shmt,Gabat,CG11899,Sply | | 2.593e-02 | -3.65 | GO\_ORGANOPHOSPHATE\_BIOSYNTHETIC\_PROCESS | MSigDB lists | GO\_ORGANOPHOSPHATE\_BIOSYNTHETIC\_PROCESS | 192 | 40 | 4161 | 644 | kri,Shmt,CG9391,Scsalpha1,CG7840,PAPLA1,Pi4KIIalpha,Stoml2,PIG-L,ACC,CG4407,Gk2,CG11089,csw,CG33156,AMPdeam,nmdyn-D7,Prip,mino,LysRS,Nadsyn,CG7322,Ac3,r-l,Pdxk,nmdyn-D6,PIG-C,Pfas,Ppcdc,sgll,Pi3K92E,Ts,r,scramb1,Synj,CG7718,Dhfr,CG2201,mal,CG6767 | | 2.593e-02 | -3.65 | GO\_RIBOSOME\_BIOGENESIS | MSigDB lists | GO\_RIBOSOME\_BIOGENESIS | 192 | 40 | 4161 | 644 | RpS30,RpS13,RpLP0,RpL13A,Rrp40,l(1)G0004,RpS27A,RpL9,RpS23,RpS28b,RpL13,RpL30,RpL23,Non1,RpS24,dco,CG11858,RpS16,RpS2,RpL38,RpS4,NHP2,mtTFB1,RpS7,mbo,RpL40,CG7488,CG7414,CG7728,RpL11,Mat89Ba,RpL37a,mRpL10,RpL26,RpL24,RpS27,RpS18,CG8611,bys,RpL12 | | 2.595e-02 | -3.65 | Peptidase\_C19\_UCH | interpro domains | IPR001394 | 22 | 6 | 11383 | 1233 | Usp16-45,not,faf,scny,Usp10,Usp39 | | 2.597e-02 | -3.65 | nuclear lumen | cellular component | GO:0031981 | 1016 | 112 | 11940 | 1105 | CG12301,MED27,hb,CG31053,trr,Mybbp1A,Snm1,fs(1)Ya,sbr,Clp,Cenp-C,mre11,mad2,CG11164,MED24,piwi,Sgf11,vers,dl,Sld5,rept,Utx,RpS2,Non1,pont,CG4021,Topors,RpS13,l(1)G0004,mute,RpS15,RYBP,CG10336,Sgf29,Elba2,Spt20,ix,gus,RpL11,Mat89Ba,DNApol-epsilon255,escl,IntS11,lin-52,Rbfox1,Ptip,RpS18,lid,Bin1,lwr,Tlk,mbo,nbs,Sirt7,CG1602,Rrp40,Su(H),bru1,d4,bys,CG46385,Top1,ph-d,CG9143,Psf3,YL-1,Cse1,Trf2,stil,gw,Nurf-38,Tfb4,PAPLA1,Orc6,Cdk7,pps,Prosbeta5,CG10055,Hat1,MED20,NHP2,TfIIFalpha,Taf7,not,scny,esc,e(y)1,Orc1,CG12018,mxc,CG12288,Tip60,SuUR,CycC,MED4,Chrac-14,CG17328,EloC,Rcd5,Bap170,HmgD,SelR,RpII140,Trf4-2,nej,Nap1,Rel,Atu,Ubc6,Rpb8,mts,HP1c | | 2.605e-02 | -3.65 | GSE411\_WT\_VS\_SOCS3\_KO\_MACROPHAGE\_IL6\_STIM\_100MIN\_UP | MSigDB lists | GSE411\_WT\_VS\_SOCS3\_KO\_MACROPHAGE\_IL6\_STIM\_100MIN\_UP | 99 | 23 | 4161 | 644 | Rpn2,RpLP0,ari-1,shv,CG30291,Dhfr,TyrRS,CG11164,Naa20A,Tspo,Grx1,Etf-QO,Nmt,CG9773,Abi,obe,CG11134,Mitofilin,kcc,CG3376,CG14715,mRpS10,CG8004 | | 2.606e-02 | -3.65 | U4/U6 x U5 tri-snRNP complex | cellular component | GO:0046540 | 19 | 5 | 11940 | 1105 | CG17768,CG6610,CG10418,SmE,LSm3 | | 2.606e-02 | -3.65 | spliceosomal tri-snRNP complex | cellular component | GO:0097526 | 19 | 5 | 11940 | 1105 | CG10418,SmE,LSm3,CG17768,CG6610 | | 2.607e-02 | -3.65 | regulation of intracellular signal transduction | biological process | GO:1902531 | 339 | 42 | 12345 | 1129 | Pp2A-29B,Rel,Herc4,Myd88,CG4853,Crag,CG10188,Cka,Lkb1,RhoGAP54D,alph,Traf6,Pura,Rbf,mts,Pdk1,Ubc6,tor,Tak1,Tao,Mos,CG17746,pn,ATPsynD,Ptp61F,chrb,Mekk1,NFAT,Iml1,PpV,dnc,ben,RhoGAP5A,Cul1,E2f1,dco,SkpA,PEK,Cbl,Pi3K92E,park,Nprl2 | | 2.611e-02 | -3.65 | GO\_MITOTIC\_CELL\_CYCLE\_CHECKPOINT | MSigDB lists | GO\_MITOTIC\_CELL\_CYCLE\_CHECKPOINT | 43 | 12 | 4161 | 644 | mad2,RpL40,Cdk1,CycB,RpS27A,Not10,CG1812,APP-BP1,Rbf,CkIIalpha,twin,CG30291 | | 2.611e-02 | -3.65 | GSE26488\_WT\_VS\_VP16\_TRANSGENIC\_HDAC7\_KO\_DOUBLE\_POSITIVE\_THYMOCYTE\_DN | MSigDB lists | GSE26488\_WT\_VS\_VP16\_TRANSGENIC\_HDAC7\_KO\_DOUBLE\_POSITIVE\_THYMOCYTE\_DN | 43 | 12 | 4161 | 644 | RpS4,eEF1beta,RpL13,dnc,RpL12,Cbp80,Hsc70-3,APP-BP1,RpL37a,RpL24,RpS13,Dph5 | | 2.611e-02 | -3.65 | GO\_NEGATIVE\_REGULATION\_OF\_CELL\_CYCLE\_PHASE\_TRANSITION | MSigDB lists | GO\_NEGATIVE\_REGULATION\_OF\_CELL\_CYCLE\_PHASE\_TRANSITION | 43 | 12 | 4161 | 644 | mad2,RpL40,Cdk1,CycB,Not10,RpS27A,CG1812,Rbf,APP-BP1,CG30291,CkIIalpha,twin | | 2.611e-02 | -3.65 | NING\_CHRONIC\_OBSTRUCTIVE\_PULMONARY\_DISEASE\_DN | MSigDB lists | NING\_CHRONIC\_OBSTRUCTIVE\_PULMONARY\_DISEASE\_DN | 43 | 12 | 4161 | 644 | RpL12,CG4332,Nca,Chd64,Gdi,RpS18,SkpA,RpS28b,RpS16,faf,Pfdn5,RpS24 | | 2.611e-02 | -3.65 | GO\_REGULATION\_OF\_CELLULAR\_AMINO\_ACID\_METABOLIC\_PROCESS | MSigDB lists | GO\_REGULATION\_OF\_CELLULAR\_AMINO\_ACID\_METABOLIC\_PROCESS | 43 | 12 | 4161 | 644 | Prosbeta5,Prosbeta6,mol,Rpn3,Prosalpha3,Prosbeta3,Prosbeta4,Prosalpha4,Rpt2,Rpn12,Prosalpha5,Rpn2 | | 2.619e-02 | -3.64 | RNA interference | biological process | GO:0016246 | 40 | 8 | 12345 | 1129 | Dcr-1,spn-E,Cbp80,r2d2,trsn,Hsc70-3,armi,piwi | | 2.619e-02 | -3.64 | glucose homeostasis | biological process | GO:0042593 | 40 | 8 | 12345 | 1129 | Pi3K92E,Hnf4,foxo,Desat1,PyK,Pfk,Lkb1,Eno | | 2.626e-02 | -3.64 | nucleoplasm part | cellular component | GO:0044451 | 401 | 49 | 11940 | 1105 | nej,lid,Bin1,Atu,Rpb8,d4,bru1,Su(H),Spt20,CycC,Sgf29,Elba2,Tip60,RYBP,Chrac-14,ix,gus,MED4,lin-52,Rcd5,IntS11,EloC,escl,RpII140,Ptip,Sgf11,MED24,rept,MED20,Utx,Sld5,vers,Taf7,pont,TfIIFalpha,CG4021,mxc,mute,esc,not,e(y)1,Psf3,MED27,Trf2,YL-1,trr,Tfb4,Clp,Cdk7 | | 2.632e-02 | -3.64 | L7ARS6FAMILY | prints domains | PR00881 | 3 | 2 | 2185 | 212 | NHP2,RpL7A | | 2.632e-02 | -3.64 | NFKBTNSCPFCT | prints domains | PR00057 | 3 | 2 | 2185 | 212 | dl,Rel | | 2.632e-02 | -3.64 | CUZNDISMTASE | prints domains | PR00068 | 3 | 2 | 2185 | 212 | Sod1,Sod3 | | 2.632e-02 | -3.64 | GLHYDRLASE20 | prints domains | PR00738 | 3 | 2 | 2185 | 212 | Hexo1,fdl | | 2.634e-02 | -3.64 | GO\_NEGATIVE\_REGULATION\_OF\_PROTEIN\_MODIFICATION\_PROCESS | MSigDB lists | GO\_NEGATIVE\_REGULATION\_OF\_PROTEIN\_MODIFICATION\_PROCESS | 181 | 38 | 4161 | 644 | Pdk1,CG4709,Gclc,Cdk5,CycB,Prosbeta5,Rpn3,CG3376,Prosbeta4,Prosalpha3,Pp2A-29B,PpD3,Smox,Prosbeta6,Ptp61F,Pka-R1,Snx6,Tspo,RpL40,Non1,CG30291,APC7,Cdk1,mad2,CG30496,Pka-R2,snz,RpS27A,Prosalpha5,lid,regucalcin,dnc,CG7530,Rpt2,Prosbeta3,mts,Rpn2,Rpn12 | | 2.639e-02 | -3.63 | regulation of JNK cascade | biological process | GO:0046328 | 63 | 11 | 12345 | 1129 | Cka,Lkb1,alph,SkpA,PEK,Mekk1,Ptp61F,park,PpV,Tak1,ben | | 2.643e-02 | -3.63 | outer mitochondrial membrane protein complex | cellular component | GO:0098799 | 13 | 4 | 11940 | 1105 | Tom70,Tom40,CG8004,CG9393 | | 2.647e-02 | -3.63 | GO\_RESPONSE\_TO\_HYDROGEN\_PEROXIDE | MSigDB lists | GO\_RESPONSE\_TO\_HYDROGEN\_PEROXIDE | 29 | 9 | 4161 | 644 | Cdk1,AIF,Grx1,Cyp1,Kap-alpha3,HDAC6,Sod1,PpD3,Prip | | 2.647e-02 | -3.63 | ATTACAT\_MIR3803P | MSigDB lists | ATTACAT\_MIR3803P | 29 | 9 | 4161 | 644 | mtgo,CG5292,Herc4,eIF4E1,Utx,Cul5,CHMP2B,CG12713,PrBP | | 2.647e-02 | -3.63 | GO\_CELL\_REDOX\_HOMEOSTASIS | MSigDB lists | GO\_CELL\_REDOX\_HOMEOSTASIS | 29 | 9 | 4161 | 644 | CG9911,ERp60,Grx1,CG4511,AIF,Jafrac2,Gclc,Trxr-1,CG14407 | | 2.647e-02 | -3.63 | GO\_FERTILIZATION | MSigDB lists | GO\_FERTILIZATION | 29 | 9 | 4161 | 644 | spn-E,Med,Cdk1,Ac3,CCT3,CCT5,Aladin,mael,mtgo | | 2.647e-02 | -3.63 | GO\_POSITIVE\_REGULATION\_OF\_PROTEIN\_IMPORT | MSigDB lists | GO\_POSITIVE\_REGULATION\_OF\_PROTEIN\_IMPORT | 29 | 9 | 4161 | 644 | CanA-14F,Smox,PDCD-5,Pp2B-14D,CanB2,p38a,Cdk1,Karybeta3,Med | | 2.647e-02 | -3.63 | GO\_NUCLEOSIDE\_DIPHOSPHATE\_METABOLIC\_PROCESS | MSigDB lists | GO\_NUCLEOSIDE\_DIPHOSPHATE\_METABOLIC\_PROCESS | 29 | 9 | 4161 | 644 | nmdyn-D7,Nc73EF,Pgm1,Gapdh1,Pgk,nmdyn-D6,Pglym78,Eno,Pfk | | 2.647e-02 | -3.63 | GO\_LIPID\_HOMEOSTASIS | MSigDB lists | GO\_LIPID\_HOMEOSTASIS | 29 | 9 | 4161 | 644 | EcR,mino,Mcad,Npc2a,Got1,CG8112,ACC,wal,crq | | 2.653e-02 | -3.63 | DNA Repair | REACTOME pathways | R-DME-73894 | 116 | 25 | 3923 | 574 | mre11,tos,Xpc,CG6923,Xpac,Pen,grp,CSN6,Cdk7,mus81,nbs,mus205,RfC38,ben,nej,Tfb4,Tip60,RpL40,RpS27A,Usp16-45,CG5285,Rpb8,Snm1,Uev1A,Usp10 | | 2.653e-02 | -3.63 | Ldsdh1 (Lipid droplet subset dehydrogenase 1) | protein interactions | 31726 | 6 | 3 | 9204 | 1112 | CG10237,Jabba,CG9186 | | 2.653e-02 | -3.63 | papi (papi) | protein interactions | 33401 | 6 | 3 | 9204 | 1112 | AGO3,piwi,tral | | 2.653e-02 | -3.63 | Rtca (RNA 3'-terminal phosphate cyclase) | protein interactions | 31175 | 6 | 3 | 9204 | 1112 | CG2678,Archease,CG4853 | | 2.653e-02 | -3.63 | ECSIT (ECSIT) | protein interactions | 40732 | 6 | 3 | 9204 | 1112 | Mekk1,Traf6,Pdrg1 | | 2.653e-02 | -3.63 | CG6550 (uncharacterized protein) | protein interactions | 36954 | 6 | 3 | 9204 | 1112 | Sld5,SERCA,alphaTub84B | | 2.653e-02 | -3.63 | CG14073 (uncharacterized protein) | protein interactions | 40046 | 6 | 3 | 9204 | 1112 | RpS24,l(3)73Ah,RYBP | | 2.653e-02 | -3.63 | mEFTs (mitochondrial translation elongation factor Ts) | protein interactions | 35060 | 6 | 3 | 9204 | 1112 | CG2162,mEFTu2,Gdi | | 2.653e-02 | -3.63 | Usp39 (Ubiquitin specific protease 39) | protein interactions | 32881 | 6 | 3 | 9204 | 1112 | AnxB10,Chd64,PHGPx | | 2.653e-02 | -3.63 | Gdap1 (Gdap1) | protein interactions | 38597 | 6 | 3 | 9204 | 1112 | Marf,Vha36-1,ix | | 2.653e-02 | -3.63 | Arc42 (Arc42) | protein interactions | 42364 | 6 | 3 | 9204 | 1112 | CG4860,SelD,r-l | | 2.653e-02 | -3.63 | lili (lilipod) | protein interactions | 42956 | 6 | 3 | 9204 | 1112 | tkv,bam,NaPi-III | | 2.653e-02 | -3.63 | bai (baiser) | protein interactions | 42996 | 6 | 3 | 9204 | 1112 | dl,tkv,CHOp24 | | 2.653e-02 | -3.63 | Picot (picot) | protein interactions | 36865 | 6 | 3 | 9204 | 1112 | capt,Chd64,CG1602 | | 2.653e-02 | -3.63 | Lhr (Lethal hybrid rescue) | protein interactions | 36957 | 6 | 3 | 9204 | 1112 | Cenp-C,RpS13,Nf-YA | | 2.653e-02 | -3.63 | Sccpdh1 (Saccheropin dehydrogenase 1) | protein interactions | 40623 | 6 | 3 | 9204 | 1112 | Bx,Ssdp,CG7997 | | 2.653e-02 | -3.63 | Yippee (yippee) | protein interactions | 32295 | 6 | 3 | 9204 | 1112 | Larp7,CG6617,cDIP | | 2.653e-02 | -3.63 | Snoo (Sno oncogene) | protein interactions | 5740414 | 6 | 3 | 9204 | 1112 | tkv,Med,Smox | | 2.653e-02 | -3.63 | CG5726 (uncharacterized protein) | protein interactions | 37082 | 6 | 3 | 9204 | 1112 | cnn,tyf,osk | | 2.664e-02 | -3.63 | aminopeptidase activity | molecular function | GO:0004177 | 55 | 10 | 12176 | 1119 | grsm,DppIII,CG32473,Dip-B,CG13630,CG2493,CG9953,CG9581,SP1029,CG14516 | | 2.664e-02 | -3.63 | UCH | pfam domains | PF00443 | 22 | 6 | 10480 | 1142 | faf,scny,Usp10,Usp16-45,Usp39,not | | 2.666e-02 | -3.62 | RHEIN\_ALL\_GLUCOCORTICOID\_THERAPY\_DN | MSigDB lists | RHEIN\_ALL\_GLUCOCORTICOID\_THERAPY\_DN | 170 | 36 | 4161 | 644 | Sccpdh1,Dph5,yip2,Prosbeta4,Jafrac2,CCT3,Pfk,Eno,CG1890,CG11858,CG4390,RpL23,Dhfr,Ts,NHP2,AGBE,Nap1,CSN6,mad2,Pgk,mEFTu1,CCT6,Cdk1,Bin1,wal,Nurf-38,grp,ND-SGDH,CG6610,Gapdh1,alphaTub84B,CG3226,Mcad,CG10932,CG11089,Stoml2 | | 2.667e-02 | -3.62 | slgA (sluggish A) | protein interactions | 33117 | 20 | 6 | 9204 | 1112 | eEF1beta,CkIIalpha,cact,RpS27A,CG3309,Arl6IP1 | | 2.667e-02 | -3.62 | Med (Medea) | protein interactions | 43725 | 20 | 6 | 9204 | 1112 | tkv,Sld5,Spx,ND-24,lwr,Smox | | 2.667e-02 | -3.62 | CG15440 (uncharacterized protein) | protein interactions | 33652 | 20 | 6 | 9204 | 1112 | alphaTub84B,RpS24,RpS27A,26-29-p,RpS30,RpS29 | | 2.667e-02 | -3.62 | Kap-alpha1 (karyopherin alpha1) | protein interactions | 40160 | 20 | 6 | 9204 | 1112 | Cse1,AMPdeam,RpLP1,GCC88,Kap-alpha3,Pen | | 2.671e-02 | -3.62 | Act5C (Actin 5C) | protein interactions | 31521 | 86 | 17 | 9204 | 1112 | PGRP-LE,CG6707,CG9911,Rpn13,glob1,Not10,COX5A,CHOp24,RhoGAP5A,Girdin,Pi3K92E,capt,osk,CG17746,Larp4B,foxo,CG11807 | | 2.678e-02 | -3.62 | REACTOME\_ASSEMBLY\_OF\_THE\_PRE\_REPLICATIVE\_COMPLEX | MSigDB lists | REACTOME\_ASSEMBLY\_OF\_THE\_PRE\_REPLICATIVE\_COMPLEX | 48 | 13 | 4161 | 644 | RpL40,Rpn3,Prosbeta5,Prosbeta6,Rpn12,Prosalpha5,RpS27A,Rpn2,Prosbeta3,Prosalpha3,Prosbeta4,Prosalpha4,Rpt2 | | 2.678e-02 | -3.62 | GO\_REGULATION\_OF\_MUSCLE\_SYSTEM\_PROCESS | MSigDB lists | GO\_REGULATION\_OF\_MUSCLE\_SYSTEM\_PROCESS | 48 | 13 | 4161 | 644 | Irk1,CG31715,mAChR-A,dnc,Calx,Tm1,CanA-14F,SERCA,AnxB10,Sod1,Gabat,Pp2B-14D,nAChRalpha4 | | 2.678e-02 | -3.62 | KEGG\_UBIQUITIN\_MEDIATED\_PROTEOLYSIS | MSigDB lists | KEGG\_UBIQUITIN\_MEDIATED\_PROTEOLYSIS | 68 | 17 | 4161 | 644 | APC7,Traf6,CG8188,Cul2,UbcE2H,CG11070,Aos1,Ubc4,CG40045,SkpA,CG4502,CG2924,Herc4,lwr,Cul1,STUB1,Cul5 | | 2.678e-02 | -3.62 | YAO\_TEMPORAL\_RESPONSE\_TO\_PROGESTERONE\_CLUSTER\_14 | MSigDB lists | YAO\_TEMPORAL\_RESPONSE\_TO\_PROGESTERONE\_CLUSTER\_14 | 68 | 17 | 4161 | 644 | CCT6,CG13630,ERp60,mRpS10,Prosbeta5,CCT3,Rpn2,Nurf-38,Prosbeta3,Gapdh1,Arts,RhoGDI,CG4332,cathD,r,NHP2,Karybeta3 | | 2.678e-02 | -3.62 | GSE37533\_PPARG1\_FOXP3\_VS\_FOXP3\_TRANSDUCED\_CD4\_TCELL\_PIOGLITAZONE\_TREATED\_DN | MSigDB lists | GSE37533\_PPARG1\_FOXP3\_VS\_FOXP3\_TRANSDUCED\_CD4\_TCELL\_PIOGLITAZONE\_TREATED\_DN | 68 | 17 | 4161 | 644 | CG14894,SelD,Rpb8,not,CG3226,Pp2B-14D,RpII140,Stip1,Klp10A,Ssdp,Top1,CCT6,Shmt,CanA-14F,yip2,Hmg-2,Irp-1B | | 2.686e-02 | -3.62 | GSE15930\_NAIVE\_VS\_48H\_IN\_VITRO\_STIM\_CD8\_TCELL\_DN | MSigDB lists | GSE15930\_NAIVE\_VS\_48H\_IN\_VITRO\_STIM\_CD8\_TCELL\_DN | 137 | 30 | 4161 | 644 | Prosalpha5,asf1,Clp,CG1774,LSm3,Prosalpha3,mad2,Pen,Hat1,CG13630,Pgk,ND-B18,Jafrac2,Coq7,Tim8,Rpn2,mRpL13,CG8188,Rpt2,CycC,Cul2,CG11899,CG12018,CG10418,Jafrac1,rept,tko,Arl5,DppIII,Stoml2 | | 2.689e-02 | -3.62 | BURTON\_ADIPOGENESIS\_PEAK\_AT\_24HR | MSigDB lists | BURTON\_ADIPOGENESIS\_PEAK\_AT\_24HR | 16 | 6 | 4161 | 644 | CycB,r,Cdk1,Karybeta3,Prip,Pen | | 2.689e-02 | -3.62 | GO\_REGULATION\_OF\_MITOCHONDRIAL\_MEMBRANE\_POTENTIAL | MSigDB lists | GO\_REGULATION\_OF\_MITOCHONDRIAL\_MEMBRANE\_POTENTIAL | 16 | 6 | 4161 | 644 | Sod1,Tspo,Stoml2,ND-75,Gclc,P5cr | | 2.689e-02 | -3.62 | MODULE\_233 | MSigDB lists | MODULE\_233 | 16 | 6 | 4161 | 644 | Ubc4,CG40045,UbcE2H,Uev1A,lwr,CG8188 | | 2.689e-02 | -3.62 | GO\_ANDROGEN\_RECEPTOR\_SIGNALING\_PATHWAY | MSigDB lists | GO\_ANDROGEN\_RECEPTOR\_SIGNALING\_PATHWAY | 16 | 6 | 4161 | 644 | CycE,MED4,Tip60,MED24,Cdk7,Fkbp59 | | 2.689e-02 | -3.62 | BIOCARTA\_NFAT\_PATHWAY | MSigDB lists | BIOCARTA\_NFAT\_PATHWAY | 16 | 6 | 4161 | 644 | Pp2B-14D,Pka-R2,Pka-R1,CanA-14F,p38a,rl | | 2.689e-02 | -3.62 | MODULE\_221 | MSigDB lists | MODULE\_221 | 16 | 6 | 4161 | 644 | Gabat,Echs1,Scsalpha1,Mcad,CG9471,Aldh | | 2.689e-02 | -3.62 | TTGCWCAAY\_CEBPB\_02 | MSigDB lists | TTGCWCAAY\_CEBPB\_02 | 16 | 6 | 4161 | 644 | Irk1,Idh,SRPK,Unr,p38a,eIF4A | | 2.689e-02 | -3.62 | ZHAN\_VARIABLE\_EARLY\_DIFFERENTIATION\_GENES\_DN | MSigDB lists | ZHAN\_VARIABLE\_EARLY\_DIFFERENTIATION\_GENES\_DN | 16 | 6 | 4161 | 644 | CCT5,anne,CycB,Mtpalpha,CG11089,Nap1 | | 2.689e-02 | -3.62 | GO\_CELLULAR\_RESPONSE\_TO\_HYDROGEN\_PEROXIDE | MSigDB lists | GO\_CELLULAR\_RESPONSE\_TO\_HYDROGEN\_PEROXIDE | 16 | 6 | 4161 | 644 | Cdk1,AIF,Prip,PpD3,HDAC6,Cyp1 | | 2.689e-02 | -3.62 | GO\_NEGATIVE\_REGULATION\_OF\_NUCLEAR\_DIVISION | MSigDB lists | GO\_NEGATIVE\_REGULATION\_OF\_NUCLEAR\_DIVISION | 16 | 6 | 4161 | 644 | Pka-R1,CkIIalpha,CG1812,nmdyn-D6,grp,mad2 | | 2.690e-02 | -3.62 | GSE17721\_POLYIC\_VS\_GARDIQUIMOD\_6H\_BMDC\_DN | MSigDB lists | GSE17721\_POLYIC\_VS\_GARDIQUIMOD\_6H\_BMDC\_DN | 94 | 22 | 4161 | 644 | CG7718,CG3631,bor,APC7,AMPdeam,CG17593,mtTFB1,CG40045,l(1)G0004,Cbp80,Atg4b,opm,ND-39,Acph-1,Irp-1B,alpha-Man-Ia,Stip1,CCT3,SdhA,Uev1A,Top1,UQCR-Q | | 2.693e-02 | -3.61 | GO\_GLUCOSE\_CATABOLIC\_PROCESS | MSigDB lists | GO\_GLUCOSE\_CATABOLIC\_PROCESS | 12 | 5 | 4161 | 644 | Pfk,Gapdh1,Eno,Pglym78,Pgk | | 2.693e-02 | -3.61 | IIZUKA\_LIVER\_CANCER\_PROGRESSION\_L0\_L1\_DN | MSigDB lists | IIZUKA\_LIVER\_CANCER\_PROGRESSION\_L0\_L1\_DN | 12 | 5 | 4161 | 644 | cathD,LysRS,Tspo,Arpc2,CG2493 | | 2.693e-02 | -3.61 | GO\_REGULATION\_OF\_NITRIC\_OXIDE\_SYNTHASE\_ACTIVITY | MSigDB lists | GO\_REGULATION\_OF\_NITRIC\_OXIDE\_SYNTHASE\_ACTIVITY | 12 | 5 | 4161 | 644 | CG7997,Dhfr,Sptr,pr,crq | | 2.693e-02 | -3.61 | GO\_PCG\_PROTEIN\_COMPLEX | MSigDB lists | GO\_PCG\_PROTEIN\_COMPLEX | 12 | 5 | 4161 | 644 | SkpA,esc,CkIIalpha,escl,l(3)73Ah | | 2.693e-02 | -3.61 | GO\_PTERIDINE\_CONTAINING\_COMPOUND\_BIOSYNTHETIC\_PROCESS | MSigDB lists | GO\_PTERIDINE\_CONTAINING\_COMPOUND\_BIOSYNTHETIC\_PROCESS | 12 | 5 | 4161 | 644 | pr,Dhfr,Sptr,Pcd,CG11089 | | 2.693e-02 | -3.61 | KYNG\_ENVIRONMENTAL\_STRESS\_RESPONSE\_NOT\_BY\_4NQO\_IN\_WS | MSigDB lists | KYNG\_ENVIRONMENTAL\_STRESS\_RESPONSE\_NOT\_BY\_4NQO\_IN\_WS | 12 | 5 | 4161 | 644 | Jafrac2,Sply,Trxr-1,scu,PDCD-5 | | 2.693e-02 | -3.61 | chr14q13 | MSigDB lists | chr14q13 | 12 | 5 | 4161 | 644 | Cka,Snx6,Trs33,cact,Sec23 | | 2.693e-02 | -3.61 | REACTOME\_AMINO\_ACID\_SYNTHESIS\_AND\_INTERCONVERSION\_TRANSAMINATION | MSigDB lists | REACTOME\_AMINO\_ACID\_SYNTHESIS\_AND\_INTERCONVERSION\_TRANSAMINATION | 12 | 5 | 4161 | 644 | aay,Got1,CG11899,CG1640,P5cr | | 2.699e-02 | -3.61 | positive regulation of gene expression | biological process | GO:0010628 | 571 | 66 | 12345 | 1129 | pont,nej,Pi3K92E,rept,e(y)1,CG3198,BigH1,cad,Trf2,ovo,TfIIFalpha,EcR,CG1602,bcd,Rbfox1,Tip60,lid,CG11710,trr,dl,foxo,not,Bx,d4,Atu,REPTOR,Myd88,Cdk7,Rel,Cf2,CG2199,lin,Sox102F,Mitf,Stat92E,Sgf11,apt,tyf,E2f1,vers,EloC,Bap170,CG17328,osk,NFAT,CycC,Rcd5,RYBP,esc,hb,Exd2,SREBP,pygo,CG2129,Tak1,Nf-YA,Wbp2,cnc,Nup358,cyc,S2P,CG8963,Mybbp1A,Su(H),Atf6,Utx | | 2.702e-02 | -3.61 | centriole replication | biological process | GO:0007099 | 26 | 6 | 12345 | 1129 | Rcd5,mts,CCT6,Pp2A-29B,cnn,tws | | 2.702e-02 | -3.61 | negative regulation of autophagy | biological process | GO:0010507 | 26 | 6 | 12345 | 1129 | scny,Pi3K92E,Smox,rl,Nprl2,Fkbp39 | | 2.702e-02 | -3.61 | ubiquitin-dependent ERAD pathway | biological process | GO:0030433 | 26 | 6 | 12345 | 1129 | Edem1,CG32640,CG6766,sip3,Hsc70-3,CG11070 | | 2.707e-02 | -3.61 | GSE39022\_LN\_VS\_SPLEEN\_DC\_DN | MSigDB lists | GSE39022\_LN\_VS\_SPLEEN\_DC\_DN | 63 | 16 | 4161 | 644 | Hmg-2,CG14407,CG11807,Sccpdh1,sws,Rnmt,CG30010,RpL13,mEFTu1,Aldh,p38a,slif,regucalcin,CG7550,GluRS-m,SkpA | | 2.707e-02 | -3.61 | GSE9006\_TYPE\_1\_DIABETES\_AT\_DX\_VS\_4MONTH\_POST\_DX\_PBMC\_DN | MSigDB lists | GSE9006\_TYPE\_1\_DIABETES\_AT\_DX\_VS\_4MONTH\_POST\_DX\_PBMC\_DN | 63 | 16 | 4161 | 644 | RpL24,tkv,mRpL17,eEF1beta,RpL12,Bin1,Tim8,CCT5,Pfdn5,Coq6,RpS13,RpS16,mRpL18,RpL40,mtTFB1,CG14894 | | 2.707e-02 | -3.61 | GSE29618\_PRE\_VS\_DAY7\_POST\_LAIV\_FLU\_VACCINE\_BCELL\_UP | MSigDB lists | GSE29618\_PRE\_VS\_DAY7\_POST\_LAIV\_FLU\_VACCINE\_BCELL\_UP | 63 | 16 | 4161 | 644 | Karybeta3,Su(H),mal,RpS30,Traf6,ste24a,RpLP0,AIF,thoc6,pr,Echs1,UQCR-Q,Pmi,Prosalpha3,Sply,PpD3 | | 2.710e-02 | -3.61 | CDC50 (CDC50) | protein interactions | 32615 | 15 | 5 | 9204 | 1112 | Fer1HCH,CG1092,CG10425,SERCA,Droj2 | | 2.710e-02 | -3.61 | CG14011 (uncharacterized protein) | protein interactions | 33790 | 15 | 5 | 9204 | 1112 | CG4730,CG9911,eEF1beta,eIF4A,CG10654 | | 2.710e-02 | -3.61 | MED25 (Mediator complex subunit 25) | protein interactions | 42385 | 15 | 5 | 9204 | 1112 | CG17002,Tlk,Usp10,Lsm12,CG3342 | | 2.710e-02 | -3.61 | TpnC4 (Troponin C isoform 4) | protein interactions | 35498 | 15 | 5 | 9204 | 1112 | up,Mf,COX5A,Blos2,CG5028 | | 2.710e-02 | -3.61 | Pop2 (Pop2) | protein interactions | 39366 | 15 | 5 | 9204 | 1112 | Pink1,cu,cup,twin,nos | | 2.712e-02 | -3.61 | REACTOME\_APC\_C\_CDH1\_MEDIATED\_DEGRADATION\_OF\_CDC20\_AND\_OTHER\_APC\_C\_CDH1\_TARGETED\_PROTEINS\_IN\_LATE\_MITOSIS\_EARLY\_G1 | MSigDB lists | REACTOME\_APC\_C\_CDH1\_MEDIATED\_DEGRADATION\_OF\_CDC20\_AND\_OTHER\_APC\_C\_CDH1\_TARGETED\_PROTEINS\_IN\_LATE\_MITOSIS\_EARLY\_G1 | 53 | 14 | 4161 | 644 | Prosalpha4,Prosbeta4,Rpt2,Prosbeta3,Prosalpha3,Prosalpha5,Rpn2,RpS27A,Rpn12,APC7,Prosbeta6,Prosbeta5,Rpn3,RpL40 | | 2.712e-02 | -3.61 | GO\_DICARBOXYLIC\_ACID\_METABOLIC\_PROCESS | MSigDB lists | GO\_DICARBOXYLIC\_ACID\_METABOLIC\_PROCESS | 53 | 14 | 4161 | 644 | ScsbetaA,Idh3b,Scsalpha1,Idh,Nc73EF,Dhfr,P5CDh1,Got1,CG11089,Shmt,CG5028,Gclc,CG1640,SdhA | | 2.712e-02 | -3.61 | GSE360\_L\_DONOVANI\_VS\_T\_GONDII\_MAC\_UP | MSigDB lists | GSE360\_L\_DONOVANI\_VS\_T\_GONDII\_MAC\_UP | 53 | 14 | 4161 | 644 | CG31729,Cyp1,CG14715,tho2,S2P,asf1,Acph-1,Tlk,Idh,CG2493,ND-49,Hsc70-3,CG44422,CG1532 | | 2.712e-02 | -3.61 | GSE1460\_NAIVE\_CD4\_TCELL\_ADULT\_BLOOD\_VS\_THYMIC\_STROMAL\_CELL\_UP | MSigDB lists | GSE1460\_NAIVE\_CD4\_TCELL\_ADULT\_BLOOD\_VS\_THYMIC\_STROMAL\_CELL\_UP | 53 | 14 | 4161 | 644 | p38a,GlyP,Tfb4,Cul1,Nca,eEF1beta,Tao,lt,RpII140,faf,Mo25,dco,RpS13,Grip75 | | 2.712e-02 | -3.61 | GSE37532\_WT\_VS\_PPARG\_KO\_VISCERAL\_ADIPOSE\_TISSUE\_TREG\_UP | MSigDB lists | GSE37532\_WT\_VS\_PPARG\_KO\_VISCERAL\_ADIPOSE\_TISSUE\_TREG\_UP | 53 | 14 | 4161 | 644 | Pen,Aos1,CG7655,SelD,Ts,PpV,CG8475,CycE,Oseg1,colt,CG32549,RhoGAP54D,alphaTub84B,LSm3 | | 2.713e-02 | -3.61 | regulation of DNA-binding transcription factor activity | biological process | GO:0051090 | 33 | 7 | 12345 | 1129 | S2P,lwr,Traf6,Aos1,rl,cact,CG13344 | | 2.720e-02 | -3.60 | GSE16385\_MONOCYTE\_VS\_12H\_ROSIGLITAZONE\_IL4\_TREATED\_MACROPHAGE\_DN | MSigDB lists | GSE16385\_MONOCYTE\_VS\_12H\_ROSIGLITAZONE\_IL4\_TREATED\_MACROPHAGE\_DN | 58 | 15 | 4161 | 644 | CG2493,CanA-14F,CG7789,Sec15,Unr,shv,Traf6,Smox,Mo25,Pp2B-14D,Prosbeta5,Tfb4,Arpc4,Plap,CG10418 | | 2.720e-02 | -3.60 | GSE8921\_UNSTIM\_0H\_VS\_TLR1\_2\_STIM\_MONOCYTE\_6H\_UP | MSigDB lists | GSE8921\_UNSTIM\_0H\_VS\_TLR1\_2\_STIM\_MONOCYTE\_6H\_UP | 58 | 15 | 4161 | 644 | CG8728,Gyg,CG11899,CG12018,Rpn2,AMPdeam,Dph5,levy,Prosbeta6,ND-B22,Stoml2,DppIII,Tim8,mRpS10,rept | | 2.720e-02 | -3.60 | MODULE\_36 | MSigDB lists | MODULE\_36 | 58 | 15 | 4161 | 644 | ari-1,trsn,Gabat,Arf51F,Uch,CG1827,mbo,dnc,milt,Hat1,CG4407,escl,alph,esc,SRPK | | 2.720e-02 | -3.60 | GSE26030\_UNSTIM\_VS\_RESTIM\_TH1\_DAY5\_POST\_POLARIZATION\_DN | MSigDB lists | GSE26030\_UNSTIM\_VS\_RESTIM\_TH1\_DAY5\_POST\_POLARIZATION\_DN | 58 | 15 | 4161 | 644 | Hmg-2,AMPdeam,isoQC,Rnmt,mal,CG30291,Idh,Prosalpha3,CG7840,CG7414,CG10932,CG42542,CG7322,CG17333,Jafrac2 | | 2.720e-02 | -3.60 | GSE37301\_COMMON\_LYMPHOID\_PROGENITOR\_VS\_CD4\_TCELL\_DN | MSigDB lists | GSE37301\_COMMON\_LYMPHOID\_PROGENITOR\_VS\_CD4\_TCELL\_DN | 58 | 15 | 4161 | 644 | CG8004,Cdk1,CG4511,CG15141,Ptp61F,Ts,Jafrac1,Thiolase,CG31729,Idh,CG3021,CSN6,CG1890,lt,Exo70 | | 2.720e-02 | -3.60 | GSE37416\_12H\_VS\_24H\_F\_TULARENSIS\_LVS\_NEUTROPHIL\_DN | MSigDB lists | GSE37416\_12H\_VS\_24H\_F\_TULARENSIS\_LVS\_NEUTROPHIL\_DN | 58 | 15 | 4161 | 644 | AGBE,Gdi,CHMP2B,Ntan1,Su(H),not,mtgo,Npc2a,Herc4,CG11807,CG1890,HDAC6,Vsp37A,Nap1,COX5A | | 2.720e-02 | -3.60 | GSE26030\_TH1\_VS\_TH17\_RESTIMULATED\_DAY5\_POST\_POLARIZATION\_DN | MSigDB lists | GSE26030\_TH1\_VS\_TH17\_RESTIMULATED\_DAY5\_POST\_POLARIZATION\_DN | 58 | 15 | 4161 | 644 | PIG-L,AcCoAS,Pdk1,shop,CG11089,ACC,CG15111,SREBP,Clp,Tsp66E,CG7728,APC7,CG11807,AMPdeam,Mekk1 | | 2.720e-02 | -3.60 | GSE21033\_CTRL\_VS\_POLYIC\_STIM\_DC\_24H\_DN | MSigDB lists | GSE21033\_CTRL\_VS\_POLYIC\_STIM\_DC\_24H\_DN | 58 | 15 | 4161 | 644 | Jafrac2,SRPK,Coq7,Aldh,AIF,RpL13,CG31759,Wnt10,CG1847,eIF3m,Prosbeta3,CG7789,Prosalpha5,Oseg1,CG4390 | | 2.721e-02 | -3.60 | csw (corkscrew) | protein interactions | 45278 | 67 | 14 | 9204 | 1112 | tor,Su(H),Stat92E,Larp4B,wal,eIF4A,RpS27,RpL37a,PyK,Fkbp12,tkv,rl,CG2747,ATPsynbeta | | 2.739e-02 | -3.60 | protein kinase regulator activity | molecular function | GO:0019887 | 71 | 12 | 12176 | 1119 | ASPP,Pka-R2,CycB,Pdk1,CycE,CycJ,Pka-R1,CG14408,RpLP1,Lkb1,Parp16,CycC | | 2.744e-02 | -3.60 | COX4 (Cytochrome c oxidase subunit 4) | protein interactions | 35279 | 31 | 8 | 9204 | 1112 | ATPsynO,Adh,ATPsynF,Tspo,COX5A,ATPsynCF6,levy,cype | | 2.744e-02 | -3.60 | MED27 (Mediator complex subunit 27) | protein interactions | 40696 | 31 | 8 | 9204 | 1112 | ix,Imp,EcR,MED4,MED20,enc,Ptip,Larp4B | | 2.744e-02 | -3.60 | apt (apontic) | protein interactions | 37734 | 31 | 8 | 9204 | 1112 | Stat92E,apt,Cenp-C,osk,CG6766,cnn,bru1,CG33156 | | 2.744e-02 | -3.60 | Invadolysin (invadolysin) | protein interactions | 49580 | 31 | 8 | 9204 | 1112 | cnc,ATPsynD,blw,CG9911,SNF4Agamma,Ady43A,ATPsynbeta,Vsp37A | | 2.762e-02 | -3.59 | GO\_CELLULAR\_HOMEOSTASIS | MSigDB lists | GO\_CELLULAR\_HOMEOSTASIS | 176 | 37 | 4161 | 644 | SERCA,Irp-1B,Trxr-1,AIF,GlyP,kcc,ERp60,CG9911,Calx,Prip,adp,Sod1,Stoml2,CG42542,zyd,regucalcin,Grx1,Mitofilin,CanA-14F,CG14407,Scox,Jafrac2,CHMP2B,Pfk,anne,Gclc,AnxB10,Vha36-1,CG1275,Pp2B-14D,nrv1,Vha13,CG4511,Irk1,Atpalpha,Pdk,Med | | 2.768e-02 | -3.59 | LysM | smart domains | SM00257 | 3 | 2 | 5086 | 506 | CG17985,red | | 2.768e-02 | -3.59 | PreSET | smart domains | SM00468 | 3 | 2 | 5086 | 506 | G9a,egg | | 2.768e-02 | -3.59 | Longin | smart domains | SM01270 | 3 | 2 | 5086 | 506 | Sec22,Vamp7 | | 2.768e-02 | -3.59 | Ribosomal\_S10 | smart domains | SM01403 | 3 | 2 | 5086 | 506 | mRpS10,mRpL48 | | 2.768e-02 | -3.59 | BEN | smart domains | SM01025 | 3 | 2 | 5086 | 506 | insv,Elba2 | | 2.768e-02 | -3.59 | Cir\_N | smart domains | SM01083 | 3 | 2 | 5086 | 506 | Cwc25,CG12818 | | 2.774e-02 | -3.58 | GSE28726\_NAIVE\_VS\_ACTIVATED\_CD4\_TCELL\_DN | MSigDB lists | GSE28726\_NAIVE\_VS\_ACTIVATED\_CD4\_TCELL\_DN | 89 | 21 | 4161 | 644 | Prosalpha5,gammaTub37C,Pdxk,r-l,Pfk,Eno,Tom40,CCT5,CycB,Pgk,Dhfr,CG12018,Prosbeta3,scu,CG8188,Ts,levy,Karybeta3,CG10932,tko,rept | | 2.774e-02 | -3.58 | GSE36476\_CTRL\_VS\_TSST\_ACT\_72H\_MEMORY\_CD4\_TCELL\_YOUNG\_DN | MSigDB lists | GSE36476\_CTRL\_VS\_TSST\_ACT\_72H\_MEMORY\_CD4\_TCELL\_YOUNG\_DN | 89 | 21 | 4161 | 644 | Prosbeta4,Ppcdc,Mitofilin,Pfas,Irp-1B,gammaTub37C,yip2,r-l,CG7997,CycB,Pglym78,msps,CG15715,alphaTub84B,Dhfr,grp,DppIII,Ts,Rpb8,Grx1,Gk2 | | 2.780e-02 | -3.58 | regulation of cellular localization | biological process | GO:0060341 | 139 | 20 | 12345 | 1129 | wapl,wash,exu,Rbf,swa,cnc,Nup358,sbr,mael,Mctp,Crag,spn-E,Sec23,tral,Ranbp21,NFAT,enc,mbo,cup,Herc4 | | 2.797e-02 | -3.58 | TIEN\_INTESTINE\_PROBIOTICS\_24HR\_UP | MSigDB lists | TIEN\_INTESTINE\_PROBIOTICS\_24HR\_UP | 267 | 53 | 4161 | 644 | CG11858,MED4,ScsbetaA,MED24,Psf3,CG7718,CG8963,CG1416,Cyp1,Adi1,Tm1,CG14894,Ts,RfC38,yip2,gammaTub37C,CG14407,Gabat,PDCD-5,CG9773,Dcr-1,LSm3,APP-BP1,ND-24,Ntan1,Pglym78,Tim8,CycB,esc,pr,anne,CG7322,CG5510,r-l,trsn,CG6610,eIF4E1,Rpn2,CG3226,Cul2,Rpt2,Jafrac1,CG11070,ste14,escl,DppIII,Mekk1,Pcd,CG8507,ND-39,Fkbp59,mad2,Echs1 | | 2.802e-02 | -3.57 | regulation of lipid localization | biological process | GO:1905952 | 48 | 9 | 12345 | 1129 | adp,Pi3K92E,foxo,Atf3,SERCA,Rab32,LpR2,dco,Seipin | | 2.802e-02 | -3.57 | response to insulin | biological process | GO:0032868 | 48 | 9 | 12345 | 1129 | raptor,Hnf4,Pi3K92E,PyK,foxo,Rab10,rl,Pdk1,CG7328 | | 2.805e-02 | -3.57 | GSE28726\_ACT\_CD4\_TCELL\_VS\_ACT\_NKTCELL\_UP | MSigDB lists | GSE28726\_ACT\_CD4\_TCELL\_VS\_ACT\_NKTCELL\_UP | 105 | 24 | 4161 | 644 | CG7550,CG4511,CG8204,Karybeta3,NHP2,eIF3i,RfC38,rept,tko,mbo,mts,Gmer,muskelin,CG8728,Gapdh1,CG33156,ScsbetaA,Rpn2,Eno,SRPK,Pgk,Cul1,ERp60,CG5116 | | 2.810e-02 | -3.57 | Larp4B (La-related protein 4B) | protein interactions | 38399 | 61 | 13 | 9204 | 1112 | csw,tws,ix,MED4,CycB,Chd64,tyf,MED20,MED27,Usp10,Rga,Marf1,CG46385 | | 2.813e-02 | -3.57 | BORCZUK\_MALIGNANT\_MESOTHELIOMA\_UP | MSigDB lists | BORCZUK\_MALIGNANT\_MESOTHELIOMA\_UP | 154 | 33 | 4161 | 644 | CG40045,CG17454,NHP2,Med,P5cr,faf,Kap3,Stip1,Jafrac2,SRPK,CCT3,CycB,CCT5,PIG-C,Arpc4,CG9248,Sccpdh1,Vamp7,SkpA,eIF3i,parvin,l(1)G0004,ND-49,Cdk7,CG8188,Rpn12,Klp10A,Pen,mad2,Pgk,Fkbp59,Nap1,Rab32 | | 2.824e-02 | -3.57 | NAD(P)-bd\_dom\_sf | interpro domains | IPR036291 | 148 | 24 | 11383 | 1233 | Scsalpha1,CG3609,Gapdh1,CG7322,Gmer,CG10425,CG31549,CG31548,CG9471,Sptr,scu,CG9265,CG3842,CG4872,Adh,ND-39,P5cr,CG4842,CG31674,CG12171,Sccpdh1,sni,Mtpalpha,Fdh | | 2.824e-02 | -3.57 | establishment of organelle localization | biological process | GO:0051656 | 148 | 21 | 12345 | 1129 | l(3)mbt,CycB,milt,Sec10,Cdk5,faf,Klp10A,alphaTub67C,cana,Pink1,mbo,Sec15,ssh,gammaTub37C,X11L,msps,wisp,Cenp-C,Exo70,Marf,jvl | | 2.855e-02 | -3.56 | GO\_ANTIGEN\_PROCESSING\_AND\_PRESENTATION\_OF\_PEPTIDE\_ANTIGEN | MSigDB lists | GO\_ANTIGEN\_PROCESSING\_AND\_PRESENTATION\_OF\_PEPTIDE\_ANTIGEN | 84 | 20 | 4161 | 644 | ERp60,Rpn3,Prosbeta5,Klp10A,Clc,Prosalpha5,Prosalpha3,Sec23,Prosbeta4,cathD,AP-1-2beta,Prosbeta6,Rpn12,Traf6,CG3860,Kap3,Rpn2,Prosbeta3,Rpt2,Prosalpha4 | | 2.855e-02 | -3.56 | GSE41978\_KLRG1\_HIGH\_VS\_LOW\_EFFECTOR\_CD8\_TCELL\_DN | MSigDB lists | GSE41978\_KLRG1\_HIGH\_VS\_LOW\_EFFECTOR\_CD8\_TCELL\_DN | 84 | 20 | 4161 | 644 | RpS18,UQCR-Q,Calx,capt,RpL12,RpL11,RpL37a,RpS23,RpL9,RpL24,Grip75,RpL38,RpS4,Pka-R1,RpS7,Atg4b,RpL23,RpLP0,RpS2,RpS16 | | 2.855e-02 | -3.56 | GSE12845\_PRE\_GC\_VS\_DARKZONE\_GC\_TONSIL\_BCELL\_DN | MSigDB lists | GSE12845\_PRE\_GC\_VS\_DARKZONE\_GC\_TONSIL\_BCELL\_DN | 84 | 20 | 4161 | 644 | Arpc1,MED20,Grip75,CG14407,CG9911,lqfR,milt,Klp10A,Aldh,isoQC,HDAC6,RpS30,Kap3,eIF4E1,CG8728,Cdk7,CG11070,CG40045,r,Crag | | 2.867e-02 | -3.55 | GO\_NUCLEOSIDE\_MONOPHOSPHATE\_BIOSYNTHETIC\_PROCESS | MSigDB lists | GO\_NUCLEOSIDE\_MONOPHOSPHATE\_BIOSYNTHETIC\_PROCESS | 34 | 10 | 4161 | 644 | r-l,kri,Ts,Stoml2,r,Shmt,CG11089,Pfas,CG6767,AMPdeam | | 2.867e-02 | -3.55 | GO\_REGULATION\_OF\_MUSCLE\_CONTRACTION | MSigDB lists | GO\_REGULATION\_OF\_MUSCLE\_CONTRACTION | 34 | 10 | 4161 | 644 | nAChRalpha4,Gabat,Sod1,SERCA,AnxB10,Tm1,dnc,Calx,mAChR-A,Irk1 | | 2.877e-02 | -3.55 | chico (chico) | protein interactions | 64880 | 37 | 9 | 9204 | 1112 | Pi3K92E,Pink1,RpL30,tws,Pp2A-29B,Hsc70-3,RpL17,RpL7A,foxo | | 2.877e-02 | -3.55 | CCT1 (Chaperonin containing TCP1 subunit 1) | protein interactions | 42649 | 37 | 9 | 9204 | 1112 | PyK,CCT6,csw,PGRP-LE,Pdk1,wal,CCT5,CCT3,foxo | | 2.880e-02 | -3.55 | Smn (survival motor neuron) | protein interactions | 39844 | 55 | 12 | 9204 | 1112 | CG10561,Moe,SkpA,Sod1,eIF3m,Actn,Atf6,CG15525,Cul1,shrb,CG2199,SmE | | 2.891e-02 | -3.54 | REACTOME\_ERKS\_ARE\_INACTIVATED | MSigDB lists | REACTOME\_ERKS\_ARE\_INACTIVATED | 5 | 3 | 4161 | 644 | mts,rl,Pp2A-29B | | 2.891e-02 | -3.54 | GO\_RECEPTOR\_CATABOLIC\_PROCESS | MSigDB lists | GO\_RECEPTOR\_CATABOLIC\_PROCESS | 5 | 3 | 4161 | 644 | snz,Cdk5,Atg6 | | 2.891e-02 | -3.54 | GO\_MACROLIDE\_BINDING | MSigDB lists | GO\_MACROLIDE\_BINDING | 5 | 3 | 4161 | 644 | shu,Fkbp59,CG14715 | | 2.891e-02 | -3.54 | BIOCARTA\_SKP2E2F\_PATHWAY | MSigDB lists | BIOCARTA\_SKP2E2F\_PATHWAY | 5 | 3 | 4161 | 644 | SkpA,CycE,Cul1 | | 2.891e-02 | -3.54 | DAZARD\_UV\_RESPONSE\_CLUSTER\_G2 | MSigDB lists | DAZARD\_UV\_RESPONSE\_CLUSTER\_G2 | 5 | 3 | 4161 | 644 | CG8188,Rpn12,Gmer | | 2.891e-02 | -3.54 | LINDSTEDT\_DENDRITIC\_CELL\_MATURATION\_B | MSigDB lists | LINDSTEDT\_DENDRITIC\_CELL\_MATURATION\_B | 5 | 3 | 4161 | 644 | escl,esc,cact | | 2.891e-02 | -3.54 | REACTOME\_REGULATION\_OF\_AMPK\_ACTIVITY\_VIA\_LKB1 | MSigDB lists | REACTOME\_REGULATION\_OF\_AMPK\_ACTIVITY\_VIA\_LKB1 | 5 | 3 | 4161 | 644 | raptor,alc,Mo25 | | 2.891e-02 | -3.54 | GO\_CAMP\_BINDING | MSigDB lists | GO\_CAMP\_BINDING | 5 | 3 | 4161 | 644 | Pka-R2,Pka-R1,dnc | | 2.891e-02 | -3.54 | REACTOME\_G2\_M\_DNA\_DAMAGE\_CHECKPOINT | MSigDB lists | REACTOME\_G2\_M\_DNA\_DAMAGE\_CHECKPOINT | 5 | 3 | 4161 | 644 | grp,Cdk1,CycB | | 2.891e-02 | -3.54 | GO\_MISFOLDED\_PROTEIN\_BINDING | MSigDB lists | GO\_MISFOLDED\_PROTEIN\_BINDING | 5 | 3 | 4161 | 644 | HDAC6,STUB1,Hsc70-3 | | 2.891e-02 | -3.54 | REACTOME\_TETRAHYDROBIOPTERIN\_BH4\_SYNTHESIS\_RECYCLING\_SALVAGE\_AND\_REGULATION | MSigDB lists | REACTOME\_TETRAHYDROBIOPTERIN\_BH4\_SYNTHESIS\_RECYCLING\_SALVAGE\_AND\_REGULATION | 5 | 3 | 4161 | 644 | Sptr,pr,Dhfr | | 2.891e-02 | -3.54 | MCDOWELL\_ACUTE\_LUNG\_INJURY\_UP | MSigDB lists | MCDOWELL\_ACUTE\_LUNG\_INJURY\_UP | 5 | 3 | 4161 | 644 | Tm1,cact,ND-75 | | 2.891e-02 | -3.54 | GO\_MYOTUBE\_CELL\_DEVELOPMENT | MSigDB lists | GO\_MYOTUBE\_CELL\_DEVELOPMENT | 5 | 3 | 4161 | 644 | CanA-14F,Wnt10,Pp2B-14D | | 2.891e-02 | -3.54 | GO\_CELLULAR\_RESPONSE\_TO\_EPIDERMAL\_GROWTH\_FACTOR\_STIMULUS | MSigDB lists | GO\_CELLULAR\_RESPONSE\_TO\_EPIDERMAL\_GROWTH\_FACTOR\_STIMULUS | 5 | 3 | 4161 | 644 | Atg6,r,Pdk1 | | 2.891e-02 | -3.54 | CHEOK\_RESPONSE\_TO\_MERCAPTOPURINE\_AND\_HD\_MTX\_DN | MSigDB lists | CHEOK\_RESPONSE\_TO\_MERCAPTOPURINE\_AND\_HD\_MTX\_DN | 5 | 3 | 4161 | 644 | Bin1,Uch,CG40045 | | 2.891e-02 | -3.54 | GO\_REGULATION\_OF\_TYPE\_I\_INTERFERON\_MEDIATED\_SIGNALING\_PATHWAY | MSigDB lists | GO\_REGULATION\_OF\_TYPE\_I\_INTERFERON\_MEDIATED\_SIGNALING\_PATHWAY | 5 | 3 | 4161 | 644 | Ptp61F,csw,Ubc4 | | 2.891e-02 | -3.54 | REACTOME\_REGULATION\_OF\_HYPOXIA\_INDUCIBLE\_FACTOR\_HIF\_BY\_OXYGEN | MSigDB lists | REACTOME\_REGULATION\_OF\_HYPOXIA\_INDUCIBLE\_FACTOR\_HIF\_BY\_OXYGEN | 5 | 3 | 4161 | 644 | Cul2,RpS27A,RpL40 | | 2.891e-02 | -3.54 | GO\_REGULATION\_OF\_HISTONE\_H3\_K9\_ACETYLATION | MSigDB lists | GO\_REGULATION\_OF\_HISTONE\_H3\_K9\_ACETYLATION | 5 | 3 | 4161 | 644 | Med,grp,AGO3 | | 2.891e-02 | -3.54 | PARK\_APL\_PATHOGENESIS\_UP | MSigDB lists | PARK\_APL\_PATHOGENESIS\_UP | 5 | 3 | 4161 | 644 | Stip1,Scsalpha1,CG11070 | | 2.891e-02 | -3.54 | GO\_LEUKOCYTE\_DEGRANULATION | MSigDB lists | GO\_LEUKOCYTE\_DEGRANULATION | 5 | 3 | 4161 | 644 | Pi4KIIalpha,Vamp7,Pi3K92E | | 2.891e-02 | -3.54 | GO\_LABYRINTHINE\_LAYER\_BLOOD\_VESSEL\_DEVELOPMENT | MSigDB lists | GO\_LABYRINTHINE\_LAYER\_BLOOD\_VESSEL\_DEVELOPMENT | 5 | 3 | 4161 | 644 | CHOp24,rl,Su(H) | | 2.891e-02 | -3.54 | PID\_CIRCADIAN\_PATHWAY | MSigDB lists | PID\_CIRCADIAN\_PATHWAY | 5 | 3 | 4161 | 644 | dco,cyc,grp | | 2.891e-02 | -3.54 | REACTOME\_OXYGEN\_DEPENDENT\_PROLINE\_HYDROXYLATION\_OF\_HYPOXIA\_INDUCIBLE\_FACTOR\_ALPHA | MSigDB lists | REACTOME\_OXYGEN\_DEPENDENT\_PROLINE\_HYDROXYLATION\_OF\_HYPOXIA\_INDUCIBLE\_FACTOR\_ALPHA | 5 | 3 | 4161 | 644 | RpS27A,RpL40,Cul2 | | 2.891e-02 | -3.54 | GO\_PYRIMIDINE\_RIBONUCLEOSIDE\_TRIPHOSPHATE\_METABOLIC\_PROCESS | MSigDB lists | GO\_PYRIMIDINE\_RIBONUCLEOSIDE\_TRIPHOSPHATE\_METABOLIC\_PROCESS | 5 | 3 | 4161 | 644 | r,nmdyn-D6,nmdyn-D7 | | 2.891e-02 | -3.54 | GO\_MYD88\_DEPENDENT\_TOLL\_LIKE\_RECEPTOR\_SIGNALING\_PATHWAY | MSigDB lists | GO\_MYD88\_DEPENDENT\_TOLL\_LIKE\_RECEPTOR\_SIGNALING\_PATHWAY | 5 | 3 | 4161 | 644 | RpS27A,RpL40,Traf6 | | 2.891e-02 | -3.54 | chrxq21 | MSigDB lists | chrxq21 | 5 | 3 | 4161 | 644 | CG6767,CG7997,Rep | | 2.891e-02 | -3.54 | KEGG\_LIMONENE\_AND\_PINENE\_DEGRADATION | MSigDB lists | KEGG\_LIMONENE\_AND\_PINENE\_DEGRADATION | 5 | 3 | 4161 | 644 | Aldh,Mtpalpha,Echs1 | | 2.891e-02 | -3.54 | GO\_NEGATIVE\_REGULATION\_OF\_VIRAL\_GENOME\_REPLICATION | MSigDB lists | GO\_NEGATIVE\_REGULATION\_OF\_VIRAL\_GENOME\_REPLICATION | 5 | 3 | 4161 | 644 | Tango10,scramb1,SRPK | | 2.891e-02 | -3.54 | REACTOME\_DOWNSTREAM\_TCR\_SIGNALING | MSigDB lists | REACTOME\_DOWNSTREAM\_TCR\_SIGNALING | 5 | 3 | 4161 | 644 | Pdk1,cact,Traf6 | | 2.900e-02 | -3.54 | regulation of multicellular organismal process | biological process | GO:0051239 | 781 | 87 | 12345 | 1129 | Desat1,Ork1,ben,Tao,png,ssh,Smox,G9a,Cul2,tws,ari-1,Sox102F,Cbl,lwr,park,Nurf-38,Stam,Cul1,swa,E2f1,dco,p38a,trsn,Atg7,csw,Su(H),Seipin,eIF4E1,mus304,opm,shv,hb,MED4,mts,exu,tor,nos,Gabat,ago,scny,grp,cnc,Wbp2,cyc,enc,bif,twe,EcR,Archease,CHOp24,cact,MED27,Pink1,Cdk5,Mtl,hwt,Pp2B-14D,Pi3K92E,rl,mael,tkv,l(3)mbt,PGRP-LE,SkpA,COX5A,MAGE,Med,Atu,tral,spn-E,Sec23,sbb,bun,mop,cup,raptor,Cul5,Top1,foxo,dl,Pdk1,Imp,COX4,bru1,Ptpmeg,up,alph | | 2.906e-02 | -3.54 | Malonate semialdehyde pathway, propanoyl-CoA => acetyl-CoA | KEGG pathways | M00013 | 5 | 3 | 3198 | 496 | Mtpalpha,Mcad,Echs1 | | 2.906e-02 | -3.54 | Malonate semialdehyde pathway, propanoyl-CoA => acetyl-CoA | KEGG pathways | dme\_M00013 | 5 | 3 | 3198 | 496 | Mcad,Mtpalpha,Echs1 | | 2.908e-02 | -3.54 | HOSHIDA\_LIVER\_CANCER\_SUBCLASS\_S3 | MSigDB lists | HOSHIDA\_LIVER\_CANCER\_SUBCLASS\_S3 | 100 | 23 | 4161 | 644 | Jheh2,Irp-1B,yip2,SREBP,CG4872,CG9391,Shmt,CG8112,ND-24,cact,CG9471,Aldh,Echs1,pr,CG14715,scu,Sod1,AnxB10,P5CDh1,Pgm1,Mcad,regucalcin,CG9485 | | 2.908e-02 | -3.54 | GSE15930\_NAIVE\_VS\_24H\_IN\_VITRO\_STIM\_INFAB\_CD8\_TCELL\_DN | MSigDB lists | GSE15930\_NAIVE\_VS\_24H\_IN\_VITRO\_STIM\_INFAB\_CD8\_TCELL\_DN | 100 | 23 | 4161 | 644 | mRpL17,Sptr,gammaTub37C,Cdk1,CCT3,CG14715,CG8004,CG5880,mRpS10,msps,CG5026,mad2,Pglym78,CCT6,CG17294,CG11164,mRpL18,Nurf-38,aay,CysRS,r,CG10418,Su(H) | | 2.924e-02 | -3.53 | eIF4E1 (eukaryotic translation initiation factor 4E1) | protein interactions | 45525 | 49 | 11 | 9204 | 1112 | Pink1,tyf,yps,cup,bam,nos,park,bru1,smg,bcd,exu | | 2.929e-02 | -3.53 | Hey (Hairy/E(spl)-related with YRPW motif) | protein interactions | 35764 | 43 | 10 | 9204 | 1112 | CG42565,Ssdp,CG9436,Hsp26,dx,cnc,Fur2,Sp7,RpL17,Marf1 | | 2.929e-02 | -3.53 | MED18 (Mediator complex subunit 18) | protein interactions | 31140 | 43 | 10 | 9204 | 1112 | CG17002,Larp4B,Ptip,MED27,Usp10,enc,MED4,MED20,ix,Ssdp | | 2.929e-02 | -3.53 | E(z) (Enhancer of zeste) | protein interactions | 39203 | 43 | 10 | 9204 | 1112 | nos,mxc,escl,Utx,Bin1,nej,esc,Rbf,piwi,Rbf2 | | 2.931e-02 | -3.53 | KAECH\_DAY8\_EFF\_VS\_MEMORY\_CD8\_TCELL\_UP | MSigDB lists | KAECH\_DAY8\_EFF\_VS\_MEMORY\_CD8\_TCELL\_UP | 79 | 19 | 4161 | 644 | Rpn12,CG11164,CG12375,cyc,Cbp80,Gdap2,Ts,CHOp24,CG42542,Prosalpha5,mRpS21,cl,Prosalpha3,Idh,mad2,Pen,ND-24,Cdk1,CG15525 | | 2.931e-02 | -3.53 | GO\_LYASE\_ACTIVITY | MSigDB lists | GO\_LYASE\_ACTIVITY | 79 | 19 | 4161 | 644 | Sply,Ppcdc,Got1,CG11134,Irp-1B,Pcd,Echs1,Eno,pr,Mtpalpha,r-l,Ac3,CG8199,Thiolase,Shmt,CG5285,mal,CG17691,Uba5 | | 2.931e-02 | -3.53 | GSE15215\_CD2\_POS\_VS\_NEG\_PDC\_DN | MSigDB lists | GSE15215\_CD2\_POS\_VS\_NEG\_PDC\_DN | 79 | 19 | 4161 | 644 | CG10344,SdhA,p38a,AIF,Eno,pen-2,APP-BP1,SERCA,Got1,Npc2a,stas,mtTFB1,Usp39,Snx6,UbcE2H,RhoGAP54D,Coq6,RpII140,RpS16 | | 2.934e-02 | -3.53 | meiosis I | biological process | GO:0007127 | 64 | 11 | 12345 | 1129 | CG31053,mus304,Klp10A,cort,tef,gammaTub37C,mtrm,Grip71,Xpc,c(2)M,mus81 | | 2.934e-02 | -3.53 | TwdlE (TweedleE) | protein interactions | 34075 | 74 | 15 | 9204 | 1112 | CG13630,Fer1HCH,plu,CG12018,nej,wal,Cdk5,PPYR1,CG9581,Atpalpha,CG13373,RpS2,mad2,Cul2,Imp | | 2.948e-02 | -3.52 | fly chr19E7-19E7\|1-65 cM | chromosome location | fly chr19E7-19E7\|1-65 cM | 13 | 3 | 25010 | 1343 | CG1532,snoRNA:Me28S-U3344b,CG32512 | | 2.948e-02 | -3.52 | fly chr19E7-19E7\|1-65 | chromosome location | fly chr19E7-19E7\|1-65 | 13 | 3 | 25010 | 1343 | CG1532,CG32512,snoRNA:Me28S-U3344b | | 2.948e-02 | -3.52 | Deadenylation-dependent mRNA decay | REACTOME pathways | R-DME-429914 | 41 | 11 | 3923 | 574 | CG11486,CG17768,Edc3,Rrp40,cN-IIIB,eIF4A,LSm3,CG8963,CG6610,CG10418,eIF4E1 | | 2.957e-02 | -3.52 | Beta-catenin independent WNT signaling | REACTOME pathways | R-DME-3858494 | 78 | 18 | 3923 | 574 | NFAT,Prosbeta3,Prosbeta6,Rpn3,Rpn12,Prosalpha5,RpS27A,Prosbeta4,Prosbeta5,Rpn2,Tak1,Prosalpha4,CanA-14F,RpL40,Rpt2,Pp2B-14D,Prosalpha3,CanB2 | | 2.962e-02 | -3.52 | regulation of cell differentiation | biological process | GO:0045595 | 428 | 51 | 12345 | 1129 | E2f1,swa,l(3)mbt,PGRP-LE,Cul1,mael,tkv,Stat92E,Pax,rl,ari-1,Cbl,lwr,Pi3K92E,G9a,lid,tws,ssh,MED27,cact,enc,bif,EcR,Cenp-C,Archease,vers,Ptpmeg,cnc,bru1,alph,COX4,Cka,exu,Imp,ago,hb,MED4,mts,dl,Seipin,raptor,mop,bun,cup,Su(H),spn-E,Sec23,csw,COX5A,MAGE,CycE | | 2.963e-02 | -3.52 | epithelial cell development | biological process | GO:0002064 | 361 | 44 | 12345 | 1129 | cact,jvl,EcR,fs(1)N,colt,Sec15,Tao,gus,Pax,e(y)1,rl,stai,Cbl,nej,Pi3K92E,E2f1,BicC,apt,toc,mael,Stat92E,SERCA,tkv,Med,spn-E,csw,Abi,CycE,r2d2,Cf2,Ptpmeg2,Pxt,fs(1)M3,armi,mop,bun,Rab5,cup,tor,Orc1,squ,Rbf,Lkb1,Cdk1 | | 2.967e-02 | -3.52 | amide biosynthetic process | biological process | GO:0043604 | 732 | 82 | 12345 | 1129 | RpL23A,eEF1beta,RpS13,mRpS11,RpL30,Ggt-1,GluRS-m,RpS2,RpS15,RpS29,eIF3i,RpLP1,Gclc,LysRS,tko,CG6094,RpS4,eIF3m,GatB,RpL31,mRpL18,CG7414,RpL34b,mRpS5,RpL29,RpL40,RpS7,mRpL36,Pym,RpL26,CG14817,RpS16,TrpRS-m,mRpL35,RpS27A,mRpL16,AsnS,RpL9,GAPsec,RpL34a,GatA,RpL37a,RpL13,RpS27,eIF4A,RpL11,RpS14b,RpS18,RpL23,RpS14a,Dhfr,eIF4E1,CG6178,RpS28b,RpL13A,SerRS-m,RpS24,AcCoAS,mRpS10,mRpS21,RpL7A,CG17691,RpS23,mRpL17,RpS30,mEFTu1,mEFTu2,CysRS,RpL17,RpS5b,CG8199,CG7009,RpL24,mRpL48,mRpL13,RpLP0,RpL38,RpL12,RpL10,TyrRS,mRpL10,mRpS23 | | 2.975e-02 | -3.51 | regulation of neurotransmitter levels | biological process | GO:0001505 | 140 | 20 | 12345 | 1129 | Gdi,Rep,Tomosyn,Fdh,Mctp,Shmt,Gabat,nej,Sec10,HDAC6,Dhfr,Sec15,Syx13,X11L,Rel,CanB2,NFAT,lqfR,Arf51F,Exo70 | | 2.979e-02 | -3.51 | GSE12963\_UNINF\_VS\_ENV\_AND\_NEF\_AND\_VPR\_DEFICIENT\_HIV1\_INF\_CD4\_TCELL\_DN | MSigDB lists | GSE12963\_UNINF\_VS\_ENV\_AND\_NEF\_AND\_VPR\_DEFICIENT\_HIV1\_INF\_CD4\_TCELL\_DN | 25 | 8 | 4161 | 644 | Gapdh1,Arts,UQCR-Q,Pen,Pglym78,r,Tom40,mRpS10 | | 2.979e-02 | -3.51 | REACTOME\_SIGNALING\_BY\_NOTCH1 | MSigDB lists | REACTOME\_SIGNALING\_BY\_NOTCH1 | 25 | 8 | 4161 | 644 | pen-2,RpL40,Su(H),Cul1,SkpA,HDAC6,RpS27A,CycC | | 2.979e-02 | -3.51 | GO\_PYRIDOXAL\_PHOSPHATE\_BINDING | MSigDB lists | GO\_PYRIDOXAL\_PHOSPHATE\_BINDING | 25 | 8 | 4161 | 644 | Got1,Sply,mal,Gabat,Pdxk,GlyP,CG1640,Shmt | | 2.979e-02 | -3.51 | MODULE\_273 | MSigDB lists | MODULE\_273 | 25 | 8 | 4161 | 644 | ND-SGDH,Nurf-38,ND-30,Su(H),ND-23,ND-51,ND-24,Tip60 | | 2.979e-02 | -3.51 | REACTOME\_PYRUVATE\_METABOLISM\_AND\_CITRIC\_ACID\_TCA\_CYCLE | MSigDB lists | REACTOME\_PYRUVATE\_METABOLISM\_AND\_CITRIC\_ACID\_TCA\_CYCLE | 25 | 8 | 4161 | 644 | CG5028,Pdk,SdhA,ScsbetaA,Idh3b,Scsalpha1,Idh,Nc73EF | | 2.979e-02 | -3.51 | GO\_ACTIVATION\_OF\_MAPK\_ACTIVITY | MSigDB lists | GO\_ACTIVATION\_OF\_MAPK\_ACTIVITY | 25 | 8 | 4161 | 644 | rl,Cdk1,p38a,RpL40,csw,Traf6,Sod1,RpS27A | | 2.996e-02 | -3.51 | outer membrane | cellular component | GO:0019867 | 48 | 9 | 11940 | 1105 | mino,Tom70,Debcl,Tspo,CG9393,Tom40,CG8004,Axs,Marf | | 2.996e-02 | -3.51 | organelle outer membrane | cellular component | GO:0031968 | 48 | 9 | 11940 | 1105 | CG9393,Tspo,Tom70,mino,Debcl,CG8004,Axs,Marf,Tom40 | | 2.998e-02 | -3.51 | GO\_REGULATION\_OF\_INNATE\_IMMUNE\_RESPONSE | MSigDB lists | GO\_REGULATION\_OF\_INNATE\_IMMUNE\_RESPONSE | 111 | 25 | 4161 | 644 | scramb1,RpL40,Ptp61F,SkpA,Prosbeta6,Ubc4,Rpn2,Traf6,Rpn12,EcR,Prosalpha4,Rpt2,csw,Prosbeta3,Rpn3,Cul1,Ect4,Uev1A,Pdk1,cact,Prosbeta5,Prosalpha5,RpS27A,Prosbeta4,Prosalpha3 | | 3.000e-02 | -3.51 | PURBEY\_TARGETS\_OF\_CTBP1\_NOT\_SATB1\_UP | MSigDB lists | PURBEY\_TARGETS\_OF\_CTBP1\_NOT\_SATB1\_UP | 74 | 18 | 4161 | 644 | CG3021,RpS30,Ptp10D,mino,CG11110,AGBE,Crag,Xpac,SkpA,Cyp1,l(1)G0004,Scsalpha1,Mitofilin,Arf51F,tkv,PDCD-5,SerRS-m,Pen | | 3.000e-02 | -3.51 | GSE21670\_UNTREATED\_VS\_TGFB\_IL6\_TREATED\_STAT3\_KO\_CD4\_TCELL\_DN | MSigDB lists | GSE21670\_UNTREATED\_VS\_TGFB\_IL6\_TREATED\_STAT3\_KO\_CD4\_TCELL\_DN | 74 | 18 | 4161 | 644 | RpS27,Sec22,sws,RpL11,Npc2a,Sply,pen-2,Moe,Pfdn2,AcCoAS,CG8188,Nadsyn,CG6144,RpL23,AnxB10,Droj2,Nprl2,CG40045 | | 3.000e-02 | -3.51 | GSE32986\_UNSTIM\_VS\_GMCSF\_AND\_CURDLAN\_HIGHDOSE\_STIM\_DC\_UP | MSigDB lists | GSE32986\_UNSTIM\_VS\_GMCSF\_AND\_CURDLAN\_HIGHDOSE\_STIM\_DC\_UP | 74 | 18 | 4161 | 644 | CG7488,Sec23,Got1,Arpc2,Pcd,Clc,MED20,LpR2,CG10425,anne,mRpL16,Pgm1,alphaTub84B,Rpn2,nmdyn-D7,RhoGAP54D,DppIII,Ranbp21 | | 3.000e-02 | -3.51 | GOTZMANN\_EPITHELIAL\_TO\_MESENCHYMAL\_TRANSITION\_DN | MSigDB lists | GOTZMANN\_EPITHELIAL\_TO\_MESENCHYMAL\_TRANSITION\_DN | 74 | 18 | 4161 | 644 | Tsp66E,Rpt2,UbcE2H,CG1847,Mcad,scu,Sod1,CG42542,Sec23,CG9143,Sec22,Exo70,p38a,cact,CCT3,Stip1,CG14715,Ntan1 | | 3.000e-02 | -3.51 | GSE27786\_LIN\_NEG\_VS\_ERYTHROBLAST\_UP | MSigDB lists | GSE27786\_LIN\_NEG\_VS\_ERYTHROBLAST\_UP | 74 | 18 | 4161 | 644 | ND-30,AIF,Tim8,Tom40,sws,CG11807,Gyg,Prosalpha3,scramb1,Aos1,Arl5,Etf-QO,Karybeta3,CG8204,r,CG11110,RpL13A,Sfxn2 | | 3.000e-02 | -3.51 | YAO\_TEMPORAL\_RESPONSE\_TO\_PROGESTERONE\_CLUSTER\_13 | MSigDB lists | YAO\_TEMPORAL\_RESPONSE\_TO\_PROGESTERONE\_CLUSTER\_13 | 74 | 18 | 4161 | 644 | Nmt,PDCD-5,Prosalpha5,LSm3,cl,ND-39,CG5026,msps,Chrac-14,Tim8,CG4390,mRpL13,ND-SGDH,ND-49,CG11899,mRpL18,Jafrac1,levy | | 3.000e-02 | -3.51 | GSE360\_DC\_VS\_MAC\_B\_MALAYI\_LOW\_DOSE\_UP | MSigDB lists | GSE360\_DC\_VS\_MAC\_B\_MALAYI\_LOW\_DOSE\_UP | 74 | 18 | 4161 | 644 | CSN6,CycE,wal,Pfk,Arpc4,capt,msps,Chrac-14,Vha36-1,TyrRS,AnxB10,LysRS,tho2,eIF3i,CysRS,RpS4,Nprl2,Tm1 | | 3.000e-02 | -3.51 | GSE15330\_HSC\_VS\_GRANULOCYTE\_MONOCYTE\_PROGENITOR\_IKAROS\_KO\_DN | MSigDB lists | GSE15330\_HSC\_VS\_GRANULOCYTE\_MONOCYTE\_PROGENITOR\_IKAROS\_KO\_DN | 74 | 18 | 4161 | 644 | Scsalpha1,CG30496,Ppcdc,Mitofilin,Jheh2,Hmg-2,Cdk1,CCT5,thoc6,Nca,CG1532,CG2201,nmdyn-D7,aay,scu,eIF4E1,mtTFB1,Fdh | | 3.002e-02 | -3.51 | sulfur compound biosynthetic process | biological process | GO:0044272 | 41 | 8 | 12345 | 1129 | oxt,CG11134,AcCoAS,Adi1,CG6178,Gclc,CG17691,CG8199 | | 3.009e-02 | -3.50 | intra-S DNA damage checkpoint | biological process | GO:0031573 | 8 | 3 | 12345 | 1129 | mus81,CG10336,mre11 | | 3.009e-02 | -3.50 | vitamin biosynthetic process | biological process | GO:0009110 | 8 | 3 | 12345 | 1129 | sgll,regucalcin,Pdxk | | 3.009e-02 | -3.50 | regulation of mitochondrial fission | biological process | GO:0090140 | 8 | 3 | 12345 | 1129 | Marf,Pink1,park | | 3.009e-02 | -3.50 | Wnt protein secretion | biological process | GO:0061355 | 8 | 3 | 12345 | 1129 | opm,Sec22,CHOp24 | | 3.009e-02 | -3.50 | mitochondrial ATP synthesis coupled proton transport | biological process | GO:0042776 | 8 | 3 | 12345 | 1129 | ATPsynbeta,ATPsynO,COX5B | | 3.009e-02 | -3.50 | positive regulation of neuron projection regeneration | biological process | GO:0070572 | 8 | 3 | 12345 | 1129 | Archease,Seipin,Imp | | 3.009e-02 | -3.50 | regulation of syncytial blastoderm mitotic cell cycle | biological process | GO:0007348 | 8 | 3 | 12345 | 1129 | mus304,grp,twe | | 3.009e-02 | -3.50 | peptidyl-histidine modification | biological process | GO:0018202 | 8 | 3 | 12345 | 1129 | CG2911,CG1578,Dph5 | | 3.009e-02 | -3.50 | ER-nucleus signaling pathway | biological process | GO:0006984 | 8 | 3 | 12345 | 1129 | PEK,SREBP,S2P | | 3.009e-02 | -3.50 | cytoplasmic mRNA processing body assembly | biological process | GO:0033962 | 8 | 3 | 12345 | 1129 | Edc3,CG17768,LSm3 | | 3.009e-02 | -3.50 | skeletal myofibril assembly | biological process | GO:0014866 | 8 | 3 | 12345 | 1129 | up,CanB2,SelR | | 3.009e-02 | -3.50 | centromere complex assembly | biological process | GO:0034508 | 8 | 3 | 12345 | 1129 | Chrac-14,Cenp-C,Hat1 | | 3.009e-02 | -3.50 | positive regulation of axon regeneration | biological process | GO:0048680 | 8 | 3 | 12345 | 1129 | Archease,Seipin,Imp | | 3.009e-02 | -3.50 | hemocyte proliferation | biological process | GO:0035172 | 8 | 3 | 12345 | 1129 | Stat92E,tub,mxc | | 3.010e-02 | -3.50 | GSE17721\_CPG\_VS\_GARDIQUIMOD\_1H\_BMDC\_DN | MSigDB lists | GSE17721\_CPG\_VS\_GARDIQUIMOD\_1H\_BMDC\_DN | 95 | 22 | 4161 | 644 | PpV,Stoml2,Rpb8,Nprl2,sgll,CG11070,Snx6,CG7718,CG9393,Traf6,Cse1,grsm,SdhA,f-cup,Fkbp59,ND-39,CG7789,CG30496,CG7488,CG14516,PpD3,Sec22 | | 3.014e-02 | -3.50 | GSE41867\_DAY15\_EFFECTOR\_VS\_DAY30\_EXHAUSTED\_CD8\_TCELL\_LCMV\_CLONE13\_UP | MSigDB lists | GSE41867\_DAY15\_EFFECTOR\_VS\_DAY30\_EXHAUSTED\_CD8\_TCELL\_LCMV\_CLONE13\_UP | 39 | 11 | 4161 | 644 | CG9471,Pi3K92E,ATP8A,Jafrac1,CG7840,Efr,mal,Pcd,Dph5,CG9393,Mctp | | 3.015e-02 | -3.50 | positive regulation of phosphorus metabolic process | biological process | GO:0010562 | 149 | 21 | 12345 | 1129 | ASPP,ben,Abi,fray,Mekk1,raptor,Scox,Mos,RpLP1,Tao,Tak1,Pgk,Parp16,CG8009,tor,Pdk1,Pi3K92E,PEK,Ankle2,Lkb1,Cka | | 3.015e-02 | -3.50 | positive regulation of phosphate metabolic process | biological process | GO:0045937 | 149 | 21 | 12345 | 1129 | fray,Abi,Mekk1,ben,ASPP,Scox,Mos,Tao,RpLP1,raptor,Pi3K92E,Tak1,Pgk,Parp16,CG8009,tor,Pdk1,Lkb1,Cka,PEK,Ankle2 | | 3.046e-02 | -3.49 | transcription factor binding | molecular function | GO:0008134 | 184 | 25 | 12176 | 1119 | Rpt2,dl,ix,E2f1,NFAT,Mybbp1A,Rbf2,rl,Utx,Bin1,Trf2,Sgf29,nej,EcR,Su(H),TfIIFalpha,asf1,Taf7,e(y)1,foxo,cact,Rbfox1,HmgD,Rbf,CG31875 | | 3.048e-02 | -3.49 | MAP kinase kinase kinase activity | molecular function | GO:0004709 | 8 | 3 | 12176 | 1119 | Tak1,Mekk1,Tao | | 3.048e-02 | -3.49 | oxidoreductase activity, acting on the CH-CH group of donors, oxygen as acceptor | molecular function | GO:0016634 | 8 | 3 | 12176 | 1119 | Ppox,CG4586,Acox57D-p | | 3.048e-02 | -3.49 | alcohol dehydrogenase (NAD) activity | molecular function | GO:0004022 | 8 | 3 | 12176 | 1119 | Adh,Fdh,CG4842 | | 3.056e-02 | -3.49 | ARM-like | interpro domains | IPR011989 | 97 | 17 | 11383 | 1233 | Ect4,wapl,CG2747,Plap,Pp2A-29B,Pen,msps,AP-1-2beta,Kap-alpha3,Kap3,raptor,CG14881,Cse1,CG1344,Ranbp21,Mo25,Rpn2 | | 3.061e-02 | -3.49 | GSE11864\_UNTREATED\_VS\_CSF1\_IFNG\_IN\_MAC\_DN | MSigDB lists | GSE11864\_UNTREATED\_VS\_CSF1\_IFNG\_IN\_MAC\_DN | 69 | 17 | 4161 | 644 | Scox,Pi4KIIalpha,Mitofilin,Rpn3,msps,CCT3,AIF,CG7997,TyrRS,CG3226,Prosbeta3,ACC,CG4511,Rpb8,GluRS-m,CG42542,Ubc4 | | 3.061e-02 | -3.49 | GSE21670\_STAT3\_KO\_VS\_WT\_CD4\_TCELL\_DN | MSigDB lists | GSE21670\_STAT3\_KO\_VS\_WT\_CD4\_TCELL\_DN | 69 | 17 | 4161 | 644 | Pgm1,TyrRS,Non1,slif,r,Karybeta3,Pi3K92E,CG11089,l(1)G0004,SREBP,Mitofilin,Pus7,Eno,CG10425,CCT5,CCT3,bys | | 3.061e-02 | -3.49 | GSE4748\_CTRL\_VS\_CYANOBACTERIUM\_LPSLIKE\_STIM\_DC\_3H\_DN | MSigDB lists | GSE4748\_CTRL\_VS\_CYANOBACTERIUM\_LPSLIKE\_STIM\_DC\_3H\_DN | 69 | 17 | 4161 | 644 | Gmer,CG11899,Rpn2,Tsp74F,CG10932,Jafrac1,Naa20A,Arts,Got1,Mitofilin,PDCD-5,gammaTub37C,Pfk,Coq7,Jafrac2,CG7997,CG9911 | | 3.061e-02 | -3.49 | GSE46606\_UNSTIM\_VS\_CD40L\_IL2\_IL5\_DAY1\_STIMULATED\_BCELL\_DN | MSigDB lists | GSE46606\_UNSTIM\_VS\_CD40L\_IL2\_IL5\_DAY1\_STIMULATED\_BCELL\_DN | 69 | 17 | 4161 | 644 | bys,r-l,Ac3,esc,Lsm12,Pfas,snz,escl,Naa20A,Cbp80,Roe1,Ranbp21,scu,ste24a,tun,CG2100,Psf3 | | 3.064e-02 | -3.49 | CG9986 (uncharacterized protein) | protein interactions | 43386 | 141 | 25 | 9204 | 1112 | CG11486,CG7328,CG46338,Fkbp12,CG42565,CG32267,LSm3,l(2)k01209,CG3376,Cenp-C,APC7,CG2678,CG1847,Sod3,Gdap2,EMC8-9,Rab5,boi,CG2061,Su(H),Scsalpha1,RpL17,Rtnl1,O-fut1,CG32512 | | 3.067e-02 | -3.48 | CG14868 (uncharacterized protein) | protein interactions | 41875 | 26 | 7 | 9204 | 1112 | vers,CG4882,CG6617,CycB,Shmt,Snx6,Prosbeta4 | | 3.067e-02 | -3.48 | Sam-S (S-adenosylmethionine Synthetase) | protein interactions | 48552 | 26 | 7 | 9204 | 1112 | foxo,CCT3,ago,Ssdp,CG4880,Pdk1,Bx | | 3.067e-02 | -3.48 | CG31343 (uncharacterized protein) | protein interactions | 326133 | 26 | 7 | 9204 | 1112 | GlyP,Adh,CG32473,Pgk,Pglym78,Scp1,Argk | | 3.067e-02 | -3.48 | CG15141 (uncharacterized protein) | protein interactions | 35044 | 26 | 7 | 9204 | 1112 | CG8435,RpS23,sol,CG6707,ATPsynB,Chd64,awd | | 3.067e-02 | -3.48 | ple (pale) | protein interactions | 38746 | 26 | 7 | 9204 | 1112 | RpS30,Plap,CG6175,hb,CG7824,CG3342,smg | | 3.067e-02 | -3.48 | CG9173 (uncharacterized protein) | protein interactions | 38165 | 26 | 7 | 9204 | 1112 | RpLP1,CG7414,RpL17,CG14464,RpS7,Neos,CG6707 | | 3.067e-02 | -3.48 | slam (slow as molasses) | protein interactions | 33890 | 26 | 7 | 9204 | 1112 | slam,RpL12,Cyt-c-p,osk,RpL7A,CG2678,RpS23 | | 3.068e-02 | -3.48 | inorganic cation transmembrane transport | biological process | GO:0098662 | 176 | 24 | 12345 | 1129 | ATPsynbeta,ATPsynE,ATPsynD,ZnT35C,fwe,CG12773,nrv1,Vha68-2,ATPsynO,ATPsyndelta,blw,ATPsynG,Fer1HCH,ATPsynB,Ork1,EMRE,zyd,ATPsynF,SERCA,Atpalpha,Vha36-1,Vha13,COX5B,ATPsynCF6 | | 3.074e-02 | -3.48 | STPHPHTASE | prints domains | PR00114 | 19 | 5 | 2185 | 212 | mts,PpV,PpD3,CanA-14F,Pp2B-14D | | 3.075e-02 | -3.48 | regulation of defense response to virus | biological process | GO:0050688 | 20 | 5 | 12345 | 1129 | G9a,Atg7,Stat92E,Rel,Traf6 | | 3.075e-02 | -3.48 | positive regulation of response to wounding | biological process | GO:1903036 | 20 | 5 | 12345 | 1129 | rl,Imp,Seipin,Archease,Mtl | | 3.075e-02 | -3.48 | protein acetylation | biological process | GO:0006473 | 89 | 14 | 12345 | 1129 | e(y)1,Hat1,YL-1,nej,pont,rept,Chrac-14,lid,d4,Spt20,Naa20A,Sgf29,Rcd5,Tip60 | | 3.076e-02 | -3.48 | Glycogen breakdown (glycogenolysis) | REACTOME pathways | R-DME-70221 | 13 | 5 | 3923 | 574 | Pgm1,CG8475,CG9485,Gyg,GlyP | | 3.076e-02 | -3.48 | mRNA decay by 5' to 3' exoribonuclease | REACTOME pathways | R-DME-430039 | 13 | 5 | 3923 | 574 | Edc3,LSm3,CG6610,CG10418,CG17768 | | 3.079e-02 | -3.48 | DNA helicase activity | molecular function | GO:0003678 | 41 | 8 | 12176 | 1119 | rept,pont,Psf3,armi,RecQ5,nbs,obe,Sld5 | | 3.079e-02 | -3.48 | acetyltransferase complex | cellular component | GO:1902493 | 88 | 14 | 11940 | 1105 | not,e(y)1,d4,pont,Rcd5,rept,Chrac-14,Naa20A,YL-1,Sgf11,Spt20,Sgf29,Tip60,nej | | 3.079e-02 | -3.48 | protein acetyltransferase complex | cellular component | GO:0031248 | 88 | 14 | 11940 | 1105 | Chrac-14,rept,Naa20A,YL-1,Sgf11,Sgf29,Spt20,Tip60,nej,d4,e(y)1,not,pont,Rcd5 | | 3.103e-02 | -3.47 | GSE10325\_BCELL\_VS\_MYELOID\_DN | MSigDB lists | GSE10325\_BCELL\_VS\_MYELOID\_DN | 44 | 12 | 4161 | 644 | Cyp6a22,CG3860,Rab32,isoQC,CG1275,Cyp6a19,shop,CG4572,Pdk,Pglym78,Aldh,CG9471 | | 3.103e-02 | -3.47 | GSE41867\_DAY6\_EFFECTOR\_VS\_DAY30\_MEMORY\_CD8\_TCELL\_LCMV\_ARMSTRONG\_DN | MSigDB lists | GSE41867\_DAY6\_EFFECTOR\_VS\_DAY30\_MEMORY\_CD8\_TCELL\_LCMV\_ARMSTRONG\_DN | 44 | 12 | 4161 | 644 | Nca,lqfR,Ntan1,Exd2,sbr,lid,CG7655,Mo25,RpS30,yip2,RpS27,Arf51F | | 3.103e-02 | -3.47 | GSE10325\_LUPUS\_CD4\_TCELL\_VS\_LUPUS\_MYELOID\_DN | MSigDB lists | GSE10325\_LUPUS\_CD4\_TCELL\_VS\_LUPUS\_MYELOID\_DN | 44 | 12 | 4161 | 644 | Rbcn-3A,CG7997,Irk1,Aldh,CG9471,Cyp6a19,cathD,CG9911,CG14516,Cyp6a22,isoQC,Rab32 | | 3.104e-02 | -3.47 | Arp2/3 protein complex | cellular component | GO:0005885 | 8 | 3 | 11940 | 1105 | Arpc4,Arpc1,Arpc2 | | 3.104e-02 | -3.47 | prefoldin complex | cellular component | GO:0016272 | 8 | 3 | 11940 | 1105 | Pfdn5,Pfdn2,Uxt | | 3.104e-02 | -3.47 | chaperonin-containing T-complex | cellular component | GO:0005832 | 8 | 3 | 11940 | 1105 | CCT3,CCT6,CCT5 | | 3.104e-02 | -3.47 | dihydrolipoyl dehydrogenase complex | cellular component | GO:0045240 | 8 | 3 | 11940 | 1105 | CG8199,CG17691,Nc73EF | | 3.106e-02 | -3.47 | Inhibition of replication initiation of damaged DNA by RB1/E2F1 | REACTOME pathways | R-DME-113501 | 9 | 4 | 3923 | 574 | Pp2A-29B,E2f1,Rbf,Rbf2 | | 3.111e-02 | -3.47 | GSE22886\_IGM\_MEMORY\_BCELL\_VS\_BM\_PLASMA\_CELL\_DN | MSigDB lists | GSE22886\_IGM\_MEMORY\_BCELL\_VS\_BM\_PLASMA\_CELL\_DN | 64 | 16 | 4161 | 644 | CG1827,CSN6,ERp60,CG15715,Pdxk,ND-B18,alpha-Man-Ia,CG14715,Cdk5,isoQC,CG10470,Pex10,CG2493,P5cr,opm,Uba5 | | 3.111e-02 | -3.47 | GSE17721\_PAM3CSK4\_VS\_CPG\_24H\_BMDC\_UP | MSigDB lists | GSE17721\_PAM3CSK4\_VS\_CPG\_24H\_BMDC\_UP | 64 | 16 | 4161 | 644 | eIF3m,Idh,mRpS21,Sccpdh1,CG15525,ERp60,eEF1beta,CG2493,Archease,CG33156,RpLP0,CG10470,slif,Irk1,CG42542,GluRS-m | | 3.111e-02 | -3.47 | GSE2128\_C57BL6\_VS\_NOD\_THYMOCYTE\_UP | MSigDB lists | GSE2128\_C57BL6\_VS\_NOD\_THYMOCYTE\_UP | 64 | 16 | 4161 | 644 | Karybeta3,Gdap2,CG10418,CkIIalpha,Cse1,aay,Eno,Caf1-105,CCT6,Uch-L5,kcc,Chrac-14,Pen,yip2,CycE,Clc | | 3.111e-02 | -3.47 | COUP\_DR1\_Q6 | MSigDB lists | COUP\_DR1\_Q6 | 64 | 16 | 4161 | 644 | Etf-QO,Rpb8,Stoml2,CG12713,Ubc4,Arl6IP1,AP-1-2beta,Phb2,GCS2beta,CG5510,Bin1,ND-75,eEF1beta,Arf51F,Sec22,Smox | | 3.111e-02 | -3.47 | GSE41087\_WT\_VS\_FOXP3\_MUT\_ANTI\_CD3\_CD28\_STIM\_CD4\_TCELL\_DN | MSigDB lists | GSE41087\_WT\_VS\_FOXP3\_MUT\_ANTI\_CD3\_CD28\_STIM\_CD4\_TCELL\_DN | 64 | 16 | 4161 | 644 | AIF,CG9471,CG8004,Prosbeta5,UQCR-Q,Nca,Got1,Gyg,SREBP,Axs,Clc,Irp-1B,Acph-1,CG11807,ACC,Vha36-1 | | 3.111e-02 | -3.47 | GSE32034\_UNTREATED\_VS\_ROSIGLIZATONE\_TREATED\_LY6C\_HIGH\_MONOCYTE\_DN | MSigDB lists | GSE32034\_UNTREATED\_VS\_ROSIGLIZATONE\_TREATED\_LY6C\_HIGH\_MONOCYTE\_DN | 64 | 16 | 4161 | 644 | RpS7,Arl5,tko,RpL30,CG3226,CG1890,RpL13A,RpS18,CG9911,CG11134,Sptr,Scsalpha1,spag,PDCD-5,RpS23,RpL24 | | 3.114e-02 | -3.47 | Deubiquitination | REACTOME pathways | R-DME-5688426 | 164 | 33 | 3923 | 574 | twe,Prosbeta5,Rpn2,Prosbeta4,RpS27A,Med,Prosalpha5,Stam,e(y)1,Traf6,not,pont,park,scny,Prosalpha3,RpL40,Rpt2,Uch-L5,Prosalpha4,Usp10,Rcd5,Usp16-45,foxo,trbd,Rpn12,Rpn3,Prosbeta6,Uch,Rpn13,nej,Atg6,faf,Prosbeta3 | | 3.117e-02 | -3.47 | GSE15930\_NAIVE\_VS\_24H\_IN\_VITRO\_STIM\_CD8\_TCELL\_DN | MSigDB lists | GSE15930\_NAIVE\_VS\_24H\_IN\_VITRO\_STIM\_CD8\_TCELL\_DN | 106 | 24 | 4161 | 644 | aay,Nurf-38,CG11164,mRpL18,CG12018,CG17294,CG10418,Su(H),CysRS,r,gammaTub37C,Sptr,mRpL17,mad2,Pglym78,CG5026,msps,CCT6,CCT3,Cdk1,CG5880,mRpS10,CG8004,CG14715 | | 3.122e-02 | -3.47 | cell communication | biological process | GO:0007154 | 1239 | 132 | 12345 | 1129 | pen-2,Stat92E,Hr78,p38a,Npc2a,Exo70,lt,CG7115,dnc,Iml1,ben,Rab10,SNF4Agamma,Tao,G9a,tor,Tak1,pygo,Pgk,grp,CG14408,Nup358,fray,S2P,CanB2,Atf6,CG6927,Su(H),HDAC6,Cdk5,Edem1,nej,spen,Nprl2,rl,tkv,Pka-R1,PGRP-LE,Gdi,Rep,bif,CG13933,EcR,fs(1)N,Hnf4,Debcl,X11L,Ac3,Syx13,gus,Sec15,nbs,Wnt10,Traf6,Pld,Rbf,Mo25,mAChR-A,Arf51F,Sec22,trk,Chd64,Myd88,Pka-R2,Rab5,CG17698,raptor,Cbl,CG17272,Nurf-38,Mitf,Rab32,Tomosyn,dco,PEK,apt,fz4,Mekk1,zyd,Smox,Uev1A,Hsc70-3,SREBP,Jafrac2,CG9391,sip3,RhoGDI,scramb1,lqfR,Atg7,Sap47,csw,Atg6,CkIIalpha,opm,Mtl,Pi3K92E,Sec10,Pp2B-14D,boi,Calx,Fkbp39,SERCA,CG3894,bcd,tub,crq,CHOp24,cact,Liprin-beta,spz,Pink1,foxo,dl,not,Pdk1,Parp16,Tlk,Cka,O-fut1,Atpalpha,Lkb1,nAChRalpha4,RhoGAP54D,sbb,REPTOR,Med,fs(1)M3,bun,Rel,disp,CanA-14F | | 3.126e-02 | -3.47 | Fatty acid metabolism | KEGG pathways | dme01212 | 44 | 12 | 3198 | 496 | Mtpalpha,whd,Thiolase,CG10932,CG4586,CG4860,Mcad,ACC,yip2,Acox57D-p,Desat1,Echs1 | | 3.126e-02 | -3.47 | Fatty acid metabolism | KEGG pathways | ko01212 | 44 | 12 | 3198 | 496 | CG4860,CG10932,Thiolase,CG4586,whd,Mtpalpha,Echs1,Desat1,Acox57D-p,Mcad,ACC,yip2 | | 3.127e-02 | -3.47 | Major pathway of rRNA processing in the nucleolus and cytosol | REACTOME pathways | R-DME-6791226 | 112 | 24 | 3923 | 574 | RpS14b,Rrp40,RpS30,RpS23,RpS29,RpS27,RpS7,RpS28b,dco,RpS24,RpS18,RpS27A,RpS16,bys,RpS14a,CG12301,RpS4,RpS5b,RpS15,RpS2,CG7728,l(1)G0004,RpS13,Mat89Ba | | 3.127e-02 | -3.47 | rRNA processing in the nucleus and cytosol | REACTOME pathways | R-DME-8868773 | 112 | 24 | 3923 | 574 | RpS29,RpS27,RpS23,RpS30,Rrp40,RpS14b,RpS16,RpS18,RpS24,RpS27A,dco,RpS28b,RpS7,RpS15,RpS5b,RpS4,CG12301,RpS14a,bys,Mat89Ba,RpS13,RpS2,CG7728,l(1)G0004 | | 3.127e-02 | -3.47 | rRNA processing | REACTOME pathways | R-DME-72312 | 112 | 24 | 3923 | 574 | RpS14b,Rrp40,RpS23,RpS30,RpS29,RpS27,RpS7,RpS28b,dco,RpS24,RpS27A,RpS18,RpS16,bys,RpS14a,CG12301,RpS4,RpS5b,RpS15,RpS2,CG7728,l(1)G0004,RpS13,Mat89Ba | | 3.133e-02 | -3.46 | ubiquitin conjugating enzyme binding | molecular function | GO:0031624 | 20 | 5 | 12176 | 1119 | park,ari-1,Aos1,Uev1A,ben | | 3.145e-02 | -3.46 | GSE21546\_SAP1A\_KO\_VS\_SAP1A\_KO\_AND\_ELK1\_KO\_ANTI\_CD3\_STIM\_DP\_THYMOCYTES\_UP | MSigDB lists | GSE21546\_SAP1A\_KO\_VS\_SAP1A\_KO\_AND\_ELK1\_KO\_ANTI\_CD3\_STIM\_DP\_THYMOCYTES\_UP | 59 | 15 | 4161 | 644 | CG2493,Sfxn2,CG11858,Sod1,mino,nrv1,CG14407,anne,SdhA,CG12713,Ublcp1,Jafrac1,Grx1,ATP8A,kcc | | 3.145e-02 | -3.46 | GO\_CELLULAR\_RESPONSE\_TO\_EXTERNAL\_STIMULUS | MSigDB lists | GO\_CELLULAR\_RESPONSE\_TO\_EXTERNAL\_STIMULUS | 59 | 15 | 4161 | 644 | SREBP,Hsc70-3,raptor,Atg6,CycE,Prip,grp,CG31715,Irk1,AIF,LpR2,Nprl2,Grx1,Iml1,Atg7 | | 3.148e-02 | -3.46 | BROWNE\_HCMV\_INFECTION\_24HR\_UP | MSigDB lists | BROWNE\_HCMV\_INFECTION\_24HR\_UP | 49 | 13 | 4161 | 644 | cact,Roe1,CG9485,Tom40,CanB2,Rab5,ATP8A,Rpn3,Fkbp59,Sptr,Gabat,nrv1,l(3)73Ah | | 3.148e-02 | -3.46 | GO\_CELLULAR\_AMINO\_ACID\_BIOSYNTHETIC\_PROCESS | MSigDB lists | GO\_CELLULAR\_AMINO\_ACID\_BIOSYNTHETIC\_PROCESS | 49 | 13 | 4161 | 644 | Gabat,aay,Dhfr,P5cr,CG11899,P5CDh1,Got1,CG11134,CG11089,Adi1,Shmt,CG1640,r | | 3.148e-02 | -3.46 | GSE30971\_CTRL\_VS\_LPS\_STIM\_MACROPHAGE\_WBP7\_KO\_4H\_DN | MSigDB lists | GSE30971\_CTRL\_VS\_LPS\_STIM\_MACROPHAGE\_WBP7\_KO\_4H\_DN | 49 | 13 | 4161 | 644 | CG7550,alpha-Man-Ia,CG7997,Nca,disp,bru1,Su(H),Hat1,Gapdh1,CG7840,CG2493,Jheh2,yip2 | | 3.148e-02 | -3.46 | GSE11924\_TFH\_VS\_TH17\_CD4\_TCELL\_UP | MSigDB lists | GSE11924\_TFH\_VS\_TH17\_CD4\_TCELL\_UP | 49 | 13 | 4161 | 644 | Iml1,shop,Moe,CG11710,fray,Grip75,adp,Ppox,MED24,CG15111,CG9143,csw,UbcE2H | | 3.156e-02 | -3.46 | ERAD pathway | biological process | GO:0036503 | 34 | 7 | 12345 | 1129 | Edem1,CG32640,CG6766,sip3,CG11070,Hsc70-3,CG8974 | | 3.156e-02 | -3.46 | salivary gland histolysis | biological process | GO:0035070 | 34 | 7 | 12345 | 1129 | DCAF12,Trf2,Utx,MED24,EcR,Idh3b,rept | | 3.160e-02 | -3.45 | GSE360\_CTRL\_VS\_T\_GONDII\_MAC\_DN | MSigDB lists | GSE360\_CTRL\_VS\_T\_GONDII\_MAC\_DN | 54 | 14 | 4161 | 644 | CG4407,l(1)G0004,GlyP,pont,CCT5,Sec22,Clc,stas,CycE,l(3)73Ah,nrv1,Pgm1,SREBP,CG12018 | | 3.160e-02 | -3.45 | GSE5679\_PPARG\_LIGAND\_ROSIGLITAZONE\_VS\_ROSIGLITAZONE\_AND\_RARA\_AGONIST\_AM580\_TREATED\_DC\_UP | MSigDB lists | GSE5679\_PPARG\_LIGAND\_ROSIGLITAZONE\_VS\_ROSIGLITAZONE\_AND\_RARA\_AGONIST\_AM580\_TREATED\_DC\_UP | 54 | 14 | 4161 | 644 | CG1640,CG7997,CG7322,lwr,PrBP,Pgk,Uev1A,Liprin-beta,ERp60,Pglym78,cl,csw,CG1532,shv | | 3.160e-02 | -3.45 | GSE26156\_DOUBLE\_POSITIVE\_VS\_CD4\_SINGLE\_POSITIVE\_THYMOCYTE\_UP | MSigDB lists | GSE26156\_DOUBLE\_POSITIVE\_VS\_CD4\_SINGLE\_POSITIVE\_THYMOCYTE\_UP | 54 | 14 | 4161 | 644 | RpL38,RpL13,RpL37a,RpL11,Rbf,eIF4A,RpL30,trsn,RpS28b,RpS13,CG5116,RpS23,Pfdn5,RpS27A | | 3.160e-02 | -3.45 | KEGG\_OOCYTE\_MEIOSIS | MSigDB lists | KEGG\_OOCYTE\_MEIOSIS | 54 | 14 | 4161 | 644 | Cul1,mad2,CycB,CanB2,Cdk1,Ac3,rl,SkpA,APC7,CycE,Pp2A-29B,Pp2B-14D,CanA-14F,mts | | 3.164e-02 | -3.45 | positive regulation of cellular protein catabolic process | biological process | GO:1903364 | 49 | 9 | 12345 | 1129 | Rpt2,Cul1,SkpA,cort,dco,Pomp,ari-1,park,Ubc6 | | 3.164e-02 | -3.45 | peptidyl-lysine methylation | biological process | GO:0018022 | 49 | 9 | 12345 | 1129 | Utx,CG9643,G9a,escl,nej,esc,trr,Ptip,egg | | 3.164e-02 | -3.45 | regulation of neuroblast proliferation | biological process | GO:1902692 | 49 | 9 | 12345 | 1129 | COX5A,mts,MAGE,EcR,MED4,bun,COX4,MED27,tws | | 3.168e-02 | -3.45 | negative regulation of developmental process | biological process | GO:0051093 | 222 | 29 | 12345 | 1129 | foxo,lwr,MED4,mts,nos,tor,park,ago,COX4,Stat92E,tkv,Ptpmeg,alph,PGRP-LE,SkpA,COX5A,bif,EcR,MAGE,Med,cact,Tao,bun,Su(H),MED27,Pink1,raptor,lid,tws | | 3.175e-02 | -3.45 | PAZ\_dom | interpro domains | IPR003100 | 7 | 3 | 11383 | 1233 | AGO3,piwi,Dcr-1 | | 3.175e-02 | -3.45 | Cullin\_homology | interpro domains | IPR016158 | 7 | 3 | 11383 | 1233 | Cul5,Cul1,Cul2 | | 3.175e-02 | -3.45 | ATPase\_F1/V1/A1\_a/bsu\_N | interpro domains | IPR004100 | 7 | 3 | 11383 | 1233 | Vha68-2,ATPsynbeta,blw | | 3.175e-02 | -3.45 | ATPase\_a/bsu\_AS | interpro domains | IPR020003 | 7 | 3 | 11383 | 1233 | blw,ATPsynbeta,Vha68-2 | | 3.175e-02 | -3.45 | IsoPropMal-DH-like\_dom | interpro domains | IPR024084 | 7 | 3 | 11383 | 1233 | Idh3b,Idh,CG5028 | | 3.175e-02 | -3.45 | Cyclin | interpro domains | IPR039361 | 7 | 3 | 11383 | 1233 | CycJ,CycE,CycB | | 3.175e-02 | -3.45 | Cullin\_N | interpro domains | IPR001373 | 7 | 3 | 11383 | 1233 | Cul5,Cul2,Cul1 | | 3.175e-02 | -3.45 | Ribosomal\_L7Ae/L30e/S12e/Gad45 | interpro domains | IPR004038 | 7 | 3 | 11383 | 1233 | RpL7A,RpL30,NHP2 | | 3.175e-02 | -3.45 | ATPase\_F1/V1/A1\_a/bsu\_nucl-bd | interpro domains | IPR000194 | 7 | 3 | 11383 | 1233 | blw,Vha68-2,ATPsynbeta | | 3.175e-02 | -3.45 | Cullin\_homology\_sf | interpro domains | IPR036317 | 7 | 3 | 11383 | 1233 | Cul5,Cul1,Cul2 | | 3.177e-02 | -3.45 | Tyr\_Pase\_AS | interpro domains | IPR016130 | 29 | 7 | 11383 | 1233 | ssh,Ptp69D,Ptpmeg2,Ptp10D,Ptp61F,Ptpmeg,csw | | 3.182e-02 | -3.45 | MORF\_SKP1A | MSigDB lists | MORF\_SKP1A | 117 | 26 | 4161 | 644 | Prosbeta4,LSm3,Clc,COX5A,Pp2A-29B,wal,ND-75,Klp10A,ND-24,Hat1,Thiolase,ND-30,Chrac-14,Pen,Rpt2,Sod1,Rpn12,LysRS,CG1890,CHOp24,levy,Prosbeta6,PpV,SkpA,CG11070,ND-51 | | 3.188e-02 | -3.45 | translation regulator activity, nucleic acid binding | molecular function | GO:0090079 | 89 | 14 | 12176 | 1119 | CG6094,CG8963,mEFTu2,eIF3m,eIF3i,Shmt,eIF4E1,heph,bam,mEFTu1,eEF1beta,eIF4A,smg,CG7414 | | 3.199e-02 | -3.44 | USP\_dom | interpro domains | IPR028889 | 23 | 6 | 11383 | 1233 | Usp39,scny,not,faf,Usp10,Usp16-45 | | 3.212e-02 | -3.44 | GSE12392\_WT\_VS\_IFNB\_KO\_CD8A\_POS\_SPLEEN\_DC\_UP | MSigDB lists | GSE12392\_WT\_VS\_IFNB\_KO\_CD8A\_POS\_SPLEEN\_DC\_UP | 85 | 20 | 4161 | 644 | Phb2,Sod1,not,CG4511,CysRS,CHOp24,Mekk1,Pi4KIIalpha,Cyp6a22,CG33181,mRpS21,ND-39,Scsalpha1,Thiolase,Cyp6a19,f-cup,crq,AcCoAS,CCT5,CG14715 | | 3.214e-02 | -3.44 | positive regulation of cell cycle phase transition | biological process | GO:1901989 | 27 | 6 | 12345 | 1129 | twe,CycE,Su(H),bru1,Sld5,APC7 | | 3.217e-02 | -3.44 | lipid droplet | cellular component | GO:0005811 | 20 | 5 | 11940 | 1105 | Sccpdh1,CG9265,dob,CG9186,Jabba | | 3.219e-02 | -3.44 | lipid homeostasis | biological process | GO:0055088 | 73 | 12 | 12345 | 1129 | dob,SNF4Agamma,Acox57D-p,slif,Npc2a,Pi3K92E,EcR,SREBP,Desat1,CG4586,CG8112,Atf3 | | 3.219e-02 | -3.44 | phospholipid biosynthetic process | biological process | GO:0008654 | 73 | 12 | 12345 | 1129 | PIG-P,Pi3K92E,Pld,PIG-L,CG7718,CG2201,mino,PIG-C,Agpat2,CG8311,CG33977,PIG-H | | 3.222e-02 | -3.44 | regulation of signaling | biological process | GO:0023051 | 855 | 94 | 12345 | 1129 | shv,Jafrac2,mts,tor,Ubc6,Tak1,Usp10,scny,ea,Ssdp,STUB1,Sap47,Su(H),CkIIalpha,Ptp10D,lwr,Cbl,Tsp86D,park,Sod1,Cul1,stas,Mctp,Aos1,Tomosyn,EndoGI,dco,E2f1,PEK,apt,Mekk1,Bap170,Ect4,PpV,dnc,NFAT,Iml1,ben,stet,Mos,CG17746,Tao,Snx6,gd,ATPsynD,tws,Cul2,Pura,foxo,Traf6,crok,Rbf,Pdk1,Imp,Dd,Cka,O-fut1,piwi,Lkb1,Ptpmeg,RhoGAP54D,alph,CycE,Crag,Nlg1,CG10188,mop,Rel,Herc4,Pp2A-29B,Myd88,CG4853,disp,rept,Pi3K92E,nej,pont,spen,Nprl2,rl,insv,RhoGAP5A,tkv,SkpA,chrb,Ptp61F,cact,Tip60,trbd,pn,snk,Pink1,spz,dx | | 3.222e-02 | -3.44 | regulation of cell communication | biological process | GO:0010646 | 855 | 94 | 12345 | 1129 | Cka,Dd,Lkb1,O-fut1,piwi,Ptpmeg,RhoGAP54D,alph,Traf6,foxo,Pura,Rbf,crok,Pdk1,Imp,Pp2A-29B,Herc4,Rel,mop,Myd88,disp,CG4853,CycE,Nlg1,Crag,CG10188,RhoGAP5A,insv,tkv,SkpA,rept,pont,nej,Pi3K92E,spen,rl,Nprl2,Tip60,trbd,pn,Pink1,dx,spz,snk,chrb,Ptp61F,cact,scny,ea,Ssdp,shv,Jafrac2,mts,Ubc6,tor,Usp10,Tak1,Su(H),CkIIalpha,Ptp10D,STUB1,Sap47,Mctp,Aos1,Cul1,stas,Tomosyn,EndoGI,E2f1,dco,apt,PEK,lwr,Cbl,Tsp86D,park,Sod1,Tao,CG17746,Mos,gd,Snx6,ATPsynD,Cul2,tws,Bap170,Mekk1,Iml1,NFAT,PpV,dnc,Ect4,stet,ben | | 3.224e-02 | -3.43 | DNA Double-Strand Break Repair | REACTOME pathways | R-DME-5693532 | 22 | 7 | 3923 | 574 | nbs,mus81,mre11,grp,Pen,Tip60,Snm1 | | 3.225e-02 | -3.43 | PAZ | pfam domains | PF02170 | 7 | 3 | 10480 | 1142 | AGO3,piwi,Dcr-1 | | 3.225e-02 | -3.43 | ATP-synt\_ab | pfam domains | PF00006 | 7 | 3 | 10480 | 1142 | blw,Vha68-2,ATPsynbeta | | 3.225e-02 | -3.43 | Cullin | pfam domains | PF00888 | 7 | 3 | 10480 | 1142 | Cul1,Cul2,Cul5 | | 3.225e-02 | -3.43 | ATP-synt\_ab\_N | pfam domains | PF02874 | 7 | 3 | 10480 | 1142 | blw,Vha68-2,ATPsynbeta | | 3.225e-02 | -3.43 | Iso\_dh | pfam domains | PF00180 | 7 | 3 | 10480 | 1142 | Idh,CG5028,Idh3b | | 3.225e-02 | -3.43 | Cyclin\_C | pfam domains | PF02984 | 7 | 3 | 10480 | 1142 | CycB,CycJ,CycE | | 3.225e-02 | -3.43 | Ribosomal\_L7Ae | pfam domains | PF01248 | 7 | 3 | 10480 | 1142 | RpL7A,NHP2,RpL30 | | 3.231e-02 | -3.43 | phospholipid binding | molecular function | GO:0005543 | 106 | 16 | 12176 | 1119 | CG8726,Exo70,CG11807,Snx6,CG5734,Hip1,CG3077,Myd88,Pld,Nup62,AnxB10,Moe,snz,Pex23,sau,crok | | 3.231e-02 | -3.43 | negative regulation of protein modification process | biological process | GO:0031400 | 98 | 15 | 12345 | 1129 | PpV,sws,park,Cbl,Ptp61F,CycE,mts,dco,CG14408,ATPsynD,alph,Utx,SkpA,CG17746,CG30496 | | 3.232e-02 | -3.43 | regulation of signal transduction | biological process | GO:0009966 | 745 | 83 | 12345 | 1129 | Pdk1,Imp,foxo,Pura,Traf6,Rbf,Ptpmeg,RhoGAP54D,alph,Dd,Cka,piwi,O-fut1,Lkb1,Crag,CG10188,CycE,disp,CG4853,mop,Pp2A-29B,Herc4,Rel,Myd88,spen,Nprl2,rept,Pi3K92E,pont,nej,SkpA,RhoGAP5A,insv,tkv,cact,chrb,Ptp61F,snk,dx,spz,pn,trbd,tor,Ubc6,Tak1,Usp10,shv,Jafrac2,mts,Ssdp,scny,ea,STUB1,CkIIalpha,Ptp10D,Su(H),park,lwr,Cbl,Tsp86D,dco,E2f1,PEK,apt,Cul1,Aos1,EndoGI,Ect4,dnc,PpV,Iml1,NFAT,ben,stet,Mekk1,Bap170,ATPsynD,tws,Cul2,Mos,CG17746,Tao,gd | | 3.238e-02 | -3.43 | regulation of proteolysis involved in cellular protein catabolic process | biological process | GO:1903050 | 57 | 10 | 12345 | 1129 | Rpt2,Cul1,SkpA,CkIIalpha,cort,Pomp,dco,ari-1,Ubc6,park | | 3.239e-02 | -3.43 | fly chr86C7-86C7\|3-50 cM | chromosome location | fly chr86C7-86C7\|3-50 cM | 23 | 4 | 25010 | 1343 | Tsp86D,SelR,CG4511,Fdh | | 3.239e-02 | -3.43 | fly chr86C7-86C7\|3-50 | chromosome location | fly chr86C7-86C7\|3-50 | 23 | 4 | 25010 | 1343 | Fdh,CG4511,Tsp86D,SelR | | 3.254e-02 | -3.43 | cellular macromolecule biosynthetic process | biological process | GO:0034645 | 1200 | 128 | 12345 | 1129 | RpL12,RpL38,RpS5b,RpLP0,RfC38,mRpL48,Efr,mEFTu2,CysRS,Nurf-38,RpL7A,fs(1)Ya,CG33977,RpL13A,RpS24,Atg7,RpL23,RpS18,RpS14b,RpS14a,Sld5,MED4,ago,eIF4A,Orc1,RpL37a,oxt,Rga,RpL34a,PIG-H,mRpL35,RpS7,mRpL36,Pym,TrpRS-m,Nmt,PIG-L,e(y)1,mtTFB1,CG10344,mRpS5,Trf2,GalT1,Gyg,RpL29,RpL31,CycE,PIG-P,LysRS,RpS4,CG6094,Tfb4,Top1,eIF3i,RpS15,O-fut1,RpL23A,Ggt-1,RpL10,TyrRS,mRpL10,mRpS23,CG7009,mRpL13,RpL24,mRpL17,RpS30,RecQ5,DNApol-epsilon255,CG12018,RpL17,mEFTu1,MED24,RpS23,E2f1,SerRS-m,RpS28b,mRpS10,mRpS21,CG8314,RpL11,eIF4E1,AGBE,Rpb8,RpS27,pygo,GatA,RpL13,mus205,PIG-C,RpL9,GAPsec,Pgm1,RpS27A,mRpL16,Orc6,RpL26,CG14817,RpS16,CG9485,spen,SuUR,CG7414,RpL34b,RpL40,Taf7,SkpA,TfIIFalpha,CG7840,GatB,eIF3m,mRpL18,tko,RpLP1,Cdk7,Rbf,CG10336,CG5880,RpS2,RpS29,Sgf29,mRpS11,eEF1beta,RpS13,pps,GluRS-m,RpII140,RpL30 | | 3.256e-02 | -3.42 | GRADE\_COLON\_CANCER\_UP | MSigDB lists | GRADE\_COLON\_CANCER\_UP | 351 | 67 | 4161 | 644 | Gmer,Tsp66E,Rpn12,mRpL13,RpS13,eIF3i,levy,O-fut1,CG11089,RhoGDI,Arpc2,RpL9,Bin1,mEFTu1,CG12018,Pp2B-14D,CG1890,SelD,Adi1,Cyp1,CG8726,CG7728,CanA-14F,Pp2A-29B,PDCD-5,RpL24,CG7322,Prosbeta5,Gapdh1,Cdk7,Prosbeta3,AMPdeam,Nurf-38,CG8188,CG42542,CG5028,ND-51,Jafrac1,CG8372,Prosalpha5,Utx,Tom40,Nca,Fbw5,RpS24,dco,GCS2beta,RpL38,RpS7,heph,PrBP,RpL40,Tspo,Tomosyn,Pmi,RpL11,sws,Lsm12,RpL26,RpS27,Pdxk,Eno,CCT5,Pfdn2,anne,CG14715,CG8314 | | 3.264e-02 | -3.42 | Ribosomal\_S10\_dom | interpro domains | IPR027486 | 3 | 2 | 11383 | 1233 | mRpL48,mRpS10 | | 3.264e-02 | -3.42 | Beta\_hexosaminidase\_sua/sub | interpro domains | IPR025705 | 3 | 2 | 11383 | 1233 | Hexo1,fdl | | 3.264e-02 | -3.42 | PurM-like\_N | interpro domains | IPR016188 | 3 | 2 | 11383 | 1233 | Pfas,SelD | | 3.264e-02 | -3.42 | Longin\_dom | interpro domains | IPR010908 | 3 | 2 | 11383 | 1233 | Vamp7,Sec22 | | 3.264e-02 | -3.42 | PLD-like\_dom | interpro domains | IPR025202 | 3 | 2 | 11383 | 1233 | Pld,CG7718 | | 3.264e-02 | -3.42 | Arm\_3 | interpro domains | IPR032413 | 3 | 2 | 11383 | 1233 | Kap-alpha3,Pen | | 3.264e-02 | -3.42 | SPOC\_C | interpro domains | IPR012921 | 3 | 2 | 11383 | 1233 | pps,spen | | 3.264e-02 | -3.42 | CSD\_1 | interpro domains | IPR019844 | 3 | 2 | 11383 | 1233 | yps,Unr | | 3.264e-02 | -3.42 | Ribosomal\_L13\_sf | interpro domains | IPR036899 | 3 | 2 | 11383 | 1233 | mRpL13,RpL13A | | 3.264e-02 | -3.42 | Importin-a\_IBB | interpro domains | IPR002652 | 3 | 2 | 11383 | 1233 | Pen,Kap-alpha3 | | 3.264e-02 | -3.42 | Pex\_N | interpro domains | IPR006845 | 3 | 2 | 11383 | 1233 | Pex10,Pex12 | | 3.264e-02 | -3.42 | Ribosomal\_L10P | interpro domains | IPR001790 | 3 | 2 | 11383 | 1233 | mRpL10,RpLP0 | | 3.264e-02 | -3.42 | NFkB/Dor | interpro domains | IPR000451 | 3 | 2 | 11383 | 1233 | Rel,dl | | 3.264e-02 | -3.42 | HEX\_eukaryotic\_N | interpro domains | IPR029019 | 3 | 2 | 11383 | 1233 | fdl,Hexo1 | | 3.264e-02 | -3.42 | GINS\_A | interpro domains | IPR021151 | 3 | 2 | 11383 | 1233 | Psf3,Sld5 | | 3.264e-02 | -3.42 | IPT\_NFkappaB | interpro domains | IPR033926 | 3 | 2 | 11383 | 1233 | Rel,dl | | 3.264e-02 | -3.42 | Glyco\_hydro\_20\_cat | interpro domains | IPR015883 | 3 | 2 | 11383 | 1233 | Hexo1,fdl | | 3.264e-02 | -3.42 | Uncharacterised\_SERF\_N | interpro domains | IPR007513 | 3 | 2 | 11383 | 1233 | CG17931,CG15715 | | 3.264e-02 | -3.42 | Sam37/metaxin\_N | interpro domains | IPR019564 | 3 | 2 | 11383 | 1233 | CG9393,CG8004 | | 3.264e-02 | -3.42 | CIR\_N\_dom | interpro domains | IPR019339 | 3 | 2 | 11383 | 1233 | CG12818,Cwc25 | | 3.264e-02 | -3.42 | Importin-a\_IBB\_sf | interpro domains | IPR036975 | 3 | 2 | 11383 | 1233 | Kap-alpha3,Pen | | 3.264e-02 | -3.42 | Ribosomal\_L7Ae/L8/Nhp2 | interpro domains | IPR018492 | 3 | 2 | 11383 | 1233 | NHP2,RpL7A | | 3.264e-02 | -3.42 | Ribosomal\_S10\_dom\_sf | interpro domains | IPR036838 | 3 | 2 | 11383 | 1233 | mRpL48,mRpS10 | | 3.264e-02 | -3.42 | SOD\_Cu/Zn\_BS | interpro domains | IPR018152 | 3 | 2 | 11383 | 1233 | Sod3,Sod1 | | 3.264e-02 | -3.42 | MPP\_PP2B | interpro domains | IPR041751 | 3 | 2 | 11383 | 1233 | Pp2B-14D,CanA-14F | | 3.264e-02 | -3.42 | Isocitrate\_DH\_NAD | interpro domains | IPR004434 | 3 | 2 | 11383 | 1233 | CG5028,Idh3b | | 3.264e-02 | -3.42 | Ribosomal\_L13 | interpro domains | IPR005822 | 3 | 2 | 11383 | 1233 | RpL13A,mRpL13 | | 3.264e-02 | -3.42 | Man6P\_isomerase\_rcpt-bd\_dom\_sf | interpro domains | IPR009011 | 3 | 2 | 11383 | 1233 | GCS2beta,CG6766 | | 3.264e-02 | -3.42 | Thiolase\_AS | interpro domains | IPR020610 | 3 | 2 | 11383 | 1233 | Thiolase,yip2 | | 3.264e-02 | -3.42 | RibuloseP-bd\_barrel | interpro domains | IPR011060 | 3 | 2 | 11383 | 1233 | r-l,CG8607 | | 3.278e-02 | -3.42 | GO\_RNA\_BINDING | MSigDB lists | GO\_RNA\_BINDING | 669 | 120 | 4161 | 644 | Thiolase,RpL12,sbr,mRpS5,Moe,bys,Eno,thoc6,SRPK,RpS18,Stip1,yip2,RpS27,CG9773,PpD3,RpL26,mtgo,eIF3m,Dcr-1,RpL11,CG7414,obe,Synj,heph,CG17593,RpS7,mtTFB1,RpL38,RpS4,RpS16,dco,mRpS23,Dhfr,RpS24,CG17187,CG10463,RpL23,Non1,Uch-L5,CCT6,Tfb4,Pus7,Irp-1B,RpS23,Fkbp59,Scsalpha1,Unr,l(1)G0004,CysRS,Karybeta3,GluRS-m,trsn,scu,RpII140,mRpL18,Cisd2,CG2100,eIF4A,CG3021,lwr,spn-E,CG8611,CCT3,RpL24,Rnmt,Mitofilin,mRpL10,RpL37a,Mat89Ba,Rcd5,CG9143,CG7728,LSm3,AGO3,CG7488,Cbp80,tko,Imp,CG10418,tho2,Ts,NHP2,Ranbp21,CG17454,RpS2,CG1890,CG11858,Edc3,TyrRS,RpL30,CG8963,mEFTu1,Top1,RpL13,Pen,eEF1beta,ERp60,mRpL16,Bin1,SerRS-m,RpS28b,Nap1,RpS27A,RpL9,Clp,mRpS21,bru1,eIF3i,Rrp40,eIF4E1,CG6610,RpL13A,LysRS,mRpL13,RpS30,RpLP0,RpS13,CG10189,piwi,Rpt2 | | 3.280e-02 | -3.42 | REACTOME\_PROTEIN\_FOLDING | MSigDB lists | REACTOME\_PROTEIN\_FOLDING | 30 | 9 | 4161 | 644 | Fbw5,CCT6,CCT5,Pfdn2,CCT3,CG1890,Pfdn5,alphaTub84B,TBCC | | 3.280e-02 | -3.42 | GO\_TRNA\_BINDING | MSigDB lists | GO\_TRNA\_BINDING | 30 | 9 | 4161 | 644 | TyrRS,CG2100,CG10189,CG7414,CG3021,LysRS,Ranbp21,CysRS,GluRS-m | | 3.280e-02 | -3.42 | GO\_MITOTIC\_DNA\_INTEGRITY\_CHECKPOINT | MSigDB lists | GO\_MITOTIC\_DNA\_INTEGRITY\_CHECKPOINT | 30 | 9 | 4161 | 644 | RpS27A,Not10,CG30291,twin,Rbf,APP-BP1,RpL40,CycB,Cdk1 | | 3.280e-02 | -3.42 | GO\_STRUCTURAL\_CONSTITUENT\_OF\_CYTOSKELETON | MSigDB lists | GO\_STRUCTURAL\_CONSTITUENT\_OF\_CYTOSKELETON | 30 | 9 | 4161 | 644 | Grip84,Grip75,gammaTub37C,Arpc2,alphaTub84B,Grip128,Tm1,Moe,Arpc4 | | 3.280e-02 | -3.42 | REACTOME\_MITOTIC\_G2\_G2\_M\_PHASES | MSigDB lists | REACTOME\_MITOTIC\_G2\_G2\_M\_PHASES | 30 | 9 | 4161 | 644 | CycB,Cdk1,Grip128,msps,Cdk7,Grip84,gammaTub37C,dco,Pp2A-29B | | 3.284e-02 | -3.42 | mEFTu1 (mitochondrial translation elongation factor Tu 1) | protein interactions | 44438 | 32 | 8 | 9204 | 1112 | CCT6,Kap-alpha3,ScsbetaA,Mcad,r,Cp1,CCT3,wal | | 3.284e-02 | -3.42 | Mlp84B (Muscle LIM protein at 84B) | protein interactions | 40849 | 32 | 8 | 9204 | 1112 | Actn,Cdk7,COX5B,CG11164,Rab10,Pfk,trbd,CG2258 | | 3.284e-02 | -3.42 | Utx (Utx histone demethylase) | protein interactions | 34377 | 32 | 8 | 9204 | 1112 | CG4853,RpL23A,Rbf,nej,Ptip,CG17068,trr,EcR | | 3.284e-02 | -3.42 | jagn (jagunal) | protein interactions | 40714 | 32 | 8 | 9204 | 1112 | Surf4,ATPsynCF6,CG31729,Karybeta3,ATPsynB,SERCA,CG10425,ATPsynO | | 3.293e-02 | -3.41 | Signaling by FGFR2 | REACTOME pathways | R-DME-5654738 | 52 | 13 | 3923 | 574 | Rbfox1,Cbp80,rl,RpS27A,Rpb8,RpL40,CG13398,TfIIFalpha,heph,mts,csw,Cbl,Pp2A-29B | | 3.301e-02 | -3.41 | PLDc\_2 | pfam domains | PF13091 | 3 | 2 | 10480 | 1142 | CG7718,Pld | | 3.301e-02 | -3.41 | PLDc | pfam domains | PF00614 | 3 | 2 | 10480 | 1142 | Pld,CG9248 | | 3.301e-02 | -3.41 | SPOC | pfam domains | PF07744 | 3 | 2 | 10480 | 1142 | spen,pps | | 3.301e-02 | -3.41 | Ribosomal\_L13 | pfam domains | PF00572 | 3 | 2 | 10480 | 1142 | mRpL13,RpL13A | | 3.301e-02 | -3.41 | Sld5 | pfam domains | PF05916 | 3 | 2 | 10480 | 1142 | Sld5,Psf3 | | 3.301e-02 | -3.41 | Glycohydro\_20b2 | pfam domains | PF14845 | 3 | 2 | 10480 | 1142 | fdl,Hexo1 | | 3.301e-02 | -3.41 | BEN | pfam domains | PF10523 | 3 | 2 | 10480 | 1142 | insv,Elba2 | | 3.301e-02 | -3.41 | Pex2\_Pex12 | pfam domains | PF04757 | 3 | 2 | 10480 | 1142 | Pex10,Pex12 | | 3.301e-02 | -3.41 | Cir\_N | pfam domains | PF10197 | 3 | 2 | 10480 | 1142 | CG12818,Cwc25 | | 3.301e-02 | -3.41 | Ribosomal\_L10 | pfam domains | PF00466 | 3 | 2 | 10480 | 1142 | mRpL10,RpLP0 | | 3.301e-02 | -3.41 | Tom37 | pfam domains | PF10568 | 3 | 2 | 10480 | 1142 | CG9393,CG8004 | | 3.301e-02 | -3.41 | Arm\_3 | pfam domains | PF16186 | 3 | 2 | 10480 | 1142 | Kap-alpha3,Pen | | 3.301e-02 | -3.41 | IBB | pfam domains | PF01749 | 3 | 2 | 10480 | 1142 | Pen,Kap-alpha3 | | 3.301e-02 | -3.41 | Longin | pfam domains | PF13774 | 3 | 2 | 10480 | 1142 | Vamp7,Sec22 | | 3.301e-02 | -3.41 | DnaJ\_CXXCXGXG | pfam domains | PF00684 | 3 | 2 | 10480 | 1142 | Droj2,DnaJ-H | | 3.301e-02 | -3.41 | Glyco\_hydro\_20 | pfam domains | PF00728 | 3 | 2 | 10480 | 1142 | fdl,Hexo1 | | 3.301e-02 | -3.41 | AIRS | pfam domains | PF00586 | 3 | 2 | 10480 | 1142 | SelD,Pfas | | 3.301e-02 | -3.41 | Ribosomal\_S10 | pfam domains | PF00338 | 3 | 2 | 10480 | 1142 | mRpS10,mRpL48 | | 3.301e-02 | -3.41 | 4F5 | pfam domains | PF04419 | 3 | 2 | 10480 | 1142 | CG17931,CG15715 | | 3.302e-02 | -3.41 | membrane | cellular component | GO:0016020 | 2722 | 277 | 11940 | 1105 | E23,Sec23,Fkbp59,tor,Roe1,alt,Pka-R2,GalT1,Synj,nAChRalpha4,Tsp74F,Arl6IP1,CG6836,CG5510,Vps16B,Arf51F,COX5A,CG9399,ND-B15,Drep2,CG12773,CG31195,AnxB10,slam,Pi4KIIalpha,stet,CG9132,UQCR-11,cype,Cyt-c-p,X11L,Tomosyn,Vha13,CG11077,CG31729,Coq8,alphaTub67C,CG6293,PIG-C,blw,Dd,Ugt35A1,Ggt-1,mol,DppIII,NaPi-III,CG31523,Tim8,Rab5,disp,Stoml2,ZnT35C,sip3,CG7091,RhoGDI,ND-B22,Hip1,ND-18,Ppox,zpg,CG8009,Tspo,Myd88,Tsp42Ef,CHMP2B,COX5B,Surf4,CG8112,Vsp37A,Eno,Ac3,EMC3,awd,Pdk1,mAChR-A,Mctp,Syx13,sbr,NP15.6,Sfxn2,Gmap,sau,CG3803,ste14,Sod3,bif,CG6707,fs(1)M3,CG5903,CG4293,CG31342,Pax,Sec15,Pex13,ND-MWFE,SREBP,tub,Phb2,UQCR-Q,zyd,ste24a,Rab10,bru1,ND-MNLL,CG2061,fdl,nSMase,kcc,PIG-H,Ho,Crag,CG10425,Calx,Jabba,toc,ND-SGDH,Tom70,Debcl,AP-1-2beta,Mitofilin,CG8728,SERCA,rpk,shrb,Clc,CG4476,Stat92E,CG8974,ATPsynbeta,Sply,Mtl,Nap1,CG10470,CG42258,CG7458,ATPsynE,Hsc70-3,hoe1,ND-39,snz,opm,CG4332,Tengl4,CG15098,Sccpdh1,CG31689,glob1,Ork1,Tao,Cchl,CHOp24,pen-2,ATPsynO,Moe,Vha36-1,sws,Topors,blot,Pex10,Adi1,Nlg1,CG33181,CG8004,CG17270,CG5447,Sec22,Hexo1,ATPsynD,crq,CG9393,Vha68-2,CG8311,fs(1)N,oxt,CG9981,nrv1,Snx6,CG16868,CG40472,CG7806,Sry-alpha,CG8726,CG7530,Vamp7,CG33977,Argk,Pi3K92E,Orc6,Ugalt,Tsp86D,Atf6,COX8,Herc4,levy,Desat1,lt,dco,Atg6,CAH2,mino,Cbl,ABCD,ATPsynF,Pmi,dob,fz4,CG5789,Marf,ND-30,Neb-cGP,tkv,mtgo,EMC8-9,Cen,Axs,GILT1,dpr6,Tsp66E,Ir41a,ND-51,SdhA,anne,ATPsynG,CG14671,EMRE,CG6927,alpha-Man-Ia,CG5969,Edem1,CG9773,yin,CG7655,Abi,Ptp69D,CG32441,ATPsynB,Pex23,ND-23,Scox,ATP8A,Agpat2,Pex12,CG31262,CG10960,SdhD,Etf-QO,Afti,CG32640,Efr,ATPsynCF6,fwe,Atpalpha,colt,ATPsyndelta,Irk1,Seipin,ND-B18,CG4562,ND-49,DIP-lambda,Fkbp12,CG11110,CG42748,Rep,Nmt,ND-75,S2P,Coq7,COX4,LpR2,Fur2,rost,Tom40,ND-24,slif,Iml1,Prip | | 3.307e-02 | -3.41 | cellular response to amino acid starvation | biological process | GO:0034198 | 14 | 4 | 12345 | 1129 | Rel,foxo,Iml1,Nprl2 | | 3.307e-02 | -3.41 | heme metabolic process | biological process | GO:0042168 | 14 | 4 | 12345 | 1129 | Ppox,Ho,CG13551,CG3803 | | 3.307e-02 | -3.41 | regulation of membrane lipid distribution | biological process | GO:0097035 | 14 | 4 | 12345 | 1129 | CG9981,scramb1,CG31729,ATP8A | | 3.307e-02 | -3.41 | neuromuscular process | biological process | GO:0050905 | 14 | 4 | 12345 | 1129 | Atpalpha,Uba5,Calx,Ufm1 | | 3.307e-02 | -3.41 | actin nucleation | biological process | GO:0045010 | 14 | 4 | 12345 | 1129 | Arpc2,Arpc4,wash,Arpc1 | | 3.307e-02 | -3.41 | porphyrin-containing compound metabolic process | biological process | GO:0006778 | 14 | 4 | 12345 | 1129 | CG3803,CG13551,Ho,Ppox | | 3.307e-02 | -3.41 | sterol homeostasis | biological process | GO:0055092 | 14 | 4 | 12345 | 1129 | Npc2a,CG8112,SNF4Agamma,EcR | | 3.307e-02 | -3.41 | female sex determination | biological process | GO:0030237 | 14 | 4 | 12345 | 1129 | ovo,ix,stil,otu | | 3.307e-02 | -3.41 | response to molecule of bacterial origin | biological process | GO:0002237 | 14 | 4 | 12345 | 1129 | Stat92E,EcR,Tak1,Vps16B | | 3.309e-02 | -3.41 | GSE27859\_DC\_VS\_CD11C\_INT\_F480\_HI\_MACROPHAGE\_UP | MSigDB lists | GSE27859\_DC\_VS\_CD11C\_INT\_F480\_HI\_MACROPHAGE\_UP | 80 | 19 | 4161 | 644 | Coq6,CG6767,mts,csw,muskelin,CG31759,l(1)G0004,ND-23,Ubc4,CG40045,DppIII,NHP2,Dph5,spag,CG3587,Gclc,Bin1,Pus7,Cdk1 | | 3.309e-02 | -3.41 | GSE17974\_CTRL\_VS\_ACT\_IL4\_AND\_ANTI\_IL12\_2H\_CD4\_TCELL\_DN | MSigDB lists | GSE17974\_CTRL\_VS\_ACT\_IL4\_AND\_ANTI\_IL12\_2H\_CD4\_TCELL\_DN | 80 | 19 | 4161 | 644 | Mitofilin,cl,mRpL17,MED20,Prosbeta5,CCT5,esc,CCT3,Top1,bys,CG17187,Cyt-c-p,Rpn12,slif,isoQC,CysRS,mbo,Cbp80,escl | | 3.309e-02 | -3.41 | GSE41867\_LCMV\_ARMSTRONG\_VS\_CLONE13\_DAY6\_EFFECTOR\_CD8\_TCELL\_UP | MSigDB lists | GSE41867\_LCMV\_ARMSTRONG\_VS\_CLONE13\_DAY6\_EFFECTOR\_CD8\_TCELL\_UP | 80 | 19 | 4161 | 644 | CG33156,Rpn12,levy,Stoml2,Ts,tko,CG10932,Prosalpha3,CG9143,Prosbeta4,Tim8,pont,Pfdn2,AIF,Cdk1,r-l,Thiolase,Shmt,mad2 | | 3.316e-02 | -3.41 | MODULE\_52 | MSigDB lists | MODULE\_52 | 112 | 25 | 4161 | 644 | isoQC,Cse1,ste24a,CkIIalpha,CG12018,scramb1,Tm1,Tsp74F,AGBE,CG14516,Got1,Rcd5,Gyg,Prosalpha3,msps,Chd64,mad2,CCT6,Hat1,Cdk1,SRPK,r-l,CycB,Echs1,pr | | 3.318e-02 | -3.41 | GO\_PEPTIDYL\_THREONINE\_MODIFICATION | MSigDB lists | GO\_PEPTIDYL\_THREONINE\_MODIFICATION | 21 | 7 | 4161 | 644 | grp,fray,Cdk1,Pdk1,rl,r,Cdk5 | | 3.318e-02 | -3.41 | KEGG\_AMYOTROPHIC\_LATERAL\_SCLEROSIS\_ALS | MSigDB lists | KEGG\_AMYOTROPHIC\_LATERAL\_SCLEROSIS\_ALS | 21 | 7 | 4161 | 644 | Pp2B-14D,Sod1,Cyt-c-p,p38a,CanB2,CanA-14F,Tom40 | | 3.318e-02 | -3.41 | REACTOME\_E2F\_MEDIATED\_REGULATION\_OF\_DNA\_REPLICATION | MSigDB lists | REACTOME\_E2F\_MEDIATED\_REGULATION\_OF\_DNA\_REPLICATION | 21 | 7 | 4161 | 644 | Pp2A-29B,CycE,Ts,Cdk1,mts,Dhfr,CycB | | 3.318e-02 | -3.41 | LIN\_MELANOMA\_COPY\_NUMBER\_DN | MSigDB lists | LIN\_MELANOMA\_COPY\_NUMBER\_DN | 21 | 7 | 4161 | 644 | mtgo,Bin1,nmd,ERp60,ScsbetaA,MED4,CG4390 | | 3.318e-02 | -3.41 | ARGGGTTAA\_UNKNOWN | MSigDB lists | ARGGGTTAA\_UNKNOWN | 21 | 7 | 4161 | 644 | O-fut1,CG9471,Mtpalpha,Thiolase,Tlk,Pex13,Abi | | 3.328e-02 | -3.40 | AcylCoA\_DH/ox\_N\_sf | interpro domains | IPR037069 | 12 | 4 | 11383 | 1233 | CG4860,CG4586,Mcad,Acox57D-p | | 3.328e-02 | -3.40 | Proteasome\_suB-type | interpro domains | IPR023333 | 12 | 4 | 11383 | 1233 | Prosbeta6,Prosbeta4,Prosbeta3,Prosbeta5 | | 3.336e-02 | -3.40 | MORF\_HDAC2 | MSigDB lists | MORF\_HDAC2 | 156 | 33 | 4161 | 644 | Chrac-14,mEFTu1,SdhA,CG5028,rept,ND-51,eIF3i,Sod1,eIF4E1,LysRS,mts,Cul1,Hat1,CG13630,Thiolase,CCT5,CG8004,Iru,Prosbeta4,CG7728,APP-BP1,eIF3m,Mitofilin,CG10418,CG1416,CG17454,tho2,r,CG4390,ScsbetaA,Cyt-c-p,Dd,TyrRS | | 3.354e-02 | -3.39 | Tspo (Translocator protein) | protein interactions | 33231 | 50 | 11 | 9204 | 1112 | ScsbetaA,Mcad,Rab5,SERCA,CG1416,CCT6,CG15098,COX4,GCS2beta,wal,Pp2A-29B | | 3.359e-02 | -3.39 | YAGI\_AML\_WITH\_INV\_16\_TRANSLOCATION | MSigDB lists | YAGI\_AML\_WITH\_INV\_16\_TRANSLOCATION | 123 | 27 | 4161 | 644 | Arf51F,RpL37a,Mitofilin,YL-1,MED20,shu,CG14516,yip2,RpS27,AIF,RpS18,CG14715,Cul1,RpL13,Thiolase,CG8314,CG11164,Dhfr,faf,dco,ste24a,RpLP0,RpS13,RpS16,ari-1,RpS4,tho2 | | 3.360e-02 | -3.39 | adenylyltransferase activity | molecular function | GO:0070566 | 14 | 4 | 12176 | 1119 | CG46385,wisp,Trf4-2,CG4407 | | 3.360e-02 | -3.39 | peptide disulfide oxidoreductase activity | molecular function | GO:0015037 | 14 | 4 | 12176 | 1119 | Trxr-1,Grx1,ERp60,CG14407 | | 3.363e-02 | -3.39 | Mtor (Megator) | protein interactions | 36264 | 21 | 6 | 9204 | 1112 | Exo70,mad2,EcR,apt,Lkb1,CG11248 | | 3.363e-02 | -3.39 | Sfmbt (Scm-related gene containing four mbt domains) | protein interactions | 34709 | 21 | 6 | 9204 | 1112 | Marf1,Pura,ph-d,UbcE2H,rept,Nap1 | | 3.363e-02 | -3.39 | CG14667 (uncharacterized protein) | protein interactions | 40643 | 21 | 6 | 9204 | 1112 | RpL37a,SelD,RpS2,mts,tral,zf30C | | 3.363e-02 | -3.39 | CG1513 (uncharacterized protein) | protein interactions | 36028 | 21 | 6 | 9204 | 1112 | AIF,Rab5,foxo,Lkb1,l(2)k01209,CG9205 | | 3.366e-02 | -3.39 | GO\_INTRACELLULAR\_SIGNAL\_TRANSDUCTION | MSigDB lists | GO\_INTRACELLULAR\_SIGNAL\_TRANSDUCTION | 387 | 73 | 4161 | 644 | Tlk,GCS2beta,Rbf,Med,RpL40,Pi3K92E,Prosbeta6,Pka-R1,Ubc4,PpD3,Prosbeta4,Arf51F,Cul1,Rpn3,p38a,Ac3,SRPK,Pdk1,CycB,Gdi,Rpn2,raptor,Prosbeta3,Tip60,dnc,Iml1,Xpac,SkpA,Prosalpha5,Mo25,Rab10,SERCA,Pka-R2,Uev1A,Cdk1,cact,rl,RhoGAP5A,Oseg1,Pp2B-14D,Mctp,Traf6,Cyt-c-p,Rab5,gus,Arl5,fray,nSMase,CanB2,Pp2A-29B,CG11807,CanA-14F,Prosalpha3,Jafrac2,Prosbeta5,Not10,CG3894,Rpn12,csw,Rpt2,CG7530,Prosalpha4,Cul2,CG7115,mts,Mekk1,RpS27A,Abi,Rab32,mael,twin,AIF,Cul5 | | 3.380e-02 | -3.39 | lilli (lilliputian) | protein interactions | 33496 | 38 | 9 | 9204 | 1112 | Su(H),Bx,tor,Rtnl1,Rbf,nej,Pi3K92E,gus,rl | | 3.380e-02 | -3.39 | CG6707 (uncharacterized protein) | protein interactions | 39168 | 38 | 9 | 9204 | 1112 | dl,Cyp1,Cse1,TORIP,mbo,CG15141,Eno,Mes2,Rcd5 | | 3.381e-02 | -3.39 | clathrin-coated vesicle | cellular component | GO:0030136 | 27 | 6 | 11940 | 1105 | Hip1,Afti,AP-1-2beta,Gga,CG9132,Clc | | 3.397e-02 | -3.38 | CULLIN\_2 | prosite domains | PS50069 | 7 | 3 | 6243 | 694 | Cul2,Cul1,Cul5 | | 3.397e-02 | -3.38 | ATPASE\_ALPHA\_BETA | prosite domains | PS00152 | 7 | 3 | 6243 | 694 | Vha68-2,ATPsynbeta,blw | | 3.397e-02 | -3.38 | PAZ | prosite domains | PS50821 | 7 | 3 | 6243 | 694 | Dcr-1,piwi,AGO3 | | 3.400e-02 | -3.38 | REACTOME\_SIGNALLING\_BY\_NGF | MSigDB lists | REACTOME\_SIGNALLING\_BY\_NGF | 75 | 18 | 4161 | 644 | nSMase,Fur2,Pka-R1,RpL40,RhoGDI,mts,Traf6,Pdk1,cact,Ac3,rl,p38a,Cdk1,pen-2,Pka-R2,Pp2A-29B,Clc,RpS27A | | 3.400e-02 | -3.38 | GSE22886\_NAIVE\_BCELL\_VS\_DC\_DN | MSigDB lists | GSE22886\_NAIVE\_BCELL\_VS\_DC\_DN | 75 | 18 | 4161 | 644 | CG8112,Top1,Prosbeta5,Cdk5,CG7997,Gclc,AIF,Synj,Grx1,Cbp80,Cyp1,CG42542,CG17593,DppIII,isoQC,Rpn2,CG10470,Rpt2 | | 3.400e-02 | -3.38 | GO\_LIPID\_CATABOLIC\_PROCESS | MSigDB lists | GO\_LIPID\_CATABOLIC\_PROCESS | 75 | 18 | 4161 | 644 | ABCD,Pex13,Mcad,CG5734,CG5966,Etf-QO,PAPLA1,CG15111,CG7840,Sply,CG9248,CG3376,Thiolase,CG7997,wal,Echs1,Mtpalpha,crq | | 3.413e-02 | -3.38 | transcription regulator activity | molecular function | GO:0140110 | 605 | 69 | 12176 | 1119 | bun,insv,Smox,Nf-YA,Uxt,CG2199,CG4730,Pax,e(y)1,Med,Trf2,cnc,trr,Taf7,apt,nej,CG10654,Atf3,Bin1,l(3)73Ah,mtTFB1,dl,E2f1,SREBP,CG2678,hb,MED24,REPTOR,foxo,CycC,CG4872,hng3,cad,Wbp2,Cf2,CG6813,EcR,Rbf2,Mybbp1A,NFAT,Rel,CG10631,CG1602,rept,pont,sbb,ovo,CG11902,ix,cyc,CG31875,Atf6,Hnf4,CG11710,Sgf29,Hr78,MED20,Su(H),CG17202,bcd,Sgf11,CG2129,CG17328,Stat92E,MED4,CG10959,MED27,pygo,Spt20 | | 3.420e-02 | -3.38 | cation transmembrane transport | biological process | GO:0098655 | 196 | 26 | 12345 | 1129 | ATPsynF,slif,Vha36-1,SERCA,Atpalpha,Vha13,CG33181,COX5B,ATPsynCF6,ATPsynD,ATPsynE,ATPsynbeta,CG12773,fwe,nrv1,ZnT35C,blw,ATPsyndelta,ATPsynO,Vha68-2,Ork1,EMRE,zyd,ATPsynG,ATPsynB,Fer1HCH | | 3.421e-02 | -3.38 | polyol metabolic process | biological process | GO:0019751 | 42 | 8 | 12345 | 1129 | l(1)G0196,CG9391,Gk1,Synj,Sptr,Pcd,pr,Gk2 | | 3.421e-02 | -3.38 | tRNA aminoacylation | biological process | GO:0043039 | 42 | 8 | 12345 | 1129 | TrpRS-m,GluRS-m,LysRS,GatA,CysRS,GatB,TyrRS,SerRS-m | | 3.421e-02 | -3.38 | cellular response to organic cyclic compound | biological process | GO:0071407 | 42 | 8 | 12345 | 1129 | Shmt,MED27,Hr78,G9a,Utx,Hnf4,EcR,Nurf-38 | | 3.423e-02 | -3.37 | PP2Ac | smart domains | SM00156 | 19 | 5 | 5086 | 506 | mts,Pp2B-14D,PpD3,CanA-14F,PpV | | 3.429e-02 | -3.37 | SOD\_CU\_ZN\_2 | prosite domains | PS00332 | 3 | 2 | 6243 | 694 | Sod1,Sod3 | | 3.429e-02 | -3.37 | LONGIN | prosite domains | PS50859 | 3 | 2 | 6243 | 694 | Sec22,Vamp7 | | 3.429e-02 | -3.37 | THIOLASE\_3 | prosite domains | PS00099 | 3 | 2 | 6243 | 694 | yip2,Thiolase | | 3.429e-02 | -3.37 | CSD\_1 | prosite domains | PS00352 | 3 | 2 | 6243 | 694 | yps,Unr | | 3.429e-02 | -3.37 | IBB | prosite domains | PS51214 | 3 | 2 | 6243 | 694 | Pen,Kap-alpha3 | | 3.436e-02 | -3.37 | CCR4-NOT complex | cellular component | GO:0030014 | 14 | 4 | 11940 | 1105 | cu,Not10,twin,Rga | | 3.437e-02 | -3.37 | SPIELMAN\_LYMPHOBLAST\_EUROPEAN\_VS\_ASIAN\_DN | MSigDB lists | SPIELMAN\_LYMPHOBLAST\_EUROPEAN\_VS\_ASIAN\_DN | 276 | 54 | 4161 | 644 | muskelin,Pgm1,RpII140,Nurf-38,Stoml2,Karybeta3,PIG-L,Fdh,Su(H),Unr,Clp,Idh3b,Prosalpha5,CG5116,Grip75,Klp10A,Cul5,Nca,Chrac-14,Uch-L5,CycC,Non1,CG1847,Vha36-1,CG2201,TyrRS,CG2493,CG6767,Sec10,aay,alc,lt,Kap3,Atg6,fray,Prosbeta6,CG4511,CG14894,Ubc4,Pdk,Snx6,Arl5,Med,CG9143,Pex12,CG11710,Mtpalpha,Jafrac2,SRPK,Ac3,nmdyn-D6,Cul1,capt,Thiolase | | 3.437e-02 | -3.37 | - | gene3d domains | 3.60.20.10 | 31 | 8 | 4929 | 622 | Prosalpha5,AsnS,Prosbeta4,Prosbeta3,Prosbeta6,Prosbeta5,Prosalpha3,Prosalpha4 | | 3.448e-02 | -3.37 | regulation of cellular component size | biological process | GO:0032535 | 178 | 24 | 12345 | 1129 | rl,Nprl2,wash,Girdin,Pdk1,Pi3K92E,Rbf,foxo,Mtl,Nup62,dco,p38a,bru1,Iml1,Arpc1,Arpc2,Mekk1,bif,Desat1,raptor,Seipin,ssh,CG12773,Arpc4 | | 3.453e-02 | -3.37 | REACTOME\_DNA\_REPLICATION | MSigDB lists | REACTOME\_DNA\_REPLICATION | 107 | 24 | 4161 | 644 | Prosbeta4,Prosalpha3,RpS27A,DNApol-epsilon255,Prosalpha5,Pp2A-29B,RpS27,Klp10A,Prosbeta5,Rpn3,msps,mad2,Rpt2,Prosalpha4,Prosbeta3,mts,CG12018,Rpn2,Rpn12,Sld5,Prosbeta6,RfC38,RpL40,Tao | | 3.453e-02 | -3.37 | GO\_REGULATION\_OF\_SYSTEM\_PROCESS | MSigDB lists | GO\_REGULATION\_OF\_SYSTEM\_PROCESS | 107 | 24 | 4161 | 644 | Tm1,bru1,Atpalpha,Med,dnc,Irk1,CG31715,Pp2B-14D,Prip,Sod1,EcR,nrv1,csw,AnxB10,Calx,mAChR-A,nAChRalpha4,Gabat,CG14516,CG44422,SERCA,Dcr-1,CanA-14F,SREBP | | 3.453e-02 | -3.37 | GO\_LYTIC\_VACUOLE\_MEMBRANE | MSigDB lists | GO\_LYTIC\_VACUOLE\_MEMBRANE | 107 | 24 | 4161 | 644 | Rab5,Iml1,RpL40,Gdap2,CG3842,AP-1-2beta,CG42542,lt,CG3860,raptor,AnxB10,Vha36-1,kcc,crq,sws,Clc,Vamp7,CG14516,Acph-1,Pi4KIIalpha,ZnT35C,CG7806,CG32473,Pex23 | | 3.454e-02 | -3.37 | Kap-alpha3 (karyopherin alpha3) | protein interactions | 41158 | 82 | 16 | 9204 | 1112 | CCT6,CG10425,Tip60,Pp2A-29B,CCT3,Pen,CG1416,dx,Smox,Atpalpha,Mcad,wal,mEFTu1,GCS2beta,Karybeta3,cnc | | 3.467e-02 | -3.36 | Oxygen-dependent proline hydroxylation of Hypoxia-inducible Factor Alpha | REACTOME pathways | R-DME-1234176 | 63 | 15 | 3923 | 574 | Prosbeta6,Rpn12,Prosalpha3,Rpn3,Rpt2,RpL40,EloC,Cul2,RpS27A,Prosalpha5,Prosbeta3,Prosbeta4,Rpn2,Prosbeta5,Prosalpha4 | | 3.467e-02 | -3.36 | Hedgehog ligand biogenesis | REACTOME pathways | R-DME-5358346 | 63 | 15 | 3923 | 574 | RpL40,Rpt2,Rpn3,Prosalpha3,Rpn12,Prosbeta6,disp,Prosalpha4,CG6766,Rpn2,Prosbeta5,Prosbeta4,Prosalpha5,Prosbeta3,RpS27A | | 3.470e-02 | -3.36 | WD40\_repeat | interpro domains | IPR001680 | 174 | 27 | 11383 | 1233 | Oseg1,Plap,Wdfy2,lt,escl,Cka,thoc6,Tomosyn,rig,Arpc1,cort,sw,DCAF12,CG8001,raptor,Aladin,Caf1-105,CG10132,ago,eIF3i,Fbw5,tws,Rbcn-3A,esc,CG3511,adp,Grip71 | | 3.478e-02 | -3.36 | GSE22886\_NAIVE\_BCELL\_VS\_BLOOD\_PLASMA\_CELL\_UP | MSigDB lists | GSE22886\_NAIVE\_BCELL\_VS\_BLOOD\_PLASMA\_CELL\_UP | 35 | 10 | 4161 | 644 | RpS23,Cka,RhoGAP54D,GAPsec,RpS27,Smox,ari-1,CG5026,Pi3K92E,oxt | | 3.478e-02 | -3.36 | GO\_CHAPERONE\_BINDING | MSigDB lists | GO\_CHAPERONE\_BINDING | 35 | 10 | 4161 | 644 | CG1890,Sod1,TBCC,Nc73EF,Hsc70-3,CG1416,Droj2,Atpalpha,Stip1,Roe1 | | 3.478e-02 | -3.36 | GO\_PYRIMIDINE\_CONTAINING\_COMPOUND\_METABOLIC\_PROCESS | MSigDB lists | GO\_PYRIMIDINE\_CONTAINING\_COMPOUND\_METABOLIC\_PROCESS | 35 | 10 | 4161 | 644 | CG5285,CG6767,nmdyn-D7,r,Ts,kri,rl,r-l,Shmt,nmdyn-D6 | | 3.478e-02 | -3.36 | REACTOME\_SIGNALING\_BY\_ERBB2 | MSigDB lists | REACTOME\_SIGNALING\_BY\_ERBB2 | 35 | 10 | 4161 | 644 | Cdk1,rl,Pdk1,Ac3,Pka-R1,STUB1,Cul5,RpL40,Pka-R2,RpS27A | | 3.478e-02 | -3.36 | MORF\_SART1 | MSigDB lists | MORF\_SART1 | 35 | 10 | 4161 | 644 | Prosbeta6,not,lwr,Prosbeta3,eIF3m,RpL30,RpL24,RpS30,RpS27A,Idh3b | | 3.478e-02 | -3.36 | Mabi (Mabiki) | protein interactions | 34727 | 11 | 4 | 9204 | 1112 | RpL38,RpLP1,RpS27A,bcd | | 3.478e-02 | -3.36 | DIP-zeta (Dpr-interacting protein zeta) | protein interactions | 34231 | 11 | 4 | 9204 | 1112 | dpr6,CG6767,cDIP,RpS27 | | 3.478e-02 | -3.36 | roq (roquin) | protein interactions | 39818 | 11 | 4 | 9204 | 1112 | Not10,Rga,zf30C,CG11486 | | 3.478e-02 | -3.36 | ND-24 (NADH dehydrogenase (ubiquinone) 24 kDa subunit) | protein interactions | 32740 | 11 | 4 | 9204 | 1112 | ND-75,CG14894,smg,Med | | 3.478e-02 | -3.36 | CG8677 (uncharacterized protein) | protein interactions | 35397 | 11 | 4 | 9204 | 1112 | rept,Tip60,Spx,cup | | 3.478e-02 | -3.36 | Sln (Silnoon) | protein interactions | 36263 | 11 | 4 | 9204 | 1112 | Lkb1,Marf,SREBP,CG2678 | | 3.478e-02 | -3.36 | TwdlW (TweedleW) | protein interactions | 42075 | 11 | 4 | 9204 | 1112 | alph,CG4991,Pdk1,E23 | | 3.478e-02 | -3.36 | His4r (Histone H4 replacement) | protein interactions | 41773 | 11 | 4 | 9204 | 1112 | Fkbp39,wisp,CG15098,e(y)1 | | 3.478e-02 | -3.36 | Z600 (Z600) | protein interactions | 39673 | 11 | 4 | 9204 | 1112 | CycE,CycB,Cdk1,Nup358 | | 3.478e-02 | -3.36 | CG6664 (uncharacterized protein) | protein interactions | 39903 | 11 | 4 | 9204 | 1112 | psq,CG2199,CG11164,Cdk1 | | 3.478e-02 | -3.36 | Cdc23 (Cell division cycle 23) | protein interactions | 35315 | 11 | 4 | 9204 | 1112 | Cdk1,cort,APC7,CycB | | 3.478e-02 | -3.36 | Pka-R1 (Protein kinase, cAMP-dependent, regulatory subunit type 1) | protein interactions | 40305 | 11 | 4 | 9204 | 1112 | dnc,capt,Pka-R1,nej | | 3.478e-02 | -3.36 | CG13124 (uncharacterized protein) | protein interactions | 34310 | 11 | 4 | 9204 | 1112 | CG2199,CG3342,dgt4,Nup58 | | 3.478e-02 | -3.36 | cic (capicua) | protein interactions | 53560 | 11 | 4 | 9204 | 1112 | rl,Rtnl1,spz,Bx | | 3.481e-02 | -3.36 | MULLIGHAN\_NPM1\_SIGNATURE\_3\_UP | MSigDB lists | MULLIGHAN\_NPM1\_SIGNATURE\_3\_UP | 91 | 21 | 4161 | 644 | Ubc4,Etf-QO,CG4572,Gk2,Dhfr,csw,CG11858,Traf6,CG6767,alc,CG10470,CG8004,Pdk1,Ptpmeg,CHMP2B,Top1,Kap-alpha3,CG9911,Got1,Acph-1,Irp-1B | | 3.484e-02 | -3.36 | GSE17721\_0.5H\_VS\_24H\_PAM3CSK4\_BMDC\_DN | MSigDB lists | GSE17721\_0.5H\_VS\_24H\_PAM3CSK4\_BMDC\_DN | 70 | 17 | 4161 | 644 | nSMase,CG8475,CG3842,opm,CG12375,Gk2,CG1416,Tlk,EcR,Mtpalpha,STUB1,Gdi,f-cup,Rab10,CG4293,CG7789,asf1 | | 3.484e-02 | -3.36 | GSE15330\_WT\_VS\_IKAROS\_KO\_GRANULOCYTE\_MONOCYTE\_PROGENITOR\_UP | MSigDB lists | GSE15330\_WT\_VS\_IKAROS\_KO\_GRANULOCYTE\_MONOCYTE\_PROGENITOR\_UP | 70 | 17 | 4161 | 644 | CG1532,csw,LysRS,Pi3K92E,CG17691,bru1,ND-23,CG31729,Herc4,stet,CG32473,asf1,pont,Bin1,pr,mRpL16,CG30010 | | 3.484e-02 | -3.36 | RICKMAN\_TUMOR\_DIFFERENTIATED\_WELL\_VS\_POORLY\_UP | MSigDB lists | RICKMAN\_TUMOR\_DIFFERENTIATED\_WELL\_VS\_POORLY\_UP | 70 | 17 | 4161 | 644 | CG15111,Mitofilin,RpS27,Abi,Nap1,alph,Aldh,CCT3,CG3226,RpL23,Sfxn2,Oseg1,RpII140,RhoGAP54D,RpS7,SelD,GluRS-m | | 3.492e-02 | -3.35 | TCOMPLEXTCP1 | prints domains | PR00304 | 8 | 3 | 2185 | 212 | CCT5,CCT6,CCT3 | | 3.526e-02 | -3.34 | Orc1 removal from chromatin | REACTOME pathways | R-DME-68949 | 74 | 17 | 3923 | 574 | Rbf,Prosalpha3,Rpt2,RpL40,Prosbeta4,RpS27A,Prosalpha5,Prosalpha4,Rbf2,Rpn2,Prosbeta5,Prosbeta6,Rpn12,Rpn3,Orc1,Prosbeta3,Orc6 | | 3.526e-02 | -3.34 | Switching of origins to a post-replicative state | REACTOME pathways | R-DME-69052 | 74 | 17 | 3923 | 574 | Prosbeta6,Rpn12,Rpn3,Orc1,Orc6,Prosbeta3,Rbf,Prosalpha3,RpL40,Rpt2,Prosbeta4,RpS27A,Prosalpha5,Prosalpha4,Rbf2,Rpn2,Prosbeta5 | | 3.529e-02 | -3.34 | regulation of MAPK cascade | biological process | GO:0043408 | 125 | 18 | 12345 | 1129 | Cbl,Tak1,tor,park,Lkb1,RhoGAP5A,Cka,PEK,alph,SkpA,Mekk1,Ptp61F,ben,PpV,CG17746,Mos,Tao,ATPsynD | | 3.533e-02 | -3.34 | DNA metabolic process | biological process | GO:0006259 | 280 | 35 | 12345 | 1129 | tos,mus205,Ubc6,Xpc,mre11,Orc1,CG10336,Exd2,Top1,Rbf,mus81,Tfb4,Sld5,mus304,MAGE,CycE,Snm1,E2f1,CG5285,obe,Dcr-1,Drep1,CG12018,Nap1,DNApol-epsilon255,c(2)M,RecQ5,Uev1A,nbs,agt,Tengl4,RfC38,Orc6,Xpac,ben | | 3.544e-02 | -3.34 | fly chr3-47 cM | chromosome location | fly chr3-47 cM | 46 | 6 | 25010 | 1343 | mRpS5,CG40472,CG40045,lncRNA:CR45159,UQCR-11,asRNA:CR45182 | | 3.544e-02 | -3.34 | fly chr3-47 | chromosome location | fly chr3-47 | 46 | 6 | 25010 | 1343 | mRpS5,UQCR-11,lncRNA:CR45159,CG40045,CG40472,asRNA:CR45182 | | 3.548e-02 | -3.34 | tubulin binding | molecular function | GO:0015631 | 142 | 20 | 12176 | 1119 | bif,Moe,Klp10A,Grip128,Grip75,Girdin,cana,TBCC,CG7716,stai,wash,Uxt,CG3226,CG1890,Grip84,Blos2,Grip71,awd,msps,cnn | | 3.557e-02 | -3.34 | regulation of lipid metabolic process | biological process | GO:0019216 | 50 | 9 | 12345 | 1129 | Desat1,foxo,SREBP,Nprl2,CG8009,Lkb1,CG13601,Dd,Seipin | | 3.557e-02 | -3.34 | response to peptide | biological process | GO:1901652 | 50 | 9 | 12345 | 1129 | raptor,Pdk1,CG7328,Rab10,rl,PyK,foxo,Hnf4,Pi3K92E | | 3.557e-02 | -3.34 | glycosyl compound metabolic process | biological process | GO:1901657 | 50 | 9 | 12345 | 1129 | awd,kri,nmdyn-D6,CG6767,CG7997,cN-IIIB,r-l,l(2)k01209,nmdyn-D7 | | 3.557e-02 | -3.34 | response to peptide hormone | biological process | GO:0043434 | 50 | 9 | 12345 | 1129 | raptor,foxo,PyK,Pi3K92E,Hnf4,CG7328,Pdk1,Rab10,rl | | 3.558e-02 | -3.34 | GSE20152\_SPHK1\_KO\_VS\_WT\_HTNFA\_OVERXPRESS\_ANKLE\_DN | MSigDB lists | GSE20152\_SPHK1\_KO\_VS\_WT\_HTNFA\_OVERXPRESS\_ANKLE\_DN | 65 | 16 | 4161 | 644 | esc,Mtpalpha,Mat89Ba,CG30496,gammaTub37C,Ppox,Prosbeta6,CG15141,Etf-QO,Cbp80,CG11089,escl,cyc,Gmer,Cse1,GAPsec | | 3.558e-02 | -3.34 | GSE1112\_OT1\_VS\_HY\_CD8AB\_THYMOCYTE\_RTOC\_CULTURE\_UP | MSigDB lists | GSE1112\_OT1\_VS\_HY\_CD8AB\_THYMOCYTE\_RTOC\_CULTURE\_UP | 65 | 16 | 4161 | 644 | nmdyn-D7,colt,CG6767,CG5734,ND-51,Nprl2,Stoml2,AP-1-2beta,Jheh2,Cyp6a22,CG11807,Pi4KIIalpha,Cyp6a19,AIF,CG9471,Eno | | 3.560e-02 | -3.34 | CG10581 (uncharacterized protein) | protein interactions | 40326 | 16 | 5 | 9204 | 1112 | Adh,Mf,up,Argk,Scp1 | | 3.560e-02 | -3.34 | CDC45L (CDC45L) | protein interactions | 31052 | 16 | 5 | 9204 | 1112 | grp,SuUR,Sld5,CG3511,Psf3 | | 3.560e-02 | -3.34 | Zpr1 (Zpr1) | protein interactions | 31851 | 16 | 5 | 9204 | 1112 | Cyt-c-p,Tspo,Atpalpha,Sod1,Top1 | | 3.560e-02 | -3.34 | Sec13 (Secretory 13) | protein interactions | 44437 | 16 | 5 | 9204 | 1112 | wal,CG1416,Tspo,Mcad,tral | | 3.560e-02 | -3.34 | spz (spatzle) | protein interactions | 43256 | 16 | 5 | 9204 | 1112 | mad2,spz,ea,Rel,cact | | 3.560e-02 | -3.34 | Cnot4 (CCR4-NOT transcription complex subunit 4) | protein interactions | 34416 | 16 | 5 | 9204 | 1112 | nos,Stat92E,Tak1,PGRP-LE,Rel | | 3.560e-02 | -3.34 | CG17855 (uncharacterized protein) | protein interactions | 34269 | 16 | 5 | 9204 | 1112 | CG15514,CG10654,RpLP1,RpL40,CG4730 | | 3.560e-02 | -3.34 | tud (tudor) | protein interactions | 37417 | 16 | 5 | 9204 | 1112 | AGO3,eIF4A,Eno,PyK,osk | | 3.560e-02 | -3.34 | Seipin (seipin) | protein interactions | 31245 | 16 | 5 | 9204 | 1112 | Seipin,mino,insv,SERCA,regucalcin | | 3.560e-02 | -3.34 | CG9248 (uncharacterized protein) | protein interactions | 35384 | 16 | 5 | 9204 | 1112 | gus,EcR,bcd,Ntan1,Stam | | 3.560e-02 | -3.34 | Cln3 (Cln3) | protein interactions | 39981 | 16 | 5 | 9204 | 1112 | Mtpalpha,Cisd2,foxo,Not10,Sod1 | | 3.560e-02 | -3.34 | PEK (pancreatic eIF-2alpha kinase) | protein interactions | 40653 | 16 | 5 | 9204 | 1112 | Pink1,CycE,grp,PEK,park | | 3.566e-02 | -3.33 | USP\_3 | prosite domains | PS50235 | 23 | 6 | 6243 | 694 | not,faf,scny,Usp16-45,Usp39,Usp10 | | 3.567e-02 | -3.33 | Fatty Acid Beta Oxidation 1 | WikiPathways | WP39 | 18 | 8 | 499 | 116 | colt,Mtpalpha,Gk1,AcCoAS,Echs1,Thiolase,whd,Gk2 | | 3.581e-02 | -3.33 | TYR\_PHOSPHATASE\_1 | prosite domains | PS00383 | 29 | 7 | 6243 | 694 | csw,Ptp61F,Ptp69D,Ptpmeg2,Ptp10D,Ptpmeg,ssh | | 3.597e-02 | -3.32 | TAKAO\_RESPONSE\_TO\_UVB\_RADIATION\_UP | MSigDB lists | TAKAO\_RESPONSE\_TO\_UVB\_RADIATION\_UP | 40 | 11 | 4161 | 644 | levy,Bin1,Moe,Jafrac1,sbr,COX5A,Pfdn5,Nap1,RpLP0,Sod1,RpS28b | | 3.597e-02 | -3.32 | GSE557\_WT\_VS\_CIITA\_KO\_DC\_DN | MSigDB lists | GSE557\_WT\_VS\_CIITA\_KO\_DC\_DN | 40 | 11 | 4161 | 644 | CG9911,Ubc4,CG7997,CG40045,ari-1,CG33181,Cse1,Cul2,Idh,Nc73EF,Tsp86D | | 3.597e-02 | -3.32 | MORF\_IKBKG | MSigDB lists | MORF\_IKBKG | 40 | 11 | 4161 | 644 | kcc,Arpc4,CSN6,P5CDh1,Gmer,Rcd5,CkIIalpha,Lsm12,Grip84,GCS2beta,CG9393 | | 3.597e-02 | -3.32 | GSE22025\_UNTREATED\_VS\_TGFB1\_AND\_PROGESTERONE\_TREATED\_CD4\_TCELL\_DN | MSigDB lists | GSE22025\_UNTREATED\_VS\_TGFB1\_AND\_PROGESTERONE\_TREATED\_CD4\_TCELL\_DN | 40 | 11 | 4161 | 644 | Atpalpha,ND-24,Gk2,Caf1-105,Klp10A,ND-75,Arpc1,lt,nrv1,CG9248,mtgo | | 3.597e-02 | -3.32 | DITTMER\_PTHLH\_TARGETS\_UP | MSigDB lists | DITTMER\_PTHLH\_TARGETS\_UP | 40 | 11 | 4161 | 644 | Ts,Mtpalpha,Cdk1,alph,pr,Marf,CG8963,Sply,Mitofilin,CG6610,Cyt-c-p | | 3.597e-02 | -3.32 | GSE22611\_NOD2\_TRANSD\_VS\_CTRL\_TRANSD\_HEK293\_MDP\_STIM\_6H\_UP | MSigDB lists | GSE22611\_NOD2\_TRANSD\_VS\_CTRL\_TRANSD\_HEK293\_MDP\_STIM\_6H\_UP | 40 | 11 | 4161 | 644 | RpS18,Mtpalpha,r-l,Pfk,Treh,bys,Uch,Prosalpha5,Edc3,nAChRalpha4,ND-SGDH | | 3.602e-02 | -3.32 | MOREAUX\_MULTIPLE\_MYELOMA\_BY\_TACI\_DN | MSigDB lists | MOREAUX\_MULTIPLE\_MYELOMA\_BY\_TACI\_DN | 86 | 20 | 4161 | 644 | Nadsyn,CG6610,Rpn12,grp,CG8963,tko,rept,Stam,Crag,CG42542,Ts,CG12713,asf1,Nmt,eIF3m,f-cup,CCT5,CG8611,Cdk1,l(1)G0196 | | 3.602e-02 | -3.32 | GSE25123\_ROSIGLITAZONE\_VS\_IL4\_AND\_ROSIGLITAZONE\_STIM\_PPARG\_KO\_MACROPHAGE\_DAY10\_UP | MSigDB lists | GSE25123\_ROSIGLITAZONE\_VS\_IL4\_AND\_ROSIGLITAZONE\_STIM\_PPARG\_KO\_MACROPHAGE\_DAY10\_UP | 86 | 20 | 4161 | 644 | Karybeta3,CG6766,CG12713,CG8726,rept,PrBP,escl,bru1,CG33156,Coq6,CG3511,esc,SelR,ND-24,mRpS5,Arf51F,APP-BP1,shu,CG14407,DNApol-epsilon255 | | 3.602e-02 | -3.32 | GSE17721\_LPS\_VS\_POLYIC\_12H\_BMDC\_UP | MSigDB lists | GSE17721\_LPS\_VS\_POLYIC\_12H\_BMDC\_UP | 86 | 20 | 4161 | 644 | l(1)G0004,tko,Cbp80,Ublcp1,Atg7,CG10470,CG8728,CG33156,CG3631,shop,Thiolase,CG8314,Echs1,Gdi,Gclc,AIF,CG33181,Iru,CG30496,Unr | | 3.605e-02 | -3.32 | regulation of stem cell proliferation | biological process | GO:0072091 | 58 | 10 | 12345 | 1129 | tws,bun,Rel,COX4,MED27,COX5A,mts,EcR,MED4,MAGE | | 3.605e-02 | -3.32 | fly chr67C5-67C5\|3-31 | chromosome location | fly chr67C5-67C5\|3-31 | 14 | 3 | 25010 | 1343 | ATPsynB,nbs,Ubc4 | | 3.605e-02 | -3.32 | fly chr67C5-67C5\|3-31 cM | chromosome location | fly chr67C5-67C5\|3-31 cM | 14 | 3 | 25010 | 1343 | ATPsynB,nbs,Ubc4 | | 3.615e-02 | -3.32 | PROTEASOME\_BETA\_1 | prosite domains | PS00854 | 12 | 4 | 6243 | 694 | Prosbeta6,Prosbeta4,Prosbeta3,Prosbeta5 | | 3.615e-02 | -3.32 | ZF\_RANBP2\_2 | prosite domains | PS50199 | 12 | 4 | 6243 | 694 | Nup358,sol,RYBP,trbd | | 3.615e-02 | -3.32 | PROTEASOME\_BETA\_2 | prosite domains | PS51476 | 12 | 4 | 6243 | 694 | Prosbeta3,Prosbeta5,Prosbeta4,Prosbeta6 | | 3.618e-02 | -3.32 | BENPORATH\_PROLIFERATION | MSigDB lists | BENPORATH\_PROLIFERATION | 60 | 15 | 4161 | 644 | Non1,Psf3,COX5A,PDCD-5,Cse1,ND-SGDH,asf1,Caf1-105,Cdk1,SRPK,Ts,pr,Pfdn2,Pen,mad2 | | 3.618e-02 | -3.32 | GSE24102\_GRANULOCYSTIC\_MDSC\_VS\_NEUTROPHIL\_UP | MSigDB lists | GSE24102\_GRANULOCYSTIC\_MDSC\_VS\_NEUTROPHIL\_UP | 60 | 15 | 4161 | 644 | ND-51,f-cup,CG1416,CG30010,Atg7,CG3376,Xpac,Aldh,Vha13,sgll,Utx,Pex10,CG1532,mRpL10,CG10038 | | 3.618e-02 | -3.32 | GSE39820\_CTRL\_VS\_TGFBETA1\_IL6\_IL23A\_CD4\_TCELL\_DN | MSigDB lists | GSE39820\_CTRL\_VS\_TGFBETA1\_IL6\_IL23A\_CD4\_TCELL\_DN | 60 | 15 | 4161 | 644 | CG14715,CG7997,Rep,RpS18,Stip1,rept,not,CG3376,Droj2,cathD,nmdyn-D6,Tip60,TyrRS,AGO3,CG2924 | | 3.618e-02 | -3.32 | GSE17721\_POLYIC\_VS\_PAM3CSK4\_24H\_BMDC\_UP | MSigDB lists | GSE17721\_POLYIC\_VS\_PAM3CSK4\_24H\_BMDC\_UP | 60 | 15 | 4161 | 644 | Pex23,Got1,Pi4KIIalpha,RpS30,CycE,MED20,CG8372,CG3803,pr,Arl6IP1,DhpD,Pi3K92E,Tsp74F,PrBP,Naa20A | | 3.618e-02 | -3.32 | CEBP\_Q2 | MSigDB lists | CEBP\_Q2 | 60 | 15 | 4161 | 644 | dco,faf,Cyp6a22,Arf51F,Nca,regucalcin,Synj,Cyp6a19,rept,dnc,cact,CG12713,p38a,Rep,CG14715 | | 3.625e-02 | -3.32 | TGF-beta signaling pathway | KEGG pathways | dme04350 | 40 | 11 | 3198 | 496 | Rbf2,Smox,Pp2A-29B,Cul1,rl,SkpA,Rbf,mts,nej,Med,tkv | | 3.625e-02 | -3.32 | TGF-beta signaling pathway | KEGG pathways | ko04350 | 40 | 11 | 3198 | 496 | rl,SkpA,tkv,Med,nej,mts,Rbf,Pp2A-29B,Rbf2,Smox,Cul1 | | 3.628e-02 | -3.32 | PID\_ERA\_GENOMIC\_PATHWAY | MSigDB lists | PID\_ERA\_GENOMIC\_PATHWAY | 17 | 6 | 4161 | 644 | Med,Sod1,cathD,Phb2,AP-1-2beta,Tsp66E | | 3.628e-02 | -3.32 | KEGG\_TRYPTOPHAN\_METABOLISM | MSigDB lists | KEGG\_TRYPTOPHAN\_METABOLISM | 17 | 6 | 4161 | 644 | Mtpalpha,Aldh,Echs1,TrpRS-m,Nc73EF,CG10932 | | 3.628e-02 | -3.32 | HUMMERICH\_SKIN\_CANCER\_PROGRESSION\_UP | MSigDB lists | HUMMERICH\_SKIN\_CANCER\_PROGRESSION\_UP | 17 | 6 | 4161 | 644 | Eno,Hsc70-3,GCS2beta,Pgk,CG3860,cathD | | 3.628e-02 | -3.32 | GTATGAT\_MIR154\_MIR487 | MSigDB lists | GTATGAT\_MIR154\_MIR487 | 17 | 6 | 4161 | 644 | DhpD,ACC,Snx6,Utx,dnc,Top1 | | 3.628e-02 | -3.32 | JOHNSTONE\_PARVB\_TARGETS\_1\_DN | MSigDB lists | JOHNSTONE\_PARVB\_TARGETS\_1\_DN | 17 | 6 | 4161 | 644 | CanA-14F,Ubc4,Mctp,Su(H),Fdh,Pp2B-14D | | 3.628e-02 | -3.32 | NADLER\_OBESITY\_DN | MSigDB lists | NADLER\_OBESITY\_DN | 17 | 6 | 4161 | 644 | SREBP,Echs1,AGBE,Aldh,Sod1,Nurf-38 | | 3.628e-02 | -3.32 | KEGG\_B\_CELL\_RECEPTOR\_SIGNALING\_PATHWAY | MSigDB lists | KEGG\_B\_CELL\_RECEPTOR\_SIGNALING\_PATHWAY | 17 | 6 | 4161 | 644 | CanA-14F,CanB2,rl,cact,Pi3K92E,Pp2B-14D | | 3.637e-02 | -3.31 | protein import | biological process | GO:0017038 | 91 | 14 | 12345 | 1129 | Pen,Nup62,Pex13,Tom40,Arts,Cse1,Pex12,Pex10,Karybeta3,Kap-alpha3,mbo,Nup358,Roe1,Topors | | 3.643e-02 | -3.31 | catalytic activity, acting on RNA | molecular function | GO:0140098 | 270 | 34 | 12176 | 1119 | GatB,CG11164,Rga,twin,TrpRS-m,CG7009,CysRS,Rpb8,Dcr-1,eIF4A,SerRS-m,CG30343,CG14100,Clp,LysRS,CG11486,Rnmt,RpII140,spn-E,Exd2,scu,mtTFB1,CG8611,CG10463,GatA,trsn,CG6967,TyrRS,GluRS-m,CG12375,Tengl4,CG31759,CG9143,cu | | 3.645e-02 | -3.31 | response to reactive oxygen species | biological process | GO:0000302 | 35 | 7 | 12345 | 1129 | ND-SGDH,spz,Sod3,p38a,Jafrac1,Sod1,rl | | 3.645e-02 | -3.31 | nuclear migration | biological process | GO:0007097 | 35 | 7 | 12345 | 1129 | gammaTub37C,ssh,faf,jvl,l(3)mbt,alphaTub67C,wisp | | 3.645e-02 | -3.31 | protein polymerization | biological process | GO:0051258 | 35 | 7 | 12345 | 1129 | msps,CG7716,SelR,gammaTub37C,Grip84,Grip128,Grip75 | | 3.645e-02 | -3.31 | synaptic growth at neuromuscular junction | biological process | GO:0051124 | 35 | 7 | 12345 | 1129 | Imp,ben,stai,Nlg1,Pdk1,Atpalpha,Liprin-beta | | 3.654e-02 | -3.31 | chromosomal region | cellular component | GO:0098687 | 204 | 27 | 11940 | 1105 | Zwilch,HP1c,nbs,Sirt7,egg,Klp10A,Moe,lwr,yem,Utx,mre11,Cenp-C,mad2,msps,piwi,cnc,Snm1,CycB,trr,EloC,Rcd5,G9a,RpL11,Top1,Tip60,SuUR,Elba2 | | 3.658e-02 | -3.31 | GO\_HEAT\_SHOCK\_PROTEIN\_BINDING | MSigDB lists | GO\_HEAT\_SHOCK\_PROTEIN\_BINDING | 45 | 12 | 4161 | 644 | Droj2,cyc,STUB1,Cdk1,Stip1,cact,shu,HDAC6,PpD3,Nc73EF,CkIIalpha,Fkbp59 | | 3.658e-02 | -3.31 | GO\_HORMONE\_RECEPTOR\_BINDING | MSigDB lists | GO\_HORMONE\_RECEPTOR\_BINDING | 45 | 12 | 4161 | 644 | CG11710,CG8204,Tip60,csw,Fkbp59,Cdk7,MED24,scu,Phb2,MED4,CycE,EcR | | 3.658e-02 | -3.31 | GSE21546\_WT\_VS\_ELK1\_KO\_DP\_THYMOCYTES\_DN | MSigDB lists | GSE21546\_WT\_VS\_ELK1\_KO\_DP\_THYMOCYTES\_DN | 45 | 12 | 4161 | 644 | Dcr-1,CG1847,Hmg-2,nAChRalpha4,ND-75,CG40045,SdhA,AIF,CG3376,CG5734,CG12773,CG17698 | | 3.658e-02 | -3.31 | GSE17721\_CTRL\_VS\_LPS\_1H\_BMDC\_UP | MSigDB lists | GSE17721\_CTRL\_VS\_LPS\_1H\_BMDC\_UP | 45 | 12 | 4161 | 644 | ND-49,CG7728,CanA-14F,RpII140,Arpc1,Pp2B-14D,Vsp37A,CycE,LysRS,fray,alpha-Man-Ia,GCC88 | | 3.658e-02 | -3.31 | GO\_REGULATION\_OF\_REACTIVE\_OXYGEN\_SPECIES\_METABOLIC\_PROCESS | MSigDB lists | GO\_REGULATION\_OF\_REACTIVE\_OXYGEN\_SPECIES\_METABOLIC\_PROCESS | 45 | 12 | 4161 | 644 | Atg6,HDAC6,Sod1,Dhfr,CG2493,regucalcin,Pdk,Tspo,mol,Coq7,p38a,CG7997 | | 3.659e-02 | -3.31 | secretion by cell | biological process | GO:0032940 | 170 | 23 | 12345 | 1129 | Tomosyn,Rep,Gdi,Sec10,nej,Tango10,Syx13,CanB2,X11L,opm,HDAC6,disp,Atg6,Sec15,Vamp7,Arf51F,Exo70,eIF3m,Rab10,Sec22,CHOp24,Crag,lqfR | | 3.659e-02 | -3.31 | GSE4590\_SMALL\_VS\_VPREB\_POS\_LARGE\_PRE\_BCELL\_UP | MSigDB lists | GSE4590\_SMALL\_VS\_VPREB\_POS\_LARGE\_PRE\_BCELL\_UP | 55 | 14 | 4161 | 644 | mtgo,Pka-R2,Kap3,CG3860,CG32549,PpD3,sws,Vamp7,Rep,Tom40,AGBE,CG14894,CG6766,Cul1 | | 3.659e-02 | -3.31 | GO\_SEX\_DIFFERENTIATION | MSigDB lists | GO\_SEX\_DIFFERENTIATION | 55 | 14 | 4161 | 644 | scu,CG11110,Sod1,CycE,tkv,csw,Unr,Idh,Sply,Fkbp59,mtgo,Med,lid,Jafrac2 | | 3.659e-02 | -3.31 | GSE5589\_LPS\_VS\_LPS\_AND\_IL10\_STIM\_MACROPHAGE\_45MIN\_DN | MSigDB lists | GSE5589\_LPS\_VS\_LPS\_AND\_IL10\_STIM\_MACROPHAGE\_45MIN\_DN | 55 | 14 | 4161 | 644 | Rab5,Nca,ND-51,Thiolase,Rbcn-3A,anne,Trxr-1,CG3894,Irp-1B,yip2,Tsp66E,CG15111,SREBP,stet | | 3.659e-02 | -3.31 | GSE2585\_CTEC\_VS\_MTEC\_THYMUS\_UP | MSigDB lists | GSE2585\_CTEC\_VS\_MTEC\_THYMUS\_UP | 55 | 14 | 4161 | 644 | CG3609,kl-2,pr,Tim8,Chd64,CG17698,zyd,raptor,Archease,scu,alc,RpL26,Smox,CG1105 | | 3.659e-02 | -3.31 | GSE6674\_ANTI\_IGM\_VS\_PL2\_3\_STIM\_BCELL\_UP | MSigDB lists | GSE6674\_ANTI\_IGM\_VS\_PL2\_3\_STIM\_BCELL\_UP | 55 | 14 | 4161 | 644 | mad2,Maf1,CG8199,Etf-QO,Sld5,CG6766,SkpA,pr,nmdyn-D7,colt,APC7,UbcE2H,CG11134,CG1774 | | 3.659e-02 | -3.31 | GSE14769\_20MIN\_VS\_360MIN\_LPS\_BMDM\_DN | MSigDB lists | GSE14769\_20MIN\_VS\_360MIN\_LPS\_BMDM\_DN | 55 | 14 | 4161 | 644 | opm,Liprin-beta,GCC88,Kap-alpha3,Cul1,Gdap2,CG4511,Vsp37A,CG5116,CG12818,LysRS,Hsc70-3,CG8728,Rab10 | | 3.667e-02 | -3.31 | Znf\_RING/FYVE/PHD | interpro domains | IPR013083 | 214 | 32 | 11383 | 1233 | Cbl,park,CG31053,d4,not,lid,CG9941,sip3,CG14435,CG17260,nej,HDAC6,CG11070,Iru,CG13344,l(3)73Ah,STUB1,CG8974,Topors,ari-1,Wdfy2,Pex12,Traf6,CG15141,pygo,Usp39,Pex10,pps,CG6923,Usp16-45,dx,trr | | 3.675e-02 | -3.30 | GO\_MEIOTIC\_CELL\_CYCLE | MSigDB lists | GO\_MEIOTIC\_CELL\_CYCLE | 50 | 13 | 4161 | 644 | spn-E,Grip128,Ac3,Pka-R1,Pp2A-29B,Grip75,gammaTub37C,shu,Grip84,piwi,mael,AGO3,mts | | 3.675e-02 | -3.30 | REACTOME\_ORC1\_REMOVAL\_FROM\_CHROMATIN | MSigDB lists | REACTOME\_ORC1\_REMOVAL\_FROM\_CHROMATIN | 50 | 13 | 4161 | 644 | Rpn12,Prosalpha5,Rpn2,RpS27A,Prosbeta3,Prosalpha3,Prosbeta4,Prosalpha4,Rpt2,RpL40,Rpn3,Prosbeta5,Prosbeta6 | | 3.675e-02 | -3.30 | GSE10422\_WT\_VS\_BAFF\_TRANSGENIC\_LN\_BCELL\_UP | MSigDB lists | GSE10422\_WT\_VS\_BAFF\_TRANSGENIC\_LN\_BCELL\_UP | 50 | 13 | 4161 | 644 | Not10,trbd,Mat89Ba,ACC,Synj,Iml1,CG3376,dnc,CG11070,CG17691,Pdk1,O-fut1,Gclc | | 3.675e-02 | -3.30 | RASHI\_RESPONSE\_TO\_IONIZING\_RADIATION\_5 | MSigDB lists | RASHI\_RESPONSE\_TO\_IONIZING\_RADIATION\_5 | 50 | 13 | 4161 | 644 | mRpL17,Gyg,twin,CkIIalpha,Mekk1,Trxr-1,CG14894,anne,Fur2,Caf1-105,Uch-L5,CG5734,Shmt | | 3.675e-02 | -3.30 | GSE40274\_CTRL\_VS\_IRF4\_TRANSDUCED\_ACTIVATED\_CD4\_TCELL\_DN | MSigDB lists | GSE40274\_CTRL\_VS\_IRF4\_TRANSDUCED\_ACTIVATED\_CD4\_TCELL\_DN | 50 | 13 | 4161 | 644 | Fkbp59,Pgm1,CG10465,CG13601,CG8188,CG32549,CCT5,Crag,Pka-R1,r-l,sgll,CG1416,CCT6 | | 3.675e-02 | -3.30 | GSE17721\_PAM3CSK4\_VS\_GADIQUIMOD\_8H\_BMDC\_DN | MSigDB lists | GSE17721\_PAM3CSK4\_VS\_GADIQUIMOD\_8H\_BMDC\_DN | 50 | 13 | 4161 | 644 | CG17691,wal,Arl6IP1,Naa20A,CG9132,shrb,CG9143,CG10465,Sfxn2,CG11899,Tlk,GAPsec,mRpL13 | | 3.675e-02 | -3.30 | GSE7768\_OVA\_WITH\_LPS\_VS\_OVA\_WITH\_MPL\_IMMUNIZED\_MOUSE\_WHOLE\_SPLEEN\_6H\_UP | MSigDB lists | GSE7768\_OVA\_WITH\_LPS\_VS\_OVA\_WITH\_MPL\_IMMUNIZED\_MOUSE\_WHOLE\_SPLEEN\_6H\_UP | 50 | 13 | 4161 | 644 | Tsp74F,r-l,dnc,shop,Tomosyn,Rbf,Pfas,Prosbeta3,Ppcdc,CG32549,Exo70,HDAC6,Smox | | 3.675e-02 | -3.30 | GSE46606\_DAY1\_VS\_DAY3\_CD40L\_IL2\_IL5\_STIMULATED\_IRF4\_KO\_BCELL\_DN | MSigDB lists | GSE46606\_DAY1\_VS\_DAY3\_CD40L\_IL2\_IL5\_STIMULATED\_IRF4\_KO\_BCELL\_DN | 50 | 13 | 4161 | 644 | CG8475,pr,Chd64,Jafrac1,PrBP,CG7718,CG3631,mRpS21,tun,l(3)mbt,CG7789,mRpL10,sau | | 3.688e-02 | -3.30 | GO\_RESPONSE\_TO\_CYTOKINE | MSigDB lists | GO\_RESPONSE\_TO\_CYTOKINE | 124 | 27 | 4161 | 644 | Prosbeta5,cact,rl,LpR2,Rpn3,bys,Prosalpha3,cl,Hsc70-3,Prosbeta4,CycE,Prosalpha5,RpS27A,Ubc4,Ts,Prosbeta6,RpL40,Med,scramb1,Prosbeta3,Gapdh1,Prosalpha4,Rpt2,RpL13A,Traf6,Rpn12,Rpn2 | | 3.712e-02 | -3.29 | RASSF8 (Ras association domain family member 8) | protein interactions | 43096 | 57 | 12 | 9204 | 1112 | cbs,CG14408,RpS14b,RpL7A,CG17002,CG40045,tos,X11L,ASPP,PyK,CG8188,RpS14a | | 3.722e-02 | -3.29 | MARKEY\_RB1\_ACUTE\_LOF\_UP | MSigDB lists | MARKEY\_RB1\_ACUTE\_LOF\_UP | 81 | 19 | 4161 | 644 | Gclc,CycB,Klp10A,Caf1-105,Cdk1,Hat1,Pen,mad2,ERp60,LSm3,Jheh2,Nap1,PpD3,Ts,Karybeta3,Jafrac1,Aos1,CG8188,Cse1 | | 3.722e-02 | -3.29 | GSE24142\_ADULT\_VS\_FETAL\_EARLY\_THYMIC\_PROGENITOR\_DN | MSigDB lists | GSE24142\_ADULT\_VS\_FETAL\_EARLY\_THYMIC\_PROGENITOR\_DN | 81 | 19 | 4161 | 644 | Imp,cyc,CG42542,Karybeta3,sgll,DppIII,Prip,CG3226,Pgm1,mts,capt,CG9911,mad2,Gclc,ND-75,Coq7,CycE,CG4293,Got1 | | 3.722e-02 | -3.29 | GSE39820\_TGFBETA1\_VS\_TGFBETA3\_IN\_IL6\_TREATED\_CD4\_TCELL\_UP | MSigDB lists | GSE39820\_TGFBETA1\_VS\_TGFBETA3\_IN\_IL6\_TREATED\_CD4\_TCELL\_UP | 81 | 19 | 4161 | 644 | Herc4,Smox,Mekk1,CG16865,alpha-Man-Ia,Thiolase,CG8112,CG8974,Vha36-1,CG8963,CG3631,UbcE2H,mino,Sec10,Crag,CG7550,O-fut1,CG2938,regucalcin | | 3.722e-02 | -3.29 | hiw (highwire) | protein interactions | 32429 | 27 | 7 | 9204 | 1112 | p38a,Atg6,raptor,SkpA,Med,faf,tkv | | 3.722e-02 | -3.29 | Mhcl (Myosin heavy chain-like) | protein interactions | 41955 | 27 | 7 | 9204 | 1112 | Trs33,vers,Pxt,26-29-p,Rab5,Cka,mad2 | | 3.722e-02 | -3.29 | CG4887 (uncharacterized protein) | protein interactions | 33304 | 27 | 7 | 9204 | 1112 | RpS28b,Dd,RpS27A,CG4586,CycE,blot,CkIIalpha | | 3.726e-02 | -3.29 | EGFR downregulation | REACTOME pathways | R-DME-182971 | 18 | 6 | 3923 | 574 | Stam,Ptpmeg,RpL40,yl,RpS27A,Cbl | | 3.726e-02 | -3.29 | TGF-beta receptor signaling activates SMADs | REACTOME pathways | R-DME-2173789 | 18 | 6 | 3923 | 574 | RpL40,Med,STUB1,Smox,Fur2,RpS27A | | 3.736e-02 | -3.29 | GSE28726\_ACT\_CD4\_TCELL\_VS\_ACT\_VA24NEG\_NKTCELL\_UP | MSigDB lists | GSE28726\_ACT\_CD4\_TCELL\_VS\_ACT\_VA24NEG\_NKTCELL\_UP | 97 | 22 | 4161 | 644 | tko,mbo,NHP2,CG4511,CG7550,AGBE,CG10470,Rpn2,ScsbetaA,muskelin,mts,Prosbeta3,ERp60,Pgk,Eno,Prosbeta5,CG8004,CG5116,Nmt,Clp,Fkbp59,Mitofilin | | 3.740e-02 | -3.29 | GRAHAM\_CML\_QUIESCENT\_VS\_NORMAL\_QUIESCENT\_UP | MSigDB lists | GRAHAM\_CML\_QUIESCENT\_VS\_NORMAL\_QUIESCENT\_UP | 26 | 8 | 4161 | 644 | Jafrac1,Cyp1,Tm1,Cdk1,CG9471,asf1,grp,Dhfr | | 3.740e-02 | -3.29 | KEGG\_CITRATE\_CYCLE\_TCA\_CYCLE | MSigDB lists | KEGG\_CITRATE\_CYCLE\_TCA\_CYCLE | 26 | 8 | 4161 | 644 | Irp-1B,Idh3b,ScsbetaA,Idh,Nc73EF,Scsalpha1,CG5028,SdhA | | 3.740e-02 | -3.29 | GSE13484\_UNSTIM\_VS\_YF17D\_VACCINE\_STIM\_PBMC\_DN | MSigDB lists | GSE13484\_UNSTIM\_VS\_YF17D\_VACCINE\_STIM\_PBMC\_DN | 26 | 8 | 4161 | 644 | mRpL17,Prosalpha3,CG8188,Irk1,Jafrac2,CG14194,ND-24,scramb1 | | 3.740e-02 | -3.29 | REACTOME\_SIGNALING\_BY\_ERBB4 | MSigDB lists | REACTOME\_SIGNALING\_BY\_ERBB4 | 26 | 8 | 4161 | 644 | RpS27A,Pdk1,rl,SkpA,Cdk1,RpL40,pen-2,Cul1 | | 3.740e-02 | -3.29 | KAMMINGA\_EZH2\_TARGETS | MSigDB lists | KAMMINGA\_EZH2\_TARGETS | 26 | 8 | 4161 | 644 | Cul2,Nap1,Karybeta3,esc,mad2,Pen,escl,Hat1 | | 3.740e-02 | -3.29 | MODULE\_177 | MSigDB lists | MODULE\_177 | 26 | 8 | 4161 | 644 | CG8188,Vamp7,CkIIalpha,Arpc4,lwr,CHOp24,crq,cact | | 3.740e-02 | -3.29 | YAMASHITA\_LIVER\_CANCER\_WITH\_EPCAM\_UP | MSigDB lists | YAMASHITA\_LIVER\_CANCER\_WITH\_EPCAM\_UP | 26 | 8 | 4161 | 644 | RpL13A,RpLP0,RpS27A,RpL9,RpS24,RpS7,SRPK,RpS4 | | 3.740e-02 | -3.29 | fly chr4A5-4A5\|1-6 | chromosome location | fly chr4A5-4A5\|1-6 | 6 | 2 | 25010 | 1343 | cib,Xpac | | 3.740e-02 | -3.29 | fly chr85D25-85D25\|3-48 | chromosome location | fly chr85D25-85D25\|3-48 | 6 | 2 | 25010 | 1343 | Kap-alpha3,CG9399 | | 3.740e-02 | -3.29 | fly chr30B12-30B12\|2-35 cM | chromosome location | fly chr30B12-30B12\|2-35 cM | 6 | 2 | 25010 | 1343 | Mtpalpha,brwl | | 3.740e-02 | -3.29 | fly chr30B12-30B12\|2-35 | chromosome location | fly chr30B12-30B12\|2-35 | 6 | 2 | 25010 | 1343 | brwl,Mtpalpha | | 3.740e-02 | -3.29 | fly chr4A5-4A5\|1-6 cM | chromosome location | fly chr4A5-4A5\|1-6 cM | 6 | 2 | 25010 | 1343 | Xpac,cib | | 3.740e-02 | -3.29 | fly chr38A2-38A2\|2-54 cM | chromosome location | fly chr38A2-38A2\|2-54 cM | 6 | 2 | 25010 | 1343 | CG10631,CG10463 | | 3.740e-02 | -3.29 | fly chr39E3-39E3\|2-54 cM | chromosome location | fly chr39E3-39E3\|2-54 cM | 6 | 2 | 25010 | 1343 | Ac3,CG2201 | | 3.740e-02 | -3.29 | fly chr77C4-77C4\|3-47 | chromosome location | fly chr77C4-77C4\|3-47 | 6 | 2 | 25010 | 1343 | CG5969,isoQC | | 3.740e-02 | -3.29 | fly chr57A8-57A8\|2-91 cM | chromosome location | fly chr57A8-57A8\|2-91 cM | 6 | 2 | 25010 | 1343 | rig,CG11110 | | 3.740e-02 | -3.29 | fly chr57A8-57A8\|2-91 | chromosome location | fly chr57A8-57A8\|2-91 | 6 | 2 | 25010 | 1343 | rig,CG11110 | | 3.740e-02 | -3.29 | fly chr60A9-60A9\|2-105 cM | chromosome location | fly chr60A9-60A9\|2-105 cM | 6 | 2 | 25010 | 1343 | CG17658,Nap1 | | 3.740e-02 | -3.29 | fly chr1C5-1C5\|1-0 | chromosome location | fly chr1C5-1C5\|1-0 | 6 | 2 | 25010 | 1343 | Sec22,Rbf | | 3.740e-02 | -3.29 | fly chr92B4-92B4\|3-67 | chromosome location | fly chr92B4-92B4\|3-67 | 6 | 2 | 25010 | 1343 | CG4686,CG4562 | | 3.740e-02 | -3.29 | fly chr13C4-13C4\|1-50 cM | chromosome location | fly chr13C4-13C4\|1-50 cM | 6 | 2 | 25010 | 1343 | CG6294,Grip128 | | 3.740e-02 | -3.29 | fly chr85D25-85D25\|3-48 cM | chromosome location | fly chr85D25-85D25\|3-48 cM | 6 | 2 | 25010 | 1343 | Kap-alpha3,CG9399 | | 3.740e-02 | -3.29 | fly chr77C4-77C4\|3-47 cM | chromosome location | fly chr77C4-77C4\|3-47 cM | 6 | 2 | 25010 | 1343 | isoQC,CG5969 | | 3.740e-02 | -3.29 | fly chr13C4-13C4\|1-50 | chromosome location | fly chr13C4-13C4\|1-50 | 6 | 2 | 25010 | 1343 | Grip128,CG6294 | | 3.740e-02 | -3.29 | fly chr39E3-39E3\|2-54 | chromosome location | fly chr39E3-39E3\|2-54 | 6 | 2 | 25010 | 1343 | CG2201,Ac3 | | 3.740e-02 | -3.29 | fly chr60A9-60A9\|2-105 | chromosome location | fly chr60A9-60A9\|2-105 | 6 | 2 | 25010 | 1343 | CG17658,Nap1 | | 3.740e-02 | -3.29 | fly chr38A2-38A2\|2-54 | chromosome location | fly chr38A2-38A2\|2-54 | 6 | 2 | 25010 | 1343 | CG10463,CG10631 | | 3.740e-02 | -3.29 | fly chr1C5-1C5\|1-0 cM | chromosome location | fly chr1C5-1C5\|1-0 cM | 6 | 2 | 25010 | 1343 | Sec22,Rbf | | 3.740e-02 | -3.29 | fly chr92B4-92B4\|3-67 cM | chromosome location | fly chr92B4-92B4\|3-67 cM | 6 | 2 | 25010 | 1343 | CG4686,CG4562 | | 3.746e-02 | -3.28 | Cyclin\_C | smart domains | SM01332 | 8 | 3 | 5086 | 506 | CycB,CycE,CycJ | | 3.747e-02 | -3.28 | GO\_REPRODUCTION | MSigDB lists | GO\_REPRODUCTION | 260 | 51 | 4161 | 644 | mts,piwi,csw,Sod1,scu,CHOp24,Fur2,lid,Su(H),cyc,Grip128,regucalcin,Fkbp59,Sply,Idh,Unr,mael,Herc4,Grip75,tkv,shu,Grip84,Mekk1,kri,rl,Cdk1,Dd,faf,Aladin,CG11110,Phb2,r,Pka-R1,Med,mtgo,AGO3,gammaTub37C,CycE,Pp2A-29B,Gabat,CCT5,pont,CycB,r-l,Jafrac2,Ac3,SRPK,CCT3,p38a,bys,spn-E | | 3.748e-02 | -3.28 | MODULE\_272 | MSigDB lists | MODULE\_272 | 9 | 4 | 4161 | 644 | r-l,P5CDh1,CG9471,P5cr | | 3.748e-02 | -3.28 | GO\_RESPONSE\_TO\_OXYGEN\_RADICAL | MSigDB lists | GO\_RESPONSE\_TO\_OXYGEN\_RADICAL | 9 | 4 | 4161 | 644 | Trxr-1,Jafrac1,Sod1,Grx1 | | 3.748e-02 | -3.28 | GO\_C\_ACYLTRANSFERASE\_ACTIVITY | MSigDB lists | GO\_C\_ACYLTRANSFERASE\_ACTIVITY | 9 | 4 | 4161 | 644 | Mtpalpha,CG10932,Thiolase,yip2 | | 3.748e-02 | -3.28 | REACTOME\_PRE\_NOTCH\_EXPRESSION\_AND\_PROCESSING | MSigDB lists | REACTOME\_PRE\_NOTCH\_EXPRESSION\_AND\_PROCESSING | 9 | 4 | 4161 | 644 | CHOp24,O-fut1,SERCA,Su(H) | | 3.748e-02 | -3.28 | BIOCARTA\_NO1\_PATHWAY | MSigDB lists | BIOCARTA\_NO1\_PATHWAY | 9 | 4 | 4161 | 644 | slif,mAChR-A,Pka-R2,Pka-R1 | | 3.748e-02 | -3.28 | GO\_SPINDLE\_CHECKPOINT | MSigDB lists | GO\_SPINDLE\_CHECKPOINT | 9 | 4 | 4161 | 644 | CkIIalpha,Tao,CG1812,mad2 | | 3.748e-02 | -3.28 | GO\_GTP\_METABOLIC\_PROCESS | MSigDB lists | GO\_GTP\_METABOLIC\_PROCESS | 9 | 4 | 4161 | 644 | mal,nmdyn-D7,AMPdeam,nmdyn-D6 | | 3.748e-02 | -3.28 | HEDENFALK\_BREAST\_CANCER\_BRACX\_UP | MSigDB lists | HEDENFALK\_BREAST\_CANCER\_BRACX\_UP | 9 | 4 | 4161 | 644 | RpS4,RpS13,RpL40,Utx | | 3.748e-02 | -3.28 | GO\_MODULATION\_BY\_SYMBIONT\_OF\_HOST\_CELLULAR\_PROCESS | MSigDB lists | GO\_MODULATION\_BY\_SYMBIONT\_OF\_HOST\_CELLULAR\_PROCESS | 9 | 4 | 4161 | 644 | Clp,Atg7,Pen,Kap-alpha3 | | 3.748e-02 | -3.28 | GO\_MICROTUBULE\_NUCLEATION | MSigDB lists | GO\_MICROTUBULE\_NUCLEATION | 9 | 4 | 4161 | 644 | Grip128,Grip84,gammaTub37C,Grip75 | | 3.748e-02 | -3.28 | GO\_REGULATION\_OF\_TRANSCRIPTION\_FROM\_RNA\_POLYMERASE\_II\_PROMOTER\_IN\_RESPONSE\_TO\_HYPOXIA | MSigDB lists | GO\_REGULATION\_OF\_TRANSCRIPTION\_FROM\_RNA\_POLYMERASE\_II\_PROMOTER\_IN\_RESPONSE\_TO\_HYPOXIA | 9 | 4 | 4161 | 644 | Cul2,RpL40,Su(H),RpS27A | | 3.748e-02 | -3.28 | GO\_MITOCHONDRIAL\_CALCIUM\_ION\_HOMEOSTASIS | MSigDB lists | GO\_MITOCHONDRIAL\_CALCIUM\_ION\_HOMEOSTASIS | 9 | 4 | 4161 | 644 | SERCA,AnxB10,Mitofilin,Calx | | 3.748e-02 | -3.28 | GO\_EXIT\_FROM\_MITOSIS | MSigDB lists | GO\_EXIT\_FROM\_MITOSIS | 9 | 4 | 4161 | 644 | shrb,tws,Sirt7,CG8188 | | 3.748e-02 | -3.28 | PID\_P38\_MKK3\_6PATHWAY | MSigDB lists | PID\_P38\_MKK3\_6PATHWAY | 9 | 4 | 4161 | 644 | Mekk1,Tao,Traf6,p38a | | 3.748e-02 | -3.28 | LEE\_LIVER\_CANCER\_ACOX1\_UP | MSigDB lists | LEE\_LIVER\_CANCER\_ACOX1\_UP | 9 | 4 | 4161 | 644 | isoQC,zyd,scramb1,RpL12 | | 3.748e-02 | -3.28 | GNF2\_RAP1B | MSigDB lists | GNF2\_RAP1B | 9 | 4 | 4161 | 644 | Mo25,Snx6,Moe,lid | | 3.748e-02 | -3.28 | GO\_G2\_DNA\_DAMAGE\_CHECKPOINT | MSigDB lists | GO\_G2\_DNA\_DAMAGE\_CHECKPOINT | 9 | 4 | 4161 | 644 | CG30291,Cdk1,Tao,grp | | 3.770e-02 | -3.28 | TYR\_PHOSPHATASE\_dom | interpro domains | IPR000387 | 30 | 7 | 11383 | 1233 | Ptp69D,Ptp10D,Ptpmeg2,ssh,csw,Ptpmeg,Ptp61F | | 3.785e-02 | -3.27 | regulation of stem cell differentiation | biological process | GO:2000736 | 28 | 6 | 12345 | 1129 | vers,Stat92E,Cenp-C,Cka,lid,bru1 | | 3.785e-02 | -3.27 | cellular response to alcohol | biological process | GO:0097306 | 28 | 6 | 12345 | 1129 | Pka-R2,MED27,EcR,Nurf-38,G9a,Utx | | 3.785e-02 | -3.27 | RNA polyadenylation | biological process | GO:0043631 | 28 | 6 | 12345 | 1129 | sbr,IntS11,Clp,Trf4-2,nos,wisp | | 3.811e-02 | -3.27 | steroid dehydrogenase activity | molecular function | GO:0016229 | 21 | 5 | 12176 | 1119 | CG12171,CG31548,CG7840,scu,CG31549 | | 3.811e-02 | -3.27 | Cyclin-like | interpro domains | IPR013763 | 18 | 5 | 11383 | 1233 | CycE,CycJ,Rbf,CycC,CycB | | 3.811e-02 | -3.27 | LSM\_dom\_euk/arc | interpro domains | IPR001163 | 18 | 5 | 11383 | 1233 | SmE,CG6610,LSm3,CG10418,CG17768 | | 3.816e-02 | -3.27 | GO\_RESPONSE\_TO\_OXIDATIVE\_STRESS | MSigDB lists | GO\_RESPONSE\_TO\_OXIDATIVE\_STRESS | 108 | 24 | 4161 | 644 | Prip,HDAC6,Sod1,Cyt-c-p,ND-49,P5cr,Tm1,cyc,Jafrac1,Cyp1,ND-23,Grx1,Etf-QO,Xpac,fray,Trxr-1,PpD3,Idh,Kap-alpha3,Cdk1,AIF,Gclc,SelR,Prosbeta5 | | 3.825e-02 | -3.26 | peroxisome | cellular component | GO:0005777 | 99 | 15 | 11940 | 1105 | Mtpalpha,CG10932,Sod1,CG4586,Marf1,Idh,CG11077,Got1,Acox57D-p,Pex10,Pex13,CG6178,ABCD,Pex12,Topors | | 3.825e-02 | -3.26 | microbody | cellular component | GO:0042579 | 99 | 15 | 11940 | 1105 | CG6178,Pex13,Pex12,ABCD,Topors,Acox57D-p,Pex10,Got1,CG11077,Idh,Marf1,CG4586,Sod1,CG10932,Mtpalpha | | 3.829e-02 | -3.26 | negative regulation of cell differentiation | biological process | GO:0045596 | 135 | 19 | 12345 | 1129 | Stat92E,COX4,alph,PGRP-LE,Ptpmeg,mts,MED4,lwr,ago,MED27,Su(H),bun,tws,lid,EcR,MAGE,bif,COX5A,cact | | 3.833e-02 | -3.26 | GCAAGAC\_MIR431 | MSigDB lists | GCAAGAC\_MIR431 | 13 | 5 | 4161 | 644 | bru1,Med,Ptp61F,lid,Irk1 | | 3.833e-02 | -3.26 | GO\_NUCLEAR\_LOCALIZATION\_SEQUENCE\_BINDING | MSigDB lists | GO\_NUCLEAR\_LOCALIZATION\_SEQUENCE\_BINDING | 13 | 5 | 4161 | 644 | Pen,Kap-alpha3,Arts,cact,Karybeta3 | | 3.833e-02 | -3.26 | DIRMEIER\_LMP1\_RESPONSE\_LATE\_DN | MSigDB lists | DIRMEIER\_LMP1\_RESPONSE\_LATE\_DN | 13 | 5 | 4161 | 644 | Gapdh1,RpS30,RpLP0,CG10418,Nca | | 3.833e-02 | -3.26 | HU\_GENOTOXIN\_ACTION\_DIRECT\_VS\_INDIRECT\_4HR | MSigDB lists | HU\_GENOTOXIN\_ACTION\_DIRECT\_VS\_INDIRECT\_4HR | 13 | 5 | 4161 | 644 | Pfk,cact,Irk1,yip2,EcR | | 3.833e-02 | -3.26 | GO\_STEROL\_BINDING | MSigDB lists | GO\_STEROL\_BINDING | 13 | 5 | 4161 | 644 | Npc2a,AnxB10,Tspo,CG8112,CG3860 | | 3.833e-02 | -3.26 | WALLACE\_PROSTATE\_CANCER\_UP | MSigDB lists | WALLACE\_PROSTATE\_CANCER\_UP | 13 | 5 | 4161 | 644 | CanA-14F,CG12018,Gmer,eIF3i,Pp2B-14D | | 3.833e-02 | -3.26 | KUROKAWA\_LIVER\_CANCER\_CHEMOTHERAPY\_DN | MSigDB lists | KUROKAWA\_LIVER\_CANCER\_CHEMOTHERAPY\_DN | 13 | 5 | 4161 | 644 | Vha36-1,CG32473,Rpb8,mad2,eEF1beta | | 3.833e-02 | -3.26 | JOSEPH\_RESPONSE\_TO\_SODIUM\_BUTYRATE\_UP | MSigDB lists | JOSEPH\_RESPONSE\_TO\_SODIUM\_BUTYRATE\_UP | 13 | 5 | 4161 | 644 | UQCR-Q,Thiolase,levy,Tsp66E,Gdi | | 3.833e-02 | -3.26 | GO\_REGULATION\_OF\_INTRACELLULAR\_ESTROGEN\_RECEPTOR\_SIGNALING\_PATHWAY | MSigDB lists | GO\_REGULATION\_OF\_INTRACELLULAR\_ESTROGEN\_RECEPTOR\_SIGNALING\_PATHWAY | 13 | 5 | 4161 | 644 | Phb2,Uba5,Cka,CG16979,Ufm1 | | 3.833e-02 | -3.26 | KEGG\_BETA\_ALANINE\_METABOLISM | MSigDB lists | KEGG\_BETA\_ALANINE\_METABOLISM | 13 | 5 | 4161 | 644 | Gabat,Echs1,Mcad,Mtpalpha,Aldh | | 3.833e-02 | -3.26 | GO\_PYRIMIDINE\_NUCLEOBASE\_METABOLIC\_PROCESS | MSigDB lists | GO\_PYRIMIDINE\_NUCLEOBASE\_METABOLIC\_PROCESS | 13 | 5 | 4161 | 644 | r,r-l,rl,kri,Ts | | 3.833e-02 | -3.26 | REACTOME\_DOWNREGULATION\_OF\_TGF\_BETA\_RECEPTOR\_SIGNALING | MSigDB lists | REACTOME\_DOWNREGULATION\_OF\_TGF\_BETA\_RECEPTOR\_SIGNALING | 13 | 5 | 4161 | 644 | STUB1,RpS27A,RpL40,Uch-L5,Smox | | 3.833e-02 | -3.26 | MOREAUX\_B\_LYMPHOCYTE\_MATURATION\_BY\_TACI\_UP | MSigDB lists | MOREAUX\_B\_LYMPHOCYTE\_MATURATION\_BY\_TACI\_UP | 13 | 5 | 4161 | 644 | grsm,Gip,Pi3K92E,RpL38,rl | | 3.833e-02 | -3.26 | MODULE\_305 | MSigDB lists | MODULE\_305 | 13 | 5 | 4161 | 644 | Aldh,Echs1,Irp-1B,CG10932,Shmt | | 3.833e-02 | -3.26 | GO\_NEGATIVE\_REGULATION\_OF\_MITOTIC\_NUCLEAR\_DIVISION | MSigDB lists | GO\_NEGATIVE\_REGULATION\_OF\_MITOTIC\_NUCLEAR\_DIVISION | 13 | 5 | 4161 | 644 | CG1812,nmdyn-D6,grp,mad2,CkIIalpha | | 3.833e-02 | -3.26 | chr5q12 | MSigDB lists | chr5q12 | 13 | 5 | 4161 | 644 | dnc,CG3511,CycB,Cdk7,Klp10A | | 3.833e-02 | -3.26 | GO\_NEGATIVE\_REGULATION\_OF\_REACTIVE\_OXYGEN\_SPECIES\_METABOLIC\_PROCESS | MSigDB lists | GO\_NEGATIVE\_REGULATION\_OF\_REACTIVE\_OXYGEN\_SPECIES\_METABOLIC\_PROCESS | 13 | 5 | 4161 | 644 | CG7997,Tspo,HDAC6,Atg6,regucalcin | | 3.838e-02 | -3.26 | HALLMARK\_FATTY\_ACID\_METABOLISM | MSigDB lists | HALLMARK\_FATTY\_ACID\_METABOLISM | 76 | 18 | 4161 | 644 | Mcad,ScsbetaA,scu,Etf-QO,CG17691,CG5028,Idh,Scsalpha1,tkv,yip2,Pcd,Jheh2,Idh3b,pr,Echs1,SdhA,CG13630,Thiolase | | 3.851e-02 | -3.26 | integral component of organelle membrane | cellular component | GO:0031301 | 160 | 22 | 11940 | 1105 | Pex10,CG9399,Pex12,ste24a,EMRE,Seipin,CG10470,Pmi,CG32441,Ugalt,CG8311,Fur2,fwe,stet,CG31523,Efr,alt,CG4293,CG5903,EMC8-9,EMC3,CG9773 | | 3.852e-02 | -3.26 | mitotic spindle | cellular component | GO:0072686 | 35 | 7 | 11940 | 1105 | Clc,Cdk1,Grip84,CycB,Moe,Klp10A,mad2 | | 3.860e-02 | -3.25 | SAM | smart domains | SM00454 | 26 | 6 | 5086 | 506 | mtrm,ph-d,Ect4,Liprin-beta,BicC,smg | | 3.861e-02 | -3.25 | GO\_LYTIC\_VACUOLE | MSigDB lists | GO\_LYTIC\_VACUOLE | 169 | 35 | 4161 | 644 | AP-1-2beta,CG42542,CG3842,Iml1,raptor,CG3860,Sod1,CG7997,crq,kcc,Npc2a,Pex23,CG7806,Got1,Pi4KIIalpha,CG14516,Pi3K92E,Gdap2,RpL40,Rab5,cathD,Vha36-1,CG2493,AnxB10,lt,CHMP2B,CG3376,CG32473,Gyg,CG1827,ZnT35C,Acph-1,Vamp7,Clc,sws | | 3.879e-02 | -3.25 | alcohol biosynthetic process | biological process | GO:0046165 | 43 | 8 | 12345 | 1129 | foxo,Marf,CG17270,l(1)G0196,pr,Sptr,CG7840,Pcd | | 3.879e-02 | -3.25 | positive regulation of Wnt signaling pathway | biological process | GO:0030177 | 43 | 8 | 12345 | 1129 | dco,Ssdp,tws,trbd,spen,mts,pont,nej | | 3.879e-02 | -3.25 | amino acid activation | biological process | GO:0043038 | 43 | 8 | 12345 | 1129 | TrpRS-m,GluRS-m,LysRS,GatA,CysRS,GatB,SerRS-m,TyrRS | | 3.879e-02 | -3.25 | positive regulation of cell size | biological process | GO:0045793 | 43 | 8 | 12345 | 1129 | p38a,raptor,Mekk1,Desat1,Pi3K92E,Pdk1,Girdin,rl | | 3.897e-02 | -3.24 | LSM | pfam domains | PF01423 | 18 | 5 | 10480 | 1142 | SmE,CG17768,CG10418,CG6610,LSm3 | | 3.923e-02 | -3.24 | isomerase activity | molecular function | GO:0016853 | 100 | 15 | 12176 | 1119 | Fkbp12,CG3511,Pgm1,Gmer,ERp60,Pglym78,Top1,CG4594,Pus7,Naxe,Fkbp59,CG4598,Gip,Cyp1,CG11858 | | 3.933e-02 | -3.24 | GO\_NEGATIVE\_REGULATION\_OF\_GENE\_EXPRESSION | MSigDB lists | GO\_NEGATIVE\_REGULATION\_OF\_GENE\_EXPRESSION | 325 | 62 | 4161 | 644 | Wnt10,AGO3,SREBP,CG4872,Dcr-1,Cka,Smox,Hmg-2,CG4709,Coq7,Eno,esc,CycB,Gclc,Cdk5,spn-E,lwr,Hat1,CG15715,CG17187,CG30291,Aladin,Sirt7,faf,Pfdn5,Phb2,Traf6,Ranbp21,heph,Maf1,Snx6,RpL40,Cbp80,Imp,mbo,Med,mael,twin,YL-1,RpS27A,shu,cact,LpR2,Bin1,piwi,Gapdh1,CG11164,RpII140,Not10,RpL13A,eIF4E1,trsn,RpS13,EcR,adp,HDAC6,Rpb8,lid,cyc,Tip60,escl,Su(H) | | 3.941e-02 | -3.23 | Sirt1 (Sirtuin 1) | protein interactions | 34708 | 39 | 9 | 9204 | 1112 | Su(H),Kcmf1,nej,foxo,Hnf4,Nap1,Atpalpha,Pink1,DnaJ-H | | 3.941e-02 | -3.23 | Kr (Kruppel) | protein interactions | 38012 | 39 | 9 | 9204 | 1112 | nej,Bin1,hb,piwi,rept,bun,Bx,ovo,Stip1 | | 3.946e-02 | -3.23 | Golgi apparatus | cellular component | GO:0005794 | 364 | 44 | 11940 | 1105 | cbs,CG8314,Rab10,ATP8A,CG31729,opm,CG5880,Rab32,CG32485,CG9981,CG3631,Sec22,stet,Pi4KIIalpha,CG10344,eIF3m,Fur2,Gga,CG9773,CG2938,Fer1HCH,CG4557,CG4293,Clc,O-fut1,CG5447,PHGPx,AP-1-2beta,alpha-Man-Ia,Ugalt,CG5510,Gmap,sau,Trs33,Afti,GalT1,CHOp24,Efr,GCC88,park,Arl5,Sec23,fdl,lqfR | | 3.948e-02 | -3.23 | GSE3982\_NEUTROPHIL\_VS\_EFF\_MEMORY\_CD4\_TCELL\_DN | MSigDB lists | GSE3982\_NEUTROPHIL\_VS\_EFF\_MEMORY\_CD4\_TCELL\_DN | 71 | 17 | 4161 | 644 | STUB1,Pus7,esc,wal,CG10344,Ptpmeg,Klp10A,Ssdp,CG7806,Ppox,Stoml2,escl,TyrRS,CycC,RpL13A,RpS16,CG4390 | | 3.955e-02 | -3.23 | SCF-dependent proteasomal ubiquitin-dependent protein catabolic process | biological process | GO:0031146 | 67 | 11 | 12345 | 1129 | RYBP,CG31633,Cul1,CG5003,SkpA,Cul2,Cul5,CG9316,ago,CG32221,Fbw5 | | 3.957e-02 | -3.23 | mRNA processing | biological process | GO:0006397 | 292 | 36 | 12345 | 1129 | obe,Rnmt,CG5986,yps,sbr,CG9684,Cbp80,CG11486,CG3198,CG10418,Bin1,SmE,Son,CG1622,Usp39,wisp,CG16865,Spx,snRNA:U12,IntS11,CG17768,bru1,rig,snRNA:U4:38AB,tho2,Clp,CG6610,Imp,LSm3,CG17187,exu,nos,heph,snRNA:U4atac,CG17454,Neos | | 3.966e-02 | -3.23 | Removal of licensing factors from origins | REACTOME pathways | R-DME-69300 | 75 | 17 | 3923 | 574 | Rbf2,Prosbeta5,Rpn2,Prosalpha4,RpS27A,Prosalpha5,Prosbeta4,Prosalpha3,Rpt2,RpL40,Rbf,Prosbeta3,Orc6,Rpn12,Rpn3,Orc1,Prosbeta6 | | 3.966e-02 | -3.23 | APC/C:Cdc20 mediated degradation of mitotic proteins | REACTOME pathways | R-DME-176409 | 75 | 17 | 3923 | 574 | Prosbeta3,Rpn3,Rpn12,Cdk1,CycB,Prosbeta6,APC7,Rpn2,Prosbeta5,Prosalpha4,Prosalpha5,mad2,RpS27A,Prosbeta4,RpL40,Rpt2,Prosalpha3 | | 3.966e-02 | -3.23 | 14-3-3zeta (14-3-3zeta) | protein interactions | 36059 | 90 | 17 | 9204 | 1112 | Cenp-C,Idh,Sec10,RpLP1,REPTOR,tral,sol,jvl,RpS27A,awd,tor,osk,foxo,faf,AnxB10,regucalcin,csw | | 3.978e-02 | -3.22 | euchromatin | cellular component | GO:0000791 | 28 | 6 | 11940 | 1105 | HP1c,lin-52,piwi,Rel,G9a,Top1 | | 3.978e-02 | -3.22 | proteasome regulatory particle | cellular component | GO:0005838 | 28 | 6 | 11940 | 1105 | Rpn13,Rpt2,Rpn3,Uch-L5,Rpn2,Rpn12 | | 3.998e-02 | -3.22 | TPR | smart domains | SM00028 | 62 | 11 | 5086 | 506 | Stip1,Tom70,adp,Utx,PpD3,CG14894,APC7,STUB1,spag,Fkbp59,CG1847 | | 3.999e-02 | -3.22 | glycerophospholipid biosynthetic process | biological process | GO:0046474 | 59 | 10 | 12345 | 1129 | CG33977,PIG-H,PIG-C,Agpat2,CG2201,PIG-L,CG7718,PIG-P,Pld,Pi3K92E | | 3.999e-02 | -3.22 | organic hydroxy compound biosynthetic process | biological process | GO:1901617 | 59 | 10 | 12345 | 1129 | Pcd,Sptr,CG7840,pr,l(1)G0196,Pdxk,CG17270,sgll,foxo,Marf | | 3.999e-02 | -3.22 | cell cycle checkpoint | biological process | GO:0000075 | 59 | 10 | 12345 | 1129 | Zwilch,nbs,mus304,grp,Cdk1,mre11,CG10336,nej,mad2,mus81 | | 4.011e-02 | -3.22 | GSE27786\_BCELL\_VS\_ERYTHROBLAST\_DN | MSigDB lists | GSE27786\_BCELL\_VS\_ERYTHROBLAST\_DN | 31 | 9 | 4161 | 644 | CG12713,CysRS,sgll,AP-1-2beta,CG11134,CG7115,Nc73EF,CG14407,HDAC6 | | 4.011e-02 | -3.22 | GO\_CELLULAR\_RESPONSE\_TO\_ACID\_CHEMICAL | MSigDB lists | GO\_CELLULAR\_RESPONSE\_TO\_ACID\_CHEMICAL | 31 | 9 | 4161 | 644 | Shmt,Pdk,ACC,CycB,Karybeta3,Prip,SREBP,raptor,Wnt10 | | 4.011e-02 | -3.22 | DARWICHE\_SKIN\_TUMOR\_PROMOTER\_UP | MSigDB lists | DARWICHE\_SKIN\_TUMOR\_PROMOTER\_UP | 31 | 9 | 4161 | 644 | Pgk,Pglym78,eEF1beta,Eno,NHP2,CG4390,Sod1,Vsp37A,PDCD-5 | | 4.011e-02 | -3.22 | REACTOME\_TOLL\_RECEPTOR\_CASCADES | MSigDB lists | REACTOME\_TOLL\_RECEPTOR\_CASCADES | 31 | 9 | 4161 | 644 | mts,Traf6,Pp2A-29B,RpS27A,rl,cact,Cdk1,p38a,RpL40 | | 4.012e-02 | -3.22 | KINSEY\_TARGETS\_OF\_EWSR1\_FLII\_FUSION\_UP | MSigDB lists | KINSEY\_TARGETS\_OF\_EWSR1\_FLII\_FUSION\_UP | 480 | 88 | 4161 | 644 | CG7530,csw,Rrp40,Sld5,CG15141,opm,ste14,Arpc2,CG7840,Nmt,GatA,Caf1-105,mRpS10,Chrac-14,Pen,CycC,Psf3,CkIIalpha,CG6767,RpS2,Cyt-c-p,ScsbetaA,CG40045,SelD,PpV,Ranbp21,Ts,scramb1,Aos1,Plap,Cyp1,Imp,Cbp80,mbo,Iru,CG6665,r-l,CG10425,CG8112,CG33156,CG3226,CG11899,grp,Rpn2,Nurf-38,CG8188,mino,trsn,GluRS-m,CHOp24,zyd,Fdh,l(1)G0004,Got1,Grip75,asf1,Cdk1,Klp10A,Pus7,Shmt,mad2,Uch-L5,Pgk,CG1532,tun,tws,Dhfr,RhoGAP54D,Sec10,CG4390,Roe1,RfC38,Usp39,heph,Med,CG11134,CG7414,CG9581,Dcr-1,eIF3m,PpD3,gammaTub37C,Stip1,CycB,pont,Eno,Cul1,msps | | 4.021e-02 | -3.21 | nucleic acid binding | molecular function | GO:0003676 | 2082 | 213 | 12176 | 1119 | Med,Rbfox1,psq,enc,CG7414,Top1,CG10492,RpL12,Nxf3,Taf7,mus205,Trf2,Rnmt,exu,pre-lola-G,CG4021,spen,CG6610,mRpS11,RpS13,CG2199,Nf-YA,CG4730,mus81,egg,CG12018,insv,Smox,CG17454,RpL9,hng3,RpL26,RfC38,CG8611,cad,CG10189,hb,Bap170,Larp4B,CG6967,Synj,Cbp80,Cf2,Tengl4,DNApol-epsilon255,CG6813,CG12288,GluRS-m,CG9143,Rcd5,CG8963,RpS2,tho2,zf30C,l(3)73Ah,Mitf,Nup62,CG3198,CG10654,Unr,RpS15,mRpL16,dl,RpL7A,cup,RpS14b,CG4709,CG31875,CG10038,del,Hnf4,RpL30,Maf1,CG4404,CG7488,PPYR1,Irp-1B,nbs,CG5986,Rbf2,RpLP0,smg,Chd64,Orc6,brwl,CG11902,mael,sbb,CG1602,CG15514,Stat92E,tef,Ssdp,gw,mRpS5,eIF3i,mRpL13,RpS5b,bru1,CG2129,CG10418,RYBP,jigr1,yem,Larp7,sbr,l(3)mbt,Elba2,Rbf,HmgD,CG9425,AIF,e(y)1,RpS16,TfIIFalpha,mEFTu1,l(1)G0004,apt,CG14100,Clp,RpL13A,cnc,CG10336,BicC,Son,Fkbp39,obe,Mat89Ba,CG7728,mEFTu2,Marf1,trsn,tos,RpL24,Wbp2,foxo,CG17187,Spx,NHP2,CG12375,EcR,LSm3,CG6276,Exd2,spn-E,armi,RpL23,Atf3,CG2162,qkr58E-2,mRpL18,E2f1,CG2678,Pus7,SREBP,wdn,eIF3m,r2d2,swa,Xpc,Atf6,CysRS,eIF4A,Dcr-1,cyc,Su(H),lid,bcd,RpS4,CG11486,eIF4E1,SLIRP1,elg1,Hr78,LysRS,Xpac,CG17768,Edc3,Utx,shep,NFAT,bys,BigH1,Mybbp1A,PDCD-5,Snm1,Rel,Neos,CG2100,RpS18,Rrp40,ovo,crm,CG10631,yps,nos,Imp,Pym,CG10959,RpS14a,eEF1beta,TyrRS,bam,RpII140,CG34116,piwi,CG4266,AGO3,osk,heph,CG17328,Shmt,CG6094 | | 4.024e-02 | -3.21 | CG16935 (uncharacterized protein) | protein interactions | 36540 | 3 | 2 | 9204 | 1112 | p38a,Rpn2 | | 4.024e-02 | -3.21 | l(3)mbn (lethal (3) malignant blood neoplasm) | protein interactions | 38701 | 3 | 2 | 9204 | 1112 | Ssdp,Bx | | 4.024e-02 | -3.21 | Nmdmc (NAD-dependent methylenetetrahydrofolate dehydrogenase) | protein interactions | 47895 | 3 | 2 | 9204 | 1112 | park,Pink1 | | 4.024e-02 | -3.21 | CG8209 (uncharacterized protein) | protein interactions | 38888 | 3 | 2 | 9204 | 1112 | Usp39,RpS27A | | 4.024e-02 | -3.21 | Sufu (SUFU negative regulator of hedgehog signaling) | protein interactions | 24069 | 3 | 2 | 9204 | 1112 | fdl,Bin1 | | 4.024e-02 | -3.21 | CG13599 (uncharacterized protein) | protein interactions | 42829 | 3 | 2 | 9204 | 1112 | GILT1,Gip | | 4.024e-02 | -3.21 | Ehbp1 (Eps15 homology domain containing protein-binding protein 1) | protein interactions | 36912 | 3 | 2 | 9204 | 1112 | mre11,Sec15 | | 4.024e-02 | -3.21 | CG14341 (uncharacterized protein) | protein interactions | 33301 | 3 | 2 | 9204 | 1112 | CG2199,Hsp26 | | 4.024e-02 | -3.21 | mdlc (midlife crisis) | protein interactions | 42397 | 3 | 2 | 9204 | 1112 | Pfdn2,CkIIalpha | | 4.024e-02 | -3.21 | Hen1 (Hen1 methyltransferase) | protein interactions | 36301 | 3 | 2 | 9204 | 1112 | piwi,CG9684 | | 4.024e-02 | -3.21 | CG31549 (uncharacterized protein) | protein interactions | 40689 | 3 | 2 | 9204 | 1112 | CG12171,CG31548 | | 4.024e-02 | -3.21 | CG8223 (uncharacterized protein) | protein interactions | 41045 | 3 | 2 | 9204 | 1112 | RpS27A,gus | | 4.024e-02 | -3.21 | Vps15 (Vacuolar protein sorting 15) | protein interactions | 41096 | 3 | 2 | 9204 | 1112 | Lkb1,Rab5 | | 4.024e-02 | -3.21 | galectin (galectin) | protein interactions | 33162 | 3 | 2 | 9204 | 1112 | Pp2B-14D,Pura | | 4.024e-02 | -3.21 | GstT3 (Glutathione S transferase T3) | protein interactions | 33047 | 3 | 2 | 9204 | 1112 | CycE,Cchl | | 4.024e-02 | -3.21 | Rnp4F (RNA-binding protein 4F) | protein interactions | 31448 | 3 | 2 | 9204 | 1112 | CG17768,LSm3 | | 4.024e-02 | -3.21 | snk (snake) | protein interactions | 41607 | 3 | 2 | 9204 | 1112 | gd,ea | | 4.024e-02 | -3.21 | CG8353 (uncharacterized protein) | protein interactions | 34123 | 3 | 2 | 9204 | 1112 | Pglym78,Gapdh1 | | 4.024e-02 | -3.21 | SCaMC (Short Calcium-binding Mitochondrial Carrier) | protein interactions | 39415 | 3 | 2 | 9204 | 1112 | grsm,Abi | | 4.024e-02 | -3.21 | P5CDh1 (delta-1-Pyrroline-5-carboxylate dehydrogenase 1) | protein interactions | 40434 | 3 | 2 | 9204 | 1112 | Plap,plu | | 4.024e-02 | -3.21 | CG8180 (uncharacterized protein) | protein interactions | 36713 | 3 | 2 | 9204 | 1112 | wisp,smg | | 4.024e-02 | -3.21 | SpdS (Spermidine Synthase) | protein interactions | 41167 | 3 | 2 | 9204 | 1112 | Rtnl1,Drep1 | | 4.024e-02 | -3.21 | CG11399 (uncharacterized protein) | protein interactions | 40295 | 3 | 2 | 9204 | 1112 | dx,Bx | | 4.024e-02 | -3.21 | Rhp (Rhophilin) | protein interactions | 32531 | 3 | 2 | 9204 | 1112 | DppIII,CG32536 | | 4.024e-02 | -3.21 | Srr (Serine racemase) | protein interactions | 41117 | 3 | 2 | 9204 | 1112 | Smox,RpL34a | | 4.024e-02 | -3.21 | AdenoK (Adenosine Kinase) | protein interactions | 39479 | 3 | 2 | 9204 | 1112 | Gyg,CG17187 | | 4.024e-02 | -3.21 | Tom7 (Translocase of outer membrane 7) | protein interactions | 35899 | 3 | 2 | 9204 | 1112 | Kyat,Pink1 | | 4.024e-02 | -3.21 | CG12219 (uncharacterized protein) | protein interactions | 31572 | 3 | 2 | 9204 | 1112 | tos,Sec10 | | 4.024e-02 | -3.21 | mbm (mushroom body miniature) | protein interactions | 33194 | 3 | 2 | 9204 | 1112 | alpha-Man-Ia,CkIIalpha | | 4.024e-02 | -3.21 | CG5793 (uncharacterized protein) | protein interactions | 42505 | 3 | 2 | 9204 | 1112 | Pfdn5,CG14881 | | 4.024e-02 | -3.21 | Rpp20 (Rpp20) | protein interactions | 3772007 | 3 | 2 | 9204 | 1112 | Ptp10D,CG9902 | | 4.025e-02 | -3.21 | MASSARWEH\_TAMOXIFEN\_RESISTANCE\_UP | MSigDB lists | MASSARWEH\_TAMOXIFEN\_RESISTANCE\_UP | 136 | 29 | 4161 | 644 | dco,CG6767,CG7115,CG11899,CG10932,cyc,ACC,ATP8A,Synj,Droj2,dnc,Crag,CG32549,Mo25,stas,CG8507,Sccpdh1,CG33181,CycE,Hsc70-3,Sply,CG4872,Idh,l(3)mbt,CG3376,milt,Pdxk,Pdk1,CG46338 | | 4.026e-02 | -3.21 | GSE1460\_INTRATHYMIC\_T\_PROGENITOR\_VS\_NAIVE\_CD4\_TCELL\_ADULT\_BLOOD\_UP | MSigDB lists | GSE1460\_INTRATHYMIC\_T\_PROGENITOR\_VS\_NAIVE\_CD4\_TCELL\_ADULT\_BLOOD\_UP | 87 | 20 | 4161 | 644 | CCT6,mad2,CCT5,Aldh,Caf1-105,Sccpdh1,Irp-1B,CG30496,ste14,CG11089,CG15141,NHP2,DppIII,Ptp61F,CG8188,grp,RhoGAP54D,CkIIalpha,CG11164,TyrRS | | 4.028e-02 | -3.21 | GSE15930\_NAIVE\_VS\_48H\_IN\_VITRO\_STIM\_IL12\_CD8\_TCELL\_DN | MSigDB lists | GSE15930\_NAIVE\_VS\_48H\_IN\_VITRO\_STIM\_IL12\_CD8\_TCELL\_DN | 114 | 25 | 4161 | 644 | CCT6,CG5026,mad2,Pen,mRpS10,CG8004,Tom40,Prosbeta5,Coq7,Jafrac2,Gk2,rept,Jafrac1,CG12375,scramb1,CG10418,NHP2,DppIII,CG8188,Nurf-38,Rpn2,Pgm1,Cul2,CG12018,CycC | | 4.041e-02 | -3.21 | GO\_PROTEIN\_FOLDING | MSigDB lists | GO\_PROTEIN\_FOLDING | 125 | 27 | 4161 | 644 | CG14715,Pfdn2,CCT5,CG5510,Jafrac2,CCT3,CCT6,CG9911,ERp60,TBCC,Fkbp59,Hsc70-3,shu,CG8507,CG14894,Roe1,Droj2,rept,Cyp1,Grx1,CkIIalpha,CG1847,CG1890,shv,GCS2beta,Pfdn5,CG3511 | | 4.043e-02 | -3.21 | positive regulation of transport | biological process | GO:0051050 | 127 | 18 | 12345 | 1129 | Hip1,Sec23,Crag,tral,Moe,mop,Rel,Tao,Snx6,Cbl,crok,Pld,Nup358,PGRP-LE,LpR2,sbr,O-fut1,SERCA | | 4.050e-02 | -3.21 | GSE12392\_WT\_VS\_IFNB\_KO\_CD8A\_NEG\_SPLEEN\_DC\_UP | MSigDB lists | GSE12392\_WT\_VS\_IFNB\_KO\_CD8A\_NEG\_SPLEEN\_DC\_UP | 66 | 16 | 4161 | 644 | not,Jafrac1,CG11089,ACC,CG1532,Gmer,Phb2,Nurf-38,pont,alpha-Man-Ia,Aldh,Top1,CG9911,Fkbp59,Mekk1,Pp2A-29B | | 4.050e-02 | -3.21 | GSE22886\_NAIVE\_BCELL\_VS\_MONOCYTE\_DN | MSigDB lists | GSE22886\_NAIVE\_BCELL\_VS\_MONOCYTE\_DN | 66 | 16 | 4161 | 644 | Top1,CCT6,Thiolase,Nca,ND-75,Tim8,Clc,CG8507,Gyg,Mitofilin,Pmi,CG1774,Grx1,cathD,isoQC,Gapdh1 | | 4.050e-02 | -3.21 | REACTOME\_FATTY\_ACID\_TRIACYLGLYCEROL\_AND\_KETONE\_BODY\_METABOLISM | MSigDB lists | REACTOME\_FATTY\_ACID\_TRIACYLGLYCEROL\_AND\_KETONE\_BODY\_METABOLISM | 66 | 16 | 4161 | 644 | CycC,MED24,Mcad,colt,MED4,mino,ACC,cyc,CG10932,Gk2,SREBP,MED20,Mtpalpha,MED27,Echs1,Thiolase | | 4.050e-02 | -3.21 | PAX4\_01 | MSigDB lists | PAX4\_01 | 66 | 16 | 4161 | 644 | APP-BP1,Arf51F,cact,esc,shop,CG2201,CG8475,Pi3K92E,AP-1-2beta,CHOp24,O-fut1,Tm1,escl,Usp39,Gk2,dnc | | 4.050e-02 | -3.21 | MORF\_PPP2CA | MSigDB lists | MORF\_PPP2CA | 66 | 16 | 4161 | 644 | eIF3m,Prosalpha3,RpS27A,Idh3b,Chrac-14,Cul1,lwr,CG13630,Thiolase,Pgk,Dd,mts,LysRS,Atg6,CG17454,Prosbeta6 | | 4.054e-02 | -3.21 | GO\_MULTI\_ORGANISM\_REPRODUCTIVE\_PROCESS | MSigDB lists | GO\_MULTI\_ORGANISM\_REPRODUCTIVE\_PROCESS | 164 | 34 | 4161 | 644 | spn-E,bys,pont,CycB,CCT5,Ac3,CCT3,SRPK,Jafrac2,r-l,Gabat,mtgo,AGO3,Med,r,faf,CG11110,Aladin,Dd,Cdk1,kri,shu,tkv,Sply,Fkbp59,mael,Herc4,regucalcin,cyc,CHOp24,lid,Fur2,Sod1,piwi | | 4.075e-02 | -3.20 | ENK\_UV\_RESPONSE\_KERATINOCYTE\_DN | MSigDB lists | ENK\_UV\_RESPONSE\_KERATINOCYTE\_DN | 181 | 37 | 4161 | 644 | Ac3,r-l,Pfk,pont,Chd64,nmdyn-D6,Cul1,CG8314,APP-BP1,Sec23,Cka,Ts,CG9485,AGBE,Pka-R1,scramb1,Med,Tlk,aay,Kap3,l(3)73Ah,Klp10A,ND-75,CG9391,mad2,Got1,Clp,Grip84,Xpac,lid,Karybeta3,Fdh,CG11070,Gapdh1,Mcad,RpII140,Cse1 | | 4.095e-02 | -3.20 | cellular response to starvation | biological process | GO:0009267 | 84 | 13 | 12345 | 1129 | SNF4Agamma,Rel,Atg6,raptor,foxo,lt,Pi3K92E,Debcl,Iml1,Atg7,Mitf,Nprl2,rl | | 4.095e-02 | -3.20 | somatic muscle development | biological process | GO:0007525 | 84 | 13 | 12345 | 1129 | Pfk,tub,Pgk,Gapdh1,Sply,PyK,foxo,spz,lid,Cf2,CG13366,Pglym78,Stat92E | | 4.132e-02 | -3.19 | cellular response to DNA damage stimulus | biological process | GO:0006974 | 218 | 28 | 12345 | 1129 | Debcl,ben,Chrac-14,MAGE,Tfb4,Sld5,nbs,Uev1A,Utx,mus304,agt,Rel,Xpac,Xpc,Ubc6,mre11,CG10336,DNApol-epsilon255,mus81,RecQ5,Rbf,tos,Snm1,grp,mus205,CG5285,Cdk1,obe | | 4.140e-02 | -3.18 | GSE1460\_DP\_THYMOCYTE\_VS\_NAIVE\_CD4\_TCELL\_ADULT\_BLOOD\_DN | MSigDB lists | GSE1460\_DP\_THYMOCYTE\_VS\_NAIVE\_CD4\_TCELL\_ADULT\_BLOOD\_DN | 61 | 15 | 4161 | 644 | dco,Pfdn5,RpL26,RpS27,RpS30,RpS13,Unr,Rbf,Adi1,RpL12,RpL13,Pdxk,RpS18,Pi3K92E,RpS7 | | 4.140e-02 | -3.18 | GSE7852\_LN\_VS\_THYMUS\_TREG\_DN | MSigDB lists | GSE7852\_LN\_VS\_THYMUS\_TREG\_DN | 61 | 15 | 4161 | 644 | CG4872,CG1275,Utx,Nap1,dco,Kap3,Pfk,SelD,Eno,Pka-R1,Pglym78,Pen,capt,Top1,ste14 | | 4.140e-02 | -3.18 | MODULE\_126 | MSigDB lists | MODULE\_126 | 61 | 15 | 4161 | 644 | Hsc70-3,Arpc2,tun,CkIIalpha,PpD3,Phb2,Cse1,r-l,RfC38,Tsp74F,Cdk1,CycB,CG7997,Cdk5,Cyp1 | | 4.140e-02 | -3.18 | REACTOME\_REGULATION\_OF\_MRNA\_STABILITY\_BY\_PROTEINS\_THAT\_BIND\_AU\_RICH\_ELEMENTS | MSigDB lists | REACTOME\_REGULATION\_OF\_MRNA\_STABILITY\_BY\_PROTEINS\_THAT\_BIND\_AU\_RICH\_ELEMENTS | 61 | 15 | 4161 | 644 | Rrp40,Rpn12,Prosalpha5,RpS27A,Rpn2,Prosbeta3,Prosalpha3,Prosbeta4,Prosalpha4,Rpt2,RpL40,Rpn3,Prosbeta5,Prosbeta6,p38a | | 4.140e-02 | -3.18 | GSE32986\_CURDLAN\_LOWDOSE\_VS\_CURDLAN\_HIGHDOSE\_STIM\_DC\_UP | MSigDB lists | GSE32986\_CURDLAN\_LOWDOSE\_VS\_CURDLAN\_HIGHDOSE\_STIM\_DC\_UP | 61 | 15 | 4161 | 644 | Nca,Cyp6a19,Tomosyn,tun,Got1,CG15111,CG10470,mal,CG3894,aay,CG9773,ABCD,Hmg-2,Cyp6a22,Mctp | | 4.167e-02 | -3.18 | CG32354 (uncharacterized protein) | protein interactions | 50302 | 22 | 6 | 9204 | 1112 | Rcd5,RpS14b,Sod3,RpS14a,CG9911,CG9941 | | 4.167e-02 | -3.18 | cdc14 (cell division cycle 14) | protein interactions | 34067 | 22 | 6 | 9204 | 1112 | bcd,heph,CG11486,Ntan1,gus,EcR | | 4.171e-02 | -3.18 | MORF\_RFC4 | MSigDB lists | MORF\_RFC4 | 82 | 19 | 4161 | 644 | Prosbeta4,APP-BP1,Mitofilin,Hat1,CG13630,SRPK,SdhA,CCT5,CG8004,Cyt-c-p,CG4390,CG8188,TyrRS,CG10418,rept,eIF3i,Ts,RfC38,r | | 4.171e-02 | -3.18 | GSE15930\_NAIVE\_VS\_72H\_IN\_VITRO\_STIM\_TRICHOSTATINA\_CD8\_TCELL\_DN | MSigDB lists | GSE15930\_NAIVE\_VS\_72H\_IN\_VITRO\_STIM\_TRICHOSTATINA\_CD8\_TCELL\_DN | 82 | 19 | 4161 | 644 | gammaTub37C,mRpL17,msps,mad2,Pglym78,milt,Hat1,Cdk1,Caf1-105,CG14715,CG8004,CG5880,CG11164,mRpL18,CG12018,ND-51,Su(H),CysRS,r | | 4.171e-02 | -3.18 | GO\_PROTEIN\_MATURATION | MSigDB lists | GO\_PROTEIN\_MATURATION | 82 | 19 | 4161 | 644 | STUB1,AIF,Jafrac2,pen-2,CG9581,CG32473,stet,CG1827,Smox,Fur2,Stoml2,Atg4b,disp,Tsp86D,CG1847,CG8728,ste24a,Cyt-c-p,CG11110 | | 4.171e-02 | -3.18 | DUTERTRE\_ESTRADIOL\_RESPONSE\_6HR\_UP | MSigDB lists | DUTERTRE\_ESTRADIOL\_RESPONSE\_6HR\_UP | 82 | 19 | 4161 | 644 | Cyp1,CG8726,Tm1,AP-1-2beta,RfC38,grp,slif,mRpS23,mal,TyrRS,Psf3,Tsp66E,Sfxn2,pont,CG7997,CG9471,crq,Mat89Ba,Fkbp59 | | 4.171e-02 | -3.18 | GSE339\_CD4POS\_VS\_CD8POS\_DC\_DN | MSigDB lists | GSE339\_CD4POS\_VS\_CD8POS\_DC\_DN | 82 | 19 | 4161 | 644 | Jafrac1,Prosbeta6,Nurf-38,RpL13A,Sod1,Pglym78,Nca,ERp60,CG9911,CG15715,r-l,Klp10A,CG15525,Gclc,CG46338,CG9773,Pi4KIIalpha,Hsc70-3,cl | | 4.171e-02 | -3.18 | GSE22935\_WT\_VS\_MYD88\_KO\_MACROPHAGE\_24H\_MBOVIS\_BCG\_STIM\_UP | MSigDB lists | GSE22935\_WT\_VS\_MYD88\_KO\_MACROPHAGE\_24H\_MBOVIS\_BCG\_STIM\_UP | 82 | 19 | 4161 | 644 | RpL38,CG5412,CG5028,tko,MED4,mino,Klp10A,Cdk1,Chrac-14,Pen,f-cup,CG3077,Pex12,Herc4,CG7728,RpL11,stet,RpL26,Ppox | | 4.173e-02 | -3.18 | GSE29618\_MONOCYTE\_VS\_PDC\_UP | MSigDB lists | GSE29618\_MONOCYTE\_VS\_PDC\_UP | 36 | 10 | 4161 | 644 | Rab32,isoQC,Cyp6a22,Gapdh1,Moe,Tspo,Cyp1,Cyp6a19,Fur2,CG9471 | | 4.173e-02 | -3.18 | GSE1432\_CTRL\_VS\_IFNG\_1H\_MICROGLIA\_UP | MSigDB lists | GSE1432\_CTRL\_VS\_IFNG\_1H\_MICROGLIA\_UP | 36 | 10 | 4161 | 644 | GCC88,Grip128,r,rl,Sld5,SdhA,PpD3,Cdk7,Arpc2,Gmer | | 4.173e-02 | -3.18 | CHNG\_MULTIPLE\_MYELOMA\_HYPERPLOID\_UP | MSigDB lists | CHNG\_MULTIPLE\_MYELOMA\_HYPERPLOID\_UP | 36 | 10 | 4161 | 644 | RpL40,eEF1beta,RpL37a,eIF4A,RpL24,RpS13,RpL13A,RpS16,RpS23,Clc | | 4.173e-02 | -3.18 | REACTOME\_SIGNALING\_BY\_FGFR | MSigDB lists | REACTOME\_SIGNALING\_BY\_FGFR | 36 | 10 | 4161 | 644 | Cdk1,Pdk1,rl,Ac3,Pka-R1,RpL40,Pka-R2,mts,RpS27A,Pp2A-29B | | 4.173e-02 | -3.18 | MOREAUX\_B\_LYMPHOCYTE\_MATURATION\_BY\_TACI\_DN | MSigDB lists | MOREAUX\_B\_LYMPHOCYTE\_MATURATION\_BY\_TACI\_DN | 36 | 10 | 4161 | 644 | Nmt,grp,asf1,CG6610,Rpn12,Ts,Cdk1,CG42542,CCT5,tko | | 4.175e-02 | -3.18 | GO\_REGULATION\_OF\_CELL\_CYCLE\_PROCESS | MSigDB lists | GO\_REGULATION\_OF\_CELL\_CYCLE\_PROCESS | 187 | 38 | 4161 | 644 | RpS27A,Prosalpha5,twin,shrb,mad2,Cdk1,CG1812,Rpn12,Not10,Rpn2,grp,Prosbeta3,CG11164,Rpt2,Prosalpha4,SkpA,Pp2A-29B,Prosalpha3,AGO3,Wnt10,Prosbeta4,APP-BP1,nmdyn-D6,Cul1,Rpn3,CycB,Cdk5,Prosbeta5,CHMP2B,APC7,Atg6,Phb2,CG30291,CkIIalpha,Rbf,RpL40,Pka-R1,Prosbeta6 | | 4.181e-02 | -3.17 | stress-activated protein kinase signaling cascade | biological process | GO:0031098 | 36 | 7 | 12345 | 1129 | Tak1,fray,Traf6,Cka,Tao,Mekk1,Mtl | | 4.181e-02 | -3.17 | regulation of insulin receptor signaling pathway | biological process | GO:0046626 | 36 | 7 | 12345 | 1129 | mts,foxo,Cul1,STUB1,Pp2A-29B,Ptp61F,SkpA | | 4.181e-02 | -3.17 | regulation of cellular response to insulin stimulus | biological process | GO:1900076 | 36 | 7 | 12345 | 1129 | foxo,Cul1,Pp2A-29B,Ptp61F,STUB1,mts,SkpA | | 4.186e-02 | -3.17 | rux (roughex) | protein interactions | 31535 | 58 | 12 | 9204 | 1112 | Pura,CycE,Sod3,enc,CycB,twe,ScsbetaA,CycJ,CG5390,Cdk1,Aladin,bcd | | 4.190e-02 | -3.17 | USP\_2 | prosite domains | PS00973 | 18 | 5 | 6243 | 694 | Usp16-45,Usp10,not,faf,scny | | 4.192e-02 | -3.17 | cellular polysaccharide biosynthetic process | biological process | GO:0033692 | 15 | 4 | 12345 | 1129 | AGBE,Pgm1,CG9485,Gyg | | 4.192e-02 | -3.17 | aspartate family amino acid biosynthetic process | biological process | GO:0009067 | 15 | 4 | 12345 | 1129 | Adi1,Got1,CG11134,AsnS | | 4.192e-02 | -3.17 | male germ-line stem cell population maintenance | biological process | GO:0036098 | 15 | 4 | 12345 | 1129 | zpg,shv,stet,lid | | 4.192e-02 | -3.17 | polysaccharide biosynthetic process | biological process | GO:0000271 | 15 | 4 | 12345 | 1129 | AGBE,Pgm1,Gyg,CG9485 | | 4.192e-02 | -3.17 | Notch receptor processing | biological process | GO:0007220 | 15 | 4 | 12345 | 1129 | Rab5,lt,pen-2,dx | | 4.192e-02 | -3.17 | negative regulation of humoral immune response | biological process | GO:0002921 | 15 | 4 | 12345 | 1129 | cact,p38a,Stat92E,scny | | 4.192e-02 | -3.17 | response to amino acid starvation | biological process | GO:1990928 | 15 | 4 | 12345 | 1129 | Nprl2,Iml1,foxo,Rel | | 4.192e-02 | -3.17 | oocyte localization involved in germarium-derived egg chamber formation | biological process | GO:0030720 | 15 | 4 | 12345 | 1129 | jvl,del,Stat92E,spn-E | | 4.192e-02 | -3.17 | egg activation | biological process | GO:0007343 | 15 | 4 | 12345 | 1129 | png,wisp,cort,plu | | 4.192e-02 | -3.17 | rhodopsin biosynthetic process | biological process | GO:0016063 | 15 | 4 | 12345 | 1129 | PAPLA1,EMC3,Hexo1,fdl | | 4.193e-02 | -3.17 | Signaling by Type 1 Insulin-like Growth Factor 1 Receptor (IGF1R) | REACTOME pathways | R-DME-2404192 | 156 | 31 | 3923 | 574 | Prosalpha3,Rpt2,RpL40,Lkb1,CG7530,PpD3,RpS27A,Pdk1,yl,Prosalpha5,Pi3K92E,Prosbeta4,Prosbeta5,Rpn2,SNF4Agamma,Prosalpha4,csw,Pp2A-29B,Prosbeta6,CG13398,Rpn12,Rpn3,eIF4E1,alph,rl,Prosbeta3,Mo25,CG4853,tor,raptor,alc | | 4.193e-02 | -3.17 | IRS-related events triggered by IGF1R | REACTOME pathways | R-DME-2428928 | 156 | 31 | 3923 | 574 | csw,Pp2A-29B,Prosbeta6,Rpn3,CG13398,Rpn12,alph,eIF4E1,Prosbeta3,rl,raptor,tor,Mo25,CG4853,alc,Lkb1,Rpt2,RpL40,Prosalpha3,CG7530,PpD3,Prosalpha5,RpS27A,yl,Pdk1,Prosbeta4,Pi3K92E,Rpn2,Prosbeta5,SNF4Agamma,Prosalpha4 | | 4.193e-02 | -3.17 | Insulin receptor signalling cascade | REACTOME pathways | R-DME-74751 | 156 | 31 | 3923 | 574 | Prosalpha4,SNF4Agamma,Rpn2,Prosbeta5,Prosbeta4,Pi3K92E,Prosalpha5,yl,Pdk1,RpS27A,PpD3,CG7530,RpL40,Rpt2,Lkb1,Prosalpha3,alc,raptor,tor,CG4853,Mo25,Prosbeta3,rl,eIF4E1,alph,Rpn3,CG13398,Rpn12,Prosbeta6,Pp2A-29B,csw | | 4.193e-02 | -3.17 | IRS-mediated signalling | REACTOME pathways | R-DME-112399 | 156 | 31 | 3923 | 574 | Rpn2,Prosbeta5,Prosalpha4,SNF4Agamma,Prosalpha5,yl,RpS27A,Pdk1,Prosbeta4,Pi3K92E,Lkb1,Rpt2,RpL40,Prosalpha3,CG7530,PpD3,alc,Prosbeta3,rl,raptor,tor,Mo25,CG4853,Rpn3,Rpn12,CG13398,alph,eIF4E1,csw,Prosbeta6,Pp2A-29B | | 4.193e-02 | -3.17 | IGF1R signaling cascade | REACTOME pathways | R-DME-2428924 | 156 | 31 | 3923 | 574 | alc,tor,raptor,Mo25,CG4853,Prosbeta3,rl,eIF4E1,alph,Rpn3,CG13398,Rpn12,Pp2A-29B,Prosbeta6,csw,Prosalpha4,SNF4Agamma,Rpn2,Prosbeta5,Prosbeta4,Pi3K92E,Prosalpha5,yl,Pdk1,RpS27A,PpD3,CG7530,Rpt2,Lkb1,RpL40,Prosalpha3 | | 4.212e-02 | -3.17 | positive regulation of receptor signaling pathway via STAT | biological process | GO:1904894 | 9 | 3 | 12345 | 1129 | Imp,Jafrac2,CycE | | 4.212e-02 | -3.17 | response to arsenic-containing substance | biological process | GO:0046685 | 9 | 3 | 12345 | 1129 | rl,p38a,Mekk1 | | 4.212e-02 | -3.17 | regulation of translational elongation | biological process | GO:0006448 | 9 | 3 | 12345 | 1129 | Dph5,CG1578,CG2911 | | 4.212e-02 | -3.17 | pronuclear fusion | biological process | GO:0007344 | 9 | 3 | 12345 | 1129 | msps,wisp,fs(1)Ya | | 4.212e-02 | -3.17 | response to cadmium ion | biological process | GO:0046686 | 9 | 3 | 12345 | 1129 | p38a,rl,Mekk1 | | 4.212e-02 | -3.17 | regulation of DNA endoreduplication | biological process | GO:0032875 | 9 | 3 | 12345 | 1129 | otu,E2f1,SuUR | | 4.212e-02 | -3.17 | regulation of triglyceride catabolic process | biological process | GO:0010896 | 9 | 3 | 12345 | 1129 | foxo,Lkb1,Nprl2 | | 4.212e-02 | -3.17 | regulation of extracellular matrix organization | biological process | GO:1903053 | 9 | 3 | 12345 | 1129 | Rab10,Sec23,Crag | | 4.212e-02 | -3.17 | positive regulation of receptor signaling pathway via JAK-STAT | biological process | GO:0046427 | 9 | 3 | 12345 | 1129 | Jafrac2,CycE,Imp | | 4.212e-02 | -3.17 | karyogamy | biological process | GO:0000741 | 9 | 3 | 12345 | 1129 | fs(1)Ya,wisp,msps | | 4.212e-02 | -3.17 | histone H3-K9 methylation | biological process | GO:0051567 | 9 | 3 | 12345 | 1129 | G9a,egg,esc | | 4.212e-02 | -3.17 | activation of MAPKK activity | biological process | GO:0000186 | 9 | 3 | 12345 | 1129 | Tao,Mekk1,Tak1 | | 4.212e-02 | -3.17 | histone deubiquitination | biological process | GO:0016578 | 9 | 3 | 12345 | 1129 | scny,Sgf11,not | | 4.212e-02 | -3.17 | behavioral response to cocaine | biological process | GO:0048148 | 9 | 3 | 12345 | 1129 | Pka-R2,cyc,dco | | 4.212e-02 | -3.17 | positive regulation of G1/S transition of mitotic cell cycle | biological process | GO:1900087 | 9 | 3 | 12345 | 1129 | CycE,Su(H),Sld5 | | 4.212e-02 | -3.17 | regulation of protein tyrosine kinase activity | biological process | GO:0061097 | 9 | 3 | 12345 | 1129 | Cbl,CG14408,ASPP | | 4.213e-02 | -3.17 | GSE360\_DC\_VS\_MAC\_L\_DONOVANI\_UP | MSigDB lists | GSE360\_DC\_VS\_MAC\_L\_DONOVANI\_UP | 56 | 14 | 4161 | 644 | Dcr-1,CG9393,CG14516,CG8188,isoQC,YL-1,slif,Jheh2,wal,cact,Klp10A,ND-24,Pen,Rpn3 | | 4.213e-02 | -3.17 | GSE5589\_LPS\_VS\_LPS\_AND\_IL10\_STIM\_MACROPHAGE\_45MIN\_UP | MSigDB lists | GSE5589\_LPS\_VS\_LPS\_AND\_IL10\_STIM\_MACROPHAGE\_45MIN\_UP | 56 | 14 | 4161 | 644 | Nprl2,Caf1-105,CG8611,ERp60,Kap-alpha3,Jafrac1,Med,Pgk,shop,CG7728,grp,isoQC,CycE,CG4390 | | 4.213e-02 | -3.17 | GSE20715\_0H\_VS\_24H\_OZONE\_TLR4\_KO\_LUNG\_DN | MSigDB lists | GSE20715\_0H\_VS\_24H\_OZONE\_TLR4\_KO\_LUNG\_DN | 56 | 14 | 4161 | 644 | Pen,cyc,alpha-Man-Ia,Echs1,fray,rl,eIF4E1,Nurf-38,dco,Pka-R2,csw,Hsc70-3,Got1,CG10465 | | 4.213e-02 | -3.17 | GSE5099\_UNSTIM\_VS\_MCSF\_TREATED\_MONOCYTE\_DAY3\_DN | MSigDB lists | GSE5099\_UNSTIM\_VS\_MCSF\_TREATED\_MONOCYTE\_DAY3\_DN | 56 | 14 | 4161 | 644 | Tspo,Arpc4,Pka-R1,PIG-L,CG31715,tzn,eIF3i,Pdxk,Pp2A-29B,CG6144,TBCC,CG12018,CG13601,AnxB10 | | 4.213e-02 | -3.17 | GSE45365\_CD8A\_DC\_VS\_CD11B\_DC\_IFNAR\_KO\_DN | MSigDB lists | GSE45365\_CD8A\_DC\_VS\_CD11B\_DC\_IFNAR\_KO\_DN | 56 | 14 | 4161 | 644 | Adi1,CG1416,CG11089,RhoGAP5A,ND-B22,CG7997,Mo25,Vsp37A,mRpL13,trsn,obe,Arpc2,CG32473,CG5292 | | 4.213e-02 | -3.17 | GSE3691\_IFN\_PRODUCING\_KILLER\_DC\_VS\_CONVENTIONAL\_DC\_SPLEEN\_UP | MSigDB lists | GSE3691\_IFN\_PRODUCING\_KILLER\_DC\_VS\_CONVENTIONAL\_DC\_SPLEEN\_UP | 56 | 14 | 4161 | 644 | Jafrac1,not,Tspo,milt,CG16865,CG46338,Etf-QO,CG8199,Pi4KIIalpha,Ppox,Pex23,Mcad,CG1774,tun | | 4.213e-02 | -3.17 | GSE8621\_LPS\_STIM\_VS\_LPS\_PRIMED\_AND\_LPS\_STIM\_MACROPHAGE\_DN | MSigDB lists | GSE8621\_LPS\_STIM\_VS\_LPS\_PRIMED\_AND\_LPS\_STIM\_MACROPHAGE\_DN | 56 | 14 | 4161 | 644 | AGBE,esc,CG17333,f-cup,Ublcp1,Fdh,escl,Kap-alpha3,Sply,ND-49,Tsp66E,Axs,Mekk1,Pfdn5 | | 4.213e-02 | -3.17 | STAT\_01 | MSigDB lists | STAT\_01 | 56 | 14 | 4161 | 644 | Rab10,ND-49,MED24,RpII140,nmdyn-D7,CG32549,Sod1,Pi4KIIalpha,ari-1,eIF4E1,scramb1,Rpn3,Gk2,Ect4 | | 4.213e-02 | -3.17 | GSE6674\_CPG\_VS\_CPG\_AND\_ANTI\_IGM\_STIM\_BCELL\_UP | MSigDB lists | GSE6674\_CPG\_VS\_CPG\_AND\_ANTI\_IGM\_STIM\_BCELL\_UP | 56 | 14 | 4161 | 644 | mad2,cathD,Jafrac2,Ts,RhoGAP54D,Oseg1,galectin,Atg6,CG9143,shrb,CG11134,mRpL18,Idh,Prosalpha3 | | 4.219e-02 | -3.17 | Signaling by FGFR1 | REACTOME pathways | R-DME-5654736 | 28 | 8 | 3923 | 574 | mts,CG13398,RpL40,Pp2A-29B,Cbl,csw,RpS27A,rl | | 4.225e-02 | -3.16 | TP53 Regulates Transcription of Cell Cycle Genes | REACTOME pathways | R-DME-6791312 | 14 | 5 | 3923 | 574 | Not10,Cdk1,Rga,CycB,twin | | 4.232e-02 | -3.16 | CG3678 (uncharacterized protein) | protein interactions | 42045 | 7 | 3 | 9204 | 1112 | EMC3,CG9399,EMC8-9 | | 4.232e-02 | -3.16 | Fsn (F-box synaptic protein) | protein interactions | 36460 | 7 | 3 | 9204 | 1112 | gus,SkpA,Cul1 | | 4.232e-02 | -3.16 | CG3902 (uncharacterized protein) | protein interactions | 40059 | 7 | 3 | 9204 | 1112 | Pen,Moe,CG5028 | | 4.232e-02 | -3.16 | Ugt36D1 (UDP-glycosyltransferase family 36 member D1) | protein interactions | 35138 | 7 | 3 | 9204 | 1112 | CG4407,CG15098,Echs1 | | 4.232e-02 | -3.16 | CG14104 (uncharacterized protein) | protein interactions | 50259 | 7 | 3 | 9204 | 1112 | Usp10,ERp60,RpL40 | | 4.232e-02 | -3.16 | dah (discontinuous actin hexagon) | protein interactions | 32459 | 7 | 3 | 9204 | 1112 | dgt4,DppIII,CG32536 | | 4.232e-02 | -3.16 | sra (sarah) | protein interactions | 47384 | 7 | 3 | 9204 | 1112 | Pp2B-14D,CanA-14F,CanB2 | | 4.232e-02 | -3.16 | phol (pleiohomeotic like) | protein interactions | 39095 | 7 | 3 | 9204 | 1112 | Sgf11,esc,Fkbp12 | | 4.232e-02 | -3.16 | LSm1 (Like Sm 1) | protein interactions | 37413 | 7 | 3 | 9204 | 1112 | CG6610,CG10418,Prosbeta5 | | 4.232e-02 | -3.16 | Zfrp8 (Zinc finger protein RP-8) | protein interactions | 45021 | 7 | 3 | 9204 | 1112 | RpS2,tral,mael | | 4.232e-02 | -3.16 | CG14589 (uncharacterized protein) | protein interactions | 35541 | 7 | 3 | 9204 | 1112 | gus,RpL38,Dip-B | | 4.232e-02 | -3.16 | toc (toucan) | protein interactions | 33526 | 7 | 3 | 9204 | 1112 | Rtnl1,Gyg,SkpA | | 4.232e-02 | -3.16 | ND-B22 (NADH dehydrogenase (ubiquinone) B22 subunit) | protein interactions | 34747 | 7 | 3 | 9204 | 1112 | AGBE,Jafrac2,Rcd5 | | 4.238e-02 | -3.16 | GO\_ERBB\_SIGNALING\_PATHWAY | MSigDB lists | GO\_ERBB\_SIGNALING\_PATHWAY | 22 | 7 | 4161 | 644 | STUB1,Cul5,csw,Pdk1,rl,RpL40,RpS27A | | 4.238e-02 | -3.16 | GO\_ORGAN\_REGENERATION | MSigDB lists | GO\_ORGAN\_REGENERATION | 22 | 7 | 4161 | 644 | Ts,Cdk1,r,CG6767,regucalcin,CG11089,CycE | | 4.238e-02 | -3.16 | GO\_MODIFICATION\_OF\_MORPHOLOGY\_OR\_PHYSIOLOGY\_OF\_OTHER\_ORGANISM | MSigDB lists | GO\_MODIFICATION\_OF\_MORPHOLOGY\_OR\_PHYSIOLOGY\_OF\_OTHER\_ORGANISM | 22 | 7 | 4161 | 644 | Ts,Clp,Smox,Atg7,Pen,Prip,Kap-alpha3 | | 4.238e-02 | -3.16 | MODULE\_278 | MSigDB lists | MODULE\_278 | 22 | 7 | 4161 | 644 | Gclc,r,Ubc4,UbcE2H,CG40045,Uev1A,CG8188 | | 4.238e-02 | -3.16 | GO\_FEMALE\_SEX\_DIFFERENTIATION | MSigDB lists | GO\_FEMALE\_SEX\_DIFFERENTIATION | 22 | 7 | 4161 | 644 | Sply,Idh,tkv,Med,CycE,Sod1,CG11110 | | 4.239e-02 | -3.16 | TYR\_PHOSPHATASE\_2 | prosite domains | PS50056 | 30 | 7 | 6243 | 694 | Ptp61F,csw,ssh,Ptp10D,Ptpmeg,Ptpmeg2,Ptp69D | | 4.247e-02 | -3.16 | organellar large ribosomal subunit | cellular component | GO:0000315 | 51 | 9 | 11940 | 1105 | mRpL13,mRpL16,mRpL36,CG6094,mRpL10,mRpL18,mRpL48,mRpL35,mRpL17 | | 4.247e-02 | -3.16 | mitochondrial large ribosomal subunit | cellular component | GO:0005762 | 51 | 9 | 11940 | 1105 | mRpL35,mRpL48,mRpL18,mRpL10,mRpL17,mRpL13,CG6094,mRpL16,mRpL36 | | 4.256e-02 | -3.16 | GSE17721\_POLYIC\_VS\_GARDIQUIMOD\_24H\_BMDC\_UP | MSigDB lists | GSE17721\_POLYIC\_VS\_GARDIQUIMOD\_24H\_BMDC\_UP | 41 | 11 | 4161 | 644 | gus,sbr,pr,DhpD,CysRS,RpL13A,yip2,colt,Utx,Sfxn2,UbcE2H | | 4.256e-02 | -3.16 | GSE21774\_CD62L\_POS\_CD56\_DIM\_VS\_CD62L\_NEG\_CD56\_DIM\_NK\_CELL\_DN | MSigDB lists | GSE21774\_CD62L\_POS\_CD56\_DIM\_VS\_CD62L\_NEG\_CD56\_DIM\_NK\_CELL\_DN | 41 | 11 | 4161 | 644 | Eno,CG7997,Pglym78,CG8974,Shmt,spn-E,CG4572,Pgk,Tspo,gus,Idh | | 4.256e-02 | -3.16 | GO\_ALPHA\_AMINO\_ACID\_BIOSYNTHETIC\_PROCESS | MSigDB lists | GO\_ALPHA\_AMINO\_ACID\_BIOSYNTHETIC\_PROCESS | 41 | 11 | 4161 | 644 | aay,Dhfr,CG11899,P5cr,P5CDh1,Got1,CG11134,CG11089,Shmt,Adi1,r | | 4.256e-02 | -3.16 | GSE24634\_NAIVE\_CD4\_TCELL\_VS\_DAY3\_IL4\_CONV\_TREG\_UP | MSigDB lists | GSE24634\_NAIVE\_CD4\_TCELL\_VS\_DAY3\_IL4\_CONV\_TREG\_UP | 41 | 11 | 4161 | 644 | RpL11,tws,CG10038,RpS27,Pfdn5,GlyP,Vha13,CG3376,CG14194,Snx6,cyc | | 4.256e-02 | -3.16 | GSE37532\_TREG\_VS\_TCONV\_PPARG\_KO\_CD4\_TCELL\_FROM\_VISCERAL\_ADIPOSE\_TISSUE\_DN | MSigDB lists | GSE37532\_TREG\_VS\_TCONV\_PPARG\_KO\_CD4\_TCELL\_FROM\_VISCERAL\_ADIPOSE\_TISSUE\_DN | 41 | 11 | 4161 | 644 | ATP8A,Moe,CG46338,Gdi,nmd,CG33181,DCAF12,Clc,CG2201,Idh,UbcE2H | | 4.256e-02 | -3.16 | GSE19198\_6H\_VS\_24H\_IL21\_TREATED\_TCELL\_UP | MSigDB lists | GSE19198\_6H\_VS\_24H\_IL21\_TREATED\_TCELL\_UP | 41 | 11 | 4161 | 644 | ERp60,CG8314,ste14,ND-B22,CG6766,CG7550,CG1640,Tsp86D,Rab10,Tsp66E,CG7840 | | 4.256e-02 | -3.16 | ZHU\_CMV\_ALL\_UP | MSigDB lists | ZHU\_CMV\_ALL\_UP | 41 | 11 | 4161 | 644 | Eno,Stip1,cact,Roe1,CG9391,Fkbp59,Prosbeta4,Rpt2,eIF4E1,Cse1,CG8188 | | 4.257e-02 | -3.16 | exodeoxyribonuclease activity, producing 5'-phosphomonoesters | molecular function | GO:0016895 | 15 | 4 | 12176 | 1119 | mre11,Snm1,tos,DNApol-epsilon255 | | 4.258e-02 | -3.16 | park (parkin) | protein interactions | 40336 | 71 | 14 | 9204 | 1112 | STUB1,foxo,PEK,ari-1,eIF4E1,Pink1,Marf,HDAC6,Shmt,ATPsynbeta,Stoml2,park,Sod1,ben | | 4.262e-02 | -3.16 | GO\_ANTIGEN\_PROCESSING\_AND\_PRESENTATION\_OF\_PEPTIDE\_ANTIGEN\_VIA\_MHC\_CLASS\_I | MSigDB lists | GO\_ANTIGEN\_PROCESSING\_AND\_PRESENTATION\_OF\_PEPTIDE\_ANTIGEN\_VIA\_MHC\_CLASS\_I | 51 | 13 | 4161 | 644 | Rpn12,Prosalpha5,Rpn2,Prosalpha3,Prosbeta3,Prosalpha4,Prosbeta4,Sec23,Rpt2,ERp60,Rpn3,Prosbeta5,Prosbeta6 | | 4.262e-02 | -3.16 | RB\_P107\_DN.V1\_UP | MSigDB lists | RB\_P107\_DN.V1\_UP | 51 | 13 | 4161 | 644 | CG7655,Sld5,CG15141,Caf1-105,RfC38,CG9471,Jafrac2,mad2,Dhfr,CycE,gammaTub37C,DNApol-epsilon255,Gabat | | 4.262e-02 | -3.16 | HMGIY\_Q6 | MSigDB lists | HMGIY\_Q6 | 51 | 13 | 4161 | 644 | eEF1beta,dnc,Imp,Su(H),Stip1,cact,O-fut1,ND-75,alph,mal,slif,Pi4KIIalpha,Sfxn2 | | 4.262e-02 | -3.16 | GSE17580\_TREG\_VS\_TEFF\_DN | MSigDB lists | GSE17580\_TREG\_VS\_TEFF\_DN | 51 | 13 | 4161 | 644 | sgll,cathD,Nca,CG3376,Marf,Atg4b,mRpS5,CG9248,Unr,stet,sws,Axs,dco | | 4.265e-02 | -3.15 | sulfurtransferase activity | molecular function | GO:0016783 | 9 | 3 | 12176 | 1119 | CG10189,CG3021,mal | | 4.265e-02 | -3.15 | glutathione disulfide oxidoreductase activity | molecular function | GO:0015038 | 9 | 3 | 12176 | 1119 | Trxr-1,Grx1,CG14407 | | 4.265e-02 | -3.15 | quinone binding | molecular function | GO:0048038 | 9 | 3 | 12176 | 1119 | UQCR-Q,SdhD,ND-49 | | 4.265e-02 | -3.15 | protein phosphorylated amino acid binding | molecular function | GO:0045309 | 9 | 3 | 12176 | 1119 | Atu,Cbl,hwt | | 4.275e-02 | -3.15 | developmental growth | biological process | GO:0048589 | 191 | 25 | 12345 | 1129 | Abi,Vha68-2,Rab10,ben,Nlg1,dnc,Pex10,Rab5,cup,Liprin-beta,bun,Cul2,Pi3K92E,Cul5,Mtl,DNApol-epsilon255,Imp,eIF4A,pygo,stai,Pdk1,Atpalpha,tkv,dco,Ptpmeg | | 4.275e-02 | -3.15 | growth | biological process | GO:0040007 | 191 | 25 | 12345 | 1129 | bun,Liprin-beta,Pex10,cup,Rab5,Cul2,Abi,dnc,Nlg1,Vha68-2,Rab10,ben,tkv,Atpalpha,dco,Ptpmeg,Cul5,Mtl,DNApol-epsilon255,Pi3K92E,Pdk1,eIF4A,stai,pygo,Imp | | 4.276e-02 | -3.15 | peroxidase activity | molecular function | GO:0004601 | 36 | 7 | 12176 | 1119 | PHGPx,Jafrac1,Jafrac2,glob1,Pxt,GstT1,GstT2 | | 4.281e-02 | -3.15 | GO\_GLUCOSE\_METABOLIC\_PROCESS | MSigDB lists | GO\_GLUCOSE\_METABOLIC\_PROCESS | 46 | 12 | 4161 | 644 | adp,CG11807,Got1,Gapdh1,Pgm1,Pdk,Pglym78,Pgk,Pfk,CG7322,p38a,Eno | | 4.281e-02 | -3.15 | GSE35685\_CD34POS\_CD38NEG\_VS\_CD34POS\_CD10POS\_BONE\_MARROW\_DN | MSigDB lists | GSE35685\_CD34POS\_CD38NEG\_VS\_CD34POS\_CD10POS\_BONE\_MARROW\_DN | 46 | 12 | 4161 | 644 | Traf6,galectin,shrb,CG9143,Herc4,Cul2,mtgo,stet,scramb1,Rab5,Gk2,cact | | 4.281e-02 | -3.15 | GSE41176\_UNSTIM\_VS\_ANTI\_IGM\_STIM\_TAK1\_KO\_BCELL\_6H\_UP | MSigDB lists | GSE41176\_UNSTIM\_VS\_ANTI\_IGM\_STIM\_TAK1\_KO\_BCELL\_6H\_UP | 46 | 12 | 4161 | 644 | ND-23,Exd2,Fbw5,spn-E,O-fut1,sgll,S2P,Ppox,Sccpdh1,CG5285,tun,ZnT35C | | 4.281e-02 | -3.15 | GSE3039\_B2\_VS\_B1\_BCELL\_DN | MSigDB lists | GSE3039\_B2\_VS\_B1\_BCELL\_DN | 46 | 12 | 4161 | 644 | TyrRS,Scsalpha1,HDAC6,CycE,grp,DNApol-epsilon255,Nurf-38,CG4502,ND-B18,Uev1A,RhoGDI,Pen | | 4.281e-02 | -3.15 | GO\_DEVELOPMENT\_OF\_PRIMARY\_SEXUAL\_CHARACTERISTICS | MSigDB lists | GO\_DEVELOPMENT\_OF\_PRIMARY\_SEXUAL\_CHARACTERISTICS | 46 | 12 | 4161 | 644 | scu,CG11110,tkv,Sod1,CycE,Unr,Sply,mtgo,Idh,Med,Jafrac2,lid | | 4.313e-02 | -3.14 | HALLMARK\_E2F\_TARGETS | MSigDB lists | HALLMARK\_E2F\_TARGETS | 93 | 21 | 4161 | 644 | Jafrac2,Cdk1,esc,Pen,mad2,Shmt,Nap1,DNApol-epsilon255,asf1,gammaTub37C,CycE,RfC38,CG15141,Sld5,escl,CG12018,Psf3,CG6767,grp,CG8188,Cse1 | | 4.313e-02 | -3.14 | chr19q13 | MSigDB lists | chr19q13 | 93 | 21 | 4161 | 644 | CG9248,CG12818,Pp2A-29B,PpD3,Tom40,CG8199,SerRS-m,CG3609,CG9471,pen-2,CG6144,CG13601,EcR,Rpn12,CG8188,RpL13A,RpS16,rept,tko,Atpalpha,Aos1 | | 4.313e-02 | -3.14 | REACTOME\_MITOTIC\_M\_M\_G1\_PHASES | MSigDB lists | REACTOME\_MITOTIC\_M\_M\_G1\_PHASES | 93 | 21 | 4161 | 644 | Tao,RpL40,Prosbeta6,Rpn12,Rpn2,Prosbeta3,mts,Rpt2,Prosalpha4,msps,Rpn3,mad2,Prosbeta5,Klp10A,RpS27,DNApol-epsilon255,RpS27A,Pp2A-29B,Prosalpha5,Prosalpha3,Prosbeta4 | | 4.313e-02 | -3.14 | GSE17974\_0H\_VS\_6H\_IN\_VITRO\_ACT\_CD4\_TCELL\_DN | MSigDB lists | GSE17974\_0H\_VS\_6H\_IN\_VITRO\_ACT\_CD4\_TCELL\_DN | 93 | 21 | 4161 | 644 | escl,CG4407,rept,Rpb8,CG7550,Irk1,scu,slif,Rpn12,CG17187,CG3226,CG12018,CG9911,Pen,CCT6,Jafrac2,esc,pont,Uch,Prosbeta4,mRpL17 | | 4.315e-02 | -3.14 | ALFANO\_MYC\_TARGETS | MSigDB lists | ALFANO\_MYC\_TARGETS | 77 | 18 | 4161 | 644 | CG13630,CG3376,mEFTu1,Ac3,pont,Tom40,Fkbp59,CG32473,Npc2a,ATP8A,Cyp1,PpV,Ubc4,Arl6IP1,Cse1,LysRS,CG1532,P5cr | | 4.315e-02 | -3.14 | GSE15139\_GMCSF\_TREATED\_VS\_UNTREATED\_NEUTROPHILS\_DN | MSigDB lists | GSE15139\_GMCSF\_TREATED\_VS\_UNTREATED\_NEUTROPHILS\_DN | 77 | 18 | 4161 | 644 | P5cr,shv,mino,Kap3,slif,CG42542,CysRS,lid,Droj2,escl,Scox,Trxr-1,DNApol-epsilon255,pont,esc,SdhA,CG5510,milt | | 4.315e-02 | -3.14 | REACTOME\_HOST\_INTERACTIONS\_OF\_HIV\_FACTORS | MSigDB lists | REACTOME\_HOST\_INTERACTIONS\_OF\_HIV\_FACTORS | 77 | 18 | 4161 | 644 | RpS27A,Prosalpha5,Prosalpha3,Prosbeta4,Rpn3,Cul5,Prosbeta5,Rpn12,Rpn2,Aladin,Prosbeta3,Rpt2,Prosalpha4,mbo,RpL40,AP-1-2beta,Prosbeta6,SkpA | | 4.315e-02 | -3.14 | GSE5589\_LPS\_AND\_IL10\_VS\_LPS\_AND\_IL6\_STIM\_IL6\_KO\_MACROPHAGE\_45MIN\_DN | MSigDB lists | GSE5589\_LPS\_AND\_IL10\_VS\_LPS\_AND\_IL6\_STIM\_IL6\_KO\_MACROPHAGE\_45MIN\_DN | 77 | 18 | 4161 | 644 | CG33156,Tsp66E,CG3894,CG17593,CG17333,Arts,TBCC,Pka-R2,CG15111,CG14407,yip2,anne,Rbcn-3A,Mtpalpha,CCT3,CG9471,Thiolase,CG15715 | | 4.315e-02 | -3.14 | GSE369\_PRE\_VS\_POST\_IL6\_INJECTION\_SOCS3\_KO\_LIVER\_DN | MSigDB lists | GSE369\_PRE\_VS\_POST\_IL6\_INJECTION\_SOCS3\_KO\_LIVER\_DN | 77 | 18 | 4161 | 644 | HDAC6,GCS2beta,CG4407,Rab5,CG10418,CG31729,CG4572,CG17691,DNApol-epsilon255,Jheh2,Cyp6a22,CG11807,CG7488,CG9248,Cyp6a19,CG9471,Mtpalpha,Gdi | | 4.315e-02 | -3.14 | GSE42021\_CD24LO\_TREG\_VS\_CD24LO\_TCONV\_THYMUS\_DN | MSigDB lists | GSE42021\_CD24LO\_TREG\_VS\_CD24LO\_TCONV\_THYMUS\_DN | 77 | 18 | 4161 | 644 | YL-1,Arpc1,Clc,eEF1beta,Rpn3,ND-30,crq,Echs1,Eno,Cdk5,Tim8,grsm,RpS16,GCS2beta,Maf1,tko,RhoGDI,Stoml2 | | 4.315e-02 | -3.14 | GSE3982\_NEUTROPHIL\_VS\_TH2\_DN | MSigDB lists | GSE3982\_NEUTROPHIL\_VS\_TH2\_DN | 77 | 18 | 4161 | 644 | Cul2,CycC,Cse1,not,ATP8A,CG12375,escl,Pfas,Prosalpha3,Prosbeta4,RpS27,Ppox,Mekk1,Tim8,Pus7,esc,Coq7,Chrac-14 | | 4.315e-02 | -3.14 | GSE32986\_UNSTIM\_VS\_GMCSF\_AND\_CURDLAN\_LOWDOSE\_STIM\_DC\_UP | MSigDB lists | GSE32986\_UNSTIM\_VS\_GMCSF\_AND\_CURDLAN\_LOWDOSE\_STIM\_DC\_UP | 77 | 18 | 4161 | 644 | LpR2,Stip1,AcCoAS,SRPK,Eno,Pfdn2,Got1,mRpL17,snz,gammaTub37C,Surf4,Rpb8,Karybeta3,AGBE,PrBP,Prosbeta3,CG6610,Cyt-c-p | | 4.319e-02 | -3.14 | ZnF\_RBZ | smart domains | SM00547 | 14 | 4 | 5086 | 506 | Nup358,RYBP,trbd,sol | | 4.331e-02 | -3.14 | fly chr89A8-89A8\|3-58 cM | chromosome location | fly chr89A8-89A8\|3-58 cM | 15 | 3 | 25010 | 1343 | Rbf2,Sap47,glob1 | | 4.331e-02 | -3.14 | fly chr21C2-21C2\|2-0.7 | chromosome location | fly chr21C2-21C2\|2-0.7 | 15 | 3 | 25010 | 1343 | crq,shv,RpLP1 | | 4.331e-02 | -3.14 | fly chr31E1-31E1\|2-42 | chromosome location | fly chr31E1-31E1\|2-42 | 15 | 3 | 25010 | 1343 | RpS27A,Wdfy2,CG17768 | | 4.331e-02 | -3.14 | fly chr21C2-21C2\|2-0.7 cM | chromosome location | fly chr21C2-21C2\|2-0.7 cM | 15 | 3 | 25010 | 1343 | shv,crq,RpLP1 | | 4.331e-02 | -3.14 | fly chr31E1-31E1\|2-42 cM | chromosome location | fly chr31E1-31E1\|2-42 cM | 15 | 3 | 25010 | 1343 | Wdfy2,CG17768,RpS27A | | 4.331e-02 | -3.14 | fly chr89A8-89A8\|3-58 | chromosome location | fly chr89A8-89A8\|3-58 | 15 | 3 | 25010 | 1343 | Rbf2,Sap47,glob1 | | 4.339e-02 | -3.14 | microtubule organizing center | cellular component | GO:0005815 | 171 | 23 | 11940 | 1105 | Uxt,Blos2,del,CycB,CG7716,gammaTub37C,Grip75,Girdin,Pp2A-29B,Cen,Grip84,toc,CycE,CG3437,mts,cnn,Grip128,spd-2,CG13366,msps,Grip71,Klp10A,alphaTub84B | | 4.342e-02 | -3.14 | chaperone complex | cellular component | GO:0101031 | 9 | 3 | 11940 | 1105 | CCT5,CCT6,CCT3 | | 4.342e-02 | -3.14 | endocytic vesicle membrane | cellular component | GO:0030666 | 9 | 3 | 11940 | 1105 | yin,AP-1-2beta,Hip1 | | 4.342e-02 | -3.14 | clathrin coat of coated pit | cellular component | GO:0030132 | 9 | 3 | 11940 | 1105 | Hip1,Clc,AP-1-2beta | | 4.342e-02 | -3.14 | replication fork protection complex | cellular component | GO:0031298 | 9 | 3 | 11940 | 1105 | Sld5,Top1,CG10336 | | 4.342e-02 | -3.14 | exocyst | cellular component | GO:0000145 | 9 | 3 | 11940 | 1105 | Sec10,Sec15,Exo70 | | 4.342e-02 | -3.14 | RNAi effector complex | cellular component | GO:0031332 | 9 | 3 | 11940 | 1105 | Dcr-1,r2d2,gw | | 4.342e-02 | -3.14 | autolysosome | cellular component | GO:0044754 | 9 | 3 | 11940 | 1105 | Synj,Vamp7,Nprl2 | | 4.342e-02 | -3.14 | RISC complex | cellular component | GO:0016442 | 9 | 3 | 11940 | 1105 | gw,r2d2,Dcr-1 | | 4.367e-02 | -3.13 | kinase regulator activity | molecular function | GO:0019207 | 76 | 12 | 12176 | 1119 | CG14408,Lkb1,CycC,Parp16,RpLP1,CycJ,CycE,Pka-R1,CycB,Pdk1,ASPP,Pka-R2 | | 4.370e-02 | -3.13 | - | gene3d domains | 2.70.130.10 | 3 | 2 | 4929 | 622 | GCS2beta,CG6766 | | 4.370e-02 | -3.13 | - | gene3d domains | 3.30.70.600 | 3 | 2 | 4929 | 622 | mRpS10,mRpL48 | | 4.370e-02 | -3.13 | - | gene3d domains | 3.90.1180.10 | 3 | 2 | 4929 | 622 | RpL13A,mRpL13 | | 4.370e-02 | -3.13 | - | gene3d domains | 1.20.5.690 | 3 | 2 | 4929 | 622 | Kap-alpha3,Pen | | 4.376e-02 | -3.13 | negative regulation of response to biotic stimulus | biological process | GO:0002832 | 44 | 8 | 12345 | 1129 | slif,p38a,Atg6,scny,GlyP,Stat92E,cact,faf | | 4.376e-02 | -3.13 | neuron remodeling | biological process | GO:0016322 | 44 | 8 | 12345 | 1129 | mop,SkpA,Atg6,shrb,EcR,shep,Stam,Imp | | 4.376e-02 | -3.13 | cellular response to insulin stimulus | biological process | GO:0032869 | 44 | 8 | 12345 | 1129 | raptor,Pdk1,rl,Rab10,PyK,foxo,Hnf4,Pi3K92E | | 4.381e-02 | -3.13 | LEE\_BMP2\_TARGETS\_DN | MSigDB lists | LEE\_BMP2\_TARGETS\_DN | 410 | 76 | 4161 | 644 | Sec10,aay,mRpS23,RpS24,Dhfr,Non1,CG31759,CG1416,Droj2,Usp39,CycE,CG33181,Lsm12,Pfas,ZnT35C,Hat1,RpL12,Kap-alpha3,bys,Pglym78,pont,Stip1,SRPK,grp,Nurf-38,CG3226,CG2100,CG11899,Cdk7,mRpL18,l(1)G0004,Fkbp59,Uch-L5,CCT6,Pgk,Shmt,mad2,RhoGAP5A,Cyt-c-p,bor,CG6767,CG17294,CG9132,CG8728,Cyp1,Cbp80,mbo,Adi1,Plap,r,Ranbp21,NHP2,CG4293,Hsc70-3,LSm3,Chd64,CCT3,Coq7,r-l,Cse1,eIF4E1,shv,slif,CG7530,rept,Naa20A,CG11089,opm,Fur2,GatA,CG3587,Rab32,Pen,CG7997,Caf1-105,AIF | | 4.386e-02 | -3.13 | GO\_PROTEIN\_LOCALIZATION | MSigDB lists | GO\_PROTEIN\_LOCALIZATION | 664 | 118 | 4161 | 644 | Dd,CG1847,RpL23,RpS24,Sec15,lt,Aladin,GAPsec,Sec10,Pex13,dco,RpS16,Kap3,RpS4,RpL38,Vha13,Rep,RpS7,Tspo,RpL40,Sec23,Tomosyn,RpL11,Arf51F,RpL26,Clc,Vamp7,CG9773,RpS27,RpS18,Gdi,Cdk5,CG8004,Kap-alpha3,Marf,CG8314,RpL12,Tsp86D,CG2100,sau,Vsp37A,HDAC6,mino,CG9393,Karybeta3,CHOp24,AP-1-2beta,GCC88,Stam,Arts,CG5734,Su(H),shrb,Rab10,Fkbp59,RpS23,Sec22,Cdk1,rl,cact,Tom40,milt,mol,CG9132,RpL30,CG8728,Pex10,Vha36-1,Phb2,Oseg1,Pp2B-14D,CG11110,Atg6,RpS2,Ranbp21,Snx6,Rab5,Atg4b,Atg7,mbo,Pex12,Hsc70-3,CanA-14F,RpL37a,RpL24,Syx13,CG5510,CHMP2B,CG46338,Tim8,CG3077,CG7530,Cse1,RpS13,RpLP0,RpS30,RpL13A,Fur2,Arl6IP1,opm,cyc,ste14,CG4041,rept,CG31729,CG44422,snz,RpL9,RpS27A,Rab32,Exo70,RpS28b,Surf4,Faf2,LpR2,ERp60,Pen,RpL13 | | 4.392e-02 | -3.13 | GO\_COFACTOR\_BIOSYNTHETIC\_PROCESS | MSigDB lists | GO\_COFACTOR\_BIOSYNTHETIC\_PROCESS | 104 | 23 | 4161 | 644 | CG1774,Sptr,ND-39,Ppcdc,Desat1,Ppox,Pcd,Coq7,AcCoAS,Pdxk,CG3803,pr,CG33156,Dhfr,mal,Coq6,ScsbetaA,Nadsyn,sgll,CG4407,ACC,CG11089,Tspo | | 4.406e-02 | -3.12 | Ribokinase-like | interpro domains | IPR029056 | 13 | 4 | 11383 | 1233 | Ady43A,CG6650,Pdxk,CG7328 | | 4.406e-02 | -3.12 | Acyl-CoA\_Oxase/DH\_cen-dom | interpro domains | IPR006091 | 13 | 4 | 11383 | 1233 | CG4860,CG4586,Mcad,Acox57D-p | | 4.406e-02 | -3.12 | Znf\_RanBP2\_sf | interpro domains | IPR036443 | 13 | 4 | 11383 | 1233 | Nup358,RYBP,sol,trbd | | 4.406e-02 | -3.12 | NADP\_OxRdtase\_dom | interpro domains | IPR023210 | 13 | 4 | 11383 | 1233 | CG10863,CG2767,CG10638,CG9436 | | 4.406e-02 | -3.12 | NADP\_OxRdtase\_dom\_sf | interpro domains | IPR036812 | 13 | 4 | 11383 | 1233 | CG2767,CG10863,CG10638,CG9436 | | 4.415e-02 | -3.12 | coated vesicle membrane | cellular component | GO:0030662 | 36 | 7 | 11940 | 1105 | Hip1,Sec22,Afti,Sec23,AP-1-2beta,Clc,CG9132 | | 4.416e-02 | -3.12 | negative regulation of stress-activated MAPK cascade | biological process | GO:0032873 | 29 | 6 | 12345 | 1129 | CG17746,Ptp61F,alph,SkpA,park,PpV | | 4.416e-02 | -3.12 | negative regulation of stress-activated protein kinase signaling cascade | biological process | GO:0070303 | 29 | 6 | 12345 | 1129 | alph,SkpA,park,PpV,CG17746,Ptp61F | | 4.416e-02 | -3.12 | vesicle-mediated transport to the plasma membrane | biological process | GO:0098876 | 29 | 6 | 12345 | 1129 | Sec15,sau,CG5734,Sec10,Tomosyn,CG5447 | | 4.418e-02 | -3.12 | WEST\_ADRENOCORTICAL\_TUMOR\_UP | MSigDB lists | WEST\_ADRENOCORTICAL\_TUMOR\_UP | 126 | 27 | 4161 | 644 | Snx6,ste14,Grx1,CG4511,DppIII,r,Ubc4,lt,GAPsec,ste24a,CG8188,CG6610,Gapdh1,CG33156,mRpL18,alphaTub84B,Chrac-14,lqfR,CCT6,Cdk1,CycB,MED27,Eno,spag,stas,CycE,Dcr-1 | | 4.421e-02 | -3.12 | IRAK1 recruits IKK complex | REACTOME pathways | R-DME-937039 | 6 | 3 | 3923 | 574 | Uev1A,Traf6,ben | | 4.421e-02 | -3.12 | IRAK1 recruits IKK complex upon TLR7/8 or 9 stimulation | REACTOME pathways | R-DME-975144 | 6 | 3 | 3923 | 574 | ben,Traf6,Uev1A | | 4.421e-02 | -3.12 | Interleukin-1 signaling | REACTOME pathways | R-DME-446652 | 6 | 3 | 3923 | 574 | Uev1A,Traf6,ben | | 4.421e-02 | -3.12 | DEx/H-box helicases activate type I IFN and inflammatory cytokines production | REACTOME pathways | R-DME-3134963 | 6 | 3 | 3923 | 574 | dl,Rel,Myd88 | | 4.421e-02 | -3.12 | Beta-oxidation of very long chain fatty acids | REACTOME pathways | R-DME-390247 | 6 | 3 | 3923 | 574 | CG4586,ABCD,Acox57D-p | | 4.421e-02 | -3.12 | PTK6 Regulates RTKs and Their Effectors AKT1 and DOK1 | REACTOME pathways | R-DME-8849469 | 6 | 3 | 3923 | 574 | RpL40,RpS27A,Cbl | | 4.421e-02 | -3.12 | Pyrimidine biosynthesis | REACTOME pathways | R-DME-500753 | 6 | 3 | 3923 | 574 | r-l,Ts,r | | 4.421e-02 | -3.12 | regulation of cellular protein catabolic process | biological process | GO:1903362 | 60 | 10 | 12345 | 1129 | park,Ubc6,ari-1,SkpA,Pomp,cort,dco,CkIIalpha,Rpt2,Cul1 | | 4.421e-02 | -3.12 | regulation of neural precursor cell proliferation | biological process | GO:2000177 | 60 | 10 | 12345 | 1129 | COX4,bun,MED27,tws,COX5A,hb,EcR,MAGE,MED4,mts | | 4.424e-02 | -3.12 | MADF | smart domains | SM00595 | 48 | 9 | 5086 | 506 | CG1602,Mes2,CG6175,CG6276,CG42526,hng3,jigr1,CG4404,brwl | | 4.425e-02 | -3.12 | peptidyl-glutamine modification | biological process | GO:0018199 | 4 | 2 | 12345 | 1129 | isoQC,QC | | 4.425e-02 | -3.12 | mitotic chromosome movement towards spindle pole | biological process | GO:0007079 | 4 | 2 | 12345 | 1129 | CycB,Klp10A | | 4.425e-02 | -3.12 | pyrimidine ribonucleoside monophosphate biosynthetic process | biological process | GO:0009174 | 4 | 2 | 12345 | 1129 | l(2)k01209,r-l | | 4.425e-02 | -3.12 | spermatogonial cell division | biological process | GO:0007284 | 4 | 2 | 12345 | 1129 | bam,Cdk1 | | 4.425e-02 | -3.12 | regulation of cohesin loading | biological process | GO:0071922 | 4 | 2 | 12345 | 1129 | wapl,Rbf | | 4.425e-02 | -3.12 | 'de novo' IMP biosynthetic process | biological process | GO:0006189 | 4 | 2 | 12345 | 1129 | CG11089,Pfas | | 4.425e-02 | -3.12 | visual learning | biological process | GO:0008542 | 4 | 2 | 12345 | 1129 | Fdh,osk | | 4.425e-02 | -3.12 | tRNA wobble position uridine thiolation | biological process | GO:0002143 | 4 | 2 | 12345 | 1129 | CG10189,CG3021 | | 4.425e-02 | -3.12 | positive regulation of histone H3-K9 methylation | biological process | GO:0051574 | 4 | 2 | 12345 | 1129 | G9a,trr | | 4.425e-02 | -3.12 | skeletal muscle thin filament assembly | biological process | GO:0030240 | 4 | 2 | 12345 | 1129 | SelR,up | | 4.425e-02 | -3.12 | negative regulation of response to DNA damage stimulus | biological process | GO:2001021 | 4 | 2 | 12345 | 1129 | Ubc6,armi | | 4.425e-02 | -3.12 | calcineurin-mediated signaling | biological process | GO:0097720 | 4 | 2 | 12345 | 1129 | Pp2B-14D,CanA-14F | | 4.425e-02 | -3.12 | mitochondrial translational elongation | biological process | GO:0070125 | 4 | 2 | 12345 | 1129 | mEFTu2,mEFTu1 | | 4.425e-02 | -3.12 | siRNA loading onto RISC involved in RNA interference | biological process | GO:0035087 | 4 | 2 | 12345 | 1129 | r2d2,Dcr-1 | | 4.425e-02 | -3.12 | female germ-line sex determination | biological process | GO:0019099 | 4 | 2 | 12345 | 1129 | ovo,otu | | 4.425e-02 | -3.12 | pyrimidine ribonucleoside monophosphate metabolic process | biological process | GO:0009173 | 4 | 2 | 12345 | 1129 | r-l,l(2)k01209 | | 4.425e-02 | -3.12 | positive regulation of protein dephosphorylation | biological process | GO:0035307 | 4 | 2 | 12345 | 1129 | Ankle2,CG8009 | | 4.425e-02 | -3.12 | vesicle tethering | biological process | GO:0099022 | 4 | 2 | 12345 | 1129 | Sec10,Sec15 | | 4.425e-02 | -3.12 | CENP-A containing chromatin organization | biological process | GO:0061641 | 4 | 2 | 12345 | 1129 | Hat1,Chrac-14 | | 4.425e-02 | -3.12 | positive regulation of dephosphorylation | biological process | GO:0035306 | 4 | 2 | 12345 | 1129 | CG8009,Ankle2 | | 4.425e-02 | -3.12 | regulation of meiotic cell cycle phase transition | biological process | GO:1901993 | 4 | 2 | 12345 | 1129 | Mos,twe | | 4.425e-02 | -3.12 | negative regulation of nuclease activity | biological process | GO:0032074 | 4 | 2 | 12345 | 1129 | EndoGI,Drep1 | | 4.425e-02 | -3.12 | UDP-xylose transmembrane transport | biological process | GO:0015790 | 4 | 2 | 12345 | 1129 | Ugalt,Efr | | 4.425e-02 | -3.12 | protein ufmylation | biological process | GO:0071569 | 4 | 2 | 12345 | 1129 | Ufm1,Uba5 | | 4.425e-02 | -3.12 | negative regulation of mitochondrion organization | biological process | GO:0010823 | 4 | 2 | 12345 | 1129 | Pink1,park | | 4.425e-02 | -3.12 | spindle assembly involved in female meiosis II | biological process | GO:0007058 | 4 | 2 | 12345 | 1129 | Grip128,Grip75 | | 4.425e-02 | -3.12 | pronuclear migration | biological process | GO:0035046 | 4 | 2 | 12345 | 1129 | alphaTub67C,wisp | | 4.425e-02 | -3.12 | germline cell cycle switching, mitotic to meiotic cell cycle | biological process | GO:0051729 | 4 | 2 | 12345 | 1129 | Iml1,Nprl2 | | 4.425e-02 | -3.12 | vesicle tethering involved in exocytosis | biological process | GO:0090522 | 4 | 2 | 12345 | 1129 | Sec10,Sec15 | | 4.425e-02 | -3.12 | CENP-A containing nucleosome assembly | biological process | GO:0034080 | 4 | 2 | 12345 | 1129 | Hat1,Chrac-14 | | 4.425e-02 | -3.12 | UDP-glucuronic acid transmembrane transport | biological process | GO:0015787 | 4 | 2 | 12345 | 1129 | Efr,Ugalt | | 4.425e-02 | -3.12 | cell cycle switching, mitotic to meiotic cell cycle | biological process | GO:0051728 | 4 | 2 | 12345 | 1129 | Nprl2,Iml1 | | 4.425e-02 | -3.12 | succinyl-CoA metabolic process | biological process | GO:0006104 | 4 | 2 | 12345 | 1129 | ScsbetaA,Scsalpha1 | | 4.425e-02 | -3.12 | astral microtubule organization | biological process | GO:0030953 | 4 | 2 | 12345 | 1129 | spd-2,Grip84 | | 4.425e-02 | -3.12 | leukocyte activation | biological process | GO:0045321 | 4 | 2 | 12345 | 1129 | Jafrac1,crq | | 4.425e-02 | -3.12 | chromatin remodeling at centromere | biological process | GO:0031055 | 4 | 2 | 12345 | 1129 | Hat1,Chrac-14 | | 4.425e-02 | -3.12 | cell cycle switching | biological process | GO:0060184 | 4 | 2 | 12345 | 1129 | Nprl2,Iml1 | | 4.425e-02 | -3.12 | regulation of I-kappaB kinase/NF-kappaB signaling | biological process | GO:0043122 | 4 | 2 | 12345 | 1129 | Traf6,Myd88 | | 4.425e-02 | -3.12 | phosphatidic acid biosynthetic process | biological process | GO:0006654 | 4 | 2 | 12345 | 1129 | Agpat2,Pld | | 4.425e-02 | -3.12 | phagosome-lysosome fusion | biological process | GO:0090385 | 4 | 2 | 12345 | 1129 | Vps16B,Snx6 | | 4.425e-02 | -3.12 | glycine biosynthetic process | biological process | GO:0006545 | 4 | 2 | 12345 | 1129 | Shmt,Dhfr | | 4.425e-02 | -3.12 | tRNA aminoacylation for mitochondrial protein translation | biological process | GO:0070127 | 4 | 2 | 12345 | 1129 | TrpRS-m,SerRS-m | | 4.425e-02 | -3.12 | regulation of nuclease activity | biological process | GO:0032069 | 4 | 2 | 12345 | 1129 | Drep1,EndoGI | | 4.425e-02 | -3.12 | glycerol-3-phosphate biosynthetic process | biological process | GO:0046167 | 4 | 2 | 12345 | 1129 | Gk2,Gk1 | | 4.425e-02 | -3.12 | regulation of protein localization to chromatin | biological process | GO:1905634 | 4 | 2 | 12345 | 1129 | wapl,Rbf | | 4.425e-02 | -3.12 | positive regulation of mitotic cell cycle, embryonic | biological process | GO:0045977 | 4 | 2 | 12345 | 1129 | png,ben | | 4.425e-02 | -3.12 | vitamin B6 biosynthetic process | biological process | GO:0042819 | 4 | 2 | 12345 | 1129 | Pdxk,sgll | | 4.425e-02 | -3.12 | pyridoxal phosphate metabolic process | biological process | GO:0042822 | 4 | 2 | 12345 | 1129 | sgll,Pdxk | | 4.425e-02 | -3.12 | peptidyl-pyroglutamic acid biosynthetic process, using glutaminyl-peptide cyclotransferase | biological process | GO:0017186 | 4 | 2 | 12345 | 1129 | isoQC,QC | | 4.425e-02 | -3.12 | pyridoxal phosphate biosynthetic process | biological process | GO:0042823 | 4 | 2 | 12345 | 1129 | Pdxk,sgll | | 4.441e-02 | -3.11 | positive regulation of protein complex assembly | biological process | GO:0031334 | 52 | 9 | 12345 | 1129 | msps,wash,Arpc1,Arpc2,CG11486,Mtl,dco,spz,Arpc4 | | 4.443e-02 | -3.11 | Activation of APC/C and APC/C:Cdc20 mediated degradation of mitotic proteins | REACTOME pathways | R-DME-176814 | 76 | 17 | 3923 | 574 | Prosalpha3,Rpt2,RpL40,Prosbeta4,mad2,RpS27A,Prosalpha5,Prosalpha4,Prosbeta5,Rpn2,APC7,Prosbeta6,CycB,Cdk1,Rpn12,Rpn3,Prosbeta3 | | 4.446e-02 | -3.11 | protein serine/threonine kinase activity | molecular function | GO:0004674 | 200 | 26 | 12176 | 1119 | alc,tkv,rl,png,CG8475,tub,Pdk1,Tfb4,p38a,Mekk1,grp,CkIIalpha,Tao,Cdk5,SRPK,Cdk1,Lkb1,Tlk,fray,Tak1,Cdk7,dco,Mo25,Pink1,CG17698,PEK | | 4.455e-02 | -3.11 | GSE43955\_1H\_VS\_20H\_ACT\_CD4\_TCELL\_WITH\_TGFB\_IL6\_UP | MSigDB lists | GSE43955\_1H\_VS\_20H\_ACT\_CD4\_TCELL\_WITH\_TGFB\_IL6\_UP | 72 | 17 | 4161 | 644 | CG15141,Arts,Tsp66E,CG10465,CG30291,CG12018,Phb2,Caf1-105,CG8199,RpS18,Gdi,CG15525,eEF1beta,mRpS5,PpD3,CG8372,Smox | | 4.455e-02 | -3.11 | REACTOME\_SYNTHESIS\_OF\_DNA | MSigDB lists | REACTOME\_SYNTHESIS\_OF\_DNA | 72 | 17 | 4161 | 644 | RfC38,Prosbeta6,Sld5,RpL40,CG12018,Prosbeta3,Prosalpha4,Rpt2,Rpn12,Rpn2,Prosbeta5,Rpn3,Prosalpha3,Prosbeta4,Prosalpha5,DNApol-epsilon255,RpS27A | | 4.462e-02 | -3.11 | FBXL7 down-regulates AURKA during mitotic entry and in early mitosis | REACTOME pathways | R-DME-8854050 | 65 | 15 | 3923 | 574 | Prosbeta4,RpS27A,Prosbeta3,Prosalpha5,Prosalpha4,Rpn2,Prosbeta5,Prosbeta6,SkpA,Rpn12,Prosalpha3,RpL40,Rpn3,Rpt2,Cul1 | | 4.464e-02 | -3.11 | tacc (transforming acidic coiled-coil protein) | protein interactions | 40589 | 28 | 7 | 9204 | 1112 | mael,Rel,PGRP-LE,cnn,msps,Tak1,Rab5 | | 4.464e-02 | -3.11 | H2.0 (Homeodomain protein 2.0) | protein interactions | 33841 | 28 | 7 | 9204 | 1112 | Ts,yl,CG15141,Chd64,RpL13A,CG11164,scramb1 | | 4.465e-02 | -3.11 | succinate-CoA ligase activity | molecular function | GO:0004774 | 4 | 2 | 12176 | 1119 | ScsbetaA,Scsalpha1 | | 4.465e-02 | -3.11 | nuclear export signal receptor activity | molecular function | GO:0005049 | 4 | 2 | 12176 | 1119 | Cse1,Ranbp21 | | 4.465e-02 | -3.11 | carbon-sulfur lyase activity | molecular function | GO:0016846 | 4 | 2 | 12176 | 1119 | Cchl,Kyat | | 4.465e-02 | -3.11 | galactoside binding | molecular function | GO:0016936 | 4 | 2 | 12176 | 1119 | galectin,CG5335 | | 4.465e-02 | -3.11 | dodecenoyl-CoA delta-isomerase activity | molecular function | GO:0004165 | 4 | 2 | 12176 | 1119 | CG4594,CG4598 | | 4.465e-02 | -3.11 | glutaminyl-peptide cyclotransferase activity | molecular function | GO:0016603 | 4 | 2 | 12176 | 1119 | QC,isoQC | | 4.465e-02 | -3.11 | phosphotyrosine residue binding | molecular function | GO:0001784 | 4 | 2 | 12176 | 1119 | Cbl,hwt | | 4.465e-02 | -3.11 | beta-N-acetylhexosaminidase activity | molecular function | GO:0004563 | 4 | 2 | 12176 | 1119 | Hexo1,fdl | | 4.465e-02 | -3.11 | ADP binding | molecular function | GO:0043531 | 4 | 2 | 12176 | 1119 | blw,Pgk | | 4.465e-02 | -3.11 | aldehyde dehydrogenase (NAD) activity | molecular function | GO:0004029 | 4 | 2 | 12176 | 1119 | Aldh,Fdh | | 4.465e-02 | -3.11 | lamin binding | molecular function | GO:0005521 | 4 | 2 | 12176 | 1119 | Topors,wash | | 4.465e-02 | -3.11 | beta-N-acetylglucosaminidase activity | molecular function | GO:0016231 | 4 | 2 | 12176 | 1119 | fdl,Hexo1 | | 4.465e-02 | -3.11 | clathrin heavy chain binding | molecular function | GO:0032050 | 4 | 2 | 12176 | 1119 | Gga,Clc | | 4.465e-02 | -3.11 | C-acetyltransferase activity | molecular function | GO:0016453 | 4 | 2 | 12176 | 1119 | yip2,CG10932 | | 4.465e-02 | -3.11 | piRNA binding | molecular function | GO:0034584 | 4 | 2 | 12176 | 1119 | piwi,AGO3 | | 4.465e-02 | -3.11 | UDP-glucuronic acid transmembrane transporter activity | molecular function | GO:0005461 | 4 | 2 | 12176 | 1119 | Ugalt,Efr | | 4.465e-02 | -3.11 | UDP-xylose transmembrane transporter activity | molecular function | GO:0005464 | 4 | 2 | 12176 | 1119 | Ugalt,Efr | | 4.485e-02 | -3.10 | GSE15330\_MEGAKARYOCYTE\_ERYTHROID\_VS\_GRANULOCYTE\_MONOCYTE\_PROGENITOR\_IKAROS\_KO\_UP | MSigDB lists | GSE15330\_MEGAKARYOCYTE\_ERYTHROID\_VS\_GRANULOCYTE\_MONOCYTE\_PROGENITOR\_IKAROS\_KO\_UP | 88 | 20 | 4161 | 644 | mbo,PrBP,Pdk,CG1640,CG15141,ste24a,AMPdeam,CG6767,P5cr,MED24,CG3631,CG30010,mRpS5,Cul1,nmdyn-D6,Tfb4,SdhA,Acph-1,Trxr-1,mRpS21 | | 4.485e-02 | -3.10 | GSE3982\_EOSINOPHIL\_VS\_TH1\_DN | MSigDB lists | GSE3982\_EOSINOPHIL\_VS\_TH1\_DN | 88 | 20 | 4161 | 644 | ND-39,Pfas,CG4293,mRpL17,Prosbeta4,CSN6,Idh3b,Axs,CG8004,Tom40,Eno,esc,Pus7,pont,CG15715,CG9393,Ts,NHP2,eIF3i,escl | | 4.490e-02 | -3.10 | Acyl-CoA\_dh\_M | pfam domains | PF02770 | 13 | 4 | 10480 | 1142 | Mcad,Acox57D-p,CG4860,CG4586 | | 4.490e-02 | -3.10 | Aldo\_ket\_red | pfam domains | PF00248 | 13 | 4 | 10480 | 1142 | CG9436,CG10638,CG2767,CG10863 | | 4.490e-02 | -3.10 | intrinsic apoptotic signaling pathway | biological process | GO:0097193 | 22 | 5 | 12345 | 1129 | Cdk5,Mekk1,Rbf,nbs,Debcl | | 4.490e-02 | -3.10 | triglyceride metabolic process | biological process | GO:0006641 | 22 | 5 | 12345 | 1129 | Gk2,ACC,dob,mino,Gk1 | | 4.490e-02 | -3.10 | actin filament bundle assembly | biological process | GO:0051017 | 22 | 5 | 12345 | 1129 | wash,Actn,Tm1,jvl,Mtl | | 4.490e-02 | -3.10 | positive regulation of chromatin silencing | biological process | GO:0031937 | 22 | 5 | 12345 | 1129 | HP1e,HP1c,piwi,egg,scny | | 4.491e-02 | -3.10 | YOSHIMURA\_MAPK8\_TARGETS\_UP | MSigDB lists | YOSHIMURA\_MAPK8\_TARGETS\_UP | 298 | 57 | 4161 | 644 | RpS27A,Abi,Prosalpha5,Exo70,Nap1,CG30496,Fkbp59,Sptr,Fbw5,Calx,Cyp6a19,Chrac-14,Treh,CG9391,GlyP,Echs1,RhoGAP5A,Sod1,RpS30,mino,ND-SGDH,Rpn2,isoQC,csw,dnc,ND-23,kl-2,Rpb8,Cyp6a22,Clc,Gabat,Pp2A-29B,CG7789,RpL11,APP-BP1,Prosbeta4,sbr,nmdyn-D6,CycB,pont,Prosbeta5,p38a,Pfk,mAChR-A,Mtpalpha,Coq7,nrv1,CG2201,Dhfr,CG1532,Cyp1,Atpalpha,Irk1,CG5966,Etf-QO,Pi3K92E,Ts | | 4.495e-02 | -3.10 | transferase activity, transferring one-carbon groups | molecular function | GO:0016741 | 128 | 18 | 12176 | 1119 | CG14100,ste14,CG9643,trr,Rnmt,G9a,escl,CG7009,CG11089,esc,agt,CG8067,egg,mtTFB1,Shmt,Ts,Dph5,r | | 4.524e-02 | -3.10 | endoplasmic reticulum tubular network | cellular component | GO:0071782 | 4 | 2 | 11940 | 1105 | Lnpk,Parp16 | | 4.524e-02 | -3.10 | epsilon DNA polymerase complex | cellular component | GO:0008622 | 4 | 2 | 11940 | 1105 | Chrac-14,DNApol-epsilon255 | | 4.524e-02 | -3.10 | GINS complex | cellular component | GO:0000811 | 4 | 2 | 11940 | 1105 | Psf3,Sld5 | | 4.524e-02 | -3.10 | glycosylphosphatidylinositol-N-acetylglucosaminyltransferase (GPI-GnT) complex | cellular component | GO:0000506 | 4 | 2 | 11940 | 1105 | PIG-C,PIG-H | | 4.524e-02 | -3.10 | DUBm complex | cellular component | GO:0071819 | 4 | 2 | 11940 | 1105 | Sgf11,not | | 4.524e-02 | -3.10 | mRNA cap binding complex | cellular component | GO:0005845 | 4 | 2 | 11940 | 1105 | eIF4E1,Cbp80 | | 4.524e-02 | -3.10 | SAM complex | cellular component | GO:0001401 | 4 | 2 | 11940 | 1105 | CG8004,CG9393 | | 4.524e-02 | -3.10 | elongin complex | cellular component | GO:0070449 | 4 | 2 | 11940 | 1105 | gus,EloC | | 4.524e-02 | -3.10 | cortical microtubule cytoskeleton | cellular component | GO:0030981 | 4 | 2 | 11940 | 1105 | alphaTub67C,Klp10A | | 4.543e-02 | -3.09 | GO\_CELLULAR\_RESPONSE\_TO\_HORMONE\_STIMULUS | MSigDB lists | GO\_CELLULAR\_RESPONSE\_TO\_HORMONE\_STIMULUS | 143 | 30 | 4161 | 644 | Moe,LpR2,AIF,CG11710,Ac3,Pdk1,rl,CycE,Pka-R2,SREBP,Fkbp59,Rab10,Got1,Wnt10,rept,Atpalpha,Tip60,ACC,Pka-R1,Vha13,Ptp61F,MED4,adp,EcR,mino,Prip,Cdk7,MED24,Vha36-1,csw | | 4.551e-02 | -3.09 | response to radiation | biological process | GO:0009314 | 183 | 24 | 12345 | 1129 | grp,Kap-alpha3,Stat92E,Fdh,Ggt-1,dco,alph,cu,RecQ5,Pld,mts,tor,Calx,CG4476,Xpac,Tfb4,Sec15,Sod3,Utx,mus304,Chrac-14,osk,Debcl,Fkbp59 | | 4.557e-02 | -3.09 | CG3815 (uncharacterized protein) | protein interactions | 31571 | 17 | 5 | 9204 | 1112 | lid,Su(H),l(3)mbt,Nap1,asf1 | | 4.557e-02 | -3.09 | CG1894 (uncharacterized protein) | protein interactions | 43378 | 17 | 5 | 9204 | 1112 | eIF4A,RpS2,CG31549,UbcE2H,GlyP | | 4.557e-02 | -3.09 | dgt5 (dim gamma-tubulin 5) | protein interactions | 36364 | 17 | 5 | 9204 | 1112 | PGRP-LE,Rel,dgt4,Tak1,Grip71 | | 4.557e-02 | -3.09 | Dera (Deoxyribose-phosphate aldolase) | protein interactions | 36358 | 17 | 5 | 9204 | 1112 | Naa20A,Prosbeta5,AnxB10,Prosalpha4,Prosbeta3 | | 4.561e-02 | -3.09 | fd59A (forkhead domain 59A) | protein interactions | 37631 | 40 | 9 | 9204 | 1112 | Rcd5,CG46385,mus205,Tim8,tws,26-29-p,dx,CycB,CG11486 | | 4.561e-02 | -3.09 | Epac (Exchange protein directly activated by cAMP) | protein interactions | 35588 | 40 | 9 | 9204 | 1112 | AOX3,twe,Arts,CG42542,scu,Cyp1,cnc,RpS7,Dhfr | | 4.561e-02 | -3.09 | Vps4 (Vacuolar protein sorting 4) | protein interactions | 32777 | 40 | 9 | 9204 | 1112 | shrb,CCT3,r,wal,Pp2A-29B,CHMP2B,Mcad,Pdk1,Kap-alpha3 | | 4.561e-02 | -3.09 | CG5568 (uncharacterized protein) | protein interactions | 38658 | 40 | 9 | 9204 | 1112 | Cul2,mad2,r-l,Cdk5,nej,CG9248,plu,png,Hexo1 | | 4.561e-02 | -3.09 | MED10 (Mediator complex subunit 10) | protein interactions | 39777 | 40 | 9 | 9204 | 1112 | Ptip,CG17002,Larp4B,ix,Ssdp,MED27,MED4,MED20,enc | | 4.571e-02 | -3.09 | kinase activator activity | molecular function | GO:0019209 | 22 | 5 | 12176 | 1119 | ASPP,Pdk1,Lkb1,RpLP1,Parp16 | | 4.571e-02 | -3.09 | protein kinase activator activity | molecular function | GO:0030295 | 22 | 5 | 12176 | 1119 | ASPP,RpLP1,Lkb1,Parp16,Pdk1 | | 4.571e-02 | -3.09 | ix (intersex) | protein interactions | 45881 | 34 | 8 | 9204 | 1112 | Ssdp,MED27,enc,MED4,MED20,Larp4B,CG17002,Ptip | | 4.571e-02 | -3.09 | CG34205 (uncharacterized protein) | protein interactions | 37526 | 34 | 8 | 9204 | 1112 | Scp1,Fer1HCH,Nf-YA,CG4390,CG10237,Adh,CG32473,Sod3 | | 4.571e-02 | -3.09 | MED30 (Mediator complex subunit 30) | protein interactions | 38078 | 34 | 8 | 9204 | 1112 | Ssdp,ix,MED20,MED4,MED27,Ptip,CG17002,Larp4B | | 4.571e-02 | -3.09 | Cul2 (Cullin 2) | protein interactions | 35420 | 34 | 8 | 9204 | 1112 | Cul5,Xpac,CG6923,tkv,Pura,EcR,lwr,CG17508 | | 4.571e-02 | -3.09 | Cp38 (Chorion protein 38) | protein interactions | 31788 | 34 | 8 | 9204 | 1112 | ssh,RpS2,CG11674,CG8679,Tsp86D,SP1029,Cul1,CkIIalpha | | 4.571e-02 | -3.09 | Gp210 (Glycoprotein 210 kDa) | protein interactions | 35512 | 34 | 8 | 9204 | 1112 | bor,cnn,CCT3,GCS2beta,wal,CG1092,RpLP0,SERCA | | 4.579e-02 | -3.08 | regulation of transport | biological process | GO:0051049 | 267 | 33 | 12345 | 1129 | Fur2,Cbl,wash,SERCA,sbr,Tomosyn,Mctp,PGRP-LE,LpR2,dco,CG8726,lt,Moe,Rab10,Fkbp59,crq,NFAT,Snx6,Tao,crok,Pld,O-fut1,Vps16B,Nup358,Sec23,Crag,Hip1,Ranbp21,tral,Rab5,mop,Rel,mbo | | 4.582e-02 | -3.08 | MORF\_RAD23A | MSigDB lists | MORF\_RAD23A | 194 | 39 | 4161 | 644 | Idh3b,SdhA,Chrac-14,mEFTu1,csw,ND-49,Sod1,LysRS,SkpA,eIF3i,CG5028,rept,ND-51,ND-23,Prosbeta4,Mitofilin,Clc,Pp2A-29B,Iru,Mtpalpha,CCT5,Hat1,ND-24,Thiolase,ND-30,capt,Dd,TyrRS,Phb2,CG4390,Cyt-c-p,CG17454,Ts,r,CG14894,CG10418,CG1416,Cyp1,tko | | 4.589e-02 | -3.08 | GSE17721\_LPS\_VS\_PAM3CSK4\_6H\_BMDC\_UP | MSigDB lists | GSE17721\_LPS\_VS\_PAM3CSK4\_6H\_BMDC\_UP | 67 | 16 | 4161 | 644 | Caf1-105,sbr,CG1774,CG9773,Pcd,nmd,Tspo,Cyp1,Jafrac1,Atg4b,Vha36-1,Sfxn2,LysRS,shv,mRpL13,Pex13 | | 4.589e-02 | -3.08 | GSE42021\_TCONV\_PLN\_VS\_CD24INT\_TCONV\_THYMUS\_UP | MSigDB lists | GSE42021\_TCONV\_PLN\_VS\_CD24INT\_TCONV\_THYMUS\_UP | 67 | 16 | 4161 | 644 | Tsp66E,Mcad,CG3226,Pex13,Kap3,Ptp10D,RfC38,tzn,Pdk,Ublcp1,APP-BP1,CG4293,Dcr-1,Cdk1,pont,pen-2 | | 4.591e-02 | -3.08 | FCERI mediated Ca+2 mobilization | REACTOME pathways | R-DME-2871809 | 10 | 4 | 3923 | 574 | CanA-14F,NFAT,Pp2B-14D,CanB2 | | 4.591e-02 | -3.08 | CLEC7A (Dectin-1) induces NFAT activation | REACTOME pathways | R-DME-5607763 | 10 | 4 | 3923 | 574 | CanB2,Pp2B-14D,CanA-14F,NFAT | | 4.591e-02 | -3.08 | TRAF6 mediated IRF7 activation in TLR7/8 or 9 signaling | REACTOME pathways | R-DME-975110 | 10 | 4 | 3923 | 574 | Uev1A,ben,Traf6,Myd88 | | 4.591e-02 | -3.08 | Insulin processing | REACTOME pathways | R-DME-264876 | 10 | 4 | 3923 | 574 | Exo70,Sec10,Sec15,ZnT35C | | 4.591e-02 | -3.08 | Depolymerisation of the Nuclear Lamina | REACTOME pathways | R-DME-4419969 | 10 | 4 | 3923 | 574 | Dd,CG8009,CycB,Cdk1 | | 4.598e-02 | -3.08 | MODULE\_16 | MSigDB lists | MODULE\_16 | 132 | 28 | 4161 | 644 | CG8199,Tfb4,CG7997,Tom40,mad2,Hat1,CCT6,CG13630,Arpc4,PIG-C,LSm3,Gyg,Prosalpha3,Vamp7,Sec22,PpD3,CG14516,CycE,NHP2,Ts,r,bru1,Rab5,CkIIalpha,CG12018,scu,ste24a,nrv1 | | 4.614e-02 | -3.08 | regulation of nuclear division | biological process | GO:0051783 | 77 | 12 | 12345 | 1129 | dgt4,ago,Moe,mad2,nej,CycE,bru1,cana,Zwilch,tws,APC7,Mos | | 4.616e-02 | -3.08 | establishment of protein localization to organelle | biological process | GO:0072594 | 165 | 22 | 12345 | 1129 | Pen,Tom40,Arts,CG14894,Tom70,Cse1,CG11110,Karybeta3,Pex10,Pex12,CG8004,mbo,CG8728,CG9393,Nup62,Pex13,Tim8,Kap-alpha3,Vsp37A,Roe1,Topors,Nup358 | | 4.620e-02 | -3.07 | GO\_SIGNAL\_TRANSDUCTION\_BY\_P53\_CLASS\_MEDIATOR | MSigDB lists | GO\_SIGNAL\_TRANSDUCTION\_BY\_P53\_CLASS\_MEDIATOR | 27 | 8 | 4161 | 644 | CycB,Cdk1,RpL40,Tip60,twin,Rbf,RpS27A,Not10 | | 4.620e-02 | -3.07 | GO\_PYRUVATE\_METABOLIC\_PROCESS | MSigDB lists | GO\_PYRUVATE\_METABOLIC\_PROCESS | 27 | 8 | 4161 | 644 | Pglym78,Pgk,Pfk,tzn,Eno,Gapdh1,Nc73EF,Pgm1 | | 4.620e-02 | -3.07 | GO\_TRICARBOXYLIC\_ACID\_METABOLIC\_PROCESS | MSigDB lists | GO\_TRICARBOXYLIC\_ACID\_METABOLIC\_PROCESS | 27 | 8 | 4161 | 644 | CG5028,SdhA,ScsbetaA,Idh3b,Irp-1B,Nc73EF,Idh,Scsalpha1 | | 4.620e-02 | -3.07 | GO\_LIPASE\_ACTIVITY | MSigDB lists | GO\_LIPASE\_ACTIVITY | 27 | 8 | 4161 | 644 | CG5966,mAChR-A,nSMase,CG3376,ERp60,CG15111,CG9248,sws | | 4.620e-02 | -3.07 | GKCGCNNNNNNNTGAYG\_UNKNOWN | MSigDB lists | GKCGCNNNNNNNTGAYG\_UNKNOWN | 27 | 8 | 4161 | 644 | Vsp37A,RpS30,shv,mino,Prosbeta4,CG11070,Marf,Coq7 | | 4.627e-02 | -3.07 | GSE21360\_SECONDARY\_VS\_TERTIARY\_MEMORY\_CD8\_TCELL\_UP | MSigDB lists | GSE21360\_SECONDARY\_VS\_TERTIARY\_MEMORY\_CD8\_TCELL\_UP | 110 | 24 | 4161 | 644 | Lsm12,Prosalpha3,CG30496,Pmi,CG7728,Got1,Hat1,CG15715,mEFTu1,ND-30,mRpL16,Tim8,Tom40,CG8004,r-l,CG9393,scu,Cdk7,Gmer,heph,Plap,CG42542,levy,Sld5 | | 4.638e-02 | -3.07 | vesicle coat | cellular component | GO:0030120 | 29 | 6 | 11940 | 1105 | AP-1-2beta,Clc,CG9132,Hip1,Afti,Sec23 | | 4.638e-02 | -3.07 | proteasome accessory complex | cellular component | GO:0022624 | 29 | 6 | 11940 | 1105 | Rpn13,Rpt2,Rpn3,Uch-L5,Rpn12,Rpn2 | | 4.642e-02 | -3.07 | mitochondrial outer membrane | cellular component | GO:0005741 | 44 | 8 | 11940 | 1105 | mino,Tom70,Debcl,Tspo,CG9393,Tom40,CG8004,Marf | | 4.644e-02 | -3.07 | RING | smart domains | SM00184 | 79 | 13 | 5086 | 506 | sip3,Iru,Cbl,Pex10,CG9941,CG8974,CG14435,CG13344,dx,Topors,Traf6,l(3)73Ah,CG6923 | | 4.659e-02 | -3.07 | GSE1740\_MCSF\_VS\_MCSF\_AND\_IFNG\_DAY2\_DERIVED\_MACROPHAGE\_WITH\_IFNA\_STIM\_UP | MSigDB lists | GSE1740\_MCSF\_VS\_MCSF\_AND\_IFNG\_DAY2\_DERIVED\_MACROPHAGE\_WITH\_IFNA\_STIM\_UP | 83 | 19 | 4161 | 644 | CG6767,mRpL13,Cse1,ScsbetaA,trsn,Adi1,ACC,Synj,Pi3K92E,Jheh2,Idh,twin,lwr,CG8314,shop,PIG-C,r-l,rl,Cdk5 | | 4.659e-02 | -3.07 | GSE17721\_POLYIC\_VS\_CPG\_16H\_BMDC\_DN | MSigDB lists | GSE17721\_POLYIC\_VS\_CPG\_16H\_BMDC\_DN | 83 | 19 | 4161 | 644 | Unr,CG7789,PpD3,Vamp7,Surf4,Coq7,CG7322,RpL12,CkIIalpha,Dhfr,CG6767,Aladin,AMPdeam,CG10470,HDAC6,CG40045,CG42542,ACC,Cbp80 | | 4.669e-02 | -3.06 | SCF ubiquitin ligase complex | cellular component | GO:0019005 | 68 | 11 | 11940 | 1105 | gus,Fbw5,CG31633,Cul1,ago,SkpA,CG5003,Cul5,Cul2,CG32221,CG9316 | | 4.678e-02 | -3.06 | Cyclin\_C-dom | interpro domains | IPR004367 | 8 | 3 | 11383 | 1233 | CycE,CycJ,CycB | | 4.678e-02 | -3.06 | Gln\_synth/guanido\_kin\_cat\_dom | interpro domains | IPR014746 | 8 | 3 | 11383 | 1233 | Gclc,Argk,GatB | | 4.678e-02 | -3.06 | Chaperone\_TCP-1 | interpro domains | IPR017998 | 8 | 3 | 11383 | 1233 | CCT5,CCT6,CCT3 | | 4.678e-02 | -3.06 | Chaperonin\_TCP-1\_CS | interpro domains | IPR002194 | 8 | 3 | 11383 | 1233 | CCT6,CCT3,CCT5 | | 4.678e-02 | -3.06 | PP2C\_BS | interpro domains | IPR000222 | 8 | 3 | 11383 | 1233 | CG17746,CG7115,alph | | 4.678e-02 | -3.06 | Ribosomal\_zn-bd | interpro domains | IPR011332 | 8 | 3 | 11383 | 1233 | RpL37a,RpS27A,RpS27 | | 4.679e-02 | -3.06 | arm (armadillo) | protein interactions | 31151 | 119 | 21 | 9204 | 1112 | Su(H),ovo,trbd,rept,foxo,Cdk1,ACC,pygo,E2f1,pont,Sec10,Exo70,lqfR,Tak1,Rbf,nej,vret,CycE,spen,tws,tkv | | 4.691e-02 | -3.06 | microtubule cytoskeleton | cellular component | GO:0015630 | 359 | 43 | 11940 | 1105 | msps,cana,mad2,CG13366,Klp10A,Moe,Grip71,CG3437,toc,spd-2,Zwilch,twe,stai,Girdin,Grip75,Tao,Cen,Axs,awd,Pp2A-29B,sw,CG1890,Kap3,CycB,alphaTub84B,Cdk1,cbs,mts,CycE,alphaTub67C,kl-2,Grip128,jvl,cnn,dgt4,Grip84,Clc,Rep,Blos2,Uxt,gammaTub37C,CG7716,del | | 4.701e-02 | -3.06 | Sec5 (Secretory 5) | protein interactions | 33563 | 59 | 12 | 9204 | 1112 | Pgk,RpL37a,Sry-alpha,CycB,Tsp66E,MED4,RpL13A,Sec15,Pax,lid,Sec10,Ect4 | | 4.701e-02 | -3.06 | Hsp60A (Heat shock protein 60A) | protein interactions | 32045 | 59 | 12 | 9204 | 1112 | csw,regucalcin,CG10638,CG4407,rl,awd,Rel,CG4949,PGRP-LE,CG1532,Tak1,mtTFB1 | | 4.701e-02 | -3.06 | wts (warts) | protein interactions | 43651 | 59 | 12 | 9204 | 1112 | LysRS,SERCA,CCT6,Wbp2,Rab5,Mcad,esc,dco,Cdk1,Cp1,Tao,Mtpalpha | | 4.701e-02 | -3.06 | sav (salvador) | protein interactions | 252554 | 59 | 12 | 9204 | 1112 | alphaTub84B,ATPsynB,cnn,Herc4,RpS7,Pdk,Cp1,RpL34b,CCT3,slif,CG9911,CG9581 | | 4.701e-02 | -3.06 | CG33017 (uncharacterized protein) | protein interactions | 36811 | 59 | 12 | 9204 | 1112 | Ssdp,mEFTu2,scu,CG42542,CG11164,spen,dx,Vha36-1,Ugalt,RpL17,eIF4A,RpL23 | | 4.701e-02 | -3.06 | CG11327 (uncharacterized protein) | protein interactions | 33940 | 59 | 12 | 9204 | 1112 | Gasz,shrb,CG42360,RpL7A,CG2678,CG17746,trsn,Hsc70-3,tral,CG31523,ND-30,trk | | 4.713e-02 | -3.05 | MORF\_FANCG | MSigDB lists | MORF\_FANCG | 62 | 15 | 4161 | 644 | Tfb4,CG9391,Rpn3,kcc,Marf,Gmer,CG7728,CG7488,Pmi,Clp,CkIIalpha,PpD3,Lsm12,CG9393,GCS2beta | | 4.713e-02 | -3.05 | GSE44732\_UNSTIM\_VS\_IL27\_STIM\_IMATURE\_DC\_DN | MSigDB lists | GSE44732\_UNSTIM\_VS\_IL27\_STIM\_IMATURE\_DC\_DN | 62 | 15 | 4161 | 644 | Idh,Tsp66E,CG7718,Cyp6a22,shv,Sccpdh1,CG14894,LpR2,CG5510,CG9471,Naa20A,Cyp6a19,cathD,disp,Nca | | 4.713e-02 | -3.05 | GSE360\_L\_DONOVANI\_VS\_B\_MALAYI\_HIGH\_DOSE\_DC\_DN | MSigDB lists | GSE360\_L\_DONOVANI\_VS\_B\_MALAYI\_HIGH\_DOSE\_DC\_DN | 62 | 15 | 4161 | 644 | oxt,Rpn3,Nca,CG5028,Rab5,eEF1beta,CG3376,CCT6,ND-51,CG4572,mEFTu1,Arf51F,AMPdeam,Rpn12,asf1 | | 4.713e-02 | -3.05 | GO\_NEGATIVE\_REGULATION\_OF\_CANONICAL\_WNT\_SIGNALING\_PATHWAY | MSigDB lists | GO\_NEGATIVE\_REGULATION\_OF\_CANONICAL\_WNT\_SIGNALING\_PATHWAY | 62 | 15 | 4161 | 644 | RpL40,Rpn3,Prosbeta5,p38a,Prosbeta6,Rpn12,Rpn2,Pfdn5,RpS27A,Prosalpha5,Prosalpha3,Prosbeta3,Rpt2,Prosalpha4,Prosbeta4 | | 4.713e-02 | -3.05 | GSE360\_CTRL\_VS\_L\_MAJOR\_DC\_UP | MSigDB lists | GSE360\_CTRL\_VS\_L\_MAJOR\_DC\_UP | 62 | 15 | 4161 | 644 | Cdk5,Stip1,Roe1,CG4572,Tip60,Idh,CanA-14F,ND-49,Pex10,Clp,S2P,RpS13,scu,Pp2B-14D,Ppox | | 4.715e-02 | -3.05 | intracellular signal transduction | biological process | GO:0035556 | 373 | 44 | 12345 | 1129 | Traf6,Pld,dl,Rbf,Pdk1,Tak1,grp,Tlk,Cka,Lkb1,CG14408,RhoGAP54D,RhoGDI,fray,REPTOR,Pka-R2,Rab5,CG17698,CanA-14F,raptor,Mtl,Cdk5,Pi3K92E,CG17272,Pp2B-14D,rl,Nprl2,Rab32,Pka-R1,p38a,Gdi,Rep,Mekk1,zyd,CG3894,CG13933,tub,dnc,Debcl,Rab10,Tao,Ac3,gus,nbs | | 4.717e-02 | -3.05 | TGCGCANK\_UNKNOWN | MSigDB lists | TGCGCANK\_UNKNOWN | 183 | 37 | 4161 | 644 | cact,AIF,SdhA,Mo25,Herc4,Rab10,escl,Su(H),ste14,dnc,PIG-L,CG31715,RpII140,adp,Sfxn2,eIF4A,sbr,SRPK,CCT3,CHMP2B,pr,esc,CG14715,Lsm12,gammaTub37C,APP-BP1,Mitofilin,Rcd5,Prosalpha3,Aos1,RpL40,Nprl2,Ubc4,RpS16,bor,Kap3,CG1847 | | 4.726e-02 | -3.05 | USP\_CS | interpro domains | IPR018200 | 19 | 5 | 11383 | 1233 | not,Usp10,faf,scny,Usp16-45 | | 4.726e-02 | -3.05 | Ser/Thr-sp\_prot-phosphatase | interpro domains | IPR006186 | 19 | 5 | 11383 | 1233 | Pp2B-14D,PpV,mts,CanA-14F,PpD3 | | 4.731e-02 | -3.05 | MED28 (Mediator complex subunit 28) | protein interactions | 43079 | 12 | 4 | 9204 | 1112 | Fdh,Grip128,png,MED20 | | 4.731e-02 | -3.05 | Mat1 (Menage a trois 1) | protein interactions | 36130 | 12 | 4 | 9204 | 1112 | CG11486,EcR,Cdk7,Mes2 | | 4.731e-02 | -3.05 | Hsl (Hormone-sensitive lipase) | protein interactions | 37289 | 12 | 4 | 9204 | 1112 | Marf1,CG42565,CG6766,Gk2 | | 4.731e-02 | -3.05 | VhaSFD (Vacuolar H[+]-ATPase SFD subunit) | protein interactions | 34997 | 12 | 4 | 9204 | 1112 | Vha36-1,Vha68-2,Mitf,Rtnl1 | | 4.731e-02 | -3.05 | l(1)G0320 (lethal (1) G0320) | protein interactions | 326234 | 12 | 4 | 9204 | 1112 | wal,CG1416,Pdk1,Tspo | | 4.731e-02 | -3.05 | CG14499 (uncharacterized protein) | protein interactions | 37086 | 12 | 4 | 9204 | 1112 | RpLP1,eIF4A,CG11164,CG9425 | | 4.731e-02 | -3.05 | CG18273 (uncharacterized protein) | protein interactions | 30991 | 12 | 4 | 9204 | 1112 | CycJ,plu,Cul2,SkpA | | 4.731e-02 | -3.05 | Sep4 (Septin 4) | protein interactions | 32646 | 12 | 4 | 9204 | 1112 | park,Ady43A,Sry-alpha,Droj2 | | 4.731e-02 | -3.05 | CG8142 (uncharacterized protein) | protein interactions | 32763 | 12 | 4 | 9204 | 1112 | Nap1,CG15047,RfC38,elg1 | | 4.731e-02 | -3.05 | SmydA-7 (SET and MYND domain containing, arthropod-specific, member 7) | protein interactions | 36916 | 12 | 4 | 9204 | 1112 | sol,Mf,CG6707,ND-24 | | 4.731e-02 | -3.05 | pelo (pelota) | protein interactions | 34286 | 12 | 4 | 9204 | 1112 | bam,park,RpS30,Pink1 | | 4.731e-02 | -3.05 | Arpc5 (Actin-related protein 2/3 complex, subunit 5) | protein interactions | 33444 | 12 | 4 | 9204 | 1112 | RpL13A,Arpc4,Arpc2,Arpc1 | | 4.731e-02 | -3.05 | ben (bendless) | protein interactions | 32358 | 12 | 4 | 9204 | 1112 | park,Uev1A,Traf6,Tak1 | | 4.742e-02 | -3.05 | GO\_KINASE\_ACTIVITY | MSigDB lists | GO\_KINASE\_ACTIVITY | 229 | 45 | 4161 | 644 | r,SelD,Nprl2,fray,Pi3K92E,Tao,CG8726,Pdk,CG17698,CkIIalpha,CG2201,CycC,dco,alc,Tlk,CG6767,CycB,Cdk5,SRPK,Pdxk,Pdk1,Pfk,p38a,nmdyn-D6,CycE,CG32549,CG8475,Gk2,CG4041,Cdk7,CG33156,csw,CG3631,grp,nmdyn-D7,Tfb4,kri,rl,l(1)G0196,Cdk1,Pgk,Pi4KIIalpha,tkv,Mo25,Mekk1 | | 4.753e-02 | -3.05 | GCM\_DENR | MSigDB lists | GCM\_DENR | 18 | 6 | 4161 | 644 | Clp,Karybeta3,CG15141,Crag,Aladin,CG1309 | | 4.753e-02 | -3.05 | GNF2\_PPP6C | MSigDB lists | GNF2\_PPP6C | 18 | 6 | 4161 | 644 | RpL40,PpV,CG17454,Prosbeta6,Arf51F,Bin1 | | 4.753e-02 | -3.05 | BACOLOD\_RESISTANCE\_TO\_ALKYLATING\_AGENTS\_DN | MSigDB lists | BACOLOD\_RESISTANCE\_TO\_ALKYLATING\_AGENTS\_DN | 18 | 6 | 4161 | 644 | RpL24,Grip75,ATP8A,r,TyrRS,CCT5 | | 4.753e-02 | -3.05 | KEGG\_CHRONIC\_MYELOID\_LEUKEMIA | MSigDB lists | KEGG\_CHRONIC\_MYELOID\_LEUKEMIA | 18 | 6 | 4161 | 644 | Pi3K92E,csw,cact,rl,Smox,Med | | 4.753e-02 | -3.05 | PID\_PLK1\_PATHWAY | MSigDB lists | PID\_PLK1\_PATHWAY | 18 | 6 | 4161 | 644 | gammaTub37C,Pp2A-29B,mts,CycB,Cdk1,Klp10A | | 4.753e-02 | -3.05 | GO\_CEREBELLAR\_CORTEX\_DEVELOPMENT | MSigDB lists | GO\_CEREBELLAR\_CORTEX\_DEVELOPMENT | 18 | 6 | 4161 | 644 | CG31715,Nc73EF,Cdk5,Hsc70-3,csw,UQCR-Q | | 4.753e-02 | -3.05 | ELVIDGE\_HIF1A\_AND\_HIF2A\_TARGETS\_DN | MSigDB lists | ELVIDGE\_HIF1A\_AND\_HIF2A\_TARGETS\_DN | 18 | 6 | 4161 | 644 | LpR2,CG7840,AGBE,Pgm1,Su(H),Pgk | | 4.753e-02 | -3.05 | KEGG\_P53\_SIGNALING\_PATHWAY | MSigDB lists | KEGG\_P53\_SIGNALING\_PATHWAY | 18 | 6 | 4161 | 644 | grp,CycE,Cyt-c-p,Tsp66E,Cdk1,CycB | | 4.753e-02 | -3.05 | REACTOME\_MAP\_KINASE\_ACTIVATION\_IN\_TLR\_CASCADE | MSigDB lists | REACTOME\_MAP\_KINASE\_ACTIVATION\_IN\_TLR\_CASCADE | 18 | 6 | 4161 | 644 | Traf6,Pp2A-29B,mts,Cdk1,p38a,rl | | 4.761e-02 | -3.04 | rpr (reaper) | protein interactions | 40015 | 92 | 17 | 9204 | 1112 | Cdk7,Imp,SkpA,RYBP,RpS27A,msps,Hsc70-3,Rbf,velo,Scox,Idh,rl,foxo,Prosbeta6,scny,Karybeta3,DCAF12 | | 4.780e-02 | -3.04 | GSE7509\_UNSTIM\_VS\_TNFA\_IL1B\_IL6\_PGE\_STIM\_DC\_UP | MSigDB lists | GSE7509\_UNSTIM\_VS\_TNFA\_IL1B\_IL6\_PGE\_STIM\_DC\_UP | 94 | 21 | 4161 | 644 | AnxB10,ND-49,Gapdh1,CG7530,Archease,CG17294,RpII140,nmdyn-D7,Ranbp21,RfC38,r,CG5734,AGO3,CG9248,Mat89Ba,PpD3,Pi4KIIalpha,CCT3,Caf1-105,Echs1,Edem1 | | 4.819e-02 | -3.03 | wound healing | biological process | GO:0042060 | 86 | 13 | 12345 | 1129 | CG30010,Gs1l,Stam,CG11089,Idgf6,Idgf4,EMC3,Utx,Arpc4,bun,grp,cold,Mtpalpha | | 4.821e-02 | -3.03 | MORF\_DDB1 | MSigDB lists | MORF\_DDB1 | 127 | 27 | 4161 | 644 | Grip84,Idh3b,CG7728,Clp,CSN6,Marf,CG13630,Cul1,Rpn3,crq,Jafrac2,SdhA,GCS2beta,CG9393,Atg6,ND-SGDH,Phb2,CkIIalpha,P5CDh1,Gmer,Dd,PrBP,rept,Crag,r,CG8204,CG17454 | | 4.823e-02 | -3.03 | GSE19888\_CTRL\_VS\_A3R\_ACT\_TREATED\_MAST\_CELL\_PRETREATED\_WITH\_A3R\_INH\_DN | MSigDB lists | GSE19888\_CTRL\_VS\_A3R\_ACT\_TREATED\_MAST\_CELL\_PRETREATED\_WITH\_A3R\_INH\_DN | 57 | 14 | 4161 | 644 | Ts,CG12713,ND-24,ste14,Uev1A,CG1847,Gapdh1,Efr,ND-39,Exo70,alc,Sod1,CycE,gammaTub37C | | 4.823e-02 | -3.03 | GSE40274\_CTRL\_VS\_FOXP3\_AND\_HELIOS\_TRANSDUCED\_ACTIVATED\_CD4\_TCELL\_UP | MSigDB lists | GSE40274\_CTRL\_VS\_FOXP3\_AND\_HELIOS\_TRANSDUCED\_ACTIVATED\_CD4\_TCELL\_UP | 57 | 14 | 4161 | 644 | Ublcp1,CG8112,CG7655,p38a,fray,CG1812,lt,sws,TrpRS-m,twin,Cisd2,CG10038,CG10465,CG10189 | | 4.823e-02 | -3.03 | GSE22611\_UNSTIM\_VS\_2H\_MDP\_STIM\_NOD2\_TRANSDUCED\_HEK293T\_CELL\_UP | MSigDB lists | GSE22611\_UNSTIM\_VS\_2H\_MDP\_STIM\_NOD2\_TRANSDUCED\_HEK293T\_CELL\_UP | 57 | 14 | 4161 | 644 | CanA-14F,Pfas,Gyg,Uch,GatA,Pp2B-14D,slif,Lsm12,sau,nAChRalpha4,CanB2,RhoGAP5A,Jafrac2,RpS18 | | 4.823e-02 | -3.03 | GSE17721\_LPS\_VS\_GARDIQUIMOD\_0.5H\_BMDC\_DN | MSigDB lists | GSE17721\_LPS\_VS\_GARDIQUIMOD\_0.5H\_BMDC\_DN | 57 | 14 | 4161 | 644 | SRPK,RpS7,AP-1-2beta,Rab5,Rpn3,Moe,CG31729,CG5734,Tspo,SERCA,shrb,Npc2a,spag,CG33181 | | 4.823e-02 | -3.03 | GSE43955\_1H\_VS\_10H\_ACT\_CD4\_TCELL\_WITH\_TGFB\_IL6\_DN | MSigDB lists | GSE43955\_1H\_VS\_10H\_ACT\_CD4\_TCELL\_WITH\_TGFB\_IL6\_DN | 57 | 14 | 4161 | 644 | NHP2,CG9485,MED27,Pka-R1,Chd64,CG9391,Pdk,CG8726,trbd,CG33156,Non1,CG9132,Aladin,tkv | | 4.844e-02 | -3.03 | GSE36078\_WT\_VS\_IL1R\_KO\_LUNG\_DC\_AFTER\_AD5\_INF\_UP | MSigDB lists | GSE36078\_WT\_VS\_IL1R\_KO\_LUNG\_DC\_AFTER\_AD5\_INF\_UP | 32 | 9 | 4161 | 644 | Caf1-105,CG3376,Rbf,AnxB10,CanA-14F,DNApol-epsilon255,Pp2B-14D,CG33181,Smox | | 4.844e-02 | -3.03 | WHITFIELD\_CELL\_CYCLE\_G1\_S | MSigDB lists | WHITFIELD\_CELL\_CYCLE\_G1\_S | 32 | 9 | 4161 | 644 | CG7414,Psf3,YL-1,CycE,CG4390,CG15141,Caf1-105,cathD,CG4407 | | 4.844e-02 | -3.03 | GO\_REGULATION\_OF\_BLOOD\_PRESSURE | MSigDB lists | GO\_REGULATION\_OF\_BLOOD\_PRESSURE | 32 | 9 | 4161 | 644 | Fur2,dnc,Tm1,Fdh,CG2493,CG32473,Sod1,CG11807,Gabat | | 4.844e-02 | -3.03 | GO\_RESPONSE\_TO\_PURINE\_CONTAINING\_COMPOUND | MSigDB lists | GO\_RESPONSE\_TO\_PURINE\_CONTAINING\_COMPOUND | 32 | 9 | 4161 | 644 | Wnt10,SREBP,Prip,CycE,Sod1,GlyP,r,dnc,Calx | | 4.844e-02 | -3.03 | MORF\_BUB1B | MSigDB lists | MORF\_BUB1B | 32 | 9 | 4161 | 644 | SRPK,RfC38,Karybeta3,Cdk1,CCT5,CG4390,Mitofilin,TyrRS,CG30496 | | 4.844e-02 | -3.03 | DARWICHE\_SQUAMOUS\_CELL\_CARCINOMA\_UP | MSigDB lists | DARWICHE\_SQUAMOUS\_CELL\_CARCINOMA\_UP | 32 | 9 | 4161 | 644 | Eno,NHP2,Pgk,eEF1beta,Pglym78,CG3226,Sod1,CG4390,PDCD-5 | | 4.844e-02 | -3.03 | GO\_NEGATIVE\_REGULATION\_OF\_BINDING | MSigDB lists | GO\_NEGATIVE\_REGULATION\_OF\_BINDING | 32 | 9 | 4161 | 644 | STUB1,cact,rept,CanA-14F,Hsc70-3,Non1,dco,Pp2B-14D,CG8507 | | 4.844e-02 | -3.03 | GSE2128\_CTRL\_VS\_MIMETOPE\_NEGATIVE\_SELECTION\_DP\_THYMOCYTE\_C57BL6\_UP | MSigDB lists | GSE2128\_CTRL\_VS\_MIMETOPE\_NEGATIVE\_SELECTION\_DP\_THYMOCYTE\_C57BL6\_UP | 32 | 9 | 4161 | 644 | Cyp1,Gclc,Gdi,Crag,CG8188,CG12818,Tsp86D,Got1,tun | | 4.846e-02 | -3.03 | response to drug | biological process | GO:0042493 | 240 | 30 | 12345 | 1129 | CG17734,EcR,MAGE,Arf51F,dnc,Aldh,AcCoAS,ben,whd,Tao,Agpat2,Cyp6v1,Pka-R2,aay,spz,nbs,Adh,Pdk1,alpha-Man-Ia,rl,Shmt,Bx,Pka-R1,mAChR-A,p38a,cyc,slif,dco,apt,glob1 | | 4.872e-02 | -3.02 | structural constituent of cytoskeleton | molecular function | GO:0005200 | 37 | 7 | 12176 | 1119 | Arpc1,gammaTub37C,Arpc2,Arpc4,Grip75,alphaTub84B,alphaTub67C | | 4.873e-02 | -3.02 | negative regulation of immune response | biological process | GO:0050777 | 61 | 10 | 12345 | 1129 | faf,cact,GlyP,scny,Stat92E,RYBP,p38a,slif,Atg6,PGRP-LE | | 4.873e-02 | -3.02 | anatomical structure regression | biological process | GO:0060033 | 61 | 10 | 12345 | 1129 | Idh3b,Trf2,DCAF12,raptor,Utx,Atg6,EcR,rept,Atg7,MED24 | | 4.909e-02 | -3.01 | CG13510 (uncharacterized protein) | protein interactions | 37596 | 53 | 11 | 9204 | 1112 | nej,squ,Cdk5,plu,boi,pen-2,CycJ,snRNA:U12,Vha36-1,Mf,grp | | 4.912e-02 | -3.01 | SHEDDEN\_LUNG\_CANCER\_POOR\_SURVIVAL\_A6 | MSigDB lists | SHEDDEN\_LUNG\_CANCER\_POOR\_SURVIVAL\_A6 | 195 | 39 | 4161 | 644 | rept,Karybeta3,Sld5,grp,CG8188,mRpL13,Cse1,Gapdh1,CG11899,mad2,Pen,Pgk,Cdk1,Klp10A,Tom40,Prosalpha5,CG30496,ND-39,Fkbp59,Snx6,Cyp1,Cbp80,CG17454,SelD,Ts,RfC38,Ubc4,r,CG6767,tun,Psf3,Hat1,CycB,CCT5,Rnmt,CycE,gammaTub37C,Sec23,Prosbeta4 | | 4.912e-02 | -3.01 | GSE17721\_LPS\_VS\_POLYIC\_2H\_BMDC\_UP | MSigDB lists | GSE17721\_LPS\_VS\_POLYIC\_2H\_BMDC\_UP | 52 | 13 | 4161 | 644 | CG44422,UbcE2H,Arf51F,mad2,opm,kcc,CG16868,cact,Ts,Stip1,Ufm1,tzn,nmd | | 4.912e-02 | -3.01 | KEGG\_INSULIN\_SIGNALING\_PATHWAY | MSigDB lists | KEGG\_INSULIN\_SIGNALING\_PATHWAY | 52 | 13 | 4161 | 644 | ACC,Pi3K92E,rl,CG8475,Pdk1,Pka-R1,GlyP,alc,Exo70,eIF4E1,raptor,Pka-R2,SREBP | | 4.912e-02 | -3.01 | LI\_DCP2\_BOUND\_MRNA | MSigDB lists | LI\_DCP2\_BOUND\_MRNA | 52 | 13 | 4161 | 644 | CG7322,ND-B18,Pdxk,Eno,nmdyn-D6,CG5028,rept,ND-23,tko,CG6028,LSm3,scu,CG9393 | | 4.912e-02 | -3.01 | GSE19941\_UNSTIM\_VS\_LPS\_STIM\_IL10\_KO\_NFKBP50\_KO\_MACROPHAGE\_UP | MSigDB lists | GSE19941\_UNSTIM\_VS\_LPS\_STIM\_IL10\_KO\_NFKBP50\_KO\_MACROPHAGE\_UP | 52 | 13 | 4161 | 644 | CG9393,CG3631,CG11134,CG15111,Efr,bru1,CG10418,mad2,Sld5,Cdk1,CG8204,Ts,mtTFB1 | | 4.912e-02 | -3.01 | GO\_RESPONSE\_TO\_REACTIVE\_OXYGEN\_SPECIES | MSigDB lists | GO\_RESPONSE\_TO\_REACTIVE\_OXYGEN\_SPECIES | 52 | 13 | 4161 | 644 | Kap-alpha3,Tm1,Jafrac1,Cyp1,Grx1,Cdk1,AIF,Trxr-1,PpD3,Prip,Sod1,HDAC6,Cyt-c-p | | 4.913e-02 | -3.01 | histone lysine methylation | biological process | GO:0034968 | 45 | 8 | 12345 | 1129 | Utx,G9a,escl,nej,trr,esc,Ptip,egg | | 4.913e-02 | -3.01 | nucleoside metabolic process | biological process | GO:0009116 | 45 | 8 | 12345 | 1129 | CG6767,nmdyn-D6,kri,awd,nmdyn-D7,l(2)k01209,cN-IIIB,r-l | | 4.913e-02 | -3.01 | negative regulation of cellular catabolic process | biological process | GO:0031330 | 45 | 8 | 12345 | 1129 | Smox,Lkb1,scny,CkIIalpha,Pi3K92E,Nprl2,rl,Fkbp39 | | 4.918e-02 | -3.01 | Cdk4 (Cyclin-dependent kinase 4) | protein interactions | 36854 | 79 | 15 | 9204 | 1112 | CG6923,bcd,Prosbeta5,CycC,CG13344,Stat92E,Rbf,ScsbetaA,CG5390,CycJ,CycE,CG11486,tws,gus,EcR | | 4.918e-02 | -3.01 | Pvr (PDGF- and VEGF-receptor related) | protein interactions | 34127 | 79 | 15 | 9204 | 1112 | NFAT,csw,Tlk,CanA-14F,Pp2B-14D,Pi3K92E,Ptpmeg,dco,Tak1,CG4266,Cbl,ago,CG5447,Tao,HmgD | | 4.927e-02 | -3.01 | ACEVEDO\_LIVER\_CANCER\_DN | MSigDB lists | ACEVEDO\_LIVER\_CANCER\_DN | 144 | 30 | 4161 | 644 | Fkbp59,Hsc70-3,Acph-1,CG14407,Grip84,CG14516,Nmt,DNApol-epsilon255,sws,Eno,Echs1,Bin1,Mtpalpha,Pdxk,cact,SerRS-m,Aldh,RpL12,Marf,Shmt,CG17294,CG33156,CG8728,Rpt2,Sod1,HDAC6,tho2,Arl6IP1,CG10932,Adi1 | | 4.929e-02 | -3.01 | UBCTHYDRLASE | prints domains | PR00707 | 4 | 2 | 2185 | 212 | Uch-L5,Uch | | 4.929e-02 | -3.01 | MAPEPTIDASE | prints domains | PR00599 | 4 | 2 | 2185 | 212 | CG9581,CG13630 | | 4.932e-02 | -3.01 | Gene Silencing by RNA | REACTOME pathways | R-DME-211000 | 39 | 10 | 3923 | 574 | gw,Dcr-1,Nup62,Ranbp21,mbo,Nup358,Ndc1,Aladin,Nup58,trsn | | 4.934e-02 | -3.01 | response to topologically incorrect protein | biological process | GO:0035966 | 53 | 9 | 12345 | 1129 | Edem1,Parp16,REPTOR,CG32640,Atf6,CG8974,PEK,Hsc70-3,sip3 | | 4.934e-02 | -3.01 | positive regulation of cytoskeleton organization | biological process | GO:0051495 | 53 | 9 | 12345 | 1129 | dco,Arpc4,Arpc1,wash,Moe,msps,Mtl,Abi,Arpc2 | | 4.951e-02 | -3.01 | Proteasome | KEGG pathways | dme03050 | 52 | 13 | 3198 | 496 | Prosbeta5,Rpn12,Prosalpha3,Rpt2,Rpn13,Rpn3,Pomp,Prosbeta3,Rpn2,Prosalpha4,Prosbeta6,Prosalpha5,Prosbeta4 | | 4.951e-02 | -3.01 | Proteasome | KEGG pathways | ko03050 | 52 | 13 | 3198 | 496 | Prosbeta5,Rpn12,Rpt2,Prosalpha3,Rpn13,Rpn3,Pomp,Rpn2,Prosbeta3,Prosalpha4,Prosbeta6,Prosalpha5,Prosbeta4 | | 4.958e-02 | -3.00 | GSE1460\_INTRATHYMIC\_T\_PROGENITOR\_VS\_DP\_THYMOCYTE\_DN | MSigDB lists | GSE1460\_INTRATHYMIC\_T\_PROGENITOR\_VS\_DP\_THYMOCYTE\_DN | 37 | 10 | 4161 | 644 | EcR,S2P,nAChRalpha4,CG7718,dnc,CG31729,CG9911,kcc,cyc,lid | | 4.958e-02 | -3.00 | GO\_PROTEIN\_KINASE\_COMPLEX | MSigDB lists | GO\_PROTEIN\_KINASE\_COMPLEX | 37 | 10 | 4161 | 644 | CG8475,lid,Tfb4,Pka-R1,alc,Mo25,CycE,CycC,Cdk7,Pka-R2 | | 4.958e-02 | -3.00 | GSE360\_CTRL\_VS\_B\_MALAYI\_LOW\_DOSE\_DC\_DN | MSigDB lists | GSE360\_CTRL\_VS\_B\_MALAYI\_LOW\_DOSE\_DC\_DN | 37 | 10 | 4161 | 644 | isoQC,CG32549,Phb2,CG8188,CycE,Prosbeta4,Pka-R2,dnc,Nprl2,GlyP | | 4.958e-02 | -3.00 | GSE37301\_HEMATOPOIETIC\_STEM\_CELL\_VS\_PRO\_BCELL\_UP | MSigDB lists | GSE37301\_HEMATOPOIETIC\_STEM\_CELL\_VS\_PRO\_BCELL\_UP | 37 | 10 | 4161 | 644 | Pp2B-14D,Abi,Tlk,CanA-14F,Gyg,Rpn3,CG5734,Synj,Med,CG17691 | | 4.967e-02 | -3.00 | ACTAYRNNNCCCR\_UNKNOWN | MSigDB lists | ACTAYRNNNCCCR\_UNKNOWN | 161 | 33 | 4161 | 644 | Tomosyn,Wdfy2,Mitofilin,Hmg-2,thoc6,Marf,Ntan1,CG10463,Rbf,CG17187,CG5285,lt,Etf-QO,Nprl2,PpV,mus81,Maf1,Rab5,PrBP,Unr,Fkbp59,RpS27A,Nmt,CG16979,lqfR,pen-2,UbcE2H,Efr,adp,CG3842,CG7550,opm,CG5734 | | 4.972e-02 | -3.00 | male meiotic nuclear division | biological process | GO:0007140 | 95 | 14 | 12345 | 1129 | Topors,eIF4E1,tef,bam,Pex10,Grip75,Cdk1,Syx13,mael,Arf51F,larp,Grip128,twe,sau | | 4.973e-02 | -3.00 | GSE34179\_THPOK\_KO\_VS\_WT\_VA14I\_NKTCELL\_DN | MSigDB lists | GSE34179\_THPOK\_KO\_VS\_WT\_VA14I\_NKTCELL\_DN | 47 | 12 | 4161 | 644 | Tsp86D,CG11899,Cul2,CG14407,GCS2beta,LysRS,Rrp40,crq,Karybeta3,kl-2,Pus7,ND-30 | | 4.973e-02 | -3.00 | WAKABAYASHI\_ADIPOGENESIS\_PPARG\_RXRA\_BOUND\_36HR | MSigDB lists | WAKABAYASHI\_ADIPOGENESIS\_PPARG\_RXRA\_BOUND\_36HR | 47 | 12 | 4161 | 644 | CG3587,CG1105,yip2,CG31759,Idh,Mcad,opm,Etf-QO,fray,Roe1,wal,thoc6 | | 4.973e-02 | -3.00 | GNF2\_ST13 | MSigDB lists | GNF2\_ST13 | 47 | 12 | 4161 | 644 | RpS4,RpL38,RpS7,eEF1beta,RpL12,RpL23,RpS24,RpL37a,RpS23,RpL9,RpL24,RpS13 | | 4.973e-02 | -3.00 | EVI1\_04 | MSigDB lists | EVI1\_04 | 47 | 12 | 4161 | 644 | Tm1,ACC,dnc,Droj2,Arts,Stoml2,Gdi,Ppox,dco,mRpS23,CG9132,stet | | 4.981e-02 | -3.00 | GSE22935\_UNSTIM\_VS\_12H\_MBOVIS\_BCG\_STIM\_MACROPHAGE\_DN | MSigDB lists | GSE22935\_UNSTIM\_VS\_12H\_MBOVIS\_BCG\_STIM\_MACROPHAGE\_DN | 89 | 20 | 4161 | 644 | ND-SGDH,lt,colt,Pgm1,Vha36-1,tko,Xpac,DppIII,Vamp7,CG32549,CSN6,mRpS21,CG3077,Ssdp,STUB1,pr,CG7322,SdhA,r-l,Mtpalpha | | 4.986e-02 | -3.00 | Signaling by FGFR | REACTOME pathways | R-DME-190236 | 55 | 13 | 3923 | 574 | Rbfox1,Cbp80,Rpb8,rl,RpS27A,TfIIFalpha,heph,mts,RpL40,CG13398,Pp2A-29B,csw,Cbl | | 4.989e-02 | -3.00 | ebi (ebi) | protein interactions | 33212 | 66 | 13 | 9204 | 1112 | ix,Ssdp,MED27,Sec23,Usp10,Pi3K92E,enc,MED4,MED20,Larp4B,CG17002,Ptip,Rbf | | 4.993e-02 | -3.00 | GSE6269\_FLU\_VS\_STAPH\_AUREUS\_INF\_PBMC\_DN | MSigDB lists | GSE6269\_FLU\_VS\_STAPH\_AUREUS\_INF\_PBMC\_DN | 42 | 11 | 4161 | 644 | Gk2,Cyp1,CG8314,cathD,CG9471,CG14516,isoQC,CG1275,Npc2a,CG9248,Gapdh1 | | 4.993e-02 | -3.00 | PELLICCIOTTA\_HDAC\_IN\_ANTIGEN\_PRESENTATION\_UP | MSigDB lists | PELLICCIOTTA\_HDAC\_IN\_ANTIGEN\_PRESENTATION\_UP | 42 | 11 | 4161 | 644 | Rpn3,ERp60,Prosbeta6,Ac3,Trs33,Sec22,Rpn2,Prosalpha5,Prosalpha4,Hsc70-3,Prosbeta3 | | 4.993e-02 | -3.00 | GSE7460\_CTRL\_VS\_TGFB\_TREATED\_ACT\_TCONV\_DN | MSigDB lists | GSE7460\_CTRL\_VS\_TGFB\_TREATED\_ACT\_TCONV\_DN | 42 | 11 | 4161 | 644 | tun,CG8963,CG7115,galectin,Grip75,RpLP0,CG11710,Vha13,CG2938,shop,Ublcp1 | | 4.993e-02 | -3.00 | GSE11057\_NAIVE\_CD4\_VS\_PBMC\_CD4\_TCELL\_UP | MSigDB lists | GSE11057\_NAIVE\_CD4\_VS\_PBMC\_CD4\_TCELL\_UP | 42 | 11 | 4161 | 644 | trbd,CG8728,Clp,Pex23,twin,tws,eEF1beta,Arts,Jafrac1,Pi3K92E,Prosbeta5 | | 4.993e-02 | -3.00 | GSE15330\_HSC\_VS\_LYMPHOID\_PRIMED\_MULTIPOTENT\_PROGENITOR\_DN | MSigDB lists | GSE15330\_HSC\_VS\_LYMPHOID\_PRIMED\_MULTIPOTENT\_PROGENITOR\_DN | 42 | 11 | 4161 | 644 | COX5A,shv,Rpn12,mad2,cathD,ND-B22,RfC38,Jafrac2,ND-B18,CG31715,CG14715 | | 4.995e-02 | -3.00 | TCP1\_3 | prosite domains | PS00995 | 8 | 3 | 6243 | 694 | CCT5,CCT6,CCT3 | | 4.995e-02 | -3.00 | TCP1\_1 | prosite domains | PS00750 | 8 | 3 | 6243 | 694 | CCT3,CCT6,CCT5 | | 4.995e-02 | -3.00 | CLIP | prosite domains | PS51888 | 8 | 3 | 6243 | 694 | ea,snk,Sp7 | | 4.995e-02 | -3.00 | PPM\_1 | prosite domains | PS01032 | 8 | 3 | 6243 | 694 | alph,CG7115,CG17746 |
