## Supplementary material for "A single DPE core promoter motif contributes to *in vivo* transcriptional regulation and affects cardiac function": SupFile2: knownResults.html

./cluster.allRlog.c300.t8//1573.cluster.txt.motifs/ - Homer Known Motif Enrichment Results


### Homer Known Motif Enrichment Results (./cluster.allRlog.c300.t8//1573.cluster.txt.motifs/)

Homer *de novo* Motif Results  
Gene Ontology Enrichment Results  
Known Motif Enrichment Results (txt file)  
Total Target Sequences = 1436, Total Background Sequences = 45578

|  |  |  |  |  |  |  |  |  |  |  |  |
| --- | --- | --- | --- | --- | --- | --- | --- | --- | --- | --- | --- |
| Rank | Motif | Name | P-value | log P-pvalue | q-value (Benjamini) | # Target Sequences with Motif | % of Targets Sequences with Motif | # Background Sequences with Motif | % of Background Sequences with Motif | Motif File | SVG |
| 1 | T G C A A G T C C T A G A C T G A G C T G T A C T C G A A G T C C G T A A G T C G A C T C T A G | Unknown1(NR/Ini-like)/Drosophila-Promoters/Homer | 1e-134 | -3.106e+02 | 0.0000 | 262.0 | 18.25% | 1175.9 | 2.58% | motif file (matrix) | svg |
| 2 | G A T C C T G A C T A G G C A T C T A G C A G T C A T G T C G A G T A C A G T C T C A G A G C T | M1BP(Zf)/S2R+-M1BP-ChIP-Seq(GSE49842)/Homer | 1e-132 | -3.044e+02 | 0.0000 | 261.0 | 18.18% | 1196.2 | 2.62% | motif file (matrix) | svg |
| 3 | A G T C C G T A C G A T A G T C G T C A A G T C A C G T C T G A | Unknown2/Drosophila-Promoters/Homer | 1e-103 | -2.375e+02 | 0.0000 | 335.0 | 23.33% | 2703.0 | 5.92% | motif file (matrix) | svg |
| 4 | C G T A T C G A G A T C G C A T C G T A A C G T G T A C T C A G G T C A G A C T C G T A C T A G | DREF/Drosophila-Promoters/Homer | 1e-98 | -2.258e+02 | 0.0000 | 232.0 | 16.16% | 1319.3 | 2.89% | motif file (matrix) | svg |
| 5 | T C G A G C T A T G A C G C T A C T A G G A T C C G A T A C T G C G A T A G C T G A C T C T A G | E-box/Drosophila-Promoters/Homer | 1e-62 | -1.435e+02 | 0.0000 | 191.0 | 13.30% | 1409.7 | 3.09% | motif file (matrix) | svg |
| 6 | T C G A G T A C T C G A T C G A C A T G A T G C A C G T A C T G A C T G A G T C C G T A C T A G A G T C A T C G A G T C | Unknown3/Drosophila-Promoters/Homer | 1e-47 | -1.102e+02 | 0.0000 | 121.0 | 8.43% | 725.5 | 1.59% | motif file (matrix) | svg |
| 7 | C T G A C G T A C G T A G C T A C T G A A C G T G T C A A G T C A G T C C T G A G T C A G C T A | Unknown4/Drosophila-Promoters/Homer | 1e-28 | -6.532e+01 | 0.0000 | 125.0 | 8.70% | 1238.2 | 2.71% | motif file (matrix) | svg |
| 8 | C T A G A G T C A G C T A C T G C G T A C A G T C G T A C T G A T A G C T G A C | Unknown5/Drosophila-Promoters/Homer | 1e-17 | -3.935e+01 | 0.0000 | 238.0 | 16.57% | 4275.7 | 9.37% | motif file (matrix) | svg |
