## Supplementary material for "A single DPE core promoter motif contributes to *in vivo* transcriptional regulation and affects cardiac function": SupFile2: geneOntology.html

Gene Ontology Results

### Gene Ontology Enrichment Results

Homer *de novo* Motif Enrichment Results  
Known Motif Enrichment Results  

###### Text file version of complete results (i.e. open with Excel) - biological process: Functional groupings of proteins (Gene Ontology) - molecular function: Mechanistic actions of proteins (Gene Ontology) - cellular component: Protein localization (Gene Ontology) - chromosome location: Genes with similar chromosome localization (NCBI Gene) - KEGG pathways: Groups of proteins in the same pathways (KEGG) - protein interactions: "Proteins interacting with a common protein (BIND, EcoCyc, HPRD)" (NCBI Gene) - interpro domains: Proteins with similar domains and features (Interpro) - pfam domains: Proteins with similar domains and features (Pfam) - smart domains: Proteins with similar domains and features (SMART) - gene3d domains: Proteins with similar domains and features (Gene3D) - prosite domains: Proteins with similar domains and features (Prosite) - prints domains: Proteins with similar domains and features (PRINTS) - MSigDB lists: "Genes sets for pathways, factor/miRNA target predictions, expression patterns, etc." (MSigDB) - BIOCYC pathways: Groups of proteins in the same pathways (BIOCYC) - COSMIC cancer mutations: Genes mutated in similar cancers (COSMIC) - GWAS genes: Genes mutated in similar diseases (GWAS Catalog) - Lipid Maps pathways: Groups of proteins in the same lipid pathways (Lipid Maps/Biosystems) - Pathway Interaction DB: Groups of proteins in the same pathways (Pathway Interaction Database) - REACTOME pathways: Groups of proteins in the same pathways (REACTOME) - SMPDB pathways: Groups of proteins in the same pathways (SMPDB) - WikiPathways: Groups of proteins in the same pathways (Wikipathways) Enriched Categories | | | | | | | | | | | | --- | --- | --- | --- | --- | --- | --- | --- | --- | --- | | P-value | ln(P) | Term | GO Tree | GO ID | # of Genes in Term | # of Target Genes in Term | # of Total Genes | # of Target Genes | Common Genes | | 5.044e-31 | -69.76 | intracellular | cellular component | GO:0005622 | 6762 | 260 | 11940 | 298 | AGO3,Arp8,CG6244,rb,Nopp140,Dbp80,CG31460,CG30423,CG11752,mon2,DCTN3-p24,CG8915,14-3-3zeta,c12.1,Cul4,CG12659,sqd,Rpt6,TfIIFbeta,g,BEAF-32,Fign,Acf,CycA,Fmr1,Mps1,Ythdf,Cdc42,RpS8,RpL36A,ver,CG17385,Orc3,Vps35,CG32473,His2Av,alpha-Man-Ib,RpS26,SF2,CG11586,ema,smt3,CIA30,PGAP1,dpp,inc,RpS3A,l(1)10Bb,Amun,CG2938,flw,Upf3,RpS19a,l(2)37Bb,Mlf,Lrr47,Cdk8,CG30105,rhi,Stlk,rhea,conu,Bmcp,CG1218,CG12877,vsg,Tctp,eIF3c,Ubqn,ND-ASHI,Rab7,Eaf6,peng,vnc,eEF5,Pex3,Ino80,smg,CG31301,Tgi,Stt3B,Mppe,eRF1,Fs(2)Ket,Tom20,Zpr1,CG6254,CG4935,krz,Dbp21E2,CG4743,CG6171,snRNA:U4:39B,CG6066,Rbm13,pll,PPP1R15,Jarid2,snf,Rop,RpL35A,E(z),ND-AGGG,Rab14,Wdr82,CG12321,CG15534,d4,Arpc5,CG11137,vap,CG8635,pav,LIMK1,HtrA2,RpS9,x16,MED21,xit,janA,Dref,noi,tsu,mRpL34,RpL15,Tcs3,E(bx),Exo84,pins,cal1,Hrb27C,Prosbeta1,ECSIT,mRpS28,sws,RpL5,Graf,MED22,Sod2,Rpn10,bai,Mdh1,RpL14,fzy,Usp32,CG14805,CG9246,PIG-K,CG4866,flr,Ncoa6,CG11920,Pten,cta,CG8223,Nulp1,Prosalpha2,mRpL30,Btk29A,eIF3b,nom,Skp2,Oxp,Stim,CG5986,CG9853,MICU1,heix,RPA3,Rcd5,mnb,Tim17b,Parp,eIF4B,viaf,CG17883,Girdin,Coq5,spoon,Phf5a,Pnn,CG6907,CG32576,DNAlig1,MED26,eEF1delta,HP4,RagA-B,EloB,Pp1-87B,NO66,CG11456,His4r,ubl,eIF3g1,mRpS9,grau,sun,Vps13D,mRpS33,mbf1,Pbp49,CCT4,Map60,su(Hw),RecQ4,Aps,Pep,lic,sm,Dek,tsr,Bub1,Arpc3A,Efa6,tkv,dgrn,me31B,Ufd4,CG4612,BthD,Pisd,Rat1,mRpS18A,CSN1b,Hsp27,Cpsf6,Cp190,CG17493,CG10672,Sgt,Nup188,Roc2,CtBP,CNBP,Not3,Rbbp5,Vap33,CG7766,GstE13,Art1,CG15168,sigmar,Rpb11,twin,CG44194,RAF2,Lst8,Rbp1,CG10365,spel1,CG12299,Nph,Rab8,Wdr33,CG5590,nsl1,VhaPPA1-1,Spn,TM9SF2 | | 5.044e-31 | -69.76 | intracellular part | cellular component | GO:0044424 | 6762 | 260 | 11940 | 298 | TM9SF2,Spn,CG5590,VhaPPA1-1,nsl1,Rab8,Wdr33,CG12299,spel1,CG10365,Nph,Rbp1,Lst8,RAF2,Rpb11,twin,CG44194,CG15168,Art1,sigmar,Vap33,GstE13,CG7766,CNBP,Not3,Rbbp5,Nup188,Sgt,Roc2,CtBP,CG17493,CG10672,Cp190,Cpsf6,CSN1b,Hsp27,Rat1,mRpS18A,me31B,Ufd4,BthD,CG4612,Pisd,tkv,dgrn,Efa6,Bub1,Arpc3A,tsr,sm,lic,Dek,Pep,su(Hw),RecQ4,Aps,CCT4,Map60,Pbp49,Vps13D,mRpS33,mbf1,mRpS9,grau,sun,eIF3g1,His4r,ubl,NO66,CG11456,EloB,Pp1-87B,MED26,eEF1delta,HP4,RagA-B,DNAlig1,Pnn,Phf5a,CG6907,CG32576,spoon,viaf,CG17883,Coq5,Girdin,eIF4B,Parp,mnb,Tim17b,RPA3,Rcd5,Stim,CG5986,heix,CG9853,MICU1,nom,Skp2,Oxp,eIF3b,mRpL30,Prosalpha2,Btk29A,CG8223,cta,Nulp1,Pten,Ncoa6,CG11920,flr,PIG-K,CG4866,CG9246,fzy,Usp32,CG14805,RpL14,bai,Mdh1,MED22,Rpn10,Sod2,sws,mRpS28,RpL5,Graf,Prosbeta1,ECSIT,cal1,Hrb27C,Exo84,pins,E(bx),mRpL34,Tcs3,RpL15,janA,Dref,tsu,noi,xit,MED21,HtrA2,RpS9,x16,pav,CG8635,LIMK1,vap,d4,CG15534,CG11137,Arpc5,CG12321,ND-AGGG,Rab14,Wdr82,E(z),Jarid2,PPP1R15,RpL35A,snf,Rop,CG6171,snRNA:U4:39B,pll,CG6066,Rbm13,Dbp21E2,CG4743,CG4935,krz,eRF1,Fs(2)Ket,Tom20,CG6254,Zpr1,Tgi,Stt3B,Mppe,CG31301,Ino80,smg,vnc,eEF5,Pex3,Eaf6,peng,Rab7,Ubqn,ND-ASHI,Tctp,eIF3c,vsg,CG12877,conu,Bmcp,CG1218,Stlk,rhea,rhi,Lrr47,CG30105,Cdk8,Mlf,l(2)37Bb,flw,Upf3,RpS19a,Amun,CG2938,l(1)10Bb,RpS3A,CIA30,smt3,inc,dpp,PGAP1,SF2,CG11586,ema,His2Av,RpS26,alpha-Man-Ib,Vps35,CG32473,ver,CG17385,Orc3,RpS8,RpL36A,Cdc42,Ythdf,Mps1,CycA,Fmr1,Acf,TfIIFbeta,g,Fign,BEAF-32,sqd,Rpt6,c12.1,Cul4,CG12659,DCTN3-p24,14-3-3zeta,CG8915,CG30423,CG11752,mon2,Dbp80,CG31460,rb,Nopp140,Arp8,AGO3,CG6244 | | 5.238e-26 | -58.21 | intracellular organelle | cellular component | GO:0043229 | 5391 | 223 | 11940 | 298 | Arp8,AGO3,CG6244,cal1,Hrb27C,Nopp140,Dbp80,CG31460,Prosbeta1,ECSIT,CG30423,mRpS28,sws,RpL5,CG11752,mon2,Graf,MED22,DCTN3-p24,Sod2,Rpn10,14-3-3zeta,CG8915,bai,c12.1,Cul4,CG12659,Mdh1,sqd,RpL14,Rpt6,TfIIFbeta,g,BEAF-32,fzy,Fign,Usp32,CG14805,CG9246,PIG-K,CG4866,Acf,CycA,flr,Fmr1,Ncoa6,CG11920,Mps1,Pten,CG8223,Nulp1,Cdc42,RpS8,mRpL30,Prosalpha2,RpL36A,ver,CG17385,Orc3,Vps35,Skp2,nom,Oxp,Stim,CG5986,His2Av,MICU1,heix,alpha-Man-Ib,RpS26,RPA3,ema,SF2,CG11586,Rcd5,mnb,smt3,Tim17b,CIA30,dpp,PGAP1,Parp,RpS3A,l(1)10Bb,CG17883,Girdin,Coq5,Amun,spoon,CG2938,flw,Pnn,Phf5a,CG6907,CG32576,RpS19a,Upf3,l(2)37Bb,DNAlig1,MED26,HP4,RagA-B,EloB,Mlf,Pp1-87B,NO66,CG11456,Lrr47,Cdk8,His4r,ubl,rhi,Stlk,rhea,mRpS9,grau,sun,Vps13D,mRpS33,mbf1,Bmcp,CG1218,Pbp49,CG12877,vsg,Tctp,Map60,su(Hw),RecQ4,ND-ASHI,Aps,Pep,Rab7,sm,Eaf6,peng,Dek,tsr,Pex3,Ino80,Bub1,Arpc3A,smg,Efa6,CG31301,tkv,Tgi,Stt3B,Mppe,dgrn,Ufd4,me31B,Fs(2)Ket,Tom20,CG6254,Zpr1,BthD,CG4612,Pisd,CG4935,Rat1,mRpS18A,Dbp21E2,CSN1b,Hsp27,CG4743,snRNA:U4:39B,CG6171,Rbm13,pll,Cpsf6,CG6066,PPP1R15,Jarid2,snf,RpL35A,Cp190,CG17493,CG10672,Nup188,E(z),Roc2,CtBP,Not3,Rab14,Rbbp5,Wdr82,ND-AGGG,Vap33,CG12321,CG15534,d4,CG11137,Arpc5,CG15168,Art1,Rpb11,pav,twin,CG44194,RpS9,HtrA2,RAF2,x16,MED21,Rbp1,spel1,CG12299,CG10365,xit,Nph,Rab8,Dref,Wdr33,tsu,noi,CG5590,VhaPPA1-1,mRpL34,nsl1,RpL15,E(bx),Spn | | 1.035e-25 | -57.53 | intracellular membrane-bounded organelle | cellular component | GO:0043231 | 4479 | 200 | 11940 | 298 | Skp2,nom,MICU1,heix,alpha-Man-Ib,RpS26,Stim,CG5986,His2Av,CG17385,Orc3,Vps35,Parp,l(1)10Bb,Rcd5,RPA3,SF2,CG11586,ema,smt3,Tim17b,CIA30,PGAP1,mnb,CG32576,Upf3,Pnn,flw,Phf5a,CG6907,DNAlig1,l(2)37Bb,CG17883,Coq5,Amun,spoon,CG2938,Pp1-87B,EloB,Mlf,NO66,CG11456,HP4,RagA-B,MED26,ECSIT,Dbp80,CG31460,Prosbeta1,CG11752,mon2,CG30423,mRpS28,sws,CG6244,Arp8,AGO3,Hrb27C,Nopp140,cal1,sqd,Rpt6,Usp32,CG14805,TfIIFbeta,g,BEAF-32,fzy,Fign,CG8915,14-3-3zeta,Rpn10,Sod2,MED22,CG12659,Mdh1,bai,c12.1,Cul4,Fmr1,CycA,Ncoa6,CG11920,CG9246,Acf,PIG-K,CG4866,Cdc42,CG8223,Nulp1,Prosalpha2,mRpL30,Mps1,Pten,snf,Cp190,PPP1R15,Jarid2,CG10672,CG4743,Hsp27,Dbp21E2,CSN1b,CG6066,pll,Rbm13,Cpsf6,CG6171,snRNA:U4:39B,CG12321,Vap33,CG11137,CG15534,d4,Roc2,CtBP,Nup188,E(z),Wdr82,Rbbp5,Rab14,ND-AGGG,Not3,RAF2,x16,RpS9,HtrA2,MED21,Rbp1,Art1,CG15168,CG44194,pav,Rpb11,twin,CG5590,VhaPPA1-1,mRpL34,nsl1,E(bx),Nph,xit,spel1,CG12299,CG10365,Wdr33,tsu,noi,Rab8,Dref,Stlk,mRpS9,grau,sun,His4r,ubl,Lrr47,Cdk8,rhi,Tctp,Map60,RecQ4,ND-ASHI,Aps,su(Hw),Bmcp,CG1218,Vps13D,mRpS33,mbf1,CG12877,Pbp49,Pex3,tsr,Bub1,Ino80,Pep,Rab7,peng,Dek,sm,Eaf6,CG4612,Pisd,Zpr1,BthD,CG6254,me31B,Ufd4,Tom20,Fs(2)Ket,Rat1,mRpS18A,CG4935,CG31301,Efa6,Stt3B,dgrn,Mppe,Tgi | | 1.586e-25 | -57.10 | organelle | cellular component | GO:0043226 | 5470 | 224 | 11940 | 298 | CG4935,mRpS18A,Rat1,Tom20,Fs(2)Ket,Ufd4,me31B,CG4612,Pisd,CG6254,Zpr1,BthD,Tgi,tkv,dgrn,Mppe,Stt3B,Efa6,CG31301,smg,Arpc3A,Ino80,Bub1,tsr,Pex3,Eaf6,sm,Dek,peng,Rab7,Pep,su(Hw),ND-ASHI,Aps,RecQ4,Map60,Tctp,Pbp49,vsg,CG12877,mbf1,mRpS33,Vps13D,CG1218,Bmcp,sun,mRpS9,grau,rhea,Stlk,rhi,Cdk8,Lrr47,ubl,His4r,Spn,E(bx),nsl1,mRpL34,VhaPPA1-1,CG5590,RpL15,Dref,Rab8,tsu,noi,Wdr33,CG10365,CG12299,spel1,xit,Nph,Rbp1,MED21,HtrA2,RpS9,RAF2,x16,RabX6,twin,Rpb11,pav,CG44194,CG15168,Art1,d4,CG15534,CG11137,Arpc5,Vap33,CG12321,Not3,ND-AGGG,Rbbp5,Rab14,Wdr82,E(z),Nup188,CtBP,Roc2,CG10672,CG17493,PPP1R15,Jarid2,Cp190,RpL35A,snf,snRNA:U4:39B,CG6171,pll,CG6066,Rbm13,Cpsf6,CSN1b,Dbp21E2,CG4743,Hsp27,RpL36A,Prosalpha2,RpS8,mRpL30,Nulp1,CG8223,Cdc42,Pten,Mps1,CG11920,Ncoa6,CycA,Fmr1,flr,CG4866,PIG-K,Acf,CG9246,Fign,BEAF-32,fzy,TfIIFbeta,g,CG14805,Usp32,Rpt6,RpL14,sqd,Cul4,c12.1,bai,CG12659,Mdh1,DCTN3-p24,MED22,CG8915,Rpn10,14-3-3zeta,Sod2,CG30423,sws,mRpS28,Graf,CG11752,mon2,RpL5,Prosbeta1,CG31460,Dbp80,ECSIT,cal1,Nopp140,Hrb27C,Arp8,AGO3,CG6244,CG11456,NO66,Mlf,EloB,Pp1-87B,MED26,RagA-B,HP4,l(2)37Bb,DNAlig1,CG6907,Pnn,flw,Phf5a,Upf3,RpS19a,CG32576,CG2938,Amun,spoon,Coq5,Girdin,CG17883,l(1)10Bb,Parp,RpS3A,mnb,dpp,PGAP1,Tim17b,smt3,CIA30,CG11586,SF2,ema,RPA3,Rcd5,CG5986,His2Av,Stim,alpha-Man-Ib,RpS26,heix,MICU1,Skp2,nom,Oxp,Vps35,ver,Orc3,CG17385 | | 3.916e-25 | -56.20 | membrane-bounded organelle | cellular component | GO:0043227 | 4716 | 205 | 11940 | 298 | Amun,spoon,CG2938,CG17883,Coq5,DNAlig1,l(2)37Bb,CG32576,Upf3,Pnn,flw,Phf5a,CG6907,HP4,RagA-B,MED26,NO66,CG11456,Pp1-87B,EloB,Mlf,Vps35,CG17385,Orc3,MICU1,heix,RpS26,alpha-Man-Ib,Stim,CG5986,His2Av,nom,Skp2,Tim17b,CIA30,smt3,dpp,PGAP1,mnb,Rcd5,RPA3,CG11586,SF2,ema,l(1)10Bb,Parp,Acf,PIG-K,CG4866,CG9246,Ncoa6,CG11920,Fmr1,CycA,Pten,Mps1,Prosalpha2,mRpL30,Cdc42,CG8223,Nulp1,Hrb27C,Nopp140,cal1,CG6244,Arp8,AGO3,CG11752,mon2,Graf,CG30423,sws,mRpS28,ECSIT,Dbp80,CG31460,Prosbeta1,Mdh1,CG12659,bai,c12.1,Cul4,14-3-3zeta,Rpn10,Sod2,CG8915,MED22,Usp32,CG14805,TfIIFbeta,g,fzy,Fign,BEAF-32,sqd,Rpt6,CG44194,Rpb11,pav,twin,Art1,CG15168,MED21,Rbp1,RabX6,x16,RAF2,HtrA2,RpS9,Wdr33,noi,tsu,Rab8,Dref,xit,Nph,CG10365,CG12299,spel1,E(bx),CG5590,mRpL34,VhaPPA1-1,nsl1,Cpsf6,pll,CG6066,Rbm13,snRNA:U4:39B,CG6171,CG4743,Hsp27,Dbp21E2,CSN1b,CG10672,snf,Cp190,Jarid2,PPP1R15,ND-AGGG,Wdr82,Rbbp5,Rab14,Not3,Roc2,CtBP,Nup188,E(z),CG11137,d4,CG15534,CG12321,Vap33,peng,Dek,Eaf6,sm,Rab7,Pep,Ino80,Bub1,Pex3,tsr,Stt3B,Mppe,dgrn,tkv,Tgi,CG31301,Efa6,mRpS18A,Rat1,CG4935,Zpr1,Pisd,CG6254,BthD,CG4612,Ufd4,me31B,Tom20,Fs(2)Ket,rhi,His4r,ubl,Lrr47,Cdk8,grau,mRpS9,sun,Stlk,vsg,CG12877,Pbp49,Bmcp,CG1218,Vps13D,mRpS33,mbf1,RecQ4,Aps,ND-ASHI,su(Hw),Tctp,Map60 | | 1.879e-21 | -47.72 | intracellular organelle part | cellular component | GO:0044446 | 3091 | 153 | 11940 | 298 | l(1)10Bb,Parp,RpS3A,CIA30,smt3,Tim17b,dpp,RPA3,ema,CG11586,SF2,Rcd5,His2Av,CG5986,heix,MICU1,RpS26,Oxp,ver,Orc3,NO66,EloB,Mlf,MED26,HP4,RagA-B,l(2)37Bb,DNAlig1,Phf5a,Pnn,CG6907,Upf3,RpS19a,spoon,CG17883,Girdin,TfIIFbeta,g,fzy,Fign,BEAF-32,CG14805,sqd,Rpt6,RpL14,c12.1,CG12659,MED22,DCTN3-p24,Sod2,14-3-3zeta,mRpS28,sws,RpL5,CG11752,Dbp80,CG31460,cal1,Hrb27C,Nopp140,Arp8,CG6244,mRpL30,RpS8,RpL36A,CG8223,Mps1,Ncoa6,CG11920,flr,PIG-K,CG4866,Acf,CG9246,d4,Arpc5,CG11137,Vap33,Rbbp5,Rab14,Wdr82,ND-AGGG,Nup188,E(z),Roc2,CtBP,CG17493,Jarid2,RpL35A,snf,Cp190,CG6171,snRNA:U4:39B,CG6066,Rbm13,CSN1b,CG4743,E(bx),nsl1,mRpL34,VhaPPA1-1,RpL15,Wdr33,tsu,noi,CG10365,spel1,xit,Nph,MED21,Rbp1,RpS9,HtrA2,x16,RAF2,pav,Rpb11,CG44194,CG15168,su(Hw),RecQ4,ND-ASHI,Tctp,Map60,Pbp49,mRpS33,Bmcp,mRpS9,sun,rhi,Cdk8,His4r,CG4935,mRpS18A,me31B,Tom20,Fs(2)Ket,CG6254,Stt3B,dgrn,Efa6,CG31301,Ino80,Bub1,Arpc3A,tsr,Pex3,sm,Eaf6,peng,Dek,Pep,Rab7 | | 3.680e-21 | -47.05 | cell | cellular component | GO:0005623 | 7811 | 265 | 11940 | 298 | Dbp21E2,CG4743,snRNA:U4:39B,CG6171,CG6066,Rbm13,pll,PPP1R15,Jarid2,snf,Rop,RpL35A,E(z),Wdr82,Rab14,ND-AGGG,CG12321,d4,CG15534,CG11137,Arpc5,vap,CG8635,pav,LIMK1,HtrA2,RpS9,RabX6,x16,MED21,xit,Dref,janA,noi,tsu,mRpL34,Tcs3,RpL15,E(bx),Lrr47,CG30105,Cdk8,rhi,Stlk,rhea,conu,Bmcp,CG1218,vsg,CG12877,Tctp,eIF3c,Ubqn,ND-ASHI,Rab7,Eaf6,peng,vnc,eEF5,Pex3,Ino80,smg,CG31301,Tgi,Stt3B,Mppe,eRF1,Tom20,Fs(2)Ket,CG6254,Zpr1,CG4935,krz,ver,CG17385,Orc3,Vps35,CG32473,His2Av,alpha-Man-Ib,RpS26,CG11586,SF2,ema,CIA30,smt3,inc,dpp,PGAP1,RpS3A,l(1)10Bb,Amun,CG2938,flw,Upf3,RpS19a,l(2)37Bb,Mlf,Arp8,AGO3,CG6244,rb,Nopp140,Dbp80,CG31460,CG30423,mon2,CG11752,DCTN3-p24,CG8915,14-3-3zeta,c12.1,Cul4,CG12659,sqd,Rpt6,g,TfIIFbeta,Fign,BEAF-32,Acf,CycA,Fmr1,Mps1,Ythdf,Cdc42,RpS8,RpL36A,CSN1b,Hsp27,Cpsf6,Cp190,CG17493,CG10672,Nup188,Sgt,Roc2,CtBP,CNBP,Not3,Rbbp5,Vap33,CG7766,GstE13,Art1,CG15168,sigmar,Rpb11,twin,CG44194,Lst8,RAF2,Rbp1,CG10365,CG12299,spel1,Nph,Rab8,Wdr33,CG5590,VhaPPA1-1,nsl1,Spn,TM9SF2,His4r,ubl,eIF3g1,mRpS9,grau,sun,Vps13D,mRpS33,mbf1,Pbp49,CCT4,Map60,su(Hw),RecQ4,Aps,Pep,lic,sm,Dek,tsr,Bub1,Arpc3A,Efa6,tkv,dgrn,Ufd4,me31B,BthD,Pisd,CG4612,mRpS18A,Rat1,eIF3b,nom,Skp2,Oxp,Stim,CG5986,MICU1,heix,CG9853,RPA3,Rcd5,mnb,Tim17b,Parp,eIF4B,viaf,CG17883,Coq5,Girdin,spoon,Phf5a,Pnn,CG6907,CG32576,Rap1,DNAlig1,MED26,eEF1delta,HP4,RagA-B,PlexA,EloB,Pp1-87B,NO66,CG11456,pins,Exo84,cal1,Hrb27C,Prosbeta1,ECSIT,mRpS28,sws,RpL5,Graf,MED22,Sod2,Rpn10,bai,Mdh1,RpL14,fzy,Usp32,Rheb,CG14805,CG9246,PIG-K,CG4866,Tsp97E,flr,Ncoa6,CG11920,Pten,CG8223,cta,Nulp1,Prosalpha2,mRpL30,Btk29A | | 3.680e-21 | -47.05 | cell part | cellular component | GO:0044464 | 7811 | 265 | 11940 | 298 | TM9SF2,Spn,nsl1,VhaPPA1-1,CG5590,Wdr33,Rab8,Nph,CG12299,spel1,CG10365,Rbp1,Lst8,RAF2,CG44194,twin,Rpb11,sigmar,CG15168,Art1,CG7766,GstE13,Vap33,Rbbp5,Not3,CNBP,CtBP,Roc2,Nup188,Sgt,CG10672,CG17493,Cp190,Cpsf6,Hsp27,CSN1b,mRpS18A,Rat1,CG4612,Pisd,BthD,Ufd4,me31B,dgrn,tkv,Efa6,Arpc3A,Bub1,tsr,Dek,lic,sm,Pep,Aps,RecQ4,su(Hw),Map60,CCT4,Pbp49,mbf1,mRpS33,Vps13D,sun,grau,mRpS9,eIF3g1,ubl,His4r,CG11456,NO66,Pp1-87B,EloB,RagA-B,PlexA,HP4,eEF1delta,MED26,DNAlig1,Rap1,CG32576,CG6907,Pnn,Phf5a,spoon,Girdin,Coq5,CG17883,viaf,eIF4B,Parp,Tim17b,mnb,Rcd5,RPA3,CG9853,MICU1,heix,CG5986,Stim,Oxp,Skp2,nom,eIF3b,Btk29A,Prosalpha2,mRpL30,Nulp1,cta,CG8223,Pten,CG11920,Ncoa6,flr,Tsp97E,CG4866,PIG-K,CG9246,Rheb,CG14805,Usp32,fzy,RpL14,Mdh1,bai,Sod2,Rpn10,MED22,Graf,RpL5,sws,mRpS28,ECSIT,Prosbeta1,Hrb27C,cal1,Exo84,pins,E(bx),RpL15,Tcs3,mRpL34,noi,tsu,janA,Dref,xit,MED21,x16,RabX6,HtrA2,RpS9,LIMK1,pav,CG8635,vap,Arpc5,CG11137,d4,CG15534,CG12321,Wdr82,ND-AGGG,Rab14,E(z),RpL35A,Rop,snf,PPP1R15,Jarid2,Rbm13,pll,CG6066,CG6171,snRNA:U4:39B,CG4743,Dbp21E2,krz,CG4935,CG6254,Zpr1,Fs(2)Ket,Tom20,eRF1,Mppe,Stt3B,Tgi,CG31301,smg,Ino80,Pex3,eEF5,vnc,peng,Eaf6,Rab7,ND-ASHI,Ubqn,eIF3c,Tctp,CG12877,vsg,CG1218,Bmcp,conu,rhea,Stlk,rhi,Cdk8,CG30105,Lrr47,Mlf,l(2)37Bb,RpS19a,Upf3,flw,CG2938,Amun,l(1)10Bb,RpS3A,inc,dpp,PGAP1,smt3,CIA30,SF2,ema,CG11586,alpha-Man-Ib,RpS26,His2Av,CG32473,Vps35,Orc3,CG17385,ver,RpL36A,RpS8,Cdc42,Ythdf,Mps1,Fmr1,CycA,Acf,BEAF-32,Fign,g,TfIIFbeta,Rpt6,sqd,CG12659,Cul4,c12.1,CG8915,14-3-3zeta,DCTN3-p24,CG11752,mon2,CG30423,CG31460,Dbp80,Nopp140,rb,CG6244,Arp8,AGO3 | | 4.359e-21 | -46.88 | protein-containing complex | cellular component | GO:0032991 | 3016 | 150 | 11940 | 298 | Rcd5,CG11586,SF2,RPA3,inc,Tim17b,smt3,CIA30,RpS3A,eIF4B,l(1)10Bb,Orc3,ver,eIF3b,Vps35,Skp2,RpS26,MICU1,CG9853,His2Av,CG5986,RagA-B,PlexA,eEF1delta,MED26,Pp1-87B,EloB,glo,RpS19a,CG6907,flw,Pnn,Phf5a,l(2)37Bb,Rpn10,DCTN3-p24,MED22,CG12659,Cul4,c12.1,RpL14,Rpt6,sqd,CG14805,fzy,g,TfIIFbeta,CG6244,Exo84,Arp8,AGO3,rb,Hrb27C,cal1,CG31460,Prosbeta1,Dbp80,CG11752,RpL5,mRpS28,Mps1,cta,RpL36A,Prosalpha2,mRpL30,RpS8,CG9246,CG32075,Acf,CG4866,PIG-K,Fmr1,CycA,CG11920,Ncoa6,CtBP,Roc2,E(z),Nup188,Sgt,ND-AGGG,Rbbp5,Wdr82,Not3,CG7766,CG11137,Arpc5,d4,CSN1b,Rbm13,CG6066,snRNA:U4:39B,Cp190,snf,RpL35A,Jarid2,PPP1R15,Fgop2,spel1,noi,tsu,Wdr33,Dref,Tcs3,RpL15,VhaPPA1-1,mRpL34,nsl1,E(bx),CG15168,twin,pav,Rpb11,x16,Lst8,RAF2,RpS9,MED21,mRpS33,Pbp49,eIF3c,Map60,CCT4,ND-ASHI,His4r,CG30105,Cdk8,eIF3g1,sun,mRpS9,Slmap,dgrn,Stt3B,tkv,CG4612,CG6254,Zpr1,Fs(2)Ket,Tom20,me31B,eRF1,mRpS18A,CG4935,Pep,Eaf6,vnc,smg,Arpc3A,Bub1,Ino80 | | 5.513e-21 | -46.65 | organelle part | cellular component | GO:0044422 | 3156 | 154 | 11940 | 298 | Rbp1,MED21,RpS9,HtrA2,RAF2,x16,pav,Rpb11,CG44194,CG15168,E(bx),VhaPPA1-1,nsl1,mRpL34,RpL15,noi,tsu,Wdr33,CG10365,spel1,xit,Nph,CG17493,Jarid2,Cp190,snf,RpL35A,snRNA:U4:39B,CG6171,CG6066,Rbm13,CSN1b,CG4743,d4,Arpc5,CG11137,Vap33,ND-AGGG,Rab14,Wdr82,Rbbp5,E(z),Nup188,CtBP,Roc2,Arpc3A,Bub1,Ino80,tsr,Pex3,sm,Eaf6,Dek,peng,Rab7,Pep,CG4935,mRpS18A,Tom20,Fs(2)Ket,me31B,CG6254,dgrn,Stt3B,Efa6,CG31301,sun,mRpS9,rhea,rhi,Cdk8,His4r,su(Hw),ND-ASHI,RecQ4,Map60,Tctp,Pbp49,mRpS33,Bmcp,l(2)37Bb,DNAlig1,CG6907,Pnn,Phf5a,RpS19a,Upf3,spoon,Girdin,CG17883,NO66,Mlf,EloB,MED26,RagA-B,HP4,His2Av,CG5986,RpS26,MICU1,heix,Oxp,ver,Orc3,l(1)10Bb,Parp,RpS3A,dpp,Tim17b,CIA30,smt3,ema,CG11586,SF2,RPA3,Rcd5,CG11920,Ncoa6,flr,CG4866,PIG-K,Acf,CG9246,RpL36A,RpS8,mRpL30,CG8223,Mps1,sws,mRpS28,RpL5,CG11752,CG31460,Dbp80,cal1,Nopp140,Hrb27C,Arp8,CG6244,fzy,BEAF-32,Fign,g,TfIIFbeta,CG14805,RpL14,Rpt6,sqd,c12.1,CG12659,DCTN3-p24,MED22,Sod2,14-3-3zeta | | 4.917e-19 | -42.16 | binding | molecular function | GO:0005488 | 5743 | 210 | 12176 | 289 | ema,Mps1,CG30423,eEF5,Zpr1,CG5589,sqd,Vps13D,nsl1,Ino80,dgrn,CG1218,CG5986,x16,Rab8,Ythdf,CG31957,Ncoa6,Dbp80,eIF3b,CG17493,CG34261,NO66,conu,grau,PlexA,DNAlig1,Graf,cta,CG4866,dah,Ubqn,Cdc42,fzy,sm,TfIIFbeta,Dlip2,CG9330,snf,Stim,CG6961,Fmr1,Hsp27,Slmap,Lst8,CG12877,mon2,RhoGAPp190,Map60,tkv,LIMK1,Girdin,I-2,CG12299,CG1815,inc,CNBP,CG7766,Tom20,CG10467,CG10209,nom,RpL5,cal1,mnb,lic,Fs(2)Ket,mbf1,Rpn10,Efa6,smg,Rab14,CG32473,Rab7,CG4935,eIF4B,Dref,CG6254,Pep,His2Av,Not3,unc-119,rhea,SF2,CG6907,Arpc5,Jarid2,pins,eEF1delta,His4r,RAF2,pav,Orc3,Nph,CG8223,c12.1,dpp,spel1,Oxp,eIF3c,PPP1R15,BEAF-32,CG10365,Cul4,CG11586,Dek,mav,Pten,14-3-3zeta,CG6066,twin,Btk29A,Rat1,Wdr82,RagA-B,CG17883,Parp,Rbp1,noi,me31B,HtrA2,mRpS18A,RpL15,Rpb11,Bub1,Tgi,RPA3,Rpt6,Rcd5,krz,flr,spoon,CtBP,sws,Rop,Rap1,CG11920,CG31301,CycA,Cdk8,CG8915,E(z),MED21,CG8635,rhi,Usp16-45,peng,Vap33,Vps35,glo,RabX6,Acf,Pbp49,Phf5a,Acbp1,E(bx),Stlk,CG9951,RpS26,smt3,CG1637,AGO3,Pex3,CG6055,Mlf,Spn,sun,ver,vap,Dbp21E2,CG7646,MED22,su(Hw),CG17385,tsu,Fign,Usp32,RecQ4,alpha-Man-Ib,CCT4,Upf3,Roc2,Sod2,eRF1,GEFmeso,Cp190,d4,Cpsf6,eIF3g1,MICU1,CG5439,CG9246,RpS9,Rheb,Ufd4,CG11456,ECSIT,tsr,RpL14,Arpc3A,CIA30,mus301,Tctp,mRpS9,CG5516,Pp1-87B,CG4612,pll,CG6244,Hrb27C | | 1.919e-17 | -38.49 | nucleus | cellular component | GO:0005634 | 2809 | 136 | 11940 | 298 | Tctp,Map60,RecQ4,Aps,su(Hw),CG1218,mbf1,CG12877,Pbp49,Stlk,grau,His4r,ubl,Lrr47,Cdk8,rhi,CG6254,Zpr1,CG4612,Ufd4,Fs(2)Ket,Rat1,CG4935,CG31301,dgrn,Tgi,tsr,Ino80,Bub1,Pep,peng,Dek,Eaf6,sm,CG12321,d4,Roc2,CtBP,Nup188,E(z),Rbbp5,Wdr82,Not3,snf,Cp190,Jarid2,Hsp27,CSN1b,Cpsf6,CG6066,Rbm13,pll,snRNA:U4:39B,CG6171,nsl1,E(bx),Nph,CG12299,spel1,Wdr33,noi,tsu,Dref,x16,RAF2,RpS9,MED21,Rbp1,Art1,Rpb11,pav,twin,sqd,Rpt6,CG14805,TfIIFbeta,Fign,BEAF-32,fzy,Rpn10,CG8915,14-3-3zeta,MED22,CG12659,c12.1,Cul4,ECSIT,Dbp80,Prosbeta1,CG6244,Arp8,Hrb27C,Nopp140,cal1,Cdc42,CG8223,Nulp1,Prosalpha2,Mps1,Pten,Fmr1,CycA,Ncoa6,CG11920,CG9246,Acf,CG4866,Parp,l(1)10Bb,Rcd5,RPA3,CG11586,SF2,smt3,mnb,nom,Skp2,CG5986,His2Av,CG17385,Orc3,Pp1-87B,EloB,Mlf,NO66,CG11456,HP4,RagA-B,MED26,Upf3,flw,Pnn,Phf5a,CG6907,DNAlig1,Amun | | 1.684e-15 | -34.02 | regulation of nitrogen compound metabolic process | biological process | GO:0051171 | 1870 | 101 | 12345 | 305 | smt3,Cpsf6,CG12299,Rbp1,PPP1R15,Not3,Parp,su(Hw),eIF3b,Pex3,Jarid2,Rcd5,sws,viaf,Fign,tkv,Tcs3,Dref,Fmr1,CG9951,MED21,Nulp1,sqd,E(z),CG9246,mbf1,nsl1,Acf,dpp,rhea,Tgi,CG6907,fzy,Ino80,Cdc42,Art1,snf,Mlf,Arp8,me31B,Cp190,MED26,mnb,CG6244,tsu,krz,flw,CG14805,NO66,CtBP,14-3-3zeta,CG6254,CG1791,spel1,Ncoa6,BEAF-32,Rheb,twin,nom,rb,CG17385,d4,Cul4,SF2,CG11456,CG4612,Dek,grau,I-2,E(bx),Upf3,TfIIFbeta,MED22,CNBP,smg,rhi,mav,x16,CycA,Pep,HtrA2,eEF5,RpS9,mus301,RAF2,CG6066,Rap1,Pten,CG1218,dgrn,AGO3,Hrb27C,Rpt6,lic,Tctp,CG1815,sigmar,PlexA,glo,Lst8,Stlk | | 3.977e-15 | -33.16 | regulation of primary metabolic process | biological process | GO:0080090 | 1894 | 101 | 12345 | 305 | krz,tsu,CG6244,mnb,Cp190,MED26,me31B,Arp8,Art1,snf,Mlf,fzy,Ino80,Cdc42,CG6907,Tgi,Acf,dpp,rhea,CG9246,mbf1,nsl1,Nulp1,sqd,E(z),MED21,CG9951,Tcs3,Dref,Fmr1,tkv,viaf,Fign,sws,Pex3,Jarid2,Rcd5,eIF3b,su(Hw),Not3,Parp,PPP1R15,Rbp1,CG12299,smt3,Cpsf6,Stlk,glo,Lst8,PlexA,sigmar,CG1815,Tctp,lic,Rpt6,AGO3,Hrb27C,dgrn,CG1218,Rap1,Pten,CG6066,RAF2,Pep,HtrA2,RpS9,eEF5,mus301,x16,CycA,rhi,mav,Upf3,CNBP,TfIIFbeta,MED22,smg,E(bx),I-2,Dek,grau,SF2,CG4612,CG11456,Cul4,nom,rb,CG17385,d4,twin,BEAF-32,Rheb,Ncoa6,spel1,CtBP,14-3-3zeta,CG6254,CG1791,flw,CG14805,NO66 | | 4.483e-15 | -33.04 | regulation of cellular metabolic process | biological process | GO:0031323 | 1987 | 104 | 12345 | 305 | Hrb27C,AGO3,lic,Rpt6,Sod2,Tctp,CG1815,PlexA,sigmar,Lst8,glo,Stlk,mav,rhi,CycA,x16,RpS9,mus301,eEF5,HtrA2,Pep,CG6066,RAF2,Pten,Rap1,CG1218,dgrn,CG11456,CG4612,SF2,grau,Dek,I-2,E(bx),CNBP,smg,TfIIFbeta,MED22,Upf3,NO66,flw,CG14805,CG1791,CtBP,CG6254,14-3-3zeta,spel1,RecQ4,Ncoa6,BEAF-32,Rheb,twin,d4,CG17385,nom,rb,Arp8,me31B,MED26,Cp190,mnb,CG6244,tsu,krz,E(z),RagA-B,sqd,Nulp1,nsl1,CG9246,mbf1,rhea,Cdk8,dpp,Acf,Tgi,CG6907,Ino80,Cdc42,fzy,snf,Mlf,Art1,eIF3b,Rcd5,Pex3,Jarid2,Graf,sws,Fign,tkv,Fmr1,Tcs3,Dref,CG9951,MED21,Cpsf6,smt3,Rbp1,CG12299,PPP1R15,Not3,Parp,su(Hw) | | 1.061e-14 | -32.18 | regulation of metabolic process | biological process | GO:0019222 | 2165 | 109 | 12345 | 305 | su(Hw),Parp,Not3,PPP1R15,CG12299,Rbp1,Cpsf6,smt3,CG9951,MED21,Tcs3,Dref,Fmr1,tkv,Fign,viaf,sws,Graf,Rcd5,g,Pex3,Jarid2,eIF3b,snf,Mlf,Art1,Cdc42,Ino80,fzy,CG6907,Tgi,Bmcp,Cdk8,rhea,Acf,dpp,nsl1,mbf1,CG9246,RagA-B,E(z),Nulp1,sqd,krz,tsu,CG6244,mnb,MED26,Cp190,me31B,Arp8,Cul4,CG17385,d4,nom,rb,twin,Rheb,BEAF-32,Ncoa6,spel1,RecQ4,CG1791,14-3-3zeta,CG6254,CtBP,NO66,CG14805,flw,smg,MED22,TfIIFbeta,CNBP,Upf3,E(bx),I-2,grau,Dek,CG11456,CG4612,SF2,CG1218,dgrn,Pten,Rap1,CG6066,RAF2,RpS9,mus301,eEF5,Pep,HtrA2,CycA,x16,mav,rhi,Stlk,Lst8,ECSIT,glo,sigmar,PlexA,CG1815,Tctp,Sod2,lic,Rpt6,Hrb27C,AGO3 | | 1.107e-14 | -32.13 | regulation of cellular process | biological process | GO:0050794 | 3560 | 151 | 12345 | 305 | Dref,Tcs3,CG9951,viaf,tkv,Graf,mthl8,sws,Parp,su(Hw),PPP1R15,Rbp1,cal1,Orc3,Cpsf6,tsr,heix,krz,Pp1-87B,mnb,Vps35,tsu,CG6244,me31B,Slmap,Rab14,Cp190,Arpc3A,RabX6,Ino80,Cdc42,DCTN3-p24,Lrr47,sqd,E(z),mbf1,Btk29A,Arpc5,CNBP,smg,MED22,E(bx),RpL14,pll,CG30423,vap,flr,SF2,CG17385,d4,Mps1,BEAF-32,Rab8,sun,spel1,RhoGAPp190,flw,BthD,glo,Fgop2,Stlk,Rab7,spoon,PlexA,Efa6,Bub1,GEFmeso,Tctp,AGO3,Hrb27C,Rpt6,lic,Pten,CG1218,Txl,CG6066,Spn,CycA,mus301,eEF5,mav,Fmr1,MED21,Fign,pav,eIF3b,Pex3,Jarid2,Rcd5,g,Not3,CG12299,smt3,Nf1,MED26,vsg,Arp8,Skp2,fzy,Sgt,Art1,Mlf,snf,CG6907,Acf,dpp,Cdk8,rhea,Tgi,LIMK1,Nulp1,RagA-B,CG9246,nsl1,conu,Upf3,TfIIFbeta,I-2,TM9SF2,Dek,grau,CG11456,CG4612,nom,rb,Cul4,Rheb,twin,RecQ4,Ncoa6,CG14805,NO66,14-3-3zeta,CtBP,CG6254,CG1791,Lst8,CG1815,sigmar,Sod2,Rap1,dgrn,RAF2,Eaf6,x16,Pep,HtrA2,RpS9,rhi,Rop,cta | | 1.470e-14 | -31.85 | protein binding | molecular function | GO:0005515 | 2410 | 113 | 12176 | 289 | Acf,RhoGAPp190,Map60,Pbp49,mon2,Hsp27,Fmr1,Lst8,Slmap,glo,Vap33,Vps35,E(bx),I-2,Girdin,tkv,Pex3,Mlf,Spn,CG7766,inc,CG9951,smt3,RpL5,cal1,sun,vap,Rpn10,mbf1,Roc2,Cp190,d4,Efa6,GEFmeso,tsu,su(Hw),MED22,Fs(2)Ket,CCT4,lic,tsr,ECSIT,CG9246,smg,CG6907,rhea,SF2,Tctp,Arpc5,CIA30,Arpc3A,Not3,His2Av,Hrb27C,pins,pav,His4r,eEF1delta,Pp1-87B,CG6244,pll,CG30423,CG8223,dpp,Nph,ema,CG10365,Cul4,Dek,BEAF-32,Zpr1,eIF3c,PPP1R15,twin,nsl1,mav,14-3-3zeta,Vps13D,Pten,me31B,HtrA2,CG17883,RagA-B,Ino80,CG1218,dgrn,Parp,Tgi,Ncoa6,Rpt6,Rpb11,PlexA,conu,krz,Rcd5,cta,Graf,flr,eIF3b,Rap1,Ubqn,TfIIFbeta,Cdc42,spoon,Rop,sws,CtBP,E(z),snf,MED21,Cdk8,Dlip2,CycA | | 2.545e-14 | -31.30 | nuclear part | cellular component | GO:0044428 | 1416 | 83 | 11940 | 298 | cal1,Hrb27C,Nopp140,Arp8,Cdk8,CG6244,Dbp80,Pbp49,c12.1,CG12659,MED22,14-3-3zeta,TfIIFbeta,fzy,BEAF-32,CG14805,sqd,Rpt6,sm,Eaf6,CG4866,peng,Acf,Dek,CG9246,Pep,Ino80,Bub1,Ncoa6,CG11920,tsr,dgrn,CG31301,Mps1,CG4935,CG8223,Fs(2)Ket,CG6254,snRNA:U4:39B,CG6066,Rbm13,CSN1b,Orc3,His2Av,CG5986,Jarid2,snf,Rbbp5,Wdr82,smt3,Nup188,RPA3,CG11586,E(z),SF2,Roc2,Rcd5,CtBP,l(1)10Bb,d4,Parp,Rpb11,MED21,Rbp1,DNAlig1,RpS9,Phf5a,Pnn,CG6907,RAF2,Upf3,x16,MED26,HP4,Wdr33,tsu,noi,spel1,Nph,E(bx),NO66,EloB,nsl1 | | 3.653e-14 | -30.94 | regulation of macromolecule metabolic process | biological process | GO:0060255 | 2019 | 103 | 12345 | 305 | Dek,grau,SF2,CG11456,CG4612,Upf3,TfIIFbeta,smg,CNBP,MED22,I-2,E(bx),RecQ4,spel1,Ncoa6,CG14805,flw,NO66,CtBP,CG6254,14-3-3zeta,CG1791,nom,rb,d4,CG17385,Cul4,Rheb,BEAF-32,twin,Tctp,AGO3,Hrb27C,lic,Rpt6,glo,Lst8,Stlk,CG1815,sigmar,x16,CycA,Pep,HtrA2,RpS9,mus301,eEF5,rhi,mav,Rap1,Pten,dgrn,CG1218,RAF2,CG6066,Graf,sws,eIF3b,Jarid2,Pex3,Rcd5,g,Fmr1,Dref,Tcs3,MED21,CG9951,viaf,Fign,tkv,Rbp1,CG12299,smt3,Cpsf6,Not3,Parp,su(Hw),PPP1R15,me31B,Cp190,MED26,Arp8,krz,mnb,CG6244,tsu,dpp,Acf,rhea,Tgi,Nulp1,sqd,E(z),CG9246,mbf1,nsl1,fzy,Cdc42,Ino80,Art1,Mlf,snf,CG6907 | | 2.313e-13 | -29.09 | regulation of biological process | biological process | GO:0050789 | 3862 | 156 | 12345 | 305 | CtBP,bai,14-3-3zeta,CG6254,CG1791,CG14805,NO66,Ncoa6,RecQ4,twin,Rheb,Cul4,nom,rb,CG11456,CG4612,Dek,grau,TM9SF2,I-2,Upf3,TfIIFbeta,conu,rhi,cta,Rop,HtrA2,Pep,RpS9,x16,Eaf6,RAF2,dgrn,Rap1,Sod2,sigmar,inc,CG1815,Lst8,smt3,CG12299,Not3,Pex3,Jarid2,g,Rcd5,pav,eIF3b,Fign,MED21,Fmr1,CG9246,nsl1,Nulp1,RagA-B,Tgi,LIMK1,Acf,dpp,Cdk8,rhea,CG6907,Art1,Mlf,snf,fzy,Sgt,Arp8,Skp2,vsg,MED26,Nf1,flw,BthD,RhoGAPp190,spel1,sun,BEAF-32,Rab8,Mps1,d4,CG17385,SF2,CG30423,vap,flr,pll,RpL14,E(bx),smg,CNBP,MED22,Btk29A,Arpc5,mav,mus301,eEF5,Spn,CycA,Txl,CG6066,CG1218,Pten,lic,Rpt6,AGO3,Hrb27C,GEFmeso,Tctp,Bub1,Efa6,PlexA,spoon,Rab7,Stlk,glo,ECSIT,Fgop2,Cpsf6,Orc3,cal1,Rbp1,PPP1R15,su(Hw),Parp,sws,Graf,mthl8,tkv,viaf,CG9951,Dref,Tcs3,mbf1,sqd,E(z),Lrr47,Bmcp,DCTN3-p24,Cdc42,Ino80,Arpc3A,RabX6,Stim,Cp190,Slmap,me31B,Rab14,tsu,CG6244,mnb,Vps35,krz,tsr,heix,Pp1-87B | | 2.674e-13 | -28.95 | biological regulation | biological process | GO:0065007 | 4350 | 169 | 12345 | 305 | Rpt6,lic,Hrb27C,AGO3,Tctp,GEFmeso,Bub1,PlexA,Efa6,spoon,Rab7,Stlk,Fgop2,ECSIT,glo,mav,eEF5,mus301,CycA,Spn,Txl,CG6066,CG1218,Pten,SF2,CG3842,flr,vap,CG30423,pll,RpL14,Girdin,E(bx),MED22,CNBP,smg,Arpc5,Btk29A,BthD,flw,RhoGAPp190,spel1,sun,Rab8,BEAF-32,Mps1,CG17385,d4,RabX6,Arpc3A,Stim,Cp190,Rab14,Slmap,me31B,tsu,CG6244,Vps35,mnb,Pp1-87B,tsr,krz,heix,mbf1,E(z),sqd,Lrr47,Bmcp,DCTN3-p24,Cdc42,Ino80,sws,Zpr1,mthl8,Graf,tkv,viaf,Vps13D,CG9951,Dref,Tcs3,Cpsf6,Orc3,Vap33,cal1,Rbp1,PPP1R15,su(Hw),Parp,Sod2,sigmar,CG1815,inc,Lst8,cta,Rop,rhi,RpS9,HtrA2,Pep,x16,Eaf6,RAF2,dgrn,Rap1,CG11456,CG4612,CSN1b,grau,Dek,CG17883,TM9SF2,I-2,TfIIFbeta,Upf3,conu,CG1791,CG6254,CtBP,14-3-3zeta,bai,NO66,CG14805,Ncoa6,RecQ4,twin,Rheb,Cul4,nom,rb,Skp2,Arp8,vsg,MED26,CG32473,Nf1,nsl1,CG9246,RagA-B,Exo84,Nulp1,LIMK1,Tgi,ver,Cdk8,rhea,dpp,Acf,CG6907,Aps,snf,Mlf,Art1,Sgt,fzy,g,Rcd5,Pex3,Jarid2,eIF3b,pav,Fign,MED21,pins,Fmr1,smt3,MICU1,CG12299,Not3 | | 1.089e-12 | -27.55 | cellular component organization or biogenesis | biological process | GO:0071840 | 2975 | 129 | 12345 | 305 | Orc3,Nopp140,Vap33,cal1,eRF1,Parp,CG4866,l(2)37Bb,sws,tkv,Dref,Vps13D,sqd,His2Av,E(z),mon2,msd5,Wdr82,Ino80,Cdc42,RpL35A,CG12659,Arpc3A,me31B,sm,Rab14,Cp190,mRpS9,His4r,CG6244,tsu,CG17493,heix,tsr,Pp1-87B,flw,Rpn10,spel1,RhoGAPp190,Nup188,Rab8,BEAF-32,d4,Mps1,CG44194,flr,Girdin,RpL14,E(bx),Btk29A,Arpc5,Spn,CycA,mus301,Pten,eIF4B,Hrb27C,CG11920,Pbp49,lic,Bub1,Tctp,Nph,Rab7,spoon,PlexA,glo,Stlk,smt3,Rbbp5,pav,Pex3,Jarid2,g,Rcd5,vnc,Fign,Fmr1,pins,Oxp,Exo84,CG9853,CG9246,nsl1,Acf,dpp,rhea,Cdk8,ver,LIMK1,CIA30,xit,fzy,Art1,snf,Arp8,RpS8,Tom20,Tim17b,CG32075,RpS19a,RpL5,HP4,CG14805,NO66,14-3-3zeta,bai,CtBP,CG1791,RecQ4,CG8223,Ncoa6,CG5589,unc-119,Rheb,rb,Cul4,grau,conu,rhi,cta,Rop,Dbp21E2,HtrA2,Eaf6,CG12321,Rap1,inc,Lst8 | | 1.257e-12 | -27.40 | cellular process | biological process | GO:0009987 | 7485 | 242 | 12345 | 305 | RpS3A,glo,Stlk,GstE13,mRpS18A,Rab7,DNAlig1,mRpL34,spoon,PlexA,Bub1,alpha-Man-Ib,Usp16-45,Nph,Tctp,Hrb27C,CG11920,AGO3,lic,Rpt6,Pbp49,RPA3,Pten,CG7766,eIF4B,CG1218,CG6066,Txl,CycA,Spn,mus301,eEF5,Hsp27,mav,Arpc5,CG6171,Btk29A,mRpL30,MED22,smg,Pisd,E(bx),Girdin,RpL14,pll,flr,CG44194,Rat1,SF2,CG3842,d4,Mps1,Rab8,BEAF-32,Nup188,sun,spel1,RhoGAPp190,BthD,Rpn10,Pect,flw,Rpb11,c12.1,CG17493,Pp1-87B,noi,tsr,krz,heix,mnb,CG6244,tsu,Rab14,me31B,sm,His4r,Cp190,mRpS9,Stim,CG12659,RabX6,snRNA:U4:39B,Arpc3A,Cdc42,Ino80,Wdr82,RpL35A,eIF3g1,Ubqn,DCTN3-p24,CG31460,CG11334,Roc2,msd5,Lrr47,CG5056,E(z),sqd,His2Av,RpS26,mon2,Prosbeta1,PIG-K,Tcs3,Dref,CG11137,janA,Vps13D,Ufd4,tkv,mthl8,l(2)37Bb,CG8635,Graf,sws,Zpr1,CG4866,CG10209,Prosalpha2,Parp,eRF1,l(1)10Bb,su(Hw),PPP1R15,Nopp140,Rbp1,Vap33,cal1,Orc3,CG15534,RpL15,Lst8,inc,Stt3B,sigmar,Sod2,CG9005,Rap1,dgrn,eEF1delta,CG12321,Eaf6,Dbp21E2,x16,RpS9,Usp32,HtrA2,Pep,Rop,cta,CCT4,rhi,conu,TfIIFbeta,Upf3,Mppe,grau,CSN1b,CG30105,RpL36A,CG34261,ubl,rb,Cul4,unc-119,CG5589,Rheb,twin,eIF3c,ND-ASHI,CG8223,RecQ4,Ncoa6,CG4935,CG10365,NO66,CG14805,mRpS33,CG1791,bai,CtBP,14-3-3zeta,HP4,CG5986,Nf1,RpS19a,CG32473,RpL5,CG32075,Tim17b,CG31957,CG11453,Skp2,Tom20,Arp8,RpS8,Sgt,fzy,snf,Art1,PGAP1,Fs(2)Ket,Aps,CG6907,Mdh1,xit,CIA30,Cdk8,rhea,dpp,Acf,LIMK1,ver,CG9853,RagA-B,Oxp,Exo84,nsl1,Fmr1,pins,Fign,Phf5a,vnc,mRpS28,eIF3b,pav,Rcd5,g,Jarid2,Pex3,Not3,Map60,Pnn,EloB,Wdr33,Rbbp5,Coq5,CG11586,MICU1,CG8132,smt3 | | 4.871e-12 | -26.05 | intracellular organelle lumen | cellular component | GO:0070013 | 1220 | 71 | 11940 | 298 | mRpS33,MED22,Sod2,CG12659,sqd,Rpt6,TfIIFbeta,BEAF-32,CG14805,Arp8,Cdk8,CG6244,cal1,Hrb27C,Nopp140,mRpS28,mRpS9,CG31301,Mps1,dgrn,CG8223,Fs(2)Ket,CG6254,mRpL30,mRpS18A,CG9246,sm,Eaf6,CG4866,peng,Acf,Dek,tsr,Bub1,Ino80,Ncoa6,CG11920,RPA3,SF2,E(z),Rcd5,CtBP,Wdr82,Rbbp5,smt3,dpp,Parp,d4,Orc3,Rbm13,Jarid2,His2Av,Nph,MED26,Wdr33,HP4,EloB,mRpL34,nsl1,NO66,E(bx),Rpb11,CG44194,HtrA2,RpS9,CG6907,x16,Upf3,MED21,Rbp1,DNAlig1 | | 4.871e-12 | -26.05 | membrane-enclosed lumen | cellular component | GO:0031974 | 1220 | 71 | 11940 | 298 | Nopp140,Hrb27C,cal1,CG6244,Cdk8,Arp8,mRpS9,mRpS28,CG12659,Sod2,MED22,mRpS33,CG14805,BEAF-32,TfIIFbeta,Rpt6,sqd,Dek,Acf,peng,CG4866,sm,Eaf6,CG9246,CG11920,Ncoa6,Bub1,Ino80,tsr,dgrn,Mps1,CG31301,mRpS18A,mRpL30,CG6254,Fs(2)Ket,CG8223,Rbm13,Orc3,His2Av,Jarid2,dpp,Rbbp5,smt3,Wdr82,Rcd5,CtBP,SF2,E(z),RPA3,d4,Parp,CG44194,Rpb11,DNAlig1,Rbp1,MED21,x16,Upf3,CG6907,HtrA2,RpS9,HP4,Wdr33,MED26,Nph,E(bx),NO66,nsl1,mRpL34,EloB | | 4.871e-12 | -26.05 | organelle lumen | cellular component | GO:0043233 | 1220 | 71 | 11940 | 298 | d4,Parp,dpp,Wdr82,smt3,Rbbp5,E(z),SF2,RPA3,CtBP,Rcd5,His2Av,Jarid2,Rbm13,Orc3,NO66,E(bx),mRpL34,nsl1,EloB,MED26,HP4,Wdr33,Nph,Rbp1,MED21,DNAlig1,CG6907,RpS9,HtrA2,Upf3,x16,Rpb11,CG44194,BEAF-32,TfIIFbeta,CG14805,Rpt6,sqd,CG12659,MED22,mRpS33,Sod2,mRpS28,mRpS9,cal1,Nopp140,Hrb27C,Cdk8,Arp8,CG6244,mRpS18A,mRpL30,Fs(2)Ket,CG8223,CG6254,dgrn,Mps1,CG31301,Bub1,Ino80,CG11920,Ncoa6,tsr,Eaf6,sm,CG4866,Acf,Dek,peng,CG9246 | | 8.776e-12 | -25.46 | intracellular non-membrane-bounded organelle | cellular component | GO:0043232 | 1752 | 89 | 11940 | 298 | CG4612,Fs(2)Ket,me31B,mRpS18A,RpL36A,RpS8,mRpL30,Mps1,CG31301,Fmr1,flr,CycA,tsr,CG11920,Arpc3A,smg,Ino80,Bub1,Pep,CG9246,Acf,peng,Eaf6,CG4866,RpL14,sqd,Map60,Tctp,RecQ4,BEAF-32,Fign,fzy,su(Hw),DCTN3-p24,mRpS33,MED22,CG12659,rhea,Dbp80,mRpS9,RpL5,mRpS28,His4r,Arp8,AGO3,Nopp140,Hrb27C,rhi,cal1,Pp1-87B,RpL15,nsl1,Mlf,mRpL34,Spn,NO66,E(bx),Nph,HP4,Dref,MED26,Upf3,RpS19a,RpS9,DNAlig1,Rbp1,Girdin,pav,RpS3A,Parp,Arpc5,d4,CtBP,Rcd5,E(z),RPA3,smt3,Not3,Cp190,Oxp,RpL35A,Jarid2,RpS26,His2Av,CG17493,Orc3,ver,Rbm13,CG6171 | | 8.776e-12 | -25.46 | non-membrane-bounded organelle | cellular component | GO:0043228 | 1752 | 89 | 11940 | 298 | CG17493,His2Av,RpS26,Jarid2,RpL35A,Oxp,Cp190,CG6171,Rbm13,ver,Orc3,d4,Arpc5,Parp,RpS3A,Not3,smt3,RPA3,E(z),Rcd5,CtBP,Rbp1,DNAlig1,RpS9,RpS19a,Upf3,pav,Girdin,E(bx),NO66,Spn,mRpL34,Mlf,nsl1,Pp1-87B,RpL15,MED26,Dref,HP4,Nph,mRpS28,mRpS9,RpL5,Dbp80,rhea,rhi,cal1,Hrb27C,Nopp140,AGO3,Arp8,His4r,su(Hw),fzy,Fign,BEAF-32,RecQ4,Tctp,Map60,sqd,RpL14,CG12659,MED22,mRpS33,DCTN3-p24,Bub1,Ino80,Arpc3A,smg,CG11920,tsr,CycA,flr,Fmr1,Eaf6,CG4866,peng,Acf,CG9246,Pep,RpS8,mRpL30,mRpS18A,RpL36A,me31B,Fs(2)Ket,CG4612,CG31301,Mps1 | | 1.939e-11 | -24.67 | regulation of cellular component organization | biological process | GO:0051128 | 791 | 53 | 12345 | 305 | Cdc42,Pten,Sgt,fzy,Rap1,E(z),RagA-B,Rop,nsl1,mav,cta,CG9246,CycA,Acf,dpp,Spn,eEF5,LIMK1,mnb,tsu,spoon,PlexA,sigmar,Lst8,krz,tsr,Nf1,Sod2,Rpt6,Arpc3A,Bub1,me31B,Cp190,Tctp,Rheb,Rab8,Mps1,su(Hw),flw,RhoGAPp190,Ncoa6,cal1,Fign,tkv,Arpc5,conu,Fmr1,Btk29A,flr,vap,pav,Jarid2,Graf | | 1.992e-11 | -24.64 | positive regulation of biological process | biological process | GO:0048518 | 1642 | 84 | 12345 | 305 | Rpt6,lic,Hrb27C,AGO3,Tctp,PlexA,Efa6,sigmar,Stlk,Lst8,Fgop2,mav,Rop,rhi,eEF5,RpS9,HtrA2,CycA,dgrn,Pten,Rap1,CG4612,flr,pll,grau,TfIIFbeta,CNBP,smg,Upf3,conu,Arpc5,CG1791,14-3-3zeta,CtBP,flw,Ncoa6,RecQ4,sun,Rheb,Cul4,d4,rb,Skp2,Arpc3A,Stim,MED26,Cp190,me31B,Slmap,CG6244,Vps35,mnb,Pp1-87B,Nf1,krz,tsr,nsl1,mbf1,RagA-B,E(z),Nulp1,Cdk8,dpp,Mlf,Sgt,Cdc42,fzy,Rcd5,g,Pex3,Graf,tkv,MED21,CG9951,Tcs3,Fmr1,Dref,Cpsf6,smt3,Orc3,cal1,PPP1R15,su(Hw),Parp | | 2.082e-11 | -24.60 | organelle organization | biological process | GO:0006996 | 1885 | 92 | 12345 | 305 | Bub1,Nph,glo,Rab7,spoon,CycA,Dbp21E2,Spn,mus301,HtrA2,cta,rhi,Pten,Eaf6,grau,Girdin,flr,CG44194,Arpc5,Btk29A,E(bx),spel1,RecQ4,CG8223,Ncoa6,NO66,Rpn10,bai,14-3-3zeta,CtBP,d4,rb,Mps1,unc-119,BEAF-32,Rab8,Nup188,Rab14,Tim17b,me31B,His4r,Cp190,CG12659,Tom20,Arp8,Arpc3A,HP4,CG17493,Pp1-87B,heix,tsr,RpS19a,CG6244,tsu,RpL5,rhea,Acf,dpp,LIMK1,ver,msd5,Oxp,E(z),Exo84,CG9853,sqd,His2Av,mon2,Cdc42,Ino80,Wdr82,fzy,snf,Art1,CIA30,l(2)37Bb,vnc,pav,g,Rcd5,Pex3,Jarid2,Fmr1,Vps13D,pins,Fign,tkv,Rbbp5,Vap33,cal1,smt3,Parp | | 2.149e-11 | -24.56 | regulation of protein metabolic process | biological process | GO:0051246 | 648 | 47 | 12345 | 305 | Ncoa6,flw,smt3,CG1791,rb,Not3,Cul4,PPP1R15,Rheb,twin,sws,eIF3b,Jarid2,CG4612,Fmr1,Tcs3,CG9951,Upf3,CNBP,smg,viaf,I-2,dpp,CycA,HtrA2,eEF5,mus301,RpS9,sqd,E(z),CG9246,mav,fzy,Rap1,Pten,Cdc42,CG1218,me31B,Tctp,Hrb27C,lic,Rpt6,glo,krz,Stlk,mnb,sigmar | | 4.912e-11 | -23.74 | positive regulation of cellular process | biological process | GO:0048522 | 1464 | 77 | 12345 | 305 | su(Hw),Parp,PPP1R15,cal1,Cpsf6,smt3,Orc3,MED21,CG9951,Fmr1,Tcs3,Dref,tkv,Graf,Rcd5,Pex3,Mlf,Cdc42,Sgt,fzy,Cdk8,dpp,nsl1,mbf1,RagA-B,E(z),Nulp1,Pp1-87B,krz,tsr,CG6244,Vps35,mnb,Cp190,Slmap,me31B,Skp2,Arpc3A,d4,sun,Rheb,Ncoa6,RecQ4,CG1791,CtBP,14-3-3zeta,flw,TfIIFbeta,smg,CNBP,Upf3,conu,Arpc5,grau,CG4612,flr,dgrn,Pten,Rap1,eEF5,RpS9,HtrA2,CycA,Rop,mav,rhi,Stlk,Lst8,Fgop2,Efa6,sigmar,PlexA,Tctp,lic,Rpt6,Hrb27C,AGO3 | | 5.957e-11 | -23.54 | regulation of macromolecule biosynthetic process | biological process | GO:0010556 | 1326 | 72 | 12345 | 305 | dgrn,CG6066,RAF2,x16,eEF5,RpS9,mus301,mav,rhi,Lst8,glo,CG1815,Hrb27C,Rpt6,d4,CG17385,nom,BEAF-32,Rheb,twin,RecQ4,Ncoa6,NO66,CG14805,CG6254,14-3-3zeta,CtBP,smg,CNBP,MED22,TfIIFbeta,Upf3,E(bx),grau,CG11456,CG4612,SF2,Ino80,snf,Mlf,Art1,CG6907,rhea,dpp,Acf,Tgi,E(z),Nulp1,sqd,nsl1,mbf1,CG9246,mnb,CG6244,me31B,MED26,Cp190,Arp8,Not3,Parp,su(Hw),Rbp1,CG12299,smt3,Fmr1,Dref,Tcs3,MED21,tkv,eIF3b,Rcd5,Pex3 | | 5.984e-11 | -23.54 | cytoplasm | cellular component | GO:0005737 | 4500 | 167 | 11940 | 298 | Fmr1,flr,CycA,PIG-K,RpL36A,Btk29A,mRpL30,RpS8,Prosalpha2,Cdc42,Pten,Ythdf,Graf,RpL5,CG11752,mon2,mRpS28,CG30423,sws,ECSIT,CG31460,Prosbeta1,Dbp80,Hrb27C,rb,Exo84,pins,AGO3,Usp32,fzy,g,RpL14,Rpt6,sqd,Mdh1,bai,CG8915,14-3-3zeta,Rpn10,Sod2,DNAlig1,l(2)37Bb,Upf3,RpS19a,CG32576,CG6907,flw,CG2938,spoon,Coq5,Girdin,viaf,CG17883,Pp1-87B,Mlf,EloB,RagA-B,eEF1delta,alpha-Man-Ib,RpS26,CG9853,MICU1,heix,Stim,Skp2,CG32473,eIF3b,Vps35,eIF4B,RpS3A,dpp,PGAP1,inc,CIA30,Tim17b,ema,smg,Pex3,eEF5,tsr,vnc,lic,Rab7,mRpS18A,krz,Zpr1,CG4612,BthD,Pisd,Fs(2)Ket,Tom20,eRF1,me31B,dgrn,Mppe,Stt3B,tkv,Efa6,sun,mRpS9,rhea,Stlk,eIF3g1,ubl,Lrr47,Aps,ND-ASHI,RecQ4,Ubqn,eIF3c,CCT4,Tctp,vsg,Bmcp,mbf1,mRpS33,conu,Vps13D,Lst8,HtrA2,RpS9,LIMK1,CG44194,twin,CG8635,pav,vap,sigmar,CG15168,Art1,TM9SF2,Spn,RpL15,VhaPPA1-1,mRpL34,nsl1,CG5590,tsu,janA,Rab8,Nph,xit,CG10365,CG10672,CG17493,Cp190,Rop,RpL35A,PPP1R15,Jarid2,pll,CG4743,Hsp27,Dbp21E2,Arpc5,CG11137,CG15534,CG7766,GstE13,CG12321,Vap33,Rab14,ND-AGGG,Not3,CNBP,Roc2,Sgt | | 6.414e-11 | -23.47 | nuclear lumen | cellular component | GO:0031981 | 1016 | 61 | 11940 | 298 | DNAlig1,Ncoa6,CG11920,Ino80,MED21,Bub1,Rbp1,x16,Upf3,tsr,RpS9,CG6907,peng,Dek,Acf,Rpb11,CG4866,Eaf6,sm,CG9246,E(bx),NO66,CG6254,CG8223,EloB,nsl1,Fs(2)Ket,HP4,Wdr33,dgrn,MED26,Nph,CG31301,Mps1,His2Av,Jarid2,Rbm13,Hrb27C,Nopp140,cal1,Orc3,CG6244,Arp8,Cdk8,CG14805,TfIIFbeta,BEAF-32,d4,sqd,Rpt6,Parp,Wdr82,smt3,Rbbp5,CG12659,CtBP,Rcd5,RPA3,MED22,SF2,E(z) | | 6.984e-11 | -23.38 | regulation of cellular protein metabolic process | biological process | GO:0032268 | 623 | 45 | 12345 | 305 | Tcs3,Fmr1,Upf3,CG9951,smg,CNBP,I-2,sws,eIF3b,Jarid2,CG4612,rb,Not3,PPP1R15,Rheb,twin,Ncoa6,flw,smt3,CG1791,glo,krz,Stlk,mnb,sigmar,me31B,Tctp,Hrb27C,Rpt6,lic,Rap1,fzy,Pten,Cdc42,CG1218,dpp,CycA,HtrA2,mus301,eEF5,RpS9,sqd,E(z),CG9246,mav | | 8.269e-11 | -23.22 | regulation of cellular macromolecule biosynthetic process | biological process | GO:2000112 | 1307 | 71 | 12345 | 305 | CG6907,snf,Mlf,Art1,Ino80,nsl1,CG9246,mbf1,E(z),sqd,Nulp1,Tgi,rhea,dpp,Acf,CG6244,mnb,Arp8,MED26,Cp190,me31B,su(Hw),Parp,Not3,smt3,Rbp1,CG12299,tkv,MED21,Dref,Fmr1,Tcs3,Rcd5,eIF3b,CG6066,RAF2,dgrn,mav,rhi,eEF5,RpS9,mus301,x16,CG1815,Lst8,glo,Rpt6,Hrb27C,twin,BEAF-32,Rheb,CG17385,d4,nom,CG6254,14-3-3zeta,CtBP,NO66,CG14805,Ncoa6,RecQ4,E(bx),MED22,smg,CNBP,TfIIFbeta,Upf3,CG4612,CG11456,SF2,grau | | 1.048e-10 | -22.98 | regulation of cellular biosynthetic process | biological process | GO:0031326 | 1371 | 73 | 12345 | 305 | tkv,MED21,Dref,Tcs3,Fmr1,Rcd5,Pex3,eIF3b,su(Hw),Not3,Parp,smt3,CG12299,Rbp1,CG6244,mnb,Arp8,MED26,Cp190,me31B,CG6907,Mlf,snf,Art1,Ino80,nsl1,mbf1,CG9246,E(z),Nulp1,sqd,Tgi,rhea,Acf,dpp,E(bx),CNBP,smg,MED22,TfIIFbeta,Upf3,CG11456,CG4612,SF2,grau,twin,BEAF-32,Rheb,CG17385,d4,nom,14-3-3zeta,CG6254,CtBP,NO66,CG14805,Ncoa6,RecQ4,PlexA,CG1815,Lst8,glo,Rpt6,Hrb27C,CG6066,RAF2,dgrn,mav,rhi,mus301,RpS9,eEF5,x16 | | 1.160e-10 | -22.88 | regulation of biosynthetic process | biological process | GO:0009889 | 1374 | 73 | 12345 | 305 | Lst8,glo,CG1815,PlexA,Hrb27C,Rpt6,dgrn,CG6066,RAF2,x16,eEF5,mus301,RpS9,mav,rhi,MED22,CNBP,TfIIFbeta,smg,Upf3,E(bx),grau,CG4612,CG11456,SF2,d4,CG17385,nom,Rheb,BEAF-32,twin,RecQ4,Ncoa6,NO66,CG14805,CtBP,CG6254,14-3-3zeta,mnb,CG6244,me31B,MED26,Cp190,Arp8,Ino80,snf,Mlf,Art1,CG6907,rhea,dpp,Acf,Tgi,E(z),sqd,Nulp1,nsl1,mbf1,CG9246,Fmr1,Dref,Tcs3,MED21,tkv,eIF3b,Rcd5,Pex3,Not3,Parp,su(Hw),Rbp1,CG12299,smt3 | | 2.927e-10 | -21.95 | cellular component organization | biological process | GO:0016043 | 2830 | 118 | 12345 | 305 | RpL5,tsu,CG6244,RpS19a,heix,tsr,Pp1-87B,CG17493,HP4,Arpc3A,Arp8,Tom20,CG12659,Cp190,His4r,me31B,sm,Tim17b,Rab14,CIA30,xit,Art1,snf,fzy,Wdr82,Ino80,Cdc42,mon2,nsl1,His2Av,sqd,Exo84,CG9853,E(z),Oxp,LIMK1,ver,msd5,dpp,Acf,Cdk8,rhea,tkv,Fign,pins,Vps13D,Fmr1,Dref,Jarid2,Pex3,Rcd5,g,pav,sws,vnc,l(2)37Bb,eRF1,Parp,smt3,Orc3,Vap33,cal1,Rbbp5,spoon,PlexA,inc,Rab7,Stlk,glo,Lst8,lic,Pbp49,Hrb27C,Tctp,Nph,Bub1,Eaf6,CG12321,eIF4B,Rap1,Pten,rhi,Rop,cta,HtrA2,mus301,Dbp21E2,Spn,CycA,E(bx),Btk29A,Arpc5,conu,CG44194,flr,Girdin,grau,Nup188,BEAF-32,unc-119,Rheb,Rab8,Cul4,Mps1,rb,d4,14-3-3zeta,bai,CtBP,CG1791,flw,Rpn10,NO66,RhoGAPp190,Ncoa6,CG8223,RecQ4,spel1 | | 3.642e-10 | -21.73 | cellular metabolic process | biological process | GO:0044237 | 4466 | 163 | 12345 | 305 | tkv,Ufd4,janA,Vps13D,Tcs3,Dref,Prosalpha2,CG4866,sws,Zpr1,CG8635,l(1)10Bb,su(Hw),Parp,eRF1,CG15534,RpL15,Orc3,Vap33,Nopp140,Rbp1,CG6244,tsu,mnb,Pp1-87B,noi,heix,krz,c12.1,snRNA:U4:39B,His4r,mRpS9,CG31460,CG11334,eIF3g1,RpL35A,Ubqn,Cdc42,Ino80,Wdr82,Prosbeta1,PIG-K,E(z),RpS26,sqd,Roc2,E(bx),Pisd,mRpL30,smg,CG6171,Btk29A,Rat1,SF2,CG3842,pll,RpL14,sun,BEAF-32,Mps1,d4,Rpb11,Pect,Rpn10,flw,spel1,mRpS18A,DNAlig1,mRpL34,Stlk,GstE13,RpS3A,Pbp49,lic,Rpt6,RPA3,CG11920,Usp16-45,Tctp,Bub1,alpha-Man-Ib,CG6066,CG7766,CG1218,eIF4B,Pten,eEF5,mus301,CycA,Rcd5,g,Jarid2,Pex3,eIF3b,vnc,mRpS28,Phf5a,Wdr33,EloB,Pnn,Not3,smt3,CG8132,CG11586,Coq5,Rbbp5,RpL5,CG32473,RpS19a,CG5986,Skp2,RpS8,Arp8,CG11453,CG31957,CG32075,Mdh1,CG6907,CIA30,xit,PGAP1,Aps,snf,Art1,Sgt,fzy,ver,LIMK1,Cdk8,Acf,Mppe,TfIIFbeta,Upf3,ubl,CG30105,RpL36A,CG34261,CSN1b,grau,twin,CG5589,Cul4,rb,NO66,CG10365,CG14805,mRpS33,CG4935,Ncoa6,ND-ASHI,eIF3c,RecQ4,Stt3B,inc,Sod2,Eaf6,eEF1delta,dgrn,rhi,Usp32,RpS9,Pep,x16 | | 5.546e-10 | -21.31 | cellular macromolecule metabolic process | biological process | GO:0044260 | 2688 | 113 | 12345 | 305 | RpS8,Arp8,Skp2,CG31957,mRpS9,His4r,RpS19a,mnb,RpL5,CG6244,tsu,krz,Pp1-87B,sqd,RpS26,E(z),PIG-K,Prosbeta1,Acf,Cdk8,Roc2,ver,LIMK1,xit,CG31460,fzy,Wdr82,Sgt,Ino80,Ubqn,Art1,RpL35A,eIF3g1,eIF3b,Prosalpha2,Jarid2,Rcd5,g,CG8635,Zpr1,mRpS28,vnc,tkv,Ufd4,Dref,janA,RpL15,Orc3,smt3,Rbbp5,Nopp140,EloB,eRF1,Parp,Not3,su(Hw),RPA3,lic,Pbp49,Rpt6,alpha-Man-Ib,Bub1,Tctp,Usp16-45,inc,mRpL34,DNAlig1,mRpS18A,Stt3B,RpS3A,Stlk,CycA,mus301,RpS9,eEF5,Usp32,Eaf6,Pten,eIF4B,eEF1delta,dgrn,CG1218,CG7766,RpL36A,CG30105,Rat1,grau,CSN1b,pll,RpL14,E(bx),Mppe,Btk29A,CG6171,Upf3,smg,TfIIFbeta,mRpL30,mRpS33,flw,NO66,Rpn10,Rpb11,RecQ4,eIF3c,spel1,Ncoa6,BEAF-32,twin,rb,d4,Cul4,Mps1 | | 1.084e-09 | -20.64 | macromolecule metabolic process | biological process | GO:0043170 | 3775 | 143 | 12345 | 305 | eIF3c,RecQ4,Ncoa6,CG4935,NO66,mRpS33,CG14805,rb,Cul4,CG5589,twin,grau,CSN1b,CG30105,RpL36A,ubl,TfIIFbeta,Upf3,Mppe,x16,Usp32,RpS9,HtrA2,Pep,rhi,dgrn,eEF1delta,Eaf6,inc,Stt3B,Rbbp5,CG11586,smt3,Not3,Pnn,EloB,Wdr33,Phf5a,vnc,mRpS28,eIF3b,Rcd5,g,Jarid2,Cdk8,Acf,ver,LIMK1,Sgt,fzy,snf,Art1,PGAP1,CG6907,xit,CG31957,CG32075,Skp2,RpS8,Arp8,CG5986,RpS19a,CG32473,RpL5,spel1,Rpn10,flw,Rpb11,d4,Mps1,BEAF-32,pll,RpL14,Dbp80,Rat1,SF2,CG6171,Btk29A,smg,mRpL30,E(bx),CycA,mus301,eEF5,Pten,CG7766,CG1218,eIF4B,CG6066,Bub1,alpha-Man-Ib,Usp16-45,Tctp,CG11920,Pbp49,lic,Rpt6,RPA3,RpS3A,Stlk,DNAlig1,mRpS18A,mRpL34,Rbp1,Nopp140,Orc3,RpL15,Parp,eRF1,l(1)10Bb,su(Hw),CG8635,Zpr1,CG4866,Prosalpha2,Tcs3,Dref,janA,tkv,Ufd4,Roc2,E(z),RpS26,sqd,Prosbeta1,PIG-K,Ino80,Wdr82,RpL35A,eIF3g1,Ubqn,CG31460,CG6961,His4r,mRpS9,snRNA:U4:39B,c12.1,Pp1-87B,krz,noi,mnb,CG6244,tsu | | 1.143e-09 | -20.59 | positive regulation of metabolic process | biological process | GO:0009893 | 937 | 55 | 12345 | 305 | CG6244,sigmar,PlexA,mnb,Stlk,krz,Lst8,lic,Rpt6,Hrb27C,AGO3,MED26,Tctp,dgrn,Cdc42,Rap1,fzy,mav,nsl1,rhi,mbf1,E(z),eEF5,RpS9,HtrA2,Cdk8,dpp,TfIIFbeta,smg,CNBP,Upf3,MED21,CG9951,Dref,Tcs3,Fmr1,g,Rcd5,CG4612,Pex3,grau,Graf,Rheb,PPP1R15,Cul4,su(Hw),Parp,d4,rb,CG1791,Cpsf6,smt3,CtBP,Ncoa6,RecQ4 | | 1.812e-09 | -20.13 | Gene Expression | REACTOME pathways | R-DME-74160 | 752 | 60 | 3923 | 151 | sm,CG11920,RpS9,Rpt6,RpL36A,snf,CG6961,twin,l(1)10Bb,eRF1,Cpsf6,Wdr33,Lst8,Rheb,eIF4B,Dek,Parp,Phf5a,Rpb11,MED26,RpS8,RpS19a,tsu,Eaf6,Cdk8,eIF3b,SF2,eEF1delta,me31B,Not3,Prosalpha2,EloB,Upf3,Cwc25,noi,Rat1,eIF3g1,RpL35A,lic,x16,eIF3c,c12.1,Prosbeta1,CG14805,RpL15,Nup188,RpS26,CG5589,RpL14,CG4866,Pbp49,RpL5,14-3-3zeta,TfIIFbeta,glo,CG32075,RagA-B,sqd,RpS3A,Rpn10 | | 2.132e-09 | -19.97 | catalytic complex | cellular component | GO:1902494 | 999 | 57 | 11940 | 298 | Acf,Eaf6,PIG-K,Rpb11,Pep,Ncoa6,l(2)37Bb,Ino80,x16,CycA,Pnn,Phf5a,flw,vnc,dgrn,noi,tsu,Stt3B,E(bx),Prosalpha2,CG4935,Pp1-87B,CG6254,nsl1,EloB,cta,CG6066,Arp8,CG11752,snf,CG31460,PPP1R15,Skp2,Prosbeta1,Jarid2,inc,Wdr82,CIA30,CG12659,Rbbp5,ND-AGGG,Cul4,c12.1,Rcd5,CtBP,Rpn10,Roc2,E(z),SF2,ND-ASHI,fzy,d4,l(1)10Bb,TfIIFbeta,Rpt6,CG7766,sqd | | 2.321e-09 | -19.88 | nucleic acid metabolic process | biological process | GO:0090304 | 1149 | 62 | 12345 | 305 | CG32075,mRpS9,Tctp,His4r,CG11920,Arp8,RpS8,snRNA:U4:39B,RPA3,Pbp49,CG5986,c12.1,noi,DNAlig1,tsu,CG6244,x16,Cdk8,Pep,ver,mus301,sqd,rhi,Ino80,dgrn,snf,PGAP1,CG6066,CG6907,Phf5a,grau,Zpr1,CG4866,CG30105,ubl,Rat1,SF2,CG6171,Tcs3,Upf3,TfIIFbeta,smg,E(bx),RecQ4,Nopp140,Rbp1,spel1,CG11586,CG4935,CG14805,Orc3,Rpb11,Parp,Not3,su(Hw),Pnn,Cul4,l(1)10Bb,EloB,CG5589,Wdr33,twin | | 4.020e-09 | -19.33 | regulation of gene expression | biological process | GO:0010468 | 1547 | 75 | 12345 | 305 | Upf3,MED22,smg,CNBP,TfIIFbeta,E(bx),grau,SF2,CG4612,CG11456,rb,nom,d4,CG17385,BEAF-32,Rheb,twin,Ncoa6,NO66,CG6254,14-3-3zeta,CtBP,glo,Lst8,CG1815,AGO3,Hrb27C,Rpt6,dgrn,CG6066,RAF2,x16,Pep,mus301,RpS9,eEF5,rhi,mav,Dref,Fmr1,Tcs3,MED21,tkv,eIF3b,g,Rcd5,Not3,Parp,su(Hw),Rbp1,CG12299,smt3,Cpsf6,mnb,tsu,CG6244,me31B,Cp190,MED26,Arp8,Ino80,Art1,snf,Mlf,CG6907,Acf,dpp,rhea,Tgi,Nulp1,sqd,E(z),mbf1,CG9246,nsl1 | | 4.508e-09 | -19.22 | ribonucleoprotein complex | cellular component | GO:1990904 | 754 | 47 | 11940 | 298 | tsu,noi,CG4935,mRpS18A,RpL36A,mRpL30,RpS8,Fs(2)Ket,mRpL34,me31B,RpL15,CG4612,CG4866,CG32075,CG9246,Pep,smg,CG11920,RpS9,Phf5a,Pnn,x16,Fmr1,RpS19a,c12.1,Not3,SF2,CG11586,mRpS33,l(1)10Bb,eIF4B,RpL14,RpS3A,sqd,snRNA:U4:39B,CG6066,Hrb27C,Rbm13,AGO3,CG5986,mRpS28,RpS26,mRpS9,RpL5,Dbp80,RpL35A,snf | | 5.305e-09 | -19.05 | positive regulation of protein metabolic process | biological process | GO:0051247 | 318 | 28 | 12345 | 305 | dpp,RpS9,eEF5,Ncoa6,HtrA2,E(z),mav,CG1791,smt3,Cdc42,Rap1,fzy,Cul4,Rheb,PPP1R15,Tctp,Hrb27C,Rpt6,lic,CG4612,Fmr1,Tcs3,Stlk,CNBP,Upf3,CG9951,krz,sigmar | | 6.515e-09 | -18.85 | positive regulation of cellular metabolic process | biological process | GO:0031325 | 875 | 51 | 12345 | 305 | lic,Rpt6,AGO3,Hrb27C,Tctp,sigmar,PlexA,CG6244,mnb,krz,Stlk,Lst8,rhi,mbf1,nsl1,mav,E(z),HtrA2,eEF5,RpS9,dpp,Cdk8,dgrn,Rap1,fzy,Cdc42,Pex3,Rcd5,CG4612,Graf,grau,Upf3,MED21,CG9951,smg,TfIIFbeta,CNBP,Fmr1,Dref,Tcs3,smt3,CtBP,Cpsf6,CG1791,Ncoa6,RecQ4,PPP1R15,Rheb,su(Hw),Parp,d4 | | 7.813e-09 | -18.67 | positive regulation of cellular protein metabolic process | biological process | GO:0032270 | 303 | 27 | 12345 | 305 | sigmar,Stlk,CNBP,CG9951,Upf3,krz,Fmr1,Tcs3,Rpt6,CG4612,lic,Hrb27C,Tctp,Rheb,PPP1R15,Cdc42,fzy,Rap1,mav,CG1791,smt3,E(z),eEF5,Ncoa6,RpS9,HtrA2,dpp | | 9.465e-09 | -18.48 | positive regulation of macromolecule metabolic process | biological process | GO:0010604 | 858 | 50 | 12345 | 305 | Upf3,MED21,CG9951,CNBP,TfIIFbeta,smg,Fmr1,Dref,Tcs3,Graf,grau,CG4612,g,Rcd5,su(Hw),Cul4,rb,d4,PPP1R15,Rheb,Ncoa6,RecQ4,smt3,CtBP,CG1791,krz,Stlk,Lst8,sigmar,CG6244,mnb,Tctp,MED26,lic,Rpt6,AGO3,Hrb27C,dgrn,fzy,Rap1,Cdc42,HtrA2,RpS9,eEF5,dpp,rhi,mbf1,nsl1,mav,E(z) | | 1.138e-08 | -18.29 | chromosome organization | biological process | GO:0051276 | 680 | 43 | 12345 | 305 | spoon,CG6244,Pp1-87B,glo,HP4,Arp8,CG12659,Cp190,His4r,Nph,Bub1,Eaf6,Art1,snf,Wdr82,Ino80,rhi,His2Av,E(z),Oxp,ver,mus301,Acf,CycA,E(bx),Fign,Btk29A,Fmr1,Jarid2,Rcd5,pav,vnc,BEAF-32,Parp,d4,CtBP,Rpn10,NO66,Ncoa6,CG8223,Rbbp5,RecQ4,spel1 | | 1.178e-08 | -18.26 | regulation of nucleobase-containing compound metabolic process | biological process | GO:0019219 | 1315 | 66 | 12345 | 305 | Pep,x16,rhi,mav,dgrn,CG6066,RAF2,Rpt6,AGO3,Hrb27C,Lst8,PlexA,CG1815,Ncoa6,spel1,CtBP,14-3-3zeta,CG6254,CG14805,NO66,nom,CG17385,d4,BEAF-32,Dek,grau,SF2,CG11456,MED22,smg,TfIIFbeta,E(bx),Tgi,dpp,Acf,rhea,CG9246,mbf1,nsl1,sqd,Nulp1,E(z),Art1,snf,Mlf,Ino80,CG6907,Cp190,MED26,Arp8,tsu,CG6244,mnb,Rbp1,CG12299,smt3,Cpsf6,su(Hw),Not3,Parp,Rcd5,MED21,Fmr1,Dref,tkv,Fign | | 2.230e-08 | -17.62 | positive regulation of nitrogen compound metabolic process | biological process | GO:0051173 | 827 | 48 | 12345 | 305 | dpp,eEF5,RpS9,HtrA2,E(z),nsl1,mav,rhi,mbf1,Cdc42,fzy,Rap1,dgrn,Tctp,Hrb27C,AGO3,lic,Rpt6,Lst8,Stlk,krz,mnb,CG6244,sigmar,PlexA,Ncoa6,CG1791,CtBP,smt3,d4,Parp,Cul4,su(Hw),Rheb,PPP1R15,grau,Rcd5,CG4612,Pex3,Tcs3,Dref,Fmr1,smg,TfIIFbeta,CNBP,Upf3,CG9951,MED21 | | 2.840e-08 | -17.38 | cellular nitrogen compound metabolic process | biological process | GO:0034641 | 2226 | 94 | 12345 | 305 | RecQ4,spel1,eIF3c,CG4935,CG14805,mRpS33,Pect,CG10365,Rpb11,Cul4,CG5589,twin,sun,grau,RpL14,CG34261,RpL36A,CG30105,SF2,ubl,Rat1,CG6171,Upf3,TfIIFbeta,mRpL30,smg,E(bx),x16,Pep,mus301,eEF5,RpS9,rhi,eEF1delta,eIF4B,dgrn,CG6066,Tctp,CG11920,RPA3,Pbp49,RpS3A,GstE13,mRpL34,mRpS18A,DNAlig1,Rbp1,Nopp140,CG11586,RpL15,CG8132,CG15534,Orc3,eRF1,Not3,Parp,su(Hw),Pnn,l(1)10Bb,EloB,Wdr33,Phf5a,CG8635,Zpr1,mRpS28,CG4866,eIF3b,Tcs3,Cdk8,ver,sqd,RpS26,Ino80,eIF3g1,RpL35A,snf,Aps,PGAP1,CG6907,CG32075,CG31957,mRpS9,His4r,Arp8,snRNA:U4:39B,RpS8,c12.1,CG5986,noi,RpS19a,RpL5,CG32473,CG6244,tsu | | 3.342e-08 | -17.21 | negative regulation of cellular process | biological process | GO:0048523 | 1291 | 64 | 12345 | 305 | tsu,mnb,krz,heix,Nf1,me31B,Slmap,snf,fzy,Sgt,Ino80,CG9246,mbf1,Nulp1,sqd,E(z),Tgi,dpp,Acf,rhea,Fmr1,g,pav,sws,Graf,su(Hw),Not3,CG12299,sigmar,PlexA,spoon,Fgop2,lic,AGO3,Hrb27C,Tctp,Bub1,CG6066,Rap1,Pten,rhi,Spn,CycA,E(bx),TM9SF2,I-2,smg,MED22,CG30423,flr,vap,RpL14,twin,Rheb,Mps1,Cul4,rb,d4,CtBP,CG6254,CG1791,flw,NO66,spel1 | | 4.783e-08 | -16.86 | nucleoplasm | cellular component | GO:0005654 | 557 | 37 | 11940 | 298 | Jarid2,Hrb27C,Cdk8,Orc3,CG6244,TfIIFbeta,d4,CG14805,Parp,sqd,Rbbp5,Wdr82,MED22,E(z),SF2,Rcd5,MED21,Rbp1,Ncoa6,CG11920,tsr,CG6907,x16,Rpb11,sm,Eaf6,Dek,CG9246,EloB,CG8223,nsl1,Fs(2)Ket,CG6254,MED26,Wdr33,CG31301,Nph | | 5.046e-08 | -16.80 | primary metabolic process | biological process | GO:0044238 | 4472 | 156 | 12345 | 305 | Roc2,Prosbeta1,PIG-K,E(z),sqd,RpS26,RpL35A,eIF3g1,Ubqn,Ino80,Wdr82,CG31460,CG11334,His4r,mRpS9,snRNA:U4:39B,Pp1-87B,noi,c12.1,tsu,CG6244,mnb,Vap33,Rbp1,Nopp140,Orc3,CG15534,RpL15,l(1)10Bb,su(Hw),Parp,eRF1,sws,Zpr1,CG8635,Prosalpha2,CG4866,janA,Dref,Tcs3,tkv,Ufd4,mus301,eEF5,CycA,CG7766,eIF4B,CG1218,Pten,CG6066,Usp16-45,Tctp,Bub1,alpha-Man-Ib,Rpt6,Pbp49,lic,RPA3,CG11920,Stlk,RpS3A,mRpS18A,DNAlig1,mRpL34,spel1,Rpb11,Rpn10,flw,Mps1,d4,sun,BEAF-32,RpL14,pll,CG3842,SF2,Rat1,mRpL30,smg,CG6171,Btk29A,E(bx),Pisd,LIMK1,ver,Cdk8,Acf,snf,Art1,Sgt,fzy,CG6907,Mdh1,xit,PGAP1,Aps,CG31957,CG32075,Skp2,Arp8,RpS8,CG11453,CG5986,RpL5,CG32473,RpS19a,CG2938,CG11586,Rbbp5,smt3,CG8132,Pnn,Not3,Wdr33,EloB,vnc,mRpS28,Phf5a,g,Rcd5,Pex3,Jarid2,eIF3b,RpS9,Usp32,Pep,HtrA2,x16,rhi,dgrn,eEF1delta,Eaf6,CG10467,Stt3B,inc,Ncoa6,CG4935,eIF3c,RecQ4,NO66,mRpS33,CG14805,Cul4,rb,twin,Rheb,CG5589,CSN1b,grau,ubl,RpL36A,CG30105,CG34261,TfIIFbeta,Upf3,Mppe | | 5.440e-08 | -16.73 | organic substance metabolic process | biological process | GO:0071704 | 4824 | 165 | 12345 | 305 | Cdk8,Acf,ver,LIMK1,PGAP1,Aps,Mdh1,CG6907,xit,Sgt,fzy,snf,Art1,CG11453,Skp2,Arp8,RpS8,CG31957,CG32075,RpS19a,CG2938,RpL5,CG32473,CG5986,CG8132,smt3,Rbbp5,Coq5,CG11586,EloB,Wdr33,Not3,Pnn,eIF3b,g,Rcd5,Jarid2,Pex3,Phf5a,vnc,mRpS28,rhi,x16,RpS9,Usp32,Pep,HtrA2,Eaf6,dgrn,eEF1delta,CG10467,inc,Stt3B,CG10365,NO66,CG14805,mRpS33,eIF3c,RecQ4,CG4935,Ncoa6,CG5589,Rheb,twin,rb,Cul4,CG30105,RpL36A,CG34261,ubl,grau,CSN1b,Mppe,TfIIFbeta,Upf3,E(z),RpS26,sqd,Prosbeta1,PIG-K,Roc2,CG31460,CG11334,Cdc42,Ino80,Wdr82,eIF3g1,RpL35A,Ubqn,snRNA:U4:39B,CG6961,His4r,mRpS9,mnb,CG6244,tsu,c12.1,Pp1-87B,krz,noi,heix,Orc3,CG15534,RpL15,Rbp1,Nopp140,Vap33,Parp,eRF1,l(1)10Bb,su(Hw),CG4866,Prosalpha2,CG8635,Zpr1,sws,Ufd4,tkv,Tcs3,Dref,janA,CycA,eEF5,mus301,CG6066,Pten,CG7766,eIF4B,CG1218,CG11920,lic,Rpt6,Pbp49,RPA3,Bub1,alpha-Man-Ib,Usp16-45,Tctp,DNAlig1,mRpS18A,mRpL34,RpS3A,Stlk,GstE13,Rpn10,Pect,flw,Rpb11,spel1,BEAF-32,sun,d4,Mps1,Dbp80,SF2,Rat1,CG3842,RpL14,pll,Pisd,E(bx),CG6171,Btk29A,mRpL30,smg | | 7.968e-08 | -16.35 | nitrogen compound metabolic process | biological process | GO:0006807 | 4233 | 149 | 12345 | 305 | twin,CG5589,Cul4,rb,CG10365,NO66,mRpS33,CG14805,Ncoa6,CG4935,eIF3c,RecQ4,Mppe,TfIIFbeta,Upf3,ubl,CG30105,RpL36A,CG34261,CSN1b,grau,Eaf6,eEF1delta,dgrn,rhi,Usp32,RpS9,HtrA2,Pep,x16,Stt3B,inc,Wdr33,EloB,Pnn,Not3,smt3,CG8132,CG11586,Rbbp5,Rcd5,g,Jarid2,eIF3b,vnc,mRpS28,Phf5a,CG6907,xit,PGAP1,Aps,snf,Art1,Sgt,fzy,LIMK1,ver,Cdk8,Acf,RpL5,CG32473,RpS19a,CG5986,Skp2,Arp8,RpS8,CG31957,CG32075,sun,BEAF-32,Mps1,d4,Rpb11,Pect,Rpn10,flw,spel1,E(bx),smg,mRpL30,CG6171,Btk29A,SF2,Rat1,pll,RpL14,CG6066,eIF4B,CG1218,Pten,eEF5,mus301,CycA,DNAlig1,mRpS18A,mRpL34,Stlk,GstE13,RpS3A,Rpt6,Pbp49,lic,RPA3,CG11920,Usp16-45,Tctp,Bub1,alpha-Man-Ib,l(1)10Bb,su(Hw),Parp,eRF1,CG15534,RpL15,Orc3,Rbp1,Nopp140,Ufd4,tkv,janA,Tcs3,Dref,Prosalpha2,CG4866,Zpr1,sws,CG8635,CG31460,CG11334,RpL35A,eIF3g1,Ubqn,Ino80,Wdr82,Prosbeta1,PIG-K,E(z),sqd,RpS26,Roc2,CG6244,tsu,mnb,Pp1-87B,noi,c12.1,snRNA:U4:39B,His4r,mRpS9 | | 8.338e-08 | -16.30 | chromosome | cellular component | GO:0005694 | 696 | 42 | 11940 | 298 | cal1,CG6171,rhi,Arp8,ver,His4r,Orc3,His2Av,Jarid2,Oxp,Cp190,CG12659,smt3,RPA3,MED22,E(z),Rcd5,CtBP,su(Hw),BEAF-32,d4,fzy,Fign,RecQ4,Tctp,Parp,sqd,Acf,Pep,Ino80,Bub1,Rbp1,DNAlig1,RpS9,MED26,Dref,HP4,Mps1,E(bx),nsl1,Mlf,Pp1-87B | | 8.373e-08 | -16.30 | RNA binding | molecular function | GO:0003723 | 929 | 49 | 12176 | 289 | CG6961,peng,Fmr1,glo,c12.1,Phf5a,eEF5,eIF3c,CG5589,sqd,RpS26,AGO3,CNBP,Rbp1,noi,Dbp21E2,CG5986,me31B,RpL5,x16,mRpS18A,RpL15,Ythdf,tsu,CG31957,mbf1,Upf3,Dbp80,eRF1,Cpsf6,eIF3g1,smg,eIF3b,RpS9,Rcd5,eIF4B,CG4866,Pep,spoon,CG11920,SF2,RpL14,sm,mRpS9,CG4612,snf,CG8915,Hrb27C,eEF1delta | | 9.313e-08 | -16.19 | regulation of translation | biological process | GO:0006417 | 144 | 17 | 12345 | 305 | me31B,eIF3b,Hrb27C,CG4612,Tcs3,Fmr1,glo,CNBP,smg,Upf3,mus301,RpS9,eEF5,sqd,Not3,Rheb,twin | | 1.045e-07 | -16.07 | female gamete generation | biological process | GO:0007292 | 761 | 44 | 12345 | 305 | pav,flr,eIF3b,grau,CSN1b,vnc,tkv,Btk29A,Fmr1,flw,bai,14-3-3zeta,smt3,RecQ4,cal1,BEAF-32,twin,Parp,Mps1,Cul4,Hrb27C,lic,me31B,Rab7,spoon,tsu,glo,tsr,Pp1-87B,sqd,rhi,nsl1,Rop,mon2,dpp,rhea,LIMK1,mus301,Fs(2)Ket,Rap1,fzy,Cdc42,Pten,snf | | 1.342e-07 | -15.82 | cytoplasmic part | cellular component | GO:0044444 | 3495 | 129 | 11940 | 298 | eEF1delta,RagA-B,Mlf,EloB,Pp1-87B,CG2938,spoon,Coq5,CG17883,l(2)37Bb,DNAlig1,flw,RpS19a,CG32576,dpp,PGAP1,CIA30,Tim17b,ema,eIF4B,RpS3A,eIF3b,Vps35,Stim,alpha-Man-Ib,RpS26,heix,MICU1,CG9853,Skp2,RpL36A,RpS8,mRpL30,Cdc42,PIG-K,CycA,Fmr1,flr,bai,Mdh1,Rpn10,Sod2,g,Usp32,RpL14,Rpt6,rb,Hrb27C,Exo84,pins,AGO3,sws,mRpS28,CG30423,Graf,RpL5,CG11752,mon2,CG31460,Dbp80,ECSIT,janA,Rab8,CG10365,xit,Spn,mRpL34,VhaPPA1-1,CG5590,RpL15,twin,pav,CG8635,CG44194,Art1,CG15168,vap,RpS9,HtrA2,Not3,Rab14,ND-AGGG,Sgt,Roc2,CG15534,CG11137,Vap33,CG7766,CG12321,Dbp21E2,CG4743,CG10672,CG17493,PPP1R15,Cp190,Rop,RpL35A,tkv,Mppe,Stt3B,Efa6,mRpS18A,Tom20,Fs(2)Ket,eRF1,me31B,Pisd,CG4612,BthD,Rab7,smg,tsr,vnc,Pex3,vsg,mbf1,Vps13D,mRpS33,conu,Bmcp,Ubqn,ND-ASHI,eIF3c,CCT4,Tctp,eIF3g1,sun,mRpS9,rhea | | 1.678e-07 | -15.60 | regulation of cellular amide metabolic process | biological process | GO:0034248 | 186 | 19 | 12345 | 305 | glo,Fmr1,Tcs3,Upf3,CNBP,smg,me31B,Hrb27C,eIF3b,Pex3,CG4612,Not3,Parp,Rheb,twin,RpS9,eEF5,mus301,sqd | | 1.723e-07 | -15.57 | negative regulation of biological process | biological process | GO:0048519 | 1502 | 69 | 12345 | 305 | CG1791,14-3-3zeta,CtBP,CG6254,NO66,flw,spel1,twin,Rheb,Mps1,Cul4,d4,rb,vap,flr,CG30423,RpL14,E(bx),TM9SF2,I-2,MED22,smg,Upf3,conu,rhi,CycA,Spn,CG6066,Pten,Rap1,lic,Hrb27C,AGO3,Tctp,Bub1,sigmar,PlexA,spoon,Rab7,Fgop2,CG12299,su(Hw),Not3,g,pav,sws,Graf,tkv,Fmr1,CG9246,mbf1,E(z),sqd,Nulp1,Tgi,rhea,dpp,Acf,snf,Sgt,Ino80,fzy,me31B,Slmap,tsu,mnb,heix,Nf1,krz | | 1.946e-07 | -15.45 | heterocyclic compound binding | molecular function | GO:1901363 | 3069 | 112 | 12176 | 289 | Rab7,Rab14,smg,eIF3g1,CG6254,Dref,CG11456,eIF4B,Rheb,RpS9,RecQ4,CCT4,lic,Fign,tsu,CG17385,su(Hw),Cp190,Cpsf6,eRF1,Upf3,mbf1,Jarid2,pll,CG4612,mRpS9,CG5516,pav,His4r,eEF1delta,Hrb27C,His2Av,Pep,RpL14,mus301,SF2,LIMK1,Acbp1,Phf5a,tkv,Stlk,CG12299,RabX6,glo,Fmr1,CG6961,peng,rhi,RhoGAPp190,Pbp49,CG12877,Acf,Dbp21E2,CG10209,ver,mnb,nom,RpL5,RpS26,CNBP,Mlf,AGO3,CG34261,eIF3b,cta,DNAlig1,Rcd5,grau,Rpb11,CG31957,Rab8,Ythdf,Rpt6,RPA3,Bub1,Dbp80,Cdk8,CG9330,Dlip2,E(z),CG8915,snf,CtBP,spoon,CG4866,sm,Cdc42,CG31301,CG11920,Rap1,BEAF-32,eIF3c,eEF5,CG5589,CG11586,Orc3,spel1,c12.1,Mps1,noi,Parp,Rbp1,RagA-B,Ino80,RpL15,mRpS18A,x16,me31B,CG5986,sqd,Rat1,Btk29A | | 2.372e-07 | -15.25 | nucleic acid binding | molecular function | GO:0003676 | 2082 | 84 | 12176 | 289 | His4r,eEF1delta,Hrb27C,CG4612,Jarid2,mRpS9,CG5516,RpL14,mus301,SF2,Pep,His2Av,CG11456,eIF4B,Dref,CG6254,RpS9,eIF3g1,smg,Cpsf6,Cp190,mbf1,Upf3,eRF1,RecQ4,su(Hw),tsu,CG17385,nom,RpL5,CG10209,Dbp21E2,ver,Mlf,CNBP,AGO3,RpS26,CG12299,Phf5a,Pbp49,Acf,CG12877,glo,rhi,CG6961,peng,Fmr1,snf,E(z),CG8915,Dlip2,sm,CG31301,CG11920,CG4866,spoon,Rcd5,grau,DNAlig1,eIF3b,RPA3,Dbp80,CG31957,Rpb11,Ythdf,x16,RpL15,mRpS18A,CG5986,me31B,Parp,Rbp1,noi,Ino80,Rat1,sqd,CG5589,CG11586,BEAF-32,eIF3c,eEF5,c12.1,spel1,Orc3 | | 2.637e-07 | -15.15 | organic cyclic compound binding | molecular function | GO:0097159 | 3086 | 112 | 12176 | 289 | RPA3,Rpt6,Bub1,Dbp80,CG31957,Rpb11,Ythdf,Rab8,cta,DNAlig1,grau,Rcd5,CG34261,eIF3b,sm,Cdc42,Rap1,CG11920,CG31301,CtBP,spoon,CG4866,CG8915,E(z),snf,Cdk8,CG9330,Dlip2,spel1,c12.1,Mps1,Orc3,CG5589,CG11586,eEF5,BEAF-32,eIF3c,Rat1,Btk29A,sqd,RpL15,mRpS18A,x16,me31B,CG5986,noi,Parp,Rbp1,RagA-B,Ino80,Cp190,Cpsf6,eRF1,Upf3,mbf1,CCT4,RecQ4,lic,Fign,tsu,CG17385,su(Hw),CG6254,Dref,CG11456,eIF4B,Rheb,RpS9,Rab7,Rab14,smg,eIF3g1,mus301,RpL14,SF2,Pep,His2Av,His4r,eEF1delta,pav,Hrb27C,Jarid2,CG4612,pll,mRpS9,CG5516,RhoGAPp190,Pbp49,CG12877,Acf,RabX6,glo,Fmr1,peng,CG6961,rhi,Stlk,CG12299,Acbp1,LIMK1,Phf5a,tkv,CNBP,Mlf,AGO3,RpS26,mnb,RpL5,nom,Dbp21E2,CG10209,ver | | 3.168e-07 | -14.97 | regulation of RNA metabolic process | biological process | GO:0051252 | 1281 | 61 | 12345 | 305 | Ncoa6,Rbp1,CG12299,Cpsf6,CtBP,CG6254,14-3-3zeta,smt3,NO66,su(Hw),CG17385,Not3,Parp,d4,nom,BEAF-32,grau,CG11456,Rcd5,SF2,TfIIFbeta,smg,MED22,MED21,Dref,Fmr1,tkv,E(bx),Tgi,Pep,rhea,x16,dpp,Acf,nsl1,mav,CG9246,rhi,mbf1,E(z),sqd,Nulp1,snf,Mlf,dgrn,Art1,Ino80,CG6907,RAF2,CG6066,MED26,Cp190,Rpt6,Arp8,Hrb27C,AGO3,Lst8,CG6244,tsu,CG1815,mnb | | 3.471e-07 | -14.87 | metabolic process | biological process | GO:0008152 | 5257 | 173 | 12345 | 305 | CycA,mus301,eEF5,CG6066,Pten,CG7766,CG1218,eIF4B,CG11920,Pbp49,lic,Rpt6,RPA3,Bub1,alpha-Man-Ib,Usp16-45,Tctp,DNAlig1,mRpS18A,mRpL34,RpS3A,Stlk,GstE13,Pect,Rpn10,flw,Rpb11,spel1,BEAF-32,sun,d4,Mps1,Dbp80,CG3842,Rat1,SF2,pll,RpL14,Pisd,E(bx),CG6171,Btk29A,smg,mRpL30,E(z),sqd,RpS26,Prosbeta1,PIG-K,Roc2,CG31460,CG11334,Cdc42,Ino80,Wdr82,eIF3g1,RpL35A,Ubqn,snRNA:U4:39B,CG6961,His4r,mRpS9,mnb,tsu,CG6244,c12.1,Pp1-87B,heix,krz,noi,Orc3,RpL15,CG15534,Nopp140,Rbp1,Vap33,Parp,eRF1,l(1)10Bb,su(Hw),CG4866,Prosalpha2,l(2)37Bb,CG8635,Zpr1,sws,tkv,Ufd4,Tcs3,Dref,janA,Vps13D,rhi,x16,RpS9,Usp32,HtrA2,Pep,Eaf6,eEF1delta,dgrn,Sod2,CG10467,inc,Stt3B,NO66,CG10365,mRpS33,CG14805,CtBP,eIF3c,ND-ASHI,RecQ4,CG4935,Ncoa6,Rheb,CG5589,twin,rb,Cul4,RpL36A,CG30105,CG34261,ubl,grau,CSN1b,GILT3,Mppe,TfIIFbeta,Upf3,Cdk8,Acf,ver,LIMK1,PGAP1,Aps,Mdh1,CG6907,CIA30,xit,Sgt,fzy,snf,Art1,CG11453,Skp2,RpS8,Arp8,CG32075,CG31957,RpS19a,CG2938,RpL5,CG32473,CG5986,CG8132,smt3,Coq5,Rbbp5,CG11586,EloB,Wdr33,Not3,Pnn,eIF3b,Rcd5,g,Pex3,Jarid2,Phf5a,vnc,mRpS28,Fign | | 3.701e-07 | -14.81 | chromosomal part | cellular component | GO:0044427 | 604 | 37 | 11940 | 298 | HP4,MED26,Mps1,E(bx),nsl1,Mlf,Acf,Pep,DNAlig1,Bub1,Ino80,Rbp1,smt3,CG12659,CtBP,Rcd5,RPA3,MED22,E(z),RecQ4,su(Hw),d4,fzy,BEAF-32,sqd,Parp,rhi,CG6171,cal1,His4r,Orc3,ver,Arp8,His2Av,Oxp,Cp190,Jarid2 | | 3.818e-07 | -14.78 | RNA metabolic process | biological process | GO:0016070 | 908 | 48 | 12345 | 305 | CG6066,CG6907,snf,sqd,rhi,Cdk8,x16,Pep,CG6244,tsu,CG5986,c12.1,noi,CG11920,Pbp49,RpS8,Arp8,snRNA:U4:39B,CG32075,His4r,mRpS9,CG5589,EloB,Wdr33,twin,Parp,Not3,l(1)10Bb,su(Hw),Pnn,CG14805,Rpb11,Rbp1,Nopp140,CG11586,CG4935,E(bx),Tcs3,TfIIFbeta,smg,Upf3,CG30105,CG4866,SF2,ubl,grau,Phf5a,Zpr1 | | 4.212e-07 | -14.68 | Ras protein signal transduction | biological process | GO:0007265 | 78 | 12 | 12345 | 305 | Cdc42,Rap1,Nf1,Rab7,Rab8,PlexA,Rab14,Graf,RhoGAPp190,Lrr47,cta,RabX6 | | 4.365e-07 | -14.64 | small GTPase mediated signal transduction | biological process | GO:0007264 | 93 | 13 | 12345 | 305 | Rab7,Rab8,Rheb,PlexA,Rap1,Cdc42,Nf1,RabX6,cta,Graf,Rab14,Lrr47,RhoGAPp190 | | 4.592e-07 | -14.59 | nucleobase-containing compound metabolic process | biological process | GO:0006139 | 1417 | 65 | 12345 | 305 | Ino80,snf,PGAP1,Aps,CG6907,Cdk8,ver,sqd,c12.1,CG5986,noi,CG6244,tsu,CG32075,His4r,mRpS9,snRNA:U4:39B,RpS8,Arp8,Parp,Not3,l(1)10Bb,Pnn,su(Hw),EloB,Wdr33,Nopp140,Rbp1,CG11586,Orc3,Tcs3,Phf5a,Zpr1,CG4866,dgrn,CG6066,x16,mus301,Pep,rhi,DNAlig1,Tctp,CG11920,Pbp49,RPA3,Cul4,CG5589,sun,twin,spel1,RecQ4,CG4935,CG14805,Rpb11,CG6171,smg,TfIIFbeta,Upf3,E(bx),grau,CG30105,CG34261,ubl,Rat1,SF2 | | 4.692e-07 | -14.57 | cellular localization | biological process | GO:0051641 | 1000 | 51 | 12345 | 305 | Parp,rb,Mps1,Rab8,Nup188,cal1,Vap33,flw,bai,Fmr1,Vps13D,pins,TM9SF2,Girdin,CG17883,sws,Dbp80,CG30423,pav,g,Pex3,SF2,Cdc42,Pten,Sgt,PGAP1,Fs(2)Ket,rhea,ema,mus301,CG9853,Exo84,CG8860,sqd,His2Av,Rop,PIG-K,Lst8,glo,Pp1-87B,Vps35,Rab7,tsu,spoon,Rab14,Tim17b,CG6961,Cp190,Hrb27C,Tom20,RabX6 | | 4.913e-07 | -14.53 | cellular aromatic compound metabolic process | biological process | GO:0006725 | 1544 | 69 | 12345 | 305 | Wdr33,EloB,l(1)10Bb,su(Hw),Pnn,Not3,Parp,Orc3,CG11586,Rbp1,Nopp140,Tcs3,g,CG4866,Zpr1,Phf5a,CG6907,PGAP1,Aps,snf,Ino80,Cdc42,sqd,ver,Cdk8,CG6244,tsu,noi,c12.1,CG5986,snRNA:U4:39B,Arp8,RpS8,His4r,mRpS9,CG32075,sun,twin,CG5589,Cul4,rb,Rpb11,CG14805,CG4935,spel1,RecQ4,E(bx),TfIIFbeta,smg,Upf3,CG6171,SF2,ubl,Rat1,CG30105,CG34261,grau,CG6066,dgrn,Pten,rhi,mus301,Pep,x16,DNAlig1,Pbp49,RPA3,CG11920,Tctp | | 6.031e-07 | -14.32 | pyrophosphatase activity | molecular function | GO:0016462 | 464 | 30 | 12176 | 289 | Arp8,Ino80,sun,RagA-B,CG9330,Dbp21E2,me31B,CG8915,pav,Aps,Rap1,Cdc42,mus301,CG12567,Rab14,Apf,Rab7,E(bx),Rheb,CG5589,cta,Rab8,Fign,RecQ4,RabX6,Acf,Dbp80,spel1,RhoGAPp190,Rpt6 | | 7.345e-07 | -14.12 | heterocycle metabolic process | biological process | GO:0046483 | 1498 | 67 | 12345 | 305 | Tcs3,Phf5a,Zpr1,CG4866,g,Not3,Parp,Pnn,su(Hw),l(1)10Bb,EloB,Wdr33,Rbp1,Nopp140,CG11586,Orc3,c12.1,CG5986,noi,tsu,CG6244,CG32075,mRpS9,His4r,snRNA:U4:39B,RpS8,Arp8,Ino80,snf,Aps,PGAP1,CG6907,Cdk8,ver,sqd,CG6171,Upf3,smg,TfIIFbeta,E(bx),grau,CG34261,CG30105,ubl,Rat1,SF2,rb,Cul4,CG5589,sun,twin,RecQ4,spel1,CG4935,CG14805,Rpb11,DNAlig1,Tctp,CG11920,RPA3,Pbp49,dgrn,CG6066,x16,Pep,mus301,rhi | | 7.959e-07 | -14.04 | oogenesis | biological process | GO:0048477 | 682 | 39 | 12345 | 305 | snf,Rap1,Pten,Cdc42,Fs(2)Ket,LIMK1,mus301,dpp,rhea,rhi,Rop,nsl1,mon2,sqd,tsr,Pp1-87B,glo,spoon,tsu,Rab7,me31B,lic,Hrb27C,Cul4,Parp,twin,BEAF-32,RecQ4,14-3-3zeta,smt3,bai,flw,Btk29A,Fmr1,tkv,CSN1b,vnc,eIF3b,flr | | 8.261e-07 | -14.01 | hydrolase activity, acting on acid anhydrides, in phosphorus-containing anhydrides | molecular function | GO:0016818 | 471 | 30 | 12176 | 289 | RecQ4,RabX6,Fign,Rab8,spel1,Rpt6,RhoGAPp190,Dbp80,Acf,Rab7,Apf,Rab14,cta,CG5589,Rheb,E(bx),Aps,CG12567,Cdc42,mus301,Rap1,CG9330,Dbp21E2,RagA-B,sun,Arp8,Ino80,pav,CG8915,me31B | | 9.426e-07 | -13.87 | nucleolus | cellular component | GO:0005730 | 226 | 20 | 11940 | 298 | MED26,HP4,smt3,MED22,CG31301,Nph,NO66,Parp,sqd,cal1,Eaf6,CG4866,peng,Hrb27C,Rbm13,Nopp140,CG9246,CG11920,RpS9,Upf3 | | 9.432e-07 | -13.87 | hydrolase activity, acting on acid anhydrides | molecular function | GO:0016817 | 474 | 30 | 12176 | 289 | E(bx),Rheb,CG5589,cta,Rab14,Apf,Rab7,Dbp80,Acf,Rpt6,spel1,RhoGAPp190,Rab8,Fign,RecQ4,RabX6,CG8915,me31B,pav,sun,Ino80,Arp8,RagA-B,Dbp21E2,CG9330,Rap1,mus301,Cdc42,CG12567,Aps | | 9.481e-07 | -13.87 | polytene chromosome | cellular component | GO:0005700 | 206 | 19 | 11940 | 298 | ver,Pep,Jarid2,Cp190,Ino80,His2Av,CtBP,Rcd5,MED26,Dref,HP4,Tctp,nsl1,Parp,Mlf,sqd,Pp1-87B,su(Hw),BEAF-32 | | 9.533e-07 | -13.86 | posttranscriptional regulation of gene expression | biological process | GO:0010608 | 208 | 19 | 12345 | 305 | Not3,twin,Rheb,RpS9,eEF5,mus301,mbf1,sqd,CNBP,smg,Upf3,Tcs3,Fmr1,glo,me31B,CG4612,Hrb27C,eIF3b,AGO3 | | 1.003e-06 | -13.81 | cellular macromolecule localization | biological process | GO:0070727 | 635 | 37 | 12345 | 305 | TM9SF2,Vps13D,g,Pex3,Dbp80,pav,sws,Rab8,Parp,rb,bai,cal1,Vap33,tsu,spoon,Rab7,Vps35,Lst8,glo,Tom20,RabX6,Hrb27C,CG6961,Rab14,Tim17b,Fs(2)Ket,Sgt,Pten,Cdc42,PIG-K,CG9853,Exo84,sqd,His2Av,CG8860,mus301,rhea | | 1.010e-06 | -13.81 | positive regulation of translation | biological process | GO:0045727 | 44 | 9 | 12345 | 305 | Rheb,CNBP,Upf3,Tcs3,Fmr1,CG4612,Hrb27C,RpS9,eEF5 | | 1.315e-06 | -13.54 | gene expression | biological process | GO:0010467 | 1554 | 68 | 12345 | 305 | tsu,CG6244,RpL5,CG32473,RpS19a,noi,c12.1,CG5986,RpS8,Arp8,snRNA:U4:39B,CG6961,His4r,mRpS9,CG32075,CG31957,CG6907,snf,RpL35A,eIF3g1,sqd,RpS26,Cdk8,Tcs3,eIF3b,CG4866,vnc,mRpS28,CG8635,Phf5a,Wdr33,EloB,l(1)10Bb,su(Hw),Pnn,Parp,eRF1,RpL15,CG11586,Nopp140,Rbp1,mRpS18A,mRpL34,RpS3A,Pbp49,CG11920,CG6066,eIF4B,eEF1delta,RpS9,eEF5,Pep,x16,E(bx),TfIIFbeta,mRpL30,SF2,ubl,RpL36A,Dbp80,RpL14,grau,CG5589,Rpb11,CG14805,mRpS33,CG4935,eIF3c | | 1.585e-06 | -13.36 | macromolecule localization | biological process | GO:0033036 | 868 | 45 | 12345 | 305 | rb,Parp,Rheb,Rab8,unc-119,cal1,Vap33,bai,14-3-3zeta,Fmr1,pins,smg,Vps13D,TM9SF2,sws,pav,Dbp80,SF2,Pex3,g,Pten,Sgt,Cdc42,Fs(2)Ket,rhea,mus301,sqd,CG8860,His2Av,CG9853,Exo84,PIG-K,mon2,glo,Lst8,Vps35,Rab7,spoon,tsu,Tim17b,Rab14,CG6961,Hrb27C,RabX6,Tom20 | | 1.644e-06 | -13.32 | regulation of protein modification process | biological process | GO:0031399 | 347 | 25 | 12345 | 305 | Tctp,sws,lic,Jarid2,Stlk,krz,sigmar,I-2,mnb,Ncoa6,CycA,dpp,CG1791,mav,CG9246,smt3,E(z),flw,CG1218,Pten,Cdc42,fzy,rb,Rap1,PPP1R15 | | 1.780e-06 | -13.24 | organic cyclic compound metabolic process | biological process | GO:1901360 | 1599 | 69 | 12345 | 305 | CG6244,tsu,c12.1,CG5986,noi,RpS8,Arp8,snRNA:U4:39B,CG32075,His4r,mRpS9,PGAP1,Aps,CG6907,Ino80,Cdc42,snf,sqd,Cdk8,ver,Tcs3,CG4866,g,Phf5a,Zpr1,EloB,Wdr33,Not3,Parp,l(1)10Bb,Pnn,su(Hw),Orc3,Nopp140,Rbp1,CG11586,DNAlig1,CG11920,Pbp49,RPA3,Tctp,CG6066,Pten,dgrn,rhi,x16,mus301,Pep,E(bx),CG6171,TfIIFbeta,smg,Upf3,CG30105,CG34261,Rat1,ubl,SF2,grau,CG5589,sun,twin,rb,Cul4,CG14805,Rpb11,spel1,RecQ4,CG4935 | | 2.010e-06 | -13.12 | regulation of cellular component biogenesis | biological process | GO:0044087 | 305 | 23 | 12345 | 305 | tsu,tkv,spoon,mnb,tsr,Nf1,Fmr1,conu,Arpc5,Btk29A,Rpt6,Pex3,Arpc3A,flr,Rheb,Rab8,Cdc42,Sgt,Pten,Rop,cal1,Acf,Spn | | 2.279e-06 | -12.99 | regulation of transcription, DNA-templated | biological process | GO:0006355 | 1144 | 54 | 12345 | 305 | BEAF-32,Not3,d4,Parp,CG17385,nom,su(Hw),NO66,CtBP,14-3-3zeta,smt3,CG6254,Rbp1,CG12299,Ncoa6,tkv,E(bx),Dref,smg,MED22,TfIIFbeta,MED21,CG11456,Rcd5,SF2,grau,CG6066,RAF2,CG6907,Ino80,Mlf,snf,dgrn,Art1,E(z),Nulp1,mav,nsl1,CG9246,mbf1,rhi,rhea,Acf,x16,dpp,Tgi,CG1815,mnb,CG6244,Lst8,Rpt6,Arp8,MED26,Cp190 | | 2.398e-06 | -12.94 | dorsal/ventral axis specification | biological process | GO:0009950 | 76 | 11 | 12345 | 305 | flw,sqd,bai,dpp,pll,mus301,spoon,tsu,glo,Parp,Fmr1 | | 2.557e-06 | -12.88 | cellular component biogenesis | biological process | GO:0044085 | 1270 | 58 | 12345 | 305 | CIA30,CG12321,Rap1,Cdc42,eIF4B,RpL35A,sqd,CG9246,Rop,mon2,Acf,Dbp21E2,dpp,rhea,msd5,LIMK1,inc,RpS19a,Rab7,spoon,RpL5,tsu,CG17493,glo,tsr,Pp1-87B,CG11920,Hrb27C,RpS8,Pbp49,Tom20,me31B,CG32075,Rab14,mRpS9,Nph,His4r,unc-119,CG5589,Cul4,Orc3,CG14805,Rpn10,smt3,CG1791,RecQ4,CG8223,Nopp140,Vap33,tkv,Fmr1,pav,CG4866,CG44194,flr,Rcd5,l(2)37Bb,RpL14 | | 2.727e-06 | -12.81 | nucleoplasm part | cellular component | GO:0044451 | 401 | 27 | 11940 | 298 | MED21,Rbp1,Ncoa6,Jarid2,CG6907,x16,Rpb11,Eaf6,Cdk8,Orc3,CG6244,TfIIFbeta,d4,CG14805,EloB,nsl1,Parp,sqd,CG6254,MED26,Wdr82,Rbbp5,Wdr33,MED22,E(z),SF2,Rcd5 | | 2.870e-06 | -12.76 | GO\_RIBONUCLEOPROTEIN\_COMPLEX | MSigDB lists | GO\_RIBONUCLEOPROTEIN\_COMPLEX | 353 | 35 | 4161 | 184 | Phf5a,l(1)10Bb,Dbp21E2,CG4935,tsu,RpL5,eIF3b,RpS26,RpL14,sm,RpL35A,c12.1,Rcd5,SF2,RpL36A,CG11920,AGO3,mRpS28,snf,CG4866,mRpS33,RpS9,vnc,CG11586,me31B,Fmr1,RpS8,mRpS9,CG9246,Cdc42,eIF3g1,noi,eIF3c,Prosalpha2,RpL15 | | 3.172e-06 | -12.66 | regulation of organelle organization | biological process | GO:0033043 | 408 | 27 | 12345 | 305 | Tctp,Cp190,Bub1,Sod2,Arpc3A,Jarid2,flr,pav,Arpc5,conu,Lst8,Btk29A,sigmar,spoon,Fign,cal1,Ncoa6,CycA,Acf,cta,CG9246,E(z),flw,Mps1,su(Hw),Cdc42,fzy | | 3.287e-06 | -12.63 | regulation of RNA biosynthetic process | biological process | GO:2001141 | 1158 | 54 | 12345 | 305 | Rpt6,Arp8,MED26,Cp190,CG1815,mnb,CG6244,Lst8,E(z),Nulp1,nsl1,mav,mbf1,rhi,CG9246,rhea,dpp,x16,Acf,Tgi,CG6066,RAF2,CG6907,Ino80,Mlf,snf,dgrn,Art1,Rcd5,CG11456,SF2,grau,tkv,E(bx),Dref,smg,TfIIFbeta,MED22,MED21,NO66,14-3-3zeta,CG6254,smt3,CtBP,CG12299,Rbp1,Ncoa6,BEAF-32,CG17385,Not3,Parp,d4,nom,su(Hw) | | 3.287e-06 | -12.63 | regulation of nucleic acid-templated transcription | biological process | GO:1903506 | 1158 | 54 | 12345 | 305 | mnb,CG1815,CG6244,Lst8,Arp8,Rpt6,Cp190,MED26,CG6066,RAF2,CG6907,Ino80,Art1,dgrn,Mlf,snf,Nulp1,E(z),CG9246,mbf1,rhi,nsl1,mav,Acf,dpp,x16,rhea,Tgi,E(bx),tkv,Dref,MED21,smg,TfIIFbeta,MED22,SF2,Rcd5,CG11456,grau,BEAF-32,nom,CG17385,d4,Not3,Parp,su(Hw),NO66,smt3,CtBP,14-3-3zeta,CG6254,Rbp1,CG12299,Ncoa6 | | 3.542e-06 | -12.55 | positive regulation of cellular amide metabolic process | biological process | GO:0034250 | 79 | 11 | 12345 | 305 | Tcs3,Parp,Fmr1,CNBP,Upf3,Rheb,eEF5,RpS9,Hrb27C,CG4612,Pex3 | | 3.621e-06 | -12.53 | regulation of intracellular signal transduction | biological process | GO:1902531 | 339 | 24 | 12345 | 305 | Slmap,GEFmeso,vap,lic,Lst8,Fgop2,conu,krz,Nf1,Pp1-87B,mnb,spoon,sigmar,Efa6,dpp,flw,RagA-B,14-3-3zeta,smt3,mav,Rap1,Pten,Cdc42,Rheb | | 4.039e-06 | -12.42 | developmental process involved in reproduction | biological process | GO:0003006 | 928 | 46 | 12345 | 305 | grau,CSN1b,vnc,eIF3b,flr,Btk29A,Fmr1,janA,E(bx),tkv,RecQ4,flw,smt3,14-3-3zeta,bai,Parp,Cul4,Mps1,BEAF-32,twin,me31B,Hrb27C,lic,glo,tsr,Pp1-87B,Rab7,spoon,tsu,dpp,rhea,HtrA2,LIMK1,mus301,sqd,Exo84,rhi,mon2,Rop,nsl1,Rap1,Pten,Cdc42,dgrn,snf,Fs(2)Ket | | 5.608e-06 | -12.09 | histone binding | molecular function | GO:0042393 | 86 | 11 | 12176 | 289 | CG8223,d4,Nph,CG9246,Ino80,Parp,CG1218,E(bx),Dek,cal1,His4r | | 5.863e-06 | -12.05 | RRM | smart domains | SM00360 | 125 | 15 | 5086 | 158 | Rbp1,Hrb27C,CG6961,glo,sqd,eIF3g1,SF2,snf,CG4612,sm,eIF4B,x16,Cpsf6,eIF3b,tsu | | 6.574e-06 | -11.93 | nucleoside-triphosphatase activity | molecular function | GO:0017111 | 442 | 27 | 12176 | 289 | Rap1,mus301,Cdc42,CG8915,me31B,pav,RagA-B,Arp8,sun,Ino80,Dbp21E2,CG9330,Acf,Dbp80,spel1,Rpt6,RhoGAPp190,Rab8,RecQ4,RabX6,Fign,Rheb,E(bx),cta,CG5589,Rab14,Rab7 | | 7.691e-06 | -11.78 | protein localization | biological process | GO:0008104 | 693 | 37 | 12345 | 305 | Cdc42,Pten,Sgt,Fs(2)Ket,rhea,PIG-K,mon2,CG8860,sqd,His2Av,Exo84,CG9853,Lst8,Vps35,Rab7,CG6961,Tim17b,Rab14,RabX6,Tom20,rb,Parp,Rheb,unc-119,Rab8,cal1,Vap33,14-3-3zeta,bai,pins,Vps13D,TM9SF2,sws,Pex3,g,pav,Dbp80 | | 7.934e-06 | -11.74 | cellular protein localization | biological process | GO:0034613 | 558 | 32 | 12345 | 305 | Exo84,CG9853,sqd,CG8860,His2Av,PIG-K,rhea,Fs(2)Ket,Cdc42,Sgt,Pten,RabX6,Tom20,Rab14,Tim17b,CG6961,Vps35,Rab7,Lst8,bai,cal1,Vap33,Rab8,Parp,rb,Dbp80,pav,g,Pex3,sws,TM9SF2,Vps13D | | 8.107e-06 | -11.72 | cytoplasmic translation | biological process | GO:0002181 | 120 | 13 | 12345 | 305 | RpS19a,RpL5,RpS3A,eRF1,RpL35A,eIF4B,RpL36A,RpS26,RpL15,RpS8,CG8635,RpS9,RpL14 | | 8.563e-06 | -11.67 | RRM\_dom | interpro domains | IPR000504 | 135 | 15 | 11383 | 331 | CG6961,tsu,glo,eIF3b,Hrb27C,snf,SF2,CG4612,Cpsf6,x16,eIF3g1,eIF4B,sqd,sm,Rbp1 | | 8.652e-06 | -11.66 | Nucleotide-bd\_a/b\_plait\_sf | interpro domains | IPR012677 | 152 | 16 | 11383 | 331 | CG4612,Upf3,x16,Rbp1,tsu,Cpsf6,sqd,eIF4B,eIF3g1,sm,CG6961,glo,eIF3b,Hrb27C,snf,SF2 | | 9.132e-06 | -11.60 | mitotic cell cycle | biological process | GO:0000278 | 432 | 27 | 12345 | 305 | Cdk8,CycA,cal1,msd5,Rpn10,E(z),Rop,14-3-3zeta,smt3,fzy,Mps1,eIF3g1,Rheb,Fs(2)Ket,Bub1,Tctp,pav,Skp2,Fmr1,CG17493,Pp1-87B,MED22,pins,noi,tsr,Fign,DNAlig1 | | 9.393e-06 | -11.58 | RRM | prosite domains | PS50102 | 131 | 15 | 6243 | 191 | Cpsf6,tsu,glo,eIF3g1,Rbp1,sqd,sm,snf,eIF3b,x16,Hrb27C,SF2,CG6961,CG4612,eIF4B | | 1.111e-05 | -11.41 | ribosomal subunit | cellular component | GO:0044391 | 180 | 16 | 11940 | 298 | mRpS33,RpS8,mRpL30,mRpS18A,RpL36A,mRpL34,RpL15,RpS3A,RpL14,mRpS28,RpL5,mRpS9,RpS26,RpS9,RpL35A,RpS19a | | 1.173e-05 | -11.35 | germ cell development | biological process | GO:0007281 | 820 | 41 | 12345 | 305 | snf,Rap1,Cdc42,Pten,Fs(2)Ket,LIMK1,mus301,dpp,rhea,rhi,nsl1,mon2,Rop,sqd,Exo84,tsr,Pp1-87B,glo,spoon,tsu,Rab7,me31B,lic,Hrb27C,Mps1,Cul4,Parp,twin,BEAF-32,RecQ4,bai,smt3,14-3-3zeta,flw,Btk29A,Fmr1,tkv,CSN1b,vnc,flr,eIF3b | | 1.287e-05 | -11.26 | - | gene3d domains | 3.30.70.330 | 152 | 16 | 4929 | 151 | CG4612,CG6961,tsu,glo,Cpsf6,sqd,sm,Upf3,x16,Rbp1,eIF3b,snf,Hrb27C,eIF3g1,eIF4B,SF2 | | 1.308e-05 | -11.24 | RBD\_domain\_sf | interpro domains | IPR035979 | 157 | 16 | 11383 | 331 | glo,CG6961,snf,SF2,Hrb27C,eIF3b,eIF4B,sqd,eIF3g1,Cpsf6,sm,tsu,x16,Upf3,CG4612,Rbp1 | | 1.390e-05 | -11.18 | chromosomal region | cellular component | GO:0098687 | 204 | 17 | 11940 | 298 | ver,Pep,cal1,Cp190,Bub1,His2Av,Mps1,Rcd5,MED26,smt3,HP4,Mlf,nsl1,Parp,sqd,fzy,BEAF-32 | | 1.416e-05 | -11.17 | RpS6 (Ribosomal protein S6) | protein interactions | 31700 | 112 | 14 | 9204 | 298 | Parp,RpL36A,RpS8,RpS3A,RpL14,RpL15,RpS19a,RpS9,RpS26,RpL5,Rcd5,GEFmeso,Rheb,RpL35A | | 1.446e-05 | -11.14 | cellular protein metabolic process | biological process | GO:0044267 | 2099 | 81 | 12345 | 305 | CG31957,mRpS9,RpS8,Skp2,Pp1-87B,mnb,RpS19a,RpL5,Acf,Roc2,Cdk8,LIMK1,RpS26,E(z),PIG-K,Prosbeta1,fzy,Wdr82,Sgt,Art1,Ubqn,RpL35A,eIF3g1,xit,CG31460,CG8635,mRpS28,vnc,eIF3b,Jarid2,Prosalpha2,g,Rcd5,Dref,janA,Ufd4,tkv,Rbbp5,RpL15,smt3,eRF1,Parp,alpha-Man-Ib,Bub1,Usp16-45,Rpt6,lic,RpS3A,Stlk,inc,mRpL34,mRpS18A,Stt3B,CycA,RpS9,Usp32,eEF5,Pten,CG1218,eEF1delta,dgrn,eIF4B,Eaf6,pll,RpL14,CSN1b,RpL36A,Btk29A,mRpL30,Mppe,eIF3c,Ncoa6,mRpS33,flw,NO66,Rpn10,rb,d4,Cul4,Mps1,BEAF-32 | | 1.534e-05 | -11.09 | catalytic step 2 spliceosome | cellular component | GO:0071013 | 126 | 13 | 11940 | 298 | noi,tsu,CG6066,c12.1,Pep,SF2,CG4935,l(1)10Bb,x16,snf,sqd,Pnn,Phf5a | | 1.625e-05 | -11.03 | regulation of signal transduction | biological process | GO:0009966 | 745 | 38 | 12345 | 305 | dpp,Nulp1,RagA-B,mav,Rap1,Cdc42,Pten,dgrn,Slmap,GEFmeso,lic,Fgop2,Lst8,Nf1,krz,Pp1-87B,mnb,Vps35,sigmar,spoon,Efa6,flw,14-3-3zeta,smt3,CtBP,Cul4,Rheb,sun,Graf,pav,CG30423,vap,conu,Fmr1,TM9SF2,I-2,E(bx),tkv | | 1.686e-05 | -10.99 | Fkbp39 (FK506-binding protein 39kD) | protein interactions | 41860 | 85 | 12 | 9204 | 298 | CG5589,CG32075,CG4866,CG14805,GEFmeso,CG11920,CG9246,His4r,Pp1-87B,His2Av,NO66,CG8108 | | 1.810e-05 | -10.92 | translation regulator activity | molecular function | GO:0045182 | 97 | 11 | 12176 | 289 | eEF5,eIF3c,eIF3g1,smg,eIF3b,eEF1delta,eIF4B,CG31957,Fmr1,CNBP,eRF1 | | 2.288e-05 | -10.69 | regulation of mRNA metabolic process | biological process | GO:1903311 | 132 | 13 | 12345 | 305 | Hrb27C,sqd,AGO3,Cpsf6,mbf1,SF2,Rbp1,x16,Pep,tsu,Fmr1,smg,snf | | 2.547e-05 | -10.58 | Tsc1 (Tsc1) | protein interactions | 42862 | 183 | 18 | 9204 | 298 | peng,Pten,TfIIFbeta,eIF3c,Nopp140,sqd,eIF3g1,Tctp,E(bx),Rheb,Lst8,14-3-3zeta,CG11920,eIF3b,spoon,Vap33,RpS9,pins | | 2.559e-05 | -10.57 | mRNA splicing, via spliceosome | biological process | GO:0000398 | 238 | 18 | 12345 | 305 | snRNA:U4:39B,SF2,ubl,Phf5a,tsu,c12.1,CG5986,noi,sqd,x16,Rbp1,Pep,CG11586,CG4935,CG6066,Pnn,snf,l(1)10Bb | | 2.559e-05 | -10.57 | RNA splicing, via transesterification reactions with bulged adenosine as nucleophile | biological process | GO:0000377 | 238 | 18 | 12345 | 305 | tsu,CG5986,c12.1,noi,ubl,snRNA:U4:39B,SF2,Phf5a,CG6066,snf,l(1)10Bb,Pnn,sqd,Rbp1,x16,CG11586,CG4935,Pep | | 2.563e-05 | -10.57 | RRM\_1 | pfam domains | PF00076 | 125 | 14 | 10480 | 319 | glo,tsu,eIF3b,CG4612,SF2,Cpsf6,x16,eIF3g1,Rbp1,CG6961,Hrb27C,sqd,snf,eIF4B | | 2.579e-05 | -10.57 | cellular process involved in reproduction in multicellular organism | biological process | GO:0022412 | 997 | 46 | 12345 | 305 | mus301,LIMK1,rhea,dpp,Rop,mon2,nsl1,rhi,Exo84,sqd,snf,Pten,Cdc42,Rap1,fzy,Fs(2)Ket,me31B,lic,Hrb27C,Pp1-87B,tsr,glo,tsu,spoon,Rab7,cal1,RecQ4,bai,smt3,14-3-3zeta,flw,Mps1,Cul4,Parp,twin,BEAF-32,vnc,CSN1b,grau,eIF3b,flr,pav,Fmr1,Btk29A,tkv,E(bx) | | 2.668e-05 | -10.53 | mRNA metabolic process | biological process | GO:0016071 | 357 | 23 | 12345 | 305 | tsu,CG5986,c12.1,Upf3,noi,smg,ubl,snRNA:U4:39B,SF2,Phf5a,CG6066,Wdr33,twin,Not3,Pnn,l(1)10Bb,snf,sqd,Rbp1,x16,Pep,CG11586,CG4935 | | 2.760e-05 | -10.50 | dorsal/ventral pattern formation | biological process | GO:0009953 | 154 | 14 | 12345 | 305 | Rab7,spoon,tkv,tsu,glo,Parp,Fmr1,CG8149,sqd,flw,bai,dpp,pll,mus301 | | 2.812e-05 | -10.48 | cell cycle | biological process | GO:0007049 | 678 | 35 | 12345 | 305 | Fign,Btk29A,Dref,Fmr1,pins,MED22,pav,Rcd5,grau,Rheb,Mps1,Orc3,flw,Rpn10,14-3-3zeta,smt3,cal1,DNAlig1,CG17493,noi,tsr,Pp1-87B,Skp2,Bub1,Tctp,Fs(2)Ket,fzy,eIF3g1,E(z),Exo84,Rop,Cdk8,CycA,msd5,mus301 | | 2.862e-05 | -10.46 | RNA splicing, via transesterification reactions | biological process | GO:0000375 | 240 | 18 | 12345 | 305 | sqd,Pep,CG11586,CG4935,x16,Rbp1,CG6066,Pnn,snf,l(1)10Bb,snRNA:U4:39B,SF2,ubl,Phf5a,tsu,noi,c12.1,CG5986 | | 3.075e-05 | -10.39 | positive regulation of protein modification process | biological process | GO:0031401 | 176 | 15 | 12345 | 305 | Tctp,Ncoa6,dpp,smt3,lic,CG1791,mav,E(z),krz,Stlk,fzy,Rap1,Cdc42,sigmar,PPP1R15 | | 3.092e-05 | -10.38 | developmental process | biological process | GO:0032502 | 2988 | 105 | 12345 | 305 | EloB,Parp,smt3,Vap33,tkv,Dref,Fmr1,janA,pins,eIF3b,pav,g,mthl8,vnc,Zpr1,sws,Fs(2)Ket,xit,peng,Cdc42,fzy,snf,E(z),Exo84,sqd,His2Av,mon2,nsl1,mbf1,Cdk8,rhea,dpp,Acf,LIMK1,mnb,tsu,Pp1-87B,CG8149,heix,krz,tsr,Nf1,Stim,Skp2,RpS8,me31B,sm,Cp190,BEAF-32,Rheb,unc-119,twin,sun,d4,Cpr73D,nom,rb,Mps1,Cul4,CG10365,BthD,flw,dah,bai,14-3-3zeta,CtBP,RecQ4,RhoGAPp190,E(bx),Arpc5,Btk29A,MED22,vap,flr,Amun,grau,Girdin,pll,CSN1b,Pten,Rap1,CG12935,dgrn,Hsp27,mav,cta,Rop,rhi,CycA,Spn,mus301,HtrA2,Rab7,Efa6,PlexA,spoon,sigmar,glo,Stlk,Hrb27C,VhaPPA1-1,Sod2,lic,RPA3,GEFmeso | | 3.095e-05 | -10.38 | negative regulation of nitrogen compound metabolic process | biological process | GO:0051172 | 625 | 33 | 12345 | 305 | me31B,Hrb27C,krz,mnb,tsu,sigmar,rhea,Acf,Tgi,E(z),sqd,Nulp1,rhi,mbf1,CG9246,Pten,Ino80,snf,CG6066,sws,Fmr1,smg,I-2,spel1,CG12299,NO66,flw,CtBP,CG6254,d4,Not3,su(Hw),twin | | 3.200e-05 | -10.35 | RpS21 (Ribosomal protein S21) | protein interactions | 33487 | 136 | 15 | 9204 | 298 | CG1218,CG6171,BthD,RpL35A,Wdr82,RpL36A,RpL14,RpL15,Bub1,RpS3A,RpS8,RpS19a,RpS9,RpS26,RpL5 | | 3.433e-05 | -10.28 | RpL23A (Ribosomal protein L23A) | protein interactions | 38208 | 121 | 14 | 9204 | 298 | RpL35A,GEFmeso,RpL5,Sgt,RpS26,RpS8,RpL15,RpL14,RpS3A,RpS19a,RpS9,Parp,RpL36A,CG11334 | | 3.451e-05 | -10.27 | regulation of biological quality | biological process | GO:0065008 | 1352 | 57 | 12345 | 305 | RhoGAPp190,Vap33,14-3-3zeta,smt3,MICU1,BthD,rb,Parp,Rheb,Rab8,Zpr1,sws,CSN1b,Girdin,CG3842,pav,flr,pins,smg,Vps13D,Btk29A,conu,Arpc5,Fmr1,Dref,tkv,ver,RpS9,Spn,dpp,rhea,mbf1,cta,Rop,Exo84,Mlf,Rap1,Pten,Cdc42,Aps,Txl,Tctp,Rab14,Arpc3A,lic,Sod2,Skp2,Stim,krz,Nf1,tsr,Lst8,spoon,PlexA,CG32473,mnb,Vps35 | | 3.514e-05 | -10.26 | chromatin organization | biological process | GO:0006325 | 466 | 27 | 12345 | 305 | E(bx),CG6244,Fmr1,HP4,Arp8,Jarid2,Rcd5,CG12659,Cp190,Nph,His4r,vnc,Eaf6,BEAF-32,Art1,snf,Wdr82,Ino80,d4,CtBP,His2Av,E(z),NO66,Ncoa6,Rbbp5,Acf,CG8223 | | 4.033e-05 | -10.12 | GSE7509\_FCGRIIB\_VS\_TNFA\_IL1B\_IL6\_PGE\_STIM\_DC\_UP | MSigDB lists | GSE7509\_FCGRIIB\_VS\_TNFA\_IL1B\_IL6\_PGE\_STIM\_DC\_UP | 92 | 14 | 4161 | 184 | CG8132,CG15168,His2Av,flr,CG5861,MICU1,inc,Stlk,Fs(2)Ket,Tim17b,Pp1-87B,CG12321,CG31957,Txl | | 4.226e-05 | -10.07 | regulation of cytoskeleton organization | biological process | GO:0051493 | 160 | 14 | 12345 | 305 | Fign,spoon,sigmar,Btk29A,conu,Cdc42,Arpc5,Lst8,flw,pav,flr,Arpc3A,cta,Cp190 | | 4.407e-05 | -10.03 | regulation of response to stimulus | biological process | GO:0048583 | 958 | 44 | 12345 | 305 | CtBP,14-3-3zeta,smt3,flw,Cul4,Parp,sun,Rheb,pll,Graf,Dek,vap,pav,CG30423,Fmr1,conu,tkv,E(bx),TM9SF2,I-2,Fign,dpp,mav,RagA-B,Nulp1,dgrn,Pten,Cdc42,Rap1,GEFmeso,Slmap,me31B,lic,Pp1-87B,Nf1,krz,Lst8,Fgop2,spoon,Efa6,sigmar,PlexA,Vps35,mnb | | 4.748e-05 | -9.96 | RpL10Ab (Ribosomal protein L10Ab) | protein interactions | 39338 | 109 | 13 | 9204 | 298 | eEF1delta,Rheb,RpL35A,RpS9,RpS19a,RpL15,RpS3A,RpL14,RpS8,RpL36A,RpL5,RpS26,me31B | | 4.905e-05 | -9.92 | RNA splicing | biological process | GO:0008380 | 250 | 18 | 12345 | 305 | snRNA:U4:39B,ubl,SF2,Phf5a,tsu,noi,c12.1,CG5986,sqd,Pep,CG4935,CG11586,x16,Rbp1,CG6066,Pnn,snf,l(1)10Bb | | 5.241e-05 | -9.86 | CAIRO\_HEPATOBLASTOMA\_CLASSES\_UP | MSigDB lists | CAIRO\_HEPATOBLASTOMA\_CLASSES\_UP | 321 | 30 | 4161 | 184 | CG8635,Skp2,CCT4,CycA,SF2,Tom20,Fs(2)Ket,CG11920,DNAlig1,mRpS33,CG8860,RpL15,CSN1b,eEF1delta,RpS9,E(z),14-3-3zeta,Art1,Lst8,eIF3b,RpL5,Parp,RpL14,RpS8,HtrA2,alpha-Man-Ib,fzy,Pp1-87B,pav,spel1 | | 5.401e-05 | -9.83 | cellular macromolecule catabolic process | biological process | GO:0044265 | 425 | 25 | 12345 | 305 | inc,tsu,Ufd4,CG6171,Upf3,smg,CG30105,Rat1,Prosalpha2,Skp2,Rpt6,alpha-Man-Ib,Zpr1,Usp16-45,twin,fzy,Sgt,Not3,Ubqn,Cul4,sqd,Rpn10,Prosbeta1,Roc2,Usp32 | | 5.767e-05 | -9.76 | nuclear chromosome part | cellular component | GO:0044454 | 322 | 21 | 11940 | 298 | Jarid2,Ino80,Bub1,His2Av,DNAlig1,Arp8,Orc3,Acf,Parp,BEAF-32,d4,E(bx),RPA3,E(z),CtBP,Rcd5,Mps1,MED26,smt3,HP4,CG12659 | | 5.931e-05 | -9.73 | multicellular organism development | biological process | GO:0007275 | 2496 | 90 | 12345 | 305 | EloB,Parp,smt3,Vap33,tkv,pins,Dref,Fmr1,g,pav,sws,Zpr1,mthl8,xit,snf,peng,Cdc42,mbf1,mon2,nsl1,sqd,E(z),LIMK1,dpp,Acf,rhea,Cdk8,tsu,mnb,Nf1,CG8149,krz,tsr,heix,Pp1-87B,RpS8,Skp2,Stim,Cp190,sm,me31B,sun,Rheb,unc-119,rb,nom,Cpr73D,d4,CtBP,bai,14-3-3zeta,dah,flw,CG10365,BthD,RhoGAPp190,RecQ4,E(bx),MED22,Btk29A,vap,flr,Amun,pll,Girdin,grau,dgrn,CG12935,Rap1,Pten,rhi,mav,Rop,cta,Hsp27,mus301,Spn,CycA,Efa6,PlexA,spoon,sigmar,Rab7,Stlk,glo,VhaPPA1-1,lic,Sod2,Hrb27C,GEFmeso | | 6.019e-05 | -9.72 | SWI/SNF superfamily-type complex | cellular component | GO:0070603 | 55 | 8 | 11940 | 298 | Acf,CG12659,Arp8,CtBP,Rcd5,d4,Ino80,E(bx) | | 6.322e-05 | -9.67 | RNA processing | biological process | GO:0006396 | 564 | 30 | 12345 | 305 | Pnn,l(1)10Bb,snf,Wdr33,CG6907,CG5589,CG6066,Pep,CG11586,CG4935,x16,Nopp140,Rbp1,Cdk8,CG14805,sqd,noi,Tcs3,c12.1,CG5986,tsu,mRpS9,Phf5a,CG32075,RpS8,ubl,SF2,snRNA:U4:39B,CG4866,CG11920 | | 6.466e-05 | -9.65 | gamete generation | biological process | GO:0007276 | 1097 | 48 | 12345 | 305 | me31B,Hrb27C,lic,glo,tsr,Pp1-87B,Rab7,spoon,tsu,dpp,rhea,HtrA2,LIMK1,mus301,sqd,Exo84,rhi,mon2,nsl1,Rop,fzy,Rap1,Pten,Cdc42,snf,Fs(2)Ket,grau,CSN1b,vnc,pav,flr,eIF3b,Btk29A,Fmr1,MED22,E(bx),tkv,RecQ4,cal1,flw,smt3,bai,14-3-3zeta,Parp,Mps1,Cul4,BEAF-32,twin | | 6.810e-05 | -9.59 | regulation of cell communication | biological process | GO:0010646 | 855 | 40 | 12345 | 305 | CG30423,pav,vap,Graf,I-2,TM9SF2,E(bx),tkv,Fmr1,conu,flw,14-3-3zeta,smt3,CtBP,Rheb,sun,Cul4,lic,Sod2,Rpt6,Slmap,GEFmeso,mnb,Vps35,sigmar,Efa6,spoon,Fgop2,Lst8,krz,Nf1,Pp1-87B,Nulp1,RagA-B,mav,dpp,Rap1,Pten,Cdc42,dgrn | | 6.810e-05 | -9.59 | regulation of signaling | biological process | GO:0023051 | 855 | 40 | 12345 | 305 | mav,RagA-B,Nulp1,dpp,dgrn,Pten,Cdc42,Rap1,Sod2,Rpt6,lic,GEFmeso,Slmap,Efa6,sigmar,spoon,Vps35,mnb,Pp1-87B,krz,Nf1,Lst8,Fgop2,14-3-3zeta,smt3,CtBP,flw,sun,Rheb,Cul4,vap,CG30423,pav,Graf,tkv,E(bx),I-2,TM9SF2,Fmr1,conu | | 7.053e-05 | -9.56 | His4:CG31611 (histone H4) | protein interactions | 318846 | 44 | 8 | 9204 | 298 | Dek,His2Av,cal1,E(bx),Acf,Art1,Nph,Parp | | 7.174e-05 | -9.54 | negative regulation of cellular metabolic process | biological process | GO:0031324 | 681 | 34 | 12345 | 305 | sws,I-2,Fmr1,smg,flw,NO66,CG6254,CtBP,CG12299,spel1,Rheb,twin,Not3,d4,su(Hw),Hrb27C,me31B,mnb,sigmar,tsu,krz,sqd,Nulp1,E(z),rhi,CG9246,mbf1,Acf,rhea,Tgi,CG6066,Ino80,Pten,snf | | 7.246e-05 | -9.53 | nuclear chromosome | cellular component | GO:0000228 | 352 | 22 | 11940 | 298 | E(bx),BEAF-32,d4,Parp,CG12659,smt3,HP4,MED26,Rcd5,Mps1,CtBP,RPA3,E(z),DNAlig1,Bub1,Ino80,His2Av,RpS9,Jarid2,Acf,Orc3,Arp8 | | 7.356e-05 | -9.52 | spliceosomal complex | cellular component | GO:0005681 | 188 | 15 | 11940 | 298 | CG6066,tsu,noi,c12.1,Pep,SF2,CG11586,CG4935,l(1)10Bb,CG5986,snf,sqd,x16,Pnn,Phf5a | | 7.633e-05 | -9.48 | RpS7 (Ribosomal protein S7) | protein interactions | 43594 | 130 | 14 | 9204 | 298 | RpS26,RpL5,RpL36A,RpS3A,RpS8,RpL14,RpL15,RpS19a,RpS9,Pten,RpL35A,noi,Rheb,Pnn | | 7.706e-05 | -9.47 | 5SrRNA:CR33353 (5S ribosomal RNA) | protein interactions | 3771903 | 84 | 11 | 9204 | 298 | RpL36A,RpS19a,RpS9,RpL14,RpS8,RpL15,RpS3A,RpS26,RpL5,Fmr1,RpL35A | | 7.791e-05 | -9.46 | response to transforming growth factor beta | biological process | GO:0071559 | 19 | 5 | 12345 | 305 | dpp,Cdc42,smt3,tkv,mav | | 7.791e-05 | -9.46 | cellular response to transforming growth factor beta stimulus | biological process | GO:0071560 | 19 | 5 | 12345 | 305 | smt3,mav,tkv,dpp,Cdc42 | | 7.905e-05 | -9.45 | positive regulation of gene expression | biological process | GO:0010628 | 571 | 30 | 12345 | 305 | Rheb,su(Hw),dgrn,d4,rb,nsl1,mbf1,CtBP,rhi,RpS9,eEF5,Ncoa6,dpp,CG6244,mnb,TfIIFbeta,CNBP,Upf3,MED21,Tcs3,Lst8,Fmr1,Dref,g,Rpt6,Rcd5,CG4612,Hrb27C,MED26,grau | | 8.219e-05 | -9.41 | foxo (forkhead box, sub-group O) | protein interactions | 41709 | 182 | 17 | 9204 | 298 | Rheb,eIF3b,14-3-3zeta,glo,RpL35A,RpS9,mnb,RAF2,pav,Pten,eIF3c,TfIIFbeta,sqd,pll,RpL14,RpL15,RpS26 | | 8.835e-05 | -9.33 | GSE23568\_CTRL\_VS\_ID3\_TRANSDUCED\_CD8\_TCELL\_DN | MSigDB lists | GSE23568\_CTRL\_VS\_ID3\_TRANSDUCED\_CD8\_TCELL\_DN | 75 | 12 | 4161 | 184 | RpS26,unc-119,mRpS28,CG12935,fzy,Sod2,pav,ubl,CycA,CG31460,CG5590,CG10343 | | 9.095e-05 | -9.31 | negative regulation of macromolecule metabolic process | biological process | GO:0010605 | 777 | 37 | 12345 | 305 | NO66,flw,CG6254,CtBP,spel1,CG12299,twin,d4,Not3,su(Hw),sws,I-2,tkv,Fmr1,smg,Upf3,E(z),Nulp1,sqd,rhi,CG9246,mbf1,rhea,Acf,dpp,Tgi,CG6066,Pten,Ino80,snf,Hrb27C,AGO3,me31B,mnb,tsu,sigmar,krz | | 9.134e-05 | -9.30 | epithelial cell development | biological process | GO:0002064 | 361 | 22 | 12345 | 305 | tsr,spoon,tkv,Rab7,me31B,flr,eIF3b,Cul4,Rap1,Parp,Cdc42,Pten,xit,Fs(2)Ket,RecQ4,dpp,rhea,smt3,rhi,Rop,cta,sqd | | 9.168e-05 | -9.30 | maintenance of protein location | biological process | GO:0045185 | 44 | 7 | 12345 | 305 | pins,Vps13D,Parp,rhea,Cdc42,14-3-3zeta,pav | | 9.384e-05 | -9.27 | Translation | REACTOME pathways | R-DME-72766 | 143 | 16 | 3923 | 151 | RpL15,eRF1,RpS26,RpS9,eIF3g1,eIF3c,RpL35A,RpL36A,RpS8,RpS19a,RpS3A,eIF3b,eEF1delta,eIF4B,RpL14,RpL5 | | 9.832e-05 | -9.23 | anatomical structure development | biological process | GO:0048856 | 2924 | 101 | 12345 | 305 | mus301,HtrA2,CycA,Spn,Rop,cta,mav,rhi,Hsp27,dgrn,Pten,Rap1,CG12935,GEFmeso,VhaPPA1-1,lic,Sod2,Hrb27C,Stlk,glo,PlexA,Efa6,spoon,sigmar,Rab7,RhoGAPp190,RecQ4,dah,CtBP,bai,14-3-3zeta,BthD,CG10365,flw,Mps1,Cul4,d4,nom,Cpr73D,rb,twin,sun,BEAF-32,Rheb,unc-119,CSN1b,pll,Girdin,grau,flr,Amun,vap,MED22,Arpc5,Btk29A,E(bx),LIMK1,Cdk8,rhea,Acf,dpp,mon2,nsl1,mbf1,Exo84,E(z),sqd,snf,peng,Cdc42,xit,Fs(2)Ket,Cp190,sm,me31B,Skp2,RpS8,Stim,Pp1-87B,Nf1,CG8149,krz,tsr,heix,tsu,mnb,Vap33,smt3,Parp,EloB,vnc,Zpr1,sws,mthl8,g,eIF3b,pav,pins,Fmr1,Dref,tkv | | 1.071e-04 | -9.14 | His2A:CG31618 (histone H2A) | protein interactions | 318855 | 35 | 7 | 9204 | 298 | Acf,E(z),His2Av,Jarid2,Dek,Parp,Nph | | 1.183e-04 | -9.04 | mRNA processing | biological process | GO:0006397 | 292 | 19 | 12345 | 305 | x16,Rbp1,CG11586,CG4935,Pep,sqd,l(1)10Bb,snf,Pnn,CG6066,Wdr33,Phf5a,SF2,ubl,snRNA:U4:39B,CG5986,c12.1,noi,tsu | | 1.209e-04 | -9.02 | S6kII (Ribosomal protein S6 kinase II) | protein interactions | 33139 | 206 | 18 | 9204 | 298 | RAF2,RpS9,RpL35A,CG9246,spoon,Fmr1,eIF3b,Pnn,SF2,sqd,RpL14,RpL15,Nph,Nopp140,TfIIFbeta,eIF3c,Cpsf6,peng | | 1.213e-04 | -9.02 | small ribosomal subunit | cellular component | GO:0015935 | 77 | 9 | 11940 | 298 | mRpS28,mRpS18A,RpS26,mRpS9,RpS8,RpS9,RpS19a,RpS3A,mRpS33 | | 1.255e-04 | -8.98 | positive regulation of signal transduction | biological process | GO:0009967 | 369 | 22 | 12345 | 305 | Pp1-87B,Lst8,Fgop2,tkv,sigmar,Efa6,Vps35,Slmap,lic,dgrn,Cdc42,Rap1,sun,Rheb,dpp,mav,CtBP,14-3-3zeta,smt3,RagA-B,flw,Nulp1 | | 1.293e-04 | -8.95 | RpS14a (Ribosomal protein S14a) | protein interactions | 47218 | 120 | 13 | 9204 | 298 | Rheb,GEFmeso,RpL35A,Pten,RpS19a,RpS9,RpS3A,RpL15,RpS8,RpL14,RpL36A,RpL5,RpS26 | | 1.297e-04 | -8.95 | GO\_RNA\_BINDING | MSigDB lists | GO\_RNA\_BINDING | 669 | 49 | 4161 | 184 | RpL15,eEF5,Rat1,Fs(2)Ket,eIF3c,Rop,noi,CG8635,CCT4,eIF3g1,RpS8,mRpS9,CG31957,Pp1-87B,CG9246,Fmr1,me31B,CG11586,His4r,Rpn10,RpS9,CG4866,snf,AGO3,mRpS28,CG11920,Tctp,RpL36A,SF2,Rcd5,c12.1,eRF1,RpL35A,CG5589,sm,RpL14,Parp,RpS26,eIF3b,RpL5,tsu,Dbp21E2,mbf1,14-3-3zeta,E(z),Art1,eEF1delta,Phf5a,Dbp80 | | 1.342e-04 | -8.92 | regulation of cellular component size | biological process | GO:0032535 | 178 | 14 | 12345 | 305 | Arpc5,conu,Cdc42,Pten,Lst8,Btk29A,spoon,Rheb,RhoGAPp190,Tctp,Girdin,lic,Arpc3A,flr | | 1.367e-04 | -8.90 | nos (nanos) | protein interactions | 42297 | 61 | 9 | 9204 | 298 | glo,smg,twin,tkv,nsl1,dpp,E(z),Not3,me31B | | 1.367e-04 | -8.90 | rictor (rapamycin-insensitive companion of Tor) | protein interactions | 32919 | 208 | 18 | 9204 | 298 | Vap33,c12.1,CG10343,Map60,Rheb,Lst8,eEF1delta,14-3-3zeta,eIF3b,Fmr1,Spn,RpL35A,RpL15,RpL14,sqd,x16,Pten,eIF3c | | 1.379e-04 | -8.89 | negative regulation of metabolic process | biological process | GO:0009892 | 823 | 38 | 12345 | 305 | me31B,AGO3,Hrb27C,krz,mnb,sigmar,tsu,Acf,dpp,rhea,Tgi,Nulp1,sqd,E(z),mbf1,CG9246,rhi,Ino80,Pten,snf,CG6066,sws,Fmr1,Upf3,smg,I-2,tkv,CG12299,spel1,flw,NO66,CtBP,CG6254,d4,Not3,su(Hw),Rheb,twin | | 1.404e-04 | -8.87 | RpL13 (Ribosomal protein L13) | protein interactions | 34329 | 105 | 12 | 9204 | 298 | RpS26,Rcd5,RpL5,RpL36A,RpS9,RpS19a,RpL14,RpL15,RpS8,RpS3A,RpL35A,Rheb | | 1.404e-04 | -8.87 | vig2 (vig2) | protein interactions | 43016 | 105 | 12 | 9204 | 298 | RpS26,RpL5,RpL36A,RpL14,RpS3A,RpS8,RpL15,RpS9,RpS19a,RpL35A,Rheb,14-3-3zeta | | 1.408e-04 | -8.87 | RpL5 (Ribosomal protein L5) | protein interactions | 3355124 | 121 | 13 | 9204 | 298 | Fmr1,noi,RpL35A,rb,RpS26,CG11586,RpL36A,RpS19a,RpS9,RpL14,RpS8,RpL15,RpS3A | | 1.437e-04 | -8.85 | ATPase complex | cellular component | GO:1904949 | 62 | 8 | 11940 | 298 | Rcd5,CtBP,Arp8,CG12659,Acf,E(bx),Ino80,d4 | | 1.500e-04 | -8.81 | regulation of actin cytoskeleton organization | biological process | GO:0032956 | 98 | 10 | 12345 | 305 | cta,Arpc3A,flr,flw,spoon,conu,Cdc42,Lst8,Arpc5,Btk29A | | 1.538e-04 | -8.78 | RpL37a (Ribosomal protein L37a) | protein interactions | 32446 | 106 | 12 | 9204 | 298 | RpL36A,RpS19a,RpS9,RpS8,RpL15,RpL14,Bub1,RpS3A,RpS26,RpL5,krz,RpL35A | | 1.538e-04 | -8.78 | RpL8 (Ribosomal protein L8) | protein interactions | 44251 | 106 | 12 | 9204 | 298 | Rheb,14-3-3zeta,RpL35A,RpL36A,RpS3A,RpS8,RpL15,RpL14,RpS9,RpS19a,RpS26,RpL5 | | 1.538e-04 | -8.78 | RpL30 (Ribosomal protein L30) | protein interactions | 44059 | 106 | 12 | 9204 | 298 | RpS3A,RpL15,RpS8,RpL14,RpS19a,RpS9,Parp,RpL36A,RpL5,RpS26,Rheb,RpL35A | | 1.544e-04 | -8.78 | fly chr40F7-40F7\|2-54 cM | chromosome location | fly chr40F7-40F7\|2-54 cM | 8 | 3 | 25010 | 358 | Slmap,RpL5,CG17493 | | 1.544e-04 | -8.78 | fly chr40F7-40F7\|2-54 | chromosome location | fly chr40F7-40F7\|2-54 | 8 | 3 | 25010 | 358 | CG17493,Slmap,RpL5 | | 1.546e-04 | -8.77 | regulation of mRNA 3'-end processing | biological process | GO:0031440 | 12 | 4 | 12345 | 305 | Fmr1,Rbp1,x16,SF2 | | 1.554e-04 | -8.77 | cup (cup) | protein interactions | 33934 | 62 | 9 | 9204 | 298 | spoon,twin,smg,Nph,Hrb27C,sqd,RpS26,me31B,Not3 | | 1.558e-04 | -8.77 | Rho\_GTPase\_activation\_prot | interpro domains | IPR008936 | 29 | 6 | 11383 | 331 | Graf,Nf1,PlexA,conu,RhoGAPp190,vap | | 1.578e-04 | -8.75 | larval development | biological process | GO:0002164 | 159 | 13 | 12345 | 305 | pll,RecQ4,dpp,rhea,RpS8,nsl1,BthD,heix,krz,CG8149,nom,Pten,EloB | | 1.581e-04 | -8.75 | Formation of a pool of free 40S subunits | REACTOME pathways | R-DME-72689 | 105 | 13 | 3923 | 151 | RpS26,eIF3b,RpS3A,RpS8,RpS19a,RpL15,RpL36A,RpL35A,eIF3c,eIF3g1,RpS9,RpL5,RpL14 | | 1.585e-04 | -8.75 | mRNA Splicing - Major Pathway | REACTOME pathways | R-DME-72163 | 165 | 17 | 3923 | 151 | snf,c12.1,x16,sm,Cwc25,noi,Wdr33,Cpsf6,l(1)10Bb,glo,TfIIFbeta,Phf5a,Rpb11,Upf3,SF2,sqd,tsu | | 1.594e-04 | -8.74 | positive regulation of cellular biosynthetic process | biological process | GO:0031328 | 594 | 30 | 12345 | 305 | CNBP,TfIIFbeta,Upf3,MED21,Fmr1,Tcs3,Lst8,Dref,CG6244,PlexA,mnb,grau,Rcd5,CG4612,Rpt6,Pex3,Hrb27C,dgrn,su(Hw),Parp,d4,Rheb,eEF5,RpS9,Ncoa6,RecQ4,nsl1,rhi,CtBP,mbf1 | | 1.594e-04 | -8.74 | positive regulation of biosynthetic process | biological process | GO:0009891 | 594 | 30 | 12345 | 305 | Rheb,d4,Parp,su(Hw),dgrn,CtBP,rhi,mbf1,nsl1,RecQ4,Ncoa6,eEF5,RpS9,mnb,PlexA,CG6244,Fmr1,Tcs3,Dref,Lst8,Upf3,MED21,TfIIFbeta,CNBP,Hrb27C,Pex3,CG4612,Rcd5,Rpt6,grau | | 1.666e-04 | -8.70 | cellular component assembly | biological process | GO:0022607 | 1109 | 47 | 12345 | 305 | Rcd5,CG44194,pav,flr,l(2)37Bb,tkv,Fmr1,smt3,CG1791,Orc3,Rpn10,Vap33,CG8223,RecQ4,unc-119,Cul4,Pbp49,Tom20,Hrb27C,Nph,His4r,me31B,Rab14,RpL5,spoon,tsu,inc,RpS19a,Rab7,tsr,Pp1-87B,CG17493,glo,mon2,Rop,sqd,msd5,LIMK1,Dbp21E2,Acf,dpp,rhea,CIA30,CG12321,eIF4B,Rap1,Cdc42 | | 1.678e-04 | -8.69 | L13a-mediated translational silencing of Ceruloplasmin expression | REACTOME pathways | R-DME-156827 | 120 | 14 | 3923 | 151 | RpL5,RpS9,eIF3g1,eIF4B,RpL14,RpL36A,eIF3c,RpL35A,eIF3b,RpL15,RpS19a,RpS8,RpS3A,RpS26 | | 1.728e-04 | -8.66 | ion binding | molecular function | GO:0043167 | 2058 | 73 | 12176 | 289 | Mps1,spel1,dpp,CG11586,CG5589,Zpr1,Btk29A,me31B,x16,RagA-B,Ino80,noi,CG1218,Parp,dgrn,Bub1,Dbp80,Rpt6,Rab8,DNAlig1,PlexA,grau,Graf,NO66,CG17493,Rap1,Cdc42,dah,CG8915,Stim,CG8635,Cdk8,CG9330,RhoGAPp190,Usp16-45,RabX6,Stlk,tkv,Acbp1,LIMK1,CG1815,CG1637,CNBP,mnb,CG7646,nom,Dbp21E2,Rpn10,Sod2,Roc2,d4,Efa6,CCT4,alpha-Man-Ib,RecQ4,Fign,lic,Usp32,Rheb,Ufd4,Dref,CG6254,MICU1,CG5439,Rab14,CG32473,Rab7,Tctp,mus301,pav,RAF2,CG5516,pll | | 1.775e-04 | -8.64 | regulation of protein complex assembly | biological process | GO:0043254 | 100 | 10 | 12345 | 305 | spoon,Btk29A,Sgt,Cdc42,Arpc5,conu,flr,Arpc3A,Rpt6,cal1 | | 1.775e-04 | -8.64 | regulation of actin filament-based process | biological process | GO:0032970 | 100 | 10 | 12345 | 305 | Arpc3A,cta,flw,flr,Btk29A,Cdc42,conu,Arpc5,Lst8,spoon | | 1.816e-04 | -8.61 | ovarian follicle cell development | biological process | GO:0030707 | 327 | 20 | 12345 | 305 | tkv,spoon,Rab7,tsr,flr,eIF3b,me31B,Fs(2)Ket,Cul4,Parp,Pten,Cdc42,Rap1,Rop,smt3,rhi,sqd,rhea,dpp,RecQ4 | | 1.832e-04 | -8.60 | regulation of phosphorus metabolic process | biological process | GO:0051174 | 302 | 19 | 12345 | 305 | PPP1R15,Cdc42,Pten,Rap1,CG1791,mav,smt3,flw,CycA,dpp,sigmar,PlexA,I-2,mnb,Stlk,krz,lic,Tctp,sws | | 1.832e-04 | -8.60 | regulation of phosphate metabolic process | biological process | GO:0019220 | 302 | 19 | 12345 | 305 | Rap1,Pten,Cdc42,PPP1R15,dpp,CycA,smt3,mav,CG1791,flw,krz,Stlk,PlexA,sigmar,mnb,I-2,Tctp,sws,lic | | 1.840e-04 | -8.60 | RpL19 (Ribosomal protein L19) | protein interactions | 37995 | 108 | 12 | 9204 | 298 | RpS9,RpS19a,RpS8,RpL14,RpL15,RpS3A,RpL36A,RpL5,CG11586,RpS26,CG5174,RpL35A | | 1.891e-04 | -8.57 | columnar/cuboidal epithelial cell development | biological process | GO:0002066 | 328 | 20 | 12345 | 305 | tkv,spoon,Rab7,tsr,eIF3b,flr,me31B,Fs(2)Ket,Cul4,Pten,Cdc42,Parp,Rap1,Rop,smt3,rhi,sqd,rhea,dpp,RecQ4 | | 1.938e-04 | -8.55 | chromatin | cellular component | GO:0000785 | 403 | 23 | 11940 | 298 | His4r,Arp8,Acf,rhi,Oxp,Jarid2,His2Av,Rbp1,Ino80,CtBP,Rcd5,E(z),MED22,CG12659,HP4,MED26,sqd,Parp,RecQ4,E(bx),BEAF-32,d4,su(Hw) | | 1.939e-04 | -8.55 | axis specification | biological process | GO:0009798 | 207 | 15 | 12345 | 305 | Fmr1,Parp,glo,tsu,spoon,mus301,pll,dpp,mon2,lic,bai,14-3-3zeta,Hrb27C,flw,sqd | | 1.953e-04 | -8.54 | RpL29 (Ribosomal protein L29) | protein interactions | 37430 | 93 | 11 | 9204 | 298 | RpL5,CG17385,RpS26,RpL15,RpL14,RpS8,RpS3A,RpS19a,RpS9,RpL36A,RpL35A | | 1.979e-04 | -8.53 | mRNA Splicing | REACTOME pathways | R-DME-72172 | 168 | 17 | 3923 | 151 | Cpsf6,Wdr33,l(1)10Bb,snf,c12.1,x16,sm,Cwc25,noi,Upf3,SF2,tsu,sqd,glo,TfIIFbeta,Phf5a,Rpb11 | | 2.005e-04 | -8.51 | GTP hydrolysis and joining of the 60S ribosomal subunit | REACTOME pathways | R-DME-72706 | 122 | 14 | 3923 | 151 | RpL35A,eIF3c,RpL36A,RpL14,eIF4B,RpL5,eIF3g1,RpS9,RpS26,RpS3A,RpS8,RpS19a,RpL15,eIF3b | | 2.009e-04 | -8.51 | RpS5a (Ribosomal protein S5a) | protein interactions | 32700 | 109 | 12 | 9204 | 298 | Rheb,lic,RpL35A,RpL36A,RpL14,RpL15,RpS8,RpS3A,RpS9,RpS19a,RpS26,RpL5 | | 2.043e-04 | -8.50 | fly chr67C10-67C10\|3-31 | chromosome location | fly chr67C10-67C10\|3-31 | 2 | 2 | 25010 | 358 | Cdk8,I-2 | | 2.043e-04 | -8.50 | fly chr67C10-67C10\|3-31 cM | chromosome location | fly chr67C10-67C10\|3-31 cM | 2 | 2 | 25010 | 358 | Cdk8,I-2 | | 2.043e-04 | -8.50 | fly chr54D2-54D2\|2-83 | chromosome location | fly chr54D2-54D2\|2-83 | 2 | 2 | 25010 | 358 | Oxp,rhi | | 2.043e-04 | -8.50 | fly chr54D2-54D2\|2-83 cM | chromosome location | fly chr54D2-54D2\|2-83 cM | 2 | 2 | 25010 | 358 | Oxp,rhi | | 2.082e-04 | -8.48 | p38b (p38b MAP kinase) | protein interactions | 34780 | 51 | 8 | 9204 | 298 | Prosbeta1,dpp,Sod2,lic,tkv,smt3,CG8108,eRF1 | | 2.119e-04 | -8.46 | RpS20 (Ribosomal protein S20) | protein interactions | 42464 | 126 | 13 | 9204 | 298 | Dref,RpL35A,GEFmeso,RpS26,RpL5,RpL36A,RpS8,RpL14,RpL15,RpS3A,Bub1,RpS19a,RpS9 | | 2.124e-04 | -8.46 | GSE32901\_TH1\_VS\_TH17\_ENRICHED\_CD4\_TCELL\_DN | MSigDB lists | GSE32901\_TH1\_VS\_TH17\_ENRICHED\_CD4\_TCELL\_DN | 59 | 10 | 4161 | 184 | janA,Rcd5,Art1,Pect,Rpn10,CG12935,eIF3c,CG32576,sigmar,Tcs3 | | 2.150e-04 | -8.45 | RpL11 (Ribosomal protein L11) | protein interactions | 37235 | 94 | 11 | 9204 | 298 | RpL35A,Fmr1,RpL5,RpS26,RpL15,RpL14,RpS8,RpS3A,RpS19a,RpS9,RpL36A | | 2.270e-04 | -8.39 | cellular developmental process | biological process | GO:0048869 | 1818 | 68 | 12345 | 305 | pav,eIF3b,vnc,tkv,Fmr1,Dref,pins,smt3,Parp,sm,me31B,mnb,tsu,Nf1,tsr,heix,Pp1-87B,sqd,Exo84,E(z),mon2,nsl1,dpp,Cdk8,rhea,LIMK1,Fs(2)Ket,xit,Cdc42,snf,flr,grau,Girdin,CSN1b,E(bx),Btk29A,Arpc5,MED22,flw,CG10365,bai,CtBP,14-3-3zeta,RecQ4,RhoGAPp190,Rheb,BEAF-32,twin,Mps1,Cul4,Hrb27C,lic,Rab7,spoon,PlexA,glo,Stlk,rhi,Rop,cta,mav,Spn,CycA,HtrA2,mus301,Rap1,Pten,dgrn | | 2.275e-04 | -8.39 | Ino80 complex | cellular component | GO:0031011 | 13 | 4 | 11940 | 298 | CG12659,Ino80,Rcd5,Arp8 | | 2.275e-04 | -8.39 | DNA helicase complex | cellular component | GO:0033202 | 13 | 4 | 11940 | 298 | Ino80,CG12659,Arp8,Rcd5 | | 2.292e-04 | -8.38 | fly chr27C4-27C4\|2-23 | chromosome location | fly chr27C4-27C4\|2-23 | 9 | 3 | 25010 | 358 | x16,Hrb27C,Rat1 | | 2.292e-04 | -8.38 | fly chr27C4-27C4\|2-23 cM | chromosome location | fly chr27C4-27C4\|2-23 cM | 9 | 3 | 25010 | 358 | x16,Rat1,Hrb27C | | 2.362e-04 | -8.35 | RpS4 (Ribosomal protein S4) | protein interactions | 39484 | 95 | 11 | 9204 | 298 | RpS9,RpS19a,RpS8,RpL14,RpS3A,RpL15,RpL36A,Parp,RpL5,RpS26,RpL35A | | 2.362e-04 | -8.35 | RpS14b (Ribosomal protein S14b) | protein interactions | 47219 | 95 | 11 | 9204 | 298 | GEFmeso,RpL35A,RpL36A,RpS19a,RpS9,RpS8,RpL15,RpS3A,RpL14,RpS26,RpL5 | | 2.387e-04 | -8.34 | RpL10 (Ribosomal protein L10) | protein interactions | 43864 | 111 | 12 | 9204 | 298 | RpL36A,RpS19a,RpS9,RpS8,RpL15,RpS3A,RpL14,RpS26,RpL5,Lst8,Rheb,RpL35A | | 2.413e-04 | -8.33 | regulation of molecular function | biological process | GO:0065009 | 523 | 27 | 12345 | 305 | HtrA2,RhoGAPp190,Acf,CycA,14-3-3zeta,smt3,CG1791,dgrn,fzy,Pten,Cdc42,PPP1R15,sws,CG17883,Graf,lic,vap,Stim,Nf1,Stlk,ECSIT,conu,sigmar,PlexA,mnb,viaf,I-2 | | 2.443e-04 | -8.32 | GO\_MRNA\_METABOLIC\_PROCESS | MSigDB lists | GO\_MRNA\_METABOLIC\_PROCESS | 268 | 25 | 4161 | 184 | TfIIFbeta,tsu,CG4935,RpL5,Phf5a,l(1)10Bb,Rpb11,twin,RpS9,RpS8,RpL35A,eRF1,ubl,RpS26,CG11586,RpL14,me31B,sm,RpL36A,SF2,c12.1,noi,Rat1,RpL15,snf | | 2.454e-04 | -8.31 | chromatin remodeling | biological process | GO:0006338 | 104 | 10 | 12345 | 305 | Arp8,Jarid2,CtBP,CG12659,Nph,CG8223,Acf,E(bx),Ino80,Fmr1 | | 2.454e-04 | -8.31 | regulation of small GTPase mediated signal transduction | biological process | GO:0051056 | 104 | 10 | 12345 | 305 | Fgop2,conu,Nf1,Pp1-87B,Efa6,Slmap,GEFmeso,vap,14-3-3zeta,smt3 | | 2.472e-04 | -8.31 | single-stranded RNA binding | molecular function | GO:0003727 | 38 | 6 | 12176 | 289 | eIF4B,CNBP,snf,Hrb27C,CG4612,Pep | | 2.480e-04 | -8.30 | RpL22 (Ribosomal protein L22) | protein interactions | 31022 | 128 | 13 | 9204 | 298 | Parp,RpL36A,RpS8,RpL15,RpL14,RpS3A,RpS19a,RpS9,Efa6,RpS26,RpL5,RpL35A,smt3 | | 2.540e-04 | -8.28 | Nonsense Mediated Decay (NMD) enhanced by the Exon Junction Complex (EJC) | REACTOME pathways | R-DME-975957 | 110 | 13 | 3923 | 151 | RpS26,eRF1,Upf3,tsu,RpS3A,RpS8,RpS19a,RpL15,RpL35A,RpL36A,RpL14,RpL5,RpS9 | | 2.540e-04 | -8.28 | Nonsense-Mediated Decay (NMD) | REACTOME pathways | R-DME-927802 | 110 | 13 | 3923 | 151 | RpL14,RpS9,RpL5,RpL35A,RpL36A,RpS19a,RpS8,tsu,RpS3A,RpL15,RpS26,eRF1,Upf3 | | 2.546e-04 | -8.28 | translation regulator activity, nucleic acid binding | molecular function | GO:0090079 | 89 | 9 | 12176 | 289 | eEF5,eIF3c,smg,eIF3g1,eIF3b,eIF4B,eEF1delta,CG31957,eRF1 | | 2.589e-04 | -8.26 | mRNA surveillance pathway | KEGG pathways | ko03015 | 73 | 10 | 3198 | 117 | Wdr33,Pnn,flw,Wdr82,Cpsf6,eRF1,Pp1-87B,Upf3,tsu,Hrb27C | | 2.589e-04 | -8.26 | mRNA surveillance pathway | KEGG pathways | dme03015 | 73 | 10 | 3198 | 117 | flw,Pnn,Wdr33,eRF1,Cpsf6,Wdr82,Hrb27C,tsu,Upf3,Pp1-87B | | 2.593e-04 | -8.26 | intracellular transport | biological process | GO:0046907 | 640 | 31 | 12345 | 305 | bai,sqd,CG8860,flw,Exo84,Vap33,ema,Nup188,Fs(2)Ket,Rab8,PGAP1,rb,Sgt,SF2,Pex3,Tom20,RabX6,g,pav,CG30423,Dbp80,Cp190,CG6961,CG17883,Tim17b,Girdin,Rab14,Vps35,Rab7,Vps13D,Fmr1 | | 2.597e-04 | -8.26 | RpS13 (Ribosomal protein S13) | protein interactions | 34149 | 112 | 12 | 9204 | 298 | RpL35A,RpS19a,RpS9,RpS8,RpL15,RpL14,RpS3A,CG11334,RpL36A,Parp,RpL5,RpS26 | | 2.597e-04 | -8.26 | RpL13A (Ribosomal protein L13A) | protein interactions | 40687 | 112 | 12 | 9204 | 298 | CG5174,Arpc5,RpL35A,RpL36A,RpS8,RpL14,RpL15,RpS3A,RpS19a,RpS9,RpS26,RpL5 | | 2.669e-04 | -8.23 | GSE16450\_CTRL\_VS\_IFNA\_6H\_STIM\_MATURE\_NEURON\_CELL\_LINE\_DN | MSigDB lists | GSE16450\_CTRL\_VS\_IFNA\_6H\_STIM\_MATURE\_NEURON\_CELL\_LINE\_DN | 72 | 11 | 4161 | 184 | CG5589,Rab8,Bmcp,sm,CG8860,CG32576,14-3-3zeta,sws,rb,Nulp1,flr | | 2.772e-04 | -8.19 | eIF3m (eukaryotic translation initiation factor 3 subunit m) | protein interactions | 36565 | 29 | 6 | 9204 | 298 | Pten,smt3,eIF3c,Rheb,eIF3b,eIF3g1 | | 2.962e-04 | -8.12 | epithelial cell differentiation | biological process | GO:0030855 | 419 | 23 | 12345 | 305 | LIMK1,rhea,RecQ4,dpp,Rop,cta,smt3,rhi,sqd,Cul4,Cdc42,Parp,Pten,Rap1,xit,Fs(2)Ket,me31B,flr,eIF3b,tsr,tkv,spoon,Rab7 | | 3.032e-04 | -8.10 | AstC-R1 (Allatostatin C receptor 1) | protein interactions | 40020 | 41 | 7 | 9204 | 298 | RpL15,RpL35A,Hsp27,RpS26,CG42676,Pp1-87B,CtBP | | 3.069e-04 | -8.09 | oocyte dorsal/ventral axis specification | biological process | GO:0007310 | 38 | 6 | 12345 | 305 | Fmr1,dpp,glo,tsu,bai,flw | | 3.111e-04 | -8.08 | RpS15 (Ribosomal protein S15) | protein interactions | 36851 | 98 | 11 | 9204 | 298 | RpL35A,RpL5,RpS26,RpS9,RpS19a,Bub1,RpS8,RpL15,RpL14,RpS3A,RpL36A | | 3.111e-04 | -8.08 | RpL6 (Ribosomal protein L6) | protein interactions | 43723 | 98 | 11 | 9204 | 298 | RpL35A,Rheb,RpL5,RpS26,RpS3A,RpL15,RpL14,RpS8,RpS9,RpS19a,RpL36A | | 3.161e-04 | -8.06 | learning | biological process | GO:0007612 | 70 | 8 | 12345 | 305 | Btk29A,Fmr1,Nf1,Pp1-87B,mnb,me31B,Orc3,14-3-3zeta | | 3.172e-04 | -8.06 | GO\_POLY\_A\_RNA\_BINDING | MSigDB lists | GO\_POLY\_A\_RNA\_BINDING | 510 | 39 | 4161 | 184 | Fmr1,me31B,eIF3g1,CG9246,RpS8,Pp1-87B,mRpS9,RpS9,Rpn10,His4r,eIF3c,Fs(2)Ket,RpL15,Rat1,noi,CCT4,CG8635,Rop,sm,CG5589,RpS26,Parp,RpL14,eRF1,RpL35A,Art1,14-3-3zeta,mbf1,Phf5a,tsu,RpL5,Dbp21E2,snf,mRpS28,AGO3,CG4866,Tctp,RpL36A,SF2 | | 3.182e-04 | -8.05 | positive regulation of cellular component organization | biological process | GO:0051130 | 290 | 18 | 12345 | 305 | Rop,E(z),cal1,Ncoa6,eEF5,Rheb,Cdc42,Arpc3A,Rpt6,flr,Tctp,Cp190,Graf,PlexA,tkv,mnb,krz,Arpc5 | | 3.229e-04 | -8.04 | positive regulation of phosphorus metabolic process | biological process | GO:0010562 | 149 | 12 | 12345 | 305 | dpp,Tctp,mav,lic,CG1791,smt3,Cdc42,Rap1,Stlk,PPP1R15,sigmar,PlexA | | 3.229e-04 | -8.04 | positive regulation of phosphate metabolic process | biological process | GO:0045937 | 149 | 12 | 12345 | 305 | PlexA,sigmar,PPP1R15,Stlk,Cdc42,Rap1,CG1791,mav,lic,smt3,Tctp,dpp | | 3.267e-04 | -8.03 | 18SrRNA:CR41548 (18S ribosomal RNA:CR41548) | protein interactions | 5740812 | 83 | 10 | 9204 | 298 | RpL5,RpS26,RpS9,RpS19a,RpS8,RpS3A,RpL15,RpL14,RpL36A,RpL35A | | 3.267e-04 | -8.03 | 5.8SrRNA:CR40454 (5.8S ribosomal RNA) | protein interactions | 5740577 | 83 | 10 | 9204 | 298 | RpL35A,RpL5,RpS26,RpL15,RpL14,RpS8,RpS3A,RpS9,RpS19a,RpL36A | | 3.281e-04 | -8.02 | cell development | biological process | GO:0048468 | 1436 | 56 | 12345 | 305 | Girdin,grau,vnc,CSN1b,eIF3b,flr,Fmr1,Dref,Btk29A,tkv,RecQ4,RhoGAPp190,flw,14-3-3zeta,bai,smt3,Parp,Cul4,Mps1,BEAF-32,twin,me31B,sm,Hrb27C,lic,glo,Pp1-87B,Stlk,Nf1,tsr,Rab7,tsu,spoon,PlexA,Cdk8,rhea,Spn,dpp,mus301,LIMK1,HtrA2,E(z),Exo84,sqd,mon2,Rop,cta,mav,nsl1,rhi,Pten,Cdc42,Rap1,snf,Fs(2)Ket,xit | | 3.299e-04 | -8.02 | cell differentiation | biological process | GO:0030154 | 1773 | 66 | 12345 | 305 | Exo84,E(z),sqd,nsl1,mon2,Cdk8,rhea,dpp,LIMK1,Fs(2)Ket,xit,Cdc42,snf,sm,me31B,mnb,tsu,Pp1-87B,tsr,Nf1,smt3,Parp,eIF3b,pav,vnc,tkv,Fmr1,Dref,pins,cta,mav,Rop,rhi,CycA,Spn,mus301,HtrA2,Pten,Rap1,dgrn,Hrb27C,lic,Rab7,PlexA,spoon,glo,Stlk,CG10365,flw,CtBP,bai,14-3-3zeta,RecQ4,RhoGAPp190,BEAF-32,Rheb,twin,Mps1,Cul4,flr,Girdin,grau,CSN1b,E(bx),Btk29A,MED22 | | 3.337e-04 | -8.01 | Cap-dependent Translation Initiation | REACTOME pathways | R-DME-72737 | 128 | 14 | 3923 | 151 | eIF3b,RpS19a,RpS3A,RpS8,RpL15,RpS26,RpL5,RpS9,eIF3g1,RpL14,eIF4B,RpL36A,RpL35A,eIF3c | | 3.337e-04 | -8.01 | Eukaryotic Translation Initiation | REACTOME pathways | R-DME-72613 | 128 | 14 | 3923 | 151 | RpS26,RpS19a,RpS3A,RpS8,RpL15,eIF3b,RpL35A,eIF3c,RpL36A,RpL14,eIF4B,eIF3g1,RpS9,RpL5 | | 3.349e-04 | -8.00 | regulation of catalytic activity | biological process | GO:0050790 | 450 | 24 | 12345 | 305 | Pten,Cdc42,fzy,PPP1R15,CycA,Acf,RhoGAPp190,HtrA2,CG1791,smt3,conu,ECSIT,Stlk,Nf1,I-2,viaf,mnb,sigmar,PlexA,Graf,CG17883,sws,vap,lic | | 3.373e-04 | -7.99 | PAL\_PRMT5\_TARGETS\_UP | MSigDB lists | PAL\_PRMT5\_TARGETS\_UP | 86 | 12 | 4161 | 184 | His2Av,noi,CG17493,SF2,vap,Fign,ND-ASHI,Fmr1,sm,snf,fzy,pav | | 3.381e-04 | -7.99 | cytoplasmic ribonucleoprotein granule | cellular component | GO:0036464 | 88 | 9 | 11940 | 298 | Not3,Hrb27C,AGO3,smg,me31B,Dbp80,Fs(2)Ket,CG4612,Fmr1 | | 3.395e-04 | -7.99 | negative regulation of cellular protein metabolic process | biological process | GO:0032269 | 218 | 15 | 12345 | 305 | Hrb27C,flw,sqd,CG9246,me31B,sws,I-2,mnb,twin,sigmar,Not3,Fmr1,Pten,smg,krz | | 3.431e-04 | -7.98 | regulation of protein stability | biological process | GO:0031647 | 54 | 7 | 12345 | 305 | Skp2,14-3-3zeta,Vps35,Mlf,CSN1b,krz,Pten | | 3.501e-04 | -7.96 | cell cycle process | biological process | GO:0022402 | 564 | 28 | 12345 | 305 | Fign,DNAlig1,Dref,CG17493,Btk29A,Pp1-87B,pins,tsr,pav,Rcd5,Skp2,Bub1,grau,Tctp,Rheb,fzy,Mps1,Exo84,Rpn10,flw,Orc3,Rop,smt3,CycA,Cdk8,cal1,mus301,msd5 | | 3.514e-04 | -7.95 | MODULE\_83 | MSigDB lists | MODULE\_83 | 182 | 19 | 4161 | 184 | ubl,eIF3g1,RpL35A,RpS8,Pp1-87B,RpL14,CG5861,RpL5,CG4935,RpS9,eEF1delta,Mdh1,Prosalpha2,RpL15,CG4866,eIF3c,Tctp,RpL36A,Arpc3A | | 3.525e-04 | -7.95 | REACTOME\_TRANSLATION | MSigDB lists | REACTOME\_TRANSLATION | 112 | 14 | 4161 | 184 | eEF1delta,RpS9,RpL36A,RpL5,eIF3b,RpS26,RpL14,eIF3c,RpS8,RpL35A,eRF1,RpL15,eIF3g1,CG8860 | | 3.529e-04 | -7.95 | columnar/cuboidal epithelial cell differentiation | biological process | GO:0002065 | 344 | 20 | 12345 | 305 | dpp,RecQ4,rhea,smt3,rhi,Rop,sqd,Cul4,Rap1,Cdc42,Parp,Pten,Fs(2)Ket,me31B,eIF3b,flr,tsr,spoon,tkv,Rab7 | | 3.539e-04 | -7.95 | Rpt3 (Regulatory particle triple-A ATPase 3) | protein interactions | 32047 | 42 | 7 | 9204 | 298 | Rpn10,Prosbeta1,Ubqn,CG12321,Prosalpha2,Rpt6,Rheb | | 3.564e-04 | -7.94 | negative regulation of protein metabolic process | biological process | GO:0051248 | 219 | 15 | 12345 | 305 | sws,me31B,CG9246,sqd,flw,Hrb27C,krz,smg,Pten,Not3,Fmr1,sigmar,twin,mnb,I-2 | | 3.596e-04 | -7.93 | regulation of mRNA processing | biological process | GO:0050684 | 109 | 10 | 12345 | 305 | Hrb27C,sqd,Cpsf6,SF2,x16,Rbp1,Pep,tsu,Fmr1,snf | | 3.617e-04 | -7.92 | regulation of multicellular organismal development | biological process | GO:2000026 | 594 | 29 | 12345 | 305 | E(z),Orc3,mav,Rop,cta,CycA,Spn,dpp,RhoGAPp190,LIMK1,Rab8,Rheb,Cdc42,Pten,Rap1,Mlf,dgrn,Hrb27C,lic,me31B,mnb,tkv,tsu,PlexA,Fmr1,Dref,MED22,Nf1,heix | | 3.681e-04 | -7.91 | ribonucleoprotein granule | cellular component | GO:0035770 | 89 | 9 | 11940 | 298 | AGO3,Not3,Hrb27C,Fs(2)Ket,me31B,Dbp80,Fmr1,CG4612,smg | | 3.713e-04 | -7.90 | RpS24 (Ribosomal protein S24) | protein interactions | 37589 | 100 | 11 | 9204 | 298 | Rpb11,RpL35A,RpS19a,RpS9,RpS3A,RpL15,RpS8,RpL14,RpL36A,RpL5,RpS26 | | 3.806e-04 | -7.87 | protein-containing complex subunit organization | biological process | GO:0043933 | 625 | 30 | 12345 | 305 | Rop,CG1791,Rpn10,flw,Orc3,Vap33,Dbp21E2,CG8223,Acf,CG12321,CIA30,Nup188,eIF4B,Cdc42,Ino80,eRF1,Pbp49,Tom20,flr,CG44194,His4r,l(2)37Bb,me31B,tkv,E(bx),RpL5,RpS19a,inc,Pp1-87B,tsr | | 3.871e-04 | -7.86 | regulation of RNA splicing | biological process | GO:0043484 | 110 | 10 | 12345 | 305 | snf,Parp,tsu,Pep,Rbp1,x16,Cpsf6,SF2,Hrb27C,sqd | | 3.878e-04 | -7.86 | intracellular signal transduction | biological process | GO:0035556 | 373 | 21 | 12345 | 305 | pll,Rab14,Graf,RabX6,lic,Stlk,Nf1,Lst8,Btk29A,PlexA,Rab7,RhoGAPp190,Lrr47,spel1,Spn,cta,Cdc42,Pten,Rap1,Rheb,Rab8 | | 3.887e-04 | -7.85 | translation factor activity, RNA binding | molecular function | GO:0008135 | 75 | 8 | 12176 | 289 | eEF5,eIF3c,eIF3b,eIF3g1,eEF1delta,eIF4B,CG31957,eRF1 | | 3.895e-04 | -7.85 | regulation of protein phosphorylation | biological process | GO:0001932 | 245 | 16 | 12345 | 305 | Rap1,Pten,Cdc42,dpp,CycA,flw,smt3,mav,CG1791,krz,Stlk,mnb,sigmar,sws,Tctp,lic | | 3.903e-04 | -7.85 | RpL27 (Ribosomal protein L27) | protein interactions | 43103 | 117 | 12 | 9204 | 298 | Rheb,RpL35A,RpL36A,RpS19a,LIMK1,RpS9,RpL14,RpS3A,RpL15,RpS8,RpS26,RpL5 | | 3.970e-04 | -7.83 | RpS10a (Ribosomal protein S10a) | protein interactions | 43321 | 85 | 10 | 9204 | 298 | RpL5,RpS26,RpS19a,RpS9,RpS8,RpL15,RpS3A,RpL14,RpL36A,RpL35A | | 4.046e-04 | -7.81 | establishment of RNA localization | biological process | GO:0051236 | 91 | 9 | 12345 | 305 | SF2,Tom20,sqd,Dbp80,CG6961,tsu,Rab7,smg,Fmr1 | | 4.049e-04 | -7.81 | RpL36 (Ribosomal protein L36) | protein interactions | 31009 | 101 | 11 | 9204 | 298 | RpL5,RpS26,RpS8,RpL15,RpL14,RpS3A,RpS19a,RpS9,RpL36A,RpL35A,Rheb | | 4.057e-04 | -7.81 | Processing of Capped Intron-Containing Pre-mRNA | REACTOME pathways | R-DME-72203 | 212 | 19 | 3923 | 151 | sm,noi,Cwc25,snf,c12.1,x16,Nup188,CG6961,Cpsf6,Wdr33,l(1)10Bb,Rpb11,Phf5a,glo,TfIIFbeta,tsu,sqd,Upf3,SF2 | | 4.153e-04 | -7.79 | PILON\_KLF1\_TARGETS\_DN | MSigDB lists | PILON\_KLF1\_TARGETS\_DN | 701 | 49 | 4161 | 184 | spel1,Pten,Fign,Parp,CG5590,tsu,flr,Rap1,sws,E(z),CIA30,Roc2,CG15168,Fgop2,Mdh1,CG12321,Tsp97E,CG11920,Exo84,Lrr47,xit,MED21,Ino80,c12.1,CG5339,Pp1-87B,CG9246,Cdc42,mRpS9,Bmcp,me31B,Vps35,CtBP,TfIIFbeta,Stt3B,twin,RpS9,cta,smt3,Rat1,eIF3c,Fs(2)Ket,CG7766,flw,CG17493,bai,Pex3,noi,Skp2 | | 4.263e-04 | -7.76 | ATPase activity | molecular function | GO:0016887 | 283 | 17 | 12176 | 289 | E(bx),CG5589,Fign,RecQ4,Dbp80,Acf,spel1,Rpt6,sun,Arp8,Ino80,Dbp21E2,CG9330,CG8915,me31B,pav,mus301 | | 4.308e-04 | -7.75 | regulation of actin filament polymerization | biological process | GO:0030833 | 56 | 7 | 12345 | 305 | Arpc3A,spoon,flr,conu,Cdc42,Arpc5,Btk29A | | 4.308e-04 | -7.75 | negative regulation of translation | biological process | GO:0017148 | 56 | 7 | 12345 | 305 | Hrb27C,sqd,twin,Not3,Fmr1,me31B,smg | | 4.317e-04 | -7.75 | positive regulation of signaling | biological process | GO:0023056 | 403 | 22 | 12345 | 305 | Lst8,Fgop2,Pp1-87B,Vps35,tkv,sigmar,Efa6,Slmap,lic,Cdc42,Rap1,dgrn,Rheb,sun,dpp,RagA-B,flw,Nulp1,mav,CtBP,smt3,14-3-3zeta | | 4.317e-04 | -7.75 | positive regulation of cell communication | biological process | GO:0010647 | 403 | 22 | 12345 | 305 | dpp,14-3-3zeta,CtBP,smt3,mav,Nulp1,flw,RagA-B,dgrn,Rap1,Cdc42,sun,Rheb,Slmap,lic,Pp1-87B,Fgop2,Lst8,Efa6,sigmar,tkv,Vps35 | | 4.411e-04 | -7.73 | RpL21 (Ribosomal protein L21) | protein interactions | 35453 | 102 | 11 | 9204 | 298 | RpL35A,RpL5,RpS26,RpS9,RpS19a,RpL14,RpS8,RpS3A,RpL15,RpL36A,Parp | | 4.411e-04 | -7.73 | RpL18 (Ribosomal protein L18) | protein interactions | 38794 | 102 | 11 | 9204 | 298 | RpL5,RpS26,RpS19a,RpS9,RpL15,RpS8,RpL14,RpS3A,RpL36A,RpL35A,Rheb | | 4.520e-04 | -7.70 | Mer (Merlin) | protein interactions | 32979 | 172 | 15 | 9204 | 298 | CG12299,RAF2,CG9005,Tctp,dpp,CG5589,RpS9,Rop,flw,RpS8,Spn,CG5439,Rab7,rhea,Pnn | | 4.520e-04 | -7.70 | IKKbeta (I-kappaB kinase beta) | protein interactions | 44432 | 172 | 15 | 9204 | 298 | Pep,RpS19a,Stim,tsr,RpL5,Map60,CtBP,CG5174,CNBP,Cp190,rhea,smt3,nsl1,Pect,Graf | | 4.697e-04 | -7.66 | olfactory learning | biological process | GO:0008355 | 41 | 6 | 12345 | 305 | Nf1,Pp1-87B,Fmr1,14-3-3zeta,Orc3,mnb | | 4.763e-04 | -7.65 | positive regulation of intracellular signal transduction | biological process | GO:1902533 | 178 | 13 | 12345 | 305 | Pp1-87B,Fgop2,Lst8,Cdc42,Rap1,sigmar,Efa6,Rheb,Slmap,lic,smt3,14-3-3zeta,RagA-B | | 4.763e-04 | -7.65 | RNA localization | biological process | GO:0006403 | 178 | 13 | 12345 | 305 | Dbp80,Hrb27C,sqd,Tom20,SF2,mus301,CG6961,Rab7,tsu,spoon,Fmr1,glo,smg | | 4.795e-04 | -7.64 | RpL34a (Ribosomal protein L34a) | protein interactions | 43169 | 87 | 10 | 9204 | 298 | RpS26,RpL5,RpL36A,RpS9,RpS19a,RpS8,RpL15,RpL14,RpS3A,RpL35A | | 4.799e-04 | -7.64 | RpS25 (Ribosomal protein S25) | protein interactions | 41343 | 103 | 11 | 9204 | 298 | RpL35A,RpL5,pins,RpS26,RpS9,RpS19a,RpS3A,RpL14,RpL15,RpS8,RpL36A | | 4.803e-04 | -7.64 | ZHANG\_TLX\_TARGETS\_36HR\_DN | MSigDB lists | ZHANG\_TLX\_TARGETS\_36HR\_DN | 65 | 10 | 4161 | 184 | CycA,SF2,vap,Cul4,E(z),pav,DNAlig1,Fign,Lrr47,Fmr1 | | 4.806e-04 | -7.64 | GSE20366\_EX\_VIVO\_VS\_HOMEOSTATIC\_CONVERSION\_TREG\_DN | MSigDB lists | GSE20366\_EX\_VIVO\_VS\_HOMEOSTATIC\_CONVERSION\_TREG\_DN | 43 | 8 | 4161 | 184 | CycA,CG41378,Stt3B,14-3-3zeta,CG5339,pins,fzy,Lrr47 | | 4.813e-04 | -7.64 | establishment of localization in cell | biological process | GO:0051649 | 784 | 35 | 12345 | 305 | CG8860,sqd,Exo84,Rop,ema,Fs(2)Ket,PGAP1,Sgt,Tom20,RabX6,Tim17b,Rab14,Cp190,CG6961,Rab7,Vps35,Pp1-87B,flw,bai,Vap33,Nup188,Rab8,rb,Mps1,pav,CG30423,Dbp80,SF2,Pex3,g,CG17883,Girdin,Fmr1,pins,Vps13D | | 4.841e-04 | -7.63 | nucleobase-containing compound catabolic process | biological process | GO:0034655 | 134 | 11 | 12345 | 305 | tsu,twin,Aps,Upf3,smg,Not3,CG6171,Rat1,sqd,CG30105,Zpr1 | | 4.854e-04 | -7.63 | helicase activity | molecular function | GO:0004386 | 97 | 9 | 12176 | 289 | RecQ4,Dbp80,mus301,Arp8,Ino80,Dbp21E2,CG8915,me31B,CG5589 | | 4.886e-04 | -7.62 | macromolecule catabolic process | biological process | GO:0009057 | 490 | 25 | 12345 | 305 | Cul4,Ubqn,Sgt,Not3,fzy,twin,Usp32,Roc2,Prosbeta1,Rpn10,sqd,smg,Upf3,CG6171,Ufd4,tsu,inc,Usp16-45,Zpr1,alpha-Man-Ib,Skp2,Rpt6,Rat1,Prosalpha2,CG30105 | | 5.152e-04 | -7.57 | regulation of neuron projection development | biological process | GO:0010975 | 114 | 10 | 12345 | 305 | LIMK1,RhoGAPp190,me31B,mav,Rop,Fmr1,Pten,Cdc42,PlexA,Rheb | | 5.157e-04 | -7.57 | oocyte axis specification | biological process | GO:0007309 | 135 | 11 | 12345 | 305 | tsu,glo,Fmr1,14-3-3zeta,bai,lic,mon2,flw,sqd,Hrb27C,dpp | | 5.266e-04 | -7.55 | purine nucleotide binding | molecular function | GO:0017076 | 821 | 35 | 12176 | 289 | Dbp21E2,RagA-B,Ino80,mnb,me31B,Btk29A,LIMK1,Acbp1,tkv,Stlk,CG5589,RabX6,RhoGAPp190,spel1,Mps1,CG9330,Cdk8,pll,pav,CG8915,Cdc42,mus301,Rap1,Rab7,Rab14,cta,DNAlig1,Rheb,CCT4,lic,Fign,Rab8,Rpt6,Dbp80,Bub1 | | 5.447e-04 | -7.52 | GSE21063\_3H\_VS\_16H\_ANTI\_IGM\_STIM\_NFATC1\_KOBCELL\_DN | MSigDB lists | GSE21063\_3H\_VS\_16H\_ANTI\_IGM\_STIM\_NFATC1\_KOBCELL\_DN | 66 | 10 | 4161 | 184 | eIF3b,xit,Rbbp5,CG5535,Lst8,Aps,fzy,eIF3c,CG6907,RpL14 | | 5.448e-04 | -7.52 | His3:CG31613 (histone H3) | protein interactions | 318847 | 73 | 9 | 9204 | 298 | Pten,Dek,rhi,14-3-3zeta,Art1,Acf,E(z),E(bx),Nph | | 5.509e-04 | -7.50 | sexual reproduction | biological process | GO:0019953 | 1300 | 51 | 12345 | 305 | pav,lncRNA:sphinx,flr,eIF3b,CSN1b,vnc,grau,E(bx),tkv,MED22,Btk29A,Fmr1,14-3-3zeta,bai,smt3,flw,cal1,RecQ4,twin,BEAF-32,Cul4,Mps1,Parp,lic,Hrb27C,Nph,me31B,spoon,tsu,Rab7,tsr,noi,Pp1-87B,glo,rhi,nsl1,Rop,mon2,sqd,Exo84,HtrA2,LIMK1,mus301,dpp,rhea,Fs(2)Ket,snf,Rap1,fzy,Pten,Cdc42 | | 5.509e-04 | -7.50 | multi-organism reproductive process | biological process | GO:0044703 | 1300 | 51 | 12345 | 305 | Exo84,sqd,nsl1,Rop,mon2,rhi,rhea,dpp,mus301,HtrA2,LIMK1,Fs(2)Ket,Pten,Cdc42,fzy,Rap1,snf,Hrb27C,lic,me31B,Nph,Rab7,tsu,spoon,glo,Pp1-87B,tsr,noi,flw,14-3-3zeta,smt3,bai,RecQ4,cal1,BEAF-32,twin,Parp,Cul4,Mps1,flr,eIF3b,lncRNA:sphinx,pav,grau,vnc,CSN1b,tkv,E(bx),Fmr1,Btk29A,MED22 | | 5.522e-04 | -7.50 | negative regulation of cellular macromolecule biosynthetic process | biological process | GO:2000113 | 438 | 23 | 12345 | 305 | twin,CG6066,su(Hw),snf,d4,Ino80,Not3,CG6254,CG9246,CtBP,rhi,Nulp1,sqd,E(z),NO66,Tgi,Acf,CG12299,rhea,smg,Fmr1,Hrb27C,me31B | | 5.615e-04 | -7.48 | myosin phosphatase activity | molecular function | GO:0017018 | 2 | 2 | 12176 | 289 | Pp1-87B,flw | | 5.617e-04 | -7.48 | positive regulation of macromolecule biosynthetic process | biological process | GO:0010557 | 552 | 27 | 12345 | 305 | grau,Rcd5,Rpt6,CG4612,Hrb27C,Upf3,MED21,TfIIFbeta,CNBP,Dref,Lst8,Tcs3,Fmr1,CG6244,mnb,eEF5,Ncoa6,RpS9,RecQ4,CtBP,rhi,mbf1,nsl1,dgrn,su(Hw),d4,Rheb | | 5.621e-04 | -7.48 | DEAD/DEAH\_box\_helicase\_dom | interpro domains | IPR011545 | 50 | 7 | 11383 | 331 | mus301,CG5589,RecQ4,Dbp80,me31B,Dbp21E2,CG8915 | | 5.733e-04 | -7.46 | JIANG\_HYPOXIA\_NORMAL | MSigDB lists | JIANG\_HYPOXIA\_NORMAL | 91 | 12 | 4161 | 184 | 14-3-3zeta,CG41378,Dbp80,cta,Cdk8,Vap33,Fs(2)Ket,Tom20,CG4866,CG3842,Cdc42,Sod2 | | 5.764e-04 | -7.46 | negative regulation of gene expression | biological process | GO:0010629 | 582 | 28 | 12345 | 305 | twin,CG6066,su(Hw),snf,Ino80,d4,Not3,rhi,CG6254,CtBP,CG9246,Nulp1,sqd,E(z),NO66,Tgi,dpp,CG12299,Acf,rhea,tsu,tkv,Upf3,smg,Fmr1,AGO3,Hrb27C,me31B | | 5.813e-04 | -7.45 | fly chr86C6-86C6\|3-50 cM | chromosome location | fly chr86C6-86C6\|3-50 cM | 12 | 3 | 25010 | 358 | Rbp1,TfIIFbeta,Art1 | | 5.813e-04 | -7.45 | fly chr86C6-86C6\|3-50 | chromosome location | fly chr86C6-86C6\|3-50 | 12 | 3 | 25010 | 358 | Art1,TfIIFbeta,Rbp1 | | 6.011e-04 | -7.42 | regulation of Ras protein signal transduction | biological process | GO:0046578 | 96 | 9 | 12345 | 305 | vap,14-3-3zeta,smt3,Slmap,GEFmeso,Efa6,Fgop2,Nf1,Pp1-87B | | 6.030e-04 | -7.41 | dl (dorsal) | protein interactions | 35047 | 74 | 9 | 9204 | 298 | bai,Dlip2,RpL15,pll,MED21,CtBP,RpS26,Cdk8,smt3 | | 6.072e-04 | -7.41 | fly chr80F-80F\|3-47 | chromosome location | fly chr80F-80F\|3-47 | 3 | 2 | 25010 | 358 | RpL15,Dbp80 | | 6.072e-04 | -7.41 | fly chr95D10-95D10\|3-82 | chromosome location | fly chr95D10-95D10\|3-82 | 3 | 2 | 25010 | 358 | Pisd,CG5986 | | 6.072e-04 | -7.41 | fly chr80F-80F\|3-47 cM | chromosome location | fly chr80F-80F\|3-47 cM | 3 | 2 | 25010 | 358 | Dbp80,RpL15 | | 6.072e-04 | -7.41 | fly chr95D10-95D10\|3-82 cM | chromosome location | fly chr95D10-95D10\|3-82 cM | 3 | 2 | 25010 | 358 | CG5986,Pisd | | 6.085e-04 | -7.40 | oxaloacetate metabolic process | biological process | GO:0006107 | 2 | 2 | 12345 | 305 | CG8132,Mdh1 | | 6.085e-04 | -7.40 | histone H4-K5 acetylation | biological process | GO:0043981 | 2 | 2 | 12345 | 305 | Eaf6,Rcd5 | | 6.085e-04 | -7.40 | zygotic specification of dorsal/ventral axis | biological process | GO:0007352 | 2 | 2 | 12345 | 305 | pll,dpp | | 6.138e-04 | -7.40 | sta (stubarista) | protein interactions | 31106 | 106 | 11 | 9204 | 298 | RpL35A,Pten,RpL5,RpS26,RpS19a,RpS9,RpS3A,RpL14,RpS8,RpL15,RpL36A | | 6.208e-04 | -7.38 | Small\_GTPase | interpro domains | IPR001806 | 66 | 8 | 11383 | 331 | Rab7,RabX6,Rab14,Rab8,Rap1,Rheb,RhoGAPp190,Cdc42 | | 6.209e-04 | -7.38 | somatic ring canal | cellular component | GO:0098549 | 2 | 2 | 11940 | 298 | vsg,pav | | 6.209e-04 | -7.38 | PTW/PP1 phosphatase complex | cellular component | GO:0072357 | 2 | 2 | 11940 | 298 | Wdr82,flw | | 6.211e-04 | -7.38 | oocyte construction | biological process | GO:0007308 | 138 | 11 | 12345 | 305 | 14-3-3zeta,bai,lic,mon2,sqd,flw,Hrb27C,dpp,tsu,glo,Fmr1 | | 6.295e-04 | -7.37 | RpL37A (Ribosomal protein L37A) | protein interactions | 44783 | 90 | 10 | 9204 | 298 | RpL5,RpS26,RpS9,RpS19a,RpS8,RpL15,RpS3A,RpL14,RpL36A,RpL35A | | 6.295e-04 | -7.37 | RpL27A (Ribosomal protein L27A) | protein interactions | 33654 | 90 | 10 | 9204 | 298 | RpL35A,RpL5,RpS26,RpS9,RpS19a,RpL14,RpS8,RpL15,RpS3A,RpL36A | | 6.417e-04 | -7.35 | GARGALOVIC\_RESPONSE\_TO\_OXIDIZED\_PHOSPHOLIPIDS\_TURQUOISE\_DN | MSigDB lists | GARGALOVIC\_RESPONSE\_TO\_OXIDIZED\_PHOSPHOLIPIDS\_TURQUOISE\_DN | 17 | 5 | 4161 | 184 | fzy,Dbp21E2,CycA,CG12935,Skp2 | | 6.500e-04 | -7.34 | smo (smoothened) | protein interactions | 33196 | 60 | 8 | 9204 | 298 | flw,smt3,eRF1,Pp1-87B,tkv,krz,CG17493,pav | | 6.596e-04 | -7.32 | regulation of actin polymerization or depolymerization | biological process | GO:0008064 | 60 | 7 | 12345 | 305 | conu,Arpc5,Cdc42,Btk29A,flr,Arpc3A,spoon | | 6.596e-04 | -7.32 | regulation of actin filament length | biological process | GO:0030832 | 60 | 7 | 12345 | 305 | flr,spoon,Arpc3A,Btk29A,Arpc5,conu,Cdc42 | | 6.641e-04 | -7.32 | GCM\_CSNK2B | MSigDB lists | GCM\_CSNK2B | 56 | 9 | 4161 | 184 | RpL14,snf,eIF3c,sm,Pp1-87B,RpS8,14-3-3zeta,Art1,l(1)10Bb | | 6.649e-04 | -7.32 | RpLP2 (Ribosomal protein LP2) | protein interactions | 36855 | 107 | 11 | 9204 | 298 | RpS26,RpL5,RpL36A,RpS3A,RpL14,RpL15,RpS8,RpS19a,RpS9,Rap1,RpL35A | | 6.741e-04 | -7.30 | transforming growth factor beta receptor signaling pathway | biological process | GO:0007179 | 17 | 4 | 12345 | 305 | Cdc42,dpp,mav,tkv | | 6.874e-04 | -7.28 | Rho1 (Rho1) | protein interactions | 36775 | 91 | 10 | 9204 | 298 | cta,PlexA,tkv,RhoGAPp190,Cdc42,flw,LIMK1,Parp,tsr,conu | | 6.874e-04 | -7.28 | RpL35 (Ribosomal protein L35) | protein interactions | 31483 | 91 | 10 | 9204 | 298 | RpL35A,RpS3A,RpL15,RpS8,RpL14,RpS19a,RpS9,RpL36A,RpL5,RpS26 | | 7.080e-04 | -7.25 | negative regulation of macromolecule biosynthetic process | biological process | GO:0010558 | 446 | 23 | 12345 | 305 | Hrb27C,me31B,Fmr1,smg,E(z),NO66,Nulp1,sqd,CtBP,CG9246,rhi,CG6254,rhea,Acf,CG12299,Tgi,CG6066,twin,d4,Not3,Ino80,snf,su(Hw) | | 7.161e-04 | -7.24 | His2B:CG17949 (histone H2B) | protein interactions | 326273 | 23 | 5 | 9204 | 298 | Dek,Acf,His2Av,Parp,Nph | | 7.161e-04 | -7.24 | CG3294 (uncharacterized protein) | protein interactions | 33677 | 23 | 5 | 9204 | 298 | Cwc25,RAF2,Rbp1,grau,rb | | 7.161e-04 | -7.24 | dup (double parked) | protein interactions | 47121 | 23 | 5 | 9204 | 298 | Fgop2,Cul4,Slmap,RAF2,Ubqn | | 7.296e-04 | -7.22 | negative regulation of cellular amide metabolic process | biological process | GO:0034249 | 61 | 7 | 12345 | 305 | twin,sqd,Hrb27C,smg,me31B,Not3,Fmr1 | | 7.349e-04 | -7.22 | DEAD | pfam domains | PF00270 | 50 | 7 | 10480 | 319 | CG5589,RecQ4,Dbp21E2,CG8915,me31B,mus301,Dbp80 | | 7.376e-04 | -7.21 | GO\_REGULATION\_OF\_ORGANELLE\_ORGANIZATION | MSigDB lists | GO\_REGULATION\_OF\_ORGANELLE\_ORGANIZATION | 355 | 29 | 4161 | 184 | Rop,Rcd5,janA,krz,Arpc3A,CCT4,Ino80,CG12321,Fs(2)Ket,AGO3,vap,CG17883,flr,14-3-3zeta,Lst8,twin,Arpc5,LIMK1,Ubqn,pav,spel1,ubl,fzy,CG9246,Cdc42,HtrA2,pins,Pten,Parp | | 7.450e-04 | -7.20 | modification-dependent protein binding | molecular function | GO:0140030 | 31 | 5 | 12176 | 289 | Graf,E(bx),Ubqn,Rpn10,Vps13D | | 7.497e-04 | -7.20 | RpL32 (Ribosomal protein L32) | protein interactions | 43573 | 92 | 10 | 9204 | 298 | RpS26,RpL5,RpL36A,RpS3A,RpL14,RpL15,RpS8,RpS9,RpS19a,RpL35A | | 7.525e-04 | -7.19 | regulation of mRNA splicing, via spliceosome | biological process | GO:0048024 | 99 | 9 | 12345 | 305 | snf,tsu,Rbp1,x16,Pep,sqd,Hrb27C,SF2,Cpsf6 | | 7.544e-04 | -7.19 | structural constituent of ribosome | molecular function | GO:0003735 | 271 | 16 | 12176 | 289 | mRpS33,RpS9,RpS3A,mRpS28,RpS8,RpL35A,mRpL34,mRpS9,RpL36A,RpL5,RpL15,mRpS18A,RpS26,RpL14,mRpL30,RpS19a | | 7.563e-04 | -7.19 | KIM\_WT1\_TARGETS\_8HR\_DN | MSigDB lists | KIM\_WT1\_TARGETS\_8HR\_DN | 26 | 6 | 4161 | 184 | RpS9,mbf1,CtBP,Sod2,TfIIFbeta,PGAP1 | | 7.626e-04 | -7.18 | regulation of transcriptional start site selection at RNA polymerase II promoter | biological process | GO:0001178 | 8 | 3 | 12345 | 305 | Rbp1,x16,SF2 | | 7.627e-04 | -7.18 | ribosome biogenesis | biological process | GO:0042254 | 211 | 14 | 12345 | 305 | CG5589,RpS19a,RpL5,RpL35A,CG11920,CG14805,CG4866,CG9246,RpS8,Nopp140,CG32075,Dbp21E2,mRpS9,RpL14 | | 7.641e-04 | -7.18 | mitotic cell cycle, embryonic | biological process | GO:0045448 | 30 | 5 | 12345 | 305 | smt3,14-3-3zeta,E(z),CycA,Fmr1 | | 7.674e-04 | -7.17 | cognition | biological process | GO:0050890 | 164 | 12 | 12345 | 305 | Pp1-87B,Nf1,Fmr1,Btk29A,spoon,mnb,me31B,CG4612,14-3-3zeta,MICU1,vsg,Orc3 | | 7.674e-04 | -7.17 | learning or memory | biological process | GO:0007611 | 164 | 12 | 12345 | 305 | me31B,CG4612,14-3-3zeta,vsg,MICU1,Orc3,Pp1-87B,Nf1,Fmr1,Btk29A,spoon,mnb | | 7.674e-04 | -7.17 | negative regulation of cellular component organization | biological process | GO:0051129 | 164 | 12 | 12345 | 305 | spoon,Mps1,su(Hw),Nf1,Pten,Fmr1,Sgt,CG9246,flr,CycA,Bub1,Spn | | 7.823e-04 | -7.15 | GSE46606\_DAY1\_VS\_DAY3\_CD40L\_IL2\_IL5\_STIMULATED\_IRF4HIGH\_BCELL\_UP | MSigDB lists | GSE46606\_DAY1\_VS\_DAY3\_CD40L\_IL2\_IL5\_STIMULATED\_IRF4HIGH\_BCELL\_UP | 69 | 10 | 4161 | 184 | CG2938,CG12321,mus301,Coq5,Mdh1,RpS26,Parp,Vap33,CCT4,CG34148 | | 7.853e-04 | -7.15 | protein phosphatase type 1 complex | cellular component | GO:0000164 | 8 | 3 | 11940 | 298 | Pp1-87B,PPP1R15,flw | | 7.966e-04 | -7.14 | GO\_NEGATIVE\_REGULATION\_OF\_NEUROTRANSMITTER\_SECRETION | MSigDB lists | GO\_NEGATIVE\_REGULATION\_OF\_NEUROTRANSMITTER\_SECRETION | 5 | 3 | 4161 | 184 | Fmr1,Rap1,Nf1 | | 7.966e-04 | -7.14 | GO\_POSITIVE\_REGULATION\_OF\_EXTRINSIC\_APOPTOTIC\_SIGNALING\_PATHWAY\_VIA\_DEATH\_DOMAIN\_RECEPTORS | MSigDB lists | GO\_POSITIVE\_REGULATION\_OF\_EXTRINSIC\_APOPTOTIC\_SIGNALING\_PATHWAY\_VIA\_DEATH\_DOMAIN\_RECEPTORS | 5 | 3 | 4161 | 184 | Nf1,tkv,Pten | | 7.966e-04 | -7.14 | LIU\_IL13\_MEMORY\_MODEL\_UP | MSigDB lists | LIU\_IL13\_MEMORY\_MODEL\_UP | 5 | 3 | 4161 | 184 | unc-119,krz,Cdc42 | | 8.084e-04 | -7.12 | actin filament-based process | biological process | GO:0030029 | 262 | 16 | 12345 | 305 | flr,pav,Arpc3A,Girdin,Arpc5,Btk29A,tsr,Exo84,mon2,cta,rhea,Spn,LIMK1,xit,Pten,Cdc42 | | 8.094e-04 | -7.12 | germarium-derived egg chamber formation | biological process | GO:0007293 | 100 | 9 | 12345 | 305 | Btk29A,Fmr1,tsu,CSN1b,mus301,14-3-3zeta,nsl1,sqd,flw | | 8.122e-04 | -7.12 | pAbp (poly(A) binding protein) | protein interactions | 37070 | 62 | 8 | 9204 | 298 | Fmr1,twin,me31B,CtBP,Art1,Not3,Hrb27C,sqd | | 8.165e-04 | -7.11 | regucalcin (regucalcin) | protein interactions | 32165 | 93 | 10 | 9204 | 298 | viaf,eEF5,Tctp,flr,eEF1delta,CG5174,eIF3b,14-3-3zeta,Graf,smt3 | | 8.336e-04 | -7.09 | ssh (slingshot) | protein interactions | 42986 | 14 | 4 | 9204 | 298 | LIMK1,Vap33,tsr,Cdc42 | | 8.337e-04 | -7.09 | Ras | pfam domains | PF00071 | 66 | 8 | 10480 | 319 | Rap1,RabX6,Rab8,Rheb,Rab14,Cdc42,RhoGAPp190,Rab7 | | 8.396e-04 | -7.08 | RpL9 (Ribosomal protein L9) | protein interactions | 34526 | 110 | 11 | 9204 | 298 | RpL5,RpS26,RpS3A,RpS8,RpL15,RpL14,RpS19a,RpS9,RpL36A,RpL35A,Rheb | | 8.430e-04 | -7.08 | fly chr35F1-35F1\|2-52 | chromosome location | fly chr35F1-35F1\|2-52 | 30 | 4 | 25010 | 358 | CG5861,heix,CG4935,fzy | | 8.430e-04 | -7.08 | fly chr35F1-35F1\|2-52 cM | chromosome location | fly chr35F1-35F1\|2-52 cM | 30 | 4 | 25010 | 358 | CG4935,heix,CG5861,fzy | | 8.431e-04 | -7.08 | CIZ1 | interpro domains | IPR026811 | 2 | 2 | 11383 | 331 | Pep,CG8108 | | 8.431e-04 | -7.08 | Ribosomal\_S4/S9 | interpro domains | IPR022801 | 2 | 2 | 11383 | 331 | CG4866,RpS9 | | 8.431e-04 | -7.08 | Ribosomal\_S4/S9\_N | interpro domains | IPR001912 | 2 | 2 | 11383 | 331 | RpS9,CG4866 | | 8.508e-04 | -7.07 | purine ribonucleotide binding | molecular function | GO:0032555 | 811 | 34 | 12176 | 289 | CG9330,Cdk8,pll,pav,CG8915,Cdc42,mus301,Rap1,Rab7,Rab14,Rheb,DNAlig1,CCT4,lic,Fign,Rab8,Rpt6,Dbp80,Bub1,Dbp21E2,RagA-B,Ino80,mnb,me31B,Btk29A,LIMK1,Acbp1,tkv,Stlk,CG5589,RabX6,spel1,RhoGAPp190,Mps1 | | 8.521e-04 | -7.07 | RNA catabolic process | biological process | GO:0006401 | 81 | 8 | 12345 | 305 | CG30105,sqd,Zpr1,twin,tsu,smg,Upf3,Not3 | | 8.535e-04 | -7.07 | histone methyltransferase complex | cellular component | GO:0035097 | 62 | 7 | 11940 | 298 | Rbbp5,Ncoa6,Wdr82,CG6254,Rcd5,E(z),Jarid2 | | 8.637e-04 | -7.05 | positive regulation of response to stimulus | biological process | GO:0048584 | 481 | 24 | 12345 | 305 | lic,pll,Slmap,sigmar,Efa6,tkv,Vps35,Pp1-87B,Fgop2,Lst8,smt3,14-3-3zeta,CtBP,mav,flw,Nulp1,RagA-B,dpp,sun,Rheb,dgrn,Rap1,Cdc42,Parp | | 8.643e-04 | -7.05 | gig (gigas) | protein interactions | 40201 | 222 | 17 | 9204 | 298 | Rheb,14-3-3zeta,eIF3b,Pp1-87B,Fmr1,CG9246,c12.1,RpS9,pins,RagA-B,pav,Dref,eIF3c,TfIIFbeta,RpL14,eIF3g1,sqd | | 8.783e-04 | -7.04 | REACTOME\_NONSENSE\_MEDIATED\_DECAY\_ENHANCED\_BY\_THE\_EXON\_JUNCTION\_COMPLEX | MSigDB lists | REACTOME\_NONSENSE\_MEDIATED\_DECAY\_ENHANCED\_BY\_THE\_EXON\_JUNCTION\_COMPLEX | 70 | 10 | 4161 | 184 | RpL15,eRF1,RpL35A,RpS8,RpL14,RpS26,RpL5,tsu,RpL36A,RpS9 | | 8.825e-04 | -7.03 | positive regulation of Ras protein signal transduction | biological process | GO:0046579 | 46 | 6 | 12345 | 305 | Pp1-87B,Slmap,Fgop2,Efa6,smt3,14-3-3zeta | | 8.825e-04 | -7.03 | positive regulation of small GTPase mediated signal transduction | biological process | GO:0051057 | 46 | 6 | 12345 | 305 | Slmap,Fgop2,Pp1-87B,Efa6,14-3-3zeta,smt3 | | 8.844e-04 | -7.03 | mitotic cell cycle process | biological process | GO:1903047 | 290 | 17 | 12345 | 305 | Rheb,Mps1,Rpn10,smt3,Rop,CycA,Cdk8,msd5,Fign,DNAlig1,pins,tsr,Pp1-87B,pav,Skp2,Bub1,Tctp | | 8.859e-04 | -7.03 | regulation of cell morphogenesis | biological process | GO:0022604 | 144 | 11 | 12345 | 305 | Cdc42,Pten,Fmr1,Rap1,PlexA,Rheb,RhoGAPp190,LIMK1,dpp,me31B,cta | | 8.881e-04 | -7.03 | RpL26 (Ribosomal protein L26) | protein interactions | 40060 | 94 | 10 | 9204 | 298 | RpL35A,RpS26,RpL5,RpL36A,RpL15,RpS3A,RpS8,RpL14,RpS9,RpS19a | | 8.929e-04 | -7.02 | larval somatic muscle development | biological process | GO:0007526 | 31 | 5 | 12345 | 305 | EloB,nsl1,nom,rhea,pll | | 9.002e-04 | -7.01 | regulation of chromosome organization | biological process | GO:0033044 | 167 | 12 | 12345 | 305 | Mps1,su(Hw),CG9246,Jarid2,E(z),Ncoa6,cal1,Cp190,Tctp,Bub1,CycA,Acf | | 9.028e-04 | -7.01 | regulation of multicellular organismal process | biological process | GO:0051239 | 781 | 34 | 12345 | 305 | Rab8,Rheb,RhoGAPp190,Orc3,bai,Dref,Fmr1,MED22,tkv,Pten,Cdc42,Rap1,Mlf,DCTN3-p24,dgrn,CycA,dpp,Spn,LIMK1,E(z),Rop,cta,mav,Nf1,heix,inc,mnb,tsu,PlexA,me31B,Tctp,Hrb27C,lic,Sod2 | | 9.057e-04 | -7.01 | RpS23 (Ribosomal protein S23) | protein interactions | 36576 | 111 | 11 | 9204 | 298 | RpS26,RpL5,RpL36A,RpS3A,RpS8,RpL14,RpL15,RpS19a,RpS9,RpL35A,Rheb | | 9.057e-04 | -7.01 | RpS27 (Ribosomal protein S27) | protein interactions | 43039 | 111 | 11 | 9204 | 298 | RpS19a,RpS9,RpS8,RpL15,RpL14,RpS3A,RpL36A,RpL5,RpS26,rb,RpL35A | | 9.139e-04 | -7.00 | Nonsense Mediated Decay (NMD) independent of the Exon Junction Complex (EJC) | REACTOME pathways | R-DME-975956 | 95 | 11 | 3923 | 151 | RpS26,eRF1,RpS19a,RpS3A,RpS8,RpL15,RpL35A,RpL36A,RpL14,RpS9,RpL5 | | 9.237e-04 | -6.99 | Ribosomal\_S4 | pfam domains | PF00163 | 2 | 2 | 10480 | 319 | RpS9,CG4866 | | 9.318e-04 | -6.98 | REACTOME\_3\_UTR\_MEDIATED\_TRANSLATIONAL\_REGULATION | MSigDB lists | REACTOME\_3\_UTR\_MEDIATED\_TRANSLATIONAL\_REGULATION | 83 | 11 | 4161 | 184 | RpS9,RpL36A,RpL5,eIF3b,RpL14,RpS26,eIF3c,RpL35A,RpS8,eIF3g1,RpL15 | | 9.486e-04 | -6.96 | system development | biological process | GO:0048731 | 1808 | 65 | 12345 | 305 | sm,me31B,Skp2,RpS8,Stim,Pp1-87B,CG8149,Nf1,tsr,heix,mnb,LIMK1,rhea,Cdk8,Acf,dpp,nsl1,mbf1,E(z),Cdc42,peng,sws,Zpr1,pav,pins,Dref,Fmr1,tkv,Vap33,smt3,EloB,GEFmeso,Sod2,VhaPPA1-1,Hrb27C,Stlk,Efa6,PlexA,sigmar,Rab7,CycA,Spn,Rop,mav,dgrn,Pten,CG12935,Rap1,pll,Girdin,flr,Amun,vap,MED22,Btk29A,E(bx),RhoGAPp190,CtBP,CG10365,BthD,flw,d4,nom,Rheb,unc-119 | | 9.564e-04 | -6.95 | GO\_POSITIVE\_REGULATION\_OF\_EXTRINSIC\_APOPTOTIC\_SIGNALING\_PATHWAY | MSigDB lists | GO\_POSITIVE\_REGULATION\_OF\_EXTRINSIC\_APOPTOTIC\_SIGNALING\_PATHWAY | 11 | 4 | 4161 | 184 | Pten,Nf1,tkv,HtrA2 | | 9.592e-04 | -6.95 | S4 | smart domains | SM00363 | 2 | 2 | 5086 | 158 | CG4866,RpS9 | | 9.592e-04 | -6.95 | Ribosomal\_S4 | smart domains | SM01390 | 2 | 2 | 5086 | 158 | CG4866,RpS9 | | 9.594e-04 | -6.95 | precatalytic spliceosome | cellular component | GO:0071011 | 144 | 11 | 11940 | 298 | c12.1,noi,SF2,Pep,CG4935,l(1)10Bb,Phf5a,Pnn,x16,sqd,snf | | 9.618e-04 | -6.95 | regulation of cell differentiation | biological process | GO:0045595 | 428 | 22 | 12345 | 305 | Hrb27C,me31B,Tctp,mnb,PlexA,tsu,tkv,Fmr1,heix,MED22,Orc3,E(z),mav,Rop,dpp,LIMK1,RhoGAPp190,Rheb,Rap1,Cdc42,Pten,Mlf | | 9.647e-04 | -6.94 | RpL39 (Ribosomal protein L39) | protein interactions | 37849 | 95 | 10 | 9204 | 298 | RpS26,RpL5,RpL36A,RpS8,RpL15,RpS3A,RpL14,RpS19a,RpS9,RpL35A | | 9.647e-04 | -6.94 | Rpt1 (Regulatory particle triple-A ATPase 1) | protein interactions | 35701 | 95 | 10 | 9204 | 298 | Zpr1,Prosbeta1,Rpt6,CCT4,Rap1,Hsp27,Ubqn,CG12321,Prosalpha2,Rheb | | 9.655e-04 | -6.94 | cellular macromolecule biosynthetic process | biological process | GO:0034645 | 1200 | 47 | 12345 | 305 | Parp,eRF1,Cul4,su(Hw),EloB,eIF3c,RecQ4,mRpS33,RpL15,Orc3,Rpb11,Dref,Btk29A,TfIIFbeta,mRpL30,E(bx),grau,CG8635,RpL14,mRpS28,RpL36A,eIF3b,RpL35A,eIF3g1,eIF4B,eEF1delta,CG31460,xit,eEF5,RpS9,RpS26,PIG-K,RpS3A,mRpS18A,DNAlig1,mRpL34,RpS19a,CG6244,Stt3B,RpL5,CG31957,His4r,mRpS9,Pbp49,Arp8,RpS8,RPA3 | | 9.692e-04 | -6.94 | histone modification | biological process | GO:0016570 | 192 | 13 | 12345 | 305 | Acf,Rbbp5,vnc,Ncoa6,E(z),NO66,Jarid2,Rcd5,Wdr82,d4,Art1,BEAF-32,Eaf6 | | 9.692e-04 | -6.94 | covalent chromatin modification | biological process | GO:0016569 | 192 | 13 | 12345 | 305 | BEAF-32,Eaf6,Wdr82,d4,Art1,NO66,E(z),Jarid2,Rcd5,Acf,Rbbp5,vnc,Ncoa6 | | 9.755e-04 | -6.93 | regulation of protein polymerization | biological process | GO:0032271 | 64 | 7 | 12345 | 305 | spoon,Arpc3A,flr,conu,Arpc5,Cdc42,Btk29A | | 9.761e-04 | -6.93 | RpL18A (Ribosomal protein L18A) | protein interactions | 36985 | 112 | 11 | 9204 | 298 | RpS26,RpL5,Parp,RpL36A,RpL14,RpS8,RpS3A,RpL15,RpS19a,RpS9,RpL35A | | 9.761e-04 | -6.93 | RpL38 (Ribosomal protein L38) | protein interactions | 3355144 | 112 | 11 | 9204 | 298 | RpL35A,su(Hw),RpL36A,RpS8,RpL14,RpS3A,RpL15,RpS9,RpS19a,RpS26,RpL5 | | 9.761e-04 | -6.93 | Pi3K21B (Pi3K21B) | protein interactions | 33203 | 112 | 11 | 9204 | 298 | pav,Rheb,14-3-3zeta,lic,tkv,Pten,glo,Nopp140,sqd,RpL14,SF2 | | 9.838e-04 | -6.92 | negative regulation of biosynthetic process | biological process | GO:0009890 | 457 | 23 | 12345 | 305 | Fmr1,smg,me31B,Hrb27C,Not3,Ino80,d4,snf,su(Hw),CG6066,twin,rhea,CG12299,Acf,Tgi,E(z),NO66,sqd,Nulp1,rhi,CG9246,CG6254,CtBP | | 9.838e-04 | -6.92 | negative regulation of cellular biosynthetic process | biological process | GO:0031327 | 457 | 23 | 12345 | 305 | Hrb27C,me31B,smg,Fmr1,CG6254,rhi,CG9246,CtBP,Nulp1,sqd,E(z),NO66,Tgi,CG12299,Acf,rhea,twin,CG6066,su(Hw),snf,Ino80,Not3,d4 | | 9.910e-04 | -6.92 | regulation of DNA metabolic process | biological process | GO:0051052 | 47 | 6 | 12345 | 305 | Dek,spel1,Dref,tsu,CG14805,Fign | | 9.961e-04 | -6.91 | pattern specification process | biological process | GO:0007389 | 544 | 26 | 12345 | 305 | EloB,Cdc42,Parp,mon2,CtBP,rhi,14-3-3zeta,bai,BthD,sqd,flw,mus301,Cdk8,dpp,tkv,tsu,spoon,Rab7,tsr,CG8149,Fmr1,glo,lic,Hrb27C,GEFmeso,pll | | 1.013e-03 | -6.90 | reproductive process | biological process | GO:0022414 | 1436 | 54 | 12345 | 305 | E(bx),tkv,Btk29A,Fmr1,janA,MED22,pav,lncRNA:sphinx,eIF3b,flr,grau,CSN1b,vnc,BEAF-32,twin,Parp,Mps1,Cul4,flw,14-3-3zeta,bai,smt3,RecQ4,cal1,Rab7,spoon,tsu,glo,noi,tsr,Pp1-87B,Hrb27C,lic,me31B,Bub1,Nph,Fs(2)Ket,fzy,Rap1,Pten,Cdc42,dgrn,snf,sqd,Exo84,rhi,nsl1,mon2,Rop,dpp,rhea,LIMK1,HtrA2,mus301 | | 1.032e-03 | -6.88 | GSE23568\_ID3\_KO\_VS\_WT\_CD8\_TCELL\_UP | MSigDB lists | GSE23568\_ID3\_KO\_VS\_WT\_CD8\_TCELL\_UP | 84 | 11 | 4161 | 184 | unc-119,RpS26,mRpS28,Sod2,fzy,Txl,DNAlig1,CG8860,Arpc3A,RpL36A,eIF3b | | 1.045e-03 | -6.86 | CG10555 (uncharacterized protein) | protein interactions | 31771 | 2 | 2 | 9204 | 298 | su(Hw),Cp190 | | 1.045e-03 | -6.86 | CG43338 (uncharacterized protein) | protein interactions | 35117 | 2 | 2 | 9204 | 298 | Cp190,su(Hw) | | 1.047e-03 | -6.86 | RpS30 (Ribosomal protein S30) | protein interactions | 42470 | 96 | 10 | 9204 | 298 | RpL35A,RpL36A,RpS19a,RpS9,RpS8,RpS3A,RpL14,RpL15,RpS26,RpL5 | | 1.051e-03 | -6.86 | RpL4 (Ribosomal protein L4) | protein interactions | 43349 | 113 | 11 | 9204 | 298 | RpL35A,RpS9,RpS19a,LIMK1,RpL14,RpS3A,RpL15,RpS8,RpL36A,RpL5,RpS26 | | 1.051e-03 | -6.86 | regulation of developmental process | biological process | GO:0050793 | 757 | 33 | 12345 | 305 | tkv,Fmr1,Dref,MED22,Orc3,RhoGAPp190,Rab8,Rheb,Hrb27C,Sod2,lic,me31B,Tctp,mnb,tsu,PlexA,heix,Nf1,E(z),cta,mav,Rop,CycA,Spn,dpp,LIMK1,Pten,Ino80,Cdc42,Rap1,Mlf,dgrn,DCTN3-p24 | | 1.056e-03 | -6.85 | lytic vacuole organization | biological process | GO:0080171 | 19 | 4 | 12345 | 305 | rb,Rab14,g,Rab7 | | 1.056e-03 | -6.85 | lysosome organization | biological process | GO:0007040 | 19 | 4 | 12345 | 305 | g,Rab7,Rab14,rb | | 1.073e-03 | -6.84 | ribonucleotide binding | molecular function | GO:0032553 | 822 | 34 | 12176 | 289 | me31B,mnb,RagA-B,Ino80,Dbp21E2,Btk29A,Stlk,CG5589,tkv,Acbp1,LIMK1,Mps1,spel1,RhoGAPp190,RabX6,CG8915,pav,Cdk8,CG9330,pll,Rap1,mus301,Cdc42,Rheb,DNAlig1,Rab14,Rab7,Bub1,Dbp80,Rpt6,Rab8,CCT4,Fign,lic | | 1.082e-03 | -6.83 | fly chr31B1-31B1\|2-40 cM | chromosome location | fly chr31B1-31B1\|2-40 cM | 32 | 4 | 25010 | 358 | eEF1delta,me31B,Pten,CG34159 | | 1.082e-03 | -6.83 | fly chr31B1-31B1\|2-40 | chromosome location | fly chr31B1-31B1\|2-40 | 32 | 4 | 25010 | 358 | Pten,eEF1delta,me31B,CG34159 | | 1.093e-03 | -6.82 | protein-containing complex binding | molecular function | GO:0044877 | 254 | 15 | 12176 | 289 | cta,eIF4B,flr,eEF5,Rab7,rhea,Ubqn,dpp,spel1,TfIIFbeta,Arpc5,Arpc3A,Spn,fzy,His2Av | | 1.096e-03 | -6.82 | INO80-type complex | cellular component | GO:0097346 | 19 | 4 | 11940 | 298 | Arp8,Rcd5,Ino80,CG12659 | | 1.103e-03 | -6.81 | purine ribonucleoside triphosphate binding | molecular function | GO:0035639 | 791 | 33 | 12176 | 289 | CG8915,pav,CG9330,Cdk8,pll,Rap1,mus301,Cdc42,DNAlig1,Rheb,Rab14,Rab7,Dbp80,Bub1,Rpt6,Rab8,CCT4,Fign,lic,me31B,mnb,RagA-B,Ino80,Dbp21E2,Btk29A,Stlk,CG5589,tkv,LIMK1,Mps1,RhoGAPp190,spel1,RabX6 | | 1.108e-03 | -6.81 | rad50 (rad50) | protein interactions | 37564 | 15 | 4 | 9204 | 298 | Tctp,His2Av,His4r,CG5174 | | 1.110e-03 | -6.80 | regulation of plasma membrane bounded cell projection organization | biological process | GO:0120035 | 148 | 11 | 12345 | 305 | Rop,mav,me31B,RhoGAPp190,LIMK1,Rheb,PlexA,Pten,Cdc42,Fmr1,tsr | | 1.123e-03 | -6.79 | regulation of torso signaling pathway | biological process | GO:0120175 | 9 | 3 | 12345 | 305 | krz,vap,14-3-3zeta | | 1.123e-03 | -6.79 | nuclear-transcribed mRNA poly(A) tail shortening | biological process | GO:0000289 | 9 | 3 | 12345 | 305 | twin,smg,Not3 | | 1.155e-03 | -6.76 | regulation of phosphorylation | biological process | GO:0042325 | 271 | 16 | 12345 | 305 | lic,sws,Tctp,sigmar,mnb,krz,Stlk,smt3,mav,CG1791,flw,dpp,CycA,Rap1,Pten,Cdc42 | | 1.156e-03 | -6.76 | cytoplasmic stress granule | cellular component | GO:0010494 | 9 | 3 | 11940 | 298 | CG4612,me31B,Dbp80 | | 1.160e-03 | -6.76 | GSE43863\_NAIVE\_VS\_MEMORY\_TFH\_CD4\_TCELL\_D150\_LCMV\_UP | MSigDB lists | GSE43863\_NAIVE\_VS\_MEMORY\_TFH\_CD4\_TCELL\_D150\_LCMV\_UP | 38 | 7 | 4161 | 184 | flw,rb,Rab8,Tsp97E,flr,PGAP1,Tctp | | 1.173e-03 | -6.75 | regulation of cell projection organization | biological process | GO:0031344 | 149 | 11 | 12345 | 305 | RhoGAPp190,LIMK1,me31B,mav,Rop,tsr,Fmr1,Cdc42,Pten,PlexA,Rheb | | 1.179e-03 | -6.74 | cytoskeleton organization | biological process | GO:0007010 | 580 | 27 | 12345 | 305 | Cdc42,Pten,14-3-3zeta,smt3,mon2,cta,sqd,Exo84,LIMK1,msd5,Vap33,Spn,rhea,tsu,Fign,tsr,pins,Pp1-87B,Btk29A,CG17493,Arpc5,Fmr1,Arpc3A,Rcd5,pav,flr,Girdin | | 1.203e-03 | -6.72 | fly chr96B17-96B17\|3-86 | chromosome location | fly chr96B17-96B17\|3-86 | 4 | 2 | 25010 | 358 | Ythdf,bai | | 1.203e-03 | -6.72 | fly chr96B17-96B17\|3-86 cM | chromosome location | fly chr96B17-96B17\|3-86 cM | 4 | 2 | 25010 | 358 | Ythdf,bai | | 1.210e-03 | -6.72 | GO\_REGULATION\_OF\_CELL\_CYCLE | MSigDB lists | GO\_REGULATION\_OF\_CELL\_CYCLE | 298 | 25 | 4161 | 184 | Fign,dpp,Pten,fzy,Cdc42,HtrA2,spel1,pav,E(z),Lst8,Art1,twin,Cdk8,Rpn10,RagA-B,Stlk,AGO3,CG12321,Prosalpha2,Rpt6,Ino80,Skp2,CG17493,CycA,Cul4 | | 1.215e-03 | -6.71 | RpL14 (Ribosomal protein L14) | protein interactions | 38983 | 115 | 11 | 9204 | 298 | RpS19a,RpS9,RpL15,RpS3A,RpS8,RpL36A,Parp,RpL5,RpS26,Rheb,RpL35A | | 1.215e-03 | -6.71 | Su(var)205 (Suppressor of variegation 205) | protein interactions | 34119 | 115 | 11 | 9204 | 298 | Pep,RpL15,ver,MED26,E(z),Acf,HP4,CG1815,fzy,His2Av,CG8108 | | 1.223e-03 | -6.71 | GTPase regulator activity | molecular function | GO:0030695 | 89 | 8 | 12176 | 289 | conu,pins,PlexA,Graf,CG17883,vap,RhoGAPp190,Nf1 | | 1.234e-03 | -6.70 | Rack1 (Receptor of activated protein kinase C 1) | protein interactions | 34070 | 133 | 12 | 9204 | 298 | Prosalpha2,Cpr73D,RpL35A,RpS3A,RpS8,RpL15,RpL14,RpS9,RpS19a,RpL36A,RpL5,RpS26 | | 1.234e-03 | -6.70 | Caf1-55 (Chromatin assembly factor 1, p55 subunit) | protein interactions | 41836 | 66 | 8 | 9204 | 298 | E(bx),E(z),Dref,Pten,Jarid2,Cp190,smg,pav | | 1.256e-03 | -6.68 | GSE30962\_PRIMARY\_VS\_SECONDARY\_ACUTE\_LCMV\_INF\_CD8\_TCELL\_UP | MSigDB lists | GSE30962\_PRIMARY\_VS\_SECONDARY\_ACUTE\_LCMV\_INF\_CD8\_TCELL\_UP | 61 | 9 | 4161 | 184 | CycA,CG5535,E(z),Stt3B,pav,DNAlig1,Fign,eIF3c,snf | | 1.261e-03 | -6.68 | anatomical structure formation involved in morphogenesis | biological process | GO:0048646 | 553 | 26 | 12345 | 305 | Fs(2)Ket,Cul4,Cdc42,nsl1,dah,smt3,rhi,14-3-3zeta,CtBP,BthD,flw,sqd,mus301,rhea,dpp,RecQ4,tsu,tkv,spoon,Fmr1,Btk29A,flr,CSN1b,Zpr1,grau,me31B | | 1.272e-03 | -6.67 | Translation initiation complex formation | REACTOME pathways | R-DME-72649 | 70 | 9 | 3923 | 151 | RpS9,eIF3g1,eIF4B,eIF3c,eIF3b,RpS19a,RpS3A,RpS8,RpS26 | | 1.282e-03 | -6.66 | maintenance of location | biological process | GO:0051235 | 67 | 7 | 12345 | 305 | 14-3-3zeta,pav,Vps13D,pins,Cdc42,rhea,Parp | | 1.292e-03 | -6.65 | Transcriptional Regulation by TP53 | REACTOME pathways | R-DME-3700989 | 99 | 11 | 3923 | 151 | Rpb11,Rheb,TfIIFbeta,lic,14-3-3zeta,twin,RagA-B,Eaf6,Lst8,EloB,Not3 | | 1.293e-03 | -6.65 | eIF3a (eukaryotic translation initiation factor 3 subunit a) | protein interactions | 40563 | 26 | 5 | 9204 | 298 | eIF3b,eIF3g1,Rheb,eIF3c,Pten | | 1.295e-03 | -6.65 | ATPase activity, coupled | molecular function | GO:0042623 | 232 | 14 | 12176 | 289 | Acf,Dbp80,mus301,Rpt6,spel1,Fign,RecQ4,E(bx),me31B,CG8915,CG5589,Arp8,Ino80,Dbp21E2 | | 1.298e-03 | -6.65 | mRNA binding | molecular function | GO:0003729 | 486 | 23 | 12176 | 289 | CG4612,Rbp1,x16,snf,Hrb27C,me31B,RpS26,sqd,CNBP,sm,SF2,smg,eIF3g1,eIF3b,eIF4B,glo,tsu,Fmr1,CG6961,Cpsf6,mbf1,Upf3,eRF1 | | 1.301e-03 | -6.64 | GO\_RIBONUCLEOPROTEIN\_COMPLEX\_BIOGENESIS | MSigDB lists | GO\_RIBONUCLEOPROTEIN\_COMPLEX\_BIOGENESIS | 250 | 22 | 4161 | 184 | CG11920,eIF3c,RpL15,CG4866,Rat1,noi,SF2,RpL36A,CG5589,RpS26,me31B,RpL14,eIF3g1,CG9246,RpS8,mRpS9,RpL35A,RpS9,RpL5,eIF3b,CG14805,Dbp21E2 | | 1.328e-03 | -6.62 | RpL28 (Ribosomal protein L28) | protein interactions | 38397 | 99 | 10 | 9204 | 298 | RpS26,RpL5,RpL36A,RpS8,RpL15,RpL14,RpS3A,RpS19a,RpS9,RpL35A | | 1.330e-03 | -6.62 | multicellular organismal reproductive process | biological process | GO:0048609 | 1285 | 49 | 12345 | 305 | Btk29A,Fmr1,MED22,E(bx),tkv,grau,CSN1b,vnc,lncRNA:sphinx,pav,flr,eIF3b,Parp,Mps1,Cul4,BEAF-32,twin,RecQ4,cal1,flw,smt3,14-3-3zeta,bai,glo,tsr,Pp1-87B,Rab7,spoon,tsu,me31B,Hrb27C,lic,fzy,Rap1,Cdc42,Pten,snf,Fs(2)Ket,dpp,rhea,HtrA2,LIMK1,mus301,sqd,Exo84,rhi,mon2,nsl1,Rop | | 1.332e-03 | -6.62 | Rpt5 (Regulatory particle triple-A ATPase 5) | protein interactions | 42805 | 52 | 7 | 9204 | 298 | Prosalpha2,Rpt6,Rop,Ubqn,CG12321,Prosbeta1,Hsp27 | | 1.339e-03 | -6.62 | regulation of anatomical structure size | biological process | GO:0090066 | 249 | 15 | 12345 | 305 | spoon,Rheb,Arpc5,Pten,conu,Cdc42,Lst8,Btk29A,lic,Arpc3A,flr,RhoGAPp190,Tctp,Girdin,dpp | | 1.363e-03 | -6.60 | aub (aubergine) | protein interactions | 34524 | 67 | 8 | 9204 | 298 | tkv,Fmr1,twin,smg,CycA,me31B,Not3,AGO3 | | 1.364e-03 | -6.60 | GO\_NUCLEAR\_TRANSCRIBED\_MRNA\_CATABOLIC\_PROCESS\_NONSENSE\_MEDIATED\_DECAY | MSigDB lists | GO\_NUCLEAR\_TRANSCRIBED\_MRNA\_CATABOLIC\_PROCESS\_NONSENSE\_MEDIATED\_DECAY | 74 | 10 | 4161 | 184 | RpL14,RpS26,eRF1,RpL35A,RpS8,RpL15,RpS9,RpL36A,RpL5,tsu | | 1.364e-03 | -6.60 | ATF\_01 | MSigDB lists | ATF\_01 | 74 | 10 | 4161 | 184 | CG5861,CycA,SF2,CG31460,l(2)37Bb,CG15168,Nf1,CG13016,Cdc42,eRF1 | | 1.365e-03 | -6.60 | GSE13522\_CTRL\_VS\_T\_CRUZI\_Y\_STRAIN\_INF\_SKIN\_BALBC\_MOUSE\_UP | MSigDB lists | GSE13522\_CTRL\_VS\_T\_CRUZI\_Y\_STRAIN\_INF\_SKIN\_BALBC\_MOUSE\_UP | 50 | 8 | 4161 | 184 | CG11586,RpS26,RpL14,RpS8,mRpS33,Phf5a,Lst8,CIA30 | | 1.397e-03 | -6.57 | LY\_AGING\_OLD\_DN | MSigDB lists | LY\_AGING\_OLD\_DN | 29 | 6 | 4161 | 184 | Parp,pav,Prosalpha2,fzy,Pp1-87B,CycA | | 1.399e-03 | -6.57 | RpL17 (Ribosomal protein L17) | protein interactions | 31613 | 117 | 11 | 9204 | 298 | RpL36A,RpS19a,RpS9,RpS3A,RpL14,RpS8,RpL15,RpS26,RpL5,Rheb,RpL35A | | 1.403e-03 | -6.57 | protein serine/threonine/tyrosine kinase activity | molecular function | GO:0004712 | 10 | 3 | 12176 | 289 | Mps1,mnb,lic | | 1.410e-03 | -6.56 | Activation of the mRNA upon binding of the cap-binding complex and eIFs, and subsequent binding to 43S | REACTOME pathways | R-DME-72662 | 71 | 9 | 3923 | 151 | RpS19a,RpS3A,RpS8,eIF3b,RpS26,eIF4B,eIF3g1,RpS9,eIF3c | | 1.410e-03 | -6.56 | Ribosomal scanning and start codon recognition | REACTOME pathways | R-DME-72702 | 71 | 9 | 3923 | 151 | eIF3b,RpS19a,RpS8,RpS3A,RpS26,RpS9,eIF3g1,eIF4B,eIF3c | | 1.414e-03 | -6.56 | GSE17186\_CD21LOW\_VS\_CD21HIGH\_TRANSITIONAL\_BCELL\_DN | MSigDB lists | GSE17186\_CD21LOW\_VS\_CD21HIGH\_TRANSITIONAL\_BCELL\_DN | 62 | 9 | 4161 | 184 | Rap1,CycA,Vps35,Skp2,DNAlig1,dpp,Fign,Stlk,CG7766 | | 1.414e-03 | -6.56 | GSE4748\_CTRL\_VS\_CYANOBACTERIUM\_LPSLIKE\_STIM\_DC\_1H\_DN | MSigDB lists | GSE4748\_CTRL\_VS\_CYANOBACTERIUM\_LPSLIKE\_STIM\_DC\_1H\_DN | 62 | 9 | 4161 | 184 | Tsp97E,Pten,CG7766,Sgt,Parp,Tctp,CG5590,flr,SF2 | | 1.427e-03 | -6.55 | positive regulation of protein phosphorylation | biological process | GO:0001934 | 130 | 10 | 12345 | 305 | smt3,mav,lic,CG1791,Tctp,dpp,sigmar,Stlk,Rap1,Cdc42 | | 1.434e-03 | -6.55 | RpS18 (Ribosomal protein S18) | protein interactions | 37292 | 100 | 10 | 9204 | 298 | RpL35A,RpS26,RpL5,RpL36A,RpS19a,RpS9,RpL15,RpS3A,RpS8,RpL14 | | 1.453e-03 | -6.53 | peptidyl-lysine modification | biological process | GO:0018205 | 153 | 11 | 12345 | 305 | d4,Wdr82,BEAF-32,Eaf6,Acf,Rbbp5,vnc,Ncoa6,E(z),Rcd5,smt3 | | 1.453e-03 | -6.53 | cellular nitrogen compound catabolic process | biological process | GO:0044270 | 153 | 11 | 12345 | 305 | CG6171,Not3,Upf3,smg,Aps,twin,tsu,Zpr1,sqd,CG30105,Rat1 | | 1.455e-03 | -6.53 | Helicase\_C | interpro domains | IPR001650 | 75 | 8 | 11383 | 331 | CG5589,Ino80,me31B,CG8915,mus301,RecQ4,Dbp80,Dbp21E2 | | 1.455e-03 | -6.53 | Helicase\_ATP-bd | interpro domains | IPR014001 | 75 | 8 | 11383 | 331 | Ino80,CG5589,me31B,CG8915,RecQ4,mus301,Dbp80,Dbp21E2 | | 1.460e-03 | -6.53 | macromolecule biosynthetic process | biological process | GO:0009059 | 1224 | 47 | 12345 | 305 | Dref,Btk29A,TfIIFbeta,mRpL30,E(bx),grau,CG8635,RpL14,mRpS28,eIF3b,RpL36A,Parp,eRF1,Cul4,su(Hw),EloB,eIF3c,RecQ4,mRpS33,RpL15,Orc3,Rpb11,RpS3A,DNAlig1,mRpS18A,RpS19a,mRpL34,CG6244,Stt3B,RpL5,CG31957,His4r,mRpS9,Pbp49,Arp8,RpS8,RPA3,RpL35A,eIF3g1,eIF4B,eEF1delta,CG31460,xit,RpS9,eEF5,RpS26,PIG-K | | 1.489e-03 | -6.51 | oocyte differentiation | biological process | GO:0009994 | 177 | 12 | 12345 | 305 | glo,Fmr1,tsu,mus301,dpp,bai,14-3-3zeta,lic,mon2,sqd,flw,Hrb27C | | 1.492e-03 | -6.51 | Iswi (Imitation SWI) | protein interactions | 36390 | 53 | 7 | 9204 | 298 | E(bx),Cp190,Acf,mbf1,CtBP,Dref,Rheb | | 1.493e-03 | -6.51 | Formation of the ternary complex, and subsequently, the 43S complex | REACTOME pathways | R-DME-72695 | 58 | 8 | 3923 | 151 | RpS26,eIF3b,RpS3A,RpS19a,RpS8,eIF3c,eIF3g1,RpS9 | | 1.495e-03 | -6.51 | transferase complex | cellular component | GO:1990234 | 496 | 24 | 11940 | 298 | vnc,CG31460,Jarid2,Skp2,CycA,Ncoa6,Rpb11,Eaf6,CG6254,CG7766,EloB,nsl1,TfIIFbeta,fzy,d4,Roc2,Rcd5,E(z),Stt3B,Wdr82,Rbbp5,dgrn,inc,Cul4 | | 1.496e-03 | -6.51 | GO\_RNA\_SPLICING\_VIA\_TRANSESTERIFICATION\_REACTIONS | MSigDB lists | GO\_RNA\_SPLICING\_VIA\_TRANSESTERIFICATION\_REACTIONS | 115 | 13 | 4161 | 184 | snf,sm,CG11586,ubl,noi,Rpb11,l(1)10Bb,Phf5a,c12.1,CG4935,tsu,SF2,TfIIFbeta | | 1.508e-03 | -6.50 | positive regulation of catalytic activity | biological process | GO:0043085 | 252 | 15 | 12345 | 305 | Graf,CG17883,RhoGAPp190,HtrA2,vap,lic,CG1791,smt3,Cdc42,conu,fzy,Stlk,Nf1,PPP1R15,PlexA | | 1.516e-03 | -6.49 | mTORC1-mediated signalling | REACTOME pathways | R-DME-166208 | 23 | 5 | 3923 | 151 | RagA-B,Rheb,eIF4B,Lst8,14-3-3zeta | | 1.536e-03 | -6.48 | PUJANA\_BRCA1\_PCC\_NETWORK | MSigDB lists | PUJANA\_BRCA1\_PCC\_NETWORK | 703 | 47 | 4161 | 184 | sigmar,DNAlig1,CG11920,snf,xit,RpL36A,SF2,CycA,His2Av,MED21,Arpc3A,pav,spel1,fzy,eRF1,sm,CG5589,RpL14,Parp,RpL5,eIF3b,E(z),Rpb11,l(1)10Bb,Art1,Dbp80,eEF5,RpL15,Prosalpha2,CG8860,Txl,Fs(2)Ket,Rbbp5,CG5535,rb,noi,CCT4,CG8635,Skp2,CG9246,HtrA2,Pp1-87B,Orc3,His4r,TfIIFbeta,vnc,cta | | 1.541e-03 | -6.48 | regulation of catabolic process | biological process | GO:0009894 | 227 | 14 | 12345 | 305 | Cdk8,Cpsf6,Rpt6,Sod2,mbf1,RagA-B,AGO3,Cul4,smg,CG9951,Fmr1,Pten,sigmar,Rheb | | 1.547e-03 | -6.47 | RpL41 (Ribosomal protein L41) | protein interactions | 251466 | 101 | 10 | 9204 | 298 | RpL36A,RpL14,RpS8,RpL15,RpS3A,RpS9,RpS19a,RpS26,RpL5,RpL35A | | 1.549e-03 | -6.47 | GO\_MACROMOLECULE\_CATABOLIC\_PROCESS | MSigDB lists | GO\_MACROMOLECULE\_CATABOLIC\_PROCESS | 407 | 31 | 4161 | 184 | Rpn10,tsu,RpL5,Usp32,Roc2,twin,RpS9,Stt3B,CG32473,Ubqn,RpS8,CG9853,fzy,HtrA2,alpha-Man-Ib,eRF1,RpL35A,RpS26,me31B,RpL14,Cul4,RpL36A,krz,Skp2,Prosalpha2,RpL15,Rpt6,Rat1,Ufd4,CG7766,Fs(2)Ket | | 1.559e-03 | -6.46 | KEGG\_REGULATION\_OF\_ACTIN\_CYTOSKELETON | MSigDB lists | KEGG\_REGULATION\_OF\_ACTIN\_CYTOSKELETON | 51 | 8 | 4161 | 184 | flw,Pp1-87B,Cdc42,Arpc3A,LIMK1,cta,Arpc5,RhoGAPp190 | | 1.568e-03 | -6.46 | mRNA cis splicing, via spliceosome | biological process | GO:0045292 | 21 | 4 | 12345 | 305 | x16,Rbp1,c12.1,SF2 | | 1.587e-03 | -6.45 | regionalization | biological process | GO:0003002 | 503 | 24 | 12345 | 305 | GEFmeso,pll,Hrb27C,lic,glo,Fmr1,CG8149,tsr,Rab7,spoon,tkv,tsu,dpp,Cdk8,mus301,sqd,flw,BthD,CtBP,bai,rhi,mon2,Parp,EloB | | 1.594e-03 | -6.44 | RpL3 (Ribosomal protein L3) | protein interactions | 41347 | 137 | 12 | 9204 | 298 | RpL15,RpS3A,RpL14,RpS8,Vlet,RpS9,RpS19a,RpL36A,RpL5,RpS26,GEFmeso,RpL35A | | 1.594e-03 | -6.44 | Snr1 (Snf5-related 1) | protein interactions | 40657 | 137 | 12 | 9204 | 298 | Cpr73D,Cpsf6,NO66,Jarid2,tkv,RpS3A,CCT4,PPP1R15,dpp,Zpr1,E(z),mbf1 | | 1.609e-03 | -6.43 | negative regulation of nucleobase-containing compound metabolic process | biological process | GO:0045934 | 417 | 21 | 12345 | 305 | Fmr1,tsu,Tgi,rhea,spel1,CG12299,Acf,rhi,mbf1,CG6254,CG9246,CtBP,NO66,E(z),sqd,Nulp1,snf,su(Hw),Ino80,d4,CG6066 | | 1.613e-03 | -6.43 | oocyte development | biological process | GO:0048599 | 155 | 11 | 12345 | 305 | glo,Fmr1,tsu,dpp,bai,14-3-3zeta,mon2,lic,flw,sqd,Hrb27C | | 1.620e-03 | -6.43 | PKB-mediated events | REACTOME pathways | R-DME-109703 | 34 | 6 | 3923 | 151 | eIF4B,Rheb,RagA-B,Lst8,Stlk,14-3-3zeta | | 1.620e-03 | -6.43 | mTOR signalling | REACTOME pathways | R-DME-165159 | 34 | 6 | 3923 | 151 | eIF4B,RagA-B,Rheb,Lst8,Stlk,14-3-3zeta | | 1.621e-03 | -6.42 | site of DNA damage | cellular component | GO:0090734 | 10 | 3 | 11940 | 298 | RPA3,CG6171,His2Av | | 1.621e-03 | -6.42 | site of double-strand break | cellular component | GO:0035861 | 10 | 3 | 11940 | 298 | His2Av,CG6171,RPA3 | | 1.624e-03 | -6.42 | Helicase\_C | pfam domains | PF00271 | 73 | 8 | 10480 | 319 | mus301,Dbp80,Ino80,CG8915,me31B,Dbp21E2,RecQ4,CG5589 | | 1.648e-03 | -6.41 | RNA Polymerase II Transcription Termination | REACTOME pathways | R-DME-73856 | 46 | 7 | 3923 | 151 | tsu,CG6961,Cpsf6,Wdr33,x16,SF2,Upf3 | | 1.648e-03 | -6.41 | mRNA 3'-end processing | REACTOME pathways | R-DME-72187 | 46 | 7 | 3923 | 151 | CG6961,tsu,Upf3,x16,SF2,Cpsf6,Wdr33 | | 1.648e-03 | -6.41 | Cleavage of Growing Transcript in the Termination Region | REACTOME pathways | R-DME-109688 | 46 | 7 | 3923 | 151 | CG6961,tsu,Upf3,Cpsf6,Wdr33,x16,SF2 | | 1.658e-03 | -6.40 | N6-methyladenosine-containing RNA binding | molecular function | GO:1990247 | 3 | 2 | 12176 | 289 | Ythdf,CG8915 | | 1.670e-03 | -6.39 | Ribosome, eukaryotes | KEGG pathways | dme\_M00177 | 92 | 10 | 3198 | 117 | RpS9,RpS8,RpL36A,RpL5,RpL35A,RpL15,RpL14,RpS26,RpS3A,RpS19a | | 1.670e-03 | -6.39 | Ribosome, eukaryotes | KEGG pathways | M00177 | 92 | 10 | 3198 | 117 | RpL15,RpL14,RpS26,RpS3A,RpS19a,RpS9,RpS8,RpL36A,RpL5,RpL35A | | 1.679e-03 | -6.39 | CREB\_01 | MSigDB lists | CREB\_01 | 76 | 10 | 4161 | 184 | Dbp21E2,CycA,CG5861,14-3-3zeta,CG15168,Roc2,Cdc42,CG13016,Rat1,Nf1 | | 1.679e-03 | -6.39 | REACTOME\_INFLUENZA\_VIRAL\_RNA\_TRANSCRIPTION\_AND\_REPLICATION | MSigDB lists | REACTOME\_INFLUENZA\_VIRAL\_RNA\_TRANSCRIPTION\_AND\_REPLICATION | 76 | 10 | 4161 | 184 | Rpb11,RpS9,RpL36A,TfIIFbeta,RpL5,RpS26,RpL14,RpS8,RpL35A,RpL15 | | 1.680e-03 | -6.39 | regulation of cellular catabolic process | biological process | GO:0031329 | 204 | 13 | 12345 | 305 | Pten,Fmr1,CG9951,smg,Rheb,sigmar,Cdk8,AGO3,RagA-B,mbf1,Rpt6,Cpsf6,Sod2 | | 1.683e-03 | -6.39 | GO\_HISTONE\_METHYLTRANSFERASE\_COMPLEX | MSigDB lists | GO\_HISTONE\_METHYLTRANSFERASE\_COMPLEX | 30 | 6 | 4161 | 184 | Rbbp5,Pp1-87B,E(z),Rcd5,Wdr82,flw | | 1.683e-03 | -6.39 | PID\_CXCR4\_PATHWAY | MSigDB lists | PID\_CXCR4\_PATHWAY | 30 | 6 | 4161 | 184 | Cdc42,Lst8,Pten,LIMK1,cta,Ubqn | | 1.695e-03 | -6.38 | associative learning | biological process | GO:0008306 | 52 | 6 | 12345 | 305 | 14-3-3zeta,Orc3,mnb,Pp1-87B,Nf1,Fmr1 | | 1.717e-03 | -6.37 | regulation of neuron differentiation | biological process | GO:0045664 | 180 | 12 | 12345 | 305 | Fmr1,Cdc42,Pten,Rap1,Rheb,mnb,PlexA,me31B,RhoGAPp190,LIMK1,mav,Rop | | 1.756e-03 | -6.34 | DEXDc | smart domains | SM00487 | 73 | 8 | 5086 | 158 | CG5589,RecQ4,me31B,Ino80,Dbp21E2,mus301,Dbp80,CG8915 | | 1.758e-03 | -6.34 | REACTOME\_METABOLISM\_OF\_MRNA | MSigDB lists | REACTOME\_METABOLISM\_OF\_MRNA | 146 | 15 | 4161 | 184 | Rpt6,Prosalpha2,RpL15,eRF1,RpL35A,RpS8,RpL14,me31B,RpS26,RpL5,Rpn10,tsu,RpL36A,RpS9,14-3-3zeta | | 1.787e-03 | -6.33 | heterocycle catabolic process | biological process | GO:0046700 | 157 | 11 | 12345 | 305 | Zpr1,Rat1,CG30105,sqd,smg,Upf3,Not3,CG6171,twin,tsu,Aps | | 1.795e-03 | -6.32 | RpL24 (Ribosomal protein L24) | protein interactions | 34754 | 103 | 10 | 9204 | 298 | RpL5,RpS26,RpS3A,RpL14,RpL15,RpS8,RpS19a,RpS9,RpL36A,RpL35A | | 1.795e-03 | -6.32 | RpL40 (Ribosomal protein L40) | protein interactions | 33629 | 103 | 10 | 9204 | 298 | RpL35A,RpS26,RpL5,RpL36A,RpS9,RpS19a,RpL15,RpS3A,RpL14,RpS8 | | 1.795e-03 | -6.32 | spectrosome organization | biological process | GO:0030721 | 3 | 2 | 12345 | 305 | tkv,dpp | | 1.795e-03 | -6.32 | beta-catenin destruction complex disassembly | biological process | GO:1904886 | 3 | 2 | 12345 | 305 | flw,Pp1-87B | | 1.795e-03 | -6.32 | positive regulation of cyclase activity | biological process | GO:0031281 | 3 | 2 | 12345 | 305 | Nf1,PlexA | | 1.795e-03 | -6.32 | Golgi to endosome transport | biological process | GO:0006895 | 3 | 2 | 12345 | 305 | g,Rab14 | | 1.795e-03 | -6.32 | histone H4-K8 acetylation | biological process | GO:0043982 | 3 | 2 | 12345 | 305 | Rcd5,Eaf6 | | 1.814e-03 | -6.31 | Thor (thor) | protein interactions | 33569 | 70 | 8 | 9204 | 298 | RAF2,Vap33,eIF3g1,RpS9,sqd,RpL14,RpL35A,Lst8 | | 1.821e-03 | -6.31 | double-strand break repair | biological process | GO:0006302 | 91 | 8 | 12345 | 305 | Tctp,mus301,RecQ4,RPA3,Arp8,dgrn,Parp,CG6171 | | 1.832e-03 | -6.30 | Mpp10 complex | cellular component | GO:0034457 | 3 | 2 | 11940 | 298 | CG4866,CG11920 | | 1.832e-03 | -6.30 | ER membrane insertion complex | cellular component | GO:0072379 | 3 | 2 | 11940 | 298 | Sgt,CG9853 | | 1.835e-03 | -6.30 | eIF3e (eukaryotic translation initiation factor 3 subunit e) | protein interactions | 39877 | 17 | 4 | 9204 | 298 | eIF3b,eIF3g1,eIF3c,Pten | | 1.847e-03 | -6.29 | GO\_METHYLTRANSFERASE\_COMPLEX | MSigDB lists | GO\_METHYLTRANSFERASE\_COMPLEX | 41 | 7 | 4161 | 184 | Pp1-87B,Rbbp5,Wdr82,flw,E(z),Rcd5,Art1 | | 1.847e-03 | -6.29 | GSE17721\_POLYIC\_VS\_PAM3CSK4\_0.5H\_BMDC\_UP | MSigDB lists | GSE17721\_POLYIC\_VS\_PAM3CSK4\_0.5H\_BMDC\_UP | 41 | 7 | 4161 | 184 | Art1,twin,Ino80,RpL14,MED22,Phf5a,Mdh1 | | 1.847e-03 | -6.29 | GSE22886\_DC\_VS\_MONOCYTE\_DN | MSigDB lists | GSE22886\_DC\_VS\_MONOCYTE\_DN | 41 | 7 | 4161 | 184 | CtBP,lic,Graf,Pisd,RpL14,CG5339,eEF1delta | | 1.880e-03 | -6.28 | Rho protein signal transduction | biological process | GO:0007266 | 22 | 4 | 12345 | 305 | cta,RhoGAPp190,Graf,Cdc42 | | 1.896e-03 | -6.27 | poly(U) RNA binding | molecular function | GO:0008266 | 11 | 3 | 12176 | 289 | CG4612,snf,Hrb27C | | 1.898e-03 | -6.27 | GO\_TRANSLATIONAL\_INITIATION | MSigDB lists | GO\_TRANSLATIONAL\_INITIATION | 104 | 12 | 4161 | 184 | RpL5,eIF3b,RpL36A,RpS9,eIF3g1,RpL15,RpS8,CG31957,RpL35A,eIF3c,RpS26,RpL14 | | 1.898e-03 | -6.27 | GO\_RIBOSOMAL\_SUBUNIT | MSigDB lists | GO\_RIBOSOMAL\_SUBUNIT | 104 | 12 | 4161 | 184 | RpS8,mRpS9,RpL35A,RpL15,CG4866,mRpS33,RpS26,RpL14,mRpS28,RpL36A,RpL5,RpS9 | | 1.918e-03 | -6.26 | HELICc | smart domains | SM00490 | 74 | 8 | 5086 | 158 | CG5589,Ino80,me31B,RecQ4,Dbp21E2,CG8915,Dbp80,mus301 | | 1.922e-03 | -6.25 | mRNA 3'-UTR binding | molecular function | GO:0003730 | 38 | 5 | 12176 | 289 | CG4612,Fmr1,sqd,smg,Hrb27C | | 1.934e-03 | -6.25 | HELICASE\_CTER | prosite domains | PS51194 | 75 | 8 | 6243 | 191 | CG5589,RecQ4,CG8915,mus301,Dbp21E2,me31B,Dbp80,Ino80 | | 1.934e-03 | -6.25 | HELICASE\_ATP\_BIND\_1 | prosite domains | PS51192 | 75 | 8 | 6243 | 191 | Ino80,me31B,Dbp80,Dbp21E2,CG8915,mus301,RecQ4,CG5589 | | 1.936e-03 | -6.25 | nuclear chromatin | cellular component | GO:0000790 | 256 | 15 | 11940 | 298 | E(z),Arp8,CtBP,Rcd5,MED26,Acf,CG12659,HP4,Parp,Jarid2,BEAF-32,His2Av,d4,Ino80,E(bx) | | 1.945e-03 | -6.24 | GO\_REGULATION\_OF\_B\_CELL\_APOPTOTIC\_PROCESS | MSigDB lists | GO\_REGULATION\_OF\_B\_CELL\_APOPTOTIC\_PROCESS | 2 | 2 | 4161 | 184 | CG9246,Pten | | 1.945e-03 | -6.24 | COURTOIS\_SENESCENCE\_TRIGGERS | MSigDB lists | COURTOIS\_SENESCENCE\_TRIGGERS | 2 | 2 | 4161 | 184 | Pten,Nf1 | | 1.945e-03 | -6.24 | TCGA\_GLIOBLASTOMA\_MUTATED | MSigDB lists | TCGA\_GLIOBLASTOMA\_MUTATED | 2 | 2 | 4161 | 184 | Nf1,Pten | | 1.956e-03 | -6.24 | regulation of actin filament organization | biological process | GO:0110053 | 72 | 7 | 12345 | 305 | spoon,Arpc3A,flr,Btk29A,Arpc5,Cdc42,conu | | 1.962e-03 | -6.23 | RpS3A (Ribosomal protein S3A) | protein interactions | 43768 | 122 | 11 | 9204 | 298 | RpL36A,RpS8,RpL14,RpL15,RpS19a,RpS9,RpS26,RpL5,Lst8,RpL35A,Rpb11 | | 1.988e-03 | -6.22 | GSE9509\_LPS\_VS\_LPS\_AND\_IL10\_STIM\_IL10\_KO\_MACROPHAGE\_10MIN\_DN | MSigDB lists | GSE9509\_LPS\_VS\_LPS\_AND\_IL10\_STIM\_IL10\_KO\_MACROPHAGE\_10MIN\_DN | 65 | 9 | 4161 | 184 | Ubqn,l(2)37Bb,Parp,eIF3c,Exo84,Arp8,CG9330,Coq5,spel1 | | 1.994e-03 | -6.22 | RpS27A (Ribosomal protein S27A) | protein interactions | 34420 | 179 | 14 | 9204 | 298 | RpL35A,Hsp27,Rpn10,14-3-3zeta,RpL5,RpS26,RpL14,RpS3A,RpL15,RpS8,RpS9,CG8223,RpS19a,RpL36A | | 2.012e-03 | -6.21 | GO\_NUCLEOLUS | MSigDB lists | GO\_NUCLEOLUS | 326 | 26 | 4161 | 184 | Rbbp5,Rcd5,CG11334,Skp2,CG8635,CG4866,Rat1,mus301,CG11920,snf,flw,RpL5,Dbp21E2,vnc,mbf1,RpS9,eEF1delta,Nf1,mRpS9,CG9246,Pp1-87B,Fmr1,Rab8,CG5589,RpS26,Parp | | 2.063e-03 | -6.18 | ribosomal large subunit biogenesis | biological process | GO:0042273 | 54 | 6 | 12345 | 305 | RpL35A,RpL14,Dbp21E2,CG32075,RpL5,CG9246 | | 2.073e-03 | -6.18 | nucleotide binding | molecular function | GO:0000166 | 955 | 37 | 12176 | 289 | CG5589,Stlk,tkv,Acbp1,LIMK1,Mps1,RhoGAPp190,spel1,RabX6,me31B,mnb,Ino80,RagA-B,Parp,Dbp21E2,Btk29A,DNAlig1,Rheb,cta,Rab14,Rab7,Bub1,Dbp80,Rpt6,Rab8,lic,Fign,CCT4,CG8915,pav,pll,Cdk8,CG9330,Rap1,Cdc42,mus301,CtBP | | 2.073e-03 | -6.18 | nucleoside phosphate binding | molecular function | GO:1901265 | 955 | 37 | 12176 | 289 | Fign,lic,CCT4,Rab8,Rpt6,Dbp80,Bub1,Rab7,Rab14,cta,Rheb,DNAlig1,CtBP,mus301,Cdc42,Rap1,pll,CG9330,Cdk8,pav,CG8915,RabX6,spel1,RhoGAPp190,Mps1,LIMK1,Acbp1,tkv,CG5589,Stlk,Btk29A,Parp,Dbp21E2,Ino80,RagA-B,mnb,me31B | | 2.074e-03 | -6.18 | RpL7 (Ribosomal protein L7) | protein interactions | 34352 | 105 | 10 | 9204 | 298 | RpL35A,RpS26,RpL5,RpL36A,RpS9,RpS19a,RpS3A,RpL15,RpS8,RpL14 | | 2.076e-03 | -6.18 | aromatic compound catabolic process | biological process | GO:0019439 | 160 | 11 | 12345 | 305 | Aps,tsu,twin,Not3,CG6171,Upf3,smg,sqd,CG30105,Rat1,Zpr1 | | 2.085e-03 | -6.17 | positive regulation of molecular function | biological process | GO:0044093 | 287 | 16 | 12345 | 305 | PPP1R15,Cdc42,fzy,CG1791,smt3,RhoGAPp190,HtrA2,PlexA,Stlk,Nf1,conu,lic,vap,Stim,Graf,CG17883 | | 2.126e-03 | -6.15 | nucleosome mobilization | biological process | GO:0042766 | 11 | 3 | 12345 | 305 | Acf,Ino80,E(bx) | | 2.156e-03 | -6.14 | calypso (calypso) | protein interactions | 36794 | 29 | 5 | 9204 | 298 | RpS9,RpS19a,Pep,RpS3A,RpL15 | | 2.190e-03 | -6.12 | GO\_RNA\_PROCESSING | MSigDB lists | GO\_RNA\_PROCESSING | 398 | 30 | 4161 | 184 | RpL14,CG11586,RpS26,CG5589,sm,RpL35A,RpS8,mRpS9,ubl,Phf5a,RpS9,twin,Rpb11,l(1)10Bb,TfIIFbeta,RpL5,CG14805,CG4935,tsu,snf,AGO3,CG11920,Rat1,Tcs3,RpL15,CG4866,c12.1,noi,RpL36A,SF2 | | 2.211e-03 | -6.11 | GO\_POSITIVE\_REGULATION\_OF\_CELLULAR\_COMPONENT\_ORGANIZATION | MSigDB lists | GO\_POSITIVE\_REGULATION\_OF\_CELLULAR\_COMPONENT\_ORGANIZATION | 311 | 25 | 4161 | 184 | AGO3,Rpt6,eEF5,Arpc3A,krz,CCT4,Cul4,RhoGAPp190,dpp,Parp,Pten,Fmr1,Cdc42,HtrA2,fzy,ubl,LIMK1,Arpc5,twin,Lst8,14-3-3zeta,E(z),Rap1,flr,TfIIFbeta | | 2.216e-03 | -6.11 | GSE21670\_TGFB\_VS\_TGFB\_AND\_IL6\_TREATED\_CD4\_TCELL\_DN | MSigDB lists | GSE21670\_TGFB\_VS\_TGFB\_AND\_IL6\_TREATED\_CD4\_TCELL\_DN | 66 | 9 | 4161 | 184 | sws,Stim,flr,Nulp1,sm,Fmr1,Rab8,Tsp97E,CG8860 | | 2.220e-03 | -6.11 | Metabolism of proteins | REACTOME pathways | R-DME-392499 | 740 | 43 | 3923 | 151 | RpS19a,RpS8,alpha-Man-Ib,CycA,eIF3b,eEF1delta,Prosalpha2,Pten,bai,Arp8,xit,Parp,eIF4B,Rcd5,Ino80,CG12659,Rab8,krz,CCT4,eRF1,Skp2,CG17883,CG8860,Rpt6,RpS9,RpL36A,RpS3A,CG32473,Rpn10,Tom20,RpL14,Usp16-45,eEF5,smt3,RpL5,Exo84,RpL15,Nup188,RpS26,eIF3g1,RpL35A,eIF3c,Prosbeta1 | | 2.237e-03 | -6.10 | MORF\_NPM1 | MSigDB lists | MORF\_NPM1 | 106 | 12 | 4161 | 184 | RpL14,RpS8,smt3,RpL15,eEF1delta,Arpc3A,RpS9,14-3-3zeta,His2Av,RpL36A,RpL5,Tctp | | 2.274e-03 | -6.09 | MISSIAGLIA\_REGULATED\_BY\_METHYLATION\_DN | MSigDB lists | MISSIAGLIA\_REGULATED\_BY\_METHYLATION\_DN | 54 | 8 | 4161 | 184 | DNAlig1,pav,snf,sm,MICU1,CycA,E(z),His2Av | | 2.280e-03 | -6.08 | GO\_NEGATIVE\_REGULATION\_OF\_SYNAPTIC\_TRANSMISSION | MSigDB lists | GO\_NEGATIVE\_REGULATION\_OF\_SYNAPTIC\_TRANSMISSION | 22 | 5 | 4161 | 184 | Rop,Nf1,Rap1,Pten,Fmr1 | | 2.298e-03 | -6.08 | ribosome | cellular component | GO:0005840 | 287 | 16 | 11940 | 298 | mRpS33,RpL36A,mRpS18A,mRpL30,RpS8,mRpL34,RpL14,RpL15,RpS3A,mRpS28,RpS26,mRpS9,RpL5,RpS9,RpS19a,RpL35A | | 2.300e-03 | -6.07 | CG18789 (uncharacterized protein) | protein interactions | 317813 | 18 | 4 | 9204 | 298 | CG6171,BthD,Wdr82,CG1218 | | 2.303e-03 | -6.07 | regulation of chromatin organization | biological process | GO:1902275 | 116 | 9 | 12345 | 305 | su(Hw),CG9246,Jarid2,E(z),Ncoa6,cal1,Cp190,Tctp,Acf | | 2.314e-03 | -6.07 | germline ring canal | cellular component | GO:0045172 | 23 | 4 | 11940 | 298 | pav,Btk29A,vsg,14-3-3zeta | | 2.351e-03 | -6.05 | positive regulation of phosphorylation | biological process | GO:0042327 | 139 | 10 | 12345 | 305 | Tctp,dpp,lic,mav,CG1791,smt3,Stlk,Cdc42,Rap1,sigmar | | 2.364e-03 | -6.05 | chromosome segregation | biological process | GO:0007059 | 187 | 12 | 12345 | 305 | Rpn10,flw,pav,14-3-3zeta,CycA,Bub1,grau,cal1,mus301,Fign,Mps1,Pp1-87B | | 2.380e-03 | -6.04 | RpL7A (Ribosomal protein L7A) | protein interactions | 31588 | 125 | 11 | 9204 | 298 | RpL35A,Rheb,RpS26,RpL5,RpL36A,RpS9,RpS19a,RpS8,RpS3A,RpL14,RpL15 | | 2.380e-03 | -6.04 | RpS26 (Ribosomal protein S26) | protein interactions | 35098 | 125 | 11 | 9204 | 298 | RpL35A,Dref,RpS8,RpL15,RpL14,Vlet,RpS3A,RpS19a,RpS9,RpL36A,RpL5 | | 2.385e-03 | -6.04 | REACTOME\_METABOLISM\_OF\_PROTEINS | MSigDB lists | REACTOME\_METABOLISM\_OF\_PROTEINS | 262 | 22 | 4161 | 184 | xit,RpL36A,CCT4,CG8860,RpL15,PGAP1,Rat1,eIF3c,PIG-K,Tom20,eIF3b,RpL5,RpS9,eEF1delta,eIF3g1,RpL35A,eRF1,alpha-Man-Ib,RpS8,Tim17b,RpL14,RpS26 | | 2.387e-03 | -6.04 | RpS28b (Ribosomal protein S28b) | protein interactions | 31897 | 107 | 10 | 9204 | 298 | RpL35A,RpL36A,RpL14,RpS8,RpL15,RpS3A,RpS19a,RpS9,RpS26,RpL5 | | 2.393e-03 | -6.04 | regulation of supramolecular fiber organization | biological process | GO:1902903 | 95 | 8 | 12345 | 305 | Cdc42,conu,Arpc5,Btk29A,Fign,spoon,flr,Arpc3A | | 2.395e-03 | -6.03 | Signaling by Type 1 Insulin-like Growth Factor 1 Receptor (IGF1R) | REACTOME pathways | R-DME-2404192 | 156 | 14 | 3923 | 151 | 14-3-3zeta,CG7054,Prosbeta1,Rheb,eIF4B,vap,Nf1,Rpt6,CG4935,Rpn10,Prosalpha2,Stlk,Lst8,RagA-B | | 2.395e-03 | -6.03 | IRS-mediated signalling | REACTOME pathways | R-DME-112399 | 156 | 14 | 3923 | 151 | CG4935,Prosalpha2,Rpn10,Lst8,Stlk,RagA-B,CG7054,14-3-3zeta,Prosbeta1,eIF4B,Rheb,Nf1,Rpt6,vap | | 2.395e-03 | -6.03 | Insulin receptor signalling cascade | REACTOME pathways | R-DME-74751 | 156 | 14 | 3923 | 151 | Prosbeta1,CG7054,14-3-3zeta,vap,Rpt6,Nf1,Rheb,eIF4B,Stlk,Lst8,Rpn10,Prosalpha2,CG4935,RagA-B | | 2.395e-03 | -6.03 | IRS-related events triggered by IGF1R | REACTOME pathways | R-DME-2428928 | 156 | 14 | 3923 | 151 | eIF4B,Rheb,Nf1,Rpt6,vap,14-3-3zeta,CG7054,Prosbeta1,RagA-B,Rpn10,Prosalpha2,CG4935,Lst8,Stlk | | 2.395e-03 | -6.03 | IGF1R signaling cascade | REACTOME pathways | R-DME-2428924 | 156 | 14 | 3923 | 151 | Rheb,eIF4B,vap,Nf1,Rpt6,CG7054,14-3-3zeta,Prosbeta1,RagA-B,CG4935,Rpn10,Prosalpha2,Stlk,Lst8 | | 2.396e-03 | -6.03 | ribonucleoprotein complex biogenesis | biological process | GO:0022613 | 291 | 16 | 12345 | 305 | CG9246,CG14805,Dbp21E2,Nopp140,CG5589,eIF4B,RpL35A,RpS8,CG4866,CG11920,RpL14,mRpS9,me31B,CG32075,RpL5,RpS19a | | 2.423e-03 | -6.02 | methyltransferase complex | cellular component | GO:0034708 | 74 | 7 | 11940 | 298 | Rbbp5,Ncoa6,Wdr82,Rcd5,CG6254,Jarid2,E(z) | | 2.424e-03 | -6.02 | GO\_CYTOSOLIC\_PART | MSigDB lists | GO\_CYTOSOLIC\_PART | 107 | 12 | 4161 | 184 | vnc,RpL36A,RpL5,CCT4,RpS9,CG9853,RpS8,RpL35A,RpL15,Rpt6,RpS26,RpL14 | | 2.441e-03 | -6.02 | PNUTS (PNUTS) | protein interactions | 33270 | 9 | 3 | 9204 | 298 | Wdr82,Pp1-87B,flw | | 2.441e-03 | -6.02 | CG5516 (uncharacterized protein) | protein interactions | 41940 | 9 | 3 | 9204 | 298 | DCTN3-p24,Rheb,dgrn | | 2.450e-03 | -6.01 | Tab2 (TAK1-associated binding protein 2) | protein interactions | 37216 | 183 | 14 | 9204 | 298 | rhea,Cp190,smt3,nsl1,Graf,CNBP,CG5174,Map60,RpL5,CtBP,Stim,RpL14,RpS3A,tsr | | 2.450e-03 | -6.01 | Calcineurin-like\_PHP\_ApaH | interpro domains | IPR004843 | 33 | 5 | 11383 | 331 | flw,Mppe,Pp1-87B,CG1637,CG15534 | | 2.480e-03 | -6.00 | APLF\_PBZ | interpro domains | IPR019406 | 3 | 2 | 11383 | 331 | CG6171,CG1218 | | 2.480e-03 | -6.00 | S4\_RNA-bd\_sf | interpro domains | IPR036986 | 3 | 2 | 11383 | 331 | RpS9,CG4866 | | 2.484e-03 | -6.00 | GSE23321\_CD8\_STEM\_CELL\_MEMORY\_VS\_EFFECTOR\_MEMORY\_CD8\_TCELL\_DN | MSigDB lists | GSE23321\_CD8\_STEM\_CELL\_MEMORY\_VS\_EFFECTOR\_MEMORY\_CD8\_TCELL\_DN | 80 | 10 | 4161 | 184 | pav,Mdh1,CG11920,Fs(2)Ket,CG6907,CycA,Pex3,CG5535,CIA30,cta | | 2.515e-03 | -5.99 | enzyme binding | molecular function | GO:0019899 | 481 | 22 | 12176 | 289 | Efa6,GEFmeso,Rpn10,Fs(2)Ket,Slmap,lic,Vps35,Fmr1,ema,I-2,Cul4,PPP1R15,nsl1,Tctp,Cdc42,CG6907,smt3,sws,eEF1delta,CG6244,CycA,CG17883 | | 2.517e-03 | -5.98 | GTPase activity | molecular function | GO:0003924 | 146 | 10 | 12176 | 289 | Rap1,Cdc42,RhoGAPp190,Rab8,RabX6,Rheb,cta,Rab14,RagA-B,Rab7 | | 2.551e-03 | -5.97 | cell division | biological process | GO:0051301 | 266 | 15 | 12345 | 305 | CSN1b,CycA,dpp,RecQ4,Rop,Exo84,flw,pav,Mps1,pins,tsr,Cdc42,Dref,twin,tkv | | 2.557e-03 | -5.97 | RpLP0 (Ribosomal protein LP0) | protein interactions | 40451 | 108 | 10 | 9204 | 298 | RpS3A,RpL14,RpL15,RpS8,RpS9,RpS19a,RpL36A,RpL5,RpS26,RpL35A | | 2.563e-03 | -5.97 | REACTOME\_MRNA\_SPLICING | MSigDB lists | REACTOME\_MRNA\_SPLICING | 55 | 8 | 4161 | 184 | snf,sm,noi,Rpb11,Phf5a,tsu,SF2,TfIIFbeta | | 2.563e-03 | -5.97 | GSE17721\_CTRL\_VS\_LPS\_12H\_BMDC\_UP | MSigDB lists | GSE17721\_CTRL\_VS\_LPS\_12H\_BMDC\_UP | 55 | 8 | 4161 | 184 | Tom20,RpS8,CG41378,janA,MED22,Vps35,CG5861,Rap1 | | 2.610e-03 | -5.95 | MODULE\_315 | MSigDB lists | MODULE\_315 | 7 | 3 | 4161 | 184 | pav,CycA,fzy | | 2.610e-03 | -5.95 | GO\_NEGATIVE\_REGULATION\_OF\_NEUROTRANSMITTER\_TRANSPORT | MSigDB lists | GO\_NEGATIVE\_REGULATION\_OF\_NEUROTRANSMITTER\_TRANSPORT | 7 | 3 | 4161 | 184 | Fmr1,Nf1,Rap1 | | 2.610e-03 | -5.95 | GO\_REGULATION\_OF\_SYNAPTIC\_VESICLE\_EXOCYTOSIS | MSigDB lists | GO\_REGULATION\_OF\_SYNAPTIC\_VESICLE\_EXOCYTOSIS | 7 | 3 | 4161 | 184 | Rap1,Rop,Fmr1 | | 2.610e-03 | -5.95 | LY\_AGING\_MIDDLE\_DN | MSigDB lists | LY\_AGING\_MIDDLE\_DN | 7 | 3 | 4161 | 184 | pav,CycA,fzy | | 2.610e-03 | -5.95 | GO\_PRONUCLEUS | MSigDB lists | GO\_PRONUCLEUS | 7 | 3 | 4161 | 184 | CycA,E(z),sm | | 2.613e-03 | -5.95 | PID\_RHOA\_PATHWAY | MSigDB lists | PID\_RHOA\_PATHWAY | 14 | 4 | 4161 | 184 | Pten,lic,LIMK1,Cdc42 | | 2.613e-03 | -5.95 | GO\_REGULATION\_OF\_NEUROTRANSMITTER\_SECRETION | MSigDB lists | GO\_REGULATION\_OF\_NEUROTRANSMITTER\_SECRETION | 14 | 4 | 4161 | 184 | Fmr1,Rap1,Nf1,Rop | | 2.624e-03 | -5.94 | GO\_RIBOSOME | MSigDB lists | GO\_RIBOSOME | 137 | 14 | 4161 | 184 | RpS8,mRpS9,RpL35A,CG4866,RpL15,mRpS33,RpS26,RpL14,Fmr1,mRpS28,RpL36A,vnc,RpL5,RpS9 | | 2.626e-03 | -5.94 | regulation of DNA-templated transcription, initiation | biological process | GO:2000142 | 24 | 4 | 12345 | 305 | Rpt6,SF2,Rbp1,x16 | | 2.626e-03 | -5.94 | regulation of transcription initiation from RNA polymerase II promoter | biological process | GO:0060260 | 24 | 4 | 12345 | 305 | x16,Rbp1,Rpt6,SF2 | | 2.649e-03 | -5.93 | TGCGCANK\_UNKNOWN | MSigDB lists | TGCGCANK\_UNKNOWN | 183 | 17 | 4161 | 184 | Vap33,CG31460,CtBP,vap,RpL5,RpL14,sm,spel1,Rcd5,Pect,CG11334,SF2,Nulp1,Stlk,ND-ASHI,CG13016,Prosalpha2 | | 2.649e-03 | -5.93 | RpS19a (Ribosomal protein S19a) | protein interactions | 32635 | 91 | 9 | 9204 | 298 | RpL36A,RpS9,RpS8,RpS3A,RpL14,RpL15,RpS26,RpL5,RpL35A | | 2.649e-03 | -5.93 | RpL36A (Ribosomal protein L36A) | protein interactions | 34098 | 91 | 9 | 9204 | 298 | RpL35A,RpL15,RpL14,RpS8,RpS3A,RpS19a,RpS9,RpS26,RpL5 | | 2.709e-03 | -5.91 | rRNA binding | molecular function | GO:0019843 | 41 | 5 | 12176 | 289 | mRpS18A,RpL5,RpS9,Dbp21E2,CG4866 | | 2.715e-03 | -5.91 | zf-CCHH | pfam domains | PF10283 | 3 | 2 | 10480 | 319 | CG6171,CG1218 | | 2.715e-03 | -5.91 | S4 | pfam domains | PF01479 | 3 | 2 | 10480 | 319 | CG4866,RpS9 | | 2.727e-03 | -5.90 | GSE11386\_NAIVE\_VS\_MEMORY\_BCELL\_UP | MSigDB lists | GSE11386\_NAIVE\_VS\_MEMORY\_BCELL\_UP | 81 | 10 | 4161 | 184 | lic,MED21,tsu,CG17493,eIF3c,ND-ASHI,Lrr47,Fign,pav,Prosalpha2 | | 2.734e-03 | -5.90 | GSE9960\_GRAM\_POS\_VS\_GRAM\_NEG\_AND\_POS\_SEPSIS\_PBMC\_UP | MSigDB lists | GSE9960\_GRAM\_POS\_VS\_GRAM\_NEG\_AND\_POS\_SEPSIS\_PBMC\_UP | 68 | 9 | 4161 | 184 | Parp,smt3,eIF3g1,Rpb11,Pisd,l(2)37Bb,Art1,eEF1delta,CtBP | | 2.737e-03 | -5.90 | RpL35A (Ribosomal protein L35A) | protein interactions | 40654 | 109 | 10 | 9204 | 298 | RpS26,RpL5,RpL36A,RpS3A,RpS8,RpL15,RpL14,RpS19a,RpS9,Rheb | | 2.737e-03 | -5.90 | S4 | prosite domains | PS50889 | 3 | 2 | 6243 | 191 | RpS9,CG4866 | | 2.741e-03 | -5.90 | - | gene3d domains | 3.10.290.10 | 3 | 2 | 4929 | 151 | RpS9,CG4866 | | 2.773e-03 | -5.89 | FEVR\_CTNNB1\_TARGETS\_DN | MSigDB lists | FEVR\_CTNNB1\_TARGETS\_DN | 232 | 20 | 4161 | 184 | flw,AGO3,Rat1,DNAlig1,CCT4,Skp2,rb,noi,CycA,SF2,Fign,Pten,fzy,spel1,pav,eEF1delta,14-3-3zeta,E(z),CIA30,CtBP | | 2.786e-03 | -5.88 | actin cytoskeleton organization | biological process | GO:0030036 | 242 | 14 | 12345 | 305 | tsr,Btk29A,Arpc5,Pten,Cdc42,LIMK1,Spn,rhea,Girdin,Arpc3A,mon2,pav,Exo84,flr | | 2.801e-03 | -5.88 | Rpt2 (Regulatory particle triple-A ATPase 2) | protein interactions | 42828 | 59 | 7 | 9204 | 298 | CG12321,Ubqn,Prosbeta1,Pten,Prosalpha2,Rpt6,Rop | | 2.801e-03 | -5.88 | osa (osa) | protein interactions | 42130 | 59 | 7 | 9204 | 298 | su(Hw),Cdk8,Cpsf6,Exo84,Dref,dpp,smt3 | | 2.801e-03 | -5.88 | Doa (Darkener of apricot) | protein interactions | 43415 | 59 | 7 | 9204 | 298 | SF2,x16,Cwc25,Pp1-87B,Rbp1,Cdk8,Cpsf6 | | 2.809e-03 | -5.87 | negative regulation of cell communication | biological process | GO:0010648 | 351 | 18 | 12345 | 305 | E(bx),TM9SF2,mnb,krz,Fgop2,Fmr1,lic,vap,CG30423,pav,Slmap,Graf,Rheb,Cul4,Pten,Rap1,CtBP,flw | | 2.811e-03 | -5.87 | SANSOM\_APC\_TARGETS\_UP | MSigDB lists | SANSOM\_APC\_TARGETS\_UP | 33 | 6 | 4161 | 184 | CG12321,RpL14,Skp2,Parp,AGO3,CG11334 | | 2.811e-03 | -5.87 | GO\_REGULATION\_OF\_SYNAPTIC\_PLASTICITY | MSigDB lists | GO\_REGULATION\_OF\_SYNAPTIC\_PLASTICITY | 33 | 6 | 4161 | 184 | Fmr1,Rab8,Pten,Nf1,Rop,fzy | | 2.819e-03 | -5.87 | DDT | smart domains | SM00571 | 3 | 2 | 5086 | 158 | Acf,E(bx) | | 2.838e-03 | -5.86 | GO\_NEGATIVE\_REGULATION\_OF\_INTRACELLULAR\_SIGNAL\_TRANSDUCTION | MSigDB lists | GO\_NEGATIVE\_REGULATION\_OF\_INTRACELLULAR\_SIGNAL\_TRANSDUCTION | 95 | 11 | 4161 | 184 | Tctp,RhoGAPp190,vap,krz,Nf1,CSN1b,Sod2,HtrA2,CG9246,Pten,Fign | | 2.840e-03 | -5.86 | poe (purity of essence) | protein interactions | 46243 | 19 | 4 | 9204 | 298 | flw,Nf1,Pten,Cdc42 | | 2.840e-03 | -5.86 | otu (ovarian tumor) | protein interactions | 31789 | 19 | 4 | 9204 | 298 | glo,Hrb27C,CycA,sqd | | 2.864e-03 | -5.86 | condensed nuclear chromosome, centromeric region | cellular component | GO:0000780 | 12 | 3 | 11940 | 298 | smt3,Bub1,Mps1 | | 2.896e-03 | -5.84 | negative regulation of signaling | biological process | GO:0023057 | 352 | 18 | 12345 | 305 | CtBP,flw,Cul4,Pten,Rap1,Rheb,Slmap,Graf,lic,vap,CG30423,pav,krz,Fgop2,Fmr1,E(bx),TM9SF2,mnb | | 2.926e-03 | -5.83 | tral (trailer hitch) | protein interactions | 39430 | 31 | 5 | 9204 | 298 | AGO3,14-3-3zeta,me31B,Dref,Fmr1 | | 2.927e-03 | -5.83 | TBPH (TAR DNA-binding protein-43 homolog) | protein interactions | 37781 | 110 | 10 | 9204 | 298 | RhoGAPp190,Ubqn,Fmr1,sm,glo,CG32576,Hrb27C,Vap33,sqd,CG1637 | | 2.927e-03 | -5.83 | RpS11 (Ribosomal protein S11) | protein interactions | 36321 | 110 | 10 | 9204 | 298 | RpL5,RpS26,RpS3A,RpL15,RpL14,RpS8,RpS9,RpS19a,RpL36A,RpL35A | | 2.927e-03 | -5.83 | RpL31 (Ribosomal protein L31) | protein interactions | 35988 | 110 | 10 | 9204 | 298 | RpL5,RpS26,RpL14,RpS8,RpL15,RpS3A,RpS9,RpS19a,RpL36A,RpL35A | | 2.927e-03 | -5.83 | RpS12 (Ribosomal protein S12) | protein interactions | 39480 | 110 | 10 | 9204 | 298 | RpS26,RpL5,RpL36A,RpL15,RpS3A,RpS8,RpL14,RpS19a,RpS9,RpL35A | | 2.927e-03 | -5.83 | RpL12 (Ribosomal protein L12) | protein interactions | 45329 | 110 | 10 | 9204 | 298 | RpL35A,RpL36A,RpS9,RpS19a,RpS3A,RpS8,RpL15,RpL14,RpS26,RpL5 | | 2.927e-03 | -5.83 | RpS15Aa (Ribosomal protein S15Aa) | protein interactions | 44150 | 110 | 10 | 9204 | 298 | RpS19a,RpS9,RpS8,RpL14,RpS3A,RpL15,RpL36A,RpL5,RpS26,RpL35A | | 2.951e-03 | -5.83 | fly chr18E1-18E1\|1-64 cM | chromosome location | fly chr18E1-18E1\|1-64 cM | 6 | 2 | 25010 | 358 | Ubqn,Cdc42 | | 2.951e-03 | -5.83 | fly chr18E1-18E1\|1-64 | chromosome location | fly chr18E1-18E1\|1-64 | 6 | 2 | 25010 | 358 | Ubqn,Cdc42 | | 2.957e-03 | -5.82 | Generic Transcription Pathway | REACTOME pathways | R-DME-212436 | 177 | 15 | 3923 | 151 | Not3,Lst8,EloB,Eaf6,Cdk8,RagA-B,twin,14-3-3zeta,lic,MED26,TfIIFbeta,Parp,Dek,Rheb,Rpb11 | | 2.962e-03 | -5.82 | BAELDE\_DIABETIC\_NEPHROPATHY\_DN | MSigDB lists | BAELDE\_DIABETIC\_NEPHROPATHY\_DN | 124 | 13 | 4161 | 184 | bai,CG32473,MED21,rb,14-3-3zeta,ema,spel1,CG8860,Prosalpha2,Parp,dpp,Vps13D,Tom20 | | 2.969e-03 | -5.82 | SRP-dependent cotranslational protein targeting to membrane | REACTOME pathways | R-DME-1799339 | 94 | 10 | 3923 | 151 | RpL35A,RpL36A,RpL14,RpL5,RpS9,RpS26,RpS8,RpS19a,RpS3A,RpL15 | | 2.979e-03 | -5.82 | nuclear-transcribed mRNA catabolic process | biological process | GO:0000956 | 58 | 6 | 12345 | 305 | smg,Upf3,Not3,tsu,twin,sqd | | 2.982e-03 | -5.82 | Metallophos | pfam domains | PF00149 | 33 | 5 | 10480 | 319 | CG15534,flw,Mppe,CG1637,Pp1-87B | | 2.982e-03 | -5.82 | GO\_KINASE\_BINDING | MSigDB lists | GO\_KINASE\_BINDING | 154 | 15 | 4161 | 184 | krz,Ubqn,14-3-3zeta,Art1,lic,CycA,Rop,flw,Stlk,Parp,Rab8,Pten,Pp1-87B,Cdc42,spel1 | | 3.008e-03 | -5.81 | cellular response to DNA damage stimulus | biological process | GO:0006974 | 218 | 13 | 12345 | 305 | DNAlig1,dgrn,CG1218,Cul4,Ino80,Parp,CG6171,RPA3,Arp8,Tctp,mus301,RecQ4,spel1 | | 3.008e-03 | -5.81 | organelle localization | biological process | GO:0051640 | 218 | 13 | 12345 | 305 | Vap33,Cp190,Rop,g,Exo84,pav,flw,Mps1,Pp1-87B,pins,Fmr1,tsu,Rab8 | | 3.012e-03 | -5.81 | P granule | cellular component | GO:0043186 | 40 | 5 | 11940 | 298 | Hrb27C,AGO3,me31B,Fs(2)Ket,Fmr1 | | 3.052e-03 | -5.79 | carbohydrate derivative binding | molecular function | GO:0097367 | 1011 | 38 | 12176 | 289 | pav,CG8915,Cdk8,CG9330,pll,mus301,Cdc42,Rap1,PlexA,DNAlig1,Rheb,CG34261,Rab7,Rab14,Rpt6,Bub1,Dbp80,CCT4,Fign,lic,Rab8,mnb,me31B,Dbp21E2,CG1218,RagA-B,Ino80,Btk29A,Stlk,CG5589,LIMK1,Acbp1,tkv,spel1,dpp,RhoGAPp190,Mps1,RabX6 | | 3.065e-03 | -5.79 | GO\_CELLULAR\_RESPONSE\_TO\_STRESS | MSigDB lists | GO\_CELLULAR\_RESPONSE\_TO\_STRESS | 498 | 35 | 4161 | 184 | DNAlig1,Mdh1,CSN1b,mus301,Rcd5,Ino80,CycA,Pten,Parp,spel1,Nf1,alpha-Man-Ib,CG9853,Art1,Rpb11,E(z),Ubqn,RagA-B,Txl,Rpt6,smt3,CG8860,Arp8,CG13016,Pex3,Cul4,Rbbp5,CG17493,Sod2,mRpS9,HtrA2,twin,Stt3B,His4r,Vap33 | | 3.067e-03 | -5.79 | regulation of mRNA catabolic process | biological process | GO:0061013 | 25 | 4 | 12345 | 305 | smg,Fmr1,mbf1,AGO3 | | 3.067e-03 | -5.79 | CG7099 (uncharacterized protein) | protein interactions | 34753 | 3 | 2 | 9204 | 298 | BEAF-32,Cp190 | | 3.067e-03 | -5.79 | phm (phantom) | protein interactions | 32857 | 3 | 2 | 9204 | 298 | flw,CNBP | | 3.067e-03 | -5.79 | TAF1C-like (TATA box-binding protein-associated factor RNA polymerase I subunit C-like) | protein interactions | 37450 | 3 | 2 | 9204 | 298 | Dref,pins | | 3.067e-03 | -5.79 | Tes (Testin) | protein interactions | 36965 | 3 | 2 | 9204 | 298 | RpL14,eEF5 | | 3.067e-03 | -5.79 | Msh6 (Msh6) | protein interactions | 39654 | 3 | 2 | 9204 | 298 | Cp190,su(Hw) | | 3.067e-03 | -5.79 | CG15116 (uncharacterized protein) | protein interactions | 37194 | 3 | 2 | 9204 | 298 | rhi,Oxp | | 3.067e-03 | -5.79 | prominin-like (prominin-like) | protein interactions | 38372 | 3 | 2 | 9204 | 298 | Hrb27C,flw | | 3.075e-03 | -5.78 | negative regulation of signal transduction | biological process | GO:0009968 | 326 | 17 | 12345 | 305 | krz,Fgop2,Fmr1,E(bx),TM9SF2,mnb,Graf,Slmap,lic,vap,pav,CG30423,Cul4,Pten,Rheb,CtBP,flw | | 3.095e-03 | -5.78 | germ-line stem cell population maintenance | biological process | GO:0030718 | 78 | 7 | 12345 | 305 | CSN1b,dpp,fzy,Rap1,nsl1,tkv,His2Av | | 3.097e-03 | -5.78 | stem cell population maintenance | biological process | GO:0019827 | 99 | 8 | 12345 | 305 | His2Av,nsl1,RPA3,dpp,CSN1b,tkv,fzy,Rap1 | | 3.097e-03 | -5.78 | maintenance of cell number | biological process | GO:0098727 | 99 | 8 | 12345 | 305 | tkv,fzy,Rap1,His2Av,nsl1,RPA3,dpp,CSN1b | | 3.127e-03 | -5.77 | Sxl (Sex lethal) | protein interactions | 3772180 | 111 | 10 | 9204 | 298 | inc,Hrb27C,Map60,me31B,Rat1,Lrr47,Stt3B,Ythdf,snf,CG5439 | | 3.172e-03 | -5.75 | polyubiquitin modification-dependent protein binding | molecular function | GO:0031593 | 13 | 3 | 12176 | 289 | Rpn10,Ubqn,Vps13D | | 3.190e-03 | -5.75 | cellular protein-containing complex assembly | biological process | GO:0034622 | 471 | 22 | 12345 | 305 | CIA30,CG12321,Cdc42,eIF4B,Orc3,Rpn10,Rop,CG1791,Acf,CG8223,Dbp21E2,Vap33,RpS19a,RpL5,tkv,tsr,CG44194,Pbp49,Tom20,me31B,l(2)37Bb,His4r | | 3.196e-03 | -5.75 | Metallo-depent\_PP-like | interpro domains | IPR029052 | 35 | 5 | 11383 | 331 | flw,CG15534,CG1637,Pp1-87B,Mppe | | 3.222e-03 | -5.74 | mitochondrial protein complex | cellular component | GO:0098798 | 270 | 15 | 11940 | 298 | CG11752,mRpS9,MICU1,mRpL30,ND-ASHI,sun,mRpS18A,l(2)37Bb,mRpS28,Tom20,mRpL34,Tim17b,CIA30,ND-AGGG,mRpS33 | | 3.225e-03 | -5.74 | GSE29949\_MICROGLIA\_BRAIN\_VS\_MONOCYTE\_BONE\_MARROW\_DN | MSigDB lists | GSE29949\_MICROGLIA\_BRAIN\_VS\_MONOCYTE\_BONE\_MARROW\_DN | 57 | 8 | 4161 | 184 | Tcs3,smt3,PGAP1,Pp1-87B,CG5589,Vps13D,vap,Pex3 | | 3.264e-03 | -5.72 | nucleotide diphosphatase activity | molecular function | GO:0004551 | 4 | 2 | 12176 | 289 | Aps,Apf | | 3.265e-03 | -5.72 | regulation of nervous system development | biological process | GO:0051960 | 356 | 18 | 12345 | 305 | LIMK1,RhoGAPp190,Spn,Rop,mav,E(z),Rap1,Pten,Cdc42,Rheb,Rab8,me31B,Nf1,MED22,Fmr1,PlexA,tkv,mnb | | 3.280e-03 | -5.72 | RODRIGUES\_THYROID\_CARCINOMA\_POORLY\_DIFFERENTIATED\_DN | MSigDB lists | RODRIGUES\_THYROID\_CARCINOMA\_POORLY\_DIFFERENTIATED\_DN | 171 | 16 | 4161 | 184 | bai,Stim,CG11334,CG5339,RpL15,Txl,sigmar,Ufd4,flw,tsu,Vap33,Rap1,Stt3B,Nf1,Aps,Pten | | 3.328e-03 | -5.71 | GO\_REGULATION\_OF\_CELL\_CYCLE\_PROCESS | MSigDB lists | GO\_REGULATION\_OF\_CELL\_CYCLE\_PROCESS | 187 | 17 | 4161 | 184 | AGO3,Prosalpha2,Rpt6,CG12321,Ino80,Cul4,CG17493,CycA,Pten,pav,spel1,Cdc42,fzy,E(z),Art1,twin,Rpn10 | | 3.328e-03 | -5.71 | upd1 (unpaired 1) | protein interactions | 32813 | 77 | 8 | 9204 | 298 | tkv,His2Av,CycA,CtBP,mthl8,dpp,E(bx),flw | | 3.338e-03 | -5.70 | RpL23 (Ribosomal protein L23) | protein interactions | 37628 | 112 | 10 | 9204 | 298 | RpL36A,RpS19a,RpS9,RpL15,RpL14,RpS3A,RpS8,RpS26,RpL5,RpL35A | | 3.338e-03 | -5.70 | RpS2 (Ribosomal protein S2) | protein interactions | 34309 | 112 | 10 | 9204 | 298 | RpS26,RpL5,RpL36A,RpS3A,RpL14,RpL15,RpS8,RpS19a,RpS9,RpL35A | | 3.379e-03 | -5.69 | Znf\_CCHC\_sf | interpro domains | IPR036875 | 11 | 3 | 11383 | 331 | x16,CG5516,CNBP | | 3.404e-03 | -5.68 | Dif (Dorsal-related immunity factor) | protein interactions | 35045 | 10 | 3 | 9204 | 298 | krz,Cdk8,MED21 | | 3.404e-03 | -5.68 | Tnpo-SR (Transportin-Serine/Arginine rich) | protein interactions | 33465 | 10 | 3 | 9204 | 298 | Rbp1,SF2,CycA | | 3.404e-03 | -5.68 | CG31457 (uncharacterized protein) | protein interactions | 326142 | 10 | 3 | 9204 | 298 | Pp1-87B,flw,RpS3A | | 3.404e-03 | -5.68 | Rab9 (Rab9) | protein interactions | 35221 | 10 | 3 | 9204 | 298 | Vps35,Nf1,Vps13D | | 3.404e-03 | -5.68 | NiPp1 (Nuclear inhibitor of Protein phosphatase 1) | protein interactions | 44748 | 10 | 3 | 9204 | 298 | flw,I-2,Pp1-87B | | 3.425e-03 | -5.68 | PEART\_HDAC\_PROLIFERATION\_CLUSTER\_UP | MSigDB lists | PEART\_HDAC\_PROLIFERATION\_CLUSTER\_UP | 24 | 5 | 4161 | 184 | RhoGAPp190,Rap1,CG17493,Pp1-87B,Stim | | 3.425e-03 | -5.68 | PUJANA\_BREAST\_CANCER\_WITH\_BRCA1\_MUTATED\_UP | MSigDB lists | PUJANA\_BREAST\_CANCER\_WITH\_BRCA1\_MUTATED\_UP | 24 | 5 | 4161 | 184 | spel1,fzy,snf,E(z),Skp2 | | 3.461e-03 | -5.67 | Tlk (Tousled-like kinase) | protein interactions | 318206 | 20 | 4 | 9204 | 298 | Ubqn,Cdc42,Slmap,Fgop2 | | 3.480e-03 | -5.66 | organic cyclic compound catabolic process | biological process | GO:1901361 | 171 | 11 | 12345 | 305 | CG30105,sqd,Rat1,Zpr1,Aps,tsu,twin,CG6171,Not3,smg,Upf3 | | 3.497e-03 | -5.66 | rRNA processing | REACTOME pathways | R-DME-72312 | 112 | 11 | 3923 | 151 | CG32075,CG14805,Rat1,CG4866,RpS9,CG11920,CG5589,RpS26,RpS3A,RpS19a,RpS8 | | 3.497e-03 | -5.66 | Major pathway of rRNA processing in the nucleolus and cytosol | REACTOME pathways | R-DME-6791226 | 112 | 11 | 3923 | 151 | RpS3A,RpS19a,RpS8,RpS26,CG5589,CG4866,Rat1,CG11920,RpS9,CG14805,CG32075 | | 3.497e-03 | -5.66 | rRNA processing in the nucleus and cytosol | REACTOME pathways | R-DME-8868773 | 112 | 11 | 3923 | 151 | CG32075,CG14805,RpS9,CG11920,Rat1,CG4866,CG5589,RpS26,RpS19a,RpS8,RpS3A | | 3.532e-03 | -5.65 | positive regulation of lyase activity | biological process | GO:0051349 | 4 | 2 | 12345 | 305 | Nf1,PlexA | | 3.532e-03 | -5.65 | regulation of cell proliferation involved in compound eye morphogenesis | biological process | GO:2000495 | 4 | 2 | 12345 | 305 | Mlf,Dref | | 3.536e-03 | -5.64 | histone methylation | biological process | GO:0016571 | 60 | 6 | 12345 | 305 | Rbbp5,Wdr82,Art1,Ncoa6,E(z),BEAF-32 | | 3.569e-03 | -5.64 | hemopoiesis | biological process | GO:0030097 | 80 | 7 | 12345 | 305 | RpS8,E(bx),CG8149,heix,Rap1,dpp,Cdc42 | | 3.583e-03 | -5.63 | establishment of vesicle localization | biological process | GO:0051650 | 42 | 5 | 12345 | 305 | Fmr1,pav,Exo84,g,Rop | | 3.603e-03 | -5.63 | condensed nuclear chromosome kinetochore | cellular component | GO:0000778 | 4 | 2 | 11940 | 298 | Mps1,Bub1 | | 3.603e-03 | -5.63 | AP-3 adaptor complex | cellular component | GO:0030123 | 4 | 2 | 11940 | 298 | g,rb | | 3.603e-03 | -5.63 | nuclear pre-replicative complex | cellular component | GO:0005656 | 4 | 2 | 11940 | 298 | Orc3,CG14805 | | 3.603e-03 | -5.63 | pre-replicative complex | cellular component | GO:0036387 | 4 | 2 | 11940 | 298 | CG14805,Orc3 | | 3.628e-03 | -5.62 | GO\_MULTI\_ORGANISM\_METABOLIC\_PROCESS | MSigDB lists | GO\_MULTI\_ORGANISM\_METABOLIC\_PROCESS | 98 | 11 | 4161 | 184 | RpL14,RpS26,RpL15,eIF3g1,RpL35A,RpS8,lic,RpS9,RpL5,eIF3b,RpL36A | | 3.649e-03 | -5.61 | PID\_PDGFRB\_PATHWAY | MSigDB lists | PID\_PDGFRB\_PATHWAY | 46 | 7 | 4161 | 184 | Pten,Arpc5,14-3-3zeta,Arpc3A,RhoGAPp190,vap,Rap1 | | 3.649e-03 | -5.61 | GO\_SMALL\_RIBOSOMAL\_SUBUNIT | MSigDB lists | GO\_SMALL\_RIBOSOMAL\_SUBUNIT | 46 | 7 | 4161 | 184 | RpS8,mRpS9,CG4866,mRpS33,RpS26,mRpS28,RpS9 | | 3.660e-03 | -5.61 | fly chr2-54 cM | chromosome location | fly chr2-54 cM | 22 | 3 | 25010 | 358 | cta,CG17715,CG12567 | | 3.660e-03 | -5.61 | fly chr2-54 | chromosome location | fly chr2-54 | 22 | 3 | 25010 | 358 | cta,CG12567,CG17715 | | 3.684e-03 | -5.60 | intercellular bridge | cellular component | GO:0045171 | 26 | 4 | 11940 | 298 | pav,Btk29A,vsg,14-3-3zeta | | 3.686e-03 | -5.60 | GSE32164\_ALTERNATIVELY\_ACT\_M2\_VS\_CMYC\_INHIBITED\_MACROPHAGE\_DN | MSigDB lists | GSE32164\_ALTERNATIVELY\_ACT\_M2\_VS\_CMYC\_INHIBITED\_MACROPHAGE\_DN | 71 | 9 | 4161 | 184 | CG10672,E(z),Art1,Phf5a,pav,DNAlig1,fzy,mRpS28,RpS26 | | 3.701e-03 | -5.60 | cellular response to stress | biological process | GO:0033554 | 537 | 24 | 12345 | 305 | RagA-B,CG5056,spel1,RecQ4,mus301,PPP1R15,Ino80,Sgt,Pten,Parp,Cul4,dgrn,CG1218,lic,Sod2,CG9005,RPA3,Arp8,alpha-Man-Ib,Tctp,DNAlig1,CG6171,Btk29A,Stlk | | 3.703e-03 | -5.60 | establishment of organelle localization | biological process | GO:0051656 | 148 | 10 | 12345 | 305 | g,Rop,flw,pav,Exo84,Cp190,pins,Mps1,Pp1-87B,Fmr1 | | 3.710e-03 | -5.60 | protein modification process | biological process | GO:0036211 | 1215 | 45 | 12345 | 305 | Jarid2,Rcd5,pll,CSN1b,vnc,Mppe,tkv,Ufd4,Btk29A,Dref,janA,flw,NO66,smt3,Rbbp5,Ncoa6,BEAF-32,d4,Parp,Mps1,Cul4,lic,Skp2,Bub1,Usp16-45,mnb,Stt3B,Pp1-87B,Stlk,E(z),PIG-K,Acf,Roc2,CycA,Cdk8,LIMK1,Usp32,xit,CG31460,Eaf6,Wdr82,Pten,Art1,dgrn,CG1218 | | 3.710e-03 | -5.60 | cellular protein modification process | biological process | GO:0006464 | 1215 | 45 | 12345 | 305 | lic,Skp2,Bub1,Usp16-45,mnb,Stt3B,Stlk,Pp1-87B,E(z),PIG-K,Acf,Cdk8,CycA,Roc2,LIMK1,Usp32,xit,Eaf6,CG31460,Wdr82,Pten,dgrn,Art1,CG1218,Jarid2,Rcd5,CSN1b,pll,vnc,Mppe,Ufd4,tkv,Btk29A,Dref,janA,flw,NO66,smt3,Rbbp5,Ncoa6,BEAF-32,Parp,d4,Mps1,Cul4 | | 3.733e-03 | -5.59 | regulation of anatomical structure morphogenesis | biological process | GO:0022603 | 250 | 14 | 12345 | 305 | Rap1,Pten,Cdc42,Fmr1,Dref,Mlf,Rheb,PlexA,tkv,dpp,me31B,LIMK1,RhoGAPp190,cta | | 3.739e-03 | -5.59 | germ plasm | cellular component | GO:0060293 | 42 | 5 | 11940 | 298 | Hrb27C,Fmr1,Fs(2)Ket,AGO3,me31B | | 3.761e-03 | -5.58 | GTPase activator activity | molecular function | GO:0005096 | 84 | 7 | 12176 | 289 | PlexA,conu,RhoGAPp190,Graf,Nf1,CG17883,vap | | 3.794e-03 | -5.57 | RpS16 (Ribosomal protein S16) | protein interactions | 37580 | 114 | 10 | 9204 | 298 | RpL35A,RpS8,RpS3A,RpL14,RpL15,RpS19a,RpS9,RpL36A,RpL5,RpS26 | | 3.804e-03 | -5.57 | regulation of localization | biological process | GO:0032879 | 478 | 22 | 12345 | 305 | PlexA,tsu,mnb,Vps35,krz,tsr,CG30423,Hrb27C,vap,Stim,Graf,Rab8,Rheb,dgrn,rb,Rap1,Parp,Pten,smt3,Rop,nsl1,Orc3 | | 3.806e-03 | -5.57 | Deubiquitination | REACTOME pathways | R-DME-5688426 | 164 | 14 | 3923 | 151 | Rpt6,CG12659,Ino80,Usp16-45,Rcd5,Prosbeta1,krz,CycA,Skp2,Arp8,Tom20,Prosalpha2,Pten,Rpn10 | | 3.809e-03 | -5.57 | osk (oskar) | protein interactions | 41066 | 96 | 9 | 9204 | 298 | sqd,RpL15,Hrb27C,SF2,CG16974,14-3-3zeta,smg,tkv,tsu | | 3.812e-03 | -5.57 | regulation of MAPK cascade | biological process | GO:0043408 | 125 | 9 | 12345 | 305 | krz,Cdc42,Rap1,sigmar,dpp,mav,lic,smt3,flw | | 3.842e-03 | -5.56 | cellular response to growth factor stimulus | biological process | GO:0071363 | 61 | 6 | 12345 | 305 | Zpr1,dpp,Cdc42,smt3,tkv,mav | | 3.843e-03 | -5.56 | NUDIX | pfam domains | PF00293 | 11 | 3 | 10480 | 319 | Aps,Apf,CG12567 | | 3.847e-03 | -5.56 | mys (myospheroid) | protein interactions | 44885 | 47 | 6 | 9204 | 298 | RhoGAPp190,sws,Rap1,rhea,Dref,tkv | | 3.847e-03 | -5.56 | sqh (spaghetti squash) | protein interactions | 31554 | 47 | 6 | 9204 | 298 | Cp190,Pp1-87B,Rop,flw,RhoGAPp190,Rheb | | 3.877e-03 | -5.55 | Rpn11 (Regulatory particle non-ATPase 11) | protein interactions | 33738 | 33 | 5 | 9204 | 298 | Txl,Rpt6,Prosalpha2,Prosbeta1,Ubqn | | 3.888e-03 | -5.55 | macromolecule modification | biological process | GO:0043412 | 1320 | 48 | 12345 | 305 | Jarid2,Rcd5,CSN1b,pll,vnc,Mppe,tkv,Ufd4,Btk29A,Dref,Tcs3,janA,flw,NO66,smt3,Rbbp5,Nopp140,Ncoa6,BEAF-32,Parp,d4,Cul4,Mps1,Skp2,lic,Bub1,Usp16-45,mnb,Stt3B,Pp1-87B,Stlk,E(z),PIG-K,Acf,CycA,Cdk8,Roc2,LIMK1,Usp32,xit,Eaf6,CG6907,CG31460,Wdr82,Pten,Art1,dgrn,CG1218 | | 3.901e-03 | -5.55 | - | gene3d domains | 3.60.21.10 | 35 | 5 | 4929 | 151 | Mppe,CG1637,Pp1-87B,CG15534,flw | | 3.945e-03 | -5.54 | Signal Transduction | REACTOME pathways | R-DME-162582 | 717 | 41 | 3923 | 151 | Prosbeta1,CG10672,tkv,rhea,Stlk,glo,TfIIFbeta,14-3-3zeta,LIMK1,pins,RhoGAPp190,Nf1,Rap1,Rpn10,CG4935,Rbbp5,RagA-B,CtBP,Arpc3A,Rpt6,Vps35,cta,vap,Lst8,Cdc42,Graf,CG7054,krz,Rpb11,flw,Parp,eIF4B,conu,Rheb,dpp,GEFmeso,Prosalpha2,Pten,Cdk8,Arpc5,VhaPPA1-1 | | 3.958e-03 | -5.53 | Translocation of GLUT4 to the plasma membrane | REACTOME pathways | R-DME-1445148 | 9 | 3 | 3923 | 151 | 14-3-3zeta,Rab8,Rab14 | | 3.974e-03 | -5.53 | positive regulation of cell size | biological process | GO:0045793 | 43 | 5 | 12345 | 305 | Rheb,lic,Lst8,Girdin,Tctp | | 3.974e-03 | -5.53 | positive regulation of Wnt signaling pathway | biological process | GO:0030177 | 43 | 5 | 12345 | 305 | flw,Nulp1,Vps35,CtBP,Pp1-87B | | 4.014e-03 | -5.52 | Signaling by Rho GTPases | REACTOME pathways | R-DME-194315 | 114 | 11 | 3923 | 151 | GEFmeso,Cdc42,Arpc5,Graf,Arpc3A,14-3-3zeta,LIMK1,conu,RhoGAPp190,flw,cta | | 4.014e-03 | -5.52 | GSE2405\_0H\_VS\_6H\_A\_PHAGOCYTOPHILUM\_STIM\_NEUTROPHIL\_UP | MSigDB lists | GSE2405\_0H\_VS\_6H\_A\_PHAGOCYTOPHILUM\_STIM\_NEUTROPHIL\_UP | 59 | 8 | 4161 | 184 | Roc2,Tom20,CG12935,CG11586,heix,smt3,Txl,Sod2 | | 4.020e-03 | -5.52 | molecular function regulator | molecular function | GO:0098772 | 563 | 24 | 12176 | 289 | dpp,RhoGAPp190,GEFmeso,Efa6,Nf1,I-2,Graf,PlexA,conu,CG1791,PPP1R15,Tctp,fzy,CG6907,14-3-3zeta,mav,mnb,eEF1delta,sigmar,Stim,pins,CycA,vap,CG17883 | | 4.039e-03 | -5.51 | key (kenny) | protein interactions | 37967 | 115 | 10 | 9204 | 298 | Graf,nsl1,Cp190,smt3,CNBP,CG5174,14-3-3zeta,Map60,tsr,Stim | | 4.039e-03 | -5.51 | ZnF\_C2HC | smart domains | SM00343 | 11 | 3 | 5086 | 158 | x16,CNBP,RecQ4 | | 4.041e-03 | -5.51 | GO\_REGULATION\_OF\_OXIDATIVE\_STRESS\_INDUCED\_INTRINSIC\_APOPTOTIC\_SIGNALING\_PATHWAY | MSigDB lists | GO\_REGULATION\_OF\_OXIDATIVE\_STRESS\_INDUCED\_INTRINSIC\_APOPTOTIC\_SIGNALING\_PATHWAY | 8 | 3 | 4161 | 184 | Sod2,HtrA2,Ubqn | | 4.041e-03 | -5.51 | GNF2\_MYD88 | MSigDB lists | GNF2\_MYD88 | 8 | 3 | 4161 | 184 | Arpc3A,Arpc5,Rab8 | | 4.041e-03 | -5.51 | GO\_DNA\_HELICASE\_COMPLEX | MSigDB lists | GO\_DNA\_HELICASE\_COMPLEX | 8 | 3 | 4161 | 184 | Rcd5,Ino80,Arp8 | | 4.066e-03 | -5.51 | centrosome | cellular component | GO:0005813 | 125 | 9 | 11940 | 298 | pav,Mps1,Girdin,Bub1,CG17493,Fign,fzy,Cp190,Map60 | | 4.089e-03 | -5.50 | protein localization to organelle | biological process | GO:0033365 | 226 | 13 | 12345 | 305 | Vap33,cal1,Tim17b,Pex3,g,Tom20,His2Av,CG9853,Vps13D,Parp,Sgt,Lst8,Fs(2)Ket | | 4.092e-03 | -5.50 | fly chr8C14-8C14\|1-26 | chromosome location | fly chr8C14-8C14\|1-26 | 7 | 2 | 25010 | 358 | Zpr1,CG9034 | | 4.092e-03 | -5.50 | fly chr34A8-34A8\|2-47 cM | chromosome location | fly chr34A8-34A8\|2-47 cM | 7 | 2 | 25010 | 358 | CG16974,CG5439 | | 4.092e-03 | -5.50 | fly chr34A8-34A8\|2-47 | chromosome location | fly chr34A8-34A8\|2-47 | 7 | 2 | 25010 | 358 | CG16974,CG5439 | | 4.092e-03 | -5.50 | fly chr8C14-8C14\|1-26 cM | chromosome location | fly chr8C14-8C14\|1-26 cM | 7 | 2 | 25010 | 358 | Zpr1,CG9034 | | 4.131e-03 | -5.49 | GO\_NEGATIVE\_REGULATION\_OF\_INTRINSIC\_APOPTOTIC\_SIGNALING\_PATHWAY | MSigDB lists | GO\_NEGATIVE\_REGULATION\_OF\_INTRINSIC\_APOPTOTIC\_SIGNALING\_PATHWAY | 25 | 5 | 4161 | 184 | Tctp,HtrA2,CG9246,Sod2,Fign | | 4.131e-03 | -5.49 | DNA metabolic process | biological process | GO:0006259 | 280 | 15 | 12345 | 305 | DNAlig1,Cul4,dgrn,Parp,CG6171,Ino80,Arp8,Rat1,RPA3,Orc3,mus301,ver,Tctp,spel1,RecQ4 | | 4.148e-03 | -5.49 | DNA repair | biological process | GO:0006281 | 175 | 11 | 12345 | 305 | DNAlig1,Ino80,Parp,CG6171,dgrn,Arp8,RPA3,spel1,RecQ4,mus301,Tctp | | 4.160e-03 | -5.48 | localization | biological process | GO:0051179 | 2206 | 73 | 12345 | 305 | Rap1,Pten,mus301,ema,Rop,cta,glo,Lst8,spoon,Rab7,VhaPPA1-1,CG4743,Hrb27C,Mps1,rb,sun,Nup188,Rheb,unc-119,Rab8,bai,14-3-3zeta,flw,smg,Btk29A,Arpc5,TM9SF2,CG17883,Girdin,SF2,CG30423,Dbp80,flr,Cdc42,Sgt,Fs(2)Ket,PGAP1,dpp,Bmcp,rhea,PIG-K,mon2,CG32576,sqd,CG8860,His2Av,CG9853,Exo84,tsr,krz,Pp1-87B,tsu,Vps35,Cp190,CG6961,Tim17b,Rab14,RabX6,Tom20,Stim,Parp,Vap33,cal1,MICU1,pins,Vps13D,Fmr1,tkv,sws,CG5535,Pex3,g,pav | | 4.168e-03 | -5.48 | response to growth factor | biological process | GO:0070848 | 62 | 6 | 12345 | 305 | smt3,mav,tkv,dpp,Cdc42,Zpr1 | | 4.168e-03 | -5.48 | regulation of cell shape | biological process | GO:0008360 | 62 | 6 | 12345 | 305 | cta,PlexA,Cdc42,Pten,dpp,Rap1 | | 4.168e-03 | -5.48 | Zn72D (Zinc-finger protein at 72D) | protein interactions | 39764 | 21 | 4 | 9204 | 298 | Rheb,Pten,His4r,Fmr1 | | 4.168e-03 | -5.48 | shn (schnurri) | protein interactions | 36171 | 21 | 4 | 9204 | 298 | eRF1,dpp,tkv,CtBP | | 4.168e-03 | -5.48 | Cen (Centrocortin) | protein interactions | 35330 | 21 | 4 | 9204 | 298 | Fgop2,Slmap,RecQ4,Ubqn | | 4.168e-03 | -5.48 | cmet (CENP-meta) | protein interactions | 53561 | 21 | 4 | 9204 | 298 | Fgop2,Slmap,RAF2,Ubqn | | 4.168e-03 | -5.48 | twin (twin) | protein interactions | 42880 | 21 | 4 | 9204 | 298 | CycA,AGO3,smg,Not3 | | 4.187e-03 | -5.48 | GO\_RNA\_CATABOLIC\_PROCESS | MSigDB lists | GO\_RNA\_CATABOLIC\_PROCESS | 129 | 13 | 4161 | 184 | RpL14,me31B,RpS26,RpL35A,eRF1,Rat1,RpS8,RpL15,twin,RpS9,RpL36A,RpL5,tsu | | 4.243e-03 | -5.46 | purine nucleoside binding | molecular function | GO:0001883 | 157 | 10 | 12176 | 289 | Rap1,Cdc42,RhoGAPp190,Rab8,RabX6,Rheb,RagA-B,Rab14,Rab7,CG34261 | | 4.245e-03 | -5.46 | GSE17974\_CTRL\_VS\_ACT\_IL4\_AND\_ANTI\_IL12\_4H\_CD4\_TCELL\_DN | MSigDB lists | GSE17974\_CTRL\_VS\_ACT\_IL4\_AND\_ANTI\_IL12\_4H\_CD4\_TCELL\_DN | 86 | 10 | 4161 | 184 | mRpS28,CG12935,ubl,Art1,CG5535,MED21,lic,Rap1,CG5861,Vps35 | | 4.245e-03 | -5.46 | MORF\_ACTG1 | MSigDB lists | MORF\_ACTG1 | 86 | 10 | 4161 | 184 | RpS8,smt3,RpL15,RpL14,RpL36A,RpL5,Tctp,Arpc3A,14-3-3zeta,RpS9 | | 4.270e-03 | -5.46 | regulation of transferase activity | biological process | GO:0051338 | 151 | 10 | 12345 | 305 | smt3,lic,CG1791,sws,CycA,mnb,Stlk,fzy,Cdc42,Pten | | 4.280e-03 | -5.45 | cnk (connector enhancer of ksr) | protein interactions | 36952 | 48 | 6 | 9204 | 298 | sqd,14-3-3zeta,Btk29A,Rap1,Nf1,pins | | 4.425e-03 | -5.42 | cellular catabolic process | biological process | GO:0044248 | 764 | 31 | 12345 | 305 | Vps13D,smg,Upf3,CG6171,Ufd4,tsu,CG32473,inc,Usp16-45,Zpr1,alpha-Man-Ib,Skp2,Rpt6,Rat1,Prosalpha2,CG30105,Cul4,Ubqn,Pten,Not3,Sgt,fzy,twin,Aps,Usp32,Roc2,Prosbeta1,CG10365,Rpn10,sqd,CG15534 | | 4.428e-03 | -5.42 | Cka (Connector of kinase to AP-1) | protein interactions | 34096 | 34 | 5 | 9204 | 298 | tkv,Fgop2,Cpr73D,pll,Slmap | | 4.429e-03 | -5.42 | CHNG\_MULTIPLE\_MYELOMA\_HYPERPLOID\_UP | MSigDB lists | CHNG\_MULTIPLE\_MYELOMA\_HYPERPLOID\_UP | 36 | 6 | 4161 | 184 | RpS9,RpL14,eIF3g1,CG4866,RpL15,RpL35A | | 4.429e-03 | -5.42 | GSE22025\_UNTREATED\_VS\_PROGESTERONE\_TREATED\_CD4\_TCELL\_UP | MSigDB lists | GSE22025\_UNTREATED\_VS\_PROGESTERONE\_TREATED\_CD4\_TCELL\_UP | 36 | 6 | 4161 | 184 | ND-ASHI,Pex3,MED22,CG5590,smt3,PGAP1 | | 4.432e-03 | -5.42 | GO\_POSITIVE\_REGULATION\_OF\_ERK1\_AND\_ERK2\_CASCADE | MSigDB lists | GO\_POSITIVE\_REGULATION\_OF\_ERK1\_AND\_ERK2\_CASCADE | 16 | 4 | 4161 | 184 | dpp,krz,Pten,Rap1 | | 4.440e-03 | -5.42 | positive regulation of mRNA metabolic process | biological process | GO:1903313 | 14 | 3 | 12345 | 305 | smg,Hrb27C,AGO3 | | 4.460e-03 | -5.41 | GSE25147\_UNSTIM\_VS\_HELIOBACTER\_PYLORI\_LPS\_STIM\_MKN45\_CELL\_UP | MSigDB lists | GSE25147\_UNSTIM\_VS\_HELIOBACTER\_PYLORI\_LPS\_STIM\_MKN45\_CELL\_UP | 60 | 8 | 4161 | 184 | Prosalpha2,viaf,RpL35A,unc-119,CG10343,Stt3B,CCT4,CG5339 | | 4.460e-03 | -5.41 | GSE17721\_12H\_VS\_24H\_POLYIC\_BMDC\_DN | MSigDB lists | GSE17721\_12H\_VS\_24H\_POLYIC\_BMDC\_DN | 60 | 8 | 4161 | 184 | CG9330,mRpS33,Parp,Pten,CG17883,eIF3b,CCT4,krz | | 4.460e-03 | -5.41 | PENG\_GLUCOSE\_DEPRIVATION\_DN | MSigDB lists | PENG\_GLUCOSE\_DEPRIVATION\_DN | 60 | 8 | 4161 | 184 | CCT4,Arpc5,noi,His2Av,Rap1,SF2,spel1,VhaPPA1-1 | | 4.460e-03 | -5.41 | GO\_POSITIVE\_REGULATION\_OF\_CELL\_CYCLE\_PROCESS | MSigDB lists | GO\_POSITIVE\_REGULATION\_OF\_CELL\_CYCLE\_PROCESS | 60 | 8 | 4161 | 184 | Pten,AGO3,pav,Cdc42,Art1,twin,Ino80,Cul4 | | 4.460e-03 | -5.41 | GSE34156\_NOD2\_LIGAND\_VS\_TLR1\_TLR2\_LIGAND\_24H\_TREATED\_MONOCYTE\_DN | MSigDB lists | GSE34156\_NOD2\_LIGAND\_VS\_TLR1\_TLR2\_LIGAND\_24H\_TREATED\_MONOCYTE\_DN | 60 | 8 | 4161 | 184 | RpL36A,Vap33,c12.1,Coq5,Fgop2,CG9246,mus301,Tom20 | | 4.476e-03 | -5.41 | GO\_REGULATION\_OF\_CELLULAR\_AMIDE\_METABOLIC\_PROCESS | MSigDB lists | GO\_REGULATION\_OF\_CELLULAR\_AMIDE\_METABOLIC\_PROCESS | 130 | 13 | 4161 | 184 | twin,RpS9,bai,eIF3b,tsu,me31B,eIF3c,AGO3,Fmr1,eRF1,CG4866,eEF5,eIF3g1 | | 4.501e-03 | -5.40 | single-stranded DNA binding | molecular function | GO:0003697 | 46 | 5 | 12176 | 289 | RPA3,Hrb27C,RecQ4,Pep,ver | | 4.513e-03 | -5.40 | Signaling by Insulin receptor | REACTOME pathways | R-DME-74752 | 185 | 15 | 3923 | 151 | Stlk,Lst8,CG4935,Rpn10,Prosalpha2,VhaPPA1-1,RagA-B,Prosbeta1,CG7054,14-3-3zeta,vap,Rpt6,Nf1,Rheb,eIF4B | | 4.567e-03 | -5.39 | ISWI-type complex | cellular component | GO:0031010 | 14 | 3 | 11940 | 298 | Acf,E(bx),CtBP | | 4.567e-03 | -5.39 | condensed chromosome kinetochore | cellular component | GO:0000777 | 14 | 3 | 11940 | 298 | Bub1,smt3,Mps1 | | 4.568e-03 | -5.39 | regulation of cell population proliferation | biological process | GO:0042127 | 229 | 13 | 12345 | 305 | DCTN3-p24,heix,Mlf,MED22,conu,Dref,Pten,Fmr1,dpp,RecQ4,Sod2,E(z),vsg | | 4.568e-03 | -5.39 | Pi3K92E (Pi3K92E) | protein interactions | 42446 | 117 | 10 | 9204 | 298 | RpS26,RpL5,RpS3A,RpL14,RpS9,Pten,RpL35A,eIF3c,Rheb,eIF3b | | 4.569e-03 | -5.39 | neb (nebbish) | protein interactions | 35293 | 11 | 3 | 9204 | 298 | Pp1-87B,pav,flw | | 4.569e-03 | -5.39 | pico (pico) | protein interactions | 33003 | 11 | 3 | 9204 | 298 | flw,Pp1-87B,Rap1 | | 4.569e-03 | -5.39 | SmD3 (Small ribonucleoprotein particle protein SmD3) | protein interactions | 36306 | 11 | 3 | 9204 | 298 | smt3,Fmr1,snf | | 4.569e-03 | -5.39 | CG12725 (uncharacterized protein) | protein interactions | 32298 | 11 | 3 | 9204 | 298 | Nopp140,krz,CtBP | | 4.569e-03 | -5.39 | Tnpo (Transportin) | protein interactions | 38721 | 11 | 3 | 9204 | 298 | sqd,Pep,CG9330 | | 4.569e-03 | -5.39 | RluA-2 (RluA pseudouridine synthase 2) | protein interactions | 34440 | 11 | 3 | 9204 | 298 | Ubqn,Fgop2,Slmap | | 4.601e-03 | -5.38 | protein-containing complex assembly | biological process | GO:0065003 | 516 | 23 | 12345 | 305 | eIF4B,Cdc42,CG12321,CIA30,Vap33,Dbp21E2,CG8223,Acf,CG1791,Rop,Rpn10,Orc3,tsr,tkv,RpL5,inc,RpS19a,His4r,l(2)37Bb,me31B,Pbp49,Tom20,CG44194 | | 4.633e-03 | -5.37 | nucleoside-triphosphatase regulator activity | molecular function | GO:0060589 | 110 | 8 | 12176 | 289 | RhoGAPp190,Nf1,PlexA,conu,pins,Graf,CG17883,vap | | 4.659e-03 | -5.37 | GO\_REGULATION\_OF\_SYNAPSE\_STRUCTURE\_OR\_ACTIVITY | MSigDB lists | GO\_REGULATION\_OF\_SYNAPSE\_STRUCTURE\_OR\_ACTIVITY | 48 | 7 | 4161 | 184 | fzy,Cdc42,Rop,Nf1,Pten,Rab8,Fmr1 | | 4.682e-03 | -5.36 | somatic muscle development | biological process | GO:0007525 | 84 | 7 | 12345 | 305 | rhea,Pten,nom,pll,pav,EloB,nsl1 | | 4.684e-03 | -5.36 | terminal branching, open tracheal system | biological process | GO:0007430 | 28 | 4 | 12345 | 305 | Zpr1,rhea,VhaPPA1-1,Rheb | | 4.684e-03 | -5.36 | germarium-derived oocyte fate determination | biological process | GO:0007294 | 28 | 4 | 12345 | 305 | Fmr1,mus301,tsu,14-3-3zeta | | 4.740e-03 | -5.35 | negative regulation of RNA metabolic process | biological process | GO:0051253 | 398 | 19 | 12345 | 305 | Fmr1,rhi,mbf1,CG6254,CG9246,CtBP,E(z),NO66,Nulp1,sqd,Tgi,rhea,Acf,CG12299,CG6066,snf,su(Hw),Ino80,d4 | | 4.780e-03 | -5.34 | GO\_STRUCTURAL\_CONSTITUENT\_OF\_RIBOSOME | MSigDB lists | GO\_STRUCTURAL\_CONSTITUENT\_OF\_RIBOSOME | 131 | 13 | 4161 | 184 | mRpS33,RpL15,CG4866,RpL35A,RpS8,mRpS9,Bmcp,RpL14,RpS26,RpL5,RpL36A,CG4743,RpS9 | | 4.839e-03 | -5.33 | ZHANG\_BREAST\_CANCER\_PROGENITORS\_UP | MSigDB lists | ZHANG\_BREAST\_CANCER\_PROGENITORS\_UP | 162 | 15 | 4161 | 184 | Rbbp5,CycA,Roc2,CG5535,E(z),Skp2,CG5339,mRpS33,Mdh1,Fgop2,RpL15,pins,Tsp97E,Parp,Fign | | 4.846e-03 | -5.33 | histone lysine methylation | biological process | GO:0034968 | 45 | 5 | 12345 | 305 | E(z),BEAF-32,Rbbp5,Wdr82,Ncoa6 | | 4.849e-03 | -5.33 | lwr (lesswright) | protein interactions | 33226 | 65 | 7 | 9204 | 298 | inc,Dlip2,Map60,Hsp27,dpp,smt3,Rpn10 | | 4.866e-03 | -5.33 | Ras\_GTPase | interpro domains | IPR039360 | 4 | 2 | 11383 | 331 | vap,Nf1 | | 4.866e-03 | -5.33 | DDT\_dom | interpro domains | IPR018501 | 4 | 2 | 11383 | 331 | Acf,E(bx) | | 4.866e-03 | -5.33 | S4\_RNA-bd | interpro domains | IPR002942 | 4 | 2 | 11383 | 331 | CG4866,RpS9 | | 4.872e-03 | -5.32 | chromatin insulator sequence binding | molecular function | GO:0043035 | 15 | 3 | 12176 | 289 | BEAF-32,su(Hw),Cp190 | | 4.881e-03 | -5.32 | GO\_CYTOSOLIC\_RIBOSOME | MSigDB lists | GO\_CYTOSOLIC\_RIBOSOME | 74 | 9 | 4161 | 184 | RpS9,vnc,RpL36A,RpL5,RpS26,RpL14,RpS8,RpL35A,RpL15 | | 4.881e-03 | -5.32 | GSE1925\_CTRL\_VS\_IFNG\_PRIMED\_MACROPHAGE\_3H\_IFNG\_STIM\_UP | MSigDB lists | GSE1925\_CTRL\_VS\_IFNG\_PRIMED\_MACROPHAGE\_3H\_IFNG\_STIM\_UP | 74 | 9 | 4161 | 184 | mRpS9,Tom20,RpS26,RagA-B,E(z),twin,Stt3B,Ino80,CCT4 | | 4.947e-03 | -5.31 | MORF\_ACP1 | MSigDB lists | MORF\_ACP1 | 102 | 11 | 4161 | 184 | Rab8,Tom20,RpL14,eIF3g1,smt3,14-3-3zeta,His2Av,Arpc3A,CG8635,RpL5,flr | | 4.977e-03 | -5.30 | GO\_POSITIVE\_REGULATION\_OF\_RESPONSE\_TO\_STIMULUS | MSigDB lists | GO\_POSITIVE\_REGULATION\_OF\_RESPONSE\_TO\_STIMULUS | 402 | 29 | 4161 | 184 | lic,janA,Skp2,Arpc3A,krz,Rpt6,Prosalpha2,Arpc5,Art1,l(1)10Bb,Lst8,14-3-3zeta,E(z),Ubqn,eEF1delta,LIMK1,Vps35,RagA-B,Rpn10,CtBP,Rap1,tkv,Pten,Parp,dpp,Nf1,Cdc42,CG32576,HtrA2 | | 5.012e-03 | -5.30 | cytosolic part | cellular component | GO:0044445 | 256 | 14 | 11940 | 298 | RpS26,RpL36A,RpL5,CG9853,RpS8,RpS9,CCT4,Rpt6,RpL14,RpS19a,RpL15,RpS3A,RpL35A,Sgt | | 5.018e-03 | -5.29 | ubiquitin-dependent protein catabolic process | biological process | GO:0006511 | 286 | 15 | 12345 | 305 | Sgt,fzy,Cul4,Ubqn,inc,Ufd4,Roc2,alpha-Man-Ib,Usp32,Usp16-45,Rpn10,Prosbeta1,Skp2,Rpt6,Prosalpha2 | | 5.031e-03 | -5.29 | row (relative of woc) | protein interactions | 36685 | 35 | 5 | 9204 | 298 | Ubqn,Fgop2,Cpr73D,Slmap,BEAF-32 | | 5.054e-03 | -5.29 | cytosolic small ribosomal subunit | cellular component | GO:0022627 | 45 | 5 | 11940 | 298 | RpS26,RpS8,RpS19a,RpS3A,RpS9 | | 5.063e-03 | -5.29 | nucleoside binding | molecular function | GO:0001882 | 161 | 10 | 12176 | 289 | Cdc42,RhoGAPp190,Rap1,RabX6,Rab8,Rheb,Rab7,CG34261,RagA-B,Rab14 | | 5.094e-03 | -5.28 | BLUM\_RESPONSE\_TO\_SALIRASIB\_DN | MSigDB lists | BLUM\_RESPONSE\_TO\_SALIRASIB\_DN | 117 | 12 | 4161 | 184 | Skp2,His2Av,E(z),SF2,CycA,Parp,Fs(2)Ket,pins,fzy,DNAlig1,pav,spel1 | | 5.096e-03 | -5.28 | positive regulation of developmental process | biological process | GO:0051094 | 259 | 14 | 12345 | 305 | Rheb,mnb,tkv,PlexA,Fmr1,Rap1,Mlf,Nf1,Hrb27C,Orc3,lic,Rop,dpp,Tctp | | 5.096e-03 | -5.28 | GSE40274\_CTRL\_VS\_FOXP3\_AND\_GATA1\_TRANSDUCED\_ACTIVATED\_CD4\_TCELL\_DN | MSigDB lists | GSE40274\_CTRL\_VS\_FOXP3\_AND\_GATA1\_TRANSDUCED\_ACTIVATED\_CD4\_TCELL\_DN | 37 | 6 | 4161 | 184 | CG14805,tsu,eIF3g1,CycA,CG41378,mRpS28 | | 5.096e-03 | -5.28 | GSE360\_CTRL\_VS\_B\_MALAYI\_LOW\_DOSE\_DC\_DN | MSigDB lists | GSE360\_CTRL\_VS\_B\_MALAYI\_LOW\_DOSE\_DC\_DN | 37 | 6 | 4161 | 184 | Cdk8,CG7766,unc-119,dpp,Parp,cta | | 5.096e-03 | -5.28 | GO\_RAS\_PROTEIN\_SIGNAL\_TRANSDUCTION | MSigDB lists | GO\_RAS\_PROTEIN\_SIGNAL\_TRANSDUCTION | 37 | 6 | 4161 | 184 | Nf1,CycA,Rap1,Fs(2)Ket,cta,LIMK1 | | 5.150e-03 | -5.27 | RpS29 (Ribosomal protein S29) | protein interactions | 41200 | 119 | 10 | 9204 | 298 | RpL5,RpS26,RpS3A,RpL15,RpL14,RpS8,RpS9,RpS19a,RpL36A,RpL35A | | 5.150e-03 | -5.27 | RpS17 (Ribosomal protein S17) | protein interactions | 39088 | 119 | 10 | 9204 | 298 | RpL36A,RpS19a,RpS9,RpL14,RpL15,RpS8,RpS3A,RpS26,RpL5,RpL35A | | 5.168e-03 | -5.27 | developmental maturation | biological process | GO:0021700 | 206 | 12 | 12345 | 305 | glo,Fmr1,tsu,Vap33,dpp,14-3-3zeta,bai,lic,mon2,flw,sqd,Hrb27C | | 5.173e-03 | -5.26 | multi-organism process | biological process | GO:0051704 | 1653 | 57 | 12345 | 305 | Cul4,Mps1,Parp,twin,BEAF-32,cal1,Vap33,RecQ4,smt3,14-3-3zeta,bai,flw,MED22,Fmr1,Btk29A,tkv,E(bx),vnc,CSN1b,pll,grau,flr,eIF3b,pav,lncRNA:sphinx,snf,dgrn,Pten,Cdc42,fzy,Rap1,Fs(2)Ket,mus301,HtrA2,LIMK1,rhea,dpp,Rop,mon2,nsl1,rhi,Exo84,sqd,Hsp27,Pp1-87B,noi,tsr,ECSIT,glo,tsu,spoon,Rab7,Nph,Rab14,me31B,lic,Hrb27C | | 5.187e-03 | -5.26 | chorion-containing eggshell formation | biological process | GO:0007304 | 131 | 9 | 12345 | 305 | Cul4,Cdc42,tkv,spoon,Fs(2)Ket,dpp,RecQ4,smt3,rhi | | 5.187e-03 | -5.26 | regulation of protein kinase activity | biological process | GO:0045859 | 131 | 9 | 12345 | 305 | Stlk,Cdc42,Pten,mnb,sws,CycA,smt3,lic,CG1791 | | 5.237e-03 | -5.25 | GSE16385\_UNTREATED\_VS\_12H\_ROSIGLITAZONE\_IL4\_TREATED\_MACROPHAGE\_UP | MSigDB lists | GSE16385\_UNTREATED\_VS\_12H\_ROSIGLITAZONE\_IL4\_TREATED\_MACROPHAGE\_UP | 49 | 7 | 4161 | 184 | janA,CG11334,flr,eRF1,alpha-Man-Ib,Fgop2,Nf1 | | 5.247e-03 | -5.25 | enzyme regulator activity | molecular function | GO:0030234 | 358 | 17 | 12176 | 289 | CG6907,fzy,14-3-3zeta,pins,mnb,sigmar,CG17883,vap,CycA,RhoGAPp190,Nf1,PlexA,conu,I-2,Graf,CG1791,PPP1R15 | | 5.251e-03 | -5.25 | regulation of transcription by RNA polymerase II | biological process | GO:0006357 | 741 | 30 | 12345 | 305 | Ino80,CG17385,d4,dgrn,BEAF-32,RAF2,CG6907,CG12299,Rbp1,dpp,x16,Ncoa6,Nulp1,mav,CG9246,rhi,mbf1,CtBP,Dref,TfIIFbeta,MED22,MED21,tkv,E(bx),grau,MED26,Rpt6,CG11456,Rcd5,SF2 | | 5.251e-03 | -5.25 | Prosalpha1 (Proteasome alpha1 subunit) | protein interactions | 45780 | 50 | 6 | 9204 | 298 | eEF1delta,Rpt6,Prosalpha2,CG12321,Ubqn,Prosbeta1 | | 5.271e-03 | -5.25 | regulation of cell development | biological process | GO:0060284 | 344 | 17 | 12345 | 305 | LIMK1,RhoGAPp190,Orc3,E(z),mav,Rop,Rap1,Pten,Cdc42,Rheb,me31B,Hrb27C,Fmr1,MED22,mnb,PlexA,tsu | | 5.330e-03 | -5.23 | vesicle localization | biological process | GO:0051648 | 46 | 5 | 12345 | 305 | pav,Exo84,g,Rop,Fmr1 | | 5.337e-03 | -5.23 | cellular response to endogenous stimulus | biological process | GO:0071495 | 181 | 11 | 12345 | 305 | smt3,mav,RagA-B,Zpr1,dpp,E(bx),tkv,Rheb,Rab8,Rap1,Cdc42 | | 5.340e-03 | -5.23 | MORF\_EIF3S6 | MSigDB lists | MORF\_EIF3S6 | 75 | 9 | 4161 | 184 | RpL5,His2Av,14-3-3zeta,eEF1delta,Mdh1,eIF3g1,snf,Rab8,RpL14 | | 5.354e-03 | -5.23 | histone demethylase activity (H3-K4 specific) | molecular function | GO:0032453 | 5 | 2 | 12176 | 289 | Jarid2,NO66 | | 5.364e-03 | -5.23 | DDT | prosite domains | PS50827 | 4 | 2 | 6243 | 191 | Acf,E(bx) | | 5.372e-03 | -5.23 | Diap2 (Death-associated inhibitor of apoptosis 2) | protein interactions | 36748 | 179 | 13 | 9204 | 298 | CG5174,CNBP,rhea,Cp190,Graf,nsl1,Dref,CCT4,Stim,HtrA2,tsr,Map60,CtBP | | 5.404e-03 | -5.22 | fly chr10B3-10B3\|1-34 | chromosome location | fly chr10B3-10B3\|1-34 | 8 | 2 | 25010 | 358 | l(1)10Bb,CG11752 | | 5.404e-03 | -5.22 | fly chr10B3-10B3\|1-34 cM | chromosome location | fly chr10B3-10B3\|1-34 cM | 8 | 2 | 25010 | 358 | CG11752,l(1)10Bb | | 5.436e-03 | -5.21 | GSE17721\_0.5H\_VS\_12H\_CPG\_BMDC\_UP | MSigDB lists | GSE17721\_0.5H\_VS\_12H\_CPG\_BMDC\_UP | 89 | 10 | 4161 | 184 | CG5861,vnc,CG41378,Stt3B,sws,eEF1delta,krz,Mdh1,Tcs3,unc-119 | | 5.448e-03 | -5.21 | regulation of neurogenesis | biological process | GO:0050767 | 261 | 14 | 12345 | 305 | RhoGAPp190,LIMK1,me31B,Rop,mav,E(z),MED22,Fmr1,Cdc42,Pten,Rap1,PlexA,Rheb,mnb | | 5.450e-03 | -5.21 | nonassociative learning | biological process | GO:0046958 | 15 | 3 | 12345 | 305 | me31B,Btk29A,Fmr1 | | 5.450e-03 | -5.21 | regulation of mRNA stability | biological process | GO:0043488 | 15 | 3 | 12345 | 305 | smg,Fmr1,mbf1 | | 5.450e-03 | -5.21 | dorsal/ventral axis specification, ovarian follicular epithelium | biological process | GO:0008069 | 15 | 3 | 12345 | 305 | Parp,sqd,spoon | | 5.467e-03 | -5.21 | KAECH\_NAIVE\_VS\_DAY8\_EFF\_CD8\_TCELL\_UP | MSigDB lists | KAECH\_NAIVE\_VS\_DAY8\_EFF\_CD8\_TCELL\_UP | 62 | 8 | 4161 | 184 | RpL14,Tsp97E,RpS8,Pisd,CG8132,CG5339,krz,RpL5 | | 5.517e-03 | -5.20 | guanyl nucleotide binding | molecular function | GO:0019001 | 163 | 10 | 12176 | 289 | Rap1,RhoGAPp190,Cdc42,Rab8,RabX6,Rheb,cta,RagA-B,Rab14,Rab7 | | 5.522e-03 | -5.20 | RasGAP | smart domains | SM00323 | 4 | 2 | 5086 | 158 | vap,Nf1 | | 5.522e-03 | -5.20 | R3H | smart domains | SM00393 | 4 | 2 | 5086 | 158 | CG8915,CG31301 | | 5.598e-03 | -5.19 | MODULE\_115 | MSigDB lists | MODULE\_115 | 17 | 4 | 4161 | 184 | eIF3g1,eIF3b,eEF1delta,eIF3c | | 5.598e-03 | -5.19 | GO\_NUCLEOTIDE\_EXCISION\_REPAIR\_DNA\_DAMAGE\_RECOGNITION | MSigDB lists | GO\_NUCLEOTIDE\_EXCISION\_REPAIR\_DNA\_DAMAGE\_RECOGNITION | 17 | 4 | 4161 | 184 | CG17493,Cul4,CSN1b,Parp | | 5.598e-03 | -5.19 | GO\_REGULATION\_OF\_NEUROTRANSMITTER\_TRANSPORT | MSigDB lists | GO\_REGULATION\_OF\_NEUROTRANSMITTER\_TRANSPORT | 17 | 4 | 4161 | 184 | Rop,Nf1,Rap1,Fmr1 | | 5.666e-03 | -5.17 | GO\_ANGIOTENSIN\_RECEPTOR\_BINDING | MSigDB lists | GO\_ANGIOTENSIN\_RECEPTOR\_BINDING | 3 | 2 | 4161 | 184 | cta,krz | | 5.666e-03 | -5.17 | GO\_PROTEIN\_AUTOPROCESSING | MSigDB lists | GO\_PROTEIN\_AUTOPROCESSING | 3 | 2 | 4161 | 184 | Parp,HtrA2 | | 5.666e-03 | -5.17 | GO\_INTRINSIC\_COMPONENT\_OF\_THE\_CYTOPLASMIC\_SIDE\_OF\_THE\_PLASMA\_MEMBRANE | MSigDB lists | GO\_INTRINSIC\_COMPONENT\_OF\_THE\_CYTOPLASMIC\_SIDE\_OF\_THE\_PLASMA\_MEMBRANE | 3 | 2 | 4161 | 184 | vap,Nf1 | | 5.666e-03 | -5.17 | GO\_MACROPHAGE\_DIFFERENTIATION | MSigDB lists | GO\_MACROPHAGE\_DIFFERENTIATION | 3 | 2 | 4161 | 184 | Parp,Cdc42 | | 5.675e-03 | -5.17 | protein metabolic process | biological process | GO:0019538 | 2600 | 83 | 12345 | 305 | eIF3c,Ncoa6,Rpn10,NO66,mRpS33,flw,d4,rb,Mps1,Cul4,BEAF-32,RpL14,CSN1b,pll,RpL36A,Btk29A,mRpL30,Mppe,CycA,Usp32,eEF5,RpS9,HtrA2,Pten,CG1218,eIF4B,eEF1delta,dgrn,Eaf6,Bub1,alpha-Man-Ib,Usp16-45,lic,Rpt6,RpS3A,Stlk,mRpS18A,mRpL34,inc,Stt3B,Rbbp5,RpL15,smt3,Parp,eRF1,CG8635,vnc,mRpS28,eIF3b,Rcd5,g,Prosalpha2,Jarid2,Dref,janA,tkv,Ufd4,Cdk8,Roc2,Acf,LIMK1,E(z),RpS26,Prosbeta1,PIG-K,Sgt,fzy,Wdr82,eIF3g1,RpL35A,Ubqn,Art1,CG31460,xit,CG31957,mRpS9,Skp2,RpS8,Pp1-87B,mnb,RpS19a,CG32473,RpL5 | | 5.675e-03 | -5.17 | regulation of alternative mRNA splicing, via spliceosome | biological process | GO:0000381 | 87 | 7 | 12345 | 305 | Rbp1,x16,snf,sqd,SF2,Cpsf6,tsu | | 5.682e-03 | -5.17 | RpLP1 (Ribosomal protein LP1) | protein interactions | 33214 | 140 | 11 | 9204 | 298 | RpS26,RpL5,RpL36A,RpS8,RpL14,RpS3A,RpL15,RpS9,RpS19a,RpL35A,14-3-3zeta | | 5.691e-03 | -5.17 | modification-dependent protein catabolic process | biological process | GO:0019941 | 290 | 15 | 12345 | 305 | inc,Ufd4,fzy,Sgt,Ubqn,Cul4,Rpn10,Prosalpha2,Prosbeta1,Skp2,Rpt6,alpha-Man-Ib,Roc2,Usp32,Usp16-45 | | 5.691e-03 | -5.17 | Rpn7 (Regulatory particle non-ATPase 7) | protein interactions | 42641 | 36 | 5 | 9204 | 298 | Prosbeta1,Rpn10,Ubqn,Prosalpha2,Rpt6 | | 5.791e-03 | -5.15 | regulation of cyclase activity | biological process | GO:0031279 | 5 | 2 | 12345 | 305 | Nf1,PlexA | | 5.791e-03 | -5.15 | histone H3-K4 demethylation | biological process | GO:0034720 | 5 | 2 | 12345 | 305 | NO66,Jarid2 | | 5.791e-03 | -5.15 | maternal specification of dorsal/ventral axis, oocyte, germ-line encoded | biological process | GO:0007311 | 5 | 2 | 12345 | 305 | glo,bai | | 5.791e-03 | -5.15 | phagolysosome assembly | biological process | GO:0001845 | 5 | 2 | 12345 | 305 | Rab7,Rab14 | | 5.791e-03 | -5.15 | synaptic vesicle budding | biological process | GO:0070142 | 5 | 2 | 12345 | 305 | g,Fmr1 | | 5.794e-03 | -5.15 | Prosalpha7 (Proteasome alpha7 subunit) | protein interactions | 36018 | 51 | 6 | 9204 | 298 | Rpt6,Prosalpha2,CG12321,Ubqn,Prosbeta1,Rpn10 | | 5.834e-03 | -5.14 | GO\_REGULATION\_OF\_CELL\_CYCLE\_G1\_S\_PHASE\_TRANSITION | MSigDB lists | GO\_REGULATION\_OF\_CELL\_CYCLE\_G1\_S\_PHASE\_TRANSITION | 38 | 6 | 4161 | 184 | Cul4,Ino80,Art1,E(z),twin,Pten | | 5.842e-03 | -5.14 | TSAI\_RESPONSE\_TO\_IONIZING\_RADIATION | MSigDB lists | TSAI\_RESPONSE\_TO\_IONIZING\_RADIATION | 27 | 5 | 4161 | 184 | sigmar,spel1,MED21,Art1,CCT4 | | 5.842e-03 | -5.14 | GCM\_PFN1 | MSigDB lists | GCM\_PFN1 | 27 | 5 | 4161 | 184 | RpL14,sm,Art1,snf,RpS8 | | 5.866e-03 | -5.14 | SCHRAMM\_INHBA\_TARGETS\_DN | MSigDB lists | SCHRAMM\_INHBA\_TARGETS\_DN | 9 | 3 | 4161 | 184 | cta,vap,Rap1 | | 5.866e-03 | -5.14 | GO\_HISTONE\_LYSINE\_N\_METHYLTRANSFERASE\_ACTIVITY | MSigDB lists | GO\_HISTONE\_LYSINE\_N\_METHYLTRANSFERASE\_ACTIVITY | 9 | 3 | 4161 | 184 | Rbbp5,Wdr82,E(z) | | 5.866e-03 | -5.14 | GO\_REGULATION\_OF\_ALTERNATIVE\_MRNA\_SPLICING\_VIA\_SPLICEOSOME | MSigDB lists | GO\_REGULATION\_OF\_ALTERNATIVE\_MRNA\_SPLICING\_VIA\_SPLICEOSOME | 9 | 3 | 4161 | 184 | tsu,Fmr1,sm | | 5.868e-03 | -5.14 | BILANGES\_SERUM\_AND\_RAPAMYCIN\_SENSITIVE\_GENES | MSigDB lists | BILANGES\_SERUM\_AND\_RAPAMYCIN\_SENSITIVE\_GENES | 50 | 7 | 4161 | 184 | RpS9,RpL14,RpL5,Tctp,RpL15,RpL36A,RpS8 | | 5.892e-03 | -5.13 | poly-pyrimidine tract binding | molecular function | GO:0008187 | 16 | 3 | 12176 | 289 | snf,Hrb27C,CG4612 | | 5.906e-03 | -5.13 | FAR/SIN/STRIPAK complex | cellular component | GO:0090443 | 5 | 2 | 11940 | 298 | Fgop2,Slmap | | 5.948e-03 | -5.12 | CG2224 (uncharacterized protein) | protein interactions | 43617 | 12 | 3 | 9204 | 298 | Acf,Spn,RpS3A | | 5.948e-03 | -5.12 | prd (paired) | protein interactions | 34629 | 12 | 3 | 9204 | 298 | CG42676,Mlf,LIMK1 | | 5.973e-03 | -5.12 | HSIAO\_HOUSEKEEPING\_GENES | MSigDB lists | HSIAO\_HOUSEKEEPING\_GENES | 198 | 17 | 4161 | 184 | RpS9,l(1)10Bb,14-3-3zeta,eEF1delta,RpL35A,RpS8,sm,RpL14,Parp,RpS26,Tctp,bai,RpL36A,CG17493,CCT4,Mdh1,CSN1b | | 6.004e-03 | -5.12 | shep (alan shepard) | protein interactions | 38605 | 4 | 2 | 9204 | 298 | su(Hw),Cp190 | | 6.004e-03 | -5.12 | CG30031 (uncharacterized protein) | protein interactions | 246404 | 4 | 2 | 9204 | 298 | Rpn10,Ubqn | | 6.004e-03 | -5.12 | Kebab (Kinetochore and EB1 associated basic protein) | protein interactions | 48440 | 4 | 2 | 9204 | 298 | BEAF-32,GEFmeso | | 6.004e-03 | -5.12 | Vm34Ca (Vitelline membrane 34Ca) | protein interactions | 34758 | 4 | 2 | 9204 | 298 | Sgt,Ubqn | | 6.004e-03 | -5.12 | CG13994 (uncharacterized protein) | protein interactions | 33853 | 4 | 2 | 9204 | 298 | Pp1-87B,I-2 | | 6.004e-03 | -5.12 | Gbs-76A (Glycogen binding subunit 76A) | protein interactions | 40102 | 4 | 2 | 9204 | 298 | flw,Pp1-87B | | 6.004e-03 | -5.12 | Irk2 (Inwardly rectifying potassium channel 2) | protein interactions | 42770 | 4 | 2 | 9204 | 298 | dpp,tkv | | 6.004e-03 | -5.12 | I-3 (Inhibitor-3) | protein interactions | 33068 | 4 | 2 | 9204 | 298 | Pp1-87B,flw | | 6.004e-03 | -5.12 | Ppi1 (Protein phosphatase 1c interacting protein 1) | protein interactions | 43581 | 4 | 2 | 9204 | 298 | Pp1-87B,flw | | 6.004e-03 | -5.12 | deltaTry (deltaTrypsin) | protein interactions | 48343 | 4 | 2 | 9204 | 298 | Ubqn,Rpn10 | | 6.004e-03 | -5.12 | CG4461 (uncharacterized protein) | protein interactions | 39074 | 4 | 2 | 9204 | 298 | krz,HtrA2 | | 6.004e-03 | -5.12 | CG31244 (uncharacterized protein) | protein interactions | 326129 | 4 | 2 | 9204 | 298 | flw,Pp1-87B | | 6.004e-03 | -5.12 | gammaTry (gammaTrypsin) | protein interactions | 36221 | 4 | 2 | 9204 | 298 | Rpn10,Ubqn | | 6.004e-03 | -5.12 | eggshell formation | biological process | GO:0030703 | 134 | 9 | 12345 | 305 | tkv,spoon,Fs(2)Ket,Cul4,Cdc42,rhi,smt3,dpp,RecQ4 | | 6.025e-03 | -5.11 | cell maturation | biological process | GO:0048469 | 184 | 11 | 12345 | 305 | 14-3-3zeta,bai,lic,mon2,flw,sqd,Hrb27C,dpp,tsu,glo,Fmr1 | | 6.031e-03 | -5.11 | GSE44649\_NAIVE\_VS\_ACTIVATED\_CD8\_TCELL\_DN | MSigDB lists | GSE44649\_NAIVE\_VS\_ACTIVATED\_CD8\_TCELL\_DN | 63 | 8 | 4161 | 184 | ND-ASHI,Pten,Lrr47,Fign,CG10343,CG4743,SF2,Skp2 | | 6.031e-03 | -5.11 | REACTOME\_PEPTIDE\_CHAIN\_ELONGATION | MSigDB lists | REACTOME\_PEPTIDE\_CHAIN\_ELONGATION | 63 | 8 | 4161 | 184 | RpL36A,RpL5,RpS9,RpS8,RpL35A,RpL15,RpS26,RpL14 | | 6.039e-03 | -5.11 | regulation of embryonic development | biological process | GO:0045995 | 88 | 7 | 12345 | 305 | Rop,tsu,cta,Fmr1,CycA,Rap1,dgrn | | 6.062e-03 | -5.11 | S6k (Ribosomal protein S6 kinase) | protein interactions | 38654 | 103 | 9 | 9204 | 298 | Rheb,14-3-3zeta,lic,Pten,Dref,Pp1-87B,Vap33,sqd,Tctp | | 6.120e-03 | -5.10 | GO\_REGULATION\_OF\_TRANSFERASE\_ACTIVITY | MSigDB lists | GO\_REGULATION\_OF\_TRANSFERASE\_ACTIVITY | 283 | 22 | 4161 | 184 | dpp,Pten,Cdc42,fzy,Nf1,LIMK1,E(z),Vap33,Rap1,Rpn10,Stlk,unc-119,CG6907,Prosalpha2,CSN1b,Rpt6,krz,CCT4,Rcd5,lic,janA,CycA | | 6.130e-03 | -5.09 | RpS8 (Ribosomal protein S8) | protein interactions | 43532 | 122 | 10 | 9204 | 298 | RpL35A,RpL5,RpS26,RpL14,RpS3A,RpL15,flw,RpS19a,RpS9,RpL36A | | 6.145e-03 | -5.09 | cellular biosynthetic process | biological process | GO:0044249 | 1702 | 58 | 12345 | 305 | CG11334,xit,CG31460,Aps,eIF4B,eEF1delta,eIF3g1,RpL35A,PIG-K,RpS26,eEF5,RpS9,Stt3B,RpL5,CG6244,RpS19a,mRpL34,mRpS18A,DNAlig1,heix,RpS3A,Arp8,RPA3,RpS8,Pbp49,CG11453,mRpS9,His4r,CG31957,sun,EloB,su(Hw),Cul4,rb,eRF1,Parp,Rpb11,Orc3,mRpS33,RpL15,CG15534,RecQ4,Coq5,eIF3c,E(bx),Pisd,mRpL30,TfIIFbeta,Btk29A,Dref,Pex3,g,eIF3b,RpL36A,mRpS28,RpL14,CG8635,grau | | 6.202e-03 | -5.08 | negative regulation of intracellular signal transduction | biological process | GO:1902532 | 111 | 8 | 12345 | 305 | vap,flw,Slmap,Rheb,mnb,Pten,Fgop2,krz | | 6.311e-03 | -5.07 | Energy dependent regulation of mTOR by LKB1-AMPK | REACTOME pathways | R-DME-380972 | 20 | 4 | 3923 | 151 | Rheb,RagA-B,Lst8,Stlk | | 6.358e-03 | -5.06 | GCM\_NPM1 | MSigDB lists | GCM\_NPM1 | 77 | 9 | 4161 | 184 | RpL14,Prosalpha2,Pp1-87B,RpS8,RpL35A,Art1,RpS9,Tctp,RpL36A | | 6.366e-03 | -5.06 | GO\_REGULATION\_OF\_APOPTOTIC\_SIGNALING\_PATHWAY | MSigDB lists | GO\_REGULATION\_OF\_APOPTOTIC\_SIGNALING\_PATHWAY | 91 | 10 | 4161 | 184 | Pten,Fign,Nf1,HtrA2,CG9246,Sod2,14-3-3zeta,Ubqn,Tctp,tkv | | 6.397e-03 | -5.05 | regulation of histone modification | biological process | GO:0031056 | 48 | 5 | 12345 | 305 | E(z),Jarid2,CG9246,Tctp,Ncoa6 | | 6.410e-03 | -5.05 | Syp (Syncrip) | protein interactions | 42460 | 37 | 5 | 9204 | 298 | Pten,Hrb27C,CG4612,Lst8,sqd | | 6.410e-03 | -5.05 | Xbp1 (X box binding protein-1) | protein interactions | 44226 | 37 | 5 | 9204 | 298 | Pep,Dref,RpS26,CG8108,His2Av | | 6.416e-03 | -5.05 | GO\_MRNA\_PROCESSING | MSigDB lists | GO\_MRNA\_PROCESSING | 167 | 15 | 4161 | 184 | snf,sm,CG11586,ubl,Rat1,noi,twin,l(1)10Bb,Rpb11,c12.1,Phf5a,tsu,CG4935,TfIIFbeta,SF2 | | 6.420e-03 | -5.05 | RNA transport | biological process | GO:0050658 | 89 | 7 | 12345 | 305 | CG6961,Fmr1,SF2,Tom20,sqd,Rab7,Dbp80 | | 6.420e-03 | -5.05 | nucleic acid transport | biological process | GO:0050657 | 89 | 7 | 12345 | 305 | CG6961,Fmr1,Tom20,SF2,Rab7,Dbp80,sqd | | 6.427e-03 | -5.05 | ZF\_CCHC | prosite domains | PS50158 | 13 | 3 | 6243 | 191 | CNBP,x16,RecQ4 | | 6.448e-03 | -5.04 | GO\_POSITIVE\_REGULATION\_OF\_CELL\_COMMUNICATION | MSigDB lists | GO\_POSITIVE\_REGULATION\_OF\_CELL\_COMMUNICATION | 355 | 26 | 4161 | 184 | Parp,dpp,Fmr1,Pten,CG32576,HtrA2,Cdc42,Nf1,eEF1delta,14-3-3zeta,l(1)10Bb,E(z),Lst8,Art1,tkv,CtBP,Rap1,Rpn10,RagA-B,Vps35,Prosalpha2,Rpt6,krz,Skp2,janA,lic | | 6.464e-03 | -5.04 | response to endogenous stimulus | biological process | GO:0009719 | 212 | 12 | 12345 | 305 | Zpr1,dpp,smt3,mav,RagA-B,su(Hw),Rap1,Cdc42,E(bx),tkv,Rheb,Rab8 | | 6.487e-03 | -5.04 | Rel (Relish) | protein interactions | 41087 | 123 | 10 | 9204 | 298 | Graf,nsl1,Cp190,CG5174,CNBP,CtBP,Map60,tsr,Vap33,Stim | | 6.501e-03 | -5.04 | GO\_NUCLEOPLASM\_PART | MSigDB lists | GO\_NUCLEOPLASM\_PART | 250 | 20 | 4161 | 184 | Wdr82,Fmr1,Pten,Pp1-87B,Phf5a,E(z),Rpb11,mbf1,Cdk8,TfIIFbeta,tsu,flw,CG6907,smt3,MED22,Rcd5,MED21,noi,SF2,Rbbp5 | | 6.539e-03 | -5.03 | Pink1 (PTEN-induced putative kinase 1) | protein interactions | 31607 | 86 | 8 | 9204 | 298 | glo,Tom20,Vps35,RpS9,Parp,heix,HtrA2,Sod2 | | 6.553e-03 | -5.03 | GSE14000\_TRANSLATED\_RNA\_VS\_MRNA\_4H\_LPS\_DC\_DN | MSigDB lists | GSE14000\_TRANSLATED\_RNA\_VS\_MRNA\_4H\_LPS\_DC\_DN | 51 | 7 | 4161 | 184 | His2Av,janA,c12.1,CG5861,Tctp,Vap33,Rap1 | | 6.553e-03 | -5.03 | KEGG\_FOCAL\_ADHESION | MSigDB lists | KEGG\_FOCAL\_ADHESION | 51 | 7 | 4161 | 184 | RhoGAPp190,Pp1-87B,Cdc42,Rap1,Pten,rhea,flw | | 6.563e-03 | -5.03 | mRNA catabolic process | biological process | GO:0006402 | 68 | 6 | 12345 | 305 | Upf3,smg,Not3,twin,tsu,sqd | | 6.586e-03 | -5.02 | negative regulation of mitotic sister chromatid segregation | biological process | GO:0033048 | 16 | 3 | 12345 | 305 | CycA,Bub1,Mps1 | | 6.586e-03 | -5.02 | negative regulation of mitotic metaphase/anaphase transition | biological process | GO:0045841 | 16 | 3 | 12345 | 305 | Mps1,Bub1,CycA | | 6.586e-03 | -5.02 | actin polymerization or depolymerization | biological process | GO:0008154 | 16 | 3 | 12345 | 305 | flr,tsr,Cdc42 | | 6.586e-03 | -5.02 | negative regulation of mitotic sister chromatid separation | biological process | GO:2000816 | 16 | 3 | 12345 | 305 | CycA,Bub1,Mps1 | | 6.602e-03 | -5.02 | regulation of kinase activity | biological process | GO:0043549 | 136 | 9 | 12345 | 305 | Stlk,Cdc42,Pten,mnb,sws,CycA,smt3,lic,CG1791 | | 6.638e-03 | -5.01 | KEGG\_RIBOSOME | MSigDB lists | KEGG\_RIBOSOME | 64 | 8 | 4161 | 184 | RpS9,RpL36A,RpL5,RpL14,RpS26,RpL35A,RpS8,RpL15 | | 6.638e-03 | -5.01 | GSE3720\_UNSTIM\_VS\_LPS\_STIM\_VD2\_GAMMADELTA\_TCELL\_UP | MSigDB lists | GSE3720\_UNSTIM\_VS\_LPS\_STIM\_VD2\_GAMMADELTA\_TCELL\_UP | 64 | 8 | 4161 | 184 | RpS8,DNAlig1,RpL14,RpL36A,RpL5,tsu,RpS9,MED21 | | 6.638e-03 | -5.01 | PECE\_MAMMARY\_STEM\_CELL\_UP | MSigDB lists | PECE\_MAMMARY\_STEM\_CELL\_UP | 64 | 8 | 4161 | 184 | eEF1delta,mbf1,RpS26,RpL35A,Sod2,ubl,RpL15,CG8860 | | 6.647e-03 | -5.01 | ZHANG\_TLX\_TARGETS\_DN | MSigDB lists | ZHANG\_TLX\_TARGETS\_DN | 39 | 6 | 4161 | 184 | SF2,CycA,DNAlig1,Lrr47,Fign,E(z) | | 6.647e-03 | -5.01 | GO\_REGULATION\_OF\_INTRINSIC\_APOPTOTIC\_SIGNALING\_PATHWAY | MSigDB lists | GO\_REGULATION\_OF\_INTRINSIC\_APOPTOTIC\_SIGNALING\_PATHWAY | 39 | 6 | 4161 | 184 | Fign,Ubqn,Tctp,HtrA2,CG9246,Sod2 | | 6.669e-03 | -5.01 | GO\_CELLULAR\_MACROMOLECULAR\_COMPLEX\_ASSEMBLY | MSigDB lists | GO\_CELLULAR\_MACROMOLECULAR\_COMPLEX\_ASSEMBLY | 338 | 25 | 4161 | 184 | Cul4,bai,SF2,CG17493,noi,CSN1b,CG12321,Tom20,Fs(2)Ket,eIF3c,Exo84,ND-ASHI,ECSIT,RpL5,eIF3b,His4r,CG12341,Rpn10,CG17883,Dbp21E2,CIA30,l(2)37Bb,eIF3g1,Parp,me31B | | 6.721e-03 | -5.00 | synapse part | cellular component | GO:0044456 | 211 | 12 | 11940 | 298 | Spn,g,Rpt6,Fmr1,Rop,Vap33,me31B,Vps35,mnb,tkv,Rab8,mav | | 6.771e-03 | -5.00 | biosynthetic process | biological process | GO:0009058 | 1781 | 60 | 12345 | 305 | Rpb11,Pect,Orc3,CG15534,mRpS33,RpL15,eIF3c,Coq5,RecQ4,sun,EloB,Cul4,su(Hw),Parp,eRF1,rb,g,Pex3,RpL36A,eIF3b,mRpS28,RpL14,grau,CG8635,E(bx),Pisd,mRpL30,TfIIFbeta,Dref,Btk29A,PIG-K,E(z),RpS26,RpS9,eEF5,CG31460,xit,CG11334,Aps,RpL35A,eIF3g1,eEF1delta,eIF4B,Pbp49,RpS8,Arp8,RPA3,CG11453,His4r,mRpS9,CG31957,CG6244,RpL5,Stt3B,mRpS18A,DNAlig1,mRpL34,RpS19a,heix,RpS3A | | 6.772e-03 | -5.00 | eukaryotic translation initiation factor 3 complex | cellular component | GO:0005852 | 16 | 3 | 11940 | 298 | eIF3c,eIF3g1,eIF3b | | 6.776e-03 | -4.99 | REACTOME\_METABOLISM\_OF\_RNA | MSigDB lists | REACTOME\_METABOLISM\_OF\_RNA | 168 | 15 | 4161 | 184 | Rpn10,tsu,RpL5,RpL36A,14-3-3zeta,RpS9,RpL15,Prosalpha2,Rpt6,RpS8,eRF1,RpL35A,RpS26,me31B,RpL14 | | 6.857e-03 | -4.98 | ZAP3 (ZAP3) | protein interactions | 31942 | 24 | 4 | 9204 | 298 | CG34159,Pp1-87B,Cpsf6,flw | | 6.857e-03 | -4.98 | His3.3A (Histone H3.3A) | protein interactions | 33736 | 24 | 4 | 9204 | 298 | CycA,Parp,Mps1,Dek | | 6.857e-03 | -4.98 | BicC (Bicaudal C) | protein interactions | 34946 | 24 | 4 | 9204 | 298 | twin,smg,Not3,me31B | | 6.860e-03 | -4.98 | Rac1 (Rac1) | protein interactions | 38146 | 124 | 10 | 9204 | 298 | LIMK1,Vps35,conu,RecQ4,GEFmeso,rhea,Graf,Pten,tkv,Fmr1 | | 6.862e-03 | -4.98 | GO\_PEPTIDE\_N\_ACETYLTRANSFERASE\_ACTIVITY | MSigDB lists | GO\_PEPTIDE\_N\_ACETYLTRANSFERASE\_ACTIVITY | 28 | 5 | 4161 | 184 | vnc,CG6907,mbf1,Rcd5,krz | | 6.883e-03 | -4.98 | fly chr85A3-85A3\|3-48 cM | chromosome location | fly chr85A3-85A3\|3-48 cM | 9 | 2 | 25010 | 358 | mRpS18A,CG31460 | | 6.883e-03 | -4.98 | fly chr85A3-85A3\|3-48 | chromosome location | fly chr85A3-85A3\|3-48 | 9 | 2 | 25010 | 358 | CG31460,mRpS18A | | 6.920e-03 | -4.97 | GO\_ESTABLISHMENT\_OF\_PROTEIN\_LOCALIZATION\_TO\_ENDOPLASMIC\_RETICULUM | MSigDB lists | GO\_ESTABLISHMENT\_OF\_PROTEIN\_LOCALIZATION\_TO\_ENDOPLASMIC\_RETICULUM | 78 | 9 | 4161 | 184 | RpL36A,RpL5,RpS9,RpS8,RpL35A,CG8860,RpL15,RpS26,RpL14 | | 6.920e-03 | -4.97 | PUJANA\_BRCA\_CENTERED\_NETWORK | MSigDB lists | PUJANA\_BRCA\_CENTERED\_NETWORK | 78 | 9 | 4161 | 184 | snf,fzy,Pp1-87B,spel1,Skp2,E(z),His2Av,CG5535,CycA | | 6.945e-03 | -4.97 | epithelium development | biological process | GO:0060429 | 1020 | 38 | 12345 | 305 | flr,eIF3b,Amun,CSN1b,Zpr1,tkv,Btk29A,flw,CtBP,smt3,RecQ4,Rheb,EloB,Parp,Cul4,Stim,VhaPPA1-1,Skp2,me31B,GEFmeso,Rab7,spoon,CG8149,tsr,sqd,Rop,cta,rhi,rhea,Cdk8,dpp,LIMK1,Fs(2)Ket,xit,Cdc42,Pten,peng,Rap1 | | 6.952e-03 | -4.97 | IKEDA\_MIR133\_TARGETS\_UP | MSigDB lists | IKEDA\_MIR133\_TARGETS\_UP | 18 | 4 | 4161 | 184 | Cdc42,CtBP,Usp32,Arpc5 | | 6.952e-03 | -4.97 | PID\_TCR\_PATHWAY | MSigDB lists | PID\_TCR\_PATHWAY | 18 | 4 | 4161 | 184 | Rap1,Cdc42,Pten,Stim | | 6.952e-03 | -4.97 | REACTOME\_SEMAPHORIN\_INTERACTIONS | MSigDB lists | REACTOME\_SEMAPHORIN\_INTERACTIONS | 18 | 4 | 4161 | 184 | PlexA,LIMK1,RhoGAPp190,Cdc42 | | 6.952e-03 | -4.97 | AATGGAG\_MIR136 | MSigDB lists | AATGGAG\_MIR136 | 18 | 4 | 4161 | 184 | Fmr1,Ino80,SF2,eRF1 | | 6.970e-03 | -4.97 | endosome | cellular component | GO:0005768 | 212 | 12 | 11940 | 298 | CG30423,g,mon2,Vps35,tkv,Rab8,Rab14,dpp,RagA-B,CG15168,ema,Rab7 | | 6.975e-03 | -4.97 | PENG\_RAPAMYCIN\_RESPONSE\_DN | MSigDB lists | PENG\_RAPAMYCIN\_RESPONSE\_DN | 137 | 13 | 4161 | 184 | vnc,eIF3b,CG14805,Dbp80,CCT4,Art1,Rpb11,CG41378,noi,eRF1,Txl,RpS26,CG11920 | | 6.983e-03 | -4.96 | positive regulation of chromatin organization | biological process | GO:1905269 | 49 | 5 | 12345 | 305 | E(z),cal1,Ncoa6,Cp190,Tctp | | 6.983e-03 | -4.96 | peptidyl-lysine methylation | biological process | GO:0018022 | 49 | 5 | 12345 | 305 | BEAF-32,E(z),Rbbp5,Wdr82,Ncoa6 | | 7.025e-03 | -4.96 | sima (similar) | protein interactions | 43580 | 206 | 14 | 9204 | 298 | Dref,peng,Txl,pav,Mlf,lic,14-3-3zeta,eIF3b,RAF2,RpL14,RpL15,c12.1,Rop,CG5516 | | 7.031e-03 | -4.96 | cullin family protein binding | molecular function | GO:0097602 | 17 | 3 | 12176 | 289 | CG9951,Roc2,inc | | 7.042e-03 | -4.96 | modification-dependent macromolecule catabolic process | biological process | GO:0043632 | 297 | 15 | 12345 | 305 | Roc2,alpha-Man-Ib,Usp32,Usp16-45,Rpn10,Rpt6,Prosbeta1,Skp2,Prosalpha2,Sgt,fzy,Cul4,Ubqn,inc,Ufd4 | | 7.068e-03 | -4.95 | anion binding | molecular function | GO:0043168 | 1099 | 39 | 12176 | 289 | Efa6,Rpt6,Bub1,Dbp80,lic,Fign,CCT4,Rab8,Graf,PlexA,Rheb,DNAlig1,Rab7,CG5439,Rab14,Cdc42,mus301,Rap1,pav,CG8915,pll,Cdk8,CG9330,RhoGAPp190,dpp,spel1,Mps1,RabX6,CG5589,Stlk,Acbp1,LIMK1,tkv,Btk29A,mnb,me31B,Dbp21E2,Ino80,RagA-B | | 7.077e-03 | -4.95 | cellular response to stimulus | biological process | GO:0051716 | 1467 | 51 | 12345 | 305 | CG6171,Btk29A,tkv,E(bx),pll,Zpr1,mthl8,Graf,Cul4,Parp,Rab8,Rheb,PPP1R15,RhoGAPp190,spel1,RecQ4,CG1791,smt3,flw,Pp1-87B,Stlk,Nf1,Lst8,PlexA,DNAlig1,Rab7,Tctp,Rab14,alpha-Man-Ib,lic,CG9005,Sod2,RabX6,RPA3,Arp8,dgrn,CG1218,Ino80,Cdc42,Sgt,Pten,Rap1,CG6066,mus301,Lrr47,dpp,Spn,mav,cta,RagA-B,CG5056 | | 7.081e-03 | -4.95 | GO\_POSTTRANSCRIPTIONAL\_REGULATION\_OF\_GENE\_EXPRESSION | MSigDB lists | GO\_POSTTRANSCRIPTIONAL\_REGULATION\_OF\_GENE\_EXPRESSION | 185 | 16 | 4161 | 184 | eIF3c,AGO3,eEF5,CG4866,Prosalpha2,Rpt6,14-3-3zeta,twin,RpS9,tsu,Rpn10,eIF3b,Fmr1,me31B,eIF3g1,eRF1 | | 7.114e-03 | -4.95 | GO\_NEGATIVE\_REGULATION\_OF\_CELL\_CYCLE | MSigDB lists | GO\_NEGATIVE\_REGULATION\_OF\_CELL\_CYCLE | 122 | 12 | 4161 | 184 | CycA,RagA-B,twin,Lst8,Art1,E(z),CG12321,HtrA2,spel1,dpp,Stlk,Pten | | 7.190e-03 | -4.94 | Prosalpha5 (Proteasome alpha5 subunit) | protein interactions | 36951 | 38 | 5 | 9204 | 298 | Prosbeta1,Ubqn,CG12321,Rpt6,Prosalpha2 | | 7.277e-03 | -4.92 | kinetochore | cellular component | GO:0000776 | 49 | 5 | 11940 | 298 | Mps1,Bub1,cal1,fzy,smt3 | | 7.277e-03 | -4.92 | pole plasm | cellular component | GO:0045495 | 49 | 5 | 11940 | 298 | Fmr1,me31B,AGO3,Fs(2)Ket,Hrb27C | | 7.282e-03 | -4.92 | chic (chickadee) | protein interactions | 33834 | 70 | 7 | 9204 | 298 | smt3,Tctp,Btk29A,spoon,Fmr1,eEF5,Cdc42 | | 7.291e-03 | -4.92 | GSE360\_CTRL\_VS\_B\_MALAYI\_LOW\_DOSE\_MAC\_UP | MSigDB lists | GSE360\_CTRL\_VS\_B\_MALAYI\_LOW\_DOSE\_MAC\_UP | 65 | 8 | 4161 | 184 | CG5589,sm,CG11920,CG9330,Rpb11,Pisd,MED21,Dbp21E2 | | 7.291e-03 | -4.92 | CREB\_Q4\_01 | MSigDB lists | CREB\_Q4\_01 | 65 | 8 | 4161 | 184 | CG13016,Cdc42,Nf1,CG15168,Roc2,14-3-3zeta,Tctp,CG5861 | | 7.297e-03 | -4.92 | GO\_PROTEASOME\_COMPLEX | MSigDB lists | GO\_PROTEASOME\_COMPLEX | 52 | 7 | 4161 | 184 | Ubqn,Txl,Rpt6,Rpn10,Prosalpha2,CG12321,CG9853 | | 7.297e-03 | -4.92 | GSE23308\_CTRL\_VS\_CORTICOSTERONE\_TREATED\_MACROPHAGE\_DN | MSigDB lists | GSE23308\_CTRL\_VS\_CORTICOSTERONE\_TREATED\_MACROPHAGE\_DN | 52 | 7 | 4161 | 184 | Rap1,sigmar,Fgop2,CG8635,Arpc5,Fs(2)Ket,Pect | | 7.540e-03 | -4.89 | MORF\_CCNI | MSigDB lists | MORF\_CCNI | 40 | 6 | 4161 | 184 | flr,RpL5,Arpc3A,RpL14,Rab8,14-3-3zeta | | 7.540e-03 | -4.89 | GSE3982\_EOSINOPHIL\_VS\_MAST\_CELL\_UP | MSigDB lists | GSE3982\_EOSINOPHIL\_VS\_MAST\_CELL\_UP | 40 | 6 | 4161 | 184 | Nulp1,ubl,Dbp21E2,Pten,me31B,Skp2 | | 7.540e-03 | -4.89 | MORI\_LARGE\_PRE\_BII\_LYMPHOCYTE\_UP | MSigDB lists | MORI\_LARGE\_PRE\_BII\_LYMPHOCYTE\_UP | 40 | 6 | 4161 | 184 | fzy,CycA,pav,DNAlig1,CG32473,His2Av | | 7.549e-03 | -4.89 | nxf2 (nuclear RNA export factor 2) | protein interactions | 39843 | 13 | 3 | 9204 | 298 | Ubqn,Slmap,Fgop2 | | 7.549e-03 | -4.89 | Rilpl (Rab interacting lysosomal protein like) | protein interactions | 31090 | 13 | 3 | 9204 | 298 | nom,Pnn,CG7766 | | 7.606e-03 | -4.88 | negative regulation of chromosome organization | biological process | GO:2001251 | 50 | 5 | 12345 | 305 | CG9246,Bub1,CycA,Mps1,su(Hw) | | 7.606e-03 | -4.88 | translation initiation factor activity | molecular function | GO:0003743 | 52 | 5 | 12176 | 289 | eIF3c,CG31957,eIF3g1,eIF3b,eIF4B | | 7.609e-03 | -4.88 | GO\_POSITIVE\_REGULATION\_OF\_CELLULAR\_COMPONENT\_BIOGENESIS | MSigDB lists | GO\_POSITIVE\_REGULATION\_OF\_CELLULAR\_COMPONENT\_BIOGENESIS | 108 | 11 | 4161 | 184 | TfIIFbeta,Cul4,Arpc3A,LIMK1,Lst8,Arpc5,twin,Cdc42,Rpt6,Parp,Fmr1 | | 7.615e-03 | -4.88 | maintenance of protein location in cell | biological process | GO:0032507 | 32 | 4 | 12345 | 305 | Vps13D,Parp,rhea,pav | | 7.666e-03 | -4.87 | Prosbeta5 (Proteasome beta5 subunit) | protein interactions | 45269 | 54 | 6 | 9204 | 298 | Prosbeta1,Rpn10,CG12321,Ubqn,Rpt6,Prosalpha2 | | 7.669e-03 | -4.87 | exocytosis | biological process | GO:0006887 | 92 | 7 | 12345 | 305 | Rab8,Rab7,Exo84,Rop,g,rb,Vap33 | | 7.832e-03 | -4.85 | reproduction | biological process | GO:0000003 | 1651 | 56 | 12345 | 305 | me31B,Bub1,Nph,Hrb27C,vsg,lic,glo,tsr,noi,Pp1-87B,Rab7,spoon,tsu,dpp,rhea,LIMK1,HtrA2,mus301,eEF5,sqd,Exo84,rhi,mon2,nsl1,Rop,fzy,Rap1,Cdc42,Pten,dgrn,snf,Fs(2)Ket,grau,CSN1b,vnc,lncRNA:sphinx,pav,flr,eIF3b,Btk29A,Fmr1,janA,MED22,E(bx),tkv,RecQ4,cal1,flw,bai,smt3,14-3-3zeta,Parp,Mps1,Cul4,BEAF-32,twin | | 7.853e-03 | -4.85 | negative regulation of metaphase/anaphase transition of cell cycle | biological process | GO:1902100 | 17 | 3 | 12345 | 305 | CycA,Bub1,Mps1 | | 7.853e-03 | -4.85 | syncytial blastoderm mitotic cell cycle | biological process | GO:0035186 | 17 | 3 | 12345 | 305 | smt3,E(z),CycA | | 7.853e-03 | -4.85 | regulation of RNA stability | biological process | GO:0043487 | 17 | 3 | 12345 | 305 | smg,Fmr1,mbf1 | | 7.853e-03 | -4.85 | negative regulation of chromosome separation | biological process | GO:1905819 | 17 | 3 | 12345 | 305 | CycA,Bub1,Mps1 | | 7.884e-03 | -4.84 | organellar small ribosomal subunit | cellular component | GO:0000314 | 32 | 4 | 11940 | 298 | mRpS18A,mRpS9,mRpS28,mRpS33 | | 7.884e-03 | -4.84 | mitochondrial small ribosomal subunit | cellular component | GO:0005763 | 32 | 4 | 11940 | 298 | mRpS9,mRpS18A,mRpS28,mRpS33 | | 7.884e-03 | -4.84 | mediator complex | cellular component | GO:0016592 | 32 | 4 | 11940 | 298 | Cdk8,MED22,MED21,MED26 | | 7.954e-03 | -4.83 | Interferon-induced\_GILT | interpro domains | IPR004911 | 5 | 2 | 11383 | 331 | GILT3,CG41378 | | 7.954e-03 | -4.83 | NUDIX\_hydrolase\_CS | interpro domains | IPR020084 | 5 | 2 | 11383 | 331 | Aps,Apf | | 7.954e-03 | -4.83 | TGF-b\_propeptide | interpro domains | IPR001111 | 5 | 2 | 11383 | 331 | dpp,mav | | 7.954e-03 | -4.83 | RasGAP\_dom | interpro domains | IPR001936 | 5 | 2 | 11383 | 331 | Nf1,vap | | 7.956e-03 | -4.83 | ROME\_INSULIN\_TARGETS\_IN\_MUSCLE\_UP | MSigDB lists | ROME\_INSULIN\_TARGETS\_IN\_MUSCLE\_UP | 171 | 15 | 4161 | 184 | Vap33,CG10343,CG5339,Ubqn,CCT4,Arpc3A,CG34148,His2Av,Ufd4,Pp1-87B,mRpS33,VhaPPA1-1,Wdr82,Fs(2)Ket,Pten | | 7.959e-03 | -4.83 | TFAM (mitochondrial transcription factor A) | protein interactions | 42433 | 25 | 4 | 9204 | 298 | RPA3,GILT3,Parp,PPP1R15 | | 7.959e-03 | -4.83 | geminin (geminin) | protein interactions | 35563 | 25 | 4 | 9204 | 298 | Slmap,grau,Fgop2,Ubqn | | 7.959e-03 | -4.83 | CG2926 (uncharacterized protein) | protein interactions | 40676 | 25 | 4 | 9204 | 298 | Rcd5,CG32075,Jarid2,flw | | 7.959e-03 | -4.83 | Not1 (Not1) | protein interactions | 5740252 | 25 | 4 | 9204 | 298 | Not3,twin,Zpr1,me31B | | 7.974e-03 | -4.83 | GO\_RRNA\_METABOLIC\_PROCESS | MSigDB lists | GO\_RRNA\_METABOLIC\_PROCESS | 155 | 14 | 4161 | 184 | RpS9,RpL36A,RpL5,CG14805,RpL14,RpS26,CG5589,CG11920,RpL35A,mRpS9,Rat1,RpS8,CG4866,RpL15 | | 7.987e-03 | -4.83 | GSE17721\_POLYIC\_VS\_GARDIQUIMOD\_6H\_BMDC\_DN | MSigDB lists | GSE17721\_POLYIC\_VS\_GARDIQUIMOD\_6H\_BMDC\_DN | 94 | 10 | 4161 | 184 | Cdk8,CG4743,eIF3b,flw,Bmcp,smt3,CG4866,CG8860,ubl,Rpt6 | | 7.991e-03 | -4.83 | GSE17721\_CTRL\_VS\_PAM3CSK4\_0.5H\_BMDC\_UP | MSigDB lists | GSE17721\_CTRL\_VS\_PAM3CSK4\_0.5H\_BMDC\_UP | 66 | 8 | 4161 | 184 | CG8860,Mdh1,Ino80,twin,CIA30,SF2,CG5590,Vps35 | | 7.991e-03 | -4.83 | GSE15930\_NAIVE\_VS\_72H\_IN\_VITRO\_STIM\_CD8\_TCELL\_DN | MSigDB lists | GSE15930\_NAIVE\_VS\_72H\_IN\_VITRO\_STIM\_CD8\_TCELL\_DN | 66 | 8 | 4161 | 184 | CG5589,unc-119,pav,fzy,Pect,E(z),CG41378,CG5590 | | 8.001e-03 | -4.83 | JUBAN\_TARGETS\_OF\_SPI1\_AND\_FLI1\_DN | MSigDB lists | JUBAN\_TARGETS\_OF\_SPI1\_AND\_FLI1\_DN | 29 | 5 | 4161 | 184 | tkv,sm,CG11920,l(1)10Bb,eIF3c | | 8.001e-03 | -4.83 | ZHANG\_RESPONSE\_TO\_IKK\_INHIBITOR\_AND\_TNF\_UP | MSigDB lists | ZHANG\_RESPONSE\_TO\_IKK\_INHIBITOR\_AND\_TNF\_UP | 29 | 5 | 4161 | 184 | sigmar,Sod2,CG5589,lic,dpp | | 8.034e-03 | -4.82 | regulation of synapse assembly | biological process | GO:0051963 | 116 | 8 | 12345 | 305 | Nf1,Fmr1,Cdc42,tkv,mnb,Rab8,Rheb,Spn | | 8.034e-03 | -4.82 | Rpn2 (Regulatory particle non-ATPase 2) | protein interactions | 43449 | 39 | 5 | 9204 | 298 | CG12321,Ubqn,Prosbeta1,Rpt6,Prosalpha2 | | 8.034e-03 | -4.82 | Cul3 (Cullin 3) | protein interactions | 34896 | 39 | 5 | 9204 | 298 | inc,flw,eIF3b,Nf1,Dref | | 8.101e-03 | -4.82 | MODULE\_239 | MSigDB lists | MODULE\_239 | 53 | 7 | 4161 | 184 | bai,Pp1-87B,CG32576,Orc3,Tom20,Arpc5,rb | | 8.101e-03 | -4.82 | GSE22313\_HEALTHY\_VS\_SLE\_MOUSE\_CD4\_TCELL\_DN | MSigDB lists | GSE22313\_HEALTHY\_VS\_SLE\_MOUSE\_CD4\_TCELL\_DN | 53 | 7 | 4161 | 184 | CycA,Aps,RhoGAPp190,pav,Parp,His2Av,rhea | | 8.101e-03 | -4.82 | OSWALD\_HEMATOPOIETIC\_STEM\_CELL\_IN\_COLLAGEN\_GEL\_UP | MSigDB lists | OSWALD\_HEMATOPOIETIC\_STEM\_CELL\_IN\_COLLAGEN\_GEL\_UP | 53 | 7 | 4161 | 184 | Sod2,Nf1,Skp2,eEF1delta,noi,Fs(2)Ket,CG5535 | | 8.101e-03 | -4.82 | GSE29949\_CD8\_NEG\_DC\_SPLEEN\_VS\_DC\_BRAIN\_DN | MSigDB lists | GSE29949\_CD8\_NEG\_DC\_SPLEEN\_VS\_DC\_BRAIN\_DN | 53 | 7 | 4161 | 184 | His4r,Tcs3,vap,Pp1-87B,CG5589,Pex3,Vps13D | | 8.109e-03 | -4.81 | GO\_REGULATION\_OF\_MEIOTIC\_NUCLEAR\_DIVISION | MSigDB lists | GO\_REGULATION\_OF\_MEIOTIC\_NUCLEAR\_DIVISION | 10 | 3 | 4161 | 184 | AGO3,fzy,spel1 | | 8.109e-03 | -4.81 | GO\_MACROMOLECULE\_TRANSMEMBRANE\_TRANSPORTER\_ACTIVITY | MSigDB lists | GO\_MACROMOLECULE\_TRANSMEMBRANE\_TRANSPORTER\_ACTIVITY | 10 | 3 | 4161 | 184 | CG8860,Tom20,Tim17b | | 8.109e-03 | -4.81 | GO\_POSITIVE\_REGULATION\_OF\_ENDOTHELIAL\_CELL\_PROLIFERATION | MSigDB lists | GO\_POSITIVE\_REGULATION\_OF\_ENDOTHELIAL\_CELL\_PROLIFERATION | 10 | 3 | 4161 | 184 | viaf,Nf1,dpp | | 8.109e-03 | -4.81 | BIOCARTA\_SALMONELLA\_PATHWAY | MSigDB lists | BIOCARTA\_SALMONELLA\_PATHWAY | 10 | 3 | 4161 | 184 | Arpc5,Arpc3A,Cdc42 | | 8.109e-03 | -4.81 | GO\_REGULATION\_OF\_EXTRINSIC\_APOPTOTIC\_SIGNALING\_PATHWAY\_VIA\_DEATH\_DOMAIN\_RECEPTORS | MSigDB lists | GO\_REGULATION\_OF\_EXTRINSIC\_APOPTOTIC\_SIGNALING\_PATHWAY\_VIA\_DEATH\_DOMAIN\_RECEPTORS | 10 | 3 | 4161 | 184 | Pten,Nf1,tkv | | 8.109e-03 | -4.81 | GO\_CALCIUM\_CHANNEL\_REGULATOR\_ACTIVITY | MSigDB lists | GO\_CALCIUM\_CHANNEL\_REGULATOR\_ACTIVITY | 10 | 3 | 4161 | 184 | Tsp97E,janA,Stim | | 8.121e-03 | -4.81 | GO\_CELLULAR\_RESPONSE\_TO\_DNA\_DAMAGE\_STIMULUS | MSigDB lists | GO\_CELLULAR\_RESPONSE\_TO\_DNA\_DAMAGE\_STIMULUS | 238 | 19 | 4161 | 184 | Parp,spel1,HtrA2,mRpS9,twin,Art1,Rpb11,His4r,DNAlig1,CSN1b,smt3,Arp8,mus301,Rcd5,Ino80,Cul4,Rbbp5,CycA,CG17493 | | 8.123e-03 | -4.81 | regulation of Wnt signaling pathway | biological process | GO:0030111 | 93 | 7 | 12345 | 305 | Pp1-87B,pav,flw,Nulp1,Vps35,CtBP,lic | | 8.128e-03 | -4.81 | PGRP-LE (Peptidoglycan recognition protein LE) | protein interactions | 32534 | 147 | 11 | 9204 | 298 | Map60,CtBP,CCT4,Stim,tsr,rhea,Cp190,nsl1,Graf,CG5174,CNBP | | 8.145e-03 | -4.81 | REACTOME\_INFLUENZA\_LIFE\_CYCLE | MSigDB lists | REACTOME\_INFLUENZA\_LIFE\_CYCLE | 109 | 11 | 4161 | 184 | Fs(2)Ket,RpL14,RpS26,RpL15,RpL35A,RpS8,RpS9,Rpb11,RpL5,TfIIFbeta,RpL36A | | 8.158e-03 | -4.81 | GSE5589\_LPS\_VS\_LPS\_AND\_IL10\_STIM\_IL6\_KO\_MACROPHAGE\_180MIN\_DN | MSigDB lists | GSE5589\_LPS\_VS\_LPS\_AND\_IL10\_STIM\_IL6\_KO\_MACROPHAGE\_180MIN\_DN | 80 | 9 | 4161 | 184 | TfIIFbeta,eIF3b,CG5590,Rpn10,CG41378,CG15168,CG5535,Sgt,PIG-K | | 8.240e-03 | -4.80 | RNA helicase activity | molecular function | GO:0003724 | 53 | 5 | 12176 | 289 | Dbp21E2,CG8915,me31B,Dbp80,CG5589 | | 8.298e-03 | -4.79 | cell fate determination | biological process | GO:0001709 | 141 | 9 | 12345 | 305 | Pten,Fmr1,pins,tsu,dpp,mus301,pav,14-3-3zeta,CtBP | | 8.474e-03 | -4.77 | Rpt6 (Regulatory particle triple-A ATPase 6) | protein interactions | 33105 | 72 | 7 | 9204 | 298 | Rpn10,Prosbeta1,Ubqn,CG12321,CCT4,Rpt6,Prosalpha2 | | 8.500e-03 | -4.77 | transmembrane receptor protein serine/threonine kinase signaling pathway | biological process | GO:0007178 | 33 | 4 | 12345 | 305 | tkv,mav,Cdc42,dpp | | 8.500e-03 | -4.77 | cell cycle DNA replication | biological process | GO:0044786 | 33 | 4 | 12345 | 305 | Btk29A,Dref,Orc3,DNAlig1 | | 8.500e-03 | -4.77 | germarium-derived oocyte differentiation | biological process | GO:0030706 | 33 | 4 | 12345 | 305 | tsu,14-3-3zeta,Fmr1,mus301 | | 8.507e-03 | -4.77 | TANG\_SENESCENCE\_TP53\_TARGETS\_DN | MSigDB lists | TANG\_SENESCENCE\_TP53\_TARGETS\_DN | 19 | 4 | 4161 | 184 | CycA,fzy,pav,E(z) | | 8.507e-03 | -4.77 | GO\_HISTONE\_METHYLTRANSFERASE\_ACTIVITY | MSigDB lists | GO\_HISTONE\_METHYLTRANSFERASE\_ACTIVITY | 19 | 4 | 4161 | 184 | Rbbp5,Art1,E(z),Wdr82 | | 8.507e-03 | -4.77 | GSE2706\_UNSTIM\_VS\_2H\_LPS\_DC\_DN | MSigDB lists | GSE2706\_UNSTIM\_VS\_2H\_LPS\_DC\_DN | 19 | 4 | 4161 | 184 | lic,E(z),sigmar,Sod2 | | 8.507e-03 | -4.77 | GO\_REGULATION\_OF\_GLIOGENESIS | MSigDB lists | GO\_REGULATION\_OF\_GLIOGENESIS | 19 | 4 | 4161 | 184 | Nf1,flr,E(z),dpp | | 8.507e-03 | -4.77 | HINATA\_NFKB\_TARGETS\_KERATINOCYTE\_UP | MSigDB lists | HINATA\_NFKB\_TARGETS\_KERATINOCYTE\_UP | 19 | 4 | 4161 | 184 | lic,dpp,sigmar,Sod2 | | 8.516e-03 | -4.77 | RHODES\_UNDIFFERENTIATED\_CANCER | MSigDB lists | RHODES\_UNDIFFERENTIATED\_CANCER | 41 | 6 | 4161 | 184 | CG41378,E(z),pav,pins,CycA,fzy | | 8.522e-03 | -4.77 | fly chr89B16-89B16\|3-58 | chromosome location | fly chr89B16-89B16\|3-58 | 10 | 2 | 25010 | 358 | mRpS33,ema | | 8.522e-03 | -4.77 | fly chr89B16-89B16\|3-58 cM | chromosome location | fly chr89B16-89B16\|3-58 cM | 10 | 2 | 25010 | 358 | ema,mRpS33 | | 8.546e-03 | -4.76 | protein ADP-ribosylation | biological process | GO:0006471 | 6 | 2 | 12345 | 305 | CG1218,Parp | | 8.546e-03 | -4.76 | regulation of lyase activity | biological process | GO:0051339 | 6 | 2 | 12345 | 305 | Nf1,PlexA | | 8.546e-03 | -4.76 | actin filament depolymerization | biological process | GO:0030042 | 6 | 2 | 12345 | 305 | tsr,flr | | 8.549e-03 | -4.76 | determination of adult lifespan | biological process | GO:0008340 | 167 | 10 | 12345 | 305 | sun,Nf1,rb,Btk29A,Sod2,g,BthD,Hsp27,mthl8,sm | | 8.549e-03 | -4.76 | Znf\_CCHC | interpro domains | IPR001878 | 15 | 3 | 11383 | 331 | RecQ4,x16,CNBP | | 8.549e-03 | -4.76 | NUDIX\_hydrolase\_dom | interpro domains | IPR000086 | 15 | 3 | 11383 | 331 | CG12567,Apf,Aps | | 8.597e-03 | -4.76 | positive regulation of developmental growth | biological process | GO:0048639 | 94 | 7 | 12345 | 305 | Rheb,mnb,lic,tkv,dpp,Nf1,Tctp | | 8.686e-03 | -4.75 | small molecule binding | molecular function | GO:0036094 | 1078 | 38 | 12176 | 289 | CG9330,Cdk8,pll,CG8915,pav,CtBP,Rap1,mus301,Cdc42,Rab14,CG34261,Rab7,DNAlig1,Rheb,cta,Rab8,CCT4,Fign,lic,Bub1,Dbp80,Rpt6,RagA-B,Ino80,Dbp21E2,Parp,me31B,mnb,Btk29A,tkv,Acbp1,LIMK1,Stlk,CG5589,RabX6,Mps1,spel1,RhoGAPp190 | | 8.691e-03 | -4.75 | GILT | pfam domains | PF03227 | 5 | 2 | 10480 | 319 | CG41378,GILT3 | | 8.691e-03 | -4.75 | RasGAP | pfam domains | PF00616 | 5 | 2 | 10480 | 319 | Nf1,vap | | 8.691e-03 | -4.75 | TGFb\_propeptide | pfam domains | PF00688 | 5 | 2 | 10480 | 319 | mav,dpp | | 8.709e-03 | -4.74 | AnxB10 (Annexin B10) | protein interactions | 33019 | 109 | 9 | 9204 | 298 | CG7054,eEF5,Tctp,Nph,smt3,eEF1delta,CG5174,14-3-3zeta,Prosalpha2 | | 8.710e-03 | -4.74 | organic substance biosynthetic process | biological process | GO:1901576 | 1731 | 58 | 12345 | 305 | g,Pex3,eIF3b,RpL36A,RpL14,mRpS28,grau,CG8635,E(bx),Pisd,TfIIFbeta,mRpL30,Dref,Btk29A,Rpb11,mRpS33,RpL15,CG15534,Orc3,eIF3c,Coq5,RecQ4,sun,EloB,Cul4,su(Hw),Parp,eRF1,rb,Pbp49,Arp8,RPA3,RpS8,CG11453,His4r,mRpS9,CG31957,CG6244,Stt3B,RpL5,mRpS18A,DNAlig1,mRpL34,RpS19a,heix,RpS3A,PIG-K,RpS26,eEF5,RpS9,CG31460,xit,CG11334,Aps,eIF3g1,RpL35A,eIF4B,eEF1delta | | 8.714e-03 | -4.74 | NURF complex | cellular component | GO:0016589 | 6 | 2 | 11940 | 298 | Acf,E(bx) | | 8.714e-03 | -4.74 | phagocytic vesicle | cellular component | GO:0045335 | 6 | 2 | 11940 | 298 | Rab14,Rab7 | | 8.740e-03 | -4.74 | TGACGTCA\_ATF3\_Q6 | MSigDB lists | TGACGTCA\_ATF3\_Q6 | 67 | 8 | 4161 | 184 | 14-3-3zeta,Roc2,CG15168,CG5861,Dbp21E2,CycA,Nf1,CG13016 | | 8.740e-03 | -4.74 | GSE3337\_CTRL\_VS\_16H\_IFNG\_IN\_CD8POS\_DC\_DN | MSigDB lists | GSE3337\_CTRL\_VS\_16H\_IFNG\_IN\_CD8POS\_DC\_DN | 67 | 8 | 4161 | 184 | l(1)10Bb,Rop,Rpn10,unc-119,Aps,Sod2,fzy,CSN1b | | 8.761e-03 | -4.74 | NUDIX\_BOX | prosite domains | PS00893 | 5 | 2 | 6243 | 191 | Aps,Apf | | 8.761e-03 | -4.74 | RAS\_GTPASE\_ACTIV\_2 | prosite domains | PS50018 | 5 | 2 | 6243 | 191 | vap,Nf1 | | 8.799e-03 | -4.73 | nuclear speck | cellular component | GO:0016607 | 33 | 4 | 11940 | 298 | Rbp1,x16,sqd,SF2 | | 8.837e-03 | -4.73 | GO\_RNA\_SPLICING | MSigDB lists | GO\_RNA\_SPLICING | 141 | 13 | 4161 | 184 | SF2,TfIIFbeta,CG4935,tsu,Phf5a,c12.1,noi,Rpb11,l(1)10Bb,ubl,CG11586,snf,sm | | 8.837e-03 | -4.73 | KIM\_WT1\_TARGETS\_DN | MSigDB lists | KIM\_WT1\_TARGETS\_DN | 141 | 13 | 4161 | 184 | Cdk8,CycA,His2Av,MED21,mbf1,RpS9,Skp2,CG4866,pav,sigmar,fzy,pins,Sod2 | | 8.837e-03 | -4.73 | GSE10239\_MEMORY\_VS\_KLRG1HIGH\_EFF\_CD8\_TCELL\_DN | MSigDB lists | GSE10239\_MEMORY\_VS\_KLRG1HIGH\_EFF\_CD8\_TCELL\_DN | 81 | 9 | 4161 | 184 | Lrr47,pav,pins,Sod2,fzy,14-3-3zeta,CG5339,Cul4,bai | | 8.837e-03 | -4.73 | REACTOME\_SRP\_DEPENDENT\_COTRANSLATIONAL\_PROTEIN\_TARGETING\_TO\_MEMBRANE | MSigDB lists | REACTOME\_SRP\_DEPENDENT\_COTRANSLATIONAL\_PROTEIN\_TARGETING\_TO\_MEMBRANE | 81 | 9 | 4161 | 184 | RpS9,RpL5,RpL36A,RpS26,RpL14,CG8860,RpL15,RpS8,RpL35A | | 8.837e-03 | -4.73 | GSE22432\_MULTIPOTENT\_PROGENITOR\_VS\_CDC\_DN | MSigDB lists | GSE22432\_MULTIPOTENT\_PROGENITOR\_VS\_CDC\_DN | 81 | 9 | 4161 | 184 | Rap1,Pect,CG5535,CCT4,eIF3g1,pav,DNAlig1,CG12321,Fign | | 8.837e-03 | -4.73 | MORF\_FBL | MSigDB lists | MORF\_FBL | 81 | 9 | 4161 | 184 | smt3,eIF3g1,RpL14,eIF3c,snf,Tom20,eIF3b,CG5861,Art1 | | 8.870e-03 | -4.73 | positive regulation of growth | biological process | GO:0045927 | 118 | 8 | 12345 | 305 | Nf1,Rheb,mnb,tkv,dpp,Tctp,RagA-B,lic | | 8.898e-03 | -4.72 | nuclear chromosome segregation | biological process | GO:0098813 | 168 | 10 | 12345 | 305 | pav,Rpn10,cal1,mus301,Bub1,CycA,grau,Fign,Pp1-87B,Mps1 | | 8.900e-03 | -4.72 | GO\_VIRAL\_LIFE\_CYCLE | MSigDB lists | GO\_VIRAL\_LIFE\_CYCLE | 157 | 14 | 4161 | 184 | CG17883,CtBP,RpL36A,RpL5,eIF3b,RpS9,RpS8,RpL35A,eIF3g1,RpL15,RpS26,me31B,RpL14,Fs(2)Ket | | 8.969e-03 | -4.71 | GSE2770\_TGFB\_AND\_IL4\_VS\_IL12\_TREATED\_ACT\_CD4\_TCELL\_48H\_DN | MSigDB lists | GSE2770\_TGFB\_AND\_IL4\_VS\_IL12\_TREATED\_ACT\_CD4\_TCELL\_48H\_DN | 54 | 7 | 4161 | 184 | Ufd4,Sod2,xit,me31B,CG5535,ND-ASHI,Roc2 | | 9.045e-03 | -4.71 | late endosome | cellular component | GO:0005770 | 72 | 6 | 11940 | 298 | g,tkv,Vps35,Rab14,ema,Rab7 | | 9.047e-03 | -4.71 | mitochondrial part | cellular component | GO:0044429 | 480 | 21 | 11940 | 298 | spoon,CG44194,CG4743,mRpS28,l(2)37Bb,CG11752,mRpS9,MICU1,sun,HtrA2,Tim17b,ND-AGGG,CIA30,mRpS33,Sod2,Bmcp,mRpL30,ND-ASHI,mRpS18A,mRpL34,Tom20 | | 9.090e-03 | -4.70 | immune system development | biological process | GO:0002520 | 95 | 7 | 12345 | 305 | E(bx),RpS8,heix,CG8149,dpp,Rap1,Cdc42 | | 9.090e-03 | -4.70 | hematopoietic or lymphoid organ development | biological process | GO:0048534 | 95 | 7 | 12345 | 305 | Cdc42,Rap1,dpp,CG8149,heix,RpS8,E(bx) | | 9.098e-03 | -4.70 | post-embryonic development | biological process | GO:0009791 | 642 | 26 | 12345 | 305 | LIMK1,RecQ4,dpp,Cdk8,rhea,smt3,nsl1,flw,BthD,nom,peng,Cdc42,Pten,EloB,pll,GEFmeso,RpS8,Stim,flr,CG8149,tsr,heix,krz,Btk29A,E(bx),tkv | | 9.109e-03 | -4.70 | establishment of protein localization | biological process | GO:0045184 | 485 | 21 | 12345 | 305 | Dbp80,g,RabX6,Tom20,Pex3,Rab14,Tim17b,CG6961,Rab7,Vps35,Vps13D,CG9853,sqd,CG8860,mon2,bai,Rab8,unc-119,Fs(2)Ket,Sgt,rb | | 9.123e-03 | -4.70 | GO\_CELL\_CYCLE\_PROCESS | MSigDB lists | GO\_CELL\_CYCLE\_PROCESS | 383 | 27 | 4161 | 184 | Rpt6,DNAlig1,Prosalpha2,Arp8,CG12321,Fs(2)Ket,flw,Stlk,CycA,CG17493,Skp2,Ino80,spel1,pav,pins,fzy,Pp1-87B,Pten,Rab8,RagA-B,vap,Orc3,Rpn10,twin,Lst8,E(z),Art1 | | 9.134e-03 | -4.70 | Rpn1 (Regulatory particle non-ATPase 1) | protein interactions | 40174 | 56 | 6 | 9204 | 298 | Prosalpha2,Rpt6,Ubqn,CG12321,Prosbeta1,Rab7 | | 9.141e-03 | -4.69 | GO\_NEGATIVE\_REGULATION\_OF\_CELL\_COMMUNICATION | MSigDB lists | GO\_NEGATIVE\_REGULATION\_OF\_CELL\_COMMUNICATION | 293 | 22 | 4161 | 184 | Rop,Tctp,RhoGAPp190,krz,janA,Rpt6,CSN1b,Prosalpha2,Rap1,Rpn10,vap,Ubqn,E(z),Sod2,HtrA2,CG9246,Cdc42,Nf1,dpp,Fign,Pten,Fmr1 | | 9.148e-03 | -4.69 | GO\_REGULATION\_OF\_CELL\_CYCLE\_PHASE\_TRANSITION | MSigDB lists | GO\_REGULATION\_OF\_CELL\_CYCLE\_PHASE\_TRANSITION | 126 | 12 | 4161 | 184 | CG12321,fzy,Prosalpha2,Rpt6,Pten,CycA,Cul4,Rpn10,Ino80,E(z),Art1,twin | | 9.171e-03 | -4.69 | RhoGEF2 (Rho guanine nucleotide exchange factor 2) | protein interactions | 36915 | 26 | 4 | 9204 | 298 | cta,Cdc42,flw,LIMK1 | | 9.171e-03 | -4.69 | Lis-1 (Lissencephaly-1) | protein interactions | 36791 | 26 | 4 | 9204 | 298 | Dlip2,smg,Hsp27,tkv | | 9.171e-03 | -4.69 | fs(1)h (female sterile (1) homeotic) | protein interactions | 31722 | 26 | 4 | 9204 | 298 | su(Hw),BEAF-32,Dref,Cp190 | | 9.221e-03 | -4.69 | cellular nitrogen compound biosynthetic process | biological process | GO:0044271 | 1175 | 42 | 12345 | 305 | Rpb11,CG15534,mRpS33,RpL15,eIF3c,RecQ4,sun,EloB,Cul4,su(Hw),Parp,eRF1,eIF3b,RpL36A,RpL14,mRpS28,grau,CG8635,E(bx),TfIIFbeta,mRpL30,RpS26,RpS9,eEF5,Aps,eIF3g1,RpL35A,eEF1delta,eIF4B,Pbp49,Arp8,RpS8,His4r,mRpS9,CG31957,CG6244,RpL5,mRpS18A,DNAlig1,RpS19a,mRpL34,RpS3A | | 9.265e-03 | -4.68 | PID\_CDC42\_PATHWAY | MSigDB lists | PID\_CDC42\_PATHWAY | 30 | 5 | 4161 | 184 | Arpc5,lic,LIMK1,Arpc3A,Cdc42 | | 9.265e-03 | -4.68 | GSE3982\_DC\_VS\_NEUTROPHIL\_DN | MSigDB lists | GSE3982\_DC\_VS\_NEUTROPHIL\_DN | 30 | 5 | 4161 | 184 | Arpc5,lic,Pten,eEF1delta,RhoGAPp190 | | 9.304e-03 | -4.68 | GSE2405\_0H\_VS\_9H\_A\_PHAGOCYTOPHILUM\_STIM\_NEUTROPHIL\_DN | MSigDB lists | GSE2405\_0H\_VS\_9H\_A\_PHAGOCYTOPHILUM\_STIM\_NEUTROPHIL\_DN | 111 | 11 | 4161 | 184 | Tctp,RpL5,14-3-3zeta,RpS9,Arpc3A,c12.1,Pp1-87B,RpS8,Aps,RpS26,RpL14 | | 9.359e-03 | -4.67 | MORF\_DEK | MSigDB lists | MORF\_DEK | 142 | 13 | 4161 | 184 | CG8635,Arpc5,Art1,14-3-3zeta,His2Av,SF2,snf,eRF1,Mdh1,Prosalpha2,CG8860,smt3,spel1 | | 9.381e-03 | -4.67 | RhoGAP18B (Rho GTPase activating protein at 18B) | protein interactions | 32898 | 14 | 3 | 9204 | 298 | Cdc42,pins,LIMK1 | | 9.381e-03 | -4.67 | CG12768 (uncharacterized protein) | protein interactions | 40513 | 14 | 3 | 9204 | 298 | eIF4B,Fmr1,CtBP | | 9.381e-03 | -4.67 | kto (kohtalo) | protein interactions | 44830 | 14 | 3 | 9204 | 298 | BEAF-32,flw,Cdk8 | | 9.381e-03 | -4.67 | Prosbeta4R2 (Proteasome beta4 subunit-related 2) | protein interactions | 33451 | 14 | 3 | 9204 | 298 | Prosbeta1,Ubqn,Prosalpha2 | | 9.381e-03 | -4.67 | PPP1R15 (Protein phosphatase 1 regulatory subunit 15) | protein interactions | 37820 | 14 | 3 | 9204 | 298 | Hrb27C,flw,Pp1-87B | | 9.450e-03 | -4.66 | mitotic cell cycle phase transition | biological process | GO:0044772 | 34 | 4 | 12345 | 305 | Rheb,Skp2,Cdk8,CycA | | 9.511e-03 | -4.66 | polysome | cellular component | GO:0005844 | 18 | 3 | 11940 | 298 | RpS26,eIF4B,Fmr1 | | 9.521e-03 | -4.65 | PTK6 Regulates RHO GTPases, RAS GTPase and MAP kinases | REACTOME pathways | R-DME-8849471 | 12 | 3 | 3923 | 151 | Nf1,RhoGAPp190,vap | | 9.541e-03 | -4.65 | GSE24634\_IL4\_VS\_CTRL\_TREATED\_NAIVE\_CD4\_TCELL\_DAY7\_UP | MSigDB lists | GSE24634\_IL4\_VS\_CTRL\_TREATED\_NAIVE\_CD4\_TCELL\_DAY7\_UP | 68 | 8 | 4161 | 184 | Arpc3A,CycA,ND-ASHI,HtrA2,fzy,CG2938,pav,Tcs3 | | 9.541e-03 | -4.65 | E2F1DP1RB\_01 | MSigDB lists | E2F1DP1RB\_01 | 68 | 8 | 4161 | 184 | Pp1-87B,spel1,DNAlig1,Fs(2)Ket,SF2,E(z),Pect,Stt3B | | 9.541e-03 | -4.65 | GSE34515\_CD16\_NEG\_MONOCYTE\_VS\_DC\_DN | MSigDB lists | GSE34515\_CD16\_NEG\_MONOCYTE\_VS\_DC\_DN | 68 | 8 | 4161 | 184 | eIF3g1,fzy,Nulp1,Rop,l(1)10Bb,noi,janA,CG41378 | | 9.541e-03 | -4.65 | ZHOU\_INFLAMMATORY\_RESPONSE\_LIVE\_UP | MSigDB lists | ZHOU\_INFLAMMATORY\_RESPONSE\_LIVE\_UP | 68 | 8 | 4161 | 184 | Fgop2,Sod2,mus301,Vps35,bai,Vap33,lic,CG11334 | | 9.544e-03 | -4.65 | organic substance catabolic process | biological process | GO:1901575 | 807 | 31 | 12345 | 305 | Skp2,Rpt6,Rat1,Prosalpha2,CG30105,Usp16-45,Zpr1,CG10467,alpha-Man-Ib,tsu,Ufd4,CG32473,inc,smg,Upf3,CG6171,Prosbeta1,Rpn10,CG10365,sqd,CG15534,Usp32,Roc2,twin,Aps,Cul4,Ubqn,Not3,Sgt,Pten,fzy | | 9.558e-03 | -4.65 | NMYC\_01 | MSigDB lists | NMYC\_01 | 82 | 9 | 4161 | 184 | cta,Art1,CG31460,CtBP,bai,Sgt,Rat1,VhaPPA1-1,CSN1b | | 9.579e-03 | -4.65 | REACTOME\_FORMATION\_OF\_THE\_TERNARY\_COMPLEX\_AND\_SUBSEQUENTLY\_THE\_43S\_COMPLEX | MSigDB lists | REACTOME\_FORMATION\_OF\_THE\_TERNARY\_COMPLEX\_AND\_SUBSEQUENTLY\_THE\_43S\_COMPLEX | 42 | 6 | 4161 | 184 | RpS26,RpS9,eIF3c,RpS8,eIF3g1,eIF3b | | 9.604e-03 | -4.65 | negative regulation of organelle organization | biological process | GO:0010639 | 96 | 7 | 12345 | 305 | su(Hw),Mps1,Bub1,CycA,CG9246,spoon,flr | | 9.628e-03 | -4.64 | multicellular organism aging | biological process | GO:0010259 | 170 | 10 | 12345 | 305 | sm,mthl8,Hsp27,BthD,g,Sod2,rb,Btk29A,Nf1,sun | | 9.762e-03 | -4.63 | RpS9 (Ribosomal protein S9) | protein interactions | 39108 | 111 | 9 | 9204 | 298 | RpS19a,RpL15,RpS3A,RpL14,RpS8,RpL36A,RpL5,RpS26,RpL35A | | 9.775e-03 | -4.63 | NUDIX | prosite domains | PS51462 | 15 | 3 | 6243 | 191 | Apf,Aps,CG12567 | | 9.780e-03 | -4.63 | protein serine/threonine phosphatase complex | cellular component | GO:0008287 | 34 | 4 | 11940 | 298 | Pp1-87B,flw,PPP1R15,Wdr82 | | 9.780e-03 | -4.63 | phosphatase complex | cellular component | GO:1903293 | 34 | 4 | 11940 | 298 | Wdr82,flw,PPP1R15,Pp1-87B | | 9.793e-03 | -4.63 | Fen1 (Flap endonuclease 1) | protein interactions | 36887 | 5 | 2 | 9204 | 298 | fzy,viaf | | 9.793e-03 | -4.63 | p24-2 (p24-related-2) | protein interactions | 318890 | 5 | 2 | 9204 | 298 | CG5174,bai | | 9.793e-03 | -4.63 | CG7029 (uncharacterized protein) | protein interactions | 42703 | 5 | 2 | 9204 | 298 | Pp1-87B,flw | | 9.793e-03 | -4.63 | Sf3a1 (Splicing factor 3a subunit 1) | protein interactions | 42048 | 5 | 2 | 9204 | 298 | noi,snf | | 9.793e-03 | -4.63 | CG13749 (uncharacterized protein) | protein interactions | 35894 | 5 | 2 | 9204 | 298 | Rcd5,RpL14 | | 9.793e-03 | -4.63 | CG9674 (uncharacterized protein) | protein interactions | 39878 | 5 | 2 | 9204 | 298 | flw,CG9951 | | 9.793e-03 | -4.63 | Slu7 (Slu7) | protein interactions | 43427 | 5 | 2 | 9204 | 298 | CG1218,CG6171 | | 9.793e-03 | -4.63 | Ide (Insulin degrading metalloproteinase) | protein interactions | 40248 | 5 | 2 | 9204 | 298 | mbf1,Pten | | 9.903e-03 | -4.61 | GO\_RIBONUCLEOPROTEIN\_GRANULE | MSigDB lists | GO\_RIBONUCLEOPROTEIN\_GRANULE | 55 | 7 | 4161 | 184 | me31B,AGO3,Fmr1,sm,Dbp21E2,Cdc42,Prosalpha2 | | 9.909e-03 | -4.61 | ONKEN\_UVEAL\_MELANOMA\_DN | MSigDB lists | ONKEN\_UVEAL\_MELANOMA\_DN | 159 | 14 | 4161 | 184 | RpL14,ECSIT,Pten,Tom20,RpL35A,RpS8,eIF3g1,RpL15,Roc2,RpS9,MED21,flr,CG17493,RpL5 | | 9.910e-03 | -4.61 | GOLDRATH\_ANTIGEN\_RESPONSE | MSigDB lists | GOLDRATH\_ANTIGEN\_RESPONSE | 97 | 10 | 4161 | 184 | unc-119,Fign,DNAlig1,fzy,CG41378,lic,His2Av,E(z),CycA,CG31460 | | 9.924e-03 | -4.61 | Rala (Ras-like protein A) | protein interactions | 31332 | 41 | 5 | 9204 | 298 | Rap1,Exo84,eIF3b,Stim,GEFmeso | | 1.001e-02 | -4.60 | GO\_RIBOSOME\_BIOGENESIS | MSigDB lists | GO\_RIBOSOME\_BIOGENESIS | 192 | 16 | 4161 | 184 | RpL36A,CG4866,RpL15,Rat1,CG11920,RpL5,CG14805,Dbp21E2,RpS9,RpS8,CG9246,mRpS9,RpL35A,CG5589,RpS26,RpL14 | | 1.007e-02 | -4.60 | GO\_PROTEIN\_COMPLEX\_BINDING | MSigDB lists | GO\_PROTEIN\_COMPLEX\_BINDING | 243 | 19 | 4161 | 184 | krz,Arpc3A,Stim,CG17493,bai,inc,CG6907,cta,rhea,Arpc5,Vap33,Rap1,Fmr1,Pten,Pp1-87B,fzy,CG9246,spel1,pav | | 1.010e-02 | -4.59 | mitochondrial respiratory chain complex I | cellular component | GO:0005747 | 53 | 5 | 11940 | 298 | l(2)37Bb,ND-ASHI,CIA30,ND-AGGG,CG11752 | | 1.010e-02 | -4.59 | respiratory chain complex I | cellular component | GO:0045271 | 53 | 5 | 11940 | 298 | l(2)37Bb,ND-ASHI,CG11752,CIA30,ND-AGGG | | 1.010e-02 | -4.59 | NADH dehydrogenase complex | cellular component | GO:0030964 | 53 | 5 | 11940 | 298 | ND-ASHI,ND-AGGG,CG11752,CIA30,l(2)37Bb | | 1.014e-02 | -4.59 | stem cell division | biological process | GO:0017145 | 97 | 7 | 12345 | 305 | tkv,twin,dpp,Cdc42,CycA,pins,Mps1 | | 1.027e-02 | -4.58 | GO\_POSITIVE\_REGULATION\_OF\_EPITHELIAL\_CELL\_PROLIFERATION | MSigDB lists | GO\_POSITIVE\_REGULATION\_OF\_EPITHELIAL\_CELL\_PROLIFERATION | 20 | 4 | 4161 | 184 | dpp,Nf1,viaf,Cdc42 | | 1.030e-02 | -4.58 | NUDIX\_hydrolase-like\_dom\_sf | interpro domains | IPR015797 | 16 | 3 | 11383 | 331 | CG12567,Apf,Aps | | 1.032e-02 | -4.57 | fly chr90C1-90C1\|3-61 cM | chromosome location | fly chr90C1-90C1\|3-61 cM | 11 | 2 | 25010 | 358 | lncRNA:CR45045,Mps1 | | 1.032e-02 | -4.57 | fly chr90C1-90C1\|3-61 | chromosome location | fly chr90C1-90C1\|3-61 | 11 | 2 | 25010 | 358 | lncRNA:CR45045,Mps1 | | 1.032e-02 | -4.57 | PUJANA\_XPRSS\_INT\_NETWORK | MSigDB lists | PUJANA\_XPRSS\_INT\_NETWORK | 83 | 9 | 4161 | 184 | E(z),His2Av,CG5535,Skp2,CycA,snf,spel1,Pp1-87B,fzy | | 1.032e-02 | -4.57 | GO\_RESPONSE\_TO\_LIGHT\_STIMULUS | MSigDB lists | GO\_RESPONSE\_TO\_LIGHT\_STIMULUS | 83 | 9 | 4161 | 184 | Ino80,krz,Cul4,flw,unc-119,CG9246,Pp1-87B,spel1,Nf1 | | 1.032e-02 | -4.57 | GSE3337\_4H\_VS\_16H\_IFNG\_IN\_CD8POS\_DC\_UP | MSigDB lists | GSE3337\_4H\_VS\_16H\_IFNG\_IN\_CD8POS\_DC\_UP | 83 | 9 | 4161 | 184 | Rpt6,CG11920,CG5861,Rpn10,mbf1,CG41378,l(1)10Bb,CG5339,Dbp80 | | 1.032e-02 | -4.57 | GSE39556\_CD8A\_DC\_VS\_NK\_CELL\_MOUSE\_3H\_POST\_POLYIC\_INJ\_UP | MSigDB lists | GSE39556\_CD8A\_DC\_VS\_NK\_CELL\_MOUSE\_3H\_POST\_POLYIC\_INJ\_UP | 83 | 9 | 4161 | 184 | pins,fzy,DNAlig1,pav,CycA,His4r,Skp2,CG5535,rb | | 1.036e-02 | -4.57 | actin filament binding | molecular function | GO:0051015 | 56 | 5 | 12176 | 289 | Spn,Arpc3A,flr,Arpc5,rhea | | 1.040e-02 | -4.57 | KIM\_ALL\_DISORDERS\_DURATION\_CORR\_DN | MSigDB lists | KIM\_ALL\_DISORDERS\_DURATION\_CORR\_DN | 69 | 8 | 4161 | 184 | smt3,eIF3g1,VhaPPA1-1,RpS8,snf,Parp,ECSIT,CG5861 | | 1.045e-02 | -4.56 | GO\_CELL\_SUBSTRATE\_JUNCTION | MSigDB lists | GO\_CELL\_SUBSTRATE\_JUNCTION | 160 | 14 | 4161 | 184 | RpL5,14-3-3zeta,Graf,RpS9,rhea,Arpc5,Arpc3A,LIMK1,cta,pav,Cdc42,Pp1-87B,RpS8,flw | | 1.045e-02 | -4.56 | protein domain specific binding | molecular function | GO:0019904 | 78 | 6 | 12176 | 289 | Pten,14-3-3zeta,pll,Vap33,Cp190,pins | | 1.047e-02 | -4.56 | oocyte fate determination | biological process | GO:0030716 | 35 | 4 | 12345 | 305 | tsu,14-3-3zeta,mus301,Fmr1 | | 1.047e-02 | -4.56 | protein alkylation | biological process | GO:0008213 | 75 | 6 | 12345 | 305 | Wdr82,Rbbp5,Art1,Ncoa6,BEAF-32,E(z) | | 1.047e-02 | -4.56 | protein methylation | biological process | GO:0006479 | 75 | 6 | 12345 | 305 | BEAF-32,E(z),Art1,Ncoa6,Wdr82,Rbbp5 | | 1.048e-02 | -4.56 | vacuolar transport | biological process | GO:0007034 | 54 | 5 | 12345 | 305 | ema,Vps13D,CG30423,Rab7,g | | 1.050e-02 | -4.56 | alt (aluminum tubes) | protein interactions | 42127 | 27 | 4 | 9204 | 298 | eEF1delta,Mps1,Rheb,Pten | | 1.056e-02 | -4.55 | intracellular protein transport | biological process | GO:0006886 | 400 | 18 | 12345 | 305 | Fs(2)Ket,Rab8,rb,Sgt,CG8860,sqd,bai,Rab7,Vps35,Vps13D,Dbp80,Pex3,RabX6,g,Tom20,Tim17b,Rab14,CG6961 | | 1.059e-02 | -4.55 | GO\_RESPONSE\_TO\_RADIATION | MSigDB lists | GO\_RESPONSE\_TO\_RADIATION | 113 | 11 | 4161 | 184 | unc-119,flw,Fign,Pp1-87B,CG9246,Nf1,spel1,eEF1delta,Ino80,krz,Cul4 | | 1.062e-02 | -4.54 | GSE22886\_NAIVE\_TCELL\_VS\_NEUTROPHIL\_UP | MSigDB lists | GSE22886\_NAIVE\_TCELL\_VS\_NEUTROPHIL\_UP | 98 | 10 | 4161 | 184 | CG5861,Art1,noi,CCT4,smt3,Rpt6,CG12321,RpL35A,CG11920,snf | | 1.066e-02 | -4.54 | ST\_INTEGRIN\_SIGNALING\_PATHWAY | MSigDB lists | ST\_INTEGRIN\_SIGNALING\_PATHWAY | 31 | 5 | 4161 | 184 | Graf,rhea,Pten,RhoGAPp190,Cdc42 | | 1.066e-02 | -4.54 | GO\_POSITIVE\_REGULATION\_OF\_BINDING | MSigDB lists | GO\_POSITIVE\_REGULATION\_OF\_BINDING | 31 | 5 | 4161 | 184 | CtBP,mbf1,Parp,dpp,krz | | 1.070e-02 | -4.54 | negative regulation of protein modification process | biological process | GO:0031400 | 98 | 7 | 12345 | 305 | I-2,flw,mnb,CG9246,Pten,krz,sws | | 1.073e-02 | -4.53 | microtubule cytoskeleton organization | biological process | GO:0000226 | 341 | 16 | 12345 | 305 | Rcd5,pav,Girdin,tsu,Fign,Pp1-87B,pins,tsr,Fmr1,CG17493,14-3-3zeta,smt3,Exo84,sqd,Vap33,msd5 | | 1.073e-02 | -4.53 | GO\_CYTOPLASMIC\_SIDE\_OF\_MEMBRANE | MSigDB lists | GO\_CYTOPLASMIC\_SIDE\_OF\_MEMBRANE | 43 | 6 | 4161 | 184 | HtrA2,vap,Nf1,cta,krz,Pten | | 1.073e-02 | -4.53 | GO\_NEGATIVE\_REGULATION\_OF\_CELL\_CYCLE\_PHASE\_TRANSITION | MSigDB lists | GO\_NEGATIVE\_REGULATION\_OF\_CELL\_CYCLE\_PHASE\_TRANSITION | 43 | 6 | 4161 | 184 | Art1,E(z),twin,Pten,CG12321,CycA | | 1.076e-02 | -4.53 | regulation of response to stress | biological process | GO:0080134 | 283 | 14 | 12345 | 305 | flw,Dek,me31B,pll,TM9SF2,Fign,E(bx),sigmar,Rap1,Cdc42,Parp,Pten,krz,dgrn | | 1.079e-02 | -4.53 | negative regulation of sister chromatid segregation | biological process | GO:0033046 | 19 | 3 | 12345 | 305 | Mps1,Bub1,CycA | | 1.079e-02 | -4.53 | negative regulation of mitotic nuclear division | biological process | GO:0045839 | 19 | 3 | 12345 | 305 | Bub1,CycA,Mps1 | | 1.079e-02 | -4.53 | GO\_LYSINE\_N\_METHYLTRANSFERASE\_ACTIVITY | MSigDB lists | GO\_LYSINE\_N\_METHYLTRANSFERASE\_ACTIVITY | 11 | 3 | 4161 | 184 | Rbbp5,E(z),Wdr82 | | 1.079e-02 | -4.53 | GO\_INO80\_TYPE\_COMPLEX | MSigDB lists | GO\_INO80\_TYPE\_COMPLEX | 11 | 3 | 4161 | 184 | Rcd5,Ino80,Arp8 | | 1.079e-02 | -4.53 | BCAT\_GDS748\_DN | MSigDB lists | BCAT\_GDS748\_DN | 11 | 3 | 4161 | 184 | CG5590,Nulp1,Skp2 | | 1.079e-02 | -4.53 | BIOCARTA\_CDC42RAC\_PATHWAY | MSigDB lists | BIOCARTA\_CDC42RAC\_PATHWAY | 11 | 3 | 4161 | 184 | Cdc42,Arpc5,Arpc3A | | 1.079e-02 | -4.53 | BASSO\_CD40\_SIGNALING\_UP | MSigDB lists | BASSO\_CD40\_SIGNALING\_UP | 11 | 3 | 4161 | 184 | sigmar,unc-119,CG41378 | | 1.079e-02 | -4.53 | SCIAN\_CELL\_CYCLE\_TARGETS\_OF\_TP53\_AND\_TP73\_DN | MSigDB lists | SCIAN\_CELL\_CYCLE\_TARGETS\_OF\_TP53\_AND\_TP73\_DN | 11 | 3 | 4161 | 184 | pav,CycA,fzy | | 1.079e-02 | -4.53 | SCHAEFFER\_SOX9\_TARGETS\_IN\_PROSTATE\_DEVELOPMENT\_DN | MSigDB lists | SCHAEFFER\_SOX9\_TARGETS\_IN\_PROSTATE\_DEVELOPMENT\_DN | 11 | 3 | 4161 | 184 | Aps,14-3-3zeta,me31B | | 1.079e-02 | -4.53 | REACTOME\_SEMA4D\_IN\_SEMAPHORIN\_SIGNALING | MSigDB lists | REACTOME\_SEMA4D\_IN\_SEMAPHORIN\_SIGNALING | 11 | 3 | 4161 | 184 | LIMK1,Cdc42,RhoGAPp190 | | 1.079e-02 | -4.53 | DAVICIONI\_PAX\_FOXO1\_SIGNATURE\_IN\_ARMS\_UP | MSigDB lists | DAVICIONI\_PAX\_FOXO1\_SIGNATURE\_IN\_ARMS\_UP | 11 | 3 | 4161 | 184 | Fmr1,Skp2,krz | | 1.079e-02 | -4.53 | BCAT\_BILD\_ET\_AL\_DN | MSigDB lists | BCAT\_BILD\_ET\_AL\_DN | 11 | 3 | 4161 | 184 | CtBP,SF2,Rap1 | | 1.081e-02 | -4.53 | MODULE\_114 | MSigDB lists | MODULE\_114 | 177 | 15 | 4161 | 184 | RpL35A,RpL15,eIF3g1,Prosalpha2,CG4866,VhaPPA1-1,ubl,Mdh1,RpL14,RpL36A,CG5861,Tctp,Arpc3A,eEF1delta,RpS9 | | 1.090e-02 | -4.52 | transforming growth factor beta receptor binding | molecular function | GO:0005160 | 7 | 2 | 12176 | 289 | dpp,mav | | 1.091e-02 | -4.52 | GO\_NEGATIVE\_REGULATION\_OF\_MITOTIC\_CELL\_CYCLE | MSigDB lists | GO\_NEGATIVE\_REGULATION\_OF\_MITOTIC\_CELL\_CYCLE | 56 | 7 | 4161 | 184 | E(z),Art1,twin,Pten,spel1,CG12321,CycA | | 1.091e-02 | -4.52 | GSE1460\_INTRATHYMIC\_T\_PROGENITOR\_VS\_THYMIC\_STROMAL\_CELL\_UP | MSigDB lists | GSE1460\_INTRATHYMIC\_T\_PROGENITOR\_VS\_THYMIC\_STROMAL\_CELL\_UP | 56 | 7 | 4161 | 184 | MICU1,noi,E(z),Stim,RpS8,sigmar,spel1 | | 1.091e-02 | -4.52 | GSE3982\_CTRL\_VS\_IGE\_STIM\_MAST\_CELL\_DN | MSigDB lists | GSE3982\_CTRL\_VS\_IGE\_STIM\_MAST\_CELL\_DN | 56 | 7 | 4161 | 184 | flw,eEF1delta,CG8635,Rcd5,CG5589,bai,ubl | | 1.095e-02 | -4.51 | proteasome-mediated ubiquitin-dependent protein catabolic process | biological process | GO:0043161 | 200 | 11 | 12345 | 305 | inc,Cul4,fzy,Sgt,Prosalpha2,Prosbeta1,Rpt6,Skp2,Rpn10,alpha-Man-Ib,Roc2 | | 1.097e-02 | -4.51 | Prosbeta7 (Proteasome beta7 subunit) | protein interactions | 40639 | 42 | 5 | 9204 | 298 | Rpt6,Prosalpha2,Prosbeta1,CG12321,Ubqn | | 1.097e-02 | -4.51 | ftz-f1 (ftz transcription factor 1) | protein interactions | 40045 | 42 | 5 | 9204 | 298 | smt3,Ino80,eIF4B,14-3-3zeta,RpS8 | | 1.097e-02 | -4.51 | su(Hw) (suppressor of Hairy wing) | protein interactions | 41740 | 42 | 5 | 9204 | 298 | Cp190,E(bx),Orc3,BEAF-32,su(Hw) | | 1.100e-02 | -4.51 | GO\_REGULATION\_OF\_LYMPHOCYTE\_APOPTOTIC\_PROCESS | MSigDB lists | GO\_REGULATION\_OF\_LYMPHOCYTE\_APOPTOTIC\_PROCESS | 4 | 2 | 4161 | 184 | CG9246,Pten | | 1.100e-02 | -4.51 | REICHERT\_G1S\_REGULATORS\_AS\_PI3K\_TARGETS | MSigDB lists | REICHERT\_G1S\_REGULATORS\_AS\_PI3K\_TARGETS | 4 | 2 | 4161 | 184 | Skp2,CycA | | 1.100e-02 | -4.51 | GOERING\_BLOOD\_HDL\_CHOLESTEROL\_QTL\_CIS | MSigDB lists | GOERING\_BLOOD\_HDL\_CHOLESTEROL\_QTL\_CIS | 4 | 2 | 4161 | 184 | RpS26,RpL14 | | 1.100e-02 | -4.51 | THUM\_MIR21\_TARGETS\_HEART\_DISEASE\_DN | MSigDB lists | THUM\_MIR21\_TARGETS\_HEART\_DISEASE\_DN | 4 | 2 | 4161 | 184 | Graf,CG5535 | | 1.100e-02 | -4.51 | GO\_POSITIVE\_REGULATION\_OF\_P38MAPK\_CASCADE | MSigDB lists | GO\_POSITIVE\_REGULATION\_OF\_P38MAPK\_CASCADE | 4 | 2 | 4161 | 184 | dpp,Art1 | | 1.100e-02 | -4.51 | KUMAMOTO\_RESPONSE\_TO\_NUTLIN\_3A\_DN | MSigDB lists | KUMAMOTO\_RESPONSE\_TO\_NUTLIN\_3A\_DN | 4 | 2 | 4161 | 184 | CycA,fzy | | 1.100e-02 | -4.51 | GO\_GOLGI\_LOCALIZATION | MSigDB lists | GO\_GOLGI\_LOCALIZATION | 4 | 2 | 4161 | 184 | Cdc42,14-3-3zeta | | 1.100e-02 | -4.51 | ONO\_FOXP3\_TARGETS\_DN | MSigDB lists | ONO\_FOXP3\_TARGETS\_DN | 4 | 2 | 4161 | 184 | Parp,CycA | | 1.103e-02 | -4.51 | Small\_GTP-bd\_dom | interpro domains | IPR005225 | 84 | 7 | 11383 | 331 | Rab14,Rab8,RabX6,Rab7,Rheb,Cdc42,Rap1 | | 1.108e-02 | -4.50 | UBIQUITIN\_2 | prosite domains | PS50053 | 29 | 4 | 6243 | 191 | Ubqn,smt3,EloB,ubl | | 1.112e-02 | -4.50 | multicellular organism reproduction | biological process | GO:0032504 | 1503 | 51 | 12345 | 305 | snf,Cdc42,Pten,Rap1,fzy,Fs(2)Ket,eEF5,mus301,LIMK1,HtrA2,rhea,dpp,nsl1,Rop,mon2,rhi,Exo84,sqd,Pp1-87B,tsr,glo,tsu,spoon,Rab7,me31B,lic,Hrb27C,vsg,Mps1,Cul4,Parp,twin,BEAF-32,cal1,RecQ4,smt3,bai,14-3-3zeta,flw,MED22,Fmr1,Btk29A,tkv,E(bx),vnc,CSN1b,grau,flr,eIF3b,lncRNA:sphinx,pav | | 1.119e-02 | -4.49 | purine ribonucleoside binding | molecular function | GO:0032550 | 154 | 9 | 12176 | 289 | Rheb,RagA-B,Rab14,Rab7,Rap1,RhoGAPp190,Cdc42,Rab8,RabX6 | | 1.119e-02 | -4.49 | GTP binding | molecular function | GO:0005525 | 154 | 9 | 12176 | 289 | Rab7,Rab14,RagA-B,Rheb,RabX6,Rab8,Cdc42,RhoGAPp190,Rap1 | | 1.131e-02 | -4.48 | GSE9006\_HEALTHY\_VS\_TYPE\_2\_DIABETES\_PBMC\_AT\_DX\_UP | MSigDB lists | GSE9006\_HEALTHY\_VS\_TYPE\_2\_DIABETES\_PBMC\_AT\_DX\_UP | 70 | 8 | 4161 | 184 | CG3842,CG8860,smt3,ECSIT,ND-ASHI,CG14805,LIMK1,mbf1 | | 1.131e-02 | -4.48 | RICKMAN\_TUMOR\_DIFFERENTIATED\_WELL\_VS\_POORLY\_UP | MSigDB lists | RICKMAN\_TUMOR\_DIFFERENTIATED\_WELL\_VS\_POORLY\_UP | 70 | 8 | 4161 | 184 | SF2,RpL5,eEF1delta,RpL35A,Mdh1,Parp,Tom20,CG7766 | | 1.131e-02 | -4.48 | GSE27786\_LSK\_VS\_CD4\_TCELL\_UP | MSigDB lists | GSE27786\_LSK\_VS\_CD4\_TCELL\_UP | 70 | 8 | 4161 | 184 | HtrA2,Parp,ND-ASHI,CG10672,MED22,CG5535,Lst8,CG15168 | | 1.131e-02 | -4.48 | GSE13411\_NAIVE\_VS\_MEMORY\_BCELL\_DN | MSigDB lists | GSE13411\_NAIVE\_VS\_MEMORY\_BCELL\_DN | 70 | 8 | 4161 | 184 | Vap33,RpL36A,Vps35,CG8132,Lst8,Rpb11,eIF3g1,VhaPPA1-1 | | 1.142e-02 | -4.47 | zinc ion binding | molecular function | GO:0008270 | 516 | 21 | 12176 | 289 | d4,Roc2,Rpn10,RecQ4,Usp16-45,CG6254,grau,CG11586,Zpr1,CG32473,CNBP,CG1815,dah,RAF2,nom,x16,Parp,dgrn,CG1218,noi,CG5516 | | 1.145e-02 | -4.47 | kra (krasavietz) | protein interactions | 40680 | 15 | 3 | 9204 | 298 | RpS9,RpS8,RpL5 | | 1.145e-02 | -4.47 | A16 (A16) | protein interactions | 34700 | 15 | 3 | 9204 | 298 | Pp1-87B,Fmr1,flw | | 1.151e-02 | -4.46 | GO\_CELL\_CYCLE | MSigDB lists | GO\_CELL\_CYCLE | 446 | 30 | 4161 | 184 | AGO3,Fs(2)Ket,flw,Stlk,Prosalpha2,CSN1b,DNAlig1,Rpt6,CG12321,Arp8,Ino80,Skp2,Cul4,CG17493,CycA,Rab8,Pten,spel1,pav,Pp1-87B,fzy,pins,Art1,E(z),Lst8,twin,Orc3,vap,Rpn10,RagA-B | | 1.155e-02 | -4.46 | maintenance of location in cell | biological process | GO:0051651 | 36 | 4 | 12345 | 305 | pav,Vps13D,rhea,Parp | | 1.155e-02 | -4.46 | Rab protein signal transduction | biological process | GO:0032482 | 36 | 4 | 12345 | 305 | Rab14,RabX6,Rab7,Rab8 | | 1.156e-02 | -4.46 | regulation of cell death | biological process | GO:0010941 | 257 | 13 | 12345 | 305 | mav,CG1791,dpp,eEF5,RpL14,HtrA2,Rheb,viaf,sigmar,Fmr1,Dref,Pten,fzy | | 1.165e-02 | -4.45 | aging | biological process | GO:0007568 | 175 | 10 | 12345 | 305 | rb,Btk29A,Nf1,sun,mthl8,sm,BthD,Hsp27,g,Sod2 | | 1.170e-02 | -4.45 | R3H\_dom\_sf | interpro domains | IPR036867 | 6 | 2 | 11383 | 331 | CG8915,CG31301 | | 1.170e-02 | -4.45 | SAP\_dom | interpro domains | IPR003034 | 6 | 2 | 11383 | 331 | CG8149,Dek | | 1.170e-02 | -4.45 | R3H\_dom | interpro domains | IPR001374 | 6 | 2 | 11383 | 331 | CG31301,CG8915 | | 1.172e-02 | -4.45 | proteolysis involved in cellular protein catabolic process | biological process | GO:0051603 | 315 | 15 | 12345 | 305 | Usp16-45,Usp32,alpha-Man-Ib,Roc2,Prosalpha2,Prosbeta1,Rpt6,Skp2,Rpn10,Ubqn,Cul4,fzy,Sgt,Ufd4,inc | | 1.174e-02 | -4.44 | endomembrane system | cellular component | GO:0012505 | 1113 | 40 | 11940 | 298 | CG32576,CG2938,spoon,PIG-K,Rab7,CG17883,CG15168,BthD,Fs(2)Ket,me31B,Mppe,RagA-B,Stt3B,tkv,Rab8,xit,CG10365,Efa6,RpS26,alpha-Man-Ib,heix,mon2,sws,CG30423,Stim,PPP1R15,CG31460,Dbp80,Vps35,CG11137,Usp32,g,Vap33,PGAP1,dpp,Rab14,bai,14-3-3zeta,ema,Nup188 | | 1.177e-02 | -4.44 | phagosome maturation | biological process | GO:0090382 | 7 | 2 | 12345 | 305 | Rab14,Rab7 | | 1.177e-02 | -4.44 | negative regulation of torso signaling pathway | biological process | GO:0120177 | 7 | 2 | 12345 | 305 | vap,krz | | 1.177e-02 | -4.44 | positive regulation of muscle organ development | biological process | GO:0048636 | 7 | 2 | 12345 | 305 | dpp,tkv | | 1.177e-02 | -4.44 | SMAD protein signal transduction | biological process | GO:0060395 | 7 | 2 | 12345 | 305 | dpp,mav | | 1.181e-02 | -4.44 | microtubule-based process | biological process | GO:0007017 | 466 | 20 | 12345 | 305 | 14-3-3zeta,smt3,sqd,Exo84,msd5,Vap33,DCTN3-p24,Map60,Rcd5,g,pav,Cp190,Girdin,tsu,Fign,tsr,pins,Pp1-87B,CG17493,Fmr1 | | 1.182e-02 | -4.44 | anatomical structure morphogenesis | biological process | GO:0009653 | 1579 | 53 | 12345 | 305 | BthD,flw,dah,smt3,14-3-3zeta,CtBP,RecQ4,Rheb,EloB,Parp,Cul4,flr,grau,Girdin,CSN1b,Zpr1,tkv,Arpc5,Dref,Fmr1,Btk29A,E(z),sqd,cta,nsl1,rhi,rhea,Cdk8,dpp,mus301,LIMK1,Fs(2)Ket,xit,Pten,peng,Cdc42,Rap1,Hrb27C,Stim,Sod2,VhaPPA1-1,sm,me31B,GEFmeso,tsu,sigmar,Efa6,PlexA,spoon,Stlk,Pp1-87B,heix,tsr | | 1.184e-02 | -4.44 | cytoplasm organization | biological process | GO:0007028 | 77 | 6 | 12345 | 305 | glo,Hrb27C,Rab7,sqd,tsu,mon2 | | 1.187e-02 | -4.43 | positive regulation of GTPase activity | biological process | GO:0043547 | 100 | 7 | 12345 | 305 | RhoGAPp190,Nf1,conu,Graf,CG17883,PlexA,vap | | 1.189e-02 | -4.43 | Ubiquitin\_dom | interpro domains | IPR000626 | 31 | 4 | 11383 | 331 | ubl,EloB,Ubqn,smt3 | | 1.194e-02 | -4.43 | Msp300 (Muscle-specific protein 300 kDa) | protein interactions | 3771968 | 28 | 4 | 9204 | 298 | Tctp,Efa6,Exo84,flw | | 1.194e-02 | -4.43 | CG18787 (uncharacterized protein) | protein interactions | 59163 | 28 | 4 | 9204 | 298 | Wdr82,BthD,CG6171,CG1218 | | 1.194e-02 | -4.43 | Wdr82 (WD repeat domain 82) | protein interactions | 34153 | 28 | 4 | 9204 | 298 | CG1218,CG9330,BthD,Rbbp5 | | 1.194e-02 | -4.43 | Rpn8 (Regulatory particle non-ATPase 8) | protein interactions | 37894 | 28 | 4 | 9204 | 298 | Prosalpha2,Rpt6,Ubqn,Prosbeta1 | | 1.194e-02 | -4.43 | Cpr92F (Cuticular protein 92F) | protein interactions | 42450 | 28 | 4 | 9204 | 298 | 14-3-3zeta,RpS3A,eIF3g1,CG8108 | | 1.198e-02 | -4.42 | CHICAS\_RB1\_TARGETS\_GROWING | MSigDB lists | CHICAS\_RB1\_TARGETS\_GROWING | 57 | 7 | 4161 | 184 | fzy,CycA,PGAP1,spel1,sigmar,Fign,E(z) | | 1.198e-02 | -4.42 | GO\_TRANSLATION\_FACTOR\_ACTIVITY\_RNA\_BINDING | MSigDB lists | GO\_TRANSLATION\_FACTOR\_ACTIVITY\_RNA\_BINDING | 57 | 7 | 4161 | 184 | eIF3c,eEF1delta,eIF3g1,eEF5,eIF3b,CG31957,eRF1 | | 1.198e-02 | -4.42 | GSE6259\_33D1\_POS\_DC\_VS\_BCELL\_DN | MSigDB lists | GSE6259\_33D1\_POS\_DC\_VS\_BCELL\_DN | 57 | 7 | 4161 | 184 | His2Av,CG41378,RpS26,Fign,RpL15,eIF3b,viaf | | 1.198e-02 | -4.42 | GSE39110\_DAY3\_VS\_DAY6\_POST\_IMMUNIZATION\_CD8\_TCELL\_DN | MSigDB lists | GSE39110\_DAY3\_VS\_DAY6\_POST\_IMMUNIZATION\_CD8\_TCELL\_DN | 57 | 7 | 4161 | 184 | Fign,Lrr47,rb,snf,fzy,CycA,DNAlig1 | | 1.198e-02 | -4.42 | GSE19198\_CTRL\_VS\_IL21\_TREATED\_TCELL\_24H\_DN | MSigDB lists | GSE19198\_CTRL\_VS\_IL21\_TREATED\_TCELL\_24H\_DN | 44 | 6 | 4161 | 184 | smt3,spel1,Pp1-87B,Tsp97E,sws,unc-119 | | 1.198e-02 | -4.42 | GSE9960\_GRAM\_NEG\_VS\_GRAM\_POS\_SEPSIS\_PBMC\_UP | MSigDB lists | GSE9960\_GRAM\_NEG\_VS\_GRAM\_POS\_SEPSIS\_PBMC\_UP | 44 | 6 | 4161 | 184 | E(z),Rab8,CG32473,sigmar,Prosalpha2,CG13016 | | 1.199e-02 | -4.42 | NRF1\_Q6 | MSigDB lists | NRF1\_Q6 | 85 | 9 | 4161 | 184 | Nf1,Fgop2,ND-ASHI,sm,Fmr1,SF2,xit,Rcd5,CG11334 | | 1.200e-02 | -4.42 | NSL complex | cellular component | GO:0044545 | 7 | 2 | 11940 | 298 | Rcd5,nsl1 | | 1.210e-02 | -4.41 | png (pan gu) | protein interactions | 31084 | 43 | 5 | 9204 | 298 | CG34159,eEF5,me31B,Pp1-87B,CycA | | 1.217e-02 | -4.41 | MORF\_G22P1 | MSigDB lists | MORF\_G22P1 | 100 | 10 | 4161 | 184 | Art1,His2Av,flr,eIF3c,Tom20,snf,RpL14,smt3,eIF3g1,Mdh1 | | 1.217e-02 | -4.41 | regulation of endocytosis | biological process | GO:0030100 | 56 | 5 | 12345 | 305 | mnb,vap,nsl1,Graf,krz | | 1.217e-02 | -4.41 | translational initiation | biological process | GO:0006413 | 56 | 5 | 12345 | 305 | CG31957,eIF3c,eIF4B,eIF3g1,eIF3b | | 1.219e-02 | -4.41 | GSE14769\_UNSTIM\_VS\_120MIN\_LPS\_BMDM\_DN | MSigDB lists | GSE14769\_UNSTIM\_VS\_120MIN\_LPS\_BMDM\_DN | 32 | 5 | 4161 | 184 | MED21,lic,dpp,cta,CG13016 | | 1.226e-02 | -4.40 | HALLMARK\_TNFA\_SIGNALING\_VIA\_NFKB | MSigDB lists | HALLMARK\_TNFA\_SIGNALING\_VIA\_NFKB | 21 | 4 | 4161 | 184 | Sod2,sigmar,dpp,lic | | 1.226e-02 | -4.40 | MAYBURD\_RESPONSE\_TO\_L663536\_DN | MSigDB lists | MAYBURD\_RESPONSE\_TO\_L663536\_DN | 21 | 4 | 4161 | 184 | MED21,spel1,RpL15,CG5590 | | 1.226e-02 | -4.40 | PUJANA\_ATM\_PCC\_NETWORK | MSigDB lists | PUJANA\_ATM\_PCC\_NETWORK | 467 | 31 | 4161 | 184 | RpL36A,SF2,CycA,RhoGAPp190,Arpc3A,CCT4,Skp2,rb,CG5535,His2Av,eEF5,sigmar,snf,Dbp21E2,Rap1,TfIIFbeta,tsu,His4r,RpL5,cta,E(z),Arpc5,fzy,HtrA2,Pp1-87B,spel1,RpL14,Parp,sm,Pten,CG5589 | | 1.227e-02 | -4.40 | fly chr83C4-83C4\|3-47.5 cM | chromosome location | fly chr83C4-83C4\|3-47.5 cM | 12 | 2 | 25010 | 358 | CG1218,dgrn | | 1.227e-02 | -4.40 | fly chr47C1-47C1\|2-63 cM | chromosome location | fly chr47C1-47C1\|2-63 cM | 12 | 2 | 25010 | 358 | CG12341,CG12935 | | 1.227e-02 | -4.40 | fly chr47C1-47C1\|2-63 | chromosome location | fly chr47C1-47C1\|2-63 | 12 | 2 | 25010 | 358 | CG12935,CG12341 | | 1.227e-02 | -4.40 | fly chr68E1-68E1\|3-37 cM | chromosome location | fly chr68E1-68E1\|3-37 cM | 12 | 2 | 25010 | 358 | CG32095,CycA | | 1.227e-02 | -4.40 | fly chr83C4-83C4\|3-47.5 | chromosome location | fly chr83C4-83C4\|3-47.5 | 12 | 2 | 25010 | 358 | dgrn,CG1218 | | 1.227e-02 | -4.40 | fly chr68E1-68E1\|3-37 | chromosome location | fly chr68E1-68E1\|3-37 | 12 | 2 | 25010 | 358 | CG32095,CycA | | 1.227e-02 | -4.40 | GSE22886\_NAIVE\_BCELL\_VS\_NEUTROPHIL\_UP | MSigDB lists | GSE22886\_NAIVE\_BCELL\_VS\_NEUTROPHIL\_UP | 71 | 8 | 4161 | 184 | Parp,RpL35A,Tsp97E,Mdh1,RpL15,smt3,noi,RpL36A | | 1.227e-02 | -4.40 | GSE17721\_LPS\_VS\_POLYIC\_8H\_BMDC\_UP | MSigDB lists | GSE17721\_LPS\_VS\_POLYIC\_8H\_BMDC\_UP | 71 | 8 | 4161 | 184 | Cdk8,Rpn10,tsu,RhoGAPp190,Arpc3A,noi,Sod2,ubl | | 1.233e-02 | -4.40 | synapse | cellular component | GO:0045202 | 285 | 14 | 11940 | 298 | g,Spn,Vap33,me31B,Rop,Rpt6,Fmr1,Rab8,Vps35,mnb,tkv,Rab14,mav,Rab7 | | 1.236e-02 | -4.39 | cellular protein catabolic process | biological process | GO:0044257 | 317 | 15 | 12345 | 305 | Prosalpha2,Skp2,Prosbeta1,Rpt6,Rpn10,Usp32,Usp16-45,alpha-Man-Ib,Roc2,Ufd4,inc,Ubqn,Cul4,fzy,Sgt | | 1.245e-02 | -4.39 | transcription factor binding | molecular function | GO:0008134 | 184 | 10 | 12176 | 289 | 14-3-3zeta,CtBP,TfIIFbeta,Rpt6,Mlf,Tgi,Ncoa6,dgrn,CG9246,E(bx) | | 1.245e-02 | -4.39 | wing disc development | biological process | GO:0035220 | 407 | 18 | 12345 | 305 | peng,Cdc42,EloB,LIMK1,rhea,dpp,CtBP,flw,CG8149,tsr,Btk29A,tkv,Rab7,GEFmeso,Skp2,Stim,Amun,flr | | 1.247e-02 | -4.38 | positive regulation of histone modification | biological process | GO:0031058 | 20 | 3 | 12345 | 305 | E(z),Ncoa6,Tctp | | 1.247e-02 | -4.38 | negative regulation of hippo signaling | biological process | GO:0035331 | 20 | 3 | 12345 | 305 | Fgop2,Slmap,mnb | | 1.247e-02 | -4.38 | negative regulation of chromosome segregation | biological process | GO:0051985 | 20 | 3 | 12345 | 305 | CycA,Bub1,Mps1 | | 1.247e-02 | -4.38 | cell cycle comprising mitosis without cytokinesis | biological process | GO:0033301 | 20 | 3 | 12345 | 305 | CycA,E(z),smt3 | | 1.256e-02 | -4.38 | positive regulation of transferase activity | biological process | GO:0051347 | 78 | 6 | 12345 | 305 | Cdc42,fzy,Stlk,CG1791,lic,smt3 | | 1.259e-02 | -4.38 | ribonucleoside binding | molecular function | GO:0032549 | 157 | 9 | 12176 | 289 | Rheb,Rab7,RagA-B,Rab14,RhoGAPp190,Cdc42,Rap1,RabX6,Rab8 | | 1.260e-02 | -4.37 | protein transport | biological process | GO:0015031 | 469 | 20 | 12345 | 305 | bai,mon2,CG8860,sqd,rb,Sgt,Fs(2)Ket,Rab8,unc-119,CG6961,Tim17b,Rab14,Pex3,RabX6,Tom20,g,Dbp80,Vps13D,Vps35,Rab7 | | 1.270e-02 | -4.37 | female meiosis chromosome segregation | biological process | GO:0016321 | 37 | 4 | 12345 | 305 | pav,mus301,Mps1,cal1 | | 1.270e-02 | -4.37 | cell cycle phase transition | biological process | GO:0044770 | 37 | 4 | 12345 | 305 | Rheb,Skp2,CycA,Cdk8 | | 1.270e-02 | -4.37 | regulation of hippo signaling | biological process | GO:0035330 | 37 | 4 | 12345 | 305 | 14-3-3zeta,mnb,Fgop2,Slmap | | 1.277e-02 | -4.36 | BLALOCK\_ALZHEIMERS\_DISEASE\_DN | MSigDB lists | BLALOCK\_ALZHEIMERS\_DISEASE\_DN | 545 | 35 | 4161 | 184 | ND-ASHI,CG11920,PIG-K,mRpS28,Mdh1,Tsp97E,CG12321,His2Av,Arpc3A,xit,Pten,RpS26,Parp,MICU1,RpL14,spel1,CG32576,RpL35A,Art1,14-3-3zeta,Arpc5,CG5861,Fs(2)Ket,Tom20,RpL15,CG2938,Pex3,CCT4,CG8635,Rop,ubl,Cdc42,HtrA2,CG8132,Orc3 | | 1.278e-02 | -4.36 | R3H | pfam domains | PF01424 | 6 | 2 | 10480 | 319 | CG8915,CG31301 | | 1.281e-02 | -4.36 | preribosome, large subunit precursor | cellular component | GO:0030687 | 20 | 3 | 11940 | 298 | Rbm13,CG9246,CG32075 | | 1.281e-02 | -4.36 | polytene chromosome chromocenter | cellular component | GO:0005701 | 20 | 3 | 11940 | 298 | HP4,MED26,His2Av | | 1.281e-02 | -4.36 | GO\_MITOTIC\_CELL\_CYCLE | MSigDB lists | GO\_MITOTIC\_CELL\_CYCLE | 302 | 22 | 4161 | 184 | Orc3,vap,Rpn10,Art1,twin,spel1,pav,fzy,pins,Rab8,Pten,CG17493,CycA,Ino80,Skp2,Prosalpha2,DNAlig1,Rpt6,CG12321,Arp8,Fs(2)Ket,flw | | 1.282e-02 | -4.36 | exoribonuclease activity | molecular function | GO:0004532 | 21 | 3 | 12176 | 289 | Not3,Rat1,twin | | 1.282e-02 | -4.36 | 3'-5' DNA helicase activity | molecular function | GO:0043138 | 21 | 3 | 12176 | 289 | RecQ4,Ino80,Arp8 | | 1.282e-02 | -4.36 | exoribonuclease activity, producing 5'-phosphomonoesters | molecular function | GO:0016896 | 21 | 3 | 12176 | 289 | Not3,Rat1,twin | | 1.282e-02 | -4.36 | repressing transcription factor binding | molecular function | GO:0070491 | 21 | 3 | 12176 | 289 | dgrn,CtBP,CG9246 | | 1.288e-02 | -4.35 | R3H | prosite domains | PS51061 | 6 | 2 | 6243 | 191 | CG31301,CG8915 | | 1.289e-02 | -4.35 | GO\_PROTEIN\_LOCALIZATION\_TO\_ENDOPLASMIC\_RETICULUM | MSigDB lists | GO\_PROTEIN\_LOCALIZATION\_TO\_ENDOPLASMIC\_RETICULUM | 86 | 9 | 4161 | 184 | RpL36A,RpL5,RpS9,RpL35A,RpS8,RpL15,CG8860,RpL14,RpS26 | | 1.290e-02 | -4.35 | - | gene3d domains | 3.30.1370.50 | 6 | 2 | 4929 | 151 | CG31301,CG8915 | | 1.297e-02 | -4.35 | adenyl ribonucleotide binding | molecular function | GO:0032559 | 655 | 25 | 12176 | 289 | me31B,CG8915,pav,mnb,Ino80,pll,Cdk8,CG9330,Dbp21E2,mus301,Btk29A,DNAlig1,CG5589,Stlk,tkv,LIMK1,Acbp1,Mps1,Dbp80,Bub1,Rpt6,spel1,lic,Fign,CCT4 | | 1.300e-02 | -4.34 | GSE2405\_0H\_VS\_24H\_A\_PHAGOCYTOPHILUM\_STIM\_NEUTROPHIL\_UP | MSigDB lists | GSE2405\_0H\_VS\_24H\_A\_PHAGOCYTOPHILUM\_STIM\_NEUTROPHIL\_UP | 101 | 10 | 4161 | 184 | RpS26,RpL14,RpS8,Pp1-87B,RpS9,Arpc3A,CCT4,Rpn10,RpL5,Tctp | | 1.302e-02 | -4.34 | GO\_REGULATION\_OF\_CYTOSKELETON\_ORGANIZATION | MSigDB lists | GO\_REGULATION\_OF\_CYTOSKELETON\_ORGANIZATION | 132 | 12 | 4161 | 184 | Arpc3A,LIMK1,Ino80,Lst8,Arpc5,janA,flr,vap,Fs(2)Ket,Cdc42,pins,pav | | 1.302e-02 | -4.34 | GO\_CELLULAR\_PROTEIN\_COMPLEX\_ASSEMBLY | MSigDB lists | GO\_CELLULAR\_PROTEIN\_COMPLEX\_ASSEMBLY | 148 | 13 | 4161 | 184 | l(2)37Bb,CIA30,Rpn10,CG12341,CG17883,bai,Exo84,ND-ASHI,Fs(2)Ket,Tom20,ECSIT,CSN1b,CG12321 | | 1.308e-02 | -4.34 | guanyl ribonucleotide binding | molecular function | GO:0032561 | 158 | 9 | 12176 | 289 | Rheb,Rab7,Rab14,RagA-B,Cdc42,RhoGAPp190,Rap1,RabX6,Rab8 | | 1.313e-02 | -4.33 | GSE16385\_UNTREATED\_VS\_12H\_ROSIGLITAZONE\_IL4\_TREATED\_MACROPHAGE\_DN | MSigDB lists | GSE16385\_UNTREATED\_VS\_12H\_ROSIGLITAZONE\_IL4\_TREATED\_MACROPHAGE\_DN | 58 | 7 | 4161 | 184 | mRpS33,sigmar,Prosalpha2,Pp1-87B,CG6907,Rpb11,Lrr47 | | 1.313e-02 | -4.33 | GSE17721\_POLYIC\_VS\_PAM3CSK4\_2H\_BMDC\_UP | MSigDB lists | GSE17721\_POLYIC\_VS\_PAM3CSK4\_2H\_BMDC\_UP | 58 | 7 | 4161 | 184 | CG4866,vap,Fign,Skp2,MED22,Phf5a,Pisd | | 1.325e-02 | -4.32 | SAP | smart domains | SM00513 | 6 | 2 | 5086 | 158 | CG8149,Dek | | 1.328e-02 | -4.32 | RNA\_helicase\_DEAD\_Q\_motif | interpro domains | IPR014014 | 32 | 4 | 11383 | 331 | Dbp21E2,me31B,Dbp80,CG5589 | | 1.330e-02 | -4.32 | sqd (squid) | protein interactions | 41666 | 44 | 5 | 9204 | 298 | Art1,Pep,Hrb27C,Parp,Rheb | | 1.330e-02 | -4.32 | Prosbeta4 (Proteasome beta4 subunit) | protein interactions | 34999 | 44 | 5 | 9204 | 298 | Rpt6,Prosalpha2,smt3,Prosbeta1,CG12321 | | 1.330e-02 | -4.32 | CG2046 (uncharacterized protein) | protein interactions | 40719 | 44 | 5 | 9204 | 298 | Prosalpha2,Rpt6,Prosbeta1,CG12321,DCTN3-p24 | | 1.333e-02 | -4.32 | GSE36891\_UNSTIM\_VS\_PAM\_TLR2\_STIM\_PERITONEAL\_MACROPHAGE\_DN | MSigDB lists | GSE36891\_UNSTIM\_VS\_PAM\_TLR2\_STIM\_PERITONEAL\_MACROPHAGE\_DN | 45 | 6 | 4161 | 184 | eIF3b,Tcs3,CG4866,CG41378,mbf1,CG12935 | | 1.333e-02 | -4.32 | GSE36888\_STAT5\_AB\_KNOCKIN\_VS\_WT\_TCELL\_IL2\_TREATED\_17H\_DN | MSigDB lists | GSE36888\_STAT5\_AB\_KNOCKIN\_VS\_WT\_TCELL\_IL2\_TREATED\_17H\_DN | 45 | 6 | 4161 | 184 | Fgop2,Sod2,Fmr1,CG15168,CG5339,MICU1 | | 1.333e-02 | -4.32 | REACTOME\_PLATELET\_ACTIVATION\_SIGNALING\_AND\_AGGREGATION | MSigDB lists | REACTOME\_PLATELET\_ACTIVATION\_SIGNALING\_AND\_AGGREGATION | 45 | 6 | 4161 | 184 | cta,krz,14-3-3zeta,Rap1,flr,Cdc42 | | 1.341e-02 | -4.31 | RpS3 (Ribosomal protein S3) | protein interactions | 42761 | 179 | 12 | 9204 | 298 | RpL5,RpS26,RpL15,RpS8,RpS3A,RpL14,Usp16-45,RpS19a,RpS9,RpL36A,RpL35A,Cpr73D | | 1.342e-02 | -4.31 | organelle membrane | cellular component | GO:0031090 | 787 | 30 | 11940 | 298 | ND-ASHI,CG11137,g,Tom20,Vap33,RagA-B,Tim17b,ND-AGGG,CIA30,Stt3B,14-3-3zeta,Bmcp,xit,ema,Efa6,RpS26,sun,MICU1,CG11752,heix,l(2)37Bb,sws,Pex3,CG31460,spoon,PIG-K,CG4743,Rab7,CG15168,CG17883 | | 1.351e-02 | -4.30 | Rpn10 (Regulatory particle non-ATPase 10) | protein interactions | 40388 | 29 | 4 | 9204 | 298 | Rpt6,Ubqn,smg,smt3 | | 1.351e-02 | -4.30 | gammaTub37C (gamma-Tubulin at 37C) | protein interactions | 35199 | 29 | 4 | 9204 | 298 | Map60,Cp190,CG7766,GILT3 | | 1.351e-02 | -4.30 | bru1 (bruno 1) | protein interactions | 34648 | 29 | 4 | 9204 | 298 | sqd,smg,Cwc25,me31B | | 1.354e-02 | -4.30 | eEF2 (eukaryotic translation elongation factor 2) | protein interactions | 35422 | 158 | 11 | 9204 | 298 | RpL35A,RpS3A,RpL15,RpL14,RpS8,RpS9,RpS19a,RpL36A,RpL5,RAF2,RpS26 | | 1.371e-02 | -4.29 | peptide transport | biological process | GO:0015833 | 473 | 20 | 12345 | 305 | Rab14,Tim17b,CG6961,Dbp80,RabX6,Tom20,g,Pex3,Vps13D,Vps35,Rab7,CG8860,sqd,mon2,bai,Sgt,rb,Rab8,unc-119,Fs(2)Ket | | 1.376e-02 | -4.29 | moon (moonshiner) | protein interactions | 32176 | 16 | 3 | 9204 | 298 | Rpb11,CG8860,Slmap | | 1.376e-02 | -4.29 | TwdlM (TweedleM) | protein interactions | 43200 | 16 | 3 | 9204 | 298 | Sgt,Ubqn,flw | | 1.376e-02 | -4.29 | MAPK family signaling cascades | REACTOME pathways | R-DME-5683057 | 117 | 10 | 3923 | 151 | Rpt6,Nf1,vap,Prosbeta1,CG7054,14-3-3zeta,Cdc42,Rpn10,CG4935,Prosalpha2 | | 1.380e-02 | -4.28 | GO\_REGULATION\_OF\_CELLULAR\_COMPONENT\_BIOGENESIS | MSigDB lists | GO\_REGULATION\_OF\_CELLULAR\_COMPONENT\_BIOGENESIS | 216 | 17 | 4161 | 184 | Fmr1,Pten,Parp,Cdc42,Lst8,twin,Arpc5,LIMK1,Ubqn,vap,CG17883,TfIIFbeta,Rap1,Rpt6,Arpc3A,Cul4,Rop | | 1.389e-02 | -4.28 | PI3K Cascade | REACTOME pathways | R-DME-109704 | 52 | 6 | 3923 | 151 | eIF4B,Rheb,RagA-B,14-3-3zeta,Lst8,Stlk | | 1.390e-02 | -4.28 | adenyl nucleotide binding | molecular function | GO:0030554 | 659 | 25 | 12176 | 289 | Btk29A,mus301,Ino80,Cdk8,Dbp21E2,CG9330,pll,me31B,CG8915,mnb,pav,CCT4,Fign,lic,Bub1,Dbp80,Mps1,Rpt6,spel1,tkv,LIMK1,Acbp1,DNAlig1,Stlk,CG5589 | | 1.393e-02 | -4.27 | BIOCARTA\_RHO\_PATHWAY | MSigDB lists | BIOCARTA\_RHO\_PATHWAY | 12 | 3 | 4161 | 184 | LIMK1,Arpc3A,Arpc5 | | 1.393e-02 | -4.27 | ST\_GA13\_PATHWAY | MSigDB lists | ST\_GA13\_PATHWAY | 12 | 3 | 4161 | 184 | Cdc42,CG7766,cta | | 1.393e-02 | -4.27 | MOOTHA\_GLYCOGEN\_METABOLISM | MSigDB lists | MOOTHA\_GLYCOGEN\_METABOLISM | 12 | 3 | 4161 | 184 | Pp1-87B,CG7766,flw | | 1.393e-02 | -4.27 | ZHAN\_EARLY\_DIFFERENTIATION\_GENES\_DN | MSigDB lists | ZHAN\_EARLY\_DIFFERENTIATION\_GENES\_DN | 12 | 3 | 4161 | 184 | fzy,DNAlig1,spel1 | | 1.393e-02 | -4.27 | GTGTGAG\_MIR342 | MSigDB lists | GTGTGAG\_MIR342 | 12 | 3 | 4161 | 184 | vap,Ino80,Fs(2)Ket | | 1.393e-02 | -4.27 | GO\_REGULATION\_OF\_SYNAPTIC\_VESICLE\_TRANSPORT | MSigDB lists | GO\_REGULATION\_OF\_SYNAPTIC\_VESICLE\_TRANSPORT | 12 | 3 | 4161 | 184 | Fmr1,Rop,Rap1 | | 1.393e-02 | -4.27 | GO\_NEGATIVE\_REGULATION\_OF\_CELL\_MATRIX\_ADHESION | MSigDB lists | GO\_NEGATIVE\_REGULATION\_OF\_CELL\_MATRIX\_ADHESION | 12 | 3 | 4161 | 184 | Pten,Nf1,vap | | 1.393e-02 | -4.27 | protein autophosphorylation | biological process | GO:0046777 | 38 | 4 | 12345 | 305 | Btk29A,Dref,pll,mnb | | 1.394e-02 | -4.27 | GATTGGY\_NFY\_Q6\_01 | MSigDB lists | GATTGGY\_NFY\_Q6\_01 | 359 | 25 | 4161 | 184 | CG17493,janA,lic,Stim,Graf,Pect,MED21,c12.1,Tsp97E,Fs(2)Ket,CG6907,snf,CG11920,Vps35,Rpn10,RpS9,Arpc5,14-3-3zeta,Usp32,Phf5a,Apf,spel1,pav,eRF1,RpL14 | | 1.403e-02 | -4.27 | protein-containing complex disassembly | biological process | GO:0032984 | 58 | 5 | 12345 | 305 | Pp1-87B,tsr,eRF1,flr,flw | | 1.406e-02 | -4.26 | protein localization to membrane | biological process | GO:0072657 | 128 | 8 | 12345 | 305 | sws,Pex3,CG9853,Exo84,Cdc42,Sgt,TM9SF2,Rab8 | | 1.428e-02 | -4.25 | nuclear-transcribed mRNA catabolic process, nonsense-mediated decay | biological process | GO:0000184 | 21 | 3 | 12345 | 305 | sqd,tsu,Upf3 | | 1.428e-02 | -4.25 | cellular response to BMP stimulus | biological process | GO:0071773 | 21 | 3 | 12345 | 305 | tkv,mav,dpp | | 1.428e-02 | -4.25 | nuclear-transcribed mRNA catabolic process, deadenylation-dependent decay | biological process | GO:0000288 | 21 | 3 | 12345 | 305 | Not3,smg,twin | | 1.428e-02 | -4.25 | BMP signaling pathway | biological process | GO:0030509 | 21 | 3 | 12345 | 305 | mav,tkv,dpp | | 1.430e-02 | -4.25 | poly(A)-specific ribonuclease activity | molecular function | GO:0004535 | 8 | 2 | 12176 | 289 | Not3,twin | | 1.431e-02 | -4.25 | fly chr13F12-13F12\|1-52 | chromosome location | fly chr13F12-13F12\|1-52 | 1 | 1 | 25010 | 358 | sun | | 1.431e-02 | -4.25 | fly chr73D7-73E1\|3-45 cM | chromosome location | fly chr73D7-73E1\|3-45 cM | 1 | 1 | 25010 | 358 | Cpr73D | | 1.431e-02 | -4.25 | fly chr52E4-52E4\|2-77 | chromosome location | fly chr52E4-52E4\|2-77 | 1 | 1 | 25010 | 358 | mRpL34 | | 1.431e-02 | -4.25 | fly chr55D3-55E1\|2-86 | chromosome location | fly chr55D3-55E1\|2-86 | 1 | 1 | 25010 | 358 | GEFmeso | | 1.431e-02 | -4.25 | fly chr58C7-58C7\|2-98 cM | chromosome location | fly chr58C7-58C7\|2-98 cM | 1 | 1 | 25010 | 358 | Vps35 | | 1.431e-02 | -4.25 | fly chr98A8-98A8\|3-94 | chromosome location | fly chr98A8-98A8\|3-94 | 1 | 1 | 25010 | 358 | pins | | 1.431e-02 | -4.25 | fly chr52D2-52D3\|2-76 cM | chromosome location | fly chr52D2-52D3\|2-76 cM | 1 | 1 | 25010 | 358 | Mlf | | 1.431e-02 | -4.25 | fly chr64A10-64A11\|3-10 cM | chromosome location | fly chr64A10-64A11\|3-10 cM | 1 | 1 | 25010 | 358 | pav | | 1.431e-02 | -4.25 | fly chr52D13-52D14\|2-76 | chromosome location | fly chr52D13-52D14\|2-76 | 1 | 1 | 25010 | 358 | Prosbeta1 | | 1.431e-02 | -4.25 | fly chr59C2-59C3\|2-102 cM | chromosome location | fly chr59C2-59C3\|2-102 cM | 1 | 1 | 25010 | 358 | DCTN3-p24 | | 1.431e-02 | -4.25 | fly chr95F1-95F2\|3-84 | chromosome location | fly chr95F1-95F2\|3-84 | 1 | 1 | 25010 | 358 | twin | | 1.431e-02 | -4.25 | fly chr70B2-70B3\|3-40 | chromosome location | fly chr70B2-70B3\|3-40 | 1 | 1 | 25010 | 358 | Tgi | | 1.431e-02 | -4.25 | fly chr46E6-46E8\|2-62 cM | chromosome location | fly chr46E6-46E8\|2-62 cM | 1 | 1 | 25010 | 358 | 14-3-3zeta | | 1.431e-02 | -4.25 | fly chr88E9-88E11\|3-56 cM | chromosome location | fly chr88E9-88E11\|3-56 cM | 1 | 1 | 25010 | 358 | Cp190 | | 1.431e-02 | -4.25 | fly chr29A1-29A3\|2-31 | chromosome location | fly chr29A1-29A3\|2-31 | 1 | 1 | 25010 | 358 | Btk29A | | 1.431e-02 | -4.25 | fly chr28E6-28E7\|2-30 | chromosome location | fly chr28E6-28E7\|2-30 | 1 | 1 | 25010 | 358 | mon2 | | 1.431e-02 | -4.25 | fly chr58C7-58C7\|2-98 | chromosome location | fly chr58C7-58C7\|2-98 | 1 | 1 | 25010 | 358 | Vps35 | | 1.431e-02 | -4.25 | fly chr51A1-51A2\|2-71 cM | chromosome location | fly chr51A1-51A2\|2-71 cM | 1 | 1 | 25010 | 358 | CG17385 | | 1.431e-02 | -4.25 | fly chr60A15-60A16\|2-105 | chromosome location | fly chr60A15-60A16\|2-105 | 1 | 1 | 25010 | 358 | PPP1R15 | | 1.431e-02 | -4.25 | fly chr45A9-45A9\|2-60 cM | chromosome location | fly chr45A9-45A9\|2-60 cM | 1 | 1 | 25010 | 358 | GstE13 | | 1.431e-02 | -4.25 | fly chr67B9-67B10\|3-29 | chromosome location | fly chr67B9-67B10\|3-29 | 1 | 1 | 25010 | 358 | Jarid2 | | 1.431e-02 | -4.25 | fly chr35A4-35B1\|2-50 | chromosome location | fly chr35A4-35B1\|2-50 | 1 | 1 | 25010 | 358 | spel1 | | 1.431e-02 | -4.25 | fly chr67B1-67D13\|3-32 cM | chromosome location | fly chr67B1-67D13\|3-32 cM | 1 | 1 | 25010 | 358 | vnc | | 1.431e-02 | -4.25 | fly chr86F8-86F9\|3-51 cM | chromosome location | fly chr86F8-86F9\|3-51 cM | 1 | 1 | 25010 | 358 | glo | | 1.431e-02 | -4.25 | fly chr55C8-55C9\|2-85 | chromosome location | fly chr55C8-55C9\|2-85 | 1 | 1 | 25010 | 358 | CG5174 | | 1.431e-02 | -4.25 | fly chr30F2-30F3\|2-38 cM | chromosome location | fly chr30F2-30F3\|2-38 cM | 1 | 1 | 25010 | 358 | Dref | | 1.431e-02 | -4.25 | fly chr87B9-87B10\|3-52 | chromosome location | fly chr87B9-87B10\|3-52 | 1 | 1 | 25010 | 358 | Pp1-87B | | 1.431e-02 | -4.25 | fly chr8C8-8C9\|1-26 cM | chromosome location | fly chr8C8-8C9\|1-26 cM | 1 | 1 | 25010 | 358 | CG7766 | | 1.431e-02 | -4.25 | fly chr60A8-60A9\|2-105 cM | chromosome location | fly chr60A8-60A9\|2-105 cM | 1 | 1 | 25010 | 358 | Upf3 | | 1.431e-02 | -4.25 | fly chr60A8-60A9\|2-105 | chromosome location | fly chr60A8-60A9\|2-105 | 1 | 1 | 25010 | 358 | Upf3 | | 1.431e-02 | -4.25 | fly chr61A5-61A5\|3-1 | chromosome location | fly chr61A5-61A5\|3-1 | 1 | 1 | 25010 | 358 | mthl8 | | 1.431e-02 | -4.25 | fly chr13F12-13F12\|1-52 cM | chromosome location | fly chr13F12-13F12\|1-52 cM | 1 | 1 | 25010 | 358 | sun | | 1.431e-02 | -4.25 | fly chr29B2-29B3\|2-31 cM | chromosome location | fly chr29B2-29B3\|2-31 cM | 1 | 1 | 25010 | 358 | Wdr82 | | 1.431e-02 | -4.25 | fly chr4F9-4F10\|1-11 | chromosome location | fly chr4F9-4F10\|1-11 | 1 | 1 | 25010 | 358 | spoon | | 1.431e-02 | -4.25 | fly chr38E4-38E5\|2-54 cM | chromosome location | fly chr38E4-38E5\|2-54 cM | 1 | 1 | 25010 | 358 | TM9SF2 | | 1.431e-02 | -4.25 | fly chr25D1-25D2\|2-16 | chromosome location | fly chr25D1-25D2\|2-16 | 1 | 1 | 25010 | 358 | tkv | | 1.431e-02 | -4.25 | fly chr66B8-66B8\|3-22 cM | chromosome location | fly chr66B8-66B8\|3-22 cM | 1 | 1 | 25010 | 358 | mus301 | | 1.431e-02 | -4.25 | fly chr2C1-2C1\|1-0.5 | chromosome location | fly chr2C1-2C1\|1-0.5 | 1 | 1 | 25010 | 358 | PIG-K | | 1.431e-02 | -4.25 | fly chr70A8-70B1\|3-40 | chromosome location | fly chr70A8-70B1\|3-40 | 1 | 1 | 25010 | 358 | flr | | 1.431e-02 | -4.25 | fly chr45A7-45A7\|2-60 cM | chromosome location | fly chr45A7-45A7\|2-60 cM | 1 | 1 | 25010 | 358 | tsu | | 1.431e-02 | -4.25 | fly chr94B5-94B6\|3-75 | chromosome location | fly chr94B5-94B6\|3-75 | 1 | 1 | 25010 | 358 | Efa6 | | 1.431e-02 | -4.25 | fly chr72E3-72F4\|3-44 | chromosome location | fly chr72E3-72F4\|3-44 | 1 | 1 | 25010 | 358 | Tcs3 | | 1.431e-02 | -4.25 | fly chr64A10-64A11\|3-10 | chromosome location | fly chr64A10-64A11\|3-10 | 1 | 1 | 25010 | 358 | pav | | 1.431e-02 | -4.25 | fly chr60A15-60A16\|2-105 cM | chromosome location | fly chr60A15-60A16\|2-105 cM | 1 | 1 | 25010 | 358 | PPP1R15 | | 1.431e-02 | -4.25 | fly chr14A5-14A6\|1-52 cM | chromosome location | fly chr14A5-14A6\|1-52 cM | 1 | 1 | 25010 | 358 | vap | | 1.431e-02 | -4.25 | fly chr35A4-35B1\|2-50 cM | chromosome location | fly chr35A4-35B1\|2-50 cM | 1 | 1 | 25010 | 358 | spel1 | | 1.431e-02 | -4.25 | fly chr10D4-10D4\|1-35 | chromosome location | fly chr10D4-10D4\|1-35 | 1 | 1 | 25010 | 358 | Vlet | | 1.431e-02 | -4.25 | fly chr87F5-87F6\|3-54 | chromosome location | fly chr87F5-87F6\|3-54 | 1 | 1 | 25010 | 358 | sqd | | 1.431e-02 | -4.25 | fly chr10D1-10D3\|1-35 cM | chromosome location | fly chr10D1-10D3\|1-35 cM | 1 | 1 | 25010 | 358 | RPA3 | | 1.431e-02 | -4.25 | fly chr14A5-14A6\|1-52 | chromosome location | fly chr14A5-14A6\|1-52 | 1 | 1 | 25010 | 358 | vap | | 1.431e-02 | -4.25 | fly chr8C1-8C4\|1-26 | chromosome location | fly chr8C1-8C4\|1-26 | 1 | 1 | 25010 | 358 | lncRNA:CR43836 | | 1.431e-02 | -4.25 | fly chr28A5-28A5\|2-27 cM | chromosome location | fly chr28A5-28A5\|2-27 cM | 1 | 1 | 25010 | 358 | lncRNA:CR44082 | | 1.431e-02 | -4.25 | fly chr41B3-41B3\|2-55 cM | chromosome location | fly chr41B3-41B3\|2-55 cM | 1 | 1 | 25010 | 358 | CG17883 | | 1.431e-02 | -4.25 | fly chr22F1-22F3\|2-6 cM | chromosome location | fly chr22F1-22F3\|2-6 cM | 1 | 1 | 25010 | 358 | dpp | | 1.431e-02 | -4.25 | fly chr87D8-87D9\|3-53 | chromosome location | fly chr87D8-87D9\|3-53 | 1 | 1 | 25010 | 358 | CtBP | | 1.431e-02 | -4.25 | fly chr14B7-14B8\|1-53 cM | chromosome location | fly chr14B7-14B8\|1-53 cM | 1 | 1 | 25010 | 358 | CG32576 | | 1.431e-02 | -4.25 | fly chr68A3-68A4\|3-34 | chromosome location | fly chr68A3-68A4\|3-34 | 1 | 1 | 25010 | 358 | CG32075 | | 1.431e-02 | -4.25 | fly chr25D3-25D3\|2-16 | chromosome location | fly chr25D3-25D3\|2-16 | 1 | 1 | 25010 | 358 | Bub1 | | 1.431e-02 | -4.25 | fly chr11A8-11A8\|1-39 | chromosome location | fly chr11A8-11A8\|1-39 | 1 | 1 | 25010 | 358 | LIMK1 | | 1.431e-02 | -4.25 | fly chr62E4-62E5\|3-3 | chromosome location | fly chr62E4-62E5\|3-3 | 1 | 1 | 25010 | 358 | Spn | | 1.431e-02 | -4.25 | fly chr61A5-61A5\|3-1 cM | chromosome location | fly chr61A5-61A5\|3-1 cM | 1 | 1 | 25010 | 358 | mthl8 | | 1.431e-02 | -4.25 | fly chr92A2-92A3\|3-66 cM | chromosome location | fly chr92A2-92A3\|3-66 cM | 1 | 1 | 25010 | 358 | Ino80 | | 1.431e-02 | -4.25 | fly chr61B2-61B3\|3-1 | chromosome location | fly chr61B2-61B3\|3-1 | 1 | 1 | 25010 | 358 | E(bx) | | 1.431e-02 | -4.25 | fly chr89A4-89A5\|3-58 cM | chromosome location | fly chr89A4-89A5\|3-58 cM | 1 | 1 | 25010 | 358 | nsl1 | | 1.431e-02 | -4.25 | fly chr52D13-52D14\|2-76 cM | chromosome location | fly chr52D13-52D14\|2-76 cM | 1 | 1 | 25010 | 358 | Prosbeta1 | | 1.431e-02 | -4.25 | fly chr12E10-12E10\|1-48 | chromosome location | fly chr12E10-12E10\|1-48 | 1 | 1 | 25010 | 358 | CG9411 | | 1.431e-02 | -4.25 | fly chr94B5-94B6\|3-75 cM | chromosome location | fly chr94B5-94B6\|3-75 cM | 1 | 1 | 25010 | 358 | Efa6 | | 1.431e-02 | -4.25 | fly chr10D6-10D7\|1-35 cM | chromosome location | fly chr10D6-10D7\|1-35 cM | 1 | 1 | 25010 | 358 | Amun | | 1.431e-02 | -4.25 | fly chr38E4-38E5\|2-54 | chromosome location | fly chr38E4-38E5\|2-54 | 1 | 1 | 25010 | 358 | TM9SF2 | | 1.431e-02 | -4.25 | fly chr69E1-69E2\|3-39 cM | chromosome location | fly chr69E1-69E2\|3-39 cM | 1 | 1 | 25010 | 358 | Vps13D | | 1.431e-02 | -4.25 | fly chr30F2-30F3\|2-38 | chromosome location | fly chr30F2-30F3\|2-38 | 1 | 1 | 25010 | 358 | Dref | | 1.431e-02 | -4.25 | fly chr89A4-89A5\|3-58 | chromosome location | fly chr89A4-89A5\|3-58 | 1 | 1 | 25010 | 358 | nsl1 | | 1.431e-02 | -4.25 | fly chr79E1-79E3\|3-47 cM | chromosome location | fly chr79E1-79E3\|3-47 cM | 1 | 1 | 25010 | 358 | lncRNA:CR45962 | | 1.431e-02 | -4.25 | fly chr95F1-95F2\|3-84 cM | chromosome location | fly chr95F1-95F2\|3-84 cM | 1 | 1 | 25010 | 358 | twin | | 1.431e-02 | -4.25 | fly chr63B9-63B10\|3-6 | chromosome location | fly chr63B9-63B10\|3-6 | 1 | 1 | 25010 | 358 | Girdin | | 1.431e-02 | -4.25 | fly chr77B4-77B5\|3-47 cM | chromosome location | fly chr77B4-77B5\|3-47 cM | 1 | 1 | 25010 | 358 | eRF1 | | 1.431e-02 | -4.25 | fly chr5A12-5A13\|1-12 cM | chromosome location | fly chr5A12-5A13\|1-12 cM | 1 | 1 | 25010 | 358 | PGAP1 | | 1.431e-02 | -4.25 | fly chr12E10-12E10\|1-48 cM | chromosome location | fly chr12E10-12E10\|1-48 cM | 1 | 1 | 25010 | 358 | CG9411 | | 1.431e-02 | -4.25 | fly chr25D3-25D3\|2-16 cM | chromosome location | fly chr25D3-25D3\|2-16 cM | 1 | 1 | 25010 | 358 | Bub1 | | 1.431e-02 | -4.25 | fly chr96B19-96B20\|3-86 cM | chromosome location | fly chr96B19-96B20\|3-86 cM | 1 | 1 | 25010 | 358 | Stt3B | | 1.431e-02 | -4.25 | fly chr56D15-56E1\|2-90 | chromosome location | fly chr56D15-56E1\|2-90 | 1 | 1 | 25010 | 358 | sm | | 1.431e-02 | -4.25 | fly chr41E3-41E4\|2-55 cM | chromosome location | fly chr41E3-41E4\|2-55 cM | 1 | 1 | 25010 | 358 | d4 | | 1.431e-02 | -4.25 | fly chr70B2-70B3\|3-40 cM | chromosome location | fly chr70B2-70B3\|3-40 cM | 1 | 1 | 25010 | 358 | Tgi | | 1.431e-02 | -4.25 | fly chr66D6-66D7\|3-26 cM | chromosome location | fly chr66D6-66D7\|3-26 cM | 1 | 1 | 25010 | 358 | rhea | | 1.431e-02 | -4.25 | fly chr45F5-45F6\|2-61 | chromosome location | fly chr45F5-45F6\|2-61 | 1 | 1 | 25010 | 358 | Map60 | | 1.431e-02 | -4.25 | fly chr11A8-11A8\|1-39 cM | chromosome location | fly chr11A8-11A8\|1-39 cM | 1 | 1 | 25010 | 358 | LIMK1 | | 1.431e-02 | -4.25 | fly chr63B9-63B10\|3-6 cM | chromosome location | fly chr63B9-63B10\|3-6 cM | 1 | 1 | 25010 | 358 | Girdin | | 1.431e-02 | -4.25 | fly chr96B19-96B20\|3-86 | chromosome location | fly chr96B19-96B20\|3-86 | 1 | 1 | 25010 | 358 | Stt3B | | 1.431e-02 | -4.25 | fly chr55C8-55C9\|2-85 cM | chromosome location | fly chr55C8-55C9\|2-85 cM | 1 | 1 | 25010 | 358 | CG5174 | | 1.431e-02 | -4.25 | fly chr17D5-17D6\|1-60 | chromosome location | fly chr17D5-17D6\|1-60 | 1 | 1 | 25010 | 358 | CG6961 | | 1.431e-02 | -4.25 | fly chr61B2-61B3\|3-1 cM | chromosome location | fly chr61B2-61B3\|3-1 cM | 1 | 1 | 25010 | 358 | E(bx) | | 1.431e-02 | -4.25 | fly chr67B9-67B10\|3-29 cM | chromosome location | fly chr67B9-67B10\|3-29 cM | 1 | 1 | 25010 | 358 | Jarid2 | | 1.431e-02 | -4.25 | fly chr70A8-70B1\|3-40 cM | chromosome location | fly chr70A8-70B1\|3-40 cM | 1 | 1 | 25010 | 358 | flr | | 1.431e-02 | -4.25 | fly chr8C1-8C4\|1-26 cM | chromosome location | fly chr8C1-8C4\|1-26 cM | 1 | 1 | 25010 | 358 | lncRNA:CR43836 | | 1.431e-02 | -4.25 | fly chr98A8-98A8\|3-94 cM | chromosome location | fly chr98A8-98A8\|3-94 cM | 1 | 1 | 25010 | 358 | pins | | 1.431e-02 | -4.25 | fly chr45A7-45A7\|2-60 | chromosome location | fly chr45A7-45A7\|2-60 | 1 | 1 | 25010 | 358 | tsu | | 1.431e-02 | -4.25 | fly chr38E5-38E6\|2-54 cM | chromosome location | fly chr38E5-38E6\|2-54 cM | 1 | 1 | 25010 | 358 | Fs(2)Ket | | 1.431e-02 | -4.25 | fly chr64E6-64E7\|3-15 | chromosome location | fly chr64E6-64E7\|3-15 | 1 | 1 | 25010 | 358 | Txl | | 1.431e-02 | -4.25 | fly chr48A3-48B2\|2-64 | chromosome location | fly chr48A3-48B2\|2-64 | 1 | 1 | 25010 | 358 | CG9005 | | 1.431e-02 | -4.25 | fly chr60D3-60D4\|2-107 | chromosome location | fly chr60D3-60D4\|2-107 | 1 | 1 | 25010 | 358 | CG4612 | | 1.431e-02 | -4.25 | fly chr87B9-87B10\|3-52 cM | chromosome location | fly chr87B9-87B10\|3-52 cM | 1 | 1 | 25010 | 358 | Pp1-87B | | 1.431e-02 | -4.25 | fly chr8C8-8C9\|1-26 | chromosome location | fly chr8C8-8C9\|1-26 | 1 | 1 | 25010 | 358 | CG7766 | | 1.431e-02 | -4.25 | fly chr77B4-77B5\|3-47 | chromosome location | fly chr77B4-77B5\|3-47 | 1 | 1 | 25010 | 358 | eRF1 | | 1.431e-02 | -4.25 | fly chr88E9-88E11\|3-56 | chromosome location | fly chr88E9-88E11\|3-56 | 1 | 1 | 25010 | 358 | Cp190 | | 1.431e-02 | -4.25 | fly chr72E3-72F4\|3-44 cM | chromosome location | fly chr72E3-72F4\|3-44 cM | 1 | 1 | 25010 | 358 | Tcs3 | | 1.431e-02 | -4.25 | fly chr79E1-79E3\|3-47 | chromosome location | fly chr79E1-79E3\|3-47 | 1 | 1 | 25010 | 358 | lncRNA:CR45962 | | 1.431e-02 | -4.25 | fly chr66C5-66C6\|3-24 cM | chromosome location | fly chr66C5-66C6\|3-24 cM | 1 | 1 | 25010 | 358 | Cpsf6 | | 1.431e-02 | -4.25 | fly chr76D2-76D3\|3-46 cM | chromosome location | fly chr76D2-76D3\|3-46 cM | 1 | 1 | 25010 | 358 | Rab8 | | 1.431e-02 | -4.25 | fly chr38E5-38E6\|2-54 | chromosome location | fly chr38E5-38E6\|2-54 | 1 | 1 | 25010 | 358 | Fs(2)Ket | | 1.431e-02 | -4.25 | fly chr66B8-66B8\|3-22 | chromosome location | fly chr66B8-66B8\|3-22 | 1 | 1 | 25010 | 358 | mus301 | | 1.431e-02 | -4.25 | fly chr9F5-9F7\|1-33 cM | chromosome location | fly chr9F5-9F7\|1-33 cM | 1 | 1 | 25010 | 358 | CG1637 | | 1.431e-02 | -4.25 | fly chr9B14-9C1\|1-31 | chromosome location | fly chr9B14-9C1\|1-31 | 1 | 1 | 25010 | 358 | flw | | 1.431e-02 | -4.25 | fly chr51A1-51A2\|2-71 | chromosome location | fly chr51A1-51A2\|2-71 | 1 | 1 | 25010 | 358 | CG17385 | | 1.431e-02 | -4.25 | fly chr10D4-10D4\|1-35 cM | chromosome location | fly chr10D4-10D4\|1-35 cM | 1 | 1 | 25010 | 358 | Vlet | | 1.431e-02 | -4.25 | fly chr60D3-60D4\|2-107 cM | chromosome location | fly chr60D3-60D4\|2-107 cM | 1 | 1 | 25010 | 358 | CG4612 | | 1.431e-02 | -4.25 | fly chr52E4-52E4\|2-77 cM | chromosome location | fly chr52E4-52E4\|2-77 cM | 1 | 1 | 25010 | 358 | mRpL34 | | 1.431e-02 | -4.25 | fly chr14B7-14B8\|1-53 | chromosome location | fly chr14B7-14B8\|1-53 | 1 | 1 | 25010 | 358 | CG32576 | | 1.431e-02 | -4.25 | fly chr41E3-41E4\|2-55 | chromosome location | fly chr41E3-41E4\|2-55 | 1 | 1 | 25010 | 358 | d4 | | 1.431e-02 | -4.25 | fly chr67C11-67D1\|3-31 cM | chromosome location | fly chr67C11-67D1\|3-31 cM | 1 | 1 | 25010 | 358 | CG8108 | | 1.431e-02 | -4.25 | fly chr87D8-87D9\|3-53 cM | chromosome location | fly chr87D8-87D9\|3-53 cM | 1 | 1 | 25010 | 358 | CtBP | | 1.431e-02 | -4.25 | fly chr41B3-41B3\|2-55 | chromosome location | fly chr41B3-41B3\|2-55 | 1 | 1 | 25010 | 358 | CG17883 | | 1.431e-02 | -4.25 | fly chr41A3-41A3\|2-55 cM | chromosome location | fly chr41A3-41A3\|2-55 cM | 1 | 1 | 25010 | 358 | Stlk | | 1.431e-02 | -4.25 | fly chr22F1-22F3\|2-6 | chromosome location | fly chr22F1-22F3\|2-6 | 1 | 1 | 25010 | 358 | dpp | | 1.431e-02 | -4.25 | fly chr16E4-16E4\|1-59 | chromosome location | fly chr16E4-16E4\|1-59 | 1 | 1 | 25010 | 358 | mnb | | 1.431e-02 | -4.25 | fly chr46E6-46E8\|2-62 | chromosome location | fly chr46E6-46E8\|2-62 | 1 | 1 | 25010 | 358 | 14-3-3zeta | | 1.431e-02 | -4.25 | fly chr85F10-85F12\|3-49 cM | chromosome location | fly chr85F10-85F12\|3-49 cM | 1 | 1 | 25010 | 358 | Fmr1 | | 1.431e-02 | -4.25 | fly chr9F5-9F7\|1-33 | chromosome location | fly chr9F5-9F7\|1-33 | 1 | 1 | 25010 | 358 | CG1637 | | 1.431e-02 | -4.25 | fly chr2C1-2C1\|1-0.5 cM | chromosome location | fly chr2C1-2C1\|1-0.5 cM | 1 | 1 | 25010 | 358 | PIG-K | | 1.431e-02 | -4.25 | fly chr29A1-29A3\|2-31 cM | chromosome location | fly chr29A1-29A3\|2-31 cM | 1 | 1 | 25010 | 358 | Btk29A | | 1.431e-02 | -4.25 | fly chr16E4-16E4\|1-59 cM | chromosome location | fly chr16E4-16E4\|1-59 cM | 1 | 1 | 25010 | 358 | mnb | | 1.431e-02 | -4.25 | fly chr4F9-4F10\|1-11 cM | chromosome location | fly chr4F9-4F10\|1-11 cM | 1 | 1 | 25010 | 358 | spoon | | 1.431e-02 | -4.25 | fly chr29B2-29B3\|2-31 | chromosome location | fly chr29B2-29B3\|2-31 | 1 | 1 | 25010 | 358 | Wdr82 | | 1.431e-02 | -4.25 | fly chr68A3-68A4\|3-34 cM | chromosome location | fly chr68A3-68A4\|3-34 cM | 1 | 1 | 25010 | 358 | CG32075 | | 1.431e-02 | -4.25 | fly chr67B1-67D13\|3-32 | chromosome location | fly chr67B1-67D13\|3-32 | 1 | 1 | 25010 | 358 | vnc | | 1.431e-02 | -4.25 | fly chr48A3-48B2\|2-64 cM | chromosome location | fly chr48A3-48B2\|2-64 cM | 1 | 1 | 25010 | 358 | CG9005 | | 1.431e-02 | -4.25 | fly chr28E6-28E7\|2-30 cM | chromosome location | fly chr28E6-28E7\|2-30 cM | 1 | 1 | 25010 | 358 | mon2 | | 1.431e-02 | -4.25 | fly chr69E1-69E2\|3-39 | chromosome location | fly chr69E1-69E2\|3-39 | 1 | 1 | 25010 | 358 | Vps13D | | 1.431e-02 | -4.25 | fly chr9B14-9C1\|1-31 cM | chromosome location | fly chr9B14-9C1\|1-31 cM | 1 | 1 | 25010 | 358 | flw | | 1.431e-02 | -4.25 | fly chr102F5-102F6\|4-0 cM | chromosome location | fly chr102F5-102F6\|4-0 cM | 1 | 1 | 25010 | 358 | PlexA | | 1.431e-02 | -4.25 | fly chr66D6-66D7\|3-26 | chromosome location | fly chr66D6-66D7\|3-26 | 1 | 1 | 25010 | 358 | rhea | | 1.431e-02 | -4.25 | fly chr86F8-86F9\|3-51 | chromosome location | fly chr86F8-86F9\|3-51 | 1 | 1 | 25010 | 358 | glo | | 1.431e-02 | -4.25 | fly chr102F5-102F6\|4-0 | chromosome location | fly chr102F5-102F6\|4-0 | 1 | 1 | 25010 | 358 | PlexA | | 1.431e-02 | -4.25 | fly chr64E6-64E7\|3-15 cM | chromosome location | fly chr64E6-64E7\|3-15 cM | 1 | 1 | 25010 | 358 | Txl | | 1.431e-02 | -4.25 | fly chr85F10-85F12\|3-49 | chromosome location | fly chr85F10-85F12\|3-49 | 1 | 1 | 25010 | 358 | Fmr1 | | 1.431e-02 | -4.25 | fly chr10D1-10D3\|1-35 | chromosome location | fly chr10D1-10D3\|1-35 | 1 | 1 | 25010 | 358 | RPA3 | | 1.431e-02 | -4.25 | fly chr67C11-67D1\|3-31 | chromosome location | fly chr67C11-67D1\|3-31 | 1 | 1 | 25010 | 358 | CG8108 | | 1.431e-02 | -4.25 | fly chr45A9-45A9\|2-60 | chromosome location | fly chr45A9-45A9\|2-60 | 1 | 1 | 25010 | 358 | GstE13 | | 1.431e-02 | -4.25 | fly chr92A2-92A3\|3-66 | chromosome location | fly chr92A2-92A3\|3-66 | 1 | 1 | 25010 | 358 | Ino80 | | 1.431e-02 | -4.25 | fly chr48B4-48C1\|2-64 cM | chromosome location | fly chr48B4-48C1\|2-64 cM | 1 | 1 | 25010 | 358 | Roc2 | | 1.431e-02 | -4.25 | fly chr56D15-56E1\|2-90 cM | chromosome location | fly chr56D15-56E1\|2-90 cM | 1 | 1 | 25010 | 358 | sm | | 1.431e-02 | -4.25 | fly chr17D5-17D6\|1-60 cM | chromosome location | fly chr17D5-17D6\|1-60 cM | 1 | 1 | 25010 | 358 | CG6961 | | 1.431e-02 | -4.25 | fly chr10D6-10D7\|1-35 | chromosome location | fly chr10D6-10D7\|1-35 | 1 | 1 | 25010 | 358 | Amun | | 1.431e-02 | -4.25 | fly chr87F5-87F6\|3-54 cM | chromosome location | fly chr87F5-87F6\|3-54 cM | 1 | 1 | 25010 | 358 | sqd | | 1.431e-02 | -4.25 | fly chr36F4-36F6\|2-53 cM | chromosome location | fly chr36F4-36F6\|2-53 cM | 1 | 1 | 25010 | 358 | RpS26 | | 1.431e-02 | -4.25 | fly chr31C4-31C6\|2-40 | chromosome location | fly chr31C4-31C6\|2-40 | 1 | 1 | 25010 | 358 | Ufd4 | | 1.431e-02 | -4.25 | fly chr25D5-25D6\|2-16 cM | chromosome location | fly chr25D5-25D6\|2-16 cM | 1 | 1 | 25010 | 358 | Ncoa6 | | 1.431e-02 | -4.25 | fly chr36F4-36F6\|2-53 | chromosome location | fly chr36F4-36F6\|2-53 | 1 | 1 | 25010 | 358 | RpS26 | | 1.431e-02 | -4.25 | fly chr31C4-31C6\|2-40 cM | chromosome location | fly chr31C4-31C6\|2-40 cM | 1 | 1 | 25010 | 358 | Ufd4 | | 1.431e-02 | -4.25 | fly chr25D1-25D2\|2-16 cM | chromosome location | fly chr25D1-25D2\|2-16 cM | 1 | 1 | 25010 | 358 | tkv | | 1.431e-02 | -4.25 | fly chr41A3-41A3\|2-55 | chromosome location | fly chr41A3-41A3\|2-55 | 1 | 1 | 25010 | 358 | Stlk | | 1.431e-02 | -4.25 | fly chr59C2-59C3\|2-102 | chromosome location | fly chr59C2-59C3\|2-102 | 1 | 1 | 25010 | 358 | DCTN3-p24 | | 1.431e-02 | -4.25 | fly chr76D2-76D3\|3-46 | chromosome location | fly chr76D2-76D3\|3-46 | 1 | 1 | 25010 | 358 | Rab8 | | 1.431e-02 | -4.25 | fly chr25D5-25D6\|2-16 | chromosome location | fly chr25D5-25D6\|2-16 | 1 | 1 | 25010 | 358 | Ncoa6 | | 1.431e-02 | -4.25 | fly chr66C5-66C6\|3-24 | chromosome location | fly chr66C5-66C6\|3-24 | 1 | 1 | 25010 | 358 | Cpsf6 | | 1.431e-02 | -4.25 | fly chr62E4-62E5\|3-3 cM | chromosome location | fly chr62E4-62E5\|3-3 cM | 1 | 1 | 25010 | 358 | Spn | | 1.431e-02 | -4.25 | fly chr52D2-52D3\|2-76 | chromosome location | fly chr52D2-52D3\|2-76 | 1 | 1 | 25010 | 358 | Mlf | | 1.431e-02 | -4.25 | fly chr45F5-45F6\|2-61 cM | chromosome location | fly chr45F5-45F6\|2-61 cM | 1 | 1 | 25010 | 358 | Map60 | | 1.431e-02 | -4.25 | fly chr5A12-5A13\|1-12 | chromosome location | fly chr5A12-5A13\|1-12 | 1 | 1 | 25010 | 358 | PGAP1 | | 1.431e-02 | -4.25 | fly chr73D7-73E1\|3-45 | chromosome location | fly chr73D7-73E1\|3-45 | 1 | 1 | 25010 | 358 | Cpr73D | | 1.431e-02 | -4.25 | fly chr28A5-28A5\|2-27 | chromosome location | fly chr28A5-28A5\|2-27 | 1 | 1 | 25010 | 358 | lncRNA:CR44082 | | 1.431e-02 | -4.25 | fly chr55D3-55E1\|2-86 cM | chromosome location | fly chr55D3-55E1\|2-86 cM | 1 | 1 | 25010 | 358 | GEFmeso | | 1.431e-02 | -4.25 | fly chr48B4-48C1\|2-64 | chromosome location | fly chr48B4-48C1\|2-64 | 1 | 1 | 25010 | 358 | Roc2 | | 1.436e-02 | -4.24 | GSE15733\_BM\_VS\_SPLEEN\_MEMORY\_CD4\_TCELL\_DN | MSigDB lists | GSE15733\_BM\_VS\_SPLEEN\_MEMORY\_CD4\_TCELL\_DN | 59 | 7 | 4161 | 184 | CG12935,CG15168,14-3-3zeta,RpL36A,eRF1,vnc,Tsp97E | | 1.438e-02 | -4.24 | SMC5 (Structural maintenance of chromosomes 5) | protein interactions | 326215 | 6 | 2 | 9204 | 298 | dgrn,E(z) | | 1.438e-02 | -4.24 | SdhD (Succinate dehydrogenase, subunit D) | protein interactions | 42808 | 6 | 2 | 9204 | 298 | Art1,smg | | 1.438e-02 | -4.24 | CG15705 (uncharacterized protein) | protein interactions | 36815 | 6 | 2 | 9204 | 298 | DCTN3-p24,Pp1-87B | | 1.438e-02 | -4.24 | CG5866 (uncharacterized protein) | protein interactions | 42097 | 6 | 2 | 9204 | 298 | PlexA,CG43367 | | 1.438e-02 | -4.24 | papi (papi) | protein interactions | 33401 | 6 | 2 | 9204 | 298 | AGO3,me31B | | 1.438e-02 | -4.24 | CG6254 (uncharacterized protein) | protein interactions | 41244 | 6 | 2 | 9204 | 298 | eEF5,krz | | 1.438e-02 | -4.24 | mute (muscle wasted) | protein interactions | 2768848 | 6 | 2 | 9204 | 298 | Cp190,su(Hw) | | 1.438e-02 | -4.24 | CG7956 (uncharacterized protein) | protein interactions | 42545 | 6 | 2 | 9204 | 298 | tsu,Rab14 | | 1.438e-02 | -4.24 | dbr (debra) | protein interactions | 33161 | 6 | 2 | 9204 | 298 | Cp190,su(Hw) | | 1.438e-02 | -4.24 | MYPT-75D (MYPT-75D) | protein interactions | 40032 | 6 | 2 | 9204 | 298 | flw,Pp1-87B | | 1.438e-02 | -4.24 | Nop17l (Nop17 like) | protein interactions | 35730 | 6 | 2 | 9204 | 298 | flw,Pp1-87B | | 1.438e-02 | -4.24 | Snoo (Sno oncogene) | protein interactions | 5740414 | 6 | 2 | 9204 | 298 | tkv,dpp | | 1.438e-02 | -4.24 | CG42748 (uncharacterized protein) | protein interactions | 35442 | 6 | 2 | 9204 | 298 | Pp1-87B,flw | | 1.438e-02 | -4.24 | slif (slimfast) | protein interactions | 40510 | 6 | 2 | 9204 | 298 | CG42676,Pten | | 1.438e-02 | -4.24 | Sara (Smad anchor for receptor activation) | protein interactions | 44263 | 6 | 2 | 9204 | 298 | flw,Pp1-87B | | 1.438e-02 | -4.24 | Sf3a2 (Splicing factor 3a subunit 2) | protein interactions | 39456 | 6 | 2 | 9204 | 298 | noi,snf | | 1.438e-02 | -4.24 | cuff (cutoff) | protein interactions | 36269 | 6 | 2 | 9204 | 298 | rhi,Rat1 | | 1.438e-02 | -4.24 | Rbpn-5 (Rabaptin-5) | protein interactions | 37395 | 6 | 2 | 9204 | 298 | Exo84,Rab14 | | 1.439e-02 | -4.24 | GSE1925\_CTRL\_VS\_IFNG\_PRIMED\_MACROPHAGE\_DN | MSigDB lists | GSE1925\_CTRL\_VS\_IFNG\_PRIMED\_MACROPHAGE\_DN | 73 | 8 | 4161 | 184 | mRpS28,Stlk,pav,eRF1,RpS8,cta,Vps35,CG31460 | | 1.448e-02 | -4.24 | REACTOME\_SIGNALING\_BY\_TGF\_BETA\_RECEPTOR\_COMPLEX | MSigDB lists | REACTOME\_SIGNALING\_BY\_TGF\_BETA\_RECEPTOR\_COMPLEX | 22 | 4 | 4161 | 184 | Parp,flw,Cdk8,Pp1-87B | | 1.448e-02 | -4.24 | GO\_GLOBAL\_GENOME\_NUCLEOTIDE\_EXCISION\_REPAIR | MSigDB lists | GO\_GLOBAL\_GENOME\_NUCLEOTIDE\_EXCISION\_REPAIR | 22 | 4 | 4161 | 184 | Cul4,smt3,CG17493,Parp | | 1.451e-02 | -4.23 | WD40/YVTN\_repeat-like\_dom\_sf | interpro domains | IPR015943 | 201 | 12 | 11383 | 331 | fzy,eIF3b,CG4935,CG14805,Rbbp5,CG43367,Wdr33,Lst8,flr,CG11141,Wdr82,PlexA | | 1.460e-02 | -4.23 | cytosolic large ribosomal subunit | cellular component | GO:0022625 | 58 | 5 | 11940 | 298 | RpL15,RpL35A,RpL14,RpL5,RpL36A | | 1.473e-02 | -4.22 | Rpt4 (Regulatory particle triple-A ATPase 4) | protein interactions | 31567 | 62 | 6 | 9204 | 298 | mRpL30,Prosalpha2,Rpt6,Prosbeta1,CG12321,Ubqn | | 1.479e-02 | -4.21 | MORI\_IMMATURE\_B\_LYMPHOCYTE\_DN | MSigDB lists | MORI\_IMMATURE\_B\_LYMPHOCYTE\_DN | 46 | 6 | 4161 | 184 | E(z),His2Av,CG32473,DNAlig1,fzy,CycA | | 1.479e-02 | -4.21 | MATTIOLI\_MGUS\_VS\_PCL | MSigDB lists | MATTIOLI\_MGUS\_VS\_PCL | 46 | 6 | 4161 | 184 | Rpn10,Tctp,CG14805,SF2,CCT4,Parp | | 1.479e-02 | -4.21 | GSE5589\_UNSTIM\_VS\_45MIN\_LPS\_AND\_IL10\_STIM\_MACROPHAGE\_UP | MSigDB lists | GSE5589\_UNSTIM\_VS\_45MIN\_LPS\_AND\_IL10\_STIM\_MACROPHAGE\_UP | 46 | 6 | 4161 | 184 | RpS9,Pisd,Pex3,rb,unc-119,HtrA2 | | 1.483e-02 | -4.21 | proteasomal protein catabolic process | biological process | GO:0010498 | 209 | 11 | 12345 | 305 | Rpn10,Rpt6,Prosbeta1,Skp2,Prosalpha2,Roc2,alpha-Man-Ib,inc,Sgt,fzy,Cul4 | | 1.488e-02 | -4.21 | GO\_MACROMOLECULAR\_COMPLEX\_BINDING | MSigDB lists | GO\_MACROMOLECULAR\_COMPLEX\_BINDING | 361 | 25 | 4161 | 184 | pav,spel1,CG9246,Pp1-87B,fzy,eRF1,Fmr1,Pten,Wdr82,Vap33,CtBP,vnc,Rap1,E(z),rhea,Arpc5,cta,eEF5,CG6907,inc,CG17493,bai,Stim,krz,Arpc3A | | 1.489e-02 | -4.21 | TP53 Regulates Transcription of Cell Cycle Genes | REACTOME pathways | R-DME-6791312 | 14 | 3 | 3923 | 151 | Not3,14-3-3zeta,twin | | 1.496e-02 | -4.20 | arm (armadillo) | protein interactions | 31151 | 119 | 9 | 9204 | 298 | tkv,eRF1,pav,CycA,Cdk8,CtBP,dpp,Cul4,MED21 | | 1.502e-02 | -4.20 | multicellular organismal process | biological process | GO:0032501 | 3844 | 113 | 12345 | 305 | tsu,CG32473,mnb,Pp1-87B,heix,krz,Nf1,tsr,CG8149,Skp2,RpS8,Stim,vsg,Cp190,me31B,sm,xit,Fs(2)Ket,snf,Cdc42,peng,fzy,mon2,nsl1,mbf1,Exo84,E(z),His2Av,sqd,LIMK1,Bmcp,Cdk8,rhea,dpp,Acf,tkv,pins,Fmr1,Dref,g,eIF3b,lncRNA:sphinx,pav,vnc,sws,Zpr1,mthl8,EloB,Parp,smt3,MICU1,Orc3,Vap33,cal1,PlexA,Efa6,sigmar,spoon,Rab7,inc,Stlk,glo,VhaPPA1-1,lic,Sod2,RPA3,Hrb27C,GEFmeso,dgrn,Pten,Rap1,CG12935,mav,Rop,cta,rhi,Hsp27,mus301,eEF5,HtrA2,CycA,Spn,E(bx),MED22,Btk29A,CG4612,Amun,flr,vap,pll,CSN1b,grau,Girdin,twin,sun,unc-119,BEAF-32,Rheb,Mps1,Cul4,d4,nom,Cpr73D,rb,dah,bai,CtBP,14-3-3zeta,BthD,CG10365,flw,RhoGAPp190,RecQ4 | | 1.503e-02 | -4.20 | branching involved in open tracheal system development | biological process | GO:0060446 | 59 | 5 | 12345 | 305 | Rheb,VhaPPA1-1,Cdc42,rhea,Zpr1 | | 1.503e-02 | -4.20 | morphogenesis of a branching epithelium | biological process | GO:0061138 | 59 | 5 | 12345 | 305 | VhaPPA1-1,Rheb,Zpr1,rhea,Cdc42 | | 1.503e-02 | -4.20 | morphogenesis of a branching structure | biological process | GO:0001763 | 59 | 5 | 12345 | 305 | Zpr1,Cdc42,rhea,VhaPPA1-1,Rheb | | 1.503e-02 | -4.20 | branching morphogenesis of an epithelial tube | biological process | GO:0048754 | 59 | 5 | 12345 | 305 | Zpr1,rhea,Cdc42,VhaPPA1-1,Rheb | | 1.503e-02 | -4.20 | PH\_DOMAIN | prosite domains | PS50003 | 66 | 6 | 6243 | 191 | Btk29A,GEFmeso,Graf,Exo84,Efa6,vap | | 1.504e-02 | -4.20 | positive regulation of multicellular organismal process | biological process | GO:0051240 | 266 | 13 | 12345 | 305 | dpp,Tctp,Orc3,Hrb27C,lic,Rop,Rap1,Fmr1,Nf1,mnb,Rheb,PlexA,tkv | | 1.509e-02 | -4.19 | SPIELMAN\_LYMPHOBLAST\_EUROPEAN\_VS\_ASIAN\_UP | MSigDB lists | SPIELMAN\_LYMPHOBLAST\_EUROPEAN\_VS\_ASIAN\_UP | 184 | 15 | 4161 | 184 | His2Av,E(z),Stim,Rpn10,TfIIFbeta,CG7766,Rab8,ND-ASHI,sm,Parp,ubl,Aps,fzy,Cdc42,RpS8 | | 1.514e-02 | -4.19 | nervous system development | biological process | GO:0007399 | 1108 | 39 | 12345 | 305 | d4,Rheb,unc-119,Vap33,RhoGAPp190,smt3,CG10365,pins,MED22,Dref,Fmr1,sws,Girdin,pav,vap,flr,dgrn,CG12935,Rap1,Pten,Cdc42,LIMK1,Acf,Spn,CycA,Cdk8,mbf1,Rop,mav,E(z),tsr,Nf1,Stlk,Pp1-87B,PlexA,mnb,sm,me31B,Hrb27C | | 1.516e-02 | -4.19 | metal ion binding | molecular function | GO:0046872 | 1047 | 36 | 12176 | 289 | dah,Tctp,CG5516,CG8635,RAF2,Stim,alpha-Man-Ib,RecQ4,Fign,Usp32,d4,Sod2,Rpn10,Roc2,NO66,CG17493,MICU1,CG32473,Dref,CG6254,Ufd4,grau,CNBP,CG1637,CG1815,noi,Parp,dgrn,CG1218,CG7646,x16,nom,Usp16-45,Zpr1,tkv,CG11586 | | 1.518e-02 | -4.19 | Rho GTPase cycle | REACTOME pathways | R-DME-194840 | 53 | 6 | 3923 | 151 | conu,RhoGAPp190,cta,Graf,GEFmeso,Cdc42 | | 1.520e-02 | -4.19 | RNASEK (Ribonuclease kappa) | protein interactions | 3355016 | 30 | 4 | 9204 | 298 | noi,14-3-3zeta,msd5,su(Hw) | | 1.520e-02 | -4.19 | ND-75 (NADH dehydrogenase (ubiquinone) 75 kDa subunit) | protein interactions | 31762 | 30 | 4 | 9204 | 298 | Lst8,Sod2,smg,CG11752 | | 1.520e-02 | -4.19 | BEAF-32 (Boundary element-associated factor of 32kD) | protein interactions | 36645 | 30 | 4 | 9204 | 298 | su(Hw),BEAF-32,Cp190,Dref | | 1.520e-02 | -4.19 | Chro (Chromator) | protein interactions | 40508 | 30 | 4 | 9204 | 298 | Cp190,nsl1,BEAF-32,Rheb | | 1.539e-02 | -4.17 | positive regulation of cell differentiation | biological process | GO:0045597 | 156 | 9 | 12345 | 305 | Mlf,Rap1,Fmr1,PlexA,mnb,Rheb,Rop,Orc3,Hrb27C | | 1.544e-02 | -4.17 | negative regulation of salivary gland boundary specification | biological process | GO:0045705 | 8 | 2 | 12345 | 305 | tkv,dpp | | 1.544e-02 | -4.17 | positive regulation of protein complex disassembly | biological process | GO:0043243 | 8 | 2 | 12345 | 305 | flr,eEF5 | | 1.544e-02 | -4.17 | regulation of salivary gland boundary specification | biological process | GO:0045704 | 8 | 2 | 12345 | 305 | tkv,dpp | | 1.544e-02 | -4.17 | regulation of pathway-restricted SMAD protein phosphorylation | biological process | GO:0060393 | 8 | 2 | 12345 | 305 | dpp,mav | | 1.544e-02 | -4.17 | actin filament polymerization | biological process | GO:0030041 | 8 | 2 | 12345 | 305 | Cdc42,tsr | | 1.544e-02 | -4.17 | positive regulation of pathway-restricted SMAD protein phosphorylation | biological process | GO:0010862 | 8 | 2 | 12345 | 305 | mav,dpp | | 1.553e-02 | -4.16 | amide transport | biological process | GO:0042886 | 479 | 20 | 12345 | 305 | Tim17b,Rab14,CG6961,Dbp80,Pex3,Tom20,g,RabX6,Vps13D,Rab7,Vps35,sqd,CG8860,bai,mon2,rb,Sgt,Fs(2)Ket,unc-119,Rab8 | | 1.554e-02 | -4.16 | CREBP1CJUN\_01 | MSigDB lists | CREBP1CJUN\_01 | 74 | 8 | 4161 | 184 | CG5861,Dbp21E2,CycA,14-3-3zeta,Roc2,CG15168,Nf1,CG13016 | | 1.562e-02 | -4.16 | organellar ribosome | cellular component | GO:0000313 | 81 | 6 | 11940 | 298 | mRpS33,mRpL34,mRpS28,mRpS9,mRpL30,mRpS18A | | 1.562e-02 | -4.16 | mitochondrial ribosome | cellular component | GO:0005761 | 81 | 6 | 11940 | 298 | mRpS28,mRpS18A,mRpL30,mRpS9,mRpL34,mRpS33 | | 1.562e-02 | -4.16 | Q\_MOTIF | prosite domains | PS51195 | 32 | 4 | 6243 | 191 | Dbp80,CG5589,me31B,Dbp21E2 | | 1.567e-02 | -4.16 | GSE33424\_CD161\_HIGH\_VS\_INT\_CD8\_TCELL\_DN | MSigDB lists | GSE33424\_CD161\_HIGH\_VS\_INT\_CD8\_TCELL\_DN | 60 | 7 | 4161 | 184 | fzy,CycA,CG10343,RpS26,CG5339,E(z),mRpS28 | | 1.567e-02 | -4.16 | GO\_REGULATION\_OF\_NEURON\_DEATH | MSigDB lists | GO\_REGULATION\_OF\_NEURON\_DEATH | 60 | 7 | 4161 | 184 | Sod2,Cdc42,HtrA2,Rop,Nf1,vap,spel1 | | 1.569e-02 | -4.15 | GO\_RRNA\_BINDING | MSigDB lists | GO\_RRNA\_BINDING | 34 | 5 | 4161 | 184 | RpS9,CG11920,Dbp21E2,RpL5,CG4866 | | 1.569e-02 | -4.15 | GSE40068\_BCL6\_POS\_VS\_NEG\_CXCR5\_POS\_TFH\_DN | MSigDB lists | GSE40068\_BCL6\_POS\_VS\_NEG\_CXCR5\_POS\_TFH\_DN | 34 | 5 | 4161 | 184 | janA,Usp32,RpL5,RhoGAPp190,RpS8 | | 1.569e-02 | -4.15 | GSE21670\_TGFB\_VS\_IL6\_TREATED\_STAT3\_KO\_CD4\_TCELL\_DN | MSigDB lists | GSE21670\_TGFB\_VS\_IL6\_TREATED\_STAT3\_KO\_CD4\_TCELL\_DN | 34 | 5 | 4161 | 184 | PGAP1,Tsp97E,sws,Usp32,Graf | | 1.569e-02 | -4.15 | GO\_N\_ACETYLTRANSFERASE\_ACTIVITY | MSigDB lists | GO\_N\_ACETYLTRANSFERASE\_ACTIVITY | 34 | 5 | 4161 | 184 | vnc,Rcd5,CG6907,mbf1,krz | | 1.569e-02 | -4.15 | GO\_POSITIVE\_REGULATION\_OF\_CELLULAR\_AMIDE\_METABOLIC\_PROCESS | MSigDB lists | GO\_POSITIVE\_REGULATION\_OF\_CELLULAR\_AMIDE\_METABOLIC\_PROCESS | 34 | 5 | 4161 | 184 | AGO3,RpS9,Fmr1,CG4866,eEF5 | | 1.574e-02 | -4.15 | Arp2/3 protein complex | cellular component | GO:0005885 | 8 | 2 | 11940 | 298 | Arpc5,Arpc3A | | 1.574e-02 | -4.15 | exon-exon junction complex | cellular component | GO:0035145 | 8 | 2 | 11940 | 298 | Pnn,tsu | | 1.580e-02 | -4.15 | DNA helicase activity | molecular function | GO:0003678 | 41 | 4 | 12176 | 289 | Arp8,Ino80,RecQ4,mus301 | | 1.593e-02 | -4.14 | Prosbeta2 (Proteasome beta2 subunit) | protein interactions | 39628 | 46 | 5 | 9204 | 298 | Rpt6,Prosalpha2,Prosbeta1,CG12321,Ubqn | | 1.597e-02 | -4.14 | regulation of autophagy | biological process | GO:0010506 | 106 | 7 | 12345 | 305 | sigmar,Sod2,Cpsf6,RagA-B,Rheb,Cdk8,Pten | | 1.597e-02 | -4.14 | asymmetric cell division | biological process | GO:0008356 | 106 | 7 | 12345 | 305 | CycA,Dref,Cdc42,pins,CSN1b,Mps1,twin | | 1.607e-02 | -4.13 | TGF-beta-rel | interpro domains | IPR015615 | 7 | 2 | 11383 | 331 | mav,dpp | | 1.607e-02 | -4.13 | Clathrin/coatomer\_adapt-like\_N | interpro domains | IPR002553 | 7 | 2 | 11383 | 331 | g,rb | | 1.607e-02 | -4.13 | TGF-b\_C | interpro domains | IPR001839 | 7 | 2 | 11383 | 331 | mav,dpp | | 1.607e-02 | -4.13 | TGFb\_CS | interpro domains | IPR017948 | 7 | 2 | 11383 | 331 | mav,dpp | | 1.611e-02 | -4.13 | Axon guidance | REACTOME pathways | R-DME-422475 | 194 | 14 | 3923 | 151 | Prosbeta1,14-3-3zeta,CG7054,Arpc3A,LIMK1,Rpt6,Nf1,vap,rhea,Cdc42,Rpn10,CG4935,Prosalpha2,Arpc5 | | 1.618e-02 | -4.12 | post-embryonic animal organ development | biological process | GO:0048569 | 481 | 20 | 12345 | 305 | Stim,flr,RpS8,pll,GEFmeso,tkv,Btk29A,heix,tsr,CG8149,flw,nsl1,dpp,rhea,Cdk8,LIMK1,EloB,nom,peng,Cdc42 | | 1.623e-02 | -4.12 | RNA Polymerase II Transcription | REACTOME pathways | R-DME-73857 | 120 | 10 | 3923 | 151 | x16,TfIIFbeta,Rpb11,SF2,Wdr33,Cpsf6,EloB,Upf3,tsu,CG6961 | | 1.624e-02 | -4.12 | synaptic vesicle transport | biological process | GO:0048489 | 22 | 3 | 12345 | 305 | Fmr1,g,Rop | | 1.624e-02 | -4.12 | response to BMP | biological process | GO:0071772 | 22 | 3 | 12345 | 305 | tkv,mav,dpp | | 1.624e-02 | -4.12 | regulation of DNA recombination | biological process | GO:0000018 | 22 | 3 | 12345 | 305 | spel1,tsu,Fign | | 1.629e-02 | -4.12 | TP53 Regulates Metabolic Genes | REACTOME pathways | R-DME-5628897 | 26 | 4 | 3923 | 151 | Lst8,14-3-3zeta,RagA-B,Rheb | | 1.631e-02 | -4.12 | CG12081 (uncharacterized protein) | protein interactions | 31799 | 17 | 3 | 9204 | 298 | E(z),RPA3,EloB | | 1.631e-02 | -4.12 | CG15525 (uncharacterized protein) | protein interactions | 43556 | 17 | 3 | 9204 | 298 | Rcd5,l(1)10Bb,CG10209 | | 1.631e-02 | -4.12 | eIF3k (eukaryotic translation initiation factor 3 subunit k) | protein interactions | 37475 | 17 | 3 | 9204 | 298 | eIF3c,eIF3g1,eIF3b | | 1.631e-02 | -4.12 | Arpc4 (Actin-related protein 2/3 complex, subunit 4) | protein interactions | 33864 | 17 | 3 | 9204 | 298 | Arpc3A,Rbp1,Arpc5 | | 1.631e-02 | -4.12 | east (enhanced adult sensory threshold) | protein interactions | 46006 | 17 | 3 | 9204 | 298 | vap,Cp190,su(Hw) | | 1.631e-02 | -4.12 | y (yellow) | protein interactions | 30980 | 17 | 3 | 9204 | 298 | su(Hw),BEAF-32,Cp190 | | 1.633e-02 | -4.11 | GO\_STRUCTURAL\_MOLECULE\_ACTIVITY | MSigDB lists | GO\_STRUCTURAL\_MOLECULE\_ACTIVITY | 220 | 17 | 4161 | 184 | Arpc3A,CG4743,RpL36A,CG4866,RpL15,mRpS33,rhea,RpS9,Arpc5,eIF3b,RpL5,RpS26,RpL14,Bmcp,mRpS9,RpS8,RpL35A | | 1.635e-02 | -4.11 | GO\_REGULATION\_OF\_NEURON\_APOPTOTIC\_PROCESS | MSigDB lists | GO\_REGULATION\_OF\_NEURON\_APOPTOTIC\_PROCESS | 47 | 6 | 4161 | 184 | Sod2,Cdc42,Rop,vap,spel1,Nf1 | | 1.635e-02 | -4.11 | GSE21927\_UNTREATED\_VS\_GMCSF\_GCSF\_TREATED\_BONE\_MARROW\_DN | MSigDB lists | GSE21927\_UNTREATED\_VS\_GMCSF\_GCSF\_TREATED\_BONE\_MARROW\_DN | 47 | 6 | 4161 | 184 | Pten,Stt3B,Parp,Skp2,Cul4,Pp1-87B | | 1.659e-02 | -4.10 | negative regulation of response to stimulus | biological process | GO:0048585 | 389 | 17 | 12345 | 305 | flw,CtBP,Rheb,Pten,Cul4,CG30423,pav,vap,lic,Graf,Slmap,mnb,TM9SF2,E(bx),Fgop2,Fmr1,krz | | 1.668e-02 | -4.09 | DNA-dependent ATPase activity | molecular function | GO:0008094 | 63 | 5 | 12176 | 289 | mus301,spel1,Arp8,Ino80,RecQ4 | | 1.671e-02 | -4.09 | Bromodomain | interpro domains | IPR001487 | 19 | 3 | 11383 | 331 | Acf,E(bx),CG1815 | | 1.685e-02 | -4.08 | nucleic acid-templated transcription | biological process | GO:0097659 | 213 | 11 | 12345 | 305 | EloB,CG6244,E(bx),Parp,TfIIFbeta,su(Hw),Pbp49,Rpb11,Arp8,grau,His4r | | 1.685e-02 | -4.08 | transcription, DNA-templated | biological process | GO:0006351 | 213 | 11 | 12345 | 305 | Arp8,Pbp49,Rpb11,grau,His4r,EloB,E(bx),CG6244,Parp,su(Hw),TfIIFbeta | | 1.694e-02 | -4.08 | KEGG\_PATHOGENIC\_ESCHERICHIA\_COLI\_INFECTION | MSigDB lists | KEGG\_PATHOGENIC\_ESCHERICHIA\_COLI\_INFECTION | 23 | 4 | 4161 | 184 | Arpc5,14-3-3zeta,Arpc3A,Cdc42 | | 1.694e-02 | -4.08 | KTGGYRSGAA\_UNKNOWN | MSigDB lists | KTGGYRSGAA\_UNKNOWN | 23 | 4 | 4161 | 184 | SF2,DNAlig1,E(z),Usp32 | | 1.694e-02 | -4.08 | MODULE\_87 | MSigDB lists | MODULE\_87 | 23 | 4 | 4161 | 184 | Cdc42,Rap1,Rab8,cta | | 1.694e-02 | -4.08 | GO\_RIBOSOME\_BINDING | MSigDB lists | GO\_RIBOSOME\_BINDING | 23 | 4 | 4161 | 184 | Fmr1,eRF1,vnc,eEF5 | | 1.694e-02 | -4.08 | GNF2\_MSH6 | MSigDB lists | GNF2\_MSH6 | 23 | 4 | 4161 | 184 | CCT4,Pp1-87B,spel1,DNAlig1 | | 1.694e-02 | -4.08 | MODULE\_149 | MSigDB lists | MODULE\_149 | 23 | 4 | 4161 | 184 | eIF3g1,eIF3b,eIF3c,eEF1delta | | 1.698e-02 | -4.08 | PH\_domain | interpro domains | IPR001849 | 71 | 6 | 11383 | 331 | Efa6,GEFmeso,Graf,Exo84,vap,Btk29A | | 1.702e-02 | -4.07 | Hrb98DE (Heterogeneous nuclear ribonucleoprotein at 98DE) | protein interactions | 43385 | 31 | 4 | 9204 | 298 | glo,Parp,Hrb27C,sqd | | 1.702e-02 | -4.07 | HP1b (Heterochromatin Protein 1b) | protein interactions | 31834 | 31 | 4 | 9204 | 298 | Tctp,Ubqn,mbf1,Rpn10 | | 1.702e-02 | -4.07 | mwh (multiple wing hairs) | protein interactions | 38131 | 31 | 4 | 9204 | 298 | smg,CG1636,RhoGAPp190,Rpt6 | | 1.702e-02 | -4.07 | Invadolysin (invadolysin) | protein interactions | 49580 | 31 | 4 | 9204 | 298 | CG7766,RpS9,CG17883,Pnn | | 1.703e-02 | -4.07 | GO\_REGULATION\_OF\_PROTEIN\_STABILITY | MSigDB lists | GO\_REGULATION\_OF\_PROTEIN\_STABILITY | 90 | 9 | 4161 | 184 | CG9853,Cdc42,Pten,dpp,Vps35,CG5861,Rop,CG41378,CCT4 | | 1.706e-02 | -4.07 | GSE19941\_UNSTIM\_VS\_LPS\_STIM\_IL10\_KO\_MACROPHAGE\_DN | MSigDB lists | GSE19941\_UNSTIM\_VS\_LPS\_STIM\_IL10\_KO\_MACROPHAGE\_DN | 61 | 7 | 4161 | 184 | Parp,snf,Stt3B,CG17493,Mdh1,vap,Tcs3 | | 1.712e-02 | -4.07 | Interleukin-3, 5 and GM-CSF signaling | REACTOME pathways | R-DME-512988 | 121 | 10 | 3923 | 151 | Prosalpha2,CG4935,Rpn10,Btk29A,14-3-3zeta,CG7054,Prosbeta1,vap,Rpt6,Nf1 | | 1.717e-02 | -4.06 | regulation of axonogenesis | biological process | GO:0050770 | 61 | 5 | 12345 | 305 | PlexA,Rheb,RhoGAPp190,LIMK1,Cdc42 | | 1.717e-02 | -4.06 | positive regulation of chromosome organization | biological process | GO:2001252 | 61 | 5 | 12345 | 305 | Tctp,Cp190,cal1,Ncoa6,E(z) | | 1.717e-02 | -4.06 | GO\_REGULATION\_OF\_MITOTIC\_CELL\_CYCLE | MSigDB lists | GO\_REGULATION\_OF\_MITOTIC\_CELL\_CYCLE | 170 | 14 | 4161 | 184 | Pten,Rpt6,spel1,Prosalpha2,fzy,Cdc42,CG12321,twin,Art1,E(z),Ino80,Rpn10,Cul4,CycA | | 1.717e-02 | -4.06 | GO\_ANCHORING\_JUNCTION | MSigDB lists | GO\_ANCHORING\_JUNCTION | 170 | 14 | 4161 | 184 | cta,Arpc3A,LIMK1,RpS9,Arpc5,rhea,Graf,14-3-3zeta,RpL5,flw,Cdc42,Pp1-87B,RpS8,pav | | 1.725e-02 | -4.06 | DODD\_NASOPHARYNGEAL\_CARCINOMA\_DN | MSigDB lists | DODD\_NASOPHARYNGEAL\_CARCINOMA\_DN | 479 | 31 | 4161 | 184 | CG13016,PGAP1,Fgop2,Prosalpha2,Lrr47,Fs(2)Ket,snf,CycA,SF2,CCT4,CG8635,Skp2,MED21,noi,CG32576,pins,RpL35A,Aps,Nf1,pav,spel1,Fign,Pten,CG5589,TfIIFbeta,Orc3,eIF3b,Phf5a,E(z),Art1,CG34148 | | 1.735e-02 | -4.05 | imd (immune deficiency) | protein interactions | 44339 | 122 | 9 | 9204 | 298 | Rpt6,Stim,tsr,Nopp140,CtBP,CNBP,CG5174,Cp190,Graf | | 1.735e-02 | -4.05 | RpL15 (Ribosomal protein L15) | protein interactions | 3354918 | 122 | 9 | 9204 | 298 | RpL35A,RpS26,RpL5,RpL36A,RpS19a,RpS9,RpL14,RpS8,RpS3A | | 1.735e-02 | -4.05 | positive regulation of cellular component biogenesis | biological process | GO:0044089 | 133 | 8 | 12345 | 305 | tkv,mnb,Rheb,Arpc5,Cdc42,Arpc3A,Rpt6,cal1 | | 1.745e-02 | -4.05 | GO\_POSITIVE\_REGULATION\_OF\_PROTEIN\_METABOLIC\_PROCESS | MSigDB lists | GO\_POSITIVE\_REGULATION\_OF\_PROTEIN\_METABOLIC\_PROCESS | 366 | 25 | 4161 | 184 | Skp2,krz,lic,Cul4,unc-119,Stlk,AGO3,Rpt6,CG4866,eEF5,Prosalpha2,Ubqn,RpS9,Lst8,E(z),Art1,Rap1,Vap33,Rpn10,dpp,Pten,Fmr1,fzy,Cdc42,HtrA2 | | 1.753e-02 | -4.04 | LIEN\_BREAST\_CARCINOMA\_METAPLASTIC\_VS\_DUCTAL\_UP | MSigDB lists | LIEN\_BREAST\_CARCINOMA\_METAPLASTIC\_VS\_DUCTAL\_UP | 13 | 3 | 4161 | 184 | CG32576,Sod2,dpp | | 1.753e-02 | -4.04 | GO\_REGULATION\_OF\_GLIAL\_CELL\_DIFFERENTIATION | MSigDB lists | GO\_REGULATION\_OF\_GLIAL\_CELL\_DIFFERENTIATION | 13 | 3 | 4161 | 184 | Nf1,flr,dpp | | 1.753e-02 | -4.04 | GO\_REGULATION\_OF\_MEIOTIC\_CELL\_CYCLE | MSigDB lists | GO\_REGULATION\_OF\_MEIOTIC\_CELL\_CYCLE | 13 | 3 | 4161 | 184 | spel1,fzy,AGO3 | | 1.753e-02 | -4.04 | GO\_REGULATION\_OF\_OXIDATIVE\_STRESS\_INDUCED\_CELL\_DEATH | MSigDB lists | GO\_REGULATION\_OF\_OXIDATIVE\_STRESS\_INDUCED\_CELL\_DEATH | 13 | 3 | 4161 | 184 | Sod2,HtrA2,Ubqn | | 1.753e-02 | -4.04 | TGF\_beta | pfam domains | PF00019 | 7 | 2 | 10480 | 319 | mav,dpp | | 1.753e-02 | -4.04 | Adaptin\_N | pfam domains | PF01602 | 7 | 2 | 10480 | 319 | rb,g | | 1.753e-02 | -4.04 | endosomal transport | biological process | GO:0016197 | 108 | 7 | 12345 | 305 | ema,Rab14,CG30423,Rab7,Vps35,Exo84,g | | 1.753e-02 | -4.04 | regulation of cell size | biological process | GO:0008361 | 108 | 7 | 12345 | 305 | Rheb,lic,Pten,Girdin,Lst8,Tctp,RhoGAPp190 | | 1.754e-02 | -4.04 | PI Metabolism | REACTOME pathways | R-DME-1483255 | 40 | 5 | 3923 | 151 | Rab14,sws,Btk29A,Pten,sigmar | | 1.762e-02 | -4.04 | ATP binding | molecular function | GO:0005524 | 639 | 24 | 12176 | 289 | CCT4,Fign,lic,spel1,Rpt6,Bub1,Dbp80,Mps1,LIMK1,tkv,Stlk,CG5589,DNAlig1,Btk29A,mus301,Dbp21E2,Cdk8,CG9330,pll,Ino80,mnb,pav,CG8915,me31B | | 1.762e-02 | -4.04 | Moe (Moesin) | protein interactions | 31816 | 143 | 10 | 9204 | 298 | eEF5,nom,flw,Nulp1,conu,smt3,Pisd,Pp1-87B,flr,eEF1delta | | 1.767e-02 | -4.04 | GSE37301\_HEMATOPOIETIC\_STEM\_CELL\_VS\_CD4\_TCELL\_UP | MSigDB lists | GSE37301\_HEMATOPOIETIC\_STEM\_CELL\_VS\_CD4\_TCELL\_UP | 35 | 5 | 4161 | 184 | Graf,Fign,CG8860,Nf1,CycA | | 1.767e-02 | -4.04 | GO\_RIBOSOMAL\_LARGE\_SUBUNIT\_BIOGENESIS | MSigDB lists | GO\_RIBOSOMAL\_LARGE\_SUBUNIT\_BIOGENESIS | 35 | 5 | 4161 | 184 | RpL14,RpL5,RpL35A,Dbp21E2,CG9246 | | 1.767e-02 | -4.04 | GSE9006\_HEALTHY\_VS\_TYPE\_1\_DIABETES\_PBMC\_AT\_DX\_DN | MSigDB lists | GSE9006\_HEALTHY\_VS\_TYPE\_1\_DIABETES\_PBMC\_AT\_DX\_DN | 35 | 5 | 4161 | 184 | DNAlig1,CycA,fzy,E(z),Rcd5 | | 1.767e-02 | -4.04 | TGF\_BETA\_2 | prosite domains | PS51362 | 7 | 2 | 6243 | 191 | mav,dpp | | 1.767e-02 | -4.04 | TGF\_BETA\_1 | prosite domains | PS00250 | 7 | 2 | 6243 | 191 | dpp,mav | | 1.769e-02 | -4.03 | WD40 | pfam domains | PF00400 | 130 | 9 | 10480 | 319 | Wdr33,flr,CG43367,CG14805,Rbbp5,CG4935,Lst8,fzy,Wdr82 | | 1.775e-02 | -4.03 | peptide N-acetyltransferase activity | molecular function | GO:0034212 | 64 | 5 | 12176 | 289 | Eaf6,vnc,Rcd5,Acf,d4 | | 1.781e-02 | -4.03 | GO\_REGULATION\_OF\_P38MAPK\_CASCADE | MSigDB lists | GO\_REGULATION\_OF\_P38MAPK\_CASCADE | 5 | 2 | 4161 | 184 | Art1,dpp | | 1.781e-02 | -4.03 | GO\_REGULATION\_BY\_VIRUS\_OF\_VIRAL\_PROTEIN\_LEVELS\_IN\_HOST\_CELL | MSigDB lists | GO\_REGULATION\_BY\_VIRUS\_OF\_VIRAL\_PROTEIN\_LEVELS\_IN\_HOST\_CELL | 5 | 2 | 4161 | 184 | Vap33,CG17883 | | 1.781e-02 | -4.03 | GO\_FOREBRAIN\_MORPHOGENESIS | MSigDB lists | GO\_FOREBRAIN\_MORPHOGENESIS | 5 | 2 | 4161 | 184 | Pten,Nf1 | | 1.781e-02 | -4.03 | REACTOME\_HORMONE\_SENSITIVE\_LIPASE\_HSL\_MEDIATED\_TRIACYLGLYCEROL\_HYDROLYSIS | MSigDB lists | REACTOME\_HORMONE\_SENSITIVE\_LIPASE\_HSL\_MEDIATED\_TRIACYLGLYCEROL\_HYDROLYSIS | 5 | 2 | 4161 | 184 | Pp1-87B,flw | | 1.781e-02 | -4.03 | GO\_LONG\_TERM\_SYNAPTIC\_DEPRESSION | MSigDB lists | GO\_LONG\_TERM\_SYNAPTIC\_DEPRESSION | 5 | 2 | 4161 | 184 | Pten,Rop | | 1.781e-02 | -4.03 | GO\_POSITIVE\_REGULATION\_BY\_HOST\_OF\_VIRAL\_PROCESS | MSigDB lists | GO\_POSITIVE\_REGULATION\_BY\_HOST\_OF\_VIRAL\_PROCESS | 5 | 2 | 4161 | 184 | Vap33,CG17883 | | 1.781e-02 | -4.03 | GO\_GLYCOPROTEIN\_CATABOLIC\_PROCESS | MSigDB lists | GO\_GLYCOPROTEIN\_CATABOLIC\_PROCESS | 5 | 2 | 4161 | 184 | Stt3B,alpha-Man-Ib | | 1.781e-02 | -4.03 | chr14q12 | MSigDB lists | chr14q12 | 5 | 2 | 4161 | 184 | Tcs3,Ufd4 | | 1.781e-02 | -4.03 | GO\_G\_PROTEIN\_BETA\_GAMMA\_SUBUNIT\_COMPLEX\_BINDING | MSigDB lists | GO\_G\_PROTEIN\_BETA\_GAMMA\_SUBUNIT\_COMPLEX\_BINDING | 5 | 2 | 4161 | 184 | cta,CG17493 | | 1.781e-02 | -4.03 | GO\_POSITIVE\_REGULATION\_OF\_PROTEIN\_COMPLEX\_DISASSEMBLY | MSigDB lists | GO\_POSITIVE\_REGULATION\_OF\_PROTEIN\_COMPLEX\_DISASSEMBLY | 5 | 2 | 4161 | 184 | eEF5,flr | | 1.781e-02 | -4.03 | GO\_NEGATIVE\_REGULATION\_OF\_STRIATED\_MUSCLE\_CELL\_DIFFERENTIATION | MSigDB lists | GO\_NEGATIVE\_REGULATION\_OF\_STRIATED\_MUSCLE\_CELL\_DIFFERENTIATION | 5 | 2 | 4161 | 184 | dpp,E(z) | | 1.781e-02 | -4.03 | LIANG\_SILENCED\_BY\_METHYLATION\_DN | MSigDB lists | LIANG\_SILENCED\_BY\_METHYLATION\_DN | 5 | 2 | 4161 | 184 | CycA,E(z) | | 1.781e-02 | -4.03 | GO\_ENTRAINMENT\_OF\_CIRCADIAN\_CLOCK\_BY\_PHOTOPERIOD | MSigDB lists | GO\_ENTRAINMENT\_OF\_CIRCADIAN\_CLOCK\_BY\_PHOTOPERIOD | 5 | 2 | 4161 | 184 | flw,Pp1-87B | | 1.781e-02 | -4.03 | GO\_PHOTOPERIODISM | MSigDB lists | GO\_PHOTOPERIODISM | 5 | 2 | 4161 | 184 | flw,Pp1-87B | | 1.781e-02 | -4.03 | GO\_POSITIVE\_REGULATION\_OF\_EXTRINSIC\_APOPTOTIC\_SIGNALING\_PATHWAY\_IN\_ABSENCE\_OF\_LIGAND | MSigDB lists | GO\_POSITIVE\_REGULATION\_OF\_EXTRINSIC\_APOPTOTIC\_SIGNALING\_PATHWAY\_IN\_ABSENCE\_OF\_LIGAND | 5 | 2 | 4161 | 184 | Nf1,HtrA2 | | 1.781e-02 | -4.03 | GO\_NEGATIVE\_REGULATION\_OF\_OXIDATIVE\_STRESS\_INDUCED\_INTRINSIC\_APOPTOTIC\_SIGNALING\_PATHWAY | MSigDB lists | GO\_NEGATIVE\_REGULATION\_OF\_OXIDATIVE\_STRESS\_INDUCED\_INTRINSIC\_APOPTOTIC\_SIGNALING\_PATHWAY | 5 | 2 | 4161 | 184 | HtrA2,Sod2 | | 1.781e-02 | -4.03 | GO\_ENTRAINMENT\_OF\_CIRCADIAN\_CLOCK | MSigDB lists | GO\_ENTRAINMENT\_OF\_CIRCADIAN\_CLOCK | 5 | 2 | 4161 | 184 | flw,Pp1-87B | | 1.781e-02 | -4.03 | GO\_TRANS\_GOLGI\_NETWORK\_TRANSPORT\_VESICLE | MSigDB lists | GO\_TRANS\_GOLGI\_NETWORK\_TRANSPORT\_VESICLE | 5 | 2 | 4161 | 184 | bai,Rab8 | | 1.781e-02 | -4.03 | LE\_NEURONAL\_DIFFERENTIATION\_DN | MSigDB lists | LE\_NEURONAL\_DIFFERENTIATION\_DN | 5 | 2 | 4161 | 184 | pav,fzy | | 1.781e-02 | -4.03 | GO\_REGULATION\_OF\_ASTROCYTE\_DIFFERENTIATION | MSigDB lists | GO\_REGULATION\_OF\_ASTROCYTE\_DIFFERENTIATION | 5 | 2 | 4161 | 184 | Nf1,dpp | | 1.781e-02 | -4.03 | GO\_REGULATION\_OF\_RETINOIC\_ACID\_RECEPTOR\_SIGNALING\_PATHWAY | MSigDB lists | GO\_REGULATION\_OF\_RETINOIC\_ACID\_RECEPTOR\_SIGNALING\_PATHWAY | 5 | 2 | 4161 | 184 | E(z),CtBP | | 1.781e-02 | -4.03 | REACTOME\_RAP1\_SIGNALLING | MSigDB lists | REACTOME\_RAP1\_SIGNALLING | 5 | 2 | 4161 | 184 | Rap1,14-3-3zeta | | 1.781e-02 | -4.03 | GO\_DNA\_LIGATION\_INVOLVED\_IN\_DNA\_REPAIR | MSigDB lists | GO\_DNA\_LIGATION\_INVOLVED\_IN\_DNA\_REPAIR | 5 | 2 | 4161 | 184 | DNAlig1,Parp | | 1.781e-02 | -4.03 | GO\_SET1C\_COMPASS\_COMPLEX | MSigDB lists | GO\_SET1C\_COMPASS\_COMPLEX | 5 | 2 | 4161 | 184 | Rbbp5,Wdr82 | | 1.781e-02 | -4.03 | WILLIAMS\_ESR1\_TARGETS\_UP | MSigDB lists | WILLIAMS\_ESR1\_TARGETS\_UP | 5 | 2 | 4161 | 184 | fzy,CycA | | 1.781e-02 | -4.03 | GO\_NEGATIVE\_REGULATION\_OF\_FIBROBLAST\_PROLIFERATION | MSigDB lists | GO\_NEGATIVE\_REGULATION\_OF\_FIBROBLAST\_PROLIFERATION | 5 | 2 | 4161 | 184 | CG41378,Nf1 | | 1.781e-02 | -4.03 | GO\_MRNA\_SPLICE\_SITE\_SELECTION | MSigDB lists | GO\_MRNA\_SPLICE\_SITE\_SELECTION | 5 | 2 | 4161 | 184 | SF2,noi | | 1.783e-02 | -4.03 | GO\_REGULATION\_OF\_PROTEIN\_SERINE\_THREONINE\_KINASE\_ACTIVITY | MSigDB lists | GO\_REGULATION\_OF\_PROTEIN\_SERINE\_THREONINE\_KINASE\_ACTIVITY | 106 | 10 | 4161 | 184 | Nf1,CSN1b,Pten,Stlk,dpp,CycA,Rap1,E(z),lic,krz | | 1.800e-02 | -4.02 | Regulation of RAS by GAPs | REACTOME pathways | R-DME-5658442 | 55 | 6 | 3923 | 151 | Prosbeta1,Rpn10,Prosalpha2,vap,Rpt6,Nf1 | | 1.802e-02 | -4.02 | GSE3982\_BCELL\_VS\_NKCELL\_UP | MSigDB lists | GSE3982\_BCELL\_VS\_NKCELL\_UP | 48 | 6 | 4161 | 184 | CIA30,Arpc3A,Apf,Prosalpha2,RpL15,TfIIFbeta | | 1.802e-02 | -4.02 | GCM\_TPT1 | MSigDB lists | GCM\_TPT1 | 48 | 6 | 4161 | 184 | RpL36A,RpL35A,RpS8,Tctp,RpL14,RpS9 | | 1.802e-02 | -4.02 | REACTOME\_ACTIVATION\_OF\_THE\_MRNA\_UPON\_BINDING\_OF\_THE\_CAP\_BINDING\_COMPLEX\_AND\_EIFS\_AND\_SUBSEQUENT\_BINDING\_TO\_43S | MSigDB lists | REACTOME\_ACTIVATION\_OF\_THE\_MRNA\_UPON\_BINDING\_OF\_THE\_CAP\_BINDING\_COMPLEX\_AND\_EIFS\_AND\_SUBSEQUENT\_BINDING\_TO\_43S | 48 | 6 | 4161 | 184 | RpS8,eIF3g1,eIF3b,RpS26,RpS9,eIF3c | | 1.805e-02 | -4.01 | regulation of hemocyte proliferation | biological process | GO:0035206 | 41 | 4 | 12345 | 305 | Sod2,DCTN3-p24,heix,Pten | | 1.805e-02 | -4.01 | E2F\_Q6 | MSigDB lists | E2F\_Q6 | 76 | 8 | 4161 | 184 | SF2,Stt3B,Pect,E(z),Phf5a,spel1,Pp1-87B,Fs(2)Ket | | 1.805e-02 | -4.01 | GSE33162\_UNTREATED\_VS\_4H\_LPS\_STIM\_HDAC3\_KO\_MACROPHAGE\_UP | MSigDB lists | GSE33162\_UNTREATED\_VS\_4H\_LPS\_STIM\_HDAC3\_KO\_MACROPHAGE\_UP | 76 | 8 | 4161 | 184 | snf,fzy,HtrA2,pins,smt3,DNAlig1,Rpt6,Arpc5 | | 1.805e-02 | -4.01 | GO\_REGULATION\_OF\_CELLULAR\_COMPONENT\_SIZE | MSigDB lists | GO\_REGULATION\_OF\_CELLULAR\_COMPONENT\_SIZE | 76 | 8 | 4161 | 184 | flr,vap,LIMK1,Arpc3A,Lst8,Arpc5,PlexA,Pten | | 1.806e-02 | -4.01 | RNA transport | KEGG pathways | dme03013 | 148 | 11 | 3198 | 117 | eIF3b,Fmr1,tsu,Upf3,eIF3g1,Nup188,Fs(2)Ket,eIF3c,eIF4B,Pnn,smt3 | | 1.806e-02 | -4.01 | RNA transport | KEGG pathways | ko03013 | 148 | 11 | 3198 | 117 | eIF4B,Pnn,smt3,eIF3c,Fs(2)Ket,eIF3b,Fmr1,tsu,eIF3g1,Upf3,Nup188 | | 1.810e-02 | -4.01 | protein kinase A binding | molecular function | GO:0051018 | 9 | 2 | 12176 | 289 | spoon,sws | | 1.810e-02 | -4.01 | poly-purine tract binding | molecular function | GO:0070717 | 9 | 2 | 12176 | 289 | Hrb27C,CG4612 | | 1.817e-02 | -4.01 | TGFB | smart domains | SM00204 | 7 | 2 | 5086 | 158 | mav,dpp | | 1.834e-02 | -4.00 | pole cell formation | biological process | GO:0007279 | 23 | 3 | 12345 | 305 | me31B,grau,Fmr1 | | 1.836e-02 | -4.00 | regulation of synaptic growth at neuromuscular junction | biological process | GO:0008582 | 109 | 7 | 12345 | 305 | Nf1,Cdc42,Fmr1,tkv,mnb,Rheb,Rab8 | | 1.841e-02 | -3.99 | large ribosomal subunit | cellular component | GO:0015934 | 108 | 7 | 11940 | 298 | RpL36A,RpL5,mRpL30,RpL14,RpL15,RpL35A,mRpL34 | | 1.843e-02 | -3.99 | supramolecular fiber organization | biological process | GO:0097435 | 188 | 10 | 12345 | 305 | Fign,Cdc42,Arpc5,tsr,flr,Arpc3A,rhea,Girdin,Spn,Vap33 | | 1.850e-02 | -3.99 | RNA biosynthetic process | biological process | GO:0032774 | 216 | 11 | 12345 | 305 | His4r,grau,Arp8,Rpb11,Pbp49,su(Hw),TfIIFbeta,Parp,E(bx),CG6244,EloB | | 1.853e-02 | -3.99 | GSE7460\_CTRL\_VS\_TGFB\_TREATED\_ACT\_CD8\_TCELL\_UP | MSigDB lists | GSE7460\_CTRL\_VS\_TGFB\_TREATED\_ACT\_CD8\_TCELL\_UP | 62 | 7 | 4161 | 184 | CG11334,MED21,Graf,eIF3g1,pav,Dbp21E2,CG9853 | | 1.855e-02 | -3.99 | peptidyl-amino acid modification | biological process | GO:0018193 | 394 | 17 | 12345 | 305 | CG31460,Eaf6,BEAF-32,CG1218,Art1,Wdr82,d4,smt3,E(z),Ncoa6,Rbbp5,Acf,Stt3B,mnb,Btk29A,Rcd5,vnc | | 1.856e-02 | -3.99 | Regulation of actin dynamics for phagocytic cup formation | REACTOME pathways | R-DME-2029482 | 27 | 4 | 3923 | 151 | Arpc5,Cdc42,Arpc3A,LIMK1 | | 1.856e-02 | -3.99 | EPHB-mediated forward signaling | REACTOME pathways | R-DME-3928662 | 27 | 4 | 3923 | 151 | vap,Arpc5,Cdc42,Arpc3A | | 1.876e-02 | -3.98 | G10 | prints domains | PR00322 | 1 | 1 | 2185 | 41 | l(1)10Bb | | 1.876e-02 | -3.98 | MNSODISMTASE | prints domains | PR01703 | 1 | 1 | 2185 | 41 | Sod2 | | 1.876e-02 | -3.98 | HISTONEH4 | prints domains | PR00623 | 1 | 1 | 2185 | 41 | His4r | | 1.876e-02 | -3.98 | PROTEASES2C | prints domains | PR00834 | 1 | 1 | 2185 | 41 | HtrA2 | | 1.876e-02 | -3.98 | GPROTEINA12 | prints domains | PR00440 | 1 | 1 | 2185 | 41 | cta | | 1.876e-02 | -3.98 | HEMOGLOBNASE | prints domains | PR00776 | 1 | 1 | 2185 | 41 | PIG-K | | 1.876e-02 | -3.98 | TCTPROTEIN | prints domains | PR01653 | 1 | 1 | 2185 | 41 | Tctp | | 1.876e-02 | -3.98 | TETRPHPHTASE | prints domains | PR01405 | 1 | 1 | 2185 | 41 | Apf | | 1.876e-02 | -3.98 | RNABINDINGM8 | prints domains | PR01738 | 1 | 1 | 2185 | 41 | tsu | | 1.876e-02 | -3.98 | RIBOSOMALL5 | prints domains | PR00058 | 1 | 1 | 2185 | 41 | RpL5 | | 1.887e-02 | -3.97 | exonuclease activity | molecular function | GO:0004527 | 65 | 5 | 12176 | 289 | Not3,twin,CG12877,Rat1,CG6171 | | 1.896e-02 | -3.97 | CG7049 (uncharacterized protein) | protein interactions | 38022 | 32 | 4 | 9204 | 298 | Wdr82,CG6171,BthD,CG1218 | | 1.896e-02 | -3.97 | par-6 (par-6) | protein interactions | 32752 | 32 | 4 | 9204 | 298 | Cdc42,Fmr1,pins,Rap1 | | 1.897e-02 | -3.96 | fly chr89A8-89A8\|3-58 cM | chromosome location | fly chr89A8-89A8\|3-58 cM | 15 | 2 | 25010 | 358 | CG4287,CG5516 | | 1.897e-02 | -3.96 | fly chr89A8-89A8\|3-58 | chromosome location | fly chr89A8-89A8\|3-58 | 15 | 2 | 25010 | 358 | CG5516,CG4287 | | 1.903e-02 | -3.96 | BROMODOMAIN\_2 | prosite domains | PS50014 | 19 | 3 | 6243 | 191 | CG1815,E(bx),Acf | | 1.910e-02 | -3.96 | GGGCGGR\_SP1\_Q6 | MSigDB lists | GGGCGGR\_SP1\_Q6 | 882 | 51 | 4161 | 184 | Graf,CG5535,Pect,lic,Rop,bai,Fs(2)Ket,inc,smt3,VhaPPA1-1,CG8860,Rpt6,Rat1,Stt3B,vap,Vps35,Vap33,CG31460,Rab8,Fmr1,heix,HtrA2,CG9246,Stim,Rcd5,His2Av,CG11334,c12.1,Tctp,PIG-K,mRpS28,Stlk,Fgop2,DNAlig1,sigmar,mRpS33,Tsp97E,Art1,Usp32,Pisd,E(z),14-3-3zeta,Arpc5,Phf5a,sm,Tim17b,RpL14,dpp,viaf,alpha-Man-Ib,RpL35A | | 1.912e-02 | -3.96 | Hsf (Heat shock factor) | protein interactions | 37068 | 18 | 3 | 9204 | 298 | Dref,Cdk8,MED21 | | 1.912e-02 | -3.96 | pix (pixie) | protein interactions | 39027 | 18 | 3 | 9204 | 298 | RpL14,eIF3b,eIF3c | | 1.912e-02 | -3.96 | eIF3h (eukaryotic translation initiation factor 3 subunit h) | protein interactions | 45682 | 18 | 3 | 9204 | 298 | eIF3c,eIF3b,eIF3g1 | | 1.912e-02 | -3.96 | mars (mars) | protein interactions | 36498 | 18 | 3 | 9204 | 298 | flw,Fs(2)Ket,Pp1-87B | | 1.912e-02 | -3.96 | sds22 (sds22) | protein interactions | 42091 | 18 | 3 | 9204 | 298 | I-2,Pp1-87B,flw | | 1.912e-02 | -3.96 | Unr (Upstream of N-ras) | protein interactions | 38963 | 18 | 3 | 9204 | 298 | sqd,Hrb27C,smg | | 1.920e-02 | -3.95 | cellular component disassembly | biological process | GO:0022411 | 110 | 7 | 12345 | 305 | flw,flr,tsr,Pp1-87B,Vps13D,eRF1,Fmr1 | | 1.928e-02 | -3.95 | signaling receptor binding | molecular function | GO:0005102 | 256 | 12 | 12176 | 289 | E(bx),CG10365,krz,cta,sun,vap,rhea,CG30423,Spn,dpp,RhoGAPp190,mav | | 1.940e-02 | -3.94 | E2F\_Q4 | MSigDB lists | E2F\_Q4 | 77 | 8 | 4161 | 184 | SF2,Pect,E(z),Stt3B,Phf5a,spel1,Pp1-87B,Fs(2)Ket | | 1.940e-02 | -3.94 | GSE7568\_IL4\_VS\_IL4\_AND\_DEXAMETHASONE\_TREATED\_MACROPHAGE\_UP | MSigDB lists | GSE7568\_IL4\_VS\_IL4\_AND\_DEXAMETHASONE\_TREATED\_MACROPHAGE\_UP | 77 | 8 | 4161 | 184 | xit,flr,noi,c12.1,DNAlig1,eIF3c,Fign,Lrr47 | | 1.953e-02 | -3.94 | histone H3-K9 methylation | biological process | GO:0051567 | 9 | 2 | 12345 | 305 | E(z),BEAF-32 | | 1.953e-02 | -3.94 | regulation of double-strand break repair | biological process | GO:2000779 | 9 | 2 | 12345 | 305 | Dek,Fign | | 1.953e-02 | -3.94 | regulation of animal organ formation | biological process | GO:0003156 | 9 | 2 | 12345 | 305 | tkv,dpp | | 1.953e-02 | -3.94 | GO\_PROTEIN\_TARGETING | MSigDB lists | GO\_PROTEIN\_TARGETING | 207 | 16 | 4161 | 184 | Tim17b,RpL14,RpS26,Vps13D,RpL35A,RpS8,RpS9,14-3-3zeta,RpL5,Tom20,Fs(2)Ket,CG8860,RpL15,Pex3,rb,RpL36A | | 1.955e-02 | -3.93 | female meiotic nuclear division | biological process | GO:0007143 | 86 | 6 | 12345 | 305 | Mps1,cal1,mus301,fzy,grau,pav | | 1.955e-02 | -3.93 | olfactory behavior | biological process | GO:0042048 | 86 | 6 | 12345 | 305 | Fmr1,Nf1,Pp1-87B,Orc3,mnb,14-3-3zeta | | 1.957e-02 | -3.93 | regulation of imaginal disc growth | biological process | GO:0045570 | 42 | 4 | 12345 | 305 | Fmr1,Pten,dpp,tkv | | 1.957e-02 | -3.93 | cytokinetic process | biological process | GO:0032506 | 42 | 4 | 12345 | 305 | tsr,Exo84,pav,flw | | 1.958e-02 | -3.93 | embryonic axis specification | biological process | GO:0000578 | 136 | 8 | 12345 | 305 | dpp,pll,sqd,Hrb27C,mon2,lic,glo,tsu | | 1.960e-02 | -3.93 | - | gene3d domains | 2.130.10.10 | 201 | 12 | 4929 | 151 | CG4935,eIF3b,CG43367,fzy,Lst8,CG11141,Wdr33,CG14805,Rbbp5,PlexA,flr,Wdr82 | | 1.964e-02 | -3.93 | YAO\_TEMPORAL\_RESPONSE\_TO\_PROGESTERONE\_CLUSTER\_2 | MSigDB lists | YAO\_TEMPORAL\_RESPONSE\_TO\_PROGESTERONE\_CLUSTER\_2 | 24 | 4 | 4161 | 184 | Fs(2)Ket,MED22,dpp,CG32473 | | 1.964e-02 | -3.93 | WANG\_TARGETS\_OF\_MLL\_CBP\_FUSION\_DN | MSigDB lists | WANG\_TARGETS\_OF\_MLL\_CBP\_FUSION\_DN | 24 | 4 | 4161 | 184 | Fs(2)Ket,bai,vnc,Rpt6 | | 1.970e-02 | -3.93 | CG15545 (uncharacterized protein) | protein interactions | 43658 | 7 | 2 | 9204 | 298 | Dek,Tsp97E | | 1.970e-02 | -3.93 | CG42684 (uncharacterized protein) | protein interactions | 32754 | 7 | 2 | 9204 | 298 | Vlet,Rcd5 | | 1.970e-02 | -3.93 | Pitslre (pitslre) | protein interactions | 40292 | 7 | 2 | 9204 | 298 | Rcd5,flw | | 1.970e-02 | -3.93 | scw (screw) | protein interactions | 46000 | 7 | 2 | 9204 | 298 | dpp,tkv | | 1.970e-02 | -3.93 | CG7168 (uncharacterized protein) | protein interactions | 42184 | 7 | 2 | 9204 | 298 | CG9951,Vlet | | 1.970e-02 | -3.93 | ric8a (ric8a) | protein interactions | 31874 | 7 | 2 | 9204 | 298 | pins,cta | | 1.970e-02 | -3.93 | Vps16A (Vacuolar protein sorting 16A) | protein interactions | 41107 | 7 | 2 | 9204 | 298 | ema,Prosbeta1 | | 1.970e-02 | -3.93 | CG15443 (uncharacterized protein) | protein interactions | 33656 | 7 | 2 | 9204 | 298 | RpS3A,eEF5 | | 1.970e-02 | -3.93 | CG7542 (uncharacterized protein) | protein interactions | 39960 | 7 | 2 | 9204 | 298 | CG34159,Cpsf6 | | 1.970e-02 | -3.93 | Nab2 (Nuclear polyadenosine RNA-binding 2) | protein interactions | 42897 | 7 | 2 | 9204 | 298 | Fmr1,CG7646 | | 1.970e-02 | -3.93 | tld (tolloid) | protein interactions | 42945 | 7 | 2 | 9204 | 298 | dpp,LIMK1 | | 1.970e-02 | -3.93 | CG4393 (uncharacterized protein) | protein interactions | 42764 | 7 | 2 | 9204 | 298 | RpS9,RpL35A | | 1.970e-02 | -3.93 | Mvd (Mevalonate diphosphate decarboxylase) | protein interactions | 32520 | 7 | 2 | 9204 | 298 | RpS9,Pisd | | 1.970e-02 | -3.93 | Fis1 (Fission, mitochondrial 1) | protein interactions | 49892 | 7 | 2 | 9204 | 298 | Rab14,Vlet | | 1.970e-02 | -3.93 | ems (empty spiracles) | protein interactions | 41697 | 7 | 2 | 9204 | 298 | su(Hw),Cp190 | | 1.970e-02 | -3.93 | prd1 (pruning defect 1) | protein interactions | 41425 | 7 | 2 | 9204 | 298 | Pp1-87B,RpL15 | | 1.970e-02 | -3.93 | Synj (Synaptojanin) | protein interactions | 37517 | 7 | 2 | 9204 | 298 | flw,mnb | | 1.975e-02 | -3.92 | Signaling by SCF-KIT | REACTOME pathways | R-DME-1433557 | 142 | 11 | 3923 | 151 | Prosbeta1,CG7054,14-3-3zeta,vap,Nf1,Rpt6,Lst8,Pten,Rpn10,Prosalpha2,CG4935 | | 1.976e-02 | -3.92 | positive regulation of nucleobase-containing compound metabolic process | biological process | GO:0045935 | 523 | 21 | 12345 | 305 | rhi,CtBP,mbf1,nsl1,Ncoa6,d4,dgrn,su(Hw),AGO3,Hrb27C,Rpt6,Rcd5,grau,mnb,PlexA,CG6244,Lst8,Dref,MED21,TfIIFbeta,smg | | 1.980e-02 | -3.92 | GSE42021\_CD24HI\_TREG\_VS\_CD24HI\_TCONV\_THYMUS\_DN | MSigDB lists | GSE42021\_CD24HI\_TREG\_VS\_CD24HI\_TCONV\_THYMUS\_DN | 49 | 6 | 4161 | 184 | Stt3B,Pisd,Nf1,eRF1,viaf,Cdk8 | | 1.980e-02 | -3.92 | GSE27241\_WT\_VS\_RORGT\_KO\_TH17\_POLARIZED\_CD4\_TCELL\_UP | MSigDB lists | GSE27241\_WT\_VS\_RORGT\_KO\_TH17\_POLARIZED\_CD4\_TCELL\_UP | 49 | 6 | 4161 | 184 | His2Av,E(z),DNAlig1,CG8860,pav,CycA | | 1.980e-02 | -3.92 | MORF\_UNG | MSigDB lists | MORF\_UNG | 49 | 6 | 4161 | 184 | spel1,xit,eIF3b,vnc,SF2,Art1 | | 1.980e-02 | -3.92 | GSE43955\_10H\_VS\_60H\_ACT\_CD4\_TCELL\_DN | MSigDB lists | GSE43955\_10H\_VS\_60H\_ACT\_CD4\_TCELL\_DN | 49 | 6 | 4161 | 184 | sigmar,Nf1,cta,Wdr82,twin,Pisd | | 1.980e-02 | -3.92 | GSE360\_L\_MAJOR\_VS\_M\_TUBERCULOSIS\_MAC\_UP | MSigDB lists | GSE360\_L\_MAJOR\_VS\_M\_TUBERCULOSIS\_MAC\_UP | 49 | 6 | 4161 | 184 | CG5535,rhea,me31B,Nulp1,RagA-B,Aps | | 1.980e-02 | -3.92 | GSE1925\_CTRL\_VS\_24H\_IFNG\_STIM\_MACROPHAGE\_UP | MSigDB lists | GSE1925\_CTRL\_VS\_24H\_IFNG\_STIM\_MACROPHAGE\_UP | 36 | 5 | 4161 | 184 | inc,MICU1,Rab8,Arp8,Tcs3 | | 1.980e-02 | -3.92 | GSE29949\_CD8\_NEG\_DC\_SPLEEN\_VS\_CD8\_POS\_DC\_SPLEEN\_UP | MSigDB lists | GSE29949\_CD8\_NEG\_DC\_SPLEEN\_VS\_CD8\_POS\_DC\_SPLEEN\_UP | 36 | 5 | 4161 | 184 | HtrA2,RhoGAPp190,Graf,Tim17b,CG41378 | | 1.980e-02 | -3.92 | GSE29615\_DAY3\_VS\_DAY7\_LAIV\_FLU\_VACCINE\_PBMC\_DN | MSigDB lists | GSE29615\_DAY3\_VS\_DAY7\_LAIV\_FLU\_VACCINE\_PBMC\_DN | 36 | 5 | 4161 | 184 | Ubqn,twin,CG2938,eRF1,CG12341 | | 1.988e-02 | -3.92 | Znf\_PHD-finger | interpro domains | IPR019787 | 36 | 4 | 11383 | 331 | E(bx),Acf,d4,CG1815 | | 1.990e-02 | -3.92 | cytoneme | cellular component | GO:0035230 | 9 | 2 | 11940 | 298 | dpp,tkv | | 1.990e-02 | -3.92 | oligosaccharyltransferase complex | cellular component | GO:0008250 | 9 | 2 | 11940 | 298 | Stt3B,CG31460 | | 2.008e-02 | -3.91 | actin filament organization | biological process | GO:0007015 | 111 | 7 | 12345 | 305 | Spn,Cdc42,Arpc5,Girdin,tsr,flr,Arpc3A | | 2.009e-02 | -3.91 | GO\_REGULATION\_OF\_PROTEIN\_COMPLEX\_DISASSEMBLY | MSigDB lists | GO\_REGULATION\_OF\_PROTEIN\_COMPLEX\_DISASSEMBLY | 63 | 7 | 4161 | 184 | pav,eEF5,eRF1,flr,pins,Fs(2)Ket,Ino80 | | 2.009e-02 | -3.91 | GSE6259\_FLT3L\_INDUCED\_33D1\_POS\_DC\_VS\_CD4\_TCELL\_DN | MSigDB lists | GSE6259\_FLT3L\_INDUCED\_33D1\_POS\_DC\_VS\_CD4\_TCELL\_DN | 63 | 7 | 4161 | 184 | Dbp80,mbf1,E(z),viaf,eIF3b,DNAlig1,Fgop2 | | 2.009e-02 | -3.91 | LI\_WILMS\_TUMOR\_VS\_FETAL\_KIDNEY\_1\_DN | MSigDB lists | LI\_WILMS\_TUMOR\_VS\_FETAL\_KIDNEY\_1\_DN | 63 | 7 | 4161 | 184 | pav,spel1,fzy,CG5589,E(z),14-3-3zeta,Parp | | 2.009e-02 | -3.91 | GO\_NEGATIVE\_REGULATION\_OF\_CELL\_CYCLE\_PROCESS | MSigDB lists | GO\_NEGATIVE\_REGULATION\_OF\_CELL\_CYCLE\_PROCESS | 63 | 7 | 4161 | 184 | Pten,twin,Art1,E(z),spel1,CycA,CG12321 | | 2.009e-02 | -3.91 | GSE9006\_TYPE\_1\_DIABETES\_AT\_DX\_VS\_4MONTH\_POST\_DX\_PBMC\_DN | MSigDB lists | GSE9006\_TYPE\_1\_DIABETES\_AT\_DX\_VS\_4MONTH\_POST\_DX\_PBMC\_DN | 63 | 7 | 4161 | 184 | Rpb11,RpL5,Mdh1,mRpS33,RpL15,RhoGAPp190,tkv | | 2.036e-02 | -3.89 | cnn (centrosomin) | protein interactions | 36491 | 105 | 8 | 9204 | 298 | rhi,Cp190,eIF3c,Hsp27,DCTN3-p24,Rpt6,MED26,CG9951 | | 2.036e-02 | -3.89 | Dsor1 (Downstream of raf1) | protein interactions | 31872 | 105 | 8 | 9204 | 298 | sqd,Rpt6,RpL15,Map60,14-3-3zeta,Nf1,glo,krz | | 2.045e-02 | -3.89 | Golgi apparatus | cellular component | GO:0005794 | 364 | 16 | 11940 | 298 | CG15168,spoon,CG2938,mon2,heix,CG30423,Efa6,ema,CG10365,Rab14,Mppe,bai,Rab8,BthD,Usp32,g | | 2.050e-02 | -3.89 | Pen (Pendulin) | protein interactions | 34338 | 49 | 5 | 9204 | 298 | Cp190,Map60,Dlip2,Fs(2)Ket,BEAF-32 | | 2.050e-02 | -3.89 | eIF4E1 (eukaryotic translation initiation factor 4E1) | protein interactions | 45525 | 49 | 5 | 9204 | 298 | me31B,snf,Rbp1,smt3,smg | | 2.056e-02 | -3.88 | GPROTEINBRPT | prints domains | PR00320 | 57 | 4 | 2185 | 41 | Wdr82,Wdr33,CG4935,Lst8 | | 2.058e-02 | -3.88 | DNA replication | biological process | GO:0006260 | 87 | 6 | 12345 | 305 | Orc3,DNAlig1,RPA3,Btk29A,RecQ4,Dref | | 2.059e-02 | -3.88 | GO\_POSITIVE\_REGULATION\_OF\_TRANSFERASE\_ACTIVITY | MSigDB lists | GO\_POSITIVE\_REGULATION\_OF\_TRANSFERASE\_ACTIVITY | 191 | 15 | 4161 | 184 | Stlk,unc-119,dpp,Pten,fzy,Cdc42,Prosalpha2,Rpt6,krz,CCT4,E(z),lic,Vap33,Rap1,Rpn10 | | 2.059e-02 | -3.88 | transcription elongation from RNA polymerase II promoter | biological process | GO:0006368 | 24 | 3 | 12345 | 305 | TfIIFbeta,EloB,CG6244 | | 2.059e-02 | -3.88 | cellular protein complex disassembly | biological process | GO:0043624 | 24 | 3 | 12345 | 305 | flr,tsr,eRF1 | | 2.059e-02 | -3.88 | negative regulation of nuclear division | biological process | GO:0051784 | 24 | 3 | 12345 | 305 | CycA,Bub1,Mps1 | | 2.069e-02 | -3.88 | enzyme activator activity | molecular function | GO:0008047 | 143 | 8 | 12176 | 289 | RhoGAPp190,Nf1,Graf,conu,PlexA,vap,CG1791,CG17883 | | 2.082e-02 | -3.87 | vesicle-mediated transport | biological process | GO:0016192 | 462 | 19 | 12345 | 305 | PGAP1,Rab8,Cdc42,rb,CG32576,Rop,bai,Exo84,Vap33,ema,Rab7,Vps35,krz,Fmr1,g,RabX6,CG30423,Rab14,CG17883 | | 2.083e-02 | -3.87 | GSE12845\_IGD\_NEG\_BLOOD\_VS\_DARKZONE\_GC\_TONSIL\_BCELL\_DN | MSigDB lists | GSE12845\_IGD\_NEG\_BLOOD\_VS\_DARKZONE\_GC\_TONSIL\_BCELL\_DN | 78 | 8 | 4161 | 184 | eIF3b,Graf,Pisd,spel1,Mdh1,DNAlig1,Parp,CG5589 | | 2.083e-02 | -3.87 | GO\_SPLICEOSOMAL\_COMPLEX | MSigDB lists | GO\_SPLICEOSOMAL\_COMPLEX | 78 | 8 | 4161 | 184 | snf,noi,l(1)10Bb,Phf5a,c12.1,tsu,CG4935,SF2 | | 2.083e-02 | -3.87 | E2F1\_Q4 | MSigDB lists | E2F1\_Q4 | 78 | 8 | 4161 | 184 | Skp2,Art1,Pect,E(z),SF2,Rop,Fs(2)Ket,Pp1-87B | | 2.083e-02 | -3.87 | REACTOME\_PROCESSING\_OF\_CAPPED\_INTRON\_CONTAINING\_PRE\_MRNA | MSigDB lists | REACTOME\_PROCESSING\_OF\_CAPPED\_INTRON\_CONTAINING\_PRE\_MRNA | 78 | 8 | 4161 | 184 | snf,sm,Phf5a,noi,Rpb11,TfIIFbeta,SF2,tsu | | 2.083e-02 | -3.87 | GSE17721\_CTRL\_VS\_POLYIC\_8H\_BMDC\_UP | MSigDB lists | GSE17721\_CTRL\_VS\_POLYIC\_8H\_BMDC\_UP | 78 | 8 | 4161 | 184 | CycA,CG17493,CG41378,Roc2,MED22,LIMK1,Mdh1,sm | | 2.090e-02 | -3.87 | protein catabolic process | biological process | GO:0030163 | 338 | 15 | 12345 | 305 | Prosalpha2,Skp2,Prosbeta1,Rpt6,Rpn10,Usp32,Usp16-45,alpha-Man-Ib,Roc2,Ufd4,inc,Ubqn,Cul4,fzy,Sgt | | 2.098e-02 | -3.86 | nucleobase-containing compound transport | biological process | GO:0015931 | 112 | 7 | 12345 | 305 | CG6961,Fmr1,SF2,Tom20,sqd,Rab7,Dbp80 | | 2.102e-02 | -3.86 | BOULE/DAZ | interpro domains | IPR037366 | 8 | 2 | 11383 | 331 | sqd,Hrb27C | | 2.103e-02 | -3.86 | cation binding | molecular function | GO:0043169 | 1072 | 36 | 12176 | 289 | tkv,Zpr1,CG11586,Usp16-45,noi,dgrn,Parp,CG1218,CG7646,x16,nom,CG1815,CG1637,CNBP,CG32473,MICU1,NO66,CG17493,Ufd4,grau,Dref,CG6254,RecQ4,alpha-Man-Ib,Fign,Usp32,Sod2,Rpn10,Roc2,d4,CG5516,Stim,RAF2,CG8635,dah,Tctp | | 2.105e-02 | -3.86 | BubR1 (Bub1-related kinase) | protein interactions | 35522 | 33 | 4 | 9204 | 298 | fzy,vap,Mps1,Bub1 | | 2.105e-02 | -3.86 | mei-41 (meiotic 41) | protein interactions | 32608 | 33 | 4 | 9204 | 298 | mus301,His4r,CycA,rhi | | 2.112e-02 | -3.86 | BYSTRYKH\_HEMATOPOIESIS\_STEM\_CELL\_QTL\_TRANS | MSigDB lists | BYSTRYKH\_HEMATOPOIESIS\_STEM\_CELL\_QTL\_TRANS | 280 | 20 | 4161 | 184 | Cdk8,Vps35,eIF3b,cta,CG32473,LIMK1,Arpc5,CIA30,l(2)37Bb,eRF1,HtrA2,viaf,fzy,CG32576,Fign,CG5589,MED22,CG8635,Mdh1,PlexA | | 2.114e-02 | -3.86 | MARTENS\_TRETINOIN\_RESPONSE\_DN | MSigDB lists | MARTENS\_TRETINOIN\_RESPONSE\_DN | 262 | 19 | 4161 | 184 | RpL14,sm,Tim17b,RpS8,CG32576,eIF3g1,Nf1,RpS9,CG34148,Art1,RpL5,Stlk,CG11920,CG12321,eEF5,RpL15,CCT4,SF2,CycA | | 2.114e-02 | -3.86 | condensed chromosome, centromeric region | cellular component | GO:0000779 | 24 | 3 | 11940 | 298 | Mps1,Bub1,smt3 | | 2.119e-02 | -3.85 | GRAESSMANN\_APOPTOSIS\_BY\_DOXORUBICIN\_DN | MSigDB lists | GRAESSMANN\_APOPTOSIS\_BY\_DOXORUBICIN\_DN | 604 | 37 | 4161 | 184 | Cul4,Skp2,CG2938,CG13016,CG3842,Arp8,Coq5,CG7766,CG6907,Fs(2)Ket,vnc,CG31460,Orc3,twin,Pp1-87B,Sod2,Wdr82,Sgt,me31B,Fmr1,CG10672,SF2,Nulp1,Rcd5,Ufd4,Tcs3,Stlk,snf,mRpS28,Apf,Dbp80,LIMK1,14-3-3zeta,eRF1,Nf1,Fign,sm | | 2.119e-02 | -3.85 | ZF\_PHD\_2 | prosite domains | PS50016 | 35 | 4 | 6243 | 191 | CG1815,d4,Acf,E(bx) | | 2.137e-02 | -3.85 | positive regulation of hydrolase activity | biological process | GO:0051345 | 165 | 9 | 12345 | 305 | HtrA2,RhoGAPp190,CG17883,Graf,vap,Nf1,conu,PlexA,PPP1R15 | | 2.150e-02 | -3.84 | chromosome, centromeric region | cellular component | GO:0000775 | 87 | 6 | 11940 | 298 | Mps1,Bub1,cal1,MED26,fzy,smt3 | | 2.159e-02 | -3.84 | GO\_VESICLE\_DOCKING\_INVOLVED\_IN\_EXOCYTOSIS | MSigDB lists | GO\_VESICLE\_DOCKING\_INVOLVED\_IN\_EXOCYTOSIS | 14 | 3 | 4161 | 184 | Rab8,Exo84,Rop | | 2.159e-02 | -3.84 | GO\_ENDOPLASMIC\_RETICULUM\_ORGANIZATION | MSigDB lists | GO\_ENDOPLASMIC\_RETICULUM\_ORGANIZATION | 14 | 3 | 4161 | 184 | CG12341,CG9853,Vap33 | | 2.159e-02 | -3.84 | GO\_REGULATION\_OF\_ENDOTHELIAL\_CELL\_PROLIFERATION | MSigDB lists | GO\_REGULATION\_OF\_ENDOTHELIAL\_CELL\_PROLIFERATION | 14 | 3 | 4161 | 184 | dpp,viaf,Nf1 | | 2.159e-02 | -3.84 | GO\_NEGATIVE\_REGULATION\_OF\_SMALL\_GTPASE\_MEDIATED\_SIGNAL\_TRANSDUCTION | MSigDB lists | GO\_NEGATIVE\_REGULATION\_OF\_SMALL\_GTPASE\_MEDIATED\_SIGNAL\_TRANSDUCTION | 14 | 3 | 4161 | 184 | Nf1,vap,RhoGAPp190 | | 2.159e-02 | -3.84 | GO\_REGULATION\_OF\_SYNAPTIC\_TRANSMISSION\_GABAERGIC | MSigDB lists | GO\_REGULATION\_OF\_SYNAPTIC\_TRANSMISSION\_GABAERGIC | 14 | 3 | 4161 | 184 | Pten,Rop,Nf1 | | 2.161e-02 | -3.83 | regulation of multi-organism process | biological process | GO:0043900 | 250 | 12 | 12345 | 305 | Parp,dgrn,TM9SF2,Rab7,E(bx),tsu,me31B,pll,Orc3,Hrb27C,smt3,Rop | | 2.165e-02 | -3.83 | meiotic chromosome segregation | biological process | GO:0045132 | 88 | 6 | 12345 | 305 | cal1,mus301,Mps1,grau,Bub1,pav | | 2.168e-02 | -3.83 | regulation of apoptotic process | biological process | GO:0042981 | 193 | 10 | 12345 | 305 | RpL14,HtrA2,eEF5,dpp,CG1791,mav,Dref,Pten,sigmar,viaf | | 2.169e-02 | -3.83 | PHD | pfam domains | PF00628 | 20 | 3 | 10480 | 319 | E(bx),d4,Acf | | 2.170e-02 | -3.83 | GSE21927\_SPLEEN\_VS\_4T1\_TUMOR\_MONOCYTE\_BALBC\_UP | MSigDB lists | GSE21927\_SPLEEN\_VS\_4T1\_TUMOR\_MONOCYTE\_BALBC\_UP | 50 | 6 | 4161 | 184 | CG5861,vnc,CtBP,Dbp21E2,Pp1-87B,CG41378 | | 2.170e-02 | -3.83 | GSE39820\_TGFBETA3\_IL6\_VS\_TGFBETA3\_IL6\_IL23A\_TREATED\_CD4\_TCELL\_DN | MSigDB lists | GSE39820\_TGFBETA3\_IL6\_VS\_TGFBETA3\_IL6\_IL23A\_TREATED\_CD4\_TCELL\_DN | 50 | 6 | 4161 | 184 | mRpS33,Coq5,CG17493,mRpS9,CG13016,CIA30 | | 2.170e-02 | -3.83 | GSE369\_PRE\_VS\_POST\_IL6\_INJECTION\_IFNG\_KO\_LIVER\_UP | MSigDB lists | GSE369\_PRE\_VS\_POST\_IL6\_INJECTION\_IFNG\_KO\_LIVER\_UP | 50 | 6 | 4161 | 184 | Arpc5,eIF3c,rb,Stlk,krz,CG9853 | | 2.175e-02 | -3.83 | GSE339\_CD4POS\_VS\_CD4CD8DN\_DC\_UP | MSigDB lists | GSE339\_CD4POS\_VS\_CD4CD8DN\_DC\_UP | 64 | 7 | 4161 | 184 | janA,RpL14,Arpc3A,RhoGAPp190,Aps,TfIIFbeta,viaf | | 2.175e-02 | -3.83 | GSE20152\_SPHK1\_KO\_VS\_HTNFA\_OVEREXPRESS\_ANKLE\_DN | MSigDB lists | GSE20152\_SPHK1\_KO\_VS\_HTNFA\_OVEREXPRESS\_ANKLE\_DN | 64 | 7 | 4161 | 184 | Pex3,Pect,Parp,RpL5,CycA,Dbp21E2,Pp1-87B | | 2.175e-02 | -3.83 | GCM\_APEX1 | MSigDB lists | GCM\_APEX1 | 64 | 7 | 4161 | 184 | Pp1-87B,RpS8,CCT4,RpL14,Art1,14-3-3zeta,snf | | 2.175e-02 | -3.83 | GSE9960\_HEALTHY\_VS\_GRAM\_POS\_SEPSIS\_PBMC\_UP | MSigDB lists | GSE9960\_HEALTHY\_VS\_GRAM\_POS\_SEPSIS\_PBMC\_UP | 64 | 7 | 4161 | 184 | His2Av,eEF1delta,Cul4,RpL15,ubl,RpS8,eRF1 | | 2.176e-02 | -3.83 | response to stress | biological process | GO:0006950 | 1136 | 39 | 12345 | 305 | mus301,RagA-B,CG5056,Hsp27,dgrn,CG1218,Pten,Cdc42,Sgt,Ino80,CIA30,Aps,Tctp,Rab14,alpha-Man-Ib,lic,CG9005,Sod2,RPA3,Arp8,Stlk,Nf1,ECSIT,glo,DNAlig1,Vap33,spel1,RecQ4,Cul4,Mps1,Parp,sun,Rheb,PPP1R15,pll,mthl8,CG6171,Btk29A,Fign | | 2.191e-02 | -3.82 | regulation of GTPase activity | biological process | GO:0043087 | 113 | 7 | 12345 | 305 | vap,PlexA,Graf,CG17883,conu,Nf1,RhoGAPp190 | | 2.191e-02 | -3.82 | regulation of neuromuscular junction development | biological process | GO:1904396 | 113 | 7 | 12345 | 305 | Fmr1,Cdc42,Nf1,mnb,Rheb,Rab8,tkv | | 2.207e-02 | -3.81 | negative regulation of transcription, DNA-templated | biological process | GO:0045892 | 371 | 16 | 12345 | 305 | d4,Ino80,snf,su(Hw),CG6066,rhea,CG12299,Acf,Tgi,NO66,E(z),Nulp1,CG6254,CtBP,rhi,CG9246 | | 2.207e-02 | -3.81 | GO\_REGULATION\_OF\_ANATOMICAL\_STRUCTURE\_MORPHOGENESIS | MSigDB lists | GO\_REGULATION\_OF\_ANATOMICAL\_STRUCTURE\_MORPHOGENESIS | 210 | 16 | 4161 | 184 | PlexA,Prosalpha2,Rpt6,RhoGAPp190,dpp,Pten,viaf,Cdc42,Nf1,LIMK1,cta,E(z),CG17883,Rap1,vap,Rpn10 | | 2.210e-02 | -3.81 | GO\_N\_ACYLTRANSFERASE\_ACTIVITY | MSigDB lists | GO\_N\_ACYLTRANSFERASE\_ACTIVITY | 37 | 5 | 4161 | 184 | vnc,krz,Rcd5,mbf1,CG6907 | | 2.210e-02 | -3.81 | KONG\_E2F3\_TARGETS | MSigDB lists | KONG\_E2F3\_TARGETS | 37 | 5 | 4161 | 184 | Fign,Skp2,E(z),fzy,CycA | | 2.210e-02 | -3.81 | GO\_PROTEIN\_ALKYLATION | MSigDB lists | GO\_PROTEIN\_ALKYLATION | 37 | 5 | 4161 | 184 | Wdr82,E(z),Art1,eRF1,Rbbp5 | | 2.215e-02 | -3.81 | regulation of programmed cell death | biological process | GO:0043067 | 222 | 11 | 12345 | 305 | eEF5,HtrA2,RpL14,dpp,mav,CG1791,Pten,Fmr1,Dref,sigmar,viaf | | 2.217e-02 | -3.81 | put (punt) | protein interactions | 41772 | 19 | 3 | 9204 | 298 | I-2,tkv,Pp1-87B | | 2.217e-02 | -3.81 | Dscam1 (Down syndrome cell adhesion molecule 1) | protein interactions | 35652 | 19 | 3 | 9204 | 298 | Vap33,vap,Fmr1 | | 2.217e-02 | -3.81 | CG32100 (uncharacterized protein) | protein interactions | 39400 | 19 | 3 | 9204 | 298 | pins,krz,Pnn | | 2.217e-02 | -3.81 | CG3548 (uncharacterized protein) | protein interactions | 37932 | 19 | 3 | 9204 | 298 | Parp,CG42676,RPA3 | | 2.217e-02 | -3.81 | Rpn5 (Regulatory particle non-ATPase 5) | protein interactions | 40717 | 19 | 3 | 9204 | 298 | Ubqn,Rpt6,Amun | | 2.217e-02 | -3.81 | Ras64B (Ras oncogene at 64B) | protein interactions | 38494 | 19 | 3 | 9204 | 298 | PlexA,tkv,dpp | | 2.228e-02 | -3.80 | histone demethylase activity | molecular function | GO:0032452 | 10 | 2 | 12176 | 289 | Jarid2,NO66 | | 2.228e-02 | -3.80 | protein kinase inhibitor activity | molecular function | GO:0004860 | 10 | 2 | 12176 | 289 | 14-3-3zeta,mnb | | 2.228e-02 | -3.80 | kinase inhibitor activity | molecular function | GO:0019210 | 10 | 2 | 12176 | 289 | 14-3-3zeta,mnb | | 2.228e-02 | -3.80 | nucleosome-dependent ATPase activity | molecular function | GO:0070615 | 10 | 2 | 12176 | 289 | E(bx),Acf | | 2.231e-02 | -3.80 | Exon junction complex (EJC) | KEGG pathways | dme\_M00430 | 17 | 3 | 3198 | 117 | Pnn,tsu,Upf3 | | 2.231e-02 | -3.80 | Exon junction complex (EJC) | KEGG pathways | M00430 | 17 | 3 | 3198 | 117 | Pnn,tsu,Upf3 | | 2.233e-02 | -3.80 | GO\_POSITIVE\_REGULATION\_OF\_MAPK\_CASCADE | MSigDB lists | GO\_POSITIVE\_REGULATION\_OF\_MAPK\_CASCADE | 79 | 8 | 4161 | 184 | Rap1,krz,lic,Art1,E(z),Cdc42,dpp,Pten | | 2.233e-02 | -3.80 | HALLMARK\_G2M\_CHECKPOINT | MSigDB lists | HALLMARK\_G2M\_CHECKPOINT | 79 | 8 | 4161 | 184 | E(z),His2Av,CG5535,SF2,CycA,Fs(2)Ket,fzy,pav | | 2.233e-02 | -3.80 | ZHOU\_INFLAMMATORY\_RESPONSE\_FIMA\_UP | MSigDB lists | ZHOU\_INFLAMMATORY\_RESPONSE\_FIMA\_UP | 79 | 8 | 4161 | 184 | Fgop2,mus301,me31B,CG5590,eIF3b,lic,CG11334,CG32473 | | 2.233e-02 | -3.80 | GO\_NUCLEOSIDE\_TRIPHOSPHATASE\_REGULATOR\_ACTIVITY | MSigDB lists | GO\_NUCLEOSIDE\_TRIPHOSPHATASE\_REGULATOR\_ACTIVITY | 79 | 8 | 4161 | 184 | vap,RhoGAPp190,CG17883,Graf,krz,Nf1,CSN1b,pins | | 2.237e-02 | -3.80 | PGRP-LC (Peptidoglycan recognition protein LC) | protein interactions | 39063 | 87 | 7 | 9204 | 298 | smt3,Graf,Stim,TM9SF2,CNBP,CG5174,tsr | | 2.254e-02 | -3.79 | regulation of hydrolase activity | biological process | GO:0051336 | 281 | 13 | 12345 | 305 | conu,Nf1,I-2,viaf,PPP1R15,PlexA,sigmar,CG17883,Graf,Acf,RhoGAPp190,HtrA2,vap | | 2.255e-02 | -3.79 | GSE31082\_DN\_VS\_DP\_THYMOCYTE\_UP | MSigDB lists | GSE31082\_DN\_VS\_DP\_THYMOCYTE\_UP | 110 | 10 | 4161 | 184 | Roc2,Rpb11,CG5339,CG10343,CG11920,ECSIT,mRpS33,CG8860,Sod2,Aps | | 2.255e-02 | -3.79 | GO\_REGULATION\_OF\_CELL\_MORPHOGENESIS | MSigDB lists | GO\_REGULATION\_OF\_CELL\_MORPHOGENESIS | 110 | 10 | 4161 | 184 | CG17883,vap,RhoGAPp190,cta,LIMK1,E(z),Cdc42,dpp,PlexA,Pten | | 2.259e-02 | -3.79 | ESC\_V6.5\_UP\_EARLY.V1\_UP | MSigDB lists | ESC\_V6.5\_UP\_EARLY.V1\_UP | 25 | 4 | 4161 | 184 | Sgt,Stim,Pect,HtrA2 | | 2.259e-02 | -3.79 | GO\_HISTONE\_METHYLATION | MSigDB lists | GO\_HISTONE\_METHYLATION | 25 | 4 | 4161 | 184 | Wdr82,Art1,E(z),Rbbp5 | | 2.259e-02 | -3.79 | GO\_NEGATIVE\_REGULATION\_OF\_CELL\_CYCLE\_G1\_S\_PHASE\_TRANSITION | MSigDB lists | GO\_NEGATIVE\_REGULATION\_OF\_CELL\_CYCLE\_G1\_S\_PHASE\_TRANSITION | 25 | 4 | 4161 | 184 | twin,Pten,Art1,E(z) | | 2.259e-02 | -3.79 | GO\_CYTOPLASMIC\_TRANSLATION | MSigDB lists | GO\_CYTOPLASMIC\_TRANSLATION | 25 | 4 | 4161 | 184 | RpL36A,RpL35A,RpL15,eIF3b | | 2.259e-02 | -3.79 | GAVIN\_FOXP3\_TARGETS\_CLUSTER\_P6 | MSigDB lists | GAVIN\_FOXP3\_TARGETS\_CLUSTER\_P6 | 25 | 4 | 4161 | 184 | Fign,unc-119,fzy,CycA | | 2.259e-02 | -3.79 | YAO\_TEMPORAL\_RESPONSE\_TO\_PROGESTERONE\_CLUSTER\_7 | MSigDB lists | YAO\_TEMPORAL\_RESPONSE\_TO\_PROGESTERONE\_CLUSTER\_7 | 25 | 4 | 4161 | 184 | Sod2,eRF1,Prosalpha2,RhoGAPp190 | | 2.259e-02 | -3.79 | CHENG\_IMPRINTED\_BY\_ESTRADIOL | MSigDB lists | CHENG\_IMPRINTED\_BY\_ESTRADIOL | 25 | 4 | 4161 | 184 | flr,CG32576,Lrr47,sm | | 2.259e-02 | -3.79 | GSE25123\_ROSIGLITAZONE\_VS\_IL4\_AND\_ROSIGLITAZONE\_STIM\_MACROPHAGE\_DAY10\_DN | MSigDB lists | GSE25123\_ROSIGLITAZONE\_VS\_IL4\_AND\_ROSIGLITAZONE\_STIM\_MACROPHAGE\_DAY10\_DN | 25 | 4 | 4161 | 184 | RpL15,cta,MED21,Tom20 | | 2.259e-02 | -3.79 | PID\_RAC1\_PATHWAY | MSigDB lists | PID\_RAC1\_PATHWAY | 25 | 4 | 4161 | 184 | Arpc3A,LIMK1,Arpc5,lic | | 2.259e-02 | -3.79 | JAZAERI\_BREAST\_CANCER\_BRCA1\_VS\_BRCA2\_UP | MSigDB lists | JAZAERI\_BREAST\_CANCER\_BRCA1\_VS\_BRCA2\_UP | 25 | 4 | 4161 | 184 | Pisd,lic,flw,CG9246 | | 2.275e-02 | -3.78 | negative regulation of phosphate metabolic process | biological process | GO:0045936 | 89 | 6 | 12345 | 305 | Pten,sws,krz,I-2,mnb,flw | | 2.275e-02 | -3.78 | negative regulation of phosphorus metabolic process | biological process | GO:0010563 | 89 | 6 | 12345 | 305 | sws,krz,Pten,mnb,flw,I-2 | | 2.284e-02 | -3.78 | larval lymph gland hemopoiesis | biological process | GO:0035167 | 44 | 4 | 12345 | 305 | heix,CG8149,dpp,RpS8 | | 2.284e-02 | -3.78 | post-embryonic hemopoiesis | biological process | GO:0035166 | 44 | 4 | 12345 | 305 | RpS8,dpp,CG8149,heix | | 2.291e-02 | -3.78 | zf-BED | pfam domains | PF02892 | 8 | 2 | 10480 | 319 | BEAF-32,Dref | | 2.298e-02 | -3.77 | synaptic vesicle localization | biological process | GO:0097479 | 25 | 3 | 12345 | 305 | Rop,g,Fmr1 | | 2.299e-02 | -3.77 | translation initiation factor binding | molecular function | GO:0031369 | 26 | 3 | 12176 | 289 | eIF3b,eIF3c,Zpr1 | | 2.317e-02 | -3.76 | acetyltransferase activity | molecular function | GO:0016407 | 93 | 6 | 12176 | 289 | CG2938,Eaf6,vnc,Rcd5,Acf,d4 | | 2.326e-02 | -3.76 | ix (intersex) | protein interactions | 45881 | 34 | 4 | 9204 | 298 | Cdk8,Cpsf6,dpp,CG34159 | | 2.326e-02 | -3.76 | MED20 (Mediator complex subunit 20) | protein interactions | 43952 | 34 | 4 | 9204 | 298 | CG34159,MED22,Cdk8,Cpsf6 | | 2.326e-02 | -3.76 | CG6843 (uncharacterized protein) | protein interactions | 40062 | 34 | 4 | 9204 | 298 | Cwc25,BthD,CG6066,RpS3A | | 2.326e-02 | -3.76 | eEF1gamma (eukaryotic translation elongation factor 1 gamma) | protein interactions | 44791 | 34 | 4 | 9204 | 298 | Exo84,CG12935,Tcs3,eEF1delta | | 2.336e-02 | -3.76 | GO\_ESTABLISHMENT\_OF\_PROTEIN\_LOCALIZATION\_TO\_ORGANELLE | MSigDB lists | GO\_ESTABLISHMENT\_OF\_PROTEIN\_LOCALIZATION\_TO\_ORGANELLE | 194 | 15 | 4161 | 184 | RpS8,RpL35A,RpL15,CG8860,Vps13D,RpS26,RpL14,Tim17b,Fs(2)Ket,Tom20,RpL36A,RpL5,Pex3,rb,RpS9 | | 2.342e-02 | -3.75 | organelle envelope | cellular component | GO:0031967 | 432 | 18 | 11940 | 298 | Nup188,14-3-3zeta,Bmcp,CIA30,Tim17b,ND-AGGG,Tom20,Fs(2)Ket,ND-ASHI,CG4743,spoon,CG44194,Dbp80,HtrA2,l(2)37Bb,MICU1,CG11752,sun | | 2.342e-02 | -3.75 | envelope | cellular component | GO:0031975 | 432 | 18 | 11940 | 298 | Tom20,Fs(2)Ket,ND-ASHI,Bmcp,14-3-3zeta,Nup188,Tim17b,CIA30,ND-AGGG,HtrA2,Dbp80,CG11752,MICU1,sun,l(2)37Bb,CG4743,CG44194,spoon | | 2.349e-02 | -3.75 | GSE19923\_WT\_VS\_E2A\_KO\_DP\_THYMOCYTE\_UP | MSigDB lists | GSE19923\_WT\_VS\_E2A\_KO\_DP\_THYMOCYTE\_UP | 65 | 7 | 4161 | 184 | CG17493,RpS8,Vap33,Wdr82,RpL14,MED22,CG8132 | | 2.349e-02 | -3.75 | GO\_POSITIVE\_REGULATION\_OF\_PROTEIN\_COMPLEX\_ASSEMBLY | MSigDB lists | GO\_POSITIVE\_REGULATION\_OF\_PROTEIN\_COMPLEX\_ASSEMBLY | 65 | 7 | 4161 | 184 | Arpc5,Lst8,Parp,Arpc3A,Rpt6,Cul4,TfIIFbeta | | 2.349e-02 | -3.75 | MORF\_ERH | MSigDB lists | MORF\_ERH | 65 | 7 | 4161 | 184 | Mdh1,Prosalpha2,smt3,eIF3c,Art1,14-3-3zeta,His2Av | | 2.360e-02 | -3.75 | polytene chromosome interband | cellular component | GO:0005705 | 44 | 4 | 11940 | 298 | BEAF-32,nsl1,Mlf,Rcd5 | | 2.372e-02 | -3.74 | GOLDRATH\_NAIVE\_VS\_EFF\_CD8\_TCELL\_DN | MSigDB lists | GOLDRATH\_NAIVE\_VS\_EFF\_CD8\_TCELL\_DN | 51 | 6 | 4161 | 184 | CG31460,CycA,DNAlig1,unc-119,CG41378,E(z) | | 2.372e-02 | -3.74 | GSE6259\_CD4\_TCELL\_VS\_CD8\_TCELL\_UP | MSigDB lists | GSE6259\_CD4\_TCELL\_VS\_CD8\_TCELL\_UP | 51 | 6 | 4161 | 184 | dpp,E(z),CycA,RpL36A,CG8860,pav | | 2.372e-02 | -3.74 | GSE9006\_HEALTHY\_VS\_TYPE\_2\_DIABETES\_PBMC\_AT\_DX\_DN | MSigDB lists | GSE9006\_HEALTHY\_VS\_TYPE\_2\_DIABETES\_PBMC\_AT\_DX\_DN | 51 | 6 | 4161 | 184 | Vps35,mon2,rb,Pex3,Vps13D,Stlk | | 2.372e-02 | -3.74 | GSE17580\_TREG\_VS\_TEFF\_DN | MSigDB lists | GSE17580\_TREG\_VS\_TEFF\_DN | 51 | 6 | 4161 | 184 | VhaPPA1-1,Aps,Rab8,sws,Lst8,RpL14 | | 2.372e-02 | -3.74 | GO\_CATALYTIC\_STEP\_2\_SPLICEOSOME | MSigDB lists | GO\_CATALYTIC\_STEP\_2\_SPLICEOSOME | 51 | 6 | 4161 | 184 | Phf5a,c12.1,noi,SF2,CG4935,tsu | | 2.372e-02 | -3.74 | GO\_PROTEIN\_TRANSPORTER\_ACTIVITY | MSigDB lists | GO\_PROTEIN\_TRANSPORTER\_ACTIVITY | 51 | 6 | 4161 | 184 | Rap1,CG8860,Vps35,Tim17b,Fs(2)Ket,Tom20 | | 2.373e-02 | -3.74 | rRNA processing | biological process | GO:0006364 | 141 | 8 | 12345 | 305 | CG5589,RpS8,CG11920,CG14805,CG4866,mRpS9,Nopp140,CG32075 | | 2.374e-02 | -3.74 | phosphatidylinositol-3,4,5-trisphosphate 3-phosphatase activity | molecular function | GO:0016314 | 1 | 1 | 12176 | 289 | Pten | | 2.374e-02 | -3.74 | histone acetyltransferase activity (H4-K5 specific) | molecular function | GO:0043995 | 1 | 1 | 12176 | 289 | Rcd5 | | 2.374e-02 | -3.74 | pre-mRNA intronic pyrimidine-rich binding | molecular function | GO:0097158 | 1 | 1 | 12176 | 289 | snf | | 2.374e-02 | -3.74 | dinucleotide insertion or deletion binding | molecular function | GO:0032139 | 1 | 1 | 12176 | 289 | spel1 | | 2.374e-02 | -3.74 | cytoskeletal regulatory protein binding | molecular function | GO:0005519 | 1 | 1 | 12176 | 289 | mon2 | | 2.374e-02 | -3.74 | bis(5'-nucleosyl)-tetraphosphatase (asymmetrical) activity | molecular function | GO:0004081 | 1 | 1 | 12176 | 289 | Apf | | 2.374e-02 | -3.74 | bis(5'-adenosyl)-hexaphosphatase activity | molecular function | GO:0034431 | 1 | 1 | 12176 | 289 | Aps | | 2.374e-02 | -3.74 | protein ADP-ribosylase activity | molecular function | GO:1990404 | 1 | 1 | 12176 | 289 | Parp | | 2.374e-02 | -3.74 | acetylation-dependent protein binding | molecular function | GO:0140033 | 1 | 1 | 12176 | 289 | E(bx) | | 2.374e-02 | -3.74 | guanine/thymine mispair binding | molecular function | GO:0032137 | 1 | 1 | 12176 | 289 | spel1 | | 2.374e-02 | -3.74 | transforming growth factor beta binding | molecular function | GO:0050431 | 1 | 1 | 12176 | 289 | tkv | | 2.374e-02 | -3.74 | single guanine insertion binding | molecular function | GO:0032142 | 1 | 1 | 12176 | 289 | spel1 | | 2.374e-02 | -3.74 | protein kinase C inhibitor activity | molecular function | GO:0008426 | 1 | 1 | 12176 | 289 | 14-3-3zeta | | 2.374e-02 | -3.74 | phosphatidylinositol trisphosphate phosphatase activity | molecular function | GO:0034594 | 1 | 1 | 12176 | 289 | Pten | | 2.374e-02 | -3.74 | endopolyphosphatase activity | molecular function | GO:0000298 | 1 | 1 | 12176 | 289 | Aps | | 2.374e-02 | -3.74 | ubiquitin-dependent protein binding | molecular function | GO:0140036 | 1 | 1 | 12176 | 289 | Graf | | 2.374e-02 | -3.74 | protein kinase A catalytic subunit binding | molecular function | GO:0034236 | 1 | 1 | 12176 | 289 | sws | | 2.374e-02 | -3.74 | mitochondrial ribosomal large subunit rRNA binding | molecular function | GO:1990400 | 1 | 1 | 12176 | 289 | Dbp21E2 | | 2.374e-02 | -3.74 | inositol-1,3,4,5-tetrakisphosphate 3-phosphatase activity | molecular function | GO:0051717 | 1 | 1 | 12176 | 289 | Pten | | 2.374e-02 | -3.74 | diphosphoinositol-polyphosphate diphosphatase activity | molecular function | GO:0008486 | 1 | 1 | 12176 | 289 | Aps | | 2.374e-02 | -3.74 | dinucleotide repeat insertion binding | molecular function | GO:0032181 | 1 | 1 | 12176 | 289 | spel1 | | 2.374e-02 | -3.74 | lysine-acetylated histone binding | molecular function | GO:0070577 | 1 | 1 | 12176 | 289 | E(bx) | | 2.374e-02 | -3.74 | bis(5'-adenosyl)-pentaphosphatase activity | molecular function | GO:0034432 | 1 | 1 | 12176 | 289 | Aps | | 2.374e-02 | -3.74 | phosphatidylinositol deacylase activity | molecular function | GO:0050185 | 1 | 1 | 12176 | 289 | PGAP1 | | 2.374e-02 | -3.74 | histone methyltransferase activity (H4-R3 specific) | molecular function | GO:0044020 | 1 | 1 | 12176 | 289 | Art1 | | 2.374e-02 | -3.74 | dolichyl pyrophosphate Glc1Man9GlcNAc2 alpha-1,3-glucosyltransferase activity | molecular function | GO:0042283 | 1 | 1 | 12176 | 289 | xit | | 2.374e-02 | -3.74 | inositol tetrakisphosphate phosphatase activity | molecular function | GO:0052743 | 1 | 1 | 12176 | 289 | Pten | | 2.374e-02 | -3.74 | SUMO binding | molecular function | GO:0032183 | 1 | 1 | 12176 | 289 | dgrn | | 2.374e-02 | -3.74 | phosphatidylinositol-3,4-bisphosphate 3-phosphatase activity | molecular function | GO:0051800 | 1 | 1 | 12176 | 289 | Pten | | 2.374e-02 | -3.74 | peptide-glutamate-N-acetyltransferase activity | molecular function | GO:1990190 | 1 | 1 | 12176 | 289 | vnc | | 2.374e-02 | -3.74 | lysophospholipase activity | molecular function | GO:0004622 | 1 | 1 | 12176 | 289 | sws | | 2.374e-02 | -3.74 | poly(G) binding | molecular function | GO:0034046 | 1 | 1 | 12176 | 289 | Hrb27C | | 2.374e-02 | -3.74 | dopamine receptor binding | molecular function | GO:0050780 | 1 | 1 | 12176 | 289 | cta | | 2.374e-02 | -3.74 | 2-hexaprenyl-6-methoxy-1,4-benzoquinone methyltransferase activity | molecular function | GO:0043334 | 1 | 1 | 12176 | 289 | Coq5 | | 2.374e-02 | -3.74 | S-adenosyl-L-methionine transmembrane transporter activity | molecular function | GO:0000095 | 1 | 1 | 12176 | 289 | CG4743 | | 2.374e-02 | -3.74 | 5'-3' exoribonuclease activity | molecular function | GO:0004534 | 1 | 1 | 12176 | 289 | Rat1 | | 2.374e-02 | -3.74 | ubiquitination-like modification-dependent protein binding | molecular function | GO:0140035 | 1 | 1 | 12176 | 289 | Graf | | 2.374e-02 | -3.74 | snRNA stem-loop binding | molecular function | GO:0035614 | 1 | 1 | 12176 | 289 | snf | | 2.374e-02 | -3.74 | D5 dopamine receptor binding | molecular function | GO:0031752 | 1 | 1 | 12176 | 289 | cta | | 2.374e-02 | -3.74 | histone acetyltransferase activity (H4-K8 specific) | molecular function | GO:0043996 | 1 | 1 | 12176 | 289 | Rcd5 | | 2.374e-02 | -3.74 | S-methyl-5-thioribose-1-phosphate isomerase activity | molecular function | GO:0046523 | 1 | 1 | 12176 | 289 | CG11334 | | 2.374e-02 | -3.74 | omega-amidase activity | molecular function | GO:0050152 | 1 | 1 | 12176 | 289 | CG8132 | | 2.374e-02 | -3.74 | bis(5'-nucleosyl)-tetraphosphatase activity | molecular function | GO:0008796 | 1 | 1 | 12176 | 289 | Apf | | 2.374e-02 | -3.74 | phosphatidylserine decarboxylase activity | molecular function | GO:0004609 | 1 | 1 | 12176 | 289 | Pisd | | 2.374e-02 | -3.74 | double-strand/single-strand DNA junction binding | molecular function | GO:0000406 | 1 | 1 | 12176 | 289 | spel1 | | 2.374e-02 | -3.74 | 8-oxo-dGDP phosphatase activity | molecular function | GO:0044715 | 1 | 1 | 12176 | 289 | CG12567 | | 2.374e-02 | -3.74 | ZnF\_BED | smart domains | SM00614 | 8 | 2 | 5086 | 158 | Dref,BEAF-32 | | 2.386e-02 | -3.74 | MORF\_PSMC1 | MSigDB lists | MORF\_PSMC1 | 111 | 10 | 4161 | 184 | bai,CG5861,eIF3b,eEF1delta,l(1)10Bb,Art1,smt3,Prosalpha2,Rpt6,eIF3c | | 2.390e-02 | -3.73 | KEGG\_SPLICEOSOME | MSigDB lists | KEGG\_SPLICEOSOME | 80 | 8 | 4161 | 184 | CG11586,snf,SF2,tsu,c12.1,Phf5a,noi,l(1)10Bb | | 2.390e-02 | -3.73 | GSE17721\_CTRL\_VS\_LPS\_24H\_BMDC\_DN | MSigDB lists | GSE17721\_CTRL\_VS\_LPS\_24H\_BMDC\_DN | 80 | 8 | 4161 | 184 | flw,CG13016,Prosalpha2,smt3,VhaPPA1-1,Phf5a,tsu,CG10343 | | 2.390e-02 | -3.73 | GSE17974\_CTRL\_VS\_ACT\_IL4\_AND\_ANTI\_IL12\_2H\_CD4\_TCELL\_DN | MSigDB lists | GSE17974\_CTRL\_VS\_ACT\_IL4\_AND\_ANTI\_IL12\_2H\_CD4\_TCELL\_DN | 80 | 8 | 4161 | 184 | CCT4,E(z),CG5535,MED21,SF2,Vps35,mRpS28,CG4866 | | 2.395e-02 | -3.73 | catabolic process | biological process | GO:0009056 | 901 | 32 | 12345 | 305 | Cul4,Not3,twin,CG15534,Rpn10,CG10365,Upf3,smg,Vps13D,CG6171,Ufd4,Zpr1,Prosalpha2,Rat1,CG30105,Ubqn,fzy,Pten,Sgt,Aps,Usp32,Roc2,Prosbeta1,sqd,CG32473,tsu,inc,CG10467,Usp16-45,alpha-Man-Ib,Skp2,Rpt6 | | 2.399e-02 | -3.73 | LOPEZ\_MBD\_TARGETS | MSigDB lists | LOPEZ\_MBD\_TARGETS | 339 | 23 | 4161 | 184 | VhaPPA1-1,Prosalpha2,Fs(2)Ket,flw,Tctp,CG17493,CCT4,Arpc3A,MED22,Skp2,spel1,Cdc42,Pp1-87B,RpS8,RpL35A,Rab8,sm,Wdr82,CG14805,Rap1,CtBP,CG41378,eEF1delta | | 2.402e-02 | -3.73 | apical constriction involved in gastrulation | biological process | GO:0003384 | 10 | 2 | 12345 | 305 | xit,cta | | 2.402e-02 | -3.73 | regulation of nuclear-transcribed mRNA poly(A) tail shortening | biological process | GO:0060211 | 10 | 2 | 12345 | 305 | smg,AGO3 | | 2.402e-02 | -3.73 | histone demethylation | biological process | GO:0016577 | 10 | 2 | 12345 | 305 | Jarid2,NO66 | | 2.402e-02 | -3.73 | negative regulation of gene silencing by RNA | biological process | GO:0060967 | 10 | 2 | 12345 | 305 | g,rb | | 2.402e-02 | -3.73 | positive regulation of nuclear-transcribed mRNA poly(A) tail shortening | biological process | GO:0060213 | 10 | 2 | 12345 | 305 | smg,AGO3 | | 2.402e-02 | -3.73 | cell cycle G1/S phase transition | biological process | GO:0044843 | 10 | 2 | 12345 | 305 | Rheb,Cdk8 | | 2.402e-02 | -3.73 | histone lysine demethylation | biological process | GO:0070076 | 10 | 2 | 12345 | 305 | Jarid2,NO66 | | 2.402e-02 | -3.73 | epithelial cell morphogenesis involved in gastrulation | biological process | GO:0003381 | 10 | 2 | 12345 | 305 | xit,cta | | 2.402e-02 | -3.73 | neuronal stem cell population maintenance | biological process | GO:0097150 | 10 | 2 | 12345 | 305 | fzy,RPA3 | | 2.402e-02 | -3.73 | G1/S transition of mitotic cell cycle | biological process | GO:0000082 | 10 | 2 | 12345 | 305 | Cdk8,Rheb | | 2.402e-02 | -3.73 | regulation of olfactory learning | biological process | GO:0090328 | 10 | 2 | 12345 | 305 | Fmr1,me31B | | 2.441e-02 | -3.71 | mRNA processing | WikiPathways | WP142 | 84 | 9 | 499 | 27 | bai,x16,SF2,Rat1,Rbp1,snf,noi,sm,Phf5a | | 2.447e-02 | -3.71 | ESC/E(Z) complex | cellular component | GO:0035098 | 10 | 2 | 11940 | 298 | E(z),Jarid2 | | 2.447e-02 | -3.71 | Cul3-RING ubiquitin ligase complex | cellular component | GO:0031463 | 10 | 2 | 11940 | 298 | inc,Roc2 | | 2.447e-02 | -3.71 | polytene chromosome, telomeric region | cellular component | GO:0035012 | 10 | 2 | 11940 | 298 | HP4,ver | | 2.447e-02 | -3.71 | condensed chromosome outer kinetochore | cellular component | GO:0000940 | 10 | 2 | 11940 | 298 | Mps1,smt3 | | 2.447e-02 | -3.71 | lamellipodium | cellular component | GO:0030027 | 10 | 2 | 11940 | 298 | Arpc5,Cdc42 | | 2.447e-02 | -3.71 | ER membrane protein complex | cellular component | GO:0072546 | 10 | 2 | 11940 | 298 | CG11137,CG15168 | | 2.456e-02 | -3.71 | GSE6259\_FLT3L\_INDUCED\_33D1\_POS\_DC\_VS\_CD8\_TCELL\_DN | MSigDB lists | GSE6259\_FLT3L\_INDUCED\_33D1\_POS\_DC\_VS\_CD8\_TCELL\_DN | 38 | 5 | 4161 | 184 | RpL14,CG41378,TfIIFbeta,Tsp97E,viaf | | 2.456e-02 | -3.71 | GO\_POSITIVE\_REGULATION\_OF\_APOPTOTIC\_SIGNALING\_PATHWAY | MSigDB lists | GO\_POSITIVE\_REGULATION\_OF\_APOPTOTIC\_SIGNALING\_PATHWAY | 38 | 5 | 4161 | 184 | HtrA2,tkv,Nf1,14-3-3zeta,Pten | | 2.456e-02 | -3.71 | LE\_EGR2\_TARGETS\_UP | MSigDB lists | LE\_EGR2\_TARGETS\_UP | 38 | 5 | 4161 | 184 | fzy,CycA,pav,Fign,E(z) | | 2.456e-02 | -3.71 | GO\_CYTOSOLIC\_LARGE\_RIBOSOMAL\_SUBUNIT | MSigDB lists | GO\_CYTOSOLIC\_LARGE\_RIBOSOMAL\_SUBUNIT | 38 | 5 | 4161 | 184 | RpL5,RpL15,RpL35A,RpL36A,RpL14 | | 2.456e-02 | -3.71 | SAKAI\_TUMOR\_INFILTRATING\_MONOCYTES\_DN | MSigDB lists | SAKAI\_TUMOR\_INFILTRATING\_MONOCYTES\_DN | 38 | 5 | 4161 | 184 | CCT4,CG41378,SF2,Cdc42,tsu | | 2.460e-02 | -3.71 | germ-line stem cell division | biological process | GO:0042078 | 45 | 4 | 12345 | 305 | dpp,Mps1,twin,tkv | | 2.460e-02 | -3.71 | mRNA transport | biological process | GO:0051028 | 45 | 4 | 12345 | 305 | sqd,Dbp80,Fmr1,CG6961 | | 2.463e-02 | -3.70 | positive regulation of organelle organization | biological process | GO:0010638 | 142 | 8 | 12345 | 305 | Arpc5,Cp190,Tctp,Ncoa6,cal1,Arpc3A,flr,E(z) | | 2.471e-02 | -3.70 | synaptic vesicle budding from endosome | biological process | GO:0016182 | 1 | 1 | 12345 | 305 | g | | 2.471e-02 | -3.70 | positive regulation of ceramide biosynthetic process | biological process | GO:2000304 | 1 | 1 | 12345 | 305 | Pex3 | | 2.471e-02 | -3.70 | positive regulation of histone exchange | biological process | GO:1900051 | 1 | 1 | 12345 | 305 | cal1 | | 2.471e-02 | -3.70 | negative regulation of protein localization to cell surface | biological process | GO:2000009 | 1 | 1 | 12345 | 305 | CG30423 | | 2.471e-02 | -3.70 | ectodermal cell fate specification | biological process | GO:0001715 | 1 | 1 | 12345 | 305 | dpp | | 2.471e-02 | -3.70 | positive regulation of actin filament depolymerization | biological process | GO:0030836 | 1 | 1 | 12345 | 305 | flr | | 2.471e-02 | -3.70 | pre-mRNA catabolic process | biological process | GO:1990261 | 1 | 1 | 12345 | 305 | Zpr1 | | 2.471e-02 | -3.70 | negative regulation of protein oligomerization | biological process | GO:0032460 | 1 | 1 | 12345 | 305 | Sgt | | 2.471e-02 | -3.70 | cardiac muscle cell-cardiac muscle cell adhesion | biological process | GO:0086042 | 1 | 1 | 12345 | 305 | rhea | | 2.471e-02 | -3.70 | histone H4-R3 methylation | biological process | GO:0043985 | 1 | 1 | 12345 | 305 | Art1 | | 2.471e-02 | -3.70 | negative regulation of presynaptic active zone assembly | biological process | GO:1905519 | 1 | 1 | 12345 | 305 | Spn | | 2.471e-02 | -3.70 | regulation of kinetochore assembly | biological process | GO:0090234 | 1 | 1 | 12345 | 305 | cal1 | | 2.471e-02 | -3.70 | myo-inositol transport | biological process | GO:0015798 | 1 | 1 | 12345 | 305 | PGAP1 | | 2.471e-02 | -3.70 | translational frameshifting | biological process | GO:0006452 | 1 | 1 | 12345 | 305 | eEF5 | | 2.471e-02 | -3.70 | regulation of chaperone-mediated protein folding | biological process | GO:1903644 | 1 | 1 | 12345 | 305 | Sgt | | 2.471e-02 | -3.70 | meiotic spindle assembly checkpoint | biological process | GO:0033316 | 1 | 1 | 12345 | 305 | Mps1 | | 2.471e-02 | -3.70 | sensitization | biological process | GO:0046960 | 1 | 1 | 12345 | 305 | Btk29A | | 2.471e-02 | -3.70 | regulation of protein desumoylation | biological process | GO:0060188 | 1 | 1 | 12345 | 305 | krz | | 2.471e-02 | -3.70 | lipoprotein transport | biological process | GO:0042953 | 1 | 1 | 12345 | 305 | unc-119 | | 2.471e-02 | -3.70 | regulation of translation at synapse | biological process | GO:0140243 | 1 | 1 | 12345 | 305 | Fmr1 | | 2.471e-02 | -3.70 | S-adenosyl-L-methionine transport | biological process | GO:0015805 | 1 | 1 | 12345 | 305 | CG4743 | | 2.471e-02 | -3.70 | negative regulation of presynapse assembly | biological process | GO:1905607 | 1 | 1 | 12345 | 305 | Spn | | 2.471e-02 | -3.70 | Cdc42 protein signal transduction | biological process | GO:0032488 | 1 | 1 | 12345 | 305 | Cdc42 | | 2.471e-02 | -3.70 | diphosphoinositol polyphosphate metabolic process | biological process | GO:0071543 | 1 | 1 | 12345 | 305 | Aps | | 2.471e-02 | -3.70 | regulation of modification of synaptic structure | biological process | GO:1905244 | 1 | 1 | 12345 | 305 | Fmr1 | | 2.471e-02 | -3.70 | menaquinone metabolic process | biological process | GO:0009233 | 1 | 1 | 12345 | 305 | heix | | 2.471e-02 | -3.70 | homotypic cell-cell adhesion | biological process | GO:0034109 | 1 | 1 | 12345 | 305 | rhea | | 2.471e-02 | -3.70 | ubiquinone biosynthetic process via 3,4-dihydroxy-5-polyprenylbenzoate | biological process | GO:0032194 | 1 | 1 | 12345 | 305 | heix | | 2.471e-02 | -3.70 | regulation of focal adhesion assembly | biological process | GO:0051893 | 1 | 1 | 12345 | 305 | Pten | | 2.471e-02 | -3.70 | diadenosine hexaphosphate catabolic process | biological process | GO:1901909 | 1 | 1 | 12345 | 305 | Aps | | 2.471e-02 | -3.70 | synapse pruning | biological process | GO:0098883 | 1 | 1 | 12345 | 305 | Fmr1 | | 2.471e-02 | -3.70 | regulation of presynaptic active zone assembly | biological process | GO:1905518 | 1 | 1 | 12345 | 305 | Spn | | 2.471e-02 | -3.70 | messenger ribonucleoprotein complex assembly | biological process | GO:1990120 | 1 | 1 | 12345 | 305 | me31B | | 2.471e-02 | -3.70 | positive regulation of CENP-A containing nucleosome assembly | biological process | GO:1903099 | 1 | 1 | 12345 | 305 | cal1 | | 2.471e-02 | -3.70 | regulation of protein folding | biological process | GO:1903332 | 1 | 1 | 12345 | 305 | Sgt | | 2.471e-02 | -3.70 | positive regulation of mitochondrial calcium ion concentration | biological process | GO:0051561 | 1 | 1 | 12345 | 305 | MICU1 | | 2.471e-02 | -3.70 | positive regulation of translational elongation | biological process | GO:0045901 | 1 | 1 | 12345 | 305 | eEF5 | | 2.471e-02 | -3.70 | positive regulation of histone phosphorylation | biological process | GO:0033129 | 1 | 1 | 12345 | 305 | Tctp | | 2.471e-02 | -3.70 | peptidyl-serine ADP-ribosylation | biological process | GO:0018312 | 1 | 1 | 12345 | 305 | CG1218 | | 2.471e-02 | -3.70 | postsynaptic modulation of chemical synaptic transmission | biological process | GO:0099170 | 1 | 1 | 12345 | 305 | Fmr1 | | 2.471e-02 | -3.70 | locomotory exploration behavior | biological process | GO:0035641 | 1 | 1 | 12345 | 305 | krz | | 2.471e-02 | -3.70 | menaquinone biosynthetic process | biological process | GO:0009234 | 1 | 1 | 12345 | 305 | heix | | 2.471e-02 | -3.70 | negative regulation of cell-substrate adhesion | biological process | GO:0010812 | 1 | 1 | 12345 | 305 | Pten | | 2.471e-02 | -3.70 | diadenosine polyphosphate catabolic process | biological process | GO:0015961 | 1 | 1 | 12345 | 305 | Aps | | 2.471e-02 | -3.70 | meiotic metaphase I plate congression | biological process | GO:0043060 | 1 | 1 | 12345 | 305 | Mps1 | | 2.471e-02 | -3.70 | positive regulation of phosphatase activity | biological process | GO:0010922 | 1 | 1 | 12345 | 305 | PPP1R15 | | 2.471e-02 | -3.70 | positive regulation of adenylate cyclase activity | biological process | GO:0045762 | 1 | 1 | 12345 | 305 | Nf1 | | 2.471e-02 | -3.70 | negative regulation of TORC2 signaling | biological process | GO:1903940 | 1 | 1 | 12345 | 305 | Rheb | | 2.471e-02 | -3.70 | exploration behavior | biological process | GO:0035640 | 1 | 1 | 12345 | 305 | krz | | 2.471e-02 | -3.70 | vitamin K biosynthetic process | biological process | GO:0042371 | 1 | 1 | 12345 | 305 | heix | | 2.471e-02 | -3.70 | protein N-linked glycosylation via arginine | biological process | GO:0042543 | 1 | 1 | 12345 | 305 | CG31460 | | 2.471e-02 | -3.70 | regulation of translation at postsynapse | biological process | GO:0140245 | 1 | 1 | 12345 | 305 | Fmr1 | | 2.471e-02 | -3.70 | S-adenosyl-L-methionine transmembrane transport | biological process | GO:1901962 | 1 | 1 | 12345 | 305 | CG4743 | | 2.471e-02 | -3.70 | DNA replication-dependent nucleosome assembly | biological process | GO:0006335 | 1 | 1 | 12345 | 305 | CG8223 | | 2.471e-02 | -3.70 | activation of store-operated calcium channel activity | biological process | GO:0032237 | 1 | 1 | 12345 | 305 | Stim | | 2.471e-02 | -3.70 | positive regulation of chaperone-mediated protein folding | biological process | GO:1903646 | 1 | 1 | 12345 | 305 | Sgt | | 2.471e-02 | -3.70 | regulation of translation at postsynapse, modulating synaptic transmission | biological process | GO:0099578 | 1 | 1 | 12345 | 305 | Fmr1 | | 2.471e-02 | -3.70 | imaginal disc-derived wing hair outgrowth | biological process | GO:0035318 | 1 | 1 | 12345 | 305 | Cdc42 | | 2.471e-02 | -3.70 | positive regulation of protein desumoylation | biological process | GO:0060189 | 1 | 1 | 12345 | 305 | krz | | 2.471e-02 | -3.70 | regulation of Cdc42 protein signal transduction | biological process | GO:0032489 | 1 | 1 | 12345 | 305 | GEFmeso | | 2.471e-02 | -3.70 | diadenosine hexaphosphate metabolic process | biological process | GO:1901908 | 1 | 1 | 12345 | 305 | Aps | | 2.471e-02 | -3.70 | diadenosine pentaphosphate catabolic process | biological process | GO:1901907 | 1 | 1 | 12345 | 305 | Aps | | 2.471e-02 | -3.70 | negative regulation of cAMP-dependent protein kinase activity | biological process | GO:2000480 | 1 | 1 | 12345 | 305 | sws | | 2.471e-02 | -3.70 | negative regulation of adherens junction organization | biological process | GO:1903392 | 1 | 1 | 12345 | 305 | Pten | | 2.471e-02 | -3.70 | positive regulation of miRNA mediated inhibition of translation | biological process | GO:1905618 | 1 | 1 | 12345 | 305 | me31B | | 2.471e-02 | -3.70 | transforming growth factor beta receptor complex assembly | biological process | GO:0007181 | 1 | 1 | 12345 | 305 | tkv | | 2.471e-02 | -3.70 | DNA replication-dependent nucleosome organization | biological process | GO:0034723 | 1 | 1 | 12345 | 305 | CG8223 | | 2.471e-02 | -3.70 | regulation of store-operated calcium channel activity | biological process | GO:1901339 | 1 | 1 | 12345 | 305 | Stim | | 2.471e-02 | -3.70 | negative regulation of cell-matrix adhesion | biological process | GO:0001953 | 1 | 1 | 12345 | 305 | Pten | | 2.471e-02 | -3.70 | polyol transport | biological process | GO:0015791 | 1 | 1 | 12345 | 305 | PGAP1 | | 2.471e-02 | -3.70 | regulation of histone exchange | biological process | GO:1900049 | 1 | 1 | 12345 | 305 | cal1 | | 2.471e-02 | -3.70 | response to redox state | biological process | GO:0051775 | 1 | 1 | 12345 | 305 | BthD | | 2.471e-02 | -3.70 | lipoprotein localization | biological process | GO:0044872 | 1 | 1 | 12345 | 305 | unc-119 | | 2.471e-02 | -3.70 | peptidyl-serine acetylation | biological process | GO:0030920 | 1 | 1 | 12345 | 305 | vnc | | 2.471e-02 | -3.70 | positive regulation of phosphoprotein phosphatase activity | biological process | GO:0032516 | 1 | 1 | 12345 | 305 | PPP1R15 | | 2.471e-02 | -3.70 | adenosine 5'-(hexahydrogen pentaphosphate) metabolic process | biological process | GO:1901910 | 1 | 1 | 12345 | 305 | Aps | | 2.471e-02 | -3.70 | N-terminal peptidyl-serine acetylation | biological process | GO:0017198 | 1 | 1 | 12345 | 305 | vnc | | 2.471e-02 | -3.70 | diadenosine polyphosphate metabolic process | biological process | GO:0015959 | 1 | 1 | 12345 | 305 | Aps | | 2.471e-02 | -3.70 | negative regulation of necrotic cell death | biological process | GO:0060547 | 1 | 1 | 12345 | 305 | fzy | | 2.471e-02 | -3.70 | diadenosine pentaphosphate metabolic process | biological process | GO:1901906 | 1 | 1 | 12345 | 305 | Aps | | 2.471e-02 | -3.70 | phagosome acidification | biological process | GO:0090383 | 1 | 1 | 12345 | 305 | Rab14 | | 2.471e-02 | -3.70 | regulation of translation at synapse, modulating synaptic transmission | biological process | GO:0099547 | 1 | 1 | 12345 | 305 | Fmr1 | | 2.471e-02 | -3.70 | Okazaki fragment processing involved in mitotic DNA replication | biological process | GO:1903461 | 1 | 1 | 12345 | 305 | DNAlig1 | | 2.471e-02 | -3.70 | positive regulation of store-operated calcium channel activity | biological process | GO:1901341 | 1 | 1 | 12345 | 305 | Stim | | 2.471e-02 | -3.70 | negative regulation of focal adhesion assembly | biological process | GO:0051895 | 1 | 1 | 12345 | 305 | Pten | | 2.471e-02 | -3.70 | adenosine 5'-(hexahydrogen pentaphosphate) catabolic process | biological process | GO:1901911 | 1 | 1 | 12345 | 305 | Aps | | 2.471e-02 | -3.70 | regulation of CENP-A containing nucleosome assembly | biological process | GO:1903097 | 1 | 1 | 12345 | 305 | cal1 | | 2.471e-02 | -3.70 | regulation of presynapse assembly | biological process | GO:1905606 | 1 | 1 | 12345 | 305 | Spn | | 2.471e-02 | -3.70 | mitochondrial large ribosomal subunit assembly | biological process | GO:1902775 | 1 | 1 | 12345 | 305 | Dbp21E2 | | 2.471e-02 | -3.70 | meiotic spindle checkpoint | biological process | GO:0044779 | 1 | 1 | 12345 | 305 | Mps1 | | 2.471e-02 | -3.70 | detection of calcium ion | biological process | GO:0005513 | 1 | 1 | 12345 | 305 | Stim | | 2.471e-02 | -3.70 | microtubule bundle formation involved in mitotic spindle midzone assembly | biological process | GO:1903562 | 1 | 1 | 12345 | 305 | pav | | 2.471e-02 | -3.70 | negative regulation of cell junction assembly | biological process | GO:1901889 | 1 | 1 | 12345 | 305 | Pten | | 2.471e-02 | -3.70 | regulation of miRNA mediated inhibition of translation | biological process | GO:1905616 | 1 | 1 | 12345 | 305 | me31B | | 2.471e-02 | -3.70 | negative regulation of lipophagy | biological process | GO:1904503 | 1 | 1 | 12345 | 305 | Rheb | | 2.471e-02 | -3.70 | establishment of pole plasm mRNA localization | biological process | GO:0046595 | 1 | 1 | 12345 | 305 | tsu | | 2.471e-02 | -3.70 | regulation of protein ADP-ribosylation | biological process | GO:0010835 | 1 | 1 | 12345 | 305 | CG1218 | | 2.471e-02 | -3.70 | regulation of cell-substrate junction assembly | biological process | GO:0090109 | 1 | 1 | 12345 | 305 | Pten | | 2.471e-02 | -3.70 | positive regulation of protein folding | biological process | GO:1903334 | 1 | 1 | 12345 | 305 | Sgt | | 2.471e-02 | -3.70 | regulation of neuron projection arborization | biological process | GO:0150011 | 1 | 1 | 12345 | 305 | me31B | | 2.471e-02 | -3.70 | regulation of actomyosin contractile ring contraction | biological process | GO:0031991 | 1 | 1 | 12345 | 305 | flw | | 2.471e-02 | -3.70 | ECS complex | KEGG pathways | dme\_M00388 | 7 | 2 | 3198 | 117 | Roc2,EloB | | 2.471e-02 | -3.70 | ECS complex | KEGG pathways | M00388 | 7 | 2 | 3198 | 117 | Roc2,EloB | | 2.475e-02 | -3.70 | RhoGAP | pfam domains | PF00620 | 21 | 3 | 10480 | 319 | Graf,RhoGAPp190,conu | | 2.475e-02 | -3.70 | asymmetric stem cell division | biological process | GO:0098722 | 67 | 5 | 12345 | 305 | Cdc42,CycA,Mps1,pins,twin | | 2.475e-02 | -3.70 | positive regulation of protein kinase activity | biological process | GO:0045860 | 67 | 5 | 12345 | 305 | smt3,lic,CG1791,Cdc42,Stlk | | 2.483e-02 | -3.70 | embryo development | biological process | GO:0009790 | 568 | 22 | 12345 | 305 | Rap1,Cdc42,xit,dpp,CycA,rhea,14-3-3zeta,smt3,mon2,cta,sqd,BthD,E(z),Btk29A,glo,Fmr1,tsu,tkv,Cp190,pll,lic,Hrb27C | | 2.486e-02 | -3.69 | cellular component morphogenesis | biological process | GO:0032989 | 667 | 25 | 12345 | 305 | E(z),smt3,cta,dpp,RecQ4,Cdk8,rhea,xit,Rap1,Cdc42,Cul4,flr,Hrb27C,sm,Girdin,spoon,PlexA,tkv,Dref,Fmr1,Arpc5,heix,tsr,Pp1-87B,Stlk | | 2.487e-02 | -3.69 | yki (yorkie) | protein interactions | 37851 | 109 | 8 | 9204 | 298 | Tgi,Ncoa6,Fs(2)Ket,tkv,RPA3,NO66,14-3-3zeta,Cpsf6 | | 2.487e-02 | -3.69 | Rheb (Ras homolog enriched in brain) | protein interactions | 117332 | 109 | 8 | 9204 | 298 | sqd,RpL14,CG5516,Tctp,14-3-3zeta,Pten,CG9246,RpL35A | | 2.491e-02 | -3.69 | Bromodomain-like\_sf | interpro domains | IPR036427 | 22 | 3 | 11383 | 331 | CG1815,Acf,E(bx) | | 2.491e-02 | -3.69 | RhoGAP\_dom | interpro domains | IPR000198 | 22 | 3 | 11383 | 331 | RhoGAPp190,conu,Graf | | 2.495e-02 | -3.69 | GO\_INTRINSIC\_COMPONENT\_OF\_ORGANELLE\_MEMBRANE | MSigDB lists | GO\_INTRINSIC\_COMPONENT\_OF\_ORGANELLE\_MEMBRANE | 96 | 9 | 4161 | 184 | CG17883,Stim,Pex3,CG15168,Tim17b,Tom20,PIG-K,heix,MICU1 | | 2.495e-02 | -3.69 | GO\_PROTEIN\_TARGETING\_TO\_MEMBRANE | MSigDB lists | GO\_PROTEIN\_TARGETING\_TO\_MEMBRANE | 96 | 9 | 4161 | 184 | RpL15,RpS8,RpL35A,RpS26,RpL14,RpL5,RpL36A,Pex3,RpS9 | | 2.496e-02 | -3.69 | TRC complex | cellular component | GO:0072380 | 1 | 1 | 11940 | 298 | Sgt | | 2.496e-02 | -3.69 | cytoplasmic vesicle lumen | cellular component | GO:0060205 | 1 | 1 | 11940 | 298 | dpp | | 2.496e-02 | -3.69 | NuA3b histone acetyltransferase complex | cellular component | GO:1990468 | 1 | 1 | 11940 | 298 | Eaf6 | | 2.496e-02 | -3.69 | MutSbeta complex | cellular component | GO:0032302 | 1 | 1 | 11940 | 298 | spel1 | | 2.496e-02 | -3.69 | cytoplasmic side of rough endoplasmic reticulum membrane | cellular component | GO:0098556 | 1 | 1 | 11940 | 298 | RpS26 | | 2.496e-02 | -3.69 | presynaptic endosome | cellular component | GO:0098830 | 1 | 1 | 11940 | 298 | g | | 2.496e-02 | -3.69 | Noc2p-Noc3p complex | cellular component | GO:0030691 | 1 | 1 | 11940 | 298 | CG9246 | | 2.496e-02 | -3.69 | secretory granule lumen | cellular component | GO:0034774 | 1 | 1 | 11940 | 298 | dpp | | 2.496e-02 | -3.69 | NuA3a histone acetyltransferase complex | cellular component | GO:1990467 | 1 | 1 | 11940 | 298 | Eaf6 | | 2.496e-02 | -3.69 | condensed nuclear chromosome outer kinetochore | cellular component | GO:0000942 | 1 | 1 | 11940 | 298 | Mps1 | | 2.496e-02 | -3.69 | vesicle lumen | cellular component | GO:0031983 | 1 | 1 | 11940 | 298 | dpp | | 2.496e-02 | -3.69 | dense core granule lumen | cellular component | GO:0098898 | 1 | 1 | 11940 | 298 | dpp | | 2.496e-02 | -3.69 | Noc1p-Noc2p complex | cellular component | GO:0030690 | 1 | 1 | 11940 | 298 | CG9246 | | 2.496e-02 | -3.69 | actomyosin, actin portion | cellular component | GO:0042643 | 1 | 1 | 11940 | 298 | flr | | 2.496e-02 | -3.69 | SUMO-targeted ubiquitin ligase complex | cellular component | GO:0033768 | 1 | 1 | 11940 | 298 | dgrn | | 2.496e-02 | -3.69 | NuA3 histone acetyltransferase complex | cellular component | GO:0033100 | 1 | 1 | 11940 | 298 | Eaf6 | | 2.499e-02 | -3.69 | RHOGAP | prosite domains | PS50238 | 21 | 3 | 6243 | 191 | RhoGAPp190,conu,Graf | | 2.501e-02 | -3.69 | structural molecule activity | molecular function | GO:0005198 | 558 | 21 | 12176 | 289 | RpS3A,Cpr73D,RpL35A,RpS8,mRpS28,RpS9,mRpS33,RpS26,Nup188,rhea,RpS19a,mRpL30,Arpc5,Arpc3A,RpL14,mRpS9,RpL36A,mRpL34,mRpS18A,RpL15,RpL5 | | 2.507e-02 | -3.69 | P-loop\_NTPase | interpro domains | IPR027417 | 511 | 23 | 11383 | 331 | me31B,pav,Ino80,Fign,RhoGAPp190,cta,Rheb,RagA-B,Dbp80,Rap1,mus301,RecQ4,Rab14,Rab7,CG9330,Dbp21E2,Cdc42,CG5589,CG8915,Rpt6,RabX6,Rab8,spel1 | | 2.523e-02 | -3.68 | GO\_REGULATION\_OF\_NEURON\_DIFFERENTIATION | MSigDB lists | GO\_REGULATION\_OF\_NEURON\_DIFFERENTIATION | 112 | 10 | 4161 | 184 | fzy,PlexA,dpp,me31B,Fmr1,Pten,Rap1,RhoGAPp190,LIMK1,E(z) | | 2.525e-02 | -3.68 | tissue development | biological process | GO:0009888 | 1113 | 38 | 12345 | 305 | Cdk8,rhea,dpp,LIMK1,sqd,Rop,cta,rhi,Cdc42,peng,Pten,Rap1,Fs(2)Ket,xit,me31B,GEFmeso,Stim,VhaPPA1-1,Skp2,CG8149,tsr,Rab7,spoon,RecQ4,flw,smt3,CtBP,Parp,Cul4,Rheb,EloB,CSN1b,Zpr1,Amun,flr,eIF3b,Btk29A,tkv | | 2.533e-02 | -3.68 | GSE34205\_HEALTHY\_VS\_FLU\_INF\_INFANT\_PBMC\_UP | MSigDB lists | GSE34205\_HEALTHY\_VS\_FLU\_INF\_INFANT\_PBMC\_UP | 66 | 7 | 4161 | 184 | bai,Tsp97E,RpL5,Tctp,RpL15,eEF1delta,CG15168 | | 2.533e-02 | -3.68 | GSE21033\_CTRL\_VS\_POLYIC\_STIM\_DC\_3H\_DN | MSigDB lists | GSE21033\_CTRL\_VS\_POLYIC\_STIM\_DC\_3H\_DN | 66 | 7 | 4161 | 184 | CG12321,HtrA2,Pp1-87B,CycA,Arp8,Pect,CG12935 | | 2.533e-02 | -3.68 | GSE360\_L\_MAJOR\_VS\_B\_MALAYI\_LOW\_DOSE\_DC\_DN | MSigDB lists | GSE360\_L\_MAJOR\_VS\_B\_MALAYI\_LOW\_DOSE\_DC\_DN | 66 | 7 | 4161 | 184 | Stim,Pisd,Vps13D,MED22,eEF1delta,Mdh1,flr | | 2.537e-02 | -3.67 | GO\_CYTOSKELETON\_ORGANIZATION | MSigDB lists | GO\_CYTOSKELETON\_ORGANIZATION | 196 | 15 | 4161 | 184 | CG17493,flr,krz,Arpc3A,Ino80,LIMK1,Graf,rhea,Arpc5,Cdc42,pins,Nf1,pav,Fign,Fs(2)Ket | | 2.542e-02 | -3.67 | macromolecule transmembrane transporter activity | molecular function | GO:0022884 | 27 | 3 | 12176 | 289 | Tom20,CG8860,Tim17b | | 2.542e-02 | -3.67 | protein transmembrane transporter activity | molecular function | GO:0008320 | 27 | 3 | 12176 | 289 | Tom20,CG8860,Tim17b | | 2.542e-02 | -3.67 | protein transporter activity | molecular function | GO:0140318 | 27 | 3 | 12176 | 289 | Tim17b,CG8860,Tom20 | | 2.547e-02 | -3.67 | Mes2 (Mesoderm-expressed 2) | protein interactions | 40514 | 20 | 3 | 9204 | 298 | eIF4B,CtBP,RecQ4 | | 2.547e-02 | -3.67 | lncRNA:Hsromega (Heat shock RNA omega) | protein interactions | 3772547 | 20 | 3 | 9204 | 298 | sqd,Pep,Sgt | | 2.547e-02 | -3.67 | CG1129 (uncharacterized protein) | protein interactions | 40599 | 20 | 3 | 9204 | 298 | BthD,Wdr82,CG1218 | | 2.547e-02 | -3.67 | Nurf-38 (Nucleosome remodeling factor - 38kD) | protein interactions | 37922 | 20 | 3 | 9204 | 298 | Dref,Pisd,E(bx) | | 2.547e-02 | -3.67 | Got1 (Glutamate oxaloacetate transaminase 1) | protein interactions | 36782 | 20 | 3 | 9204 | 298 | CG1218,BthD,Wdr82 | | 2.547e-02 | -3.67 | Nup98-96 (Nucleoporin 98-96kD) | protein interactions | 42816 | 20 | 3 | 9204 | 298 | su(Hw),Pten,Cp190 | | 2.547e-02 | -3.67 | woc (without children) | protein interactions | 47249 | 20 | 3 | 9204 | 298 | Ubqn,Slmap,Fgop2 | | 2.549e-02 | -3.67 | Adaptive Immune System | REACTOME pathways | R-DME-1280218 | 186 | 13 | 3923 | 151 | 14-3-3zeta,Prosbeta1,Rap1,Rpt6,Prosalpha2,Rpn10,Pten,Cdc42,Skp2,GILT3,CG41378,Lst8,Stim | | 2.551e-02 | -3.67 | histone H3-K4 methylation | biological process | GO:0051568 | 26 | 3 | 12345 | 305 | Ncoa6,Rbbp5,Wdr82 | | 2.551e-02 | -3.67 | drug transmembrane transport | biological process | GO:0006855 | 26 | 3 | 12345 | 305 | CG4743,CG5535,Bmcp | | 2.551e-02 | -3.67 | negative regulation of protein kinase activity | biological process | GO:0006469 | 26 | 3 | 12345 | 305 | Pten,sws,mnb | | 2.551e-02 | -3.67 | DNA-templated transcription, elongation | biological process | GO:0006354 | 26 | 3 | 12345 | 305 | CG6244,EloB,TfIIFbeta | | 2.551e-02 | -3.67 | mitochondrial envelope | cellular component | GO:0005740 | 313 | 14 | 11940 | 298 | HtrA2,Tom20,MICU1,CG11752,ND-ASHI,sun,l(2)37Bb,Bmcp,CG4743,CIA30,Tim17b,ND-AGGG,CG44194,spoon | | 2.555e-02 | -3.67 | GSE32986\_CURDLAN\_LOWDOSE\_VS\_GMCSF\_AND\_CURDLAN\_LOWDOSE\_STIM\_DC\_DN | MSigDB lists | GSE32986\_CURDLAN\_LOWDOSE\_VS\_GMCSF\_AND\_CURDLAN\_LOWDOSE\_STIM\_DC\_DN | 81 | 8 | 4161 | 184 | CG31460,eIF3b,Ino80,Phf5a,CG5535,lic,Aps,eRF1 | | 2.555e-02 | -3.67 | GO\_REGULATION\_OF\_NEURON\_PROJECTION\_DEVELOPMENT | MSigDB lists | GO\_REGULATION\_OF\_NEURON\_PROJECTION\_DEVELOPMENT | 81 | 8 | 4161 | 184 | RhoGAPp190,Rap1,E(z),LIMK1,fzy,Pten,Fmr1,PlexA | | 2.555e-02 | -3.67 | GO\_REGULATION\_OF\_CELL\_DIVISION | MSigDB lists | GO\_REGULATION\_OF\_CELL\_DIVISION | 81 | 8 | 4161 | 184 | Cdc42,fzy,CG12321,spel1,pav,AGO3,Pten,CG17493 | | 2.557e-02 | -3.67 | RHO GTPases activate PAKs | REACTOME pathways | R-DME-5627123 | 17 | 3 | 3923 | 151 | flw,LIMK1,Cdc42 | | 2.561e-02 | -3.66 | tor (torso) | protein interactions | 35717 | 35 | 4 | 9204 | 298 | Btk29A,dpp,tkv,14-3-3zeta | | 2.561e-02 | -3.66 | l(2)gl (lethal (2) giant larvae) | protein interactions | 33156 | 35 | 4 | 9204 | 298 | Rheb,Exo84,Fmr1,dpp | | 2.561e-02 | -3.66 | h (hairy) | protein interactions | 38995 | 35 | 4 | 9204 | 298 | dgrn,CG12659,CtBP,DCTN3-p24 | | 2.571e-02 | -3.66 | CG4707 (uncharacterized protein) | protein interactions | 37935 | 8 | 2 | 9204 | 298 | CG6171,BthD | | 2.571e-02 | -3.66 | Ibf1 (Insulator binding factor 1) | protein interactions | 41103 | 8 | 2 | 9204 | 298 | su(Hw),Cp190 | | 2.571e-02 | -3.66 | CG14079 (uncharacterized protein) | protein interactions | 40086 | 8 | 2 | 9204 | 298 | eEF1delta,krz | | 2.571e-02 | -3.66 | Tollo (tollo) | protein interactions | 44497 | 8 | 2 | 9204 | 298 | dpp,tkv | | 2.571e-02 | -3.66 | CG7564 (uncharacterized protein) | protein interactions | 39956 | 8 | 2 | 9204 | 298 | Vap33,x16 | | 2.571e-02 | -3.66 | Trpml (Transient receptor potential cation channel, mucolipin) | protein interactions | 40152 | 8 | 2 | 9204 | 298 | Rheb,RagA-B | | 2.571e-02 | -3.66 | CG42674 (uncharacterized protein) | protein interactions | 40205 | 8 | 2 | 9204 | 298 | pins,RecQ4 | | 2.571e-02 | -3.66 | BOD1 (Biorientation Defective 1) | protein interactions | 43352 | 8 | 2 | 9204 | 298 | CG4743,RpL15 | | 2.571e-02 | -3.66 | unk (unkempt) | protein interactions | 42738 | 8 | 2 | 9204 | 298 | Lst8,Rheb | | 2.571e-02 | -3.66 | nesd (nessun dorma) | protein interactions | 35298 | 8 | 2 | 9204 | 298 | 14-3-3zeta,pav | | 2.571e-02 | -3.66 | CG31802 (uncharacterized protein) | protein interactions | 318949 | 8 | 2 | 9204 | 298 | CG9246,smg | | 2.571e-02 | -3.66 | dgt1 (dim gamma-tubulin 1) | protein interactions | 43529 | 8 | 2 | 9204 | 298 | nsl1,Rcd5 | | 2.571e-02 | -3.66 | Ocrl (Oculocerebrorenal syndrome of Lowe) | protein interactions | 31157 | 8 | 2 | 9204 | 298 | Rab7,Rab14 | | 2.571e-02 | -3.66 | CG7744 (uncharacterized protein) | protein interactions | 37239 | 8 | 2 | 9204 | 298 | sun,Acbp1 | | 2.580e-02 | -3.66 | PID\_FAK\_PATHWAY | MSigDB lists | PID\_FAK\_PATHWAY | 26 | 4 | 4161 | 184 | Graf,Rap1,RhoGAPp190,vap | | 2.580e-02 | -3.66 | GSE9988\_ANTI\_TREM1\_VS\_LOW\_LPS\_MONOCYTE\_DN | MSigDB lists | GSE9988\_ANTI\_TREM1\_VS\_LOW\_LPS\_MONOCYTE\_DN | 26 | 4 | 4161 | 184 | Sod2,sigmar,krz,E(z) | | 2.580e-02 | -3.66 | GSE40274\_CTRL\_VS\_FOXP3\_AND\_GATA1\_TRANSDUCED\_ACTIVATED\_CD4\_TCELL\_UP | MSigDB lists | GSE40274\_CTRL\_VS\_FOXP3\_AND\_GATA1\_TRANSDUCED\_ACTIVATED\_CD4\_TCELL\_UP | 26 | 4 | 4161 | 184 | dpp,Arpc5,Rap1,RpL5 | | 2.580e-02 | -3.66 | GO\_REGULATION\_OF\_RNA\_SPLICING | MSigDB lists | GO\_REGULATION\_OF\_RNA\_SPLICING | 26 | 4 | 4161 | 184 | sm,Fmr1,RpS26,tsu | | 2.580e-02 | -3.66 | MODULE\_123 | MSigDB lists | MODULE\_123 | 26 | 4 | 4161 | 184 | pins,TfIIFbeta,E(z),cta | | 2.586e-02 | -3.66 | GSE5099\_UNSTIM\_VS\_MCSF\_TREATED\_MONOCYTE\_DAY3\_UP | MSigDB lists | GSE5099\_UNSTIM\_VS\_MCSF\_TREATED\_MONOCYTE\_DAY3\_UP | 52 | 6 | 4161 | 184 | PlexA,Rpb11,CG6907,Tsp97E,CG3842,CG8860 | | 2.586e-02 | -3.66 | KAECH\_DAY8\_EFF\_VS\_DAY15\_EFF\_CD8\_TCELL\_DN | MSigDB lists | KAECH\_DAY8\_EFF\_VS\_DAY15\_EFF\_CD8\_TCELL\_DN | 52 | 6 | 4161 | 184 | HtrA2,RpL5,CG8132,RpL14,krz,Pisd | | 2.586e-02 | -3.66 | GSE5589\_LPS\_VS\_LPS\_AND\_IL10\_STIM\_MACROPHAGE\_180MIN\_UP | MSigDB lists | GSE5589\_LPS\_VS\_LPS\_AND\_IL10\_STIM\_MACROPHAGE\_180MIN\_UP | 52 | 6 | 4161 | 184 | Pex3,CG41378,dpp,sigmar,HtrA2,CG10672 | | 2.586e-02 | -3.66 | GSE17721\_CTRL\_VS\_PAM3CSK4\_4H\_BMDC\_DN | MSigDB lists | GSE17721\_CTRL\_VS\_PAM3CSK4\_4H\_BMDC\_DN | 52 | 6 | 4161 | 184 | Arpc5,flw,cta,Rop,mus301,CG13016 | | 2.586e-02 | -3.66 | CEBPDELTA\_Q6 | MSigDB lists | CEBPDELTA\_Q6 | 52 | 6 | 4161 | 184 | SF2,Cdc42,lic,CG15168,CG5589,Pten | | 2.586e-02 | -3.66 | KIM\_WT1\_TARGETS\_12HR\_DN | MSigDB lists | KIM\_WT1\_TARGETS\_12HR\_DN | 52 | 6 | 4161 | 184 | CtBP,Rap1,Sod2,TfIIFbeta,CG8635,flw | | 2.594e-02 | -3.65 | HONRADO\_BREAST\_CANCER\_BRCA1\_VS\_BRCA2 | MSigDB lists | HONRADO\_BREAST\_CANCER\_BRCA1\_VS\_BRCA2 | 6 | 2 | 4161 | 184 | CycA,Skp2 | | 2.594e-02 | -3.65 | REACTOME\_SIGNALING\_BY\_BMP | MSigDB lists | REACTOME\_SIGNALING\_BY\_BMP | 6 | 2 | 4161 | 184 | dpp,tkv | | 2.594e-02 | -3.65 | KORKOLA\_YOLK\_SAC\_TUMOR | MSigDB lists | KORKOLA\_YOLK\_SAC\_TUMOR | 6 | 2 | 4161 | 184 | PlexA,dpp | | 2.594e-02 | -3.65 | GO\_NEGATIVE\_REGULATION\_OF\_CALCIUM\_ION\_DEPENDENT\_EXOCYTOSIS | MSigDB lists | GO\_NEGATIVE\_REGULATION\_OF\_CALCIUM\_ION\_DEPENDENT\_EXOCYTOSIS | 6 | 2 | 4161 | 184 | Fmr1,Rap1 | | 2.594e-02 | -3.65 | ABDELMOHSEN\_ELAVL4\_TARGETS | MSigDB lists | ABDELMOHSEN\_ELAVL4\_TARGETS | 6 | 2 | 4161 | 184 | Parp,Cdc42 | | 2.594e-02 | -3.65 | GO\_NEGATIVE\_REGULATION\_OF\_TELOMERE\_MAINTENANCE\_VIA\_TELOMERE\_LENGTHENING | MSigDB lists | GO\_NEGATIVE\_REGULATION\_OF\_TELOMERE\_MAINTENANCE\_VIA\_TELOMERE\_LENGTHENING | 6 | 2 | 4161 | 184 | Parp,Rcd5 | | 2.594e-02 | -3.65 | GO\_POSITIVE\_REGULATION\_OF\_CARTILAGE\_DEVELOPMENT | MSigDB lists | GO\_POSITIVE\_REGULATION\_OF\_CARTILAGE\_DEVELOPMENT | 6 | 2 | 4161 | 184 | dpp,tkv | | 2.594e-02 | -3.65 | CHEOK\_RESPONSE\_TO\_MERCAPTOPURINE\_DN | MSigDB lists | CHEOK\_RESPONSE\_TO\_MERCAPTOPURINE\_DN | 6 | 2 | 4161 | 184 | Vap33,Cul4 | | 2.594e-02 | -3.65 | PID\_S1P\_S1P4\_PATHWAY | MSigDB lists | PID\_S1P\_S1P4\_PATHWAY | 6 | 2 | 4161 | 184 | Cdc42,cta | | 2.594e-02 | -3.65 | CASORELLI\_APL\_SECONDARY\_VS\_DE\_NOVO\_DN | MSigDB lists | CASORELLI\_APL\_SECONDARY\_VS\_DE\_NOVO\_DN | 6 | 2 | 4161 | 184 | eEF1delta,RpL14 | | 2.594e-02 | -3.65 | GO\_POSITIVE\_REGULATION\_OF\_STEM\_CELL\_DIFFERENTIATION | MSigDB lists | GO\_POSITIVE\_REGULATION\_OF\_STEM\_CELL\_DIFFERENTIATION | 6 | 2 | 4161 | 184 | E(z),dpp | | 2.594e-02 | -3.65 | GO\_MODULATION\_BY\_HOST\_OF\_VIRAL\_PROCESS | MSigDB lists | GO\_MODULATION\_BY\_HOST\_OF\_VIRAL\_PROCESS | 6 | 2 | 4161 | 184 | CG17883,Vap33 | | 2.594e-02 | -3.65 | GO\_AGGRESOME | MSigDB lists | GO\_AGGRESOME | 6 | 2 | 4161 | 184 | Rat1,Ubqn | | 2.594e-02 | -3.65 | GO\_ENDOPLASMIC\_RETICULUM\_SUBCOMPARTMENT | MSigDB lists | GO\_ENDOPLASMIC\_RETICULUM\_SUBCOMPARTMENT | 6 | 2 | 4161 | 184 | Fs(2)Ket,Stim | | 2.594e-02 | -3.65 | GO\_ENDOPLASMIC\_RETICULUM\_TUBULAR\_NETWORK | MSigDB lists | GO\_ENDOPLASMIC\_RETICULUM\_TUBULAR\_NETWORK | 6 | 2 | 4161 | 184 | Stim,Fs(2)Ket | | 2.594e-02 | -3.65 | GO\_POSITIVE\_REGULATION\_OF\_INSULIN\_SECRETION\_INVOLVED\_IN\_CELLULAR\_RESPONSE\_TO\_GLUCOSE\_STIMULUS | MSigDB lists | GO\_POSITIVE\_REGULATION\_OF\_INSULIN\_SECRETION\_INVOLVED\_IN\_CELLULAR\_RESPONSE\_TO\_GLUCOSE\_STIMULUS | 6 | 2 | 4161 | 184 | janA,krz | | 2.594e-02 | -3.65 | GO\_TRANSLATION\_REGULATOR\_ACTIVITY | MSigDB lists | GO\_TRANSLATION\_REGULATOR\_ACTIVITY | 6 | 2 | 4161 | 184 | RpS9,Fmr1 | | 2.594e-02 | -3.65 | WANG\_LSD1\_TARGETS\_DN | MSigDB lists | WANG\_LSD1\_TARGETS\_DN | 6 | 2 | 4161 | 184 | Exo84,Rop | | 2.594e-02 | -3.65 | GO\_HISTONE\_METHYLTRANSFERASE\_ACTIVITY\_H3\_K4\_SPECIFIC\_ | MSigDB lists | GO\_HISTONE\_METHYLTRANSFERASE\_ACTIVITY\_H3\_K4\_SPECIFIC\_ | 6 | 2 | 4161 | 184 | Rbbp5,Wdr82 | | 2.594e-02 | -3.65 | HAHTOLA\_MYCOSIS\_FUNGOIDES\_DN | MSigDB lists | HAHTOLA\_MYCOSIS\_FUNGOIDES\_DN | 6 | 2 | 4161 | 184 | Cdc42,Art1 | | 2.594e-02 | -3.65 | GO\_POSITIVE\_REGULATION\_OF\_EPITHELIAL\_TO\_MESENCHYMAL\_TRANSITION | MSigDB lists | GO\_POSITIVE\_REGULATION\_OF\_EPITHELIAL\_TO\_MESENCHYMAL\_TRANSITION | 6 | 2 | 4161 | 184 | E(z),dpp | | 2.594e-02 | -3.65 | REACTOME\_THROMBIN\_SIGNALLING\_THROUGH\_PROTEINASE\_ACTIVATED\_RECEPTORS\_PARS | MSigDB lists | REACTOME\_THROMBIN\_SIGNALLING\_THROUGH\_PROTEINASE\_ACTIVATED\_RECEPTORS\_PARS | 6 | 2 | 4161 | 184 | cta,krz | | 2.605e-02 | -3.65 | cytoskeletal protein binding | molecular function | GO:0008092 | 330 | 14 | 12176 | 289 | tsr,flr,Girdin,pav,smg,Map60,Cp190,Arpc5,mon2,Spn,Arpc3A,inc,rhea,Pten | | 2.613e-02 | -3.64 | GO\_REGULATION\_OF\_TRANSLATIONAL\_ELONGATION | MSigDB lists | GO\_REGULATION\_OF\_TRANSLATIONAL\_ELONGATION | 15 | 3 | 4161 | 184 | RpS9,eEF5,CG4866 | | 2.613e-02 | -3.64 | GO\_TRANSLATION\_PREINITIATION\_COMPLEX | MSigDB lists | GO\_TRANSLATION\_PREINITIATION\_COMPLEX | 15 | 3 | 4161 | 184 | eIF3c,eIF3b,eIF3g1 | | 2.613e-02 | -3.64 | GO\_ORGANELLAR\_SMALL\_RIBOSOMAL\_SUBUNIT | MSigDB lists | GO\_ORGANELLAR\_SMALL\_RIBOSOMAL\_SUBUNIT | 15 | 3 | 4161 | 184 | mRpS28,mRpS9,mRpS33 | | 2.613e-02 | -3.64 | PID\_LKB1\_PATHWAY | MSigDB lists | PID\_LKB1\_PATHWAY | 15 | 3 | 4161 | 184 | Lst8,14-3-3zeta,Stlk | | 2.613e-02 | -3.64 | BIOCARTA\_MET\_PATHWAY | MSigDB lists | BIOCARTA\_MET\_PATHWAY | 15 | 3 | 4161 | 184 | Rap1,vap,Pten | | 2.613e-02 | -3.64 | SIG\_INSULIN\_RECEPTOR\_PATHWAY\_IN\_CARDIAC\_MYOCYTES | MSigDB lists | SIG\_INSULIN\_RECEPTOR\_PATHWAY\_IN\_CARDIAC\_MYOCYTES | 15 | 3 | 4161 | 184 | 14-3-3zeta,Pten,Cdc42 | | 2.613e-02 | -3.64 | chr13q14 | MSigDB lists | chr13q14 | 15 | 3 | 4161 | 184 | Bmcp,TfIIFbeta,Tctp | | 2.620e-02 | -3.64 | negative regulation of protein phosphorylation | biological process | GO:0001933 | 68 | 5 | 12345 | 305 | mnb,flw,krz,sws,Pten | | 2.625e-02 | -3.64 | negative regulation of nucleic acid-templated transcription | biological process | GO:1903507 | 379 | 16 | 12345 | 305 | CG6254,CtBP,rhi,CG9246,Nulp1,NO66,E(z),Tgi,CG12299,Acf,rhea,CG6066,su(Hw),snf,Ino80,d4 | | 2.625e-02 | -3.64 | negative regulation of RNA biosynthetic process | biological process | GO:1902679 | 379 | 16 | 12345 | 305 | CG9246,CtBP,CG6254,rhi,Nulp1,E(z),NO66,Tgi,CG12299,Acf,rhea,CG6066,su(Hw),snf,Ino80,d4 | | 2.627e-02 | -3.64 | intracellular mRNA localization | biological process | GO:0008298 | 92 | 6 | 12345 | 305 | glo,mus301,sqd,Hrb27C,spoon,tsu | | 2.627e-02 | -3.64 | regulation of cell morphogenesis involved in differentiation | biological process | GO:0010769 | 92 | 6 | 12345 | 305 | RhoGAPp190,LIMK1,Cdc42,Fmr1,PlexA,Rheb | | 2.631e-02 | -3.64 | regulation of response to external stimulus | biological process | GO:0032101 | 228 | 11 | 12345 | 305 | pll,me31B,smt3,krz,dgrn,Pten,Parp,PlexA,E(bx),TM9SF2,Rheb | | 2.631e-02 | -3.64 | regulation of developmental growth | biological process | GO:0048638 | 228 | 11 | 12345 | 305 | lic,dpp,Tctp,Rheb,Rab8,mnb,tkv,Pten,Cdc42,Fmr1,Nf1 | | 2.638e-02 | -3.64 | 14-3-3zeta (14-3-3zeta) | protein interactions | 36059 | 90 | 7 | 9204 | 298 | Tctp,ver,Pten,14-3-3zeta,mon2,CG5174,Rheb | | 2.643e-02 | -3.63 | membrane docking | biological process | GO:0022406 | 46 | 4 | 12345 | 305 | Rop,Rab8,Exo84,Vap33 | | 2.643e-02 | -3.63 | organelle localization by membrane tethering | biological process | GO:0140056 | 46 | 4 | 12345 | 305 | Exo84,Rab8,Rop,Vap33 | | 2.649e-02 | -3.63 | GSE22432\_CONVENTIONAL\_CDC\_VS\_PLASMACYTOID\_PDC\_UP | MSigDB lists | GSE22432\_CONVENTIONAL\_CDC\_VS\_PLASMACYTOID\_PDC\_UP | 97 | 9 | 4161 | 184 | Rap1,CG14805,CCT4,CG12321,eIF3g1,spel1,pav,Tcs3,DNAlig1 | | 2.650e-02 | -3.63 | anterior/posterior axis specification | biological process | GO:0009948 | 144 | 8 | 12345 | 305 | mus301,dpp,mon2,lic,Hrb27C,sqd,glo,tsu | | 2.651e-02 | -3.63 | STPPase\_N | interpro domains | IPR031675 | 9 | 2 | 11383 | 331 | flw,Pp1-87B | | 2.656e-02 | -3.63 | protein kinase regulator activity | molecular function | GO:0019887 | 71 | 5 | 12176 | 289 | mnb,CG6907,CycA,14-3-3zeta,CG1791 | | 2.664e-02 | -3.63 | nitrogen compound transport | biological process | GO:0071705 | 605 | 23 | 12345 | 305 | CG8860,sqd,mon2,bai,unc-119,Rab8,Fs(2)Ket,Sgt,rb,Dbp80,RabX6,g,Tom20,Pex3,SF2,Rab14,Tim17b,CG5535,CG6961,Rab7,Vps35,Fmr1,Vps13D | | 2.665e-02 | -3.62 | GO\_NEGATIVE\_REGULATION\_OF\_ORGANELLE\_ORGANIZATION | MSigDB lists | GO\_NEGATIVE\_REGULATION\_OF\_ORGANELLE\_ORGANIZATION | 113 | 10 | 4161 | 184 | Rcd5,Ubqn,Ino80,spel1,pav,pins,CG9246,CG12321,Fs(2)Ket,Parp | | 2.677e-02 | -3.62 | Tak1 (TGF-beta activated kinase 1) | protein interactions | 39659 | 153 | 10 | 9204 | 298 | tsr,Stim,Rab8,Map60,CNBP,CG5174,nsl1,Graf,rhea,Cp190 | | 2.681e-02 | -3.62 | protein tag | molecular function | GO:0031386 | 11 | 2 | 12176 | 289 | ubl,smt3 | | 2.681e-02 | -3.62 | GDP binding | molecular function | GO:0019003 | 11 | 2 | 12176 | 289 | Rap1,Rheb | | 2.681e-02 | -3.62 | GO\_REGULATION\_OF\_KINASE\_ACTIVITY | MSigDB lists | GO\_REGULATION\_OF\_KINASE\_ACTIVITY | 180 | 14 | 4161 | 184 | Cdc42,CSN1b,Nf1,unc-119,dpp,Stlk,Pten,CG6907,CycA,Rap1,Vap33,krz,lic,E(z) | | 2.687e-02 | -3.62 | Ub-specific processing proteases | REACTOME pathways | R-DME-5689880 | 130 | 10 | 3923 | 151 | Usp16-45,Rpt6,krz,Prosbeta1,CycA,Prosalpha2,Pten,Rpn10,Tom20,Skp2 | | 2.697e-02 | -3.61 | GO\_REGULATION\_OF\_CELL\_DEVELOPMENT | MSigDB lists | GO\_REGULATION\_OF\_CELL\_DEVELOPMENT | 163 | 13 | 4161 | 184 | Rap1,flr,RhoGAPp190,LIMK1,E(z),Cdc42,fzy,Nf1,dpp,me31B,PlexA,Pten,Fmr1 | | 2.701e-02 | -3.61 | park (parkin) | protein interactions | 40336 | 71 | 6 | 9204 | 298 | Usp32,Sod2,Parp,heix,HtrA2,Vps35 | | 2.711e-02 | -3.61 | GO\_REGULATION\_OF\_INTRACELLULAR\_SIGNAL\_TRANSDUCTION | MSigDB lists | GO\_REGULATION\_OF\_INTRACELLULAR\_SIGNAL\_TRANSDUCTION | 343 | 23 | 4161 | 184 | CSN1b,krz,lic,Graf,Tctp,RhoGAPp190,dpp,Fign,Pten,Sod2,CG9246,HtrA2,Cdc42,CG32576,Nf1,eEF1delta,Ubqn,Lst8,Art1,E(z),Rap1,RagA-B,vap | | 2.720e-02 | -3.60 | GSE34156\_NOD2\_LIGAND\_VS\_TLR1\_TLR2\_LIGAND\_6H\_TREATED\_MONOCYTE\_UP | MSigDB lists | GSE34156\_NOD2\_LIGAND\_VS\_TLR1\_TLR2\_LIGAND\_6H\_TREATED\_MONOCYTE\_UP | 39 | 5 | 4161 | 184 | CG5590,sigmar,Rop,Fmr1,cta | | 2.721e-02 | -3.60 | Rap1 signalling | REACTOME pathways | R-DME-392517 | 7 | 2 | 3923 | 151 | Rap1,14-3-3zeta | | 2.726e-02 | -3.60 | E2F\_Q3 | MSigDB lists | E2F\_Q3 | 67 | 7 | 4161 | 184 | SF2,Pp1-87B,DNAlig1,Fs(2)Ket,Stt3B,E(z),Pect | | 2.726e-02 | -3.60 | GSE42021\_TCONV\_PLN\_VS\_CD24INT\_TCONV\_THYMUS\_UP | MSigDB lists | GSE42021\_TCONV\_PLN\_VS\_CD24INT\_TCONV\_THYMUS\_UP | 67 | 7 | 4161 | 184 | Coq5,tsu,VhaPPA1-1,CG4866,flr,SF2,Rap1 | | 2.726e-02 | -3.60 | KEGG\_MAPK\_SIGNALING\_PATHWAY | MSigDB lists | KEGG\_MAPK\_SIGNALING\_PATHWAY | 67 | 7 | 4161 | 184 | lic,ECSIT,krz,vap,Nf1,Rap1,Cdc42 | | 2.726e-02 | -3.60 | GSE25846\_IL10\_POS\_VS\_NEG\_CD8\_TCELL\_DAY7\_POST\_CORONAVIRUS\_BRAIN\_DN | MSigDB lists | GSE25846\_IL10\_POS\_VS\_NEG\_CD8\_TCELL\_DAY7\_POST\_CORONAVIRUS\_BRAIN\_DN | 67 | 7 | 4161 | 184 | Tom20,noi,CG3842,alpha-Man-Ib,sigmar,vap,CG5590 | | 2.726e-02 | -3.60 | GO\_BLOOD\_VESSEL\_MORPHOGENESIS | MSigDB lists | GO\_BLOOD\_VESSEL\_MORPHOGENESIS | 67 | 7 | 4161 | 184 | Nf1,vap,Cdc42,viaf,Pten,cta,CG32473 | | 2.753e-02 | -3.59 | DNA recombination | biological process | GO:0006310 | 93 | 6 | 12345 | 305 | DNAlig1,RPA3,spel1,RecQ4,mus301,dgrn | | 2.753e-02 | -3.59 | eggshell chorion assembly | biological process | GO:0007306 | 93 | 6 | 12345 | 305 | smt3,spoon,Cul4,Cdc42,dpp,RecQ4 | | 2.768e-02 | -3.59 | microtubule organizing center | cellular component | GO:0005815 | 171 | 9 | 11940 | 298 | Cp190,Map60,Fign,fzy,CG17493,Bub1,Mps1,Girdin,pav | | 2.769e-02 | -3.59 | positive regulation of kinase activity | biological process | GO:0033674 | 69 | 5 | 12345 | 305 | Cdc42,Stlk,lic,CG1791,smt3 | | 2.781e-02 | -3.58 | cytoplasmic vesicle | cellular component | GO:0031410 | 378 | 16 | 11940 | 298 | g,Vap33,bai,tkv,Rab8,Rab14,vsg,dpp,RagA-B,ema,CG30423,mon2,Graf,Vps35,CG15168,Rab7 | | 2.781e-02 | -3.58 | intracellular vesicle | cellular component | GO:0097708 | 378 | 16 | 11940 | 298 | Rab8,tkv,bai,vsg,Rab14,dpp,RagA-B,ema,g,Vap33,Vps35,CG15168,Rab7,CG30423,mon2,Graf | | 2.781e-02 | -3.58 | regulation of immune system process | biological process | GO:0002682 | 230 | 11 | 12345 | 305 | heix,DCTN3-p24,dgrn,Mlf,Pten,Parp,E(bx),pll,me31B,smt3,Sod2 | | 2.806e-02 | -3.57 | fly chr3-47 cM | chromosome location | fly chr3-47 cM | 46 | 3 | 25010 | 358 | ND-AGGG,MED21,eIF4B | | 2.806e-02 | -3.57 | fly chr3-47 | chromosome location | fly chr3-47 | 46 | 3 | 25010 | 358 | eIF4B,ND-AGGG,MED21 | | 2.810e-02 | -3.57 | regulation of cellular response to stress | biological process | GO:0080135 | 119 | 7 | 12345 | 305 | Cdc42,Pten,Dek,Rap1,sigmar,Fign,flw | | 2.810e-02 | -3.57 | regulation of vesicle-mediated transport | biological process | GO:0060627 | 119 | 7 | 12345 | 305 | mnb,Rab8,vap,nsl1,Rap1,Graf,krz | | 2.810e-02 | -3.57 | Rpn9 (Regulatory particle non-ATPase 9) | protein interactions | 42802 | 36 | 4 | 9204 | 298 | Rpt6,Prosbeta1,Rpn10,Ubqn | | 2.810e-02 | -3.57 | Irbp (Inverted repeat-binding protein) | protein interactions | 117419 | 36 | 4 | 9204 | 298 | CG43367,Parp,RhoGAPp190,RPA3 | | 2.811e-02 | -3.57 | GO\_POSITIVE\_REGULATION\_OF\_BIOSYNTHETIC\_PROCESS | MSigDB lists | GO\_POSITIVE\_REGULATION\_OF\_BIOSYNTHETIC\_PROCESS | 401 | 26 | 4161 | 184 | tkv,Cdk8,CtBP,TfIIFbeta,Rpb11,Art1,l(1)10Bb,mbf1,RpS9,Phf5a,Nf1,Cdc42,Fmr1,Pten,Parp,dpp,CycA,MED21,lic,krz,Ino80,CCT4,eEF5,CG4866,Rpt6,AGO3 | | 2.813e-02 | -3.57 | GO\_POSITIVE\_REGULATION\_OF\_NEURON\_DIFFERENTIATION | MSigDB lists | GO\_POSITIVE\_REGULATION\_OF\_NEURON\_DIFFERENTIATION | 53 | 6 | 4161 | 184 | E(z),Fmr1,dpp,LIMK1,RhoGAPp190,Rap1 | | 2.813e-02 | -3.57 | OUELLET\_OVARIAN\_CANCER\_INVASIVE\_VS\_LMP\_UP | MSigDB lists | OUELLET\_OVARIAN\_CANCER\_INVASIVE\_VS\_LMP\_UP | 53 | 6 | 4161 | 184 | Mdh1,Prosalpha2,bai,CG41378,14-3-3zeta,Parp | | 2.813e-02 | -3.57 | GSE32164\_RESTING\_DIFFERENTIATED\_VS\_ALTERNATIVELY\_ACT\_M2\_MACROPHAGE\_UP | MSigDB lists | GSE32164\_RESTING\_DIFFERENTIATED\_VS\_ALTERNATIVELY\_ACT\_M2\_MACROPHAGE\_UP | 53 | 6 | 4161 | 184 | RpS26,E(z),CycA,fzy,pav,CG5590 | | 2.813e-02 | -3.57 | GSE9006\_TYPE\_1\_DIABETES\_AT\_DX\_VS\_1MONTH\_POST\_DX\_PBMC\_DN | MSigDB lists | GSE9006\_TYPE\_1\_DIABETES\_AT\_DX\_VS\_1MONTH\_POST\_DX\_PBMC\_DN | 53 | 6 | 4161 | 184 | RpL15,Mdh1,tkv,CG12935,CG5339,RpL14 | | 2.813e-02 | -3.57 | GSE18791\_UNSTIM\_VS\_NEWCATSLE\_VIRUS\_DC\_6H\_UP | MSigDB lists | GSE18791\_UNSTIM\_VS\_NEWCATSLE\_VIRUS\_DC\_6H\_UP | 53 | 6 | 4161 | 184 | me31B,Lrr47,Wdr82,CG34148,CG32576,spel1 | | 2.819e-02 | -3.57 | regulation of histone methylation | biological process | GO:0031060 | 27 | 3 | 12345 | 305 | Ncoa6,E(z),Jarid2 | | 2.819e-02 | -3.57 | regulation of gene silencing by RNA | biological process | GO:0060966 | 27 | 3 | 12345 | 305 | rb,me31B,g | | 2.819e-02 | -3.57 | negative regulation of kinase activity | biological process | GO:0033673 | 27 | 3 | 12345 | 305 | mnb,sws,Pten | | 2.819e-02 | -3.57 | negative regulation of gene silencing | biological process | GO:0060969 | 27 | 3 | 12345 | 305 | g,rb,su(Hw) | | 2.819e-02 | -3.57 | lysosomal transport | biological process | GO:0007041 | 27 | 3 | 12345 | 305 | g,Rab7,ema | | 2.819e-02 | -3.57 | Znf\_PHD | interpro domains | IPR001965 | 40 | 4 | 11383 | 331 | CG1815,d4,E(bx),Acf | | 2.820e-02 | -3.57 | Akt1 (Akt1) | protein interactions | 41957 | 199 | 12 | 9204 | 298 | Nph,RpS9,sqd,Girdin,CCT4,Rpt6,Rop,Tctp,lic,Rpn10,Pten,Dref | | 2.832e-02 | -3.56 | - | gene3d domains | 1.10.555.10 | 22 | 3 | 4929 | 151 | conu,Graf,RhoGAPp190 | | 2.832e-02 | -3.56 | - | gene3d domains | 1.20.920.10 | 22 | 3 | 4929 | 151 | Acf,E(bx),CG1815 | | 2.842e-02 | -3.56 | fly chr65D2-65D2\|3-18 | chromosome location | fly chr65D2-65D2\|3-18 | 2 | 1 | 25010 | 358 | lncRNA:CR45416 | | 2.842e-02 | -3.56 | fly chr3F7-3F7\|1-5 | chromosome location | fly chr3F7-3F7\|1-5 | 2 | 1 | 25010 | 358 | CG2938 | | 2.842e-02 | -3.56 | fly chr48E6-48E6\|2-66 | chromosome location | fly chr48E6-48E6\|2-66 | 2 | 1 | 25010 | 358 | CG8860 | | 2.842e-02 | -3.56 | fly chr81F-81F\|3-47.1 | chromosome location | fly chr81F-81F\|3-47.1 | 2 | 1 | 25010 | 358 | Parp | | 2.842e-02 | -3.56 | fly chr85F15-85F15\|3-49 | chromosome location | fly chr85F15-85F15\|3-49 | 2 | 1 | 25010 | 358 | CG6254 | | 2.842e-02 | -3.56 | fly chr48D7-48D7\|2-65 | chromosome location | fly chr48D7-48D7\|2-65 | 2 | 1 | 25010 | 358 | Mppe | | 2.842e-02 | -3.56 | fly chr3F9-3F9\|1-5 cM | chromosome location | fly chr3F9-3F9\|1-5 cM | 2 | 1 | 25010 | 358 | Vap33 | | 2.842e-02 | -3.56 | fly chr8D7-8D7\|1-27 cM | chromosome location | fly chr8D7-8D7\|1-27 cM | 2 | 1 | 25010 | 358 | c12.1 | | 2.842e-02 | -3.56 | fly chr3F7-3F7\|1-5 cM | chromosome location | fly chr3F7-3F7\|1-5 cM | 2 | 1 | 25010 | 358 | CG2938 | | 2.842e-02 | -3.56 | fly chr14F4-14F4\|1-54 cM | chromosome location | fly chr14F4-14F4\|1-54 cM | 2 | 1 | 25010 | 358 | RpS19a | | 2.842e-02 | -3.56 | fly chr49F7-49F8\|2-68 cM | chromosome location | fly chr49F7-49F8\|2-68 cM | 2 | 1 | 25010 | 358 | Orc3 | | 2.842e-02 | -3.56 | fly chr6C5-6C5\|1-18 | chromosome location | fly chr6C5-6C5\|1-18 | 2 | 1 | 25010 | 358 | ND-ASHI | | 2.842e-02 | -3.56 | fly chr97D4-97D5\|3-92 | chromosome location | fly chr97D4-97D5\|3-92 | 2 | 1 | 25010 | 358 | CG5500 | | 2.842e-02 | -3.56 | fly chr64D6-64D6\|3-14 | chromosome location | fly chr64D6-64D6\|3-14 | 2 | 1 | 25010 | 358 | Eaf6 | | 2.842e-02 | -3.56 | fly chr48E6-48E6\|2-66 cM | chromosome location | fly chr48E6-48E6\|2-66 cM | 2 | 1 | 25010 | 358 | CG8860 | | 2.842e-02 | -3.56 | fly chr6C5-6C5\|1-18 cM | chromosome location | fly chr6C5-6C5\|1-18 cM | 2 | 1 | 25010 | 358 | ND-ASHI | | 2.842e-02 | -3.56 | fly chr28E9-28E9\|2-30 cM | chromosome location | fly chr28E9-28E9\|2-30 cM | 2 | 1 | 25010 | 358 | Acbp1 | | 2.842e-02 | -3.56 | fly chr88D9-88D9\|3-56 | chromosome location | fly chr88D9-88D9\|3-56 | 2 | 1 | 25010 | 358 | lncRNA:CR45599 | | 2.842e-02 | -3.56 | fly chr8B7-8B7\|1-25 cM | chromosome location | fly chr8B7-8B7\|1-25 cM | 2 | 1 | 25010 | 358 | Rbm13 | | 2.842e-02 | -3.56 | fly chr14F4-14F4\|1-54 | chromosome location | fly chr14F4-14F4\|1-54 | 2 | 1 | 25010 | 358 | RpS19a | | 2.842e-02 | -3.56 | fly chr48D7-48D7\|2-65 cM | chromosome location | fly chr48D7-48D7\|2-65 cM | 2 | 1 | 25010 | 358 | Mppe | | 2.842e-02 | -3.56 | fly chr3F9-3F9\|1-5 | chromosome location | fly chr3F9-3F9\|1-5 | 2 | 1 | 25010 | 358 | Vap33 | | 2.842e-02 | -3.56 | fly chr102E1-102E1\|4-0 cM | chromosome location | fly chr102E1-102E1\|4-0 cM | 2 | 1 | 25010 | 358 | MED26 | | 2.842e-02 | -3.56 | fly chr64D6-64D6\|3-14 cM | chromosome location | fly chr64D6-64D6\|3-14 cM | 2 | 1 | 25010 | 358 | Eaf6 | | 2.842e-02 | -3.56 | fly chr102E1-102E1\|4-0 | chromosome location | fly chr102E1-102E1\|4-0 | 2 | 1 | 25010 | 358 | MED26 | | 2.842e-02 | -3.56 | fly chr85F15-85F15\|3-49 cM | chromosome location | fly chr85F15-85F15\|3-49 cM | 2 | 1 | 25010 | 358 | CG6254 | | 2.842e-02 | -3.56 | fly chr76D3-76D3\|3-46 cM | chromosome location | fly chr76D3-76D3\|3-46 cM | 2 | 1 | 25010 | 358 | Usp32 | | 2.842e-02 | -3.56 | fly chr8D7-8D7\|1-27 | chromosome location | fly chr8D7-8D7\|1-27 | 2 | 1 | 25010 | 358 | c12.1 | | 2.842e-02 | -3.56 | fly chr65D2-65D2\|3-18 cM | chromosome location | fly chr65D2-65D2\|3-18 cM | 2 | 1 | 25010 | 358 | lncRNA:CR45416 | | 2.842e-02 | -3.56 | fly chr28E9-28E9\|2-30 | chromosome location | fly chr28E9-28E9\|2-30 | 2 | 1 | 25010 | 358 | Acbp1 | | 2.842e-02 | -3.56 | fly chr7F7-7F7\|1-23 cM | chromosome location | fly chr7F7-7F7\|1-23 cM | 2 | 1 | 25010 | 358 | CG12659 | | 2.842e-02 | -3.56 | fly chr88D9-88D9\|3-56 cM | chromosome location | fly chr88D9-88D9\|3-56 cM | 2 | 1 | 25010 | 358 | lncRNA:CR45599 | | 2.842e-02 | -3.56 | fly chr97D4-97D5\|3-92 cM | chromosome location | fly chr97D4-97D5\|3-92 cM | 2 | 1 | 25010 | 358 | CG5500 | | 2.842e-02 | -3.56 | fly chr7F7-7F7\|1-23 | chromosome location | fly chr7F7-7F7\|1-23 | 2 | 1 | 25010 | 358 | CG12659 | | 2.842e-02 | -3.56 | fly chr49F7-49F8\|2-68 | chromosome location | fly chr49F7-49F8\|2-68 | 2 | 1 | 25010 | 358 | Orc3 | | 2.842e-02 | -3.56 | fly chr81F-81F\|3-47.1 cM | chromosome location | fly chr81F-81F\|3-47.1 cM | 2 | 1 | 25010 | 358 | Parp | | 2.842e-02 | -3.56 | fly chr76D3-76D3\|3-46 | chromosome location | fly chr76D3-76D3\|3-46 | 2 | 1 | 25010 | 358 | Usp32 | | 2.842e-02 | -3.56 | fly chr8B7-8B7\|1-25 | chromosome location | fly chr8B7-8B7\|1-25 | 2 | 1 | 25010 | 358 | Rbm13 | | 2.855e-02 | -3.56 | cytosolic ribosome | cellular component | GO:0022626 | 200 | 10 | 11940 | 298 | RpS8,RpL5,RpL36A,RpS26,RpS9,RpS3A,RpL15,RpL35A,RpL14,RpS19a | | 2.861e-02 | -3.55 | GO\_ORGANIC\_CYCLIC\_COMPOUND\_CATABOLIC\_PROCESS | MSigDB lists | GO\_ORGANIC\_CYCLIC\_COMPOUND\_CATABOLIC\_PROCESS | 199 | 15 | 4161 | 184 | RpL15,RpS8,Rat1,Aps,RpL35A,eRF1,Fs(2)Ket,RpS26,RpL14,me31B,tsu,RpL5,RpL36A,RpS9,twin | | 2.873e-02 | -3.55 | Frs2-mediated activation | REACTOME pathways | R-DME-170968 | 113 | 9 | 3923 | 151 | Rpn10,CG4935,Prosalpha2,vap,Rpt6,Nf1,CG7054,14-3-3zeta,Prosbeta1 | | 2.873e-02 | -3.55 | RAF/MAP kinase cascade | REACTOME pathways | R-DME-5673001 | 113 | 9 | 3923 | 151 | Rpn10,Prosalpha2,CG4935,Prosbeta1,14-3-3zeta,CG7054,vap,Nf1,Rpt6 | | 2.873e-02 | -3.55 | SOS-mediated signalling | REACTOME pathways | R-DME-112412 | 113 | 9 | 3923 | 151 | vap,Rpt6,Nf1,Prosbeta1,CG7054,14-3-3zeta,CG4935,Rpn10,Prosalpha2 | | 2.873e-02 | -3.55 | Signalling to p38 via RIT and RIN | REACTOME pathways | R-DME-187706 | 113 | 9 | 3923 | 151 | Prosalpha2,Rpn10,CG4935,14-3-3zeta,CG7054,Prosbeta1,vap,Rpt6,Nf1 | | 2.873e-02 | -3.55 | Prolonged ERK activation events | REACTOME pathways | R-DME-169893 | 113 | 9 | 3923 | 151 | vap,Nf1,Rpt6,Prosbeta1,14-3-3zeta,CG7054,CG4935,Prosalpha2,Rpn10 | | 2.873e-02 | -3.55 | Interleukin receptor SHC signaling | REACTOME pathways | R-DME-912526 | 113 | 9 | 3923 | 151 | Rpt6,Nf1,vap,14-3-3zeta,CG7054,Prosbeta1,CG4935,Rpn10,Prosalpha2 | | 2.873e-02 | -3.55 | GRB2 events in EGFR signaling | REACTOME pathways | R-DME-179812 | 113 | 9 | 3923 | 151 | CG4935,Rpn10,Prosalpha2,14-3-3zeta,CG7054,Prosbeta1,vap,Nf1,Rpt6 | | 2.873e-02 | -3.55 | SHC1 events in EGFR signaling | REACTOME pathways | R-DME-180336 | 113 | 9 | 3923 | 151 | CG4935,Prosalpha2,Rpn10,CG7054,14-3-3zeta,Prosbeta1,vap,Nf1,Rpt6 | | 2.873e-02 | -3.55 | ARMS-mediated activation | REACTOME pathways | R-DME-170984 | 113 | 9 | 3923 | 151 | Prosbeta1,CG7054,14-3-3zeta,vap,Rpt6,Nf1,Prosalpha2,Rpn10,CG4935 | | 2.873e-02 | -3.55 | RET signaling | REACTOME pathways | R-DME-8853659 | 113 | 9 | 3923 | 151 | vap,Rpt6,Nf1,Prosbeta1,14-3-3zeta,CG7054,Rpn10,Prosalpha2,CG4935 | | 2.882e-02 | -3.55 | regulation of animal organ morphogenesis | biological process | GO:2000027 | 94 | 6 | 12345 | 305 | tkv,Mlf,dpp,Rap1,Dref,Pten | | 2.886e-02 | -3.55 | STPPase\_N | pfam domains | PF16891 | 9 | 2 | 10480 | 319 | Pp1-87B,flw | | 2.888e-02 | -3.54 | oocyte nucleus localization involved in oocyte dorsal/ventral axis specification | biological process | GO:0051663 | 11 | 2 | 12345 | 305 | tsu,flw | | 2.888e-02 | -3.54 | protein demethylation | biological process | GO:0006482 | 11 | 2 | 12345 | 305 | NO66,Jarid2 | | 2.888e-02 | -3.54 | vesicle transport along microtubule | biological process | GO:0047496 | 11 | 2 | 12345 | 305 | pav,g | | 2.888e-02 | -3.54 | base-excision repair | biological process | GO:0006284 | 11 | 2 | 12345 | 305 | RPA3,RecQ4 | | 2.888e-02 | -3.54 | Arp2/3 complex-mediated actin nucleation | biological process | GO:0034314 | 11 | 2 | 12345 | 305 | Arpc5,Arpc3A | | 2.888e-02 | -3.54 | regulation of nuclear-transcribed mRNA catabolic process, deadenylation-dependent decay | biological process | GO:1900151 | 11 | 2 | 12345 | 305 | smg,AGO3 | | 2.888e-02 | -3.54 | negative regulation of mRNA metabolic process | biological process | GO:1903312 | 11 | 2 | 12345 | 305 | mbf1,Fmr1 | | 2.888e-02 | -3.54 | vesicle cytoskeletal trafficking | biological process | GO:0099518 | 11 | 2 | 12345 | 305 | pav,g | | 2.888e-02 | -3.54 | positive regulation of nuclear-transcribed mRNA catabolic process, deadenylation-dependent decay | biological process | GO:1900153 | 11 | 2 | 12345 | 305 | smg,AGO3 | | 2.888e-02 | -3.54 | protein dealkylation | biological process | GO:0008214 | 11 | 2 | 12345 | 305 | Jarid2,NO66 | | 2.893e-02 | -3.54 | U2 snRNP | cellular component | GO:0005686 | 27 | 3 | 11940 | 298 | snf,Phf5a,noi | | 2.894e-02 | -3.54 | PUJANA\_CHEK2\_PCC\_NETWORK | MSigDB lists | PUJANA\_CHEK2\_PCC\_NETWORK | 383 | 25 | 4161 | 184 | CycA,SF2,CG8635,Skp2,CCT4,His2Av,MED21,CG5535,DNAlig1,eEF5,CG8860,Prosalpha2,snf,vnc,cta,Rpb11,Art1,E(z),Pp1-87B,fzy,spel1,Nf1,Parp,CG5589,sm | | 2.902e-02 | -3.54 | eIF3f1 (eukaryotic translation initiation factor 3 subunit f1) | protein interactions | 40587 | 21 | 3 | 9204 | 298 | eIF3c,eIF3g1,eIF3b | | 2.902e-02 | -3.54 | tut (tumorous testis) | protein interactions | 317996 | 21 | 3 | 9204 | 298 | Wdr33,CG4612,twin | | 2.902e-02 | -3.54 | Art8 (Arginine methyltransferase 8) | protein interactions | 34528 | 21 | 3 | 9204 | 298 | Wdr82,EloB,CG1218 | | 2.902e-02 | -3.54 | Patr-1 (Protein associated with topo II related - 1) | protein interactions | 42058 | 21 | 3 | 9204 | 298 | me31B,Not3,twin | | 2.902e-02 | -3.54 | coro (coro) | protein interactions | 35596 | 21 | 3 | 9204 | 298 | tkv,CG7766,dpp | | 2.908e-02 | -3.54 | DAZAP1\_RRM2 | interpro domains | IPR034131 | 1 | 1 | 11383 | 331 | Hrb27C | | 2.908e-02 | -3.54 | Ribosomal\_S26e | interpro domains | IPR000892 | 1 | 1 | 11383 | 331 | RpS26 | | 2.908e-02 | -3.54 | PRC2\_HTH\_1 | interpro domains | IPR041343 | 1 | 1 | 11383 | 331 | E(z) | | 2.908e-02 | -3.54 | TMEM18 | interpro domains | IPR026721 | 1 | 1 | 11383 | 331 | Tmem18 | | 2.908e-02 | -3.54 | Myeloid\_leukemia\_factor | interpro domains | IPR019376 | 1 | 1 | 11383 | 331 | Mlf | | 2.908e-02 | -3.54 | DCTN3 | interpro domains | IPR009991 | 1 | 1 | 11383 | 331 | DCTN3-p24 | | 2.908e-02 | -3.54 | Nucleoporin\_Nup188 | interpro domains | IPR018864 | 1 | 1 | 11383 | 331 | Nup188 | | 2.908e-02 | -3.54 | Rab14 | interpro domains | IPR030702 | 1 | 1 | 11383 | 331 | Rab14 | | 2.908e-02 | -3.54 | PS\_Dcarbxylase | interpro domains | IPR003817 | 1 | 1 | 11383 | 331 | Pisd | | 2.908e-02 | -3.54 | WSTF\_Acf1\_Cbp146 | interpro domains | IPR013136 | 1 | 1 | 11383 | 331 | Acf | | 2.908e-02 | -3.54 | Peptide\_chain-rel\_eRF1/aRF1 | interpro domains | IPR004403 | 1 | 1 | 11383 | 331 | eRF1 | | 2.908e-02 | -3.54 | Peptidase\_M22\_CS | interpro domains | IPR017860 | 1 | 1 | 11383 | 331 | Tcs3 | | 2.908e-02 | -3.54 | NDUFB8 | interpro domains | IPR008699 | 1 | 1 | 11383 | 331 | ND-ASHI | | 2.908e-02 | -3.54 | Peroxin-3 | interpro domains | IPR006966 | 1 | 1 | 11383 | 331 | Pex3 | | 2.908e-02 | -3.54 | CPL\_dom | interpro domains | IPR012959 | 1 | 1 | 11383 | 331 | peng | | 2.908e-02 | -3.54 | Ribosomal\_L35A | interpro domains | IPR001780 | 1 | 1 | 11383 | 331 | RpL35A | | 2.908e-02 | -3.54 | GET4 | interpro domains | IPR007317 | 1 | 1 | 11383 | 331 | CG9853 | | 2.908e-02 | -3.54 | Kif23\_Arf-bd | interpro domains | IPR032384 | 1 | 1 | 11383 | 331 | pav | | 2.908e-02 | -3.54 | INO80C/Ies6 | interpro domains | IPR029525 | 1 | 1 | 11383 | 331 | CG12659 | | 2.908e-02 | -3.54 | Rbsml\_L5\_euk/L18\_arc | interpro domains | IPR005485 | 1 | 1 | 11383 | 331 | RpL5 | | 2.908e-02 | -3.54 | Ribosomal\_S4\_CS | interpro domains | IPR018079 | 1 | 1 | 11383 | 331 | RpS9 | | 2.908e-02 | -3.54 | Transl\_elong\_IF5A\_C | interpro domains | IPR020189 | 1 | 1 | 11383 | 331 | eEF5 | | 2.908e-02 | -3.54 | Vps35\_C | interpro domains | IPR042491 | 1 | 1 | 11383 | 331 | Vps35 | | 2.908e-02 | -3.54 | ARPC5\_sf | interpro domains | IPR036743 | 1 | 1 | 11383 | 331 | Arpc5 | | 2.908e-02 | -3.54 | NDUFAF1 | interpro domains | IPR039131 | 1 | 1 | 11383 | 331 | CIA30 | | 2.908e-02 | -3.54 | COMMD10 | interpro domains | IPR037361 | 1 | 1 | 11383 | 331 | Vlet | | 2.908e-02 | -3.54 | ZPR1\_ZnF1/2 | interpro domains | IPR042452 | 1 | 1 | 11383 | 331 | Zpr1 | | 2.908e-02 | -3.54 | RRM\_RBM8 | interpro domains | IPR033744 | 1 | 1 | 11383 | 331 | tsu | | 2.908e-02 | -3.54 | Arp8/plant\_Arp9 | interpro domains | IPR027668 | 1 | 1 | 11383 | 331 | Arp8 | | 2.908e-02 | -3.54 | CENP-T/H4\_C | interpro domains | IPR035425 | 1 | 1 | 11383 | 331 | His4r | | 2.908e-02 | -3.54 | Exo84 | interpro domains | IPR033961 | 1 | 1 | 11383 | 331 | Exo84 | | 2.908e-02 | -3.54 | CCR4-NOT\_su3/5 | interpro domains | IPR012270 | 1 | 1 | 11383 | 331 | Not3 | | 2.908e-02 | -3.54 | TFIIF\_beta\_HTH | interpro domains | IPR040450 | 1 | 1 | 11383 | 331 | TfIIFbeta | | 2.908e-02 | -3.54 | Ribosomal\_L15e | interpro domains | IPR000439 | 1 | 1 | 11383 | 331 | RpL15 | | 2.908e-02 | -3.54 | RasGAP\_C\_SH2 | interpro domains | IPR035842 | 1 | 1 | 11383 | 331 | vap | | 2.908e-02 | -3.54 | AP3B\_C | interpro domains | IPR029390 | 1 | 1 | 11383 | 331 | rb | | 2.908e-02 | -3.54 | GPI\_transamidase | interpro domains | IPR028361 | 1 | 1 | 11383 | 331 | PIG-K | | 2.908e-02 | -3.54 | ARPC5 | interpro domains | IPR006789 | 1 | 1 | 11383 | 331 | Arpc5 | | 2.908e-02 | -3.54 | pG1 | interpro domains | IPR039007 | 1 | 1 | 11383 | 331 | RhoGAPp190 | | 2.908e-02 | -3.54 | NADH-UbQ\_OxRdtase-assoc\_prot30 | interpro domains | IPR013857 | 1 | 1 | 11383 | 331 | CIA30 | | 2.908e-02 | -3.54 | CXC\_dom | interpro domains | IPR026489 | 1 | 1 | 11383 | 331 | E(z) | | 2.908e-02 | -3.54 | Vps55/LEPROT | interpro domains | IPR007262 | 1 | 1 | 11383 | 331 | CG30423 | | 2.908e-02 | -3.54 | Purple\_acid\_Pase\_N | interpro domains | IPR015914 | 1 | 1 | 11383 | 331 | CG1637 | | 2.908e-02 | -3.54 | Cdc42 | interpro domains | IPR037874 | 1 | 1 | 11383 | 331 | Cdc42 | | 2.908e-02 | -3.54 | SOD\_C\_sf | interpro domains | IPR036314 | 1 | 1 | 11383 | 331 | Sod2 | | 2.908e-02 | -3.54 | TNFAIP8-like | interpro domains | IPR008477 | 1 | 1 | 11383 | 331 | sigmar | | 2.908e-02 | -3.54 | EloA-BP1 | interpro domains | IPR031736 | 1 | 1 | 11383 | 331 | CG12877 | | 2.908e-02 | -3.54 | Agenet-like\_dom | interpro domains | IPR008395 | 1 | 1 | 11383 | 331 | Fmr1 | | 2.908e-02 | -3.54 | Coa4-like | interpro domains | IPR039870 | 1 | 1 | 11383 | 331 | CG44194 | | 2.908e-02 | -3.54 | Ribosomal\_protein\_S8e\_euk | interpro domains | IPR042563 | 1 | 1 | 11383 | 331 | RpS8 | | 2.908e-02 | -3.54 | BPTF | interpro domains | IPR038028 | 1 | 1 | 11383 | 331 | E(bx) | | 2.908e-02 | -3.54 | CG7185-like | interpro domains | IPR034771 | 1 | 1 | 11383 | 331 | Cpsf6 | | 2.908e-02 | -3.54 | Prot\_Pase1\_reg-su15A/B\_C | interpro domains | IPR019523 | 1 | 1 | 11383 | 331 | PPP1R15 | | 2.908e-02 | -3.54 | ZPR1\_A/B\_dom | interpro domains | IPR042451 | 1 | 1 | 11383 | 331 | Zpr1 | | 2.908e-02 | -3.54 | SNX29\_PX | interpro domains | IPR037916 | 1 | 1 | 11383 | 331 | CG5439 | | 2.908e-02 | -3.54 | Rap1 | interpro domains | IPR038851 | 1 | 1 | 11383 | 331 | Rap1 | | 2.908e-02 | -3.54 | PHAT\_dom\_sf | interpro domains | IPR037093 | 1 | 1 | 11383 | 331 | smg | | 2.908e-02 | -3.54 | Malate\_DH\_NAD-dep\_euk | interpro domains | IPR011274 | 1 | 1 | 11383 | 331 | Mdh1 | | 2.908e-02 | -3.54 | CD164\_MGC24 | interpro domains | IPR007947 | 1 | 1 | 11383 | 331 | vsg | | 2.908e-02 | -3.54 | VGLL4 | interpro domains | IPR028184 | 1 | 1 | 11383 | 331 | Tgi | | 2.908e-02 | -3.54 | Mediator\_Med21 | interpro domains | IPR021384 | 1 | 1 | 11383 | 331 | MED21 | | 2.908e-02 | -3.54 | PSMG2\_sf | interpro domains | IPR038389 | 1 | 1 | 11383 | 331 | CG12321 | | 2.908e-02 | -3.54 | Las1 | interpro domains | IPR007174 | 1 | 1 | 11383 | 331 | CG32075 | | 2.908e-02 | -3.54 | RPL14\_KOW | interpro domains | IPR041985 | 1 | 1 | 11383 | 331 | RpL14 | | 2.908e-02 | -3.54 | Ribosomal\_S8e | interpro domains | IPR001047 | 1 | 1 | 11383 | 331 | RpS8 | | 2.908e-02 | -3.54 | Kae1/OSGEP | interpro domains | IPR034680 | 1 | 1 | 11383 | 331 | Tcs3 | | 2.908e-02 | -3.54 | L35A\_sf | interpro domains | IPR038661 | 1 | 1 | 11383 | 331 | RpL35A | | 2.908e-02 | -3.54 | ZPR1 | interpro domains | IPR040141 | 1 | 1 | 11383 | 331 | Zpr1 | | 2.908e-02 | -3.54 | DUF3449 | interpro domains | IPR024598 | 1 | 1 | 11383 | 331 | noi | | 2.908e-02 | -3.54 | Pinin\_SDK\_MemA | interpro domains | IPR006786 | 1 | 1 | 11383 | 331 | Pnn | | 2.908e-02 | -3.54 | Hub1/Ubl5 | interpro domains | IPR039732 | 1 | 1 | 11383 | 331 | ubl | | 2.908e-02 | -3.54 | Histone\_H4 | interpro domains | IPR001951 | 1 | 1 | 11383 | 331 | His4r | | 2.908e-02 | -3.54 | Ribosomal\_S26e\_sf | interpro domains | IPR038551 | 1 | 1 | 11383 | 331 | RpS26 | | 2.908e-02 | -3.54 | SF3A3\_dom | interpro domains | IPR031774 | 1 | 1 | 11383 | 331 | noi | | 2.908e-02 | -3.54 | DUF4516 | interpro domains | IPR027858 | 1 | 1 | 11383 | 331 | CG42308 | | 2.908e-02 | -3.54 | Snu23 | interpro domains | IPR040107 | 1 | 1 | 11383 | 331 | CG11586 | | 2.908e-02 | -3.54 | DUF4528 | interpro domains | IPR029245 | 1 | 1 | 11383 | 331 | CG34148 | | 2.908e-02 | -3.54 | Akap1 | interpro domains | IPR033104 | 1 | 1 | 11383 | 331 | spoon | | 2.908e-02 | -3.54 | Cwf\_Cwc\_15 | interpro domains | IPR006973 | 1 | 1 | 11383 | 331 | c12.1 | | 2.908e-02 | -3.54 | PADR1 | interpro domains | IPR012982 | 1 | 1 | 11383 | 331 | Parp | | 2.908e-02 | -3.54 | KIF23 | interpro domains | IPR032924 | 1 | 1 | 11383 | 331 | pav | | 2.908e-02 | -3.54 | snRNA-activating\_su3 | interpro domains | IPR022042 | 1 | 1 | 11383 | 331 | Pbp49 | | 2.908e-02 | -3.54 | Talin\_cent | interpro domains | IPR015224 | 1 | 1 | 11383 | 331 | rhea | | 2.908e-02 | -3.54 | Pelle\_death | interpro domains | IPR037924 | 1 | 1 | 11383 | 331 | pll | | 2.908e-02 | -3.54 | FAM214/SPAC3H8 | interpro domains | IPR033473 | 1 | 1 | 11383 | 331 | CG9005 | | 2.908e-02 | -3.54 | Malate\_DH\_type2 | interpro domains | IPR010945 | 1 | 1 | 11383 | 331 | Mdh1 | | 2.908e-02 | -3.54 | Elongator\_complex\_protein\_4 | interpro domains | IPR008728 | 1 | 1 | 11383 | 331 | CG6907 | | 2.908e-02 | -3.54 | TMEM85/Emc4 | interpro domains | IPR009445 | 1 | 1 | 11383 | 331 | CG11137 | | 2.908e-02 | -3.54 | Oxidoreductase-like\_N | interpro domains | IPR019180 | 1 | 1 | 11383 | 331 | CG5500 | | 2.908e-02 | -3.54 | PSD | interpro domains | IPR033177 | 1 | 1 | 11383 | 331 | Pisd | | 2.908e-02 | -3.54 | TECPR2 | interpro domains | IPR029772 | 1 | 1 | 11383 | 331 | CG11141 | | 2.908e-02 | -3.54 | DREV\_MeTrfase | interpro domains | IPR007884 | 1 | 1 | 11383 | 331 | CG5339 | | 2.908e-02 | -3.54 | Proteasome\_assmbl\_chp\_2\_euk | interpro domains | IPR016562 | 1 | 1 | 11383 | 331 | CG12321 | | 2.908e-02 | -3.54 | TFIIF\_beta | interpro domains | IPR003196 | 1 | 1 | 11383 | 331 | TfIIFbeta | | 2.908e-02 | -3.54 | Peptidase\_S1C | interpro domains | IPR001940 | 1 | 1 | 11383 | 331 | HtrA2 | | 2.908e-02 | -3.54 | Ribosomal\_S1/3\_euk | interpro domains | IPR027500 | 1 | 1 | 11383 | 331 | RpS3A | | 2.908e-02 | -3.54 | CLEC16A/TT9 | interpro domains | IPR039272 | 1 | 1 | 11383 | 331 | ema | | 2.908e-02 | -3.54 | Xrn2/3/4 | interpro domains | IPR017151 | 1 | 1 | 11383 | 331 | Rat1 | | 2.908e-02 | -3.54 | RasGAP\_SH3 | interpro domains | IPR035652 | 1 | 1 | 11383 | 331 | vap | | 2.908e-02 | -3.54 | DUF4743 | interpro domains | IPR031804 | 1 | 1 | 11383 | 331 | CG12567 | | 2.908e-02 | -3.54 | M1Pi\_N | interpro domains | IPR027363 | 1 | 1 | 11383 | 331 | CG11334 | | 2.908e-02 | -3.54 | Pinin | interpro domains | IPR039853 | 1 | 1 | 11383 | 331 | Pnn | | 2.908e-02 | -3.54 | Ribosomal\_L15e\_CS | interpro domains | IPR020925 | 1 | 1 | 11383 | 331 | RpL15 | | 2.908e-02 | -3.54 | STIM | interpro domains | IPR037608 | 1 | 1 | 11383 | 331 | Stim | | 2.908e-02 | -3.54 | Tmem223 | interpro domains | IPR026100 | 1 | 1 | 11383 | 331 | CG12935 | | 2.908e-02 | -3.54 | SF3a60\_bindingd | interpro domains | IPR021966 | 1 | 1 | 11383 | 331 | noi | | 2.908e-02 | -3.54 | Mn/Fe\_SOD\_N\_sf | interpro domains | IPR036324 | 1 | 1 | 11383 | 331 | Sod2 | | 2.908e-02 | -3.54 | Talin\_cent\_sf | interpro domains | IPR036476 | 1 | 1 | 11383 | 331 | rhea | | 2.908e-02 | -3.54 | Scm3/HJURP | interpro domains | IPR018465 | 1 | 1 | 11383 | 331 | cal1 | | 2.908e-02 | -3.54 | FXMRP1\_C\_core | interpro domains | IPR022034 | 1 | 1 | 11383 | 331 | Fmr1 | | 2.908e-02 | -3.54 | SRSF1\_RRM1 | interpro domains | IPR034520 | 1 | 1 | 11383 | 331 | SF2 | | 2.908e-02 | -3.54 | Talin1/2-RS | interpro domains | IPR037438 | 1 | 1 | 11383 | 331 | rhea | | 2.908e-02 | -3.54 | HPF1 | interpro domains | IPR019361 | 1 | 1 | 11383 | 331 | CG1218 | | 2.908e-02 | -3.54 | NSL1 | interpro domains | IPR026180 | 1 | 1 | 11383 | 331 | nsl1 | | 2.908e-02 | -3.54 | Ribosomal\_S3Ae | interpro domains | IPR001593 | 1 | 1 | 11383 | 331 | RpS3A | | 2.908e-02 | -3.54 | Ribosomal\_protein\_L14 | interpro domains | IPR039660 | 1 | 1 | 11383 | 331 | RpL14 | | 2.908e-02 | -3.54 | Ndufb8\_metazoa | interpro domains | IPR016551 | 1 | 1 | 11383 | 331 | ND-ASHI | | 2.908e-02 | -3.54 | ALG8 | interpro domains | IPR039487 | 1 | 1 | 11383 | 331 | xit | | 2.908e-02 | -3.54 | AP3\_complex\_dsu | interpro domains | IPR017105 | 1 | 1 | 11383 | 331 | g | | 2.908e-02 | -3.54 | TNFAIP8\_sf | interpro domains | IPR038355 | 1 | 1 | 11383 | 331 | sigmar | | 2.908e-02 | -3.54 | Ribosomal\_L44e | interpro domains | IPR000552 | 1 | 1 | 11383 | 331 | RpL36A | | 2.908e-02 | -3.54 | WGR\_domain | interpro domains | IPR008893 | 1 | 1 | 11383 | 331 | Parp | | 2.908e-02 | -3.54 | RBP11 | interpro domains | IPR037685 | 1 | 1 | 11383 | 331 | Rpb11 | | 2.908e-02 | -3.54 | FMRP\_KH0 | interpro domains | IPR040472 | 1 | 1 | 11383 | 331 | Fmr1 | | 2.908e-02 | -3.54 | MCRS\_N | interpro domains | IPR025999 | 1 | 1 | 11383 | 331 | Rcd5 | | 2.908e-02 | -3.54 | RBBP5/Swd1 | interpro domains | IPR037850 | 1 | 1 | 11383 | 331 | Rbbp5 | | 2.908e-02 | -3.54 | IF-M1Pi | interpro domains | IPR005251 | 1 | 1 | 11383 | 331 | CG11334 | | 2.908e-02 | -3.54 | FMR1 | interpro domains | IPR040148 | 1 | 1 | 11383 | 331 | Fmr1 | | 2.908e-02 | -3.54 | PARP | interpro domains | IPR008288 | 1 | 1 | 11383 | 331 | Parp | | 2.908e-02 | -3.54 | Mon2-like | interpro domains | IPR026829 | 1 | 1 | 11383 | 331 | mon2 | | 2.908e-02 | -3.54 | VP13D\_UBA | interpro domains | IPR041969 | 1 | 1 | 11383 | 331 | Vps13D | | 2.908e-02 | -3.54 | CLEC16A/TT9\_N | interpro domains | IPR019155 | 1 | 1 | 11383 | 331 | ema | | 2.908e-02 | -3.54 | Magnesium\_transport | interpro domains | IPR018937 | 1 | 1 | 11383 | 331 | CG15168 | | 2.908e-02 | -3.54 | Sde2\_N | interpro domains | IPR024974 | 1 | 1 | 11383 | 331 | CG5986 | | 2.908e-02 | -3.54 | Not\_N | interpro domains | IPR007207 | 1 | 1 | 11383 | 331 | Not3 | | 2.908e-02 | -3.54 | NKAP | interpro domains | IPR040466 | 1 | 1 | 11383 | 331 | CG6066 | | 2.908e-02 | -3.54 | Bifunc\_PIno\_P3\_Pase/Pase\_PTEN | interpro domains | IPR017361 | 1 | 1 | 11383 | 331 | Pten | | 2.908e-02 | -3.54 | Vps35 | interpro domains | IPR005378 | 1 | 1 | 11383 | 331 | Vps35 | | 2.908e-02 | -3.54 | CPSF6/7 | interpro domains | IPR034772 | 1 | 1 | 11383 | 331 | Cpsf6 | | 2.908e-02 | -3.54 | Acf1\_Bromo | interpro domains | IPR037325 | 1 | 1 | 11383 | 331 | Acf | | 2.908e-02 | -3.54 | UBIAD1 | interpro domains | IPR026046 | 1 | 1 | 11383 | 331 | heix | | 2.908e-02 | -3.54 | R3H\_NRF | interpro domains | IPR034071 | 1 | 1 | 11383 | 331 | CG31301 | | 2.908e-02 | -3.54 | SGTA\_dimer | interpro domains | IPR032374 | 1 | 1 | 11383 | 331 | Sgt | | 2.908e-02 | -3.54 | UbiE/COQ5\_MeTrFase | interpro domains | IPR004033 | 1 | 1 | 11383 | 331 | Coq5 | | 2.908e-02 | -3.54 | Rbsml\_L5e/L18P\_C | interpro domains | IPR025607 | 1 | 1 | 11383 | 331 | RpL5 | | 2.908e-02 | -3.54 | TCTP | interpro domains | IPR034737 | 1 | 1 | 11383 | 331 | Tctp | | 2.908e-02 | -3.54 | Srp40/Nopp140 | interpro domains | IPR039191 | 1 | 1 | 11383 | 331 | Nopp140 | | 2.908e-02 | -3.54 | Mn/Fe\_SOD\_C | interpro domains | IPR019832 | 1 | 1 | 11383 | 331 | Sod2 | | 2.908e-02 | -3.54 | APLF | interpro domains | IPR039253 | 1 | 1 | 11383 | 331 | CG6171 | | 2.908e-02 | -3.54 | Mn/Fe\_SOD | interpro domains | IPR001189 | 1 | 1 | 11383 | 331 | Sod2 | | 2.908e-02 | -3.54 | eIF3B\_RRM | interpro domains | IPR034363 | 1 | 1 | 11383 | 331 | eIF3b | | 2.908e-02 | -3.54 | Proteasome\_subunit\_alpha2 | interpro domains | IPR034644 | 1 | 1 | 11383 | 331 | Prosalpha2 | | 2.908e-02 | -3.54 | ECSIT | interpro domains | IPR010418 | 1 | 1 | 11383 | 331 | ECSIT | | 2.908e-02 | -3.54 | Gprotein\_alpha\_12/13 | interpro domains | IPR000469 | 1 | 1 | 11383 | 331 | cta | | 2.908e-02 | -3.54 | Histone\_H4\_CS | interpro domains | IPR019809 | 1 | 1 | 11383 | 331 | His4r | | 2.908e-02 | -3.54 | Ino80 | interpro domains | IPR031047 | 1 | 1 | 11383 | 331 | Ino80 | | 2.908e-02 | -3.54 | Srp40\_C | interpro domains | IPR007718 | 1 | 1 | 11383 | 331 | Nopp140 | | 2.908e-02 | -3.54 | Ribosomal\_L34 | interpro domains | IPR000271 | 1 | 1 | 11383 | 331 | mRpL34 | | 2.908e-02 | -3.54 | TMEM65 | interpro domains | IPR019537 | 1 | 1 | 11383 | 331 | CG17715 | | 2.908e-02 | -3.54 | G10 | interpro domains | IPR001748 | 1 | 1 | 11383 | 331 | l(1)10Bb | | 2.908e-02 | -3.54 | Msh2 | interpro domains | IPR032642 | 1 | 1 | 11383 | 331 | spel1 | | 2.908e-02 | -3.54 | Cas1\_AcylTrans\_dom | interpro domains | IPR012419 | 1 | 1 | 11383 | 331 | CG2938 | | 2.908e-02 | -3.54 | ELOB | interpro domains | IPR039049 | 1 | 1 | 11383 | 331 | EloB | | 2.908e-02 | -3.54 | EIF3C | interpro domains | IPR027516 | 1 | 1 | 11383 | 331 | eIF3c | | 2.908e-02 | -3.54 | Translational\_control\_tumour\_p | interpro domains | IPR018105 | 1 | 1 | 11383 | 331 | Tctp | | 2.908e-02 | -3.54 | CUL4B | interpro domains | IPR033044 | 1 | 1 | 11383 | 331 | Cul4 | | 2.908e-02 | -3.54 | NKAP\_C | interpro domains | IPR009269 | 1 | 1 | 11383 | 331 | CG6066 | | 2.908e-02 | -3.54 | Mon2\_C | interpro domains | IPR032817 | 1 | 1 | 11383 | 331 | mon2 | | 2.908e-02 | -3.54 | UbiE/COQ5\_MeTrFase\_CS | interpro domains | IPR023576 | 1 | 1 | 11383 | 331 | Coq5 | | 2.908e-02 | -3.54 | EIF3C\_N\_dom | interpro domains | IPR008905 | 1 | 1 | 11383 | 331 | eIF3c | | 2.908e-02 | -3.54 | DNA\_rep\_checkpnt\_protein | interpro domains | IPR021110 | 1 | 1 | 11383 | 331 | RecQ4 | | 2.908e-02 | -3.54 | Neurabin-1/2\_PDZ | interpro domains | IPR040645 | 1 | 1 | 11383 | 331 | Spn | | 2.908e-02 | -3.54 | Poly(ADP-ribose)pol\_reg\_dom\_sf | interpro domains | IPR036616 | 1 | 1 | 11383 | 331 | Parp | | 2.908e-02 | -3.54 | Znf\_PARP\_sf | interpro domains | IPR036957 | 1 | 1 | 11383 | 331 | Parp | | 2.908e-02 | -3.54 | p120-RasGAP | interpro domains | IPR028554 | 1 | 1 | 11383 | 331 | vap | | 2.908e-02 | -3.54 | Ribosomal\_S3Ae\_CS | interpro domains | IPR018281 | 1 | 1 | 11383 | 331 | RpS3A | | 2.908e-02 | -3.54 | RACK7\_Bromo\_dom | interpro domains | IPR037967 | 1 | 1 | 11383 | 331 | CG1815 | | 2.908e-02 | -3.54 | JmjC\_protein | interpro domains | IPR039994 | 1 | 1 | 11383 | 331 | NO66 | | 2.908e-02 | -3.54 | DUF4210 | interpro domains | IPR025261 | 1 | 1 | 11383 | 331 | CG9005 | | 2.908e-02 | -3.54 | Tetra\_PHTase | interpro domains | IPR003565 | 1 | 1 | 11383 | 331 | Apf | | 2.908e-02 | -3.54 | PHF5 | interpro domains | IPR005345 | 1 | 1 | 11383 | 331 | Phf5a | | 2.908e-02 | -3.54 | Exo84\_C | interpro domains | IPR032403 | 1 | 1 | 11383 | 331 | Exo84 | | 2.908e-02 | -3.54 | Pre-SET\_CXC | interpro domains | IPR041355 | 1 | 1 | 11383 | 331 | E(z) | | 2.908e-02 | -3.54 | TMEM147 | interpro domains | IPR019164 | 1 | 1 | 11383 | 331 | CG5861 | | 2.908e-02 | -3.54 | Release\_factor\_eRF1/aRF1\_N | interpro domains | IPR024049 | 1 | 1 | 11383 | 331 | eRF1 | | 2.908e-02 | -3.54 | Mn/Fe\_SOD\_BS | interpro domains | IPR019833 | 1 | 1 | 11383 | 331 | Sod2 | | 2.908e-02 | -3.54 | Kif23\_Arf-bd\_sf | interpro domains | IPR038105 | 1 | 1 | 11383 | 331 | pav | | 2.908e-02 | -3.54 | Purple\_acid\_Pase-like\_N | interpro domains | IPR008963 | 1 | 1 | 11383 | 331 | CG1637 | | 2.908e-02 | -3.54 | DAZAP1\_RRM1 | interpro domains | IPR034134 | 1 | 1 | 11383 | 331 | Hrb27C | | 2.908e-02 | -3.54 | Peptidase\_C13 | interpro domains | IPR001096 | 1 | 1 | 11383 | 331 | PIG-K | | 2.908e-02 | -3.54 | MED26 | interpro domains | IPR042376 | 1 | 1 | 11383 | 331 | MED26 | | 2.908e-02 | -3.54 | Rfa2 | interpro domains | IPR013970 | 1 | 1 | 11383 | 331 | RPA3 | | 2.908e-02 | -3.54 | Ribosomal\_S4/S9\_euk/arc | interpro domains | IPR005710 | 1 | 1 | 11383 | 331 | RpS9 | | 2.908e-02 | -3.54 | CWC25 | interpro domains | IPR022209 | 1 | 1 | 11383 | 331 | Cwc25 | | 2.908e-02 | -3.54 | Prot\_kin\_Mps1\_fam | interpro domains | IPR027084 | 1 | 1 | 11383 | 331 | Mps1 | | 2.908e-02 | -3.54 | Proteasome\_assmbl\_chaperone\_2 | interpro domains | IPR019151 | 1 | 1 | 11383 | 331 | CG12321 | | 2.908e-02 | -3.54 | Proteasome\_beta\_Pre3 | interpro domains | IPR037559 | 1 | 1 | 11383 | 331 | Prosbeta1 | | 2.908e-02 | -3.54 | Ribosomal\_L35Ae\_CS | interpro domains | IPR018266 | 1 | 1 | 11383 | 331 | RpL35A | | 2.908e-02 | -3.54 | PSD\_type1\_euk | interpro domains | IPR033661 | 1 | 1 | 11383 | 331 | Pisd | | 2.908e-02 | -3.54 | Purple\_acid\_PPase\_C\_dom | interpro domains | IPR025733 | 1 | 1 | 11383 | 331 | CG1637 | | 2.908e-02 | -3.54 | PHAT\_dom | interpro domains | IPR015327 | 1 | 1 | 11383 | 331 | smg | | 2.908e-02 | -3.54 | Translation\_control\_tumour\_CS | interpro domains | IPR018103 | 1 | 1 | 11383 | 331 | Tctp | | 2.908e-02 | -3.54 | Ribosomal\_S28\_mit | interpro domains | IPR019375 | 1 | 1 | 11383 | 331 | mRpS28 | | 2.908e-02 | -3.54 | SF3A3 | interpro domains | IPR031776 | 1 | 1 | 11383 | 331 | noi | | 2.908e-02 | -3.54 | Eaf6 | interpro domains | IPR015418 | 1 | 1 | 11383 | 331 | Eaf6 | | 2.908e-02 | -3.54 | VAPA | interpro domains | IPR030229 | 1 | 1 | 11383 | 331 | Vap33 | | 2.908e-02 | -3.54 | TFIIF\_beta\_N | interpro domains | IPR040504 | 1 | 1 | 11383 | 331 | TfIIFbeta | | 2.908e-02 | -3.54 | Trans\_elong\_IF5A\_hypusine\_site | interpro domains | IPR019769 | 1 | 1 | 11383 | 331 | eEF5 | | 2.908e-02 | -3.54 | Uncharacterised\_TMEM39 | interpro domains | IPR019397 | 1 | 1 | 11383 | 331 | CG13016 | | 2.908e-02 | -3.54 | OXLD1 | interpro domains | IPR039251 | 1 | 1 | 11383 | 331 | CG5500 | | 2.908e-02 | -3.54 | XTBD | interpro domains | IPR021859 | 1 | 1 | 11383 | 331 | CG31301 | | 2.908e-02 | -3.54 | Ribosomal\_S27/S33\_mit | interpro domains | IPR013219 | 1 | 1 | 11383 | 331 | mRpS33 | | 2.908e-02 | -3.54 | AP3\_beta | interpro domains | IPR026740 | 1 | 1 | 11383 | 331 | rb | | 2.908e-02 | -3.54 | PGAP1-like | interpro domains | IPR012908 | 1 | 1 | 11383 | 331 | PGAP1 | | 2.908e-02 | -3.54 | RhoGAP\_pG2 | interpro domains | IPR039006 | 1 | 1 | 11383 | 331 | RhoGAPp190 | | 2.908e-02 | -3.54 | KRTCAP2 | interpro domains | IPR018614 | 1 | 1 | 11383 | 331 | CG31460 | | 2.908e-02 | -3.54 | Znf\_PARP | interpro domains | IPR001510 | 1 | 1 | 11383 | 331 | Parp | | 2.908e-02 | -3.54 | RNA-bd\_8 | interpro domains | IPR008111 | 1 | 1 | 11383 | 331 | tsu | | 2.908e-02 | -3.54 | EIF1AD | interpro domains | IPR039294 | 1 | 1 | 11383 | 331 | CG31957 | | 2.908e-02 | -3.54 | Rbsml\_L15e\_core\_dom\_sf | interpro domains | IPR024794 | 1 | 1 | 11383 | 331 | RpL15 | | 2.908e-02 | -3.54 | MPP\_PAP | interpro domains | IPR041792 | 1 | 1 | 11383 | 331 | CG1637 | | 2.908e-02 | -3.54 | RNase\_H2\_suC | interpro domains | IPR013924 | 1 | 1 | 11383 | 331 | CG30105 | | 2.908e-02 | -3.54 | BUD31/G10-rel\_CS | interpro domains | IPR018230 | 1 | 1 | 11383 | 331 | l(1)10Bb | | 2.908e-02 | -3.54 | Exo84\_C\_2 | interpro domains | IPR042560 | 1 | 1 | 11383 | 331 | Exo84 | | 2.908e-02 | -3.54 | Smaug\_SAM | interpro domains | IPR037634 | 1 | 1 | 11383 | 331 | smg | | 2.908e-02 | -3.54 | UPF3\_dom | interpro domains | IPR005120 | 1 | 1 | 11383 | 331 | Upf3 | | 2.908e-02 | -3.54 | RagA/B | interpro domains | IPR039397 | 1 | 1 | 11383 | 331 | RagA-B | | 2.908e-02 | -3.54 | MLST8 | interpro domains | IPR037588 | 1 | 1 | 11383 | 331 | Lst8 | | 2.908e-02 | -3.54 | RasGAP\_N\_SH2 | interpro domains | IPR035841 | 1 | 1 | 11383 | 331 | vap | | 2.908e-02 | -3.54 | Initiation\_fac\_2B\_a/b/d | interpro domains | IPR011559 | 1 | 1 | 11383 | 331 | CG11334 | | 2.908e-02 | -3.54 | Ribosomal\_L14e\_dom | interpro domains | IPR002784 | 1 | 1 | 11383 | 331 | RpL14 | | 2.908e-02 | -3.54 | MS4A | interpro domains | IPR030417 | 1 | 1 | 11383 | 331 | CG17600 | | 2.908e-02 | -3.54 | ORC\_WH\_C | interpro domains | IPR040855 | 1 | 1 | 11383 | 331 | Orc3 | | 2.908e-02 | -3.54 | UNC119\_homologue\_A | interpro domains | IPR032977 | 1 | 1 | 11383 | 331 | unc-119 | | 2.908e-02 | -3.54 | Upf3 | interpro domains | IPR039722 | 1 | 1 | 11383 | 331 | Upf3 | | 2.908e-02 | -3.54 | Ald1\_epimerase\_CS | interpro domains | IPR018052 | 1 | 1 | 11383 | 331 | CG10467 | | 2.908e-02 | -3.54 | CCDC22 | interpro domains | IPR008530 | 1 | 1 | 11383 | 331 | CG9951 | | 2.908e-02 | -3.54 | Ribosomal\_L30\_bac-type | interpro domains | IPR005996 | 1 | 1 | 11383 | 331 | mRpL30 | | 2.908e-02 | -3.54 | PGAP1/BST1 | interpro domains | IPR039529 | 1 | 1 | 11383 | 331 | PGAP1 | | 2.908e-02 | -3.54 | Mak16 | interpro domains | IPR006958 | 1 | 1 | 11383 | 331 | Rbm13 | | 2.908e-02 | -3.54 | Chap\_CCT\_delta | interpro domains | IPR012717 | 1 | 1 | 11383 | 331 | CCT4 | | 2.908e-02 | -3.54 | TPD52 | interpro domains | IPR007327 | 1 | 1 | 11383 | 331 | CG5174 | | 2.908e-02 | -3.54 | DYRK1A/B\_MNB | interpro domains | IPR028318 | 1 | 1 | 11383 | 331 | mnb | | 2.908e-02 | -3.54 | Znf\_ZPR1 | interpro domains | IPR004457 | 1 | 1 | 11383 | 331 | Zpr1 | | 2.908e-02 | -3.54 | Poly(ADP-ribose)pol\_reg\_dom | interpro domains | IPR004102 | 1 | 1 | 11383 | 331 | Parp | | 2.908e-02 | -3.54 | EIF3B | interpro domains | IPR011400 | 1 | 1 | 11383 | 331 | eIF3b | | 2.908e-02 | -3.54 | IF5A-like | interpro domains | IPR001884 | 1 | 1 | 11383 | 331 | eEF5 | | 2.908e-02 | -3.54 | LysoPLipase\_patatin\_CS | interpro domains | IPR001423 | 1 | 1 | 11383 | 331 | sws | | 2.908e-02 | -3.54 | PUM3 | interpro domains | IPR040059 | 1 | 1 | 11383 | 331 | peng | | 2.908e-02 | -3.54 | ECIST\_C | interpro domains | IPR029342 | 1 | 1 | 11383 | 331 | ECSIT | | 2.908e-02 | -3.54 | Noc2 | interpro domains | IPR005343 | 1 | 1 | 11383 | 331 | CG9246 | | 2.908e-02 | -3.54 | MCRS1 | interpro domains | IPR037912 | 1 | 1 | 11383 | 331 | Rcd5 | | 2.908e-02 | -3.54 | Ribosomal\_S8e\_CS | interpro domains | IPR018283 | 1 | 1 | 11383 | 331 | RpS8 | | 2.908e-02 | -3.54 | MBF1\_N | interpro domains | IPR013729 | 1 | 1 | 11383 | 331 | mbf1 | | 2.908e-02 | -3.54 | Mn/Fe\_SOD\_N | interpro domains | IPR019831 | 1 | 1 | 11383 | 331 | Sod2 | | 2.908e-02 | -3.54 | CD20-like | interpro domains | IPR007237 | 1 | 1 | 11383 | 331 | CG17600 | | 2.908e-02 | -3.54 | PADR1\_dom\_sf | interpro domains | IPR038650 | 1 | 1 | 11383 | 331 | Parp | | 2.908e-02 | -3.54 | RhoGAP-FF1 | interpro domains | IPR032835 | 1 | 1 | 11383 | 331 | RhoGAPp190 | | 2.908e-02 | -3.54 | ORC3 | interpro domains | IPR020795 | 1 | 1 | 11383 | 331 | Orc3 | | 2.908e-02 | -3.54 | DNA\_mismatch\_repair\_Msh2 | interpro domains | IPR011184 | 1 | 1 | 11383 | 331 | spel1 | | 2.908e-02 | -3.54 | Transmembrane\_pr\_170 | interpro domains | IPR019334 | 1 | 1 | 11383 | 331 | CG12341 | | 2.908e-02 | -3.54 | TCF25/Rqc1 | interpro domains | IPR006994 | 1 | 1 | 11383 | 331 | Nulp1 | | 2.908e-02 | -3.54 | WGR\_dom\_sf | interpro domains | IPR036930 | 1 | 1 | 11383 | 331 | Parp | | 2.908e-02 | -3.54 | SOAR | interpro domains | IPR032393 | 1 | 1 | 11383 | 331 | Stim | | 2.908e-02 | -3.54 | Tudor\_FRX1 | interpro domains | IPR041560 | 1 | 1 | 11383 | 331 | Fmr1 | | 2.908e-02 | -3.54 | Vinculin-bd\_dom | interpro domains | IPR015009 | 1 | 1 | 11383 | 331 | rhea | | 2.909e-02 | -3.54 | GSE15930\_NAIVE\_VS\_72H\_IN\_VITRO\_STIM\_IFNAB\_CD8\_TCELL\_DN | MSigDB lists | GSE15930\_NAIVE\_VS\_72H\_IN\_VITRO\_STIM\_IFNAB\_CD8\_TCELL\_DN | 83 | 8 | 4161 | 184 | unc-119,CG5589,fzy,DNAlig1,pav,CG41378,E(z),CG5590 | | 2.921e-02 | -3.53 | endoplasmic reticulum | cellular component | GO:0005783 | 475 | 19 | 11940 | 298 | PPP1R15,CG31460,CG32576,Stim,sws,alpha-Man-Ib,RpS26,heix,CG17883,CG15168,PIG-K,Vap33,me31B,g,CG11137,xit,bai,PGAP1,Stt3B | | 2.924e-02 | -3.53 | pole plasm assembly | biological process | GO:0007315 | 70 | 5 | 12345 | 305 | mon2,tsu,sqd,Hrb27C,glo | | 2.924e-02 | -3.53 | negative regulation of phosphorylation | biological process | GO:0042326 | 70 | 5 | 12345 | 305 | sws,krz,Pten,flw,mnb | | 2.927e-02 | -3.53 | GSE13484\_3H\_UNSTIM\_VS\_YF17D\_VACCINE\_STIM\_PBMC\_DN | MSigDB lists | GSE13484\_3H\_UNSTIM\_VS\_YF17D\_VACCINE\_STIM\_PBMC\_DN | 27 | 4 | 4161 | 184 | E(z),Sod2,Cul4,sigmar | | 2.927e-02 | -3.53 | GO\_RNA\_SECONDARY\_STRUCTURE\_UNWINDING | MSigDB lists | GO\_RNA\_SECONDARY\_STRUCTURE\_UNWINDING | 27 | 4 | 4161 | 184 | me31B,Dbp80,CG5589,Dbp21E2 | | 2.927e-02 | -3.53 | GO\_MAINTENANCE\_OF\_LOCATION\_IN\_CELL | MSigDB lists | GO\_MAINTENANCE\_OF\_LOCATION\_IN\_CELL | 27 | 4 | 4161 | 184 | pins,CG9853,rhea,Vps13D | | 2.930e-02 | -3.53 | GO\_NUCLEOTIDE\_EXCISION\_REPAIR | MSigDB lists | GO\_NUCLEOTIDE\_EXCISION\_REPAIR | 68 | 7 | 4161 | 184 | CSN1b,Cul4,smt3,DNAlig1,CG17493,Rpb11,Parp | | 2.930e-02 | -3.53 | E2F4DP1\_01 | MSigDB lists | E2F4DP1\_01 | 68 | 7 | 4161 | 184 | Phf5a,E(z),His2Av,Stt3B,SF2,spel1,DNAlig1 | | 2.930e-02 | -3.53 | GSE9006\_TYPE\_1\_VS\_TYPE\_2\_DIABETES\_PBMC\_AT\_DX\_DN | MSigDB lists | GSE9006\_TYPE\_1\_VS\_TYPE\_2\_DIABETES\_PBMC\_AT\_DX\_DN | 68 | 7 | 4161 | 184 | Mdh1,Vps35,CG5339,CG8132,MICU1,Stlk,rb | | 2.930e-02 | -3.53 | GSE17721\_CTRL\_VS\_GARDIQUIMOD\_8H\_BMDC\_UP | MSigDB lists | GSE17721\_CTRL\_VS\_GARDIQUIMOD\_8H\_BMDC\_UP | 68 | 7 | 4161 | 184 | CG4866,Tsp97E,CG3842,RpS8,Rap1,CycA,RpS26 | | 2.937e-02 | -3.53 | RhoGAP | smart domains | SM00324 | 22 | 3 | 5086 | 158 | RhoGAPp190,conu,Graf | | 2.943e-02 | -3.53 | MLL1/2 complex | cellular component | GO:0044665 | 11 | 2 | 11940 | 298 | Rbbp5,Rcd5 | | 2.948e-02 | -3.52 | rRNA metabolic process | biological process | GO:0016072 | 147 | 8 | 12345 | 305 | mRpS9,CG32075,Nopp140,RpS8,CG4866,CG14805,CG11920,CG5589 | | 2.953e-02 | -3.52 | GO\_AMIDE\_BIOSYNTHETIC\_PROCESS | MSigDB lists | GO\_AMIDE\_BIOSYNTHETIC\_PROCESS | 290 | 20 | 4161 | 184 | Bmcp,RpS26,RpL14,eIF3g1,RpS8,mRpS9,CG31957,RpL35A,eRF1,RpS9,eEF1delta,RpL5,eIF3b,eIF3c,mRpS28,eEF5,RpL15,mRpS33,RpL36A,CG4743 | | 2.961e-02 | -3.52 | VEGFA-VEGFR2 Pathway | REACTOME pathways | R-DME-4420097 | 151 | 11 | 3923 | 151 | vap,Rpt6,Nf1,Prosbeta1,CG7054,14-3-3zeta,Cdc42,Lst8,Rpn10,Prosalpha2,CG4935 | | 2.961e-02 | -3.52 | Signaling by VEGF | REACTOME pathways | R-DME-194138 | 151 | 11 | 3923 | 151 | Prosbeta1,14-3-3zeta,CG7054,vap,Rpt6,Nf1,Cdc42,Lst8,CG4935,Rpn10,Prosalpha2 | | 2.967e-02 | -3.52 | GSE22432\_PDC\_VS\_TGFB1\_TREATEDCOMMON\_DC\_PROGENITOR\_UP | MSigDB lists | GSE22432\_PDC\_VS\_TGFB1\_TREATEDCOMMON\_DC\_PROGENITOR\_UP | 115 | 10 | 4161 | 184 | Tim17b,Mdh1,CG15168,CG5535,Phf5a,CG8635,CG4935,CG10343,TfIIFbeta,Rap1 | | 2.969e-02 | -3.52 | positive regulation of RNA metabolic process | biological process | GO:0051254 | 513 | 20 | 12345 | 305 | Dref,Lst8,smg,TfIIFbeta,MED21,mnb,CG6244,grau,Hrb27C,AGO3,Rpt6,Rcd5,d4,su(Hw),dgrn,Ncoa6,nsl1,rhi,mbf1,CtBP | | 2.976e-02 | -3.51 | organelle assembly | biological process | GO:0070925 | 385 | 16 | 12345 | 305 | Pp1-87B,CG17493,Fmr1,RpL5,RpS19a,Rab7,Nph,me31B,Rab14,Rcd5,pav,flr,unc-119,msd5,Dbp21E2,rhea | | 2.979e-02 | -3.51 | cell population proliferation | biological process | GO:0008283 | 175 | 9 | 12345 | 305 | CSN1b,pll,CycA,nsl1,pins,eIF4B,Fmr1,Pten,Cdc42 | | 2.983e-02 | -3.51 | MAP2K and MAPK activation | REACTOME pathways | R-DME-5674135 | 18 | 3 | 3923 | 151 | CG7054,14-3-3zeta,CG4935 | | 2.987e-02 | -3.51 | dos (daughter of sevenless) | protein interactions | 38321 | 54 | 5 | 9204 | 298 | 14-3-3zeta,RAF2,pins,Ythdf,glo | | 3.001e-02 | -3.51 | MORF\_IKBKG | MSigDB lists | MORF\_IKBKG | 40 | 5 | 4161 | 184 | rb,sm,Rcd5,tsu,Aps | | 3.001e-02 | -3.51 | GSE26351\_WNT\_VS\_BMP\_PATHWAY\_STIM\_HEMATOPOIETIC\_PROGENITORS\_UP | MSigDB lists | GSE26351\_WNT\_VS\_BMP\_PATHWAY\_STIM\_HEMATOPOIETIC\_PROGENITORS\_UP | 40 | 5 | 4161 | 184 | CG8860,Rat1,Rap1,RpL36A,LIMK1 | | 3.001e-02 | -3.51 | GO\_ERAD\_PATHWAY | MSigDB lists | GO\_ERAD\_PATHWAY | 40 | 5 | 4161 | 184 | alpha-Man-Ib,CG9853,Rpt6,Ubqn,Stt3B | | 3.001e-02 | -3.51 | STEARMAN\_LUNG\_CANCER\_EARLY\_VS\_LATE\_UP | MSigDB lists | STEARMAN\_LUNG\_CANCER\_EARLY\_VS\_LATE\_UP | 40 | 5 | 4161 | 184 | His2Av,Art1,Arpc3A,vap,SF2 | | 3.001e-02 | -3.51 | GSE32986\_UNSTIM\_VS\_GMCSF\_STIM\_DC\_UP | MSigDB lists | GSE32986\_UNSTIM\_VS\_GMCSF\_STIM\_DC\_UP | 40 | 5 | 4161 | 184 | Sgt,CG7766,Pex3,RpL35A,RpL5 | | 3.015e-02 | -3.50 | oocyte anterior/posterior axis specification | biological process | GO:0007314 | 95 | 6 | 12345 | 305 | sqd,Hrb27C,tsu,lic,mon2,glo | | 3.020e-02 | -3.50 | Interleukin-2 signaling | REACTOME pathways | R-DME-451927 | 114 | 9 | 3923 | 151 | CG4935,Prosalpha2,Rpn10,vap,Rpt6,Nf1,Prosbeta1,CG7054,14-3-3zeta | | 3.020e-02 | -3.50 | MAPK1/MAPK3 signaling | REACTOME pathways | R-DME-5684996 | 114 | 9 | 3923 | 151 | Nf1,Rpt6,vap,14-3-3zeta,CG7054,Prosbeta1,CG4935,Rpn10,Prosalpha2 | | 3.020e-02 | -3.50 | Signaling by Leptin | REACTOME pathways | R-DME-2586552 | 114 | 9 | 3923 | 151 | Rpn10,CG4935,Prosalpha2,14-3-3zeta,CG7054,Prosbeta1,Nf1,Rpt6,vap | | 3.035e-02 | -3.50 | regulation of multicellular organism growth | biological process | GO:0040014 | 48 | 4 | 12345 | 305 | lic,Nf1,Tctp,Pten | | 3.035e-02 | -3.50 | exocytic process | biological process | GO:0140029 | 48 | 4 | 12345 | 305 | Rab8,Exo84,Rop,Vap33 | | 3.035e-02 | -3.49 | preribosome | cellular component | GO:0030684 | 70 | 5 | 11940 | 298 | CG9246,CG32075,Rbm13,CG11920,CG4866 | | 3.044e-02 | -3.49 | Ribosomal\_S3Ae | pfam domains | PF01015 | 1 | 1 | 10480 | 319 | RpS3A | | 3.044e-02 | -3.49 | EloA-BP1 | pfam domains | PF15870 | 1 | 1 | 10480 | 319 | CG12877 | | 3.044e-02 | -3.49 | Sod\_Fe\_N | pfam domains | PF00081 | 1 | 1 | 10480 | 319 | Sod2 | | 3.044e-02 | -3.49 | G10 | pfam domains | PF01125 | 1 | 1 | 10480 | 319 | l(1)10Bb | | 3.044e-02 | -3.49 | DUF2053 | pfam domains | PF09767 | 1 | 1 | 10480 | 319 | CG5861 | | 3.044e-02 | -3.49 | ORC3\_N | pfam domains | PF07034 | 1 | 1 | 10480 | 319 | Orc3 | | 3.044e-02 | -3.49 | ECSIT\_C | pfam domains | PF14784 | 1 | 1 | 10480 | 319 | ECSIT | | 3.044e-02 | -3.49 | MCRS\_N | pfam domains | PF13325 | 1 | 1 | 10480 | 319 | Rcd5 | | 3.044e-02 | -3.49 | zf-SNAP50\_C | pfam domains | PF12251 | 1 | 1 | 10480 | 319 | Pbp49 | | 3.044e-02 | -3.49 | MKLP1\_Arf\_bdg | pfam domains | PF16540 | 1 | 1 | 10480 | 319 | pav | | 3.044e-02 | -3.49 | PDZ\_5 | pfam domains | PF17817 | 1 | 1 | 10480 | 319 | Spn | | 3.044e-02 | -3.49 | SF3a60\_bindingd | pfam domains | PF12108 | 1 | 1 | 10480 | 319 | noi | | 3.044e-02 | -3.49 | KH\_9 | pfam domains | PF17904 | 1 | 1 | 10480 | 319 | Fmr1 | | 3.044e-02 | -3.49 | Metallophos\_C | pfam domains | PF14008 | 1 | 1 | 10480 | 319 | CG1637 | | 3.044e-02 | -3.49 | VBS | pfam domains | PF08913 | 1 | 1 | 10480 | 319 | rhea | | 3.044e-02 | -3.49 | Ribosomal\_L35Ae | pfam domains | PF01247 | 1 | 1 | 10480 | 319 | RpL35A | | 3.044e-02 | -3.49 | PAC2 | pfam domains | PF09754 | 1 | 1 | 10480 | 319 | CG12321 | | 3.044e-02 | -3.49 | RNase\_H2\_suC | pfam domains | PF08615 | 1 | 1 | 10480 | 319 | CG30105 | | 3.044e-02 | -3.49 | MMgT | pfam domains | PF10270 | 1 | 1 | 10480 | 319 | CG15168 | | 3.044e-02 | -3.49 | Cwf\_Cwc\_15 | pfam domains | PF04889 | 1 | 1 | 10480 | 319 | c12.1 | | 3.044e-02 | -3.49 | Talin\_middle | pfam domains | PF09141 | 1 | 1 | 10480 | 319 | rhea | | 3.044e-02 | -3.49 | CPL | pfam domains | PF08144 | 1 | 1 | 10480 | 319 | peng | | 3.044e-02 | -3.49 | DUF3449 | pfam domains | PF11931 | 1 | 1 | 10480 | 319 | noi | | 3.044e-02 | -3.49 | DUF758 | pfam domains | PF05527 | 1 | 1 | 10480 | 319 | sigmar | | 3.044e-02 | -3.49 | FXMRP1\_C\_core | pfam domains | PF12235 | 1 | 1 | 10480 | 319 | Fmr1 | | 3.044e-02 | -3.49 | Peroxin-3 | pfam domains | PF04882 | 1 | 1 | 10480 | 319 | Pex3 | | 3.044e-02 | -3.49 | Tmemb\_170 | pfam domains | PF10190 | 1 | 1 | 10480 | 319 | CG12341 | | 3.044e-02 | -3.49 | DUF4210 | pfam domains | PF13915 | 1 | 1 | 10480 | 319 | CG9005 | | 3.044e-02 | -3.49 | Tmp39 | pfam domains | PF10271 | 1 | 1 | 10480 | 319 | CG13016 | | 3.044e-02 | -3.49 | PRC2\_HTH\_1 | pfam domains | PF18118 | 1 | 1 | 10480 | 319 | E(z) | | 3.044e-02 | -3.49 | TMEM65 | pfam domains | PF10507 | 1 | 1 | 10480 | 319 | CG17715 | | 3.044e-02 | -3.49 | Exo84\_C | pfam domains | PF16528 | 1 | 1 | 10480 | 319 | Exo84 | | 3.044e-02 | -3.49 | MBF1 | pfam domains | PF08523 | 1 | 1 | 10480 | 319 | mbf1 | | 3.044e-02 | -3.49 | Tcf25 | pfam domains | PF04910 | 1 | 1 | 10480 | 319 | Nulp1 | | 3.044e-02 | -3.49 | Ribosomal\_L5e | pfam domains | PF17144 | 1 | 1 | 10480 | 319 | RpL5 | | 3.044e-02 | -3.49 | ORC\_WH\_C | pfam domains | PF18137 | 1 | 1 | 10480 | 319 | Orc3 | | 3.044e-02 | -3.49 | Ribosomal\_L15e | pfam domains | PF00827 | 1 | 1 | 10480 | 319 | RpL15 | | 3.044e-02 | -3.49 | preSET\_CXC | pfam domains | PF18264 | 1 | 1 | 10480 | 319 | E(z) | | 3.044e-02 | -3.49 | HTH\_3 | pfam domains | PF01381 | 1 | 1 | 10480 | 319 | mbf1 | | 3.044e-02 | -3.49 | TFIIF\_beta\_N | pfam domains | PF17683 | 1 | 1 | 10480 | 319 | TfIIFbeta | | 3.044e-02 | -3.49 | zf-ZPR1 | pfam domains | PF03367 | 1 | 1 | 10480 | 319 | Zpr1 | | 3.044e-02 | -3.49 | DREV | pfam domains | PF05219 | 1 | 1 | 10480 | 319 | CG5339 | | 3.044e-02 | -3.49 | CENP-T\_C | pfam domains | PF15511 | 1 | 1 | 10480 | 319 | His4r | | 3.044e-02 | -3.49 | PARP\_reg | pfam domains | PF02877 | 1 | 1 | 10480 | 319 | Parp | | 3.044e-02 | -3.49 | PHAT | pfam domains | PF09246 | 1 | 1 | 10480 | 319 | smg | | 3.044e-02 | -3.49 | DUF4528 | pfam domains | PF15031 | 1 | 1 | 10480 | 319 | CG34148 | | 3.044e-02 | -3.49 | Cas1\_AcylT | pfam domains | PF07779 | 1 | 1 | 10480 | 319 | CG2938 | | 3.044e-02 | -3.49 | Dynactin\_p22 | pfam domains | PF07426 | 1 | 1 | 10480 | 319 | DCTN3-p24 | | 3.044e-02 | -3.49 | MGC-24 | pfam domains | PF05283 | 1 | 1 | 10480 | 319 | vsg | | 3.044e-02 | -3.49 | DUF4516 | pfam domains | PF14990 | 1 | 1 | 10480 | 319 | CG42308 | | 3.044e-02 | -3.49 | Nup188 | pfam domains | PF10487 | 1 | 1 | 10480 | 319 | Nup188 | | 3.044e-02 | -3.49 | WAC\_Acf1\_DNA\_bd | pfam domains | PF10537 | 1 | 1 | 10480 | 319 | Acf | | 3.044e-02 | -3.49 | RhoGAP-FF1 | pfam domains | PF16512 | 1 | 1 | 10480 | 319 | RhoGAPp190 | | 3.044e-02 | -3.49 | Noc2 | pfam domains | PF03715 | 1 | 1 | 10480 | 319 | CG9246 | | 3.044e-02 | -3.49 | PS\_Dcarbxylase | pfam domains | PF02666 | 1 | 1 | 10480 | 319 | Pisd | | 3.044e-02 | -3.49 | Not3 | pfam domains | PF04065 | 1 | 1 | 10480 | 319 | Not3 | | 3.044e-02 | -3.49 | SRP40\_C | pfam domains | PF05022 | 1 | 1 | 10480 | 319 | Nopp140 | | 3.044e-02 | -3.49 | MRP-S33 | pfam domains | PF08293 | 1 | 1 | 10480 | 319 | mRpS33 | | 3.044e-02 | -3.49 | CD20 | pfam domains | PF04103 | 1 | 1 | 10480 | 319 | CG17600 | | 3.044e-02 | -3.49 | TCTP | pfam domains | PF00838 | 1 | 1 | 10480 | 319 | Tctp | | 3.044e-02 | -3.49 | MRP-S35 | pfam domains | PF10246 | 1 | 1 | 10480 | 319 | mRpS28 | | 3.044e-02 | -3.49 | Ribosomal\_L44 | pfam domains | PF00935 | 1 | 1 | 10480 | 319 | RpL36A | | 3.044e-02 | -3.49 | Ribosomal\_L18\_c | pfam domains | PF14204 | 1 | 1 | 10480 | 319 | RpL5 | | 3.044e-02 | -3.49 | PGAP1 | pfam domains | PF07819 | 1 | 1 | 10480 | 319 | PGAP1 | | 3.044e-02 | -3.49 | NDUF\_B8 | pfam domains | PF05821 | 1 | 1 | 10480 | 319 | ND-ASHI | | 3.044e-02 | -3.49 | Oxidored-like | pfam domains | PF09791 | 1 | 1 | 10480 | 319 | CG5500 | | 3.044e-02 | -3.49 | DUF410 | pfam domains | PF04190 | 1 | 1 | 10480 | 319 | CG9853 | | 3.044e-02 | -3.49 | Ubie\_methyltran | pfam domains | PF01209 | 1 | 1 | 10480 | 319 | Coq5 | | 3.044e-02 | -3.49 | FPL | pfam domains | PF09758 | 1 | 1 | 10480 | 319 | ema | | 3.044e-02 | -3.49 | Peptidase\_C13 | pfam domains | PF01650 | 1 | 1 | 10480 | 319 | PIG-K | | 3.044e-02 | -3.49 | NuA4 | pfam domains | PF09340 | 1 | 1 | 10480 | 319 | Eaf6 | | 3.044e-02 | -3.49 | TMEM223 | pfam domains | PF14640 | 1 | 1 | 10480 | 319 | CG12935 | | 3.044e-02 | -3.49 | CIA30 | pfam domains | PF08547 | 1 | 1 | 10480 | 319 | CIA30 | | 3.044e-02 | -3.49 | Med21 | pfam domains | PF11221 | 1 | 1 | 10480 | 319 | MED21 | | 3.044e-02 | -3.49 | XTBD | pfam domains | PF11952 | 1 | 1 | 10480 | 319 | CG31301 | | 3.044e-02 | -3.49 | WGR | pfam domains | PF05406 | 1 | 1 | 10480 | 319 | Parp | | 3.044e-02 | -3.49 | TMEM18 | pfam domains | PF14770 | 1 | 1 | 10480 | 319 | Tmem18 | | 3.044e-02 | -3.49 | Rep\_fac-A\_3 | pfam domains | PF08661 | 1 | 1 | 10480 | 319 | RPA3 | | 3.044e-02 | -3.49 | Mak16 | pfam domains | PF04874 | 1 | 1 | 10480 | 319 | Rbm13 | | 3.044e-02 | -3.49 | CWC25 | pfam domains | PF12542 | 1 | 1 | 10480 | 319 | Cwc25 | | 3.044e-02 | -3.49 | Nkap\_C | pfam domains | PF06047 | 1 | 1 | 10480 | 319 | CG6066 | | 3.044e-02 | -3.49 | eIF-3c\_N | pfam domains | PF05470 | 1 | 1 | 10480 | 319 | eIF3c | | 3.044e-02 | -3.49 | Cupin\_4 | pfam domains | PF08007 | 1 | 1 | 10480 | 319 | NO66 | | 3.044e-02 | -3.49 | TPD52 | pfam domains | PF04201 | 1 | 1 | 10480 | 319 | CG5174 | | 3.044e-02 | -3.49 | Ribosomal\_S26e | pfam domains | PF01283 | 1 | 1 | 10480 | 319 | RpS26 | | 3.044e-02 | -3.49 | zf-PARP | pfam domains | PF00645 | 1 | 1 | 10480 | 319 | Parp | | 3.044e-02 | -3.49 | Pinin\_SDK\_memA | pfam domains | PF04696 | 1 | 1 | 10480 | 319 | Pnn | | 3.044e-02 | -3.49 | PADR1 | pfam domains | PF08063 | 1 | 1 | 10480 | 319 | Parp | | 3.044e-02 | -3.49 | SGTA\_dimer | pfam domains | PF16546 | 1 | 1 | 10480 | 319 | Sgt | | 3.044e-02 | -3.49 | Ribosomal\_L14e | pfam domains | PF01929 | 1 | 1 | 10480 | 319 | RpL14 | | 3.044e-02 | -3.49 | Ribosomal\_L34 | pfam domains | PF00468 | 1 | 1 | 10480 | 319 | mRpL34 | | 3.044e-02 | -3.49 | Sod\_Fe\_C | pfam domains | PF02777 | 1 | 1 | 10480 | 319 | Sod2 | | 3.044e-02 | -3.49 | SOAR | pfam domains | PF16533 | 1 | 1 | 10480 | 319 | Stim | | 3.044e-02 | -3.49 | PAXNEB | pfam domains | PF05625 | 1 | 1 | 10480 | 319 | CG6907 | | 3.044e-02 | -3.49 | Mon2\_C | pfam domains | PF16206 | 1 | 1 | 10480 | 319 | mon2 | | 3.044e-02 | -3.49 | AP3B1\_C | pfam domains | PF14796 | 1 | 1 | 10480 | 319 | rb | | 3.044e-02 | -3.49 | PHF5 | pfam domains | PF03660 | 1 | 1 | 10480 | 319 | Phf5a | | 3.044e-02 | -3.49 | Keratin\_assoc | pfam domains | PF09775 | 1 | 1 | 10480 | 319 | CG31460 | | 3.044e-02 | -3.49 | DUF4743 | pfam domains | PF15916 | 1 | 1 | 10480 | 319 | CG12567 | | 3.044e-02 | -3.49 | DUF812 | pfam domains | PF05667 | 1 | 1 | 10480 | 319 | CG9951 | | 3.044e-02 | -3.49 | DUF1077 | pfam domains | PF06417 | 1 | 1 | 10480 | 319 | CG11137 | | 3.044e-02 | -3.49 | Chromosome\_seg | pfam domains | PF13889 | 1 | 1 | 10480 | 319 | CG9005 | | 3.044e-02 | -3.49 | Sde2\_N\_Ubi | pfam domains | PF13019 | 1 | 1 | 10480 | 319 | CG5986 | | 3.044e-02 | -3.49 | Pur\_ac\_phosph\_N | pfam domains | PF16656 | 1 | 1 | 10480 | 319 | CG1637 | | 3.044e-02 | -3.49 | eIF-5a | pfam domains | PF01287 | 1 | 1 | 10480 | 319 | eEF5 | | 3.044e-02 | -3.49 | Vps35 | pfam domains | PF03635 | 1 | 1 | 10480 | 319 | Vps35 | | 3.044e-02 | -3.49 | TFIIF\_beta | pfam domains | PF02270 | 1 | 1 | 10480 | 319 | TfIIFbeta | | 3.044e-02 | -3.49 | SF3A3 | pfam domains | PF16837 | 1 | 1 | 10480 | 319 | noi | | 3.044e-02 | -3.49 | DUF2228 | pfam domains | PF10228 | 1 | 1 | 10480 | 319 | CG1218 | | 3.044e-02 | -3.49 | Tudor\_FRX1 | pfam domains | PF18336 | 1 | 1 | 10480 | 319 | Fmr1 | | 3.044e-02 | -3.49 | Mlf1IP | pfam domains | PF10248 | 1 | 1 | 10480 | 319 | Mlf | | 3.044e-02 | -3.49 | Drc1-Sld2 | pfam domains | PF11719 | 1 | 1 | 10480 | 319 | RecQ4 | | 3.044e-02 | -3.49 | PDZ\_2 | pfam domains | PF13180 | 1 | 1 | 10480 | 319 | HtrA2 | | 3.044e-02 | -3.49 | P16-Arc | pfam domains | PF04699 | 1 | 1 | 10480 | 319 | Arpc5 | | 3.044e-02 | -3.49 | Vps55 | pfam domains | PF04133 | 1 | 1 | 10480 | 319 | CG30423 | | 3.044e-02 | -3.49 | Smg4\_UPF3 | pfam domains | PF03467 | 1 | 1 | 10480 | 319 | Upf3 | | 3.044e-02 | -3.49 | ECSIT | pfam domains | PF06239 | 1 | 1 | 10480 | 319 | ECSIT | | 3.044e-02 | -3.49 | PP1c\_bdg | pfam domains | PF10488 | 1 | 1 | 10480 | 319 | PPP1R15 | | 3.044e-02 | -3.49 | Scm3 | pfam domains | PF10384 | 1 | 1 | 10480 | 319 | cal1 | | 3.044e-02 | -3.49 | Las1 | pfam domains | PF04031 | 1 | 1 | 10480 | 319 | CG32075 | | 3.048e-02 | -3.49 | histone acetyltransferase activity | molecular function | GO:0004402 | 50 | 4 | 12176 | 289 | Acf,Rcd5,d4,Eaf6 | | 3.052e-02 | -3.49 | GSE37416\_0H\_VS\_12H\_F\_TULARENSIS\_LVS\_NEUTROPHIL\_UP | MSigDB lists | GSE37416\_0H\_VS\_12H\_F\_TULARENSIS\_LVS\_NEUTROPHIL\_UP | 54 | 6 | 4161 | 184 | Usp32,Graf,CG7766,CG15168,Rat1,CG5590 | | 3.052e-02 | -3.49 | GSE22886\_NAIVE\_TCELL\_VS\_MONOCYTE\_DN | MSigDB lists | GSE22886\_NAIVE\_TCELL\_VS\_MONOCYTE\_DN | 54 | 6 | 4161 | 184 | CG5339,Pisd,Graf,CG41378,CtBP,VhaPPA1-1 | | 3.052e-02 | -3.49 | GSE22886\_NAIVE\_CD4\_TCELL\_VS\_MONOCYTE\_DN | MSigDB lists | GSE22886\_NAIVE\_CD4\_TCELL\_VS\_MONOCYTE\_DN | 54 | 6 | 4161 | 184 | CtBP,VhaPPA1-1,CG5339,CG41378,Pisd,Graf | | 3.052e-02 | -3.49 | GSE3982\_BASOPHIL\_VS\_NKCELL\_UP | MSigDB lists | GSE3982\_BASOPHIL\_VS\_NKCELL\_UP | 54 | 6 | 4161 | 184 | sigmar,Tctp,CIA30,14-3-3zeta,Skp2,Apf | | 3.052e-02 | -3.49 | GSE15330\_MEGAKARYOCYTE\_ERYTHROID\_VS\_GRANULOCYTE\_MONOCYTE\_PROGENITOR\_DN | MSigDB lists | GSE15330\_MEGAKARYOCYTE\_ERYTHROID\_VS\_GRANULOCYTE\_MONOCYTE\_PROGENITOR\_DN | 54 | 6 | 4161 | 184 | Ufd4,Ubqn,lic,Bmcp,Exo84,MED21 | | 3.052e-02 | -3.49 | GSE30971\_WBP7\_HET\_VS\_KO\_MACROPHAGE\_UP | MSigDB lists | GSE30971\_WBP7\_HET\_VS\_KO\_MACROPHAGE\_UP | 54 | 6 | 4161 | 184 | pav,Arpc3A,cta,rb,Arpc5,CG5589 | | 3.052e-02 | -3.49 | GO\_REGULATION\_OF\_NUCLEAR\_DIVISION | MSigDB lists | GO\_REGULATION\_OF\_NUCLEAR\_DIVISION | 54 | 6 | 4161 | 184 | spel1,CG12321,Cdc42,fzy,Pten,AGO3 | | 3.053e-02 | -3.49 | positive regulation of cell development | biological process | GO:0010720 | 148 | 8 | 12345 | 305 | Rop,Orc3,Hrb27C,Rap1,Fmr1,PlexA,mnb,Rheb | | 3.059e-02 | -3.49 | G10\_1 | prosite domains | PS00997 | 1 | 1 | 6243 | 191 | l(1)10Bb | | 3.059e-02 | -3.49 | HISTONE\_H4 | prosite domains | PS00047 | 1 | 1 | 6243 | 191 | His4r | | 3.059e-02 | -3.49 | SOD\_MN | prosite domains | PS00088 | 1 | 1 | 6243 | 191 | Sod2 | | 3.059e-02 | -3.49 | SAM\_MT\_UBIE | prosite domains | PS51608 | 1 | 1 | 6243 | 191 | Coq5 | | 3.059e-02 | -3.49 | RIBOSOMAL\_S3AE | prosite domains | PS01191 | 1 | 1 | 6243 | 191 | RpS3A | | 3.059e-02 | -3.49 | PARP\_ZN\_FINGER\_1 | prosite domains | PS00347 | 1 | 1 | 6243 | 191 | Parp | | 3.059e-02 | -3.49 | G10\_2 | prosite domains | PS00998 | 1 | 1 | 6243 | 191 | l(1)10Bb | | 3.059e-02 | -3.49 | PG1 | prosite domains | PS51852 | 1 | 1 | 6243 | 191 | RhoGAPp190 | | 3.059e-02 | -3.49 | PARP\_ALPHA\_HD | prosite domains | PS51060 | 1 | 1 | 6243 | 191 | Parp | | 3.059e-02 | -3.49 | PARP\_ZN\_FINGER\_2 | prosite domains | PS50064 | 1 | 1 | 6243 | 191 | Parp | | 3.059e-02 | -3.49 | RIBOSOMAL\_S4 | prosite domains | PS00632 | 1 | 1 | 6243 | 191 | RpS9 | | 3.059e-02 | -3.49 | RIBOSOMAL\_L15E | prosite domains | PS01194 | 1 | 1 | 6243 | 191 | RpL15 | | 3.059e-02 | -3.49 | UPF0028 | prosite domains | PS01237 | 1 | 1 | 6243 | 191 | sws | | 3.059e-02 | -3.49 | TCTP\_2 | prosite domains | PS01003 | 1 | 1 | 6243 | 191 | Tctp | | 3.059e-02 | -3.49 | UBIE\_1 | prosite domains | PS01183 | 1 | 1 | 6243 | 191 | Coq5 | | 3.059e-02 | -3.49 | PG2 | prosite domains | PS51853 | 1 | 1 | 6243 | 191 | RhoGAPp190 | | 3.059e-02 | -3.49 | RIBOSOMAL\_L35AE | prosite domains | PS01105 | 1 | 1 | 6243 | 191 | RpL35A | | 3.059e-02 | -3.49 | CXC | prosite domains | PS51633 | 1 | 1 | 6243 | 191 | E(z) | | 3.059e-02 | -3.49 | DBINO | prosite domains | PS51413 | 1 | 1 | 6243 | 191 | Ino80 | | 3.059e-02 | -3.49 | TCTP\_1 | prosite domains | PS01002 | 1 | 1 | 6243 | 191 | Tctp | | 3.059e-02 | -3.49 | HTH\_CROC1 | prosite domains | PS50943 | 1 | 1 | 6243 | 191 | mbf1 | | 3.059e-02 | -3.49 | RIBOSOMAL\_S26E | prosite domains | PS00733 | 1 | 1 | 6243 | 191 | RpS26 | | 3.059e-02 | -3.49 | IF5A\_HYPUSINE | prosite domains | PS00302 | 1 | 1 | 6243 | 191 | eEF5 | | 3.059e-02 | -3.49 | UBIE\_2 | prosite domains | PS01184 | 1 | 1 | 6243 | 191 | Coq5 | | 3.059e-02 | -3.49 | WAC | prosite domains | PS51136 | 1 | 1 | 6243 | 191 | Acf | | 3.059e-02 | -3.49 | ALDOSE\_1\_EPIMERASE | prosite domains | PS00545 | 1 | 1 | 6243 | 191 | CG10467 | | 3.059e-02 | -3.49 | GLYCOPROTEASE | prosite domains | PS01016 | 1 | 1 | 6243 | 191 | Tcs3 | | 3.059e-02 | -3.49 | AGENET\_LIKE | prosite domains | PS51641 | 1 | 1 | 6243 | 191 | Fmr1 | | 3.059e-02 | -3.49 | TCTP\_3 | prosite domains | PS51797 | 1 | 1 | 6243 | 191 | Tctp | | 3.059e-02 | -3.49 | RIBOSOMAL\_S8E | prosite domains | PS01193 | 1 | 1 | 6243 | 191 | RpS8 | | 3.059e-02 | -3.49 | XTBD | prosite domains | PS51827 | 1 | 1 | 6243 | 191 | CG31301 | | 3.059e-02 | -3.49 | RIBOSOMAL\_L44E | prosite domains | PS01172 | 1 | 1 | 6243 | 191 | RpL36A | | 3.060e-02 | -3.49 | protein kinase activity | molecular function | GO:0004672 | 274 | 12 | 12176 | 289 | lic,Mps1,Bub1,Btk29A,CG7766,tkv,pll,CycA,Cdk8,LIMK1,mnb,Stlk | | 3.064e-02 | -3.49 | - | gene3d domains | 3.30.1740.20 | 1 | 1 | 4929 | 151 | RpS26 | | 3.064e-02 | -3.49 | - | gene3d domains | 1.20.58.1220 | 1 | 1 | 4929 | 151 | Exo84 | | 3.064e-02 | -3.49 | - | gene3d domains | 2.20.25.420 | 1 | 1 | 4929 | 151 | Zpr1 | | 3.064e-02 | -3.49 | - | gene3d domains | 1.20.1440.160 | 1 | 1 | 4929 | 151 | sigmar | | 3.064e-02 | -3.49 | - | gene3d domains | 1.25.40.190 | 1 | 1 | 4929 | 151 | Arpc5 | | 3.064e-02 | -3.49 | - | gene3d domains | 2.40.10.190 | 1 | 1 | 4929 | 151 | RpL35A | | 3.064e-02 | -3.49 | - | gene3d domains | 3.30.960.10 | 1 | 1 | 4929 | 151 | eRF1 | | 3.064e-02 | -3.49 | - | gene3d domains | 2.40.500.20 | 1 | 1 | 4929 | 151 | Sod2 | | 3.064e-02 | -3.49 | - | gene3d domains | 1.10.168.20 | 1 | 1 | 4929 | 151 | RpS8 | | 3.064e-02 | -3.49 | - | gene3d domains | 3.40.50.10900 | 1 | 1 | 4929 | 151 | CG12321 | | 3.064e-02 | -3.49 | - | gene3d domains | 3.30.1740.10 | 1 | 1 | 4929 | 151 | Parp | | 3.064e-02 | -3.49 | - | gene3d domains | 1.20.142.10 | 1 | 1 | 4929 | 151 | Parp | | 3.064e-02 | -3.49 | - | gene3d domains | 2.60.120.1040 | 1 | 1 | 4929 | 151 | Zpr1 | | 3.064e-02 | -3.49 | - | gene3d domains | 1.25.40.660 | 1 | 1 | 4929 | 151 | Vps35 | | 3.064e-02 | -3.49 | - | gene3d domains | 3.40.50.12000 | 1 | 1 | 4929 | 151 | pav | | 3.064e-02 | -3.49 | - | gene3d domains | 2.60.40.380 | 1 | 1 | 4929 | 151 | CG1637 | | 3.064e-02 | -3.49 | - | gene3d domains | 3.40.1120.10 | 1 | 1 | 4929 | 151 | RpL15 | | 3.064e-02 | -3.49 | - | gene3d domains | 1.25.40.170 | 1 | 1 | 4929 | 151 | smg | | 3.064e-02 | -3.49 | - | gene3d domains | 2.20.140.10 | 1 | 1 | 4929 | 151 | Parp | | 3.064e-02 | -3.49 | - | gene3d domains | 1.10.287.990 | 1 | 1 | 4929 | 151 | Sod2 | | 3.064e-02 | -3.49 | - | gene3d domains | 2.20.25.630 | 1 | 1 | 4929 | 151 | Parp | | 3.064e-02 | -3.49 | - | gene3d domains | 1.20.120.420 | 1 | 1 | 4929 | 151 | CG11334 | | 3.068e-02 | -3.48 | Transport of Mature mRNA derived from an Intron-Containing Transcript | REACTOME pathways | R-DME-159236 | 62 | 6 | 3923 | 151 | tsu,CG6961,Nup188,SF2,x16,Upf3 | | 3.068e-02 | -3.48 | Transport of Mature Transcript to Cytoplasm | REACTOME pathways | R-DME-72202 | 62 | 6 | 3923 | 151 | SF2,x16,Upf3,tsu,CG6961,Nup188 | | 3.073e-02 | -3.48 | MED6 (Mediator complex subunit 6) | protein interactions | 44728 | 37 | 4 | 9204 | 298 | MED22,Cdk8,Cpsf6,CG34159 | | 3.073e-02 | -3.48 | Imp (IGF-II mRNA-binding protein) | protein interactions | 32009 | 37 | 4 | 9204 | 298 | sqd,Rheb,Hrb27C,His2Av | | 3.073e-02 | -3.48 | ana2 (anastral spindle 2) | protein interactions | 35878 | 37 | 4 | 9204 | 298 | Ubqn,Pten,Slmap,Fgop2 | | 3.073e-02 | -3.48 | Spindly (spindly) | protein interactions | 33578 | 37 | 4 | 9204 | 298 | lic,flw,nsl1,Pp1-87B | | 3.073e-02 | -3.48 | MED11 (Mediator complex subunit 11) | protein interactions | 40042 | 37 | 4 | 9204 | 298 | MED22,Cpsf6,Cdk8,CG34159 | | 3.073e-02 | -3.48 | chico (chico) | protein interactions | 64880 | 37 | 4 | 9204 | 298 | Pten,RpS26,14-3-3zeta,RpL14 | | 3.085e-02 | -3.48 | Calr (Calreticulin) | protein interactions | 41166 | 93 | 7 | 9204 | 298 | eIF4B,flr,eEF1delta,Tctp,smt3,eEF5,CG7054 | | 3.089e-02 | -3.48 | Developmental Biology | REACTOME pathways | R-DME-1266738 | 211 | 14 | 3923 | 151 | LIMK1,CG7054,14-3-3zeta,Arpc3A,Prosbeta1,vap,Nf1,Rpt6,CG4935,Rpn10,Prosalpha2,Cdc42,rhea,Arpc5 | | 3.098e-02 | -3.47 | CREB\_02 | MSigDB lists | CREB\_02 | 84 | 8 | 4161 | 184 | CG15168,14-3-3zeta,SF2,Dbp21E2,Tctp,CG13016,Cdc42,Nf1 | | 3.098e-02 | -3.47 | GSE17721\_0.5H\_VS\_12H\_POLYIC\_BMDC\_UP | MSigDB lists | GSE17721\_0.5H\_VS\_12H\_POLYIC\_BMDC\_UP | 84 | 8 | 4161 | 184 | Phf5a,krz,LIMK1,CIA30,janA,Rap1,CG31460,CSN1b | | 3.101e-02 | -3.47 | positive regulation of transmembrane receptor protein serine/threonine kinase signaling pathway | biological process | GO:0090100 | 28 | 3 | 12345 | 305 | dpp,tkv,mav | | 3.101e-02 | -3.47 | actomyosin contractile ring organization | biological process | GO:0044837 | 28 | 3 | 12345 | 305 | pav,Exo84,tsr | | 3.101e-02 | -3.47 | positive regulation of multicellular organism growth | biological process | GO:0040018 | 28 | 3 | 12345 | 305 | Nf1,Tctp,lic | | 3.103e-02 | -3.47 | WD40\_repeat\_dom | interpro domains | IPR017986 | 150 | 9 | 11383 | 331 | flr,Lst8,Wdr82,CG43367,Rbbp5,Wdr33,fzy,CG14805,CG4935 | | 3.107e-02 | -3.47 | Ribosomal\_L15e | smart domains | SM01384 | 1 | 1 | 5086 | 158 | RpL15 | | 3.107e-02 | -3.47 | ECSIT\_Cterm | smart domains | SM01284 | 1 | 1 | 5086 | 158 | ECSIT | | 3.107e-02 | -3.47 | H4 | smart domains | SM00417 | 1 | 1 | 5086 | 158 | His4r | | 3.107e-02 | -3.47 | WGR | smart domains | SM00773 | 1 | 1 | 5086 | 158 | Parp | | 3.107e-02 | -3.47 | eIF-5a | smart domains | SM01376 | 1 | 1 | 5086 | 158 | eEF5 | | 3.107e-02 | -3.47 | zf-PARP | smart domains | SM01336 | 1 | 1 | 5086 | 158 | Parp | | 3.107e-02 | -3.47 | Zpr1 | smart domains | SM00709 | 1 | 1 | 5086 | 158 | Zpr1 | | 3.107e-02 | -3.47 | Ribosomal\_S3Ae | smart domains | SM01397 | 1 | 1 | 5086 | 158 | RpS3A | | 3.107e-02 | -3.47 | AP3B1\_C | smart domains | SM01355 | 1 | 1 | 5086 | 158 | rb | | 3.107e-02 | -3.47 | DUF4210 | smart domains | SM01177 | 1 | 1 | 5086 | 158 | CG9005 | | 3.107e-02 | -3.47 | PADR1 | smart domains | SM01335 | 1 | 1 | 5086 | 158 | Parp | | 3.107e-02 | -3.47 | HTH\_XRE | smart domains | SM00530 | 1 | 1 | 5086 | 158 | mbf1 | | 3.114e-02 | -3.47 | GO\_NEGATIVE\_REGULATION\_OF\_CELL\_SUBSTRATE\_ADHESION | MSigDB lists | GO\_NEGATIVE\_REGULATION\_OF\_CELL\_SUBSTRATE\_ADHESION | 16 | 3 | 4161 | 184 | Pten,vap,Nf1 | | 3.114e-02 | -3.47 | GO\_EUKARYOTIC\_TRANSLATION\_INITIATION\_FACTOR\_3\_COMPLEX | MSigDB lists | GO\_EUKARYOTIC\_TRANSLATION\_INITIATION\_FACTOR\_3\_COMPLEX | 16 | 3 | 4161 | 184 | eIF3c,eIF3b,eIF3g1 | | 3.114e-02 | -3.47 | KYNG\_RESPONSE\_TO\_H2O2 | MSigDB lists | KYNG\_RESPONSE\_TO\_H2O2 | 16 | 3 | 4161 | 184 | CG5589,CycA,RpL5 | | 3.114e-02 | -3.47 | HU\_ANGIOGENESIS\_DN | MSigDB lists | HU\_ANGIOGENESIS\_DN | 16 | 3 | 4161 | 184 | Mdh1,His2Av,PlexA | | 3.114e-02 | -3.47 | GO\_REGULATION\_OF\_MRNA\_SPLICING\_VIA\_SPLICEOSOME | MSigDB lists | GO\_REGULATION\_OF\_MRNA\_SPLICING\_VIA\_SPLICEOSOME | 16 | 3 | 4161 | 184 | sm,Fmr1,tsu | | 3.114e-02 | -3.47 | BURTON\_ADIPOGENESIS\_PEAK\_AT\_24HR | MSigDB lists | BURTON\_ADIPOGENESIS\_PEAK\_AT\_24HR | 16 | 3 | 4161 | 184 | fzy,CycA,SF2 | | 3.114e-02 | -3.47 | GSE7509\_UNSTIM\_VS\_IFNA\_STIM\_IMMATURE\_DC\_DN | MSigDB lists | GSE7509\_UNSTIM\_VS\_IFNA\_STIM\_IMMATURE\_DC\_DN | 16 | 3 | 4161 | 184 | sws,Nulp1,Rop | | 3.114e-02 | -3.47 | ZHAN\_VARIABLE\_EARLY\_DIFFERENTIATION\_GENES\_DN | MSigDB lists | ZHAN\_VARIABLE\_EARLY\_DIFFERENTIATION\_GENES\_DN | 16 | 3 | 4161 | 184 | noi,Rpn10,Rpt6 | | 3.114e-02 | -3.47 | XU\_RESPONSE\_TO\_TRETINOIN\_AND\_NSC682994\_DN | MSigDB lists | XU\_RESPONSE\_TO\_TRETINOIN\_AND\_NSC682994\_DN | 16 | 3 | 4161 | 184 | eIF3b,Art1,eIF3c | | 3.114e-02 | -3.47 | GO\_SMAD\_BINDING | MSigDB lists | GO\_SMAD\_BINDING | 16 | 3 | 4161 | 184 | tkv,dpp,Parp | | 3.114e-02 | -3.47 | JAZAG\_TGFB1\_SIGNALING\_VIA\_SMAD4\_DN | MSigDB lists | JAZAG\_TGFB1\_SIGNALING\_VIA\_SMAD4\_DN | 16 | 3 | 4161 | 184 | unc-119,dpp,Vap33 | | 3.143e-02 | -3.46 | GSE22589\_HEALTHY\_VS\_HIV\_AND\_SIV\_INFECTED\_DC\_DN | MSigDB lists | GSE22589\_HEALTHY\_VS\_HIV\_AND\_SIV\_INFECTED\_DC\_DN | 69 | 7 | 4161 | 184 | lic,Fs(2)Ket,CG15168,Skp2,RhoGAPp190,Nf1,CG2938 | | 3.143e-02 | -3.46 | GSE6259\_DEC205\_POS\_DC\_VS\_CD8\_TCELL\_UP | MSigDB lists | GSE6259\_DEC205\_POS\_DC\_VS\_CD8\_TCELL\_UP | 69 | 7 | 4161 | 184 | Tom20,Tim17b,CG5339,mRpS33,CG5590,pins,Tsp97E | | 3.143e-02 | -3.46 | GO\_REGULATION\_OF\_ACTIN\_FILAMENT\_BASED\_PROCESS | MSigDB lists | GO\_REGULATION\_OF\_ACTIN\_FILAMENT\_BASED\_PROCESS | 69 | 7 | 4161 | 184 | vap,flr,Arpc5,janA,Lst8,LIMK1,Arpc3A | | 3.143e-02 | -3.46 | GSE5589\_UNSTIM\_VS\_45MIN\_LPS\_AND\_IL10\_STIM\_MACROPHAGE\_DN | MSigDB lists | GSE5589\_UNSTIM\_VS\_45MIN\_LPS\_AND\_IL10\_STIM\_MACROPHAGE\_DN | 69 | 7 | 4161 | 184 | Sod2,Mdh1,CCT4,CG11334,Fs(2)Ket,Stt3B,Tom20 | | 3.143e-02 | -3.46 | E2F1\_Q6\_01 | MSigDB lists | E2F1\_Q6\_01 | 69 | 7 | 4161 | 184 | dpp,Phf5a,Stt3B,E(z),Pect,SF2,spel1 | | 3.143e-02 | -3.46 | GO\_REGULATION\_OF\_MAP\_KINASE\_ACTIVITY | MSigDB lists | GO\_REGULATION\_OF\_MAP\_KINASE\_ACTIVITY | 69 | 7 | 4161 | 184 | CSN1b,Nf1,Rap1,lic,E(z),dpp,krz | | 3.150e-02 | -3.46 | GSE14000\_TRANSLATED\_RNA\_VS\_MRNA\_4H\_LPS\_DC\_UP | MSigDB lists | GSE14000\_TRANSLATED\_RNA\_VS\_MRNA\_4H\_LPS\_DC\_UP | 100 | 9 | 4161 | 184 | CG5589,CG6907,mRpS28,Ufd4,CIA30,Pex3,tsu,vap,CG17493 | | 3.152e-02 | -3.46 | positive regulation of neurogenesis | biological process | GO:0050769 | 96 | 6 | 12345 | 305 | Rap1,Fmr1,mnb,Rheb,PlexA,Rop | | 3.172e-02 | -3.45 | NCAM signaling for neurite out-growth | REACTOME pathways | R-DME-375165 | 115 | 9 | 3923 | 151 | Nf1,Rpt6,vap,Prosbeta1,14-3-3zeta,CG7054,Prosalpha2,CG4935,Rpn10 | | 3.181e-02 | -3.45 | chromocenter | cellular component | GO:0010369 | 28 | 3 | 11940 | 298 | His2Av,MED26,HP4 | | 3.181e-02 | -3.45 | PcG protein complex | cellular component | GO:0031519 | 28 | 3 | 11940 | 298 | RAF2,E(z),Jarid2 | | 3.181e-02 | -3.45 | nuclear ubiquitin ligase complex | cellular component | GO:0000152 | 28 | 3 | 11940 | 298 | Roc2,fzy,dgrn | | 3.181e-02 | -3.45 | 90S preribosome | cellular component | GO:0030686 | 28 | 3 | 11940 | 298 | CG11920,CG4866,CG9246 | | 3.194e-02 | -3.44 | WD\_REPEATS\_2 | prosite domains | PS50082 | 144 | 9 | 6243 | 191 | Rbbp5,Lst8,Wdr33,CG14805,flr,Wdr82,CG43367,CG4935,fzy | | 3.202e-02 | -3.44 | 4E-T (eIF4E-Transporter) | protein interactions | 43836 | 55 | 5 | 9204 | 298 | RpL15,RpS8,CCT4,RpL5,me31B | | 3.216e-02 | -3.44 | BMP signaling | KEGG pathways | dme\_M00679 | 8 | 2 | 3198 | 117 | dpp,tkv | | 3.216e-02 | -3.44 | BMP signaling | KEGG pathways | M00679 | 8 | 2 | 3198 | 117 | dpp,tkv | | 3.226e-02 | -3.43 | GO\_REGULATION\_OF\_NERVOUS\_SYSTEM\_DEVELOPMENT | MSigDB lists | GO\_REGULATION\_OF\_NERVOUS\_SYSTEM\_DEVELOPMENT | 150 | 12 | 4161 | 184 | Nf1,fzy,Fmr1,Pten,PlexA,dpp,me31B,RhoGAPp190,flr,Rap1,E(z),LIMK1 | | 3.236e-02 | -3.43 | Rcd-1 (Required for cell differentiation 1) | protein interactions | 32966 | 9 | 2 | 9204 | 298 | twin,Not3 | | 3.236e-02 | -3.43 | tobi (target of brain insulin) | protein interactions | 43072 | 9 | 2 | 9204 | 298 | tsr,I-2 | | 3.236e-02 | -3.43 | cort (cortex) | protein interactions | 33937 | 9 | 2 | 9204 | 298 | CycA,grau | | 3.236e-02 | -3.43 | RagC-D (Ras-related GTP binding C/D) | protein interactions | 35793 | 9 | 2 | 9204 | 298 | Rheb,RagA-B | | 3.236e-02 | -3.43 | CG2258 (uncharacterized protein) | protein interactions | 31729 | 9 | 2 | 9204 | 298 | CG12877,Ythdf | | 3.236e-02 | -3.43 | Gyc76C (Guanylyl cyclase at 76C) | protein interactions | 8674026 | 9 | 2 | 9204 | 298 | dpp,PlexA | | 3.236e-02 | -3.43 | CG31198 (uncharacterized protein) | protein interactions | 318622 | 9 | 2 | 9204 | 298 | CG5500,CG32473 | | 3.236e-02 | -3.43 | Cyp6t3 (Cyp6t3) | protein interactions | 36318 | 9 | 2 | 9204 | 298 | Acbp1,sun | | 3.236e-02 | -3.43 | Sec3 (Secretory 3) | protein interactions | 39940 | 9 | 2 | 9204 | 298 | CG9951,Exo84 | | 3.236e-02 | -3.43 | Cks30A (Cyclin-dependent kinase subunit 30A) | protein interactions | 34250 | 9 | 2 | 9204 | 298 | CycA,Skp2 | | 3.236e-02 | -3.43 | Edem1 (ER degradation enhancer, mannosidase alpha-like 1) | protein interactions | 31154 | 9 | 2 | 9204 | 298 | krz,Fgop2 | | 3.236e-02 | -3.43 | CG14718 (uncharacterized protein) | protein interactions | 41413 | 9 | 2 | 9204 | 298 | CNBP,RpL36A | | 3.238e-02 | -3.43 | CG14357 (uncharacterized protein) | protein interactions | 41716 | 1 | 1 | 9204 | 298 | CtBP | | 3.238e-02 | -3.43 | Pex1 (Peroxin 1) | protein interactions | 45460 | 1 | 1 | 9204 | 298 | MED22 | | 3.238e-02 | -3.43 | CG12506 (uncharacterized protein) | protein interactions | 33249 | 1 | 1 | 9204 | 298 | Sgt | | 3.238e-02 | -3.43 | CG13471 (uncharacterized protein) | protein interactions | 39604 | 1 | 1 | 9204 | 298 | Rcd5 | | 3.238e-02 | -3.43 | CG9101 (uncharacterized protein) | protein interactions | 32452 | 1 | 1 | 9204 | 298 | CG1636 | | 3.238e-02 | -3.43 | CG7840 (uncharacterized protein) | protein interactions | 34136 | 1 | 1 | 9204 | 298 | unc-119 | | 3.238e-02 | -3.43 | mir-304 (mir-304 stem loop) | protein interactions | 12797982 | 1 | 1 | 9204 | 298 | Fmr1 | | 3.238e-02 | -3.43 | ND-B8 (NADH dehydrogenase (ubiquinone) B8 subunit) | protein interactions | 50178 | 1 | 1 | 9204 | 298 | smg | | 3.238e-02 | -3.43 | CG4520 (uncharacterized protein) | protein interactions | 41902 | 1 | 1 | 9204 | 298 | RpL5 | | 3.238e-02 | -3.43 | CG7142 (uncharacterized protein) | protein interactions | 42194 | 1 | 1 | 9204 | 298 | Sgt | | 3.238e-02 | -3.43 | Nhe3 (Na[+]/H[+] hydrogen exchanger 3) | protein interactions | 33939 | 1 | 1 | 9204 | 298 | Cpr73D | | 3.238e-02 | -3.43 | mir-1 (mir-1 stem loop) | protein interactions | 12798026 | 1 | 1 | 9204 | 298 | Fmr1 | | 3.238e-02 | -3.43 | CG13847 (uncharacterized protein) | protein interactions | 42636 | 1 | 1 | 9204 | 298 | Exo84 | | 3.238e-02 | -3.43 | PARP1 (poly(ADP-ribose) polymerase 1) | protein interactions | 142 | 1 | 1 | 9204 | 298 | CG1218 | | 3.238e-02 | -3.43 | CG14926 (uncharacterized protein) | protein interactions | 34567 | 1 | 1 | 9204 | 298 | Zpr1 | | 3.238e-02 | -3.43 | CG13054 (uncharacterized protein) | protein interactions | 39780 | 1 | 1 | 9204 | 298 | DNAlig1 | | 3.238e-02 | -3.43 | CG15617 (uncharacterized protein) | protein interactions | 36871 | 1 | 1 | 9204 | 298 | Cpr73D | | 3.238e-02 | -3.43 | mir-79 (mir-79 stem loop) | protein interactions | 12798238 | 1 | 1 | 9204 | 298 | Fmr1 | | 3.238e-02 | -3.43 | CG8549 (uncharacterized protein) | protein interactions | 38749 | 1 | 1 | 9204 | 298 | Vlet | | 3.238e-02 | -3.43 | CG8952 (uncharacterized protein) | protein interactions | 32525 | 1 | 1 | 9204 | 298 | smt3 | | 3.238e-02 | -3.43 | CG2574 (uncharacterized protein) | protein interactions | 32190 | 1 | 1 | 9204 | 298 | Art1 | | 3.238e-02 | -3.43 | Obp57a (Odorant-binding protein 57a) | protein interactions | 246670 | 1 | 1 | 9204 | 298 | Sgt | | 3.238e-02 | -3.43 | CG2811 (uncharacterized protein) | protein interactions | 37988 | 1 | 1 | 9204 | 298 | RpL15 | | 3.238e-02 | -3.43 | CG6324 (uncharacterized protein) | protein interactions | 32481 | 1 | 1 | 9204 | 298 | Not3 | | 3.238e-02 | -3.43 | PML (promyelocytic leukemia) | protein interactions | 5371 | 1 | 1 | 9204 | 298 | smt3 | | 3.238e-02 | -3.43 | Cpr11B (Cuticular protein 11B) | protein interactions | 32203 | 1 | 1 | 9204 | 298 | BthD | | 3.238e-02 | -3.43 | Pask (PAS kinase) | protein interactions | 37824 | 1 | 1 | 9204 | 298 | CtBP | | 3.238e-02 | -3.43 | CG15343 (uncharacterized protein) | protein interactions | 31765 | 1 | 1 | 9204 | 298 | Art1 | | 3.238e-02 | -3.43 | mir-276b (mir-276b stem loop) | protein interactions | 12798170 | 1 | 1 | 9204 | 298 | Fmr1 | | 3.238e-02 | -3.43 | NAAT1 (Nutrient Amino Acid Transporter 1) | protein interactions | 31457 | 1 | 1 | 9204 | 298 | Nf1 | | 3.238e-02 | -3.43 | CG9125 (uncharacterized protein) | protein interactions | 2768882 | 1 | 1 | 9204 | 298 | Rat1 | | 3.238e-02 | -3.43 | SSH1 (slingshot protein phosphatase 1) | protein interactions | 54434 | 1 | 1 | 9204 | 298 | LIMK1 | | 3.238e-02 | -3.43 | Drip (drip) | protein interactions | 36236 | 1 | 1 | 9204 | 298 | Sgt | | 3.238e-02 | -3.43 | CG6310 (uncharacterized protein) | protein interactions | 39237 | 1 | 1 | 9204 | 298 | RpS26 | | 3.238e-02 | -3.43 | mir-9c (mir-9c stem loop) | protein interactions | 12798236 | 1 | 1 | 9204 | 298 | Fmr1 | | 3.238e-02 | -3.43 | CG2990 (uncharacterized protein) | protein interactions | 31934 | 1 | 1 | 9204 | 298 | flw | | 3.238e-02 | -3.43 | Dak1 (Dak1) | protein interactions | 43165 | 1 | 1 | 9204 | 298 | CG8132 | | 3.238e-02 | -3.43 | Orcokinin (orcokinin) | protein interactions | 37806 | 1 | 1 | 9204 | 298 | pins | | 3.238e-02 | -3.43 | fzr2 (fizzy-related 2) | protein interactions | 37808 | 1 | 1 | 9204 | 298 | fzy | | 3.238e-02 | -3.43 | mir-12 (mir-12 stem loop) | protein interactions | 12797983 | 1 | 1 | 9204 | 298 | Fmr1 | | 3.238e-02 | -3.43 | CG14182 (uncharacterized protein) | protein interactions | 40181 | 1 | 1 | 9204 | 298 | Sgt | | 3.238e-02 | -3.43 | CG15172 (uncharacterized protein) | protein interactions | 35167 | 1 | 1 | 9204 | 298 | krz | | 3.238e-02 | -3.43 | Cfl1 (cofilin 1, non-muscle) | protein interactions | 12631 | 1 | 1 | 9204 | 298 | LIMK1 | | 3.238e-02 | -3.43 | CG11755 (uncharacterized protein) | protein interactions | 41031 | 1 | 1 | 9204 | 298 | Cpsf6 | | 3.238e-02 | -3.43 | CG5381 (uncharacterized protein) | protein interactions | 34389 | 1 | 1 | 9204 | 298 | nsl1 | | 3.238e-02 | -3.43 | mir-125 (mir-125 stem loop) | protein interactions | 12798013 | 1 | 1 | 9204 | 298 | Fmr1 | | 3.238e-02 | -3.43 | CG5877 (uncharacterized protein) | protein interactions | 32455 | 1 | 1 | 9204 | 298 | krz | | 3.238e-02 | -3.43 | SBF1 (SET binding factor 1) | protein interactions | 6305 | 1 | 1 | 9204 | 298 | E(z) | | 3.238e-02 | -3.43 | CG8260 (uncharacterized protein) | protein interactions | 32521 | 1 | 1 | 9204 | 298 | MED26 | | 3.238e-02 | -3.43 | CG1750 (uncharacterized protein) | protein interactions | 43699 | 1 | 1 | 9204 | 298 | noi | | 3.238e-02 | -3.43 | Ogg1 (8-oxoguanine DNA glycosylase) | protein interactions | 31806 | 1 | 1 | 9204 | 298 | Sod2 | | 3.238e-02 | -3.43 | TotC (Turandot C) | protein interactions | 117462 | 1 | 1 | 9204 | 298 | Ubqn | | 3.238e-02 | -3.43 | yellow-d (yellow-d) | protein interactions | 37703 | 1 | 1 | 9204 | 298 | Mps1 | | 3.238e-02 | -3.43 | CG16854 (uncharacterized protein) | protein interactions | 34539 | 1 | 1 | 9204 | 298 | 14-3-3zeta | | 3.238e-02 | -3.43 | CG14374 (uncharacterized protein) | protein interactions | 50026 | 1 | 1 | 9204 | 298 | CG10209 | | 3.238e-02 | -3.43 | Srp54k (Signal recognition particle protein 54k) | protein interactions | 38593 | 1 | 1 | 9204 | 298 | CG5174 | | 3.238e-02 | -3.43 | ND-13A (NADH dehydrogenase (ubiquinone) 13 kDa A subunit) | protein interactions | 33744 | 1 | 1 | 9204 | 298 | Pect | | 3.238e-02 | -3.43 | CG7341 (uncharacterized protein) | protein interactions | 40004 | 1 | 1 | 9204 | 298 | ver | | 3.238e-02 | -3.43 | CG6048 (uncharacterized protein) | protein interactions | 31528 | 1 | 1 | 9204 | 298 | Vap33 | | 3.238e-02 | -3.43 | jtb (jetboil) | protein interactions | 36838 | 1 | 1 | 9204 | 298 | Arpc5 | | 3.238e-02 | -3.43 | Prosalpha3T (Proteasome alpha3 subunit, Testis-specific) | protein interactions | 43679 | 1 | 1 | 9204 | 298 | Ubqn | | 3.238e-02 | -3.43 | Sid (Stress induced DNase) | protein interactions | 43390 | 1 | 1 | 9204 | 298 | Pep | | 3.238e-02 | -3.43 | CG10413 (uncharacterized protein) | protein interactions | 35111 | 1 | 1 | 9204 | 298 | Bub1 | | 3.238e-02 | -3.43 | Epg5 (ectopic P-granules autophagy protein 5) | protein interactions | 42256 | 1 | 1 | 9204 | 298 | Rab7 | | 3.238e-02 | -3.43 | ppk30 (pickpocket 30) | protein interactions | 43487 | 1 | 1 | 9204 | 298 | Vap33 | | 3.238e-02 | -3.43 | pre-mod(mdg4)-T (pre-mod(mdg4)-T) | protein interactions | 10178861 | 1 | 1 | 9204 | 298 | su(Hw) | | 3.238e-02 | -3.43 | CG30275 (uncharacterized protein) | protein interactions | 246522 | 1 | 1 | 9204 | 298 | flw | | 3.238e-02 | -3.43 | RpIIIC160 (RNA polymerase III subunit C160) | protein interactions | 32524 | 1 | 1 | 9204 | 298 | Pp1-87B | | 3.238e-02 | -3.43 | mth (methuselah) | protein interactions | 38058 | 1 | 1 | 9204 | 298 | sun | | 3.238e-02 | -3.43 | CG15760 (uncharacterized protein) | protein interactions | 32324 | 1 | 1 | 9204 | 298 | RpS9 | | 3.238e-02 | -3.43 | Naa80 (N(alpha)-acetyltransferase 80) | protein interactions | 41195 | 1 | 1 | 9204 | 298 | Arpc5 | | 3.242e-02 | -3.43 | imaginal disc-derived wing hair organization | biological process | GO:0035317 | 49 | 4 | 12345 | 305 | Cdc42,LIMK1,tsr,flr | | 3.242e-02 | -3.43 | pole plasm oskar mRNA localization | biological process | GO:0045451 | 49 | 4 | 12345 | 305 | tsu,sqd,Hrb27C,glo | | 3.242e-02 | -3.43 | RNA export from nucleus | biological process | GO:0006405 | 49 | 4 | 12345 | 305 | CG6961,Dbp80,sqd,SF2 | | 3.242e-02 | -3.43 | negative regulation of mitotic cell cycle phase transition | biological process | GO:1901991 | 49 | 4 | 12345 | 305 | Bub1,CycA,Mps1,Tctp | | 3.251e-02 | -3.43 | Znf\_C2H2\_jaz | interpro domains | IPR022755 | 10 | 2 | 11383 | 331 | Pep,CG11586 | | 3.251e-02 | -3.43 | Cullin\_repeat-like\_dom\_sf | interpro domains | IPR016159 | 10 | 2 | 11383 | 331 | Cul4,Exo84 | | 3.281e-02 | -3.42 | Fcgamma receptor (FCGR) dependent phagocytosis | REACTOME pathways | R-DME-2029480 | 32 | 4 | 3923 | 151 | Arpc3A,LIMK1,Cdc42,Arpc5 | | 3.282e-02 | -3.42 | NaCP60E (Na channel protein 60E) | protein interactions | 37981 | 22 | 3 | 9204 | 298 | CycA,msd5,RpL5 | | 3.282e-02 | -3.42 | Hrb87F (Heterogeneous nuclear ribonucleoprotein at 87F) | protein interactions | 48535 | 22 | 3 | 9204 | 298 | sqd,Hrb27C,glo | | 3.282e-02 | -3.42 | Rca1 (Regulator of cyclin A1) | protein interactions | 33959 | 22 | 3 | 9204 | 298 | CycA,ver,GILT3 | | 3.282e-02 | -3.42 | corolla (corolla) | protein interactions | 32739 | 22 | 3 | 9204 | 298 | flw,eIF3c,Pp1-87B | | 3.282e-02 | -3.42 | CG13319 (uncharacterized protein) | protein interactions | 36423 | 22 | 3 | 9204 | 298 | Prosbeta1,CG12321,Prosalpha2 | | 3.282e-02 | -3.42 | Pka-C1 (Protein kinase, cAMP-dependent, catalytic subunit 1) | protein interactions | 34284 | 22 | 3 | 9204 | 298 | PlexA,dpp,Nf1 | | 3.287e-02 | -3.42 | RHO GTPase Effectors | REACTOME pathways | R-DME-195258 | 63 | 6 | 3923 | 151 | Arpc3A,14-3-3zeta,LIMK1,Cdc42,Arpc5,flw | | 3.290e-02 | -3.41 | GO\_PEPTIDE\_METABOLIC\_PROCESS | MSigDB lists | GO\_PEPTIDE\_METABOLIC\_PROCESS | 312 | 21 | 4161 | 184 | Bmcp,RpS26,RpL14,eIF3g1,CG31957,mRpS9,RpS8,RpL35A,eRF1,RpS9,CG32473,eEF1delta,eIF3b,RpL5,eIF3c,mRpS28,eEF5,RpL15,mRpS33,RpL36A,CG4743 | | 3.292e-02 | -3.41 | organic substance transport | biological process | GO:0071702 | 719 | 26 | 12345 | 305 | Rab7,Vps35,Fmr1,Vps13D,CG4743,Dbp80,Pex3,SF2,Tom20,g,RabX6,CG5535,Tim17b,Rab14,CG6961,Fs(2)Ket,unc-119,PGAP1,Rab8,rb,Sgt,sqd,CG8860,bai,mon2,Bmcp | | 3.292e-02 | -3.41 | regulation of mitotic cell cycle phase transition | biological process | GO:1901990 | 97 | 6 | 12345 | 305 | Mps1,Tctp,CycA,Bub1,fzy,RecQ4 | | 3.295e-02 | -3.41 | GSE557\_CIITA\_KO\_VS\_I\_AB\_KO\_DC\_DN | MSigDB lists | GSE557\_CIITA\_KO\_VS\_I\_AB\_KO\_DC\_DN | 85 | 8 | 4161 | 184 | CG5339,Pect,Vps35,Rbbp5,CG11586,Tim17b,CG9853,smt3 | | 3.299e-02 | -3.41 | CHIANG\_LIVER\_CANCER\_SUBCLASS\_PROLIFERATION\_UP | MSigDB lists | CHIANG\_LIVER\_CANCER\_SUBCLASS\_PROLIFERATION\_UP | 41 | 5 | 4161 | 184 | pav,fzy,CtBP,CycA,E(z) | | 3.299e-02 | -3.41 | GSE27786\_LSK\_VS\_ERYTHROBLAST\_DN | MSigDB lists | GSE27786\_LSK\_VS\_ERYTHROBLAST\_DN | 41 | 5 | 4161 | 184 | flw,CG11586,Lst8,CG9853,Rap1 | | 3.299e-02 | -3.41 | PID\_ERBB1\_DOWNSTREAM\_PATHWAY | MSigDB lists | PID\_ERBB1\_DOWNSTREAM\_PATHWAY | 41 | 5 | 4161 | 184 | Cdc42,Lst8,14-3-3zeta,Arpc5,Arpc3A | | 3.301e-02 | -3.41 | ABRAMSON\_INTERACT\_WITH\_AIRE | MSigDB lists | ABRAMSON\_INTERACT\_WITH\_AIRE | 28 | 4 | 4161 | 184 | SF2,spel1,Parp,Fs(2)Ket | | 3.301e-02 | -3.41 | GO\_PROTEIN\_METHYLTRANSFERASE\_ACTIVITY | MSigDB lists | GO\_PROTEIN\_METHYLTRANSFERASE\_ACTIVITY | 28 | 4 | 4161 | 184 | Rbbp5,E(z),Art1,Wdr82 | | 3.301e-02 | -3.41 | GO\_CHANNEL\_REGULATOR\_ACTIVITY | MSigDB lists | GO\_CHANNEL\_REGULATOR\_ACTIVITY | 28 | 4 | 4161 | 184 | vap,Tsp97E,Stim,janA | | 3.301e-02 | -3.41 | GO\_REGULATION\_OF\_CALCIUM\_ION\_TRANSMEMBRANE\_TRANSPORT | MSigDB lists | GO\_REGULATION\_OF\_CALCIUM\_ION\_TRANSMEMBRANE\_TRANSPORT | 28 | 4 | 4161 | 184 | Tsp97E,Fmr1,Stim,Ubqn | | 3.301e-02 | -3.41 | ISHIDA\_E2F\_TARGETS | MSigDB lists | ISHIDA\_E2F\_TARGETS | 28 | 4 | 4161 | 184 | E(z),CycA,fzy,DNAlig1 | | 3.301e-02 | -3.41 | DE\_YY1\_TARGETS\_DN | MSigDB lists | DE\_YY1\_TARGETS\_DN | 28 | 4 | 4161 | 184 | Pten,cta,vap,flr | | 3.301e-02 | -3.41 | GNF2\_NS | MSigDB lists | GNF2\_NS | 28 | 4 | 4161 | 184 | mRpS33,Prosalpha2,Parp,CCT4 | | 3.301e-02 | -3.41 | AGGGCAG\_MIR18A | MSigDB lists | AGGGCAG\_MIR18A | 28 | 4 | 4161 | 184 | Stlk,rhea,Fmr1,14-3-3zeta | | 3.301e-02 | -3.41 | GSE16385\_IFNG\_TNF\_VS\_IL4\_STIM\_MACROPHAGE\_ROSIGLITAZONE\_TREATED\_UP | MSigDB lists | GSE16385\_IFNG\_TNF\_VS\_IL4\_STIM\_MACROPHAGE\_ROSIGLITAZONE\_TREATED\_UP | 28 | 4 | 4161 | 184 | Skp2,Tom20,twin,Usp32 | | 3.301e-02 | -3.41 | GSE9946\_LISTERIA\_INF\_MATURE\_VS\_PROSTAGLANDINE2\_TREATED\_MATURE\_DC\_UP | MSigDB lists | GSE9946\_LISTERIA\_INF\_MATURE\_VS\_PROSTAGLANDINE2\_TREATED\_MATURE\_DC\_UP | 28 | 4 | 4161 | 184 | Ubqn,Rbbp5,Prosalpha2,Nf1 | | 3.302e-02 | -3.41 | Regulation of mitotic cell cycle | REACTOME pathways | R-DME-453276 | 80 | 7 | 3923 | 151 | Prosalpha2,Rpn10,Skp2,fzy,Prosbeta1,Rpt6,CycA | | 3.302e-02 | -3.41 | APC/C-mediated degradation of cell cycle proteins | REACTOME pathways | R-DME-174143 | 80 | 7 | 3923 | 151 | CycA,Rpt6,Prosalpha2,Rpn10,Prosbeta1,fzy,Skp2 | | 3.303e-02 | -3.41 | UBQ | smart domains | SM00213 | 23 | 3 | 5086 | 158 | Ubqn,smt3,EloB | | 3.305e-02 | -3.41 | GSE32034\_UNTREATED\_VS\_ROSIGLIZATONE\_TREATED\_LY6C\_HIGH\_MONOCYTE\_UP | MSigDB lists | GSE32034\_UNTREATED\_VS\_ROSIGLIZATONE\_TREATED\_LY6C\_HIGH\_MONOCYTE\_UP | 55 | 6 | 4161 | 184 | Ubqn,dpp,PlexA,lic,Tim17b,RpL15 | | 3.305e-02 | -3.41 | MORF\_DDX11 | MSigDB lists | MORF\_DDX11 | 55 | 6 | 4161 | 184 | tsu,Aps,Rcd5,rb,sm,rhea | | 3.305e-02 | -3.41 | GSE2405\_S\_AUREUS\_VS\_UNTREATED\_NEUTROPHIL\_DN | MSigDB lists | GSE2405\_S\_AUREUS\_VS\_UNTREATED\_NEUTROPHIL\_DN | 55 | 6 | 4161 | 184 | E(z),Skp2,Fign,DNAlig1,pins,CycA | | 3.305e-02 | -3.41 | GSE22196\_HEALTHY\_VS\_OBESE\_MOUSE\_SKIN\_GAMMADELTA\_TCELL\_DN | MSigDB lists | GSE22196\_HEALTHY\_VS\_OBESE\_MOUSE\_SKIN\_GAMMADELTA\_TCELL\_DN | 55 | 6 | 4161 | 184 | Vap33,RpL36A,ND-ASHI,14-3-3zeta,sm,eEF1delta | | 3.305e-02 | -3.41 | GSE1791\_CTRL\_VS\_NEUROMEDINU\_IN\_T\_CELL\_LINE\_3H\_DN | MSigDB lists | GSE1791\_CTRL\_VS\_NEUROMEDINU\_IN\_T\_CELL\_LINE\_3H\_DN | 55 | 6 | 4161 | 184 | Coq5,ubl,Rpn10,CG8132,Wdr82,Roc2 | | 3.332e-02 | -3.40 | GO\_SINGLE\_ORGANISM\_CELLULAR\_LOCALIZATION | MSigDB lists | GO\_SINGLE\_ORGANISM\_CELLULAR\_LOCALIZATION | 369 | 24 | 4161 | 184 | Pex3,rb,Rop,bai,RpL36A,Tom20,Fs(2)Ket,RpL15,CG8860,RpS9,RpL5,Vap33,CG17883,Rab8,Tim17b,Vps13D,RpS26,RpL14,pav,RpS8,CG9853,Cdc42,pins,RpL35A | | 3.335e-02 | -3.40 | peptide metabolic process | biological process | GO:0006518 | 788 | 28 | 12345 | 305 | CG31957,CG8635,mRpS28,mRpS9,RpL14,eIF3b,RpL36A,RpS8,RpS3A,GstE13,mRpL30,mRpL34,RpS19a,mRpS18A,CG32473,RpL5,eIF3c,eEF5,RpS9,RpS26,RpL15,mRpS33,CG10365,eRF1,eEF1delta,eIF4B,eIF3g1,RpL35A | | 3.344e-02 | -3.40 | immune system process | biological process | GO:0002376 | 328 | 14 | 12345 | 305 | E(bx),Parp,Pten,Cdc42,ECSIT,Rap1,heix,CG8149,dgrn,lic,RpS8,dpp,alpha-Man-Ib,pll | | 3.350e-02 | -3.40 | peptide transmembrane transporter activity | molecular function | GO:1904680 | 30 | 3 | 12176 | 289 | Tim17b,CG8860,Tom20 | | 3.350e-02 | -3.40 | Rpn6 (Regulatory particle non-ATPase 6) | protein interactions | 36638 | 38 | 4 | 9204 | 298 | Prosalpha2,Rpt6,Ubqn,Prosbeta1 | | 3.350e-02 | -3.40 | Pomp (pomp) | protein interactions | 35342 | 38 | 4 | 9204 | 298 | Rpt6,Prosalpha2,CG12321,Prosbeta1 | | 3.367e-02 | -3.39 | GSE17721\_CTRL\_VS\_CPG\_1H\_BMDC\_UP | MSigDB lists | GSE17721\_CTRL\_VS\_CPG\_1H\_BMDC\_UP | 70 | 7 | 4161 | 184 | smt3,fzy,CG4743,bai,janA,LIMK1,Dbp80 | | 3.397e-02 | -3.38 | establishment or maintenance of polarity of follicular epithelium | biological process | GO:0016334 | 29 | 3 | 12345 | 305 | Parp,sqd,spoon | | 3.397e-02 | -3.38 | regulation of protein complex disassembly | biological process | GO:0043244 | 29 | 3 | 12345 | 305 | eEF5,flr,Fign | | 3.397e-02 | -3.38 | negative regulation of transferase activity | biological process | GO:0051348 | 29 | 3 | 12345 | 305 | mnb,sws,Pten | | 3.407e-02 | -3.38 | GO\_POSITIVE\_REGULATION\_OF\_PROTEIN\_MODIFICATION\_PROCESS | MSigDB lists | GO\_POSITIVE\_REGULATION\_OF\_PROTEIN\_MODIFICATION\_PROCESS | 276 | 19 | 4161 | 184 | Vap33,Rap1,Rpn10,Ubqn,E(z),Lst8,Art1,Cdc42,fzy,dpp,Pten,krz,Skp2,lic,Prosalpha2,Rpt6,Stlk,unc-119,AGO3 | | 3.408e-02 | -3.38 | imaginal disc-derived wing morphogenesis | biological process | GO:0007476 | 298 | 13 | 12345 | 305 | tkv,EloB,tsr,Cdc42,peng,Btk29A,flr,Stim,flw,GEFmeso,LIMK1,rhea,dpp | | 3.410e-02 | -3.38 | hpo (hippo) | protein interactions | 37247 | 95 | 7 | 9204 | 298 | Fgop2,Slmap,Ncoa6,eEF1delta,Rap1,Tgi,RpS26 | | 3.410e-02 | -3.38 | quinone biosynthetic process | biological process | GO:1901663 | 12 | 2 | 12345 | 305 | Coq5,heix | | 3.410e-02 | -3.38 | female germline ring canal formation | biological process | GO:0007301 | 12 | 2 | 12345 | 305 | flw,Btk29A | | 3.410e-02 | -3.38 | habituation | biological process | GO:0046959 | 12 | 2 | 12345 | 305 | Fmr1,me31B | | 3.410e-02 | -3.38 | quinone metabolic process | biological process | GO:1901661 | 12 | 2 | 12345 | 305 | heix,Coq5 | | 3.410e-02 | -3.38 | anterograde axonal transport | biological process | GO:0008089 | 12 | 2 | 12345 | 305 | g,Fmr1 | | 3.410e-02 | -3.38 | histone H3-K9 modification | biological process | GO:0061647 | 12 | 2 | 12345 | 305 | E(z),BEAF-32 | | 3.410e-02 | -3.38 | ubiquinone metabolic process | biological process | GO:0006743 | 12 | 2 | 12345 | 305 | Coq5,heix | | 3.410e-02 | -3.38 | regulation of exit from mitosis | biological process | GO:0007096 | 12 | 2 | 12345 | 305 | CycA,fzy | | 3.410e-02 | -3.38 | ubiquinone biosynthetic process | biological process | GO:0006744 | 12 | 2 | 12345 | 305 | Coq5,heix | | 3.410e-02 | -3.38 | positive regulation of mRNA catabolic process | biological process | GO:0061014 | 12 | 2 | 12345 | 305 | AGO3,smg | | 3.410e-02 | -3.38 | anterior Malpighian tubule development | biological process | GO:0061327 | 12 | 2 | 12345 | 305 | dpp,tkv | | 3.422e-02 | -3.38 | positive regulation of MAPK cascade | biological process | GO:0043410 | 73 | 5 | 12345 | 305 | smt3,sigmar,lic,Rap1,Cdc42 | | 3.426e-02 | -3.37 | CCT7 (Chaperonin containing TCP1 subunit 7) | protein interactions | 41054 | 56 | 5 | 9204 | 298 | Fmr1,Lst8,Rheb,viaf,CCT4 | | 3.426e-02 | -3.37 | MED17 (Mediator complex subunit 17) | protein interactions | 42175 | 56 | 5 | 9204 | 298 | MED22,Cdk8,Cpsf6,CG34159,MED26 | | 3.429e-02 | -3.37 | kinase regulator activity | molecular function | GO:0019207 | 76 | 5 | 12176 | 289 | CG6907,mnb,CycA,14-3-3zeta,CG1791 | | 3.437e-02 | -3.37 | maternal determination of anterior/posterior axis, embryo | biological process | GO:0008358 | 98 | 6 | 12345 | 305 | lic,tsu,mon2,Hrb27C,sqd,glo | | 3.437e-02 | -3.37 | mitochondrial translation | biological process | GO:0032543 | 98 | 6 | 12345 | 305 | mRpL34,mRpS33,mRpS18A,mRpS9,mRpS28,mRpL30 | | 3.441e-02 | -3.37 | PHD | smart domains | SM00249 | 40 | 4 | 5086 | 158 | CG1815,Acf,E(bx),d4 | | 3.455e-02 | -3.37 | peptide-lysine-N-acetyltransferase activity | molecular function | GO:0061733 | 52 | 4 | 12176 | 289 | d4,Acf,Rcd5,Eaf6 | | 3.455e-02 | -3.37 | Mismatch repair | KEGG pathways | dme03430 | 20 | 3 | 3198 | 117 | spel1,DNAlig1,RPA3 | | 3.455e-02 | -3.37 | Mismatch repair | KEGG pathways | ko03430 | 20 | 3 | 3198 | 117 | spel1,DNAlig1,RPA3 | | 3.458e-02 | -3.36 | non-sensory hair organization | biological process | GO:0035316 | 50 | 4 | 12345 | 305 | Cdc42,LIMK1,tsr,flr | | 3.458e-02 | -3.36 | regulation of cell adhesion | biological process | GO:0030155 | 50 | 4 | 12345 | 305 | tkv,PlexA,Pten,Rap1 | | 3.458e-02 | -3.36 | negative regulation of cell cycle phase transition | biological process | GO:1901988 | 50 | 4 | 12345 | 305 | Tctp,Mps1,Bub1,CycA | | 3.458e-02 | -3.36 | hair cell differentiation | biological process | GO:0035315 | 50 | 4 | 12345 | 305 | LIMK1,tsr,Cdc42,flr | | 3.501e-02 | -3.35 | GSE17721\_0.5H\_VS\_24H\_POLYIC\_BMDC\_UP | MSigDB lists | GSE17721\_0.5H\_VS\_24H\_POLYIC\_BMDC\_UP | 86 | 8 | 4161 | 184 | sm,sigmar,CG4866,fzy,Phf5a,CycA,Rap1,CG17493 | | 3.516e-02 | -3.35 | JOHNSTONE\_PARVB\_TARGETS\_3\_DN | MSigDB lists | JOHNSTONE\_PARVB\_TARGETS\_3\_DN | 333 | 22 | 4161 | 184 | mRpS9,CG32576,fzy,pins,spel1,pav,Fign,Sgt,RpL14,Fmr1,Pten,eEF1delta,E(z),twin,Stt3B,Lrr47,flw,CycA,RhoGAPp190,Skp2,CG5535,His2Av | | 3.519e-02 | -3.35 | GO\_REGULATION\_OF\_ANATOMICAL\_STRUCTURE\_SIZE | MSigDB lists | GO\_REGULATION\_OF\_ANATOMICAL\_STRUCTURE\_SIZE | 102 | 9 | 4161 | 184 | Lst8,Arpc5,Arpc3A,LIMK1,vap,flr,Pten,PlexA,Sod2 | | 3.519e-02 | -3.35 | GO\_ACTIN\_FILAMENT\_BASED\_PROCESS | MSigDB lists | GO\_ACTIN\_FILAMENT\_BASED\_PROCESS | 102 | 9 | 4161 | 184 | Cdc42,Nf1,pav,krz,Arpc3A,LIMK1,Arpc5,Graf,flr | | 3.527e-02 | -3.34 | GO\_CYSTEINE\_TYPE\_ENDOPEPTIDASE\_INHIBITOR\_ACTIVITY\_INVOLVED\_IN\_APOPTOTIC\_PROCESS | MSigDB lists | GO\_CYSTEINE\_TYPE\_ENDOPEPTIDASE\_INHIBITOR\_ACTIVITY\_INVOLVED\_IN\_APOPTOTIC\_PROCESS | 7 | 2 | 4161 | 184 | krz,sigmar | | 3.527e-02 | -3.34 | HASLINGER\_B\_CLL\_WITH\_CHROMOSOME\_12\_TRISOMY | MSigDB lists | HASLINGER\_B\_CLL\_WITH\_CHROMOSOME\_12\_TRISOMY | 7 | 2 | 4161 | 184 | Arpc3A,Rcd5 | | 3.527e-02 | -3.34 | GO\_INOSITOL\_PHOSPHATE\_CATABOLIC\_PROCESS | MSigDB lists | GO\_INOSITOL\_PHOSPHATE\_CATABOLIC\_PROCESS | 7 | 2 | 4161 | 184 | Aps,Pten | | 3.527e-02 | -3.34 | GO\_CYSTEINE\_TYPE\_ENDOPEPTIDASE\_REGULATOR\_ACTIVITY\_INVOLVED\_IN\_APOPTOTIC\_PROCESS | MSigDB lists | GO\_CYSTEINE\_TYPE\_ENDOPEPTIDASE\_REGULATOR\_ACTIVITY\_INVOLVED\_IN\_APOPTOTIC\_PROCESS | 7 | 2 | 4161 | 184 | sigmar,krz | | 3.527e-02 | -3.34 | GO\_ARP2\_3\_PROTEIN\_COMPLEX | MSigDB lists | GO\_ARP2\_3\_PROTEIN\_COMPLEX | 7 | 2 | 4161 | 184 | Arpc3A,Arpc5 | | 3.527e-02 | -3.34 | GO\_POLY\_PYRIMIDINE\_TRACT\_BINDING | MSigDB lists | GO\_POLY\_PYRIMIDINE\_TRACT\_BINDING | 7 | 2 | 4161 | 184 | Rcd5,Fmr1 | | 3.527e-02 | -3.34 | GO\_ALCOHOL\_DEHYDROGENASE\_NADP\_ACTIVITY | MSigDB lists | GO\_ALCOHOL\_DEHYDROGENASE\_NADP\_ACTIVITY | 7 | 2 | 4161 | 184 | CG10672,CG3842 | | 3.527e-02 | -3.34 | GO\_DNA\_LIGATION | MSigDB lists | GO\_DNA\_LIGATION | 7 | 2 | 4161 | 184 | DNAlig1,Parp | | 3.527e-02 | -3.34 | REACTOME\_SEMA3A\_PAK\_DEPENDENT\_AXON\_REPULSION | MSigDB lists | REACTOME\_SEMA3A\_PAK\_DEPENDENT\_AXON\_REPULSION | 7 | 2 | 4161 | 184 | LIMK1,PlexA | | 3.527e-02 | -3.34 | DORN\_ADENOVIRUS\_INFECTION\_48HR\_DN | MSigDB lists | DORN\_ADENOVIRUS\_INFECTION\_48HR\_DN | 7 | 2 | 4161 | 184 | SF2,lic | | 3.527e-02 | -3.34 | REACTOME\_PI3K\_EVENTS\_IN\_ERBB4\_SIGNALING | MSigDB lists | REACTOME\_PI3K\_EVENTS\_IN\_ERBB4\_SIGNALING | 7 | 2 | 4161 | 184 | Lst8,Pten | | 3.527e-02 | -3.34 | GO\_POLY\_PURINE\_TRACT\_BINDING | MSigDB lists | GO\_POLY\_PURINE\_TRACT\_BINDING | 7 | 2 | 4161 | 184 | Fmr1,Rcd5 | | 3.527e-02 | -3.34 | GO\_ACTIN\_NUCLEATION | MSigDB lists | GO\_ACTIN\_NUCLEATION | 7 | 2 | 4161 | 184 | Arpc3A,Arpc5 | | 3.527e-02 | -3.34 | BIOCARTA\_IL1R\_PATHWAY | MSigDB lists | BIOCARTA\_IL1R\_PATHWAY | 7 | 2 | 4161 | 184 | lic,ECSIT | | 3.527e-02 | -3.34 | GO\_PROTEIN\_CHANNEL\_ACTIVITY | MSigDB lists | GO\_PROTEIN\_CHANNEL\_ACTIVITY | 7 | 2 | 4161 | 184 | Tim17b,Tom20 | | 3.527e-02 | -3.34 | GO\_ARP2\_3\_COMPLEX\_MEDIATED\_ACTIN\_NUCLEATION | MSigDB lists | GO\_ARP2\_3\_COMPLEX\_MEDIATED\_ACTIN\_NUCLEATION | 7 | 2 | 4161 | 184 | Arpc3A,Arpc5 | | 3.527e-02 | -3.34 | GO\_EXON\_EXON\_JUNCTION\_COMPLEX | MSigDB lists | GO\_EXON\_EXON\_JUNCTION\_COMPLEX | 7 | 2 | 4161 | 184 | SF2,tsu | | 3.527e-02 | -3.34 | WANG\_RECURRENT\_LIVER\_CANCER\_UP | MSigDB lists | WANG\_RECURRENT\_LIVER\_CANCER\_UP | 7 | 2 | 4161 | 184 | CG17493,Cul4 | | 3.527e-02 | -3.34 | GO\_REGULATION\_OF\_PHAGOCYTOSIS | MSigDB lists | GO\_REGULATION\_OF\_PHAGOCYTOSIS | 7 | 2 | 4161 | 184 | Pten,Rap1 | | 3.527e-02 | -3.34 | GO\_PROTEASOME\_ASSEMBLY | MSigDB lists | GO\_PROTEASOME\_ASSEMBLY | 7 | 2 | 4161 | 184 | Rpn10,CG12321 | | 3.527e-02 | -3.34 | PENG\_GLUTAMINE\_DEPRIVATION\_UP | MSigDB lists | PENG\_GLUTAMINE\_DEPRIVATION\_UP | 7 | 2 | 4161 | 184 | vap,lic | | 3.527e-02 | -3.34 | KEGG\_CYTOKINE\_CYTOKINE\_RECEPTOR\_INTERACTION | MSigDB lists | KEGG\_CYTOKINE\_CYTOKINE\_RECEPTOR\_INTERACTION | 7 | 2 | 4161 | 184 | dpp,tkv | | 3.527e-02 | -3.34 | GO\_HETEROTRIMERIC\_G\_PROTEIN\_COMPLEX | MSigDB lists | GO\_HETEROTRIMERIC\_G\_PROTEIN\_COMPLEX | 7 | 2 | 4161 | 184 | cta,krz | | 3.527e-02 | -3.34 | GO\_REGULATION\_OF\_LONG\_TERM\_NEURONAL\_SYNAPTIC\_PLASTICITY | MSigDB lists | GO\_REGULATION\_OF\_LONG\_TERM\_NEURONAL\_SYNAPTIC\_PLASTICITY | 7 | 2 | 4161 | 184 | Nf1,Rab8 | | 3.527e-02 | -3.34 | REACTOME\_PIP3\_ACTIVATES\_AKT\_SIGNALING | MSigDB lists | REACTOME\_PIP3\_ACTIVATES\_AKT\_SIGNALING | 7 | 2 | 4161 | 184 | Pten,Lst8 | | 3.527e-02 | -3.34 | DASU\_IL6\_SIGNALING\_UP | MSigDB lists | DASU\_IL6\_SIGNALING\_UP | 7 | 2 | 4161 | 184 | Sod2,CG41378 | | 3.527e-02 | -3.34 | REACTOME\_PI3K\_AKT\_ACTIVATION | MSigDB lists | REACTOME\_PI3K\_AKT\_ACTIVATION | 7 | 2 | 4161 | 184 | Lst8,Pten | | 3.527e-02 | -3.34 | GO\_POSITIVE\_REGULATION\_OF\_INTRACELLULAR\_STEROID\_HORMONE\_RECEPTOR\_SIGNALING\_PATHWAY | MSigDB lists | GO\_POSITIVE\_REGULATION\_OF\_INTRACELLULAR\_STEROID\_HORMONE\_RECEPTOR\_SIGNALING\_PATHWAY | 7 | 2 | 4161 | 184 | l(1)10Bb,Skp2 | | 3.527e-02 | -3.34 | GHANDHI\_DIRECT\_IRRADIATION\_UP | MSigDB lists | GHANDHI\_DIRECT\_IRRADIATION\_UP | 7 | 2 | 4161 | 184 | dpp,Sod2 | | 3.533e-02 | -3.34 | N (Notch) | protein interactions | 31293 | 301 | 16 | 9204 | 298 | tkv,krz,RPA3,dgrn,Amun,Cdk8,CtBP,dpp,MED21,RpL15,flw,Stim,sm,Btk29A,Mlf,Cdc42 | | 3.537e-02 | -3.34 | zf-C2H2\_jaz | pfam domains | PF12171 | 10 | 2 | 10480 | 319 | CG11586,Pep | | 3.538e-02 | -3.34 | positive regulation of catabolic process | biological process | GO:0009896 | 125 | 7 | 12345 | 305 | Cpsf6,Rpt6,AGO3,smg,Cul4,CG9951,Cdk8 | | 3.538e-02 | -3.34 | SUMOylation of DNA damage response and repair proteins | REACTOME pathways | R-DME-3108214 | 8 | 2 | 3923 | 151 | smt3,Parp | | 3.538e-02 | -3.34 | activated TAK1 mediates p38 MAPK activation | REACTOME pathways | R-DME-450302 | 8 | 2 | 3923 | 151 | pll,lic | | 3.555e-02 | -3.34 | inner mitochondrial membrane protein complex | cellular component | GO:0098800 | 151 | 8 | 11940 | 298 | MICU1,CG11752,ND-ASHI,sun,l(2)37Bb,CIA30,Tim17b,ND-AGGG | | 3.571e-02 | -3.33 | GSE37416\_CTRL\_VS\_6H\_F\_TULARENSIS\_LVS\_NEUTROPHIL\_UP | MSigDB lists | GSE37416\_CTRL\_VS\_6H\_F\_TULARENSIS\_LVS\_NEUTROPHIL\_UP | 56 | 6 | 4161 | 184 | Dbp21E2,vnc,Aps,VhaPPA1-1,Cul4,CG11586 | | 3.571e-02 | -3.33 | GSE10500\_ARTHRITIC\_SYNOVIAL\_FLUID\_VS\_HEALTHY\_MACROPHAGE\_UP | MSigDB lists | GSE10500\_ARTHRITIC\_SYNOVIAL\_FLUID\_VS\_HEALTHY\_MACROPHAGE\_UP | 56 | 6 | 4161 | 184 | Tsp97E,His2Av,inc,Phf5a,cta,eEF1delta | | 3.571e-02 | -3.33 | GSE17721\_CTRL\_VS\_LPS\_0.5H\_BMDC\_DN | MSigDB lists | GSE17721\_CTRL\_VS\_LPS\_0.5H\_BMDC\_DN | 56 | 6 | 4161 | 184 | CG13016,CG17883,Rpn10,ubl,Stlk,lic | | 3.571e-02 | -3.33 | GSE37301\_RAG2\_KO\_VS\_RAG2\_AND\_ETS1\_KO\_NK\_CELL\_UP | MSigDB lists | GSE37301\_RAG2\_KO\_VS\_RAG2\_AND\_ETS1\_KO\_NK\_CELL\_UP | 56 | 6 | 4161 | 184 | CG9330,fzy,Phf5a,flw,CG32473,Exo84 | | 3.571e-02 | -3.33 | GO\_REGULATION\_OF\_SMALL\_GTPASE\_MEDIATED\_SIGNAL\_TRANSDUCTION | MSigDB lists | GO\_REGULATION\_OF\_SMALL\_GTPASE\_MEDIATED\_SIGNAL\_TRANSDUCTION | 56 | 6 | 4161 | 184 | Nf1,vap,RhoGAPp190,Cdc42,Graf,krz | | 3.571e-02 | -3.33 | GSE5589\_LPS\_VS\_LPS\_AND\_IL6\_STIM\_MACROPHAGE\_45MIN\_UP | MSigDB lists | GSE5589\_LPS\_VS\_LPS\_AND\_IL6\_STIM\_MACROPHAGE\_45MIN\_UP | 56 | 6 | 4161 | 184 | mon2,CG11586,me31B,MED22,sws,Roc2 | | 3.571e-02 | -3.33 | REACTOME\_SIGNALING\_BY\_THE\_B\_CELL\_RECEPTOR\_BCR | MSigDB lists | REACTOME\_SIGNALING\_BY\_THE\_B\_CELL\_RECEPTOR\_BCR | 56 | 6 | 4161 | 184 | Stim,Lst8,Pten,Rpn10,Prosalpha2,Rpt6 | | 3.571e-02 | -3.33 | GSE7831\_UNSTIM\_VS\_CPG\_STIM\_PDC\_1H\_DN | MSigDB lists | GSE7831\_UNSTIM\_VS\_CPG\_STIM\_PDC\_1H\_DN | 56 | 6 | 4161 | 184 | VhaPPA1-1,vap,Sod2,bai,Ufd4,E(z) | | 3.571e-02 | -3.33 | GSE17721\_PAM3CSK4\_VS\_GADIQUIMOD\_24H\_BMDC\_UP | MSigDB lists | GSE17721\_PAM3CSK4\_VS\_GADIQUIMOD\_24H\_BMDC\_UP | 56 | 6 | 4161 | 184 | Pect,RpS9,krz,eEF1delta,ubl,vnc | | 3.578e-02 | -3.33 | vesicle | cellular component | GO:0031982 | 422 | 17 | 11940 | 298 | Vap33,g,ema,dpp,RagA-B,Rab14,vsg,bai,tkv,Rab8,RabX6,Graf,mon2,CG30423,Rab7,CG15168,Vps35 | | 3.581e-02 | -3.33 | LEE\_BMP2\_TARGETS\_DN | MSigDB lists | LEE\_BMP2\_TARGETS\_DN | 410 | 26 | 4161 | 184 | SF2,CG10343,CCT4,Skp2,CG5535,CSN1b,Txl,Lrr47,Tom20,Fs(2)Ket,CG14805,LIMK1,Phf5a,Ubqn,Art1,l(2)37Bb,E(z),CG15168,RpS9,CG32576,HtrA2,CG9246,eRF1,Fign,dpp,Parp | | 3.598e-02 | -3.32 | long-term memory | biological process | GO:0007616 | 74 | 5 | 12345 | 305 | vsg,CG4612,spoon,Fmr1,Nf1 | | 3.600e-02 | -3.32 | FLECHNER\_BIOPSY\_KIDNEY\_TRANSPLANT\_OK\_VS\_DONOR\_UP | MSigDB lists | FLECHNER\_BIOPSY\_KIDNEY\_TRANSPLANT\_OK\_VS\_DONOR\_UP | 205 | 15 | 4161 | 184 | Rpt6,Mdh1,eRF1,Rab8,Parp,xit,SF2,bai,His2Av,Pex3,Arpc5,CIA30,Dbp80,CG8635,eEF1delta | | 3.601e-02 | -3.32 | GSE13547\_CTRL\_VS\_ANTI\_IGM\_STIM\_BCELL\_12H\_DN | MSigDB lists | GSE13547\_CTRL\_VS\_ANTI\_IGM\_STIM\_BCELL\_12H\_DN | 71 | 7 | 4161 | 184 | mRpS28,CG11586,eEF1delta,RpL14,eIF3g1,RpL5,RpL36A | | 3.601e-02 | -3.32 | GSE33425\_CD161\_INT\_VS\_NEG\_CD8\_TCELL\_UP | MSigDB lists | GSE33425\_CD161\_INT\_VS\_NEG\_CD8\_TCELL\_UP | 71 | 7 | 4161 | 184 | CG10672,Cdk8,CG5861,flw,krz,Roc2,Rab8 | | 3.601e-02 | -3.32 | GSE27786\_LIN\_NEG\_VS\_NKCELL\_UP | MSigDB lists | GSE27786\_LIN\_NEG\_VS\_NKCELL\_UP | 71 | 7 | 4161 | 184 | CG11334,CG12321,mus301,vnc,pins,eIF3g1,Rpt6 | | 3.601e-02 | -3.32 | LU\_EZH2\_TARGETS\_UP | MSigDB lists | LU\_EZH2\_TARGETS\_UP | 71 | 7 | 4161 | 184 | Phf5a,CG31460,CG4743,Sod2,Rpn10,CG5590,CG4935 | | 3.613e-02 | -3.32 | InR (Insulin-like receptor) | protein interactions | 42549 | 76 | 6 | 9204 | 298 | glo,SF2,Pten,RpL15,14-3-3zeta,CNBP | | 3.613e-02 | -3.32 | CG6084 (uncharacterized protein) | protein interactions | 39304 | 76 | 6 | 9204 | 298 | Sgt,eEF5,Sod2,smt3,eEF1delta,flw | | 3.613e-02 | -3.32 | regulation of synapse organization | biological process | GO:0050807 | 153 | 8 | 12345 | 305 | Spn,mnb,Rab8,Rheb,tkv,Cdc42,Fmr1,Nf1 | | 3.616e-02 | -3.32 | GSE46606\_DAY1\_VS\_DAY3\_CD40L\_IL2\_IL5\_STIMULATED\_BCELL\_DN | MSigDB lists | GSE46606\_DAY1\_VS\_DAY3\_CD40L\_IL2\_IL5\_STIMULATED\_BCELL\_DN | 42 | 5 | 4161 | 184 | sigmar,dpp,me31B,mbf1,CG5535 | | 3.616e-02 | -3.32 | GSE360\_L\_DONOVANI\_VS\_B\_MALAYI\_LOW\_DOSE\_DC\_UP | MSigDB lists | GSE360\_L\_DONOVANI\_VS\_B\_MALAYI\_LOW\_DOSE\_DC\_UP | 42 | 5 | 4161 | 184 | E(z),lic,CG17493,Sod2,RagA-B | | 3.616e-02 | -3.32 | GSE2770\_TGFB\_AND\_IL4\_VS\_TGFB\_AND\_IL12\_TREATED\_ACT\_CD4\_TCELL\_6H\_UP | MSigDB lists | GSE2770\_TGFB\_AND\_IL4\_VS\_TGFB\_AND\_IL12\_TREATED\_ACT\_CD4\_TCELL\_6H\_UP | 42 | 5 | 4161 | 184 | unc-119,E(z),CG17883,CG31460,CG14805 | | 3.616e-02 | -3.32 | REACTOME\_SCFSKP2\_MEDIATED\_DEGRADATION\_OF\_P27\_P21 | MSigDB lists | REACTOME\_SCFSKP2\_MEDIATED\_DEGRADATION\_OF\_P27\_P21 | 42 | 5 | 4161 | 184 | Skp2,CycA,Rpt6,Prosalpha2,Rpn10 | | 3.616e-02 | -3.32 | GO\_NEGATIVE\_REGULATION\_OF\_NEURON\_DEATH | MSigDB lists | GO\_NEGATIVE\_REGULATION\_OF\_NEURON\_DEATH | 42 | 5 | 4161 | 184 | Rop,vap,spel1,Sod2,HtrA2 | | 3.616e-02 | -3.32 | GSE16385\_UNTREATED\_VS\_12H\_ROSIGLITAZONE\_IFNG\_TNF\_TREATED\_MACROPHAGE\_DN | MSigDB lists | GSE16385\_UNTREATED\_VS\_12H\_ROSIGLITAZONE\_IFNG\_TNF\_TREATED\_MACROPHAGE\_DN | 42 | 5 | 4161 | 184 | Nf1,Fgop2,Orc3,tkv,Stlk | | 3.616e-02 | -3.32 | GSE32901\_NAIVE\_VS\_TH17\_NEG\_CD4\_TCELL\_UP | MSigDB lists | GSE32901\_NAIVE\_VS\_TH17\_NEG\_CD4\_TCELL\_UP | 42 | 5 | 4161 | 184 | pav,DNAlig1,CycA,Rcd5,CG41378 | | 3.616e-02 | -3.32 | GO\_REGULATION\_OF\_ACTIN\_FILAMENT\_LENGTH | MSigDB lists | GO\_REGULATION\_OF\_ACTIN\_FILAMENT\_LENGTH | 42 | 5 | 4161 | 184 | Arpc3A,Arpc5,Lst8,flr,vap | | 3.636e-02 | -3.31 | response to stimulus | biological process | GO:0050896 | 2655 | 79 | 12345 | 305 | Cul4,Mps1,Rheb,Rab8,sun,spel1,RecQ4,RhoGAPp190,BthD,flw,CG1791,14-3-3zeta,CG6171,Btk29A,E(bx),pll,vap,Pten,Rap1,dgrn,CG1218,CG6066,Spn,mus301,Hsp27,mav,cta,Rop,Lst8,glo,ECSIT,Stlk,Rab7,DNAlig1,PlexA,spoon,alpha-Man-Ib,Tctp,Hrb27C,CG4743,lic,CG9005,Sod2,RPA3,Parp,su(Hw),PPP1R15,Vap33,Orc3,smt3,Fmr1,Fign,tkv,mthl8,Graf,CG5535,Zpr1,Ino80,Sgt,Cdc42,Aps,CIA30,Bmcp,Cdk8,dpp,Lrr47,CG5056,E(z),RagA-B,Pp1-87B,Nf1,Vps35,mnb,Rab14,sm,vsg,Stim,RabX6,Arp8 | | 3.641e-02 | -3.31 | GO\_REGULATION\_OF\_CELL\_PROJECTION\_ORGANIZATION | MSigDB lists | GO\_REGULATION\_OF\_CELL\_PROJECTION\_ORGANIZATION | 119 | 10 | 4161 | 184 | Pten,Fmr1,PlexA,Cdc42,fzy,E(z),LIMK1,RhoGAPp190,Rap1,CG17883 | | 3.641e-02 | -3.31 | Signaling by Interleukins | REACTOME pathways | R-DME-449147 | 156 | 11 | 3923 | 151 | 14-3-3zeta,CG7054,Prosbeta1,Nf1,Rpt6,vap,Prosalpha2,CG4935,Rpn10,pll,Btk29A | | 3.649e-02 | -3.31 | cell fate commitment | biological process | GO:0045165 | 301 | 13 | 12345 | 305 | mus301,dpp,CycA,14-3-3zeta,CtBP,pav,pins,Rap1,Fmr1,Cdc42,Pten,tsu,tkv | | 3.661e-02 | -3.31 | GO\_TRANSLATION\_INITIATION\_FACTOR\_BINDING | MSigDB lists | GO\_TRANSLATION\_INITIATION\_FACTOR\_BINDING | 17 | 3 | 4161 | 184 | Fmr1,eIF3c,eIF3b | | 3.661e-02 | -3.31 | GO\_NUCLEOTIDE\_EXCISION\_REPAIR\_DNA\_DUPLEX\_UNWINDING | MSigDB lists | GO\_NUCLEOTIDE\_EXCISION\_REPAIR\_DNA\_DUPLEX\_UNWINDING | 17 | 3 | 4161 | 184 | Parp,CG17493,Cul4 | | 3.661e-02 | -3.31 | GO\_PEPTIDYL\_LYSINE\_METHYLATION | MSigDB lists | GO\_PEPTIDYL\_LYSINE\_METHYLATION | 17 | 3 | 4161 | 184 | Wdr82,E(z),Rbbp5 | | 3.661e-02 | -3.31 | DER\_IFN\_GAMMA\_RESPONSE\_UP | MSigDB lists | DER\_IFN\_GAMMA\_RESPONSE\_UP | 17 | 3 | 4161 | 184 | VhaPPA1-1,Parp,CG41378 | | 3.661e-02 | -3.31 | GO\_REGULATION\_OF\_CALCIUM\_ION\_DEPENDENT\_EXOCYTOSIS | MSigDB lists | GO\_REGULATION\_OF\_CALCIUM\_ION\_DEPENDENT\_EXOCYTOSIS | 17 | 3 | 4161 | 184 | Rop,Rap1,Fmr1 | | 3.661e-02 | -3.31 | GUO\_HEX\_TARGETS\_DN | MSigDB lists | GUO\_HEX\_TARGETS\_DN | 17 | 3 | 4161 | 184 | Rpb11,Sod2,DNAlig1 | | 3.661e-02 | -3.31 | ZHAN\_MULTIPLE\_MYELOMA\_SUBGROUPS | MSigDB lists | ZHAN\_MULTIPLE\_MYELOMA\_SUBGROUPS | 17 | 3 | 4161 | 184 | E(z),CCT4,spel1 | | 3.661e-02 | -3.31 | GGCCAGT\_MIR193A\_MIR193B | MSigDB lists | GGCCAGT\_MIR193A\_MIR193B | 17 | 3 | 4161 | 184 | Arpc5,Pten,14-3-3zeta | | 3.661e-02 | -3.31 | PID\_CXCR3\_PATHWAY | MSigDB lists | PID\_CXCR3\_PATHWAY | 17 | 3 | 4161 | 184 | krz,lic,Lst8 | | 3.661e-02 | -3.31 | GSE32901\_TH1\_VS\_TH17\_NEG\_CD4\_TCELL\_DN | MSigDB lists | GSE32901\_TH1\_VS\_TH17\_NEG\_CD4\_TCELL\_DN | 17 | 3 | 4161 | 184 | sigmar,fzy,Fign | | 3.661e-02 | -3.31 | SIG\_PIP3\_SIGNALING\_IN\_CARDIAC\_MYOCTES | MSigDB lists | SIG\_PIP3\_SIGNALING\_IN\_CARDIAC\_MYOCTES | 17 | 3 | 4161 | 184 | Cdc42,14-3-3zeta,Pten | | 3.661e-02 | -3.31 | BROWNE\_HCMV\_INFECTION\_8HR\_UP | MSigDB lists | BROWNE\_HCMV\_INFECTION\_8HR\_UP | 17 | 3 | 4161 | 184 | cta,Rbbp5,Sod2 | | 3.661e-02 | -3.31 | Signalling to ERKs | REACTOME pathways | R-DME-187687 | 118 | 9 | 3923 | 151 | CG4935,Rpn10,Prosalpha2,Nf1,Rpt6,vap,CG7054,14-3-3zeta,Prosbeta1 | | 3.661e-02 | -3.31 | Signalling to RAS | REACTOME pathways | R-DME-167044 | 118 | 9 | 3923 | 151 | Rpn10,CG4935,Prosalpha2,14-3-3zeta,CG7054,Prosbeta1,vap,Nf1,Rpt6 | | 3.661e-02 | -3.31 | FCERI mediated MAPK activation | REACTOME pathways | R-DME-2871796 | 118 | 9 | 3923 | 151 | vap,Nf1,Rpt6,Prosbeta1,14-3-3zeta,CG7054,CG4935,Rpn10,Prosalpha2 | | 3.665e-02 | -3.31 | morphogenesis of an epithelium | biological process | GO:0002009 | 625 | 23 | 12345 | 305 | Zpr1,CSN1b,GEFmeso,Stim,flr,VhaPPA1-1,Btk29A,tsr,tkv,spoon,rhea,Cdk8,dpp,LIMK1,sqd,flw,cta,peng,Parp,Cdc42,Rap1,Rheb,EloB | | 3.682e-02 | -3.30 | regulation of dendrite development | biological process | GO:0050773 | 51 | 4 | 12345 | 305 | Rop,mav,Pten,Fmr1 | | 3.686e-02 | -3.30 | fand (fandango) | protein interactions | 36514 | 23 | 3 | 9204 | 298 | CG12659,RAF2,l(1)10Bb | | 3.686e-02 | -3.30 | mbt (mushroom bodies tiny) | protein interactions | 32631 | 23 | 3 | 9204 | 298 | LIMK1,tsr,Cdc42 | | 3.686e-02 | -3.30 | fs(1)K10 (female sterile (1) K10) | protein interactions | 47781 | 23 | 3 | 9204 | 298 | sqd,Hrb27C,DCTN3-p24 | | 3.686e-02 | -3.30 | Arp6 (Actin-related protein 6) | protein interactions | 32514 | 23 | 3 | 9204 | 298 | Rpt6,RpS3A,me31B | | 3.686e-02 | -3.30 | Ccs (Copper chaperone for superoxide dismutase) | protein interactions | 46035 | 23 | 3 | 9204 | 298 | CG1218,Wdr82,BthD | | 3.686e-02 | -3.30 | CG1218 (uncharacterized protein) | protein interactions | 40718 | 23 | 3 | 9204 | 298 | CG6171,BthD,Wdr82 | | 3.686e-02 | -3.30 | Arp3 (Actin-related protein 3) | protein interactions | 38898 | 23 | 3 | 9204 | 298 | Arpc3A,RpL35A,Arpc5 | | 3.686e-02 | -3.30 | CG7231 (uncharacterized protein) | protein interactions | 34086 | 23 | 3 | 9204 | 298 | Cdc42,RpS3A,CG5174 | | 3.701e-02 | -3.30 | GO\_REGULATION\_OF\_ERK1\_AND\_ERK2\_CASCADE | MSigDB lists | GO\_REGULATION\_OF\_ERK1\_AND\_ERK2\_CASCADE | 29 | 4 | 4161 | 184 | krz,dpp,Pten,Rap1 | | 3.701e-02 | -3.30 | GO\_REGULATION\_OF\_DENDRITE\_DEVELOPMENT | MSigDB lists | GO\_REGULATION\_OF\_DENDRITE\_DEVELOPMENT | 29 | 4 | 4161 | 184 | E(z),Fmr1,Pten,fzy | | 3.701e-02 | -3.30 | GO\_N\_METHYLTRANSFERASE\_ACTIVITY | MSigDB lists | GO\_N\_METHYLTRANSFERASE\_ACTIVITY | 29 | 4 | 4161 | 184 | Wdr82,E(z),Art1,Rbbp5 | | 3.701e-02 | -3.30 | FLOTHO\_PEDIATRIC\_ALL\_THERAPY\_RESPONSE\_UP | MSigDB lists | FLOTHO\_PEDIATRIC\_ALL\_THERAPY\_RESPONSE\_UP | 29 | 4 | 4161 | 184 | Tctp,RpL5,Tom20,RpS9 | | 3.701e-02 | -3.30 | GSE10147\_IL3\_AND\_HIVP17\_VS\_IL3\_AND\_CPG\_STIM\_PDC\_UP | MSigDB lists | GSE10147\_IL3\_AND\_HIVP17\_VS\_IL3\_AND\_CPG\_STIM\_PDC\_UP | 29 | 4 | 4161 | 184 | RpL36A,CG31460,CG32473,RpS26 | | 3.701e-02 | -3.30 | ATAGGAA\_MIR202 | MSigDB lists | ATAGGAA\_MIR202 | 29 | 4 | 4161 | 184 | PGAP1,CtBP,vap,Pten | | 3.701e-02 | -3.30 | GSE13522\_CTRL\_VS\_T\_CRUZI\_Y\_STRAIN\_INF\_SKIN\_IFNAR\_KO\_UP | MSigDB lists | GSE13522\_CTRL\_VS\_T\_CRUZI\_Y\_STRAIN\_INF\_SKIN\_IFNAR\_KO\_UP | 29 | 4 | 4161 | 184 | RpS8,Stim,RpL14,CG5339 | | 3.703e-02 | -3.30 | REACTOME\_ADAPTIVE\_IMMUNE\_SYSTEM | MSigDB lists | REACTOME\_ADAPTIVE\_IMMUNE\_SYSTEM | 188 | 14 | 4161 | 184 | Cdc42,fzy,Rpt6,CG8860,pav,Prosalpha2,Lrr47,Pten,Rap1,Rpn10,Skp2,14-3-3zeta,Stim,Lst8 | | 3.707e-02 | -3.29 | cytoplasmic microtubule organization | biological process | GO:0031122 | 30 | 3 | 12345 | 305 | Fign,Girdin,Vap33 | | 3.707e-02 | -3.29 | regulation of mitotic metaphase/anaphase transition | biological process | GO:0030071 | 30 | 3 | 12345 | 305 | CycA,Bub1,Mps1 | | 3.714e-02 | -3.29 | GO\_WOUND\_HEALING | MSigDB lists | GO\_WOUND\_HEALING | 103 | 9 | 4161 | 184 | Fgop2,Nf1,Cdc42,Rop,14-3-3zeta,rb,E(z),krz,cta | | 3.714e-02 | -3.29 | GO\_SMALL\_GTPASE\_MEDIATED\_SIGNAL\_TRANSDUCTION | MSigDB lists | GO\_SMALL\_GTPASE\_MEDIATED\_SIGNAL\_TRANSDUCTION | 103 | 9 | 4161 | 184 | Nf1,Cdc42,Rab8,Fs(2)Ket,RhoGAPp190,Rap1,CycA,LIMK1,cta | | 3.715e-02 | -3.29 | GO\_POSITIVE\_REGULATION\_OF\_CELL\_CYCLE | MSigDB lists | GO\_POSITIVE\_REGULATION\_OF\_CELL\_CYCLE | 87 | 8 | 4161 | 184 | Cul4,Ino80,twin,Art1,Cdc42,pav,AGO3,Pten | | 3.715e-02 | -3.29 | FUJII\_YBX1\_TARGETS\_DN | MSigDB lists | FUJII\_YBX1\_TARGETS\_DN | 87 | 8 | 4161 | 184 | pav,DNAlig1,sm,Fign,Vap33,Cdk8,His2Av,Skp2 | | 3.715e-02 | -3.29 | GSE6674\_ANTI\_IGM\_VS\_ANTI\_IGM\_AND\_CPG\_STIM\_BCELL\_DN | MSigDB lists | GSE6674\_ANTI\_IGM\_VS\_ANTI\_IGM\_AND\_CPG\_STIM\_BCELL\_DN | 87 | 8 | 4161 | 184 | eIF3b,Ubqn,Art1,CG9246,Sod2,CSN1b,eIF3g1,snf | | 3.715e-02 | -3.29 | GSE17721\_CTRL\_VS\_POLYIC\_24H\_BMDC\_UP | MSigDB lists | GSE17721\_CTRL\_VS\_POLYIC\_24H\_BMDC\_UP | 87 | 8 | 4161 | 184 | sigmar,Pp1-87B,Tom20,CG5861,Cul4,CycA,janA,Stim | | 3.715e-02 | -3.29 | RIZKI\_TUMOR\_INVASIVENESS\_3D\_DN | MSigDB lists | RIZKI\_TUMOR\_INVASIVENESS\_3D\_DN | 87 | 8 | 4161 | 184 | CCT4,CG41378,Art1,Orc3,dpp,Tim17b,Cdc42,Txl | | 3.718e-02 | -3.29 | OSIALOPTASE | prints domains | PR00789 | 2 | 1 | 2185 | 41 | Tcs3 | | 3.718e-02 | -3.29 | 1433ZETA | prints domains | PR00305 | 2 | 1 | 2185 | 41 | 14-3-3zeta | | 3.718e-02 | -3.29 | OM20RECEPTOR | prints domains | PR00351 | 2 | 1 | 2185 | 41 | Tom20 | | 3.718e-02 | -3.29 | EUOM20RECPTR | prints domains | PR01989 | 2 | 1 | 2185 | 41 | Tom20 | | 3.733e-02 | -3.29 | cellular component assembly involved in morphogenesis | biological process | GO:0010927 | 154 | 8 | 12345 | 305 | smt3,flr,rhea,RecQ4,dpp,spoon,Cul4,Cdc42 | | 3.746e-02 | -3.28 | Ubx (Ultrabithorax) | protein interactions | 42034 | 118 | 8 | 9204 | 298 | tkv,eIF4B,E(z),E(bx),dpp,CG32576,Hrb27C,su(Hw) | | 3.781e-02 | -3.28 | positive regulation of neuron differentiation | biological process | GO:0045666 | 75 | 5 | 12345 | 305 | PlexA,Rop,mnb,Rheb,Rap1 | | 3.787e-02 | -3.27 | MODULE\_151 | MSigDB lists | MODULE\_151 | 171 | 13 | 4161 | 184 | RpL14,RpL35A,RpL15,eIF3g1,VhaPPA1-1,CG4866,Prosalpha2,ubl,Mdh1,Arpc3A,eEF1delta,RpS9,RpL36A | | 3.846e-02 | -3.26 | GSE26030\_TH1\_VS\_TH17\_RESTIMULATED\_DAY15\_POST\_POLARIZATION\_UP | MSigDB lists | GSE26030\_TH1\_VS\_TH17\_RESTIMULATED\_DAY15\_POST\_POLARIZATION\_UP | 72 | 7 | 4161 | 184 | Skp2,CG5339,His2Av,rhea,CG12321,CycA,RhoGAPp190 | | 3.850e-02 | -3.26 | GSE17721\_LPS\_VS\_GARDIQUIMOD\_0.5H\_BMDC\_DN | MSigDB lists | GSE17721\_LPS\_VS\_GARDIQUIMOD\_0.5H\_BMDC\_DN | 57 | 6 | 4161 | 184 | Rap1,Aps,CG32576,smt3,RpS26,Art1 | | 3.850e-02 | -3.26 | GSE29618\_PRE\_VS\_DAY7\_POST\_TIV\_FLU\_VACCINE\_PDC\_DN | MSigDB lists | GSE29618\_PRE\_VS\_DAY7\_POST\_TIV\_FLU\_VACCINE\_PDC\_DN | 57 | 6 | 4161 | 184 | CG9246,CtBP,Rbbp5,Graf,Rab8,Fs(2)Ket | | 3.850e-02 | -3.26 | GSE33425\_CD8\_ALPHAALPHA\_VS\_ALPHABETA\_CD161\_HIGH\_TCELL\_DN | MSigDB lists | GSE33425\_CD8\_ALPHAALPHA\_VS\_ALPHABETA\_CD161\_HIGH\_TCELL\_DN | 57 | 6 | 4161 | 184 | CG10672,pav,Phf5a,mRpS28,Rab8,E(z) | | 3.850e-02 | -3.26 | GSE339\_EX\_VIVO\_VS\_IN\_CULTURE\_CD8POS\_DC\_DN | MSigDB lists | GSE339\_EX\_VIVO\_VS\_IN\_CULTURE\_CD8POS\_DC\_DN | 57 | 6 | 4161 | 184 | cta,lic,Tim17b,eRF1,Vap33,Mdh1 | | 3.850e-02 | -3.26 | PURBEY\_TARGETS\_OF\_CTBP1\_AND\_SATB1\_DN | MSigDB lists | PURBEY\_TARGETS\_OF\_CTBP1\_AND\_SATB1\_DN | 57 | 6 | 4161 | 184 | Rap1,Dbp21E2,Phf5a,rhea,lic,sm | | 3.870e-02 | -3.25 | GO\_REGULATION\_OF\_PROTEIN\_MODIFICATION\_PROCESS | MSigDB lists | GO\_REGULATION\_OF\_PROTEIN\_MODIFICATION\_PROCESS | 413 | 26 | 4161 | 184 | Rpn10,Vap33,Rap1,Lst8,E(z),Art1,LIMK1,Ubqn,Nf1,viaf,Cdc42,CG9246,fzy,Pten,dpp,CycA,lic,krz,Skp2,CSN1b,Prosalpha2,Rpt6,AGO3,CG6907,Stlk,unc-119 | | 3.899e-02 | -3.24 | Rib\_L2\_dom2 | interpro domains | IPR014722 | 11 | 2 | 11383 | 331 | eEF5,RpL14 | | 3.899e-02 | -3.24 | Translation\_prot\_SH3-like\_sf | interpro domains | IPR008991 | 11 | 2 | 11383 | 331 | eEF5,RpL14 | | 3.903e-02 | -3.24 | eIF4A (eukaryotic translation initiation factor 4A) | protein interactions | 33835 | 58 | 5 | 9204 | 298 | Dref,Pten,dpp,Rheb,LIMK1 | | 3.911e-02 | -3.24 | BROMODOMAIN | prints domains | PR00503 | 17 | 2 | 2185 | 41 | Acf,E(bx) | | 3.914e-02 | -3.24 | positive regulation of protein complex assembly | biological process | GO:0031334 | 52 | 4 | 12345 | 305 | Arpc5,cal1,Rpt6,Arpc3A | | 3.914e-02 | -3.24 | morphogenesis of follicular epithelium | biological process | GO:0016333 | 52 | 4 | 12345 | 305 | Parp,CSN1b,sqd,spoon | | 3.916e-02 | -3.24 | actin binding | molecular function | GO:0003779 | 133 | 7 | 12176 | 289 | Arpc5,tsr,flr,Spn,Arpc3A,rhea,Pten | | 3.929e-02 | -3.24 | UCH proteinases | REACTOME pathways | R-DME-5689603 | 101 | 8 | 3923 | 151 | Prosbeta1,Rpt6,Ino80,CG12659,Rcd5,Arp8,Rpn10,Prosalpha2 | | 3.938e-02 | -3.23 | CATTTCA\_MIR203 | MSigDB lists | CATTTCA\_MIR203 | 88 | 8 | 4161 | 184 | Rap1,SF2,Rat1,me31B,flw,Fs(2)Ket,sm,Fmr1 | | 3.938e-02 | -3.23 | GSE17721\_CTRL\_VS\_CPG\_4H\_BMDC\_UP | MSigDB lists | GSE17721\_CTRL\_VS\_CPG\_4H\_BMDC\_UP | 88 | 8 | 4161 | 184 | CG41378,janA,Rap1,eIF3b,CG5861,Aps,RpS8,CSN1b | | 3.946e-02 | -3.23 | Epac (Exchange protein directly activated by cAMP) | protein interactions | 35588 | 40 | 4 | 9204 | 298 | Pten,Rap1,RpS9,janA | | 3.946e-02 | -3.23 | homer (homer) | protein interactions | 33944 | 40 | 4 | 9204 | 298 | EloB,CG1637,RpL35A,Roc2 | | 3.950e-02 | -3.23 | GSE2706\_2H\_VS\_8H\_R848\_AND\_LPS\_STIM\_DC\_DN | MSigDB lists | GSE2706\_2H\_VS\_8H\_R848\_AND\_LPS\_STIM\_DC\_DN | 43 | 5 | 4161 | 184 | Sod2,CG32576,Rbbp5,CG8860,Prosalpha2 | | 3.950e-02 | -3.23 | GSE22886\_NAIVE\_CD4\_TCELL\_VS\_12H\_ACT\_TH1\_UP | MSigDB lists | GSE22886\_NAIVE\_CD4\_TCELL\_VS\_12H\_ACT\_TH1\_UP | 43 | 5 | 4161 | 184 | Stim,RpS9,CG7766,Vps13D,krz | | 3.950e-02 | -3.23 | GSE3982\_MAST\_CELL\_VS\_BASOPHIL\_DN | MSigDB lists | GSE3982\_MAST\_CELL\_VS\_BASOPHIL\_DN | 43 | 5 | 4161 | 184 | HtrA2,Prosalpha2,Nf1,CG5590,Skp2 | | 3.950e-02 | -3.23 | GO\_CELLULAR\_RESPONSE\_TO\_RADIATION | MSigDB lists | GO\_CELLULAR\_RESPONSE\_TO\_RADIATION | 43 | 5 | 4161 | 184 | eEF1delta,Fign,Ino80,CG9246,Cul4 | | 3.950e-02 | -3.23 | GO\_MITOTIC\_CELL\_CYCLE\_CHECKPOINT | MSigDB lists | GO\_MITOTIC\_CELL\_CYCLE\_CHECKPOINT | 43 | 5 | 4161 | 184 | CycA,CG12321,spel1,twin,Art1 | | 3.950e-02 | -3.23 | WATANABE\_RECTAL\_CANCER\_RADIOTHERAPY\_RESPONSIVE\_UP | MSigDB lists | WATANABE\_RECTAL\_CANCER\_RADIOTHERAPY\_RESPONSIVE\_UP | 43 | 5 | 4161 | 184 | Cul4,Nulp1,RpL36A,flr,Rap1 | | 3.950e-02 | -3.23 | GSE34156\_TLR1\_TLR2\_LIGAND\_VS\_NOD2\_AND\_TLR1\_TLR2\_LIGAND\_6H\_TREATED\_MONOCYTE\_UP | MSigDB lists | GSE34156\_TLR1\_TLR2\_LIGAND\_VS\_NOD2\_AND\_TLR1\_TLR2\_LIGAND\_6H\_TREATED\_MONOCYTE\_UP | 43 | 5 | 4161 | 184 | Aps,Nf1,Skp2,Rab8,Bmcp | | 3.954e-02 | -3.23 | N-acetyltransferase activity | molecular function | GO:0008080 | 79 | 5 | 12176 | 289 | Eaf6,vnc,Rcd5,Acf,d4 | | 3.954e-02 | -3.23 | core promoter binding | molecular function | GO:0001047 | 32 | 3 | 12176 | 289 | Dref,CG6254,Pbp49 | | 3.960e-02 | -3.23 | Ube3a (Ubiquitin protein ligase E3A) | protein interactions | 39266 | 10 | 2 | 9204 | 298 | tkv,Rpn10 | | 3.960e-02 | -3.23 | CG13097 (uncharacterized protein) | protein interactions | 34184 | 10 | 2 | 9204 | 298 | CG11920,CG4866 | | 3.960e-02 | -3.23 | Dpy-30L1 (Dpy-30-like 1) | protein interactions | 34480 | 10 | 2 | 9204 | 298 | Rbbp5,Wdr82 | | 3.960e-02 | -3.23 | Roc1a (Regulator of cullins 1a) | protein interactions | 31014 | 10 | 2 | 9204 | 298 | Zpr1,Cul4 | | 3.960e-02 | -3.23 | wech (wech) | protein interactions | 44653 | 10 | 2 | 9204 | 298 | noi,rhea | | 3.960e-02 | -3.23 | mRpL23 (mitochondrial ribosomal protein L23) | protein interactions | 38302 | 10 | 2 | 9204 | 298 | mRpL30,CG7766 | | 3.960e-02 | -3.23 | stet (stem cell tumor) | protein interactions | 38169 | 10 | 2 | 9204 | 298 | tkv,dpp | | 3.960e-02 | -3.23 | mei-P22 (meiotic P22) | protein interactions | 38797 | 10 | 2 | 9204 | 298 | RecQ4,14-3-3zeta | | 3.960e-02 | -3.23 | ken (ken and barbie) | protein interactions | 37785 | 10 | 2 | 9204 | 298 | dpp,E(bx) | | 3.960e-02 | -3.23 | l(1)G0004 (lethal (1) G0004) | protein interactions | 33033 | 10 | 2 | 9204 | 298 | Cpsf6,Dref | | 3.960e-02 | -3.23 | DCTN6-p27 (Dynactin 6, p27 subunit) | protein interactions | 35195 | 10 | 2 | 9204 | 298 | DCTN3-p24,Ufd4 | | 3.960e-02 | -3.23 | YME1L (YME1 like ATPase) | protein interactions | 37636 | 10 | 2 | 9204 | 298 | HtrA2,Sod2 | | 3.960e-02 | -3.23 | CG12576 (uncharacterized protein) | protein interactions | 33135 | 10 | 2 | 9204 | 298 | eEF5,cal1 | | 3.960e-02 | -3.23 | Topors (Topoisomerase I-interacting protein) | protein interactions | 37188 | 10 | 2 | 9204 | 298 | Cp190,su(Hw) | | 3.960e-02 | -3.23 | CG15152 (uncharacterized protein) | protein interactions | 35082 | 10 | 2 | 9204 | 298 | Sgt,DCTN3-p24 | | 3.960e-02 | -3.23 | wac (wee Augmin) | protein interactions | 38044 | 10 | 2 | 9204 | 298 | msd5,Cdk8 | | 3.960e-02 | -3.23 | cv-2 (crossveinless 2) | protein interactions | 45280 | 10 | 2 | 9204 | 298 | dpp,tkv | | 3.960e-02 | -3.23 | C3G (C3G guanyl-nucleotide exchange factor) | protein interactions | 31618 | 10 | 2 | 9204 | 298 | Rap1,Cdc42 | | 3.960e-02 | -3.23 | ec (echinus) | protein interactions | 31339 | 10 | 2 | 9204 | 298 | flw,Pp1-87B | | 3.962e-02 | -3.23 | amide biosynthetic process | biological process | GO:0043604 | 732 | 26 | 12345 | 305 | mRpL30,RpS3A,RpL5,RpS19a,mRpL34,mRpS18A,mRpS28,mRpS9,RpL14,CG8635,CG31957,RpS8,RpL36A,eIF3b,eEF1delta,eIF4B,eIF3g1,RpL35A,eRF1,eEF5,RpS9,eIF3c,RpS26,mRpS33,RpL15,CG15534 | | 3.966e-02 | -3.23 | protein localization to chromosome | biological process | GO:0034502 | 13 | 2 | 12345 | 305 | His2Av,cal1 | | 3.966e-02 | -3.23 | actomyosin contractile ring contraction | biological process | GO:0000916 | 13 | 2 | 12345 | 305 | pav,Exo84 | | 3.966e-02 | -3.23 | mismatch repair | biological process | GO:0006298 | 13 | 2 | 12345 | 305 | RPA3,spel1 | | 3.966e-02 | -3.23 | contractile ring contraction | biological process | GO:0036213 | 13 | 2 | 12345 | 305 | Exo84,pav | | 3.966e-02 | -3.23 | positive regulation of histone methylation | biological process | GO:0031062 | 13 | 2 | 12345 | 305 | Ncoa6,E(z) | | 3.966e-02 | -3.23 | larval salivary gland morphogenesis | biological process | GO:0007436 | 13 | 2 | 12345 | 305 | BthD,Pten | | 3.966e-02 | -3.23 | sensory perception of touch | biological process | GO:0050975 | 13 | 2 | 12345 | 305 | Cdc42,Fmr1 | | 3.966e-02 | -3.23 | peptidyl-arginine modification | biological process | GO:0018195 | 13 | 2 | 12345 | 305 | CG31460,Art1 | | 3.969e-02 | -3.23 | translation | biological process | GO:0006412 | 698 | 25 | 12345 | 305 | RpL5,RpS19a,mRpL34,mRpS18A,mRpL30,RpS3A,RpS8,eIF3b,RpL36A,mRpS28,mRpS9,RpL14,CG8635,CG31957,eIF4B,eEF1delta,RpL35A,eIF3g1,eRF1,RpS26,RpL15,mRpS33,RpS9,eEF5,eIF3c | | 3.975e-02 | -3.23 | GO\_CATALYTIC\_COMPLEX | MSigDB lists | GO\_CATALYTIC\_COMPLEX | 453 | 28 | 4161 | 184 | vnc,CG31460,TfIIFbeta,l(2)37Bb,E(z),Art1,Rpb11,mbf1,Roc2,CIA30,Stt3B,cta,fzy,Pp1-87B,Wdr82,Cul4,Rbbp5,MED21,Rcd5,Ino80,krz,Skp2,Arp8,ND-ASHI,CG6907,PIG-K,CG7766,flw | | 3.979e-02 | -3.22 | Signaling by EGFR | REACTOME pathways | R-DME-177929 | 178 | 12 | 3923 | 151 | Lst8,Cdc42,Prosalpha2,Pten,Rpn10,CG4935,Prosbeta1,CG7054,14-3-3zeta,Rpt6,Nf1,vap | | 3.981e-02 | -3.22 | epithelium migration | biological process | GO:0090132 | 156 | 8 | 12345 | 305 | tsr,Rab7,spoon,Girdin,rhea,dpp,flr,sqd | | 3.989e-02 | -3.22 | Formation of Incision Complex in GG-NER | REACTOME pathways | R-DME-5696395 | 34 | 4 | 3923 | 151 | Usp16-45,Parp,Cul4,smt3 | | 3.994e-02 | -3.22 | WD\_REPEATS\_REGION | prosite domains | PS50294 | 150 | 9 | 6243 | 191 | fzy,CG43367,CG4935,Wdr82,flr,Wdr33,CG14805,Lst8,Rbbp5 | | 4.004e-02 | -3.22 | SCF(Skp2)-mediated degradation of p27/p21 | REACTOME pathways | R-DME-187577 | 66 | 6 | 3923 | 151 | Rpt6,CycA,Prosalpha2,Rpn10,Skp2,Prosbeta1 | | 4.014e-02 | -3.22 | VEGFR2 mediated cell proliferation | REACTOME pathways | R-DME-5218921 | 120 | 9 | 3923 | 151 | Prosbeta1,14-3-3zeta,CG7054,Nf1,Rpt6,vap,Prosalpha2,Rpn10,CG4935 | | 4.032e-02 | -3.21 | regulation of metaphase/anaphase transition of cell cycle | biological process | GO:1902099 | 31 | 3 | 12345 | 305 | CycA,Bub1,Mps1 | | 4.032e-02 | -3.21 | meiosis II cell cycle process | biological process | GO:0061983 | 31 | 3 | 12345 | 305 | Bub1,fzy,Exo84 | | 4.039e-02 | -3.21 | postsynaptic density | cellular component | GO:0014069 | 13 | 2 | 11940 | 298 | Spn,Fmr1 | | 4.039e-02 | -3.21 | female germline ring canal | cellular component | GO:0035324 | 13 | 2 | 11940 | 298 | vsg,pav | | 4.039e-02 | -3.21 | neuron to neuron synapse | cellular component | GO:0098984 | 13 | 2 | 11940 | 298 | Spn,Fmr1 | | 4.039e-02 | -3.21 | asymmetric synapse | cellular component | GO:0032279 | 13 | 2 | 11940 | 298 | Spn,Fmr1 | | 4.058e-02 | -3.20 | regulation of cell cycle phase transition | biological process | GO:1901987 | 102 | 6 | 12345 | 305 | RecQ4,fzy,CycA,Bub1,Tctp,Mps1 | | 4.058e-02 | -3.20 | sister chromatid segregation | biological process | GO:0000819 | 102 | 6 | 12345 | 305 | pav,Fign,Rpn10,Bub1,CycA,Pp1-87B | | 4.069e-02 | -3.20 | cellular amide metabolic process | biological process | GO:0043603 | 872 | 30 | 12345 | 305 | eIF3b,RpL36A,RpS8,CG31957,CG8635,mRpS9,mRpS28,RpL14,mRpS18A,mRpL34,RpS19a,RpL5,CG32473,RpS3A,mRpL30,GstE13,CG10365,CG8132,CG15534,RpS26,RpL15,mRpS33,eIF3c,eEF5,RpS9,eRF1,eIF3g1,RpL35A,eIF4B,eEF1delta | | 4.087e-02 | -3.20 | macromolecule methylation | biological process | GO:0043414 | 129 | 7 | 12345 | 305 | E(z),BEAF-32,Nopp140,Rbbp5,Wdr82,Art1,Ncoa6 | | 4.101e-02 | -3.19 | GSE9006\_HEALTHY\_VS\_TYPE\_1\_DIABETES\_PBMC\_4MONTH\_POST\_DX\_UP | MSigDB lists | GSE9006\_HEALTHY\_VS\_TYPE\_1\_DIABETES\_PBMC\_4MONTH\_POST\_DX\_UP | 73 | 7 | 4161 | 184 | smt3,Graf,14-3-3zeta,Pten,Fs(2)Ket,ECSIT,flw | | 4.101e-02 | -3.19 | GSE17721\_CTRL\_VS\_GARDIQUIMOD\_6H\_BMDC\_UP | MSigDB lists | GSE17721\_CTRL\_VS\_GARDIQUIMOD\_6H\_BMDC\_UP | 73 | 7 | 4161 | 184 | twin,sws,krz,LIMK1,Mdh1,Pp1-87B,RpS8 | | 4.101e-02 | -3.19 | E2F1\_Q6 | MSigDB lists | E2F1\_Q6 | 73 | 7 | 4161 | 184 | SF2,spel1,Phf5a,Fs(2)Ket,Stt3B,His2Av,E(z) | | 4.101e-02 | -3.19 | GO\_POSITIVE\_REGULATION\_OF\_NERVOUS\_SYSTEM\_DEVELOPMENT | MSigDB lists | GO\_POSITIVE\_REGULATION\_OF\_NERVOUS\_SYSTEM\_DEVELOPMENT | 73 | 7 | 4161 | 184 | RhoGAPp190,Rap1,fzy,E(z),Fmr1,dpp,LIMK1 | | 4.101e-02 | -3.19 | GSE22886\_NAIVE\_CD4\_TCELL\_VS\_NEUTROPHIL\_UP | MSigDB lists | GSE22886\_NAIVE\_CD4\_TCELL\_VS\_NEUTROPHIL\_UP | 73 | 7 | 4161 | 184 | E(z),CG11920,noi,smt3,CG4866,Mdh1,Dbp21E2 | | 4.101e-02 | -3.19 | GO\_DNA\_RECOMBINATION | MSigDB lists | GO\_DNA\_RECOMBINATION | 73 | 7 | 4161 | 184 | Ino80,Parp,Rcd5,mus301,Arp8,spel1,DNAlig1 | | 4.101e-02 | -3.19 | GO\_SIDE\_OF\_MEMBRANE | MSigDB lists | GO\_SIDE\_OF\_MEMBRANE | 73 | 7 | 4161 | 184 | Nf1,vap,HtrA2,Pten,CG32473,krz,cta | | 4.114e-02 | -3.19 | extrinsic component of membrane | cellular component | GO:0019898 | 76 | 5 | 11940 | 298 | dah,Rop,Lrr47,cta,Vps13D | | 4.115e-02 | -3.19 | GEFmeso (Guanine nucleotide exchange factor in mesoderm) | protein interactions | 37134 | 24 | 3 | 9204 | 298 | CG8860,Cdc42,nsl1 | | 4.115e-02 | -3.19 | SCAR (SCAR) | protein interactions | 34519 | 24 | 3 | 9204 | 298 | LIMK1,Cdc42,Fmr1 | | 4.115e-02 | -3.19 | elfless (elfless) | protein interactions | 35076 | 24 | 3 | 9204 | 298 | RpS26,CG10209,RpL36A | | 4.115e-02 | -3.19 | Lim1 (LIM homeobox 1) | protein interactions | 31813 | 24 | 3 | 9204 | 298 | Mdh1,Hsp27,CG5516 | | 4.127e-02 | -3.19 | GO\_POSITIVE\_REGULATION\_OF\_KINASE\_ACTIVITY | MSigDB lists | GO\_POSITIVE\_REGULATION\_OF\_KINASE\_ACTIVITY | 105 | 9 | 4161 | 184 | E(z),lic,krz,Vap33,Rap1,Stlk,dpp,unc-119,Cdc42 | | 4.128e-02 | -3.19 | GO\_POSITIVE\_REGULATION\_OF\_CELL\_DIVISION | MSigDB lists | GO\_POSITIVE\_REGULATION\_OF\_CELL\_DIVISION | 30 | 4 | 4161 | 184 | pav,Cdc42,Pten,AGO3 | | 4.128e-02 | -3.19 | GO\_MITOTIC\_DNA\_INTEGRITY\_CHECKPOINT | MSigDB lists | GO\_MITOTIC\_DNA\_INTEGRITY\_CHECKPOINT | 30 | 4 | 4161 | 184 | twin,Art1,CycA,spel1 | | 4.128e-02 | -3.19 | GO\_NEGATIVE\_REGULATION\_OF\_CHROMOSOME\_ORGANIZATION | MSigDB lists | GO\_NEGATIVE\_REGULATION\_OF\_CHROMOSOME\_ORGANIZATION | 30 | 4 | 4161 | 184 | CG12321,CG9246,Parp,Rcd5 | | 4.142e-02 | -3.18 | ATF3\_Q6 | MSigDB lists | ATF3\_Q6 | 58 | 6 | 4161 | 184 | 14-3-3zeta,l(2)37Bb,Cdc42,CG13016,CycA,CG5861 | | 4.142e-02 | -3.18 | GOLDRATH\_EFF\_VS\_MEMORY\_CD8\_TCELL\_UP | MSigDB lists | GOLDRATH\_EFF\_VS\_MEMORY\_CD8\_TCELL\_UP | 58 | 6 | 4161 | 184 | pav,DNAlig1,fzy,CycA,E(z),Stt3B | | 4.142e-02 | -3.18 | GSE7768\_OVA\_ALONE\_VS\_OVA\_WITH\_MPL\_IMMUNIZED\_MOUSE\_WHOLE\_SPLEEN\_6H\_UP | MSigDB lists | GSE7768\_OVA\_ALONE\_VS\_OVA\_WITH\_MPL\_IMMUNIZED\_MOUSE\_WHOLE\_SPLEEN\_6H\_UP | 58 | 6 | 4161 | 184 | pav,fzy,CycA,E(z),CG7766,Skp2 | | 4.142e-02 | -3.18 | GSE37301\_LYMPHOID\_PRIMED\_MPP\_VS\_COMMON\_LYMPHOID\_PROGENITOR\_DN | MSigDB lists | GSE37301\_LYMPHOID\_PRIMED\_MPP\_VS\_COMMON\_LYMPHOID\_PROGENITOR\_DN | 58 | 6 | 4161 | 184 | CG5339,Arpc3A,Nulp1,spel1,Sod2,eRF1 | | 4.142e-02 | -3.18 | GSE3720\_LPS\_VS\_PMA\_STIM\_VD1\_GAMMADELTA\_TCELL\_DN | MSigDB lists | GSE3720\_LPS\_VS\_PMA\_STIM\_VD1\_GAMMADELTA\_TCELL\_DN | 58 | 6 | 4161 | 184 | viaf,RpL36A,eIF3g1,RpS26,His2Av,CG41378 | | 4.142e-02 | -3.18 | GSE1925\_CTRL\_VS\_24H\_IFNG\_STIM\_IFNG\_PRIMED\_MACROPHAGE\_UP | MSigDB lists | GSE1925\_CTRL\_VS\_24H\_IFNG\_STIM\_IFNG\_PRIMED\_MACROPHAGE\_UP | 58 | 6 | 4161 | 184 | Arpc3A,PIG-K,Roc2,ema,ubl,CG10343 | | 4.154e-02 | -3.18 | positive regulation of antimicrobial humoral response | biological process | GO:0002760 | 53 | 4 | 12345 | 305 | pll,dgrn,Parp,smt3 | | 4.154e-02 | -3.18 | positive regulation of production of molecular mediator of immune response | biological process | GO:0002702 | 53 | 4 | 12345 | 305 | smt3,pll,dgrn,Parp | | 4.154e-02 | -3.18 | positive regulation of antimicrobial peptide production | biological process | GO:0002225 | 53 | 4 | 12345 | 305 | smt3,Parp,pll,dgrn | | 4.154e-02 | -3.18 | positive regulation of immune effector process | biological process | GO:0002699 | 53 | 4 | 12345 | 305 | smt3,Parp,dgrn,pll | | 4.154e-02 | -3.18 | ATP-dependent chromatin remodeling | biological process | GO:0043044 | 53 | 4 | 12345 | 305 | CG8223,Ino80,CtBP,Arp8 | | 4.154e-02 | -3.18 | epidermal cell differentiation | biological process | GO:0009913 | 53 | 4 | 12345 | 305 | LIMK1,tsr,Cdc42,flr | | 4.156e-02 | -3.18 | wts (warts) | protein interactions | 43651 | 59 | 5 | 9204 | 298 | CycA,Ncoa6,lic,mnb,Tgi | | 4.156e-02 | -3.18 | Prosbeta3 (Proteasome beta3 subunit) | protein interactions | 41079 | 59 | 5 | 9204 | 298 | Prosbeta1,Ubqn,CG12321,Prosalpha2,Rpt6 | | 4.156e-02 | -3.18 | Stip1 (Stress induced phosphoprotein 1) | protein interactions | 33202 | 59 | 5 | 9204 | 298 | eEF5,smt3,Lst8,eEF1delta,Nph | | 4.162e-02 | -3.18 | DNA-dependent DNA replication | biological process | GO:0006261 | 77 | 5 | 12345 | 305 | Dref,RecQ4,Btk29A,DNAlig1,Orc3 | | 4.164e-02 | -3.18 | AUF1 (hnRNP D0) binds and destabilizes mRNA | REACTOME pathways | R-DME-450408 | 50 | 5 | 3923 | 151 | Rpt6,sqd,Prosbeta1,Rpn10,Prosalpha2 | | 4.174e-02 | -3.18 | GO\_MACROMOLECULAR\_COMPLEX\_ASSEMBLY | MSigDB lists | GO\_MACROMOLECULAR\_COMPLEX\_ASSEMBLY | 554 | 33 | 4161 | 184 | inc,Fs(2)Ket,eIF3c,Tom20,noi,bai,CG17493,Rbbp5,Cul4,me31B,Sod2,CG9246,eIF3g1,His4r,Rpn10,ECSIT,Exo84,ND-ASHI,CG12321,CSN1b,Ino80,CG10672,SF2,Parp,MICU1,pav,CIA30,l(2)37Bb,Dbp21E2,CG17883,eIF3b,RpL5,CG12341 | | 4.181e-02 | -3.17 | TARTE\_PLASMA\_CELL\_VS\_PLASMABLAST\_DN | MSigDB lists | TARTE\_PLASMA\_CELL\_VS\_PLASMABLAST\_DN | 156 | 12 | 4161 | 184 | fzy,eRF1,Sod2,pav,Prosalpha2,Rpt6,CG11586,CycA,Rpn10,CCT4,E(z),CG41378 | | 4.218e-02 | -3.17 | DNA binding | molecular function | GO:0003677 | 984 | 32 | 12176 | 289 | Mlf,sqd,nom,Ino80,ver,Parp,CG10209,Acf,Pbp49,spel1,rhi,Orc3,CG12299,BEAF-32,SF2,His2Av,Pep,E(z),Hrb27C,His4r,Jarid2,mbf1,Cp190,RPA3,su(Hw),Rpb11,RecQ4,grau,DNAlig1,CG11456,Dref,CG6254 | | 4.224e-02 | -3.16 | apoptotic process | biological process | GO:0006915 | 103 | 6 | 12345 | 305 | dpp,spel1,pll,sws,CG10209,sigmar | | 4.232e-02 | -3.16 | Golgi vesicle transport | biological process | GO:0048193 | 130 | 7 | 12345 | 305 | CG17883,Rab14,bai,g,Rab8,Exo84,PGAP1 | | 4.233e-02 | -3.16 | fly chr73A9-73A9\|3-44 | chromosome location | fly chr73A9-73A9\|3-44 | 3 | 1 | 25010 | 358 | mbf1 | | 4.233e-02 | -3.16 | fly chr43C3-43C3\|2-56 cM | chromosome location | fly chr43C3-43C3\|2-56 cM | 3 | 1 | 25010 | 358 | CG11141 | | 4.233e-02 | -3.16 | fly chr73D3-73D3\|3-45 cM | chromosome location | fly chr73D3-73D3\|3-45 cM | 3 | 1 | 25010 | 358 | CG9951 | | 4.233e-02 | -3.16 | fly chr57B9-57B9\|2-92 cM | chromosome location | fly chr57B9-57B9\|2-92 cM | 3 | 1 | 25010 | 358 | grau | | 4.233e-02 | -3.16 | fly chr23E3-23E3\|2-9 cM | chromosome location | fly chr23E3-23E3\|2-9 cM | 3 | 1 | 25010 | 358 | Fign | | 4.233e-02 | -3.16 | fly chr14A8-14A8\|1-52 cM | chromosome location | fly chr14A8-14A8\|1-52 cM | 3 | 1 | 25010 | 358 | CG8958 | | 4.233e-02 | -3.16 | fly chr5F2-5F2\|1-16 | chromosome location | fly chr5F2-5F2\|1-16 | 3 | 1 | 25010 | 358 | CG3842 | | 4.233e-02 | -3.16 | fly chr88C9-88C9\|3-55 cM | chromosome location | fly chr88C9-88C9\|3-55 cM | 3 | 1 | 25010 | 358 | His4r | | 4.233e-02 | -3.16 | fly chr88C9-88C9\|3-55 | chromosome location | fly chr88C9-88C9\|3-55 | 3 | 1 | 25010 | 358 | His4r | | 4.233e-02 | -3.16 | fly chr27C6-27C6\|2-23 | chromosome location | fly chr27C6-27C6\|2-23 | 3 | 1 | 25010 | 358 | Fgop2 | | 4.233e-02 | -3.16 | fly chr98B2-98B2\|3-94 | chromosome location | fly chr98B2-98B2\|3-94 | 3 | 1 | 25010 | 358 | CG12877 | | 4.233e-02 | -3.16 | fly chr8F1-8F1\|1-29 cM | chromosome location | fly chr8F1-8F1\|1-29 cM | 3 | 1 | 25010 | 358 | Lst8 | | 4.233e-02 | -3.16 | fly chr23E3-23E3\|2-9 | chromosome location | fly chr23E3-23E3\|2-9 | 3 | 1 | 25010 | 358 | Fign | | 4.233e-02 | -3.16 | fly chr94A16-94A16\|3-74 cM | chromosome location | fly chr94A16-94A16\|3-74 cM | 3 | 1 | 25010 | 358 | CG7054 | | 4.233e-02 | -3.16 | fly chr27C6-27C6\|2-23 cM | chromosome location | fly chr27C6-27C6\|2-23 cM | 3 | 1 | 25010 | 358 | Fgop2 | | 4.233e-02 | -3.16 | fly chr8F1-8F1\|1-29 | chromosome location | fly chr8F1-8F1\|1-29 | 3 | 1 | 25010 | 358 | Lst8 | | 4.233e-02 | -3.16 | fly chr3A7-3A7\|1-1 | chromosome location | fly chr3A7-3A7\|1-1 | 3 | 1 | 25010 | 358 | eIF3g1 | | 4.233e-02 | -3.16 | fly chr11D10-11D10\|1-41 | chromosome location | fly chr11D10-11D10\|1-41 | 3 | 1 | 25010 | 358 | lic | | 4.233e-02 | -3.16 | fly chr14A8-14A8\|1-52 | chromosome location | fly chr14A8-14A8\|1-52 | 3 | 1 | 25010 | 358 | CG8958 | | 4.233e-02 | -3.16 | fly chr76B8-76B8\|3-45 | chromosome location | fly chr76B8-76B8\|3-45 | 3 | 1 | 25010 | 358 | CG9330 | | 4.233e-02 | -3.16 | fly chr85A10-85A10\|3-48 | chromosome location | fly chr85A10-85A10\|3-48 | 3 | 1 | 25010 | 358 | CG8223 | | 4.233e-02 | -3.16 | fly chr98B2-98B2\|3-94 cM | chromosome location | fly chr98B2-98B2\|3-94 cM | 3 | 1 | 25010 | 358 | CG12877 | | 4.233e-02 | -3.16 | fly chr73A9-73A9\|3-44 cM | chromosome location | fly chr73A9-73A9\|3-44 cM | 3 | 1 | 25010 | 358 | mbf1 | | 4.233e-02 | -3.16 | fly chr94A16-94A16\|3-74 | chromosome location | fly chr94A16-94A16\|3-74 | 3 | 1 | 25010 | 358 | CG7054 | | 4.233e-02 | -3.16 | fly chr36B5-36B5\|2-53 | chromosome location | fly chr36B5-36B5\|2-53 | 3 | 1 | 25010 | 358 | lncRNA:CR45285 | | 4.233e-02 | -3.16 | fly chr5F2-5F2\|1-16 cM | chromosome location | fly chr5F2-5F2\|1-16 cM | 3 | 1 | 25010 | 358 | CG3842 | | 4.233e-02 | -3.16 | fly chr43C3-43C3\|2-56 | chromosome location | fly chr43C3-43C3\|2-56 | 3 | 1 | 25010 | 358 | CG11141 | | 4.233e-02 | -3.16 | fly chr11D10-11D10\|1-41 cM | chromosome location | fly chr11D10-11D10\|1-41 cM | 3 | 1 | 25010 | 358 | lic | | 4.233e-02 | -3.16 | fly chr49A6-49A6\|2-67 | chromosome location | fly chr49A6-49A6\|2-67 | 3 | 1 | 25010 | 358 | Tmem18 | | 4.233e-02 | -3.16 | fly chr73D3-73D3\|3-45 | chromosome location | fly chr73D3-73D3\|3-45 | 3 | 1 | 25010 | 358 | CG9951 | | 4.233e-02 | -3.16 | fly chr97D11-97D11\|3-92 | chromosome location | fly chr97D11-97D11\|3-92 | 3 | 1 | 25010 | 358 | Tsp97E | | 4.233e-02 | -3.16 | fly chr85D18-85D18\|3-48 cM | chromosome location | fly chr85D18-85D18\|3-48 cM | 3 | 1 | 25010 | 358 | CG8149 | | 4.233e-02 | -3.16 | fly chr76B8-76B8\|3-45 cM | chromosome location | fly chr76B8-76B8\|3-45 cM | 3 | 1 | 25010 | 358 | CG9330 | | 4.233e-02 | -3.16 | fly chr3A7-3A7\|1-1 cM | chromosome location | fly chr3A7-3A7\|1-1 cM | 3 | 1 | 25010 | 358 | eIF3g1 | | 4.233e-02 | -3.16 | fly chr60A8-60A8\|2-105 | chromosome location | fly chr60A8-60A8\|2-105 | 3 | 1 | 25010 | 358 | DNAlig1 | | 4.233e-02 | -3.16 | fly chr49A6-49A6\|2-67 cM | chromosome location | fly chr49A6-49A6\|2-67 cM | 3 | 1 | 25010 | 358 | Tmem18 | | 4.233e-02 | -3.16 | fly chr89E12-89E12\|3-59 | chromosome location | fly chr89E12-89E12\|3-59 | 3 | 1 | 25010 | 358 | cal1 | | 4.233e-02 | -3.16 | fly chr85D18-85D18\|3-48 | chromosome location | fly chr85D18-85D18\|3-48 | 3 | 1 | 25010 | 358 | CG8149 | | 4.233e-02 | -3.16 | fly chr89E12-89E12\|3-59 cM | chromosome location | fly chr89E12-89E12\|3-59 cM | 3 | 1 | 25010 | 358 | cal1 | | 4.233e-02 | -3.16 | fly chr85A10-85A10\|3-48 cM | chromosome location | fly chr85A10-85A10\|3-48 cM | 3 | 1 | 25010 | 358 | CG8223 | | 4.233e-02 | -3.16 | fly chr57B9-57B9\|2-92 | chromosome location | fly chr57B9-57B9\|2-92 | 3 | 1 | 25010 | 358 | grau | | 4.233e-02 | -3.16 | fly chr36B5-36B5\|2-53 cM | chromosome location | fly chr36B5-36B5\|2-53 cM | 3 | 1 | 25010 | 358 | lncRNA:CR45285 | | 4.233e-02 | -3.16 | fly chr97D11-97D11\|3-92 cM | chromosome location | fly chr97D11-97D11\|3-92 cM | 3 | 1 | 25010 | 358 | Tsp97E | | 4.233e-02 | -3.16 | fly chr60A8-60A8\|2-105 cM | chromosome location | fly chr60A8-60A8\|2-105 cM | 3 | 1 | 25010 | 358 | DNAlig1 | | 4.233e-02 | -3.16 | demethylase activity | molecular function | GO:0032451 | 14 | 2 | 12176 | 289 | Jarid2,NO66 | | 4.242e-02 | -3.16 | fly chr88F1-88F1\|3-57 | chromosome location | fly chr88F1-88F1\|3-57 | 23 | 2 | 25010 | 358 | CG6171,CG31301 | | 4.242e-02 | -3.16 | fly chr88F1-88F1\|3-57 cM | chromosome location | fly chr88F1-88F1\|3-57 cM | 23 | 2 | 25010 | 358 | CG31301,CG6171 | | 4.254e-02 | -3.16 | PID\_PLK1\_PATHWAY | MSigDB lists | PID\_PLK1\_PATHWAY | 18 | 3 | 4161 | 184 | flw,Tctp,fzy | | 4.254e-02 | -3.16 | SIG\_CHEMOTAXIS | MSigDB lists | SIG\_CHEMOTAXIS | 18 | 3 | 4161 | 184 | Pten,LIMK1,Cdc42 | | 4.254e-02 | -3.16 | PID\_EPHB\_FWD\_PATHWAY | MSigDB lists | PID\_EPHB\_FWD\_PATHWAY | 18 | 3 | 4161 | 184 | vap,Rap1,Cdc42 | | 4.254e-02 | -3.16 | MCLACHLAN\_DENTAL\_CARIES\_UP | MSigDB lists | MCLACHLAN\_DENTAL\_CARIES\_UP | 18 | 3 | 4161 | 184 | sigmar,Sod2,CG41378 | | 4.254e-02 | -3.16 | RASHI\_RESPONSE\_TO\_IONIZING\_RADIATION\_2 | MSigDB lists | RASHI\_RESPONSE\_TO\_IONIZING\_RADIATION\_2 | 18 | 3 | 4161 | 184 | Nulp1,Rop,cta | | 4.254e-02 | -3.16 | GO\_FORMATION\_OF\_TRANSLATION\_PREINITIATION\_COMPLEX | MSigDB lists | GO\_FORMATION\_OF\_TRANSLATION\_PREINITIATION\_COMPLEX | 18 | 3 | 4161 | 184 | eIF3c,eIF3b,eIF3g1 | | 4.254e-02 | -3.16 | STONER\_ESOPHAGEAL\_CARCINOGENESIS\_UP | MSigDB lists | STONER\_ESOPHAGEAL\_CARCINOGENESIS\_UP | 18 | 3 | 4161 | 184 | RagA-B,Txl,lic | | 4.254e-02 | -3.16 | KYNG\_DNA\_DAMAGE\_BY\_4NQO\_OR\_UV | MSigDB lists | KYNG\_DNA\_DAMAGE\_BY\_4NQO\_OR\_UV | 18 | 3 | 4161 | 184 | Pect,Art1,VhaPPA1-1 | | 4.254e-02 | -3.16 | GSE2706\_UNSTIM\_VS\_2H\_LPS\_AND\_R848\_DC\_DN | MSigDB lists | GSE2706\_UNSTIM\_VS\_2H\_LPS\_AND\_R848\_DC\_DN | 18 | 3 | 4161 | 184 | CG13016,sigmar,E(z) | | 4.258e-02 | -3.16 | wing disc morphogenesis | biological process | GO:0007472 | 308 | 13 | 12345 | 305 | Cdc42,peng,Btk29A,tsr,EloB,tkv,rhea,dpp,LIMK1,GEFmeso,Stim,flr,flw | | 4.266e-02 | -3.15 | CCT2 (Chaperonin containing TCP1 subunit 2) | protein interactions | 31838 | 41 | 4 | 9204 | 298 | fzy,Rop,Rpt6,CCT4 | | 4.266e-02 | -3.15 | Prosbeta1 (Proteasome beta1 subunit) | protein interactions | 46058 | 41 | 4 | 9204 | 298 | Prosalpha2,Rpt6,Ubqn,CG12321 | | 4.266e-02 | -3.15 | Prosalpha3 (Proteasome alpha3 subunit) | protein interactions | 37378 | 41 | 4 | 9204 | 298 | Prosbeta1,CG12321,Prosalpha2,Rpt6 | | 4.271e-02 | -3.15 | ZF\_BED | prosite domains | PS50808 | 11 | 2 | 6243 | 191 | BEAF-32,Dref | | 4.276e-02 | -3.15 | amide transmembrane transporter activity | molecular function | GO:0042887 | 33 | 3 | 12176 | 289 | Tim17b,CG8860,Tom20 | | 4.277e-02 | -3.15 | - | gene3d domains | 2.30.30.30 | 11 | 2 | 4929 | 151 | eEF5,RpL14 | | 4.303e-02 | -3.15 | GSE45365\_NK\_CELL\_VS\_CD8\_TCELL\_UP | MSigDB lists | GSE45365\_NK\_CELL\_VS\_CD8\_TCELL\_UP | 44 | 5 | 4161 | 184 | Arpc3A,CG11334,bai,CycA,Pp1-87B | | 4.303e-02 | -3.15 | GSE21546\_WT\_VS\_SAP1A\_KO\_DP\_THYMOCYTES\_DN | MSigDB lists | GSE21546\_WT\_VS\_SAP1A\_KO\_DP\_THYMOCYTES\_DN | 44 | 5 | 4161 | 184 | flr,eRF1,DNAlig1,eIF3b,RpL5 | | 4.303e-02 | -3.15 | GO\_S\_ADENOSYLMETHIONINE\_DEPENDENT\_METHYLTRANSFERASE\_ACTIVITY | MSigDB lists | GO\_S\_ADENOSYLMETHIONINE\_DEPENDENT\_METHYLTRANSFERASE\_ACTIVITY | 44 | 5 | 4161 | 184 | Wdr82,E(z),Art1,Coq5,Rbbp5 | | 4.303e-02 | -3.15 | GSE25677\_MPL\_VS\_R848\_STIM\_BCELL\_DN | MSigDB lists | GSE25677\_MPL\_VS\_R848\_STIM\_BCELL\_DN | 44 | 5 | 4161 | 184 | PGAP1,Graf,Rab8,Parp,cta | | 4.303e-02 | -3.15 | GSE3039\_CD4\_TCELL\_VS\_B1\_BCELL\_DN | MSigDB lists | GSE3039\_CD4\_TCELL\_VS\_B1\_BCELL\_DN | 44 | 5 | 4161 | 184 | Sod2,sigmar,Pisd,AGO3,CG34148 | | 4.303e-02 | -3.15 | MODULE\_124 | MSigDB lists | MODULE\_124 | 44 | 5 | 4161 | 184 | Rpb11,sm,CycA,TfIIFbeta,pav | | 4.303e-02 | -3.15 | GSE15324\_ELF4\_KO\_VS\_WT\_ACTIVATED\_CD8\_TCELL\_DN | MSigDB lists | GSE15324\_ELF4\_KO\_VS\_WT\_ACTIVATED\_CD8\_TCELL\_DN | 44 | 5 | 4161 | 184 | pav,Fgop2,ubl,Dbp21E2,Pten | | 4.310e-02 | -3.14 | aromatic compound biosynthetic process | biological process | GO:0019438 | 469 | 18 | 12345 | 305 | grau,His4r,Pbp49,g,Arp8,TfIIFbeta,DNAlig1,CG6244,E(bx),RecQ4,Rpb11,Parp,rb,Cul4,su(Hw),Aps,EloB,sun | | 4.314e-02 | -3.14 | spliceosomal snRNP complex | cellular component | GO:0097525 | 77 | 5 | 11940 | 298 | Phf5a,CG11586,snf,snRNA:U4:39B,noi | | 4.317e-02 | -3.14 | hydrolase activity | molecular function | GO:0016787 | 1775 | 53 | 12176 | 289 | me31B,HtrA2,CG8132,RagA-B,Ino80,sun,Dbp21E2,twin,CG1637,Rat1,CG12567,Mppe,Pten,Aps,Prosbeta1,E(bx),CG6171,CG5589,Apf,Acf,CG12877,spel1,RhoGAPp190,Usp16-45,RabX6,CG8915,pav,Pp1-87B,Arp8,CG9330,PIG-K,Rap1,PGAP1,Cdc42,mus301,Not3,sws,Rheb,janA,cta,Rab14,CG32473,Rab7,flw,Dbp80,CG15534,Prosalpha2,Rpt6,Rab8,RecQ4,alpha-Man-Ib,Usp32,Fign | | 4.317e-02 | -3.14 | nucleobase-containing compound biosynthetic process | biological process | GO:0034654 | 404 | 16 | 12345 | 305 | TfIIFbeta,E(bx),CG6244,DNAlig1,His4r,grau,Arp8,Pbp49,su(Hw),Cul4,Parp,sun,EloB,Aps,RecQ4,Rpb11 | | 4.320e-02 | -3.14 | peptide biosynthetic process | biological process | GO:0043043 | 704 | 25 | 12345 | 305 | eEF1delta,eIF4B,eIF3g1,RpL35A,eRF1,RpS9,eEF5,eIF3c,RpL15,RpS26,mRpS33,mRpL30,RpS3A,RpL5,RpS19a,mRpL34,mRpS18A,mRpS28,mRpS9,RpL14,CG31957,CG8635,RpS8,eIF3b,RpL36A | | 4.356e-02 | -3.13 | mitochondrion | cellular component | GO:0005739 | 800 | 28 | 11940 | 298 | Pisd,mRpL34,Tom20,mRpL30,mRpS18A,ND-ASHI,Bmcp,Sod2,mRpS33,ND-AGGG,Tim17b,CIA30,ECSIT,HtrA2,DNAlig1,MICU1,CG11752,heix,mRpS9,sun,mRpS28,l(2)37Bb,CG4743,AGO3,Dbp21E2,Coq5,CG44194,spoon | | 4.367e-02 | -3.13 | GSE27786\_LIN\_NEG\_VS\_ERYTHROBLAST\_UP | MSigDB lists | GSE27786\_LIN\_NEG\_VS\_ERYTHROBLAST\_UP | 74 | 7 | 4161 | 184 | CG4743,TfIIFbeta,HtrA2,PIG-K,sws,CG11920,Art1 | | 4.367e-02 | -3.13 | GSE17721\_LPS\_VS\_POLYIC\_4H\_BMDC\_DN | MSigDB lists | GSE17721\_LPS\_VS\_POLYIC\_4H\_BMDC\_DN | 74 | 7 | 4161 | 184 | Parp,unc-119,CG41378,Pisd,mRpS33,Rpn10,Fgop2 | | 4.367e-02 | -3.13 | GSE20727\_CTRL\_VS\_ROS\_INHIBITOR\_TREATED\_DC\_UP | MSigDB lists | GSE20727\_CTRL\_VS\_ROS\_INHIBITOR\_TREATED\_DC\_UP | 74 | 7 | 4161 | 184 | CG31460,Mdh1,CG5861,xit,Lst8,CG12935,CIA30 | | 4.381e-02 | -3.13 | protein targeting | biological process | GO:0006605 | 131 | 7 | 12345 | 305 | Tom20,g,Pex3,CG8860,Vps13D,Sgt,Tim17b | | 4.381e-02 | -3.13 | positive regulation of nervous system development | biological process | GO:0051962 | 131 | 7 | 12345 | 305 | Rop,tkv,PlexA,Rheb,mnb,Fmr1,Rap1 | | 4.394e-02 | -3.13 | innate immune response | biological process | GO:0045087 | 104 | 6 | 12345 | 305 | Cdc42,Pten,Parp,ECSIT,pll,dgrn | | 4.403e-02 | -3.12 | positive regulation of humoral immune response | biological process | GO:0002922 | 54 | 4 | 12345 | 305 | smt3,Parp,pll,dgrn | | 4.403e-02 | -3.12 | vacuole organization | biological process | GO:0007033 | 54 | 4 | 12345 | 305 | g,Rab7,Rab14,rb | | 4.415e-02 | -3.12 | GO\_REGULATION\_OF\_MAPK\_CASCADE | MSigDB lists | GO\_REGULATION\_OF\_MAPK\_CASCADE | 123 | 10 | 4161 | 184 | E(z),Art1,lic,krz,Rap1,Pten,dpp,CSN1b,Nf1,Cdc42 | | 4.415e-02 | -3.12 | MODULE\_32 | MSigDB lists | MODULE\_32 | 123 | 10 | 4161 | 184 | RpS8,eRF1,RpL15,eIF3g1,sm,Fmr1,eIF3c,snf,eEF1delta,noi | | 4.419e-02 | -3.12 | 14-3-3epsilon (14-3-3epsilon) | protein interactions | 42186 | 60 | 5 | 9204 | 298 | Rheb,14-3-3zeta,PlexA,smt3,Tctp | | 4.419e-02 | -3.12 | grk (gurken) | protein interactions | 34171 | 60 | 5 | 9204 | 298 | me31B,RpS19a,sqd,RpL14,Hrb27C | | 4.422e-02 | -3.12 | GO\_PLATELET\_DERIVED\_GROWTH\_FACTOR\_RECEPTOR\_BINDING | MSigDB lists | GO\_PLATELET\_DERIVED\_GROWTH\_FACTOR\_RECEPTOR\_BINDING | 1 | 1 | 4161 | 184 | Pten | | 4.422e-02 | -3.12 | UROSEVIC\_RESPONSE\_TO\_IMIQUIMOD | MSigDB lists | UROSEVIC\_RESPONSE\_TO\_IMIQUIMOD | 1 | 1 | 4161 | 184 | CG41378 | | 4.422e-02 | -3.12 | MATZUK\_CUMULUS\_EXPANSION | MSigDB lists | MATZUK\_CUMULUS\_EXPANSION | 1 | 1 | 4161 | 184 | tkv | | 4.422e-02 | -3.12 | RUAN\_RESPONSE\_TO\_TNF\_UP | MSigDB lists | RUAN\_RESPONSE\_TO\_TNF\_UP | 1 | 1 | 4161 | 184 | Sod2 | | 4.422e-02 | -3.12 | ZHU\_SKIL\_TARGETS\_DN | MSigDB lists | ZHU\_SKIL\_TARGETS\_DN | 1 | 1 | 4161 | 184 | CycA | | 4.422e-02 | -3.12 | GO\_POSITIVE\_REGULATION\_OF\_ACTIN\_FILAMENT\_DEPOLYMERIZATION | MSigDB lists | GO\_POSITIVE\_REGULATION\_OF\_ACTIN\_FILAMENT\_DEPOLYMERIZATION | 1 | 1 | 4161 | 184 | flr | | 4.422e-02 | -3.12 | GO\_REGULATION\_OF\_ODONTOGENESIS\_OF\_DENTIN\_CONTAINING\_TOOTH | MSigDB lists | GO\_REGULATION\_OF\_ODONTOGENESIS\_OF\_DENTIN\_CONTAINING\_TOOTH | 1 | 1 | 4161 | 184 | dpp | | 4.422e-02 | -3.12 | GO\_ENAMEL\_MINERALIZATION | MSigDB lists | GO\_ENAMEL\_MINERALIZATION | 1 | 1 | 4161 | 184 | Stim | | 4.422e-02 | -3.12 | TESAR\_JAK\_TARGETS\_MOUSE\_ES\_D3\_DN | MSigDB lists | TESAR\_JAK\_TARGETS\_MOUSE\_ES\_D3\_DN | 1 | 1 | 4161 | 184 | AGO3 | | 4.422e-02 | -3.12 | GO\_ACROSOME\_ASSEMBLY | MSigDB lists | GO\_ACROSOME\_ASSEMBLY | 1 | 1 | 4161 | 184 | CG17883 | | 4.422e-02 | -3.12 | HINATA\_NFKB\_TARGETS\_KERATINOCYTE\_DN | MSigDB lists | HINATA\_NFKB\_TARGETS\_KERATINOCYTE\_DN | 1 | 1 | 4161 | 184 | ema | | 4.422e-02 | -3.12 | WANG\_THOC1\_TARGETS\_UP | MSigDB lists | WANG\_THOC1\_TARGETS\_UP | 1 | 1 | 4161 | 184 | CycA | | 4.422e-02 | -3.12 | GO\_NEGATIVE\_REGULATION\_OF\_STEM\_CELL\_PROLIFERATION | MSigDB lists | GO\_NEGATIVE\_REGULATION\_OF\_STEM\_CELL\_PROLIFERATION | 1 | 1 | 4161 | 184 | Nf1 | | 4.422e-02 | -3.12 | MYLLYKANGAS\_AMPLIFICATION\_HOT\_SPOT\_27 | MSigDB lists | MYLLYKANGAS\_AMPLIFICATION\_HOT\_SPOT\_27 | 1 | 1 | 4161 | 184 | Graf | | 4.422e-02 | -3.12 | chr1q | MSigDB lists | chr1q | 1 | 1 | 4161 | 184 | Arpc5 | | 4.422e-02 | -3.12 | SIMBULAN\_UV\_RESPONSE\_NORMAL\_UP | MSigDB lists | SIMBULAN\_UV\_RESPONSE\_NORMAL\_UP | 1 | 1 | 4161 | 184 | Rpb11 | | 4.422e-02 | -3.12 | GO\_NEGATIVE\_REGULATION\_OF\_B\_CELL\_APOPTOTIC\_PROCESS | MSigDB lists | GO\_NEGATIVE\_REGULATION\_OF\_B\_CELL\_APOPTOTIC\_PROCESS | 1 | 1 | 4161 | 184 | CG9246 | | 4.422e-02 | -3.12 | SCHLINGEMANN\_SKIN\_CARCINOGENESIS\_TPA\_DN | MSigDB lists | SCHLINGEMANN\_SKIN\_CARCINOGENESIS\_TPA\_DN | 1 | 1 | 4161 | 184 | Mdh1 | | 4.422e-02 | -3.12 | KUROKAWA\_LIVER\_CANCER\_EARLY\_RECURRENCE\_DN | MSigDB lists | KUROKAWA\_LIVER\_CANCER\_EARLY\_RECURRENCE\_DN | 1 | 1 | 4161 | 184 | Vps35 | | 4.422e-02 | -3.12 | PID\_P38\_GAMMA\_DELTA\_PATHWAY | MSigDB lists | PID\_P38\_GAMMA\_DELTA\_PATHWAY | 1 | 1 | 4161 | 184 | lic | | 4.422e-02 | -3.12 | chr1p | MSigDB lists | chr1p | 1 | 1 | 4161 | 184 | noi | | 4.422e-02 | -3.12 | PETRETTO\_BLOOD\_PRESSURE\_DN | MSigDB lists | PETRETTO\_BLOOD\_PRESSURE\_DN | 1 | 1 | 4161 | 184 | tsu | | 4.422e-02 | -3.12 | GO\_WHITE\_FAT\_CELL\_DIFFERENTIATION | MSigDB lists | GO\_WHITE\_FAT\_CELL\_DIFFERENTIATION | 1 | 1 | 4161 | 184 | CtBP | | 4.422e-02 | -3.12 | BASSO\_HAIRY\_CELL\_LEUKEMIA\_DN | MSigDB lists | BASSO\_HAIRY\_CELL\_LEUKEMIA\_DN | 1 | 1 | 4161 | 184 | sigmar | | 4.422e-02 | -3.12 | GO\_DENDRITIC\_CELL\_MIGRATION | MSigDB lists | GO\_DENDRITIC\_CELL\_MIGRATION | 1 | 1 | 4161 | 184 | Cdc42 | | 4.422e-02 | -3.12 | WATANABE\_ULCERATIVE\_COLITIS\_WITH\_CANCER\_DN | MSigDB lists | WATANABE\_ULCERATIVE\_COLITIS\_WITH\_CANCER\_DN | 1 | 1 | 4161 | 184 | twin | | 4.422e-02 | -3.12 | GO\_NEGATIVE\_REGULATION\_OF\_INTERLEUKIN\_8\_PRODUCTION | MSigDB lists | GO\_NEGATIVE\_REGULATION\_OF\_INTERLEUKIN\_8\_PRODUCTION | 1 | 1 | 4161 | 184 | krz | | 4.422e-02 | -3.12 | chr6q | MSigDB lists | chr6q | 1 | 1 | 4161 | 184 | Orc3 | | 4.422e-02 | -3.12 | GO\_NEGATIVE\_REGULATION\_OF\_CARDIAC\_MUSCLE\_TISSUE\_DEVELOPMENT | MSigDB lists | GO\_NEGATIVE\_REGULATION\_OF\_CARDIAC\_MUSCLE\_TISSUE\_DEVELOPMENT | 1 | 1 | 4161 | 184 | dpp | | 4.422e-02 | -3.12 | chrxq23 | MSigDB lists | chrxq23 | 1 | 1 | 4161 | 184 | Cul4 | | 4.422e-02 | -3.12 | GO\_REGULATION\_OF\_PSEUDOPODIUM\_ASSEMBLY | MSigDB lists | GO\_REGULATION\_OF\_PSEUDOPODIUM\_ASSEMBLY | 1 | 1 | 4161 | 184 | Cdc42 | | 4.422e-02 | -3.12 | GO\_EPITHELIAL\_CELL\_CELL\_ADHESION | MSigDB lists | GO\_EPITHELIAL\_CELL\_CELL\_ADHESION | 1 | 1 | 4161 | 184 | Cdc42 | | 4.422e-02 | -3.12 | REACTOME\_ELEVATION\_OF\_CYTOSOLIC\_CA2\_LEVELS | MSigDB lists | REACTOME\_ELEVATION\_OF\_CYTOSOLIC\_CA2\_LEVELS | 1 | 1 | 4161 | 184 | Stim | | 4.422e-02 | -3.12 | CHANG\_POU5F1\_TARGETS\_DN | MSigDB lists | CHANG\_POU5F1\_TARGETS\_DN | 1 | 1 | 4161 | 184 | Txl | | 4.422e-02 | -3.12 | GO\_POSITIVE\_REGULATION\_OF\_OSTEOBLAST\_PROLIFERATION | MSigDB lists | GO\_POSITIVE\_REGULATION\_OF\_OSTEOBLAST\_PROLIFERATION | 1 | 1 | 4161 | 184 | dpp | | 4.422e-02 | -3.12 | GO\_NEGATIVE\_REGULATION\_OF\_MULTICELLULAR\_ORGANISMAL\_METABOLIC\_PROCESS | MSigDB lists | GO\_NEGATIVE\_REGULATION\_OF\_MULTICELLULAR\_ORGANISMAL\_METABOLIC\_PROCESS | 1 | 1 | 4161 | 184 | Rap1 | | 4.422e-02 | -3.12 | PID\_FRA\_PATHWAY | MSigDB lists | PID\_FRA\_PATHWAY | 1 | 1 | 4161 | 184 | CycA | | 4.436e-02 | -3.12 | CD28 co-stimulation | REACTOME pathways | R-DME-389356 | 9 | 2 | 3923 | 151 | Lst8,Cdc42 | | 4.436e-02 | -3.12 | MHC class II antigen presentation | REACTOME pathways | R-DME-2132295 | 9 | 2 | 3923 | 151 | CG41378,GILT3 | | 4.444e-02 | -3.11 | regulation of growth | biological process | GO:0040008 | 310 | 13 | 12345 | 305 | Rab7,Rab8,Rheb,mnb,tkv,Cdc42,Pten,Fmr1,Nf1,RagA-B,lic,dpp,Tctp | | 4.449e-02 | -3.11 | GO\_DETECTION\_OF\_STIMULUS | MSigDB lists | GO\_DETECTION\_OF\_STIMULUS | 59 | 6 | 4161 | 184 | mRpS9,Cul4,krz,Parp,unc-119,Stim | | 4.449e-02 | -3.11 | USF\_Q6\_01 | MSigDB lists | USF\_Q6\_01 | 59 | 6 | 4161 | 184 | eIF3b,flr,CG31460,bai,Stt3B,cta | | 4.449e-02 | -3.11 | GSE26669\_CD4\_VS\_CD8\_TCELL\_IN\_MLR\_DN | MSigDB lists | GSE26669\_CD4\_VS\_CD8\_TCELL\_IN\_MLR\_DN | 59 | 6 | 4161 | 184 | RpL14,Stlk,LIMK1,Pten,Sod2,xit | | 4.469e-02 | -3.11 | w (white) | protein interactions | 31271 | 80 | 6 | 9204 | 298 | Cp190,E(z),g,rb,su(Hw),BEAF-32 | | 4.518e-02 | -3.10 | EcR (Ecdysone receptor) | protein interactions | 35540 | 101 | 7 | 9204 | 298 | CycA,Cdk8,MED26,Pten,Dek,RecQ4,E(bx) | | 4.525e-02 | -3.10 | protein kinase binding | molecular function | GO:0019901 | 82 | 5 | 12176 | 289 | Cdc42,Slmap,CycA,lic,sws | | 4.534e-02 | -3.09 | SAM | smart domains | SM00454 | 26 | 3 | 5086 | 158 | Spn,Stim,smg | | 4.535e-02 | -3.09 | membrane protein complex | cellular component | GO:0098796 | 533 | 20 | 11940 | 298 | CG15168,rb,PIG-K,Vps35,CG31460,CG11752,MICU1,sun,l(2)37Bb,CIA30,ND-AGGG,Tim17b,Stt3B,RagA-B,cta,Tom20,VhaPPA1-1,ND-ASHI,CG11137,g | | 4.536e-02 | -3.09 | GO\_CELLULAR\_MACROMOLECULE\_LOCALIZATION | MSigDB lists | GO\_CELLULAR\_MACROMOLECULE\_LOCALIZATION | 498 | 30 | 4161 | 184 | Rap1,RpL5,Vps35,RagA-B,E(z),14-3-3zeta,RpS9,rhea,RpS8,Cdc42,CG9853,RpL35A,RpS26,Vps13D,RpL14,Tim17b,Rab8,bai,RpL36A,Rop,rb,Pex3,PGAP1,RpL15,CG8860,Stlk,Exo84,Tom20,Fs(2)Ket,PIG-K | | 4.552e-02 | -3.09 | demethylation | biological process | GO:0070988 | 14 | 2 | 12345 | 305 | Jarid2,NO66 | | 4.552e-02 | -3.09 | dorsal closure, amnioserosa morphology change | biological process | GO:0046664 | 14 | 2 | 12345 | 305 | Cdc42,tkv | | 4.552e-02 | -3.09 | homologous chromosome segregation | biological process | GO:0045143 | 14 | 2 | 12345 | 305 | grau,Mps1 | | 4.552e-02 | -3.09 | negative regulation of receptor signaling pathway via STAT | biological process | GO:1904893 | 14 | 2 | 12345 | 305 | CG30423,E(bx) | | 4.552e-02 | -3.09 | protein targeting to vacuole | biological process | GO:0006623 | 14 | 2 | 12345 | 305 | Vps13D,g | | 4.552e-02 | -3.09 | pupariation | biological process | GO:0035073 | 14 | 2 | 12345 | 305 | E(bx),smt3 | | 4.552e-02 | -3.09 | regulation of cytoplasmic translation | biological process | GO:2000765 | 14 | 2 | 12345 | 305 | me31B,CNBP | | 4.552e-02 | -3.09 | terminal button organization | biological process | GO:0072553 | 14 | 2 | 12345 | 305 | Vap33,Fmr1 | | 4.552e-02 | -3.09 | meiosis II | biological process | GO:0007135 | 14 | 2 | 12345 | 305 | fzy,Bub1 | | 4.552e-02 | -3.09 | vesicle budding from membrane | biological process | GO:0006900 | 14 | 2 | 12345 | 305 | Fmr1,g | | 4.552e-02 | -3.09 | actin nucleation | biological process | GO:0045010 | 14 | 2 | 12345 | 305 | Arpc5,Arpc3A | | 4.552e-02 | -3.09 | protein depolymerization | biological process | GO:0051261 | 14 | 2 | 12345 | 305 | tsr,flr | | 4.552e-02 | -3.09 | negative regulation of receptor signaling pathway via JAK-STAT | biological process | GO:0046426 | 14 | 2 | 12345 | 305 | E(bx),CG30423 | | 4.552e-02 | -3.09 | regulation of peptidoglycan recognition protein signaling pathway | biological process | GO:0061058 | 14 | 2 | 12345 | 305 | dgrn,TM9SF2 | | 4.552e-02 | -3.09 | endosome to lysosome transport | biological process | GO:0008333 | 14 | 2 | 12345 | 305 | Rab7,ema | | 4.552e-02 | -3.09 | regulation of chromatin assembly | biological process | GO:0010847 | 14 | 2 | 12345 | 305 | cal1,Acf | | 4.560e-02 | -3.09 | Translation Factors | WikiPathways | WP537 | 38 | 5 | 499 | 27 | eIF4B,eRF1,eEF5,eIF3b,eEF1delta | | 4.566e-02 | -3.09 | cytoskeleton-dependent intracellular transport | biological process | GO:0030705 | 79 | 5 | 12345 | 305 | pav,g,Girdin,Fmr1,Vap33 | | 4.566e-02 | -3.09 | regulation of canonical Wnt signaling pathway | biological process | GO:0060828 | 79 | 5 | 12345 | 305 | Pp1-87B,pav,flw,CtBP,lic | | 4.566e-02 | -3.09 | histone acetylation | biological process | GO:0016573 | 79 | 5 | 12345 | 305 | vnc,Acf,d4,Rcd5,Eaf6 | | 4.567e-02 | -3.09 | Pa1 (PTIP associated 1) | protein interactions | 32079 | 25 | 3 | 9204 | 298 | CG34159,Cpsf6,Ncoa6 | | 4.567e-02 | -3.09 | NC2alpha (Negative Cofactor 2alpha) | protein interactions | 37471 | 25 | 3 | 9204 | 298 | CG9951,CtBP,Nopp140 | | 4.567e-02 | -3.09 | CG1635 (uncharacterized protein) | protein interactions | 43720 | 25 | 3 | 9204 | 298 | Sod2,RpL35A,dah | | 4.568e-02 | -3.09 | GNF2\_PAK2 | MSigDB lists | GNF2\_PAK2 | 8 | 2 | 4161 | 184 | Arpc5,Arpc3A | | 4.568e-02 | -3.09 | HALLMARK\_PANCREAS\_BETA\_CELLS | MSigDB lists | HALLMARK\_PANCREAS\_BETA\_CELLS | 8 | 2 | 4161 | 184 | Rop,CG6907 | | 4.568e-02 | -3.09 | GCM\_MSN | MSigDB lists | GCM\_MSN | 8 | 2 | 4161 | 184 | sm,14-3-3zeta | | 4.568e-02 | -3.09 | GO\_OXALOACETATE\_METABOLIC\_PROCESS | MSigDB lists | GO\_OXALOACETATE\_METABOLIC\_PROCESS | 8 | 2 | 4161 | 184 | Mdh1,CG8132 | | 4.568e-02 | -3.09 | GO\_GTP\_DEPENDENT\_PROTEIN\_BINDING | MSigDB lists | GO\_GTP\_DEPENDENT\_PROTEIN\_BINDING | 8 | 2 | 4161 | 184 | rb,Cdc42 | | 4.568e-02 | -3.09 | TCGA\_GLIOBLASTOMA\_COPY\_NUMBER\_DN | MSigDB lists | TCGA\_GLIOBLASTOMA\_COPY\_NUMBER\_DN | 8 | 2 | 4161 | 184 | Pten,Nf1 | | 4.568e-02 | -3.09 | GO\_REGULATION\_OF\_OLIGODENDROCYTE\_DIFFERENTIATION | MSigDB lists | GO\_REGULATION\_OF\_OLIGODENDROCYTE\_DIFFERENTIATION | 8 | 2 | 4161 | 184 | flr,Nf1 | | 4.568e-02 | -3.09 | GO\_NEGATIVE\_REGULATION\_OF\_REGULATED\_SECRETORY\_PATHWAY | MSigDB lists | GO\_NEGATIVE\_REGULATION\_OF\_REGULATED\_SECRETORY\_PATHWAY | 8 | 2 | 4161 | 184 | Fmr1,Rap1 | | 4.568e-02 | -3.09 | GO\_ESTABLISHMENT\_OF\_MITOTIC\_SPINDLE\_LOCALIZATION | MSigDB lists | GO\_ESTABLISHMENT\_OF\_MITOTIC\_SPINDLE\_LOCALIZATION | 8 | 2 | 4161 | 184 | pins,Fs(2)Ket | | 4.568e-02 | -3.09 | REACTOME\_PI3K\_EVENTS\_IN\_ERBB2\_SIGNALING | MSigDB lists | REACTOME\_PI3K\_EVENTS\_IN\_ERBB2\_SIGNALING | 8 | 2 | 4161 | 184 | Pten,Lst8 | | 4.568e-02 | -3.09 | DORN\_ADENOVIRUS\_INFECTION\_32HR\_DN | MSigDB lists | DORN\_ADENOVIRUS\_INFECTION\_32HR\_DN | 8 | 2 | 4161 | 184 | lic,SF2 | | 4.568e-02 | -3.09 | PID\_RAS\_PATHWAY | MSigDB lists | PID\_RAS\_PATHWAY | 8 | 2 | 4161 | 184 | vap,Nf1 | | 4.568e-02 | -3.09 | GO\_REGULATION\_OF\_LEUKOCYTE\_APOPTOTIC\_PROCESS | MSigDB lists | GO\_REGULATION\_OF\_LEUKOCYTE\_APOPTOTIC\_PROCESS | 8 | 2 | 4161 | 184 | Pten,CG9246 | | 4.568e-02 | -3.09 | GO\_CENTROSOME\_LOCALIZATION | MSigDB lists | GO\_CENTROSOME\_LOCALIZATION | 8 | 2 | 4161 | 184 | pins,PlexA | | 4.568e-02 | -3.09 | GO\_NEGATIVE\_REGULATION\_OF\_RESPONSE\_TO\_OXIDATIVE\_STRESS | MSigDB lists | GO\_NEGATIVE\_REGULATION\_OF\_RESPONSE\_TO\_OXIDATIVE\_STRESS | 8 | 2 | 4161 | 184 | Sod2,HtrA2 | | 4.568e-02 | -3.09 | GO\_FOREBRAIN\_REGIONALIZATION | MSigDB lists | GO\_FOREBRAIN\_REGIONALIZATION | 8 | 2 | 4161 | 184 | dpp,PGAP1 | | 4.568e-02 | -3.09 | GO\_BRAIN\_MORPHOGENESIS | MSigDB lists | GO\_BRAIN\_MORPHOGENESIS | 8 | 2 | 4161 | 184 | Pten,Nf1 | | 4.568e-02 | -3.09 | GO\_CYSTEINE\_TYPE\_ENDOPEPTIDASE\_INHIBITOR\_ACTIVITY | MSigDB lists | GO\_CYSTEINE\_TYPE\_ENDOPEPTIDASE\_INHIBITOR\_ACTIVITY | 8 | 2 | 4161 | 184 | krz,sigmar | | 4.568e-02 | -3.09 | GO\_ACTIVATION\_OF\_ANAPHASE\_PROMOTING\_COMPLEX\_ACTIVITY | MSigDB lists | GO\_ACTIVATION\_OF\_ANAPHASE\_PROMOTING\_COMPLEX\_ACTIVITY | 8 | 2 | 4161 | 184 | Pten,fzy | | 4.568e-02 | -3.09 | GO\_NEGATIVE\_REGULATION\_OF\_EXOCYTOSIS | MSigDB lists | GO\_NEGATIVE\_REGULATION\_OF\_EXOCYTOSIS | 8 | 2 | 4161 | 184 | Rap1,Fmr1 | | 4.568e-02 | -3.09 | PID\_PI3KCI\_AKT\_PATHWAY | MSigDB lists | PID\_PI3KCI\_AKT\_PATHWAY | 8 | 2 | 4161 | 184 | 14-3-3zeta,Lst8 | | 4.568e-02 | -3.09 | GNF2\_MSN | MSigDB lists | GNF2\_MSN | 8 | 2 | 4161 | 184 | Cdc42,Arpc5 | | 4.568e-02 | -3.09 | GO\_NEGATIVE\_REGULATION\_OF\_CALCIUM\_ION\_TRANSMEMBRANE\_TRANSPORT | MSigDB lists | GO\_NEGATIVE\_REGULATION\_OF\_CALCIUM\_ION\_TRANSMEMBRANE\_TRANSPORT | 8 | 2 | 4161 | 184 | Ubqn,Fmr1 | | 4.568e-02 | -3.09 | STAMBOLSKY\_BOUND\_BY\_MUTATED\_TP53 | MSigDB lists | STAMBOLSKY\_BOUND\_BY\_MUTATED\_TP53 | 8 | 2 | 4161 | 184 | DNAlig1,Skp2 | | 4.568e-02 | -3.09 | GO\_RHO\_PROTEIN\_SIGNAL\_TRANSDUCTION | MSigDB lists | GO\_RHO\_PROTEIN\_SIGNAL\_TRANSDUCTION | 8 | 2 | 4161 | 184 | cta,LIMK1 | | 4.568e-02 | -3.09 | BIOCARTA\_MAL\_PATHWAY | MSigDB lists | BIOCARTA\_MAL\_PATHWAY | 8 | 2 | 4161 | 184 | Cdc42,LIMK1 | | 4.570e-02 | -3.09 | BOYAULT\_LIVER\_CANCER\_SUBCLASS\_G3\_UP | MSigDB lists | BOYAULT\_LIVER\_CANCER\_SUBCLASS\_G3\_UP | 107 | 9 | 4161 | 184 | CycA,eIF3b,CCT4,E(z),CG5535,CG32576,Pp1-87B,CG8860,Fs(2)Ket | | 4.570e-02 | -3.09 | REACTOME\_DNA\_REPLICATION | MSigDB lists | REACTOME\_DNA\_REPLICATION | 107 | 9 | 4161 | 184 | Prosalpha2,pav,Rpt6,DNAlig1,fzy,Pp1-87B,Rpn10,Orc3,CycA | | 4.574e-02 | -3.08 | endoplasmic reticulum membrane | cellular component | GO:0005789 | 217 | 10 | 11940 | 298 | Stt3B,PIG-K,xit,CG17883,CG15168,CG11137,RpS26,sws,Vap33,CG31460 | | 4.581e-02 | -3.08 | KEGG\_FC\_GAMMA\_R\_MEDIATED\_PHAGOCYTOSIS | MSigDB lists | KEGG\_FC\_GAMMA\_R\_MEDIATED\_PHAGOCYTOSIS | 31 | 4 | 4161 | 184 | Cdc42,LIMK1,Arpc3A,Arpc5 | | 4.581e-02 | -3.08 | GSE19888\_ADENOSINE\_A3R\_INH\_VS\_ACT\_WITH\_INHIBITOR\_PRETREATMENT\_IN\_MAST\_CELL\_DN | MSigDB lists | GSE19888\_ADENOSINE\_A3R\_INH\_VS\_ACT\_WITH\_INHIBITOR\_PRETREATMENT\_IN\_MAST\_CELL\_DN | 31 | 4 | 4161 | 184 | Tsp97E,CG9330,Aps,Parp | | 4.581e-02 | -3.08 | MORF\_PPP2R4 | MSigDB lists | MORF\_PPP2R4 | 31 | 4 | 4161 | 184 | Art1,l(1)10Bb,CG5861,bai | | 4.581e-02 | -3.08 | GO\_MITOCHONDRIAL\_RESPIRATORY\_CHAIN\_COMPLEX\_I\_BIOGENESIS | MSigDB lists | GO\_MITOCHONDRIAL\_RESPIRATORY\_CHAIN\_COMPLEX\_I\_BIOGENESIS | 31 | 4 | 4161 | 184 | ND-ASHI,l(2)37Bb,CIA30,ECSIT | | 4.581e-02 | -3.08 | GSE13547\_CTRL\_VS\_ANTI\_IGM\_STIM\_BCELL\_2H\_DN | MSigDB lists | GSE13547\_CTRL\_VS\_ANTI\_IGM\_STIM\_BCELL\_2H\_DN | 31 | 4 | 4161 | 184 | Pisd,MED21,CCT4,CG31460 | | 4.581e-02 | -3.08 | GSE35825\_UNTREATED\_VS\_IFNA\_STIM\_MACROPHAGE\_DN | MSigDB lists | GSE35825\_UNTREATED\_VS\_IFNA\_STIM\_MACROPHAGE\_DN | 31 | 4 | 4161 | 184 | rb,cta,Rop,Sod2 | | 4.581e-02 | -3.08 | GO\_PROTEIN\_TRANSMEMBRANE\_TRANSPORT | MSigDB lists | GO\_PROTEIN\_TRANSMEMBRANE\_TRANSPORT | 31 | 4 | 4161 | 184 | Tim17b,Pex3,Tom20,CG8860 | | 4.581e-02 | -3.08 | GSE29617\_CTRL\_VS\_DAY7\_TIV\_FLU\_VACCINE\_PBMC\_2008\_UP | MSigDB lists | GSE29617\_CTRL\_VS\_DAY7\_TIV\_FLU\_VACCINE\_PBMC\_2008\_UP | 31 | 4 | 4161 | 184 | eRF1,cta,inc,MED21 | | 4.581e-02 | -3.08 | GSE6269\_HEALTHY\_VS\_STAPH\_AUREUS\_INF\_PBMC\_UP | MSigDB lists | GSE6269\_HEALTHY\_VS\_STAPH\_AUREUS\_INF\_PBMC\_UP | 31 | 4 | 4161 | 184 | Fs(2)Ket,CG12935,PGAP1,Arp8 | | 4.590e-02 | -3.08 | Znf\_BED | interpro domains | IPR003656 | 12 | 2 | 11383 | 331 | Dref,BEAF-32 | | 4.599e-02 | -3.08 | bam (bag of marbles) | protein interactions | 43038 | 42 | 4 | 9204 | 298 | CycA,tkv,twin,Not3 | | 4.599e-02 | -3.08 | Parp (Poly-(ADP-ribose) polymerase) | protein interactions | 3355109 | 42 | 4 | 9204 | 298 | His2Av,Parp,RpL14,sqd | | 4.599e-02 | -3.08 | CG1532 (uncharacterized protein) | protein interactions | 33076 | 42 | 4 | 9204 | 298 | Wdr82,CG6171,BthD,CG1218 | | 4.625e-02 | -3.07 | GO\_REGULATION\_OF\_PROTEIN\_COMPLEX\_ASSEMBLY | MSigDB lists | GO\_REGULATION\_OF\_PROTEIN\_COMPLEX\_ASSEMBLY | 124 | 10 | 4161 | 184 | Lst8,Arpc5,Arpc3A,Cul4,vap,Rop,TfIIFbeta,Parp,Rpt6,Cdc42 | | 4.636e-02 | -3.07 | polytene chromosome band | cellular component | GO:0005704 | 14 | 2 | 11940 | 298 | Cp190,HP4 | | 4.636e-02 | -3.07 | Set1C/COMPASS complex | cellular component | GO:0048188 | 14 | 2 | 11940 | 298 | Wdr82,Rbbp5 | | 4.636e-02 | -3.07 | MLL3/4 complex | cellular component | GO:0044666 | 14 | 2 | 11940 | 298 | Rbbp5,Ncoa6 | | 4.636e-02 | -3.07 | Cul4-RING E3 ubiquitin ligase complex | cellular component | GO:0080008 | 14 | 2 | 11940 | 298 | Cul4,Roc2 | | 4.636e-02 | -3.07 | proteasome regulatory particle, base subcomplex | cellular component | GO:0008540 | 14 | 2 | 11940 | 298 | Rpt6,Rpn10 | | 4.636e-02 | -3.07 | neuronal ribonucleoprotein granule | cellular component | GO:0071598 | 14 | 2 | 11940 | 298 | me31B,Fmr1 | | 4.636e-02 | -3.07 | CCR4-NOT complex | cellular component | GO:0030014 | 14 | 2 | 11940 | 298 | Not3,twin | | 4.644e-02 | -3.07 | GSE27786\_CD4\_TCELL\_VS\_NKTCELL\_UP | MSigDB lists | GSE27786\_CD4\_TCELL\_VS\_NKTCELL\_UP | 75 | 7 | 4161 | 184 | CG4866,mon2,CG10672,vnc,l(1)10Bb,CG15168,Wdr82 | | 4.644e-02 | -3.07 | GSE17721\_0.5H\_VS\_4H\_CPG\_BMDC\_UP | MSigDB lists | GSE17721\_0.5H\_VS\_4H\_CPG\_BMDC\_UP | 75 | 7 | 4161 | 184 | unc-119,krz,Stt3B,sws,CG12321,fzy,CG17883 | | 4.644e-02 | -3.07 | GSE2770\_IL12\_AND\_TGFB\_VS\_IL4\_TREATED\_ACT\_CD4\_TCELL\_6H\_UP | MSigDB lists | GSE2770\_IL12\_AND\_TGFB\_VS\_IL4\_TREATED\_ACT\_CD4\_TCELL\_6H\_UP | 75 | 7 | 4161 | 184 | Tim17b,Fign,Rbbp5,Cul4,Mdh1,CycA,pins | | 4.660e-02 | -3.07 | negative regulation of response to external stimulus | biological process | GO:0032102 | 55 | 4 | 12345 | 305 | krz,Pten,E(bx),TM9SF2 | | 4.660e-02 | -3.07 | cortical actin cytoskeleton organization | biological process | GO:0030866 | 55 | 4 | 12345 | 305 | Cdc42,tsr,pav,Exo84 | | 4.660e-02 | -3.07 | REACTOME\_MRNA\_PROCESSING | MSigDB lists | REACTOME\_MRNA\_PROCESSING | 91 | 8 | 4161 | 184 | sm,snf,Phf5a,Rpb11,noi,TfIIFbeta,SF2,tsu | | 4.674e-02 | -3.06 | GSE7831\_UNSTIM\_VS\_INFLUENZA\_STIM\_PDC\_1H\_UP | MSigDB lists | GSE7831\_UNSTIM\_VS\_INFLUENZA\_STIM\_PDC\_1H\_UP | 45 | 5 | 4161 | 184 | CG41378,Graf,sws,Mdh1,CG9330 | | 4.674e-02 | -3.06 | GSE6674\_PL2\_3\_VS\_ANTI\_IGM\_AND\_CPG\_STIM\_BCELL\_DN | MSigDB lists | GSE6674\_PL2\_3\_VS\_ANTI\_IGM\_AND\_CPG\_STIM\_BCELL\_DN | 45 | 5 | 4161 | 184 | E(z),rb,Wdr82,vap,Ufd4 | | 4.674e-02 | -3.06 | GSE25123\_IL4\_VS\_IL4\_AND\_ROSIGLITAZONE\_STIM\_MACROPHAGE\_DAY10\_DN | MSigDB lists | GSE25123\_IL4\_VS\_IL4\_AND\_ROSIGLITAZONE\_STIM\_MACROPHAGE\_DAY10\_DN | 45 | 5 | 4161 | 184 | ubl,CG9853,Vap33,CG5535,CG32473 | | 4.674e-02 | -3.06 | GO\_POSITIVE\_REGULATION\_OF\_NEURON\_PROJECTION\_DEVELOPMENT | MSigDB lists | GO\_POSITIVE\_REGULATION\_OF\_NEURON\_PROJECTION\_DEVELOPMENT | 45 | 5 | 4161 | 184 | Rap1,RhoGAPp190,LIMK1,E(z),Fmr1 | | 4.674e-02 | -3.06 | GSE33162\_UNTREATED\_VS\_4H\_LPS\_STIM\_HDAC3\_HET\_MACROPHAGE\_DN | MSigDB lists | GSE33162\_UNTREATED\_VS\_4H\_LPS\_STIM\_HDAC3\_HET\_MACROPHAGE\_DN | 45 | 5 | 4161 | 184 | krz,Sgt,Roc2,RpS9,CG4866 | | 4.689e-02 | -3.06 | cellular response to oxygen-containing compound | biological process | GO:1901701 | 133 | 7 | 12345 | 305 | Sod2,E(bx),RagA-B,CG5056,Rab8,Rheb,Rap1 | | 4.691e-02 | -3.06 | phosphatidylinositol-3,4-bisphosphate phosphatase activity | molecular function | GO:0106017 | 2 | 1 | 12176 | 289 | Pten | | 4.691e-02 | -3.06 | dolichyl pyrophosphate Man9GlcNAc2 alpha-1,3-glucosyltransferase activity | molecular function | GO:0042281 | 2 | 1 | 12176 | 289 | xit | | 4.691e-02 | -3.06 | MutLalpha complex binding | molecular function | GO:0032405 | 2 | 1 | 12176 | 289 | spel1 | | 4.691e-02 | -3.06 | 2-polyprenyl-6-methoxy-1,4-benzoquinone methyltransferase activity | molecular function | GO:0008425 | 2 | 1 | 12176 | 289 | Coq5 | | 4.691e-02 | -3.06 | cyclase regulator activity | molecular function | GO:0010851 | 2 | 1 | 12176 | 289 | PlexA | | 4.691e-02 | -3.06 | mitogen-activated protein kinase kinase kinase binding | molecular function | GO:0031435 | 2 | 1 | 12176 | 289 | lic | | 4.691e-02 | -3.06 | sequence-specific mRNA binding | molecular function | GO:1990825 | 2 | 1 | 12176 | 289 | eRF1 | | 4.691e-02 | -3.06 | mismatch repair complex binding | molecular function | GO:0032404 | 2 | 1 | 12176 | 289 | spel1 | | 4.691e-02 | -3.06 | poly-ADP-D-ribose binding | molecular function | GO:0072572 | 2 | 1 | 12176 | 289 | CG1218 | | 4.691e-02 | -3.06 | single thymine insertion binding | molecular function | GO:0032143 | 2 | 1 | 12176 | 289 | spel1 | | 4.691e-02 | -3.06 | single base insertion or deletion binding | molecular function | GO:0032138 | 2 | 1 | 12176 | 289 | spel1 | | 4.691e-02 | -3.06 | peptide-serine-N-acetyltransferase activity | molecular function | GO:1990189 | 2 | 1 | 12176 | 289 | vnc | | 4.691e-02 | -3.06 | DNA insertion or deletion binding | molecular function | GO:0032135 | 2 | 1 | 12176 | 289 | spel1 | | 4.691e-02 | -3.06 | G-quadruplex RNA binding | molecular function | GO:0002151 | 2 | 1 | 12176 | 289 | Rcd5 | | 4.691e-02 | -3.06 | translation release factor activity, codon specific | molecular function | GO:0016149 | 2 | 1 | 12176 | 289 | eRF1 | | 4.691e-02 | -3.06 | TFIIF-class transcription factor complex binding | molecular function | GO:0001096 | 2 | 1 | 12176 | 289 | TfIIFbeta | | 4.691e-02 | -3.06 | semaphorin receptor activity | molecular function | GO:0017154 | 2 | 1 | 12176 | 289 | PlexA | | 4.691e-02 | -3.06 | calcium-induced calcium release activity | molecular function | GO:0048763 | 2 | 1 | 12176 | 289 | Stim | | 4.691e-02 | -3.06 | mRNA 3'-UTR AU-rich region binding | molecular function | GO:0035925 | 2 | 1 | 12176 | 289 | sqd | | 4.691e-02 | -3.06 | quinone cofactor methyltransferase activity | molecular function | GO:0030580 | 2 | 1 | 12176 | 289 | Coq5 | | 4.691e-02 | -3.06 | guanylate cyclase regulator activity | molecular function | GO:0030249 | 2 | 1 | 12176 | 289 | PlexA | | 4.691e-02 | -3.06 | guanylate cyclase activator activity | molecular function | GO:0030250 | 2 | 1 | 12176 | 289 | PlexA | | 4.691e-02 | -3.06 | cyclase activator activity | molecular function | GO:0010853 | 2 | 1 | 12176 | 289 | PlexA | | 4.691e-02 | -3.06 | acid sphingomyelin phosphodiesterase activity | molecular function | GO:0061750 | 2 | 1 | 12176 | 289 | CG15534 | | 4.692e-02 | -3.06 | Cdc42 (Cdc42) | protein interactions | 32981 | 61 | 5 | 9204 | 298 | GEFmeso,LIMK1,CG12299,tkv,smt3 | | 4.701e-02 | -3.06 | Lkb1 (Lkb1 kinase) | protein interactions | 41673 | 81 | 6 | 9204 | 298 | RpL14,eIF3b,pav,pins,RpL35A,Stlk | | 4.722e-02 | -3.05 | protein stabilization | biological process | GO:0050821 | 33 | 3 | 12345 | 305 | CSN1b,Mlf,14-3-3zeta | | 4.722e-02 | -3.05 | pole cell development | biological process | GO:0007277 | 33 | 3 | 12345 | 305 | me31B,grau,Fmr1 | | 4.722e-02 | -3.05 | germ-line cyst formation | biological process | GO:0048134 | 33 | 3 | 12345 | 305 | CSN1b,Fmr1,nsl1 | | 4.726e-02 | -3.05 | N-acyltransferase activity | molecular function | GO:0016410 | 83 | 5 | 12176 | 289 | d4,Rcd5,Acf,Eaf6,vnc | | 4.738e-02 | -3.05 | eIF3d1 (eukaryotic translation initiation factor 3 subunit d1) | protein interactions | 42789 | 11 | 2 | 9204 | 298 | Hrb27C,eIF3c | | 4.738e-02 | -3.05 | CG13005 (uncharacterized protein) | protein interactions | 32670 | 11 | 2 | 9204 | 298 | CG11456,Nph | | 4.738e-02 | -3.05 | I-2 (Inhibitor-2) | protein interactions | 39156 | 11 | 2 | 9204 | 298 | Pp1-87B,flw | | 4.738e-02 | -3.05 | TwdlD (TweedleD) | protein interactions | 43210 | 11 | 2 | 9204 | 298 | Ubqn,Sgt | | 4.738e-02 | -3.05 | spin (spinster) | protein interactions | 45380 | 11 | 2 | 9204 | 298 | dpp,tkv | | 4.738e-02 | -3.05 | Ulp1 (Ulp1) | protein interactions | 32910 | 11 | 2 | 9204 | 298 | krz,smt3 | | 4.738e-02 | -3.05 | CG31204 (uncharacterized protein) | protein interactions | 318625 | 11 | 2 | 9204 | 298 | Mps1,CycA | | 4.738e-02 | -3.05 | CG9723 (uncharacterized protein) | protein interactions | 32639 | 11 | 2 | 9204 | 298 | Fs(2)Ket,Vap33 | | 4.738e-02 | -3.05 | Sec15 (Secretory 15) | protein interactions | 42499 | 11 | 2 | 9204 | 298 | dpp,Rop | | 4.738e-02 | -3.05 | CG6712 (uncharacterized protein) | protein interactions | 34631 | 11 | 2 | 9204 | 298 | Lst8,Rbm13 | | 4.738e-02 | -3.05 | CG14881 (uncharacterized protein) | protein interactions | 41990 | 11 | 2 | 9204 | 298 | CycA,Map60 | | 4.738e-02 | -3.05 | ver (verrocchio) | protein interactions | 39432 | 11 | 2 | 9204 | 298 | ver,14-3-3zeta | | 4.738e-02 | -3.05 | CG7546 (uncharacterized protein) | protein interactions | 38844 | 11 | 2 | 9204 | 298 | CG9853,Rbp1 | | 4.765e-02 | -3.04 | post-embryonic appendage morphogenesis | biological process | GO:0035120 | 345 | 14 | 12345 | 305 | tsr,Cdc42,peng,Btk29A,tkv,EloB,LIMK1,GEFmeso,Cdk8,rhea,dpp,flr,Stim,flw | | 4.769e-02 | -3.04 | KARLSSON\_TGFB1\_TARGETS\_UP | MSigDB lists | KARLSSON\_TGFB1\_TARGETS\_UP | 60 | 6 | 4161 | 184 | TfIIFbeta,eIF3b,CG8635,Fs(2)Ket,Tom20,Graf | | 4.769e-02 | -3.04 | THEILGAARD\_NEUTROPHIL\_AT\_SKIN\_WOUND\_DN | MSigDB lists | THEILGAARD\_NEUTROPHIL\_AT\_SKIN\_WOUND\_DN | 60 | 6 | 4161 | 184 | Arpc3A,CG7766,Rpb11,SF2,Pp1-87B,vap | | 4.769e-02 | -3.04 | GO\_CELLULAR\_RESPONSE\_TO\_OXIDATIVE\_STRESS | MSigDB lists | GO\_CELLULAR\_RESPONSE\_TO\_OXIDATIVE\_STRESS | 60 | 6 | 4161 | 184 | E(z),Parp,Txl,Sod2,CycA,HtrA2 | | 4.769e-02 | -3.04 | GSE2826\_XID\_VS\_BTK\_KO\_BCELL\_UP | MSigDB lists | GSE2826\_XID\_VS\_BTK\_KO\_BCELL\_UP | 60 | 6 | 4161 | 184 | TfIIFbeta,eIF3g1,cta,CCT4,Stt3B,Pisd | | 4.769e-02 | -3.04 | BENPORATH\_PROLIFERATION | MSigDB lists | BENPORATH\_PROLIFERATION | 60 | 6 | 4161 | 184 | tsu,spel1,pins,His2Av,E(z),Parp | | 4.789e-02 | -3.04 | Pdk1 (Phosphoinositide-dependent kinase 1) | protein interactions | 38017 | 124 | 8 | 9204 | 298 | Rpt6,RpS9,sqd,RpL36A,RpS26,lic,eIF3c,Dref | | 4.798e-02 | -3.04 | GO\_REGULATION\_OF\_MULTICELLULAR\_ORGANISMAL\_DEVELOPMENT | MSigDB lists | GO\_REGULATION\_OF\_MULTICELLULAR\_ORGANISMAL\_DEVELOPMENT | 325 | 21 | 4161 | 184 | His4r,Rpn10,Rap1,flr,tkv,Art1,E(z),LIMK1,Nf1,Cdc42,fzy,viaf,Pten,Fmr1,dpp,me31B,RhoGAPp190,rb,Rpt6,Prosalpha2,PlexA | | 4.803e-02 | -3.04 | STPHPHTASE | prints domains | PR00114 | 19 | 2 | 2185 | 41 | flw,Pp1-87B | | 4.803e-02 | -3.04 | SH2DOMAIN | prints domains | PR00401 | 19 | 2 | 2185 | 41 | Btk29A,vap | | 4.847e-02 | -3.03 | anterior/posterior axis specification, embryo | biological process | GO:0008595 | 134 | 7 | 12345 | 305 | Hrb27C,sqd,tsu,mon2,lic,dpp,glo | | 4.847e-02 | -3.03 | tripartite regional subdivision | biological process | GO:0007351 | 134 | 7 | 12345 | 305 | dpp,glo,lic,tsu,mon2,sqd,Hrb27C | | 4.852e-02 | -3.03 | heterocycle biosynthetic process | biological process | GO:0018130 | 476 | 18 | 12345 | 305 | DNAlig1,E(bx),CG6244,TfIIFbeta,Arp8,Pbp49,g,grau,His4r,Aps,EloB,sun,rb,Parp,su(Hw),Cul4,Rpb11,RecQ4 | | 4.854e-02 | -3.03 | tissue morphogenesis | biological process | GO:0048729 | 644 | 23 | 12345 | 305 | LIMK1,dpp,Cdk8,rhea,cta,flw,sqd,Rap1,Cdc42,peng,Parp,EloB,Rheb,GEFmeso,CSN1b,Zpr1,VhaPPA1-1,Stim,flr,tsr,Btk29A,spoon,tkv | | 4.880e-02 | -3.02 | neurotransmitter receptor transport, postsynaptic endosome to lysosome | biological process | GO:0098943 | 2 | 1 | 12345 | 305 | g | | 4.880e-02 | -3.02 | phagolysosome assembly involved in apoptotic cell clearance | biological process | GO:0090387 | 2 | 1 | 12345 | 305 | Rab14 | | 4.880e-02 | -3.02 | cytoplasmic translational termination | biological process | GO:0002184 | 2 | 1 | 12345 | 305 | eRF1 | | 4.880e-02 | -3.02 | positive regulation of translational termination | biological process | GO:0045905 | 2 | 1 | 12345 | 305 | eEF5 | | 4.880e-02 | -3.02 | regulation of protein localization to cell surface | biological process | GO:2000008 | 2 | 1 | 12345 | 305 | CG30423 | | 4.880e-02 | -3.02 | neuron death | biological process | GO:0070997 | 2 | 1 | 12345 | 305 | sws | | 4.880e-02 | -3.02 | negative regulation of phosphatidylinositol 3-kinase activity | biological process | GO:0043553 | 2 | 1 | 12345 | 305 | Pten | | 4.880e-02 | -3.02 | DNA rewinding | biological process | GO:0036292 | 2 | 1 | 12345 | 305 | RecQ4 | | 4.880e-02 | -3.02 | positive regulation of torso signaling pathway | biological process | GO:0120176 | 2 | 1 | 12345 | 305 | 14-3-3zeta | | 4.880e-02 | -3.02 | regulation of guanylate cyclase activity | biological process | GO:0031282 | 2 | 1 | 12345 | 305 | PlexA | | 4.880e-02 | -3.02 | positive regulation of cytoplasmic translation | biological process | GO:2000767 | 2 | 1 | 12345 | 305 | CNBP | | 4.880e-02 | -3.02 | mitotic spindle midzone assembly | biological process | GO:0051256 | 2 | 1 | 12345 | 305 | pav | | 4.880e-02 | -3.02 | vitamin K metabolic process | biological process | GO:0042373 | 2 | 1 | 12345 | 305 | heix | | 4.880e-02 | -3.02 | tRNA threonylcarbamoyladenosine modification | biological process | GO:0002949 | 2 | 1 | 12345 | 305 | Tcs3 | | 4.880e-02 | -3.02 | negative regulation of inflammatory response | biological process | GO:0050728 | 2 | 1 | 12345 | 305 | krz | | 4.880e-02 | -3.02 | regulation of mitochondrial depolarization | biological process | GO:0051900 | 2 | 1 | 12345 | 305 | Dref | | 4.880e-02 | -3.02 | dendrite regeneration | biological process | GO:0031104 | 2 | 1 | 12345 | 305 | Pten | | 4.880e-02 | -3.02 | glial cell apoptotic process | biological process | GO:0034349 | 2 | 1 | 12345 | 305 | sws | | 4.880e-02 | -3.02 | positive regulation of calcium ion transmembrane transporter activity | biological process | GO:1901021 | 2 | 1 | 12345 | 305 | Stim | | 4.880e-02 | -3.02 | negative regulation of lipid kinase activity | biological process | GO:0090219 | 2 | 1 | 12345 | 305 | Pten | | 4.880e-02 | -3.02 | cellular response to epidermal growth factor stimulus | biological process | GO:0071364 | 2 | 1 | 12345 | 305 | Zpr1 | | 4.880e-02 | -3.02 | protein kinase B signaling | biological process | GO:0043491 | 2 | 1 | 12345 | 305 | Pten | | 4.880e-02 | -3.02 | transforming growth factor beta receptor signaling pathway involved in endodermal cell fate specification | biological process | GO:0060799 | 2 | 1 | 12345 | 305 | tkv | | 4.880e-02 | -3.02 | positive regulation of translational fidelity | biological process | GO:0045903 | 2 | 1 | 12345 | 305 | RpS9 | | 4.880e-02 | -3.02 | regulation of inflammatory response | biological process | GO:0050727 | 2 | 1 | 12345 | 305 | krz | | 4.880e-02 | -3.02 | positive regulation of mRNA processing | biological process | GO:0050685 | 2 | 1 | 12345 | 305 | Hrb27C | | 4.880e-02 | -3.02 | RNA import into mitochondrion | biological process | GO:0035927 | 2 | 1 | 12345 | 305 | Tom20 | | 4.880e-02 | -3.02 | synaptic vesicle clustering | biological process | GO:0097091 | 2 | 1 | 12345 | 305 | Fmr1 | | 4.880e-02 | -3.02 | negative regulation of axon regeneration | biological process | GO:0048681 | 2 | 1 | 12345 | 305 | Pten | | 4.880e-02 | -3.02 | Golgi vesicle fusion to target membrane | biological process | GO:0048210 | 2 | 1 | 12345 | 305 | Rab8 | | 4.880e-02 | -3.02 | negative regulation of meiotic chromosome separation | biological process | GO:1905133 | 2 | 1 | 12345 | 305 | Mps1 | | 4.880e-02 | -3.02 | negative regulation of metaphase/anaphase transition of meiotic cell cycle | biological process | GO:1902103 | 2 | 1 | 12345 | 305 | Mps1 | | 4.880e-02 | -3.02 | maternal determination of dorsal/ventral axis, ovarian follicular epithelium, germ-line encoded | biological process | GO:0008070 | 2 | 1 | 12345 | 305 | spoon | | 4.880e-02 | -3.02 | positive regulation of mRNA splicing, via spliceosome | biological process | GO:0048026 | 2 | 1 | 12345 | 305 | Hrb27C | | 4.880e-02 | -3.02 | N-terminal peptidyl-glutamic acid acetylation | biological process | GO:0018002 | 2 | 1 | 12345 | 305 | vnc | | 4.880e-02 | -3.02 | tRNA import into mitochondrion | biological process | GO:0016031 | 2 | 1 | 12345 | 305 | Tom20 | | 4.880e-02 | -3.02 | phagosome maturation involved in apoptotic cell clearance | biological process | GO:0090386 | 2 | 1 | 12345 | 305 | Rab14 | | 4.880e-02 | -3.02 | positive regulation of sphingolipid biosynthetic process | biological process | GO:0090154 | 2 | 1 | 12345 | 305 | Pex3 | | 4.880e-02 | -3.02 | negative regulation of GTPase activity | biological process | GO:0034260 | 2 | 1 | 12345 | 305 | conu | | 4.880e-02 | -3.02 | transforming growth factor beta receptor signaling pathway involved in primary germ layer cell fate commitment | biological process | GO:0060797 | 2 | 1 | 12345 | 305 | tkv | | 4.880e-02 | -3.02 | regulation of metaphase/anaphase transition of meiotic cell cycle | biological process | GO:1902102 | 2 | 1 | 12345 | 305 | Mps1 | | 4.880e-02 | -3.02 | endodermal cell fate specification | biological process | GO:0001714 | 2 | 1 | 12345 | 305 | tkv | | 4.880e-02 | -3.02 | polarized secretion of basement membrane proteins in epithelium | biological process | GO:0061865 | 2 | 1 | 12345 | 305 | Rab8 | | 4.880e-02 | -3.02 | meiotic metaphase plate congression | biological process | GO:0051311 | 2 | 1 | 12345 | 305 | Mps1 | | 4.880e-02 | -3.02 | Rap protein signal transduction | biological process | GO:0032486 | 2 | 1 | 12345 | 305 | Rap1 | | 4.880e-02 | -3.02 | protein poly-ADP-ribosylation | biological process | GO:0070212 | 2 | 1 | 12345 | 305 | Parp | | 4.880e-02 | -3.02 | fat-soluble vitamin biosynthetic process | biological process | GO:0042362 | 2 | 1 | 12345 | 305 | heix | | 4.880e-02 | -3.02 | negative regulation of neuron projection regeneration | biological process | GO:0070571 | 2 | 1 | 12345 | 305 | Pten | | 4.880e-02 | -3.02 | response to epidermal growth factor | biological process | GO:0070849 | 2 | 1 | 12345 | 305 | Zpr1 | | 4.880e-02 | -3.02 | plus-end-directed vesicle transport along microtubule | biological process | GO:0072383 | 2 | 1 | 12345 | 305 | pav | | 4.880e-02 | -3.02 | semaphorin-plexin signaling pathway involved in neuron projection guidance | biological process | GO:1902285 | 2 | 1 | 12345 | 305 | PlexA | | 4.880e-02 | -3.02 | achiasmate meiosis I | biological process | GO:0000705 | 2 | 1 | 12345 | 305 | Mps1 | | 4.880e-02 | -3.02 | negative regulation of Arp2/3 complex-mediated actin nucleation | biological process | GO:0034316 | 2 | 1 | 12345 | 305 | flr | | 4.880e-02 | -3.02 | regulation of centromere complex assembly | biological process | GO:0090230 | 2 | 1 | 12345 | 305 | cal1 | | 4.880e-02 | -3.02 | negative regulation of meiotic cell cycle phase transition | biological process | GO:1901994 | 2 | 1 | 12345 | 305 | Mps1 | | 4.880e-02 | -3.02 | regulation of meiotic chromosome separation | biological process | GO:1905132 | 2 | 1 | 12345 | 305 | Mps1 | | 4.880e-02 | -3.02 | neuron apoptotic process | biological process | GO:0051402 | 2 | 1 | 12345 | 305 | sws | | 4.880e-02 | -3.02 | positive regulation of transcription by transcription factor localization | biological process | GO:0061586 | 2 | 1 | 12345 | 305 | Lst8 | | 4.880e-02 | -3.02 | DNA replication, Okazaki fragment processing | biological process | GO:0033567 | 2 | 1 | 12345 | 305 | DNAlig1 | | 4.880e-02 | -3.02 | regulation of cell junction assembly | biological process | GO:1901888 | 2 | 1 | 12345 | 305 | Pten | | 4.880e-02 | -3.02 | mitochondrial outer membrane translocase complex assembly | biological process | GO:0070096 | 2 | 1 | 12345 | 305 | Tom20 | | 4.880e-02 | -3.02 | positive regulation of calcium ion transmembrane transport | biological process | GO:1904427 | 2 | 1 | 12345 | 305 | Stim | | 4.880e-02 | -3.02 | semaphorin-plexin signaling pathway involved in axon guidance | biological process | GO:1902287 | 2 | 1 | 12345 | 305 | PlexA | | 4.880e-02 | -3.02 | semaphorin-plexin signaling pathway involved in regulation of photoreceptor cell axon guidance | biological process | GO:2000305 | 2 | 1 | 12345 | 305 | PlexA | | 4.880e-02 | -3.02 | regulation of transcription from RNA polymerase II promoter involved in spermatogenesis | biological process | GO:1902064 | 2 | 1 | 12345 | 305 | MED22 | | 4.880e-02 | -3.02 | plus-end-directed organelle transport along microtubule | biological process | GO:0072386 | 2 | 1 | 12345 | 305 | pav | | 4.880e-02 | -3.02 | protein import into peroxisome membrane | biological process | GO:0045046 | 2 | 1 | 12345 | 305 | Pex3 | | 4.880e-02 | -3.02 | cellular response to cAMP | biological process | GO:0071320 | 2 | 1 | 12345 | 305 | Rap1 | | 4.880e-02 | -3.02 | regulation of cAMP-dependent protein kinase activity | biological process | GO:2000479 | 2 | 1 | 12345 | 305 | sws | | 4.880e-02 | -3.02 | response to rotenone | biological process | GO:1904647 | 2 | 1 | 12345 | 305 | Vps35 | | 4.880e-02 | -3.02 | regulation of protein oligomerization | biological process | GO:0032459 | 2 | 1 | 12345 | 305 | Sgt | | 4.880e-02 | -3.02 | fusion cell fate specification | biological process | GO:0035156 | 2 | 1 | 12345 | 305 | dpp | | 4.880e-02 | -3.02 | positive regulation of guanylate cyclase activity | biological process | GO:0031284 | 2 | 1 | 12345 | 305 | PlexA | | 4.892e-02 | -3.02 | GO\_POSITIVE\_REGULATION\_OF\_ACTIN\_FILAMENT\_POLYMERIZATION | MSigDB lists | GO\_POSITIVE\_REGULATION\_OF\_ACTIN\_FILAMENT\_POLYMERIZATION | 19 | 3 | 4161 | 184 | Lst8,Arpc5,Arpc3A | | 4.892e-02 | -3.02 | AAGGGAT\_MIR188 | MSigDB lists | AAGGGAT\_MIR188 | 19 | 3 | 4161 | 184 | Pten,l(1)10Bb,CG13016 | | 4.892e-02 | -3.02 | ZHOU\_TNF\_SIGNALING\_4HR | MSigDB lists | ZHOU\_TNF\_SIGNALING\_4HR | 19 | 3 | 4161 | 184 | noi,Sod2,RpS8 | | 4.892e-02 | -3.02 | RODRIGUES\_NTN1\_TARGETS\_DN | MSigDB lists | RODRIGUES\_NTN1\_TARGETS\_DN | 19 | 3 | 4161 | 184 | Ufd4,Graf,LIMK1 | | 4.892e-02 | -3.02 | PID\_AURORA\_B\_PATHWAY | MSigDB lists | PID\_AURORA\_B\_PATHWAY | 19 | 3 | 4161 | 184 | Pp1-87B,pav,vap | | 4.892e-02 | -3.02 | GSE14413\_UNSTIM\_VS\_IFNB\_STIM\_L929\_CELLS\_UP | MSigDB lists | GSE14413\_UNSTIM\_VS\_IFNB\_STIM\_L929\_CELLS\_UP | 19 | 3 | 4161 | 184 | Prosalpha2,Acbp1,Dbp21E2 | | 4.892e-02 | -3.02 | AGCTCCT\_MIR28 | MSigDB lists | AGCTCCT\_MIR28 | 19 | 3 | 4161 | 184 | Txl,HtrA2,rhea | | 4.892e-02 | -3.02 | MORI\_MATURE\_B\_LYMPHOCYTE\_UP | MSigDB lists | MORI\_MATURE\_B\_LYMPHOCYTE\_UP | 19 | 3 | 4161 | 184 | twin,Rab8,Pisd | | 4.892e-02 | -3.02 | GUTIERREZ\_MULTIPLE\_MYELOMA\_UP | MSigDB lists | GUTIERREZ\_MULTIPLE\_MYELOMA\_UP | 19 | 3 | 4161 | 184 | 14-3-3zeta,eRF1,xit | | 4.892e-02 | -3.02 | GO\_CELLULAR\_RESPONSE\_TO\_IONIZING\_RADIATION | MSigDB lists | GO\_CELLULAR\_RESPONSE\_TO\_IONIZING\_RADIATION | 19 | 3 | 4161 | 184 | Ino80,Fign,eEF1delta | | 4.892e-02 | -3.02 | GO\_CELLULAR\_RESPONSE\_TO\_UV | MSigDB lists | GO\_CELLULAR\_RESPONSE\_TO\_UV | 19 | 3 | 4161 | 184 | Ino80,Cul4,CG9246 | | 4.892e-02 | -3.02 | GO\_REGULATION\_OF\_FIBROBLAST\_PROLIFERATION | MSigDB lists | GO\_REGULATION\_OF\_FIBROBLAST\_PROLIFERATION | 19 | 3 | 4161 | 184 | CG41378,Nf1,CycA | | 4.892e-02 | -3.02 | MODULE\_447 | MSigDB lists | MODULE\_447 | 19 | 3 | 4161 | 184 | snf,eIF3c,CtBP | | 4.892e-02 | -3.02 | PID\_FOXO\_PATHWAY | MSigDB lists | PID\_FOXO\_PATHWAY | 19 | 3 | 4161 | 184 | Skp2,14-3-3zeta,Sod2 | | 4.912e-02 | -3.01 | pbl (pebble) | protein interactions | 38879 | 147 | 9 | 9204 | 298 | LIMK1,Rop,Cdc42,flr,pav,RhoGAPp190,PlexA,His2Av,Rap1 | | 4.917e-02 | -3.01 | organonitrogen compound catabolic process | biological process | GO:1901565 | 510 | 19 | 12345 | 305 | Roc2,Usp32,Rpn10,CG10365,CG15534,Prosbeta1,Sgt,fzy,Cul4,Ubqn,Aps,alpha-Man-Ib,Usp16-45,Skp2,Rpt6,Prosalpha2,inc,Ufd4,CG32473 | | 4.919e-02 | -3.01 | TTGCACT\_MIR130A\_MIR301\_MIR130B | MSigDB lists | TTGCACT\_MIR130A\_MIR301\_MIR130B | 92 | 8 | 4161 | 184 | Fmr1,me31B,CG32576,Usp32,CG15168,Ino80,RhoGAPp190,flr | | 4.924e-02 | -3.01 | dorsal appendage formation | biological process | GO:0046843 | 56 | 4 | 12345 | 305 | smt3,spoon,Cdc42,dpp | | 4.924e-02 | -3.01 | pole plasm mRNA localization | biological process | GO:0019094 | 56 | 4 | 12345 | 305 | glo,tsu,Hrb27C,sqd | | 4.925e-02 | -3.01 | organonitrogen compound metabolic process | biological process | GO:1901564 | 3232 | 93 | 12345 | 305 | Stt3B,inc,mRpL34,mRpS18A,GstE13,Stlk,RpS3A,lic,Rpt6,Usp16-45,alpha-Man-Ib,Bub1,Eaf6,dgrn,eEF1delta,eIF4B,CG1218,Pten,HtrA2,eEF5,Usp32,RpS9,CycA,Mppe,mRpL30,Btk29A,CG34261,RpL36A,pll,CSN1b,RpL14,sun,BEAF-32,Cul4,Mps1,rb,d4,flw,mRpS33,Rpn10,Pect,NO66,CG10365,Ncoa6,eIF3c,RpL5,CG32473,RpS19a,mnb,Pp1-87B,RpS8,Skp2,mRpS9,CG31957,xit,CG11334,CG31460,Aps,Ubqn,Art1,RpL35A,eIF3g1,fzy,Wdr82,Sgt,PIG-K,Prosbeta1,RpS26,E(z),LIMK1,Acf,Roc2,Cdk8,Ufd4,tkv,janA,Dref,Prosalpha2,Jarid2,Rcd5,g,eIF3b,mRpS28,sws,vnc,CG8635,eRF1,Parp,smt3,CG15534,CG8132,RpL15,Rbbp5 | | 4.928e-02 | -3.01 | regulation of protein localization | biological process | GO:0032880 | 107 | 6 | 12345 | 305 | Parp,tsr,dgrn,Vps35,CG30423,smt3 | | 4.928e-02 | -3.01 | nuclear transport | biological process | GO:0051169 | 107 | 6 | 12345 | 305 | CG6961,SF2,Dbp80,Fs(2)Ket,sqd,Nup188 | | 4.928e-02 | -3.01 | nucleocytoplasmic transport | biological process | GO:0006913 | 107 | 6 | 12345 | 305 | CG6961,Fs(2)Ket,Nup188,sqd,Dbp80,SF2 | | 4.930e-02 | -3.01 | VCB complex | cellular component | GO:0030891 | 2 | 1 | 11940 | 298 | EloB | | 4.930e-02 | -3.01 | endolysosome membrane | cellular component | GO:0036020 | 2 | 1 | 11940 | 298 | ema | | 4.930e-02 | -3.01 | Gtr1-Gtr2 GTPase complex | cellular component | GO:1990131 | 2 | 1 | 11940 | 298 | RagA-B | | 4.930e-02 | -3.01 | semaphorin receptor complex | cellular component | GO:0002116 | 2 | 1 | 11940 | 298 | PlexA | | 4.930e-02 | -3.01 | translation release factor complex | cellular component | GO:0018444 | 2 | 1 | 11940 | 298 | eRF1 | | 4.930e-02 | -3.01 | BAT3 complex | cellular component | GO:0071818 | 2 | 1 | 11940 | 298 | CG9853 | | 4.930e-02 | -3.01 | MutSalpha complex | cellular component | GO:0032301 | 2 | 1 | 11940 | 298 | spel1 | | 4.932e-02 | -3.01 | GSE9006\_TYPE\_1\_VS\_TYPE\_2\_DIABETES\_PBMC\_AT\_DX\_UP | MSigDB lists | GSE9006\_TYPE\_1\_VS\_TYPE\_2\_DIABETES\_PBMC\_AT\_DX\_UP | 76 | 7 | 4161 | 184 | CG14805,CG9330,HtrA2,CG9246,mbf1,eEF1delta,MED22 | | 4.932e-02 | -3.01 | GSE22886\_IL2\_VS\_IL15\_STIM\_NKCELL\_DN | MSigDB lists | GSE22886\_IL2\_VS\_IL15\_STIM\_NKCELL\_DN | 76 | 7 | 4161 | 184 | CIA30,E(z),CG8635,Orc3,CG4866,RpS8,Vap33 | | 4.932e-02 | -3.01 | GSE10239\_NAIVE\_VS\_MEMORY\_CD8\_TCELL\_UP | MSigDB lists | GSE10239\_NAIVE\_VS\_MEMORY\_CD8\_TCELL\_UP | 76 | 7 | 4161 | 184 | CG5339,E(z),Tom20,CG9853,CG2938,Pp1-87B,Sod2 | | 4.933e-02 | -3.01 | Fadd (Fas-associated death domain) | protein interactions | 42594 | 103 | 7 | 9204 | 298 | Cp190,Graf,nsl1,Stim,CG5174,CNBP,tsr | | 4.941e-02 | -3.01 | capt (capulet) | protein interactions | 45233 | 82 | 6 | 9204 | 298 | eEF5,smt3,CG5174,tsr,eEF1delta,eIF4B | | 4.945e-02 | -3.01 | Ras85D (Ras oncogene at 85D) | protein interactions | 41140 | 241 | 13 | 9204 | 298 | vap,Cdc42,dpp,CG6907,RpL15,Pten,Graf,fzy,tsu,Nf1,glo,smt3,Btk29A | | 4.947e-02 | -3.01 | Rm62 (Rm62) | protein interactions | 40739 | 43 | 4 | 9204 | 298 | Rheb,Fmr1,Pten,Cp190 | | 4.947e-02 | -3.01 | E(z) (Enhancer of zeste) | protein interactions | 39203 | 43 | 4 | 9204 | 298 | E(z),Hsp27,Jarid2,mbf1 | | 4.947e-02 | -3.01 | Poxm (Pox meso) | protein interactions | 40990 | 43 | 4 | 9204 | 298 | conu,His2Av,RpL35A,smt3 | | 4.948e-02 | -3.01 | DAP12 signaling | REACTOME pathways | R-DME-2424491 | 164 | 11 | 3923 | 151 | Lst8,CG4935,Rpn10,Pten,Prosalpha2,Prosbeta1,CG7054,14-3-3zeta,Rpt6,Nf1,vap | | 4.948e-02 | -3.01 | DAP12 interactions | REACTOME pathways | R-DME-2172127 | 164 | 11 | 3923 | 151 | CG7054,14-3-3zeta,Prosbeta1,vap,Nf1,Rpt6,Pten,Prosalpha2,Rpn10,CG4935,Lst8 | | 4.948e-02 | -3.01 | Downstream signal transduction | REACTOME pathways | R-DME-186763 | 164 | 11 | 3923 | 151 | Lst8,CG4935,Pten,Prosalpha2,Rpn10,Prosbeta1,14-3-3zeta,CG7054,vap,Nf1,Rpt6 | | 4.948e-02 | -3.01 | Signaling by PDGF | REACTOME pathways | R-DME-186797 | 164 | 11 | 3923 | 151 | Lst8,Prosalpha2,Pten,Rpn10,CG4935,Nf1,Rpt6,vap,Prosbeta1,CG7054,14-3-3zeta | | 4.956e-02 | -3.00 | exonuclease activity, active with either ribo- or deoxyribonucleic acids and producing 5'-phosphomonoesters | molecular function | GO:0016796 | 35 | 3 | 12176 | 289 | Not3,twin,Rat1 | | 4.978e-02 | -3.00 | protein phosphorylation | biological process | GO:0006468 | 284 | 12 | 12345 | 305 | lic,LIMK1,pll,CycA,Bub1,Cdk8,tkv,mnb,Mps1,Stlk,Dref,Btk29A | | 4.984e-02 | -3.00 | EF-hand\_5 | pfam domains | PF13202 | 12 | 2 | 10480 | 319 | Usp32,MICU1 | | 4.992e-02 | -3.00 | peptidyl-lysine acetylation | biological process | GO:0018394 | 81 | 5 | 12345 | 305 | vnc,Acf,d4,Eaf6,Rcd5 | | 4.992e-02 | -3.00 | internal peptidyl-lysine acetylation | biological process | GO:0018393 | 81 | 5 | 12345 | 305 | Eaf6,Rcd5,vnc,Acf,d4 | | 4.992e-02 | -3.00 | chaeta development | biological process | GO:0022416 | 81 | 5 | 12345 | 305 | CtBP,Amun,Stim,Cdk8,Dref | | 4.992e-02 | -3.00 | internal protein amino acid acetylation | biological process | GO:0006475 | 81 | 5 | 12345 | 305 | Eaf6,Rcd5,vnc,Acf,d4 |
