## Supplementary material for "A single DPE core promoter motif contributes to *in vivo* transcriptional regulation and affects cardiac function": SupFile2: knownResults.html

./cluster.allRlog.c300.t8//1596.cluster.txt.motifs/ - Homer Known Motif Enrichment Results


### Homer Known Motif Enrichment Results (./cluster.allRlog.c300.t8//1596.cluster.txt.motifs/)

Homer *de novo* Motif Results  
Gene Ontology Enrichment Results  
Known Motif Enrichment Results (txt file)  
Total Target Sequences = 361, Total Background Sequences = 26798

|  |  |  |  |  |  |  |  |  |  |  |  |
| --- | --- | --- | --- | --- | --- | --- | --- | --- | --- | --- | --- |
| Rank | Motif | Name | P-value | log P-pvalue | q-value (Benjamini) | # Target Sequences with Motif | % of Targets Sequences with Motif | # Background Sequences with Motif | % of Background Sequences with Motif | Motif File | SVG |
| 1 | C G T A T C G A G A T C G C A T C G T A A C G T G T A C T C A G G T C A G A C T C G T A C T A G | DREF/Drosophila-Promoters/Homer | 1e-47 | -1.102e+02 | 0.0000 | 91.0 | 25.21% | 980.8 | 3.66% | motif file (matrix) | svg |
| 2 | T G C A A G T C C T A G A C T G A G C T G T A C T C G A A G T C C G T A A G T C G A C T C T A G | Unknown1(NR/Ini-like)/Drosophila-Promoters/Homer | 1e-37 | -8.521e+01 | 0.0000 | 74.0 | 20.50% | 838.3 | 3.13% | motif file (matrix) | svg |
| 3 | G A T C C T G A C T A G G C A T C T A G C A G T C A T G T C G A G T A C A G T C T C A G A G C T | M1BP(Zf)/S2R+-M1BP-ChIP-Seq(GSE49842)/Homer | 1e-36 | -8.400e+01 | 0.0000 | 72.0 | 19.94% | 800.5 | 2.99% | motif file (matrix) | svg |
| 4 | A G T C C G T A C G A T A G T C G T C A A G T C A C G T C T G A | Unknown2/Drosophila-Promoters/Homer | 1e-34 | -8.053e+01 | 0.0000 | 103.0 | 28.53% | 1855.7 | 6.93% | motif file (matrix) | svg |
| 5 | T C G A G T A C T C G A T C G A C A T G A T G C A C G T A C T G A C T G A G T C C G T A C T A G A G T C A T C G A G T C | Unknown3/Drosophila-Promoters/Homer | 1e-17 | -4.003e+01 | 0.0000 | 36.0 | 9.97% | 422.0 | 1.58% | motif file (matrix) | svg |
| 6 | T C G A G C T A T G A C G C T A C T A G G A T C C G A T A C T G C G A T A G C T G A C T C T A G | E-box/Drosophila-Promoters/Homer | 1e-12 | -2.781e+01 | 0.0000 | 45.0 | 12.47% | 966.5 | 3.61% | motif file (matrix) | svg |
| 7 | C T G A C G T A C G T A G C T A C T G A A C G T G T C A A G T C A G T C C T G A G T C A G C T A | Unknown4/Drosophila-Promoters/Homer | 1e-11 | -2.689e+01 | 0.0000 | 41.0 | 11.36% | 838.4 | 3.13% | motif file (matrix) | svg |
| 8 | C T A G A G T C A G C T A C T G C G T A C A G T C G T A C T G A T A G C T G A C | Unknown5/Drosophila-Promoters/Homer | 1e-7 | -1.753e+01 | 0.0000 | 74.0 | 20.50% | 2836.7 | 10.59% | motif file (matrix) | svg |
