## Supplementary material for "A single DPE core promoter motif contributes to *in vivo* transcriptional regulation and affects cardiac function": SupFile3: meme.html


Help poup.

[
close ]

The statistical significance of the motif. MEME usually finds the most
statistically significant (low E-value) motifs first. It is unusual to
consider a motif with an E-value larger than 0.05 significant so, as an
additional indicator, MEME displays these partially transparent.

The E-value of a motif is based on its log likelihood ratio, width,
sites, the background letter frequencies (given in the command line
summary), and the size of the training set.

The E-value is an estimate of the expected number of motifs with the
given log likelihood ratio (or higher), and with the same width and site
count, that one would find in a similarly sized set of random
sequences (sequences where each position is independent and letters are
chosen according to the background letter frequencies).

[
close ]

The number of sites contributing to the construction of the motif.

[
close ]

The width of the motif. Each motif describes a pattern of a fixed
width, as no gaps are allowed in MEME motifs.

[
close ]

Show more information on the motif.

[
close ]

Submit your motif to another MEME Suite program or download in various
text formats or as a logo.

###### Supported Programs

Tomtom
:   Tomtom is a tool for searching for similar known motifs.
    [manual]

MAST
:   MAST is a tool for searching biological sequence databases for
    sequences that contain one or more of a group of known motifs.
    [manual]

FIMO
:   FIMO is a tool for searching biological sequence databases for
    sequences that contain one or more known motifs.
    [manual]

GOMO
:   GOMO is a tool for identifying possible roles (Gene Ontology
    terms) for DNA binding motifs.
    [manual]

SpaMo
:   SpaMo is a tool for inferring possible transcription factor
    complexes by finding motifs with enriched spacings.
    [manual]

[
close ]

The log likelihood ratio of the motif.The log likelihood ratio is the
logarithm of the ratio of the probability of the occurrences of the motif
given the motif model (likelihood given the motif) versus their
probability given the background model (likelihood given the null model).
(Normally the background model is a 0-order Markov model using the
background letter frequencies, but higher order Markov models may be
specified via the -bfile option to MEME.).

[
close ]

The information content of the motif in bits. It is equal to the sum
of the uncorrected information content, R(), in the columns of the pwm.
This is equal relative entropy of the motif relative to a uniform
background frequency model.

[
close ]

The relative entropy of the motif.

re = llr / (sites \* ln(2))

[
close ]

The Bayes Threshold.

[
close ]

The strand used for the motif site.

+
:   The motif site was found in the sequence as it was supplied.

-
:   The motif site was found in the reverse complement of the supplied sequence.

[
close ]

The position in the sequence where the motif site starts. If a motif
started right at the begining of a sequence it would be described as
starting at position 1.

[
close ]

The probability that an equal or better site would be found in a
random sequence of the same length conforming to the background letter
frequencies.

[
close ]

A motif site with the 10 flanking letters on either side.

When the site is not on the given strand then the site
and both flanks are reverse complemented so they align.

[
close ]

The name of the sequences as given in the FASTA file.

The number to the left of the sequence name is the ordinal
of the sequence.

[
close ]

This is the combined match *p*-value.

The combined match *p*-value is defined as the probability that a
random sequence (with the same length and conforming to the background)
would have motif-sequence match p-values such that the product is smaller
or equal to the value calulated for the sequence under test.

The motif-sequence match p-value is defined as the probability that a
random sequence (with the same length and conforming to the background)
would have a match to the motif under test with a score greater or equal
to the largest found in the sequence under test.

[
close ]

This diagram shows the location of motif sites.

[
close ]

|  |  |
| --- | --- |
| Motif | 1 |
| *p*-value | 8.23e-7 |
| Start | 23 |
| End | 33 |

Scanned Site

|  |  |
| --- | --- |
| Motif | 1 |
| *p*-value | 8.23e-7 |
| Start | 23 |
| End | 33 |

.

↥

⇢

*E*-value:

Site Count:

Width:

StandardReverse
Complement

Log Likelihood Ratio:

Information Content:

Relative Entropy:

Bayes Threshold:

x

#### Submit or Download

⇧⬆

⇩⬇

Submit MotifDownload MotifDownload Logo

###### Submit to program

|  |  |  |
| --- | --- | --- |
|  | Tomtom | Find similar motifs in published libraries or a library you supply. |
|  | FIMO | Find motif occurrences in sequence data. |
|  | MAST | Rank sequences by affinity to groups of motifs. |
|  | GOMo | Identify possible roles (Gene Ontology terms) for motifs. |
|  | SpaMo | Find other motifs that are enriched at specific close spacings which might imply the existance of a complex. |

Format:

Count Matrix
Probability Matrix
Minimal MEME
FASTA
Raw

|  |  |
| --- | --- |
| Format: | PNG (for web) EPS (for publication) |
| Orientation: | Normal Reverse Complement |
| Small Sample Correction: | Off On |
| Width: | cm |
| Height: | cm |

### MEME

#### Multiple Em for Motif Elicitation

For further information on how to interpret these results or to get a
copy of the MEME software please access
http://meme-suite.org.

If you use MEME in your research, please cite the following paper:  

Timothy L. Bailey and Charles Elkan,
"Fitting a mixture model by expectation maximization to discover motifs in biopolymers",
*Proceedings of the Second International Conference on Intelligent Systems
for Molecular Biology*, pp. 28-36, AAAI Press, Menlo Park, California, 1994.
[pdf]

Discovered Motifs
  |  
Motif Locations
  |  
Program information


### Javascript is required to view these results!

### Your browser does not support canvas!

#### Discovered Motifs

Please wait... Loading...

If the page has fully loaded and this message does not disappear then an error may have occurred.

#### Motif Locations

Please wait... Loading...

If the page has fully loaded and this message does not disappear then an error may have occurred.

###### MEME version

(Release date: )

###### Reference

Timothy L. Bailey and Charles Elkan,
"Fitting a mixture model by expectation maximization to discover motifs in biopolymers",
*Proceedings of the Second International Conference on Intelligent Systems
for Molecular Biology*, pp. 28-36, AAAI Press, Menlo Park, California, 1994.

###### Command line summary
