## Supplementary figures and images for "A single DPE core promoter motif contributes to *in vivo* transcriptional regulation and affects cardiac function"

### logo1.png

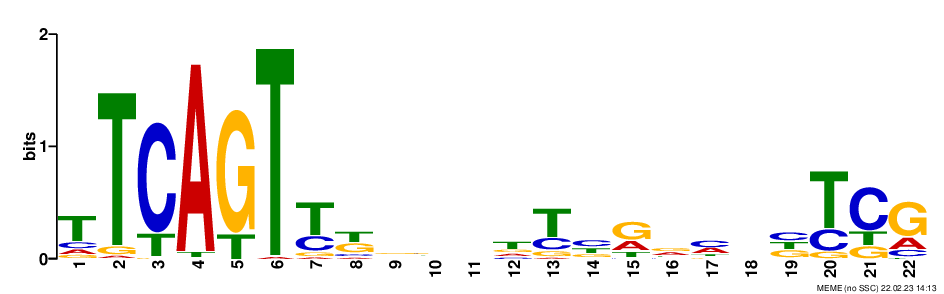

### logo1.png

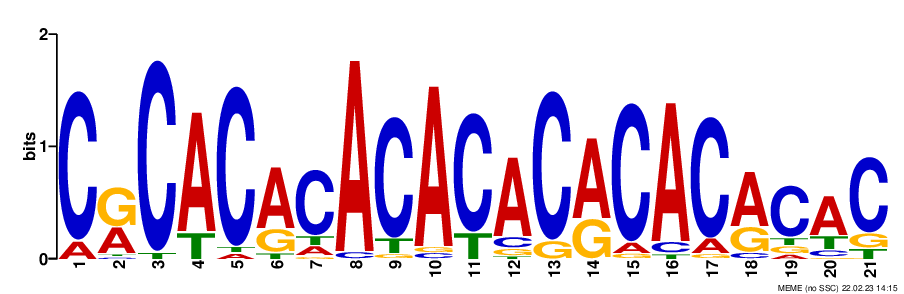

### logo1.png

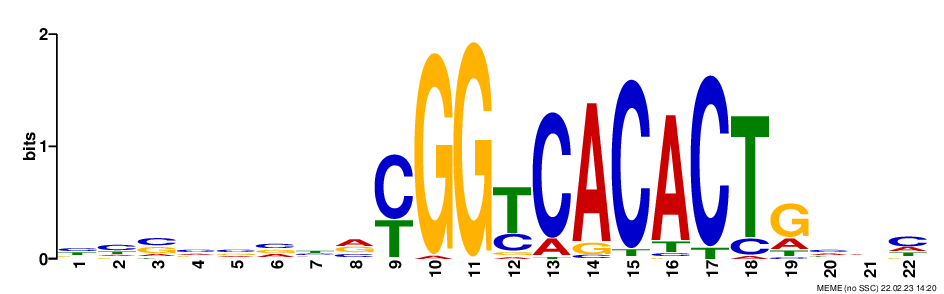

### logo1.png

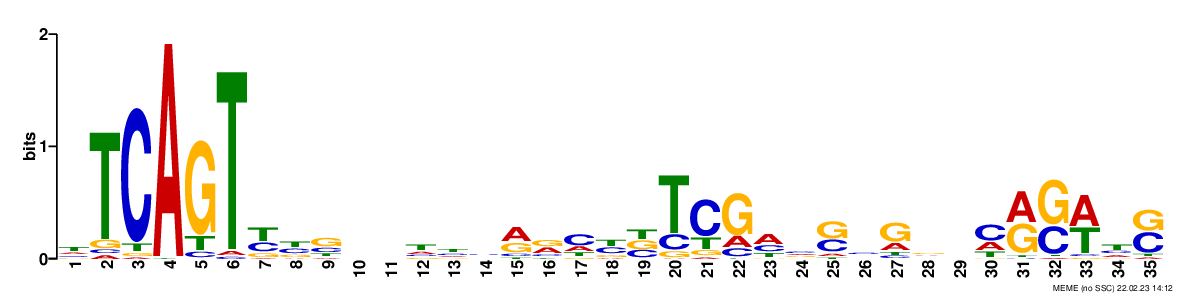

### logo1.png

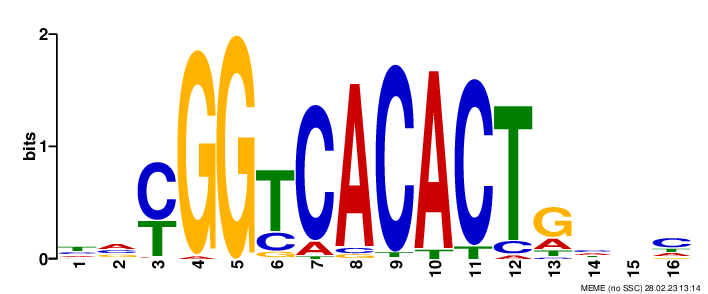

### logo2.png

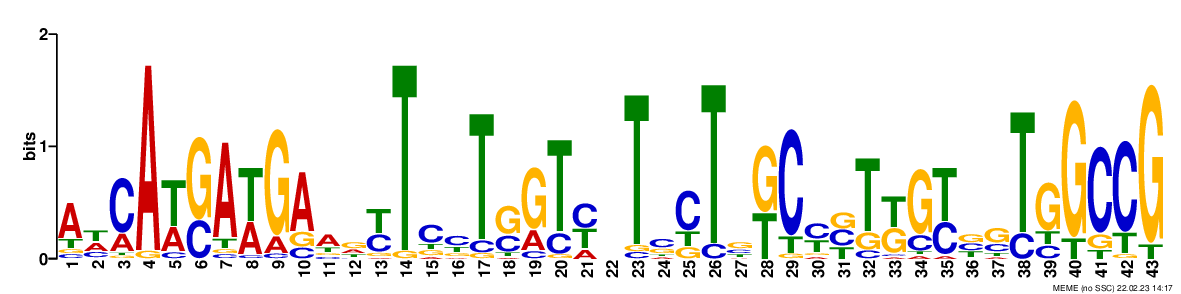

### logo2.png

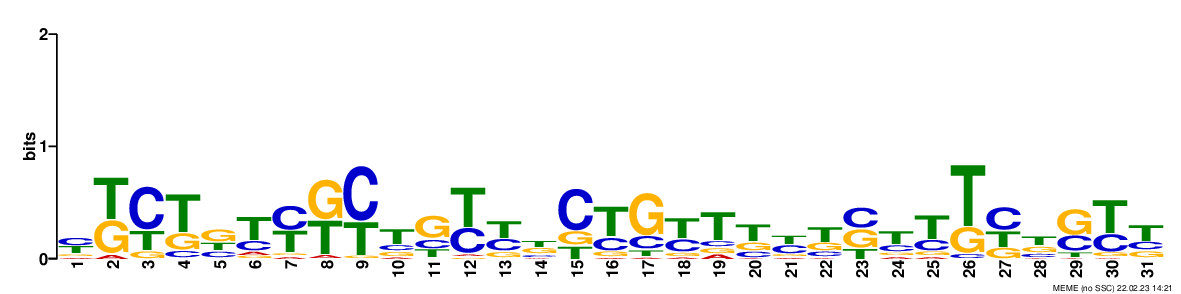

### logo2.png

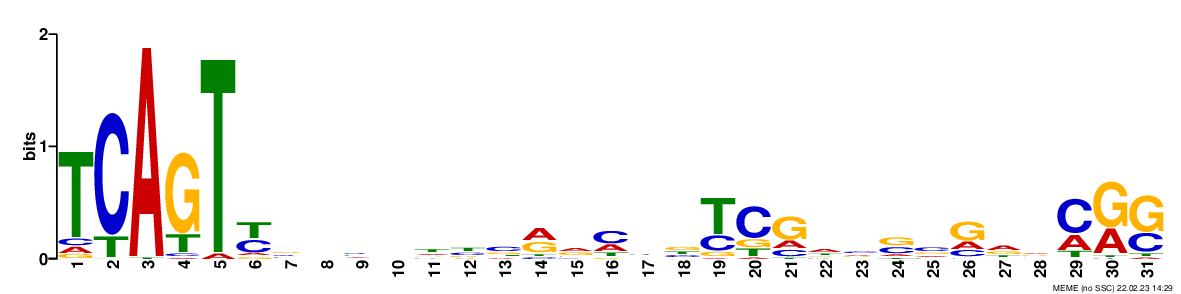

### logo2.png

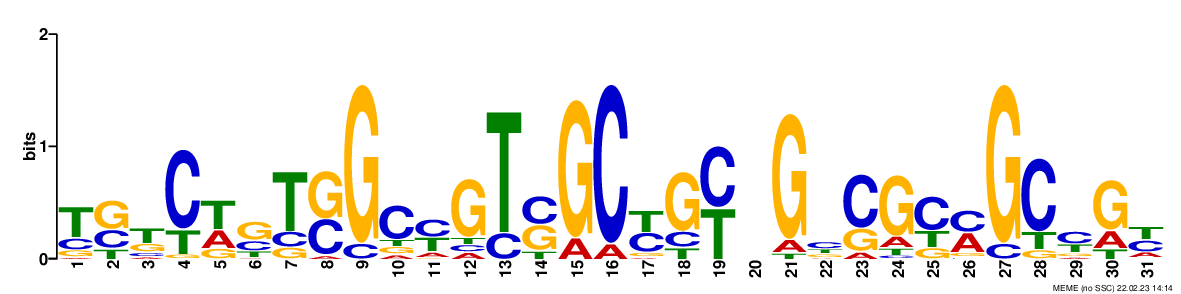

### logo2.png

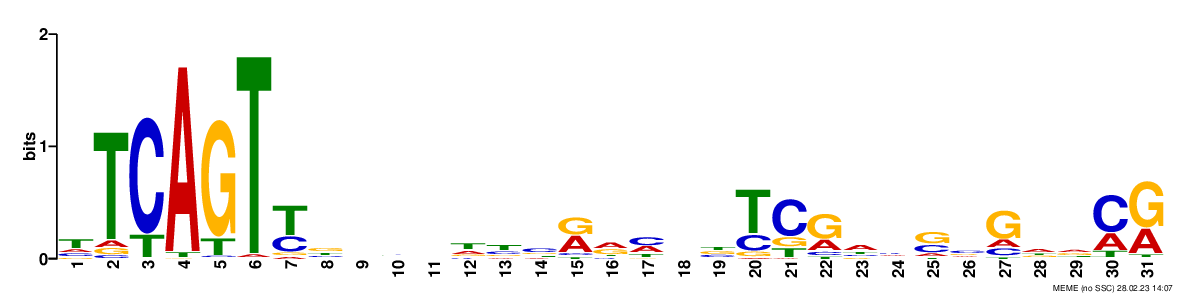

### logo3.png

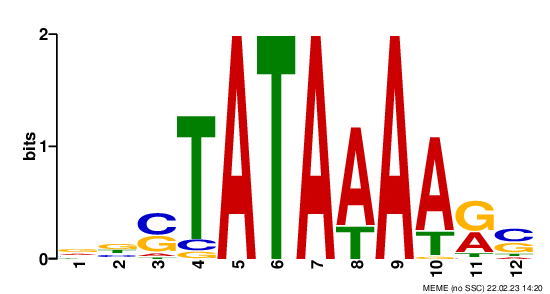

### logo3.png

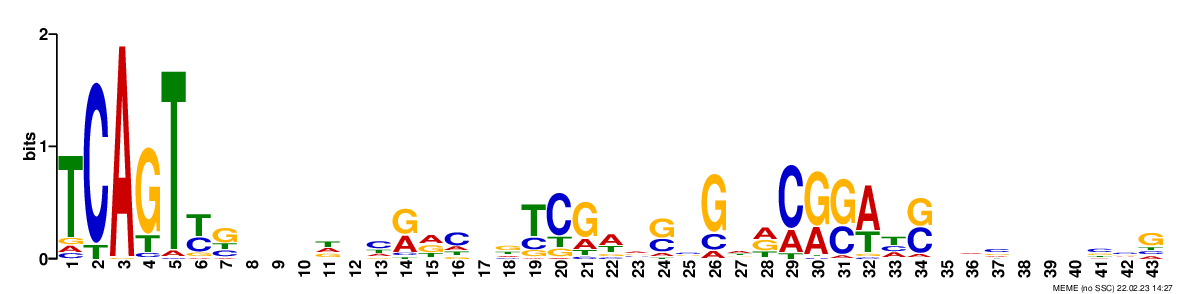

### logo3.png

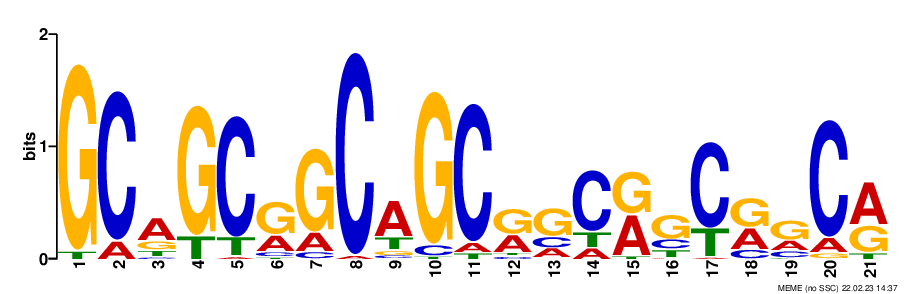

### logo3.png

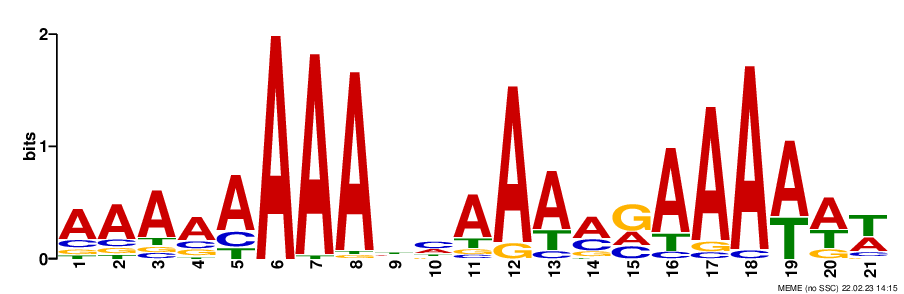

### logo3.png

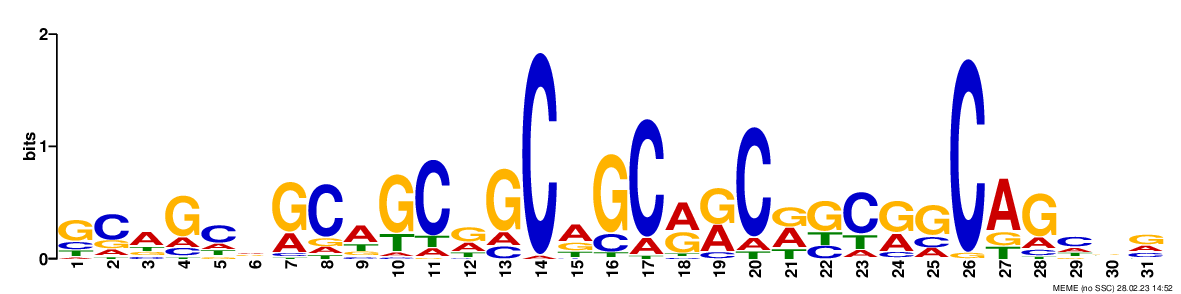

### logo4.png

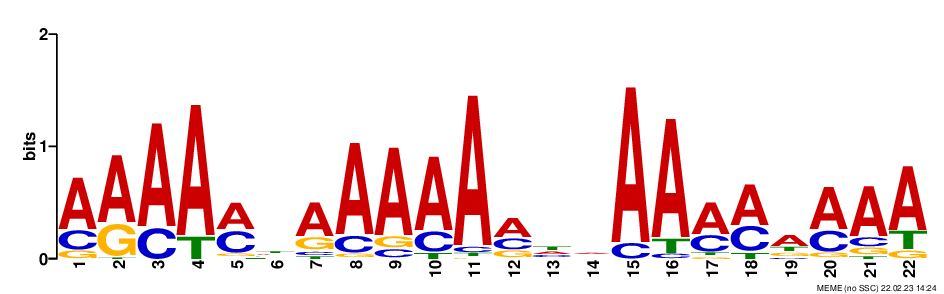

### logo4.png

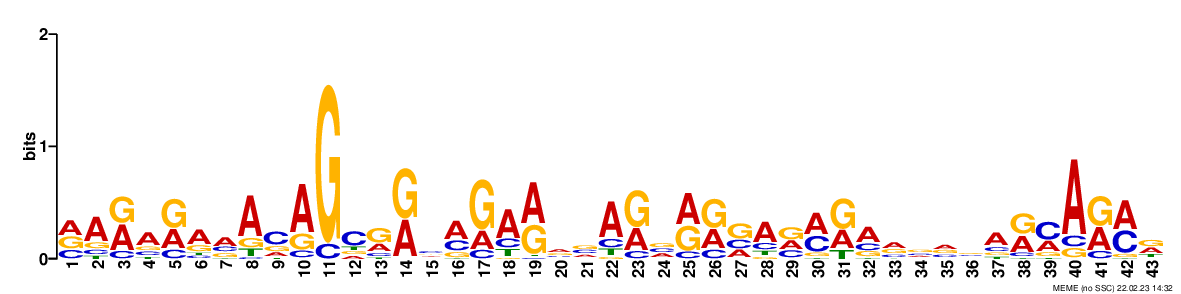

### logo4.png

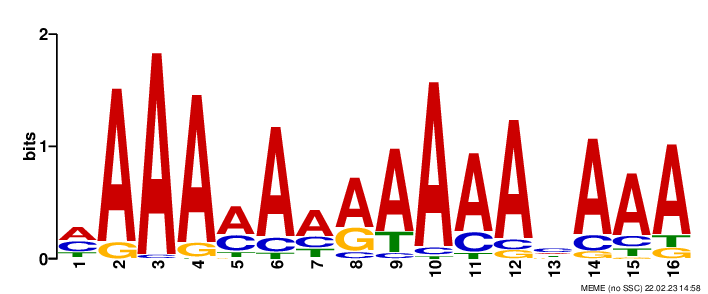

### logo4.png

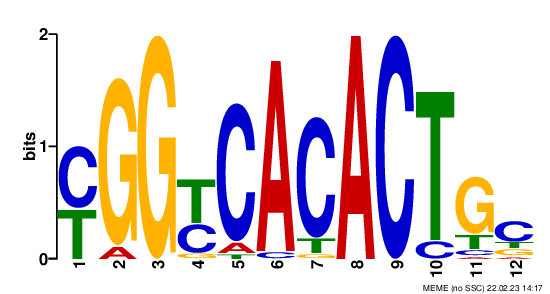

### logo4.png

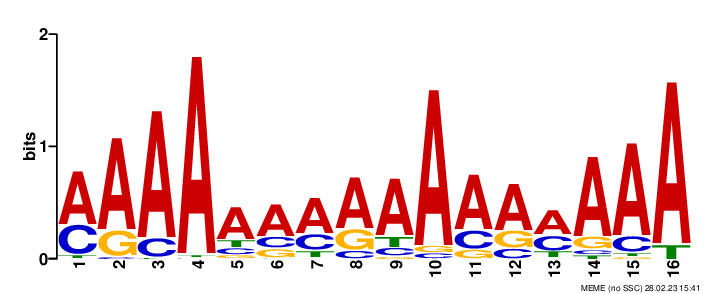

### logo5.png

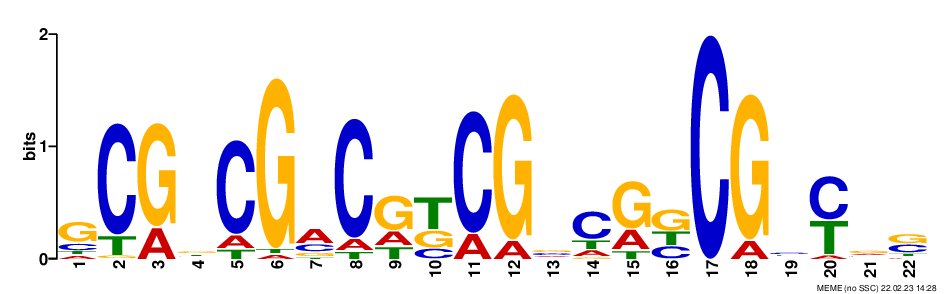

### logo5.png

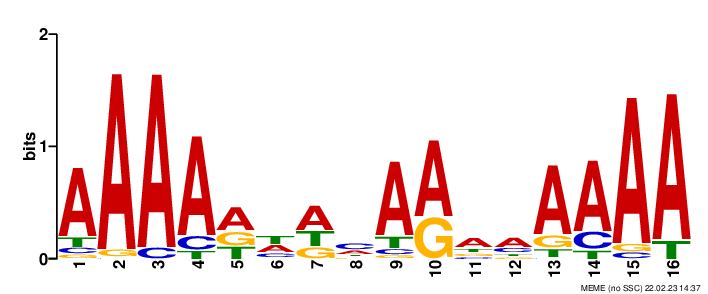

### logo5.png

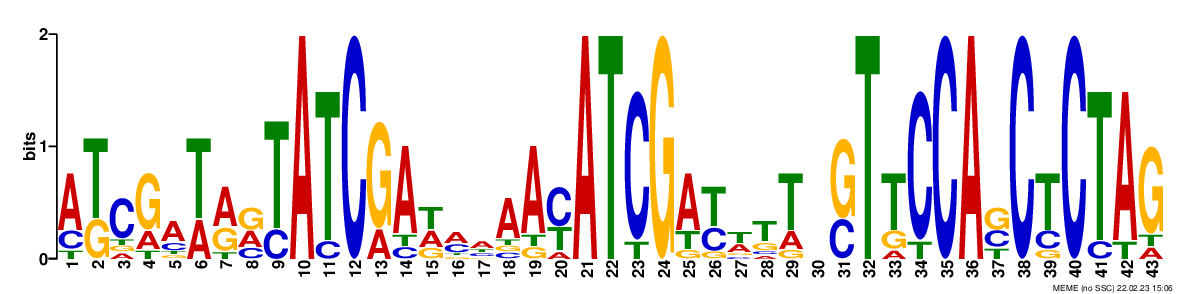

### logo5.png

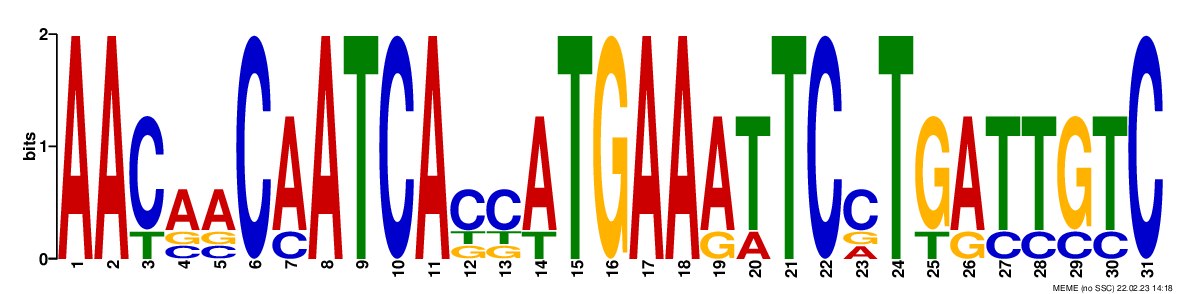

### logo5.png

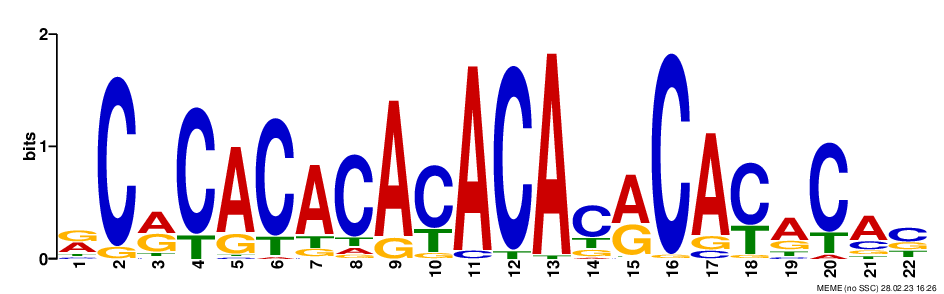

### logo6.png

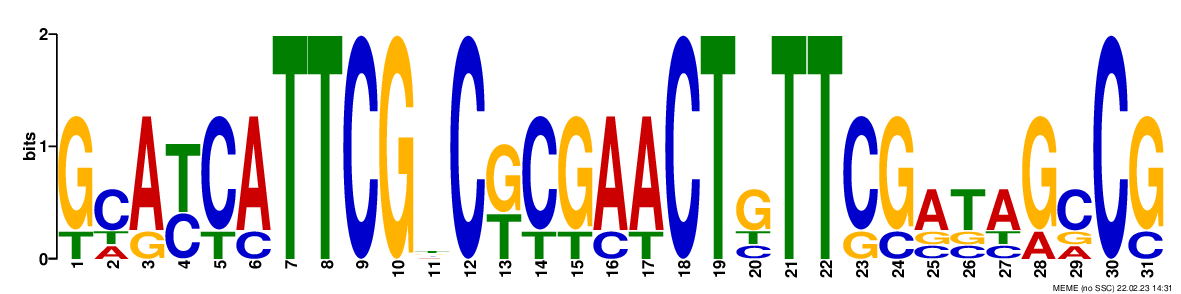

### logo6.png

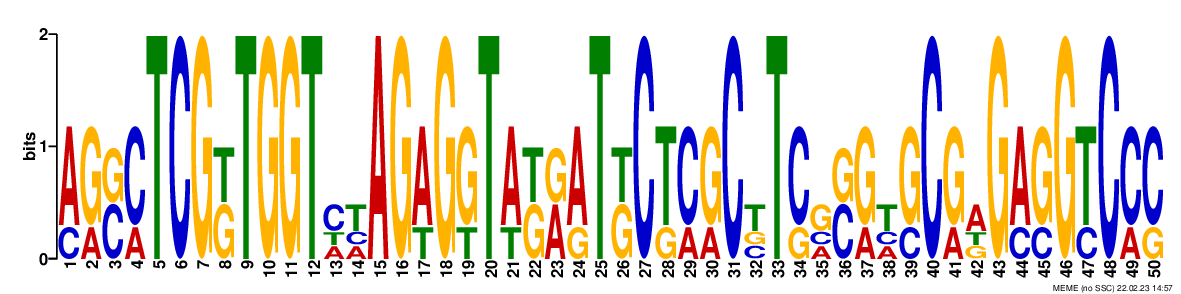

### logo6.png

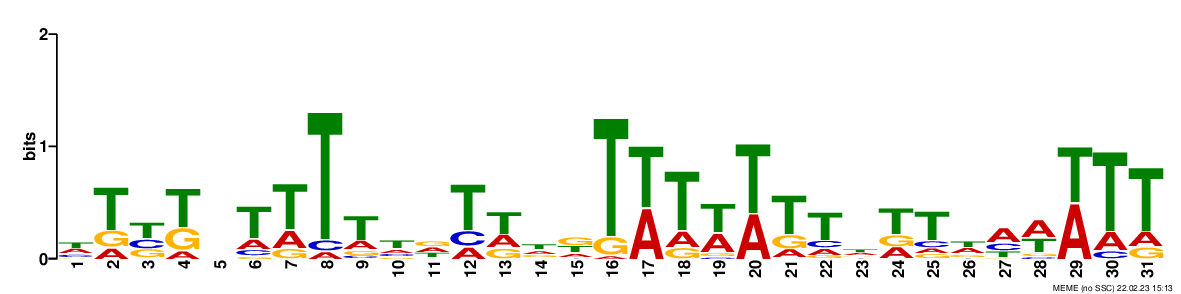

### logo6.png

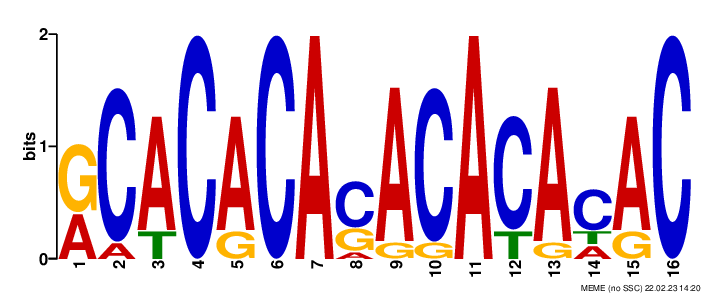

### logo6.png

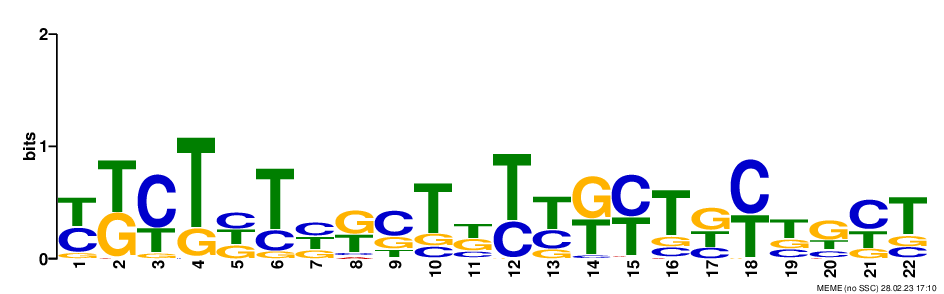

### logo7.png

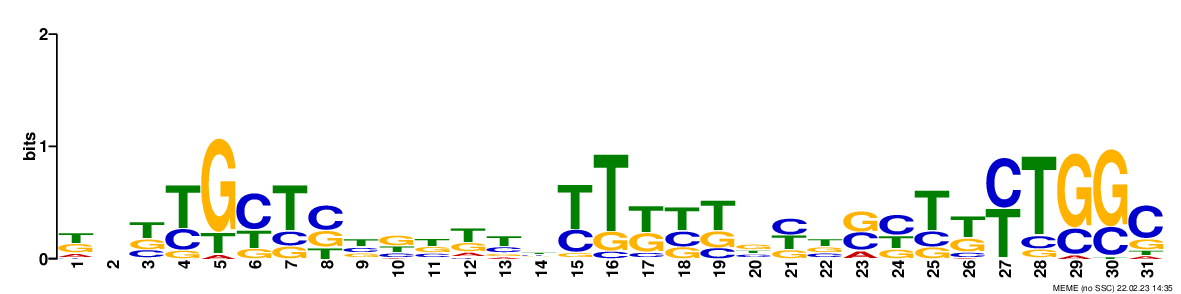

### logo7.png

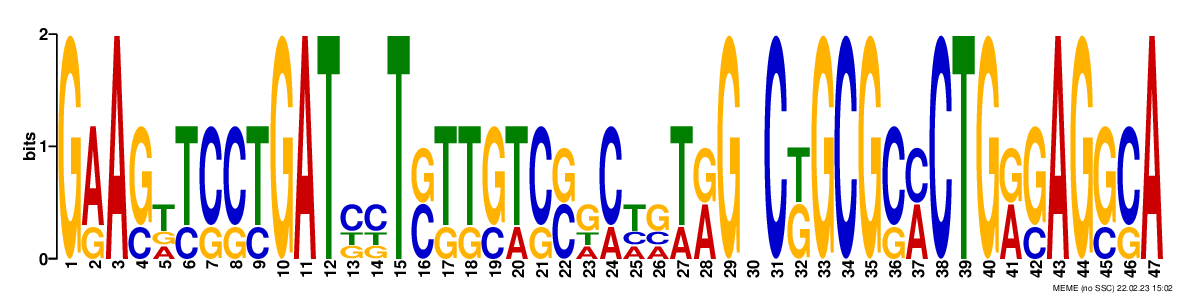

### logo7.png

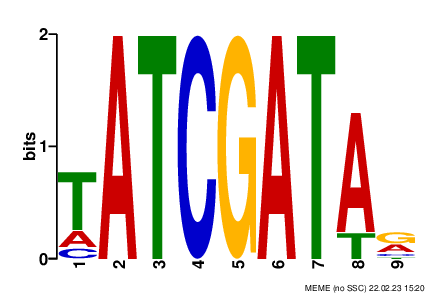

### logo7.png

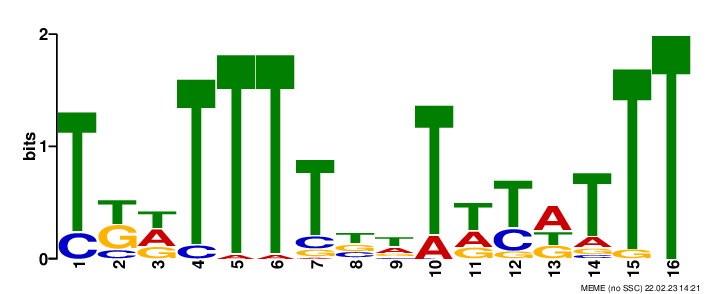

### logo7.png

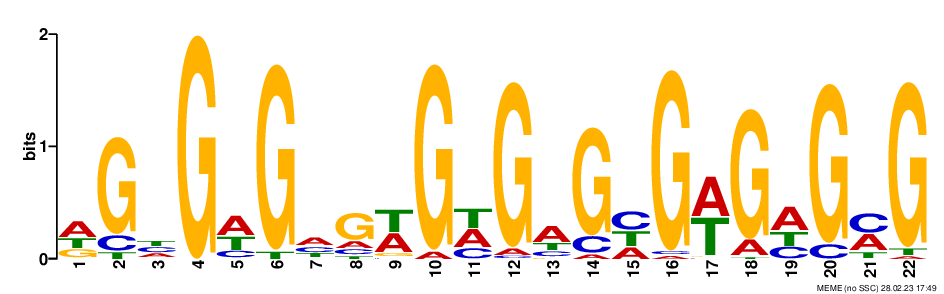

### logo8.png

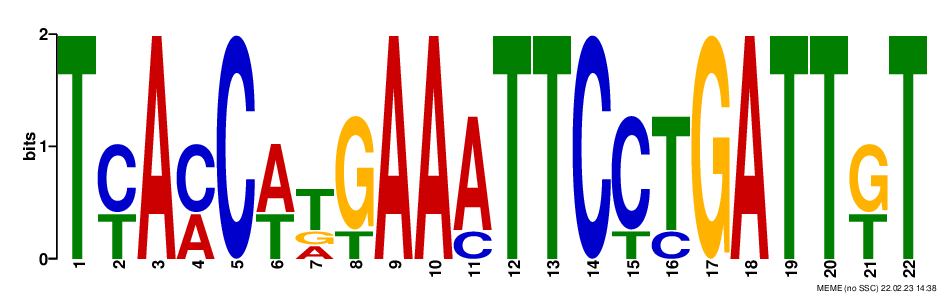

### logo8.png

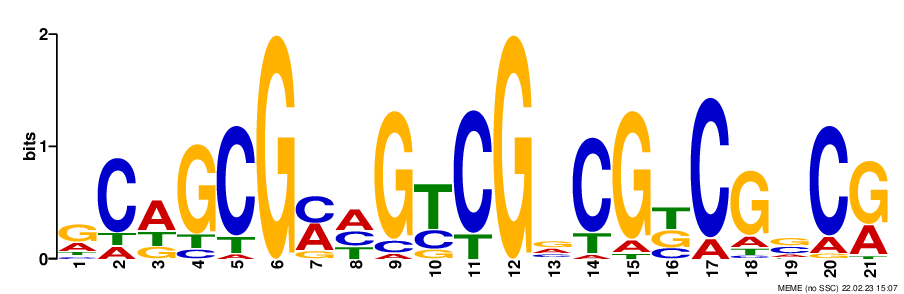

### logo8.png

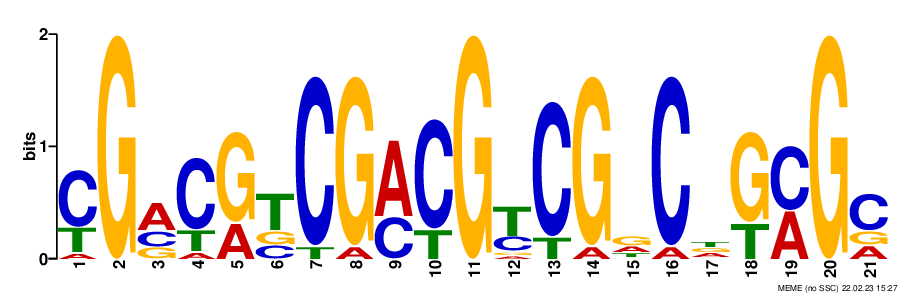

### logo8.png

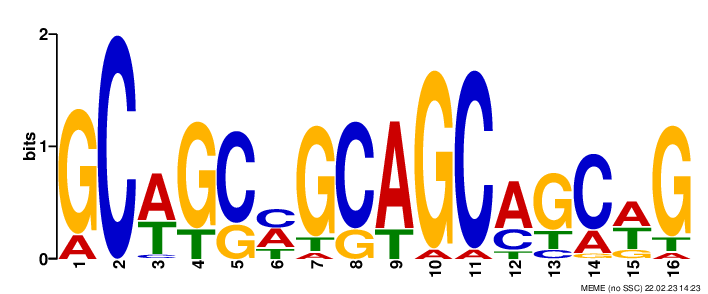

### logo8.png

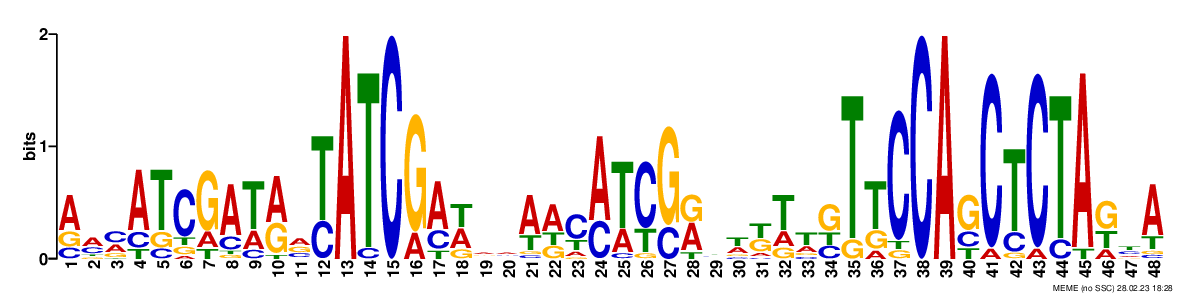

### logo9.png

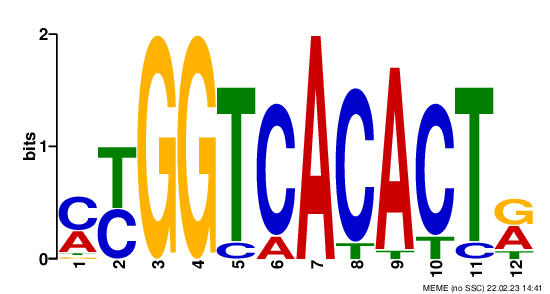

### logo9.png

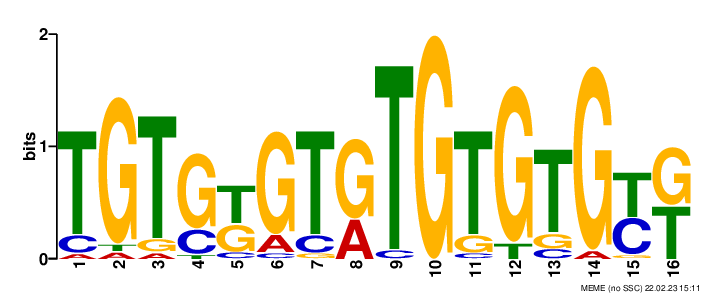

### logo9.png

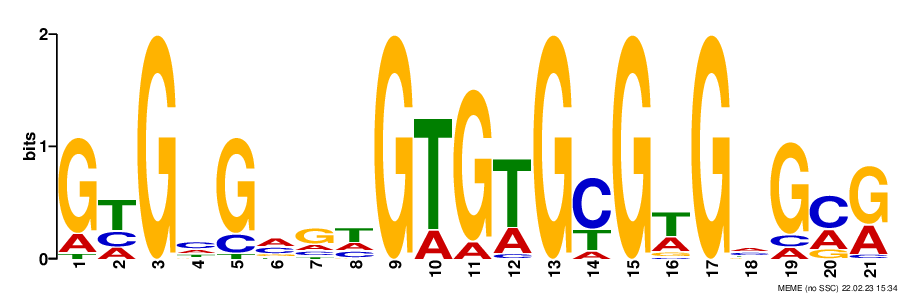

### logo9.png

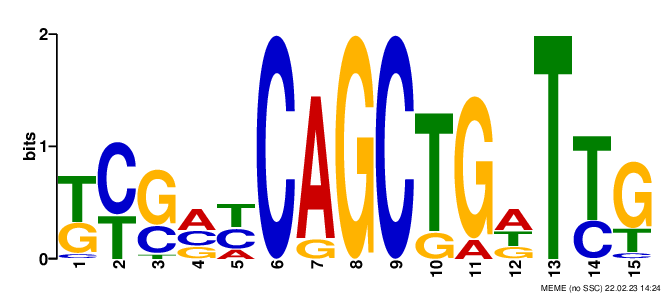

### logo9.png

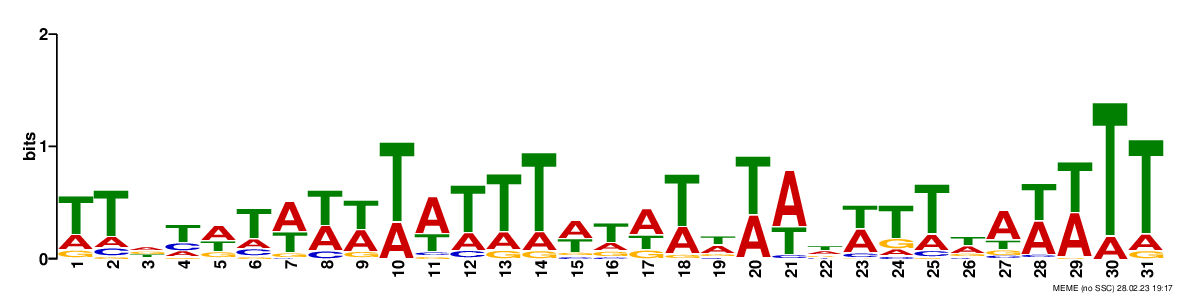

### logo10.png

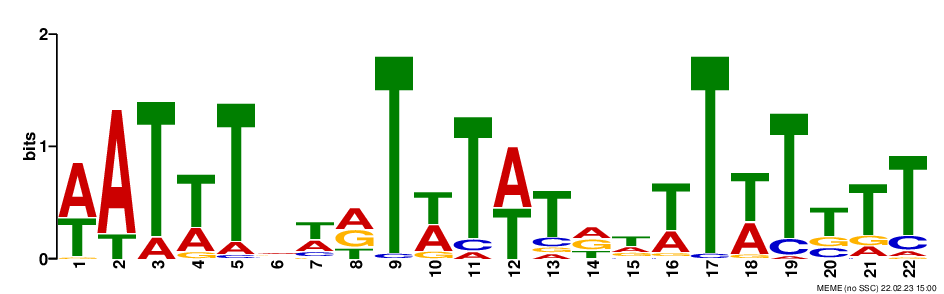

### logo10.png

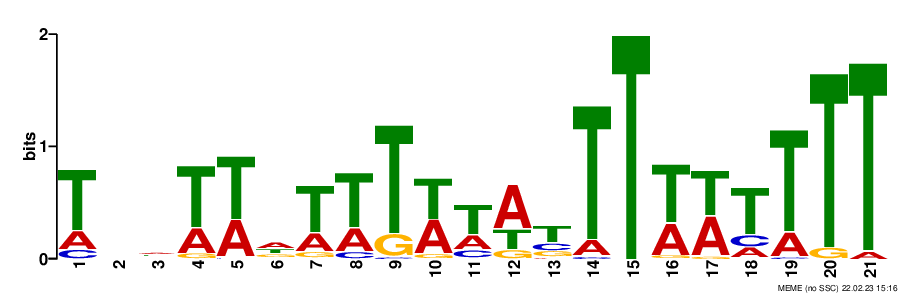

### logo10.png

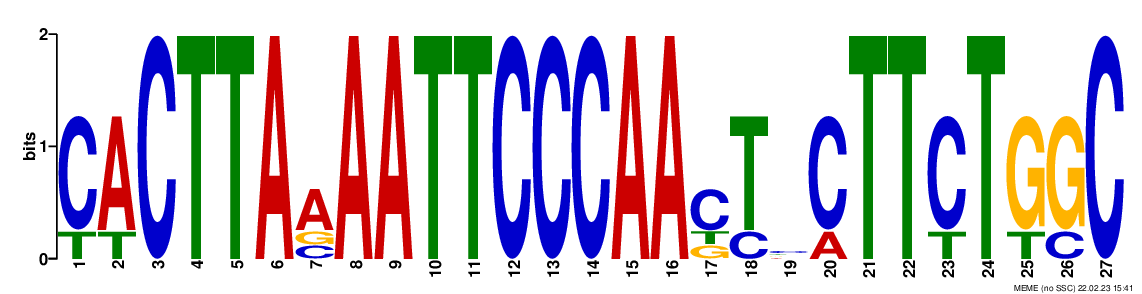

### logo10.png

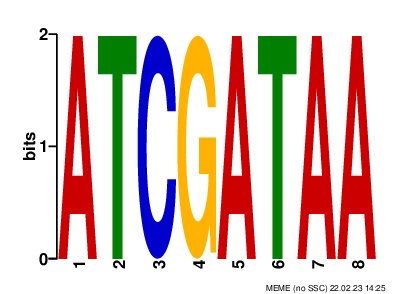

### logo10.png

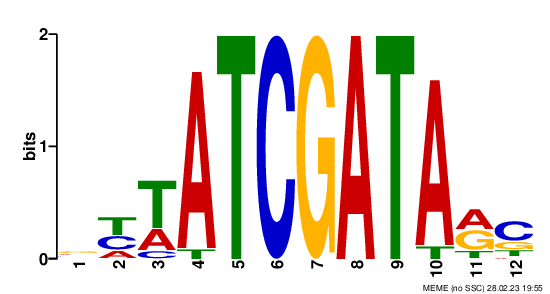

### logo_rc1.png

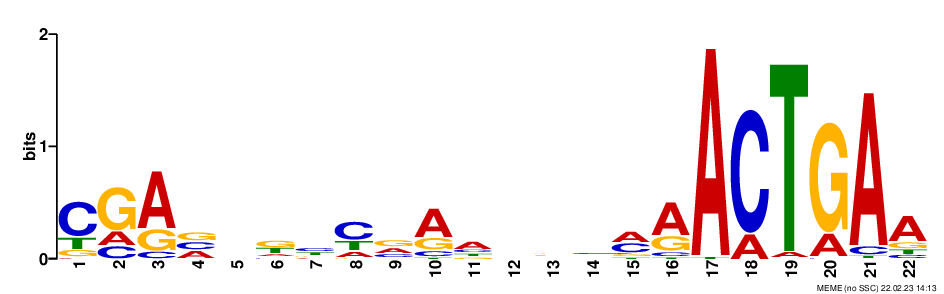

### logo_rc1.png

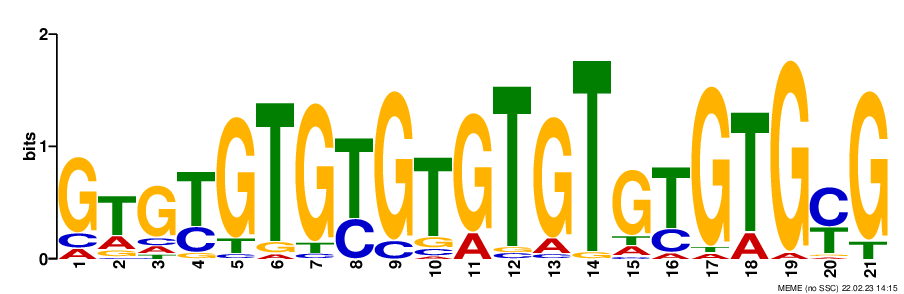

### logo_rc1.png

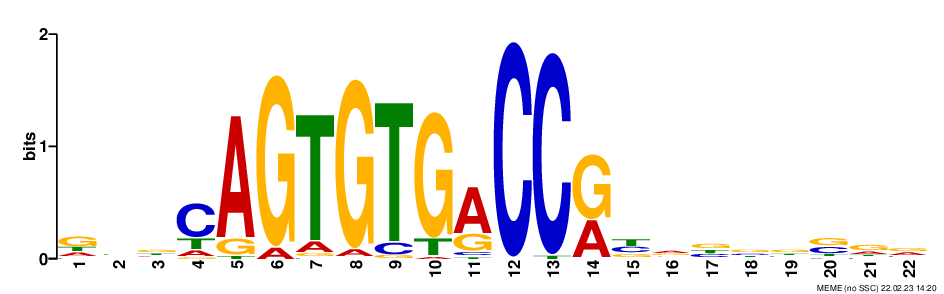

### logo_rc1.png

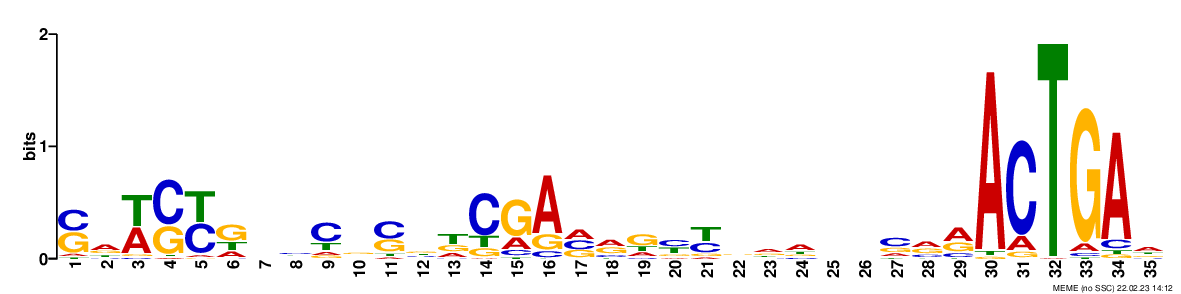

### logo_rc1.png

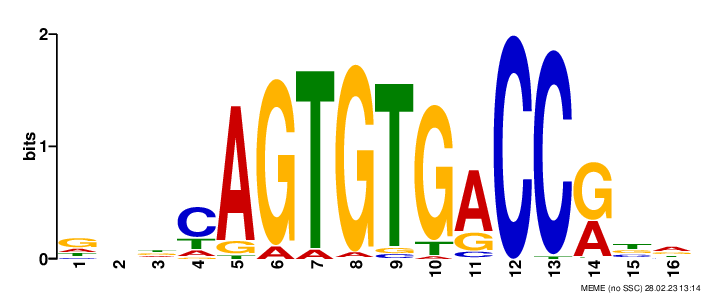

### logo_rc2.png

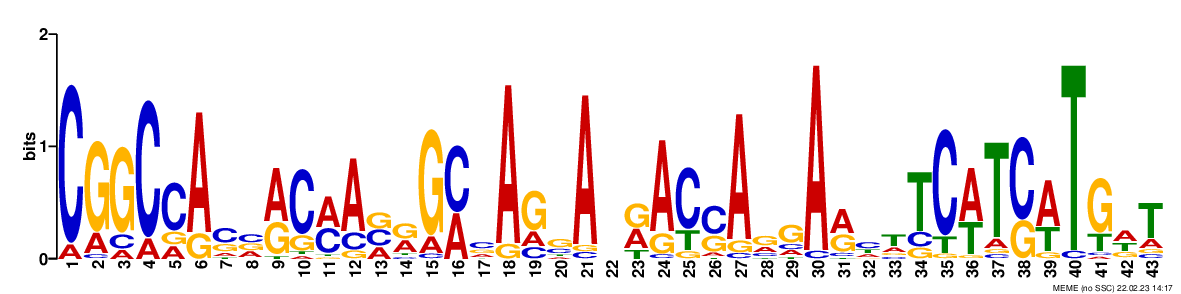

### logo_rc2.png

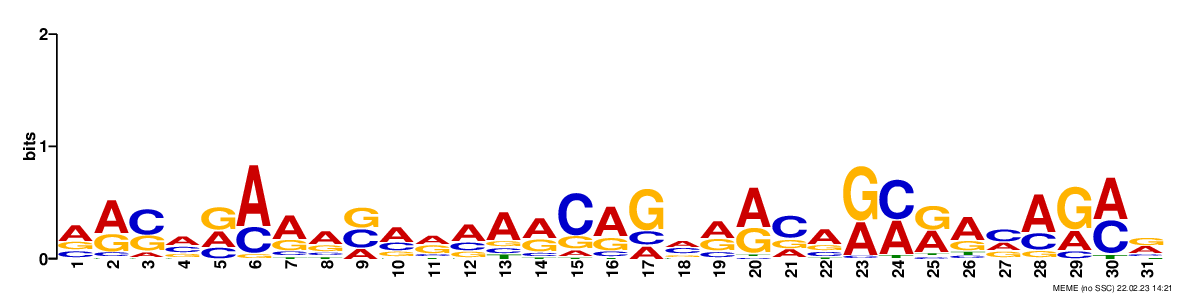

### logo_rc2.png

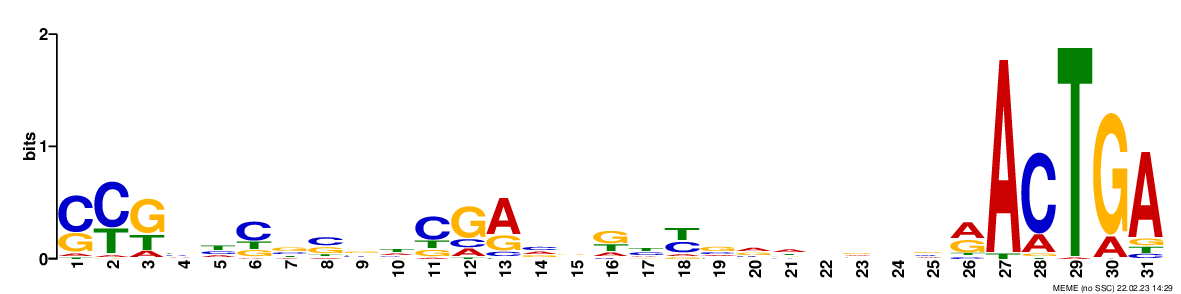

### logo_rc2.png

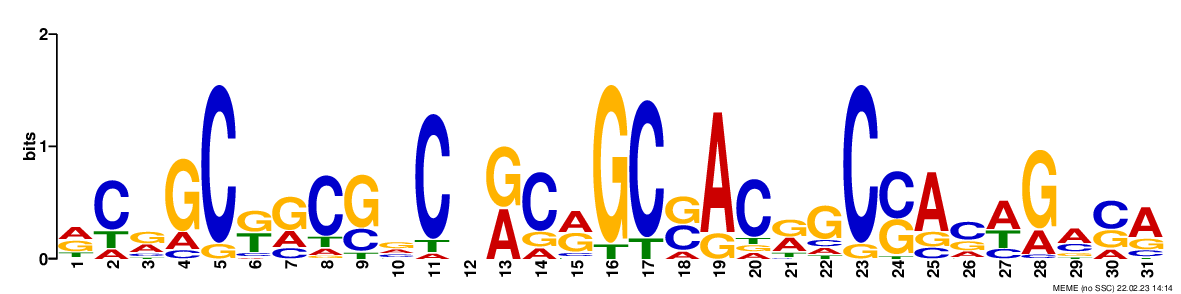

### logo_rc2.png

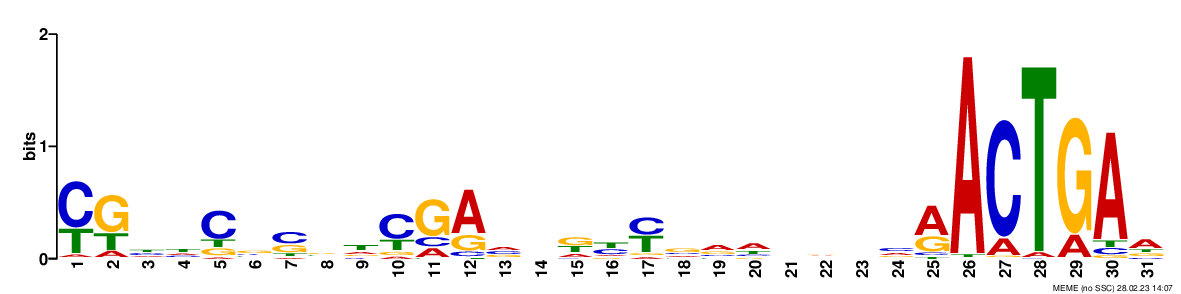

### logo_rc3.png

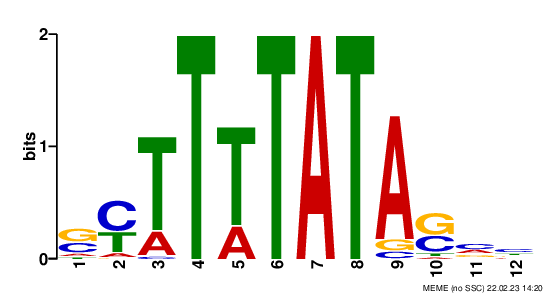

### logo_rc3.png

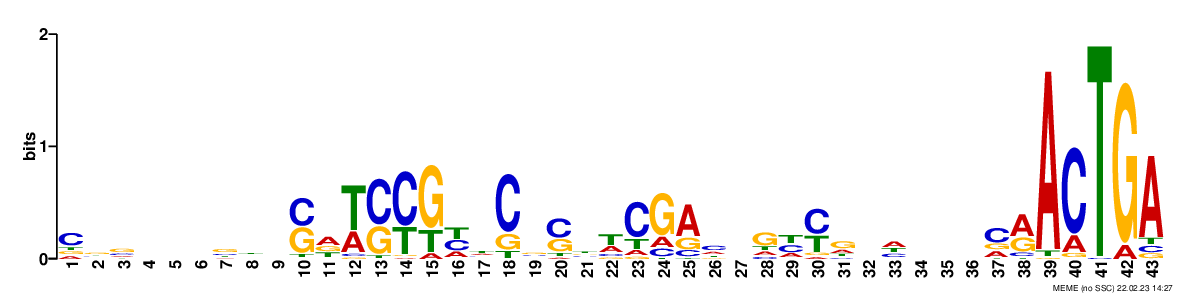

### logo_rc3.png

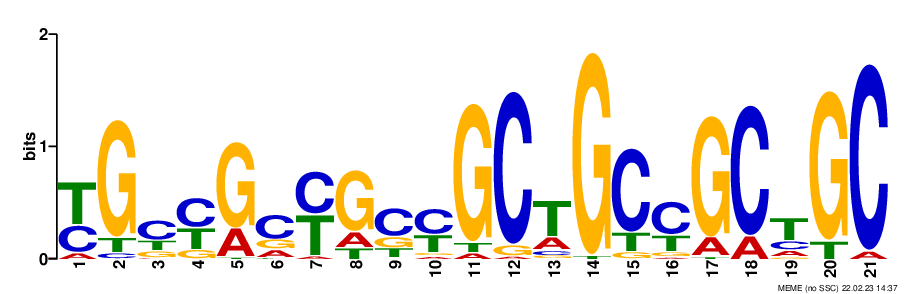

### logo_rc3.png

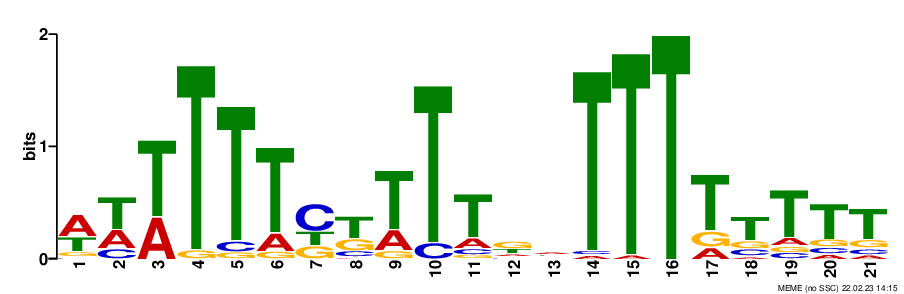

### logo_rc3.png

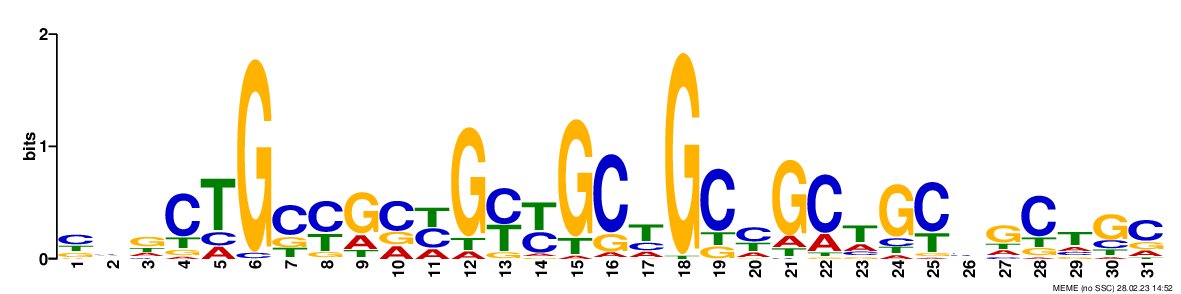

### logo_rc4.png

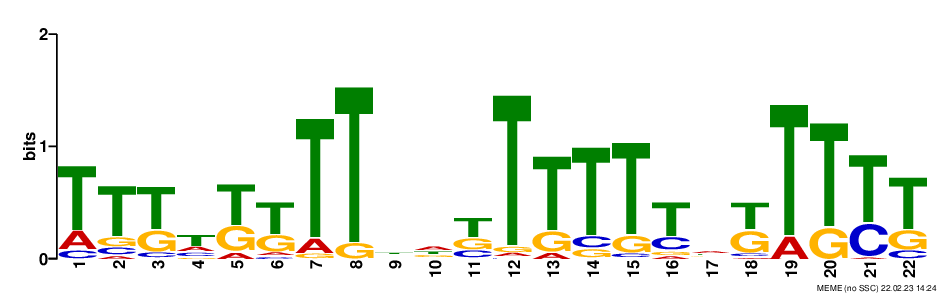

### logo_rc4.png

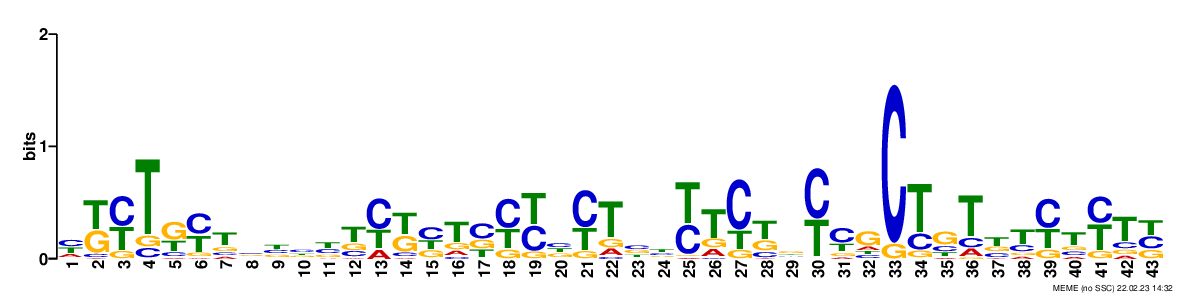

### logo_rc4.png

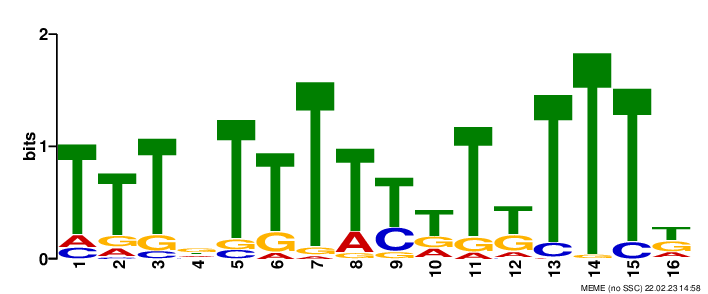

### logo_rc4.png

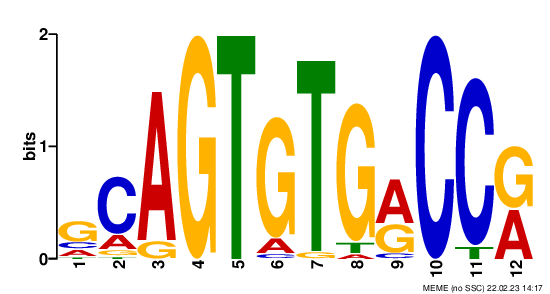

### logo_rc4.png

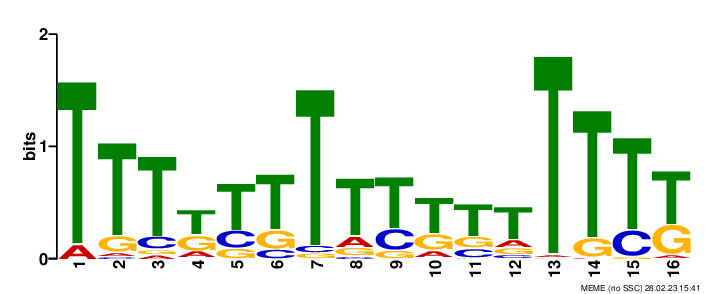

### logo_rc5.png

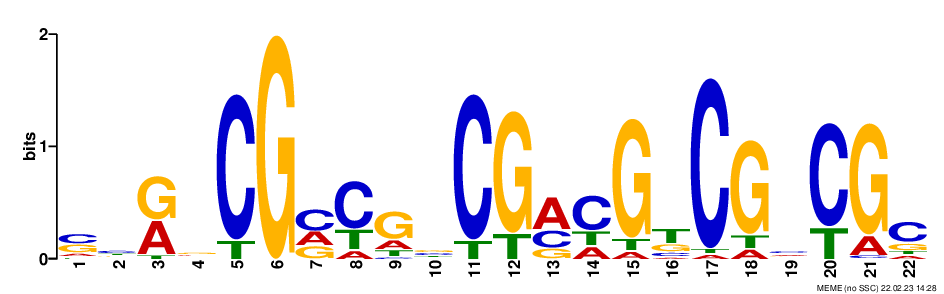

### logo_rc5.png

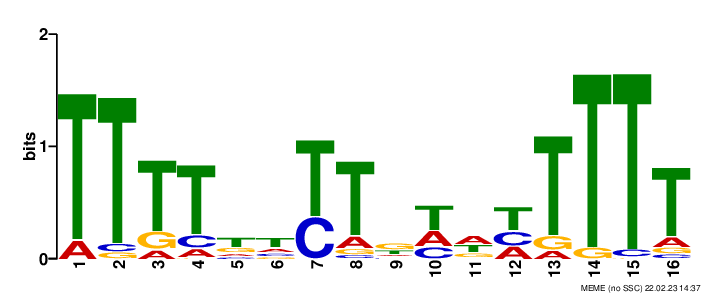

### logo_rc5.png

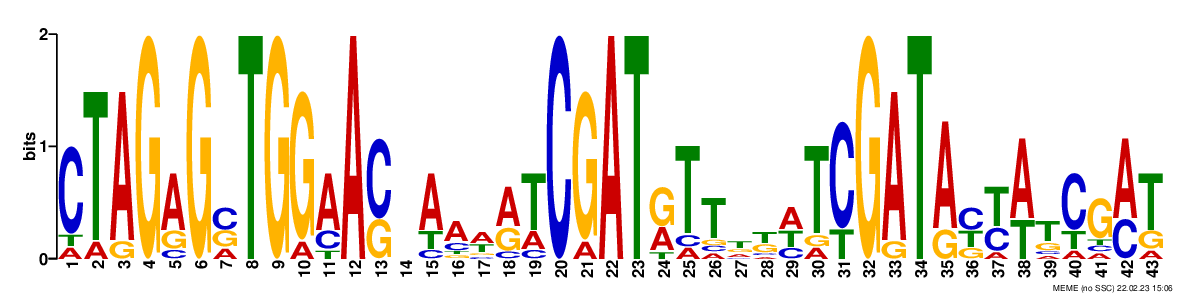

### logo_rc5.png

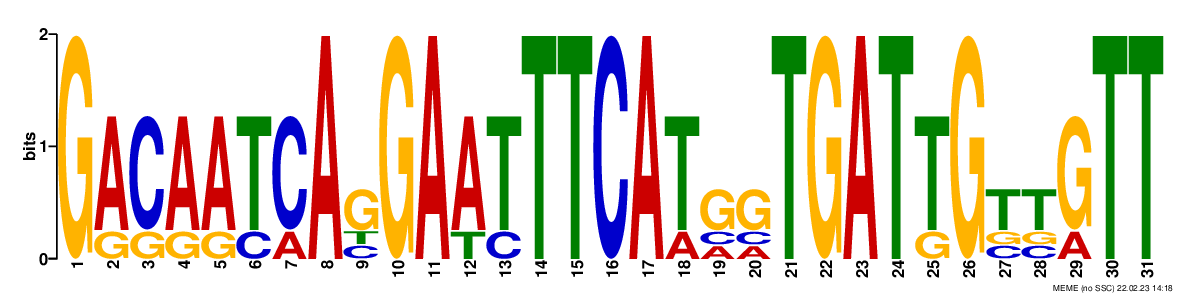

### logo_rc5.png

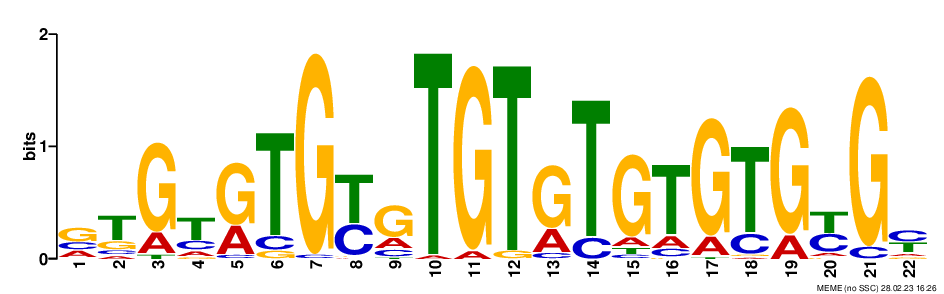

### logo_rc6.png

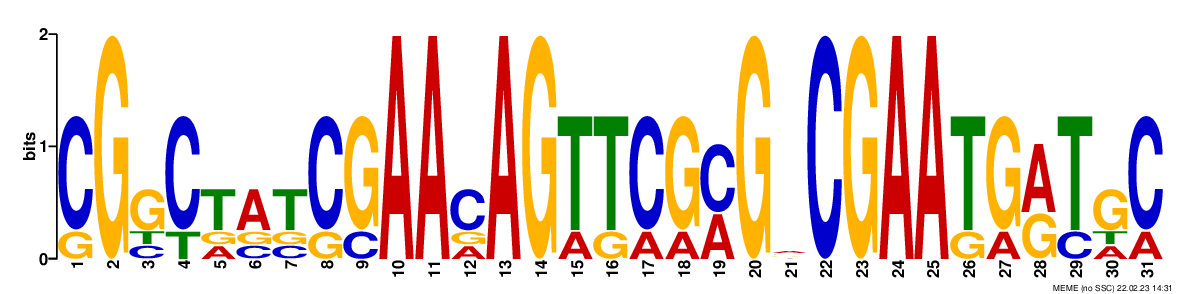

### logo_rc6.png

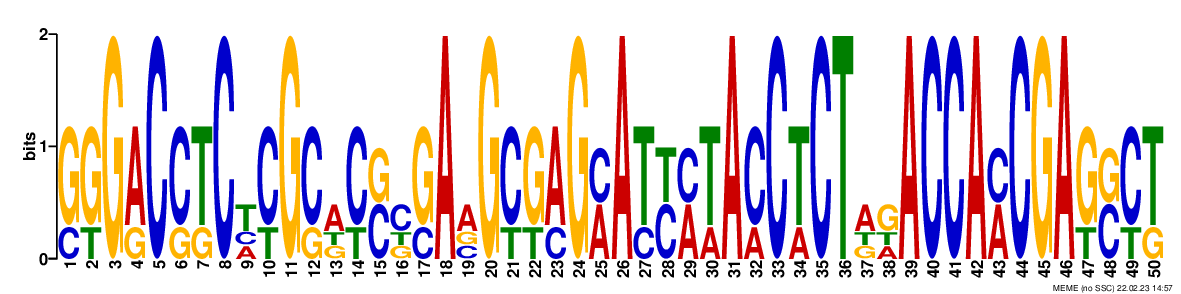

### logo_rc6.png

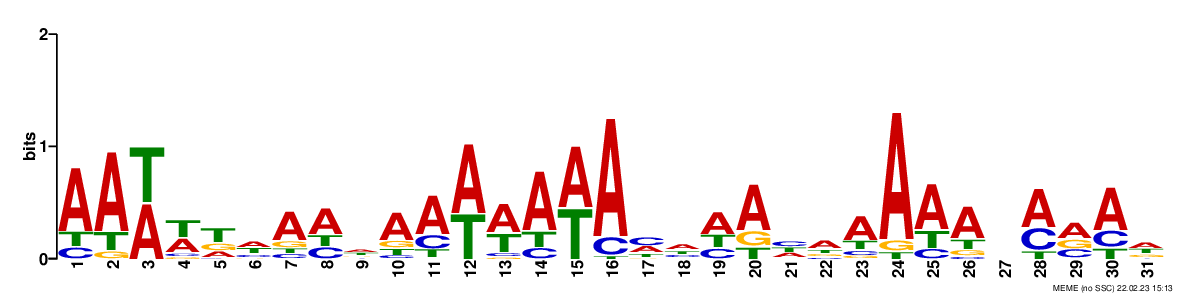

### logo_rc6.png

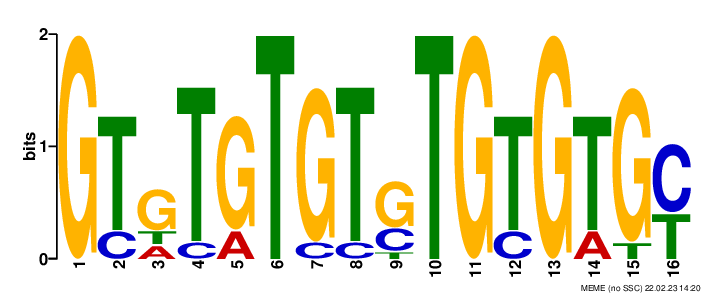

### logo_rc6.png

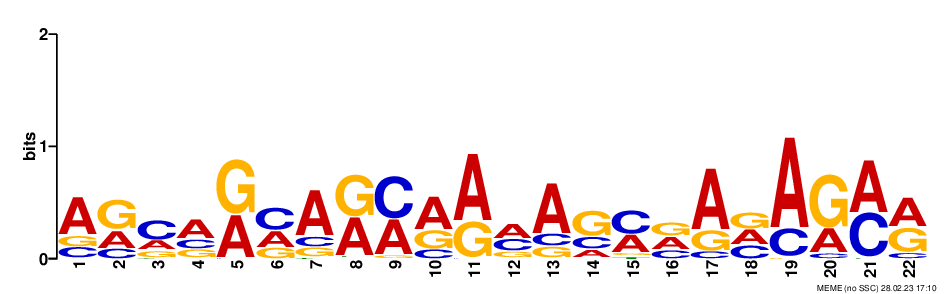

### logo_rc7.png

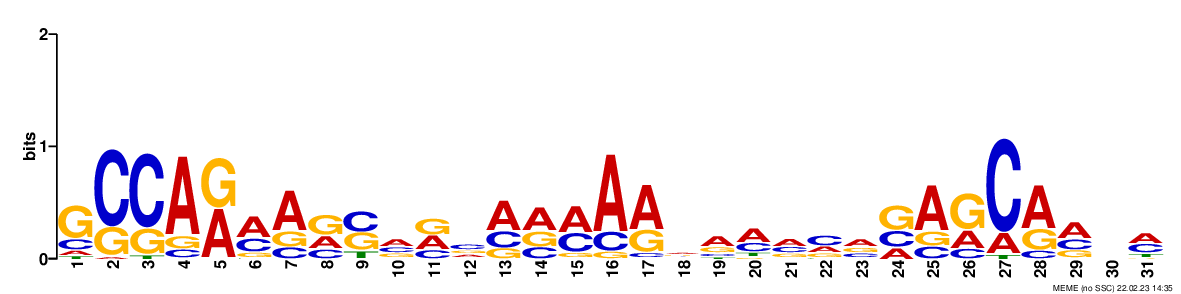

### logo_rc7.png

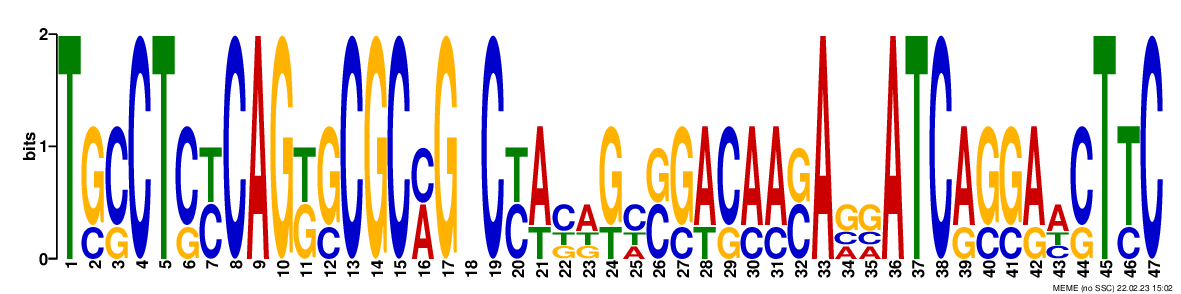

### logo_rc7.png

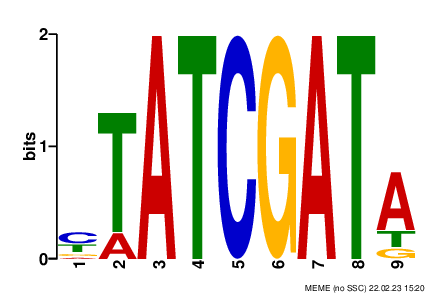

### logo_rc7.png

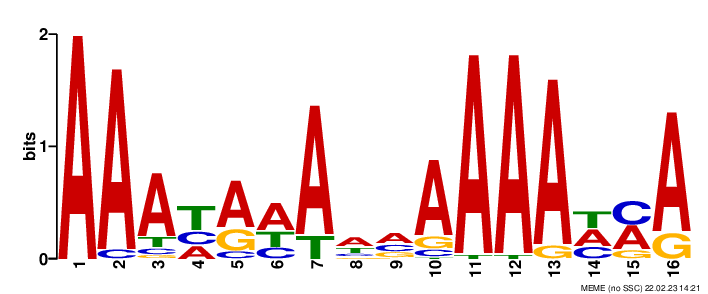

### logo_rc7.png

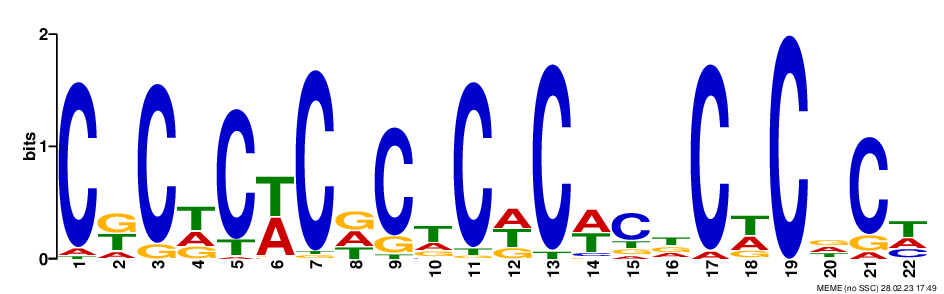

### logo_rc8.png

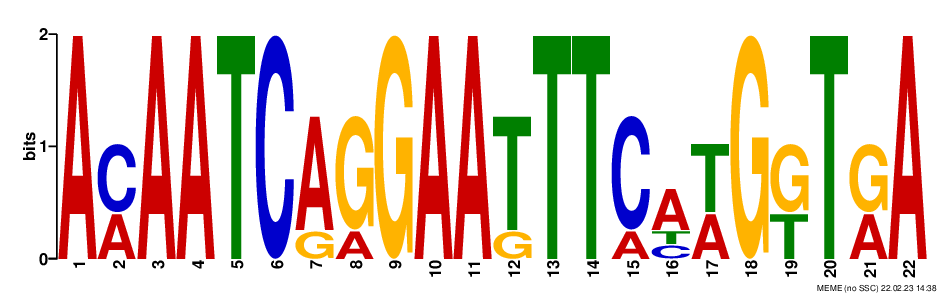

### logo_rc8.png

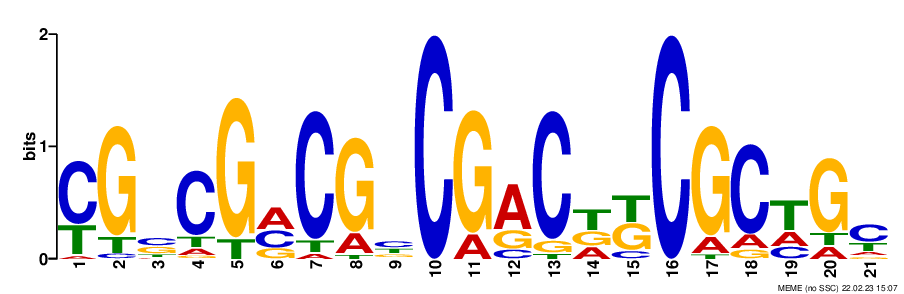

### logo_rc8.png

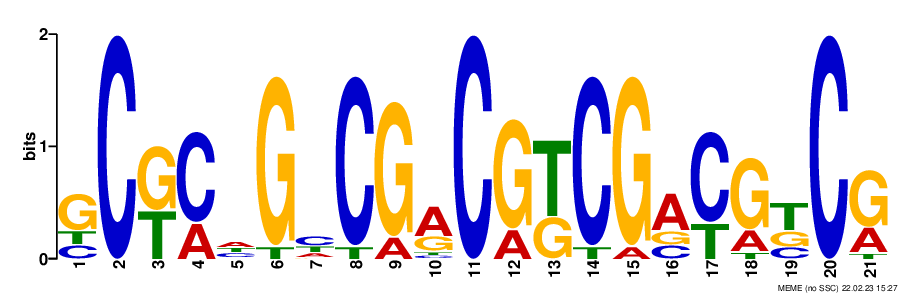

### logo_rc8.png

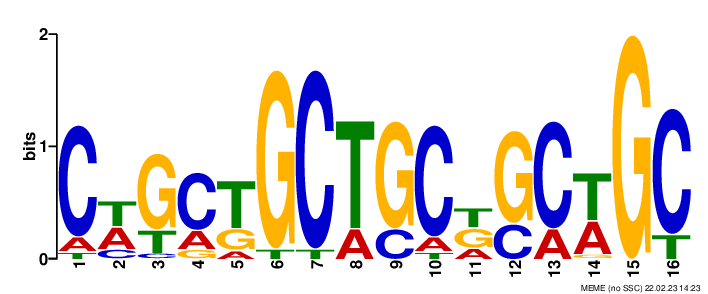

### logo_rc8.png

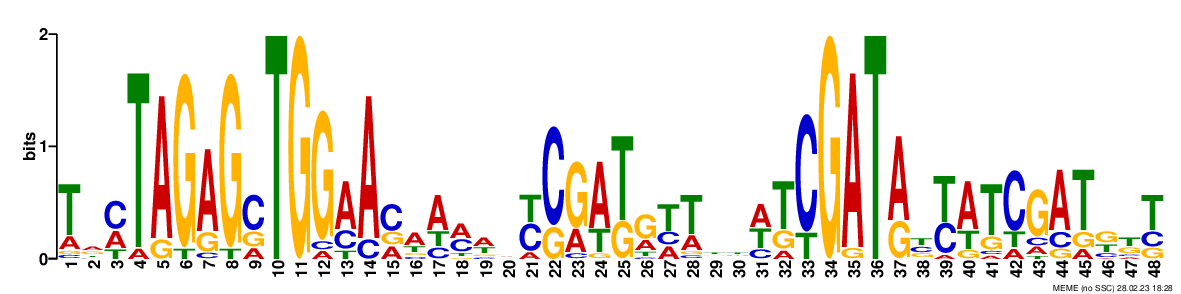

### logo_rc9.png

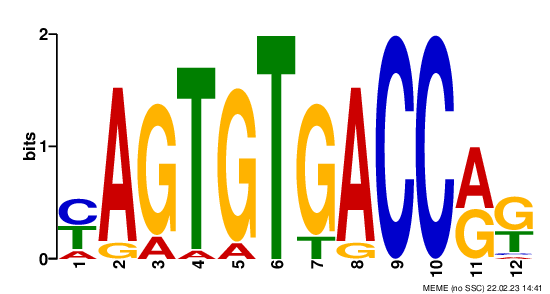

### logo_rc9.png

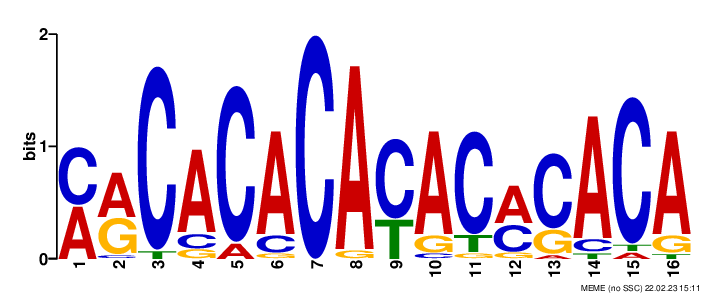

### logo_rc9.png

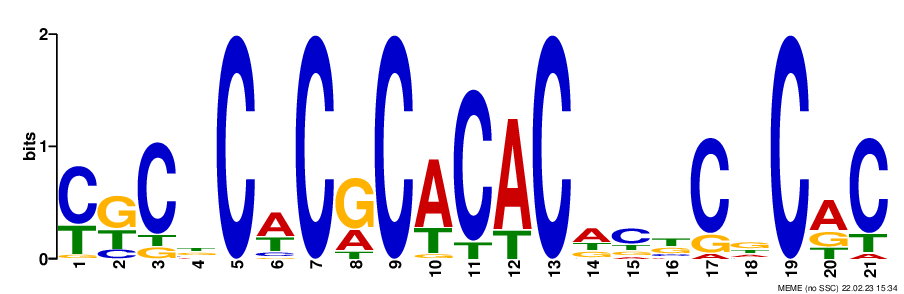

### logo_rc9.png

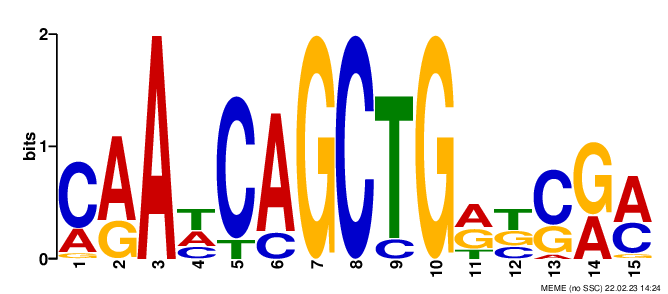

### logo_rc9.png

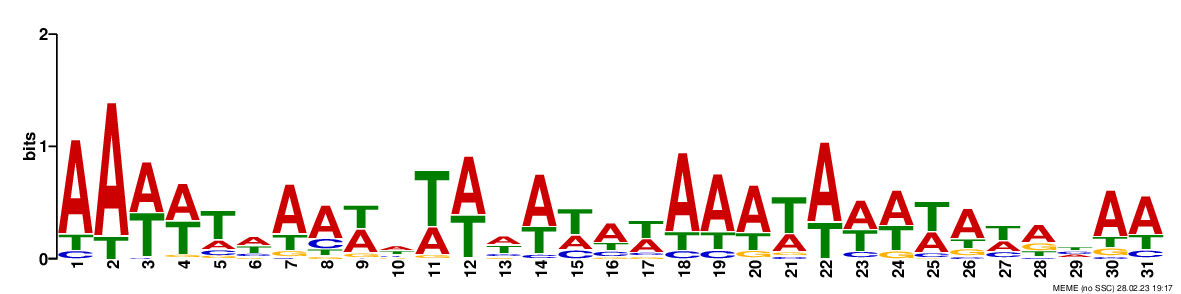

### logo_rc10.png

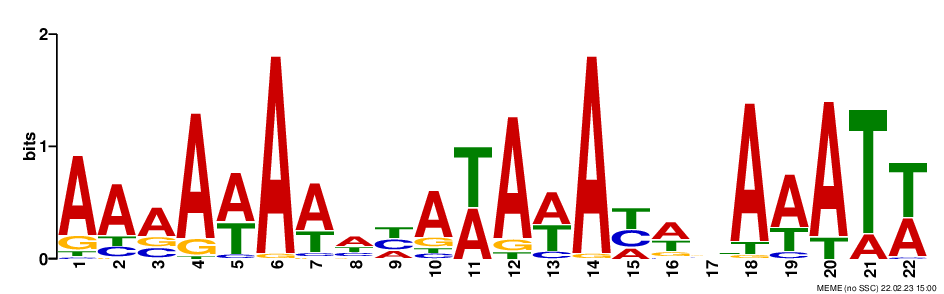

### logo_rc10.png

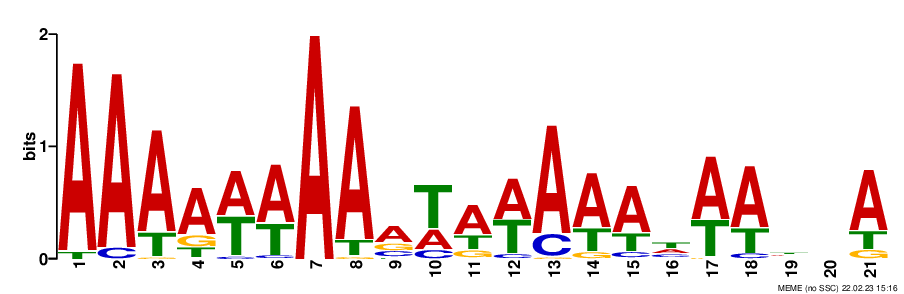

### logo_rc10.png

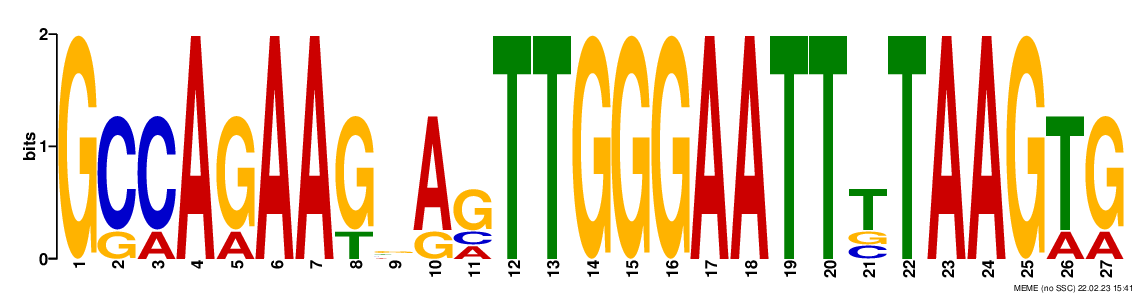

### logo_rc10.png

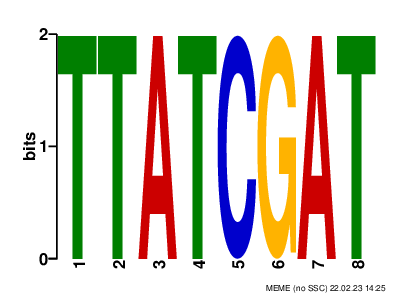

### logo_rc10.png

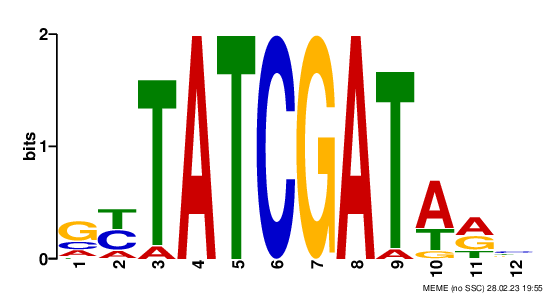
